## Supplementary material for "High-quality carnivore genomes from roadkill samples enable species delimitation in aardwolf and bat-eared fox": Table S6

| Species | Common name | Collection number | Origin | Sample type | Raw reads | Mito reads | Mean coverage | SRA Accession | Genbank Accession |
|---|---|---|---|---|---|---|---|---|---|
| *Proteles cristata cristata* | Southern aardwolf | NMB (TS307) | South Africa | Tissue | 4,779,321 | 6,665 | 39x | Pending | Pending |
| *Proteles cristata septentrionalis* | Eastern aardwolf | NMS Z.2018.54 | Tanzania | Tissue | 7,597,504 | 2,709 | 19x | Pending | Pending |
| *Otocyon megalotis megalotis* | Southern bat-eared fox | NMB (TS305) | South Africa | Tissue | 6,719,302 | 4,559 | 27x | Pending | Pending |
| *Otocyon megalotis virgatus* | Eastern bat-eared fox | FMNH 158128 | Tanzania | Tissue | 6,838,309 | 37,719 | 219x | Pending | Pending |
| *Speothos venaticus* | Bush dog | ISEM T1624 | French Guiana | Tissue | 5,271,782 | 4,763 | 27x | Pending | Pending |
| *Vulpes vulpes* | Red fox | ISEM T3611 | France | Tissue | 8,355,398 | 3,050 | 18x | Pending | Pending |
| *Parahyaena brunnea* | Brown hyaena | ISEM FD126 | Toulon Zoo | Feces | 11,739,905 | 3,243 | 17x | Pending | Pending |
| *Bdeogale nigripes* | Black-footed mongoose | FMNH 167685 | Gabon | Tissue | 6,839,178 | 8,215 | 61x | SRR6053065* | Pending |
| *Fossa fossana* | Malagasy civet | FMNH 156648 | Madagascar | Tissue | 4,914,258 | 17,689 | 131x | SRR6053060* | Pending |
| *Viverra tangalunga* | Malayan civet | FMNH 146957 | Philippines | Tissue | 5,648,442 | 23,116 | 205x | SRR6053069* | Pending |

*: UCE capture reads from Esselstyn et al. (2017); NMB: National Museum, Bloemfontein, South Africa; ISEM: Institut des Sciences de l'Evolution, Montpellier, France; FMNH: Field Museum of Natural History, Chicago, IL, USA.
