## Supplementary material for "High-quality carnivore genomes from roadkill samples enable species delimitation in aardwolf and bat-eared fox": Table S10

| Species | Order | Family | Accession | AssemblyName | Database | Assembly | Busco scores | | | | | OMM | Annotation |
|---|---|---|---|---|---|---|---|---|---|---|---|---|---|
| | | | | | | | Complete | Fragmented | Missing | Single | Duplicated | | |
| *Acinonyx jubatus* | Carnivora | Felidae | GCF_001443585.1 | aciJub1 | NCBI | Acinonyx jubatus [aciJub1] | 90.5 | 6.0 | 3.5 | 90.2 | 0.3 | Yes | Yes |
| *Ailuropoda melanoleuca* | Carnivora | Ursidae | GCF_000004335.2 | AilMel_1.0 | NCBI | Ailuropoda melanoleuca [AilMel_1.0] | 92.9 | 4.6 | 2.5 | 92.6 | 0.3 | Yes | Yes |
| *Ailurus fulgens* | Carnivora | Ailuridae | ASM200746v1_DNAZoo_HiC | ASM200746v1_DNAZoo_HiC | DNAZoo | Ailurus fulgens [ASM200746v1_DNAZoo_HiC] | 95.4 | 2.3 | 2.3 | 94.6 | 0.8 | No | Yes |
| *Arctocephalus gazella* | Carnivora | Otariidae | GCA_900500725.1 | ArcGazv1.4 | NCBI | Arctocephalus gazella [ArcGazv1.4] | 81.6 | 12.4 | 6.0 | 80.8 | 0.8 | No | No |
| *Callorhinus ursinus* | Carnivora | Otariidae | GCF_003265705.1 | ASM326570v1 | NCBI | Callorhinus ursinus [ASM326570v1] | 93.0 | 3.6 | 3.4 | 81.2 | 11.8 | No | Yes |
| *Canis lupus* | Carnivora | Canidae | GCF_000002285.3 | CanFam3.1 | NCBI | Canis lupus [CanFam3.1] | 95.3 | 2.4 | 2.3 | 94.0 | 1.3 | Yes | Yes |
| *Crocuta crocuta* | Carnivora | Hyaenidae | GCA_008692635.1 | BGI_CrCroc_1.0 | NCBI | Crocuta crocuta [BGI_CrCroc_1.0] | 95.5 | 2.5 | 2.0 | 95.0 | 0.5 | No | No |
| *Cryptoprocta ferox* | Carnivora | Eupleridae | GCA_004023885.1 | CryFer_v1_BIUU | NCBI | Cryptoprocta ferox [CryFer_v1_BIUU] | 85.9 | 10.6 | 3.5 | 84.9 | 1.0 | No | No |
| *Enhydra lutris* | Carnivora | Mustelidae | GCF_002288905.1 | ASM228890v2 | NCBI | Enhydra lutris [ASM228890v2] | 94.2 | 3.5 | 2.3 | 93.2 | 1.0 | Yes | Yes |
| *Eumetopias jubatus* | Carnivora | Otariidae | GCF_004028035.1 | ASM402803v1 | NCBI | Eumetopias jubatus [ASM402803v1] | 93.4 | 3.7 | 2.9 | 91.2 | 2.2 | No | Yes |
| *Felis catus* | Carnivora | Felidae | GCF_000181335.3 | Felis_catus_9.0 | NCBI | Felis catus [Felis_catus_9] | 93.1 | 3.8 | 3.1 | 92.7 | 0.4 | Yes | Yes |
| *Gulo gulo* | Carnivora | Mustelidae | GCA_900006375.2 | Gulo_2.2_annotated | NCBI | Gulo gulo [Gulo_2.2] | 85.4 | 10.6 | 4.0 | 85.0 | 0.4 | No | Yes |
| *Helogale parvula* | Carnivora | Herpestidae | GCA_004023845.1 | HelPar_v1_BIUU | NCBI | Helogale parvula [HelPar_v1_BIUU] | 83.9 | 11.8 | 4.3 | 83.2 | 0.7 | No | No |
| *Hyaena hyaena* | Carnivora | Hyaenidae | GCA_003009895.1 | ASM300989v1 | NCBI | Hyaena hyaena [ASM300989v1] | 93.5 | 4.0 | 2.5 | 92.8 | 0.7 | No | No |
| *Leptonychotes weddellii* | Carnivora | Phocidae | GCF_000349705.1 | LepWed1.0 | NCBI | Leptonychotes weddellii [LepWed1.0] | 78.8 | 15.3 | 5.9 | 77.8 | 1.0 | Yes | Yes |
| *Lutra lutra* | Carnivora | Mustelidae | mLutLut1_DNAZoo_HiC | mLutLut1_DNAZoo_HiC | DNAZoo | Lutra lutra [mLutLut1_DNAZoo_HiC] | 95.5 | 2.2 | 2.3 | 94.1 | 1.4 | No | Yes |
| *Lycaon pictus* | Carnivora | Canidae | sis2-181106_DNAZoo_HiC | sis2-181106_DNAZoo_HiC | DNAZoo | Lycaon pictus [sis2-181106_DNAZoo_HiC] | 94.7 | 2.5 | 2.8 | 93.5 | 1.2 | No | Yes |
| *Lynx canadensis* | Carnivora | Felidae | GCA_007474595.1 | mLynCan4_v1.p | NCBI | Lynx canadensis [mLynCan4_v1.p_VGP] | 94.6 | 2.7 | 2.7 | 94.2 | 0.4 | No | Yes |
| *Lynx pardinus* | Carnivora | Felidae | GCA_900661375.1 | LYPA1.0 | NCBI | Lynx pardinus [LYPA1.0] | 94.4 | 3.1 | 2.5 | 93.9 | 0.5 | No | Yes |
| *Mellivora capensis* | Carnivora | Mustelidae | GCA_004024625.1 | MelCap_v1_BIUU | NCBI | Mellivora capensis [MelCap_v1_BIUU] | 78.2 | 16.1 | 5.7 | 77.7 | 0.5 | No | No |
| *Mirounga angustirostris* | Carnivora | Phocidae | Mirounga_angustirostris_DNAZoo_HiC | Mirounga_angustirostris_DNAZoo_HiC | DNAZoo | Mirounga angustirostris [Mirounga_angustirostris_DNAZoo_HiC] | 94.7 | 2.8 | 2.5 | 93.7 | 1.0 | No | Yes |
| *Mungos mungo* | Carnivora | Herpestidae | GCA_004023785.1 | MunMun_v1_BIUU | NCBI | Mungos mungo [MunMun_v1_BIUU] | 87.4 | 9.1 | 3.5 | 86.6 | 0.8 | No | No |
| *Mustela putorius* | Carnivora | Mustelidae | GCF_000215625.1 | MusPutFur1.0 | NCBI | Mustela putorius [MusPutFur1.0] | 93.5 | 3.9 | 2.6 | 92.9 | 0.6 | Yes | Yes |
| *Nasua narica* | Carnivora | Procyonidae | Nasua_narica_DNAZoo_HiC | Nasua_narica_DNAZoo_HiC | DNAZoo | Nasua narica [Nasua_narica_DNAZoo_HiC] | 90.7 | 5.0 | 4.3 | 90.1 | 0.6 | No | No |
| *Neofelis nebulosa* | Carnivora | Felidae | Neofelis_nebulosa_DNAZoo_HiC | Neofelis_nebulosa_DNAZoo_HiC | DNAZoo | Neofelis nebulosa [Neofelis_nebulosa_DNAZoo_HiC] | 95.8 | 1.9 | 2.3 | 95.1 | 0.7 | No | Yes |
| *Neomonachus schauinslandi* | Carnivora | Phocidae | GCF_002201575.1 | ASM220157v1 | NCBI | Neomonachus schauinslandi [ASM220157v1] | 93.0 | 4.3 | 2.7 | 91.9 | 1.1 | Yes | Yes |
| *Neovison vison* | Carnivora | Mustelidae | GCA_900108605.1 | NNQGG.v01 | NCBI | Neovison vison [NNQGG.v01] | 94.0 | 3.8 | 2.2 | 93.4 | 0.6 | No | No |
| *Odobenus rosmarus* | Carnivora | Odobenidae | GCF_000321225.1 | Oros_1.0 | NCBI | Odobenus rosmarus [Oros_1.0] | 93.4 | 3.8 | 2.8 | 91.9 | 1.5 | Yes | Yes |
| *Otocyon megalotis* | Carnivora | Canidae | ISEM_TS305_MaSuRCA | Otoc_meg_TS305_MaSuRCA | ISEM | Otocyon megalotis [ISEM_TS305_MaSuRCA] | 92.9 | 4.2 | 2.9 | 91.7 | 1.2 | No | Yes |
| *Panthera leo* | Carnivora | Felidae | GCA_008795835.1 | PanLeo1.0 | NCBI | Panthera leo [PanLeo1.0] | 96.2 | 1.9 | 1.9 | 95.6 | 0.6 | No | No |
| *Panthera onca* | Carnivora | Felidae | GCA_004023805.1 | PanOnc_v1_BIUU | NCBI | Panthera onca [PanOnc_v1_BIUU] | 78.4 | 15.7 | 5.9 | 78.0 | 0.4 | No | No |
| *Panthera pardus* | Carnivora | Felidae | GCF_001857705.1 | PanPar1.0 | NCBI | Panthera pardus [PanPar1.0] | 93.5 | 3.8 | 2.7 | 92.7 | 0.8 | Yes | Yes |
| *Panthera tigris* | Carnivora | Felidae | GCF_000464555.1 | PanTig1.0 | NCBI | Panthera tigris [PanTig1.0] | 91.5 | 5.3 | 3.2 | 91.2 | 0.3 | Yes | Yes |
| *Paradoxurus hermaphroditus* | Carnivora | Viverridae | GCA_004024585.1 | ParHer_v1_BIUU | NCBI | Paradoxurus hermaphroditus [ParHer_v1_BIUU] | 74.8 | 18.4 | 6.8 | 74.2 | 0.6 | No | No |
| *Phoca vitulina* | Carnivora | Phocidae | GSC_HSeal_1.0_DNAZoo_HiC | GSC_HSeal_1.0_DNAZoo_HiC | DNAZoo | Phoca vitulina [GSC_HSeal_1.0_DNAZoo_HiC] | 95.2 | 2.3 | 2.5 | 94.1 | 1.1 | No | Yes |
| *Potos flavus* | Carnivora | Procyonidae | Potos_flavus_DNAZoo_HiC | Potos_flavus_DNAZoo_HiC | DNAZoo | Potos flavus [Potos_flavus_DNAZoo_HiC] | 92.7 | 4.4 | 2.9 | 92.2 | 0.5 | No | Yes |
| *Prionailurus bengalensis* | Carnivora | Felidae | GCA_005406085.1 | Prionailurus_bengalensis_euptilurus_v01 | NCBI | Prionailurus bengalensis [Prionailurus_bengalensis_euptilurus_v01] | 84.2 | 11.0 | 4.8 | 83.7 | 0.5 | No | No |
| *Procyon lotor* | Carnivora | Procyonidae | pl-1k_DNAZoo_HiC | pl-1k_DNAZoo_HiC | DNAZoo | Procyon lotor [pl-1k_DNAZoo_HiC] | 87.8 | 7.2 | 5.0 | 87.2 | 0.6 | No | Yes |
| *Proteles cristatus* | Carnivora | Hyaenidae | ISEM_TS307_MaSuRCA | Prot_cri_TS307_MaSuRCA | ISEM | Proteles cristatus [ISEM_TS307_MaSuRCA] | 92.8 | 3.8 | 3.4 | 92.3 | 0.5 | No | Yes |
| *Pteronura brasiliensis* | Carnivora | Mustelidae | GCA_004024605.1 | PteBra_v1_BIUU | NCBI | Pteronura brasiliensis [PteBra_v1_BIUU] | 82.0 | 13.3 | 4.7 | 81.4 | 0.6 | No | No |
| *Puma concolor* | Carnivora | Felidae | GCF_003327715.1 | PumCon1.0 | NCBI | Puma concolor [PumCon1] | 93.4 | 3.4 | 3.2 | 93.0 | 0.4 | No | Yes |
| *Spilogale gracilis* | Carnivora | Mephitidae | GCA_004023965.1 | SpiGra_v1_BIUU | NCBI | Spilogale gracilis [SpiGra_v1_BIUU] | 76.7 | 17.1 | 6.2 | 76.1 | 0.6 | No | No |
| *Suricata suricatta* | Carnivora | Herpestidae | GCF_006229205.1 | meerkat_22Aug2017_6uvM2_HiC | DNAZoo | Suricata suricatta [meerkat_22Aug2017_6uvM2_DNAZoo_HiC] | 94.7 | 2.6 | 2.7 | 93.5 | 1.2 | No | Yes |
| *Taxidea taxus* | Carnivora | Mustelidae | GCA_003697995.1 | ASM369799v1 | NCBI | Taxidea taxus [ASM369799v1] | 58.5 | 28.4 | 13.1 | 57.5 | 1.0 | No | No |
| *Ursus americanus* | Carnivora | Ursidae | ASM334442v1_DNAZoo_HiC | ASM334442v1_DNAZoo_HiC | DNAZoo | Ursus americanus [ASM334442v1_DNAZoo_HiC] | 93.9 | 3.3 | 2.8 | 93.3 | 0.6 | No | Yes |
| *Ursus arctos* | Carnivora | Ursidae | ASM358476v1_DNAZoo_HiC | ASM358476v1_DNAZoo_HiC | DNAZoo | Ursus arctos [ASM358476v1_DNAZoo_HiC] | 95.7 | 2.0 | 2.3 | 94.8 | 0.9 | No | Yes |
| *Ursus maritimus* | Carnivora | Ursidae | GCF_000687225.1 | UrsMar_1.0 | NCBI | Ursus maritimus [UrsMar_1] | 91.8 | 5.5 | 2.7 | 91.4 | 0.4 | Yes | Yes |
| *Ursus thibetanus* | Carnivora | Ursidae | GCA_009660055.1 | ASM966005v1 | NCBI | Ursus thibetanus [ASM966005v1] | 95.8 | 1.8 | 2.4 | 94.4 | 1.4 | No | No |
| *Vulpes vulpes* | Carnivora | Canidae | VulVul2.2_DNAZoo_HiC | VulVul2.2_DNAZoo_HiC | NCBI | Vulpes vulpes [VulVul2.2_DNAZoo_HiC] | 94.7 | 2.8 | 2.5 | 93.3 | 1.4 | No | Yes |
| *Zalophus californianus* | Carnivora | Otariidae | GCF_900631625.1 | zalCal2.2 | NCBI | Zalophus californianus [zalCal2.2] | 93.3 | 3.7 | 3.0 | 91.8 | 1.5 | No | Yes |
