## Supplementary material for "High-quality carnivore genomes from roadkill samples enable species delimitation in aardwolf and bat-eared fox": Table S9

| Contigs name | length | GC | N | bam0 | superkingdom.t. | superkingdom.s. | superkingdom.c. | phylum.t. | phylum.s. | phylum.c. | order.t. | order.s. | order.c. | family.t. | family.s. | family.c. | genus.t. | genus.s. | genus.c | species.t. | species.s. | species.c. |
|---|---|---|---|---|---|---|---|---|---|---|---|---|---|---|---|---|---|---|---|---|---|---|
| scf7180000052649 | 5588 | 0.5104 | 0 | 98.094 | Eukaryota | 7866.0 | 0 | Chordata | 7866.0 | 0 | Carnivora | 7866.0 | 0 | Canidae | 7866.0 | 0 | Canis | 6251.0 | 1 | Canis lupus | 5162.0 | 2 |
| scf7180000045803 | 160177 | 0.5209 | 0 | 56.7306 | Eukaryota | 30832.0 | 0 | Chordata | 30832.0 | 0 | Carnivora | 20790.0 | 1 | Canidae | 15792.0 | 2 | Canis | 11588.0 | 4 | Canis lupus | 11588.0 | 4 |
| scf7180000042293 | 774332 | 0.3674 | 0 | 59.0354 | Eukaryota | 41722.0 | 0 | Chordata | 41722.0 | 0 | Carnivora | 41722.0 | 0 | Canidae | 41722.0 | 0 | Canis | 41722.0 | 0 | Canis lupus | 41722.0 | 0 |
| scf7180000051360 | 2695 | 0.4505 | 0 | 561.424 | no-hit | 0.0 | 0 | no-hit | 0.0 | 0 | no-hit | 0.0 | 0 | no-hit | 0.0 | 0 | no-hit | 0.0 | 0 | no-hit | 0.0 | 0 |
| scf7180000044612 | 96519 | 0.3763 | 0 | 58.5429 | Eukaryota | 61940.0 | 0 | Chordata | 61940.0 | 0 | Carnivora | 61940.0 | 0 | Canidae | 61940.0 | 0 | Canis | 61940.0 | 0 | Canis lupus | 61940.0 | 0 |
| scf7180000045736 | 172290 | 0.4424 | 0 | 58.5426 | Eukaryota | 40370.0 | 0 | Chordata | 40370.0 | 0 | Carnivora | 40370.0 | 0 | Canidae | 40370.0 | 0 | Canis | 40370.0 | 0 | Canis lupus | 40370.0 | 0 |
| scf7180000041819 | 797790 | 0.4822 | 0 | 59.4729 | Eukaryota | 122794.0 | 0 | Chordata | 122794.0 | 0 | Carnivora | 122794.0 | 0 | Canidae | 122794.0 | 0 | Canis | 122794.0 | 0 | Canis lupus | 122794.0 | 0 |
| scf7180000041440 | 152689 | 0.3624 | 0 | 56.9334 | Eukaryota | 19383.0 | 0 | Chordata | 19383.0 | 0 | Carnivora | 17430.0 | 1 | Canidae | 17430.0 | 1 | Canis | 17430.0 | 1 | Canis lupus | 17430.0 | 1 |
| scf7180000054322 | 40679 | 0.445 | 0 | 616.2034 | Eukaryota | 73991.0 | 0 | Chordata | 52502.0 | 2 | Gadiformes | 47903.0 | 4 | Gadidae | 47903.0 | 4 | Gadus | 47903.0 | 4 | Gadus morhua | 47903.0 | 4 |
| scf7180000051699 | 9099 | 0.4497 | 0 | 1611.4698 | no-hit | 0.0 | 0 | no-hit | 0.0 | 0 | no-hit | 0.0 | 0 | no-hit | 0.0 | 0 | no-hit | 0.0 | 0 | no-hit | 0.0 | 0 |
| scf7180000051945 | 6388 | 0.44 | 0 | 570.561 | Eukaryota | 11985.0 | 0 | Chordata | 11399.0 | 1 | Gadiformes | 4864.0 | 6 | Gadidae | 4864.0 | 6 | Gadus | 4864.0 | 6 | Gadus morhua | 4864.0 | 6 |
| scf7180000049347 | 25489 | 0.4345 | 0 | 136.6632 | Eukaryota | 12040.0 | 0 | Chordata | 10471.0 | 2 | Synbranchiformes | 3748.0 | 6 | Mastacembelidae | 3748.0 | 6 | Mastacembelus | 3748.0 | 7 | Mastacembelus armatus | 3748.0 | 7 |
| scf7180000050473 | 19576 | 0.4432 | 0 | 273.3977 | no-hit | 0.0 | 0 | no-hit | 0.0 | 0 | no-hit | 0.0 | 0 | no-hit | 0.0 | 0 | no-hit | 0.0 | 0 | no-hit | 0.0 | 0 |
| scf7180000043676 | 3615 | 0.4415 | 0 | 52.9794 | Eukaryota | 987.0 | 0 | Chordata | 987.0 | 0 | Syngnathiformes | 767.0 | 1 | Syngnathidae | 767.0 | 1 | Syngnathus | 767.0 | 1 | Syngnathus acus | 767.0 | 1 |
| scf7180000051192 | 2842 | 0.4514 | 0 | 239.754 | no-hit | 0.0 | 0 | no-hit | 0.0 | 0 | no-hit | 0.0 | 0 | no-hit | 0.0 | 0 | no-hit | 0.0 | 0 | no-hit | 0.0 | 0 |
| scf7180000048756 | 41459 | 0.4353 | 0 | 122.9914 | Eukaryota | 27711.0 | 0 | Chordata | 19372.0 | 2 | Gadiformes | 6787.0 | 6 | Gadidae | 6787.0 | 6 | Gadus | 6787.0 | 7 | Gadus morhua | 6787.0 | 8 |
| scf7180000040866 | 105305 | 0.4231 | 0 | 59.2827 | Eukaryota | 33747.0 | 0 | Chordata | 33747.0 | 0 | Carnivora | 33747.0 | 0 | Canidae | 33747.0 | 0 | Canis | 33747.0 | 0 | Canis lupus | 33747.0 | 0 |
| scf7180000052688 | 18477 | 0.4437 | 0 | 58.6501 | Eukaryota | 24213.0 | 0 | Chordata | 24213.0 | 0 | Carnivora | 20079.0 | 1 | Felidae | 12803.0 | 3 | Zalophus | 4888.0 | 5 | Zalophus californianus | 4888.0 | 5 |
| scf7180000042822 | 612949 | 0.4106 | 0 | 59.1118 | Eukaryota | 17205.0 | 0 | Chordata | 17205.0 | 0 | Carnivora | 15428.0 | 1 | Canidae | 15428.0 | 1 | Canis | 15428.0 | 1 | Canis lupus | 15428.0 | 1 |
| scf7180000049084 | 4718 | 0.4487 | 0 | 210.1655 | Eukaryota | 1202.0 | 0 | Chordata | 1202.0 | 0 | Tetraodontiformes | 952.0 | 1 | Tetraodontidae | 952.0 | 1 | Takifugu | 952.0 | 1 | Takifugu rubripes | 952.0 | 1 |
| scf7180000045678 | 195035 | 0.5218 | 0 | 56.9615 | Eukaryota | 72011.0 | 0 | Chordata | 72011.0 | 0 | Carnivora | 72011.0 | 0 | Canidae | 72011.0 | 0 | Canis | 72011.0 | 0 | Canis lupus | 72011.0 | 0 |
| scf7180000051070 | 22834 | 0.4473 | 0 | 41.9888 | Eukaryota | 9788.0 | 0 | Chordata | 8979.0 | 1 | Carnivora | 8979.0 | 1 | Canidae | 8979.0 | 1 | Canis | 4565.0 | 2 | Canis lupus | 4565.0 | 2 |
| scf7180000041332 | 6475 | 0.348 | 0 | 35.0239 | Eukaryota | 3379.0 | 0 | Chordata | 3379.0 | 0 | Carnivora | 3379.0 | 0 | Canidae | 3379.0 | 0 | Canis | 3379.0 | 0 | Canis lupus | 3379.0 | 0 |
| scf7180000050133 | 3154 | 0.455 | 0 | 700.0062 | no-hit | 0.0 | 0 | no-hit | 0.0 | 0 | no-hit | 0.0 | 0 | no-hit | 0.0 | 0 | no-hit | 0.0 | 0 | no-hit | 0.0 | 0 |
| scf7180000051130 | 4853 | 0.434 | 0 | 81.4756 | no-hit | 0.0 | 0 | no-hit | 0.0 | 0 | no-hit | 0.0 | 0 | no-hit | 0.0 | 0 | no-hit | 0.0 | 0 | no-hit | 0.0 | 0 |
| scf7180000043045 | 15567 | 0.4395 | 0 | 87.7674 | Eukaryota | 1041.0 | 0 | Platyhelminthes | 728.0 | 1 | Polystomatidea | 728.0 | 1 | Polystomatidae | 728.0 | 1 | Protopolystoma | 728.0 | 1 | Protopolystoma xenopodis | 728.0 | 1 |
| scf7180000055007 | 69838 | 0.3807 | 0 | 64.3969 | Eukaryota | 74215.0 | 0 | Chordata | 74215.0 | 0 | Carnivora | 74215.0 | 0 | Canidae | 74215.0 | 0 | Canis | 74215.0 | 0 | Canis lupus | 74215.0 | 0 |
| scf7180000046152 | 127840 | 0.4462 | 0 | 57.7918 | Eukaryota | 6016.0 | 0 | Chordata | 6016.0 | 0 | Primates | 3059.0 | 3 | Hominidae | 3059.0 | 3 | Homo | 2453.0 | 5 | Homo sapiens | 2453.0 | 5 |
| scf7180000054944 | 32529 | 0.4025 | 0 | 833.2832 | Eukaryota | 89820.0 | 0 | Chordata | 89820.0 | 0 | Carnivora | 89820.0 | 0 | Canidae | 89820.0 | 0 | Canis | 89820.0 | 0 | Canis lupus | 89820.0 | 0 |
| scf7180000043725 | 448867 | 0.377 | 0 | 59.5412 | Eukaryota | 53169.0 | 0 | Chordata | 53169.0 | 0 | Carnivora | 53169.0 | 0 | Canidae | 53169.0 | 0 | Canis | 53169.0 | 0 | Canis lupus | 53169.0 | 0 |
| scf7180000047433 | 75873 | 0.4526 | 0 | 59.5233 | Eukaryota | 25724.0 | 0 | Chordata | 25724.0 | 0 | Carnivora | 25724.0 | 0 | Canidae | 15240.0 | 1 | Canis | 15240.0 | 1 | Canis lupus | 15240.0 | 1 |
| scf7180000052509 | 3178 | 0.4487 | 0 | 297.8833 | no-hit | 0.0 | 0 | no-hit | 0.0 | 0 | no-hit | 0.0 | 0 | no-hit | 0.0 | 0 | no-hit | 0.0 | 0 | no-hit | 0.0 | 0 |
| scf7180000048715 | 39033 | 0.4075 | 0 | 59.5795 | Eukaryota | 6184.0 | 0 | Chordata | 6184.0 | 0 | Primates | 2935.0 | 4 | Hominidae | 2319.0 | 5 | Homo | 2319.0 | 5 | Homo sapiens | 2319.0 | 5 |
| scf7180000047876 | 1776 | 0.4291 | 0 | 67.5453 | Eukaryota | 335.0 | 0 | Chordata | 335.0 | 0 | Kurtiformes | 335.0 | 0 | Apogonidae | 335.0 | 0 | Sphaeramia | 335.0 | 0 | Sphaeramia orbicularis | 335.0 | 0 |
| scf7180000040613 | 284341 | 0.4107 | 0 | 62.3239 | Eukaryota | 93570.0 | 0 | Chordata | 93570.0 | 0 | Carnivora | 93570.0 | 0 | Canidae | 93570.0 | 0 | Canis | 93570.0 | 0 | Canis lupus | 93570.0 | 0 |
| scf7180000048314 | 5337 | 0.5183 | 0 | 203.0012 | Eukaryota | 15534.0 | 0 | Chordata | 15534.0 | 0 | Carnivora | 15534.0 | 0 | Canidae | 15534.0 | 0 | Canis | 15534.0 | 0 | Canis lupus | 15534.0 | 0 |
| scf7180000049665 | 48087 | 0.4432 | 0 | 57.8857 | Eukaryota | 26802.0 | 0 | Chordata | 26802.0 | 0 | Carnivora | 26802.0 | 0 | Otariidae | 15318.0 | 1 | Zalophus | 15318.0 | 2 | Zalophus californianus | 15318.0 | 2 |
| scf7180000051669 | 31489 | 0.5226 | 0 | 3150.6261 | Eukaryota | 34992.0 | 0 | Chordata | 34992.0 | 0 | Carnivora | 34992.0 | 0 | Canidae | 34992.0 | 0 | Canis | 34992.0 | 0 | Canis lupus | 34992.0 | 0 |
| scf7180000048505 | 11527 | 0.4808 | 0 | 603.0479 | Eukaryota | 63060.0 | 0 | Chordata | 63060.0 | 0 | Carnivora | 63060.0 | 0 | Canidae | 63060.0 | 0 | Canis | 63060.0 | 0 | Canis lupus | 63060.0 | 0 |
| scf7180000050656 | 5937 | 0.4418 | 0 | 97.8283 | no-hit | 0.0 | 0 | no-hit | 0.0 | 0 | no-hit | 0.0 | 0 | no-hit | 0.0 | 0 | no-hit | 0.0 | 0 | no-hit | 0.0 | 0 |
| scf7180000043893 | 54907 | 0.4025 | 0 | 55.0606 | Eukaryota | 63003.0 | 0 | Chordata | 63003.0 | 0 | Carnivora | 63003.0 | 0 | Canidae | 63003.0 | 0 | Canis | 63003.0 | 0 | Canis lupus | 63003.0 | 0 |
| scf7180000045356 | 169837 | 0.4651 | 0 | 59.4862 | Eukaryota | 59044.0 | 0 | Chordata | 59044.0 | 0 | Carnivora | 59044.0 | 0 | Canidae | 59044.0 | 0 | Canis | 59044.0 | 0 | Canis lupus | 59044.0 | 0 |
| scf7180000050595 | 6212 | 0.4363 | 0 | 104.1794 | Eukaryota | 6428.0 | 0 | Chordata | 5260.0 | 1 | Tetraodontiformes | 4484.0 | 2 | Tetraodontidae | 4484.0 | 2 | Takifugu | 4484.0 | 2 | Takifugu rubripes | 4484.0 | 2 |
| scf7180000044867 | 23710 | 0.369 | 0 | 48.5974 | Eukaryota | 4737.0 | 0 | Chordata | 4737.0 | 0 | Carnivora | 2485.0 | 2 | Canidae | 2485.0 | 2 | Canis | 2485.0 | 3 | Canis lupus | 2485.0 | 3 |
| scf7180000051202 | 27484 | 0.4137 | 0 | 58.9903 | Eukaryota | 18197.0 | 0 | Chordata | 18197.0 | 0 | Primates | 10953.0 | 2 | Hominidae | 8312.0 | 4 | Homo | 8312.0 | 4 | Homo sapiens | 8312.0 | 4 |
| scf7180000055962 | 14584 | 0.443 | 0 | 86.4301 | Eukaryota | 466.0 | 0 | unresolved | 233.0 | 1 | unresolved | 233.0 | 1 | unresolved | 233.0 | 1 | unresolved | 233.0 | 1 | unresolved | 233.0 | 1 |
| scf7180000046759 | 2245 | 0.4463 | 0 | 142.5114 | no-hit | 0.0 | 0 | no-hit | 0.0 | 0 | no-hit | 0.0 | 0 | no-hit | 0.0 | 0 | no-hit | 0.0 | 0 | no-hit | 0.0 | 0 |
| scf7180000049217 | 8261 | 0.4381 | 0 | 188.1155 | Eukaryota | 6805.0 | 0 | Chordata | 5649.0 | 1 | Tetraodontiformes | 3099.0 | 2 | Tetraodontidae | 3099.0 | 2 | Takifugu | 3099.0 | 3 | Takifugu rubripes | 3099.0 | 3 |
| scf7180000047225 | 76643 | 0.3922 | 0 | 57.6233 | Eukaryota | 6673.0 | 0 | Chordata | 6673.0 | 0 | Primates | 4193.0 | 2 | Hominidae | 2480.0 | 4 | Homo | 2480.0 | 4 | Homo sapiens | 2480.0 | 4 |
| scf7180000052425 | 2045 | 0.4357 | 0 | 28.3122 | Eukaryota | 4633.0 | 0 | Chordata | 4633.0 | 0 | Tetraodontiformes | 4633.0 | 0 | Tetraodontidae | 4633.0 | 0 | Takifugu | 4633.0 | 0 | Takifugu rubripes | 4633.0 | 0 |
| scf7180000044120 | 118492 | 0.4378 | 0 | 55.1941 | Eukaryota | 11769.0 | 1 | Chordata | 11769.0 | 1 | Carnivora | 7317.0 | 2 | Canidae | 6548.0 | 3 | Canis | 5667.0 | 4 | Canis lupus | 5667.0 | 4 |
| scf7180000050074 | 10926 | 0.4619 | 0 | 70.4846 | Eukaryota | 32038.0 | 0 | Chordata | 32038.0 | 0 | Carnivora | 32038.0 | 0 | Canidae | 32038.0 | 0 | Canis | 32038.0 | 0 | Canis lupus | 32038.0 | 0 |
| scf7180000052458 | 4002 | 0.452 | 0 | 895.6779 | Eukaryota | 459.0 | 0 | Platyhelminthes | 459.0 | 0 | Polystomatidea | 459.0 | 0 | Polystomatidae | 459.0 | 0 | Protopolystoma | 459.0 | 0 | Protopolystoma xenopodis | 459.0 | 0 |
| scf7180000056491 | 237047 | 0.3933 | 0 | 60.5622 | Eukaryota | 55674.0 | 0 | Chordata | 55674.0 | 0 | Carnivora | 36254.0 | 2 | Canidae | 36254.0 | 2 | Canis | 36254.0 | 3 | Canis lupus | 36254.0 | 3 |
| scf7180000053728 | 2610 | 0.508 | 0 | 30.608 | Eukaryota | 6655.0 | 0 | Chordata | 6655.0 | 0 | Carnivora | 6655.0 | 0 | Canidae | 6655.0 | 0 | Canis | 4848.0 | 1 | Canis lupus | 3888.0 | 2 |
| scf7180000045050 | 249934 | 0.3852 | 0 | 59.6837 | Eukaryota | 60550.0 | 0 | Chordata | 60550.0 | 0 | Carnivora | 60550.0 | 0 | Canidae | 60550.0 | 0 | Canis | 60550.0 | 0 | Canis lupus | 60550.0 | 0 |
| scf7180000056175 | 2743569 | 0.3639 | 0 | 60.9344 | Eukaryota | 93840.0 | 0 | Chordata | 93840.0 | 0 | Carnivora | 93840.0 | 0 | Canidae | 93840.0 | 0 | Canis | 93840.0 | 0 | Canis lupus | 93840.0 | 0 |
| scf7180000047711 | 95095 | 0.3379 | 0 | 60.0093 | Eukaryota | 13898.0 | 0 | Chordata | 13898.0 | 0 | Carnivora | 13898.0 | 0 | Canidae | 13898.0 | 0 | Canis | 13898.0 | 0 | Canis lupus | 13898.0 | 0 |
| scf7180000042249 | 468380 | 0.3641 | 0 | 60.7688 | Eukaryota | 91361.0 | 0 | Chordata | 91361.0 | 0 | Carnivora | 91361.0 | 0 | Canidae | 91361.0 | 0 | Canis | 91361.0 | 0 | Canis lupus | 91361.0 | 0 |
| scf7180000049514 | 44759 | 0.3578 | 0 | 60.309 | Eukaryota | 15587.0 | 0 | Chordata | 15587.0 | 0 | Primates | 9551.0 | 6 | Hominidae | 9551.0 | 6 | Homo | 9551.0 | 6 | Homo sapiens | 9551.0 | 6 |
| scf7180000054670 | 43920 | 0.4458 | 0 | 152.5966 | Eukaryota | 96296.0 | 0 | Chordata | 96296.0 | 0 | Carnivora | 96296.0 | 0 | Canidae | 96296.0 | 0 | Canis | 96296.0 | 0 | Canis lupus | 96296.0 | 0 |
| scf7180000045454 | 12845 | 0.4434 | 0 | 334.5052 | Eukaryota | 93882.0 | 0 | Chordata | 93882.0 | 0 | Carnivora | 93882.0 | 0 | Canidae | 93882.0 | 0 | Canis | 93882.0 | 0 | Canis lupus | 93882.0 | 0 |
| scf7180000055738 | 7280 | 0.4438 | 0 | 656.4785 | no-hit | 0.0 | 0 | no-hit | 0.0 | 0 | no-hit | 0.0 | 0 | no-hit | 0.0 | 0 | no-hit | 0.0 | 0 | no-hit | 0.0 | 0 |
| scf7180000054432 | 20343 | 0.4318 | 0 | 357.1074 | Eukaryota | 26193.0 | 0 | Chordata | 26193.0 | 0 | Carnivora | 26193.0 | 0 | Canidae | 26193.0 | 0 | Canis | 26193.0 | 0 | Canis lupus | 26193.0 | 0 |
| scf7180000046933 | 80129 | 0.3894 | 0 | 60.2961 | Eukaryota | 8784.0 | 0 | Chordata | 8784.0 | 0 | Carnivora | 7496.0 | 1 | Felidae | 6674.0 | 2 | Felis | 5813.0 | 3 | Felis catus | 5813.0 | 3 |
| scf7180000052257 | 7838 | 0.4468 | 0 | 671.0125 | Eukaryota | 5688.0 | 0 | Chordata | 2975.0 | 2 | Tetraodontiformes | 2064.0 | 4 | Tetraodontidae | 2064.0 | 4 | Takifugu | 2064.0 | 4 | Takifugu rubripes | 2064.0 | 4 |
| scf7180000046038 | 178430 | 0.3956 | 0 | 60.7034 | Eukaryota | 80618.0 | 0 | Chordata | 80618.0 | 0 | Carnivora | 80618.0 | 0 | Canidae | 80618.0 | 0 | Canis | 80618.0 | 0 | Canis lupus | 80618.0 | 0 |
| scf7180000055095 | 16491 | 0.4625 | 0 | 156.9813 | Eukaryota | 9390.0 | 0 | Chordata | 9390.0 | 0 | Carnivora | 9390.0 | 0 | Canidae | 9390.0 | 0 | Canis | 5675.0 | 1 | Canis lupus | 5675.0 | 1 |
| scf7180000049139 | 7572 | 0.4414 | 0 | 57.3909 | Eukaryota | 9276.0 | 0 | Chordata | 9276.0 | 0 | Carnivora | 9276.0 | 0 | Canidae | 9276.0 | 0 | Canis | 9276.0 | 0 | Canis lupus | 9276.0 | 0 |
| scf7180000042028 | 142110 | 0.4199 | 0 | 55.0118 | Eukaryota | 134983.0 | 0 | Chordata | 134983.0 | 0 | Carnivora | 134983.0 | 0 | Canidae | 134983.0 | 0 | Canis | 134983.0 | 0 | Canis lupus | 134983.0 | 0 |
| scf7180000046526 | 92836 | 0.5216 | 0 | 55.7878 | Eukaryota | 14078.0 | 0 | Chordata | 14078.0 | 0 | Carnivora | 14078.0 | 0 | Canidae | 14078.0 | 0 | Canis | 9508.0 | 1 | Canis lupus | 9508.0 | 1 |
| scf7180000051442 | 5540 | 0.4542 | 0 | 2116.4554 | Eukaryota | 4952.0 | 0 | Chordata | 4952.0 | 0 | Kurtiformes | 2286.0 | 2 | Apogonidae | 2286.0 | 2 | Sphaeramia | 2286.0 | 2 | Sphaeramia orbicularis | 2286.0 | 2 |
| scf7180000046955 | 19256 | 0.4871 | 0 | 1238.6977 | Eukaryota | 80529.0 | 0 | Chordata | 80529.0 | 0 | Carnivora | 80529.0 | 0 | Canidae | 80529.0 | 0 | Canis | 80529.0 | 0 | Canis lupus | 80529.0 | 0 |
| scf7180000051526 | 4987 | 0.4562 | 0 | 74.6164 | no-hit | 0.0 | 0 | no-hit | 0.0 | 0 | no-hit | 0.0 | 0 | no-hit | 0.0 | 0 | no-hit | 0.0 | 0 | no-hit | 0.0 | 0 |
| scf7180000048688 | 4943 | 0.4457 | 0 | 251.716 | Eukaryota | 1057.0 | 0 | Echinodermata | 481.0 | 2 | Forcipulatida | 481.0 | 2 | Asteriidae | 481.0 | 2 | Asterias | 481.0 | 2 | Asterias rubens | 481.0 | 2 |
| scf7180000041714 | 939748 | 0.4026 | 0 | 59.7081 | Eukaryota | 93454.0 | 0 | Chordata | 93454.0 | 0 | Carnivora | 93454.0 | 0 | Canidae | 93454.0 | 0 | Canis | 93454.0 | 0 | Canis lupus | 93454.0 | 0 |
| scf7180000056369 | 573852 | 0.4574 | 0 | 59.1285 | Eukaryota | 33583.0 | 0 | Chordata | 33583.0 | 0 | Carnivora | 33583.0 | 0 | Canidae | 33583.0 | 0 | Canis | 23588.0 | 1 | Canis lupus | 23588.0 | 1 |
| scf7180000045911 | 72720 | 0.3883 | 0 | 63.3226 | Eukaryota | 80615.0 | 0 | Chordata | 80615.0 | 0 | Carnivora | 80615.0 | 0 | Canidae | 80615.0 | 0 | Canis | 80615.0 | 0 | Canis lupus | 80615.0 | 0 |
| scf7180000052199 | 9019 | 0.4468 | 0 | 842.9026 | no-hit | 0.0 | 0 | no-hit | 0.0 | 0 | no-hit | 0.0 | 0 | no-hit | 0.0 | 0 | no-hit | 0.0 | 0 | no-hit | 0.0 | 0 |
| scf7180000056273 | 1569792 | 0.3665 | 0 | 60.1465 | Eukaryota | 117356.0 | 0 | Chordata | 117356.0 | 0 | Carnivora | 117356.0 | 0 | Canidae | 117356.0 | 0 | Canis | 103942.0 | 1 | Canis lupus | 103942.0 | 1 |
| scf7180000048479 | 22802 | 0.4411 | 0 | 107.1137 | no-hit | 0.0 | 0 | no-hit | 0.0 | 0 | no-hit | 0.0 | 0 | no-hit | 0.0 | 0 | no-hit | 0.0 | 0 | no-hit | 0.0 | 0 |
| scf7180000054052 | 11880 | 0.5105 | 0 | 244.0748 | Eukaryota | 43645.0 | 0 | Chordata | 43645.0 | 0 | Carnivora | 43645.0 | 0 | Canidae | 43645.0 | 0 | Canis | 43645.0 | 0 | Canis lupus | 43645.0 | 0 |
| scf7180000051047 | 6711 | 0.4509 | 0 | 1240.5043 | Eukaryota | 2071.0 | 0 | Platyhelminthes | 826.0 | 2 | Polystomatidea | 826.0 | 2 | Polystomatidae | 826.0 | 2 | Protopolystoma | 826.0 | 2 | Protopolystoma xenopodis | 826.0 | 2 |
| scf7180000047351 | 2703494 | 0.3834 | 0 | 59.969 | Eukaryota | 74059.0 | 0 | Chordata | 74059.0 | 0 | Carnivora | 74059.0 | 0 | Canidae | 74059.0 | 0 | Canis | 74059.0 | 0 | Canis lupus | 74059.0 | 0 |
| scf7180000053423 | 6096 | 0.5033 | 0 | 72.2899 | Eukaryota | 8317.0 | 0 | Chordata | 8317.0 | 0 | Carnivora | 8317.0 | 0 | Canidae | 8317.0 | 0 | Canis | 6758.0 | 1 | Canis lupus | 5629.0 | 2 |
| scf7180000050284 | 12714 | 0.4417 | 0 | 29.841 | Eukaryota | 501.0 | 0 | Chordata | 501.0 | 0 | Columbiformes | 501.0 | 0 | Columbidae | 501.0 | 0 | Streptopelia | 501.0 | 0 | Streptopelia turtur | 501.0 | 0 |
| scf7180000053202 | 2416 | 0.4553 | 0 | 286.1752 | no-hit | 0.0 | 0 | no-hit | 0.0 | 0 | no-hit | 0.0 | 0 | no-hit | 0.0 | 0 | no-hit | 0.0 | 0 | no-hit | 0.0 | 0 |
| scf7180000046360 | 131410 | 0.4199 | 0 | 59.2238 | Eukaryota | 13191.0 | 0 | Chordata | 13191.0 | 0 | Carnivora | 13191.0 | 0 | Canidae | 13191.0 | 0 | Canis | 13191.0 | 0 | Canis lupus | 13191.0 | 0 |
| scf7180000041864 | 270569 | 0.4379 | 0 | 59.8041 | Eukaryota | 23376.0 | 0 | Chordata | 23376.0 | 0 | Carnivora | 23376.0 | 0 | Canidae | 23376.0 | 0 | Canis | 23376.0 | 0 | Canis lupus | 23376.0 | 0 |
| scf7180000040937 | 791976 | 0.4198 | 0 | 59.7422 | Eukaryota | 61398.0 | 0 | Chordata | 61398.0 | 0 | Carnivora | 61398.0 | 0 | Canidae | 61398.0 | 0 | Canis | 51126.0 | 1 | Canis lupus | 51126.0 | 1 |
| scf7180000044236 | 159435 | 0.3978 | 0 | 58.8863 | Eukaryota | 80862.0 | 0 | Chordata | 80862.0 | 0 | Carnivora | 80862.0 | 0 | Canidae | 80862.0 | 0 | Canis | 80862.0 | 0 | Canis lupus | 80862.0 | 0 |
| scf7180000052434 | 4173 | 0.4443 | 0 | 94.0975 | Eukaryota | 1141.0 | 0 | Chordata | 760.0 | 1 | Tetraodontiformes | 760.0 | 1 | Tetraodontidae | 760.0 | 1 | Takifugu | 760.0 | 1 | Takifugu rubripes | 760.0 | 1 |
| scf7180000042793 | 113932 | 0.4394 | 0 | 59.6655 | Eukaryota | 37134.0 | 0 | Chordata | 37134.0 | 0 | Carnivora | 37134.0 | 0 | Canidae | 37134.0 | 0 | Canis | 37134.0 | 0 | Canis lupus | 37134.0 | 0 |
| scf7180000041021 | 624615 | 0.4355 | 0 | 59.4186 | Eukaryota | 69286.0 | 0 | Chordata | 69286.0 | 0 | Carnivora | 69286.0 | 0 | Canidae | 69286.0 | 0 | Canis | 57149.0 | 1 | Canis lupus | 57149.0 | 1 |
| scf7180000045159 | 216851 | 0.4131 | 0 | 60.8783 | Eukaryota | 18732.0 | 0 | Chordata | 18732.0 | 0 | Carnivora | 18732.0 | 0 | Canidae | 18732.0 | 0 | Canis | 18732.0 | 0 | Canis lupus | 18732.0 | 0 |
| scf7180000052048 | 22750 | 0.586 | 0 | 555.1565 | Eukaryota | 18591.0 | 0 | Chordata | 18591.0 | 0 | Carnivora | 16773.0 | 1 | Felidae | 7248.0 | 4 | Acinonyx | 5436.0 | 6 | Acinonyx jubatus | 5436.0 | 6 |
| scf7180000042728 | 3541 | 0.4002 | 0 | 289.9436 | Eukaryota | 2892.0 | 0 | Chordata | 2892.0 | 0 | Carnivora | 2892.0 | 0 | Canidae | 2892.0 | 0 | Canis | 2892.0 | 0 | Canis lupus | 2892.0 | 0 |
| scf7180000044253 | 3641 | 0.4444 | 0 | 254.6077 | Eukaryota | 6303.0 | 0 | Chordata | 6049.0 | 1 | Kurtiformes | 4698.0 | 2 | Apogonidae | 4698.0 | 2 | Sphaeramia | 4698.0 | 2 | Sphaeramia orbicularis | 4698.0 | 2 |
| scf7180000043917 | 28759 | 0.4025 | 0 | 228.3547 | Eukaryota | 90181.0 | 0 | Chordata | 90181.0 | 0 | Carnivora | 90181.0 | 0 | Canidae | 90181.0 | 0 | Canis | 90181.0 | 0 | Canis lupus | 90181.0 | 0 |
| scf7180000044489 | 98387 | 0.4229 | 0 | 57.8334 | Eukaryota | 11687.0 | 0 | Chordata | 11687.0 | 0 | Primates | 5076.0 | 2 | Hominidae | 5076.0 | 3 | Homo | 5076.0 | 5 | Homo sapiens | 5076.0 | 5 |
| scf7180000046765 | 11453 | 0.4432 | 0 | 145.593 | Eukaryota | 7740.0 | 0 | Chordata | 7740.0 | 0 | Tetraodontiformes | 4056.0 | 3 | Tetraodontidae | 4056.0 | 3 | Takifugu | 4056.0 | 3 | Takifugu rubripes | 4056.0 | 3 |
| scf7180000045137 | 261934 | 0.4424 | 0 | 58.3855 | Eukaryota | 40137.0 | 0 | Chordata | 40137.0 | 0 | Carnivora | 40137.0 | 0 | Herpestidae | 20478.0 | 2 | Suricata | 20478.0 | 3 | Suricata suricatta | 20478.0 | 3 |
| scf7180000051389 | 10866 | 0.446 | 0 | 60.3727 | Eukaryota | 1330.0 | 0 | Platyhelminthes | 1015.0 | 1 | Polystomatidea | 1015.0 | 2 | Polystomatidae | 1015.0 | 2 | Protopolystoma | 1015.0 | 2 | Protopolystoma xenopodis | 1015.0 | 2 |
| scf7180000045801 | 9807 | 0.4225 | 0 | 161.1537 | Eukaryota | 44855.0 | 0 | Chordata | 44855.0 | 0 | Carnivora | 44855.0 | 0 | Canidae | 44855.0 | 0 | Canis | 44855.0 | 0 | Canis lupus | 44855.0 | 0 |
| scf7180000047119 | 85491 | 0.4167 | 0 | 56.2434 | Eukaryota | 6839.0 | 0 | Chordata | 6839.0 | 0 | Primates | 3521.0 | 2 | Hominidae | 2636.0 | 3 | Homo | 2636.0 | 4 | Homo sapiens | 2636.0 | 4 |
| scf7180000049124 | 26385 | 0.4489 | 0 | 119.7881 | Eukaryota | 9172.0 | 0 | Chordata | 9172.0 | 0 | Carnivora | 8403.0 | 1 | Canidae | 8403.0 | 1 | Canis | 6351.0 | 2 | Canis lupus | 6351.0 | 2 |
| scf7180000047379 | 83064 | 0.4045 | 0 | 62.127 | Eukaryota | 15040.0 | 0 | Chordata | 15040.0 | 0 | Perissodactyla | 5752.0 | 2 | Equidae | 5752.0 | 3 | Equus | 5752.0 | 3 | Equus przewalskii | 5752.0 | 3 |
| scf7180000041691 | 1230393 | 0.3777 | 0 | 60.17 | Eukaryota | 86232.0 | 0 | Chordata | 86232.0 | 0 | Carnivora | 86232.0 | 0 | Canidae | 86232.0 | 0 | Canis | 86232.0 | 0 | Canis lupus | 86232.0 | 0 |
| scf7180000049493 | 7252 | 0.4359 | 0 | 50.5401 | Eukaryota | 278.0 | 0 | Platyhelminthes | 278.0 | 0 | Polystomatidea | 278.0 | 0 | Polystomatidae | 278.0 | 0 | Protopolystoma | 278.0 | 0 | Protopolystoma xenopodis | 278.0 | 0 |
| scf7180000056125 | 16130 | 0.4435 | 0 | 1021.8356 | Eukaryota | 1525.0 | 0 | Platyhelminthes | 580.0 | 2 | Polystomatidea | 580.0 | 2 | Polystomatidae | 580.0 | 2 | Protopolystoma | 580.0 | 2 | Protopolystoma xenopodis | 580.0 | 2 |
| scf7180000046699 | 90576 | 0.5056 | 0 | 57.7544 | Eukaryota | 22663.0 | 0 | Chordata | 22663.0 | 0 | Carnivora | 22663.0 | 0 | Canidae | 22663.0 | 0 | Canis | 15058.0 | 1 | Canis lupus | 15058.0 | 1 |
| scf7180000051319 | 19850 | 0.5042 | 0 | 1138.8044 | Eukaryota | 9525.0 | 0 | Chordata | 9525.0 | 0 | Carnivora | 9525.0 | 0 | Canidae | 9525.0 | 0 | Canis | 8281.0 | 1 | Canis lupus | 6993.0 | 2 |
| scf7180000048912 | 12891 | 0.4327 | 0 | 124.0936 | Eukaryota | 2588.0 | 0 | Chordata | 2588.0 | 0 | Carnivora | 2588.0 | 0 | Canidae | 2588.0 | 0 | Canis | 2588.0 | 0 | Canis lupus | 2588.0 | 0 |
| scf7180000044536 | 211552 | 0.4523 | 0 | 59.1355 | Eukaryota | 54868.0 | 0 | Chordata | 54868.0 | 0 | Carnivora | 54868.0 | 0 | Canidae | 54868.0 | 0 | Canis | 54868.0 | 0 | Canis lupus | 54868.0 | 0 |
| scf7180000041690 | 282193 | 0.3663 | 0 | 60.5174 | Eukaryota | 81362.0 | 0 | Chordata | 81362.0 | 0 | Carnivora | 81362.0 | 0 | Canidae | 81362.0 | 0 | Canis | 81362.0 | 0 | Canis lupus | 81362.0 | 0 |
| scf7180000055813 | 11815 | 0.4409 | 0 | 578.3161 | no-hit | 0.0 | 0 | no-hit | 0.0 | 0 | no-hit | 0.0 | 0 | no-hit | 0.0 | 0 | no-hit | 0.0 | 0 | no-hit | 0.0 | 0 |
| scf7180000040924 | 119247 | 0.3664 | 0 | 57.8582 | Eukaryota | 11509.0 | 0 | Chordata | 11509.0 | 0 | Primates | 5882.0 | 2 | Hominidae | 5882.0 | 2 | Homo | 5198.0 | 3 | Homo sapiens | 5198.0 | 3 |
| scf7180000042153 | 624082 | 0.3543 | 100 | 59.2726 | Eukaryota | 23253.0 | 0 | Chordata | 23253.0 | 0 | Carnivora | 23253.0 | 0 | Canidae | 23253.0 | 0 | Canis | 23253.0 | 0 | Canis lupus | 23253.0 | 0 |
| scf7180000045516 | 210403 | 0.414 | 0 | 60.007 | Eukaryota | 62217.0 | 0 | Chordata | 62217.0 | 0 | Carnivora | 62217.0 | 0 | Canidae | 58394.0 | 1 | Canis | 47722.0 | 2 | Canis lupus | 47722.0 | 2 |
| scf7180000056492 | 104910 | 0.4024 | 0 | 60.6245 | Eukaryota | 53356.0 | 0 | Chordata | 53356.0 | 0 | Carnivora | 53356.0 | 0 | Canidae | 53356.0 | 0 | Canis | 53356.0 | 0 | Canis lupus | 53356.0 | 0 |
| scf7180000054660 | 23431 | 0.4353 | 0 | 221.3758 | Eukaryota | 1239.0 | 0 | Platyhelminthes | 673.0 | 1 | Polystomatidea | 673.0 | 1 | Polystomatidae | 673.0 | 1 | Protopolystoma | 673.0 | 1 | Protopolystoma xenopodis | 673.0 | 1 |
| scf7180000050044 | 9510 | 0.5002 | 0 | 98.0948 | Eukaryota | 7665.0 | 0 | Chordata | 7665.0 | 0 | Carnivora | 7665.0 | 0 | Canidae | 7665.0 | 0 | Canis | 5680.0 | 1 | Canis lupus | 4540.0 | 2 |
| scf7180000042618 | 129383 | 0.4064 | 0 | 56.6732 | Eukaryota | 39634.0 | 0 | Chordata | 39634.0 | 0 | Carnivora | 39634.0 | 0 | Canidae | 39634.0 | 0 | Canis | 39634.0 | 0 | Canis lupus | 39634.0 | 0 |
| scf7180000045964 | 29740 | 0.4252 | 0 | 51.1205 | Eukaryota | 5936.0 | 0 | Chordata | 5936.0 | 0 | Carnivora | 5936.0 | 0 | Otariidae | 2971.0 | 1 | Callorhinus | 2383.0 | 3 | Callorhinus ursinus | 2383.0 | 3 |
| scf7180000054815 | 22378 | 0.3878 | 0 | 173.593 | Eukaryota | 5842.0 | 0 | Chordata | 5014.0 | 1 | Carnivora | 3542.0 | 2 | Canidae | 2764.0 | 3 | Canis | 2764.0 | 3 | Canis lupus | 2764.0 | 3 |
| scf7180000040813 | 1676055 | 0.4098 | 0 | 60.8035 | Eukaryota | 97267.0 | 0 | Chordata | 97267.0 | 0 | Carnivora | 97267.0 | 0 | Canidae | 97267.0 | 0 | Canis | 65278.0 | 1 | Canis lupus | 65278.0 | 1 |
| scf7180000049194 | 26332 | 0.3938 | 0 | 180.4835 | Eukaryota | 74331.0 | 0 | Chordata | 74331.0 | 0 | Carnivora | 74331.0 | 0 | Canidae | 74331.0 | 0 | Canis | 74331.0 | 0 | Canis lupus | 74331.0 | 0 |
| scf7180000047981 | 65206 | 0.3453 | 0 | 58.4798 | Eukaryota | 55319.0 | 0 | Chordata | 55319.0 | 0 | Carnivora | 55319.0 | 0 | Canidae | 55319.0 | 0 | Canis | 55319.0 | 0 | Canis lupus | 55319.0 | 0 |
| scf7180000042270 | 109261 | 0.4745 | 0 | 57.8174 | Eukaryota | 50021.0 | 0 | Chordata | 50021.0 | 0 | Carnivora | 50021.0 | 0 | Canidae | 50021.0 | 0 | Canis | 42191.0 | 1 | Canis lupus | 42191.0 | 1 |
| scf7180000042704 | 685084 | 0.4001 | 0 | 59.8417 | Eukaryota | 64934.0 | 0 | Chordata | 64934.0 | 0 | Carnivora | 64934.0 | 0 | Canidae | 39109.0 | 1 | Canis | 39109.0 | 4 | Canis lupus | 39109.0 | 4 |
| scf7180000054639 | 26244 | 0.5081 | 0 | 323.1754 | Eukaryota | 4968.0 | 0 | Chordata | 4968.0 | 0 | Carnivora | 4244.0 | 1 | Canidae | 3907.0 | 3 | Canis | 3360.0 | 4 | Canis lupus | 3360.0 | 4 |
| scf7180000043122 | 553804 | 0.3942 | 0 | 58.9701 | Eukaryota | 37750.0 | 0 | Chordata | 37750.0 | 0 | Carnivora | 21761.0 | 2 | Canidae | 18436.0 | 5 | Canis | 14319.0 | 7 | Canis lupus | 14319.0 | 7 |
| scf7180000049442 | 28415 | 0.5061 | 0 | 799.052 | Eukaryota | 9958.0 | 0 | Chordata | 9958.0 | 0 | Carnivora | 9958.0 | 0 | Canidae | 9958.0 | 0 | Canis | 7932.0 | 2 | Canis lupus | 6622.0 | 3 |
| scf7180000044953 | 163840 | 0.4677 | 0 | 59.1534 | Eukaryota | 54663.0 | 0 | Chordata | 54663.0 | 0 | Carnivora | 54663.0 | 0 | Canidae | 54663.0 | 0 | Canis | 54663.0 | 0 | Canis lupus | 54663.0 | 0 |
| scf7180000040539 | 2429456 | 0.3986 | 0 | 60.5078 | Eukaryota | 59989.0 | 0 | Chordata | 59989.0 | 0 | Carnivora | 59989.0 | 0 | Canidae | 59989.0 | 0 | Canis | 51991.0 | 1 | Canis lupus | 51991.0 | 1 |
| scf7180000055557 | 8483 | 0.4399 | 0 | 153.5602 | Eukaryota | 718.0 | 0 | Platyhelminthes | 518.0 | 1 | Polystomatidea | 518.0 | 1 | Polystomatidae | 518.0 | 1 | Protopolystoma | 518.0 | 1 | Protopolystoma xenopodis | 518.0 | 1 |
| scf7180000045738 | 140624 | 0.4224 | 0 | 58.4271 | Eukaryota | 27676.0 | 0 | Chordata | 27676.0 | 0 | Carnivora | 27676.0 | 0 | Canidae | 27676.0 | 0 | Canis | 27676.0 | 0 | Canis lupus | 27676.0 | 0 |
| scf7180000055251 | 65155 | 0.4441 | 0 | 139.6862 | Eukaryota | 9552.0 | 0 | Chordata | 9552.0 | 0 | Carnivora | 9552.0 | 0 | Canidae | 9552.0 | 0 | Canis | 5678.0 | 1 | Canis lupus | 5678.0 | 1 |
| scf7180000054363 | 767058 | 0.4183 | 0 | 58.6617 | Eukaryota | 92013.0 | 0 | Chordata | 92013.0 | 0 | Carnivora | 92013.0 | 0 | Canidae | 92013.0 | 0 | Canis | 92013.0 | 0 | Canis lupus | 92013.0 | 0 |
| scf7180000045538 | 528791 | 0.454 | 0 | 58.6846 | Eukaryota | 95815.0 | 0 | Chordata | 95815.0 | 0 | Carnivora | 95815.0 | 0 | Canidae | 95815.0 | 0 | Canis | 95815.0 | 0 | Canis lupus | 95815.0 | 0 |
| scf7180000048899 | 2797 | 0.453 | 0 | 20.0025 | Eukaryota | 23823.0 | 0 | Chordata | 23823.0 | 0 | Carnivora | 23823.0 | 0 | Canidae | 23823.0 | 0 | Canis | 23823.0 | 0 | Canis lupus | 23823.0 | 0 |
| scf7180000050550 | 30643 | 0.5027 | 0 | 383.3764 | Eukaryota | 9174.0 | 0 | Chordata | 9174.0 | 0 | Carnivora | 9174.0 | 0 | Canidae | 9174.0 | 0 | Canis | 7958.0 | 1 | Canis lupus | 6698.0 | 2 |
| scf7180000052619 | 4829 | 0.4413 | 0 | 1461.7796 | Eukaryota | 1991.0 | 0 | Platyhelminthes | 791.0 | 2 | Polystomatidea | 791.0 | 2 | Polystomatidae | 791.0 | 2 | Protopolystoma | 791.0 | 2 | Protopolystoma xenopodis | 791.0 | 2 |
| scf7180000048736 | 21170 | 0.5058 | 0 | 661.6138 | Eukaryota | 9275.0 | 0 | Chordata | 9275.0 | 0 | Carnivora | 9275.0 | 0 | Canidae | 9275.0 | 0 | Canis | 7537.0 | 1 | Canis lupus | 6315.0 | 2 |
| scf7180000051431 | 5419 | 0.4388 | 0 | 73.1818 | Eukaryota | 557.0 | 0 | Platyhelminthes | 557.0 | 0 | Polystomatidea | 368.0 | 1 | Polystomatidae | 368.0 | 1 | Protopolystoma | 368.0 | 1 | Protopolystoma xenopodis | 368.0 | 1 |
| scf7180000043974 | 374428 | 0.3975 | 0 | 59.658 | Eukaryota | 33288.0 | 0 | Chordata | 33288.0 | 0 | Carnivora | 33288.0 | 0 | Canidae | 33288.0 | 0 | Canis | 33288.0 | 0 | Canis lupus | 33288.0 | 0 |
| scf7180000042931 | 74220 | 0.35 | 0 | 56.058 | Eukaryota | 25915.0 | 0 | Chordata | 25915.0 | 0 | Carnivora | 25915.0 | 0 | Phocidae | 5242.0 | 5 | Odobenus | 5127.0 | 8 | Odobenus rosmarus | 5127.0 | 8 |
| scf7180000044354 | 317173 | 0.4998 | 0 | 57.3447 | Eukaryota | 28332.0 | 0 | Chordata | 28332.0 | 0 | Carnivora | 28332.0 | 0 | Canidae | 28332.0 | 0 | Canis | 19644.0 | 1 | Canis lupus | 19644.0 | 1 |
| scf7180000048493 | 47474 | 0.3276 | 0 | 58.4194 | Eukaryota | 38789.0 | 0 | Chordata | 38789.0 | 0 | Carnivora | 38789.0 | 0 | Canidae | 38789.0 | 0 | Canis | 38789.0 | 0 | Canis lupus | 38789.0 | 0 |
| scf7180000056287 | 983555 | 0.3938 | 0 | 61.606 | Eukaryota | 98355.0 | 0 | Chordata | 98355.0 | 0 | Carnivora | 98355.0 | 0 | Canidae | 98355.0 | 0 | Canis | 98355.0 | 0 | Canis lupus | 98355.0 | 0 |
| scf7180000050474 | 8998 | 0.446 | 0 | 82.681 | Eukaryota | 5811.0 | 0 | Echinodermata | 2063.0 | 2 | Forcipulatida | 2063.0 | 4 | Asteriidae | 2063.0 | 4 | Asterias | 2063.0 | 4 | Asterias rubens | 2063.0 | 4 |
| scf7180000054383 | 372228 | 0.4227 | 0 | 60.3637 | Eukaryota | 44794.0 | 0 | Chordata | 44794.0 | 0 | Carnivora | 44794.0 | 0 | Canidae | 44794.0 | 0 | Canis | 26850.0 | 1 | Canis lupus | 26850.0 | 1 |
| scf7180000053339 | 4568 | 0.442 | 0 | 1742.9992 | Eukaryota | 2470.0 | 0 | Platyhelminthes | 1074.0 | 2 | Polystomatidea | 1074.0 | 2 | Polystomatidae | 1074.0 | 2 | Protopolystoma | 1074.0 | 2 | Protopolystoma xenopodis | 1074.0 | 2 |
| scf7180000049806 | 4455 | 0.4496 | 0 | 181.4517 | no-hit | 0.0 | 0 | no-hit | 0.0 | 0 | no-hit | 0.0 | 0 | no-hit | 0.0 | 0 | no-hit | 0.0 | 0 | no-hit | 0.0 | 0 |
| scf7180000048980 | 7221 | 0.454 | 0 | 54.7031 | no-hit | 0.0 | 0 | no-hit | 0.0 | 0 | no-hit | 0.0 | 0 | no-hit | 0.0 | 0 | no-hit | 0.0 | 0 | no-hit | 0.0 | 0 |
| scf7180000046181 | 180374 | 0.4404 | 0 | 55.6996 | Eukaryota | 33243.0 | 0 | Chordata | 33243.0 | 0 | Carnivora | 27624.0 | 2 | Canidae | 8450.0 | 6 | Canis | 8450.0 | 7 | Canis lupus | 8450.0 | 8 |
| scf7180000055612 | 12489 | 0.4398 | 0 | 279.3869 | Eukaryota | 6847.0 | 0 | Chordata | 4955.0 | 2 | Kurtiformes | 4744.0 | 3 | Apogonidae | 4744.0 | 3 | Sphaeramia | 4744.0 | 3 | Sphaeramia orbicularis | 4744.0 | 3 |
| scf7180000041806 | 38200 | 0.4724 | 0 | 61.3386 | Eukaryota | 15299.0 | 0 | Chordata | 15299.0 | 0 | Carnivora | 15299.0 | 0 | Canidae | 15299.0 | 0 | Canis | 15299.0 | 0 | Canis lupus | 15299.0 | 0 |
| scf7180000048603 | 31286 | 0.4386 | 0 | 140.7594 | Eukaryota | 3515.0 | 0 | Platyhelminthes | 2799.0 | 1 | Polystomatidea | 2799.0 | 2 | Polystomatidae | 2799.0 | 2 | Protopolystoma | 2799.0 | 2 | Protopolystoma xenopodis | 2799.0 | 2 |
| scf7180000050971 | 19469 | 0.3381 | 0 | 55.9624 | Eukaryota | 5273.0 | 0 | Chordata | 5273.0 | 0 | Carnivora | 2250.0 | 2 | Canidae | 2250.0 | 2 | Canis | 2250.0 | 3 | Canis lupus | 2250.0 | 3 |
| scf7180000052306 | 2494 | 0.4539 | 0 | 325.6899 | Eukaryota | 1904.0 | 0 | Chordata | 1532.0 | 1 | Columbiformes | 948.0 | 2 | Columbidae | 948.0 | 2 | Streptopelia | 948.0 | 2 | Streptopelia turtur | 948.0 | 2 |
| scf7180000051589 | 6134 | 0.4405 | 0 | 1849.4823 | no-hit | 0.0 | 0 | no-hit | 0.0 | 0 | no-hit | 0.0 | 0 | no-hit | 0.0 | 0 | no-hit | 0.0 | 0 | no-hit | 0.0 | 0 |
| scf7180000047595 | 97153 | 0.4138 | 0 | 70.9022 | Eukaryota | 53683.0 | 0 | Chordata | 53683.0 | 0 | Carnivora | 53683.0 | 0 | Canidae | 53683.0 | 0 | Canis | 53683.0 | 0 | Canis lupus | 53683.0 | 0 |
| scf7180000052679 | 14227 | 0.5215 | 0 | 58.2276 | Eukaryota | 11398.0 | 0 | Chordata | 11398.0 | 0 | Carnivora | 11398.0 | 0 | Canidae | 11063.0 | 1 | Canis | 11063.0 | 1 | Canis lupus | 11063.0 | 1 |
| scf7180000048341 | 2593 | 0.447 | 0 | 259.8166 | no-hit | 0.0 | 0 | no-hit | 0.0 | 0 | no-hit | 0.0 | 0 | no-hit | 0.0 | 0 | no-hit | 0.0 | 0 | no-hit | 0.0 | 0 |
| scf7180000041804 | 349862 | 0.4231 | 0 | 58.8589 | Eukaryota | 77749.0 | 0 | Chordata | 77749.0 | 0 | Carnivora | 77749.0 | 0 | Canidae | 77749.0 | 0 | Canis | 77749.0 | 0 | Canis lupus | 77749.0 | 0 |
| scf7180000049717 | 16826 | 0.4588 | 0 | 226.0166 | Eukaryota | 1200.0 | 0 | Chordata | 1200.0 | 0 | Syngnathiformes | 821.0 | 1 | Syngnathidae | 821.0 | 1 | Syngnathus | 821.0 | 1 | Syngnathus acus | 821.0 | 1 |
| scf7180000048443 | 54835 | 0.368 | 0 | 59.4537 | Eukaryota | 69471.0 | 0 | Chordata | 69471.0 | 0 | Carnivora | 69471.0 | 0 | Canidae | 69471.0 | 0 | Canis | 69471.0 | 0 | Canis lupus | 69471.0 | 0 |
| scf7180000041418 | 877605 | 0.4169 | 0 | 59.8643 | Eukaryota | 56525.0 | 0 | Chordata | 56525.0 | 0 | Carnivora | 56525.0 | 0 | Canidae | 56525.0 | 0 | Canis | 38813.0 | 1 | Canis lupus | 38813.0 | 1 |
| scf7180000050729 | 6463 | 0.4425 | 0 | 68.7486 | no-hit | 0.0 | 0 | no-hit | 0.0 | 0 | no-hit | 0.0 | 0 | no-hit | 0.0 | 0 | no-hit | 0.0 | 0 | no-hit | 0.0 | 0 |
| scf7180000054729 | 24069 | 0.4689 | 0 | 203.8369 | Eukaryota | 85825.0 | 0 | Chordata | 85825.0 | 0 | Carnivora | 85825.0 | 0 | Canidae | 85825.0 | 0 | Canis | 85825.0 | 0 | Canis lupus | 85825.0 | 0 |
| scf7180000055248 | 10134 | 0.4417 | 0 | 130.8023 | Eukaryota | 3090.0 | 0 | Echinodermata | 1875.0 | 2 | Forcipulatida | 1875.0 | 3 | Asteriidae | 1875.0 | 3 | Asterias | 1875.0 | 3 | Asterias rubens | 1875.0 | 3 |
| scf7180000040645 | 126756 | 0.5551 | 0 | 55.5998 | Eukaryota | 23440.0 | 0 | Chordata | 23440.0 | 0 | Carnivora | 23440.0 | 0 | Canidae | 23440.0 | 0 | Canis | 21151.0 | 1 | Canis lupus | 21151.0 | 1 |
| scf7180000044360 | 49060 | 0.369 | 0 | 51.1207 | Eukaryota | 6115.0 | 0 | Chordata | 6115.0 | 0 | Carnivora | 4925.0 | 2 | Canidae | 4925.0 | 2 | Canis | 4925.0 | 2 | Canis lupus | 4925.0 | 2 |
| scf7180000045203 | 204174 | 0.4595 | 0 | 60.2554 | Eukaryota | 80372.0 | 0 | Chordata | 80372.0 | 0 | Carnivora | 80372.0 | 0 | Canidae | 80372.0 | 0 | Canis | 80372.0 | 0 | Canis lupus | 80372.0 | 0 |
| scf7180000052251 | 14233 | 0.4519 | 0 | 80.7456 | Eukaryota | 55143.0 | 0 | Chordata | 55143.0 | 0 | Carnivora | 55143.0 | 0 | Canidae | 55143.0 | 0 | Canis | 55143.0 | 0 | Canis lupus | 55143.0 | 0 |
| scf7180000040611 | 236869 | 0.3883 | 0 | 59.1703 | Eukaryota | 42875.0 | 0 | Chordata | 42875.0 | 0 | Carnivora | 42875.0 | 0 | Canidae | 42875.0 | 0 | Canis | 42875.0 | 0 | Canis lupus | 42875.0 | 0 |

| | | | | | | | | | | | | | | | | | | | | | | |
|---|---|---|---|---|---|---|---|---|---|---|---|---|---|---|---|---|---|---|---|---|---|---|
| scf7180000056579 | 89855 | 0.3866 | 0 | 60.4332 | Eukaryota | 7491.0 | 0 | Chordata | 7491.0 | 0 | Primates | 5391.0 | 1 | Hominidae | 4115.0 | 3 | Homo | 4115.0 | 3 | Homo sapiens | 4115.0 | 3 |
| scf7180000042351 | 715664 | 0.3557 | 0 | 60.5802 | Eukaryota | 87714.0 | 0 | Chordata | 87714.0 | 0 | Carnivora | 87714.0 | 0 | Canidae | 87714.0 | 0 | Canis | 87714.0 | 0 | Canis lupus | 87714.0 | 0 |
| scf7180000044347 | 319600 | 0.3647 | 0 | 60.8777 | Eukaryota | 29771.0 | 0 | Chordata | 29771.0 | 0 | Carnivora | 29771.0 | 0 | Canidae | 29771.0 | 0 | Vulpes | 22434.0 | 1 | Vulpes vulpes | 22434.0 | 1 |
| scf7180000045731 | 218745 | 0.3567 | 0 | 58.8231 | Eukaryota | 88212.0 | 0 | Chordata | 88212.0 | 0 | Carnivora | 88212.0 | 0 | Canidae | 88212.0 | 0 | Canis | 88212.0 | 0 | Canis lupus | 88212.0 | 0 |
| scf7180000043839 | 256014 | 0.3421 | 0 | 58.9138 | Eukaryota | 91655.0 | 0 | Chordata | 91655.0 | 0 | Carnivora | 91655.0 | 0 | Canidae | 91655.0 | 0 | Canis | 91655.0 | 0 | Canis lupus | 91655.0 | 0 |
| scf7180000056460 | 157830 | 0.343 | 0 | 64.2598 | Eukaryota | 90003.0 | 0 | Chordata | 90003.0 | 0 | Carnivora | 90003.0 | 0 | Canidae | 90003.0 | 0 | Canis | 90003.0 | 0 | Canis lupus | 90003.0 | 0 |
| scf7180000043977 | 3210 | 0.4333 | 0 | 46.896 | Eukaryota | 20213.0 | 0 | Chordata | 20213.0 | 0 | Carnivora | 20213.0 | 0 | Canidae | 20213.0 | 0 | Canis | 20213.0 | 0 | Canis lupus | 20213.0 | 0 |
| scf7180000044423 | 318504 | 0.3757 | 0 | 60.9054 | Eukaryota | 89629.0 | 0 | Chordata | 89629.0 | 0 | Carnivora | 89629.0 | 0 | Canidae | 89629.0 | 0 | Canis | 89629.0 | 0 | Canis lupus | 89629.0 | 0 |
| scf7180000046987 | 102867 | 0.4421 | 0 | 55.4399 | Eukaryota | 77581.0 | 0 | Chordata | 77581.0 | 0 | Carnivora | 77581.0 | 0 | Canidae | 77581.0 | 0 | Canis | 77581.0 | 0 | Canis lupus | 77581.0 | 0 |
| scf7180000040684 | 1587697 | 0.4389 | 0 | 58.8343 | Eukaryota | 163887.0 | 0 | Chordata | 163887.0 | 0 | Carnivora | 163887.0 | 0 | Canidae | 163887.0 | 0 | Canis | 163887.0 | 0 | Canis lupus | 163887.0 | 0 |
| scf7180000052601 | 4647 | 0.4407 | 0 | 116.4737 | Eukaryota | 1556.0 | 0 | Chordata | 1275.0 | 1 | Columbiformes | 907.0 | 2 | Columbidae | 907.0 | 2 | Streptopelia | 907.0 | 2 | Streptopelia turtur | 907.0 | 2 |
| scf7180000046551 | 54930 | 0.4086 | 0 | 50.9967 | Eukaryota | 12624.0 | 0 | Chordata | 12624.0 | 0 | Carnivora | 12624.0 | 0 | Canidae | 12624.0 | 0 | Canis | 12624.0 | 0 | Canis lupus | 12624.0 | 0 |
| scf7180000055356 | 28182 | 0.5386 | 0 | 622.5075 | Eukaryota | 15984.0 | 0 | Chordata | 15984.0 | 0 | Carnivora | 15984.0 | 0 | Canidae | 15984.0 | 0 | Canis | 15984.0 | 0 | Canis lupus | 15984.0 | 0 |
| scf7180000044973 | 250998 | 0.3571 | 0 | 60.3643 | Eukaryota | 25317.0 | 0 | Chordata | 25317.0 | 0 | Carnivora | 25317.0 | 0 | Canidae | 25317.0 | 0 | Canis | 25317.0 | 0 | Canis lupus | 25317.0 | 0 |
| scf7180000044785 | 207294 | 0.395 | 0 | 59.5588 | Eukaryota | 36539.0 | 0 | Chordata | 36539.0 | 0 | Carnivora | 36539.0 | 0 | Canidae | 36539.0 | 0 | Canis | 36539.0 | 0 | Canis lupus | 36539.0 | 0 |
| scf7180000048830 | 44195 | 0.4546 | 0 | 85.9435 | Eukaryota | 87490.0 | 0 | Chordata | 87490.0 | 0 | Carnivora | 87490.0 | 0 | Canidae | 87490.0 | 0 | Canis | 87490.0 | 0 | Canis lupus | 87490.0 | 0 |
| scf7180000051848 | 2153 | 0.4552 | 0 | 246.9835 | no-hit | 0.0 | 0 | no-hit | 0.0 | 0 | no-hit | 0.0 | 0 | no-hit | 0.0 | 0 | no-hit | 0.0 | 0 | no-hit | 0.0 | 0 |
| scf7180000052463 | 3547 | 0.4452 | 0 | 258.4573 | Eukaryota | 16835.0 | 0 | Chordata | 12387.0 | 2 | Gadiformes | 8265.0 | 5 | Gadidae | 8265.0 | 5 | Gadus | 8265.0 | 5 | Gadus morhua | 8265.0 | 5 |
| scf7180000049369 | 6950 | 0.4393 | 0 | 98.7705 | no-hit | 0.0 | 0 | no-hit | 0.0 | 0 | no-hit | 0.0 | 0 | no-hit | 0.0 | 0 | no-hit | 0.0 | 0 | no-hit | 0.0 | 0 |
| scf7180000052852 | 7137 | 0.4377 | 0 | 101.8522 | Eukaryota | 3379.0 | 0 | Chordata | 3379.0 | 0 | Kurtiformes | 3379.0 | 0 | Apogonidae | 3379.0 | 0 | Sphaeramia | 3379.0 | 0 | Sphaeramia orbicularis | 3379.0 | 0 |
| scf7180000055358 | 23352 | 0.4396 | 0 | 177.1337 | Eukaryota | 326.0 | 0 | Platyhelminthes | 326.0 | 0 | Polystomatidea | 326.0 | 0 | Polystomatidae | 326.0 | 0 | Protopolystoma | 326.0 | 0 | Protopolystoma xenopodis | 326.0 | 0 |
| scf7180000043333 | 466181 | 0.4231 | 0 | 58.9291 | Eukaryota | 32276.0 | 0 | Chordata | 32276.0 | 0 | Carnivora | 32276.0 | 0 | Canidae | 13278.0 | 3 | Canis | 13278.0 | 3 | Canis lupus | 13278.0 | 3 |
| scf7180000049947 | 4825 | 0.4311 | 0 | 78.8474 | no-hit | 0.0 | 0 | no-hit | 0.0 | 0 | no-hit | 0.0 | 0 | no-hit | 0.0 | 0 | no-hit | 0.0 | 0 | no-hit | 0.0 | 0 |
| scf7180000041756 | 120764 | 0.3693 | 0 | 54.6631 | Eukaryota | 6347.0 | 1 | Chordata | 6347.0 | 1 | Primates | 2963.0 | 3 | Hominidae | 2963.0 | 3 | Homo | 2963.0 | 4 | Homo sapiens | 2963.0 | 4 |
| scf7180000053916 | 3194 | 0.449 | 0 | 560.5197 | Eukaryota | 2496.0 | 0 | Echinodermata | 926.0 | 3 | Forcipulatida | 926.0 | 3 | Asteriidae | 926.0 | 3 | Asterias | 926.0 | 3 | Asterias rubens | 926.0 | 3 |
| scf7180000046904 | 10010 | 0.4354 | 0 | 239.2914 | no-hit | 0.0 | 0 | no-hit | 0.0 | 0 | no-hit | 0.0 | 0 | no-hit | 0.0 | 0 | no-hit | 0.0 | 0 | no-hit | 0.0 | 0 |
| scf7180000043842 | 113502 | 0.4046 | 0 | 83.1938 | Eukaryota | 49215.0 | 0 | Chordata | 49215.0 | 0 | Carnivora | 49215.0 | 0 | Canidae | 43908.0 | 2 | Canis | 35395.0 | 3 | Canis lupus | 35395.0 | 3 |
| scf7180000052482 | 18394 | 0.3421 | 0 | 57.6981 | Eukaryota | 2974.0 | 1 | Chordata | 2974.0 | 1 | Carnivora | 1797.0 | 2 | Canidae | 1797.0 | 2 | Canis | 1797.0 | 3 | Canis lupus | 1797.0 | 3 |
| scf7180000050844 | 29170 | 0.5177 | 0 | 59.769 | Eukaryota | 6211.0 | 0 | Chordata | 6211.0 | 0 | Carnivora | 6211.0 | 0 | Canidae | 5712.0 | 1 | Canis | 5712.0 | 1 | Canis lupus | 5712.0 | 1 |
| scf7180000046374 | 40927 | 0.3939 | 0 | 54.384 | Eukaryota | 34588.0 | 0 | Chordata | 34588.0 | 0 | Carnivora | 34588.0 | 0 | Canidae | 34588.0 | 0 | Canis | 34588.0 | 0 | Canis lupus | 34588.0 | 0 |
| scf7180000050538 | 4806 | 0.4476 | 0 | 239.2641 | Eukaryota | 2683.0 | 0 | Chordata | 2253.0 | 1 | Tetraodontiformes | 1719.0 | 3 | Tetraodontidae | 1719.0 | 3 | Takifugu | 1719.0 | 3 | Takifugu rubripes | 1719.0 | 3 |
| scf7180000051159 | 4611 | 0.4407 | 0 | 145.7131 | no-hit | 0.0 | 0 | no-hit | 0.0 | 0 | no-hit | 0.0 | 0 | no-hit | 0.0 | 0 | no-hit | 0.0 | 0 | no-hit | 0.0 | 0 |
| scf7180000053530 | 2078 | 0.449 | 0 | 197.8558 | no-hit | 0.0 | 0 | no-hit | 0.0 | 0 | no-hit | 0.0 | 0 | no-hit | 0.0 | 0 | no-hit | 0.0 | 0 | no-hit | 0.0 | 0 |
| scf7180000052771 | 29612 | 0.3764 | 0 | 59.9445 | Eukaryota | 20122.0 | 0 | Chordata | 20122.0 | 0 | Carnivora | 20122.0 | 0 | Canidae | 20122.0 | 0 | Canis | 17440.0 | 1 | Canis lupus | 17440.0 | 1 |
| scf7180000046262 | 33398 | 0.4422 | 0 | 128.0599 | Eukaryota | 92386.0 | 0 | Chordata | 92386.0 | 0 | Carnivora | 92386.0 | 0 | Canidae | 92386.0 | 0 | Canis | 92386.0 | 0 | Canis lupus | 92386.0 | 0 |
| scf7180000048844 | 2214 | 0.4476 | 0 | 181.0008 | no-hit | 0.0 | 0 | no-hit | 0.0 | 0 | no-hit | 0.0 | 0 | no-hit | 0.0 | 0 | no-hit | 0.0 | 0 | no-hit | 0.0 | 0 |
| scf7180000043266 | 528395 | 0.5487 | 0 | 57.1062 | Eukaryota | 59955.0 | 0 | Chordata | 59955.0 | 0 | Carnivora | 59955.0 | 0 | Canidae | 59955.0 | 0 | Canis | 59955.0 | 0 | Canis lupus | 59955.0 | 0 |
| scf7180000050034 | 4070 | 0.4437 | 0 | 790.7239 | no-hit | 0.0 | 0 | no-hit | 0.0 | 0 | no-hit | 0.0 | 0 | no-hit | 0.0 | 0 | no-hit | 0.0 | 0 | no-hit | 0.0 | 0 |
| scf7180000050492 | 8048 | 0.4459 | 0 | 51.5392 | Eukaryota | 14850.0 | 0 | Chordata | 12483.0 | 3 | Gadiformes | 10045.0 | 5 | Gadidae | 10045.0 | 5 | Gadus | 10045.0 | 5 | Gadus morhua | 10045.0 | 5 |
| scf7180000048104 | 23002 | 0.4637 | 0 | 56.0551 | Eukaryota | 9650.0 | 0 | Chordata | 9650.0 | 0 | Carnivora | 9650.0 | 0 | Canidae | 9650.0 | 0 | Canis | 7844.0 | 1 | Canis lupus | 6563.0 | 2 |
| scf7180000044366 | 3358 | 0.436 | 0 | 187.608 | Eukaryota | 764.0 | 0 | Chordata | 764.0 | 0 | Kurtiformes | 764.0 | 0 | Apogonidae | 764.0 | 0 | Sphaeramia | 764.0 | 0 | Sphaeramia orbicularis | 764.0 | 0 |
| scf7180000040903 | 33051 | 0.3582 | 0 | 56.1479 | Eukaryota | 93045.0 | 0 | Chordata | 93045.0 | 0 | Carnivora | 93045.0 | 0 | Canidae | 93045.0 | 0 | Canis | 93045.0 | 0 | Canis lupus | 93045.0 | 0 |
| scf7180000041851 | 46808 | 0.4369 | 0 | 109.1367 | Eukaryota | 65548.0 | 0 | Chordata | 65548.0 | 0 | Carnivora | 65548.0 | 0 | Canidae | 65548.0 | 0 | Canis | 65548.0 | 0 | Canis lupus | 65548.0 | 0 |
| scf7180000040483 | 3619195 | 0.4076 | 0 | 60.2561 | Eukaryota | 99065.0 | 0 | Chordata | 99065.0 | 0 | Carnivora | 99065.0 | 0 | Canidae | 99065.0 | 0 | Canis | 99065.0 | 0 | Canis lupus | 99065.0 | 0 |
| scf7180000041754 | 280962 | 0.3832 | 0 | 59.2201 | Eukaryota | 34976.0 | 0 | Chordata | 34976.0 | 0 | Carnivora | 34976.0 | 0 | Canidae | 34976.0 | 0 | Canis | 34976.0 | 0 | Canis lupus | 34976.0 | 0 |
| scf7180000053548 | 6345 | 0.5039 | 0 | 151.394 | Eukaryota | 8567.0 | 0 | Chordata | 8567.0 | 0 | Carnivora | 8567.0 | 0 | Canidae | 8567.0 | 0 | Canis | 6923.0 | 1 | Canis lupus | 5732.0 | 2 |
| scf7180000055447 | 19884 | 0.4459 | 0 | 55.7629 | Eukaryota | 2410.0 | 0 | Chordata | 1513.0 | 1 | Tetraodontiformes | 1283.0 | 2 | Tetraodontidae | 1283.0 | 2 | Takifugu | 1283.0 | 2 | Takifugu rubripes | 1283.0 | 2 |
| scf7180000047804 | 24506 | 0.4377 | 0 | 95.1952 | Eukaryota | 11531.0 | 0 | Chordata | 6438.0 | 2 | Strigeidida | 3711.0 | 6 | Schistosomatidae | 3711.0 | 6 | Schistosoma | 3711.0 | 6 | Schistosoma mansoni | 3711.0 | 6 |
| scf7180000046482 | 11681 | 0.4434 | 0 | 82.3359 | Eukaryota | 13349.0 | 0 | Chordata | 13349.0 | 0 | Kurtiformes | 6240.0 | 2 | Apogonidae | 6240.0 | 2 | Sphaeramia | 6240.0 | 2 | Sphaeramia orbicularis | 6240.0 | 2 |
| scf7180000043995 | 21899 | 0.3286 | 0 | 40.8525 | Eukaryota | 3270.0 | 0 | Chordata | 3270.0 | 0 | Carnivora | 3270.0 | 0 | Canidae | 3270.0 | 0 | Canis | 2943.0 | 1 | Canis lupus | 2943.0 | 1 |
| scf7180000044362 | 214278 | 0.4362 | 0 | 59.4652 | Eukaryota | 44845.0 | 0 | Chordata | 44845.0 | 0 | Carnivora | 44845.0 | 0 | Canidae | 44845.0 | 0 | Canis | 44845.0 | 0 | Canis lupus | 44845.0 | 0 |
| scf7180000041845 | 896661 | 0.3974 | 0 | 59.9369 | Eukaryota | 83737.0 | 0 | Chordata | 83737.0 | 0 | Carnivora | 83737.0 | 0 | Canidae | 83737.0 | 0 | Canis | 83737.0 | 0 | Canis lupus | 83737.0 | 0 |
| scf7180000049533 | 4099 | 0.436 | 0 | 80.6873 | Eukaryota | 362.0 | 0 | Chordata | 362.0 | 0 | Beloniformes | 362.0 | 0 | Adrianichthyidae | 362.0 | 0 | Oryzias | 362.0 | 0 | Oryzias latipes | 362.0 | 0 |
| scf7180000049619 | 5464 | 0.4286 | 0 | 111.9642 | no-hit | 0.0 | 0 | no-hit | 0.0 | 0 | no-hit | 0.0 | 0 | no-hit | 0.0 | 0 | no-hit | 0.0 | 0 | no-hit | 0.0 | 0 |
| scf7180000047517 | 3762 | 0.4508 | 0 | 362.8319 | Eukaryota | 791.0 | 0 | Chordata | 791.0 | 0 | Salmoniformes | 791.0 | 0 | Salmonidae | 791.0 | 0 | Coregonus | 791.0 | 0 | Coregonus sp. 'balchen' | 791.0 | 0 |
| scf7180000042122 | 805253 | 0.4266 | 0 | 59.6906 | Eukaryota | 91714.0 | 0 | Chordata | 91714.0 | 0 | Carnivora | 91714.0 | 0 | Canidae | 91714.0 | 0 | Canis | 91714.0 | 0 | Canis lupus | 91714.0 | 0 |
| scf7180000050710 | 4179 | 0.4503 | 0 | 296.3646 | Eukaryota | 299.0 | 0 | Chordata | 299.0 | 0 | Salmoniformes | 299.0 | 0 | Salmonidae | 299.0 | 0 | Salmo | 152.0 | 1 | Salmo trutta | 152.0 | 1 |
| scf7180000052730 | 3142 | 0.4478 | 0 | 139.6332 | Eukaryota | 354.0 | 0 | Platyhelminthes | 354.0 | 0 | Polystomatidea | 354.0 | 0 | Polystomatidae | 354.0 | 0 | Protopolystoma | 354.0 | 0 | Protopolystoma xenopodis | 354.0 | 0 |
| scf7180000042573 | 624076 | 0.4126 | 0 | 59.2124 | Eukaryota | 95405.0 | 0 | Chordata | 95405.0 | 0 | Carnivora | 95405.0 | 0 | Canidae | 95405.0 | 0 | Canis | 95405.0 | 0 | Canis lupus | 95405.0 | 0 |
| scf7180000046835 | 28505 | 0.439 | 0 | 752.9707 | Eukaryota | 55500.0 | 0 | Chordata | 55500.0 | 0 | Carnivora | 55500.0 | 0 | Canidae | 55500.0 | 0 | Canis | 55500.0 | 0 | Canis lupus | 55500.0 | 0 |
| scf7180000055045 | 22619 | 0.3956 | 0 | 668.3826 | Eukaryota | 80268.0 | 0 | Chordata | 80268.0 | 0 | Carnivora | 80268.0 | 0 | Canidae | 80268.0 | 0 | Canis | 80268.0 | 0 | Canis lupus | 80268.0 | 0 |
| scf7180000040994 | 21838 | 0.4378 | 0 | 188.157 | Eukaryota | 239.0 | 0 | Platyhelminthes | 239.0 | 0 | Polystomatidea | 239.0 | 0 | Polystomatidae | 239.0 | 0 | Protopolystoma | 239.0 | 0 | Protopolystoma xenopodis | 239.0 | 0 |
| scf7180000053437 | 3993 | 0.4533 | 0 | 997.3394 | Eukaryota | 641.0 | 0 | Chordata | 641.0 | 0 | Salmoniformes | 641.0 | 0 | Salmonidae | 641.0 | 0 | Salmo | 641.0 | 0 | Salmo trutta | 641.0 | 0 |
| scf7180000041098 | 1227713 | 0.4103 | 0 | 59.8796 | Eukaryota | 89989.0 | 0 | Chordata | 89989.0 | 0 | Carnivora | 89989.0 | 0 | Canidae | 89989.0 | 0 | Canis | 89989.0 | 0 | Canis lupus | 89989.0 | 0 |
| scf7180000042656 | 686672 | 0.3966 | 0 | 59.2212 | Eukaryota | 74949.0 | 0 | Chordata | 74949.0 | 0 | Carnivora | 74949.0 | 0 | Canidae | 40183.0 | 1 | Canis | 28667.0 | 4 | Canis lupus | 28667.0 | 4 |
| scf7180000041937 | 848310 | 0.3778 | 0 | 59.9177 | Eukaryota | 50119.0 | 0 | Chordata | 50119.0 | 0 | Carnivora | 50119.0 | 0 | Canidae | 45886.0 | 1 | Canis | 39132.0 | 2 | Canis lupus | 39132.0 | 2 |
| scf7180000046499 | 108666 | 0.4398 | 0 | 59.4402 | Eukaryota | 8112.0 | 0 | Chordata | 8112.0 | 0 | Carnivora | 5301.0 | 1 | Canidae | 5301.0 | 2 | Canis | 5301.0 | 2 | Canis lupus | 5301.0 | 2 |
| scf7180000052150 | 9589 | 0.4514 | 0 | 1970.9159 | Eukaryota | 1915.0 | 0 | Echinodermata | 889.0 | 2 | Forcipulatida | 889.0 | 2 | Asteriidae | 889.0 | 2 | Asterias | 889.0 | 2 | Asterias rubens | 889.0 | 2 |
| scf7180000045712 | 3853 | 0.4487 | 0 | 166.2072 | Eukaryota | 15504.0 | 0 | Chordata | 11215.0 | 2 | Gadiformes | 7450.0 | 6 | Gadidae | 7450.0 | 6 | Gadus | 7450.0 | 6 | Gadus morhua | 7450.0 | 6 |
| scf7180000047863 | 3310 | 0.4444 | 0 | 72.9554 | no-hit | 0.0 | 0 | no-hit | 0.0 | 0 | no-hit | 0.0 | 0 | no-hit | 0.0 | 0 | no-hit | 0.0 | 0 | no-hit | 0.0 | 0 |
| scf7180000053867 | 1903 | 0.444 | 0 | 159.7818 | Eukaryota | 656.0 | 0 | Platyhelminthes | 656.0 | 0 | Polystomatidea | 656.0 | 0 | Polystomatidae | 656.0 | 0 | Protopolystoma | 656.0 | 0 | Protopolystoma xenopodis | 656.0 | 0 |
| scf7180000054936 | 17620 | 0.4447 | 0 | 139.0955 | Eukaryota | 1050.0 | 0 | Chordata | 1050.0 | 0 | Carnivora | 1050.0 | 0 | Canidae | 1050.0 | 0 | Canis | 738.0 | 1 | Canis lupus | 738.0 | 1 |
| scf7180000046811 | 46640 | 0.438 | 0 | 81.2428 | Eukaryota | 56958.0 | 0 | Chordata | 56958.0 | 0 | Carnivora | 56958.0 | 0 | Canidae | 56958.0 | 0 | Canis | 56958.0 | 0 | Canis lupus | 56958.0 | 0 |
| scf7180000045828 | 87639 | 0.4625 | 0 | 62.0979 | Eukaryota | 15695.0 | 0 | Chordata | 15695.0 | 0 | Carnivora | 15695.0 | 0 | Canidae | 15695.0 | 0 | Canis | 15695.0 | 0 | Canis lupus | 15695.0 | 0 |
| scf7180000051348 | 9481 | 0.5055 | 0 | 312.6815 | Eukaryota | 8989.0 | 0 | Chordata | 8989.0 | 0 | Carnivora | 8989.0 | 0 | Canidae | 8989.0 | 0 | Canis | 7810.0 | 1 | Canis lupus | 6577.0 | 2 |
| scf7180000053153 | 2238 | 0.4446 | 0 | 168.7696 | Eukaryota | 1221.0 | 0 | Streptophyta | 457.0 | 2 | Fabales | 457.0 | 2 | Fabaceae | 457.0 | 2 | Cicer | 457.0 | 2 | Cicer arietinum | 457.0 | 2 |
| scf7180000042262 | 724491 | 0.3654 | 0 | 61.0954 | Eukaryota | 46570.0 | 0 | Chordata | 46570.0 | 0 | Carnivora | 46570.0 | 0 | Canidae | 46570.0 | 0 | Canis | 27779.0 | 1 | Canis lupus | 27779.0 | 1 |
| scf7180000044600 | 317826 | 0.4291 | 0 | 59.3638 | Eukaryota | 29598.0 | 0 | Chordata | 29598.0 | 0 | Carnivora | 29598.0 | 0 | Canidae | 29598.0 | 0 | Canis | 29598.0 | 0 | Canis lupus | 29598.0 | 0 |
| scf7180000047052 | 28123 | 0.441 | 0 | 229.7946 | Eukaryota | 3540.0 | 0 | Chordata | 3540.0 | 0 | Kurtiformes | 3540.0 | 0 | Apogonidae | 3540.0 | 0 | Sphaeramia | 3540.0 | 0 | Sphaeramia orbicularis | 3540.0 | 0 |
| scf7180000048444 | 38268 | 0.3692 | 0 | 61.3962 | Eukaryota | 8333.0 | 1 | Chordata | 8333.0 | 1 | Carnivora | 6195.0 | 4 | Ursidae | 4046.0 | 6 | Ursus | 4046.0 | 6 | Ursus arctos | 4046.0 | 6 |
| scf7180000045798 | 36595 | 0.386 | 0 | 52.1574 | Eukaryota | 6006.0 | 0 | Chordata | 6006.0 | 0 | Carnivora | 3170.0 | 1 | Hominidae | 2836.0 | 3 | Canis | 1939.0 | 4 | Canis lupus | 1939.0 | 4 |
| scf7180000041007 | 1616640 | 0.4062 | 0 | 60.2276 | Eukaryota | 100347.0 | 0 | Chordata | 100347.0 | 0 | Carnivora | 100347.0 | 0 | Canidae | 100347.0 | 0 | Canis | 100347.0 | 0 | Canis lupus | 100347.0 | 0 |
| scf7180000048738 | 77651 | 0.4142 | 0 | 58.085 | Eukaryota | 10194.0 | 0 | Chordata | 10194.0 | 0 | Carnivora | 4958.0 | 3 | Ursidae | 2974.0 | 5 | Ursus | 2974.0 | 6 | Canis lupus | 1984.0 | 8 |
| scf7180000042271 | 718865 | 0.3365 | 0 | 58.5865 | Eukaryota | 91175.0 | 0 | Chordata | 91175.0 | 0 | Carnivora | 91175.0 | 0 | Canidae | 91175.0 | 0 | Canis | 91175.0 | 0 | Canis lupus | 91175.0 | 0 |
| scf7180000055589 | 5586 | 0.4456 | 0 | 99.9226 | no-hit | 0.0 | 0 | no-hit | 0.0 | 0 | no-hit | 0.0 | 0 | no-hit | 0.0 | 0 | no-hit | 0.0 | 0 | no-hit | 0.0 | 0 |
| scf7180000052707 | 4673 | 0.4376 | 0 | 225.3311 | Eukaryota | 623.0 | 0 | Chordata | 345.0 | 1 | Polystomatidea | 278.0 | 2 | Polystomatidae | 278.0 | 2 | Protopolystoma | 278.0 | 2 | Protopolystoma xenopodis | 278.0 | 2 |
| scf7180000046815 | 98879 | 0.4161 | 0 | 55.4035 | Eukaryota | 14656.0 | 0 | Chordata | 14656.0 | 0 | Carnivora | 14656.0 | 0 | Canidae | 6214.0 | 2 | Canis | 4714.0 | 5 | Canis lupus | 4714.0 | 5 |
| scf7180000041762 | 110636 | 0.3458 | 0 | 59.8895 | Eukaryota | 71302.0 | 0 | Chordata | 71302.0 | 0 | Carnivora | 71302.0 | 0 | Canidae | 71302.0 | 0 | Canis | 71302.0 | 0 | Canis lupus | 71302.0 | 0 |
| scf7180000046437 | 100397 | 0.3544 | 0 | 60.7422 | Eukaryota | 35504.0 | 0 | Chordata | 35504.0 | 0 | Carnivora | 35504.0 | 0 | Canidae | 25765.0 | 2 | Canis | 18459.0 | 3 | Canis lupus | 18459.0 | 3 |
| scf7180000045581 | 197827 | 0.328 | 0 | 58.716 | Eukaryota | 20989.0 | 0 | Chordata | 20989.0 | 0 | Carnivora | 20989.0 | 0 | Felidae | 20989.0 | 0 | Felis | 20989.0 | 0 | Felis catus | 20989.0 | 0 |
| scf7180000047548 | 68152 | 0.459 | 0 | 59.7625 | Eukaryota | 33645.0 | 0 | Chordata | 33645.0 | 0 | Rodentia | 15313.0 | 3 | Muridae | 15313.0 | 3 | Mus | 15313.0 | 4 | Mus musculus | 15313.0 | 4 |
| scf7180000041787 | 198883 | 0.476 | 0 | 58.7771 | Eukaryota | 16965.0 | 1 | Chordata | 16965.0 | 1 | Carnivora | 14033.0 | 3 | Canidae | 10884.0 | 4 | Canis | 9931.0 | 5 | Canis lupus | 9931.0 | 5 |
| scf7180000043107 | 252393 | 0.3968 | 0 | 63.3 | Eukaryota | 37396.0 | 0 | Chordata | 37396.0 | 0 | Carnivora | 37396.0 | 0 | Canidae | 37396.0 | 0 | Canis | 37396.0 | 0 | Canis lupus | 37396.0 | 0 |
| scf7180000055171 | 37911 | 0.4459 | 0 | 149.0225 | Eukaryota | 761.0 | 0 | Chordata | 761.0 | 0 | Kurtiformes | 454.0 | 1 | Apogonidae | 454.0 | 1 | Sphaeramia | 454.0 | 1 | Sphaeramia orbicularis | 454.0 | 1 |
| scf7180000043088 | 560785 | 0.4122 | 0 | 59.0829 | Eukaryota | 91726.0 | 0 | Chordata | 91726.0 | 0 | Carnivora | 91726.0 | 0 | Canidae | 91726.0 | 0 | Canis | 91726.0 | 0 | Canis lupus | 91726.0 | 0 |
| scf7180000045015 | 304896 | 0.4597 | 0 | 59.7707 | Eukaryota | 80098.0 | 0 | Chordata | 80098.0 | 0 | Carnivora | 80098.0 | 0 | Canidae | 80098.0 | 0 | Canis | 80098.0 | 0 | Canis lupus | 80098.0 | 0 |
| scf7180000053857 | 4266 | 0.4543 | 0 | 489.205 | Eukaryota | 1964.0 | 0 | Chordata | 1655.0 | 1 | Gonorynchiformes | 902.0 | 3 | Chanidae | 902.0 | 3 | Chanos | 902.0 | 3 | Chanos chanos | 902.0 | 3 |
| scf7180000041067 | 208506 | 0.4304 | 0 | 58.168 | Eukaryota | 24381.0 | 0 | Chordata | 24381.0 | 0 | Carnivora | 24381.0 | 0 | Canidae | 16456.0 | 3 | Vulpes | 12431.0 | 4 | Vulpes vulpes | 12431.0 | 4 |
| scf7180000041515 | 1008408 | 0.3874 | 0 | 58.8994 | Eukaryota | 46825.0 | 0 | Chordata | 46825.0 | 0 | Carnivora | 46825.0 | 0 | Canidae | 46825.0 | 0 | Canis | 46825.0 | 0 | Canis lupus | 46825.0 | 0 |
| scf7180000047033 | 107904 | 0.3955 | 0 | 59.1992 | Eukaryota | 32499.0 | 0 | Chordata | 32499.0 | 0 | Carnivora | 32499.0 | 0 | Canidae | 32499.0 | 0 | Canis | 32499.0 | 0 | Canis lupus | 32499.0 | 0 |
| scf7180000050374 | 34774 | 0.3651 | 0 | 61.1221 | Eukaryota | 12473.0 | 0 | Chordata | 12473.0 | 0 | Carnivora | 12473.0 | 0 | Canidae | 4895.0 | 2 | Odobenus | 3525.0 | 4 | Odobenus rosmarus | 3525.0 | 4 |
| scf7180000045968 | 150993 | 0.4001 | 0 | 57.8159 | Eukaryota | 93347.0 | 0 | Chordata | 93347.0 | 0 | Carnivora | 93347.0 | 0 | Canidae | 40196.0 | 2 | Canis | 30078.0 | 4 | Canis lupus | 30078.0 | 5 |
| scf7180000045663 | 50409 | 0.3835 | 0 | 57.7023 | Eukaryota | 32417.0 | 0 | Chordata | 32417.0 | 0 | Carnivora | 32417.0 | 0 | Canidae | 32417.0 | 0 | Canis | 32417.0 | 0 | Canis lupus | 32417.0 | 0 |
| scf7180000054954 | 22149 | 0.391 | 0 | 527.1444 | Eukaryota | 36058.0 | 0 | Chordata | 36058.0 | 0 | Carnivora | 36058.0 | 0 | Canidae | 36058.0 | 0 | Canis | 36058.0 | 0 | Canis lupus | 36058.0 | 0 |
| scf7180000045115 | 22563 | 0.4101 | 0 | 53.3976 | Eukaryota | 4608.0 | 1 | Chordata | 4608.0 | 1 | Primates | 2226.0 | 4 | Hominidae | 2226.0 | 5 | Homo | 2226.0 | 6 | Homo sapiens | 2226.0 | 6 |
| scf7180000054038 | 4390 | 0.4599 | 0 | 72.1972 | no-hit | 0.0 | 0 | no-hit | 0.0 | 0 | no-hit | 0.0 | 0 | no-hit | 0.0 | 0 | no-hit | 0.0 | 0 | no-hit | 0.0 | 0 |
| scf7180000056555 | 106415 | 0.4219 | 0 | 56.554 | Eukaryota | 34329.0 | 0 | Chordata | 34329.0 | 0 | Carnivora | 34329.0 | 0 | Canidae | 34329.0 | 0 | Canis | 34329.0 | 0 | Canis lupus | 34329.0 | 0 |
| scf7180000046388 | 12578 | 0.422 | 0 | 65.9338 | Eukaryota | 1687.0 | 0 | Chordata | 1687.0 | 0 | Carnivora | 1687.0 | 0 | Canidae | 1687.0 | 0 | Canis | 1687.0 | 0 | Canis lupus | 1687.0 | 0 |
| scf7180000042870 | 782297 | 0.4985 | 0 | 59.0659 | Eukaryota | 65639.0 | 0 | Chordata | 65639.0 | 0 | Carnivora | 65639.0 | 0 | Canidae | 65639.0 | 0 | Canis | 61201.0 | 1 | Canis lupus | 61201.0 | 1 |
| scf7180000044534 | 317520 | 0.6099 | 0 | 55.2217 | Eukaryota | 71183.0 | 0 | Chordata | 71183.0 | 0 | Carnivora | 71183.0 | 0 | Canidae | 71183.0 | 0 | Canis | 52304.0 | 1 | Canis lupus | 52304.0 | 1 |
| scf7180000050316 | 20035 | 0.4559 | 0 | 2622.2337 | Eukaryota | 44569.0 | 0 | Chordata | 44569.0 | 0 | Carnivora | 44569.0 | 0 | Canidae | 44569.0 | 0 | Canis | 44569.0 | 0 | Canis lupus | 44569.0 | 0 |
| scf7180000046521 | 43533 | 0.4472 | 0 | 250.7168 | Eukaryota | 3304.0 | 0 | Chordata | 2018.0 | 1 | Polystomatidea | 1286.0 | 3 | Polystomatidae | 1286.0 | 3 | Protopolystoma | 1286.0 | 3 | Protopolystoma xenopodis | 1286.0 | 3 |
| scf7180000051392 | 7456 | 0.4476 | 0 | 1243.7136 | no-hit | 0.0 | 0 | no-hit | 0.0 | 0 | no-hit | 0.0 | 0 | no-hit | 0.0 | 0 | no-hit | 0.0 | 0 | no-hit | 0.0 | 0 |
| scf7180000048412 | 48710 | 0.4389 | 0 | 236.8593 | Eukaryota | 629.0 | 0 | Chordata | 629.0 | 0 | Kurtiformes | 425.0 | 1 | Apogonidae | 425.0 | 1 | Sphaeramia | 425.0 | 1 | Sphaeramia orbicularis | 425.0 | 1 |
| scf7180000048866 | 15643 | 0.4467 | 0 | 113.3578 | Eukaryota | 12924.0 | 0 | Platyhelminthes | 12737.0 | 1 | Strigeidida | 11287.0 | 2 | Schistosomatidae | 11287.0 | 2 | Schistosoma | 11287.0 | 2 | Schistosoma mansoni | 11287.0 | 2 |
| scf7180000043315 | 48540 | 0.367 | 0 | 56.6942 | Eukaryota | 69389.0 | 0 | Chordata | 69389.0 | 0 | Carnivora | 69389.0 | 0 | Canidae | 69389.0 | 0 | Canis | 69389.0 | 0 | Canis lupus | 69389.0 | 0 |
| scf7180000056548 | 229454 | 0.3875 | 0 | 60.2897 | Eukaryota | 22585.0 | 0 | Chordata | 22585.0 | 0 | Carnivora | 22585.0 | 0 | Canidae | 21026.0 | 1 | Canis | 18577.0 | 2 | Canis lupus | 18577.0 | 2 |
| scf7180000048455 | 26386 | 0.443 | 0 | 148.9158 | Eukaryota | 6294.0 | 0 | Chordata | 4647.0 | 1 | Polystomatidea | 1647.0 | 5 | Polystomatidae | 1647.0 | 5 | Protopolystoma | 1647.0 | 5 | Protopolystoma xenopodis | 1647.0 | 5 |
| scf7180000048065 | 47782 | 0.4595 | 0 | 118.9176 | Eukaryota | 14121.0 | 0 | Chordata | 14121.0 | 0 | Carnivora | 14121.0 | 0 | Canidae | 14121.0 | 0 | Canis | 8319.0 | 1 | Canis lupus | 8319.0 | 1 |
| scf7180000046753 | 43419 | 0.4396 | 0 | 126.3453 | Eukaryota | 9545.0 | 0 | Chordata | 8343.0 | 1 | Kurtiformes | 3984.0 | 3 | Apogonidae | 3984.0 | 3 | Sphaeramia | 3984.0 | 3 | Sphaeramia orbicularis | 3984.0 | 4 |
| scf7180000054809 | 23449 | 0.4539 | 0 | 196.6089 | no-hit | 0.0 | 0 | no-hit | 0.0 | 0 | no-hit | 0.0 | 0 | no-hit | 0.0 | 0 | no-hit | 0.0 | 0 | no-hit | 0.0 | 0 |
| scf7180000053690 | 2078 | 0.4533 | 0 | 610.1599 | no-hit | 0.0 | 0 | no-hit | 0.0 | 0 | no-hit | 0.0 | 0 | no-hit | 0.0 | 0 | no-hit | 0.0 | 0 | no-hit | 0.0 | 0 |
| scf7180000049034 | 48092 | 0.4702 | 0 | 56.4026 | Eukaryota | 40256.0 | 0 | Chordata | 40256.0 | 0 | Carnivora | 40256.0 | 0 | Otariidae | 15119.0 | 2 | Zalophus | 15119.0 | 4 | Zalophus californianus | 15119.0 | 4 |
| scf7180000044486 | 12354 | 0.4088 | 0 | 39.0411 | Eukaryota | 21531.0 | 0 | Chordata | 21531.0 | 0 | Carnivora | 21531.0 | 0 | Canidae | 18683.0 | 2 | Canis | 17598.0 | 4 | Canis lupus | 17598.0 | 4 |
| scf7180000050117 | 12817 | 0.4477 | 0 | 2147.5345 | Eukaryota | 2654.0 | 0 | Platyhelminthes | 960.0 | 3 | Polystomatidea | 960.0 | 3 | Polystomatidae | 960.0 | 3 | Protopolystoma | 960.0 | 3 | Protopolystoma xenopodis | 960.0 | 3 |
| scf7180000040674 | 1819777 | 0.4285 | 0 | 59.5658 | Eukaryota | 92150.0 | 0 | Chordata | 92150.0 | 0 | Carnivora | 92150.0 | 0 | Canidae | 92150.0 | 0 | Canis | 92150.0 | 0 | Canis lupus | 92150.0 | 0 |
| scf7180000045268 | 235111 | 0.3549 | 0 | 60.2851 | Eukaryota | 19458.0 | 0 | Chordata | 19458.0 | 0 | Carnivora | 9021.0 | 2 | Hominidae | 8231.0 | 3 | Homo | 8231.0 | 3 | Homo sapiens | 8231.0 | 3 |
| scf7180000041881 | 20412 | 0.4454 | 0 | 392.3476 | Eukaryota | 92390.0 | 0 | Chordata | 92390.0 | 0 | Carnivora | 92390.0 | 0 | Canidae | 92390.0 | 0 | Canis | 92390.0 | 0 | Canis lupus | 92390.0 | 0 |
| scf7180000052226 | 10347 | 0.5028 | 0 | 1357.3486 | Eukaryota | 8874.0 | 0 | Chordata | 8874.0 | 0 | Carnivora | 8874.0 | 0 | Canidae | 8874.0 | 0 | Canis | 7697.0 | 1 | Canis lupus | 6521.0 | 2 |
| scf7180000042496 | 24999 | 0.4153 | 0 | 52.4881 | Eukaryota | 13705.0 | 0 | Chordata | 13705.0 | 0 | Carnivora | 10790.0 | 1 | Canidae | 10790.0 | 1 | Canis | 10790.0 | 1 | Canis lupus | 10790.0 | 1 |
| scf7180000047171 | 89000 | 0.5093 | 0 | 57.7114 | Eukaryota | 12088.0 | 0 | Chordata | 12088.0 | 0 | Carnivora | 10737.0 | 1 | Canidae | 10737.0 | 2 | Canis | 10737.0 | 2 | Canis lupus | 10737.0 | 2 |
| scf7180000050925 | 3672 | 0.4493 | 0 | 563.0261 | no-hit | 0.0 | 0 | no-hit | 0.0 | 0 | no-hit | 0.0 | 0 | no-hit | 0.0 | 0 | no-hit | 0.0 | 0 | no-hit | 0.0 | 0 |
| scf7180000050156 | 4409 | 0.4507 | 0 | 182.7841 | no-hit | 0.0 | 0 | no-hit | 0.0 | 0 | no-hit | 0.0 | 0 | no-hit | 0.0 | 0 | no-hit | 0.0 | 0 | no-hit | 0.0 | 0 |
| scf7180000044717 | 29773 | 0.6615 | 0 | 48.4331 | Eukaryota | 4856.0 | 0 | Chordata | 4856.0 | 0 | Carnivora | 4856.0 | 0 | Canidae | 4856.0 | 0 | Canis | 4368.0 | 1 | Canis lupus | 4368.0 | 1 |
| scf7180000046986 | 126011 | 0.3777 | 0 | 59.1387 | Eukaryota | 5218.0 | 0 | Chordata | 5218.0 | 0 | Carnivora | 5218.0 | 0 | Canidae | 5218.0 | 0 | Canis | 5218.0 | 0 | Canis lupus | 5218.0 | 0 |
| scf7180000049133 | 12670 | 0.4422 | 0 | 75.7241 | Eukaryota | 9335.0 | 0 | Chordata | 7401.0 | 1 | Tetraodontiformes | 4260.0 | 3 | Tetraodontidae | 4260.0 | 3 | Takifugu | 4260.0 | 3 | Takifugu rubripes | 4260.0 | 3 |
| scf7180000042286 | 700693 | 0.428 | 0 | 59.8769 | Eukaryota | 34232.0 | 0 | Chordata | 34232.0 | 0 | Carnivora | 34232.0 | 0 | Canidae | 34232.0 | 0 | Canis | 20582.0 | 1 | Canis lupus | 20582.0 | 1 |
| scf7180000044983 | 248860 | 0.3957 | 0 | 57.7167 | Eukaryota | 46080.0 | 0 | Chordata | 46080.0 | 0 | Carnivora | 46080.0 | 0 | Canidae | 46080.0 | 0 | Canis | 40072.0 | 1 | Canis lupus | 40072.0 | 1 |
| scf7180000055339 | 11432 | 0.4401 | 0 | 299.4829 | Eukaryota | 19646.0 | 0 | Chordata | 11930.0 | 1 | Strigeidida | 7716.0 | 5 | Schistosomatidae | 7716.0 | 5 | Schistosoma | 7716.0 | 5 | Schistosoma mansoni | 7716.0 | 6 |
| scf7180000051483 | 20548 | 0.4413 | 0 | 58.4313 | Eukaryota | 4247.0 | 0 | Chordata | 4028.0 | 1 | Salmoniformes | 4028.0 | 1 | Salmonidae | 4028.0 | 1 | Salmo | 3506.0 | 2 | Salmo trutta | 3506.0 | 2 |
| scf7180000050891 | 16424 | 0.5359 | 0 | 3724.1958 | Eukaryota | 33372.0 | 0 | Chordata | 33372.0 | 0 | Carnivora | 33372.0 | 0 | Canidae | 33372.0 | 0 | Canis | 33372.0 | 0 | Canis lupus | 33372.0 | 0 |
| scf7180000053755 | 2436 | 0.4241 | 0 | 356.0906 | no-hit | 0.0 | 0 | no-hit | 0.0 | 0 | no-hit | 0.0 | 0 | no-hit | 0.0 | 0 | no-hit | 0.0 | 0 | no-hit | 0.0 | 0 |
| scf7180000041749 | 245849 | 0.4157 | 0 | 63.9751 | Eukaryota | 91203.0 | 0 | Chordata | 91203.0 | 0 | Carnivora | 91203.0 | 0 | Canidae | 91203.0 | 0 | Canis | 91203.0 | 0 | Canis lupus | 91203.0 | 0 |
| scf7180000055001 | 23009 | 0.4444 | 0 | 159.7283 | Eukaryota | 91504.0 | 0 | Chordata | 91504.0 | 0 | Carnivora | 91504.0 | 0 | Canidae | 91504.0 | 0 | Canis | 91504.0 | 0 | Canis lupus | 91504.0 | 0 |
| scf7180000055904 | 9475 | 0.4421 | 0 | 267.18 | Eukaryota | 721.0 | 0 | Chordata | 721.0 | 0 | Tetraodontiformes | 534.0 | 1 | Tetraodontidae | 534.0 | 1 | Takifugu | 534.0 | 1 | Takifugu rubripes | 534.0 | 1 |
| scf7180000041651 | 775806 | 0.3545 | 0 | 60.2738 | Eukaryota | 40607.0 | 0 | Chordata | 40607.0 | 0 | Carnivora | 40607.0 | 0 | Canidae | 40607.0 | 0 | Canis | 37761.0 | 1 | Canis lupus | 37761.0 | 1 |
| scf7180000045225 | 71091 | 0.3798 | 0 | 65.1103 | Eukaryota | 115755.0 | 0 | Chordata | 115755.0 | 0 | Carnivora | 115755.0 | 0 | Canidae | 115755.0 | 0 | Canis | 115755.0 | 0 | Canis lupus | 115755.0 | 0 |
| scf7180000044823 | 4329 | 0.4019 | 0 | 24.2546 | Eukaryota | 1976.0 | 0 | Chordata | 1976.0 | 0 | Carnivora | 1976.0 | 0 | Canidae | 1976.0 | 0 | Canis | 1789.0 | 1 | Canis lupus | 1789.0 | 1 |
| scf7180000055484 | 23654 | 0.378 | 0 | 93.5581 | Eukaryota | 7404.0 | 0 | Chordata | 7404.0 | 0 | Carnivora | 7404.0 | 0 | Canidae | 7404.0 | 0 | Canis | 7404.0 | 0 | Canis lupus | 7404.0 | 0 |
| scf7180000041559 | 901685 | 0.4973 | 0 | 59.2258 | Eukaryota | 127122.0 | 0 | Chordata | 127122.0 | 0 | Carnivora | 127122.0 | 0 | Canidae | 127122.0 | 0 | Canis | 92206.0 | 1 | Canis lupus | 92206.0 | 1 |
| scf7180000042105 | 564892 | 0.4051 | 0 | 59.8558 | Eukaryota | 52809.0 | 0 | Chordata | 52809.0 | 0 | Carnivora | 52809.0 | 0 | Canidae | 52809.0 | 0 | Canis | 52809.0 | 0 | Canis lupus | 52809.0 | 0 |
| scf7180000048683 | 2249 | 0.4526 | 0 | 202.4239 | Eukaryota | 3780.0 | 0 | Chordata | 3780.0 | 0 | Carnivora | 2159.0 | 2 | Canidae | 2159.0 | 2 | Canis | 2159.0 | 2 | Canis lupus | 2159.0 | 2 |
| scf7180000050200 | 33637 | 0.6236 | 0 | 53.4175 | Eukaryota | 6211.0 | 0 | Chordata | 6211.0 | 0 | Carnivora | 6211.0 | 0 | Canidae | 6211.0 | 0 | Canis | 6211.0 | 0 | Canis lupus | 6211.0 | 0 |
| scf7180000053020 | 6613 | 0.453 | 0 | 1420.7911 | Eukaryota | 1566.0 | 0 | Chordata | 1566.0 | 0 | Columbiformes | 568.0 | 3 | Columbidae | 568.0 | 3 | Streptopelia | 568.0 | 3 | Streptopelia turtur | 568.0 | 3 |
| scf7180000048597 | 23193 | 0.4392 | 0 | 175.7002 | Eukaryota | 4215.0 | 0 | Chordata | 3057.0 | 2 | Salmoniformes | 3057.0 | 2 | Salmonidae | 3057.0 | 2 | Salmo | 2667.0 | 3 | Salmo trutta | 2667.0 | 3 |
| scf7180000051643 | 17246 | 0.4407 | 0 | 54.8848 | Eukaryota | 450.0 | 0 | Chordata | 450.0 | 0 | Tetraodontiformes | 278.0 | 1 | Tetraodontidae | 278.0 | 1 | Takifugu | 278.0 | 1 | Takifugu rubripes | 278.0 | 1 |
| scf7180000053502 | 1541 | 0.4588 | 0 | 98.616 | no-hit | 0.0 | 0 | no-hit | 0.0 | 0 | no-hit | 0.0 | 0 | no-hit | 0.0 | 0 | no-hit | 0.0 | 0 | no-hit | 0.0 | 0 |
| scf7180000046278 | 209123 | 0.3367 | 0 | 60.0402 | Eukaryota | 33998.0 | 0 | Chordata | 33998.0 | 0 | Carnivora | 33998.0 | 0 | Otariidae | 13424.0 | 3 | Zalophus | 10068.0 | 5 | Zalophus californianus | 10068.0 | 6 |
| scf7180000047813 | 5574 | 0.4433 | 0 | 164.8888 | no-hit | 0.0 | 0 | no-hit | 0.0 | 0 | no-hit | 0.0 | 0 | no-hit | 0.0 | 0 | no-hit | 0.0 | 0 | no-hit | 0.0 | 0 |
| scf7180000044037 | 255617 | 0.4625 | 0 | 56.9942 | Eukaryota | 26036.0 | 0 | Chordata | 26036.0 | 0 | Carnivora | 26036.0 | 0 | Canidae | 26036.0 | 0 | Canis | 15742.0 | 1 | Canis lupus | 15742.0 | 1 |
| scf7180000040985 | 1499298 | 0.3958 | 0 | 59.3596 | Eukaryota | 69172.0 | 0 | Chordata | 69172.0 | 0 | Carnivora | 69172.0 | 0 | Canidae | 69172.0 | 0 | Canis | 69172.0 | 0 | Canis lupus | 69172.0 | 0 |
| scf7180000044535 | 12146 | 0.3198 | 0 | 29.2976 | Eukaryota | 1730.0 | 0 | Chordata | 1730.0 | 0 | Carnivora | 1730.0 | 0 | Canidae | 1730.0 | 0 | Canis | 1730.0 | 0 | Canis lupus | 1730.0 | 0 |
| scf7180000042300 | 691604 | 0.3785 | 0 | 59.7497 | Eukaryota | 78519.0 | 0 | Chordata | 78519.0 | 0 | Carnivora | 78519.0 | 0 | Canidae | 78519.0 | 0 | Canis | 67706.0 | 1 | Canis lupus | 67706.0 | 1 |
| scf7180000042952 | 340494 | 0.3646 | 0 | 58.758 | Eukaryota | 94327.0 | 0 | Chordata | 94327.0 | 0 | Carnivora | 94327.0 | 0 | Canidae | 94327.0 | 0 | Canis | 94327.0 | 0 | Canis lupus | 94327.0 | 0 |
| scf7180000051922 | 10337 | 0.4457 | 0 | 252.7163 | Eukaryota | 43049.0 | 0 | Chordata | 33732.0 | 1 | Gadiformes | 25896.0 | 4 | Gadidae | 25896.0 | 4 | Gadus | 25896.0 | 4 | Gadus morhua | 25896.0 | 4 |
| scf7180000049273 | 3676 | 0.4483 | 0 | 608.7799 | no-hit | 0.0 | 0 | no-hit | 0.0 | 0 | no-hit | 0.0 | 0 | no-hit | 0.0 | 0 | no-hit | 0.0 | 0 | no-hit | 0.0 | 0 |
| scf7180000043448 | 457023 | 0.4437 | 0 | 60.5873 | Eukaryota | 79785.0 | 0 | Chordata | 79785.0 | 0 | Carnivora | 79785.0 | 0 | Canidae | 79785.0 | 0 | Canis | 79785.0 | 0 | Canis lupus | 79785.0 | 0 |
| scf7180000049221 | 23108 | 0.4543 | 0 | 311.7706 | Eukaryota | 11503.0 | 0 | Chordata | 11503.0 | 0 | Tetraodontiformes | 4798.0 | 3 | Tetraodontidae | 4798.0 | 3 | Takifugu | 4798.0 | 3 | Takifugu rubripes | 4085.0 | 4 |
| scf7180000051467 | 14357 | 0.4089 | 0 | 58.9872 | Eukaryota | 3747.0 | 0 | Chordata | 3747.0 | 0 | Carnivora | 3747.0 | 0 | Canidae | 3747.0 | 0 | Canis | 3747.0 | 0 | Canis lupus | 3747.0 | 0 |
| scf7180000041466 | 305403 | 0.3483 | 0 | 58.4641 | Eukaryota | 92989.0 | 0 | Chordata | 92989.0 | 0 | Carnivora | 92989.0 | 0 | Canidae | 92989.0 | 0 | Canis | 92989.0 | 0 | Canis lupus | 92989.0 | 0 |
| scf7180000045721 | 48210 | 0.4428 | 0 | 112.1458 | Eukaryota | 2759.0 | 0 | Chordata | 2208.0 | 1 | Kurtiformes | 990.0 | 3 | Apogonidae | 990.0 | 3 | Sphaeramia | 990.0 | 3 | Sphaeramia orbicularis | 990.0 | 3 |
| scf7180000041755 | 421476 | 0.5304 | 0 | 59.3278 | Eukaryota | 71000.0 | 0 | Chordata | 71000.0 | 0 | Carnivora | 71000.0 | 0 | Canidae | 71000.0 | 0 | Canis | 71000.0 | 0 | Canis lupus | 71000.0 | 0 |
| scf7180000052087 | 6386 | 0.4541 | 0 | 1521.008 | no-hit | 0.0 | 0 | no-hit | 0.0 | 0 | no-hit | 0.0 | 0 | no-hit | 0.0 | 0 | no-hit | 0.0 | 0 | no-hit | 0.0 | 0 |
| scf7180000043662 | 569933 | 0.3949 | 0 | 58.826 | Eukaryota | 56012.0 | 0 | Chordata | 56012.0 | 0 | Carnivora | 56012.0 | 0 | Canidae | 56012.0 | 0 | Canis | 56012.0 | 0 | Canis lupus | 56012.0 | 0 |
| scf7180000049360 | 11595 | 0.4423 | 0 | 60.1789 | Eukaryota | 20686.0 | 0 | Chordata | 12829.0 | 1 | Strigeidida | 7857.0 | 5 | Schistosomatidae | 7857.0 | 5 | Schistosoma | 7857.0 | 5 | Schistosoma mansoni | 7857.0 | 6 |
| scf7180000045146 | 28161 | 0.4102 | 0 | 54.2179 | Eukaryota | 85942.0 | 0 | Chordata | 85942.0 | 0 | Carnivora | 85942.0 | 0 | Canidae | 85942.0 | 0 | Canis | 85942.0 | 0 | Canis lupus | 85942.0 | 0 |
| scf7180000053004 | 5557 | 0.4555 | 0 | 2518.5937 | Eukaryota | 3916.0 | 0 | Platyhelminthes | 2167.0 | 3 | Polystomatidea | 2167.0 | 3 | Polystomatidae | 2167.0 | 3 | Protopolystoma | 2167.0 | 3 | Protopolystoma xenopodis | 2167.0 | 3 |
| scf7180000053134 | 7582 | 0.4401 | 0 | 153.5327 | Eukaryota | 259.0 | 0 | Platyhelminthes | 259.0 | 0 | Polystomatidea | 259.0 | 0 | Polystomatidae | 259.0 | 0 | Protopolystoma | 259.0 | 0 | Protopolystoma xenopodis | 259.0 | 0 |

| | | | | | | | | | | | | | | | | | | | | | | |
|---|---|---|---|---|---|---|---|---|---|---|---|---|---|---|---|---|---|---|---|---|---|---|
| scf7180000055945 | 18205 | 0.4455 | 0 | 769.1833 | Eukaryota | 3259.0 | 0 | Platyhelminthes | 1756.0 | 2 | Polystomatidea | 1756.0 | 2 | Polystomatidae | 1756.0 | 2 | Protopolystoma | 1756.0 | 2 | Protopolystoma xenopodis | 1756.0 | 2 |
| scf7180000045569 | 195911 | 0.5461 | 0 | 95.5587 | Eukaryota | 91453.0 | 0 | Chordata | 91453.0 | 0 | Carnivora | 91453.0 | 0 | Canidae | 91453.0 | 0 | Canis | 91453.0 | 0 | Canis lupus | 91453.0 | 0 |
| scf7180000043754 | 99887 | 0.4153 | 0 | 57.9523 | Eukaryota | 90246.0 | 0 | Chordata | 90246.0 | 0 | Carnivora | 90246.0 | 0 | Canidae | 90246.0 | 0 | Canis | 90246.0 | 0 | Canis lupus | 90246.0 | 0 |
| scf7180000045676 | 187700 | 0.3835 | 0 | 60.3638 | Eukaryota | 47296.0 | 0 | Chordata | 47296.0 | 0 | Carnivora | 47296.0 | 0 | Canidae | 35454.0 | 1 | Canis | 30432.0 | 3 | Canis lupus | 30432.0 | 3 |
| scf7180000046509 | 28896 | 0.4001 | 0 | 52.7847 | Eukaryota | 8574.0 | 0 | Chordata | 8574.0 | 0 | Carnivora | 8574.0 | 0 | Canidae | 8574.0 | 0 | Canis | 7698.0 | 1 | Canis lupus | 7698.0 | 1 |
| scf7180000048275 | 43116 | 0.3647 | 0 | 55.4584 | Eukaryota | 78121.0 | 0 | Chordata | 78121.0 | 0 | Carnivora | 78121.0 | 0 | Canidae | 78121.0 | 0 | Canis | 78121.0 | 0 | Canis lupus | 78121.0 | 0 |
| scf7180000046209 | 29347 | 0.434 | 0 | 269.8386 | Eukaryota | 42598.0 | 0 | Chordata | 42598.0 | 0 | Carnivora | 42598.0 | 0 | Canidae | 42598.0 | 0 | Canis | 42598.0 | 0 | Canis lupus | 42598.0 | 0 |
| scf7180000048647 | 59314 | 0.3871 | 0 | 58.1316 | Eukaryota | 21712.0 | 0 | Chordata | 21712.0 | 0 | Carnivora | 21712.0 | 0 | Canidae | 21712.0 | 0 | Canis | 21712.0 | 0 | Canis lupus | 21712.0 | 0 |
| scf7180000053866 | 2973 | 0.4649 | 0 | 109.1463 | Eukaryota | 3753.0 | 0 | Chordata | 3570.0 | 1 | Salmoniformes | 3182.0 | 2 | Salmonidae | 3182.0 | 2 | Coregonus | 1772.0 | 3 | Coregonus sp. 'balchen' | 1772.0 | 3 |
| scf7180000043711 | 129523 | 0.369 | 0 | 59.6908 | Eukaryota | 84624.0 | 0 | Chordata | 84624.0 | 0 | Carnivora | 84624.0 | 0 | Canidae | 84624.0 | 0 | Canis | 84624.0 | 0 | Canis lupus | 84624.0 | 0 |
| scf7180000047879 | 2416 | 0.4507 | 0 | 113.1991 | no-hit | 0.0 | 0 | no-hit | 0.0 | 0 | no-hit | 0.0 | 0 | no-hit | 0.0 | 0 | no-hit | 0.0 | 0 | no-hit | 0.0 | 0 |
| scf7180000040478 | 5068101 | 0.4467 | 0 | 59.495 | Eukaryota | 93719.0 | 0 | Chordata | 93719.0 | 0 | Carnivora | 93719.0 | 0 | Canidae | 93719.0 | 0 | Canis | 63737.0 | 1 | Canis lupus | 63737.0 | 1 |
| scf7180000049135 | 24420 | 0.4438 | 0 | 57.947 | Eukaryota | 38135.0 | 0 | Chordata | 31807.0 | 2 | Gadiformes | 25083.0 | 6 | Gadidae | 25083.0 | 6 | Gadus | 25083.0 | 6 | Gadus morhua | 25083.0 | 6 |
| scf7180000043853 | 472812 | 0.3851 | 0 | 59.9725 | Eukaryota | 58800.0 | 0 | Chordata | 58800.0 | 0 | Carnivora | 58800.0 | 0 | Canidae | 58800.0 | 0 | Canis | 58800.0 | 0 | Canis lupus | 58800.0 | 0 |
| scf7180000054208 | 25901 | 0.4787 | 0 | 204.419 | Eukaryota | 12176.0 | 0 | Chordata | 12176.0 | 0 | Carnivora | 10220.0 | 1 | Canidae | 10220.0 | 1 | Canis | 8809.0 | 2 | Canis lupus | 7933.0 | 3 |
| scf7180000056140 | 2061 | 0.4386 | 0 | 113.6954 | no-hit | 0.0 | 0 | no-hit | 0.0 | 0 | no-hit | 0.0 | 0 | no-hit | 0.0 | 0 | no-hit | 0.0 | 0 | no-hit | 0.0 | 0 |
| scf7180000054373 | 380480 | 0.3459 | 0 | 59.8464 | Eukaryota | 93635.0 | 0 | Chordata | 93635.0 | 0 | Carnivora | 93635.0 | 0 | Canidae | 93635.0 | 0 | Canis | 93635.0 | 0 | Canis lupus | 93635.0 | 0 |
| scf7180000043805 | 38143 | 0.475 | 0 | 131.9476 | Eukaryota | 9993.0 | 0 | Chordata | 9993.0 | 0 | Carnivora | 9993.0 | 0 | Canidae | 9040.0 | 1 | Canis | 6502.0 | 2 | Canis lupus | 6502.0 | 2 |
| scf7180000041289 | 123149 | 0.3764 | 0 | 57.1618 | Eukaryota | 91854.0 | 0 | Chordata | 91854.0 | 0 | Carnivora | 91854.0 | 0 | Canidae | 91854.0 | 0 | Canis | 91854.0 | 0 | Canis lupus | 91854.0 | 0 |
| scf7180000044008 | 92342 | 0.4192 | 0 | 57.0965 | Eukaryota | 45633.0 | 0 | Chordata | 45633.0 | 0 | Carnivora | 45633.0 | 0 | Canidae | 45633.0 | 0 | Canis | 30132.0 | 1 | Canis lupus | 30132.0 | 1 |
| scf7180000055255 | 22183 | 0.5161 | 0 | 620.1278 | Eukaryota | 15562.0 | 0 | Chordata | 15562.0 | 0 | Carnivora | 15562.0 | 0 | Canidae | 15562.0 | 0 | Canis | 15562.0 | 0 | Canis lupus | 15562.0 | 0 |
| scf7180000051200 | 17637 | 0.4158 | 0 | 57.3049 | Eukaryota | 9600.0 | 0 | Chordata | 9600.0 | 0 | Carnivora | 9600.0 | 0 | Canidae | 9600.0 | 0 | Canis | 9600.0 | 0 | Canis lupus | 9600.0 | 0 |
| scf7180000053124 | 4122 | 0.4391 | 0 | 159.1484 | no-hit | 0.0 | 0 | no-hit | 0.0 | 0 | no-hit | 0.0 | 0 | no-hit | 0.0 | 0 | no-hit | 0.0 | 0 | no-hit | 0.0 | 0 |
| scf7180000054943 | 23933 | 0.4129 | 0 | 557.697 | Eukaryota | 92407.0 | 0 | Chordata | 92407.0 | 0 | Carnivora | 92407.0 | 0 | Canidae | 92407.0 | 0 | Canis | 92407.0 | 0 | Canis lupus | 92407.0 | 0 |
| scf7180000045080 | 281458 | 0.3811 | 0 | 61.4382 | Eukaryota | 91057.0 | 0 | Chordata | 91057.0 | 0 | Carnivora | 91057.0 | 0 | Canidae | 91057.0 | 0 | Canis | 91057.0 | 0 | Canis lupus | 91057.0 | 0 |
| scf7180000054497 | 51318 | 0.5207 | 0 | 323.9161 | Eukaryota | 33345.0 | 0 | Chordata | 33345.0 | 0 | Carnivora | 33345.0 | 0 | Canidae | 33345.0 | 0 | Canis | 33345.0 | 0 | Canis lupus | 33345.0 | 0 |
| scf7180000047282 | 84685 | 0.4266 | 0 | 59.1555 | Eukaryota | 35906.0 | 0 | Chordata | 35906.0 | 0 | Carnivora | 35906.0 | 0 | Otariidae | 23456.0 | 1 | Zalophus | 10068.0 | 4 | Zalophus californianus | 10068.0 | 4 |
| scf7180000049908 | 6958 | 0.4477 | 0 | 1137.2477 | Eukaryota | 496.0 | 0 | Streptophyta | 496.0 | 0 | Fabales | 496.0 | 0 | Fabaceae | 496.0 | 0 | Cicer | 496.0 | 0 | Cicer arietinum | 496.0 | 0 |
| scf7180000048457 | 80510 | 0.3269 | 0 | 59.4225 | Eukaryota | 28841.0 | 0 | Chordata | 28841.0 | 0 | Carnivora | 28841.0 | 0 | Canidae | 28841.0 | 0 | Canis | 25947.0 | 1 | Canis lupus | 25947.0 | 1 |
| scf7180000049844 | 3069 | 0.4529 | 0 | 501.528 | Eukaryota | 217.0 | 0 | Streptophyta | 217.0 | 0 | Fabales | 217.0 | 0 | Fabaceae | 217.0 | 0 | Cicer | 217.0 | 0 | Cicer arietinum | 217.0 | 0 |
| scf7180000046057 | 4224 | 0.5509 | 0 | 26.798 | Eukaryota | 20425.0 | 0 | Chordata | 20425.0 | 0 | Carnivora | 20425.0 | 0 | Canidae | 12801.0 | 2 | Canis | 9920.0 | 4 | Canis lupus | 9920.0 | 5 |
| scf7180000049692 | 8734 | 0.4452 | 0 | 1068.5652 | Eukaryota | 174.0 | 0 | Euglenozoa | 174.0 | 0 | Kinetoplastida | 174.0 | 0 | Trypanosomatidae | 174.0 | 0 | Leishmania | 174.0 | 0 | Leishmania donovani | 174.0 | 0 |
| scf7180000041672 | 1004026 | 0.4257 | 0 | 60.2512 | Eukaryota | 77566.0 | 0 | Chordata | 77566.0 | 0 | Carnivora | 77566.0 | 0 | Canidae | 77566.0 | 0 | Canis | 49998.0 | 1 | Canis lupus | 49998.0 | 1 |
| scf7180000041999 | 964404 | 0.352 | 100 | 59.335 | Eukaryota | 92776.0 | 0 | Chordata | 92776.0 | 0 | Carnivora | 92776.0 | 0 | Canidae | 92776.0 | 0 | Canis | 92776.0 | 0 | Canis lupus | 92776.0 | 0 |
| scf7180000040844 | 1626617 | 0.4092 | 0 | 57.6663 | Eukaryota | 56980.0 | 0 | Chordata | 56980.0 | 0 | Carnivora | 56980.0 | 0 | Canidae | 56980.0 | 0 | Canis | 56980.0 | 0 | Canis lupus | 56980.0 | 0 |
| scf7180000042442 | 357401 | 0.4706 | 0 | 57.3242 | Eukaryota | 43531.0 | 0 | Chordata | 43531.0 | 0 | Carnivora | 43531.0 | 0 | Canidae | 43531.0 | 0 | Canis | 43531.0 | 0 | Canis lupus | 43531.0 | 0 |
| scf7180000044743 | 264677 | 0.3671 | 0 | 60.9122 | Eukaryota | 32750.0 | 0 | Chordata | 32750.0 | 0 | Carnivora | 32750.0 | 0 | Canidae | 32750.0 | 0 | Canis | 32750.0 | 0 | Canis lupus | 32750.0 | 0 |
| scf7180000050071 | 12489 | 0.4493 | 0 | 1607.2092 | Eukaryota | 1570.0 | 0 | Chordata | 1011.0 | 1 | Kurtiformes | 756.0 | 2 | Apogonidae | 756.0 | 2 | Sphaeramia | 756.0 | 2 | Sphaeramia orbicularis | 756.0 | 2 |
| scf7180000056441 | 416819 | 0.3883 | 0 | 59.9164 | Eukaryota | 79905.0 | 0 | Chordata | 79905.0 | 0 | Carnivora | 79905.0 | 0 | Canidae | 79905.0 | 0 | Canis | 79905.0 | 0 | Canis lupus | 79905.0 | 0 |
| scf7180000048932 | 5330 | 0.4548 | 0 | 71.4013 | no-hit | 0.0 | 0 | no-hit | 0.0 | 0 | no-hit | 0.0 | 0 | no-hit | 0.0 | 0 | no-hit | 0.0 | 0 | no-hit | 0.0 | 0 |
| scf7180000055134 | 15228 | 0.4332 | 0 | 178.7589 | no-hit | 0.0 | 0 | no-hit | 0.0 | 0 | no-hit | 0.0 | 0 | no-hit | 0.0 | 0 | no-hit | 0.0 | 0 | no-hit | 0.0 | 0 |
| scf7180000050109 | 21476 | 0.4386 | 0 | 63.8792 | Eukaryota | 2345.0 | 0 | Chordata | 1925.0 | 1 | Kurtiformes | 1623.0 | 2 | Apogonidae | 1623.0 | 2 | Sphaeramia | 1623.0 | 2 | Sphaeramia orbicularis | 1623.0 | 2 |
| scf7180000043479 | 24360 | 0.355 | 0 | 47.1351 | Eukaryota | 35093.0 | 0 | Chordata | 35093.0 | 0 | Carnivora | 35093.0 | 0 | Canidae | 35093.0 | 0 | Canis | 35093.0 | 0 | Canis lupus | 35093.0 | 0 |
| scf7180000051919 | 3852 | 0.4499 | 0 | 381.2072 | Eukaryota | 3084.0 | 0 | Platyhelminthes | 1658.0 | 2 | Polystomatidea | 1658.0 | 2 | Polystomatidae | 1658.0 | 2 | Protopolystoma | 1658.0 | 2 | Protopolystoma xenopodis | 1658.0 | 2 |
| scf7180000042569 | 606206 | 0.3368 | 0 | 59.5778 | Eukaryota | 59302.0 | 0 | Chordata | 59302.0 | 0 | Carnivora | 59302.0 | 0 | Canidae | 59302.0 | 0 | Canis | 59302.0 | 0 | Canis lupus | 59302.0 | 0 |
| scf7180000048584 | 22693 | 0.4402 | 0 | 205.3604 | Eukaryota | 7881.0 | 0 | Chordata | 5901.0 | 1 | Carnivora | 4395.0 | 2 | Canidae | 4395.0 | 2 | Canis | 4395.0 | 2 | Canis lupus | 4395.0 | 2 |
| scf7180000045313 | 58644 | 0.4377 | 0 | 111.5325 | Eukaryota | 15956.0 | 0 | Chordata | 10266.0 | 2 | Gadiformes | 4407.0 | 7 | Gadidae | 4407.0 | 7 | Gadus | 4407.0 | 7 | Gadus morhua | 4407.0 | 7 |
| scf7180000050905 | 16119 | 0.5184 | 0 | 224.6618 | Eukaryota | 15787.0 | 0 | Chordata | 15787.0 | 0 | Carnivora | 15787.0 | 0 | Canidae | 15787.0 | 0 | Canis | 15787.0 | 0 | Canis lupus | 15787.0 | 0 |
| scf7180000047068 | 28305 | 0.4361 | 0 | 266.6396 | Eukaryota | 24109.0 | 0 | Chordata | 19421.0 | 2 | Gadiformes | 6791.0 | 6 | Gadidae | 6791.0 | 6 | Gadus | 6791.0 | 7 | Gadus morhua | 6791.0 | 7 |
| scf7180000045877 | 164727 | 0.4538 | 0 | 62.2296 | Eukaryota | 78860.0 | 0 | Chordata | 78860.0 | 0 | Carnivora | 78860.0 | 0 | Canidae | 78860.0 | 0 | Canis | 70175.0 | 1 | Canis lupus | 70175.0 | 1 |
| scf7180000052853 | 13397 | 0.4353 | 0 | 52.4631 | Eukaryota | 12717.0 | 0 | Chordata | 9674.0 | 2 | Chordata-undef | 3446.0 | 7 | Ambassidae | 3446.0 | 7 | Parambassis | 3446.0 | 7 | Parambassis ranga | 3446.0 | 7 |
| scf7180000045771 | 195523 | 0.3787 | 0 | 58.5967 | Eukaryota | 32247.0 | 0 | Chordata | 32247.0 | 0 | Carnivora | 32247.0 | 0 | Canidae | 32247.0 | 0 | Canis | 29028.0 | 1 | Canis lupus | 29028.0 | 1 |
| scf7180000048016 | 15123 | 0.4376 | 0 | 446.2662 | no-hit | 0.0 | 0 | no-hit | 0.0 | 0 | no-hit | 0.0 | 0 | no-hit | 0.0 | 0 | no-hit | 0.0 | 0 | no-hit | 0.0 | 0 |
| scf7180000043549 | 1034091 | 0.4595 | 0 | 58.5415 | Eukaryota | 87907.0 | 0 | Chordata | 87907.0 | 0 | Carnivora | 87907.0 | 0 | Canidae | 87907.0 | 0 | Canis | 79216.0 | 1 | Canis lupus | 79216.0 | 1 |
| scf7180000043749 | 418902 | 0.3896 | 0 | 58.5655 | Eukaryota | 66733.0 | 0 | Chordata | 66733.0 | 0 | Carnivora | 40477.0 | 1 | Suidae | 26256.0 | 3 | Sus | 26256.0 | 4 | Sus scrofa | 26256.0 | 4 |
| scf7180000049352 | 17566 | 0.4411 | 0 | 344.2489 | Eukaryota | 1343.0 | 0 | Chordata | 715.0 | 2 | Fabales | 487.0 | 4 | Fabaceae | 487.0 | 4 | Cicer | 487.0 | 4 | Cicer arietinum | 487.0 | 4 |
| scf7180000040703 | 1867329 | 0.3987 | 0 | 58.7843 | Eukaryota | 50612.0 | 0 | Chordata | 50612.0 | 0 | Carnivora | 50612.0 | 0 | Canidae | 50612.0 | 0 | Canis | 50612.0 | 0 | Canis lupus | 50612.0 | 0 |
| scf7180000049452 | 4769 | 0.455 | 0 | 487.9974 | Eukaryota | 1915.0 | 0 | Platyhelminthes | 1005.0 | 2 | Polystomatidea | 1005.0 | 2 | Polystomatidae | 1005.0 | 2 | Protopolystoma | 1005.0 | 2 | Protopolystoma xenopodis | 1005.0 | 2 |
| scf7180000045135 | 238825 | 0.3974 | 0 | 59.8657 | Eukaryota | 105520.0 | 0 | Chordata | 105520.0 | 0 | Carnivora | 105520.0 | 0 | Canidae | 67109.0 | 3 | Vulpes | 45713.0 | 4 | Vulpes vulpes | 45713.0 | 4 |
| scf7180000042037 | 153018 | 0.3624 | 0 | 59.6977 | Eukaryota | 42724.0 | 0 | Chordata | 42724.0 | 0 | Carnivora | 42724.0 | 0 | Canidae | 42724.0 | 0 | Canis | 42724.0 | 0 | Canis lupus | 42724.0 | 0 |
| scf7180000049204 | 46063 | 0.4447 | 0 | 79.4076 | no-hit | 0.0 | 0 | no-hit | 0.0 | 0 | no-hit | 0.0 | 0 | no-hit | 0.0 | 0 | no-hit | 0.0 | 0 | no-hit | 0.0 | 0 |
| scf7180000050347 | 30360 | 0.4492 | 0 | 56.8091 | Eukaryota | 81972.0 | 0 | Chordata | 81972.0 | 0 | Carnivora | 81972.0 | 0 | Canidae | 81972.0 | 0 | Canis | 81972.0 | 0 | Canis lupus | 81972.0 | 0 |
| scf7180000043051 | 486134 | 0.4043 | 0 | 59.1833 | Eukaryota | 46543.0 | 0 | Chordata | 46543.0 | 0 | Carnivora | 46543.0 | 0 | Canidae | 46543.0 | 0 | Canis | 40421.0 | 1 | Canis lupus | 40421.0 | 1 |
| scf7180000045576 | 30878 | 0.4907 | 0 | 47.5342 | Eukaryota | 13648.0 | 0 | Chordata | 13648.0 | 0 | Carnivora | 10679.0 | 1 | Canidae | 8572.0 | 3 | Canis | 4748.0 | 4 | Canis lupus | 4748.0 | 4 |
| scf7180000048512 | 10190 | 0.4393 | 0 | 64.4161 | Eukaryota | 11326.0 | 0 | Chordata | 11326.0 | 0 | Carnivora | 11326.0 | 0 | Otariidae | 4288.0 | 3 | Canis | 2585.0 | 5 | Canis lupus | 2585.0 | 5 |
| scf7180000046196 | 157561 | 0.3411 | 0 | 59.5594 | Eukaryota | 56063.0 | 0 | Chordata | 56063.0 | 0 | Carnivora | 56063.0 | 0 | Canidae | 56063.0 | 0 | Canis | 56063.0 | 0 | Canis lupus | 56063.0 | 0 |
| scf7180000050683 | 35938 | 0.3562 | 0 | 68.1608 | Eukaryota | 67991.0 | 0 | Chordata | 67991.0 | 0 | Carnivora | 67991.0 | 0 | Canidae | 67991.0 | 0 | Canis | 67991.0 | 0 | Canis lupus | 67991.0 | 0 |
| scf7180000048108 | 38530 | 0.4428 | 0 | 149.4751 | Eukaryota | 39640.0 | 0 | Chordata | 31011.0 | 2 | Gadiformes | 23263.0 | 5 | Gadidae | 23263.0 | 5 | Gadus | 23263.0 | 5 | Gadus morhua | 23263.0 | 5 |
| scf7180000045925 | 157924 | 0.3778 | 0 | 60.6064 | Eukaryota | 28629.0 | 0 | Chordata | 28629.0 | 0 | Carnivora | 28629.0 | 0 | Canidae | 28629.0 | 0 | Canis | 28629.0 | 0 | Canis lupus | 28629.0 | 0 |
| scf7180000050357 | 24374 | 0.4458 | 0 | 61.1738 | Eukaryota | 55505.0 | 0 | Chordata | 55505.0 | 0 | Carnivora | 55505.0 | 0 | Canidae | 47527.0 | 1 | Canis | 31643.0 | 4 | Canis lupus | 31643.0 | 4 |
| scf7180000051892 | 6829 | 0.4561 | 0 | 589.7487 | Eukaryota | 2218.0 | 0 | Platyhelminthes | 1232.0 | 2 | Polystomatidea | 1232.0 | 2 | Polystomatidae | 1232.0 | 2 | Protopolystoma | 1232.0 | 2 | Protopolystoma xenopodis | 1232.0 | 2 |
| scf7180000051109 | 17868 | 0.4345 | 0 | 121.1671 | Eukaryota | 2490.0 | 0 | Chordata | 1498.0 | 1 | Polystomatidea | 853.0 | 4 | Polystomatidae | 853.0 | 4 | Protopolystoma | 853.0 | 5 | Protopolystoma xenopodis | 853.0 | 5 |
| scf7180000042323 | 576182 | 0.464 | 0 | 58.7126 | Eukaryota | 44780.0 | 0 | Chordata | 44780.0 | 0 | Carnivora | 44780.0 | 0 | Canidae | 44780.0 | 0 | Canis | 40384.0 | 1 | Canis lupus | 40384.0 | 1 |
| scf7180000044323 | 75369 | 0.3907 | 0 | 68.1627 | Eukaryota | 77604.0 | 0 | Chordata | 77604.0 | 0 | Carnivora | 77604.0 | 0 | Canidae | 77604.0 | 0 | Canis | 77604.0 | 0 | Canis lupus | 77604.0 | 0 |
| scf7180000042459 | 25620 | 0.3955 | 0 | 55.2836 | Eukaryota | 92258.0 | 0 | Chordata | 92258.0 | 0 | Carnivora | 92258.0 | 0 | Canidae | 92258.0 | 0 | Canis | 92258.0 | 0 | Canis lupus | 92258.0 | 0 |
| scf7180000041334 | 1475675 | 0.3669 | 100 | 59.2589 | Eukaryota | 124104.0 | 0 | Chordata | 124104.0 | 0 | Carnivora | 124104.0 | 0 | Canidae | 124104.0 | 0 | Canis | 124104.0 | 0 | Canis lupus | 124104.0 | 0 |
| scf7180000041658 | 368001 | 0.4092 | 0 | 59.7361 | Eukaryota | 62774.0 | 0 | Chordata | 62774.0 | 0 | Carnivora | 62774.0 | 0 | Canidae | 62774.0 | 0 | Canis | 62774.0 | 0 | Canis lupus | 62774.0 | 0 |
| scf7180000052594 | 3751 | 0.4441 | 0 | 777.116 | no-hit | 0.0 | 0 | no-hit | 0.0 | 0 | no-hit | 0.0 | 0 | no-hit | 0.0 | 0 | no-hit | 0.0 | 0 | no-hit | 0.0 | 0 |
| scf7180000041155 | 1324525 | 0.3948 | 0 | 60.9642 | Eukaryota | 109648.0 | 0 | Chordata | 109648.0 | 0 | Carnivora | 109648.0 | 0 | Canidae | 109648.0 | 0 | Canis | 109648.0 | 0 | Canis lupus | 109648.0 | 0 |
| scf7180000042224 | 201762 | 0.3618 | 0 | 59.0889 | Eukaryota | 55444.0 | 0 | Chordata | 55444.0 | 0 | Carnivora | 55444.0 | 0 | Canidae | 55444.0 | 0 | Canis | 55444.0 | 0 | Canis lupus | 55444.0 | 0 |
| scf7180000048380 | 10842 | 0.4535 | 0 | 804.1029 | Eukaryota | 182.0 | 0 | Streptophyta | 182.0 | 0 | Fabales | 182.0 | 0 | Fabaceae | 182.0 | 0 | Cicer | 182.0 | 0 | Cicer arietinum | 182.0 | 0 |
| scf7180000056444 | 273100 | 0.4286 | 0 | 60.7082 | Eukaryota | 56554.0 | 0 | Chordata | 56554.0 | 0 | Carnivora | 56554.0 | 0 | Canidae | 56554.0 | 0 | Canis | 56554.0 | 0 | Canis lupus | 56554.0 | 0 |
| scf7180000052350 | 5386 | 0.4543 | 0 | 776.4507 | Eukaryota | 2542.0 | 0 | Chordata | 2542.0 | 0 | Carnivora | 2542.0 | 0 | Canidae | 2542.0 | 0 | Canis | 2542.0 | 0 | Canis lupus | 2542.0 | 0 |
| scf7180000056589 | 323361 | 0.3603 | 0 | 60.7941 | Eukaryota | 93549.0 | 0 | Chordata | 93549.0 | 0 | Carnivora | 93549.0 | 0 | Canidae | 93549.0 | 0 | Canis | 65510.0 | 1 | Canis lupus | 65510.0 | 1 |
| scf7180000045748 | 162652 | 0.3847 | 0 | 60.7749 | Eukaryota | 11719.0 | 0 | Chordata | 11719.0 | 0 | Carnivora | 11719.0 | 0 | Canidae | 11719.0 | 0 | Canis | 11719.0 | 0 | Canis lupus | 11719.0 | 0 |
| scf7180000042198 | 78109 | 0.4136 | 0 | 162.4959 | Eukaryota | 133990.0 | 0 | Chordata | 133990.0 | 0 | Carnivora | 133990.0 | 0 | Canidae | 133990.0 | 0 | Canis | 133990.0 | 0 | Canis lupus | 133990.0 | 0 |
| scf7180000048766 | 41639 | 0.395 | 0 | 61.3502 | Eukaryota | 65647.0 | 0 | Chordata | 65647.0 | 0 | Carnivora | 65647.0 | 0 | Canidae | 22876.0 | 3 | Leptonychotes | 18294.0 | 6 | Leptonychotes weddellii | 18294.0 | 6 |
| scf7180000056407 | 495417 | 0.3561 | 0 | 59.9282 | Eukaryota | 91534.0 | 0 | Chordata | 91534.0 | 0 | Carnivora | 91534.0 | 0 | Canidae | 91534.0 | 0 | Canis | 91534.0 | 0 | Canis lupus | 91534.0 | 0 |
| scf7180000046133 | 133031 | 0.5429 | 0 | 59.0587 | Eukaryota | 16490.0 | 0 | Chordata | 16490.0 | 0 | Carnivora | 16490.0 | 0 | Canidae | 16490.0 | 0 | Canis | 9810.0 | 1 | Canis lupus | 9810.0 | 1 |
| scf7180000041567 | 1319221 | 0.4908 | 100 | 58.8536 | Eukaryota | 156544.0 | 0 | Chordata | 156544.0 | 0 | Carnivora | 156544.0 | 0 | Canidae | 156544.0 | 0 | Canis | 156544.0 | 0 | Canis lupus | 156544.0 | 0 |
| scf7180000052872 | 8637 | 0.4387 | 0 | 105.5394 | Eukaryota | 7597.0 | 0 | Chordata | 7597.0 | 0 | Carnivora | 7597.0 | 0 | Canidae | 7597.0 | 0 | Canis | 7597.0 | 0 | Canis lupus | 7597.0 | 0 |
| scf7180000041276 | 1539792 | 0.3652 | 100 | 59.1808 | Eukaryota | 92924.0 | 0 | Chordata | 92924.0 | 0 | Carnivora | 92924.0 | 0 | Canidae | 92924.0 | 0 | Canis | 92924.0 | 0 | Canis lupus | 92924.0 | 0 |
| scf7180000050136 | 37525 | 0.4551 | 0 | 59.6022 | Eukaryota | 33655.0 | 1 | Chordata | 30986.0 | 2 | Tetraodontiformes | 11263.0 | 5 | Tetraodontidae | 11263.0 | 5 | Takifugu | 11263.0 | 5 | Takifugu rubripes | 11263.0 | 5 |
| scf7180000048827 | 24821 | 0.4525 | 0 | 1111.8892 | Eukaryota | 3551.0 | 0 | Platyhelminthes | 1937.0 | 3 | Polystomatidea | 1937.0 | 3 | Polystomatidae | 1937.0 | 3 | Protopolystoma | 1937.0 | 3 | Protopolystoma xenopodis | 1937.0 | 3 |
| scf7180000052887 | 5966 | 0.4309 | 0 | 48.9817 | Eukaryota | 10000.0 | 0 | Chordata | 9663.0 | 1 | Kurtiformes | 4016.0 | 3 | Apogonidae | 4016.0 | 3 | Sphaeramia | 4016.0 | 4 | Sphaeramia orbicularis | 4016.0 | 4 |
| scf7180000051248 | 4702 | 0.4462 | 0 | 811.5072 | no-hit | 0.0 | 0 | no-hit | 0.0 | 0 | no-hit | 0.0 | 0 | no-hit | 0.0 | 0 | no-hit | 0.0 | 0 | no-hit | 0.0 | 0 |
| scf7180000044802 | 297793 | 0.408 | 0 | 61.0434 | Eukaryota | 89004.0 | 0 | Chordata | 89004.0 | 0 | Carnivora | 89004.0 | 0 | Canidae | 89004.0 | 0 | Canis | 89004.0 | 0 | Canis lupus | 89004.0 | 0 |
| scf7180000056191 | 1499030 | 0.4659 | 0 | 58.4614 | Eukaryota | 86669.0 | 0 | Chordata | 86669.0 | 0 | Carnivora | 86669.0 | 0 | Canidae | 86669.0 | 0 | Canis | 78111.0 | 1 | Canis lupus | 78111.0 | 1 |
| scf7180000043685 | 746305 | 0.462 | 0 | 57.9829 | Eukaryota | 87141.0 | 0 | Chordata | 87141.0 | 0 | Carnivora | 87141.0 | 0 | Canidae | 87141.0 | 0 | Canis | 87141.0 | 0 | Canis lupus | 87141.0 | 0 |
| scf7180000043085 | 102018 | 0.3728 | 0 | 58.8984 | Eukaryota | 26165.0 | 0 | Chordata | 26165.0 | 0 | Carnivora | 26165.0 | 0 | Felidae | 12933.0 | 3 | Leptonychotes | 5253.0 | 7 | Leptonychotes weddellii | 5253.0 | 8 |
| scf7180000051720 | 8922 | 0.4378 | 0 | 128.432 | no-hit | 0.0 | 0 | no-hit | 0.0 | 0 | no-hit | 0.0 | 0 | no-hit | 0.0 | 0 | no-hit | 0.0 | 0 | no-hit | 0.0 | 0 |
| scf7180000051981 | 8560 | 0.4464 | 0 | 169.8409 | Eukaryota | 13272.0 | 0 | Platyhelminthes | 6738.0 | 3 | Strigeidida | 6738.0 | 7 | Schistosomatidae | 6738.0 | 7 | Schistosoma | 6738.0 | 7 | Schistosoma mansoni | 6738.0 | 7 |
| scf7180000054864 | 27869 | 0.3992 | 0 | 330.8398 | Eukaryota | 69879.0 | 0 | Chordata | 69879.0 | 0 | Carnivora | 69879.0 | 0 | Canidae | 69879.0 | 0 | Canis | 69879.0 | 0 | Canis lupus | 69879.0 | 0 |
| scf7180000045611 | 90845 | 0.437 | 0 | 200.4262 | Eukaryota | 4061.0 | 0 | Platyhelminthes | 2271.0 | 1 | Polystomatidea | 2271.0 | 2 | Polystomatidae | 2271.0 | 2 | Protopolystoma | 2271.0 | 2 | Protopolystoma xenopodis | 2271.0 | 2 |
| scf7180000049314 | 652551 | 0.3978 | 0 | 60.9297 | Eukaryota | 107629.0 | 0 | Chordata | 107629.0 | 0 | Carnivora | 107629.0 | 0 | Canidae | 107629.0 | 0 | Canis | 92597.0 | 1 | Canis lupus | 92597.0 | 1 |
| scf7180000052166 | 5885 | 0.4464 | 0 | 1158.6002 | no-hit | 0.0 | 0 | no-hit | 0.0 | 0 | no-hit | 0.0 | 0 | no-hit | 0.0 | 0 | no-hit | 0.0 | 0 | no-hit | 0.0 | 0 |
| scf7180000050112 | 14918 | 0.4541 | 0 | 3173.7595 | Eukaryota | 3829.0 | 0 | Chordata | 2720.0 | 1 | Gonorynchiformes | 1146.0 | 3 | Chanidae | 1146.0 | 3 | Chanos | 1146.0 | 3 | Chanos chanos | 1146.0 | 3 |
| scf7180000048399 | 60040 | 0.4996 | 0 | 224.9343 | Eukaryota | 8445.0 | 0 | Chordata | 8445.0 | 0 | Carnivora | 8445.0 | 0 | Canidae | 8445.0 | 0 | Canis | 6352.0 | 1 | Canis lupus | 5231.0 | 2 |
| scf7180000041943 | 797359 | 0.3744 | 0 | 104.7683 | Eukaryota | 91165.0 | 0 | Chordata | 91165.0 | 0 | Carnivora | 91165.0 | 0 | Canidae | 91165.0 | 0 | Canis | 91165.0 | 0 | Canis lupus | 91165.0 | 0 |
| scf7180000046746 | 87302 | 0.3962 | 0 | 59.2079 | Eukaryota | 7910.0 | 0 | Chordata | 7910.0 | 0 | Carnivora | 5911.0 | 1 | Canidae | 5911.0 | 1 | Canis | 5911.0 | 2 | Canis lupus | 5911.0 | 2 |
| scf7180000041854 | 282739 | 0.373 | 0 | 59.3743 | Eukaryota | 55566.0 | 0 | Chordata | 55566.0 | 0 | Carnivora | 55566.0 | 0 | Canidae | 55566.0 | 0 | Canis | 55566.0 | 0 | Canis lupus | 55566.0 | 0 |
| scf7180000043859 | 374821 | 0.4273 | 100 | 58.8638 | Eukaryota | 92983.0 | 0 | Chordata | 92983.0 | 0 | Carnivora | 92983.0 | 0 | Canidae | 92983.0 | 0 | Canis | 92983.0 | 0 | Canis lupus | 92983.0 | 0 |
| scf7180000045545 | 29034 | 0.375 | 100 | 39.45 | Eukaryota | 78531.0 | 0 | Chordata | 78531.0 | 0 | Carnivora | 78531.0 | 0 | Canidae | 78531.0 | 0 | Canis | 78531.0 | 0 | Canis lupus | 78531.0 | 0 |
| scf7180000055448 | 28682 | 0.4422 | 0 | 134.0646 | Eukaryota | 25790.0 | 0 | Chordata | 22444.0 | 3 | Gadiformes | 18473.0 | 6 | Gadidae | 18473.0 | 6 | Gadus | 18473.0 | 6 | Gadus morhua | 18473.0 | 6 |
| scf7180000041484 | 529407 | 0.3647 | 0 | 59.9467 | Eukaryota | 90810.0 | 0 | Chordata | 90810.0 | 0 | Carnivora | 90810.0 | 0 | Canidae | 90810.0 | 0 | Canis | 90810.0 | 0 | Canis lupus | 90810.0 | 0 |
| scf7180000050956 | 6227 | 0.4468 | 0 | 995.4895 | no-hit | 0.0 | 0 | no-hit | 0.0 | 0 | no-hit | 0.0 | 0 | no-hit | 0.0 | 0 | no-hit | 0.0 | 0 | no-hit | 0.0 | 0 |
| scf7180000041018 | 1423377 | 0.3962 | 0 | 59.558 | Eukaryota | 59439.0 | 0 | Chordata | 59439.0 | 0 | Carnivora | 59439.0 | 0 | Canidae | 59439.0 | 0 | Canis | 59439.0 | 0 | Canis lupus | 59439.0 | 0 |
| scf7180000040669 | 2039414 | 0.3981 | 0 | 60.0049 | Eukaryota | 93153.0 | 0 | Chordata | 93153.0 | 0 | Carnivora | 93153.0 | 0 | Canidae | 93153.0 | 0 | Canis | 93153.0 | 0 | Canis lupus | 93153.0 | 0 |
| scf7180000041722 | 319934 | 0.4148 | 0 | 59.3895 | Eukaryota | 59342.0 | 0 | Chordata | 59342.0 | 0 | Carnivora | 59342.0 | 0 | Canidae | 59342.0 | 0 | Canis | 59342.0 | 0 | Canis lupus | 59342.0 | 0 |
| scf7180000044947 | 160008 | 0.351 | 0 | 58.834 | Eukaryota | 89458.0 | 0 | Chordata | 89458.0 | 0 | Carnivora | 89458.0 | 0 | Canidae | 89458.0 | 0 | Canis | 89458.0 | 0 | Canis lupus | 89458.0 | 0 |
| scf7180000041617 | 458824 | 0.4285 | 0 | 62.2029 | Eukaryota | 102917.0 | 0 | Chordata | 102917.0 | 0 | Carnivora | 102917.0 | 0 | Canidae | 102917.0 | 0 | Canis | 102917.0 | 0 | Canis lupus | 102917.0 | 0 |
| scf7180000041923 | 589352 | 0.3466 | 0 | 58.9368 | Eukaryota | 89656.0 | 0 | Chordata | 89656.0 | 0 | Carnivora | 89656.0 | 0 | Canidae | 89656.0 | 0 | Canis | 89656.0 | 0 | Canis lupus | 89656.0 | 0 |
| scf7180000054592 | 142259 | 0.3954 | 0 | 93.6652 | Eukaryota | 67025.0 | 0 | Chordata | 67025.0 | 0 | Carnivora | 67025.0 | 0 | Canidae | 67025.0 | 0 | Canis | 67025.0 | 0 | Canis lupus | 67025.0 | 0 |
| scf7180000049470 | 3227 | 0.4465 | 0 | 491.5274 | no-hit | 0.0 | 0 | no-hit | 0.0 | 0 | no-hit | 0.0 | 0 | no-hit | 0.0 | 0 | no-hit | 0.0 | 0 | no-hit | 0.0 | 0 |
| scf7180000046756 | 40672 | 0.4342 | 0 | 227.4124 | Eukaryota | 803.0 | 0 | Platyhelminthes | 803.0 | 0 | Polystomatidea | 803.0 | 0 | Polystomatidae | 803.0 | 0 | Protopolystoma | 803.0 | 0 | Protopolystoma xenopodis | 803.0 | 0 |
| scf7180000044492 | 319372 | 0.4621 | 0 | 59.0801 | Eukaryota | 92712.0 | 0 | Chordata | 92712.0 | 0 | Carnivora | 92712.0 | 0 | Canidae | 92712.0 | 0 | Canis | 92712.0 | 0 | Canis lupus | 92712.0 | 0 |
| scf7180000046930 | 121799 | 0.3575 | 0 | 57.712 | Eukaryota | 37527.0 | 0 | Chordata | 37527.0 | 0 | Carnivora | 37527.0 | 0 | Canidae | 37527.0 | 0 | Canis | 37527.0 | 0 | Canis lupus | 37527.0 | 0 |
| scf7180000045052 | 16888 | 0.4447 | 0 | 112.8978 | no-hit | 0.0 | 0 | no-hit | 0.0 | 0 | no-hit | 0.0 | 0 | no-hit | 0.0 | 0 | no-hit | 0.0 | 0 | no-hit | 0.0 | 0 |
| scf7180000042477 | 340639 | 0.4111 | 0 | 63.9738 | Eukaryota | 91845.0 | 0 | Chordata | 91845.0 | 0 | Carnivora | 91845.0 | 0 | Canidae | 91845.0 | 0 | Canis | 91845.0 | 0 | Canis lupus | 91845.0 | 0 |
| scf7180000048004 | 4563 | 0.613 | 0 | 26.1187 | no-hit | 0.0 | 0 | no-hit | 0.0 | 0 | no-hit | 0.0 | 0 | no-hit | 0.0 | 0 | no-hit | 0.0 | 0 | no-hit | 0.0 | 0 |
| scf7180000042166 | 122708 | 0.4409 | 0 | 59.3977 | Eukaryota | 55600.0 | 0 | Chordata | 55600.0 | 0 | Carnivora | 55600.0 | 0 | Canidae | 55600.0 | 0 | Canis | 35770.0 | 1 | Canis lupus | 35770.0 | 1 |
| scf7180000056377 | 204226 | 0.4373 | 0 | 58.7181 | Eukaryota | 13991.0 | 0 | Chordata | 13991.0 | 0 | Carnivora | 13991.0 | 0 | Canidae | 13991.0 | 0 | Canis | 13991.0 | 0 | Canis lupus | 13991.0 | 0 |
| scf7180000047441 | 24883 | 0.4355 | 0 | 217.0279 | Eukaryota | 440.0 | 0 | Chordata | 440.0 | 0 | Tetraodontiformes | 440.0 | 0 | Tetraodontidae | 440.0 | 0 | Takifugu | 440.0 | 0 | Takifugu rubripes | 440.0 | 0 |
| scf7180000042519 | 326269 | 0.3621 | 0 | 58.7827 | Eukaryota | 85925.0 | 0 | Chordata | 85925.0 | 0 | Carnivora | 85925.0 | 0 | Canidae | 85925.0 | 0 | Canis | 85925.0 | 0 | Canis lupus | 85925.0 | 0 |
| scf7180000052571 | 3321 | 0.4399 | 0 | 232.4019 | Eukaryota | 2481.0 | 0 | Platyhelminthes | 1193.0 | 3 | Polystomatidea | 1193.0 | 3 | Polystomatidae | 1193.0 | 3 | Protopolystoma | 1193.0 | 3 | Protopolystoma xenopodis | 1193.0 | 3 |
| scf7180000047135 | 6305 | 0.453 | 0 | 43.0464 | Eukaryota | 8950.0 | 0 | Chordata | 8950.0 | 0 | Carnivora | 8950.0 | 0 | Canidae | 8950.0 | 0 | Canis | 5456.0 | 1 | Canis lupus | 5456.0 | 1 |
| scf7180000052131 | 7881 | 0.4392 | 0 | 54.0346 | Eukaryota | 1117.0 | 0 | Platyhelminthes | 869.0 | 1 | Polystomatidea | 869.0 | 1 | Polystomatidae | 869.0 | 1 | Protopolystoma | 869.0 | 1 | Protopolystoma xenopodis | 869.0 | 1 |
| scf7180000047693 | 53439 | 0.3937 | 0 | 60.812 | Eukaryota | 90079.0 | 0 | Chordata | 90079.0 | 0 | Carnivora | 90079.0 | 0 | Canidae | 90079.0 | 0 | Canis | 90079.0 | 0 | Canis lupus | 90079.0 | 0 |
| scf7180000043880 | 466360 | 0.3591 | 0 | 59.7643 | Eukaryota | 62676.0 | 0 | Chordata | 62676.0 | 0 | Carnivora | 62676.0 | 0 | Canidae | 45060.0 | 3 | Canis | 45060.0 | 3 | Canis lupus | 45060.0 | 3 |
| scf7180000042437 | 636538 | 0.4401 | 0 | 60.936 | Eukaryota | 45325.0 | 1 | Chordata | 45325.0 | 1 | Carnivora | 31744.0 | 3 | Canidae | 26456.0 | 4 | Canis | 20343.0 | 5 | Canis lupus | 20343.0 | 5 |
| scf7180000046690 | 126662 | 0.3885 | 0 | 58.9878 | Eukaryota | 59585.0 | 0 | Chordata | 59585.0 | 0 | Carnivora | 47946.0 | 1 | Canidae | 25834.0 | 4 | Canis | 20262.0 | 5 | Canis lupus | 20262.0 | 6 |
| scf7180000044388 | 456222 | 0.4343 | 0 | 58.8893 | Eukaryota | 37802.0 | 0 | Chordata | 37802.0 | 0 | Carnivora | 37802.0 | 0 | Canidae | 37802.0 | 0 | Canis | 37802.0 | 0 | Canis lupus | 37802.0 | 0 |
| scf7180000052271 | 2080 | 0.4361 | 0 | 539.2017 | Eukaryota | 351.0 | 0 | Platyhelminthes | 351.0 | 0 | Polystomatidea | 351.0 | 0 | Polystomatidae | 351.0 | 0 | Protopolystoma | 351.0 | 0 | Protopolystoma xenopodis | 351.0 | 0 |
| scf7180000041291 | 9438 | 0.4339 | 0 | 39.1099 | Eukaryota | 30261.0 | 0 | Chordata | 30261.0 | 0 | Carnivora | 30261.0 | 0 | Canidae | 30261.0 | 0 | Canis | 30261.0 | 0 | Canis lupus | 30261.0 | 0 |
| scf7180000042313 | 289797 | 0.3996 | 0 | 58.6201 | Eukaryota | 16956.0 | 1 | Chordata | 16956.0 | 1 | Primates | 7747.0 | 3 | Hominidae | 7747.0 | 3 | Homo | 5283.0 | 7 | Homo sapiens | 5283.0 | 7 |
| scf7180000054726 | 24183 | 0.4459 | 0 | 252.6158 | Eukaryota | 19647.0 | 0 | Chordata | 19647.0 | 0 | Carnivora | 19647.0 | 0 | Canidae | 11177.0 | 1 | Canis | 9040.0 | 3 | Canis lupus | 9040.0 | 3 |
| scf7180000042601 | 260018 | 0.3933 | 0 | 60.5615 | Eukaryota | 31966.0 | 0 | Chordata | 31966.0 | 0 | Carnivora | 31966.0 | 0 | Canidae | 17999.0 | 4 | Canis | 10633.0 | 6 | Canis lupus | 10633.0 | 6 |
| scf7180000045558 | 157213 | 0.6387 | 0 | 52.8462 | Eukaryota | 22673.0 | 0 | Chordata | 22673.0 | 0 | Carnivora | 22673.0 | 0 | Canidae | 22673.0 | 0 | Canis | 12812.0 | 1 | Canis lupus | 12812.0 | 1 |
| scf7180000056457 | 303913 | 0.5602 | 0 | 55.2843 | Eukaryota | 16461.0 | 0 | Chordata | 16461.0 | 0 | Carnivora | 16461.0 | 0 | Canidae | 16461.0 | 0 | Canis | 16461.0 | 0 | Canis lupus | 16461.0 | 0 |
| scf7180000049424 | 4243 | 0.4476 | 0 | 197.356 | no-hit | 0.0 | 0 | no-hit | 0.0 | 0 | no-hit | 0.0 | 0 | no-hit | 0.0 | 0 | no-hit | 0.0 | 0 | no-hit | 0.0 | 0 |
| scf7180000052343 | 10182 | 0.5173 | 0 | 393.2149 | Eukaryota | 15759.0 | 0 | Chordata | 15759.0 | 0 | Carnivora | 15759.0 | 0 | Canidae | 15759.0 | 0 | Canis | 15759.0 | 0 | Canis lupus | 15759.0 | 0 |
| scf7180000042736 | 68115 | 0.4118 | 0 | 60.0643 | Eukaryota | 87410.0 | 0 | Chordata | 87410.0 | 0 | Carnivora | 87410.0 | 0 | Canidae | 87410.0 | 0 | Canis | 87410.0 | 0 | Canis lupus | 87410.0 | 0 |
| scf7180000045503 | 214126 | 0.3702 | 0 | 60.9366 | Eukaryota | 54483.0 | 0 | Chordata | 54483.0 | 0 | Carnivora | 54483.0 | 0 | Canidae | 54483.0 | 0 | Canis | 54483.0 | 0 | Canis lupus | 54483.0 | 0 |
| scf7180000053354 | 3726 | 0.4369 | 0 | 452.134 | no-hit | 0.0 | 0 | no-hit | 0.0 | 0 | no-hit | 0.0 | 0 | no-hit | 0.0 | 0 | no-hit | 0.0 | 0 | no-hit | 0.0 | 0 |
| scf7180000044370 | 4802 | 0.444 | 0 | 75.7298 | Eukaryota | 515.0 | 0 | Chordata | 515.0 | 0 | Kurtiformes | 348.0 | 1 | Apogonidae | 348.0 | 1 | Sphaeramia | 348.0 | 1 | Sphaeramia orbicularis | 348.0 | 1 |
| scf7180000041439 | 41954 | 0.4285 | 0 | 102.1266 | Eukaryota | 2757.0 | 1 | Chordata | 2757.0 | 1 | Carnivora | 2757.0 | 1 | Canidae | 2757.0 | 1 | Canis | 2757.0 | 1 | Canis lupus | 2757.0 | 1 |
| scf7180000048406 | 11991 | 0.44 | 0 | 216.2733 | Eukaryota | 1649.0 | 0 | Chordata | 1065.0 | 1 | Kurtiformes | 898.0 | 2 | Apogonidae | 898.0 | 2 | Sphaeramia | 898.0 | 2 | Sphaeramia orbicularis | 898.0 | 2 |
| scf7180000040722 | 117895 | 0.3885 | 0 | 68.4354 | Eukaryota | 19538.0 | 0 | Chordata | 19538.0 | 0 | Carnivora | 17637.0 | 1 | Canidae | 15309.0 | 2 | Canis | 12444.0 | 4 | Canis lupus | 12444.0 | 4 |
| scf7180000043249 | 260066 | 0.3596 | 0 | 58.8934 | Eukaryota | 84867.0 | 0 | Chordata | 84867.0 | 0 | Carnivora | 84867.0 | 0 | Canidae | 84867.0 | 0 | Canis | 84867.0 | 0 | Canis lupus | 84867.0 | 0 |
| scf7180000048403 | 6918 | 0.4351 | 0 | 202.5924 | Eukaryota | 594.0 | 0 | Platyhelminthes | 594.0 | 0 | Polystomatidea | 594.0 | 0 | Polystomatidae | 594.0 | 0 | Protopolystoma | 594.0 | 0 | Protopolystoma xenopodis | 594.0 | 0 |
| scf7180000048220 | 37572 | 0.4463 | 0 | 145.8134 | Eukaryota | 42048.0 | 0 | Chordata | 21837.0 | 2 | Cyclophyllidea | 10994.0 | 9 | Hymenolepididae | 10994.0 | 9 | Hymenolepis | 10994.0 | 9 | Hymenolepis microstoma | 10994.0 | 9 |
| scf7180000041213 | 1222923 | 0.4438 | 0 | 59.2777 | Eukaryota | 111272.0 | 0 | Chordata | 111272.0 | 0 | Carnivora | 111272.0 | 0 | Canidae | 111272.0 | 0 | Canis | 111272.0 | 0 | Canis lupus | 111272.0 | 0 |
| scf7180000051695 | 5109 | 0.4339 | 0 | 173.2749 | no-hit | 0.0 | 0 | no-hit | 0.0 | 0 | no-hit | 0.0 | 0 | no-hit | 0.0 | 0 | no-hit | 0.0 | 0 | no-hit | 0.0 | 0 |
| scf7180000048133 | 15903 | 0.4348 | 0 | 75.3917 | Eukaryota | 9594.0 | 0 | Chordata | 9594.0 | 0 | Tetraodontiformes | 4554.0 | 3 | Tetraodontidae | 4554.0 | 3 | Takifugu | 4554.0 | 4 | Takifugu rubripes | 4554.0 | 4 |
| scf7180000055139 | 4586 | 0.4389 | 0 | 374.2691 | no-hit | 0.0 | 0 | no-hit | 0.0 | 0 | no-hit | 0.0 | 0 | no-hit | 0.0 | 0 | no-hit | 0.0 | 0 | no-hit | 0.0 | 0 |
| scf7180000045887 | 3185 | 0.444 | 0 | 532.6095 | Eukaryota | 4983.0 | 0 | Chordata | 4983.0 | 0 | Carnivora | 4983.0 | 0 | Canidae | 4983.0 | 0 | Canis | 4983.0 | 0 | Canis lupus | 4983.0 | 0 |
| scf7180000045926 | 182853 | 0.3995 | 0 | 61.1613 | Eukaryota | 45185.0 | 0 | Chordata | 45185.0 | 0 | Carnivora | 45185.0 | 0 | Otariidae | 12705.0 | 3 | Zalophus | 12705.0 | 4 | Zalophus californianus | 12705.0 | 5 |
| scf7180000042299 | 713443 | 0.4023 | 0 | 60.3103 | Eukaryota | 89266.0 | 0 | Chordata | 89266.0 | 0 | Carnivora | 89266.0 | 0 | Canidae | 89266.0 | 0 | Canis | 89266.0 | 0 | Canis lupus | 89266.0 | 0 |
| scf7180000044278 | 115115 | 0.3922 | 0 | 83.0611 | Eukaryota | 94088.0 | 0 | Chordata | 94088.0 | 0 | Carnivora | 94088.0 | 0 | Canidae | 94088.0 | 0 | Canis | 94088.0 | 0 | Canis lupus | 94088.0 | 0 |
| scf7180000049437 | 5897 | 0.4328 | 0 | 197.0279 | no-hit | 0.0 | 0 | no-hit | 0.0 | 0 | no-hit | 0.0 | 0 | no-hit | 0.0 | 0 | no-hit | 0.0 | 0 | no-hit | 0.0 | 0 |
| scf7180000050842 | 25331 | 0.5019 | 0 | 358.0347 | Eukaryota | 9471.0 | 0 | Chordata | 9471.0 | 0 | Carnivora | 9471.0 | 0 | Canidae | 9471.0 | 0 | Canis | 8265.0 | 1 | Canis lupus | 6978.0 | 2 |
| scf7180000054834 | 4232 | 0.4362 | 0 | 325.3647 | Eukaryota | 2561.0 | 0 | Chordata | 2561.0 | 0 | Carnivora | 2561.0 | 0 | Canidae | 2561.0 | 0 | Canis | 2561.0 | 0 | Canis lupus | 2561.0 | 0 |
| scf7180000056611 | 438005 | 0.3802 | 0 | 60.5594 | Eukaryota | 68753.0 | 0 | Chordata | 68753.0 | 0 | Carnivora | 68753.0 | 0 | Canidae | 68753.0 | 0 | Canis | 51047.0 | 1 | Canis lupus | 51047.0 | 1 |
| scf7180000054547 | 26072 | 0.4412 | 0 | 311.0749 | Eukaryota | 3027.0 | 0 | Platyhelminthes | 1641.0 | 2 | Polystomatidea | 1641.0 | 2 | Polystomatidae | 1641.0 | 2 | Protopolystoma | 1641.0 | 2 | Protopolystoma xenopodis | 1641.0 | 2 |
| scf7180000051555 | 11127 | 0.4418 | 0 | 1512.1619 | Eukaryota | 470.0 | 0 | Chordata | 470.0 | 0 | Kurtiformes | 470.0 | 0 | Apogonidae | 470.0 | 0 | Sphaeramia | 470.0 | 0 | Sphaeramia orbicularis | 470.0 | 0 |
| scf7180000048081 | 51511 | 0.5324 | 0 | 57.7257 | Eukaryota | 14184.0 | 0 | Chordata | 14184.0 | 0 | Carnivora | 12436.0 | 1 | Canidae | 7399.0 | 5 | Canis | 4603.0 | 7 | Canis lupus | 4603.0 | 7 |
| scf7180000041892 | 333563 | 0.4117 | 0 | 59.4718 | Eukaryota | 107430.0 | 0 | Chordata | 107430.0 | 0 | Carnivora | 107430.0 | 0 | Canidae | 57884.0 | 2 | Canis | 45010.0 | 5 | Canis lupus | 45010.0 | 6 |
| scf7180000043909 | 399560 | 0.3389 | 0 | 58.6077 | Eukaryota | 89752.0 | 0 | Chordata | 89752.0 | 0 | Carnivora | 89752.0 | 0 | Canidae | 89752.0 | 0 | Canis | 89752.0 | 0 | Canis lupus | 89752.0 | 0 |

| | | | | | | | | | | | | | | | | | | | |
|---|---|---|---|---|---|---|---|---|---|---|---|---|---|---|---|---|---|---|---|
| scf7180000050687 | 19201 | 0.4474 | 0 | 57.9497 | Eukaryota | 1376.0 | 0 | Platyhelminthes | 712.0 | 2 | Polystomatidea | 712.0 | 3 | Polystomatidae | 712.0 | 3 | Protopolystoma | 712.0 | 3 | Protopolystoma xenopodis | 712.0 | 3 |
| scf7180000049827 | 7367 | 0.4372 | 0 | 153.1743 | no-hit | 0.0 | 0 | no-hit | 0.0 | 0 | no-hit | 0.0 | 0 | no-hit | 0.0 | 0 | no-hit | 0.0 | 0 | no-hit | 0.0 | 0 |
| scf7180000044722 | 95697 | 0.461 | 0 | 56.4218 | Eukaryota | 23654.0 | 0 | Chordata | 23654.0 | 0 | Carnivora | 23654.0 | 0 | Canidae | 23654.0 | 0 | Canis | 23654.0 | 0 | Canis lupus | 23654.0 | 0 |
| scf7180000042327 | 706921 | 0.478 | 0 | 59.8513 | Eukaryota | 179203.0 | 0 | Chordata | 179203.0 | 0 | Carnivora | 179203.0 | 0 | Canidae | 174817.0 | 1 | Canis | 174817.0 | 1 | Canis lupus | 174817.0 | 1 |
| scf7180000045143 | 68716 | 0.4166 | 0 | 57.9049 | Eukaryota | 61361.0 | 0 | Chordata | 61361.0 | 0 | Carnivora | 61361.0 | 0 | Canidae | 61361.0 | 0 | Canis | 52286.0 | 1 | Canis lupus | 52286.0 | 1 |
| scf7180000052960 | 5663 | 0.4478 | 0 | 1769.1424 | no-hit | 0.0 | 0 | no-hit | 0.0 | 0 | no-hit | 0.0 | 0 | no-hit | 0.0 | 0 | no-hit | 0.0 | 0 | no-hit | 0.0 | 0 |
| scf7180000048482 | 76210 | 0.4074 | 0 | 60.1854 | Eukaryota | 49583.0 | 0 | Chordata | 49583.0 | 0 | Carnivora | 49583.0 | 0 | Canidae | 49583.0 | 0 | Canis | 49583.0 | 0 | Canis lupus | 49583.0 | 0 |
| scf7180000050454 | 9646 | 0.4494 | 0 | 1602.0294 | Eukaryota | 3376.0 | 0 | Platyhelminthes | 1892.0 | 2 | Polystomatidea | 1892.0 | 2 | Polystomatidae | 1892.0 | 2 | Protopolystoma | 1892.0 | 2 | Protopolystoma xenopodis | 1892.0 | 2 |
| scf7180000041157 | 365486 | 0.3451 | 0 | 58.8137 | Eukaryota | 76073.0 | 0 | Chordata | 76073.0 | 0 | Carnivora | 76073.0 | 0 | Canidae | 76073.0 | 0 | Canis | 76073.0 | 0 | Canis lupus | 76073.0 | 0 |
| scf7180000049305 | 30562 | 0.4223 | 0 | 58.9539 | Eukaryota | 5605.0 | 0 | Chordata | 5605.0 | 0 | Carnivora | 3673.0 | 1 | Otariidae | 2236.0 | 2 | Homo | 1932.0 | 4 | Homo sapiens | 1932.0 | 4 |
| scf7180000052374 | 14319 | 0.6464 | 0 | 334.3726 | Eukaryota | 34370.0 | 0 | Chordata | 34370.0 | 0 | Artiodactyla | 24033.0 | 2 | Bovidae | 24033.0 | 2 | Ovis | 15654.0 | 3 | Ovis aries | 15654.0 | 3 |
| scf7180000044445 | 15726 | 0.4119 | 0 | 54.299 | Eukaryota | 26820.0 | 0 | Chordata | 26820.0 | 0 | Carnivora | 22175.0 | 2 | Canidae | 18468.0 | 3 | Canis | 18468.0 | 6 | Canis lupus | 18468.0 | 6 |
| scf7180000047383 | 85806 | 0.375 | 0 | 61.6185 | Eukaryota | 9989.0 | 0 | Chordata | 9989.0 | 0 | Carnivora | 9989.0 | 0 | Canidae | 7112.0 | 2 | Canis | 6090.0 | 3 | Canis lupus | 6090.0 | 4 |
| scf7180000053800 | 3554 | 0.5059 | 0 | 26.7247 | Eukaryota | 5802.0 | 0 | Chordata | 5802.0 | 0 | Carnivora | 5802.0 | 0 | Canidae | 5802.0 | 0 | Canis | 3864.0 | 1 | Canis lupus | 3136.0 | 2 |
| scf7180000044719 | 217252 | 0.3721 | 0 | 60.1371 | Eukaryota | 22024.0 | 0 | Chordata | 22024.0 | 0 | Carnivora | 22024.0 | 0 | Canidae | 22024.0 | 0 | Canis | 22024.0 | 0 | Canis lupus | 22024.0 | 0 |
| scf7180000043813 | 113426 | 0.3494 | 0 | 54.951 | Eukaryota | 85679.0 | 0 | Chordata | 85679.0 | 0 | Carnivora | 85679.0 | 0 | Canidae | 85679.0 | 0 | Canis | 85679.0 | 0 | Canis lupus | 85679.0 | 0 |
| scf7180000050519 | 2990 | 0.4331 | 0 | 86.6378 | Eukaryota | 139.0 | 0 | Chordata | 139.0 | 0 | Salmoniformes | 139.0 | 0 | Salmonidae | 139.0 | 0 | Coregonus | 139.0 | 0 | Coregonus sp. 'balchen' | 139.0 | 0 |
| scf7180000043850 | 34135 | 0.5346 | 0 | 50.2865 | Eukaryota | 34981.0 | 0 | Chordata | 34981.0 | 0 | Carnivora | 34981.0 | 0 | Canidae | 34981.0 | 0 | Canis | 34981.0 | 0 | Canis lupus | 34981.0 | 0 |
| scf7180000042426 | 54569 | 0.3862 | 0 | 62.0062 | Eukaryota | 92162.0 | 0 | Chordata | 92162.0 | 0 | Carnivora | 92162.0 | 0 | Canidae | 92162.0 | 0 | Canis | 92162.0 | 0 | Canis lupus | 92162.0 | 0 |
| scf7180000051553 | 10950 | 0.4509 | 0 | 1368.01 | no-hit | 0.0 | 0 | no-hit | 0.0 | 0 | no-hit | 0.0 | 0 | no-hit | 0.0 | 0 | no-hit | 0.0 | 0 | no-hit | 0.0 | 0 |
| scf7180000047392 | 20207 | 0.4392 | 0 | 310.2475 | no-hit | 0.0 | 0 | no-hit | 0.0 | 0 | no-hit | 0.0 | 0 | no-hit | 0.0 | 0 | no-hit | 0.0 | 0 | no-hit | 0.0 | 0 |
| scf7180000043659 | 399433 | 0.4517 | 0 | 57.7159 | Eukaryota | 80187.0 | 0 | Chordata | 80187.0 | 0 | Carnivora | 80187.0 | 0 | Canidae | 80187.0 | 0 | Canis | 64965.0 | 1 | Canis lupus | 64965.0 | 1 |
| scf7180000052721 | 5689 | 0.4959 | 0 | 82.0814 | Eukaryota | 8198.0 | 0 | Chordata | 8198.0 | 0 | Carnivora | 8198.0 | 0 | Canidae | 8198.0 | 0 | Canis | 6708.0 | 1 | Canis lupus | 5732.0 | 2 |
| scf7180000051626 | 21942 | 0.5035 | 0 | 418.5938 | Eukaryota | 9288.0 | 0 | Chordata | 9288.0 | 0 | Carnivora | 9288.0 | 0 | Canidae | 9288.0 | 0 | Canis | 7523.0 | 1 | Canis lupus | 6235.0 | 2 |
| scf7180000051152 | 17781 | 0.4347 | 0 | 61.4434 | Eukaryota | 3387.0 | 0 | Chordata | 3387.0 | 0 | Carnivora | 1510.0 | 2 | Canidae | 1510.0 | 3 | Canis | 1510.0 | 3 | Canis lupus | 1510.0 | 3 |
| scf7180000046464 | 119534 | 0.4392 | 0 | 80.6044 | Eukaryota | 91677.0 | 0 | Chordata | 91677.0 | 0 | Carnivora | 91677.0 | 0 | Canidae | 91677.0 | 0 | Canis | 91677.0 | 0 | Canis lupus | 91677.0 | 0 |
| scf7180000049606 | 9004 | 0.4499 | 0 | 184.1669 | Eukaryota | 571.0 | 0 | Chordata | 571.0 | 0 | Kurtiformes | 571.0 | 0 | Apogonidae | 571.0 | 0 | Sphaeramia | 571.0 | 0 | Sphaeramia orbicularis | 571.0 | 0 |
| scf7180000053325 | 3179 | 0.4281 | 0 | 108.9494 | no-hit | 0.0 | 0 | no-hit | 0.0 | 0 | no-hit | 0.0 | 0 | no-hit | 0.0 | 0 | no-hit | 0.0 | 0 | no-hit | 0.0 | 0 |
| scf7180000052006 | 9722 | 0.4425 | 0 | 88.7289 | Eukaryota | 7422.0 | 0 | Chordata | 6995.0 | 1 | Tetraodontiformes | 5037.0 | 2 | Tetraodontidae | 5037.0 | 2 | Takifugu | 5037.0 | 2 | Takifugu rubripes | 4392.0 | 3 |
| scf7180000049406 | 11683 | 0.4418 | 0 | 573.3126 | Eukaryota | 4594.0 | 0 | Platyhelminthes | 2477.0 | 3 | Polystomatidea | 2477.0 | 3 | Polystomatidae | 2477.0 | 3 | Protopolystoma | 2477.0 | 3 | Protopolystoma xenopodis | 2477.0 | 3 |
| scf7180000053222 | 1738 | 0.454 | 0 | 188.7254 | no-hit | 0.0 | 0 | no-hit | 0.0 | 0 | no-hit | 0.0 | 0 | no-hit | 0.0 | 0 | no-hit | 0.0 | 0 | no-hit | 0.0 | 0 |
| scf7180000045409 | 31790 | 0.4436 | 0 | 1037.4586 | Eukaryota | 3713.0 | 0 | Chordata | 1969.0 | 1 | Polystomatidea | 1744.0 | 2 | Polystomatidae | 1744.0 | 2 | Protopolystoma | 1744.0 | 2 | Protopolystoma xenopodis | 1744.0 | 2 |
| scf7180000043014 | 566799 | 0.4038 | 0 | 60.3672 | Eukaryota | 94986.0 | 0 | Chordata | 94986.0 | 0 | Carnivora | 94986.0 | 0 | Canidae | 94986.0 | 0 | Canis | 94986.0 | 0 | Canis lupus | 94986.0 | 0 |
| scf7180000046606 | 120292 | 0.3487 | 0 | 58.8493 | Eukaryota | 12954.0 | 0 | Chordata | 12954.0 | 0 | Carnivora | 11006.0 | 1 | Canidae | 11006.0 | 1 | Canis | 11006.0 | 2 | Canis lupus | 11006.0 | 2 |
| scf7180000041282 | 145330 | 0.4843 | 0 | 57.0956 | Eukaryota | 32180.0 | 0 | Chordata | 32180.0 | 0 | Carnivora | 32180.0 | 0 | Canidae | 32180.0 | 0 | Canis | 29832.0 | 1 | Canis lupus | 29832.0 | 1 |
| scf7180000046318 | 5179 | 0.4931 | 0 | 32.7848 | Eukaryota | 3294.0 | 0 | Chordata | 3294.0 | 0 | Carnivora | 2848.0 | 1 | Canidae | 2848.0 | 1 | Canis | 2848.0 | 1 | Canis lupus | 2848.0 | 1 |
| scf7180000041479 | 1017698 | 0.4007 | 0 | 60.6625 | Eukaryota | 195141.0 | 0 | Chordata | 195141.0 | 0 | Carnivora | 195141.0 | 0 | Canidae | 195141.0 | 0 | Canis | 195141.0 | 0 | Canis lupus | 195141.0 | 0 |
| scf7180000042954 | 285968 | 0.3687 | 0 | 60.6526 | Eukaryota | 26108.0 | 0 | Chordata | 26108.0 | 0 | Carnivora | 26108.0 | 0 | Canidae | 26108.0 | 0 | Canis | 23070.0 | 1 | Canis lupus | 23070.0 | 1 |
| scf7180000044070 | 344914 | 0.4304 | 0 | 59.9171 | Eukaryota | 72388.0 | 0 | Chordata | 72388.0 | 0 | Carnivora | 72388.0 | 0 | Canidae | 66881.0 | 1 | Canis | 66881.0 | 1 | Canis lupus | 66881.0 | 1 |
| scf7180000042605 | 14852 | 0.3861 | 0 | 39.4041 | Eukaryota | 4468.0 | 0 | Chordata | 4468.0 | 0 | Carnivora | 4468.0 | 0 | Canidae | 4468.0 | 0 | Canis | 4468.0 | 0 | Canis lupus | 4468.0 | 0 |
| scf7180000043841 | 393922 | 0.4222 | 0 | 59.2688 | Eukaryota | 25872.0 | 0 | Chordata | 25872.0 | 0 | Carnivora | 25872.0 | 0 | Canidae | 25872.0 | 0 | Canis | 25872.0 | 0 | Canis lupus | 25872.0 | 0 |
| scf7180000054879 | 17975 | 0.4222 | 0 | 231.8003 | Eukaryota | 63957.0 | 0 | Chordata | 63957.0 | 0 | Carnivora | 63957.0 | 0 | Canidae | 63957.0 | 0 | Canis | 63957.0 | 0 | Canis lupus | 63957.0 | 0 |
| scf7180000045904 | 44365 | 0.4443 | 0 | 72.6428 | Eukaryota | 81706.0 | 0 | Platyhelminthes | 46512.0 | 1 | Strigeidida | 34401.0 | 4 | Schistosomatidae | 34401.0 | 4 | Schistosoma | 34401.0 | 4 | Schistosoma mansoni | 34401.0 | 4 |
| scf7180000044353 | 65414 | 0.5091 | 0 | 57.8046 | Eukaryota | 36924.0 | 0 | Chordata | 36924.0 | 0 | Carnivora | 36924.0 | 0 | Canidae | 36924.0 | 0 | Canis | 31726.0 | 1 | Canis lupus | 31726.0 | 1 |
| scf7180000046533 | 42203 | 0.4428 | 0 | 106.2132 | Eukaryota | 15265.0 | 0 | Chordata | 12225.0 | 3 | Gadiformes | 6464.0 | 7 | Gadidae | 6464.0 | 7 | Gadus | 6464.0 | 7 | Gadus morhua | 6464.0 | 7 |
| scf7180000052110 | 3905 | 0.4551 | 0 | 445.5359 | Eukaryota | 747.0 | 0 | Streptophyta | 484.0 | 1 | Fabales | 484.0 | 1 | Fabaceae | 484.0 | 1 | Cicer | 484.0 | 1 | Cicer arietinum | 484.0 | 1 |
| scf7180000056304 | 264315 | 0.3587 | 0 | 58.4325 | Eukaryota | 54204.0 | 0 | Chordata | 54204.0 | 0 | Carnivora | 54204.0 | 0 | Canidae | 54204.0 | 0 | Canis | 54204.0 | 0 | Canis lupus | 54204.0 | 0 |
| scf7180000044005 | 41005 | 0.3965 | 0 | 54.4995 | Eukaryota | 94900.0 | 0 | Chordata | 94900.0 | 0 | Carnivora | 94900.0 | 0 | Canidae | 94900.0 | 0 | Canis | 94900.0 | 0 | Canis lupus | 94900.0 | 0 |
| scf7180000042818 | 583377 | 0.4121 | 0 | 60.8195 | Eukaryota | 61816.0 | 0 | Chordata | 61816.0 | 0 | Carnivora | 52867.0 | 1 | Canidae | 39692.0 | 2 | Canis | 25014.0 | 4 | Canis lupus | 25014.0 | 5 |
| scf7180000056545 | 204819 | 0.3802 | 0 | 59.8306 | Eukaryota | 11457.0 | 0 | Chordata | 11457.0 | 0 | Carnivora | 11457.0 | 0 | Canidae | 11457.0 | 0 | Canis | 8906.0 | 1 | Canis lupus | 8906.0 | 1 |
| scf7180000056610 | 594025 | 0.3458 | 0 | 59.889 | Eukaryota | 119776.0 | 0 | Chordata | 119776.0 | 0 | Carnivora | 46893.0 | 2 | Bovidae | 41279.0 | 4 | Ovis | 41279.0 | 4 | Rousettus aegyptiacus | 31604.0 | 5 |
| scf7180000046688 | 90214 | 0.3372 | 0 | 60.1971 | Eukaryota | 43568.0 | 0 | Chordata | 43568.0 | 0 | Carnivora | 43568.0 | 0 | Canidae | 31689.0 | 2 | Canis | 25449.0 | 3 | Canis lupus | 25449.0 | 3 |
| scf7180000053498 | 14148 | 0.4167 | 0 | 51.4716 | Eukaryota | 8726.0 | 0 | Chordata | 8726.0 | 0 | Primates | 3697.0 | 4 | Hominidae | 2838.0 | 5 | Homo | 2838.0 | 6 | Homo sapiens | 2838.0 | 6 |
| scf7180000048185 | 52861 | 0.4479 | 0 | 93.2922 | Eukaryota | 11013.0 | 0 | Chordata | 10185.0 | 1 | Carnivora | 9387.0 | 2 | Canidae | 9387.0 | 2 | Canis | 8419.0 | 3 | Canis lupus | 7739.0 | 4 |
| scf7180000047357 | 64068 | 0.4428 | 0 | 215.8532 | Eukaryota | 96793.0 | 0 | Platyhelminthes | 51052.0 | 1 | Strigeidida | 37680.0 | 4 | Schistosomatidae | 37680.0 | 4 | Schistosoma | 37680.0 | 4 | Schistosoma mansoni | 37680.0 | 4 |
| scf7180000045099 | 230565 | 0.4294 | 0 | 60.5398 | Eukaryota | 72043.0 | 0 | Chordata | 72043.0 | 0 | Carnivora | 72043.0 | 0 | Canidae | 72043.0 | 0 | Canis | 72043.0 | 0 | Canis lupus | 72043.0 | 0 |
| scf7180000054787 | 20607 | 0.3983 | 0 | 93.2994 | Eukaryota | 89700.0 | 0 | Chordata | 89700.0 | 0 | Carnivora | 89700.0 | 0 | Canidae | 89700.0 | 0 | Canis | 89700.0 | 0 | Canis lupus | 89700.0 | 0 |
| scf7180000045492 | 126714 | 0.3599 | 0 | 57.5353 | Eukaryota | 91520.0 | 0 | Chordata | 91520.0 | 0 | Carnivora | 91520.0 | 0 | Canidae | 91520.0 | 0 | Canis | 91520.0 | 0 | Canis lupus | 91520.0 | 0 |
| scf7180000048696 | 26001 | 0.4446 | 0 | 250.7905 | Eukaryota | 20797.0 | 0 | Chordata | 11420.0 | 3 | Gadiformes | 7836.0 | 6 | Gadidae | 7836.0 | 6 | Gadus | 7836.0 | 6 | Gadus morhua | 7836.0 | 6 |
| scf7180000040637 | 222576 | 0.3579 | 0 | 57.9316 | Eukaryota | 16234.0 | 0 | Chordata | 16234.0 | 0 | Carnivora | 16234.0 | 0 | Otariidae | 11302.0 | 1 | Canis | 4932.0 | 3 | Canis lupus | 4932.0 | 3 |
| scf7180000053158 | 3647 | 0.4513 | 0 | 310.699 | Eukaryota | 1626.0 | 0 | Echinodermata | 584.0 | 3 | Forcipulatida | 584.0 | 3 | Asteriidae | 584.0 | 3 | Asterias | 584.0 | 3 | Asterias rubens | 584.0 | 3 |
| scf7180000048145 | 96473 | 0.3445 | 0 | 60.2661 | Eukaryota | 8338.0 | 0 | Chordata | 8338.0 | 0 | Carnivora | 8338.0 | 0 | Canidae | 3340.0 | 2 | Canis | 3340.0 | 3 | Canis lupus | 3340.0 | 3 |
| scf7180000050804 | 5698 | 0.4456 | 0 | 104.0096 | Eukaryota | 1018.0 | 0 | Chordata | 1018.0 | 0 | Salmoniformes | 1018.0 | 0 | Salmonidae | 1018.0 | 0 | Salmo | 1018.0 | 0 | Salmo trutta | 1018.0 | 0 |
| scf7180000056572 | 151455 | 0.5283 | 0 | 58.6764 | Eukaryota | 62405.0 | 0 | Chordata | 62405.0 | 0 | Carnivora | 62405.0 | 0 | Canidae | 62405.0 | 0 | Canis | 57511.0 | 1 | Canis lupus | 57511.0 | 1 |
| scf7180000041961 | 839563 | 0.4499 | 0 | 60.6997 | Eukaryota | 66958.0 | 0 | Chordata | 66958.0 | 0 | Carnivora | 66958.0 | 0 | Canidae | 21585.0 | 4 | Canis | 14427.0 | 8 | Canis lupus | 14427.0 | 8 |
| scf7180000051628 | 6039 | 0.4338 | 0 | 63.4494 | no-hit | 0.0 | 0 | no-hit | 0.0 | 0 | no-hit | 0.0 | 0 | no-hit | 0.0 | 0 | no-hit | 0.0 | 0 | no-hit | 0.0 | 0 |
| scf7180000045449 | 250600 | 0.3722 | 0 | 60.0677 | Eukaryota | 22199.0 | 0 | Chordata | 22199.0 | 0 | Carnivora | 19774.0 | 1 | Phocidae | 8772.0 | 3 | Leptonychotes | 8772.0 | 3 | Leptonychotes weddellii | 8772.0 | 3 |
| scf7180000048701 | 5974 | 0.4381 | 0 | 66.5137 | no-hit | 0.0 | 0 | no-hit | 0.0 | 0 | no-hit | 0.0 | 0 | no-hit | 0.0 | 0 | no-hit | 0.0 | 0 | no-hit | 0.0 | 0 |
| scf7180000047065 | 7739 | 0.455 | 0 | 26.6304 | Eukaryota | 4035.0 | 0 | Chordata | 4035.0 | 0 | Carnivora | 4035.0 | 0 | Canidae | 4035.0 | 0 | Canis | 4035.0 | 0 | Canis lupus | 4035.0 | 0 |
| scf7180000041826 | 559468 | 0.3527 | 0 | 60.0703 | Eukaryota | 91663.0 | 0 | Chordata | 91663.0 | 0 | Carnivora | 91663.0 | 0 | Canidae | 91663.0 | 0 | Canis | 91663.0 | 0 | Canis lupus | 91663.0 | 0 |
| scf7180000054467 | 161556 | 0.4621 | 0 | 67.5076 | Eukaryota | 82220.0 | 0 | Chordata | 82220.0 | 0 | Carnivora | 82220.0 | 0 | Canidae | 82220.0 | 0 | Canis | 82220.0 | 0 | Canis lupus | 82220.0 | 0 |
| scf7180000044248 | 313499 | 0.4248 | 0 | 59.4656 | Eukaryota | 55724.0 | 0 | Chordata | 55724.0 | 0 | Carnivora | 55724.0 | 0 | Mustelidae | 39568.0 | 1 | Mustela | 39568.0 | 1 | Mustela putorius | 39568.0 | 1 |
| scf7180000044628 | 226307 | 0.4572 | 0 | 59.6626 | Eukaryota | 57845.0 | 0 | Chordata | 57845.0 | 0 | Carnivora | 57845.0 | 0 | Canidae | 43465.0 | 2 | Canis | 32386.0 | 3 | Canis lupus | 32386.0 | 3 |
| scf7180000045972 | 4330 | 0.4178 | 0 | 29.8379 | Eukaryota | 31543.0 | 0 | Chordata | 31543.0 | 0 | Carnivora | 31543.0 | 0 | Canidae | 31543.0 | 0 | Canis | 31543.0 | 0 | Canis lupus | 31543.0 | 0 |
| scf7180000051469 | 16660 | 0.3924 | 0 | 58.2054 | Eukaryota | 11801.0 | 0 | Chordata | 11801.0 | 0 | Carnivora | 11801.0 | 0 | Canidae | 11801.0 | 0 | Canis | 11801.0 | 0 | Canis lupus | 11801.0 | 0 |
| scf7180000043477 | 543233 | 0.3966 | 0 | 60.6431 | Eukaryota | 75704.0 | 0 | Chordata | 75704.0 | 0 | Carnivora | 75704.0 | 0 | Canidae | 48371.0 | 2 | Canis | 29089.0 | 3 | Canis lupus | 29089.0 | 4 |
| scf7180000047489 | 20633 | 0.4441 | 0 | 53.7828 | Eukaryota | 4814.0 | 0 | Chordata | 3369.0 | 1 | Kurtiformes | 1790.0 | 3 | Apogonidae | 1790.0 | 3 | Sphaeramia | 1790.0 | 4 | Sphaeramia orbicularis | 1790.0 | 4 |
| scf7180000042901 | 7525 | 0.4918 | 0 | 54.7927 | Eukaryota | 31691.0 | 0 | Chordata | 31691.0 | 0 | Carnivora | 31691.0 | 0 | Canidae | 31691.0 | 0 | Canis | 31691.0 | 0 | Canis lupus | 31691.0 | 0 |
| scf7180000040540 | 2539794 | 0.3839 | 0 | 59.8817 | Eukaryota | 167920.0 | 0 | Chordata | 167920.0 | 0 | Carnivora | 167920.0 | 0 | Canidae | 167920.0 | 0 | Canis | 167920.0 | 0 | Canis lupus | 167920.0 | 0 |
| scf7180000050664 | 12371 | 0.4461 | 0 | 74.2729 | no-hit | 0.0 | 0 | no-hit | 0.0 | 0 | no-hit | 0.0 | 0 | no-hit | 0.0 | 0 | no-hit | 0.0 | 0 | no-hit | 0.0 | 0 |
| scf7180000047589 | 23044 | 0.4515 | 0 | 1685.178 | no-hit | 0.0 | 0 | no-hit | 0.0 | 0 | no-hit | 0.0 | 0 | no-hit | 0.0 | 0 | no-hit | 0.0 | 0 | no-hit | 0.0 | 0 |
| scf7180000042336 | 276018 | 0.3382 | 0 | 58.153 | Eukaryota | 50075.0 | 0 | Chordata | 50075.0 | 0 | Carnivora | 50075.0 | 0 | Canidae | 50075.0 | 0 | Canis | 50075.0 | 0 | Canis lupus | 50075.0 | 0 |
| scf7180000048202 | 76090 | 0.4356 | 0 | 62.216 | Eukaryota | 70803.0 | 0 | Chordata | 70803.0 | 0 | Carnivora | 70803.0 | 0 | Canidae | 70803.0 | 0 | Canis | 70803.0 | 0 | Canis lupus | 70803.0 | 0 |
| scf7180000043412 | 458687 | 0.4529 | 0 | 57.5284 | Eukaryota | 64415.0 | 0 | Chordata | 64415.0 | 0 | Carnivora | 64415.0 | 0 | Canidae | 40746.0 | 2 | Canis | 33745.0 | 4 | Canis lupus | 33745.0 | 4 |
| scf7180000045856 | 89071 | 0.3875 | 0 | 56.6326 | Eukaryota | 58589.0 | 0 | Chordata | 58589.0 | 0 | Carnivora | 58589.0 | 0 | Canidae | 58589.0 | 0 | Canis | 58589.0 | 0 | Canis lupus | 58589.0 | 0 |
| scf7180000043520 | 446640 | 0.3516 | 0 | 61.6098 | Eukaryota | 52032.0 | 0 | Chordata | 52032.0 | 0 | Carnivora | 42042.0 | 1 | Canidae | 42042.0 | 1 | Canis | 42042.0 | 1 | Canis lupus | 42042.0 | 1 |
| scf7180000047947 | 12058 | 0.4426 | 0 | 62.2706 | Eukaryota | 9543.0 | 0 | Chordata | 8090.0 | 1 | Kurtiformes | 3893.0 | 3 | Apogonidae | 3893.0 | 3 | Sphaeramia | 3893.0 | 3 | Sphaeramia orbicularis | 3893.0 | 3 |
| scf7180000048302 | 13734 | 0.6625 | 0 | 409.8992 | Eukaryota | 66958.0 | 0 | Chordata | 66958.0 | 0 | Artiodactyla | 27700.0 | 3 | Bovidae | 27700.0 | 3 | Ovis | 27700.0 | 4 | Ovis aries | 27700.0 | 4 |
| scf7180000051105 | 4829 | 0.4543 | 0 | 105.8331 | no-hit | 0.0 | 0 | no-hit | 0.0 | 0 | no-hit | 0.0 | 0 | no-hit | 0.0 | 0 | no-hit | 0.0 | 0 | no-hit | 0.0 | 0 |
| scf7180000045830 | 155411 | 0.4079 | 0 | 59.9522 | Eukaryota | 63393.0 | 0 | Chordata | 63393.0 | 0 | Carnivora | 63393.0 | 0 | Canidae | 63393.0 | 0 | Canis | 63393.0 | 0 | Canis lupus | 63393.0 | 0 |
| scf7180000043785 | 401167 | 0.3564 | 0 | 59.1477 | Eukaryota | 50814.0 | 0 | Chordata | 50814.0 | 0 | Carnivora | 50814.0 | 0 | Canidae | 50814.0 | 0 | Canis | 50814.0 | 0 | Canis lupus | 50814.0 | 0 |
| scf7180000051851 | 2802 | 0.4465 | 0 | 541.9815 | no-hit | 0.0 | 0 | no-hit | 0.0 | 0 | no-hit | 0.0 | 0 | no-hit | 0.0 | 0 | no-hit | 0.0 | 0 | no-hit | 0.0 | 0 |
| scf7180000051901 | 4496 | 0.4448 | 0 | 258.2502 | Eukaryota | 1421.0 | 0 | Platyhelminthes | 1421.0 | 0 | Polystomatidea | 1085.0 | 2 | Polystomatidae | 1085.0 | 2 | Protopolystoma | 1085.0 | 2 | Protopolystoma xenopodis | 1085.0 | 2 |
| scf7180000041032 | 665380 | 0.3888 | 0 | 60.3492 | Eukaryota | 93722.0 | 0 | Chordata | 93722.0 | 0 | Carnivora | 93722.0 | 0 | Canidae | 93722.0 | 0 | Canis | 93722.0 | 0 | Canis lupus | 93722.0 | 0 |
| scf7180000046475 | 18394 | 0.3753 | 0 | 49.6202 | Eukaryota | 3838.0 | 1 | Chordata | 3838.0 | 1 | Carnivora | 2693.0 | 2 | Canidae | 2693.0 | 3 | Canis | 1956.0 | 4 | Canis lupus | 1956.0 | 4 |
| scf7180000047615 | 8524 | 0.4503 | 0 | 159.553 | Eukaryota | 2217.0 | 0 | Chordata | 1500.0 | 2 | Kurtiformes | 1269.0 | 3 | Apogonidae | 1269.0 | 3 | Sphaeramia | 1269.0 | 3 | Sphaeramia orbicularis | 1269.0 | 3 |
| scf7180000046741 | 15811 | 0.4344 | 0 | 241.3653 | no-hit | 0.0 | 0 | no-hit | 0.0 | 0 | no-hit | 0.0 | 0 | no-hit | 0.0 | 0 | no-hit | 0.0 | 0 | no-hit | 0.0 | 0 |
| scf7180000045314 | 15022 | 0.4317 | 0 | 383.7704 | Eukaryota | 89893.0 | 0 | Chordata | 89893.0 | 0 | Carnivora | 89893.0 | 0 | Canidae | 89893.0 | 0 | Canis | 89893.0 | 0 | Canis lupus | 89893.0 | 0 |
| scf7180000042020 | 4640 | 0.4011 | 0 | 26.9471 | Eukaryota | 3366.0 | 0 | Chordata | 3366.0 | 0 | Carnivora | 3366.0 | 0 | Canidae | 3366.0 | 0 | Canis | 3366.0 | 0 | Canis lupus | 3366.0 | 0 |
| scf7180000055641 | 13661 | 0.4372 | 0 | 292.2762 | no-hit | 0.0 | 0 | no-hit | 0.0 | 0 | no-hit | 0.0 | 0 | no-hit | 0.0 | 0 | no-hit | 0.0 | 0 | no-hit | 0.0 | 0 |
| scf7180000055588 | 17547 | 0.4436 | 0 | 107.4862 | Eukaryota | 1348.0 | 0 | Platyhelminthes | 873.0 | 1 | Polystomatidea | 873.0 | 1 | Polystomatidae | 873.0 | 1 | Protopolystoma | 873.0 | 1 | Protopolystoma xenopodis | 873.0 | 1 |
| scf7180000047599 | 100802 | 0.4168 | 0 | 59.3598 | Eukaryota | 9464.0 | 1 | Chordata | 9464.0 | 1 | Primates | 6234.0 | 3 | Hominidae | 4987.0 | 4 | Homo | 4987.0 | 6 | Homo sapiens | 4987.0 | 6 |
| scf7180000051771 | 17654 | 0.4457 | 0 | 57.3382 | no-hit | 0.0 | 0 | no-hit | 0.0 | 0 | no-hit | 0.0 | 0 | no-hit | 0.0 | 0 | no-hit | 0.0 | 0 | no-hit | 0.0 | 0 |
| scf7180000040730 | 1837418 | 0.3589 | 0 | 59.7063 | Eukaryota | 94641.0 | 0 | Chordata | 94641.0 | 0 | Carnivora | 94641.0 | 0 | Canidae | 94641.0 | 0 | Canis | 94641.0 | 0 | Canis lupus | 94641.0 | 0 |
| scf7180000054541 | 68833 | 0.4236 | 0 | 141.3028 | Eukaryota | 116859.0 | 0 | Chordata | 116859.0 | 0 | Carnivora | 116859.0 | 0 | Canidae | 92530.0 | 2 | Canis | 92530.0 | 2 | Canis lupus | 92530.0 | 2 |
| scf7180000041481 | 1043299 | 0.439 | 0 | 60.8768 | Eukaryota | 44126.0 | 0 | Chordata | 44126.0 | 0 | Carnivora | 44126.0 | 0 | Canidae | 44126.0 | 0 | Canis | 38851.0 | 1 | Canis lupus | 38851.0 | 1 |
| scf7180000043006 | 550973 | 0.4199 | 0 | 60.2748 | Eukaryota | 47091.0 | 0 | Chordata | 47091.0 | 0 | Carnivora | 26977.0 | 1 | Canidae | 26977.0 | 2 | Canis | 18594.0 | 4 | Canis lupus | 18594.0 | 4 |
| scf7180000040969 | 281245 | 0.4147 | 0 | 58.5855 | Eukaryota | 53283.0 | 0 | Chordata | 53283.0 | 0 | Carnivora | 53283.0 | 0 | Canidae | 53283.0 | 0 | Canis | 53283.0 | 0 | Canis lupus | 53283.0 | 0 |
| scf7180000050597 | 32930 | 0.4533 | 0 | 65.8566 | Eukaryota | 30300.0 | 0 | Chordata | 30300.0 | 0 | Carnivora | 30300.0 | 0 | Canidae | 30300.0 | 0 | Canis | 30300.0 | 0 | Canis lupus | 30300.0 | 0 |
| scf7180000042161 | 754319 | 0.372 | 0 | 58.8905 | Eukaryota | 57417.0 | 0 | Chordata | 57417.0 | 0 | Carnivora | 57417.0 | 0 | Canidae | 57417.0 | 0 | Canis | 57417.0 | 0 | Canis lupus | 57417.0 | 0 |
| scf7180000050367 | 10594 | 0.4531 | 0 | 892.0093 | Eukaryota | 20115.0 | 0 | Chordata | 19487.0 | 1 | Gadiformes | 12991.0 | 6 | Gadidae | 12991.0 | 6 | Gadus | 12991.0 | 6 | Gadus morhua | 12991.0 | 6 |
| scf7180000045983 | 162744 | 0.6776 | 0 | 54.2465 | Eukaryota | 32217.0 | 0 | Chordata | 32217.0 | 0 | Carnivora | 32217.0 | 0 | Canidae | 32217.0 | 0 | Canis | 24376.0 | 1 | Canis lupus | 24376.0 | 1 |
| scf7180000055831 | 12584 | 0.4361 | 0 | 66.0884 | Eukaryota | 3252.0 | 0 | Platyhelminthes | 2020.0 | 1 | Polystomatidea | 2020.0 | 3 | Polystomatidae | 2020.0 | 3 | Protopolystoma | 2020.0 | 3 | Protopolystoma xenopodis | 2020.0 | 3 |
| scf7180000043450 | 133320 | 0.5354 | 0 | 54.7293 | Eukaryota | 35760.0 | 0 | Chordata | 35760.0 | 0 | Carnivora | 35760.0 | 0 | Canidae | 35760.0 | 0 | Canis | 35760.0 | 0 | Canis lupus | 35760.0 | 0 |
| scf7180000045417 | 221165 | 0.3951 | 0 | 59.1184 | Eukaryota | 95123.0 | 0 | Chordata | 95123.0 | 0 | Carnivora | 95123.0 | 0 | Canidae | 95123.0 | 0 | Canis | 95123.0 | 0 | Canis lupus | 95123.0 | 0 |
| scf7180000050023 | 17105 | 0.4427 | 0 | 174.4244 | Eukaryota | 3148.0 | 0 | Chordata | 3148.0 | 0 | Carnivora | 3148.0 | 0 | Canidae | 3148.0 | 0 | Canis | 3148.0 | 0 | Canis lupus | 3148.0 | 0 |
| scf7180000050078 | 7409 | 0.4478 | 0 | 1775.4637 | Eukaryota | 1179.0 | 0 | Echinodermata | 688.0 | 2 | Forcipulatida | 688.0 | 2 | Asteriidae | 688.0 | 2 | Asterias | 688.0 | 2 | Asterias rubens | 688.0 | 2 |
| scf7180000045271 | 262762 | 0.463 | 100 | 58.8151 | Eukaryota | 41333.0 | 0 | Chordata | 41333.0 | 0 | Carnivora | 41333.0 | 0 | Canidae | 41333.0 | 0 | Canis | 41333.0 | 0 | Canis lupus | 41333.0 | 0 |
| scf7180000045496 | 208012 | 0.339 | 0 | 58.9335 | Eukaryota | 71056.0 | 0 | Chordata | 71056.0 | 0 | Carnivora | 71056.0 | 0 | Canidae | 71056.0 | 0 | Canis | 71056.0 | 0 | Canis lupus | 71056.0 | 0 |
| scf7180000040728 | 1947400 | 0.3715 | 0 | 61.1768 | Eukaryota | 130102.0 | 0 | Chordata | 130102.0 | 0 | Carnivora | 130102.0 | 0 | Canidae | 130102.0 | 0 | Vulpes | 84210.0 | 1 | Vulpes vulpes | 84210.0 | 1 |
| scf7180000047448 | 176604 | 0.4465 | 0 | 58.4365 | Eukaryota | 11550.0 | 0 | Chordata | 11550.0 | 0 | Carnivora | 11550.0 | 0 | Canidae | 11550.0 | 0 | Canis | 11550.0 | 0 | Canis lupus | 11550.0 | 0 |
| scf7180000048648 | 5661 | 0.4393 | 0 | 58.7685 | Eukaryota | 11268.0 | 0 | Chordata | 11268.0 | 0 | Carnivora | 11268.0 | 0 | Otariidae | 4264.0 | 3 | Canis | 2574.0 | 5 | Canis lupus | 2574.0 | 5 |
| scf7180000042042 | 19751 | 0.4389 | 0 | 59.4387 | Eukaryota | 49767.0 | 0 | Chordata | 49767.0 | 0 | Carnivora | 49767.0 | 0 | Canidae | 49767.0 | 0 | Canis | 49767.0 | 0 | Canis lupus | 49767.0 | 0 |
| scf7180000042812 | 77704 | 0.4192 | 0 | 58.1877 | Eukaryota | 75187.0 | 0 | Chordata | 75187.0 | 0 | Carnivora | 75187.0 | 0 | Canidae | 75187.0 | 0 | Canis | 75187.0 | 0 | Canis lupus | 75187.0 | 0 |
| scf7180000044442 | 302104 | 0.385 | 0 | 75.6801 | Eukaryota | 89328.0 | 0 | Chordata | 89328.0 | 0 | Carnivora | 89328.0 | 0 | Canidae | 89328.0 | 0 | Canis | 89328.0 | 0 | Canis lupus | 89328.0 | 0 |
| scf7180000041133 | 183872 | 0.4705 | 0 | 72.0018 | Eukaryota | 115246.0 | 0 | Chordata | 115246.0 | 0 | Carnivora | 115246.0 | 0 | Canidae | 115246.0 | 0 | Canis | 115246.0 | 0 | Canis lupus | 115246.0 | 0 |
| scf7180000042516 | 688181 | 0.4562 | 0 | 59.3922 | Eukaryota | 41818.0 | 0 | Chordata | 41818.0 | 0 | Carnivora | 33554.0 | 1 | Canidae | 33554.0 | 1 | Canis | 33554.0 | 1 | Canis lupus | 33554.0 | 1 |
| scf7180000046493 | 10902 | 0.4329 | 0 | 78.1585 | Eukaryota | 3115.0 | 0 | Platyhelminthes | 1187.0 | 3 | Polystomatidea | 1187.0 | 3 | Polystomatidae | 1187.0 | 3 | Protopolystoma | 1187.0 | 3 | Protopolystoma xenopodis | 1187.0 | 3 |
| scf7180000046429 | 7801 | 0.4571 | 0 | 149.0057 | Eukaryota | 3412.0 | 0 | Chordata | 3412.0 | 0 | Carnivora | 1942.0 | 2 | Canidae | 1942.0 | 2 | Canis | 1942.0 | 2 | Canis lupus | 1942.0 | 2 |
| scf7180000051310 | 6655 | 0.5077 | 0 | 124.9477 | Eukaryota | 7910.0 | 0 | Chordata | 7910.0 | 0 | Carnivora | 7910.0 | 0 | Canidae | 7910.0 | 0 | Canis | 6469.0 | 1 | Canis lupus | 5417.0 | 2 |
| scf7180000052190 | 3533 | 0.4464 | 0 | 355.4152 | no-hit | 0.0 | 0 | no-hit | 0.0 | 0 | no-hit | 0.0 | 0 | no-hit | 0.0 | 0 | no-hit | 0.0 | 0 | no-hit | 0.0 | 0 |
| scf7180000048511 | 3860 | 0.435 | 0 | 93.4021 | no-hit | 0.0 | 0 | no-hit | 0.0 | 0 | no-hit | 0.0 | 0 | no-hit | 0.0 | 0 | no-hit | 0.0 | 0 | no-hit | 0.0 | 0 |
| scf7180000056466 | 548057 | 0.4257 | 0 | 60.0001 | Eukaryota | 80265.0 | 0 | Chordata | 80265.0 | 0 | Carnivora | 80265.0 | 0 | Canidae | 80265.0 | 0 | Canis | 80265.0 | 0 | Canis lupus | 80265.0 | 0 |
| scf7180000046247 | 61850 | 0.5377 | 0 | 68.3908 | Eukaryota | 23858.0 | 0 | Chordata | 23858.0 | 0 | Carnivora | 22471.0 | 1 | Canidae | 22471.0 | 1 | Canis | 22471.0 | 1 | Canis lupus | 22471.0 | 1 |
| scf7180000051586 | 1410 | 0.4369 | 0 | 449.199 | no-hit | 0.0 | 0 | no-hit | 0.0 | 0 | no-hit | 0.0 | 0 | no-hit | 0.0 | 0 | no-hit | 0.0 | 0 | no-hit | 0.0 | 0 |
| scf7180000047704 | 4824 | 0.4442 | 0 | 107.7499 | no-hit | 0.0 | 0 | no-hit | 0.0 | 0 | no-hit | 0.0 | 0 | no-hit | 0.0 | 0 | no-hit | 0.0 | 0 | no-hit | 0.0 | 0 |
| scf7180000043276 | 172144 | 0.3567 | 0 | 56.0013 | Eukaryota | 92041.0 | 0 | Chordata | 92041.0 | 0 | Carnivora | 92041.0 | 0 | Canidae | 92041.0 | 0 | Canis | 92041.0 | 0 | Canis lupus | 92041.0 | 0 |
| scf7180000049216 | 52220 | 0.4411 | 0 | 65.0196 | Eukaryota | 10057.0 | 0 | Chordata | 7944.0 | 1 | Tetraodontiformes | 4845.0 | 4 | Tetraodontidae | 4845.0 | 4 | Takifugu | 4845.0 | 4 | Takifugu rubripes | 4845.0 | 4 |
| scf7180000040720 | 9086 | 0.3903 | 0 | 31.3888 | Eukaryota | 2356.0 | 1 | Chordata | 2356.0 | 1 | Primates | 1254.0 | 2 | Hominidae | 1254.0 | 3 | Canis | 708.0 | 4 | Canis lupus | 708.0 | 4 |
| scf7180000048051 | 10632 | 0.4402 | 0 | 62.0318 | Eukaryota | 3587.0 | 0 | Chordata | 3195.0 | 1 | Kurtiformes | 2862.0 | 2 | Apogonidae | 2862.0 | 2 | Sphaeramia | 2862.0 | 2 | Sphaeramia orbicularis | 2862.0 | 2 |
| scf7180000056398 | 1027455 | 0.4097 | 0 | 59.9898 | Eukaryota | 75776.0 | 0 | Chordata | 75776.0 | 0 | Carnivora | 75776.0 | 0 | Canidae | 75776.0 | 0 | Canis | 70184.0 | 1 | Canis lupus | 70184.0 | 1 |
| scf7180000045822 | 13968 | 0.4466 | 0 | 163.4839 | Eukaryota | 38273.0 | 0 | Chordata | 26851.0 | 1 | Cyclophyllidea | 7649.0 | 8 | Hymenolepididae | 7649.0 | 8 | Hymenolepis | 7649.0 | 8 | Hymenolepis microstoma | 7649.0 | 8 |
| scf7180000043835 | 302992 | 0.4151 | 0 | 59.4013 | Eukaryota | 55479.0 | 0 | Chordata | 55479.0 | 0 | Carnivora | 55479.0 | 0 | Canidae | 55479.0 | 0 | Canis | 55479.0 | 0 | Canis lupus | 55479.0 | 0 |
| scf7180000054452 | 16038 | 0.4431 | 0 | 166.3072 | no-hit | 0.0 | 0 | no-hit | 0.0 | 0 | no-hit | 0.0 | 0 | no-hit | 0.0 | 0 | no-hit | 0.0 | 0 | no-hit | 0.0 | 0 |
| scf7180000043587 | 423168 | 0.4779 | 0 | 58.1329 | Eukaryota | 39507.0 | 0 | Chordata | 39507.0 | 0 | Carnivora | 28404.0 | 3 | Canidae | 28404.0 | 3 | Canis | 28404.0 | 3 | Canis lupus | 28404.0 | 3 |
| scf7180000044714 | 70590 | 0.3941 | 0 | 88.4074 | Eukaryota | 47683.0 | 0 | Chordata | 47683.0 | 0 | Carnivora | 47683.0 | 0 | Canidae | 47683.0 | 0 | Canis | 47683.0 | 0 | Canis lupus | 47683.0 | 0 |
| scf7180000051011 | 11931 | 0.503 | 0 | 734.1218 | Eukaryota | 9196.0 | 0 | Chordata | 9196.0 | 0 | Carnivora | 9196.0 | 0 | Canidae | 9196.0 | 0 | Canis | 7464.0 | 1 | Canis lupus | 6236.0 | 2 |
| scf7180000056541 | 174741 | 0.3672 | 0 | 59.787 | Eukaryota | 59066.0 | 0 | Chordata | 59066.0 | 0 | Carnivora | 59066.0 | 0 | Canidae | 59066.0 | 0 | Canis | 59066.0 | 0 | Canis lupus | 59066.0 | 0 |
| scf7180000048582 | 41620 | 0.454 | 0 | 56.8797 | Eukaryota | 3304.0 | 0 | Chordata | 3304.0 | 0 | Carnivora | 2105.0 | 2 | Canidae | 2105.0 | 2 | Canis | 2105.0 | 3 | Canis lupus | 2105.0 | 3 |
| scf7180000041898 | 889545 | 0.4652 | 0 | 58.322 | Eukaryota | 54648.0 | 0 | Chordata | 54648.0 | 0 | Carnivora | 54648.0 | 0 | Canidae | 54648.0 | 0 | Canis | 54648.0 | 0 | Canis lupus | 54648.0 | 0 |
| scf7180000056409 | 547395 | 0.491 | 0 | 58.7568 | Eukaryota | 96558.0 | 0 | Chordata | 96558.0 | 0 | Carnivora | 96558.0 | 0 | Canidae | 96558.0 | 0 | Canis | 96558.0 | 0 | Canis lupus | 96558.0 | 0 |
| scf7180000044410 | 10794 | 0.4493 | 0 | 253.1062 | Eukaryota | 7381.0 | 0 | Chordata | 6563.0 | 1 | Kurtiformes | 5239.0 | 2 | Apogonidae | 5239.0 | 2 | Sphaeramia | 5239.0 | 2 | Sphaeramia orbicularis | 5239.0 | 2 |
| scf7180000046335 | 129604 | 0.3602 | 0 | 59.8513 | Eukaryota | 62653.0 | 0 | Chordata | 62653.0 | 0 | Carnivora | 62653.0 | 0 | Canidae | 62653.0 | 0 | Canis | 62653.0 | 0 | Canis lupus | 62653.0 | 0 |
| scf7180000040933 | 609218 | 0.3852 | 0 | 60.9143 | Eukaryota | 60422.0 | 0 | Chordata | 60422.0 | 0 | Carnivora | 60422.0 | 0 | Canidae | 60422.0 | 0 | Canis | 36234.0 | 1 | Canis lupus | 36234.0 | 1 |
| scf7180000042287 | 556625 | 0.3941 | 0 | 60.3386 | Eukaryota | 51859.0 | 0 | Chordata | 51859.0 | 0 | Carnivora | 51859.0 | 0 | Canidae | 51859.0 | 0 | Canis | 51859.0 | 0 | Canis lupus | 51859.0 | 0 |
| scf7180000045745 | 190263 | 0.4214 | 0 | 59.0275 | Eukaryota | 47112.0 | 0 | Chordata | 47112.0 | 0 | Carnivora | 47112.0 | 0 | Canidae | 47112.0 | 0 | Canis | 42275.0 | 1 | Canis lupus | 42275.0 | 1 |
| scf7180000055039 | 16832 | 0.4755 | 0 | 266.6106 | Eukaryota | 9447.0 | 0 | Chordata | 9447.0 | 0 | Carnivora | 9447.0 | 0 | Canidae | 9447.0 | 0 | Canis | 5691.0 | 1 | Canis lupus | 5691.0 | 1 |
| scf7180000041685 | 74749 | 0.3975 | 0 | 57.9589 | Eukaryota | 15825.0 | 1 | Chordata | 15825.0 | 1 | Carnivora | 14116.0 | 2 | Canidae | 14116.0 | 2 | Canis | 14116.0 | 2 | Canis lupus | 14116.0 | 2 |
| scf7180000052588 | 14707 | 0.3778 | 0 | 60.954 | Eukaryota | 3532.0 | 1 | Chordata | 3532.0 | 1 | Carnivora | 1313.0 | 4 | Canidae | 1313.0 | 4 | Canis | 1313.0 | 4 | Canis lupus | 1313.0 | 4 |
| scf7180000042894 | 617733 | 0.3649 | 0 | 58.6561 | Eukaryota | 66246.0 | 0 | Chordata | 66246.0 | 0 | Carnivora | 66246.0 | 0 | Canidae | 66246.0 | 0 | Canis | 66246.0 | 0 | Canis lupus | 66246.0 | 0 |
| scf7180000053791 | 3452 | 0.5041 | 0 | 303.4872 | Eukaryota | 6249.0 | 0 | Chordata | 6249.0 | 0 | Carnivora | 6249.0 | 0 | Canidae | 6249.0 | 0 | Canis | 3915.0 | 1 | Canis lupus | 2911.0 | 2 |
| scf7180000051410 | 8885 | 0.4506 | 0 | 1631.334 | Eukaryota | 17598.0 | 0 | Chordata | 15294.0 | 2 | Gadiformes | 9592.0 | 7 | Gadidae | 9592.0 | 7 | Gadus | 9592.0 | 7 | Gadus morhua | 9592.0 | 7 |
| scf7180000051875 | 7540 | 0.4497 | 0 | 1771.6739 | Eukaryota | 1645.0 | 0 | Chordata | 1340.0 | 2 | Salmoniformes | 1060.0 | 4 | Salmonidae | 1060.0 | 4 | Salmo | 810.0 | 5 | Salmo trutta | 810.0 | 5 |
| scf7180000054679 | 142354 | 0.44 | 0 | 148.3548 | Eukaryota | 72870.0 | 0 | Platyhelminthes | 47103.0 | 1 | Strigeidida | 35369.0 | 5 | Schistosomatidae | 35369.0 | 5 | Schistosoma | 35369.0 | 5 | Schistosoma mansoni | 35369.0 | 5 |
| scf7180000052094 | 3167 | 0.4373 | 0 | 253.4393 | no-hit | 0.0 | 0 | no-hit | 0.0 | 0 | no-hit | 0.0 | 0 | no-hit | 0.0 | 0 | no-hit | 0.0 | 0 | no-hit | 0.0 | 0 |
| scf7180000056479 | 330377 | 0.4061 | 0 | 60.104 | Eukaryota | 77724.0 | 0 | Chordata | 77724.0 | 0 | Carnivora | 77724.0 | 0 | Canidae | 77724.0 | 0 | Canis | 77724.0 | 0 | Canis lupus | 77724.0 | 0 |
| scf7180000042201 | 683473 | 0.3985 | 0 | 59.8809 | Eukaryota | 67646.0 | 0 | Chordata | 67646.0 | 0 | Carnivora | 67646.0 | 0 | Canidae | 67646.0 | 0 | Canis | 57978.0 | 1 | Canis lupus | 57978.0 | 1 |
| scf7180000049478 | 5532 | 0.4349 | 0 | 257.2781 | no-hit | 0.0 | 0 | no-hit | 0.0 | 0 | no-hit | 0.0 | 0 | no-hit | 0.0 | 0 | no-hit | 0.0 | 0 | no-hit | 0.0 | 0 |
| scf7180000043827 | 340883 | 0.3933 | 0 | 62.2928 | Eukaryota | 94157.0 | 0 | Chordata | 94157.0 | 0 | Carnivora | 94157.0 | 0 | Canidae | 94157.0 | 0 | Canis | 94157.0 | 0 | Canis lupus | 94157.0 | 0 |
| scf7180000048895 | 11187 | 0.4403 | 0 | 86.3904 | no-hit | 0.0 | 0 | no-hit | 0.0 | 0 | no-hit | 0.0 | 0 | no-hit | 0.0 | 0 | no-hit | 0.0 | 0 | no-hit | 0.0 | 0 |
| scf7180000053944 | 6385 | 0.5045 | 0 | 56.9749 | Eukaryota | 6753.0 | 0 | Chordata | 6753.0 | 0 | Carnivora | 6753.0 | 0 | Canidae | 6753.0 | 0 | Canis | 6276.0 | 1 | Canis lupus | 6276.0 | 1 |
| scf7180000049456 | 4755 | 0.437 | 0 | 107.5622 | Eukaryota | 2707.0 | 0 | Chordata | 1536.0 | 1 | Salmoniformes | 1536.0 | 1 | Salmonidae | 1536.0 | 1 | Protopolystoma | 1171.0 | 2 | Protopolystoma xenopodis | 1171.0 | 2 |
| scf7180000049150 | 6969 | 0.4391 | 0 | 99.6961 | Eukaryota | 19127.0 | 1 | Chordata | 16914.0 | 2 | Gadiformes | 6834.0 | 6 | Gadidae | 6834.0 | 6 | Gadus | 6834.0 | 6 | Gadus morhua | 6834.0 | 6 |
| scf7180000043341 | 292851 | 0.3629 | 0 | 58.1234 | Eukaryota | 79941.0 | 0 | Chordata | 79941.0 | 0 | Carnivora | 79941.0 | 0 | Canidae | 79941.0 | 0 | Canis | 79941.0 | 0 | Canis lupus | 79941.0 | 0 |
| scf7180000050211 | 15042 | 0.4556 | 0 | 986.3442 | Eukaryota | 33117.0 | 0 | Chordata | 33117.0 | 0 | Carnivora | 33117.0 | 0 | Canidae | 33117.0 | 0 | Canis | 33117.0 | 0 | Canis lupus | 33117.0 | 0 |
| scf7180000041394 | 1125386 | 0.3911 | 0 | 60.205 | Eukaryota | 101802.0 | 0 | Chordata | 101802.0 | 0 | Carnivora | 101802.0 | 0 | Canidae | 101802.0 | 0 | Canis | 101802.0 | 0 | Canis lupus | 101802.0 | 0 |
| scf7180000040558 | 109335 | 0.4584 | 0 | 57.0333 | Eukaryota | 42119.0 | 0 | Chordata | 42119.0 | 0 | Carnivora | 42119.0 | 0 | Canidae | 14404.0 | 4 | Canis | 9617.0 | 8 | Canis lupus | 9617.0 | 8 |
| scf7180000052189 | 5849 | 0.4491 | 0 | 675.1476 | no-hit | 0.0 | 0 | no-hit | 0.0 | 0 | no-hit | 0.0 | 0 | no-hit | 0.0 | 0 | no-hit | 0.0 | 0 | no-hit | 0.0 | 0 |

| | | | | | | | | | | | | | | | | | | | | | | |
|---|---|---|---|---|---|---|---|---|---|---|---|---|---|---|---|---|---|---|---|---|---|---|
| scf7180000053203 | 3713 | 0.5009 | 0 | 60.9371 | Eukaryota | 5721.0 | 0 | Chordata | 5721.0 | 0 | Carnivora | 5721.0 | 0 | Canidae | 5721.0 | 0 | Canis | 4996.0 | 1 | Canis lupus | 4996.0 | 1 |
| scf7180000048834 | 13535 | 0.4454 | 0 | 1680.8429 | Eukaryota | 565.0 | 0 | Platyhelminthes | 235.0 | 2 | Polystomatidea | 235.0 | 2 | Polystomatidae | 235.0 | 2 | Protopolystoma | 235.0 | 2 | Protopolystoma xenopodis | 235.0 | 2 |
| scf7180000047185 | 5635 | 0.3681 | 0 | 37.3818 | Eukaryota | 12517.0 | 0 | Chordata | 12517.0 | 0 | Carnivora | 12517.0 | 0 | Canidae | 12517.0 | 0 | Canis | 12517.0 | 0 | Canis lupus | 12517.0 | 0 |
| scf7180000043220 | 66890 | 0.3934 | 0 | 59.4681 | Eukaryota | 92807.0 | 0 | Chordata | 92807.0 | 0 | Carnivora | 92807.0 | 0 | Canidae | 92807.0 | 0 | Canis | 92807.0 | 0 | Canis lupus | 92807.0 | 0 |
| scf7180000041710 | 26297 | 0.4269 | 0 | 52.1683 | Eukaryota | 41031.0 | 0 | Chordata | 41031.0 | 0 | Carnivora | 41031.0 | 0 | Canidae | 41031.0 | 0 | Canis | 41031.0 | 0 | Canis lupus | 41031.0 | 0 |
| scf7180000053096 | 3206 | 0.4451 | 0 | 345.8481 | Eukaryota | 2821.0 | 0 | Platyhelminthes | 1380.0 | 3 | Polystomatidea | 1380.0 | 3 | Polystomatidae | 1380.0 | 3 | Protopolystoma | 1380.0 | 3 | Protopolystoma xenopodis | 1380.0 | 3 |
| scf7180000045837 | 208646 | 0.4556 | 0 | 60.5248 | Eukaryota | 60250.0 | 0 | Chordata | 60250.0 | 0 | Carnivora | 60250.0 | 0 | Canidae | 60250.0 | 0 | Canis | 60250.0 | 0 | Canis lupus | 60250.0 | 0 |
| scf7180000049808 | 30541 | 0.4539 | 0 | 56.6897 | Eukaryota | 31143.0 | 0 | Chordata | 31143.0 | 0 | Carnivora | 31143.0 | 0 | Canidae | 31143.0 | 0 | Canis | 31143.0 | 0 | Canis lupus | 31143.0 | 0 |
| scf7180000045802 | 170200 | 0.5504 | 0 | 57.715 | Eukaryota | 35610.0 | 0 | Chordata | 35610.0 | 0 | Carnivora | 35610.0 | 0 | Canidae | 35610.0 | 0 | Canis | 32725.0 | 1 | Canis lupus | 32725.0 | 1 |
| scf7180000049012 | 12753 | 0.3829 | 0 | 91.8813 | Eukaryota | 55582.0 | 0 | Chordata | 55582.0 | 0 | Carnivora | 55582.0 | 0 | Canidae | 55582.0 | 0 | Canis | 55582.0 | 0 | Canis lupus | 55582.0 | 0 |
| scf7180000040930 | 636201 | 0.3821 | 0 | 58.4708 | Eukaryota | 41731.0 | 0 | Chordata | 41731.0 | 0 | Carnivora | 41731.0 | 0 | Canidae | 34981.0 | 1 | Canis | 34981.0 | 1 | Canis lupus | 34981.0 | 1 |
| scf7180000055745 | 6029 | 0.4404 | 0 | 230.8469 | Eukaryota | 974.0 | 0 | Platyhelminthes | 568.0 | 1 | Polystomatidea | 568.0 | 1 | Polystomatidae | 568.0 | 1 | Protopolystoma | 568.0 | 2 | Protopolystoma xenopodis | 568.0 | 2 |
| scf7180000048608 | 4139 | 0.4501 | 0 | 753.1485 | Eukaryota | 1184.0 | 0 | Platyhelminthes | 419.0 | 2 | Polystomatidea | 419.0 | 2 | Polystomatidae | 419.0 | 2 | Protopolystoma | 419.0 | 2 | Protopolystoma xenopodis | 419.0 | 2 |
| scf7180000044512 | 239033 | 0.3947 | 0 | 59.7853 | Eukaryota | 17646.0 | 0 | Chordata | 17646.0 | 0 | Carnivora | 17646.0 | 0 | Canidae | 17646.0 | 0 | Canis | 17646.0 | 0 | Canis lupus | 17646.0 | 0 |
| scf7180000042441 | 687727 | 0.3997 | 0 | 60.9604 | Eukaryota | 57455.0 | 0 | Chordata | 57455.0 | 0 | Carnivora | 57455.0 | 0 | Canidae | 52899.0 | 1 | Canis | 41031.0 | 2 | Canis lupus | 41031.0 | 2 |
| scf7180000055459 | 29399 | 0.3456 | 0 | 139.8631 | Eukaryota | 7040.0 | 1 | Chordata | 7040.0 | 1 | Carnivora | 3559.0 | 3 | Canidae | 3559.0 | 3 | Vulpes | 2228.0 | 5 | Vulpes vulpes | 2228.0 | 5 |
| scf7180000054750 | 23962 | 0.4346 | 0 | 134.8601 | Eukaryota | 23727.0 | 0 | Chordata | 23727.0 | 0 | Carnivora | 23727.0 | 0 | Canidae | 23727.0 | 0 | Canis | 23727.0 | 0 | Canis lupus | 23727.0 | 0 |
| scf7180000044961 | 20545 | 0.4102 | 0 | 389.5417 | Eukaryota | 91122.0 | 0 | Chordata | 91122.0 | 0 | Carnivora | 91122.0 | 0 | Canidae | 91122.0 | 0 | Canis | 91122.0 | 0 | Canis lupus | 91122.0 | 0 |
| scf7180000046230 | 38389 | 0.4434 | 0 | 147.6076 | Eukaryota | 6466.0 | 0 | Chordata | 4635.0 | 1 | Tetraodontiformes | 2645.0 | 4 | Tetraodontidae | 2645.0 | 4 | Takifugu | 2645.0 | 4 | Takifugu rubripes | 2645.0 | 4 |
| scf7180000045044 | 269184 | 0.3389 | 0 | 59.6715 | Eukaryota | 57189.0 | 0 | Chordata | 57189.0 | 0 | Carnivora | 43843.0 | 1 | Canidae | 26437.0 | 3 | Canis | 26437.0 | 3 | Canis lupus | 26437.0 | 3 |
| scf7180000055391 | 15165 | 0.4196 | 0 | 138.2939 | Eukaryota | 6049.0 | 0 | Chordata | 6049.0 | 0 | Carnivora | 6049.0 | 0 | Felidae | 1971.0 | 5 | Felis | 1971.0 | 7 | Felis catus | 1971.0 | 7 |
| scf7180000052969 | 4643 | 0.4491 | 0 | 1091.0885 | Eukaryota | 1374.0 | 0 | Echinodermata | 817.0 | 2 | Forcipulatida | 817.0 | 2 | Asteriidae | 817.0 | 2 | Asterias | 817.0 | 2 | Asterias rubens | 817.0 | 2 |
| scf7180000046316 | 5547 | 0.4584 | 0 | 85.6135 | Eukaryota | 1931.0 | 0 | Chordata | 1313.0 | 1 | Tetraodontiformes | 1100.0 | 2 | Tetraodontidae | 1100.0 | 2 | Takifugu | 1100.0 | 2 | Takifugu rubripes | 1100.0 | 2 |
| scf7180000049568 | 8941 | 0.383 | 0 | 99.6372 | Eukaryota | 68547.0 | 0 | Chordata | 68547.0 | 0 | Carnivora | 68547.0 | 0 | Canidae | 68547.0 | 0 | Canis | 68547.0 | 0 | Canis lupus | 68547.0 | 0 |
| scf7180000045045 | 76008 | 0.3896 | 0 | 58.6336 | Eukaryota | 8703.0 | 0 | Chordata | 8703.0 | 0 | Carnivora | 8703.0 | 0 | Canidae | 8703.0 | 0 | Canis | 8703.0 | 0 | Canis lupus | 8703.0 | 0 |
| scf7180000046468 | 138580 | 0.4068 | 0 | 66.4074 | Eukaryota | 50094.0 | 0 | Chordata | 50094.0 | 0 | Carnivora | 50094.0 | 0 | Canidae | 50094.0 | 0 | Canis | 50094.0 | 0 | Canis lupus | 50094.0 | 0 |
| scf7180000051825 | 17658 | 0.3705 | 0 | 30.9173 | Eukaryota | 19139.0 | 0 | Chordata | 19139.0 | 0 | Carnivora | 19139.0 | 0 | Canidae | 19139.0 | 0 | Canis | 19139.0 | 0 | Canis lupus | 19139.0 | 0 |
| scf7180000053829 | 3027 | 0.4536 | 0 | 366.0417 | no-hit | 0.0 | 0 | no-hit | 0.0 | 0 | no-hit | 0.0 | 0 | no-hit | 0.0 | 0 | no-hit | 0.0 | 0 | no-hit | 0.0 | 0 |
| scf7180000047604 | 26751 | 0.4467 | 0 | 164.4857 | Eukaryota | 4652.0 | 0 | Chordata | 3675.0 | 1 | Kurtiformes | 1802.0 | 5 | Apogonidae | 1802.0 | 5 | Sphaeramia | 1802.0 | 5 | Sphaeramia orbicularis | 1802.0 | 5 |
| scf7180000054935 | 20773 | 0.4453 | 0 | 301.2807 | Eukaryota | 7031.0 | 0 | Chordata | 7031.0 | 0 | Tetraodontiformes | 4741.0 | 2 | Tetraodontidae | 4741.0 | 2 | Takifugu | 4741.0 | 2 | Takifugu rubripes | 4098.0 | 3 |
| scf7180000045432 | 6262 | 0.4487 | 0 | 46.8326 | Eukaryota | 7221.0 | 0 | Chordata | 7221.0 | 0 | Carnivora | 5813.0 | 1 | Canidae | 5813.0 | 1 | Canis | 5813.0 | 1 | Canis lupus | 5813.0 | 1 |
| scf7180000052911 | 6050 | 0.4506 | 0 | 1009.2802 | Eukaryota | 2212.0 | 0 | Platyhelminthes | 1427.0 | 1 | Polystomatidea | 1427.0 | 1 | Polystomatidae | 1427.0 | 1 | Protopolystoma | 1427.0 | 1 | Protopolystoma xenopodis | 1427.0 | 1 |
| scf7180000050135 | 4400 | 0.4377 | 0 | 58.8743 | Eukaryota | 244.0 | 0 | Chordata | 244.0 | 0 | Kurtiformes | 244.0 | 0 | Apogonidae | 244.0 | 0 | Sphaeramia | 244.0 | 0 | Sphaeramia orbicularis | 244.0 | 0 |
| scf7180000042074 | 86190 | 0.4385 | 0 | 56.6281 | Eukaryota | 84489.0 | 0 | Chordata | 84489.0 | 0 | Carnivora | 84489.0 | 0 | Canidae | 82082.0 | 1 | Canis | 82082.0 | 1 | Canis lupus | 82082.0 | 1 |
| scf7180000048368 | 6429 | 0.4365 | 0 | 215.3912 | Eukaryota | 3138.0 | 0 | Chordata | 3138.0 | 0 | Carnivora | 3138.0 | 0 | Canidae | 3138.0 | 0 | Canis | 3138.0 | 0 | Canis lupus | 3138.0 | 0 |
| scf7180000049165 | 9706 | 0.436 | 0 | 102.5751 | no-hit | 0.0 | 0 | no-hit | 0.0 | 0 | no-hit | 0.0 | 0 | no-hit | 0.0 | 0 | no-hit | 0.0 | 0 | no-hit | 0.0 | 0 |
| scf7180000050976 | 19977 | 0.5034 | 0 | 495.6933 | Eukaryota | 9322.0 | 0 | Chordata | 9322.0 | 0 | Carnivora | 9322.0 | 0 | Canidae | 9322.0 | 0 | Canis | 8120.0 | 1 | Canis lupus | 6868.0 | 2 |
| scf7180000043608 | 75899 | 0.3438 | 0 | 59.1344 | Eukaryota | 93527.0 | 0 | Chordata | 93527.0 | 0 | Carnivora | 93527.0 | 0 | Canidae | 93527.0 | 0 | Canis | 93527.0 | 0 | Canis lupus | 93527.0 | 0 |
| scf7180000042638 | 491471 | 0.3791 | 0 | 58.6115 | Eukaryota | 25763.0 | 0 | Chordata | 25763.0 | 0 | Carnivora | 25763.0 | 0 | Canidae | 25763.0 | 0 | Canis | 25763.0 | 0 | Canis lupus | 25763.0 | 0 |
| scf7180000047927 | 28934 | 0.4168 | 0 | 57.4595 | Eukaryota | 33068.0 | 0 | Chordata | 33068.0 | 0 | Carnivora | 33068.0 | 0 | Canidae | 33068.0 | 0 | Canis | 33068.0 | 0 | Canis lupus | 33068.0 | 0 |
| scf7180000043727 | 3532 | 0.4499 | 0 | 35.069 | Eukaryota | 34584.0 | 0 | Chordata | 34584.0 | 0 | Carnivora | 34584.0 | 0 | Canidae | 34584.0 | 0 | Canis | 34584.0 | 0 | Canis lupus | 34584.0 | 0 |
| scf7180000049762 | 5559 | 0.4402 | 0 | 102.1867 | Eukaryota | 664.0 | 0 | Chordata | 664.0 | 0 | Kurtiformes | 664.0 | 0 | Apogonidae | 664.0 | 0 | Sphaeramia | 664.0 | 0 | Sphaeramia orbicularis | 664.0 | 0 |
| scf7180000054411 | 28549 | 0.4436 | 0 | 243.2053 | Eukaryota | 4627.0 | 0 | Chordata | 3597.0 | 1 | Kurtiformes | 3177.0 | 2 | Apogonidae | 3177.0 | 2 | Sphaeramia | 3177.0 | 2 | Sphaeramia orbicularis | 3177.0 | 2 |
| scf7180000046107 | 11022 | 0.4416 | 0 | 43.9481 | Eukaryota | 88477.0 | 0 | Chordata | 88477.0 | 0 | Carnivora | 88477.0 | 0 | Canidae | 88477.0 | 0 | Canis | 88477.0 | 0 | Canis lupus | 88477.0 | 0 |
| scf7180000041677 | 1104405 | 0.3822 | 100 | 60.4753 | Eukaryota | 131824.0 | 0 | Chordata | 131824.0 | 0 | Carnivora | 131824.0 | 0 | Canidae | 131824.0 | 0 | Canis | 109349.0 | 1 | Canis lupus | 109349.0 | 1 |
| scf7180000047862 | 55176 | 0.488 | 0 | 60.3218 | Eukaryota | 26081.0 | 0 | Chordata | 26081.0 | 0 | Carnivora | 26081.0 | 0 | Canidae | 26081.0 | 0 | Canis | 17073.0 | 1 | Canis lupus | 17073.0 | 1 |
| scf7180000044814 | 279315 | 0.3824 | 0 | 60.3286 | Eukaryota | 50757.0 | 0 | Chordata | 50757.0 | 0 | Carnivora | 29735.0 | 2 | Canidae | 29735.0 | 2 | Canis | 29735.0 | 2 | Canis lupus | 29735.0 | 2 |
| scf7180000047683 | 67893 | 0.5788 | 0 | 58.283 | Eukaryota | 3161.0 | 0 | Chordata | 3161.0 | 0 | Carnivora | 3161.0 | 0 | Canidae | 3161.0 | 0 | Canis | 3161.0 | 0 | Canis lupus | 3161.0 | 0 |
| scf7180000048627 | 46696 | 0.4471 | 0 | 236.4704 | Eukaryota | 8675.0 | 0 | Chordata | 6312.0 | 2 | Tetraodontiformes | 4911.0 | 4 | Tetraodontidae | 4911.0 | 4 | Takifugu | 4911.0 | 4 | Takifugu rubripes | 4911.0 | 4 |
| scf7180000047163 | 85014 | 0.5183 | 0 | 57.5768 | Eukaryota | 11958.0 | 0 | Chordata | 11958.0 | 0 | Carnivora | 11958.0 | 0 | Canidae | 11958.0 | 0 | Canis | 11958.0 | 0 | Canis lupus | 11958.0 | 0 |
| scf7180000041299 | 403866 | 0.3818 | 0 | 60.0485 | Eukaryota | 86176.0 | 0 | Chordata | 86176.0 | 0 | Carnivora | 86176.0 | 0 | Canidae | 86176.0 | 0 | Canis | 86176.0 | 0 | Canis lupus | 86176.0 | 0 |
| scf7180000049284 | 36282 | 0.372 | 0 | 59.7723 | Eukaryota | 5036.0 | 0 | Chordata | 5036.0 | 0 | Primates | 2808.0 | 2 | Hominidae | 2808.0 | 3 | Homo | 1990.0 | 4 | Homo sapiens | 1990.0 | 4 |
| scf7180000042342 | 517757 | 0.3895 | 0 | 58.1337 | Eukaryota | 91958.0 | 0 | Chordata | 91958.0 | 0 | Carnivora | 91958.0 | 0 | Canidae | 91958.0 | 0 | Canis | 91958.0 | 0 | Canis lupus | 91958.0 | 0 |
| scf7180000043089 | 281728 | 0.5321 | 0 | 59.0771 | Eukaryota | 55017.0 | 0 | Chordata | 55017.0 | 0 | Carnivora | 55017.0 | 0 | Canidae | 28255.0 | 2 | Panthera | 23301.0 | 3 | Panthera pardus | 23301.0 | 3 |
| scf7180000041343 | 1140675 | 0.394 | 0 | 59.2296 | Eukaryota | 107945.0 | 0 | Chordata | 107945.0 | 0 | Carnivora | 107945.0 | 0 | Canidae | 107945.0 | 0 | Canis | 107945.0 | 0 | Canis lupus | 107945.0 | 0 |
| scf7180000046176 | 125774 | 0.3516 | 0 | 58.8187 | Eukaryota | 86021.0 | 0 | Chordata | 86021.0 | 0 | Carnivora | 86021.0 | 0 | Canidae | 86021.0 | 0 | Canis | 86021.0 | 0 | Canis lupus | 86021.0 | 0 |
| scf7180000044027 | 377753 | 0.4989 | 0 | 56.5697 | Eukaryota | 30164.0 | 0 | Chordata | 30164.0 | 0 | Carnivora | 30164.0 | 0 | Canidae | 30164.0 | 0 | Canis | 27026.0 | 1 | Canis lupus | 27026.0 | 1 |
| scf7180000052108 | 10375 | 0.4376 | 0 | 349.8835 | Eukaryota | 1966.0 | 0 | Platyhelminthes | 755.0 | 2 | Polystomatidea | 755.0 | 2 | Polystomatidae | 755.0 | 2 | Protopolystoma | 755.0 | 2 | Protopolystoma xenopodis | 755.0 | 2 |
| scf7180000043746 | 32064 | 0.4206 | 0 | 64.1821 | Eukaryota | 8021.0 | 0 | Chordata | 8021.0 | 0 | Carnivora | 8021.0 | 0 | Canidae | 8021.0 | 0 | Canis | 8021.0 | 0 | Canis lupus | 8021.0 | 0 |
| scf7180000051770 | 10850 | 0.4377 | 0 | 246.5236 | Eukaryota | 58264.0 | 0 | Chordata | 58264.0 | 0 | Carnivora | 58264.0 | 0 | Canidae | 58264.0 | 0 | Canis | 58264.0 | 0 | Canis lupus | 58264.0 | 0 |
| scf7180000049623 | 19924 | 0.4448 | 0 | 167.2074 | Eukaryota | 5786.0 | 0 | Chordata | 4225.0 | 2 | Blenniiformes | 2030.0 | 7 | Gobiesocidae | 2030.0 | 7 | Gouania | 2030.0 | 7 | Gouania willdenowi | 2030.0 | 7 |
| scf7180000042507 | 339342 | 0.4265 | 0 | 63.7402 | Eukaryota | 95026.0 | 0 | Chordata | 95026.0 | 0 | Carnivora | 95026.0 | 0 | Canidae | 95026.0 | 0 | Canis | 95026.0 | 0 | Canis lupus | 95026.0 | 0 |
| scf7180000053272 | 2813 | 0.4486 | 0 | 361.9278 | no-hit | 0.0 | 0 | no-hit | 0.0 | 0 | no-hit | 0.0 | 0 | no-hit | 0.0 | 0 | no-hit | 0.0 | 0 | no-hit | 0.0 | 0 |
| scf7180000043382 | 247438 | 0.4055 | 0 | 60.1165 | Eukaryota | 106188.0 | 0 | Chordata | 106188.0 | 0 | Carnivora | 106188.0 | 0 | Canidae | 106188.0 | 0 | Canis | 106188.0 | 0 | Canis lupus | 106188.0 | 0 |
| scf7180000053044 | 13632 | 0.4448 | 0 | 90.7575 | no-hit | 0.0 | 0 | no-hit | 0.0 | 0 | no-hit | 0.0 | 0 | no-hit | 0.0 | 0 | no-hit | 0.0 | 0 | no-hit | 0.0 | 0 |
| scf7180000041589 | 536317 | 0.4177 | 0 | 59.5887 | Eukaryota | 56668.0 | 0 | Chordata | 56668.0 | 0 | Carnivora | 51923.0 | 1 | Canidae | 43390.0 | 2 | Canis | 33295.0 | 4 | Canis lupus | 33295.0 | 4 |
| scf7180000043648 | 34516 | 0.4327 | 0 | 60.418 | Eukaryota | 87277.0 | 0 | Chordata | 87277.0 | 0 | Carnivora | 87277.0 | 0 | Canidae | 87277.0 | 0 | Canis | 87277.0 | 0 | Canis lupus | 87277.0 | 0 |
| scf7180000052801 | 2270 | 0.452 | 0 | 518.8884 | no-hit | 0.0 | 0 | no-hit | 0.0 | 0 | no-hit | 0.0 | 0 | no-hit | 0.0 | 0 | no-hit | 0.0 | 0 | no-hit | 0.0 | 0 |
| scf7180000054922 | 25008 | 0.3852 | 0 | 307.0631 | Eukaryota | 13897.0 | 0 | Chordata | 13897.0 | 0 | Carnivora | 13897.0 | 0 | Canidae | 13897.0 | 0 | Canis | 13897.0 | 0 | Canis lupus | 13897.0 | 0 |
| scf7180000051733 | 29882 | 0.5097 | 0 | 48.4132 | Eukaryota | 26341.0 | 0 | Chordata | 26341.0 | 0 | Carnivora | 26341.0 | 0 | Canidae | 26341.0 | 0 | Canis | 23724.0 | 1 | Canis lupus | 23724.0 | 1 |
| scf7180000049739 | 19687 | 0.4458 | 0 | 56.9487 | Eukaryota | 24217.0 | 0 | Platyhelminthes | 12970.0 | 2 | Cyclophyllidea | 7699.0 | 8 | Hymenolepididae | 7699.0 | 8 | Hymenolepis | 7699.0 | 8 | Hymenolepis microstoma | 7699.0 | 8 |
| scf7180000043639 | 481457 | 0.4766 | 0 | 58.093 | Eukaryota | 33777.0 | 0 | Chordata | 33777.0 | 0 | Carnivora | 33777.0 | 0 | Canidae | 15372.0 | 2 | Canis | 11534.0 | 5 | Canis lupus | 11534.0 | 5 |
| scf7180000051544 | 19952 | 0.6097 | 0 | 59.4659 | Eukaryota | 15239.0 | 0 | Chordata | 15239.0 | 0 | Carnivora | 15239.0 | 0 | Canidae | 15239.0 | 0 | Canis | 14003.0 | 1 | Canis lupus | 14003.0 | 1 |
| scf7180000043812 | 430036 | 0.4241 | 0 | 60.1259 | Eukaryota | 37747.0 | 0 | Chordata | 37747.0 | 0 | Carnivora | 37747.0 | 0 | Canidae | 37747.0 | 0 | Canis | 34775.0 | 1 | Canis lupus | 34775.0 | 1 |
| scf7180000046337 | 53351 | 0.4388 | 0 | 134.5207 | Eukaryota | 9685.0 | 0 | Chordata | 7445.0 | 1 | Kurtiformes | 2586.0 | 4 | Apogonidae | 2586.0 | 4 | Sphaeramia | 2586.0 | 4 | Sphaeramia orbicularis | 2586.0 | 4 |
| scf7180000052818 | 9466 | 0.4394 | 0 | 558.7155 | no-hit | 0.0 | 0 | no-hit | 0.0 | 0 | no-hit | 0.0 | 0 | no-hit | 0.0 | 0 | no-hit | 0.0 | 0 | no-hit | 0.0 | 0 |
| scf7180000047084 | 9782 | 0.4386 | 0 | 80.5367 | Eukaryota | 4067.0 | 1 | Chordata | 2458.0 | 2 | Polystomatidea | 1609.0 | 6 | Polystomatidae | 1609.0 | 6 | Protopolystoma | 1609.0 | 6 | Protopolystoma xenopodis | 1609.0 | 6 |
| scf7180000048560 | 71292 | 0.402 | 0 | 62.4466 | Eukaryota | 34164.0 | 0 | Chordata | 34164.0 | 0 | Carnivora | 34164.0 | 0 | Canidae | 22565.0 | 3 | Canis | 18821.0 | 4 | Canis lupus | 18821.0 | 5 |
| scf7180000045102 | 272976 | 0.3911 | 0 | 59.1428 | Eukaryota | 61021.0 | 0 | Chordata | 61021.0 | 0 | Carnivora | 61021.0 | 0 | Canidae | 61021.0 | 0 | Canis | 51745.0 | 1 | Canis lupus | 51745.0 | 1 |
| scf7180000043130 | 233714 | 0.4272 | 0 | 60.0275 | Eukaryota | 32359.0 | 0 | Chordata | 32359.0 | 0 | Carnivora | 32359.0 | 0 | Canidae | 29363.0 | 1 | Canis | 25399.0 | 2 | Canis lupus | 25399.0 | 2 |
| scf7180000042467 | 115152 | 0.3409 | 0 | 57.804 | Eukaryota | 83040.0 | 0 | Chordata | 83040.0 | 0 | Carnivora | 83040.0 | 0 | Canidae | 83040.0 | 0 | Canis | 83040.0 | 0 | Canis lupus | 83040.0 | 0 |
| scf7180000050928 | 29360 | 0.5039 | 0 | 724.786 | Eukaryota | 9587.0 | 0 | Chordata | 9587.0 | 0 | Carnivora | 9587.0 | 0 | Canidae | 9587.0 | 0 | Canis | 7774.0 | 1 | Canis lupus | 6464.0 | 2 |
| scf7180000042673 | 87504 | 0.3423 | 0 | 60.4392 | Eukaryota | 93271.0 | 0 | Chordata | 93271.0 | 0 | Carnivora | 93271.0 | 0 | Canidae | 93271.0 | 0 | Canis | 93271.0 | 0 | Canis lupus | 93271.0 | 0 |
| scf7180000042424 | 339428 | 0.3448 | 0 | 58.8371 | Eukaryota | 85704.0 | 0 | Chordata | 85704.0 | 0 | Carnivora | 85704.0 | 0 | Canidae | 85704.0 | 0 | Canis | 85704.0 | 0 | Canis lupus | 85704.0 | 0 |
| scf7180000053085 | 3658 | 0.4497 | 0 | 712.3628 | no-hit | 0.0 | 0 | no-hit | 0.0 | 0 | no-hit | 0.0 | 0 | no-hit | 0.0 | 0 | no-hit | 0.0 | 0 | no-hit | 0.0 | 0 |
| scf7180000042030 | 675756 | 0.3943 | 0 | 59.369 | Eukaryota | 50768.0 | 0 | Chordata | 50768.0 | 0 | Carnivora | 50768.0 | 0 | Canidae | 50768.0 | 0 | Canis | 50768.0 | 0 | Canis lupus | 50768.0 | 0 |
| scf7180000043092 | 472369 | 0.3714 | 0 | 58.9154 | Eukaryota | 40779.0 | 0 | Chordata | 40779.0 | 0 | Carnivora | 40779.0 | 0 | Canidae | 40779.0 | 0 | Canis | 40779.0 | 0 | Canis lupus | 40779.0 | 0 |
| scf7180000053711 | 3478 | 0.4948 | 0 | 254.1203 | Eukaryota | 5182.0 | 0 | Chordata | 5182.0 | 0 | Carnivora | 5182.0 | 0 | Canidae | 5182.0 | 0 | Canis | 4478.0 | 1 | Canis lupus | 4478.0 | 1 |
| scf7180000056234 | 1262501 | 0.3493 | 0 | 59.1829 | Eukaryota | 94082.0 | 0 | Chordata | 94082.0 | 0 | Carnivora | 94082.0 | 0 | Canidae | 94082.0 | 0 | Canis | 94082.0 | 0 | Canis lupus | 94082.0 | 0 |
| scf7180000050027 | 7862 | 0.4359 | 0 | 134.9579 | Eukaryota | 4156.0 | 0 | Chordata | 2458.0 | 1 | Polystomatidea | 1100.0 | 6 | Polystomatidae | 1100.0 | 6 | Protopolystoma | 1100.0 | 6 | Protopolystoma xenopodis | 1100.0 | 6 |
| scf7180000047625 | 18314 | 0.4356 | 0 | 157.3542 | Eukaryota | 2367.0 | 0 | Chordata | 1650.0 | 1 | Tetraodontiformes | 880.0 | 4 | Tetraodontidae | 880.0 | 4 | Takifugu | 880.0 | 4 | Takifugu rubripes | 880.0 | 4 |
| scf7180000042529 | 687292 | 0.4785 | 0 | 58.1495 | Eukaryota | 42816.0 | 0 | Chordata | 42816.0 | 0 | Carnivora | 42816.0 | 0 | Canidae | 39366.0 | 1 | Canis | 39366.0 | 1 | Canis lupus | 39366.0 | 1 |
| scf7180000052945 | 2880 | 0.4524 | 0 | 660.0321 | no-hit | 0.0 | 0 | no-hit | 0.0 | 0 | no-hit | 0.0 | 0 | no-hit | 0.0 | 0 | no-hit | 0.0 | 0 | no-hit | 0.0 | 0 |
| scf7180000041259 | 1113931 | 0.3779 | 0 | 60.261 | Eukaryota | 63729.0 | 0 | Chordata | 63729.0 | 0 | Carnivora | 63729.0 | 0 | Canidae | 51645.0 | 2 | Canis | 45068.0 | 3 | Canis lupus | 45068.0 | 3 |
| scf7180000051686 | 2987 | 0.4412 | 0 | 575.7552 | Eukaryota | 1428.0 | 0 | Chordata | 1428.0 | 0 | Columbiformes | 462.0 | 3 | Columbidae | 462.0 | 3 | Streptopelia | 462.0 | 3 | Streptopelia turtur | 462.0 | 3 |
| scf7180000055190 | 18937 | 0.5191 | 0 | 1155.3709 | Eukaryota | 16134.0 | 0 | Chordata | 16134.0 | 0 | Carnivora | 16134.0 | 0 | Canidae | 16134.0 | 0 | Canis | 16134.0 | 0 | Canis lupus | 16134.0 | 0 |
| scf7180000045884 | 155721 | 0.4628 | 0 | 61.5875 | Eukaryota | 17214.0 | 0 | Chordata | 17214.0 | 0 | Carnivora | 6351.0 | 3 | Canidae | 6351.0 | 4 | Homo | 5162.0 | 5 | Homo sapiens | 5162.0 | 5 |
| scf7180000041873 | 899579 | 0.3634 | 0 | 60.1498 | Eukaryota | 82577.0 | 0 | Chordata | 82577.0 | 0 | Carnivora | 82577.0 | 0 | Canidae | 82577.0 | 0 | Canis | 82577.0 | 0 | Canis lupus | 82577.0 | 0 |
| scf7180000042047 | 911440 | 0.4943 | 0 | 59.396 | Eukaryota | 74730.0 | 0 | Chordata | 74730.0 | 0 | Carnivora | 74730.0 | 0 | Canidae | 74730.0 | 0 | Canis | 74730.0 | 0 | Canis lupus | 74730.0 | 0 |
| scf7180000048477 | 69942 | 0.4443 | 0 | 59.6449 | Eukaryota | 56384.0 | 0 | Chordata | 56384.0 | 0 | Carnivora | 56384.0 | 0 | Canidae | 56384.0 | 0 | Vulpes | 34124.0 | 1 | Vulpes vulpes | 34124.0 | 1 |
| scf7180000040619 | 2032426 | 0.4362 | 0 | 60.0711 | Eukaryota | 89981.0 | 0 | Chordata | 89981.0 | 0 | Carnivora | 89981.0 | 0 | Canidae | 89981.0 | 0 | Canis | 89981.0 | 0 | Canis lupus | 89981.0 | 0 |
| scf7180000055298 | 24076 | 0.4389 | 0 | 324.2511 | Eukaryota | 1513.0 | 0 | Platyhelminthes | 913.0 | 1 | Polystomatidea | 913.0 | 3 | Polystomatidae | 913.0 | 3 | Protopolystoma | 913.0 | 3 | Protopolystoma xenopodis | 913.0 | 3 |
| scf7180000045337 | 3851 | 0.4536 | 0 | 188.8962 | no-hit | 0.0 | 0 | no-hit | 0.0 | 0 | no-hit | 0.0 | 0 | no-hit | 0.0 | 0 | no-hit | 0.0 | 0 | no-hit | 0.0 | 0 |
| scf7180000045286 | 54794 | 0.4454 | 0 | 100.1902 | Eukaryota | 67488.0 | 0 | Chordata | 67488.0 | 0 | Carnivora | 67488.0 | 0 | Canidae | 67488.0 | 0 | Canis | 67488.0 | 0 | Canis lupus | 67488.0 | 0 |
| scf7180000047018 | 99169 | 0.3682 | 0 | 60.7361 | Eukaryota | 7191.0 | 0 | Chordata | 7191.0 | 0 | Carnivora | 6437.0 | 1 | Felidae | 5616.0 | 2 | Felis | 5616.0 | 2 | Felis catus | 5616.0 | 2 |
| scf7180000042358 | 768813 | 0.3736 | 0 | 59.2367 | Eukaryota | 57440.0 | 0 | Chordata | 57440.0 | 0 | Carnivora | 57440.0 | 0 | Canidae | 57440.0 | 0 | Canis | 57440.0 | 0 | Canis lupus | 57440.0 | 0 |
| scf7180000041750 | 895872 | 0.3713 | 0 | 60.9588 | Eukaryota | 59707.0 | 0 | Chordata | 59707.0 | 0 | Carnivora | 59707.0 | 0 | Canidae | 59707.0 | 0 | Canis | 59707.0 | 0 | Canis lupus | 59707.0 | 0 |
| scf7180000054288 | 22826 | 0.4255 | 0 | 433.0602 | Eukaryota | 8191.0 | 0 | Chordata | 8191.0 | 0 | Primates | 4863.0 | 2 | Hominidae | 3692.0 | 3 | Homo | 3692.0 | 4 | Homo sapiens | 3692.0 | 4 |
| scf7180000056583 | 33614 | 0.3875 | 0 | 74.3944 | Eukaryota | 61907.0 | 0 | Chordata | 61907.0 | 0 | Carnivora | 61907.0 | 0 | Canidae | 61907.0 | 0 | Canis | 61907.0 | 0 | Canis lupus | 61907.0 | 0 |
| scf7180000056557 | 125908 | 0.5342 | 0 | 57.0449 | Eukaryota | 19602.0 | 0 | Chordata | 19602.0 | 0 | Carnivora | 19602.0 | 0 | Canidae | 19602.0 | 0 | Canis | 19602.0 | 0 | Canis lupus | 19602.0 | 0 |
| scf7180000050391 | 3763 | 0.4459 | 0 | 389.1514 | no-hit | 0.0 | 0 | no-hit | 0.0 | 0 | no-hit | 0.0 | 0 | no-hit | 0.0 | 0 | no-hit | 0.0 | 0 | no-hit | 0.0 | 0 |
| scf7180000041168 | 1195133 | 0.4583 | 0 | 58.4406 | Eukaryota | 90746.0 | 0 | Chordata | 90746.0 | 0 | Carnivora | 90746.0 | 0 | Canidae | 90746.0 | 0 | Canis | 90746.0 | 0 | Canis lupus | 90746.0 | 0 |
| scf7180000044669 | 257233 | 0.3544 | 0 | 59.1581 | Eukaryota | 19911.0 | 0 | Chordata | 19911.0 | 0 | Carnivora | 19911.0 | 0 | Canidae | 19911.0 | 0 | Canis | 19911.0 | 0 | Canis lupus | 19911.0 | 0 |
| scf7180000051028 | 4305 | 0.5094 | 0 | 58.1486 | Eukaryota | 7965.0 | 0 | Chordata | 7965.0 | 0 | Carnivora | 7965.0 | 0 | Canidae | 7965.0 | 0 | Canis | 6358.0 | 1 | Canis lupus | 5181.0 | 2 |
| scf7180000055919 | 16328 | 0.4331 | 0 | 172.6287 | Eukaryota | 14883.0 | 0 | Chordata | 8242.0 | 2 | Gadiformes | 4471.0 | 6 | Gadidae | 4471.0 | 6 | Gadus | 4471.0 | 6 | Gadus morhua | 4471.0 | 6 |
| scf7180000054728 | 249604 | 0.3657 | 0 | 59.7658 | Eukaryota | 89017.0 | 0 | Chordata | 89017.0 | 0 | Carnivora | 89017.0 | 0 | Canidae | 89017.0 | 0 | Canis | 89017.0 | 0 | Canis lupus | 89017.0 | 0 |
| scf7180000054450 | 14334 | 0.4399 | 0 | 109.7948 | Eukaryota | 7295.0 | 0 | Platyhelminthes | 2783.0 | 2 | Polystomatidea | 2783.0 | 4 | Polystomatidae | 2783.0 | 4 | Protopolystoma | 2783.0 | 4 | Protopolystoma xenopodis | 2783.0 | 4 |
| scf7180000045493 | 5878 | 0.4388 | 0 | 103.9455 | no-hit | 0.0 | 0 | no-hit | 0.0 | 0 | no-hit | 0.0 | 0 | no-hit | 0.0 | 0 | no-hit | 0.0 | 0 | no-hit | 0.0 | 0 |
| scf7180000043226 | 39267 | 0.557 | 0 | 141.7624 | Eukaryota | 83823.0 | 0 | Chordata | 83823.0 | 0 | Carnivora | 83823.0 | 0 | Canidae | 83823.0 | 0 | Canis | 83823.0 | 0 | Canis lupus | 83823.0 | 0 |
| scf7180000053787 | 2678 | 0.4473 | 0 | 296.9964 | no-hit | 0.0 | 0 | no-hit | 0.0 | 0 | no-hit | 0.0 | 0 | no-hit | 0.0 | 0 | no-hit | 0.0 | 0 | no-hit | 0.0 | 0 |
| scf7180000051477 | 9219 | 0.4443 | 0 | 68.5475 | Eukaryota | 337.0 | 0 | Chordata | 337.0 | 0 | Tetraodontiformes | 337.0 | 0 | Tetraodontidae | 337.0 | 0 | Takifugu | 337.0 | 0 | Takifugu rubripes | 337.0 | 0 |
| scf7180000049994 | 7333 | 0.4346 | 0 | 256.7404 | Eukaryota | 1416.0 | 1 | Chordata | 1229.0 | 2 | Columbiformes | 1042.0 | 3 | Columbidae | 1042.0 | 3 | Streptopelia | 1042.0 | 3 | Streptopelia turtur | 1042.0 | 3 |
| scf7180000056000 | 5828 | 0.4384 | 0 | 297.2995 | no-hit | 0.0 | 0 | no-hit | 0.0 | 0 | no-hit | 0.0 | 0 | no-hit | 0.0 | 0 | no-hit | 0.0 | 0 | no-hit | 0.0 | 0 |
| scf7180000042536 | 286782 | 0.4115 | 0 | 60.1751 | Eukaryota | 20499.0 | 0 | Chordata | 20499.0 | 0 | Carnivora | 20499.0 | 0 | Canidae | 20499.0 | 0 | Canis | 20499.0 | 0 | Canis lupus | 20499.0 | 0 |
| scf7180000041957 | 834972 | 0.4372 | 0 | 61.3387 | Eukaryota | 56848.0 | 0 | Chordata | 56848.0 | 0 | Carnivora | 56848.0 | 0 | Canidae | 56848.0 | 0 | Canis | 56848.0 | 0 | Canis lupus | 56848.0 | 0 |
| scf7180000041764 | 5406 | 0.4145 | 0 | 28.5922 | Eukaryota | 6572.0 | 0 | Chordata | 6572.0 | 0 | Carnivora | 6572.0 | 0 | Canidae | 6572.0 | 0 | Canis | 6263.0 | 1 | Canis lupus | 6263.0 | 1 |
| scf7180000040883 | 782447 | 0.6327 | 0 | 55.1234 | Eukaryota | 68021.0 | 0 | Chordata | 68021.0 | 0 | Carnivora | 68021.0 | 0 | Canidae | 68021.0 | 0 | Canis | 48211.0 | 1 | Canis lupus | 48211.0 | 1 |
| scf7180000044545 | 671654 | 0.346 | 100 | 59.503 | Eukaryota | 75653.0 | 1 | Chordata | 75653.0 | 1 | Primates | 27463.0 | 4 | Bovidae | 22750.0 | 5 | unresolved | 19826.0 | 6 | unresolved | 19826.0 | 7 |
| scf7180000046368 | 5367 | 0.4399 | 0 | 145.9393 | Eukaryota | 390.0 | 0 | Platyhelminthes | 390.0 | 0 | Polystomatidea | 390.0 | 0 | Polystomatidae | 390.0 | 0 | Protopolystoma | 390.0 | 0 | Protopolystoma xenopodis | 390.0 | 0 |
| scf7180000047734 | 34382 | 0.445 | 0 | 217.0552 | Eukaryota | 3074.0 | 0 | Chordata | 3074.0 | 0 | Carnivora | 3074.0 | 0 | Canidae | 3074.0 | 0 | Canis | 3074.0 | 0 | Canis lupus | 3074.0 | 0 |
| scf7180000041676 | 869360 | 0.4878 | 0 | 59.7284 | Eukaryota | 56238.0 | 0 | Chordata | 56238.0 | 0 | Carnivora | 56238.0 | 0 | Canidae | 56238.0 | 0 | Canis | 41869.0 | 1 | Canis lupus | 41869.0 | 1 |
| scf7180000040881 | 829585 | 0.4175 | 0 | 59.1647 | Eukaryota | 90244.0 | 0 | Chordata | 90244.0 | 0 | Carnivora | 90244.0 | 0 | Canidae | 90244.0 | 0 | Canis | 90244.0 | 0 | Canis lupus | 90244.0 | 0 |
| scf7180000043308 | 498520 | 0.3846 | 0 | 59.2665 | Eukaryota | 35299.0 | 0 | Chordata | 35299.0 | 0 | Carnivora | 35299.0 | 0 | Canidae | 35299.0 | 0 | Canis | 35299.0 | 0 | Canis lupus | 35299.0 | 0 |
| scf7180000041743 | 284947 | 0.3736 | 0 | 57.8285 | Eukaryota | 93312.0 | 0 | Chordata | 93312.0 | 0 | Carnivora | 93312.0 | 0 | Canidae | 93312.0 | 0 | Canis | 93312.0 | 0 | Canis lupus | 93312.0 | 0 |
| scf7180000051356 | 7921 | 0.445 | 0 | 331.079 | no-hit | 0.0 | 0 | no-hit | 0.0 | 0 | no-hit | 0.0 | 0 | no-hit | 0.0 | 0 | no-hit | 0.0 | 0 | no-hit | 0.0 | 0 |
| scf7180000056577 | 81779 | 0.4213 | 0 | 59.8416 | Eukaryota | 28265.0 | 0 | Chordata | 28265.0 | 0 | Carnivora | 28265.0 | 0 | Canidae | 11781.0 | 3 | Canis | 7823.0 | 6 | Canis lupus | 7823.0 | 6 |
| scf7180000042392 | 228834 | 0.3985 | 0 | 60.0835 | Eukaryota | 39742.0 | 1 | Chordata | 39742.0 | 1 | Carnivora | 33373.0 | 3 | Canidae | 33373.0 | 3 | Canis | 30109.0 | 4 | Canis lupus | 30109.0 | 4 |
| scf7180000042598 | 19566 | 0.4279 | 0 | 50.8506 | Eukaryota | 55806.0 | 0 | Chordata | 55806.0 | 0 | Carnivora | 55806.0 | 0 | Canidae | 55806.0 | 0 | Canis | 55806.0 | 0 | Canis lupus | 55806.0 | 0 |
| scf7180000046760 | 106326 | 0.3549 | 0 | 58.0722 | Eukaryota | 27087.0 | 0 | Chordata | 27087.0 | 0 | Carnivora | 27087.0 | 0 | Canidae | 24388.0 | 1 | Canis | 24388.0 | 1 | Canis lupus | 24388.0 | 1 |
| scf7180000050187 | 2974 | 0.4475 | 0 | 480.7996 | Eukaryota | 1197.0 | 0 | Platyhelminthes | 709.0 | 1 | Polystomatidea | 709.0 | 1 | Polystomatidae | 709.0 | 1 | Protopolystoma | 709.0 | 1 | Protopolystoma xenopodis | 709.0 | 1 |
| scf7180000041042 | 102823 | 0.3669 | 0 | 59.8532 | Eukaryota | 69965.0 | 0 | Chordata | 69965.0 | 0 | Carnivora | 69965.0 | 0 | Canidae | 69965.0 | 0 | Canis | 69965.0 | 0 | Canis lupus | 69965.0 | 0 |
| scf7180000056196 | 1454336 | 0.3869 | 0 | 60.964 | Eukaryota | 86980.0 | 0 | Chordata | 86980.0 | 0 | Carnivora | 86980.0 | 0 | Canidae | 86980.0 | 0 | Canis | 86980.0 | 0 | Canis lupus | 86980.0 | 0 |
| scf7180000044187 | 5576 | 0.4437 | 0 | 222.534 | Eukaryota | 13034.0 | 0 | Chordata | 13034.0 | 0 | Carnivora | 13034.0 | 0 | Canidae | 13034.0 | 0 | Canis | 13034.0 | 0 | Canis lupus | 13034.0 | 0 |
| scf7180000045447 | 29160 | 0.4341 | 0 | 49.4111 | Eukaryota | 2778.0 | 0 | Chordata | 2778.0 | 0 | Carnivora | 2778.0 | 0 | Canidae | 2778.0 | 0 | Canis | 2526.0 | 1 | Canis lupus | 2526.0 | 1 |
| scf7180000043664 | 397286 | 0.3817 | 0 | 59.8545 | Eukaryota | 67506.0 | 0 | Chordata | 67506.0 | 0 | Carnivora | 67506.0 | 0 | Canidae | 67506.0 | 0 | Vulpes | 40734.0 | 1 | Vulpes vulpes | 40734.0 | 1 |
| scf7180000044588 | 35707 | 0.4886 | 0 | 50.6378 | Eukaryota | 23125.0 | 0 | Chordata | 23125.0 | 0 | Carnivora | 23125.0 | 0 | Canidae | 23125.0 | 0 | Canis | 19858.0 | 1 | Canis lupus | 19858.0 | 1 |
| scf7180000049142 | 14500 | 0.4458 | 0 | 150.7829 | no-hit | 0.0 | 0 | no-hit | 0.0 | 0 | no-hit | 0.0 | 0 | no-hit | 0.0 | 0 | no-hit | 0.0 | 0 | no-hit | 0.0 | 0 |
| scf7180000046302 | 19889 | 0.4381 | 0 | 68.4791 | Eukaryota | 46097.0 | 0 | Chordata | 31791.0 | 2 | Gadiformes | 25613.0 | 5 | Gadidae | 25613.0 | 5 | Gadus | 25613.0 | 5 | Gadus morhua | 25613.0 | 5 |
| scf7180000043956 | 182602 | 0.5309 | 0 | 55.9577 | Eukaryota | 27178.0 | 0 | Chordata | 27178.0 | 0 | Carnivora | 27178.0 | 0 | Canidae | 27178.0 | 0 | Canis | 27178.0 | 0 | Canis lupus | 27178.0 | 0 |
| scf7180000046798 | 17238 | 0.4434 | 0 | 130.5694 | Eukaryota | 91663.0 | 0 | Chordata | 91663.0 | 0 | Carnivora | 91663.0 | 0 | Canidae | 91663.0 | 0 | Canis | 91663.0 | 0 | Canis lupus | 91663.0 | 0 |
| scf7180000053393 | 18967 | 0.505 | 0 | 123.3247 | Eukaryota | 8809.0 | 0 | Chordata | 8809.0 | 0 | Carnivora | 8809.0 | 0 | Canidae | 8809.0 | 0 | Canis | 7048.0 | 1 | Canis lupus | 5830.0 | 2 |
| scf7180000053209 | 5723 | 0.4529 | 0 | 519.9358 | no-hit | 0.0 | 0 | no-hit | 0.0 | 0 | no-hit | 0.0 | 0 | no-hit | 0.0 | 0 | no-hit | 0.0 | 0 | no-hit | 0.0 | 0 |
| scf7180000047499 | 96698 | 0.576 | 0 | 56.9033 | Eukaryota | 33023.0 | 0 | Chordata | 33023.0 | 0 | Carnivora | 33023.0 | 0 | Canidae | 31113.0 | 1 | Canis | 17344.0 | 2 | Canis lupus | 17344.0 | 2 |
| scf7180000043039 | 509369 | 0.4308 | 0 | 59.053 | Eukaryota | 153515.0 | 0 | Chordata | 153515.0 | 0 | Carnivora | 153515.0 | 0 | Canidae | 153515.0 | 0 | Canis | 121463.0 | 1 | Canis lupus | 121463.0 | 1 |
| scf7180000041587 | 453061 | 0.3838 | 0 | 59.2982 | Eukaryota | 36368.0 | 0 | Chordata | 36368.0 | 0 | Carnivora | 36368.0 | 0 | Canidae | 36368.0 | 0 | Canis | 36368.0 | 0 | Canis lupus | 36368.0 | 0 |
| scf7180000050616 | 8717 | 0.4511 | 0 | 561.7956 | no-hit | 0.0 | 0 | no-hit | 0.0 | 0 | no-hit | 0.0 | 0 | no-hit | 0.0 | 0 | no-hit | 0.0 | 0 | no-hit | 0.0 | 0 |
| scf7180000044815 | 8192 | 0.4023 | 0 | 29.2237 | Eukaryota | 3310.0 | 0 | Chordata | 3310.0 | 0 | Carnivora | 3310.0 | 0 | Canidae | 3310.0 | 0 | Canis | 3310.0 | 0 | Canis lupus | 3310.0 | 0 |
| scf7180000044271 | 12418 | 0.4146 | 0 | 45.8493 | Eukaryota | 30758.0 | 0 | Chordata | 30758.0 | 0 | Carnivora | 30758.0 | 0 | Canidae | 30758.0 | 0 | Canis | 30758.0 | 0 | Canis lupus | 30758.0 | 0 |
| scf7180000047312 | 37024 | 0.4401 | 0 | 101.9112 | no-hit | 0.0 | 0 | no-hit | 0.0 | 0 | no-hit | 0.0 | 0 | no-hit | 0.0 | 0 | no-hit | 0.0 | 0 | no-hit | 0.0 | 0 |
| scf7180000044805 | 37236 | 0.4377 | 0 | 177.497 | Eukaryota | 28491.0 | 0 | Chordata | 28491.0 | 0 | Gadiformes | 18623.0 | 5 | Gadidae | 18623.0 | 5 | Gadus | 18623.0 | 5 | Gadus morhua | 18623.0 | 5 |
| scf7180000052560 | 3815 | 0.4495 | 0 | 200.0575 | Eukaryota | 1491.0 | 0 | Echinodermata | 797.0 | 2 | Forcipulatida | 797.0 | 2 | Asteriidae | 797.0 | 2 | Asterias | 797.0 | 2 | Asterias rubens | 797.0 | 2 |
| scf7180000041503 | 477327 | 0.3848 | 0 | 58.9015 | Eukaryota | 55407.0 | 0 | Chordata | 55407.0 | 0 | Carnivora | 55407.0 | 0 | Canidae | 55407.0 | 0 | Canis | 55407.0 | 0 | Canis lupus | 55407.0 | 0 |
| scf7180000041802 | 31486 | 0.415 | 0 | 57.947 | Eukaryota | 81273.0 | 0 | Chordata | 81273.0 | 0 | Carnivora | 81273.0 | 0 | Canidae | 81273.0 | 0 | Canis | 81273.0 | 0 | Canis lupus | 81273.0 | 0 |
| scf7180000042600 | 36400 | 0.4364 | 0 | 55.1712 | Eukaryota | 57719.0 | 0 | Chordata | 57719.0 | 0 | Carnivora | 57719.0 | 0 | Canidae | 57719.0 | 0 | Canis | 57719.0 | 0 | Canis lupus | 57719.0 | 0 |
| scf7180000056420 | 405694 | 0.4242 | 0 | 61.1483 | Eukaryota | 88056.0 | 0 | Chordata | 88056.0 | 0 | Carnivora | 88056.0 | 0 | Canidae | 88056.0 | 0 | Canis | 67598.0 | 1 | Canis lupus | 67598.0 | 1 |
| scf7180000054957 | 23208 | 0.4454 | 0 | 239.6208 | Eukaryota | 2058.0 | 0 | Chordata | 2058.0 | 0 | Anabantiformes | 976.0 | 3 | Anabantidae | 976.0 | 3 | Anabas | 976.0 | 3 | Anabas testudineus | 976.0 | 3 |
| scf7180000044411 | 370124 | 0.3433 | 0 | 59.4385 | Eukaryota | 22675.0 | 0 | Chordata | 22675.0 | 0 | Carnivora | 22675.0 | 0 | Canidae | 22675.0 | 0 | Canis | 22675.0 | 0 | Canis lupus | 22675.0 | 0 |
| scf7180000041902 | 543076 | 0.4737 | 0 | 57.8264 | Eukaryota | 37072.0 | 0 | Chordata | 37072.0 | 0 | Carnivora | 37072.0 | 0 | Canidae | 37072.0 | 0 | Canis | 37072.0 | 0 | Canis lupus | 37072.0 | 0 |
| scf7180000050295 | 9812 | 0.4409 | 0 | 70.5044 | no-hit | 0.0 | 0 | no-hit | 0.0 | 0 | no-hit | 0.0 | 0 | no-hit | 0.0 | 0 | no-hit | 0.0 | 0 | no-hit | 0.0 | 0 |
| scf7180000040598 | 2136866 | 0.4093 | 0 | 59.9941 | Eukaryota | 64832.0 | 0 | Chordata | 64832.0 | 0 | Carnivora | 64832.0 | 0 | Canidae | 64832.0 | 0 | Canis | 52283.0 | 1 | Canis lupus | 52283.0 | 1 |
| scf7180000051594 | 35140 | 0.4247 | 0 | 54.1319 | Eukaryota | 39073.0 | 0 | Chordata | 39073.0 | 0 | Carnivora | 39073.0 | 0 | Canidae | 39073.0 | 0 | Canis | 39073.0 | 0 | Canis lupus | 39073.0 | 0 |
| scf7180000045817 | 162869 | 0.4011 | 0 | 59.4981 | Eukaryota | 57557.0 | 0 | Chordata | 57557.0 | 0 | Carnivora | 57557.0 | 0 | Canidae | 57557.0 | 0 | Canis | 57557.0 | 0 | Canis lupus | 57557.0 | 0 |
| scf7180000050459 | 10388 | 0.4396 | 0 | 197.2176 | Eukaryota | 4700.0 | 0 | Chordata | 4259.0 | 2 | Kurtiformes | 2444.0 | 4 | Apogonidae | 2444.0 | 4 | Sphaeramia | 2444.0 | 4 | Sphaeramia orbicularis | 2444.0 | 4 |
| scf7180000046872 | 17490 | 0.493 | 0 | 158.5485 | Eukaryota | 90725.0 | 0 | Chordata | 90725.0 | 0 | Carnivora | 90725.0 | 0 | Canidae | 90725.0 | 0 | Canis | 90725.0 | 0 | Canis lupus | 90725.0 | 0 |
| scf7180000053973 | 5376 | 0.3093 | 0 | 62.6585 | Eukaryota | 143.0 | 0 | Chordata | 143.0 | 0 | Carnivora | 143.0 | 0 | Canidae | 143.0 | 0 | Canis | 143.0 | 0 | Canis lupus | 143.0 | 0 |
| scf7180000051554 | 3489 | 0.4494 | 0 | 397.0767 | no-hit | 0.0 | 0 | no-hit | 0.0 | 0 | no-hit | 0.0 | 0 | no-hit | 0.0 | 0 | no-hit | 0.0 | 0 | no-hit | 0.0 | 0 |
| scf7180000044466 | 282423 | 0.5219 | 0 | 60.2111 | Eukaryota | 43530.0 | 0 | Chordata | 43530.0 | 0 | Carnivora | 43530.0 | 0 | Canidae | 43530.0 | 0 | Canis | 34780.0 | 1 | Canis lupus | 34780.0 | 1 |
| scf7180000041183 | 1265344 | 0.3756 | 0 | 59.7498 | Eukaryota | 93211.0 | 0 | Chordata | 93211.0 | 0 | Carnivora | 93211.0 | 0 | Canidae | 93211.0 | 0 | Canis | 93211.0 | 0 | Canis lupus | 93211.0 | 0 |

| | | | | | | | | | | | | | | | | | | | | | | |
|---|---|---|---|---|---|---|---|---|---|---|---|---|---|---|---|---|---|---|---|---|---|---|
| scf7180000045436 | 722299 | 0.3818 | 0 | 61.0398 | Eukaryota | 94993.0 | 0 | Chordata | 94993.0 | 0 | Carnivora | 94993.0 | 0 | Canidae | 94993.0 | 0 | Canis | 94993.0 | 0 | Canis lupus | 94993.0 | 0 |
| scf7180000041932 | 119309 | 0.3871 | 0 | 58.8238 | Eukaryota | 24325.0 | 0 | Chordata | 24325.0 | 0 | Carnivora | 21734.0 | 1 | Canidae | 21734.0 | 1 | Canis | 15420.0 | 2 | Canis lupus | 15420.0 | 2 |
| scf7180000051914 | 2655 | 0.4554 | 0 | 889.0537 | no-hit | 0.0 | 0 | no-hit | 0.0 | 0 | no-hit | 0.0 | 0 | no-hit | 0.0 | 0 | no-hit | 0.0 | 0 | no-hit | 0.0 | 0 |
| scf7180000056606 | 2199252 | 0.3625 | 0 | 60.3855 | Eukaryota | 92312.0 | 0 | Chordata | 92312.0 | 0 | Carnivora | 92312.0 | 0 | Canidae | 92312.0 | 0 | Canis | 92312.0 | 0 | Canis lupus | 92312.0 | 0 |
| scf7180000045836 | 5920 | 0.5404 | 0 | 34.8349 | Eukaryota | 13670.0 | 0 | Chordata | 13670.0 | 0 | Carnivora | 13341.0 | 1 | Canidae | 13341.0 | 1 | Canis | 13341.0 | 1 | Canis lupus | 13341.0 | 1 |
| scf7180000044571 | 297392 | 0.4899 | 0 | 59.164 | Eukaryota | 46637.0 | 0 | Chordata | 46637.0 | 0 | Carnivora | 46637.0 | 0 | Canidae | 24818.0 | 3 | Canis | 18507.0 | 6 | Canis lupus | 18507.0 | 6 |
| scf7180000042032 | 100209 | 0.3623 | 0 | 60.3083 | Eukaryota | 21608.0 | 0 | Chordata | 21608.0 | 0 | Primates | 8174.0 | 3 | Hominidae | 6812.0 | 4 | Homo | 6812.0 | 6 | Homo sapiens | 6812.0 | 6 |
| scf7180000055418 | 15458 | 0.4462 | 0 | 392.8605 | no-hit | 0.0 | 0 | no-hit | 0.0 | 0 | no-hit | 0.0 | 0 | no-hit | 0.0 | 0 | no-hit | 0.0 | 0 | no-hit | 0.0 | 0 |
| scf7180000046066 | 159778 | 0.4013 | 0 | 59.3038 | Eukaryota | 41128.0 | 0 | Chordata | 41128.0 | 0 | Carnivora | 41128.0 | 0 | Canidae | 41128.0 | 0 | Canis | 24684.0 | 1 | Canis lupus | 24684.0 | 1 |
| scf7180000046892 | 49006 | 0.4542 | 0 | 102.7523 | Eukaryota | 409.0 | 0 | Platyhelminthes | 409.0 | 0 | Polystomatidea | 409.0 | 0 | Polystomatidae | 409.0 | 0 | Protopolystoma | 409.0 | 0 | Protopolystoma xenopodis | 409.0 | 0 |
| scf7180000055033 | 41683 | 0.4384 | 0 | 218.3851 | Eukaryota | 7159.0 | 0 | Chordata | 5402.0 | 1 | Tetraodontiformes | 4764.0 | 2 | Tetraodontidae | 4764.0 | 2 | Takifugu | 4764.0 | 2 | Takifugu rubripes | 4764.0 | 2 |
| scf7180000049480 | 12460 | 0.4476 | 0 | 996.1797 | Eukaryota | 502.0 | 0 | Streptophyta | 502.0 | 0 | Fabales | 502.0 | 0 | Fabaceae | 502.0 | 0 | Cicer | 502.0 | 0 | Cicer arietinum | 502.0 | 0 |
| scf7180000041090 | 57754 | 0.3601 | 0 | 57.832 | Eukaryota | 82151.0 | 0 | Chordata | 82151.0 | 0 | Carnivora | 82151.0 | 0 | Canidae | 82151.0 | 0 | Canis | 82151.0 | 0 | Canis lupus | 82151.0 | 0 |
| scf7180000047580 | 80242 | 0.4834 | 0 | 57.7304 | Eukaryota | 16007.0 | 0 | Chordata | 16007.0 | 0 | Carnivora | 16007.0 | 0 | Canidae | 16007.0 | 0 | Canis | 16007.0 | 0 | Canis lupus | 16007.0 | 0 |
| scf7180000040803 | 598600 | 0.4129 | 0 | 60.1973 | Eukaryota | 63596.0 | 0 | Chordata | 63596.0 | 0 | Carnivora | 63596.0 | 0 | Canidae | 56049.0 | 1 | Canis | 50263.0 | 3 | Canis lupus | 50263.0 | 3 |
| scf7180000040845 | 1880481 | 0.4021 | 0 | 61.3066 | Eukaryota | 75585.0 | 0 | Chordata | 75585.0 | 0 | Carnivora | 75585.0 | 0 | Canidae | 75585.0 | 0 | Canis | 69538.0 | 1 | Canis lupus | 69538.0 | 1 |
| scf7180000047030 | 118432 | 0.4281 | 0 | 59.4044 | Eukaryota | 18725.0 | 0 | Chordata | 18725.0 | 0 | Primates | 9853.0 | 4 | Hominidae | 9853.0 | 4 | Pan | 5481.0 | 5 | Pan troglodytes | 5481.0 | 5 |
| scf7180000054768 | 14389 | 0.4454 | 0 | 326.136 | Eukaryota | 9032.0 | 0 | Chordata | 9032.0 | 0 | Tetraodontiformes | 5040.0 | 2 | Tetraodontidae | 5040.0 | 2 | Takifugu | 5040.0 | 2 | Takifugu rubripes | 4364.0 | 3 |
| scf7180000055130 | 102359 | 0.4492 | 0 | 74.5579 | Eukaryota | 46306.0 | 0 | Chordata | 46306.0 | 0 | Carnivora | 46306.0 | 0 | Canidae | 46306.0 | 0 | Canis | 41968.0 | 1 | Canis lupus | 41968.0 | 1 |
| scf7180000055862 | 6556 | 0.4524 | 0 | 1956.0011 | Eukaryota | 2878.0 | 0 | Chordata | 2303.0 | 1 | Carnivora | 2303.0 | 1 | Canidae | 2303.0 | 1 | Canis | 2303.0 | 1 | Canis lupus | 2303.0 | 1 |
| scf7180000056426 | 346120 | 0.3882 | 0 | 61.159 | Eukaryota | 43637.0 | 0 | Chordata | 43637.0 | 0 | Carnivora | 43637.0 | 0 | Canidae | 28101.0 | 1 | Canis | 28101.0 | 1 | Canis lupus | 28101.0 | 1 |
| scf7180000056346 | 541934 | 0.3832 | 0 | 59.5652 | Eukaryota | 28102.0 | 0 | Chordata | 28102.0 | 0 | Carnivora | 28102.0 | 0 | Canidae | 28102.0 | 0 | Canis | 28102.0 | 0 | Canis lupus | 28102.0 | 0 |
| scf7180000044562 | 137488 | 0.3579 | 0 | 58.6545 | Eukaryota | 88395.0 | 0 | Chordata | 88395.0 | 0 | Carnivora | 88395.0 | 0 | Canidae | 88395.0 | 0 | Canis | 88395.0 | 0 | Canis lupus | 88395.0 | 0 |
| scf7180000046502 | 59741 | 0.4121 | 0 | 54.6213 | Eukaryota | 26280.0 | 0 | Chordata | 26280.0 | 0 | Carnivora | 26280.0 | 0 | Canidae | 26280.0 | 0 | Canis | 26280.0 | 0 | Canis lupus | 26280.0 | 0 |
| scf7180000042025 | 975346 | 0.4143 | 0 | 60.9545 | Eukaryota | 91502.0 | 0 | Chordata | 91502.0 | 0 | Carnivora | 91502.0 | 0 | Canidae | 91502.0 | 0 | Canis | 91502.0 | 0 | Canis lupus | 91502.0 | 0 |
| scf7180000047778 | 44640 | 0.4406 | 0 | 112.1345 | Eukaryota | 5544.0 | 0 | Chordata | 5211.0 | 1 | Synbranchiformes | 2756.0 | 3 | Mastacembelidae | 2756.0 | 3 | Mastacembelus | 2756.0 | 3 | Mastacembelus armatus | 2756.0 | 3 |
| scf7180000050339 | 8484 | 0.4371 | 0 | 38.0328 | Eukaryota | 12819.0 | 0 | Chordata | 11792.0 | 1 | Primates | 8756.0 | 3 | Hominidae | 8163.0 | 4 | Homo | 8163.0 | 4 | Homo sapiens | 8163.0 | 4 |
| scf7180000044720 | 31966 | 0.4331 | 0 | 45.4767 | Eukaryota | 11797.0 | 0 | Chordata | 11797.0 | 0 | Carnivora | 11797.0 | 0 | Canidae | 4265.0 | 3 | Canis | 2850.0 | 7 | Canis lupus | 2850.0 | 8 |
| scf7180000048447 | 25105 | 0.44 | 0 | 72.6029 | Eukaryota | 2928.0 | 0 | Chordata | 1591.0 | 1 | Polystomatidea | 1337.0 | 3 | Polystomatidae | 1337.0 | 3 | Protopolystoma | 1337.0 | 3 | Protopolystoma xenopodis | 1337.0 | 3 |
| scf7180000054711 | 23895 | 0.4412 | 0 | 268.2766 | Eukaryota | 11065.0 | 0 | Chordata | 9253.0 | 1 | Blenniiformes | 5888.0 | 3 | Gobiesocidae | 5888.0 | 3 | Gouania | 5888.0 | 3 | Gouania willdenowi | 5888.0 | 3 |
| scf7180000056182 | 2417454 | 0.4099 | 0 | 60.2079 | Eukaryota | 57843.0 | 0 | Chordata | 57843.0 | 0 | Carnivora | 57843.0 | 0 | Canidae | 57843.0 | 0 | Canis | 57843.0 | 0 | Canis lupus | 57843.0 | 0 |
| scf7180000053238 | 6094 | 0.4526 | 0 | 1184.1115 | Eukaryota | 20786.0 | 0 | Chordata | 18791.0 | 1 | Gadiformes | 9870.0 | 6 | Gadidae | 9870.0 | 6 | Gadus | 9870.0 | 6 | Gadus morhua | 9870.0 | 6 |
| scf7180000044318 | 61321 | 0.4233 | 0 | 57.9519 | Eukaryota | 26187.0 | 0 | Chordata | 26187.0 | 0 | Carnivora | 26187.0 | 0 | Canidae | 26187.0 | 0 | Canis | 26187.0 | 0 | Canis lupus | 26187.0 | 0 |
| scf7180000045639 | 149278 | 0.3934 | 0 | 58.8732 | Eukaryota | 42240.0 | 0 | Chordata | 42240.0 | 0 | Carnivora | 42240.0 | 0 | Canidae | 42240.0 | 0 | Canis | 42240.0 | 0 | Canis lupus | 42240.0 | 0 |
| scf7180000047672 | 15816 | 0.448 | 0 | 2737.6362 | Eukaryota | 4626.0 | 0 | Platyhelminthes | 2561.0 | 3 | Polystomatidea | 2561.0 | 3 | Polystomatidae | 2561.0 | 3 | Protopolystoma | 2561.0 | 3 | Protopolystoma xenopodis | 2561.0 | 3 |
| scf7180000041235 | 1287159 | 0.4158 | 0 | 59.8267 | Eukaryota | 77055.0 | 0 | Chordata | 77055.0 | 0 | Carnivora | 77055.0 | 0 | Canidae | 77055.0 | 0 | Canis | 77055.0 | 0 | Canis lupus | 77055.0 | 0 |
| scf7180000055353 | 33314 | 0.4405 | 0 | 111.4288 | Eukaryota | 492.0 | 0 | Platyhelminthes | 296.0 | 1 | Polystomatidea | 296.0 | 1 | Polystomatidae | 296.0 | 1 | Protopolystoma | 296.0 | 1 | Protopolystoma xenopodis | 296.0 | 1 |
| scf7180000040950 | 1104390 | 0.3808 | 0 | 59.5435 | Eukaryota | 55539.0 | 0 | Chordata | 55539.0 | 0 | Carnivora | 55539.0 | 0 | Canidae | 55539.0 | 0 | Canis | 55539.0 | 0 | Canis lupus | 55539.0 | 0 |
| scf7180000050952 | 18672 | 0.4416 | 0 | 158.2115 | Eukaryota | 29979.0 | 0 | Chordata | 18362.0 | 2 | Gadiformes | 15796.0 | 5 | Gadidae | 15796.0 | 5 | Gadus | 15796.0 | 5 | Gadus morhua | 15796.0 | 5 |
| scf7180000050603 | 4671 | 0.4312 | 0 | 107.5467 | no-hit | 0.0 | 0 | no-hit | 0.0 | 0 | no-hit | 0.0 | 0 | no-hit | 0.0 | 0 | no-hit | 0.0 | 0 | no-hit | 0.0 | 0 |
| scf7180000043108 | 86946 | 0.3955 | 0 | 54.2053 | Eukaryota | 25198.0 | 0 | Chordata | 25198.0 | 0 | Carnivora | 25198.0 | 0 | Canidae | 25198.0 | 0 | Canis | 25198.0 | 0 | Canis lupus | 25198.0 | 0 |
| scf7180000054780 | 135402 | 0.439 | 0 | 233.253 | Eukaryota | 60982.0 | 0 | Platyhelminthes | 35388.0 | 3 | Strigeidida | 23500.0 | 7 | Schistosomatidae | 23500.0 | 7 | Schistosoma | 23500.0 | 7 | Schistosoma mansoni | 23500.0 | 7 |
| scf7180000051804 | 2359 | 0.4544 | 0 | 427.2252 | no-hit | 0.0 | 0 | no-hit | 0.0 | 0 | no-hit | 0.0 | 0 | no-hit | 0.0 | 0 | no-hit | 0.0 | 0 | no-hit | 0.0 | 0 |
| scf7180000042880 | 629095 | 0.3752 | 0 | 59.4268 | Eukaryota | 23455.0 | 0 | Chordata | 23455.0 | 0 | Primates | 17077.0 | 2 | Hominidae | 17077.0 | 2 | Homo | 17077.0 | 3 | Homo sapiens | 17077.0 | 3 |
| scf7180000053872 | 2509 | 0.5026 | 0 | 89.0902 | Eukaryota | 6352.0 | 0 | Chordata | 6352.0 | 0 | Carnivora | 6352.0 | 0 | Canidae | 6352.0 | 0 | Canis | 4411.0 | 1 | Canis lupus | 3381.0 | 2 |
| scf7180000049471 | 28647 | 0.4332 | 0 | 193.1129 | Eukaryota | 2959.0 | 0 | Chordata | 1493.0 | 1 | Synbranchiformes | 1493.0 | 1 | Mastacembelidae | 1493.0 | 1 | Mastacembelus | 1493.0 | 1 | Mastacembelus armatus | 1493.0 | 1 |
| scf7180000040799 | 3948 | 0.4397 | 0 | 524.918 | no-hit | 0.0 | 0 | no-hit | 0.0 | 0 | no-hit | 0.0 | 0 | no-hit | 0.0 | 0 | no-hit | 0.0 | 0 | no-hit | 0.0 | 0 |
| scf7180000043928 | 14172 | 0.415 | 0 | 205.7121 | Eukaryota | 91525.0 | 0 | Chordata | 91525.0 | 0 | Carnivora | 91525.0 | 0 | Canidae | 91525.0 | 0 | Canis | 91525.0 | 0 | Canis lupus | 91525.0 | 0 |
| scf7180000049063 | 45472 | 0.4404 | 0 | 92.4772 | Eukaryota | 4831.0 | 0 | Chordata | 4831.0 | 0 | Carnivora | 2642.0 | 2 | Canidae | 2642.0 | 2 | Canis | 2642.0 | 2 | Canis lupus | 2642.0 | 2 |
| scf7180000046155 | 76234 | 0.3815 | 0 | 57.7566 | Eukaryota | 82189.0 | 0 | Chordata | 82189.0 | 0 | Carnivora | 82189.0 | 0 | Canidae | 82189.0 | 0 | Canis | 82189.0 | 0 | Canis lupus | 82189.0 | 0 |
| scf7180000052844 | 4673 | 0.4502 | 0 | 1151.7974 | no-hit | 0.0 | 0 | no-hit | 0.0 | 0 | no-hit | 0.0 | 0 | no-hit | 0.0 | 0 | no-hit | 0.0 | 0 | no-hit | 0.0 | 0 |
| scf7180000044179 | 340811 | 0.4271 | 0 | 60.3365 | Eukaryota | 76369.0 | 0 | Chordata | 76369.0 | 0 | Carnivora | 76369.0 | 0 | Canidae | 52537.0 | 2 | Canis | 43416.0 | 4 | Canis lupus | 43416.0 | 4 |
| scf7180000043432 | 271250 | 0.391 | 0 | 56.6244 | Eukaryota | 38359.0 | 0 | Chordata | 38359.0 | 0 | Carnivora | 38359.0 | 0 | Canidae | 38359.0 | 0 | Canis | 38359.0 | 0 | Canis lupus | 38359.0 | 0 |
| scf7180000055465 | 6679 | 0.4469 | 0 | 93.6419 | Eukaryota | 2635.0 | 0 | Chordata | 2635.0 | 0 | Carnivora | 2635.0 | 0 | Canidae | 2635.0 | 0 | Canis | 2635.0 | 0 | Canis lupus | 2635.0 | 0 |
| scf7180000040733 | 19804 | 0.3738 | 0 | 49.7205 | Eukaryota | 76689.0 | 0 | Chordata | 76689.0 | 0 | Carnivora | 76689.0 | 0 | Canidae | 76689.0 | 0 | Canis | 76689.0 | 0 | Canis lupus | 76689.0 | 0 |
| scf7180000042257 | 243338 | 0.4224 | 0 | 57.4866 | Eukaryota | 61283.0 | 0 | Chordata | 61283.0 | 0 | Carnivora | 61283.0 | 0 | Canidae | 61283.0 | 0 | Canis | 61283.0 | 0 | Canis lupus | 61283.0 | 0 |
| scf7180000043916 | 24873 | 0.4472 | 0 | 205.4501 | Eukaryota | 51448.0 | 0 | Chordata | 51448.0 | 0 | Carnivora | 51448.0 | 0 | Canidae | 51448.0 | 0 | Canis | 51448.0 | 0 | Canis lupus | 51448.0 | 0 |
| scf7180000054355 | 33159 | 0.4415 | 0 | 554.1335 | Eukaryota | 55224.0 | 0 | Chordata | 55224.0 | 0 | Carnivora | 55224.0 | 0 | Canidae | 55224.0 | 0 | Canis | 55224.0 | 0 | Canis lupus | 55224.0 | 0 |
| scf7180000050890 | 2669 | 0.4384 | 0 | 141.3698 | no-hit | 0.0 | 0 | no-hit | 0.0 | 0 | no-hit | 0.0 | 0 | no-hit | 0.0 | 0 | no-hit | 0.0 | 0 | no-hit | 0.0 | 0 |
| scf7180000052641 | 2443 | 0.4507 | 0 | 209.3567 | Eukaryota | 432.0 | 0 | Streptophyta | 432.0 | 0 | Fabales | 432.0 | 0 | Fabaceae | 432.0 | 0 | Cicer | 432.0 | 0 | Cicer arietinum | 432.0 | 0 |
| scf7180000042657 | 31511 | 0.3458 | 0 | 48.8874 | Eukaryota | 38235.0 | 0 | Chordata | 38235.0 | 0 | Carnivora | 38235.0 | 0 | Canidae | 38235.0 | 0 | Canis | 38235.0 | 0 | Canis lupus | 38235.0 | 0 |
| scf7180000046546 | 142678 | 0.3904 | 0 | 57.6143 | Eukaryota | 16288.0 | 0 | Chordata | 16288.0 | 0 | Carnivora | 16288.0 | 0 | Canidae | 16288.0 | 0 | Canis | 16288.0 | 0 | Canis lupus | 16288.0 | 0 |
| scf7180000041421 | 251729 | 0.4195 | 0 | 59.7316 | Eukaryota | 49158.0 | 0 | Chordata | 49158.0 | 0 | Carnivora | 49158.0 | 0 | Canidae | 49158.0 | 0 | Canis | 49158.0 | 0 | Canis lupus | 49158.0 | 0 |
| scf7180000045465 | 67585 | 0.3729 | 0 | 57.1047 | Eukaryota | 35163.0 | 0 | Chordata | 35163.0 | 0 | Carnivora | 35163.0 | 0 | Canidae | 35163.0 | 0 | Canis | 35163.0 | 0 | Canis lupus | 35163.0 | 0 |
| scf7180000041084 | 72699 | 0.407 | 0 | 58.2546 | Eukaryota | 67964.0 | 0 | Chordata | 67964.0 | 0 | Carnivora | 67964.0 | 0 | Canidae | 67964.0 | 0 | Canis | 67964.0 | 0 | Canis lupus | 67964.0 | 0 |
| scf7180000050564 | 30970 | 0.4342 | 0 | 57.4336 | Eukaryota | 19981.0 | 0 | Chordata | 19981.0 | 0 | Carnivora | 14705.0 | 3 | Canidae | 7616.0 | 4 | Vulpes | 5233.0 | 8 | Vulpes vulpes | 5233.0 | 8 |
| scf7180000055462 | 15935 | 0.442 | 0 | 533.9082 | Eukaryota | 1344.0 | 0 | Platyhelminthes | 993.0 | 1 | Polystomatidea | 993.0 | 1 | Polystomatidae | 993.0 | 1 | Protopolystoma | 993.0 | 1 | Protopolystoma xenopodis | 993.0 | 1 |
| scf7180000054702 | 21620 | 0.5244 | 0 | 223.1111 | Eukaryota | 3487.0 | 0 | Chordata | 3487.0 | 0 | Carnivora | 3487.0 | 0 | Canidae | 3487.0 | 0 | Canis | 2764.0 | 1 | Canis lupus | 2764.0 | 1 |
| scf7180000056286 | 969266 | 0.4091 | 0 | 59.5448 | Eukaryota | 46382.0 | 0 | Chordata | 46382.0 | 0 | Carnivora | 46382.0 | 0 | Canidae | 42394.0 | 1 | Vulpes | 22189.0 | 2 | Vulpes vulpes | 22189.0 | 2 |
| scf7180000047269 | 12659 | 0.4644 | 0 | 78.0716 | Eukaryota | 8242.0 | 0 | Chordata | 8242.0 | 0 | Carnivora | 8242.0 | 0 | Canidae | 8242.0 | 0 | Canis | 7126.0 | 1 | Canis lupus | 6015.0 | 2 |
| scf7180000052291 | 7876 | 0.4518 | 0 | 1757.3966 | Eukaryota | 3142.0 | 0 | Platyhelminthes | 1171.0 | 3 | Polystomatidea | 1171.0 | 3 | Polystomatidae | 1171.0 | 3 | Protopolystoma | 1171.0 | 4 | Protopolystoma xenopodis | 1171.0 | 4 |
| scf7180000045957 | 58730 | 0.4399 | 0 | 180.5021 | Eukaryota | 70623.0 | 0 | Chordata | 70623.0 | 0 | Carnivora | 70623.0 | 0 | Canidae | 70623.0 | 0 | Canis | 70623.0 | 0 | Canis lupus | 70623.0 | 0 |
| scf7180000042853 | 1337542 | 0.4116 | 0 | 59.7665 | Eukaryota | 47969.0 | 0 | Chordata | 47969.0 | 0 | Carnivora | 47969.0 | 0 | Canidae | 47969.0 | 0 | Canis | 47969.0 | 0 | Canis lupus | 47969.0 | 0 |
| scf7180000052207 | 3471 | 0.4587 | 0 | 300.0266 | Eukaryota | 1317.0 | 0 | Echinodermata | 569.0 | 2 | Forcipulatida | 569.0 | 2 | Asteriidae | 569.0 | 2 | Asterias | 569.0 | 2 | Asterias rubens | 569.0 | 2 |
| scf7180000052610 | 3814 | 0.4518 | 0 | 643.1083 | no-hit | 0.0 | 0 | no-hit | 0.0 | 0 | no-hit | 0.0 | 0 | no-hit | 0.0 | 0 | no-hit | 0.0 | 0 | no-hit | 0.0 | 0 |
| scf7180000052339 | 9328 | 0.507 | 0 | 334.0599 | Eukaryota | 8992.0 | 0 | Chordata | 8992.0 | 0 | Carnivora | 8992.0 | 0 | Canidae | 8992.0 | 0 | Canis | 7229.0 | 1 | Canis lupus | 5941.0 | 2 |
| scf7180000044483 | 458126 | 0.3723 | 0 | 61.1231 | Eukaryota | 111852.0 | 0 | Chordata | 111852.0 | 0 | Carnivora | 111852.0 | 0 | Canidae | 102022.0 | 1 | Canis | 65938.0 | 2 | Canis lupus | 65938.0 | 2 |
| scf7180000049976 | 36121 | 0.3662 | 0 | 59.1238 | Eukaryota | 7263.0 | 0 | Chordata | 6703.0 | 1 | Primates | 3065.0 | 6 | Hominidae | 3065.0 | 6 | Homo | 3065.0 | 6 | Homo sapiens | 3065.0 | 6 |
| scf7180000048978 | 19972 | 0.4413 | 0 | 62.2467 | Eukaryota | 6014.0 | 0 | Chordata | 5472.0 | 1 | Tetraodontiformes | 3235.0 | 4 | Tetraodontidae | 3235.0 | 4 | Takifugu | 3235.0 | 4 | Takifugu rubripes | 3235.0 | 4 |
| scf7180000047789 | 6933 | 0.4406 | 0 | 89.3104 | Eukaryota | 5894.0 | 0 | Chordata | 4736.0 | 1 | Kurtiformes | 4442.0 | 2 | Apogonidae | 4442.0 | 2 | Sphaeramia | 4442.0 | 2 | Sphaeramia orbicularis | 4442.0 | 2 |
| scf7180000048142 | 7653 | 0.4397 | 0 | 118.7693 | no-hit | 0.0 | 0 | no-hit | 0.0 | 0 | no-hit | 0.0 | 0 | no-hit | 0.0 | 0 | no-hit | 0.0 | 0 | no-hit | 0.0 | 0 |
| scf7180000055313 | 61401 | 0.4421 | 0 | 186.0253 | Eukaryota | 7202.0 | 0 | Chordata | 7202.0 | 0 | Carnivora | 6274.0 | 1 | Canidae | 6274.0 | 1 | Canis | 6274.0 | 1 | Canis lupus | 6274.0 | 1 |
| scf7180000049269 | 3707 | 0.4427 | 0 | 346.9807 | Eukaryota | 411.0 | 0 | Platyhelminthes | 239.0 | 1 | Polystomatidea | 239.0 | 1 | Polystomatidae | 239.0 | 1 | Protopolystoma | 239.0 | 1 | Protopolystoma xenopodis | 239.0 | 1 |
| scf7180000045689 | 32238 | 0.4354 | 0 | 117.9045 | Eukaryota | 88358.0 | 0 | Chordata | 88358.0 | 0 | Carnivora | 88358.0 | 0 | Canidae | 88358.0 | 0 | Canis | 88358.0 | 0 | Canis lupus | 88358.0 | 0 |
| scf7180000055723 | 22418 | 0.4331 | 0 | 190.1142 | no-hit | 0.0 | 0 | no-hit | 0.0 | 0 | no-hit | 0.0 | 0 | no-hit | 0.0 | 0 | no-hit | 0.0 | 0 | no-hit | 0.0 | 0 |
| scf7180000042029 | 236472 | 0.354 | 0 | 63.9134 | Eukaryota | 34938.0 | 0 | Chordata | 34938.0 | 0 | Carnivora | 34938.0 | 0 | Canidae | 34938.0 | 0 | Canis | 34938.0 | 0 | Canis lupus | 34938.0 | 0 |
| scf7180000045399 | 195262 | 0.3587 | 0 | 58.2 | Eukaryota | 64208.0 | 0 | Chordata | 64208.0 | 0 | Carnivora | 64208.0 | 0 | Canidae | 64208.0 | 0 | Canis | 64208.0 | 0 | Canis lupus | 64208.0 | 0 |
| scf7180000052941 | 6482 | 0.4426 | 0 | 56.6241 | Eukaryota | 2907.0 | 0 | Chordata | 2907.0 | 0 | Carnivora | 2907.0 | 0 | Canidae | 2907.0 | 0 | Canis | 2907.0 | 0 | Canis lupus | 2907.0 | 0 |
| scf7180000043715 | 565619 | 0.4262 | 0 | 58.447 | Eukaryota | 43196.0 | 0 | Chordata | 43196.0 | 0 | Carnivora | 43196.0 | 0 | Canidae | 43196.0 | 0 | Canis | 43196.0 | 0 | Canis lupus | 43196.0 | 0 |
| scf7180000043596 | 411608 | 0.4511 | 0 | 58.3257 | Eukaryota | 66083.0 | 0 | Chordata | 66083.0 | 0 | Carnivora | 66083.0 | 0 | Canidae | 66083.0 | 0 | Vulpes | 53683.0 | 1 | Vulpes vulpes | 53683.0 | 1 |
| scf7180000054585 | 16260 | 0.4689 | 0 | 214.4558 | Eukaryota | 15946.0 | 0 | Chordata | 15946.0 | 0 | Carnivora | 15946.0 | 0 | Canidae | 14324.0 | 1 | Canis | 14324.0 | 1 | Canis lupus | 14324.0 | 1 |
| scf7180000044818 | 245110 | 0.3446 | 0 | 61.7146 | Eukaryota | 23014.0 | 1 | Chordata | 23014.0 | 1 | Primates | 17434.0 | 3 | Hominidae | 17434.0 | 4 | Homo | 11614.0 | 5 | Homo sapiens | 11614.0 | 5 |
| scf7180000048071 | 7778 | 0.4479 | 0 | 104.1474 | no-hit | 0.0 | 0 | no-hit | 0.0 | 0 | no-hit | 0.0 | 0 | no-hit | 0.0 | 0 | no-hit | 0.0 | 0 | no-hit | 0.0 | 0 |
| scf7180000056449 | 496046 | 0.3629 | 0 | 59.3003 | Eukaryota | 56538.0 | 0 | Chordata | 56538.0 | 0 | Carnivora | 56538.0 | 0 | Canidae | 56538.0 | 0 | Canis | 56538.0 | 0 | Canis lupus | 56538.0 | 0 |
| scf7180000042891 | 599677 | 0.4786 | 0 | 58.6053 | Eukaryota | 43545.0 | 0 | Chordata | 43545.0 | 0 | Carnivora | 43545.0 | 0 | Canidae | 43545.0 | 0 | Canis | 34242.0 | 1 | Canis lupus | 34242.0 | 1 |
| scf7180000048872 | 36286 | 0.4497 | 0 | 84.0513 | Eukaryota | 85905.0 | 0 | Chordata | 85905.0 | 0 | Carnivora | 85905.0 | 0 | Canidae | 85905.0 | 0 | Canis | 85905.0 | 0 | Canis lupus | 85905.0 | 0 |
| scf7180000047014 | 5275 | 0.4351 | 0 | 62.0083 | no-hit | 0.0 | 0 | no-hit | 0.0 | 0 | no-hit | 0.0 | 0 | no-hit | 0.0 | 0 | no-hit | 0.0 | 0 | no-hit | 0.0 | 0 |
| scf7180000043655 | 392971 | 0.3377 | 0 | 59.1536 | Eukaryota | 13871.0 | 0 | Chordata | 13871.0 | 0 | Carnivora | 13871.0 | 0 | Canidae | 13871.0 | 0 | Canis | 13871.0 | 0 | Canis lupus | 13871.0 | 0 |
| scf7180000054154 | 8976 | 0.44 | 0 | 838.5189 | no-hit | 0.0 | 0 | no-hit | 0.0 | 0 | no-hit | 0.0 | 0 | no-hit | 0.0 | 0 | no-hit | 0.0 | 0 | no-hit | 0.0 | 0 |
| scf7180000048080 | 66048 | 0.5551 | 0 | 57.8525 | Eukaryota | 17494.0 | 0 | Chordata | 17494.0 | 0 | Carnivora | 17494.0 | 0 | Canidae | 17494.0 | 0 | Vulpes | 11037.0 | 1 | Vulpes vulpes | 11037.0 | 1 |
| scf7180000048533 | 24819 | 0.4414 | 0 | 76.0293 | Eukaryota | 26421.0 | 1 | Chordata | 26421.0 | 1 | Gadiformes | 10054.0 | 5 | Gadidae | 10054.0 | 5 | Gadus | 10054.0 | 5 | Gadus morhua | 10054.0 | 6 |
| scf7180000042607 | 681347 | 0.3877 | 0 | 60.8285 | Eukaryota | 108541.0 | 0 | Chordata | 108541.0 | 0 | Carnivora | 108541.0 | 0 | Canidae | 108541.0 | 0 | Canis | 108541.0 | 0 | Canis lupus | 108541.0 | 0 |
| scf7180000041209 | 1455834 | 0.4181 | 0 | 60.6831 | Eukaryota | 68793.0 | 0 | Chordata | 68793.0 | 0 | Carnivora | 68793.0 | 0 | Canidae | 59357.0 | 1 | Canis | 37660.0 | 3 | Canis lupus | 37660.0 | 3 |
| scf7180000048009 | 11465 | 0.4495 | 0 | 251.9337 | Eukaryota | 3291.0 | 0 | Chordata | 3291.0 | 0 | Carnivora | 2500.0 | 1 | Canidae | 2500.0 | 1 | Canis | 2500.0 | 1 | Canis lupus | 2500.0 | 1 |
| scf7180000046178 | 116012 | 0.4157 | 0 | 52.1515 | Eukaryota | 22326.0 | 0 | Chordata | 22326.0 | 0 | Carnivora | 22326.0 | 0 | Canidae | 22326.0 | 0 | Canis | 15582.0 | 1 | Canis lupus | 15582.0 | 1 |
| scf7180000045329 | 36597 | 0.4831 | 0 | 55.116 | Eukaryota | 6574.0 | 0 | Chordata | 6574.0 | 0 | Carnivora | 6154.0 | 1 | Mustelidae | 3031.0 | 4 | Mustela | 3031.0 | 6 | Mustela putorius | 3031.0 | 6 |
| scf7180000048106 | 73210 | 0.3757 | 0 | 60.0055 | Eukaryota | 19078.0 | 0 | Chordata | 19078.0 | 0 | Carnivora | 19078.0 | 0 | Canidae | 11963.0 | 3 | Canis | 11963.0 | 3 | Canis lupus | 11963.0 | 3 |
| scf7180000045633 | 161753 | 0.5156 | 0 | 56.3066 | Eukaryota | 38905.0 | 0 | Chordata | 38905.0 | 0 | Carnivora | 38905.0 | 0 | Canidae | 38905.0 | 0 | Canis | 31268.0 | 1 | Canis lupus | 31268.0 | 1 |
| scf7180000055405 | 13857 | 0.4295 | 0 | 117.5366 | Eukaryota | 3974.0 | 0 | Echinodermata | 1991.0 | 2 | Forcipulatida | 1991.0 | 2 | Asteriidae | 1991.0 | 2 | Asterias | 1991.0 | 2 | Asterias rubens | 1991.0 | 2 |
| scf7180000052005 | 6416 | 0.4353 | 0 | 21.1103 | Eukaryota | 200.0 | 0 | Chordata | 200.0 | 0 | Salmoniformes | 200.0 | 0 | Salmonidae | 200.0 | 0 | Coregonus | 200.0 | 0 | Coregonus sp. 'balchen' | 200.0 | 0 |
| scf7180000044297 | 60691 | 0.3875 | 0 | 55.5114 | Eukaryota | 20123.0 | 0 | Chordata | 20123.0 | 0 | Carnivora | 20123.0 | 0 | Canidae | 20123.0 | 0 | Canis | 20123.0 | 0 | Canis lupus | 20123.0 | 0 |
| scf7180000053685 | 5155 | 0.4433 | 0 | 98.4588 | Eukaryota | 15006.0 | 0 | Chordata | 12617.0 | 2 | Chordata-undef | 6166.0 | 4 | Ambassidae | 6166.0 | 4 | Parambassis | 6166.0 | 4 | Parambassis ranga | 6166.0 | 4 |
| scf7180000041192 | 1302102 | 0.354 | 0 | 59.7225 | Eukaryota | 198516.0 | 0 | Chordata | 198516.0 | 0 | Carnivora | 189598.0 | 1 | Canidae | 150717.0 | 3 | Canis | 140128.0 | 4 | Canis lupus | 140128.0 | 4 |
| scf7180000055997 | 5718 | 0.4388 | 0 | 63.1886 | Eukaryota | 23228.0 | 0 | Chordata | 18754.0 | 1 | Gadiformes | 10985.0 | 5 | Gadidae | 10985.0 | 5 | Gadus | 10985.0 | 5 | Gadus morhua | 10985.0 | 5 |
| scf7180000044863 | 257069 | 0.4178 | 0 | 59.2996 | Eukaryota | 105028.0 | 0 | Chordata | 105028.0 | 0 | Carnivora | 105028.0 | 0 | Canidae | 105028.0 | 0 | Canis | 105028.0 | 0 | Canis lupus | 105028.0 | 0 |
| scf7180000050634 | 8265 | 0.4479 | 0 | 1183.7479 | Eukaryota | 1654.0 | 0 | Echinodermata | 933.0 | 2 | Forcipulatida | 933.0 | 2 | Asteriidae | 933.0 | 2 | Asterias | 933.0 | 2 | Asterias rubens | 933.0 | 2 |
| scf7180000056204 | 1050047 | 0.402 | 0 | 60.7234 | Eukaryota | 92052.0 | 0 | Chordata | 92052.0 | 0 | Carnivora | 92052.0 | 0 | Canidae | 92052.0 | 0 | Canis | 92052.0 | 0 | Canis lupus | 92052.0 | 0 |
| scf7180000040580 | 1182998 | 0.4433 | 0 | 59.1426 | Eukaryota | 130648.0 | 0 | Chordata | 130648.0 | 0 | Carnivora | 130648.0 | 0 | Canidae | 130648.0 | 0 | Canis | 130648.0 | 0 | Canis lupus | 130648.0 | 0 |
| scf7180000046386 | 9369 | 0.4205 | 0 | 115.2775 | Eukaryota | 81894.0 | 0 | Chordata | 81894.0 | 0 | Carnivora | 81894.0 | 0 | Canidae | 81894.0 | 0 | Canis | 81894.0 | 0 | Canis lupus | 81894.0 | 0 |
| scf7180000055151 | 27141 | 0.4377 | 0 | 517.7754 | Eukaryota | 45211.0 | 0 | Chordata | 37117.0 | 2 | Gadiformes | 25909.0 | 6 | Gadidae | 25909.0 | 6 | Gadus | 25909.0 | 6 | Gadus morhua | 25909.0 | 6 |
| scf7180000053075 | 2737 | 0.4593 | 0 | 754.4611 | no-hit | 0.0 | 0 | no-hit | 0.0 | 0 | no-hit | 0.0 | 0 | no-hit | 0.0 | 0 | no-hit | 0.0 | 0 | no-hit | 0.0 | 0 |
| scf7180000041415 | 588807 | 0.4405 | 0 | 60.7236 | Eukaryota | 58891.0 | 0 | Chordata | 58891.0 | 0 | Carnivora | 58891.0 | 0 | Canidae | 49915.0 | 1 | Canis | 38366.0 | 3 | Canis lupus | 38366.0 | 3 |
| scf7180000045242 | 198295 | 0.4687 | 0 | 62.1685 | Eukaryota | 52003.0 | 0 | Chordata | 52003.0 | 0 | Carnivora | 52003.0 | 0 | Canidae | 52003.0 | 0 | Canis | 52003.0 | 0 | Canis lupus | 52003.0 | 0 |
| scf7180000041629 | 796615 | 0.4189 | 0 | 60.5005 | Eukaryota | 102701.0 | 0 | Chordata | 102701.0 | 0 | Carnivora | 102701.0 | 0 | Otariidae | 55488.0 | 2 | Ailuropoda | 26781.0 | 4 | Ailuropoda melanoleuca | 26781.0 | 4 |
| scf7180000041296 | 174966 | 0.3539 | 100 | 58.8023 | Eukaryota | 84722.0 | 0 | Chordata | 84722.0 | 0 | Carnivora | 84722.0 | 0 | Canidae | 84722.0 | 0 | Canis | 84722.0 | 0 | Canis lupus | 84722.0 | 0 |
| scf7180000043161 | 461223 | 0.4153 | 0 | 61.9885 | Eukaryota | 89488.0 | 0 | Chordata | 89488.0 | 0 | Carnivora | 89488.0 | 0 | Canidae | 89488.0 | 0 | Canis | 89488.0 | 0 | Canis lupus | 89488.0 | 0 |
| scf7180000055514 | 17099 | 0.4316 | 0 | 119.7073 | no-hit | 0.0 | 0 | no-hit | 0.0 | 0 | no-hit | 0.0 | 0 | no-hit | 0.0 | 0 | no-hit | 0.0 | 0 | no-hit | 0.0 | 0 |
| scf7180000051272 | 21592 | 0.4743 | 0 | 57.3415 | Eukaryota | 16365.0 | 0 | Chordata | 16365.0 | 0 | Carnivora | 16365.0 | 0 | Canidae | 16365.0 | 0 | Canis | 15775.0 | 1 | Canis lupus | 15775.0 | 1 |
| scf7180000049109 | 8765 | 0.5045 | 0 | 168.5999 | Eukaryota | 8443.0 | 0 | Chordata | 8443.0 | 0 | Carnivora | 8443.0 | 0 | Canidae | 8443.0 | 0 | Canis | 7361.0 | 1 | Canis lupus | 6229.0 | 2 |
| scf7180000056568 | 152634 | 0.5278 | 0 | 56.0691 | Eukaryota | 46726.0 | 0 | Chordata | 46726.0 | 0 | Carnivora | 46726.0 | 0 | Canidae | 25263.0 | 4 | Canis | 19722.0 | 8 | Canis lupus | 19722.0 | 8 |
| scf7180000041560 | 994379 | 0.4287 | 0 | 59.6775 | Eukaryota | 140303.0 | 0 | Chordata | 140303.0 | 0 | Carnivora | 140303.0 | 0 | Canidae | 140303.0 | 0 | Canis | 132155.0 | 1 | Canis lupus | 132155.0 | 1 |
| scf7180000052584 | 2161 | 0.4489 | 0 | 217.7113 | no-hit | 0.0 | 0 | no-hit | 0.0 | 0 | no-hit | 0.0 | 0 | no-hit | 0.0 | 0 | no-hit | 0.0 | 0 | no-hit | 0.0 | 0 |
| scf7180000050847 | 3630 | 0.441 | 0 | 767.3564 | no-hit | 0.0 | 0 | no-hit | 0.0 | 0 | no-hit | 0.0 | 0 | no-hit | 0.0 | 0 | no-hit | 0.0 | 0 | no-hit | 0.0 | 0 |
| scf7180000054555 | 24415 | 0.4403 | 0 | 696.0605 | Eukaryota | 6701.0 | 0 | Platyhelminthes | 4636.0 | 1 | Cyclophyllidea | 4366.0 | 2 | Hymenolepididae | 4366.0 | 2 | Hymenolepis | 4366.0 | 2 | Hymenolepis microstoma | 4366.0 | 2 |
| scf7180000042375 | 666955 | 0.4214 | 0 | 60.5297 | Eukaryota | 58625.0 | 0 | Chordata | 58625.0 | 0 | Carnivora | 58625.0 | 0 | Canidae | 58625.0 | 0 | Vulpes | 53775.0 | 1 | Vulpes vulpes | 53775.0 | 1 |
| scf7180000048183 | 52607 | 0.4454 | 0 | 58.6056 | Eukaryota | 14805.0 | 0 | Chordata | 14805.0 | 0 | Carnivora | 13419.0 | 1 | Canidae | 5029.0 | 4 | Canis | 5029.0 | 7 | Canis lupus | 5029.0 | 7 |
| scf7180000051663 | 13447 | 0.4439 | 0 | 58.0251 | Eukaryota | 8718.0 | 0 | Chordata | 8718.0 | 0 | Carnivora | 8718.0 | 0 | Felidae | 7859.0 | 1 | Felis | 7859.0 | 1 | Felis catus | 7859.0 | 1 |
| scf7180000045732 | 236418 | 0.3905 | 0 | 61.6782 | Eukaryota | 68890.0 | 0 | Chordata | 68890.0 | 0 | Carnivora | 68890.0 | 0 | Canidae | 68890.0 | 0 | Canis | 68890.0 | 0 | Canis lupus | 68890.0 | 0 |
| scf7180000048446 | 68894 | 0.4441 | 0 | 175.8116 | Eukaryota | 22150.0 | 0 | Chordata | 17978.0 | 1 | Carnivora | 14242.0 | 2 | Canidae | 14242.0 | 2 | Canis | 14242.0 | 2 | Canis lupus | 14242.0 | 2 |
| scf7180000056518 | 244104 | 0.4423 | 0 | 60.42 | Eukaryota | 67486.0 | 0 | Chordata | 67486.0 | 0 | Carnivora | 67486.0 | 0 | Canidae | 67486.0 | 0 | Vulpes | 40590.0 | 1 | Vulpes vulpes | 40590.0 | 1 |
| scf7180000049403 | 8042 | 0.4549 | 0 | 194.759 | Eukaryota | 475.0 | 0 | Chordata | 475.0 | 0 | Syngnathiformes | 475.0 | 0 | Syngnathidae | 475.0 | 0 | Syngnathus | 475.0 | 0 | Syngnathus acus | 475.0 | 0 |
| scf7180000056603 | 321957 | 0.3708 | 100 | 58.3984 | Eukaryota | 107907.0 | 0 | Chordata | 107907.0 | 0 | Carnivora | 107907.0 | 0 | Canidae | 107907.0 | 0 | Canis | 107907.0 | 0 | Canis lupus | 107907.0 | 0 |
| scf7180000045078 | 73504 | 0.4385 | 0 | 292.4266 | Eukaryota | 101416.0 | 0 | Chordata | 57671.0 | 1 | Gadiformes | 57671.0 | 2 | Gadidae | 57671.0 | 2 | Gadus | 57671.0 | 2 | Gadus morhua | 57671.0 | 2 |
| scf7180000050628 | 4422 | 0.4435 | 0 | 458.265 | no-hit | 0.0 | 0 | no-hit | 0.0 | 0 | no-hit | 0.0 | 0 | no-hit | 0.0 | 0 | no-hit | 0.0 | 0 | no-hit | 0.0 | 0 |
| scf7180000042932 | 617814 | 0.4014 | 0 | 60.6794 | Eukaryota | 88827.0 | 0 | Chordata | 88827.0 | 0 | Carnivora | 88827.0 | 0 | Canidae | 88827.0 | 0 | Canis | 88827.0 | 0 | Canis lupus | 88827.0 | 0 |
| scf7180000056243 | 226473 | 0.3587 | 0 | 64.3441 | Eukaryota | 95254.0 | 0 | Chordata | 95254.0 | 0 | Carnivora | 95254.0 | 0 | Canidae | 95254.0 | 0 | Canis | 95254.0 | 0 | Canis lupus | 95254.0 | 0 |
| scf7180000048907 | 22563 | 0.4394 | 0 | 285.6597 | Eukaryota | 68860.0 | 0 | Chordata | 43483.0 | 1 | Gadiformes | 34885.0 | 4 | Gadidae | 34885.0 | 4 | Gadus | 34885.0 | 4 | Gadus morhua | 34885.0 | 4 |
| scf7180000049402 | 12616 | 0.5744 | 0 | 353.8189 | Eukaryota | 56140.0 | 0 | Chordata | 56140.0 | 0 | Primates | 21929.0 | 3 | Bovidae | 16695.0 | 4 | Ovis | 16695.0 | 5 | Ovis aries | 16695.0 | 5 |
| scf7180000054179 | 208882 | 0.4318 | 0 | 57.456 | Eukaryota | 29156.0 | 0 | Chordata | 29156.0 | 0 | Carnivora | 29156.0 | 0 | Canidae | 29156.0 | 0 | Canis | 29156.0 | 0 | Canis lupus | 29156.0 | 0 |
| scf7180000042452 | 656030 | 0.363 | 0 | 60.0513 | Eukaryota | 104534.0 | 0 | Chordata | 104534.0 | 0 | Carnivora | 104534.0 | 0 | Canidae | 104534.0 | 0 | Canis | 84912.0 | 1 | Canis lupus | 84912.0 | 1 |
| scf7180000044343 | 10614 | 0.3522 | 0 | 29.9837 | Eukaryota | 2319.0 | 0 | Chordata | 2319.0 | 0 | Carnivora | 2319.0 | 0 | Canidae | 2319.0 | 0 | Canis | 2319.0 | 0 | Canis lupus | 2319.0 | 0 |
| scf7180000052321 | 3214 | 0.4437 | 0 | 597.9994 | no-hit | 0.0 | 0 | no-hit | 0.0 | 0 | no-hit | 0.0 | 0 | no-hit | 0.0 | 0 | no-hit | 0.0 | 0 | no-hit | 0.0 | 0 |
| scf7180000048288 | 48834 | 0.3934 | 0 | 61.1699 | Eukaryota | 53109.0 | 0 | Chordata | 53109.0 | 0 | Carnivora | 53109.0 | 0 | Canidae | 53109.0 | 0 | Canis | 53109.0 | 0 | Canis lupus | 53109.0 | 0 |
| scf7180000052733 | 3121 | 0.5107 | 0 | 124.8146 | Eukaryota | 4508.0 | 0 | Chordata | 4508.0 | 0 | Carnivora | 4508.0 | 0 | Canidae | 4508.0 | 0 | Nyctereutes | 2382.0 | 1 | Nyctereutes procyonoides | 2382.0 | 2 |
| scf7180000047538 | 100108 | 0.3821 | 0 | 61.3777 | Eukaryota | 48145.0 | 0 | Chordata | 48145.0 | 0 | Carnivora | 48145.0 | 0 | Canidae | 25973.0 | 6 | Canis | 14911.0 | 8 | Canis lupus | 14911.0 | 8 |
| scf7180000053194 | 2204 | 0.446 | 0 | 1337.5725 | no-hit | 0.0 | 0 | no-hit | 0.0 | 0 | no-hit | 0.0 | 0 | no-hit | 0.0 | 0 | no-hit | 0.0 | 0 | no-hit | 0.0 | 0 |
| scf7180000042645 | 606288 | 0.4312 | 0 | 59.855 | Eukaryota | 64075.0 | 0 | Chordata | 64075.0 | 0 | Carnivora | 64075.0 | 0 | Canidae | 64075.0 | 0 | Canis | 64075.0 | 0 | Canis lupus | 64075.0 | 0 |
| scf7180000047794 | 83272 | 0.3848 | 0 | 57.1141 | Eukaryota | 30448.0 | 0 | Chordata | 30448.0 | 0 | Carnivora | 30448.0 | 0 | Canidae | 30448.0 | 0 | Canis | 30448.0 | 0 | Canis lupus | 30448.0 | 0 |
| scf7180000055090 | 10149 | 0.4372 | 0 | 95.5851 | Eukaryota | 377.0 | 0 | Chordata | 377.0 | 0 | Tetraodontiformes | 377.0 | 0 | Tetraodontidae | 377.0 | 0 | Takifugu | 377.0 | 0 | Takifugu rubripes | 377.0 | 0 |
| scf7180000054829 | 25072 | 0.4415 | 0 | 320.5034 | Eukaryota | 5465.0 | 0 | Chordata | 3311.0 | 1 | Kurtiformes | 2690.0 | 2 | Apogonidae | 2690.0 | 2 | Sphaeramia | 2690.0 | 2 | Sphaeramia orbicularis | 2690.0 | 2 |
| scf7180000047058 | 20678 | 0.3845 | 0 | 48.525 | Eukaryota | 82676.0 | 0 | Chordata | 82676.0 | 0 | Carnivora | 82676.0 | 0 | Canidae | 82676.0 | 0 | Canis | 82676.0 | 0 | Canis lupus | 82676.0 | 0 |
| scf7180000047936 | 7910 | 0.4067 | 0 | 25.5119 | Eukaryota | 1593.0 | 0 | Chordata | 1593.0 | 0 | Carnivora | 1593.0 | 0 | Canidae | 1593.0 | 0 | Vulpes | 1593.0 | 0 | Vulpes vulpes | 1593.0 | 0 |
| scf7180000046942 | 56008 | 0.5425 | 0 | 56.1015 | Eukaryota | 10208.0 | 0 | Chordata | 10208.0 | 0 | Carnivora | 10208.0 | 0 | Canidae | 8777.0 | 2 | Canis | 5628.0 | 3 | Canis lupus | 5628.0 | 3 |
| scf7180000049326 | 22811 | 0.4494 | 0 | 1872.4095 | Eukaryota | 239.0 | 0 | Streptophyta | 239.0 | 0 | Fabales | 239.0 | 0 | Fabaceae | 239.0 | 0 | Cicer | 239.0 | 0 | Cicer arietinum | 239.0 | 0 |
| scf7180000045019 | 564430 | 0.428 | 0 | 60.6372 | Eukaryota | 61030.0 | 0 | Chordata | 61030.0 | 0 | Carnivora | 61030.0 | 0 | Canidae | 55549.0 | 1 | Vulpes | 55549.0 | 1 | Vulpes vulpes | 55549.0 | 1 |
| scf7180000043305 | 479835 | 0.3576 | 0 | 60.5079 | Eukaryota | 34006.0 | 0 | Chordata | 34006.0 | 0 | Carnivora | 34006.0 | 0 | Canidae | 34006.0 | 0 | Canis | 34006.0 | 0 | Canis lupus | 34006.0 | 0 |
| scf7180000043777 | 50233 | 0.3627 | 0 | 64.1518 | Eukaryota | 92262.0 | 0 | Chordata | 92262.0 | 0 | Carnivora | 92262.0 | 0 | Canidae | 92262.0 | 0 | Canis | 92262.0 | 0 | Canis lupus | 92262.0 | 0 |
| scf7180000044397 | 327745 | 0.5773 | 0 | 57.7355 | Eukaryota | 31741.0 | 0 | Chordata | 31741.0 | 0 | Carnivora | 31741.0 | 0 | Canidae | 31741.0 | 0 | Canis | 25325.0 | 1 | Canis lupus | 22420.0 | 2 |
| scf7180000044068 | 493962 | 0.4158 | 100 | 59.7881 | Eukaryota | 30101.0 | 0 | Chordata | 30101.0 | 0 | Carnivora | 30101.0 | 0 | Canidae | 30101.0 | 0 | Canis | 30101.0 | 0 | Canis lupus | 30101.0 | 0 |
| scf7180000052883 | 11963 | 0.5072 | 0 | 795.4453 | Eukaryota | 8902.0 | 0 | Chordata | 8902.0 | 0 | Carnivora | 8902.0 | 0 | Canidae | 8902.0 | 0 | Canis | 7234.0 | 1 | Canis lupus | 6044.0 | 2 |
| scf7180000053563 | 7661 | 0.5067 | 0 | 41.8808 | Eukaryota | 8368.0 | 0 | Chordata | 8368.0 | 0 | Carnivora | 8368.0 | 0 | Canidae | 8368.0 | 0 | Canis | 6755.0 | 1 | Canis lupus | 5597.0 | 2 |
| scf7180000043131 | 259255 | 0.4182 | 0 | 58.9707 | Eukaryota | 39654.0 | 0 | Chordata | 39654.0 | 0 | Carnivora | 39654.0 | 0 | Canidae | 39654.0 | 0 | Canis | 39654.0 | 0 | Canis lupus | 39654.0 | 0 |
| scf7180000044457 | 125874 | 0.3876 | 0 | 57.4986 | Eukaryota | 63468.0 | 0 | Chordata | 63468.0 | 0 | Carnivora | 63468.0 | 0 | Canidae | 63468.0 | 0 | Canis | 63468.0 | 0 | Canis lupus | 63468.0 | 0 |
| scf7180000053772 | 2079 | 0.4483 | 0 | 48.0634 | Eukaryota | 178.0 | 0 | Chordata | 178.0 | 0 | Osteoglossiformes | 178.0 | 0 | Osteoglossidae | 178.0 | 0 | Scleropages | 178.0 | 0 | Scleropages formosus | 178.0 | 0 |
| scf7180000044351 | 347237 | 0.4646 | 0 | 58.6522 | Eukaryota | 57189.0 | 0 | Chordata | 57189.0 | 0 | Carnivora | 57189.0 | 0 | Canidae | 40061.0 | 2 | Vulpes | 40061.0 | 2 | Vulpes vulpes | 40061.0 | 2 |
| scf7180000045747 | 193751 | 0.3962 | 0 | 59.0856 | Eukaryota | 26712.0 | 0 | Chordata | 26712.0 | 0 | Carnivora | 26712.0 | 0 | Canidae | 26712.0 | 0 | Canis | 26712.0 | 0 | Canis lupus | 26712.0 | 0 |
| scf7180000044435 | 14137 | 0.5822 | 0 | 48.4377 | Eukaryota | 21287.0 | 0 | Chordata | 21287.0 | 0 | Carnivora | 20875.0 | 1 | Canidae | 19979.0 | 2 | Canis | 19979.0 | 2 | Canis lupus | 19979.0 | 2 |
| scf7180000040847 | 30141 | 0.3661 | 0 | 61.8062 | Eukaryota | 63727.0 | 0 | Chordata | 63727.0 | 0 | Carnivora | 63727.0 | 0 | Canidae | 63727.0 | 0 | Canis | 63727.0 | 0 | Canis lupus | 63727.0 | 0 |

| | | | | | | | | | | | | | | | | | | | | | |
|---|---|---|---|---|---|---|---|---|---|---|---|---|---|---|---|---|---|---|---|---|---|
| scf7180000045374 | 179822 | 0.3792 | 0 | 60.35 | Eukaryota | 62397.0 | 0 | Chordata | 62397.0 | 0 | Carnivora | 62397.0 | 0 | Canidae | 62397.0 | 0 | Canis | 62397.0 | 0 | Canis lupus | 62397.0 | 0 |
| scf7180000051461 | 35051 | 0.3777 | 0 | 61.8009 | Eukaryota | 5711.0 | 0 | Chordata | 5711.0 | 0 | Carnivora | 4900.0 | 1 | Canidae | 4272.0 | 2 | Canis | 4272.0 | 2 | Canis lupus | 4272.0 | 2 |
| scf7180000050793 | 3039 | 0.4442 | 0 | 80.2317 | no-hit | 0.0 | 0 | no-hit | 0.0 | 0 | no-hit | 0.0 | 0 | no-hit | 0.0 | 0 | no-hit | 0.0 | 0 | no-hit | 0.0 | 0 |
| scf7180000052971 | 3010 | 0.4435 | 0 | 331.4538 | Eukaryota | 10528.0 | 0 | Chordata | 10047.0 | 1 | Gadiformes | 4677.0 | 6 | Gadidae | 4677.0 | 6 | Gadus | 4677.0 | 6 | Gadus morhua | 4677.0 | 6 |
| scf7180000044208 | 26309 | 0.3712 | 0 | 56.9415 | Eukaryota | 7974.0 | 0 | Chordata | 7974.0 | 0 | Carnivora | 6044.0 | 1 | Otariidae | 3352.0 | 2 | Canis | 2692.0 | 4 | Canis lupus | 2692.0 | 4 |
| scf7180000056337 | 535861 | 0.4063 | 0 | 56.294 | Eukaryota | 38320.0 | 0 | Chordata | 38320.0 | 0 | Carnivora | 38320.0 | 0 | Canidae | 38320.0 | 0 | Canis | 33173.0 | 1 | Canis lupus | 33173.0 | 1 |
| scf7180000045535 | 223299 | 0.3891 | 0 | 59.4064 | Eukaryota | 19916.0 | 0 | Chordata | 19916.0 | 0 | Primates | 14610.0 | 3 | Hominidae | 10883.0 | 4 | Homo | 8949.0 | 5 | Homo sapiens | 8949.0 | 5 |
| scf7180000054871 | 21618 | 0.4407 | 0 | 159.1124 | Eukaryota | 9553.0 | 0 | Chordata | 8150.0 | 1 | Kurtiformes | 4464.0 | 3 | Apogonidae | 4464.0 | 3 | Sphaeramia | 4464.0 | 3 | Sphaeramia orbicularis | 4464.0 | 3 |
| scf7180000043004 | 11234 | 0.361 | 0 | 52.0367 | Eukaryota | 32261.0 | 0 | Chordata | 32261.0 | 0 | Carnivora | 32261.0 | 0 | Canidae | 32261.0 | 0 | Canis | 32261.0 | 0 | Canis lupus | 32261.0 | 0 |
| scf7180000041679 | 1718340 | 0.521 | 0 | 58.7198 | Eukaryota | 69479.0 | 0 | Chordata | 69479.0 | 0 | Carnivora | 69479.0 | 0 | Canidae | 69479.0 | 0 | Vulpes | 49047.0 | 1 | Vulpes vulpes | 49047.0 | 1 |
| scf7180000045940 | 40011 | 0.4142 | 0 | 216.0126 | Eukaryota | 48056.0 | 0 | Chordata | 48056.0 | 0 | Carnivora | 48056.0 | 0 | Canidae | 48056.0 | 0 | Canis | 48056.0 | 0 | Canis lupus | 48056.0 | 0 |
| scf7180000046323 | 32326 | 0.4392 | 0 | 124.8163 | Eukaryota | 10216.0 | 0 | Chordata | 10216.0 | 0 | Tetraodontiformes | 6313.0 | 2 | Tetraodontidae | 6313.0 | 2 | Takifugu | 6313.0 | 2 | Takifugu rubripes | 6313.0 | 2 |
| scf7180000045371 | 205827 | 0.3869 | 0 | 60.5199 | Eukaryota | 27307.0 | 0 | Chordata | 27307.0 | 0 | Carnivora | 27307.0 | 0 | Canidae | 27307.0 | 0 | Canis | 27307.0 | 0 | Canis lupus | 27307.0 | 0 |
| scf7180000042391 | 60110 | 0.5135 | 0 | 54.9679 | Eukaryota | 6497.0 | 0 | Chordata | 6497.0 | 0 | Carnivora | 2565.0 | 2 | Canidae | 2565.0 | 2 | Canis | 2565.0 | 3 | Canis lupus | 2565.0 | 3 |
| scf7180000041977 | 115966 | 0.3924 | 0 | 57.9312 | Eukaryota | 9271.0 | 0 | Chordata | 9271.0 | 0 | Carnivora | 7510.0 | 1 | Canidae | 4173.0 | 3 | Canis | 3009.0 | 6 | Canis lupus | 3009.0 | 6 |
| scf7180000042349 | 654934 | 0.4313 | 0 | 59.2665 | Eukaryota | 45451.0 | 0 | Chordata | 45451.0 | 0 | Carnivora | 45451.0 | 0 | Canidae | 45451.0 | 0 | Canis | 28029.0 | 1 | Canis lupus | 28029.0 | 1 |
| scf7180000049901 | 3148 | 0.4527 | 0 | 353.708 | no-hit | 0.0 | 0 | no-hit | 0.0 | 0 | no-hit | 0.0 | 0 | no-hit | 0.0 | 0 | no-hit | 0.0 | 0 | no-hit | 0.0 | 0 |
| scf7180000042482 | 258992 | 0.345 | 0 | 57.9254 | Eukaryota | 85586.0 | 0 | Chordata | 85586.0 | 0 | Carnivora | 85586.0 | 0 | Canidae | 85586.0 | 0 | Canis | 85586.0 | 0 | Canis lupus | 85586.0 | 0 |
| scf7180000045677 | 184722 | 0.4006 | 0 | 59.2316 | Eukaryota | 91080.0 | 0 | Chordata | 91080.0 | 0 | Carnivora | 91080.0 | 0 | Canidae | 91080.0 | 0 | Canis | 79392.0 | 1 | Canis lupus | 79392.0 | 1 |
| scf7180000051472 | 7356 | 0.4474 | 0 | 1364.0397 | Eukaryota | 2840.0 | 1 | Echinodermata | 1700.0 | 3 | Forcipulatida | 1700.0 | 3 | Asteriidae | 1700.0 | 3 | Asterias | 1700.0 | 3 | Asterias rubens | 1700.0 | 3 |
| scf7180000049924 | 9774 | 0.4504 | 0 | 1602.6543 | Eukaryota | 18240.0 | 0 | Chordata | 17031.0 | 2 | Gadiformes | 11968.0 | 6 | Gadidae | 11968.0 | 6 | Gadus | 11968.0 | 6 | Gadus morhua | 11968.0 | 6 |
| scf7180000047025 | 83609 | 0.3585 | 0 | 58.5817 | Eukaryota | 28524.0 | 0 | Chordata | 28524.0 | 0 | Carnivora | 28524.0 | 0 | Canidae | 11549.0 | 3 | Canis | 7571.0 | 4 | Canis lupus | 7571.0 | 5 |
| scf7180000047194 | 90038 | 0.4621 | 0 | 62.7612 | Eukaryota | 57035.0 | 0 | Chordata | 57035.0 | 0 | Carnivora | 57035.0 | 0 | Canidae | 57035.0 | 0 | Vulpes | 28885.0 | 1 | Vulpes vulpes | 28885.0 | 1 |
| scf7180000049463 | 44285 | 0.5078 | 0 | 329.1316 | Eukaryota | 9676.0 | 0 | Chordata | 9676.0 | 0 | Carnivora | 9676.0 | 0 | Canidae | 9676.0 | 0 | Nyctereutes | 5278.0 | 1 | Nyctereutes procyonoides | 5278.0 | 1 |
| scf7180000046910 | 80313 | 0.3798 | 0 | 56.9195 | Eukaryota | 12764.0 | 0 | Chordata | 12764.0 | 0 | Primates | 10031.0 | 1 | Hominidae | 6879.0 | 3 | Homo | 6879.0 | 4 | Homo sapiens | 6879.0 | 4 |
| scf7180000049798 | 44565 | 0.5026 | 0 | 625.2794 | Eukaryota | 9424.0 | 0 | Chordata | 9424.0 | 0 | Carnivora | 9424.0 | 0 | Canidae | 9424.0 | 0 | Canis | 8184.0 | 1 | Canis lupus | 6896.0 | 2 |
| scf7180000052931 | 3009 | 0.4543 | 0 | 83.9287 | Eukaryota | 243.0 | 0 | Platyhelminthes | 243.0 | 0 | Polystomatidea | 243.0 | 0 | Polystomatidae | 243.0 | 0 | Protopolystoma | 243.0 | 0 | Protopolystoma xenopodis | 243.0 | 0 |
| scf7180000050312 | 2744 | 0.4417 | 0 | 133.2384 | no-hit | 0.0 | 0 | no-hit | 0.0 | 0 | no-hit | 0.0 | 0 | no-hit | 0.0 | 0 | no-hit | 0.0 | 0 | no-hit | 0.0 | 0 |
| scf7180000054486 | 493490 | 0.4407 | 0 | 91.428 | Eukaryota | 91896.0 | 0 | Chordata | 91896.0 | 0 | Carnivora | 91896.0 | 0 | Canidae | 91896.0 | 0 | Canis | 91896.0 | 0 | Canis lupus | 91896.0 | 0 |
| scf7180000056522 | 212487 | 0.358 | 0 | 60.329 | Eukaryota | 31270.0 | 0 | Chordata | 31270.0 | 0 | Carnivora | 31270.0 | 0 | Canidae | 31270.0 | 0 | Canis | 31270.0 | 0 | Canis lupus | 31270.0 | 0 |
| scf7180000044234 | 204116 | 0.3796 | 0 | 59.5631 | Eukaryota | 49409.0 | 0 | Chordata | 49409.0 | 0 | Carnivora | 49409.0 | 0 | Canidae | 49409.0 | 0 | Canis | 49409.0 | 0 | Canis lupus | 49409.0 | 0 |
| scf7180000042245 | 2410 | 0.4017 | 0 | 553.7461 | no-hit | 0.0 | 0 | no-hit | 0.0 | 0 | no-hit | 0.0 | 0 | no-hit | 0.0 | 0 | no-hit | 0.0 | 0 | no-hit | 0.0 | 0 |
| scf7180000040625 | 2004520 | 0.4244 | 0 | 59.5558 | Eukaryota | 75706.0 | 0 | Chordata | 75706.0 | 0 | Carnivora | 75706.0 | 0 | Canidae | 69835.0 | 1 | Canis | 55795.0 | 2 | Canis lupus | 55795.0 | 2 |
| scf7180000044407 | 66258 | 0.4333 | 0 | 107.8056 | Eukaryota | 94475.0 | 0 | Chordata | 94475.0 | 0 | Carnivora | 94475.0 | 0 | Canidae | 94475.0 | 0 | Canis | 94475.0 | 0 | Canis lupus | 94475.0 | 0 |
| scf7180000056271 | 429632 | 0.3888 | 0 | 60.359 | Eukaryota | 108788.0 | 0 | Chordata | 108788.0 | 0 | Carnivora | 108788.0 | 0 | Canidae | 108788.0 | 0 | Canis | 108788.0 | 0 | Canis lupus | 108788.0 | 0 |
| scf7180000048174 | 61867 | 0.4382 | 0 | 76.6707 | Eukaryota | 92424.0 | 0 | Chordata | 92424.0 | 0 | Carnivora | 92424.0 | 0 | Canidae | 92424.0 | 0 | Canis | 92424.0 | 0 | Canis lupus | 92424.0 | 0 |
| scf7180000048078 | 80927 | 0.3925 | 0 | 61.2643 | Eukaryota | 83374.0 | 0 | Chordata | 83374.0 | 0 | Carnivora | 83374.0 | 0 | Canidae | 83374.0 | 0 | Canis | 50045.0 | 1 | Canis lupus | 50045.0 | 1 |
| scf7180000041008 | 79364 | 0.5073 | 0 | 56.0985 | Eukaryota | 39041.0 | 0 | Chordata | 39041.0 | 0 | Carnivora | 32996.0 | 1 | Canidae | 32996.0 | 1 | Vulpes | 18256.0 | 2 | Vulpes vulpes | 18256.0 | 2 |
| scf7180000053376 | 2523 | 0.4423 | 0 | 44.3495 | no-hit | 0.0 | 0 | no-hit | 0.0 | 0 | no-hit | 0.0 | 0 | no-hit | 0.0 | 0 | no-hit | 0.0 | 0 | no-hit | 0.0 | 0 |
| scf7180000049990 | 9718 | 0.5118 | 0 | 1421.6798 | Eukaryota | 8499.0 | 0 | Chordata | 8499.0 | 0 | Carnivora | 8499.0 | 0 | Canidae | 8499.0 | 0 | Canis | 6817.0 | 1 | Canis lupus | 5605.0 | 2 |
| scf7180000051422 | 14794 | 0.4693 | 0 | 58.8542 | Eukaryota | 2576.0 | 1 | Chordata | 2576.0 | 1 | Carnivora | 2110.0 | 2 | Canidae | 2110.0 | 2 | Canis | 2110.0 | 2 | Canis lupus | 2110.0 | 2 |
| scf7180000047131 | 19432 | 0.3767 | 0 | 48.0075 | Eukaryota | 28697.0 | 0 | Chordata | 28697.0 | 0 | Carnivora | 28697.0 | 0 | Canidae | 15484.0 | 3 | Canis | 10398.0 | 7 | Canis lupus | 10398.0 | 7 |
| scf7180000045298 | 190415 | 0.4634 | 0 | 59.2325 | Eukaryota | 26256.0 | 0 | Chordata | 26256.0 | 0 | Carnivora | 26256.0 | 0 | Canidae | 26256.0 | 0 | Canis | 23443.0 | 1 | Canis lupus | 23443.0 | 1 |
| scf7180000048179 | 61700 | 0.3835 | 0 | 73.9302 | Eukaryota | 92977.0 | 0 | Chordata | 92977.0 | 0 | Carnivora | 92977.0 | 0 | Canidae | 92977.0 | 0 | Canis | 92977.0 | 0 | Canis lupus | 92977.0 | 0 |
| scf7180000045171 | 96303 | 0.384 | 0 | 56.7787 | Eukaryota | 40475.0 | 0 | Chordata | 40475.0 | 0 | Carnivora | 40475.0 | 0 | Canidae | 40475.0 | 0 | Canis | 40475.0 | 0 | Canis lupus | 40475.0 | 0 |
| scf7180000043675 | 459502 | 0.4207 | 0 | 59.1743 | Eukaryota | 18582.0 | 0 | Chordata | 18582.0 | 0 | Carnivora | 16903.0 | 1 | Canidae | 16903.0 | 1 | Canis | 16903.0 | 1 | Canis lupus | 16903.0 | 1 |
| scf7180000048060 | 15194 | 0.4398 | 0 | 80.4768 | no-hit | 0.0 | 0 | no-hit | 0.0 | 0 | no-hit | 0.0 | 0 | no-hit | 0.0 | 0 | no-hit | 0.0 | 0 | no-hit | 0.0 | 0 |
| scf7180000055406 | 40738 | 0.4407 | 0 | 162.9572 | Eukaryota | 11783.0 | 0 | Chordata | 7878.0 | 2 | Tetraodontiformes | 6303.0 | 4 | Tetraodontidae | 6303.0 | 4 | Takifugu | 6303.0 | 4 | Takifugu rubripes | 6303.0 | 4 |
| scf7180000052595 | 6847 | 0.4441 | 0 | 324.5979 | no-hit | 0.0 | 0 | no-hit | 0.0 | 0 | no-hit | 0.0 | 0 | no-hit | 0.0 | 0 | no-hit | 0.0 | 0 | no-hit | 0.0 | 0 |
| scf7180000050718 | 14825 | 0.5029 | 0 | 781.1718 | Eukaryota | 8841.0 | 0 | Chordata | 8841.0 | 0 | Carnivora | 8841.0 | 0 | Canidae | 8841.0 | 0 | Canis | 7137.0 | 1 | Canis lupus | 5921.0 | 2 |
| scf7180000045154 | 208117 | 0.3938 | 0 | 59.2907 | Eukaryota | 15639.0 | 0 | Chordata | 15639.0 | 0 | Carnivora | 14610.0 | 1 | Canidae | 14610.0 | 1 | Canis | 14610.0 | 1 | Canis lupus | 14610.0 | 1 |
| scf7180000041877 | 355859 | 0.3471 | 0 | 58.6472 | Eukaryota | 84634.0 | 0 | Chordata | 84634.0 | 0 | Carnivora | 84634.0 | 0 | Canidae | 84634.0 | 0 | Canis | 84634.0 | 0 | Canis lupus | 84634.0 | 0 |
| scf7180000053677 | 4232 | 0.4428 | 0 | 30.16 | Eukaryota | 3652.0 | 0 | Platyhelminthes | 2343.0 | 1 | Polystomatidea | 2343.0 | 3 | Polystomatidae | 2343.0 | 3 | Protopolystoma | 2343.0 | 3 | Protopolystoma xenopodis | 2343.0 | 3 |
| scf7180000042172 | 154528 | 0.3826 | 0 | 58.9485 | Eukaryota | 11836.0 | 0 | Chordata | 11836.0 | 0 | Carnivora | 11836.0 | 0 | Canidae | 11836.0 | 0 | Canis | 11836.0 | 0 | Canis lupus | 11836.0 | 0 |
| scf7180000051396 | 5349 | 0.4577 | 0 | 36.9725 | Eukaryota | 2938.0 | 0 | Chordata | 2596.0 | 1 | Tetraodontiformes | 2596.0 | 1 | Tetraodontidae | 2596.0 | 1 | Takifugu | 2596.0 | 1 | Takifugu rubripes | 2596.0 | 1 |
| scf7180000055894 | 36978 | 0.4382 | 0 | 306.8634 | Eukaryota | 7913.0 | 0 | Chordata | 5010.0 | 1 | Polystomatidea | 2903.0 | 3 | Polystomatidae | 2903.0 | 3 | Protopolystoma | 2903.0 | 3 | Protopolystoma xenopodis | 2903.0 | 3 |
| scf7180000050272 | 33006 | 0.44 | 0 | 86.0495 | Eukaryota | 8155.0 | 0 | Chordata | 5107.0 | 2 | Tetraodontiformes | 5107.0 | 2 | Tetraodontidae | 5107.0 | 2 | Takifugu | 5107.0 | 2 | Takifugu rubripes | 5107.0 | 2 |
| scf7180000041150 | 1224808 | 0.4282 | 0 | 59.7289 | Eukaryota | 102228.0 | 0 | Chordata | 102228.0 | 0 | Carnivora | 102228.0 | 0 | Canidae | 102228.0 | 0 | Canis | 78463.0 | 1 | Canis lupus | 78463.0 | 1 |
| scf7180000043732 | 82149 | 0.4424 | 0 | 56.0196 | Eukaryota | 56480.0 | 0 | Chordata | 56480.0 | 0 | Carnivora | 56480.0 | 0 | Canidae | 56480.0 | 0 | Canis | 56480.0 | 0 | Canis lupus | 56480.0 | 0 |
| scf7180000055379 | 14979 | 0.4355 | 0 | 426.3395 | Eukaryota | 2849.0 | 0 | Chordata | 2849.0 | 0 | Carnivora | 2849.0 | 0 | Canidae | 2849.0 | 0 | Canis | 2849.0 | 0 | Canis lupus | 2849.0 | 0 |
| scf7180000056241 | 83236 | 0.3822 | 0 | 58.8504 | Eukaryota | 57379.0 | 0 | Chordata | 57379.0 | 0 | Carnivora | 57379.0 | 0 | Canidae | 57379.0 | 0 | Canis | 57379.0 | 0 | Canis lupus | 57379.0 | 0 |
| scf7180000042053 | 795995 | 0.4638 | 0 | 60.4883 | Eukaryota | 64785.0 | 0 | Chordata | 64785.0 | 0 | Carnivora | 64785.0 | 0 | Canidae | 64785.0 | 0 | Canis | 52232.0 | 1 | Canis lupus | 52232.0 | 1 |
| scf7180000055337 | 22684 | 0.4519 | 0 | 284.7038 | Eukaryota | 12451.0 | 0 | Chordata | 12451.0 | 0 | Carnivora | 12451.0 | 0 | Canidae | 12451.0 | 0 | Canis | 10127.0 | 1 | Canis lupus | 9159.0 | 2 |
| scf7180000043862 | 412203 | 0.3711 | 0 | 59.9902 | Eukaryota | 34067.0 | 0 | Chordata | 34067.0 | 0 | Carnivora | 34067.0 | 0 | Canidae | 34067.0 | 0 | Canis | 25869.0 | 1 | Canis lupus | 25869.0 | 1 |
| scf7180000045123 | 179431 | 0.4404 | 0 | 58.5527 | Eukaryota | 59419.0 | 0 | Chordata | 59419.0 | 0 | Carnivora | 59419.0 | 0 | Canidae | 59419.0 | 0 | Canis | 59419.0 | 0 | Canis lupus | 59419.0 | 0 |
| scf7180000044811 | 23368 | 0.4084 | 0 | 67.2761 | Eukaryota | 65405.0 | 0 | Chordata | 65405.0 | 0 | Carnivora | 65405.0 | 0 | Canidae | 65405.0 | 0 | Canis | 65405.0 | 0 | Canis lupus | 65405.0 | 0 |
| scf7180000054560 | 23209 | 0.4077 | 0 | 922.8551 | Eukaryota | 88965.0 | 0 | Chordata | 88965.0 | 0 | Carnivora | 88965.0 | 0 | Canidae | 88965.0 | 0 | Canis | 88965.0 | 0 | Canis lupus | 88965.0 | 0 |
| scf7180000056439 | 386415 | 0.3829 | 0 | 60.3993 | Eukaryota | 58737.0 | 0 | Chordata | 58737.0 | 0 | Carnivora | 50849.0 | 1 | Canidae | 29540.0 | 4 | Canis | 23750.0 | 6 | Canis lupus | 23750.0 | 6 |
| scf7180000053934 | 4207 | 0.4493 | 0 | 339.4834 | Eukaryota | 561.0 | 0 | Chordata | 333.0 | 1 | Tetraodontiformes | 333.0 | 1 | Tetraodontidae | 333.0 | 1 | Takifugu | 333.0 | 1 | Takifugu rubripes | 333.0 | 1 |
| scf7180000049683 | 32222 | 0.4399 | 0 | 57.3421 | Eukaryota | 11659.0 | 0 | Chordata | 11659.0 | 0 | Carnivora | 11659.0 | 0 | Canidae | 7708.0 | 2 | Canis | 6389.0 | 5 | Canis lupus | 6389.0 | 5 |
| scf7180000042402 | 398176 | 0.4321 | 0 | 59.0973 | Eukaryota | 86997.0 | 0 | Chordata | 86997.0 | 0 | Carnivora | 86997.0 | 0 | Canidae | 86997.0 | 0 | Canis | 86997.0 | 0 | Canis lupus | 86997.0 | 0 |
| scf7180000043152 | 72820 | 0.4139 | 0 | 59.9266 | Eukaryota | 63744.0 | 0 | Chordata | 63744.0 | 0 | Carnivora | 63744.0 | 0 | Canidae | 63744.0 | 0 | Canis | 63744.0 | 0 | Canis lupus | 63744.0 | 0 |
| scf7180000042938 | 64383 | 0.4271 | 0 | 58.8878 | Eukaryota | 23290.0 | 0 | Chordata | 23290.0 | 0 | Carnivora | 23290.0 | 0 | Canidae | 23290.0 | 0 | Canis | 23290.0 | 0 | Canis lupus | 23290.0 | 0 |
| scf7180000047361 | 6128 | 0.7244 | 0 | 32.9307 | Eukaryota | 50598.0 | 0 | Chordata | 50598.0 | 0 | Carnivora | 50598.0 | 0 | Canidae | 50598.0 | 0 | Canis | 50598.0 | 0 | Canis lupus | 50598.0 | 0 |
| scf7180000046923 | 92332 | 0.4567 | 0 | 59.4137 | Eukaryota | 19579.0 | 0 | Chordata | 19579.0 | 0 | Carnivora | 18288.0 | 1 | Canidae | 11395.0 | 3 | Canis | 11395.0 | 3 | Canis lupus | 11395.0 | 4 |
| scf7180000051114 | 13638 | 0.4278 | 0 | 198.8556 | no-hit | 0.0 | 0 | no-hit | 0.0 | 0 | no-hit | 0.0 | 0 | no-hit | 0.0 | 0 | no-hit | 0.0 | 0 | no-hit | 0.0 | 0 |
| scf7180000043134 | 574967 | 0.4856 | 0 | 59.1159 | Eukaryota | 116963.0 | 0 | Chordata | 116963.0 | 0 | Carnivora | 116963.0 | 0 | Canidae | 107271.0 | 1 | Canis | 71463.0 | 2 | Canis lupus | 71463.0 | 2 |
| scf7180000044022 | 58716 | 0.4404 | 0 | 404.9365 | Eukaryota | 13879.0 | 0 | Chordata | 13879.0 | 0 | Tetraodontiformes | 8132.0 | 3 | Tetraodontidae | 8132.0 | 3 | Takifugu | 8132.0 | 3 | Takifugu rubripes | 8132.0 | 3 |
| scf7180000043248 | 4401 | 0.4269 | 0 | 25.5724 | Eukaryota | 5465.0 | 0 | Chordata | 5465.0 | 0 | Carnivora | 5465.0 | 0 | Canidae | 5465.0 | 0 | Canis | 5465.0 | 0 | Canis lupus | 5465.0 | 0 |
| scf7180000045226 | 214524 | 0.3641 | 0 | 61.7127 | Eukaryota | 16821.0 | 0 | Chordata | 16821.0 | 0 | Primates | 6105.0 | 3 | Hominidae | 4725.0 | 6 | Homo | 4725.0 | 6 | Homo sapiens | 4725.0 | 6 |
| scf7180000049746 | 34296 | 0.6434 | 0 | 55.6082 | Eukaryota | 21982.0 | 0 | Chordata | 21982.0 | 0 | Carnivora | 21982.0 | 0 | Canidae | 21982.0 | 0 | Canis | 15747.0 | 1 | Canis lupus | 15747.0 | 1 |
| scf7180000044558 | 265772 | 0.464 | 0 | 59.7071 | Eukaryota | 25728.0 | 0 | Chordata | 25728.0 | 0 | Carnivora | 23753.0 | 1 | Ursidae | 19711.0 | 2 | Ursus | 19711.0 | 2 | Ursus maritimus | 11233.0 | 3 |
| scf7180000044509 | 281339 | 0.4595 | 0 | 59.458 | Eukaryota | 68568.0 | 0 | Chordata | 68568.0 | 0 | Carnivora | 68568.0 | 0 | Canidae | 56844.0 | 1 | Canis | 56844.0 | 1 | Canis lupus | 56844.0 | 1 |
| scf7180000052344 | 6888 | 0.4698 | 0 | 55.4966 | Eukaryota | 4758.0 | 0 | Chordata | 4758.0 | 0 | Carnivora | 4758.0 | 0 | Canidae | 4758.0 | 0 | Canis | 4758.0 | 0 | Canis lupus | 4758.0 | 0 |
| scf7180000053199 | 9452 | 0.445 | 0 | 911.3824 | Eukaryota | 1994.0 | 0 | Echinodermata | 808.0 | 2 | Forcipulatida | 808.0 | 2 | Asteriidae | 808.0 | 2 | Asterias | 808.0 | 2 | Asterias rubens | 808.0 | 2 |
| scf7180000051182 | 26457 | 0.503 | 0 | 482.7463 | Eukaryota | 9361.0 | 0 | Chordata | 9361.0 | 0 | Carnivora | 9361.0 | 0 | Canidae | 9361.0 | 0 | Canis | 7643.0 | 1 | Canis lupus | 6412.0 | 2 |
| scf7180000040717 | 725488 | 0.4026 | 0 | 59.4648 | Eukaryota | 43464.0 | 0 | Chordata | 43464.0 | 0 | Carnivora | 43464.0 | 0 | Canidae | 43464.0 | 0 | Canis | 34336.0 | 1 | Canis lupus | 34336.0 | 1 |
| scf7180000045445 | 73778 | 0.4414 | 0 | 77.5261 | Eukaryota | 89533.0 | 0 | Chordata | 89533.0 | 0 | Carnivora | 89533.0 | 0 | Canidae | 89533.0 | 0 | Canis | 89533.0 | 0 | Canis lupus | 89533.0 | 0 |
| scf7180000050371 | 30123 | 0.3826 | 0 | 59.9902 | Eukaryota | 16818.0 | 0 | Chordata | 16818.0 | 0 | Carnivora | 16818.0 | 0 | Canidae | 16818.0 | 0 | Canis | 16818.0 | 0 | Canis lupus | 16818.0 | 0 |
| scf7180000043199 | 527794 | 0.3368 | 0 | 58.0863 | Eukaryota | 75419.0 | 0 | Chordata | 75419.0 | 0 | Carnivora | 75419.0 | 0 | Canidae | 75419.0 | 0 | Canis | 75419.0 | 0 | Canis lupus | 75419.0 | 0 |
| scf7180000053689 | 6786 | 0.4524 | 0 | 1193.0301 | Eukaryota | 425.0 | 0 | Platyhelminthes | 425.0 | 0 | Polystomatidea | 425.0 | 0 | Polystomatidae | 425.0 | 0 | Protopolystoma | 425.0 | 0 | Protopolystoma xenopodis | 425.0 | 0 |
| scf7180000043151 | 435094 | 0.4144 | 0 | 58.4225 | Eukaryota | 42176.0 | 0 | Chordata | 42176.0 | 0 | Carnivora | 42176.0 | 0 | Canidae | 42176.0 | 0 | Canis | 42176.0 | 0 | Canis lupus | 42176.0 | 0 |
| scf7180000048753 | 4785 | 0.4568 | 0 | 226.9634 | Eukaryota | 25697.0 | 0 | Platyhelminthes | 12982.0 | 1 | Strigeidida | 10398.0 | 4 | Schistosomatidae | 10398.0 | 4 | Schistosoma | 10398.0 | 4 | Schistosoma mansoni | 10398.0 | 4 |
| scf7180000055159 | 11870 | 0.4486 | 0 | 168.484 | Eukaryota | 2929.0 | 0 | Chordata | 2929.0 | 0 | Carnivora | 2929.0 | 0 | Canidae | 2929.0 | 0 | Canis | 2929.0 | 0 | Canis lupus | 2929.0 | 0 |
| scf7180000054790 | 14429 | 0.4784 | 0 | 274.2938 | Eukaryota | 18723.0 | 0 | Chordata | 18723.0 | 0 | Carnivora | 18723.0 | 0 | Canidae | 18723.0 | 0 | Canis | 18723.0 | 0 | Canis lupus | 18723.0 | 0 |
| scf7180000048053 | 14728 | 0.4405 | 0 | 90.9032 | Eukaryota | 3434.0 | 0 | Platyhelminthes | 1533.0 | 2 | Polystomatidea | 1533.0 | 2 | Polystomatidae | 1533.0 | 2 | Protopolystoma | 1533.0 | 2 | Protopolystoma xenopodis | 1533.0 | 2 |
| scf7180000043993 | 28235 | 0.4476 | 0 | 55.0204 | Eukaryota | 7473.0 | 0 | Chordata | 7473.0 | 0 | Carnivora | 7473.0 | 0 | Felidae | 5983.0 | 2 | Lynx | 2390.0 | 4 | Lynx canadensis | 2390.0 | 4 |
| scf7180000042319 | 65396 | 0.399 | 0 | 52.9202 | Eukaryota | 90178.0 | 0 | Chordata | 90178.0 | 0 | Carnivora | 90178.0 | 0 | Canidae | 90178.0 | 0 | Canis | 90178.0 | 0 | Canis lupus | 90178.0 | 0 |
| scf7180000043508 | 415812 | 0.5225 | 0 | 58.9473 | Eukaryota | 38703.0 | 0 | Chordata | 38703.0 | 0 | Carnivora | 38703.0 | 0 | Canidae | 38703.0 | 0 | Canis | 27909.0 | 1 | Canis lupus | 27909.0 | 1 |
| scf7180000042933 | 589870 | 0.4141 | 0 | 57.5087 | Eukaryota | 36911.0 | 0 | Chordata | 36911.0 | 0 | Carnivora | 36911.0 | 0 | Canidae | 36911.0 | 0 | Canis | 36911.0 | 0 | Canis lupus | 36911.0 | 0 |
| scf7180000055730 | 26967 | 0.4295 | 0 | 245.3674 | Eukaryota | 13938.0 | 0 | Chordata | 13938.0 | 0 | Carnivora | 13938.0 | 0 | Canidae | 13938.0 | 0 | Canis | 13938.0 | 0 | Canis lupus | 13938.0 | 0 |
| scf7180000048565 | 5777 | 0.4464 | 0 | 513.0263 | no-hit | 0.0 | 0 | no-hit | 0.0 | 0 | no-hit | 0.0 | 0 | no-hit | 0.0 | 0 | no-hit | 0.0 | 0 | no-hit | 0.0 | 0 |
| scf7180000056343 | 870764 | 0.4362 | 0 | 58.8248 | Eukaryota | 63525.0 | 0 | Chordata | 63525.0 | 0 | Carnivora | 63525.0 | 0 | Canidae | 58459.0 | 1 | Canis | 39364.0 | 2 | Canis lupus | 39364.0 | 2 |
| scf7180000044728 | 4663 | 0.3841 | 0 | 44.2811 | Eukaryota | 3004.0 | 0 | Chordata | 3004.0 | 0 | Carnivora | 3004.0 | 0 | Canidae | 3004.0 | 0 | Canis | 3004.0 | 0 | Canis lupus | 3004.0 | 0 |
| scf7180000056510 | 257898 | 0.4383 | 0 | 58.7402 | Eukaryota | 82392.0 | 0 | Chordata | 82392.0 | 0 | Carnivora | 82392.0 | 0 | Canidae | 82392.0 | 0 | Canis | 82392.0 | 0 | Canis lupus | 82392.0 | 0 |
| scf7180000055587 | 20550 | 0.4478 | 0 | 138.1561 | Eukaryota | 492.0 | 0 | Chordata | 492.0 | 0 | Beloniformes | 492.0 | 0 | Adrianichthyidae | 492.0 | 0 | Oryzias | 492.0 | 0 | Oryzias latipes | 492.0 | 0 |
| scf7180000052363 | 13366 | 0.4515 | 0 | 118.4473 | Eukaryota | 3459.0 | 0 | Platyhelminthes | 2607.0 | 1 | Polystomatidea | 2607.0 | 2 | Polystomatidae | 2607.0 | 2 | Protopolystoma | 2607.0 | 2 | Protopolystoma xenopodis | 2607.0 | 2 |
| scf7180000041237 | 48826 | 0.3694 | 0 | 55.4694 | Eukaryota | 22803.0 | 0 | Chordata | 22803.0 | 0 | Carnivora | 22803.0 | 0 | Canidae | 22803.0 | 0 | Canis | 22803.0 | 0 | Canis lupus | 22803.0 | 0 |
| scf7180000043059 | 389517 | 0.4105 | 0 | 57.9726 | Eukaryota | 56560.0 | 0 | Chordata | 56560.0 | 0 | Carnivora | 56560.0 | 0 | Phocidae | 17979.0 | 3 | Leptonychotes | 17979.0 | 3 | Leptonychotes weddellii | 17979.0 | 3 |
| scf7180000047123 | 3205 | 0.4393 | 0 | 89.554 | no-hit | 0.0 | 0 | no-hit | 0.0 | 0 | no-hit | 0.0 | 0 | no-hit | 0.0 | 0 | no-hit | 0.0 | 0 | no-hit | 0.0 | 0 |
| scf7180000050753 | 10376 | 0.4446 | 0 | 96.5034 | Eukaryota | 285.0 | 0 | Chordata | 285.0 | 0 | Tetraodontiformes | 285.0 | 0 | Tetraodontidae | 285.0 | 0 | Takifugu | 285.0 | 0 | Takifugu rubripes | 285.0 | 0 |
| scf7180000056438 | 363507 | 0.3399 | 0 | 59.0057 | Eukaryota | 71845.0 | 0 | Chordata | 71845.0 | 0 | Carnivora | 71845.0 | 0 | Canidae | 71845.0 | 0 | Canis | 71845.0 | 0 | Canis lupus | 71845.0 | 0 |
| scf7180000044518 | 278195 | 0.509 | 0 | 58.5936 | Eukaryota | 69303.0 | 0 | Chordata | 69303.0 | 0 | Carnivora | 69303.0 | 0 | Canidae | 69303.0 | 0 | Canis | 69303.0 | 0 | Canis lupus | 69303.0 | 0 |
| scf7180000040593 | 629467 | 0.3716 | 0 | 59.7183 | Eukaryota | 94555.0 | 0 | Chordata | 94555.0 | 0 | Carnivora | 94555.0 | 0 | Canidae | 94555.0 | 0 | Canis | 94555.0 | 0 | Canis lupus | 94555.0 | 0 |
| scf7180000049542 | 2451 | 0.43 | 0 | 64.6033 | no-hit | 0.0 | 0 | no-hit | 0.0 | 0 | no-hit | 0.0 | 0 | no-hit | 0.0 | 0 | no-hit | 0.0 | 0 | no-hit | 0.0 | 0 |
| scf7180000055410 | 19258 | 0.4394 | 0 | 297.3698 | Eukaryota | 998.0 | 0 | Platyhelminthes | 998.0 | 0 | Polystomatidea | 998.0 | 0 | Polystomatidae | 998.0 | 0 | Protopolystoma | 998.0 | 0 | Protopolystoma xenopodis | 998.0 | 0 |
| scf7180000042115 | 713214 | 0.3957 | 0 | 59.5551 | Eukaryota | 37046.0 | 0 | Chordata | 37046.0 | 0 | Carnivora | 37046.0 | 0 | Canidae | 37046.0 | 0 | Canis | 28926.0 | 1 | Canis lupus | 28926.0 | 1 |
| scf7180000056181 | 923259 | 0.4011 | 0 | 59.8042 | Eukaryota | 77116.0 | 0 | Chordata | 77116.0 | 0 | Carnivora | 77116.0 | 0 | Canidae | 38663.0 | 2 | Ailuropoda | 30584.0 | 3 | Ailuropoda melanoleuca | 30584.0 | 3 |
| scf7180000056330 | 796453 | 0.4122 | 0 | 59.6982 | Eukaryota | 55170.0 | 0 | Chordata | 55170.0 | 0 | Carnivora | 55170.0 | 0 | Canidae | 55170.0 | 0 | Canis | 55170.0 | 0 | Canis lupus | 55170.0 | 0 |
| scf7180000042330 | 426167 | 0.4862 | 0 | 55.8743 | Eukaryota | 50082.0 | 0 | Chordata | 50082.0 | 0 | Carnivora | 50082.0 | 0 | Canidae | 50082.0 | 0 | Canis | 50082.0 | 0 | Canis lupus | 50082.0 | 0 |
| scf7180000049482 | 2109 | 0.4457 | 0 | 356.8898 | Eukaryota | 789.0 | 0 | Chordata | 789.0 | 0 | Kurtiformes | 789.0 | 0 | Apogonidae | 789.0 | 0 | Sphaeramia | 789.0 | 0 | Sphaeramia orbicularis | 789.0 | 0 |
| scf7180000042194 | 28010 | 0.4318 | 0 | 55.6621 | Eukaryota | 40162.0 | 0 | Chordata | 40162.0 | 0 | Carnivora | 40162.0 | 0 | Canidae | 40162.0 | 0 | Canis | 36226.0 | 1 | Canis lupus | 36226.0 | 1 |
| scf7180000048927 | 4619 | 0.5025 | 0 | 123.1233 | Eukaryota | 8572.0 | 0 | Chordata | 8572.0 | 0 | Carnivora | 8572.0 | 0 | Canidae | 8572.0 | 0 | Canis | 6395.0 | 1 | Canis lupus | 5201.0 | 2 |
| scf7180000041924 | 784677 | 0.475 | 0 | 59.4123 | Eukaryota | 64658.0 | 0 | Chordata | 64658.0 | 0 | Carnivora | 64658.0 | 0 | Canidae | 64658.0 | 0 | Canis | 55736.0 | 1 | Canis lupus | 55736.0 | 1 |
| scf7180000044552 | 327334 | 0.3803 | 0 | 60.6297 | Eukaryota | 92603.0 | 0 | Chordata | 92603.0 | 0 | Carnivora | 92603.0 | 0 | Canidae | 92603.0 | 0 | Canis | 92603.0 | 0 | Canis lupus | 92603.0 | 0 |
| scf7180000044359 | 281496 | 0.3789 | 0 | 61.3166 | Eukaryota | 35103.0 | 0 | Chordata | 35103.0 | 0 | Carnivora | 35103.0 | 0 | Canidae | 35103.0 | 0 | Canis | 35103.0 | 0 | Canis lupus | 35103.0 | 0 |
| scf7180000044843 | 237178 | 0.6067 | 0 | 55.7398 | Eukaryota | 39500.0 | 0 | Chordata | 39500.0 | 0 | Carnivora | 39500.0 | 0 | Canidae | 39500.0 | 0 | Vulpes | 27319.0 | 1 | Vulpes vulpes | 27319.0 | 1 |
| scf7180000044795 | 256941 | 0.5204 | 0 | 58.2278 | Eukaryota | 48475.0 | 0 | Chordata | 48475.0 | 0 | Carnivora | 48475.0 | 0 | Canidae | 48475.0 | 0 | Canis | 42831.0 | 1 | Canis lupus | 42831.0 | 1 |
| scf7180000042729 | 634634 | 0.3843 | 0 | 58.9707 | Eukaryota | 66041.0 | 0 | Chordata | 66041.0 | 0 | Carnivora | 66041.0 | 0 | Canidae | 66041.0 | 0 | Canis | 66041.0 | 0 | Canis lupus | 66041.0 | 0 |
| scf7180000046660 | 109265 | 0.3494 | 0 | 58.6774 | Eukaryota | 14268.0 | 0 | Chordata | 14268.0 | 0 | Carnivora | 14268.0 | 0 | Otariidae | 4278.0 | 5 | Leptonychotes | 1469.0 | 9 | Leptonychotes weddellii | 1469.0 | 9 |
| scf7180000045413 | 44569 | 0.3667 | 0 | 55.6238 | Eukaryota | 54300.0 | 0 | Chordata | 54300.0 | 0 | Carnivora | 54300.0 | 0 | Canidae | 54300.0 | 0 | Canis | 54300.0 | 0 | Canis lupus | 54300.0 | 0 |
| scf7180000050394 | 27742 | 0.3998 | 0 | 59.647 | Eukaryota | 3396.0 | 0 | Chordata | 3396.0 | 0 | Carnivora | 3396.0 | 0 | Canidae | 3045.0 | 1 | Canis | 3045.0 | 1 | Canis lupus | 3045.0 | 1 |
| scf7180000049122 | 37379 | 0.4982 | 0 | 59.299 | Eukaryota | 14711.0 | 0 | Chordata | 14711.0 | 0 | Carnivora | 14711.0 | 0 | Canidae | 10577.0 | 1 | Canis | 10577.0 | 1 | Canis lupus | 10577.0 | 1 |
| scf7180000056544 | 211582 | 0.3414 | 0 | 58.9132 | Eukaryota | 91260.0 | 0 | Chordata | 91260.0 | 0 | Carnivora | 91260.0 | 0 | Canidae | 91260.0 | 0 | Canis | 91260.0 | 0 | Canis lupus | 91260.0 | 0 |
| scf7180000047968 | 53946 | 0.5001 | 0 | 57.86 | Eukaryota | 65779.0 | 0 | Chordata | 65779.0 | 0 | Carnivora | 65779.0 | 0 | Felidae | 36855.0 | 1 | Felis | 36855.0 | 2 | Felis catus | 36855.0 | 2 |
| scf7180000053521 | 6699 | 0.4878 | 0 | 91.6531 | Eukaryota | 31725.0 | 0 | Chordata | 31725.0 | 0 | Carnivora | 31725.0 | 0 | Canidae | 31725.0 | 0 | Canis | 31725.0 | 0 | Canis lupus | 31725.0 | 0 |
| scf7180000055510 | 3948 | 0.4354 | 0 | 82.2903 | Eukaryota | 13455.0 | 0 | Chordata | 9694.0 | 1 | Strigeidida | 3761.0 | 4 | Schistosomatidae | 3761.0 | 4 | Schistosoma | 3761.0 | 5 | Schistosoma mansoni | 3761.0 | 6 |
| scf7180000047417 | 31973 | 0.4612 | 0 | 179.816 | Eukaryota | 1468.0 | 0 | Platyhelminthes | 856.0 | 1 | Polystomatidea | 856.0 | 1 | Polystomatidae | 856.0 | 1 | Protopolystoma | 856.0 | 1 | Protopolystoma xenopodis | 856.0 | 1 |
| scf7180000048682 | 18936 | 0.4432 | 0 | 141.0682 | Eukaryota | 5991.0 | 1 | Chordata | 4983.0 | 2 | Salmoniformes | 3202.0 | 5 | Salmonidae | 3202.0 | 5 | Salmo | 2935.0 | 6 | Salmo trutta | 2935.0 | 6 |
| scf7180000040574 | 2240840 | 0.3567 | 0 | 60.6311 | Eukaryota | 94499.0 | 0 | Chordata | 94499.0 | 0 | Carnivora | 94499.0 | 0 | Canidae | 94499.0 | 0 | Canis | 94499.0 | 0 | Canis lupus | 94499.0 | 0 |
| scf7180000050435 | 24147 | 0.4142 | 0 | 57.1581 | Eukaryota | 4392.0 | 0 | Chordata | 4392.0 | 0 | Carnivora | 4392.0 | 0 | Canidae | 4392.0 | 0 | Canis | 4392.0 | 0 | Canis lupus | 4392.0 | 0 |
| scf7180000046158 | 20062 | 0.4539 | 0 | 209.5869 | Eukaryota | 88342.0 | 0 | Chordata | 88342.0 | 0 | Carnivora | 88342.0 | 0 | Canidae | 88342.0 | 0 | Canis | 88342.0 | 0 | Canis lupus | 88342.0 | 0 |
| scf7180000054933 | 8677 | 0.4385 | 0 | 144.8938 | no-hit | 0.0 | 0 | no-hit | 0.0 | 0 | no-hit | 0.0 | 0 | no-hit | 0.0 | 0 | no-hit | 0.0 | 0 | no-hit | 0.0 | 0 |
| scf7180000043537 | 410383 | 0.4255 | 0 | 60.2334 | Eukaryota | 96330.0 | 0 | Chordata | 96330.0 | 0 | Carnivora | 96330.0 | 0 | Canidae | 96330.0 | 0 | Canis | 96330.0 | 0 | Canis lupus | 96330.0 | 0 |
| scf7180000052414 | 11975 | 0.5061 | 0 | 581.1173 | Eukaryota | 9138.0 | 0 | Chordata | 9138.0 | 0 | Carnivora | 9138.0 | 0 | Canidae | 9138.0 | 0 | Canis | 7411.0 | 1 | Canis lupus | 6156.0 | 2 |
| scf7180000051529 | 9044 | 0.5036 | 0 | 35.7066 | Eukaryota | 8141.0 | 0 | Chordata | 8141.0 | 0 | Carnivora | 8141.0 | 0 | Canidae | 8141.0 | 0 | Canis | 6108.0 | 1 | Canis lupus | 4964.0 | 2 |
| scf7180000056324 | 785823 | 0.3474 | 0 | 59.5386 | Eukaryota | 76088.0 | 0 | Chordata | 76088.0 | 0 | Carnivora | 76088.0 | 0 | Canidae | 76088.0 | 0 | Canis | 76088.0 | 0 | Canis lupus | 76088.0 | 0 |
| scf7180000047245 | 89817 | 0.472 | 0 | 59.1754 | Eukaryota | 41786.0 | 0 | Chordata | 41786.0 | 0 | Carnivora | 41786.0 | 0 | Canidae | 41786.0 | 0 | Canis | 36269.0 | 1 | Canis lupus | 36269.0 | 1 |
| scf7180000048585 | 46593 | 0.6774 | 0 | 52.1659 | Eukaryota | 26466.0 | 0 | Chordata | 26466.0 | 0 | Carnivora | 26466.0 | 0 | Canidae | 22990.0 | 1 | Canis | 22990.0 | 2 | Canis lupus | 22990.0 | 2 |
| scf7180000053983 | 5365 | 0.4578 | 0 | 64.9939 | no-hit | 0.0 | 0 | no-hit | 0.0 | 0 | no-hit | 0.0 | 0 | no-hit | 0.0 | 0 | no-hit | 0.0 | 0 | no-hit | 0.0 | 0 |
| scf7180000055544 | 8225 | 0.4398 | 0 | 134.1111 | Eukaryota | 36493.0 | 0 | Chordata | 24584.0 | 2 | Gadiformes | 19787.0 | 5 | Gadidae | 19787.0 | 5 | Gadus | 19787.0 | 5 | Gadus morhua | 19787.0 | 5 |
| scf7180000048787 | 21928 | 0.4475 | 0 | 271.0347 | Eukaryota | 709.0 | 0 | Platyhelminthes | 435.0 | 1 | Polystomatidea | 435.0 | 1 | Polystomatidae | 435.0 | 1 | Protopolystoma | 435.0 | 1 | Protopolystoma xenopodis | 435.0 | 1 |
| scf7180000050811 | 14976 | 0.4062 | 0 | 386.3219 | Eukaryota | 34604.0 | 0 | Chordata | 34604.0 | 0 | Carnivora | 34604.0 | 0 | Canidae | 34604.0 | 0 | Canis | 34604.0 | 0 | Canis lupus | 34604.0 | 0 |
| scf7180000041838 | 470162 | 0.3498 | 0 | 59.0165 | Eukaryota | 130769.0 | 0 | Chordata | 130769.0 | 0 | Carnivora | 130769.0 | 0 | Canidae | 130769.0 | 0 | Canis | 130769.0 | 0 | Canis lupus | 130769.0 | 0 |
| scf7180000044609 | 159250 | 0.406 | 0 | 58.1794 | Eukaryota | 79743.0 | 0 | Chordata | 79743.0 | 0 | Carnivora | 79743.0 | 0 | Canidae | 79743.0 | 0 | Canis | 79743.0 | 0 | Canis lupus | 79743.0 | 0 |
| scf7180000044412 | 22882 | 0.4441 | 0 | 279.9658 | Eukaryota | 972.0 | 0 | Chordata | 620.0 | 1 | Salmoniformes | 414.0 | 2 | Salmonidae | 414.0 | 2 | Coregonus | 414.0 | 2 | Coregonus sp. 'balchen' | 414.0 | 2 |
| scf7180000056173 | 3445228 | 0.3852 | 0 | 60.4835 | Eukaryota | 101760.0 | 0 | Chordata | 101760.0 | 0 | Carnivora | 101760.0 | 0 | Canidae | 101760.0 | 0 | Canis | 101760.0 | 0 | Canis lupus | 101760.0 | 0 |
| scf7180000044914 | 111316 | 0.3907 | 0 | 61.7093 | Eukaryota | 47567.0 | 0 | Chordata | 47567.0 | 0 | Carnivora | 47567.0 | 0 | Canidae | 47567.0 | 0 | Canis | 47567.0 | 0 | Canis lupus | 47567.0 | 0 |
| scf7180000047212 | 43808 | 0.4598 | 0 | 56.8633 | Eukaryota | 38337.0 | 0 | Chordata | 38337.0 | 0 | Carnivora | 38337.0 | 0 | Canidae | 38337.0 | 0 | Canis | 38337.0 | 0 | Canis lupus | 38337.0 | 0 |
| scf7180000052253 | 13344 | 0.5311 | 0 | 56.5229 | Eukaryota | 2002.0 | 0 | Chordata | 2002.0 | 0 | Carnivora | 1177.0 | 2 | Canidae | 1177.0 | 2 | Canis | 1177.0 | 2 | Canis lupus | 1177.0 | 2 |
| scf7180000053514 | 1829 | 0.4511 | 0 | 369.2705 | no-hit | 0.0 | 0 | no-hit | 0.0 | 0 | no-hit | 0.0 | 0 | no-hit | 0.0 | 0 | no-hit | 0.0 | 0 | no-hit | 0.0 | 0 |
| scf7180000040850 | 1810380 | 0.3693 | 100 | 59.5031 | Eukaryota | 68373.0 | 0 | Chordata | 68373.0 | 0 | Carnivora | 68373.0 | 0 | Canidae | 68373.0 | 0 | Canis | 68373.0 | 0 | Canis lupus | 68373.0 | 0 |
| scf7180000047308 | 95278 | 0.4499 | 0 | 58.6404 | Eukaryota | 29300.0 | 0 | Chordata | 29300.0 | 0 | Carnivora | 21236.0 | 1 | Canidae | 9257.0 | 4 | Canis | 6121.0 | 6 | Canis lupus | 6121.0 | 7 |
| scf7180000047675 | 5709 | 0.4276 | 0 | 57.5679 | Eukaryota | 1815.0 | 0 | Platyhelminthes | 1345.0 | 1 | Polystomatidea | 1345.0 | 1 | Polystomatidae | 1345.0 | 1 | Protopolystoma | 1345.0 | 1 | Protopolystoma xenopodis | 1345.0 | 1 |
| scf7180000052259 | 7789 | 0.5066 | 0 | 346.3616 | Eukaryota | 9080.0 | 0 | Chordata | 9080.0 | 0 | Carnivora | 9080.0 | 0 | Canidae | 9080.0 | 0 | Canis | 7348.0 | 1 | Canis lupus | 6078.0 | 2 |
| scf7180000042625 | 258796 | 0.428 | 0 | 60.4443 | Eukaryota | 96813.0 | 0 | Chordata | 96813.0 | 0 | Carnivora | 96813.0 | 0 | Canidae | 96813.0 | 0 | Canis | 96813.0 | 0 | Canis lupus | 96813.0 | 0 |
| scf7180000054440 | 46279 | 0.4431 | 0 | 460.0808 | Eukaryota | 66419.0 | 0 | Chordata | 46021.0 | 2 | Gadiformes | 39425.0 | 5 | Gadidae | 39425.0 | 5 | Gadus | 39425.0 | 5 | Gadus morhua | 39425.0 | 5 |
| scf7180000040767 | 112265 | 0.3842 | 100 | 56.6255 | Eukaryota | 61153.0 | 0 | Chordata | 61153.0 | 0 | Carnivora | 61153.0 | 0 | Canidae | 61153.0 | 0 | Canis | 61153.0 | 0 | Canis lupus | 61153.0 | 0 |
| scf7180000052128 | 10319 | 0.441 | 0 | 190.8829 | Eukaryota | 8868.0 | 0 | Chordata | 7951.0 | 1 | Tetraodontiformes | 5003.0 | 2 | Tetraodontidae | 5003.0 | 2 | Takifugu | 4620.0 | 3 | Takifugu rubripes | 4620.0 | 3 |
| scf7180000046495 | 23434 | 0.4412 | 0 | 96.3075 | Eukaryota | 1608.0 | 0 | Chordata | 1608.0 | 0 | Tetraodontiformes | 1002.0 | 1 | Tetraodontidae | 1002.0 | 1 | Takifugu | 1002.0 | 1 | Takifugu rubripes | 1002.0 | 1 |
| scf7180000053875 | 4673 | 0.4415 | 0 | 311.0866 | no-hit | 0.0 | 0 | no-hit | 0.0 | 0 | no-hit | 0.0 | 0 | no-hit | 0.0 | 0 | no-hit | 0.0 | 0 | no-hit | 0.0 | 0 |
| scf7180000048986 | 29760 | 0.4527 | 0 | 1289.5218 | Eukaryota | 7555.0 | 0 | Chordata | 4384.0 | 2 | Salmoniformes | 3367.0 | 4 | Salmonidae | 3367.0 | 4 | Salmo | 3367.0 | 4 | Salmo trutta | 3367.0 | 4 |
| scf7180000045192 | 98382 | 0.476 | 0 | 60.4708 | Eukaryota | 37482.0 | 0 | Chordata | 37482.0 | 0 | Carnivora | 37482.0 | 0 | Canidae | 24430.0 | 3 | Canis | 21532.0 | 6 | Canis lupus | 21532.0 | 6 |
| scf7180000047100 | 23121 | 0.4343 | 0 | 272.041 | Eukaryota | 53973.0 | 0 | Chordata | 53973.0 | 0 | Carnivora | 53973.0 | 0 | Canidae | 53973.0 | 0 | Canis | 53973.0 | 0 | Canis lupus | 53973.0 | 0 |
| scf7180000050214 | 8335 | 0.3677 | 0 | 25.7252 | Eukaryota | 40853.0 | 0 | Chordata | 40853.0 | 0 | Carnivora | 40853.0 | 0 | Canidae | 40853.0 | 0 | Canis | 40853.0 | 0 | Canis lupus | 40853.0 | 0 |
| scf7180000042449 | 448670 | 0.4198 | 0 | 60.1015 | Eukaryota | 32916.0 | 0 | Chordata | 32916.0 | 0 | Carnivora | 32916.0 | 0 | Canidae | 32916.0 | 0 | Canis | 32916.0 | 0 | Canis lupus | 32916.0 | 0 |
| scf7180000049079 | 54403 | 0.4397 | 0 | 41.3623 | Eukaryota | 90648.0 | 0 | Chordata | 90648.0 | 0 | Carnivora | 90648.0 | 0 | Canidae | 90648.0 | 0 | Canis | 90648.0 | 0 | Canis lupus | 90648.0 | 0 |
| scf7180000047717 | 7072 | 0.4429 | 0 | 38.249 | Eukaryota | 29268.0 | 0 | Chordata | 24129.0 | 2 | Gadiformes | 14558.0 | 6 | Gadidae | 14558.0 | 6 | Gadus | 14558.0 | 6 | Gadus morhua | 14558.0 | 6 |

| | | | | | | | | | | | | | | | | | | | | | | |
|---|---|---|---|---|---|---|---|---|---|---|---|---|---|---|---|---|---|---|---|---|---|---|
| scf7180000041766 | 801770 | 0.4031 | 0 | 59.6781 | Eukaryota | 53266.0 | 0 | Chordata | 53266.0 | 0 | Carnivora | 53266.0 | 0 | Canidae | 53266.0 | 0 | Canis | 53266.0 | 0 | Canis lupus | 53266.0 | 0 |
| scf7180000051763 | 5676 | 0.4468 | 0 | 478.0749 | no-hit | 0.0 | 0 | no-hit | 0.0 | 0 | no-hit | 0.0 | 0 | no-hit | 0.0 | 0 | no-hit | 0.0 | 0 | no-hit | 0.0 | 0 |
| scf7180000047723 | 62051 | 0.5026 | 0 | 58.6204 | Eukaryota | 12767.0 | 0 | Chordata | 12767.0 | 0 | Carnivora | 12767.0 | 0 | Canidae | 6051.0 | 3 | Canis | 4019.0 | 8 | Canis lupus | 4019.0 | 8 |
| scf7180000050578 | 13381 | 0.453 | 0 | 1223.9833 | Eukaryota | 10029.0 | 0 | Chordata | 8222.0 | 2 | Synbranchiformes | 3407.0 | 7 | Mastacembelidae | 3407.0 | 7 | Mastacembelus | 3407.0 | 8 | Mastacembelus armatus | 3407.0 | 8 |
| scf7180000044458 | 325080 | 0.3584 | 0 | 59.7606 | Eukaryota | 83429.0 | 0 | Chordata | 83429.0 | 0 | Carnivora | 83429.0 | 0 | Canidae | 83429.0 | 0 | Canis | 83429.0 | 0 | Canis lupus | 83429.0 | 0 |
| scf7180000042350 | 136609 | 0.3799 | 0 | 57.4604 | Eukaryota | 51182.0 | 0 | Chordata | 51182.0 | 0 | Carnivora | 51182.0 | 0 | Canidae | 41921.0 | 3 | Canis | 41921.0 | 3 | Canis lupus | 41921.0 | 3 |
| scf7180000050231 | 5437 | 0.4455 | 0 | 86.1969 | Eukaryota | 198.0 | 0 | Platyhelminthes | 198.0 | 0 | Polystomatidea | 198.0 | 0 | Polystomatidae | 198.0 | 0 | Protopolystoma | 198.0 | 0 | Protopolystoma xenopodis | 198.0 | 0 |
| scf7180000045031 | 27078 | 0.4315 | 0 | 44.8304 | Eukaryota | 3659.0 | 0 | Chordata | 3659.0 | 0 | Carnivora | 2402.0 | 2 | Canidae | 2402.0 | 2 | Canis | 2402.0 | 2 | Canis lupus | 2402.0 | 2 |
| scf7180000045574 | 7140 | 0.4339 | 0 | 34.6228 | Eukaryota | 6179.0 | 0 | Chordata | 5280.0 | 1 | Kurtiformes | 3142.0 | 3 | Apogonidae | 3142.0 | 3 | Sphaeramia | 3142.0 | 3 | Sphaeramia orbicularis | 3142.0 | 3 |
| scf7180000045570 | 197500 | 0.4195 | 0 | 58.7954 | Eukaryota | 42149.0 | 0 | Chordata | 42149.0 | 0 | Carnivora | 42149.0 | 0 | Canidae | 42149.0 | 0 | Canis | 42149.0 | 0 | Canis lupus | 42149.0 | 0 |
| scf7180000044549 | 232989 | 0.5149 | 0 | 57.4148 | Eukaryota | 28265.0 | 0 | Chordata | 28265.0 | 0 | Carnivora | 28265.0 | 0 | Canidae | 28265.0 | 0 | Canis | 25461.0 | 1 | Canis lupus | 25461.0 | 1 |
| scf7180000047405 | 8209 | 0.4427 | 0 | 253.3234 | Eukaryota | 1509.0 | 0 | Platyhelminthes | 1359.0 | 1 | Polystomatidea | 1359.0 | 1 | Polystomatidae | 1359.0 | 1 | Protopolystoma | 1359.0 | 1 | Protopolystoma xenopodis | 1359.0 | 1 |
| scf7180000050236 | 19978 | 0.5062 | 0 | 135.4038 | Eukaryota | 32856.0 | 0 | Chordata | 32856.0 | 0 | Carnivora | 32856.0 | 0 | Canidae | 32856.0 | 0 | Canis | 32856.0 | 0 | Canis lupus | 32856.0 | 0 |
| scf7180000041245 | 222356 | 0.4409 | 0 | 58.4249 | Eukaryota | 50660.0 | 0 | Chordata | 50660.0 | 0 | Carnivora | 47033.0 | 1 | Canidae | 31076.0 | 4 | Canis | 25980.0 | 5 | Canis lupus | 25980.0 | 5 |
| scf7180000052063 | 16301 | 0.4458 | 0 | 149.6641 | Eukaryota | 29395.0 | 0 | Chordata | 25326.0 | 1 | Gadiformes | 13566.0 | 5 | Gadidae | 13566.0 | 5 | Gadus | 13566.0 | 5 | Gadus morhua | 13566.0 | 5 |
| scf7180000051745 | 5945 | 0.4486 | 0 | 61.52 | Eukaryota | 846.0 | 0 | Platyhelminthes | 846.0 | 0 | Polystomatidea | 846.0 | 0 | Polystomatidae | 846.0 | 0 | Protopolystoma | 846.0 | 0 | Protopolystoma xenopodis | 846.0 | 0 |
| scf7180000051826 | 19723 | 0.437 | 0 | 97.303 | Eukaryota | 39024.0 | 0 | Platyhelminthes | 22468.0 | 2 | Cyclophyllidea | 11651.0 | 5 | Hymenolepididae | 11651.0 | 5 | Hymenolepis | 11651.0 | 5 | Hymenolepis microstoma | 11651.0 | 5 |
| scf7180000055614 | 14213 | 0.443 | 0 | 241.0948 | Eukaryota | 17641.0 | 1 | Chordata | 15356.0 | 3 | Gadiformes | 9747.0 | 7 | Gadidae | 9747.0 | 7 | Gadus | 9747.0 | 7 | Gadus morhua | 9747.0 | 7 |
| scf7180000049444 | 4643 | 0.4495 | 0 | 48.4511 | Eukaryota | 1724.0 | 0 | Platyhelminthes | 1030.0 | 1 | Polystomatidea | 1030.0 | 2 | Polystomatidae | 1030.0 | 2 | Protopolystoma | 1030.0 | 2 | Protopolystoma xenopodis | 1030.0 | 2 |
| scf7180000041292 | 175742 | 0.3728 | 0 | 61.1736 | Eukaryota | 16073.0 | 0 | Chordata | 16073.0 | 0 | Primates | 8493.0 | 2 | Hominidae | 8493.0 | 3 | Homo | 8493.0 | 3 | Homo sapiens | 8493.0 | 3 |
| scf7180000048845 | 46699 | 0.3619 | 0 | 54.4157 | Eukaryota | 59798.0 | 0 | Chordata | 59798.0 | 0 | Carnivora | 59798.0 | 0 | Canidae | 59798.0 | 0 | Canis | 59798.0 | 0 | Canis lupus | 59798.0 | 0 |
| scf7180000046200 | 155499 | 0.5554 | 0 | 57.8022 | Eukaryota | 58533.0 | 0 | Chordata | 58533.0 | 0 | Carnivora | 58533.0 | 0 | Canidae | 54464.0 | 2 | Canis | 47640.0 | 3 | Canis lupus | 47640.0 | 3 |
| scf7180000046407 | 74596 | 0.4403 | 0 | 122.2096 | Eukaryota | 79191.0 | 0 | Chordata | 79191.0 | 0 | Carnivora | 79191.0 | 0 | Canidae | 79191.0 | 0 | Canis | 79191.0 | 0 | Canis lupus | 79191.0 | 0 |
| scf7180000056483 | 16582 | 0.4384 | 0 | 119.3437 | Eukaryota | 37365.0 | 0 | Chordata | 27078.0 | 2 | Gadiformes | 24903.0 | 5 | Gadidae | 24903.0 | 5 | Gadus | 24903.0 | 5 | Gadus morhua | 24903.0 | 5 |
| scf7180000047582 | 11636 | 0.4377 | 0 | 79.2887 | no-hit | 0.0 | 0 | no-hit | 0.0 | 0 | no-hit | 0.0 | 0 | no-hit | 0.0 | 0 | no-hit | 0.0 | 0 | no-hit | 0.0 | 0 |
| scf7180000046883 | 7102 | 0.3662 | 0 | 32.9395 | Eukaryota | 6055.0 | 0 | Chordata | 6055.0 | 0 | Carnivora | 5321.0 | 1 | Canidae | 5321.0 | 1 | Canis | 5321.0 | 1 | Canis lupus | 5321.0 | 1 |
| scf7180000046346 | 135192 | 0.3574 | 0 | 59.77 | Eukaryota | 23635.0 | 0 | Chordata | 23635.0 | 0 | Carnivora | 23635.0 | 0 | Canidae | 23635.0 | 0 | Canis | 23635.0 | 0 | Canis lupus | 23635.0 | 0 |
| scf7180000046122 | 37805 | 0.5631 | 0 | 54.6936 | Eukaryota | 8995.0 | 0 | Chordata | 8995.0 | 0 | Carnivora | 8995.0 | 0 | Felidae | 4487.0 | 2 | Panthera | 2103.0 | 5 | Panthera pardus | 2103.0 | 5 |
| scf7180000045666 | 180047 | 0.6105 | 0 | 55.5348 | Eukaryota | 9417.0 | 0 | Chordata | 9417.0 | 0 | Carnivora | 9417.0 | 0 | Canidae | 7942.0 | 1 | Canis | 7942.0 | 2 | Canis lupus | 7942.0 | 2 |
| scf7180000042191 | 738073 | 0.463 | 0 | 59.0034 | Eukaryota | 74071.0 | 0 | Chordata | 74071.0 | 0 | Carnivora | 74071.0 | 0 | Canidae | 74071.0 | 0 | Canis | 51793.0 | 1 | Canis lupus | 51793.0 | 1 |
| scf7180000046638 | 122074 | 0.3493 | 0 | 60.9095 | Eukaryota | 90619.0 | 0 | Chordata | 90619.0 | 0 | Carnivora | 90619.0 | 0 | Canidae | 90619.0 | 0 | Canis | 90619.0 | 0 | Canis lupus | 90619.0 | 0 |
| scf7180000046378 | 145371 | 0.4257 | 0 | 59.2982 | Eukaryota | 23232.0 | 0 | Chordata | 23232.0 | 0 | Carnivora | 23232.0 | 0 | Felidae | 8530.0 | 3 | Mustela | 4526.0 | 6 | Mustela putorius | 4526.0 | 6 |
| scf7180000045834 | 169006 | 0.4526 | 0 | 56.1258 | Eukaryota | 91465.0 | 0 | Chordata | 91465.0 | 0 | Carnivora | 91465.0 | 0 | Canidae | 91465.0 | 0 | Canis | 91465.0 | 0 | Canis lupus | 91465.0 | 0 |
| scf7180000053703 | 8325 | 0.504 | 0 | 81.9413 | Eukaryota | 7432.0 | 0 | Chordata | 7432.0 | 0 | Carnivora | 7432.0 | 0 | Canidae | 7432.0 | 0 | Canis | 5380.0 | 1 | Canis lupus | 4333.0 | 2 |
| scf7180000051756 | 7784 | 0.4451 | 0 | 66.3415 | Eukaryota | 3202.0 | 0 | Chordata | 2668.0 | 1 | Carnivora | 2668.0 | 1 | Canidae | 2668.0 | 1 | Canis | 2668.0 | 1 | Canis lupus | 2668.0 | 1 |
| scf7180000053722 | 2129 | 0.4528 | 0 | 270.8516 | no-hit | 0.0 | 0 | no-hit | 0.0 | 0 | no-hit | 0.0 | 0 | no-hit | 0.0 | 0 | no-hit | 0.0 | 0 | no-hit | 0.0 | 0 |
| scf7180000053000 | 4160 | 0.4397 | 0 | 99.4583 | Eukaryota | 2806.0 | 0 | Chordata | 2806.0 | 0 | Carnivora | 2806.0 | 0 | Canidae | 2806.0 | 0 | Canis | 2806.0 | 0 | Canis lupus | 2806.0 | 0 |
| scf7180000049473 | 13648 | 0.5188 | 0 | 398.0298 | Eukaryota | 15758.0 | 0 | Chordata | 15758.0 | 0 | Carnivora | 15758.0 | 0 | Canidae | 15758.0 | 0 | Canis | 15758.0 | 0 | Canis lupus | 15758.0 | 0 |
| scf7180000044691 | 272564 | 0.4032 | 0 | 61.7369 | Eukaryota | 22578.0 | 0 | Chordata | 22578.0 | 0 | Carnivora | 22578.0 | 0 | Canidae | 22578.0 | 0 | Canis | 22578.0 | 0 | Canis lupus | 22578.0 | 0 |
| scf7180000042802 | 570777 | 0.481 | 0 | 58.5904 | Eukaryota | 98750.0 | 0 | Chordata | 98750.0 | 0 | Carnivora | 98750.0 | 0 | Canidae | 98750.0 | 0 | Canis | 69125.0 | 1 | Canis lupus | 69125.0 | 1 |
| scf7180000054554 | 16982 | 0.4337 | 0 | 427.3478 | Eukaryota | 384.0 | 0 | Chordata | 384.0 | 0 | Kurtiformes | 230.0 | 1 | Apogonidae | 230.0 | 1 | Sphaeramia | 230.0 | 1 | Sphaeramia orbicularis | 230.0 | 1 |
| scf7180000049942 | 29420 | 0.4443 | 0 | 150.3462 | Eukaryota | 3227.0 | 0 | Platyhelminthes | 1629.0 | 1 | Polystomatidea | 1629.0 | 2 | Polystomatidae | 1629.0 | 2 | Protopolystoma | 1629.0 | 2 | Protopolystoma xenopodis | 1629.0 | 2 |
| scf7180000044156 | 221598 | 0.4077 | 0 | 62.9637 | Eukaryota | 99981.0 | 0 | Chordata | 99981.0 | 0 | Carnivora | 99981.0 | 0 | Canidae | 99981.0 | 0 | Canis | 99981.0 | 0 | Canis lupus | 99981.0 | 0 |
| scf7180000040982 | 1486522 | 0.4023 | 0 | 59.6964 | Eukaryota | 72731.0 | 0 | Chordata | 72731.0 | 0 | Carnivora | 72731.0 | 0 | Canidae | 72731.0 | 0 | Canis | 64648.0 | 1 | Canis lupus | 64648.0 | 1 |
| scf7180000048474 | 36223 | 0.4378 | 0 | 109.919 | Eukaryota | 4610.0 | 0 | Chordata | 2902.0 | 1 | Tetraodontiformes | 1864.0 | 3 | Tetraodontidae | 1864.0 | 3 | Takifugu | 1864.0 | 3 | Takifugu rubripes | 1864.0 | 3 |
| scf7180000043775 | 672888 | 0.4425 | 0 | 57.5214 | Eukaryota | 46120.0 | 0 | Chordata | 46120.0 | 0 | Carnivora | 46120.0 | 0 | Canidae | 46120.0 | 0 | Canis | 46120.0 | 0 | Canis lupus | 46120.0 | 0 |
| scf7180000056160 | 6108 | 0.4473 | 0 | 101.203 | Eukaryota | 272.0 | 0 | Chordata | 272.0 | 0 | Salmoniformes | 272.0 | 0 | Salmonidae | 272.0 | 0 | Coregonus | 272.0 | 0 | Coregonus sp. 'balchen' | 272.0 | 0 |
| scf7180000045386 | 10420 | 0.4012 | 0 | 28.5191 | Eukaryota | 15919.0 | 0 | Chordata | 15919.0 | 0 | Carnivora | 15919.0 | 0 | Canidae | 7839.0 | 2 | Canis | 7839.0 | 3 | Canis lupus | 7839.0 | 3 |
| scf7180000050722 | 18143 | 0.4515 | 0 | 61.243 | Eukaryota | 3365.0 | 1 | Chordata | 3365.0 | 1 | Primates | 2216.0 | 3 | Hominidae | 2216.0 | 3 | Homo | 1931.0 | 4 | Homo sapiens | 1931.0 | 4 |
| scf7180000046099 | 32956 | 0.4328 | 0 | 63.345 | Eukaryota | 8698.0 | 0 | Chordata | 8166.0 | 1 | Chordata-undef | 3308.0 | 4 | Ambassidae | 3308.0 | 4 | Parambassis | 3308.0 | 5 | Parambassis ranga | 3308.0 | 5 |
| scf7180000044380 | 344053 | 0.3735 | 0 | 60.421 | Eukaryota | 26131.0 | 0 | Chordata | 26131.0 | 0 | Carnivora | 26131.0 | 0 | Canidae | 21291.0 | 1 | Canis | 21291.0 | 1 | Canis lupus | 21291.0 | 1 |
| scf7180000051074 | 12744 | 0.4436 | 0 | 88.2299 | Eukaryota | 40482.0 | 0 | Platyhelminthes | 22433.0 | 2 | Strigeidida | 13679.0 | 7 | Schistosomatidae | 13679.0 | 7 | Schistosoma | 13679.0 | 7 | Schistosoma mansoni | 13679.0 | 7 |
| scf7180000044873 | 95352 | 0.3847 | 0 | 57.6152 | Eukaryota | 12271.0 | 0 | Chordata | 12271.0 | 0 | Carnivora | 12271.0 | 0 | Canidae | 12271.0 | 0 | Canis | 12271.0 | 0 | Canis lupus | 12271.0 | 0 |
| scf7180000040542 | 838162 | 0.4161 | 0 | 58.3692 | Eukaryota | 36335.0 | 0 | Chordata | 36335.0 | 0 | Carnivora | 32772.0 | 1 | Canidae | 30221.0 | 2 | Canis | 20234.0 | 3 | Canis lupus | 20234.0 | 3 |
| scf7180000045111 | 267346 | 0.3439 | 0 | 60.9514 | Eukaryota | 47000.0 | 0 | Chordata | 47000.0 | 0 | Carnivora | 47000.0 | 0 | Canidae | 47000.0 | 0 | Canis | 47000.0 | 0 | Canis lupus | 47000.0 | 0 |
| scf7180000048722 | 47636 | 0.3757 | 0 | 60.6926 | Eukaryota | 17683.0 | 0 | Chordata | 17683.0 | 0 | Carnivora | 17683.0 | 0 | Canidae | 15325.0 | 1 | Canis | 13884.0 | 2 | Canis lupus | 13884.0 | 2 |
| scf7180000045457 | 208504 | 0.3692 | 0 | 74.4621 | Eukaryota | 94998.0 | 0 | Chordata | 94998.0 | 0 | Carnivora | 94998.0 | 0 | Canidae | 94998.0 | 0 | Canis | 94998.0 | 0 | Canis lupus | 94998.0 | 0 |
| scf7180000052953 | 6312 | 0.4426 | 0 | 326.736 | Eukaryota | 263.0 | 0 | Platyhelminthes | 263.0 | 0 | Polystomatidea | 263.0 | 0 | Polystomatidae | 263.0 | 0 | Protopolystoma | 263.0 | 0 | Protopolystoma xenopodis | 263.0 | 0 |
| scf7180000052800 | 2347 | 0.444 | 0 | 192.1977 | Eukaryota | 862.0 | 0 | Platyhelminthes | 862.0 | 0 | Polystomatidea | 862.0 | 0 | Polystomatidae | 862.0 | 0 | Protopolystoma | 862.0 | 0 | Protopolystoma xenopodis | 862.0 | 0 |
| scf7180000048334 | 9438 | 0.4408 | 0 | 89.0829 | Eukaryota | 3301.0 | 0 | Chordata | 2277.0 | 2 | Kurtiformes | 1554.0 | 3 | Apogonidae | 1554.0 | 3 | Sphaeramia | 1554.0 | 3 | Sphaeramia orbicularis | 1554.0 | 3 |
| scf7180000044296 | 344949 | 0.3757 | 0 | 59.8088 | Eukaryota | 51586.0 | 0 | Chordata | 51586.0 | 0 | Carnivora | 51586.0 | 0 | Canidae | 51586.0 | 0 | Canis | 51586.0 | 0 | Canis lupus | 51586.0 | 0 |
| scf7180000051112 | 5861 | 0.4424 | 0 | 64.3108 | Eukaryota | 6445.0 | 0 | Chordata | 3343.0 | 1 | Polystomatidea | 3102.0 | 3 | Polystomatidae | 3102.0 | 3 | Protopolystoma | 3102.0 | 3 | Protopolystoma xenopodis | 3102.0 | 3 |
| scf7180000042135 | 208376 | 0.4599 | 0 | 61.5891 | Eukaryota | 93162.0 | 0 | Chordata | 93162.0 | 0 | Carnivora | 93162.0 | 0 | Canidae | 93162.0 | 0 | Canis | 93162.0 | 0 | Canis lupus | 93162.0 | 0 |
| scf7180000045792 | 154616 | 0.6709 | 0 | 53.7061 | Eukaryota | 56090.0 | 0 | Chordata | 56090.0 | 0 | Carnivora | 56090.0 | 0 | Canidae | 56090.0 | 0 | Canis | 56090.0 | 0 | Canis lupus | 56090.0 | 0 |
| scf7180000046065 | 181339 | 0.3896 | 0 | 60.2003 | Eukaryota | 63112.0 | 0 | Chordata | 63112.0 | 0 | Carnivora | 63112.0 | 0 | Canidae | 45693.0 | 2 | Canis | 25632.0 | 4 | Canis lupus | 25632.0 | 4 |
| scf7180000045240 | 242025 | 0.4216 | 0 | 59.9509 | Eukaryota | 61338.0 | 0 | Chordata | 61338.0 | 0 | Carnivora | 61338.0 | 0 | Canidae | 61338.0 | 0 | Canis | 61338.0 | 0 | Canis lupus | 61338.0 | 0 |
| scf7180000040964 | 44422 | 0.3912 | 0 | 55.4095 | Eukaryota | 3993.0 | 0 | Chordata | 3993.0 | 0 | Carnivora | 2064.0 | 2 | Canidae | 2064.0 | 2 | Canis | 2064.0 | 2 | Canis lupus | 2064.0 | 2 |
| scf7180000040880 | 1776684 | 0.3576 | 100 | 60.263 | Eukaryota | 91285.0 | 0 | Chordata | 91285.0 | 0 | Carnivora | 91285.0 | 0 | Canidae | 91285.0 | 0 | Canis | 91285.0 | 0 | Canis lupus | 91285.0 | 0 |
| scf7180000042373 | 484462 | 0.4001 | 0 | 59.0639 | Eukaryota | 93777.0 | 0 | Chordata | 93777.0 | 0 | Carnivora | 93777.0 | 0 | Canidae | 93777.0 | 0 | Canis | 93777.0 | 0 | Canis lupus | 93777.0 | 0 |
| scf7180000050732 | 6783 | 0.4426 | 0 | 97.27 | Eukaryota | 461.0 | 0 | Platyhelminthes | 461.0 | 0 | Polystomatidea | 461.0 | 0 | Polystomatidae | 461.0 | 0 | Protopolystoma | 461.0 | 0 | Protopolystoma xenopodis | 461.0 | 0 |
| scf7180000053024 | 6612 | 0.4527 | 0 | 1029.6951 | Eukaryota | 2488.0 | 0 | Chordata | 2488.0 | 0 | Carnivora | 2220.0 | 1 | Canidae | 2220.0 | 1 | Canis | 1974.0 | 2 | Canis lupus | 1974.0 | 2 |
| scf7180000053635 | 2304 | 0.4384 | 0 | 1455.1427 | no-hit | 0.0 | 0 | no-hit | 0.0 | 0 | no-hit | 0.0 | 0 | no-hit | 0.0 | 0 | no-hit | 0.0 | 0 | no-hit | 0.0 | 0 |
| scf7180000046890 | 16143 | 0.3974 | 0 | 51.0748 | Eukaryota | 89680.0 | 0 | Chordata | 89680.0 | 0 | Carnivora | 89680.0 | 0 | Canidae | 89680.0 | 0 | Canis | 89680.0 | 0 | Canis lupus | 89680.0 | 0 |
| scf7180000054107 | 3218 | 0.4198 | 0 | 161.2414 | Eukaryota | 141.0 | 0 | Platyhelminthes | 141.0 | 0 | Polystomatidea | 141.0 | 0 | Polystomatidae | 141.0 | 0 | Protopolystoma | 141.0 | 0 | Protopolystoma xenopodis | 141.0 | 0 |
| scf7180000046072 | 11831 | 0.3711 | 0 | 48.4761 | Eukaryota | 79292.0 | 0 | Chordata | 79292.0 | 0 | Carnivora | 79292.0 | 0 | Canidae | 79292.0 | 0 | Canis | 79292.0 | 0 | Canis lupus | 79292.0 | 0 |
| scf7180000040533 | 2948507 | 0.413 | 100 | 60.2374 | Eukaryota | 166229.0 | 0 | Chordata | 166229.0 | 0 | Carnivora | 166229.0 | 0 | Canidae | 166229.0 | 0 | Canis | 166229.0 | 0 | Canis lupus | 166229.0 | 0 |
| scf7180000040569 | 2119176 | 0.3912 | 0 | 59.7145 | Eukaryota | 93777.0 | 0 | Chordata | 93777.0 | 0 | Carnivora | 93777.0 | 0 | Canidae | 93777.0 | 0 | Canis | 93777.0 | 0 | Canis lupus | 93777.0 | 0 |
| scf7180000043028 | 62978 | 0.3823 | 0 | 58.9889 | Eukaryota | 69778.0 | 0 | Chordata | 69778.0 | 0 | Carnivora | 69778.0 | 0 | Canidae | 69778.0 | 0 | Canis | 47835.0 | 1 | Canis lupus | 47835.0 | 1 |
| scf7180000048969 | 10386 | 0.4487 | 0 | 76.5797 | Eukaryota | 2684.0 | 0 | Chordata | 2684.0 | 0 | Carnivora | 2684.0 | 0 | Canidae | 2684.0 | 0 | Canis | 2684.0 | 0 | Canis lupus | 2684.0 | 0 |
| scf7180000044385 | 1988 | 0.4502 | 0 | 466.9002 | Eukaryota | 1810.0 | 0 | Platyhelminthes | 672.0 | 3 | Polystomatidea | 672.0 | 3 | Polystomatidae | 672.0 | 3 | Protopolystoma | 672.0 | 3 | Protopolystoma xenopodis | 672.0 | 3 |
| scf7180000054299 | 22775 | 0.4426 | 0 | 118.055 | Eukaryota | 2049.0 | 0 | Chordata | 1814.0 | 1 | Kurtiformes | 1369.0 | 3 | Apogonidae | 1369.0 | 3 | Sphaeramia | 1369.0 | 3 | Sphaeramia orbicularis | 1369.0 | 3 |
| scf7180000050805 | 4855 | 0.4453 | 0 | 202.7878 | no-hit | 0.0 | 0 | no-hit | 0.0 | 0 | no-hit | 0.0 | 0 | no-hit | 0.0 | 0 | no-hit | 0.0 | 0 | no-hit | 0.0 | 0 |
| scf7180000055597 | 25168 | 0.4385 | 0 | 203.6003 | Eukaryota | 86398.0 | 0 | Platyhelminthes | 44842.0 | 1 | Strigeidida | 32742.0 | 4 | Schistosomatidae | 32742.0 | 4 | Schistosoma | 32742.0 | 4 | Schistosoma mansoni | 32742.0 | 4 |
| scf7180000042276 | 383520 | 0.4411 | 0 | 58.2868 | Eukaryota | 57007.0 | 0 | Chordata | 57007.0 | 0 | Carnivora | 57007.0 | 0 | Canidae | 57007.0 | 0 | Canis | 57007.0 | 0 | Canis lupus | 57007.0 | 0 |
| scf7180000043102 | 28625 | 0.3871 | 0 | 63.8977 | Eukaryota | 48700.0 | 0 | Chordata | 48700.0 | 0 | Carnivora | 48700.0 | 0 | Canidae | 48700.0 | 0 | Canis | 48700.0 | 0 | Canis lupus | 48700.0 | 0 |
| scf7180000043263 | 677440 | 0.3529 | 100 | 60.3274 | Eukaryota | 93768.0 | 0 | Chordata | 93768.0 | 0 | Carnivora | 93768.0 | 0 | Canidae | 93768.0 | 0 | Canis | 93768.0 | 0 | Canis lupus | 93768.0 | 0 |
| scf7180000049026 | 7577 | 0.442 | 0 | 183.4439 | no-hit | 0.0 | 0 | no-hit | 0.0 | 0 | no-hit | 0.0 | 0 | no-hit | 0.0 | 0 | no-hit | 0.0 | 0 | no-hit | 0.0 | 0 |
| scf7180000048975 | 12927 | 0.4545 | 0 | 65.4219 | Eukaryota | 4490.0 | 0 | Chordata | 2605.0 | 1 | Polystomatidea | 1885.0 | 3 | Polystomatidae | 1885.0 | 3 | Protopolystoma | 1885.0 | 3 | Protopolystoma xenopodis | 1885.0 | 3 |
| scf7180000048669 | 12862 | 0.4429 | 0 | 116.392 | no-hit | 0.0 | 0 | no-hit | 0.0 | 0 | no-hit | 0.0 | 0 | no-hit | 0.0 | 0 | no-hit | 0.0 | 0 | no-hit | 0.0 | 0 |
| scf7180000044664 | 299760 | 0.4321 | 0 | 59.546 | Eukaryota | 95574.0 | 0 | Chordata | 95574.0 | 0 | Carnivora | 95574.0 | 0 | Canidae | 95574.0 | 0 | Canis | 86067.0 | 1 | Canis lupus | 86067.0 | 1 |
| scf7180000050727 | 50804 | 0.3617 | 0 | 53.7861 | Eukaryota | 28615.0 | 0 | Chordata | 28615.0 | 0 | Carnivora | 28615.0 | 0 | Canidae | 28615.0 | 0 | Canis | 28615.0 | 0 | Canis lupus | 28615.0 | 0 |
| scf7180000044721 | 302911 | 0.3807 | 0 | 62.0514 | Eukaryota | 20394.0 | 0 | Chordata | 20394.0 | 0 | Carnivora | 20394.0 | 0 | Canidae | 18443.0 | 1 | Canis | 18443.0 | 1 | Canis lupus | 18443.0 | 1 |
| scf7180000056252 | 388872 | 0.4394 | 0 | 57.9617 | Eukaryota | 35291.0 | 0 | Chordata | 35291.0 | 0 | Carnivora | 35291.0 | 0 | Canidae | 35291.0 | 0 | Canis | 30391.0 | 1 | Canis lupus | 30391.0 | 1 |
| scf7180000051689 | 31158 | 0.5026 | 0 | 1231.225 | Eukaryota | 9680.0 | 0 | Chordata | 9680.0 | 0 | Carnivora | 9680.0 | 0 | Canidae | 9680.0 | 0 | Canis | 8991.0 | 1 | Canis lupus | 7699.0 | 2 |
| scf7180000047845 | 68245 | 0.3971 | 0 | 61.2863 | Eukaryota | 26210.0 | 0 | Chordata | 26210.0 | 0 | Carnivora | 26210.0 | 0 | Canidae | 26210.0 | 0 | Canis | 26210.0 | 0 | Canis lupus | 26210.0 | 0 |
| scf7180000040944 | 90050 | 0.4388 | 0 | 58.0672 | Eukaryota | 49338.0 | 0 | Chordata | 49338.0 | 0 | Carnivora | 49338.0 | 0 | Canidae | 49338.0 | 0 | Canis | 42606.0 | 1 | Canis lupus | 42606.0 | 1 |
| scf7180000043701 | 395389 | 0.4566 | 0 | 58.1999 | Eukaryota | 34219.0 | 0 | Chordata | 34219.0 | 0 | Carnivora | 34219.0 | 0 | Canidae | 29360.0 | 1 | Canis | 29360.0 | 1 | Canis lupus | 29360.0 | 1 |
| scf7180000053552 | 3174 | 0.4543 | 0 | 874.7309 | no-hit | 0.0 | 0 | no-hit | 0.0 | 0 | no-hit | 0.0 | 0 | no-hit | 0.0 | 0 | no-hit | 0.0 | 0 | no-hit | 0.0 | 0 |
| scf7180000044881 | 303621 | 0.3881 | 0 | 59.3296 | Eukaryota | 17592.0 | 0 | Chordata | 17592.0 | 0 | Carnivora | 17592.0 | 0 | Canidae | 17592.0 | 0 | Canis | 17592.0 | 0 | Canis lupus | 17592.0 | 0 |
| scf7180000048143 | 23290 | 0.4443 | 0 | 223.5819 | Eukaryota | 3068.0 | 0 | Chordata | 3068.0 | 0 | Carnivora | 3068.0 | 0 | Canidae | 3068.0 | 0 | Canis | 3068.0 | 0 | Canis lupus | 3068.0 | 0 |
| scf7180000043430 | 447224 | 0.3543 | 0 | 58.8895 | Eukaryota | 104565.0 | 0 | Chordata | 104565.0 | 0 | Carnivora | 104565.0 | 0 | Canidae | 104565.0 | 0 | Canis | 104565.0 | 0 | Canis lupus | 104565.0 | 0 |
| scf7180000044994 | 274619 | 0.338 | 0 | 59.1368 | Eukaryota | 24453.0 | 0 | Chordata | 24453.0 | 0 | Carnivora | 24453.0 | 0 | Canidae | 24453.0 | 0 | Canis | 24453.0 | 0 | Canis lupus | 24453.0 | 0 |
| scf7180000041442 | 1103222 | 0.3702 | 0 | 59.9909 | Eukaryota | 52949.0 | 0 | Chordata | 52949.0 | 0 | Carnivora | 52949.0 | 0 | Canidae | 52949.0 | 0 | Canis | 52949.0 | 0 | Canis lupus | 52949.0 | 0 |
| scf7180000054992 | 11733 | 0.4378 | 0 | 95.2891 | no-hit | 0.0 | 0 | no-hit | 0.0 | 0 | no-hit | 0.0 | 0 | no-hit | 0.0 | 0 | no-hit | 0.0 | 0 | no-hit | 0.0 | 0 |
| scf7180000042228 | 739559 | 0.3719 | 0 | 60.5016 | Eukaryota | 49424.0 | 0 | Chordata | 49424.0 | 0 | Carnivora | 49424.0 | 0 | Canidae | 26069.0 | 2 | Canis | 21788.0 | 6 | Canis lupus | 21788.0 | 6 |
| scf7180000055176 | 15425 | 0.4433 | 0 | 123.7069 | Eukaryota | 12223.0 | 0 | Chordata | 11879.0 | 1 | Tetraodontiformes | 11879.0 | 1 | Tetraodontidae | 11879.0 | 1 | Takifugu | 11879.0 | 1 | Takifugu rubripes | 11879.0 | 1 |
| scf7180000043814 | 265067 | 0.3681 | 0 | 59.5749 | Eukaryota | 14839.0 | 0 | Chordata | 14839.0 | 0 | Primates | 9512.0 | 2 | Hominidae | 7155.0 | 4 | Homo | 7155.0 | 4 | Homo sapiens | 7155.0 | 4 |
| scf7180000045657 | 7751 | 0.3698 | 0 | 53.4973 | Eukaryota | 6348.0 | 0 | Chordata | 6348.0 | 0 | Primates | 2628.0 | 2 | Hominidae | 2628.0 | 2 | Homo | 2628.0 | 3 | Homo sapiens | 2628.0 | 3 |
| scf7180000048944 | 8003 | 0.4521 | 0 | 127.5078 | Eukaryota | 4610.0 | 0 | Chordata | 2139.0 | 2 | Polystomatidea | 1772.0 | 4 | Polystomatidae | 1772.0 | 4 | Protopolystoma | 1772.0 | 5 | Protopolystoma xenopodis | 1772.0 | 5 |
| scf7180000041790 | 427680 | 0.4015 | 0 | 58.7918 | Eukaryota | 95766.0 | 0 | Chordata | 95766.0 | 0 | Carnivora | 95766.0 | 0 | Canidae | 95766.0 | 0 | Canis | 95766.0 | 0 | Canis lupus | 95766.0 | 0 |
| scf7180000051206 | 18466 | 0.38 | 0 | 60.7283 | Eukaryota | 7360.0 | 0 | Chordata | 7360.0 | 0 | Carnivora | 7360.0 | 0 | Canidae | 3023.0 | 3 | Canis | 2269.0 | 6 | Canis lupus | 2269.0 | 6 |
| scf7180000041758 | 1064549 | 0.3736 | 0 | 60.7816 | Eukaryota | 62529.0 | 0 | Chordata | 62529.0 | 0 | Carnivora | 62529.0 | 0 | Canidae | 62529.0 | 0 | Canis | 62529.0 | 0 | Canis lupus | 62529.0 | 0 |
| scf7180000052753 | 9115 | 0.4286 | 0 | 60.0657 | Eukaryota | 5166.0 | 0 | Chordata | 5166.0 | 0 | Carnivora | 2385.0 | 2 | Canidae | 2385.0 | 2 | Canis | 1967.0 | 4 | Canis lupus | 1967.0 | 4 |
| scf7180000046532 | 15177 | 0.6205 | 0 | 32.4308 | Eukaryota | 4614.0 | 0 | Chordata | 4614.0 | 0 | Carnivora | 4614.0 | 0 | Canidae | 3483.0 | 1 | Canis | 1986.0 | 2 | Canis lupus | 1986.0 | 2 |
| scf7180000050365 | 17264 | 0.4505 | 0 | 1355.9779 | Eukaryota | 14330.0 | 0 | Chordata | 14330.0 | 0 | Tetraodontiformes | 9601.0 | 2 | Tetraodontidae | 9601.0 | 2 | Takifugu | 9601.0 | 2 | Takifugu rubripes | 9601.0 | 2 |
| scf7180000041988 | 879974 | 0.4197 | 0 | 59.4135 | Eukaryota | 58881.0 | 0 | Chordata | 58881.0 | 0 | Carnivora | 49713.0 | 1 | Canidae | 44599.0 | 2 | Canis | 24197.0 | 4 | Canis lupus | 24197.0 | 4 |
| scf7180000052829 | 5402 | 0.4415 | 0 | 122.209 | Eukaryota | 532.0 | 0 | Platyhelminthes | 532.0 | 0 | Polystomatidea | 532.0 | 0 | Polystomatidae | 532.0 | 0 | Protopolystoma | 532.0 | 0 | Protopolystoma xenopodis | 532.0 | 0 |
| scf7180000056542 | 250090 | 0.3732 | 0 | 59.4707 | Eukaryota | 48388.0 | 0 | Chordata | 48388.0 | 0 | Carnivora | 48388.0 | 0 | Canidae | 48388.0 | 0 | Canis | 39882.0 | 1 | Canis lupus | 39882.0 | 1 |
| scf7180000047436 | 5069 | 0.4549 | 0 | 37.218 | Eukaryota | 1192.0 | 0 | Chordata | 920.0 | 1 | Rodentia | 422.0 | 3 | Muridae | 422.0 | 3 | Mus | 422.0 | 3 | Mus musculus | 422.0 | 3 |
| scf7180000045800 | 14925 | 0.4074 | 0 | 51.2154 | Eukaryota | 62449.0 | 0 | Chordata | 62449.0 | 0 | Carnivora | 62449.0 | 0 | Canidae | 62449.0 | 0 | Canis | 62449.0 | 0 | Canis lupus | 62449.0 | 0 |
| scf7180000052603 | 7374 | 0.4471 | 0 | 409.4181 | Eukaryota | 158.0 | 0 | Streptophyta | 158.0 | 0 | Fabales | 158.0 | 0 | Fabaceae | 158.0 | 0 | Cicer | 158.0 | 0 | Cicer arietinum | 158.0 | 0 |
| scf7180000056035 | 8813 | 0.4384 | 0 | 115.9362 | Eukaryota | 1486.0 | 0 | Platyhelminthes | 952.0 | 1 | Polystomatidea | 952.0 | 1 | Polystomatidae | 952.0 | 1 | Protopolystoma | 952.0 | 1 | Protopolystoma xenopodis | 952.0 | 1 |
| scf7180000042934 | 83836 | 0.4407 | 0 | 67.2241 | Eukaryota | 95259.0 | 0 | Chordata | 95259.0 | 0 | Carnivora | 95259.0 | 0 | Canidae | 95259.0 | 0 | Canis | 95259.0 | 0 | Canis lupus | 95259.0 | 0 |
| scf7180000044393 | 80830 | 0.4463 | 0 | 57.7772 | Eukaryota | 80536.0 | 0 | Chordata | 80536.0 | 0 | Carnivora | 80536.0 | 0 | Canidae | 80536.0 | 0 | Canis | 80536.0 | 0 | Canis lupus | 80536.0 | 0 |
| scf7180000040951 | 401705 | 0.421 | 0 | 61.4416 | Eukaryota | 34370.0 | 0 | Chordata | 34370.0 | 0 | Carnivora | 34370.0 | 0 | Canidae | 34370.0 | 0 | Canis | 34370.0 | 0 | Canis lupus | 34370.0 | 0 |
| scf7180000048343 | 3714 | 0.4467 | 0 | 54.0544 | Eukaryota | 261.0 | 0 | Platyhelminthes | 261.0 | 0 | Polystomatidea | 261.0 | 0 | Polystomatidae | 261.0 | 0 | Protopolystoma | 261.0 | 0 | Protopolystoma xenopodis | 261.0 | 0 |
| scf7180000040671 | 2020234 | 0.4302 | 0 | 59.5509 | Eukaryota | 149363.0 | 0 | Chordata | 149363.0 | 0 | Carnivora | 149363.0 | 0 | Canidae | 149363.0 | 0 | Canis | 133921.0 | 1 | Canis lupus | 133921.0 | 1 |
| scf7180000043347 | 120385 | 0.4035 | 0 | 76.6561 | Eukaryota | 66021.0 | 0 | Chordata | 66021.0 | 0 | Carnivora | 66021.0 | 0 | Canidae | 66021.0 | 0 | Canis | 66021.0 | 0 | Canis lupus | 66021.0 | 0 |
| scf7180000053405 | 5081 | 0.4389 | 0 | 255.3527 | no-hit | 0.0 | 0 | no-hit | 0.0 | 0 | no-hit | 0.0 | 0 | no-hit | 0.0 | 0 | no-hit | 0.0 | 0 | no-hit | 0.0 | 0 |
| scf7180000046259 | 63630 | 0.4436 | 0 | 193.9285 | Eukaryota | 58812.0 | 0 | Chordata | 42238.0 | 2 | Gadiformes | 33805.0 | 5 | Gadidae | 33805.0 | 5 | Gadus | 33805.0 | 5 | Gadus morhua | 33805.0 | 5 |
| scf7180000049985 | 3656 | 0.4303 | 0 | 48.7553 | Eukaryota | 12845.0 | 0 | Chordata | 12845.0 | 0 | Carnivora | 12845.0 | 0 | Canidae | 12845.0 | 0 | Canis | 12845.0 | 0 | Canis lupus | 12845.0 | 0 |
| scf7180000044149 | 87267 | 0.3596 | 0 | 54.978 | Eukaryota | 27290.0 | 0 | Chordata | 27290.0 | 0 | Carnivora | 27290.0 | 0 | Canidae | 27290.0 | 0 | Canis | 27290.0 | 0 | Canis lupus | 27290.0 | 0 |
| scf7180000049116 | 25591 | 0.3776 | 0 | 57.8019 | Eukaryota | 23944.0 | 0 | Chordata | 23944.0 | 0 | Carnivora | 23944.0 | 0 | Canidae | 23944.0 | 0 | Canis | 23944.0 | 0 | Canis lupus | 23944.0 | 0 |
| scf7180000055802 | 31216 | 0.4495 | 0 | 106.2262 | Eukaryota | 189.0 | 0 | Platyhelminthes | 189.0 | 0 | Diphyllobothriidea | 189.0 | 0 | Diphyllobothriidae | 189.0 | 0 | Dibothriocephalus | 189.0 | 0 | Dibothriocephalus latus | 189.0 | 0 |
| scf7180000051954 | 6748 | 0.4342 | 0 | 409.4924 | Eukaryota | 35780.0 | 0 | Chordata | 24788.0 | 2 | Gadiformes | 20202.0 | 5 | Gadidae | 20202.0 | 5 | Gadus | 20202.0 | 5 | Gadus morhua | 20202.0 | 5 |
| scf7180000056229 | 883850 | 0.4167 | 0 | 59.6 | Eukaryota | 93414.0 | 0 | Chordata | 93414.0 | 0 | Carnivora | 93414.0 | 0 | Canidae | 93414.0 | 0 | Canis | 93414.0 | 0 | Canis lupus | 93414.0 | 0 |
| scf7180000045944 | 147741 | 0.561 | 0 | 56.1321 | Eukaryota | 28138.0 | 0 | Chordata | 28138.0 | 0 | Carnivora | 28138.0 | 0 | Canidae | 28138.0 | 0 | Canis | 17105.0 | 1 | Canis lupus | 17105.0 | 1 |
| scf7180000044078 | 413377 | 0.3899 | 0 | 59.8403 | Eukaryota | 45363.0 | 0 | Chordata | 45363.0 | 0 | Carnivora | 45363.0 | 0 | Canidae | 45363.0 | 0 | Canis | 45363.0 | 0 | Canis lupus | 45363.0 | 0 |
| scf7180000051435 | 7840 | 0.4457 | 0 | 399.1558 | Eukaryota | 169.0 | 0 | Chordata | 169.0 | 0 | Salmoniformes | 169.0 | 0 | Salmonidae | 169.0 | 0 | Salmo | 169.0 | 0 | Salmo trutta | 169.0 | 0 |
| scf7180000050164 | 34476 | 0.5969 | 0 | 52.5138 | Eukaryota | 13302.0 | 0 | Chordata | 13302.0 | 0 | Carnivora | 13302.0 | 0 | Canidae | 13302.0 | 0 | Canis | 8077.0 | 1 | Canis lupus | 8077.0 | 1 |
| scf7180000056219 | 126983 | 0.4198 | 0 | 57.312 | Eukaryota | 76593.0 | 0 | Chordata | 76593.0 | 0 | Carnivora | 76593.0 | 0 | Canidae | 76593.0 | 0 | Canis | 76593.0 | 0 | Canis lupus | 76593.0 | 0 |
| scf7180000048617 | 54925 | 0.4531 | 0 | 61.2584 | Eukaryota | 6763.0 | 0 | Chordata | 6763.0 | 0 | Carnivora | 2874.0 | 2 | Hominidae | 1894.0 | 5 | Homo | 1894.0 | 6 | Homo sapiens | 1894.0 | 6 |
| scf7180000052107 | 3249 | 0.4506 | 0 | 1199.0932 | no-hit | 0.0 | 0 | no-hit | 0.0 | 0 | no-hit | 0.0 | 0 | no-hit | 0.0 | 0 | no-hit | 0.0 | 0 | no-hit | 0.0 | 0 |
| scf7180000051659 | 6896 | 0.451 | 0 | 1133.6992 | Eukaryota | 174.0 | 0 | Platyhelminthes | 174.0 | 0 | Polystomatidea | 174.0 | 0 | Polystomatidae | 174.0 | 0 | Protopolystoma | 174.0 | 0 | Protopolystoma xenopodis | 174.0 | 0 |
| scf7180000046573 | 74283 | 0.4351 | 0 | 309.718 | Eukaryota | 46722.0 | 0 | Chordata | 46722.0 | 0 | Carnivora | 46722.0 | 0 | Canidae | 46722.0 | 0 | Canis | 46722.0 | 0 | Canis lupus | 46722.0 | 0 |
| scf7180000049877 | 22829 | 0.5293 | 0 | 54.2572 | Eukaryota | 10647.0 | 0 | Chordata | 10647.0 | 0 | Carnivora | 5879.0 | 3 | Canidae | 5879.0 | 4 | Canis | 3219.0 | 6 | Canis lupus | 3219.0 | 6 |
| scf7180000043158 | 110170 | 0.3394 | 0 | 63.5724 | Eukaryota | 90419.0 | 0 | Chordata | 90419.0 | 0 | Carnivora | 90419.0 | 0 | Canidae | 90419.0 | 0 | Canis | 90419.0 | 0 | Canis lupus | 90419.0 | 0 |
| scf7180000051803 | 9484 | 0.5147 | 0 | 412.5514 | Eukaryota | 32618.0 | 0 | Chordata | 32618.0 | 0 | Carnivora | 32618.0 | 0 | Canidae | 32618.0 | 0 | Canis | 32618.0 | 0 | Canis lupus | 32618.0 | 0 |
| scf7180000046549 | 4142 | 0.3317 | 0 | 48.8144 | no-hit | 0.0 | 0 | no-hit | 0.0 | 0 | no-hit | 0.0 | 0 | no-hit | 0.0 | 0 | no-hit | 0.0 | 0 | no-hit | 0.0 | 0 |
| scf7180000042368 | 86409 | 0.3809 | 0 | 55.0225 | Eukaryota | 23386.0 | 0 | Chordata | 23386.0 | 0 | Carnivora | 23386.0 | 0 | Canidae | 23386.0 | 0 | Canis | 23386.0 | 0 | Canis lupus | 23386.0 | 0 |
| scf7180000041844 | 834861 | 0.5481 | 0 | 58.2485 | Eukaryota | 53301.0 | 0 | Chordata | 53301.0 | 0 | Carnivora | 53301.0 | 0 | Canidae | 53301.0 | 0 | Canis | 53301.0 | 0 | Canis lupus | 53301.0 | 0 |
| scf7180000041941 | 880231 | 0.3917 | 0 | 58.4639 | Eukaryota | 39094.0 | 0 | Chordata | 39094.0 | 0 | Carnivora | 39094.0 | 0 | Canidae | 39094.0 | 0 | Canis | 39094.0 | 0 | Canis lupus | 39094.0 | 0 |
| scf7180000049738 | 3514 | 0.4368 | 0 | 42.6168 | no-hit | 0.0 | 0 | no-hit | 0.0 | 0 | no-hit | 0.0 | 0 | no-hit | 0.0 | 0 | no-hit | 0.0 | 0 | no-hit | 0.0 | 0 |
| scf7180000044191 | 46237 | 0.4518 | 0 | 66.0098 | Eukaryota | 40459.0 | 0 | Chordata | 40459.0 | 0 | Carnivora | 28307.0 | 1 | Canidae | 28307.0 | 1 | Canis | 28307.0 | 1 | Canis lupus | 28307.0 | 1 |
| scf7180000049462 | 42198 | 0.4461 | 0 | 71.6651 | Eukaryota | 937.0 | 0 | Platyhelminthes | 492.0 | 1 | Polystomatidea | 492.0 | 2 | Polystomatidae | 492.0 | 2 | Protopolystoma | 492.0 | 2 | Protopolystoma xenopodis | 492.0 | 2 |
| scf7180000046027 | 153052 | 0.4747 | 0 | 58.7757 | Eukaryota | 33900.0 | 0 | Chordata | 33900.0 | 0 | Carnivora | 33900.0 | 0 | Canidae | 33900.0 | 0 | Canis | 33900.0 | 0 | Canis lupus | 33900.0 | 0 |
| scf7180000048277 | 3797 | 0.4459 | 0 | 61.3023 | no-hit | 0.0 | 0 | no-hit | 0.0 | 0 | no-hit | 0.0 | 0 | no-hit | 0.0 | 0 | no-hit | 0.0 | 0 | no-hit | 0.0 | 0 |
| scf7180000054692 | 21481 | 0.4406 | 0 | 201.4476 | Eukaryota | 1110.0 | 0 | Streptophyta | 847.0 | 1 | Fabales | 847.0 | 1 | Fabaceae | 847.0 | 1 | Cicer | 847.0 | 1 | Cicer arietinum | 847.0 | 1 |
| scf7180000043313 | 14934 | 0.3903 | 0 | 191.9051 | Eukaryota | 82204.0 | 0 | Chordata | 82204.0 | 0 | Carnivora | 82204.0 | 0 | Canidae | 82204.0 | 0 | Canis | 82204.0 | 0 | Canis lupus | 82204.0 | 0 |
| scf7180000055077 | 34561 | 0.4423 | 0 | 154.9141 | Eukaryota | 847.0 | 0 | Platyhelminthes | 492.0 | 1 | Polystomatidea | 492.0 | 1 | Polystomatidae | 492.0 | 1 | Protopolystoma | 492.0 | 1 | Protopolystoma xenopodis | 492.0 | 1 |
| scf7180000047445 | 48563 | 0.5514 | 0 | 53.8944 | Eukaryota | 8748.0 | 0 | Chordata | 8748.0 | 0 | Carnivora | 8748.0 | 0 | Canidae | 7166.0 | 1 | Canis | 5526.0 | 2 | Canis lupus | 5526.0 | 2 |
| scf7180000050642 | 12755 | 0.4471 | 0 | 856.1488 | Eukaryota | 3288.0 | 0 | Platyhelminthes | 1726.0 | 2 | Polystomatidea | 1726.0 | 2 | Polystomatidae | 1726.0 | 2 | Protopolystoma | 1726.0 | 2 | Protopolystoma xenopodis | 1726.0 | 2 |
| scf7180000053336 | 4382 | 0.4169 | 0 | 21.7343 | Eukaryota | 15701.0 | 0 | Chordata | 15701.0 | 0 | Carnivora | 15701.0 | 0 | Canidae | 15701.0 | 0 | Canis | 15701.0 | 0 | Canis lupus | 15701.0 | 0 |
| scf7180000040556 | 2344992 | 0.4017 | 0 | 59.9424 | Eukaryota | 88082.0 | 0 | Chordata | 88082.0 | 0 | Carnivora | 88082.0 | 0 | Canidae | 88082.0 | 0 | Canis | 88082.0 | 0 | Canis lupus | 88082.0 | 0 |
| scf7180000052184 | 6883 | 0.452 | 0 | 455.5044 | Eukaryota | 19172.0 | 0 | Chordata | 17219.0 | 1 | Gadiformes | 9077.0 | 6 | Gadidae | 9077.0 | 6 | Gadus | 9077.0 | 6 | Gadus morhua | 9077.0 | 6 |
| scf7180000055993 | 10638 | 0.4461 | 0 | 153.2126 | Eukaryota | 18641.0 | 0 | Chordata | 14073.0 | 2 | Gadiformes | 6606.0 | 7 | Gadidae | 6606.0 | 7 | Gadus | 6606.0 | 7 | Gadus morhua | 6606.0 | 7 |
| scf7180000054011 | 10000 | 0.3767 | 0 | 63.3761 | Eukaryota | 3285.0 | 0 | Chordata | 2686.0 | 1 | Carnivora | 1523.0 | 2 | Canidae | 1523.0 | 3 | Canis | 1308.0 | 4 | Canis lupus | 1308.0 | 4 |
| scf7180000046416 | 11038 | 0.5262 | 0 | 30.0106 | Eukaryota | 47398.0 | 0 | Chordata | 47398.0 | 0 | Carnivora | 47398.0 | 0 | Canidae | 47398.0 | 0 | Canis | 47398.0 | 0 | Canis lupus | 47398.0 | 0 |
| scf7180000042219 | 682745 | 0.3808 | 0 | 61.6236 | Eukaryota | 61485.0 | 0 | Chordata | 61485.0 | 0 | Carnivora | 61485.0 | 0 | Canidae | 61485.0 | 0 | Canis | 61485.0 | 0 | Canis lupus | 61485.0 | 0 |
| scf7180000045546 | 163230 | 0.3875 | 0 | 58.9584 | Eukaryota | 65088.0 | 0 | Chordata | 65088.0 | 0 | Carnivora | 65088.0 | 0 | Canidae | 65088.0 | 0 | Canis | 65088.0 | 0 | Canis lupus | 65088.0 | 0 |
| scf7180000056442 | 560018 | 0.4259 | 0 | 59.6447 | Eukaryota | 61243.0 | 0 | Chordata | 61243.0 | 0 | Carnivora | 61243.0 | 0 | Canidae | 61243.0 | 0 | Canis | 61243.0 | 0 | Canis lupus | 61243.0 | 0 |
| scf7180000048882 | 13695 | 0.4429 | 0 | 82.5306 | Eukaryota | 3410.0 | 0 | Chordata | 3410.0 | 0 | Carnivora | 1626.0 | 3 | Canidae | 1626.0 | 3 | Canis | 1626.0 | 3 | Canis lupus | 1626.0 | 3 |
| scf7180000042723 | 3114 | 0.5122 | 0 | 30.9604 | no-hit | 0.0 | 0 | no-hit | 0.0 | 0 | no-hit | 0.0 | 0 | no-hit | 0.0 | 0 | no-hit | 0.0 | 0 | no-hit | 0.0 | 0 |
| scf7180000046226 | 131774 | 0.3643 | 0 | 60.3191 | Eukaryota | 51711.0 | 0 | Chordata | 51711.0 | 0 | Carnivora | 51711.0 | 0 | Canidae | 51711.0 | 0 | Canis | 51711.0 | 0 | Canis lupus | 51711.0 | 0 |
| scf7180000053054 | 4611 | 0.4487 | 0 | 344.8128 | Eukaryota | 1027.0 | 0 | Platyhelminthes | 1027.0 | 0 | Polystomatidea | 1027.0 | 0 | Polystomatidae | 1027.0 | 0 | Protopolystoma | 1027.0 | 0 | Protopolystoma xenopodis | 1027.0 | 0 |
| scf7180000043633 | 174560 | 0.3753 | 0 | 58.7786 | Eukaryota | 107921.0 | 0 | Chordata | 107921.0 | 0 | Carnivora | 107921.0 | 0 | Canidae | 107921.0 | 0 | Canis | 107921.0 | 0 | Canis lupus | 107921.0 | 0 |
| scf7180000053816 | 3574 | 0.4524 | 0 | 248.3138 | no-hit | 0.0 | 0 | no-hit | 0.0 | 0 | no-hit | 0.0 | 0 | no-hit | 0.0 | 0 | no-hit | 0.0 | 0 | no-hit | 0.0 | 0 |
| scf7180000049091 | 10311 | 0.4418 | 0 | 1258.3008 | no-hit | 0.0 | 0 | no-hit | 0.0 | 0 | no-hit | 0.0 | 0 | no-hit | 0.0 | 0 | no-hit | 0.0 | 0 | no-hit | 0.0 | 0 |

| | | | | | | | | | | | | | | | | | | | | | |
|---|---|---|---|---|---|---|---|---|---|---|---|---|---|---|---|---|---|---|---|---|---|
| scf7180000051521 | 10344 | 0.4461 | 0 | 169.6423 | Eukaryota | 515.0 | 0 | Chordata | 263.0 | 1 | Kurtiformes | 263.0 | 1 | Apogonidae | 263.0 | 1 | Sphaeramia | 263.0 | 1 | Sphaeramia orbicularis | 263.0 | 1 |
| scf7180000050539 | 2611 | 0.455 | 0 | 457.3997 | Eukaryota | 2448.0 | 0 | Platyhelminthes | 1458.0 | 2 | Polystomatidea | 1458.0 | 2 | Polystomatidae | 1458.0 | 2 | Protopolystoma | 1458.0 | 2 | Protopolystoma xenopodis | 1458.0 | 2 |
| scf7180000047089 | 82721 | 0.4115 | 0 | 60.6811 | Eukaryota | 11414.0 | 0 | Chordata | 11414.0 | 0 | Apterygiformes | 10278.0 | 1 | Apterygidae | 10278.0 | 1 | Apteryx | 10278.0 | 1 | Apteryx mantelli | 10278.0 | 1 |
| scf7180000055768 | 4315 | 0.4382 | 0 | 122.6613 | no-hit | 0.0 | 0 | no-hit | 0.0 | 0 | no-hit | 0.0 | 0 | no-hit | 0.0 | 0 | no-hit | 0.0 | 0 | no-hit | 0.0 | 0 |
| scf7180000049316 | 2742 | 0.45 | 0 | 592.0387 | no-hit | 0.0 | 0 | no-hit | 0.0 | 0 | no-hit | 0.0 | 0 | no-hit | 0.0 | 0 | no-hit | 0.0 | 0 | no-hit | 0.0 | 0 |
| scf7180000049335 | 18685 | 0.446 | 0 | 620.0806 | Eukaryota | 3454.0 | 0 | Platyhelminthes | 2026.0 | 2 | Polystomatidea | 2026.0 | 2 | Polystomatidae | 2026.0 | 2 | Protopolystoma | 2026.0 | 2 | Protopolystoma xenopodis | 2026.0 | 2 |
| scf7180000046299 | 145772 | 0.4717 | 0 | 59.2145 | Eukaryota | 27295.0 | 0 | Chordata | 27295.0 | 0 | Carnivora | 27295.0 | 0 | Canidae | 27295.0 | 0 | Canis | 27295.0 | 0 | Canis lupus | 27295.0 | 0 |
| scf7180000044167 | 15194 | 0.4195 | 0 | 80.1343 | Eukaryota | 2559.0 | 0 | Chordata | 2559.0 | 0 | Carnivora | 2559.0 | 0 | Canidae | 2559.0 | 0 | Canis | 2559.0 | 0 | Canis lupus | 2559.0 | 0 |
| scf7180000049879 | 3787 | 0.5382 | 0 | 99.5659 | Eukaryota | 31527.0 | 0 | Chordata | 31527.0 | 0 | Carnivora | 31527.0 | 0 | Canidae | 31527.0 | 0 | Canis | 31527.0 | 0 | Canis lupus | 31527.0 | 0 |
| scf7180000044398 | 332354 | 0.4954 | 0 | 56.3794 | Eukaryota | 12037.0 | 0 | Chordata | 12037.0 | 0 | Carnivora | 7507.0 | 1 | Canidae | 7507.0 | 1 | Canis | 7507.0 | 1 | Canis lupus | 7507.0 | 1 |
| scf7180000046249 | 145408 | 0.4056 | 0 | 59.5883 | Eukaryota | 15728.0 | 0 | Chordata | 15728.0 | 0 | Carnivora | 7895.0 | 3 | Canidae | 7895.0 | 3 | Canis | 7895.0 | 3 | Canis lupus | 7895.0 | 3 |
| scf7180000041731 | 86638 | 0.3672 | 0 | 55.3682 | Eukaryota | 102637.0 | 0 | Chordata | 102637.0 | 0 | Carnivora | 102637.0 | 0 | Canidae | 102637.0 | 0 | Canis | 102637.0 | 0 | Canis lupus | 102637.0 | 0 |
| scf7180000050368 | 8629 | 0.5052 | 0 | 110.4924 | Eukaryota | 7741.0 | 0 | Chordata | 7741.0 | 0 | Carnivora | 7741.0 | 0 | Canidae | 7741.0 | 0 | Canis | 5766.0 | 1 | Canis lupus | 4743.0 | 2 |
| scf7180000044715 | 25551 | 0.4504 | 0 | 231.1827 | Eukaryota | 10103.0 | 0 | Chordata | 8458.0 | 1 | Carnivora | 5530.0 | 2 | Canidae | 5530.0 | 2 | Canis | 4890.0 | 3 | Canis lupus | 4265.0 | 4 |
| scf7180000046918 | 102315 | 0.4152 | 0 | 60.3106 | Eukaryota | 8596.0 | 0 | Chordata | 8596.0 | 0 | Carnivora | 5425.0 | 1 | Felidae | 5425.0 | 1 | Felis | 5425.0 | 1 | Felis catus | 5425.0 | 1 |
| scf7180000051813 | 4235 | 0.4524 | 0 | 1732.6156 | no-hit | 0.0 | 0 | no-hit | 0.0 | 0 | no-hit | 0.0 | 0 | no-hit | 0.0 | 0 | no-hit | 0.0 | 0 | no-hit | 0.0 | 0 |
| scf7180000052928 | 7807 | 0.5043 | 0 | 102.394 | Eukaryota | 8513.0 | 0 | Chordata | 8513.0 | 0 | Carnivora | 8513.0 | 0 | Canidae | 8513.0 | 0 | Canis | 6908.0 | 1 | Canis lupus | 5745.0 | 2 |
| scf7180000049564 | 8515 | 0.5446 | 0 | 544.6089 | Eukaryota | 31245.0 | 0 | Chordata | 31245.0 | 0 | Carnivora | 31245.0 | 0 | Canidae | 31245.0 | 0 | Canis | 31245.0 | 0 | Canis lupus | 31245.0 | 0 |
| scf7180000045514 | 3162 | 0.3934 | 0 | 28.6799 | Eukaryota | 2322.0 | 0 | Chordata | 2322.0 | 0 | Carnivora | 2322.0 | 0 | Canidae | 2322.0 | 0 | Canis | 2322.0 | 0 | Canis lupus | 2322.0 | 0 |
| scf7180000042054 | 801805 | 0.4187 | 0 | 60.081 | Eukaryota | 42024.0 | 0 | Chordata | 42024.0 | 0 | Carnivora | 42024.0 | 0 | Canidae | 42024.0 | 0 | Canis | 42024.0 | 0 | Canis lupus | 42024.0 | 0 |
| scf7180000052504 | 3706 | 0.445 | 0 | 191.6176 | no-hit | 0.0 | 0 | no-hit | 0.0 | 0 | no-hit | 0.0 | 0 | no-hit | 0.0 | 0 | no-hit | 0.0 | 0 | no-hit | 0.0 | 0 |
| scf7180000050428 | 26736 | 0.4051 | 0 | 60.8304 | Eukaryota | 4004.0 | 0 | Chordata | 4004.0 | 0 | Carnivora | 1847.0 | 2 | Canidae | 1847.0 | 2 | Canis | 1847.0 | 2 | Canis lupus | 1847.0 | 2 |
| scf7180000053260 | 21064 | 0.4261 | 0 | 59.5509 | Eukaryota | 21684.0 | 0 | Chordata | 21200.0 | 1 | Carnivora | 17092.0 | 3 | Canidae | 12792.0 | 5 | Canis | 12792.0 | 6 | Canis lupus | 12792.0 | 6 |
| scf7180000056576 | 120212 | 0.3413 | 0 | 57.1254 | Eukaryota | 55677.0 | 0 | Chordata | 55677.0 | 0 | Carnivora | 55677.0 | 0 | Canidae | 55677.0 | 0 | Canis | 55677.0 | 0 | Canis lupus | 55677.0 | 0 |
| scf7180000041147 | 31882 | 0.3731 | 0 | 145.9921 | Eukaryota | 20889.0 | 0 | Chordata | 20889.0 | 0 | Carnivora | 20889.0 | 0 | Canidae | 20889.0 | 0 | Canis | 20889.0 | 0 | Canis lupus | 20889.0 | 0 |
| scf7180000056209 | 427634 | 0.4081 | 0 | 62.8331 | Eukaryota | 87276.0 | 0 | Chordata | 87276.0 | 0 | Carnivora | 87276.0 | 0 | Canidae | 87276.0 | 0 | Canis | 87276.0 | 0 | Canis lupus | 87276.0 | 0 |
| scf7180000042387 | 489786 | 0.351 | 0 | 60.5279 | Eukaryota | 92255.0 | 0 | Chordata | 92255.0 | 0 | Carnivora | 92255.0 | 0 | Canidae | 92255.0 | 0 | Canis | 92255.0 | 0 | Canis lupus | 92255.0 | 0 |
| scf7180000041580 | 78262 | 0.3676 | 0 | 55.6332 | Eukaryota | 11849.0 | 0 | Chordata | 11849.0 | 0 | Artiodactyla | 6469.0 | 2 | Bovidae | 6469.0 | 2 | Ovis | 6469.0 | 2 | Ovis canadensis | 6469.0 | 2 |
| scf7180000048424 | 8040 | 0.431 | 0 | 128.1185 | no-hit | 0.0 | 0 | no-hit | 0.0 | 0 | no-hit | 0.0 | 0 | no-hit | 0.0 | 0 | no-hit | 0.0 | 0 | no-hit | 0.0 | 0 |
| scf7180000048815 | 42505 | 0.4434 | 0 | 163.7864 | Eukaryota | 2754.0 | 0 | Chordata | 2754.0 | 0 | Carnivora | 2754.0 | 0 | Canidae | 2754.0 | 0 | Canis | 2754.0 | 0 | Canis lupus | 2754.0 | 0 |
| scf7180000041966 | 854310 | 0.4666 | 0 | 59.6106 | Eukaryota | 67478.0 | 0 | Chordata | 67478.0 | 0 | Carnivora | 67478.0 | 0 | Canidae | 67478.0 | 0 | Canis | 67478.0 | 0 | Canis lupus | 67478.0 | 0 |
| scf7180000056299 | 659323 | 0.402 | 0 | 59.4972 | Eukaryota | 49205.0 | 0 | Chordata | 49205.0 | 0 | Carnivora | 49205.0 | 0 | Otariidae | 32728.0 | 2 | Zalophus | 32728.0 | 2 | Zalophus californianus | 32728.0 | 2 |
| scf7180000053617 | 3163 | 0.4518 | 0 | 255.3641 | no-hit | 0.0 | 0 | no-hit | 0.0 | 0 | no-hit | 0.0 | 0 | no-hit | 0.0 | 0 | no-hit | 0.0 | 0 | no-hit | 0.0 | 0 |
| scf7180000053576 | 4834 | 0.5056 | 0 | 176.4573 | Eukaryota | 7288.0 | 0 | Chordata | 7288.0 | 0 | Carnivora | 7288.0 | 0 | Canidae | 7288.0 | 0 | Canis | 5813.0 | 1 | Canis lupus | 4753.0 | 2 |
| scf7180000044312 | 80711 | 0.4169 | 0 | 56.4615 | Eukaryota | 31868.0 | 0 | Chordata | 31868.0 | 0 | Carnivora | 31868.0 | 0 | Canidae | 31868.0 | 0 | Canis | 31868.0 | 0 | Canis lupus | 31868.0 | 0 |
| scf7180000049308 | 3612 | 0.4485 | 0 | 359.7809 | Eukaryota | 734.0 | 0 | Chordata | 734.0 | 0 | Tetraodontiformes | 734.0 | 0 | Tetraodontidae | 734.0 | 0 | Takifugu | 734.0 | 0 | Takifugu rubripes | 734.0 | 0 |
| scf7180000054869 | 26600 | 0.4493 | 0 | 447.8244 | Eukaryota | 89197.0 | 0 | Chordata | 89197.0 | 0 | Carnivora | 89197.0 | 0 | Canidae | 89197.0 | 0 | Canis | 89197.0 | 0 | Canis lupus | 89197.0 | 0 |
| scf7180000051837 | 12708 | 0.4628 | 0 | 52.5125 | Eukaryota | 3161.0 | 0 | Chordata | 2643.0 | 1 | Carnivora | 2643.0 | 1 | Canidae | 2643.0 | 1 | Canis | 2643.0 | 1 | Canis lupus | 2643.0 | 1 |
| scf7180000053032 | 2878 | 0.4375 | 0 | 150.7691 | no-hit | 0.0 | 0 | no-hit | 0.0 | 0 | no-hit | 0.0 | 0 | no-hit | 0.0 | 0 | no-hit | 0.0 | 0 | no-hit | 0.0 | 0 |
| scf7180000049999 | 4533 | 0.4542 | 0 | 900.0856 | Eukaryota | 15613.0 | 0 | Chordata | 13041.0 | 2 | Gadiformes | 7261.0 | 7 | Gadidae | 7261.0 | 7 | Gadus | 7261.0 | 7 | Gadus morhua | 7261.0 | 7 |
| scf7180000055449 | 12593 | 0.4414 | 0 | 133.4565 | Eukaryota | 2936.0 | 0 | Chordata | 2799.0 | 1 | Columbiformes | 1748.0 | 4 | Columbidae | 1748.0 | 4 | Streptopelia | 1748.0 | 4 | Streptopelia turtur | 1748.0 | 4 |
| scf7180000051522 | 16468 | 0.4429 | 0 | 279.3298 | Eukaryota | 40212.0 | 0 | Chordata | 26677.0 | 2 | Gadiformes | 19523.0 | 5 | Gadidae | 19523.0 | 5 | Gadus | 19523.0 | 5 | Gadus morhua | 19523.0 | 5 |
| scf7180000053796 | 2358 | 0.4559 | 0 | 275.9436 | no-hit | 0.0 | 0 | no-hit | 0.0 | 0 | no-hit | 0.0 | 0 | no-hit | 0.0 | 0 | no-hit | 0.0 | 0 | no-hit | 0.0 | 0 |
| scf7180000040967 | 1521082 | 0.3601 | 0 | 60.8013 | Eukaryota | 112040.0 | 0 | Chordata | 112040.0 | 0 | Carnivora | 112040.0 | 0 | Canidae | 112040.0 | 0 | Canis | 112040.0 | 0 | Canis lupus | 112040.0 | 0 |
| scf7180000040955 | 1410742 | 0.5006 | 0 | 59.7396 | Eukaryota | 97318.0 | 0 | Chordata | 97318.0 | 0 | Carnivora | 97318.0 | 0 | Canidae | 97318.0 | 0 | Canis | 87558.0 | 1 | Canis lupus | 87558.0 | 1 |
| scf7180000045174 | 238952 | 0.3896 | 0 | 59.8409 | Eukaryota | 52328.0 | 0 | Chordata | 52328.0 | 0 | Carnivora | 52328.0 | 0 | Canidae | 27498.0 | 1 | Felis | 24830.0 | 2 | Felis catus | 24830.0 | 2 |
| scf7180000054069 | 3959 | 0.4489 | 0 | 689.5547 | no-hit | 0.0 | 0 | no-hit | 0.0 | 0 | no-hit | 0.0 | 0 | no-hit | 0.0 | 0 | no-hit | 0.0 | 0 | no-hit | 0.0 | 0 |
| scf7180000050572 | 9801 | 0.501 | 0 | 46.1043 | Eukaryota | 8437.0 | 0 | Chordata | 8437.0 | 0 | Carnivora | 8437.0 | 0 | Canidae | 8437.0 | 0 | Canis | 6325.0 | 2 | Canis lupus | 5183.0 | 3 |
| scf7180000048372 | 7705 | 0.44 | 0 | 192.791 | Eukaryota | 492.0 | 0 | Platyhelminthes | 492.0 | 0 | Polystomatidea | 492.0 | 0 | Polystomatidae | 492.0 | 0 | Protopolystoma | 492.0 | 0 | Protopolystoma xenopodis | 492.0 | 0 |
| scf7180000049681 | 6846 | 0.4549 | 0 | 73.3217 | Eukaryota | 675.0 | 0 | Chordata | 534.0 | 1 | Tetraodontiformes | 534.0 | 1 | Tetraodontidae | 534.0 | 1 | Takifugu | 534.0 | 1 | Takifugu rubripes | 534.0 | 1 |
| scf7180000052281 | 6500 | 0.4528 | 0 | 880.1574 | no-hit | 0.0 | 0 | no-hit | 0.0 | 0 | no-hit | 0.0 | 0 | no-hit | 0.0 | 0 | no-hit | 0.0 | 0 | no-hit | 0.0 | 0 |
| scf7180000054941 | 27366 | 0.4988 | 0 | 64.3081 | Eukaryota | 19642.0 | 0 | Chordata | 19642.0 | 0 | Carnivora | 19642.0 | 0 | Canidae | 19642.0 | 0 | Canis | 19642.0 | 0 | Canis lupus | 19642.0 | 0 |
| scf7180000051258 | 41622 | 0.476 | 0 | 53.4853 | Eukaryota | 4768.0 | 0 | Chordata | 4768.0 | 0 | Carnivora | 2045.0 | 3 | Hominidae | 1347.0 | 4 | Homo | 1347.0 | 5 | Homo sapiens | 1347.0 | 5 |
| scf7180000052925 | 6289 | 0.4368 | 0 | 63.2236 | no-hit | 0.0 | 0 | no-hit | 0.0 | 0 | no-hit | 0.0 | 0 | no-hit | 0.0 | 0 | no-hit | 0.0 | 0 | no-hit | 0.0 | 0 |
| scf7180000043043 | 124084 | 0.3877 | 0 | 64.7724 | Eukaryota | 93107.0 | 0 | Chordata | 93107.0 | 0 | Carnivora | 93107.0 | 0 | Canidae | 93107.0 | 0 | Canis | 93107.0 | 0 | Canis lupus | 93107.0 | 0 |
| scf7180000043822 | 427886 | 0.3687 | 0 | 60.7312 | Eukaryota | 77290.0 | 0 | Chordata | 77290.0 | 0 | Carnivora | 77290.0 | 0 | Phocidae | 36128.0 | 2 | Leptonychotes | 28964.0 | 5 | Leptonychotes weddellii | 28964.0 | 5 |
| scf7180000055256 | 13543 | 0.519 | 0 | 452.5906 | Eukaryota | 15489.0 | 0 | Chordata | 15489.0 | 0 | Carnivora | 15489.0 | 0 | Canidae | 15489.0 | 0 | Canis | 15489.0 | 0 | Canis lupus | 15489.0 | 0 |
| scf7180000056318 | 767899 | 0.407 | 0 | 60.2011 | Eukaryota | 75116.0 | 0 | Chordata | 75116.0 | 0 | Carnivora | 75116.0 | 0 | Canidae | 75116.0 | 0 | Canis | 75116.0 | 0 | Canis lupus | 75116.0 | 0 |
| scf7180000051912 | 3418 | 0.4623 | 0 | 400.186 | Eukaryota | 2872.0 | 0 | Chordata | 2872.0 | 0 | Carnivora | 2872.0 | 0 | Canidae | 2872.0 | 0 | Canis | 2872.0 | 0 | Canis lupus | 2872.0 | 0 |
| scf7180000044701 | 240707 | 0.3827 | 0 | 58.9176 | Eukaryota | 93267.0 | 0 | Chordata | 93267.0 | 0 | Carnivora | 93267.0 | 0 | Canidae | 93267.0 | 0 | Canis | 93267.0 | 0 | Canis lupus | 93267.0 | 0 |
| scf7180000042642 | 332541 | 0.3824 | 0 | 59.4562 | Eukaryota | 79043.0 | 0 | Chordata | 79043.0 | 0 | Carnivora | 79043.0 | 0 | Canidae | 36219.0 | 3 | Canis | 27209.0 | 6 | Canis lupus | 27209.0 | 7 |
| scf7180000056581 | 36245 | 0.4444 | 0 | 57.8769 | Eukaryota | 59402.0 | 0 | Chordata | 59402.0 | 0 | Carnivora | 59402.0 | 0 | Canidae | 59402.0 | 0 | Canis | 59402.0 | 0 | Canis lupus | 59402.0 | 0 |
| scf7180000043966 | 302621 | 0.5482 | 0 | 56.9121 | Eukaryota | 52779.0 | 0 | Chordata | 52779.0 | 0 | Carnivora | 52779.0 | 0 | Canidae | 35502.0 | 3 | Canis | 35502.0 | 3 | Canis lupus | 35502.0 | 3 |
| scf7180000045637 | 188193 | 0.3443 | 0 | 57.6614 | Eukaryota | 72642.0 | 0 | Chordata | 72642.0 | 0 | Carnivora | 72642.0 | 0 | Canidae | 72642.0 | 0 | Canis | 72642.0 | 0 | Canis lupus | 72642.0 | 0 |
| scf7180000046750 | 90391 | 0.5007 | 0 | 66.3924 | Eukaryota | 9791.0 | 0 | Chordata | 9791.0 | 0 | Carnivora | 9791.0 | 0 | Canidae | 8490.0 | 1 | Canis | 8490.0 | 1 | Canis lupus | 8490.0 | 1 |
| scf7180000052967 | 3476 | 0.5014 | 0 | 119.2541 | Eukaryota | 5675.0 | 0 | Chordata | 5675.0 | 0 | Carnivora | 5675.0 | 0 | Canidae | 5675.0 | 0 | Canis | 3251.0 | 1 | Nyctereutes procyonoides | 2424.0 | 2 |
| scf7180000047784 | 56709 | 0.4782 | 0 | 60.9924 | Eukaryota | 31045.0 | 0 | Chordata | 31045.0 | 0 | Carnivora | 28121.0 | 1 | Canidae | 25216.0 | 2 | Canis | 25216.0 | 2 | Canis lupus | 25216.0 | 2 |
| scf7180000043386 | 471235 | 0.4202 | 0 | 59.5224 | Eukaryota | 30633.0 | 0 | Chordata | 30633.0 | 0 | Primates | 21080.0 | 2 | Hominidae | 12053.0 | 6 | Homo | 12053.0 | 6 | Homo sapiens | 12053.0 | 6 |
| scf7180000040700 | 220890 | 0.4251 | 0 | 59.0359 | Eukaryota | 36806.0 | 0 | Chordata | 36806.0 | 0 | Carnivora | 36806.0 | 0 | Canidae | 36806.0 | 0 | Canis | 36806.0 | 0 | Canis lupus | 36806.0 | 0 |
| scf7180000040975 | 1600418 | 0.4068 | 0 | 59.3221 | Eukaryota | 104391.0 | 0 | Chordata | 104391.0 | 0 | Carnivora | 104391.0 | 0 | Canidae | 88323.0 | 1 | Canis | 51291.0 | 2 | Canis lupus | 51291.0 | 2 |
| scf7180000049965 | 4270 | 0.4525 | 0 | 1119.0593 | no-hit | 0.0 | 0 | no-hit | 0.0 | 0 | no-hit | 0.0 | 0 | no-hit | 0.0 | 0 | no-hit | 0.0 | 0 | no-hit | 0.0 | 0 |
| scf7180000054131 | 25449 | 0.4414 | 0 | 209.4761 | Eukaryota | 7499.0 | 0 | Chordata | 6844.0 | 1 | Salmoniformes | 3861.0 | 3 | Salmonidae | 3861.0 | 3 | Salmo | 3678.0 | 4 | Salmo trutta | 3678.0 | 4 |
| scf7180000050532 | 19051 | 0.4395 | 0 | 64.5184 | Eukaryota | 44540.0 | 0 | Chordata | 27400.0 | 2 | Gadiformes | 14134.0 | 6 | Gadidae | 14134.0 | 6 | Gadus | 14134.0 | 6 | Gadus morhua | 14134.0 | 6 |
| scf7180000054772 | 11656 | 0.5178 | 0 | 270.186 | Eukaryota | 15722.0 | 0 | Chordata | 15722.0 | 0 | Carnivora | 15722.0 | 0 | Canidae | 15722.0 | 0 | Canis | 15722.0 | 0 | Canis lupus | 15722.0 | 0 |
| scf7180000043205 | 4306 | 0.343 | 0 | 27.2244 | Eukaryota | 3235.0 | 0 | Chordata | 3235.0 | 0 | Carnivora | 2322.0 | 1 | Canidae | 2322.0 | 1 | Canis | 2322.0 | 1 | Canis lupus | 2322.0 | 1 |
| scf7180000052508 | 4462 | 0.4496 | 0 | 887.2897 | no-hit | 0.0 | 0 | no-hit | 0.0 | 0 | no-hit | 0.0 | 0 | no-hit | 0.0 | 0 | no-hit | 0.0 | 0 | no-hit | 0.0 | 0 |
| scf7180000048219 | 10541 | 0.3819 | 0 | 319.9835 | Eukaryota | 22834.0 | 0 | Chordata | 22834.0 | 0 | Carnivora | 22834.0 | 0 | Canidae | 22834.0 | 0 | Canis | 22834.0 | 0 | Canis lupus | 22834.0 | 0 |
| scf7180000041536 | 716605 | 0.3482 | 0 | 58.4531 | Eukaryota | 65740.0 | 0 | Chordata | 65740.0 | 0 | Carnivora | 65740.0 | 0 | Canidae | 65740.0 | 0 | Canis | 65740.0 | 0 | Canis lupus | 65740.0 | 0 |
| scf7180000051948 | 21901 | 0.3385 | 0 | 58.8137 | Eukaryota | 7140.0 | 0 | Chordata | 7140.0 | 0 | Carnivora | 6465.0 | 1 | Canidae | 3300.0 | 2 | Canis | 3300.0 | 2 | Canis lupus | 3300.0 | 2 |
| scf7180000047978 | 16789 | 0.4405 | 0 | 161.3944 | Eukaryota | 7646.0 | 0 | Chordata | 6812.0 | 1 | Kurtiformes | 3041.0 | 3 | Apogonidae | 3041.0 | 3 | Sphaeramia | 3041.0 | 3 | Sphaeramia orbicularis | 3041.0 | 3 |
| scf7180000042841 | 49933 | 0.364 | 0 | 54.0547 | Eukaryota | 4796.0 | 0 | Chordata | 4352.0 | 1 | Perissodactyla | 2750.0 | 3 | Equidae | 2750.0 | 4 | Equus | 2750.0 | 4 | Equus caballus | 2750.0 | 4 |
| scf7180000042703 | 528564 | 0.4487 | 0 | 58.4817 | Eukaryota | 74563.0 | 0 | Chordata | 74563.0 | 0 | Carnivora | 74563.0 | 0 | Canidae | 74563.0 | 0 | Canis | 43677.0 | 1 | Canis lupus | 43677.0 | 1 |
| scf7180000047692 | 73788 | 0.4068 | 0 | 60.0891 | Eukaryota | 35344.0 | 0 | Chordata | 35344.0 | 0 | Carnivora | 35344.0 | 0 | Canidae | 35344.0 | 0 | Canis | 35344.0 | 0 | Canis lupus | 35344.0 | 0 |
| scf7180000040738 | 1771471 | 0.4321 | 0 | 60.4723 | Eukaryota | 91487.0 | 0 | Chordata | 91487.0 | 0 | Carnivora | 91487.0 | 0 | Canidae | 91487.0 | 0 | Canis | 81502.0 | 1 | Canis lupus | 81502.0 | 1 |
| scf7180000054039 | 2692 | 0.4502 | 0 | 285.6888 | Eukaryota | 243.0 | 0 | Streptophyta | 243.0 | 0 | Fabales | 243.0 | 0 | Fabaceae | 243.0 | 0 | Cicer | 243.0 | 0 | Cicer arietinum | 243.0 | 0 |
| scf7180000041377 | 243953 | 0.3466 | 0 | 72.4223 | Eukaryota | 92395.0 | 0 | Chordata | 92395.0 | 0 | Carnivora | 92395.0 | 0 | Canidae | 92395.0 | 0 | Canis | 92395.0 | 0 | Canis lupus | 92395.0 | 0 |
| scf7180000041460 | 938627 | 0.4462 | 0 | 59.0668 | Eukaryota | 88040.0 | 0 | Chordata | 88040.0 | 0 | Carnivora | 88040.0 | 0 | Canidae | 88040.0 | 0 | Canis | 88040.0 | 0 | Canis lupus | 88040.0 | 0 |
| scf7180000045296 | 209590 | 0.4081 | 0 | 58.8206 | Eukaryota | 35487.0 | 0 | Chordata | 35487.0 | 0 | Carnivora | 35487.0 | 0 | Otariidae | 22085.0 | 1 | Zalophus | 22085.0 | 2 | Zalophus californianus | 22085.0 | 2 |
| scf7180000046129 | 179008 | 0.4371 | 0 | 59.8322 | Eukaryota | 14896.0 | 0 | Chordata | 14896.0 | 0 | Carnivora | 12021.0 | 1 | Canidae | 11042.0 | 2 | Canis | 9245.0 | 3 | Canis lupus | 9245.0 | 3 |
| scf7180000055534 | 24499 | 0.5147 | 0 | 604.1924 | Eukaryota | 16158.0 | 0 | Chordata | 16158.0 | 0 | Carnivora | 16158.0 | 0 | Canidae | 16158.0 | 0 | Canis | 16158.0 | 0 | Canis lupus | 16158.0 | 0 |
| scf7180000040878 | 1647557 | 0.4498 | 0 | 57.9941 | Eukaryota | 74368.0 | 0 | Chordata | 74368.0 | 0 | Carnivora | 74368.0 | 0 | Canidae | 72039.0 | 1 | Canis | 53979.0 | 2 | Canis lupus | 53979.0 | 2 |
| scf7180000052376 | 11867 | 0.5019 | 0 | 180.5979 | Eukaryota | 8450.0 | 0 | Chordata | 8450.0 | 0 | Carnivora | 8450.0 | 0 | Canidae | 8450.0 | 0 | Canis | 6921.0 | 1 | Canis lupus | 5849.0 | 2 |
| scf7180000048337 | 7251 | 0.4499 | 0 | 1249.4842 | Eukaryota | 22164.0 | 0 | Chordata | 19208.0 | 2 | Gadiformes | 11503.0 | 7 | Gadidae | 11503.0 | 7 | Gadus | 11503.0 | 7 | Gadus morhua | 11503.0 | 7 |
| scf7180000053545 | 3289 | 0.4542 | 0 | 1405.2088 | Eukaryota | 699.0 | 0 | Platyhelminthes | 699.0 | 0 | Polystomatidea | 699.0 | 0 | Polystomatidae | 699.0 | 0 | Protopolystoma | 699.0 | 0 | Protopolystoma xenopodis | 699.0 | 0 |
| scf7180000051928 | 19906 | 0.5206 | 0 | 663.4735 | Eukaryota | 16598.0 | 0 | Chordata | 16598.0 | 0 | Carnivora | 16598.0 | 0 | Canidae | 16598.0 | 0 | Canis | 16598.0 | 0 | Canis lupus | 16598.0 | 0 |
| scf7180000051989 | 8313 | 0.4975 | 0 | 70.3838 | Eukaryota | 8726.0 | 0 | Chordata | 8726.0 | 0 | Carnivora | 8726.0 | 0 | Canidae | 8726.0 | 0 | Canis | 7111.0 | 1 | Canis lupus | 5973.0 | 2 |
| scf7180000048622 | 59762 | 0.3905 | 0 | 60.8135 | Eukaryota | 6990.0 | 0 | Chordata | 6990.0 | 0 | Carnivora | 4314.0 | 2 | Canidae | 4314.0 | 2 | Canis | 4314.0 | 2 | Canis lupus | 4314.0 | 2 |
| scf7180000052947 | 2438 | 0.4541 | 0 | 1329.0544 | no-hit | 0.0 | 0 | no-hit | 0.0 | 0 | no-hit | 0.0 | 0 | no-hit | 0.0 | 0 | no-hit | 0.0 | 0 | no-hit | 0.0 | 0 |
| scf7180000049172 | 3447 | 0.4444 | 0 | 45.8231 | Eukaryota | 5339.0 | 0 | Chordata | 5078.0 | 1 | Kurtiformes | 4797.0 | 2 | Apogonidae | 4797.0 | 2 | Sphaeramia | 4797.0 | 2 | Sphaeramia orbicularis | 4797.0 | 2 |
| scf7180000050789 | 22919 | 0.4806 | 0 | 385.8536 | Eukaryota | 85707.0 | 0 | Chordata | 85707.0 | 0 | Carnivora | 85707.0 | 0 | Canidae | 85707.0 | 0 | Canis | 85707.0 | 0 | Canis lupus | 85707.0 | 0 |
| scf7180000048415 | 17918 | 0.4396 | 0 | 325.2362 | Eukaryota | 1059.0 | 0 | Chordata | 796.0 | 1 | Columbiformes | 566.0 | 2 | Columbidae | 566.0 | 2 | Streptopelia | 566.0 | 2 | Streptopelia turtur | 566.0 | 2 |
| scf7180000046095 | 16133 | 0.49 | 0 | 69.9297 | Eukaryota | 24561.0 | 0 | Chordata | 24561.0 | 0 | Carnivora | 24561.0 | 0 | Canidae | 24561.0 | 0 | Canis | 24561.0 | 0 | Canis lupus | 24561.0 | 0 |
| scf7180000047493 | 65688 | 0.4889 | 0 | 59.0569 | Eukaryota | 19425.0 | 0 | Chordata | 19425.0 | 0 | Carnivora | 19425.0 | 0 | Canidae | 12804.0 | 2 | Canis | 8552.0 | 3 | Canis lupus | 8552.0 | 3 |
| scf7180000055434 | 23742 | 0.4482 | 0 | 345.9545 | Eukaryota | 9349.0 | 0 | Chordata | 9349.0 | 0 | Carnivora | 9349.0 | 0 | Canidae | 9349.0 | 0 | Canis | 5551.0 | 1 | Canis lupus | 5551.0 | 1 |
| scf7180000048311 | 10430 | 0.4357 | 0 | 102.0164 | Eukaryota | 4647.0 | 0 | Chordata | 4647.0 | 0 | Tetraodontiformes | 1581.0 | 5 | Tetraodontidae | 1581.0 | 5 | Takifugu | 1581.0 | 6 | Takifugu rubripes | 1581.0 | 6 |
| scf7180000044331 | 35382 | 0.6099 | 0 | 66.9777 | Eukaryota | 32430.0 | 0 | Chordata | 32430.0 | 0 | Carnivora | 32430.0 | 0 | Canidae | 32430.0 | 0 | Canis | 32430.0 | 0 | Canis lupus | 32430.0 | 0 |
| scf7180000046856 | 75271 | 0.4002 | 0 | 60.5128 | Eukaryota | 13371.0 | 0 | Chordata | 13371.0 | 0 | Carnivora | 13371.0 | 0 | Canidae | 13371.0 | 0 | Canis | 9402.0 | 3 | Canis aureus | 3942.0 | 6 |
| scf7180000043643 | 64862 | 0.3823 | 0 | 58.1461 | Eukaryota | 61963.0 | 0 | Chordata | 61963.0 | 0 | Carnivora | 61963.0 | 0 | Canidae | 61963.0 | 0 | Canis | 61963.0 | 0 | Canis lupus | 61963.0 | 0 |
| scf7180000051095 | 4744 | 0.4313 | 0 | 55.428 | no-hit | 0.0 | 0 | no-hit | 0.0 | 0 | no-hit | 0.0 | 0 | no-hit | 0.0 | 0 | no-hit | 0.0 | 0 | no-hit | 0.0 | 0 |
| scf7180000054473 | 17916 | 0.4606 | 0 | 63.6529 | Eukaryota | 47206.0 | 0 | Chordata | 47206.0 | 0 | Carnivora | 47206.0 | 0 | Canidae | 47206.0 | 0 | Canis | 47206.0 | 0 | Canis lupus | 47206.0 | 0 |
| scf7180000047601 | 82027 | 0.4099 | 0 | 55.4066 | Eukaryota | 13724.0 | 0 | Chordata | 13724.0 | 0 | Carnivora | 12538.0 | 1 | Canidae | 5219.0 | 4 | Canis | 5219.0 | 5 | Canis lupus | 5219.0 | 5 |
| scf7180000050853 | 2046 | 0.4545 | 0 | 170.3627 | no-hit | 0.0 | 0 | no-hit | 0.0 | 0 | no-hit | 0.0 | 0 | no-hit | 0.0 | 0 | no-hit | 0.0 | 0 | no-hit | 0.0 | 0 |
| scf7180000050033 | 8766 | 0.439 | 0 | 163.371 | no-hit | 0.0 | 0 | no-hit | 0.0 | 0 | no-hit | 0.0 | 0 | no-hit | 0.0 | 0 | no-hit | 0.0 | 0 | no-hit | 0.0 | 0 |
| scf7180000045706 | 90544 | 0.3777 | 0 | 57.1828 | Eukaryota | 88698.0 | 0 | Chordata | 88698.0 | 0 | Carnivora | 88698.0 | 0 | Canidae | 88698.0 | 0 | Canis | 88698.0 | 0 | Canis lupus | 88698.0 | 0 |
| scf7180000052274 | 3275 | 0.4388 | 0 | 112.3903 | no-hit | 0.0 | 0 | no-hit | 0.0 | 0 | no-hit | 0.0 | 0 | no-hit | 0.0 | 0 | no-hit | 0.0 | 0 | no-hit | 0.0 | 0 |
| scf7180000044125 | 209167 | 0.372 | 0 | 60.5376 | Eukaryota | 18716.0 | 0 | Chordata | 18716.0 | 0 | Primates | 18716.0 | 0 | Hominidae | 16913.0 | 1 | Homo | 9220.0 | 2 | Homo sapiens | 9220.0 | 2 |
| scf7180000042794 | 537714 | 0.4702 | 0 | 60.0223 | Eukaryota | 29013.0 | 0 | Chordata | 29013.0 | 0 | Carnivora | 29013.0 | 0 | Canidae | 29013.0 | 0 | Canis | 26023.0 | 1 | Canis lupus | 26023.0 | 1 |
| scf7180000044000 | 359496 | 0.479 | 0 | 57.9022 | Eukaryota | 59075.0 | 0 | Chordata | 59075.0 | 0 | Carnivora | 59075.0 | 0 | Canidae | 47547.0 | 1 | Canis | 34761.0 | 3 | Canis lupus | 34761.0 | 3 |
| scf7180000044709 | 337235 | 0.357 | 0 | 59.9207 | Eukaryota | 57721.0 | 0 | Chordata | 57721.0 | 0 | Carnivora | 57721.0 | 0 | Canidae | 57721.0 | 0 | Canis | 57721.0 | 0 | Canis lupus | 57721.0 | 0 |
| scf7180000040509 | 2312918 | 0.3923 | 0 | 60.6448 | Eukaryota | 94150.0 | 0 | Chordata | 94150.0 | 0 | Carnivora | 94150.0 | 0 | Canidae | 94150.0 | 0 | Canis | 94150.0 | 0 | Canis lupus | 94150.0 | 0 |
| scf7180000043969 | 345473 | 0.5484 | 0 | 58.2373 | Eukaryota | 28055.0 | 0 | Chordata | 28055.0 | 0 | Carnivora | 28055.0 | 0 | Canidae | 28055.0 | 0 | Canis | 18152.0 | 1 | Canis lupus | 18152.0 | 1 |
| scf7180000041445 | 666017 | 0.5168 | 0 | 58.6525 | Eukaryota | 83150.0 | 0 | Chordata | 83150.0 | 0 | Carnivora | 83150.0 | 0 | Canidae | 83150.0 | 0 | Canis | 78891.0 | 1 | Canis lupus | 78891.0 | 1 |
| scf7180000046827 | 8784 | 0.4323 | 0 | 47.8281 | no-hit | 0.0 | 0 | no-hit | 0.0 | 0 | no-hit | 0.0 | 0 | no-hit | 0.0 | 0 | no-hit | 0.0 | 0 | no-hit | 0.0 | 0 |
| scf7180000054071 | 4145 | 0.4548 | 0 | 1133.3451 | no-hit | 0.0 | 0 | no-hit | 0.0 | 0 | no-hit | 0.0 | 0 | no-hit | 0.0 | 0 | no-hit | 0.0 | 0 | no-hit | 0.0 | 0 |
| scf7180000045367 | 187890 | 0.3872 | 0 | 59.3437 | Eukaryota | 127187.0 | 0 | Chordata | 127187.0 | 0 | Carnivora | 127187.0 | 0 | Canidae | 127187.0 | 0 | Canis | 121272.0 | 1 | Canis lupus | 121272.0 | 1 |
| scf7180000046187 | 170905 | 0.4689 | 0 | 58.0692 | Eukaryota | 15373.0 | 1 | Chordata | 15373.0 | 1 | Primates | 10920.0 | 2 | Hominidae | 4615.0 | 5 | Ursus | 4453.0 | 6 | Ursus arctos | 4453.0 | 6 |
| scf7180000051173 | 4119 | 0.4462 | 0 | 213.1842 | no-hit | 0.0 | 0 | no-hit | 0.0 | 0 | no-hit | 0.0 | 0 | no-hit | 0.0 | 0 | no-hit | 0.0 | 0 | no-hit | 0.0 | 0 |
| scf7180000052499 | 8269 | 0.4575 | 0 | 62.6707 | Eukaryota | 6502.0 | 0 | Echinodermata | 2307.0 | 2 | Forcipulatida | 2307.0 | 3 | Asteriidae | 2307.0 | 3 | Asterias | 2307.0 | 4 | Asterias rubens | 2307.0 | 4 |
| scf7180000046615 | 9116 | 0.4115 | 0 | 35.4389 | Eukaryota | 34003.0 | 0 | Chordata | 34003.0 | 0 | Carnivora | 34003.0 | 0 | Canidae | 34003.0 | 0 | Canis | 34003.0 | 0 | Canis lupus | 34003.0 | 0 |
| scf7180000050788 | 2998 | 0.449 | 0 | 395.5348 | Eukaryota | 1998.0 | 0 | Platyhelminthes | 777.0 | 2 | Polystomatidea | 777.0 | 2 | Polystomatidae | 777.0 | 2 | Protopolystoma | 777.0 | 2 | Protopolystoma xenopodis | 777.0 | 2 |
| scf7180000049149 | 10745 | 0.4457 | 0 | 193.0505 | Eukaryota | 7711.0 | 0 | Chordata | 7711.0 | 0 | Tetraodontiformes | 5913.0 | 1 | Tetraodontidae | 5913.0 | 1 | Takifugu | 5913.0 | 1 | Takifugu rubripes | 5913.0 | 1 |
| scf7180000050463 | 29210 | 0.4405 | 0 | 65.9765 | Eukaryota | 789.0 | 0 | Chordata | 789.0 | 0 | Columbiformes | 414.0 | 1 | Columbidae | 414.0 | 1 | Streptopelia | 414.0 | 1 | Streptopelia turtur | 414.0 | 1 |
| scf7180000051223 | 25016 | 0.5026 | 0 | 200.0629 | Eukaryota | 9202.0 | 0 | Chordata | 9202.0 | 0 | Carnivora | 9202.0 | 0 | Canidae | 9202.0 | 0 | Canis | 8056.0 | 1 | Canis lupus | 6860.0 | 2 |
| scf7180000051835 | 15305 | 0.4427 | 0 | 79.9101 | Eukaryota | 17021.0 | 0 | Platyhelminthes | 9231.0 | 1 | Cyclophyllidea | 4662.0 | 7 | Hymenolepididae | 4662.0 | 7 | Hymenolepis | 4662.0 | 7 | Hymenolepis microstoma | 4662.0 | 7 |
| scf7180000047388 | 54278 | 0.4356 | 0 | 101.6772 | Eukaryota | 7005.0 | 0 | Chordata | 6458.0 | 1 | Kurtiformes | 3028.0 | 4 | Apogonidae | 3028.0 | 4 | Sphaeramia | 3028.0 | 4 | Sphaeramia orbicularis | 3028.0 | 4 |
| scf7180000052097 | 10124 | 0.4455 | 0 | 112.0397 | Eukaryota | 666.0 | 0 | Platyhelminthes | 409.0 | 1 | Polystomatidea | 409.0 | 1 | Polystomatidae | 409.0 | 1 | Protopolystoma | 409.0 | 1 | Protopolystoma xenopodis | 409.0 | 1 |
| scf7180000047859 | 98222 | 0.3503 | 0 | 59.0896 | Eukaryota | 12060.0 | 1 | Chordata | 12060.0 | 1 | Carnivora | 5274.0 | 3 | Canidae | 5274.0 | 3 | Canis | 5274.0 | 4 | Canis lupus | 5274.0 | 4 |
| scf7180000051568 | 2912 | 0.4571 | 0 | 249.1373 | Eukaryota | 1225.0 | 0 | Chordata | 1225.0 | 0 | Tetraodontiformes | 988.0 | 1 | Tetraodontidae | 988.0 | 1 | Takifugu | 988.0 | 1 | Takifugu rubripes | 988.0 | 1 |
| scf7180000056192 | 1681947 | 0.3863 | 0 | 59.8767 | Eukaryota | 93723.0 | 0 | Chordata | 93723.0 | 0 | Carnivora | 93723.0 | 0 | Canidae | 93723.0 | 0 | Canis | 93723.0 | 0 | Canis lupus | 93723.0 | 0 |
| scf7180000041972 | 281767 | 0.3708 | 0 | 57.8079 | Eukaryota | 85123.0 | 0 | Chordata | 85123.0 | 0 | Carnivora | 85123.0 | 0 | Canidae | 85123.0 | 0 | Canis | 85123.0 | 0 | Canis lupus | 85123.0 | 0 |
| scf7180000048350 | 33099 | 0.4453 | 0 | 129.8378 | Eukaryota | 10080.0 | 0 | Chordata | 6539.0 | 1 | Polystomatidea | 3541.0 | 3 | Polystomatidae | 3541.0 | 3 | Protopolystoma | 3541.0 | 3 | Protopolystoma xenopodis | 3541.0 | 3 |
| scf7180000044104 | 3548 | 0.3771 | 0 | 26.1098 | Eukaryota | 5198.0 | 0 | Chordata | 5198.0 | 0 | Primates | 4034.0 | 1 | Hominidae | 4034.0 | 1 | Pan | 2326.0 | 2 | Pan troglodytes | 2326.0 | 2 |
| scf7180000045671 | 156195 | 0.5789 | 0 | 55.3656 | Eukaryota | 39117.0 | 0 | Chordata | 39117.0 | 0 | Carnivora | 39117.0 | 0 | Canidae | 37628.0 | 1 | Canis | 25674.0 | 2 | Canis lupus | 25674.0 | 2 |
| scf7180000045193 | 170563 | 0.3873 | 0 | 60.7823 | Eukaryota | 23800.0 | 0 | Chordata | 23800.0 | 0 | Carnivora | 23800.0 | 0 | Canidae | 23800.0 | 0 | Canis | 21083.0 | 1 | Canis lupus | 21083.0 | 1 |
| scf7180000041959 | 792607 | 0.5345 | 0 | 58.6894 | Eukaryota | 61745.0 | 0 | Chordata | 61745.0 | 0 | Carnivora | 61745.0 | 0 | Canidae | 29921.0 | 2 | Leptonychotes | 21552.0 | 4 | Leptonychotes weddellii | 21552.0 | 4 |
| scf7180000049539 | 24039 | 0.4398 | 0 | 559.2159 | Eukaryota | 2752.0 | 0 | Chordata | 2752.0 | 0 | Carnivora | 2752.0 | 0 | Canidae | 2752.0 | 0 | Canis | 2752.0 | 0 | Canis lupus | 2752.0 | 0 |
| scf7180000053412 | 4697 | 0.4486 | 0 | 368.0004 | Eukaryota | 2264.0 | 0 | Platyhelminthes | 869.0 | 2 | Polystomatidea | 869.0 | 2 | Polystomatidae | 869.0 | 2 | Protopolystoma | 869.0 | 2 | Protopolystoma xenopodis | 869.0 | 2 |
| scf7180000053708 | 3744 | 0.4479 | 0 | 133.2714 | no-hit | 0.0 | 0 | no-hit | 0.0 | 0 | no-hit | 0.0 | 0 | no-hit | 0.0 | 0 | no-hit | 0.0 | 0 | no-hit | 0.0 | 0 |
| scf7180000048700 | 44857 | 0.668 | 0 | 55.7046 | Eukaryota | 38452.0 | 0 | Chordata | 38452.0 | 0 | Carnivora | 38452.0 | 0 | Canidae | 34948.0 | 2 | Canis | 33623.0 | 4 | Canis lupus | 33623.0 | 4 |
| scf7180000053105 | 7733 | 0.4469 | 0 | 530.9013 | Eukaryota | 2831.0 | 0 | Chordata | 2831.0 | 0 | Tetraodontiformes | 2564.0 | 1 | Tetraodontidae | 2564.0 | 1 | Takifugu | 2564.0 | 1 | Takifugu rubripes | 2564.0 | 1 |
| scf7180000047247 | 15706 | 0.5194 | 0 | 962.6459 | Eukaryota | 32403.0 | 0 | Chordata | 32403.0 | 0 | Carnivora | 32403.0 | 0 | Canidae | 32403.0 | 0 | Canis | 32403.0 | 0 | Canis lupus | 32403.0 | 0 |
| scf7180000055722 | 9075 | 0.4334 | 0 | 108.1733 | no-hit | 0.0 | 0 | no-hit | 0.0 | 0 | no-hit | 0.0 | 0 | no-hit | 0.0 | 0 | no-hit | 0.0 | 0 | no-hit | 0.0 | 0 |
| scf7180000046695 | 98153 | 0.3654 | 0 | 56.6471 | Eukaryota | 76240.0 | 0 | Chordata | 76240.0 | 0 | Carnivora | 76240.0 | 0 | Canidae | 76240.0 | 0 | Canis | 76240.0 | 0 | Canis lupus | 76240.0 | 0 |
| scf7180000051713 | 7967 | 0.445 | 0 | 2558.8212 | Eukaryota | 398.0 | 0 | Platyhelminthes | 398.0 | 0 | Polystomatidea | 398.0 | 0 | Polystomatidae | 398.0 | 0 | Protopolystoma | 398.0 | 0 | Protopolystoma xenopodis | 398.0 | 0 |
| scf7180000053216 | 6005 | 0.5204 | 0 | 84.3921 | Eukaryota | 3228.0 | 0 | Chordata | 3228.0 | 0 | Carnivora | 3228.0 | 0 | Canidae | 3228.0 | 0 | Canis | 2406.0 | 1 | Canis lupus | 2406.0 | 1 |
| scf7180000041769 | 213730 | 0.3462 | 0 | 57.3446 | Eukaryota | 125511.0 | 0 | Chordata | 125511.0 | 0 | Carnivora | 125511.0 | 0 | Canidae | 125511.0 | 0 | Canis | 111476.0 | 1 | Canis lupus | 111476.0 | 1 |
| scf7180000045814 | 8485 | 0.4354 | 0 | 227.8479 | Eukaryota | 577.0 | 0 | Platyhelminthes | 577.0 | 0 | Polystomatidea | 577.0 | 0 | Polystomatidae | 577.0 | 0 | Protopolystoma | 577.0 | 0 | Protopolystoma xenopodis | 577.0 | 0 |
| scf7180000041230 | 216595 | 0.4049 | 0 | 59.8828 | Eukaryota | 64443.0 | 0 | Chordata | 64443.0 | 0 | Carnivora | 64443.0 | 0 | Canidae | 64443.0 | 0 | Canis | 64443.0 | 0 | Canis lupus | 64443.0 | 0 |
| scf7180000041978 | 547796 | 0.3865 | 0 | 60.1256 | Eukaryota | 71347.0 | 0 | Chordata | 71347.0 | 0 | Carnivora | 71347.0 | 0 | Canidae | 71347.0 | 0 | Canis | 71347.0 | 0 | Canis lupus | 71347.0 | 0 |
| scf7180000040968 | 8407 | 0.4708 | 0 | 66.3158 | Eukaryota | 25149.0 | 0 | Chordata | 25149.0 | 0 | Carnivora | 25149.0 | 0 | Canidae | 25149.0 | 0 | Canis | 25149.0 | 0 | Canis lupus | 25149.0 | 0 |
| scf7180000044363 | 324601 | 0.4139 | 0 | 59.0659 | Eukaryota | 44640.0 | 0 | Chordata | 44640.0 | 0 | Carnivora | 44640.0 | 0 | Canidae | 23635.0 | 2 | Canis | 14223.0 | 4 | Canis lupus | 14223.0 | 4 |
| scf7180000055626 | 17933 | 0.4434 | 0 | 112.7471 | Eukaryota | 4792.0 | 0 | Platyhelminthes | 3957.0 | 1 | Cyclophyllidea | 3666.0 | 3 | Hymenolepididae | 3666.0 | 3 | Hymenolepis | 3666.0 | 3 | Hymenolepis microstoma | 3666.0 | 3 |
| scf7180000046867 | 109218 | 0.3714 | 0 | 59.9963 | Eukaryota | 7208.0 | 0 | Chordata | 7208.0 | 0 | Carnivora | 6037.0 | 1 | Felidae | 4592.0 | 2 | Felis | 3967.0 | 3 | Felis catus | 3967.0 | 3 |
| scf7180000045392 | 229112 | 0.4822 | 0 | 57.7816 | Eukaryota | 51543.0 | 0 | Chordata | 51543.0 | 0 | Carnivora | 51543.0 | 0 | Canidae | 31799.0 | 2 | Canis | 25656.0 | 4 | Canis lupus | 25656.0 | 4 |
| scf7180000047459 | 70892 | 0.4138 | 0 | 56.8451 | Eukaryota | 20954.0 | 0 | Chordata | 20954.0 | 0 | Carnivora | 20954.0 | 0 | Canidae | 19788.0 | 1 | Canis | 18979.0 | 2 | Canis lupus | 18979.0 | 2 |
| scf7180000046117 | 134508 | 0.4335 | 0 | 59.4906 | Eukaryota | 11866.0 | 0 | Chordata | 11866.0 | 0 | Carnivora | 11866.0 | 0 | Canidae | 11866.0 | 0 | Canis | 11866.0 | 0 | Canis lupus | 11866.0 | 0 |
| scf7180000051447 | 10300 | 0.4549 | 0 | 1049.4491 | Eukaryota | 17501.0 | 0 | Chordata | 14648.0 | 3 | Gadiformes | 10394.0 | 7 | Gadidae | 10394.0 | 7 | Gadus | 10394.0 | 7 | Gadus morhua | 10394.0 | 7 |
| scf7180000044782 | 10251 | 0.4272 | 0 | 36.4397 | Eukaryota | 33013.0 | 0 | Chordata | 33013.0 | 0 | Carnivora | 33013.0 | 0 | Canidae | 33013.0 | 0 | Canis | 23128.0 | 1 | Canis lupus | 23128.0 | 1 |
| scf7180000052068 | 5074 | 0.4511 | 0 | 246.9609 | no-hit | 0.0 | 0 | no-hit | 0.0 | 0 | no-hit | 0.0 | 0 | no-hit | 0.0 | 0 | no-hit | 0.0 | 0 | no-hit | 0.0 | 0 |
| scf7180000041635 | 954394 | 0.4958 | 0 | 58.9842 | Eukaryota | 51088.0 | 0 | Chordata | 51088.0 | 0 | Carnivora | 51088.0 | 0 | Canidae | 51088.0 | 0 | Canis | 51088.0 | 0 | Canis lupus | 51088.0 | 0 |
| scf7180000048015 | 51179 | 0.4497 | 0 | 80.6625 | Eukaryota | 27015.0 | 0 | Chordata | 27015.0 | 0 | Carnivora | 27015.0 | 0 | Canidae | 27015.0 | 0 | Canis | 27015.0 | 0 | Canis lupus | 27015.0 | 0 |
| scf7180000050083 | 5154 | 0.4562 | 0 | 1098.3027 | Eukaryota | 3219.0 | 0 | Platyhelminthes | 1941.0 | 2 | Polystomatidea | 1941.0 | 2 | Polystomatidae | 1941.0 | 2 | Protopolystoma | 1941.0 | 2 | Protopolystoma xenopodis | 1941.0 | 2 |
| scf7180000046880 | 3990 | 0.3862 | 0 | 32.7718 | Eukaryota | 3196.0 | 0 | Chordata | 3196.0 | 0 | Carnivora | 3196.0 | 0 | Canidae | 3196.0 | 0 | Canis | 3196.0 | 0 | Canis lupus | 3196.0 | 0 |
| scf7180000040479 | 1115550 | 0.3809 | 0 | 61.0257 | Eukaryota | 92159.0 | 0 | Chordata | 92159.0 | 0 | Carnivora | 92159.0 | 0 | Canidae | 92159.0 | 0 | Canis | 83036.0 | 1 | Canis lupus | 83036.0 | 1 |
| scf7180000047488 | 82937 | 0.4386 | 0 | 115.8677 | Eukaryota | 85071.0 | 0 | Chordata | 85071.0 | 0 | Carnivora | 85071.0 | 0 | Canidae | 85071.0 | 0 | Canis | 85071.0 | 0 | Canis lupus | 85071.0 | 0 |
| scf7180000054202 | 23218 | 0.4403 | 0 | 165.6259 | Eukaryota | 29098.0 | 1 | Chordata | 29098.0 | 1 | Tetraodontiformes | 10078.0 | 5 | Tetraodontidae | 10078.0 | 5 | Takifugu | 10078.0 | 5 | Takifugu rubripes | 10078.0 | 5 |
| scf7180000047653 | 18749 | 0.4192 | 0 | 56.4738 | Eukaryota | 34750.0 | 0 | Chordata | 34750.0 | 0 | Carnivora | 34750.0 | 0 | Canidae | 34750.0 | 0 | Canis | 34750.0 | 0 | Canis lupus | 34750.0 | 0 |
| scf7180000043895 | 56320 | 0.3584 | 0 | 51.1449 | Eukaryota | 56034.0 | 0 | Chordata | 56034.0 | 0 | Carnivora | 56034.0 | 0 | Canidae | 56034.0 | 0 | Canis | 56034.0 | 0 | Canis lupus | 56034.0 | 0 |

| | | | | | | | | | | | | | | | | | | | | | | |
|---|---|---|---|---|---|---|---|---|---|---|---|---|---|---|---|---|---|---|---|---|---|---|
| scf7180000056211 | 142936 | 0.4178 | 0 | 160.3997 | Eukaryota | 46377.0 | 0 | Chordata | 46377.0 | 0 | Carnivora | 41691.0 | 1 | Canidae | 41691.0 | 1 | Canis | 41691.0 | 1 | Canis lupus | 41691.0 | 1 |
| scf7180000051823 | 5700 | 0.4514 | 0 | 1084.0362 | Eukaryota | 2939.0 | 0 | Platyhelminthes | 1655.0 | 2 | Polystomatidea | 1655.0 | 2 | Polystomatidae | 1655.0 | 2 | Protopolystoma | 1655.0 | 2 | Protopolystoma xenopodis | 1655.0 | 2 |
| scf7180000053488 | 5355 | 0.504 | 0 | 208.3202 | Eukaryota | 7913.0 | 0 | Chordata | 7913.0 | 0 | Carnivora | 7913.0 | 0 | Canidae | 7913.0 | 0 | Canis | 5922.0 | 1 | Canis lupus | 4835.0 | 2 |
| scf7180000053764 | 6490 | 0.516 | 0 | 98.534 | Eukaryota | 12228.0 | 0 | Chordata | 12228.0 | 0 | Carnivora | 12228.0 | 0 | Canidae | 12228.0 | 0 | Canis | 12228.0 | 0 | Canis lupus | 12228.0 | 0 |
| scf7180000045120 | 89987 | 0.4197 | 0 | 54.5843 | Eukaryota | 55413.0 | 0 | Chordata | 55413.0 | 0 | Carnivora | 55413.0 | 0 | Canidae | 55413.0 | 0 | Canis | 55413.0 | 0 | Canis lupus | 55413.0 | 0 |
| scf7180000046135 | 195584 | 0.4027 | 0 | 58.77 | Eukaryota | 47526.0 | 0 | Chordata | 47526.0 | 0 | Carnivora | 47526.0 | 0 | Canidae | 47526.0 | 0 | Canis | 43551.0 | 1 | Canis lupus | 43551.0 | 1 |
| scf7180000052939 | 2861 | 0.4474 | 0 | 753.5044 | no-hit | 0.0 | 0 | no-hit | 0.0 | 0 | no-hit | 0.0 | 0 | no-hit | 0.0 | 0 | no-hit | 0.0 | 0 | no-hit | 0.0 | 0 |
| scf7180000050837 | 12777 | 0.4564 | 0 | 104.4782 | Eukaryota | 58971.0 | 0 | Chordata | 58971.0 | 0 | Carnivora | 58971.0 | 0 | Canidae | 58971.0 | 0 | Canis | 58971.0 | 0 | Canis lupus | 58971.0 | 0 |
| scf7180000041352 | 1071387 | 0.4679 | 0 | 59.4827 | Eukaryota | 67466.0 | 0 | Chordata | 67466.0 | 0 | Carnivora | 67466.0 | 0 | Canidae | 67466.0 | 0 | Canis | 62202.0 | 1 | Canis lupus | 62202.0 | 1 |
| scf7180000050290 | 8240 | 0.4521 | 0 | 684.2565 | Eukaryota | 3459.0 | 0 | Chordata | 2649.0 | 1 | Gonorynchiformes | 994.0 | 3 | Chanidae | 994.0 | 3 | Chanos | 994.0 | 3 | Chanos chanos | 994.0 | 3 |
| scf7180000041019 | 1472992 | 0.3817 | 0 | 59.8921 | Eukaryota | 33887.0 | 0 | Chordata | 33887.0 | 0 | Carnivora | 33887.0 | 0 | Canidae | 33887.0 | 0 | Canis | 30518.0 | 1 | Canis lupus | 30518.0 | 1 |
| scf7180000046225 | 17433 | 0.4451 | 0 | 107.5696 | Eukaryota | 88494.0 | 0 | Chordata | 88494.0 | 0 | Carnivora | 88494.0 | 0 | Canidae | 88494.0 | 0 | Canis | 88494.0 | 0 | Canis lupus | 88494.0 | 0 |
| scf7180000042073 | 806391 | 0.3452 | 0 | 59.1243 | Eukaryota | 74932.0 | 0 | Chordata | 74932.0 | 0 | Carnivora | 74932.0 | 0 | Canidae | 74932.0 | 0 | Canis | 74932.0 | 0 | Canis lupus | 74932.0 | 0 |
| scf7180000056528 | 267853 | 0.4515 | 0 | 58.0187 | Eukaryota | 35360.0 | 0 | Chordata | 35360.0 | 0 | Carnivora | 35360.0 | 0 | Canidae | 35360.0 | 0 | Canis | 35360.0 | 0 | Canis lupus | 35360.0 | 0 |
| scf7180000049722 | 10327 | 0.4541 | 0 | 2225.1575 | Eukaryota | 4296.0 | 0 | Platyhelminthes | 2488.0 | 3 | Polystomatidea | 2488.0 | 3 | Polystomatidae | 2488.0 | 3 | Protopolystoma | 2488.0 | 3 | Protopolystoma xenopodis | 2488.0 | 3 |
| scf7180000056005 | 17993 | 0.5499 | 0 | 438.8765 | Eukaryota | 5842.0 | 0 | Chordata | 5842.0 | 0 | Carnivora | 5842.0 | 0 | Canidae | 5842.0 | 0 | Canis | 5186.0 | 1 | Canis lupus | 5186.0 | 1 |
| scf7180000054952 | 22110 | 0.441 | 0 | 219.3375 | Eukaryota | 3280.0 | 0 | Platyhelminthes | 2036.0 | 1 | Polystomatidea | 2036.0 | 2 | Polystomatidae | 2036.0 | 2 | Protopolystoma | 2036.0 | 2 | Protopolystoma xenopodis | 2036.0 | 2 |
| scf7180000049662 | 15794 | 0.4238 | 0 | 54.543 | Eukaryota | 49469.0 | 0 | Chordata | 49469.0 | 0 | Carnivora | 49469.0 | 0 | Canidae | 31518.0 | 2 | Vulpes | 27874.0 | 3 | Vulpes vulpes | 27874.0 | 3 |
| scf7180000055765 | 26742 | 0.4418 | 0 | 107.4949 | Eukaryota | 287.0 | 0 | Chordata | 287.0 | 0 | Kurtiformes | 287.0 | 0 | Apogonidae | 287.0 | 0 | Sphaeramia | 287.0 | 0 | Sphaeramia orbicularis | 287.0 | 0 |
| scf7180000045204 | 249969 | 0.4521 | 0 | 60.2313 | Eukaryota | 43384.0 | 0 | Chordata | 43384.0 | 0 | Carnivora | 43384.0 | 0 | Canidae | 43384.0 | 0 | Canis | 39195.0 | 1 | Canis lupus | 39195.0 | 1 |
| scf7180000047184 | 17297 | 0.4011 | 0 | 38.9116 | Eukaryota | 3903.0 | 0 | Chordata | 3903.0 | 0 | Carnivora | 3903.0 | 0 | Canidae | 3090.0 | 1 | Canis | 3090.0 | 1 | Canis lupus | 3090.0 | 1 |
| scf7180000042995 | 310275 | 0.3986 | 0 | 59.7392 | Eukaryota | 90231.0 | 0 | Chordata | 90231.0 | 0 | Carnivora | 90231.0 | 0 | Canidae | 90231.0 | 0 | Canis | 90231.0 | 0 | Canis lupus | 90231.0 | 0 |
| scf7180000044045 | 46681 | 0.4085 | 0 | 59.1554 | Eukaryota | 3708.0 | 0 | Chordata | 3708.0 | 0 | Artiodactyla | 1860.0 | 2 | Canidae | 1393.0 | 3 | Canis | 1393.0 | 3 | Canis lupus | 1393.0 | 3 |
| scf7180000055485 | 53274 | 0.4488 | 0 | 64.2119 | Eukaryota | 81258.0 | 0 | Chordata | 81258.0 | 0 | Carnivora | 81258.0 | 0 | Canidae | 81258.0 | 0 | Canis | 81258.0 | 0 | Canis lupus | 81258.0 | 0 |
| scf7180000043736 | 22797 | 0.4562 | 0 | 49.973 | Eukaryota | 48635.0 | 0 | Chordata | 48635.0 | 0 | Carnivora | 48635.0 | 0 | Canidae | 48635.0 | 0 | Canis | 48635.0 | 0 | Canis lupus | 48635.0 | 0 |
| scf7180000054776 | 29629 | 0.4376 | 0 | 204.9275 | Eukaryota | 7106.0 | 0 | Chordata | 7106.0 | 0 | Carnivora | 7106.0 | 0 | Canidae | 7106.0 | 0 | Canis | 7106.0 | 0 | Canis lupus | 7106.0 | 0 |
| scf7180000044547 | 244678 | 0.3662 | 0 | 58.7707 | Eukaryota | 84138.0 | 0 | Chordata | 84138.0 | 0 | Carnivora | 84138.0 | 0 | Canidae | 84138.0 | 0 | Canis | 84138.0 | 0 | Canis lupus | 84138.0 | 0 |
| scf7180000045680 | 162082 | 0.4847 | 0 | 58.4608 | Eukaryota | 38533.0 | 0 | Chordata | 38533.0 | 0 | Carnivora | 38533.0 | 0 | Canidae | 38533.0 | 0 | Canis | 38533.0 | 0 | Canis lupus | 38533.0 | 0 |
| scf7180000051680 | 6919 | 0.4386 | 0 | 129.432 | Eukaryota | 3100.0 | 0 | Chordata | 3100.0 | 0 | Carnivora | 3100.0 | 0 | Canidae | 3100.0 | 0 | Canis | 3100.0 | 0 | Canis lupus | 3100.0 | 0 |
| scf7180000048877 | 61027 | 0.485 | 0 | 59.6083 | Eukaryota | 96754.0 | 0 | Chordata | 96754.0 | 0 | Carnivora | 96754.0 | 0 | Canidae | 87302.0 | 1 | Canis | 67099.0 | 2 | Canis lupus | 67099.0 | 2 |
| scf7180000049504 | 15632 | 0.4539 | 0 | 2539.3561 | Eukaryota | 1562.0 | 0 | Echinodermata | 577.0 | 3 | Forcipulatida | 577.0 | 3 | Asteriidae | 577.0 | 3 | Asterias | 577.0 | 3 | Asterias rubens | 577.0 | 3 |
| scf7180000047235 | 22056 | 0.3751 | 0 | 57.2147 | Eukaryota | 52248.0 | 0 | Chordata | 52248.0 | 0 | Carnivora | 52248.0 | 0 | Canidae | 52248.0 | 0 | Canis | 52248.0 | 0 | Canis lupus | 52248.0 | 0 |
| scf7180000042283 | 901230 | 0.5022 | 0 | 56.4182 | Eukaryota | 71019.0 | 0 | Chordata | 71019.0 | 0 | Carnivora | 71019.0 | 0 | Canidae | 71019.0 | 0 | Canis | 60405.0 | 1 | Canis lupus | 60405.0 | 1 |
| scf7180000047749 | 78449 | 0.3647 | 0 | 59.6021 | Eukaryota | 17380.0 | 0 | Chordata | 17380.0 | 0 | Carnivora | 17380.0 | 0 | Canidae | 17380.0 | 0 | Canis | 17380.0 | 0 | Canis lupus | 17380.0 | 0 |
| scf7180000049303 | 8509 | 0.4529 | 0 | 2700.8359 | Eukaryota | 3330.0 | 0 | Platyhelminthes | 1642.0 | 3 | Polystomatidea | 1642.0 | 3 | Polystomatidae | 1642.0 | 3 | Protopolystoma | 1642.0 | 3 | Protopolystoma xenopodis | 1642.0 | 3 |
| scf7180000043414 | 140222 | 0.4049 | 0 | 59.2386 | Eukaryota | 60788.0 | 0 | Chordata | 60788.0 | 0 | Carnivora | 60788.0 | 0 | Canidae | 60788.0 | 0 | Canis | 60788.0 | 0 | Canis lupus | 60788.0 | 0 |
| scf7180000047785 | 26280 | 0.4409 | 0 | 259.7873 | Eukaryota | 3401.0 | 0 | Chordata | 2695.0 | 1 | Carnivora | 2695.0 | 1 | Canidae | 2695.0 | 1 | Canis | 2695.0 | 1 | Canis lupus | 2695.0 | 1 |
| scf7180000047043 | 88467 | 0.3875 | 0 | 60.5343 | Eukaryota | 9338.0 | 0 | Chordata | 9338.0 | 0 | Primates | 5656.0 | 2 | Hominidae | 5656.0 | 3 | Homo | 5656.0 | 4 | Homo sapiens | 5656.0 | 4 |
| scf7180000055999 | 3798 | 0.4344 | 0 | 374.9455 | no-hit | 0.0 | 0 | no-hit | 0.0 | 0 | no-hit | 0.0 | 0 | no-hit | 0.0 | 0 | no-hit | 0.0 | 0 | no-hit | 0.0 | 0 |
| scf7180000051606 | 10631 | 0.4581 | 0 | 60.4589 | Eukaryota | 12528.0 | 0 | Chordata | 12528.0 | 0 | Carnivora | 12528.0 | 0 | Canidae | 12528.0 | 0 | Canis | 12528.0 | 0 | Canis lupus | 12528.0 | 0 |
| scf7180000049905 | 15659 | 0.4454 | 0 | 117.2884 | Eukaryota | 7633.0 | 0 | Chordata | 6287.0 | 1 | Salmoniformes | 3150.0 | 2 | Salmonidae | 3150.0 | 2 | Sphaeramia | 3137.0 | 3 | Sphaeramia orbicularis | 3137.0 | 3 |
| scf7180000046530 | 116600 | 0.6208 | 0 | 54.9426 | Eukaryota | 36632.0 | 0 | Chordata | 36632.0 | 0 | Carnivora | 36632.0 | 0 | Canidae | 36632.0 | 0 | Canis | 28074.0 | 1 | Canis lupus | 28074.0 | 1 |
| scf7180000052690 | 3466 | 0.4553 | 0 | 216.7858 | Eukaryota | 502.0 | 0 | Streptophyta | 502.0 | 0 | Fabales | 502.0 | 0 | Fabaceae | 502.0 | 0 | Cicer | 502.0 | 0 | Cicer arietinum | 502.0 | 0 |
| scf7180000041591 | 154146 | 0.3654 | 0 | 56.6004 | Eukaryota | 55651.0 | 0 | Chordata | 55651.0 | 0 | Carnivora | 55651.0 | 0 | Canidae | 55651.0 | 0 | Canis | 30408.0 | 1 | Canis lupus | 30408.0 | 1 |
| scf7180000048213 | 32649 | 0.4425 | 0 | 79.8531 | Eukaryota | 6623.0 | 0 | Chordata | 5431.0 | 1 | Kurtiformes | 1651.0 | 4 | Apogonidae | 1651.0 | 4 | Sphaeramia | 1651.0 | 4 | Sphaeramia orbicularis | 1651.0 | 4 |
| scf7180000056381 | 73215 | 0.3687 | 0 | 56.0225 | Eukaryota | 8852.0 | 0 | Chordata | 8852.0 | 0 | Carnivora | 6895.0 | 1 | Canidae | 6895.0 | 1 | Canis | 6895.0 | 2 | Canis lupus | 6895.0 | 2 |
| scf7180000046394 | 18447 | 0.4357 | 0 | 594.1129 | no-hit | 0.0 | 0 | no-hit | 0.0 | 0 | no-hit | 0.0 | 0 | no-hit | 0.0 | 0 | no-hit | 0.0 | 0 | no-hit | 0.0 | 0 |
| scf7180000044708 | 267894 | 0.4077 | 0 | 58.426 | Eukaryota | 63303.0 | 0 | Chordata | 63303.0 | 0 | Carnivora | 63303.0 | 0 | Canidae | 63303.0 | 0 | Canis | 63303.0 | 0 | Canis lupus | 63303.0 | 0 |
| scf7180000044039 | 528148 | 0.4403 | 0 | 58.4435 | Eukaryota | 21602.0 | 0 | Chordata | 21602.0 | 0 | Carnivora | 21602.0 | 0 | Canidae | 21602.0 | 0 | Canis | 19066.0 | 1 | Canis lupus | 19066.0 | 1 |
| scf7180000041989 | 181634 | 0.5423 | 0 | 56.854 | Eukaryota | 48529.0 | 0 | Chordata | 48529.0 | 0 | Carnivora | 48529.0 | 0 | Canidae | 48529.0 | 0 | Canis | 40214.0 | 1 | Canis lupus | 40214.0 | 1 |
| scf7180000042096 | 9432 | 0.4078 | 0 | 44.1835 | Eukaryota | 3365.0 | 0 | Chordata | 3365.0 | 0 | Carnivora | 2659.0 | 1 | Canidae | 2659.0 | 1 | Canis | 2659.0 | 1 | Canis lupus | 2659.0 | 1 |
| scf7180000046036 | 16725 | 0.4519 | 0 | 50.69 | Eukaryota | 13971.0 | 0 | Chordata | 13971.0 | 0 | Carnivora | 13971.0 | 0 | Canidae | 13971.0 | 0 | Canis | 13971.0 | 0 | Canis lupus | 13971.0 | 0 |
| scf7180000043881 | 218754 | 0.4455 | 0 | 58.8486 | Eukaryota | 68153.0 | 0 | Chordata | 68153.0 | 0 | Carnivora | 68153.0 | 0 | Canidae | 56071.0 | 2 | Canis | 49675.0 | 3 | Canis lupus | 49675.0 | 4 |
| scf7180000053416 | 5089 | 0.4496 | 0 | 1124.424 | Eukaryota | 3782.0 | 0 | Platyhelminthes | 2353.0 | 2 | Polystomatidea | 2353.0 | 2 | Polystomatidae | 2353.0 | 2 | Protopolystoma | 2353.0 | 2 | Protopolystoma xenopodis | 2353.0 | 2 |
| scf7180000042602 | 625047 | 0.3746 | 0 | 59.6096 | Eukaryota | 45519.0 | 0 | Chordata | 45519.0 | 0 | Carnivora | 45519.0 | 0 | Canidae | 45519.0 | 0 | Canis | 45519.0 | 0 | Canis lupus | 45519.0 | 0 |
| scf7180000051197 | 3738 | 0.4588 | 0 | 360.0796 | Eukaryota | 878.0 | 0 | Chordata | 449.0 | 2 | Columbiformes | 449.0 | 2 | Columbidae | 449.0 | 2 | Streptopelia | 449.0 | 2 | Streptopelia turtur | 449.0 | 2 |
| scf7180000050134 | 2660 | 0.4526 | 0 | 249.0658 | Eukaryota | 1722.0 | 0 | Platyhelminthes | 674.0 | 3 | Polystomatidea | 674.0 | 3 | Polystomatidae | 674.0 | 3 | Protopolystoma | 674.0 | 3 | Protopolystoma xenopodis | 674.0 | 3 |
| scf7180000046622 | 79233 | 0.3239 | 0 | 53.2814 | Eukaryota | 4814.0 | 1 | Chordata | 4814.0 | 1 | Artiodactyla | 2123.0 | 4 | Bovidae | 2123.0 | 4 | undef | 1984.0 | 5 | eukaryotic synthetic construct | 1984.0 | 5 |
| scf7180000043377 | 42503 | 0.4123 | 0 | 61.8418 | Eukaryota | 42258.0 | 0 | Chordata | 42258.0 | 0 | Carnivora | 42258.0 | 0 | Canidae | 42258.0 | 0 | Canis | 42258.0 | 0 | Canis lupus | 42258.0 | 0 |
| scf7180000053803 | 4261 | 0.5095 | 0 | 162.7813 | Eukaryota | 8093.0 | 0 | Chordata | 8093.0 | 0 | Carnivora | 8093.0 | 0 | Canidae | 8093.0 | 0 | Canis | 6418.0 | 1 | Canis lupus | 5174.0 | 2 |
| scf7180000045216 | 246061 | 0.4134 | 0 | 59.7782 | Eukaryota | 135146.0 | 0 | Chordata | 135146.0 | 0 | Carnivora | 135146.0 | 0 | Canidae | 135146.0 | 0 | Canis | 80874.0 | 1 | Canis lupus | 80874.0 | 1 |
| scf7180000051971 | 3143 | 0.4327 | 0 | 65.9859 | Eukaryota | 2147.0 | 0 | Platyhelminthes | 1644.0 | 1 | Polystomatidea | 1644.0 | 1 | Polystomatidae | 1644.0 | 1 | Protopolystoma | 1644.0 | 1 | Protopolystoma xenopodis | 1644.0 | 1 |
| scf7180000049888 | 8504 | 0.4392 | 0 | 248.2842 | no-hit | 0.0 | 0 | no-hit | 0.0 | 0 | no-hit | 0.0 | 0 | no-hit | 0.0 | 0 | no-hit | 0.0 | 0 | no-hit | 0.0 | 0 |
| scf7180000052399 | 6697 | 0.659 | 0 | 39.769 | Eukaryota | 13731.0 | 0 | Chordata | 13731.0 | 0 | Carnivora | 13731.0 | 0 | Canidae | 10238.0 | 1 | Vulpes | 10238.0 | 1 | Vulpes vulpes | 10238.0 | 1 |
| scf7180000041319 | 271947 | 0.3547 | 0 | 62.3626 | Eukaryota | 59870.0 | 0 | Chordata | 59870.0 | 0 | Carnivora | 59870.0 | 0 | Canidae | 59870.0 | 0 | Canis | 59870.0 | 0 | Canis lupus | 59870.0 | 0 |
| scf7180000049525 | 4482 | 0.4984 | 0 | 48.0434 | Eukaryota | 8042.0 | 0 | Chordata | 8042.0 | 0 | Carnivora | 8042.0 | 0 | Canidae | 8042.0 | 0 | Canis | 6484.0 | 1 | Canis lupus | 5334.0 | 2 |
| scf7180000052247 | 1944 | 0.4408 | 0 | 325.0048 | Eukaryota | 607.0 | 0 | Platyhelminthes | 607.0 | 0 | Polystomatidea | 422.0 | 1 | Polystomatidae | 422.0 | 1 | Protopolystoma | 422.0 | 1 | Protopolystoma xenopodis | 422.0 | 1 |
| scf7180000054080 | 9221 | 0.5043 | 0 | 73.2595 | Eukaryota | 32331.0 | 0 | Chordata | 32331.0 | 0 | Carnivora | 32331.0 | 0 | Canidae | 32331.0 | 0 | Canis | 32331.0 | 0 | Canis lupus | 32331.0 | 0 |
| scf7180000046479 | 100121 | 0.5113 | 0 | 57.8142 | Eukaryota | 22618.0 | 0 | Chordata | 22618.0 | 0 | Carnivora | 22618.0 | 0 | Canidae | 22618.0 | 0 | Canis | 18283.0 | 1 | Canis lupus | 18283.0 | 1 |
| scf7180000040570 | 2381792 | 0.3758 | 0 | 60.0806 | Eukaryota | 64213.0 | 0 | Chordata | 64213.0 | 0 | Carnivora | 64213.0 | 0 | Canidae | 64213.0 | 0 | Canis | 64213.0 | 0 | Canis lupus | 64213.0 | 0 |
| scf7180000044077 | 319425 | 0.3429 | 0 | 58.4748 | Eukaryota | 25131.0 | 0 | Chordata | 25131.0 | 0 | Carnivora | 25131.0 | 0 | Phocidae | 7641.0 | 4 | Leptonychotes | 5068.0 | 8 | Leptonychotes weddellii | 5068.0 | 8 |
| scf7180000056520 | 300146 | 0.3914 | 0 | 56.9527 | Eukaryota | 25707.0 | 0 | Chordata | 25707.0 | 0 | Carnivora | 25707.0 | 0 | Canidae | 25707.0 | 0 | Canis | 25707.0 | 0 | Canis lupus | 25707.0 | 0 |
| scf7180000055627 | 23403 | 0.4412 | 0 | 281.4386 | Eukaryota | 7142.0 | 0 | Chordata | 7142.0 | 0 | Carnivora | 7142.0 | 0 | Canidae | 7142.0 | 0 | Canis | 7142.0 | 0 | Canis lupus | 7142.0 | 0 |
| scf7180000045494 | 213195 | 0.5138 | 0 | 59.865 | Eukaryota | 30300.0 | 0 | Chordata | 30300.0 | 0 | Carnivora | 30300.0 | 0 | Canidae | 30300.0 | 0 | Canis | 24879.0 | 1 | Canis lupus | 24879.0 | 1 |
| scf7180000047654 | 68527 | 0.4871 | 0 | 57.869 | Eukaryota | 66353.0 | 0 | Chordata | 66353.0 | 0 | Carnivora | 57822.0 | 3 | Canidae | 48637.0 | 5 | Canis | 48637.0 | 6 | Canis lupus | 48637.0 | 6 |
| scf7180000048995 | 35941 | 0.4692 | 0 | 53.0884 | Eukaryota | 16924.0 | 0 | Chordata | 16924.0 | 0 | Carnivora | 15918.0 | 1 | Canidae | 15918.0 | 1 | Canis | 9339.0 | 2 | Canis lupus | 9339.0 | 2 |
| scf7180000054042 | 3145 | 0.4518 | 0 | 779.3302 | Eukaryota | 387.0 | 0 | Echinodermata | 207.0 | 1 | Forcipulatida | 207.0 | 1 | Asteriidae | 207.0 | 1 | Asterias | 207.0 | 1 | Asterias rubens | 207.0 | 1 |
| scf7180000049654 | 43261 | 0.3857 | 0 | 62.4515 | Eukaryota | 10696.0 | 0 | Chordata | 10696.0 | 0 | Artiodactyla | 6351.0 | 1 | Bovidae | 6351.0 | 1 | Bison | 4722.0 | 2 | Bison bison | 4722.0 | 2 |
| scf7180000044420 | 41538 | 0.4355 | 0 | 53.8346 | Eukaryota | 4177.0 | 0 | Chordata | 4177.0 | 0 | Carnivora | 4177.0 | 0 | Canidae | 4177.0 | 0 | Canis | 4177.0 | 0 | Canis lupus | 4177.0 | 0 |
| scf7180000048541 | 16870 | 0.4392 | 0 | 102.5524 | Eukaryota | 2744.0 | 0 | Chordata | 2522.0 | 1 | Tetraodontiformes | 1953.0 | 2 | Tetraodontidae | 1953.0 | 2 | Takifugu | 1953.0 | 2 | Takifugu rubripes | 1953.0 | 2 |
| scf7180000056585 | 2046682 | 0.4599 | 0 | 60.2956 | Eukaryota | 105547.0 | 0 | Chordata | 105547.0 | 0 | Carnivora | 105547.0 | 0 | Canidae | 105547.0 | 0 | Canis | 84571.0 | 1 | Canis lupus | 84571.0 | 1 |
| scf7180000049097 | 16064 | 0.4325 | 0 | 185.995 | Eukaryota | 88237.0 | 0 | Chordata | 88237.0 | 0 | Carnivora | 88237.0 | 0 | Canidae | 88237.0 | 0 | Canis | 88237.0 | 0 | Canis lupus | 88237.0 | 0 |
| scf7180000042763 | 6915 | 0.333 | 0 | 37.6498 | Eukaryota | 5118.0 | 0 | Chordata | 5118.0 | 0 | Carnivora | 5118.0 | 0 | Canidae | 5118.0 | 0 | Canis | 5118.0 | 0 | Canis lupus | 5118.0 | 0 |
| scf7180000044514 | 322517 | 0.4061 | 0 | 62.3772 | Eukaryota | 90319.0 | 0 | Chordata | 90319.0 | 0 | Carnivora | 90319.0 | 0 | Canidae | 90319.0 | 0 | Canis | 90319.0 | 0 | Canis lupus | 90319.0 | 0 |
| scf7180000046596 | 213806 | 0.6458 | 0 | 55.8011 | Eukaryota | 44520.0 | 0 | Chordata | 44520.0 | 0 | Carnivora | 44520.0 | 0 | Canidae | 44520.0 | 0 | Vulpes | 28648.0 | 1 | Vulpes vulpes | 28648.0 | 1 |
| scf7180000041293 | 30294 | 0.4524 | 0 | 133.94 | Eukaryota | 35127.0 | 0 | Chordata | 35127.0 | 0 | Carnivora | 35127.0 | 0 | Canidae | 35127.0 | 0 | Canis | 35127.0 | 0 | Canis lupus | 35127.0 | 0 |
| scf7180000043848 | 345809 | 0.3693 | 0 | 60.1947 | Eukaryota | 20718.0 | 0 | Chordata | 20718.0 | 0 | Carnivora | 20718.0 | 0 | Canidae | 20718.0 | 0 | Canis | 18477.0 | 1 | Canis lupus | 18477.0 | 1 |
| scf7180000042726 | 144728 | 0.3685 | 0 | 60.7591 | Eukaryota | 45944.0 | 0 | Chordata | 45944.0 | 0 | Carnivora | 45944.0 | 0 | Canidae | 45944.0 | 0 | Canis | 45944.0 | 0 | Canis lupus | 45944.0 | 0 |
| scf7180000040854 | 19988 | 0.4499 | 0 | 36.9601 | Eukaryota | 21571.0 | 0 | Chordata | 21571.0 | 0 | Carnivora | 21571.0 | 0 | Canidae | 21571.0 | 0 | Canis | 21571.0 | 0 | Canis lupus | 21571.0 | 0 |
| scf7180000047228 | 19239 | 0.4411 | 0 | 152.4464 | Eukaryota | 1013.0 | 0 | Chordata | 1013.0 | 0 | Tetraodontiformes | 713.0 | 1 | Tetraodontidae | 713.0 | 1 | Takifugu | 713.0 | 1 | Takifugu rubripes | 713.0 | 1 |
| scf7180000044998 | 237946 | 0.4348 | 0 | 59.3022 | Eukaryota | 51706.0 | 0 | Chordata | 51706.0 | 0 | Carnivora | 51706.0 | 0 | Canidae | 42265.0 | 1 | Canis | 42265.0 | 1 | Canis lupus | 42265.0 | 1 |
| scf7180000045838 | 162622 | 0.4065 | 0 | 60.7925 | Eukaryota | 36715.0 | 0 | Chordata | 36715.0 | 0 | Carnivora | 36715.0 | 0 | Canidae | 27616.0 | 1 | Canis | 20612.0 | 2 | Canis lupus | 20612.0 | 2 |
| scf7180000052459 | 22924 | 0.5069 | 0 | 389.2202 | Eukaryota | 33449.0 | 0 | Chordata | 33449.0 | 0 | Carnivora | 33449.0 | 0 | Canidae | 33449.0 | 0 | Canis | 33449.0 | 0 | Canis lupus | 33449.0 | 0 |
| scf7180000050609 | 2190 | 0.4516 | 0 | 454.3715 | Eukaryota | 2742.0 | 0 | Chordata | 2205.0 | 1 | Tetraodontiformes | 1259.0 | 2 | Tetraodontidae | 1259.0 | 2 | Takifugu | 1259.0 | 2 | Takifugu rubripes | 1259.0 | 2 |
| scf7180000044633 | 173954 | 0.481 | 0 | 57.9041 | Eukaryota | 32513.0 | 0 | Chordata | 32513.0 | 0 | Carnivora | 32513.0 | 0 | Canidae | 32513.0 | 0 | Vulpes | 20532.0 | 1 | Vulpes vulpes | 20532.0 | 1 |
| scf7180000045737 | 91608 | 0.3859 | 0 | 60.3548 | Eukaryota | 51269.0 | 0 | Chordata | 51269.0 | 0 | Carnivora | 51269.0 | 0 | Canidae | 51269.0 | 0 | Canis | 46159.0 | 1 | Canis lupus | 46159.0 | 1 |
| scf7180000054932 | 16549 | 0.4416 | 0 | 346.3074 | Eukaryota | 1828.0 | 0 | Chordata | 1828.0 | 0 | Tetraodontiformes | 1328.0 | 2 | Tetraodontidae | 1328.0 | 2 | Takifugu | 1328.0 | 2 | Takifugu rubripes | 1328.0 | 2 |
| scf7180000052056 | 4091 | 0.452 | 0 | 1775.3152 | Eukaryota | 3517.0 | 0 | Platyhelminthes | 1910.0 | 3 | Polystomatidea | 1910.0 | 3 | Polystomatidae | 1910.0 | 3 | Protopolystoma | 1910.0 | 3 | Protopolystoma xenopodis | 1910.0 | 3 |
| scf7180000045250 | 210993 | 0.5309 | 0 | 56.4821 | Eukaryota | 50005.0 | 0 | Chordata | 50005.0 | 0 | Carnivora | 50005.0 | 0 | Canidae | 18488.0 | 4 | Canis | 12414.0 | 8 | Canis lupus | 12414.0 | 8 |
| scf7180000046321 | 20950 | 0.4398 | 0 | 276.2037 | no-hit | 0.0 | 0 | no-hit | 0.0 | 0 | no-hit | 0.0 | 0 | no-hit | 0.0 | 0 | no-hit | 0.0 | 0 | no-hit | 0.0 | 0 |
| scf7180000040872 | 1396885 | 0.3723 | 0 | 59.8286 | Eukaryota | 90335.0 | 0 | Chordata | 90335.0 | 0 | Carnivora | 90335.0 | 0 | Canidae | 90335.0 | 0 | Canis | 81607.0 | 1 | Canis lupus | 81607.0 | 1 |
| scf7180000046476 | 27115 | 0.442 | 0 | 93.6698 | Eukaryota | 652.0 | 0 | Streptophyta | 652.0 | 0 | Fabales | 652.0 | 0 | Fabaceae | 652.0 | 0 | Cicer | 652.0 | 0 | Cicer arietinum | 652.0 | 0 |
| scf7180000041253 | 1181536 | 0.4368 | 0 | 59.2035 | Eukaryota | 161172.0 | 0 | Chordata | 161172.0 | 0 | Carnivora | 161172.0 | 0 | Canidae | 161172.0 | 0 | Canis | 161172.0 | 0 | Canis lupus | 161172.0 | 0 |
| scf7180000049940 | 44790 | 0.3439 | 0 | 59.8519 | Eukaryota | 6691.0 | 0 | Chordata | 6691.0 | 0 | Primates | 3369.0 | 4 | Hominidae | 2798.0 | 5 | Homo | 2798.0 | 5 | Homo sapiens | 2798.0 | 5 |
| scf7180000052897 | 4904 | 0.4533 | 0 | 503.0581 | Eukaryota | 4490.0 | 0 | Chordata | 2323.0 | 2 | Polystomatidea | 1618.0 | 4 | Polystomatidae | 1618.0 | 4 | Protopolystoma | 1618.0 | 4 | Protopolystoma xenopodis | 1618.0 | 4 |
| scf7180000051958 | 15311 | 0.4153 | 0 | 50.7825 | Eukaryota | 26433.0 | 0 | Chordata | 26433.0 | 0 | Carnivora | 26433.0 | 0 | Canidae | 14184.0 | 2 | Canis | 11340.0 | 4 | Canis lupus | 11340.0 | 4 |
| scf7180000054831 | 22471 | 0.4413 | 0 | 326.225 | Eukaryota | 2640.0 | 0 | Chordata | 2640.0 | 0 | Carnivora | 2640.0 | 0 | Canidae | 2640.0 | 0 | Canis | 2640.0 | 0 | Canis lupus | 2640.0 | 0 |
| scf7180000054465 | 25253 | 0.4444 | 0 | 487.3691 | Eukaryota | 2756.0 | 0 | Chordata | 2756.0 | 0 | Carnivora | 2308.0 | 1 | Canidae | 2308.0 | 1 | Canis | 2308.0 | 1 | Canis lupus | 2308.0 | 1 |
| scf7180000050276 | 13239 | 0.454 | 0 | 47.2811 | Eukaryota | 10686.0 | 0 | Chordata | 5421.0 | 1 | Strigeidida | 4819.0 | 6 | Schistosomatidae | 4819.0 | 6 | Schistosoma | 4819.0 | 6 | Schistosoma mansoni | 4819.0 | 6 |
| scf7180000041931 | 201664 | 0.3668 | 0 | 61.3273 | Eukaryota | 53834.0 | 0 | Chordata | 53834.0 | 0 | Carnivora | 53834.0 | 0 | Canidae | 53834.0 | 0 | Canis | 43160.0 | 1 | Canis lupus | 43160.0 | 1 |
| scf7180000041626 | 1082811 | 0.3663 | 0 | 60.0924 | Eukaryota | 102700.0 | 0 | Chordata | 102700.0 | 0 | Carnivora | 102700.0 | 0 | Canidae | 102700.0 | 0 | Canis | 102700.0 | 0 | Canis lupus | 102700.0 | 0 |
| scf7180000045598 | 12126 | 0.4714 | 0 | 42.5327 | Eukaryota | 2764.0 | 0 | Chordata | 2764.0 | 0 | Carnivora | 2468.0 | 1 | Canidae | 2468.0 | 1 | Canis | 2468.0 | 1 | Canis lupus | 2468.0 | 1 |
| scf7180000041930 | 952825 | 0.5605 | 100 | 57.1337 | Eukaryota | 91590.0 | 0 | Chordata | 91590.0 | 0 | Carnivora | 91590.0 | 0 | Canidae | 91590.0 | 0 | Canis | 91590.0 | 0 | Canis lupus | 91590.0 | 0 |
| scf7180000046136 | 148781 | 0.3787 | 0 | 59.6129 | Eukaryota | 51007.0 | 0 | Chordata | 51007.0 | 0 | Carnivora | 51007.0 | 0 | Canidae | 51007.0 | 0 | Canis | 51007.0 | 0 | Canis lupus | 51007.0 | 0 |
| scf7180000045723 | 192966 | 0.4499 | 0 | 60.0731 | Eukaryota | 44134.0 | 0 | Chordata | 44134.0 | 0 | Carnivora | 44134.0 | 0 | Canidae | 44134.0 | 0 | Canis | 33118.0 | 1 | Canis lupus | 33118.0 | 1 |
| scf7180000046762 | 66609 | 0.4583 | 0 | 58.2872 | Eukaryota | 14637.0 | 0 | Chordata | 14637.0 | 0 | Carnivora | 14637.0 | 0 | Felidae | 5754.0 | 3 | Neomonachus | 2984.0 | 7 | Neomonachus schauinslandi | 2984.0 | 7 |
| scf7180000051592 | 35680 | 0.5896 | 0 | 59.3367 | Eukaryota | 27298.0 | 0 | Chordata | 27298.0 | 0 | Carnivora | 27298.0 | 0 | Canidae | 27298.0 | 0 | Canis | 24284.0 | 1 | Canis lupus | 24284.0 | 1 |
| scf7180000055275 | 35817 | 0.4421 | 0 | 91.3346 | Eukaryota | 964.0 | 0 | Platyhelminthes | 773.0 | 1 | Polystomatidea | 773.0 | 1 | Polystomatidae | 773.0 | 1 | Protopolystoma | 773.0 | 1 | Protopolystoma xenopodis | 773.0 | 1 |
| scf7180000040792 | 1815415 | 0.4108 | 0 | 83.3109 | Eukaryota | 125858.0 | 0 | Chordata | 125858.0 | 0 | Carnivora | 125858.0 | 0 | Canidae | 125858.0 | 0 | Vulpes | 102241.0 | 1 | Vulpes vulpes | 102241.0 | 1 |
| scf7180000054340 | 15944 | 0.5172 | 0 | 442.023 | Eukaryota | 15930.0 | 0 | Chordata | 15930.0 | 0 | Carnivora | 15930.0 | 0 | Canidae | 15930.0 | 0 | Canis | 15930.0 | 0 | Canis lupus | 15930.0 | 0 |
| scf7180000051762 | 4817 | 0.4536 | 0 | 1302.0354 | no-hit | 0.0 | 0 | no-hit | 0.0 | 0 | no-hit | 0.0 | 0 | no-hit | 0.0 | 0 | no-hit | 0.0 | 0 | no-hit | 0.0 | 0 |
| scf7180000045791 | 181499 | 0.3977 | 0 | 60.6032 | Eukaryota | 52505.0 | 0 | Chordata | 52505.0 | 0 | Carnivora | 52505.0 | 0 | Canidae | 42638.0 | 1 | Canis | 31122.0 | 3 | Canis lupus | 31122.0 | 3 |
| scf7180000054302 | 16881 | 0.4026 | 0 | 154.0405 | Eukaryota | 6013.0 | 0 | Chordata | 6013.0 | 0 | Carnivora | 5455.0 | 1 | Felidae | 1971.0 | 5 | Felis | 1971.0 | 7 | Felis catus | 1971.0 | 7 |
| scf7180000055219 | 27427 | 0.441 | 0 | 157.9932 | Eukaryota | 53827.0 | 0 | Chordata | 33299.0 | 2 | Gadiformes | 15439.0 | 7 | Gadidae | 15439.0 | 7 | Gadus | 15439.0 | 7 | Gadus morhua | 15439.0 | 7 |
| scf7180000048096 | 15668 | 0.5203 | 0 | 1114.2804 | Eukaryota | 16082.0 | 0 | Chordata | 16082.0 | 0 | Carnivora | 16082.0 | 0 | Canidae | 16082.0 | 0 | Canis | 16082.0 | 0 | Canis lupus | 16082.0 | 0 |
| scf7180000053984 | 6343 | 0.4585 | 0 | 57.8239 | Eukaryota | 250.0 | 0 | Chordata | 250.0 | 0 | Kurtiformes | 250.0 | 0 | Apogonidae | 250.0 | 0 | Sphaeramia | 250.0 | 0 | Sphaeramia orbicularis | 250.0 | 0 |
| scf7180000050776 | 3429 | 0.4369 | 0 | 138.7222 | no-hit | 0.0 | 0 | no-hit | 0.0 | 0 | no-hit | 0.0 | 0 | no-hit | 0.0 | 0 | no-hit | 0.0 | 0 | no-hit | 0.0 | 0 |
| scf7180000051792 | 15359 | 0.4425 | 0 | 158.1571 | Eukaryota | 8534.0 | 0 | Chordata | 6882.0 | 3 | Tetraodontiformes | 3367.0 | 5 | Tetraodontidae | 3367.0 | 5 | Takifugu | 3367.0 | 5 | Takifugu rubripes | 3367.0 | 5 |
| scf7180000052173 | 15308 | 0.5031 | 0 | 54.2517 | Eukaryota | 8355.0 | 0 | Chordata | 8355.0 | 0 | Carnivora | 8355.0 | 0 | Canidae | 8355.0 | 0 | Canis | 6789.0 | 1 | Canis lupus | 5614.0 | 2 |
| scf7180000056364 | 865024 | 0.3697 | 0 | 61.0891 | Eukaryota | 59149.0 | 0 | Chordata | 59149.0 | 0 | Carnivora | 59149.0 | 0 | Canidae | 28191.0 | 2 | Canis | 15542.0 | 5 | Canis lupus | 15542.0 | 5 |
| scf7180000049449 | 35131 | 0.4445 | 0 | 69.2562 | Eukaryota | 85535.0 | 0 | Chordata | 85535.0 | 0 | Carnivora | 85535.0 | 0 | Canidae | 85535.0 | 0 | Canis | 85535.0 | 0 | Canis lupus | 85535.0 | 0 |
| scf7180000046635 | 18626 | 0.424 | 0 | 216.0287 | Eukaryota | 1926.0 | 0 | Chordata | 1719.0 | 1 | Carnivora | 1719.0 | 1 | Herpestidae | 756.0 | 3 | Suricata | 756.0 | 4 | Suricata suricatta | 756.0 | 4 |
| scf7180000040769 | 11597 | 0.4069 | 0 | 47.5245 | Eukaryota | 40447.0 | 0 | Chordata | 40447.0 | 0 | Carnivora | 40447.0 | 0 | Canidae | 40447.0 | 0 | Canis | 40447.0 | 0 | Canis lupus | 40447.0 | 0 |
| scf7180000043531 | 383679 | 0.3909 | 0 | 60.316 | Eukaryota | 65103.0 | 0 | Chordata | 65103.0 | 0 | Carnivora | 65103.0 | 0 | Canidae | 65103.0 | 0 | Canis | 45402.0 | 1 | Canis lupus | 45402.0 | 1 |
| scf7180000050267 | 9605 | 0.4382 | 0 | 65.4947 | Eukaryota | 11104.0 | 0 | Chordata | 11104.0 | 0 | Tetraodontiformes | 11104.0 | 0 | Tetraodontidae | 11104.0 | 0 | Takifugu | 11104.0 | 0 | Takifugu rubripes | 11104.0 | 0 |
| scf7180000055078 | 54171 | 0.4496 | 0 | 217.2783 | Eukaryota | 10788.0 | 0 | Chordata | 7863.0 | 2 | Kurtiformes | 4958.0 | 4 | Apogonidae | 4958.0 | 4 | Sphaeramia | 4958.0 | 4 | Sphaeramia orbicularis | 4958.0 | 4 |
| scf7180000050601 | 2621 | 0.4456 | 0 | 246.2192 | Eukaryota | 1696.0 | 0 | Chordata | 1285.0 | 1 | Carnivora | 1285.0 | 1 | Canidae | 1285.0 | 1 | Canis | 1144.0 | 2 | Canis lupus | 1144.0 | 2 |
| scf7180000050012 | 4519 | 0.4415 | 0 | 80.7603 | Eukaryota | 2060.0 | 0 | Chordata | 1494.0 | 1 | Salmoniformes | 716.0 | 3 | Salmonidae | 716.0 | 3 | Coregonus | 716.0 | 3 | Coregonus sp. 'balchen' | 716.0 | 3 |
| scf7180000049035 | 54373 | 0.5298 | 0 | 58.5207 | Eukaryota | 17590.0 | 0 | Chordata | 17590.0 | 0 | Carnivora | 17590.0 | 0 | Canidae | 14280.0 | 1 | Canis | 14280.0 | 1 | Canis lupus | 14280.0 | 1 |
| scf7180000051446 | 6214 | 0.4551 | 0 | 1355.6755 | Eukaryota | 135.0 | 0 | Chordata | 135.0 | 0 | Salmoniformes | 135.0 | 0 | Salmonidae | 135.0 | 0 | Salmo | 135.0 | 0 | Salmo trutta | 135.0 | 0 |
| scf7180000049331 | 2962 | 0.4382 | 0 | 260.6573 | no-hit | 0.0 | 0 | no-hit | 0.0 | 0 | no-hit | 0.0 | 0 | no-hit | 0.0 | 0 | no-hit | 0.0 | 0 | no-hit | 0.0 | 0 |
| scf7180000043982 | 579787 | 0.4066 | 0 | 59.2319 | Eukaryota | 62793.0 | 0 | Chordata | 62793.0 | 0 | Carnivora | 62793.0 | 0 | Canidae | 62793.0 | 0 | Canis | 51598.0 | 1 | Canis lupus | 51598.0 | 1 |
| scf7180000050425 | 3217 | 0.4358 | 0 | 527.3115 | Eukaryota | 1292.0 | 0 | Chordata | 990.0 | 1 | Kurtiformes | 819.0 | 2 | Apogonidae | 819.0 | 2 | Sphaeramia | 819.0 | 2 | Sphaeramia orbicularis | 819.0 | 2 |
| scf7180000040794 | 22715 | 0.4952 | 0 | 145.5972 | Eukaryota | 9469.0 | 0 | Chordata | 9469.0 | 0 | Carnivora | 9469.0 | 0 | Canidae | 9469.0 | 0 | Canis | 6971.0 | 1 | Canis lupus | 6971.0 | 1 |
| scf7180000052916 | 5600 | 0.5059 | 0 | 97.6282 | Eukaryota | 8155.0 | 0 | Chordata | 8155.0 | 0 | Carnivora | 8155.0 | 0 | Canidae | 8155.0 | 0 | Canis | 6077.0 | 1 | Canis lupus | 4946.0 | 2 |
| scf7180000045619 | 29061 | 0.4169 | 0 | 66.4267 | Eukaryota | 89031.0 | 0 | Chordata | 89031.0 | 0 | Carnivora | 89031.0 | 0 | Canidae | 89031.0 | 0 | Canis | 89031.0 | 0 | Canis lupus | 89031.0 | 0 |
| scf7180000050953 | 3814 | 0.4491 | 0 | 531.6252 | no-hit | 0.0 | 0 | no-hit | 0.0 | 0 | no-hit | 0.0 | 0 | no-hit | 0.0 | 0 | no-hit | 0.0 | 0 | no-hit | 0.0 | 0 |
| scf7180000054341 | 20486 | 0.4497 | 0 | 244.8356 | Eukaryota | 81509.0 | 0 | Chordata | 81509.0 | 0 | Carnivora | 81509.0 | 0 | Canidae | 81509.0 | 0 | Canis | 81509.0 | 0 | Canis lupus | 81509.0 | 0 |
| scf7180000043445 | 421373 | 0.4939 | 0 | 59.7632 | Eukaryota | 28178.0 | 0 | Chordata | 28178.0 | 0 | Carnivora | 28178.0 | 0 | Canidae | 17726.0 | 1 | Canis | 13353.0 | 3 | Canis lupus | 13353.0 | 3 |
| scf7180000042407 | 86319 | 0.4371 | 0 | 58.6929 | Eukaryota | 47437.0 | 0 | Chordata | 47437.0 | 0 | Carnivora | 47437.0 | 0 | Canidae | 47437.0 | 0 | Canis | 47437.0 | 0 | Canis lupus | 47437.0 | 0 |
| scf7180000041990 | 98804 | 0.3976 | 0 | 56.6636 | Eukaryota | 89828.0 | 0 | Chordata | 89828.0 | 0 | Carnivora | 89828.0 | 0 | Canidae | 89828.0 | 0 | Canis | 89828.0 | 0 | Canis lupus | 89828.0 | 0 |
| scf7180000044468 | 270665 | 0.5234 | 0 | 58.9306 | Eukaryota | 94219.0 | 0 | Chordata | 94219.0 | 0 | Carnivora | 94219.0 | 0 | Canidae | 94219.0 | 0 | Canis | 85958.0 | 1 | Canis lupus | 85958.0 | 1 |
| scf7180000055530 | 39326 | 0.4437 | 0 | 152.5735 | Eukaryota | 34358.0 | 0 | Platyhelminthes | 21173.0 | 1 | Strigeidida | 20186.0 | 2 | Schistosomatidae | 20186.0 | 2 | Schistosoma | 20186.0 | 2 | Schistosoma mansoni | 20186.0 | 2 |
| scf7180000050332 | 15207 | 0.4391 | 0 | 96.4502 | Eukaryota | 89063.0 | 0 | Chordata | 89063.0 | 0 | Carnivora | 89063.0 | 0 | Canidae | 89063.0 | 0 | Canis | 89063.0 | 0 | Canis lupus | 89063.0 | 0 |
| scf7180000049740 | 8114 | 0.4493 | 0 | 1485.9108 | Eukaryota | 4341.0 | 0 | Platyhelminthes | 2183.0 | 3 | Polystomatidea | 2183.0 | 3 | Polystomatidae | 2183.0 | 3 | Protopolystoma | 2183.0 | 4 | Protopolystoma xenopodis | 2183.0 | 4 |
| scf7180000048385 | 58328 | 0.6221 | 0 | 54.9854 | Eukaryota | 16701.0 | 0 | Chordata | 16701.0 | 0 | Carnivora | 16701.0 | 0 | Canidae | 16701.0 | 0 | Canis | 16701.0 | 0 | Canis lupus | 16701.0 | 0 |
| scf7180000044181 | 50450 | 0.4433 | 0 | 56.7064 | Eukaryota | 8352.0 | 0 | Chordata | 8352.0 | 0 | Carnivora | 8352.0 | 0 | Canidae | 7191.0 | 1 | Canis | 6070.0 | 2 | Canis lupus | 6070.0 | 3 |
| scf7180000055281 | 28174 | 0.4398 | 0 | 104.4788 | Eukaryota | 15146.0 | 0 | Chordata | 9491.0 | 2 | Gadiformes | 4471.0 | 6 | Gadidae | 4471.0 | 6 | Gadus | 4471.0 | 6 | Gadus morhua | 4471.0 | 6 |
| scf7180000043728 | 434922 | 0.367 | 0 | 56.9905 | Eukaryota | 50678.0 | 0 | Chordata | 50678.0 | 0 | Carnivora | 50678.0 | 0 | Canidae | 50678.0 | 0 | Canis | 50678.0 | 0 | Canis lupus | 50678.0 | 0 |
| scf7180000046561 | 111874 | 0.4617 | 0 | 58.8139 | Eukaryota | 18588.0 | 0 | Chordata | 18588.0 | 0 | Carnivora | 18588.0 | 0 | Canidae | 18588.0 | 0 | Canis | 18588.0 | 0 | Canis lupus | 18588.0 | 0 |
| scf7180000042690 | 287532 | 0.5015 | 0 | 58.0805 | Eukaryota | 43784.0 | 0 | Chordata | 43784.0 | 0 | Carnivora | 43784.0 | 0 | Canidae | 43784.0 | 0 | Canis | 43784.0 | 0 | Canis lupus | 43784.0 | 0 |
| scf7180000048418 | 6946 | 0.4411 | 0 | 121.6798 | Eukaryota | 1025.0 | 0 | Platyhelminthes | 559.0 | 1 | Polystomatidea | 559.0 | 2 | Polystomatidae | 559.0 | 2 | Protopolystoma | 559.0 | 2 | Protopolystoma xenopodis | 559.0 | 2 |
| scf7180000045274 | 37614 | 0.4423 | 0 | 216.627 | Eukaryota | 7380.0 | 0 | Chordata | 7380.0 | 0 | Carnivora | 3052.0 | 3 | Canidae | 3052.0 | 3 | Canis | 3052.0 | 3 | Canis lupus | 3052.0 | 3 |
| scf7180000054644 | 121933 | 0.4181 | 0 | 152.5449 | Eukaryota | 37067.0 | 0 | Chordata | 37067.0 | 0 | Carnivora | 37067.0 | 0 | Canidae | 37067.0 | 0 | Canis | 37067.0 | 0 | Canis lupus | 37067.0 | 0 |
| scf7180000041682 | 388134 | 0.3952 | 0 | 58.6438 | Eukaryota | 63134.0 | 0 | Chordata | 63134.0 | 0 | Carnivora | 63134.0 | 0 | Canidae | 63134.0 | 0 | Canis | 63134.0 | 0 | Canis lupus | 63134.0 | 0 |
| scf7180000043575 | 441928 | 0.3712 | 0 | 58.5321 | Eukaryota | 32010.0 | 0 | Chordata | 32010.0 | 0 | Carnivora | 32010.0 | 0 | Canidae | 32010.0 | 0 | Canis | 32010.0 | 0 | Canis lupus | 32010.0 | 0 |
| scf7180000045381 | 197084 | 0.3942 | 0 | 60.364 | Eukaryota | 37399.0 | 0 | Chordata | 37399.0 | 0 | Carnivora | 37399.0 | 0 | Canidae | 37399.0 | 0 | Canis | 37399.0 | 0 | Canis lupus | 37399.0 | 0 |
| scf7180000046276 | 117466 | 0.379 | 0 | 60.6029 | Eukaryota | 18986.0 | 0 | Chordata | 18986.0 | 0 | Carnivora | 18986.0 | 0 | Canidae | 6005.0 | 4 | Canis | 4010.0 | 8 | Canis lupus | 4010.0 | 8 |
| scf7180000049089 | 16615 | 0.4403 | 0 | 198.4172 | Eukaryota | 676.0 | 0 | Platyhelminthes | 344.0 | 1 | Polystomatidea | 344.0 | 2 | Polystomatidae | 344.0 | 2 | Protopolystoma | 344.0 | 2 | Protopolystoma xenopodis | 344.0 | 2 |
| scf7180000048414 | 61622 | 0.469 | 0 | 55.9185 | Eukaryota | 30650.0 | 0 | Chordata | 30650.0 | 0 | Carnivora | 30650.0 | 0 | Canidae | 30650.0 | 0 | Canis | 30650.0 | 0 | Canis lupus | 30650.0 | 0 |
| scf7180000044937 | 240781 | 0.514 | 0 | 56.9877 | Eukaryota | 49232.0 | 0 | Chordata | 49232.0 | 0 | Carnivora | 49232.0 | 0 | Canidae | 39242.0 | 1 | Canis | 29385.0 | 3 | Canis lupus | 29385.0 | 4 |
| scf7180000040989 | 1454440 | 0.4246 | 0 | 59.8934 | Eukaryota | 121842.0 | 0 | Chordata | 121842.0 | 0 | Carnivora | 121842.0 | 0 | Canidae | 121842.0 | 0 | Canis | 108889.0 | 1 | Canis lupus | 108889.0 | 1 |
| scf7180000056218 | 567237 | 0.4192 | 0 | 60.5642 | Eukaryota | 55453.0 | 0 | Chordata | 55453.0 | 0 | Carnivora | 55453.0 | 0 | Canidae | 36048.0 | 2 | Canis | 21475.0 | 3 | Canis lupus | 21475.0 | 3 |
| scf7180000041888 | 842252 | 0.3989 | 0 | 60.2122 | Eukaryota | 95524.0 | 0 | Chordata | 95524.0 | 0 | Carnivora | 95524.0 | 0 | Canidae | 95524.0 | 0 | Canis | 95524.0 | 0 | Canis lupus | 95524.0 | 0 |
| scf7180000048991 | 45072 | 0.6523 | 0 | 54.0504 | Eukaryota | 31259.0 | 0 | Chordata | 31259.0 | 0 | Carnivora | 31259.0 | 0 | Canidae | 27517.0 | 2 | Canis | 24842.0 | 3 | Canis lupus | 24842.0 | 3 |
| scf7180000042996 | 579998 | 0.3774 | 0 | 60.1314 | Eukaryota | 90080.0 | 0 | Chordata | 90080.0 | 0 | Carnivora | 90080.0 | 0 | Canidae | 90080.0 | 0 | Canis | 90080.0 | 0 | Canis lupus | 90080.0 | 0 |
| scf7180000056253 | 224375 | 0.4515 | 0 | 57.4045 | Eukaryota | 40296.0 | 0 | Chordata | 40296.0 | 0 | Carnivora | 37125.0 | 1 | Canidae | 29113.0 | 3 | Canis | 25179.0 | 4 | Canis lupus | 25179.0 | 4 |
| scf7180000048889 | 22808 | 0.4339 | 0 | 106.7033 | Eukaryota | 1719.0 | 0 | Platyhelminthes | 879.0 | 1 | Polystomatidea | 879.0 | 2 | Polystomatidae | 879.0 | 2 | Protopolystoma | 879.0 | 2 | Protopolystoma xenopodis | 879.0 | 2 |

| | | | | | | | | | | | | | | | | | | | | | | |
|---|---|---|---|---|---|---|---|---|---|---|---|---|---|---|---|---|---|---|---|---|---|---|
| scf7180000055576 | 24585 | 0.441 | 0 | 175.9395 | Eukaryota | 924.0 | 0 | Platyhelminthes | 924.0 | 0 | Polystomatidea | 924.0 | 0 | Polystomatidae | 924.0 | 0 | Protopolystoma | 924.0 | 0 | Protopolystoma xenopodis | 924.0 | 0 |
| scf7180000053462 | 4575 | 0.5143 | 0 | 65.8935 | Eukaryota | 30120.0 | 0 | Chordata | 30120.0 | 0 | Carnivora | 30120.0 | 0 | Canidae | 30120.0 | 0 | Canis | 30120.0 | 0 | Canis lupus | 30120.0 | 0 |
| scf7180000055784 | 9302 | 0.4366 | 0 | 449.8426 | Eukaryota | 1213.0 | 0 | Platyhelminthes | 1213.0 | 0 | Polystomatidea | 1213.0 | 0 | Polystomatidae | 1213.0 | 0 | Protopolystoma | 1213.0 | 0 | Protopolystoma xenopodis | 1213.0 | 0 |
| scf7180000044426 | 314491 | 0.3929 | 0 | 60.5825 | Eukaryota | 56715.0 | 0 | Chordata | 56715.0 | 0 | Carnivora | 56715.0 | 0 | Canidae | 56715.0 | 0 | Canis | 40491.0 | 1 | Canis lupus | 40491.0 | 1 |
| scf7180000042039 | 840592 | 0.376 | 0 | 60.1467 | Eukaryota | 88150.0 | 0 | Chordata | 88150.0 | 0 | Carnivora | 88150.0 | 0 | Canidae | 88150.0 | 0 | Canis | 88150.0 | 0 | Canis lupus | 88150.0 | 0 |
| scf7180000043499 | 81815 | 0.3793 | 0 | 56.816 | Eukaryota | 38324.0 | 0 | Chordata | 38324.0 | 0 | Carnivora | 38324.0 | 0 | Canidae | 31838.0 | 1 | Canis | 19847.0 | 2 | Canis lupus | 19847.0 | 2 |
| scf7180000056301 | 781219 | 0.3945 | 0 | 59.5061 | Eukaryota | 47556.0 | 0 | Chordata | 47556.0 | 0 | Carnivora | 47556.0 | 0 | Canidae | 39980.0 | 1 | Canis | 23204.0 | 2 | Canis lupus | 23204.0 | 2 |
| scf7180000041808 | 954035 | 0.3873 | 0 | 60.4086 | Eukaryota | 28992.0 | 0 | Chordata | 28992.0 | 0 | Carnivora | 28992.0 | 0 | Canidae | 28992.0 | 0 | Canis | 25461.0 | 1 | Canis lupus | 25461.0 | 1 |
| scf7180000046219 | 178076 | 0.5256 | 0 | 56.3943 | Eukaryota | 27328.0 | 0 | Chordata | 27328.0 | 0 | Carnivora | 27328.0 | 0 | Canidae | 27328.0 | 0 | Canis | 27328.0 | 0 | Canis lupus | 27328.0 | 0 |
| scf7180000056455 | 316675 | 0.4184 | 0 | 62.1668 | Eukaryota | 32058.0 | 0 | Chordata | 32058.0 | 0 | Carnivora | 32058.0 | 0 | Canidae | 32058.0 | 0 | Canis | 32058.0 | 0 | Canis lupus | 32058.0 | 0 |
| scf7180000040965 | 395158 | 0.3815 | 0 | 59.707 | Eukaryota | 44922.0 | 0 | Chordata | 44922.0 | 0 | Carnivora | 44922.0 | 0 | Canidae | 44922.0 | 0 | Canis | 44922.0 | 0 | Canis lupus | 44922.0 | 0 |
| scf7180000056400 | 258370 | 0.488 | 0 | 56.5597 | Eukaryota | 43878.0 | 0 | Chordata | 43878.0 | 0 | Carnivora | 43878.0 | 0 | Canidae | 33510.0 | 1 | Canis | 24582.0 | 3 | Canis lupus | 24582.0 | 3 |
| scf7180000048562 | 4194 | 0.432 | 0 | 102.6229 | no-hit | 0.0 | 0 | no-hit | 0.0 | 0 | no-hit | 0.0 | 0 | no-hit | 0.0 | 0 | no-hit | 0.0 | 0 | no-hit | 0.0 | 0 |
| scf7180000047316 | 88539 | 0.4109 | 0 | 62.6129 | Eukaryota | 77418.0 | 0 | Chordata | 77418.0 | 0 | Carnivora | 77418.0 | 0 | Canidae | 77418.0 | 0 | Canis | 77418.0 | 0 | Canis lupus | 77418.0 | 0 |
| scf7180000045021 | 4303 | 0.3667 | 0 | 56.3614 | Eukaryota | 3062.0 | 0 | Chordata | 3062.0 | 0 | Carnivora | 3062.0 | 0 | Canidae | 3062.0 | 0 | Canis | 3062.0 | 0 | Canis lupus | 3062.0 | 0 |
| scf7180000053225 | 3021 | 0.4479 | 0 | 433.6105 | no-hit | 0.0 | 0 | no-hit | 0.0 | 0 | no-hit | 0.0 | 0 | no-hit | 0.0 | 0 | no-hit | 0.0 | 0 | no-hit | 0.0 | 0 |
| scf7180000056484 | 372950 | 0.4071 | 0 | 58.9731 | Eukaryota | 53966.0 | 0 | Chordata | 53966.0 | 0 | Carnivora | 53966.0 | 0 | Canidae | 53966.0 | 0 | Canis | 48294.0 | 1 | Canis lupus | 48294.0 | 1 |
| scf7180000041091 | 368932 | 0.4178 | 0 | 57.2719 | Eukaryota | 60864.0 | 0 | Chordata | 60864.0 | 0 | Carnivora | 60864.0 | 0 | Canidae | 60864.0 | 0 | Canis | 48324.0 | 1 | Canis lupus | 48324.0 | 1 |
| scf7180000049466 | 5756 | 0.4375 | 0 | 124.1415 | no-hit | 0.0 | 0 | no-hit | 0.0 | 0 | no-hit | 0.0 | 0 | no-hit | 0.0 | 0 | no-hit | 0.0 | 0 | no-hit | 0.0 | 0 |
| scf7180000045410 | 205237 | 0.4017 | 0 | 61.0781 | Eukaryota | 93761.0 | 0 | Chordata | 93761.0 | 0 | Carnivora | 93761.0 | 0 | Canidae | 93761.0 | 0 | Canis | 93761.0 | 0 | Canis lupus | 93761.0 | 0 |
| scf7180000042156 | 773899 | 0.4229 | 0 | 59.0871 | Eukaryota | 58802.0 | 0 | Chordata | 58802.0 | 0 | Carnivora | 58802.0 | 0 | Canidae | 58802.0 | 0 | Canis | 52297.0 | 1 | Canis lupus | 52297.0 | 1 |
| scf7180000050610 | 25704 | 0.4388 | 0 | 62.7666 | Eukaryota | 14852.0 | 0 | Chordata | 9327.0 | 2 | Gadiformes | 4455.0 | 6 | Gadidae | 4455.0 | 6 | Gadus | 4455.0 | 6 | Gadus morhua | 4455.0 | 6 |
| scf7180000049624 | 16069 | 0.443 | 100 | 59.4204 | Eukaryota | 41617.0 | 0 | Chordata | 23658.0 | 2 | Gadiformes | 10737.0 | 7 | Gadidae | 10737.0 | 7 | Gadus | 10737.0 | 7 | Gadus morhua | 10737.0 | 7 |
| scf7180000046417 | 22610 | 0.4364 | 0 | 143.5918 | Eukaryota | 32985.0 | 0 | Chordata | 32985.0 | 0 | Carnivora | 32985.0 | 0 | Canidae | 32985.0 | 0 | Canis | 32985.0 | 0 | Canis lupus | 32985.0 | 0 |
| scf7180000045474 | 228120 | 0.3783 | 0 | 59.4645 | Eukaryota | 90686.0 | 0 | Chordata | 90686.0 | 0 | Carnivora | 90686.0 | 0 | Canidae | 90686.0 | 0 | Canis | 90686.0 | 0 | Canis lupus | 90686.0 | 0 |
| scf7180000044414 | 17497 | 0.4434 | 0 | 91.3447 | Eukaryota | 16343.0 | 0 | Chordata | 13090.0 | 1 | Tetraodontiformes | 4608.0 | 5 | Tetraodontidae | 4608.0 | 5 | Takifugu | 4608.0 | 5 | Takifugu rubripes | 4608.0 | 5 |
| scf7180000044447 | 174048 | 0.3639 | 0 | 60.558 | Eukaryota | 44861.0 | 0 | Chordata | 44861.0 | 0 | Carnivora | 44861.0 | 0 | Canidae | 44861.0 | 0 | Canis | 44861.0 | 0 | Canis lupus | 44861.0 | 0 |
| scf7180000048083 | 16474 | 0.4434 | 0 | 57.4173 | Eukaryota | 4704.0 | 0 | Chordata | 4704.0 | 0 | Tetraodontiformes | 4704.0 | 0 | Tetraodontidae | 4704.0 | 0 | Takifugu | 4704.0 | 0 | Takifugu rubripes | 4704.0 | 0 |
| scf7180000049293 | 51790 | 0.3625 | 0 | 59.9346 | Eukaryota | 29830.0 | 0 | Chordata | 29830.0 | 0 | Carnivora | 29830.0 | 0 | Canidae | 19775.0 | 3 | Canis | 15533.0 | 4 | Canis lupus | 15533.0 | 5 |
| scf7180000047422 | 86132 | 0.4201 | 0 | 60.6542 | Eukaryota | 29765.0 | 0 | Chordata | 29765.0 | 0 | Carnivora | 29765.0 | 0 | Canidae | 29765.0 | 0 | Canis | 29765.0 | 0 | Canis lupus | 29765.0 | 0 |
| scf7180000043593 | 162394 | 0.3896 | 0 | 58.7804 | Eukaryota | 32755.0 | 0 | Chordata | 32755.0 | 0 | Carnivora | 32755.0 | 0 | Canidae | 16453.0 | 1 | Ursus | 16302.0 | 2 | Ursus arctos | 16302.0 | 2 |
| scf7180000046349 | 16411 | 0.4427 | 0 | 283.5981 | no-hit | 0.0 | 0 | no-hit | 0.0 | 0 | no-hit | 0.0 | 0 | no-hit | 0.0 | 0 | no-hit | 0.0 | 0 | no-hit | 0.0 | 0 |
| scf7180000045411 | 125265 | 0.4364 | 0 | 58.0648 | Eukaryota | 14235.0 | 0 | Chordata | 14235.0 | 0 | Carnivora | 14235.0 | 0 | Canidae | 14235.0 | 0 | Canis | 11111.0 | 1 | Canis lupus | 11111.0 | 1 |
| scf7180000056249 | 1090297 | 0.4618 | 0 | 59.3356 | Eukaryota | 88030.0 | 0 | Chordata | 88030.0 | 0 | Carnivora | 88030.0 | 0 | Canidae | 88030.0 | 0 | Canis | 88030.0 | 0 | Canis lupus | 88030.0 | 0 |
| scf7180000054515 | 105040 | 0.4445 | 0 | 166.2395 | Eukaryota | 46408.0 | 0 | Chordata | 37065.0 | 1 | Carnivora | 37065.0 | 1 | Canidae | 37065.0 | 1 | Canis | 37065.0 | 1 | Canis lupus | 37065.0 | 1 |
| scf7180000055838 | 9751 | 0.4439 | 0 | 112.9897 | Eukaryota | 3152.0 | 0 | Chordata | 3152.0 | 0 | Carnivora | 3152.0 | 0 | Canidae | 3152.0 | 0 | Canis | 3152.0 | 0 | Canis lupus | 3152.0 | 0 |
| scf7180000048469 | 3051 | 0.3789 | 0 | 28.6572 | Eukaryota | 2592.0 | 0 | Chordata | 2592.0 | 0 | Primates | 1902.0 | 1 | Hominidae | 1902.0 | 1 | Pan | 1575.0 | 2 | Pan troglodytes | 1575.0 | 2 |
| scf7180000043144 | 108370 | 0.4284 | 0 | 55.9972 | Eukaryota | 52143.0 | 0 | Chordata | 52143.0 | 0 | Carnivora | 52143.0 | 0 | Canidae | 52143.0 | 0 | Canis | 52143.0 | 0 | Canis lupus | 52143.0 | 0 |
| scf7180000054160 | 76004 | 0.4407 | 0 | 389.4129 | Eukaryota | 93528.0 | 0 | Chordata | 93528.0 | 0 | Carnivora | 93528.0 | 0 | Canidae | 93528.0 | 0 | Canis | 93528.0 | 0 | Canis lupus | 93528.0 | 0 |
| scf7180000052587 | 4434 | 0.4477 | 0 | 1188.7632 | no-hit | 0.0 | 0 | no-hit | 0.0 | 0 | no-hit | 0.0 | 0 | no-hit | 0.0 | 0 | no-hit | 0.0 | 0 | no-hit | 0.0 | 0 |
| scf7180000044885 | 27374 | 0.3779 | 0 | 45.5339 | Eukaryota | 7721.0 | 0 | Chordata | 7721.0 | 0 | Primates | 4887.0 | 2 | Hominidae | 4887.0 | 2 | Homo | 3274.0 | 3 | Homo sapiens | 3274.0 | 3 |
| scf7180000049968 | 9896 | 0.4434 | 0 | 92.0992 | no-hit | 0.0 | 0 | no-hit | 0.0 | 0 | no-hit | 0.0 | 0 | no-hit | 0.0 | 0 | no-hit | 0.0 | 0 | no-hit | 0.0 | 0 |
| scf7180000048601 | 5485 | 0.4397 | 0 | 100.2681 | Eukaryota | 5301.0 | 0 | Chordata | 4447.0 | 2 | Kurtiformes | 2609.0 | 4 | Apogonidae | 2609.0 | 4 | Sphaeramia | 2609.0 | 4 | Sphaeramia orbicularis | 2609.0 | 4 |
| scf7180000048079 | 49308 | 0.3558 | 0 | 58.7032 | Eukaryota | 10354.0 | 0 | Chordata | 10354.0 | 0 | Carnivora | 5885.0 | 2 | Felidae | 5184.0 | 3 | Felis | 4450.0 | 5 | Felis catus | 4450.0 | 5 |
| scf7180000046092 | 112779 | 0.5016 | 0 | 55.7254 | Eukaryota | 16166.0 | 0 | Chordata | 16166.0 | 0 | Carnivora | 16166.0 | 0 | Canidae | 16166.0 | 0 | Canis | 16166.0 | 0 | Canis lupus | 16166.0 | 0 |
| scf7180000041780 | 541541 | 0.3964 | 0 | 61.5406 | Eukaryota | 84747.0 | 0 | Chordata | 84747.0 | 0 | Carnivora | 84747.0 | 0 | Canidae | 84747.0 | 0 | Canis | 84747.0 | 0 | Canis lupus | 84747.0 | 0 |
| scf7180000056467 | 349046 | 0.406 | 0 | 60.3734 | Eukaryota | 55673.0 | 0 | Chordata | 55673.0 | 0 | Carnivora | 55673.0 | 0 | Canidae | 55673.0 | 0 | Canis | 55673.0 | 0 | Canis lupus | 55673.0 | 0 |
| scf7180000041997 | 116999 | 0.3498 | 0 | 56.8792 | Eukaryota | 30881.0 | 0 | Chordata | 30881.0 | 0 | Carnivora | 30881.0 | 0 | Canidae | 30881.0 | 0 | Canis | 30881.0 | 0 | Canis lupus | 30881.0 | 0 |
| scf7180000056162 | 4731 | 0.446 | 0 | 158.8479 | no-hit | 0.0 | 0 | no-hit | 0.0 | 0 | no-hit | 0.0 | 0 | no-hit | 0.0 | 0 | no-hit | 0.0 | 0 | no-hit | 0.0 | 0 |
| scf7180000042874 | 711047 | 0.4969 | 0 | 58.2766 | Eukaryota | 70175.0 | 0 | Chordata | 70175.0 | 0 | Carnivora | 70175.0 | 0 | Canidae | 70175.0 | 0 | Canis | 55201.0 | 1 | Canis lupus | 55201.0 | 1 |
| scf7180000046980 | 20579 | 0.4472 | 0 | 52.2358 | Eukaryota | 73144.0 | 0 | Chordata | 62254.0 | 1 | Carnivora | 44573.0 | 2 | Canidae | 44573.0 | 2 | Canis | 44573.0 | 2 | Canis lupus | 44573.0 | 2 |
| scf7180000051930 | 10649 | 0.5066 | 0 | 147.4951 | Eukaryota | 8396.0 | 0 | Chordata | 8396.0 | 0 | Carnivora | 8396.0 | 0 | Canidae | 8396.0 | 0 | Canis | 7272.0 | 1 | Canis lupus | 6135.0 | 2 |
| scf7180000054218 | 29523 | 0.3869 | 0 | 601.7282 | Eukaryota | 39015.0 | 0 | Chordata | 39015.0 | 0 | Carnivora | 39015.0 | 0 | Canidae | 39015.0 | 0 | Canis | 39015.0 | 0 | Canis lupus | 39015.0 | 0 |
| scf7180000042016 | 290388 | 0.3764 | 0 | 61.1641 | Eukaryota | 32823.0 | 0 | Chordata | 32823.0 | 0 | Carnivora | 32823.0 | 0 | Canidae | 32823.0 | 0 | Canis | 32823.0 | 0 | Canis lupus | 32823.0 | 0 |
| scf7180000046522 | 11081 | 0.4447 | 0 | 82.5084 | Eukaryota | 3170.0 | 0 | Chordata | 3170.0 | 0 | Carnivora | 3170.0 | 0 | Canidae | 3170.0 | 0 | Canis | 3170.0 | 0 | Canis lupus | 3170.0 | 0 |
| scf7180000042044 | 196903 | 0.3832 | 0 | 58.1372 | Eukaryota | 77180.0 | 0 | Chordata | 77180.0 | 0 | Carnivora | 77180.0 | 0 | Canidae | 77180.0 | 0 | Canis | 77180.0 | 0 | Canis lupus | 77180.0 | 0 |
| scf7180000045901 | 9100 | 0.3836 | 0 | 26.222 | Eukaryota | 66792.0 | 0 | Chordata | 66792.0 | 0 | Carnivora | 66792.0 | 0 | Canidae | 66792.0 | 0 | Canis | 66792.0 | 0 | Canis lupus | 66792.0 | 0 |
| scf7180000051032 | 9115 | 0.3954 | 0 | 32.645 | Eukaryota | 1964.0 | 0 | Chordata | 1964.0 | 0 | Carnivora | 1964.0 | 0 | Canidae | 1964.0 | 0 | Canis | 1964.0 | 0 | Canis lupus | 1964.0 | 0 |
| scf7180000044827 | 149507 | 0.3881 | 0 | 57.9693 | Eukaryota | 57389.0 | 0 | Chordata | 57389.0 | 0 | Carnivora | 57389.0 | 0 | Canidae | 57389.0 | 0 | Canis | 57389.0 | 0 | Canis lupus | 57389.0 | 0 |
| scf7180000043932 | 394982 | 0.394 | 0 | 59.3775 | Eukaryota | 80718.0 | 0 | Chordata | 80718.0 | 0 | Carnivora | 80718.0 | 0 | Canidae | 80718.0 | 0 | Canis | 63606.0 | 1 | Canis lupus | 63606.0 | 1 |
| scf7180000045789 | 227983 | 0.3967 | 0 | 72.2866 | Eukaryota | 43652.0 | 0 | Chordata | 43652.0 | 0 | Carnivora | 43652.0 | 0 | Canidae | 43652.0 | 0 | Canis | 43652.0 | 0 | Canis lupus | 43652.0 | 0 |
| scf7180000041397 | 155441 | 0.4832 | 0 | 90.2722 | Eukaryota | 89641.0 | 0 | Chordata | 89641.0 | 0 | Carnivora | 89641.0 | 0 | Canidae | 89641.0 | 0 | Canis | 89641.0 | 0 | Canis lupus | 89641.0 | 0 |
| scf7180000050800 | 5012 | 0.4366 | 0 | 92.4158 | Eukaryota | 279.0 | 0 | Platyhelminthes | 279.0 | 0 | Polystomatidea | 279.0 | 0 | Polystomatidae | 279.0 | 0 | Protopolystoma | 279.0 | 0 | Protopolystoma xenopodis | 279.0 | 0 |
| scf7180000055625 | 17925 | 0.4412 | 0 | 313.256 | Eukaryota | 11332.0 | 0 | Platyhelminthes | 10934.0 | 1 | Cyclophyllidea | 9588.0 | 2 | Hymenolepididae | 9588.0 | 2 | Hymenolepis | 9588.0 | 2 | Hymenolepis microstoma | 9588.0 | 2 |
| scf7180000042339 | 806826 | 0.3952 | 0 | 60.541 | Eukaryota | 109820.0 | 0 | Chordata | 109820.0 | 0 | Carnivora | 109820.0 | 0 | Canidae | 109820.0 | 0 | Canis | 95212.0 | 1 | Canis lupus | 95212.0 | 1 |
| scf7180000048829 | 55181 | 0.4572 | 0 | 57.6709 | Eukaryota | 33976.0 | 0 | Chordata | 33976.0 | 0 | Carnivora | 33976.0 | 0 | Canidae | 33976.0 | 0 | Canis | 33976.0 | 0 | Canis lupus | 33976.0 | 0 |
| scf7180000041621 | 334165 | 0.543 | 0 | 53.0302 | Eukaryota | 33031.0 | 0 | Chordata | 33031.0 | 0 | Carnivora | 33031.0 | 0 | Canidae | 33031.0 | 0 | Canis | 30480.0 | 1 | Canis lupus | 30480.0 | 1 |
| scf7180000047763 | 60029 | 0.5917 | 0 | 53.4815 | Eukaryota | 25758.0 | 1 | Chordata | 25758.0 | 1 | Rodentia | 6812.0 | 5 | Muridae | 6812.0 | 5 | Mus | 6812.0 | 6 | Mus musculus | 6812.0 | 6 |
| scf7180000052781 | 4107 | 0.4336 | 0 | 125.4204 | Eukaryota | 1823.0 | 0 | Chordata | 1823.0 | 0 | Kurtiformes | 1349.0 | 2 | Apogonidae | 1349.0 | 2 | Sphaeramia | 1349.0 | 2 | Sphaeramia orbicularis | 1349.0 | 2 |
| scf7180000043686 | 164532 | 0.3908 | 0 | 62.2045 | Eukaryota | 56459.0 | 0 | Chordata | 56459.0 | 0 | Carnivora | 56459.0 | 0 | Canidae | 56459.0 | 0 | Canis | 56459.0 | 0 | Canis lupus | 56459.0 | 0 |
| scf7180000044602 | 303859 | 0.3453 | 0 | 58.4161 | Eukaryota | 87465.0 | 0 | Chordata | 87465.0 | 0 | Carnivora | 87465.0 | 0 | Canidae | 87465.0 | 0 | Canis | 87465.0 | 0 | Canis lupus | 87465.0 | 0 |
| scf7180000049535 | 5327 | 0.4564 | 0 | 57.3974 | no-hit | 0.0 | 0 | no-hit | 0.0 | 0 | no-hit | 0.0 | 0 | no-hit | 0.0 | 0 | no-hit | 0.0 | 0 | no-hit | 0.0 | 0 |
| scf7180000047147 | 95896 | 0.5013 | 0 | 55.6331 | Eukaryota | 20675.0 | 0 | Chordata | 20675.0 | 0 | Carnivora | 20675.0 | 0 | Canidae | 20675.0 | 0 | Canis | 20675.0 | 0 | Canis lupus | 20675.0 | 0 |
| scf7180000046789 | 13398 | 0.4385 | 0 | 129.8662 | Eukaryota | 51253.0 | 0 | Chordata | 36981.0 | 2 | Gadiformes | 31730.0 | 5 | Gadidae | 31730.0 | 5 | Gadus | 31730.0 | 5 | Gadus morhua | 31730.0 | 5 |
| scf7180000043027 | 607736 | 0.3585 | 0 | 61.2223 | Eukaryota | 45274.0 | 0 | Chordata | 45274.0 | 0 | Carnivora | 45274.0 | 0 | Canidae | 45274.0 | 0 | Canis | 45274.0 | 0 | Canis lupus | 45274.0 | 0 |
| scf7180000051142 | 7580 | 0.4497 | 0 | 2187.1701 | no-hit | 0.0 | 0 | no-hit | 0.0 | 0 | no-hit | 0.0 | 0 | no-hit | 0.0 | 0 | no-hit | 0.0 | 0 | no-hit | 0.0 | 0 |
| scf7180000049379 | 12314 | 0.4199 | 0 | 278.7507 | Eukaryota | 66944.0 | 0 | Chordata | 66944.0 | 0 | Carnivora | 66944.0 | 0 | Canidae | 66944.0 | 0 | Canis | 66944.0 | 0 | Canis lupus | 66944.0 | 0 |
| scf7180000055254 | 14653 | 0.4913 | 0 | 259.4277 | Eukaryota | 37921.0 | 0 | Chordata | 37921.0 | 0 | Carnivora | 37921.0 | 0 | Canidae | 37921.0 | 0 | Canis | 37921.0 | 0 | Canis lupus | 37921.0 | 0 |
| scf7180000046423 | 31771 | 0.439 | 0 | 126.3045 | Eukaryota | 26394.0 | 1 | Chordata | 26394.0 | 1 | Gadiformes | 10981.0 | 6 | Gadidae | 10981.0 | 6 | Gadus | 10981.0 | 6 | Gadus morhua | 10981.0 | 6 |
| scf7180000044686 | 341713 | 0.3492 | 0 | 59.0757 | Eukaryota | 25849.0 | 0 | Chordata | 25849.0 | 0 | Carnivora | 25849.0 | 0 | Canidae | 25849.0 | 0 | Canis | 25849.0 | 0 | Canis lupus | 25849.0 | 0 |
| scf7180000040915 | 1925740 | 0.3499 | 100 | 59.4826 | Eukaryota | 92500.0 | 0 | Chordata | 92500.0 | 0 | Carnivora | 92500.0 | 0 | Canidae | 92500.0 | 0 | Canis | 55583.0 | 1 | Canis lupus | 55583.0 | 1 |
| scf7180000055013 | 59947 | 0.4392 | 0 | 182.7108 | Eukaryota | 5432.0 | 0 | Chordata | 3888.0 | 1 | Tetraodontiformes | 2736.0 | 3 | Tetraodontidae | 2736.0 | 3 | Takifugu | 2736.0 | 3 | Takifugu rubripes | 2736.0 | 3 |
| scf7180000042840 | 608418 | 0.4518 | 0 | 58.6153 | Eukaryota | 53321.0 | 0 | Chordata | 53321.0 | 0 | Carnivora | 53321.0 | 0 | Canidae | 53321.0 | 0 | Canis | 53321.0 | 0 | Canis lupus | 53321.0 | 0 |
| scf7180000044786 | 175148 | 0.5014 | 0 | 61.6329 | Eukaryota | 89614.0 | 0 | Chordata | 89614.0 | 0 | Carnivora | 89614.0 | 0 | Canidae | 89614.0 | 0 | Canis | 89614.0 | 0 | Canis lupus | 89614.0 | 0 |
| scf7180000041340 | 102586 | 0.365 | 0 | 53.669 | Eukaryota | 88736.0 | 0 | Chordata | 88736.0 | 0 | Carnivora | 88736.0 | 0 | Canidae | 88736.0 | 0 | Canis | 88736.0 | 0 | Canis lupus | 88736.0 | 0 |
| scf7180000050223 | 41402 | 0.4012 | 0 | 58.7725 | Eukaryota | 6918.0 | 0 | Chordata | 6918.0 | 0 | Perissodactyla | 4445.0 | 2 | Equidae | 3392.0 | 3 | Equus | 3392.0 | 3 | Equus przewalskii | 2317.0 | 4 |
| scf7180000043563 | 83130 | 0.5042 | 0 | 68.5385 | Eukaryota | 16877.0 | 0 | Chordata | 16877.0 | 0 | Carnivora | 16877.0 | 0 | Canidae | 13262.0 | 2 | Vulpes | 6999.0 | 4 | Vulpes vulpes | 6999.0 | 4 |
| scf7180000045705 | 32419 | 0.3617 | 0 | 92.5079 | Eukaryota | 32816.0 | 0 | Chordata | 32816.0 | 0 | Carnivora | 32816.0 | 0 | Canidae | 32816.0 | 0 | Canis | 32816.0 | 0 | Canis lupus | 32816.0 | 0 |
| scf7180000044421 | 368147 | 0.3394 | 0 | 58.5094 | Eukaryota | 30314.0 | 0 | Chordata | 30314.0 | 0 | Carnivora | 30314.0 | 0 | Canidae | 30314.0 | 0 | Canis | 15177.0 | 1 | Canis lupus | 15177.0 | 1 |
| scf7180000052400 | 23515 | 0.4364 | 0 | 145.3248 | Eukaryota | 4813.0 | 0 | Chordata | 3098.0 | 4 | Columbiformes | 2192.0 | 7 | Columbidae | 2192.0 | 7 | Streptopelia | 2192.0 | 8 | Streptopelia turtur | 2192.0 | 8 |
| scf7180000048131 | 24276 | 0.4354 | 0 | 110.5037 | Eukaryota | 19170.0 | 0 | Chordata | 17133.0 | 2 | Gadiformes | 9782.0 | 7 | Gadidae | 9782.0 | 7 | Gadus | 9782.0 | 7 | Gadus morhua | 9782.0 | 7 |
| scf7180000040626 | 2158011 | 0.3853 | 0 | 60.1413 | Eukaryota | 94985.0 | 0 | Chordata | 94985.0 | 0 | Carnivora | 94985.0 | 0 | Canidae | 94985.0 | 0 | Canis | 94985.0 | 0 | Canis lupus | 94985.0 | 0 |
| scf7180000046951 | 73267 | 0.3636 | 0 | 64.322 | Eukaryota | 86746.0 | 0 | Chordata | 86746.0 | 0 | Carnivora | 86746.0 | 0 | Canidae | 86746.0 | 0 | Canis | 86746.0 | 0 | Canis lupus | 86746.0 | 0 |
| scf7180000042199 | 754370 | 0.3958 | 0 | 60.3941 | Eukaryota | 74011.0 | 0 | Chordata | 74011.0 | 0 | Carnivora | 74011.0 | 0 | Canidae | 74011.0 | 0 | Canis | 74011.0 | 0 | Canis lupus | 74011.0 | 0 |
| scf7180000046736 | 76974 | 0.3709 | 0 | 60.3301 | Eukaryota | 20898.0 | 0 | Chordata | 20898.0 | 0 | Carnivora | 20898.0 | 0 | Canidae | 20898.0 | 0 | Canis | 20898.0 | 0 | Canis lupus | 20898.0 | 0 |
| scf7180000051390 | 4213 | 0.4498 | 0 | 397.3991 | no-hit | 0.0 | 0 | no-hit | 0.0 | 0 | no-hit | 0.0 | 0 | no-hit | 0.0 | 0 | no-hit | 0.0 | 0 | no-hit | 0.0 | 0 |
| scf7180000045807 | 3815 | 0.5151 | 0 | 37.3445 | Eukaryota | 23815.0 | 0 | Chordata | 23815.0 | 0 | Carnivora | 23815.0 | 0 | Canidae | 23815.0 | 0 | Canis | 21624.0 | 1 | Canis lupus | 21624.0 | 1 |
| scf7180000052572 | 3985 | 0.4487 | 0 | 373.0227 | Eukaryota | 165.0 | 0 | Streptophyta | 165.0 | 0 | Fabales | 165.0 | 0 | Fabaceae | 165.0 | 0 | Cicer | 165.0 | 0 | Cicer arietinum | 165.0 | 0 |
| scf7180000040758 | 41685 | 0.3809 | 0 | 56.9471 | Eukaryota | 4182.0 | 0 | Chordata | 4182.0 | 0 | Carnivora | 2345.0 | 2 | Canidae | 1317.0 | 4 | Canis | 1317.0 | 4 | Canis lupus | 1317.0 | 4 |
| scf7180000045515 | 166110 | 0.4164 | 0 | 58.0317 | Eukaryota | 56191.0 | 0 | Chordata | 56191.0 | 0 | Carnivora | 56191.0 | 0 | Canidae | 44661.0 | 4 | Canis | 35282.0 | 5 | Canis lupus | 35282.0 | 5 |
| scf7180000052768 | 14740 | 0.4218 | 0 | 3446.6351 | Eukaryota | 103801.0 | 0 | Chordata | 103801.0 | 0 | Carnivora | 103801.0 | 0 | Canidae | 103801.0 | 0 | Canis | 87659.0 | 1 | Canis latrans | 58527.0 | 2 |
| scf7180000048234 | 54684 | 0.3729 | 0 | 57.417 | Eukaryota | 26692.0 | 0 | Chordata | 26692.0 | 0 | Carnivora | 26692.0 | 0 | Canidae | 26692.0 | 0 | Canis | 26692.0 | 0 | Canis lupus | 26692.0 | 0 |
| scf7180000046109 | 150551 | 0.4226 | 0 | 60.3086 | Eukaryota | 14816.0 | 0 | Chordata | 14816.0 | 0 | Carnivora | 14816.0 | 0 | Canidae | 14816.0 | 0 | Canis | 14816.0 | 0 | Canis lupus | 14816.0 | 0 |
| scf7180000048568 | 11745 | 0.4393 | 0 | 86.1079 | no-hit | 0.0 | 0 | no-hit | 0.0 | 0 | no-hit | 0.0 | 0 | no-hit | 0.0 | 0 | no-hit | 0.0 | 0 | no-hit | 0.0 | 0 |
| scf7180000048915 | 27063 | 0.4429 | 0 | 41.3841 | Eukaryota | 9877.0 | 0 | Chordata | 8115.0 | 1 | Salmoniformes | 4031.0 | 3 | Salmonidae | 4031.0 | 3 | Salmo | 4031.0 | 3 | Salmo trutta | 4031.0 | 3 |
| scf7180000056474 | 207444 | 0.423 | 0 | 63.0757 | Eukaryota | 94646.0 | 0 | Chordata | 94646.0 | 0 | Carnivora | 94646.0 | 0 | Canidae | 94646.0 | 0 | Canis | 94646.0 | 0 | Canis lupus | 94646.0 | 0 |
| scf7180000043808 | 228545 | 0.371 | 0 | 58.6195 | Eukaryota | 39152.0 | 0 | Chordata | 39152.0 | 0 | Carnivora | 39152.0 | 0 | Canidae | 18522.0 | 4 | Vulpes | 9378.0 | 7 | Vulpes vulpes | 9378.0 | 7 |
| scf7180000049841 | 5467 | 0.4458 | 0 | 42.981 | Eukaryota | 2770.0 | 0 | Chordata | 2770.0 | 0 | Carnivora | 2770.0 | 0 | Canidae | 2770.0 | 0 | Canis | 2770.0 | 0 | Canis lupus | 2770.0 | 0 |
| scf7180000042838 | 9736 | 0.4908 | 0 | 57.2234 | Eukaryota | 2652.0 | 0 | Chordata | 2652.0 | 0 | Carnivora | 2652.0 | 0 | Canidae | 2652.0 | 0 | Canis | 2319.0 | 1 | Canis lupus | 2319.0 | 1 |
| scf7180000055377 | 42184 | 0.4413 | 0 | 120.927 | Eukaryota | 3095.0 | 0 | Chordata | 2771.0 | 1 | Carnivora | 2771.0 | 1 | Canidae | 2771.0 | 1 | Canis | 2771.0 | 1 | Canis lupus | 2771.0 | 1 |
| scf7180000046793 | 100642 | 0.589 | 0 | 57.5705 | Eukaryota | 53211.0 | 0 | Chordata | 53211.0 | 0 | Carnivora | 53211.0 | 0 | Canidae | 53211.0 | 0 | Canis | 37313.0 | 1 | Canis lupus | 37313.0 | 1 |
| scf7180000044607 | 8691 | 0.465 | 0 | 56.8881 | Eukaryota | 9200.0 | 0 | Chordata | 9200.0 | 0 | Carnivora | 9200.0 | 0 | Canidae | 9200.0 | 0 | Canis | 9200.0 | 0 | Canis lupus | 9200.0 | 0 |
| scf7180000043421 | 397343 | 0.4385 | 0 | 61.1083 | Eukaryota | 54976.0 | 0 | Chordata | 54976.0 | 0 | Carnivora | 54976.0 | 0 | Canidae | 54976.0 | 0 | Vulpes | 29818.0 | 1 | Vulpes vulpes | 29818.0 | 1 |
| scf7180000045022 | 266804 | 0.3789 | 0 | 59.7468 | Eukaryota | 21957.0 | 0 | Chordata | 21957.0 | 0 | Carnivora | 21957.0 | 0 | Canidae | 21957.0 | 0 | Canis | 21957.0 | 0 | Canis lupus | 21957.0 | 0 |
| scf7180000048841 | 80595 | 0.4045 | 0 | 58.2734 | Eukaryota | 8000.0 | 0 | Chordata | 8000.0 | 0 | Apterygiformes | 7172.0 | 1 | Apterygidae | 7172.0 | 1 | Apteryx | 7172.0 | 1 | Apteryx mantelli | 7172.0 | 1 |
| scf7180000052557 | 5035 | 0.4564 | 0 | 1083.6387 | Eukaryota | 3545.0 | 0 | Platyhelminthes | 1977.0 | 2 | Polystomatidea | 1977.0 | 2 | Polystomatidae | 1977.0 | 2 | Protopolystoma | 1977.0 | 2 | Protopolystoma xenopodis | 1977.0 | 2 |
| scf7180000044787 | 288035 | 0.3888 | 0 | 59.991 | Eukaryota | 32579.0 | 0 | Chordata | 32579.0 | 0 | Carnivora | 32579.0 | 0 | Canidae | 32579.0 | 0 | Canis | 27655.0 | 1 | Canis lupus | 27655.0 | 1 |
| scf7180000041735 | 900703 | 0.4398 | 0 | 59.1967 | Eukaryota | 122448.0 | 0 | Chordata | 122448.0 | 0 | Carnivora | 122448.0 | 0 | Canidae | 122448.0 | 0 | Vulpes | 83172.0 | 1 | Vulpes vulpes | 83172.0 | 1 |
| scf7180000045694 | 198431 | 0.3971 | 0 | 61.6675 | Eukaryota | 31216.0 | 0 | Chordata | 31216.0 | 0 | Carnivora | 31216.0 | 0 | Canidae | 31216.0 | 0 | Canis | 15982.0 | 1 | Canis lupus | 15982.0 | 1 |
| scf7180000053626 | 3414 | 0.4549 | 0 | 501.0378 | Eukaryota | 13724.0 | 0 | Chordata | 10176.0 | 2 | Synbranchiformes | 2938.0 | 6 | Mastacembelidae | 2938.0 | 6 | Mastacembelus | 2938.0 | 7 | Mastacembelus armatus | 2938.0 | 8 |
| scf7180000044537 | 289032 | 0.3765 | 0 | 58.6299 | Eukaryota | 90576.0 | 0 | Chordata | 90576.0 | 0 | Carnivora | 90576.0 | 0 | Canidae | 90576.0 | 0 | Canis | 90576.0 | 0 | Canis lupus | 90576.0 | 0 |
| scf7180000045916 | 9830 | 0.3642 | 0 | 50.9715 | Eukaryota | 3229.0 | 0 | Chordata | 3229.0 | 0 | Carnivora | 2492.0 | 1 | Canidae | 2492.0 | 1 | Canis | 2492.0 | 1 | Canis lupus | 2492.0 | 1 |
| scf7180000043374 | 430117 | 0.3433 | 0 | 58.9984 | Eukaryota | 86018.0 | 0 | Chordata | 86018.0 | 0 | Carnivora | 86018.0 | 0 | Canidae | 86018.0 | 0 | Canis | 86018.0 | 0 | Canis lupus | 86018.0 | 0 |
| scf7180000040641 | 104020 | 0.4233 | 0 | 60.5566 | Eukaryota | 54332.0 | 0 | Chordata | 54332.0 | 0 | Carnivora | 54332.0 | 0 | Felidae | 28742.0 | 1 | Canis | 12832.0 | 4 | Canis lupus | 12832.0 | 4 |
| scf7180000050066 | 3841 | 0.4514 | 0 | 254.3578 | no-hit | 0.0 | 0 | no-hit | 0.0 | 0 | no-hit | 0.0 | 0 | no-hit | 0.0 | 0 | no-hit | 0.0 | 0 | no-hit | 0.0 | 0 |
| scf7180000045453 | 191766 | 0.4414 | 0 | 61.7062 | Eukaryota | 112552.0 | 0 | Chordata | 112552.0 | 0 | Carnivora | 112552.0 | 0 | Canidae | 78954.0 | 3 | Canis | 72119.0 | 6 | Canis lupus | 72119.0 | 7 |
| scf7180000050526 | 12847 | 0.4449 | 0 | 266.5882 | Eukaryota | 278.0 | 0 | Chordata | 278.0 | 0 | Kurtiformes | 278.0 | 0 | Apogonidae | 278.0 | 0 | Sphaeramia | 278.0 | 0 | Sphaeramia orbicularis | 278.0 | 0 |
| scf7180000044652 | 212579 | 0.349 | 0 | 59.0898 | Eukaryota | 74751.0 | 0 | Chordata | 74751.0 | 0 | Carnivora | 74751.0 | 0 | Canidae | 30156.0 | 2 | Vulpes | 30156.0 | 2 | Vulpes vulpes | 30156.0 | 2 |
| scf7180000043861 | 177329 | 0.3579 | 0 | 59.3011 | Eukaryota | 52711.0 | 0 | Chordata | 52711.0 | 0 | Carnivora | 52711.0 | 0 | Canidae | 52711.0 | 0 | Canis | 52711.0 | 0 | Canis lupus | 52711.0 | 0 |
| scf7180000055146 | 35214 | 0.4461 | 0 | 167.6889 | Eukaryota | 4132.0 | 0 | Chordata | 3691.0 | 1 | Columbiformes | 2357.0 | 5 | Columbidae | 2357.0 | 5 | Streptopelia | 2357.0 | 5 | Streptopelia turtur | 2357.0 | 5 |
| scf7180000042064 | 475061 | 0.4478 | 0 | 59.7209 | Eukaryota | 52177.0 | 1 | Chordata | 52177.0 | 1 | Carnivora | 47979.0 | 2 | Canidae | 47979.0 | 2 | Canis | 34831.0 | 3 | Canis lupus | 34831.0 | 3 |
| scf7180000044195 | 89643 | 0.6084 | 0 | 53.801 | Eukaryota | 10484.0 | 0 | Chordata | 10484.0 | 0 | Carnivora | 7114.0 | 3 | Canidae | 7114.0 | 4 | Canis | 4602.0 | 5 | Canis lupus | 4602.0 | 5 |
| scf7180000054168 | 1341089 | 0.3604 | 0 | 60.879 | Eukaryota | 90958.0 | 0 | Chordata | 90958.0 | 0 | Carnivora | 90958.0 | 0 | Canidae | 90958.0 | 0 | Canis | 90958.0 | 0 | Canis lupus | 90958.0 | 0 |
| scf7180000050583 | 3619 | 0.4482 | 0 | 53.4601 | Eukaryota | 189.0 | 0 | Platyhelminthes | 189.0 | 0 | Polystomatidea | 189.0 | 0 | Polystomatidae | 189.0 | 0 | Protopolystoma | 189.0 | 0 | Protopolystoma xenopodis | 189.0 | 0 |
| scf7180000042297 | 711004 | 0.37 | 0 | 60.0412 | Eukaryota | 93106.0 | 0 | Chordata | 93106.0 | 0 | Carnivora | 93106.0 | 0 | Canidae | 93106.0 | 0 | Canis | 93106.0 | 0 | Canis lupus | 93106.0 | 0 |
| scf7180000043802 | 366551 | 0.4325 | 0 | 58.3095 | Eukaryota | 44346.0 | 0 | Chordata | 44346.0 | 0 | Carnivora | 44346.0 | 0 | Canidae | 23426.0 | 4 | Canis | 23426.0 | 5 | Canis lupus | 23426.0 | 5 |
| scf7180000056339 | 630609 | 0.3602 | 0 | 60.1317 | Eukaryota | 90840.0 | 0 | Chordata | 90840.0 | 0 | Carnivora | 90840.0 | 0 | Canidae | 90840.0 | 0 | Canis | 90840.0 | 0 | Canis lupus | 90840.0 | 0 |
| scf7180000044158 | 7895 | 0.4348 | 0 | 358.9492 | Eukaryota | 134.0 | 0 | Chordata | 134.0 | 0 | Salmoniformes | 134.0 | 0 | Salmonidae | 134.0 | 0 | Coregonus | 134.0 | 0 | Coregonus sp. 'balchen' | 134.0 | 0 |
| scf7180000047196 | 21985 | 0.4453 | 0 | 75.7573 | Eukaryota | 2035.0 | 0 | Chordata | 2035.0 | 0 | Salmoniformes | 1484.0 | 2 | Salmonidae | 1484.0 | 2 | Coregonus | 929.0 | 3 | Coregonus sp. 'balchen' | 929.0 | 3 |
| scf7180000044496 | 307383 | 0.3982 | 0 | 58.4613 | Eukaryota | 17437.0 | 0 | Chordata | 17437.0 | 0 | Carnivora | 17437.0 | 0 | Canidae | 17437.0 | 0 | Canis | 17437.0 | 0 | Canis lupus | 17437.0 | 0 |
| scf7180000041413 | 598657 | 0.4486 | 0 | 58.4166 | Eukaryota | 101047.0 | 0 | Chordata | 101047.0 | 0 | Carnivora | 101047.0 | 0 | Canidae | 69388.0 | 2 | Canis | 69388.0 | 2 | Canis lupus | 69388.0 | 2 |
| scf7180000042346 | 92098 | 0.396 | 0 | 57.7931 | Eukaryota | 12637.0 | 0 | Chordata | 12637.0 | 0 | Carnivora | 12637.0 | 0 | Canidae | 12637.0 | 0 | Canis | 12637.0 | 0 | Canis lupus | 12637.0 | 0 |
| scf7180000044338 | 30267 | 0.3761 | 0 | 55.2384 | Eukaryota | 29785.0 | 0 | Chordata | 29785.0 | 0 | Carnivora | 29785.0 | 0 | Canidae | 29785.0 | 0 | Canis | 29785.0 | 0 | Canis lupus | 29785.0 | 0 |
| scf7180000054451 | 47595 | 0.4414 | 0 | 154.291 | Eukaryota | 11267.0 | 0 | Chordata | 9316.0 | 1 | Carnivora | 6759.0 | 3 | Canidae | 6759.0 | 3 | Canis | 4296.0 | 5 | Canis lupus | 4296.0 | 5 |
| scf7180000051812 | 3157 | 0.4495 | 0 | 276.9916 | Eukaryota | 1711.0 | 0 | Echinodermata | 763.0 | 2 | Forcipulatida | 763.0 | 2 | Asteriidae | 763.0 | 2 | Asterias | 763.0 | 2 | Asterias rubens | 763.0 | 2 |
| scf7180000041358 | 1204649 | 0.3902 | 0 | 59.9593 | Eukaryota | 92010.0 | 0 | Chordata | 92010.0 | 0 | Carnivora | 92010.0 | 0 | Canidae | 92010.0 | 0 | Canis | 92010.0 | 0 | Canis lupus | 92010.0 | 0 |
| scf7180000053196 | 2209 | 0.5079 | 0 | 20.4381 | Eukaryota | 5734.0 | 0 | Chordata | 5734.0 | 0 | Carnivora | 5734.0 | 0 | Canidae | 5734.0 | 0 | Canis | 4021.0 | 1 | Canis lupus | 3182.0 | 2 |
| scf7180000051135 | 16539 | 0.4561 | 0 | 58.1671 | Eukaryota | 8108.0 | 0 | Chordata | 4923.0 | 2 | Salmoniformes | 4077.0 | 5 | Salmonidae | 4077.0 | 5 | Salmo | 4077.0 | 5 | Salmo trutta | 4077.0 | 5 |
| scf7180000051217 | 13898 | 0.446 | 0 | 575.4173 | Eukaryota | 3392.0 | 0 | Platyhelminthes | 2088.0 | 2 | Polystomatidea | 2088.0 | 2 | Polystomatidae | 2088.0 | 2 | Protopolystoma | 2088.0 | 2 | Protopolystoma xenopodis | 2088.0 | 2 |
| scf7180000044845 | 338873 | 0.5592 | 0 | 56.5818 | Eukaryota | 78674.0 | 0 | Chordata | 78674.0 | 0 | Carnivora | 78674.0 | 0 | Canidae | 78674.0 | 0 | Canis | 70027.0 | 1 | Canis lupus | 70027.0 | 1 |
| scf7180000043104 | 4417 | 0.5196 | 0 | 23.4768 | Eukaryota | 3836.0 | 0 | Chordata | 3836.0 | 0 | Primates | 2033.0 | 2 | Hominidae | 1414.0 | 3 | Homo | 1414.0 | 3 | Homo sapiens | 1414.0 | 3 |
| scf7180000053048 | 6998 | 0.4417 | 0 | 33.0385 | no-hit | 0.0 | 0 | no-hit | 0.0 | 0 | no-hit | 0.0 | 0 | no-hit | 0.0 | 0 | no-hit | 0.0 | 0 | no-hit | 0.0 | 0 |
| scf7180000047040 | 28770 | 0.4387 | 0 | 177.7026 | Eukaryota | 1405.0 | 0 | Chordata | 835.0 | 2 | Polystomatidea | 351.0 | 4 | Polystomatidae | 351.0 | 4 | Protopolystoma | 351.0 | 4 | Protopolystoma xenopodis | 351.0 | 4 |
| scf7180000046002 | 104272 | 0.4281 | 0 | 57.171 | Eukaryota | 7233.0 | 0 | Chordata | 7233.0 | 0 | Carnivora | 6309.0 | 1 | Canidae | 6309.0 | 1 | Vulpes | 6309.0 | 1 | Vulpes vulpes | 6309.0 | 1 |
| scf7180000049312 | 9876 | 0.4417 | 0 | 1327.2936 | Eukaryota | 3757.0 | 0 | Chordata | 3757.0 | 0 | Salmoniformes | 3376.0 | 1 | Salmonidae | 3376.0 | 1 | Coregonus | 1692.0 | 2 | Coregonus sp. 'balchen' | 1692.0 | 2 |
| scf7180000042090 | 1343300 | 0.3903 | 0 | 60.9459 | Eukaryota | 61871.0 | 0 | Chordata | 61871.0 | 0 | Carnivora | 61871.0 | 0 | Canidae | 38115.0 | 1 | Zalophus | 23756.0 | 2 | Zalophus californianus | 23756.0 | 2 |
| scf7180000050155 | 38144 | 0.4396 | 0 | 58.6896 | Eukaryota | 2534.0 | 0 | Platyhelminthes | 2093.0 | 1 | Polystomatidea | 2093.0 | 2 | Polystomatidae | 2093.0 | 2 | Protopolystoma | 2093.0 | 2 | Protopolystoma xenopodis | 2093.0 | 2 |
| scf7180000041348 | 1102732 | 0.4304 | 0 | 58.7674 | Eukaryota | 94454.0 | 0 | Chordata | 94454.0 | 0 | Carnivora | 94454.0 | 0 | Canidae | 94454.0 | 0 | Canis | 94454.0 | 0 | Canis lupus | 94454.0 | 0 |
| scf7180000046347 | 28624 | 0.4456 | 0 | 219.3862 | Eukaryota | 2744.0 | 0 | Chordata | 2744.0 | 0 | Carnivora | 2744.0 | 0 | Canidae | 2744.0 | 0 | Canis | 2744.0 | 0 | Canis lupus | 2744.0 | 0 |
| scf7180000048892 | 43103 | 0.4965 | 0 | 80.022 | Eukaryota | 9597.0 | 0 | Chordata | 9597.0 | 0 | Carnivora | 9597.0 | 0 | Canidae | 9597.0 | 0 | Canis | 8965.0 | 1 | Canis lupus | 7819.0 | 2 |
| scf7180000044835 | 275130 | 0.3573 | 0 | 60.2967 | Eukaryota | 37808.0 | 0 | Chordata | 37808.0 | 0 | Carnivora | 37808.0 | 0 | Canidae | 27611.0 | 2 | Canis | 23690.0 | 4 | Canis lupus | 23690.0 | 4 |
| scf7180000042491 | 23777 | 0.4109 | 0 | 858.6034 | Eukaryota | 88145.0 | 0 | Chordata | 88145.0 | 0 | Carnivora | 88145.0 | 0 | Canidae | 88145.0 | 0 | Canis | 88145.0 | 0 | Canis lupus | 88145.0 | 0 |
| scf7180000044453 | 299234 | 0.4516 | 0 | 64.3801 | Eukaryota | 86474.0 | 0 | Chordata | 86474.0 | 0 | Carnivora | 86474.0 | 0 | Canidae | 86474.0 | 0 | Canis | 86474.0 | 0 | Canis lupus | 86474.0 | 0 |
| scf7180000042764 | 561124 | 0.4075 | 0 | 60.1652 | Eukaryota | 50736.0 | 0 | Chordata | 50736.0 | 0 | Carnivora | 50736.0 | 0 | Canidae | 50736.0 | 0 | Canis | 38672.0 | 1 | Canis lupus | 38672.0 | 1 |
| scf7180000048686 | 30332 | 0.5014 | 0 | 667.2869 | Eukaryota | 9214.0 | 0 | Chordata | 9214.0 | 0 | Carnivora | 9214.0 | 0 | Canidae | 9214.0 | 0 | Canis | 7465.0 | 1 | Canis lupus | 6199.0 | 2 |
| scf7180000048660 | 2760 | 0.4518 | 0 | 78.584 | no-hit | 0.0 | 0 | no-hit | 0.0 | 0 | no-hit | 0.0 | 0 | no-hit | 0.0 | 0 | no-hit | 0.0 | 0 | no-hit | 0.0 | 0 |
| scf7180000046326 | 6273 | 0.4398 | 0 | 88.6475 | no-hit | 0.0 | 0 | no-hit | 0.0 | 0 | no-hit | 0.0 | 0 | no-hit | 0.0 | 0 | no-hit | 0.0 | 0 | no-hit | 0.0 | 0 |
| scf7180000055233 | 2870 | 0.4575 | 0 | 97.6498 | no-hit | 0.0 | 0 | no-hit | 0.0 | 0 | no-hit | 0.0 | 0 | no-hit | 0.0 | 0 | no-hit | 0.0 | 0 | no-hit | 0.0 | 0 |
| scf7180000053978 | 5004 | 0.442 | 0 | 31.7472 | no-hit | 0.0 | 0 | no-hit | 0.0 | 0 | no-hit | 0.0 | 0 | no-hit | 0.0 | 0 | no-hit | 0.0 | 0 | no-hit | 0.0 | 0 |
| scf7180000045561 | 194490 | 0.4615 | 0 | 60.2111 | Eukaryota | 26886.0 | 0 | Chordata | 26886.0 | 0 | Carnivora | 26886.0 | 0 | Canidae | 26886.0 | 0 | Canis | 22911.0 | 1 | Canis lupus | 22911.0 | 1 |
| scf7180000041085 | 128087 | 0.415 | 0 | 60.2067 | Eukaryota | 9118.0 | 0 | Chordata | 9118.0 | 0 | Carnivora | 6550.0 | 1 | Otariidae | 3880.0 | 2 | Homo | 2568.0 | 4 | Homo sapiens | 2568.0 | 4 |
| scf7180000042921 | 563794 | 0.3747 | 0 | 58.7673 | Eukaryota | 16014.0 | 0 | Chordata | 16014.0 | 0 | Carnivora | 14200.0 | 1 | Canidae | 14200.0 | 1 | Canis | 14200.0 | 1 | Canis lupus | 14200.0 | 1 |
| scf7180000042778 | 39765 | 0.4591 | 0 | 53.5618 | Eukaryota | 11780.0 | 0 | Chordata | 11780.0 | 0 | Carnivora | 11780.0 | 0 | Canidae | 7647.0 | 1 | Canis | 7647.0 | 2 | Canis lupus | 7647.0 | 2 |
| scf7180000051033 | 7903 | 0.35 | 0 | 34.7045 | Eukaryota | 11698.0 | 0 | Chordata | 11698.0 | 0 | Carnivora | 11698.0 | 0 | Canidae | 8088.0 | 1 | Canis | 8088.0 | 2 | Canis lupus | 8088.0 | 2 |
| scf7180000052789 | 14518 | 0.4244 | 0 | 57.876 | Eukaryota | 3256.0 | 0 | Chordata | 3256.0 | 0 | Carnivora | 3256.0 | 0 | Canidae | 3256.0 | 0 | Canis | 3256.0 | 0 | Canis lupus | 3256.0 | 0 |
| scf7180000043324 | 499449 | 0.3685 | 0 | 59.012 | Eukaryota | 93494.0 | 0 | Chordata | 93494.0 | 0 | Carnivora | 93494.0 | 0 | Canidae | 93494.0 | 0 | Canis | 93494.0 | 0 | Canis lupus | 93494.0 | 0 |
| scf7180000040521 | 2606080 | 0.4025 | 0 | 61.924 | Eukaryota | 120754.0 | 0 | Chordata | 120754.0 | 0 | Carnivora | 120754.0 | 0 | Canidae | 120754.0 | 0 | Canis | 120754.0 | 0 | Canis lupus | 120754.0 | 0 |
| scf7180000040870 | 1472792 | 0.4347 | 100 | 59.4325 | Eukaryota | 54391.0 | 0 | Chordata | 54391.0 | 0 | Carnivora | 54391.0 | 0 | Canidae | 54391.0 | 0 | Canis | 46511.0 | 1 | Canis lupus | 46511.0 | 1 |

| | | | | | | | | | | | | | | | | | | | | | | |
|---|---|---|---|---|---|---|---|---|---|---|---|---|---|---|---|---|---|---|---|---|---|---|
| scf7180000041636 | 944068 | 0.3574 | 0 | 61.0561 | Eukaryota | 93662.0 | 0 | Chordata | 93662.0 | 0 | Carnivora | 93662.0 | 0 | Canidae | 93662.0 | 0 | Canis | 93662.0 | 0 | Canis lupus | 93662.0 | 0 |
| scf7180000044781 | 197778 | 0.3623 | 0 | 58.8049 | Eukaryota | 13866.0 | 0 | Chordata | 13866.0 | 0 | Carnivora | 13866.0 | 0 | Canidae | 13866.0 | 0 | Canis | 13866.0 | 0 | Canis lupus | 13866.0 | 0 |
| scf7180000050565 | 28833 | 0.4171 | 0 | 61.1399 | Eukaryota | 59620.0 | 0 | Chordata | 59620.0 | 0 | Carnivora | 59620.0 | 0 | Canidae | 49481.0 | 2 | Canis | 28484.0 | 3 | Canis lupus | 28484.0 | 4 |
| scf7180000048573 | 14555 | 0.4456 | 0 | 259.2705 | no-hit | 0.0 | 0 | no-hit | 0.0 | 0 | no-hit | 0.0 | 0 | no-hit | 0.0 | 0 | no-hit | 0.0 | 0 | no-hit | 0.0 | 0 |
| scf7180000050720 | 10592 | 0.5789 | 0 | 305.029 | Eukaryota | 42938.0 | 0 | Chordata | 42938.0 | 0 | Artiodactyla | 30119.0 | 2 | Bovidae | 30119.0 | 2 | Ovis | 17048.0 | 3 | Ovis aries | 17048.0 | 3 |
| scf7180000050868 | 37183 | 0.504 | 0 | 1165.9912 | Eukaryota | 9425.0 | 0 | Chordata | 9425.0 | 0 | Carnivora | 9425.0 | 0 | Canidae | 9425.0 | 0 | Canis | 8182.0 | 1 | Canis lupus | 6882.0 | 2 |
| scf7180000051995 | 10668 | 0.5061 | 0 | 208.8964 | Eukaryota | 8807.0 | 0 | Chordata | 8807.0 | 0 | Carnivora | 8807.0 | 0 | Canidae | 8807.0 | 0 | Canis | 7091.0 | 1 | Canis lupus | 5851.0 | 2 |
| scf7180000042881 | 15267 | 0.3789 | 0 | 48.4404 | Eukaryota | 3916.0 | 0 | Chordata | 3916.0 | 0 | Carnivora | 3916.0 | 0 | Canidae | 3916.0 | 0 | Canis | 3916.0 | 0 | Canis lupus | 3916.0 | 0 |
| scf7180000041913 | 784089 | 0.3656 | 0 | 59.6671 | Eukaryota | 145637.0 | 0 | Chordata | 145637.0 | 0 | Carnivora | 145637.0 | 0 | Canidae | 81322.0 | 1 | Canis | 65045.0 | 3 | Canis lupus | 65045.0 | 3 |
| scf7180000051292 | 10648 | 0.4423 | 0 | 242.9646 | Eukaryota | 206.0 | 0 | Chordata | 206.0 | 0 | Tetraodontiformes | 206.0 | 0 | Tetraodontidae | 206.0 | 0 | Takifugu | 206.0 | 0 | Takifugu rubripes | 206.0 | 0 |
| scf7180000051018 | 13796 | 0.4611 | 0 | 147.0752 | Eukaryota | 10157.0 | 0 | Chordata | 10157.0 | 0 | Carnivora | 8884.0 | 1 | Canidae | 8884.0 | 1 | Canis | 5011.0 | 2 | Canis lupus | 5011.0 | 2 |
| scf7180000049472 | 16922 | 0.4475 | 0 | 1421.0806 | Eukaryota | 3447.0 | 0 | Platyhelminthes | 1900.0 | 2 | Polystomatidea | 1900.0 | 2 | Polystomatidae | 1900.0 | 2 | Protopolystoma | 1900.0 | 2 | Protopolystoma xenopodis | 1900.0 | 2 |
| scf7180000050803 | 19103 | 0.5159 | 0 | 921.9329 | Eukaryota | 17128.0 | 0 | Chordata | 17128.0 | 0 | Carnivora | 17128.0 | 0 | Canidae | 17128.0 | 0 | Canis | 17128.0 | 0 | Canis lupus | 17128.0 | 0 |
| scf7180000044099 | 61002 | 0.5418 | 0 | 52.8929 | Eukaryota | 4559.0 | 0 | Chordata | 4559.0 | 0 | Primates | 2949.0 | 1 | Hominidae | 2064.0 | 2 | Homo | 2064.0 | 2 | Homo sapiens | 2064.0 | 3 |
| scf7180000047679 | 61647 | 0.3922 | 0 | 60.3554 | Eukaryota | 14214.0 | 0 | Chordata | 14214.0 | 0 | Carnivora | 14214.0 | 0 | Canidae | 14214.0 | 0 | Canis | 14214.0 | 0 | Canis lupus | 14214.0 | 0 |
| scf7180000042582 | 342289 | 0.4643 | 0 | 58.7577 | Eukaryota | 64357.0 | 0 | Chordata | 64357.0 | 0 | Carnivora | 64357.0 | 0 | Canidae | 64357.0 | 0 | Canis | 64357.0 | 0 | Canis lupus | 64357.0 | 0 |
| scf7180000042746 | 371501 | 0.4472 | 0 | 59.0326 | Eukaryota | 58285.0 | 0 | Chordata | 58285.0 | 0 | Carnivora | 58285.0 | 0 | Canidae | 58285.0 | 0 | Canis | 58285.0 | 0 | Canis lupus | 58285.0 | 0 |
| scf7180000041088 | 63380 | 0.4204 | 0 | 50.7359 | Eukaryota | 11102.0 | 0 | Chordata | 11102.0 | 0 | Primates | 4688.0 | 2 | Hominidae | 4688.0 | 3 | Mus | 3948.0 | 4 | Mus musculus | 3948.0 | 4 |
| scf7180000052212 | 4470 | 0.451 | 0 | 782.2258 | no-hit | 0.0 | 0 | no-hit | 0.0 | 0 | no-hit | 0.0 | 0 | no-hit | 0.0 | 0 | no-hit | 0.0 | 0 | no-hit | 0.0 | 0 |
| scf7180000044345 | 335057 | 0.3646 | 0 | 58.6043 | Eukaryota | 33715.0 | 0 | Chordata | 33715.0 | 0 | Carnivora | 33715.0 | 0 | Canidae | 24090.0 | 2 | Canis | 20559.0 | 4 | Canis lupus | 20559.0 | 4 |
| scf7180000051250 | 12240 | 0.4528 | 0 | 908.0949 | Eukaryota | 424.0 | 0 | Platyhelminthes | 424.0 | 0 | Polystomatidea | 424.0 | 0 | Polystomatidae | 424.0 | 0 | Protopolystoma | 424.0 | 0 | Protopolystoma xenopodis | 424.0 | 0 |
| scf7180000047932 | 8185 | 0.4468 | 0 | 453.9747 | Eukaryota | 267.0 | 0 | Platyhelminthes | 267.0 | 0 | Polystomatidea | 267.0 | 0 | Polystomatidae | 267.0 | 0 | Protopolystoma | 267.0 | 0 | Protopolystoma xenopodis | 267.0 | 0 |
| scf7180000047890 | 79697 | 0.3756 | 0 | 61.2352 | Eukaryota | 25819.0 | 0 | Chordata | 25819.0 | 0 | Carnivora | 25819.0 | 0 | Canidae | 25819.0 | 0 | Canis | 25819.0 | 0 | Canis lupus | 25819.0 | 0 |
| scf7180000050719 | 10949 | 0.5019 | 0 | 364.8851 | Eukaryota | 8232.0 | 0 | Chordata | 8232.0 | 0 | Carnivora | 8232.0 | 0 | Canidae | 8232.0 | 0 | Canis | 5909.0 | 1 | Canis lupus | 4680.0 | 2 |
| scf7180000044991 | 100486 | 0.3752 | 0 | 55.2438 | Eukaryota | 18324.0 | 0 | Chordata | 18324.0 | 0 | Carnivora | 18324.0 | 0 | Canidae | 18324.0 | 0 | Canis | 18324.0 | 0 | Canis lupus | 18324.0 | 0 |
| scf7180000050611 | 3140 | 0.4452 | 0 | 69.281 | no-hit | 0.0 | 0 | no-hit | 0.0 | 0 | no-hit | 0.0 | 0 | no-hit | 0.0 | 0 | no-hit | 0.0 | 0 | no-hit | 0.0 | 0 |
| scf7180000053503 | 1803 | 0.4515 | 0 | 150.531 | no-hit | 0.0 | 0 | no-hit | 0.0 | 0 | no-hit | 0.0 | 0 | no-hit | 0.0 | 0 | no-hit | 0.0 | 0 | no-hit | 0.0 | 0 |
| scf7180000051013 | 14180 | 0.437 | 0 | 34.8303 | Eukaryota | 7254.0 | 0 | Chordata | 2820.0 | 3 | Forcipulatida | 1967.0 | 5 | Asteriidae | 1967.0 | 5 | Asterias | 1967.0 | 6 | Asterias rubens | 1967.0 | 6 |
| scf7180000049187 | 45733 | 0.4387 | 0 | 74.7544 | Eukaryota | 1841.0 | 0 | Platyhelminthes | 1393.0 | 1 | Polystomatidea | 1393.0 | 1 | Polystomatidae | 1393.0 | 1 | Protopolystoma | 1393.0 | 1 | Protopolystoma xenopodis | 1393.0 | 1 |
| scf7180000044903 | 247641 | 0.4169 | 0 | 59.0852 | Eukaryota | 57954.0 | 0 | Chordata | 57954.0 | 0 | Carnivora | 57954.0 | 0 | Otariidae | 42562.0 | 1 | Zalophus | 32020.0 | 2 | Zalophus californianus | 32020.0 | 2 |
| scf7180000056145 | 12478 | 0.523 | 0 | 612.4028 | Eukaryota | 16050.0 | 0 | Chordata | 16050.0 | 0 | Carnivora | 16050.0 | 0 | Canidae | 16050.0 | 0 | Canis | 16050.0 | 0 | Canis lupus | 16050.0 | 0 |
| scf7180000049992 | 8531 | 0.4327 | 0 | 399.2795 | Eukaryota | 46559.0 | 0 | Chordata | 46559.0 | 0 | Carnivora | 46559.0 | 0 | Canidae | 46559.0 | 0 | Canis | 46559.0 | 0 | Canis lupus | 46559.0 | 0 |
| scf7180000053119 | 5791 | 0.4462 | 0 | 231.6843 | Eukaryota | 1982.0 | 0 | Platyhelminthes | 1107.0 | 1 | Polystomatidea | 1107.0 | 2 | Polystomatidae | 1107.0 | 2 | Protopolystoma | 1107.0 | 2 | Protopolystoma xenopodis | 1107.0 | 2 |
| scf7180000041427 | 253716 | 0.4266 | 0 | 56.5936 | Eukaryota | 37182.0 | 0 | Chordata | 37182.0 | 0 | Carnivora | 37182.0 | 0 | Canidae | 37182.0 | 0 | Canis | 29027.0 | 1 | Canis lupus | 29027.0 | 1 |
| scf7180000056214 | 1366355 | 0.3928 | 0 | 65.106 | Eukaryota | 91490.0 | 0 | Chordata | 91490.0 | 0 | Carnivora | 91490.0 | 0 | Canidae | 91490.0 | 0 | Canis | 91490.0 | 0 | Canis lupus | 91490.0 | 0 |
| scf7180000052561 | 2989 | 0.458 | 0 | 393.5917 | Eukaryota | 1406.0 | 0 | Echinodermata | 610.0 | 2 | Forcipulatida | 610.0 | 2 | Asteriidae | 610.0 | 2 | Asterias | 610.0 | 2 | Asterias rubens | 610.0 | 2 |
| scf7180000050495 | 4225 | 0.4469 | 0 | 319.1905 | Eukaryota | 1753.0 | 0 | Platyhelminthes | 938.0 | 2 | Polystomatidea | 938.0 | 2 | Polystomatidae | 938.0 | 2 | Protopolystoma | 938.0 | 2 | Protopolystoma xenopodis | 938.0 | 2 |
| scf7180000050946 | 24295 | 0.4487 | 0 | 79.2168 | Eukaryota | 3856.0 | 0 | Chordata | 3856.0 | 0 | Carnivora | 2513.0 | 1 | Canidae | 2513.0 | 1 | Canis | 2513.0 | 1 | Canis lupus | 2513.0 | 1 |
| scf7180000045778 | 150305 | 0.4312 | 0 | 58.4415 | Eukaryota | 14388.0 | 0 | Chordata | 14388.0 | 0 | Carnivora | 10445.0 | 2 | Canidae | 7764.0 | 3 | Canis | 5802.0 | 6 | Canis lupus | 5802.0 | 6 |
| scf7180000050581 | 2285 | 0.4363 | 0 | 1184.0773 | Eukaryota | 741.0 | 0 | Chordata | 741.0 | 0 | Columbiformes | 741.0 | 0 | Columbidae | 741.0 | 0 | Streptopelia | 741.0 | 0 | Streptopelia turtur | 741.0 | 0 |
| scf7180000047944 | 76945 | 0.4619 | 0 | 59.9927 | Eukaryota | 23145.0 | 0 | Chordata | 23145.0 | 0 | Carnivora | 20185.0 | 2 | Canidae | 18210.0 | 3 | Canis | 18210.0 | 3 | Canis lupus | 18210.0 | 4 |
| scf7180000047158 | 18297 | 0.4355 | 0 | 73.1788 | Eukaryota | 8725.0 | 0 | Chordata | 7058.0 | 1 | Carnivora | 6265.0 | 2 | Canidae | 6265.0 | 2 | Canis | 5799.0 | 3 | Canis lupus | 5799.0 | 3 |
| scf7180000042538 | 4696 | 0.3822 | 0 | 49.7883 | Eukaryota | 32670.0 | 0 | Chordata | 32670.0 | 0 | Carnivora | 32670.0 | 0 | Canidae | 32670.0 | 0 | Canis | 32670.0 | 0 | Canis lupus | 32670.0 | 0 |
| scf7180000054819 | 62880 | 0.4363 | 0 | 130.5063 | Eukaryota | 47610.0 | 0 | Chordata | 29187.0 | 3 | Gadiformes | 24899.0 | 5 | Gadidae | 24899.0 | 5 | Gadus | 24899.0 | 5 | Gadus morhua | 24899.0 | 5 |
| scf7180000048442 | 46717 | 0.3933 | 0 | 58.9371 | Eukaryota | 90727.0 | 0 | Chordata | 90727.0 | 0 | Carnivora | 90727.0 | 0 | Canidae | 90727.0 | 0 | Canis | 90727.0 | 0 | Canis lupus | 90727.0 | 0 |
| scf7180000055232 | 24016 | 0.441 | 0 | 268.7453 | Eukaryota | 9207.0 | 0 | Chordata | 6380.0 | 1 | Kurtiformes | 3512.0 | 2 | Apogonidae | 3512.0 | 2 | Sphaeramia | 3512.0 | 2 | Sphaeramia orbicularis | 3512.0 | 2 |
| scf7180000043927 | 349699 | 0.4679 | 0 | 59.6598 | Eukaryota | 86277.0 | 0 | Chordata | 86277.0 | 0 | Carnivora | 86277.0 | 0 | Canidae | 72282.0 | 1 | Canis | 59510.0 | 2 | Canis lupus | 59510.0 | 2 |
| scf7180000048876 | 52082 | 0.4192 | 0 | 74.6474 | Eukaryota | 44675.0 | 0 | Chordata | 44675.0 | 0 | Carnivora | 44675.0 | 0 | Canidae | 44675.0 | 0 | Canis | 44675.0 | 0 | Canis lupus | 44675.0 | 0 |
| scf7180000042809 | 80853 | 0.3502 | 0 | 56.4847 | Eukaryota | 51009.0 | 0 | Chordata | 51009.0 | 0 | Carnivora | 51009.0 | 0 | Canidae | 51009.0 | 0 | Canis | 51009.0 | 0 | Canis lupus | 51009.0 | 0 |
| scf7180000050916 | 29621 | 0.5021 | 0 | 791.9426 | Eukaryota | 9285.0 | 0 | Chordata | 9285.0 | 0 | Carnivora | 9285.0 | 0 | Canidae | 9285.0 | 0 | Canis | 8063.0 | 1 | Canis lupus | 6782.0 | 2 |
| scf7180000052177 | 11821 | 0.4387 | 0 | 130.1971 | Eukaryota | 1748.0 | 0 | Platyhelminthes | 1036.0 | 1 | Polystomatidea | 899.0 | 2 | Polystomatidae | 899.0 | 2 | Protopolystoma | 899.0 | 2 | Protopolystoma xenopodis | 899.0 | 2 |
| scf7180000049515 | 20337 | 0.5052 | 0 | 449.3069 | Eukaryota | 9274.0 | 0 | Chordata | 9274.0 | 0 | Carnivora | 9274.0 | 0 | Canidae | 9274.0 | 0 | Canis | 7516.0 | 1 | Canis lupus | 6242.0 | 2 |
| scf7180000048395 | 72394 | 0.4381 | 0 | 55.3854 | Eukaryota | 16217.0 | 0 | Chordata | 16217.0 | 0 | Carnivora | 16217.0 | 0 | Canidae | 6386.0 | 2 | Panthera | 4929.0 | 5 | Canis lupus | 4710.0 | 6 |
| scf7180000043404 | 438166 | 0.3531 | 0 | 60.4377 | Eukaryota | 65684.0 | 0 | Chordata | 65684.0 | 0 | Carnivora | 65684.0 | 0 | Canidae | 65684.0 | 0 | Canis | 65684.0 | 0 | Canis lupus | 65684.0 | 0 |
| scf7180000050235 | 7408 | 0.4414 | 0 | 68.936 | Eukaryota | 372.0 | 0 | Chordata | 372.0 | 0 | Salmoniformes | 372.0 | 0 | Salmonidae | 372.0 | 0 | Coregonus | 372.0 | 0 | Coregonus sp. 'balchen' | 372.0 | 0 |
| scf7180000042359 | 741538 | 0.3568 | 0 | 59.5314 | Eukaryota | 59906.0 | 0 | Chordata | 59906.0 | 0 | Carnivora | 59906.0 | 0 | Canidae | 59906.0 | 0 | Canis | 59906.0 | 0 | Canis lupus | 59906.0 | 0 |
| scf7180000052030 | 20786 | 0.5234 | 0 | 2590.9318 | Eukaryota | 34273.0 | 0 | Chordata | 34273.0 | 0 | Carnivora | 34273.0 | 0 | Canidae | 34273.0 | 0 | Canis | 34273.0 | 0 | Canis lupus | 34273.0 | 0 |
| scf7180000044069 | 75144 | 0.4585 | 0 | 128.3429 | Eukaryota | 15636.0 | 0 | Chordata | 15636.0 | 0 | Carnivora | 15636.0 | 0 | Canidae | 15636.0 | 0 | Canis | 9670.0 | 1 | Canis lupus | 9670.0 | 1 |
| scf7180000056505 | 219587 | 0.4571 | 0 | 58.165 | Eukaryota | 35528.0 | 0 | Chordata | 35528.0 | 0 | Carnivora | 35528.0 | 0 | Canidae | 14110.0 | 2 | Canis | 14110.0 | 3 | Canis lupus | 14110.0 | 3 |
| scf7180000044262 | 3849 | 0.3879 | 0 | 22.3778 | Eukaryota | 54352.0 | 0 | Chordata | 54352.0 | 0 | Carnivora | 54352.0 | 0 | Canidae | 54352.0 | 0 | Canis | 54352.0 | 0 | Canis lupus | 54352.0 | 0 |
| scf7180000048697 | 67766 | 0.4248 | 0 | 59.6939 | Eukaryota | 29996.0 | 0 | Chordata | 29996.0 | 0 | Carnivora | 29996.0 | 0 | Canidae | 29996.0 | 0 | Canis | 17932.0 | 1 | Canis lupus | 17932.0 | 1 |
| scf7180000041620 | 225066 | 0.3728 | 0 | 58.9816 | Eukaryota | 8126.0 | 0 | Chordata | 8126.0 | 0 | Primates | 3578.0 | 2 | Hominidae | 3578.0 | 2 | Homo | 3578.0 | 3 | Homo sapiens | 3578.0 | 3 |
| scf7180000049440 | 35793 | 0.4726 | 0 | 52.9371 | Eukaryota | 15027.0 | 0 | Chordata | 15027.0 | 0 | Carnivora | 15027.0 | 0 | Canidae | 15027.0 | 0 | Canis | 15027.0 | 0 | Canis lupus | 15027.0 | 0 |
| scf7180000051775 | 5906 | 0.446 | 0 | 255.5036 | Eukaryota | 686.0 | 0 | Chordata | 686.0 | 0 | Kurtiformes | 538.0 | 1 | Apogonidae | 538.0 | 1 | Sphaeramia | 538.0 | 1 | Sphaeramia orbicularis | 538.0 | 1 |
| scf7180000056119 | 29883 | 0.4403 | 0 | 173.5657 | Eukaryota | 561.0 | 0 | Platyhelminthes | 289.0 | 1 | Polystomatidea | 289.0 | 1 | Polystomatidae | 289.0 | 1 | Protopolystoma | 289.0 | 1 | Protopolystoma xenopodis | 289.0 | 1 |
| scf7180000053720 | 3709 | 0.4335 | 0 | 94.6439 | no-hit | 0.0 | 0 | no-hit | 0.0 | 0 | no-hit | 0.0 | 0 | no-hit | 0.0 | 0 | no-hit | 0.0 | 0 | no-hit | 0.0 | 0 |
| scf7180000053648 | 1785 | 0.5384 | 0 | 43.151 | Eukaryota | 17927.0 | 0 | Chordata | 17927.0 | 0 | Carnivora | 17927.0 | 0 | Canidae | 17927.0 | 0 | Canis | 17927.0 | 0 | Canis lupus | 17927.0 | 0 |
| scf7180000052574 | 2685 | 0.4566 | 0 | 461.8506 | Eukaryota | 10351.0 | 0 | Chordata | 9117.0 | 2 | Gadiformes | 3938.0 | 7 | Gadidae | 3938.0 | 7 | Gadus | 3938.0 | 7 | Gadus morhua | 3938.0 | 7 |
| scf7180000040715 | 81313 | 0.3586 | 0 | 58.8645 | Eukaryota | 21161.0 | 0 | Chordata | 21161.0 | 0 | Carnivora | 21161.0 | 0 | Canidae | 21161.0 | 0 | Canis | 21161.0 | 0 | Canis lupus | 21161.0 | 0 |
| scf7180000054026 | 3056 | 0.3953 | 0 | 44.9591 | Eukaryota | 2835.0 | 0 | Chordata | 2835.0 | 0 | Carnivora | 2835.0 | 0 | Canidae | 2835.0 | 0 | Canis | 2835.0 | 0 | Canis lupus | 2835.0 | 0 |
| scf7180000045470 | 138041 | 0.3447 | 100 | 56.5301 | Eukaryota | 83026.0 | 0 | Chordata | 83026.0 | 0 | Carnivora | 83026.0 | 0 | Canidae | 83026.0 | 0 | Canis | 83026.0 | 0 | Canis lupus | 83026.0 | 0 |
| scf7180000042734 | 100872 | 0.3828 | 0 | 57.6224 | Eukaryota | 9295.0 | 0 | Chordata | 9295.0 | 0 | Carnivora | 4532.0 | 2 | Felidae | 2980.0 | 4 | Panthera | 2980.0 | 5 | Panthera pardus | 2980.0 | 5 |
| scf7180000049481 | 21922 | 0.4391 | 0 | 301.9298 | no-hit | 0.0 | 0 | no-hit | 0.0 | 0 | no-hit | 0.0 | 0 | no-hit | 0.0 | 0 | no-hit | 0.0 | 0 | no-hit | 0.0 | 0 |
| scf7180000043753 | 79999 | 0.375 | 0 | 65.3578 | Eukaryota | 96293.0 | 0 | Chordata | 96293.0 | 0 | Carnivora | 96293.0 | 0 | Canidae | 96293.0 | 0 | Canis | 96293.0 | 0 | Canis lupus | 96293.0 | 0 |
| scf7180000044642 | 14043 | 0.4414 | 0 | 121.8816 | Eukaryota | 682.0 | 0 | Platyhelminthes | 682.0 | 0 | Polystomatidea | 682.0 | 0 | Polystomatidae | 682.0 | 0 | Protopolystoma | 682.0 | 0 | Protopolystoma xenopodis | 682.0 | 0 |
| scf7180000050843 | 29719 | 0.4466 | 0 | 55.6878 | Eukaryota | 49150.0 | 0 | Platyhelminthes | 28533.0 | 2 | Strigeidida | 15796.0 | 6 | Schistosomatidae | 15796.0 | 6 | Schistosoma | 15796.0 | 6 | Schistosoma mansoni | 15796.0 | 6 |
| scf7180000045967 | 6613 | 0.4434 | 0 | 95.5011 | Eukaryota | 27017.0 | 0 | Chordata | 20161.0 | 2 | Gadiformes | 14110.0 | 4 | Gadidae | 14110.0 | 4 | Gadus | 14110.0 | 4 | Gadus morhua | 14110.0 | 4 |
| scf7180000054573 | 17174 | 0.4513 | 0 | 249.7026 | Eukaryota | 35390.0 | 0 | Chordata | 35390.0 | 0 | Carnivora | 35390.0 | 0 | Canidae | 35390.0 | 0 | Canis | 32924.0 | 1 | Canis lupus | 32924.0 | 1 |
| scf7180000051685 | 2438 | 0.4516 | 0 | 397.6383 | no-hit | 0.0 | 0 | no-hit | 0.0 | 0 | no-hit | 0.0 | 0 | no-hit | 0.0 | 0 | no-hit | 0.0 | 0 | no-hit | 0.0 | 0 |
| scf7180000052373 | 10314 | 0.515 | 0 | 495.1686 | Eukaryota | 16091.0 | 0 | Chordata | 16091.0 | 0 | Carnivora | 16091.0 | 0 | Canidae | 16091.0 | 0 | Canis | 16091.0 | 0 | Canis lupus | 16091.0 | 0 |
| scf7180000044754 | 13078 | 0.352 | 0 | 39.9341 | Eukaryota | 25236.0 | 0 | Chordata | 25236.0 | 0 | Carnivora | 25236.0 | 0 | Canidae | 10964.0 | 2 | Canis | 8240.0 | 6 | Canis lupus | 8240.0 | 6 |
| scf7180000046728 | 36158 | 0.4315 | 0 | 271.6823 | Eukaryota | 29661.0 | 0 | Chordata | 21585.0 | 2 | Gadiformes | 6946.0 | 6 | Gadidae | 6946.0 | 6 | Gadus | 6946.0 | 7 | Gadus morhua | 6946.0 | 7 |
| scf7180000043610 | 434569 | 0.5345 | 0 | 57.0137 | Eukaryota | 48176.0 | 0 | Chordata | 48176.0 | 0 | Carnivora | 48176.0 | 0 | Canidae | 45016.0 | 1 | Canis | 41091.0 | 2 | Canis lupus | 41091.0 | 2 |
| scf7180000042246 | 618054 | 0.406 | 0 | 60.6116 | Eukaryota | 89019.0 | 0 | Chordata | 89019.0 | 0 | Carnivora | 89019.0 | 0 | Canidae | 89019.0 | 0 | Canis | 89019.0 | 0 | Canis lupus | 89019.0 | 0 |
| scf7180000043759 | 343778 | 0.3793 | 0 | 61.1726 | Eukaryota | 77645.0 | 0 | Chordata | 77645.0 | 0 | Carnivora | 77645.0 | 0 | Canidae | 77645.0 | 0 | Canis | 77645.0 | 0 | Canis lupus | 77645.0 | 0 |
| scf7180000048577 | 27478 | 0.4344 | 0 | 57.4027 | Eukaryota | 50011.0 | 0 | Chordata | 50011.0 | 0 | Carnivora | 50011.0 | 0 | Canidae | 50011.0 | 0 | Canis | 50011.0 | 0 | Canis lupus | 50011.0 | 0 |
| scf7180000043141 | 534816 | 0.3573 | 0 | 59.9873 | Eukaryota | 92592.0 | 0 | Chordata | 92592.0 | 0 | Carnivora | 92592.0 | 0 | Canidae | 92592.0 | 0 | Canis | 92592.0 | 0 | Canis lupus | 92592.0 | 0 |
| scf7180000045544 | 44255 | 0.3394 | 0 | 53.1814 | Eukaryota | 55474.0 | 0 | Chordata | 55474.0 | 0 | Carnivora | 55474.0 | 0 | Otariidae | 18779.0 | 3 | Canis | 11484.0 | 7 | Canis lupus | 11484.0 | 7 |
| scf7180000054496 | 86804 | 0.3799 | 0 | 503.4357 | Eukaryota | 90291.0 | 0 | Chordata | 90291.0 | 0 | Carnivora | 90291.0 | 0 | Canidae | 90291.0 | 0 | Canis | 90291.0 | 0 | Canis lupus | 90291.0 | 0 |
| scf7180000053558 | 2845 | 0.4457 | 0 | 351.3172 | no-hit | 0.0 | 0 | no-hit | 0.0 | 0 | no-hit | 0.0 | 0 | no-hit | 0.0 | 0 | no-hit | 0.0 | 0 | no-hit | 0.0 | 0 |
| scf7180000045403 | 374090 | 0.4187 | 0 | 59.4142 | Eukaryota | 79953.0 | 0 | Chordata | 79953.0 | 0 | Carnivora | 79953.0 | 0 | Canidae | 79953.0 | 0 | Canis | 79953.0 | 0 | Canis lupus | 79953.0 | 0 |
| scf7180000055497 | 22510 | 0.4427 | 0 | 147.0446 | Eukaryota | 75265.0 | 0 | Platyhelminthes | 48984.0 | 1 | Strigeidida | 37080.0 | 5 | Schistosomatidae | 37080.0 | 5 | Schistosoma | 37080.0 | 5 | Schistosoma mansoni | 37080.0 | 5 |
| scf7180000053941 | 1980 | 0.4742 | 0 | 119.7923 | no-hit | 0.0 | 0 | no-hit | 0.0 | 0 | no-hit | 0.0 | 0 | no-hit | 0.0 | 0 | no-hit | 0.0 | 0 | no-hit | 0.0 | 0 |
| scf7180000045141 | 30916 | 0.4416 | 0 | 286.845 | Eukaryota | 66276.0 | 0 | Chordata | 46487.0 | 2 | Gadiformes | 38282.0 | 5 | Gadidae | 38282.0 | 5 | Gadus | 38282.0 | 5 | Gadus morhua | 38282.0 | 5 |
| scf7180000044707 | 280975 | 0.4139 | 0 | 59.7786 | Eukaryota | 48613.0 | 0 | Chordata | 48613.0 | 0 | Carnivora | 48613.0 | 0 | Canidae | 48613.0 | 0 | Canis | 38563.0 | 1 | Canis lupus | 38563.0 | 1 |
| scf7180000040973 | 128222 | 0.37 | 0 | 59.3792 | Eukaryota | 23337.0 | 0 | Chordata | 23337.0 | 0 | Carnivora | 23337.0 | 0 | Canidae | 10725.0 | 2 | Ursus | 7110.0 | 3 | Ursus arctos | 7110.0 | 3 |
| scf7180000050307 | 19134 | 0.5029 | 0 | 469.7973 | Eukaryota | 9064.0 | 0 | Chordata | 9064.0 | 0 | Carnivora | 9064.0 | 0 | Canidae | 9064.0 | 0 | Canis | 7857.0 | 1 | Canis lupus | 6587.0 | 2 |
| scf7180000042056 | 986908 | 0.3634 | 0 | 59.8134 | Eukaryota | 93302.0 | 0 | Chordata | 93302.0 | 0 | Carnivora | 93302.0 | 0 | Canidae | 93302.0 | 0 | Canis | 93302.0 | 0 | Canis lupus | 93302.0 | 0 |
| scf7180000043370 | 262258 | 0.4428 | 0 | 58.3218 | Eukaryota | 48974.0 | 0 | Chordata | 48974.0 | 0 | Carnivora | 45164.0 | 1 | Canidae | 29760.0 | 3 | Canis | 29760.0 | 5 | Canis lupus | 29760.0 | 6 |
| scf7180000053289 | 7331 | 0.3526 | 0 | 28.9111 | Eukaryota | 4238.0 | 0 | Chordata | 4238.0 | 0 | Artiodactyla | 2493.0 | 2 | Suidae | 2493.0 | 2 | Sus | 2493.0 | 2 | Sus scrofa | 2493.0 | 2 |
| scf7180000051504 | 2180 | 0.4564 | 0 | 161.2635 | Eukaryota | 7883.0 | 0 | Chordata | 7883.0 | 0 | Tetraodontiformes | 5190.0 | 2 | Tetraodontidae | 5190.0 | 2 | Takifugu | 5190.0 | 2 | Takifugu rubripes | 5190.0 | 2 |
| scf7180000040622 | 13476 | 0.4633 | 0 | 282.1539 | Eukaryota | 73516.0 | 0 | Chordata | 73516.0 | 0 | Carnivora | 73516.0 | 0 | Canidae | 73516.0 | 0 | Canis | 73516.0 | 0 | Canis lupus | 73516.0 | 0 |
| scf7180000045004 | 30033 | 0.3674 | 0 | 52.6146 | Eukaryota | 12666.0 | 0 | Chordata | 12666.0 | 0 | Apterygiformes | 8925.0 | 2 | Apterygidae | 8925.0 | 2 | Apteryx | 8925.0 | 2 | Apteryx mantelli | 8925.0 | 2 |
| scf7180000043544 | 282342 | 0.4862 | 0 | 59.515 | Eukaryota | 50733.0 | 0 | Chordata | 50733.0 | 0 | Carnivora | 50733.0 | 0 | Canidae | 50733.0 | 0 | Canis | 50733.0 | 0 | Canis lupus | 50733.0 | 0 |
| scf7180000046544 | 9193 | 0.4412 | 0 | 205.4032 | Eukaryota | 311.0 | 0 | Chordata | 311.0 | 0 | Kurtiformes | 311.0 | 0 | Apogonidae | 311.0 | 0 | Sphaeramia | 311.0 | 0 | Sphaeramia orbicularis | 311.0 | 0 |
| scf7180000043086 | 13160 | 0.4249 | 0 | 27.6136 | Eukaryota | 64417.0 | 0 | Chordata | 64417.0 | 0 | Carnivora | 64417.0 | 0 | Canidae | 64417.0 | 0 | Canis | 64417.0 | 0 | Canis lupus | 64417.0 | 0 |
| scf7180000046787 | 87676 | 0.4407 | 0 | 131.1953 | Eukaryota | 87677.0 | 0 | Chordata | 87677.0 | 0 | Carnivora | 87677.0 | 0 | Canidae | 87677.0 | 0 | Canis | 87677.0 | 0 | Canis lupus | 87677.0 | 0 |
| scf7180000054512 | 197071 | 0.3781 | 0 | 435.0109 | Eukaryota | 44745.0 | 0 | Chordata | 44745.0 | 0 | Carnivora | 44745.0 | 0 | Canidae | 44745.0 | 0 | Canis | 44745.0 | 0 | Canis lupus | 44745.0 | 0 |
| scf7180000041301 | 1135541 | 0.4902 | 0 | 56.3376 | Eukaryota | 77442.0 | 0 | Chordata | 77442.0 | 0 | Carnivora | 77442.0 | 0 | Canidae | 43891.0 | 3 | Canis | 35196.0 | 5 | Canis lupus | 35196.0 | 6 |
| scf7180000044115 | 339313 | 0.3854 | 0 | 60.8088 | Eukaryota | 38860.0 | 0 | Chordata | 38860.0 | 0 | Carnivora | 30304.0 | 1 | Canidae | 30304.0 | 1 | Canis | 30304.0 | 1 | Canis lupus | 30304.0 | 1 |
| scf7180000055315 | 30316 | 0.4412 | 0 | 119.1092 | Eukaryota | 39133.0 | 0 | Chordata | 30267.0 | 2 | Gadiformes | 25765.0 | 6 | Gadidae | 25765.0 | 6 | Gadus | 25765.0 | 6 | Gadus morhua | 25765.0 | 6 |
| scf7180000045531 | 352016 | 0.4753 | 0 | 59.0565 | Eukaryota | 16590.0 | 0 | Chordata | 16590.0 | 0 | Carnivora | 9889.0 | 1 | Canidae | 9889.0 | 1 | Canis | 9889.0 | 1 | Canis lupus | 9889.0 | 1 |
| scf7180000042203 | 55556 | 0.4546 | 0 | 54.3565 | Eukaryota | 63407.0 | 0 | Chordata | 63407.0 | 0 | Carnivora | 63407.0 | 0 | Canidae | 63407.0 | 0 | Canis | 63407.0 | 0 | Canis lupus | 63407.0 | 0 |
| scf7180000053554 | 3019 | 0.4525 | 0 | 367.141 | no-hit | 0.0 | 0 | no-hit | 0.0 | 0 | no-hit | 0.0 | 0 | no-hit | 0.0 | 0 | no-hit | 0.0 | 0 | no-hit | 0.0 | 0 |
| scf7180000042370 | 5328 | 0.4084 | 0 | 19.6824 | Eukaryota | 48061.0 | 0 | Chordata | 48061.0 | 0 | Carnivora | 48061.0 | 0 | Canidae | 48061.0 | 0 | Canis | 48061.0 | 0 | Canis lupus | 48061.0 | 0 |
| scf7180000052617 | 3870 | 0.4496 | 0 | 382.628 | no-hit | 0.0 | 0 | no-hit | 0.0 | 0 | no-hit | 0.0 | 0 | no-hit | 0.0 | 0 | no-hit | 0.0 | 0 | no-hit | 0.0 | 0 |
| scf7180000041031 | 1495835 | 0.3435 | 0 | 59.7646 | Eukaryota | 83492.0 | 0 | Chordata | 83492.0 | 0 | Carnivora | 83492.0 | 0 | Canidae | 83492.0 | 0 | Canis | 83492.0 | 0 | Canis lupus | 83492.0 | 0 |
| scf7180000051341 | 23420 | 0.4381 | 0 | 71.2877 | Eukaryota | 616.0 | 0 | Platyhelminthes | 616.0 | 0 | Polystomatidea | 616.0 | 0 | Polystomatidae | 616.0 | 0 | Protopolystoma | 616.0 | 0 | Protopolystoma xenopodis | 616.0 | 0 |
| scf7180000048516 | 23546 | 0.4373 | 0 | 316.0058 | no-hit | 0.0 | 0 | no-hit | 0.0 | 0 | no-hit | 0.0 | 0 | no-hit | 0.0 | 0 | no-hit | 0.0 | 0 | no-hit | 0.0 | 0 |
| scf7180000042422 | 711184 | 0.427 | 0 | 57.9344 | Eukaryota | 40576.0 | 0 | Chordata | 40576.0 | 0 | Carnivora | 40576.0 | 0 | Canidae | 40576.0 | 0 | Canis | 40576.0 | 0 | Canis lupus | 40576.0 | 0 |
| scf7180000046080 | 174830 | 0.3743 | 0 | 62.4937 | Eukaryota | 87995.0 | 0 | Chordata | 87995.0 | 0 | Carnivora | 87995.0 | 0 | Canidae | 87995.0 | 0 | Canis | 87995.0 | 0 | Canis lupus | 87995.0 | 0 |
| scf7180000054577 | 28088 | 0.3486 | 0 | 494.769 | Eukaryota | 5166.0 | 0 | Chordata | 5166.0 | 0 | Primates | 3588.0 | 1 | Hominidae | 2810.0 | 2 | Homo | 2810.0 | 2 | Homo sapiens | 2810.0 | 2 |
| scf7180000047279 | 68732 | 0.4119 | 0 | 57.9747 | Eukaryota | 77970.0 | 0 | Chordata | 77970.0 | 0 | Carnivora | 77970.0 | 0 | Canidae | 77970.0 | 0 | Canis | 77970.0 | 0 | Canis lupus | 77970.0 | 0 |
| scf7180000050770 | 2812 | 0.4477 | 0 | 167.4945 | no-hit | 0.0 | 0 | no-hit | 0.0 | 0 | no-hit | 0.0 | 0 | no-hit | 0.0 | 0 | no-hit | 0.0 | 0 | no-hit | 0.0 | 0 |
| scf7180000042151 | 768191 | 0.4583 | 0 | 58.8043 | Eukaryota | 126839.0 | 0 | Chordata | 126839.0 | 0 | Carnivora | 126839.0 | 0 | Canidae | 126839.0 | 0 | Canis | 91781.0 | 1 | Canis lupus | 91781.0 | 1 |
| scf7180000053434 | 2351 | 0.4547 | 0 | 222.1981 | Eukaryota | 294.0 | 0 | Streptophyta | 294.0 | 0 | Fabales | 294.0 | 0 | Fabaceae | 294.0 | 0 | Cicer | 294.0 | 0 | Cicer arietinum | 294.0 | 0 |
| scf7180000041745 | 3028 | 0.4174 | 0 | 35.8358 | Eukaryota | 23795.0 | 0 | Chordata | 23795.0 | 0 | Carnivora | 23795.0 | 0 | Canidae | 23795.0 | 0 | Canis | 23795.0 | 0 | Canis lupus | 23795.0 | 0 |
| scf7180000050392 | 10572 | 0.4464 | 0 | 85.6932 | no-hit | 0.0 | 0 | no-hit | 0.0 | 0 | no-hit | 0.0 | 0 | no-hit | 0.0 | 0 | no-hit | 0.0 | 0 | no-hit | 0.0 | 0 |
| scf7180000043056 | 46956 | 0.3923 | 0 | 47.5592 | Eukaryota | 64915.0 | 0 | Chordata | 64915.0 | 0 | Carnivora | 64915.0 | 0 | Canidae | 64915.0 | 0 | Canis | 64915.0 | 0 | Canis lupus | 64915.0 | 0 |
| scf7180000046534 | 8025 | 0.4373 | 0 | 127.8068 | Eukaryota | 4307.0 | 0 | Chordata | 3896.0 | 1 | Kurtiformes | 2826.0 | 4 | Apogonidae | 2826.0 | 4 | Sphaeramia | 2826.0 | 4 | Sphaeramia orbicularis | 2826.0 | 4 |
| scf7180000042092 | 180951 | 0.3916 | 0 | 56.4912 | Eukaryota | 92873.0 | 0 | Chordata | 92873.0 | 0 | Carnivora | 92873.0 | 0 | Canidae | 92873.0 | 0 | Canis | 92873.0 | 0 | Canis lupus | 92873.0 | 0 |
| scf7180000042139 | 731411 | 0.4623 | 0 | 58.6254 | Eukaryota | 51962.0 | 0 | Chordata | 51962.0 | 0 | Carnivora | 51962.0 | 0 | Canidae | 51962.0 | 0 | Canis | 45361.0 | 1 | Canis lupus | 45361.0 | 1 |
| scf7180000050961 | 10718 | 0.4605 | 0 | 456.8139 | Eukaryota | 3456.0 | 0 | Echinodermata | 1704.0 | 2 | Forcipulatida | 1704.0 | 3 | Asteriidae | 1704.0 | 3 | Asterias | 1704.0 | 3 | Asterias rubens | 1704.0 | 3 |
| scf7180000046857 | 8266 | 0.4106 | 0 | 137.5384 | Eukaryota | 3423.0 | 0 | Chordata | 3423.0 | 0 | Primates | 1880.0 | 1 | Hominidae | 1880.0 | 1 | Pan | 1880.0 | 1 | Pan troglodytes | 1880.0 | 1 |
| scf7180000053311 | 6485 | 0.4319 | 0 | 49.8768 | no-hit | 0.0 | 0 | no-hit | 0.0 | 0 | no-hit | 0.0 | 0 | no-hit | 0.0 | 0 | no-hit | 0.0 | 0 | no-hit | 0.0 | 0 |
| scf7180000052246 | 9979 | 0.4462 | 0 | 64.6568 | Eukaryota | 2559.0 | 0 | Chordata | 2559.0 | 0 | Carnivora | 2559.0 | 0 | Canidae | 2559.0 | 0 | Canis | 2559.0 | 0 | Canis lupus | 2559.0 | 0 |
| scf7180000041599 | 37941 | 0.4375 | 0 | 104.6777 | Eukaryota | 4479.0 | 0 | Chordata | 4159.0 | 1 | Tetraodontiformes | 3751.0 | 3 | Tetraodontidae | 3751.0 | 3 | Takifugu | 3751.0 | 3 | Takifugu rubripes | 3751.0 | 3 |
| scf7180000056068 | 722598 | 0.3416 | 0 | 58.9443 | Eukaryota | 43898.0 | 0 | Chordata | 43898.0 | 0 | Carnivora | 43898.0 | 0 | Canidae | 43898.0 | 0 | Canis | 43898.0 | 0 | Canis lupus | 43898.0 | 0 |
| scf7180000051815 | 20878 | 0.6082 | 0 | 59.1155 | Eukaryota | 5880.0 | 0 | Chordata | 5880.0 | 0 | Carnivora | 5880.0 | 0 | Canidae | 5880.0 | 0 | Canis | 3105.0 | 1 | Canis lupus | 3105.0 | 1 |
| scf7180000043997 | 117077 | 0.4599 | 0 | 56.2101 | Eukaryota | 6796.0 | 1 | Chordata | 6796.0 | 1 | Primates | 4137.0 | 4 | Hominidae | 3252.0 | 5 | Homo | 3252.0 | 6 | Homo sapiens | 3252.0 | 6 |
| scf7180000045660 | 3785 | 0.4544 | 0 | 662.9887 | Eukaryota | 2558.0 | 0 | Chordata | 2558.0 | 0 | Carnivora | 2558.0 | 0 | Canidae | 2558.0 | 0 | Canis | 2558.0 | 0 | Canis lupus | 2558.0 | 0 |
| scf7180000043934 | 115522 | 0.4138 | 0 | 57.0985 | Eukaryota | 81078.0 | 0 | Chordata | 81078.0 | 0 | Carnivora | 81078.0 | 0 | Canidae | 81078.0 | 0 | Canis | 81078.0 | 0 | Canis lupus | 81078.0 | 0 |
| scf7180000042470 | 13047 | 0.4672 | 0 | 57.7797 | Eukaryota | 51378.0 | 0 | Chordata | 51378.0 | 0 | Carnivora | 51378.0 | 0 | Canidae | 51378.0 | 0 | Canis | 51378.0 | 0 | Canis lupus | 51378.0 | 0 |
| scf7180000046199 | 23657 | 0.4363 | 0 | 127.7276 | Eukaryota | 7544.0 | 0 | Chordata | 6322.0 | 1 | Anabantiformes | 3088.0 | 3 | Anabantidae | 3088.0 | 3 | Anabas | 3088.0 | 3 | Anabas testudineus | 3088.0 | 3 |
| scf7180000054454 | 16138 | 0.4244 | 0 | 458.0824 | Eukaryota | 45768.0 | 0 | Chordata | 45768.0 | 0 | Carnivora | 45768.0 | 0 | Canidae | 45768.0 | 0 | Canis | 45768.0 | 0 | Canis lupus | 45768.0 | 0 |
| scf7180000048635 | 55881 | 0.3732 | 0 | 57.463 | Eukaryota | 53631.0 | 0 | Chordata | 53631.0 | 0 | Carnivora | 31692.0 | 2 | Mustelidae | 31692.0 | 2 | Mustela | 31692.0 | 2 | Mustela putorius | 31692.0 | 2 |
| scf7180000056169 | 3175353 | 0.3751 | 0 | 60.638 | Eukaryota | 95074.0 | 0 | Chordata | 95074.0 | 0 | Carnivora | 95074.0 | 0 | Canidae | 95074.0 | 0 | Canis | 95074.0 | 0 | Canis lupus | 95074.0 | 0 |
| scf7180000045922 | 161450 | 0.4012 | 0 | 58.9943 | Eukaryota | 88076.0 | 0 | Chordata | 88076.0 | 0 | Carnivora | 88076.0 | 0 | Canidae | 88076.0 | 0 | Canis | 88076.0 | 0 | Canis lupus | 88076.0 | 0 |
| scf7180000055492 | 18176 | 0.5254 | 0 | 597.4949 | Eukaryota | 2968.0 | 0 | Chordata | 2968.0 | 0 | Carnivora | 2968.0 | 0 | Canidae | 2968.0 | 0 | Canis | 2260.0 | 1 | Canis lupus | 2260.0 | 1 |
| scf7180000045991 | 5525 | 0.4259 | 0 | 152.8575 | no-hit | 0.0 | 0 | no-hit | 0.0 | 0 | no-hit | 0.0 | 0 | no-hit | 0.0 | 0 | no-hit | 0.0 | 0 | no-hit | 0.0 | 0 |
| scf7180000050011 | 9813 | 0.4657 | 0 | 28.4932 | Eukaryota | 12750.0 | 0 | Chordata | 12750.0 | 0 | Carnivora | 12750.0 | 0 | Canidae | 12750.0 | 0 | Canis | 12750.0 | 0 | Canis lupus | 12750.0 | 0 |
| scf7180000053625 | 1701 | 0.4274 | 0 | 312.0015 | Eukaryota | 390.0 | 0 | Platyhelminthes | 390.0 | 0 | Polystomatidea | 390.0 | 0 | Polystomatidae | 390.0 | 0 | Protopolystoma | 390.0 | 0 | Protopolystoma xenopodis | 390.0 | 0 |
| scf7180000042534 | 127428 | 0.3864 | 0 | 53.7516 | Eukaryota | 82556.0 | 0 | Chordata | 82556.0 | 0 | Carnivora | 82556.0 | 0 | Canidae | 82556.0 | 0 | Canis | 82556.0 | 0 | Canis lupus | 82556.0 | 0 |
| scf7180000049064 | 14387 | 0.451 | 0 | 1263.9302 | Eukaryota | 2702.0 | 0 | Platyhelminthes | 1387.0 | 2 | Polystomatidea | 1387.0 | 2 | Polystomatidae | 1387.0 | 2 | Protopolystoma | 1387.0 | 2 | Protopolystoma xenopodis | 1387.0 | 2 |
| scf7180000044332 | 27793 | 0.4547 | 0 | 113.7149 | Eukaryota | 92033.0 | 0 | Chordata | 92033.0 | 0 | Carnivora | 92033.0 | 0 | Canidae | 92033.0 | 0 | Canis | 92033.0 | 0 | Canis lupus | 92033.0 | 0 |
| scf7180000044531 | 47038 | 0.3648 | 0 | 59.7852 | Eukaryota | 121341.0 | 0 | Chordata | 121341.0 | 0 | Carnivora | 121341.0 | 0 | Canidae | 121341.0 | 0 | Canis | 121341.0 | 0 | Canis lupus | 121341.0 | 0 |
| scf7180000054083 | 4783 | 0.4399 | 0 | 427.5399 | no-hit | 0.0 | 0 | no-hit | 0.0 | 0 | no-hit | 0.0 | 0 | no-hit | 0.0 | 0 | no-hit | 0.0 | 0 | no-hit | 0.0 | 0 |
| scf7180000049641 | 25150 | 0.5101 | 0 | 424.6054 | Eukaryota | 35728.0 | 0 | Chordata | 35728.0 | 0 | Carnivora | 35728.0 | 0 | Canidae | 35728.0 | 0 | Canis | 35728.0 | 0 | Canis lupus | 35728.0 | 0 |
| scf7180000042236 | 743372 | 0.4332 | 0 | 59.1133 | Eukaryota | 65780.0 | 0 | Chordata | 65780.0 | 0 | Carnivora | 65780.0 | 0 | Canidae | 65780.0 | 0 | Canis | 52516.0 | 1 | Canis lupus | 52516.0 | 1 |
| scf7180000045108 | 68116 | 0.394 | 0 | 55.7542 | Eukaryota | 71114.0 | 0 | Chordata | 71114.0 | 0 | Carnivora | 71114.0 | 0 | Canidae | 37745.0 | 3 | Canis | 25097.0 | 6 | Canis lupus | 25097.0 | 7 |
| scf7180000041523 | 17088 | 0.413 | 0 | 56.8998 | Eukaryota | 86503.0 | 0 | Chordata | 86503.0 | 0 | Carnivora | 86503.0 | 0 | Canidae | 86503.0 | 0 | Canis | 86503.0 | 0 | Canis lupus | 86503.0 | 0 |
| scf7180000050319 | 24461 | 0.3981 | 0 | 61.4349 | Eukaryota | 5154.0 | 0 | Chordata | 5154.0 | 0 | Primates | 2963.0 | 2 | Hominidae | 2298.0 | 3 | Homo | 2298.0 | 3 | Homo sapiens | 2298.0 | 3 |
| scf7180000047628 | 7511 | 0.458 | 0 | 83.4943 | Eukaryota | 8097.0 | 0 | Chordata | 6508.0 | 1 | Kurtiformes | 4154.0 | 3 | Apogonidae | 4154.0 | 3 | Sphaeramia | 4154.0 | 3 | Sphaeramia orbicularis | 4154.0 | 3 |
| scf7180000051405 | 8570 | 0.4536 | 0 | 2067.2605 | Eukaryota | 2841.0 | 0 | Chordata | 2331.0 | 1 | Carnivora | 2331.0 | 1 | Canidae | 2331.0 | 1 | Canis | 2331.0 | 1 | Canis lupus | 2331.0 | 1 |
| scf7180000045533 | 69628 | 0.4449 | 0 | 156.4118 | Eukaryota | 15784.0 | 0 | Chordata | 9135.0 | 2 | Gadiformes | 4802.0 | 6 | Gadidae | 4802.0 | 6 | Gadus | 4802.0 | 6 | Gadus morhua | 4802.0 | 6 |
| scf7180000041528 | 1056440 | 0.4006 | 0 | 68.741 | Eukaryota | 93649.0 | 0 | Chordata | 93649.0 | 0 | Carnivora | 93649.0 | 0 | Canidae | 93649.0 | 0 | Canis | 93649.0 | 0 | Canis lupus | 93649.0 | 0 |
| scf7180000049028 | 23835 | 0.4447 | 0 | 125.4036 | Eukaryota | 512.0 | 0 | Platyhelminthes | 512.0 | 0 | Polystomatidea | 512.0 | 0 | Polystomatidae | 512.0 | 0 | Protopolystoma | 512.0 | 0 | Protopolystoma xenopodis | 512.0 | 0 |
| scf7180000051920 | 1510 | 0.3934 | 0 | 39.2912 | Eukaryota | 3781.0 | 0 | Chordata | 3781.0 | 0 | Primates | 1468.0 | 3 | Hominidae | 1468.0 | 3 | Homo | 1468.0 | 3 | Homo sapiens | 1468.0 | 3 |
| scf7180000042063 | 582406 | 0.3837 | 0 | 58.9376 | Eukaryota | 87507.0 | 0 | Chordata | 87507.0 | 0 | Carnivora | 87507.0 | 0 | Canidae | 87507.0 | 0 | Canis | 87507.0 | 0 | Canis lupus | 87507.0 | 0 |
| scf7180000051211 | 5418 | 0.4526 | 0 | 766.9179 | no-hit | 0.0 | 0 | no-hit | 0.0 | 0 | no-hit | 0.0 | 0 | no-hit | 0.0 | 0 | no-hit | 0.0 | 0 | no-hit | 0.0 | 0 |
| scf7180000045477 | 155518 | 0.4104 | 0 | 60.7944 | Eukaryota | 46887.0 | 0 | Chordata | 46887.0 | 0 | Carnivora | 46887.0 | 0 | Canidae | 46887.0 | 0 | Canis | 46887.0 | 0 | Canis lupus | 46887.0 | 0 |
| scf7180000056519 | 240009 | 0.4165 | 0 | 57.84 | Eukaryota | 18079.0 | 0 | Chordata | 18079.0 | 0 | Carnivora | 18079.0 | 0 | Canidae | 16426.0 | 1 | Vulpes | 9453.0 | 2 | Vulpes vulpes | 9453.0 | 2 |
| scf7180000051079 | 20834 | 0.4386 | 0 | 62.9437 | Eukaryota | 3573.0 | 0 | Chordata | 2586.0 | 1 | Polystomatidea | 987.0 | 4 | Polystomatidae | 987.0 | 4 | Protopolystoma | 987.0 | 4 | Protopolystoma xenopodis | 987.0 | 4 |
| scf7180000040882 | 1674926 | 0.3702 | 0 | 59.2868 | Eukaryota | 93034.0 | 0 | Chordata | 93034.0 | 0 | Carnivora | 93034.0 | 0 | Canidae | 93034.0 | 0 | Canis | 93034.0 | 0 | Canis lupus | 93034.0 | 0 |
| scf7180000048742 | 5096 | 0.4555 | 0 | 811.7004 | no-hit | 0.0 | 0 | no-hit | 0.0 | 0 | no-hit | 0.0 | 0 | no-hit | 0.0 | 0 | no-hit | 0.0 | 0 | no-hit | 0.0 | 0 |
| scf7180000042814 | 614622 | 0.3644 | 0 | 59.207 | Eukaryota | 32443.0 | 0 | Chordata | 32443.0 | 0 | Carnivora | 32443.0 | 0 | Canidae | 32443.0 | 0 | Canis | 32443.0 | 0 | Canis lupus | 32443.0 | 0 |
| scf7180000052786 | 8071 | 0.431 | 0 | 180.518 | no-hit | 0.0 | 0 | no-hit | 0.0 | 0 | no-hit | 0.0 | 0 | no-hit | 0.0 | 0 | no-hit | 0.0 | 0 | no-hit | 0.0 | 0 |
| scf7180000044494 | 284718 | 0.5036 | 0 | 57.5683 | Eukaryota | 22319.0 | 0 | Chordata | 22319.0 | 0 | Carnivora | 22319.0 | 0 | Canidae | 14259.0 | 1 | Zalophus | 8060.0 | 2 | Zalophus californianus | 8060.0 | 2 |

| | | | | | | | | | | | | | | | | | | | | |
|---|---|---|---|---|---|---|---|---|---|---|---|---|---|---|---|---|---|---|---|---|
| scf7180000043748 | 212307 | 0.4426 | 0 | 58.6139 | Eukaryota | 42433.0 | 0 | Chordata | 42433.0 | 0 | Carnivora | 42433.0 | 0 | Canidae | 42433.0 | 0 | Canis | 34089.0 | 1 | Canis lupus | 34089.0 | 1 |
| scf7180000053088 | 7836 | 0.4798 | 0 | 1501.5004 | Eukaryota | 19650.0 | 0 | Chordata | 19650.0 | 0 | Carnivora | 19650.0 | 0 | Canidae | 19650.0 | 0 | Canis | 19650.0 | 0 | Canis lupus | 19650.0 | 0 |
| scf7180000042099 | 89197 | 0.3978 | 0 | 59.5881 | Eukaryota | 29762.0 | 0 | Chordata | 29762.0 | 0 | Carnivora | 29762.0 | 0 | Canidae | 29762.0 | 0 | Canis | 29762.0 | 0 | Canis lupus | 29762.0 | 0 |
| scf7180000055158 | 15292 | 0.4465 | 0 | 252.016 | Eukaryota | 7457.0 | 0 | Chordata | 7457.0 | 0 | Tetraodontiformes | 5322.0 | 1 | Tetraodontidae | 5322.0 | 1 | Takifugu | 5322.0 | 1 | Takifugu rubripes | 5322.0 | 1 |
| scf7180000041622 | 46839 | 0.4204 | 0 | 57.7336 | Eukaryota | 63825.0 | 0 | Chordata | 63825.0 | 0 | Carnivora | 63825.0 | 0 | Canidae | 63825.0 | 0 | Canis | 63825.0 | 0 | Canis lupus | 63825.0 | 0 |
| scf7180000043457 | 442219 | 0.4182 | 0 | 58.7475 | Eukaryota | 32773.0 | 0 | Chordata | 32773.0 | 0 | Carnivora | 32773.0 | 0 | Canidae | 32773.0 | 0 | Canis | 26367.0 | 1 | Canis lupus | 26367.0 | 1 |
| scf7180000048905 | 22803 | 0.4517 | 0 | 197.694 | Eukaryota | 93546.0 | 0 | Chordata | 93546.0 | 0 | Carnivora | 93546.0 | 0 | Canidae | 93546.0 | 0 | Canis | 93546.0 | 0 | Canis lupus | 93546.0 | 0 |
| scf7180000054456 | 55912 | 0.4407 | 0 | 652.3787 | Eukaryota | 5829.0 | 0 | Platyhelminthes | 3955.0 | 2 | Polystomatidea | 3955.0 | 2 | Polystomatidae | 3955.0 | 2 | Protopolystoma | 3955.0 | 2 | Protopolystoma xenopodis | 3955.0 | 2 |
| scf7180000055613 | 11745 | 0.4388 | 0 | 166.5772 | Eukaryota | 6858.0 | 0 | Chordata | 4576.0 | 2 | Kurtiformes | 4420.0 | 3 | Apogonidae | 4420.0 | 3 | Sphaeramia | 4420.0 | 3 | Sphaeramia orbicularis | 4420.0 | 3 |
| scf7180000053041 | 12693 | 0.3873 | 0 | 56.8285 | Eukaryota | 35572.0 | 0 | Chordata | 35572.0 | 0 | Carnivora | 35572.0 | 0 | Canidae | 35572.0 | 0 | Canis | 35572.0 | 0 | Canis lupus | 35572.0 | 0 |
| scf7180000041980 | 36818 | 0.415 | 0 | 58.1988 | Eukaryota | 49362.0 | 0 | Chordata | 49362.0 | 0 | Carnivora | 49362.0 | 0 | Canidae | 49362.0 | 0 | Canis | 49362.0 | 0 | Canis lupus | 49362.0 | 0 |
| scf7180000041009 | 1505688 | 0.4926 | 0 | 58.5185 | Eukaryota | 66845.0 | 0 | Chordata | 66845.0 | 0 | Carnivora | 66845.0 | 0 | Canidae | 66845.0 | 0 | Canis | 66845.0 | 0 | Canis lupus | 66845.0 | 0 |
| scf7180000049751 | 33015 | 0.5029 | 0 | 443.675 | Eukaryota | 8830.0 | 0 | Chordata | 8830.0 | 0 | Carnivora | 8830.0 | 0 | Canidae | 8830.0 | 0 | Canis | 7145.0 | 1 | Canis lupus | 5984.0 | 2 |
| scf7180000044882 | 33261 | 0.4926 | 0 | 498.0451 | Eukaryota | 9564.0 | 0 | Chordata | 9564.0 | 0 | Carnivora | 9564.0 | 0 | Canidae | 9564.0 | 0 | Canis | 8891.0 | 1 | Canis lupus | 7630.0 | 2 |
| scf7180000042306 | 137069 | 0.3944 | 0 | 62.5756 | Eukaryota | 17810.0 | 1 | Chordata | 17810.0 | 1 | Carnivora | 9583.0 | 3 | Canidae | 7832.0 | 4 | Canis | 5844.0 | 5 | Canis lupus | 5844.0 | 5 |
| scf7180000049589 | 36950 | 0.4493 | 0 | 61.0121 | Eukaryota | 11033.0 | 0 | Chordata | 11033.0 | 0 | Carnivora | 11033.0 | 0 | Canidae | 11033.0 | 0 | Canis | 8462.0 | 1 | Canis lupus | 8462.0 | 1 |
| scf7180000045244 | 199928 | 0.4523 | 0 | 60.3305 | Eukaryota | 49613.0 | 0 | Chordata | 49613.0 | 0 | Carnivora | 49613.0 | 0 | Canidae | 49613.0 | 0 | Canis | 49613.0 | 0 | Canis lupus | 49613.0 | 0 |
| scf7180000043615 | 4751 | 0.3502 | 0 | 20.1832 | Eukaryota | 27304.0 | 0 | Chordata | 27304.0 | 0 | Carnivora | 27304.0 | 0 | Canidae | 27304.0 | 0 | Canis | 27304.0 | 0 | Canis lupus | 27304.0 | 0 |
| scf7180000056131 | 4681 | 0.4399 | 0 | 145.98 | Eukaryota | 16640.0 | 0 | Chordata | 13933.0 | 2 | Gadiformes | 3389.0 | 6 | Gadidae | 3389.0 | 6 | Gadus | 3389.0 | 7 | Gadus morhua | 3389.0 | 7 |
| scf7180000056575 | 92873 | 0.3723 | 0 | 61.1018 | Eukaryota | 15862.0 | 0 | Chordata | 15862.0 | 0 | Artiodactyla | 7820.0 | 2 | Suidae | 7820.0 | 2 | Sus | 7820.0 | 2 | Sus scrofa | 7820.0 | 2 |
| scf7180000050809 | 2961 | 0.4448 | 0 | 690.663 | no-hit | 0.0 | 0 | no-hit | 0.0 | 0 | no-hit | 0.0 | 0 | no-hit | 0.0 | 0 | no-hit | 0.0 | 0 | no-hit | 0.0 | 0 |
| scf7180000052359 | 6202 | 0.5044 | 0 | 231.876 | Eukaryota | 7962.0 | 0 | Chordata | 7962.0 | 0 | Carnivora | 7962.0 | 0 | Canidae | 7962.0 | 0 | Canis | 5786.0 | 1 | Canis lupus | 4629.0 | 2 |
| scf7180000055450 | 2764 | 0.4345 | 0 | 85.0133 | Eukaryota | 2142.0 | 0 | Chordata | 2142.0 | 0 | Columbiformes | 2142.0 | 0 | Columbidae | 2142.0 | 0 | Streptopelia | 2142.0 | 0 | Streptopelia turtur | 2142.0 | 0 |
| scf7180000050907 | 18798 | 0.4464 | 0 | 576.6034 | Eukaryota | 2011.0 | 0 | Platyhelminthes | 983.0 | 2 | Polystomatidea | 983.0 | 2 | Polystomatidae | 983.0 | 2 | Protopolystoma | 983.0 | 2 | Protopolystoma xenopodis | 983.0 | 2 |
| scf7180000051291 | 20475 | 0.4802 | 0 | 93.5959 | Eukaryota | 8994.0 | 0 | Chordata | 8994.0 | 0 | Carnivora | 8994.0 | 0 | Canidae | 8994.0 | 0 | Canis | 7796.0 | 1 | Canis lupus | 6567.0 | 2 |
| scf7180000046738 | 100672 | 0.5584 | 0 | 54.0161 | Eukaryota | 64172.0 | 0 | Chordata | 64172.0 | 0 | Carnivora | 64172.0 | 0 | Ursidae | 17580.0 | 5 | Canis | 16790.0 | 7 | Canis lupus | 16790.0 | 8 |
| scf7180000051675 | 6982 | 0.4529 | 0 | 546.0963 | Eukaryota | 492.0 | 0 | Streptophyta | 492.0 | 0 | Fabales | 492.0 | 0 | Fabaceae | 492.0 | 0 | Cicer | 492.0 | 0 | Cicer arietinum | 492.0 | 0 |
| scf7180000041829 | 26802 | 0.3639 | 0 | 54.9864 | Eukaryota | 3311.0 | 0 | Chordata | 3311.0 | 0 | Primates | 1517.0 | 2 | Hominidae | 1517.0 | 2 | Homo | 1517.0 | 2 | Homo sapiens | 1517.0 | 2 |
| scf7180000045103 | 219960 | 0.3878 | 0 | 58.9578 | Eukaryota | 9058.0 | 0 | Chordata | 9058.0 | 0 | Carnivora | 6597.0 | 1 | Canidae | 6597.0 | 1 | Canis | 6597.0 | 1 | Canis lupus | 6597.0 | 1 |
| scf7180000054066 | 3593 | 0.4448 | 0 | 325.6527 | no-hit | 0.0 | 0 | no-hit | 0.0 | 0 | no-hit | 0.0 | 0 | no-hit | 0.0 | 0 | no-hit | 0.0 | 0 | no-hit | 0.0 | 0 |
| scf7180000047376 | 78566 | 0.4176 | 0 | 59.823 | Eukaryota | 61738.0 | 0 | Chordata | 61738.0 | 0 | Carnivora | 61738.0 | 0 | Canidae | 61738.0 | 0 | Canis | 61738.0 | 0 | Canis lupus | 61738.0 | 0 |
| scf7180000040550 | 2331643 | 0.5263 | 0 | 58.4533 | Eukaryota | 84591.0 | 0 | Chordata | 84591.0 | 0 | Carnivora | 84591.0 | 0 | Canidae | 84591.0 | 0 | Canis | 75385.0 | 1 | Canis lupus | 75385.0 | 1 |
| scf7180000045414 | 267387 | 0.3605 | 0 | 59.4633 | Eukaryota | 93052.0 | 0 | Chordata | 93052.0 | 0 | Carnivora | 93052.0 | 0 | Canidae | 93052.0 | 0 | Canis | 93052.0 | 0 | Canis lupus | 93052.0 | 0 |
| scf7180000056232 | 211332 | 0.3705 | 0 | 59.423 | Eukaryota | 72701.0 | 0 | Chordata | 72701.0 | 0 | Carnivora | 72701.0 | 0 | Canidae | 39147.0 | 3 | Ursus | 24805.0 | 4 | Ursus arctos | 24805.0 | 4 |
| scf7180000049417 | 16545 | 0.4032 | 0 | 424.4723 | Eukaryota | 29661.0 | 0 | Chordata | 29661.0 | 0 | Carnivora | 29661.0 | 0 | Canidae | 16959.0 | 4 | Canis | 14205.0 | 6 | Canis lupus | 14205.0 | 6 |
| scf7180000040754 | 37536 | 0.4355 | 0 | 176.8375 | Eukaryota | 2281.0 | 0 | Platyhelminthes | 1643.0 | 1 | Polystomatidea | 1167.0 | 2 | Polystomatidae | 1167.0 | 2 | Protopolystoma | 1167.0 | 2 | Protopolystoma xenopodis | 1167.0 | 2 |
| scf7180000054172 | 42854 | 0.4445 | 0 | 467.2193 | Eukaryota | 72603.0 | 0 | Chordata | 54931.0 | 1 | Gadiformes | 45204.0 | 4 | Gadidae | 45204.0 | 4 | Gadus | 45204.0 | 4 | Gadus morhua | 45204.0 | 4 |
| scf7180000055173 | 32565 | 0.4405 | 0 | 137.7122 | Eukaryota | 7635.0 | 0 | Chordata | 6846.0 | 1 | Kurtiformes | 5391.0 | 2 | Apogonidae | 5391.0 | 2 | Sphaeramia | 5391.0 | 2 | Sphaeramia orbicularis | 5391.0 | 2 |
| scf7180000055891 | 9388 | 0.757 | 0 | 177.7288 | Eukaryota | 2449.0 | 0 | Chordata | 2449.0 | 0 | Rodentia | 1221.0 | 4 | Caviidae | 723.0 | 6 | Cavia | 723.0 | 6 | Cavia porcellus | 723.0 | 6 |
| scf7180000049791 | 12229 | 0.4408 | 0 | 54.9524 | no-hit | 0.0 | 0 | no-hit | 0.0 | 0 | no-hit | 0.0 | 0 | no-hit | 0.0 | 0 | no-hit | 0.0 | 0 | no-hit | 0.0 | 0 |
| scf7180000049526 | 43900 | 0.5344 | 0 | 58.2853 | Eukaryota | 30647.0 | 0 | Chordata | 30647.0 | 0 | Carnivora | 30647.0 | 0 | Canidae | 30647.0 | 0 | Canis | 30647.0 | 0 | Canis lupus | 30647.0 | 0 |
| scf7180000041322 | 509146 | 0.3671 | 0 | 60.7683 | Eukaryota | 47858.0 | 0 | Chordata | 47858.0 | 0 | Carnivora | 47858.0 | 0 | Canidae | 47858.0 | 0 | Canis | 47858.0 | 0 | Canis lupus | 47858.0 | 0 |
| scf7180000041384 | 1945478 | 0.4704 | 100 | 58.7849 | Eukaryota | 110476.0 | 0 | Chordata | 110476.0 | 0 | Carnivora | 110476.0 | 0 | Canidae | 110476.0 | 0 | Canis | 83675.0 | 1 | Canis lupus | 83675.0 | 1 |
| scf7180000056227 | 1557230 | 0.3697 | 0 | 60.3706 | Eukaryota | 82938.0 | 0 | Chordata | 82938.0 | 0 | Carnivora | 82938.0 | 0 | Canidae | 62902.0 | 2 | Canis | 37259.0 | 3 | Canis lupus | 37259.0 | 3 |
| scf7180000047177 | 3856 | 0.4157 | 0 | 26.285 | Eukaryota | 8714.0 | 0 | Chordata | 8714.0 | 0 | Carnivora | 8714.0 | 0 | Canidae | 8714.0 | 0 | Canis | 8714.0 | 0 | Canis lupus | 8714.0 | 0 |
| scf7180000052626 | 5750 | 0.4471 | 0 | 883.2704 | Eukaryota | 2709.0 | 0 | Platyhelminthes | 1487.0 | 3 | Polystomatidea | 1487.0 | 3 | Polystomatidae | 1487.0 | 3 | Protopolystoma | 1487.0 | 3 | Protopolystoma xenopodis | 1487.0 | 3 |
| scf7180000043458 | 480961 | 0.3577 | 0 | 60.669 | Eukaryota | 139354.0 | 0 | Chordata | 139354.0 | 0 | Carnivora | 70600.0 | 2 | Canidae | 62511.0 | 3 | Canis | 62511.0 | 5 | Canis lupus | 62511.0 | 5 |
| scf7180000044226 | 54671 | 0.439 | 0 | 370.0126 | Eukaryota | 78029.0 | 0 | Chordata | 78029.0 | 0 | Carnivora | 78029.0 | 0 | Canidae | 78029.0 | 0 | Canis | 78029.0 | 0 | Canis lupus | 78029.0 | 0 |
| scf7180000042121 | 609227 | 0.3924 | 0 | 58.8407 | Eukaryota | 77081.0 | 0 | Chordata | 77081.0 | 0 | Carnivora | 77081.0 | 0 | Canidae | 77081.0 | 0 | Canis | 77081.0 | 0 | Canis lupus | 77081.0 | 0 |
| scf7180000042055 | 230458 | 0.3809 | 0 | 60.0976 | Eukaryota | 48450.0 | 0 | Chordata | 48450.0 | 0 | Carnivora | 48450.0 | 0 | Canidae | 48450.0 | 0 | Canis | 45840.0 | 1 | Canis lupus | 45840.0 | 1 |
| scf7180000043207 | 145898 | 0.4495 | 0 | 55.7399 | Eukaryota | 46864.0 | 0 | Chordata | 46864.0 | 0 | Carnivora | 46864.0 | 0 | Canidae | 28141.0 | 3 | Canis | 25446.0 | 6 | Canis lupus | 25446.0 | 6 |
| scf7180000040504 | 2474121 | 0.4015 | 0 | 58.7286 | Eukaryota | 95401.0 | 0 | Chordata | 95401.0 | 0 | Carnivora | 95401.0 | 0 | Canidae | 95401.0 | 0 | Canis | 95401.0 | 0 | Canis lupus | 95401.0 | 0 |
| scf7180000041383 | 76051 | 0.376 | 0 | 53.9607 | Eukaryota | 40919.0 | 0 | Chordata | 40919.0 | 0 | Carnivora | 40919.0 | 0 | Canidae | 40919.0 | 0 | Canis | 24344.0 | 1 | Canis lupus | 24344.0 | 1 |
| scf7180000046922 | 25825 | 0.4381 | 0 | 81.5467 | Eukaryota | 6505.0 | 0 | Chordata | 4831.0 | 1 | Synbranchiformes | 2959.0 | 3 | Mastacembelidae | 2959.0 | 3 | Mastacembelus | 2959.0 | 3 | Mastacembelus armatus | 2959.0 | 3 |
| scf7180000045079 | 205295 | 0.3842 | 0 | 60.9613 | Eukaryota | 40278.0 | 0 | Chordata | 40278.0 | 0 | Carnivora | 40278.0 | 0 | Canidae | 40278.0 | 0 | Canis | 40278.0 | 0 | Canis lupus | 40278.0 | 0 |
| scf7180000046891 | 8842 | 0.4414 | 0 | 62.5235 | Eukaryota | 532.0 | 0 | Platyhelminthes | 532.0 | 0 | Polystomatidea | 532.0 | 0 | Polystomatidae | 532.0 | 0 | Protopolystoma | 532.0 | 0 | Protopolystoma xenopodis | 532.0 | 0 |
| scf7180000042485 | 645556 | 0.366 | 0 | 60.7019 | Eukaryota | 77299.0 | 0 | Chordata | 77299.0 | 0 | Carnivora | 77299.0 | 0 | Canidae | 77299.0 | 0 | Canis | 77299.0 | 0 | Canis lupus | 77299.0 | 0 |
| scf7180000043335 | 560932 | 0.4311 | 0 | 60.293 | Eukaryota | 98646.0 | 0 | Chordata | 98646.0 | 0 | Carnivora | 98646.0 | 0 | Canidae | 98646.0 | 0 | Canis | 49898.0 | 1 | Canis lupus | 49898.0 | 1 |
| scf7180000043670 | 232653 | 0.3653 | 0 | 58.6151 | Eukaryota | 89152.0 | 0 | Chordata | 89152.0 | 0 | Carnivora | 89152.0 | 0 | Canidae | 89152.0 | 0 | Canis | 89152.0 | 0 | Canis lupus | 89152.0 | 0 |
| scf7180000052409 | 3378 | 0.4408 | 0 | 51.8818 | Eukaryota | 5697.0 | 0 | Chordata | 3807.0 | 1 | Kurtiformes | 1976.0 | 2 | Apogonidae | 1976.0 | 2 | Sphaeramia | 1976.0 | 2 | Sphaeramia orbicularis | 1976.0 | 2 |
| scf7180000056072 | 13167 | 0.4333 | 0 | 190.6625 | Eukaryota | 17508.0 | 0 | Chordata | 17508.0 | 0 | Carnivora | 17508.0 | 0 | Canidae | 17508.0 | 0 | Canis | 17508.0 | 0 | Canis lupus | 17508.0 | 0 |
| scf7180000046458 | 57160 | 0.3523 | 0 | 62.2343 | Eukaryota | 25605.0 | 0 | Chordata | 25605.0 | 0 | Carnivora | 25605.0 | 0 | Canidae | 25605.0 | 0 | Canis | 25605.0 | 0 | Canis lupus | 25605.0 | 0 |
| scf7180000041775 | 95393 | 0.4559 | 0 | 57.2535 | Eukaryota | 18813.0 | 0 | Chordata | 18813.0 | 0 | Carnivora | 18813.0 | 0 | Canidae | 18813.0 | 0 | Canis | 18813.0 | 0 | Canis lupus | 18813.0 | 0 |
| scf7180000046097 | 27423 | 0.5072 | 0 | 166.8031 | Eukaryota | 8626.0 | 0 | Chordata | 8626.0 | 0 | Carnivora | 8626.0 | 0 | Canidae | 8626.0 | 0 | Canis | 7005.0 | 1 | Canis lupus | 5822.0 | 2 |
| scf7180000040731 | 18035 | 0.4389 | 0 | 170.2491 | Eukaryota | 2593.0 | 0 | Platyhelminthes | 1506.0 | 1 | Polystomatidea | 992.0 | 3 | Polystomatidae | 992.0 | 3 | Protopolystoma | 992.0 | 3 | Protopolystoma xenopodis | 992.0 | 3 |
| scf7180000044583 | 257757 | 0.4981 | 100 | 57.9963 | Eukaryota | 67196.0 | 0 | Chordata | 67196.0 | 0 | Carnivora | 67196.0 | 0 | Canidae | 61815.0 | 2 | Canis | 34134.0 | 4 | Canis lupus | 34134.0 | 4 |
| scf7180000048022 | 4799 | 0.4447 | 0 | 130.0006 | Eukaryota | 17100.0 | 1 | Chordata | 17100.0 | 1 | Tetraodontiformes | 4858.0 | 5 | Tetraodontidae | 4858.0 | 5 | Takifugu | 4858.0 | 5 | Takifugu rubripes | 4858.0 | 5 |
| scf7180000044216 | 16918 | 0.3916 | 0 | 128.8093 | Eukaryota | 90903.0 | 0 | Chordata | 90903.0 | 0 | Carnivora | 90903.0 | 0 | Canidae | 90903.0 | 0 | Canis | 90903.0 | 0 | Canis lupus | 90903.0 | 0 |
| scf7180000052403 | 4165 | 0.443 | 0 | 121.6433 | Eukaryota | 10178.0 | 0 | Chordata | 6200.0 | 3 | Gadiformes | 2730.0 | 7 | Gadidae | 2730.0 | 7 | Gadus | 2730.0 | 7 | Gadus morhua | 2730.0 | 7 |
| scf7180000046424 | 12976 | 0.3855 | 0 | 52.8417 | Eukaryota | 8731.0 | 0 | Chordata | 8731.0 | 0 | Carnivora | 8731.0 | 0 | Canidae | 6448.0 | 1 | Canis | 4630.0 | 2 | Canis lupus | 4630.0 | 2 |
| scf7180000041687 | 1028500 | 0.3689 | 0 | 60.9296 | Eukaryota | 101967.0 | 0 | Chordata | 101967.0 | 0 | Carnivora | 101967.0 | 0 | Canidae | 101967.0 | 0 | Canis | 101967.0 | 0 | Canis lupus | 101967.0 | 0 |
| scf7180000046220 | 23126 | 0.3969 | 0 | 125.5462 | Eukaryota | 45091.0 | 0 | Chordata | 31884.0 | 1 | Gadiformes | 17555.0 | 7 | Gadidae | 17555.0 | 7 | Gadus | 17555.0 | 7 | Gadus morhua | 17555.0 | 7 |
| scf7180000042015 | 808313 | 0.4753 | 0 | 60.302 | Eukaryota | 61534.0 | 0 | Chordata | 61534.0 | 0 | Carnivora | 61534.0 | 0 | Canidae | 61534.0 | 0 | Canis | 45592.0 | 1 | Canis lupus | 45592.0 | 1 |
| scf7180000051836 | 8973 | 0.4468 | 0 | 843.0477 | Eukaryota | 167.0 | 0 | Platyhelminthes | 167.0 | 0 | Polystomatidea | 167.0 | 0 | Polystomatidae | 167.0 | 0 | Protopolystoma | 167.0 | 0 | Protopolystoma xenopodis | 167.0 | 0 |
| scf7180000055702 | 16485 | 0.4401 | 0 | 229.1765 | Eukaryota | 34786.0 | 0 | Chordata | 28544.0 | 2 | Gadiformes | 17678.0 | 6 | Gadidae | 17678.0 | 6 | Gadus | 17678.0 | 6 | Gadus morhua | 17678.0 | 6 |
| scf7180000047765 | 62496 | 0.366 | 0 | 58.0198 | Eukaryota | 9875.0 | 1 | Chordata | 9875.0 | 1 | Carnivora | 5054.0 | 2 | Canidae | 5054.0 | 2 | Canis | 5054.0 | 3 | Canis lupus | 5054.0 | 3 |
| scf7180000047458 | 82178 | 0.3674 | 0 | 58.8798 | Eukaryota | 25804.0 | 0 | Chordata | 25804.0 | 0 | Carnivora | 25804.0 | 0 | Canidae | 25804.0 | 0 | Canis | 23209.0 | 1 | Canis lupus | 23209.0 | 1 |
| scf7180000040918 | 1153658 | 0.3912 | 0 | 59.1207 | Eukaryota | 81725.0 | 0 | Chordata | 81725.0 | 0 | Carnivora | 81725.0 | 0 | Canidae | 81725.0 | 0 | Canis | 81725.0 | 0 | Canis lupus | 81725.0 | 0 |
| scf7180000042050 | 847605 | 0.4466 | 0 | 60.5243 | Eukaryota | 94409.0 | 0 | Chordata | 94409.0 | 0 | Carnivora | 94409.0 | 0 | Canidae | 94409.0 | 0 | Canis | 94409.0 | 0 | Canis lupus | 94409.0 | 0 |
| scf7180000044603 | 359844 | 0.3464 | 0 | 59.8432 | Eukaryota | 41322.0 | 0 | Chordata | 41322.0 | 0 | Carnivora | 41322.0 | 0 | Canidae | 41322.0 | 0 | Canis | 41322.0 | 0 | Canis lupus | 41322.0 | 0 |
| scf7180000053790 | 2290 | 0.5017 | 0 | 40.9644 | Eukaryota | 4148.0 | 0 | Chordata | 4148.0 | 0 | Carnivora | 4148.0 | 0 | Canidae | 4148.0 | 0 | Canis | 3397.0 | 1 | Canis lupus | 3397.0 | 1 |
| scf7180000048941 | 7303 | 0.5017 | 0 | 1402.8184 | Eukaryota | 34031.0 | 0 | Chordata | 34031.0 | 0 | Carnivora | 34031.0 | 0 | Canidae | 34031.0 | 0 | Canis | 34031.0 | 0 | Canis lupus | 34031.0 | 0 |
| scf7180000042587 | 63887 | 0.3914 | 0 | 57.3899 | Eukaryota | 17900.0 | 0 | Chordata | 17900.0 | 0 | Carnivora | 17900.0 | 0 | Canidae | 17900.0 | 0 | Canis | 17900.0 | 0 | Canis lupus | 17900.0 | 0 |
| scf7180000048307 | 9052 | 0.442 | 0 | 78.5369 | Eukaryota | 748.0 | 0 | Chordata | 600.0 | 1 | Columbiformes | 394.0 | 2 | Columbidae | 394.0 | 2 | Streptopelia | 394.0 | 2 | Streptopelia turtur | 394.0 | 2 |
| scf7180000044939 | 226848 | 0.4225 | 0 | 59.982 | Eukaryota | 31974.0 | 0 | Chordata | 31974.0 | 0 | Carnivora | 31974.0 | 0 | Felidae | 16628.0 | 2 | Panthera | 11116.0 | 4 | Panthera pardus | 11116.0 | 4 |
| scf7180000046160 | 23754 | 0.4363 | 0 | 236.9612 | Eukaryota | 95639.0 | 0 | Chordata | 95639.0 | 0 | Carnivora | 95639.0 | 0 | Canidae | 95639.0 | 0 | Canis | 95639.0 | 0 | Canis lupus | 95639.0 | 0 |
| scf7180000041024 | 874519 | 0.4228 | 0 | 60.1757 | Eukaryota | 26504.0 | 0 | Chordata | 26504.0 | 0 | Carnivora | 26504.0 | 0 | Canidae | 26504.0 | 0 | Canis | 26504.0 | 0 | Canis lupus | 26504.0 | 0 |
| scf7180000050540 | 9447 | 0.4445 | 0 | 97.4819 | no-hit | 0.0 | 0 | no-hit | 0.0 | 0 | no-hit | 0.0 | 0 | no-hit | 0.0 | 0 | no-hit | 0.0 | 0 | no-hit | 0.0 | 0 |
| scf7180000054814 | 25653 | 0.4426 | 0 | 333.4053 | no-hit | 0.0 | 0 | no-hit | 0.0 | 0 | no-hit | 0.0 | 0 | no-hit | 0.0 | 0 | no-hit | 0.0 | 0 | no-hit | 0.0 | 0 |
| scf7180000049494 | 20701 | 0.4508 | 0 | 631.325 | Eukaryota | 9730.0 | 0 | Chordata | 8808.0 | 1 | Tetraodontiformes | 5750.0 | 3 | Tetraodontidae | 5750.0 | 3 | Takifugu | 5750.0 | 3 | Takifugu rubripes | 5750.0 | 3 |
| scf7180000052793 | 4652 | 0.4518 | 0 | 1276.5337 | unresolved | 183.0 | 1 | unresolved | 183.0 | 1 | unresolved | 183.0 | 1 | unresolved | 183.0 | 1 | unresolved | 183.0 | 1 | unresolved | 183.0 | 1 |
| scf7180000054715 | 30437 | 0.4392 | 0 | 223.7463 | Eukaryota | 12606.0 | 1 | Chordata | 10095.0 | 2 | Tetraodontiformes | 8500.0 | 4 | Tetraodontidae | 8500.0 | 4 | Takifugu | 8500.0 | 4 | Takifugu rubripes | 8500.0 | 4 |
| scf7180000054991 | 30090 | 0.4362 | 0 | 213.8518 | Eukaryota | 28246.0 | 0 | Chordata | 16807.0 | 1 | Strigeidida | 8526.0 | 6 | Schistosomatidae | 8526.0 | 6 | Schistosoma | 8526.0 | 6 | Schistosoma mansoni | 8526.0 | 6 |
| scf7180000046032 | 142741 | 0.3875 | 0 | 60.0946 | Eukaryota | 37618.0 | 0 | Chordata | 37618.0 | 0 | Carnivora | 37618.0 | 0 | Canidae | 37618.0 | 0 | Canis | 37618.0 | 0 | Canis lupus | 37618.0 | 0 |
| scf7180000042505 | 662840 | 0.409 | 0 | 61.0042 | Eukaryota | 88906.0 | 0 | Chordata | 88906.0 | 0 | Carnivora | 88906.0 | 0 | Canidae | 88906.0 | 0 | Canis | 88906.0 | 0 | Canis lupus | 88906.0 | 0 |
| scf7180000044692 | 311974 | 0.4606 | 0 | 58.898 | Eukaryota | 58087.0 | 0 | Chordata | 58087.0 | 0 | Carnivora | 58087.0 | 0 | Canidae | 58087.0 | 0 | Canis | 58087.0 | 0 | Canis lupus | 58087.0 | 0 |
| scf7180000049350 | 11546 | 0.4535 | 0 | 86.4989 | no-hit | 0.0 | 0 | no-hit | 0.0 | 0 | no-hit | 0.0 | 0 | no-hit | 0.0 | 0 | no-hit | 0.0 | 0 | no-hit | 0.0 | 0 |
| scf7180000048184 | 54368 | 0.4637 | 0 | 60.017 | Eukaryota | 35526.0 | 0 | Chordata | 35526.0 | 0 | Carnivora | 35526.0 | 0 | Canidae | 35526.0 | 0 | Canis | 28700.0 | 1 | Canis lupus | 28700.0 | 1 |
| scf7180000050964 | 11547 | 0.5196 | 0 | 654.6014 | Eukaryota | 13858.0 | 0 | Chordata | 13858.0 | 0 | Carnivora | 13858.0 | 0 | Canidae | 13858.0 | 0 | Canis | 13858.0 | 0 | Canis lupus | 13858.0 | 0 |
| scf7180000047338 | 6639 | 0.4903 | 0 | 122.3875 | Eukaryota | 25676.0 | 0 | Chordata | 25676.0 | 0 | Carnivora | 25676.0 | 0 | Canidae | 25676.0 | 0 | Canis | 25676.0 | 0 | Canis lupus | 25676.0 | 0 |
| scf7180000045248 | 129793 | 0.4654 | 0 | 58.2004 | Eukaryota | 60754.0 | 0 | Chordata | 60754.0 | 0 | Carnivora | 57518.0 | 1 | Canidae | 33736.0 | 4 | Canis | 19649.0 | 6 | Canis lupus | 19649.0 | 6 |
| scf7180000040827 | 1588581 | 0.4415 | 0 | 58.8417 | Eukaryota | 72892.0 | 0 | Chordata | 72892.0 | 0 | Carnivora | 72892.0 | 0 | Canidae | 72892.0 | 0 | Canis | 43352.0 | 1 | Canis lupus | 43352.0 | 1 |
| scf7180000047738 | 72000 | 0.4133 | 0 | 55.5475 | Eukaryota | 40273.0 | 0 | Chordata | 40273.0 | 0 | Carnivora | 40273.0 | 0 | Canidae | 40273.0 | 0 | Canis | 33404.0 | 1 | Canis lupus | 33404.0 | 1 |
| scf7180000043353 | 440735 | 0.4743 | 0 | 57.5974 | Eukaryota | 58562.0 | 0 | Chordata | 58562.0 | 0 | Carnivora | 58562.0 | 0 | Canidae | 58562.0 | 0 | Canis | 58562.0 | 0 | Canis lupus | 58562.0 | 0 |
| scf7180000050949 | 6835 | 0.4462 | 0 | 136.2575 | Eukaryota | 1338.0 | 0 | Chordata | 1338.0 | 0 | Synbranchiformes | 966.0 | 1 | Mastacembelidae | 966.0 | 1 | Mastacembelus | 966.0 | 1 | Mastacembelus armatus | 966.0 | 1 |
| scf7180000046682 | 22552 | 0.4386 | 0 | 141.1456 | Eukaryota | 11824.0 | 0 | Chordata | 11111.0 | 1 | Tetraodontiformes | 7064.0 | 3 | Tetraodontidae | 7064.0 | 3 | Takifugu | 7064.0 | 4 | Takifugu rubripes | 7064.0 | 4 |
| scf7180000043953 | 37391 | 0.4763 | 0 | 92.1943 | Eukaryota | 56453.0 | 0 | Chordata | 56453.0 | 0 | Carnivora | 56453.0 | 0 | Canidae | 56453.0 | 0 | Canis | 56453.0 | 0 | Canis lupus | 56453.0 | 0 |
| scf7180000056608 | 447971 | 0.3954 | 0 | 62.5296 | Eukaryota | 91623.0 | 0 | Chordata | 91623.0 | 0 | Carnivora | 91623.0 | 0 | Canidae | 91623.0 | 0 | Canis | 91623.0 | 0 | Canis lupus | 91623.0 | 0 |
| scf7180000050378 | 15666 | 0.4489 | 0 | 58.0781 | Eukaryota | 73743.0 | 0 | Chordata | 73743.0 | 0 | Carnivora | 73743.0 | 0 | Canidae | 73743.0 | 0 | Canis | 73743.0 | 0 | Canis lupus | 73743.0 | 0 |
| scf7180000048551 | 10112 | 0.4393 | 0 | 179.8134 | Eukaryota | 7330.0 | 0 | Chordata | 7330.0 | 0 | Carnivora | 7330.0 | 0 | Canidae | 7330.0 | 0 | Canis | 7330.0 | 0 | Canis lupus | 7330.0 | 0 |
| scf7180000040981 | 56988 | 0.3913 | 0 | 59.4061 | Eukaryota | 105337.0 | 0 | Chordata | 105337.0 | 0 | Carnivora | 105337.0 | 0 | Canidae | 105337.0 | 0 | Canis | 105337.0 | 0 | Canis lupus | 105337.0 | 0 |
| scf7180000049302 | 5070 | 0.4481 | 0 | 359.0187 | no-hit | 0.0 | 0 | no-hit | 0.0 | 0 | no-hit | 0.0 | 0 | no-hit | 0.0 | 0 | no-hit | 0.0 | 0 | no-hit | 0.0 | 0 |
| scf7180000043634 | 16122 | 0.438 | 0 | 91.0004 | no-hit | 0.0 | 0 | no-hit | 0.0 | 0 | no-hit | 0.0 | 0 | no-hit | 0.0 | 0 | no-hit | 0.0 | 0 | no-hit | 0.0 | 0 |
| scf7180000052926 | 10427 | 0.4402 | 0 | 67.2943 | Eukaryota | 34582.0 | 0 | Chordata | 34582.0 | 0 | Carnivora | 34582.0 | 0 | Canidae | 34582.0 | 0 | Canis | 34582.0 | 0 | Canis lupus | 34582.0 | 0 |
| scf7180000052511 | 5783 | 0.4473 | 0 | 381.8001 | Eukaryota | 808.0 | 0 | Chordata | 808.0 | 0 | Tetraodontiformes | 808.0 | 0 | Tetraodontidae | 808.0 | 0 | Takifugu | 808.0 | 0 | Takifugu rubripes | 808.0 | 0 |
| scf7180000044951 | 207537 | 0.4208 | 0 | 59.9216 | Eukaryota | 14447.0 | 0 | Chordata | 14447.0 | 0 | Carnivora | 14447.0 | 0 | Canidae | 14447.0 | 0 | Canis | 11972.0 | 1 | Canis lupus | 11972.0 | 1 |
| scf7180000042577 | 571585 | 0.39 | 0 | 60.6643 | Eukaryota | 63818.0 | 0 | Chordata | 63818.0 | 0 | Carnivora | 63818.0 | 0 | Canidae | 63818.0 | 0 | Canis | 63818.0 | 0 | Canis lupus | 63818.0 | 0 |
| scf7180000047674 | 83804 | 0.3615 | 0 | 56.9347 | Eukaryota | 71452.0 | 0 | Chordata | 71452.0 | 0 | Carnivora | 71452.0 | 0 | Canidae | 71452.0 | 0 | Canis | 71452.0 | 0 | Canis lupus | 71452.0 | 0 |
| scf7180000043348 | 647345 | 0.3945 | 0 | 59.5889 | Eukaryota | 37708.0 | 0 | Chordata | 37708.0 | 0 | Carnivora | 37708.0 | 0 | Canidae | 37708.0 | 0 | Canis | 37708.0 | 0 | Canis lupus | 37708.0 | 0 |
| scf7180000046101 | 149829 | 0.589 | 0 | 57.2855 | Eukaryota | 42233.0 | 0 | Chordata | 42233.0 | 0 | Carnivora | 42233.0 | 0 | Canidae | 42233.0 | 0 | Canis | 35440.0 | 1 | Canis lupus | 35440.0 | 1 |
| scf7180000046188 | 12977 | 0.3552 | 0 | 44.3355 | Eukaryota | 3186.0 | 0 | Chordata | 3186.0 | 0 | Carnivora | 1323.0 | 2 | Canidae | 1323.0 | 4 | Canis | 1323.0 | 4 | Canis lupus | 1323.0 | 4 |
| scf7180000043035 | 70887 | 0.405 | 0 | 61.2652 | Eukaryota | 86176.0 | 0 | Chordata | 86176.0 | 0 | Carnivora | 86176.0 | 0 | Canidae | 83071.0 | 1 | Canis | 83071.0 | 1 | Canis lupus | 83071.0 | 1 |
| scf7180000041273 | 256146 | 0.4093 | 0 | 57.6354 | Eukaryota | 54698.0 | 0 | Chordata | 54698.0 | 0 | Carnivora | 54698.0 | 0 | Canidae | 54698.0 | 0 | Canis | 54698.0 | 0 | Canis lupus | 54698.0 | 0 |
| scf7180000049979 | 28274 | 0.4207 | 0 | 61.1292 | Eukaryota | 5950.0 | 0 | Chordata | 5950.0 | 0 | Carnivora | 5950.0 | 0 | Canidae | 5950.0 | 0 | Canis | 5950.0 | 0 | Canis lupus | 5950.0 | 0 |
| scf7180000055691 | 10888 | 0.4463 | 0 | 470.3785 | Eukaryota | 713.0 | 0 | Platyhelminthes | 713.0 | 0 | Polystomatidea | 713.0 | 0 | Polystomatidae | 713.0 | 0 | Protopolystoma | 713.0 | 0 | Protopolystoma xenopodis | 713.0 | 0 |
| scf7180000045902 | 126218 | 0.4456 | 0 | 57.9593 | Eukaryota | 14243.0 | 0 | Chordata | 14243.0 | 0 | Primates | 9400.0 | 2 | Hominidae | 6626.0 | 3 | Homo | 6626.0 | 4 | Homo sapiens | 6626.0 | 4 |
| scf7180000053777 | 3281 | 0.441 | 0 | 109.1314 | no-hit | 0.0 | 0 | no-hit | 0.0 | 0 | no-hit | 0.0 | 0 | no-hit | 0.0 | 0 | no-hit | 0.0 | 0 | no-hit | 0.0 | 0 |
| scf7180000043178 | 325715 | 0.3599 | 0 | 57.5233 | Eukaryota | 13199.0 | 0 | Chordata | 13199.0 | 0 | Primates | 9821.0 | 1 | Hominidae | 9821.0 | 1 | Homo | 9821.0 | 2 | Homo sapiens | 9821.0 | 2 |
| scf7180000049780 | 28262 | 0.3623 | 0 | 125.9841 | Eukaryota | 4618.0 | 0 | Chordata | 4618.0 | 0 | Carnivora | 4618.0 | 0 | Canidae | 2524.0 | 1 | Canis | 2524.0 | 1 | Canis lupus | 2524.0 | 1 |
| scf7180000041502 | 265914 | 0.4385 | 0 | 58.6683 | Eukaryota | 42977.0 | 0 | Chordata | 42977.0 | 0 | Carnivora | 42977.0 | 0 | Canidae | 33789.0 | 2 | Canis | 23803.0 | 3 | Canis lupus | 23803.0 | 3 |
| scf7180000042076 | 629924 | 0.4233 | 0 | 58.8944 | Eukaryota | 87786.0 | 0 | Chordata | 87786.0 | 0 | Carnivora | 87786.0 | 0 | Canidae | 87786.0 | 0 | Canis | 87786.0 | 0 | Canis lupus | 87786.0 | 0 |
| scf7180000050839 | 13144 | 0.5052 | 0 | 216.1389 | Eukaryota | 8730.0 | 0 | Chordata | 8730.0 | 0 | Carnivora | 8730.0 | 0 | Canidae | 8730.0 | 0 | Canis | 7042.0 | 1 | Canis lupus | 5846.0 | 2 |
| scf7180000047665 | 81768 | 0.4251 | 0 | 61.643 | Eukaryota | 22457.0 | 0 | Chordata | 22457.0 | 0 | Carnivora | 22457.0 | 0 | Canidae | 15172.0 | 1 | Canis | 15172.0 | 1 | Canis lupus | 15172.0 | 1 |
| scf7180000048020 | 14129 | 0.4426 | 0 | 86.4416 | Eukaryota | 5094.0 | 0 | Platyhelminthes | 2569.0 | 2 | Polystomatidea | 2569.0 | 3 | Polystomatidae | 2569.0 | 3 | Protopolystoma | 2569.0 | 4 | Protopolystoma xenopodis | 2569.0 | 4 |
| scf7180000047797 | 23853 | 0.4433 | 0 | 125.3471 | Eukaryota | 89795.0 | 0 | Chordata | 89795.0 | 0 | Carnivora | 89795.0 | 0 | Canidae | 89795.0 | 0 | Canis | 89795.0 | 0 | Canis lupus | 89795.0 | 0 |
| scf7180000044576 | 324601 | 0.4036 | 0 | 60.1622 | Eukaryota | 34752.0 | 0 | Chordata | 34752.0 | 0 | Carnivora | 34752.0 | 0 | Canidae | 34752.0 | 0 | Canis | 34752.0 | 0 | Canis lupus | 34752.0 | 0 |
| scf7180000046273 | 12470 | 0.4342 | 0 | 629.8027 | Eukaryota | 2575.0 | 0 | Chordata | 2575.0 | 0 | Carnivora | 2575.0 | 0 | Canidae | 2575.0 | 0 | Canis | 2575.0 | 0 | Canis lupus | 2575.0 | 0 |
| scf7180000049948 | 6550 | 0.454 | 0 | 1362.9663 | Eukaryota | 2517.0 | 0 | Echinodermata | 950.0 | 3 | Forcipulatida | 950.0 | 3 | Asteriidae | 950.0 | 3 | Asterias | 950.0 | 3 | Asterias rubens | 950.0 | 3 |
| scf7180000048856 | 41724 | 0.4383 | 0 | 79.2522 | Eukaryota | 10941.0 | 0 | Chordata | 10941.0 | 0 | Carnivora | 8976.0 | 1 | Canidae | 8976.0 | 1 | Canis | 8976.0 | 1 | Canis lupus | 8976.0 | 1 |
| scf7180000043771 | 349585 | 0.6078 | 0 | 56.8522 | Eukaryota | 66535.0 | 0 | Chordata | 66535.0 | 0 | Carnivora | 66535.0 | 0 | Canidae | 66535.0 | 0 | Canis | 66535.0 | 0 | Canis lupus | 66535.0 | 0 |
| scf7180000051754 | 8288 | 0.4486 | 0 | 984.3104 | Eukaryota | 21909.0 | 0 | Chordata | 19822.0 | 1 | Gadiformes | 11836.0 | 6 | Gadidae | 11836.0 | 6 | Gadus | 11836.0 | 6 | Gadus morhua | 11836.0 | 6 |
| scf7180000050036 | 21679 | 0.4405 | 0 | 100.0229 | Eukaryota | 8875.0 | 0 | Chordata | 8875.0 | 0 | Kurtiformes | 3734.0 | 3 | Apogonidae | 3734.0 | 3 | Sphaeramia | 3734.0 | 3 | Sphaeramia orbicularis | 3734.0 | 3 |
| scf7180000048452 | 27164 | 0.4528 | 0 | 109.4891 | Eukaryota | 6498.0 | 0 | Chordata | 5702.0 | 1 | Kurtiformes | 2886.0 | 2 | Apogonidae | 2886.0 | 2 | Sphaeramia | 2886.0 | 2 | Sphaeramia orbicularis | 2886.0 | 2 |
| scf7180000044773 | 87991 | 0.3722 | 0 | 59.8341 | Eukaryota | 21596.0 | 0 | Chordata | 21596.0 | 0 | Carnivora | 21596.0 | 0 | Canidae | 21596.0 | 0 | Canis | 21596.0 | 0 | Canis lupus | 21596.0 | 0 |
| scf7180000040595 | 2051362 | 0.5038 | 0 | 58.4804 | Eukaryota | 60248.0 | 0 | Chordata | 60248.0 | 0 | Carnivora | 60248.0 | 0 | Canidae | 60248.0 | 0 | Canis | 40607.0 | 1 | Canis lupus | 40607.0 | 1 |
| scf7180000053347 | 13057 | 0.4436 | 0 | 310.6194 | Eukaryota | 2892.0 | 0 | Chordata | 1798.0 | 1 | Tetraodontiformes | 1258.0 | 2 | Tetraodontidae | 1258.0 | 2 | Takifugu | 1258.0 | 2 | Takifugu rubripes | 1258.0 | 2 |
| scf7180000047719 | 21105 | 0.515 | 0 | 345.2065 | Eukaryota | 36371.0 | 0 | Chordata | 36371.0 | 0 | Carnivora | 36371.0 | 0 | Canidae | 36371.0 | 0 | Canis | 36371.0 | 0 | Canis lupus | 36371.0 | 0 |
| scf7180000042331 | 794652 | 0.4183 | 0 | 59.4711 | Eukaryota | 54771.0 | 0 | Chordata | 54771.0 | 0 | Carnivora | 54771.0 | 0 | Canidae | 54771.0 | 0 | Canis | 42989.0 | 1 | Canis lupus | 42989.0 | 1 |
| scf7180000049876 | 38624 | 0.4156 | 0 | 52.2269 | Eukaryota | 12319.0 | 0 | Chordata | 12319.0 | 0 | Carnivora | 12319.0 | 0 | Canidae | 12319.0 | 0 | Canis | 12319.0 | 0 | Canis lupus | 12319.0 | 0 |
| scf7180000041137 | 802100 | 0.3911 | 0 | 59.4477 | Eukaryota | 50185.0 | 0 | Chordata | 50185.0 | 0 | Carnivora | 50185.0 | 0 | Canidae | 43118.0 | 1 | Canis | 43118.0 | 2 | Canis lupus | 43118.0 | 2 |
| scf7180000049853 | 4883 | 0.4417 | 0 | 104.9558 | Eukaryota | 609.0 | 0 | Chordata | 407.0 | 1 | Blenniiformes | 407.0 | 1 | Gobiesocidae | 407.0 | 1 | Gouania | 407.0 | 1 | Gouania willdenowi | 407.0 | 1 |
| scf7180000045179 | 26442 | 0.4416 | 0 | 152.4532 | Eukaryota | 261.0 | 0 | Platyhelminthes | 261.0 | 0 | Polystomatidea | 261.0 | 0 | Polystomatidae | 261.0 | 0 | Protopolystoma | 261.0 | 0 | Protopolystoma xenopodis | 261.0 | 0 |
| scf7180000043010 | 408511 | 0.3626 | 0 | 57.6184 | Eukaryota | 18889.0 | 0 | Chordata | 18889.0 | 0 | Carnivora | 18889.0 | 0 | Canidae | 18889.0 | 0 | Canis | 18889.0 | 0 | Canis lupus | 18889.0 | 0 |
| scf7180000056284 | 957577 | 0.4315 | 0 | 60.9153 | Eukaryota | 63731.0 | 0 | Chordata | 63731.0 | 0 | Carnivora | 63731.0 | 0 | Canidae | 26136.0 | 4 | Canis | 19844.0 | 5 | Canis lupus | 19844.0 | 5 |
| scf7180000041659 | 166535 | 0.376 | 0 | 58.2899 | Eukaryota | 80774.0 | 0 | Chordata | 80774.0 | 0 | Carnivora | 80774.0 | 0 | Canidae | 80774.0 | 0 | Canis | 80774.0 | 0 | Canis lupus | 80774.0 | 0 |
| scf7180000046879 | 38336 | 0.4356 | 0 | 146.7324 | Eukaryota | 6173.0 | 0 | Chordata | 4917.0 | 2 | Kurtiformes | 3437.0 | 4 | Apogonidae | 3437.0 | 4 | Sphaeramia | 3437.0 | 4 | Sphaeramia orbicularis | 3437.0 | 4 |
| scf7180000045291 | 175392 | 0.3798 | 0 | 59.2535 | Eukaryota | 18478.0 | 0 | Chordata | 18478.0 | 0 | Carnivora | 18478.0 | 0 | Felidae | 18478.0 | 0 | Felis | 18478.0 | 0 | Felis catus | 18478.0 | 0 |
| scf7180000050830 | 27923 | 0.4471 | 0 | 137.8579 | Eukaryota | 1743.0 | 0 | Platyhelminthes | 914.0 | 1 | Polystomatidea | 914.0 | 1 | Polystomatidae | 914.0 | 1 | Protopolystoma | 914.0 | 1 | Protopolystoma xenopodis | 914.0 | 1 |
| scf7180000053493 | 4162 | 0.4524 | 0 | 508.1504 | no-hit | 0.0 | 0 | no-hit | 0.0 | 0 | no-hit | 0.0 | 0 | no-hit | 0.0 | 0 | no-hit | 0.0 | 0 | no-hit | 0.0 | 0 |
| scf7180000041566 | 321999 | 0.418 | 100 | 58.2109 | Eukaryota | 52143.0 | 0 | Chordata | 52143.0 | 0 | Carnivora | 52143.0 | 0 | Canidae | 48601.0 | 1 | Vulpes | 29250.0 | 2 | Vulpes vulpes | 29250.0 | 2 |
| scf7180000042906 | 598733 | 0.3584 | 0 | 59.7432 | Eukaryota | 86703.0 | 0 | Chordata | 86703.0 | 0 | Carnivora | 86703.0 | 0 | Canidae | 86703.0 | 0 | Canis | 86703.0 | 0 | Canis lupus | 86703.0 | 0 |
| scf7180000046292 | 65008 | 0.4382 | 0 | 59.8168 | Eukaryota | 39515.0 | 0 | Chordata | 39515.0 | 0 | Carnivora | 39515.0 | 0 | Canidae | 39515.0 | 0 | Canis | 39515.0 | 0 | Canis lupus | 39515.0 | 0 |
| scf7180000047214 | 18022 | 0.4101 | 0 | 52.2761 | Eukaryota | 20825.0 | 0 | Chordata | 20825.0 | 0 | Carnivora | 20825.0 | 0 | Canidae | 20825.0 | 0 | Canis | 20825.0 | 0 | Canis lupus | 20825.0 | 0 |
| scf7180000050882 | 5674 | 0.4468 | 0 | 971.5753 | no-hit | 0.0 | 0 | no-hit | 0.0 | 0 | no-hit | 0.0 | 0 | no-hit | 0.0 | 0 | no-hit | 0.0 | 0 | no-hit | 0.0 | 0 |
| scf7180000051315 | 17213 | 0.5033 | 0 | 460.0264 | Eukaryota | 8946.0 | 0 | Chordata | 8946.0 | 0 | Carnivora | 8946.0 | 0 | Canidae | 8946.0 | 0 | Canis | 7269.0 | 1 | Canis lupus | 6047.0 | 2 |
| scf7180000047519 | 67849 | 0.438 | 0 | 113.2914 | Eukaryota | 41873.0 | 0 | Platyhelminthes | 23105.0 | 2 | Strigeidida | 23105.0 | 5 | Schistosomatidae | 23105.0 | 5 | Schistosoma | 23105.0 | 5 | Schistosoma mansoni | 23105.0 | 5 |
| scf7180000050887 | 20594 | 0.4525 | 0 | 65.0292 | Eukaryota | 12753.0 | 0 | Chordata | 11749.0 | 1 | Anabantiformes | 4161.0 | 5 | Anabantidae | 4161.0 | 5 | Anabas | 4161.0 | 5 | Anabas testudineus | 4161.0 | 6 |
| scf7180000043206 | 518943 | 0.3611 | 0 | 59.689 | Eukaryota | 36367.0 | 0 | Chordata | 36367.0 | 0 | Carnivora | 36367.0 | 0 | Canidae | 36367.0 | 0 | Canis | 36367.0 | 0 | Canis lupus | 36367.0 | 0 |
| scf7180000041858 | 500825 | 0.3963 | 0 | 60.9866 | Eukaryota | 91581.0 | 0 | Chordata | 91581.0 | 0 | Carnivora | 91581.0 | 0 | Canidae | 91581.0 | 0 | Canis | 91581.0 | 0 | Canis lupus | 91581.0 | 0 |
| scf7180000052858 | 6611 | 0.4397 | 0 | 181.2197 | Eukaryota | 1169.0 | 0 | Chordata | 768.0 | 1 | Tetraodontiformes | 768.0 | 1 | Tetraodontidae | 768.0 | 1 | Takifugu | 768.0 | 1 | Takifugu rubripes | 768.0 | 1 |
| scf7180000043429 | 3351 | 0.3859 | 0 | 27.9554 | Eukaryota | 3288.0 | 0 | Chordata | 3288.0 | 0 | Carnivora | 3288.0 | 0 | Canidae | 3288.0 | 0 | Canis | 3288.0 | 0 | Canis lupus | 3288.0 | 0 |
| scf7180000043230 | 401049 | 0.5609 | 0 | 56.0536 | Eukaryota | 41563.0 | 0 | Chordata | 41563.0 | 0 | Carnivora | 41563.0 | 0 | Canidae | 41563.0 | 0 | Canis | 29125.0 | 1 | Canis lupus | 29125.0 | 1 |
| scf7180000054894 | 13252 | 0.367 | 0 | 658.8707 | Eukaryota | 31093.0 | 0 | Chordata | 31093.0 | 0 | Carnivora | 31093.0 | 0 | Canidae | 31093.0 | 0 | Canis | 31093.0 | 0 | Canis lupus | 31093.0 | 0 |
| scf7180000052657 | 16562 | 0.5125 | 0 | 364.7404 | Eukaryota | 33394.0 | 0 | Chordata | 33394.0 | 0 | Carnivora | 33394.0 | 0 | Canidae | 33394.0 | 0 | Canis | 33394.0 | 0 | Canis lupus | 33394.0 | 0 |
| scf7180000041443 | 74653 | 0.4389 | 0 | 57.5594 | Eukaryota | 104100.0 | 0 | Chordata | 104100.0 | 0 | Carnivora | 104100.0 | 0 | Canidae | 104100.0 | 0 | Canis | 104100.0 | 0 | Canis lupus | 104100.0 | 0 |
| scf7180000052962 | 2344 | 0.4578 | 0 | 208.3187 | no-hit | 0.0 | 0 | no-hit | 0.0 | 0 | no-hit | 0.0 | 0 | no-hit | 0.0 | 0 | no-hit | 0.0 | 0 | no-hit | 0.0 | 0 |
| scf7180000047216 | 47807 | 0.3491 | 0 | 57.6342 | Eukaryota | 12058.0 | 0 | Chordata | 12058.0 | 0 | Eulipotyphla | 10334.0 | 1 | Soricidae | 10334.0 | 1 | Crocidura | 10334.0 | 1 | Crocidura grayi | 3452.0 | 5 |
| scf7180000043124 | 550253 | 0.5004 | 0 | 56.7433 | Eukaryota | 57209.0 | 0 | Chordata | 57209.0 | 0 | Carnivora | 57209.0 | 0 | Canidae | 57209.0 | 0 | Canis | 48493.0 | 1 | Canis lupus | 48493.0 | 1 |

| | | | | | | | | | | | | | | | | | | | | | | |
|---|---|---|---|---|---|---|---|---|---|---|---|---|---|---|---|---|---|---|---|---|---|---|
| scf7180000051125 | 7660 | 0.4419 | 0 | 79.679 | no-hit | 0.0 | 0 | no-hit | 0.0 | 0 | no-hit | 0.0 | 0 | no-hit | 0.0 | 0 | no-hit | 0.0 | 0 | no-hit | 0.0 | 0 |
| scf7180000046047 | 154434 | 0.3721 | 0 | 60.618 | Eukaryota | 11349.0 | 0 | Chordata | 11349.0 | 0 | Carnivora | 11349.0 | 0 | Canidae | 10157.0 | 1 | Canis | 10157.0 | 1 | Canis lupus | 10157.0 | 1 |
| scf7180000040947 | 393456 | 0.4072 | 0 | 59.213 | Eukaryota | 58390.0 | 0 | Chordata | 58390.0 | 0 | Carnivora | 58390.0 | 0 | Canidae | 58390.0 | 0 | Canis | 46103.0 | 1 | Canis lupus | 46103.0 | 1 |
| scf7180000046277 | 149339 | 0.391 | 0 | 60.7702 | Eukaryota | 81703.0 | 0 | Chordata | 81703.0 | 0 | Carnivora | 81703.0 | 0 | Canidae | 81703.0 | 0 | Canis | 81703.0 | 0 | Canis lupus | 81703.0 | 0 |
| scf7180000046735 | 2412 | 0.459 | 0 | 25.5147 | Eukaryota | 3910.0 | 1 | Chordata | 3910.0 | 1 | Carnivora | 2234.0 | 3 | Canidae | 2234.0 | 3 | Canis | 2234.0 | 4 | Canis lupus | 2234.0 | 4 |
| scf7180000053223 | 2317 | 0.4454 | 0 | 270.6627 | no-hit | 0.0 | 0 | no-hit | 0.0 | 0 | no-hit | 0.0 | 0 | no-hit | 0.0 | 0 | no-hit | 0.0 | 0 | no-hit | 0.0 | 0 |
| scf7180000046582 | 3558 | 0.4241 | 0 | 52.0363 | Eukaryota | 24091.0 | 0 | Chordata | 24091.0 | 0 | Carnivora | 24091.0 | 0 | Canidae | 24091.0 | 0 | Canis | 24091.0 | 0 | Canis lupus | 24091.0 | 0 |
| scf7180000040757 | 14475 | 0.4423 | 0 | 284.6938 | Eukaryota | 202.0 | 0 | Platyhelminthes | 202.0 | 0 | Polystomatidea | 202.0 | 0 | Polystomatidae | 202.0 | 0 | Protopolystoma | 202.0 | 0 | Protopolystoma xenopodis | 202.0 | 0 |
| scf7180000048266 | 2512 | 0.455 | 0 | 319.2125 | no-hit | 0.0 | 0 | no-hit | 0.0 | 0 | no-hit | 0.0 | 0 | no-hit | 0.0 | 0 | no-hit | 0.0 | 0 | no-hit | 0.0 | 0 |
| scf7180000045328 | 143117 | 0.373 | 0 | 58.7776 | Eukaryota | 26712.0 | 0 | Chordata | 26712.0 | 0 | Carnivora | 26712.0 | 0 | Canidae | 26712.0 | 0 | Canis | 26712.0 | 0 | Canis lupus | 26712.0 | 0 |
| scf7180000042100 | 12464 | 0.5182 | 0 | 1040.8094 | Eukaryota | 33332.0 | 0 | Chordata | 33332.0 | 0 | Carnivora | 33332.0 | 0 | Canidae | 33332.0 | 0 | Canis | 33332.0 | 0 | Canis lupus | 33332.0 | 0 |
| scf7180000040603 | 393647 | 0.4117 | 0 | 59.3882 | Eukaryota | 95475.0 | 0 | Chordata | 95475.0 | 0 | Carnivora | 95475.0 | 0 | Canidae | 69480.0 | 1 | Vulpes | 51200.0 | 2 | Vulpes vulpes | 51200.0 | 2 |
| scf7180000041886 | 801133 | 0.3925 | 0 | 60.285 | Eukaryota | 94441.0 | 0 | Chordata | 94441.0 | 0 | Carnivora | 94441.0 | 0 | Canidae | 94441.0 | 0 | Canis | 94441.0 | 0 | Canis lupus | 94441.0 | 0 |
| scf7180000045609 | 4766 | 0.3764 | 0 | 29.0739 | Eukaryota | 6302.0 | 0 | Chordata | 6302.0 | 0 | Carnivora | 6302.0 | 0 | Felidae | 5733.0 | 1 | Felis | 5733.0 | 1 | Felis catus | 5733.0 | 1 |
| scf7180000041066 | 83106 | 0.5431 | 0 | 55.2211 | Eukaryota | 53327.0 | 0 | Chordata | 53327.0 | 0 | Carnivora | 53327.0 | 0 | Canidae | 51428.0 | 1 | Canis | 47778.0 | 2 | Canis lupus | 47778.0 | 2 |
| scf7180000042780 | 550701 | 0.3866 | 0 | 60.2673 | Eukaryota | 38707.0 | 0 | Chordata | 38707.0 | 0 | Carnivora | 38707.0 | 0 | Canidae | 38707.0 | 0 | Vulpes | 31105.0 | 1 | Vulpes vulpes | 31105.0 | 1 |
| scf7180000045505 | 174170 | 0.3983 | 0 | 60.4118 | Eukaryota | 47738.0 | 0 | Chordata | 47738.0 | 0 | Carnivora | 47738.0 | 0 | Canidae | 47738.0 | 0 | Canis | 47738.0 | 0 | Canis lupus | 47738.0 | 0 |
| scf7180000050500 | 13952 | 0.4422 | 0 | 138.9309 | no-hit | 0.0 | 0 | no-hit | 0.0 | 0 | no-hit | 0.0 | 0 | no-hit | 0.0 | 0 | no-hit | 0.0 | 0 | no-hit | 0.0 | 0 |
| scf7180000052117 | 7725 | 0.4502 | 0 | 1067.5507 | Eukaryota | 2778.0 | 0 | Chordata | 1982.0 | 1 | Columbiformes | 922.0 | 3 | Columbidae | 922.0 | 3 | Streptopelia | 922.0 | 3 | Streptopelia turtur | 922.0 | 3 |
| scf7180000041974 | 157511 | 0.5075 | 0 | 58.3566 | Eukaryota | 79889.0 | 0 | Chordata | 79889.0 | 0 | Carnivora | 79889.0 | 0 | Canidae | 67852.0 | 1 | Canis | 56662.0 | 3 | Canis lupus | 56662.0 | 3 |
| scf7180000051796 | 4859 | 0.4554 | 0 | 1627.9022 | no-hit | 0.0 | 0 | no-hit | 0.0 | 0 | no-hit | 0.0 | 0 | no-hit | 0.0 | 0 | no-hit | 0.0 | 0 | no-hit | 0.0 | 0 |
| scf7180000047404 | 17780 | 0.4345 | 0 | 79.4484 | Eukaryota | 8089.0 | 0 | Chordata | 7730.0 | 1 | Kurtiformes | 3455.0 | 3 | Apogonidae | 3455.0 | 3 | Sphaeramia | 3455.0 | 4 | Sphaeramia orbicularis | 3455.0 | 4 |
| scf7180000046796 | 106131 | 0.3978 | 0 | 60.5547 | Eukaryota | 23814.0 | 0 | Chordata | 23814.0 | 0 | Carnivora | 23814.0 | 0 | Canidae | 23814.0 | 0 | Canis | 23814.0 | 0 | Canis lupus | 23814.0 | 0 |
| scf7180000041976 | 617182 | 0.3913 | 0 | 60.4561 | Eukaryota | 91776.0 | 0 | Chordata | 91776.0 | 0 | Carnivora | 91776.0 | 0 | Canidae | 91776.0 | 0 | Canis | 91776.0 | 0 | Canis lupus | 91776.0 | 0 |
| scf7180000043883 | 159549 | 0.4342 | 0 | 58.6868 | Eukaryota | 13493.0 | 0 | Chordata | 13493.0 | 0 | Artiodactyla | 7869.0 | 1 | Suidae | 6659.0 | 3 | Sus | 6659.0 | 4 | Sus scrofa | 6659.0 | 5 |
| scf7180000051997 | 13185 | 0.5035 | 0 | 454.4095 | Eukaryota | 8573.0 | 0 | Chordata | 8573.0 | 0 | Carnivora | 8573.0 | 0 | Canidae | 8573.0 | 0 | Canis | 7462.0 | 1 | Canis lupus | 6336.0 | 2 |
| scf7180000046412 | 7960 | 0.371 | 0 | 24.6324 | Eukaryota | 22473.0 | 0 | Chordata | 22473.0 | 0 | Carnivora | 22473.0 | 0 | Canidae | 22473.0 | 0 | Canis | 22473.0 | 0 | Canis lupus | 22473.0 | 0 |
| scf7180000044833 | 85527 | 0.3894 | 0 | 56.4875 | Eukaryota | 72680.0 | 0 | Chordata | 72680.0 | 0 | Carnivora | 72680.0 | 0 | Canidae | 72680.0 | 0 | Canis | 72680.0 | 0 | Canis lupus | 72680.0 | 0 |
| scf7180000043803 | 332926 | 0.4057 | 0 | 59.5746 | Eukaryota | 87743.0 | 0 | Chordata | 87743.0 | 0 | Carnivora | 87743.0 | 0 | Canidae | 87743.0 | 0 | Canis | 87743.0 | 0 | Canis lupus | 87743.0 | 0 |
| scf7180000052351 | 15836 | 0.5221 | 0 | 513.4277 | Eukaryota | 4743.0 | 0 | Chordata | 4743.0 | 0 | Carnivora | 4743.0 | 0 | Canidae | 4743.0 | 0 | Canis | 4059.0 | 1 | Canis lupus | 4059.0 | 1 |
| scf7180000041057 | 103525 | 0.3769 | 0 | 60.2705 | Eukaryota | 91519.0 | 0 | Chordata | 91519.0 | 0 | Carnivora | 91519.0 | 0 | Canidae | 91519.0 | 0 | Canis | 91519.0 | 0 | Canis lupus | 91519.0 | 0 |
| scf7180000053509 | 2164 | 0.4524 | 0 | 394.2952 | no-hit | 0.0 | 0 | no-hit | 0.0 | 0 | no-hit | 0.0 | 0 | no-hit | 0.0 | 0 | no-hit | 0.0 | 0 | no-hit | 0.0 | 0 |
| scf7180000049582 | 6990 | 0.4435 | 0 | 115.3856 | no-hit | 0.0 | 0 | no-hit | 0.0 | 0 | no-hit | 0.0 | 0 | no-hit | 0.0 | 0 | no-hit | 0.0 | 0 | no-hit | 0.0 | 0 |
| scf7180000043269 | 459985 | 0.4495 | 0 | 60.4161 | Eukaryota | 111189.0 | 0 | Chordata | 111189.0 | 0 | Carnivora | 111189.0 | 0 | Canidae | 99055.0 | 1 | Canis | 65843.0 | 2 | Canis lupus | 65843.0 | 2 |
| scf7180000053413 | 4241 | 0.4515 | 0 | 406.372 | Eukaryota | 14442.0 | 0 | Chordata | 14442.0 | 0 | Tetraodontiformes | 5754.0 | 4 | Tetraodontidae | 5754.0 | 4 | Takifugu | 5754.0 | 4 | Takifugu rubripes | 5754.0 | 4 |
| scf7180000043103 | 336895 | 0.3855 | 0 | 60.1206 | Eukaryota | 89849.0 | 0 | Chordata | 89849.0 | 0 | Carnivora | 89849.0 | 0 | Canidae | 89849.0 | 0 | Canis | 89849.0 | 0 | Canis lupus | 89849.0 | 0 |
| scf7180000052995 | 6406 | 0.5062 | 0 | 145.22 | Eukaryota | 8617.0 | 0 | Chordata | 8617.0 | 0 | Carnivora | 8617.0 | 0 | Canidae | 8617.0 | 0 | Canis | 7045.0 | 1 | Canis lupus | 5982.0 | 2 |
| scf7180000053151 | 5308 | 0.4452 | 0 | 86.6495 | no-hit | 0.0 | 0 | no-hit | 0.0 | 0 | no-hit | 0.0 | 0 | no-hit | 0.0 | 0 | no-hit | 0.0 | 0 | no-hit | 0.0 | 0 |
| scf7180000054260 | 48931 | 0.3708 | 0 | 853.7029 | Eukaryota | 65535.0 | 0 | Chordata | 65535.0 | 0 | Carnivora | 65535.0 | 0 | Canidae | 65535.0 | 0 | Canis | 65535.0 | 0 | Canis lupus | 65535.0 | 0 |
| scf7180000050604 | 6130 | 0.4326 | 0 | 136.2855 | Eukaryota | 3029.0 | 0 | Chordata | 2673.0 | 1 | Columbiformes | 1428.0 | 5 | Columbidae | 1428.0 | 5 | Streptopelia | 1428.0 | 5 | Streptopelia turtur | 1428.0 | 5 |
| scf7180000044975 | 216601 | 0.5707 | 0 | 56.5611 | Eukaryota | 13513.0 | 0 | Chordata | 13513.0 | 0 | Carnivora | 7834.0 | 1 | Canidae | 2959.0 | 5 | Canis | 2959.0 | 7 | Canis lupus | 2959.0 | 7 |
| scf7180000050317 | 7745 | 0.4396 | 0 | 236.9345 | Eukaryota | 537.0 | 0 | Streptophyta | 537.0 | 0 | Fabales | 537.0 | 0 | Fabaceae | 537.0 | 0 | Cicer | 537.0 | 0 | Cicer arietinum | 537.0 | 0 |
| scf7180000050313 | 6462 | 0.4426 | 0 | 578.9065 | no-hit | 0.0 | 0 | no-hit | 0.0 | 0 | no-hit | 0.0 | 0 | no-hit | 0.0 | 0 | no-hit | 0.0 | 0 | no-hit | 0.0 | 0 |
| scf7180000056506 | 239231 | 0.4144 | 0 | 63.5994 | Eukaryota | 56650.0 | 0 | Chordata | 56650.0 | 0 | Carnivora | 56650.0 | 0 | Canidae | 56650.0 | 0 | Canis | 52251.0 | 1 | Canis lupus | 52251.0 | 1 |
| scf7180000047155 | 9824 | 0.4425 | 0 | 212.9237 | no-hit | 0.0 | 0 | no-hit | 0.0 | 0 | no-hit | 0.0 | 0 | no-hit | 0.0 | 0 | no-hit | 0.0 | 0 | no-hit | 0.0 | 0 |
| scf7180000044910 | 267866 | 0.4782 | 0 | 58.0907 | Eukaryota | 73311.0 | 0 | Chordata | 73311.0 | 0 | Carnivora | 73311.0 | 0 | Canidae | 73311.0 | 0 | Canis | 73311.0 | 0 | Canis lupus | 73311.0 | 0 |
| scf7180000043211 | 497396 | 0.398 | 0 | 60.0936 | Eukaryota | 91615.0 | 0 | Chordata | 91615.0 | 0 | Carnivora | 91615.0 | 0 | Canidae | 91615.0 | 0 | Canis | 91615.0 | 0 | Canis lupus | 91615.0 | 0 |
| scf7180000043329 | 453119 | 0.4313 | 0 | 58.8211 | Eukaryota | 22473.0 | 0 | Chordata | 22473.0 | 0 | Carnivora | 22473.0 | 0 | Canidae | 22473.0 | 0 | Canis | 22473.0 | 0 | Canis lupus | 22473.0 | 0 |
| scf7180000048098 | 17265 | 0.4462 | 0 | 184.0291 | Eukaryota | 5333.0 | 0 | Chordata | 2536.0 | 2 | Forcipulatida | 1683.0 | 4 | Asteriidae | 1683.0 | 4 | Asterias | 1683.0 | 5 | Asterias rubens | 1683.0 | 5 |
| scf7180000046645 | 120312 | 0.5643 | 0 | 56.4604 | Eukaryota | 22615.0 | 0 | Chordata | 22615.0 | 0 | Carnivora | 22615.0 | 0 | Canidae | 20841.0 | 1 | Canis | 12565.0 | 3 | Canis lupus | 12565.0 | 3 |
| scf7180000043498 | 400270 | 0.3451 | 100 | 59.085 | Eukaryota | 84373.0 | 0 | Chordata | 84373.0 | 0 | Carnivora | 84373.0 | 0 | Canidae | 84373.0 | 0 | Canis | 84373.0 | 0 | Canis lupus | 84373.0 | 0 |
| scf7180000050528 | 7571 | 0.4419 | 0 | 119.0246 | no-hit | 0.0 | 0 | no-hit | 0.0 | 0 | no-hit | 0.0 | 0 | no-hit | 0.0 | 0 | no-hit | 0.0 | 0 | no-hit | 0.0 | 0 |
| scf7180000049006 | 53932 | 0.36 | 0 | 61.1367 | Eukaryota | 31251.0 | 0 | Chordata | 31251.0 | 0 | Carnivora | 31251.0 | 0 | Canidae | 17274.0 | 3 | Canis | 13824.0 | 4 | Canis lupus | 13824.0 | 4 |
| scf7180000045000 | 80499 | 0.4474 | 0 | 58.6128 | Eukaryota | 26174.0 | 0 | Chordata | 26174.0 | 0 | Carnivora | 25025.0 | 1 | Canidae | 22809.0 | 3 | Vulpes | 13755.0 | 4 | Vulpes vulpes | 13755.0 | 4 |
| scf7180000049967 | 8763 | 0.4633 | 0 | 166.6043 | Eukaryota | 2505.0 | 0 | Chordata | 1666.0 | 1 | Syngnathiformes | 859.0 | 3 | Syngnathidae | 859.0 | 3 | Syngnathus | 859.0 | 3 | Syngnathus acus | 859.0 | 3 |
| scf7180000047605 | 21622 | 0.4714 | 0 | 153.1798 | Eukaryota | 9544.0 | 0 | Chordata | 9544.0 | 0 | Carnivora | 9544.0 | 0 | Canidae | 9544.0 | 0 | Canis | 8334.0 | 1 | Canis lupus | 7113.0 | 2 |
| scf7180000041517 | 71632 | 0.3743 | 0 | 55.1172 | Eukaryota | 51184.0 | 0 | Chordata | 51184.0 | 0 | Carnivora | 51184.0 | 0 | Canidae | 51184.0 | 0 | Canis | 51184.0 | 0 | Canis lupus | 51184.0 | 0 |
| scf7180000055517 | 40498 | 0.4443 | 0 | 162.3519 | Eukaryota | 4433.0 | 0 | Chordata | 4198.0 | 1 | Salmoniformes | 2653.0 | 3 | Salmonidae | 2653.0 | 3 | Salmo | 1969.0 | 4 | Salmo trutta | 1969.0 | 4 |
| scf7180000045308 | 10198 | 0.4369 | 0 | 47.767 | Eukaryota | 36291.0 | 0 | Chordata | 36291.0 | 0 | Carnivora | 36291.0 | 0 | Canidae | 36291.0 | 0 | Canis | 36291.0 | 0 | Canis lupus | 36291.0 | 0 |
| scf7180000050070 | 1148 | 0.466 | 0 | 1103.2065 | no-hit | 0.0 | 0 | no-hit | 0.0 | 0 | no-hit | 0.0 | 0 | no-hit | 0.0 | 0 | no-hit | 0.0 | 0 | no-hit | 0.0 | 0 |
| scf7180000052460 | 4401 | 0.4472 | 0 | 544.2642 | Eukaryota | 1611.0 | 0 | Echinodermata | 595.0 | 2 | Forcipulatida | 595.0 | 2 | Asteriidae | 595.0 | 2 | Asterias | 595.0 | 2 | Asterias rubens | 595.0 | 2 |
| scf7180000044897 | 273753 | 0.438 | 0 | 59.4303 | Eukaryota | 24244.0 | 0 | Chordata | 24244.0 | 0 | Carnivora | 24244.0 | 0 | Canidae | 24244.0 | 0 | Canis | 24244.0 | 0 | Canis lupus | 24244.0 | 0 |
| scf7180000049768 | 6864 | 0.4331 | 0 | 158.7883 | Eukaryota | 846.0 | 0 | Platyhelminthes | 846.0 | 0 | Polystomatidea | 846.0 | 0 | Polystomatidae | 846.0 | 0 | Protopolystoma | 846.0 | 0 | Protopolystoma xenopodis | 846.0 | 0 |
| scf7180000056239 | 753960 | 0.4519 | 0 | 59.591 | Eukaryota | 67459.0 | 0 | Chordata | 67459.0 | 0 | Carnivora | 67459.0 | 0 | Canidae | 67459.0 | 0 | Canis | 47752.0 | 1 | Canis lupus | 47752.0 | 1 |
| scf7180000044161 | 317381 | 0.5208 | 0 | 57.3293 | Eukaryota | 34154.0 | 0 | Chordata | 34154.0 | 0 | Carnivora | 34154.0 | 0 | Canidae | 34154.0 | 0 | Vulpes | 18576.0 | 1 | Vulpes vulpes | 18576.0 | 1 |
| scf7180000045571 | 139337 | 0.4932 | 0 | 58.2879 | Eukaryota | 90380.0 | 0 | Chordata | 90380.0 | 0 | Carnivora | 90380.0 | 0 | Canidae | 90380.0 | 0 | Canis | 90380.0 | 0 | Canis lupus | 90380.0 | 0 |
| scf7180000041606 | 1014800 | 0.4891 | 0 | 58.8339 | Eukaryota | 59323.0 | 0 | Chordata | 59323.0 | 0 | Carnivora | 59323.0 | 0 | Canidae | 59323.0 | 0 | Canis | 54641.0 | 1 | Canis lupus | 54641.0 | 1 |
| scf7180000052556 | 2809 | 0.4482 | 0 | 661.9394 | no-hit | 0.0 | 0 | no-hit | 0.0 | 0 | no-hit | 0.0 | 0 | no-hit | 0.0 | 0 | no-hit | 0.0 | 0 | no-hit | 0.0 | 0 |
| scf7180000051049 | 14808 | 0.5165 | 0 | 164.0684 | Eukaryota | 15600.0 | 0 | Chordata | 15600.0 | 0 | Carnivora | 15600.0 | 0 | Canidae | 15600.0 | 0 | Canis | 15600.0 | 0 | Canis lupus | 15600.0 | 0 |
| scf7180000041512 | 1068542 | 0.3998 | 0 | 60.177 | Eukaryota | 85508.0 | 0 | Chordata | 85508.0 | 0 | Carnivora | 85508.0 | 0 | Canidae | 85508.0 | 0 | Canis | 85508.0 | 0 | Canis lupus | 85508.0 | 0 |
| scf7180000056475 | 377592 | 0.3646 | 0 | 59.9396 | Eukaryota | 52054.0 | 0 | Chordata | 52054.0 | 0 | Carnivora | 52054.0 | 0 | Canidae | 26898.0 | 3 | Canis | 21490.0 | 5 | Canis lupus | 21490.0 | 5 |
| scf7180000047328 | 102200 | 0.3992 | 0 | 61.1436 | Eukaryota | 23202.0 | 0 | Chordata | 23202.0 | 0 | Carnivora | 23202.0 | 0 | Phocidae | 6916.0 | 3 | Leptonychotes | 4587.0 | 7 | Leptonychotes weddellii | 4587.0 | 7 |
| scf7180000055846 | 3983 | 0.4409 | 0 | 124.5884 | Eukaryota | 5824.0 | 0 | Chordata | 5198.0 | 1 | Kurtiformes | 4893.0 | 2 | Apogonidae | 4893.0 | 2 | Sphaeramia | 4893.0 | 2 | Sphaeramia orbicularis | 4893.0 | 2 |
| scf7180000056609 | 1110134 | 0.3752 | 0 | 61.6438 | Eukaryota | 55492.0 | 0 | Chordata | 55492.0 | 0 | Carnivora | 55492.0 | 0 | Canidae | 23972.0 | 3 | Canis | 23972.0 | 4 | Canis lupus | 23972.0 | 5 |
| scf7180000042923 | 616011 | 0.4011 | 0 | 60.3929 | Eukaryota | 58415.0 | 0 | Chordata | 58415.0 | 0 | Carnivora | 58415.0 | 0 | Canidae | 58415.0 | 0 | Canis | 46448.0 | 1 | Canis lupus | 46448.0 | 1 |
| scf7180000052915 | 4247 | 0.454 | 0 | 1102.3268 | Eukaryota | 3449.0 | 0 | Platyhelminthes | 2020.0 | 2 | Polystomatidea | 2020.0 | 2 | Polystomatidae | 2020.0 | 2 | Protopolystoma | 2020.0 | 2 | Protopolystoma xenopodis | 2020.0 | 2 |
| scf7180000043036 | 975638 | 0.3929 | 0 | 59.8193 | Eukaryota | 62007.0 | 0 | Chordata | 62007.0 | 0 | Carnivora | 62007.0 | 0 | Canidae | 62007.0 | 0 | Canis | 62007.0 | 0 | Canis lupus | 62007.0 | 0 |
| scf7180000047359 | 57157 | 0.4484 | 0 | 58.4178 | Eukaryota | 14052.0 | 0 | Chordata | 14052.0 | 0 | Carnivora | 14052.0 | 0 | Canidae | 14052.0 | 0 | Canis | 14052.0 | 0 | Canis lupus | 14052.0 | 0 |
| scf7180000043058 | 132783 | 0.4372 | 0 | 58.0833 | Eukaryota | 59817.0 | 0 | Chordata | 59817.0 | 0 | Carnivora | 59817.0 | 0 | Canidae | 59817.0 | 0 | Canis | 43759.0 | 1 | Canis lupus | 43759.0 | 1 |
| scf7180000048869 | 12878 | 0.447 | 0 | 140.3739 | Eukaryota | 616.0 | 0 | Platyhelminthes | 616.0 | 0 | Polystomatidea | 616.0 | 0 | Polystomatidae | 616.0 | 0 | Protopolystoma | 616.0 | 0 | Protopolystoma xenopodis | 616.0 | 0 |
| scf7180000043915 | 5878 | 0.4345 | 0 | 147.9497 | no-hit | 0.0 | 0 | no-hit | 0.0 | 0 | no-hit | 0.0 | 0 | no-hit | 0.0 | 0 | no-hit | 0.0 | 0 | no-hit | 0.0 | 0 |
| scf7180000051383 | 3688 | 0.4376 | 0 | 103.8967 | no-hit | 0.0 | 0 | no-hit | 0.0 | 0 | no-hit | 0.0 | 0 | no-hit | 0.0 | 0 | no-hit | 0.0 | 0 | no-hit | 0.0 | 0 |
| scf7180000042661 | 259078 | 0.3935 | 0 | 65.5351 | Eukaryota | 92935.0 | 0 | Chordata | 92935.0 | 0 | Carnivora | 92935.0 | 0 | Canidae | 92935.0 | 0 | Canis | 92935.0 | 0 | Canis lupus | 92935.0 | 0 |
| scf7180000055333 | 42445 | 0.4444 | 0 | 470.1059 | Eukaryota | 10130.0 | 0 | Chordata | 10130.0 | 0 | Carnivora | 7928.0 | 2 | Canidae | 7928.0 | 2 | Canis | 4219.0 | 3 | Canis lupus | 4219.0 | 3 |
| scf7180000041385 | 8496 | 0.4131 | 0 | 28.2699 | Eukaryota | 4054.0 | 1 | Chordata | 4054.0 | 1 | Carnivora | 2340.0 | 3 | Canidae | 2340.0 | 3 | Canis | 2340.0 | 3 | Canis lupus | 2340.0 | 3 |
| scf7180000052329 | 13359 | 0.5955 | 0 | 64.1049 | Eukaryota | 18738.0 | 0 | Chordata | 18738.0 | 0 | Carnivora | 18738.0 | 0 | Canidae | 18738.0 | 0 | Vulpes | 11274.0 | 1 | Vulpes vulpes | 11274.0 | 1 |
| scf7180000040895 | 13900 | 0.3842 | 0 | 52.9328 | Eukaryota | 82246.0 | 0 | Chordata | 82246.0 | 0 | Carnivora | 82246.0 | 0 | Canidae | 82246.0 | 0 | Canis | 82246.0 | 0 | Canis lupus | 82246.0 | 0 |
| scf7180000041788 | 13439 | 0.4028 | 0 | 49.753 | Eukaryota | 29186.0 | 0 | Chordata | 29186.0 | 0 | Carnivora | 29186.0 | 0 | Felidae | 8561.0 | 5 | Canis | 3136.0 | 9 | Canis lupus | 3136.0 | 9 |
| scf7180000046926 | 12888 | 0.4468 | 0 | 132.2391 | Eukaryota | 4568.0 | 0 | Chordata | 4568.0 | 0 | Tetraodontiformes | 2817.0 | 3 | Tetraodontidae | 2817.0 | 3 | Takifugu | 2817.0 | 3 | Takifugu rubripes | 2817.0 | 3 |
| scf7180000043684 | 83863 | 0.4292 | 0 | 55.8071 | Eukaryota | 25189.0 | 0 | Chordata | 25189.0 | 0 | Carnivora | 25189.0 | 0 | Canidae | 25189.0 | 0 | Canis | 25189.0 | 0 | Canis lupus | 25189.0 | 0 |
| scf7180000049469 | 3367 | 0.4479 | 0 | 339.0485 | no-hit | 0.0 | 0 | no-hit | 0.0 | 0 | no-hit | 0.0 | 0 | no-hit | 0.0 | 0 | no-hit | 0.0 | 0 | no-hit | 0.0 | 0 |
| scf7180000044698 | 7133 | 0.4563 | 0 | 30.3521 | Eukaryota | 56614.0 | 0 | Chordata | 56614.0 | 0 | Carnivora | 56614.0 | 0 | Canidae | 56614.0 | 0 | Canis | 56614.0 | 0 | Canis lupus | 56614.0 | 0 |
| scf7180000047210 | 98545 | 0.3865 | 0 | 59.7145 | Eukaryota | 8364.0 | 0 | Chordata | 8364.0 | 0 | Carnivora | 4406.0 | 2 | Canidae | 4406.0 | 2 | Canis | 4406.0 | 3 | Canis lupus | 4406.0 | 3 |
| scf7180000049984 | 11619 | 0.5201 | 0 | 449.6649 | Eukaryota | 15468.0 | 0 | Chordata | 15468.0 | 0 | Carnivora | 15468.0 | 0 | Canidae | 15468.0 | 0 | Canis | 15468.0 | 0 | Canis lupus | 15468.0 | 0 |
| scf7180000053395 | 16131 | 0.5339 | 0 | 553.8364 | Eukaryota | 33732.0 | 0 | Chordata | 33732.0 | 0 | Carnivora | 33732.0 | 0 | Canidae | 33732.0 | 0 | Canis | 33732.0 | 0 | Canis lupus | 33732.0 | 0 |
| scf7180000047481 | 19554 | 0.4414 | 0 | 368.6754 | Eukaryota | 997.0 | 0 | Platyhelminthes | 997.0 | 0 | Polystomatidea | 997.0 | 0 | Polystomatidae | 997.0 | 0 | Protopolystoma | 997.0 | 0 | Protopolystoma xenopodis | 997.0 | 0 |
| scf7180000043326 | 80971 | 0.4043 | 0 | 56.5454 | Eukaryota | 34570.0 | 0 | Chordata | 34570.0 | 0 | Carnivora | 34570.0 | 0 | Canidae | 34570.0 | 0 | Canis | 34570.0 | 0 | Canis lupus | 34570.0 | 0 |
| scf7180000045236 | 198127 | 0.4196 | 0 | 57.1942 | Eukaryota | 92556.0 | 0 | Chordata | 92556.0 | 0 | Carnivora | 92556.0 | 0 | Canidae | 92556.0 | 0 | Canis | 63536.0 | 1 | Canis lupus | 63536.0 | 1 |
| scf7180000043496 | 240223 | 0.4046 | 0 | 58.9995 | Eukaryota | 59859.0 | 0 | Chordata | 59859.0 | 0 | Carnivora | 59859.0 | 0 | Canidae | 59859.0 | 0 | Canis | 59859.0 | 0 | Canis lupus | 59859.0 | 0 |
| scf7180000048123 | 10712 | 0.45 | 0 | 927.8769 | no-hit | 0.0 | 0 | no-hit | 0.0 | 0 | no-hit | 0.0 | 0 | no-hit | 0.0 | 0 | no-hit | 0.0 | 0 | no-hit | 0.0 | 0 |
| scf7180000045208 | 220317 | 0.4592 | 0 | 60.455 | Eukaryota | 23176.0 | 0 | Chordata | 23176.0 | 0 | Carnivora | 23176.0 | 0 | Canidae | 12526.0 | 1 | Canis | 12526.0 | 1 | Canis lupus | 12526.0 | 1 |
| scf7180000045841 | 36519 | 0.3842 | 0 | 50.5593 | Eukaryota | 10457.0 | 0 | Chordata | 10457.0 | 0 | Carnivora | 6193.0 | 3 | Canidae | 6193.0 | 5 | Canis | 6193.0 | 5 | Canis lupus | 6193.0 | 5 |
| scf7180000041811 | 873657 | 0.4346 | 0 | 59.4744 | Eukaryota | 158470.0 | 0 | Chordata | 158470.0 | 0 | Carnivora | 158470.0 | 0 | Canidae | 158470.0 | 0 | Canis | 110654.0 | 1 | Canis lupus | 110654.0 | 1 |
| scf7180000052472 | 14207 | 0.3828 | 0 | 48.6753 | Eukaryota | 12120.0 | 0 | Chordata | 12120.0 | 0 | Carnivora | 12120.0 | 0 | Phocidae | 4944.0 | 3 | Leptonychotes | 4944.0 | 5 | Leptonychotes weddellii | 4944.0 | 6 |
| scf7180000046540 | 130833 | 0.4 | 0 | 60.3353 | Eukaryota | 13271.0 | 0 | Chordata | 13271.0 | 0 | Carnivora | 13271.0 | 0 | Canidae | 7419.0 | 3 | Canis | 5834.0 | 5 | Canis lupus | 5834.0 | 5 |
| scf7180000042936 | 525284 | 0.4017 | 0 | 59.423 | Eukaryota | 89290.0 | 0 | Chordata | 89290.0 | 0 | Carnivora | 89290.0 | 0 | Canidae | 89290.0 | 0 | Canis | 89290.0 | 0 | Canis lupus | 89290.0 | 0 |
| scf7180000051165 | 9561 | 0.447 | 0 | 185.5901 | Eukaryota | 2901.0 | 0 | Chordata | 2901.0 | 0 | Carnivora | 2901.0 | 0 | Canidae | 2901.0 | 0 | Canis | 2901.0 | 0 | Canis lupus | 2901.0 | 0 |
| scf7180000047162 | 218544 | 0.4046 | 100 | 58.8273 | Eukaryota | 93434.0 | 0 | Chordata | 93434.0 | 0 | Carnivora | 93434.0 | 0 | Canidae | 93434.0 | 0 | Canis | 93434.0 | 0 | Canis lupus | 93434.0 | 0 |
| scf7180000049365 | 27701 | 0.3493 | 0 | 56.2924 | Eukaryota | 29280.0 | 0 | Chordata | 29280.0 | 0 | Carnivora | 29280.0 | 0 | Canidae | 29280.0 | 0 | Canis | 29280.0 | 0 | Canis lupus | 29280.0 | 0 |
| scf7180000048849 | 4257 | 0.444 | 0 | 155.4462 | Eukaryota | 15661.0 | 0 | Chordata | 13361.0 | 2 | Gadiformes | 7209.0 | 6 | Gadidae | 7209.0 | 6 | Gadus | 7209.0 | 6 | Gadus morhua | 7209.0 | 6 |
| scf7180000051427 | 1977 | 0.4426 | 0 | 748.9329 | no-hit | 0.0 | 0 | no-hit | 0.0 | 0 | no-hit | 0.0 | 0 | no-hit | 0.0 | 0 | no-hit | 0.0 | 0 | no-hit | 0.0 | 0 |
| scf7180000053136 | 7907 | 0.4376 | 0 | 35.411 | Eukaryota | 2743.0 | 0 | Chordata | 2743.0 | 0 | Carnivora | 2743.0 | 0 | Canidae | 2743.0 | 0 | Canis | 2743.0 | 0 | Canis lupus | 2743.0 | 0 |
| scf7180000047555 | 32845 | 0.5497 | 0 | 516.8233 | Eukaryota | 45895.0 | 0 | Chordata | 45895.0 | 0 | Artiodactyla | 27696.0 | 2 | Bovidae | 27696.0 | 3 | Ovis | 18373.0 | 4 | Ovis aries | 18373.0 | 4 |
| scf7180000051878 | 3936 | 0.4492 | 0 | 157.9092 | no-hit | 0.0 | 0 | no-hit | 0.0 | 0 | no-hit | 0.0 | 0 | no-hit | 0.0 | 0 | no-hit | 0.0 | 0 | no-hit | 0.0 | 0 |
| scf7180000045707 | 36443 | 0.3576 | 0 | 54.5635 | Eukaryota | 73817.0 | 0 | Chordata | 73817.0 | 0 | Carnivora | 73817.0 | 0 | Canidae | 73817.0 | 0 | Canis | 73817.0 | 0 | Canis lupus | 73817.0 | 0 |
| scf7180000056342 | 525807 | 0.3582 | 0 | 59.8672 | Eukaryota | 32133.0 | 0 | Chordata | 32133.0 | 0 | Primates | 18922.0 | 2 | Cercopithecidae | 10282.0 | 3 | Homo | 8640.0 | 5 | Homo sapiens | 8640.0 | 5 |
| scf7180000051542 | 5403 | 0.4503 | 0 | 510.7314 | no-hit | 0.0 | 0 | no-hit | 0.0 | 0 | no-hit | 0.0 | 0 | no-hit | 0.0 | 0 | no-hit | 0.0 | 0 | no-hit | 0.0 | 0 |
| scf7180000040835 | 1818279 | 0.4254 | 0 | 60.0432 | Eukaryota | 94656.0 | 0 | Chordata | 94656.0 | 0 | Carnivora | 94656.0 | 0 | Canidae | 94656.0 | 0 | Canis | 94656.0 | 0 | Canis lupus | 94656.0 | 0 |
| scf7180000049019 | 61780 | 0.4025 | 0 | 56.283 | Eukaryota | 88203.0 | 0 | Chordata | 88203.0 | 0 | Carnivora | 88203.0 | 0 | Canidae | 88203.0 | 0 | Canis | 88203.0 | 0 | Canis lupus | 88203.0 | 0 |
| scf7180000053120 | 8124 | 0.47 | 0 | 38.3088 | Eukaryota | 3484.0 | 1 | Chordata | 3484.0 | 1 | Primates | 1796.0 | 3 | Hominidae | 1796.0 | 3 | Homo | 1796.0 | 3 | Homo sapiens | 1796.0 | 3 |
| scf7180000047792 | 22393 | 0.4393 | 0 | 270.4208 | Eukaryota | 29267.0 | 0 | Chordata | 28415.0 | 1 | Gadiformes | 19216.0 | 6 | Gadidae | 19216.0 | 6 | Gadus | 19216.0 | 6 | Gadus morhua | 19216.0 | 6 |
| scf7180000053264 | 6998 | 0.5071 | 0 | 297.8302 | Eukaryota | 8431.0 | 0 | Chordata | 8431.0 | 0 | Carnivora | 8431.0 | 0 | Canidae | 8431.0 | 0 | Canis | 6697.0 | 1 | Canis lupus | 5434.0 | 2 |
| scf7180000049761 | 11706 | 0.5049 | 0 | 700.5502 | Eukaryota | 8591.0 | 0 | Chordata | 8591.0 | 0 | Carnivora | 8591.0 | 0 | Canidae | 8591.0 | 0 | Canis | 6988.0 | 1 | Canis lupus | 5829.0 | 2 |
| scf7180000045874 | 73424 | 0.4201 | 0 | 60.4366 | Eukaryota | 80309.0 | 0 | Chordata | 80309.0 | 0 | Carnivora | 80309.0 | 0 | Canidae | 80309.0 | 0 | Canis | 80309.0 | 0 | Canis lupus | 80309.0 | 0 |
| scf7180000042011 | 1062910 | 0.4344 | 0 | 60.5502 | Eukaryota | 93037.0 | 0 | Chordata | 93037.0 | 0 | Carnivora | 93037.0 | 0 | Canidae | 93037.0 | 0 | Canis | 93037.0 | 0 | Canis lupus | 93037.0 | 0 |
| scf7180000047590 | 77183 | 0.4333 | 0 | 59.5211 | Eukaryota | 49746.0 | 0 | Chordata | 49746.0 | 0 | Carnivora | 49746.0 | 0 | Canidae | 27067.0 | 2 | Canis | 27067.0 | 3 | Canis lupus | 27067.0 | 4 |
| scf7180000049356 | 35171 | 0.4463 | 0 | 1080.218 | Eukaryota | 1167.0 | 0 | Platyhelminthes | 632.0 | 1 | Polystomatidea | 632.0 | 2 | Polystomatidae | 632.0 | 2 | Protopolystoma | 632.0 | 2 | Protopolystoma xenopodis | 632.0 | 2 |
| scf7180000054437 | 218591 | 0.3709 | 0 | 161.7001 | Eukaryota | 91123.0 | 0 | Chordata | 91123.0 | 0 | Carnivora | 91123.0 | 0 | Canidae | 91123.0 | 0 | Canis | 91123.0 | 0 | Canis lupus | 91123.0 | 0 |
| scf7180000052147 | 3083 | 0.435 | 0 | 317.1486 | no-hit | 0.0 | 0 | no-hit | 0.0 | 0 | no-hit | 0.0 | 0 | no-hit | 0.0 | 0 | no-hit | 0.0 | 0 | no-hit | 0.0 | 0 |
| scf7180000045070 | 15128 | 0.5429 | 0 | 1304.857 | Eukaryota | 32799.0 | 0 | Chordata | 32799.0 | 0 | Carnivora | 32799.0 | 0 | Canidae | 32799.0 | 0 | Canis | 32799.0 | 0 | Canis lupus | 32799.0 | 0 |
| scf7180000047550 | 43721 | 0.6301 | 0 | 56.3515 | Eukaryota | 14422.0 | 0 | Chordata | 14422.0 | 0 | Carnivora | 14422.0 | 0 | Canidae | 14422.0 | 0 | Canis | 11239.0 | 1 | Canis lupus | 11239.0 | 1 |
| scf7180000048171 | 66822 | 0.349 | 0 | 60.5995 | Eukaryota | 20961.0 | 0 | Chordata | 20961.0 | 0 | Carnivora | 20961.0 | 0 | Canidae | 20961.0 | 0 | Canis | 20961.0 | 0 | Canis lupus | 20961.0 | 0 |
| scf7180000043090 | 580639 | 0.3499 | 0 | 59.3646 | Eukaryota | 86660.0 | 0 | Chordata | 86660.0 | 0 | Carnivora | 86660.0 | 0 | Canidae | 86660.0 | 0 | Canis | 76811.0 | 1 | Canis lupus | 76811.0 | 1 |
| scf7180000044866 | 116921 | 0.3458 | 0 | 57.2789 | Eukaryota | 22024.0 | 0 | Chordata | 22024.0 | 0 | Carnivora | 22024.0 | 0 | Canidae | 15916.0 | 2 | Canis | 13685.0 | 4 | Canis lupus | 11503.0 | 5 |
| scf7180000044310 | 288096 | 0.3867 | 0 | 60.6268 | Eukaryota | 99515.0 | 0 | Chordata | 99515.0 | 0 | Carnivora | 99515.0 | 0 | Canidae | 99515.0 | 0 | Canis | 68973.0 | 1 | Canis lupus | 68973.0 | 1 |
| scf7180000046675 | 64920 | 0.4048 | 0 | 60.473 | Eukaryota | 73659.0 | 0 | Chordata | 73659.0 | 0 | Carnivora | 73659.0 | 0 | Canidae | 73659.0 | 0 | Canis | 73659.0 | 0 | Canis lupus | 73659.0 | 0 |
| scf7180000052495 | 9272 | 0.4089 | 0 | 60.7185 | Eukaryota | 3442.0 | 0 | Chordata | 3442.0 | 0 | Carnivora | 2776.0 | 1 | Canidae | 2776.0 | 1 | Canis | 2776.0 | 1 | Canis lupus | 2776.0 | 1 |
| scf7180000046282 | 141203 | 0.4183 | 0 | 59.8526 | Eukaryota | 84975.0 | 0 | Chordata | 84975.0 | 0 | Carnivora | 84975.0 | 0 | Canidae | 84975.0 | 0 | Canis | 84975.0 | 0 | Canis lupus | 84975.0 | 0 |
| scf7180000056305 | 350127 | 0.4374 | 0 | 57.853 | Eukaryota | 28695.0 | 1 | Chordata | 28695.0 | 1 | Carnivora | 23609.0 | 2 | Canidae | 19057.0 | 3 | Canis | 15936.0 | 4 | Canis lupus | 15936.0 | 4 |
| scf7180000052539 | 14880 | 0.4474 | 0 | 435.734 | Eukaryota | 54048.0 | 0 | Chordata | 54048.0 | 0 | Carnivora | 54048.0 | 0 | Canidae | 54048.0 | 0 | Canis | 54048.0 | 0 | Canis lupus | 54048.0 | 0 |
| scf7180000041821 | 3317 | 0.3838 | 0 | 27.732 | Eukaryota | 3731.0 | 0 | Chordata | 3731.0 | 0 | Carnivora | 2833.0 | 1 | Canidae | 2833.0 | 1 | Canis | 2833.0 | 1 | Canis lupus | 2833.0 | 1 |
| scf7180000056265 | 994205 | 0.3782 | 0 | 60.3511 | Eukaryota | 93083.0 | 0 | Chordata | 93083.0 | 0 | Carnivora | 93083.0 | 0 | Canidae | 93083.0 | 0 | Canis | 93083.0 | 0 | Canis lupus | 93083.0 | 0 |
| scf7180000052176 | 5569 | 0.441 | 0 | 117.2359 | Eukaryota | 873.0 | 0 | Platyhelminthes | 873.0 | 0 | Polystomatidea | 873.0 | 0 | Polystomatidae | 873.0 | 0 | Protopolystoma | 873.0 | 0 | Protopolystoma xenopodis | 873.0 | 0 |
| scf7180000041578 | 1069825 | 0.3919 | 0 | 60.7007 | Eukaryota | 95000.0 | 0 | Chordata | 95000.0 | 0 | Carnivora | 95000.0 | 0 | Canidae | 95000.0 | 0 | Canis | 95000.0 | 0 | Canis lupus | 95000.0 | 0 |
| scf7180000043945 | 347698 | 0.3837 | 0 | 76.2158 | Eukaryota | 91551.0 | 0 | Chordata | 91551.0 | 0 | Carnivora | 91551.0 | 0 | Canidae | 91551.0 | 0 | Canis | 91551.0 | 0 | Canis lupus | 91551.0 | 0 |
| scf7180000055551 | 13630 | 0.4428 | 0 | 235.4467 | Eukaryota | 611.0 | 0 | Chordata | 611.0 | 0 | Kurtiformes | 350.0 | 1 | Apogonidae | 350.0 | 1 | Sphaeramia | 350.0 | 1 | Sphaeramia orbicularis | 350.0 | 1 |
| scf7180000056480 | 281148 | 0.3571 | 0 | 60.2708 | Eukaryota | 52634.0 | 0 | Chordata | 52634.0 | 0 | Carnivora | 52634.0 | 0 | Canidae | 52634.0 | 0 | Canis | 52634.0 | 0 | Canis lupus | 52634.0 | 0 |
| scf7180000052216 | 5847 | 0.4459 | 0 | 703.3719 | Eukaryota | 3185.0 | 1 | Platyhelminthes | 1475.0 | 4 | Polystomatidea | 1475.0 | 4 | Polystomatidae | 1475.0 | 4 | Protopolystoma | 1475.0 | 4 | Protopolystoma xenopodis | 1475.0 | 4 |
| scf7180000045373 | 186644 | 0.3863 | 0 | 60.4676 | Eukaryota | 26254.0 | 0 | Chordata | 26254.0 | 0 | Carnivora | 26254.0 | 0 | Canidae | 26254.0 | 0 | Canis | 26254.0 | 0 | Canis lupus | 26254.0 | 0 |
| scf7180000056370 | 18203 | 0.435 | 0 | 51.2988 | Eukaryota | 17091.0 | 0 | Chordata | 17091.0 | 0 | Carnivora | 13417.0 | 1 | Canidae | 13417.0 | 1 | Canis | 13417.0 | 1 | Canis lupus | 13417.0 | 1 |
| scf7180000042434 | 136250 | 0.4066 | 0 | 55.7841 | Eukaryota | 34241.0 | 0 | Chordata | 34241.0 | 0 | Carnivora | 34241.0 | 0 | Canidae | 34241.0 | 0 | Canis | 22734.0 | 1 | Canis lupus | 22734.0 | 1 |
| scf7180000040680 | 1911198 | 0.3806 | 0 | 59.5264 | Eukaryota | 118200.0 | 0 | Chordata | 118200.0 | 0 | Carnivora | 118200.0 | 0 | Canidae | 118200.0 | 0 | Canis | 70872.0 | 1 | Canis lupus | 70872.0 | 1 |
| scf7180000046313 | 127121 | 0.4049 | 0 | 59.2128 | Eukaryota | 28915.0 | 0 | Chordata | 28915.0 | 0 | Carnivora | 28915.0 | 0 | Canidae | 28915.0 | 0 | Canis | 28915.0 | 0 | Canis lupus | 28915.0 | 0 |
| scf7180000049071 | 20092 | 0.44 | 0 | 134.8557 | Eukaryota | 91721.0 | 0 | Chordata | 91721.0 | 0 | Carnivora | 91721.0 | 0 | Canidae | 91721.0 | 0 | Canis | 91721.0 | 0 | Canis lupus | 91721.0 | 0 |
| scf7180000040548 | 2347518 | 0.4336 | 0 | 59.7359 | Eukaryota | 94901.0 | 0 | Chordata | 94901.0 | 0 | Carnivora | 94901.0 | 0 | Canidae | 94901.0 | 0 | Canis | 94901.0 | 0 | Canis lupus | 94901.0 | 0 |
| scf7180000049279 | 64629 | 0.4674 | 0 | 61.6835 | Eukaryota | 22054.0 | 0 | Chordata | 22054.0 | 0 | Carnivora | 22054.0 | 0 | Canidae | 12418.0 | 2 | Canis | 9919.0 | 4 | Canis lupus | 9919.0 | 4 |
| scf7180000045722 | 188071 | 0.3663 | 0 | 60.3768 | Eukaryota | 10050.0 | 0 | Chordata | 10050.0 | 0 | Carnivora | 7870.0 | 1 | Canidae | 6894.0 | 2 | Canis | 6894.0 | 2 | Canis lupus | 6894.0 | 2 |
| scf7180000052175 | 3088 | 0.4401 | 0 | 579.044 | no-hit | 0.0 | 0 | no-hit | 0.0 | 0 | no-hit | 0.0 | 0 | no-hit | 0.0 | 0 | no-hit | 0.0 | 0 | no-hit | 0.0 | 0 |
| scf7180000042154 | 579323 | 0.3645 | 0 | 59.0689 | Eukaryota | 89533.0 | 0 | Chordata | 89533.0 | 0 | Carnivora | 89533.0 | 0 | Canidae | 89533.0 | 0 | Canis | 89533.0 | 0 | Canis lupus | 89533.0 | 0 |
| scf7180000042318 | 305669 | 0.384 | 0 | 59.4446 | Eukaryota | 55348.0 | 0 | Chordata | 55348.0 | 0 | Carnivora | 46740.0 | 1 | Canidae | 46740.0 | 1 | Canis | 46740.0 | 2 | Canis lupus | 46740.0 | 2 |
| scf7180000054209 | 24426 | 0.4435 | 0 | 166.4528 | Eukaryota | 8645.0 | 0 | Chordata | 7301.0 | 1 | Kurtiformes | 5064.0 | 4 | Apogonidae | 5064.0 | 4 | Sphaeramia | 5064.0 | 4 | Sphaeramia orbicularis | 5064.0 | 4 |
| scf7180000056546 | 229835 | 0.374 | 0 | 60.248 | Eukaryota | 25971.0 | 0 | Chordata | 25971.0 | 0 | Carnivora | 25971.0 | 0 | Canidae | 25971.0 | 0 | Canis | 25971.0 | 0 | Canis lupus | 25971.0 | 0 |
| scf7180000047120 | 102449 | 0.3414 | 0 | 59.1243 | Eukaryota | 10476.0 | 0 | Chordata | 10476.0 | 0 | Carnivora | 6278.0 | 2 | Canidae | 5622.0 | 3 | Canis | 5622.0 | 3 | Canis lupus | 5622.0 | 3 |
| scf7180000052618 | 13977 | 0.4487 | 0 | 65.8039 | Eukaryota | 906.0 | 0 | Platyhelminthes | 906.0 | 0 | Polystomatidea | 906.0 | 0 | Polystomatidae | 906.0 | 0 | Protopolystoma | 906.0 | 0 | Protopolystoma xenopodis | 906.0 | 0 |
| scf7180000041664 | 1032754 | 0.4096 | 0 | 59.6029 | Eukaryota | 34831.0 | 0 | Chordata | 34831.0 | 0 | Carnivora | 34831.0 | 0 | Canidae | 34831.0 | 0 | Vulpes | 19892.0 | 1 | Vulpes vulpes | 19892.0 | 1 |
| scf7180000055602 | 26195 | 0.4447 | 0 | 187.199 | Eukaryota | 94499.0 | 0 | Chordata | 94499.0 | 0 | Carnivora | 94499.0 | 0 | Canidae | 94499.0 | 0 | Canis | 94499.0 | 0 | Canis lupus | 94499.0 | 0 |
| scf7180000043255 | 130356 | 0.3721 | 0 | 57.5378 | Eukaryota | 89185.0 | 0 | Chordata | 89185.0 | 0 | Carnivora | 89185.0 | 0 | Canidae | 89185.0 | 0 | Canis | 89185.0 | 0 | Canis lupus | 89185.0 | 0 |
| scf7180000052777 | 6615 | 0.4404 | 0 | 37.6067 | Eukaryota | 1119.0 | 0 | Platyhelminthes | 943.0 | 1 | Polystomatidea | 943.0 | 1 | Polystomatidae | 943.0 | 1 | Protopolystoma | 943.0 | 1 | Protopolystoma xenopodis | 943.0 | 1 |
| scf7180000045529 | 13148 | 0.5078 | 0 | 1810.5318 | Eukaryota | 40484.0 | 0 | Chordata | 40484.0 | 0 | Carnivora | 40484.0 | 0 | Canidae | 40484.0 | 0 | Canis | 40484.0 | 0 | Canis lupus | 40484.0 | 0 |
| scf7180000044556 | 342064 | 0.3451 | 0 | 59.373 | Eukaryota | 91117.0 | 0 | Chordata | 91117.0 | 0 | Carnivora | 91117.0 | 0 | Canidae | 91117.0 | 0 | Canis | 91117.0 | 0 | Canis lupus | 91117.0 | 0 |
| scf7180000044157 | 42422 | 0.5162 | 0 | 57.574 | Eukaryota | 22999.0 | 0 | Chordata | 22999.0 | 0 | Carnivora | 22999.0 | 0 | Canidae | 19218.0 | 1 | Canis | 16804.0 | 3 | Canis lupus | 16804.0 | 3 |
| scf7180000041001 | 1538395 | 0.3892 | 0 | 59.7285 | Eukaryota | 90327.0 | 0 | Chordata | 90327.0 | 0 | Carnivora | 90327.0 | 0 | Canidae | 90327.0 | 0 | Canis | 90327.0 | 0 | Canis lupus | 90327.0 | 0 |
| scf7180000048881 | 19895 | 0.4671 | 0 | 61.9972 | Eukaryota | 6710.0 | 0 | Chordata | 6710.0 | 0 | Carnivora | 6710.0 | 0 | Canidae | 4859.0 | 1 | Canis | 2837.0 | 2 | Canis lupus | 2837.0 | 2 |
| scf7180000046846 | 6171 | 0.45 | 0 | 173.563 | Eukaryota | 3568.0 | 0 | Chordata | 3568.0 | 0 | Carnivora | 2644.0 | 1 | Canidae | 2644.0 | 1 | Canis | 2644.0 | 1 | Canis lupus | 2644.0 | 1 |
| scf7180000055461 | 30252 | 0.4405 | 0 | 342.321 | Eukaryota | 562.0 | 0 | Platyhelminthes | 562.0 | 0 | Polystomatidea | 562.0 | 0 | Polystomatidae | 562.0 | 0 | Protopolystoma | 562.0 | 0 | Protopolystoma xenopodis | 562.0 | 0 |
| scf7180000043631 | 80518 | 0.407 | 0 | 58.2063 | Eukaryota | 93588.0 | 0 | Chordata | 93588.0 | 0 | Carnivora | 93588.0 | 0 | Canidae | 93588.0 | 0 | Canis | 93588.0 | 0 | Canis lupus | 93588.0 | 0 |
| scf7180000043224 | 803240 | 0.3992 | 0 | 60.1224 | Eukaryota | 88397.0 | 0 | Chordata | 88397.0 | 0 | Carnivora | 88397.0 | 0 | Canidae | 88397.0 | 0 | Canis | 88397.0 | 0 | Canis lupus | 88397.0 | 0 |
| scf7180000044887 | 31577 | 0.5901 | 0 | 42.0649 | Eukaryota | 32443.0 | 0 | Chordata | 32443.0 | 0 | Carnivora | 32443.0 | 0 | Canidae | 32443.0 | 0 | Canis | 28604.0 | 1 | Canis lupus | 28604.0 | 1 |

| | | | | | | | | | | | | | | | | | | | | | | |
|---|---|---|---|---|---|---|---|---|---|---|---|---|---|---|---|---|---|---|---|---|---|---|
| scf7180000050095 | 36691 | 0.4533 | 0 | 166.9421 | Eukaryota | 6026.0 | 0 | Chordata | 3612.0 | 1 | Tetraodontiformes | 2496.0 | 3 | Tetraodontidae | 2496.0 | 3 | Takifugu | 2496.0 | 3 | Takifugu rubripes | 2496.0 | 3 |
| scf7180000042555 | 617492 | 0.3907 | 0 | 59.5816 | Eukaryota | 92644.0 | 0 | Chordata | 92644.0 | 0 | Carnivora | 92644.0 | 0 | Canidae | 92644.0 | 0 | Canis | 92644.0 | 0 | Canis lupus | 92644.0 | 0 |
| scf7180000044166 | 66888 | 0.3878 | 0 | 57.8184 | Eukaryota | 21934.0 | 0 | Chordata | 21934.0 | 0 | Carnivora | 19982.0 | 1 | Canidae | 17307.0 | 3 | Canis | 16372.0 | 5 | Canis lupus | 16372.0 | 5 |
| scf7180000043618 | 35811 | 0.4375 | 0 | 55.3738 | Eukaryota | 75312.0 | 0 | Chordata | 75312.0 | 0 | Carnivora | 75312.0 | 0 | Canidae | 75312.0 | 0 | Canis | 75312.0 | 0 | Canis lupus | 75312.0 | 0 |
| scf7180000044303 | 198873 | 0.4443 | 0 | 58.0107 | Eukaryota | 17718.0 | 0 | Chordata | 17718.0 | 0 | Carnivora | 17718.0 | 0 | Canidae | 17718.0 | 0 | Canis | 17718.0 | 0 | Canis lupus | 17718.0 | 0 |
| scf7180000049425 | 41828 | 0.4509 | 0 | 87.7555 | Eukaryota | 87737.0 | 0 | Chordata | 87737.0 | 0 | Carnivora | 87737.0 | 0 | Canidae | 87737.0 | 0 | Canis | 87737.0 | 0 | Canis lupus | 87737.0 | 0 |
| scf7180000043533 | 440785 | 0.3811 | 0 | 60.055 | Eukaryota | 61012.0 | 0 | Chordata | 61012.0 | 0 | Carnivora | 61012.0 | 0 | Canidae | 61012.0 | 0 | Canis | 49802.0 | 1 | Canis lupus | 49802.0 | 1 |
| scf7180000056332 | 570438 | 0.4372 | 0 | 59.1322 | Eukaryota | 59522.0 | 0 | Chordata | 59522.0 | 0 | Carnivora | 59522.0 | 0 | Canidae | 59522.0 | 0 | Canis | 59522.0 | 0 | Canis lupus | 59522.0 | 0 |
| scf7180000040607 | 721862 | 0.3546 | 0 | 58.7007 | Eukaryota | 88302.0 | 0 | Chordata | 88302.0 | 0 | Carnivora | 88302.0 | 0 | Canidae | 88302.0 | 0 | Canis | 88302.0 | 0 | Canis lupus | 88302.0 | 0 |
| scf7180000051099 | 2199 | 0.4543 | 0 | 353.8207 | Eukaryota | 281.0 | 0 | Platyhelminthes | 281.0 | 0 | Polystomatidea | 281.0 | 0 | Polystomatidae | 281.0 | 0 | Protopolystoma | 281.0 | 0 | Protopolystoma xenopodis | 281.0 | 0 |
| scf7180000049143 | 13571 | 0.4496 | 0 | 1171.8726 | Eukaryota | 243.0 | 0 | Streptophyta | 243.0 | 0 | Fabales | 243.0 | 0 | Fabaceae | 243.0 | 0 | Cicer | 243.0 | 0 | Cicer arietinum | 243.0 | 0 |
| scf7180000053647 | 7697 | 0.4443 | 0 | 196.8454 | Eukaryota | 37888.0 | 0 | Chordata | 26726.0 | 2 | Gadiformes | 20130.0 | 5 | Gadidae | 20130.0 | 5 | Gadus | 20130.0 | 5 | Gadus morhua | 20130.0 | 5 |
| scf7180000051031 | 30454 | 0.3539 | 0 | 59.6458 | Eukaryota | 13423.0 | 0 | Chordata | 13423.0 | 0 | Carnivora | 11021.0 | 1 | Canidae | 4394.0 | 3 | Canis | 2912.0 | 5 | Canis lupus | 2912.0 | 5 |
| scf7180000052515 | 13814 | 0.368 | 0 | 61.579 | Eukaryota | 15306.0 | 0 | Chordata | 15306.0 | 0 | Primates | 8040.0 | 1 | Hominidae | 8040.0 | 1 | Canis | 7266.0 | 2 | Canis lupus | 7266.0 | 2 |
| scf7180000044269 | 13501 | 0.4464 | 0 | 218.856 | Eukaryota | 86500.0 | 0 | Chordata | 86500.0 | 0 | Carnivora | 86500.0 | 0 | Canidae | 86500.0 | 0 | Canis | 86500.0 | 0 | Canis lupus | 86500.0 | 0 |
| scf7180000043472 | 386363 | 0.444 | 0 | 56.4981 | Eukaryota | 71198.0 | 0 | Chordata | 71198.0 | 0 | Carnivora | 71198.0 | 0 | Canidae | 71198.0 | 0 | Canis | 71198.0 | 0 | Canis lupus | 71198.0 | 0 |
| scf7180000044689 | 201950 | 0.4406 | 0 | 129.8043 | Eukaryota | 90634.0 | 0 | Chordata | 90634.0 | 0 | Carnivora | 90634.0 | 0 | Canidae | 90634.0 | 0 | Canis | 90634.0 | 0 | Canis lupus | 90634.0 | 0 |
| scf7180000049144 | 26555 | 0.4495 | 0 | 1197.8504 | Eukaryota | 15918.0 | 0 | Chordata | 15918.0 | 0 | Gadiformes | 10831.0 | 5 | Gadidae | 10831.0 | 5 | Gadus | 10831.0 | 5 | Gadus morhua | 10831.0 | 5 |
| scf7180000041252 | 5666 | 0.389 | 0 | 55.7315 | Eukaryota | 9191.0 | 0 | Chordata | 9191.0 | 0 | Carnivora | 8164.0 | 1 | Felidae | 5804.0 | 2 | Panthera | 2890.0 | 4 | Panthera pardus | 2890.0 | 4 |
| scf7180000040520 | 1773472 | 0.4855 | 0 | 87.8623 | Eukaryota | 110352.0 | 0 | Chordata | 110352.0 | 0 | Carnivora | 110352.0 | 0 | Canidae | 110352.0 | 0 | Vulpes | 88360.0 | 1 | Vulpes vulpes | 88360.0 | 1 |
| scf7180000047762 | 6026 | 0.4457 | 0 | 500.7016 | Eukaryota | 455.0 | 0 | Chordata | 455.0 | 0 | Kurtiformes | 455.0 | 0 | Apogonidae | 455.0 | 0 | Sphaeramia | 455.0 | 0 | Sphaeramia orbicularis | 455.0 | 0 |
| scf7180000043067 | 14216 | 0.3592 | 0 | 32.7652 | Eukaryota | 3074.0 | 0 | Chordata | 3074.0 | 0 | Carnivora | 3074.0 | 0 | Canidae | 3074.0 | 0 | Canis | 3074.0 | 0 | Canis lupus | 3074.0 | 0 |
| scf7180000053122 | 8588 | 0.5044 | 0 | 308.7892 | Eukaryota | 7889.0 | 0 | Chordata | 7889.0 | 0 | Carnivora | 7889.0 | 0 | Canidae | 7889.0 | 0 | Canis | 6786.0 | 1 | Canis lupus | 5649.0 | 2 |
| scf7180000051479 | 10353 | 0.4502 | 0 | 1234.3578 | Eukaryota | 20839.0 | 0 | Chordata | 18417.0 | 2 | Gadiformes | 12949.0 | 7 | Gadidae | 12949.0 | 7 | Gadus | 12949.0 | 7 | Gadus morhua | 12949.0 | 7 |
| scf7180000047860 | 21169 | 0.4394 | 0 | 106.6182 | Eukaryota | 15825.0 | 0 | Chordata | 15825.0 | 0 | Carnivora | 15825.0 | 0 | Canidae | 15825.0 | 0 | Canis | 15825.0 | 0 | Canis lupus | 15825.0 | 0 |
| scf7180000043439 | 279488 | 0.3919 | 0 | 60.0194 | Eukaryota | 81937.0 | 0 | Chordata | 81937.0 | 0 | Carnivora | 81937.0 | 0 | Canidae | 81937.0 | 0 | Canis | 72616.0 | 1 | Canis lupus | 72616.0 | 1 |
| scf7180000051926 | 10198 | 0.509 | 0 | 1310.3398 | Eukaryota | 8199.0 | 0 | Chordata | 8199.0 | 0 | Carnivora | 8199.0 | 0 | Canidae | 8199.0 | 0 | Canis | 5987.0 | 1 | Canis lupus | 4758.0 | 2 |
| scf7180000043169 | 337635 | 0.4878 | 0 | 57.7919 | Eukaryota | 94668.0 | 0 | Chordata | 94668.0 | 0 | Carnivora | 94668.0 | 0 | Canidae | 94668.0 | 0 | Canis | 94668.0 | 0 | Canis lupus | 94668.0 | 0 |
| scf7180000041350 | 1392149 | 0.3892 | 0 | 60.7508 | Eukaryota | 87153.0 | 0 | Chordata | 87153.0 | 0 | Carnivora | 87153.0 | 0 | Canidae | 87153.0 | 0 | Canis | 87153.0 | 0 | Canis lupus | 87153.0 | 0 |
| scf7180000042660 | 139686 | 0.4999 | 0 | 59.4128 | Eukaryota | 24904.0 | 0 | Chordata | 24904.0 | 0 | Carnivora | 24904.0 | 0 | Canidae | 24904.0 | 0 | Vulpes | 12511.0 | 1 | Vulpes vulpes | 12511.0 | 1 |
| scf7180000051648 | 2748 | 0.4472 | 0 | 186.2623 | Eukaryota | 1082.0 | 0 | Echinodermata | 634.0 | 2 | Forcipulatida | 634.0 | 2 | Asteriidae | 634.0 | 2 | Asterias | 634.0 | 2 | Asterias rubens | 634.0 | 2 |
| scf7180000046958 | 54045 | 0.4388 | 0 | 247.8415 | Eukaryota | 20073.0 | 0 | Chordata | 17147.0 | 2 | Gadiformes | 6454.0 | 6 | Gadidae | 6454.0 | 6 | Gadus | 6454.0 | 6 | Gadus morhua | 6454.0 | 6 |
| scf7180000050301 | 15027 | 0.4416 | 0 | 499.8032 | Eukaryota | 557.0 | 0 | Platyhelminthes | 557.0 | 0 | Diphyllobothriidea | 557.0 | 0 | Diphyllobothriidae | 557.0 | 0 | Spirometra | 557.0 | 0 | Spirometra erinaceieuropaei | 557.0 | 0 |
| scf7180000042052 | 8097 | 0.4067 | 0 | 49.8936 | Eukaryota | 2293.0 | 0 | Chordata | 2293.0 | 0 | Carnivora | 2293.0 | 0 | Canidae | 2293.0 | 0 | Canis | 2293.0 | 0 | Canis lupus | 2293.0 | 0 |
| scf7180000042097 | 41577 | 0.3421 | 0 | 53.5088 | Eukaryota | 10859.0 | 0 | Chordata | 10859.0 | 0 | Carnivora | 10859.0 | 0 | Canidae | 10859.0 | 0 | Canis | 10859.0 | 0 | Canis lupus | 10859.0 | 0 |
| scf7180000055389 | 182859 | 0.4452 | 0 | 256.9628 | Eukaryota | 18509.0 | 0 | Chordata | 18509.0 | 0 | Carnivora | 18509.0 | 0 | Canidae | 15677.0 | 1 | Canis | 14035.0 | 2 | Canis lupus | 14035.0 | 2 |
| scf7180000050241 | 5522 | 0.4529 | 0 | 57.067 | Eukaryota | 18889.0 | 1 | Chordata | 18889.0 | 1 | Kurtiformes | 4948.0 | 5 | Apogonidae | 4948.0 | 5 | Sphaeramia | 4948.0 | 5 | Sphaeramia orbicularis | 4948.0 | 5 |
| scf7180000045369 | 242334 | 0.4263 | 0 | 60.7759 | Eukaryota | 46836.0 | 0 | Chordata | 46836.0 | 0 | Carnivora | 46836.0 | 0 | Canidae | 28035.0 | 3 | Canis | 17320.0 | 4 | Canis lupus | 17320.0 | 4 |
| scf7180000045567 | 156984 | 0.3886 | 0 | 60.5848 | Eukaryota | 90562.0 | 0 | Chordata | 90562.0 | 0 | Carnivora | 90562.0 | 0 | Canidae | 90562.0 | 0 | Canis | 90562.0 | 0 | Canis lupus | 90562.0 | 0 |
| scf7180000043057 | 381465 | 0.3714 | 0 | 59.9024 | Eukaryota | 89295.0 | 0 | Chordata | 89295.0 | 0 | Carnivora | 89295.0 | 0 | Canidae | 89295.0 | 0 | Canis | 89295.0 | 0 | Canis lupus | 89295.0 | 0 |
| scf7180000045688 | 47371 | 0.3622 | 0 | 131.0456 | Eukaryota | 93036.0 | 0 | Chordata | 93036.0 | 0 | Carnivora | 93036.0 | 0 | Canidae | 93036.0 | 0 | Canis | 93036.0 | 0 | Canis lupus | 93036.0 | 0 |
| scf7180000050654 | 20952 | 0.4486 | 0 | 202.2179 | Eukaryota | 1418.0 | 0 | Platyhelminthes | 747.0 | 1 | Polystomatidea | 747.0 | 1 | Polystomatidae | 747.0 | 1 | Protopolystoma | 747.0 | 1 | Protopolystoma xenopodis | 747.0 | 1 |
| scf7180000043898 | 401849 | 0.3729 | 0 | 59.1079 | Eukaryota | 90195.0 | 0 | Chordata | 90195.0 | 0 | Carnivora | 90195.0 | 0 | Canidae | 90195.0 | 0 | Canis | 90195.0 | 0 | Canis lupus | 90195.0 | 0 |
| scf7180000050873 | 16413 | 0.6721 | 0 | 51.5902 | Eukaryota | 15189.0 | 0 | Chordata | 15189.0 | 0 | Carnivora | 15189.0 | 0 | Canidae | 10539.0 | 2 | Canis | 9059.0 | 4 | Canis lupus | 9059.0 | 4 |
| scf7180000055704 | 20218 | 0.434 | 0 | 129.9224 | Eukaryota | 3734.0 | 0 | Chordata | 2222.0 | 1 | Polystomatidea | 1512.0 | 4 | Polystomatidae | 1512.0 | 4 | Protopolystoma | 1512.0 | 4 | Protopolystoma xenopodis | 1512.0 | 4 |
| scf7180000052240 | 3164 | 0.4516 | 0 | 513.7743 | no-hit | 0.0 | 0 | no-hit | 0.0 | 0 | no-hit | 0.0 | 0 | no-hit | 0.0 | 0 | no-hit | 0.0 | 0 | no-hit | 0.0 | 0 |
| scf7180000055890 | 7543 | 0.4326 | 0 | 220.5281 | no-hit | 0.0 | 0 | no-hit | 0.0 | 0 | no-hit | 0.0 | 0 | no-hit | 0.0 | 0 | no-hit | 0.0 | 0 | no-hit | 0.0 | 0 |
| scf7180000044774 | 10823 | 0.4395 | 0 | 68.0779 | Eukaryota | 2827.0 | 0 | Chordata | 2827.0 | 0 | Carnivora | 2053.0 | 1 | Canidae | 2053.0 | 1 | Canis | 2053.0 | 1 | Canis lupus | 2053.0 | 1 |
| scf7180000048636 | 18289 | 0.4414 | 0 | 200.1967 | Eukaryota | 976.0 | 0 | Chordata | 976.0 | 0 | Syngnathiformes | 765.0 | 1 | Syngnathidae | 765.0 | 1 | Syngnathus | 765.0 | 1 | Syngnathus acus | 765.0 | 1 |
| scf7180000050984 | 3450 | 0.4504 | 0 | 453.8353 | no-hit | 0.0 | 0 | no-hit | 0.0 | 0 | no-hit | 0.0 | 0 | no-hit | 0.0 | 0 | no-hit | 0.0 | 0 | no-hit | 0.0 | 0 |
| scf7180000050889 | 4881 | 0.5044 | 0 | 119.1 | Eukaryota | 8315.0 | 0 | Chordata | 8315.0 | 0 | Carnivora | 8315.0 | 0 | Canidae | 8315.0 | 0 | Canis | 6734.0 | 1 | Canis lupus | 5622.0 | 2 |
| scf7180000055318 | 24289 | 0.4988 | 0 | 781.0772 | Eukaryota | 90234.0 | 0 | Chordata | 90234.0 | 0 | Carnivora | 90234.0 | 0 | Canidae | 90234.0 | 0 | Canis | 90234.0 | 0 | Canis lupus | 90234.0 | 0 |
| scf7180000043682 | 434807 | 0.4005 | 0 | 59.4006 | Eukaryota | 71445.0 | 0 | Chordata | 71445.0 | 0 | Carnivora | 71445.0 | 0 | Canidae | 59349.0 | 2 | Canis | 44989.0 | 3 | Canis lupus | 44989.0 | 3 |
| scf7180000048552 | 11399 | 0.4334 | 0 | 112.8509 | Eukaryota | 1406.0 | 0 | Platyhelminthes | 555.0 | 2 | Polystomatidea | 555.0 | 3 | Polystomatidae | 555.0 | 3 | Protopolystoma | 555.0 | 3 | Protopolystoma xenopodis | 555.0 | 3 |
| scf7180000049460 | 4208 | 0.4449 | 0 | 570.34 | no-hit | 0.0 | 0 | no-hit | 0.0 | 0 | no-hit | 0.0 | 0 | no-hit | 0.0 | 0 | no-hit | 0.0 | 0 | no-hit | 0.0 | 0 |
| scf7180000043773 | 406627 | 0.4559 | 0 | 60.1063 | Eukaryota | 91184.0 | 0 | Chordata | 91184.0 | 0 | Carnivora | 91184.0 | 0 | Canidae | 91184.0 | 0 | Canis | 91184.0 | 0 | Canis lupus | 91184.0 | 0 |
| scf7180000044737 | 289979 | 0.5263 | 0 | 56.1353 | Eukaryota | 28147.0 | 0 | Chordata | 28147.0 | 0 | Carnivora | 28147.0 | 0 | Canidae | 28147.0 | 0 | Canis | 28147.0 | 0 | Canis lupus | 28147.0 | 0 |
| scf7180000054896 | 7092 | 0.4437 | 0 | 134.9156 | Eukaryota | 518.0 | 0 | Chordata | 518.0 | 0 | Columbiformes | 518.0 | 0 | Columbidae | 518.0 | 0 | Streptopelia | 518.0 | 0 | Streptopelia turtur | 518.0 | 0 |
| scf7180000047230 | 68819 | 0.3454 | 0 | 57.1811 | Eukaryota | 12926.0 | 0 | Chordata | 12926.0 | 0 | Carnivora | 12926.0 | 0 | Canidae | 12926.0 | 0 | Canis | 12926.0 | 0 | Canis lupus | 12926.0 | 0 |
| scf7180000050363 | 4892 | 0.4526 | 0 | 561.9528 | Eukaryota | 3249.0 | 0 | Chordata | 2573.0 | 1 | Tetraodontiformes | 1822.0 | 3 | Tetraodontidae | 1822.0 | 3 | Takifugu | 1822.0 | 3 | Takifugu rubripes | 1822.0 | 3 |
| scf7180000046714 | 49750 | 0.3691 | 0 | 64.9054 | Eukaryota | 61990.0 | 0 | Chordata | 61990.0 | 0 | Carnivora | 61990.0 | 0 | Canidae | 61990.0 | 0 | Canis | 61990.0 | 0 | Canis lupus | 61990.0 | 0 |
| scf7180000048056 | 9901 | 0.472 | 0 | 70.6986 | Eukaryota | 7876.0 | 0 | Chordata | 7876.0 | 0 | Carnivora | 4542.0 | 1 | Canidae | 4542.0 | 1 | Homo | 3334.0 | 2 | Homo sapiens | 3334.0 | 2 |
| scf7180000055948 | 11270 | 0.4388 | 0 | 1401.0788 | Eukaryota | 4083.0 | 0 | Chordata | 3135.0 | 1 | Polystomatidea | 948.0 | 5 | Polystomatidae | 948.0 | 5 | Protopolystoma | 948.0 | 6 | Protopolystoma xenopodis | 948.0 | 6 |
| scf7180000047253 | 27310 | 0.4118 | 0 | 75.6119 | Eukaryota | 30309.0 | 0 | Chordata | 30309.0 | 0 | Carnivora | 30309.0 | 0 | Canidae | 30309.0 | 0 | Canis | 30309.0 | 0 | Canis lupus | 30309.0 | 0 |
| scf7180000045788 | 155661 | 0.5743 | 0 | 56.5151 | Eukaryota | 39281.0 | 0 | Chordata | 39281.0 | 0 | Carnivora | 39281.0 | 0 | Canidae | 39281.0 | 0 | Canis | 34381.0 | 1 | Canis lupus | 34381.0 | 1 |
| scf7180000045107 | 260913 | 0.5106 | 0 | 57.6371 | Eukaryota | 31595.0 | 0 | Chordata | 31595.0 | 0 | Carnivora | 21097.0 | 3 | Canidae | 21097.0 | 3 | Canis | 21097.0 | 3 | Canis lupus | 21097.0 | 3 |
| scf7180000050862 | 22029 | 0.445 | 0 | 86.0306 | Eukaryota | 6665.0 | 1 | Chordata | 6665.0 | 1 | Tetraodontiformes | 3054.0 | 4 | Tetraodontidae | 3054.0 | 4 | Takifugu | 3054.0 | 4 | Takifugu rubripes | 2621.0 | 5 |
| scf7180000048068 | 46494 | 0.4375 | 0 | 130.9065 | Eukaryota | 2326.0 | 0 | Platyhelminthes | 1856.0 | 1 | Polystomatidea | 1856.0 | 1 | Polystomatidae | 1856.0 | 1 | Protopolystoma | 1856.0 | 1 | Protopolystoma xenopodis | 1856.0 | 1 |
| scf7180000044346 | 365841 | 0.45 | 0 | 58.4909 | Eukaryota | 51560.0 | 0 | Chordata | 51560.0 | 0 | Carnivora | 51560.0 | 0 | Canidae | 34682.0 | 2 | Canis | 29634.0 | 5 | Canis lupus | 29634.0 | 5 |
| scf7180000043069 | 887236 | 0.4161 | 0 | 58.4814 | Eukaryota | 78699.0 | 0 | Chordata | 78699.0 | 0 | Carnivora | 78699.0 | 0 | Canidae | 54934.0 | 3 | Canis | 31610.0 | 4 | Canis lupus | 31610.0 | 4 |
| scf7180000046903 | 89721 | 0.4889 | 0 | 58.8109 | Eukaryota | 136434.0 | 0 | Chordata | 136434.0 | 0 | Carnivora | 136434.0 | 0 | Canidae | 136434.0 | 0 | Canis | 136434.0 | 0 | Canis lupus | 136434.0 | 0 |
| scf7180000043525 | 342175 | 0.4098 | 0 | 61.8888 | Eukaryota | 36782.0 | 0 | Chordata | 36782.0 | 0 | Carnivora | 36782.0 | 0 | Canidae | 33655.0 | 1 | Canis | 22433.0 | 2 | Canis lupus | 22433.0 | 2 |
| scf7180000048740 | 22274 | 0.4436 | 0 | 185.5254 | Eukaryota | 8735.0 | 0 | Chordata | 8735.0 | 0 | Carnivora | 8735.0 | 0 | Canidae | 8735.0 | 0 | Canis | 8735.0 | 0 | Canis lupus | 8735.0 | 0 |
| scf7180000048025 | 20557 | 0.445 | 0 | 200.3566 | Eukaryota | 176.0 | 0 | Chordata | 176.0 | 0 | Salmoniformes | 176.0 | 0 | Salmonidae | 176.0 | 0 | Coregonus | 176.0 | 0 | Coregonus sp. 'balchen' | 176.0 | 0 |
| scf7180000043857 | 72972 | 0.5185 | 0 | 57.2681 | Eukaryota | 35376.0 | 0 | Chordata | 35376.0 | 0 | Carnivora | 35376.0 | 0 | Canidae | 18647.0 | 3 | Canis | 14741.0 | 6 | Canis lupus | 14741.0 | 6 |
| scf7180000046583 | 138662 | 0.4333 | 0 | 58.5004 | Eukaryota | 25867.0 | 0 | Chordata | 25867.0 | 0 | Carnivora | 20347.0 | 3 | Canidae | 14269.0 | 5 | Canis | 14269.0 | 5 | Canis lupus | 14269.0 | 5 |
| scf7180000043967 | 32778 | 0.3589 | 0 | 46.8866 | Eukaryota | 63041.0 | 0 | Chordata | 63041.0 | 0 | Carnivora | 63041.0 | 0 | Canidae | 63041.0 | 0 | Canis | 63041.0 | 0 | Canis lupus | 63041.0 | 0 |
| scf7180000046373 | 36585 | 0.3879 | 0 | 61.1976 | Eukaryota | 62267.0 | 0 | Chordata | 62267.0 | 0 | Carnivora | 62267.0 | 0 | Canidae | 62267.0 | 0 | Canis | 62267.0 | 0 | Canis lupus | 62267.0 | 0 |
| scf7180000043384 | 186577 | 0.4632 | 0 | 60.5646 | Eukaryota | 88282.0 | 0 | Chordata | 88282.0 | 0 | Carnivora | 88282.0 | 0 | Canidae | 88282.0 | 0 | Canis | 88282.0 | 0 | Canis lupus | 88282.0 | 0 |
| scf7180000045364 | 8186 | 0.4388 | 0 | 62.7181 | Eukaryota | 975.0 | 0 | Platyhelminthes | 975.0 | 0 | Polystomatidea | 975.0 | 0 | Polystomatidae | 975.0 | 0 | Protopolystoma | 975.0 | 0 | Protopolystoma xenopodis | 975.0 | 0 |
| scf7180000056289 | 79930 | 0.4425 | 0 | 70.6232 | Eukaryota | 94958.0 | 0 | Chordata | 94958.0 | 0 | Carnivora | 94958.0 | 0 | Canidae | 94958.0 | 0 | Canis | 94958.0 | 0 | Canis lupus | 94958.0 | 0 |
| scf7180000048643 | 50748 | 0.4364 | 0 | 58.7475 | Eukaryota | 51938.0 | 0 | Chordata | 30005.0 | 2 | Cyclophyllidea | 12194.0 | 9 | Hymenolepididae | 12194.0 | 9 | Hymenolepis | 12194.0 | 9 | Hymenolepis microstoma | 12194.0 | 9 |
| scf7180000046671 | 10945 | 0.4407 | 0 | 44.0314 | Eukaryota | 18266.0 | 0 | Chordata | 18266.0 | 0 | Carnivora | 18266.0 | 0 | Canidae | 18266.0 | 0 | Canis | 18266.0 | 0 | Canis lupus | 18266.0 | 0 |
| scf7180000043921 | 176697 | 0.411 | 0 | 60.6442 | Eukaryota | 50032.0 | 0 | Chordata | 50032.0 | 0 | Carnivora | 50032.0 | 0 | Canidae | 50032.0 | 0 | Canis | 40026.0 | 1 | Canis lupus | 40026.0 | 1 |
| scf7180000047048 | 43420 | 0.4398 | 0 | 232.1136 | Eukaryota | 20842.0 | 0 | Chordata | 16790.0 | 3 | Gadiformes | 10602.0 | 7 | Gadidae | 10602.0 | 7 | Gadus | 10602.0 | 7 | Gadus morhua | 10602.0 | 7 |
| scf7180000051311 | 3388 | 0.4445 | 0 | 105.7616 | Eukaryota | 172.0 | 0 | Platyhelminthes | 172.0 | 0 | Polystomatidea | 172.0 | 0 | Polystomatidae | 172.0 | 0 | Protopolystoma | 172.0 | 0 | Protopolystoma xenopodis | 172.0 | 0 |
| scf7180000055816 | 6123 | 0.453 | 0 | 131.469 | Eukaryota | 2854.0 | 0 | Chordata | 2854.0 | 0 | Carnivora | 2854.0 | 0 | Canidae | 2854.0 | 0 | Canis | 2854.0 | 0 | Canis lupus | 2854.0 | 0 |
| scf7180000054255 | 10384 | 0.4244 | 0 | 200.7017 | Eukaryota | 23062.0 | 0 | Chordata | 23062.0 | 0 | Carnivora | 23062.0 | 0 | Canidae | 23062.0 | 0 | Canis | 23062.0 | 0 | Canis lupus | 23062.0 | 0 |
| scf7180000053641 | 2933 | 0.4514 | 0 | 439.7001 | no-hit | 0.0 | 0 | no-hit | 0.0 | 0 | no-hit | 0.0 | 0 | no-hit | 0.0 | 0 | no-hit | 0.0 | 0 | no-hit | 0.0 | 0 |
| scf7180000056225 | 300968 | 0.366 | 0 | 58.9201 | Eukaryota | 84696.0 | 0 | Chordata | 84696.0 | 0 | Carnivora | 84696.0 | 0 | Canidae | 84696.0 | 0 | Canis | 84696.0 | 0 | Canis lupus | 84696.0 | 0 |
| scf7180000042352 | 688497 | 0.391 | 0 | 61.5026 | Eukaryota | 45822.0 | 0 | Chordata | 45822.0 | 0 | Carnivora | 45822.0 | 0 | Canidae | 45822.0 | 0 | Vulpes | 36720.0 | 1 | Vulpes vulpes | 36720.0 | 1 |
| scf7180000042258 | 8373 | 0.3724 | 0 | 36.0507 | Eukaryota | 42915.0 | 0 | Chordata | 42915.0 | 0 | Carnivora | 42915.0 | 0 | Canidae | 42915.0 | 0 | Canis | 42915.0 | 0 | Canis lupus | 42915.0 | 0 |
| scf7180000043778 | 15203 | 0.4398 | 0 | 49.3119 | Eukaryota | 3549.0 | 0 | Chordata | 3024.0 | 1 | Tetraodontiformes | 1311.0 | 3 | Tetraodontidae | 1311.0 | 3 | Takifugu | 1311.0 | 3 | Takifugu rubripes | 1311.0 | 3 |
| scf7180000056077 | 39900 | 0.498 | 0 | 283.6409 | Eukaryota | 9491.0 | 0 | Chordata | 9491.0 | 0 | Carnivora | 9491.0 | 0 | Canidae | 9491.0 | 0 | Canis | 8826.0 | 1 | Canis lupus | 7569.0 | 2 |
| scf7180000044031 | 173017 | 0.4503 | 0 | 57.3393 | Eukaryota | 14711.0 | 0 | Chordata | 14711.0 | 0 | Carnivora | 14711.0 | 0 | Canidae | 14711.0 | 0 | Canis | 14711.0 | 0 | Canis lupus | 14711.0 | 0 |
| scf7180000045122 | 87969 | 0.4059 | 0 | 63.5067 | Eukaryota | 12687.0 | 0 | Chordata | 12687.0 | 0 | Primates | 5915.0 | 3 | Hominidae | 5915.0 | 4 | Homo | 5915.0 | 5 | Homo sapiens | 5915.0 | 5 |
| scf7180000041869 | 442729 | 0.4456 | 0 | 59.4316 | Eukaryota | 35598.0 | 0 | Chordata | 35598.0 | 0 | Carnivora | 35598.0 | 0 | Canidae | 20457.0 | 3 | Canis | 20457.0 | 3 | Canis lupus | 20457.0 | 3 |
| scf7180000050602 | 30312 | 0.5451 | 0 | 57.7833 | Eukaryota | 34204.0 | 0 | Chordata | 34204.0 | 0 | Carnivora | 34204.0 | 0 | Canidae | 28614.0 | 2 | Canis | 23995.0 | 4 | Canis lupus | 23995.0 | 4 |
| scf7180000056533 | 152212 | 0.383 | 0 | 59.5069 | Eukaryota | 80020.0 | 0 | Chordata | 80020.0 | 0 | Carnivora | 80020.0 | 0 | Canidae | 80020.0 | 0 | Canis | 80020.0 | 0 | Canis lupus | 80020.0 | 0 |
| scf7180000054532 | 16766 | 0.3505 | 0 | 140.3914 | Eukaryota | 2594.0 | 1 | Chordata | 2594.0 | 1 | Primates | 2594.0 | 1 | Hominidae | 2259.0 | 2 | Homo | 2033.0 | 3 | Homo sapiens | 2033.0 | 3 |
| scf7180000043987 | 336537 | 0.5275 | 0 | 60.0268 | Eukaryota | 152032.0 | 0 | Chordata | 152032.0 | 0 | Carnivora | 142118.0 | 1 | Canidae | 142118.0 | 1 | Canis | 136797.0 | 2 | Canis lupus | 136797.0 | 2 |
| scf7180000041874 | 89830 | 0.4217 | 0 | 58.3079 | Eukaryota | 13363.0 | 0 | Chordata | 13363.0 | 0 | Carnivora | 8209.0 | 2 | Otariidae | 6465.0 | 4 | Zalophus | 6465.0 | 4 | Zalophus californianus | 6465.0 | 4 |
| scf7180000053307 | 6655 | 0.4573 | 0 | 83.3636 | no-hit | 0.0 | 0 | no-hit | 0.0 | 0 | no-hit | 0.0 | 0 | no-hit | 0.0 | 0 | no-hit | 0.0 | 0 | no-hit | 0.0 | 0 |
| scf7180000044173 | 34902 | 0.4152 | 0 | 95.1046 | Eukaryota | 14413.0 | 0 | Chordata | 9329.0 | 2 | Strigeidida | 4444.0 | 6 | Schistosomatidae | 4444.0 | 6 | Schistosoma | 4444.0 | 6 | Schistosoma mansoni | 4444.0 | 6 |
| scf7180000041842 | 583929 | 0.3631 | 0 | 59.6887 | Eukaryota | 89834.0 | 0 | Chordata | 89834.0 | 0 | Carnivora | 89834.0 | 0 | Canidae | 89834.0 | 0 | Canis | 83221.0 | 1 | Canis lupus | 83221.0 | 1 |
| scf7180000044392 | 153180 | 0.4222 | 0 | 61.7258 | Eukaryota | 59483.0 | 0 | Chordata | 59483.0 | 0 | Carnivora | 59483.0 | 0 | Canidae | 59483.0 | 0 | Canis | 59483.0 | 0 | Canis lupus | 59483.0 | 0 |
| scf7180000053160 | 2404 | 0.4447 | 0 | 479.0869 | no-hit | 0.0 | 0 | no-hit | 0.0 | 0 | no-hit | 0.0 | 0 | no-hit | 0.0 | 0 | no-hit | 0.0 | 0 | no-hit | 0.0 | 0 |
| scf7180000051373 | 5285 | 0.4498 | 0 | 733.091 | Eukaryota | 2319.0 | 0 | Echinodermata | 933.0 | 2 | Forcipulatida | 933.0 | 2 | Asteriidae | 933.0 | 2 | Asterias | 933.0 | 2 | Asterias rubens | 933.0 | 2 |
| scf7180000046113 | 101516 | 0.3779 | 0 | 57.9605 | Eukaryota | 50990.0 | 0 | Chordata | 50990.0 | 0 | Carnivora | 50990.0 | 0 | Canidae | 50990.0 | 0 | Canis | 50990.0 | 0 | Canis lupus | 50990.0 | 0 |
| scf7180000041530 | 184408 | 0.4365 | 0 | 60.055 | Eukaryota | 71163.0 | 0 | Chordata | 71163.0 | 0 | Carnivora | 71163.0 | 0 | Canidae | 71163.0 | 0 | Canis | 71163.0 | 0 | Canis lupus | 71163.0 | 0 |
| scf7180000040961 | 686468 | 0.4059 | 0 | 60.9524 | Eukaryota | 149706.0 | 0 | Chordata | 149706.0 | 0 | Carnivora | 149706.0 | 0 | Canidae | 149706.0 | 0 | Canis | 119760.0 | 1 | Canis lupus | 119760.0 | 1 |
| scf7180000043505 | 420601 | 0.3481 | 0 | 59.0578 | Eukaryota | 74749.0 | 0 | Chordata | 74749.0 | 0 | Carnivora | 74749.0 | 0 | Canidae | 74749.0 | 0 | Canis | 74749.0 | 0 | Canis lupus | 74749.0 | 0 |
| scf7180000043602 | 481417 | 0.4646 | 0 | 58.9702 | Eukaryota | 41313.0 | 0 | Chordata | 41313.0 | 0 | Carnivora | 22356.0 | 5 | Canidae | 22356.0 | 5 | Canis | 22356.0 | 5 | Canis lupus | 22356.0 | 5 |
| scf7180000056509 | 202770 | 0.3672 | 0 | 61.0255 | Eukaryota | 46471.0 | 0 | Chordata | 46471.0 | 0 | Carnivora | 46471.0 | 0 | Canidae | 46471.0 | 0 | Canis | 46471.0 | 0 | Canis lupus | 46471.0 | 0 |
| scf7180000051042 | 3378 | 0.4464 | 0 | 234.4119 | Eukaryota | 547.0 | 0 | Platyhelminthes | 547.0 | 0 | Polystomatidea | 547.0 | 0 | Polystomatidae | 547.0 | 0 | Protopolystoma | 547.0 | 0 | Protopolystoma xenopodis | 547.0 | 0 |
| scf7180000046646 | 22025 | 0.6531 | 0 | 42.8104 | Eukaryota | 23677.0 | 0 | Chordata | 23677.0 | 0 | Carnivora | 23677.0 | 0 | Canidae | 23677.0 | 0 | Canis | 23677.0 | 0 | Canis lupus | 23677.0 | 0 |
| scf7180000044025 | 333901 | 0.3733 | 0 | 61.0256 | Eukaryota | 52162.0 | 0 | Chordata | 52162.0 | 0 | Carnivora | 52162.0 | 0 | Canidae | 35067.0 | 2 | Canis | 26304.0 | 4 | Canis lupus | 26304.0 | 5 |
| scf7180000054449 | 59282 | 0.4404 | 0 | 127.8901 | Eukaryota | 91579.0 | 0 | Chordata | 91579.0 | 0 | Carnivora | 91579.0 | 0 | Canidae | 91579.0 | 0 | Canis | 91579.0 | 0 | Canis lupus | 91579.0 | 0 |
| scf7180000054749 | 33759 | 0.4403 | 0 | 142.7002 | Eukaryota | 2067.0 | 0 | Chordata | 1099.0 | 2 | Polystomatidea | 781.0 | 4 | Polystomatidae | 781.0 | 4 | Protopolystoma | 781.0 | 4 | Protopolystoma xenopodis | 781.0 | 4 |
| scf7180000043943 | 73084 | 0.4066 | 0 | 54.3515 | Eukaryota | 25250.0 | 0 | Chordata | 25250.0 | 0 | Carnivora | 25250.0 | 0 | Canidae | 25250.0 | 0 | Canis | 25250.0 | 0 | Canis lupus | 25250.0 | 0 |
| scf7180000052646 | 4890 | 0.4528 | 0 | 339.188 | no-hit | 0.0 | 0 | no-hit | 0.0 | 0 | no-hit | 0.0 | 0 | no-hit | 0.0 | 0 | no-hit | 0.0 | 0 | no-hit | 0.0 | 0 |
| scf7180000043323 | 541467 | 0.3685 | 0 | 60.6429 | Eukaryota | 93036.0 | 0 | Chordata | 93036.0 | 0 | Carnivora | 93036.0 | 0 | Canidae | 93036.0 | 0 | Canis | 93036.0 | 0 | Canis lupus | 93036.0 | 0 |
| scf7180000054701 | 30753 | 0.4928 | 0 | 194.0318 | Eukaryota | 25036.0 | 0 | Chordata | 25036.0 | 0 | Artiodactyla | 17586.0 | 1 | Bovidae | 17586.0 | 1 | Ovis | 10038.0 | 3 | Ovis aries | 10038.0 | 3 |
| scf7180000044062 | 211816 | 0.4576 | 0 | 57.9943 | Eukaryota | 90025.0 | 0 | Chordata | 90025.0 | 0 | Carnivora | 90025.0 | 0 | Canidae | 90025.0 | 0 | Canis | 79434.0 | 1 | Canis lupus | 79434.0 | 1 |
| scf7180000044072 | 12452 | 0.5133 | 0 | 49.5589 | Eukaryota | 4826.0 | 0 | Chordata | 4826.0 | 0 | Carnivora | 1618.0 | 5 | Hominidae | 1190.0 | 6 | Homo | 1190.0 | 6 | Homo sapiens | 1190.0 | 6 |
| scf7180000041096 | 307649 | 0.443 | 0 | 57.5184 | Eukaryota | 121016.0 | 0 | Chordata | 121016.0 | 0 | Carnivora | 121016.0 | 0 | Canidae | 121016.0 | 0 | Canis | 121016.0 | 0 | Canis lupus | 121016.0 | 0 |
| scf7180000042316 | 711801 | 0.4086 | 0 | 60.6103 | Eukaryota | 53260.0 | 0 | Chordata | 53260.0 | 0 | Carnivora | 41955.0 | 2 | Canidae | 34519.0 | 3 | Canis | 20690.0 | 5 | Canis lupus | 20690.0 | 5 |
| scf7180000046935 | 81125 | 0.4926 | 0 | 58.717 | Eukaryota | 12180.0 | 0 | Chordata | 12180.0 | 0 | Carnivora | 12180.0 | 0 | Canidae | 12180.0 | 0 | Canis | 12180.0 | 0 | Canis lupus | 12180.0 | 0 |
| scf7180000040736 | 1664436 | 0.3955 | 0 | 59.2109 | Eukaryota | 82909.0 | 0 | Chordata | 82909.0 | 0 | Carnivora | 82909.0 | 0 | Canidae | 82909.0 | 0 | Canis | 82909.0 | 0 | Canis lupus | 82909.0 | 0 |
| scf7180000053071 | 2172 | 0.4761 | 0 | 421.5963 | no-hit | 0.0 | 0 | no-hit | 0.0 | 0 | no-hit | 0.0 | 0 | no-hit | 0.0 | 0 | no-hit | 0.0 | 0 | no-hit | 0.0 | 0 |
| scf7180000046504 | 37227 | 0.3376 | 0 | 57.9684 | Eukaryota | 4656.0 | 0 | Chordata | 4656.0 | 0 | Carnivora | 4656.0 | 0 | Felidae | 2627.0 | 2 | Panthera | 2301.0 | 3 | Panthera pardus | 1785.0 | 4 |
| scf7180000041509 | 1001481 | 0.4388 | 0 | 59.6613 | Eukaryota | 46967.0 | 0 | Chordata | 46967.0 | 0 | Carnivora | 46967.0 | 0 | Canidae | 46967.0 | 0 | Canis | 46967.0 | 0 | Canis lupus | 46967.0 | 0 |
| scf7180000054331 | 16053 | 0.4394 | 0 | 67.3279 | Eukaryota | 23866.0 | 0 | Chordata | 23866.0 | 0 | Carnivora | 23866.0 | 0 | Canidae | 23866.0 | 0 | Canis | 23866.0 | 0 | Canis lupus | 23866.0 | 0 |
| scf7180000040886 | 2138481 | 0.4934 | 100 | 59.3947 | Eukaryota | 106554.0 | 0 | Chordata | 106554.0 | 0 | Carnivora | 106554.0 | 0 | Canidae | 106554.0 | 0 | Canis | 106554.0 | 0 | Canis lupus | 106554.0 | 0 |
| scf7180000047113 | 25529 | 0.4359 | 0 | 81.1824 | Eukaryota | 91853.0 | 0 | Chordata | 91853.0 | 0 | Carnivora | 91853.0 | 0 | Canidae | 91853.0 | 0 | Canis | 91853.0 | 0 | Canis lupus | 91853.0 | 0 |
| scf7180000053902 | 3948 | 0.5079 | 0 | 114.3562 | Eukaryota | 5855.0 | 0 | Chordata | 5855.0 | 0 | Carnivora | 5855.0 | 0 | Canidae | 5855.0 | 0 | Canis | 2943.0 | 1 | Nyctereutes procyonoides | 2912.0 | 2 |
| scf7180000048261 | 51060 | 0.4413 | 0 | 613.7878 | Eukaryota | 4160.0 | 0 | Platyhelminthes | 3137.0 | 2 | Polystomatidea | 3137.0 | 2 | Polystomatidae | 3137.0 | 2 | Protopolystoma | 3137.0 | 2 | Protopolystoma xenopodis | 3137.0 | 2 |
| scf7180000044348 | 410039 | 0.381 | 0 | 58.8294 | Eukaryota | 27683.0 | 0 | Chordata | 27683.0 | 0 | Carnivora | 27683.0 | 0 | Canidae | 27683.0 | 0 | Canis | 27683.0 | 0 | Canis lupus | 27683.0 | 0 |
| scf7180000055362 | 7580 | 0.4327 | 0 | 119.278 | no-hit | 0.0 | 0 | no-hit | 0.0 | 0 | no-hit | 0.0 | 0 | no-hit | 0.0 | 0 | no-hit | 0.0 | 0 | no-hit | 0.0 | 0 |
| scf7180000053871 | 1929 | 0.45 | 0 | 239.9159 | no-hit | 0.0 | 0 | no-hit | 0.0 | 0 | no-hit | 0.0 | 0 | no-hit | 0.0 | 0 | no-hit | 0.0 | 0 | no-hit | 0.0 | 0 |
| scf7180000043336 | 466338 | 0.4372 | 0 | 58.4726 | Eukaryota | 41544.0 | 0 | Chordata | 41544.0 | 0 | Carnivora | 38688.0 | 1 | Canidae | 14852.0 | 3 | Canis | 14852.0 | 3 | Canis lupus | 14852.0 | 4 |
| scf7180000042737 | 591306 | 0.4588 | 0 | 60.1906 | Eukaryota | 59818.0 | 0 | Chordata | 59818.0 | 0 | Carnivora | 59818.0 | 0 | Canidae | 59818.0 | 0 | Canis | 49848.0 | 1 | Canis lupus | 49848.0 | 1 |
| scf7180000049575 | 12363 | 0.4442 | 0 | 86.7393 | Eukaryota | 7136.0 | 0 | Chordata | 4468.0 | 1 | Kurtiformes | 3267.0 | 3 | Apogonidae | 3267.0 | 3 | Sphaeramia | 3267.0 | 3 | Sphaeramia orbicularis | 3267.0 | 3 |
| scf7180000044085 | 364191 | 0.537 | 0 | 53.9265 | Eukaryota | 10382.0 | 0 | Chordata | 10382.0 | 0 | Artiodactyla | 4276.0 | 3 | Suidae | 4276.0 | 5 | Sus | 4276.0 | 5 | Sus scrofa | 4276.0 | 5 |
| scf7180000051093 | 7446 | 0.4384 | 0 | 575.0762 | Eukaryota | 449.0 | 0 | Platyhelminthes | 449.0 | 0 | Polystomatidea | 449.0 | 0 | Polystomatidae | 449.0 | 0 | Protopolystoma | 449.0 | 0 | Protopolystoma xenopodis | 449.0 | 0 |
| scf7180000045490 | 138360 | 0.5856 | 0 | 56.5912 | Eukaryota | 60958.0 | 0 | Chordata | 60958.0 | 0 | Carnivora | 60958.0 | 0 | Canidae | 60958.0 | 0 | Canis | 60958.0 | 0 | Canis lupus | 60958.0 | 0 |
| scf7180000051828 | 16961 | 0.4537 | 0 | 662.7267 | Eukaryota | 9536.0 | 0 | Chordata | 7392.0 | 3 | Tetraodontiformes | 7392.0 | 3 | Tetraodontidae | 7392.0 | 3 | Takifugu | 7392.0 | 3 | Takifugu rubripes | 7392.0 | 3 |
| scf7180000042627 | 14494 | 0.4356 | 0 | 284.8682 | Eukaryota | 4659.0 | 0 | Chordata | 2966.0 | 1 | Salmoniformes | 2411.0 | 3 | Salmonidae | 2411.0 | 3 | Salmo | 2411.0 | 3 | Salmo trutta | 2411.0 | 3 |
| scf7180000044389 | 179529 | 0.4682 | 0 | 57.2162 | Eukaryota | 88951.0 | 0 | Chordata | 88951.0 | 0 | Carnivora | 88951.0 | 0 | Canidae | 88951.0 | 0 | Canis | 88951.0 | 0 | Canis lupus | 88951.0 | 0 |
| scf7180000053883 | 16222 | 0.4874 | 0 | 727.0849 | Eukaryota | 9076.0 | 0 | Chordata | 9076.0 | 0 | Carnivora | 9076.0 | 0 | Canidae | 9076.0 | 0 | Canis | 7310.0 | 1 | Canis lupus | 6031.0 | 2 |
| scf7180000046961 | 86271 | 0.4033 | 0 | 61.312 | Eukaryota | 9852.0 | 0 | Chordata | 9852.0 | 0 | Carnivora | 9852.0 | 0 | Canidae | 8745.0 | 1 | Canis | 8745.0 | 1 | Canis lupus | 8745.0 | 1 |
| scf7180000042612 | 487878 | 0.4035 | 0 | 60.2702 | Eukaryota | 54957.0 | 0 | Chordata | 54957.0 | 0 | Carnivora | 54957.0 | 0 | Canidae | 44079.0 | 1 | Canis | 34228.0 | 2 | Canis lupus | 34228.0 | 2 |
| scf7180000041763 | 602629 | 0.4205 | 0 | 59.13 | Eukaryota | 97472.0 | 0 | Chordata | 97472.0 | 0 | Carnivora | 97472.0 | 0 | Canidae | 97472.0 | 0 | Canis | 97472.0 | 0 | Canis lupus | 97472.0 | 0 |
| scf7180000053103 | 5513 | 0.4564 | 0 | 300.2207 | no-hit | 0.0 | 0 | no-hit | 0.0 | 0 | no-hit | 0.0 | 0 | no-hit | 0.0 | 0 | no-hit | 0.0 | 0 | no-hit | 0.0 | 0 |
| scf7180000044273 | 257335 | 0.3578 | 0 | 59.4309 | Eukaryota | 17001.0 | 0 | Chordata | 17001.0 | 0 | Apterygiformes | 12418.0 | 2 | Apterygidae | 12418.0 | 2 | Apteryx | 12418.0 | 2 | Apteryx mantelli | 12418.0 | 2 |
| scf7180000040802 | 3133442 | 0.3852 | 100 | 59.9402 | Eukaryota | 76436.0 | 0 | Chordata | 76436.0 | 0 | Carnivora | 76436.0 | 0 | Canidae | 76436.0 | 0 | Canis | 52971.0 | 1 | Canis lupus | 52971.0 | 1 |
| scf7180000047670 | 87592 | 0.4179 | 0 | 60.3443 | Eukaryota | 36176.0 | 0 | Chordata | 36176.0 | 0 | Carnivora | 36176.0 | 0 | Canidae | 17306.0 | 3 | Canis | 13361.0 | 6 | Canis lupus | 13361.0 | 7 |
| scf7180000047473 | 8903 | 0.4348 | 0 | 94.7137 | Eukaryota | 2036.0 | 0 | Chordata | 2036.0 | 0 | Salmoniformes | 849.0 | 3 | Salmonidae | 849.0 | 3 | Salmo | 849.0 | 3 | Salmo trutta | 849.0 | 3 |
| scf7180000047387 | 111558 | 0.3903 | 0 | 58.8428 | Eukaryota | 53762.0 | 0 | Chordata | 53762.0 | 0 | Carnivora | 53762.0 | 0 | Canidae | 53762.0 | 0 | Canis | 53762.0 | 0 | Canis lupus | 53762.0 | 0 |
| scf7180000040552 | 83935 | 0.3704 | 0 | 56.6102 | Eukaryota | 35849.0 | 0 | Chordata | 35849.0 | 0 | Carnivora | 35849.0 | 0 | Canidae | 35849.0 | 0 | Canis | 35849.0 | 0 | Canis lupus | 35849.0 | 0 |
| scf7180000050544 | 9142 | 0.4424 | 0 | 278.3211 | Eukaryota | 46336.0 | 0 | Chordata | 32968.0 | 2 | Gadiformes | 27228.0 | 5 | Gadidae | 27228.0 | 5 | Gadus | 27228.0 | 5 | Gadus morhua | 27228.0 | 5 |
| scf7180000046703 | 109298 | 0.3373 | 0 | 59.6358 | Eukaryota | 14448.0 | 0 | Chordata | 14448.0 | 0 | Carnivora | 14448.0 | 0 | Canidae | 14448.0 | 0 | Canis | 14448.0 | 0 | Canis lupus | 14448.0 | 0 |
| scf7180000040614 | 388480 | 0.412 | 0 | 59.7765 | Eukaryota | 42196.0 | 0 | Chordata | 42196.0 | 0 | Carnivora | 42196.0 | 0 | Canidae | 42196.0 | 0 | Canis | 42196.0 | 0 | Canis lupus | 42196.0 | 0 |
| scf7180000048125 | 5760 | 0.43 | 0 | 134.5137 | no-hit | 0.0 | 0 | no-hit | 0.0 | 0 | no-hit | 0.0 | 0 | no-hit | 0.0 | 0 | no-hit | 0.0 | 0 | no-hit | 0.0 | 0 |
| scf7180000052160 | 5030 | 0.4423 | 0 | 1035.7763 | Eukaryota | 2948.0 | 0 | Platyhelminthes | 1572.0 | 3 | Polystomatidea | 1418.0 | 4 | Polystomatidae | 1418.0 | 4 | Protopolystoma | 1418.0 | 4 | Protopolystoma xenopodis | 1418.0 | 4 |
| scf7180000046118 | 178817 | 0.362 | 0 | 59.9456 | Eukaryota | 9104.0 | 0 | Chordata | 9104.0 | 0 | Primates | 4177.0 | 2 | Hominidae | 4177.0 | 2 | Homo | 4177.0 | 3 | Homo sapiens | 4177.0 | 3 |
| scf7180000053361 | 3712 | 0.4534 | 0 | 413.9513 | Eukaryota | 2712.0 | 0 | Platyhelminthes | 1502.0 | 2 | Polystomatidea | 1502.0 | 2 | Polystomatidae | 1502.0 | 2 | Protopolystoma | 1502.0 | 2 | Protopolystoma xenopodis | 1502.0 | 2 |
| scf7180000043076 | 6598 | 0.6723 | 0 | 27.3903 | no-hit | 0.0 | 0 | no-hit | 0.0 | 0 | no-hit | 0.0 | 0 | no-hit | 0.0 | 0 | no-hit | 0.0 | 0 | no-hit | 0.0 | 0 |
| scf7180000051968 | 12911 | 0.4424 | 0 | 194.7957 | Eukaryota | 47903.0 | 0 | Chordata | 32481.0 | 2 | Gadiformes | 24857.0 | 5 | Gadidae | 24857.0 | 5 | Gadus | 24857.0 | 5 | Gadus morhua | 24857.0 | 5 |
| scf7180000045279 | 50223 | 0.4442 | 0 | 78.141 | Eukaryota | 99545.0 | 0 | Chordata | 99545.0 | 0 | Carnivora | 99545.0 | 0 | Canidae | 99545.0 | 0 | Canis | 99545.0 | 0 | Canis lupus | 99545.0 | 0 |
| scf7180000048002 | 81895 | 0.3756 | 0 | 59.5646 | Eukaryota | 20791.0 | 0 | Chordata | 20791.0 | 0 | Carnivora | 20791.0 | 0 | Canidae | 20791.0 | 0 | Canis | 20791.0 | 0 | Canis lupus | 20791.0 | 0 |
| scf7180000043580 | 88222 | 0.4659 | 0 | 59.3884 | Eukaryota | 36839.0 | 0 | Chordata | 36839.0 | 0 | Carnivora | 36839.0 | 0 | Canidae | 36839.0 | 0 | Canis | 25845.0 | 1 | Canis lupus | 25845.0 | 1 |
| scf7180000043426 | 437946 | 0.4375 | 0 | 60.3688 | Eukaryota | 30063.0 | 0 | Chordata | 30063.0 | 0 | Carnivora | 30063.0 | 0 | Canidae | 28253.0 | 1 | Canis | 25927.0 | 2 | Canis lupus | 25927.0 | 2 |
| scf7180000048465 | 6045 | 0.4415 | 0 | 60.5238 | Eukaryota | 1945.0 | 0 | Chordata | 1945.0 | 0 | Synbranchiformes | 1417.0 | 2 | Mastacembelidae | 1417.0 | 2 | Mastacembelus | 1417.0 | 2 | Mastacembelus armatus | 1417.0 | 2 |
| scf7180000055307 | 13334 | 0.4399 | 0 | 221.4605 | Eukaryota | 337.0 | 0 | Chordata | 337.0 | 0 | Osteoglossiformes | 176.0 | 1 | Osteoglossidae | 176.0 | 1 | Scleropages | 176.0 | 1 | Scleropages formosus | 176.0 | 1 |
| scf7180000043091 | 29051 | 0.3639 | 0 | 50.2548 | Eukaryota | 28152.0 | 0 | Chordata | 28152.0 | 0 | Carnivora | 28152.0 | 0 | Canidae | 28152.0 | 0 | Canis | 28152.0 | 0 | Canis lupus | 28152.0 | 0 |
| scf7180000042463 | 729339 | 0.3875 | 0 | 60.1936 | Eukaryota | 59852.0 | 0 | Chordata | 59852.0 | 0 | Carnivora | 59852.0 | 0 | Canidae | 59852.0 | 0 | Canis | 59852.0 | 0 | Canis lupus | 59852.0 | 0 |
| scf7180000051633 | 9451 | 0.4418 | 0 | 486.9554 | no-hit | 0.0 | 0 | no-hit | 0.0 | 0 | no-hit | 0.0 | 0 | no-hit | 0.0 | 0 | no-hit | 0.0 | 0 | no-hit | 0.0 | 0 |

| | | | | | | | | | | | | | | | | | | | | | |
|---|---|---|---|---|---|---|---|---|---|---|---|---|---|---|---|---|---|---|---|---|---|
| scf7180000041274 | 567358 | 0.3811 | 0 | 59.0104 | Eukaryota | 61523.0 | 0 | Chordata | 61523.0 | 0 | Carnivora | 61523.0 | 0 | Canidae | 61523.0 | 0 | Canis | 61523.0 | 0 | Canis lupus | 61523.0 | 0 |
| scf7180000044276 | 633129 | 0.4163 | 0 | 60.4699 | Eukaryota | 114618.0 | 0 | Chordata | 114618.0 | 0 | Carnivora | 114618.0 | 0 | Canidae | 114618.0 | 0 | Canis | 114618.0 | 0 | Canis lupus | 114618.0 | 0 |
| scf7180000052647 | 8507 | 0.446 | 0 | 354.9665 | Eukaryota | 3483.0 | 0 | Platyhelminthes | 2232.0 | 1 | Polystomatidea | 2232.0 | 3 | Polystomatidae | 2232.0 | 3 | Protopolystoma | 2232.0 | 3 | Protopolystoma xenopodis | 2232.0 | 3 |
| scf7180000041612 | 9428 | 0.4244 | 0 | 26.7863 | Eukaryota | 8422.0 | 0 | Chordata | 8422.0 | 0 | Carnivora | 8000.0 | 1 | Canidae | 8000.0 | 1 | Canis | 8000.0 | 1 | Canis lupus | 8000.0 | 1 |
| scf7180000055690 | 21807 | 0.4383 | 0 | 100.1518 | Eukaryota | 996.0 | 0 | Platyhelminthes | 829.0 | 1 | Polystomatidea | 829.0 | 1 | Polystomatidae | 829.0 | 1 | Protopolystoma | 829.0 | 1 | Protopolystoma xenopodis | 829.0 | 1 |
| scf7180000050816 | 2689 | 0.4682 | 0 | 441.1319 | no-hit | 0.0 | 0 | no-hit | 0.0 | 0 | no-hit | 0.0 | 0 | no-hit | 0.0 | 0 | no-hit | 0.0 | 0 | no-hit | 0.0 | 0 |
| scf7180000041106 | 140553 | 0.3569 | 0 | 58.0317 | Eukaryota | 13787.0 | 0 | Chordata | 13787.0 | 0 | Carnivora | 8889.0 | 1 | Canidae | 8889.0 | 1 | Canis | 8889.0 | 2 | Canis lupus | 8889.0 | 2 |
| scf7180000041074 | 793936 | 0.389 | 0 | 58.3255 | Eukaryota | 91209.0 | 0 | Chordata | 91209.0 | 0 | Carnivora | 91209.0 | 0 | Canidae | 91209.0 | 0 | Canis | 91209.0 | 0 | Canis lupus | 91209.0 | 0 |
| scf7180000048163 | 3203 | 0.4667 | 0 | 56.3819 | Eukaryota | 905.0 | 0 | Chordata | 905.0 | 0 | Kurtiformes | 905.0 | 0 | Apogonidae | 905.0 | 0 | Sphaeramia | 905.0 | 0 | Sphaeramia orbicularis | 905.0 | 0 |
| scf7180000055560 | 87873 | 0.4374 | 0 | 183.0747 | Eukaryota | 42942.0 | 0 | Chordata | 35480.0 | 1 | Gadiformes | 18373.0 | 6 | Gadidae | 18373.0 | 6 | Gadus | 18373.0 | 6 | Gadus morhua | 18373.0 | 6 |
| scf7180000043965 | 335366 | 0.4726 | 0 | 57.2212 | Eukaryota | 52153.0 | 0 | Chordata | 52153.0 | 0 | Carnivora | 52153.0 | 0 | Canidae | 52153.0 | 0 | Canis | 52153.0 | 0 | Canis lupus | 52153.0 | 0 |
| scf7180000050955 | 9187 | 0.4451 | 0 | 60.8271 | Eukaryota | 3104.0 | 0 | Chordata | 3104.0 | 0 | Carnivora | 3104.0 | 0 | Canidae | 3104.0 | 0 | Canis | 3104.0 | 0 | Canis lupus | 3104.0 | 0 |
| scf7180000051188 | 9449 | 0.4476 | 0 | 1224.9303 | no-hit | 0.0 | 0 | no-hit | 0.0 | 0 | no-hit | 0.0 | 0 | no-hit | 0.0 | 0 | no-hit | 0.0 | 0 | no-hit | 0.0 | 0 |
| scf7180000043866 | 437634 | 0.3604 | 0 | 59.3766 | Eukaryota | 92789.0 | 0 | Chordata | 92789.0 | 0 | Carnivora | 92789.0 | 0 | Canidae | 92789.0 | 0 | Canis | 92789.0 | 0 | Canis lupus | 92789.0 | 0 |
| scf7180000045105 | 86226 | 0.3726 | 0 | 58.9479 | Eukaryota | 27643.0 | 0 | Chordata | 27643.0 | 0 | Carnivora | 27643.0 | 0 | Canidae | 27643.0 | 0 | Canis | 27643.0 | 0 | Canis lupus | 27643.0 | 0 |
| scf7180000043488 | 62713 | 0.4366 | 0 | 54.371 | Eukaryota | 16084.0 | 0 | Chordata | 16084.0 | 0 | Carnivora | 16084.0 | 0 | Canidae | 16084.0 | 0 | Canis | 12382.0 | 1 | Canis lupus | 12382.0 | 1 |
| scf7180000044928 | 208322 | 0.4957 | 0 | 59.3985 | Eukaryota | 35259.0 | 0 | Chordata | 35259.0 | 0 | Carnivora | 35259.0 | 0 | Canidae | 35259.0 | 0 | Canis | 28202.0 | 1 | Canis lupus | 28202.0 | 1 |
| scf7180000056321 | 766264 | 0.4731 | 0 | 60.2109 | Eukaryota | 82720.0 | 0 | Chordata | 82720.0 | 0 | Carnivora | 82720.0 | 0 | Canidae | 82720.0 | 0 | Canis | 82720.0 | 0 | Canis lupus | 82720.0 | 0 |
| scf7180000056006 | 12357 | 0.4338 | 0 | 140.3539 | Eukaryota | 38444.0 | 0 | Chordata | 34865.0 | 1 | Gadiformes | 29019.0 | 5 | Gadidae | 29019.0 | 5 | Gadus | 29019.0 | 5 | Gadus morhua | 29019.0 | 5 |
| scf7180000048244 | 9603 | 0.4453 | 0 | 149.5244 | Eukaryota | 8840.0 | 0 | Chordata | 7473.0 | 2 | Tetraodontiformes | 4301.0 | 4 | Tetraodontidae | 4301.0 | 4 | Takifugu | 4301.0 | 4 | Takifugu rubripes | 3283.0 | 5 |
| scf7180000054192 | 22598 | 0.3632 | 0 | 470.3454 | Eukaryota | 3887.0 | 0 | Chordata | 3887.0 | 0 | Carnivora | 2273.0 | 1 | Canidae | 2273.0 | 2 | Canis | 2273.0 | 2 | Canis lupus | 2273.0 | 2 |
| scf7180000041199 | 1241243 | 0.389 | 0 | 59.8435 | Eukaryota | 42840.0 | 0 | Chordata | 42840.0 | 0 | Carnivora | 42840.0 | 0 | Canidae | 42840.0 | 0 | Canis | 42840.0 | 0 | Canis lupus | 42840.0 | 0 |
| scf7180000043223 | 18297 | 0.4381 | 0 | 50.7011 | Eukaryota | 47775.0 | 0 | Chordata | 47775.0 | 0 | Carnivora | 47775.0 | 0 | Canidae | 47775.0 | 0 | Canis | 47775.0 | 0 | Canis lupus | 47775.0 | 0 |
| scf7180000052821 | 2322 | 0.4591 | 0 | 399.6845 | Eukaryota | 524.0 | 0 | Streptophyta | 278.0 | 1 | Fabales | 278.0 | 1 | Fabaceae | 278.0 | 1 | Cicer | 278.0 | 1 | Cicer arietinum | 278.0 | 1 |
| scf7180000052232 | 5735 | 0.4544 | 0 | 422.8826 | Eukaryota | 1607.0 | 0 | Echinodermata | 521.0 | 3 | Forcipulatida | 521.0 | 3 | Asteriidae | 521.0 | 3 | Asterias | 521.0 | 3 | Asterias rubens | 521.0 | 3 |
| scf7180000056430 | 460574 | 0.3631 | 0 | 60.7485 | Eukaryota | 53647.0 | 0 | Chordata | 53647.0 | 0 | Carnivora | 53647.0 | 0 | Canidae | 53647.0 | 0 | Canis | 53647.0 | 0 | Canis lupus | 53647.0 | 0 |
| scf7180000045605 | 196274 | 0.3676 | 0 | 57.4312 | Eukaryota | 20762.0 | 0 | Chordata | 20762.0 | 0 | Carnivora | 20762.0 | 0 | Canidae | 18721.0 | 1 | Canis | 14626.0 | 2 | Canis lupus | 14626.0 | 2 |
| scf7180000043975 | 539710 | 0.3484 | 0 | 59.6926 | Eukaryota | 88593.0 | 0 | Chordata | 88593.0 | 0 | Carnivora | 88593.0 | 0 | Canidae | 88593.0 | 0 | Canis | 88593.0 | 0 | Canis lupus | 88593.0 | 0 |
| scf7180000041115 | 1400042 | 0.3363 | 0 | 58.7415 | Eukaryota | 88899.0 | 0 | Chordata | 88899.0 | 0 | Carnivora | 88899.0 | 0 | Canidae | 88899.0 | 0 | Canis | 88899.0 | 0 | Canis lupus | 88899.0 | 0 |
| scf7180000042069 | 833104 | 0.4108 | 0 | 60.0074 | Eukaryota | 71773.0 | 0 | Chordata | 71773.0 | 0 | Carnivora | 71773.0 | 0 | Canidae | 71773.0 | 0 | Canis | 71773.0 | 0 | Canis lupus | 71773.0 | 0 |
| scf7180000040893 | 256071 | 0.3878 | 0 | 61.1749 | Eukaryota | 78911.0 | 0 | Chordata | 78911.0 | 0 | Carnivora | 78911.0 | 0 | Canidae | 78911.0 | 0 | Canis | 78911.0 | 0 | Canis lupus | 78911.0 | 0 |
| scf7180000048273 | 25633 | 0.4387 | 0 | 117.592 | Eukaryota | 2189.0 | 0 | Chordata | 2189.0 | 0 | Kurtiformes | 1534.0 | 1 | Apogonidae | 1534.0 | 1 | Sphaeramia | 1534.0 | 1 | Sphaeramia orbicularis | 1534.0 | 1 |
| scf7180000051902 | 9897 | 0.4374 | 0 | 135.6042 | Eukaryota | 1555.0 | 0 | Chordata | 967.0 | 1 | Columbiformes | 736.0 | 2 | Columbidae | 736.0 | 2 | Streptopelia | 736.0 | 2 | Streptopelia turtur | 736.0 | 2 |
| scf7180000056256 | 418134 | 0.4039 | 0 | 58.0651 | Eukaryota | 89575.0 | 0 | Chordata | 89575.0 | 0 | Carnivora | 89575.0 | 0 | Canidae | 89575.0 | 0 | Canis | 89575.0 | 0 | Canis lupus | 89575.0 | 0 |
| scf7180000044367 | 287641 | 0.5057 | 0 | 58.7565 | Eukaryota | 42102.0 | 0 | Chordata | 42102.0 | 0 | Carnivora | 42102.0 | 0 | Canidae | 42102.0 | 0 | Canis | 42102.0 | 0 | Canis lupus | 42102.0 | 0 |
| scf7180000041718 | 939023 | 0.4323 | 0 | 60.0291 | Eukaryota | 93062.0 | 0 | Chordata | 93062.0 | 0 | Carnivora | 93062.0 | 0 | Canidae | 93062.0 | 0 | Canis | 93062.0 | 0 | Canis lupus | 93062.0 | 0 |
| scf7180000043996 | 419847 | 0.3735 | 0 | 60.7009 | Eukaryota | 90042.0 | 0 | Chordata | 90042.0 | 0 | Carnivora | 90042.0 | 0 | Canidae | 90042.0 | 0 | Canis | 90042.0 | 0 | Canis lupus | 90042.0 | 0 |
| scf7180000043482 | 430134 | 0.4122 | 0 | 59.6582 | Eukaryota | 61552.0 | 0 | Chordata | 61552.0 | 0 | Carnivora | 61552.0 | 0 | Canidae | 42437.0 | 1 | Canis | 34808.0 | 2 | Canis lupus | 34808.0 | 2 |
| scf7180000048165 | 78907 | 0.4999 | 0 | 564.8752 | Eukaryota | 9783.0 | 0 | Chordata | 9783.0 | 0 | Carnivora | 9783.0 | 0 | Canidae | 9783.0 | 0 | Canis | 8554.0 | 1 | Canis lupus | 7251.0 | 2 |
| scf7180000045061 | 215369 | 0.4897 | 0 | 57.7147 | Eukaryota | 30626.0 | 0 | Chordata | 30626.0 | 0 | Carnivora | 30626.0 | 0 | Canidae | 30626.0 | 0 | Canis | 27748.0 | 1 | Canis lupus | 27748.0 | 1 |
| scf7180000041312 | 212601 | 0.4153 | 0 | 60.7219 | Eukaryota | 51622.0 | 0 | Chordata | 51622.0 | 0 | Carnivora | 51622.0 | 0 | Canidae | 51622.0 | 0 | Canis | 51622.0 | 0 | Canis lupus | 51622.0 | 0 |
| scf7180000053320 | 4969 | 0.45 | 0 | 131.0254 | Eukaryota | 4583.0 | 0 | Chordata | 4583.0 | 0 | Tetraodontiformes | 4180.0 | 1 | Tetraodontidae | 4180.0 | 1 | Takifugu | 4180.0 | 1 | Takifugu rubripes | 4180.0 | 1 |
| scf7180000042706 | 36780 | 0.4859 | 0 | 31.1865 | Eukaryota | 68005.0 | 0 | Chordata | 68005.0 | 0 | Carnivora | 68005.0 | 0 | Canidae | 50653.0 | 2 | Canis | 50653.0 | 4 | Canis lupus | 50653.0 | 4 |
| scf7180000045827 | 134098 | 0.4729 | 0 | 59.884 | Eukaryota | 50645.0 | 0 | Chordata | 50645.0 | 0 | Carnivora | 50645.0 | 0 | Canidae | 36269.0 | 2 | Canis | 30005.0 | 5 | Canis lupus | 30005.0 | 5 |
| scf7180000053679 | 2512 | 0.4566 | 0 | 287.6693 | no-hit | 0.0 | 0 | no-hit | 0.0 | 0 | no-hit | 0.0 | 0 | no-hit | 0.0 | 0 | no-hit | 0.0 | 0 | no-hit | 0.0 | 0 |
| scf7180000052886 | 2934 | 0.4513 | 0 | 165.6739 | Eukaryota | 2917.0 | 0 | Chordata | 2917.0 | 0 | Carnivora | 2917.0 | 0 | Canidae | 2917.0 | 0 | Canis | 2917.0 | 0 | Canis lupus | 2917.0 | 0 |
| scf7180000041899 | 958601 | 0.3687 | 0 | 60.6007 | Eukaryota | 89452.0 | 0 | Chordata | 89452.0 | 0 | Carnivora | 89452.0 | 0 | Canidae | 89452.0 | 0 | Canis | 89452.0 | 0 | Canis lupus | 89452.0 | 0 |
| scf7180000050665 | 11653 | 0.4486 | 0 | 548.5278 | Eukaryota | 568.0 | 0 | Platyhelminthes | 568.0 | 0 | Polystomatidea | 568.0 | 0 | Polystomatidae | 568.0 | 0 | Protopolystoma | 568.0 | 0 | Protopolystoma xenopodis | 568.0 | 0 |
| scf7180000041987 | 838448 | 0.3706 | 0 | 59.1591 | Eukaryota | 131710.0 | 0 | Chordata | 131710.0 | 0 | Carnivora | 131710.0 | 0 | Canidae | 131710.0 | 0 | Canis | 131710.0 | 0 | Canis lupus | 131710.0 | 0 |
| scf7180000054321 | 18405 | 0.4614 | 0 | 401.97 | Eukaryota | 11559.0 | 0 | Chordata | 11559.0 | 0 | Carnivora | 11559.0 | 0 | Canidae | 11559.0 | 0 | Canis | 8854.0 | 1 | Canis lupus | 7917.0 | 2 |
| scf7180000041839 | 328110 | 0.3737 | 0 | 60.1517 | Eukaryota | 35840.0 | 0 | Chordata | 35840.0 | 0 | Carnivora | 35840.0 | 0 | Canidae | 24671.0 | 1 | Canis | 20962.0 | 4 | Canis lupus | 20962.0 | 5 |
| scf7180000052636 | 4532 | 0.4417 | 0 | 153.4207 | Eukaryota | 3748.0 | 0 | Chordata | 2897.0 | 1 | Salmoniformes | 2436.0 | 4 | Salmonidae | 2436.0 | 4 | Salmo | 2436.0 | 4 | Salmo trutta | 2436.0 | 4 |
| scf7180000053058 | 2770 | 0.5087 | 0 | 327.9029 | Eukaryota | 6221.0 | 0 | Chordata | 6221.0 | 0 | Carnivora | 6221.0 | 0 | Canidae | 6221.0 | 0 | Canis | 4783.0 | 1 | Canis lupus | 3786.0 | 2 |
| scf7180000046983 | 100419 | 0.4735 | 0 | 59.7637 | Eukaryota | 18348.0 | 0 | Chordata | 18348.0 | 0 | Carnivora | 18348.0 | 0 | Canidae | 18348.0 | 0 | Canis | 18348.0 | 0 | Canis lupus | 18348.0 | 0 |
| scf7180000051048 | 5502 | 0.4446 | 0 | 83.9016 | Eukaryota | 285.0 | 0 | Platyhelminthes | 285.0 | 0 | Polystomatidea | 285.0 | 0 | Polystomatidae | 285.0 | 0 | Protopolystoma | 285.0 | 0 | Protopolystoma xenopodis | 285.0 | 0 |
| scf7180000042126 | 820448 | 0.3539 | 0 | 60.0314 | Eukaryota | 57083.0 | 0 | Chordata | 57083.0 | 0 | Carnivora | 57083.0 | 0 | Canidae | 57083.0 | 0 | Canis | 57083.0 | 0 | Canis lupus | 57083.0 | 0 |
| scf7180000041107 | 1350899 | 0.3626 | 0 | 59.9741 | Eukaryota | 94383.0 | 0 | Chordata | 94383.0 | 0 | Carnivora | 94383.0 | 0 | Canidae | 94383.0 | 0 | Canis | 94383.0 | 0 | Canis lupus | 94383.0 | 0 |
| scf7180000041336 | 109583 | 0.3535 | 0 | 60.0209 | Eukaryota | 8239.0 | 1 | Chordata | 8239.0 | 1 | Carnivora | 6957.0 | 2 | Canidae | 6957.0 | 2 | Canis | 6957.0 | 2 | Canis lupus | 6957.0 | 2 |
| scf7180000045408 | 191402 | 0.3455 | 0 | 58.8734 | Eukaryota | 55728.0 | 0 | Chordata | 55728.0 | 0 | Carnivora | 55728.0 | 0 | Canidae | 55728.0 | 0 | Canis | 55728.0 | 0 | Canis lupus | 55728.0 | 0 |
| scf7180000056344 | 740237 | 0.4856 | 0 | 63.0516 | Eukaryota | 73125.0 | 0 | Chordata | 73125.0 | 0 | Carnivora | 73125.0 | 0 | Canidae | 73125.0 | 0 | Canis | 41182.0 | 1 | Canis lupus | 41182.0 | 1 |
| scf7180000040934 | 160426 | 0.3659 | 0 | 60.885 | Eukaryota | 62239.0 | 0 | Chordata | 62239.0 | 0 | Carnivora | 62239.0 | 0 | Canidae | 62239.0 | 0 | Canis | 62239.0 | 0 | Canis lupus | 62239.0 | 0 |
| scf7180000050202 | 17647 | 0.3851 | 0 | 53.8514 | Eukaryota | 40001.0 | 0 | Chordata | 40001.0 | 0 | Carnivora | 40001.0 | 0 | Canidae | 40001.0 | 0 | Canis | 40001.0 | 0 | Canis lupus | 40001.0 | 0 |
| scf7180000050149 | 2319 | 0.4411 | 0 | 150.0265 | Eukaryota | 283.0 | 0 | Chordata | 283.0 | 0 | Salmoniformes | 283.0 | 0 | Salmonidae | 283.0 | 0 | Salmo | 283.0 | 0 | Salmo trutta | 283.0 | 0 |
| scf7180000049317 | 5349 | 0.4472 | 0 | 105.2256 | no-hit | 0.0 | 0 | no-hit | 0.0 | 0 | no-hit | 0.0 | 0 | no-hit | 0.0 | 0 | no-hit | 0.0 | 0 | no-hit | 0.0 | 0 |
| scf7180000049882 | 29201 | 0.5197 | 0 | 58.3911 | Eukaryota | 5067.0 | 0 | Chordata | 5067.0 | 0 | Carnivora | 4499.0 | 1 | Canidae | 4499.0 | 1 | Canis | 4499.0 | 1 | Canis lupus | 4499.0 | 1 |
| scf7180000044229 | 7262 | 0.4424 | 0 | 52.6384 | Eukaryota | 8585.0 | 0 | Chordata | 8585.0 | 0 | Carnivora | 8585.0 | 0 | Felidae | 5016.0 | 1 | Canis | 3569.0 | 3 | Canis lupus | 3569.0 | 5 |
| scf7180000053699 | 6201 | 0.5056 | 0 | 156.1686 | Eukaryota | 7737.0 | 0 | Chordata | 7737.0 | 0 | Carnivora | 7737.0 | 0 | Canidae | 7737.0 | 0 | Canis | 6166.0 | 1 | Canis lupus | 5014.0 | 2 |
| scf7180000049914 | 2799 | 0.4387 | 0 | 134.4785 | no-hit | 0.0 | 0 | no-hit | 0.0 | 0 | no-hit | 0.0 | 0 | no-hit | 0.0 | 0 | no-hit | 0.0 | 0 | no-hit | 0.0 | 0 |
| scf7180000051605 | 5227 | 0.4477 | 0 | 910.1656 | no-hit | 0.0 | 0 | no-hit | 0.0 | 0 | no-hit | 0.0 | 0 | no-hit | 0.0 | 0 | no-hit | 0.0 | 0 | no-hit | 0.0 | 0 |
| scf7180000049516 | 27970 | 0.443 | 0 | 56.1883 | Eukaryota | 41482.0 | 0 | Chordata | 27117.0 | 2 | Carnivora | 17988.0 | 4 | Canidae | 17988.0 | 4 | Canis | 17988.0 | 4 | Canis lupus | 17988.0 | 4 |
| scf7180000049224 | 32955 | 0.4419 | 0 | 71.4366 | Eukaryota | 1276.0 | 0 | Platyhelminthes | 1100.0 | 1 | Polystomatidea | 1100.0 | 1 | Polystomatidae | 1100.0 | 1 | Protopolystoma | 1100.0 | 1 | Protopolystoma xenopodis | 1100.0 | 1 |
| scf7180000042576 | 214096 | 0.4968 | 0 | 58.6541 | Eukaryota | 87214.0 | 0 | Chordata | 87214.0 | 0 | Carnivora | 87214.0 | 0 | Canidae | 87214.0 | 0 | Canis | 87214.0 | 0 | Canis lupus | 87214.0 | 0 |
| scf7180000055825 | 30991 | 0.7579 | 0 | 232.4128 | Eukaryota | 2425.0 | 0 | Chordata | 2425.0 | 0 | Rodentia | 984.0 | 4 | Caviidae | 723.0 | 6 | Cavia | 723.0 | 6 | Cavia porcellus | 723.0 | 6 |
| scf7180000044247 | 184961 | 0.347 | 0 | 58.7647 | Eukaryota | 31075.0 | 0 | Chordata | 31075.0 | 0 | Carnivora | 31075.0 | 0 | Canidae | 31075.0 | 0 | Canis | 31075.0 | 0 | Canis lupus | 31075.0 | 0 |
| scf7180000044130 | 366713 | 0.4474 | 0 | 60.3831 | Eukaryota | 44724.0 | 0 | Chordata | 44724.0 | 0 | Carnivora | 44724.0 | 0 | Canidae | 44724.0 | 0 | Canis | 44724.0 | 0 | Canis lupus | 44724.0 | 0 |
| scf7180000049385 | 9595 | 0.4485 | 0 | 130.9904 | Eukaryota | 2824.0 | 0 | Chordata | 2824.0 | 0 | Carnivora | 2824.0 | 0 | Canidae | 2824.0 | 0 | Canis | 2824.0 | 0 | Canis lupus | 2824.0 | 0 |
| scf7180000044378 | 386320 | 0.3658 | 0 | 61.0016 | Eukaryota | 49299.0 | 0 | Chordata | 49299.0 | 0 | Carnivora | 49299.0 | 0 | Canidae | 49299.0 | 0 | Canis | 49299.0 | 0 | Canis lupus | 49299.0 | 0 |
| scf7180000056360 | 279399 | 0.4159 | 0 | 58.8524 | Eukaryota | 29232.0 | 0 | Chordata | 29232.0 | 0 | Carnivora | 26450.0 | 1 | Canidae | 6084.0 | 6 | Neovison | 5614.0 | 7 | Neovison vison | 5614.0 | 7 |
| scf7180000040905 | 72690 | 0.3992 | 0 | 53.2053 | Eukaryota | 28128.0 | 0 | Chordata | 28128.0 | 0 | Carnivora | 28128.0 | 0 | Canidae | 28128.0 | 0 | Canis | 28128.0 | 0 | Canis lupus | 28128.0 | 0 |
| scf7180000041823 | 688462 | 0.3847 | 0 | 60.0808 | Eukaryota | 94323.0 | 0 | Chordata | 94323.0 | 0 | Carnivora | 94323.0 | 0 | Canidae | 94323.0 | 0 | Canis | 94323.0 | 0 | Canis lupus | 94323.0 | 0 |
| scf7180000056202 | 536663 | 0.3854 | 0 | 58.5817 | Eukaryota | 103769.0 | 0 | Chordata | 103769.0 | 0 | Carnivora | 92311.0 | 1 | Canidae | 92311.0 | 1 | Canis | 67801.0 | 2 | Canis lupus | 67801.0 | 2 |
| scf7180000043752 | 410755 | 0.372 | 0 | 60.8386 | Eukaryota | 62080.0 | 0 | Chordata | 62080.0 | 0 | Carnivora | 62080.0 | 0 | Canidae | 62080.0 | 0 | Canis | 49672.0 | 1 | Canis lupus | 49672.0 | 1 |
| scf7180000049029 | 13821 | 0.448 | 0 | 407.4791 | Eukaryota | 54234.0 | 0 | Chordata | 40733.0 | 2 | Gadiformes | 33559.0 | 5 | Gadidae | 33559.0 | 5 | Gadus | 33559.0 | 5 | Gadus morhua | 33559.0 | 5 |
| scf7180000043227 | 500246 | 0.4885 | 0 | 58.6181 | Eukaryota | 37411.0 | 0 | Chordata | 37411.0 | 0 | Carnivora | 37411.0 | 0 | Canidae | 37411.0 | 0 | Canis | 23857.0 | 1 | Canis lupus | 23857.0 | 1 |
| scf7180000051588 | 6458 | 0.4379 | 0 | 998.785 | no-hit | 0.0 | 0 | no-hit | 0.0 | 0 | no-hit | 0.0 | 0 | no-hit | 0.0 | 0 | no-hit | 0.0 | 0 | no-hit | 0.0 | 0 |
| scf7180000044999 | 253667 | 0.3909 | 0 | 58.9172 | Eukaryota | 87054.0 | 0 | Chordata | 87054.0 | 0 | Carnivora | 87054.0 | 0 | Canidae | 87054.0 | 0 | Canis | 87054.0 | 0 | Canis lupus | 87054.0 | 0 |
| scf7180000043589 | 165579 | 0.3332 | 0 | 59.2205 | Eukaryota | 92828.0 | 0 | Chordata | 92828.0 | 0 | Carnivora | 92828.0 | 0 | Canidae | 92828.0 | 0 | Canis | 92828.0 | 0 | Canis lupus | 92828.0 | 0 |
| scf7180000051285 | 2163 | 0.4401 | 0 | 123.4923 | Eukaryota | 1913.0 | 0 | Chordata | 1489.0 | 1 | Salmoniformes | 1043.0 | 3 | Salmonidae | 1043.0 | 3 | Coregonus | 725.0 | 4 | Coregonus sp. 'balchen' | 725.0 | 4 |
| scf7180000055892 | 8040 | 0.758 | 0 | 181.2464 | Eukaryota | 2425.0 | 0 | Chordata | 2425.0 | 0 | Rodentia | 984.0 | 4 | Caviidae | 723.0 | 6 | Cavia | 723.0 | 6 | Cavia porcellus | 723.0 | 6 |
| scf7180000050510 | 7115 | 0.4444 | 0 | 187.602 | Eukaryota | 2903.0 | 0 | Chordata | 2903.0 | 0 | Carnivora | 2903.0 | 0 | Canidae | 2903.0 | 0 | Canis | 2903.0 | 0 | Canis lupus | 2903.0 | 0 |
| scf7180000043003 | 183636 | 0.3861 | 0 | 59.3524 | Eukaryota | 60239.0 | 0 | Chordata | 60239.0 | 0 | Carnivora | 60239.0 | 0 | Canidae | 60239.0 | 0 | Canis | 60239.0 | 0 | Canis lupus | 60239.0 | 0 |
| scf7180000053979 | 3183 | 0.5096 | 0 | 107.5862 | Eukaryota | 4598.0 | 0 | Chordata | 4598.0 | 0 | Carnivora | 4598.0 | 0 | Canidae | 4598.0 | 0 | Canis | 3681.0 | 1 | Canis lupus | 3681.0 | 1 |
| scf7180000048192 | 3937 | 0.4516 | 0 | 52.2815 | Eukaryota | 654.0 | 0 | Chordata | 654.0 | 0 | Salmoniformes | 654.0 | 0 | Salmonidae | 654.0 | 0 | Salmo | 654.0 | 0 | Salmo trutta | 654.0 | 0 |
| scf7180000043487 | 477579 | 0.407 | 0 | 59.467 | Eukaryota | 89830.0 | 0 | Chordata | 89830.0 | 0 | Carnivora | 89830.0 | 0 | Canidae | 89830.0 | 0 | Canis | 89830.0 | 0 | Canis lupus | 89830.0 | 0 |
| scf7180000044300 | 11587 | 0.4743 | 0 | 50.2043 | Eukaryota | 1852.0 | 0 | Chordata | 1852.0 | 0 | Carnivora | 1852.0 | 0 | Canidae | 1852.0 | 0 | Canis | 1852.0 | 0 | Canis lupus | 1852.0 | 0 |
| scf7180000052262 | 2356 | 0.4567 | 0 | 290.8676 | no-hit | 0.0 | 0 | no-hit | 0.0 | 0 | no-hit | 0.0 | 0 | no-hit | 0.0 | 0 | no-hit | 0.0 | 0 | no-hit | 0.0 | 0 |
| scf7180000045251 | 251934 | 0.5326 | 0 | 57.7117 | Eukaryota | 44416.0 | 0 | Chordata | 44416.0 | 0 | Carnivora | 44416.0 | 0 | Canidae | 44416.0 | 0 | Canis | 44416.0 | 0 | Canis lupus | 44416.0 | 0 |
| scf7180000056507 | 228674 | 0.3807 | 0 | 59.5747 | Eukaryota | 100474.0 | 0 | Chordata | 100474.0 | 0 | Carnivora | 100474.0 | 0 | Canidae | 100474.0 | 0 | Canis | 100474.0 | 0 | Canis lupus | 100474.0 | 0 |
| scf7180000040584 | 2043501 | 0.3775 | 0 | 61.0807 | Eukaryota | 85142.0 | 0 | Chordata | 85142.0 | 0 | Carnivora | 85142.0 | 0 | Canidae | 75363.0 | 1 | Canis | 63773.0 | 2 | Canis lupus | 63773.0 | 2 |
| scf7180000042592 | 17303 | 0.4513 | 0 | 164.3902 | Eukaryota | 87809.0 | 0 | Chordata | 87809.0 | 0 | Carnivora | 87809.0 | 0 | Canidae | 87809.0 | 0 | Canis | 87809.0 | 0 | Canis lupus | 87809.0 | 0 |
| scf7180000044202 | 188728 | 0.3608 | 0 | 57.498 | Eukaryota | 23864.0 | 0 | Chordata | 23864.0 | 0 | Carnivora | 23864.0 | 0 | Canidae | 23864.0 | 0 | Canis | 23864.0 | 0 | Canis lupus | 23864.0 | 0 |
| scf7180000054548 | 32885 | 0.4417 | 0 | 332.2616 | Eukaryota | 3507.0 | 0 | Platyhelminthes | 2205.0 | 2 | Polystomatidea | 2205.0 | 2 | Polystomatidae | 2205.0 | 2 | Protopolystoma | 2205.0 | 2 | Protopolystoma xenopodis | 2205.0 | 2 |
| scf7180000050833 | 18892 | 0.4619 | 0 | 207.1124 | Eukaryota | 94628.0 | 0 | Chordata | 94628.0 | 0 | Carnivora | 94628.0 | 0 | Canidae | 94628.0 | 0 | Canis | 94628.0 | 0 | Canis lupus | 94628.0 | 0 |
| scf7180000055201 | 174287 | 0.4407 | 0 | 215.308 | Eukaryota | 60235.0 | 0 | Chordata | 36935.0 | 2 | Gadiformes | 33799.0 | 4 | Gadidae | 33799.0 | 4 | Gadus | 33799.0 | 4 | Gadus morhua | 33799.0 | 4 |
| scf7180000042622 | 78657 | 0.3869 | 0 | 58.5761 | Eukaryota | 90256.0 | 0 | Chordata | 90256.0 | 0 | Carnivora | 90256.0 | 0 | Canidae | 90256.0 | 0 | Canis | 90256.0 | 0 | Canis lupus | 90256.0 | 0 |
| scf7180000054837 | 4641 | 0.4331 | 0 | 128.0905 | Eukaryota | 1037.0 | 0 | Chordata | 1037.0 | 0 | Kurtiformes | 1037.0 | 0 | Apogonidae | 1037.0 | 0 | Sphaeramia | 1037.0 | 0 | Sphaeramia orbicularis | 1037.0 | 0 |
| scf7180000048117 | 3920 | 0.4263 | 0 | 105.5476 | no-hit | 0.0 | 0 | no-hit | 0.0 | 0 | no-hit | 0.0 | 0 | no-hit | 0.0 | 0 | no-hit | 0.0 | 0 | no-hit | 0.0 | 0 |
| scf7180000044028 | 110090 | 0.4315 | 0 | 59.2871 | Eukaryota | 75021.0 | 0 | Chordata | 75021.0 | 0 | Carnivora | 75021.0 | 0 | Canidae | 75021.0 | 0 | Canis | 75021.0 | 0 | Canis lupus | 75021.0 | 0 |
| scf7180000047928 | 71194 | 0.4585 | 0 | 140.9024 | Eukaryota | 32429.0 | 0 | Chordata | 23983.0 | 1 | Carnivora | 13831.0 | 3 | Canidae | 13831.0 | 3 | Canis | 13831.0 | 3 | Canis lupus | 13831.0 | 3 |
| scf7180000046564 | 37494 | 0.4529 | 0 | 103.2294 | Eukaryota | 105213.0 | 0 | Chordata | 61309.0 | 1 | Gadiformes | 61309.0 | 2 | Gadidae | 61309.0 | 2 | Gadus | 61309.0 | 2 | Gadus morhua | 61309.0 | 2 |
| scf7180000045178 | 204330 | 0.3998 | 0 | 59.7121 | Eukaryota | 54224.0 | 0 | Chordata | 54224.0 | 0 | Carnivora | 54224.0 | 0 | Canidae | 54224.0 | 0 | Canis | 54224.0 | 0 | Canis lupus | 54224.0 | 0 |
| scf7180000047290 | 5421 | 0.4385 | 0 | 178.2068 | Eukaryota | 621.0 | 0 | Chordata | 621.0 | 0 | Kurtiformes | 621.0 | 0 | Apogonidae | 621.0 | 0 | Sphaeramia | 621.0 | 0 | Sphaeramia orbicularis | 621.0 | 0 |
| scf7180000045704 | 188517 | 0.4049 | 0 | 59.6515 | Eukaryota | 93750.0 | 0 | Chordata | 93750.0 | 0 | Carnivora | 93750.0 | 0 | Canidae | 93750.0 | 0 | Canis | 93750.0 | 0 | Canis lupus | 93750.0 | 0 |
| scf7180000040841 | 275899 | 0.3405 | 0 | 58.7176 | Eukaryota | 26088.0 | 0 | Chordata | 26088.0 | 0 | Carnivora | 26088.0 | 0 | Canidae | 26088.0 | 0 | Canis | 26088.0 | 0 | Canis lupus | 26088.0 | 0 |
| scf7180000042620 | 463412 | 0.4509 | 0 | 58.109 | Eukaryota | 94128.0 | 0 | Chordata | 94128.0 | 0 | Carnivora | 94128.0 | 0 | Canidae | 94128.0 | 0 | Canis | 94128.0 | 0 | Canis lupus | 94128.0 | 0 |
| scf7180000046362 | 8231 | 0.4433 | 0 | 138.1549 | Eukaryota | 1160.0 | 0 | Platyhelminthes | 775.0 | 1 | Diphyllobothriidea | 775.0 | 1 | Diphyllobothriidae | 775.0 | 1 | Dibothriocephalus | 775.0 | 1 | Dibothriocephalus latus | 775.0 | 1 |
| scf7180000043960 | 404947 | 0.4955 | 0 | 59.6173 | Eukaryota | 62450.0 | 0 | Chordata | 62450.0 | 0 | Carnivora | 62450.0 | 0 | Canidae | 62450.0 | 0 | Canis | 62450.0 | 0 | Canis lupus | 62450.0 | 0 |
| scf7180000052126 | 10298 | 0.4479 | 0 | 46.9747 | no-hit | 0.0 | 0 | no-hit | 0.0 | 0 | no-hit | 0.0 | 0 | no-hit | 0.0 | 0 | no-hit | 0.0 | 0 | no-hit | 0.0 | 0 |
| scf7180000052602 | 10312 | 0.3889 | 0 | 58.5566 | Eukaryota | 18917.0 | 0 | Chordata | 18917.0 | 0 | Carnivora | 18917.0 | 0 | Canidae | 18917.0 | 0 | Canis | 18917.0 | 0 | Canis lupus | 18917.0 | 0 |
| scf7180000053316 | 13063 | 0.502 | 0 | 229.7301 | Eukaryota | 9274.0 | 0 | Chordata | 9274.0 | 0 | Carnivora | 9274.0 | 0 | Canidae | 9274.0 | 0 | Canis | 8043.0 | 1 | Canis lupus | 6788.0 | 2 |
| scf7180000054525 | 216901 | 0.3635 | 0 | 63.4802 | Eukaryota | 47584.0 | 0 | Chordata | 47584.0 | 0 | Carnivora | 47584.0 | 0 | Canidae | 47584.0 | 0 | Canis | 47584.0 | 0 | Canis lupus | 47584.0 | 0 |
| scf7180000042129 | 10933 | 0.4421 | 0 | 102.3807 | Eukaryota | 4213.0 | 0 | Chordata | 4213.0 | 0 | Carnivora | 4213.0 | 0 | Canidae | 4213.0 | 0 | Canis | 4213.0 | 0 | Canis lupus | 4213.0 | 0 |
| scf7180000051509 | 10484 | 0.4443 | 0 | 193.302 | Eukaryota | 1625.0 | 0 | Chordata | 1625.0 | 0 | Kurtiformes | 1264.0 | 1 | Apogonidae | 1264.0 | 1 | Sphaeramia | 1264.0 | 1 | Sphaeramia orbicularis | 1264.0 | 1 |
| scf7180000051710 | 4278 | 0.4521 | 0 | 872.4724 | Eukaryota | 2355.0 | 0 | Platyhelminthes | 1199.0 | 2 | Polystomatidea | 1199.0 | 2 | Polystomatidae | 1199.0 | 2 | Protopolystoma | 1199.0 | 2 | Protopolystoma xenopodis | 1199.0 | 2 |
| scf7180000051456 | 15941 | 0.5046 | 0 | 328.9658 | Eukaryota | 9216.0 | 0 | Chordata | 9216.0 | 0 | Carnivora | 9216.0 | 0 | Canidae | 9216.0 | 0 | Canis | 7478.0 | 1 | Canis lupus | 6201.0 | 2 |
| scf7180000048691 | 45965 | 0.4057 | 0 | 60.125 | Eukaryota | 13606.0 | 0 | Chordata | 13606.0 | 0 | Carnivora | 8919.0 | 2 | Canidae | 6046.0 | 4 | Canis | 4380.0 | 5 | Canis lupus | 4380.0 | 7 |
| scf7180000053668 | 3295 | 0.4604 | 0 | 701.404 | no-hit | 0.0 | 0 | no-hit | 0.0 | 0 | no-hit | 0.0 | 0 | no-hit | 0.0 | 0 | no-hit | 0.0 | 0 | no-hit | 0.0 | 0 |
| scf7180000044861 | 146093 | 0.5327 | 0 | 55.7572 | Eukaryota | 25772.0 | 0 | Chordata | 25772.0 | 0 | Carnivora | 23893.0 | 1 | Canidae | 18367.0 | 2 | Canis | 9275.0 | 3 | Canis lupus | 9275.0 | 3 |
| scf7180000048503 | 27764 | 0.4586 | 0 | 52.2236 | Eukaryota | 13516.0 | 0 | Chordata | 13516.0 | 0 | Carnivora | 13516.0 | 0 | Canidae | 13516.0 | 0 | Canis | 13516.0 | 0 | Canis lupus | 13516.0 | 0 |
| scf7180000047739 | 23428 | 0.4484 | 0 | 110.0828 | Eukaryota | 46686.0 | 0 | Chordata | 32754.0 | 1 | Gadiformes | 32754.0 | 2 | Gadidae | 32754.0 | 2 | Gadus | 32754.0 | 2 | Gadus morhua | 32754.0 | 2 |
| scf7180000053536 | 4125 | 0.4514 | 0 | 669.2492 | Eukaryota | 220.0 | 0 | Streptophyta | 220.0 | 0 | Fabales | 220.0 | 0 | Fabaceae | 220.0 | 0 | Cicer | 220.0 | 0 | Cicer arietinum | 220.0 | 0 |
| scf7180000049727 | 7155 | 0.4551 | 0 | 399.1646 | no-hit | 0.0 | 0 | no-hit | 0.0 | 0 | no-hit | 0.0 | 0 | no-hit | 0.0 | 0 | no-hit | 0.0 | 0 | no-hit | 0.0 | 0 |
| scf7180000056137 | 5235 | 0.4369 | 0 | 85.2233 | Eukaryota | 417.0 | 0 | Platyhelminthes | 417.0 | 0 | Polystomatidea | 417.0 | 0 | Polystomatidae | 417.0 | 0 | Protopolystoma | 417.0 | 0 | Protopolystoma xenopodis | 417.0 | 0 |
| scf7180000041341 | 1298767 | 0.3869 | 0 | 60.0861 | Eukaryota | 49828.0 | 0 | Chordata | 49828.0 | 0 | Carnivora | 49828.0 | 0 | Canidae | 28168.0 | 2 | Canis | 16574.0 | 6 | Canis lupus | 16574.0 | 6 |
| scf7180000047891 | 53451 | 0.6384 | 0 | 53.8683 | Eukaryota | 44203.0 | 0 | Chordata | 44203.0 | 0 | Carnivora | 44203.0 | 0 | Canidae | 40907.0 | 1 | Canis | 36114.0 | 2 | Canis lupus | 36114.0 | 2 |
| scf7180000055801 | 26031 | 0.4388 | 0 | 83.4155 | Eukaryota | 4716.0 | 0 | Chordata | 1927.0 | 2 | Nymphaeales | 1620.0 | 6 | Nymphaeaceae | 1620.0 | 6 | Nymphaea | 1620.0 | 6 | Nymphaea colorata | 1620.0 | 6 |
| scf7180000050106 | 9365 | 0.4325 | 0 | 94.7788 | Eukaryota | 3451.0 | 0 | Platyhelminthes | 2160.0 | 1 | Polystomatidea | 2160.0 | 2 | Polystomatidae | 2160.0 | 2 | Protopolystoma | 2160.0 | 3 | Protopolystoma xenopodis | 2160.0 | 3 |
| scf7180000043526 | 405826 | 0.3538 | 0 | 60.4128 | Eukaryota | 69425.0 | 0 | Chordata | 69425.0 | 0 | Carnivora | 69425.0 | 0 | Canidae | 69425.0 | 0 | Canis | 69425.0 | 0 | Canis lupus | 69425.0 | 0 |
| scf7180000043494 | 59652 | 0.3461 | 0 | 55.2808 | Eukaryota | 86817.0 | 0 | Chordata | 86817.0 | 0 | Carnivora | 86817.0 | 0 | Canidae | 86817.0 | 0 | Canis | 86817.0 | 0 | Canis lupus | 86817.0 | 0 |
| scf7180000043106 | 96741 | 0.5297 | 0 | 55.5116 | Eukaryota | 74493.0 | 0 | Chordata | 74493.0 | 0 | Carnivora | 74493.0 | 0 | Canidae | 74493.0 | 0 | Canis | 74493.0 | 0 | Canis lupus | 74493.0 | 0 |
| scf7180000055105 | 24704 | 0.4184 | 0 | 80.0646 | Eukaryota | 33366.0 | 0 | Chordata | 33366.0 | 0 | Carnivora | 33366.0 | 0 | Canidae | 24624.0 | 1 | Canis | 21070.0 | 3 | Canis lupus | 21070.0 | 4 |
| scf7180000049632 | 14020 | 0.4371 | 0 | 97.0064 | Eukaryota | 499.0 | 0 | Chordata | 338.0 | 1 | Rodentia | 338.0 | 1 | Muridae | 338.0 | 1 | Mus | 338.0 | 1 | Mus musculus | 338.0 | 1 |
| scf7180000044456 | 40274 | 0.4032 | 0 | 55.689 | Eukaryota | 82257.0 | 0 | Chordata | 82257.0 | 0 | Carnivora | 82257.0 | 0 | Canidae | 82257.0 | 0 | Canis | 82257.0 | 0 | Canis lupus | 82257.0 | 0 |
| scf7180000052670 | 2293 | 0.4461 | 0 | 459.0981 | no-hit | 0.0 | 0 | no-hit | 0.0 | 0 | no-hit | 0.0 | 0 | no-hit | 0.0 | 0 | no-hit | 0.0 | 0 | no-hit | 0.0 | 0 |
| scf7180000040923 | 1538508 | 0.3961 | 0 | 60.0256 | Eukaryota | 89423.0 | 0 | Chordata | 89423.0 | 0 | Carnivora | 89423.0 | 0 | Canidae | 89423.0 | 0 | Canis | 89423.0 | 0 | Canis lupus | 89423.0 | 0 |
| scf7180000052346 | 3210 | 0.4978 | 0 | 73.5663 | Eukaryota | 6719.0 | 0 | Chordata | 6719.0 | 0 | Carnivora | 6719.0 | 0 | Canidae | 6719.0 | 0 | Canis | 4607.0 | 1 | Canis lupus | 3672.0 | 2 |
| scf7180000054412 | 52533 | 0.4418 | 0 | 139.622 | Eukaryota | 93357.0 | 0 | Chordata | 93357.0 | 0 | Carnivora | 93357.0 | 0 | Canidae | 93357.0 | 0 | Canis | 93357.0 | 0 | Canis lupus | 93357.0 | 0 |
| scf7180000044151 | 31669 | 0.3477 | 0 | 45.8826 | Eukaryota | 8433.0 | 0 | Chordata | 8433.0 | 0 | Primates | 3174.0 | 3 | Hominidae | 3174.0 | 3 | Homo | 3174.0 | 4 | Homo sapiens | 3174.0 | 4 |
| scf7180000042675 | 665607 | 0.3972 | 100 | 59.986 | Eukaryota | 56492.0 | 0 | Chordata | 56492.0 | 0 | Carnivora | 56492.0 | 0 | Canidae | 56492.0 | 0 | Canis | 56492.0 | 0 | Canis lupus | 56492.0 | 0 |
| scf7180000044940 | 7804 | 0.3538 | 0 | 65.6714 | Eukaryota | 37106.0 | 0 | Chordata | 37106.0 | 0 | Carnivora | 37106.0 | 0 | Canidae | 37106.0 | 0 | Canis | 37106.0 | 0 | Canis lupus | 37106.0 | 0 |
| scf7180000047121 | 12974 | 0.3893 | 0 | 55.5175 | Eukaryota | 85953.0 | 0 | Chordata | 85953.0 | 0 | Carnivora | 85953.0 | 0 | Canidae | 85953.0 | 0 | Canis | 85953.0 | 0 | Canis lupus | 85953.0 | 0 |
| scf7180000051977 | 15545 | 0.4986 | 0 | 217.1301 | Eukaryota | 33637.0 | 0 | Chordata | 33637.0 | 0 | Carnivora | 33637.0 | 0 | Canidae | 33637.0 | 0 | Canis | 33637.0 | 0 | Canis lupus | 33637.0 | 0 |
| scf7180000048741 | 37790 | 0.4035 | 0 | 60.5374 | Eukaryota | 8038.0 | 0 | Chordata | 8038.0 | 0 | Primates | 5410.0 | 1 | Hominidae | 5410.0 | 1 | Homo | 3268.0 | 2 | Homo sapiens | 3268.0 | 3 |
| scf7180000056476 | 271756 | 0.3374 | 0 | 58.773 | Eukaryota | 17974.0 | 0 | Chordata | 17974.0 | 0 | Carnivora | 17974.0 | 0 | Canidae | 17974.0 | 0 | Canis | 17974.0 | 0 | Canis lupus | 17974.0 | 0 |
| scf7180000046653 | 98560 | 0.4438 | 0 | 69.3145 | Eukaryota | 48171.0 | 0 | Platyhelminthes | 23115.0 | 2 | Cyclophyllidea | 11910.0 | 6 | Hymenolepididae | 11910.0 | 6 | Hymenolepis | 11910.0 | 6 | Hymenolepis microstoma | 11910.0 | 6 |
| scf7180000049491 | 14512 | 0.4509 | 0 | 1119.6349 | Eukaryota | 2268.0 | 0 | Echinodermata | 915.0 | 2 | Forcipulatida | 915.0 | 2 | Asteriidae | 915.0 | 2 | Asterias | 915.0 | 2 | Asterias rubens | 915.0 | 2 |
| scf7180000041408 | 1084218 | 0.3514 | 0 | 58.9282 | Eukaryota | 85322.0 | 0 | Chordata | 85322.0 | 0 | Carnivora | 85322.0 | 0 | Canidae | 85322.0 | 0 | Vulpes | 45267.0 | 1 | Vulpes vulpes | 45267.0 | 1 |
| scf7180000043316 | 96259 | 0.3911 | 0 | 59.4809 | Eukaryota | 11210.0 | 0 | Chordata | 11210.0 | 0 | Primates | 7172.0 | 3 | Hominidae | 7172.0 | 3 | Homo | 7172.0 | 3 | Homo sapiens | 7172.0 | 3 |
| scf7180000041219 | 3590 | 0.4209 | 0 | 27.9064 | Eukaryota | 3299.0 | 0 | Chordata | 3299.0 | 0 | Carnivora | 2977.0 | 1 | Felidae | 1985.0 | 2 | Felis | 1665.0 | 3 | Felis catus | 1665.0 | 3 |
| scf7180000040705 | 1792483 | 0.4359 | 0 | 59.9419 | Eukaryota | 153742.0 | 0 | Chordata | 153742.0 | 0 | Carnivora | 153742.0 | 0 | Canidae | 142966.0 | 1 | Canis | 142966.0 | 1 | Canis lupus | 142966.0 | 1 |
| scf7180000042732 | 12624 | 0.4343 | 0 | 800.7207 | no-hit | 0.0 | 0 | no-hit | 0.0 | 0 | no-hit | 0.0 | 0 | no-hit | 0.0 | 0 | no-hit | 0.0 | 0 | no-hit | 0.0 | 0 |
| scf7180000042232 | 118634 | 0.4238 | 0 | 56.93 | Eukaryota | 41138.0 | 0 | Chordata | 41138.0 | 0 | Carnivora | 41138.0 | 0 | Canidae | 41138.0 | 0 | Canis | 41138.0 | 0 | Canis lupus | 41138.0 | 0 |
| scf7180000051494 | 7056 | 0.3798 | 0 | 55.4475 | Eukaryota | 15898.0 | 0 | Chordata | 15898.0 | 0 | Carnivora | 15898.0 | 0 | Canidae | 6886.0 | 3 | Canis | 5190.0 | 6 | Canis lupus | 5190.0 | 7 |
| scf7180000043344 | 454856 | 0.4782 | 0 | 58.8565 | Eukaryota | 30074.0 | 0 | Chordata | 30074.0 | 0 | Carnivora | 30074.0 | 0 | Canidae | 30074.0 | 0 | Canis | 30074.0 | 0 | Canis lupus | 30074.0 | 0 |
| scf7180000055907 | 18914 | 0.4215 | 0 | 234.1742 | Eukaryota | 7190.0 | 0 | Chordata | 7190.0 | 0 | Carnivora | 7190.0 | 0 | Felidae | 5776.0 | 1 | Felis | 5022.0 | 2 | Felis catus | 5022.0 | 2 |
| scf7180000041243 | 1910429 | 0.3893 | 0 | 59.2007 | Eukaryota | 76636.0 | 0 | Chordata | 76636.0 | 0 | Carnivora | 76636.0 | 0 | Canidae | 76636.0 | 0 | Canis | 58752.0 | 1 | Canis lupus | 58752.0 | 1 |
| scf7180000041649 | 537149 | 0.421 | 0 | 59.0371 | Eukaryota | 89153.0 | 0 | Chordata | 89153.0 | 0 | Carnivora | 89153.0 | 0 | Canidae | 89153.0 | 0 | Canis | 89153.0 | 0 | Canis lupus | 89153.0 | 0 |
| scf7180000056406 | 550301 | 0.3705 | 0 | 59.8699 | Eukaryota | 54294.0 | 0 | Chordata | 54294.0 | 0 | Carnivora | 54294.0 | 0 | Canidae | 54294.0 | 0 | Canis | 54294.0 | 0 | Canis lupus | 54294.0 | 0 |
| scf7180000049792 | 62349 | 0.3773 | 0 | 63.4925 | Eukaryota | 6648.0 | 1 | Chordata | 6648.0 | 1 | Carnivora | 4355.0 | 2 | Canidae | 4355.0 | 2 | Canis | 4355.0 | 2 | Canis lupus | 4355.0 | 2 |
| scf7180000049382 | 19317 | 0.4343 | 0 | 156.1887 | no-hit | 0.0 | 0 | no-hit | 0.0 | 0 | no-hit | 0.0 | 0 | no-hit | 0.0 | 0 | no-hit | 0.0 | 0 | no-hit | 0.0 | 0 |
| scf7180000050305 | 10908 | 0.449 | 0 | 988.2225 | Eukaryota | 21103.0 | 0 | Chordata | 19114.0 | 1 | Gadiformes | 13211.0 | 6 | Gadidae | 13211.0 | 6 | Gadus | 13211.0 | 6 | Gadus morhua | 13211.0 | 6 |
| scf7180000040523 | 294499 | 0.3876 | 0 | 59.048 | Eukaryota | 95165.0 | 0 | Chordata | 95165.0 | 0 | Carnivora | 95165.0 | 0 | Canidae | 95165.0 | 0 | Canis | 95165.0 | 0 | Canis lupus | 95165.0 | 0 |
| scf7180000050635 | 2952 | 0.4441 | 0 | 226.4567 | no-hit | 0.0 | 0 | no-hit | 0.0 | 0 | no-hit | 0.0 | 0 | no-hit | 0.0 | 0 | no-hit | 0.0 | 0 | no-hit | 0.0 | 0 |
| scf7180000056354 | 775690 | 0.3582 | 0 | 59.3432 | Eukaryota | 83871.0 | 0 | Chordata | 83871.0 | 0 | Carnivora | 83871.0 | 0 | Canidae | 83871.0 | 0 | Canis | 83871.0 | 0 | Canis lupus | 83871.0 | 0 |
| scf7180000045128 | 304884 | 0.4138 | 0 | 58.949 | Eukaryota | 39111.0 | 0 | Chordata | 39111.0 | 0 | Carnivora | 39111.0 | 0 | Canidae | 39111.0 | 0 | Canis | 39111.0 | 0 | Canis lupus | 39111.0 | 0 |
| scf7180000056551 | 122072 | 0.4837 | 0 | 57.7791 | Eukaryota | 17479.0 | 0 | Chordata | 17479.0 | 0 | Carnivora | 11243.0 | 2 | Canidae | 11243.0 | 3 | Canis | 11243.0 | 4 | Canis lupus | 11243.0 | 4 |
| scf7180000050992 | 12959 | 0.4455 | 0 | 456.3704 | no-hit | 0.0 | 0 | no-hit | 0.0 | 0 | no-hit | 0.0 | 0 | no-hit | 0.0 | 0 | no-hit | 0.0 | 0 | no-hit | 0.0 | 0 |
| scf7180000052909 | 2749 | 0.446 | 0 | 593.2289 | no-hit | 0.0 | 0 | no-hit | 0.0 | 0 | no-hit | 0.0 | 0 | no-hit | 0.0 | 0 | no-hit | 0.0 | 0 | no-hit | 0.0 | 0 |

| | | | | | | | | | | | | | | | | | | | | | | |
|---|---|---|---|---|---|---|---|---|---|---|---|---|---|---|---|---|---|---|---|---|---|---|
| scf7180000049275 | 40158 | 0.4564 | 0 | 77.4634 | Eukaryota | 5754.0 | 0 | Chordata | 3482.0 | 2 | Kurtiformes | 1970.0 | 4 | Apogonidae | 1970.0 | 4 | Sphaeramia | 1970.0 | 5 | Sphaeramia orbicularis | 1970.0 | 5 |
| scf7180000040997 | 14460 | 0.4369 | 0 | 580.9286 | no-hit | 0.0 | 0 | no-hit | 0.0 | 0 | no-hit | 0.0 | 0 | no-hit | 0.0 | 0 | no-hit | 0.0 | 0 | no-hit | 0.0 | 0 |
| scf7180000042143 | 752177 | 0.4691 | 0 | 58.0667 | Eukaryota | 92802.0 | 0 | Chordata | 92802.0 | 0 | Carnivora | 92802.0 | 0 | Canidae | 92802.0 | 0 | Canis | 92802.0 | 0 | Canis lupus | 92802.0 | 0 |
| scf7180000042878 | 650147 | 0.375 | 0 | 60.7306 | Eukaryota | 66898.0 | 0 | Chordata | 66898.0 | 0 | Carnivora | 66898.0 | 0 | Canidae | 66898.0 | 0 | Canis | 60781.0 | 1 | Canis lupus | 60781.0 | 1 |
| scf7180000049508 | 27748 | 0.5032 | 0 | 280.8634 | Eukaryota | 10214.0 | 0 | Chordata | 10214.0 | 0 | Carnivora | 10214.0 | 0 | Canidae | 10214.0 | 0 | Canis | 8955.0 | 1 | Canis lupus | 7659.0 | 2 |
| scf7180000045556 | 148226 | 0.3522 | 0 | 60.5041 | Eukaryota | 53503.0 | 0 | Chordata | 53503.0 | 0 | Carnivora | 53503.0 | 0 | Canidae | 53503.0 | 0 | Canis | 53503.0 | 0 | Canis lupus | 53503.0 | 0 |
| scf7180000054280 | 30105 | 0.4399 | 0 | 355.5103 | Eukaryota | 17333.0 | 0 | Chordata | 12090.0 | 2 | Gadiformes | 8072.0 | 6 | Gadidae | 8072.0 | 6 | Gadus | 8072.0 | 6 | Gadus morhua | 8072.0 | 6 |
| scf7180000053546 | 2958 | 0.4577 | 0 | 763.8516 | no-hit | 0.0 | 0 | no-hit | 0.0 | 0 | no-hit | 0.0 | 0 | no-hit | 0.0 | 0 | no-hit | 0.0 | 0 | no-hit | 0.0 | 0 |
| scf7180000040723 | 146454 | 0.3454 | 0 | 59.439 | Eukaryota | 18438.0 | 0 | Chordata | 18438.0 | 0 | Carnivora | 18438.0 | 0 | Canidae | 18438.0 | 0 | Canis | 18438.0 | 0 | Canis lupus | 18438.0 | 0 |
| scf7180000043363 | 436678 | 0.5854 | 0 | 55.5037 | Eukaryota | 28979.0 | 0 | Chordata | 28979.0 | 0 | Carnivora | 28979.0 | 0 | Canidae | 28979.0 | 0 | Canis | 24046.0 | 1 | Canis lupus | 24046.0 | 1 |
| scf7180000041543 | 224023 | 0.3806 | 0 | 59.8324 | Eukaryota | 94633.0 | 0 | Chordata | 94633.0 | 0 | Carnivora | 94633.0 | 0 | Canidae | 94633.0 | 0 | Canis | 94633.0 | 0 | Canis lupus | 94633.0 | 0 |
| scf7180000047465 | 23116 | 0.4361 | 0 | 346.0474 | Eukaryota | 17189.0 | 0 | Chordata | 17189.0 | 0 | Carnivora | 17189.0 | 0 | Canidae | 17189.0 | 0 | Canis | 17189.0 | 0 | Canis lupus | 17189.0 | 0 |
| scf7180000040644 | 665870 | 0.4095 | 0 | 59.616 | Eukaryota | 69764.0 | 0 | Chordata | 69764.0 | 0 | Carnivora | 69764.0 | 0 | Canidae | 69764.0 | 0 | Canis | 60278.0 | 1 | Canis lupus | 60278.0 | 1 |
| scf7180000048792 | 13321 | 0.4406 | 0 | 69.4937 | Eukaryota | 2275.0 | 0 | Platyhelminthes | 1501.0 | 1 | Polystomatidea | 1501.0 | 2 | Polystomatidae | 1501.0 | 2 | Protopolystoma | 1501.0 | 3 | Protopolystoma xenopodis | 1501.0 | 3 |
| scf7180000054259 | 21788 | 0.3548 | 0 | 337.6934 | Eukaryota | 13921.0 | 0 | Chordata | 13921.0 | 0 | Carnivora | 13921.0 | 0 | Canidae | 8108.0 | 2 | Canis | 6276.0 | 5 | Canis lupus | 6276.0 | 5 |
| scf7180000040724 | 970607 | 0.386 | 0 | 63.916 | Eukaryota | 58634.0 | 0 | Chordata | 58634.0 | 0 | Carnivora | 58634.0 | 0 | Canidae | 58634.0 | 0 | Canis | 58634.0 | 0 | Canis lupus | 58634.0 | 0 |
| scf7180000053256 | 12476 | 0.5487 | 0 | 352.7418 | Eukaryota | 27641.0 | 0 | Chordata | 27641.0 | 0 | Artiodactyla | 19032.0 | 2 | Bovidae | 19032.0 | 2 | Ovis | 10886.0 | 4 | Ovis aries | 10886.0 | 4 |
| scf7180000051069 | 2505 | 0.4499 | 0 | 490.5707 | no-hit | 0.0 | 0 | no-hit | 0.0 | 0 | no-hit | 0.0 | 0 | no-hit | 0.0 | 0 | no-hit | 0.0 | 0 | no-hit | 0.0 | 0 |
| scf7180000053956 | 12160 | 0.4536 | 0 | 58.2281 | Eukaryota | 5229.0 | 0 | Platyhelminthes | 3401.0 | 1 | Polystomatidea | 3401.0 | 2 | Polystomatidae | 3401.0 | 2 | Protopolystoma | 3401.0 | 2 | Protopolystoma xenopodis | 3401.0 | 2 |
| scf7180000049039 | 11562 | 0.4384 | 0 | 52.7886 | Eukaryota | 263.0 | 0 | Chordata | 263.0 | 0 | Kurtiformes | 263.0 | 0 | Apogonidae | 263.0 | 0 | Sphaeramia | 263.0 | 0 | Sphaeramia orbicularis | 263.0 | 0 |
| scf7180000040617 | 47048 | 0.3291 | 0 | 52.433 | Eukaryota | 23495.0 | 0 | Chordata | 23495.0 | 0 | Carnivora | 23495.0 | 0 | Canidae | 23495.0 | 0 | Canis | 23495.0 | 0 | Canis lupus | 23495.0 | 0 |
| scf7180000046899 | 95715 | 0.4515 | 0 | 57.5747 | Eukaryota | 5464.0 | 0 | Chordata | 5464.0 | 0 | Carnivora | 5464.0 | 0 | Canidae | 5464.0 | 0 | Canis | 5464.0 | 0 | Canis lupus | 5464.0 | 0 |
| scf7180000044277 | 250867 | 0.4222 | 0 | 58.758 | Eukaryota | 51463.0 | 0 | Chordata | 51463.0 | 0 | Carnivora | 51463.0 | 0 | Canidae | 51463.0 | 0 | Canis | 44752.0 | 1 | Canis lupus | 44752.0 | 1 |
| scf7180000056198 | 357509 | 0.3649 | 0 | 58.3099 | Eukaryota | 111863.0 | 0 | Chordata | 111863.0 | 0 | Carnivora | 111863.0 | 0 | Canidae | 111863.0 | 0 | Canis | 111863.0 | 0 | Canis lupus | 111863.0 | 0 |
| scf7180000047956 | 64896 | 0.5425 | 0 | 56.8104 | Eukaryota | 39362.0 | 0 | Chordata | 39362.0 | 0 | Carnivora | 39362.0 | 0 | Canidae | 30733.0 | 1 | Vulpes | 16930.0 | 3 | Vulpes vulpes | 16930.0 | 3 |
| scf7180000046565 | 48525 | 0.3975 | 0 | 56.5884 | Eukaryota | 11034.0 | 0 | Chordata | 11034.0 | 0 | Primates | 7566.0 | 3 | Hominidae | 7566.0 | 3 | Homo | 6101.0 | 5 | Homo sapiens | 6101.0 | 5 |
| scf7180000048031 | 5042 | 0.4344 | 0 | 159.8622 | no-hit | 0.0 | 0 | no-hit | 0.0 | 0 | no-hit | 0.0 | 0 | no-hit | 0.0 | 0 | no-hit | 0.0 | 0 | no-hit | 0.0 | 0 |
| scf7180000045434 | 108583 | 0.4306 | 0 | 56.7668 | Eukaryota | 26533.0 | 0 | Chordata | 26533.0 | 0 | Carnivora | 20564.0 | 1 | Canidae | 20564.0 | 1 | Canis | 20564.0 | 1 | Canis lupus | 20564.0 | 1 |
| scf7180000044289 | 70571 | 0.417 | 0 | 75.4279 | Eukaryota | 93919.0 | 0 | Chordata | 93919.0 | 0 | Carnivora | 93919.0 | 0 | Canidae | 93919.0 | 0 | Canis | 93919.0 | 0 | Canis lupus | 93919.0 | 0 |
| scf7180000047390 | 81686 | 0.504 | 0 | 58.5862 | Eukaryota | 19619.0 | 0 | Chordata | 19619.0 | 0 | Carnivora | 19619.0 | 0 | Canidae | 19619.0 | 0 | Canis | 15029.0 | 1 | Canis lupus | 15029.0 | 1 |
| scf7180000050366 | 42727 | 0.4426 | 0 | 61.1156 | Eukaryota | 26187.0 | 0 | Platyhelminthes | 18872.0 | 1 | Cyclophyllidea | 12643.0 | 6 | Hymenolepididae | 12643.0 | 6 | Hymenolepis | 12643.0 | 6 | Hymenolepis microstoma | 12643.0 | 6 |
| scf7180000056601 | 415418 | 0.3669 | 100 | 59.5155 | Eukaryota | 89601.0 | 0 | Chordata | 89601.0 | 0 | Carnivora | 89601.0 | 0 | Canidae | 89601.0 | 0 | Canis | 89601.0 | 0 | Canis lupus | 89601.0 | 0 |
| scf7180000053824 | 4669 | 0.4583 | 0 | 324.1718 | Eukaryota | 534.0 | 0 | Platyhelminthes | 534.0 | 0 | Polystomatidea | 534.0 | 0 | Polystomatidae | 534.0 | 0 | Protopolystoma | 534.0 | 0 | Protopolystoma xenopodis | 534.0 | 0 |
| scf7180000049451 | 6288 | 0.4467 | 0 | 1664.2282 | Eukaryota | 2320.0 | 0 | Platyhelminthes | 1395.0 | 2 | Polystomatidea | 1395.0 | 2 | Polystomatidae | 1395.0 | 2 | Protopolystoma | 1395.0 | 2 | Protopolystoma xenopodis | 1395.0 | 2 |
| scf7180000051649 | 13058 | 0.4346 | 0 | 120.3543 | Eukaryota | 21343.0 | 0 | Chordata | 13300.0 | 2 | Strigeidida | 7466.0 | 6 | Schistosomatidae | 7466.0 | 6 | Schistosoma | 7466.0 | 6 | Schistosoma mansoni | 7466.0 | 6 |
| scf7180000048951 | 18009 | 0.433 | 0 | 144.9774 | no-hit | 0.0 | 0 | no-hit | 0.0 | 0 | no-hit | 0.0 | 0 | no-hit | 0.0 | 0 | no-hit | 0.0 | 0 | no-hit | 0.0 | 0 |
| scf7180000055857 | 24295 | 0.4857 | 0 | 222.0127 | Eukaryota | 14990.0 | 0 | Chordata | 14990.0 | 0 | Carnivora | 14990.0 | 0 | Canidae | 14990.0 | 0 | Canis | 9172.0 | 1 | Canis lupus | 9172.0 | 1 |
| scf7180000043094 | 491633 | 0.4506 | 0 | 59.4883 | Eukaryota | 43884.0 | 0 | Chordata | 43884.0 | 0 | Carnivora | 43884.0 | 0 | Canidae | 43884.0 | 0 | Canis | 43884.0 | 0 | Canis lupus | 43884.0 | 0 |
| scf7180000047821 | 63718 | 0.4431 | 0 | 113.6121 | Eukaryota | 9518.0 | 0 | Chordata | 8146.0 | 1 | Tetraodontiformes | 6334.0 | 4 | Tetraodontidae | 6334.0 | 4 | Takifugu | 6334.0 | 4 | Takifugu rubripes | 6334.0 | 4 |
| scf7180000052086 | 1867 | 0.4456 | 0 | 151.7125 | no-hit | 0.0 | 0 | no-hit | 0.0 | 0 | no-hit | 0.0 | 0 | no-hit | 0.0 | 0 | no-hit | 0.0 | 0 | no-hit | 0.0 | 0 |
| scf7180000044143 | 64693 | 0.3992 | 0 | 55.7458 | Eukaryota | 25290.0 | 0 | Chordata | 25290.0 | 0 | Carnivora | 25290.0 | 0 | Canidae | 25290.0 | 0 | Canis | 25290.0 | 0 | Canis lupus | 25290.0 | 0 |
| scf7180000042953 | 10500 | 0.3785 | 0 | 41.1764 | Eukaryota | 3067.0 | 0 | Chordata | 3067.0 | 0 | Carnivora | 2531.0 | 1 | Canidae | 2531.0 | 1 | Canis | 2531.0 | 1 | Canis lupus | 2531.0 | 1 |
| scf7180000041214 | 262805 | 0.4021 | 0 | 62.266 | Eukaryota | 92927.0 | 0 | Chordata | 92927.0 | 0 | Carnivora | 92927.0 | 0 | Canidae | 92927.0 | 0 | Canis | 92927.0 | 0 | Canis lupus | 92927.0 | 0 |
| scf7180000049349 | 3478 | 0.4482 | 0 | 732.3962 | no-hit | 0.0 | 0 | no-hit | 0.0 | 0 | no-hit | 0.0 | 0 | no-hit | 0.0 | 0 | no-hit | 0.0 | 0 | no-hit | 0.0 | 0 |
| scf7180000042680 | 531769 | 0.4111 | 0 | 59.5241 | Eukaryota | 75143.0 | 0 | Chordata | 75143.0 | 0 | Carnivora | 75143.0 | 0 | Canidae | 75143.0 | 0 | Canis | 75143.0 | 0 | Canis lupus | 75143.0 | 0 |
| scf7180000052186 | 4620 | 0.4377 | 0 | 395.706 | no-hit | 0.0 | 0 | no-hit | 0.0 | 0 | no-hit | 0.0 | 0 | no-hit | 0.0 | 0 | no-hit | 0.0 | 0 | no-hit | 0.0 | 0 |
| scf7180000046039 | 6918 | 0.4644 | 0 | 748.6063 | Eukaryota | 235.0 | 0 | Chordata | 235.0 | 0 | Carnivora | 235.0 | 0 | Ursidae | 235.0 | 0 | Ursus | 235.0 | 0 | Ursus arctos | 235.0 | 0 |
| scf7180000046981 | 111576 | 0.4815 | 0 | 57.1145 | Eukaryota | 48464.0 | 0 | Chordata | 48464.0 | 0 | Carnivora | 48464.0 | 0 | Canidae | 48464.0 | 0 | Canis | 48464.0 | 0 | Canis lupus | 48464.0 | 0 |
| scf7180000047546 | 74705 | 0.4253 | 0 | 58.6183 | Eukaryota | 19250.0 | 0 | Chordata | 19250.0 | 0 | Carnivora | 19250.0 | 0 | Canidae | 19250.0 | 0 | Canis | 19250.0 | 0 | Canis lupus | 19250.0 | 0 |
| scf7180000055630 | 23620 | 0.4523 | 0 | 1772.5506 | no-hit | 0.0 | 0 | no-hit | 0.0 | 0 | no-hit | 0.0 | 0 | no-hit | 0.0 | 0 | no-hit | 0.0 | 0 | no-hit | 0.0 | 0 |
| scf7180000051639 | 5275 | 0.45 | 0 | 116.5185 | Eukaryota | 7348.0 | 0 | Chordata | 7348.0 | 0 | Carnivora | 7348.0 | 0 | Canidae | 7348.0 | 0 | Nyctereutes | 4389.0 | 1 | Nyctereutes procyonoides | 4389.0 | 1 |
| scf7180000047145 | 18283 | 0.4431 | 0 | 125.1489 | Eukaryota | 5503.0 | 0 | Chordata | 5503.0 | 0 | Carnivora | 5503.0 | 0 | Canidae | 5503.0 | 0 | Canis | 5503.0 | 0 | Canis lupus | 5503.0 | 0 |
| scf7180000052620 | 8903 | 0.4113 | 0 | 59.8007 | Eukaryota | 37026.0 | 0 | Chordata | 37026.0 | 0 | Carnivora | 37026.0 | 0 | Canidae | 15666.0 | 2 | Callorhinus | 15110.0 | 4 | Callorhinus ursinus | 15110.0 | 4 |
| scf7180000043153 | 498711 | 0.3861 | 0 | 61.0048 | Eukaryota | 91409.0 | 0 | Chordata | 91409.0 | 0 | Carnivora | 91409.0 | 0 | Canidae | 91409.0 | 0 | Canis | 91409.0 | 0 | Canis lupus | 91409.0 | 0 |
| scf7180000052544 | 16577 | 0.5035 | 0 | 373.8603 | Eukaryota | 8417.0 | 0 | Chordata | 8417.0 | 0 | Carnivora | 8417.0 | 0 | Canidae | 8417.0 | 0 | Canis | 6217.0 | 1 | Canis lupus | 5086.0 | 2 |
| scf7180000043247 | 547514 | 0.4094 | 0 | 60.2735 | Eukaryota | 65492.0 | 0 | Chordata | 65492.0 | 0 | Carnivora | 65492.0 | 0 | Mustelidae | 34260.0 | 3 | Mustela | 34260.0 | 6 | Mustela putorius | 34260.0 | 6 |
| scf7180000041250 | 1247933 | 0.4057 | 0 | 56.8957 | Eukaryota | 65082.0 | 0 | Chordata | 65082.0 | 0 | Artiodactyla | 58559.0 | 1 | Bovidae | 45192.0 | 2 | Bubalus | 19382.0 | 4 | Bubalus bubalis | 19382.0 | 4 |
| scf7180000042189 | 45766 | 0.6016 | 0 | 51.127 | Eukaryota | 16589.0 | 0 | Chordata | 16589.0 | 0 | Carnivora | 13660.0 | 3 | Canidae | 13660.0 | 3 | Canis | 13660.0 | 3 | Canis lupus | 13660.0 | 3 |
| scf7180000050379 | 5271 | 0.4506 | 0 | 94.9838 | no-hit | 0.0 | 0 | no-hit | 0.0 | 0 | no-hit | 0.0 | 0 | no-hit | 0.0 | 0 | no-hit | 0.0 | 0 | no-hit | 0.0 | 0 |
| scf7180000054054 | 6319 | 0.5034 | 0 | 471.8433 | Eukaryota | 8252.0 | 0 | Chordata | 8252.0 | 0 | Carnivora | 8252.0 | 0 | Canidae | 8252.0 | 0 | Canis | 6609.0 | 1 | Canis lupus | 5411.0 | 2 |
| scf7180000050287 | 6512 | 0.4413 | 0 | 53.9457 | Eukaryota | 18894.0 | 0 | Chordata | 12162.0 | 1 | Strigeidida | 5900.0 | 6 | Schistosomatidae | 5900.0 | 6 | Schistosoma | 5900.0 | 6 | Schistosoma mansoni | 5900.0 | 6 |
| scf7180000040911 | 85133 | 0.3883 | 0 | 56.2201 | Eukaryota | 60736.0 | 0 | Chordata | 60736.0 | 0 | Carnivora | 60736.0 | 0 | Canidae | 60736.0 | 0 | Canis | 60736.0 | 0 | Canis lupus | 60736.0 | 0 |
| scf7180000046711 | 5767 | 0.4451 | 0 | 251.4644 | no-hit | 0.0 | 0 | no-hit | 0.0 | 0 | no-hit | 0.0 | 0 | no-hit | 0.0 | 0 | no-hit | 0.0 | 0 | no-hit | 0.0 | 0 |
| scf7180000052296 | 12270 | 0.4205 | 0 | 55.1293 | Eukaryota | 4358.0 | 0 | Chordata | 4358.0 | 0 | Carnivora | 2615.0 | 2 | Canidae | 2615.0 | 2 | Canis | 2615.0 | 3 | Canis lupus | 2615.0 | 3 |
| scf7180000053741 | 10041 | 0.5029 | 0 | 866.875 | Eukaryota | 9641.0 | 0 | Chordata | 9641.0 | 0 | Carnivora | 9641.0 | 0 | Canidae | 9641.0 | 0 | Canis | 8430.0 | 1 | Canis lupus | 7192.0 | 2 |
| scf7180000055584 | 14550 | 0.4752 | 0 | 68.4257 | Eukaryota | 9453.0 | 0 | Chordata | 9453.0 | 0 | Carnivora | 9453.0 | 0 | Canidae | 9453.0 | 0 | Canis | 5539.0 | 1 | Canis lupus | 5539.0 | 1 |
| scf7180000045401 | 10161 | 0.3937 | 0 | 27.4534 | Eukaryota | 2513.0 | 0 | Chordata | 2513.0 | 0 | Carnivora | 2513.0 | 0 | Canidae | 2513.0 | 0 | Canis | 2513.0 | 0 | Canis lupus | 2513.0 | 0 |
| scf7180000049274 | 41556 | 0.5033 | 0 | 59.519 | Eukaryota | 41121.0 | 0 | Chordata | 41121.0 | 0 | Carnivora | 41121.0 | 0 | Canidae | 41121.0 | 0 | Canis | 32262.0 | 1 | Canis lupus | 32262.0 | 1 |
| scf7180000042668 | 1029998 | 0.3546 | 0 | 59.5487 | Eukaryota | 94599.0 | 0 | Chordata | 94599.0 | 0 | Carnivora | 94599.0 | 0 | Canidae | 94599.0 | 0 | Canis | 94599.0 | 0 | Canis lupus | 94599.0 | 0 |
| scf7180000050105 | 14621 | 0.4498 | 0 | 900.4493 | Eukaryota | 3999.0 | 0 | Platyhelminthes | 2015.0 | 3 | Polystomatidea | 2015.0 | 3 | Polystomatidae | 2015.0 | 3 | Protopolystoma | 2015.0 | 3 | Protopolystoma xenopodis | 2015.0 | 3 |
| scf7180000049594 | 36020 | 0.4431 | 0 | 163.16 | Eukaryota | 3187.0 | 0 | Chordata | 2750.0 | 1 | Columbiformes | 2321.0 | 2 | Columbidae | 2321.0 | 2 | Streptopelia | 2321.0 | 2 | Streptopelia turtur | 2321.0 | 2 |
| scf7180000053835 | 2300 | 0.4561 | 0 | 192.3916 | no-hit | 0.0 | 0 | no-hit | 0.0 | 0 | no-hit | 0.0 | 0 | no-hit | 0.0 | 0 | no-hit | 0.0 | 0 | no-hit | 0.0 | 0 |
| scf7180000053856 | 3893 | 0.4392 | 0 | 273.7202 | no-hit | 0.0 | 0 | no-hit | 0.0 | 0 | no-hit | 0.0 | 0 | no-hit | 0.0 | 0 | no-hit | 0.0 | 0 | no-hit | 0.0 | 0 |
| scf7180000046440 | 118215 | 0.4139 | 0 | 61.3064 | Eukaryota | 72088.0 | 0 | Chordata | 72088.0 | 0 | Carnivora | 72088.0 | 0 | Canidae | 72088.0 | 0 | Canis | 72088.0 | 0 | Canis lupus | 72088.0 | 0 |
| scf7180000048344 | 51032 | 0.3447 | 0 | 57.4364 | Eukaryota | 5677.0 | 0 | Chordata | 5677.0 | 0 | Carnivora | 4503.0 | 1 | Canidae | 2693.0 | 3 | Canis | 2693.0 | 3 | Canis lupus | 2693.0 | 3 |
| scf7180000047592 | 75276 | 0.4868 | 0 | 58.5764 | Eukaryota | 34673.0 | 0 | Chordata | 34673.0 | 0 | Carnivora | 34673.0 | 0 | Canidae | 30416.0 | 1 | Canis | 20668.0 | 2 | Canis lupus | 20668.0 | 3 |
| scf7180000048780 | 7225 | 0.3954 | 0 | 42.6073 | Eukaryota | 2846.0 | 0 | Chordata | 2846.0 | 0 | Carnivora | 2846.0 | 0 | Canidae | 2846.0 | 0 | Canis | 2846.0 | 0 | Canis lupus | 2846.0 | 0 |
| scf7180000052745 | 9463 | 0.4401 | 0 | 74.7546 | no-hit | 0.0 | 0 | no-hit | 0.0 | 0 | no-hit | 0.0 | 0 | no-hit | 0.0 | 0 | no-hit | 0.0 | 0 | no-hit | 0.0 | 0 |
| scf7180000047063 | 38340 | 0.4426 | 0 | 179.397 | Eukaryota | 9155.0 | 0 | Chordata | 9155.0 | 0 | Carnivora | 3912.0 | 3 | Canidae | 3912.0 | 3 | Canis | 3912.0 | 3 | Canis lupus | 3912.0 | 3 |
| scf7180000043864 | 292547 | 0.3674 | 0 | 59.1996 | Eukaryota | 93117.0 | 0 | Chordata | 93117.0 | 0 | Carnivora | 93117.0 | 0 | Canidae | 93117.0 | 0 | Canis | 93117.0 | 0 | Canis lupus | 93117.0 | 0 |
| scf7180000042711 | 8925 | 0.44 | 0 | 46.4019 | Eukaryota | 14953.0 | 0 | Chordata | 14953.0 | 0 | Carnivora | 14953.0 | 0 | Canidae | 14953.0 | 0 | Canis | 14953.0 | 0 | Canis lupus | 14953.0 | 0 |
| scf7180000044511 | 278874 | 0.3806 | 0 | 59.8455 | Eukaryota | 92156.0 | 0 | Chordata | 92156.0 | 0 | Carnivora | 92156.0 | 0 | Canidae | 92156.0 | 0 | Canis | 92156.0 | 0 | Canis lupus | 92156.0 | 0 |
| scf7180000041349 | 25823 | 0.3995 | 0 | 45.165 | Eukaryota | 105927.0 | 0 | Chordata | 105927.0 | 0 | Carnivora | 105927.0 | 0 | Canidae | 105927.0 | 0 | Canis | 90503.0 | 1 | Canis lupus | 90503.0 | 1 |
| scf7180000054601 | 105260 | 0.4392 | 0 | 96.5855 | Eukaryota | 94909.0 | 0 | Chordata | 94909.0 | 0 | Carnivora | 94909.0 | 0 | Canidae | 94909.0 | 0 | Canis | 94909.0 | 0 | Canis lupus | 94909.0 | 0 |
| scf7180000042479 | 172994 | 0.4286 | 0 | 58.9151 | Eukaryota | 51288.0 | 0 | Chordata | 51288.0 | 0 | Carnivora | 51288.0 | 0 | Canidae | 51288.0 | 0 | Canis | 30703.0 | 1 | Canis lupus | 30703.0 | 1 |
| scf7180000052645 | 4286 | 0.4545 | 0 | 551.6731 | no-hit | 0.0 | 0 | no-hit | 0.0 | 0 | no-hit | 0.0 | 0 | no-hit | 0.0 | 0 | no-hit | 0.0 | 0 | no-hit | 0.0 | 0 |
| scf7180000045187 | 87484 | 0.3492 | 0 | 53.1671 | Eukaryota | 34742.0 | 0 | Chordata | 34742.0 | 0 | Carnivora | 34742.0 | 0 | Canidae | 34742.0 | 0 | Canis | 34742.0 | 0 | Canis lupus | 34742.0 | 0 |
| scf7180000055743 | 4852 | 0.4435 | 0 | 151.3737 | Eukaryota | 8537.0 | 0 | Chordata | 8537.0 | 0 | Kurtiformes | 4597.0 | 3 | Apogonidae | 4597.0 | 3 | Sphaeramia | 4597.0 | 3 | Sphaeramia orbicularis | 4597.0 | 3 |
| scf7180000048309 | 8732 | 0.4512 | 0 | 149.1068 | Eukaryota | 2762.0 | 0 | Chordata | 2762.0 | 0 | Carnivora | 2762.0 | 0 | Canidae | 2762.0 | 0 | Canis | 2762.0 | 0 | Canis lupus | 2762.0 | 0 |
| scf7180000049192 | 35735 | 0.3974 | 0 | 63.3633 | Eukaryota | 21122.0 | 0 | Chordata | 21122.0 | 0 | Carnivora | 21122.0 | 0 | Canidae | 21122.0 | 0 | Canis | 21122.0 | 0 | Canis lupus | 21122.0 | 0 |
| scf7180000046491 | 16589 | 0.4333 | 0 | 71.8898 | Eukaryota | 3061.0 | 0 | Platyhelminthes | 2044.0 | 1 | Polystomatidea | 2044.0 | 1 | Polystomatidae | 2044.0 | 1 | Protopolystoma | 2044.0 | 2 | Protopolystoma xenopodis | 2044.0 | 2 |
| scf7180000042001 | 17458 | 0.4089 | 0 | 68.5125 | Eukaryota | 86586.0 | 0 | Chordata | 86586.0 | 0 | Carnivora | 86586.0 | 0 | Canidae | 86586.0 | 0 | Canis | 86586.0 | 0 | Canis lupus | 86586.0 | 0 |
| scf7180000042005 | 40636 | 0.372 | 0 | 60.7897 | Eukaryota | 87981.0 | 0 | Chordata | 87981.0 | 0 | Carnivora | 87981.0 | 0 | Canidae | 87981.0 | 0 | Canis | 87981.0 | 0 | Canis lupus | 87981.0 | 0 |
| scf7180000040564 | 392325 | 0.4742 | 0 | 59.3628 | Eukaryota | 39711.0 | 0 | Chordata | 39711.0 | 0 | Carnivora | 39711.0 | 0 | Canidae | 22461.0 | 2 | Vulpes | 14582.0 | 5 | Vulpes vulpes | 14582.0 | 5 |
| scf7180000042385 | 145888 | 0.4024 | 0 | 62.4619 | Eukaryota | 90416.0 | 0 | Chordata | 90416.0 | 0 | Carnivora | 90416.0 | 0 | Canidae | 90416.0 | 0 | Canis | 90416.0 | 0 | Canis lupus | 90416.0 | 0 |
| scf7180000041395 | 220368 | 0.4655 | 0 | 57.4529 | Eukaryota | 51418.0 | 0 | Chordata | 51418.0 | 0 | Carnivora | 51418.0 | 0 | Canidae | 51418.0 | 0 | Canis | 51418.0 | 0 | Canis lupus | 51418.0 | 0 |
| scf7180000041378 | 1327045 | 0.4044 | 0 | 58.8558 | Eukaryota | 80409.0 | 0 | Chordata | 80409.0 | 0 | Carnivora | 80409.0 | 0 | Canidae | 80409.0 | 0 | Canis | 67139.0 | 1 | Canis lupus | 67139.0 | 1 |
| scf7180000045764 | 199774 | 0.4164 | 0 | 59.3265 | Eukaryota | 49954.0 | 0 | Chordata | 49954.0 | 0 | Carnivora | 49954.0 | 0 | Otariidae | 20042.0 | 3 | Zalophus | 20042.0 | 3 | Zalophus californianus | 20042.0 | 4 |
| scf7180000045442 | 22599 | 0.4823 | 0 | 187.9122 | Eukaryota | 15049.0 | 0 | Chordata | 15049.0 | 0 | Carnivora | 15049.0 | 0 | Canidae | 15049.0 | 0 | Canis | 9163.0 | 1 | Canis lupus | 9163.0 | 1 |
| scf7180000044543 | 19242 | 0.5166 | 0 | 152.7946 | Eukaryota | 22289.0 | 0 | Chordata | 22289.0 | 0 | Carnivora | 22289.0 | 0 | Canidae | 22289.0 | 0 | Canis | 17793.0 | 1 | Canis lupus | 17793.0 | 1 |
| scf7180000043243 | 333882 | 0.4068 | 0 | 58.9193 | Eukaryota | 94152.0 | 0 | Chordata | 94152.0 | 0 | Carnivora | 94152.0 | 0 | Canidae | 94152.0 | 0 | Canis | 94152.0 | 0 | Canis lupus | 94152.0 | 0 |
| scf7180000047910 | 61742 | 0.4523 | 0 | 58.465 | Eukaryota | 6310.0 | 0 | Chordata | 6310.0 | 0 | Carnivora | 6310.0 | 0 | Canidae | 6310.0 | 0 | Canis | 6310.0 | 0 | Canis lupus | 6310.0 | 0 |
| scf7180000042935 | 18971 | 0.3986 | 0 | 54.0366 | Eukaryota | 62019.0 | 0 | Chordata | 62019.0 | 0 | Carnivora | 62019.0 | 0 | Canidae | 62019.0 | 0 | Canis | 62019.0 | 0 | Canis lupus | 62019.0 | 0 |
| scf7180000043690 | 393163 | 0.3625 | 0 | 60.6914 | Eukaryota | 54565.0 | 0 | Chordata | 54565.0 | 0 | Carnivora | 54565.0 | 0 | Canidae | 54565.0 | 0 | Canis | 54565.0 | 0 | Canis lupus | 54565.0 | 0 |
| scf7180000042114 | 742717 | 0.4225 | 0 | 59.5737 | Eukaryota | 85082.0 | 0 | Chordata | 85082.0 | 0 | Carnivora | 85082.0 | 0 | Canidae | 85082.0 | 0 | Vulpes | 42560.0 | 1 | Vulpes vulpes | 42560.0 | 1 |
| scf7180000040557 | 2377810 | 0.3999 | 0 | 60.3436 | Eukaryota | 120449.0 | 0 | Chordata | 120449.0 | 0 | Carnivora | 120449.0 | 0 | Canidae | 120449.0 | 0 | Vulpes | 60555.0 | 1 | Vulpes vulpes | 60555.0 | 1 |
| scf7180000054746 | 22837 | 0.5901 | 0 | 637.6919 | Eukaryota | 33973.0 | 0 | Chordata | 33973.0 | 0 | Primates | 33973.0 | 0 | Hominidae | 23779.0 | 1 | Homo | 23779.0 | 1 | Homo sapiens | 23779.0 | 1 |
| scf7180000045402 | 30599 | 0.5207 | 0 | 714.9511 | Eukaryota | 16155.0 | 0 | Chordata | 16155.0 | 0 | Carnivora | 16155.0 | 0 | Canidae | 16155.0 | 0 | Canis | 16155.0 | 0 | Canis lupus | 16155.0 | 0 |
| scf7180000056570 | 112526 | 0.3917 | 0 | 58.1967 | Eukaryota | 59953.0 | 0 | Chordata | 59953.0 | 0 | Carnivora | 59953.0 | 0 | Canidae | 59953.0 | 0 | Canis | 42406.0 | 1 | Canis lupus | 42406.0 | 1 |
| scf7180000041632 | 320762 | 0.3925 | 0 | 59.9006 | Eukaryota | 85661.0 | 0 | Chordata | 85661.0 | 0 | Carnivora | 85661.0 | 0 | Canidae | 85661.0 | 0 | Canis | 85661.0 | 0 | Canis lupus | 85661.0 | 0 |
| scf7180000044087 | 67226 | 0.4435 | 0 | 133.9295 | Eukaryota | 7703.0 | 0 | Chordata | 4955.0 | 2 | Kurtiformes | 4594.0 | 3 | Apogonidae | 4594.0 | 3 | Sphaeramia | 4594.0 | 3 | Sphaeramia orbicularis | 4594.0 | 3 |
| scf7180000048449 | 2872 | 0.4495 | 0 | 139.8944 | no-hit | 0.0 | 0 | no-hit | 0.0 | 0 | no-hit | 0.0 | 0 | no-hit | 0.0 | 0 | no-hit | 0.0 | 0 | no-hit | 0.0 | 0 |
| scf7180000043125 | 6022 | 0.3892 | 0 | 21.8001 | Eukaryota | 24943.0 | 0 | Chordata | 24943.0 | 0 | Carnivora | 24943.0 | 0 | Canidae | 24943.0 | 0 | Canis | 24943.0 | 0 | Canis lupus | 24943.0 | 0 |
| scf7180000041697 | 21871 | 0.4287 | 0 | 690.5039 | Eukaryota | 51502.0 | 0 | Chordata | 51502.0 | 0 | Carnivora | 51502.0 | 0 | Canidae | 51502.0 | 0 | Canis | 51502.0 | 0 | Canis lupus | 51502.0 | 0 |
| scf7180000047336 | 6479 | 0.4448 | 0 | 263.4511 | no-hit | 0.0 | 0 | no-hit | 0.0 | 0 | no-hit | 0.0 | 0 | no-hit | 0.0 | 0 | no-hit | 0.0 | 0 | no-hit | 0.0 | 0 |
| scf7180000047594 | 90811 | 0.3778 | 0 | 56.5089 | Eukaryota | 32991.0 | 0 | Chordata | 32991.0 | 0 | Carnivora | 32991.0 | 0 | Canidae | 32991.0 | 0 | Canis | 32991.0 | 0 | Canis lupus | 32991.0 | 0 |
| scf7180000048596 | 64180 | 0.3969 | 0 | 60.4141 | Eukaryota | 5815.0 | 0 | Chordata | 5815.0 | 0 | Carnivora | 4107.0 | 1 | Canidae | 3696.0 | 2 | Canis | 3696.0 | 2 | Canis lupus | 3696.0 | 2 |
| scf7180000048684 | 52771 | 0.5671 | 0 | 55.3957 | Eukaryota | 24627.0 | 0 | Chordata | 24627.0 | 0 | Carnivora | 24627.0 | 0 | Canidae | 18134.0 | 1 | Canis | 16506.0 | 3 | Canis lupus | 16506.0 | 3 |
| scf7180000050909 | 18521 | 0.4427 | 0 | 106.3366 | Eukaryota | 3336.0 | 0 | Chordata | 3336.0 | 0 | Carnivora | 3336.0 | 0 | Canidae | 3336.0 | 0 | Canis | 3336.0 | 0 | Canis lupus | 3336.0 | 0 |
| scf7180000044663 | 35434 | 0.4368 | 0 | 161.5329 | Eukaryota | 38892.0 | 0 | Chordata | 26509.0 | 2 | Chordata-undef | 9107.0 | 7 | Ambassidae | 9107.0 | 7 | Parambassis | 9107.0 | 7 | Parambassis ranga | 9107.0 | 7 |
| scf7180000044477 | 13599 | 0.4656 | 0 | 92.9965 | Eukaryota | 37455.0 | 0 | Chordata | 37455.0 | 0 | Carnivora | 37455.0 | 0 | Canidae | 37455.0 | 0 | Canis | 37455.0 | 0 | Canis lupus | 37455.0 | 0 |
| scf7180000041017 | 1467534 | 0.4016 | 0 | 60.3529 | Eukaryota | 96993.0 | 0 | Chordata | 96993.0 | 0 | Carnivora | 96993.0 | 0 | Canidae | 96993.0 | 0 | Canis | 77376.0 | 1 | Canis lupus | 77376.0 | 1 |
| scf7180000049926 | 2903 | 0.4599 | 0 | 237.3156 | Eukaryota | 778.0 | 0 | Streptophyta | 391.0 | 1 | Fabales | 391.0 | 1 | Fabaceae | 391.0 | 1 | Cicer | 391.0 | 1 | Cicer arietinum | 391.0 | 1 |
| scf7180000049256 | 29156 | 0.4385 | 0 | 159.2626 | Eukaryota | 17459.0 | 0 | Platyhelminthes | 13759.0 | 1 | Cyclophyllidea | 12285.0 | 4 | Hymenolepididae | 12285.0 | 4 | Hymenolepis | 12285.0 | 4 | Hymenolepis microstoma | 12285.0 | 4 |
| scf7180000045173 | 233967 | 0.3966 | 0 | 59.9943 | Eukaryota | 47109.0 | 0 | Chordata | 47109.0 | 0 | Carnivora | 47109.0 | 0 | Canidae | 47109.0 | 0 | Canis | 42462.0 | 1 | Canis lupus | 42462.0 | 1 |
| scf7180000043449 | 70035 | 0.3691 | 0 | 55.1572 | Eukaryota | 73164.0 | 0 | Chordata | 73164.0 | 0 | Carnivora | 73164.0 | 0 | Canidae | 73164.0 | 0 | Canis | 73164.0 | 0 | Canis lupus | 73164.0 | 0 |
| scf7180000049572 | 6093 | 0.4353 | 0 | 98.0142 | Eukaryota | 9832.0 | 0 | Chordata | 6743.0 | 2 | Gadiformes | 3892.0 | 6 | Gadidae | 3892.0 | 6 | Gadus | 3892.0 | 6 | Gadus morhua | 3892.0 | 7 |
| scf7180000050402 | 32646 | 0.5036 | 0 | 451.3961 | Eukaryota | 9293.0 | 0 | Chordata | 9293.0 | 0 | Carnivora | 9293.0 | 0 | Canidae | 9293.0 | 0 | Canis | 8086.0 | 1 | Canis lupus | 6813.0 | 2 |
| scf7180000047029 | 30511 | 0.4447 | 0 | 208.8797 | Eukaryota | 2010.0 | 0 | Platyhelminthes | 1010.0 | 1 | Polystomatidea | 1010.0 | 3 | Polystomatidae | 1010.0 | 3 | Protopolystoma | 1010.0 | 4 | Protopolystoma xenopodis | 1010.0 | 4 |
| scf7180000042609 | 661478 | 0.4102 | 0 | 59.5897 | Eukaryota | 63126.0 | 0 | Chordata | 63126.0 | 0 | Carnivora | 58352.0 | 1 | Canidae | 58352.0 | 1 | Canis | 44446.0 | 2 | Canis lupus | 44446.0 | 2 |
| scf7180000053308 | 8373 | 0.4012 | 0 | 57.6501 | Eukaryota | 4539.0 | 1 | Chordata | 4539.0 | 1 | undef | 2372.0 | 3 | undef | 2372.0 | 3 | undef | 2372.0 | 4 | eukaryotic synthetic construct | 2372.0 | 4 |
| scf7180000052531 | 5151 | 0.4514 | 0 | 703.5889 | Eukaryota | 16096.0 | 0 | Chordata | 15360.0 | 1 | Gadiformes | 7258.0 | 6 | Gadidae | 7258.0 | 6 | Gadus | 7258.0 | 6 | Gadus morhua | 7258.0 | 6 |
| scf7180000041306 | 640063 | 0.3517 | 0 | 59.0017 | Eukaryota | 56611.0 | 0 | Chordata | 56611.0 | 0 | Carnivora | 56611.0 | 0 | Canidae | 56611.0 | 0 | Canis | 56611.0 | 0 | Canis lupus | 56611.0 | 0 |
| scf7180000043704 | 419661 | 0.5812 | 0 | 58.1865 | Eukaryota | 38645.0 | 0 | Chordata | 38645.0 | 0 | Carnivora | 38645.0 | 0 | Canidae | 38645.0 | 0 | Canis | 38645.0 | 0 | Canis lupus | 38645.0 | 0 |
| scf7180000047332 | 19478 | 0.566 | 0 | 72.5626 | Eukaryota | 83122.0 | 0 | Chordata | 83122.0 | 0 | Carnivora | 83122.0 | 0 | Canidae | 83122.0 | 0 | Canis | 83122.0 | 0 | Canis lupus | 83122.0 | 0 |
| scf7180000043451 | 212603 | 0.3625 | 0 | 57.6254 | Eukaryota | 8120.0 | 0 | Chordata | 7536.0 | 1 | Carnivora | 5677.0 | 2 | Canidae | 5677.0 | 2 | Canis | 5677.0 | 2 | Canis lupus | 5677.0 | 2 |
| scf7180000052991 | 5719 | 0.4371 | 0 | 236.2619 | Viruses | 3172.0 | 1 | Viruses-undef | 3172.0 | 1 | Herpesvirales | 3172.0 | 2 | Herpesviridae | 3172.0 | 2 | Roseolovirus | 3172.0 | 2 | Human betaherpesvirus 7 | 3172.0 | 2 |
| scf7180000055936 | 8872 | 0.4496 | 0 | 439.9391 | Eukaryota | 4340.0 | 0 | Chordata | 2514.0 | 1 | Kurtiformes | 1911.0 | 2 | Apogonidae | 1911.0 | 2 | Sphaeramia | 1911.0 | 3 | Sphaeramia orbicularis | 1911.0 | 3 |
| scf7180000049170 | 14560 | 0.4373 | 0 | 112.1654 | Eukaryota | 710.0 | 0 | Platyhelminthes | 710.0 | 0 | Polystomatidea | 710.0 | 0 | Polystomatidae | 710.0 | 0 | Protopolystoma | 710.0 | 0 | Protopolystoma xenopodis | 710.0 | 0 |
| scf7180000052623 | 7085 | 0.3884 | 0 | 54.9818 | Eukaryota | 3409.0 | 0 | Chordata | 3409.0 | 0 | Carnivora | 3409.0 | 0 | Canidae | 2485.0 | 2 | Canis | 2485.0 | 2 | Canis lupus | 2485.0 | 2 |
| scf7180000051304 | 13735 | 0.4401 | 0 | 96.509 | Eukaryota | 7542.0 | 0 | Chordata | 5138.0 | 1 | Synbranchiformes | 3033.0 | 4 | Mastacembelidae | 3033.0 | 4 | Mastacembelus | 3033.0 | 4 | Mastacembelus armatus | 3033.0 | 4 |
| scf7180000043253 | 823777 | 0.426 | 0 | 59.8006 | Eukaryota | 64124.0 | 0 | Chordata | 64124.0 | 0 | Carnivora | 64124.0 | 0 | Canidae | 64124.0 | 0 | Canis | 57656.0 | 1 | Canis lupus | 57656.0 | 1 |
| scf7180000041812 | 986350 | 0.3701 | 0 | 60.2992 | Eukaryota | 152145.0 | 0 | Chordata | 152145.0 | 0 | Carnivora | 152145.0 | 0 | Canidae | 152145.0 | 0 | Canis | 139081.0 | 1 | Canis lupus | 139081.0 | 1 |
| scf7180000050437 | 24998 | 0.4703 | 0 | 1234.7447 | Eukaryota | 90454.0 | 0 | Chordata | 90454.0 | 0 | Carnivora | 90454.0 | 0 | Canidae | 90454.0 | 0 | Canis | 90454.0 | 0 | Canis lupus | 90454.0 | 0 |
| scf7180000043330 | 457821 | 0.4877 | 0 | 59.2969 | Eukaryota | 63320.0 | 0 | Chordata | 63320.0 | 0 | Carnivora | 63320.0 | 0 | Canidae | 63320.0 | 0 | Canis | 63320.0 | 0 | Canis lupus | 63320.0 | 0 |
| scf7180000043574 | 361076 | 0.4391 | 0 | 59.6073 | Eukaryota | 85962.0 | 0 | Chordata | 85962.0 | 0 | Carnivora | 85962.0 | 0 | Canidae | 85962.0 | 0 | Canis | 85962.0 | 0 | Canis lupus | 85962.0 | 0 |
| scf7180000048755 | 3159 | 0.4425 | 0 | 69.1114 | no-hit | 0.0 | 0 | no-hit | 0.0 | 0 | no-hit | 0.0 | 0 | no-hit | 0.0 | 0 | no-hit | 0.0 | 0 | no-hit | 0.0 | 0 |
| scf7180000047004 | 54100 | 0.4365 | 0 | 100.2609 | Eukaryota | 81059.0 | 0 | Chordata | 81059.0 | 0 | Carnivora | 81059.0 | 0 | Canidae | 81059.0 | 0 | Canis | 81059.0 | 0 | Canis lupus | 81059.0 | 0 |
| scf7180000047834 | 5544 | 0.4531 | 0 | 154.7743 | no-hit | 0.0 | 0 | no-hit | 0.0 | 0 | no-hit | 0.0 | 0 | no-hit | 0.0 | 0 | no-hit | 0.0 | 0 | no-hit | 0.0 | 0 |
| scf7180000043272 | 113158 | 0.3726 | 0 | 57.7717 | Eukaryota | 28458.0 | 0 | Chordata | 28458.0 | 0 | Carnivora | 28458.0 | 0 | Canidae | 28458.0 | 0 | Canis | 28458.0 | 0 | Canis lupus | 28458.0 | 0 |
| scf7180000047914 | 7020 | 0.4538 | 0 | 194.0515 | Eukaryota | 2874.0 | 0 | Chordata | 2874.0 | 0 | Carnivora | 2874.0 | 0 | Canidae | 2874.0 | 0 | Canis | 2874.0 | 0 | Canis lupus | 2874.0 | 0 |
| scf7180000049760 | 39678 | 0.3565 | 0 | 58.916 | Eukaryota | 7235.0 | 0 | Chordata | 7235.0 | 0 | Carnivora | 5475.0 | 1 | Canidae | 3456.0 | 2 | Canis | 3456.0 | 2 | Canis lupus | 3456.0 | 2 |
| scf7180000048021 | 10643 | 0.439 | 0 | 169.4119 | Eukaryota | 756.0 | 0 | Platyhelminthes | 756.0 | 0 | Polystomatidea | 756.0 | 0 | Polystomatidae | 756.0 | 0 | Protopolystoma | 756.0 | 0 | Protopolystoma xenopodis | 756.0 | 0 |
| scf7180000041455 | 467744 | 0.3415 | 0 | 58.7071 | Eukaryota | 71716.0 | 0 | Chordata | 71716.0 | 0 | Carnivora | 71716.0 | 0 | Canidae | 71716.0 | 0 | Canis | 71716.0 | 0 | Canis lupus | 71716.0 | 0 |
| scf7180000043222 | 310658 | 0.3976 | 0 | 60.3822 | Eukaryota | 49066.0 | 0 | Chordata | 49066.0 | 0 | Carnivora | 49066.0 | 0 | Canidae | 49066.0 | 0 | Canis | 49066.0 | 0 | Canis lupus | 49066.0 | 0 |
| scf7180000049422 | 9312 | 0.4381 | 0 | 50.747 | Eukaryota | 12907.0 | 1 | Platyhelminthes | 6840.0 | 4 | Strigeidida | 6840.0 | 7 | Schistosomatidae | 6840.0 | 7 | Schistosoma | 6840.0 | 7 | Schistosoma mansoni | 6840.0 | 7 |
| scf7180000043192 | 534511 | 0.3848 | 0 | 61.0918 | Eukaryota | 47320.0 | 0 | Chordata | 47320.0 | 0 | Carnivora | 47320.0 | 0 | Canidae | 47320.0 | 0 | Canis | 47320.0 | 0 | Canis lupus | 47320.0 | 0 |
| scf7180000051575 | 3912 | 0.4489 | 0 | 747.7942 | Eukaryota | 673.0 | 0 | Chordata | 673.0 | 0 | Salmoniformes | 460.0 | 1 | Salmonidae | 460.0 | 1 | Salmo | 460.0 | 1 | Salmo trutta | 460.0 | 1 |
| scf7180000052083 | 3906 | 0.4598 | 0 | 398.2881 | Eukaryota | 326.0 | 0 | Streptophyta | 326.0 | 0 | Fabales | 326.0 | 0 | Fabaceae | 326.0 | 0 | Cicer | 326.0 | 0 | Cicer arietinum | 326.0 | 0 |
| scf7180000046743 | 131882 | 0.4097 | 0 | 58.6577 | Eukaryota | 17050.0 | 0 | Chordata | 17050.0 | 0 | Carnivora | 17050.0 | 0 | Canidae | 17050.0 | 0 | Canis | 17050.0 | 0 | Canis lupus | 17050.0 | 0 |
| scf7180000043024 | 477448 | 0.4033 | 0 | 58.9568 | Eukaryota | 63349.0 | 0 | Chordata | 63349.0 | 0 | Carnivora | 63349.0 | 0 | Canidae | 63349.0 | 0 | Canis | 63349.0 | 0 | Canis lupus | 63349.0 | 0 |
| scf7180000046112 | 169160 | 0.3739 | 0 | 61.2388 | Eukaryota | 35312.0 | 1 | Chordata | 35312.0 | 1 | Primates | 16598.0 | 3 | Hominidae | 16598.0 | 4 | Homo | 10222.0 | 6 | Homo sapiens | 10222.0 | 6 |
| scf7180000053470 | 6301 | 0.4899 | 0 | 266.392 | Eukaryota | 33191.0 | 0 | Chordata | 33191.0 | 0 | Carnivora | 33191.0 | 0 | Canidae | 33191.0 | 0 | Canis | 33191.0 | 0 | Canis lupus | 33191.0 | 0 |
| scf7180000053028 | 3303 | 0.4435 | 0 | 125.1457 | Eukaryota | 592.0 | 0 | Platyhelminthes | 592.0 | 0 | Polystomatidea | 592.0 | 0 | Polystomatidae | 592.0 | 0 | Protopolystoma | 592.0 | 0 | Protopolystoma xenopodis | 592.0 | 0 |
| scf7180000044088 | 411373 | 0.4333 | 0 | 59.7908 | Eukaryota | 142615.0 | 0 | Chordata | 142615.0 | 0 | Carnivora | 133233.0 | 1 | Canidae | 113337.0 | 2 | Canis | 113337.0 | 2 | Canis lupus | 113337.0 | 2 |
| scf7180000044596 | 297972 | 0.3679 | 0 | 60.2662 | Eukaryota | 93383.0 | 0 | Chordata | 93383.0 | 0 | Carnivora | 93383.0 | 0 | Canidae | 93383.0 | 0 | Canis | 93383.0 | 0 | Canis lupus | 93383.0 | 0 |
| scf7180000053977 | 7014 | 0.5607 | 0 | 58.8774 | Eukaryota | 3420.0 | 0 | Chordata | 3420.0 | 0 | Carnivora | 3420.0 | 0 | Canidae | 2926.0 | 1 | Canis | 2274.0 | 2 | Canis lupus | 2274.0 | 2 |
| scf7180000054625 | 13009 | 0.4571 | 0 | 257.4276 | Eukaryota | 75483.0 | 0 | Chordata | 75483.0 | 0 | Carnivora | 75483.0 | 0 | Canidae | 75483.0 | 0 | Canis | 75483.0 | 0 | Canis lupus | 75483.0 | 0 |
| scf7180000042344 | 677799 | 0.4171 | 0 | 60.4088 | Eukaryota | 50955.0 | 0 | Chordata | 50955.0 | 0 | Carnivora | 50955.0 | 0 | Canidae | 50955.0 | 0 | Canis | 44787.0 | 1 | Canis lupus | 44787.0 | 1 |
| scf7180000048064 | 23772 | 0.4464 | 0 | 184.2226 | Eukaryota | 135.0 | 0 | Chordata | 135.0 | 0 | Salmoniformes | 135.0 | 0 | Salmonidae | 135.0 | 0 | Salmo | 135.0 | 0 | Salmo trutta | 135.0 | 0 |
| scf7180000044997 | 239250 | 0.3682 | 0 | 60.407 | Eukaryota | 41879.0 | 0 | Chordata | 41879.0 | 0 | Carnivora | 41879.0 | 0 | Canidae | 41879.0 | 0 | Canis | 41879.0 | 0 | Canis lupus | 41879.0 | 0 |
| scf7180000041704 | 989070 | 0.3438 | 0 | 60.1474 | Eukaryota | 96799.0 | 0 | Chordata | 96799.0 | 0 | Carnivora | 96799.0 | 0 | Canidae | 96799.0 | 0 | Canis | 96799.0 | 0 | Canis lupus | 96799.0 | 0 |
| scf7180000045642 | 153081 | 0.4039 | 0 | 58.4535 | Eukaryota | 39368.0 | 0 | Chordata | 39368.0 | 0 | Carnivora | 39368.0 | 0 | Canidae | 39368.0 | 0 | Canis | 39368.0 | 0 | Canis lupus | 39368.0 | 0 |
| scf7180000052791 | 20923 | 0.5038 | 0 | 493.6282 | Eukaryota | 68459.0 | 0 | Chordata | 68459.0 | 0 | Carnivora | 68459.0 | 0 | Canidae | 68459.0 | 0 | Canis | 68459.0 | 0 | Canis lupus | 68459.0 | 0 |
| scf7180000042471 | 8327 | 0.4013 | 0 | 27.5937 | Eukaryota | 11294.0 | 0 | Chordata | 11294.0 | 0 | Carnivora | 11294.0 | 0 | Canidae | 3658.0 | 4 | Canis | 2433.0 | 8 | Canis lupus | 2433.0 | 8 |
| scf7180000041847 | 437957 | 0.3828 | 0 | 58.638 | Eukaryota | 86120.0 | 0 | Chordata | 86120.0 | 0 | Carnivora | 86120.0 | 0 | Canidae | 86120.0 | 0 | Canis | 86120.0 | 0 | Canis lupus | 86120.0 | 0 |
| scf7180000044086 | 313684 | 0.4873 | 0 | 58.6827 | Eukaryota | 30953.0 | 0 | Chordata | 30953.0 | 0 | Carnivora | 30953.0 | 0 | Canidae | 10867.0 | 4 | Canis | 7993.0 | 7 | Canis lupus | 7993.0 | 8 |
| scf7180000054448 | 353707 | 0.432 | 0 | 66.7438 | Eukaryota | 70019.0 | 0 | Chordata | 70019.0 | 0 | Carnivora | 70019.0 | 0 | Canidae | 70019.0 | 0 | Canis | 54954.0 | 1 | Canis lupus | 54954.0 | 1 |

| | | | | | | | | | | | | | | | | | | | | | | |
|---|---|---|---|---|---|---|---|---|---|---|---|---|---|---|---|---|---|---|---|---|---|---|
| scf7180000044219 | 364871 | 0.5792 | 0 | 58.2039 | Eukaryota | 48240.0 | 0 | Chordata | 48240.0 | 0 | Carnivora | 48240.0 | 0 | Canidae | 48240.0 | 0 | Canis | 42238.0 | 1 | Canis lupus | 42238.0 | 1 |
| scf7180000043400 | 507149 | 0.438 | 0 | 58.2453 | Eukaryota | 46009.0 | 0 | Chordata | 46009.0 | 0 | Carnivora | 46009.0 | 0 | Canidae | 46009.0 | 0 | Canis | 46009.0 | 0 | Canis lupus | 46009.0 | 0 |
| scf7180000056188 | 202020 | 0.3621 | 0 | 57.4716 | Eukaryota | 10839.0 | 0 | Chordata | 10839.0 | 0 | Carnivora | 8002.0 | 2 | Canidae | 8002.0 | 2 | Canis | 8002.0 | 2 | Canis lupus | 8002.0 | 2 |
| scf7180000047138 | 99348 | 0.3761 | 0 | 55.6566 | Eukaryota | 35027.0 | 0 | Chordata | 35027.0 | 0 | Carnivora | 35027.0 | 0 | Canidae | 35027.0 | 0 | Canis | 35027.0 | 0 | Canis lupus | 35027.0 | 0 |
| scf7180000046285 | 140444 | 0.5096 | 0 | 57.6275 | Eukaryota | 126556.0 | 0 | Chordata | 126556.0 | 0 | Carnivora | 126556.0 | 0 | Canidae | 118868.0 | 1 | Canis | 92879.0 | 2 | Canis lupus | 92879.0 | 2 |
| scf7180000051676 | 9370 | 0.4442 | 0 | 137.9683 | no-hit | 0.0 | 0 | no-hit | 0.0 | 0 | no-hit | 0.0 | 0 | no-hit | 0.0 | 0 | no-hit | 0.0 | 0 | no-hit | 0.0 | 0 |
| scf7180000052881 | 19902 | 0.5391 | 0 | 305.9346 | Eukaryota | 25441.0 | 0 | Chordata | 25441.0 | 0 | Artiodactyla | 17847.0 | 1 | Bovidae | 17847.0 | 1 | Ovis | 10188.0 | 3 | Ovis aries | 10188.0 | 3 |
| scf7180000042472 | 190096 | 0.3828 | 0 | 59.8754 | Eukaryota | 84794.0 | 0 | Chordata | 84794.0 | 0 | Carnivora | 84794.0 | 0 | Canidae | 84794.0 | 0 | Canis | 55535.0 | 1 | Canis lupus | 55535.0 | 1 |
| scf7180000048416 | 19083 | 0.4325 | 0 | 108.0458 | Eukaryota | 22662.0 | 0 | Chordata | 21028.0 | 2 | Gadiformes | 18666.0 | 6 | Gadidae | 18666.0 | 6 | Gadus | 18666.0 | 6 | Gadus morhua | 18666.0 | 6 |
| scf7180000049176 | 27387 | 0.4437 | 0 | 397.3296 | Eukaryota | 41435.0 | 0 | Chordata | 23388.0 | 2 | Cyclophyllidea | 11834.0 | 9 | Hymenolepididae | 11834.0 | 9 | Hymenolepis | 11834.0 | 9 | Hymenolepis microstoma | 11834.0 | 9 |
| scf7180000056296 | 267381 | 0.4227 | 0 | 59.888 | Eukaryota | 81280.0 | 0 | Chordata | 81280.0 | 0 | Carnivora | 81280.0 | 0 | Canidae | 57556.0 | 2 | Canis | 50834.0 | 3 | Canis lupus | 50834.0 | 3 |
| scf7180000051808 | 15303 | 0.4411 | 0 | 146.6458 | Eukaryota | 13153.0 | 0 | Chordata | 7193.0 | 2 | Gadiformes | 4194.0 | 5 | Gadidae | 4194.0 | 5 | Gadus | 4194.0 | 5 | Gadus morhua | 4194.0 | 5 |
| scf7180000047514 | 55094 | 0.4376 | 0 | 132.7587 | Eukaryota | 43488.0 | 0 | Chordata | 20147.0 | 2 | Cyclophyllidea | 11635.0 | 7 | Hymenolepididae | 11635.0 | 7 | Hymenolepis | 11635.0 | 7 | Hymenolepis microstoma | 11635.0 | 7 |
| scf7180000055876 | 13019 | 0.4386 | 0 | 143.031 | Eukaryota | 929.0 | 0 | Platyhelminthes | 929.0 | 0 | Polystomatidea | 929.0 | 0 | Polystomatidae | 929.0 | 0 | Protopolystoma | 929.0 | 0 | Protopolystoma xenopodis | 929.0 | 0 |
| scf7180000050204 | 10105 | 0.4488 | 0 | 224.2433 | Eukaryota | 15200.0 | 0 | Chordata | 9983.0 | 3 | Gadiformes | 4604.0 | 6 | Gadidae | 4604.0 | 6 | Gadus | 4604.0 | 6 | Gadus morhua | 4604.0 | 6 |
| scf7180000047325 | 26466 | 0.4384 | 0 | 209.8577 | Eukaryota | 15496.0 | 0 | Chordata | 9373.0 | 3 | Gadiformes | 4416.0 | 7 | Gadidae | 4416.0 | 7 | Gadus | 4416.0 | 7 | Gadus morhua | 4416.0 | 7 |
| scf7180000050798 | 31133 | 0.45 | 0 | 70.8698 | Eukaryota | 455.0 | 0 | Platyhelminthes | 455.0 | 0 | Polystomatidea | 455.0 | 0 | Polystomatidae | 455.0 | 0 | Protopolystoma | 455.0 | 0 | Protopolystoma xenopodis | 455.0 | 0 |
| scf7180000053386 | 3589 | 0.4547 | 0 | 358.9326 | Eukaryota | 439.0 | 0 | Platyhelminthes | 265.0 | 1 | Polystomatidea | 265.0 | 1 | Polystomatidae | 265.0 | 1 | Protopolystoma | 265.0 | 1 | Protopolystoma xenopodis | 265.0 | 1 |
| scf7180000053862 | 3590 | 0.5072 | 0 | 41.4972 | Eukaryota | 6937.0 | 0 | Chordata | 6937.0 | 0 | Carnivora | 6937.0 | 0 | Canidae | 6937.0 | 0 | Canis | 4994.0 | 1 | Canis lupus | 3933.0 | 2 |
| scf7180000052436 | 3556 | 0.4409 | 0 | 336.1027 | no-hit | 0.0 | 0 | no-hit | 0.0 | 0 | no-hit | 0.0 | 0 | no-hit | 0.0 | 0 | no-hit | 0.0 | 0 | no-hit | 0.0 | 0 |
| scf7180000041564 | 15093 | 0.364 | 0 | 50.0505 | Eukaryota | 7535.0 | 0 | Chordata | 7535.0 | 0 | Carnivora | 6609.0 | 1 | Felidae | 4590.0 | 2 | Felis | 4590.0 | 2 | Felis catus | 4590.0 | 2 |
| scf7180000047215 | 62626 | 0.4002 | 0 | 229.3746 | Eukaryota | 15490.0 | 0 | Chordata | 15490.0 | 0 | Carnivora | 15490.0 | 0 | Ursidae | 9411.0 | 3 | Ursus | 9411.0 | 5 | Ursus arctos | 9411.0 | 5 |
| scf7180000041277 | 756316 | 0.3534 | 0 | 59.5458 | Eukaryota | 92076.0 | 0 | Chordata | 92076.0 | 0 | Carnivora | 92076.0 | 0 | Canidae | 92076.0 | 0 | Canis | 92076.0 | 0 | Canis lupus | 92076.0 | 0 |
| scf7180000041136 | 1404443 | 0.4764 | 0 | 59.9198 | Eukaryota | 89393.0 | 0 | Chordata | 89393.0 | 0 | Carnivora | 89393.0 | 0 | Canidae | 89393.0 | 0 | Vulpes | 53641.0 | 1 | Vulpes vulpes | 53641.0 | 1 |
| scf7180000055662 | 21262 | 0.4344 | 0 | 312.1772 | Eukaryota | 174.0 | 0 | Chordata | 174.0 | 0 | Kurtiformes | 174.0 | 0 | Apogonidae | 174.0 | 0 | Sphaeramia | 174.0 | 0 | Sphaeramia orbicularis | 174.0 | 0 |
| scf7180000055103 | 60727 | 0.436 | 0 | 107.514 | Eukaryota | 62681.0 | 0 | Chordata | 62681.0 | 0 | Carnivora | 62681.0 | 0 | Canidae | 62681.0 | 0 | Canis | 62681.0 | 0 | Canis lupus | 62681.0 | 0 |
| scf7180000042691 | 2693 | 0.4768 | 0 | 17.1031 | Eukaryota | 19152.0 | 0 | Chordata | 19152.0 | 0 | Carnivora | 19152.0 | 0 | Canidae | 19152.0 | 0 | Canis | 19152.0 | 0 | Canis lupus | 19152.0 | 0 |
| scf7180000040782 | 1361314 | 0.3443 | 0 | 59.3435 | Eukaryota | 94879.0 | 0 | Chordata | 94879.0 | 0 | Carnivora | 94879.0 | 0 | Canidae | 94879.0 | 0 | Canis | 94879.0 | 0 | Canis lupus | 94879.0 | 0 |
| scf7180000045168 | 43755 | 0.4403 | 0 | 57.1046 | Eukaryota | 48442.0 | 0 | Chordata | 48442.0 | 0 | Carnivora | 48442.0 | 0 | Canidae | 48442.0 | 0 | Canis | 48442.0 | 0 | Canis lupus | 48442.0 | 0 |
| scf7180000043671 | 447742 | 0.4231 | 0 | 58.7574 | Eukaryota | 51448.0 | 0 | Chordata | 51448.0 | 0 | Carnivora | 51448.0 | 0 | Canidae | 51448.0 | 0 | Canis | 39304.0 | 1 | Canis lupus | 39304.0 | 1 |
| scf7180000056539 | 172716 | 0.3538 | 0 | 57.7994 | Eukaryota | 8452.0 | 0 | Chordata | 8452.0 | 0 | Carnivora | 6076.0 | 2 | Canidae | 6076.0 | 2 | Canis | 6076.0 | 2 | Canis lupus | 6076.0 | 2 |
| scf7180000043677 | 114562 | 0.4081 | 0 | 90.4171 | Eukaryota | 35262.0 | 0 | Chordata | 35262.0 | 0 | Carnivora | 35262.0 | 0 | Canidae | 35262.0 | 0 | Canis | 24159.0 | 1 | Canis lupus | 24159.0 | 1 |
| scf7180000045966 | 153529 | 0.3627 | 0 | 61.7397 | Eukaryota | 76178.0 | 0 | Chordata | 76178.0 | 0 | Carnivora | 76178.0 | 0 | Canidae | 76178.0 | 0 | Canis | 76178.0 | 0 | Canis lupus | 76178.0 | 0 |
| scf7180000040773 | 15637 | 0.4539 | 0 | 44.4319 | Eukaryota | 3820.0 | 0 | Chordata | 3820.0 | 0 | Carnivora | 2292.0 | 1 | Canidae | 2292.0 | 1 | Canis | 2292.0 | 1 | Canis lupus | 2292.0 | 1 |
| scf7180000041565 | 108256 | 0.4164 | 0 | 57.2027 | Eukaryota | 45252.0 | 0 | Chordata | 45252.0 | 0 | Carnivora | 45252.0 | 0 | Canidae | 45252.0 | 0 | Canis | 45252.0 | 0 | Canis lupus | 45252.0 | 0 |
| scf7180000055698 | 19085 | 0.4431 | 0 | 173.6991 | Eukaryota | 28830.0 | 0 | Chordata | 26776.0 | 1 | Gadiformes | 15909.0 | 6 | Gadidae | 15909.0 | 6 | Gadus | 15909.0 | 6 | Gadus morhua | 15909.0 | 6 |
| scf7180000047972 | 48436 | 0.3886 | 0 | 55.6017 | Eukaryota | 12598.0 | 0 | Chordata | 12598.0 | 0 | Carnivora | 12598.0 | 0 | Canidae | 12598.0 | 0 | Canis | 12598.0 | 0 | Canis lupus | 12598.0 | 0 |
| scf7180000053555 | 3972 | 0.4484 | 0 | 392.6855 | no-hit | 0.0 | 0 | no-hit | 0.0 | 0 | no-hit | 0.0 | 0 | no-hit | 0.0 | 0 | no-hit | 0.0 | 0 | no-hit | 0.0 | 0 |
| scf7180000047798 | 100660 | 0.3532 | 0 | 60.4362 | Eukaryota | 9236.0 | 0 | Chordata | 9236.0 | 0 | Primates | 6201.0 | 2 | Hominidae | 6201.0 | 2 | Homo | 6201.0 | 2 | Homo sapiens | 6201.0 | 2 |
| scf7180000042438 | 633077 | 0.4554 | 0 | 58.7102 | Eukaryota | 28975.0 | 0 | Chordata | 28975.0 | 0 | Carnivora | 28975.0 | 0 | Canidae | 28975.0 | 0 | Canis | 28975.0 | 0 | Canis lupus | 28975.0 | 0 |
| scf7180000053642 | 3742 | 0.43 | 0 | 63.768 | Eukaryota | 586.0 | 0 | Platyhelminthes | 586.0 | 0 | Polystomatidea | 586.0 | 0 | Polystomatidae | 586.0 | 0 | Protopolystoma | 586.0 | 0 | Protopolystoma xenopodis | 586.0 | 0 |
| scf7180000045821 | 150563 | 0.4327 | 0 | 57.5697 | Eukaryota | 62853.0 | 0 | Chordata | 62853.0 | 0 | Carnivora | 62853.0 | 0 | Canidae | 62853.0 | 0 | Canis | 62853.0 | 0 | Canis lupus | 62853.0 | 0 |
| scf7180000056308 | 81274 | 0.3864 | 0 | 58.5278 | Eukaryota | 11751.0 | 0 | Chordata | 11751.0 | 0 | Carnivora | 11751.0 | 0 | Canidae | 7584.0 | 1 | Canis | 7584.0 | 1 | Canis lupus | 7584.0 | 1 |
| scf7180000047557 | 46565 | 0.4416 | 0 | 132.2884 | Eukaryota | 5366.0 | 0 | Chordata | 5366.0 | 0 | Kurtiformes | 2734.0 | 2 | Apogonidae | 2734.0 | 2 | Sphaeramia | 2734.0 | 2 | Sphaeramia orbicularis | 2734.0 | 2 |
| scf7180000053672 | 2955 | 0.443 | 0 | 1002.3016 | no-hit | 0.0 | 0 | no-hit | 0.0 | 0 | no-hit | 0.0 | 0 | no-hit | 0.0 | 0 | no-hit | 0.0 | 0 | no-hit | 0.0 | 0 |
| scf7180000042403 | 775626 | 0.3837 | 0 | 59.465 | Eukaryota | 44426.0 | 0 | Chordata | 44426.0 | 0 | Carnivora | 44426.0 | 0 | Phocidae | 31542.0 | 2 | Leptonychotes | 31542.0 | 2 | Leptonychotes weddellii | 31542.0 | 2 |
| scf7180000054375 | 23225 | 0.4437 | 0 | 344.2104 | Eukaryota | 25371.0 | 0 | Chordata | 17019.0 | 1 | Strigeidida | 8352.0 | 5 | Schistosomatidae | 8352.0 | 6 | Schistosoma | 8352.0 | 6 | Schistosoma mansoni | 8352.0 | 6 |
| scf7180000052442 | 8243 | 0.4477 | 0 | 65.0624 | no-hit | 0.0 | 0 | no-hit | 0.0 | 0 | no-hit | 0.0 | 0 | no-hit | 0.0 | 0 | no-hit | 0.0 | 0 | no-hit | 0.0 | 0 |
| scf7180000053527 | 6967 | 0.4358 | 0 | 55.4835 | Eukaryota | 7872.0 | 0 | Platyhelminthes | 4504.0 | 1 | Cyclophyllidea | 3110.0 | 4 | Hymenolepididae | 3110.0 | 4 | Hymenolepis | 3110.0 | 4 | Hymenolepis microstoma | 3110.0 | 4 |
| scf7180000042220 | 1378765 | 0.3887 | 0 | 59.9492 | Eukaryota | 87774.0 | 0 | Chordata | 87774.0 | 0 | Carnivora | 87774.0 | 0 | Canidae | 87774.0 | 0 | Canis | 87774.0 | 0 | Canis lupus | 87774.0 | 0 |
| scf7180000053874 | 4281 | 0.4506 | 0 | 468.7389 | no-hit | 0.0 | 0 | no-hit | 0.0 | 0 | no-hit | 0.0 | 0 | no-hit | 0.0 | 0 | no-hit | 0.0 | 0 | no-hit | 0.0 | 0 |
| scf7180000042791 | 629677 | 0.4746 | 0 | 59.8494 | Eukaryota | 57293.0 | 0 | Chordata | 57293.0 | 0 | Carnivora | 57293.0 | 0 | Canidae | 57293.0 | 0 | Canis | 57293.0 | 0 | Canis lupus | 57293.0 | 0 |
| scf7180000046637 | 136845 | 0.5133 | 0 | 60.6358 | Eukaryota | 212371.0 | 0 | Chordata | 212371.0 | 0 | Carnivora | 212371.0 | 0 | Canidae | 212371.0 | 0 | Canis | 145470.0 | 1 | Canis lupus | 145470.0 | 1 |
| scf7180000044172 | 337159 | 0.4044 | 0 | 60.1511 | Eukaryota | 70528.0 | 0 | Chordata | 70528.0 | 0 | Carnivora | 70528.0 | 0 | Canidae | 70528.0 | 0 | Canis | 63953.0 | 1 | Canis lupus | 63953.0 | 1 |
| scf7180000046455 | 6921 | 0.3945 | 0 | 47.2799 | Eukaryota | 10623.0 | 0 | Chordata | 10623.0 | 0 | Carnivora | 10623.0 | 0 | Canidae | 10623.0 | 0 | Canis | 10623.0 | 0 | Canis lupus | 10623.0 | 0 |
| scf7180000044415 | 306214 | 0.3473 | 0 | 60.3726 | Eukaryota | 66641.0 | 0 | Chordata | 66641.0 | 0 | Carnivora | 66641.0 | 0 | Canidae | 66641.0 | 0 | Canis | 66641.0 | 0 | Canis lupus | 66641.0 | 0 |
| scf7180000051593 | 23942 | 0.3506 | 0 | 58.4844 | Eukaryota | 7312.0 | 0 | Chordata | 7312.0 | 0 | Carnivora | 7312.0 | 0 | Canidae | 4265.0 | 2 | Canis | 4265.0 | 2 | Canis lupus | 4265.0 | 2 |
| scf7180000054136 | 22413 | 0.3784 | 0 | 327.8238 | Eukaryota | 57196.0 | 0 | Chordata | 57196.0 | 0 | Carnivora | 57196.0 | 0 | Canidae | 57196.0 | 0 | Canis | 57196.0 | 0 | Canis lupus | 57196.0 | 0 |
| scf7180000056178 | 2346210 | 0.4007 | 0 | 59.8181 | Eukaryota | 54640.0 | 0 | Chordata | 54640.0 | 0 | Carnivora | 54640.0 | 0 | Canidae | 54640.0 | 0 | Canis | 50375.0 | 1 | Canis lupus | 50375.0 | 1 |
| scf7180000048318 | 9131 | 0.4871 | 0 | 116.1306 | Eukaryota | 67319.0 | 0 | Chordata | 67319.0 | 0 | Carnivora | 67319.0 | 0 | Canidae | 67319.0 | 0 | Canis | 67319.0 | 0 | Canis lupus | 67319.0 | 0 |
| scf7180000046068 | 172506 | 0.3664 | 0 | 60.5736 | Eukaryota | 27801.0 | 0 | Chordata | 27801.0 | 0 | Carnivora | 27801.0 | 0 | Canidae | 27801.0 | 0 | Canis | 27801.0 | 0 | Canis lupus | 27801.0 | 0 |
| scf7180000042547 | 32499 | 0.341 | 0 | 53.7066 | Eukaryota | 11138.0 | 0 | Chordata | 11138.0 | 0 | Carnivora | 11138.0 | 0 | Canidae | 11138.0 | 0 | Canis | 11138.0 | 0 | Canis lupus | 11138.0 | 0 |
| scf7180000055142 | 18860 | 0.4417 | 0 | 216.7839 | Eukaryota | 19274.0 | 0 | Chordata | 18544.0 | 1 | Tetraodontiformes | 15517.0 | 2 | Tetraodontidae | 15517.0 | 2 | Takifugu | 15517.0 | 2 | Takifugu rubripes | 14423.0 | 3 |
| scf7180000043657 | 231870 | 0.3841 | 0 | 59.6912 | Eukaryota | 22535.0 | 0 | Chordata | 22535.0 | 0 | Carnivora | 22535.0 | 0 | Canidae | 22535.0 | 0 | Canis | 22535.0 | 0 | Canis lupus | 22535.0 | 0 |
| scf7180000044513 | 288389 | 0.4135 | 0 | 61.062 | Eukaryota | 44214.0 | 0 | Chordata | 44214.0 | 0 | Carnivora | 44214.0 | 0 | Canidae | 44214.0 | 0 | Canis | 34886.0 | 1 | Canis lupus | 34886.0 | 1 |
| scf7180000046757 | 13274 | 0.4427 | 0 | 179.0396 | Eukaryota | 3139.0 | 0 | Chordata | 2193.0 | 1 | Tetraodontiformes | 2058.0 | 2 | Tetraodontidae | 2058.0 | 2 | Takifugu | 2058.0 | 2 | Takifugu rubripes | 2058.0 | 2 |
| scf7180000042520 | 13791 | 0.3704 | 0 | 44.6716 | Eukaryota | 4265.0 | 0 | Chordata | 4265.0 | 0 | Carnivora | 2375.0 | 1 | Canidae | 2375.0 | 1 | Canis | 2375.0 | 1 | Canis lupus | 2375.0 | 1 |
| scf7180000055996 | 11693 | 0.4396 | 0 | 121.9528 | Eukaryota | 1602.0 | 0 | Platyhelminthes | 879.0 | 1 | Polystomatidea | 879.0 | 2 | Polystomatidae | 879.0 | 2 | Protopolystoma | 879.0 | 2 | Protopolystoma xenopodis | 879.0 | 2 |
| scf7180000046939 | 28604 | 0.3493 | 0 | 50.4261 | Eukaryota | 4027.0 | 0 | Chordata | 4027.0 | 0 | Carnivora | 2691.0 | 2 | Canidae | 2691.0 | 2 | Canis | 2691.0 | 2 | Canis lupus | 2691.0 | 2 |
| scf7180000044704 | 310457 | 0.3577 | 0 | 59.1359 | Eukaryota | 86812.0 | 0 | Chordata | 86812.0 | 0 | Carnivora | 86812.0 | 0 | Canidae | 86812.0 | 0 | Canis | 86812.0 | 0 | Canis lupus | 86812.0 | 0 |
| scf7180000046580 | 15211 | 0.4419 | 0 | 165.2643 | no-hit | 0.0 | 0 | no-hit | 0.0 | 0 | no-hit | 0.0 | 0 | no-hit | 0.0 | 0 | no-hit | 0.0 | 0 | no-hit | 0.0 | 0 |
| scf7180000053064 | 7887 | 0.4554 | 0 | 1409.3904 | Eukaryota | 5734.0 | 0 | Echinodermata | 2699.0 | 2 | Forcipulatida | 2699.0 | 2 | Asteriidae | 2699.0 | 2 | Asterias | 2699.0 | 2 | Asterias rubens | 2699.0 | 2 |
| scf7180000043905 | 338945 | 0.4231 | 0 | 82.418 | Eukaryota | 43921.0 | 0 | Chordata | 43921.0 | 0 | Carnivora | 31911.0 | 1 | Canidae | 14843.0 | 4 | Canis | 14843.0 | 5 | Canis lupus | 14843.0 | 5 |
| scf7180000054533 | 13848 | 0.4478 | 0 | 514.7987 | Eukaryota | 3522.0 | 0 | Chordata | 2834.0 | 1 | Carnivora | 1836.0 | 2 | Canidae | 1836.0 | 2 | Canis | 1836.0 | 2 | Canis lupus | 1836.0 | 2 |
| scf7180000048887 | 29101 | 0.4358 | 0 | 139.7707 | Eukaryota | 31120.0 | 0 | Chordata | 24716.0 | 2 | Gadiformes | 11251.0 | 7 | Gadidae | 11251.0 | 7 | Gadus | 11251.0 | 7 | Gadus morhua | 11251.0 | 7 |
| scf7180000046689 | 20859 | 0.4417 | 0 | 56.1566 | Eukaryota | 14646.0 | 0 | Chordata | 10725.0 | 3 | Tetraodontiformes | 10725.0 | 3 | Tetraodontidae | 10725.0 | 3 | Takifugu | 10725.0 | 3 | Takifugu rubripes | 10725.0 | 3 |
| scf7180000043712 | 579461 | 0.4043 | 200 | 59.2421 | Eukaryota | 61710.0 | 0 | Chordata | 61710.0 | 0 | Carnivora | 61710.0 | 0 | Canidae | 61710.0 | 0 | Canis | 61710.0 | 0 | Canis lupus | 61710.0 | 0 |
| scf7180000051499 | 7419 | 0.4416 | 0 | 61.1349 | Eukaryota | 1847.0 | 0 | Chordata | 1847.0 | 0 | Columbiformes | 1847.0 | 0 | Columbidae | 1847.0 | 0 | Streptopelia | 1847.0 | 0 | Streptopelia turtur | 1847.0 | 0 |
| scf7180000043120 | 99721 | 0.4496 | 0 | 60.0189 | Eukaryota | 94162.0 | 0 | Chordata | 94162.0 | 0 | Carnivora | 94162.0 | 0 | Canidae | 94162.0 | 0 | Canis | 94162.0 | 0 | Canis lupus | 94162.0 | 0 |
| scf7180000047718 | 44169 | 0.3996 | 0 | 56.0892 | Eukaryota | 15489.0 | 0 | Chordata | 15489.0 | 0 | Carnivora | 15489.0 | 0 | Canidae | 15489.0 | 0 | Canis | 15489.0 | 0 | Canis lupus | 15489.0 | 0 |
| scf7180000046828 | 16768 | 0.4373 | 0 | 106.6912 | Eukaryota | 5089.0 | 0 | Chordata | 4240.0 | 1 | Tetraodontiformes | 2352.0 | 4 | Tetraodontidae | 2352.0 | 4 | Takifugu | 2352.0 | 5 | Takifugu rubripes | 2352.0 | 5 |
| scf7180000047551 | 96335 | 0.6855 | 0 | 55.7294 | Eukaryota | 34983.0 | 0 | Chordata | 34983.0 | 0 | Carnivora | 34983.0 | 0 | Canidae | 13250.0 | 2 | Ursus | 12484.0 | 3 | Ursus arctos | 12484.0 | 3 |
| scf7180000041968 | 17039 | 0.4198 | 0 | 37.4085 | Eukaryota | 36460.0 | 0 | Chordata | 36460.0 | 0 | Carnivora | 26761.0 | 2 | Canidae | 12676.0 | 4 | Canis | 9093.0 | 7 | Canis lupus | 9093.0 | 8 |
| scf7180000051398 | 15112 | 0.504 | 0 | 890.5734 | Eukaryota | 9275.0 | 0 | Chordata | 9275.0 | 0 | Carnivora | 9275.0 | 0 | Canidae | 9275.0 | 0 | Canis | 7572.0 | 1 | Canis lupus | 6332.0 | 2 |
| scf7180000054271 | 1064568 | 0.3548 | 0 | 58.8484 | Eukaryota | 108284.0 | 0 | Chordata | 108284.0 | 0 | Carnivora | 108284.0 | 0 | Canidae | 108284.0 | 0 | Canis | 108284.0 | 0 | Canis lupus | 108284.0 | 0 |
| scf7180000041637 | 473469 | 0.3823 | 0 | 60.2157 | Eukaryota | 90452.0 | 0 | Chordata | 90452.0 | 0 | Carnivora | 90452.0 | 0 | Canidae | 90452.0 | 0 | Canis | 90452.0 | 0 | Canis lupus | 90452.0 | 0 |
| scf7180000056298 | 176518 | 0.41 | 0 | 53.6879 | Eukaryota | 13949.0 | 0 | Chordata | 13949.0 | 0 | Carnivora | 13949.0 | 0 | Canidae | 13949.0 | 0 | Canis | 13949.0 | 0 | Canis lupus | 13949.0 | 0 |
| scf7180000052580 | 2896 | 0.4523 | 0 | 534.633 | Eukaryota | 9787.0 | 0 | Chordata | 7541.0 | 3 | Gadiformes | 4624.0 | 7 | Gadidae | 4624.0 | 7 | Gadus | 4624.0 | 7 | Gadus morhua | 4624.0 | 7 |
| scf7180000046953 | 8418 | 0.438 | 0 | 86.6621 | Eukaryota | 19753.0 | 1 | Chordata | 15847.0 | 2 | Gadiformes | 9912.0 | 6 | Gadidae | 9912.0 | 6 | Gadus | 9912.0 | 6 | Gadus morhua | 9912.0 | 6 |
| scf7180000046445 | 3260 | 0.4202 | 0 | 71.081 | no-hit | 0.0 | 0 | no-hit | 0.0 | 0 | no-hit | 0.0 | 0 | no-hit | 0.0 | 0 | no-hit | 0.0 | 0 | no-hit | 0.0 | 0 |
| scf7180000051128 | 10613 | 0.4046 | 0 | 61.0886 | Eukaryota | 5306.0 | 0 | Chordata | 5306.0 | 0 | Carnivora | 5306.0 | 0 | Canidae | 5306.0 | 0 | Canis | 5306.0 | 0 | Canis lupus | 5306.0 | 0 |
| scf7180000042911 | 602842 | 0.3851 | 0 | 62.0504 | Eukaryota | 59490.0 | 0 | Chordata | 59490.0 | 0 | Carnivora | 59490.0 | 0 | Canidae | 59490.0 | 0 | Canis | 34776.0 | 1 | Canis lupus | 34776.0 | 1 |
| scf7180000043730 | 15210 | 0.424 | 0 | 70.6147 | Eukaryota | 89127.0 | 0 | Chordata | 89127.0 | 0 | Carnivora | 89127.0 | 0 | Canidae | 89127.0 | 0 | Canis | 89127.0 | 0 | Canis lupus | 89127.0 | 0 |
| scf7180000053327 | 7723 | 0.4369 | 0 | 56.7405 | Eukaryota | 359.0 | 0 | Chordata | 359.0 | 0 | Kurtiformes | 359.0 | 0 | Apogonidae | 359.0 | 0 | Sphaeramia | 359.0 | 0 | Sphaeramia orbicularis | 359.0 | 0 |
| scf7180000047759 | 90869 | 0.3848 | 0 | 60.1827 | Eukaryota | 29274.0 | 0 | Chordata | 29274.0 | 0 | Carnivora | 29274.0 | 0 | Canidae | 29274.0 | 0 | Canis | 29274.0 | 0 | Canis lupus | 29274.0 | 0 |
| scf7180000045823 | 198585 | 0.4545 | 0 | 57.7417 | Eukaryota | 15242.0 | 0 | Chordata | 15242.0 | 0 | Carnivora | 15242.0 | 0 | Canidae | 15242.0 | 0 | Canis | 15242.0 | 0 | Canis lupus | 15242.0 | 0 |
| scf7180000044009 | 113195 | 0.3415 | 0 | 56.8149 | Eukaryota | 10703.0 | 0 | Chordata | 10703.0 | 0 | Carnivora | 10703.0 | 0 | Canidae | 10703.0 | 0 | Canis | 10703.0 | 0 | Canis lupus | 10703.0 | 0 |
| scf7180000045125 | 183892 | 0.4974 | 0 | 55.8319 | Eukaryota | 60272.0 | 0 | Chordata | 60272.0 | 0 | Carnivora | 60272.0 | 0 | Canidae | 60272.0 | 0 | Canis | 35325.0 | 1 | Canis lupus | 35325.0 | 1 |
| scf7180000053168 | 5610 | 0.4424 | 0 | 596.1358 | Eukaryota | 15517.0 | 1 | Chordata | 15517.0 | 1 | Tetraodontiformes | 5475.0 | 5 | Tetraodontidae | 5475.0 | 5 | Takifugu | 5475.0 | 5 | Takifugu rubripes | 5475.0 | 5 |
| scf7180000056053 | 95455 | 0.4086 | 0 | 191.5917 | Eukaryota | 16288.0 | 0 | Chordata | 16288.0 | 0 | Carnivora | 16288.0 | 0 | Canidae | 16288.0 | 0 | Canis | 16288.0 | 0 | Canis lupus | 16288.0 | 0 |
| scf7180000044932 | 237732 | 0.4128 | 0 | 60.0977 | Eukaryota | 68501.0 | 0 | Chordata | 68501.0 | 0 | Carnivora | 68501.0 | 0 | Canidae | 68501.0 | 0 | Canis | 68501.0 | 0 | Canis lupus | 68501.0 | 0 |
| scf7180000040708 | 176365 | 0.3538 | 0 | 60.222 | Eukaryota | 92156.0 | 0 | Chordata | 92156.0 | 0 | Carnivora | 92156.0 | 0 | Canidae | 92156.0 | 0 | Canis | 92156.0 | 0 | Canis lupus | 92156.0 | 0 |
| scf7180000044739 | 309198 | 0.4037 | 0 | 60.531 | Eukaryota | 86842.0 | 0 | Chordata | 86842.0 | 0 | Carnivora | 86842.0 | 0 | Canidae | 86842.0 | 0 | Canis | 86842.0 | 0 | Canis lupus | 86842.0 | 0 |
| scf7180000047953 | 43369 | 0.4378 | 0 | 58.2369 | Eukaryota | 4785.0 | 0 | Chordata | 4785.0 | 0 | Carnivora | 3591.0 | 1 | Felidae | 2502.0 | 3 | Acinonyx | 1260.0 | 5 | Acinonyx jubatus | 1260.0 | 5 |
| scf7180000042213 | 499134 | 0.5158 | 0 | 58.4927 | Eukaryota | 47463.0 | 0 | Chordata | 47463.0 | 0 | Carnivora | 47463.0 | 0 | Canidae | 47463.0 | 0 | Canis | 47463.0 | 0 | Canis lupus | 47463.0 | 0 |
| scf7180000055264 | 17489 | 0.4375 | 0 | 177.7142 | Eukaryota | 5642.0 | 0 | Platyhelminthes | 2846.0 | 1 | Polystomatidea | 2846.0 | 3 | Polystomatidae | 2846.0 | 3 | Protopolystoma | 2846.0 | 3 | Protopolystoma xenopodis | 2846.0 | 3 |
| scf7180000043023 | 42348 | 0.4487 | 0 | 105.8895 | Eukaryota | 43671.0 | 0 | Chordata | 43671.0 | 0 | Carnivora | 43671.0 | 0 | Canidae | 43671.0 | 0 | Canis | 43671.0 | 0 | Canis lupus | 43671.0 | 0 |
| scf7180000044011 | 345922 | 0.3607 | 0 | 59.3799 | Eukaryota | 20045.0 | 0 | Chordata | 20045.0 | 0 | Carnivora | 20045.0 | 0 | Canidae | 14144.0 | 3 | Canis | 12077.0 | 4 | Canis lupus | 12077.0 | 4 |
| scf7180000046677 | 13522 | 0.4439 | 0 | 65.0353 | Eukaryota | 90481.0 | 0 | Chordata | 90481.0 | 0 | Carnivora | 90481.0 | 0 | Canidae | 90481.0 | 0 | Canis | 90481.0 | 0 | Canis lupus | 90481.0 | 0 |
| scf7180000041117 | 6323 | 0.3475 | 0 | 29.6104 | Eukaryota | 2239.0 | 0 | Chordata | 2239.0 | 0 | Carnivora | 2239.0 | 0 | Canidae | 2239.0 | 0 | Canis | 2239.0 | 0 | Canis lupus | 2239.0 | 0 |
| scf7180000041771 | 232113 | 0.3904 | 0 | 59.4185 | Eukaryota | 83079.0 | 0 | Chordata | 83079.0 | 0 | Carnivora | 83079.0 | 0 | Canidae | 83079.0 | 0 | Canis | 83079.0 | 0 | Canis lupus | 83079.0 | 0 |
| scf7180000043948 | 208755 | 0.4263 | 0 | 60.6876 | Eukaryota | 44930.0 | 0 | Chordata | 44930.0 | 0 | Carnivora | 44930.0 | 0 | Canidae | 44930.0 | 0 | Canis | 38939.0 | 1 | Canis lupus | 38939.0 | 1 |
| scf7180000045407 | 208186 | 0.583 | 0 | 55.65 | Eukaryota | 40930.0 | 0 | Chordata | 40930.0 | 0 | Carnivora | 40930.0 | 0 | Canidae | 40930.0 | 0 | Canis | 40930.0 | 0 | Canis lupus | 40930.0 | 0 |
| scf7180000042324 | 28950 | 0.4117 | 0 | 52.7986 | Eukaryota | 13480.0 | 0 | Chordata | 13480.0 | 0 | Carnivora | 13480.0 | 0 | Canidae | 13480.0 | 0 | Canis | 13480.0 | 0 | Canis lupus | 13480.0 | 0 |
| scf7180000052023 | 13989 | 0.4424 | 0 | 329.3695 | Eukaryota | 2595.0 | 0 | Platyhelminthes | 1469.0 | 2 | Polystomatidea | 1469.0 | 2 | Polystomatidae | 1469.0 | 2 | Protopolystoma | 1469.0 | 2 | Protopolystoma xenopodis | 1469.0 | 2 |
| scf7180000042102 | 13566 | 0.4449 | 0 | 36.0615 | Eukaryota | 13881.0 | 0 | Chordata | 13881.0 | 0 | Carnivora | 9694.0 | 2 | Canidae | 7281.0 | 4 | Canis | 7281.0 | 5 | Canis lupus | 7281.0 | 5 |
| scf7180000054849 | 23565 | 0.4465 | 0 | 284.4067 | Eukaryota | 11729.0 | 0 | Chordata | 11729.0 | 0 | Carnivora | 8184.0 | 2 | Canidae | 8184.0 | 2 | Canis | 4347.0 | 3 | Canis lupus | 4347.0 | 3 |
| scf7180000053682 | 10335 | 0.3469 | 0 | 58.4515 | Eukaryota | 4359.0 | 0 | Chordata | 4359.0 | 0 | Carnivora | 3880.0 | 1 | Canidae | 2060.0 | 2 | Canis | 2060.0 | 3 | Canis lupus | 2060.0 | 3 |
| scf7180000055443 | 29513 | 0.3579 | 0 | 618.9667 | Eukaryota | 5701.0 | 0 | Chordata | 5701.0 | 0 | Primates | 4719.0 | 1 | Hominidae | 3167.0 | 3 | Homo | 3167.0 | 3 | Homo sapiens | 3167.0 | 3 |
| scf7180000052806 | 2955 | 0.4362 | 0 | 167.8428 | Eukaryota | 237.0 | 0 | Chordata | 237.0 | 0 | Salmoniformes | 237.0 | 0 | Salmonidae | 237.0 | 0 | Coregonus | 237.0 | 0 | Coregonus sp. 'balchen' | 237.0 | 0 |
| scf7180000046353 | 53874 | 0.4476 | 0 | 69.2326 | Eukaryota | 75584.0 | 0 | Chordata | 75584.0 | 0 | Carnivora | 75584.0 | 0 | Canidae | 75584.0 | 0 | Canis | 75584.0 | 0 | Canis lupus | 75584.0 | 0 |
| scf7180000042347 | 709709 | 0.3905 | 0 | 60.3128 | Eukaryota | 161103.0 | 0 | Chordata | 161103.0 | 0 | Carnivora | 161103.0 | 0 | Canidae | 161103.0 | 0 | Canis | 161103.0 | 0 | Canis lupus | 161103.0 | 0 |
| scf7180000041800 | 121609 | 0.3857 | 0 | 61.5351 | Eukaryota | 90484.0 | 0 | Chordata | 90484.0 | 0 | Carnivora | 90484.0 | 0 | Canidae | 90484.0 | 0 | Canis | 90484.0 | 0 | Canis lupus | 90484.0 | 0 |
| scf7180000056502 | 174834 | 0.577 | 0 | 56.7092 | Eukaryota | 10298.0 | 0 | Chordata | 10298.0 | 0 | Carnivora | 8613.0 | 1 | Canidae | 8613.0 | 1 | Canis | 8613.0 | 1 | Canis lupus | 8613.0 | 1 |
| scf7180000042747 | 580376 | 0.3904 | 0 | 59.155 | Eukaryota | 66204.0 | 0 | Chordata | 66204.0 | 0 | Carnivora | 66204.0 | 0 | Canidae | 66204.0 | 0 | Canis | 57339.0 | 1 | Canis lupus | 57339.0 | 1 |
| scf7180000048294 | 72187 | 0.3527 | 0 | 59.7762 | Eukaryota | 9655.0 | 0 | Chordata | 9655.0 | 0 | Carnivora | 5313.0 | 2 | Canidae | 5313.0 | 2 | Canis | 5313.0 | 2 | Canis lupus | 5313.0 | 2 |
| scf7180000041647 | 972183 | 0.3944 | 0 | 60.0158 | Eukaryota | 216837.0 | 0 | Chordata | 216837.0 | 0 | Carnivora | 216837.0 | 0 | Canidae | 216837.0 | 0 | Canis | 216837.0 | 0 | Canis lupus | 216837.0 | 0 |
| scf7180000054730 | 259929 | 0.4442 | 0 | 85.107 | Eukaryota | 88000.0 | 0 | Chordata | 88000.0 | 0 | Carnivora | 88000.0 | 0 | Canidae | 88000.0 | 0 | Canis | 88000.0 | 0 | Canis lupus | 88000.0 | 0 |
| scf7180000043783 | 393119 | 0.4813 | 0 | 60.6985 | Eukaryota | 64786.0 | 0 | Chordata | 64786.0 | 0 | Carnivora | 64786.0 | 0 | Canidae | 64786.0 | 0 | Vulpes | 54016.0 | 1 | Vulpes vulpes | 54016.0 | 1 |
| scf7180000040831 | 1658909 | 0.4056 | 0 | 60.2718 | Eukaryota | 94864.0 | 0 | Chordata | 94864.0 | 0 | Carnivora | 94864.0 | 0 | Canidae | 94864.0 | 0 | Canis | 94864.0 | 0 | Canis lupus | 94864.0 | 0 |
| scf7180000046634 | 37891 | 0.4414 | 0 | 268.7131 | Eukaryota | 23048.0 | 0 | Chordata | 14886.0 | 1 | Strigeidida | 8162.0 | 5 | Schistosomatidae | 8162.0 | 5 | Schistosoma | 8162.0 | 5 | Schistosoma mansoni | 8162.0 | 5 |
| scf7180000042248 | 508161 | 0.363 | 0 | 60.0465 | Eukaryota | 93611.0 | 0 | Chordata | 93611.0 | 0 | Carnivora | 93611.0 | 0 | Canidae | 93611.0 | 0 | Canis | 93611.0 | 0 | Canis lupus | 93611.0 | 0 |
| scf7180000042852 | 128763 | 0.4042 | 0 | 57.2624 | Eukaryota | 59915.0 | 0 | Chordata | 59915.0 | 0 | Carnivora | 59915.0 | 0 | Canidae | 59915.0 | 0 | Canis | 59915.0 | 0 | Canis lupus | 59915.0 | 0 |
| scf7180000056007 | 15018 | 0.5113 | 0 | 388.9842 | Eukaryota | 15135.0 | 0 | Chordata | 15135.0 | 0 | Carnivora | 15135.0 | 0 | Canidae | 15135.0 | 0 | Canis | 15135.0 | 0 | Canis lupus | 15135.0 | 0 |
| scf7180000048593 | 57077 | 0.4452 | 0 | 59.8355 | Eukaryota | 34847.0 | 0 | Chordata | 34847.0 | 0 | Carnivora | 34847.0 | 0 | Canidae | 34847.0 | 0 | Canis | 22158.0 | 1 | Canis lupus | 22158.0 | 1 |
| scf7180000041860 | 60358 | 0.3781 | 0 | 60.3617 | Eukaryota | 88834.0 | 0 | Chordata | 88834.0 | 0 | Carnivora | 88834.0 | 0 | Canidae | 88834.0 | 0 | Canis | 88834.0 | 0 | Canis lupus | 88834.0 | 0 |
| scf7180000056208 | 1236156 | 0.4346 | 0 | 60.1086 | Eukaryota | 229165.0 | 0 | Chordata | 229165.0 | 0 | Carnivora | 229165.0 | 0 | Canidae | 229165.0 | 0 | Canis | 229165.0 | 0 | Canis lupus | 229165.0 | 0 |
| scf7180000052276 | 2353 | 0.4505 | 0 | 260.1627 | Eukaryota | 2225.0 | 0 | Platyhelminthes | 1019.0 | 2 | Polystomatidea | 1019.0 | 2 | Polystomatidae | 1019.0 | 2 | Protopolystoma | 1019.0 | 2 | Protopolystoma xenopodis | 1019.0 | 2 |
| scf7180000041828 | 73376 | 0.3939 | 0 | 61.7954 | Eukaryota | 46396.0 | 0 | Chordata | 46396.0 | 0 | Carnivora | 46396.0 | 0 | Canidae | 46396.0 | 0 | Canis | 46396.0 | 0 | Canis lupus | 46396.0 | 0 |
| scf7180000055486 | 28184 | 0.4415 | 0 | 462.26 | no-hit | 0.0 | 0 | no-hit | 0.0 | 0 | no-hit | 0.0 | 0 | no-hit | 0.0 | 0 | no-hit | 0.0 | 0 | no-hit | 0.0 | 0 |
| scf7180000056020 | 9177 | 0.4412 | 0 | 353.309 | no-hit | 0.0 | 0 | no-hit | 0.0 | 0 | no-hit | 0.0 | 0 | no-hit | 0.0 | 0 | no-hit | 0.0 | 0 | no-hit | 0.0 | 0 |
| scf7180000048422 | 21993 | 0.4432 | 0 | 144.7628 | Eukaryota | 2969.0 | 0 | Chordata | 2341.0 | 1 | Carnivora | 2341.0 | 1 | Canidae | 2341.0 | 1 | Canis | 2341.0 | 1 | Canis lupus | 2341.0 | 1 |
| scf7180000045035 | 209589 | 0.4795 | 0 | 59.1319 | Eukaryota | 55528.0 | 0 | Chordata | 55528.0 | 0 | Carnivora | 55528.0 | 0 | Canidae | 55528.0 | 0 | Canis | 49629.0 | 1 | Canis lupus | 49629.0 | 1 |
| scf7180000048621 | 4557 | 0.4349 | 0 | 155.634 | no-hit | 0.0 | 0 | no-hit | 0.0 | 0 | no-hit | 0.0 | 0 | no-hit | 0.0 | 0 | no-hit | 0.0 | 0 | no-hit | 0.0 | 0 |
| scf7180000046387 | 73748 | 0.4369 | 0 | 56.0516 | Eukaryota | 57297.0 | 0 | Chordata | 57297.0 | 0 | Carnivora | 57297.0 | 0 | Canidae | 33738.0 | 3 | Canis | 27040.0 | 5 | Canis lupus | 27040.0 | 6 |
| scf7180000045132 | 174566 | 0.3989 | 0 | 60.5974 | Eukaryota | 16414.0 | 0 | Chordata | 16414.0 | 0 | Apterygiformes | 8776.0 | 1 | Apterygidae | 8776.0 | 1 | Apteryx | 8776.0 | 2 | Apteryx mantelli | 8776.0 | 2 |
| scf7180000041848 | 930534 | 0.3866 | 0 | 60.9636 | Eukaryota | 103782.0 | 0 | Chordata | 103782.0 | 0 | Carnivora | 103782.0 | 0 | Canidae | 103782.0 | 0 | Vulpes | 62394.0 | 1 | Vulpes vulpes | 62394.0 | 1 |
| scf7180000042340 | 14564 | 0.5216 | 0 | 54.5396 | Eukaryota | 37686.0 | 0 | Chordata | 37686.0 | 0 | Carnivora | 37686.0 | 0 | Canidae | 30511.0 | 2 | Canis | 26671.0 | 3 | Canis lupus | 26671.0 | 3 |
| scf7180000055234 | 32630 | 0.4421 | 0 | 110.7741 | Eukaryota | 2388.0 | 0 | Platyhelminthes | 1347.0 | 1 | Polystomatidea | 1347.0 | 1 | Polystomatidae | 1347.0 | 1 | Protopolystoma | 1347.0 | 1 | Protopolystoma xenopodis | 1347.0 | 1 |
| scf7180000055982 | 32941 | 0.4353 | 0 | 317.7897 | Eukaryota | 910.0 | 0 | Chordata | 536.0 | 1 | Tetraodontiformes | 536.0 | 1 | Tetraodontidae | 536.0 | 1 | Takifugu | 536.0 | 1 | Takifugu rubripes | 536.0 | 1 |
| scf7180000043334 | 731399 | 0.3894 | 0 | 59.7015 | Eukaryota | 45843.0 | 0 | Chordata | 45843.0 | 0 | Carnivora | 45843.0 | 0 | Canidae | 45843.0 | 0 | Canis | 37997.0 | 1 | Canis lupus | 37997.0 | 1 |
| scf7180000048381 | 24074 | 0.4483 | 0 | 1049.2977 | Eukaryota | 20536.0 | 0 | Chordata | 18871.0 | 1 | Gadiformes | 12374.0 | 5 | Gadidae | 12374.0 | 5 | Gadus | 12374.0 | 5 | Gadus morhua | 12374.0 | 5 |
| scf7180000053242 | 5177 | 0.4458 | 0 | 498.6213 | no-hit | 0.0 | 0 | no-hit | 0.0 | 0 | no-hit | 0.0 | 0 | no-hit | 0.0 | 0 | no-hit | 0.0 | 0 | no-hit | 0.0 | 0 |
| scf7180000056390 | 454381 | 0.3708 | 0 | 59.2485 | Eukaryota | 88265.0 | 0 | Chordata | 88265.0 | 0 | Carnivora | 88265.0 | 0 | Canidae | 88265.0 | 0 | Canis | 88265.0 | 0 | Canis lupus | 88265.0 | 0 |
| scf7180000055181 | 169819 | 0.3801 | 0 | 74.4986 | Eukaryota | 90636.0 | 0 | Chordata | 90636.0 | 0 | Carnivora | 90636.0 | 0 | Canidae | 90636.0 | 0 | Canis | 90636.0 | 0 | Canis lupus | 90636.0 | 0 |
| scf7180000043320 | 938679 | 0.429 | 100 | 60.2405 | Eukaryota | 91101.0 | 0 | Chordata | 91101.0 | 0 | Carnivora | 91101.0 | 0 | Canidae | 91101.0 | 0 | Canis | 91101.0 | 0 | Canis lupus | 91101.0 | 0 |
| scf7180000043516 | 914702 | 0.3415 | 0 | 59.1313 | Eukaryota | 25554.0 | 0 | Chordata | 25554.0 | 0 | Carnivora | 25554.0 | 0 | Canidae | 20735.0 | 1 | Canis | 18260.0 | 3 | Canis lupus | 18260.0 | 3 |
| scf7180000044573 | 284129 | 0.4143 | 0 | 58.4066 | Eukaryota | 58452.0 | 0 | Chordata | 58452.0 | 0 | Carnivora | 58452.0 | 0 | Canidae | 58452.0 | 0 | Canis | 58452.0 | 0 | Canis lupus | 58452.0 | 0 |
| scf7180000043830 | 91204 | 0.5154 | 0 | 72.0418 | Eukaryota | 63792.0 | 0 | Chordata | 63792.0 | 0 | Carnivora | 63792.0 | 0 | Canidae | 63792.0 | 0 | Canis | 63792.0 | 0 | Canis lupus | 63792.0 | 0 |
| scf7180000046076 | 212356 | 0.47 | 0 | 57.9172 | Eukaryota | 28608.0 | 0 | Chordata | 28608.0 | 0 | Carnivora | 28608.0 | 0 | Canidae | 28608.0 | 0 | Canis | 28608.0 | 0 | Canis lupus | 28608.0 | 0 |
| scf7180000044286 | 373838 | 0.3958 | 0 | 59.9868 | Eukaryota | 92663.0 | 0 | Chordata | 92663.0 | 0 | Carnivora | 92663.0 | 0 | Canidae | 92663.0 | 0 | Canis | 92663.0 | 0 | Canis lupus | 92663.0 | 0 |
| scf7180000047552 | 75184 | 0.4673 | 0 | 60.2886 | Eukaryota | 31160.0 | 0 | Chordata | 31160.0 | 0 | Carnivora | 31160.0 | 0 | Canidae | 16284.0 | 1 | Vulpes | 13137.0 | 4 | Vulpes vulpes | 13137.0 | 4 |
| scf7180000044675 | 291705 | 0.4554 | 0 | 59.1894 | Eukaryota | 19508.0 | 0 | Chordata | 19508.0 | 0 | Carnivora | 19508.0 | 0 | Canidae | 19508.0 | 0 | Canis | 9992.0 | 1 | Canis lupus | 9992.0 | 1 |
| scf7180000056617 | 99369 | 0.6473 | 0 | 53.5577 | Eukaryota | 20716.0 | 0 | Chordata | 20716.0 | 0 | Carnivora | 20716.0 | 0 | Canidae | 20716.0 | 0 | Canis | 20716.0 | 0 | Canis lupus | 20716.0 | 0 |
| scf7180000047285 | 78590 | 0.4521 | 0 | 60.3892 | Eukaryota | 30512.0 | 0 | Chordata | 30512.0 | 0 | Carnivora | 30512.0 | 0 | Canidae | 15793.0 | 3 | Canis | 12596.0 | 5 | Canis lupus | 12596.0 | 6 |
| scf7180000051116 | 3710 | 0.4472 | 0 | 565.3254 | Eukaryota | 2636.0 | 0 | Platyhelminthes | 1260.0 | 2 | Polystomatidea | 1260.0 | 2 | Polystomatidae | 1260.0 | 2 | Protopolystoma | 1260.0 | 2 | Protopolystoma xenopodis | 1260.0 | 2 |
| scf7180000056462 | 396931 | 0.4372 | 0 | 58.0105 | Eukaryota | 65175.0 | 0 | Chordata | 65175.0 | 0 | Carnivora | 65175.0 | 0 | Canidae | 59407.0 | 1 | Canis | 32350.0 | 2 | Canis lupus | 32350.0 | 2 |
| scf7180000052455 | 3365 | 0.4493 | 0 | 125.7011 | Eukaryota | 3478.0 | 0 | Chordata | 2426.0 | 2 | Chordata-undef | 696.0 | 7 | Ambassidae | 696.0 | 7 | Parambassis | 696.0 | 7 | Parambassis ranga | 696.0 | 7 |
| scf7180000048916 | 43034 | 0.4434 | 0 | 61.9192 | Eukaryota | 18029.0 | 0 | Chordata | 17445.0 | 1 | Tetraodontiformes | 14813.0 | 2 | Tetraodontidae | 14813.0 | 2 | Takifugu | 14813.0 | 2 | Takifugu rubripes | 14813.0 | 2 |
| scf7180000047186 | 83985 | 0.5637 | 0 | 54.1242 | Eukaryota | 3852.0 | 0 | Chordata | 3852.0 | 0 | Carnivora | 3852.0 | 0 | Canidae | 3408.0 | 1 | Canis | 2852.0 | 2 | Canis lupus | 2852.0 | 2 |
| scf7180000048218 | 65741 | 0.5218 | 0 | 56.6937 | Eukaryota | 67608.0 | 0 | Chordata | 67608.0 | 0 | Carnivora | 67608.0 | 0 | Canidae | 67608.0 | 0 | Canis | 67608.0 | 0 | Canis lupus | 67608.0 | 0 |
| scf7180000045100 | 109216 | 0.4384 | 0 | 55.9696 | Eukaryota | 29295.0 | 0 | Chordata | 29295.0 | 0 | Carnivora | 29295.0 | 0 | Canidae | 29295.0 | 0 | Canis | 29295.0 | 0 | Canis lupus | 29295.0 | 0 |
| scf7180000052878 | 6021 | 0.4408 | 0 | 372.8235 | no-hit | 0.0 | 0 | no-hit | 0.0 | 0 | no-hit | 0.0 | 0 | no-hit | 0.0 | 0 | no-hit | 0.0 | 0 | no-hit | 0.0 | 0 |
| scf7180000040909 | 278536 | 0.4239 | 0 | 57.7039 | Eukaryota | 69613.0 | 0 | Chordata | 69613.0 | 0 | Carnivora | 69613.0 | 0 | Canidae | 69613.0 | 0 | Canis | 69613.0 | 0 | Canis lupus | 69613.0 | 0 |
| scf7180000046376 | 67212 | 0.4424 | 0 | 86.356 | Eukaryota | 57819.0 | 0 | Chordata | 57819.0 | 0 | Carnivora | 57819.0 | 0 | Canidae | 57819.0 | 0 | Canis | 57819.0 | 0 | Canis lupus | 57819.0 | 0 |
| scf7180000042758 | 202544 | 0.4906 | 0 | 56.8931 | Eukaryota | 32853.0 | 0 | Chordata | 32853.0 | 0 | Carnivora | 32853.0 | 0 | Canidae | 32853.0 | 0 | Canis | 32853.0 | 0 | Canis lupus | 32853.0 | 0 |
| scf7180000041656 | 834349 | 0.4192 | 0 | 59.9413 | Eukaryota | 63449.0 | 0 | Chordata | 63449.0 | 0 | Carnivora | 63449.0 | 0 | Canidae | 63449.0 | 0 | Canis | 59557.0 | 1 | Canis lupus | 59557.0 | 1 |
| scf7180000056604 | 62788 | 0.4631 | 0 | 58.9788 | Eukaryota | 31585.0 | 0 | Chordata | 31585.0 | 0 | Carnivora | 31585.0 | 0 | Canidae | 15906.0 | 4 | Vulpes | 12056.0 | 7 | Vulpes vulpes | 12056.0 | 7 |

| | | | | | | | | | | | | | | | | | | | | | | |
|---|---|---|---|---|---|---|---|---|---|---|---|---|---|---|---|---|---|---|---|---|---|---|
| scf7180000046146 | 174394 | 0.3507 | 0 | 59.8089 | Eukaryota | 38326.0 | 0 | Chordata | 38326.0 | 0 | Carnivora | 38326.0 | 0 | Canidae | 38326.0 | 0 | Canis | 38326.0 | 0 | Canis lupus | 38326.0 | 0 |
| scf7180000042591 | 205397 | 0.383 | 0 | 58.5024 | Eukaryota | 66015.0 | 0 | Chordata | 66015.0 | 0 | Carnivora | 66015.0 | 0 | Mustelidae | 46341.0 | 1 | Mustela | 46341.0 | 1 | Mustela putorius | 46341.0 | 1 |
| scf7180000051463 | 3515 | 0.4518 | 0 | 737.7902 | no-hit | 0.0 | 0 | no-hit | 0.0 | 0 | no-hit | 0.0 | 0 | no-hit | 0.0 | 0 | no-hit | 0.0 | 0 | no-hit | 0.0 | 0 |
| scf7180000056170 | 652810 | 0.5275 | 0 | 58.1368 | Eukaryota | 60302.0 | 0 | Chordata | 60302.0 | 0 | Carnivora | 60302.0 | 0 | Canidae | 60302.0 | 0 | Canis | 49845.0 | 1 | Canis lupus | 49845.0 | 1 |
| scf7180000050945 | 7970 | 0.4476 | 0 | 450.8595 | Eukaryota | 3570.0 | 0 | Platyhelminthes | 2018.0 | 3 | Polystomatidea | 2018.0 | 3 | Polystomatidae | 2018.0 | 3 | Protopolystoma | 2018.0 | 3 | Protopolystoma xenopodis | 2018.0 | 3 |
| scf7180000048836 | 42417 | 0.3836 | 0 | 58.5839 | Eukaryota | 32457.0 | 0 | Chordata | 32457.0 | 0 | Carnivora | 32457.0 | 0 | Canidae | 32457.0 | 0 | Canis | 32457.0 | 0 | Canis lupus | 32457.0 | 0 |
| scf7180000051493 | 6671 | 0.5385 | 0 | 31.6545 | Eukaryota | 16848.0 | 0 | Chordata | 16848.0 | 0 | Carnivora | 14391.0 | 1 | Canidae | 13979.0 | 2 | Canis | 12776.0 | 4 | Canis lupus | 12776.0 | 4 |
| scf7180000042939 | 54432 | 0.4569 | 0 | 56.1339 | Eukaryota | 54040.0 | 0 | Chordata | 44457.0 | 2 | Gadiformes | 30608.0 | 6 | Gadidae | 30608.0 | 6 | Gadus | 30608.0 | 6 | Gadus morhua | 30608.0 | 6 |
| scf7180000043879 | 110181 | 0.3778 | 0 | 57.1846 | Eukaryota | 90529.0 | 0 | Chordata | 90529.0 | 0 | Carnivora | 90529.0 | 0 | Canidae | 90529.0 | 0 | Canis | 90529.0 | 0 | Canis lupus | 90529.0 | 0 |
| scf7180000045577 | 166351 | 0.4261 | 0 | 59.3914 | Eukaryota | 68739.0 | 0 | Chordata | 68739.0 | 0 | Carnivora | 68739.0 | 0 | Canidae | 68739.0 | 0 | Canis | 40763.0 | 1 | Canis lupus | 40763.0 | 1 |
| scf7180000046332 | 18483 | 0.4444 | 0 | 89.9141 | Eukaryota | 2237.0 | 0 | Platyhelminthes | 1156.0 | 1 | Polystomatidea | 1156.0 | 2 | Polystomatidae | 1156.0 | 2 | Protopolystoma | 1156.0 | 2 | Protopolystoma xenopodis | 1156.0 | 2 |
| scf7180000051700 | 3187 | 0.4481 | 0 | 431.1927 | no-hit | 0.0 | 0 | no-hit | 0.0 | 0 | no-hit | 0.0 | 0 | no-hit | 0.0 | 0 | no-hit | 0.0 | 0 | no-hit | 0.0 | 0 |
| scf7180000051214 | 9537 | 0.4511 | 0 | 1436.2604 | no-hit | 0.0 | 0 | no-hit | 0.0 | 0 | no-hit | 0.0 | 0 | no-hit | 0.0 | 0 | no-hit | 0.0 | 0 | no-hit | 0.0 | 0 |
| scf7180000040499 | 2811361 | 0.3766 | 0 | 60.4015 | Eukaryota | 90940.0 | 0 | Chordata | 90940.0 | 0 | Carnivora | 90940.0 | 0 | Canidae | 90940.0 | 0 | Canis | 90940.0 | 0 | Canis lupus | 90940.0 | 0 |
| scf7180000046613 | 112318 | 0.5786 | 0 | 56.3352 | Eukaryota | 17746.0 | 0 | Chordata | 17746.0 | 0 | Carnivora | 17746.0 | 0 | Canidae | 7228.0 | 2 | Leptonychotes | 5259.0 | 4 | Leptonychotes weddellii | 5259.0 | 4 |
| scf7180000047827 | 19041 | 0.4456 | 0 | 139.1424 | Eukaryota | 3093.0 | 0 | Chordata | 3093.0 | 0 | Carnivora | 3093.0 | 0 | Canidae | 3093.0 | 0 | Canis | 3093.0 | 0 | Canis lupus | 3093.0 | 0 |
| scf7180000041033 | 1272327 | 0.4779 | 0 | 57.6353 | Eukaryota | 44166.0 | 0 | Chordata | 44166.0 | 0 | Carnivora | 44166.0 | 0 | Canidae | 44166.0 | 0 | Canis | 44166.0 | 0 | Canis lupus | 44166.0 | 0 |
| scf7180000046193 | 133148 | 0.3979 | 0 | 58.4117 | Eukaryota | 55304.0 | 0 | Chordata | 55304.0 | 0 | Carnivora | 55304.0 | 0 | Canidae | 55304.0 | 0 | Canis | 55304.0 | 0 | Canis lupus | 55304.0 | 0 |
| scf7180000047091 | 100412 | 0.3644 | 0 | 59.1161 | Eukaryota | 16828.0 | 0 | Chordata | 16828.0 | 0 | Carnivora | 16828.0 | 0 | Canidae | 6913.0 | 2 | Canis | 6913.0 | 5 | Canis lupus | 6913.0 | 6 |
| scf7180000042017 | 157846 | 0.3933 | 0 | 60.1014 | Eukaryota | 44812.0 | 0 | Chordata | 44812.0 | 0 | Carnivora | 44812.0 | 0 | Otariidae | 23095.0 | 1 | Zalophus | 23095.0 | 2 | Zalophus californianus | 23095.0 | 2 |
| scf7180000043401 | 6395 | 0.3403 | 0 | 29.3584 | Eukaryota | 9787.0 | 0 | Chordata | 9787.0 | 0 | Carnivora | 9787.0 | 0 | Canidae | 9787.0 | 0 | Canis | 9787.0 | 0 | Canis lupus | 9787.0 | 0 |
| scf7180000048149 | 18903 | 0.4482 | 0 | 1251.4293 | Eukaryota | 5951.0 | 0 | Chordata | 2668.0 | 3 | Polystomatidea | 2131.0 | 5 | Polystomatidae | 2131.0 | 5 | Protopolystoma | 2131.0 | 5 | Protopolystoma xenopodis | 2131.0 | 5 |
| scf7180000055014 | 33040 | 0.4415 | 0 | 139.1662 | Eukaryota | 8419.0 | 0 | Chordata | 7728.0 | 1 | Chordata-undef | 2531.0 | 6 | Ambassidae | 2531.0 | 6 | Parambassis | 2531.0 | 7 | Parambassis ranga | 2531.0 | 7 |
| scf7180000045253 | 23985 | 0.5688 | 0 | 44.7367 | Eukaryota | 24191.0 | 0 | Chordata | 24191.0 | 0 | Carnivora | 24191.0 | 0 | Canidae | 24191.0 | 0 | Canis | 21594.0 | 1 | Canis lupus | 21594.0 | 1 |
| scf7180000050940 | 14023 | 0.4088 | 0 | 59.1489 | Eukaryota | 5698.0 | 0 | Chordata | 5698.0 | 0 | Carnivora | 5698.0 | 0 | Canidae | 5698.0 | 0 | Canis | 5698.0 | 0 | Canis lupus | 5698.0 | 0 |
| scf7180000051269 | 4019 | 0.4421 | 0 | 70.2504 | Eukaryota | 6159.0 | 0 | Chordata | 6159.0 | 0 | Kurtiformes | 4671.0 | 1 | Apogonidae | 4671.0 | 1 | Sphaeramia | 4671.0 | 1 | Sphaeramia orbicularis | 4671.0 | 1 |
| scf7180000050705 | 2729 | 0.4456 | 0 | 89.1549 | no-hit | 0.0 | 0 | no-hit | 0.0 | 0 | no-hit | 0.0 | 0 | no-hit | 0.0 | 0 | no-hit | 0.0 | 0 | no-hit | 0.0 | 0 |
| scf7180000040486 | 3757373 | 0.4393 | 0 | 59.4513 | Eukaryota | 96725.0 | 0 | Chordata | 96725.0 | 0 | Carnivora | 96725.0 | 0 | Canidae | 96725.0 | 0 | Canis | 76463.0 | 1 | Canis lupus | 76463.0 | 1 |
| scf7180000050648 | 12628 | 0.4499 | 0 | 206.3116 | no-hit | 0.0 | 0 | no-hit | 0.0 | 0 | no-hit | 0.0 | 0 | no-hit | 0.0 | 0 | no-hit | 0.0 | 0 | no-hit | 0.0 | 0 |
| scf7180000047219 | 131024 | 0.4026 | 0 | 58.7518 | Eukaryota | 91330.0 | 0 | Chordata | 91330.0 | 0 | Carnivora | 91330.0 | 0 | Canidae | 91330.0 | 0 | Canis | 91330.0 | 0 | Canis lupus | 91330.0 | 0 |
| scf7180000051338 | 6703 | 0.4394 | 0 | 68.364 | no-hit | 0.0 | 0 | no-hit | 0.0 | 0 | no-hit | 0.0 | 0 | no-hit | 0.0 | 0 | no-hit | 0.0 | 0 | no-hit | 0.0 | 0 |
| scf7180000045130 | 260691 | 0.3529 | 0 | 61.0605 | Eukaryota | 34959.0 | 0 | Chordata | 34959.0 | 0 | Carnivora | 34959.0 | 0 | Canidae | 26189.0 | 1 | Canis | 26189.0 | 1 | Canis lupus | 26189.0 | 1 |
| scf7180000043416 | 48830 | 0.4329 | 0 | 92.7629 | Eukaryota | 51739.0 | 0 | Chordata | 51739.0 | 0 | Carnivora | 51739.0 | 0 | Canidae | 51739.0 | 0 | Canis | 51739.0 | 0 | Canis lupus | 51739.0 | 0 |
| scf7180000046537 | 133064 | 0.4417 | 0 | 56.2729 | Eukaryota | 34452.0 | 0 | Chordata | 34452.0 | 0 | Carnivora | 34452.0 | 0 | Canidae | 34452.0 | 0 | Canis | 34452.0 | 0 | Canis lupus | 34452.0 | 0 |
| scf7180000047858 | 3834 | 0.4541 | 0 | 1427.1267 | no-hit | 0.0 | 0 | no-hit | 0.0 | 0 | no-hit | 0.0 | 0 | no-hit | 0.0 | 0 | no-hit | 0.0 | 0 | no-hit | 0.0 | 0 |
| scf7180000053112 | 3634 | 0.5058 | 0 | 54.6 | Eukaryota | 6257.0 | 0 | Chordata | 6257.0 | 0 | Carnivora | 6257.0 | 0 | Canidae | 6257.0 | 0 | Canis | 4178.0 | 1 | Canis lupus | 3346.0 | 2 |
| scf7180000044878 | 164304 | 0.3673 | 0 | 59.2025 | Eukaryota | 56429.0 | 0 | Chordata | 56429.0 | 0 | Carnivora | 56429.0 | 0 | Canidae | 56429.0 | 0 | Canis | 56429.0 | 0 | Canis lupus | 56429.0 | 0 |
| scf7180000056562 | 129068 | 0.4813 | 0 | 55.487 | Eukaryota | 21234.0 | 0 | Chordata | 21234.0 | 0 | Carnivora | 21234.0 | 0 | Canidae | 21234.0 | 0 | Canis | 21234.0 | 0 | Canis lupus | 21234.0 | 0 |
| scf7180000044723 | 44486 | 0.3972 | 0 | 57.8626 | Eukaryota | 28524.0 | 0 | Chordata | 28524.0 | 0 | Carnivora | 28524.0 | 0 | Canidae | 28524.0 | 0 | Canis | 28524.0 | 0 | Canis lupus | 28524.0 | 0 |
| scf7180000047010 | 93312 | 0.4879 | 0 | 50.6983 | Eukaryota | 24144.0 | 0 | Chordata | 24144.0 | 0 | Carnivora | 24144.0 | 0 | Canidae | 24144.0 | 0 | Canis | 20382.0 | 1 | Canis lupus | 20382.0 | 1 |
| scf7180000043321 | 539913 | 0.3387 | 0 | 58.8339 | Eukaryota | 12609.0 | 0 | Chordata | 12609.0 | 0 | Carnivora | 12609.0 | 0 | Canidae | 12609.0 | 0 | Canis | 12609.0 | 0 | Canis lupus | 12609.0 | 0 |
| scf7180000052433 | 2411 | 0.4475 | 0 | 505.056 | no-hit | 0.0 | 0 | no-hit | 0.0 | 0 | no-hit | 0.0 | 0 | no-hit | 0.0 | 0 | no-hit | 0.0 | 0 | no-hit | 0.0 | 0 |
| scf7180000056174 | 2640449 | 0.4004 | 0 | 58.8306 | Eukaryota | 30048.0 | 0 | Chordata | 30048.0 | 0 | Carnivora | 30048.0 | 0 | Canidae | 30048.0 | 0 | Canis | 27063.0 | 1 | Canis lupus | 27063.0 | 1 |
| scf7180000042280 | 170459 | 0.4898 | 0 | 55.052 | Eukaryota | 39199.0 | 1 | Chordata | 39199.0 | 1 | Carnivora | 32419.0 | 2 | Canidae | 24427.0 | 3 | Canis | 20196.0 | 5 | Canis lupus | 20196.0 | 5 |
| scf7180000046308 | 110601 | 0.5212 | 0 | 55.977 | Eukaryota | 25064.0 | 0 | Chordata | 25064.0 | 0 | Carnivora | 25064.0 | 0 | Canidae | 20624.0 | 1 | Canis | 16659.0 | 2 | Canis lupus | 16659.0 | 2 |
| scf7180000052498 | 13920 | 0.3382 | 0 | 57.7826 | Eukaryota | 3169.0 | 0 | Chordata | 3169.0 | 0 | Carnivora | 1643.0 | 2 | Canidae | 1643.0 | 3 | Canis | 1643.0 | 3 | Canis lupus | 1643.0 | 3 |
| scf7180000051424 | 8602 | 0.4497 | 0 | 59.2917 | Eukaryota | 1267.0 | 0 | Platyhelminthes | 1047.0 | 1 | Polystomatidea | 1047.0 | 1 | Polystomatidae | 1047.0 | 1 | Protopolystoma | 1047.0 | 1 | Protopolystoma xenopodis | 1047.0 | 1 |
| scf7180000051186 | 4085 | 0.4465 | 0 | 411.9216 | Eukaryota | 220.0 | 0 | Chordata | 220.0 | 0 | Kurtiformes | 220.0 | 0 | Apogonidae | 220.0 | 0 | Sphaeramia | 220.0 | 0 | Sphaeramia orbicularis | 220.0 | 0 |
| scf7180000050384 | 1936 | 0.4329 | 0 | 108.5362 | no-hit | 0.0 | 0 | no-hit | 0.0 | 0 | no-hit | 0.0 | 0 | no-hit | 0.0 | 0 | no-hit | 0.0 | 0 | no-hit | 0.0 | 0 |
| scf7180000042860 | 527510 | 0.4026 | 0 | 57.8275 | Eukaryota | 26626.0 | 0 | Chordata | 26626.0 | 0 | Carnivora | 21757.0 | 2 | Canidae | 21757.0 | 2 | Vulpes | 17010.0 | 3 | Vulpes vulpes | 17010.0 | 3 |
| scf7180000046822 | 132758 | 0.3501 | 0 | 60.3767 | Eukaryota | 90986.0 | 0 | Chordata | 90986.0 | 0 | Carnivora | 90986.0 | 0 | Canidae | 90986.0 | 0 | Canis | 90986.0 | 0 | Canis lupus | 90986.0 | 0 |
| scf7180000049236 | 34433 | 0.4395 | 0 | 75.8897 | Eukaryota | 93170.0 | 0 | Chordata | 93170.0 | 0 | Carnivora | 93170.0 | 0 | Canidae | 93170.0 | 0 | Canis | 93170.0 | 0 | Canis lupus | 93170.0 | 0 |
| scf7180000042848 | 480566 | 0.4347 | 0 | 59.044 | Eukaryota | 89467.0 | 0 | Chordata | 89467.0 | 0 | Carnivora | 89467.0 | 0 | Canidae | 54816.0 | 3 | Canis | 36344.0 | 4 | Canis lupus | 36344.0 | 4 |
| scf7180000053021 | 13844 | 0.489 | 0 | 92.5725 | Eukaryota | 8714.0 | 0 | Chordata | 8714.0 | 0 | Carnivora | 8714.0 | 0 | Canidae | 8714.0 | 0 | Canis | 7540.0 | 1 | Canis lupus | 6344.0 | 2 |
| scf7180000052612 | 9345 | 0.4464 | 0 | 230.4927 | no-hit | 0.0 | 0 | no-hit | 0.0 | 0 | no-hit | 0.0 | 0 | no-hit | 0.0 | 0 | no-hit | 0.0 | 0 | no-hit | 0.0 | 0 |
| scf7180000054937 | 18644 | 0.4555 | 0 | 295.8298 | Eukaryota | 11749.0 | 0 | Chordata | 11749.0 | 0 | Carnivora | 11749.0 | 0 | Canidae | 11749.0 | 0 | Canis | 9740.0 | 1 | Canis lupus | 8755.0 | 2 |
| scf7180000052770 | 7550 | 0.4641 | 0 | 203.4353 | Eukaryota | 21323.0 | 0 | Chordata | 21323.0 | 0 | Carnivora | 21323.0 | 0 | Canidae | 21323.0 | 0 | Canis | 21323.0 | 0 | Canis lupus | 21323.0 | 0 |
| scf7180000046862 | 24467 | 0.4776 | 0 | 297.0486 | Eukaryota | 50507.0 | 0 | Chordata | 50507.0 | 0 | Carnivora | 50507.0 | 0 | Canidae | 50507.0 | 0 | Canis | 50507.0 | 0 | Canis lupus | 50507.0 | 0 |
| scf7180000041257 | 1091294 | 0.3677 | 0 | 59.5432 | Eukaryota | 36535.0 | 0 | Chordata | 36535.0 | 0 | Carnivora | 36535.0 | 0 | Canidae | 36535.0 | 0 | Canis | 29109.0 | 1 | Canis lupus | 29109.0 | 1 |
| scf7180000045501 | 212248 | 0.3629 | 0 | 58.0472 | Eukaryota | 132589.0 | 0 | Chordata | 132589.0 | 0 | Carnivora | 132589.0 | 0 | Canidae | 124725.0 | 1 | Canis | 107901.0 | 2 | Canis lupus | 107901.0 | 2 |
| scf7180000050444 | 19257 | 0.5015 | 0 | 546.5159 | Eukaryota | 9199.0 | 0 | Chordata | 9199.0 | 0 | Carnivora | 9199.0 | 0 | Canidae | 9199.0 | 0 | Canis | 7983.0 | 1 | Canis lupus | 6706.0 | 2 |
| scf7180000053430 | 2484 | 0.4533 | 0 | 699.3097 | no-hit | 0.0 | 0 | no-hit | 0.0 | 0 | no-hit | 0.0 | 0 | no-hit | 0.0 | 0 | no-hit | 0.0 | 0 | no-hit | 0.0 | 0 |
| scf7180000052440 | 10849 | 0.4458 | 0 | 90.1373 | Eukaryota | 6895.0 | 0 | Chordata | 3639.0 | 1 | Polystomatidea | 3256.0 | 3 | Polystomatidae | 3256.0 | 3 | Protopolystoma | 3256.0 | 3 | Protopolystoma xenopodis | 3256.0 | 3 |
| scf7180000042715 | 360300 | 0.4319 | 0 | 59.6999 | Eukaryota | 40416.0 | 0 | Chordata | 40416.0 | 0 | Carnivora | 40416.0 | 0 | Otariidae | 15360.0 | 3 | Zalophus | 11553.0 | 6 | Zalophus californianus | 11553.0 | 6 |
| scf7180000054293 | 595965 | 0.4256 | 0 | 68.4021 | Eukaryota | 119199.0 | 0 | Chordata | 119199.0 | 0 | Carnivora | 119199.0 | 0 | Canidae | 119199.0 | 0 | Vulpes | 93513.0 | 1 | Vulpes vulpes | 93513.0 | 1 |
| scf7180000054766 | 16218 | 0.44 | 0 | 316.1509 | Eukaryota | 3994.0 | 0 | Platyhelminthes | 2530.0 | 2 | Polystomatidea | 2395.0 | 3 | Polystomatidae | 2395.0 | 3 | Protopolystoma | 2395.0 | 3 | Protopolystoma xenopodis | 2395.0 | 3 |
| scf7180000044793 | 276225 | 0.3438 | 0 | 60.0605 | Eukaryota | 20554.0 | 0 | Chordata | 20554.0 | 0 | Carnivora | 20554.0 | 0 | Felidae | 18577.0 | 1 | Felis | 18577.0 | 1 | Felis catus | 18577.0 | 1 |
| scf7180000052467 | 5400 | 0.458 | 0 | 134.0364 | Eukaryota | 1884.0 | 0 | Chordata | 1884.0 | 0 | Tetraodontiformes | 1884.0 | 0 | Tetraodontidae | 1884.0 | 0 | Takifugu | 1884.0 | 0 | Takifugu rubripes | 1884.0 | 0 |
| scf7180000048962 | 33573 | 0.4423 | 0 | 57.8869 | Eukaryota | 10287.0 | 0 | Chordata | 10287.0 | 0 | Carnivora | 6970.0 | 2 | Ursidae | 4076.0 | 3 | Canis | 2894.0 | 4 | Canis lupus | 2894.0 | 5 |
| scf7180000046608 | 110113 | 0.4856 | 0 | 58.4193 | Eukaryota | 62861.0 | 0 | Chordata | 62861.0 | 0 | Carnivora | 62861.0 | 0 | Canidae | 62861.0 | 0 | Canis | 43973.0 | 1 | Canis lupus | 43973.0 | 1 |
| scf7180000052318 | 14239 | 0.4299 | 0 | 43.5494 | Eukaryota | 5458.0 | 0 | Chordata | 5458.0 | 0 | Chiroptera | 3484.0 | 2 | Pteropodidae | 3484.0 | 2 | Rousettus | 3484.0 | 3 | Rousettus aegyptiacus | 3484.0 | 3 |
| scf7180000044813 | 302950 | 0.4085 | 0 | 63.1266 | Eukaryota | 95005.0 | 0 | Chordata | 95005.0 | 0 | Carnivora | 95005.0 | 0 | Canidae | 95005.0 | 0 | Canis | 95005.0 | 0 | Canis lupus | 95005.0 | 0 |
| scf7180000053877 | 2965 | 0.457 | 0 | 447.5925 | Eukaryota | 2716.0 | 0 | Platyhelminthes | 1678.0 | 3 | Polystomatidea | 1678.0 | 3 | Polystomatidae | 1678.0 | 3 | Protopolystoma | 1678.0 | 3 | Protopolystoma xenopodis | 1678.0 | 3 |
| scf7180000055594 | 19195 | 0.4117 | 0 | 863.7624 | Eukaryota | 88535.0 | 0 | Chordata | 88535.0 | 0 | Carnivora | 88535.0 | 0 | Canidae | 88535.0 | 0 | Canis | 88535.0 | 0 | Canis lupus | 88535.0 | 0 |
| scf7180000047937 | 71195 | 0.4405 | 0 | 113.974 | Eukaryota | 42293.0 | 0 | Chordata | 26976.0 | 2 | Gadiformes | 18491.0 | 5 | Gadidae | 18491.0 | 5 | Gadus | 18491.0 | 5 | Gadus morhua | 18491.0 | 5 |
| scf7180000046702 | 98500 | 0.4237 | 0 | 59.9479 | Eukaryota | 55722.0 | 0 | Chordata | 55722.0 | 0 | Carnivora | 55722.0 | 0 | Canidae | 42329.0 | 4 | Canis | 35812.0 | 6 | Canis lupus | 35812.0 | 6 |
| scf7180000046260 | 124477 | 0.6917 | 0 | 53.0985 | Eukaryota | 59410.0 | 0 | Chordata | 59410.0 | 0 | Carnivora | 59410.0 | 0 | Canidae | 59410.0 | 0 | Vulpes | 59410.0 | 0 | Vulpes vulpes | 59410.0 | 0 |
| scf7180000045211 | 33057 | 0.4492 | 0 | 195.13 | Eukaryota | 9409.0 | 0 | Chordata | 8684.0 | 1 | Carnivora | 8684.0 | 1 | Canidae | 8684.0 | 1 | Canis | 5661.0 | 2 | Canis lupus | 5661.0 | 2 |
| scf7180000045341 | 113041 | 0.4127 | 0 | 57.2042 | Eukaryota | 90832.0 | 0 | Chordata | 90832.0 | 0 | Carnivora | 90832.0 | 0 | Canidae | 90832.0 | 0 | Canis | 90832.0 | 0 | Canis lupus | 90832.0 | 0 |
| scf7180000054332 | 1256672 | 0.3866 | 0 | 60.1387 | Eukaryota | 92523.0 | 0 | Chordata | 92523.0 | 0 | Carnivora | 92523.0 | 0 | Canidae | 92523.0 | 0 | Canis | 92523.0 | 0 | Canis lupus | 92523.0 | 0 |
| scf7180000043703 | 404455 | 0.3342 | 0 | 59.4772 | Eukaryota | 92954.0 | 0 | Chordata | 92954.0 | 0 | Carnivora | 92954.0 | 0 | Canidae | 92954.0 | 0 | Canis | 92954.0 | 0 | Canis lupus | 92954.0 | 0 |
| scf7180000044212 | 4362 | 0.4523 | 0 | 215.4166 | Eukaryota | 1259.0 | 0 | Streptophyta | 437.0 | 2 | Fabales | 437.0 | 2 | Fabaceae | 437.0 | 2 | Cicer | 437.0 | 2 | Cicer arietinum | 437.0 | 2 |
| scf7180000040655 | 146688 | 0.5116 | 0 | 54.4294 | Eukaryota | 33000.0 | 0 | Chordata | 33000.0 | 0 | Carnivora | 33000.0 | 0 | Canidae | 33000.0 | 0 | Canis | 25265.0 | 1 | Canis lupus | 25265.0 | 1 |
| scf7180000052361 | 4204 | 0.4512 | 0 | 708.4194 | no-hit | 0.0 | 0 | no-hit | 0.0 | 0 | no-hit | 0.0 | 0 | no-hit | 0.0 | 0 | no-hit | 0.0 | 0 | no-hit | 0.0 | 0 |
| scf7180000044267 | 318160 | 0.3882 | 0 | 58.7466 | Eukaryota | 44900.0 | 0 | Chordata | 44900.0 | 0 | Carnivora | 44900.0 | 0 | Canidae | 44900.0 | 0 | Canis | 44900.0 | 0 | Canis lupus | 44900.0 | 0 |
| scf7180000043397 | 476147 | 0.4237 | 0 | 60.6375 | Eukaryota | 46942.0 | 0 | Chordata | 46942.0 | 0 | Carnivora | 46942.0 | 0 | Canidae | 46942.0 | 0 | Canis | 41898.0 | 1 | Canis lupus | 41898.0 | 1 |
| scf7180000041948 | 8476 | 0.3804 | 0 | 29.4879 | Eukaryota | 12307.0 | 0 | Chordata | 12307.0 | 0 | Carnivora | 12307.0 | 0 | Canidae | 12307.0 | 0 | Canis | 12307.0 | 0 | Canis lupus | 12307.0 | 0 |
| scf7180000048630 | 6937 | 0.4358 | 0 | 112.7786 | Eukaryota | 7591.0 | 0 | Chordata | 6527.0 | 3 | Gadiformes | 2815.0 | 8 | Gadidae | 2815.0 | 8 | Gadus | 2815.0 | 8 | Gadus morhua | 2815.0 | 8 |
| scf7180000044108 | 517544 | 0.4152 | 0 | 64.5992 | Eukaryota | 95165.0 | 0 | Chordata | 95165.0 | 0 | Carnivora | 95165.0 | 0 | Canidae | 95165.0 | 0 | Canis | 95165.0 | 0 | Canis lupus | 95165.0 | 0 |
| scf7180000048883 | 19279 | 0.5051 | 0 | 771.4082 | Eukaryota | 9134.0 | 0 | Chordata | 9134.0 | 0 | Carnivora | 9134.0 | 0 | Canidae | 9134.0 | 0 | Canis | 7368.0 | 1 | Canis lupus | 6114.0 | 2 |
| scf7180000041073 | 1373840 | 0.4548 | 0 | 59.9374 | Eukaryota | 68284.0 | 0 | Chordata | 68284.0 | 0 | Carnivora | 68284.0 | 0 | Canidae | 68284.0 | 0 | Canis | 46174.0 | 1 | Canis lupus | 46174.0 | 1 |
| scf7180000046824 | 95309 | 0.3482 | 0 | 57.7422 | Eukaryota | 18182.0 | 0 | Chordata | 18182.0 | 0 | Artiodactyla | 5894.0 | 5 | Bovidae | 5894.0 | 5 | Homo | 5653.0 | 6 | Homo sapiens | 5653.0 | 6 |
| scf7180000043705 | 96900 | 0.3406 | 0 | 54.3826 | Eukaryota | 39622.0 | 0 | Chordata | 39622.0 | 0 | Carnivora | 39622.0 | 0 | Canidae | 39622.0 | 0 | Canis | 39622.0 | 0 | Canis lupus | 39622.0 | 0 |
| scf7180000048262 | 8630 | 0.4366 | 0 | 116.9624 | no-hit | 0.0 | 0 | no-hit | 0.0 | 0 | no-hit | 0.0 | 0 | no-hit | 0.0 | 0 | no-hit | 0.0 | 0 | no-hit | 0.0 | 0 |
| scf7180000056105 | 19750 | 0.4422 | 0 | 101.5575 | Eukaryota | 1687.0 | 0 | Platyhelminthes | 1138.0 | 2 | Polystomatidea | 1138.0 | 3 | Polystomatidae | 1138.0 | 3 | Protopolystoma | 1138.0 | 3 | Protopolystoma xenopodis | 1138.0 | 3 |
| scf7180000041186 | 778817 | 0.4116 | 0 | 59.0977 | Eukaryota | 90775.0 | 0 | Chordata | 90775.0 | 0 | Carnivora | 90775.0 | 0 | Canidae | 90775.0 | 0 | Canis | 90775.0 | 0 | Canis lupus | 90775.0 | 0 |
| scf7180000044395 | 316331 | 0.3864 | 0 | 60.1351 | Eukaryota | 102808.0 | 0 | Chordata | 102808.0 | 0 | Carnivora | 102808.0 | 0 | Canidae | 61922.0 | 1 | Canis | 40994.0 | 3 | Canis lupus | 40994.0 | 3 |
| scf7180000043210 | 410866 | 0.3516 | 0 | 66.7477 | Eukaryota | 87810.0 | 0 | Chordata | 87810.0 | 0 | Carnivora | 87810.0 | 0 | Canidae | 87810.0 | 0 | Canis | 87810.0 | 0 | Canis lupus | 87810.0 | 0 |
| scf7180000043462 | 309064 | 0.3862 | 0 | 61.1028 | Eukaryota | 95140.0 | 0 | Chordata | 95140.0 | 0 | Carnivora | 95140.0 | 0 | Canidae | 95140.0 | 0 | Canis | 95140.0 | 0 | Canis lupus | 95140.0 | 0 |
| scf7180000041835 | 682239 | 0.3479 | 0 | 59.5511 | Eukaryota | 55265.0 | 0 | Chordata | 55265.0 | 0 | Carnivora | 55265.0 | 0 | Canidae | 55265.0 | 0 | Canis | 55265.0 | 0 | Canis lupus | 55265.0 | 0 |
| scf7180000045878 | 152352 | 0.5194 | 0 | 58.9518 | Eukaryota | 24920.0 | 0 | Chordata | 24920.0 | 0 | Carnivora | 24920.0 | 0 | Canidae | 24920.0 | 0 | Canis | 24920.0 | 0 | Canis lupus | 24920.0 | 0 |
| scf7180000044871 | 1028994 | 0.5284 | 100 | 56.9251 | Eukaryota | 98068.0 | 0 | Chordata | 98068.0 | 0 | Carnivora | 98068.0 | 0 | Canidae | 98068.0 | 0 | Canis | 79316.0 | 1 | Canis lupus | 79316.0 | 1 |
| scf7180000049434 | 54953 | 0.4375 | 0 | 60.0813 | Eukaryota | 24460.0 | 0 | Chordata | 24460.0 | 0 | Carnivora | 24460.0 | 0 | Canidae | 18466.0 | 3 | Canis | 12396.0 | 4 | Canis lupus | 12396.0 | 5 |
| scf7180000055604 | 6608 | 0.4411 | 0 | 191.0287 | no-hit | 0.0 | 0 | no-hit | 0.0 | 0 | no-hit | 0.0 | 0 | no-hit | 0.0 | 0 | no-hit | 0.0 | 0 | no-hit | 0.0 | 0 |
| scf7180000054678 | 35398 | 0.4417 | 0 | 118.2184 | Eukaryota | 1744.0 | 0 | Platyhelminthes | 1744.0 | 0 | Polystomatidea | 1744.0 | 0 | Polystomatidae | 1744.0 | 0 | Protopolystoma | 1744.0 | 0 | Protopolystoma xenopodis | 1744.0 | 0 |
| scf7180000045415 | 251493 | 0.3766 | 0 | 60.6056 | Eukaryota | 54015.0 | 0 | Chordata | 54015.0 | 0 | Carnivora | 54015.0 | 0 | Canidae | 54015.0 | 0 | Canis | 54015.0 | 0 | Canis lupus | 54015.0 | 0 |
| scf7180000040889 | 1633562 | 0.4643 | 0 | 60.1751 | Eukaryota | 76474.0 | 0 | Chordata | 76474.0 | 0 | Carnivora | 76474.0 | 0 | Canidae | 76474.0 | 0 | Canis | 52607.0 | 1 | Canis lupus | 52607.0 | 1 |
| scf7180000041283 | 218873 | 0.4223 | 0 | 55.1685 | Eukaryota | 87139.0 | 0 | Chordata | 87139.0 | 0 | Carnivora | 87139.0 | 0 | Canidae | 87139.0 | 0 | Canis | 87139.0 | 0 | Canis lupus | 87139.0 | 0 |
| scf7180000052014 | 3009 | 0.4477 | 0 | 128.7253 | no-hit | 0.0 | 0 | no-hit | 0.0 | 0 | no-hit | 0.0 | 0 | no-hit | 0.0 | 0 | no-hit | 0.0 | 0 | no-hit | 0.0 | 0 |
| scf7180000043637 | 430337 | 0.4157 | 0 | 59.8335 | Eukaryota | 58064.0 | 0 | Chordata | 58064.0 | 0 | Carnivora | 58064.0 | 0 | Canidae | 53109.0 | 1 | Canis | 43022.0 | 2 | Canis lupus | 43022.0 | 2 |
| scf7180000041231 | 1299019 | 0.4469 | 0 | 59.4119 | Eukaryota | 57399.0 | 0 | Chordata | 57399.0 | 0 | Carnivora | 57399.0 | 0 | Canidae | 57399.0 | 0 | Canis | 52887.0 | 1 | Canis lupus | 52887.0 | 1 |
| scf7180000052217 | 3210 | 0.4573 | 0 | 832.8192 | no-hit | 0.0 | 0 | no-hit | 0.0 | 0 | no-hit | 0.0 | 0 | no-hit | 0.0 | 0 | no-hit | 0.0 | 0 | no-hit | 0.0 | 0 |
| scf7180000050904 | 16812 | 0.518 | 0 | 1017.2524 | Eukaryota | 16143.0 | 0 | Chordata | 16143.0 | 0 | Carnivora | 16143.0 | 0 | Canidae | 16143.0 | 0 | Canis | 16143.0 | 0 | Canis lupus | 16143.0 | 0 |
| scf7180000051961 | 8470 | 0.37 | 0 | 60.8302 | Eukaryota | 3086.0 | 0 | Chordata | 3086.0 | 0 | Carnivora | 2184.0 | 2 | Canidae | 1161.0 | 4 | Canis | 1161.0 | 4 | Canis lupus | 1161.0 | 4 |
| scf7180000044593 | 99218 | 0.3664 | 0 | 59.0735 | Eukaryota | 45762.0 | 0 | Chordata | 45762.0 | 0 | Carnivora | 45762.0 | 0 | Canidae | 45762.0 | 0 | Canis | 45762.0 | 0 | Canis lupus | 45762.0 | 0 |
| scf7180000051453 | 6044 | 0.4466 | 0 | 99.1747 | no-hit | 0.0 | 0 | no-hit | 0.0 | 0 | no-hit | 0.0 | 0 | no-hit | 0.0 | 0 | no-hit | 0.0 | 0 | no-hit | 0.0 | 0 |
| scf7180000049524 | 21280 | 0.4507 | 0 | 1405.9659 | Eukaryota | 25725.0 | 0 | Chordata | 22607.0 | 2 | Gadiformes | 15920.0 | 6 | Gadidae | 15920.0 | 6 | Gadus | 15920.0 | 6 | Gadus morhua | 15920.0 | 6 |
| scf7180000041279 | 1322222 | 0.3777 | 0 | 60.7965 | Eukaryota | 114786.0 | 0 | Chordata | 114786.0 | 0 | Carnivora | 114786.0 | 0 | Canidae | 114786.0 | 0 | Canis | 114786.0 | 0 | Canis lupus | 114786.0 | 0 |
| scf7180000046503 | 120104 | 0.3442 | 0 | 58.5731 | Eukaryota | 17850.0 | 0 | Chordata | 17850.0 | 0 | Carnivora | 17850.0 | 0 | Canidae | 17850.0 | 0 | Canis | 17850.0 | 0 | Canis lupus | 17850.0 | 0 |
| scf7180000051814 | 2767 | 0.4478 | 0 | 350.0764 | Eukaryota | 2775.0 | 0 | Platyhelminthes | 1750.0 | 2 | Polystomatidea | 1750.0 | 4 | Polystomatidae | 1750.0 | 4 | Protopolystoma | 1750.0 | 4 | Protopolystoma xenopodis | 1750.0 | 4 |
| scf7180000040516 | 505860 | 0.5169 | 0 | 57.4688 | Eukaryota | 88235.0 | 0 | Chordata | 88235.0 | 0 | Carnivora | 88235.0 | 0 | Canidae | 88235.0 | 0 | Canis | 88235.0 | 0 | Canis lupus | 88235.0 | 0 |
| scf7180000043625 | 71380 | 0.3782 | 0 | 59.1396 | Eukaryota | 59287.0 | 0 | Chordata | 59287.0 | 0 | Carnivora | 59287.0 | 0 | Canidae | 59287.0 | 0 | Canis | 59287.0 | 0 | Canis lupus | 59287.0 | 0 |
| scf7180000050341 | 30422 | 0.4394 | 0 | 74.4378 | Eukaryota | 4024.0 | 0 | Platyhelminthes | 2272.0 | 1 | Polystomatidea | 2272.0 | 4 | Polystomatidae | 2272.0 | 4 | Protopolystoma | 2272.0 | 4 | Protopolystoma xenopodis | 2272.0 | 4 |
| scf7180000042559 | 997114 | 0.3827 | 0 | 61.1358 | Eukaryota | 191017.0 | 0 | Chordata | 191017.0 | 0 | Carnivora | 191017.0 | 0 | Canidae | 191017.0 | 0 | Canis | 191017.0 | 0 | Canis lupus | 191017.0 | 0 |
| scf7180000042757 | 21412 | 0.4288 | 0 | 194.173 | Eukaryota | 38316.0 | 0 | Chordata | 38316.0 | 0 | Carnivora | 38316.0 | 0 | Canidae | 38316.0 | 0 | Canis | 38316.0 | 0 | Canis lupus | 38316.0 | 0 |
| scf7180000054264 | 12870 | 0.442 | 0 | 132.8992 | no-hit | 0.0 | 0 | no-hit | 0.0 | 0 | no-hit | 0.0 | 0 | no-hit | 0.0 | 0 | no-hit | 0.0 | 0 | no-hit | 0.0 | 0 |
| scf7180000047165 | 30174 | 0.446 | 0 | 68.3316 | Eukaryota | 4829.0 | 0 | Platyhelminthes | 3049.0 | 1 | Polystomatidea | 2890.0 | 4 | Polystomatidae | 2890.0 | 4 | Protopolystoma | 2890.0 | 4 | Protopolystoma xenopodis | 2890.0 | 4 |
| scf7180000043699 | 66889 | 0.4961 | 0 | 53.4313 | Eukaryota | 8497.0 | 0 | Chordata | 8497.0 | 0 | Artiodactyla | 4046.0 | 2 | Camelidae | 3225.0 | 3 | Camelus | 3225.0 | 4 | Camelus dromedarius | 3225.0 | 4 |
| scf7180000054748 | 40497 | 0.4641 | 0 | 408.9291 | Eukaryota | 53868.0 | 0 | Chordata | 53868.0 | 0 | Carnivora | 53868.0 | 0 | Canidae | 53868.0 | 0 | Canis | 53868.0 | 0 | Canis lupus | 53868.0 | 0 |
| scf7180000048515 | 52116 | 0.6217 | 0 | 53.6204 | Eukaryota | 48204.0 | 0 | Chordata | 48204.0 | 0 | Carnivora | 48204.0 | 0 | Canidae | 48204.0 | 0 | Canis | 48204.0 | 0 | Canis lupus | 48204.0 | 0 |
| scf7180000045596 | 185693 | 0.3823 | 0 | 59.844 | Eukaryota | 58743.0 | 0 | Chordata | 58743.0 | 0 | Carnivora | 58743.0 | 0 | Canidae | 58743.0 | 0 | Canis | 58743.0 | 0 | Canis lupus | 58743.0 | 0 |
| scf7180000041272 | 956201 | 0.4124 | 0 | 57.4592 | Eukaryota | 65970.0 | 0 | Chordata | 65970.0 | 0 | Carnivora | 65970.0 | 0 | Canidae | 65970.0 | 0 | Canis | 65970.0 | 0 | Canis lupus | 65970.0 | 0 |
| scf7180000044982 | 172161 | 0.3809 | 0 | 59.3679 | Eukaryota | 84351.0 | 0 | Chordata | 84351.0 | 0 | Carnivora | 84351.0 | 0 | Canidae | 84351.0 | 0 | Canis | 84351.0 | 0 | Canis lupus | 84351.0 | 0 |
| scf7180000041142 | 1283052 | 0.4089 | 0 | 59.6417 | Eukaryota | 93829.0 | 0 | Chordata | 93829.0 | 0 | Carnivora | 93829.0 | 0 | Canidae | 93829.0 | 0 | Canis | 93829.0 | 0 | Canis lupus | 93829.0 | 0 |
| scf7180000050913 | 3947 | 0.4495 | 0 | 497.8886 | no-hit | 0.0 | 0 | no-hit | 0.0 | 0 | no-hit | 0.0 | 0 | no-hit | 0.0 | 0 | no-hit | 0.0 | 0 | no-hit | 0.0 | 0 |
| scf7180000050912 | 5237 | 0.4436 | 0 | 237.1378 | Eukaryota | 2223.0 | 0 | Platyhelminthes | 1472.0 | 2 | Polystomatidea | 1472.0 | 2 | Polystomatidae | 1472.0 | 2 | Protopolystoma | 1472.0 | 2 | Protopolystoma xenopodis | 1472.0 | 2 |
| scf7180000047529 | 82050 | 0.4384 | 0 | 71.2246 | Eukaryota | 63072.0 | 0 | Chordata | 38765.0 | 1 | Gadiformes | 27092.0 | 5 | Gadidae | 27092.0 | 5 | Gadus | 27092.0 | 5 | Gadus morhua | 27092.0 | 5 |
| scf7180000056526 | 280909 | 0.5184 | 0 | 56.1503 | Eukaryota | 59481.0 | 0 | Chordata | 59481.0 | 0 | Carnivora | 59481.0 | 0 | Canidae | 55471.0 | 1 | Canis | 42604.0 | 2 | Canis lupus | 42604.0 | 2 |
| scf7180000049634 | 7410 | 0.4421 | 0 | 197.7271 | Eukaryota | 396.0 | 0 | Platyhelminthes | 396.0 | 0 | Polystomatidea | 396.0 | 0 | Polystomatidae | 396.0 | 0 | Protopolystoma | 396.0 | 0 | Protopolystoma xenopodis | 396.0 | 0 |
| scf7180000050733 | 19397 | 0.4219 | 0 | 262.7133 | Eukaryota | 33648.0 | 0 | Chordata | 33648.0 | 0 | Carnivora | 33648.0 | 0 | Canidae | 33648.0 | 0 | Canis | 33648.0 | 0 | Canis lupus | 33648.0 | 0 |
| scf7180000040709 | 1730185 | 0.3687 | 0 | 60.993 | Eukaryota | 92171.0 | 0 | Chordata | 92171.0 | 0 | Carnivora | 92171.0 | 0 | Canidae | 92171.0 | 0 | Canis | 81174.0 | 1 | Canis lupus | 81174.0 | 1 |
| scf7180000049587 | 39490 | 0.4399 | 0 | 221.8311 | Eukaryota | 16030.0 | 0 | Chordata | 16030.0 | 0 | Tetraodontiformes | 8409.0 | 4 | Tetraodontidae | 8409.0 | 4 | Takifugu | 8409.0 | 4 | Takifugu rubripes | 7389.0 | 5 |
| scf7180000048851 | 14149 | 0.4383 | 0 | 41.9229 | Eukaryota | 799.0 | 0 | Platyhelminthes | 799.0 | 0 | Polystomatidea | 799.0 | 0 | Polystomatidae | 799.0 | 0 | Protopolystoma | 799.0 | 0 | Protopolystoma xenopodis | 799.0 | 0 |
| scf7180000042160 | 247322 | 0.3745 | 0 | 61.0164 | Eukaryota | 88826.0 | 0 | Chordata | 88826.0 | 0 | Carnivora | 88826.0 | 0 | Canidae | 88826.0 | 0 | Canis | 88826.0 | 0 | Canis lupus | 88826.0 | 0 |
| scf7180000045582 | 197451 | 0.3436 | 0 | 60.827 | Eukaryota | 66094.0 | 0 | Chordata | 66094.0 | 0 | Carnivora | 66094.0 | 0 | Canidae | 66094.0 | 0 | Canis | 66094.0 | 0 | Canis lupus | 66094.0 | 0 |
| scf7180000042325 | 36440 | 0.4679 | 0 | 61.9874 | Eukaryota | 30920.0 | 0 | Chordata | 30920.0 | 0 | Carnivora | 30920.0 | 0 | Canidae | 30920.0 | 0 | Canis | 30920.0 | 0 | Canis lupus | 30920.0 | 0 |
| scf7180000044176 | 80570 | 0.3685 | 0 | 58.784 | Eukaryota | 10851.0 | 0 | Chordata | 10851.0 | 0 | Carnivora | 6279.0 | 1 | Hominidae | 4572.0 | 3 | Homo | 4572.0 | 3 | Homo sapiens | 4572.0 | 3 |
| scf7180000040654 | 220567 | 0.3894 | 0 | 58.6646 | Eukaryota | 106952.0 | 0 | Chordata | 106952.0 | 0 | Carnivora | 106952.0 | 0 | Canidae | 106952.0 | 0 | Canis | 106952.0 | 0 | Canis lupus | 106952.0 | 0 |
| scf7180000048270 | 50847 | 0.4813 | 0 | 60.217 | Eukaryota | 4709.0 | 0 | Chordata | 4709.0 | 0 | Primates | 2895.0 | 1 | Hominidae | 2895.0 | 1 | Canis | 1814.0 | 2 | Canis lupus | 1814.0 | 2 |
| scf7180000051578 | 2525 | 0.4539 | 0 | 439.7942 | no-hit | 0.0 | 0 | no-hit | 0.0 | 0 | no-hit | 0.0 | 0 | no-hit | 0.0 | 0 | no-hit | 0.0 | 0 | no-hit | 0.0 | 0 |
| scf7180000050090 | 3246 | 0.4559 | 0 | 445.6814 | Eukaryota | 1683.0 | 0 | Echinodermata | 569.0 | 3 | Forcipulatida | 569.0 | 3 | Asteriidae | 569.0 | 3 | Asterias | 569.0 | 3 | Asterias rubens | 569.0 | 3 |
| scf7180000043845 | 418448 | 0.3981 | 0 | 59.829 | Eukaryota | 63472.0 | 0 | Chordata | 63472.0 | 0 | Carnivora | 63472.0 | 0 | Canidae | 63472.0 | 0 | Canis | 63472.0 | 0 | Canis lupus | 63472.0 | 0 |
| scf7180000048581 | 5860 | 0.4514 | 0 | 194.0312 | Eukaryota | 2704.0 | 0 | Chordata | 2704.0 | 0 | Carnivora | 2704.0 | 0 | Canidae | 2704.0 | 0 | Canis | 2704.0 | 0 | Canis lupus | 2704.0 | 0 |
| scf7180000041089 | 1318954 | 0.3425 | 0 | 58.4384 | Eukaryota | 80628.0 | 0 | Chordata | 80628.0 | 0 | Carnivora | 80628.0 | 0 | Canidae | 80628.0 | 0 | Canis | 80628.0 | 0 | Canis lupus | 80628.0 | 0 |
| scf7180000041590 | 65050 | 0.4269 | 0 | 79.3495 | Eukaryota | 7481.0 | 0 | Chordata | 7481.0 | 0 | Artiodactyla | 3160.0 | 3 | Bovidae | 2520.0 | 4 | Homo | 2106.0 | 5 | Homo sapiens | 2106.0 | 6 |
| scf7180000050691 | 15454 | 0.4526 | 0 | 58.1508 | Eukaryota | 3047.0 | 0 | Chordata | 3047.0 | 0 | Carnivora | 3047.0 | 0 | Canidae | 2296.0 | 1 | Canis | 2296.0 | 2 | Canis lupus | 2296.0 | 2 |
| scf7180000049824 | 25104 | 0.4465 | 0 | 536.7515 | no-hit | 0.0 | 0 | no-hit | 0.0 | 0 | no-hit | 0.0 | 0 | no-hit | 0.0 | 0 | no-hit | 0.0 | 0 | no-hit | 0.0 | 0 |
| scf7180000050712 | 2086 | 0.4386 | 0 | 132.4792 | no-hit | 0.0 | 0 | no-hit | 0.0 | 0 | no-hit | 0.0 | 0 | no-hit | 0.0 | 0 | no-hit | 0.0 | 0 | no-hit | 0.0 | 0 |
| scf7180000051158 | 3363 | 0.5046 | 0 | 42.3638 | Eukaryota | 6942.0 | 0 | Chordata | 6942.0 | 0 | Carnivora | 6942.0 | 0 | Canidae | 6942.0 | 0 | Canis | 4536.0 | 1 | Canis lupus | 3484.0 | 2 |
| scf7180000056612 | 486176 | 0.3506 | 0 | 59.6545 | Eukaryota | 92652.0 | 0 | Chordata | 92652.0 | 0 | Carnivora | 92652.0 | 0 | Canidae | 92652.0 | 0 | Canis | 92652.0 | 0 | Canis lupus | 92652.0 | 0 |
| scf7180000048786 | 29689 | 0.4412 | 0 | 137.2172 | Eukaryota | 4928.0 | 0 | Platyhelminthes | 2560.0 | 1 | Polystomatidea | 2560.0 | 1 | Polystomatidae | 2560.0 | 1 | Protopolystoma | 2560.0 | 1 | Protopolystoma xenopodis | 2560.0 | 1 |
| scf7180000041783 | 111356 | 0.3373 | 0 | 57.8408 | Eukaryota | 7610.0 | 0 | Chordata | 7610.0 | 0 | Artiodactyla | 2329.0 | 3 | Bovidae | 2329.0 | 4 | Homo | 2099.0 | 6 | Homo sapiens | 2099.0 | 6 |
| scf7180000042348 | 536493 | 0.3722 | 0 | 62.2589 | Eukaryota | 94184.0 | 0 | Chordata | 94184.0 | 0 | Carnivora | 94184.0 | 0 | Canidae | 94184.0 | 0 | Canis | 94184.0 | 0 | Canis lupus | 94184.0 | 0 |
| scf7180000050310 | 23164 | 0.3733 | 0 | 63.903 | Eukaryota | 18045.0 | 0 | Chordata | 18045.0 | 0 | Carnivora | 16185.0 | 1 | Canidae | 8057.0 | 3 | Canis | 5218.0 | 6 | Canis lupus | 5218.0 | 6 |
| scf7180000045650 | 164780 | 0.5644 | 0 | 54.8672 | Eukaryota | 60958.0 | 0 | Chordata | 60958.0 | 0 | Carnivora | 60958.0 | 0 | Canidae | 41490.0 | 1 | Vulpes | 20811.0 | 2 | Vulpes vulpes | 20811.0 | 2 |
| scf7180000040495 | 3236734 | 0.409 | 0 | 59.7584 | Eukaryota | 79842.0 | 0 | Chordata | 79842.0 | 0 | Carnivora | 79842.0 | 0 | Canidae | 79842.0 | 0 | Canis | 72763.0 | 1 | Canis lupus | 72763.0 | 1 |
| scf7180000040567 | 20873 | 0.475 | 0 | 51.2632 | Eukaryota | 13798.0 | 0 | Chordata | 13798.0 | 0 | Carnivora | 13798.0 | 0 | Canidae | 7760.0 | 3 | Canis | 5847.0 | 5 | Canis lupus | 5847.0 | 5 |
| scf7180000056058 | 18233 | 0.4527 | 0 | 448.82 | Eukaryota | 8284.0 | 0 | Chordata | 6399.0 | 1 | Tetraodontiformes | 2485.0 | 5 | Tetraodontidae | 2485.0 | 5 | Takifugu | 2485.0 | 5 | Takifugu rubripes | 2485.0 | 5 |
| scf7180000054358 | 43843 | 0.4432 | 0 | 348.2937 | Eukaryota | 92838.0 | 0 | Chordata | 92838.0 | 0 | Carnivora | 92838.0 | 0 | Canidae | 92838.0 | 0 | Canis | 92838.0 | 0 | Canis lupus | 92838.0 | 0 |
| scf7180000049294 | 11977 | 0.4447 | 0 | 275.6547 | Eukaryota | 2354.0 | 0 | Platyhelminthes | 1485.0 | 1 | Polystomatidea | 1485.0 | 1 | Polystomatidae | 1485.0 | 1 | Protopolystoma | 1485.0 | 1 | Protopolystoma xenopodis | 1485.0 | 1 |
| scf7180000052961 | 3104 | 0.4462 | 0 | 316.47 | Eukaryota | 386.0 | 0 | Platyhelminthes | 386.0 | 0 | Polystomatidea | 386.0 | 0 | Polystomatidae | 386.0 | 0 | Protopolystoma | 386.0 | 0 | Protopolystoma xenopodis | 386.0 | 0 |
| scf7180000056257 | 1179149 | 0.4847 | 0 | 58.9091 | Eukaryota | 105190.0 | 0 | Chordata | 105190.0 | 0 | Carnivora | 105190.0 | 0 | Canidae | 105190.0 | 0 | Vulpes | 105190.0 | 0 | Vulpes vulpes | 105190.0 | 0 |
| scf7180000044021 | 276087 | 0.5043 | 0 | 58.2969 | Eukaryota | 36950.0 | 0 | Chordata | 36950.0 | 0 | Carnivora | 36950.0 | 0 | Canidae | 36950.0 | 0 | Canis | 31430.0 | 1 | Canis lupus | 31430.0 | 1 |
| scf7180000040662 | 1816344 | 0.4796 | 0 | 59.5324 | Eukaryota | 81960.0 | 0 | Chordata | 81960.0 | 0 | Carnivora | 81960.0 | 0 | Canidae | 81960.0 | 0 | Canis | 81960.0 | 0 | Canis lupus | 81960.0 | 0 |
| scf7180000049391 | 14094 | 0.4447 | 0 | 373.5626 | Eukaryota | 400.0 | 0 | Streptophyta | 211.0 | 1 | Fabales | 211.0 | 1 | Fabaceae | 211.0 | 1 | Cicer | 211.0 | 1 | Cicer arietinum | 211.0 | 1 |
| scf7180000055792 | 9285 | 0.4521 | 0 | 319.7725 | Eukaryota | 2445.0 | 0 | Chordata | 2156.0 | 1 | Carnivora | 2156.0 | 1 | Canidae | 2156.0 | 1 | Canis | 2156.0 | 1 | Canis lupus | 2156.0 | 1 |
| scf7180000046538 | 132440 | 0.4598 | 0 | 59.7368 | Eukaryota | 33304.0 | 0 | Chordata | 33304.0 | 0 | Carnivora | 33304.0 | 0 | Canidae | 33304.0 | 0 | Canis | 33304.0 | 0 | Canis lupus | 33304.0 | 0 |
| scf7180000056481 | 314345 | 0.4915 | 0 | 58.1081 | Eukaryota | 52723.0 | 0 | Chordata | 52723.0 | 0 | Carnivora | 52723.0 | 0 | Canidae | 52723.0 | 0 | Canis | 52723.0 | 0 | Canis lupus | 52723.0 | 0 |

| | | | | | | | | | | | | | | | | | | | | | | |
|---|---|---|---|---|---|---|---|---|---|---|---|---|---|---|---|---|---|---|---|---|---|---|
| scf7180000042026 | 811451 | 0.5017 | 0 | 58.0475 | Eukaryota | 43465.0 | 0 | Chordata | 43465.0 | 0 | Carnivora | 43465.0 | 0 | Canidae | 43465.0 | 0 | Canis | 29395.0 | 1 | Canis lupus | 29395.0 | 1 |
| scf7180000048260 | 23503 | 0.4342 | 0 | 125.4644 | Eukaryota | 7491.0 | 0 | Chordata | 6034.0 | 1 | Kurtiformes | 2897.0 | 4 | Apogonidae | 2897.0 | 4 | Sphaeramia | 2897.0 | 4 | Sphaeramia orbicularis | 2897.0 | 4 |
| scf7180000044960 | 285837 | 0.374 | 0 | 57.5141 | Eukaryota | 41798.0 | 0 | Chordata | 41798.0 | 0 | Carnivora | 41798.0 | 0 | Canidae | 41798.0 | 0 | Canis | 34572.0 | 1 | Canis lupus | 34572.0 | 1 |
| scf7180000047793 | 60439 | 0.3982 | 0 | 59.7771 | Eukaryota | 55855.0 | 0 | Chordata | 55855.0 | 0 | Carnivora | 55855.0 | 0 | Canidae | 55855.0 | 0 | Canis | 55855.0 | 0 | Canis lupus | 55855.0 | 0 |
| scf7180000047516 | 22631 | 0.4387 | 0 | 120.3124 | Eukaryota | 5164.0 | 0 | Chordata | 4346.0 | 1 | Kurtiformes | 2278.0 | 3 | Apogonidae | 2278.0 | 3 | Sphaeramia | 2278.0 | 3 | Sphaeramia orbicularis | 2278.0 | 3 |
| scf7180000048628 | 23869 | 0.4533 | 0 | 74.9453 | Eukaryota | 6513.0 | 0 | Platyhelminthes | 4520.0 | 1 | Polystomatidea | 4520.0 | 1 | Polystomatidae | 4520.0 | 1 | Protopolystoma | 4520.0 | 1 | Protopolystoma xenopodis | 4520.0 | 1 |
| scf7180000051148 | 3055 | 0.4478 | 0 | 149.9128 | no-hit | 0.0 | 0 | no-hit | 0.0 | 0 | no-hit | 0.0 | 0 | no-hit | 0.0 | 0 | no-hit | 0.0 | 0 | no-hit | 0.0 | 0 |
| scf7180000042366 | 6105 | 0.3894 | 0 | 82.0933 | Eukaryota | 10315.0 | 0 | Chordata | 10315.0 | 0 | Carnivora | 7830.0 | 3 | Canidae | 7830.0 | 3 | Canis | 7830.0 | 3 | Canis lupus | 7830.0 | 3 |
| scf7180000054013 | 3586 | 0.4506 | 0 | 54.2786 | Eukaryota | 385.0 | 0 | Chordata | 226.0 | 1 | Kurtiformes | 226.0 | 1 | Apogonidae | 226.0 | 1 | Sphaeramia | 226.0 | 1 | Sphaeramia orbicularis | 226.0 | 1 |
| scf7180000049883 | 9721 | 0.5007 | 0 | 99.1377 | Eukaryota | 7373.0 | 0 | Chordata | 7373.0 | 0 | Carnivora | 7373.0 | 0 | Canidae | 7373.0 | 0 | Canis | 5296.0 | 1 | Canis lupus | 4155.0 | 2 |
| scf7180000044124 | 42966 | 0.4281 | 0 | 56.4242 | Eukaryota | 59597.0 | 0 | Chordata | 59597.0 | 0 | Carnivora | 59597.0 | 0 | Canidae | 59597.0 | 0 | Canis | 59597.0 | 0 | Canis lupus | 59597.0 | 0 |
| scf7180000045361 | 195241 | 0.461 | 0 | 57.3172 | Eukaryota | 13277.0 | 0 | Chordata | 13277.0 | 0 | Carnivora | 13277.0 | 0 | Canidae | 13277.0 | 0 | Canis | 13277.0 | 0 | Canis lupus | 13277.0 | 0 |
| scf7180000041169 | 613348 | 0.4888 | 0 | 58.9611 | Eukaryota | 48197.0 | 0 | Chordata | 48197.0 | 0 | Carnivora | 48197.0 | 0 | Canidae | 37979.0 | 1 | Canis | 33883.0 | 3 | Canis lupus | 33883.0 | 3 |
| scf7180000049512 | 49015 | 0.3678 | 0 | 59.277 | Eukaryota | 48352.0 | 0 | Chordata | 48352.0 | 0 | Carnivora | 48352.0 | 0 | Canidae | 48352.0 | 0 | Canis | 48352.0 | 0 | Canis lupus | 48352.0 | 0 |
| scf7180000042621 | 177098 | 0.3494 | 0 | 58.4593 | Eukaryota | 51090.0 | 0 | Chordata | 51090.0 | 0 | Carnivora | 51090.0 | 0 | Canidae | 51090.0 | 0 | Canis | 51090.0 | 0 | Canis lupus | 51090.0 | 0 |
| scf7180000056240 | 1519465 | 0.4528 | 0 | 59.8112 | Eukaryota | 111990.0 | 0 | Chordata | 111990.0 | 0 | Carnivora | 111990.0 | 0 | Canidae | 111990.0 | 0 | Canis | 111990.0 | 0 | Canis lupus | 111990.0 | 0 |
| scf7180000041516 | 967099 | 0.5006 | 0 | 58.6413 | Eukaryota | 78091.0 | 0 | Chordata | 78091.0 | 0 | Carnivora | 78091.0 | 0 | Canidae | 78091.0 | 0 | Canis | 70828.0 | 1 | Canis lupus | 70828.0 | 1 |
| scf7180000044152 | 374255 | 0.3951 | 0 | 61.4982 | Eukaryota | 105634.0 | 0 | Chordata | 105634.0 | 0 | Carnivora | 105634.0 | 0 | Canidae | 105634.0 | 0 | Canis | 105634.0 | 0 | Canis lupus | 105634.0 | 0 |
| scf7180000042992 | 22870 | 0.5043 | 0 | 69.0198 | Eukaryota | 43758.0 | 0 | Chordata | 43758.0 | 0 | Carnivora | 43758.0 | 0 | Canidae | 43758.0 | 0 | Canis | 43758.0 | 0 | Canis lupus | 43758.0 | 0 |
| scf7180000045230 | 208199 | 0.3765 | 0 | 61.6696 | Eukaryota | 32931.0 | 0 | Chordata | 32931.0 | 0 | Carnivora | 32931.0 | 0 | Canidae | 32931.0 | 0 | Canis | 32931.0 | 0 | Canis lupus | 32931.0 | 0 |
| scf7180000051864 | 16147 | 0.3888 | 0 | 59.4664 | Eukaryota | 6183.0 | 0 | Chordata | 6183.0 | 0 | Carnivora | 6183.0 | 0 | Canidae | 6183.0 | 0 | Canis | 6183.0 | 0 | Canis lupus | 6183.0 | 0 |
| scf7180000044807 | 71263 | 0.3869 | 0 | 57.3379 | Eukaryota | 66170.0 | 0 | Chordata | 66170.0 | 0 | Carnivora | 66170.0 | 0 | Canidae | 66170.0 | 0 | Canis | 66170.0 | 0 | Canis lupus | 66170.0 | 0 |
| scf7180000045500 | 48455 | 0.5169 | 0 | 53.4318 | Eukaryota | 9694.0 | 0 | Chordata | 9694.0 | 0 | Carnivora | 7506.0 | 1 | Ursidae | 3232.0 | 4 | Ailuropoda | 3232.0 | 6 | Ailuropoda melanoleuca | 3232.0 | 6 |
| scf7180000041525 | 422933 | 0.3485 | 0 | 59.3339 | Eukaryota | 89468.0 | 0 | Chordata | 89468.0 | 0 | Carnivora | 89468.0 | 0 | Canidae | 89468.0 | 0 | Canis | 89468.0 | 0 | Canis lupus | 89468.0 | 0 |
| scf7180000052775 | 7163 | 0.4515 | 0 | 75.3989 | Eukaryota | 581.0 | 0 | Chordata | 581.0 | 0 | Kurtiformes | 427.0 | 1 | Apogonidae | 427.0 | 1 | Sphaeramia | 427.0 | 1 | Sphaeramia orbicularis | 427.0 | 1 |
| scf7180000044697 | 46740 | 0.375 | 0 | 61.3021 | Eukaryota | 75671.0 | 0 | Chordata | 75671.0 | 0 | Carnivora | 75671.0 | 0 | Canidae | 75671.0 | 0 | Canis | 75671.0 | 0 | Canis lupus | 75671.0 | 0 |
| scf7180000054684 | 20074 | 0.4406 | 0 | 185.8756 | Eukaryota | 3197.0 | 0 | Platyhelminthes | 1816.0 | 1 | Polystomatidea | 1816.0 | 3 | Polystomatidae | 1816.0 | 3 | Protopolystoma | 1816.0 | 3 | Protopolystoma xenopodis | 1816.0 | 3 |
| scf7180000043804 | 351667 | 0.4198 | 0 | 59.4018 | Eukaryota | 75258.0 | 0 | Chordata | 75258.0 | 0 | Carnivora | 75258.0 | 0 | Canidae | 75258.0 | 0 | Canis | 75258.0 | 0 | Canis lupus | 75258.0 | 0 |
| scf7180000042494 | 31492 | 0.3768 | 0 | 54.9695 | Eukaryota | 6113.0 | 0 | Chordata | 6113.0 | 0 | Carnivora | 3129.0 | 1 | Hominidae | 2984.0 | 2 | Homo | 2084.0 | 4 | Homo sapiens | 2084.0 | 4 |
| scf7180000048319 | 4616 | 0.4482 | 0 | 215.9933 | Eukaryota | 200.0 | 0 | Platyhelminthes | 200.0 | 0 | Polystomatidea | 200.0 | 0 | Polystomatidae | 200.0 | 0 | Protopolystoma | 200.0 | 0 | Protopolystoma xenopodis | 200.0 | 0 |
| scf7180000047042 | 2864 | 0.3897 | 0 | 32.5273 | Eukaryota | 36048.0 | 0 | Chordata | 36048.0 | 0 | Carnivora | 36048.0 | 0 | Canidae | 36048.0 | 0 | Canis | 36048.0 | 0 | Canis lupus | 36048.0 | 0 |
| scf7180000042332 | 597014 | 0.4541 | 0 | 59.2568 | Eukaryota | 74582.0 | 0 | Chordata | 74582.0 | 0 | Carnivora | 74582.0 | 0 | Canidae | 74582.0 | 0 | Vulpes | 46852.0 | 1 | Vulpes vulpes | 46852.0 | 1 |
| scf7180000053845 | 3261 | 0.4376 | 0 | 162.9455 | no-hit | 0.0 | 0 | no-hit | 0.0 | 0 | no-hit | 0.0 | 0 | no-hit | 0.0 | 0 | no-hit | 0.0 | 0 | no-hit | 0.0 | 0 |
| scf7180000054506 | 129121 | 0.4347 | 0 | 135.4808 | Eukaryota | 94482.0 | 0 | Chordata | 94482.0 | 0 | Carnivora | 94482.0 | 0 | Canidae | 94482.0 | 0 | Canis | 94482.0 | 0 | Canis lupus | 94482.0 | 0 |
| scf7180000051393 | 8079 | 0.4416 | 0 | 164.4601 | Eukaryota | 1321.0 | 0 | Streptophyta | 1321.0 | 0 | Poales | 1321.0 | 0 | Poaceae | 1321.0 | 0 | Oryza | 1321.0 | 0 | Oryza sativa | 1321.0 | 0 |
| scf7180000044631 | 321502 | 0.3839 | 0 | 60.8137 | Eukaryota | 32080.0 | 0 | Chordata | 32080.0 | 0 | Carnivora | 32080.0 | 0 | Canidae | 32080.0 | 0 | Vulpes | 32080.0 | 0 | Vulpes vulpes | 32080.0 | 0 |
| scf7180000046042 | 126704 | 0.4081 | 0 | 58.3942 | Eukaryota | 12390.0 | 0 | Chordata | 12390.0 | 0 | Carnivora | 12390.0 | 0 | Canidae | 12390.0 | 0 | Canis | 12390.0 | 0 | Canis lupus | 12390.0 | 0 |
| scf7180000041614 | 8168 | 0.4607 | 0 | 38.7077 | Eukaryota | 79636.0 | 0 | Chordata | 79636.0 | 0 | Carnivora | 79636.0 | 0 | Canidae | 79636.0 | 0 | Canis | 79636.0 | 0 | Canis lupus | 79636.0 | 0 |
| scf7180000051903 | 1558 | 0.4602 | 0 | 110.7346 | no-hit | 0.0 | 0 | no-hit | 0.0 | 0 | no-hit | 0.0 | 0 | no-hit | 0.0 | 0 | no-hit | 0.0 | 0 | no-hit | 0.0 | 0 |
| scf7180000041703 | 358210 | 0.3703 | 0 | 60.1514 | Eukaryota | 91487.0 | 0 | Chordata | 91487.0 | 0 | Carnivora | 91487.0 | 0 | Canidae | 91487.0 | 0 | Canis | 91487.0 | 0 | Canis lupus | 91487.0 | 0 |
| scf7180000041285 | 1223346 | 0.399 | 0 | 60.3115 | Eukaryota | 101798.0 | 0 | Chordata | 101798.0 | 0 | Carnivora | 101798.0 | 0 | Canidae | 101798.0 | 0 | Canis | 70618.0 | 1 | Canis lupus | 70618.0 | 1 |
| scf7180000050947 | 19000 | 0.4343 | 0 | 60.5966 | Eukaryota | 32594.0 | 0 | Chordata | 32594.0 | 0 | Carnivora | 32594.0 | 0 | Canidae | 20774.0 | 1 | Leptonychotes | 11820.0 | 2 | Leptonychotes weddellii | 11820.0 | 2 |
| scf7180000056365 | 50585 | 0.4321 | 0 | 55.1507 | Eukaryota | 57702.0 | 0 | Chordata | 57702.0 | 0 | Carnivora | 57702.0 | 0 | Canidae | 32870.0 | 3 | Canis | 27531.0 | 6 | Canis lupus | 27531.0 | 6 |
| scf7180000046966 | 104076 | 0.3875 | 0 | 61.5691 | Eukaryota | 43855.0 | 0 | Chordata | 43855.0 | 0 | Carnivora | 43855.0 | 0 | Canidae | 37029.0 | 1 | Canis | 28166.0 | 3 | Canis lupus | 28166.0 | 3 |
| scf7180000045185 | 202914 | 0.3993 | 0 | 58.3571 | Eukaryota | 91388.0 | 0 | Chordata | 91388.0 | 0 | Carnivora | 91388.0 | 0 | Canidae | 91388.0 | 0 | Canis | 91388.0 | 0 | Canis lupus | 91388.0 | 0 |
| scf7180000051484 | 19519 | 0.4487 | 0 | 58.3921 | Eukaryota | 7537.0 | 0 | Chordata | 7537.0 | 0 | Carnivora | 4272.0 | 1 | Canidae | 4272.0 | 2 | Canis | 4272.0 | 2 | Canis lupus | 4272.0 | 2 |
| scf7180000054323 | 40047 | 0.4402 | 0 | 603.1077 | Eukaryota | 21951.0 | 0 | Chordata | 18009.0 | 2 | Gadiformes | 5962.0 | 6 | Gadidae | 5962.0 | 6 | Gadus | 5962.0 | 7 | Gadus morhua | 5962.0 | 7 |
| scf7180000046205 | 54531 | 0.4442 | 0 | 71.7913 | Eukaryota | 37514.0 | 0 | Chordata | 37514.0 | 0 | Carnivora | 37514.0 | 0 | Canidae | 37514.0 | 0 | Canis | 37514.0 | 0 | Canis lupus | 37514.0 | 0 |
| scf7180000052893 | 19441 | 0.3873 | 0 | 57.1272 | Eukaryota | 23750.0 | 0 | Chordata | 23750.0 | 0 | Carnivora | 23750.0 | 0 | Canidae | 23750.0 | 0 | Canis | 23750.0 | 0 | Canis lupus | 23750.0 | 0 |
| scf7180000042804 | 104661 | 0.4059 | 0 | 57.0057 | Eukaryota | 63316.0 | 0 | Chordata | 63316.0 | 0 | Carnivora | 63316.0 | 0 | Canidae | 63316.0 | 0 | Canis | 63316.0 | 0 | Canis lupus | 63316.0 | 0 |
| scf7180000042393 | 356289 | 0.4465 | 0 | 57.6083 | Eukaryota | 27694.0 | 0 | Chordata | 27694.0 | 0 | Carnivora | 27694.0 | 0 | Canidae | 27694.0 | 0 | Canis | 27694.0 | 0 | Canis lupus | 27694.0 | 0 |
| scf7180000044687 | 162623 | 0.4254 | 0 | 58.0503 | Eukaryota | 92678.0 | 0 | Chordata | 92678.0 | 0 | Carnivora | 92678.0 | 0 | Canidae | 92678.0 | 0 | Canis | 92678.0 | 0 | Canis lupus | 92678.0 | 0 |
| scf7180000047391 | 4262 | 0.4338 | 0 | 97.6886 | no-hit | 0.0 | 0 | no-hit | 0.0 | 0 | no-hit | 0.0 | 0 | no-hit | 0.0 | 0 | no-hit | 0.0 | 0 | no-hit | 0.0 | 0 |
| scf7180000045416 | 182495 | 0.426 | 0 | 61.3262 | Eukaryota | 23699.0 | 0 | Chordata | 23699.0 | 0 | Carnivora | 23699.0 | 0 | Canidae | 23699.0 | 0 | Canis | 17529.0 | 1 | Canis lupus | 17529.0 | 1 |
| scf7180000045554 | 11250 | 0.48 | 0 | 102.364 | Eukaryota | 17020.0 | 0 | Chordata | 17020.0 | 0 | Carnivora | 17020.0 | 0 | Canidae | 17020.0 | 0 | Canis | 11207.0 | 1 | Canis lupus | 11207.0 | 1 |
| scf7180000042564 | 9435 | 0.4391 | 0 | 334.8635 | Eukaryota | 22243.0 | 0 | Chordata | 22243.0 | 0 | Carnivora | 22243.0 | 0 | Canidae | 22243.0 | 0 | Canis | 22243.0 | 0 | Canis lupus | 22243.0 | 0 |
| scf7180000055751 | 32749 | 0.445 | 0 | 200.4415 | Eukaryota | 49615.0 | 0 | Chordata | 41488.0 | 1 | Gadiformes | 28436.0 | 6 | Gadidae | 28436.0 | 6 | Gadus | 28436.0 | 6 | Gadus morhua | 28436.0 | 6 |
| scf7180000044632 | 226981 | 0.5193 | 0 | 60.0558 | Eukaryota | 17064.0 | 0 | Chordata | 17064.0 | 0 | Carnivora | 17064.0 | 0 | Canidae | 17064.0 | 0 | Canis | 17064.0 | 0 | Canis lupus | 17064.0 | 0 |
| scf7180000044399 | 312751 | 0.4931 | 0 | 57.482 | Eukaryota | 49586.0 | 0 | Chordata | 49586.0 | 0 | Carnivora | 49586.0 | 0 | Canidae | 29914.0 | 2 | Canis | 24192.0 | 4 | Canis lupus | 24192.0 | 4 |
| scf7180000054736 | 324441 | 0.3793 | 0 | 64.5061 | Eukaryota | 63375.0 | 0 | Chordata | 63375.0 | 0 | Carnivora | 63375.0 | 0 | Canidae | 63375.0 | 0 | Canis | 63375.0 | 0 | Canis lupus | 63375.0 | 0 |
| scf7180000055573 | 22214 | 0.4449 | 0 | 1497.2141 | Eukaryota | 11706.0 | 0 | Platyhelminthes | 11521.0 | 1 | Strigeidida | 10547.0 | 2 | Schistosomatidae | 10547.0 | 2 | Schistosoma | 10547.0 | 2 | Schistosoma mansoni | 10547.0 | 2 |
| scf7180000054113 | 3887 | 0.4492 | 0 | 746.746 | Eukaryota | 2115.0 | 0 | Platyhelminthes | 974.0 | 2 | Polystomatidea | 974.0 | 2 | Polystomatidae | 974.0 | 2 | Protopolystoma | 974.0 | 2 | Protopolystoma xenopodis | 974.0 | 2 |
| scf7180000043906 | 96477 | 0.5266 | 0 | 57.6959 | Eukaryota | 48276.0 | 0 | Chordata | 48276.0 | 0 | Carnivora | 48276.0 | 0 | Canidae | 48276.0 | 0 | Canis | 48276.0 | 0 | Canis lupus | 48276.0 | 0 |
| scf7180000051078 | 8010 | 0.4511 | 0 | 1715.9769 | no-hit | 0.0 | 0 | no-hit | 0.0 | 0 | no-hit | 0.0 | 0 | no-hit | 0.0 | 0 | no-hit | 0.0 | 0 | no-hit | 0.0 | 0 |
| scf7180000045820 | 7232 | 0.4378 | 0 | 161.125 | Eukaryota | 1409.0 | 0 | Chordata | 751.0 | 1 | Tetraodontiformes | 751.0 | 1 | Tetraodontidae | 751.0 | 1 | Takifugu | 751.0 | 1 | Takifugu rubripes | 751.0 | 1 |
| scf7180000046618 | 143754 | 0.3565 | 0 | 61.1826 | Eukaryota | 8565.0 | 0 | Chordata | 8565.0 | 0 | Carnivora | 8565.0 | 0 | Felidae | 5345.0 | 1 | Felis | 5345.0 | 2 | Felis catus | 5345.0 | 2 |
| scf7180000053357 | 6575 | 0.4412 | 0 | 271.0027 | no-hit | 0.0 | 0 | no-hit | 0.0 | 0 | no-hit | 0.0 | 0 | no-hit | 0.0 | 0 | no-hit | 0.0 | 0 | no-hit | 0.0 | 0 |
| scf7180000047110 | 10794 | 0.4351 | 0 | 48.5769 | Eukaryota | 55515.0 | 0 | Chordata | 55515.0 | 0 | Carnivora | 55515.0 | 0 | Canidae | 55515.0 | 0 | Canis | 55515.0 | 0 | Canis lupus | 55515.0 | 0 |
| scf7180000056079 | 23016 | 0.4387 | 0 | 743.9775 | Eukaryota | 4707.0 | 0 | Chordata | 4707.0 | 0 | Salmoniformes | 2377.0 | 1 | Salmonidae | 2377.0 | 1 | Salmo | 2377.0 | 1 | Salmo trutta | 2377.0 | 1 |
| scf7180000047358 | 90003 | 0.3583 | 0 | 61.6118 | Eukaryota | 21592.0 | 0 | Chordata | 21592.0 | 0 | Artiodactyla | 11990.0 | 2 | Bovidae | 6265.0 | 4 | Sus | 5725.0 | 5 | Sus scrofa | 5725.0 | 5 |
| scf7180000045620 | 53363 | 0.4298 | 0 | 53.2646 | Eukaryota | 44896.0 | 0 | Chordata | 44896.0 | 0 | Carnivora | 44896.0 | 0 | Canidae | 44896.0 | 0 | Canis | 44896.0 | 0 | Canis lupus | 44896.0 | 0 |
| scf7180000049933 | 23227 | 0.4409 | 0 | 558.7847 | Eukaryota | 56617.0 | 0 | Chordata | 37703.0 | 2 | Gadiformes | 30718.0 | 5 | Gadidae | 30718.0 | 5 | Gadus | 30718.0 | 5 | Gadus morhua | 30718.0 | 5 |
| scf7180000045473 | 47562 | 0.4396 | 0 | 185.3346 | Eukaryota | 3854.0 | 0 | Chordata | 1743.0 | 2 | Polystomatidea | 1279.0 | 5 | Polystomatidae | 1279.0 | 5 | Protopolystoma | 1279.0 | 6 | Protopolystoma xenopodis | 1279.0 | 6 |
| scf7180000043367 | 509685 | 0.3986 | 0 | 61.1306 | Eukaryota | 60970.0 | 0 | Chordata | 60970.0 | 0 | Artiodactyla | 53503.0 | 1 | Suidae | 53503.0 | 1 | Sus | 53503.0 | 1 | Sus scrofa | 53503.0 | 1 |
| scf7180000046783 | 100566 | 0.3792 | 0 | 59.7125 | Eukaryota | 20374.0 | 1 | Chordata | 20374.0 | 1 | Carnivora | 11480.0 | 3 | Canidae | 11480.0 | 3 | Canis | 11480.0 | 4 | Canis lupus | 11480.0 | 4 |
| scf7180000050139 | 22798 | 0.4395 | 0 | 67.8795 | Eukaryota | 600.0 | 0 | Platyhelminthes | 424.0 | 1 | Polystomatidea | 259.0 | 2 | Polystomatidae | 259.0 | 2 | Protopolystoma | 259.0 | 2 | Protopolystoma xenopodis | 259.0 | 2 |
| scf7180000051760 | 9542 | 0.4394 | 0 | 95.9796 | Eukaryota | 2186.0 | 0 | Platyhelminthes | 1125.0 | 2 | Polystomatidea | 1125.0 | 2 | Polystomatidae | 1125.0 | 2 | Protopolystoma | 1125.0 | 2 | Protopolystoma xenopodis | 1125.0 | 2 |
| scf7180000045932 | 31011 | 0.3605 | 0 | 31.3158 | Eukaryota | 4041.0 | 0 | Chordata | 4041.0 | 0 | Carnivora | 2141.0 | 1 | Canidae | 2141.0 | 1 | Canis | 2141.0 | 1 | Canis lupus | 2141.0 | 1 |
| scf7180000051888 | 6366 | 0.4471 | 0 | 334.6536 | Eukaryota | 315.0 | 0 | Platyhelminthes | 315.0 | 0 | Polystomatidea | 315.0 | 0 | Polystomatidae | 315.0 | 0 | Protopolystoma | 315.0 | 0 | Protopolystoma xenopodis | 315.0 | 0 |
| scf7180000041562 | 157408 | 0.3602 | 0 | 55.744 | Eukaryota | 66519.0 | 0 | Chordata | 66519.0 | 0 | Carnivora | 66519.0 | 0 | Canidae | 66519.0 | 0 | Canis | 66519.0 | 0 | Canis lupus | 66519.0 | 0 |
| scf7180000049348 | 4350 | 0.4515 | 0 | 260.218 | Eukaryota | 1725.0 | 0 | Echinodermata | 815.0 | 2 | Forcipulatida | 815.0 | 2 | Asteriidae | 815.0 | 2 | Asterias | 815.0 | 2 | Asterias rubens | 815.0 | 2 |
| scf7180000050209 | 27057 | 0.4917 | 0 | 57.3341 | Eukaryota | 11090.0 | 0 | Chordata | 11090.0 | 0 | Carnivora | 7665.0 | 1 | Canidae | 7665.0 | 2 | Canis | 6405.0 | 3 | Canis lupus | 6405.0 | 3 |
| scf7180000044888 | 231966 | 0.3847 | 0 | 58.1672 | Eukaryota | 25784.0 | 0 | Chordata | 25784.0 | 0 | Carnivora | 25784.0 | 0 | Mustelidae | 12755.0 | 2 | Mustela | 12755.0 | 2 | Mustela putorius | 12755.0 | 2 |
| scf7180000055189 | 49357 | 0.4541 | 0 | 221.1381 | Eukaryota | 17618.0 | 0 | Chordata | 17618.0 | 0 | Carnivora | 14905.0 | 1 | Canidae | 14905.0 | 1 | Canis | 14905.0 | 1 | Canis lupus | 14905.0 | 1 |
| scf7180000041584 | 9576 | 0.3841 | 0 | 37.3099 | Eukaryota | 4465.0 | 0 | Chordata | 4465.0 | 0 | Carnivora | 2450.0 | 2 | Canidae | 2450.0 | 2 | Canis | 2450.0 | 3 | Canis lupus | 2450.0 | 3 |
| scf7180000041609 | 1009344 | 0.5111 | 0 | 58.576 | Eukaryota | 48702.0 | 0 | Chordata | 48702.0 | 0 | Carnivora | 48702.0 | 0 | Canidae | 38100.0 | 2 | Canis | 38100.0 | 3 | Canis lupus | 38100.0 | 3 |
| scf7180000051378 | 3580 | 0.4534 | 0 | 1427.4193 | no-hit | 0.0 | 0 | no-hit | 0.0 | 0 | no-hit | 0.0 | 0 | no-hit | 0.0 | 0 | no-hit | 0.0 | 0 | no-hit | 0.0 | 0 |
| scf7180000041946 | 56685 | 0.3971 | 0 | 63.5362 | Eukaryota | 65752.0 | 0 | Chordata | 65752.0 | 0 | Carnivora | 65752.0 | 0 | Canidae | 65752.0 | 0 | Canis | 65752.0 | 0 | Canis lupus | 65752.0 | 0 |
| scf7180000052884 | 5693 | 0.4386 | 0 | 140.5305 | no-hit | 0.0 | 0 | no-hit | 0.0 | 0 | no-hit | 0.0 | 0 | no-hit | 0.0 | 0 | no-hit | 0.0 | 0 | no-hit | 0.0 | 0 |
| scf7180000046807 | 61378 | 0.3703 | 0 | 57.6161 | Eukaryota | 13348.0 | 0 | Chordata | 13348.0 | 0 | Carnivora | 13348.0 | 0 | Canidae | 7045.0 | 3 | Canis | 5658.0 | 6 | Canis lupus | 5658.0 | 6 |
| scf7180000050866 | 23359 | 0.3813 | 0 | 60.2448 | Eukaryota | 20562.0 | 0 | Chordata | 20562.0 | 0 | Carnivora | 14050.0 | 2 | Canidae | 14050.0 | 2 | Canis | 12504.0 | 3 | Canis lupus | 12504.0 | 3 |
| scf7180000049467 | 25107 | 0.45 | 0 | 762.4386 | Eukaryota | 4019.0 | 0 | Platyhelminthes | 2492.0 | 2 | Polystomatidea | 2492.0 | 2 | Polystomatidae | 2492.0 | 2 | Protopolystoma | 2492.0 | 2 | Protopolystoma xenopodis | 2492.0 | 2 |
| scf7180000051929 | 16911 | 0.3525 | 0 | 60.4575 | Eukaryota | 27331.0 | 0 | Chordata | 27331.0 | 0 | Carnivora | 27331.0 | 0 | Canidae | 27331.0 | 0 | Canis | 27331.0 | 0 | Canis lupus | 27331.0 | 0 |
| scf7180000049183 | 31693 | 0.4384 | 0 | 156.2967 | Eukaryota | 45518.0 | 0 | Chordata | 37371.0 | 2 | Gadiformes | 28380.0 | 6 | Gadidae | 28380.0 | 6 | Gadus | 28380.0 | 6 | Gadus morhua | 28380.0 | 6 |
| scf7180000042296 | 368069 | 0.4411 | 0 | 58.6547 | Eukaryota | 119190.0 | 0 | Chordata | 119190.0 | 0 | Carnivora | 119190.0 | 0 | Canidae | 119190.0 | 0 | Canis | 119190.0 | 0 | Canis lupus | 119190.0 | 0 |
| scf7180000049162 | 6463 | 0.4472 | 0 | 688.0699 | Eukaryota | 1483.0 | 0 | Echinodermata | 512.0 | 2 | Forcipulatida | 512.0 | 2 | Asteriidae | 512.0 | 2 | Asterias | 512.0 | 2 | Asterias rubens | 512.0 | 2 |
| scf7180000045525 | 175272 | 0.3631 | 0 | 59.9762 | Eukaryota | 48060.0 | 0 | Chordata | 48060.0 | 0 | Carnivora | 48060.0 | 0 | Canidae | 48060.0 | 0 | Canis | 48060.0 | 0 | Canis lupus | 48060.0 | 0 |
| scf7180000040566 | 2397037 | 0.3711 | 0 | 60.9286 | Eukaryota | 93035.0 | 0 | Chordata | 93035.0 | 0 | Carnivora | 93035.0 | 0 | Canidae | 93035.0 | 0 | Canis | 93035.0 | 0 | Canis lupus | 93035.0 | 0 |
| scf7180000055215 | 104034 | 0.4419 | 0 | 82.202 | Eukaryota | 6727.0 | 0 | Platyhelminthes | 3498.0 | 1 | Polystomatidea | 3498.0 | 5 | Polystomatidae | 3498.0 | 5 | Protopolystoma | 3498.0 | 5 | Protopolystoma xenopodis | 3498.0 | 5 |
| scf7180000048803 | 15561 | 0.4424 | 0 | 105.5772 | Eukaryota | 4819.0 | 0 | Chordata | 3855.0 | 1 | Kurtiformes | 3487.0 | 2 | Apogonidae | 3487.0 | 2 | Sphaeramia | 3487.0 | 2 | Sphaeramia orbicularis | 3487.0 | 2 |
| scf7180000054185 | 24070 | 0.3771 | 0 | 156.7903 | Eukaryota | 79889.0 | 0 | Chordata | 79889.0 | 0 | Carnivora | 79889.0 | 0 | Canidae | 79889.0 | 0 | Canis | 79889.0 | 0 | Canis lupus | 79889.0 | 0 |
| scf7180000049263 | 49159 | 0.5019 | 0 | 393.8074 | Eukaryota | 8953.0 | 0 | Chordata | 8953.0 | 0 | Carnivora | 8953.0 | 0 | Canidae | 8953.0 | 0 | Canis | 7176.0 | 1 | Canis lupus | 5884.0 | 2 |
| scf7180000043351 | 476028 | 0.3946 | 0 | 59.8644 | Eukaryota | 100433.0 | 0 | Chordata | 100433.0 | 0 | Carnivora | 100433.0 | 0 | Canidae | 100433.0 | 0 | Vulpes | 60333.0 | 1 | Vulpes vulpes | 60333.0 | 1 |
| scf7180000048535 | 5814 | 0.3973 | 0 | 24.3926 | Eukaryota | 23370.0 | 0 | Chordata | 23370.0 | 0 | Carnivora | 23370.0 | 0 | Canidae | 21475.0 | 1 | Canis | 21475.0 | 1 | Canis lupus | 21475.0 | 1 |
| scf7180000042365 | 792455 | 0.3442 | 0 | 58.8392 | Eukaryota | 41666.0 | 0 | Chordata | 41666.0 | 0 | Carnivora | 41666.0 | 0 | Canidae | 41666.0 | 0 | Canis | 41666.0 | 0 | Canis lupus | 41666.0 | 0 |
| scf7180000041361 | 1142786 | 0.4034 | 0 | 59.8518 | Eukaryota | 52468.0 | 0 | Chordata | 52468.0 | 0 | Carnivora | 52468.0 | 0 | Canidae | 52468.0 | 0 | Canis | 39700.0 | 1 | Canis lupus | 39700.0 | 1 |
| scf7180000050091 | 54897 | 0.4395 | 0 | 59.5761 | Eukaryota | 14548.0 | 1 | Chordata | 14548.0 | 1 | Primates | 14548.0 | 1 | Hominidae | 14548.0 | 1 | Homo | 12928.0 | 2 | Homo sapiens | 12928.0 | 2 |
| scf7180000047697 | 68714 | 0.3593 | 0 | 57.7627 | Eukaryota | 55649.0 | 0 | Chordata | 55649.0 | 0 | Carnivora | 55649.0 | 0 | Canidae | 55649.0 | 0 | Canis | 55649.0 | 0 | Canis lupus | 55649.0 | 0 |
| scf7180000056470 | 290045 | 0.5441 | 0 | 56.0136 | Eukaryota | 35722.0 | 0 | Chordata | 35722.0 | 0 | Carnivora | 35722.0 | 0 | Canidae | 23341.0 | 2 | Canis | 12914.0 | 4 | Canis lupus | 12914.0 | 4 |
| scf7180000041801 | 46286 | 0.3508 | 0 | 49.4451 | Eukaryota | 37443.0 | 0 | Chordata | 37443.0 | 0 | Carnivora | 37443.0 | 0 | Canidae | 37443.0 | 0 | Canis | 37443.0 | 0 | Canis lupus | 37443.0 | 0 |
| scf7180000041513 | 549803 | 0.3391 | 0 | 58.655 | Eukaryota | 83260.0 | 0 | Chordata | 83260.0 | 0 | Carnivora | 83260.0 | 0 | Canidae | 83260.0 | 0 | Canis | 83260.0 | 0 | Canis lupus | 83260.0 | 0 |
| scf7180000053072 | 3704 | 0.4611 | 0 | 1159.6 | no-hit | 0.0 | 0 | no-hit | 0.0 | 0 | no-hit | 0.0 | 0 | no-hit | 0.0 | 0 | no-hit | 0.0 | 0 | no-hit | 0.0 | 0 |
| scf7180000056501 | 268818 | 0.3834 | 0 | 57.2756 | Eukaryota | 24436.0 | 0 | Chordata | 24436.0 | 0 | Carnivora | 24436.0 | 0 | Canidae | 24436.0 | 0 | Canis | 24436.0 | 0 | Canis lupus | 24436.0 | 0 |
| scf7180000048685 | 55853 | 0.3897 | 0 | 60.6692 | Eukaryota | 22564.0 | 0 | Chordata | 22564.0 | 0 | Carnivora | 22564.0 | 0 | Canidae | 7541.0 | 4 | Canis | 5031.0 | 7 | Canis lupus | 5031.0 | 7 |
| scf7180000050695 | 34986 | 0.4397 | 0 | 57.9217 | Eukaryota | 8212.0 | 0 | Chordata | 6267.0 | 2 | Tetraodontiformes | 1830.0 | 6 | Tetraodontidae | 1830.0 | 6 | Takifugu | 1830.0 | 7 | Protopolystoma xenopodis | 1222.0 | 8 |
| scf7180000052362 | 15551 | 0.4878 | 0 | 259.6909 | Eukaryota | 89161.0 | 0 | Chordata | 89161.0 | 0 | Carnivora | 89161.0 | 0 | Canidae | 89161.0 | 0 | Canis | 89161.0 | 0 | Canis lupus | 89161.0 | 0 |
| scf7180000046310 | 142740 | 0.4953 | 0 | 59.3539 | Eukaryota | 45977.0 | 0 | Chordata | 45977.0 | 0 | Carnivora | 45977.0 | 0 | Canidae | 29998.0 | 2 | Canis | 17120.0 | 3 | Canis lupus | 17120.0 | 4 |
| scf7180000056117 | 4961 | 0.432 | 0 | 126.9881 | Eukaryota | 782.0 | 0 | Chordata | 782.0 | 0 | Salmoniformes | 782.0 | 0 | Salmonidae | 782.0 | 0 | Salmo | 782.0 | 0 | Salmo trutta | 782.0 | 0 |
| scf7180000055635 | 22355 | 0.4424 | 0 | 253.2047 | no-hit | 0.0 | 0 | no-hit | 0.0 | 0 | no-hit | 0.0 | 0 | no-hit | 0.0 | 0 | no-hit | 0.0 | 0 | no-hit | 0.0 | 0 |
| scf7180000047232 | 99698 | 0.3658 | 0 | 61.0285 | Eukaryota | 19159.0 | 0 | Chordata | 19159.0 | 0 | Carnivora | 19159.0 | 0 | Canidae | 19159.0 | 0 | Canis | 14222.0 | 1 | Canis lupus | 14222.0 | 1 |
| scf7180000046171 | 125102 | 0.4202 | 0 | 58.0094 | Eukaryota | 10086.0 | 1 | Chordata | 9326.0 | 2 | Primates | 3918.0 | 5 | Hominidae | 3918.0 | 5 | Homo | 3918.0 | 6 | Homo sapiens | 3918.0 | 6 |
| scf7180000042777 | 588172 | 0.4233 | 0 | 59.8313 | Eukaryota | 82311.0 | 0 | Chordata | 82311.0 | 0 | Carnivora | 82311.0 | 0 | Canidae | 82311.0 | 0 | Canis | 82311.0 | 0 | Canis lupus | 82311.0 | 0 |
| scf7180000041945 | 129191 | 0.4678 | 0 | 56.1592 | Eukaryota | 6646.0 | 0 | Chordata | 6646.0 | 0 | Carnivora | 2195.0 | 4 | Camelidae | 1626.0 | 6 | Camelus | 1626.0 | 7 | Homo sapiens | 1243.0 | 9 |
| scf7180000048919 | 5912 | 0.4482 | 0 | 127.8202 | Eukaryota | 1355.0 | 0 | Chordata | 929.0 | 1 | Tetraodontiformes | 648.0 | 2 | Tetraodontidae | 648.0 | 2 | Takifugu | 648.0 | 2 | Takifugu rubripes | 648.0 | 2 |
| scf7180000041507 | 271164 | 0.369 | 0 | 81.8826 | Eukaryota | 65849.0 | 0 | Chordata | 65849.0 | 0 | Carnivora | 65849.0 | 0 | Canidae | 65849.0 | 0 | Canis | 65849.0 | 0 | Canis lupus | 65849.0 | 0 |
| scf7180000045743 | 175211 | 0.4132 | 0 | 58.532 | Eukaryota | 38941.0 | 0 | Chordata | 38941.0 | 0 | Carnivora | 36050.0 | 1 | Canidae | 17861.0 | 4 | Canis | 13192.0 | 5 | Canis lupus | 13192.0 | 5 |
| scf7180000043136 | 498888 | 0.419 | 0 | 58.2745 | Eukaryota | 37850.0 | 0 | Chordata | 37850.0 | 0 | Carnivora | 37850.0 | 0 | Canidae | 24952.0 | 2 | Canis | 20292.0 | 4 | Canis lupus | 20292.0 | 4 |
| scf7180000043554 | 416620 | 0.3554 | 0 | 58.5016 | Eukaryota | 80642.0 | 0 | Chordata | 80642.0 | 0 | Carnivora | 80642.0 | 0 | Canidae | 80642.0 | 0 | Canis | 80642.0 | 0 | Canis lupus | 80642.0 | 0 |
| scf7180000056345 | 714654 | 0.3723 | 0 | 60.4626 | Eukaryota | 77755.0 | 0 | Chordata | 77755.0 | 0 | Carnivora | 77755.0 | 0 | Canidae | 77755.0 | 0 | Canis | 71075.0 | 1 | Canis lupus | 71075.0 | 1 |
| scf7180000050599 | 7433 | 0.4437 | 0 | 84.9131 | Eukaryota | 1496.0 | 0 | Chordata | 885.0 | 1 | Polystomatidea | 611.0 | 3 | Polystomatidae | 611.0 | 3 | Protopolystoma | 611.0 | 3 | Protopolystoma xenopodis | 611.0 | 3 |
| scf7180000049581 | 13881 | 0.4431 | 0 | 79.2408 | no-hit | 0.0 | 0 | no-hit | 0.0 | 0 | no-hit | 0.0 | 0 | no-hit | 0.0 | 0 | no-hit | 0.0 | 0 | no-hit | 0.0 | 0 |
| scf7180000053465 | 1500 | 0.452 | 0 | 842.0654 | no-hit | 0.0 | 0 | no-hit | 0.0 | 0 | no-hit | 0.0 | 0 | no-hit | 0.0 | 0 | no-hit | 0.0 | 0 | no-hit | 0.0 | 0 |
| scf7180000047394 | 81256 | 0.4053 | 0 | 62.6943 | Eukaryota | 78430.0 | 0 | Chordata | 78430.0 | 0 | Carnivora | 78430.0 | 0 | Canidae | 78430.0 | 0 | Canis | 70060.0 | 1 | Canis lupus | 70060.0 | 1 |
| scf7180000052530 | 2824 | 0.4533 | 0 | 106.7502 | no-hit | 0.0 | 0 | no-hit | 0.0 | 0 | no-hit | 0.0 | 0 | no-hit | 0.0 | 0 | no-hit | 0.0 | 0 | no-hit | 0.0 | 0 |
| scf7180000046454 | 155388 | 0.4082 | 0 | 58.9446 | Eukaryota | 63151.0 | 0 | Chordata | 63151.0 | 0 | Carnivora | 63151.0 | 0 | Canidae | 63151.0 | 0 | Canis | 63151.0 | 0 | Canis lupus | 63151.0 | 0 |
| scf7180000053865 | 8556 | 0.4472 | 0 | 284.2143 | Eukaryota | 12316.0 | 0 | Chordata | 11892.0 | 1 | Tetraodontiformes | 5529.0 | 5 | Tetraodontidae | 5529.0 | 5 | Takifugu | 5529.0 | 5 | Takifugu rubripes | 5529.0 | 5 |
| scf7180000046016 | 34368 | 0.3834 | 0 | 50.6043 | Eukaryota | 20421.0 | 0 | Chordata | 20421.0 | 0 | Apterygiformes | 18206.0 | 1 | Apterygidae | 18206.0 | 1 | Apteryx | 18206.0 | 1 | Apteryx mantelli | 18206.0 | 1 |
| scf7180000041981 | 113751 | 0.3667 | 0 | 57.8777 | Eukaryota | 25736.0 | 0 | Chordata | 25736.0 | 0 | Carnivora | 25736.0 | 0 | Canidae | 25736.0 | 0 | Canis | 25736.0 | 0 | Canis lupus | 25736.0 | 0 |
| scf7180000051119 | 34442 | 0.4399 | 0 | 206.5228 | Eukaryota | 9418.0 | 0 | Chordata | 9418.0 | 0 | Carnivora | 9418.0 | 0 | Canidae | 9418.0 | 0 | Canis | 9418.0 | 0 | Canis lupus | 9418.0 | 0 |
| scf7180000042915 | 566527 | 0.4266 | 0 | 63.7869 | Eukaryota | 91089.0 | 0 | Chordata | 91089.0 | 0 | Carnivora | 91089.0 | 0 | Canidae | 91089.0 | 0 | Canis | 91089.0 | 0 | Canis lupus | 91089.0 | 0 |
| scf7180000053402 | 16329 | 0.5251 | 0 | 175.0245 | Eukaryota | 33369.0 | 0 | Chordata | 33369.0 | 0 | Carnivora | 33369.0 | 0 | Canidae | 33369.0 | 0 | Canis | 33369.0 | 0 | Canis lupus | 33369.0 | 0 |
| scf7180000043000 | 527414 | 0.4725 | 0 | 60.3229 | Eukaryota | 30395.0 | 0 | Chordata | 30395.0 | 0 | Carnivora | 30395.0 | 0 | Canidae | 30395.0 | 0 | Canis | 27412.0 | 1 | Canis lupus | 27412.0 | 1 |
| scf7180000040702 | 1872203 | 0.4371 | 0 | 59.0081 | Eukaryota | 67120.0 | 0 | Chordata | 67120.0 | 0 | Carnivora | 67120.0 | 0 | Canidae | 67120.0 | 0 | Canis | 53485.0 | 1 | Canis lupus | 53485.0 | 1 |
| scf7180000050678 | 7508 | 0.439 | 0 | 211.0892 | Eukaryota | 382.0 | 0 | Chordata | 382.0 | 0 | Tetraodontiformes | 224.0 | 1 | Tetraodontidae | 224.0 | 1 | Takifugu | 224.0 | 1 | Takifugu rubripes | 224.0 | 1 |
| scf7180000050794 | 2430 | 0.4572 | 0 | 366.1257 | Eukaryota | 291.0 | 0 | Platyhelminthes | 291.0 | 0 | Polystomatidea | 291.0 | 0 | Polystomatidae | 291.0 | 0 | Protopolystoma | 291.0 | 0 | Protopolystoma xenopodis | 291.0 | 0 |
| scf7180000047304 | 2498 | 0.4464 | 0 | 147.4013 | no-hit | 0.0 | 0 | no-hit | 0.0 | 0 | no-hit | 0.0 | 0 | no-hit | 0.0 | 0 | no-hit | 0.0 | 0 | no-hit | 0.0 | 0 |
| scf7180000052114 | 10187 | 0.4554 | 0 | 1218.3654 | Eukaryota | 2355.0 | 0 | Platyhelminthes | 1460.0 | 2 | Polystomatidea | 1460.0 | 2 | Polystomatidae | 1460.0 | 2 | Protopolystoma | 1460.0 | 2 | Protopolystoma xenopodis | 1460.0 | 2 |
| scf7180000042169 | 81405 | 0.4584 | 0 | 66.9972 | Eukaryota | 88247.0 | 0 | Chordata | 88247.0 | 0 | Carnivora | 88247.0 | 0 | Canidae | 88247.0 | 0 | Canis | 88247.0 | 0 | Canis lupus | 88247.0 | 0 |
| scf7180000047294 | 34159 | 0.4445 | 0 | 147.6186 | Eukaryota | 4737.0 | 0 | Chordata | 4203.0 | 1 | Kurtiformes | 2550.0 | 2 | Apogonidae | 2550.0 | 2 | Sphaeramia | 2550.0 | 2 | Sphaeramia orbicularis | 2550.0 | 2 |
| scf7180000051656 | 15818 | 0.4362 | 0 | 190.4997 | Eukaryota | 787.0 | 0 | Chordata | 787.0 | 0 | Salmoniformes | 544.0 | 1 | Salmonidae | 544.0 | 1 | Salmo | 289.0 | 2 | Salmo trutta | 289.0 | 2 |
| scf7180000053365 | 22009 | 0.5036 | 0 | 106.1492 | Eukaryota | 8845.0 | 0 | Chordata | 8845.0 | 0 | Carnivora | 8845.0 | 0 | Canidae | 8845.0 | 0 | Canis | 7122.0 | 1 | Canis lupus | 5893.0 | 2 |
| scf7180000056180 | 2372338 | 0.3968 | 0 | 59.0493 | Eukaryota | 64860.0 | 0 | Chordata | 64860.0 | 0 | Carnivora | 64860.0 | 0 | Canidae | 64860.0 | 0 | Canis | 64860.0 | 0 | Canis lupus | 64860.0 | 0 |
| scf7180000054250 | 42638 | 0.3859 | 0 | 103.7778 | Eukaryota | 8438.0 | 0 | Chordata | 8438.0 | 0 | Carnivora | 8438.0 | 0 | Canidae | 8438.0 | 0 | Canis | 6631.0 | 1 | Canis lupus | 6631.0 | 1 |
| scf7180000049715 | 27435 | 0.5166 | 0 | 728.563 | Eukaryota | 34190.0 | 0 | Chordata | 34190.0 | 0 | Carnivora | 34190.0 | 0 | Canidae | 34190.0 | 0 | Canis | 34190.0 | 0 | Canis lupus | 34190.0 | 0 |
| scf7180000041706 | 218023 | 0.4304 | 0 | 55.7184 | Eukaryota | 30357.0 | 0 | Chordata | 30357.0 | 0 | Carnivora | 30357.0 | 0 | Canidae | 30357.0 | 0 | Canis | 30357.0 | 0 | Canis lupus | 30357.0 | 0 |
| scf7180000044419 | 10420 | 0.5643 | 0 | 295.4548 | Eukaryota | 30479.0 | 0 | Chordata | 30479.0 | 0 | Artiodactyla | 21458.0 | 1 | Bovidae | 21458.0 | 2 | Ovis | 12171.0 | 3 | Ovis aries | 12171.0 | 3 |
| scf7180000050814 | 2292 | 0.4263 | 0 | 48.9896 | no-hit | 0.0 | 0 | no-hit | 0.0 | 0 | no-hit | 0.0 | 0 | no-hit | 0.0 | 0 | no-hit | 0.0 | 0 | no-hit | 0.0 | 0 |
| scf7180000046805 | 107797 | 0.4084 | 0 | 60.267 | Eukaryota | 25430.0 | 0 | Chordata | 25430.0 | 0 | Carnivora | 25430.0 | 0 | Canidae | 25430.0 | 0 | Canis | 25430.0 | 0 | Canis lupus | 25430.0 | 0 |
| scf7180000052057 | 1704 | 0.4501 | 0 | 478.3644 | Eukaryota | 7968.0 | 0 | Chordata | 6785.0 | 2 | Gadiformes | 3077.0 | 6 | Gadidae | 3077.0 | 6 | Gadus | 3077.0 | 6 | Gadus morhua | 3077.0 | 6 |
| scf7180000044825 | 259851 | 0.4814 | 0 | 58.2527 | Eukaryota | 17037.0 | 0 | Chordata | 17037.0 | 0 | Primates | 9054.0 | 4 | Aotidae | 5380.0 | 5 | Aotus | 5380.0 | 5 | Aotus nancymaae | 5380.0 | 5 |
| scf7180000041069 | 1340516 | 0.3762 | 0 | 59.6981 | Eukaryota | 92455.0 | 0 | Chordata | 92455.0 | 0 | Carnivora | 92455.0 | 0 | Canidae | 92455.0 | 0 | Canis | 92455.0 | 0 | Canis lupus | 92455.0 | 0 |
| scf7180000040721 | 1774329 | 0.3689 | 0 | 60.0576 | Eukaryota | 82235.0 | 0 | Chordata | 82235.0 | 0 | Carnivora | 82235.0 | 0 | Canidae | 82235.0 | 0 | Canis | 82235.0 | 0 | Canis lupus | 82235.0 | 0 |
| scf7180000040541 | 2339580 | 0.4103 | 0 | 59.5299 | Eukaryota | 78162.0 | 0 | Chordata | 78162.0 | 0 | Carnivora | 78162.0 | 0 | Canidae | 78162.0 | 0 | Canis | 66459.0 | 1 | Canis lupus | 66459.0 | 1 |
| scf7180000040949 | 297821 | 0.377 | 0 | 59.3807 | Eukaryota | 90525.0 | 0 | Chordata | 90525.0 | 0 | Carnivora | 90525.0 | 0 | Canidae | 90525.0 | 0 | Canis | 90525.0 | 0 | Canis lupus | 90525.0 | 0 |
| scf7180000043154 | 357797 | 0.3654 | 0 | 60.2703 | Eukaryota | 58349.0 | 0 | Chordata | 58349.0 | 0 | Carnivora | 58349.0 | 0 | Canidae | 34920.0 | 2 | Canis | 30002.0 | 5 | Canis lupus | 30002.0 | 5 |
| scf7180000040526 | 30888 | 0.3816 | 0 | 50.3915 | Eukaryota | 2886.0 | 0 | Chordata | 2886.0 | 0 | Carnivora | 2886.0 | 0 | Canidae | 2886.0 | 0 | Canis | 2886.0 | 0 | Canis lupus | 2886.0 | 0 |
| scf7180000049120 | 46100 | 0.5672 | 0 | 58.233 | Eukaryota | 6370.0 | 1 | Chordata | 6370.0 | 1 | Artiodactyla | 3415.0 | 3 | Bovidae | 1826.0 | 5 | Sus | 1589.0 | 7 | Sus scrofa | 1589.0 | 7 |
| scf7180000048690 | 32875 | 0.4489 | 0 | 85.6839 | Eukaryota | 4615.0 | 0 | Chordata | 3776.0 | 1 | Columbiformes | 2407.0 | 4 | Columbidae | 2407.0 | 4 | Streptopelia | 2407.0 | 4 | Streptopelia turtur | 2407.0 | 4 |
| scf7180000048718 | 39452 | 0.455 | 0 | 57.4528 | Eukaryota | 4638.0 | 1 | Chordata | 4638.0 | 1 | Primates | 2431.0 | 2 | Hominidae | 2431.0 | 3 | Homo | 2431.0 | 3 | Homo sapiens | 2431.0 | 3 |
| scf7180000044962 | 240881 | 0.3912 | 0 | 61.5979 | Eukaryota | 68682.0 | 0 | Chordata | 68682.0 | 0 | Carnivora | 68682.0 | 0 | Canidae | 68682.0 | 0 | Canis | 43290.0 | 1 | Canis lupus | 43290.0 | 1 |
| scf7180000043665 | 424006 | 0.389 | 0 | 63.0311 | Eukaryota | 89545.0 | 0 | Chordata | 89545.0 | 0 | Carnivora | 89545.0 | 0 | Canidae | 89545.0 | 0 | Canis | 89545.0 | 0 | Canis lupus | 89545.0 | 0 |
| scf7180000043612 | 143132 | 0.3607 | 0 | 59.1532 | Eukaryota | 53631.0 | 0 | Chordata | 53631.0 | 0 | Carnivora | 53631.0 | 0 | Otariidae | 20124.0 | 2 | Odobenus | 16914.0 | 3 | Odobenus rosmarus | 16914.0 | 3 |
| scf7180000047389 | 76538 | 0.3324 | 0 | 59.6385 | Eukaryota | 73636.0 | 0 | Chordata | 73636.0 | 0 | Carnivora | 73636.0 | 0 | Canidae | 73636.0 | 0 | Canis | 73636.0 | 0 | Canis lupus | 73636.0 | 0 |
| scf7180000052406 | 6103 | 0.454 | 0 | 454.9871 | Eukaryota | 239.0 | 0 | Streptophyta | 239.0 | 0 | Fabales | 239.0 | 0 | Fabaceae | 239.0 | 0 | Cicer | 239.0 | 0 | Cicer arietinum | 239.0 | 0 |
| scf7180000043636 | 5158 | 0.4752 | 0 | 44.7738 | Eukaryota | 38688.0 | 0 | Chordata | 38688.0 | 0 | Carnivora | 38688.0 | 0 | Canidae | 38688.0 | 0 | Canis | 38688.0 | 0 | Canis lupus | 38688.0 | 0 |
| scf7180000040546 | 2375848 | 0.3718 | 0 | 60.2356 | Eukaryota | 54020.0 | 0 | Chordata | 54020.0 | 0 | Carnivora | 54020.0 | 0 | Canidae | 54020.0 | 0 | Canis | 54020.0 | 0 | Canis lupus | 54020.0 | 0 |
| scf7180000047044 | 33301 | 0.3358 | 0 | 59.3014 | Eukaryota | 3856.0 | 0 | Chordata | 3856.0 | 0 | Carnivora | 1671.0 | 3 | Bovidae | 1589.0 | 4 | Canis | 1364.0 | 5 | Canis lupus | 1364.0 | 5 |
| scf7180000042196 | 344488 | 0.3431 | 0 | 58.0473 | Eukaryota | 91258.0 | 0 | Chordata | 91258.0 | 0 | Carnivora | 91258.0 | 0 | Canidae | 91258.0 | 0 | Canis | 91258.0 | 0 | Canis lupus | 91258.0 | 0 |

| | | | | | | | | | | | | | | | | | | | | | |
|---|---|---|---|---|---|---|---|---|---|---|---|---|---|---|---|---|---|---|---|---|---|
| scf7180000041583 | 932270 | 0.4036 | 0 | 60.0465 | Eukaryota | 36395.0 | 0 | Chordata | 36395.0 | 0 | Carnivora | 36395.0 | 0 | Canidae | 36395.0 | 0 | Vulpes | 36395.0 | 0 | Vulpes vulpes | 36395.0 | 0 |
| scf7180000047889 | 65377 | 0.3367 | 0 | 58.8622 | Eukaryota | 61829.0 | 0 | Chordata | 61829.0 | 0 | Carnivora | 61829.0 | 0 | Canidae | 61829.0 | 0 | Canis | 61829.0 | 0 | Canis lupus | 61829.0 | 0 |
| scf7180000055909 | 2714 | 0.4506 | 0 | 363.7683 | no-hit | 0.0 | 0 | no-hit | 0.0 | 0 | no-hit | 0.0 | 0 | no-hit | 0.0 | 0 | no-hit | 0.0 | 0 | no-hit | 0.0 | 0 |
| scf7180000056356 | 509792 | 0.4922 | 0 | 58.0278 | Eukaryota | 17896.0 | 0 | Chordata | 17896.0 | 0 | Carnivora | 17896.0 | 0 | Canidae | 17896.0 | 0 | Canis | 17896.0 | 0 | Canis lupus | 17896.0 | 0 |
| scf7180000042667 | 460407 | 0.3656 | 0 | 61.9953 | Eukaryota | 95200.0 | 0 | Chordata | 95200.0 | 0 | Carnivora | 95200.0 | 0 | Canidae | 95200.0 | 0 | Canis | 95200.0 | 0 | Canis lupus | 95200.0 | 0 |
| scf7180000042655 | 8006 | 0.3626 | 0 | 46.5604 | Eukaryota | 3195.0 | 0 | Chordata | 3195.0 | 0 | Carnivora | 3195.0 | 0 | Canidae | 3195.0 | 0 | Canis | 3195.0 | 0 | Canis lupus | 3195.0 | 0 |
| scf7180000052732 | 17849 | 0.6029 | 0 | 546.1094 | Eukaryota | 60291.0 | 0 | Chordata | 60291.0 | 0 | Primates | 45364.0 | 2 | Hominidae | 45364.0 | 2 | Homo | 45364.0 | 2 | Homo sapiens | 45364.0 | 2 |
| scf7180000043865 | 129833 | 0.3417 | 0 | 59.7878 | Eukaryota | 57959.0 | 0 | Chordata | 57959.0 | 0 | Carnivora | 57959.0 | 0 | Canidae | 57959.0 | 0 | Canis | 57959.0 | 0 | Canis lupus | 57959.0 | 0 |
| scf7180000042588 | 45815 | 0.389 | 0 | 53.2647 | Eukaryota | 17797.0 | 0 | Chordata | 17797.0 | 0 | Carnivora | 17797.0 | 0 | Canidae | 12575.0 | 1 | Canis | 10918.0 | 2 | Canis lupus | 10918.0 | 2 |
| scf7180000053063 | 4732 | 0.4666 | 0 | 468.057 | no-hit | 0.0 | 0 | no-hit | 0.0 | 0 | no-hit | 0.0 | 0 | no-hit | 0.0 | 0 | no-hit | 0.0 | 0 | no-hit | 0.0 | 0 |
| scf7180000047202 | 27291 | 0.4487 | 0 | 75.3117 | Eukaryota | 2833.0 | 0 | Chordata | 2833.0 | 0 | Carnivora | 2833.0 | 0 | Canidae | 2833.0 | 0 | Canis | 2833.0 | 0 | Canis lupus | 2833.0 | 0 |
| scf7180000050022 | 13713 | 0.4405 | 0 | 88.2574 | Eukaryota | 316.0 | 0 | Platyhelminthes | 316.0 | 0 | Polystomatidea | 316.0 | 0 | Polystomatidae | 316.0 | 0 | Protopolystoma | 316.0 | 0 | Protopolystoma xenopodis | 316.0 | 0 |
| scf7180000046137 | 170404 | 0.346 | 0 | 61.2414 | Eukaryota | 38471.0 | 0 | Chordata | 38471.0 | 0 | Carnivora | 38471.0 | 0 | Canidae | 38471.0 | 0 | Canis | 38471.0 | 0 | Canis lupus | 38471.0 | 0 |
| scf7180000044116 | 22695 | 0.4423 | 0 | 407.7723 | Eukaryota | 4613.0 | 0 | Chordata | 2546.0 | 1 | Polystomatidea | 2067.0 | 2 | Polystomatidae | 2067.0 | 2 | Protopolystoma | 2067.0 | 3 | Protopolystoma xenopodis | 2067.0 | 3 |
| scf7180000043071 | 540263 | 0.3635 | 0 | 59.604 | Eukaryota | 47213.0 | 0 | Chordata | 47213.0 | 0 | Carnivora | 47213.0 | 0 | Canidae | 47213.0 | 0 | Canis | 27113.0 | 1 | Canis lupus | 27113.0 | 1 |
| scf7180000045116 | 261160 | 0.377 | 0 | 59.2475 | Eukaryota | 42178.0 | 0 | Chordata | 42178.0 | 0 | Carnivora | 42178.0 | 0 | Canidae | 26691.0 | 2 | Canis | 17780.0 | 4 | Canis lupus | 17780.0 | 5 |
| scf7180000051287 | 7980 | 0.5128 | 0 | 186.5874 | Eukaryota | 4086.0 | 0 | Chordata | 4086.0 | 0 | Carnivora | 4086.0 | 0 | Canidae | 4086.0 | 0 | Canis | 3391.0 | 1 | Canis lupus | 3391.0 | 1 |
| scf7180000047837 | 24381 | 0.3921 | 0 | 55.0939 | Eukaryota | 41505.0 | 0 | Chordata | 41505.0 | 0 | Carnivora | 41505.0 | 0 | Canidae | 36258.0 | 1 | Canis | 36258.0 | 2 | Canis lupus | 36258.0 | 2 |
| scf7180000054716 | 12540 | 0.4415 | 0 | 79.7877 | no-hit | 0.0 | 0 | no-hit | 0.0 | 0 | no-hit | 0.0 | 0 | no-hit | 0.0 | 0 | no-hit | 0.0 | 0 | no-hit | 0.0 | 0 |
| scf7180000052305 | 3907 | 0.4492 | 0 | 654.6411 | no-hit | 0.0 | 0 | no-hit | 0.0 | 0 | no-hit | 0.0 | 0 | no-hit | 0.0 | 0 | no-hit | 0.0 | 0 | no-hit | 0.0 | 0 |
| scf7180000047154 | 122675 | 0.3848 | 0 | 59.1857 | Eukaryota | 18773.0 | 0 | Chordata | 18773.0 | 0 | Carnivora | 18773.0 | 0 | Canidae | 17341.0 | 1 | Canis | 15176.0 | 2 | Canis lupus | 15176.0 | 2 |
| scf7180000047965 | 76925 | 0.5334 | 0 | 59.5533 | Eukaryota | 38332.0 | 0 | Chordata | 38332.0 | 0 | Carnivora | 38332.0 | 0 | Canidae | 30096.0 | 3 | Canis | 25100.0 | 4 | Canis lupus | 25100.0 | 4 |
| scf7180000052480 | 3228 | 0.4504 | 0 | 578.5175 | no-hit | 0.0 | 0 | no-hit | 0.0 | 0 | no-hit | 0.0 | 0 | no-hit | 0.0 | 0 | no-hit | 0.0 | 0 | no-hit | 0.0 | 0 |
| scf7180000043042 | 502731 | 0.3993 | 0 | 59.6756 | Eukaryota | 91868.0 | 0 | Chordata | 91868.0 | 0 | Carnivora | 91868.0 | 0 | Canidae | 91868.0 | 0 | Canis | 91868.0 | 0 | Canis lupus | 91868.0 | 0 |
| scf7180000047645 | 11877 | 0.445 | 0 | 147.034 | Eukaryota | 189.0 | 0 | Echinodermata | 189.0 | 0 | Forcipulatida | 189.0 | 0 | Asteriidae | 189.0 | 0 | Asterias | 189.0 | 0 | Asterias rubens | 189.0 | 0 |
| scf7180000043383 | 69889 | 0.5104 | 0 | 159.2755 | Eukaryota | 48891.0 | 0 | Chordata | 48891.0 | 0 | Carnivora | 48891.0 | 0 | Canidae | 48891.0 | 0 | Canis | 48891.0 | 0 | Canis lupus | 48891.0 | 0 |
| scf7180000048733 | 7628 | 0.4459 | 0 | 171.3523 | Eukaryota | 3395.0 | 0 | Platyhelminthes | 2069.0 | 2 | Polystomatidea | 2069.0 | 2 | Polystomatidae | 2069.0 | 2 | Protopolystoma | 2069.0 | 2 | Protopolystoma xenopodis | 2069.0 | 2 |
| scf7180000040877 | 1698121 | 0.3609 | 0 | 57.6472 | Eukaryota | 54298.0 | 0 | Chordata | 54298.0 | 0 | Carnivora | 54298.0 | 0 | Canidae | 54298.0 | 0 | Canis | 48822.0 | 1 | Canis lupus | 48822.0 | 1 |
| scf7180000047418 | 35100 | 0.4435 | 0 | 104.7062 | Eukaryota | 4379.0 | 0 | Chordata | 4379.0 | 0 | Carnivora | 1712.0 | 5 | Canidae | 1712.0 | 5 | Canis | 1712.0 | 5 | Canis lupus | 1712.0 | 5 |
| scf7180000045023 | 25381 | 0.374 | 0 | 45.6901 | Eukaryota | 135102.0 | 0 | Chordata | 135102.0 | 0 | Carnivora | 135102.0 | 0 | Canidae | 135102.0 | 0 | Canis | 135102.0 | 0 | Canis lupus | 135102.0 | 0 |
| scf7180000046342 | 23636 | 0.4388 | 0 | 229.9202 | Eukaryota | 4689.0 | 0 | Echinodermata | 1949.0 | 2 | Forcipulatida | 1949.0 | 3 | Asteriidae | 1949.0 | 3 | Asterias | 1949.0 | 4 | Asterias rubens | 1949.0 | 4 |
| scf7180000052842 | 5175 | 0.5063 | 0 | 128.2736 | Eukaryota | 6848.0 | 0 | Chordata | 6848.0 | 0 | Carnivora | 6848.0 | 0 | Canidae | 6848.0 | 0 | Canis | 4454.0 | 1 | Canis lupus | 3420.0 | 2 |
| scf7180000050121 | 13043 | 0.4511 | 0 | 1563.4328 | Eukaryota | 3389.0 | 0 | Platyhelminthes | 1989.0 | 2 | Polystomatidea | 1989.0 | 2 | Polystomatidae | 1989.0 | 2 | Protopolystoma | 1989.0 | 2 | Protopolystoma xenopodis | 1989.0 | 2 |
| scf7180000048805 | 5219 | 0.4461 | 0 | 68.5571 | Eukaryota | 2267.0 | 0 | Chordata | 1296.0 | 1 | Polystomatidea | 971.0 | 3 | Polystomatidae | 971.0 | 3 | Protopolystoma | 971.0 | 3 | Protopolystoma xenopodis | 971.0 | 3 |
| scf7180000042136 | 402283 | 0.4349 | 0 | 58.6922 | Eukaryota | 49283.0 | 0 | Chordata | 49283.0 | 0 | Carnivora | 49283.0 | 0 | Canidae | 49283.0 | 0 | Canis | 39259.0 | 1 | Canis lupus | 39259.0 | 1 |
| scf7180000041449 | 1141645 | 0.3441 | 0 | 58.7261 | Eukaryota | 65625.0 | 0 | Chordata | 65625.0 | 0 | Carnivora | 65625.0 | 0 | Canidae | 65625.0 | 0 | Canis | 65625.0 | 0 | Canis lupus | 65625.0 | 0 |
| scf7180000056401 | 502679 | 0.3813 | 0 | 60.3121 | Eukaryota | 47621.0 | 0 | Chordata | 47621.0 | 0 | Carnivora | 47621.0 | 0 | Canidae | 47621.0 | 0 | Canis | 34508.0 | 1 | Canis lupus | 34508.0 | 1 |
| scf7180000042443 | 193701 | 0.5497 | 0 | 59.6235 | Eukaryota | 17929.0 | 0 | Chordata | 17929.0 | 0 | Carnivora | 17929.0 | 0 | Canidae | 17929.0 | 0 | Canis | 17929.0 | 0 | Canis lupus | 17929.0 | 0 |
| scf7180000055047 | 25443 | 0.4265 | 0 | 135.7801 | Eukaryota | 9865.0 | 0 | Chordata | 9865.0 | 0 | Carnivora | 9865.0 | 0 | Canidae | 9865.0 | 0 | Canis | 9865.0 | 0 | Canis lupus | 9865.0 | 0 |
| scf7180000053180 | 8981 | 0.4409 | 0 | 91.0517 | Eukaryota | 1869.0 | 0 | Chordata | 1130.0 | 1 | Polystomatidea | 739.0 | 4 | Polystomatidae | 739.0 | 4 | Protopolystoma | 739.0 | 4 | Protopolystoma xenopodis | 739.0 | 4 |
| scf7180000046938 | 16409 | 0.3764 | 0 | 53.0617 | Eukaryota | 38523.0 | 0 | Chordata | 38523.0 | 0 | Carnivora | 38523.0 | 0 | Canidae | 38523.0 | 0 | Canis | 38523.0 | 0 | Canis lupus | 38523.0 | 0 |
| scf7180000041744 | 344376 | 0.379 | 0 | 59.6665 | Eukaryota | 62951.0 | 0 | Chordata | 62951.0 | 0 | Carnivora | 62951.0 | 0 | Canidae | 62951.0 | 0 | Canis | 56249.0 | 1 | Canis lupus | 56249.0 | 1 |
| scf7180000042214 | 775561 | 0.3742 | 0 | 59.724 | Eukaryota | 35103.0 | 0 | Chordata | 35103.0 | 0 | Carnivora | 35103.0 | 0 | Canidae | 35103.0 | 0 | Canis | 35103.0 | 0 | Canis lupus | 35103.0 | 0 |
| scf7180000048513 | 48145 | 0.44 | 0 | 60.4887 | Eukaryota | 9377.0 | 0 | Chordata | 6845.0 | 1 | Polystomatidea | 2532.0 | 4 | Polystomatidae | 2532.0 | 4 | Protopolystoma | 2532.0 | 4 | Protopolystoma xenopodis | 2532.0 | 4 |
| scf7180000042826 | 62266 | 0.3735 | 0 | 56.9115 | Eukaryota | 62975.0 | 0 | Chordata | 62975.0 | 0 | Carnivora | 62975.0 | 0 | Canidae | 62975.0 | 0 | Canis | 62975.0 | 0 | Canis lupus | 62975.0 | 0 |
| scf7180000040517 | 168738 | 0.372 | 0 | 59.4696 | Eukaryota | 46254.0 | 0 | Chordata | 46254.0 | 0 | Carnivora | 46254.0 | 0 | Canidae | 46254.0 | 0 | Canis | 46254.0 | 0 | Canis lupus | 46254.0 | 0 |
| scf7180000050589 | 7164 | 0.4463 | 0 | 1439.3717 | no-hit | 0.0 | 0 | no-hit | 0.0 | 0 | no-hit | 0.0 | 0 | no-hit | 0.0 | 0 | no-hit | 0.0 | 0 | no-hit | 0.0 | 0 |
| scf7180000044026 | 121082 | 0.6307 | 0 | 53.9039 | Eukaryota | 29390.0 | 0 | Chordata | 29390.0 | 0 | Carnivora | 29390.0 | 0 | Canidae | 29390.0 | 0 | Canis | 24386.0 | 1 | Canis lupus | 24386.0 | 1 |
| scf7180000052312 | 3741 | 0.4542 | 0 | 338.6451 | Eukaryota | 1008.0 | 0 | Platyhelminthes | 743.0 | 1 | Polystomatidea | 743.0 | 1 | Polystomatidae | 743.0 | 1 | Protopolystoma | 743.0 | 1 | Protopolystoma xenopodis | 743.0 | 1 |
| scf7180000051179 | 18796 | 0.3557 | 0 | 58.4161 | Eukaryota | 4807.0 | 0 | Chordata | 4807.0 | 0 | Carnivora | 4807.0 | 0 | Canidae | 4807.0 | 0 | Canis | 4807.0 | 0 | Canis lupus | 4807.0 | 0 |
| scf7180000043221 | 521306 | 0.4517 | 0 | 58.5726 | Eukaryota | 51278.0 | 0 | Chordata | 51278.0 | 0 | Carnivora | 51278.0 | 0 | Canidae | 51278.0 | 0 | Canis | 33315.0 | 1 | Canis lupus | 33315.0 | 1 |
| scf7180000044738 | 313950 | 0.3955 | 0 | 60.0499 | Eukaryota | 115912.0 | 0 | Chordata | 115912.0 | 0 | Carnivora | 115912.0 | 0 | Canidae | 115912.0 | 0 | Canis | 115912.0 | 0 | Canis lupus | 115912.0 | 0 |
| scf7180000051824 | 5930 | 0.4444 | 0 | 983.0141 | Eukaryota | 633.0 | 0 | Streptophyta | 498.0 | 1 | Fabales | 498.0 | 1 | Fabaceae | 498.0 | 1 | Cicer | 498.0 | 1 | Cicer arietinum | 498.0 | 1 |
| scf7180000054757 | 20018 | 0.4459 | 0 | 342.4054 | Eukaryota | 2562.0 | 0 | Platyhelminthes | 1375.0 | 1 | Polystomatidea | 1375.0 | 2 | Polystomatidae | 1375.0 | 2 | Protopolystoma | 1375.0 | 2 | Protopolystoma xenopodis | 1375.0 | 2 |
| scf7180000056293 | 317458 | 0.407 | 0 | 62.6495 | Eukaryota | 90031.0 | 0 | Chordata | 90031.0 | 0 | Carnivora | 90031.0 | 0 | Canidae | 90031.0 | 0 | Canis | 90031.0 | 0 | Canis lupus | 90031.0 | 0 |
| scf7180000042361 | 352006 | 0.4331 | 0 | 58.5229 | Eukaryota | 95344.0 | 0 | Chordata | 95344.0 | 0 | Carnivora | 95344.0 | 0 | Canidae | 45526.0 | 2 | Canis | 45526.0 | 2 | Canis lupus | 45526.0 | 2 |
| scf7180000049395 | 10384 | 0.4419 | 0 | 478.95 | Eukaryota | 1213.0 | 0 | Platyhelminthes | 775.0 | 1 | Polystomatidea | 775.0 | 1 | Polystomatidae | 775.0 | 1 | Protopolystoma | 775.0 | 1 | Protopolystoma xenopodis | 775.0 | 1 |
| scf7180000045383 | 209672 | 0.4453 | 0 | 59.0477 | Eukaryota | 41011.0 | 0 | Chordata | 41011.0 | 0 | Carnivora | 41011.0 | 0 | Canidae | 23326.0 | 1 | Vulpes | 13863.0 | 3 | Vulpes vulpes | 13863.0 | 3 |
| scf7180000041921 | 862422 | 0.4303 | 0 | 60.0604 | Eukaryota | 61931.0 | 0 | Chordata | 61931.0 | 0 | Carnivora | 61931.0 | 0 | Canidae | 61931.0 | 0 | Canis | 61931.0 | 0 | Canis lupus | 61931.0 | 0 |
| scf7180000052757 | 13243 | 0.4537 | 0 | 65.6096 | Eukaryota | 3156.0 | 0 | Chordata | 3156.0 | 0 | Carnivora | 3156.0 | 0 | Canidae | 3156.0 | 0 | Canis | 3156.0 | 0 | Canis lupus | 3156.0 | 0 |
| scf7180000041953 | 859349 | 0.4222 | 0 | 58.3487 | Eukaryota | 66271.0 | 0 | Chordata | 66271.0 | 0 | Carnivora | 66271.0 | 0 | Canidae | 66271.0 | 0 | Canis | 66271.0 | 0 | Canis lupus | 66271.0 | 0 |
| scf7180000056511 | 199067 | 0.3695 | 0 | 60.6666 | Eukaryota | 56074.0 | 0 | Chordata | 56074.0 | 0 | Carnivora | 56074.0 | 0 | Canidae | 56074.0 | 0 | Canis | 38622.0 | 1 | Canis lupus | 38622.0 | 1 |
| scf7180000043332 | 173864 | 0.402 | 0 | 59.9471 | Eukaryota | 12148.0 | 0 | Chordata | 12148.0 | 0 | Primates | 12148.0 | 0 | Hominidae | 9546.0 | 1 | Homo | 9546.0 | 1 | Homo sapiens | 9546.0 | 1 |
| scf7180000042944 | 606089 | 0.5105 | 0 | 56.7428 | Eukaryota | 25483.0 | 0 | Chordata | 25483.0 | 0 | Carnivora | 25483.0 | 0 | Felidae | 15981.0 | 1 | Felis | 15981.0 | 2 | Felis catus | 15981.0 | 2 |
| scf7180000055930 | 9948 | 0.544 | 0 | 73.7912 | Eukaryota | 15457.0 | 0 | Chordata | 15457.0 | 0 | Carnivora | 15457.0 | 0 | Canidae | 15457.0 | 0 | Canis | 14373.0 | 1 | Canis lupus | 14373.0 | 2 |
| scf7180000051597 | 5505 | 0.4425 | 0 | 134.1742 | Eukaryota | 437.0 | 0 | Chordata | 259.0 | 1 | Salmoniformes | 259.0 | 1 | Salmonidae | 259.0 | 1 | Coregonus | 259.0 | 1 | Coregonus sp. 'balchen' | 259.0 | 1 |
| scf7180000042842 | 131386 | 0.572 | 0 | 58.7217 | Eukaryota | 24880.0 | 0 | Chordata | 24880.0 | 0 | Carnivora | 18127.0 | 1 | Canidae | 15458.0 | 3 | Canis | 15458.0 | 3 | Canis lupus | 15458.0 | 3 |
| scf7180000045960 | 5612 | 0.4498 | 0 | 522.4994 | Eukaryota | 202.0 | 0 | Chordata | 202.0 | 0 | Salmoniformes | 202.0 | 0 | Salmonidae | 202.0 | 0 | Salmo | 202.0 | 0 | Salmo trutta | 202.0 | 0 |
| scf7180000051817 | 8665 | 0.4478 | 0 | 624.358 | Eukaryota | 622.0 | 0 | Chordata | 455.0 | 1 | Tetraodontiformes | 455.0 | 1 | Tetraodontidae | 455.0 | 1 | Takifugu | 455.0 | 1 | Takifugu rubripes | 455.0 | 1 |
| scf7180000053652 | 4771 | 0.4278 | 0 | 106.6561 | Eukaryota | 8367.0 | 0 | Chordata | 7228.0 | 2 | Gadiformes | 3260.0 | 6 | Gadidae | 3260.0 | 6 | Gadus | 3260.0 | 6 | Gadus morhua | 3260.0 | 6 |
| scf7180000049164 | 11010 | 0.4413 | 0 | 95.3778 | Eukaryota | 230.0 | 0 | Chordata | 230.0 | 0 | Kurtiformes | 230.0 | 0 | Apogonidae | 230.0 | 0 | Sphaeramia | 230.0 | 0 | Sphaeramia orbicularis | 230.0 | 0 |
| scf7180000051086 | 7930 | 0.4362 | 0 | 61.0676 | no-hit | 0.0 | 0 | no-hit | 0.0 | 0 | no-hit | 0.0 | 0 | no-hit | 0.0 | 0 | no-hit | 0.0 | 0 | no-hit | 0.0 | 0 |
| scf7180000050739 | 21586 | 0.4547 | 0 | 50.8056 | Eukaryota | 11401.0 | 0 | Chordata | 6909.0 | 1 | Tetraodontiformes | 3860.0 | 5 | Tetraodontidae | 3860.0 | 5 | Takifugu | 3860.0 | 5 | Takifugu rubripes | 3860.0 | 5 |
| scf7180000043764 | 85681 | 0.432 | 0 | 67.9496 | Eukaryota | 69921.0 | 0 | Chordata | 69921.0 | 0 | Carnivora | 69921.0 | 0 | Canidae | 69921.0 | 0 | Canis | 69921.0 | 0 | Canis lupus | 69921.0 | 0 |
| scf7180000056600 | 548332 | 0.3996 | 0 | 60.0768 | Eukaryota | 29961.0 | 0 | Chordata | 29961.0 | 0 | Carnivora | 29961.0 | 0 | Canidae | 29961.0 | 0 | Canis | 29961.0 | 0 | Canis lupus | 29961.0 | 0 |
| scf7180000044996 | 95714 | 0.3666 | 0 | 59.3153 | Eukaryota | 84520.0 | 0 | Chordata | 84520.0 | 0 | Carnivora | 84520.0 | 0 | Canidae | 84520.0 | 0 | Canis | 84520.0 | 0 | Canis lupus | 84520.0 | 0 |
| scf7180000045086 | 220383 | 0.3854 | 0 | 59.8742 | Eukaryota | 86465.0 | 0 | Chordata | 86465.0 | 0 | Carnivora | 86465.0 | 0 | Canidae | 86465.0 | 0 | Canis | 86465.0 | 0 | Canis lupus | 86465.0 | 0 |
| scf7180000055558 | 8075 | 0.4394 | 0 | 333.8524 | Eukaryota | 907.0 | 0 | Platyhelminthes | 529.0 | 1 | Polystomatidea | 529.0 | 1 | Polystomatidae | 529.0 | 1 | Protopolystoma | 529.0 | 1 | Protopolystoma xenopodis | 529.0 | 1 |
| scf7180000046669 | 48296 | 0.4976 | 0 | 51.5818 | Eukaryota | 60973.0 | 0 | Chordata | 60973.0 | 0 | Carnivora | 60973.0 | 0 | Canidae | 60973.0 | 0 | Canis | 60973.0 | 0 | Canis lupus | 60973.0 | 0 |
| scf7180000049199 | 10065 | 0.3488 | 0 | 38.2765 | Eukaryota | 22247.0 | 0 | Chordata | 22247.0 | 0 | Carnivora | 22247.0 | 0 | Canidae | 22247.0 | 0 | Canis | 22247.0 | 0 | Canis lupus | 22247.0 | 0 |
| scf7180000055856 | 11218 | 0.435 | 0 | 152.4124 | Eukaryota | 1301.0 | 0 | Platyhelminthes | 1301.0 | 0 | Polystomatidea | 1301.0 | 0 | Polystomatidae | 1301.0 | 0 | Protopolystoma | 1301.0 | 0 | Protopolystoma xenopodis | 1301.0 | 0 |
| scf7180000045617 | 169864 | 0.4093 | 0 | 61.0638 | Eukaryota | 36826.0 | 0 | Chordata | 36826.0 | 0 | Carnivora | 36826.0 | 0 | Canidae | 29920.0 | 2 | Canis | 26215.0 | 3 | Canis lupus | 26215.0 | 3 |
| scf7180000042416 | 740027 | 0.3648 | 0 | 60.1699 | Eukaryota | 136175.0 | 0 | Chordata | 136175.0 | 0 | Carnivora | 136175.0 | 0 | Canidae | 136175.0 | 0 | Canis | 136175.0 | 0 | Canis lupus | 136175.0 | 0 |
| scf7180000049344 | 10833 | 0.4548 | 0 | 127.1064 | no-hit | 0.0 | 0 | no-hit | 0.0 | 0 | no-hit | 0.0 | 0 | no-hit | 0.0 | 0 | no-hit | 0.0 | 0 | no-hit | 0.0 | 0 |
| scf7180000046978 | 111434 | 0.3927 | 0 | 59.9953 | Eukaryota | 21882.0 | 0 | Chordata | 21882.0 | 0 | Carnivora | 21882.0 | 0 | Canidae | 21882.0 | 0 | Canis | 21882.0 | 0 | Canis lupus | 21882.0 | 0 |
| scf7180000041371 | 177717 | 0.396 | 0 | 60.0355 | Eukaryota | 53674.0 | 0 | Chordata | 53674.0 | 0 | Carnivora | 53674.0 | 0 | Canidae | 53674.0 | 0 | Canis | 53674.0 | 0 | Canis lupus | 53674.0 | 0 |
| scf7180000050726 | 17763 | 0.4995 | 0 | 338.8928 | Eukaryota | 9078.0 | 0 | Chordata | 9078.0 | 0 | Carnivora | 9078.0 | 0 | Canidae | 9078.0 | 0 | Canis | 7904.0 | 1 | Canis lupus | 6738.0 | 2 |
| scf7180000048495 | 94621 | 0.4503 | 0 | 54.3875 | Eukaryota | 107840.0 | 0 | Chordata | 107840.0 | 0 | Carnivora | 107840.0 | 0 | Canidae | 107840.0 | 0 | Canis | 107840.0 | 0 | Canis lupus | 107840.0 | 0 |
| scf7180000045984 | 27996 | 0.4364 | 0 | 154.8983 | Eukaryota | 15069.0 | 0 | Chordata | 10186.0 | 2 | Gadiformes | 4261.0 | 7 | Gadidae | 4261.0 | 7 | Gadus | 4261.0 | 7 | Gadus morhua | 4261.0 | 7 |
| scf7180000043964 | 407495 | 0.4283 | 0 | 58.1238 | Eukaryota | 33406.0 | 0 | Chordata | 33406.0 | 0 | Carnivora | 23290.0 | 1 | Canidae | 23290.0 | 2 | Vulpes | 13336.0 | 4 | Vulpes vulpes | 13336.0 | 4 |
| scf7180000042578 | 677273 | 0.4503 | 0 | 58.5535 | Eukaryota | 51623.0 | 0 | Chordata | 51623.0 | 0 | Carnivora | 51623.0 | 0 | Canidae | 51623.0 | 0 | Canis | 51623.0 | 0 | Canis lupus | 51623.0 | 0 |
| scf7180000051917 | 5836 | 0.4486 | 0 | 1842.5494 | no-hit | 0.0 | 0 | no-hit | 0.0 | 0 | no-hit | 0.0 | 0 | no-hit | 0.0 | 0 | no-hit | 0.0 | 0 | no-hit | 0.0 | 0 |
| scf7180000042937 | 39391 | 0.3584 | 0 | 58.0538 | Eukaryota | 13934.0 | 0 | Chordata | 13934.0 | 0 | Carnivora | 8590.0 | 1 | Hominidae | 5344.0 | 4 | Pan | 3587.0 | 6 | Pan troglodytes | 3587.0 | 7 |
| scf7180000054653 | 29558 | 0.4459 | 0 | 446.7451 | Eukaryota | 92480.0 | 0 | Chordata | 92480.0 | 0 | Carnivora | 92480.0 | 0 | Canidae | 92480.0 | 0 | Canis | 92480.0 | 0 | Canis lupus | 92480.0 | 0 |
| scf7180000045246 | 305447 | 0.4556 | 0 | 58.9498 | Eukaryota | 29591.0 | 0 | Chordata | 29591.0 | 0 | Carnivora | 29591.0 | 0 | Canidae | 29591.0 | 0 | Canis | 29591.0 | 0 | Canis lupus | 29591.0 | 0 |
| scf7180000052890 | 4117 | 0.4445 | 0 | 118.8584 | no-hit | 0.0 | 0 | no-hit | 0.0 | 0 | no-hit | 0.0 | 0 | no-hit | 0.0 | 0 | no-hit | 0.0 | 0 | no-hit | 0.0 | 0 |
| scf7180000053513 | 2468 | 0.4489 | 0 | 310.5113 | no-hit | 0.0 | 0 | no-hit | 0.0 | 0 | no-hit | 0.0 | 0 | no-hit | 0.0 | 0 | no-hit | 0.0 | 0 | no-hit | 0.0 | 0 |
| scf7180000052462 | 6853 | 0.4531 | 0 | 44.8632 | no-hit | 0.0 | 0 | no-hit | 0.0 | 0 | no-hit | 0.0 | 0 | no-hit | 0.0 | 0 | no-hit | 0.0 | 0 | no-hit | 0.0 | 0 |
| scf7180000043114 | 504328 | 0.4098 | 0 | 61.0452 | Eukaryota | 85828.0 | 0 | Chordata | 85828.0 | 0 | Carnivora | 85828.0 | 0 | Canidae | 85828.0 | 0 | Canis | 80238.0 | 1 | Canis lupus | 80238.0 | 1 |
| scf7180000055674 | 12729 | 0.443 | 0 | 212.2487 | Eukaryota | 44828.0 | 0 | Chordata | 32657.0 | 2 | Gadiformes | 28553.0 | 5 | Gadidae | 28553.0 | 5 | Gadus | 28553.0 | 5 | Gadus morhua | 28553.0 | 5 |
| scf7180000046024 | 99346 | 0.4467 | 0 | 57.5109 | Eukaryota | 62173.0 | 0 | Chordata | 62173.0 | 0 | Carnivora | 62173.0 | 0 | Canidae | 62173.0 | 0 | Canis | 62173.0 | 0 | Canis lupus | 62173.0 | 0 |
| scf7180000055617 | 13190 | 0.4393 | 0 | 127.5739 | Eukaryota | 3377.0 | 0 | Chordata | 3377.0 | 0 | Carnivora | 2458.0 | 2 | Canidae | 2458.0 | 2 | Canis | 2458.0 | 2 | Canis lupus | 2458.0 | 2 |
| scf7180000054827 | 55043 | 0.4407 | 0 | 115.1771 | Eukaryota | 64656.0 | 0 | Chordata | 48739.0 | 1 | Gadiformes | 36906.0 | 5 | Gadidae | 36906.0 | 5 | Gadus | 36906.0 | 5 | Gadus morhua | 36906.0 | 5 |
| scf7180000051749 | 11226 | 0.4472 | 0 | 88.9994 | no-hit | 0.0 | 0 | no-hit | 0.0 | 0 | no-hit | 0.0 | 0 | no-hit | 0.0 | 0 | no-hit | 0.0 | 0 | no-hit | 0.0 | 0 |
| scf7180000045466 | 173004 | 0.4311 | 0 | 56.2735 | Eukaryota | 18940.0 | 0 | Chordata | 18940.0 | 0 | Carnivora | 13653.0 | 1 | Canidae | 13653.0 | 2 | Vulpes | 9482.0 | 3 | Vulpes vulpes | 9482.0 | 3 |
| scf7180000056221 | 812240 | 0.361 | 0 | 59.5092 | Eukaryota | 91650.0 | 0 | Chordata | 91650.0 | 0 | Carnivora | 91650.0 | 0 | Canidae | 91650.0 | 0 | Canis | 91650.0 | 0 | Canis lupus | 91650.0 | 0 |
| scf7180000043555 | 31295 | 0.5467 | 0 | 55.2865 | Eukaryota | 2195.0 | 0 | Chordata | 2195.0 | 0 | Carnivora | 2195.0 | 0 | Canidae | 2195.0 | 0 | Canis | 2195.0 | 0 | Canis lupus | 2195.0 | 0 |
| scf7180000041597 | 429113 | 0.4211 | 0 | 59.4144 | Eukaryota | 80877.0 | 0 | Chordata | 80877.0 | 0 | Carnivora | 80877.0 | 0 | Canidae | 80877.0 | 0 | Canis | 80877.0 | 0 | Canis lupus | 80877.0 | 0 |
| scf7180000043652 | 392571 | 0.3659 | 0 | 97.2393 | Eukaryota | 91887.0 | 0 | Chordata | 91887.0 | 0 | Carnivora | 91887.0 | 0 | Canidae | 91887.0 | 0 | Canis | 91887.0 | 0 | Canis lupus | 91887.0 | 0 |
| scf7180000045484 | 207715 | 0.4156 | 0 | 60.2686 | Eukaryota | 20260.0 | 0 | Chordata | 20260.0 | 0 | Carnivora | 20260.0 | 0 | Odobenidae | 6103.0 | 5 | Odobenus | 6103.0 | 6 | Odobenus rosmarus | 6103.0 | 7 |
| scf7180000054382 | 65275 | 0.4236 | 0 | 384.2536 | Eukaryota | 53374.0 | 0 | Chordata | 53374.0 | 0 | Carnivora | 53374.0 | 0 | Canidae | 38691.0 | 1 | Canis | 26437.0 | 2 | Canis lupus | 26437.0 | 3 |
| scf7180000053346 | 11375 | 0.5009 | 0 | 149.0849 | Eukaryota | 8902.0 | 0 | Chordata | 8902.0 | 0 | Carnivora | 8902.0 | 0 | Canidae | 8902.0 | 0 | Canis | 7704.0 | 1 | Canis lupus | 6438.0 | 2 |
| scf7180000052951 | 6455 | 0.3593 | 0 | 65.6693 | Eukaryota | 2964.0 | 0 | Chordata | 2964.0 | 0 | Carnivora | 2964.0 | 0 | Canidae | 2964.0 | 0 | Canis | 2670.0 | 1 | Canis lupus | 2670.0 | 1 |
| scf7180000047086 | 55078 | 0.5934 | 0 | 56.4039 | Eukaryota | 28350.0 | 0 | Chordata | 28350.0 | 0 | Carnivora | 28350.0 | 0 | Canidae | 28350.0 | 0 | Canis | 28350.0 | 0 | Canis lupus | 28350.0 | 0 |
| scf7180000043118 | 463221 | 0.3946 | 0 | 60.7471 | Eukaryota | 32428.0 | 0 | Chordata | 32428.0 | 0 | Carnivora | 32428.0 | 0 | Canidae | 32428.0 | 0 | Canis | 16657.0 | 1 | Canis lupus | 16657.0 | 1 |
| scf7180000052028 | 8394 | 0.4369 | 0 | 131.6435 | Eukaryota | 619.0 | 0 | Platyhelminthes | 463.0 | 1 | Polystomatidea | 463.0 | 1 | Polystomatidae | 463.0 | 1 | Protopolystoma | 463.0 | 1 | Protopolystoma xenopodis | 463.0 | 1 |
| scf7180000049867 | 31197 | 0.6782 | 0 | 53.5375 | Eukaryota | 19766.0 | 0 | Chordata | 19766.0 | 0 | Carnivora | 19766.0 | 0 | Canidae | 18866.0 | 1 | Canis | 16479.0 | 2 | Canis lupus | 16479.0 | 2 |
| scf7180000048591 | 17382 | 0.5055 | 0 | 790.4522 | Eukaryota | 9178.0 | 0 | Chordata | 9178.0 | 0 | Carnivora | 9178.0 | 0 | Canidae | 9178.0 | 0 | Canis | 7430.0 | 1 | Canis lupus | 6164.0 | 2 |
| scf7180000048694 | 47740 | 0.3935 | 0 | 61.3017 | Eukaryota | 7828.0 | 0 | Chordata | 7828.0 | 0 | Primates | 3336.0 | 2 | Hominidae | 3336.0 | 3 | Homo | 3336.0 | 4 | Homo sapiens | 3336.0 | 4 |
| scf7180000052977 | 5109 | 0.4506 | 0 | 60.2795 | no-hit | 0.0 | 0 | no-hit | 0.0 | 0 | no-hit | 0.0 | 0 | no-hit | 0.0 | 0 | no-hit | 0.0 | 0 | no-hit | 0.0 | 0 |
| scf7180000040485 | 899087 | 0.3539 | 0 | 59.6205 | Eukaryota | 40393.0 | 0 | Chordata | 40393.0 | 0 | Carnivora | 40393.0 | 0 | Canidae | 40393.0 | 0 | Canis | 40393.0 | 0 | Canis lupus | 40393.0 | 0 |
| scf7180000050410 | 28720 | 0.4414 | 0 | 50.0503 | no-hit | 0.0 | 0 | no-hit | 0.0 | 0 | no-hit | 0.0 | 0 | no-hit | 0.0 | 0 | no-hit | 0.0 | 0 | no-hit | 0.0 | 0 |
| scf7180000053769 | 7731 | 0.4513 | 0 | 793.781 | Eukaryota | 19186.0 | 0 | Chordata | 18360.0 | 1 | Gadiformes | 10806.0 | 5 | Gadidae | 10806.0 | 5 | Gadus | 10806.0 | 5 | Gadus morhua | 10806.0 | 5 |
| scf7180000044035 | 355111 | 0.4399 | 0 | 59.0393 | Eukaryota | 82762.0 | 0 | Chordata | 82762.0 | 0 | Carnivora | 82762.0 | 0 | Canidae | 82762.0 | 0 | Canis | 63764.0 | 1 | Canis lupus | 63764.0 | 1 |
| scf7180000043296 | 377819 | 0.4434 | 0 | 59.8378 | Eukaryota | 92045.0 | 0 | Chordata | 92045.0 | 0 | Carnivora | 92045.0 | 0 | Canidae | 92045.0 | 0 | Canis | 92045.0 | 0 | Canis lupus | 92045.0 | 0 |
| scf7180000041266 | 660668 | 0.3626 | 0 | 60.3574 | Eukaryota | 84534.0 | 0 | Chordata | 84534.0 | 0 | Carnivora | 84534.0 | 0 | Canidae | 84534.0 | 0 | Canis | 84534.0 | 0 | Canis lupus | 84534.0 | 0 |
| scf7180000053571 | 2708 | 0.4446 | 0 | 197.4217 | no-hit | 0.0 | 0 | no-hit | 0.0 | 0 | no-hit | 0.0 | 0 | no-hit | 0.0 | 0 | no-hit | 0.0 | 0 | no-hit | 0.0 | 0 |
| scf7180000055086 | 28651 | 0.4402 | 0 | 201.5824 | Eukaryota | 3928.0 | 0 | Echinodermata | 2209.0 | 2 | Forcipulatida | 2209.0 | 2 | Asteriidae | 2209.0 | 2 | Asterias | 2209.0 | 2 | Asterias rubens | 2209.0 | 2 |
| scf7180000040774 | 1884218 | 0.3762 | 0 | 60.1827 | Eukaryota | 82146.0 | 0 | Chordata | 82146.0 | 0 | Carnivora | 82146.0 | 0 | Canidae | 82146.0 | 0 | Canis | 82146.0 | 0 | Canis lupus | 82146.0 | 0 |
| scf7180000056385 | 515251 | 0.3634 | 0 | 59.8058 | Eukaryota | 93576.0 | 0 | Chordata | 93576.0 | 0 | Carnivora | 93576.0 | 0 | Canidae | 93576.0 | 0 | Canis | 93576.0 | 0 | Canis lupus | 93576.0 | 0 |
| scf7180000044153 | 173030 | 0.4295 | 0 | 58.5269 | Eukaryota | 24830.0 | 0 | Chordata | 24830.0 | 0 | Carnivora | 24830.0 | 0 | Canidae | 10761.0 | 4 | Vulpes | 5724.0 | 7 | Vulpes vulpes | 5724.0 | 7 |
| scf7180000052124 | 19596 | 0.4412 | 0 | 50.4607 | Eukaryota | 32834.0 | 0 | Platyhelminthes | 17430.0 | 2 | Strigeidida | 17430.0 | 5 | Schistosomatidae | 17430.0 | 5 | Schistosoma | 17430.0 | 5 | Schistosoma mansoni | 17430.0 | 6 |
| scf7180000040936 | 2840 | 0.4018 | 0 | 57.3121 | Eukaryota | 1552.0 | 1 | Chordata | 1552.0 | 1 | Primates | 1042.0 | 2 | Hominidae | 1042.0 | 2 | Homo | 686.0 | 3 | Homo sapiens | 686.0 | 3 |
| scf7180000049837 | 37125 | 0.4514 | 0 | 68.6345 | Eukaryota | 12376.0 | 0 | Chordata | 12376.0 | 0 | Carnivora | 12376.0 | 0 | Canidae | 12376.0 | 0 | Canis | 12376.0 | 0 | Canis lupus | 12376.0 | 0 |
| scf7180000049080 | 12553 | 0.4401 | 0 | 115.9714 | Eukaryota | 25998.0 | 0 | Chordata | 18035.0 | 2 | Gadiformes | 13193.0 | 7 | Gadidae | 13193.0 | 7 | Gadus | 13193.0 | 7 | Gadus morhua | 13193.0 | 7 |
| scf7180000045046 | 198903 | 0.4517 | 0 | 60.3739 | Eukaryota | 57940.0 | 0 | Chordata | 57940.0 | 0 | Carnivora | 57940.0 | 0 | Canidae | 57940.0 | 0 | Canis | 57940.0 | 0 | Canis lupus | 57940.0 | 0 |
| scf7180000044175 | 84085 | 0.3369 | 0 | 58.9278 | Eukaryota | 24889.0 | 1 | Chordata | 24889.0 | 1 | Primates | 11155.0 | 3 | Hominidae | 11155.0 | 3 | Homo | 11155.0 | 4 | Homo sapiens | 11155.0 | 4 |
| scf7180000040496 | 2882596 | 0.3909 | 0 | 60.5395 | Eukaryota | 95237.0 | 0 | Chordata | 95237.0 | 0 | Carnivora | 95237.0 | 0 | Canidae | 95237.0 | 0 | Canis | 95237.0 | 0 | Canis lupus | 95237.0 | 0 |
| scf7180000050894 | 5370 | 0.4508 | 0 | 39.6142 | Eukaryota | 987.0 | 0 | Chordata | 987.0 | 0 | Salmoniformes | 987.0 | 0 | Salmonidae | 987.0 | 0 | Salmo | 987.0 | 0 | Salmo trutta | 987.0 | 0 |
| scf7180000054056 | 2126 | 0.4586 | 0 | 470.2528 | no-hit | 0.0 | 0 | no-hit | 0.0 | 0 | no-hit | 0.0 | 0 | no-hit | 0.0 | 0 | no-hit | 0.0 | 0 | no-hit | 0.0 | 0 |
| scf7180000044288 | 89352 | 0.5984 | 0 | 54.7028 | Eukaryota | 22419.0 | 0 | Chordata | 22419.0 | 0 | Carnivora | 22419.0 | 0 | Canidae | 22419.0 | 0 | Canis | 11328.0 | 1 | Canis lupus | 11328.0 | 1 |
| scf7180000049778 | 3195 | 0.4507 | 0 | 94.7617 | Eukaryota | 2337.0 | 0 | Chordata | 2337.0 | 0 | Carnivora | 2337.0 | 0 | Canidae | 2337.0 | 0 | Canis | 2337.0 | 0 | Canis lupus | 2337.0 | 0 |
| scf7180000055102 | 67488 | 0.4422 | 0 | 239.8403 | Eukaryota | 7482.0 | 0 | Chordata | 4254.0 | 2 | Kurtiformes | 2748.0 | 3 | Apogonidae | 2748.0 | 3 | Sphaeramia | 2748.0 | 3 | Sphaeramia orbicularis | 2748.0 | 3 |
| scf7180000044684 | 284842 | 0.3632 | 0 | 60.1355 | Eukaryota | 92327.0 | 0 | Chordata | 92327.0 | 0 | Carnivora | 92327.0 | 0 | Canidae | 92327.0 | 0 | Canis | 92327.0 | 0 | Canis lupus | 92327.0 | 0 |
| scf7180000051641 | 6088 | 0.4537 | 0 | 59.9254 | no-hit | 0.0 | 0 | no-hit | 0.0 | 0 | no-hit | 0.0 | 0 | no-hit | 0.0 | 0 | no-hit | 0.0 | 0 | no-hit | 0.0 | 0 |
| scf7180000046774 | 90603 | 0.3498 | 0 | 59.1617 | Eukaryota | 4626.0 | 0 | Chordata | 4626.0 | 0 | Carnivora | 1744.0 | 2 | Canidae | 1744.0 | 2 | Canis | 1744.0 | 2 | Canis lupus | 1744.0 | 2 |
| scf7180000042418 | 141920 | 0.3584 | 0 | 58.8818 | Eukaryota | 40220.0 | 0 | Chordata | 40220.0 | 0 | Carnivora | 40220.0 | 0 | Canidae | 40220.0 | 0 | Canis | 40220.0 | 0 | Canis lupus | 40220.0 | 0 |
| scf7180000042714 | 608905 | 0.3666 | 0 | 60.0283 | Eukaryota | 63055.0 | 0 | Chordata | 63055.0 | 0 | Carnivora | 63055.0 | 0 | Canidae | 63055.0 | 0 | Canis | 63055.0 | 0 | Canis lupus | 63055.0 | 0 |
| scf7180000050632 | 5484 | 0.454 | 0 | 1049.2707 | Eukaryota | 1705.0 | 0 | Echinodermata | 774.0 | 2 | Forcipulatida | 774.0 | 2 | Asteriidae | 774.0 | 2 | Asterias | 774.0 | 2 | Asterias rubens | 774.0 | 2 |
| scf7180000040793 | 1785625 | 0.3685 | 0 | 59.7027 | Eukaryota | 82493.0 | 0 | Chordata | 82493.0 | 0 | Carnivora | 82493.0 | 0 | Canidae | 82493.0 | 0 | Canis | 82493.0 | 0 | Canis lupus | 82493.0 | 0 |
| scf7180000054685 | 137892 | 0.4387 | 0 | 84.6343 | Eukaryota | 91470.0 | 0 | Chordata | 91470.0 | 0 | Carnivora | 91470.0 | 0 | Canidae | 91470.0 | 0 | Canis | 91470.0 | 0 | Canis lupus | 91470.0 | 0 |
| scf7180000052061 | 8791 | 0.4101 | 0 | 31.6873 | Eukaryota | 64321.0 | 0 | Chordata | 64321.0 | 0 | Carnivora | 64321.0 | 0 | Canidae | 64321.0 | 0 | Canis | 64321.0 | 0 | Canis lupus | 64321.0 | 0 |
| scf7180000040606 | 30749 | 0.3766 | 0 | 47.01 | Eukaryota | 28801.0 | 0 | Chordata | 28801.0 | 0 | Carnivora | 28801.0 | 0 | Canidae | 28801.0 | 0 | Canis | 28801.0 | 0 | Canis lupus | 28801.0 | 0 |
| scf7180000042990 | 23057 | 0.4116 | 0 | 51.8924 | Eukaryota | 89468.0 | 0 | Chordata | 89468.0 | 0 | Carnivora | 89468.0 | 0 | Canidae | 89468.0 | 0 | Canis | 89468.0 | 0 | Canis lupus | 89468.0 | 0 |
| scf7180000042710 | 9462 | 0.3244 | 0 | 45.666 | Eukaryota | 2564.0 | 0 | Chordata | 2564.0 | 0 | Carnivora | 2564.0 | 0 | Canidae | 2564.0 | 0 | Canis | 2564.0 | 0 | Canis lupus | 2564.0 | 0 |
| scf7180000041820 | 869365 | 0.372 | 0 | 60.4983 | Eukaryota | 64119.0 | 0 | Chordata | 64119.0 | 0 | Carnivora | 64119.0 | 0 | Canidae | 64119.0 | 0 | Canis | 64119.0 | 0 | Canis lupus | 64119.0 | 0 |
| scf7180000043716 | 88555 | 0.5096 | 0 | 55.3905 | Eukaryota | 6107.0 | 0 | Chordata | 6107.0 | 0 | Carnivora | 2815.0 | 2 | Canidae | 2815.0 | 2 | Canis | 2815.0 | 3 | Canis lupus | 2815.0 | 3 |
| scf7180000054689 | 71884 | 0.4411 | 0 | 147.0251 | Eukaryota | 23065.0 | 1 | Chordata | 23065.0 | 1 | Gadiformes | 8503.0 | 7 | Gadidae | 8503.0 | 7 | Gadus | 8503.0 | 7 | Gadus morhua | 8503.0 | 7 |
| scf7180000051732 | 6574 | 0.436 | 0 | 68.3003 | Eukaryota | 3850.0 | 0 | Echinodermata | 1997.0 | 2 | Forcipulatida | 1997.0 | 4 | Asteriidae | 1997.0 | 4 | Asterias | 1997.0 | 4 | Asterias rubens | 1997.0 | 4 |
| scf7180000050838 | 28171 | 0.3631 | 0 | 58.9419 | Eukaryota | 5649.0 | 0 | Chordata | 5649.0 | 0 | Primates | 2381.0 | 2 | Hominidae | 2381.0 | 2 | Homo | 2381.0 | 2 | Homo sapiens | 2381.0 | 2 |
| scf7180000053005 | 3253 | 0.5091 | 0 | 54.3554 | Eukaryota | 3658.0 | 0 | Chordata | 3658.0 | 0 | Carnivora | 3658.0 | 0 | Canidae | 3658.0 | 0 | Canis | 2683.0 | 1 | Canis lupus | 2683.0 | 1 |
| scf7180000051560 | 13040 | 0.4438 | 0 | 316.6218 | Eukaryota | 20300.0 | 0 | Chordata | 18914.0 | 1 | Gadiformes | 14430.0 | 4 | Gadidae | 14430.0 | 4 | Gadus | 14430.0 | 4 | Gadus morhua | 14430.0 | 4 |
| scf7180000045455 | 150718 | 0.3729 | 0 | 61.8924 | Eukaryota | 91902.0 | 0 | Chordata | 91902.0 | 0 | Carnivora | 91902.0 | 0 | Canidae | 91902.0 | 0 | Canis | 91902.0 | 0 | Canis lupus | 91902.0 | 0 |
| scf7180000052816 | 19667 | 0.4434 | 0 | 54.8896 | Eukaryota | 48006.0 | 0 | Chordata | 48006.0 | 0 | Carnivora | 48006.0 | 0 | Canidae | 48006.0 | 0 | Canis | 48006.0 | 0 | Canis lupus | 48006.0 | 0 |
| scf7180000052663 | 7412 | 0.4498 | 0 | 1983.9516 | no-hit | 0.0 | 0 | no-hit | 0.0 | 0 | no-hit | 0.0 | 0 | no-hit | 0.0 | 0 | no-hit | 0.0 | 0 | no-hit | 0.0 | 0 |
| scf7180000052975 | 4623 | 0.4536 | 0 | 565.6913 | Eukaryota | 2425.0 | 0 | Platyhelminthes | 1243.0 | 2 | Polystomatidea | 1243.0 | 2 | Polystomatidae | 1243.0 | 2 | Protopolystoma | 1243.0 | 2 | Protopolystoma xenopodis | 1243.0 | 2 |
| scf7180000045541 | 32002 | 0.439 | 0 | 91.8065 | Eukaryota | 3056.0 | 0 | Chordata | 2485.0 | 1 | Tetraodontiformes | 1652.0 | 3 | Tetraodontidae | 1652.0 | 3 | Takifugu | 1652.0 | 3 | Takifugu rubripes | 1652.0 | 3 |
| scf7180000041194 | 486378 | 0.3502 | 0 | 59.7132 | Eukaryota | 86134.0 | 0 | Chordata | 86134.0 | 0 | Carnivora | 86134.0 | 0 | Canidae | 86134.0 | 0 | Canis | 86134.0 | 0 | Canis lupus | 86134.0 | 0 |
| scf7180000055101 | 75360 | 0.4388 | 0 | 198.2781 | Eukaryota | 99540.0 | 0 | Platyhelminthes | 51261.0 | 1 | Gadiformes | 48279.0 | 2 | Gadidae | 48279.0 | 2 | Gadus | 48279.0 | 2 | Gadus morhua | 48279.0 | 2 |
| scf7180000045032 | 280743 | 0.401 | 0 | 59.1871 | Eukaryota | 39701.0 | 0 | Chordata | 39701.0 | 0 | Carnivora | 39701.0 | 0 | Canidae | 36269.0 | 1 | Canis | 32751.0 | 2 | Canis lupus | 32751.0 | 2 |
| scf7180000049003 | 2316 | 0.4529 | 0 | 241.3421 | Eukaryota | 390.0 | 0 | Chordata | 390.0 | 0 | Kurtiformes | 390.0 | 0 | Apogonidae | 390.0 | 0 | Sphaeramia | 390.0 | 0 | Sphaeramia orbicularis | 390.0 | 0 |
| scf7180000044634 | 25932 | 0.3363 | 0 | 55.6721 | Eukaryota | 46875.0 | 0 | Chordata | 46875.0 | 0 | Carnivora | 46875.0 | 0 | Canidae | 46875.0 | 0 | Canis | 46875.0 | 0 | Canis lupus | 46875.0 | 0 |
| scf7180000053833 | 2182 | 0.4455 | 0 | 329.7446 | no-hit | 0.0 | 0 | no-hit | 0.0 | 0 | no-hit | 0.0 | 0 | no-hit | 0.0 | 0 | no-hit | 0.0 | 0 | no-hit | 0.0 | 0 |
| scf7180000047060 | 11703 | 0.5512 | 0 | 38.0332 | Eukaryota | 25219.0 | 0 | Chordata | 25219.0 | 0 | Carnivora | 25219.0 | 0 | Canidae | 25219.0 | 0 | Canis | 25219.0 | 0 | Canis lupus | 25219.0 | 0 |
| scf7180000043914 | 380515 | 0.37 | 0 | 60.4468 | Eukaryota | 98233.0 | 0 | Chordata | 98233.0 | 0 | Carnivora | 74995.0 | 1 | Canidae | 46211.0 | 3 | Canis | 34207.0 | 4 | Canis lupus | 34207.0 | 7 |
| scf7180000048026 | 26380 | 0.4512 | 0 | 96.1268 | Eukaryota | 10854.0 | 0 | Chordata | 10854.0 | 0 | Carnivora | 8240.0 | 2 | Canidae | 8240.0 | 2 | Canis | 4298.0 | 3 | Canis lupus | 4298.0 | 3 |
| scf7180000040848 | 1569736 | 0.3866 | 0 | 59.2983 | Eukaryota | 86743.0 | 0 | Chordata | 86743.0 | 0 | Carnivora | 81439.0 | 1 | Canidae | 81439.0 | 1 | Canis | 81439.0 | 1 | Canis lupus | 81439.0 | 1 |
| scf7180000054308 | 14748 | 0.4409 | 0 | 271.1663 | Eukaryota | 516.0 | 0 | Platyhelminthes | 516.0 | 0 | Polystomatidea | 516.0 | 0 | Polystomatidae | 516.0 | 0 | Protopolystoma | 516.0 | 0 | Protopolystoma xenopodis | 516.0 | 0 |

| | | | | | | | | | | | | | | | | | | | |
|---|---|---|---|---|---|---|---|---|---|---|---|---|---|---|---|---|---|---|---|
| scf7180000048445 | 50109 | 0.3834 | 0 | 59.6524 | Eukaryota | 6313.0 | 1 | Chordata | 6313.0 | 1 | Carnivora | 4203.0 | 2 | Canidae | 4203.0 | 2 | Canis | 4203.0 | 2 | Canis lupus | 4203.0 | 2 |
| scf7180000045734 | 5701 | 0.4404 | 0 | 27.5235 | Eukaryota | 6598.0 | 0 | Chordata | 6598.0 | 0 | Carnivora | 6598.0 | 0 | Canidae | 2139.0 | 3 | Canis | 1426.0 | 6 | Canis lupus | 1426.0 | 6 |
| scf7180000046433 | 66873 | 0.4547 | 0 | 121.5911 | Eukaryota | 5047.0 | 0 | Chordata | 4590.0 | 1 | Synbranchiformes | 1709.0 | 4 | Mastacembelidae | 1709.0 | 4 | Mastacembelus | 1709.0 | 5 | Mastacembelus armatus | 1709.0 | 5 |
| scf7180000052129 | 15382 | 0.441 | 0 | 53.8723 | Eukaryota | 370.0 | 0 | Chordata | 370.0 | 0 | Salmoniformes | 370.0 | 0 | Salmonidae | 370.0 | 0 | Salmo | 370.0 | 0 | Salmo trutta | 370.0 | 0 |
| scf7180000041581 | 1529491 | 0.3517 | 100 | 59.298 | Eukaryota | 40789.0 | 0 | Chordata | 40789.0 | 0 | Carnivora | 40789.0 | 0 | Canidae | 40789.0 | 0 | Canis | 40789.0 | 0 | Canis lupus | 40789.0 | 0 |
| scf7180000040820 | 1679238 | 0.4287 | 0 | 59.05 | Eukaryota | 69848.0 | 0 | Chordata | 69848.0 | 0 | Carnivora | 69848.0 | 0 | Canidae | 69848.0 | 0 | Canis | 69848.0 | 0 | Canis lupus | 69848.0 | 0 |
| scf7180000053943 | 2239 | 0.4511 | 0 | 65.7809 | Eukaryota | 392.0 | 0 | Platyhelminthes | 392.0 | 0 | Polystomatidea | 392.0 | 0 | Polystomatidae | 392.0 | 0 | Protopolystoma | 392.0 | 0 | Protopolystoma xenopodis | 392.0 | 0 |
| scf7180000053318 | 4371 | 0.5102 | 0 | 131.2997 | Eukaryota | 7772.0 | 0 | Chordata | 7772.0 | 0 | Carnivora | 7772.0 | 0 | Canidae | 7772.0 | 0 | Canis | 5825.0 | 1 | Canis lupus | 4744.0 | 2 |
| scf7180000044481 | 300857 | 0.4507 | 0 | 77.8845 | Eukaryota | 43642.0 | 0 | Chordata | 43642.0 | 0 | Carnivora | 43642.0 | 0 | Canidae | 43642.0 | 0 | Canis | 29533.0 | 1 | Canis lupus | 29533.0 | 1 |
| scf7180000046035 | 177253 | 0.3642 | 0 | 60.4562 | Eukaryota | 37549.0 | 0 | Chordata | 37549.0 | 0 | Carnivora | 37549.0 | 0 | Canidae | 37549.0 | 0 | Canis | 33944.0 | 1 | Canis lupus | 33944.0 | 1 |
| scf7180000052706 | 7224 | 0.451 | 0 | 122.7066 | Eukaryota | 2800.0 | 0 | Chordata | 2800.0 | 0 | Carnivora | 2800.0 | 0 | Canidae | 2800.0 | 0 | Canis | 2800.0 | 0 | Canis lupus | 2800.0 | 0 |
| scf7180000049042 | 26885 | 0.4415 | 0 | 95.4875 | Eukaryota | 4815.0 | 0 | Chordata | 4815.0 | 0 | Tetraodontiformes | 2072.0 | 4 | Tetraodontidae | 2072.0 | 4 | Takifugu | 2072.0 | 5 | Takifugu rubripes | 2072.0 | 5 |
| scf7180000052214 | 4765 | 0.4527 | 0 | 704.2269 | Eukaryota | 1260.0 | 0 | Echinodermata | 472.0 | 2 | Forcipulatida | 472.0 | 2 | Asteriidae | 472.0 | 2 | Asterias | 472.0 | 2 | Asterias rubens | 472.0 | 2 |
| scf7180000043476 | 412198 | 0.4782 | 0 | 58.6762 | Eukaryota | 94521.0 | 0 | Chordata | 94521.0 | 0 | Carnivora | 94521.0 | 0 | Canidae | 80181.0 | 1 | Canis | 67686.0 | 4 | Canis lupus | 67686.0 | 4 |
| scf7180000049297 | 27946 | 0.4883 | 0 | 109.5633 | Eukaryota | 15078.0 | 0 | Chordata | 15078.0 | 0 | Carnivora | 15078.0 | 0 | Canidae | 15078.0 | 0 | Canis | 15078.0 | 0 | Canis lupus | 15078.0 | 0 |
| scf7180000046664 | 6438 | 0.44 | 0 | 82.4549 | no-hit | 0.0 | 0 | no-hit | 0.0 | 0 | no-hit | 0.0 | 0 | no-hit | 0.0 | 0 | no-hit | 0.0 | 0 | no-hit | 0.0 | 0 |
| scf7180000043261 | 469374 | 0.422 | 0 | 60.4633 | Eukaryota | 44406.0 | 0 | Chordata | 44406.0 | 0 | Carnivora | 44406.0 | 0 | Canidae | 44406.0 | 0 | Canis | 32730.0 | 1 | Canis lupus | 32730.0 | 1 |
| scf7180000055472 | 19310 | 0.4003 | 0 | 651.2862 | Eukaryota | 73940.0 | 0 | Chordata | 73940.0 | 0 | Carnivora | 73940.0 | 0 | Canidae | 73940.0 | 0 | Canis | 73940.0 | 0 | Canis lupus | 73940.0 | 0 |
| scf7180000049743 | 15835 | 0.4375 | 0 | 134.8171 | Eukaryota | 10633.0 | 0 | Chordata | 10633.0 | 0 | Tetraodontiformes | 4975.0 | 3 | Tetraodontidae | 4975.0 | 3 | Takifugu | 4975.0 | 3 | Takifugu rubripes | 4975.0 | 3 |
| scf7180000056303 | 698002 | 0.3993 | 0 | 61.7377 | Eukaryota | 32068.0 | 1 | Chordata | 32068.0 | 1 | Chiroptera | 15816.0 | 3 | Pteropodidae | 15816.0 | 3 | Pteropus | 15816.0 | 4 | Pteropus vampyrus | 15816.0 | 4 |
| scf7180000054300 | 316699 | 0.4095 | 0 | 72.5992 | Eukaryota | 91001.0 | 0 | Chordata | 91001.0 | 0 | Carnivora | 91001.0 | 0 | Canidae | 91001.0 | 0 | Canis | 91001.0 | 0 | Canis lupus | 91001.0 | 0 |
| scf7180000047341 | 6822 | 0.4553 | 0 | 101.0815 | Eukaryota | 368.0 | 0 | Platyhelminthes | 368.0 | 0 | Polystomatidea | 368.0 | 0 | Polystomatidae | 368.0 | 0 | Protopolystoma | 368.0 | 0 | Protopolystoma xenopodis | 368.0 | 0 |
| scf7180000055038 | 9419 | 0.4362 | 0 | 116.8066 | no-hit | 0.0 | 0 | no-hit | 0.0 | 0 | no-hit | 0.0 | 0 | no-hit | 0.0 | 0 | no-hit | 0.0 | 0 | no-hit | 0.0 | 0 |
| scf7180000054022 | 4143 | 0.4502 | 0 | 2025.6555 | Eukaryota | 3660.0 | 0 | Platyhelminthes | 1562.0 | 3 | Polystomatidea | 1562.0 | 3 | Polystomatidae | 1562.0 | 3 | Protopolystoma | 1562.0 | 3 | Protopolystoma xenopodis | 1562.0 | 3 |
| scf7180000048906 | 9936 | 0.7088 | 0 | 35.9843 | Eukaryota | 46508.0 | 0 | Chordata | 46508.0 | 0 | Carnivora | 46508.0 | 0 | Canidae | 46508.0 | 0 | Canis | 46508.0 | 0 | Canis lupus | 46508.0 | 0 |
| scf7180000043585 | 494967 | 0.3889 | 0 | 60.2106 | Eukaryota | 94308.0 | 0 | Chordata | 94308.0 | 0 | Carnivora | 94308.0 | 0 | Canidae | 94308.0 | 0 | Canis | 94308.0 | 0 | Canis lupus | 94308.0 | 0 |
| scf7180000052851 | 3128 | 0.4479 | 0 | 247.282 | no-hit | 0.0 | 0 | no-hit | 0.0 | 0 | no-hit | 0.0 | 0 | no-hit | 0.0 | 0 | no-hit | 0.0 | 0 | no-hit | 0.0 | 0 |
| scf7180000055052 | 24253 | 0.4392 | 0 | 236.2541 | Eukaryota | 83650.0 | 0 | Platyhelminthes | 44080.0 | 1 | Strigeidida | 32143.0 | 4 | Schistosomatidae | 32143.0 | 4 | Schistosoma | 32143.0 | 4 | Schistosoma mansoni | 32143.0 | 4 |
| scf7180000055656 | 3297 | 0.438 | 0 | 157.4568 | no-hit | 0.0 | 0 | no-hit | 0.0 | 0 | no-hit | 0.0 | 0 | no-hit | 0.0 | 0 | no-hit | 0.0 | 0 | no-hit | 0.0 | 0 |
| scf7180000043604 | 27697 | 0.5236 | 0 | 54.6047 | Eukaryota | 8863.0 | 0 | Chordata | 8863.0 | 0 | Carnivora | 7037.0 | 2 | Canidae | 4051.0 | 5 | Canis | 4051.0 | 6 | Canis lupus | 4051.0 | 7 |
| scf7180000052817 | 10194 | 0.446 | 0 | 32.6102 | Eukaryota | 62718.0 | 0 | Chordata | 62718.0 | 0 | Carnivora | 62718.0 | 0 | Canidae | 62718.0 | 0 | Canis | 62718.0 | 0 | Canis lupus | 62718.0 | 0 |
| scf7180000051510 | 8429 | 0.447 | 0 | 337.3974 | Eukaryota | 2582.0 | 0 | Chordata | 2070.0 | 1 | Carnivora | 2070.0 | 1 | Canidae | 2070.0 | 1 | Canis | 2070.0 | 1 | Canis lupus | 2070.0 | 1 |
| scf7180000041059 | 1238146 | 0.379 | 0 | 66.3822 | Eukaryota | 154104.0 | 0 | Chordata | 154104.0 | 0 | Carnivora | 154104.0 | 0 | Canidae | 154104.0 | 0 | Canis | 154104.0 | 0 | Canis lupus | 154104.0 | 0 |
| scf7180000054416 | 7508 | 0.4389 | 0 | 291.3936 | Eukaryota | 864.0 | 0 | Platyhelminthes | 607.0 | 1 | Polystomatidea | 607.0 | 1 | Polystomatidae | 607.0 | 1 | Protopolystoma | 607.0 | 1 | Protopolystoma xenopodis | 607.0 | 1 |
| scf7180000045906 | 26587 | 0.4406 | 0 | 266.7658 | Eukaryota | 88879.0 | 0 | Chordata | 88879.0 | 0 | Carnivora | 88879.0 | 0 | Canidae | 88879.0 | 0 | Canis | 88879.0 | 0 | Canis lupus | 88879.0 | 0 |
| scf7180000056586 | 926494 | 0.4452 | 0 | 58.3775 | Eukaryota | 42775.0 | 0 | Chordata | 42775.0 | 0 | Carnivora | 42775.0 | 0 | Canidae | 32247.0 | 1 | Canis | 26730.0 | 2 | Canis lupus | 26730.0 | 2 |
| scf7180000043569 | 429926 | 0.5777 | 0 | 55.5153 | Eukaryota | 15050.0 | 1 | Chordata | 15050.0 | 1 | Carnivora | 12500.0 | 3 | Canidae | 12500.0 | 3 | Canis | 11103.0 | 5 | Canis lupus | 11103.0 | 5 |
| scf7180000048707 | 69045 | 0.4444 | 0 | 88.7933 | Eukaryota | 3586.0 | 0 | Chordata | 1658.0 | 2 | Polystomatidea | 1578.0 | 5 | Polystomatidae | 1578.0 | 5 | Protopolystoma | 1578.0 | 6 | Protopolystoma xenopodis | 1578.0 | 6 |
| scf7180000040689 | 192807 | 0.3661 | 0 | 61.3015 | Eukaryota | 11337.0 | 0 | Chordata | 11337.0 | 0 | Chiroptera | 3245.0 | 4 | Pteropodidae | 3245.0 | 4 | Homo | 2896.0 | 5 | Homo sapiens | 2896.0 | 6 |
| scf7180000043628 | 14751 | 0.4416 | 0 | 69.2927 | Eukaryota | 34243.0 | 0 | Chordata | 28028.0 | 2 | Gadiformes | 20324.0 | 6 | Gadidae | 20324.0 | 6 | Gadus | 20324.0 | 6 | Gadus morhua | 20324.0 | 6 |
| scf7180000044232 | 139315 | 0.3509 | 0 | 58.0646 | Eukaryota | 61135.0 | 0 | Chordata | 61135.0 | 0 | Carnivora | 61135.0 | 0 | Canidae | 61135.0 | 0 | Vulpes | 55818.0 | 1 | Vulpes vulpes | 55818.0 | 1 |
| scf7180000041803 | 876558 | 0.4165 | 0 | 60.7038 | Eukaryota | 57869.0 | 0 | Chordata | 57869.0 | 0 | Carnivora | 57869.0 | 0 | Canidae | 35250.0 | 2 | Canis | 35250.0 | 3 | Canis lupus | 35250.0 | 3 |
| scf7180000051960 | 9703 | 0.4356 | 0 | 41.4745 | Eukaryota | 87535.0 | 0 | Chordata | 87535.0 | 0 | Carnivora | 87535.0 | 0 | Canidae | 87535.0 | 0 | Canis | 87535.0 | 0 | Canis lupus | 87535.0 | 0 |
| scf7180000056071 | 323013 | 0.3973 | 0 | 63.0157 | Eukaryota | 21578.0 | 0 | Chordata | 21578.0 | 0 | Apterygiformes | 14806.0 | 2 | Apterygidae | 14806.0 | 2 | Apteryx | 14806.0 | 2 | Apteryx mantelli | 14806.0 | 2 |
| scf7180000047869 | 16398 | 0.4362 | 0 | 81.3836 | Eukaryota | 963.0 | 0 | Chordata | 963.0 | 0 | Primates | 356.0 | 2 | Hominidae | 356.0 | 2 | Homo | 356.0 | 2 | Homo sapiens | 356.0 | 2 |
| scf7180000050302 | 57087 | 0.5189 | 0 | 59.4444 | Eukaryota | 26067.0 | 0 | Chordata | 26067.0 | 0 | Carnivora | 17646.0 | 1 | Canidae | 11364.0 | 3 | Canis | 11364.0 | 3 | Canis lupus | 11364.0 | 3 |
| scf7180000047066 | 4454 | 0.4499 | 0 | 56.9044 | no-hit | 0.0 | 0 | no-hit | 0.0 | 0 | no-hit | 0.0 | 0 | no-hit | 0.0 | 0 | no-hit | 0.0 | 0 | no-hit | 0.0 | 0 |
| scf7180000047657 | 109135 | 0.4471 | 0 | 58.6221 | Eukaryota | 29619.0 | 0 | Chordata | 29619.0 | 0 | Carnivora | 15448.0 | 2 | Canidae | 13117.0 | 4 | Canis | 10457.0 | 6 | Canis lupus | 10457.0 | 6 |
| scf7180000051257 | 28976 | 0.4911 | 0 | 55.7822 | Eukaryota | 8575.0 | 0 | Chordata | 8575.0 | 0 | Carnivora | 8575.0 | 0 | Canidae | 8575.0 | 0 | Canis | 6909.0 | 1 | Canis lupus | 5724.0 | 2 |
| scf7180000048057 | 28864 | 0.4505 | 0 | 139.0747 | Eukaryota | 3576.0 | 0 | Chordata | 2522.0 | 1 | Carnivora | 1375.0 | 3 | Canidae | 1375.0 | 3 | Canis | 1375.0 | 3 | Canis lupus | 1375.0 | 3 |
| scf7180000040764 | 4284 | 0.4493 | 0 | 240.4465 | no-hit | 0.0 | 0 | no-hit | 0.0 | 0 | no-hit | 0.0 | 0 | no-hit | 0.0 | 0 | no-hit | 0.0 | 0 | no-hit | 0.0 | 0 |
| scf7180000041822 | 169308 | 0.3836 | 0 | 59.5053 | Eukaryota | 27337.0 | 0 | Chordata | 27337.0 | 0 | Carnivora | 27337.0 | 0 | Canidae | 13965.0 | 2 | Canis | 11689.0 | 3 | Canis lupus | 11689.0 | 3 |
| scf7180000043693 | 32095 | 0.3573 | 0 | 58.5947 | Eukaryota | 33833.0 | 0 | Chordata | 33833.0 | 0 | Carnivora | 33833.0 | 0 | Canidae | 33833.0 | 0 | Canis | 33833.0 | 0 | Canis lupus | 33833.0 | 0 |
| scf7180000048281 | 61239 | 0.5028 | 0 | 821.7163 | Eukaryota | 9644.0 | 0 | Chordata | 9644.0 | 0 | Carnivora | 9644.0 | 0 | Canidae | 9644.0 | 0 | Canis | 8413.0 | 1 | Canis lupus | 7121.0 | 2 |
| scf7180000045896 | 197862 | 0.3799 | 0 | 59.3954 | Eukaryota | 40675.0 | 0 | Chordata | 40675.0 | 0 | Carnivora | 40675.0 | 0 | Canidae | 40675.0 | 0 | Canis | 40675.0 | 0 | Canis lupus | 40675.0 | 0 |
| scf7180000042132 | 814156 | 0.3645 | 0 | 58.6725 | Eukaryota | 56162.0 | 0 | Chordata | 56162.0 | 0 | Carnivora | 56162.0 | 0 | Canidae | 56162.0 | 0 | Canis | 56162.0 | 0 | Canis lupus | 56162.0 | 0 |
| scf7180000056458 | 632835 | 0.3908 | 0 | 60.5286 | Eukaryota | 94987.0 | 0 | Chordata | 94987.0 | 0 | Carnivora | 94987.0 | 0 | Canidae | 94987.0 | 0 | Canis | 94987.0 | 0 | Canis lupus | 94987.0 | 0 |
| scf7180000043228 | 184524 | 0.5187 | 0 | 57.2335 | Eukaryota | 31153.0 | 0 | Chordata | 31153.0 | 0 | Carnivora | 31153.0 | 0 | Canidae | 31153.0 | 0 | Canis | 28602.0 | 1 | Canis lupus | 28602.0 | 1 |
| scf7180000047942 | 54956 | 0.4431 | 0 | 60.3289 | Eukaryota | 49494.0 | 0 | Chordata | 49494.0 | 0 | Carnivora | 49494.0 | 0 | Canidae | 49494.0 | 0 | Canis | 49494.0 | 0 | Canis lupus | 49494.0 | 0 |
| scf7180000046786 | 99282 | 0.3626 | 0 | 56.9949 | Eukaryota | 78591.0 | 0 | Chordata | 78591.0 | 0 | Carnivora | 78591.0 | 0 | Canidae | 78591.0 | 0 | Canis | 78591.0 | 0 | Canis lupus | 78591.0 | 0 |
| scf7180000049179 | 6827 | 0.4561 | 0 | 82.3306 | Eukaryota | 739.0 | 0 | Platyhelminthes | 437.0 | 1 | Polystomatidea | 437.0 | 1 | Polystomatidae | 437.0 | 1 | Protopolystoma | 437.0 | 1 | Protopolystoma xenopodis | 437.0 | 1 |
| scf7180000041901 | 997679 | 0.3437 | 0 | 59.5286 | Eukaryota | 90710.0 | 0 | Chordata | 90710.0 | 0 | Carnivora | 90710.0 | 0 | Canidae | 90710.0 | 0 | Canis | 90710.0 | 0 | Canis lupus | 90710.0 | 0 |
| scf7180000045303 | 205541 | 0.4013 | 0 | 55.3419 | Eukaryota | 85437.0 | 0 | Chordata | 85437.0 | 0 | Carnivora | 85437.0 | 0 | Canidae | 85437.0 | 0 | Canis | 85437.0 | 0 | Canis lupus | 85437.0 | 0 |
| scf7180000046851 | 117555 | 0.6458 | 0 | 54.6136 | Eukaryota | 20357.0 | 0 | Chordata | 20357.0 | 0 | Carnivora | 20357.0 | 0 | Canidae | 17635.0 | 1 | Vulpes | 11428.0 | 3 | Vulpes vulpes | 11428.0 | 3 |
| scf7180000043238 | 99360 | 0.3741 | 0 | 58.1484 | Eukaryota | 18849.0 | 0 | Chordata | 18849.0 | 0 | Carnivora | 18849.0 | 0 | Canidae | 18849.0 | 0 | Canis | 18849.0 | 0 | Canis lupus | 18849.0 | 0 |
| scf7180000056535 | 241411 | 0.4174 | 0 | 60.9323 | Eukaryota | 35890.0 | 0 | Chordata | 35890.0 | 0 | Carnivora | 35890.0 | 0 | Canidae | 35890.0 | 0 | Canis | 35890.0 | 0 | Canis lupus | 35890.0 | 0 |
| scf7180000050942 | 5273 | 0.4485 | 0 | 339.3 | Eukaryota | 481.0 | 0 | Streptophyta | 481.0 | 0 | Fabales | 481.0 | 0 | Fabaceae | 481.0 | 0 | Cicer | 481.0 | 0 | Cicer arietinum | 481.0 | 0 |
| scf7180000051850 | 19802 | 0.5103 | 0 | 1082.4238 | Eukaryota | 8831.0 | 0 | Chordata | 8831.0 | 0 | Carnivora | 8831.0 | 0 | Canidae | 8831.0 | 0 | Canis | 4936.0 | 1 | Canis lupus | 4936.0 | 1 |
| scf7180000045780 | 205604 | 0.437 | 0 | 59.7794 | Eukaryota | 16227.0 | 0 | Chordata | 16227.0 | 0 | Chiroptera | 8156.0 | 2 | Pteropodidae | 8156.0 | 3 | Pteropus | 8156.0 | 3 | Pteropus alecto | 5432.0 | 4 |
| scf7180000046128 | 31935 | 0.4408 | 0 | 210.46 | Eukaryota | 14335.0 | 0 | Chordata | 8675.0 | 3 | Gadiformes | 4639.0 | 7 | Gadidae | 4639.0 | 7 | Gadus | 4639.0 | 7 | Gadus morhua | 4639.0 | 7 |
| scf7180000052477 | 9011 | 0.4501 | 0 | 99.0798 | Eukaryota | 1081.0 | 0 | Chordata | 554.0 | 1 | Polystomatidea | 527.0 | 2 | Polystomatidae | 527.0 | 2 | Protopolystoma | 527.0 | 2 | Protopolystoma xenopodis | 527.0 | 2 |
| scf7180000054714 | 21318 | 0.4397 | 0 | 156.0173 | Eukaryota | 86876.0 | 0 | Chordata | 86876.0 | 0 | Carnivora | 86876.0 | 0 | Canidae | 86876.0 | 0 | Canis | 86876.0 | 0 | Canis lupus | 86876.0 | 0 |
| scf7180000049826 | 5591 | 0.4382 | 0 | 154.1714 | Eukaryota | 405.0 | 0 | Chordata | 405.0 | 0 | Tetraodontiformes | 405.0 | 0 | Tetraodontidae | 405.0 | 0 | Takifugu | 405.0 | 0 | Takifugu rubripes | 405.0 | 0 |
| scf7180000051844 | 6263 | 0.5103 | 0 | 91.6767 | Eukaryota | 6939.0 | 0 | Chordata | 6939.0 | 0 | Carnivora | 6939.0 | 0 | Canidae | 6939.0 | 0 | Canis | 4598.0 | 1 | Canis lupus | 3566.0 | 2 |
| scf7180000056161 | 3899 | 0.4468 | 0 | 131.3437 | no-hit | 0.0 | 0 | no-hit | 0.0 | 0 | no-hit | 0.0 | 0 | no-hit | 0.0 | 0 | no-hit | 0.0 | 0 | no-hit | 0.0 | 0 |
| scf7180000042024 | 70681 | 0.3969 | 0 | 60.3227 | Eukaryota | 54060.0 | 0 | Chordata | 54060.0 | 0 | Carnivora | 54060.0 | 0 | Canidae | 54060.0 | 0 | Canis | 54060.0 | 0 | Canis lupus | 54060.0 | 0 |
| scf7180000052233 | 5050 | 0.365 | 0 | 54.3108 | Eukaryota | 26473.0 | 0 | Chordata | 26473.0 | 0 | Carnivora | 26473.0 | 0 | Canidae | 26473.0 | 0 | Canis | 26473.0 | 0 | Canis lupus | 26473.0 | 0 |
| scf7180000053399 | 3863 | 0.4478 | 0 | 52.9799 | Eukaryota | 145.0 | 0 | Platyhelminthes | 145.0 | 0 | Polystomatidea | 145.0 | 0 | Polystomatidae | 145.0 | 0 | Protopolystoma | 145.0 | 0 | Protopolystoma xenopodis | 145.0 | 0 |
| scf7180000050975 | 25696 | 0.502 | 0 | 394.4638 | Eukaryota | 9210.0 | 0 | Chordata | 9210.0 | 0 | Carnivora | 9210.0 | 0 | Canidae | 9210.0 | 0 | Canis | 7988.0 | 1 | Canis lupus | 6703.0 | 2 |
| scf7180000042243 | 786658 | 0.4207 | 0 | 58.6011 | Eukaryota | 58554.0 | 0 | Chordata | 58554.0 | 0 | Carnivora | 58554.0 | 0 | Canidae | 46501.0 | 1 | Canis | 37684.0 | 5 | Canis lupus | 37684.0 | 5 |
| scf7180000053917 | 4504 | 0.4489 | 0 | 322.5571 | no-hit | 0.0 | 0 | no-hit | 0.0 | 0 | no-hit | 0.0 | 0 | no-hit | 0.0 | 0 | no-hit | 0.0 | 0 | no-hit | 0.0 | 0 |
| scf7180000049113 | 10205 | 0.4425 | 0 | 409.668 | Eukaryota | 1699.0 | 0 | Chordata | 1020.0 | 1 | Gonorynchiformes | 1020.0 | 1 | Chanidae | 1020.0 | 1 | Chanos | 1020.0 | 1 | Chanos chanos | 1020.0 | 1 |
| scf7180000053409 | 6708 | 0.5061 | 0 | 377.4868 | Eukaryota | 8325.0 | 0 | Chordata | 8325.0 | 0 | Carnivora | 8325.0 | 0 | Canidae | 8325.0 | 0 | Canis | 6186.0 | 1 | Canis lupus | 5007.0 | 2 |
| scf7180000051231 | 27490 | 0.3821 | 0 | 55.4324 | Eukaryota | 11998.0 | 0 | Chordata | 11998.0 | 0 | Carnivora | 11998.0 | 0 | Canidae | 11998.0 | 0 | Canis | 11998.0 | 0 | Canis lupus | 11998.0 | 0 |
| scf7180000052093 | 13280 | 0.6889 | 0 | 57.344 | Eukaryota | 55212.0 | 0 | Chordata | 55212.0 | 0 | Carnivora | 55212.0 | 0 | Canidae | 34127.0 | 2 | Canis | 27320.0 | 4 | Canis lupus | 27320.0 | 4 |
| scf7180000045372 | 185668 | 0.4489 | 0 | 60.3475 | Eukaryota | 38535.0 | 0 | Chordata | 38535.0 | 0 | Carnivora | 38535.0 | 0 | Canidae | 38535.0 | 0 | Canis | 38535.0 | 0 | Canis lupus | 38535.0 | 0 |
| scf7180000050148 | 21248 | 0.4557 | 0 | 54.4978 | Eukaryota | 39260.0 | 0 | Chordata | 39260.0 | 0 | Carnivora | 39260.0 | 0 | Canidae | 39260.0 | 0 | Canis | 39260.0 | 0 | Canis lupus | 39260.0 | 0 |
| scf7180000049040 | 47630 | 0.4463 | 0 | 71.3783 | Eukaryota | 9745.0 | 0 | Chordata | 9745.0 | 0 | Carnivora | 9745.0 | 0 | Canidae | 9745.0 | 0 | Canis | 6812.0 | 1 | Canis lupus | 6812.0 | 1 |
| scf7180000043339 | 25840 | 0.3835 | 0 | 42.8292 | Eukaryota | 83631.0 | 0 | Chordata | 83631.0 | 0 | Carnivora | 83631.0 | 0 | Canidae | 83631.0 | 0 | Canis | 83631.0 | 0 | Canis lupus | 83631.0 | 0 |
| scf7180000049547 | 18675 | 0.5017 | 0 | 310.7916 | Eukaryota | 6774.0 | 0 | Chordata | 6774.0 | 0 | Carnivora | 6774.0 | 0 | Canidae | 6774.0 | 0 | Canis | 5572.0 | 2 | Canis lupus | 5572.0 | 2 |
| scf7180000048956 | 21746 | 0.444 | 0 | 115.0794 | Eukaryota | 375.0 | 0 | Chordata | 375.0 | 0 | Salmoniformes | 375.0 | 0 | Salmonidae | 375.0 | 0 | Salmo | 375.0 | 0 | Salmo trutta | 375.0 | 0 |
| scf7180000042751 | 604854 | 0.4129 | 0 | 60.6205 | Eukaryota | 103559.0 | 0 | Chordata | 103559.0 | 0 | Carnivora | 103559.0 | 0 | Canidae | 103559.0 | 0 | Canis | 73929.0 | 1 | Canis lupus | 73929.0 | 1 |
| scf7180000040670 | 1909396 | 0.3781 | 0 | 58.4427 | Eukaryota | 91102.0 | 0 | Chordata | 91102.0 | 0 | Carnivora | 91102.0 | 0 | Canidae | 91102.0 | 0 | Canis | 91102.0 | 0 | Canis lupus | 91102.0 | 0 |
| scf7180000041908 | 11372 | 0.4861 | 0 | 93.9091 | Eukaryota | 32270.0 | 0 | Chordata | 32270.0 | 0 | Carnivora | 32270.0 | 0 | Canidae | 32270.0 | 0 | Canis | 32270.0 | 0 | Canis lupus | 32270.0 | 0 |
| scf7180000044796 | 602492 | 0.4342 | 0 | 59.8838 | Eukaryota | 109525.0 | 0 | Chordata | 109525.0 | 0 | Carnivora | 109525.0 | 0 | Canidae | 109525.0 | 0 | Canis | 98227.0 | 1 | Canis lupus | 98227.0 | 1 |
| scf7180000049604 | 12112 | 0.4596 | 0 | 141.442 | Eukaryota | 4938.0 | 0 | Chordata | 3253.0 | 1 | Salmoniformes | 2137.0 | 4 | Salmonidae | 2137.0 | 4 | Salmo | 2137.0 | 4 | Salmo trutta | 2137.0 | 4 |
| scf7180000055956 | 16603 | 0.443 | 0 | 114.248 | Eukaryota | 14826.0 | 0 | Chordata | 9273.0 | 3 | Gadiformes | 4359.0 | 7 | Gadidae | 4359.0 | 7 | Gadus | 4359.0 | 7 | Gadus morhua | 4359.0 | 7 |
| scf7180000052205 | 4663 | 0.4504 | 0 | 755.0117 | no-hit | 0.0 | 0 | no-hit | 0.0 | 0 | no-hit | 0.0 | 0 | no-hit | 0.0 | 0 | no-hit | 0.0 | 0 | no-hit | 0.0 | 0 |
| scf7180000050657 | 4394 | 0.447 | 0 | 266.229 | no-hit | 0.0 | 0 | no-hit | 0.0 | 0 | no-hit | 0.0 | 0 | no-hit | 0.0 | 0 | no-hit | 0.0 | 0 | no-hit | 0.0 | 0 |
| scf7180000043295 | 473193 | 0.3671 | 0 | 59.9676 | Eukaryota | 93038.0 | 0 | Chordata | 93038.0 | 0 | Carnivora | 93038.0 | 0 | Canidae | 93038.0 | 0 | Canis | 93038.0 | 0 | Canis lupus | 93038.0 | 0 |
| scf7180000056619 | 94701 | 0.3803 | 0 | 60.7261 | Eukaryota | 15211.0 | 0 | Chordata | 15211.0 | 0 | Carnivora | 15211.0 | 0 | Canidae | 12938.0 | 1 | Canis | 10868.0 | 2 | Canis lupus | 10868.0 | 2 |
| scf7180000041198 | 278230 | 0.4039 | 0 | 58.7639 | Eukaryota | 121500.0 | 0 | Chordata | 121500.0 | 0 | Carnivora | 121500.0 | 0 | Canidae | 121500.0 | 0 | Canis | 84945.0 | 1 | Canis lupus | 84945.0 | 1 |
| scf7180000047919 | 16307 | 0.4393 | 0 | 113.3138 | Eukaryota | 4372.0 | 0 | Chordata | 4026.0 | 1 | Tetraodontiformes | 2527.0 | 4 | Tetraodontidae | 2527.0 | 4 | Takifugu | 2527.0 | 4 | Takifugu rubripes | 2527.0 | 4 |
| scf7180000055416 | 4759 | 0.4524 | 0 | 661.6722 | no-hit | 0.0 | 0 | no-hit | 0.0 | 0 | no-hit | 0.0 | 0 | no-hit | 0.0 | 0 | no-hit | 0.0 | 0 | no-hit | 0.0 | 0 |
| scf7180000053926 | 11328 | 0.5031 | 0 | 1128.3924 | Eukaryota | 9183.0 | 0 | Chordata | 9183.0 | 0 | Carnivora | 9183.0 | 0 | Canidae | 9183.0 | 0 | Canis | 7400.0 | 1 | Canis lupus | 6093.0 | 2 |
| scf7180000045235 | 36137 | 0.5013 | 0 | 284.7226 | Eukaryota | 9076.0 | 0 | Chordata | 9076.0 | 0 | Carnivora | 9076.0 | 0 | Canidae | 9076.0 | 0 | Canis | 7389.0 | 1 | Canis lupus | 6154.0 | 2 |
| scf7180000046145 | 136671 | 0.62 | 0 | 55.3923 | Eukaryota | 52503.0 | 0 | Chordata | 52503.0 | 0 | Carnivora | 52503.0 | 0 | Canidae | 52503.0 | 0 | Canis | 43982.0 | 1 | Canis lupus | 43982.0 | 1 |
| scf7180000047851 | 4455 | 0.4386 | 0 | 69.6251 | Eukaryota | 2933.0 | 0 | Chordata | 2933.0 | 0 | Carnivora | 2933.0 | 0 | Canidae | 2933.0 | 0 | Canis | 2933.0 | 0 | Canis lupus | 2933.0 | 0 |
| scf7180000041441 | 53711 | 0.4407 | 0 | 69.1786 | Eukaryota | 25954.0 | 0 | Chordata | 25954.0 | 0 | Carnivora | 25954.0 | 0 | Canidae | 25954.0 | 0 | Canis | 25954.0 | 0 | Canis lupus | 25954.0 | 0 |
| scf7180000049094 | 17925 | 0.4327 | 0 | 220.4776 | Eukaryota | 3621.0 | 0 | Platyhelminthes | 2632.0 | 2 | Polystomatidea | 2632.0 | 2 | Polystomatidae | 2632.0 | 2 | Protopolystoma | 2632.0 | 2 | Protopolystoma xenopodis | 2632.0 | 2 |
| scf7180000050996 | 14961 | 0.4437 | 0 | 394.51 | Eukaryota | 3890.0 | 0 | Platyhelminthes | 2446.0 | 2 | Polystomatidea | 2446.0 | 2 | Polystomatidae | 2446.0 | 2 | Protopolystoma | 2446.0 | 2 | Protopolystoma xenopodis | 2446.0 | 2 |
| scf7180000056026 | 9187 | 0.446 | 0 | 1035.3629 | Eukaryota | 880.0 | 0 | Platyhelminthes | 511.0 | 1 | Polystomatidea | 511.0 | 2 | Polystomatidae | 511.0 | 2 | Protopolystoma | 511.0 | 2 | Protopolystoma xenopodis | 511.0 | 2 |
| scf7180000043772 | 60227 | 0.3542 | 0 | 52.0419 | Eukaryota | 74559.0 | 0 | Chordata | 74559.0 | 0 | Carnivora | 74559.0 | 0 | Canidae | 74559.0 | 0 | Canis | 74559.0 | 0 | Canis lupus | 74559.0 | 0 |
| scf7180000042671 | 6658 | 0.4336 | 0 | 113.9675 | no-hit | 0.0 | 0 | no-hit | 0.0 | 0 | no-hit | 0.0 | 0 | no-hit | 0.0 | 0 | no-hit | 0.0 | 0 | no-hit | 0.0 | 0 |
| scf7180000049311 | 11121 | 0.4504 | 0 | 1079.4449 | Eukaryota | 3320.0 | 0 | Platyhelminthes | 1782.0 | 2 | Polystomatidea | 1782.0 | 2 | Polystomatidae | 1782.0 | 2 | Protopolystoma | 1782.0 | 2 | Protopolystoma xenopodis | 1782.0 | 2 |
| scf7180000047816 | 34303 | 0.4606 | 0 | 68.4966 | Eukaryota | 12172.0 | 0 | Chordata | 12172.0 | 0 | Carnivora | 8409.0 | 1 | Canidae | 8409.0 | 1 | Canis | 7716.0 | 2 | Canis lupus | 6397.0 | 3 |
| scf7180000042046 | 20529 | 0.6194 | 0 | 154.1293 | Eukaryota | 18287.0 | 0 | Chordata | 18287.0 | 0 | Carnivora | 18287.0 | 0 | Canidae | 18287.0 | 0 | Canis | 16683.0 | 1 | Canis lupus | 16683.0 | 1 |
| scf7180000041185 | 466010 | 0.4396 | 0 | 58.9532 | Eukaryota | 110296.0 | 0 | Chordata | 110296.0 | 0 | Carnivora | 110296.0 | 0 | Canidae | 110296.0 | 0 | Canis | 110296.0 | 0 | Canis lupus | 110296.0 | 0 |
| scf7180000055582 | 38145 | 0.4386 | 0 | 171.8517 | Eukaryota | 962.0 | 0 | Platyhelminthes | 823.0 | 1 | Polystomatidea | 823.0 | 1 | Polystomatidae | 823.0 | 1 | Protopolystoma | 823.0 | 1 | Protopolystoma xenopodis | 823.0 | 1 |
| scf7180000055257 | 6257 | 0.4454 | 0 | 102.8271 | no-hit | 0.0 | 0 | no-hit | 0.0 | 0 | no-hit | 0.0 | 0 | no-hit | 0.0 | 0 | no-hit | 0.0 | 0 | no-hit | 0.0 | 0 |
| scf7180000044950 | 2041 | 0.4517 | 0 | 230.6391 | no-hit | 0.0 | 0 | no-hit | 0.0 | 0 | no-hit | 0.0 | 0 | no-hit | 0.0 | 0 | no-hit | 0.0 | 0 | no-hit | 0.0 | 0 |
| scf7180000051752 | 2504 | 0.4493 | 0 | 1154.8733 | Eukaryota | 1668.0 | 0 | Echinodermata | 837.0 | 2 | Forcipulatida | 837.0 | 2 | Asteriidae | 837.0 | 2 | Asterias | 837.0 | 2 | Asterias rubens | 837.0 | 2 |
| scf7180000042372 | 13109 | 0.4922 | 0 | 1141.4536 | Eukaryota | 34184.0 | 0 | Chordata | 34184.0 | 0 | Carnivora | 34184.0 | 0 | Canidae | 34184.0 | 0 | Canis | 34184.0 | 0 | Canis lupus | 34184.0 | 0 |
| scf7180000042708 | 614070 | 0.4111 | 0 | 60.2469 | Eukaryota | 90944.0 | 0 | Chordata | 90944.0 | 0 | Carnivora | 90944.0 | 0 | Canidae | 90944.0 | 0 | Canis | 90944.0 | 0 | Canis lupus | 90944.0 | 0 |
| scf7180000046820 | 104838 | 0.4704 | 0 | 58.58 | Eukaryota | 15447.0 | 0 | Chordata | 15447.0 | 0 | Carnivora | 13880.0 | 1 | Canidae | 9010.0 | 3 | Canis | 7087.0 | 5 | Canis lupus | 7087.0 | 5 |
| scf7180000047696 | 20564 | 0.5262 | 0 | 436.7104 | Eukaryota | 4266.0 | 0 | Chordata | 4266.0 | 0 | Carnivora | 4266.0 | 0 | Canidae | 4266.0 | 0 | Canis | 3619.0 | 1 | Canis lupus | 3619.0 | 1 |
| scf7180000048045 | 54493 | 0.397 | 0 | 59.6334 | Eukaryota | 58068.0 | 0 | Chordata | 58068.0 | 0 | Carnivora | 58068.0 | 0 | Canidae | 58068.0 | 0 | Canis | 58068.0 | 0 | Canis lupus | 58068.0 | 0 |
| scf7180000054059 | 5113 | 0.4358 | 0 | 134.664 | no-hit | 0.0 | 0 | no-hit | 0.0 | 0 | no-hit | 0.0 | 0 | no-hit | 0.0 | 0 | no-hit | 0.0 | 0 | no-hit | 0.0 | 0 |
| scf7180000044383 | 290469 | 0.3947 | 0 | 61.0977 | Eukaryota | 85780.0 | 0 | Chordata | 85780.0 | 0 | Carnivora | 85780.0 | 0 | Canidae | 85780.0 | 0 | Canis | 85780.0 | 0 | Canis lupus | 85780.0 | 0 |
| scf7180000047327 | 3250 | 0.42 | 0 | 21.6581 | Eukaryota | 40189.0 | 0 | Chordata | 40189.0 | 0 | Carnivora | 40189.0 | 0 | Canidae | 40189.0 | 0 | Canis | 40189.0 | 0 | Canis lupus | 40189.0 | 0 |
| scf7180000045051 | 239904 | 0.4134 | 0 | 60.6887 | Eukaryota | 56286.0 | 0 | Chordata | 56286.0 | 0 | Carnivora | 56286.0 | 0 | Canidae | 56286.0 | 0 | Vulpes | 45080.0 | 1 | Vulpes vulpes | 45080.0 | 1 |
| scf7180000046168 | 142184 | 0.4899 | 0 | 58.4175 | Eukaryota | 17088.0 | 0 | Chordata | 17088.0 | 0 | Carnivora | 17088.0 | 0 | Ursidae | 9672.0 | 2 | Ursus | 9672.0 | 2 | Ursus arctos | 5540.0 | 3 |
| scf7180000043806 | 13119 | 0.3983 | 0 | 39.7398 | Eukaryota | 79833.0 | 0 | Chordata | 79833.0 | 0 | Carnivora | 79833.0 | 0 | Canidae | 79833.0 | 0 | Canis | 79833.0 | 0 | Canis lupus | 79833.0 | 0 |
| scf7180000054928 | 98992 | 0.3378 | 0 | 58.0675 | Eukaryota | 52001.0 | 0 | Chordata | 52001.0 | 0 | Carnivora | 52001.0 | 0 | Canidae | 52001.0 | 0 | Canis | 52001.0 | 0 | Canis lupus | 52001.0 | 0 |
| scf7180000047986 | 65595 | 0.3702 | 0 | 61.2174 | Eukaryota | 12358.0 | 1 | Chordata | 12358.0 | 1 | Carnivora | 5505.0 | 4 | Felidae | 5505.0 | 4 | Felis | 5505.0 | 4 | Felis catus | 5505.0 | 4 |
| scf7180000041331 | 55732 | 0.3976 | 0 | 57.5419 | Eukaryota | 9565.0 | 0 | Chordata | 9565.0 | 0 | Carnivora | 7715.0 | 1 | Canidae | 7715.0 | 2 | Canis | 7715.0 | 2 | Canis lupus | 7715.0 | 3 |
| scf7180000053436 | 5068 | 0.5099 | 0 | 127.131 | Eukaryota | 6154.0 | 0 | Chordata | 6154.0 | 0 | Carnivora | 6154.0 | 0 | Canidae | 6154.0 | 0 | Canis | 4153.0 | 1 | Canis lupus | 3251.0 | 2 |
| scf7180000045508 | 155629 | 0.4775 | 0 | 57.7071 | Eukaryota | 37835.0 | 0 | Chordata | 37835.0 | 0 | Carnivora | 37835.0 | 0 | Canidae | 37835.0 | 0 | Canis | 33487.0 | 1 | Canis lupus | 33487.0 | 1 |
| scf7180000050431 | 10232 | 0.4348 | 0 | 48.384 | no-hit | 0.0 | 0 | no-hit | 0.0 | 0 | no-hit | 0.0 | 0 | no-hit | 0.0 | 0 | no-hit | 0.0 | 0 | no-hit | 0.0 | 0 |
| scf7180000047567 | 13708 | 0.4489 | 0 | 338.7694 | Eukaryota | 3149.0 | 0 | Chordata | 2232.0 | 1 | Carnivora | 2232.0 | 1 | Canidae | 2232.0 | 1 | Canis | 2232.0 | 1 | Canis lupus | 2232.0 | 1 |
| scf7180000040927 | 1629251 | 0.3624 | 0 | 60.626 | Eukaryota | 89181.0 | 0 | Chordata | 89181.0 | 0 | Carnivora | 89181.0 | 0 | Canidae | 89181.0 | 0 | Canis | 89181.0 | 0 | Canis lupus | 89181.0 | 0 |
| scf7180000042035 | 230076 | 0.3771 | 0 | 57.8845 | Eukaryota | 13157.0 | 0 | Chordata | 13157.0 | 0 | Carnivora | 13157.0 | 0 | Canidae | 13157.0 | 0 | Canis | 11890.0 | 1 | Canis lupus | 11890.0 | 1 |
| scf7180000043756 | 110670 | 0.3956 | 0 | 58.7871 | Eukaryota | 91369.0 | 0 | Chordata | 91369.0 | 0 | Carnivora | 91369.0 | 0 | Canidae | 91369.0 | 0 | Canis | 91369.0 | 0 | Canis lupus | 91369.0 | 0 |
| scf7180000049758 | 35949 | 0.3614 | 0 | 67.3763 | Eukaryota | 65493.0 | 0 | Chordata | 65493.0 | 0 | Carnivora | 65493.0 | 0 | Canidae | 65493.0 | 0 | Canis | 65493.0 | 0 | Canis lupus | 65493.0 | 0 |
| scf7180000055917 | 23218 | 0.4462 | 0 | 99.176 | Eukaryota | 498.0 | 0 | Platyhelminthes | 498.0 | 0 | Polystomatidea | 498.0 | 0 | Polystomatidae | 498.0 | 0 | Protopolystoma | 498.0 | 0 | Protopolystoma xenopodis | 498.0 | 0 |
| scf7180000055261 | 11503 | 0.3716 | 0 | 226.8622 | Eukaryota | 43118.0 | 0 | Chordata | 43118.0 | 0 | Carnivora | 43118.0 | 0 | Canidae | 43118.0 | 0 | Canis | 43118.0 | 0 | Canis lupus | 43118.0 | 0 |
| scf7180000045397 | 191875 | 0.4019 | 0 | 59.6647 | Eukaryota | 77110.0 | 0 | Chordata | 77110.0 | 0 | Carnivora | 77110.0 | 0 | Canidae | 77110.0 | 0 | Canis | 77110.0 | 0 | Canis lupus | 77110.0 | 0 |
| scf7180000043407 | 113789 | 0.3663 | 0 | 58.4632 | Eukaryota | 86473.0 | 0 | Chordata | 86473.0 | 0 | Carnivora | 86473.0 | 0 | Canidae | 86473.0 | 0 | Canis | 86473.0 | 0 | Canis lupus | 86473.0 | 0 |
| scf7180000041558 | 14863 | 0.4555 | 0 | 31.4177 | Eukaryota | 4243.0 | 0 | Chordata | 4243.0 | 0 | Carnivora | 4243.0 | 0 | Canidae | 4243.0 | 0 | Canis | 4243.0 | 0 | Canis lupus | 4243.0 | 0 |
| scf7180000045729 | 136167 | 0.3808 | 0 | 62.9672 | Eukaryota | 51002.0 | 0 | Chordata | 51002.0 | 0 | Carnivora | 51002.0 | 0 | Canidae | 51002.0 | 0 | Canis | 45836.0 | 1 | Canis lupus | 45836.0 | 1 |
| scf7180000049328 | 14010 | 0.4454 | 0 | 453.1898 | Eukaryota | 2944.0 | 1 | Platyhelminthes | 1101.0 | 4 | Polystomatidea | 1101.0 | 4 | Polystomatidae | 1101.0 | 4 | Protopolystoma | 1101.0 | 4 | Protopolystoma xenopodis | 1101.0 | 4 |
| scf7180000049832 | 3305 | 0.443 | 0 | 80.8211 | no-hit | 0.0 | 0 | no-hit | 0.0 | 0 | no-hit | 0.0 | 0 | no-hit | 0.0 | 0 | no-hit | 0.0 | 0 | no-hit | 0.0 | 0 |
| scf7180000053154 | 5081 | 0.445 | 0 | 268.6427 | Eukaryota | 556.0 | 0 | Platyhelminthes | 556.0 | 0 | Polystomatidea | 556.0 | 0 | Polystomatidae | 556.0 | 0 | Protopolystoma | 556.0 | 0 | Protopolystoma xenopodis | 556.0 | 0 |
| scf7180000051056 | 7519 | 0.3709 | 0 | 106.1083 | Eukaryota | 5315.0 | 0 | Chordata | 5315.0 | 0 | Carnivora | 5315.0 | 0 | Felidae | 3084.0 | 1 | Felis | 3084.0 | 1 | Felis catus | 3084.0 | 1 |
| scf7180000044470 | 90137 | 0.4188 | 0 | 58.8758 | Eukaryota | 88201.0 | 0 | Chordata | 88201.0 | 0 | Carnivora | 88201.0 | 0 | Canidae | 88201.0 | 0 | Canis | 88201.0 | 0 | Canis lupus | 88201.0 | 0 |
| scf7180000042971 | 50194 | 0.4864 | 0 | 108.5218 | Eukaryota | 45397.0 | 0 | Chordata | 45397.0 | 0 | Carnivora | 45397.0 | 0 | Canidae | 45397.0 | 0 | Canis | 45397.0 | 0 | Canis lupus | 45397.0 | 0 |
| scf7180000041053 | 1368351 | 0.3672 | 0 | 59.7551 | Eukaryota | 88684.0 | 0 | Chordata | 88684.0 | 0 | Carnivora | 88684.0 | 0 | Canidae | 88684.0 | 0 | Canis | 88684.0 | 0 | Canis lupus | 88684.0 | 0 |
| scf7180000043543 | 29968 | 0.5539 | 0 | 53.7012 | Eukaryota | 4160.0 | 0 | Chordata | 4160.0 | 0 | Carnivora | 3236.0 | 2 | Canidae | 2245.0 | 4 | Canis | 1641.0 | 6 | Canis lupus | 1641.0 | 6 |
| scf7180000042147 | 774224 | 0.408 | 0 | 58.6055 | Eukaryota | 99190.0 | 0 | Chordata | 99190.0 | 0 | Carnivora | 99190.0 | 0 | Canidae | 99190.0 | 0 | Canis | 99190.0 | 0 | Canis lupus | 99190.0 | 0 |
| scf7180000042646 | 249332 | 0.3971 | 0 | 60.5036 | Eukaryota | 93745.0 | 0 | Chordata | 93745.0 | 0 | Carnivora | 93745.0 | 0 | Canidae | 93745.0 | 0 | Canis | 93745.0 | 0 | Canis lupus | 93745.0 | 0 |
| scf7180000040642 | 2562573 | 0.4974 | 0 | 59.0193 | Eukaryota | 188052.0 | 0 | Chordata | 188052.0 | 0 | Carnivora | 188052.0 | 0 | Canidae | 188052.0 | 0 | Canis | 188052.0 | 0 | Canis lupus | 188052.0 | 0 |
| scf7180000042489 | 11687 | 0.3962 | 0 | 29.8136 | Eukaryota | 3548.0 | 0 | Chordata | 3548.0 | 0 | Carnivora | 3548.0 | 0 | Canidae | 1328.0 | 4 | Canis | 892.0 | 7 | Canis lupus | 892.0 | 8 |
| scf7180000051759 | 7047 | 0.453 | 0 | 552.8756 | Eukaryota | 3502.0 | 0 | Platyhelminthes | 2002.0 | 2 | Polystomatidea | 2002.0 | 2 | Polystomatidae | 2002.0 | 2 | Protopolystoma | 2002.0 | 2 | Protopolystoma xenopodis | 2002.0 | 2 |
| scf7180000042043 | 906575 | 0.34 | 0 | 59.0237 | Eukaryota | 68605.0 | 0 | Chordata | 68605.0 | 0 | Carnivora | 68605.0 | 0 | Canidae | 68605.0 | 0 | Canis | 68605.0 | 0 | Canis lupus | 68605.0 | 0 |
| scf7180000044693 | 1589704 | 0.4159 | 0 | 60.6328 | Eukaryota | 91690.0 | 0 | Chordata | 91690.0 | 0 | Carnivora | 91690.0 | 0 | Canidae | 91690.0 | 0 | Canis | 91690.0 | 0 | Canis lupus | 91690.0 | 0 |
| scf7180000051704 | 3915 | 0.4511 | 0 | 410.2006 | no-hit | 0.0 | 0 | no-hit | 0.0 | 0 | no-hit | 0.0 | 0 | no-hit | 0.0 | 0 | no-hit | 0.0 | 0 | no-hit | 0.0 | 0 |
| scf7180000042233 | 298045 | 0.4761 | 0 | 56.8944 | Eukaryota | 41869.0 | 0 | Chordata | 41869.0 | 0 | Carnivora | 35061.0 | 2 | Canidae | 35061.0 | 2 | Canis | 19231.0 | 3 | Canis lupus | 19231.0 | 3 |
| scf7180000042719 | 590151 | 0.3632 | 0 | 61.1076 | Eukaryota | 48180.0 | 0 | Chordata | 48180.0 | 0 | Carnivora | 29315.0 | 1 | Canidae | 29315.0 | 1 | Canis | 29315.0 | 1 | Canis lupus | 29315.0 | 1 |
| scf7180000043034 | 50139 | 0.4115 | 0 | 58.5346 | Eukaryota | 19402.0 | 0 | Chordata | 19402.0 | 0 | Carnivora | 19402.0 | 0 | Canidae | 19402.0 | 0 | Canis | 19402.0 | 0 | Canis lupus | 19402.0 | 0 |
| scf7180000052548 | 6178 | 0.4518 | 0 | 514.476 | no-hit | 0.0 | 0 | no-hit | 0.0 | 0 | no-hit | 0.0 | 0 | no-hit | 0.0 | 0 | no-hit | 0.0 | 0 | no-hit | 0.0 | 0 |
| scf7180000048468 | 7415 | 0.4415 | 0 | 85.5052 | Eukaryota | 34122.0 | 0 | Chordata | 23824.0 | 2 | Gadiformes | 17075.0 | 5 | Gadidae | 17075.0 | 5 | Gadus | 17075.0 | 5 | Gadus morhua | 17075.0 | 5 |
| scf7180000054596 | 16079 | 0.4445 | 0 | 455.9763 | no-hit | 0.0 | 0 | no-hit | 0.0 | 0 | no-hit | 0.0 | 0 | no-hit | 0.0 | 0 | no-hit | 0.0 | 0 | no-hit | 0.0 | 0 |
| scf7180000052722 | 2787 | 0.4442 | 0 | 209.9857 | no-hit | 0.0 | 0 | no-hit | 0.0 | 0 | no-hit | 0.0 | 0 | no-hit | 0.0 | 0 | no-hit | 0.0 | 0 | no-hit | 0.0 | 0 |
| scf7180000048727 | 58329 | 0.3711 | 0 | 59.8271 | Eukaryota | 30492.0 | 0 | Chordata | 30492.0 | 0 | Carnivora | 30492.0 | 0 | Canidae | 30492.0 | 0 | Canis | 30492.0 | 0 | Canis lupus | 30492.0 | 0 |
| scf7180000050963 | 8965 | 0.4495 | 0 | 924.5649 | no-hit | 0.0 | 0 | no-hit | 0.0 | 0 | no-hit | 0.0 | 0 | no-hit | 0.0 | 0 | no-hit | 0.0 | 0 | no-hit | 0.0 | 0 |
| scf7180000047437 | 114811 | 0.3758 | 0 | 59.8965 | Eukaryota | 12367.0 | 0 | Chordata | 12367.0 | 0 | Carnivora | 10146.0 | 1 | Felidae | 7870.0 | 2 | Felis | 2646.0 | 5 | Felis catus | 2646.0 | 5 |
| scf7180000042980 | 168554 | 0.3509 | 0 | 57.1756 | Eukaryota | 62420.0 | 0 | Chordata | 62420.0 | 0 | Carnivora | 62420.0 | 0 | Canidae | 62420.0 | 0 | Canis | 62420.0 | 0 | Canis lupus | 62420.0 | 0 |
| scf7180000043552 | 445336 | 0.3958 | 0 | 60.1643 | Eukaryota | 73982.0 | 0 | Chordata | 73982.0 | 0 | Carnivora | 73982.0 | 0 | Canidae | 73982.0 | 0 | Canis | 73982.0 | 0 | Canis lupus | 73982.0 | 0 |

| | | | | | | | | | | | | | | | | | | | | | | |
|---|---|---|---|---|---|---|---|---|---|---|---|---|---|---|---|---|---|---|---|---|---|---|
| scf7180000049854 | 7820 | 0.4267 | 0 | 65.2542 | Eukaryota | 1844.0 | 0 | Chordata | 1844.0 | 0 | Salmoniformes | 1844.0 | 0 | Salmonidae | 1844.0 | 0 | Salmo | 1844.0 | 0 | Salmo trutta | 1844.0 | 0 |
| scf7180000041837 | 343840 | 0.34 | 0 | 57.9762 | Eukaryota | 71437.0 | 0 | Chordata | 71437.0 | 0 | Carnivora | 71437.0 | 0 | Canidae | 71437.0 | 0 | Canis | 71437.0 | 0 | Canis lupus | 71437.0 | 0 |
| scf7180000053192 | 6422 | 0.4472 | 0 | 93.7425 | Eukaryota | 12235.0 | 0 | Chordata | 6871.0 | 2 | Strigeidida | 3890.0 | 6 | Schistosomatidae | 3890.0 | 6 | Schistosoma | 3890.0 | 6 | Schistosoma mansoni | 3890.0 | 6 |
| scf7180000040954 | 697813 | 0.3851 | 0 | 61.0119 | Eukaryota | 74352.0 | 0 | Chordata | 74352.0 | 0 | Carnivora | 74352.0 | 0 | Canidae | 74352.0 | 0 | Canis | 65753.0 | 1 | Canis lupus | 65753.0 | 1 |
| scf7180000052182 | 8730 | 0.4507 | 0 | 296.9108 | Eukaryota | 3024.0 | 0 | Platyhelminthes | 1480.0 | 3 | Polystomatidea | 1480.0 | 3 | Polystomatidae | 1480.0 | 3 | Protopolystoma | 1480.0 | 3 | Protopolystoma xenopodis | 1480.0 | 3 |
| scf7180000041013 | 66424 | 0.432 | 0 | 156.2037 | Eukaryota | 63745.0 | 0 | Chordata | 43788.0 | 2 | Gadiformes | 38226.0 | 5 | Gadidae | 38226.0 | 5 | Gadus | 38226.0 | 5 | Gadus morhua | 38226.0 | 5 |
| scf7180000041458 | 1025301 | 0.4114 | 0 | 58.4584 | Eukaryota | 43994.0 | 0 | Chordata | 43994.0 | 0 | Carnivora | 43994.0 | 0 | Canidae | 43994.0 | 0 | Canis | 33096.0 | 1 | Canis lupus | 33096.0 | 1 |
| scf7180000055817 | 22325 | 0.4404 | 0 | 479.4014 | no-hit | 0.0 | 0 | no-hit | 0.0 | 0 | no-hit | 0.0 | 0 | no-hit | 0.0 | 0 | no-hit | 0.0 | 0 | no-hit | 0.0 | 0 |
| scf7180000042899 | 612075 | 0.3389 | 0 | 59.2533 | Eukaryota | 88819.0 | 0 | Chordata | 88819.0 | 0 | Carnivora | 88819.0 | 0 | Canidae | 88819.0 | 0 | Canis | 88819.0 | 0 | Canis lupus | 88819.0 | 0 |
| scf7180000047190 | 7046 | 0.4312 | 0 | 73.1794 | Eukaryota | 32377.0 | 0 | Chordata | 17581.0 | 2 | Gadiformes | 8892.0 | 6 | Gadidae | 8892.0 | 6 | Gadus | 8892.0 | 6 | Gadus morhua | 8892.0 | 6 |
| scf7180000042948 | 260332 | 0.3682 | 0 | 59.6784 | Eukaryota | 87797.0 | 0 | Chordata | 87797.0 | 0 | Carnivora | 87797.0 | 0 | Canidae | 87797.0 | 0 | Canis | 87797.0 | 0 | Canis lupus | 87797.0 | 0 |
| scf7180000040681 | 1950912 | 0.4822 | 0 | 58.7781 | Eukaryota | 78925.0 | 0 | Chordata | 78925.0 | 0 | Carnivora | 78925.0 | 0 | Canidae | 78925.0 | 0 | Canis | 78925.0 | 0 | Canis lupus | 78925.0 | 0 |
| scf7180000050321 | 13607 | 0.3679 | 0 | 58.6719 | Eukaryota | 10131.0 | 0 | Chordata | 10131.0 | 0 | Carnivora | 10131.0 | 0 | Canidae | 10131.0 | 0 | Canis | 10131.0 | 0 | Canis lupus | 10131.0 | 0 |
| scf7180000042630 | 435665 | 0.3931 | 0 | 60.36 | Eukaryota | 89987.0 | 0 | Chordata | 89987.0 | 0 | Carnivora | 89987.0 | 0 | Canidae | 89987.0 | 0 | Canis | 89987.0 | 0 | Canis lupus | 89987.0 | 0 |
| scf7180000048374 | 3169 | 0.4336 | 0 | 203.3133 | no-hit | 0.0 | 0 | no-hit | 0.0 | 0 | no-hit | 0.0 | 0 | no-hit | 0.0 | 0 | no-hit | 0.0 | 0 | no-hit | 0.0 | 0 |
| scf7180000042456 | 524292 | 0.433 | 0 | 59.5547 | Eukaryota | 82630.0 | 0 | Chordata | 82630.0 | 0 | Carnivora | 82630.0 | 0 | Canidae | 82630.0 | 0 | Canis | 82630.0 | 0 | Canis lupus | 82630.0 | 0 |
| scf7180000053659 | 3696 | 0.4413 | 0 | 79.4172 | Eukaryota | 3521.0 | 0 | Chordata | 3521.0 | 0 | Salmoniformes | 2292.0 | 2 | Salmonidae | 2292.0 | 2 | Salmo | 1674.0 | 3 | Salmo trutta | 1674.0 | 3 |
| scf7180000055809 | 11198 | 0.4412 | 0 | 88.7688 | Eukaryota | 5626.0 | 0 | Chordata | 5103.0 | 1 | Kurtiformes | 3668.0 | 3 | Apogonidae | 3668.0 | 3 | Sphaeramia | 3668.0 | 3 | Sphaeramia orbicularis | 3668.0 | 3 |
| scf7180000044974 | 248803 | 0.3572 | 0 | 59.0033 | Eukaryota | 33325.0 | 0 | Chordata | 33325.0 | 0 | Carnivora | 33325.0 | 0 | Canidae | 33325.0 | 0 | Canis | 33325.0 | 0 | Canis lupus | 33325.0 | 0 |
| scf7180000042678 | 53922 | 0.4564 | 0 | 432.758 | Eukaryota | 69486.0 | 0 | Chordata | 69486.0 | 0 | Carnivora | 69486.0 | 0 | Canidae | 69486.0 | 0 | Canis | 69486.0 | 0 | Canis lupus | 69486.0 | 0 |
| scf7180000055538 | 21795 | 0.4468 | 0 | 185.1733 | no-hit | 0.0 | 0 | no-hit | 0.0 | 0 | no-hit | 0.0 | 0 | no-hit | 0.0 | 0 | no-hit | 0.0 | 0 | no-hit | 0.0 | 0 |
| scf7180000043720 | 96903 | 0.4329 | 0 | 58.9299 | Eukaryota | 32585.0 | 0 | Chordata | 32585.0 | 0 | Carnivora | 32585.0 | 0 | Canidae | 32585.0 | 0 | Canis | 32585.0 | 0 | Canis lupus | 32585.0 | 0 |
| scf7180000048632 | 60257 | 0.4845 | 0 | 58.1244 | Eukaryota | 7019.0 | 0 | Chordata | 7019.0 | 0 | Carnivora | 3292.0 | 2 | Canidae | 3292.0 | 2 | Homo | 3111.0 | 3 | Homo sapiens | 3111.0 | 3 |
| scf7180000056313 | 930197 | 0.4106 | 0 | 59.4195 | Eukaryota | 65385.0 | 0 | Chordata | 65385.0 | 0 | Carnivora | 65385.0 | 0 | Canidae | 65385.0 | 0 | Canis | 65385.0 | 0 | Canis lupus | 65385.0 | 0 |
| scf7180000040732 | 30542 | 0.4404 | 0 | 204.97 | Eukaryota | 2145.0 | 0 | Platyhelminthes | 1487.0 | 1 | Polystomatidea | 985.0 | 2 | Polystomatidae | 985.0 | 2 | Protopolystoma | 985.0 | 2 | Protopolystoma xenopodis | 985.0 | 2 |
| scf7180000045869 | 31122 | 0.4327 | 0 | 236.7689 | Eukaryota | 89738.0 | 0 | Chordata | 89738.0 | 0 | Carnivora | 89738.0 | 0 | Canidae | 89738.0 | 0 | Canis | 89738.0 | 0 | Canis lupus | 89738.0 | 0 |
| scf7180000048714 | 52945 | 0.4383 | 0 | 58.9343 | Eukaryota | 33544.0 | 0 | Chordata | 33544.0 | 0 | Carnivora | 33544.0 | 0 | Canidae | 33544.0 | 0 | Canis | 17676.0 | 1 | Canis lupus | 17676.0 | 1 |
| scf7180000047667 | 58109 | 0.3758 | 0 | 58.887 | Eukaryota | 69272.0 | 0 | Chordata | 69272.0 | 0 | Carnivora | 69272.0 | 0 | Canidae | 69272.0 | 0 | Canis | 69272.0 | 0 | Canis lupus | 69272.0 | 0 |
| scf7180000052106 | 4093 | 0.4478 | 0 | 402.9787 | Eukaryota | 3279.0 | 0 | Chordata | 2342.0 | 1 | Gonorynchiformes | 1088.0 | 3 | Chanidae | 1088.0 | 3 | Chanos | 1088.0 | 3 | Chanos chanos | 1088.0 | 3 |
| scf7180000040910 | 1444091 | 0.4531 | 0 | 57.9528 | Eukaryota | 45424.0 | 0 | Chordata | 45424.0 | 0 | Carnivora | 45424.0 | 0 | Canidae | 45424.0 | 0 | Canis | 45424.0 | 0 | Canis lupus | 45424.0 | 0 |
| scf7180000056340 | 297944 | 0.3747 | 0 | 58.5077 | Eukaryota | 66451.0 | 0 | Chordata | 66451.0 | 0 | Carnivora | 66451.0 | 0 | Canidae | 66451.0 | 0 | Canis | 66451.0 | 0 | Canis lupus | 66451.0 | 0 |
| scf7180000042425 | 194994 | 0.3581 | 0 | 58.7303 | Eukaryota | 69484.0 | 0 | Chordata | 69484.0 | 0 | Carnivora | 69484.0 | 0 | Canidae | 69484.0 | 0 | Canis | 69484.0 | 0 | Canis lupus | 69484.0 | 0 |
| scf7180000047810 | 10407 | 0.5139 | 0 | 406.3922 | Eukaryota | 3869.0 | 0 | Chordata | 3869.0 | 0 | Carnivora | 3869.0 | 0 | Canidae | 3869.0 | 0 | Canis | 3168.0 | 1 | Canis lupus | 3168.0 | 1 |
| scf7180000042190 | 816031 | 0.3821 | 0 | 60.0609 | Eukaryota | 64104.0 | 0 | Chordata | 64104.0 | 0 | Carnivora | 64104.0 | 0 | Canidae | 52687.0 | 2 | Canis | 45693.0 | 3 | Canis lupus | 45693.0 | 3 |
| scf7180000048148 | 55964 | 0.4102 | 0 | 59.8979 | Eukaryota | 50360.0 | 0 | Chordata | 50360.0 | 0 | Carnivora | 50360.0 | 0 | Canidae | 30642.0 | 3 | Canis | 22005.0 | 6 | Canis lupus | 22005.0 | 7 |
| scf7180000043788 | 382055 | 0.5381 | 0 | 74.9514 | Eukaryota | 52450.0 | 0 | Chordata | 52450.0 | 0 | Carnivora | 52450.0 | 0 | Canidae | 52450.0 | 0 | unresolved | 26225.0 | 1 | unresolved | 26225.0 | 1 |
| scf7180000044140 | 44205 | 0.4407 | 0 | 52.1231 | Eukaryota | 3668.0 | 0 | Chordata | 3668.0 | 0 | Carnivora | 1878.0 | 3 | Canidae | 1878.0 | 3 | Canis | 1878.0 | 4 | Canis lupus | 1878.0 | 4 |
| scf7180000044056 | 386449 | 0.545 | 0 | 56.6367 | Eukaryota | 87962.0 | 0 | Chordata | 87962.0 | 0 | Carnivora | 87962.0 | 0 | Canidae | 87962.0 | 0 | Canis | 74978.0 | 1 | Canis lupus | 74978.0 | 1 |
| scf7180000041568 | 608254 | 0.4186 | 0 | 58.5086 | Eukaryota | 113234.0 | 0 | Chordata | 113234.0 | 0 | Carnivora | 113234.0 | 0 | Canidae | 113234.0 | 0 | Canis | 102868.0 | 1 | Canis lupus | 102868.0 | 1 |
| scf7180000047909 | 36901 | 0.4442 | 0 | 81.9559 | Eukaryota | 42784.0 | 1 | Chordata | 33310.0 | 3 | Gadiformes | 18889.0 | 6 | Gadidae | 18889.0 | 6 | Gadus | 18889.0 | 6 | Gadus morhua | 18889.0 | 6 |
| scf7180000052949 | 3111 | 0.4481 | 0 | 169.4219 | no-hit | 0.0 | 0 | no-hit | 0.0 | 0 | no-hit | 0.0 | 0 | no-hit | 0.0 | 0 | no-hit | 0.0 | 0 | no-hit | 0.0 | 0 |
| scf7180000043791 | 377506 | 0.5239 | 0 | 59.234 | Eukaryota | 31092.0 | 0 | Chordata | 31092.0 | 0 | Carnivora | 31092.0 | 0 | Canidae | 31092.0 | 0 | Canis | 21645.0 | 1 | Canis lupus | 21645.0 | 1 |
| scf7180000055242 | 25587 | 0.4351 | 0 | 307.4675 | no-hit | 0.0 | 0 | no-hit | 0.0 | 0 | no-hit | 0.0 | 0 | no-hit | 0.0 | 0 | no-hit | 0.0 | 0 | no-hit | 0.0 | 0 |
| scf7180000041577 | 924985 | 0.3957 | 0 | 59.6109 | Eukaryota | 59426.0 | 0 | Chordata | 59426.0 | 0 | Carnivora | 59426.0 | 0 | Canidae | 59426.0 | 0 | Canis | 52709.0 | 1 | Canis lupus | 52709.0 | 1 |
| scf7180000047680 | 62829 | 0.4737 | 0 | 57.2298 | Eukaryota | 4623.0 | 0 | Chordata | 4623.0 | 0 | Primates | 2075.0 | 3 | Hominidae | 2075.0 | 3 | Canis | 1636.0 | 4 | Canis lupus | 1636.0 | 4 |
| scf7180000048958 | 6723 | 0.4302 | 0 | 129.0252 | no-hit | 0.0 | 0 | no-hit | 0.0 | 0 | no-hit | 0.0 | 0 | no-hit | 0.0 | 0 | no-hit | 0.0 | 0 | no-hit | 0.0 | 0 |
| scf7180000042616 | 399073 | 0.4358 | 0 | 59.2357 | Eukaryota | 80308.0 | 0 | Chordata | 80308.0 | 0 | Carnivora | 80308.0 | 0 | Canidae | 80308.0 | 0 | Canis | 80308.0 | 0 | Canis lupus | 80308.0 | 0 |
| scf7180000044369 | 373963 | 0.3847 | 0 | 58.8499 | Eukaryota | 111615.0 | 0 | Chordata | 111615.0 | 0 | Carnivora | 111615.0 | 0 | Canidae | 111615.0 | 0 | Canis | 72135.0 | 1 | Canis lupus | 72135.0 | 1 |
| scf7180000055402 | 58190 | 0.4422 | 0 | 141.6665 | Eukaryota | 2105.0 | 0 | Platyhelminthes | 1093.0 | 2 | Polystomatidea | 1093.0 | 2 | Polystomatidae | 1093.0 | 2 | Protopolystoma | 1093.0 | 3 | Protopolystoma xenopodis | 1093.0 | 3 |
| scf7180000043930 | 84240 | 0.3548 | 0 | 57.21 | Eukaryota | 20444.0 | 0 | Chordata | 20444.0 | 0 | Carnivora | 20444.0 | 0 | Canidae | 20444.0 | 0 | Canis | 20444.0 | 0 | Canis lupus | 20444.0 | 0 |
| scf7180000045026 | 13721 | 0.4371 | 0 | 51.9639 | Eukaryota | 39426.0 | 0 | Chordata | 39426.0 | 0 | Carnivora | 39426.0 | 0 | Canidae | 39426.0 | 0 | Canis | 39426.0 | 0 | Canis lupus | 39426.0 | 0 |
| scf7180000051963 | 15575 | 0.3829 | 0 | 31.2679 | Eukaryota | 6573.0 | 0 | Chordata | 6573.0 | 0 | Carnivora | 6573.0 | 0 | Canidae | 5856.0 | 1 | Canis | 5856.0 | 1 | Canis lupus | 5856.0 | 1 |
| scf7180000050120 | 12246 | 0.4443 | 0 | 466.2611 | Eukaryota | 1820.0 | 0 | Platyhelminthes | 914.0 | 2 | Polystomatidea | 914.0 | 2 | Polystomatidae | 914.0 | 2 | Protopolystoma | 914.0 | 2 | Protopolystoma xenopodis | 914.0 | 2 |
| scf7180000053067 | 5113 | 0.4403 | 0 | 261.3631 | no-hit | 0.0 | 0 | no-hit | 0.0 | 0 | no-hit | 0.0 | 0 | no-hit | 0.0 | 0 | no-hit | 0.0 | 0 | no-hit | 0.0 | 0 |
| scf7180000043763 | 182210 | 0.427 | 0 | 59.5991 | Eukaryota | 58380.0 | 0 | Chordata | 58380.0 | 0 | Carnivora | 58380.0 | 0 | Canidae | 58380.0 | 0 | Canis | 58380.0 | 0 | Canis lupus | 58380.0 | 0 |
| scf7180000043868 | 772114 | 0.3777 | 0 | 60.4127 | Eukaryota | 55666.0 | 0 | Chordata | 55666.0 | 0 | Carnivora | 55666.0 | 0 | Canidae | 55666.0 | 0 | Canis | 55666.0 | 0 | Canis lupus | 55666.0 | 0 |
| scf7180000046994 | 22476 | 0.439 | 0 | 104.1636 | Eukaryota | 41845.0 | 0 | Chordata | 35366.0 | 1 | Gadiformes | 24294.0 | 5 | Gadidae | 24294.0 | 5 | Gadus | 24294.0 | 5 | Gadus morhua | 24294.0 | 5 |
| scf7180000046718 | 56982 | 0.5291 | 0 | 56.9555 | Eukaryota | 80384.0 | 0 | Chordata | 80384.0 | 0 | Carnivora | 80384.0 | 0 | Canidae | 80384.0 | 0 | Canis | 80384.0 | 0 | Canis lupus | 80384.0 | 0 |
| scf7180000045186 | 242756 | 0.371 | 0 | 61.1977 | Eukaryota | 12757.0 | 0 | Chordata | 12757.0 | 0 | Primates | 7821.0 | 1 | Hominidae | 6760.0 | 2 | Homo | 6760.0 | 2 | Homo sapiens | 6760.0 | 2 |
| scf7180000043678 | 422334 | 0.4045 | 0 | 61.6543 | Eukaryota | 53671.0 | 0 | Chordata | 53671.0 | 0 | Carnivora | 53671.0 | 0 | Canidae | 53671.0 | 0 | Canis | 35088.0 | 1 | Canis lupus | 35088.0 | 1 |
| scf7180000053533 | 3494 | 0.5049 | 0 | 51.351 | Eukaryota | 7000.0 | 0 | Chordata | 7000.0 | 0 | Carnivora | 7000.0 | 0 | Canidae | 7000.0 | 0 | Canis | 5447.0 | 1 | Canis lupus | 4328.0 | 2 |
| scf7180000047449 | 2717 | 0.4284 | 0 | 84.1854 | no-hit | 0.0 | 0 | no-hit | 0.0 | 0 | no-hit | 0.0 | 0 | no-hit | 0.0 | 0 | no-hit | 0.0 | 0 | no-hit | 0.0 | 0 |
| scf7180000049222 | 54737 | 0.4524 | 0 | 64.0431 | Eukaryota | 84718.0 | 0 | Chordata | 84718.0 | 0 | Carnivora | 84718.0 | 0 | Canidae | 84718.0 | 0 | Canis | 84718.0 | 0 | Canis lupus | 84718.0 | 0 |
| scf7180000053345 | 8920 | 0.4178 | 0 | 55.7047 | Eukaryota | 79396.0 | 0 | Chordata | 79396.0 | 0 | Carnivora | 79396.0 | 0 | Canidae | 79396.0 | 0 | Canis | 79396.0 | 0 | Canis lupus | 79396.0 | 0 |
| scf7180000051839 | 3146 | 0.4501 | 0 | 806.6675 | Eukaryota | 440.0 | 0 | Chordata | 440.0 | 0 | Tetraodontiformes | 440.0 | 0 | Tetraodontidae | 440.0 | 0 | Takifugu | 440.0 | 0 | Takifugu rubripes | 440.0 | 0 |
| scf7180000045883 | 150494 | 0.3381 | 0 | 58.4208 | Eukaryota | 54051.0 | 0 | Chordata | 54051.0 | 0 | Carnivora | 54051.0 | 0 | Canidae | 54051.0 | 0 | Canis | 54051.0 | 0 | Canis lupus | 54051.0 | 0 |
| scf7180000046586 | 18276 | 0.4386 | 0 | 235.0448 | Eukaryota | 2535.0 | 0 | Chordata | 1533.0 | 1 | Salmoniformes | 1169.0 | 2 | Salmonidae | 1169.0 | 2 | Protopolystoma | 1002.0 | 3 | Protopolystoma xenopodis | 1002.0 | 3 |
| scf7180000040846 | 1622526 | 0.4085 | 0 | 60.578 | Eukaryota | 72275.0 | 0 | Chordata | 72275.0 | 0 | Carnivora | 72275.0 | 0 | Canidae | 72275.0 | 0 | Canis | 72275.0 | 0 | Canis lupus | 72275.0 | 0 |
| scf7180000044349 | 193386 | 0.3811 | 0 | 56.7852 | Eukaryota | 40820.0 | 0 | Chordata | 40820.0 | 0 | Carnivora | 40820.0 | 0 | Canidae | 40820.0 | 0 | Canis | 40820.0 | 0 | Canis lupus | 40820.0 | 0 |
| scf7180000046844 | 9596 | 0.4424 | 0 | 378.5445 | no-hit | 0.0 | 0 | no-hit | 0.0 | 0 | no-hit | 0.0 | 0 | no-hit | 0.0 | 0 | no-hit | 0.0 | 0 | no-hit | 0.0 | 0 |
| scf7180000042666 | 156169 | 0.3773 | 0 | 59.9687 | Eukaryota | 31837.0 | 0 | Chordata | 31837.0 | 0 | Carnivora | 31837.0 | 0 | Canidae | 19285.0 | 2 | Canis | 16060.0 | 3 | Canis lupus | 16060.0 | 3 |
| scf7180000049267 | 7600 | 0.4338 | 0 | 65.7222 | Eukaryota | 33381.0 | 0 | Chordata | 33381.0 | 0 | Carnivora | 33381.0 | 0 | Canidae | 33381.0 | 0 | Canis | 33381.0 | 0 | Canis lupus | 33381.0 | 0 |
| scf7180000046566 | 1380917 | 0.4332 | 0 | 59.1328 | Eukaryota | 110920.0 | 0 | Chordata | 110920.0 | 0 | Carnivora | 110920.0 | 0 | Canidae | 110920.0 | 0 | Canis | 110920.0 | 0 | Canis lupus | 110920.0 | 0 |
| scf7180000041537 | 55345 | 0.4344 | 0 | 54.5449 | Eukaryota | 11015.0 | 0 | Chordata | 11015.0 | 0 | Carnivora | 4694.0 | 3 | Canidae | 4694.0 | 3 | Canis | 4694.0 | 5 | Canis lupus | 4694.0 | 5 |
| scf7180000048811 | 65194 | 0.4281 | 0 | 59.8377 | Eukaryota | 51394.0 | 0 | Chordata | 51394.0 | 0 | Carnivora | 51394.0 | 0 | Canidae | 51394.0 | 0 | Canis | 51394.0 | 0 | Canis lupus | 51394.0 | 0 |
| scf7180000040525 | 2848339 | 0.3946 | 0 | 60.6454 | Eukaryota | 117955.0 | 0 | Chordata | 117955.0 | 0 | Carnivora | 117955.0 | 0 | Canidae | 117955.0 | 0 | Canis | 117955.0 | 0 | Canis lupus | 117955.0 | 0 |
| scf7180000045412 | 193475 | 0.4238 | 0 | 58.7496 | Eukaryota | 20598.0 | 0 | Chordata | 20598.0 | 0 | Carnivora | 20598.0 | 0 | Canidae | 20598.0 | 0 | Canis | 20598.0 | 0 | Canis lupus | 20598.0 | 0 |
| scf7180000050353 | 4399 | 0.4415 | 0 | 330.6999 | no-hit | 0.0 | 0 | no-hit | 0.0 | 0 | no-hit | 0.0 | 0 | no-hit | 0.0 | 0 | no-hit | 0.0 | 0 | no-hit | 0.0 | 0 |
| scf7180000048208 | 24067 | 0.4482 | 0 | 118.2426 | Eukaryota | 5732.0 | 0 | Platyhelminthes | 3405.0 | 1 | Strigeidida | 2880.0 | 3 | Schistosomatidae | 2880.0 | 3 | Schistosoma | 2880.0 | 3 | Schistosoma mansoni | 2880.0 | 3 |
| scf7180000052203 | 5297 | 0.4463 | 0 | 425.3289 | Eukaryota | 2515.0 | 0 | Platyhelminthes | 1082.0 | 2 | Polystomatidea | 1082.0 | 2 | Polystomatidae | 1082.0 | 2 | Protopolystoma | 1082.0 | 2 | Protopolystoma xenopodis | 1082.0 | 2 |
| scf7180000056016 | 19537 | 0.6782 | 0 | 470.329 | Eukaryota | 35655.0 | 0 | Chordata | 35655.0 | 0 | Primates | 27458.0 | 1 | Cercopithecidae | 13739.0 | 3 | Macaca | 13739.0 | 4 | Macaca mulatta | 13739.0 | 4 |
| scf7180000052241 | 8049 | 0.5658 | 0 | 52.9714 | Eukaryota | 25606.0 | 0 | Chordata | 25606.0 | 0 | Carnivora | 25606.0 | 0 | Canidae | 25606.0 | 0 | Canis | 25606.0 | 0 | Canis lupus | 25606.0 | 0 |
| scf7180000054818 | 17998 | 0.4014 | 0 | 172.2184 | Eukaryota | 5340.0 | 0 | Chordata | 5340.0 | 0 | Carnivora | 5340.0 | 0 | Canidae | 5340.0 | 0 | Canis | 5340.0 | 0 | Canis lupus | 5340.0 | 0 |
| scf7180000042698 | 61422 | 0.4721 | 0 | 55.3718 | Eukaryota | 31373.0 | 0 | Chordata | 31373.0 | 0 | Carnivora | 31373.0 | 0 | Canidae | 31373.0 | 0 | Canis | 18809.0 | 1 | Canis lupus | 18809.0 | 1 |
| scf7180000043894 | 26943 | 0.3595 | 0 | 52.7815 | Eukaryota | 24017.0 | 0 | Chordata | 24017.0 | 0 | Carnivora | 24017.0 | 0 | Canidae | 24017.0 | 0 | Canis | 24017.0 | 0 | Canis lupus | 24017.0 | 0 |
| scf7180000041161 | 1288511 | 0.4823 | 0 | 57.4328 | Eukaryota | 69505.0 | 0 | Chordata | 69505.0 | 0 | Carnivora | 69505.0 | 0 | Canidae | 69505.0 | 0 | Canis | 69505.0 | 0 | Canis lupus | 69505.0 | 0 |
| scf7180000053757 | 11552 | 0.447 | 0 | 334.846 | Eukaryota | 4977.0 | 0 | Platyhelminthes | 1788.0 | 3 | Polystomatidea | 1788.0 | 5 | Polystomatidae | 1788.0 | 5 | Protopolystoma | 1788.0 | 5 | Protopolystoma xenopodis | 1788.0 | 5 |
| scf7180000051281 | 4397 | 0.4408 | 0 | 831.4311 | Eukaryota | 456.0 | 0 | Platyhelminthes | 315.0 | 1 | Polystomatidea | 315.0 | 1 | Polystomatidae | 315.0 | 1 | Protopolystoma | 315.0 | 1 | Protopolystoma xenopodis | 315.0 | 1 |
| scf7180000049545 | 22668 | 0.4457 | 0 | 128.2206 | Eukaryota | 4132.0 | 0 | Chordata | 4132.0 | 0 | Carnivora | 2826.0 | 1 | Canidae | 2826.0 | 1 | Canis | 2826.0 | 1 | Canis lupus | 2826.0 | 1 |
| scf7180000044111 | 210575 | 0.3444 | 0 | 59.2116 | Eukaryota | 55742.0 | 0 | Chordata | 55742.0 | 0 | Carnivora | 55742.0 | 0 | Canidae | 55742.0 | 0 | Canis | 51293.0 | 1 | Canis lupus | 51293.0 | 1 |
| scf7180000041414 | 36467 | 0.4106 | 0 | 55.1876 | Eukaryota | 29427.0 | 0 | Chordata | 29427.0 | 0 | Carnivora | 29427.0 | 0 | Canidae | 29427.0 | 0 | Canis | 29427.0 | 0 | Canis lupus | 29427.0 | 0 |
| scf7180000041798 | 858406 | 0.373 | 0 | 60.1363 | Eukaryota | 48700.0 | 0 | Chordata | 48700.0 | 0 | Carnivora | 48700.0 | 0 | Canidae | 35998.0 | 1 | Canis | 35998.0 | 1 | Canis lupus | 35998.0 | 1 |
| scf7180000054227 | 14734 | 0.4291 | 0 | 173.7936 | Eukaryota | 81452.0 | 0 | Chordata | 81452.0 | 0 | Carnivora | 81452.0 | 0 | Canidae | 81452.0 | 0 | Canis | 81452.0 | 0 | Canis lupus | 81452.0 | 0 |
| scf7180000056186 | 1759739 | 0.3982 | 0 | 59.1037 | Eukaryota | 134737.0 | 0 | Chordata | 134737.0 | 0 | Carnivora | 134737.0 | 0 | Canidae | 134737.0 | 0 | Canis | 93159.0 | 1 | Canis lupus | 93159.0 | 1 |
| scf7180000042395 | 689602 | 0.376 | 0 | 60.2632 | Eukaryota | 105595.0 | 0 | Chordata | 105595.0 | 0 | Carnivora | 105595.0 | 0 | Canidae | 105595.0 | 0 | Canis | 105595.0 | 0 | Canis lupus | 105595.0 | 0 |
| scf7180000045297 | 185357 | 0.4171 | 0 | 60.8201 | Eukaryota | 85723.0 | 0 | Chordata | 85723.0 | 0 | Carnivora | 85723.0 | 0 | Canidae | 85723.0 | 0 | Canis | 85723.0 | 0 | Canis lupus | 85723.0 | 0 |
| scf7180000043984 | 6683 | 0.4306 | 0 | 419.5689 | Eukaryota | 2237.0 | 0 | Chordata | 2237.0 | 0 | Perissodactyla | 902.0 | 3 | Equidae | 902.0 | 4 | Equus | 902.0 | 4 | Equus caballus | 902.0 | 4 |
| scf7180000050818 | 21601 | 0.444 | 0 | 60.9858 | Eukaryota | 4069.0 | 0 | Chordata | 2793.0 | 2 | Kurtiformes | 2055.0 | 4 | Apogonidae | 2055.0 | 4 | Sphaeramia | 2055.0 | 5 | Sphaeramia orbicularis | 2055.0 | 5 |
| scf7180000052675 | 3967 | 0.4454 | 0 | 1144.1524 | Eukaryota | 233.0 | 0 | Chordata | 233.0 | 0 | Syngnathiformes | 233.0 | 0 | Syngnathidae | 233.0 | 0 | Syngnathus | 233.0 | 0 | Syngnathus acus | 233.0 | 0 |
| scf7180000048310 | 15508 | 0.4416 | 0 | 56.3039 | Eukaryota | 49524.0 | 0 | Platyhelminthes | 27639.0 | 1 | Strigeidida | 17140.0 | 7 | Schistosomatidae | 17140.0 | 7 | Schistosoma | 17140.0 | 7 | Schistosoma mansoni | 17140.0 | 7 |
| scf7180000045418 | 184582 | 0.4991 | 0 | 67.1593 | Eukaryota | 54830.0 | 0 | Chordata | 54830.0 | 0 | Carnivora | 54830.0 | 0 | Canidae | 54830.0 | 0 | Canis | 54830.0 | 0 | Canis lupus | 54830.0 | 0 |
| scf7180000042140 | 853368 | 0.467 | 0 | 58.9205 | Eukaryota | 55564.0 | 0 | Chordata | 55564.0 | 0 | Carnivora | 46248.0 | 1 | Canidae | 31820.0 | 3 | Canis | 15938.0 | 4 | Canis lupus | 15938.0 | 6 |
| scf7180000047352 | 73651 | 0.3973 | 0 | 58.853 | Eukaryota | 57804.0 | 0 | Chordata | 57804.0 | 0 | Carnivora | 57804.0 | 0 | Canidae | 57804.0 | 0 | Canis | 41499.0 | 1 | Canis lupus | 41499.0 | 1 |
| scf7180000046670 | 12449 | 0.5038 | 0 | 370.7098 | Eukaryota | 9464.0 | 0 | Chordata | 9464.0 | 0 | Carnivora | 9464.0 | 0 | Canidae | 9464.0 | 0 | Canis | 7681.0 | 1 | Canis lupus | 6374.0 | 2 |
| scf7180000052100 | 13742 | 0.4346 | 0 | 157.765 | Viruses | 403.0 | 0 | Viruses-undef | 403.0 | 0 | Herpesvirales | 403.0 | 0 | Herpesviridae | 403.0 | 0 | Mardivirus | 403.0 | 0 | Gallid alphaherpesvirus 2 | 403.0 | 0 |
| scf7180000055963 | 15873 | 0.4371 | 0 | 91.6739 | Eukaryota | 33260.0 | 0 | Chordata | 20008.0 | 2 | Gadiformes | 16983.0 | 4 | Gadidae | 16983.0 | 4 | Gadus | 16983.0 | 4 | Gadus morhua | 16983.0 | 4 |
| scf7180000040804 | 1690227 | 0.4997 | 0 | 58.8466 | Eukaryota | 82563.0 | 0 | Chordata | 82563.0 | 0 | Carnivora | 82563.0 | 0 | Canidae | 82563.0 | 0 | Canis | 67172.0 | 1 | Canis lupus | 67172.0 | 1 |
| scf7180000049073 | 18254 | 0.4223 | 0 | 791.9585 | Eukaryota | 46661.0 | 0 | Chordata | 46661.0 | 0 | Carnivora | 46661.0 | 0 | Canidae | 46661.0 | 0 | Canis | 46661.0 | 0 | Canis lupus | 46661.0 | 0 |
| scf7180000049244 | 19309 | 0.3805 | 0 | 61.9178 | Eukaryota | 4613.0 | 0 | Chordata | 4613.0 | 0 | Carnivora | 4613.0 | 0 | Canidae | 4613.0 | 0 | Canis | 4613.0 | 0 | Canis lupus | 4613.0 | 0 |
| scf7180000043170 | 513268 | 0.4686 | 0 | 60.6683 | Eukaryota | 76559.0 | 0 | Chordata | 76559.0 | 0 | Carnivora | 76559.0 | 0 | Canidae | 76559.0 | 0 | Canis | 76559.0 | 0 | Canis lupus | 76559.0 | 0 |
| scf7180000041474 | 1056317 | 0.5808 | 0 | 58.486 | Eukaryota | 117149.0 | 0 | Chordata | 117149.0 | 0 | Carnivora | 117149.0 | 0 | Canidae | 117149.0 | 0 | Canis | 101686.0 | 1 | Canis lupus | 101686.0 | 1 |
| scf7180000043949 | 355052 | 0.4095 | 0 | 60.2481 | Eukaryota | 83645.0 | 0 | Chordata | 83645.0 | 0 | Carnivora | 83645.0 | 0 | Canidae | 71123.0 | 2 | Canis | 68005.0 | 3 | Canis lupus | 68005.0 | 3 |
| scf7180000048674 | 70147 | 0.3613 | 0 | 58.1852 | Eukaryota | 50799.0 | 0 | Chordata | 50799.0 | 0 | Carnivora | 50799.0 | 0 | Canidae | 50799.0 | 0 | Canis | 49170.0 | 1 | Canis lupus | 49170.0 | 1 |
| scf7180000045631 | 227433 | 0.3329 | 0 | 58.294 | Eukaryota | 16090.0 | 0 | Chordata | 16090.0 | 0 | Carnivora | 16090.0 | 0 | Canidae | 16090.0 | 0 | Canis | 16090.0 | 0 | Canis lupus | 16090.0 | 0 |
| scf7180000055968 | 14598 | 0.3778 | 0 | 290.0937 | Eukaryota | 35469.0 | 0 | Chordata | 35469.0 | 0 | Carnivora | 35469.0 | 0 | Canidae | 35469.0 | 0 | Canis | 35469.0 | 0 | Canis lupus | 35469.0 | 0 |
| scf7180000049043 | 4874 | 0.4364 | 0 | 81.8987 | no-hit | 0.0 | 0 | no-hit | 0.0 | 0 | no-hit | 0.0 | 0 | no-hit | 0.0 | 0 | no-hit | 0.0 | 0 | no-hit | 0.0 | 0 |
| scf7180000042022 | 17512 | 0.4408 | 0 | 56.7629 | Eukaryota | 39780.0 | 0 | Chordata | 39780.0 | 0 | Carnivora | 39780.0 | 0 | Canidae | 39780.0 | 0 | Canis | 39780.0 | 0 | Canis lupus | 39780.0 | 0 |
| scf7180000044390 | 316340 | 0.516 | 0 | 60.7042 | Eukaryota | 62957.0 | 0 | Chordata | 62957.0 | 0 | Carnivora | 62957.0 | 0 | Canidae | 62957.0 | 0 | Canis | 51191.0 | 1 | Canis lupus | 51191.0 | 1 |
| scf7180000043817 | 255267 | 0.4211 | 0 | 55.5525 | Eukaryota | 39438.0 | 0 | Chordata | 39438.0 | 0 | Carnivora | 39438.0 | 0 | Canidae | 39438.0 | 0 | Vulpes | 22490.0 | 1 | Vulpes vulpes | 22490.0 | 1 |
| scf7180000042925 | 615874 | 0.4061 | 0 | 59.3135 | Eukaryota | 41066.0 | 0 | Chordata | 41066.0 | 0 | Carnivora | 41066.0 | 0 | Canidae | 41066.0 | 0 | Canis | 41066.0 | 0 | Canis lupus | 41066.0 | 0 |
| scf7180000051145 | 12293 | 0.4565 | 0 | 783.6387 | Eukaryota | 2844.0 | 0 | Platyhelminthes | 1236.0 | 3 | Polystomatidea | 1236.0 | 3 | Polystomatidae | 1236.0 | 3 | Protopolystoma | 1236.0 | 3 | Protopolystoma xenopodis | 1236.0 | 3 |
| scf7180000048767 | 35269 | 0.4389 | 0 | 194.7337 | Eukaryota | 9979.0 | 0 | Chordata | 9979.0 | 0 | Kurtiformes | 4314.0 | 4 | Apogonidae | 4314.0 | 4 | Sphaeramia | 4314.0 | 4 | Sphaeramia orbicularis | 4314.0 | 4 |
| scf7180000043346 | 25727 | 0.4433 | 0 | 70.9235 | Eukaryota | 167.0 | 0 | Platyhelminthes | 167.0 | 0 | Polystomatidea | 167.0 | 0 | Polystomatidae | 167.0 | 0 | Protopolystoma | 167.0 | 0 | Protopolystoma xenopodis | 167.0 | 0 |
| scf7180000042847 | 327795 | 0.5043 | 0 | 57.3409 | Eukaryota | 11164.0 | 0 | Chordata | 11164.0 | 0 | Carnivora | 3766.0 | 4 | Canidae | 3766.0 | 4 | Canis | 3766.0 | 4 | Canis lupus | 3766.0 | 4 |
| scf7180000048467 | 22888 | 0.4349 | 0 | 128.9948 | Eukaryota | 6387.0 | 0 | Chordata | 4277.0 | 2 | Kurtiformes | 3493.0 | 5 | Apogonidae | 3493.0 | 5 | Sphaeramia | 3493.0 | 5 | Sphaeramia orbicularis | 3493.0 | 5 |
| scf7180000052981 | 4138 | 0.4519 | 0 | 373.2223 | no-hit | 0.0 | 0 | no-hit | 0.0 | 0 | no-hit | 0.0 | 0 | no-hit | 0.0 | 0 | no-hit | 0.0 | 0 | no-hit | 0.0 | 0 |
| scf7180000048888 | 42962 | 0.5837 | 0 | 59.8486 | Eukaryota | 35958.0 | 0 | Chordata | 35958.0 | 0 | Carnivora | 35958.0 | 0 | Canidae | 16101.0 | 3 | Canis | 9261.0 | 5 | Canis lupus | 9261.0 | 5 |
| scf7180000056064 | 22384 | 0.4429 | 0 | 239.2578 | Eukaryota | 36622.0 | 0 | Chordata | 23925.0 | 2 | Gadiformes | 17530.0 | 5 | Gadidae | 17530.0 | 5 | Gadus | 17530.0 | 5 | Gadus morhua | 17530.0 | 5 |
| scf7180000045578 | 152571 | 0.3814 | 0 | 57.3364 | Eukaryota | 87089.0 | 0 | Chordata | 87089.0 | 0 | Carnivora | 87089.0 | 0 | Canidae | 87089.0 | 0 | Canis | 87089.0 | 0 | Canis lupus | 87089.0 | 0 |
| scf7180000046810 | 29593 | 0.4431 | 0 | 95.0328 | Eukaryota | 9112.0 | 0 | Chordata | 9112.0 | 0 | Carnivora | 8471.0 | 1 | Canidae | 8471.0 | 1 | Canis | 8471.0 | 1 | Canis lupus | 8471.0 | 1 |
| scf7180000040926 | 603783 | 0.4372 | 0 | 66.6277 | Eukaryota | 91277.0 | 0 | Chordata | 91277.0 | 0 | Carnivora | 91277.0 | 0 | Canidae | 91277.0 | 0 | Canis | 91277.0 | 0 | Canis lupus | 91277.0 | 0 |
| scf7180000041639 | 294080 | 0.4647 | 100 | 57.3271 | Eukaryota | 38553.0 | 0 | Chordata | 38553.0 | 0 | Carnivora | 38553.0 | 0 | Canidae | 20511.0 | 2 | Canis | 20511.0 | 2 | Canis lupus | 20511.0 | 2 |
| scf7180000044185 | 338854 | 0.3482 | 0 | 61.3108 | Eukaryota | 33125.0 | 0 | Chordata | 33125.0 | 0 | Carnivora | 33125.0 | 0 | Canidae | 33125.0 | 0 | Canis | 33125.0 | 0 | Canis lupus | 33125.0 | 0 |
| scf7180000046614 | 58716 | 0.4474 | 0 | 68.7085 | Eukaryota | 63244.0 | 0 | Chordata | 63244.0 | 0 | Carnivora | 63244.0 | 0 | Canidae | 63244.0 | 0 | Canis | 63244.0 | 0 | Canis lupus | 63244.0 | 0 |
| scf7180000042185 | 674512 | 0.3865 | 0 | 60.2268 | Eukaryota | 95391.0 | 0 | Chordata | 95391.0 | 0 | Carnivora | 95391.0 | 0 | Canidae | 95391.0 | 0 | Canis | 95391.0 | 0 | Canis lupus | 95391.0 | 0 |
| scf7180000042562 | 49424 | 0.348 | 0 | 56.7911 | Eukaryota | 60288.0 | 0 | Chordata | 60288.0 | 0 | Carnivora | 60288.0 | 0 | Canidae | 60288.0 | 0 | Canis | 60288.0 | 0 | Canis lupus | 60288.0 | 0 |
| scf7180000052069 | 5291 | 0.4451 | 0 | 256.7261 | no-hit | 0.0 | 0 | no-hit | 0.0 | 0 | no-hit | 0.0 | 0 | no-hit | 0.0 | 0 | no-hit | 0.0 | 0 | no-hit | 0.0 | 0 |
| scf7180000041094 | 337605 | 0.4072 | 0 | 58.5436 | Eukaryota | 19825.0 | 0 | Chordata | 19825.0 | 0 | Carnivora | 9640.0 | 2 | Hominidae | 6762.0 | 4 | Callorhinus | 3488.0 | 8 | Callorhinus ursinus | 3488.0 | 8 |
| scf7180000050511 | 9500 | 0.4415 | 0 | 57.5904 | Eukaryota | 3148.0 | 0 | Chordata | 1617.0 | 2 | Polystomatidea | 1202.0 | 3 | Polystomatidae | 1202.0 | 3 | Protopolystoma | 1202.0 | 3 | Protopolystoma xenopodis | 1202.0 | 3 |
| scf7180000040864 | 1652555 | 0.4005 | 0 | 60.0398 | Eukaryota | 107321.0 | 0 | Chordata | 107321.0 | 0 | Carnivora | 107321.0 | 0 | Canidae | 107321.0 | 0 | Canis | 63727.0 | 1 | Canis lupus | 63727.0 | 1 |
| scf7180000041561 | 246079 | 0.3714 | 100 | 58.5593 | Eukaryota | 47002.0 | 0 | Chordata | 47002.0 | 0 | Primates | 22881.0 | 3 | Canidae | 14484.0 | 5 | Canis | 9952.0 | 7 | Canis lupus | 9952.0 | 7 |
| scf7180000046389 | 4010 | 0.4172 | 0 | 303.5336 | Eukaryota | 6631.0 | 0 | Chordata | 3379.0 | 3 | Carnivora | 3234.0 | 4 | Canidae | 3234.0 | 4 | Canis | 3234.0 | 4 | Canis lupus | 3234.0 | 4 |
| scf7180000046772 | 20313 | 0.4355 | 0 | 121.526 | Eukaryota | 5638.0 | 0 | Chordata | 4937.0 | 1 | Synbranchiformes | 2545.0 | 2 | Mastacembelidae | 2545.0 | 2 | Mastacembelus | 2545.0 | 2 | Mastacembelus armatus | 2545.0 | 2 |
| scf7180000041108 | 512293 | 0.3466 | 0 | 59.6562 | Eukaryota | 136506.0 | 0 | Chordata | 136506.0 | 0 | Carnivora | 136506.0 | 0 | Canidae | 136506.0 | 0 | Canis | 136506.0 | 0 | Canis lupus | 136506.0 | 0 |
| scf7180000046081 | 166419 | 0.4477 | 0 | 58.5283 | Eukaryota | 33223.0 | 0 | Chordata | 33223.0 | 0 | Carnivora | 33223.0 | 0 | Canidae | 29643.0 | 1 | Canis | 25059.0 | 2 | Canis lupus | 25059.0 | 2 |
| scf7180000045784 | 15354 | 0.4463 | 0 | 268.9123 | Eukaryota | 30885.0 | 0 | Chordata | 18231.0 | 2 | Strigeidida | 8086.0 | 8 | Schistosomatidae | 8086.0 | 8 | Schistosoma | 8086.0 | 8 | Schistosoma mansoni | 8086.0 | 8 |
| scf7180000043667 | 34313 | 0.5106 | 0 | 242.6087 | Eukaryota | 9476.0 | 0 | Chordata | 9476.0 | 0 | Carnivora | 9476.0 | 0 | Canidae | 7817.0 | 1 | Canis | 6581.0 | 2 | Canis lupus | 6581.0 | 2 |
| scf7180000047880 | 67414 | 0.5272 | 0 | 58.4423 | Eukaryota | 27651.0 | 0 | Chordata | 27651.0 | 0 | Carnivora | 27651.0 | 0 | Canidae | 27651.0 | 0 | Canis | 27651.0 | 0 | Canis lupus | 27651.0 | 0 |
| scf7180000050162 | 4495 | 0.4501 | 0 | 346.3881 | Eukaryota | 463.0 | 0 | Chordata | 463.0 | 0 | Columbiformes | 254.0 | 1 | Columbidae | 254.0 | 1 | Streptopelia | 254.0 | 1 | Streptopelia turtur | 254.0 | 1 |
| scf7180000043765 | 277689 | 0.3818 | 0 | 62.7401 | Eukaryota | 156358.0 | 0 | Chordata | 156358.0 | 0 | Carnivora | 92128.0 | 1 | Canidae | 85138.0 | 2 | Canis | 85138.0 | 4 | Canis lupus | 85138.0 | 4 |
| scf7180000051351 | 9990 | 0.4534 | 0 | 293.2857 | Eukaryota | 33198.0 | 0 | Chordata | 33198.0 | 0 | Carnivora | 33198.0 | 0 | Canidae | 33198.0 | 0 | Canis | 33198.0 | 0 | Canis lupus | 33198.0 | 0 |
| scf7180000046989 | 12767 | 0.4422 | 0 | 68.8812 | Eukaryota | 80147.0 | 0 | Chordata | 80147.0 | 0 | Carnivora | 80147.0 | 0 | Canidae | 80147.0 | 0 | Canis | 80147.0 | 0 | Canis lupus | 80147.0 | 0 |
| scf7180000045039 | 240369 | 0.3461 | 0 | 58.995 | Eukaryota | 88034.0 | 0 | Chordata | 88034.0 | 0 | Carnivora | 88034.0 | 0 | Canidae | 88034.0 | 0 | Canis | 88034.0 | 0 | Canis lupus | 88034.0 | 0 |
| scf7180000055539 | 23836 | 0.4474 | 0 | 228.0237 | no-hit | 0.0 | 0 | no-hit | 0.0 | 0 | no-hit | 0.0 | 0 | no-hit | 0.0 | 0 | no-hit | 0.0 | 0 | no-hit | 0.0 | 0 |
| scf7180000048989 | 47490 | 0.4401 | 0 | 72.1668 | no-hit | 0.0 | 0 | no-hit | 0.0 | 0 | no-hit | 0.0 | 0 | no-hit | 0.0 | 0 | no-hit | 0.0 | 0 | no-hit | 0.0 | 0 |
| scf7180000043444 | 12850 | 0.4742 | 0 | 109.2004 | Eukaryota | 95487.0 | 0 | Chordata | 95487.0 | 0 | Carnivora | 95487.0 | 0 | Canidae | 95487.0 | 0 | Canis | 95487.0 | 0 | Canis lupus | 95487.0 | 0 |
| scf7180000052388 | 10907 | 0.4572 | 0 | 888.6706 | Eukaryota | 19134.0 | 0 | Chordata | 19134.0 | 0 | Carnivora | 19134.0 | 0 | Canidae | 19134.0 | 0 | Canis | 19134.0 | 0 | Canis lupus | 19134.0 | 0 |
| scf7180000050638 | 23511 | 0.4603 | 0 | 57.8022 | Eukaryota | 6029.0 | 0 | Chordata | 6029.0 | 0 | Carnivora | 6029.0 | 0 | Canidae | 6029.0 | 0 | Canis | 6029.0 | 0 | Canis lupus | 6029.0 | 0 |
| scf7180000045011 | 213964 | 0.4378 | 0 | 57.8547 | Eukaryota | 35456.0 | 0 | Chordata | 35456.0 | 0 | Carnivora | 35456.0 | 0 | Canidae | 35456.0 | 0 | Canis | 35456.0 | 0 | Canis lupus | 35456.0 | 0 |
| scf7180000056408 | 379614 | 0.4801 | 0 | 58.8931 | Eukaryota | 55974.0 | 0 | Chordata | 55974.0 | 0 | Carnivora | 55974.0 | 0 | Canidae | 55974.0 | 0 | Canis | 55974.0 | 0 | Canis lupus | 55974.0 | 0 |
| scf7180000055579 | 10426 | 0.4387 | 0 | 982.1071 | no-hit | 0.0 | 0 | no-hit | 0.0 | 0 | no-hit | 0.0 | 0 | no-hit | 0.0 | 0 | no-hit | 0.0 | 0 | no-hit | 0.0 | 0 |
| scf7180000047991 | 33203 | 0.4607 | 0 | 137.86 | Eukaryota | 47957.0 | 0 | Chordata | 47957.0 | 0 | Carnivora | 47957.0 | 0 | Canidae | 47957.0 | 0 | Canis | 47957.0 | 0 | Canis lupus | 47957.0 | 0 |
| scf7180000044758 | 207147 | 0.3542 | 0 | 57.6175 | Eukaryota | 66572.0 | 0 | Chordata | 66572.0 | 0 | Carnivora | 66572.0 | 0 | Canidae | 66572.0 | 0 | Canis | 66572.0 | 0 | Canis lupus | 66572.0 | 0 |
| scf7180000042130 | 839932 | 0.4218 | 100 | 59.2959 | Eukaryota | 55638.0 | 0 | Chordata | 55638.0 | 0 | Carnivora | 55638.0 | 0 | Canidae | 12555.0 | 5 | Ursus | 11029.0 | 8 | Vulpes vulpes | 6292.0 | 9 |
| scf7180000041996 | 4379 | 0.422 | 0 | 27.9293 | Eukaryota | 1917.0 | 0 | Chordata | 1917.0 | 0 | Carnivora | 1917.0 | 0 | Canidae | 1917.0 | 0 | Canis | 1734.0 | 1 | Canis lupus | 1734.0 | 1 |
| scf7180000052170 | 3320 | 0.4524 | 0 | 191.5973 | Eukaryota | 459.0 | 0 | Streptophyta | 459.0 | 0 | Fabales | 459.0 | 0 | Fabaceae | 459.0 | 0 | Cicer | 459.0 | 0 | Cicer arietinum | 459.0 | 0 |
| scf7180000049033 | 8221 | 0.4363 | 0 | 223.2 | Viruses | 3201.0 | 1 | Viruses-undef | 3201.0 | 2 | Herpesvirales | 3201.0 | 3 | Herpesviridae | 3201.0 | 3 | Roseolovirus | 3201.0 | 3 | Human betaherpesvirus 7 | 3201.0 | 3 |
| scf7180000051372 | 2488 | 0.4473 | 0 | 215.4538 | Eukaryota | 1055.0 | 0 | Echinodermata | 462.0 | 2 | Forcipulatida | 462.0 | 2 | Asteriidae | 462.0 | 2 | Asterias | 462.0 | 2 | Asterias rubens | 462.0 | 2 |
| scf7180000048797 | 5538 | 0.4442 | 0 | 110.4446 | Eukaryota | 3952.0 | 0 | Platyhelminthes | 1554.0 | 4 | Polystomatidea | 1554.0 | 6 | Polystomatidae | 1554.0 | 6 | Protopolystoma | 1554.0 | 6 | Protopolystoma xenopodis | 1554.0 | 6 |
| scf7180000053802 | 4723 | 0.4408 | 0 | 38.6222 | Eukaryota | 548.0 | 0 | Chordata | 548.0 | 0 | Carnivora | 548.0 | 0 | Canidae | 548.0 | 0 | Canis | 548.0 | 0 | Canis lupus | 548.0 | 0 |
| scf7180000046639 | 123137 | 0.3775 | 0 | 58.505 | Eukaryota | 95051.0 | 0 | Chordata | 95051.0 | 0 | Carnivora | 95051.0 | 0 | Canidae | 95051.0 | 0 | Canis | 95051.0 | 0 | Canis lupus | 95051.0 | 0 |
| scf7180000051874 | 7388 | 0.4518 | 0 | 571.3138 | Eukaryota | 405.0 | 0 | Platyhelminthes | 405.0 | 0 | Polystomatidea | 405.0 | 0 | Polystomatidae | 405.0 | 0 | Protopolystoma | 405.0 | 0 | Protopolystoma xenopodis | 405.0 | 0 |
| scf7180000045311 | 2444 | 0.4374 | 0 | 169.4505 | Eukaryota | 15234.0 | 0 | Chordata | 13599.0 | 1 | Gadiformes | 3454.0 | 6 | Gadidae | 3454.0 | 6 | Gadus | 3454.0 | 6 | Gadus morhua | 3454.0 | 6 |
| scf7180000042174 | 769176 | 0.4199 | 0 | 59.6445 | Eukaryota | 248892.0 | 0 | Chordata | 248892.0 | 0 | Carnivora | 216354.0 | 1 | Canidae | 216354.0 | 1 | Canis | 216354.0 | 2 | Canis lupus | 216354.0 | 2 |
| scf7180000041894 | 302523 | 0.3676 | 0 | 58.936 | Eukaryota | 14241.0 | 0 | Chordata | 14241.0 | 0 | Primates | 5615.0 | 3 | Hominidae | 5615.0 | 3 | Homo | 5615.0 | 5 | Homo sapiens | 5615.0 | 5 |
| scf7180000042500 | 90880 | 0.4001 | 0 | 63.7145 | Eukaryota | 92396.0 | 0 | Chordata | 92396.0 | 0 | Carnivora | 92396.0 | 0 | Canidae | 92396.0 | 0 | Canis | 92396.0 | 0 | Canis lupus | 92396.0 | 0 |
| scf7180000053330 | 2876 | 0.4392 | 0 | 212.0831 | no-hit | 0.0 | 0 | no-hit | 0.0 | 0 | no-hit | 0.0 | 0 | no-hit | 0.0 | 0 | no-hit | 0.0 | 0 | no-hit | 0.0 | 0 |
| scf7180000042397 | 372836 | 0.3484 | 0 | 68.0089 | Eukaryota | 14236.0 | 0 | Chordata | 14236.0 | 0 | Carnivora | 14236.0 | 0 | Canidae | 14236.0 | 0 | Canis | 14236.0 | 0 | Canis lupus | 14236.0 | 0 |

| | | | | | | | | | | | | | | | | | | | | | |
|---|---|---|---|---|---|---|---|---|---|---|---|---|---|---|---|---|---|---|---|---|---|
| scf7180000042399 | 1126582 | 0.4183 | 0 | 58.4248 | Eukaryota | 69244.0 | 0 | Chordata | 69244.0 | 0 | Carnivora | 69244.0 | 0 | Canidae | 69244.0 | 0 | Canis | 69244.0 | 0 | Canis lupus | 69244.0 | 0 |
| scf7180000044484 | 193033 | 0.4085 | 0 | 58.1528 | Eukaryota | 28544.0 | 0 | Chordata | 28544.0 | 0 | Carnivora | 28544.0 | 0 | Canidae | 20995.0 | 2 | Canis | 20995.0 | 2 | Canis lupus | 20995.0 | 2 |
| scf7180000046744 | 56767 | 0.4468 | 0 | 220.2871 | Eukaryota | 3226.0 | 0 | Chordata | 3226.0 | 0 | Carnivora | 3226.0 | 0 | Canidae | 3226.0 | 0 | Canis | 3226.0 | 0 | Canis lupus | 3226.0 | 0 |
| scf7180000044783 | 527954 | 0.4242 | 100 | 58.7642 | Eukaryota | 64985.0 | 0 | Chordata | 64985.0 | 0 | Carnivora | 64985.0 | 0 | Canidae | 64985.0 | 0 | Canis | 64985.0 | 0 | Canis lupus | 64985.0 | 0 |
| scf7180000056373 | 174563 | 0.3915 | 0 | 59.0832 | Eukaryota | 25565.0 | 0 | Chordata | 25565.0 | 0 | Carnivora | 22182.0 | 2 | Canidae | 18011.0 | 4 | Canis | 18011.0 | 4 | Canis lupus | 18011.0 | 4 |
| scf7180000046371 | 124324 | 0.3586 | 0 | 60.2614 | Eukaryota | 11818.0 | 0 | Chordata | 11818.0 | 0 | Carnivora | 11818.0 | 0 | Canidae | 10672.0 | 1 | Canis | 8362.0 | 2 | Canis lupus | 8362.0 | 2 |
| scf7180000046408 | 27967 | 0.4363 | 0 | 160.744 | Eukaryota | 6648.0 | 0 | Platyhelminthes | 2361.0 | 3 | Polystomatidea | 2361.0 | 4 | Polystomatidae | 2361.0 | 4 | Protopolystoma | 2361.0 | 5 | Protopolystoma xenopodis | 2361.0 | 5 |
| scf7180000042888 | 103668 | 0.5161 | 0 | 55.8073 | Eukaryota | 8116.0 | 0 | Chordata | 8116.0 | 0 | Carnivora | 8116.0 | 0 | Canidae | 8116.0 | 0 | Canis | 8116.0 | 0 | Canis lupus | 8116.0 | 0 |
| scf7180000042329 | 3998 | 0.4262 | 0 | 44.6172 | Eukaryota | 1119.0 | 0 | Chordata | 1119.0 | 0 | Carnivora | 1119.0 | 0 | Canidae | 566.0 | 2 | Canis | 566.0 | 2 | Canis lupus | 566.0 | 2 |
| scf7180000040920 | 796774 | 0.4027 | 0 | 59.9919 | Eukaryota | 88727.0 | 0 | Chordata | 88727.0 | 0 | Carnivora | 88727.0 | 0 | Canidae | 88727.0 | 0 | Canis | 88727.0 | 0 | Canis lupus | 88727.0 | 0 |
| scf7180000050865 | 34416 | 0.3993 | 0 | 52.3748 | Eukaryota | 6693.0 | 0 | Chordata | 6693.0 | 0 | Artiodactyla | 5307.0 | 2 | Delphinidae | 3987.0 | 3 | Lagenorhynchus | 2001.0 | 4 | Lagenorhynchus obliquidens | 2001.0 | 4 |
| scf7180000050052 | 7371 | 0.4359 | 0 | 60.1033 | Eukaryota | 233.0 | 0 | Platyhelminthes | 233.0 | 0 | Polystomatidea | 233.0 | 0 | Polystomatidae | 233.0 | 0 | Protopolystoma | 233.0 | 0 | Protopolystoma xenopodis | 233.0 | 0 |
| scf7180000050216 | 22465 | 0.3925 | 0 | 59.5032 | Eukaryota | 5088.0 | 1 | Chordata | 5088.0 | 1 | Primates | 3271.0 | 3 | Hominidae | 3271.0 | 3 | Homo | 3271.0 | 3 | Homo sapiens | 3271.0 | 3 |
| scf7180000043681 | 392127 | 0.4428 | 0 | 60.3172 | Eukaryota | 66248.0 | 0 | Chordata | 66248.0 | 0 | Carnivora | 66248.0 | 0 | Canidae | 66248.0 | 0 | Canis | 66248.0 | 0 | Canis lupus | 66248.0 | 0 |
| scf7180000053121 | 4180 | 0.51 | 0 | 86.4384 | Eukaryota | 6843.0 | 0 | Chordata | 6843.0 | 0 | Carnivora | 6843.0 | 0 | Canidae | 6843.0 | 0 | Canis | 5186.0 | 1 | Canis lupus | 4336.0 | 2 |
| scf7180000048523 | 7885 | 0.4419 | 0 | 126.8253 | no-hit | 0.0 | 0 | no-hit | 0.0 | 0 | no-hit | 0.0 | 0 | no-hit | 0.0 | 0 | no-hit | 0.0 | 0 | no-hit | 0.0 | 0 |
| scf7180000050606 | 14872 | 0.4635 | 0 | 57.0233 | Eukaryota | 3974.0 | 1 | Chordata | 3974.0 | 1 | Carnivora | 3092.0 | 3 | Canidae | 1386.0 | 5 | Vulpes | 1386.0 | 6 | Vulpes vulpes | 1386.0 | 6 |
| scf7180000047547 | 18014 | 0.5576 | 0 | 403.335 | Eukaryota | 6002.0 | 0 | Chordata | 6002.0 | 0 | Carnivora | 6002.0 | 0 | Canidae | 6002.0 | 0 | Canis | 5333.0 | 1 | Canis lupus | 5333.0 | 1 |
| scf7180000050877 | 4269 | 0.442 | 0 | 653.7452 | Eukaryota | 16214.0 | 1 | Chordata | 16214.0 | 1 | Tetraodontiformes | 6040.0 | 4 | Tetraodontidae | 6040.0 | 4 | Takifugu | 6040.0 | 4 | Takifugu rubripes | 6040.0 | 4 |
| scf7180000044154 | 160087 | 0.5013 | 0 | 57.7248 | Eukaryota | 10591.0 | 0 | Chordata | 10591.0 | 0 | Carnivora | 4966.0 | 3 | Hominidae | 4043.0 | 4 | Homo | 3006.0 | 7 | Homo sapiens | 3006.0 | 7 |
| scf7180000053285 | 2184 | 0.4478 | 0 | 318.1832 | no-hit | 0.0 | 0 | no-hit | 0.0 | 0 | no-hit | 0.0 | 0 | no-hit | 0.0 | 0 | no-hit | 0.0 | 0 | no-hit | 0.0 | 0 |
| scf7180000051620 | 13841 | 0.5016 | 0 | 333.258 | Eukaryota | 8431.0 | 0 | Chordata | 8431.0 | 0 | Carnivora | 8431.0 | 0 | Canidae | 8431.0 | 0 | Canis | 6734.0 | 1 | Canis lupus | 5542.0 | 2 |
| scf7180000047250 | 42867 | 0.4389 | 0 | 54.014 | Eukaryota | 44887.0 | 0 | Chordata | 44887.0 | 0 | Carnivora | 44887.0 | 0 | Canidae | 44887.0 | 0 | Canis | 44887.0 | 0 | Canis lupus | 44887.0 | 0 |
| scf7180000053495 | 2758 | 0.5112 | 0 | 330.4146 | Eukaryota | 5318.0 | 0 | Chordata | 5318.0 | 0 | Carnivora | 5318.0 | 0 | Canidae | 5318.0 | 0 | Canis | 2879.0 | 1 | Nyctereutes procyonoides | 2439.0 | 2 |
| scf7180000046833 | 13549 | 0.449 | 0 | 310.7002 | Eukaryota | 10061.0 | 0 | Chordata | 10061.0 | 0 | Tetraodontiformes | 3833.0 | 4 | Tetraodontidae | 3833.0 | 4 | Takifugu | 3833.0 | 4 | Takifugu rubripes | 3833.0 | 4 |
| scf7180000047137 | 7493 | 0.4478 | 0 | 123.6843 | no-hit | 0.0 | 0 | no-hit | 0.0 | 0 | no-hit | 0.0 | 0 | no-hit | 0.0 | 0 | no-hit | 0.0 | 0 | no-hit | 0.0 | 0 |
| scf7180000047842 | 55101 | 0.3414 | 0 | 54.8384 | Eukaryota | 5800.0 | 0 | Chordata | 5800.0 | 0 | Carnivora | 3986.0 | 1 | Canidae | 2908.0 | 2 | Canis | 2908.0 | 3 | Canis lupus | 2908.0 | 3 |
| scf7180000043513 | 396093 | 0.4338 | 0 | 60.3938 | Eukaryota | 30010.0 | 0 | Chordata | 30010.0 | 0 | Carnivora | 16856.0 | 2 | Felidae | 15105.0 | 4 | Felis | 15105.0 | 4 | Felis catus | 15105.0 | 4 |
| scf7180000045796 | 153524 | 0.5354 | 0 | 58.239 | Eukaryota | 40738.0 | 0 | Chordata | 40738.0 | 0 | Carnivora | 40738.0 | 0 | Canidae | 31268.0 | 2 | Vulpes | 19534.0 | 4 | Vulpes vulpes | 19534.0 | 4 |
| scf7180000043138 | 446853 | 0.4339 | 0 | 118.5732 | Eukaryota | 94672.0 | 0 | Chordata | 94672.0 | 0 | Carnivora | 94672.0 | 0 | Canidae | 94672.0 | 0 | Canis | 94672.0 | 0 | Canis lupus | 94672.0 | 0 |
| scf7180000052457 | 10733 | 0.4273 | 0 | 143.383 | Eukaryota | 1525.0 | 0 | Platyhelminthes | 1306.0 | 1 | Polystomatidea | 1306.0 | 1 | Polystomatidae | 1306.0 | 1 | Protopolystoma | 1306.0 | 1 | Protopolystoma xenopodis | 1306.0 | 1 |
| scf7180000045552 | 15555 | 0.4431 | 0 | 203.0303 | Eukaryota | 20895.0 | 0 | Chordata | 12659.0 | 2 | Strigeidida | 7713.0 | 6 | Schistosomatidae | 7713.0 | 6 | Schistosoma | 7713.0 | 6 | Schistosoma mansoni | 7713.0 | 6 |
| scf7180000053788 | 2792 | 0.447 | 0 | 196.1421 | Eukaryota | 794.0 | 0 | Chordata | 375.0 | 2 | Salmoniformes | 375.0 | 2 | Salmonidae | 375.0 | 2 | Coregonus | 375.0 | 2 | Coregonus sp. 'balchen' | 375.0 | 2 |
| scf7180000049678 | 3280 | 0.4427 | 0 | 123.0482 | Eukaryota | 620.0 | 0 | Platyhelminthes | 620.0 | 0 | Polystomatidea | 472.0 | 1 | Polystomatidae | 472.0 | 1 | Protopolystoma | 472.0 | 1 | Protopolystoma xenopodis | 472.0 | 1 |
| scf7180000040524 | 1421339 | 0.3911 | 0 | 59.6511 | Eukaryota | 74273.0 | 0 | Chordata | 74273.0 | 0 | Carnivora | 74273.0 | 0 | Canidae | 74273.0 | 0 | Canis | 74273.0 | 0 | Canis lupus | 74273.0 | 0 |
| scf7180000049922 | 5003 | 0.4501 | 0 | 308.5895 | no-hit | 0.0 | 0 | no-hit | 0.0 | 0 | no-hit | 0.0 | 0 | no-hit | 0.0 | 0 | no-hit | 0.0 | 0 | no-hit | 0.0 | 0 |
| scf7180000055111 | 28267 | 0.4331 | 0 | 692.0352 | Eukaryota | 37379.0 | 0 | Chordata | 37379.0 | 0 | Carnivora | 37379.0 | 0 | Canidae | 37379.0 | 0 | Canis | 37379.0 | 0 | Canis lupus | 37379.0 | 0 |
| scf7180000053799 | 2477 | 0.4457 | 0 | 330.0787 | no-hit | 0.0 | 0 | no-hit | 0.0 | 0 | no-hit | 0.0 | 0 | no-hit | 0.0 | 0 | no-hit | 0.0 | 0 | no-hit | 0.0 | 0 |
| scf7180000043087 | 496714 | 0.4377 | 0 | 58.9099 | Eukaryota | 68375.0 | 0 | Chordata | 68375.0 | 0 | Carnivora | 68375.0 | 0 | Canidae | 46794.0 | 1 | Canis | 46794.0 | 1 | Canis lupus | 46794.0 | 2 |
| scf7180000047626 | 34673 | 0.3922 | 0 | 57.1837 | Eukaryota | 35739.0 | 0 | Chordata | 35739.0 | 0 | Carnivora | 35739.0 | 0 | Canidae | 35739.0 | 0 | Canis | 35739.0 | 0 | Canis lupus | 35739.0 | 0 |
| scf7180000043446 | 503255 | 0.3481 | 0 | 59.9175 | Eukaryota | 55703.0 | 0 | Chordata | 55703.0 | 0 | Carnivora | 55703.0 | 0 | Canidae | 55703.0 | 0 | Canis | 55703.0 | 0 | Canis lupus | 55703.0 | 0 |
| scf7180000048996 | 19044 | 0.4413 | 0 | 99.6109 | Eukaryota | 14542.0 | 0 | Chordata | 9345.0 | 2 | Gadiformes | 4209.0 | 6 | Gadidae | 4209.0 | 6 | Gadus | 4209.0 | 6 | Gadus morhua | 4209.0 | 6 |
| scf7180000056322 | 822810 | 0.4064 | 0 | 59.6418 | Eukaryota | 115465.0 | 0 | Chordata | 115465.0 | 0 | Carnivora | 115465.0 | 0 | Canidae | 115465.0 | 0 | Canis | 115465.0 | 0 | Canis lupus | 115465.0 | 0 |
| scf7180000045312 | 44686 | 0.3809 | 0 | 58.7101 | Eukaryota | 72156.0 | 0 | Chordata | 72156.0 | 0 | Carnivora | 72156.0 | 0 | Canidae | 72156.0 | 0 | Canis | 72156.0 | 0 | Canis lupus | 72156.0 | 0 |
| scf7180000043722 | 292735 | 0.4252 | 0 | 58.5376 | Eukaryota | 69676.0 | 0 | Chordata | 69676.0 | 0 | Carnivora | 69676.0 | 0 | Canidae | 69676.0 | 0 | Canis | 69676.0 | 0 | Canis lupus | 69676.0 | 0 |
| scf7180000047411 | 26867 | 0.4023 | 0 | 80.0257 | Eukaryota | 5124.0 | 1 | Chordata | 5124.0 | 1 | Artiodactyla | 1848.0 | 3 | Bovidae | 1848.0 | 3 | Canis | 1652.0 | 4 | Canis lupus | 1652.0 | 4 |
| scf7180000046069 | 57796 | 0.6416 | 0 | 52.6775 | Eukaryota | 95440.0 | 0 | Chordata | 95440.0 | 0 | Carnivora | 95440.0 | 0 | Canidae | 95440.0 | 0 | Canis | 68667.0 | 1 | Canis lupus | 68667.0 | 1 |
| scf7180000049121 | 13675 | 0.4349 | 0 | 153.2479 | Eukaryota | 21076.0 | 0 | Chordata | 12992.0 | 1 | Strigeidida | 7568.0 | 6 | Schistosomatidae | 7568.0 | 6 | Schistosoma | 7568.0 | 6 | Schistosoma mansoni | 7568.0 | 6 |
| scf7180000049022 | 4181 | 0.4499 | 0 | 780.6851 | Eukaryota | 2454.0 | 0 | Platyhelminthes | 1037.0 | 2 | Polystomatidea | 1037.0 | 2 | Polystomatidae | 1037.0 | 2 | Protopolystoma | 1037.0 | 2 | Protopolystoma xenopodis | 1037.0 | 2 |
| scf7180000056153 | 10485 | 0.4417 | 0 | 337.0177 | Eukaryota | 3313.0 | 0 | Chordata | 1951.0 | 1 | Kurtiformes | 1705.0 | 2 | Apogonidae | 1705.0 | 2 | Sphaeramia | 1705.0 | 2 | Sphaeramia orbicularis | 1705.0 | 2 |
| scf7180000044574 | 65974 | 0.4 | 0 | 51.8018 | Eukaryota | 22265.0 | 0 | Chordata | 22265.0 | 0 | Carnivora | 17469.0 | 1 | Canidae | 17469.0 | 1 | Canis | 17469.0 | 1 | Canis lupus | 17469.0 | 1 |
| scf7180000043901 | 408014 | 0.416 | 0 | 59.9058 | Eukaryota | 47789.0 | 0 | Chordata | 47789.0 | 0 | Carnivora | 47789.0 | 0 | Canidae | 47789.0 | 0 | Canis | 39799.0 | 1 | Canis lupus | 39799.0 | 1 |
| scf7180000040587 | 2094970 | 0.4031 | 0 | 60.1644 | Eukaryota | 92516.0 | 0 | Chordata | 92516.0 | 0 | Carnivora | 92516.0 | 0 | Canidae | 92516.0 | 0 | Canis | 92516.0 | 0 | Canis lupus | 92516.0 | 0 |
| scf7180000046686 | 8380 | 0.4427 | 0 | 184.2952 | no-hit | 0.0 | 0 | no-hit | 0.0 | 0 | no-hit | 0.0 | 0 | no-hit | 0.0 | 0 | no-hit | 0.0 | 0 | no-hit | 0.0 | 0 |
| scf7180000042692 | 225709 | 0.4123 | 0 | 56.4865 | Eukaryota | 70074.0 | 0 | Chordata | 70074.0 | 0 | Carnivora | 70074.0 | 0 | Canidae | 70074.0 | 0 | Canis | 70074.0 | 0 | Canis lupus | 70074.0 | 0 |
| scf7180000053806 | 3204 | 0.4519 | 0 | 359.5865 | no-hit | 0.0 | 0 | no-hit | 0.0 | 0 | no-hit | 0.0 | 0 | no-hit | 0.0 | 0 | no-hit | 0.0 | 0 | no-hit | 0.0 | 0 |
| scf7180000045318 | 6853 | 0.4394 | 0 | 172.2193 | Eukaryota | 3042.0 | 0 | Platyhelminthes | 1872.0 | 1 | Polystomatidea | 1872.0 | 2 | Polystomatidae | 1872.0 | 2 | Protopolystoma | 1872.0 | 2 | Protopolystoma xenopodis | 1872.0 | 2 |
| scf7180000045028 | 210722 | 0.486 | 0 | 59.826 | Eukaryota | 20304.0 | 0 | Chordata | 20304.0 | 0 | Carnivora | 20304.0 | 0 | Canidae | 20304.0 | 0 | Canis | 18505.0 | 1 | Canis lupus | 18505.0 | 1 |
| scf7180000052798 | 3734 | 0.4547 | 0 | 454.3362 | Eukaryota | 2914.0 | 0 | Chordata | 2914.0 | 0 | Carnivora | 2914.0 | 0 | Canidae | 2914.0 | 0 | Canis | 2914.0 | 0 | Canis lupus | 2914.0 | 0 |
| scf7180000046912 | 91990 | 0.4395 | 0 | 124.4882 | Eukaryota | 91852.0 | 0 | Chordata | 91852.0 | 0 | Carnivora | 91852.0 | 0 | Canidae | 91852.0 | 0 | Canis | 91852.0 | 0 | Canis lupus | 91852.0 | 0 |
| scf7180000048576 | 3567 | 0.4522 | 0 | 79.3348 | Eukaryota | 319.0 | 0 | Platyhelminthes | 165.0 | 1 | Polystomatidea | 165.0 | 1 | Polystomatidae | 165.0 | 1 | Protopolystoma | 165.0 | 1 | Protopolystoma xenopodis | 165.0 | 1 |
| scf7180000054173 | 12156 | 0.4457 | 0 | 144.2052 | no-hit | 0.0 | 0 | no-hit | 0.0 | 0 | no-hit | 0.0 | 0 | no-hit | 0.0 | 0 | no-hit | 0.0 | 0 | no-hit | 0.0 | 0 |
| scf7180000050490 | 5447 | 0.4346 | 0 | 75.1704 | Eukaryota | 3569.0 | 0 | Platyhelminthes | 2818.0 | 1 | Polystomatidea | 2818.0 | 2 | Polystomatidae | 2818.0 | 2 | Protopolystoma | 2818.0 | 2 | Protopolystoma xenopodis | 2818.0 | 2 |
| scf7180000041063 | 103363 | 0.372 | 0 | 59.6047 | Eukaryota | 47907.0 | 0 | Chordata | 47907.0 | 0 | Carnivora | 47907.0 | 0 | Canidae | 47907.0 | 0 | Canis | 47907.0 | 0 | Canis lupus | 47907.0 | 0 |
| scf7180000046627 | 19391 | 0.4385 | 0 | 96.6743 | Eukaryota | 904.0 | 0 | Chordata | 904.0 | 0 | Tetraodontiformes | 704.0 | 1 | Tetraodontidae | 704.0 | 1 | Takifugu | 704.0 | 1 | Takifugu rubripes | 704.0 | 1 |
| scf7180000047896 | 13558 | 0.4611 | 0 | 115.7623 | Eukaryota | 77383.0 | 0 | Chordata | 77383.0 | 0 | Carnivora | 77383.0 | 0 | Canidae | 77383.0 | 0 | Canis | 77383.0 | 0 | Canis lupus | 77383.0 | 0 |
| scf7180000050936 | 4132 | 0.4446 | 0 | 819.3844 | no-hit | 0.0 | 0 | no-hit | 0.0 | 0 | no-hit | 0.0 | 0 | no-hit | 0.0 | 0 | no-hit | 0.0 | 0 | no-hit | 0.0 | 0 |
| scf7180000049277 | 11798 | 0.4466 | 0 | 64.062 | Eukaryota | 17831.0 | 0 | Chordata | 14607.0 | 2 | Synbranchiformes | 4496.0 | 7 | Mastacembelidae | 4496.0 | 7 | Mastacembelus | 4496.0 | 7 | Mastacembelus armatus | 4496.0 | 7 |
| scf7180000053860 | 4356 | 0.4477 | 0 | 332.7397 | Eukaryota | 3897.0 | 0 | Platyhelminthes | 1881.0 | 3 | Polystomatidea | 1881.0 | 3 | Polystomatidae | 1881.0 | 3 | Protopolystoma | 1881.0 | 3 | Protopolystoma xenopodis | 1881.0 | 3 |
| scf7180000041406 | 750325 | 0.4251 | 0 | 60.6042 | Eukaryota | 94919.0 | 0 | Chordata | 94919.0 | 0 | Carnivora | 94919.0 | 0 | Canidae | 94919.0 | 0 | Canis | 94919.0 | 0 | Canis lupus | 94919.0 | 0 |
| scf7180000046656 | 13264 | 0.4546 | 0 | 78.6795 | no-hit | 0.0 | 0 | no-hit | 0.0 | 0 | no-hit | 0.0 | 0 | no-hit | 0.0 | 0 | no-hit | 0.0 | 0 | no-hit | 0.0 | 0 |
| scf7180000042615 | 2751 | 0.3526 | 0 | 26.7218 | Eukaryota | 3547.0 | 0 | Chordata | 3547.0 | 0 | Carnivora | 3140.0 | 1 | Canidae | 3140.0 | 1 | Canis | 2100.0 | 2 | Canis lupus | 2100.0 | 2 |
| scf7180000052236 | 8086 | 0.4519 | 0 | 577.2843 | Eukaryota | 5554.0 | 0 | Chordata | 2425.0 | 3 | Polystomatidea | 2044.0 | 5 | Polystomatidae | 2044.0 | 5 | Protopolystoma | 2044.0 | 5 | Protopolystoma xenopodis | 2044.0 | 5 |
| scf7180000049911 | 5467 | 0.4478 | 0 | 615.7271 | no-hit | 0.0 | 0 | no-hit | 0.0 | 0 | no-hit | 0.0 | 0 | no-hit | 0.0 | 0 | no-hit | 0.0 | 0 | no-hit | 0.0 | 0 |
| scf7180000046261 | 7939 | 0.4381 | 0 | 106.1489 | Eukaryota | 405.0 | 0 | Chordata | 405.0 | 0 | Tetraodontiformes | 405.0 | 0 | Tetraodontidae | 405.0 | 0 | Takifugu | 405.0 | 0 | Takifugu rubripes | 405.0 | 0 |
| scf7180000049159 | 18268 | 0.4451 | 0 | 265.3603 | Eukaryota | 30370.0 | 0 | Chordata | 19531.0 | 1 | Cyclophyllidea | 10839.0 | 7 | Hymenolepididae | 10839.0 | 7 | Hymenolepis | 10839.0 | 7 | Hymenolepis microstoma | 10839.0 | 7 |
| scf7180000040582 | 2306009 | 0.3745 | 0 | 59.4056 | Eukaryota | 73136.0 | 0 | Chordata | 73136.0 | 0 | Carnivora | 73136.0 | 0 | Canidae | 73136.0 | 0 | Canis | 41150.0 | 1 | Canis lupus | 41150.0 | 1 |
| scf7180000041767 | 1128348 | 0.3904 | 0 | 59.7213 | Eukaryota | 93445.0 | 0 | Chordata | 93445.0 | 0 | Carnivora | 93445.0 | 0 | Canidae | 93445.0 | 0 | Canis | 93445.0 | 0 | Canis lupus | 93445.0 | 0 |
| scf7180000043535 | 38016 | 0.4362 | 0 | 179.2798 | Eukaryota | 25132.0 | 0 | Chordata | 12251.0 | 2 | Strigeidida | 7675.0 | 5 | Schistosomatidae | 7675.0 | 5 | Schistosoma | 7675.0 | 5 | Schistosoma mansoni | 7675.0 | 5 |
| scf7180000051464 | 7211 | 0.4547 | 0 | 824.9138 | no-hit | 0.0 | 0 | no-hit | 0.0 | 0 | no-hit | 0.0 | 0 | no-hit | 0.0 | 0 | no-hit | 0.0 | 0 | no-hit | 0.0 | 0 |
| scf7180000056613 | 21218 | 0.4377 | 0 | 148.4192 | Eukaryota | 2397.0 | 0 | Platyhelminthes | 1765.0 | 2 | Polystomatidea | 1765.0 | 3 | Polystomatidae | 1765.0 | 3 | Protopolystoma | 1765.0 | 3 | Protopolystoma xenopodis | 1765.0 | 3 |
| scf7180000049125 | 31495 | 0.3855 | 0 | 30.3223 | Eukaryota | 54410.0 | 0 | Chordata | 54410.0 | 0 | Carnivora | 54410.0 | 0 | Canidae | 54410.0 | 0 | Canis | 54410.0 | 0 | Canis lupus | 54410.0 | 0 |
| scf7180000053340 | 7069 | 0.4356 | 0 | 40.8972 | Eukaryota | 6037.0 | 0 | Platyhelminthes | 2363.0 | 3 | Polystomatidea | 2363.0 | 5 | Polystomatidae | 2363.0 | 5 | Protopolystoma | 2363.0 | 5 | Protopolystoma xenopodis | 2363.0 | 5 |
| scf7180000045769 | 118289 | 0.4128 | 0 | 59.176 | Eukaryota | 92639.0 | 0 | Chordata | 92639.0 | 0 | Carnivora | 92639.0 | 0 | Canidae | 92639.0 | 0 | Canis | 92639.0 | 0 | Canis lupus | 92639.0 | 0 |
| scf7180000045882 | 163740 | 0.4644 | 0 | 58.7191 | Eukaryota | 25068.0 | 0 | Chordata | 25068.0 | 0 | Carnivora | 25068.0 | 0 | Canidae | 25068.0 | 0 | Canis | 19544.0 | 1 | Canis lupus | 19544.0 | 1 |
| scf7180000045645 | 179189 | 0.461 | 0 | 60.8188 | Eukaryota | 67901.0 | 0 | Chordata | 67901.0 | 0 | Carnivora | 67901.0 | 0 | Canidae | 22044.0 | 3 | Canis | 14744.0 | 7 | Canis lupus | 14744.0 | 7 |
| scf7180000045615 | 23506 | 0.4181 | 0 | 55.0912 | Eukaryota | 20958.0 | 0 | Chordata | 20958.0 | 0 | Carnivora | 20958.0 | 0 | Canidae | 15894.0 | 1 | Canis | 15894.0 | 2 | Canis lupus | 15894.0 | 2 |
| scf7180000049412 | 32621 | 0.4456 | 0 | 117.2461 | Eukaryota | 75467.0 | 0 | Chordata | 48702.0 | 1 | Gadiformes | 30820.0 | 5 | Gadidae | 30820.0 | 6 | Gadus | 30820.0 | 6 | Gadus morhua | 30820.0 | 6 |
| scf7180000041381 | 702085 | 0.3403 | 0 | 58.2977 | Eukaryota | 61242.0 | 0 | Chordata | 61242.0 | 0 | Carnivora | 61242.0 | 0 | Canidae | 61242.0 | 0 | Canis | 61242.0 | 0 | Canis lupus | 61242.0 | 0 |
| scf7180000049713 | 15350 | 0.4484 | 0 | 120.6242 | no-hit | 0.0 | 0 | no-hit | 0.0 | 0 | no-hit | 0.0 | 0 | no-hit | 0.0 | 0 | no-hit | 0.0 | 0 | no-hit | 0.0 | 0 |
| scf7180000048258 | 74957 | 0.5786 | 0 | 57.0482 | Eukaryota | 10043.0 | 0 | Chordata | 10043.0 | 0 | Carnivora | 10043.0 | 0 | Canidae | 10043.0 | 0 | Canis | 10043.0 | 0 | Canis lupus | 10043.0 | 0 |
| scf7180000041430 | 1155779 | 0.353 | 0 | 59.8345 | Eukaryota | 50244.0 | 0 | Chordata | 50244.0 | 0 | Carnivora | 50244.0 | 0 | Canidae | 50244.0 | 0 | Canis | 50244.0 | 0 | Canis lupus | 50244.0 | 0 |
| scf7180000047600 | 7342 | 0.3405 | 0 | 47.7232 | Eukaryota | 30376.0 | 0 | Chordata | 30376.0 | 0 | Carnivora | 30376.0 | 0 | Canidae | 30376.0 | 0 | Canis | 30376.0 | 0 | Canis lupus | 30376.0 | 0 |
| scf7180000053692 | 2017 | 0.4462 | 0 | 478.378 | no-hit | 0.0 | 0 | no-hit | 0.0 | 0 | no-hit | 0.0 | 0 | no-hit | 0.0 | 0 | no-hit | 0.0 | 0 | no-hit | 0.0 | 0 |
| scf7180000044503 | 18388 | 0.3428 | 0 | 33.9566 | Eukaryota | 65966.0 | 0 | Chordata | 65966.0 | 0 | Carnivora | 65966.0 | 0 | Canidae | 65966.0 | 0 | Canis | 65966.0 | 0 | Canis lupus | 65966.0 | 0 |
| scf7180000040942 | 764068 | 0.3953 | 0 | 59.7879 | Eukaryota | 60286.0 | 0 | Chordata | 60286.0 | 0 | Carnivora | 60286.0 | 0 | Canidae | 60286.0 | 0 | Canis | 60286.0 | 0 | Canis lupus | 60286.0 | 0 |
| scf7180000043273 | 334820 | 0.3855 | 0 | 58.8168 | Eukaryota | 25943.0 | 0 | Chordata | 25943.0 | 0 | Carnivora | 25943.0 | 0 | Canidae | 23169.0 | 1 | Canis | 20301.0 | 2 | Canis lupus | 20301.0 | 2 |
| scf7180000047242 | 133286 | 0.472 | 0 | 59.3713 | Eukaryota | 24069.0 | 0 | Chordata | 24069.0 | 0 | Carnivora | 24069.0 | 0 | Canidae | 24069.0 | 0 | Canis | 15676.0 | 2 | Canis lupus | 15676.0 | 3 |
| scf7180000056309 | 742192 | 0.3841 | 0 | 61.0292 | Eukaryota | 93113.0 | 0 | Chordata | 93113.0 | 0 | Carnivora | 93113.0 | 0 | Canidae | 93113.0 | 0 | Canis | 93113.0 | 0 | Canis lupus | 93113.0 | 0 |
| scf7180000056561 | 118868 | 0.3943 | 0 | 59.7272 | Eukaryota | 6215.0 | 0 | Chordata | 6215.0 | 0 | Primates | 4841.0 | 1 | Hominidae | 3948.0 | 2 | Homo | 2762.0 | 3 | Homo sapiens | 2762.0 | 3 |
| scf7180000056335 | 459823 | 0.4112 | 0 | 59.7823 | Eukaryota | 88486.0 | 0 | Chordata | 88486.0 | 0 | Carnivora | 88486.0 | 0 | Canidae | 88486.0 | 0 | Canis | 88486.0 | 0 | Canis lupus | 88486.0 | 0 |
| scf7180000049361 | 15722 | 0.501 | 0 | 163.0707 | Eukaryota | 8215.0 | 0 | Chordata | 8215.0 | 0 | Carnivora | 8215.0 | 0 | Canidae | 8215.0 | 0 | Canis | 6694.0 | 1 | Canis lupus | 5707.0 | 2 |
| scf7180000054175 | 32304 | 0.4425 | 0 | 137.9105 | Eukaryota | 969.0 | 0 | Chordata | 664.0 | 1 | Salmoniformes | 372.0 | 2 | Salmonidae | 372.0 | 2 | Protopolystoma | 305.0 | 3 | Protopolystoma xenopodis | 305.0 | 3 |
| scf7180000042355 | 657183 | 0.4473 | 0 | 58.4833 | Eukaryota | 58943.0 | 0 | Chordata | 58943.0 | 0 | Carnivora | 58943.0 | 0 | Canidae | 58943.0 | 0 | Canis | 58943.0 | 0 | Canis lupus | 58943.0 | 0 |
| scf7180000050271 | 4053 | 0.4414 | 0 | 330.4891 | Eukaryota | 989.0 | 0 | Echinodermata | 494.0 | 2 | Forcipulatida | 494.0 | 2 | Asteriidae | 494.0 | 2 | Asterias | 494.0 | 2 | Asterias rubens | 494.0 | 2 |
| scf7180000053819 | 6353 | 0.4505 | 0 | 1004.9092 | no-hit | 0.0 | 0 | no-hit | 0.0 | 0 | no-hit | 0.0 | 0 | no-hit | 0.0 | 0 | no-hit | 0.0 | 0 | no-hit | 0.0 | 0 |
| scf7180000056376 | 494415 | 0.3866 | 0 | 59.3136 | Eukaryota | 37685.0 | 0 | Chordata | 37685.0 | 0 | Carnivora | 37685.0 | 0 | Canidae | 37685.0 | 0 | Canis | 37685.0 | 0 | Canis lupus | 37685.0 | 0 |
| scf7180000042457 | 672119 | 0.4519 | 0 | 56.0612 | Eukaryota | 91084.0 | 0 | Chordata | 91084.0 | 0 | Carnivora | 91084.0 | 0 | Canidae | 91084.0 | 0 | Canis | 91084.0 | 0 | Canis lupus | 91084.0 | 0 |
| scf7180000054269 | 146955 | 0.3844 | 0 | 229.6525 | Eukaryota | 89134.0 | 0 | Chordata | 89134.0 | 0 | Carnivora | 89134.0 | 0 | Canidae | 89134.0 | 0 | Canis | 89134.0 | 0 | Canis lupus | 89134.0 | 0 |
| scf7180000052742 | 1786 | 0.4524 | 0 | 208.2035 | Eukaryota | 549.0 | 0 | Echinodermata | 549.0 | 0 | Forcipulatida | 549.0 | 0 | Asteriidae | 549.0 | 0 | Asterias | 549.0 | 0 | Asterias rubens | 549.0 | 0 |
| scf7180000051314 | 4437 | 0.4503 | 0 | 125.5052 | Eukaryota | 828.0 | 0 | Platyhelminthes | 470.0 | 2 | Polystomatidea | 470.0 | 2 | Polystomatidae | 470.0 | 2 | Protopolystoma | 470.0 | 2 | Protopolystoma xenopodis | 470.0 | 2 |
| scf7180000048172 | 29050 | 0.4494 | 0 | 262.6097 | Eukaryota | 2917.0 | 0 | Chordata | 2917.0 | 0 | Carnivora | 2917.0 | 0 | Canidae | 2917.0 | 0 | Canis | 2917.0 | 0 | Canis lupus | 2917.0 | 0 |
| scf7180000055467 | 9132 | 0.4432 | 0 | 181.5571 | Eukaryota | 2919.0 | 0 | Chordata | 1972.0 | 1 | Kurtiformes | 1051.0 | 3 | Apogonidae | 1051.0 | 3 | Sphaeramia | 1051.0 | 3 | Sphaeramia orbicularis | 1051.0 | 3 |
| scf7180000053193 | 2501 | 0.4562 | 0 | 952.0691 | no-hit | 0.0 | 0 | no-hit | 0.0 | 0 | no-hit | 0.0 | 0 | no-hit | 0.0 | 0 | no-hit | 0.0 | 0 | no-hit | 0.0 | 0 |
| scf7180000045347 | 205415 | 0.4313 | 0 | 59.2909 | Eukaryota | 83748.0 | 0 | Chordata | 83748.0 | 0 | Carnivora | 83748.0 | 0 | Canidae | 53108.0 | 1 | Vulpes | 41484.0 | 2 | Vulpes vulpes | 41484.0 | 2 |
| scf7180000055283 | 14662 | 0.4482 | 0 | 118.7682 | Eukaryota | 9194.0 | 0 | Chordata | 9194.0 | 0 | Carnivora | 9194.0 | 0 | Canidae | 9194.0 | 0 | Nyctereutes | 4940.0 | 1 | Nyctereutes procyonoides | 4940.0 | 1 |
| scf7180000053146 | 8358 | 0.469 | 0 | 332.2465 | Eukaryota | 8055.0 | 0 | Chordata | 8055.0 | 0 | Carnivora | 4074.0 | 3 | Canidae | 4074.0 | 3 | Canis | 2490.0 | 4 | Nyctereutes procyonoides | 1584.0 | 5 |
| scf7180000041657 | 12927 | 0.4406 | 0 | 56.1622 | no-hit | 0.0 | 0 | no-hit | 0.0 | 0 | no-hit | 0.0 | 0 | no-hit | 0.0 | 0 | no-hit | 0.0 | 0 | no-hit | 0.0 | 0 |
| scf7180000052848 | 2783 | 0.4567 | 0 | 269.989 | no-hit | 0.0 | 0 | no-hit | 0.0 | 0 | no-hit | 0.0 | 0 | no-hit | 0.0 | 0 | no-hit | 0.0 | 0 | no-hit | 0.0 | 0 |
| scf7180000055068 | 12871 | 0.4388 | 0 | 97.7106 | Eukaryota | 8192.0 | 0 | Chordata | 7287.0 | 1 | Chordata-undef | 4135.0 | 5 | Ambassidae | 4135.0 | 5 | Parambassis | 4135.0 | 6 | Parambassis ranga | 4135.0 | 6 |
| scf7180000049068 | 21189 | 0.4415 | 0 | 362.1487 | Eukaryota | 3849.0 | 0 | Platyhelminthes | 2490.0 | 2 | Polystomatidea | 2490.0 | 2 | Polystomatidae | 2490.0 | 2 | Protopolystoma | 2490.0 | 2 | Protopolystoma xenopodis | 2490.0 | 2 |
| scf7180000048252 | 32202 | 0.6036 | 0 | 60.7838 | Eukaryota | 11266.0 | 0 | Chordata | 11266.0 | 0 | Carnivora | 11266.0 | 0 | Canidae | 9855.0 | 1 | Vulpes | 6480.0 | 2 | Vulpes vulpes | 6480.0 | 2 |
| scf7180000050020 | 5806 | 0.4492 | 0 | 1768.0123 | no-hit | 0.0 | 0 | no-hit | 0.0 | 0 | no-hit | 0.0 | 0 | no-hit | 0.0 | 0 | no-hit | 0.0 | 0 | no-hit | 0.0 | 0 |
| scf7180000049140 | 7097 | 0.445 | 0 | 155.9585 | Eukaryota | 2832.0 | 0 | Chordata | 2832.0 | 0 | Carnivora | 2832.0 | 0 | Canidae | 2832.0 | 0 | Canis | 2832.0 | 0 | Canis lupus | 2832.0 | 0 |
| scf7180000040908 | 1197023 | 0.3698 | 0 | 96.5065 | Eukaryota | 91622.0 | 0 | Chordata | 91622.0 | 0 | Carnivora | 91622.0 | 0 | Canidae | 91622.0 | 0 | Canis | 91622.0 | 0 | Canis lupus | 91622.0 | 0 |
| scf7180000043937 | 413145 | 0.3761 | 0 | 60.7701 | Eukaryota | 59417.0 | 0 | Chordata | 59417.0 | 0 | Carnivora | 59417.0 | 0 | Canidae | 59417.0 | 0 | Canis | 59417.0 | 0 | Canis lupus | 59417.0 | 0 |
| scf7180000055235 | 24433 | 0.3765 | 0 | 590.2006 | Eukaryota | 16022.0 | 0 | Chordata | 16022.0 | 0 | Carnivora | 16022.0 | 0 | Canidae | 16022.0 | 0 | Canis | 16022.0 | 0 | Canis lupus | 16022.0 | 0 |
| scf7180000044197 | 400807 | 0.4117 | 0 | 59.9186 | Eukaryota | 54078.0 | 0 | Chordata | 54078.0 | 0 | Carnivora | 54078.0 | 0 | Canidae | 54078.0 | 0 | Canis | 41664.0 | 1 | Canis lupus | 41664.0 | 1 |
| scf7180000054646 | 3663 | 0.442 | 0 | 202.2852 | no-hit | 0.0 | 0 | no-hit | 0.0 | 0 | no-hit | 0.0 | 0 | no-hit | 0.0 | 0 | no-hit | 0.0 | 0 | no-hit | 0.0 | 0 |
| scf7180000050696 | 5565 | 0.4494 | 0 | 1428.674 | Eukaryota | 289.0 | 0 | Chordata | 289.0 | 0 | Salmoniformes | 289.0 | 0 | Salmonidae | 289.0 | 0 | Salmo | 289.0 | 0 | Salmo trutta | 289.0 | 0 |
| scf7180000040851 | 30834 | 0.3703 | 0 | 59.5647 | Eukaryota | 65206.0 | 0 | Chordata | 65206.0 | 0 | Carnivora | 65206.0 | 0 | Canidae | 65206.0 | 0 | Canis | 65206.0 | 0 | Canis lupus | 65206.0 | 0 |
| scf7180000054423 | 23762 | 0.4268 | 0 | 242.2583 | Eukaryota | 80883.0 | 0 | Chordata | 80883.0 | 0 | Carnivora | 80883.0 | 0 | Canidae | 80883.0 | 0 | Canis | 80883.0 | 0 | Canis lupus | 80883.0 | 0 |
| scf7180000051583 | 3391 | 0.4468 | 0 | 335.2676 | no-hit | 0.0 | 0 | no-hit | 0.0 | 0 | no-hit | 0.0 | 0 | no-hit | 0.0 | 0 | no-hit | 0.0 | 0 | no-hit | 0.0 | 0 |
| scf7180000046576 | 130518 | 0.5237 | 0 | 57.6408 | Eukaryota | 51203.0 | 0 | Chordata | 51203.0 | 0 | Carnivora | 51203.0 | 0 | Canidae | 51203.0 | 0 | Canis | 51203.0 | 0 | Canis lupus | 51203.0 | 0 |
| scf7180000040796 | 176409 | 0.373 | 0 | 59.0306 | Eukaryota | 91814.0 | 0 | Chordata | 91814.0 | 0 | Carnivora | 91814.0 | 0 | Canidae | 91814.0 | 0 | Canis | 91814.0 | 0 | Canis lupus | 91814.0 | 0 |
| scf7180000047099 | 97047 | 0.3578 | 0 | 58.4805 | Eukaryota | 48903.0 | 0 | Chordata | 48903.0 | 0 | Carnivora | 48903.0 | 0 | Canidae | 48903.0 | 0 | Canis | 48903.0 | 0 | Canis lupus | 48903.0 | 0 |
| scf7180000043327 | 460982 | 0.3772 | 0 | 60.2115 | Eukaryota | 91858.0 | 0 | Chordata | 91858.0 | 0 | Carnivora | 91858.0 | 0 | Canidae | 91858.0 | 0 | Canis | 91858.0 | 0 | Canis lupus | 91858.0 | 0 |
| scf7180000054735 | 81686 | 0.4393 | 0 | 167.4345 | Eukaryota | 6205.0 | 0 | Platyhelminthes | 3187.0 | 1 | Polystomatidea | 3187.0 | 3 | Polystomatidae | 3187.0 | 3 | Protopolystoma | 3187.0 | 3 | Protopolystoma xenopodis | 3187.0 | 4 |
| scf7180000050340 | 7663 | 0.4496 | 0 | 107.5101 | no-hit | 0.0 | 0 | no-hit | 0.0 | 0 | no-hit | 0.0 | 0 | no-hit | 0.0 | 0 | no-hit | 0.0 | 0 | no-hit | 0.0 | 0 |
| scf7180000046194 | 99179 | 0.3812 | 0 | 58.2387 | Eukaryota | 8881.0 | 0 | Chordata | 8881.0 | 0 | Carnivora | 8881.0 | 0 | Canidae | 7297.0 | 2 | Canis | 7297.0 | 2 | Canis lupus | 7297.0 | 2 |
| scf7180000046797 | 136404 | 0.398 | 0 | 58.8125 | Eukaryota | 12331.0 | 0 | Chordata | 12331.0 | 0 | Carnivora | 6593.0 | 2 | Hominidae | 4105.0 | 4 | Homo | 2823.0 | 6 | Homo sapiens | 2823.0 | 6 |
| scf7180000046901 | 98965 | 0.3912 | 0 | 59.1219 | Eukaryota | 21627.0 | 0 | Chordata | 21627.0 | 0 | Carnivora | 21627.0 | 0 | Canidae | 21627.0 | 0 | Canis | 21627.0 | 0 | Canis lupus | 21627.0 | 0 |
| scf7180000044915 | 39420 | 0.5313 | 0 | 53.9579 | Eukaryota | 41520.0 | 0 | Chordata | 41520.0 | 0 | Carnivora | 41520.0 | 0 | Canidae | 41520.0 | 0 | Canis | 41520.0 | 0 | Canis lupus | 41520.0 | 0 |
| scf7180000047319 | 80533 | 0.444 | 0 | 137.0157 | Eukaryota | 23002.0 | 0 | Platyhelminthes | 12474.0 | 1 | Strigeidida | 7179.0 | 6 | Schistosomatidae | 7179.0 | 6 | Schistosoma | 7179.0 | 6 | Schistosoma mansoni | 7179.0 | 6 |
| scf7180000044482 | 286989 | 0.3608 | 0 | 60.0868 | Eukaryota | 14098.0 | 0 | Chordata | 14098.0 | 0 | Artiodactyla | 6440.0 | 2 | Hominidae | 5356.0 | 4 | Homo | 4026.0 | 6 | Homo sapiens | 4026.0 | 6 |
| scf7180000046430 | 95516 | 0.4356 | 0 | 61.9597 | Eukaryota | 99784.0 | 0 | Chordata | 99784.0 | 0 | Carnivora | 99784.0 | 0 | Canidae | 99784.0 | 0 | Canis | 99784.0 | 0 | Canis lupus | 99784.0 | 0 |
| scf7180000049400 | 9924 | 0.4407 | 0 | 67.9257 | Eukaryota | 420.0 | 0 | Chordata | 420.0 | 0 | Kurtiformes | 420.0 | 0 | Apogonidae | 420.0 | 0 | Sphaeramia | 420.0 | 0 | Sphaeramia orbicularis | 420.0 | 0 |
| scf7180000054820 | 22868 | 0.4351 | 0 | 197.4905 | Eukaryota | 11491.0 | 0 | Chordata | 7768.0 | 3 | Forcipulatida | 2673.0 | 7 | Asteriidae | 2673.0 | 7 | Asterias | 2673.0 | 7 | Asterias rubens | 2673.0 | 7 |
| scf7180000041163 | 1358887 | 0.4375 | 0 | 59.499 | Eukaryota | 74531.0 | 0 | Chordata | 74531.0 | 0 | Carnivora | 74531.0 | 0 | Canidae | 74531.0 | 0 | Canis | 51995.0 | 1 | Canis lupus | 51995.0 | 1 |
| scf7180000056187 | 1534934 | 0.4651 | 0 | 58.8194 | Eukaryota | 53483.0 | 0 | Chordata | 53483.0 | 0 | Carnivora | 53483.0 | 0 | Canidae | 53483.0 | 0 | Canis | 53483.0 | 0 | Canis lupus | 53483.0 | 0 |
| scf7180000050901 | 7935 | 0.4514 | 0 | 1021.6221 | Eukaryota | 1021.0 | 0 | Platyhelminthes | 535.0 | 1 | Polystomatidea | 535.0 | 1 | Polystomatidae | 535.0 | 1 | Protopolystoma | 535.0 | 1 | Protopolystoma xenopodis | 535.0 | 1 |
| scf7180000042796 | 125828 | 0.391 | 0 | 57.9704 | Eukaryota | 7366.0 | 0 | Chordata | 7366.0 | 0 | Carnivora | 7366.0 | 0 | Canidae | 7366.0 | 0 | Canis | 7366.0 | 0 | Canis lupus | 7366.0 | 0 |
| scf7180000047585 | 85026 | 0.4219 | 0 | 58.3659 | Eukaryota | 26209.0 | 0 | Chordata | 26209.0 | 0 | Carnivora | 26209.0 | 0 | Canidae | 21745.0 | 1 | Canis | 16051.0 | 3 | Canis lupus | 16051.0 | 3 |
| scf7180000046643 | 102210 | 0.3797 | 0 | 60.2809 | Eukaryota | 20952.0 | 0 | Chordata | 20952.0 | 0 | Carnivora | 20952.0 | 0 | Canidae | 20952.0 | 0 | Canis | 20952.0 | 0 | Canis lupus | 20952.0 | 0 |
| scf7180000055772 | 18873 | 0.439 | 0 | 453.9579 | no-hit | 0.0 | 0 | no-hit | 0.0 | 0 | no-hit | 0.0 | 0 | no-hit | 0.0 | 0 | no-hit | 0.0 | 0 | no-hit | 0.0 | 0 |
| scf7180000052430 | 9328 | 0.448 | 0 | 332.0792 | no-hit | 0.0 | 0 | no-hit | 0.0 | 0 | no-hit | 0.0 | 0 | no-hit | 0.0 | 0 | no-hit | 0.0 | 0 | no-hit | 0.0 | 0 |
| scf7180000041574 | 918600 | 0.3886 | 0 | 71.2869 | Eukaryota | 57973.0 | 0 | Chordata | 57973.0 | 0 | Carnivora | 57973.0 | 0 | Canidae | 57973.0 | 0 | Canis | 51751.0 | 1 | Canis lupus | 51751.0 | 1 |
| scf7180000051724 | 9870 | 0.5157 | 0 | 248.0903 | Eukaryota | 15589.0 | 0 | Chordata | 15589.0 | 0 | Carnivora | 15589.0 | 0 | Canidae | 15589.0 | 0 | Canis | 15589.0 | 0 | Canis lupus | 15589.0 | 0 |
| scf7180000052799 | 14099 | 0.5017 | 0 | 172.4482 | Eukaryota | 7901.0 | 0 | Chordata | 7901.0 | 0 | Carnivora | 7901.0 | 0 | Canidae | 7901.0 | 0 | Canis | 6785.0 | 1 | Canis lupus | 6785.0 | 1 |
| scf7180000051727 | 5451 | 0.4522 | 0 | 509.9098 | Eukaryota | 1087.0 | 0 | Chordata | 1087.0 | 0 | Rodentia | 352.0 | 3 | Muridae | 352.0 | 3 | Mus | 352.0 | 3 | Mus musculus | 352.0 | 3 |
| scf7180000041100 | 36867 | 0.4725 | 0 | 54.2218 | Eukaryota | 14428.0 | 0 | Chordata | 14428.0 | 0 | Carnivora | 14428.0 | 0 | Canidae | 14428.0 | 0 | Canis | 14428.0 | 0 | Canis lupus | 14428.0 | 0 |
| scf7180000044387 | 353592 | 0.3503 | 0 | 60.0748 | Eukaryota | 26533.0 | 0 | Chordata | 26533.0 | 0 | Carnivora | 26533.0 | 0 | Canidae | 26533.0 | 0 | Canis | 26533.0 | 0 | Canis lupus | 26533.0 | 0 |
| scf7180000051599 | 10410 | 0.4384 | 0 | 121.2101 | no-hit | 0.0 | 0 | no-hit | 0.0 | 0 | no-hit | 0.0 | 0 | no-hit | 0.0 | 0 | no-hit | 0.0 | 0 | no-hit | 0.0 | 0 |
| scf7180000046560 | 56036 | 0.3934 | 0 | 55.76 | Eukaryota | 74077.0 | 0 | Chordata | 74077.0 | 0 | Carnivora | 74077.0 | 0 | Canidae | 74077.0 | 0 | Canis | 74077.0 | 0 | Canis lupus | 74077.0 | 0 |
| scf7180000047395 | 116101 | 0.4238 | 0 | 60.9893 | Eukaryota | 79167.0 | 0 | Chordata | 79167.0 | 0 | Carnivora | 79167.0 | 0 | Canidae | 79167.0 | 0 | Canis | 79167.0 | 0 | Canis lupus | 79167.0 | 0 |
| scf7180000041080 | 1320514 | 0.3858 | 0 | 60.4829 | Eukaryota | 89930.0 | 0 | Chordata | 89930.0 | 0 | Carnivora | 89930.0 | 0 | Canidae | 89930.0 | 0 | Canis | 89930.0 | 0 | Canis lupus | 89930.0 | 0 |
| scf7180000048118 | 19468 | 0.437 | 0 | 172.1799 | Eukaryota | 10247.0 | 0 | Chordata | 8890.0 | 2 | Kurtiformes | 5951.0 | 3 | Apogonidae | 5951.0 | 3 | Sphaeramia | 5951.0 | 3 | Sphaeramia orbicularis | 5951.0 | 3 |
| scf7180000041036 | 1359600 | 0.3442 | 0 | 58.6361 | Eukaryota | 54921.0 | 0 | Chordata | 54921.0 | 0 | Carnivora | 54921.0 | 0 | Canidae | 54921.0 | 0 | Canis | 54921.0 | 0 | Canis lupus | 54921.0 | 0 |
| scf7180000044012 | 386389 | 0.3814 | 0 | 60.928 | Eukaryota | 37181.0 | 0 | Chordata | 37181.0 | 0 | Carnivora | 37181.0 | 0 | Canidae | 32207.0 | 2 | Canis | 24725.0 | 3 | Canis lupus | 24725.0 | 3 |
| scf7180000044164 | 19779 | 0.4117 | 0 | 51.3621 | Eukaryota | 60296.0 | 0 | Chordata | 60296.0 | 0 | Carnivora | 60296.0 | 0 | Canidae | 60296.0 | 0 | Canis | 60296.0 | 0 | Canis lupus | 60296.0 | 0 |
| scf7180000042851 | 497339 | 0.4746 | 0 | 58.2768 | Eukaryota | 20222.0 | 0 | Chordata | 20222.0 | 0 | Carnivora | 20222.0 | 0 | Canidae | 18751.0 | 1 | Canis | 14920.0 | 2 | Canis lupus | 14920.0 | 2 |
| scf7180000041268 | 1221536 | 0.4265 | 0 | 60.1403 | Eukaryota | 76770.0 | 0 | Chordata | 76770.0 | 0 | Carnivora | 76770.0 | 0 | Canidae | 76770.0 | 0 | Canis | 46578.0 | 1 | Canis lupus | 46578.0 | 1 |
| scf7180000043361 | 536083 | 0.3654 | 0 | 60.7255 | Eukaryota | 94488.0 | 0 | Chordata | 94488.0 | 0 | Carnivora | 94488.0 | 0 | Canidae | 94488.0 | 0 | Canis | 94488.0 | 0 | Canis lupus | 94488.0 | 0 |
| scf7180000051347 | 6390 | 0.4407 | 0 | 187.5484 | no-hit | 0.0 | 0 | no-hit | 0.0 | 0 | no-hit | 0.0 | 0 | no-hit | 0.0 | 0 | no-hit | 0.0 | 0 | no-hit | 0.0 | 0 |
| scf7180000044252 | 332527 | 0.4771 | 0 | 58.7736 | Eukaryota | 30114.0 | 0 | Chordata | 30114.0 | 0 | Carnivora | 27305.0 | 1 | Canidae | 17348.0 | 4 | Canis | 10096.0 | 5 | Canis lupus | 10096.0 | 5 |
| scf7180000041666 | 222004 | 0.4525 | 0 | 59.5789 | Eukaryota | 31760.0 | 0 | Chordata | 31760.0 | 0 | Carnivora | 31760.0 | 0 | Canidae | 31760.0 | 0 | Canis | 31760.0 | 0 | Canis lupus | 31760.0 | 0 |
| scf7180000040498 | 700543 | 0.3501 | 100 | 59.7998 | Eukaryota | 94233.0 | 0 | Chordata | 94233.0 | 0 | Carnivora | 94233.0 | 0 | Canidae | 94233.0 | 0 | Canis | 94233.0 | 0 | Canis lupus | 94233.0 | 0 |
| scf7180000042561 | 280205 | 0.4269 | 0 | 59.5208 | Eukaryota | 19078.0 | 0 | Chordata | 19078.0 | 0 | Carnivora | 10802.0 | 1 | Canidae | 8600.0 | 2 | Canis | 8600.0 | 2 | Canis lupus | 8600.0 | 2 |

| | | | | | | | | | | | | | | | | | | | | | | |
|---|---|---|---|---|---|---|---|---|---|---|---|---|---|---|---|---|---|---|---|---|---|---|
| scf7180000044585 | 32335 | 0.4723 | 0 | 213.7755 | Eukaryota | 879.0 | 0 | Chordata | 879.0 | 0 | Carnivora | 405.0 | 2 | Phocidae | 405.0 | 2 | Leptonychotes | 405.0 | 2 | Leptonychotes weddellii | 405.0 | 2 |
| scf7180000052310 | 3423 | 0.4522 | 0 | 420.2791 | Eukaryota | 4530.0 | 0 | Chordata | 4023.0 | 1 | Gadiformes | 1499.0 | 7 | Gadidae | 1499.0 | 7 | Gadus | 1499.0 | 7 | Gadus morhua | 1499.0 | 7 |
| scf7180000042473 | 15698 | 0.4508 | 0 | 30.1775 | Eukaryota | 54679.0 | 0 | Chordata | 54679.0 | 0 | Carnivora | 54679.0 | 0 | Canidae | 54679.0 | 0 | Canis | 54679.0 | 0 | Canis lupus | 54679.0 | 0 |
| scf7180000044306 | 3848 | 0.4397 | 0 | 149.179 | no-hit | 0.0 | 0 | no-hit | 0.0 | 0 | no-hit | 0.0 | 0 | no-hit | 0.0 | 0 | no-hit | 0.0 | 0 | no-hit | 0.0 | 0 |
| scf7180000049264 | 11879 | 0.4995 | 0 | 34.4029 | Eukaryota | 3864.0 | 0 | Chordata | 3864.0 | 0 | Carnivora | 3864.0 | 0 | Canidae | 2608.0 | 4 | Canis | 1410.0 | 5 | Canis lupus | 1410.0 | 5 |
| scf7180000056212 | 609000 | 0.3734 | 0 | 58.8791 | Eukaryota | 75949.0 | 0 | Chordata | 75949.0 | 0 | Carnivora | 75949.0 | 0 | Canidae | 75949.0 | 0 | Canis | 75949.0 | 0 | Canis lupus | 75949.0 | 0 |
| scf7180000046758 | 88083 | 0.4756 | 0 | 55.4614 | Eukaryota | 18868.0 | 0 | Chordata | 18868.0 | 0 | Carnivora | 18868.0 | 0 | Canidae | 18868.0 | 0 | Canis | 16409.0 | 1 | Canis lupus | 16409.0 | 1 |
| scf7180000048084 | 54146 | 0.5012 | 0 | 533.1907 | Eukaryota | 9671.0 | 0 | Chordata | 9671.0 | 0 | Carnivora | 9671.0 | 0 | Canidae | 9671.0 | 0 | Canis | 8486.0 | 1 | Canis lupus | 7266.0 | 2 |
| scf7180000041302 | 1303286 | 0.443 | 0 | 59.7743 | Eukaryota | 127452.0 | 0 | Chordata | 127452.0 | 0 | Carnivora | 127452.0 | 0 | Canidae | 127452.0 | 0 | Canis | 84652.0 | 1 | Canis lupus | 84652.0 | 1 |
| scf7180000052008 | 10440 | 0.4402 | 0 | 200.5473 | no-hit | 0.0 | 0 | no-hit | 0.0 | 0 | no-hit | 0.0 | 0 | no-hit | 0.0 | 0 | no-hit | 0.0 | 0 | no-hit | 0.0 | 0 |
| scf7180000056418 | 657163 | 0.3625 | 0 | 59.1055 | Eukaryota | 89578.0 | 0 | Chordata | 89578.0 | 0 | Carnivora | 89578.0 | 0 | Canidae | 89578.0 | 0 | Canis | 89578.0 | 0 | Canis lupus | 89578.0 | 0 |
| scf7180000054875 | 7462 | 0.5147 | 0 | 290.9406 | Eukaryota | 3002.0 | 0 | Chordata | 3002.0 | 0 | Carnivora | 3002.0 | 0 | Canidae | 1940.0 | 2 | Canis | 1940.0 | 2 | Canis lupus | 1940.0 | 2 |
| scf7180000045512 | 187840 | 0.4983 | 0 | 59.9683 | Eukaryota | 51963.0 | 0 | Chordata | 51963.0 | 0 | Carnivora | 51963.0 | 0 | Canidae | 51963.0 | 0 | Canis | 51963.0 | 0 | Canis lupus | 51963.0 | 0 |
| scf7180000050548 | 17467 | 0.4409 | 0 | 61.669 | Eukaryota | 2255.0 | 0 | Chordata | 1900.0 | 1 | Salmoniformes | 1900.0 | 1 | Salmonidae | 1900.0 | 1 | Coregonus | 1237.0 | 2 | Coregonus sp. 'balchen' | 1237.0 | 2 |
| scf7180000042674 | 294368 | 0.3728 | 0 | 59.4814 | Eukaryota | 84052.0 | 0 | Chordata | 84052.0 | 0 | Carnivora | 84052.0 | 0 | Canidae | 84052.0 | 0 | Canis | 84052.0 | 0 | Canis lupus | 84052.0 | 0 |
| scf7180000048747 | 57448 | 0.4388 | 0 | 93.7995 | Eukaryota | 94675.0 | 0 | Chordata | 49787.0 | 1 | Strigeidida | 32578.0 | 3 | Schistosomatidae | 32578.0 | 3 | Schistosoma | 32578.0 | 3 | Schistosoma mansoni | 32578.0 | 3 |
| scf7180000042731 | 573942 | 0.3535 | 0 | 59.4432 | Eukaryota | 65053.0 | 0 | Chordata | 65053.0 | 0 | Carnivora | 65053.0 | 0 | Canidae | 65053.0 | 0 | Canis | 65053.0 | 0 | Canis lupus | 65053.0 | 0 |
| scf7180000040489 | 1567649 | 0.3845 | 0 | 60.5428 | Eukaryota | 88449.0 | 0 | Chordata | 88449.0 | 0 | Carnivora | 88449.0 | 0 | Canidae | 88449.0 | 0 | Canis | 88449.0 | 0 | Canis lupus | 88449.0 | 0 |
| scf7180000044627 | 262207 | 0.6192 | 0 | 56.262 | Eukaryota | 94353.0 | 0 | Chordata | 94353.0 | 0 | Carnivora | 94353.0 | 0 | Canidae | 94353.0 | 0 | Canis | 94353.0 | 0 | Canis lupus | 94353.0 | 0 |
| scf7180000049782 | 21930 | 0.4516 | 0 | 1219.9454 | Eukaryota | 6360.0 | 0 | Chordata | 6134.0 | 1 | Kurtiformes | 4980.0 | 2 | Apogonidae | 4980.0 | 2 | Sphaeramia | 4980.0 | 2 | Sphaeramia orbicularis | 4980.0 | 2 |
| scf7180000043630 | 445236 | 0.3383 | 0 | 59.3935 | Eukaryota | 89437.0 | 0 | Chordata | 89437.0 | 0 | Carnivora | 89437.0 | 0 | Canidae | 89437.0 | 0 | Canis | 89437.0 | 0 | Canis lupus | 89437.0 | 0 |
| scf7180000042386 | 635705 | 0.4322 | 0 | 59.2181 | Eukaryota | 92447.0 | 0 | Chordata | 92447.0 | 0 | Carnivora | 92447.0 | 0 | Canidae | 92447.0 | 0 | Canis | 92447.0 | 0 | Canis lupus | 92447.0 | 0 |
| scf7180000050298 | 6693 | 0.5029 | 0 | 40.5509 | Eukaryota | 8005.0 | 0 | Chordata | 8005.0 | 0 | Carnivora | 8005.0 | 0 | Canidae | 8005.0 | 0 | Canis | 4195.0 | 1 | Canis lupus | 4195.0 | 1 |
| scf7180000055407 | 27450 | 0.4211 | 0 | 111.7752 | Eukaryota | 96995.0 | 0 | Chordata | 96995.0 | 0 | Carnivora | 96995.0 | 0 | Canidae | 96995.0 | 0 | Canis | 96995.0 | 0 | Canis lupus | 96995.0 | 0 |
| scf7180000042877 | 654090 | 0.3858 | 0 | 59.2762 | Eukaryota | 115785.0 | 0 | Chordata | 115785.0 | 0 | Carnivora | 115785.0 | 0 | Canidae | 115785.0 | 0 | Vulpes | 70260.0 | 1 | Vulpes vulpes | 70260.0 | 1 |
| scf7180000054159 | 469670 | 0.4229 | 0 | 60.6302 | Eukaryota | 61425.0 | 0 | Chordata | 61425.0 | 0 | Carnivora | 61425.0 | 0 | Canidae | 61425.0 | 0 | Canis | 61425.0 | 0 | Canis lupus | 61425.0 | 0 |
| scf7180000046841 | 6422 | 0.4041 | 0 | 32.6589 | Eukaryota | 2439.0 | 0 | Chordata | 2439.0 | 0 | Carnivora | 2439.0 | 0 | Canidae | 2439.0 | 0 | Canis | 2439.0 | 0 | Canis lupus | 2439.0 | 0 |
| scf7180000049329 | 3320 | 0.4569 | 0 | 446.2112 | Eukaryota | 636.0 | 0 | Platyhelminthes | 636.0 | 0 | Polystomatidea | 636.0 | 0 | Polystomatidae | 636.0 | 0 | Protopolystoma | 636.0 | 0 | Protopolystoma xenopodis | 636.0 | 0 |
| scf7180000052564 | 10698 | 0.5183 | 0 | 316.6983 | Eukaryota | 15874.0 | 0 | Chordata | 15874.0 | 0 | Carnivora | 15874.0 | 0 | Canidae | 15874.0 | 0 | Canis | 15874.0 | 0 | Canis lupus | 15874.0 | 0 |
| scf7180000045377 | 210400 | 0.4972 | 0 | 59.3369 | Eukaryota | 40727.0 | 0 | Chordata | 40727.0 | 0 | Carnivora | 40727.0 | 0 | Canidae | 40727.0 | 0 | Canis | 29009.0 | 1 | Canis lupus | 29009.0 | 1 |
| scf7180000047447 | 97388 | 0.4372 | 0 | 57.1254 | Eukaryota | 82072.0 | 0 | Chordata | 82072.0 | 0 | Carnivora | 82072.0 | 0 | Canidae | 82072.0 | 0 | Canis | 82072.0 | 0 | Canis lupus | 82072.0 | 0 |
| scf7180000053261 | 9839 | 0.3404 | 0 | 56.0425 | Eukaryota | 2842.0 | 0 | Chordata | 2842.0 | 0 | Carnivora | 2842.0 | 0 | Canidae | 2842.0 | 0 | Canis | 2842.0 | 0 | Canis lupus | 2842.0 | 0 |
| scf7180000051473 | 8106 | 0.451 | 0 | 1117.8845 | Eukaryota | 3279.0 | 0 | Platyhelminthes | 1727.0 | 3 | Polystomatidea | 1727.0 | 3 | Polystomatidae | 1727.0 | 3 | Protopolystoma | 1727.0 | 3 | Protopolystoma xenopodis | 1727.0 | 3 |
| scf7180000048283 | 25050 | 0.4406 | 0 | 71.173 | Eukaryota | 15478.0 | 0 | Chordata | 9693.0 | 2 | Gadiformes | 4571.0 | 7 | Gadidae | 4571.0 | 7 | Gadus | 4571.0 | 7 | Gadus morhua | 4571.0 | 7 |
| scf7180000044298 | 305476 | 0.4347 | 0 | 60.9035 | Eukaryota | 81550.0 | 0 | Chordata | 81550.0 | 0 | Carnivora | 81550.0 | 0 | Canidae | 81550.0 | 0 | Vulpes | 81550.0 | 0 | Vulpes vulpes | 81550.0 | 0 |
| scf7180000040586 | 2311472 | 0.3793 | 0 | 60.7808 | Eukaryota | 95353.0 | 0 | Chordata | 95353.0 | 0 | Carnivora | 95353.0 | 0 | Canidae | 95353.0 | 0 | Canis | 95353.0 | 0 | Canis lupus | 95353.0 | 0 |
| scf7180000055084 | 41921 | 0.5039 | 0 | 274.1814 | Eukaryota | 9324.0 | 0 | Chordata | 9324.0 | 0 | Carnivora | 9324.0 | 0 | Canidae | 9324.0 | 0 | Canis | 8122.0 | 1 | Canis lupus | 6870.0 | 2 |
| scf7180000042227 | 43994 | 0.4514 | 0 | 52.654 | Eukaryota | 32568.0 | 0 | Chordata | 32568.0 | 0 | Carnivora | 32568.0 | 0 | Canidae | 32568.0 | 0 | Canis | 32568.0 | 0 | Canis lupus | 32568.0 | 0 |
| scf7180000049266 | 9666 | 0.4447 | 0 | 175.242 | Eukaryota | 1028.0 | 0 | Chordata | 1028.0 | 0 | Kurtiformes | 1028.0 | 0 | Apogonidae | 1028.0 | 0 | Sphaeramia | 1028.0 | 0 | Sphaeramia orbicularis | 1028.0 | 0 |
| scf7180000055903 | 5118 | 0.4386 | 0 | 194.3246 | Eukaryota | 507.0 | 0 | Chordata | 507.0 | 0 | Tetraodontiformes | 320.0 | 1 | Tetraodontidae | 320.0 | 1 | Takifugu | 320.0 | 1 | Takifugu rubripes | 320.0 | 1 |
| scf7180000053780 | 5109 | 0.4574 | 0 | 422.8224 | Eukaryota | 1051.0 | 0 | Platyhelminthes | 796.0 | 1 | Polystomatidea | 796.0 | 1 | Polystomatidae | 796.0 | 1 | Protopolystoma | 796.0 | 1 | Protopolystoma xenopodis | 796.0 | 1 |
| scf7180000056311 | 976728 | 0.4427 | 0 | 58.9054 | Eukaryota | 94577.0 | 0 | Chordata | 94577.0 | 0 | Carnivora | 94577.0 | 0 | Canidae | 94577.0 | 0 | Canis | 94577.0 | 0 | Canis lupus | 94577.0 | 0 |
| scf7180000054140 | 47879 | 0.3468 | 0 | 82.3389 | Eukaryota | 5112.0 | 0 | Chordata | 5112.0 | 0 | Carnivora | 2778.0 | 3 | Canidae | 1768.0 | 4 | Canis | 1768.0 | 5 | Canis lupus | 1768.0 | 5 |
| scf7180000044649 | 271738 | 0.3781 | 0 | 59.4721 | Eukaryota | 88614.0 | 0 | Chordata | 88614.0 | 0 | Carnivora | 88614.0 | 0 | Canidae | 88614.0 | 0 | Canis | 88614.0 | 0 | Canis lupus | 88614.0 | 0 |
| scf7180000042889 | 747588 | 0.418 | 0 | 60.8288 | Eukaryota | 92823.0 | 0 | Chordata | 92823.0 | 0 | Carnivora | 92823.0 | 0 | Canidae | 92823.0 | 0 | Canis | 92823.0 | 0 | Canis lupus | 92823.0 | 0 |
| scf7180000053066 | 3124 | 0.4465 | 0 | 343.5144 | no-hit | 0.0 | 0 | no-hit | 0.0 | 0 | no-hit | 0.0 | 0 | no-hit | 0.0 | 0 | no-hit | 0.0 | 0 | no-hit | 0.0 | 0 |
| scf7180000042683 | 120230 | 0.5212 | 0 | 54.9059 | Eukaryota | 24705.0 | 0 | Chordata | 24705.0 | 0 | Carnivora | 24705.0 | 0 | Canidae | 24705.0 | 0 | Canis | 16795.0 | 1 | Canis lupus | 16795.0 | 1 |
| scf7180000052836 | 4790 | 0.4428 | 0 | 192.6645 | no-hit | 0.0 | 0 | no-hit | 0.0 | 0 | no-hit | 0.0 | 0 | no-hit | 0.0 | 0 | no-hit | 0.0 | 0 | no-hit | 0.0 | 0 |
| scf7180000056556 | 72395 | 0.4395 | 0 | 118.4972 | Eukaryota | 33086.0 | 0 | Chordata | 25057.0 | 2 | Tetraodontiformes | 8221.0 | 6 | Tetraodontidae | 8221.0 | 6 | Takifugu | 8221.0 | 6 | Takifugu rubripes | 8221.0 | 6 |
| scf7180000049711 | 7029 | 0.4028 | 0 | 46.9792 | Eukaryota | 597.0 | 0 | Chordata | 597.0 | 0 | Salmoniformes | 597.0 | 0 | Salmonidae | 597.0 | 0 | Salmo | 597.0 | 0 | Salmo trutta | 597.0 | 0 |
| scf7180000045460 | 168428 | 0.4395 | 0 | 110.8292 | Eukaryota | 100907.0 | 0 | Platyhelminthes | 50539.0 | 1 | Gadiformes | 50368.0 | 2 | Gadidae | 50368.0 | 2 | Gadus | 50368.0 | 2 | Gadus morhua | 50368.0 | 2 |
| scf7180000049598 | 14670 | 0.508 | 0 | 113.1336 | Eukaryota | 8643.0 | 0 | Chordata | 8643.0 | 0 | Carnivora | 8643.0 | 0 | Canidae | 8643.0 | 0 | Canis | 6949.0 | 1 | Canis lupus | 5740.0 | 2 |
| scf7180000044355 | 177180 | 0.4648 | 0 | 57.4006 | Eukaryota | 21890.0 | 0 | Chordata | 21890.0 | 0 | Primates | 16426.0 | 2 | Hominidae | 16426.0 | 2 | Homo | 16426.0 | 2 | Homo sapiens | 16426.0 | 2 |
| scf7180000045592 | 190099 | 0.3547 | 0 | 57.8026 | Eukaryota | 54501.0 | 0 | Chordata | 54501.0 | 0 | Carnivora | 54501.0 | 0 | Canidae | 54501.0 | 0 | Canis | 54501.0 | 0 | Canis lupus | 54501.0 | 0 |
| scf7180000049475 | 8620 | 0.4542 | 0 | 888.1222 | Eukaryota | 2368.0 | 0 | Echinodermata | 850.0 | 3 | Forcipulatida | 850.0 | 3 | Asteriidae | 850.0 | 3 | Asterias | 850.0 | 3 | Asterias rubens | 850.0 | 3 |
| scf7180000050198 | 26207 | 0.3993 | 0 | 58.3001 | Eukaryota | 7159.0 | 0 | Chordata | 6181.0 | 1 | Artiodactyla | 2459.0 | 4 | Suidae | 2459.0 | 4 | Sus | 2459.0 | 4 | Sus scrofa | 2459.0 | 4 |
| scf7180000043919 | 428200 | 0.3646 | 0 | 60.1674 | Eukaryota | 30125.0 | 0 | Chordata | 30125.0 | 0 | Carnivora | 30125.0 | 0 | Canidae | 22418.0 | 2 | Canis | 14045.0 | 3 | Canis lupus | 14045.0 | 3 |
| scf7180000052453 | 4069 | 0.4502 | 0 | 519.9564 | no-hit | 0.0 | 0 | no-hit | 0.0 | 0 | no-hit | 0.0 | 0 | no-hit | 0.0 | 0 | no-hit | 0.0 | 0 | no-hit | 0.0 | 0 |
| scf7180000053344 | 9133 | 0.4397 | 0 | 54.1492 | Eukaryota | 361.0 | 0 | Platyhelminthes | 361.0 | 0 | Polystomatidea | 361.0 | 0 | Polystomatidae | 361.0 | 0 | Protopolystoma | 361.0 | 0 | Protopolystoma xenopodis | 361.0 | 0 |
| scf7180000051832 | 4358 | 0.4461 | 0 | 463.9798 | no-hit | 0.0 | 0 | no-hit | 0.0 | 0 | no-hit | 0.0 | 0 | no-hit | 0.0 | 0 | no-hit | 0.0 | 0 | no-hit | 0.0 | 0 |
| scf7180000044425 | 98633 | 0.4633 | 0 | 56.2962 | Eukaryota | 41846.0 | 0 | Chordata | 41846.0 | 0 | Carnivora | 41846.0 | 0 | Canidae | 41846.0 | 0 | Canis | 41846.0 | 0 | Canis lupus | 41846.0 | 0 |
| scf7180000049212 | 16479 | 0.5111 | 0 | 1131.9696 | Eukaryota | 8585.0 | 0 | Chordata | 8585.0 | 0 | Carnivora | 8585.0 | 0 | Canidae | 8585.0 | 0 | Canis | 6909.0 | 1 | Canis lupus | 5687.0 | 2 |
| scf7180000051753 | 8688 | 0.4503 | 0 | 1097.9594 | Eukaryota | 3300.0 | 0 | Platyhelminthes | 1718.0 | 3 | Polystomatidea | 1718.0 | 3 | Polystomatidae | 1718.0 | 3 | Protopolystoma | 1718.0 | 3 | Protopolystoma xenopodis | 1718.0 | 3 |
| scf7180000053921 | 5558 | 0.4435 | 0 | 651.1771 | no-hit | 0.0 | 0 | no-hit | 0.0 | 0 | no-hit | 0.0 | 0 | no-hit | 0.0 | 0 | no-hit | 0.0 | 0 | no-hit | 0.0 | 0 |
| scf7180000053046 | 15431 | 0.4613 | 0 | 69.2324 | Eukaryota | 11247.0 | 0 | Chordata | 11247.0 | 0 | Carnivora | 11247.0 | 0 | Felidae | 11247.0 | 0 | Leopardus | 3399.0 | 6 | unresolved | 1136.0 | 9 |
| scf7180000043627 | 48517 | 0.3901 | 0 | 48.7054 | Eukaryota | 24409.0 | 0 | Chordata | 24409.0 | 0 | Carnivora | 24409.0 | 0 | Canidae | 24409.0 | 0 | Canis | 24409.0 | 0 | Canis lupus | 24409.0 | 0 |
| scf7180000042825 | 33176 | 0.4899 | 0 | 566.6307 | Eukaryota | 36122.0 | 0 | Chordata | 36122.0 | 0 | Carnivora | 36122.0 | 0 | Canidae | 36122.0 | 0 | Canis | 36122.0 | 0 | Canis lupus | 36122.0 | 0 |
| scf7180000051255 | 11600 | 0.44 | 0 | 252.5573 | no-hit | 0.0 | 0 | no-hit | 0.0 | 0 | no-hit | 0.0 | 0 | no-hit | 0.0 | 0 | no-hit | 0.0 | 0 | no-hit | 0.0 | 0 |
| scf7180000042134 | 162225 | 0.3696 | 0 | 153.1633 | Eukaryota | 90939.0 | 0 | Chordata | 90939.0 | 0 | Carnivora | 90939.0 | 0 | Canidae | 90939.0 | 0 | Canis | 90939.0 | 0 | Canis lupus | 90939.0 | 0 |
| scf7180000047453 | 22071 | 0.4023 | 0 | 50.4319 | Eukaryota | 27782.0 | 0 | Chordata | 27782.0 | 0 | Carnivora | 27782.0 | 0 | Canidae | 27782.0 | 0 | Canis | 27782.0 | 0 | Canis lupus | 27782.0 | 0 |
| scf7180000052841 | 2850 | 0.4414 | 0 | 349.6052 | no-hit | 0.0 | 0 | no-hit | 0.0 | 0 | no-hit | 0.0 | 0 | no-hit | 0.0 | 0 | no-hit | 0.0 | 0 | no-hit | 0.0 | 0 |
| scf7180000056513 | 290317 | 0.3557 | 0 | 61.0208 | Eukaryota | 28057.0 | 0 | Chordata | 28057.0 | 0 | Carnivora | 28057.0 | 0 | Canidae | 28057.0 | 0 | Canis | 20485.0 | 1 | Canis lupus | 20485.0 | 1 |
| scf7180000050935 | 4303 | 0.4439 | 0 | 640.5366 | no-hit | 0.0 | 0 | no-hit | 0.0 | 0 | no-hit | 0.0 | 0 | no-hit | 0.0 | 0 | no-hit | 0.0 | 0 | no-hit | 0.0 | 0 |
| scf7180000053551 | 2191 | 0.4459 | 0 | 454.3423 | no-hit | 0.0 | 0 | no-hit | 0.0 | 0 | no-hit | 0.0 | 0 | no-hit | 0.0 | 0 | no-hit | 0.0 | 0 | no-hit | 0.0 | 0 |
| scf7180000042436 | 79131 | 0.4426 | 0 | 60.1342 | Eukaryota | 34022.0 | 0 | Chordata | 34022.0 | 0 | Carnivora | 34022.0 | 0 | Canidae | 34022.0 | 0 | Canis | 34022.0 | 0 | Canis lupus | 34022.0 | 0 |
| scf7180000042091 | 18434 | 0.395 | 0 | 46.7105 | Eukaryota | 71635.0 | 0 | Chordata | 71635.0 | 0 | Carnivora | 71635.0 | 0 | Canidae | 71635.0 | 0 | Canis | 71635.0 | 0 | Canis lupus | 71635.0 | 0 |
| scf7180000056503 | 282952 | 0.349 | 0 | 59.4391 | Eukaryota | 20801.0 | 0 | Chordata | 20801.0 | 0 | Carnivora | 20801.0 | 0 | Canidae | 20801.0 | 0 | Canis | 20801.0 | 0 | Canis lupus | 20801.0 | 0 |
| scf7180000042321 | 942988 | 0.5516 | 0 | 59.0908 | Eukaryota | 128967.0 | 0 | Chordata | 128967.0 | 0 | Carnivora | 128967.0 | 0 | Canidae | 128967.0 | 0 | Canis | 98644.0 | 1 | Canis lupus | 98644.0 | 1 |
| scf7180000046367 | 162033 | 0.5066 | 0 | 58.9659 | Eukaryota | 36564.0 | 0 | Chordata | 36564.0 | 0 | Carnivora | 36564.0 | 0 | Canidae | 36564.0 | 0 | Canis | 36564.0 | 0 | Canis lupus | 36564.0 | 0 |
| scf7180000043878 | 391292 | 0.523 | 0 | 56.7716 | Eukaryota | 14145.0 | 0 | Chordata | 14145.0 | 0 | Carnivora | 14145.0 | 0 | Canidae | 14145.0 | 0 | Canis | 11472.0 | 1 | Canis lupus | 11472.0 | 1 |
| scf7180000054096 | 6307 | 0.4362 | 0 | 111.0475 | Eukaryota | 2253.0 | 0 | Chordata | 1158.0 | 1 | Polystomatidea | 1095.0 | 2 | Polystomatidae | 1095.0 | 2 | Protopolystoma | 1095.0 | 3 | Protopolystoma xenopodis | 1095.0 | 3 |
| scf7180000052258 | 11338 | 0.4518 | 0 | 636.74 | Eukaryota | 3196.0 | 0 | Chordata | 2721.0 | 1 | Carnivora | 2721.0 | 1 | Canidae | 2721.0 | 1 | Canis | 2721.0 | 1 | Canis lupus | 2721.0 | 1 |
| scf7180000048048 | 50298 | 0.4058 | 0 | 58.7661 | Eukaryota | 30766.0 | 0 | Chordata | 30766.0 | 0 | Carnivora | 30766.0 | 0 | Canidae | 30766.0 | 0 | Canis | 30766.0 | 0 | Canis lupus | 30766.0 | 0 |
| scf7180000047633 | 8542 | 0.4487 | 0 | 289.1582 | Eukaryota | 5982.0 | 0 | Platyhelminthes | 3316.0 | 1 | Polystomatidea | 3316.0 | 1 | Polystomatidae | 3316.0 | 1 | Protopolystoma | 3316.0 | 1 | Protopolystoma xenopodis | 3316.0 | 1 |
| scf7180000040507 | 441366 | 0.3839 | 0 | 67.5858 | Eukaryota | 94701.0 | 0 | Chordata | 94701.0 | 0 | Carnivora | 94701.0 | 0 | Canidae | 94701.0 | 0 | Canis | 94701.0 | 0 | Canis lupus | 94701.0 | 0 |
| scf7180000056328 | 832010 | 0.4676 | 0 | 58.8492 | Eukaryota | 48871.0 | 0 | Chordata | 48871.0 | 0 | Carnivora | 48871.0 | 0 | Canidae | 48871.0 | 0 | Canis | 48871.0 | 0 | Canis lupus | 48871.0 | 0 |
| scf7180000045584 | 34732 | 0.3815 | 0 | 53.5952 | Eukaryota | 12597.0 | 0 | Chordata | 12597.0 | 0 | Carnivora | 12597.0 | 0 | Canidae | 12597.0 | 0 | Canis | 12597.0 | 0 | Canis lupus | 12597.0 | 0 |
| scf7180000051353 | 4400 | 0.4541 | 0 | 315.8396 | Eukaryota | 1183.0 | 0 | Platyhelminthes | 689.0 | 1 | Polystomatidea | 689.0 | 1 | Polystomatidae | 689.0 | 1 | Protopolystoma | 689.0 | 1 | Protopolystoma xenopodis | 689.0 | 1 |
| scf7180000047406 | 7914 | 0.444 | 0 | 107.8981 | Eukaryota | 346.0 | 0 | Platyhelminthes | 346.0 | 0 | Polystomatidea | 346.0 | 0 | Polystomatidae | 346.0 | 0 | Protopolystoma | 346.0 | 0 | Protopolystoma xenopodis | 346.0 | 0 |
| scf7180000041062 | 1451660 | 0.3598 | 0 | 60.0015 | Eukaryota | 89228.0 | 0 | Chordata | 89228.0 | 0 | Carnivora | 89228.0 | 0 | Canidae | 89228.0 | 0 | Canis | 89228.0 | 0 | Canis lupus | 89228.0 | 0 |
| scf7180000052784 | 8868 | 0.4419 | 0 | 80.5185 | Eukaryota | 2499.0 | 0 | Chordata | 1504.0 | 1 | Columbiformes | 1199.0 | 2 | Columbidae | 1199.0 | 2 | Streptopelia | 1199.0 | 2 | Streptopelia turtur | 1199.0 | 2 |
| scf7180000056380 | 639325 | 0.3875 | 0 | 60.1706 | Eukaryota | 66983.0 | 0 | Chordata | 66983.0 | 0 | Carnivora | 66983.0 | 0 | Otariidae | 33974.0 | 1 | Zalophus | 33974.0 | 2 | Zalophus californianus | 33974.0 | 2 |
| scf7180000052715 | 5255 | 0.4426 | 0 | 93.9607 | Eukaryota | 3042.0 | 0 | Chordata | 3042.0 | 0 | Carnivora | 3042.0 | 0 | Canidae | 3042.0 | 0 | Canis | 3042.0 | 0 | Canis lupus | 3042.0 | 0 |
| scf7180000044685 | 23093 | 0.4393 | 0 | 144.8817 | Eukaryota | 3666.0 | 0 | Chordata | 2723.0 | 1 | Tetraodontiformes | 2560.0 | 2 | Tetraodontidae | 2560.0 | 2 | Takifugu | 2560.0 | 2 | Takifugu rubripes | 2560.0 | 2 |
| scf7180000055547 | 34644 | 0.4376 | 0 | 276.1117 | Eukaryota | 5996.0 | 0 | Chordata | 5246.0 | 1 | Tetraodontiformes | 2560.0 | 6 | Tetraodontidae | 2560.0 | 6 | Takifugu | 2560.0 | 6 | Takifugu rubripes | 2560.0 | 6 |
| scf7180000050504 | 36829 | 0.432 | 0 | 51.6847 | Eukaryota | 13873.0 | 0 | Chordata | 13873.0 | 0 | Carnivora | 13873.0 | 0 | Canidae | 13873.0 | 0 | Canis | 13873.0 | 0 | Canis lupus | 13873.0 | 0 |
| scf7180000044095 | 393366 | 0.3673 | 0 | 61.2987 | Eukaryota | 20192.0 | 0 | Chordata | 20192.0 | 0 | Carnivora | 20192.0 | 0 | Canidae | 12318.0 | 3 | Canis | 9989.0 | 4 | Canis lupus | 9989.0 | 4 |
| scf7180000051687 | 7010 | 0.4447 | 0 | 338.8031 | Eukaryota | 24344.0 | 0 | Chordata | 8936.0 | 3 | Strigeidida | 8363.0 | 5 | Schistosomatidae | 8363.0 | 5 | Schistosoma | 8363.0 | 5 | Schistosoma mansoni | 8363.0 | 6 |
| scf7180000040759 | 33482 | 0.3791 | 0 | 76.7279 | Eukaryota | 93767.0 | 0 | Chordata | 93767.0 | 0 | Carnivora | 93767.0 | 0 | Canidae | 93767.0 | 0 | Canis | 93767.0 | 0 | Canis lupus | 93767.0 | 0 |
| scf7180000056349 | 1115610 | 0.4327 | 0 | 59.0651 | Eukaryota | 62227.0 | 0 | Chordata | 62227.0 | 0 | Carnivora | 62227.0 | 0 | Canidae | 62227.0 | 0 | Canis | 46242.0 | 1 | Canis lupus | 46242.0 | 1 |
| scf7180000051001 | 27184 | 0.3589 | 0 | 57.1046 | Eukaryota | 5721.0 | 0 | Chordata | 5721.0 | 0 | Carnivora | 5721.0 | 0 | Canidae | 5721.0 | 0 | Canis | 5721.0 | 0 | Canis lupus | 5721.0 | 0 |
| scf7180000045586 | 4415 | 0.3803 | 0 | 25.9629 | Eukaryota | 3717.0 | 0 | Chordata | 3717.0 | 0 | Carnivora | 3717.0 | 0 | Canidae | 3717.0 | 0 | Canis | 3717.0 | 0 | Canis lupus | 3717.0 | 0 |
| scf7180000046274 | 111916 | 0.4447 | 0 | 73.1436 | Eukaryota | 9569.0 | 0 | Chordata | 9206.0 | 1 | Blenniiformes | 5355.0 | 4 | Blenniidae | 5355.0 | 4 | Salarias | 5355.0 | 4 | Salarias fasciatus | 5355.0 | 4 |
| scf7180000043020 | 56382 | 0.3989 | 0 | 62.05 | Eukaryota | 63438.0 | 0 | Chordata | 63438.0 | 0 | Carnivora | 63438.0 | 0 | Canidae | 63438.0 | 0 | Canis | 63438.0 | 0 | Canis lupus | 63438.0 | 0 |
| scf7180000050981 | 13747 | 0.4456 | 0 | 1871.5745 | no-hit | 0.0 | 0 | no-hit | 0.0 | 0 | no-hit | 0.0 | 0 | no-hit | 0.0 | 0 | no-hit | 0.0 | 0 | no-hit | 0.0 | 0 |
| scf7180000055569 | 12958 | 0.441 | 0 | 157.1088 | Eukaryota | 1402.0 | 0 | Chordata | 1402.0 | 0 | Kurtiformes | 869.0 | 2 | Apogonidae | 869.0 | 2 | Sphaeramia | 869.0 | 2 | Sphaeramia orbicularis | 869.0 | 2 |
| scf7180000054310 | 17913 | 0.4453 | 0 | 153.2429 | Eukaryota | 50513.0 | 0 | Chordata | 50513.0 | 0 | Carnivora | 50513.0 | 0 | Canidae | 50513.0 | 0 | Canis | 50513.0 | 0 | Canis lupus | 50513.0 | 0 |
| scf7180000056288 | 323820 | 0.381 | 0 | 61.6248 | Eukaryota | 43053.0 | 0 | Chordata | 43053.0 | 0 | Carnivora | 43053.0 | 0 | Canidae | 26813.0 | 1 | Canis | 22375.0 | 2 | Canis lupus | 22375.0 | 2 |
| scf7180000052738 | 5610 | 0.4102 | 0 | 61.3205 | Eukaryota | 3852.0 | 0 | Chordata | 3852.0 | 0 | Carnivora | 3201.0 | 1 | Canidae | 3201.0 | 1 | Canis | 3201.0 | 1 | Canis lupus | 3201.0 | 1 |
| scf7180000043751 | 34695 | 0.3846 | 0 | 53.2661 | Eukaryota | 4280.0 | 0 | Chordata | 4280.0 | 0 | Artiodactyla | 2200.0 | 1 | Hominidae | 2080.0 | 2 | Homo | 1406.0 | 5 | Homo sapiens | 1406.0 | 5 |
| scf7180000049066 | 40546 | 0.5027 | 0 | 584.6401 | Eukaryota | 9367.0 | 0 | Chordata | 9367.0 | 0 | Carnivora | 9367.0 | 0 | Canidae | 9367.0 | 0 | Canis | 7607.0 | 1 | Canis lupus | 6326.0 | 2 |
| scf7180000045489 | 194900 | 0.4274 | 0 | 70.4311 | Eukaryota | 112547.0 | 0 | Chordata | 112547.0 | 0 | Carnivora | 112547.0 | 0 | Canidae | 112547.0 | 0 | Canis | 112547.0 | 0 | Canis lupus | 112547.0 | 0 |
| scf7180000045928 | 134215 | 0.3894 | 0 | 57.2464 | Eukaryota | 37967.0 | 0 | Chordata | 37967.0 | 0 | Carnivora | 37967.0 | 0 | Canidae | 37967.0 | 0 | Canis | 37967.0 | 0 | Canis lupus | 37967.0 | 0 |
| scf7180000048773 | 10067 | 0.434 | 0 | 188.4235 | Eukaryota | 228.0 | 0 | Platyhelminthes | 228.0 | 0 | Polystomatidea | 228.0 | 0 | Polystomatidae | 228.0 | 0 | Protopolystoma | 228.0 | 0 | Protopolystoma xenopodis | 228.0 | 0 |
| scf7180000042548 | 590206 | 0.4441 | 0 | 61.6393 | Eukaryota | 85031.0 | 0 | Chordata | 85031.0 | 0 | Carnivora | 85031.0 | 0 | Canidae | 85031.0 | 0 | Canis | 75577.0 | 1 | Canis lupus | 75577.0 | 1 |
| scf7180000051352 | 7353 | 0.4367 | 0 | 70.6903 | no-hit | 0.0 | 0 | no-hit | 0.0 | 0 | no-hit | 0.0 | 0 | no-hit | 0.0 | 0 | no-hit | 0.0 | 0 | no-hit | 0.0 | 0 |
| scf7180000043947 | 38267 | 0.4221 | 0 | 106.404 | Eukaryota | 11839.0 | 0 | Chordata | 11839.0 | 0 | Artiodactyla | 5734.0 | 2 | Bovidae | 5734.0 | 2 | Homo | 5112.0 | 3 | Homo sapiens | 5112.0 | 3 |
| scf7180000044989 | 221968 | 0.3921 | 0 | 61.1888 | Eukaryota | 32876.0 | 0 | Chordata | 32876.0 | 0 | Carnivora | 32876.0 | 0 | Canidae | 32876.0 | 0 | Canis | 32876.0 | 0 | Canis lupus | 32876.0 | 0 |
| scf7180000048209 | 49622 | 0.3948 | 0 | 59.876 | Eukaryota | 85557.0 | 0 | Chordata | 85557.0 | 0 | Carnivora | 85557.0 | 0 | Canidae | 55032.0 | 2 | Canis | 44681.0 | 3 | Canis lupus | 44681.0 | 3 |
| scf7180000049421 | 23308 | 0.4381 | 0 | 104.699 | Eukaryota | 2218.0 | 0 | Platyhelminthes | 1185.0 | 1 | Polystomatidea | 1185.0 | 1 | Polystomatidae | 1185.0 | 1 | Protopolystoma | 1185.0 | 1 | Protopolystoma xenopodis | 1185.0 | 1 |
| scf7180000052686 | 14769 | 0.4463 | 0 | 68.9616 | Eukaryota | 768.0 | 0 | Platyhelminthes | 551.0 | 1 | Polystomatidea | 551.0 | 1 | Polystomatidae | 551.0 | 1 | Protopolystoma | 551.0 | 1 | Protopolystoma xenopodis | 551.0 | 1 |
| scf7180000043599 | 497721 | 0.3881 | 0 | 60.3409 | Eukaryota | 38855.0 | 0 | Chordata | 38855.0 | 0 | Carnivora | 38855.0 | 0 | Canidae | 38855.0 | 0 | Canis | 38855.0 | 0 | Canis lupus | 38855.0 | 0 |
| scf7180000040551 | 14639 | 0.3797 | 0 | 34.6537 | Eukaryota | 5879.0 | 0 | Chordata | 5879.0 | 0 | Rodentia | 3720.0 | 2 | Cricetidae | 2684.0 | 4 | Mesocricetus | 2684.0 | 4 | Mesocricetus auratus | 2684.0 | 4 |
| scf7180000052357 | 6387 | 0.439 | 0 | 118.6198 | no-hit | 0.0 | 0 | no-hit | 0.0 | 0 | no-hit | 0.0 | 0 | no-hit | 0.0 | 0 | no-hit | 0.0 | 0 | no-hit | 0.0 | 0 |
| scf7180000049691 | 34580 | 0.4565 | 0 | 56.4909 | Eukaryota | 8898.0 | 0 | Chordata | 8360.0 | 1 | Salmoniformes | 3710.0 | 4 | Salmonidae | 3710.0 | 4 | Salmo | 2895.0 | 5 | Salmo trutta | 2895.0 | 5 |
| scf7180000052840 | 3463 | 0.4381 | 0 | 95.0175 | Eukaryota | 346.0 | 0 | Platyhelminthes | 346.0 | 0 | Polystomatidea | 346.0 | 0 | Polystomatidae | 346.0 | 0 | Protopolystoma | 346.0 | 0 | Protopolystoma xenopodis | 346.0 | 0 |
| scf7180000046644 | 135434 | 0.6019 | 0 | 54.3266 | Eukaryota | 23389.0 | 0 | Chordata | 23389.0 | 0 | Carnivora | 23389.0 | 0 | Canidae | 8852.0 | 4 | Canis | 6333.0 | 6 | Canis lupus | 6333.0 | 6 |
| scf7180000052992 | 4146 | 0.4091 | 0 | 64.7093 | Eukaryota | 34074.0 | 0 | Chordata | 34074.0 | 0 | Carnivora | 34074.0 | 0 | Canidae | 34074.0 | 0 | Canis | 34074.0 | 0 | Canis lupus | 34074.0 | 0 |
| scf7180000041047 | 354483 | 0.3566 | 0 | 60.252 | Eukaryota | 94919.0 | 0 | Chordata | 94919.0 | 0 | Carnivora | 94919.0 | 0 | Canidae | 94919.0 | 0 | Canis | 94919.0 | 0 | Canis lupus | 94919.0 | 0 |
| scf7180000052454 | 2452 | 0.4572 | 0 | 213.417 | Eukaryota | 463.0 | 0 | Streptophyta | 463.0 | 0 | Fabales | 463.0 | 0 | Fabaceae | 463.0 | 0 | Cicer | 463.0 | 0 | Cicer arietinum | 463.0 | 0 |
| scf7180000050743 | 26542 | 0.5056 | 0 | 692.6434 | Eukaryota | 8973.0 | 0 | Chordata | 8973.0 | 0 | Carnivora | 8973.0 | 0 | Canidae | 8973.0 | 0 | Canis | 7781.0 | 1 | Canis lupus | 6540.0 | 2 |
| scf7180000053286 | 3245 | 0.4545 | 0 | 626.2874 | no-hit | 0.0 | 0 | no-hit | 0.0 | 0 | no-hit | 0.0 | 0 | no-hit | 0.0 | 0 | no-hit | 0.0 | 0 | no-hit | 0.0 | 0 |
| scf7180000056197 | 1442958 | 0.3707 | 0 | 59.5939 | Eukaryota | 122987.0 | 0 | Chordata | 122987.0 | 0 | Carnivora | 122987.0 | 0 | Canidae | 122987.0 | 0 | Vulpes | 75640.0 | 1 | Vulpes vulpes | 75640.0 | 1 |
| scf7180000046641 | 100798 | 0.438 | 0 | 159.1105 | Eukaryota | 38651.0 | 0 | Platyhelminthes | 22724.0 | 2 | Strigeidida | 22724.0 | 5 | Schistosomatidae | 22724.0 | 5 | Schistosoma | 22724.0 | 5 | Schistosoma mansoni | 22724.0 | 6 |
| scf7180000046130 | 4703 | 0.5397 | 0 | 39.8378 | Eukaryota | 5262.0 | 0 | Chordata | 5262.0 | 0 | Carnivora | 5262.0 | 0 | Canidae | 5262.0 | 0 | Canis | 5262.0 | 0 | Canis lupus | 5262.0 | 0 |
| scf7180000050032 | 5462 | 0.4513 | 0 | 397.7829 | Eukaryota | 5151.0 | 0 | Chordata | 5151.0 | 0 | Tetraodontiformes | 3401.0 | 2 | Tetraodontidae | 3401.0 | 2 | Takifugu | 3401.0 | 2 | Takifugu rubripes | 3401.0 | 2 |
| scf7180000050972 | 13789 | 0.4459 | 0 | 118.2672 | no-hit | 0.0 | 0 | no-hit | 0.0 | 0 | no-hit | 0.0 | 0 | no-hit | 0.0 | 0 | no-hit | 0.0 | 0 | no-hit | 0.0 | 0 |
| scf7180000053150 | 3446 | 0.4446 | 0 | 428.9519 | no-hit | 0.0 | 0 | no-hit | 0.0 | 0 | no-hit | 0.0 | 0 | no-hit | 0.0 | 0 | no-hit | 0.0 | 0 | no-hit | 0.0 | 0 |
| scf7180000046026 | 165819 | 0.4193 | 0 | 59.7821 | Eukaryota | 12490.0 | 0 | Chordata | 12490.0 | 0 | Carnivora | 10661.0 | 1 | Ursidae | 5592.0 | 4 | Ursus | 5592.0 | 4 | Ursus arctos | 5592.0 | 4 |
| scf7180000044586 | 26396 | 0.4438 | 0 | 273.3147 | Eukaryota | 29464.0 | 0 | Chordata | 24329.0 | 2 | Gadiformes | 13766.0 | 7 | Gadidae | 13766.0 | 7 | Gadus | 13766.0 | 7 | Gadus morhua | 13766.0 | 7 |
| scf7180000051337 | 8564 | 0.4402 | 0 | 64.6595 | Eukaryota | 2529.0 | 0 | Chordata | 2529.0 | 0 | Kurtiformes | 2529.0 | 0 | Apogonidae | 2529.0 | 0 | Sphaeramia | 2529.0 | 0 | Sphaeramia orbicularis | 2529.0 | 0 |
| scf7180000044101 | 363973 | 0.4327 | 0 | 57.827 | Eukaryota | 28547.0 | 0 | Chordata | 28547.0 | 0 | Carnivora | 28547.0 | 0 | Canidae | 28547.0 | 0 | Canis | 16950.0 | 1 | Canis lupus | 16950.0 | 1 |
| scf7180000046169 | 189329 | 0.3783 | 0 | 60.9089 | Eukaryota | 19242.0 | 0 | Chordata | 19242.0 | 0 | Carnivora | 13609.0 | 2 | Canidae | 11616.0 | 3 | Canis | 11616.0 | 3 | Canis lupus | 11616.0 | 3 |
| scf7180000046239 | 90965 | 0.5822 | 0 | 57.5646 | Eukaryota | 42377.0 | 0 | Chordata | 42377.0 | 0 | Carnivora | 42377.0 | 0 | Canidae | 42377.0 | 0 | Canis | 36648.0 | 1 | Canis lupus | 36648.0 | 1 |
| scf7180000054531 | 21433 | 0.344 | 0 | 131.2685 | Eukaryota | 5840.0 | 0 | Chordata | 5840.0 | 0 | Carnivora | 4302.0 | 1 | Felidae | 2360.0 | 2 | Acinonyx | 2360.0 | 2 | Acinonyx jubatus | 2360.0 | 2 |
| scf7180000050303 | 27478 | 0.5345 | 0 | 393.4947 | Eukaryota | 25496.0 | 0 | Chordata | 25496.0 | 0 | Artiodactyla | 17886.0 | 1 | Bovidae | 17886.0 | 1 | Ovis | 10212.0 | 3 | Ovis aries | 10212.0 | 3 |
| scf7180000048044 | 64326 | 0.4402 | 0 | 59.2798 | Eukaryota | 43977.0 | 0 | Chordata | 43977.0 | 0 | Carnivora | 43977.0 | 0 | Canidae | 43977.0 | 0 | Canis | 43977.0 | 0 | Canis lupus | 43977.0 | 0 |
| scf7180000042554 | 901686 | 0.3497 | 0 | 58.4567 | Eukaryota | 51853.0 | 0 | Chordata | 51853.0 | 0 | Carnivora | 51853.0 | 0 | Canidae | 51853.0 | 0 | Canis | 51853.0 | 0 | Canis lupus | 51853.0 | 0 |
| scf7180000047988 | 68391 | 0.5555 | 0 | 55.4063 | Eukaryota | 6130.0 | 0 | Chordata | 6130.0 | 0 | Perissodactyla | 3180.0 | 4 | Equidae | 3180.0 | 4 | Equus | 3180.0 | 5 | Equus caballus | 3180.0 | 5 |
| scf7180000041432 | 959632 | 0.3571 | 100 | 59.7303 | Eukaryota | 98793.0 | 0 | Chordata | 98793.0 | 0 | Carnivora | 98793.0 | 0 | Canidae | 98793.0 | 0 | Canis | 98793.0 | 0 | Canis lupus | 98793.0 | 0 |
| scf7180000050404 | 2824 | 0.4487 | 0 | 419.6752 | Eukaryota | 2516.0 | 0 | Platyhelminthes | 1173.0 | 3 | Polystomatidea | 1173.0 | 3 | Polystomatidae | 1173.0 | 3 | Protopolystoma | 1173.0 | 3 | Protopolystoma xenopodis | 1173.0 | 3 |
| scf7180000043390 | 1085947 | 0.4214 | 0 | 58.7468 | Eukaryota | 53461.0 | 0 | Chordata | 53461.0 | 0 | Carnivora | 53461.0 | 0 | Canidae | 53461.0 | 0 | Canis | 45085.0 | 1 | Canis lupus | 45085.0 | 1 |
| scf7180000041544 | 16278 | 0.3184 | 0 | 29.5259 | Eukaryota | 9975.0 | 0 | Chordata | 9975.0 | 0 | Carnivora | 9975.0 | 0 | Felidae | 6219.0 | 2 | Felis | 6219.0 | 2 | Felis catus | 6219.0 | 2 |
| scf7180000050629 | 9125 | 0.4876 | 0 | 70.3658 | Eukaryota | 13923.0 | 0 | Chordata | 13923.0 | 0 | Carnivora | 13923.0 | 0 | Canidae | 13923.0 | 0 | Canis | 9500.0 | 1 | Canis lupus | 8489.0 | 2 |
| scf7180000040718 | 1964055 | 0.47 | 0 | 58.6451 | Eukaryota | 61792.0 | 0 | Chordata | 61792.0 | 0 | Carnivora | 61792.0 | 0 | Canidae | 61792.0 | 0 | Canis | 61792.0 | 0 | Canis lupus | 61792.0 | 0 |
| scf7180000055041 | 22476 | 0.4429 | 0 | 94.3111 | no-hit | 0.0 | 0 | no-hit | 0.0 | 0 | no-hit | 0.0 | 0 | no-hit | 0.0 | 0 | no-hit | 0.0 | 0 | no-hit | 0.0 | 0 |
| scf7180000052609 | 6877 | 0.4531 | 0 | 1035.2718 | Eukaryota | 3223.0 | 0 | Platyhelminthes | 1825.0 | 2 | Polystomatidea | 1825.0 | 2 | Polystomatidae | 1825.0 | 2 | Protopolystoma | 1825.0 | 2 | Protopolystoma xenopodis | 1825.0 | 2 |
| scf7180000041865 | 83973 | 0.4315 | 0 | 56.1322 | Eukaryota | 18065.0 | 0 | Chordata | 18065.0 | 0 | Carnivora | 13996.0 | 2 | Canidae | 9970.0 | 3 | Canis | 9970.0 | 3 | Canis lupus | 9970.0 | 3 |
| scf7180000048212 | 16360 | 0.4441 | 0 | 105.7096 | Eukaryota | 14659.0 | 0 | Chordata | 9198.0 | 3 | Gadiformes | 4233.0 | 7 | Gadidae | 4233.0 | 7 | Gadus | 4233.0 | 7 | Gadus morhua | 4233.0 | 7 |
| scf7180000044093 | 202060 | 0.4142 | 0 | 60.5754 | Eukaryota | 48075.0 | 0 | Chordata | 48075.0 | 0 | Carnivora | 48075.0 | 0 | Canidae | 48075.0 | 0 | Canis | 48075.0 | 0 | Canis lupus | 48075.0 | 0 |
| scf7180000041309 | 10103 | 0.4114 | 0 | 47.7011 | Eukaryota | 88089.0 | 0 | Chordata | 88089.0 | 0 | Carnivora | 88089.0 | 0 | Canidae | 88089.0 | 0 | Canis | 88089.0 | 0 | Canis lupus | 88089.0 | 0 |
| scf7180000055750 | 16818 | 0.4402 | 0 | 252.1877 | no-hit | 0.0 | 0 | no-hit | 0.0 | 0 | no-hit | 0.0 | 0 | no-hit | 0.0 | 0 | no-hit | 0.0 | 0 | no-hit | 0.0 | 0 |
| scf7180000044379 | 4513 | 0.4345 | 0 | 351.3779 | Eukaryota | 40797.0 | 0 | Chordata | 40797.0 | 0 | Carnivora | 40797.0 | 0 | Canidae | 40797.0 | 0 | Canis | 40797.0 | 0 | Canis lupus | 40797.0 | 0 |
| scf7180000047975 | 48902 | 0.4407 | 0 | 179.9954 | Eukaryota | 501.0 | 0 | Chordata | 501.0 | 0 | Salmoniformes | 501.0 | 0 | Salmonidae | 501.0 | 0 | Salmo | 501.0 | 0 | Salmo trutta | 501.0 | 0 |
| scf7180000056306 | 662647 | 0.4412 | 0 | 59.7218 | Eukaryota | 171390.0 | 0 | Chordata | 171390.0 | 0 | Carnivora | 171390.0 | 0 | Canidae | 171390.0 | 0 | Canis | 171390.0 | 0 | Canis lupus | 171390.0 | 0 |
| scf7180000049355 | 52524 | 0.4429 | 0 | 62.9167 | Eukaryota | 92260.0 | 0 | Chordata | 92260.0 | 0 | Carnivora | 92260.0 | 0 | Canidae | 92260.0 | 0 | Canis | 92260.0 | 0 | Canis lupus | 92260.0 | 0 |
| scf7180000046142 | 140951 | 0.4849 | 0 | 58.4743 | Eukaryota | 12963.0 | 0 | Chordata | 12963.0 | 0 | Carnivora | 12963.0 | 0 | Canidae | 6348.0 | 2 | Canis | 4733.0 | 7 | Canis lupus | 4733.0 | 7 |
| scf7180000051400 | 8526 | 0.4281 | 0 | 76.8507 | Eukaryota | 33398.0 | 0 | Chordata | 16812.0 | 1 | Gadiformes | 13474.0 | 3 | Gadidae | 13474.0 | 3 | Gadus | 13474.0 | 3 | Gadus morhua | 13474.0 | 3 |
| scf7180000041895 | 171047 | 0.414 | 0 | 60.7822 | Eukaryota | 33447.0 | 0 | Chordata | 33447.0 | 0 | Carnivora | 33447.0 | 0 | Canidae | 33447.0 | 0 | Canis | 33447.0 | 0 | Canis lupus | 33447.0 | 0 |
| scf7180000054613 | 17775 | 0.4307 | 0 | 226.5397 | Eukaryota | 35056.0 | 0 | Chordata | 24902.0 | 2 | Gadiformes | 10993.0 | 6 | Gadidae | 10993.0 | 7 | Gadus | 10993.0 | 7 | Gadus morhua | 10993.0 | 7 |
| scf7180000042817 | 210958 | 0.3999 | 0 | 59.6813 | Eukaryota | 97548.0 | 0 | Chordata | 97548.0 | 0 | Carnivora | 97548.0 | 0 | Canidae | 97548.0 | 0 | Canis | 97548.0 | 0 | Canis lupus | 97548.0 | 0 |
| scf7180000050738 | 6224 | 0.4335 | 0 | 94.6969 | Eukaryota | 226.0 | 0 | Platyhelminthes | 226.0 | 0 | Polystomatidea | 226.0 | 0 | Polystomatidae | 226.0 | 0 | Protopolystoma | 226.0 | 0 | Protopolystoma xenopodis | 226.0 | 0 |
| scf7180000052153 | 3577 | 0.447 | 0 | 1369.9819 | no-hit | 0.0 | 0 | no-hit | 0.0 | 0 | no-hit | 0.0 | 0 | no-hit | 0.0 | 0 | no-hit | 0.0 | 0 | no-hit | 0.0 | 0 |
| scf7180000050151 | 22823 | 0.5006 | 0 | 555.4958 | Eukaryota | 9565.0 | 0 | Chordata | 9565.0 | 0 | Carnivora | 9565.0 | 0 | Canidae | 9565.0 | 0 | Canis | 8358.0 | 1 | Canis lupus | 7090.0 | 2 |
| scf7180000044032 | 273300 | 0.4079 | 0 | 59.4288 | Eukaryota | 34583.0 | 0 | Chordata | 34583.0 | 0 | Carnivora | 34583.0 | 0 | Canidae | 34583.0 | 0 | Canis | 34583.0 | 0 | Canis lupus | 34583.0 | 0 |
| scf7180000052986 | 6306 | 0.4399 | 0 | 282.5631 | no-hit | 0.0 | 0 | no-hit | 0.0 | 0 | no-hit | 0.0 | 0 | no-hit | 0.0 | 0 | no-hit | 0.0 | 0 | no-hit | 0.0 | 0 |
| scf7180000053270 | 9320 | 0.3461 | 0 | 63.826 | Eukaryota | 4938.0 | 0 | Chordata | 4938.0 | 0 | Primates | 3090.0 | 2 | Hominidae | 3090.0 | 2 | Pan | 1560.0 | 3 | Pan troglodytes | 1560.0 | 3 |
| scf7180000046734 | 7458 | 0.4362 | 0 | 116.8155 | no-hit | 0.0 | 0 | no-hit | 0.0 | 0 | no-hit | 0.0 | 0 | no-hit | 0.0 | 0 | no-hit | 0.0 | 0 | no-hit | 0.0 | 0 |

| | | | | | | | | | | | | | | | | | | | | | |
|---|---|---|---|---|---|---|---|---|---|---|---|---|---|---|---|---|---|---|---|---|---|
| scf7180000040612 | 2062372 | 0.434 | 0 | 59.3877 | Eukaryota | 80203.0 | 0 | Chordata | 80203.0 | 0 | Carnivora | 80203.0 | 0 | Canidae | 80203.0 | 0 | Canis | 80203.0 | 0 | Canis lupus | 80203.0 | 0 |
| scf7180000042040 | 924982 | 0.3489 | 0 | 59.8315 | Eukaryota | 94974.0 | 0 | Chordata | 94974.0 | 0 | Carnivora | 94974.0 | 0 | Canidae | 94974.0 | 0 | Canis | 94974.0 | 0 | Canis lupus | 94974.0 | 0 |
| scf7180000041329 | 1084190 | 0.4505 | 0 | 57.8894 | Eukaryota | 47034.0 | 0 | Chordata | 47034.0 | 0 | Carnivora | 47034.0 | 0 | Canidae | 47034.0 | 0 | Canis | 43327.0 | 1 | Canis lupus | 43327.0 | 1 |
| scf7180000040666 | 2747427 | 0.4648 | 0 | 59.5632 | Eukaryota | 107143.0 | 0 | Chordata | 107143.0 | 0 | Carnivora | 107143.0 | 0 | Canidae | 107143.0 | 0 | Canis | 57666.0 | 1 | Canis lupus | 57666.0 | 1 |
| scf7180000040781 | 56253 | 0.3803 | 0 | 60.2795 | Eukaryota | 33207.0 | 0 | Chordata | 33207.0 | 0 | Carnivora | 33207.0 | 0 | Canidae | 33207.0 | 0 | Canis | 33207.0 | 0 | Canis lupus | 33207.0 | 0 |
| scf7180000047183 | 5202 | 0.5058 | 0 | 69.6066 | Eukaryota | 7177.0 | 0 | Chordata | 7177.0 | 0 | Carnivora | 7177.0 | 0 | Canidae | 7177.0 | 0 | Canis | 6395.0 | 1 | Canis lupus | 5487.0 | 2 |
| scf7180000042768 | 129816 | 0.398 | 0 | 61.2489 | Eukaryota | 78047.0 | 0 | Chordata | 78047.0 | 0 | Carnivora | 78047.0 | 0 | Canidae | 78047.0 | 0 | Canis | 78047.0 | 0 | Canis lupus | 78047.0 | 0 |
| scf7180000052964 | 5918 | 0.5039 | 0 | 48.9173 | Eukaryota | 7701.0 | 0 | Chordata | 7701.0 | 0 | Carnivora | 7701.0 | 0 | Canidae | 7701.0 | 0 | Canis | 6190.0 | 1 | Canis lupus | 5129.0 | 2 |
| scf7180000042384 | 677282 | 0.4203 | 0 | 60.1506 | Eukaryota | 91535.0 | 0 | Chordata | 91535.0 | 0 | Carnivora | 91535.0 | 0 | Canidae | 91535.0 | 0 | Canis | 53499.0 | 1 | Canis lupus | 53499.0 | 1 |
| scf7180000054693 | 53397 | 0.438 | 0 | 163.2594 | Eukaryota | 1568.0 | 0 | Platyhelminthes | 1087.0 | 1 | Polystomatidea | 1087.0 | 2 | Polystomatidae | 1087.0 | 2 | Protopolystoma | 1087.0 | 2 | Protopolystoma xenopodis | 1087.0 | 2 |
| scf7180000048637 | 25995 | 0.5426 | 0 | 822.2068 | Eukaryota | 14081.0 | 0 | Chordata | 14081.0 | 0 | Carnivora | 14081.0 | 0 | Canidae | 14081.0 | 0 | Canis | 13366.0 | 1 | Canis lupus | 13366.0 | 1 |
| scf7180000048355 | 17811 | 0.4535 | 0 | 1675.6602 | Eukaryota | 2022.0 | 0 | Streptophyta | 2022.0 | 0 | Fabales | 2022.0 | 0 | Fabaceae | 2022.0 | 0 | Cicer | 2022.0 | 0 | Cicer arietinum | 2022.0 | 0 |
| scf7180000050522 | 10514 | 0.4453 | 0 | 96.8578 | Eukaryota | 2967.0 | 0 | Chordata | 2543.0 | 1 | Carnivora | 2543.0 | 1 | Canidae | 2543.0 | 1 | Canis | 2543.0 | 1 | Canis lupus | 2543.0 | 1 |
| scf7180000051355 | 16031 | 0.4377 | 0 | 42.504 | Eukaryota | 3562.0 | 0 | Chordata | 3373.0 | 1 | Columbiformes | 2328.0 | 3 | Columbidae | 2328.0 | 3 | Streptopelia | 2328.0 | 3 | Streptopelia turtur | 2328.0 | 3 |
| scf7180000041122 | 402662 | 0.3607 | 0 | 59.2929 | Eukaryota | 93739.0 | 0 | Chordata | 93739.0 | 0 | Carnivora | 93739.0 | 0 | Canidae | 93739.0 | 0 | Canis | 93739.0 | 0 | Canis lupus | 93739.0 | 0 |
| scf7180000046086 | 18449 | 0.4369 | 0 | 61.4966 | Eukaryota | 40234.0 | 0 | Chordata | 40234.0 | 0 | Carnivora | 40234.0 | 0 | Canidae | 40234.0 | 0 | Canis | 40234.0 | 0 | Canis lupus | 40234.0 | 0 |
| scf7180000047400 | 30821 | 0.4152 | 0 | 57.5045 | Eukaryota | 13115.0 | 0 | Chordata | 13115.0 | 0 | Carnivora | 13115.0 | 0 | Canidae | 8903.0 | 1 | Canis | 8903.0 | 1 | Canis lupus | 8903.0 | 1 |
| scf7180000048990 | 64138 | 0.4447 | 0 | 82.0499 | Eukaryota | 32200.0 | 0 | Chordata | 32200.0 | 0 | Carnivora | 27276.0 | 1 | Canidae | 24829.0 | 2 | Canis | 21687.0 | 3 | Canis lupus | 21687.0 | 3 |
| scf7180000041179 | 1282269 | 0.3538 | 0 | 60.5614 | Eukaryota | 95140.0 | 0 | Chordata | 95140.0 | 0 | Carnivora | 95140.0 | 0 | Canidae | 95140.0 | 0 | Canis | 95140.0 | 0 | Canis lupus | 95140.0 | 0 |
| scf7180000053221 | 3909 | 0.4564 | 0 | 684.7947 | no-hit | 0.0 | 0 | no-hit | 0.0 | 0 | no-hit | 0.0 | 0 | no-hit | 0.0 | 0 | no-hit | 0.0 | 0 | no-hit | 0.0 | 0 |
| scf7180000041592 | 32614 | 0.3931 | 0 | 57.359 | Eukaryota | 51592.0 | 0 | Chordata | 51592.0 | 0 | Carnivora | 51592.0 | 0 | Canidae | 51592.0 | 0 | Canis | 51592.0 | 0 | Canis lupus | 51592.0 | 0 |
| scf7180000049447 | 5201 | 0.438 | 0 | 92.4097 | no-hit | 0.0 | 0 | no-hit | 0.0 | 0 | no-hit | 0.0 | 0 | no-hit | 0.0 | 0 | no-hit | 0.0 | 0 | no-hit | 0.0 | 0 |
| scf7180000042792 | 572248 | 0.4231 | 0 | 59.0023 | Eukaryota | 68577.0 | 0 | Chordata | 68577.0 | 0 | Carnivora | 68577.0 | 0 | Canidae | 68577.0 | 0 | Canis | 34580.0 | 1 | Canis lupus | 34580.0 | 1 |
| scf7180000044131 | 644836 | 0.4309 | 0 | 59.7011 | Eukaryota | 98510.0 | 0 | Chordata | 98510.0 | 0 | Carnivora | 98510.0 | 0 | Canidae | 98510.0 | 0 | Canis | 68642.0 | 1 | Canis lupus | 68642.0 | 1 |
| scf7180000041228 | 5436 | 0.3825 | 0 | 174.46 | Eukaryota | 1122.0 | 0 | Chordata | 1122.0 | 0 | Primates | 761.0 | 1 | Hominidae | 761.0 | 1 | Homo | 761.0 | 1 | Homo sapiens | 761.0 | 1 |
| scf7180000044302 | 6874 | 0.4488 | 0 | 311.6635 | no-hit | 0.0 | 0 | no-hit | 0.0 | 0 | no-hit | 0.0 | 0 | no-hit | 0.0 | 0 | no-hit | 0.0 | 0 | no-hit | 0.0 | 0 |
| scf7180000042085 | 245412 | 0.396 | 0 | 56.5432 | Eukaryota | 90880.0 | 0 | Chordata | 90880.0 | 0 | Carnivora | 90880.0 | 0 | Canidae | 90880.0 | 0 | Canis | 90880.0 | 0 | Canis lupus | 90880.0 | 0 |
| scf7180000049185 | 30833 | 0.4227 | 0 | 62.5121 | Eukaryota | 12443.0 | 0 | Chordata | 12443.0 | 0 | Carnivora | 12443.0 | 0 | Canidae | 12443.0 | 0 | Canis | 12443.0 | 0 | Canis lupus | 12443.0 | 0 |
| scf7180000045351 | 8356 | 0.4178 | 0 | 28.1986 | Eukaryota | 14995.0 | 0 | Chordata | 14995.0 | 0 | Carnivora | 14995.0 | 0 | Canidae | 14995.0 | 0 | Canis | 14995.0 | 0 | Canis lupus | 14995.0 | 0 |
| scf7180000052630 | 4970 | 0.4437 | 0 | 426.1921 | Eukaryota | 312.0 | 0 | Platyhelminthes | 178.0 | 1 | Polystomatidea | 178.0 | 1 | Polystomatidae | 178.0 | 1 | Protopolystoma | 178.0 | 1 | Protopolystoma xenopodis | 178.0 | 1 |
| scf7180000047908 | 42407 | 0.4407 | 0 | 366.81 | Eukaryota | 100202.0 | 0 | Chordata | 56802.0 | 1 | Gadiformes | 56802.0 | 2 | Gadidae | 56802.0 | 2 | Gadus | 56802.0 | 2 | Gadus morhua | 56802.0 | 2 |
| scf7180000043550 | 369060 | 0.5017 | 0 | 57.7308 | Eukaryota | 14204.0 | 0 | Chordata | 14204.0 | 0 | Carnivora | 14204.0 | 0 | Canidae | 14204.0 | 0 | Canis | 14204.0 | 0 | Canis lupus | 14204.0 | 0 |
| scf7180000047009 | 11259 | 0.3959 | 0 | 29.8491 | Eukaryota | 2574.0 | 0 | Chordata | 2574.0 | 0 | Carnivora | 2574.0 | 0 | Canidae | 2574.0 | 0 | Canis | 1554.0 | 1 | Canis lupus | 1554.0 | 1 |
| scf7180000050977 | 10725 | 0.4255 | 0 | 150.8658 | Eukaryota | 72734.0 | 0 | Chordata | 72734.0 | 0 | Carnivora | 72734.0 | 0 | Canidae | 72734.0 | 0 | Canis | 72734.0 | 0 | Canis lupus | 72734.0 | 0 |
| scf7180000051482 | 5051 | 0.4407 | 0 | 73.6709 | no-hit | 0.0 | 0 | no-hit | 0.0 | 0 | no-hit | 0.0 | 0 | no-hit | 0.0 | 0 | no-hit | 0.0 | 0 | no-hit | 0.0 | 0 |
| scf7180000048364 | 58408 | 0.3822 | 0 | 58.7259 | Eukaryota | 88523.0 | 0 | Chordata | 88523.0 | 0 | Carnivora | 88523.0 | 0 | Canidae | 88523.0 | 0 | Canis | 88523.0 | 0 | Canis lupus | 88523.0 | 0 |
| scf7180000044246 | 327623 | 0.4321 | 0 | 60.9087 | Eukaryota | 16310.0 | 0 | Chordata | 16310.0 | 0 | Carnivora | 16310.0 | 0 | Ursidae | 11441.0 | 3 | Ursus | 9036.0 | 4 | Ursus arctos | 5339.0 | 5 |
| scf7180000042192 | 635072 | 0.3976 | 0 | 59.8938 | Eukaryota | 68260.0 | 0 | Chordata | 68260.0 | 0 | Carnivora | 68260.0 | 0 | Canidae | 49965.0 | 2 | Canis | 42181.0 | 4 | Canis lupus | 42181.0 | 4 |
| scf7180000052501 | 6144 | 0.4492 | 0 | 439.0332 | no-hit | 0.0 | 0 | no-hit | 0.0 | 0 | no-hit | 0.0 | 0 | no-hit | 0.0 | 0 | no-hit | 0.0 | 0 | no-hit | 0.0 | 0 |
| scf7180000056433 | 254324 | 0.3639 | 0 | 58.719 | Eukaryota | 42556.0 | 0 | Chordata | 42556.0 | 0 | Carnivora | 42556.0 | 0 | Canidae | 42556.0 | 0 | Canis | 42556.0 | 0 | Canis lupus | 42556.0 | 0 |
| scf7180000050649 | 7265 | 0.5039 | 0 | 230.3279 | Eukaryota | 7525.0 | 0 | Chordata | 7525.0 | 0 | Carnivora | 7525.0 | 0 | Canidae | 7525.0 | 0 | Canis | 5716.0 | 1 | Canis lupus | 4741.0 | 2 |
| scf7180000042663 | 1307740 | 0.4219 | 0 | 59.6212 | Eukaryota | 104425.0 | 0 | Chordata | 104425.0 | 0 | Carnivora | 104425.0 | 0 | Canidae | 104425.0 | 0 | Canis | 104425.0 | 0 | Canis lupus | 104425.0 | 0 |
| scf7180000040991 | 1429432 | 0.4856 | 0 | 58.9845 | Eukaryota | 121465.0 | 0 | Chordata | 121465.0 | 0 | Carnivora | 121465.0 | 0 | Canidae | 121465.0 | 0 | Canis | 100507.0 | 1 | Canis lupus | 100507.0 | 1 |
| scf7180000040639 | 1988642 | 0.3862 | 0 | 60.7643 | Eukaryota | 137830.0 | 0 | Chordata | 137830.0 | 0 | Carnivora | 109328.0 | 1 | Canidae | 109328.0 | 1 | Canis | 109328.0 | 1 | Canis lupus | 109328.0 | 1 |
| scf7180000045563 | 65555 | 0.4407 | 0 | 150.2143 | Eukaryota | 5163.0 | 0 | Chordata | 5163.0 | 0 | Columbiformes | 2266.0 | 3 | Columbidae | 2266.0 | 3 | Streptopelia | 2266.0 | 4 | Streptopelia turtur | 2266.0 | 4 |
| scf7180000041403 | 1137888 | 0.3494 | 0 | 59.8372 | Eukaryota | 93446.0 | 0 | Chordata | 93446.0 | 0 | Carnivora | 93446.0 | 0 | Canidae | 93446.0 | 0 | Canis | 93446.0 | 0 | Canis lupus | 93446.0 | 0 |
| scf7180000041120 | 137134 | 0.3664 | 0 | 60.2414 | Eukaryota | 94627.0 | 0 | Chordata | 94627.0 | 0 | Carnivora | 94627.0 | 0 | Canidae | 94627.0 | 0 | Canis | 94627.0 | 0 | Canis lupus | 94627.0 | 0 |
| scf7180000043908 | 296613 | 0.3908 | 0 | 60.2164 | Eukaryota | 14550.0 | 0 | Chordata | 14550.0 | 0 | Carnivora | 14550.0 | 0 | Canidae | 14550.0 | 0 | Canis | 14550.0 | 0 | Canis lupus | 14550.0 | 0 |
| scf7180000041610 | 991216 | 0.3774 | 0 | 60.4535 | Eukaryota | 90966.0 | 0 | Chordata | 90966.0 | 0 | Carnivora | 90966.0 | 0 | Canidae | 90966.0 | 0 | Canis | 90966.0 | 0 | Canis lupus | 90966.0 | 0 |
| scf7180000041729 | 954919 | 0.3856 | 0 | 61.2335 | Eukaryota | 58160.0 | 0 | Chordata | 58160.0 | 0 | Carnivora | 58160.0 | 0 | Canidae | 58160.0 | 0 | Canis | 58160.0 | 0 | Canis lupus | 58160.0 | 0 |
| scf7180000042427 | 582131 | 0.5007 | 0 | 59.9676 | Eukaryota | 46649.0 | 0 | Chordata | 46649.0 | 0 | Carnivora | 46649.0 | 0 | Canidae | 46649.0 | 0 | Canis | 42045.0 | 1 | Canis lupus | 42045.0 | 1 |
| scf7180000041933 | 63141 | 0.4297 | 0 | 55.9785 | Eukaryota | 92750.0 | 0 | Chordata | 92750.0 | 0 | Carnivora | 92750.0 | 0 | Canidae | 92750.0 | 0 | Canis | 92750.0 | 0 | Canis lupus | 92750.0 | 0 |
| scf7180000047412 | 9212 | 0.4454 | 0 | 258.9469 | Eukaryota | 2772.0 | 0 | Chordata | 2772.0 | 0 | Carnivora | 2772.0 | 0 | Canidae | 2772.0 | 0 | Canis | 2772.0 | 0 | Canis lupus | 2772.0 | 0 |
| scf7180000052239 | 8982 | 0.4429 | 0 | 221.2519 | Eukaryota | 8270.0 | 0 | Platyhelminthes | 3766.0 | 2 | Polystomatidea | 3766.0 | 3 | Polystomatidae | 3766.0 | 3 | Protopolystoma | 3766.0 | 3 | Protopolystoma xenopodis | 3766.0 | 3 |
| scf7180000054035 | 12410 | 0.625 | 0 | 232.1203 | Eukaryota | 65488.0 | 0 | Chordata | 65488.0 | 0 | Primates | 22687.0 | 3 | Bovidae | 20820.0 | 4 | Ovis | 20820.0 | 4 | Ovis aries | 20820.0 | 4 |
| scf7180000053523 | 2765 | 0.4456 | 0 | 296.2627 | no-hit | 0.0 | 0 | no-hit | 0.0 | 0 | no-hit | 0.0 | 0 | no-hit | 0.0 | 0 | no-hit | 0.0 | 0 | no-hit | 0.0 | 0 |
| scf7180000046496 | 140214 | 0.5066 | 0 | 60.5336 | Eukaryota | 24205.0 | 0 | Chordata | 24205.0 | 0 | Carnivora | 24205.0 | 0 | Canidae | 19978.0 | 2 | Canis | 13518.0 | 3 | Canis lupus | 13518.0 | 3 |
| scf7180000044444 | 336273 | 0.3687 | 0 | 61.1782 | Eukaryota | 82717.0 | 0 | Chordata | 82717.0 | 0 | Carnivora | 82717.0 | 0 | Canidae | 82717.0 | 0 | Canis | 82717.0 | 0 | Canis lupus | 82717.0 | 0 |
| scf7180000056115 | 6022 | 0.4495 | 0 | 258.8785 | Eukaryota | 331.0 | 0 | Chordata | 331.0 | 0 | Kurtiformes | 331.0 | 0 | Apogonidae | 331.0 | 0 | Sphaeramia | 331.0 | 0 | Sphaeramia orbicularis | 331.0 | 0 |
| scf7180000050880 | 13396 | 0.412 | 0 | 54.2833 | Eukaryota | 42009.0 | 0 | Chordata | 42009.0 | 0 | Carnivora | 42009.0 | 0 | Canidae | 42009.0 | 0 | Canis | 42009.0 | 0 | Canis lupus | 42009.0 | 0 |
| scf7180000050217 | 16383 | 0.375 | 0 | 56.5053 | Eukaryota | 51140.0 | 0 | Chordata | 51140.0 | 0 | Carnivora | 51140.0 | 0 | Canidae | 51140.0 | 0 | Canis | 51140.0 | 0 | Canis lupus | 51140.0 | 0 |
| scf7180000049076 | 15488 | 0.4386 | 0 | 89.3743 | Eukaryota | 5019.0 | 0 | Chordata | 4015.0 | 1 | Salmoniformes | 2379.0 | 2 | Salmonidae | 2379.0 | 2 | Salmo | 2379.0 | 2 | Salmo trutta | 2379.0 | 2 |
| scf7180000041190 | 1313156 | 0.3555 | 0 | 59.9481 | Eukaryota | 89655.0 | 0 | Chordata | 89655.0 | 0 | Carnivora | 89655.0 | 0 | Canidae | 89655.0 | 0 | Canis | 89655.0 | 0 | Canis lupus | 89655.0 | 0 |
| scf7180000051196 | 4603 | 0.4545 | 0 | 548.7587 | Eukaryota | 163.0 | 0 | Streptophyta | 163.0 | 0 | Fabales | 163.0 | 0 | Fabaceae | 163.0 | 0 | Cicer | 163.0 | 0 | Cicer arietinum | 163.0 | 0 |
| scf7180000051240 | 13385 | 0.603 | 0 | 56.6511 | Eukaryota | 1927.0 | 0 | Chordata | 1927.0 | 0 | Carnivora | 1927.0 | 0 | Canidae | 1927.0 | 0 | Canis | 1927.0 | 0 | Canis lupus | 1927.0 | 0 |
| scf7180000044622 | 406183 | 0.3983 | 0 | 58.2863 | Eukaryota | 41388.0 | 0 | Chordata | 41388.0 | 0 | Carnivora | 41388.0 | 0 | Canidae | 14274.0 | 4 | Canis | 9533.0 | 6 | Canis lupus | 9533.0 | 6 |
| scf7180000055031 | 26476 | 0.4463 | 0 | 146.9353 | Eukaryota | 11460.0 | 0 | Chordata | 11460.0 | 0 | Tetraodontiformes | 5735.0 | 3 | Tetraodontidae | 5735.0 | 3 | Takifugu | 5735.0 | 3 | Takifugu rubripes | 4972.0 | 4 |
| scf7180000053906 | 5521 | 0.4523 | 0 | 28.4444 | Eukaryota | 34702.0 | 0 | Chordata | 34702.0 | 0 | Carnivora | 34702.0 | 0 | Canidae | 34702.0 | 0 | Canis | 34702.0 | 0 | Canis lupus | 34702.0 | 0 |
| scf7180000048761 | 19282 | 0.4436 | 0 | 206.5941 | Eukaryota | 91196.0 | 0 | Chordata | 91196.0 | 0 | Carnivora | 91196.0 | 0 | Canidae | 91196.0 | 0 | Canis | 91196.0 | 0 | Canis lupus | 91196.0 | 0 |
| scf7180000041571 | 961449 | 0.4114 | 0 | 59.415 | Eukaryota | 89763.0 | 0 | Chordata | 89763.0 | 0 | Carnivora | 89763.0 | 0 | Canidae | 89763.0 | 0 | Canis | 89763.0 | 0 | Canis lupus | 89763.0 | 0 |
| scf7180000052718 | 5390 | 0.4432 | 0 | 413.2349 | Eukaryota | 4658.0 | 0 | Chordata | 2176.0 | 2 | Polystomatidea | 1927.0 | 3 | Polystomatidae | 1927.0 | 3 | Protopolystoma | 1927.0 | 3 | Protopolystoma xenopodis | 1927.0 | 3 |
| scf7180000042837 | 584218 | 0.4232 | 0 | 58.4807 | Eukaryota | 45424.0 | 0 | Chordata | 45424.0 | 0 | Carnivora | 45424.0 | 0 | Canidae | 45424.0 | 0 | Canis | 40576.0 | 1 | Canis lupus | 40576.0 | 1 |
| scf7180000048833 | 63290 | 0.4756 | 0 | 60.3036 | Eukaryota | 21198.0 | 0 | Chordata | 21198.0 | 0 | Carnivora | 21198.0 | 0 | Canidae | 21198.0 | 0 | Canis | 21198.0 | 0 | Canis lupus | 21198.0 | 0 |
| scf7180000041188 | 311827 | 0.3821 | 0 | 60.8801 | Eukaryota | 46782.0 | 0 | Chordata | 46782.0 | 0 | Carnivora | 46782.0 | 0 | Canidae | 46782.0 | 0 | Canis | 46782.0 | 0 | Canis lupus | 46782.0 | 0 |
| scf7180000041203 | 564953 | 0.5389 | 0 | 59.2727 | Eukaryota | 82240.0 | 0 | Chordata | 82240.0 | 0 | Carnivora | 82240.0 | 0 | Canidae | 82240.0 | 0 | Canis | 82240.0 | 0 | Canis lupus | 82240.0 | 0 |
| scf7180000046694 | 144840 | 0.4966 | 0 | 58.4754 | Eukaryota | 11844.0 | 0 | Chordata | 11844.0 | 0 | Carnivora | 7852.0 | 2 | Canidae | 3763.0 | 5 | Canis | 2810.0 | 8 | Canis lupus | 2810.0 | 8 |
| scf7180000041909 | 774638 | 0.4127 | 0 | 59.945 | Eukaryota | 89572.0 | 0 | Chordata | 89572.0 | 0 | Carnivora | 89572.0 | 0 | Canidae | 89572.0 | 0 | Canis | 89572.0 | 0 | Canis lupus | 89572.0 | 0 |
| scf7180000054463 | 25183 | 0.4459 | 0 | 348.1164 | Eukaryota | 38839.0 | 0 | Chordata | 38839.0 | 0 | Carnivora | 38839.0 | 0 | Canidae | 38839.0 | 0 | Canis | 38839.0 | 0 | Canis lupus | 38839.0 | 0 |
| scf7180000047439 | 95101 | 0.4518 | 0 | 59.1846 | Eukaryota | 20456.0 | 0 | Chordata | 20456.0 | 0 | Carnivora | 20456.0 | 0 | Canidae | 20456.0 | 0 | Canis | 15564.0 | 1 | Canis lupus | 15564.0 | 1 |
| scf7180000041863 | 28619 | 0.3801 | 0 | 52.9729 | Eukaryota | 91261.0 | 0 | Chordata | 91261.0 | 0 | Carnivora | 91261.0 | 0 | Canidae | 91261.0 | 0 | Canis | 91261.0 | 0 | Canis lupus | 91261.0 | 0 |
| scf7180000041308 | 788850 | 0.453 | 0 | 60.9466 | Eukaryota | 80347.0 | 0 | Chordata | 80347.0 | 0 | Carnivora | 80347.0 | 0 | Canidae | 80347.0 | 0 | Canis | 71641.0 | 1 | Canis lupus | 71641.0 | 1 |
| scf7180000043694 | 257237 | 0.4737 | 0 | 68.4724 | Eukaryota | 54528.0 | 0 | Chordata | 54528.0 | 0 | Carnivora | 54528.0 | 0 | Canidae | 54528.0 | 0 | Canis | 54528.0 | 0 | Canis lupus | 54528.0 | 0 |
| scf7180000044284 | 7558 | 0.4395 | 0 | 213.3251 | Eukaryota | 1806.0 | 0 | Chordata | 1806.0 | 0 | Tetraodontiformes | 1067.0 | 1 | Tetraodontidae | 1067.0 | 1 | Takifugu | 1067.0 | 1 | Takifugu rubripes | 1067.0 | 1 |
| scf7180000040698 | 1961874 | 0.3621 | 0 | 59.74 | Eukaryota | 124447.0 | 0 | Chordata | 124447.0 | 0 | Carnivora | 124447.0 | 0 | Canidae | 124447.0 | 0 | Canis | 124447.0 | 0 | Canis lupus | 124447.0 | 0 |
| scf7180000056320 | 484100 | 0.4415 | 0 | 69.8869 | Eukaryota | 51317.0 | 0 | Chordata | 51317.0 | 0 | Carnivora | 51317.0 | 0 | Canidae | 51317.0 | 0 | Canis | 46107.0 | 1 | Canis lupus | 46107.0 | 1 |
| scf7180000055091 | 19897 | 0.435 | 0 | 349.5752 | Eukaryota | 7839.0 | 0 | Chordata | 5780.0 | 1 | Salmoniformes | 3209.0 | 3 | Salmonidae | 3209.0 | 3 | Salmo | 3209.0 | 3 | Salmo trutta | 3209.0 | 3 |
| scf7180000041546 | 85780 | 0.4501 | 0 | 97.3072 | Eukaryota | 92851.0 | 0 | Chordata | 92851.0 | 0 | Carnivora | 92851.0 | 0 | Canidae | 92851.0 | 0 | Canis | 92851.0 | 0 | Canis lupus | 92851.0 | 0 |
| scf7180000042077 | 705550 | 0.4364 | 0 | 58.1353 | Eukaryota | 76011.0 | 0 | Chordata | 76011.0 | 0 | Carnivora | 76011.0 | 0 | Canidae | 51758.0 | 1 | Canis | 42821.0 | 3 | Canis lupus | 42821.0 | 3 |
| scf7180000042019 | 134765 | 0.3862 | 0 | 56.6391 | Eukaryota | 54922.0 | 0 | Chordata | 54922.0 | 0 | Carnivora | 54922.0 | 0 | Canidae | 54922.0 | 0 | Canis | 54922.0 | 0 | Canis lupus | 54922.0 | 0 |
| scf7180000050828 | 1708 | 0.4491 | 0 | 35.6505 | no-hit | 0.0 | 0 | no-hit | 0.0 | 0 | no-hit | 0.0 | 0 | no-hit | 0.0 | 0 | no-hit | 0.0 | 0 | no-hit | 0.0 | 0 |
| scf7180000054384 | 77578 | 0.4543 | 0 | 99.8954 | Eukaryota | 89256.0 | 0 | Chordata | 89256.0 | 0 | Carnivora | 89256.0 | 0 | Canidae | 89256.0 | 0 | Canis | 89256.0 | 0 | Canis lupus | 89256.0 | 0 |
| scf7180000051326 | 8074 | 0.4368 | 0 | 173.7956 | Eukaryota | 17086.0 | 0 | Chordata | 11246.0 | 2 | Gonorynchiformes | 6111.0 | 5 | Chanidae | 6111.0 | 5 | Chanos | 6111.0 | 5 | Chanos chanos | 6111.0 | 5 |
| scf7180000043049 | 586186 | 0.3789 | 0 | 60.6948 | Eukaryota | 25049.0 | 0 | Chordata | 25049.0 | 0 | Carnivora | 22410.0 | 1 | Canidae | 22410.0 | 1 | Canis | 22410.0 | 1 | Canis lupus | 22410.0 | 1 |
| scf7180000056245 | 1060542 | 0.3877 | 0 | 60.7768 | Eukaryota | 45375.0 | 0 | Chordata | 45375.0 | 0 | Carnivora | 45375.0 | 0 | Canidae | 45375.0 | 0 | Canis | 40702.0 | 1 | Canis lupus | 40702.0 | 1 |
| scf7180000053596 | 5909 | 0.451 | 0 | 455.7448 | no-hit | 0.0 | 0 | no-hit | 0.0 | 0 | no-hit | 0.0 | 0 | no-hit | 0.0 | 0 | no-hit | 0.0 | 0 | no-hit | 0.0 | 0 |
| scf7180000050163 | 4189 | 0.4426 | 0 | 70.5055 | no-hit | 0.0 | 0 | no-hit | 0.0 | 0 | no-hit | 0.0 | 0 | no-hit | 0.0 | 0 | no-hit | 0.0 | 0 | no-hit | 0.0 | 0 |
| scf7180000047791 | 76408 | 0.4662 | 0 | 59.4863 | Eukaryota | 6076.0 | 0 | Chordata | 6076.0 | 0 | Carnivora | 6076.0 | 0 | Canidae | 6076.0 | 0 | Canis | 6076.0 | 0 | Canis lupus | 6076.0 | 0 |
| scf7180000048147 | 24605 | 0.4379 | 0 | 69.3086 | Eukaryota | 3528.0 | 0 | Platyhelminthes | 2237.0 | 1 | Polystomatidea | 2237.0 | 1 | Polystomatidae | 2237.0 | 1 | Protopolystoma | 2237.0 | 1 | Protopolystoma xenopodis | 2237.0 | 1 |
| scf7180000055414 | 44906 | 0.4452 | 0 | 208.1929 | Eukaryota | 16940.0 | 0 | Chordata | 13181.0 | 2 | Gadiformes | 4638.0 | 5 | Gadidae | 4638.0 | 5 | Gadus | 4638.0 | 5 | Gadus morhua | 4638.0 | 5 |
| scf7180000041508 | 222023 | 0.3633 | 0 | 57.4518 | Eukaryota | 31420.0 | 0 | Chordata | 31420.0 | 0 | Carnivora | 31420.0 | 0 | Otariidae | 26142.0 | 1 | Callorhinus | 11442.0 | 3 | Callorhinus ursinus | 11442.0 | 3 |
| scf7180000043433 | 81139 | 0.3967 | 0 | 280.6775 | Eukaryota | 92767.0 | 0 | Chordata | 92767.0 | 0 | Carnivora | 92767.0 | 0 | Canidae | 92767.0 | 0 | Canis | 92767.0 | 0 | Canis lupus | 92767.0 | 0 |
| scf7180000052294 | 2328 | 0.4506 | 0 | 381.8878 | no-hit | 0.0 | 0 | no-hit | 0.0 | 0 | no-hit | 0.0 | 0 | no-hit | 0.0 | 0 | no-hit | 0.0 | 0 | no-hit | 0.0 | 0 |
| scf7180000045289 | 205551 | 0.4593 | 0 | 60.3202 | Eukaryota | 86107.0 | 0 | Chordata | 86107.0 | 0 | Carnivora | 86107.0 | 0 | Canidae | 86107.0 | 0 | Canis | 86107.0 | 0 | Canis lupus | 86107.0 | 0 |
| scf7180000043532 | 24113 | 0.3892 | 0 | 59.138 | Eukaryota | 93015.0 | 0 | Chordata | 93015.0 | 0 | Carnivora | 93015.0 | 0 | Canidae | 93015.0 | 0 | Canis | 93015.0 | 0 | Canis lupus | 93015.0 | 0 |
| scf7180000044015 | 407625 | 0.3976 | 0 | 61.5781 | Eukaryota | 93061.0 | 0 | Chordata | 93061.0 | 0 | Carnivora | 93061.0 | 0 | Canidae | 93061.0 | 0 | Canis | 93061.0 | 0 | Canis lupus | 93061.0 | 0 |
| scf7180000055200 | 102740 | 0.4417 | 0 | 745.0216 | Eukaryota | 21092.0 | 0 | Chordata | 20305.0 | 1 | Tetraodontiformes | 12644.0 | 4 | Tetraodontidae | 12644.0 | 4 | Takifugu | 12644.0 | 4 | Takifugu rubripes | 12644.0 | 4 |
| scf7180000041076 | 1432979 | 0.3772 | 0 | 58.3486 | Eukaryota | 58455.0 | 0 | Chordata | 58455.0 | 0 | Carnivora | 58455.0 | 0 | Canidae | 58455.0 | 0 | Canis | 58455.0 | 0 | Canis lupus | 58455.0 | 0 |
| scf7180000053280 | 15651 | 0.4506 | 0 | 60.5696 | Eukaryota | 10113.0 | 0 | Chordata | 10113.0 | 0 | Carnivora | 10113.0 | 0 | Canidae | 10113.0 | 0 | Canis | 10113.0 | 0 | Canis lupus | 10113.0 | 0 |
| scf7180000043942 | 4000 | 0.4698 | 0 | 27.2904 | Eukaryota | 4066.0 | 0 | Chordata | 4066.0 | 0 | Carnivora | 2568.0 | 1 | Canidae | 2568.0 | 1 | Canis | 2568.0 | 1 | Canis lupus | 2568.0 | 1 |
| scf7180000054602 | 87896 | 0.4432 | 0 | 123.4686 | Eukaryota | 75241.0 | 0 | Chordata | 75241.0 | 0 | Carnivora | 75241.0 | 0 | Canidae | 75241.0 | 0 | Canis | 75241.0 | 0 | Canis lupus | 75241.0 | 0 |
| scf7180000053452 | 3710 | 0.4496 | 0 | 637.2535 | no-hit | 0.0 | 0 | no-hit | 0.0 | 0 | no-hit | 0.0 | 0 | no-hit | 0.0 | 0 | no-hit | 0.0 | 0 | no-hit | 0.0 | 0 |
| scf7180000040716 | 244964 | 0.4052 | 0 | 58.478 | Eukaryota | 91525.0 | 0 | Chordata | 91525.0 | 0 | Carnivora | 91525.0 | 0 | Canidae | 91525.0 | 0 | Canis | 91525.0 | 0 | Canis lupus | 91525.0 | 0 |
| scf7180000049790 | 31523 | 0.4382 | 0 | 198.4593 | Eukaryota | 1909.0 | 0 | Chordata | 1909.0 | 0 | Tetraodontiformes | 1772.0 | 1 | Tetraodontidae | 1772.0 | 1 | Takifugu | 1772.0 | 1 | Takifugu rubripes | 1772.0 | 1 |
| scf7180000053980 | 2318 | 0.4538 | 0 | 312.6433 | Eukaryota | 714.0 | 0 | Platyhelminthes | 442.0 | 1 | Polystomatidea | 442.0 | 1 | Polystomatidae | 442.0 | 1 | Protopolystoma | 442.0 | 1 | Protopolystoma xenopodis | 442.0 | 1 |
| scf7180000047011 | 8094 | 0.4341 | 0 | 36.2807 | no-hit | 0.0 | 0 | no-hit | 0.0 | 0 | no-hit | 0.0 | 0 | no-hit | 0.0 | 0 | no-hit | 0.0 | 0 | no-hit | 0.0 | 0 |
| scf7180000056348 | 823438 | 0.4097 | 0 | 63.0307 | Eukaryota | 59109.0 | 0 | Chordata | 59109.0 | 0 | Carnivora | 59109.0 | 0 | Canidae | 59109.0 | 0 | Canis | 59109.0 | 0 | Canis lupus | 59109.0 | 0 |
| scf7180000045259 | 64370 | 0.5623 | 100 | 52.987 | Eukaryota | 24344.0 | 0 | Chordata | 24344.0 | 0 | Carnivora | 24344.0 | 0 | Canidae | 24344.0 | 0 | Canis | 24344.0 | 0 | Canis lupus | 24344.0 | 0 |
| scf7180000041598 | 979350 | 0.4945 | 0 | 57.1564 | Eukaryota | 56281.0 | 0 | Chordata | 56281.0 | 0 | Carnivora | 56281.0 | 0 | Canidae | 56281.0 | 0 | Canis | 44578.0 | 1 | Canis lupus | 44578.0 | 1 |
| scf7180000047708 | 6314 | 0.4462 | 0 | 143.9565 | Eukaryota | 813.0 | 0 | Chordata | 813.0 | 0 | Kurtiformes | 813.0 | 0 | Apogonidae | 813.0 | 0 | Sphaeramia | 813.0 | 0 | Sphaeramia orbicularis | 813.0 | 0 |
| scf7180000041667 | 397597 | 0.3644 | 0 | 59.8463 | Eukaryota | 102808.0 | 0 | Chordata | 102808.0 | 0 | Carnivora | 102808.0 | 0 | Canidae | 102808.0 | 0 | Canis | 102808.0 | 0 | Canis lupus | 102808.0 | 0 |
| scf7180000041796 | 900636 | 0.3771 | 0 | 60.7089 | Eukaryota | 31305.0 | 0 | Chordata | 31305.0 | 0 | Carnivora | 31305.0 | 0 | Canidae | 31305.0 | 0 | Canis | 28605.0 | 1 | Canis lupus | 28605.0 | 1 |
| scf7180000051412 | 3778 | 0.4494 | 0 | 771.7344 | no-hit | 0.0 | 0 | no-hit | 0.0 | 0 | no-hit | 0.0 | 0 | no-hit | 0.0 | 0 | no-hit | 0.0 | 0 | no-hit | 0.0 | 0 |
| scf7180000052300 | 3572 | 0.5006 | 0 | 168.3549 | Eukaryota | 5603.0 | 0 | Chordata | 5603.0 | 0 | Carnivora | 5603.0 | 0 | Canidae | 5603.0 | 0 | Canis | 4023.0 | 1 | Canis lupus | 3222.0 | 2 |
| scf7180000046969 | 92326 | 0.4504 | 0 | 116.7819 | Eukaryota | 78291.0 | 0 | Chordata | 78291.0 | 0 | Carnivora | 67334.0 | 1 | Canidae | 67334.0 | 1 | Canis | 67334.0 | 1 | Canis lupus | 67334.0 | 1 |
| scf7180000049477 | 29928 | 0.5181 | 0 | 56.0684 | Eukaryota | 31728.0 | 0 | Chordata | 31728.0 | 0 | Carnivora | 31728.0 | 0 | Canidae | 28818.0 | 3 | Canis | 27887.0 | 4 | Canis lupus | 27887.0 | 4 |
| scf7180000047996 | 64008 | 0.498 | 0 | 153.9322 | Eukaryota | 9418.0 | 0 | Chordata | 9418.0 | 0 | Carnivora | 9418.0 | 0 | Canidae | 9418.0 | 0 | Canis | 8801.0 | 1 | Canis lupus | 7664.0 | 2 |
| scf7180000045939 | 106225 | 0.4438 | 0 | 180.7955 | Eukaryota | 94257.0 | 0 | Chordata | 94257.0 | 0 | Carnivora | 94257.0 | 0 | Canidae | 94257.0 | 0 | Canis | 94257.0 | 0 | Canis lupus | 94257.0 | 0 |
| scf7180000050518 | 11464 | 0.5027 | 0 | 330.2662 | Eukaryota | 8526.0 | 0 | Chordata | 8526.0 | 0 | Carnivora | 8526.0 | 0 | Canidae | 8526.0 | 0 | Canis | 6944.0 | 1 | Canis lupus | 5824.0 | 2 |
| scf7180000054081 | 6404 | 0.4403 | 0 | 155.4238 | Eukaryota | 5965.0 | 0 | Platyhelminthes | 3330.0 | 2 | Polystomatidea | 3330.0 | 3 | Polystomatidae | 3330.0 | 3 | Protopolystoma | 3330.0 | 3 | Protopolystoma xenopodis | 3330.0 | 3 |
| scf7180000055115 | 13202 | 0.4503 | 0 | 123.0132 | Eukaryota | 3123.0 | 0 | Chordata | 3123.0 | 0 | Carnivora | 2507.0 | 1 | Canidae | 2507.0 | 1 | Canis | 2507.0 | 1 | Canis lupus | 2507.0 | 1 |
| scf7180000051184 | 5046 | 0.4449 | 0 | 1444.8906 | no-hit | 0.0 | 0 | no-hit | 0.0 | 0 | no-hit | 0.0 | 0 | no-hit | 0.0 | 0 | no-hit | 0.0 | 0 | no-hit | 0.0 | 0 |
| scf7180000040887 | 427261 | 0.3842 | 0 | 59.7944 | Eukaryota | 89237.0 | 0 | Chordata | 89237.0 | 0 | Carnivora | 89237.0 | 0 | Canidae | 89237.0 | 0 | Canis | 89237.0 | 0 | Canis lupus | 89237.0 | 0 |
| scf7180000042760 | 7893 | 0.4049 | 0 | 42.4301 | Eukaryota | 6248.0 | 1 | Chordata | 6248.0 | 1 | unresolved | 2742.0 | 4 | unresolved | 2742.0 | 4 | unresolved | 2742.0 | 5 | unresolved | 2742.0 | 5 |
| scf7180000049935 | 32670 | 0.4442 | 0 | 140.6388 | Eukaryota | 3189.0 | 0 | Chordata | 3189.0 | 0 | Kurtiformes | 1430.0 | 4 | Apogonidae | 1430.0 | 4 | Sphaeramia | 1430.0 | 4 | Sphaeramia orbicularis | 1430.0 | 4 |
| scf7180000049196 | 60406 | 0.3546 | 0 | 60.4538 | Eukaryota | 23475.0 | 0 | Chordata | 23475.0 | 0 | Carnivora | 23475.0 | 0 | Felidae | 8008.0 | 3 | Felis | 8008.0 | 3 | Felis catus | 8008.0 | 4 |
| scf7180000054947 | 13005 | 0.4375 | 0 | 231.9852 | no-hit | 0.0 | 0 | no-hit | 0.0 | 0 | no-hit | 0.0 | 0 | no-hit | 0.0 | 0 | no-hit | 0.0 | 0 | no-hit | 0.0 | 0 |
| scf7180000044906 | 223846 | 0.5064 | 0 | 57.0676 | Eukaryota | 20493.0 | 0 | Chordata | 20493.0 | 0 | Carnivora | 20493.0 | 0 | Canidae | 8297.0 | 2 | Canis | 5593.0 | 5 | Canis lupus | 5593.0 | 5 |
| scf7180000043173 | 493187 | 0.4259 | 0 | 59.2149 | Eukaryota | 63976.0 | 0 | Chordata | 63976.0 | 0 | Carnivora | 63976.0 | 0 | Canidae | 63976.0 | 0 | Vulpes | 39195.0 | 1 | Vulpes vulpes | 39195.0 | 1 |
| scf7180000048519 | 29125 | 0.4532 | 0 | 55.2645 | Eukaryota | 18707.0 | 0 | Chordata | 18707.0 | 0 | Carnivora | 18707.0 | 0 | Canidae | 18707.0 | 0 | Canis | 18707.0 | 0 | Canis lupus | 18707.0 | 0 |
| scf7180000046071 | 145949 | 0.6127 | 0 | 55.5037 | Eukaryota | 20696.0 | 0 | Chordata | 20696.0 | 0 | Artiodactyla | 17467.0 | 1 | Delphinidae | 11082.0 | 2 | Globicephala | 11082.0 | 3 | Globicephala melas | 11082.0 | 3 |
| scf7180000041323 | 21373 | 0.4523 | 0 | 813.265 | Eukaryota | 46230.0 | 0 | Chordata | 46230.0 | 0 | Carnivora | 46230.0 | 0 | Canidae | 46230.0 | 0 | Canis | 46230.0 | 0 | Canis lupus | 46230.0 | 0 |
| scf7180000047895 | 78059 | 0.3508 | 0 | 60.3066 | Eukaryota | 6038.0 | 1 | Chordata | 5685.0 | 2 | Artiodactyla | 2600.0 | 4 | Bovidae | 2600.0 | 4 | Homo | 2473.0 | 5 | Homo sapiens | 2473.0 | 5 |
| scf7180000042277 | 668566 | 0.3585 | 0 | 60.2899 | Eukaryota | 41866.0 | 0 | Chordata | 41866.0 | 0 | Carnivora | 41866.0 | 0 | Canidae | 30222.0 | 1 | Canis | 30222.0 | 1 | Canis lupus | 30222.0 | 1 |
| scf7180000042163 | 1144596 | 0.4083 | 0 | 61.0483 | Eukaryota | 87332.0 | 0 | Chordata | 87332.0 | 0 | Carnivora | 87332.0 | 0 | Canidae | 87332.0 | 0 | Canis | 80176.0 | 1 | Canis lupus | 80176.0 | 1 |
| scf7180000041359 | 401097 | 0.4233 | 0 | 61.0502 | Eukaryota | 87027.0 | 0 | Chordata | 87027.0 | 0 | Carnivora | 87027.0 | 0 | Canidae | 87027.0 | 0 | Canis | 87027.0 | 0 | Canis lupus | 87027.0 | 0 |
| scf7180000047111 | 10489 | 0.4561 | 0 | 91.5812 | no-hit | 0.0 | 0 | no-hit | 0.0 | 0 | no-hit | 0.0 | 0 | no-hit | 0.0 | 0 | no-hit | 0.0 | 0 | no-hit | 0.0 | 0 |
| scf7180000044757 | 262412 | 0.3794 | 0 | 60.308 | Eukaryota | 24552.0 | 0 | Chordata | 24552.0 | 0 | Carnivora | 24552.0 | 0 | Canidae | 24552.0 | 0 | Canis | 24552.0 | 0 | Canis lupus | 24552.0 | 0 |
| scf7180000041355 | 39894 | 0.3777 | 0 | 53.5093 | Eukaryota | 88782.0 | 0 | Chordata | 88782.0 | 0 | Carnivora | 88782.0 | 0 | Canidae | 88782.0 | 0 | Canis | 88782.0 | 0 | Canis lupus | 88782.0 | 0 |
| scf7180000040859 | 1565177 | 0.4542 | 0 | 59.2847 | Eukaryota | 76244.0 | 0 | Chordata | 76244.0 | 0 | Carnivora | 76244.0 | 0 | Canidae | 76244.0 | 0 | Canis | 76244.0 | 0 | Canis lupus | 76244.0 | 0 |
| scf7180000051601 | 21145 | 0.455 | 0 | 57.4338 | Eukaryota | 13956.0 | 0 | Chordata | 13956.0 | 0 | Carnivora | 9466.0 | 1 | Canidae | 9466.0 | 1 | Canis | 8380.0 | 2 | Canis lupus | 8380.0 | 2 |
| scf7180000054076 | 6611 | 0.524 | 0 | 71.0276 | Eukaryota | 29727.0 | 0 | Chordata | 29727.0 | 0 | Carnivora | 29727.0 | 0 | Canidae | 29727.0 | 0 | Canis | 29727.0 | 0 | Canis lupus | 29727.0 | 0 |
| scf7180000040790 | 96194 | 0.4038 | 0 | 56.908 | Eukaryota | 72413.0 | 0 | Chordata | 72413.0 | 0 | Carnivora | 72413.0 | 0 | Canidae | 72413.0 | 0 | Canis | 72413.0 | 0 | Canis lupus | 72413.0 | 0 |
| scf7180000046570 | 118616 | 0.3662 | 0 | 57.589 | Eukaryota | 55811.0 | 0 | Chordata | 55811.0 | 0 | Carnivora | 55811.0 | 0 | Canidae | 55811.0 | 0 | Canis | 55811.0 | 0 | Canis lupus | 55811.0 | 0 |
| scf7180000042617 | 437365 | 0.3632 | 0 | 59.0922 | Eukaryota | 88658.0 | 0 | Chordata | 88658.0 | 0 | Carnivora | 88658.0 | 0 | Canidae | 88658.0 | 0 | Canis | 88658.0 | 0 | Canis lupus | 88658.0 | 0 |
| scf7180000045985 | 280592 | 0.4219 | 0 | 58.2354 | Eukaryota | 83310.0 | 0 | Chordata | 83310.0 | 0 | Carnivora | 83310.0 | 0 | Canidae | 83310.0 | 0 | Canis | 83310.0 | 0 | Canis lupus | 83310.0 | 0 |
| scf7180000052658 | 14027 | 0.4536 | 0 | 62.5733 | Eukaryota | 9259.0 | 0 | Chordata | 7848.0 | 1 | Salmoniformes | 4046.0 | 4 | Salmonidae | 4046.0 | 4 | Salmo | 4046.0 | 4 | Salmo trutta | 4046.0 | 4 |
| scf7180000052334 | 16659 | 0.5425 | 0 | 365.0456 | Eukaryota | 24114.0 | 0 | Chordata | 24114.0 | 0 | Artiodactyla | 16938.0 | 1 | Bovidae | 16938.0 | 1 | Ovis | 9678.0 | 3 | Ovis aries | 9678.0 | 3 |
| scf7180000048712 | 32519 | 0.3762 | 0 | 62.3238 | Eukaryota | 4925.0 | 0 | Chordata | 4925.0 | 0 | Carnivora | 3798.0 | 1 | Canidae | 3297.0 | 2 | Canis | 3297.0 | 2 | Canis lupus | 3297.0 | 2 |
| scf7180000045388 | 190130 | 0.4222 | 0 | 60.6927 | Eukaryota | 22163.0 | 0 | Chordata | 22163.0 | 0 | Carnivora | 22163.0 | 0 | Canidae | 22163.0 | 0 | Canis | 22163.0 | 0 | Canis lupus | 22163.0 | 0 |
| scf7180000042942 | 5245 | 0.4553 | 0 | 31.3055 | Eukaryota | 6459.0 | 0 | Chordata | 6459.0 | 0 | Carnivora | 6459.0 | 0 | Canidae | 6459.0 | 0 | Canis | 6459.0 | 0 | Canis lupus | 6459.0 | 0 |
| scf7180000044924 | 416001 | 0.4456 | 0 | 60.0761 | Eukaryota | 80280.0 | 0 | Chordata | 80280.0 | 0 | Carnivora | 80280.0 | 0 | Canidae | 80280.0 | 0 | Canis | 70745.0 | 1 | Canis lupus | 70745.0 | 1 |
| scf7180000043576 | 29268 | 0.3632 | 0 | 54.3677 | Eukaryota | 86788.0 | 0 | Chordata | 86788.0 | 0 | Carnivora | 86788.0 | 0 | Canidae | 86788.0 | 0 | Canis | 86788.0 | 0 | Canis lupus | 86788.0 | 0 |
| scf7180000046710 | 106509 | 0.3753 | 0 | 60.3923 | Eukaryota | 10323.0 | 0 | Chordata | 10323.0 | 0 | Carnivora | 10323.0 | 0 | Canidae | 10323.0 | 0 | Canis | 10323.0 | 0 | Canis lupus | 10323.0 | 0 |
| scf7180000047176 | 31546 | 0.4374 | 0 | 114.5806 | Eukaryota | 9989.0 | 0 | Chordata | 9495.0 | 1 | Kurtiformes | 3263.0 | 4 | Apogonidae | 3263.0 | 4 | Sphaeramia | 3263.0 | 4 | Sphaeramia orbicularis | 3263.0 | 4 |
| scf7180000051440 | 21980 | 0.563 | 0 | 56.4536 | Eukaryota | 28456.0 | 0 | Chordata | 28456.0 | 0 | Carnivora | 28456.0 | 0 | Canidae | 25596.0 | 1 | Vulpes | 14524.0 | 2 | Vulpes vulpes | 14524.0 | 3 |
| scf7180000051045 | 8369 | 0.4543 | 0 | 794.6459 | no-hit | 0.0 | 0 | no-hit | 0.0 | 0 | no-hit | 0.0 | 0 | no-hit | 0.0 | 0 | no-hit | 0.0 | 0 | no-hit | 0.0 | 0 |
| scf7180000051451 | 4753 | 0.45 | 0 | 635.8451 | Eukaryota | 6256.0 | 0 | Chordata | 4116.0 | 3 | Tetraodontiformes | 4116.0 | 3 | Tetraodontidae | 4116.0 | 3 | Takifugu | 4116.0 | 3 | Takifugu rubripes | 4116.0 | 3 |
| scf7180000053141 | 11322 | 0.5006 | 0 | 150.0011 | Eukaryota | 9222.0 | 0 | Chordata | 9222.0 | 0 | Carnivora | 9222.0 | 0 | Canidae | 9222.0 | 0 | Canis | 7547.0 | 1 | Canis lupus | 6321.0 | 2 |
| scf7180000046984 | 74586 | 0.3725 | 0 | 59.6294 | Eukaryota | 8474.0 | 0 | Chordata | 8474.0 | 0 | Carnivora | 8474.0 | 0 | Canidae | 8474.0 | 0 | Canis | 8474.0 | 0 | Canis lupus | 8474.0 | 0 |
| scf7180000043710 | 302969 | 0.3986 | 0 | 60.6138 | Eukaryota | 65531.0 | 0 | Chordata | 65531.0 | 0 | Carnivora | 65531.0 | 0 | Canidae | 65531.0 | 0 | Canis | 65531.0 | 0 | Canis lupus | 65531.0 | 0 |
| scf7180000043570 | 424227 | 0.4154 | 0 | 61.1927 | Eukaryota | 67680.0 | 0 | Chordata | 67680.0 | 0 | Carnivora | 67680.0 | 0 | Canidae | 67680.0 | 0 | Canis | 52526.0 | 1 | Canis lupus | 52526.0 | 1 |
| scf7180000045112 | 17090 | 0.4421 | 0 | 30.1025 | Eukaryota | 3734.0 | 0 | Chordata | 1929.0 | 1 | Polystomatidea | 1805.0 | 4 | Polystomatidae | 1805.0 | 4 | Protopolystoma | 1805.0 | 4 | Protopolystoma xenopodis | 1805.0 | 4 |
| scf7180000052632 | 4122 | 0.4527 | 0 | 660.4878 | Eukaryota | 211.0 | 0 | Chordata | 211.0 | 0 | Kurtiformes | 211.0 | 0 | Apogonidae | 211.0 | 0 | Sphaeramia | 211.0 | 0 | Sphaeramia orbicularis | 211.0 | 0 |
| scf7180000056515 | 202929 | 0.3827 | 0 | 58.9978 | Eukaryota | 39487.0 | 0 | Chordata | 39487.0 | 0 | Carnivora | 39487.0 | 0 | Canidae | 39487.0 | 0 | Canis | 39487.0 | 0 | Canis lupus | 39487.0 | 0 |
| scf7180000045340 | 226967 | 0.4217 | 0 | 58.1821 | Eukaryota | 56616.0 | 0 | Chordata | 56616.0 | 0 | Carnivora | 56616.0 | 0 | Canidae | 56616.0 | 0 | Canis | 56616.0 | 0 | Canis lupus | 56616.0 | 0 |
| scf7180000049488 | 42386 | 0.3399 | 0 | 60.8614 | Eukaryota | 17468.0 | 0 | Chordata | 17468.0 | 0 | Carnivora | 14696.0 | 2 | Canidae | 10788.0 | 3 | Canis | 6724.0 | 4 | Canis lupus | 6724.0 | 4 |
| scf7180000050001 | 22459 | 0.4378 | 0 | 67.0271 | Eukaryota | 10881.0 | 0 | Chordata | 6593.0 | 2 | Tetraodontiformes | 6593.0 | 2 | Tetraodontidae | 6593.0 | 2 | Takifugu | 6593.0 | 2 | Takifugu rubripes | 6593.0 | 2 |
| scf7180000044256 | 171201 | 0.3573 | 0 | 59.1483 | Eukaryota | 35285.0 | 0 | Chordata | 35285.0 | 0 | Carnivora | 17218.0 | 2 | Monodontidae | 10623.0 | 3 | Monodon | 10623.0 | 3 | Monodon monoceros | 10623.0 | 3 |

| | | | | | | | | | | | | | | | | | | | | | | |
|---|---|---|---|---|---|---|---|---|---|---|---|---|---|---|---|---|---|---|---|---|---|---|
| scf7180000046436 | 12494 | 0.447 | 0 | 183.219 | Eukaryota | 17437.0 | 0 | Chordata | 8484.0 | 4 | Gadiformes | 6732.0 | 6 | Gadidae | 6732.0 | 6 | Gadus | 6732.0 | 6 | Gadus morhua | 6732.0 | 7 |
| scf7180000051931 | 13914 | 0.4648 | 0 | 58.4247 | Eukaryota | 3505.0 | 0 | Chordata | 3505.0 | 0 | Carnivora | 3505.0 | 0 | Canidae | 3505.0 | 0 | Canis | 3505.0 | 0 | Canis lupus | 3505.0 | 0 |
| scf7180000043992 | 373296 | 0.4158 | 0 | 57.8505 | Eukaryota | 25587.0 | 0 | Chordata | 25587.0 | 0 | Carnivora | 25587.0 | 0 | Canidae | 25587.0 | 0 | Canis | 23186.0 | 1 | Canis lupus | 23186.0 | 1 |
| scf7180000041911 | 396158 | 0.3976 | 0 | 59.3954 | Eukaryota | 31120.0 | 0 | Chordata | 31120.0 | 0 | Carnivora | 28874.0 | 1 | Canidae | 17523.0 | 3 | Canis | 13493.0 | 5 | Canis lupus | 13493.0 | 6 |
| scf7180000053783 | 3675 | 0.4493 | 0 | 358.6406 | no-hit | 0.0 | 0 | no-hit | 0.0 | 0 | no-hit | 0.0 | 0 | no-hit | 0.0 | 0 | no-hit | 0.0 | 0 | no-hit | 0.0 | 0 |
| scf7180000044666 | 292291 | 0.3978 | 0 | 59.1536 | Eukaryota | 27894.0 | 0 | Chordata | 27894.0 | 0 | Carnivora | 27894.0 | 0 | Canidae | 27894.0 | 0 | Canis | 24830.0 | 1 | Canis lupus | 24830.0 | 1 |
| scf7180000051677 | 4083 | 0.4489 | 0 | 698.4715 | Eukaryota | 2694.0 | 0 | Platyhelminthes | 1252.0 | 3 | Polystomatidea | 1252.0 | 3 | Polystomatidae | 1252.0 | 3 | Protopolystoma | 1252.0 | 3 | Protopolystoma xenopodis | 1252.0 | 3 |
| scf7180000042218 | 221910 | 0.3792 | 0 | 59.3497 | Eukaryota | 51712.0 | 0 | Chordata | 51712.0 | 0 | Carnivora | 51712.0 | 0 | Canidae | 35970.0 | 2 | Canis | 35970.0 | 3 | Canis lupus | 35970.0 | 3 |
| scf7180000043214 | 304362 | 0.3612 | 0 | 61.1144 | Eukaryota | 24729.0 | 0 | Chordata | 24729.0 | 0 | Primates | 17125.0 | 2 | Hominidae | 17125.0 | 2 | Homo | 15185.0 | 3 | Homo sapiens | 15185.0 | 3 |
| scf7180000044954 | 233046 | 0.4965 | 0 | 58.0384 | Eukaryota | 28692.0 | 0 | Chordata | 28692.0 | 0 | Carnivora | 28692.0 | 0 | Canidae | 28692.0 | 0 | Canis | 26409.0 | 1 | Canis lupus | 26409.0 | 1 |
| scf7180000056207 | 1423757 | 0.4889 | 0 | 60.2058 | Eukaryota | 93541.0 | 0 | Chordata | 93541.0 | 0 | Carnivora | 93541.0 | 0 | Canidae | 93541.0 | 0 | Canis | 73763.0 | 1 | Canis lupus | 73763.0 | 1 |
| scf7180000054511 | 15986 | 0.3367 | 0 | 136.6039 | Eukaryota | 4753.0 | 0 | Chordata | 4753.0 | 0 | Carnivora | 2360.0 | 2 | Felidae | 2360.0 | 4 | Acinonyx | 2360.0 | 4 | Acinonyx jubatus | 2360.0 | 4 |
| scf7180000042993 | 134360 | 0.4142 | 0 | 92.0146 | Eukaryota | 76748.0 | 0 | Chordata | 76748.0 | 0 | Carnivora | 76748.0 | 0 | Canidae | 76748.0 | 0 | Canis | 76748.0 | 0 | Canis lupus | 76748.0 | 0 |
| scf7180000048985 | 37825 | 0.6309 | 0 | 53.6053 | Eukaryota | 47131.0 | 1 | Chordata | 47131.0 | 1 | Carnivora | 43805.0 | 2 | Canidae | 33101.0 | 3 | Canis | 33101.0 | 3 | Canis lupus | 33101.0 | 3 |
| scf7180000041694 | 797241 | 0.3524 | 0 | 59.663 | Eukaryota | 64761.0 | 0 | Chordata | 64761.0 | 0 | Carnivora | 64761.0 | 0 | Canidae | 64761.0 | 0 | Canis | 64761.0 | 0 | Canis lupus | 64761.0 | 0 |
| scf7180000046289 | 115343 | 0.4614 | 0 | 56.9694 | Eukaryota | 48987.0 | 0 | Chordata | 48987.0 | 0 | Carnivora | 48987.0 | 0 | Canidae | 42831.0 | 2 | Canis | 42831.0 | 3 | Canis lupus | 42831.0 | 3 |
| scf7180000052380 | 10876 | 0.5027 | 0 | 142.1676 | Eukaryota | 8184.0 | 0 | Chordata | 8184.0 | 0 | Carnivora | 8184.0 | 0 | Canidae | 8184.0 | 0 | Canis | 6188.0 | 1 | Canis lupus | 5080.0 | 2 |
| scf7180000049904 | 26826 | 0.4366 | 0 | 134.542 | Eukaryota | 21196.0 | 1 | Chordata | 17160.0 | 2 | Gadiformes | 12895.0 | 5 | Gadidae | 12895.0 | 5 | Gadus | 12895.0 | 5 | Gadus morhua | 12895.0 | 6 |
| scf7180000052828 | 8750 | 0.4449 | 0 | 54.6904 | Eukaryota | 145.0 | 0 | Chordata | 145.0 | 0 | Salmoniformes | 145.0 | 0 | Salmonidae | 145.0 | 0 | Salmo | 145.0 | 0 | Salmo trutta | 145.0 | 0 |
| scf7180000040547 | 383557 | 0.4651 | 0 | 59.6543 | Eukaryota | 51599.0 | 0 | Chordata | 51599.0 | 0 | Carnivora | 51599.0 | 0 | Canidae | 31093.0 | 3 | Canis | 26212.0 | 4 | Canis lupus | 26212.0 | 4 |
| scf7180000053463 | 14246 | 0.4384 | 0 | 83.2475 | Eukaryota | 6847.0 | 0 | Chordata | 4293.0 | 2 | Salmoniformes | 2935.0 | 4 | Salmonidae | 2935.0 | 4 | Salmo | 2935.0 | 4 | Salmo trutta | 2935.0 | 4 |
| scf7180000056371 | 74039 | 0.3813 | 0 | 54.9017 | Eukaryota | 59120.0 | 0 | Chordata | 59120.0 | 0 | Carnivora | 59120.0 | 0 | Canidae | 59120.0 | 0 | Canis | 59120.0 | 0 | Canis lupus | 59120.0 | 0 |
| scf7180000054032 | 7746 | 0.5062 | 0 | 123.8773 | Eukaryota | 8788.0 | 0 | Chordata | 8788.0 | 0 | Carnivora | 8788.0 | 0 | Canidae | 8788.0 | 0 | Canis | 6508.0 | 1 | Canis lupus | 5264.0 | 2 |
| scf7180000045431 | 115308 | 0.5729 | 0 | 54.0304 | Eukaryota | 33488.0 | 0 | Chordata | 33488.0 | 0 | Carnivora | 33488.0 | 0 | Canidae | 33488.0 | 0 | Canis | 28168.0 | 1 | Canis lupus | 28168.0 | 1 |
| scf7180000055163 | 3325 | 0.4382 | 0 | 490.7751 | no-hit | 0.0 | 0 | no-hit | 0.0 | 0 | no-hit | 0.0 | 0 | no-hit | 0.0 | 0 | no-hit | 0.0 | 0 | no-hit | 0.0 | 0 |
| scf7180000044662 | 236284 | 0.4616 | 0 | 57.2959 | Eukaryota | 91669.0 | 0 | Chordata | 91669.0 | 0 | Carnivora | 91669.0 | 0 | Canidae | 91669.0 | 0 | Canis | 91669.0 | 0 | Canis lupus | 91669.0 | 0 |
| scf7180000047640 | 27472 | 0.4372 | 0 | 174.4906 | Eukaryota | 824.0 | 0 | Platyhelminthes | 545.0 | 1 | Polystomatidea | 545.0 | 1 | Polystomatidae | 545.0 | 1 | Protopolystoma | 545.0 | 1 | Protopolystoma xenopodis | 545.0 | 1 |
| scf7180000049205 | 28221 | 0.5064 | 0 | 217.8903 | Eukaryota | 9026.0 | 0 | Chordata | 9026.0 | 0 | Carnivora | 9026.0 | 0 | Canidae | 9026.0 | 0 | Canis | 4735.0 | 1 | Canis lupus | 4735.0 | 1 |
| scf7180000041092 | 80418 | 0.3997 | 0 | 58.5149 | Eukaryota | 31422.0 | 0 | Chordata | 31422.0 | 0 | Carnivora | 31422.0 | 0 | Felidae | 12947.0 | 4 | Panthera | 6477.0 | 6 | Zalophus californianus | 6140.0 | 8 |
| scf7180000042974 | 608287 | 0.4234 | 0 | 60.5861 | Eukaryota | 86728.0 | 0 | Chordata | 86728.0 | 0 | Carnivora | 86728.0 | 0 | Canidae | 86728.0 | 0 | Canis | 86728.0 | 0 | Canis lupus | 86728.0 | 0 |
| scf7180000052266 | 3807 | 0.4423 | 0 | 254.1005 | Eukaryota | 307.0 | 0 | Chordata | 307.0 | 0 | Kurtiformes | 307.0 | 0 | Apogonidae | 307.0 | 0 | Sphaeramia | 307.0 | 0 | Sphaeramia orbicularis | 307.0 | 0 |
| scf7180000042648 | 100628 | 0.6481 | 0 | 51.7447 | Eukaryota | 8986.0 | 0 | Chordata | 8986.0 | 0 | Carnivora | 8986.0 | 0 | Canidae | 8986.0 | 0 | Canis | 8986.0 | 0 | Canis lupus | 8986.0 | 0 |
| scf7180000047157 | 111000 | 0.4029 | 0 | 57.0698 | Eukaryota | 32760.0 | 0 | Chordata | 32760.0 | 0 | Carnivora | 32760.0 | 0 | Canidae | 32760.0 | 0 | Canis | 32760.0 | 0 | Canis lupus | 32760.0 | 0 |
| scf7180000044769 | 28690 | 0.3921 | 0 | 49.3135 | Eukaryota | 14263.0 | 0 | Chordata | 14263.0 | 0 | Artiodactyla | 5477.0 | 3 | Canidae | 4048.0 | 4 | Canis | 4048.0 | 5 | Canis lupus | 4048.0 | 5 |
| scf7180000044582 | 2871 | 0.371 | 0 | 25.5596 | Eukaryota | 3514.0 | 1 | Chordata | 3514.0 | 1 | Carnivora | 2277.0 | 2 | Canidae | 2277.0 | 3 | Canis | 1946.0 | 4 | Canis lupus | 1946.0 | 4 |
| scf7180000054205 | 23265 | 0.5065 | 0 | 152.697 | Eukaryota | 8836.0 | 0 | Chordata | 8836.0 | 0 | Carnivora | 8836.0 | 0 | Canidae | 8836.0 | 0 | Canis | 6203.0 | 1 | Canis lupus | 6203.0 | 1 |
| scf7180000050661 | 4519 | 0.4472 | 0 | 104.0527 | Eukaryota | 206.0 | 0 | Platyhelminthes | 206.0 | 0 | Polystomatidea | 206.0 | 0 | Polystomatidae | 206.0 | 0 | Protopolystoma | 206.0 | 0 | Protopolystoma xenopodis | 206.0 | 0 |
| scf7180000051273 | 10155 | 0.4373 | 0 | 49.7374 | Eukaryota | 37368.0 | 0 | Chordata | 26160.0 | 1 | Gadiformes | 12232.0 | 5 | Gadidae | 12232.0 | 6 | Gadus | 12232.0 | 6 | Gadus morhua | 12232.0 | 6 |
| scf7180000049167 | 3240 | 0.4435 | 0 | 267.6471 | no-hit | 0.0 | 0 | no-hit | 0.0 | 0 | no-hit | 0.0 | 0 | no-hit | 0.0 | 0 | no-hit | 0.0 | 0 | no-hit | 0.0 | 0 |
| scf7180000040749 | 5845 | 0.4513 | 0 | 426.0541 | Eukaryota | 9186.0 | 0 | Chordata | 8369.0 | 1 | Tetraodontiformes | 4687.0 | 4 | Tetraodontidae | 4687.0 | 4 | Takifugu | 4687.0 | 4 | Takifugu rubripes | 4687.0 | 4 |
| scf7180000044144 | 395989 | 0.3741 | 0 | 58.899 | Eukaryota | 104594.0 | 0 | Chordata | 104594.0 | 0 | Carnivora | 104594.0 | 0 | Canidae | 104594.0 | 0 | Canis | 104594.0 | 0 | Canis lupus | 104594.0 | 0 |
| scf7180000051323 | 7348 | 0.4423 | 0 | 391.4071 | no-hit | 0.0 | 0 | no-hit | 0.0 | 0 | no-hit | 0.0 | 0 | no-hit | 0.0 | 0 | no-hit | 0.0 | 0 | no-hit | 0.0 | 0 |
| scf7180000051987 | 7061 | 0.447 | 0 | 213.7834 | Eukaryota | 279.0 | 0 | Chordata | 279.0 | 0 | Salmoniformes | 279.0 | 0 | Salmonidae | 279.0 | 0 | Coregonus | 279.0 | 0 | Coregonus sp. 'balchen' | 279.0 | 0 |
| scf7180000048472 | 31735 | 0.4394 | 0 | 337.7668 | Eukaryota | 3588.0 | 0 | Chordata | 2132.0 | 1 | Kurtiformes | 1877.0 | 2 | Apogonidae | 1877.0 | 2 | Sphaeramia | 1877.0 | 2 | Sphaeramia orbicularis | 1877.0 | 2 |
| scf7180000053760 | 3726 | 0.4466 | 0 | 516.3505 | Eukaryota | 2502.0 | 0 | Platyhelminthes | 1170.0 | 2 | Polystomatidea | 1170.0 | 2 | Polystomatidae | 1170.0 | 2 | Protopolystoma | 1170.0 | 2 | Protopolystoma xenopodis | 1170.0 | 2 |
| scf7180000051671 | 10053 | 0.4442 | 0 | 282.0776 | Eukaryota | 3361.0 | 1 | Echinodermata | 2211.0 | 3 | Forcipulatida | 2211.0 | 3 | Asteriidae | 2211.0 | 3 | Asterias | 2211.0 | 3 | Asterias rubens | 2211.0 | 3 |
| scf7180000048370 | 2043 | 0.4464 | 0 | 27.1299 | Eukaryota | 2852.0 | 0 | Chordata | 2852.0 | 0 | Carnivora | 2852.0 | 0 | Canidae | 2852.0 | 0 | Canis | 2852.0 | 0 | Canis lupus | 2852.0 | 0 |
| scf7180000045757 | 89710 | 0.3641 | 0 | 56.6136 | Eukaryota | 35164.0 | 0 | Chordata | 35164.0 | 0 | Carnivora | 35164.0 | 0 | Canidae | 35164.0 | 0 | Canis | 35164.0 | 0 | Canis lupus | 35164.0 | 0 |
| scf7180000045630 | 14408 | 0.4364 | 0 | 125.4042 | no-hit | 0.0 | 0 | no-hit | 0.0 | 0 | no-hit | 0.0 | 0 | no-hit | 0.0 | 0 | no-hit | 0.0 | 0 | no-hit | 0.0 | 0 |
| scf7180000055229 | 22335 | 0.4412 | 0 | 228.2311 | Eukaryota | 19874.0 | 0 | Chordata | 15375.0 | 3 | Gadiformes | 10556.0 | 8 | Gadidae | 10556.0 | 8 | Gadus | 10556.0 | 8 | Gadus morhua | 10556.0 | 8 |
| scf7180000053230 | 8856 | 0.5556 | 0 | 550.9505 | Eukaryota | 17165.0 | 0 | Chordata | 17165.0 | 0 | Carnivora | 17165.0 | 0 | Canidae | 17165.0 | 0 | Canis | 17165.0 | 0 | Canis lupus | 17165.0 | 0 |
| scf7180000050863 | 3111 | 0.4442 | 0 | 308.3972 | no-hit | 0.0 | 0 | no-hit | 0.0 | 0 | no-hit | 0.0 | 0 | no-hit | 0.0 | 0 | no-hit | 0.0 | 0 | no-hit | 0.0 | 0 |
| scf7180000052529 | 3050 | 0.4187 | 0 | 81.2221 | Eukaryota | 1147.0 | 0 | Chordata | 1147.0 | 0 | Salmoniformes | 1147.0 | 0 | Salmonidae | 1147.0 | 0 | Salmo | 1147.0 | 0 | Salmo trutta | 1147.0 | 0 |
| scf7180000047097 | 92329 | 0.3872 | 0 | 61.9028 | Eukaryota | 10569.0 | 0 | Chordata | 10569.0 | 0 | Carnivora | 6960.0 | 2 | Canidae | 3115.0 | 4 | Canis | 2084.0 | 8 | Canis lupus | 2084.0 | 8 |
| scf7180000046140 | 22720 | 0.4383 | 0 | 121.3244 | Eukaryota | 4811.0 | 0 | Chordata | 4286.0 | 1 | Kurtiformes | 2829.0 | 2 | Apogonidae | 2829.0 | 2 | Sphaeramia | 2829.0 | 2 | Sphaeramia orbicularis | 2829.0 | 2 |
| scf7180000047549 | 21225 | 0.4573 | 0 | 167.9923 | Eukaryota | 17763.0 | 0 | Chordata | 16193.0 | 2 | Gadiformes | 7474.0 | 6 | Gadidae | 7474.0 | 6 | Gadus | 7474.0 | 6 | Gadus morhua | 7474.0 | 6 |
| scf7180000042120 | 778199 | 0.3905 | 100 | 58.1296 | Eukaryota | 41665.0 | 0 | Chordata | 41665.0 | 0 | Carnivora | 41665.0 | 0 | Canidae | 41665.0 | 0 | Vulpes | 30534.0 | 1 | Vulpes vulpes | 30534.0 | 1 |
| scf7180000054286 | 21722 | 0.4634 | 0 | 127.5274 | Eukaryota | 9236.0 | 0 | Chordata | 9236.0 | 0 | Carnivora | 9236.0 | 0 | Canidae | 9236.0 | 0 | Canis | 5604.0 | 1 | Canis lupus | 5604.0 | 1 |
| scf7180000056323 | 525979 | 0.4253 | 0 | 58.281 | Eukaryota | 42704.0 | 0 | Chordata | 42704.0 | 0 | Carnivora | 42704.0 | 0 | Canidae | 42704.0 | 0 | Canis | 39440.0 | 1 | Canis lupus | 39440.0 | 1 |
| scf7180000046595 | 77166 | 0.3722 | 0 | 56.4453 | Eukaryota | 53847.0 | 0 | Chordata | 53847.0 | 0 | Carnivora | 53847.0 | 0 | Canidae | 53847.0 | 0 | Canis | 53847.0 | 0 | Canis lupus | 53847.0 | 0 |
| scf7180000053832 | 3453 | 0.5013 | 0 | 229.5465 | Eukaryota | 7513.0 | 0 | Chordata | 7513.0 | 0 | Carnivora | 7513.0 | 0 | Canidae | 7513.0 | 0 | Canis | 4966.0 | 1 | Canis lupus | 3909.0 | 2 |
| scf7180000054575 | 22717 | 0.4166 | 0 | 883.2143 | Eukaryota | 5155.0 | 0 | Chordata | 5155.0 | 0 | Carnivora | 3385.0 | 1 | Canidae | 1828.0 | 2 | Canis | 1828.0 | 2 | Canis lupus | 1828.0 | 2 |
| scf7180000053081 | 2446 | 0.4575 | 0 | 424.7958 | no-hit | 0.0 | 0 | no-hit | 0.0 | 0 | no-hit | 0.0 | 0 | no-hit | 0.0 | 0 | no-hit | 0.0 | 0 | no-hit | 0.0 | 0 |
| scf7180000046134 | 124129 | 0.3735 | 0 | 61.3864 | Eukaryota | 47334.0 | 0 | Chordata | 47334.0 | 0 | Carnivora | 47334.0 | 0 | Canidae | 47334.0 | 0 | Canis | 47334.0 | 0 | Canis lupus | 47334.0 | 0 |
| scf7180000053904 | 2304 | 0.5013 | 0 | 840.2873 | Eukaryota | 5207.0 | 0 | Chordata | 5207.0 | 0 | Carnivora | 5207.0 | 0 | Canidae | 5207.0 | 0 | Canis | 4546.0 | 2 | Canis lupus | 4546.0 | 2 |
| scf7180000045041 | 36289 | 0.4513 | 0 | 55.4583 | Eukaryota | 38722.0 | 0 | Chordata | 38722.0 | 0 | Carnivora | 38722.0 | 0 | Canidae | 38722.0 | 0 | Canis | 36723.0 | 1 | Canis lupus | 36723.0 | 1 |
| scf7180000049092 | 4098 | 0.4505 | 0 | 126.5848 | Eukaryota | 276.0 | 0 | Platyhelminthes | 139.0 | 1 | Polystomatidea | 139.0 | 1 | Polystomatidae | 139.0 | 1 | Protopolystoma | 139.0 | 1 | Protopolystoma xenopodis | 139.0 | 1 |
| scf7180000047618 | 112126 | 0.461 | 0 | 59.3976 | Eukaryota | 9680.0 | 0 | Chordata | 9680.0 | 0 | Carnivora | 4272.0 | 2 | Hominidae | 3447.0 | 4 | Homo | 3447.0 | 4 | Homo sapiens | 3447.0 | 4 |
| scf7180000056554 | 184163 | 0.4085 | 0 | 61.4334 | Eukaryota | 16279.0 | 0 | Chordata | 16279.0 | 0 | Carnivora | 16279.0 | 0 | Canidae | 16279.0 | 0 | Canis | 16279.0 | 0 | Canis lupus | 16279.0 | 0 |
| scf7180000056464 | 429641 | 0.3475 | 0 | 58.555 | Eukaryota | 43036.0 | 0 | Chordata | 43036.0 | 0 | Carnivora | 43036.0 | 0 | Canidae | 43036.0 | 0 | Canis | 43036.0 | 0 | Canis lupus | 43036.0 | 0 |
| scf7180000054228 | 30099 | 0.4026 | 0 | 417.7545 | Eukaryota | 72024.0 | 0 | Chordata | 72024.0 | 0 | Carnivora | 72024.0 | 0 | Canidae | 72024.0 | 0 | Canis | 72024.0 | 0 | Canis lupus | 72024.0 | 0 |
| scf7180000042414 | 652883 | 0.4336 | 0 | 62.1211 | Eukaryota | 59358.0 | 0 | Chordata | 59358.0 | 0 | Carnivora | 59358.0 | 0 | Canidae | 59358.0 | 0 | Canis | 59358.0 | 0 | Canis lupus | 59358.0 | 0 |
| scf7180000044460 | 6848 | 0.4492 | 0 | 306.7859 | Eukaryota | 4373.0 | 0 | Chordata | 2888.0 | 2 | Polystomatidea | 1046.0 | 5 | Polystomatidae | 1046.0 | 5 | Protopolystoma | 1046.0 | 5 | Protopolystoma xenopodis | 1046.0 | 5 |
| scf7180000052099 | 7670 | 0.4488 | 0 | 544.3204 | Eukaryota | 387.0 | 0 | Chordata | 387.0 | 0 | Kurtiformes | 387.0 | 0 | Apogonidae | 387.0 | 0 | Sphaeramia | 387.0 | 0 | Sphaeramia orbicularis | 387.0 | 0 |
| scf7180000047259 | 60875 | 0.4046 | 0 | 58.7847 | Eukaryota | 29762.0 | 0 | Chordata | 29762.0 | 0 | Carnivora | 29762.0 | 0 | Canidae | 29762.0 | 0 | Canis | 21075.0 | 1 | Canis lupus | 21075.0 | 1 |
| scf7180000047080 | 22833 | 0.434 | 0 | 80.2832 | Eukaryota | 17205.0 | 0 | Chordata | 8543.0 | 3 | Strigeidida | 6298.0 | 7 | Schistosomatidae | 6298.0 | 7 | Schistosoma | 6298.0 | 7 | Schistosoma mansoni | 6298.0 | 7 |
| scf7180000051785 | 3087 | 0.5092 | 0 | 31.9056 | Eukaryota | 5843.0 | 0 | Chordata | 5843.0 | 0 | Carnivora | 5843.0 | 0 | Canidae | 5843.0 | 0 | Canis | 4139.0 | 1 | Canis lupus | 3246.0 | 2 |
| scf7180000046548 | 89703 | 0.3857 | 0 | 58.0888 | Eukaryota | 64671.0 | 0 | Chordata | 64671.0 | 0 | Carnivora | 64671.0 | 0 | Canidae | 64671.0 | 0 | Canis | 64671.0 | 0 | Canis lupus | 64671.0 | 0 |
| scf7180000042566 | 104685 | 0.431 | 0 | 63.5326 | Eukaryota | 80636.0 | 0 | Chordata | 80636.0 | 0 | Carnivora | 80636.0 | 0 | Canidae | 80636.0 | 0 | Canis | 80636.0 | 0 | Canis lupus | 80636.0 | 0 |
| scf7180000051238 | 2252 | 0.456 | 0 | 778.5163 | no-hit | 0.0 | 0 | no-hit | 0.0 | 0 | no-hit | 0.0 | 0 | no-hit | 0.0 | 0 | no-hit | 0.0 | 0 | no-hit | 0.0 | 0 |
| scf7180000049085 | 5482 | 0.4365 | 0 | 254.8779 | no-hit | 0.0 | 0 | no-hit | 0.0 | 0 | no-hit | 0.0 | 0 | no-hit | 0.0 | 0 | no-hit | 0.0 | 0 | no-hit | 0.0 | 0 |
| scf7180000051843 | 1909 | 0.4557 | 0 | 128.5414 | no-hit | 0.0 | 0 | no-hit | 0.0 | 0 | no-hit | 0.0 | 0 | no-hit | 0.0 | 0 | no-hit | 0.0 | 0 | no-hit | 0.0 | 0 |
| scf7180000043483 | 444157 | 0.4155 | 0 | 60.6492 | Eukaryota | 53011.0 | 0 | Chordata | 53011.0 | 0 | Carnivora | 53011.0 | 0 | Canidae | 48735.0 | 1 | Canis | 38697.0 | 2 | Canis lupus | 38697.0 | 2 |
| scf7180000042095 | 772829 | 0.3809 | 0 | 60.4255 | Eukaryota | 89745.0 | 0 | Chordata | 89745.0 | 0 | Carnivora | 89745.0 | 0 | Canidae | 89745.0 | 0 | Canis | 89745.0 | 0 | Canis lupus | 89745.0 | 0 |
| scf7180000054097 | 2686 | 0.4311 | 0 | 70.6251 | no-hit | 0.0 | 0 | no-hit | 0.0 | 0 | no-hit | 0.0 | 0 | no-hit | 0.0 | 0 | no-hit | 0.0 | 0 | no-hit | 0.0 | 0 |
| scf7180000047180 | 10576 | 0.3753 | 0 | 29.8454 | Eukaryota | 53358.0 | 0 | Chordata | 53358.0 | 0 | Carnivora | 53358.0 | 0 | Canidae | 53358.0 | 0 | Canis | 53358.0 | 0 | Canis lupus | 53358.0 | 0 |
| scf7180000045406 | 83577 | 0.5567 | 0 | 56.5258 | Eukaryota | 20520.0 | 0 | Chordata | 20520.0 | 0 | Carnivora | 20520.0 | 0 | Canidae | 13595.0 | 3 | Canis | 11990.0 | 4 | Canis lupus | 11990.0 | 4 |
| scf7180000044333 | 100686 | 0.3908 | 0 | 59.6427 | Eukaryota | 92111.0 | 0 | Chordata | 92111.0 | 0 | Carnivora | 92111.0 | 0 | Canidae | 92111.0 | 0 | Canis | 92111.0 | 0 | Canis lupus | 92111.0 | 0 |
| scf7180000053200 | 2888 | 0.4529 | 0 | 442.9022 | no-hit | 0.0 | 0 | no-hit | 0.0 | 0 | no-hit | 0.0 | 0 | no-hit | 0.0 | 0 | no-hit | 0.0 | 0 | no-hit | 0.0 | 0 |
| scf7180000044341 | 30383 | 0.351 | 0 | 68.7865 | Eukaryota | 94537.0 | 0 | Chordata | 94537.0 | 0 | Carnivora | 94537.0 | 0 | Canidae | 94537.0 | 0 | Canis | 94537.0 | 0 | Canis lupus | 94537.0 | 0 |
| scf7180000041311 | 861986 | 0.3634 | 0 | 62.0319 | Eukaryota | 129922.0 | 0 | Chordata | 129922.0 | 0 | Carnivora | 129922.0 | 0 | Canidae | 129922.0 | 0 | Canis | 129922.0 | 0 | Canis lupus | 129922.0 | 0 |
| scf7180000043972 | 104315 | 0.3531 | 0 | 61.8085 | Eukaryota | 92759.0 | 0 | Chordata | 92759.0 | 0 | Carnivora | 92759.0 | 0 | Canidae | 92759.0 | 0 | Canis | 92759.0 | 0 | Canis lupus | 92759.0 | 0 |
| scf7180000041126 | 31557 | 0.392 | 0 | 51.5249 | Eukaryota | 6111.0 | 0 | Chordata | 6111.0 | 0 | Carnivora | 6111.0 | 0 | Felidae | 2735.0 | 2 | Felis | 2735.0 | 2 | Felis catus | 2735.0 | 2 |
| scf7180000043376 | 91544 | 0.4248 | 0 | 59.7304 | Eukaryota | 23589.0 | 0 | Chordata | 23589.0 | 0 | Carnivora | 23589.0 | 0 | Canidae | 21596.0 | 1 | Canis | 17550.0 | 2 | Canis lupus | 17550.0 | 2 |
| scf7180000045283 | 74658 | 0.4555 | 0 | 57.8814 | Eukaryota | 38398.0 | 0 | Chordata | 38398.0 | 0 | Carnivora | 38398.0 | 0 | Canidae | 38398.0 | 0 | Canis | 29540.0 | 1 | Canis lupus | 29540.0 | 1 |
| scf7180000050993 | 24837 | 0.457 | 0 | 55.8635 | Eukaryota | 5867.0 | 0 | Platyhelminthes | 3751.0 | 2 | Polystomatidea | 3751.0 | 3 | Polystomatidae | 3751.0 | 3 | Protopolystoma | 3751.0 | 3 | Protopolystoma xenopodis | 3751.0 | 3 |
| scf7180000047903 | 61904 | 0.3617 | 0 | 61.4373 | Eukaryota | 46067.0 | 0 | Chordata | 46067.0 | 0 | Carnivora | 46067.0 | 0 | Canidae | 46067.0 | 0 | Canis | 46067.0 | 0 | Canis lupus | 46067.0 | 0 |
| scf7180000045126 | 21998 | 0.4407 | 0 | 94.308 | Eukaryota | 5117.0 | 0 | Chordata | 3873.0 | 1 | Kurtiformes | 3873.0 | 1 | Apogonidae | 3873.0 | 1 | Sphaeramia | 3873.0 | 1 | Sphaeramia orbicularis | 3873.0 | 1 |
| scf7180000047073 | 87144 | 0.5256 | 0 | 57.6834 | Eukaryota | 33456.0 | 0 | Chordata | 33456.0 | 0 | Carnivora | 33456.0 | 0 | Felidae | 19836.0 | 1 | Canis | 13620.0 | 3 | Canis lupus | 13620.0 | 3 |
| scf7180000050009 | 4361 | 0.4526 | 0 | 488.9306 | Eukaryota | 2448.0 | 0 | Chordata | 2448.0 | 0 | Carnivora | 2448.0 | 0 | Canidae | 2448.0 | 0 | Canis | 2448.0 | 0 | Canis lupus | 2448.0 | 0 |
| scf7180000042193 | 21186 | 0.5195 | 0 | 904.5726 | Eukaryota | 16143.0 | 0 | Chordata | 16143.0 | 0 | Carnivora | 16143.0 | 0 | Canidae | 16143.0 | 0 | Canis | 16143.0 | 0 | Canis lupus | 16143.0 | 0 |
| scf7180000044476 | 24527 | 0.4941 | 0 | 85.0666 | Eukaryota | 45688.0 | 0 | Chordata | 45688.0 | 0 | Carnivora | 45688.0 | 0 | Canidae | 45688.0 | 0 | Canis | 45688.0 | 0 | Canis lupus | 45688.0 | 0 |
| scf7180000047423 | 57135 | 0.6314 | 0 | 55.1156 | Eukaryota | 49834.0 | 0 | Chordata | 49834.0 | 0 | Carnivora | 49834.0 | 0 | Canidae | 49834.0 | 0 | Vulpes | 29545.0 | 1 | Vulpes vulpes | 29545.0 | 1 |
| scf7180000044471 | 310355 | 0.3565 | 0 | 61.6711 | Eukaryota | 59442.0 | 0 | Chordata | 59442.0 | 0 | Carnivora | 59442.0 | 0 | Canidae | 32842.0 | 3 | Vulpes | 19659.0 | 5 | Vulpes vulpes | 19659.0 | 5 |
| scf7180000054306 | 124701 | 0.3869 | 0 | 1703.037 | Eukaryota | 35304.0 | 0 | Chordata | 35304.0 | 0 | Carnivora | 35304.0 | 0 | Canidae | 35304.0 | 0 | Canis | 35304.0 | 0 | Canis lupus | 35304.0 | 0 |
| scf7180000043391 | 457556 | 0.3439 | 0 | 59.7859 | Eukaryota | 36765.0 | 0 | Chordata | 36765.0 | 0 | Primates | 17537.0 | 2 | Hominidae | 17537.0 | 2 | Homo | 17537.0 | 3 | Homo sapiens | 17537.0 | 3 |
| scf7180000048960 | 4182 | 0.4512 | 0 | 195.1946 | Eukaryota | 2780.0 | 0 | Chordata | 2780.0 | 0 | Carnivora | 2780.0 | 0 | Canidae | 2780.0 | 0 | Canis | 2780.0 | 0 | Canis lupus | 2780.0 | 0 |
| scf7180000040914 | 751730 | 0.3874 | 0 | 60.0172 | Eukaryota | 106517.0 | 0 | Chordata | 106517.0 | 0 | Carnivora | 106517.0 | 0 | Canidae | 106517.0 | 0 | Canis | 106517.0 | 0 | Canis lupus | 106517.0 | 0 |
| scf7180000047427 | 70068 | 0.3993 | 0 | 59.6621 | Eukaryota | 33180.0 | 0 | Chordata | 33180.0 | 0 | Carnivora | 33180.0 | 0 | Canidae | 24840.0 | 1 | Canis | 24840.0 | 1 | Canis lupus | 24840.0 | 2 |
| scf7180000049872 | 50987 | 0.4319 | 0 | 57.2192 | Eukaryota | 36066.0 | 0 | Chordata | 36066.0 | 0 | Carnivora | 36066.0 | 0 | Canidae | 36066.0 | 0 | Canis | 36066.0 | 0 | Canis lupus | 36066.0 | 0 |
| scf7180000054254 | 29570 | 0.4702 | 0 | 462.5096 | Eukaryota | 37482.0 | 0 | Chordata | 37482.0 | 0 | Carnivora | 37482.0 | 0 | Canidae | 37482.0 | 0 | Canis | 37482.0 | 0 | Canis lupus | 37482.0 | 0 |
| scf7180000054878 | 62344 | 0.3741 | 0 | 90.1272 | Eukaryota | 93082.0 | 0 | Chordata | 93082.0 | 0 | Carnivora | 93082.0 | 0 | Canidae | 93082.0 | 0 | Canis | 93082.0 | 0 | Canis lupus | 93082.0 | 0 |
| scf7180000050523 | 5106 | 0.4493 | 0 | 822.8006 | no-hit | 0.0 | 0 | no-hit | 0.0 | 0 | no-hit | 0.0 | 0 | no-hit | 0.0 | 0 | no-hit | 0.0 | 0 | no-hit | 0.0 | 0 |
| scf7180000047474 | 5964 | 0.4425 | 0 | 108.4024 | Eukaryota | 2690.0 | 0 | Chordata | 2690.0 | 0 | Carnivora | 2690.0 | 0 | Canidae | 2690.0 | 0 | Canis | 2690.0 | 0 | Canis lupus | 2690.0 | 0 |
| scf7180000050943 | 11707 | 0.4438 | 0 | 227.199 | Eukaryota | 4554.0 | 0 | Platyhelminthes | 2697.0 | 2 | Polystomatidea | 2697.0 | 2 | Polystomatidae | 2697.0 | 2 | Protopolystoma | 2697.0 | 2 | Protopolystoma xenopodis | 2697.0 | 2 |
| scf7180000045763 | 59548 | 0.4403 | 0 | 147.3985 | Eukaryota | 102087.0 | 0 | Chordata | 57748.0 | 1 | Gadiformes | 57748.0 | 2 | Gadidae | 57748.0 | 2 | Gadus | 57748.0 | 2 | Gadus morhua | 57748.0 | 2 |
| scf7180000053811 | 2159 | 0.4488 | 0 | 259.9167 | no-hit | 0.0 | 0 | no-hit | 0.0 | 0 | no-hit | 0.0 | 0 | no-hit | 0.0 | 0 | no-hit | 0.0 | 0 | no-hit | 0.0 | 0 |
| scf7180000041054 | 129115 | 0.3634 | 0 | 57.1375 | Eukaryota | 27291.0 | 0 | Chordata | 27291.0 | 0 | Carnivora | 27291.0 | 0 | Canidae | 27291.0 | 0 | Canis | 27291.0 | 0 | Canis lupus | 27291.0 | 0 |
| scf7180000047642 | 7128 | 0.4346 | 0 | 120.307 | no-hit | 0.0 | 0 | no-hit | 0.0 | 0 | no-hit | 0.0 | 0 | no-hit | 0.0 | 0 | no-hit | 0.0 | 0 | no-hit | 0.0 | 0 |
| scf7180000049697 | 7280 | 0.3876 | 0 | 114.6496 | Eukaryota | 2217.0 | 0 | Chordata | 2217.0 | 0 | Carnivora | 1683.0 | 1 | Canidae | 1683.0 | 1 | Canis | 1683.0 | 1 | Canis lupus | 1683.0 | 1 |
| scf7180000050081 | 26061 | 0.6184 | 0 | 55.5064 | Eukaryota | 10654.0 | 0 | Chordata | 10654.0 | 0 | Carnivora | 10654.0 | 0 | Canidae | 8719.0 | 4 | Canis | 8332.0 | 6 | Canis lupus | 8332.0 | 6 |
| scf7180000051468 | 12738 | 0.4536 | 0 | 96.4735 | no-hit | 0.0 | 0 | no-hit | 0.0 | 0 | no-hit | 0.0 | 0 | no-hit | 0.0 | 0 | no-hit | 0.0 | 0 | no-hit | 0.0 | 0 |
| scf7180000053006 | 1477 | 0.4374 | 0 | 664.5513 | no-hit | 0.0 | 0 | no-hit | 0.0 | 0 | no-hit | 0.0 | 0 | no-hit | 0.0 | 0 | no-hit | 0.0 | 0 | no-hit | 0.0 | 0 |
| scf7180000044109 | 317955 | 0.4016 | 0 | 59.9049 | Eukaryota | 44380.0 | 0 | Chordata | 44380.0 | 0 | Carnivora | 44380.0 | 0 | Canidae | 44380.0 | 0 | Canis | 44380.0 | 0 | Canis lupus | 44380.0 | 0 |
| scf7180000043365 | 593329 | 0.426 | 0 | 59.9599 | Eukaryota | 48357.0 | 0 | Chordata | 48357.0 | 0 | Carnivora | 48357.0 | 0 | Canidae | 48357.0 | 0 | Canis | 27647.0 | 1 | Canis lupus | 27647.0 | 1 |
| scf7180000044479 | 37373 | 0.3884 | 0 | 54.7027 | Eukaryota | 91983.0 | 0 | Chordata | 91983.0 | 0 | Carnivora | 91983.0 | 0 | Canidae | 91983.0 | 0 | Canis | 91983.0 | 0 | Canis lupus | 91983.0 | 0 |
| scf7180000051220 | 7456 | 0.4519 | 0 | 788.018 | Eukaryota | 2852.0 | 0 | Platyhelminthes | 1297.0 | 3 | Polystomatidea | 1297.0 | 3 | Polystomatidae | 1297.0 | 3 | Protopolystoma | 1297.0 | 3 | Protopolystoma xenopodis | 1297.0 | 3 |
| scf7180000048250 | 30374 | 0.4398 | 0 | 111.4841 | Eukaryota | 4011.0 | 0 | Platyhelminthes | 2356.0 | 1 | Polystomatidea | 1811.0 | 3 | Polystomatidae | 1811.0 | 3 | Protopolystoma | 1811.0 | 4 | Protopolystoma xenopodis | 1811.0 | 4 |
| scf7180000052096 | 12989 | 0.5033 | 0 | 161.8624 | Eukaryota | 8564.0 | 0 | Chordata | 8564.0 | 0 | Carnivora | 8564.0 | 0 | Canidae | 8564.0 | 0 | Canis | 6965.0 | 1 | Canis lupus | 5822.0 | 2 |
| scf7180000046771 | 92568 | 0.4455 | 0 | 56.7646 | Eukaryota | 50751.0 | 0 | Chordata | 50751.0 | 0 | Carnivora | 50751.0 | 0 | Canidae | 50751.0 | 0 | Canis | 50751.0 | 0 | Canis lupus | 50751.0 | 0 |
| scf7180000041346 | 68759 | 0.3855 | 0 | 56.9831 | Eukaryota | 38805.0 | 0 | Chordata | 38805.0 | 0 | Carnivora | 38805.0 | 0 | Canidae | 38805.0 | 0 | Canis | 38805.0 | 0 | Canis lupus | 38805.0 | 0 |
| scf7180000054510 | 1263620 | 0.3582 | 0 | 60.9759 | Eukaryota | 93967.0 | 0 | Chordata | 93967.0 | 0 | Carnivora | 93967.0 | 0 | Canidae | 93967.0 | 0 | Canis | 93967.0 | 0 | Canis lupus | 93967.0 | 0 |
| scf7180000044321 | 377779 | 0.358 | 0 | 60.0157 | Eukaryota | 46186.0 | 0 | Chordata | 46186.0 | 0 | Carnivora | 46186.0 | 0 | Canidae | 46186.0 | 0 | Canis | 46186.0 | 0 | Canis lupus | 46186.0 | 0 |
| scf7180000052242 | 16727 | 0.5039 | 0 | 417.8391 | Eukaryota | 9350.0 | 0 | Chordata | 9350.0 | 0 | Carnivora | 9350.0 | 0 | Canidae | 9350.0 | 0 | Canis | 7550.0 | 1 | Canis lupus | 6206.0 | 2 |
| scf7180000044699 | 21501 | 0.437 | 0 | 233.9675 | Eukaryota | 3824.0 | 0 | Chordata | 3255.0 | 1 | Kurtiformes | 3255.0 | 1 | Apogonidae | 3255.0 | 1 | Sphaeramia | 3255.0 | 1 | Sphaeramia orbicularis | 3255.0 | 1 |
| scf7180000042216 | 396740 | 0.3795 | 0 | 58.0306 | Eukaryota | 94054.0 | 0 | Chordata | 94054.0 | 0 | Carnivora | 94054.0 | 0 | Canidae | 94054.0 | 0 | Canis | 94054.0 | 0 | Canis lupus | 94054.0 | 0 |
| scf7180000041310 | 149846 | 0.4943 | 0 | 57.655 | Eukaryota | 72965.0 | 0 | Chordata | 72965.0 | 0 | Carnivora | 72965.0 | 0 | Canidae | 69729.0 | 1 | Canis | 59296.0 | 2 | Canis lupus | 59296.0 | 2 |
| scf7180000049522 | 22137 | 0.4466 | 0 | 127.3642 | Eukaryota | 433.0 | 0 | Chordata | 433.0 | 0 | Tetraodontiformes | 433.0 | 0 | Tetraodontidae | 433.0 | 0 | Takifugu | 433.0 | 0 | Takifugu rubripes | 433.0 | 0 |
| scf7180000051063 | 18565 | 0.6429 | 0 | 364.7958 | Eukaryota | 35698.0 | 0 | Chordata | 35698.0 | 0 | Primates | 27370.0 | 1 | Hominidae | 17098.0 | 2 | Macaca | 10272.0 | 3 | Macaca mulatta | 10272.0 | 3 |
| scf7180000044263 | 70728 | 0.3782 | 0 | 54.6517 | Eukaryota | 59770.0 | 0 | Chordata | 59770.0 | 0 | Carnivora | 59770.0 | 0 | Canidae | 59770.0 | 0 | Canis | 59770.0 | 0 | Canis lupus | 59770.0 | 0 |
| scf7180000056148 | 4641 | 0.4426 | 0 | 146.7616 | no-hit | 0.0 | 0 | no-hit | 0.0 | 0 | no-hit | 0.0 | 0 | no-hit | 0.0 | 0 | no-hit | 0.0 | 0 | no-hit | 0.0 | 0 |
| scf7180000054483 | 34080 | 0.3928 | 0 | 444.7758 | Eukaryota | 40655.0 | 0 | Chordata | 40655.0 | 0 | Carnivora | 40655.0 | 0 | Canidae | 40655.0 | 0 | Canis | 40655.0 | 0 | Canis lupus | 40655.0 | 0 |
| scf7180000055349 | 39742 | 0.4391 | 0 | 114.7416 | Eukaryota | 14719.0 | 0 | Chordata | 14719.0 | 0 | Carnivora | 14719.0 | 0 | Canidae | 14719.0 | 0 | Canis | 14719.0 | 0 | Canis lupus | 14719.0 | 0 |
| scf7180000049032 | 39444 | 0.405 | 0 | 57.8504 | Eukaryota | 74169.0 | 0 | Chordata | 74169.0 | 0 | Carnivora | 74169.0 | 0 | Canidae | 74169.0 | 0 | Canis | 74169.0 | 0 | Canis lupus | 74169.0 | 0 |
| scf7180000042396 | 523991 | 0.4098 | 0 | 60.0148 | Eukaryota | 85988.0 | 0 | Chordata | 85988.0 | 0 | Carnivora | 85988.0 | 0 | Canidae | 85988.0 | 0 | Canis | 85988.0 | 0 | Canis lupus | 85988.0 | 0 |
| scf7180000051703 | 4322 | 0.4387 | 0 | 207.1147 | Eukaryota | 3143.0 | 0 | Chordata | 1627.0 | 1 | Polystomatidea | 1516.0 | 2 | Polystomatidae | 1516.0 | 2 | Protopolystoma | 1516.0 | 2 | Protopolystoma xenopodis | 1516.0 | 2 |
| scf7180000055266 | 14174 | 0.4399 | 0 | 199.8358 | Eukaryota | 61673.0 | 0 | Chordata | 43375.0 | 2 | Gadiformes | 38064.0 | 5 | Gadidae | 38064.0 | 5 | Gadus | 38064.0 | 5 | Gadus morhua | 38064.0 | 5 |
| scf7180000050458 | 17177 | 0.5032 | 0 | 752.2795 | Eukaryota | 9364.0 | 0 | Chordata | 9364.0 | 0 | Carnivora | 9364.0 | 0 | Canidae | 9364.0 | 0 | Canis | 8142.0 | 1 | Canis lupus | 6863.0 | 2 |
| scf7180000048988 | 5094 | 0.4578 | 0 | 115.4992 | Eukaryota | 477.0 | 0 | Platyhelminthes | 477.0 | 0 | Polystomatidea | 477.0 | 0 | Polystomatidae | 477.0 | 0 | Protopolystoma | 477.0 | 0 | Protopolystoma xenopodis | 477.0 | 0 |
| scf7180000043168 | 77595 | 0.3379 | 0 | 55.6839 | Eukaryota | 15584.0 | 0 | Chordata | 15584.0 | 0 | Carnivora | 15584.0 | 0 | Canidae | 15584.0 | 0 | Canis | 15584.0 | 0 | Canis lupus | 15584.0 | 0 |
| scf7180000041604 | 416772 | 0.5299 | 0 | 55.611 | Eukaryota | 24207.0 | 0 | Chordata | 24207.0 | 0 | Carnivora | 24207.0 | 0 | Canidae | 24207.0 | 0 | Vulpes | 15971.0 | 1 | Vulpes vulpes | 15971.0 | 1 |
| scf7180000051471 | 2897 | 0.4474 | 0 | 579.899 | Eukaryota | 1029.0 | 0 | Echinodermata | 532.0 | 1 | Forcipulatida | 532.0 | 1 | Asteriidae | 532.0 | 1 | Asterias | 532.0 | 1 | Asterias rubens | 532.0 | 1 |
| scf7180000054907 | 21247 | 0.4403 | 0 | 140.1824 | Eukaryota | 4797.0 | 0 | Chordata | 4199.0 | 1 | Kurtiformes | 3466.0 | 4 | Apogonidae | 3466.0 | 4 | Sphaeramia | 3466.0 | 4 | Sphaeramia orbicularis | 3466.0 | 4 |
| scf7180000045482 | 220707 | 0.3511 | 0 | 59.8182 | Eukaryota | 92678.0 | 0 | Chordata | 92678.0 | 0 | Carnivora | 92678.0 | 0 | Canidae | 92678.0 | 0 | Canis | 92678.0 | 0 | Canis lupus | 92678.0 | 0 |
| scf7180000045775 | 38257 | 0.4672 | 0 | 54.146 | Eukaryota | 13151.0 | 0 | Chordata | 13151.0 | 0 | Rodentia | 7074.0 | 1 | Muridae | 7074.0 | 1 | Mus | 7074.0 | 2 | Mus musculus | 7074.0 | 2 |
| scf7180000045517 | 180232 | 0.3563 | 0 | 58.8794 | Eukaryota | 20266.0 | 1 | Chordata | 20266.0 | 1 | Carnivora | 16072.0 | 2 | Ursidae | 16072.0 | 2 | Ursus | 16072.0 | 2 | Ursus arctos | 16072.0 | 2 |
| scf7180000052227 | 13664 | 0.4409 | 0 | 382.4142 | Eukaryota | 274.0 | 0 | Platyhelminthes | 274.0 | 0 | Polystomatidea | 274.0 | 0 | Polystomatidae | 274.0 | 0 | Protopolystoma | 274.0 | 0 | Protopolystoma xenopodis | 274.0 | 0 |
| scf7180000043362 | 472460 | 0.3548 | 0 | 59.3212 | Eukaryota | 31830.0 | 0 | Chordata | 31830.0 | 0 | Carnivora | 31830.0 | 0 | Canidae | 31830.0 | 0 | Canis | 31830.0 | 0 | Canis lupus | 31830.0 | 0 |
| scf7180000046003 | 3858 | 0.5052 | 0 | 35.6805 | Eukaryota | 6725.0 | 0 | Chordata | 6725.0 | 0 | Carnivora | 6725.0 | 0 | Canidae | 6725.0 | 0 | Canis | 4298.0 | 1 | Canis lupus | 3183.0 | 2 |
| scf7180000040531 | 301019 | 0.3743 | 0 | 59.8775 | Eukaryota | 40977.0 | 0 | Chordata | 40977.0 | 0 | Carnivora | 40977.0 | 0 | Canidae | 40977.0 | 0 | Canis | 24288.0 | 1 | Canis lupus | 24288.0 | 1 |
| scf7180000053640 | 2203 | 0.4576 | 0 | 340.7167 | Eukaryota | 2738.0 | 0 | Platyhelminthes | 1475.0 | 2 | Polystomatidea | 1475.0 | 2 | Polystomatidae | 1475.0 | 2 | Protopolystoma | 1475.0 | 2 | Protopolystoma xenopodis | 1475.0 | 2 |
| scf7180000049670 | 6175 | 0.4554 | 0 | 26.5064 | Eukaryota | 5576.0 | 0 | Chordata | 5576.0 | 0 | Primates | 2928.0 | 1 | Hominidae | 2928.0 | 1 | Homo | 2928.0 | 1 | Homo sapiens | 2928.0 | 1 |
| scf7180000041671 | 902080 | 0.4266 | 0 | 58.2336 | Eukaryota | 89144.0 | 0 | Chordata | 89144.0 | 0 | Carnivora | 89144.0 | 0 | Canidae | 89144.0 | 0 | Canis | 62807.0 | 1 | Canis lupus | 62807.0 | 1 |
| scf7180000053378 | 9240 | 0.4013 | 0 | 45.2565 | Eukaryota | 3376.0 | 0 | Chordata | 3376.0 | 0 | Carnivora | 2727.0 | 1 | Canidae | 2727.0 | 1 | Canis | 2727.0 | 1 | Canis lupus | 2727.0 | 1 |
| scf7180000046328 | 229465 | 0.682 | 0 | 54.0406 | Eukaryota | 38104.0 | 0 | Chordata | 38104.0 | 0 | Carnivora | 38104.0 | 0 | Canidae | 38104.0 | 0 | Canis | 29376.0 | 1 | Canis lupus | 29376.0 | 1 |
| scf7180000049513 | 22604 | 0.4374 | 0 | 253.9382 | Eukaryota | 3867.0 | 0 | Platyhelminthes | 3253.0 | 1 | Polystomatidea | 3253.0 | 2 | Polystomatidae | 3253.0 | 2 | Protopolystoma | 3253.0 | 2 | Protopolystoma xenopodis | 3253.0 | 2 |
| scf7180000049098 | 5598 | 0.4528 | 0 | 79.4413 | Eukaryota | 15972.0 | 0 | Chordata | 10977.0 | 2 | Strigeidida | 4359.0 | 7 | Schistosomatidae | 4359.0 | 7 | Schistosoma | 4359.0 | 7 | Schistosoma mansoni | 4359.0 | 7 |
| scf7180000046393 | 4958 | 0.4397 | 0 | 127.5373 | Eukaryota | 1143.0 | 0 | Platyhelminthes | 912.0 | 1 | Polystomatidea | 777.0 | 2 | Polystomatidae | 777.0 | 2 | Protopolystoma | 777.0 | 2 | Protopolystoma xenopodis | 777.0 | 2 |
| scf7180000053849 | 11011 | 0.5099 | 0 | 375.1431 | Eukaryota | 8854.0 | 0 | Chordata | 8854.0 | 0 | Carnivora | 8854.0 | 0 | Canidae | 8854.0 | 0 | Canis | 7140.0 | 1 | Canis lupus | 5898.0 | 2 |
| scf7180000051316 | 3603 | 0.5046 | 0 | 128.4479 | Eukaryota | 7988.0 | 0 | Chordata | 7988.0 | 0 | Carnivora | 7988.0 | 0 | Canidae | 7988.0 | 0 | Canis | 5864.0 | 1 | Canis lupus | 4724.0 | 2 |
| scf7180000049181 | 48934 | 0.365 | 0 | 52.736 | Eukaryota | 36839.0 | 0 | Chordata | 36839.0 | 0 | Carnivora | 36839.0 | 0 | Canidae | 36839.0 | 0 | Canis | 36839.0 | 0 | Canis lupus | 36839.0 | 0 |
| scf7180000043166 | 13549 | 0.4445 | 0 | 43.3665 | Eukaryota | 67476.0 | 0 | Chordata | 67476.0 | 0 | Carnivora | 67476.0 | 0 | Canidae | 67476.0 | 0 | Canis | 67476.0 | 0 | Canis lupus | 67476.0 | 0 |
| scf7180000048557 | 69392 | 0.3282 | 0 | 63.247 | Eukaryota | 30361.0 | 0 | Chordata | 30361.0 | 0 | Carnivora | 30361.0 | 0 | Canidae | 30361.0 | 0 | Canis | 21322.0 | 1 | Canis lupus | 21322.0 | 1 |
| scf7180000052795 | 8971 | 0.416 | 0 | 59.7535 | Eukaryota | 4251.0 | 0 | Chordata | 4251.0 | 0 | Carnivora | 1674.0 | 2 | Muridae | 1433.0 | 4 | Homo | 1144.0 | 7 | Homo sapiens | 1144.0 | 7 |
| scf7180000047472 | 12743 | 0.4369 | 0 | 69.4398 | Eukaryota | 91694.0 | 0 | Chordata | 91694.0 | 0 | Carnivora | 91694.0 | 0 | Canidae | 91694.0 | 0 | Canis | 91694.0 | 0 | Canis lupus | 91694.0 | 0 |
| scf7180000042027 | 204057 | 0.3709 | 0 | 64.5058 | Eukaryota | 44309.0 | 0 | Chordata | 44309.0 | 0 | Carnivora | 44309.0 | 0 | Canidae | 44309.0 | 0 | Canis | 35477.0 | 1 | Canis lupus | 35477.0 | 1 |
| scf7180000054374 | 79353 | 0.4565 | 0 | 85.6047 | Eukaryota | 97935.0 | 0 | Chordata | 97935.0 | 0 | Carnivora | 97935.0 | 0 | Canidae | 97935.0 | 0 | Canis | 97935.0 | 0 | Canis lupus | 97935.0 | 0 |
| scf7180000041256 | 440893 | 0.4344 | 0 | 58.8832 | Eukaryota | 38059.0 | 0 | Chordata | 38059.0 | 0 | Carnivora | 38059.0 | 0 | Canidae | 38059.0 | 0 | Canis | 38059.0 | 0 | Canis lupus | 38059.0 | 0 |

| | | | | | | | | | | | | | | | | | | | | | | |
|---|---|---|---|---|---|---|---|---|---|---|---|---|---|---|---|---|---|---|---|---|---|---|
| scf7180000043739 | 153687 | 0.4653 | 0 | 58.5582 | Eukaryota | 48242.0 | 0 | Chordata | 48242.0 | 0 | Carnivora | 48242.0 | 0 | Canidae | 48242.0 | 0 | Canis | 48242.0 | 0 | Canis lupus | 48242.0 | 0 |
| scf7180000046312 | 137115 | 0.3739 | 0 | 59.6832 | Eukaryota | 88448.0 | 0 | Chordata | 88448.0 | 0 | Carnivora | 88448.0 | 0 | Canidae | 73365.0 | 1 | Canis | 64260.0 | 3 | Canis lupus | 64260.0 | 3 |
| scf7180000053901 | 2835 | 0.4575 | 0 | 548.3853 | Eukaryota | 427.0 | 0 | Platyhelminthes | 427.0 | 0 | Polystomatidea | 427.0 | 0 | Polystomatidae | 427.0 | 0 | Protopolystoma | 427.0 | 0 | Protopolystoma xenopodis | 427.0 | 0 |
| scf7180000047381 | 102767 | 0.4787 | 0 | 60.0276 | Eukaryota | 26968.0 | 0 | Chordata | 26968.0 | 0 | Carnivora | 26968.0 | 0 | Canidae | 22747.0 | 1 | Canis | 16720.0 | 3 | Canis lupus | 16720.0 | 3 |
| scf7180000044311 | 18671 | 0.4511 | 0 | 70.814 | Eukaryota | 3337.0 | 0 | Chordata | 2191.0 | 1 | Carnivora | 2191.0 | 1 | Canidae | 2191.0 | 1 | Canis | 2191.0 | 1 | Canis lupus | 2191.0 | 1 |
| scf7180000042725 | 630878 | 0.4558 | 0 | 57.7059 | Eukaryota | 79400.0 | 0 | Chordata | 79400.0 | 0 | Carnivora | 79400.0 | 0 | Canidae | 79400.0 | 0 | Canis | 79400.0 | 0 | Canis lupus | 79400.0 | 0 |
| scf7180000046748 | 115618 | 0.3691 | 0 | 58.1472 | Eukaryota | 54282.0 | 0 | Chordata | 54282.0 | 0 | Carnivora | 54282.0 | 0 | Canidae | 54282.0 | 0 | Canis | 43352.0 | 1 | Canis lupus | 43352.0 | 1 |
| scf7180000042285 | 718045 | 0.4773 | 0 | 58.7751 | Eukaryota | 41473.0 | 0 | Chordata | 41473.0 | 0 | Carnivora | 41473.0 | 0 | Canidae | 41473.0 | 0 | Canis | 41473.0 | 0 | Canis lupus | 41473.0 | 0 |
| scf7180000042922 | 462423 | 0.4207 | 0 | 60.0039 | Eukaryota | 91702.0 | 0 | Chordata | 91702.0 | 0 | Carnivora | 91702.0 | 0 | Canidae | 91702.0 | 0 | Canis | 91702.0 | 0 | Canis lupus | 91702.0 | 0 |
| scf7180000040940 | 1433259 | 0.4262 | 0 | 59.4776 | Eukaryota | 177873.0 | 0 | Chordata | 177873.0 | 0 | Carnivora | 177873.0 | 0 | Canidae | 177873.0 | 0 | Canis | 177873.0 | 0 | Canis lupus | 177873.0 | 0 |
| scf7180000052734 | 5732 | 0.3707 | 0 | 60.6056 | Eukaryota | 4477.0 | 0 | Chordata | 4477.0 | 0 | Carnivora | 3599.0 | 1 | Canidae | 3599.0 | 1 | Canis | 3192.0 | 2 | Canis lupus | 3192.0 | 2 |
| scf7180000052230 | 15707 | 0.5023 | 0 | 345.2565 | Eukaryota | 9023.0 | 0 | Chordata | 9023.0 | 0 | Carnivora | 9023.0 | 0 | Canidae | 9023.0 | 0 | Canis | 7304.0 | 1 | Canis lupus | 6056.0 | 2 |
| scf7180000043319 | 432269 | 0.4592 | 0 | 58.8876 | Eukaryota | 53380.0 | 0 | Chordata | 53380.0 | 0 | Carnivora | 53380.0 | 0 | Canidae | 48504.0 | 1 | Canis | 43080.0 | 2 | Canis lupus | 43080.0 | 2 |
| scf7180000045065 | 204391 | 0.4195 | 0 | 59.1786 | Eukaryota | 77490.0 | 0 | Chordata | 77490.0 | 0 | Carnivora | 77490.0 | 0 | Canidae | 77490.0 | 0 | Canis | 77490.0 | 0 | Canis lupus | 77490.0 | 0 |
| scf7180000049578 | 4914 | 0.4538 | 0 | 666.193 | Eukaryota | 391.0 | 0 | Chordata | 248.0 | 1 | Tetraodontiformes | 248.0 | 1 | Tetraodontidae | 248.0 | 1 | Takifugu | 248.0 | 1 | Takifugu rubripes | 248.0 | 1 |
| scf7180000043280 | 375369 | 0.4508 | 100 | 56.6841 | Eukaryota | 47861.0 | 0 | Chordata | 47861.0 | 0 | Carnivora | 47861.0 | 0 | Canidae | 43984.0 | 1 | Canis | 39068.0 | 2 | Canis lupus | 39068.0 | 2 |
| scf7180000051598 | 17287 | 0.527 | 0 | 481.6976 | Eukaryota | 4695.0 | 0 | Chordata | 4695.0 | 0 | Carnivora | 4695.0 | 0 | Canidae | 4695.0 | 0 | Canis | 3987.0 | 1 | Canis lupus | 3987.0 | 1 |
| scf7180000055739 | 10807 | 0.4433 | 0 | 368.3432 | no-hit | 0.0 | 0 | no-hit | 0.0 | 0 | no-hit | 0.0 | 0 | no-hit | 0.0 | 0 | no-hit | 0.0 | 0 | no-hit | 0.0 | 0 |
| scf7180000053691 | 4765 | 0.4512 | 0 | 738.4676 | Eukaryota | 2609.0 | 0 | Platyhelminthes | 1776.0 | 1 | Polystomatidea | 1776.0 | 1 | Polystomatidae | 1776.0 | 1 | Protopolystoma | 1776.0 | 1 | Protopolystoma xenopodis | 1776.0 | 1 |
| scf7180000045787 | 179728 | 0.5521 | 0 | 54.3006 | Eukaryota | 40762.0 | 0 | Chordata | 40762.0 | 0 | Carnivora | 35943.0 | 1 | Canidae | 35943.0 | 1 | Canis | 35943.0 | 1 | Canis lupus | 35943.0 | 1 |
| scf7180000052936 | 4731 | 0.3893 | 0 | 61.4381 | Eukaryota | 4825.0 | 0 | Chordata | 4825.0 | 0 | Carnivora | 2761.0 | 1 | Canidae | 2761.0 | 2 | Canis | 2761.0 | 2 | Canis lupus | 2761.0 | 2 |
| scf7180000043355 | 164564 | 0.391 | 0 | 58.7579 | Eukaryota | 90076.0 | 0 | Chordata | 90076.0 | 0 | Carnivora | 90076.0 | 0 | Canidae | 90076.0 | 0 | Canis | 90076.0 | 0 | Canis lupus | 90076.0 | 0 |
| scf7180000043357 | 493980 | 0.4215 | 0 | 61.9942 | Eukaryota | 53253.0 | 0 | Chordata | 53253.0 | 0 | Carnivora | 53253.0 | 0 | Canidae | 53253.0 | 0 | Canis | 47567.0 | 1 | Canis lupus | 47567.0 | 1 |
| scf7180000042173 | 792769 | 0.4588 | 0 | 58.5449 | Eukaryota | 54907.0 | 0 | Chordata | 54907.0 | 0 | Carnivora | 54907.0 | 0 | Canidae | 54907.0 | 0 | Canis | 33727.0 | 1 | Canis lupus | 33727.0 | 1 |
| scf7180000047432 | 86080 | 0.5363 | 0 | 55.9296 | Eukaryota | 8930.0 | 0 | Chordata | 8930.0 | 0 | Carnivora | 8930.0 | 0 | Mustelidae | 5869.0 | 1 | Mustela | 4209.0 | 3 | Mustela putorius | 4209.0 | 3 |
| scf7180000047525 | 81442 | 0.5415 | 0 | 57.5188 | Eukaryota | 28707.0 | 0 | Chordata | 28707.0 | 0 | Carnivora | 28707.0 | 0 | Canidae | 19434.0 | 3 | Canis | 14350.0 | 5 | Canis lupus | 14350.0 | 5 |
| scf7180000041473 | 596601 | 0.3544 | 0 | 61.3566 | Eukaryota | 20091.0 | 0 | Chordata | 20091.0 | 0 | Carnivora | 20091.0 | 0 | Canidae | 12287.0 | 3 | Canis | 10233.0 | 5 | Canis lupus | 10233.0 | 5 |
| scf7180000045915 | 19785 | 0.4174 | 0 | 42.2254 | Eukaryota | 3405.0 | 0 | Chordata | 3405.0 | 0 | Carnivora | 2456.0 | 1 | Canidae | 2456.0 | 1 | Canis | 2456.0 | 2 | Canis lupus | 2456.0 | 2 |
| scf7180000042141 | 216439 | 0.3907 | 0 | 59.0134 | Eukaryota | 60514.0 | 0 | Chordata | 60514.0 | 0 | Carnivora | 60514.0 | 0 | Canidae | 60514.0 | 0 | Canis | 60514.0 | 0 | Canis lupus | 60514.0 | 0 |
| scf7180000046649 | 119041 | 0.5177 | 0 | 58.1267 | Eukaryota | 9515.0 | 0 | Chordata | 9515.0 | 0 | Carnivora | 9515.0 | 0 | Canidae | 9515.0 | 0 | Canis | 7411.0 | 1 | Canis lupus | 7411.0 | 1 |
| scf7180000056437 | 480974 | 0.3691 | 0 | 60.6595 | Eukaryota | 90453.0 | 0 | Chordata | 90453.0 | 0 | Carnivora | 90453.0 | 0 | Canidae | 90453.0 | 0 | Canis | 90453.0 | 0 | Canis lupus | 90453.0 | 0 |
| scf7180000044440 | 23700 | 0.483 | 0 | 79.6245 | Eukaryota | 22211.0 | 0 | Chordata | 22211.0 | 0 | Carnivora | 22211.0 | 0 | Canidae | 22211.0 | 0 | Canis | 22211.0 | 0 | Canis lupus | 22211.0 | 0 |
| scf7180000049586 | 40634 | 0.4495 | 0 | 58.0102 | Eukaryota | 34580.0 | 0 | Chordata | 34580.0 | 0 | Carnivora | 34580.0 | 0 | Canidae | 25277.0 | 1 | Canis | 25277.0 | 1 | Canis lupus | 25277.0 | 1 |
| scf7180000043707 | 426623 | 0.445 | 0 | 59.0701 | Eukaryota | 79430.0 | 0 | Chordata | 79430.0 | 0 | Carnivora | 79430.0 | 0 | Canidae | 79430.0 | 0 | Canis | 79430.0 | 0 | Canis lupus | 79430.0 | 0 |
| scf7180000055293 | 47180 | 0.4393 | 0 | 445.7874 | Eukaryota | 3680.0 | 0 | Chordata | 2533.0 | 1 | Kurtiformes | 1775.0 | 3 | Apogonidae | 1775.0 | 3 | Sphaeramia | 1775.0 | 3 | Sphaeramia orbicularis | 1775.0 | 3 |
| scf7180000042057 | 26972 | 0.3804 | 0 | 58.5134 | Eukaryota | 92639.0 | 0 | Chordata | 92639.0 | 0 | Carnivora | 92639.0 | 0 | Canidae | 92639.0 | 0 | Canis | 92639.0 | 0 | Canis lupus | 92639.0 | 0 |
| scf7180000055815 | 9696 | 0.4495 | 0 | 1616.9291 | Eukaryota | 1457.0 | 0 | Streptophyta | 1457.0 | 0 | Fabales | 1457.0 | 0 | Fabaceae | 1457.0 | 0 | Cicer | 1457.0 | 0 | Cicer arietinum | 1457.0 | 0 |
| scf7180000042465 | 89570 | 0.4202 | 0 | 58.3624 | Eukaryota | 88387.0 | 0 | Chordata | 88387.0 | 0 | Carnivora | 88387.0 | 0 | Canidae | 88387.0 | 0 | Canis | 88387.0 | 0 | Canis lupus | 88387.0 | 0 |
| scf7180000056534 | 165437 | 0.3597 | 0 | 60.6366 | Eukaryota | 16440.0 | 0 | Chordata | 16440.0 | 0 | Carnivora | 16440.0 | 0 | Canidae | 16440.0 | 0 | Canis | 16440.0 | 0 | Canis lupus | 16440.0 | 0 |
| scf7180000054246 | 1038965 | 0.3923 | 0 | 58.1062 | Eukaryota | 21946.0 | 0 | Chordata | 21946.0 | 0 | Carnivora | 21946.0 | 0 | Canidae | 21946.0 | 0 | Canis | 17679.0 | 1 | Canis lupus | 17679.0 | 1 |
| scf7180000042338 | 40094 | 0.3898 | 0 | 56.7308 | Eukaryota | 91301.0 | 0 | Chordata | 91301.0 | 0 | Carnivora | 91301.0 | 0 | Canidae | 91301.0 | 0 | Canis | 91301.0 | 0 | Canis lupus | 91301.0 | 0 |
| scf7180000041234 | 23870 | 0.3693 | 0 | 45.0067 | Eukaryota | 14073.0 | 0 | Chordata | 14073.0 | 0 | Carnivora | 14073.0 | 0 | Canidae | 14073.0 | 0 | Canis | 14073.0 | 0 | Canis lupus | 14073.0 | 0 |
| scf7180000050999 | 7184 | 0.4431 | 0 | 101.4032 | Eukaryota | 195.0 | 0 | Chordata | 195.0 | 0 | Salmoniformes | 195.0 | 0 | Salmonidae | 195.0 | 0 | Coregonus | 195.0 | 0 | Coregonus sp. 'balchen' | 195.0 | 0 |
| scf7180000042309 | 287842 | 0.4618 | 0 | 61.7697 | Eukaryota | 93931.0 | 0 | Chordata | 93931.0 | 0 | Carnivora | 93931.0 | 0 | Canidae | 93931.0 | 0 | Canis | 93931.0 | 0 | Canis lupus | 93931.0 | 0 |
| scf7180000044789 | 286213 | 0.4367 | 0 | 59.1037 | Eukaryota | 127998.0 | 0 | Chordata | 127998.0 | 0 | Carnivora | 127998.0 | 0 | Canidae | 127998.0 | 0 | Canis | 127998.0 | 0 | Canis lupus | 127998.0 | 0 |
| scf7180000052011 | 16536 | 0.4403 | 0 | 208.1854 | Eukaryota | 3893.0 | 0 | Platyhelminthes | 2497.0 | 2 | Polystomatidea | 2497.0 | 2 | Polystomatidae | 2497.0 | 2 | Protopolystoma | 2497.0 | 2 | Protopolystoma xenopodis | 2497.0 | 2 |
| scf7180000041248 | 601277 | 0.3511 | 0 | 59.1152 | Eukaryota | 95956.0 | 0 | Chordata | 95956.0 | 0 | Carnivora | 95956.0 | 0 | Canidae | 95956.0 | 0 | Canis | 95956.0 | 0 | Canis lupus | 95956.0 | 0 |
| scf7180000051787 | 16261 | 0.5037 | 0 | 743.464 | Eukaryota | 9503.0 | 0 | Chordata | 9503.0 | 0 | Carnivora | 9503.0 | 0 | Canidae | 9503.0 | 0 | Canis | 7720.0 | 1 | Canis lupus | 6420.0 | 2 |
| scf7180000051543 | 21991 | 0.505 | 0 | 131.6473 | Eukaryota | 9270.0 | 0 | Chordata | 9270.0 | 0 | Carnivora | 9270.0 | 0 | Canidae | 9270.0 | 0 | Canis | 7493.0 | 1 | Canis lupus | 6236.0 | 2 |
| scf7180000054939 | 89844 | 0.4413 | 0 | 209.6886 | Eukaryota | 29167.0 | 0 | Chordata | 29167.0 | 0 | Gadiformes | 19272.0 | 5 | Gadidae | 19272.0 | 5 | Gadus | 19272.0 | 5 | Gadus morhua | 19272.0 | 5 |
| scf7180000043787 | 60242 | 0.4093 | 0 | 60.6507 | Eukaryota | 9912.0 | 0 | Chordata | 9912.0 | 0 | Primates | 7299.0 | 1 | Hominidae | 4650.0 | 2 | Homo | 4060.0 | 3 | Homo sapiens | 4060.0 | 3 |
| scf7180000051549 | 5028 | 0.4491 | 0 | 634.1908 | Eukaryota | 1875.0 | 0 | Platyhelminthes | 767.0 | 2 | Polystomatidea | 767.0 | 2 | Polystomatidae | 767.0 | 2 | Protopolystoma | 767.0 | 2 | Protopolystoma xenopodis | 767.0 | 2 |
| scf7180000046979 | 75936 | 0.3974 | 0 | 57.7862 | Eukaryota | 12251.0 | 0 | Chordata | 12251.0 | 0 | Carnivora | 7388.0 | 2 | Canidae | 7388.0 | 2 | Canis | 7388.0 | 3 | Canis lupus | 7388.0 | 3 |
| scf7180000046832 | 8546 | 0.4443 | 0 | 163.0093 | Eukaryota | 10390.0 | 0 | Chordata | 10390.0 | 0 | Tetraodontiformes | 4222.0 | 2 | Tetraodontidae | 4222.0 | 2 | Takifugu | 4222.0 | 2 | Sphaeramia orbicularis | 3625.0 | 3 |
| scf7180000048238 | 15264 | 0.4459 | 0 | 159.6444 | Eukaryota | 3033.0 | 0 | Chordata | 2350.0 | 1 | Carnivora | 2350.0 | 1 | Canidae | 2350.0 | 1 | Canis | 2350.0 | 1 | Canis lupus | 2350.0 | 1 |
| scf7180000046784 | 20494 | 0.4416 | 0 | 303.4713 | Eukaryota | 663.0 | 0 | Platyhelminthes | 489.0 | 1 | Polystomatidea | 489.0 | 1 | Polystomatidae | 489.0 | 1 | Protopolystoma | 489.0 | 1 | Protopolystoma xenopodis | 489.0 | 1 |
| scf7180000045284 | 18574 | 0.372 | 0 | 44.0888 | Eukaryota | 51380.0 | 0 | Chordata | 51380.0 | 0 | Carnivora | 51380.0 | 0 | Canidae | 39395.0 | 4 | Vulpes | 27655.0 | 5 | Vulpes vulpes | 27655.0 | 5 |
| scf7180000047403 | 3985 | 0.4592 | 0 | 99.7448 | no-hit | 0.0 | 0 | no-hit | 0.0 | 0 | no-hit | 0.0 | 0 | no-hit | 0.0 | 0 | no-hit | 0.0 | 0 | no-hit | 0.0 | 0 |
| scf7180000041016 | 718123 | 0.4023 | 0 | 59.5886 | Eukaryota | 74457.0 | 0 | Chordata | 74457.0 | 0 | Carnivora | 74457.0 | 0 | Canidae | 74457.0 | 0 | Canis | 67199.0 | 1 | Canis lupus | 67199.0 | 1 |
| scf7180000042892 | 321957 | 0.4009 | 0 | 58.4057 | Eukaryota | 22575.0 | 0 | Chordata | 22575.0 | 0 | Carnivora | 22575.0 | 0 | Canidae | 22575.0 | 0 | Canis | 22575.0 | 0 | Canis lupus | 22575.0 | 0 |
| scf7180000043601 | 753160 | 0.444 | 0 | 59.6037 | Eukaryota | 96427.0 | 0 | Chordata | 96427.0 | 0 | Carnivora | 96427.0 | 0 | Canidae | 96427.0 | 0 | Canis | 96427.0 | 0 | Canis lupus | 96427.0 | 0 |
| scf7180000041160 | 1340220 | 0.3771 | 0 | 60.2553 | Eukaryota | 57650.0 | 0 | Chordata | 57650.0 | 0 | Carnivora | 57650.0 | 0 | Canidae | 57650.0 | 0 | Canis | 42200.0 | 1 | Canis lupus | 42200.0 | 1 |
| scf7180000055427 | 23458 | 0.442 | 0 | 104.8005 | Eukaryota | 1736.0 | 0 | Platyhelminthes | 1204.0 | 1 | Polystomatidea | 1204.0 | 1 | Polystomatidae | 1204.0 | 1 | Protopolystoma | 1204.0 | 1 | Protopolystoma xenopodis | 1204.0 | 1 |
| scf7180000044239 | 10437 | 0.4287 | 0 | 29.6819 | Eukaryota | 4523.0 | 0 | Chordata | 4523.0 | 0 | Carnivora | 4523.0 | 0 | Canidae | 3187.0 | 1 | Canis | 3187.0 | 2 | Canis lupus | 3187.0 | 2 |
| scf7180000040818 | 265553 | 0.4184 | 0 | 59.5016 | Eukaryota | 61200.0 | 0 | Chordata | 61200.0 | 0 | Carnivora | 61200.0 | 0 | Canidae | 61200.0 | 0 | Canis | 61200.0 | 0 | Canis lupus | 61200.0 | 0 |
| scf7180000044170 | 363849 | 0.4772 | 0 | 59.3179 | Eukaryota | 24479.0 | 0 | Chordata | 24479.0 | 0 | Carnivora | 24479.0 | 0 | Canidae | 24479.0 | 0 | Canis | 24479.0 | 0 | Canis lupus | 24479.0 | 0 |
| scf7180000047661 | 95560 | 0.4194 | 0 | 60.3327 | Eukaryota | 23623.0 | 0 | Chordata | 23623.0 | 0 | Carnivora | 23623.0 | 0 | Canidae | 21600.0 | 1 | Canis | 19127.0 | 2 | Canis lupus | 19127.0 | 2 |
| scf7180000047129 | 91562 | 0.4972 | 0 | 60.2658 | Eukaryota | 24565.0 | 0 | Chordata | 24565.0 | 0 | Carnivora | 24565.0 | 0 | Canidae | 24565.0 | 0 | Canis | 16517.0 | 1 | Canis lupus | 16517.0 | 1 |
| scf7180000047872 | 6716 | 0.4626 | 0 | 36.8281 | no-hit | 0.0 | 0 | no-hit | 0.0 | 0 | no-hit | 0.0 | 0 | no-hit | 0.0 | 0 | no-hit | 0.0 | 0 | no-hit | 0.0 | 0 |
| scf7180000055732 | 10993 | 0.517 | 0 | 471.8508 | Eukaryota | 15879.0 | 0 | Chordata | 15879.0 | 0 | Carnivora | 15879.0 | 0 | Canidae | 15879.0 | 0 | Canis | 15879.0 | 0 | Canis lupus | 15879.0 | 0 |
| scf7180000045824 | 51095 | 0.5186 | 0 | 53.8312 | Eukaryota | 24229.0 | 0 | Chordata | 24229.0 | 0 | Carnivora | 24229.0 | 0 | Canidae | 23680.0 | 1 | Canis | 22112.0 | 2 | Canis lupus | 22112.0 | 2 |
| scf7180000047146 | 204766 | 0.447 | 100 | 57.3102 | Eukaryota | 15492.0 | 0 | Chordata | 15492.0 | 0 | Carnivora | 12091.0 | 3 | Canidae | 12091.0 | 3 | Canis | 12091.0 | 4 | Canis lupus | 12091.0 | 4 |
| scf7180000055884 | 29830 | 0.4477 | 0 | 454.2387 | Eukaryota | 3791.0 | 0 | Platyhelminthes | 2380.0 | 3 | Polystomatidea | 2380.0 | 3 | Polystomatidae | 2380.0 | 3 | Protopolystoma | 2380.0 | 3 | Protopolystoma xenopodis | 2380.0 | 3 |
| scf7180000044842 | 15736 | 0.3798 | 0 | 54.9838 | Eukaryota | 3936.0 | 0 | Chordata | 3936.0 | 0 | Carnivora | 2291.0 | 2 | Canidae | 2291.0 | 2 | Canis | 2291.0 | 3 | Canis lupus | 2291.0 | 3 |
| scf7180000048656 | 34140 | 0.441 | 0 | 80.3863 | no-hit | 0.0 | 0 | no-hit | 0.0 | 0 | no-hit | 0.0 | 0 | no-hit | 0.0 | 0 | no-hit | 0.0 | 0 | no-hit | 0.0 | 0 |
| scf7180000054079 | 5188 | 0.5089 | 0 | 95.9015 | Eukaryota | 7830.0 | 0 | Chordata | 7830.0 | 0 | Carnivora | 7830.0 | 0 | Canidae | 7830.0 | 0 | Canis | 5192.0 | 1 | Canis lupus | 4012.0 | 2 |
| scf7180000044854 | 272296 | 0.59 | 0 | 55.058 | Eukaryota | 21377.0 | 0 | Chordata | 21377.0 | 0 | Carnivora | 21377.0 | 0 | Canidae | 21377.0 | 0 | Canis | 21377.0 | 0 | Canis lupus | 21377.0 | 0 |
| scf7180000043654 | 394707 | 0.455 | 0 | 60.2209 | Eukaryota | 100890.0 | 0 | Chordata | 100890.0 | 0 | Carnivora | 100890.0 | 0 | Canidae | 100890.0 | 0 | Canis | 100890.0 | 0 | Canis lupus | 100890.0 | 0 |
| scf7180000050715 | 9911 | 0.5035 | 0 | 672.6521 | Eukaryota | 8937.0 | 0 | Chordata | 8937.0 | 0 | Carnivora | 8937.0 | 0 | Canidae | 8937.0 | 0 | Canis | 7279.0 | 1 | Canis lupus | 6063.0 | 2 |
| scf7180000044339 | 37003 | 0.3715 | 0 | 60.0211 | Eukaryota | 7651.0 | 0 | Chordata | 7651.0 | 0 | Carnivora | 7651.0 | 0 | Otariidae | 3849.0 | 3 | Zalophus | 1543.0 | 6 | Zalophus californianus | 1543.0 | 6 |
| scf7180000047984 | 8489 | 0.4456 | 0 | 175.7981 | Eukaryota | 215.0 | 0 | Streptophyta | 215.0 | 0 | Fabales | 215.0 | 0 | Fabaceae | 215.0 | 0 | Cicer | 215.0 | 0 | Cicer arietinum | 215.0 | 0 |
| scf7180000047938 | 26533 | 0.4336 | 0 | 197.6668 | Eukaryota | 237.0 | 0 | Platyhelminthes | 237.0 | 0 | Polystomatidea | 237.0 | 0 | Polystomatidae | 237.0 | 0 | Protopolystoma | 237.0 | 0 | Protopolystoma xenopodis | 237.0 | 0 |
| scf7180000043821 | 330599 | 0.3539 | 0 | 60.2112 | Eukaryota | 12527.0 | 0 | Chordata | 12527.0 | 0 | Primates | 8623.0 | 1 | Hominidae | 8623.0 | 1 | Homo | 8623.0 | 1 | Homo sapiens | 8623.0 | 1 |
| scf7180000055663 | 8967 | 0.4345 | 0 | 848.6903 | Eukaryota | 1788.0 | 0 | Platyhelminthes | 1352.0 | 1 | Polystomatidea | 1078.0 | 2 | Polystomatidae | 1078.0 | 2 | Protopolystoma | 1078.0 | 2 | Protopolystoma xenopodis | 1078.0 | 2 |
| scf7180000045160 | 251282 | 0.4274 | 0 | 58.4305 | Eukaryota | 29480.0 | 0 | Chordata | 29480.0 | 0 | Carnivora | 29480.0 | 0 | Canidae | 29480.0 | 0 | Canis | 29480.0 | 0 | Canis lupus | 29480.0 | 0 |
| scf7180000056582 | 638587 | 0.4123 | 0 | 61.2833 | Eukaryota | 91701.0 | 0 | Chordata | 91701.0 | 0 | Carnivora | 91701.0 | 0 | Canidae | 57313.0 | 3 | Canis | 49204.0 | 6 | Canis lupus | 49204.0 | 6 |
| scf7180000042065 | 224466 | 0.3587 | 0 | 57.3227 | Eukaryota | 77987.0 | 0 | Chordata | 77987.0 | 0 | Carnivora | 77987.0 | 0 | Canidae | 77987.0 | 0 | Canis | 77987.0 | 0 | Canis lupus | 77987.0 | 0 |
| scf7180000040657 | 486246 | 0.378 | 0 | 59.1865 | Eukaryota | 38973.0 | 0 | Chordata | 38973.0 | 0 | Carnivora | 26553.0 | 1 | Canidae | 14725.0 | 2 | Vulpes | 14725.0 | 2 | Vulpes vulpes | 14725.0 | 2 |
| scf7180000051010 | 14619 | 0.5035 | 0 | 190.184 | Eukaryota | 9099.0 | 0 | Chordata | 9099.0 | 0 | Carnivora | 9099.0 | 0 | Canidae | 9099.0 | 0 | Canis | 7303.0 | 1 | Canis lupus | 6044.0 | 2 |
| scf7180000051734 | 4074 | 0.432 | 0 | 175.2352 | no-hit | 0.0 | 0 | no-hit | 0.0 | 0 | no-hit | 0.0 | 0 | no-hit | 0.0 | 0 | no-hit | 0.0 | 0 | no-hit | 0.0 | 0 |
| scf7180000046240 | 22275 | 0.4303 | 0 | 57.2596 | Eukaryota | 28483.0 | 0 | Chordata | 28483.0 | 0 | Carnivora | 28483.0 | 0 | Canidae | 28483.0 | 0 | Canis | 28483.0 | 0 | Canis lupus | 28483.0 | 0 |
| scf7180000053411 | 5352 | 0.4411 | 0 | 66.0998 | no-hit | 0.0 | 0 | no-hit | 0.0 | 0 | no-hit | 0.0 | 0 | no-hit | 0.0 | 0 | no-hit | 0.0 | 0 | no-hit | 0.0 | 0 |
| scf7180000055598 | 12332 | 0.3956 | 0 | 512.3332 | Eukaryota | 51392.0 | 0 | Chordata | 51392.0 | 0 | Carnivora | 51392.0 | 0 | Canidae | 51392.0 | 0 | Canis | 51392.0 | 0 | Canis lupus | 51392.0 | 0 |
| scf7180000044755 | 48324 | 0.4972 | 0 | 249.0794 | Eukaryota | 14536.0 | 0 | Chordata | 14536.0 | 0 | Carnivora | 14536.0 | 0 | Canidae | 14536.0 | 0 | Canis | 14536.0 | 0 | Canis lupus | 14536.0 | 0 |
| scf7180000043907 | 301895 | 0.3676 | 0 | 59.0612 | Eukaryota | 90004.0 | 0 | Chordata | 90004.0 | 0 | Carnivora | 90004.0 | 0 | Canidae | 90004.0 | 0 | Canis | 90004.0 | 0 | Canis lupus | 90004.0 | 0 |
| scf7180000045127 | 235840 | 0.4801 | 0 | 58.0216 | Eukaryota | 8006.0 | 0 | Chordata | 8006.0 | 0 | Carnivora | 8006.0 | 0 | Canidae | 8006.0 | 0 | Canis | 8006.0 | 0 | Canis lupus | 8006.0 | 0 |
| scf7180000044706 | 13957 | 0.361 | 0 | 29.8588 | Eukaryota | 20874.0 | 0 | Chordata | 20874.0 | 0 | Primates | 14749.0 | 1 | Hominidae | 8505.0 | 4 | Homo | 4628.0 | 7 | Homo sapiens | 4628.0 | 8 |
| scf7180000047335 | 14968 | 0.4303 | 0 | 35.1698 | Eukaryota | 14716.0 | 0 | Chordata | 14716.0 | 0 | Carnivora | 12996.0 | 1 | Canidae | 12996.0 | 1 | Canis | 12996.0 | 1 | Canis lupus | 12996.0 | 1 |
| scf7180000044405 | 305110 | 0.3895 | 0 | 57.5851 | Eukaryota | 60312.0 | 0 | Chordata | 60312.0 | 0 | Carnivora | 60312.0 | 0 | Canidae | 60312.0 | 0 | Canis | 34580.0 | 1 | Canis lupus | 34580.0 | 1 |
| scf7180000046322 | 7317 | 0.4358 | 0 | 26.3881 | Eukaryota | 3913.0 | 0 | Chordata | 3913.0 | 0 | Primates | 1596.0 | 2 | Hominidae | 1596.0 | 2 | Homo | 1596.0 | 3 | Homo sapiens | 1596.0 | 3 |
| scf7180000048225 | 6877 | 0.4442 | 0 | 274.9373 | Eukaryota | 7541.0 | 0 | Chordata | 5714.0 | 1 | Kurtiformes | 2961.0 | 3 | Apogonidae | 2961.0 | 3 | Sphaeramia | 2961.0 | 3 | Sphaeramia orbicularis | 2961.0 | 3 |
| scf7180000044580 | 84346 | 0.4403 | 0 | 54.4139 | Eukaryota | 13282.0 | 0 | Chordata | 13282.0 | 0 | Carnivora | 13282.0 | 0 | Canidae | 13282.0 | 0 | Canis | 13282.0 | 0 | Canis lupus | 13282.0 | 0 |
| scf7180000049752 | 29594 | 0.5036 | 0 | 481.1931 | Eukaryota | 9340.0 | 0 | Chordata | 9340.0 | 0 | Carnivora | 9340.0 | 0 | Canidae | 9340.0 | 0 | Canis | 7590.0 | 1 | Canis lupus | 6304.0 | 2 |
| scf7180000051420 | 5337 | 0.451 | 0 | 896.1846 | no-hit | 0.0 | 0 | no-hit | 0.0 | 0 | no-hit | 0.0 | 0 | no-hit | 0.0 | 0 | no-hit | 0.0 | 0 | no-hit | 0.0 | 0 |
| scf7180000048561 | 68608 | 0.4123 | 0 | 60.2661 | Eukaryota | 13604.0 | 0 | Chordata | 13604.0 | 0 | Carnivora | 8461.0 | 2 | Canidae | 8461.0 | 3 | Canis | 8461.0 | 4 | Canis lupus | 8461.0 | 4 |
| scf7180000050264 | 20731 | 0.4416 | 0 | 52.2482 | Eukaryota | 13101.0 | 0 | Chordata | 13101.0 | 0 | Tetraodontiformes | 13101.0 | 0 | Tetraodontidae | 13101.0 | 0 | Takifugu | 13101.0 | 0 | Takifugu rubripes | 13101.0 | 0 |
| scf7180000040861 | 69954 | 0.4191 | 0 | 54.9663 | Eukaryota | 66209.0 | 0 | Chordata | 66209.0 | 0 | Carnivora | 66209.0 | 0 | Canidae | 66209.0 | 0 | Canis | 66209.0 | 0 | Canis lupus | 66209.0 | 0 |
| scf7180000042623 | 561882 | 0.4407 | 0 | 56.9423 | Eukaryota | 62181.0 | 0 | Chordata | 62181.0 | 0 | Carnivora | 62181.0 | 0 | Canidae | 47747.0 | 2 | Canis | 47747.0 | 3 | Canis lupus | 47747.0 | 3 |
| scf7180000041433 | 5160 | 0.3589 | 0 | 29.2493 | Eukaryota | 30191.0 | 0 | Chordata | 30191.0 | 0 | Carnivora | 30191.0 | 0 | Canidae | 30191.0 | 0 | Canis | 30191.0 | 0 | Canis lupus | 30191.0 | 0 |
| scf7180000043440 | 43154 | 0.3556 | 0 | 53.9322 | Eukaryota | 3368.0 | 0 | Chordata | 3368.0 | 0 | Carnivora | 2298.0 | 1 | Canidae | 1934.0 | 3 | Canis | 1934.0 | 3 | Canis lupus | 1934.0 | 3 |
| scf7180000043165 | 467148 | 0.3786 | 0 | 60.3425 | Eukaryota | 18073.0 | 0 | Chordata | 18073.0 | 0 | Carnivora | 18073.0 | 0 | Canidae | 16133.0 | 1 | Canis | 12812.0 | 2 | Canis lupus | 12812.0 | 2 |
| scf7180000053814 | 5846 | 0.5099 | 0 | 547.2991 | Eukaryota | 6545.0 | 0 | Chordata | 6545.0 | 0 | Carnivora | 6545.0 | 0 | Canidae | 6545.0 | 0 | Canis | 3651.0 | 1 | Nyctereutes procyonoides | 2894.0 | 2 |
| scf7180000049225 | 27957 | 0.4385 | 0 | 73.0243 | Eukaryota | 5460.0 | 0 | Chordata | 4000.0 | 1 | Synbranchiformes | 2747.0 | 4 | Mastacembelidae | 2747.0 | 4 | Mastacembelus | 2747.0 | 4 | Mastacembelus armatus | 2747.0 | 4 |
| scf7180000040763 | 1773253 | 0.3929 | 0 | 59.8004 | Eukaryota | 93695.0 | 0 | Chordata | 93695.0 | 0 | Carnivora | 93695.0 | 0 | Canidae | 93695.0 | 0 | Canis | 93695.0 | 0 | Canis lupus | 93695.0 | 0 |
| scf7180000053093 | 2927 | 0.4441 | 0 | 479.8519 | no-hit | 0.0 | 0 | no-hit | 0.0 | 0 | no-hit | 0.0 | 0 | no-hit | 0.0 | 0 | no-hit | 0.0 | 0 | no-hit | 0.0 | 0 |
| scf7180000042018 | 9607 | 0.4411 | 0 | 84.6988 | Eukaryota | 2171.0 | 0 | Chordata | 2171.0 | 0 | Carnivora | 1912.0 | 1 | Canidae | 1912.0 | 1 | Canis | 1912.0 | 1 | Canis lupus | 1912.0 | 1 |
| scf7180000049650 | 50178 | 0.5556 | 0 | 59.4304 | Eukaryota | 22535.0 | 0 | Chordata | 22535.0 | 0 | Carnivora | 22535.0 | 0 | Canidae | 22535.0 | 0 | Vulpes | 11545.0 | 1 | Vulpes vulpes | 11545.0 | 1 |
| scf7180000052102 | 2870 | 0.4537 | 0 | 247.2978 | no-hit | 0.0 | 0 | no-hit | 0.0 | 0 | no-hit | 0.0 | 0 | no-hit | 0.0 | 0 | no-hit | 0.0 | 0 | no-hit | 0.0 | 0 |
| scf7180000050631 | 25248 | 0.4445 | 0 | 183.9449 | Eukaryota | 1646.0 | 0 | Platyhelminthes | 1420.0 | 1 | Polystomatidea | 1420.0 | 1 | Polystomatidae | 1420.0 | 1 | Protopolystoma | 1420.0 | 1 | Protopolystoma xenopodis | 1420.0 | 1 |
| scf7180000056590 | 20695 | 0.4383 | 0 | 144.1227 | no-hit | 0.0 | 0 | no-hit | 0.0 | 0 | no-hit | 0.0 | 0 | no-hit | 0.0 | 0 | no-hit | 0.0 | 0 | no-hit | 0.0 | 0 |
| scf7180000056558 | 137229 | 0.6115 | 0 | 56.357 | Eukaryota | 33456.0 | 0 | Chordata | 33456.0 | 0 | Carnivora | 33456.0 | 0 | Canidae | 33456.0 | 0 | Canis | 28923.0 | 1 | Canis lupus | 28923.0 | 1 |
| scf7180000049929 | 9521 | 0.4514 | 0 | 522.3082 | no-hit | 0.0 | 0 | no-hit | 0.0 | 0 | no-hit | 0.0 | 0 | no-hit | 0.0 | 0 | no-hit | 0.0 | 0 | no-hit | 0.0 | 0 |
| scf7180000050017 | 3403 | 0.4426 | 0 | 97.6487 | Eukaryota | 1474.0 | 0 | Platyhelminthes | 958.0 | 1 | Polystomatidea | 958.0 | 2 | Polystomatidae | 958.0 | 2 | Protopolystoma | 958.0 | 2 | Protopolystoma xenopodis | 958.0 | 2 |
| scf7180000042439 | 8057 | 0.4204 | 0 | 30.7953 | Eukaryota | 37093.0 | 0 | Chordata | 37093.0 | 0 | Carnivora | 37093.0 | 0 | Canidae | 37093.0 | 0 | Canis | 37093.0 | 0 | Canis lupus | 37093.0 | 0 |
| scf7180000044839 | 213547 | 0.383 | 0 | 59.6566 | Eukaryota | 57936.0 | 0 | Chordata | 57936.0 | 0 | Carnivora | 57936.0 | 0 | Canidae | 57936.0 | 0 | Canis | 44938.0 | 1 | Canis lupus | 44938.0 | 1 |
| scf7180000055387 | 8618 | 0.4485 | 0 | 162.2966 | Eukaryota | 655.0 | 0 | Chordata | 344.0 | 1 | Kurtiformes | 344.0 | 1 | Apogonidae | 344.0 | 1 | Sphaeramia | 344.0 | 1 | Sphaeramia orbicularis | 344.0 | 1 |
| scf7180000054546 | 11422 | 0.4272 | 0 | 44.1063 | Eukaryota | 43466.0 | 0 | Chordata | 43466.0 | 0 | Carnivora | 43466.0 | 0 | Canidae | 43466.0 | 0 | Canis | 43466.0 | 0 | Canis lupus | 43466.0 | 0 |
| scf7180000053130 | 1655 | 0.4423 | 0 | 154.7157 | no-hit | 0.0 | 0 | no-hit | 0.0 | 0 | no-hit | 0.0 | 0 | no-hit | 0.0 | 0 | no-hit | 0.0 | 0 | no-hit | 0.0 | 0 |
| scf7180000048764 | 9636 | 0.4431 | 0 | 126.1933 | Eukaryota | 14671.0 | 0 | Chordata | 9186.0 | 2 | Gadiformes | 4403.0 | 6 | Gadidae | 4403.0 | 6 | Gadus | 4403.0 | 6 | Gadus morhua | 4403.0 | 6 |
| scf7180000045750 | 194194 | 0.3509 | 0 | 60.9267 | Eukaryota | 94500.0 | 0 | Chordata | 94500.0 | 0 | Carnivora | 94500.0 | 0 | Canidae | 94500.0 | 0 | Canis | 94500.0 | 0 | Canis lupus | 94500.0 | 0 |
| scf7180000047169 | 71498 | 0.4583 | 0 | 56.4853 | Eukaryota | 41716.0 | 0 | Chordata | 41716.0 | 0 | Carnivora | 41716.0 | 0 | Canidae | 41716.0 | 0 | Vulpes | 21144.0 | 1 | Vulpes vulpes | 21144.0 | 1 |
| scf7180000043100 | 263774 | 0.3808 | 0 | 64.909 | Eukaryota | 30880.0 | 0 | Chordata | 30880.0 | 0 | Carnivora | 30880.0 | 0 | Canidae | 30880.0 | 0 | Canis | 30880.0 | 0 | Canis lupus | 30880.0 | 0 |
| scf7180000044713 | 77336 | 0.3901 | 0 | 57.5056 | Eukaryota | 90303.0 | 0 | Chordata | 90303.0 | 0 | Carnivora | 90303.0 | 0 | Canidae | 90303.0 | 0 | Canis | 90303.0 | 0 | Canis lupus | 90303.0 | 0 |
| scf7180000052292 | 5321 | 0.4644 | 0 | 543.5187 | Eukaryota | 47093.0 | 0 | Chordata | 47093.0 | 0 | Carnivora | 47093.0 | 0 | Canidae | 47093.0 | 0 | Canis | 47093.0 | 0 | Canis lupus | 47093.0 | 0 |
| scf7180000042546 | 179085 | 0.4202 | 0 | 58.854 | Eukaryota | 77130.0 | 0 | Chordata | 77130.0 | 0 | Carnivora | 77130.0 | 0 | Canidae | 77130.0 | 0 | Canis | 77130.0 | 0 | Canis lupus | 77130.0 | 0 |
| scf7180000048356 | 44564 | 0.6864 | 0 | 53.5746 | Eukaryota | 14180.0 | 0 | Chordata | 14180.0 | 0 | Carnivora | 14180.0 | 0 | Canidae | 12354.0 | 1 | Canis | 11402.0 | 3 | Canis lupus | 11402.0 | 3 |
| scf7180000041139 | 15648 | 0.4433 | 0 | 160.3331 | Eukaryota | 1513.0 | 0 | Platyhelminthes | 893.0 | 1 | Polystomatidea | 893.0 | 2 | Polystomatidae | 893.0 | 2 | Protopolystoma | 893.0 | 2 | Protopolystoma xenopodis | 893.0 | 2 |
| scf7180000053582 | 1430 | 0.435 | 0 | 164.4105 | no-hit | 0.0 | 0 | no-hit | 0.0 | 0 | no-hit | 0.0 | 0 | no-hit | 0.0 | 0 | no-hit | 0.0 | 0 | no-hit | 0.0 | 0 |
| scf7180000050562 | 2443 | 0.447 | 0 | 624.4486 | no-hit | 0.0 | 0 | no-hit | 0.0 | 0 | no-hit | 0.0 | 0 | no-hit | 0.0 | 0 | no-hit | 0.0 | 0 | no-hit | 0.0 | 0 |
| scf7180000050352 | 7482 | 0.4368 | 0 | 138.1256 | no-hit | 0.0 | 0 | no-hit | 0.0 | 0 | no-hit | 0.0 | 0 | no-hit | 0.0 | 0 | no-hit | 0.0 | 0 | no-hit | 0.0 | 0 |
| scf7180000046366 | 30484 | 0.3672 | 0 | 70.0994 | Eukaryota | 84070.0 | 0 | Chordata | 84070.0 | 0 | Carnivora | 84070.0 | 0 | Canidae | 84070.0 | 0 | Canis | 84070.0 | 0 | Canis lupus | 84070.0 | 0 |
| scf7180000055426 | 37675 | 0.4408 | 0 | 82.1232 | Eukaryota | 7031.0 | 0 | Chordata | 5181.0 | 1 | Tetraodontiformes | 3045.0 | 3 | Tetraodontidae | 3045.0 | 3 | Takifugu | 3045.0 | 3 | Takifugu rubripes | 3045.0 | 3 |
| scf7180000042083 | 149814 | 0.413 | 0 | 61.017 | Eukaryota | 40587.0 | 0 | Chordata | 40587.0 | 0 | Carnivora | 23097.0 | 2 | Canidae | 23097.0 | 3 | Canis | 18570.0 | 4 | Canis lupus | 18570.0 | 4 |
| scf7180000052151 | 4339 | 0.4508 | 0 | 419.7132 | no-hit | 0.0 | 0 | no-hit | 0.0 | 0 | no-hit | 0.0 | 0 | no-hit | 0.0 | 0 | no-hit | 0.0 | 0 | no-hit | 0.0 | 0 |
| scf7180000041409 | 1094365 | 0.4323 | 0 | 59.6453 | Eukaryota | 76782.0 | 0 | Chordata | 76782.0 | 0 | Carnivora | 76782.0 | 0 | Canidae | 76782.0 | 0 | Canis | 64390.0 | 1 | Canis lupus | 64390.0 | 1 |
| scf7180000051596 | 22661 | 0.5016 | 0 | 1030.4109 | Eukaryota | 9281.0 | 0 | Chordata | 9281.0 | 0 | Carnivora | 9281.0 | 0 | Canidae | 9281.0 | 0 | Canis | 8037.0 | 1 | Canis lupus | 6782.0 | 2 |
| scf7180000041684 | 913013 | 0.4421 | 0 | 58.0995 | Eukaryota | 132656.0 | 0 | Chordata | 132656.0 | 0 | Carnivora | 132656.0 | 0 | Canidae | 132656.0 | 0 | Canis | 132656.0 | 0 | Canis lupus | 132656.0 | 0 |
| scf7180000042345 | 409957 | 0.3487 | 0 | 59.4283 | Eukaryota | 35903.0 | 0 | Chordata | 35903.0 | 0 | Carnivora | 35903.0 | 0 | Canidae | 26208.0 | 1 | Canis | 18678.0 | 3 | Canis lupus | 18678.0 | 3 |
| scf7180000054816 | 35765 | 0.3853 | 0 | 205.8789 | Eukaryota | 34323.0 | 0 | Chordata | 34323.0 | 0 | Carnivora | 34323.0 | 0 | Canidae | 34323.0 | 0 | Canis | 34323.0 | 0 | Canis lupus | 34323.0 | 0 |
| scf7180000048998 | 14700 | 0.4343 | 0 | 183.6219 | no-hit | 0.0 | 0 | no-hit | 0.0 | 0 | no-hit | 0.0 | 0 | no-hit | 0.0 | 0 | no-hit | 0.0 | 0 | no-hit | 0.0 | 0 |
| scf7180000048396 | 5284 | 0.4336 | 0 | 134.6643 | no-hit | 0.0 | 0 | no-hit | 0.0 | 0 | no-hit | 0.0 | 0 | no-hit | 0.0 | 0 | no-hit | 0.0 | 0 | no-hit | 0.0 | 0 |
| scf7180000041022 | 8212 | 0.4424 | 0 | 27.6157 | Eukaryota | 4456.0 | 0 | Chordata | 4456.0 | 0 | Carnivora | 4456.0 | 0 | Canidae | 4456.0 | 0 | Canis | 4456.0 | 0 | Canis lupus | 4456.0 | 0 |
| scf7180000043209 | 425785 | 0.4891 | 0 | 58.1777 | Eukaryota | 35187.0 | 0 | Chordata | 35187.0 | 0 | Carnivora | 35187.0 | 0 | Canidae | 35187.0 | 0 | Canis | 23587.0 | 1 | Canis lupus | 23587.0 | 1 |
| scf7180000046704 | 101366 | 0.4416 | 0 | 117.4544 | Eukaryota | 9399.0 | 0 | Chordata | 7040.0 | 1 | Kurtiformes | 2917.0 | 5 | Apogonidae | 2917.0 | 5 | Sphaeramia | 2917.0 | 5 | Sphaeramia orbicularis | 2917.0 | 5 |
| scf7180000049585 | 4475 | 0.444 | 0 | 158.922 | no-hit | 0.0 | 0 | no-hit | 0.0 | 0 | no-hit | 0.0 | 0 | no-hit | 0.0 | 0 | no-hit | 0.0 | 0 | no-hit | 0.0 | 0 |
| scf7180000048303 | 7983 | 0.4448 | 0 | 68.8275 | Eukaryota | 24655.0 | 0 | Chordata | 24655.0 | 0 | Gadiformes | 14842.0 | 5 | Gadidae | 14842.0 | 5 | Gadus | 14842.0 | 5 | Gadus morhua | 14842.0 | 5 |
| scf7180000048023 | 66976 | 0.3683 | 0 | 58.3505 | Eukaryota | 38733.0 | 0 | Chordata | 38733.0 | 0 | Carnivora | 38733.0 | 0 | Canidae | 38733.0 | 0 | Canis | 38733.0 | 0 | Canis lupus | 38733.0 | 0 |
| scf7180000055748 | 29151 | 0.4421 | 0 | 183.7576 | Eukaryota | 5682.0 | 0 | Chordata | 3813.0 | 2 | Tetraodontiformes | 2332.0 | 4 | Tetraodontidae | 2332.0 | 4 | Takifugu | 2332.0 | 5 | Takifugu rubripes | 2332.0 | 5 |
| scf7180000040560 | 2219923 | 0.3958 | 0 | 59.3394 | Eukaryota | 93453.0 | 0 | Chordata | 93453.0 | 0 | Carnivora | 93453.0 | 0 | Canidae | 93453.0 | 0 | Canis | 93453.0 | 0 | Canis lupus | 93453.0 | 0 |
| scf7180000053026 | 12994 | 0.5056 | 0 | 184.4095 | Eukaryota | 8819.0 | 0 | Chordata | 8819.0 | 0 | Carnivora | 8819.0 | 0 | Canidae | 8819.0 | 0 | Canis | 7713.0 | 1 | Canis lupus | 6610.0 | 2 |
| scf7180000053348 | 5915 | 0.4343 | 0 | 39.8356 | no-hit | 0.0 | 0 | no-hit | 0.0 | 0 | no-hit | 0.0 | 0 | no-hit | 0.0 | 0 | no-hit | 0.0 | 0 | no-hit | 0.0 | 0 |
| scf7180000045269 | 13240 | 0.4152 | 0 | 289.3943 | Eukaryota | 29208.0 | 0 | Chordata | 29208.0 | 0 | Carnivora | 29208.0 | 0 | Canidae | 29208.0 | 0 | Canis | 29208.0 | 0 | Canis lupus | 29208.0 | 0 |
| scf7180000041006 | 950597 | 0.397 | 0 | 70.1854 | Eukaryota | 93831.0 | 0 | Chordata | 93831.0 | 0 | Carnivora | 93831.0 | 0 | Canidae | 93831.0 | 0 | Canis | 93831.0 | 0 | Canis lupus | 93831.0 | 0 |
| scf7180000043638 | 75134 | 0.4921 | 0 | 54.3405 | Eukaryota | 30010.0 | 0 | Chordata | 30010.0 | 0 | Carnivora | 30010.0 | 0 | Canidae | 27557.0 | 1 | Canis | 17963.0 | 2 | Canis lupus | 17963.0 | 2 |
| scf7180000049744 | 40314 | 0.4632 | 0 | 60.0198 | Eukaryota | 5493.0 | 0 | Chordata | 5493.0 | 0 | Carnivora | 4473.0 | 1 | Canidae | 4473.0 | 1 | Canis | 4473.0 | 1 | Canis lupus | 4473.0 | 1 |
| scf7180000052912 | 1683 | 0.4569 | 0 | 699.2489 | Eukaryota | 143.0 | 0 | Platyhelminthes | 143.0 | 0 | Polystomatidea | 143.0 | 0 | Polystomatidae | 143.0 | 0 | Protopolystoma | 143.0 | 0 | Protopolystoma xenopodis | 143.0 | 0 |
| scf7180000052966 | 6688 | 0.4432 | 0 | 47.8182 | Eukaryota | 999.0 | 0 | Platyhelminthes | 631.0 | 1 | Polystomatidea | 631.0 | 1 | Polystomatidae | 631.0 | 1 | Protopolystoma | 631.0 | 1 | Protopolystoma xenopodis | 631.0 | 1 |
| scf7180000055422 | 43740 | 0.4406 | 0 | 85.2372 | Eukaryota | 5481.0 | 0 | Chordata | 3896.0 | 1 | Tetraodontiformes | 1581.0 | 6 | Tetraodontidae | 1581.0 | 6 | Takifugu | 1581.0 | 6 | Takifugu rubripes | 1581.0 | 6 |
| scf7180000047684 | 59469 | 0.4364 | 0 | 215.9819 | Eukaryota | 21269.0 | 0 | Chordata | 13366.0 | 1 | Strigeidida | 7903.0 | 5 | Schistosomatidae | 7903.0 | 5 | Schistosoma | 7903.0 | 5 | Schistosoma mansoni | 7903.0 | 5 |
| scf7180000052122 | 7510 | 0.4457 | 0 | 547.8784 | no-hit | 0.0 | 0 | no-hit | 0.0 | 0 | no-hit | 0.0 | 0 | no-hit | 0.0 | 0 | no-hit | 0.0 | 0 | no-hit | 0.0 | 0 |
| scf7180000043877 | 104170 | 0.3526 | 0 | 56.4389 | Eukaryota | 44380.0 | 0 | Chordata | 44380.0 | 0 | Carnivora | 44380.0 | 0 | Canidae | 44380.0 | 0 | Canis | 44380.0 | 0 | Canis lupus | 44380.0 | 0 |
| scf7180000048847 | 37554 | 0.4321 | 0 | 265.1827 | Eukaryota | 85069.0 | 0 | Chordata | 85069.0 | 0 | Carnivora | 85069.0 | 0 | Canidae | 85069.0 | 0 | Canis | 85069.0 | 0 | Canis lupus | 85069.0 | 0 |
| scf7180000055267 | 10234 | 0.4388 | 0 | 130.4523 | Eukaryota | 54266.0 | 0 | Chordata | 39072.0 | 2 | Gadiformes | 33969.0 | 5 | Gadidae | 33969.0 | 5 | Gadus | 33969.0 | 5 | Gadus morhua | 33969.0 | 5 |
| scf7180000053149 | 3129 | 0.449 | 0 | 186.5353 | no-hit | 0.0 | 0 | no-hit | 0.0 | 0 | no-hit | 0.0 | 0 | no-hit | 0.0 | 0 | no-hit | 0.0 | 0 | no-hit | 0.0 | 0 |
| scf7180000044013 | 9954 | 0.4422 | 0 | 74.0212 | Eukaryota | 32740.0 | 0 | Chordata | 32740.0 | 0 | Carnivora | 29426.0 | 1 | Canidae | 29426.0 | 1 | Canis | 29426.0 | 1 | Canis lupus | 29426.0 | 1 |
| scf7180000047873 | 18521 | 0.4477 | 0 | 721.3845 | Eukaryota | 3636.0 | 0 | Platyhelminthes | 2179.0 | 2 | Polystomatidea | 2179.0 | 2 | Polystomatidae | 2179.0 | 2 | Protopolystoma | 2179.0 | 2 | Protopolystoma xenopodis | 2179.0 | 2 |
| scf7180000050280 | 12698 | 0.4443 | 0 | 144.2798 | Eukaryota | 547.0 | 0 | Chordata | 547.0 | 0 | Salmoniformes | 547.0 | 0 | Salmonidae | 547.0 | 0 | Coregonus | 402.0 | 1 | Coregonus sp. 'balchen' | 402.0 | 1 |
| scf7180000040839 | 1664978 | 0.3877 | 0 | 59.1351 | Eukaryota | 93698.0 | 0 | Chordata | 93698.0 | 0 | Carnivora | 93698.0 | 0 | Canidae | 93698.0 | 0 | Canis | 93698.0 | 0 | Canis lupus | 93698.0 | 0 |

| | | | | | | | | | | | | | | | | | | | | | | |
|---|---|---|---|---|---|---|---|---|---|---|---|---|---|---|---|---|---|---|---|---|---|---|
| scf7180000046270 | 128509 | 0.3543 | 0 | 61.7261 | Eukaryota | 88727.0 | 0 | Chordata | 88727.0 | 0 | Carnivora | 63105.0 | 2 | Canidae | 63105.0 | 2 | Canis | 63105.0 | 4 | Canis lupus | 63105.0 | 4 |
| scf7180000050486 | 14710 | 0.5051 | 0 | 310.9349 | Eukaryota | 9212.0 | 0 | Chordata | 9212.0 | 0 | Carnivora | 9212.0 | 0 | Canidae | 9212.0 | 0 | Canis | 6828.0 | 1 | Canis lupus | 5505.0 | 2 |
| scf7180000041060 | 156116 | 0.3657 | 0 | 59.1781 | Eukaryota | 89417.0 | 0 | Chordata | 89417.0 | 0 | Carnivora | 89417.0 | 0 | Canidae | 89417.0 | 0 | Canis | 89417.0 | 0 | Canis lupus | 89417.0 | 0 |
| scf7180000046350 | 4102 | 0.4166 | 0 | 29.0541 | Eukaryota | 18758.0 | 0 | Chordata | 18758.0 | 0 | Carnivora | 18758.0 | 0 | Canidae | 18758.0 | 0 | Canis | 18758.0 | 0 | Canis lupus | 18758.0 | 0 |
| scf7180000051562 | 2610 | 0.4502 | 0 | 346.7081 | no-hit | 0.0 | 0 | no-hit | 0.0 | 0 | no-hit | 0.0 | 0 | no-hit | 0.0 | 0 | no-hit | 0.0 | 0 | no-hit | 0.0 | 0 |
| scf7180000055998 | 8162 | 0.4376 | 0 | 133.6797 | Eukaryota | 3007.0 | 0 | Platyhelminthes | 2067.0 | 1 | Polystomatidea | 2067.0 | 2 | Polystomatidae | 2067.0 | 2 | Protopolystoma | 2067.0 | 3 | Protopolystoma xenopodis | 2067.0 | 3 |
| scf7180000041906 | 2120 | 0.366 | 0 | 27.4947 | Eukaryota | 2654.0 | 0 | Chordata | 2654.0 | 0 | Carnivora | 1055.0 | 2 | Canidae | 1055.0 | 2 | Vulpes | 1055.0 | 5 | Vulpes vulpes | 1055.0 | 5 |
| scf7180000041588 | 7299 | 0.4701 | 0 | 58.8066 | Eukaryota | 15632.0 | 0 | Chordata | 15632.0 | 0 | Carnivora | 15632.0 | 0 | Canidae | 14732.0 | 1 | Canis | 12237.0 | 2 | Canis lupus | 12237.0 | 2 |
| scf7180000052161 | 4129 | 0.435 | 0 | 103.6397 | no-hit | 0.0 | 0 | no-hit | 0.0 | 0 | no-hit | 0.0 | 0 | no-hit | 0.0 | 0 | no-hit | 0.0 | 0 | no-hit | 0.0 | 0 |
| scf7180000053560 | 2856 | 0.4482 | 0 | 228.5108 | no-hit | 0.0 | 0 | no-hit | 0.0 | 0 | no-hit | 0.0 | 0 | no-hit | 0.0 | 0 | no-hit | 0.0 | 0 | no-hit | 0.0 | 0 |
| scf7180000040694 | 1778541 | 0.382 | 0 | 59.6945 | Eukaryota | 102181.0 | 0 | Chordata | 102181.0 | 0 | Carnivora | 102181.0 | 0 | Canidae | 102181.0 | 0 | Canis | 102181.0 | 0 | Canis lupus | 102181.0 | 0 |
| scf7180000047363 | 23116 | 0.4417 | 0 | 131.4373 | Eukaryota | 6334.0 | 0 | Chordata | 3830.0 | 1 | Kurtiformes | 3830.0 | 2 | Apogonidae | 3830.0 | 2 | Sphaeramia | 3830.0 | 2 | Sphaeramia orbicularis | 3830.0 | 2 |
| scf7180000050817 | 3323 | 0.4523 | 0 | 774.0761 | no-hit | 0.0 | 0 | no-hit | 0.0 | 0 | no-hit | 0.0 | 0 | no-hit | 0.0 | 0 | no-hit | 0.0 | 0 | no-hit | 0.0 | 0 |
| scf7180000043517 | 112669 | 0.3437 | 0 | 57.2626 | Eukaryota | 10246.0 | 0 | Chordata | 10246.0 | 0 | Carnivora | 10246.0 | 0 | Canidae | 10246.0 | 0 | Canis | 10246.0 | 0 | Canis lupus | 10246.0 | 0 |
| scf7180000047852 | 82713 | 0.4196 | 0 | 59.1439 | Eukaryota | 17306.0 | 0 | Chordata | 17306.0 | 0 | Carnivora | 17306.0 | 0 | Canidae | 5500.0 | 4 | Canis | 3662.0 | 8 | Canis lupus | 3662.0 | 8 |
| scf7180000042771 | 54303 | 0.3495 | 0 | 53.3082 | Eukaryota | 48298.0 | 0 | Chordata | 48298.0 | 0 | Carnivora | 48298.0 | 0 | Canidae | 48298.0 | 0 | Canis | 48298.0 | 0 | Canis lupus | 48298.0 | 0 |
| scf7180000054439 | 7422 | 0.4361 | 0 | 122.2905 | Eukaryota | 3282.0 | 0 | Chordata | 3282.0 | 0 | Tetraodontiformes | 2844.0 | 1 | Tetraodontidae | 2844.0 | 1 | Takifugu | 2844.0 | 1 | Takifugu rubripes | 2844.0 | 1 |
| scf7180000043683 | 550286 | 0.3597 | 100 | 60.1678 | Eukaryota | 76351.0 | 0 | Chordata | 76351.0 | 0 | Carnivora | 76351.0 | 0 | Canidae | 76351.0 | 0 | Canis | 76351.0 | 0 | Canis lupus | 76351.0 | 0 |
| scf7180000048633 | 3542 | 0.4531 | 0 | 62.0424 | no-hit | 0.0 | 0 | no-hit | 0.0 | 0 | no-hit | 0.0 | 0 | no-hit | 0.0 | 0 | no-hit | 0.0 | 0 | no-hit | 0.0 | 0 |
| scf7180000053062 | 3840 | 0.4451 | 0 | 245.4452 | no-hit | 0.0 | 0 | no-hit | 0.0 | 0 | no-hit | 0.0 | 0 | no-hit | 0.0 | 0 | no-hit | 0.0 | 0 | no-hit | 0.0 | 0 |
| scf7180000051941 | 22421 | 0.5046 | 0 | 653.2782 | Eukaryota | 9690.0 | 0 | Chordata | 9690.0 | 0 | Carnivora | 9690.0 | 0 | Canidae | 9690.0 | 0 | Canis | 7868.0 | 1 | Canis lupus | 6556.0 | 2 |
| scf7180000040508 | 883098 | 0.3802 | 0 | 60.7201 | Eukaryota | 49170.0 | 0 | Chordata | 49170.0 | 0 | Carnivora | 49170.0 | 0 | Canidae | 49170.0 | 0 | Canis | 44510.0 | 1 | Canis lupus | 44510.0 | 1 |
| scf7180000055773 | 10940 | 0.4474 | 0 | 68.67 | no-hit | 0.0 | 0 | no-hit | 0.0 | 0 | no-hit | 0.0 | 0 | no-hit | 0.0 | 0 | no-hit | 0.0 | 0 | no-hit | 0.0 | 0 |
| scf7180000042876 | 793788 | 0.4604 | 0 | 58.6409 | Eukaryota | 113910.0 | 0 | Chordata | 113910.0 | 0 | Carnivora | 113910.0 | 0 | Canidae | 113910.0 | 0 | Canis | 113910.0 | 0 | Canis lupus | 113910.0 | 0 |
| scf7180000044778 | 45158 | 0.4553 | 0 | 54.7859 | Eukaryota | 96018.0 | 0 | Chordata | 96018.0 | 0 | Carnivora | 96018.0 | 0 | Canidae | 96018.0 | 0 | Canis | 96018.0 | 0 | Canis lupus | 96018.0 | 0 |
| scf7180000041905 | 122472 | 0.3789 | 0 | 59.5079 | Eukaryota | 87066.0 | 0 | Chordata | 87066.0 | 0 | Carnivora | 87066.0 | 0 | Canidae | 87066.0 | 0 | Canis | 87066.0 | 0 | Canis lupus | 87066.0 | 0 |
| scf7180000053201 | 3322 | 0.4557 | 0 | 188.427 | no-hit | 0.0 | 0 | no-hit | 0.0 | 0 | no-hit | 0.0 | 0 | no-hit | 0.0 | 0 | no-hit | 0.0 | 0 | no-hit | 0.0 | 0 |
| scf7180000056614 | 305403 | 0.5981 | 0 | 57.417 | Eukaryota | 28626.0 | 0 | Chordata | 28626.0 | 0 | Carnivora | 28626.0 | 0 | Canidae | 26245.0 | 1 | Canis | 22848.0 | 2 | Canis lupus | 22848.0 | 2 |
| scf7180000043293 | 216452 | 0.3842 | 0 | 59.4126 | Eukaryota | 90040.0 | 0 | Chordata | 90040.0 | 0 | Carnivora | 90040.0 | 0 | Canidae | 90040.0 | 0 | Canis | 90040.0 | 0 | Canis lupus | 90040.0 | 0 |
| scf7180000044837 | 4818 | 0.4772 | 0 | 65.3786 | Eukaryota | 29576.0 | 0 | Chordata | 29576.0 | 0 | Carnivora | 29576.0 | 0 | Canidae | 29576.0 | 0 | Canis | 29576.0 | 0 | Canis lupus | 29576.0 | 0 |
| scf7180000044880 | 298751 | 0.4232 | 0 | 58.5918 | Eukaryota | 32977.0 | 0 | Chordata | 32977.0 | 0 | Primates | 20389.0 | 1 | Hominidae | 17426.0 | 2 | Homo | 17426.0 | 2 | Homo sapiens | 17426.0 | 2 |
| scf7180000040919 | 1631187 | 0.3829 | 0 | 61.2605 | Eukaryota | 59167.0 | 0 | Chordata | 59167.0 | 0 | Carnivora | 59167.0 | 0 | Canidae | 59167.0 | 0 | Canis | 59167.0 | 0 | Canis lupus | 59167.0 | 0 |
| scf7180000041949 | 161134 | 0.437 | 0 | 57.7712 | Eukaryota | 22523.0 | 1 | Chordata | 22523.0 | 1 | undef | 10794.0 | 5 | undef | 10794.0 | 5 | undef | 10794.0 | 7 | eukaryotic synthetic construct | 10794.0 | 7 |
| scf7180000043903 | 4108 | 0.435 | 0 | 71.5473 | no-hit | 0.0 | 0 | no-hit | 0.0 | 0 | no-hit | 0.0 | 0 | no-hit | 0.0 | 0 | no-hit | 0.0 | 0 | no-hit | 0.0 | 0 |
| scf7180000040829 | 1632512 | 0.435 | 0 | 59.8547 | Eukaryota | 101110.0 | 0 | Chordata | 101110.0 | 0 | Carnivora | 101110.0 | 0 | Canidae | 101110.0 | 0 | Canis | 70300.0 | 1 | Canis lupus | 70300.0 | 1 |
| scf7180000047897 | 40643 | 0.5412 | 0 | 59.1786 | Eukaryota | 3770.0 | 0 | Chordata | 3770.0 | 0 | Carnivora | 3770.0 | 0 | Canidae | 2203.0 | 1 | Canis | 2203.0 | 1 | Canis lupus | 2203.0 | 2 |
| scf7180000051611 | 3410 | 0.5012 | 0 | 245.7572 | Eukaryota | 5961.0 | 0 | Chordata | 5961.0 | 0 | Carnivora | 5961.0 | 0 | Canidae | 5961.0 | 0 | Canis | 4845.0 | 2 | Canis lupus | 4845.0 | 2 |
| scf7180000041305 | 265087 | 0.423 | 0 | 60.8239 | Eukaryota | 101646.0 | 0 | Chordata | 101646.0 | 0 | Carnivora | 101646.0 | 0 | Canidae | 101646.0 | 0 | Canis | 99189.0 | 1 | Canis lupus | 99189.0 | 1 |
| scf7180000044711 | 262842 | 0.3797 | 0 | 58.9431 | Eukaryota | 40389.0 | 0 | Chordata | 40389.0 | 0 | Carnivora | 40389.0 | 0 | Canidae | 40389.0 | 0 | Canis | 40389.0 | 0 | Canis lupus | 40389.0 | 0 |
| scf7180000047208 | 7055 | 0.4126 | 0 | 36.3863 | Eukaryota | 65382.0 | 0 | Chordata | 65382.0 | 0 | Carnivora | 65382.0 | 0 | Canidae | 65382.0 | 0 | Canis | 65382.0 | 0 | Canis lupus | 65382.0 | 0 |
| scf7180000044741 | 235457 | 0.3871 | 0 | 59.5781 | Eukaryota | 87724.0 | 0 | Chordata | 87724.0 | 0 | Carnivora | 87724.0 | 0 | Canidae | 87724.0 | 0 | Canis | 87724.0 | 0 | Canis lupus | 87724.0 | 0 |
| scf7180000046045 | 181558 | 0.3307 | 0 | 59.4698 | Eukaryota | 17136.0 | 0 | Chordata | 17136.0 | 0 | Carnivora | 17136.0 | 0 | Canidae | 13478.0 | 1 | Canis | 13478.0 | 2 | Canis lupus | 13478.0 | 2 |
| scf7180000041104 | 1294738 | 0.4428 | 0 | 58.6259 | Eukaryota | 92279.0 | 0 | Chordata | 92279.0 | 0 | Carnivora | 92279.0 | 0 | Canidae | 92279.0 | 0 | Canis | 92279.0 | 0 | Canis lupus | 92279.0 | 0 |
| scf7180000048825 | 12898 | 0.4443 | 0 | 108.8243 | Eukaryota | 11179.0 | 0 | Chordata | 6117.0 | 3 | Strigeidida | 3533.0 | 7 | Schistosomatidae | 3533.0 | 7 | Schistosoma | 3533.0 | 7 | Schistosoma mansoni | 3533.0 | 7 |
| scf7180000043755 | 75141 | 0.3456 | 0 | 54.7302 | Eukaryota | 8218.0 | 0 | Chordata | 8218.0 | 0 | Carnivora | 3152.0 | 3 | Canidae | 3152.0 | 3 | Canis | 3152.0 | 3 | Canis lupus | 3152.0 | 3 |
| scf7180000048542 | 3655 | 0.4402 | 0 | 56.6357 | no-hit | 0.0 | 0 | no-hit | 0.0 | 0 | no-hit | 0.0 | 0 | no-hit | 0.0 | 0 | no-hit | 0.0 | 0 | no-hit | 0.0 | 0 |
| scf7180000051905 | 5679 | 0.4342 | 0 | 94.002 | Eukaryota | 2471.0 | 0 | Platyhelminthes | 1600.0 | 2 | Polystomatidea | 1600.0 | 2 | Polystomatidae | 1600.0 | 2 | Protopolystoma | 1600.0 | 2 | Protopolystoma xenopodis | 1600.0 | 2 |
| scf7180000045499 | 95965 | 0.4484 | 0 | 58.1067 | Eukaryota | 73536.0 | 0 | Chordata | 73536.0 | 0 | Carnivora | 73536.0 | 0 | Canidae | 73536.0 | 0 | Canis | 73536.0 | 0 | Canis lupus | 73536.0 | 0 |
| scf7180000045264 | 414153 | 0.3453 | 0 | 69.6432 | Eukaryota | 96083.0 | 0 | Chordata | 96083.0 | 0 | Carnivora | 96083.0 | 0 | Canidae | 70319.0 | 2 | Canis | 52592.0 | 4 | Canis lupus | 52592.0 | 4 |
| scf7180000041388 | 1179498 | 0.3436 | 0 | 58.2042 | Eukaryota | 85457.0 | 0 | Chordata | 85457.0 | 0 | Carnivora | 85457.0 | 0 | Canidae | 85457.0 | 0 | Canis | 85457.0 | 0 | Canis lupus | 85457.0 | 0 |
| scf7180000053985 | 4124 | 0.5048 | 0 | 77.8004 | Eukaryota | 6148.0 | 0 | Chordata | 6148.0 | 0 | Carnivora | 6148.0 | 0 | Canidae | 6148.0 | 0 | Canis | 4352.0 | 1 | Canis lupus | 3330.0 | 2 |
| scf7180000044431 | 319623 | 0.4908 | 0 | 58.739 | Eukaryota | 39659.0 | 0 | Chordata | 39659.0 | 0 | Carnivora | 39659.0 | 0 | Canidae | 39659.0 | 0 | Canis | 35780.0 | 1 | Canis lupus | 35780.0 | 1 |
| scf7180000046473 | 24406 | 0.4444 | 0 | 358.2289 | Eukaryota | 716.0 | 0 | Chordata | 431.0 | 1 | Tetraodontiformes | 431.0 | 1 | Tetraodontidae | 431.0 | 1 | Takifugu | 431.0 | 1 | Takifugu rubripes | 431.0 | 1 |
| scf7180000048157 | 46300 | 0.4435 | 0 | 574.8345 | Eukaryota | 49986.0 | 0 | Chordata | 49986.0 | 0 | Carnivora | 49986.0 | 0 | Canidae | 49986.0 | 0 | Canis | 49986.0 | 0 | Canis lupus | 49986.0 | 0 |
| scf7180000054176 | 28870 | 0.444 | 0 | 307.8495 | Eukaryota | 12660.0 | 0 | Chordata | 11206.0 | 1 | Kurtiformes | 6630.0 | 3 | Apogonidae | 6630.0 | 3 | Sphaeramia | 6630.0 | 3 | Sphaeramia orbicularis | 6630.0 | 4 |
| scf7180000050659 | 3906 | 0.4427 | 0 | 329.5695 | no-hit | 0.0 | 0 | no-hit | 0.0 | 0 | no-hit | 0.0 | 0 | no-hit | 0.0 | 0 | no-hit | 0.0 | 0 | no-hit | 0.0 | 0 |
| scf7180000050160 | 3291 | 0.4427 | 0 | 74.3254 | no-hit | 0.0 | 0 | no-hit | 0.0 | 0 | no-hit | 0.0 | 0 | no-hit | 0.0 | 0 | no-hit | 0.0 | 0 | no-hit | 0.0 | 0 |
| scf7180000054579 | 61266 | 0.3488 | 0 | 1079.9603 | Eukaryota | 10823.0 | 0 | Chordata | 10823.0 | 0 | Primates | 8349.0 | 2 | Hominidae | 5640.0 | 4 | Homo | 5640.0 | 4 | Homo sapiens | 5640.0 | 4 |
| scf7180000043640 | 388485 | 0.3991 | 0 | 60.68 | Eukaryota | 41511.0 | 0 | Chordata | 41511.0 | 0 | Carnivora | 41511.0 | 0 | Canidae | 41511.0 | 0 | Canis | 41511.0 | 0 | Canis lupus | 41511.0 | 0 |
| scf7180000052407 | 3371 | 0.4373 | 0 | 242.3864 | no-hit | 0.0 | 0 | no-hit | 0.0 | 0 | no-hit | 0.0 | 0 | no-hit | 0.0 | 0 | no-hit | 0.0 | 0 | no-hit | 0.0 | 0 |
| scf7180000049138 | 12310 | 0.4376 | 0 | 328.3376 | Eukaryota | 1662.0 | 0 | Chordata | 1662.0 | 0 | Carnivora | 1662.0 | 0 | Canidae | 1662.0 | 0 | Canis | 1662.0 | 0 | Canis lupus | 1662.0 | 0 |
| scf7180000049020 | 15669 | 0.4429 | 0 | 78.9755 | Eukaryota | 418.0 | 0 | Chordata | 418.0 | 0 | Salmoniformes | 418.0 | 0 | Salmonidae | 418.0 | 0 | Salmo | 418.0 | 0 | Salmo trutta | 418.0 | 0 |
| scf7180000052416 | 5593 | 0.4513 | 0 | 1097.4007 | Eukaryota | 374.0 | 0 | Chordata | 374.0 | 0 | Kurtiformes | 374.0 | 0 | Apogonidae | 374.0 | 0 | Sphaeramia | 374.0 | 0 | Sphaeramia orbicularis | 374.0 | 0 |
| scf7180000052576 | 10808 | 0.4858 | 0 | 58.602 | Eukaryota | 2756.0 | 0 | Chordata | 2756.0 | 0 | Carnivora | 2756.0 | 0 | Canidae | 2756.0 | 0 | Canis | 2756.0 | 0 | Canis lupus | 2756.0 | 0 |
| scf7180000056497 | 183914 | 0.4895 | 0 | 55.8514 | Eukaryota | 54210.0 | 0 | Chordata | 54210.0 | 0 | Carnivora | 54210.0 | 0 | Canidae | 54210.0 | 0 | Canis | 45024.0 | 1 | Canis lupus | 45024.0 | 1 |
| scf7180000049945 | 15318 | 0.4359 | 0 | 1747.2896 | Eukaryota | 19394.0 | 0 | Chordata | 19394.0 | 0 | Carnivora | 19394.0 | 0 | Canidae | 19394.0 | 0 | Canis | 19394.0 | 0 | Canis lupus | 19394.0 | 0 |
| scf7180000054394 | 25306 | 0.4109 | 0 | 1286.3701 | Eukaryota | 88660.0 | 0 | Chordata | 88660.0 | 0 | Carnivora | 88660.0 | 0 | Canidae | 88660.0 | 0 | Canis | 88660.0 | 0 | Canis lupus | 88660.0 | 0 |
| scf7180000047676 | 48055 | 0.4442 | 0 | 284.6451 | Eukaryota | 835.0 | 0 | Chordata | 676.0 | 1 | Tetraodontiformes | 676.0 | 1 | Tetraodontidae | 676.0 | 1 | Takifugu | 676.0 | 1 | Takifugu rubripes | 676.0 | 1 |
| scf7180000047064 | 48506 | 0.5063 | 0 | 57.5306 | Eukaryota | 94664.0 | 0 | Chordata | 94664.0 | 0 | Carnivora | 94664.0 | 0 | Canidae | 94664.0 | 0 | Canis | 94664.0 | 0 | Canis lupus | 94664.0 | 0 |
| scf7180000043991 | 384876 | 0.3729 | 0 | 59.1024 | Eukaryota | 16411.0 | 0 | Chordata | 16411.0 | 0 | Carnivora | 16411.0 | 0 | Canidae | 16411.0 | 0 | Canis | 16411.0 | 0 | Canis lupus | 16411.0 | 0 |
| scf7180000044847 | 238797 | 0.4278 | 0 | 59.2456 | Eukaryota | 14388.0 | 0 | Chordata | 14388.0 | 0 | Primates | 5750.0 | 2 | Hominidae | 5750.0 | 3 | Homo | 5750.0 | 4 | Homo sapiens | 5750.0 | 4 |
| scf7180000056383 | 344173 | 0.4822 | 0 | 57.7537 | Eukaryota | 75681.0 | 0 | Chordata | 75681.0 | 0 | Carnivora | 75681.0 | 0 | Canidae | 75681.0 | 0 | Canis | 75681.0 | 0 | Canis lupus | 75681.0 | 0 |
| scf7180000050273 | 11234 | 0.4404 | 0 | 43.8252 | Eukaryota | 1984.0 | 0 | Chordata | 1826.0 | 1 | Gonorynchiformes | 1024.0 | 4 | Chanidae | 1024.0 | 4 | Chanos | 1024.0 | 4 | Chanos chanos | 1024.0 | 4 |
| scf7180000049154 | 14072 | 0.4419 | 0 | 236.345 | no-hit | 0.0 | 0 | no-hit | 0.0 | 0 | no-hit | 0.0 | 0 | no-hit | 0.0 | 0 | no-hit | 0.0 | 0 | no-hit | 0.0 | 0 |
| scf7180000047501 | 68723 | 0.4962 | 0 | 57.4636 | Eukaryota | 16619.0 | 0 | Chordata | 16619.0 | 0 | Carnivora | 16619.0 | 0 | Canidae | 6096.0 | 3 | Canis | 6096.0 | 5 | Canis lupus | 6096.0 | 6 |
| scf7180000054211 | 121032 | 0.4802 | 0 | 168.0274 | Eukaryota | 61544.0 | 0 | Chordata | 61544.0 | 0 | Carnivora | 61544.0 | 0 | Canidae | 61544.0 | 0 | Canis | 61544.0 | 0 | Canis lupus | 61544.0 | 0 |
| scf7180000052780 | 9878 | 0.4696 | 0 | 61.0899 | Eukaryota | 11788.0 | 0 | Chordata | 11788.0 | 0 | Carnivora | 11788.0 | 0 | Canidae | 11788.0 | 0 | Canis | 9058.0 | 1 | Canis lupus | 8136.0 | 2 |
| scf7180000047373 | 82823 | 0.4139 | 0 | 59.6387 | Eukaryota | 27718.0 | 0 | Chordata | 27718.0 | 0 | Carnivora | 27718.0 | 0 | Canidae | 18673.0 | 2 | Canis | 14610.0 | 3 | Canis lupus | 14610.0 | 3 |
| scf7180000051190 | 2936 | 0.4343 | 0 | 156.9196 | no-hit | 0.0 | 0 | no-hit | 0.0 | 0 | no-hit | 0.0 | 0 | no-hit | 0.0 | 0 | no-hit | 0.0 | 0 | no-hit | 0.0 | 0 |
| scf7180000044136 | 28982 | 0.367 | 0 | 66.0062 | Eukaryota | 90725.0 | 0 | Chordata | 90725.0 | 0 | Carnivora | 90725.0 | 0 | Canidae | 90725.0 | 0 | Canis | 90725.0 | 0 | Canis lupus | 90725.0 | 0 |
| scf7180000041816 | 230358 | 0.5096 | 0 | 56.9144 | Eukaryota | 70754.0 | 0 | Chordata | 70754.0 | 0 | Carnivora | 70754.0 | 0 | Canidae | 70754.0 | 0 | Canis | 46802.0 | 1 | Canis lupus | 46802.0 | 1 |
| scf7180000051690 | 9017 | 0.4454 | 0 | 220.826 | Eukaryota | 3076.0 | 0 | Platyhelminthes | 1991.0 | 2 | Polystomatidea | 1991.0 | 2 | Polystomatidae | 1991.0 | 2 | Protopolystoma | 1991.0 | 2 | Protopolystoma xenopodis | 1991.0 | 2 |
| scf7180000041410 | 590019 | 0.4388 | 0 | 58.8063 | Eukaryota | 46444.0 | 0 | Chordata | 46444.0 | 0 | Carnivora | 46444.0 | 0 | Canidae | 42535.0 | 1 | Canis | 39731.0 | 3 | Canis lupus | 39731.0 | 3 |
| scf7180000047255 | 90971 | 0.3849 | 0 | 60.2811 | Eukaryota | 33167.0 | 0 | Chordata | 33167.0 | 0 | Carnivora | 33167.0 | 0 | Canidae | 30612.0 | 1 | Canis | 20412.0 | 2 | Canis lupus | 20412.0 | 2 |
| scf7180000053372 | 4055 | 0.4481 | 0 | 318.1975 | no-hit | 0.0 | 0 | no-hit | 0.0 | 0 | no-hit | 0.0 | 0 | no-hit | 0.0 | 0 | no-hit | 0.0 | 0 | no-hit | 0.0 | 0 |
| scf7180000052978 | 5311 | 0.4468 | 0 | 205.8814 | no-hit | 0.0 | 0 | no-hit | 0.0 | 0 | no-hit | 0.0 | 0 | no-hit | 0.0 | 0 | no-hit | 0.0 | 0 | no-hit | 0.0 | 0 |
| scf7180000048567 | 11472 | 0.3831 | 0 | 42.2242 | Eukaryota | 3499.0 | 0 | Chordata | 3499.0 | 0 | Carnivora | 2761.0 | 1 | Felidae | 1831.0 | 3 | Felis | 1441.0 | 4 | Felis catus | 1441.0 | 4 |
| scf7180000045425 | 12635 | 0.3879 | 0 | 42.7655 | Eukaryota | 9042.0 | 0 | Chordata | 9042.0 | 0 | Primates | 5612.0 | 2 | Hominidae | 3384.0 | 5 | Pan | 2262.0 | 6 | Pan troglodytes | 2262.0 | 6 |
| scf7180000051172 | 21119 | 0.3995 | 0 | 59.7347 | Eukaryota | 20402.0 | 0 | Chordata | 20402.0 | 0 | Carnivora | 20402.0 | 0 | Canidae | 20402.0 | 0 | Canis | 20402.0 | 0 | Canis lupus | 20402.0 | 0 |
| scf7180000046345 | 76394 | 0.4161 | 0 | 61.1969 | Eukaryota | 60133.0 | 0 | Chordata | 60133.0 | 0 | Carnivora | 60133.0 | 0 | Canidae | 60133.0 | 0 | Canis | 60133.0 | 0 | Canis lupus | 60133.0 | 0 |
| scf7180000044841 | 282596 | 0.3383 | 0 | 59.0761 | Eukaryota | 58524.0 | 0 | Chordata | 58524.0 | 0 | Carnivora | 58524.0 | 0 | Canidae | 58524.0 | 0 | Canis | 58524.0 | 0 | Canis lupus | 58524.0 | 0 |
| scf7180000047608 | 5742 | 0.3765 | 0 | 51.0058 | no-hit | 0.0 | 0 | no-hit | 0.0 | 0 | no-hit | 0.0 | 0 | no-hit | 0.0 | 0 | no-hit | 0.0 | 0 | no-hit | 0.0 | 0 |
| scf7180000047321 | 18822 | 0.4436 | 0 | 91.4883 | no-hit | 0.0 | 0 | no-hit | 0.0 | 0 | no-hit | 0.0 | 0 | no-hit | 0.0 | 0 | no-hit | 0.0 | 0 | no-hit | 0.0 | 0 |
| scf7180000045854 | 162197 | 0.398 | 0 | 61.7307 | Eukaryota | 68848.0 | 0 | Chordata | 68848.0 | 0 | Carnivora | 68848.0 | 0 | Canidae | 68848.0 | 0 | Canis | 68848.0 | 0 | Canis lupus | 68848.0 | 0 |
| scf7180000044801 | 11318 | 0.4364 | 0 | 207.5674 | Eukaryota | 3520.0 | 0 | Platyhelminthes | 1759.0 | 3 | Polystomatidea | 1759.0 | 3 | Polystomatidae | 1759.0 | 3 | Protopolystoma | 1759.0 | 3 | Protopolystoma xenopodis | 1759.0 | 3 |
| scf7180000053507 | 2555 | 0.4481 | 0 | 371.3326 | no-hit | 0.0 | 0 | no-hit | 0.0 | 0 | no-hit | 0.0 | 0 | no-hit | 0.0 | 0 | no-hit | 0.0 | 0 | no-hit | 0.0 | 0 |
| scf7180000046406 | 146476 | 0.4951 | 0 | 60.1007 | Eukaryota | 27445.0 | 0 | Chordata | 27445.0 | 0 | Carnivora | 27445.0 | 0 | Canidae | 20762.0 | 2 | Canis | 17657.0 | 3 | Canis lupus | 17657.0 | 3 |
| scf7180000049684 | 19867 | 0.4486 | 0 | 832.8561 | Eukaryota | 8374.0 | 0 | Chordata | 5441.0 | 2 | Tetraodontiformes | 2730.0 | 5 | Tetraodontidae | 2730.0 | 5 | Takifugu | 2730.0 | 5 | Takifugu rubripes | 2730.0 | 5 |
| scf7180000047838 | 29022 | 0.4369 | 0 | 87.358 | Eukaryota | 18038.0 | 0 | Chordata | 12302.0 | 1 | Cyclophyllidea | 5736.0 | 4 | Hymenolepididae | 5736.0 | 4 | Hymenolepis | 5736.0 | 4 | Hymenolepis microstoma | 5736.0 | 5 |
| scf7180000051860 | 7723 | 0.4529 | 0 | 1277.7064 | Eukaryota | 3624.0 | 0 | Platyhelminthes | 2039.0 | 3 | Polystomatidea | 2039.0 | 3 | Polystomatidae | 2039.0 | 3 | Protopolystoma | 2039.0 | 3 | Protopolystoma xenopodis | 2039.0 | 3 |
| scf7180000053809 | 2670 | 0.4989 | 0 | 131.5222 | Eukaryota | 6721.0 | 0 | Chordata | 6721.0 | 0 | Carnivora | 6721.0 | 0 | Canidae | 6721.0 | 0 | Canis | 4359.0 | 1 | Canis lupus | 3268.0 | 2 |
| scf7180000055785 | 11671 | 0.4453 | 0 | 88.6605 | Eukaryota | 15669.0 | 0 | Chordata | 14663.0 | 1 | Gadiformes | 9992.0 | 5 | Gadidae | 9992.0 | 5 | Gadus | 9992.0 | 5 | Gadus morhua | 9992.0 | 5 |
| scf7180000056399 | 882639 | 0.4622 | 0 | 59.6171 | Eukaryota | 65742.0 | 0 | Chordata | 65742.0 | 0 | Carnivora | 65742.0 | 0 | Canidae | 65742.0 | 0 | Canis | 65742.0 | 0 | Canis lupus | 65742.0 | 0 |
| scf7180000052218 | 7302 | 0.4475 | 0 | 146.2524 | no-hit | 0.0 | 0 | no-hit | 0.0 | 0 | no-hit | 0.0 | 0 | no-hit | 0.0 | 0 | no-hit | 0.0 | 0 | no-hit | 0.0 | 0 |
| scf7180000054647 | 95219 | 0.4268 | 0 | 120.6916 | Eukaryota | 38174.0 | 0 | Chordata | 38174.0 | 0 | Carnivora | 38174.0 | 0 | Canidae | 38174.0 | 0 | Canis | 38174.0 | 0 | Canis lupus | 38174.0 | 0 |
| scf7180000044671 | 122102 | 0.3722 | 0 | 57.4228 | Eukaryota | 10373.0 | 0 | Chordata | 10373.0 | 0 | Carnivora | 10373.0 | 0 | Mustelidae | 8024.0 | 1 | Mustela | 8024.0 | 1 | Mustela putorius | 8024.0 | 1 |
| scf7180000047479 | 17730 | 0.4437 | 0 | 119.2629 | Eukaryota | 2612.0 | 0 | Chordata | 2612.0 | 0 | Carnivora | 2612.0 | 0 | Canidae | 2612.0 | 0 | Canis | 2612.0 | 0 | Canis lupus | 2612.0 | 0 |
| scf7180000041499 | 161976 | 0.3659 | 0 | 59.52 | Eukaryota | 65781.0 | 0 | Chordata | 65781.0 | 0 | Carnivora | 65781.0 | 0 | Canidae | 65781.0 | 0 | Canis | 65781.0 | 0 | Canis lupus | 65781.0 | 0 |
| scf7180000049211 | 17942 | 0.5037 | 0 | 692.9454 | Eukaryota | 9989.0 | 0 | Chordata | 9989.0 | 0 | Carnivora | 9989.0 | 0 | Canidae | 9989.0 | 0 | Canis | 8754.0 | 1 | Canis lupus | 7469.0 | 2 |
| scf7180000055278 | 31617 | 0.3749 | 0 | 153.3514 | Eukaryota | 5915.0 | 0 | Chordata | 5098.0 | 1 | Carnivora | 3582.0 | 2 | Canidae | 2798.0 | 3 | Canis | 2798.0 | 3 | Canis lupus | 2798.0 | 3 |
| scf7180000054142 | 30974 | 0.4447 | 0 | 387.0799 | Eukaryota | 49781.0 | 0 | Chordata | 30113.0 | 1 | Strigeidida | 10063.0 | 6 | Schistosomatidae | 10063.0 | 6 | Schistosoma | 10063.0 | 6 | Schistosoma mansoni | 10063.0 | 6 |
| scf7180000047565 | 15785 | 0.4359 | 0 | 99.9165 | Eukaryota | 3046.0 | 0 | Chordata | 3046.0 | 0 | Carnivora | 3046.0 | 0 | Canidae | 3046.0 | 0 | Canis | 3046.0 | 0 | Canis lupus | 3046.0 | 0 |
| scf7180000042523 | 635978 | 0.3481 | 0 | 59.3919 | Eukaryota | 89992.0 | 0 | Chordata | 89992.0 | 0 | Carnivora | 89992.0 | 0 | Canidae | 89992.0 | 0 | Canis | 89992.0 | 0 | Canis lupus | 89992.0 | 0 |
| scf7180000046676 | 106943 | 0.4099 | 0 | 60.2446 | Eukaryota | 21643.0 | 0 | Chordata | 21643.0 | 0 | Carnivora | 19120.0 | 1 | Canidae | 19120.0 | 1 | Canis | 19120.0 | 1 | Canis lupus | 19120.0 | 1 |
| scf7180000043872 | 190426 | 0.3484 | 0 | 58.7822 | Eukaryota | 13525.0 | 0 | Chordata | 13525.0 | 0 | Carnivora | 11475.0 | 1 | Canidae | 11475.0 | 1 | Canis | 11475.0 | 1 | Canis lupus | 11475.0 | 1 |
| scf7180000043600 | 2059493 | 0.4155 | 0 | 59.996 | Eukaryota | 93254.0 | 0 | Chordata | 93254.0 | 0 | Carnivora | 93254.0 | 0 | Canidae | 93254.0 | 0 | Canis | 93254.0 | 0 | Canis lupus | 93254.0 | 0 |
| scf7180000051979 | 15444 | 0.5027 | 0 | 270.3865 | Eukaryota | 9192.0 | 0 | Chordata | 9192.0 | 0 | Carnivora | 9192.0 | 0 | Canidae | 9192.0 | 0 | Canis | 7985.0 | 1 | Canis lupus | 6754.0 | 2 |
| scf7180000053912 | 3644 | 0.4536 | 0 | 435.3153 | no-hit | 0.0 | 0 | no-hit | 0.0 | 0 | no-hit | 0.0 | 0 | no-hit | 0.0 | 0 | no-hit | 0.0 | 0 | no-hit | 0.0 | 0 |
| scf7180000043193 | 462972 | 0.4003 | 0 | 61.3061 | Eukaryota | 59592.0 | 0 | Chordata | 59592.0 | 0 | Carnivora | 59592.0 | 0 | Canidae | 59592.0 | 0 | Canis | 59592.0 | 0 | Canis lupus | 59592.0 | 0 |
| scf7180000051218 | 4090 | 0.4482 | 0 | 440.4619 | Eukaryota | 315.0 | 0 | Chordata | 315.0 | 0 | Salmoniformes | 178.0 | 1 | Salmonidae | 178.0 | 1 | Coregonus | 178.0 | 1 | Coregonus sp. 'balchen' | 178.0 | 1 |
| scf7180000046341 | 28775 | 0.4341 | 0 | 97.8293 | Eukaryota | 9389.0 | 0 | Chordata | 3782.0 | 3 | Polystomatidea | 2357.0 | 6 | Polystomatidae | 2357.0 | 6 | Protopolystoma | 2357.0 | 6 | Protopolystoma xenopodis | 2357.0 | 6 |
| scf7180000050182 | 13474 | 0.4433 | 0 | 216.8033 | Eukaryota | 10728.0 | 0 | Chordata | 7758.0 | 2 | Gadiformes | 4074.0 | 6 | Gadidae | 4074.0 | 6 | Gadus | 4074.0 | 6 | Gadus morhua | 4074.0 | 6 |
| scf7180000051256 | 25960 | 0.4437 | 0 | 76.0907 | Eukaryota | 64062.0 | 0 | Chordata | 44475.0 | 2 | Gadiformes | 38003.0 | 5 | Gadidae | 38003.0 | 5 | Gadus | 38003.0 | 5 | Gadus morhua | 38003.0 | 5 |
| scf7180000054431 | 24810 | 0.4809 | 0 | 430.8071 | Eukaryota | 37548.0 | 0 | Chordata | 37548.0 | 0 | Carnivora | 37548.0 | 0 | Canidae | 37548.0 | 0 | Canis | 37548.0 | 0 | Canis lupus | 37548.0 | 0 |
| scf7180000042458 | 312497 | 0.3825 | 0 | 59.8923 | Eukaryota | 56031.0 | 0 | Chordata | 56031.0 | 0 | Carnivora | 44237.0 | 1 | Canidae | 44237.0 | 1 | Vulpes | 27710.0 | 2 | Vulpes vulpes | 27710.0 | 2 |
| scf7180000050127 | 2795 | 0.4569 | 0 | 320.2683 | no-hit | 0.0 | 0 | no-hit | 0.0 | 0 | no-hit | 0.0 | 0 | no-hit | 0.0 | 0 | no-hit | 0.0 | 0 | no-hit | 0.0 | 0 |
| scf7180000048708 | 8568 | 0.4383 | 0 | 147.9215 | no-hit | 0.0 | 0 | no-hit | 0.0 | 0 | no-hit | 0.0 | 0 | no-hit | 0.0 | 0 | no-hit | 0.0 | 0 | no-hit | 0.0 | 0 |
| scf7180000044204 | 198843 | 0.4428 | 0 | 60.979 | Eukaryota | 56407.0 | 0 | Chordata | 56407.0 | 0 | Carnivora | 56407.0 | 0 | Canidae | 23258.0 | 2 | Canis | 23258.0 | 5 | Canis lupus | 23258.0 | 6 |
| scf7180000048666 | 52408 | 0.4393 | 0 | 147.7058 | Eukaryota | 28006.0 | 0 | Chordata | 28006.0 | 0 | Gadiformes | 17918.0 | 4 | Gadidae | 17918.0 | 4 | Gadus | 17918.0 | 4 | Gadus morhua | 17918.0 | 5 |
| scf7180000050966 | 2606 | 0.4559 | 0 | 394.3211 | no-hit | 0.0 | 0 | no-hit | 0.0 | 0 | no-hit | 0.0 | 0 | no-hit | 0.0 | 0 | no-hit | 0.0 | 0 | no-hit | 0.0 | 0 |
| scf7180000041814 | 790576 | 0.3882 | 0 | 60.0358 | Eukaryota | 46511.0 | 0 | Chordata | 46511.0 | 0 | Carnivora | 46511.0 | 0 | Canidae | 46511.0 | 0 | Canis | 42045.0 | 1 | Canis lupus | 42045.0 | 1 |
| scf7180000052279 | 3544 | 0.4501 | 0 | 513.6075 | Eukaryota | 486.0 | 0 | Streptophyta | 486.0 | 0 | Fabales | 486.0 | 0 | Fabaceae | 486.0 | 0 | Cicer | 486.0 | 0 | Cicer arietinum | 486.0 | 0 |
| scf7180000046189 | 123749 | 0.4999 | 0 | 56.6008 | Eukaryota | 27420.0 | 0 | Chordata | 27420.0 | 0 | Carnivora | 18837.0 | 3 | Canidae | 17608.0 | 5 | Canis | 17608.0 | 6 | Canis lupus | 17608.0 | 6 |
| scf7180000040528 | 110680 | 0.3482 | 0 | 73.4443 | Eukaryota | 90183.0 | 0 | Chordata | 90183.0 | 0 | Carnivora | 90183.0 | 0 | Canidae | 90183.0 | 0 | Canis | 90183.0 | 0 | Canis lupus | 90183.0 | 0 |
| scf7180000044142 | 4874 | 0.517 | 0 | 26.9881 | Eukaryota | 1727.0 | 0 | Chordata | 1727.0 | 0 | Carnivora | 1404.0 | 1 | Canidae | 1404.0 | 1 | Canis | 1130.0 | 3 | Canis lupus | 1130.0 | 3 |
| scf7180000052307 | 3614 | 0.4541 | 0 | 613.9676 | Eukaryota | 415.0 | 0 | Platyhelminthes | 415.0 | 0 | Polystomatidea | 415.0 | 0 | Polystomatidae | 415.0 | 0 | Protopolystoma | 415.0 | 0 | Protopolystoma xenopodis | 415.0 | 0 |
| scf7180000054978 | 7983 | 0.4387 | 0 | 176.559 | Eukaryota | 14389.0 | 0 | Chordata | 9597.0 | 3 | Strigeidida | 3129.0 | 7 | Schistosomatidae | 3129.0 | 7 | Schistosoma | 3129.0 | 7 | Schistosoma mansoni | 3129.0 | 7 |
| scf7180000049004 | 15507 | 0.3794 | 0 | 54.1177 | Eukaryota | 76350.0 | 0 | Chordata | 76350.0 | 0 | Carnivora | 76350.0 | 0 | Canidae | 76350.0 | 0 | Canis | 76350.0 | 0 | Canis lupus | 76350.0 | 0 |
| scf7180000042145 | 505867 | 0.3626 | 0 | 115.5787 | Eukaryota | 95096.0 | 0 | Chordata | 95096.0 | 0 | Carnivora | 95096.0 | 0 | Canidae | 95096.0 | 0 | Canis | 95096.0 | 0 | Canis lupus | 95096.0 | 0 |
| scf7180000045833 | 169992 | 0.519 | 0 | 58.1622 | Eukaryota | 37344.0 | 0 | Chordata | 37344.0 | 0 | Carnivora | 37344.0 | 0 | Canidae | 37344.0 | 0 | Canis | 37344.0 | 0 | Canis lupus | 37344.0 | 0 |
| scf7180000045950 | 225477 | 0.4072 | 0 | 58.7316 | Eukaryota | 67218.0 | 0 | Chordata | 67218.0 | 0 | Carnivora | 67218.0 | 0 | Canidae | 67218.0 | 0 | Canis | 67218.0 | 0 | Canis lupus | 67218.0 | 0 |
| scf7180000054529 | 324324 | 0.3754 | 0 | 213.6253 | Eukaryota | 92094.0 | 0 | Chordata | 92094.0 | 0 | Carnivora | 92094.0 | 0 | Canidae | 92094.0 | 0 | Canis | 92094.0 | 0 | Canis lupus | 92094.0 | 0 |
| scf7180000041730 | 1289985 | 0.378 | 0 | 60.2361 | Eukaryota | 384352.0 | 0 | Chordata | 384352.0 | 0 | Carnivora | 384352.0 | 0 | Canidae | 384352.0 | 0 | Canis | 384352.0 | 0 | Canis lupus | 384352.0 | 0 |
| scf7180000050131 | 8724 | 0.4554 | 0 | 482.316 | Eukaryota | 1907.0 | 0 | Platyhelminthes | 715.0 | 2 | Polystomatidea | 715.0 | 2 | Polystomatidae | 715.0 | 2 | Protopolystoma | 715.0 | 2 | Protopolystoma xenopodis | 715.0 | 2 |
| scf7180000045880 | 178080 | 0.3738 | 0 | 59.5719 | Eukaryota | 59907.0 | 0 | Chordata | 59907.0 | 0 | Carnivora | 59907.0 | 0 | Canidae | 59907.0 | 0 | Canis | 59907.0 | 0 | Canis lupus | 59907.0 | 0 |
| scf7180000044097 | 38169 | 0.3748 | 0 | 56.0673 | Eukaryota | 61263.0 | 0 | Chordata | 61263.0 | 0 | Carnivora | 61263.0 | 0 | Canidae | 61263.0 | 0 | Canis | 61263.0 | 0 | Canis lupus | 61263.0 | 0 |
| scf7180000046244 | 22096 | 0.3637 | 0 | 57.4644 | Eukaryota | 87601.0 | 0 | Chordata | 87601.0 | 0 | Carnivora | 87601.0 | 0 | Canidae | 87601.0 | 0 | Canis | 87601.0 | 0 | Canis lupus | 87601.0 | 0 |
| scf7180000043484 | 471648 | 0.5341 | 0 | 57.3621 | Eukaryota | 26321.0 | 0 | Chordata | 26321.0 | 0 | Carnivora | 26321.0 | 0 | Canidae | 26321.0 | 0 | Canis | 23600.0 | 1 | Canis lupus | 23600.0 | 1 |
| scf7180000055312 | 31269 | 0.4426 | 0 | 218.4045 | Eukaryota | 807.0 | 0 | Chordata | 577.0 | 1 | Tetraodontiformes | 577.0 | 1 | Tetraodontidae | 577.0 | 1 | Takifugu | 577.0 | 1 | Takifugu rubripes | 577.0 | 1 |
| scf7180000043741 | 359156 | 0.3724 | 0 | 60.1458 | Eukaryota | 31180.0 | 1 | Chordata | 31180.0 | 1 | Carnivora | 24168.0 | 2 | Otariidae | 24168.0 | 2 | Callorhinus | 10416.0 | 4 | Callorhinus ursinus | 10416.0 | 4 |
| scf7180000042541 | 661643 | 0.445 | 0 | 59.4935 | Eukaryota | 69278.0 | 0 | Chordata | 69278.0 | 0 | Carnivora | 69278.0 | 0 | Canidae | 69278.0 | 0 | Canis | 61115.0 | 1 | Canis lupus | 61115.0 | 1 |
| scf7180000047081 | 84795 | 0.4164 | 0 | 61.4878 | Eukaryota | 23238.0 | 0 | Chordata | 23238.0 | 0 | Carnivora | 23238.0 | 0 | Canidae | 23238.0 | 0 | Canis | 13641.0 | 1 | Canis lupus | 13641.0 | 1 |
| scf7180000043080 | 536443 | 0.3903 | 0 | 59.7728 | Eukaryota | 91322.0 | 0 | Chordata | 91322.0 | 0 | Carnivora | 91322.0 | 0 | Canidae | 91322.0 | 0 | Canis | 91322.0 | 0 | Canis lupus | 91322.0 | 0 |
| scf7180000043060 | 17795 | 0.3766 | 0 | 46.5307 | Eukaryota | 49265.0 | 0 | Chordata | 49265.0 | 0 | Carnivora | 49265.0 | 0 | Canidae | 11168.0 | 5 | Ursus | 9790.0 | 8 | Vulpes vulpes | 5602.0 | 9 |
| scf7180000041650 | 992790 | 0.3921 | 0 | 60.0295 | Eukaryota | 35758.0 | 0 | Chordata | 35758.0 | 0 | Carnivora | 35758.0 | 0 | Canidae | 35758.0 | 0 | Canis | 35758.0 | 0 | Canis lupus | 35758.0 | 0 |
| scf7180000041624 | 86914 | 0.3914 | 0 | 60.586 | Eukaryota | 42906.0 | 0 | Chordata | 42906.0 | 0 | Carnivora | 42906.0 | 0 | Canidae | 42906.0 | 0 | Canis | 42906.0 | 0 | Canis lupus | 42906.0 | 0 |
| scf7180000052016 | 6148 | 0.5054 | 0 | 56.5259 | Eukaryota | 8360.0 | 0 | Chordata | 8360.0 | 0 | Carnivora | 8360.0 | 0 | Canidae | 8360.0 | 0 | Canis | 6142.0 | 1 | Canis lupus | 4940.0 | 2 |
| scf7180000056264 | 242384 | 0.3944 | 0 | 60.8064 | Eukaryota | 93914.0 | 0 | Chordata | 93914.0 | 0 | Carnivora | 93914.0 | 0 | Canidae | 93914.0 | 0 | Canis | 93914.0 | 0 | Canis lupus | 93914.0 | 0 |
| scf7180000048663 | 4242 | 0.4364 | 0 | 62.9688 | Eukaryota | 263.0 | 0 | Chordata | 263.0 | 0 | Beloniformes | 263.0 | 0 | Adrianichthyidae | 263.0 | 0 | Oryzias | 263.0 | 0 | Oryzias latipes | 263.0 | 0 |
| scf7180000047695 | 83195 | 0.4336 | 0 | 59.4815 | Eukaryota | 11991.0 | 0 | Chordata | 11991.0 | 0 | Carnivora | 11991.0 | 0 | Canidae | 11991.0 | 0 | Canis | 8382.0 | 1 | Canis lupus | 8382.0 | 1 |
| scf7180000045773 | 32511 | 0.4287 | 0 | 87.4557 | Eukaryota | 25692.0 | 0 | Chordata | 25692.0 | 0 | Carnivora | 25692.0 | 0 | Canidae | 25692.0 | 0 | Canis | 25692.0 | 0 | Canis lupus | 25692.0 | 0 |
| scf7180000056235 | 1167264 | 0.3481 | 0 | 59.2931 | Eukaryota | 92222.0 | 0 | Chordata | 92222.0 | 0 | Carnivora | 92222.0 | 0 | Canidae | 92222.0 | 0 | Canis | 92222.0 | 0 | Canis lupus | 92222.0 | 0 |
| scf7180000046755 | 121073 | 0.3628 | 0 | 60.4513 | Eukaryota | 58151.0 | 0 | Chordata | 58151.0 | 0 | Carnivora | 58151.0 | 0 | Canidae | 58151.0 | 0 | Canis | 52276.0 | 1 | Canis lupus | 52276.0 | 1 |
| scf7180000045811 | 158094 | 0.523 | 0 | 57.8814 | Eukaryota | 52653.0 | 0 | Chordata | 52653.0 | 0 | Carnivora | 52653.0 | 0 | Canidae | 52653.0 | 0 | Canis | 52653.0 | 0 | Canis lupus | 52653.0 | 0 |
| scf7180000056413 | 559221 | 0.3597 | 0 | 60.916 | Eukaryota | 52319.0 | 0 | Chordata | 52319.0 | 0 | Carnivora | 52319.0 | 0 | Canidae | 52319.0 | 0 | Canis | 52319.0 | 0 | Canis lupus | 52319.0 | 0 |
| scf7180000044748 | 79303 | 0.3727 | 0 | 55.5149 | Eukaryota | 5195.0 | 1 | Chordata | 5195.0 | 1 | Primates | 5195.0 | 1 | Hominidae | 5195.0 | 1 | Homo | 5195.0 | 1 | Homo sapiens | 5195.0 | 1 |
| scf7180000048005 | 15977 | 0.4862 | 0 | 631.1146 | Eukaryota | 40393.0 | 0 | Chordata | 40393.0 | 0 | Carnivora | 40393.0 | 0 | Canidae | 40393.0 | 0 | Canis | 40393.0 | 0 | Canis lupus | 40393.0 | 0 |
| scf7180000047243 | 4817 | 0.4563 | 0 | 570.0 | no-hit | 0.0 | 0 | no-hit | 0.0 | 0 | no-hit | 0.0 | 0 | no-hit | 0.0 | 0 | no-hit | 0.0 | 0 | no-hit | 0.0 | 0 |
| scf7180000048860 | 25748 | 0.4354 | 0 | 107.2636 | Eukaryota | 93808.0 | 0 | Platyhelminthes | 47841.0 | 1 | Gadiformes | 36223.0 | 4 | Gadidae | 36223.0 | 4 | Gadus | 36223.0 | 4 | Gadus morhua | 36223.0 | 4 |
| scf7180000042272 | 198660 | 0.3676 | 0 | 56.4022 | Eukaryota | 48562.0 | 0 | Chordata | 48562.0 | 0 | Carnivora | 48562.0 | 0 | Canidae | 48562.0 | 0 | Canis | 48562.0 | 0 | Canis lupus | 48562.0 | 0 |
| scf7180000044478 | 244423 | 0.5215 | 0 | 57.5481 | Eukaryota | 38467.0 | 0 | Chordata | 38467.0 | 0 | Carnivora | 38467.0 | 0 | Canidae | 35226.0 | 1 | Canis | 35226.0 | 1 | Canis lupus | 35226.0 | 1 |
| scf7180000042430 | 674076 | 0.3935 | 0 | 60.0078 | Eukaryota | 126902.0 | 0 | Chordata | 126902.0 | 0 | Carnivora | 126902.0 | 0 | Canidae | 126902.0 | 0 | Canis | 101417.0 | 1 | Canis lupus | 101417.0 | 1 |
| scf7180000045207 | 125480 | 0.4991 | 0 | 60.4061 | Eukaryota | 54608.0 | 0 | Chordata | 54608.0 | 0 | Carnivora | 54608.0 | 0 | Canidae | 54608.0 | 0 | Canis | 43712.0 | 1 | Canis lupus | 43712.0 | 1 |
| scf7180000053935 | 7446 | 0.4447 | 0 | 88.5391 | Eukaryota | 1378.0 | 0 | Chordata | 1378.0 | 0 | Kurtiformes | 1378.0 | 0 | Apogonidae | 1378.0 | 0 | Sphaeramia | 1378.0 | 0 | Sphaeramia orbicularis | 1378.0 | 0 |
| scf7180000048791 | 5958 | 0.4357 | 0 | 141.137 | no-hit | 0.0 | 0 | no-hit | 0.0 | 0 | no-hit | 0.0 | 0 | no-hit | 0.0 | 0 | no-hit | 0.0 | 0 | no-hit | 0.0 | 0 |
| scf7180000044638 | 422481 | 0.4346 | 0 | 58.4698 | Eukaryota | 39310.0 | 0 | Chordata | 39310.0 | 0 | Carnivora | 39310.0 | 0 | Canidae | 39310.0 | 0 | Canis | 33939.0 | 1 | Canis lupus | 33939.0 | 1 |
| scf7180000051305 | 8482 | 0.435 | 0 | 92.7268 | no-hit | 0.0 | 0 | no-hit | 0.0 | 0 | no-hit | 0.0 | 0 | no-hit | 0.0 | 0 | no-hit | 0.0 | 0 | no-hit | 0.0 | 0 |
| scf7180000046609 | 92473 | 0.6195 | 0 | 56.9988 | Eukaryota | 13383.0 | 0 | Chordata | 13383.0 | 0 | Carnivora | 13383.0 | 0 | Canidae | 12367.0 | 1 | Canis | 8971.0 | 2 | Canis lupus | 8971.0 | 2 |

| | | | | | | | | | | | | | | | | | | | | | | |
|---|---|---|---|---|---|---|---|---|---|---|---|---|---|---|---|---|---|---|---|---|---|---|
| scf7180000044299 | 357555 | 0.3542 | 0 | 60.4908 | Eukaryota | 91596.0 | 0 | Chordata | 91596.0 | 0 | Carnivora | 91596.0 | 0 | Canidae | 91596.0 | 0 | Canis | 91596.0 | 0 | Canis lupus | 91596.0 | 0 |
| scf7180000051550 | 47490 | 0.389 | 0 | 56.9477 | Eukaryota | 90044.0 | 0 | Chordata | 90044.0 | 0 | Carnivora | 90044.0 | 0 | Canidae | 90044.0 | 0 | Canis | 90044.0 | 0 | Canis lupus | 90044.0 | 0 |
| scf7180000056060 | 11926 | 0.454 | 0 | 1953.3587 | no-hit | 0.0 | 0 | no-hit | 0.0 | 0 | no-hit | 0.0 | 0 | no-hit | 0.0 | 0 | no-hit | 0.0 | 0 | no-hit | 0.0 | 0 |
| scf7180000041264 | 1137804 | 0.4625 | 0 | 59.9811 | Eukaryota | 74833.0 | 0 | Chordata | 74833.0 | 0 | Carnivora | 74833.0 | 0 | Canidae | 74833.0 | 0 | Canis | 67673.0 | 1 | Canis lupus | 67673.0 | 1 |
| scf7180000051434 | 5253 | 0.4533 | 0 | 1146.6088 | no-hit | 0.0 | 0 | no-hit | 0.0 | 0 | no-hit | 0.0 | 0 | no-hit | 0.0 | 0 | no-hit | 0.0 | 0 | no-hit | 0.0 | 0 |
| scf7180000052105 | 13451 | 0.5271 | 0 | 248.6493 | Eukaryota | 31999.0 | 0 | Chordata | 31999.0 | 0 | Carnivora | 31999.0 | 0 | Canidae | 31999.0 | 0 | Canis | 31999.0 | 0 | Canis lupus | 31999.0 | 0 |
| scf7180000055253 | 143374 | 0.4381 | 0 | 174.2788 | Eukaryota | 18805.0 | 0 | Chordata | 13718.0 | 2 | Kurtiformes | 5248.0 | 7 | Apogonidae | 5248.0 | 7 | Sphaeramia | 5248.0 | 7 | Sphaeramia orbicularis | 5248.0 | 7 |
| scf7180000055371 | 42125 | 0.4819 | 0 | 201.0736 | Eukaryota | 92597.0 | 0 | Chordata | 92597.0 | 0 | Carnivora | 92597.0 | 0 | Canidae | 92597.0 | 0 | Canis | 92597.0 | 0 | Canis lupus | 92597.0 | 0 |
| scf7180000045509 | 52613 | 0.4372 | 0 | 133.2521 | Eukaryota | 22685.0 | 0 | Chordata | 12286.0 | 1 | Strigeidida | 10399.0 | 6 | Schistosomatidae | 10399.0 | 6 | Schistosoma | 10399.0 | 6 | Schistosoma mansoni | 10399.0 | 6 |
| scf7180000048423 | 45190 | 0.4379 | 0 | 126.1271 | Eukaryota | 16484.0 | 0 | Chordata | 15238.0 | 1 | Tetraodontiformes | 8899.0 | 2 | Tetraodontidae | 8899.0 | 2 | Takifugu | 8899.0 | 2 | Takifugu rubripes | 8899.0 | 2 |
| scf7180000045058 | 83747 | 0.3791 | 0 | 56.5089 | Eukaryota | 56328.0 | 0 | Chordata | 56328.0 | 0 | Carnivora | 56328.0 | 0 | Canidae | 56328.0 | 0 | Canis | 56328.0 | 0 | Canis lupus | 56328.0 | 0 |
| scf7180000045427 | 15335 | 0.4489 | 0 | 166.4564 | Eukaryota | 1916.0 | 0 | Chordata | 1648.0 | 1 | Salmoniformes | 1182.0 | 2 | Salmonidae | 1182.0 | 2 | Coregonus | 856.0 | 3 | Coregonus sp. 'balchen' | 856.0 | 3 |
| scf7180000052924 | 5885 | 0.453 | 0 | 397.6228 | Eukaryota | 2014.0 | 0 | Echinodermata | 894.0 | 3 | Forcipulatida | 894.0 | 3 | Asteriidae | 894.0 | 3 | Asterias | 894.0 | 3 | Asterias rubens | 894.0 | 3 |
| scf7180000053542 | 5068 | 0.4463 | 0 | 222.6589 | Eukaryota | 294.0 | 0 | Platyhelminthes | 294.0 | 0 | Polystomatidea | 294.0 | 0 | Polystomatidae | 294.0 | 0 | Protopolystoma | 294.0 | 0 | Protopolystoma xenopodis | 294.0 | 0 |
| scf7180000048000 | 17049 | 0.5088 | 0 | 392.1891 | Eukaryota | 9113.0 | 0 | Chordata | 9113.0 | 0 | Carnivora | 9113.0 | 0 | Canidae | 9113.0 | 0 | Canis | 6796.0 | 1 | Canis lupus | 5554.0 | 2 |
| scf7180000040505 | 25237 | 0.3964 | 0 | 52.6788 | Eukaryota | 11037.0 | 0 | Chordata | 11037.0 | 0 | Carnivora | 11037.0 | 0 | Canidae | 11037.0 | 0 | Canis | 11037.0 | 0 | Canis lupus | 11037.0 | 0 |
| scf7180000047966 | 26632 | 0.4466 | 0 | 984.1227 | Eukaryota | 2488.0 | 0 | Platyhelminthes | 1019.0 | 2 | Forcipulatida | 953.0 | 3 | Asteriidae | 953.0 | 3 | Asterias | 953.0 | 3 | Asterias rubens | 953.0 | 3 |
| scf7180000041097 | 1236835 | 0.5362 | 0 | 58.2096 | Eukaryota | 88212.0 | 0 | Chordata | 88212.0 | 0 | Carnivora | 88212.0 | 0 | Canidae | 88212.0 | 0 | Canis | 77700.0 | 1 | Canis lupus | 77700.0 | 1 |
| scf7180000045093 | 78778 | 0.4122 | 0 | 58.1707 | Eukaryota | 34024.0 | 0 | Chordata | 34024.0 | 0 | Carnivora | 34024.0 | 0 | Canidae | 34024.0 | 0 | Canis | 34024.0 | 0 | Canis lupus | 34024.0 | 0 |
| scf7180000051117 | 3618 | 0.4514 | 0 | 528.3475 | Eukaryota | 2220.0 | 0 | Platyhelminthes | 1032.0 | 2 | Polystomatidea | 1032.0 | 2 | Polystomatidae | 1032.0 | 2 | Protopolystoma | 1032.0 | 2 | Protopolystoma xenopodis | 1032.0 | 2 |
| scf7180000048695 | 19793 | 0.4465 | 0 | 793.8075 | Eukaryota | 2339.0 | 0 | Platyhelminthes | 887.0 | 3 | Polystomatidea | 887.0 | 3 | Polystomatidae | 887.0 | 3 | Protopolystoma | 887.0 | 3 | Protopolystoma xenopodis | 887.0 | 3 |
| scf7180000052492 | 8170 | 0.5054 | 0 | 180.2893 | Eukaryota | 7766.0 | 0 | Chordata | 7766.0 | 0 | Carnivora | 7766.0 | 0 | Canidae | 7766.0 | 0 | Canis | 6123.0 | 1 | Canis lupus | 4951.0 | 2 |
| scf7180000042955 | 348804 | 0.3696 | 0 | 64.2037 | Eukaryota | 94393.0 | 0 | Chordata | 94393.0 | 0 | Carnivora | 94393.0 | 0 | Canidae | 94393.0 | 0 | Canis | 94393.0 | 0 | Canis lupus | 94393.0 | 0 |
| scf7180000048705 | 3815 | 0.4689 | 0 | 27.5565 | Eukaryota | 3273.0 | 0 | Chordata | 3273.0 | 0 | Carnivora | 3273.0 | 0 | Canidae | 3273.0 | 0 | Canis | 3273.0 | 0 | Canis lupus | 3273.0 | 0 |
| scf7180000048409 | 49963 | 0.4447 | 0 | 58.9622 | Eukaryota | 47267.0 | 0 | Chordata | 47267.0 | 0 | Carnivora | 32463.0 | 2 | Canidae | 32463.0 | 2 | Canis | 32463.0 | 2 | Canis lupus | 32463.0 | 2 |
| scf7180000042070 | 24794 | 0.44 | 0 | 107.219 | Eukaryota | 7399.0 | 0 | Chordata | 6567.0 | 1 | Synbranchiformes | 2743.0 | 4 | Mastacembelidae | 2743.0 | 4 | Mastacembelus | 2743.0 | 4 | Mastacembelus armatus | 2743.0 | 4 |
| scf7180000054106 | 2681 | 0.448 | 0 | 644.4302 | Eukaryota | 1556.0 | 0 | Echinodermata | 662.0 | 2 | Forcipulatida | 662.0 | 2 | Asteriidae | 662.0 | 2 | Asterias | 662.0 | 2 | Asterias rubens | 662.0 | 2 |
| scf7180000046303 | 111022 | 0.6292 | 0 | 56.1924 | Eukaryota | 33401.0 | 0 | Chordata | 33401.0 | 0 | Carnivora | 33401.0 | 0 | Canidae | 28088.0 | 2 | Canis | 25740.0 | 4 | Canis lupus | 25740.0 | 4 |
| scf7180000053550 | 2739 | 0.444 | 0 | 130.204 | no-hit | 0.0 | 0 | no-hit | 0.0 | 0 | no-hit | 0.0 | 0 | no-hit | 0.0 | 0 | no-hit | 0.0 | 0 | no-hit | 0.0 | 0 |
| scf7180000043331 | 203405 | 0.4698 | 0 | 56.0837 | Eukaryota | 11154.0 | 0 | Chordata | 11154.0 | 0 | Carnivora | 11154.0 | 0 | Canidae | 11154.0 | 0 | Canis | 11154.0 | 0 | Canis lupus | 11154.0 | 0 |
| scf7180000049868 | 9482 | 0.5024 | 0 | 107.2212 | Eukaryota | 8988.0 | 0 | Chordata | 8988.0 | 0 | Carnivora | 8988.0 | 0 | Canidae | 8988.0 | 0 | Canis | 7846.0 | 1 | Canis lupus | 6682.0 | 2 |
| scf7180000049849 | 4195 | 0.4462 | 0 | 103.5289 | no-hit | 0.0 | 0 | no-hit | 0.0 | 0 | no-hit | 0.0 | 0 | no-hit | 0.0 | 0 | no-hit | 0.0 | 0 | no-hit | 0.0 | 0 |
| scf7180000041462 | 1057615 | 0.393 | 0 | 60.0357 | Eukaryota | 66855.0 | 0 | Chordata | 66855.0 | 0 | Carnivora | 66855.0 | 0 | Canidae | 66855.0 | 0 | Canis | 66855.0 | 0 | Canis lupus | 66855.0 | 0 |
| scf7180000047492 | 74333 | 0.4409 | 0 | 125.9572 | Eukaryota | 1329.0 | 0 | Chordata | 848.0 | 1 | Kurtiformes | 670.0 | 3 | Apogonidae | 670.0 | 3 | Sphaeramia | 670.0 | 3 | Sphaeramia orbicularis | 670.0 | 3 |
| scf7180000042518 | 610951 | 0.4054 | 0 | 61.0281 | Eukaryota | 433100.0 | 0 | Chordata | 433100.0 | 0 | Carnivora | 433100.0 | 0 | Canidae | 433100.0 | 0 | Canis | 433100.0 | 0 | Canis lupus | 433100.0 | 0 |
| scf7180000049580 | 31448 | 0.3594 | 0 | 48.0296 | Eukaryota | 4121.0 | 0 | Chordata | 4121.0 | 0 | Carnivora | 4121.0 | 0 | Canidae | 4121.0 | 0 | Canis | 4121.0 | 0 | Canis lupus | 4121.0 | 0 |
| scf7180000048497 | 85870 | 0.5142 | 0 | 58.1498 | Eukaryota | 7603.0 | 1 | Chordata | 7603.0 | 1 | Carnivora | 3927.0 | 3 | Canidae | 3927.0 | 3 | Canis | 3927.0 | 4 | Canis lupus | 3927.0 | 4 |
| scf7180000043538 | 281989 | 0.355 | 0 | 59.4603 | Eukaryota | 85488.0 | 0 | Chordata | 85488.0 | 0 | Carnivora | 85488.0 | 0 | Canidae | 85488.0 | 0 | Canis | 85488.0 | 0 | Canis lupus | 85488.0 | 0 |
| scf7180000056310 | 527816 | 0.4046 | 0 | 59.2616 | Eukaryota | 87337.0 | 0 | Chordata | 87337.0 | 0 | Carnivora | 87337.0 | 0 | Canidae | 87337.0 | 0 | Canis | 87337.0 | 0 | Canis lupus | 87337.0 | 0 |
| scf7180000050323 | 18070 | 0.5056 | 0 | 993.2278 | Eukaryota | 9505.0 | 0 | Chordata | 9505.0 | 0 | Carnivora | 9505.0 | 0 | Canidae | 9505.0 | 0 | Canis | 8211.0 | 1 | Canis lupus | 6875.0 | 2 |
| scf7180000045475 | 260806 | 0.3888 | 0 | 60.5841 | Eukaryota | 49908.0 | 0 | Chordata | 49908.0 | 0 | Carnivora | 49908.0 | 0 | Canidae | 49908.0 | 0 | Canis | 49908.0 | 0 | Canis lupus | 49908.0 | 0 |
| scf7180000044656 | 266988 | 0.4018 | 0 | 187.3215 | Eukaryota | 14541.0 | 0 | Chordata | 14541.0 | 0 | Carnivora | 14541.0 | 0 | Canidae | 14541.0 | 0 | Canis | 14541.0 | 0 | Canis lupus | 14541.0 | 0 |
| scf7180000045076 | 244854 | 0.3686 | 0 | 61.0363 | Eukaryota | 52203.0 | 0 | Chordata | 52203.0 | 0 | Carnivora | 52203.0 | 0 | Canidae | 52203.0 | 0 | Canis | 52203.0 | 0 | Canis lupus | 52203.0 | 0 |
| scf7180000046073 | 15163 | 0.4605 | 0 | 191.0834 | Eukaryota | 11658.0 | 0 | Chordata | 11658.0 | 0 | Carnivora | 11658.0 | 0 | Canidae | 11658.0 | 0 | Canis | 8954.0 | 1 | Canis lupus | 8012.0 | 2 |
| scf7180000054217 | 1496866 | 0.4248 | 0 | 61.5197 | Eukaryota | 85402.0 | 0 | Chordata | 85402.0 | 0 | Carnivora | 85402.0 | 0 | Canidae | 85402.0 | 0 | Canis | 54227.0 | 1 | Canis lupus | 54227.0 | 1 |
| scf7180000041171 | 1300448 | 0.4023 | 0 | 60.1743 | Eukaryota | 88978.0 | 0 | Chordata | 88978.0 | 0 | Carnivora | 88978.0 | 0 | Canidae | 88978.0 | 0 | Canis | 88978.0 | 0 | Canis lupus | 88978.0 | 0 |
| scf7180000044169 | 366595 | 0.4337 | 0 | 58.5459 | Eukaryota | 50999.0 | 0 | Chordata | 50999.0 | 0 | Carnivora | 50999.0 | 0 | Canidae | 50999.0 | 0 | Canis | 46437.0 | 1 | Canis lupus | 46437.0 | 1 |
| scf7180000044776 | 61574 | 0.3873 | 0 | 57.2785 | Eukaryota | 23330.0 | 0 | Chordata | 23330.0 | 0 | Carnivora | 23330.0 | 0 | Canidae | 23330.0 | 0 | Canis | 23330.0 | 0 | Canis lupus | 23330.0 | 0 |
| scf7180000054915 | 106172 | 0.4944 | 0 | 55.7582 | Eukaryota | 66749.0 | 0 | Chordata | 66749.0 | 0 | Carnivora | 66749.0 | 0 | Canidae | 66749.0 | 0 | Canis | 66749.0 | 0 | Canis lupus | 66749.0 | 0 |
| scf7180000042819 | 564850 | 0.3385 | 0 | 58.6786 | Eukaryota | 56633.0 | 0 | Chordata | 56633.0 | 0 | Carnivora | 56633.0 | 0 | Canidae | 56633.0 | 0 | Canis | 56633.0 | 0 | Canis lupus | 56633.0 | 0 |
| scf7180000048233 | 23640 | 0.4553 | 0 | 138.4362 | no-hit | 0.0 | 0 | no-hit | 0.0 | 0 | no-hit | 0.0 | 0 | no-hit | 0.0 | 0 | no-hit | 0.0 | 0 | no-hit | 0.0 | 0 |
| scf7180000048291 | 19200 | 0.4529 | 0 | 51.7183 | Eukaryota | 61667.0 | 0 | Chordata | 52501.0 | 1 | Carnivora | 43958.0 | 2 | Canidae | 43958.0 | 2 | Canis | 43958.0 | 2 | Canis lupus | 43958.0 | 2 |
| scf7180000055351 | 128242 | 0.4434 | 0 | 196.0249 | Eukaryota | 97615.0 | 0 | Platyhelminthes | 50444.0 | 1 | Gadiformes | 47171.0 | 2 | Gadidae | 47171.0 | 2 | Gadus | 47171.0 | 2 | Gadus morhua | 47171.0 | 2 |
| scf7180000053497 | 5849 | 0.5098 | 0 | 464.6686 | Eukaryota | 7665.0 | 0 | Chordata | 7665.0 | 0 | Carnivora | 7665.0 | 0 | Canidae | 7665.0 | 0 | Canis | 6607.0 | 1 | Canis lupus | 5507.0 | 2 |
| scf7180000055595 | 26469 | 0.4339 | 0 | 165.7072 | Eukaryota | 9541.0 | 0 | Chordata | 7191.0 | 1 | Salmoniformes | 4849.0 | 5 | Salmonidae | 4849.0 | 5 | Salmo | 4849.0 | 5 | Salmo trutta | 4849.0 | 5 |
| scf7180000047740 | 50066 | 0.4404 | 0 | 91.2733 | Eukaryota | 5169.0 | 0 | Platyhelminthes | 4100.0 | 1 | Polystomatidea | 4100.0 | 2 | Polystomatidae | 4100.0 | 2 | Protopolystoma | 4100.0 | 2 | Protopolystoma xenopodis | 4100.0 | 2 |
| scf7180000042515 | 48775 | 0.4629 | 0 | 49.216 | Eukaryota | 32392.0 | 0 | Chordata | 32392.0 | 0 | Carnivora | 29228.0 | 1 | Felidae | 26060.0 | 2 | Lynx | 13056.0 | 4 | Lynx canadensis | 13056.0 | 4 |
| scf7180000055566 | 25971 | 0.441 | 0 | 267.1616 | no-hit | 0.0 | 0 | no-hit | 0.0 | 0 | no-hit | 0.0 | 0 | no-hit | 0.0 | 0 | no-hit | 0.0 | 0 | no-hit | 0.0 | 0 |
| scf7180000046957 | 11593 | 0.4395 | 0 | 112.1745 | no-hit | 0.0 | 0 | no-hit | 0.0 | 0 | no-hit | 0.0 | 0 | no-hit | 0.0 | 0 | no-hit | 0.0 | 0 | no-hit | 0.0 | 0 |
| scf7180000040884 | 1726715 | 0.4619 | 0 | 59.1157 | Eukaryota | 81897.0 | 0 | Chordata | 81897.0 | 0 | Carnivora | 81897.0 | 0 | Canidae | 81897.0 | 0 | Canis | 81897.0 | 0 | Canis lupus | 81897.0 | 0 |
| scf7180000049523 | 28644 | 0.4413 | 0 | 59.0042 | Eukaryota | 967.0 | 0 | Chordata | 579.0 | 1 | Tetraodontiformes | 579.0 | 1 | Tetraodontidae | 579.0 | 1 | Takifugu | 579.0 | 1 | Takifugu rubripes | 579.0 | 1 |
| scf7180000040876 | 266728 | 0.398 | 0 | 102.6636 | Eukaryota | 89582.0 | 0 | Chordata | 89582.0 | 0 | Carnivora | 89582.0 | 0 | Canidae | 89582.0 | 0 | Canis | 89582.0 | 0 | Canis lupus | 89582.0 | 0 |
| scf7180000056465 | 282073 | 0.3713 | 0 | 61.3476 | Eukaryota | 55018.0 | 0 | Chordata | 55018.0 | 0 | Carnivora | 55018.0 | 0 | Canidae | 55018.0 | 0 | Canis | 55018.0 | 0 | Canis lupus | 55018.0 | 0 |
| scf7180000049387 | 30849 | 0.3793 | 0 | 59.9406 | Eukaryota | 8704.0 | 0 | Chordata | 8704.0 | 0 | Carnivora | 8704.0 | 0 | Canidae | 8704.0 | 0 | Canis | 7333.0 | 1 | Canis lupus | 7333.0 | 1 |
| scf7180000044261 | 324276 | 0.3747 | 0 | 60.0195 | Eukaryota | 55127.0 | 0 | Chordata | 55127.0 | 0 | Carnivora | 55127.0 | 0 | Canidae | 38706.0 | 3 | Canis | 38706.0 | 3 | Canis lupus | 38706.0 | 3 |
| scf7180000043083 | 33893 | 0.3708 | 0 | 578.1718 | Eukaryota | 29778.0 | 0 | Chordata | 29778.0 | 0 | Carnivora | 29778.0 | 0 | Canidae | 29778.0 | 0 | Canis | 29778.0 | 0 | Canis lupus | 29778.0 | 0 |
| scf7180000043672 | 440315 | 0.3619 | 0 | 60.4713 | Eukaryota | 54614.0 | 0 | Chordata | 54614.0 | 0 | Carnivora | 54614.0 | 0 | Canidae | 54614.0 | 0 | Canis | 54614.0 | 0 | Canis lupus | 54614.0 | 0 |
| scf7180000042502 | 62095 | 0.3508 | 0 | 57.481 | Eukaryota | 37002.0 | 0 | Chordata | 37002.0 | 0 | Carnivora | 37002.0 | 0 | Canidae | 37002.0 | 0 | Canis | 37002.0 | 0 | Canis lupus | 37002.0 | 0 |
| scf7180000056431 | 480729 | 0.3629 | 0 | 60.4057 | Eukaryota | 97362.0 | 0 | Chordata | 97362.0 | 0 | Carnivora | 97362.0 | 0 | Canidae | 97362.0 | 0 | Canis | 97362.0 | 0 | Canis lupus | 97362.0 | 0 |
| scf7180000052741 | 9522 | 0.4327 | 0 | 204.1544 | Eukaryota | 3504.0 | 0 | Platyhelminthes | 2235.0 | 1 | Polystomatidea | 2235.0 | 2 | Polystomatidae | 2235.0 | 2 | Protopolystoma | 2235.0 | 3 | Protopolystoma xenopodis | 2235.0 | 3 |
| scf7180000047701 | 31752 | 0.3739 | 0 | 125.0402 | Eukaryota | 5622.0 | 0 | Chordata | 5622.0 | 0 | Carnivora | 5032.0 | 1 | Canidae | 5032.0 | 1 | Canis | 5032.0 | 1 | Canis lupus | 5032.0 | 1 |
| scf7180000044604 | 240202 | 0.5218 | 0 | 61.4102 | Eukaryota | 25845.0 | 0 | Chordata | 25845.0 | 0 | Carnivora | 25845.0 | 0 | Canidae | 10965.0 | 2 | Zalophus | 10760.0 | 3 | Zalophus californianus | 10760.0 | 3 |
| scf7180000042702 | 3261 | 0.3456 | 0 | 20.428 | Eukaryota | 8519.0 | 0 | Chordata | 8519.0 | 0 | Carnivora | 8519.0 | 0 | Canidae | 8519.0 | 0 | Canis | 8519.0 | 0 | Canis lupus | 8519.0 | 0 |
| scf7180000055984 | 2850 | 0.4463 | 0 | 308.1529 | no-hit | 0.0 | 0 | no-hit | 0.0 | 0 | no-hit | 0.0 | 0 | no-hit | 0.0 | 0 | no-hit | 0.0 | 0 | no-hit | 0.0 | 0 |
| scf7180000051682 | 10648 | 0.5147 | 0 | 28.8615 | Eukaryota | 5346.0 | 0 | Chordata | 5346.0 | 0 | Carnivora | 5346.0 | 0 | Canidae | 4817.0 | 1 | Canis | 4817.0 | 1 | Canis lupus | 4817.0 | 1 |
| scf7180000049636 | 26529 | 0.4354 | 0 | 226.5198 | Eukaryota | 619.0 | 0 | Chordata | 619.0 | 0 | Tetraodontiformes | 619.0 | 0 | Tetraodontidae | 619.0 | 0 | Takifugu | 619.0 | 0 | Takifugu rubripes | 619.0 | 0 |
| scf7180000042433 | 161879 | 0.6117 | 0 | 54.4343 | Eukaryota | 57712.0 | 0 | Chordata | 57712.0 | 0 | Carnivora | 57712.0 | 0 | Canidae | 57712.0 | 0 | Canis | 47842.0 | 1 | Canis lupus | 47842.0 | 1 |
| scf7180000045293 | 26499 | 0.4278 | 0 | 254.439 | Eukaryota | 47040.0 | 0 | Chordata | 34832.0 | 1 | Gadiformes | 29965.0 | 4 | Gadidae | 29965.0 | 4 | Gadus | 29965.0 | 4 | Gadus morhua | 29965.0 | 4 |
| scf7180000040836 | 1658067 | 0.3681 | 0 | 60.0232 | Eukaryota | 89053.0 | 0 | Chordata | 89053.0 | 0 | Carnivora | 89053.0 | 0 | Canidae | 89053.0 | 0 | Canis | 89053.0 | 0 | Canis lupus | 89053.0 | 0 |
| scf7180000043406 | 454670 | 0.3673 | 0 | 59.0694 | Eukaryota | 93169.0 | 0 | Chordata | 93169.0 | 0 | Carnivora | 93169.0 | 0 | Canidae | 93169.0 | 0 | Canis | 93169.0 | 0 | Canis lupus | 93169.0 | 0 |
| scf7180000046263 | 2344 | 0.4386 | 0 | 32.1387 | Eukaryota | 3252.0 | 0 | Chordata | 3252.0 | 0 | Carnivora | 3252.0 | 0 | Canidae | 3252.0 | 0 | Canis | 3252.0 | 0 | Canis lupus | 3252.0 | 0 |
| scf7180000044461 | 271813 | 0.3782 | 0 | 60.8357 | Eukaryota | 33658.0 | 0 | Chordata | 33658.0 | 0 | Carnivora | 33658.0 | 0 | Canidae | 33658.0 | 0 | Canis | 33658.0 | 0 | Canis lupus | 33658.0 | 0 |
| scf7180000054099 | 4337 | 0.5066 | 0 | 51.2096 | Eukaryota | 5865.0 | 0 | Chordata | 5865.0 | 0 | Carnivora | 5865.0 | 0 | Canidae | 5865.0 | 0 | Canis | 3822.0 | 1 | Canis lupus | 3014.0 | 2 |
| scf7180000040710 | 20231 | 0.4156 | 0 | 56.0927 | Eukaryota | 33758.0 | 0 | Chordata | 33758.0 | 0 | Carnivora | 33758.0 | 0 | Canidae | 33758.0 | 0 | Canis | 33758.0 | 0 | Canis lupus | 33758.0 | 0 |
| scf7180000041446 | 604244 | 0.4626 | 0 | 58.3786 | Eukaryota | 65580.0 | 0 | Chordata | 65580.0 | 0 | Carnivora | 65580.0 | 0 | Canidae | 65580.0 | 0 | Vulpes | 39921.0 | 1 | Vulpes vulpes | 39921.0 | 1 |
| scf7180000040530 | 540861 | 0.3522 | 0 | 58.2357 | Eukaryota | 82175.0 | 0 | Chordata | 82175.0 | 0 | Carnivora | 82175.0 | 0 | Canidae | 82175.0 | 0 | Canis | 82175.0 | 0 | Canis lupus | 82175.0 | 0 |
| scf7180000048555 | 9177 | 0.4409 | 0 | 41.9944 | no-hit | 0.0 | 0 | no-hit | 0.0 | 0 | no-hit | 0.0 | 0 | no-hit | 0.0 | 0 | no-hit | 0.0 | 0 | no-hit | 0.0 | 0 |
| scf7180000047115 | 24920 | 0.4562 | 0 | 102.7506 | Eukaryota | 9343.0 | 0 | Chordata | 9343.0 | 0 | Carnivora | 8683.0 | 1 | Canidae | 8683.0 | 1 | Canis | 4804.0 | 2 | Canis lupus | 4804.0 | 2 |
| scf7180000052164 | 6983 | 0.4498 | 0 | 748.0662 | no-hit | 0.0 | 0 | no-hit | 0.0 | 0 | no-hit | 0.0 | 0 | no-hit | 0.0 | 0 | no-hit | 0.0 | 0 | no-hit | 0.0 | 0 |
| scf7180000047790 | 43760 | 0.4382 | 0 | 167.1377 | Eukaryota | 100867.0 | 0 | Chordata | 57397.0 | 1 | Gadiformes | 57397.0 | 2 | Gadidae | 57397.0 | 2 | Gadus | 57397.0 | 2 | Gadus morhua | 57397.0 | 2 |
| scf7180000053432 | 2506 | 0.5016 | 0 | 164.8947 | Eukaryota | 520.0 | 0 | Chordata | 520.0 | 0 | Carnivora | 520.0 | 0 | Canidae | 520.0 | 0 | Canis | 520.0 | 0 | Canis lupus | 520.0 | 0 |
| scf7180000045863 | 61930 | 0.4012 | 0 | 57.8484 | Eukaryota | 52896.0 | 0 | Chordata | 52896.0 | 0 | Carnivora | 52896.0 | 0 | Canidae | 52896.0 | 0 | Canis | 52896.0 | 0 | Canis lupus | 52896.0 | 0 |
| scf7180000043666 | 648079 | 0.5049 | 0 | 57.1215 | Eukaryota | 103687.0 | 0 | Chordata | 103687.0 | 0 | Carnivora | 103687.0 | 0 | Canidae | 103687.0 | 0 | Canis | 92848.0 | 1 | Canis lupus | 92848.0 | 1 |
| scf7180000045199 | 17017 | 0.4473 | 0 | 267.558 | Eukaryota | 4594.0 | 0 | Chordata | 4283.0 | 1 | Tetraodontiformes | 1567.0 | 4 | Tetraodontidae | 1567.0 | 4 | Takifugu | 1567.0 | 6 | Takifugu rubripes | 1567.0 | 6 |
| scf7180000042281 | 742043 | 0.4308 | 0 | 59.6947 | Eukaryota | 91667.0 | 0 | Chordata | 91667.0 | 0 | Carnivora | 91667.0 | 0 | Canidae | 91667.0 | 0 | Canis | 91667.0 | 0 | Canis lupus | 91667.0 | 0 |
| scf7180000043553 | 239216 | 0.4247 | 0 | 57.0445 | Eukaryota | 9717.0 | 0 | Chordata | 9717.0 | 0 | Primates | 6175.0 | 2 | Hominidae | 6175.0 | 2 | Homo | 3218.0 | 3 | Homo sapiens | 3218.0 | 3 |
| scf7180000042634 | 35128 | 0.4999 | 0 | 49.5872 | Eukaryota | 49186.0 | 0 | Chordata | 49186.0 | 0 | Carnivora | 49186.0 | 0 | Canidae | 49186.0 | 0 | Canis | 39350.0 | 1 | Canis lupus | 39350.0 | 1 |
| scf7180000041216 | 1180752 | 0.4294 | 0 | 61.0699 | Eukaryota | 58491.0 | 0 | Chordata | 58491.0 | 0 | Carnivora | 58491.0 | 0 | Canidae | 58491.0 | 0 | Canis | 49617.0 | 1 | Canis lupus | 49617.0 | 1 |
| scf7180000052194 | 6157 | 0.452 | 0 | 848.3394 | Eukaryota | 714.0 | 0 | Echinodermata | 564.0 | 1 | Forcipulatida | 564.0 | 1 | Asteriidae | 564.0 | 1 | Asterias | 564.0 | 1 | Asterias rubens | 564.0 | 1 |
| scf7180000042208 | 546530 | 0.6 | 0 | 59.3993 | Eukaryota | 69541.0 | 0 | Chordata | 69541.0 | 0 | Carnivora | 69541.0 | 0 | Canidae | 69541.0 | 0 | Canis | 63110.0 | 1 | Canis lupus | 63110.0 | 1 |
| scf7180000051721 | 29344 | 0.4473 | 0 | 170.4799 | Eukaryota | 41970.0 | 0 | Platyhelminthes | 18670.0 | 3 | Strigeidida | 11759.0 | 7 | Schistosomatidae | 11759.0 | 7 | Schistosoma | 11759.0 | 7 | Schistosoma mansoni | 11759.0 | 7 |
| scf7180000042104 | 751805 | 0.4232 | 0 | 59.3463 | Eukaryota | 45426.0 | 0 | Chordata | 45426.0 | 0 | Carnivora | 45426.0 | 0 | Canidae | 45426.0 | 0 | Canis | 29425.0 | 1 | Canis lupus | 29425.0 | 1 |
| scf7180000044054 | 185535 | 0.4257 | 0 | 57.7827 | Eukaryota | 50710.0 | 0 | Chordata | 50710.0 | 0 | Carnivora | 50710.0 | 0 | Canidae | 50710.0 | 0 | Vulpes | 41102.0 | 1 | Vulpes vulpes | 41102.0 | 1 |
| scf7180000047460 | 13078 | 0.4382 | 0 | 106.461 | Eukaryota | 10701.0 | 0 | Chordata | 9143.0 | 1 | Kurtiformes | 4528.0 | 3 | Apogonidae | 4528.0 | 3 | Sphaeramia | 4528.0 | 4 | Sphaeramia orbicularis | 4528.0 | 4 |
| scf7180000046271 | 20994 | 0.4396 | 0 | 238.1799 | Eukaryota | 4745.0 | 0 | Chordata | 4745.0 | 0 | Carnivora | 2607.0 | 2 | Canidae | 2607.0 | 2 | Canis | 2607.0 | 2 | Canis lupus | 2607.0 | 2 |
| scf7180000053511 | 3056 | 0.4601 | 0 | 102.0521 | no-hit | 0.0 | 0 | no-hit | 0.0 | 0 | no-hit | 0.0 | 0 | no-hit | 0.0 | 0 | no-hit | 0.0 | 0 | no-hit | 0.0 | 0 |
| scf7180000051371 | 6506 | 0.5131 | 0 | 28.8376 | Eukaryota | 2331.0 | 0 | Chordata | 2331.0 | 0 | Carnivora | 2331.0 | 0 | Canidae | 2331.0 | 0 | Canis | 2331.0 | 0 | Canis lupus | 1879.0 | 1 |
| scf7180000043851 | 161116 | 0.428 | 0 | 59.7949 | Eukaryota | 19541.0 | 0 | Chordata | 19541.0 | 0 | Carnivora | 19541.0 | 0 | Canidae | 16038.0 | 1 | Canis | 16038.0 | 2 | Canis lupus | 16038.0 | 2 |
| scf7180000056276 | 910772 | 0.4865 | 0 | 60.5275 | Eukaryota | 171384.0 | 0 | Chordata | 171384.0 | 0 | Carnivora | 171384.0 | 0 | Canidae | 171384.0 | 0 | Canis | 171384.0 | 0 | Canis lupus | 171384.0 | 0 |
| scf7180000040833 | 1564427 | 0.4314 | 0 | 59.7239 | Eukaryota | 98315.0 | 0 | Chordata | 98315.0 | 0 | Carnivora | 98315.0 | 0 | Canidae | 98315.0 | 0 | Canis | 98315.0 | 0 | Canis lupus | 98315.0 | 0 |
| scf7180000040922 | 1570516 | 0.3933 | 0 | 60.452 | Eukaryota | 90471.0 | 0 | Chordata | 90471.0 | 0 | Carnivora | 90471.0 | 0 | Canidae | 90471.0 | 0 | Canis | 90471.0 | 0 | Canis lupus | 90471.0 | 0 |
| scf7180000044110 | 313389 | 0.5113 | 0 | 58.7263 | Eukaryota | 42538.0 | 0 | Chordata | 42538.0 | 0 | Carnivora | 42538.0 | 0 | Canidae | 42538.0 | 0 | Canis | 21568.0 | 1 | Canis lupus | 21568.0 | 1 |
| scf7180000041202 | 3768 | 0.4289 | 0 | 38.57 | Eukaryota | 17416.0 | 0 | Chordata | 17416.0 | 0 | Carnivora | 17416.0 | 0 | Canidae | 17416.0 | 0 | Canis | 17416.0 | 0 | Canis lupus | 17416.0 | 0 |
| scf7180000040568 | 351908 | 0.3674 | 0 | 59.086 | Eukaryota | 92152.0 | 0 | Chordata | 92152.0 | 0 | Carnivora | 92152.0 | 0 | Canidae | 92152.0 | 0 | Canis | 92152.0 | 0 | Canis lupus | 92152.0 | 0 |
| scf7180000042382 | 4804 | 0.3853 | 0 | 19.5001 | Eukaryota | 64107.0 | 0 | Chordata | 64107.0 | 0 | Carnivora | 64107.0 | 0 | Canidae | 64107.0 | 0 | Canis | 64107.0 | 0 | Canis lupus | 64107.0 | 0 |
| scf7180000042291 | 37623 | 0.4062 | 0 | 46.9178 | Eukaryota | 5830.0 | 0 | Chordata | 5830.0 | 0 | Primates | 3124.0 | 1 | Hominidae | 2569.0 | 3 | Homo | 2569.0 | 3 | Homo sapiens | 2569.0 | 3 |
| scf7180000045752 | 10388 | 0.4462 | 0 | 538.0085 | Eukaryota | 152.0 | 0 | Chordata | 152.0 | 0 | Salmoniformes | 152.0 | 0 | Salmonidae | 152.0 | 0 | Salmo | 152.0 | 0 | Salmo trutta | 152.0 | 0 |
| scf7180000045751 | 45425 | 0.4418 | 0 | 144.6881 | Eukaryota | 1973.0 | 0 | Chordata | 1208.0 | 1 | Polystomatidea | 765.0 | 3 | Polystomatidae | 765.0 | 3 | Protopolystoma | 765.0 | 3 | Protopolystoma xenopodis | 765.0 | 3 |
| scf7180000049569 | 3787 | 0.3945 | 0 | 16.079 | Eukaryota | 51611.0 | 0 | Chordata | 51611.0 | 0 | Carnivora | 51611.0 | 0 | Canidae | 51611.0 | 0 | Canis | 51611.0 | 0 | Canis lupus | 51611.0 | 0 |
| scf7180000051141 | 13019 | 0.4559 | 0 | 60.9321 | no-hit | 0.0 | 0 | no-hit | 0.0 | 0 | no-hit | 0.0 | 0 | no-hit | 0.0 | 0 | no-hit | 0.0 | 0 | no-hit | 0.0 | 0 |
| scf7180000043195 | 84963 | 0.4152 | 0 | 58.2652 | Eukaryota | 59346.0 | 0 | Chordata | 59346.0 | 0 | Carnivora | 59346.0 | 0 | Canidae | 59346.0 | 0 | Canis | 59346.0 | 0 | Canis lupus | 59346.0 | 0 |
| scf7180000047520 | 59118 | 0.4407 | 0 | 160.6324 | Eukaryota | 12634.0 | 0 | Chordata | 10884.0 | 1 | Tetraodontiformes | 6768.0 | 5 | Tetraodontidae | 6768.0 | 5 | Takifugu | 6768.0 | 5 | Takifugu rubripes | 6768.0 | 5 |
| scf7180000054446 | 298923 | 0.3644 | 0 | 58.9402 | Eukaryota | 86790.0 | 0 | Chordata | 86790.0 | 0 | Carnivora | 86790.0 | 0 | Canidae | 86790.0 | 0 | Canis | 86790.0 | 0 | Canis lupus | 86790.0 | 0 |
| scf7180000043352 | 391047 | 0.5012 | 0 | 58.3171 | Eukaryota | 70352.0 | 0 | Chordata | 70352.0 | 0 | Carnivora | 46960.0 | 1 | Canidae | 33024.0 | 4 | Canis | 26375.0 | 7 | Canis lupus | 26375.0 | 7 |
| scf7180000053707 | 2632 | 0.4441 | 0 | 224.9719 | no-hit | 0.0 | 0 | no-hit | 0.0 | 0 | no-hit | 0.0 | 0 | no-hit | 0.0 | 0 | no-hit | 0.0 | 0 | no-hit | 0.0 | 0 |
| scf7180000050177 | 8320 | 0.4424 | 0 | 571.2755 | Eukaryota | 3411.0 | 0 | Platyhelminthes | 1722.0 | 3 | Polystomatidea | 1722.0 | 3 | Polystomatidae | 1722.0 | 3 | Protopolystoma | 1722.0 | 3 | Protopolystoma xenopodis | 1722.0 | 3 |
| scf7180000041391 | 111666 | 0.3816 | 0 | 59.1029 | Eukaryota | 87573.0 | 0 | Chordata | 87573.0 | 0 | Carnivora | 87573.0 | 0 | Canidae | 87573.0 | 0 | Canis | 87573.0 | 0 | Canis lupus | 87573.0 | 0 |
| scf7180000045451 | 174645 | 0.486 | 0 | 58.6785 | Eukaryota | 37608.0 | 0 | Chordata | 37608.0 | 0 | Carnivora | 37608.0 | 0 | Canidae | 37608.0 | 0 | Canis | 22476.0 | 1 | Canis lupus | 22476.0 | 1 |
| scf7180000043566 | 432285 | 0.4039 | 0 | 60.2977 | Eukaryota | 90464.0 | 0 | Chordata | 90464.0 | 0 | Carnivora | 90464.0 | 0 | Canidae | 90464.0 | 0 | Canis | 90464.0 | 0 | Canis lupus | 90464.0 | 0 |
| scf7180000044695 | 264609 | 0.3681 | 0 | 59.1961 | Eukaryota | 31155.0 | 0 | Chordata | 31155.0 | 0 | Carnivora | 24471.0 | 2 | Canidae | 19447.0 | 3 | Canis | 11782.0 | 4 | Canis lupus | 11782.0 | 4 |
| scf7180000044123 | 208762 | 0.415 | 0 | 58.0181 | Eukaryota | 17653.0 | 0 | Chordata | 17653.0 | 0 | Carnivora | 17653.0 | 0 | Canidae | 10969.0 | 2 | Canis | 7445.0 | 5 | Canis lupus | 7445.0 | 5 |
| scf7180000056262 | 1047883 | 0.3746 | 0 | 60.1088 | Eukaryota | 74730.0 | 0 | Chordata | 74730.0 | 0 | Carnivora | 74730.0 | 0 | Canidae | 74730.0 | 0 | Canis | 43025.0 | 1 | Canis lupus | 43025.0 | 1 |
| scf7180000042885 | 666924 | 0.3441 | 0 | 58.4964 | Eukaryota | 84867.0 | 0 | Chordata | 84867.0 | 0 | Carnivora | 84867.0 | 0 | Canidae | 84867.0 | 0 | Canis | 84867.0 | 0 | Canis lupus | 84867.0 | 0 |
| scf7180000040974 | 1368513 | 0.4609 | 0 | 59.9362 | Eukaryota | 101665.0 | 0 | Chordata | 101665.0 | 0 | Carnivora | 101665.0 | 0 | Canidae | 101665.0 | 0 | Canis | 101665.0 | 0 | Canis lupus | 101665.0 | 0 |
| scf7180000042651 | 666368 | 0.4004 | 100 | 59.9987 | Eukaryota | 82559.0 | 0 | Chordata | 82559.0 | 0 | Carnivora | 82559.0 | 0 | Canidae | 52221.0 | 2 | Canis | 29739.0 | 4 | Canis lupus | 29739.0 | 4 |
| scf7180000044948 | 191727 | 0.5128 | 0 | 66.5322 | Eukaryota | 38198.0 | 0 | Chordata | 38198.0 | 0 | Carnivora | 38198.0 | 0 | Canidae | 20581.0 | 3 | Canis | 16329.0 | 6 | Canis lupus | 16329.0 | 6 |
| scf7180000046973 | 101168 | 0.4458 | 0 | 59.8768 | Eukaryota | 60780.0 | 0 | Chordata | 48445.0 | 2 | Gadiformes | 37881.0 | 5 | Gadidae | 37881.0 | 5 | Gadus | 37881.0 | 5 | Gadus morhua | 37881.0 | 5 |
| scf7180000054377 | 538137 | 0.383 | 0 | 59.1011 | Eukaryota | 34712.0 | 0 | Chordata | 34712.0 | 0 | Carnivora | 34712.0 | 0 | Canidae | 34712.0 | 0 | Canis | 34712.0 | 0 | Canis lupus | 34712.0 | 0 |
| scf7180000040497 | 2931511 | 0.4309 | 0 | 60.3507 | Eukaryota | 94314.0 | 0 | Chordata | 94314.0 | 0 | Carnivora | 94314.0 | 0 | Canidae | 94314.0 | 0 | Canis | 94314.0 | 0 | Canis lupus | 94314.0 | 0 |
| scf7180000051328 | 16125 | 0.441 | 0 | 53.9317 | Eukaryota | 6200.0 | 0 | Chordata | 2772.0 | 2 | Kurtiformes | 2772.0 | 2 | Apogonidae | 2772.0 | 2 | Sphaeramia | 2772.0 | 2 | Sphaeramia orbicularis | 2772.0 | 2 |
| scf7180000046692 | 98408 | 0.3589 | 0 | 62.323 | Eukaryota | 13138.0 | 0 | Chordata | 13138.0 | 0 | Carnivora | 7966.0 | 2 | Canidae | 7966.0 | 2 | Vulpes | 4552.0 | 3 | Vulpes vulpes | 4552.0 | 3 |
| scf7180000053068 | 5743 | 0.4484 | 0 | 927.7771 | no-hit | 0.0 | 0 | no-hit | 0.0 | 0 | no-hit | 0.0 | 0 | no-hit | 0.0 | 0 | no-hit | 0.0 | 0 | no-hit | 0.0 | 0 |
| scf7180000048450 | 12007 | 0.4501 | 0 | 1882.1973 | no-hit | 0.0 | 0 | no-hit | 0.0 | 0 | no-hit | 0.0 | 0 | no-hit | 0.0 | 0 | no-hit | 0.0 | 0 | no-hit | 0.0 | 0 |
| scf7180000047104 | 52848 | 0.3378 | 0 | 57.548 | Eukaryota | 48796.0 | 0 | Chordata | 48796.0 | 0 | Carnivora | 48796.0 | 0 | Canidae | 48796.0 | 0 | Canis | 48796.0 | 0 | Canis lupus | 48796.0 | 0 |
| scf7180000042488 | 6372 | 0.344 | 0 | 26.9121 | Eukaryota | 161.0 | 0 | Chordata | 161.0 | 0 | Carnivora | 161.0 | 0 | Canidae | 161.0 | 0 | Canis | 161.0 | 0 | Canis lupus | 161.0 | 0 |
| scf7180000046837 | 3248 | 0.4415 | 0 | 80.6772 | Eukaryota | 137.0 | 0 | Chordata | 137.0 | 0 | Osteoglossiformes | 137.0 | 0 | Osteoglossidae | 137.0 | 0 | Scleropages | 137.0 | 0 | Scleropages formosus | 137.0 | 0 |
| scf7180000051108 | 2592 | 0.4487 | 0 | 465.471 | no-hit | 0.0 | 0 | no-hit | 0.0 | 0 | no-hit | 0.0 | 0 | no-hit | 0.0 | 0 | no-hit | 0.0 | 0 | no-hit | 0.0 | 0 |
| scf7180000042893 | 130519 | 0.4082 | 0 | 57.9145 | Eukaryota | 87708.0 | 0 | Chordata | 87708.0 | 0 | Carnivora | 87708.0 | 0 | Canidae | 87708.0 | 0 | Canis | 87708.0 | 0 | Canis lupus | 87708.0 | 0 |
| scf7180000046715 | 36975 | 0.4035 | 0 | 52.406 | Eukaryota | 8618.0 | 0 | Chordata | 8618.0 | 0 | Carnivora | 6112.0 | 1 | Hominidae | 2506.0 | 4 | Homo | 2506.0 | 4 | Homo sapiens | 2506.0 | 4 |
| scf7180000051776 | 6331 | 0.4423 | 0 | 312.2739 | Eukaryota | 1972.0 | 0 | Chordata | 1972.0 | 0 | Tetraodontiformes | 1972.0 | 0 | Tetraodontidae | 1972.0 | 0 | Takifugu | 1972.0 | 0 | Takifugu rubripes | 1972.0 | 0 |
| scf7180000041313 | 637360 | 0.4355 | 0 | 59.1749 | Eukaryota | 57028.0 | 0 | Chordata | 57028.0 | 0 | Carnivora | 43261.0 | 1 | Canidae | 26621.0 | 3 | Canis | 26621.0 | 6 | Canis lupus | 26621.0 | 6 |
| scf7180000041026 | 1334859 | 0.3704 | 0 | 58.6477 | Eukaryota | 78109.0 | 0 | Chordata | 78109.0 | 0 | Carnivora | 78109.0 | 0 | Canidae | 78109.0 | 0 | Canis | 78109.0 | 0 | Canis lupus | 78109.0 | 0 |
| scf7180000050456 | 20029 | 0.4445 | 0 | 99.3243 | Eukaryota | 2471.0 | 0 | Chordata | 1911.0 | 1 | Carnivora | 1376.0 | 3 | Canidae | 1376.0 | 3 | Canis | 1376.0 | 3 | Canis lupus | 1376.0 | 3 |
| scf7180000045131 | 200919 | 0.3724 | 0 | 59.8523 | Eukaryota | 53962.0 | 0 | Chordata | 53962.0 | 0 | Carnivora | 53962.0 | 0 | Canidae | 53962.0 | 0 | Canis | 53962.0 | 0 | Canis lupus | 53962.0 | 0 |
| scf7180000053218 | 5992 | 0.5247 | 0 | 290.8562 | Eukaryota | 28646.0 | 0 | Chordata | 28646.0 | 0 | Carnivora | 28646.0 | 0 | Canidae | 28646.0 | 0 | Canis | 28646.0 | 0 | Canis lupus | 28646.0 | 0 |
| scf7180000042279 | 490599 | 0.3781 | 0 | 59.6636 | Eukaryota | 89304.0 | 0 | Chordata | 89304.0 | 0 | Carnivora | 89304.0 | 0 | Canidae | 89304.0 | 0 | Canis | 89304.0 | 0 | Canis lupus | 89304.0 | 0 |
| scf7180000048620 | 20507 | 0.5221 | 0 | 450.5628 | Eukaryota | 17844.0 | 0 | Chordata | 17844.0 | 0 | Carnivora | 17844.0 | 0 | Canidae | 17844.0 | 0 | Canis | 17844.0 | 0 | Canis lupus | 17844.0 | 0 |
| scf7180000045270 | 183621 | 0.5589 | 0 | 55.8115 | Eukaryota | 50440.0 | 0 | Chordata | 50440.0 | 0 | Carnivora | 50440.0 | 0 | Canidae | 50440.0 | 0 | Canis | 35614.0 | 1 | Canis lupus | 35614.0 | 1 |
| scf7180000050856 | 18188 | 0.5485 | 0 | 53.7876 | Eukaryota | 6371.0 | 1 | Chordata | 6371.0 | 1 | Carnivora | 4015.0 | 3 | Canidae | 4015.0 | 3 | Vulpes | 2486.0 | 5 | Vulpes vulpes | 2486.0 | 5 |
| scf7180000056599 | 630025 | 0.4499 | 100 | 58.6502 | Eukaryota | 93287.0 | 0 | Chordata | 93287.0 | 0 | Carnivora | 93287.0 | 0 | Canidae | 93287.0 | 0 | Canis | 93287.0 | 0 | Canis lupus | 93287.0 | 0 |
| scf7180000054338 | 108299 | 0.3648 | 0 | 57.2377 | Eukaryota | 5381.0 | 1 | Chordata | 5381.0 | 1 | Primates | 2935.0 | 3 | Hominidae | 2935.0 | 4 | Homo | 2935.0 | 5 | Homo sapiens | 2935.0 | 5 |
| scf7180000043299 | 129146 | 0.4026 | 0 | 59.2495 | Eukaryota | 83400.0 | 0 | Chordata | 83400.0 | 0 | Carnivora | 83400.0 | 0 | Canidae | 83400.0 | 0 | Canis | 83400.0 | 0 | Canis lupus | 83400.0 | 0 |
| scf7180000049410 | 8453 | 0.4272 | 0 | 71.8809 | Eukaryota | 82336.0 | 0 | Chordata | 82336.0 | 0 | Carnivora | 82336.0 | 0 | Canidae | 82336.0 | 0 | Canis | 82336.0 | 0 | Canis lupus | 82336.0 | 0 |
| scf7180000045404 | 208100 | 0.3771 | 0 | 59.2437 | Eukaryota | 24285.0 | 0 | Chordata | 24285.0 | 0 | Carnivora | 12914.0 | 2 | Canidae | 12914.0 | 4 | Canis | 7616.0 | 5 | Canis lupus | 7616.0 | 5 |
| scf7180000045921 | 162932 | 0.6032 | 0 | 55.7889 | Eukaryota | 36270.0 | 0 | Chordata | 36270.0 | 0 | Carnivora | 36270.0 | 0 | Canidae | 36270.0 | 0 | Canis | 29162.0 | 1 | Canis lupus | 29162.0 | 1 |
| scf7180000045339 | 8302 | 0.4195 | 0 | 26.6267 | Eukaryota | 4997.0 | 0 | Chordata | 4997.0 | 0 | Carnivora | 4997.0 | 0 | Canidae | 4997.0 | 0 | Canis | 4053.0 | 1 | Canis lupus | 4053.0 | 1 |
| scf7180000044855 | 257067 | 0.4965 | 0 | 58.137 | Eukaryota | 81937.0 | 0 | Chordata | 81937.0 | 0 | Carnivora | 81937.0 | 0 | Canidae | 81937.0 | 0 | Canis | 81937.0 | 0 | Canis lupus | 81937.0 | 0 |
| scf7180000041238 | 161314 | 0.3661 | 0 | 58.1052 | Eukaryota | 77238.0 | 0 | Chordata | 77238.0 | 0 | Carnivora | 77238.0 | 0 | Canidae | 77238.0 | 0 | Canis | 77238.0 | 0 | Canis lupus | 77238.0 | 0 |
| scf7180000045603 | 194335 | 0.4478 | 0 | 59.5669 | Eukaryota | 38702.0 | 0 | Chordata | 38702.0 | 0 | Carnivora | 38702.0 | 0 | Canidae | 31350.0 | 2 | Canis | 23603.0 | 5 | Canis lupus | 23603.0 | 5 |
| scf7180000046936 | 109334 | 0.3645 | 0 | 60.3087 | Eukaryota | 20066.0 | 0 | Chordata | 20066.0 | 0 | Carnivora | 5785.0 | 5 | Hominidae | 5311.0 | 6 | Homo | 5311.0 | 7 | Homo sapiens | 5311.0 | 8 |
| scf7180000051321 | 3393 | 0.4427 | 0 | 167.1899 | no-hit | 0.0 | 0 | no-hit | 0.0 | 0 | no-hit | 0.0 | 0 | no-hit | 0.0 | 0 | no-hit | 0.0 | 0 | no-hit | 0.0 | 0 |
| scf7180000049268 | 51456 | 0.3645 | 0 | 59.1478 | Eukaryota | 9231.0 | 1 | Chordata | 9231.0 | 1 | Apterygiformes | 3606.0 | 3 | Apterygidae | 3606.0 | 3 | Apteryx | 3606.0 | 4 | Apteryx mantelli | 3606.0 | 4 |
| scf7180000043109 | 398317 | 0.348 | 0 | 58.8392 | Eukaryota | 136218.0 | 0 | Chordata | 136218.0 | 0 | Carnivora | 136218.0 | 0 | Canidae | 136218.0 | 0 | Canis | 136218.0 | 0 | Canis lupus | 136218.0 | 0 |
| scf7180000045555 | 181673 | 0.4566 | 0 | 57.8982 | Eukaryota | 21928.0 | 0 | Chordata | 21928.0 | 0 | Carnivora | 19999.0 | 1 | Canidae | 14596.0 | 3 | Canis | 14596.0 | 3 | Canis lupus | 14596.0 | 3 |
| scf7180000056290 | 845655 | 0.5641 | 0 | 57.4324 | Eukaryota | 52513.0 | 0 | Chordata | 52513.0 | 0 | Carnivora | 52513.0 | 0 | Canidae | 52513.0 | 0 | Canis | 46481.0 | 1 | Canis lupus | 46481.0 | 1 |
| scf7180000053893 | 2818 | 0.4436 | 0 | 862.9302 | Eukaryota | 998.0 | 0 | Platyhelminthes | 857.0 | 1 | Polystomatidea | 857.0 | 1 | Polystomatidae | 857.0 | 1 | Protopolystoma | 857.0 | 1 | Protopolystoma xenopodis | 857.0 | 1 |
| scf7180000044118 | 9818 | 0.4213 | 0 | 23.6947 | Eukaryota | 91249.0 | 0 | Chordata | 91249.0 | 0 | Carnivora | 91249.0 | 0 | Canidae | 91249.0 | 0 | Canis | 91249.0 | 0 | Canis lupus | 91249.0 | 0 |
| scf7180000043918 | 243552 | 0.3596 | 0 | 57.7972 | Eukaryota | 65875.0 | 0 | Chordata | 65875.0 | 0 | Carnivora | 65875.0 | 0 | Canidae | 65875.0 | 0 | Canis | 65875.0 | 0 | Canis lupus | 65875.0 | 0 |

| | | | | | | | | | | | | | | | | | | | | | |
|---|---|---|---|---|---|---|---|---|---|---|---|---|---|---|---|---|---|---|---|---|---|
| scf7180000052621 | 3870 | 0.447 | 0 | 398.4966 | no-hit | 0.0 | 0 | no-hit | 0.0 | 0 | no-hit | 0.0 | 0 | no-hit | 0.0 | 0 | no-hit | 0.0 | 0 | no-hit | 0.0 | 0 |
| scf7180000046500 | 94150 | 0.363 | 0 | 60.0851 | Eukaryota | 37629.0 | 0 | Chordata | 37629.0 | 0 | Carnivora | 37629.0 | 0 | Canidae | 37629.0 | 0 | Canis | 37629.0 | 0 | Canis lupus | 37629.0 | 0 |
| scf7180000040636 | 666415 | 0.385 | 0 | 59.7153 | Eukaryota | 54205.0 | 0 | Chordata | 54205.0 | 0 | Carnivora | 54205.0 | 0 | Canidae | 54205.0 | 0 | Canis | 54205.0 | 0 | Canis lupus | 54205.0 | 0 |
| scf7180000053125 | 2928 | 0.5584 | 0 | 53.8122 | Eukaryota | 6382.0 | 0 | Chordata | 6382.0 | 0 | Carnivora | 6382.0 | 0 | Canidae | 6382.0 | 0 | Canis | 6382.0 | 0 | Canis lupus | 6382.0 | 0 |
| scf7180000052895 | 3135 | 0.451 | 0 | 210.9965 | Eukaryota | 4895.0 | 0 | Chordata | 2278.0 | 2 | Polystomatidea | 1850.0 | 4 | Polystomatidae | 1850.0 | 4 | Protopolystoma | 1850.0 | 4 | Protopolystoma xenopodis | 1850.0 | 4 |
| scf7180000043958 | 32641 | 0.4144 | 0 | 57.3514 | Eukaryota | 93392.0 | 0 | Chordata | 93392.0 | 0 | Carnivora | 93392.0 | 0 | Canidae | 93392.0 | 0 | Canis | 93392.0 | 0 | Canis lupus | 93392.0 | 0 |
| scf7180000045378 | 182472 | 0.3691 | 0 | 59.1285 | Eukaryota | 48251.0 | 1 | Chordata | 48251.0 | 1 | Carnivora | 44500.0 | 2 | Canidae | 25061.0 | 4 | Canis | 16773.0 | 6 | Canis lupus | 16773.0 | 6 |
| scf7180000040892 | 117256 | 0.3715 | 0 | 65.5283 | Eukaryota | 85866.0 | 0 | Chordata | 85866.0 | 0 | Carnivora | 85866.0 | 0 | Canidae | 85866.0 | 0 | Canis | 85866.0 | 0 | Canis lupus | 85866.0 | 0 |
| scf7180000055110 | 56597 | 0.405 | 0 | 705.858 | Eukaryota | 32450.0 | 0 | Chordata | 32450.0 | 0 | Carnivora | 32450.0 | 0 | Canidae | 32450.0 | 0 | Canis | 32450.0 | 0 | Canis lupus | 32450.0 | 0 |
| scf7180000041674 | 538723 | 0.4244 | 0 | 60.6315 | Eukaryota | 90526.0 | 0 | Chordata | 90526.0 | 0 | Carnivora | 90526.0 | 0 | Canidae | 90526.0 | 0 | Canis | 90526.0 | 0 | Canis lupus | 90526.0 | 0 |
| scf7180000054121 | 2291 | 0.4609 | 0 | 303.0387 | Eukaryota | 204.0 | 0 | Platyhelminthes | 204.0 | 0 | Polystomatidea | 204.0 | 0 | Polystomatidae | 204.0 | 0 | Protopolystoma | 204.0 | 0 | Protopolystoma xenopodis | 204.0 | 0 |
| scf7180000042234 | 737665 | 0.4178 | 0 | 66.9016 | Eukaryota | 91494.0 | 0 | Chordata | 91494.0 | 0 | Carnivora | 91494.0 | 0 | Canidae | 91494.0 | 0 | Canis | 91494.0 | 0 | Canis lupus | 91494.0 | 0 |
| scf7180000043041 | 258183 | 0.3721 | 0 | 61.1684 | Eukaryota | 29207.0 | 0 | Chordata | 29207.0 | 0 | Carnivora | 14306.0 | 2 | Canidae | 8093.0 | 7 | Canis | 8093.0 | 8 | Canis lupus | 8093.0 | 8 |
| scf7180000048132 | 72979 | 0.418 | 0 | 60.4931 | Eukaryota | 20949.0 | 0 | Chordata | 20949.0 | 0 | Carnivora | 13140.0 | 2 | Canidae | 12294.0 | 3 | Canis | 12294.0 | 4 | Canis lupus | 12294.0 | 4 |
| scf7180000046858 | 56603 | 0.4404 | 0 | 113.821 | Eukaryota | 25754.0 | 0 | Chordata | 25754.0 | 0 | Carnivora | 20206.0 | 1 | Canidae | 20206.0 | 1 | Canis | 20206.0 | 1 | Canis lupus | 20206.0 | 1 |
| scf7180000045602 | 196721 | 0.368 | 0 | 60.5184 | Eukaryota | 24214.0 | 0 | Chordata | 24214.0 | 0 | Carnivora | 24214.0 | 0 | Canidae | 24214.0 | 0 | Canis | 24214.0 | 0 | Canis lupus | 24214.0 | 0 |
| scf7180000044223 | 365715 | 0.4072 | 0 | 61.0488 | Eukaryota | 80861.0 | 0 | Chordata | 80861.0 | 0 | Carnivora | 80861.0 | 0 | Canidae | 80861.0 | 0 | Canis | 75306.0 | 1 | Canis lupus | 75306.0 | 1 |
| scf7180000052804 | 2418 | 0.457 | 0 | 793.03 | Eukaryota | 614.0 | 0 | Platyhelminthes | 431.0 | 1 | Polystomatidea | 431.0 | 1 | Polystomatidae | 431.0 | 1 | Protopolystoma | 431.0 | 1 | Protopolystoma xenopodis | 431.0 | 1 |
| scf7180000042884 | 673556 | 0.4324 | 0 | 56.9927 | Eukaryota | 27378.0 | 0 | Chordata | 27378.0 | 0 | Carnivora | 27378.0 | 0 | Canidae | 27378.0 | 0 | Canis | 27378.0 | 0 | Canis lupus | 27378.0 | 0 |
| scf7180000041701 | 701096 | 0.3756 | 0 | 60.3948 | Eukaryota | 61083.0 | 0 | Chordata | 61083.0 | 0 | Carnivora | 61083.0 | 0 | Canidae | 49817.0 | 1 | Canis | 38144.0 | 2 | Canis lupus | 38144.0 | 3 |
| scf7180000051619 | 17734 | 0.4565 | 0 | 75.6707 | Eukaryota | 8638.0 | 0 | Chordata | 8638.0 | 0 | Carnivora | 8638.0 | 0 | Canidae | 8638.0 | 0 | Canis | 8638.0 | 0 | Canis lupus | 8638.0 | 0 |
| scf7180000042739 | 8404 | 0.4421 | 0 | 223.7739 | Eukaryota | 719.0 | 0 | Chordata | 411.0 | 1 | Tetraodontiformes | 411.0 | 1 | Tetraodontidae | 411.0 | 1 | Takifugu | 411.0 | 1 | Takifugu rubripes | 411.0 | 1 |
| scf7180000048574 | 9507 | 0.4455 | 0 | 162.1196 | Eukaryota | 2741.0 | 0 | Chordata | 2741.0 | 0 | Carnivora | 2741.0 | 0 | Canidae | 2741.0 | 0 | Canis | 2741.0 | 0 | Canis lupus | 2741.0 | 0 |
| scf7180000051992 | 19934 | 0.5018 | 0 | 487.6836 | Eukaryota | 9125.0 | 0 | Chordata | 9125.0 | 0 | Carnivora | 9125.0 | 0 | Canidae | 9125.0 | 0 | Canis | 7923.0 | 1 | Canis lupus | 6655.0 | 2 |
| scf7180000040740 | 3697 | 0.4544 | 0 | 967.6703 | Eukaryota | 2097.0 | 0 | Platyhelminthes | 878.0 | 2 | Polystomatidea | 878.0 | 2 | Polystomatidae | 878.0 | 2 | Protopolystoma | 878.0 | 2 | Protopolystoma xenopodis | 878.0 | 2 |
| scf7180000040784 | 116448 | 0.4855 | 0 | 56.4512 | Eukaryota | 57807.0 | 0 | Chordata | 57807.0 | 0 | Carnivora | 57807.0 | 0 | Canidae | 57807.0 | 0 | Canis | 50856.0 | 1 | Canis lupus | 50856.0 | 1 |
| scf7180000053842 | 5050 | 0.4388 | 0 | 206.1133 | Eukaryota | 2306.0 | 0 | Platyhelminthes | 1378.0 | 1 | Polystomatidea | 1098.0 | 2 | Polystomatidae | 1098.0 | 2 | Protopolystoma | 1098.0 | 2 | Protopolystoma xenopodis | 1098.0 | 2 |
| scf7180000043337 | 28212 | 0.4126 | 0 | 84.3596 | Eukaryota | 67401.0 | 0 | Chordata | 67401.0 | 0 | Carnivora | 67401.0 | 0 | Canidae | 67401.0 | 0 | Canis | 67401.0 | 0 | Canis lupus | 67401.0 | 0 |
| scf7180000046256 | 28285 | 0.4502 | 0 | 241.0363 | Eukaryota | 12629.0 | 0 | Chordata | 12629.0 | 0 | Carnivora | 12629.0 | 0 | Canidae | 12629.0 | 0 | Canis | 10831.0 | 1 | Canis lupus | 9889.0 | 2 |
| scf7180000049834 | 3997 | 0.4573 | 0 | 489.7723 | Eukaryota | 8651.0 | 0 | Chordata | 4535.0 | 1 | Cyclophyllidea | 2628.0 | 3 | Hymenolepididae | 2628.0 | 3 | Hymenolepis | 2628.0 | 3 | Hymenolepis microstoma | 2628.0 | 3 |
| scf7180000043185 | 179200 | 0.4791 | 0 | 56.0224 | Eukaryota | 33702.0 | 0 | Chordata | 33702.0 | 0 | Carnivora | 33702.0 | 0 | Canidae | 33702.0 | 0 | Canis | 33702.0 | 0 | Canis lupus | 33702.0 | 0 |
| scf7180000047293 | 99397 | 0.3647 | 0 | 60.9018 | Eukaryota | 36780.0 | 0 | Chordata | 36780.0 | 0 | Carnivora | 34105.0 | 1 | Felidae | 19924.0 | 2 | Panthera | 11952.0 | 4 | Panthera pardus | 11952.0 | 4 |
| scf7180000051488 | 2834 | 0.4319 | 0 | 49.491 | no-hit | 0.0 | 0 | no-hit | 0.0 | 0 | no-hit | 0.0 | 0 | no-hit | 0.0 | 0 | no-hit | 0.0 | 0 | no-hit | 0.0 | 0 |
| scf7180000040600 | 2934344 | 0.3801 | 0 | 61.2197 | Eukaryota | 95043.0 | 0 | Chordata | 95043.0 | 0 | Carnivora | 95043.0 | 0 | Canidae | 95043.0 | 0 | Canis | 95043.0 | 0 | Canis lupus | 95043.0 | 0 |
| scf7180000051003 | 20088 | 0.4984 | 0 | 253.9807 | Eukaryota | 9273.0 | 0 | Chordata | 9273.0 | 0 | Carnivora | 9273.0 | 0 | Canidae | 9273.0 | 0 | Canis | 8077.0 | 1 | Canis lupus | 6855.0 | 2 |
| scf7180000052072 | 3685 | 0.4366 | 0 | 115.8447 | no-hit | 0.0 | 0 | no-hit | 0.0 | 0 | no-hit | 0.0 | 0 | no-hit | 0.0 | 0 | no-hit | 0.0 | 0 | no-hit | 0.0 | 0 |
| scf7180000053131 | 2174 | 0.454 | 0 | 815.1861 | no-hit | 0.0 | 0 | no-hit | 0.0 | 0 | no-hit | 0.0 | 0 | no-hit | 0.0 | 0 | no-hit | 0.0 | 0 | no-hit | 0.0 | 0 |
| scf7180000041994 | 930327 | 0.4192 | 0 | 58.636 | Eukaryota | 93593.0 | 0 | Chordata | 93593.0 | 0 | Carnivora | 93593.0 | 0 | Canidae | 93593.0 | 0 | Canis | 93593.0 | 0 | Canis lupus | 93593.0 | 0 |
| scf7180000051183 | 19808 | 0.3479 | 0 | 60.2792 | Eukaryota | 2304.0 | 0 | Chordata | 2304.0 | 0 | Carnivora | 2304.0 | 0 | Canidae | 2304.0 | 0 | Vulpes | 1808.0 | 1 | Vulpes vulpes | 1808.0 | 1 |
| scf7180000044287 | 43839 | 0.3971 | 0 | 59.6099 | Eukaryota | 94762.0 | 0 | Chordata | 94762.0 | 0 | Carnivora | 94762.0 | 0 | Canidae | 94762.0 | 0 | Canis | 94762.0 | 0 | Canis lupus | 94762.0 | 0 |
| scf7180000051938 | 13542 | 0.4536 | 0 | 1681.2519 | Eukaryota | 2664.0 | 0 | Platyhelminthes | 1278.0 | 2 | Polystomatidea | 1278.0 | 2 | Polystomatidae | 1278.0 | 2 | Protopolystoma | 1278.0 | 2 | Protopolystoma xenopodis | 1278.0 | 2 |
| scf7180000049158 | 40579 | 0.4266 | 0 | 58.7172 | Eukaryota | 41914.0 | 0 | Chordata | 41914.0 | 0 | Carnivora | 41914.0 | 0 | Canidae | 23951.0 | 2 | Canis | 19204.0 | 3 | Canis lupus | 19204.0 | 4 |
| scf7180000047931 | 4094 | 0.4399 | 0 | 180.1631 | no-hit | 0.0 | 0 | no-hit | 0.0 | 0 | no-hit | 0.0 | 0 | no-hit | 0.0 | 0 | no-hit | 0.0 | 0 | no-hit | 0.0 | 0 |
| scf7180000042603 | 208951 | 0.3744 | 0 | 133.7211 | Eukaryota | 62439.0 | 0 | Chordata | 62439.0 | 0 | Carnivora | 62439.0 | 0 | Canidae | 62439.0 | 0 | Canis | 62439.0 | 0 | Canis lupus | 62439.0 | 0 |
| scf7180000049910 | 6947 | 0.4471 | 0 | 1793.5639 | Eukaryota | 15340.0 | 0 | Chordata | 13291.0 | 3 | Gadiformes | 8507.0 | 7 | Gadidae | 8507.0 | 7 | Gadus | 8507.0 | 7 | Gadus morhua | 8507.0 | 7 |
| scf7180000046585 | 78558 | 0.3436 | 0 | 60.4966 | Eukaryota | 55968.0 | 0 | Chordata | 55968.0 | 0 | Carnivora | 55968.0 | 0 | Canidae | 55968.0 | 0 | Canis | 55968.0 | 0 | Canis lupus | 55968.0 | 0 |
| scf7180000044337 | 4584 | 0.4492 | 0 | 182.6749 | Eukaryota | 322.0 | 0 | Platyhelminthes | 322.0 | 0 | Polystomatidea | 322.0 | 0 | Polystomatidae | 322.0 | 0 | Protopolystoma | 322.0 | 0 | Protopolystoma xenopodis | 322.0 | 0 |
| scf7180000048752 | 47534 | 0.4562 | 0 | 59.8374 | Eukaryota | 111862.0 | 0 | Chordata | 111862.0 | 0 | Carnivora | 111862.0 | 0 | Canidae | 111862.0 | 0 | Canis | 89560.0 | 1 | Canis lupus | 89560.0 | 1 |
| scf7180000044280 | 339919 | 0.3675 | 0 | 60.7428 | Eukaryota | 27585.0 | 0 | Chordata | 27585.0 | 0 | Carnivora | 27585.0 | 0 | Mustelidae | 15372.0 | 2 | Mustela | 15372.0 | 3 | Mustela putorius | 15372.0 | 3 |
| scf7180000043957 | 171713 | 0.3622 | 0 | 57.5687 | Eukaryota | 7301.0 | 0 | Chordata | 6330.0 | 1 | Primates | 3483.0 | 4 | Hominidae | 3483.0 | 5 | Homo | 3483.0 | 5 | Homo sapiens | 3483.0 | 5 |
| scf7180000040692 | 1948621 | 0.378 | 0 | 60.2748 | Eukaryota | 93166.0 | 0 | Chordata | 93166.0 | 0 | Carnivora | 93166.0 | 0 | Canidae | 93166.0 | 0 | Canis | 93166.0 | 0 | Canis lupus | 93166.0 | 0 |
| scf7180000049426 | 4413 | 0.4333 | 0 | 102.4538 | Eukaryota | 222.0 | 0 | Euglenozoa | 222.0 | 0 | Kinetoplastida | 222.0 | 0 | Trypanosomatidae | 222.0 | 0 | Leishmania | 222.0 | 0 | Leishmania infantum | 222.0 | 0 |
| scf7180000053102 | 14504 | 0.5004 | 0 | 58.3714 | Eukaryota | 8713.0 | 0 | Chordata | 8713.0 | 0 | Carnivora | 8713.0 | 0 | Canidae | 8713.0 | 0 | Canis | 7055.0 | 1 | Canis lupus | 5841.0 | 2 |
| scf7180000041919 | 209561 | 0.4174 | 0 | 60.7609 | Eukaryota | 82411.0 | 0 | Chordata | 82411.0 | 0 | Carnivora | 82411.0 | 0 | Canidae | 82411.0 | 0 | Canis | 82411.0 | 0 | Canis lupus | 82411.0 | 0 |
| scf7180000040867 | 1593832 | 0.4267 | 0 | 59.5259 | Eukaryota | 159504.0 | 0 | Chordata | 159504.0 | 0 | Carnivora | 159504.0 | 0 | Canidae | 159504.0 | 0 | Canis | 134523.0 | 1 | Canis lupus | 134523.0 | 1 |
| scf7180000056312 | 665750 | 0.3403 | 0 | 59.2595 | Eukaryota | 42107.0 | 0 | Chordata | 42107.0 | 0 | Carnivora | 42107.0 | 0 | Canidae | 42107.0 | 0 | Canis | 42107.0 | 0 | Canis lupus | 42107.0 | 0 |
| scf7180000048476 | 5282 | 0.4665 | 0 | 132.8678 | Eukaryota | 46639.0 | 0 | Chordata | 46639.0 | 0 | Carnivora | 46639.0 | 0 | Canidae | 46639.0 | 0 | Canis | 46639.0 | 0 | Canis lupus | 46639.0 | 0 |
| scf7180000051897 | 11240 | 0.4459 | 0 | 203.9378 | Eukaryota | 40672.0 | 0 | Chordata | 29909.0 | 1 | Gadiformes | 24042.0 | 6 | Gadidae | 24042.0 | 6 | Gadus | 24042.0 | 6 | Gadus morhua | 24042.0 | 6 |
| scf7180000042440 | 639042 | 0.4277 | 0 | 61.2865 | Eukaryota | 99202.0 | 0 | Chordata | 99202.0 | 0 | Carnivora | 99202.0 | 0 | Canidae | 99202.0 | 0 | Canis | 89481.0 | 1 | Canis lupus | 89481.0 | 1 |
| scf7180000044799 | 10056 | 0.4149 | 0 | 43.9907 | Eukaryota | 51041.0 | 0 | Chordata | 51041.0 | 0 | Carnivora | 51041.0 | 0 | Canidae | 51041.0 | 0 | Canis | 51041.0 | 0 | Canis lupus | 51041.0 | 0 |
| scf7180000045831 | 29947 | 0.4382 | 0 | 109.0809 | Eukaryota | 50142.0 | 0 | Chordata | 50142.0 | 0 | Carnivora | 50142.0 | 0 | Canidae | 50142.0 | 0 | Canis | 50142.0 | 0 | Canis lupus | 50142.0 | 0 |
| scf7180000045139 | 43210 | 0.3468 | 0 | 51.0228 | Eukaryota | 4569.0 | 0 | Chordata | 4569.0 | 0 | Carnivora | 3826.0 | 1 | Canidae | 2130.0 | 2 | Canis | 2130.0 | 2 | Canis lupus | 2130.0 | 2 |
| scf7180000054794 | 22487 | 0.45 | 0 | 296.0389 | Eukaryota | 33083.0 | 0 | Chordata | 20044.0 | 2 | Strigeidida | 8294.0 | 8 | Schistosomatidae | 8294.0 | 8 | Schistosoma | 8294.0 | 8 | Schistosoma mansoni | 8294.0 | 8 |
| scf7180000052469 | 2920 | 0.4425 | 0 | 452.6189 | no-hit | 0.0 | 0 | no-hit | 0.0 | 0 | no-hit | 0.0 | 0 | no-hit | 0.0 | 0 | no-hit | 0.0 | 0 | no-hit | 0.0 | 0 |
| scf7180000044003 | 279469 | 0.3681 | 0 | 58.5411 | Eukaryota | 65746.0 | 0 | Chordata | 65746.0 | 0 | Carnivora | 65746.0 | 0 | Canidae | 65746.0 | 0 | Canis | 65746.0 | 0 | Canis lupus | 65746.0 | 0 |
| scf7180000046553 | 12114 | 0.4292 | 0 | 99.9822 | Eukaryota | 63556.0 | 0 | Chordata | 63556.0 | 0 | Carnivora | 63556.0 | 0 | Canidae | 63556.0 | 0 | Canis | 63556.0 | 0 | Canis lupus | 63556.0 | 0 |
| scf7180000042307 | 123967 | 0.3668 | 0 | 67.6117 | Eukaryota | 37222.0 | 0 | Chordata | 37222.0 | 0 | Carnivora | 37222.0 | 0 | Canidae | 37222.0 | 0 | Canis | 37222.0 | 0 | Canis lupus | 37222.0 | 0 |
| scf7180000054420 | 19697 | 0.3937 | 0 | 156.9076 | Eukaryota | 89996.0 | 0 | Chordata | 89996.0 | 0 | Carnivora | 89996.0 | 0 | Canidae | 89996.0 | 0 | Canis | 89996.0 | 0 | Canis lupus | 89996.0 | 0 |
| scf7180000051163 | 4077 | 0.4516 | 0 | 473.6517 | Eukaryota | 7021.0 | 0 | Chordata | 3883.0 | 3 | Chordata-undef | 2636.0 | 7 | Ambassidae | 2636.0 | 7 | Parambassis | 2636.0 | 7 | Parambassis ranga | 2636.0 | 7 |
| scf7180000046982 | 85156 | 0.3749 | 0 | 61.6274 | Eukaryota | 5924.0 | 1 | Chordata | 5924.0 | 1 | Primates | 3712.0 | 3 | Hominidae | 1925.0 | 4 | Homo | 1925.0 | 5 | Homo sapiens | 1925.0 | 6 |
| scf7180000048091 | 63394 | 0.5599 | 0 | 56.9414 | Eukaryota | 46086.0 | 0 | Chordata | 46086.0 | 0 | Carnivora | 46086.0 | 0 | Canidae | 38309.0 | 3 | Canis | 32853.0 | 4 | Canis lupus | 32853.0 | 4 |
| scf7180000049072 | 48718 | 0.4419 | 0 | 71.4534 | Eukaryota | 13724.0 | 0 | Chordata | 13724.0 | 0 | Carnivora | 13724.0 | 0 | Canidae | 13724.0 | 0 | Canis | 13724.0 | 0 | Canis lupus | 13724.0 | 0 |
| scf7180000041241 | 204396 | 0.3746 | 0 | 58.5699 | Eukaryota | 91442.0 | 0 | Chordata | 91442.0 | 0 | Carnivora | 91442.0 | 0 | Canidae | 91442.0 | 0 | Canis | 91442.0 | 0 | Canis lupus | 91442.0 | 0 |
| scf7180000043062 | 48147 | 0.3884 | 0 | 52.8792 | Eukaryota | 17150.0 | 0 | Chordata | 17150.0 | 0 | Carnivora | 17150.0 | 0 | Canidae | 17150.0 | 0 | Canis | 17150.0 | 0 | Canis lupus | 17150.0 | 0 |
| scf7180000042421 | 44985 | 0.4492 | 0 | 263.7032 | Eukaryota | 26482.0 | 1 | Chordata | 26482.0 | 1 | Carnivora | 9732.0 | 4 | Gadidae | 7923.0 | 5 | Gadus | 7923.0 | 6 | Gadus morhua | 7923.0 | 6 |
| scf7180000053807 | 3414 | 0.4455 | 0 | 874.4819 | Eukaryota | 1007.0 | 0 | Echinodermata | 520.0 | 2 | Forcipulatida | 520.0 | 2 | Asteriidae | 520.0 | 2 | Asterias | 520.0 | 2 | Asterias rubens | 520.0 | 2 |
| scf7180000049380 | 37451 | 0.3866 | 0 | 58.8758 | Eukaryota | 42530.0 | 0 | Chordata | 42530.0 | 0 | Carnivora | 42530.0 | 0 | Canidae | 42530.0 | 0 | Canis | 42530.0 | 0 | Canis lupus | 42530.0 | 0 |
| scf7180000049336 | 6284 | 0.4465 | 0 | 109.5905 | Eukaryota | 825.0 | 0 | Platyhelminthes | 825.0 | 0 | Polystomatidea | 825.0 | 0 | Polystomatidae | 825.0 | 0 | Protopolystoma | 825.0 | 0 | Protopolystoma xenopodis | 825.0 | 0 |
| scf7180000042483 | 63327 | 0.4336 | 0 | 55.923 | Eukaryota | 11810.0 | 0 | Chordata | 11810.0 | 0 | Carnivora | 11810.0 | 0 | Canidae | 10376.0 | 1 | Canis | 9011.0 | 2 | Canis lupus | 9011.0 | 2 |
| scf7180000043613 | 282371 | 0.4417 | 0 | 58.3583 | Eukaryota | 15580.0 | 0 | Chordata | 15580.0 | 0 | Carnivora | 15580.0 | 0 | Canidae | 15580.0 | 0 | Canis | 15580.0 | 0 | Canis lupus | 15580.0 | 0 |
| scf7180000042850 | 570334 | 0.4713 | 0 | 59.746 | Eukaryota | 47567.0 | 0 | Chordata | 47567.0 | 0 | Carnivora | 47567.0 | 0 | Canidae | 47567.0 | 0 | Canis | 36516.0 | 1 | Canis lupus | 36516.0 | 1 |
| scf7180000045624 | 19188 | 0.3686 | 0 | 57.989 | Eukaryota | 20723.0 | 0 | Chordata | 20723.0 | 0 | Carnivora | 20723.0 | 0 | Canidae | 20723.0 | 0 | Canis | 20723.0 | 0 | Canis lupus | 20723.0 | 0 |
| scf7180000043101 | 109864 | 0.4114 | 0 | 60.5862 | Eukaryota | 15025.0 | 0 | Chordata | 15025.0 | 0 | Carnivora | 15025.0 | 0 | Otariidae | 4470.0 | 5 | Odobenus | 2991.0 | 7 | Odobenus rosmarus | 2991.0 | 7 |
| scf7180000055153 | 63966 | 0.4369 | 0 | 211.7592 | Eukaryota | 3675.0 | 0 | Chordata | 2318.0 | 1 | Carnivora | 1933.0 | 2 | Canidae | 1933.0 | 2 | Canis | 1933.0 | 2 | Canis lupus | 1933.0 | 2 |
| scf7180000048240 | 22226 | 0.3941 | 0 | 55.6473 | Eukaryota | 87147.0 | 0 | Chordata | 87147.0 | 0 | Carnivora | 87147.0 | 0 | Canidae | 87147.0 | 0 | Canis | 87147.0 | 0 | Canis lupus | 87147.0 | 0 |
| scf7180000049562 | 45326 | 0.4981 | 0 | 47.4726 | Eukaryota | 34996.0 | 0 | Chordata | 34996.0 | 0 | Carnivora | 34996.0 | 0 | Canidae | 29309.0 | 1 | Canis | 18845.0 | 3 | Canis lupus | 18845.0 | 3 |
| scf7180000056294 | 774296 | 0.427 | 0 | 60.2149 | Eukaryota | 58574.0 | 0 | Chordata | 58574.0 | 0 | Carnivora | 58574.0 | 0 | Canidae | 53174.0 | 1 | Vulpes | 28547.0 | 2 | Vulpes vulpes | 28547.0 | 2 |
| scf7180000048120 | 4194 | 0.4204 | 0 | 29.5025 | Eukaryota | 19579.0 | 0 | Chordata | 19579.0 | 0 | Carnivora | 19579.0 | 0 | Canidae | 19579.0 | 0 | Canis | 19579.0 | 0 | Canis lupus | 19579.0 | 0 |
| scf7180000043405 | 89037 | 0.3469 | 0 | 59.5155 | Eukaryota | 24613.0 | 0 | Chordata | 24613.0 | 0 | Carnivora | 24613.0 | 0 | Canidae | 24613.0 | 0 | Canis | 24613.0 | 0 | Canis lupus | 24613.0 | 0 |
| scf7180000046806 | 60323 | 0.4526 | 0 | 1394.7565 | Eukaryota | 4960.0 | 0 | Platyhelminthes | 2282.0 | 3 | Polystomatidea | 2282.0 | 5 | Polystomatidae | 2282.0 | 5 | Protopolystoma | 2282.0 | 5 | Protopolystoma xenopodis | 2282.0 | 5 |
| scf7180000056280 | 245031 | 0.3771 | 0 | 56.4787 | Eukaryota | 49490.0 | 0 | Chordata | 49490.0 | 0 | Carnivora | 49490.0 | 0 | Canidae | 49490.0 | 0 | Canis | 49490.0 | 0 | Canis lupus | 49490.0 | 0 |
| scf7180000055850 | 16887 | 0.4416 | 0 | 469.7469 | Eukaryota | 3850.0 | 0 | Chordata | 1992.0 | 1 | Polystomatidea | 1858.0 | 2 | Polystomatidae | 1858.0 | 2 | Protopolystoma | 1858.0 | 2 | Protopolystoma xenopodis | 1858.0 | 2 |
| scf7180000042907 | 578632 | 0.444 | 0 | 59.6944 | Eukaryota | 27038.0 | 0 | Chordata | 27038.0 | 0 | Carnivora | 27038.0 | 0 | Canidae | 21456.0 | 1 | Canis | 21456.0 | 1 | Canis lupus | 21456.0 | 2 |
| scf7180000053070 | 4375 | 0.4487 | 0 | 316.6333 | no-hit | 0.0 | 0 | no-hit | 0.0 | 0 | no-hit | 0.0 | 0 | no-hit | 0.0 | 0 | no-hit | 0.0 | 0 | no-hit | 0.0 | 0 |
| scf7180000049322 | 7137 | 0.4401 | 0 | 322.8205 | no-hit | 0.0 | 0 | no-hit | 0.0 | 0 | no-hit | 0.0 | 0 | no-hit | 0.0 | 0 | no-hit | 0.0 | 0 | no-hit | 0.0 | 0 |
| scf7180000053743 | 6084 | 0.4461 | 0 | 194.3081 | Eukaryota | 936.0 | 0 | Chordata | 603.0 | 1 | Kurtiformes | 603.0 | 1 | Apogonidae | 603.0 | 1 | Sphaeramia | 603.0 | 1 | Sphaeramia orbicularis | 603.0 | 1 |
| scf7180000051865 | 4320 | 0.4576 | 0 | 143.903 | Eukaryota | 3800.0 | 0 | Chordata | 3800.0 | 0 | Carnivora | 3800.0 | 0 | Canidae | 3800.0 | 0 | Canis | 2893.0 | 1 | Canis lupus | 2893.0 | 1 |
| scf7180000050057 | 20409 | 0.3859 | 0 | 58.4722 | Eukaryota | 53335.0 | 0 | Chordata | 53335.0 | 0 | Carnivora | 53335.0 | 0 | Canidae | 53335.0 | 0 | Canis | 53335.0 | 0 | Canis lupus | 53335.0 | 0 |
| scf7180000048293 | 78644 | 0.5047 | 0 | 56.7974 | Eukaryota | 19754.0 | 0 | Chordata | 19754.0 | 0 | Carnivora | 19754.0 | 0 | Canidae | 8162.0 | 4 | Canis | 6520.0 | 8 | Canis lupus | 6520.0 | 8 |
| scf7180000051523 | 3463 | 0.4505 | 0 | 587.3127 | Eukaryota | 789.0 | 0 | Chordata | 580.0 | 1 | Kurtiformes | 580.0 | 1 | Apogonidae | 580.0 | 1 | Sphaeramia | 580.0 | 1 | Sphaeramia orbicularis | 580.0 | 1 |
| scf7180000041456 | 520539 | 0.4927 | 0 | 58.9501 | Eukaryota | 54175.0 | 0 | Chordata | 54175.0 | 0 | Carnivora | 54175.0 | 0 | Canidae | 54175.0 | 0 | Canis | 54175.0 | 0 | Canis lupus | 54175.0 | 0 |
| scf7180000055327 | 6112 | 0.4483 | 0 | 911.3803 | no-hit | 0.0 | 0 | no-hit | 0.0 | 0 | no-hit | 0.0 | 0 | no-hit | 0.0 | 0 | no-hit | 0.0 | 0 | no-hit | 0.0 | 0 |
| scf7180000041205 | 79987 | 0.5333 | 0 | 55.0207 | Eukaryota | 60809.0 | 0 | Chordata | 60809.0 | 0 | Carnivora | 60809.0 | 0 | Canidae | 60809.0 | 0 | Canis | 42581.0 | 1 | Canis lupus | 42581.0 | 1 |
| scf7180000044119 | 3239 | 0.4183 | 0 | 24.182 | Eukaryota | 25806.0 | 0 | Chordata | 25806.0 | 0 | Carnivora | 25806.0 | 0 | Canidae | 25806.0 | 0 | Canis | 25806.0 | 0 | Canis lupus | 25806.0 | 0 |
| scf7180000052827 | 20833 | 0.3467 | 0 | 48.6511 | Eukaryota | 52310.0 | 0 | Chordata | 52310.0 | 0 | Carnivora | 52310.0 | 0 | Canidae | 52310.0 | 0 | Canis | 52310.0 | 0 | Canis lupus | 52310.0 | 0 |
| scf7180000052990 | 2431 | 0.4463 | 0 | 473.6053 | no-hit | 0.0 | 0 | no-hit | 0.0 | 0 | no-hit | 0.0 | 0 | no-hit | 0.0 | 0 | no-hit | 0.0 | 0 | no-hit | 0.0 | 0 |
| scf7180000042571 | 653002 | 0.3972 | 0 | 59.638 | Eukaryota | 61648.0 | 0 | Chordata | 61648.0 | 0 | Carnivora | 61648.0 | 0 | Canidae | 61648.0 | 0 | Canis | 55781.0 | 1 | Canis lupus | 55781.0 | 1 |
| scf7180000052865 | 3669 | 0.4434 | 0 | 288.9065 | no-hit | 0.0 | 0 | no-hit | 0.0 | 0 | no-hit | 0.0 | 0 | no-hit | 0.0 | 0 | no-hit | 0.0 | 0 | no-hit | 0.0 | 0 |
| scf7180000042807 | 393823 | 0.3624 | 0 | 58.3804 | Eukaryota | 90410.0 | 0 | Chordata | 90410.0 | 0 | Carnivora | 90410.0 | 0 | Canidae | 90410.0 | 0 | Canis | 90410.0 | 0 | Canis lupus | 90410.0 | 0 |
| scf7180000045641 | 189049 | 0.4125 | 0 | 61.3219 | Eukaryota | 49536.0 | 0 | Chordata | 49536.0 | 0 | Carnivora | 49536.0 | 0 | Canidae | 45286.0 | 1 | Canis | 40268.0 | 2 | Canis lupus | 40268.0 | 2 |
| scf7180000053519 | 9094 | 0.5049 | 0 | 125.8169 | Eukaryota | 8390.0 | 0 | Chordata | 8390.0 | 0 | Carnivora | 8390.0 | 0 | Canidae | 8390.0 | 0 | Canis | 6144.0 | 1 | Canis lupus | 4900.0 | 2 |
| scf7180000054945 | 16020 | 0.4504 | 0 | 952.3784 | Eukaryota | 45834.0 | 0 | Chordata | 45834.0 | 0 | Carnivora | 45834.0 | 0 | Canidae | 45834.0 | 0 | Canis | 45834.0 | 0 | Canis lupus | 45834.0 | 0 |
| scf7180000050744 | 36650 | 0.3806 | 0 | 55.3806 | Eukaryota | 26922.0 | 0 | Chordata | 26922.0 | 0 | Carnivora | 26922.0 | 0 | Canidae | 25413.0 | 1 | Canis | 20387.0 | 2 | Canis lupus | 20387.0 | 2 |
| scf7180000053060 | 4523 | 0.4526 | 0 | 528.3179 | Eukaryota | 2941.0 | 0 | Chordata | 2253.0 | 2 | Synbranchiformes | 1103.0 | 4 | Mastacembelidae | 1103.0 | 4 | Mastacembelus | 1103.0 | 4 | Mastacembelus armatus | 1103.0 | 4 |
| scf7180000045659 | 4377 | 0.4531 | 0 | 446.657 | Eukaryota | 2535.0 | 0 | Chordata | 2535.0 | 0 | Carnivora | 2535.0 | 0 | Canidae | 2535.0 | 0 | Canis | 2535.0 | 0 | Canis lupus | 2535.0 | 0 |
| scf7180000052141 | 12231 | 0.4412 | 0 | 336.7346 | Eukaryota | 2382.0 | 0 | Platyhelminthes | 1253.0 | 2 | Polystomatidea | 1253.0 | 2 | Polystomatidae | 1253.0 | 2 | Protopolystoma | 1253.0 | 2 | Protopolystoma xenopodis | 1253.0 | 2 |
| scf7180000046791 | 4828 | 0.4377 | 0 | 79.6608 | no-hit | 0.0 | 0 | no-hit | 0.0 | 0 | no-hit | 0.0 | 0 | no-hit | 0.0 | 0 | no-hit | 0.0 | 0 | no-hit | 0.0 | 0 |
| scf7180000050571 | 8662 | 0.4367 | 0 | 56.9283 | no-hit | 0.0 | 0 | no-hit | 0.0 | 0 | no-hit | 0.0 | 0 | no-hit | 0.0 | 0 | no-hit | 0.0 | 0 | no-hit | 0.0 | 0 |
| scf7180000041154 | 181421 | 0.3763 | 0 | 59.4574 | Eukaryota | 55984.0 | 0 | Chordata | 55984.0 | 0 | Carnivora | 55984.0 | 0 | Canidae | 55984.0 | 0 | Canis | 55984.0 | 0 | Canis lupus | 55984.0 | 0 |
| scf7180000047847 | 61508 | 0.3707 | 0 | 60.8069 | Eukaryota | 8974.0 | 0 | Chordata | 8974.0 | 0 | Carnivora | 8974.0 | 0 | Canidae | 8974.0 | 0 | Canis | 8974.0 | 0 | Canis lupus | 8974.0 | 0 |
| scf7180000041486 | 628800 | 0.3381 | 0 | 59.3144 | Eukaryota | 33712.0 | 0 | Chordata | 33712.0 | 0 | Carnivora | 27534.0 | 1 | Canidae | 12171.0 | 4 | Mustela | 12018.0 | 5 | Mustela putorius | 12018.0 | 5 |
| scf7180000043961 | 359900 | 0.3893 | 0 | 59.3285 | Eukaryota | 94236.0 | 0 | Chordata | 94236.0 | 0 | Carnivora | 94236.0 | 0 | Canidae | 94236.0 | 0 | Canis | 94236.0 | 0 | Canis lupus | 94236.0 | 0 |
| scf7180000048772 | 3983 | 0.4487 | 0 | 84.4092 | no-hit | 0.0 | 0 | no-hit | 0.0 | 0 | no-hit | 0.0 | 0 | no-hit | 0.0 | 0 | no-hit | 0.0 | 0 | no-hit | 0.0 | 0 |
| scf7180000041220 | 49245 | 0.376 | 0 | 56.3396 | Eukaryota | 8660.0 | 0 | Chordata | 8660.0 | 0 | Carnivora | 6663.0 | 1 | Canidae | 6663.0 | 1 | Canis | 6663.0 | 1 | Canis lupus | 6663.0 | 1 |
| scf7180000051407 | 17921 | 0.5194 | 0 | 1198.6494 | Eukaryota | 33407.0 | 0 | Chordata | 33407.0 | 0 | Carnivora | 33407.0 | 0 | Canidae | 33407.0 | 0 | Canis | 33407.0 | 0 | Canis lupus | 33407.0 | 0 |
| scf7180000054251 | 25251 | 0.3761 | 0 | 115.9084 | Eukaryota | 20640.0 | 0 | Chordata | 20640.0 | 0 | Carnivora | 20640.0 | 0 | Canidae | 20640.0 | 0 | Canis | 20640.0 | 0 | Canis lupus | 20640.0 | 0 |
| scf7180000043200 | 169216 | 0.3782 | 0 | 63.459 | Eukaryota | 93633.0 | 0 | Chordata | 93633.0 | 0 | Carnivora | 93633.0 | 0 | Canidae | 93633.0 | 0 | Canis | 93633.0 | 0 | Canis lupus | 93633.0 | 0 |
| scf7180000052248 | 8807 | 0.4402 | 0 | 204.6111 | Eukaryota | 2568.0 | 0 | Platyhelminthes | 1626.0 | 2 | Polystomatidea | 1626.0 | 2 | Polystomatidae | 1626.0 | 2 | Protopolystoma | 1626.0 | 2 | Protopolystoma xenopodis | 1626.0 | 2 |
| scf7180000045164 | 228686 | 0.3451 | 0 | 59.8299 | Eukaryota | 63371.0 | 0 | Chordata | 63371.0 | 0 | Carnivora | 63371.0 | 0 | Canidae | 63371.0 | 0 | Canis | 35000.0 | 1 | Canis lupus | 35000.0 | 1 |
| scf7180000051225 | 12276 | 0.4437 | 0 | 60.5925 | Eukaryota | 3364.0 | 0 | Platyhelminthes | 1893.0 | 3 | Polystomatidea | 1893.0 | 4 | Polystomatidae | 1893.0 | 4 | Protopolystoma | 1893.0 | 4 | Protopolystoma xenopodis | 1893.0 | 4 |
| scf7180000048719 | 14715 | 0.4353 | 0 | 77.5526 | Eukaryota | 2710.0 | 0 | Chordata | 2710.0 | 0 | Carnivora | 2222.0 | 1 | Canidae | 2222.0 | 1 | Canis | 2222.0 | 1 | Canis lupus | 2222.0 | 1 |
| scf7180000040532 | 2499898 | 0.4429 | 0 | 59.7888 | Eukaryota | 91439.0 | 0 | Chordata | 91439.0 | 0 | Carnivora | 91439.0 | 0 | Canidae | 91439.0 | 0 | Vulpes | 61797.0 | 1 | Vulpes vulpes | 61797.0 | 1 |
| scf7180000044963 | 186571 | 0.3795 | 0 | 55.0016 | Eukaryota | 7971.0 | 0 | Chordata | 7971.0 | 0 | Carnivora | 3995.0 | 2 | Canidae | 3995.0 | 2 | Canis | 3326.0 | 4 | Canis lupus | 3326.0 | 4 |
| scf7180000051801 | 2476 | 0.4451 | 0 | 135.8893 | no-hit | 0.0 | 0 | no-hit | 0.0 | 0 | no-hit | 0.0 | 0 | no-hit | 0.0 | 0 | no-hit | 0.0 | 0 | no-hit | 0.0 | 0 |
| scf7180000050619 | 6091 | 0.4561 | 0 | 504.5222 | Eukaryota | 2930.0 | 0 | Platyhelminthes | 1672.0 | 2 | Polystomatidea | 1672.0 | 2 | Polystomatidae | 1672.0 | 2 | Protopolystoma | 1672.0 | 2 | Protopolystoma xenopodis | 1672.0 | 2 |
| scf7180000040807 | 1658909 | 0.3974 | 0 | 59.7663 | Eukaryota | 104586.0 | 0 | Chordata | 104586.0 | 0 | Carnivora | 104586.0 | 0 | Canidae | 104586.0 | 0 | Canis | 91458.0 | 1 | Canis lupus | 91458.0 | 1 |
| scf7180000048489 | 29321 | 0.4994 | 0 | 162.2668 | Eukaryota | 8112.0 | 0 | Chordata | 8112.0 | 0 | Carnivora | 8112.0 | 0 | Canidae | 8112.0 | 0 | Canis | 6701.0 | 1 | Canis lupus | 5799.0 | 2 |
| scf7180000044775 | 306925 | 0.3392 | 0 | 58.9648 | Eukaryota | 71555.0 | 0 | Chordata | 71555.0 | 0 | Carnivora | 71555.0 | 0 | Canidae | 71555.0 | 0 | Canis | 71555.0 | 0 | Canis lupus | 71555.0 | 0 |
| scf7180000051402 | 1226 | 0.4494 | 0 | 288.2349 | no-hit | 0.0 | 0 | no-hit | 0.0 | 0 | no-hit | 0.0 | 0 | no-hit | 0.0 | 0 | no-hit | 0.0 | 0 | no-hit | 0.0 | 0 |
| scf7180000045954 | 159389 | 0.4005 | 0 | 62.1456 | Eukaryota | 59893.0 | 0 | Chordata | 59893.0 | 0 | Carnivora | 46892.0 | 2 | Canidae | 46892.0 | 2 | Canis | 46892.0 | 3 | Canis lupus | 46892.0 | 3 |
| scf7180000053356 | 2990 | 0.4522 | 0 | 497.3297 | Eukaryota | 1895.0 | 0 | Echinodermata | 841.0 | 2 | Forcipulatida | 841.0 | 2 | Asteriidae | 841.0 | 2 | Asterias | 841.0 | 2 | Asterias rubens | 841.0 | 2 |
| scf7180000050420 | 24333 | 0.3702 | 0 | 63.8274 | Eukaryota | 19641.0 | 0 | Chordata | 19641.0 | 0 | Carnivora | 19641.0 | 0 | Canidae | 8160.0 | 3 | Canis | 5312.0 | 6 | Canis lupus | 5312.0 | 8 |
| scf7180000049943 | 40852 | 0.4356 | 0 | 59.1019 | Eukaryota | 22177.0 | 0 | Chordata | 22177.0 | 0 | Carnivora | 22177.0 | 0 | Canidae | 22177.0 | 0 | Canis | 22177.0 | 0 | Canis lupus | 22177.0 | 0 |
| scf7180000050824 | 13757 | 0.4537 | 0 | 158.4725 | Eukaryota | 1120.0 | 0 | Platyhelminthes | 1120.0 | 0 | Polystomatidea | 1120.0 | 0 | Polystomatidae | 1120.0 | 0 | Protopolystoma | 1120.0 | 0 | Protopolystoma xenopodis | 1120.0 | 0 |
| scf7180000047848 | 34190 | 0.4465 | 0 | 164.6551 | Eukaryota | 14945.0 | 0 | Chordata | 9506.0 | 3 | Gadiformes | 4512.0 | 6 | Gadidae | 4512.0 | 6 | Gadus | 4512.0 | 6 | Gadus morhua | 4512.0 | 7 |
| scf7180000043680 | 369384 | 0.4216 | 0 | 59.1715 | Eukaryota | 31046.0 | 0 | Chordata | 31046.0 | 0 | Carnivora | 31046.0 | 0 | Canidae | 31046.0 | 0 | Canis | 22676.0 | 1 | Canis lupus | 22676.0 | 1 |
| scf7180000040646 | 1870572 | 0.4032 | 0 | 59.6342 | Eukaryota | 91980.0 | 0 | Chordata | 91980.0 | 0 | Carnivora | 91980.0 | 0 | Canidae | 91980.0 | 0 | Canis | 91980.0 | 0 | Canis lupus | 91980.0 | 0 |
| scf7180000054116 | 4507 | 0.4418 | 0 | 80.6578 | Eukaryota | 625.0 | 0 | Platyhelminthes | 475.0 | 1 | Polystomatidea | 475.0 | 1 | Polystomatidae | 475.0 | 1 | Protopolystoma | 475.0 | 1 | Protopolystoma xenopodis | 475.0 | 1 |
| scf7180000047450 | 62818 | 0.3553 | 0 | 65.1912 | Eukaryota | 82427.0 | 0 | Chordata | 82427.0 | 0 | Carnivora | 82427.0 | 0 | Canidae | 82427.0 | 0 | Canis | 82427.0 | 0 | Canis lupus | 82427.0 | 0 |
| scf7180000049384 | 14570 | 0.4351 | 0 | 191.1718 | Eukaryota | 15933.0 | 0 | Platyhelminthes | 11074.0 | 1 | Cyclophyllidea | 8911.0 | 4 | Hymenolepididae | 8911.0 | 4 | Hymenolepis | 8911.0 | 4 | Hymenolepis microstoma | 8911.0 | 4 |
| scf7180000044636 | 790387 | 0.5242 | 0 | 58.7354 | Eukaryota | 85870.0 | 0 | Chordata | 85870.0 | 0 | Carnivora | 85870.0 | 0 | Canidae | 85870.0 | 0 | Canis | 85870.0 | 0 | Canis lupus | 85870.0 | 0 |
| scf7180000052078 | 16334 | 0.502 | 0 | 986.093 | Eukaryota | 9469.0 | 0 | Chordata | 9469.0 | 0 | Carnivora | 9469.0 | 0 | Canidae | 9469.0 | 0 | Canis | 7661.0 | 1 | Canis lupus | 6380.0 | 2 |
| scf7180000055553 | 24332 | 0.4368 | 0 | 101.3943 | Eukaryota | 78248.0 | 0 | Chordata | 78248.0 | 0 | Carnivora | 78248.0 | 0 | Canidae | 78248.0 | 0 | Canis | 78248.0 | 0 | Canis lupus | 78248.0 | 0 |
| scf7180000040590 | 2371627 | 0.3908 | 0 | 59.6536 | Eukaryota | 89661.0 | 0 | Chordata | 89661.0 | 0 | Carnivora | 89661.0 | 0 | Canidae | 89661.0 | 0 | Canis | 89661.0 | 0 | Canis lupus | 89661.0 | 0 |
| scf7180000044487 | 383281 | 0.3725 | 0 | 100.914 | Eukaryota | 128967.0 | 0 | Chordata | 128967.0 | 0 | Carnivora | 128967.0 | 0 | Canidae | 128967.0 | 0 | Canis | 128967.0 | 0 | Canis lupus | 128967.0 | 0 |
| scf7180000050089 | 4122 | 0.4262 | 0 | 181.2435 | no-hit | 0.0 | 0 | no-hit | 0.0 | 0 | no-hit | 0.0 | 0 | no-hit | 0.0 | 0 | no-hit | 0.0 | 0 | no-hit | 0.0 | 0 |
| scf7180000048704 | 27916 | 0.5024 | 0 | 699.4873 | Eukaryota | 9600.0 | 0 | Chordata | 9600.0 | 0 | Carnivora | 9600.0 | 0 | Canidae | 9600.0 | 0 | Canis | 8358.0 | 1 | Canis lupus | 7058.0 | 2 |
| scf7180000056242 | 78789 | 0.4319 | 0 | 54.9143 | Eukaryota | 18640.0 | 0 | Chordata | 18640.0 | 0 | Carnivora | 18640.0 | 0 | Canidae | 18640.0 | 0 | Canis | 11208.0 | 1 | Canis lupus | 11208.0 | 1 |
| scf7180000054438 | 16514 | 0.4433 | 0 | 304.1891 | Eukaryota | 2579.0 | 0 | Platyhelminthes | 1422.0 | 2 | Polystomatidea | 1422.0 | 4 | Polystomatidae | 1422.0 | 4 | Protopolystoma | 1422.0 | 4 | Protopolystoma xenopodis | 1422.0 | 4 |
| scf7180000051985 | 4474 | 0.4535 | 0 | 635.3646 | Eukaryota | 1131.0 | 0 | Echinodermata | 586.0 | 2 | Forcipulatida | 586.0 | 2 | Asteriidae | 586.0 | 2 | Asterias | 586.0 | 2 | Asterias rubens | 586.0 | 2 |
| scf7180000042748 | 3968 | 0.5502 | 0 | 26.4612 | Eukaryota | 4745.0 | 0 | Chordata | 4745.0 | 0 | Carnivora | 4745.0 | 0 | Canidae | 4745.0 | 0 | Canis | 4745.0 | 0 | Canis lupus | 4745.0 | 0 |
| scf7180000051965 | 10473 | 0.3916 | 0 | 55.8054 | Eukaryota | 29073.0 | 0 | Chordata | 29073.0 | 0 | Carnivora | 29073.0 | 0 | Canidae | 29073.0 | 0 | Canis | 29073.0 | 0 | Canis lupus | 29073.0 | 0 |
| scf7180000048500 | 63259 | 0.4645 | 0 | 56.6948 | Eukaryota | 13947.0 | 0 | Chordata | 13947.0 | 0 | Carnivora | 12622.0 | 1 | Canidae | 9261.0 | 3 | Canis | 6608.0 | 4 | Canis lupus | 6608.0 | 4 |
| scf7180000045654 | 22618 | 0.4447 | 0 | 218.7025 | Eukaryota | 1595.0 | 0 | Chordata | 1321.0 | 1 | Tetraodontiformes | 789.0 | 2 | Tetraodontidae | 789.0 | 2 | Takifugu | 789.0 | 2 | Takifugu rubripes | 789.0 | 2 |
| scf7180000048114 | 7988 | 0.4414 | 0 | 181.1883 | Eukaryota | 6578.0 | 0 | Chordata | 6578.0 | 0 | Tetraodontiformes | 4233.0 | 3 | Tetraodontidae | 4233.0 | 3 | Takifugu | 4233.0 | 3 | Takifugu rubripes | 4233.0 | 3 |
| scf7180000053228 | 8696 | 0.4419 | 0 | 53.0101 | Eukaryota | 17846.0 | 1 | Chordata | 17846.0 | 1 | Kurtiformes | 5161.0 | 6 | Apogonidae | 5161.0 | 6 | Sphaeramia | 5161.0 | 6 | Sphaeramia orbicularis | 5161.0 | 6 |
| scf7180000052810 | 6686 | 0.4471 | 0 | 423.6618 | Eukaryota | 2674.0 | 0 | Chordata | 1929.0 | 2 | Kurtiformes | 1794.0 | 3 | Apogonidae | 1794.0 | 3 | Sphaeramia | 1794.0 | 3 | Sphaeramia orbicularis | 1794.0 | 3 |
| scf7180000050488 | 17710 | 0.4422 | 0 | 117.0834 | no-hit | 0.0 | 0 | no-hit | 0.0 | 0 | no-hit | 0.0 | 0 | no-hit | 0.0 | 0 | no-hit | 0.0 | 0 | no-hit | 0.0 | 0 |
| scf7180000054101 | 23718 | 0.4476 | 0 | 78.0171 | Eukaryota | 28454.0 | 0 | Chordata | 28454.0 | 0 | Carnivora | 28454.0 | 0 | Canidae | 28454.0 | 0 | Canis | 28454.0 | 0 | Canis lupus | 28454.0 | 0 |
| scf7180000041112 | 606910 | 0.3521 | 0 | 58.7357 | Eukaryota | 89836.0 | 0 | Chordata | 89836.0 | 0 | Carnivora | 89836.0 | 0 | Canidae | 89836.0 | 0 | Canis | 89836.0 | 0 | Canis lupus | 89836.0 | 0 |
| scf7180000052666 | 10252 | 0.5011 | 0 | 51.3608 | Eukaryota | 8769.0 | 0 | Chordata | 8769.0 | 0 | Carnivora | 8769.0 | 0 | Canidae | 8769.0 | 0 | Canis | 7061.0 | 1 | Canis lupus | 5908.0 | 2 |
| scf7180000042635 | 25151 | 0.4354 | 0 | 206.459 | Eukaryota | 342.0 | 0 | Platyhelminthes | 342.0 | 0 | Polystomatidea | 342.0 | 0 | Polystomatidae | 342.0 | 0 | Protopolystoma | 342.0 | 0 | Protopolystoma xenopodis | 342.0 | 0 |
| scf7180000047074 | 857443 | 0.4247 | 100 | 64.3152 | Eukaryota | 54277.0 | 0 | Chordata | 54277.0 | 0 | Carnivora | 54277.0 | 0 | Canidae | 54277.0 | 0 | Canis | 43193.0 | 1 | Canis lupus | 43193.0 | 1 |
| scf7180000048676 | 5398 | 0.4437 | 0 | 84.1924 | no-hit | 0.0 | 0 | no-hit | 0.0 | 0 | no-hit | 0.0 | 0 | no-hit | 0.0 | 0 | no-hit | 0.0 | 0 | no-hit | 0.0 | 0 |
| scf7180000054889 | 11703 | 0.4461 | 0 | 140.8819 | Eukaryota | 392.0 | 0 | Chordata | 392.0 | 0 | Kurtiformes | 392.0 | 0 | Apogonidae | 392.0 | 0 | Sphaeramia | 392.0 | 0 | Sphaeramia orbicularis | 392.0 | 0 |
| scf7180000052332 | 22068 | 0.4461 | 0 | 59.3356 | Eukaryota | 56978.0 | 0 | Chordata | 56978.0 | 0 | Carnivora | 56978.0 | 0 | Canidae | 49054.0 | 3 | Canis | 47310.0 | 5 | Canis lupus | 47310.0 | 6 |
| scf7180000047229 | 20183 | 0.4491 | 0 | 156.1205 | Eukaryota | 1148.0 | 0 | Platyhelminthes | 952.0 | 1 | Polystomatidea | 952.0 | 1 | Polystomatidae | 952.0 | 1 | Protopolystoma | 952.0 | 1 | Protopolystoma xenopodis | 952.0 | 1 |
| scf7180000041631 | 568742 | 0.3915 | 0 | 59.8169 | Eukaryota | 105266.0 | 0 | Chordata | 105266.0 | 0 | Carnivora | 105266.0 | 0 | Canidae | 105266.0 | 0 | Canis | 105266.0 | 0 | Canis lupus | 105266.0 | 0 |

| | | | | | | | | | | | | | | | | | | | | | | |
|---|---|---|---|---|---|---|---|---|---|---|---|---|---|---|---|---|---|---|---|---|---|---|
| scf7180000041375 | 50690 | 0.3879 | 0 | 52.9599 | Eukaryota | 11566.0 | 0 | Chordata | 11566.0 | 0 | Carnivora | 11566.0 | 0 | Canidae | 9252.0 | 1 | Canis | 9252.0 | 1 | Canis lupus | 9252.0 | 1 |
| scf7180000053293 | 17729 | 0.3458 | 0 | 47.8488 | Eukaryota | 3484.0 | 0 | Chordata | 3484.0 | 0 | Carnivora | 2493.0 | 1 | Canidae | 2493.0 | 1 | Canis | 2493.0 | 1 | Canis lupus | 2493.0 | 1 |
| scf7180000041258 | 1174207 | 0.4517 | 0 | 58.6911 | Eukaryota | 33606.0 | 0 | Chordata | 33606.0 | 0 | Carnivora | 33606.0 | 0 | Canidae | 33606.0 | 0 | Canis | 33606.0 | 0 | Canis lupus | 33606.0 | 0 |
| scf7180000045931 | 39037 | 0.4413 | 0 | 237.6771 | Eukaryota | 6453.0 | 0 | Chordata | 6453.0 | 0 | Kurtiformes | 3531.0 | 1 | Apogonidae | 3531.0 | 1 | Sphaeramia | 3531.0 | 1 | Sphaeramia orbicularis | 3531.0 | 1 |
| scf7180000050782 | 24664 | 0.5384 | 0 | 51.4865 | Eukaryota | 6603.0 | 0 | Chordata | 6603.0 | 0 | Carnivora | 4929.0 | 2 | Canidae | 4929.0 | 2 | Canis | 2767.0 | 3 | Canis lupus | 2767.0 | 3 |
| scf7180000046517 | 19504 | 0.4437 | 0 | 276.6217 | Eukaryota | 4471.0 | 0 | Chordata | 3291.0 | 2 | Kurtiformes | 3291.0 | 2 | Apogonidae | 3291.0 | 2 | Sphaeramia | 3291.0 | 2 | Sphaeramia orbicularis | 3291.0 | 2 |
| scf7180000041618 | 117692 | 0.3925 | 0 | 58.1517 | Eukaryota | 89921.0 | 0 | Chordata | 89921.0 | 0 | Carnivora | 89921.0 | 0 | Canidae | 89921.0 | 0 | Canis | 89921.0 | 0 | Canis lupus | 89921.0 | 0 |
| scf7180000045398 | 76906 | 0.4113 | 0 | 59.925 | Eukaryota | 98561.0 | 0 | Chordata | 98561.0 | 0 | Carnivora | 98561.0 | 0 | Canidae | 98561.0 | 0 | Canis | 98561.0 | 0 | Canis lupus | 98561.0 | 0 |
| scf7180000044096 | 43887 | 0.343 | 0 | 55.836 | Eukaryota | 30854.0 | 0 | Chordata | 30854.0 | 0 | Carnivora | 30854.0 | 0 | Canidae | 30854.0 | 0 | Canis | 30854.0 | 0 | Canis lupus | 30854.0 | 0 |
| scf7180000049803 | 31124 | 0.5028 | 0 | 218.0467 | Eukaryota | 8787.0 | 0 | Chordata | 8787.0 | 0 | Carnivora | 8787.0 | 0 | Canidae | 8787.0 | 0 | Canis | 6546.0 | 1 | Canis lupus | 5309.0 | 2 |
| scf7180000047366 | 76114 | 0.3697 | 0 | 61.4562 | Eukaryota | 49008.0 | 0 | Chordata | 49008.0 | 0 | Carnivora | 49008.0 | 0 | Canidae | 49008.0 | 0 | Canis | 41724.0 | 1 | Canis lupus | 41724.0 | 1 |
| scf7180000044920 | 11009 | 0.7167 | 0 | 38.9417 | Eukaryota | 67626.0 | 0 | Chordata | 67626.0 | 0 | Carnivora | 67626.0 | 0 | Canidae | 67626.0 | 0 | Canis | 67626.0 | 0 | Canis lupus | 67626.0 | 0 |
| scf7180000041692 | 932023 | 0.4264 | 0 | 60.8467 | Eukaryota | 103062.0 | 0 | Chordata | 103062.0 | 0 | Carnivora | 103062.0 | 0 | Canidae | 103062.0 | 0 | Canis | 67116.0 | 1 | Canis lupus | 67116.0 | 1 |
| scf7180000052146 | 4274 | 0.4443 | 0 | 107.4297 | no-hit | 0.0 | 0 | no-hit | 0.0 | 0 | no-hit | 0.0 | 0 | no-hit | 0.0 | 0 | no-hit | 0.0 | 0 | no-hit | 0.0 | 0 |
| scf7180000055624 | 28695 | 0.4442 | 0 | 165.9791 | Eukaryota | 38602.0 | 0 | Chordata | 27140.0 | 2 | Gadiformes | 20932.0 | 5 | Gadidae | 20932.0 | 5 | Gadus | 20932.0 | 5 | Gadus morhua | 20932.0 | 5 |
| scf7180000053454 | 8353 | 0.4214 | 0 | 58.4735 | Eukaryota | 3039.0 | 1 | Chordata | 3039.0 | 1 | Carnivora | 2504.0 | 2 | Canidae | 2504.0 | 2 | Canis | 1317.0 | 3 | Canis lupus | 1317.0 | 3 |
| scf7180000054425 | 39457 | 0.3712 | 0 | 848.2954 | Eukaryota | 93810.0 | 0 | Chordata | 93810.0 | 0 | Carnivora | 93810.0 | 0 | Canidae | 93810.0 | 0 | Canis | 93810.0 | 0 | Canis lupus | 93810.0 | 0 |
| scf7180000042769 | 24006 | 0.4419 | 0 | 58.8433 | Eukaryota | 42633.0 | 0 | Chordata | 42633.0 | 0 | Carnivora | 42633.0 | 0 | Canidae | 33928.0 | 3 | Canis | 29486.0 | 4 | Canis lupus | 29486.0 | 4 |
| scf7180000044046 | 173781 | 0.3732 | 0 | 63.8779 | Eukaryota | 117740.0 | 0 | Chordata | 117740.0 | 0 | Carnivora | 117740.0 | 0 | Canidae | 117740.0 | 0 | Canis | 117740.0 | 0 | Canis lupus | 117740.0 | 0 |
| scf7180000052918 | 11996 | 0.4382 | 0 | 144.3715 | Eukaryota | 2554.0 | 0 | Platyhelminthes | 1843.0 | 2 | Polystomatidea | 1843.0 | 2 | Polystomatidae | 1843.0 | 2 | Protopolystoma | 1843.0 | 2 | Protopolystoma xenopodis | 1843.0 | 2 |
| scf7180000041402 | 73352 | 0.3512 | 0 | 58.6441 | Eukaryota | 49317.0 | 0 | Chordata | 49317.0 | 0 | Carnivora | 49317.0 | 0 | Canidae | 49317.0 | 0 | Canis | 49317.0 | 0 | Canis lupus | 49317.0 | 0 |
| scf7180000055501 | 10748 | 0.4392 | 0 | 67.6935 | no-hit | 0.0 | 0 | no-hit | 0.0 | 0 | no-hit | 0.0 | 0 | no-hit | 0.0 | 0 | no-hit | 0.0 | 0 | no-hit | 0.0 | 0 |
| scf7180000056374 | 427341 | 0.3503 | 0 | 59.4967 | Eukaryota | 84740.0 | 0 | Chordata | 84740.0 | 0 | Carnivora | 84740.0 | 0 | Canidae | 84740.0 | 0 | Canis | 84740.0 | 0 | Canis lupus | 84740.0 | 0 |
| scf7180000047234 | 20259 | 0.3966 | 0 | 49.3552 | Eukaryota | 11837.0 | 0 | Chordata | 11837.0 | 0 | Carnivora | 11837.0 | 0 | Canidae | 11837.0 | 0 | Canis | 11837.0 | 0 | Canis lupus | 11837.0 | 0 |
| scf7180000047298 | 90476 | 0.432 | 0 | 60.0084 | Eukaryota | 26294.0 | 0 | Chordata | 26294.0 | 0 | Carnivora | 26294.0 | 0 | Canidae | 26294.0 | 0 | Canis | 26294.0 | 0 | Canis lupus | 26294.0 | 0 |
| scf7180000053532 | 2102 | 0.4577 | 0 | 174.1183 | no-hit | 0.0 | 0 | no-hit | 0.0 | 0 | no-hit | 0.0 | 0 | no-hit | 0.0 | 0 | no-hit | 0.0 | 0 | no-hit | 0.0 | 0 |
| scf7180000053723 | 10603 | 0.5143 | 0 | 1258.1294 | Eukaryota | 32536.0 | 0 | Chordata | 32536.0 | 0 | Carnivora | 32536.0 | 0 | Canidae | 32536.0 | 0 | Canis | 32536.0 | 0 | Canis lupus | 32536.0 | 0 |
| scf7180000054231 | 11217 | 0.5032 | 0 | 414.2911 | Eukaryota | 8456.0 | 0 | Chordata | 8456.0 | 0 | Carnivora | 8456.0 | 0 | Canidae | 8456.0 | 0 | Canis | 5370.0 | 1 | Canis lupus | 5370.0 | 1 |
| scf7180000047156 | 62165 | 0.3531 | 0 | 60.1664 | Eukaryota | 61157.0 | 0 | Chordata | 61157.0 | 0 | Carnivora | 61157.0 | 0 | Canidae | 61157.0 | 0 | Canis | 61157.0 | 0 | Canis lupus | 61157.0 | 0 |
| scf7180000053237 | 4632 | 0.5199 | 0 | 414.2871 | Eukaryota | 13281.0 | 0 | Chordata | 13281.0 | 0 | Carnivora | 13281.0 | 0 | Canidae | 13281.0 | 0 | Canis | 13281.0 | 0 | Canis lupus | 13281.0 | 0 |
| scf7180000046515 | 8033 | 0.4922 | 0 | 54.0812 | Eukaryota | 15093.0 | 0 | Chordata | 15093.0 | 0 | Carnivora | 15093.0 | 0 | Canidae | 15093.0 | 0 | Canis | 15093.0 | 0 | Canis lupus | 15093.0 | 0 |
| scf7180000049777 | 12945 | 0.4457 | 0 | 103.3635 | Eukaryota | 856.0 | 0 | Chordata | 856.0 | 0 | Tetraodontiformes | 856.0 | 0 | Tetraodontidae | 856.0 | 0 | Takifugu | 856.0 | 0 | Takifugu rubripes | 856.0 | 0 |
| scf7180000049413 | 38724 | 0.4161 | 0 | 60.2129 | Eukaryota | 4857.0 | 0 | Chordata | 4857.0 | 0 | Perissodactyla | 1721.0 | 4 | Equidae | 1275.0 | 6 | Equus | 1275.0 | 6 | Mustela putorius | 992.0 | 7 |
| scf7180000054221 | 924907 | 0.4282 | 0 | 61.8634 | Eukaryota | 84825.0 | 0 | Chordata | 84825.0 | 0 | Carnivora | 84825.0 | 0 | Canidae | 84825.0 | 0 | Canis | 57096.0 | 1 | Canis lupus | 57096.0 | 1 |
| scf7180000054620 | 6222 | 0.4447 | 0 | 198.0492 | no-hit | 0.0 | 0 | no-hit | 0.0 | 0 | no-hit | 0.0 | 0 | no-hit | 0.0 | 0 | no-hit | 0.0 | 0 | no-hit | 0.0 | 0 |
| scf7180000043592 | 444040 | 0.572 | 0 | 57.0564 | Eukaryota | 46205.0 | 0 | Chordata | 46205.0 | 0 | Carnivora | 46205.0 | 0 | Canidae | 46205.0 | 0 | Canis | 37010.0 | 1 | Canis lupus | 37010.0 | 1 |
| scf7180000051029 | 11050 | 0.4458 | 0 | 645.8116 | Eukaryota | 2600.0 | 0 | Platyhelminthes | 1040.0 | 3 | Forcipulatida | 933.0 | 4 | Asteriidae | 933.0 | 4 | Asterias | 933.0 | 4 | Asterias rubens | 933.0 | 4 |
| scf7180000045317 | 222222 | 0.3659 | 0 | 60.2219 | Eukaryota | 78796.0 | 0 | Chordata | 78796.0 | 0 | Carnivora | 78796.0 | 0 | Canidae | 78796.0 | 0 | Canis | 78796.0 | 0 | Canis lupus | 78796.0 | 0 |
| scf7180000047747 | 50748 | 0.4382 | 0 | 110.5214 | Eukaryota | 18543.0 | 0 | Platyhelminthes | 14741.0 | 1 | Cyclophyllidea | 12028.0 | 3 | Hymenolepididae | 12028.0 | 3 | Hymenolepis | 12028.0 | 4 | Hymenolepis microstoma | 12028.0 | 4 |
| scf7180000046110 | 18253 | 0.5014 | 0 | 52.6267 | Eukaryota | 3801.0 | 0 | Chordata | 3801.0 | 0 | Carnivora | 3801.0 | 0 | Canidae | 3801.0 | 0 | Canis | 3411.0 | 1 | Canis lupus | 3411.0 | 1 |
| scf7180000049672 | 3575 | 0.4389 | 0 | 607.6727 | no-hit | 0.0 | 0 | no-hit | 0.0 | 0 | no-hit | 0.0 | 0 | no-hit | 0.0 | 0 | no-hit | 0.0 | 0 | no-hit | 0.0 | 0 |
| scf7180000055323 | 14397 | 0.4429 | 0 | 385.4469 | Eukaryota | 58479.0 | 0 | Chordata | 58479.0 | 0 | Carnivora | 58479.0 | 0 | Canidae | 58479.0 | 0 | Canis | 58479.0 | 0 | Canis lupus | 58479.0 | 0 |
| scf7180000050096 | 25169 | 0.3956 | 0 | 57.9233 | Eukaryota | 32124.0 | 0 | Chordata | 32124.0 | 0 | Carnivora | 32124.0 | 0 | Canidae | 32124.0 | 0 | Canis | 32124.0 | 0 | Canis lupus | 32124.0 | 0 |
| scf7180000044112 | 71516 | 0.4494 | 0 | 57.5845 | Eukaryota | 61877.0 | 0 | Chordata | 61877.0 | 0 | Carnivora | 61877.0 | 0 | Canidae | 61877.0 | 0 | Canis | 61877.0 | 0 | Canis lupus | 61877.0 | 0 |
| scf7180000050927 | 5859 | 0.441 | 0 | 186.1365 | no-hit | 0.0 | 0 | no-hit | 0.0 | 0 | no-hit | 0.0 | 0 | no-hit | 0.0 | 0 | no-hit | 0.0 | 0 | no-hit | 0.0 | 0 |
| scf7180000055213 | 28900 | 0.4296 | 0 | 193.7994 | Eukaryota | 7944.0 | 0 | Chordata | 4214.0 | 2 | Kurtiformes | 3314.0 | 3 | Apogonidae | 3314.0 | 3 | Sphaeramia | 3314.0 | 3 | Sphaeramia orbicularis | 3314.0 | 3 |
| scf7180000050508 | 6369 | 0.4434 | 0 | 64.3293 | Eukaryota | 2406.0 | 0 | Platyhelminthes | 1912.0 | 1 | Polystomatidea | 1912.0 | 1 | Polystomatidae | 1912.0 | 1 | Protopolystoma | 1912.0 | 1 | Protopolystoma xenopodis | 1912.0 | 1 |
| scf7180000041759 | 202833 | 0.3803 | 0 | 57.9712 | Eukaryota | 91295.0 | 0 | Chordata | 91295.0 | 0 | Carnivora | 91295.0 | 0 | Canidae | 91295.0 | 0 | Canis | 91295.0 | 0 | Canis lupus | 91295.0 | 0 |
| scf7180000051959 | 11552 | 0.5052 | 0 | 118.6208 | Eukaryota | 8570.0 | 0 | Chordata | 8570.0 | 0 | Carnivora | 8570.0 | 0 | Canidae | 8570.0 | 0 | Nyctereutes | 4325.0 | 1 | Nyctereutes procyonoides | 4325.0 | 1 |
| scf7180000054755 | 60580 | 0.4436 | 0 | 200.681 | Eukaryota | 27085.0 | 1 | Chordata | 26488.0 | 2 | Gadiformes | 10619.0 | 6 | Gadidae | 10619.0 | 6 | Gadus | 10619.0 | 7 | Gadus morhua | 10619.0 | 7 |
| scf7180000044292 | 51270 | 0.4287 | 0 | 105.0972 | Eukaryota | 39787.0 | 0 | Chordata | 39787.0 | 0 | Carnivora | 39787.0 | 0 | Canidae | 39787.0 | 0 | Canis | 39787.0 | 0 | Canis lupus | 39787.0 | 0 |
| scf7180000053738 | 10805 | 0.3424 | 0 | 43.8553 | Eukaryota | 7194.0 | 0 | Chordata | 7194.0 | 0 | Carnivora | 7194.0 | 0 | Canidae | 7194.0 | 0 | Canis | 7194.0 | 0 | Canis lupus | 7194.0 | 0 |
| scf7180000043502 | 402514 | 0.4278 | 0 | 58.3379 | Eukaryota | 65918.0 | 0 | Chordata | 65918.0 | 0 | Carnivora | 65918.0 | 0 | Canidae | 65918.0 | 0 | Canis | 47912.0 | 1 | Canis lupus | 47912.0 | 1 |
| scf7180000043583 | 3259749 | 0.4263 | 0 | 59.6658 | Eukaryota | 72486.0 | 0 | Chordata | 72486.0 | 0 | Carnivora | 72486.0 | 0 | Canidae | 72486.0 | 0 | Canis | 50928.0 | 1 | Canis lupus | 50928.0 | 1 |
| scf7180000045937 | 115249 | 0.3912 | 0 | 499.2199 | Eukaryota | 7278.0 | 0 | Chordata | 7278.0 | 0 | Primates | 3070.0 | 2 | Hominidae | 3070.0 | 2 | Sus | 2746.0 | 4 | Sus scrofa | 2746.0 | 4 |
| scf7180000040834 | 146613 | 0.3897 | 0 | 56.9775 | Eukaryota | 72811.0 | 0 | Chordata | 72811.0 | 0 | Carnivora | 72811.0 | 0 | Canidae | 72811.0 | 0 | Canis | 72811.0 | 0 | Canis lupus | 72811.0 | 0 |
| scf7180000046952 | 22540 | 0.4385 | 0 | 130.8652 | Eukaryota | 24631.0 | 0 | Chordata | 17673.0 | 1 | Strigeidida | 6958.0 | 7 | Schistosomatidae | 6958.0 | 7 | Schistosoma | 6958.0 | 7 | Schistosoma mansoni | 6958.0 | 7 |
| scf7180000040658 | 827175 | 0.4048 | 0 | 59.7322 | Eukaryota | 99469.0 | 0 | Chordata | 99469.0 | 0 | Carnivora | 99469.0 | 0 | Canidae | 99469.0 | 0 | Canis | 99469.0 | 0 | Canis lupus | 99469.0 | 0 |
| scf7180000049163 | 9604 | 0.4394 | 0 | 65.2984 | no-hit | 0.0 | 0 | no-hit | 0.0 | 0 | no-hit | 0.0 | 0 | no-hit | 0.0 | 0 | no-hit | 0.0 | 0 | no-hit | 0.0 | 0 |
| scf7180000044735 | 261891 | 0.3996 | 0 | 61.6904 | Eukaryota | 35321.0 | 0 | Chordata | 35321.0 | 0 | Carnivora | 35321.0 | 0 | Canidae | 35321.0 | 0 | Canis | 35321.0 | 0 | Canis lupus | 35321.0 | 0 |
| scf7180000046849 | 101017 | 0.4835 | 0 | 57.1997 | Eukaryota | 13800.0 | 0 | Chordata | 13800.0 | 0 | Carnivora | 13800.0 | 0 | Canidae | 13800.0 | 0 | Canis | 12278.0 | 1 | Canis lupus | 12278.0 | 1 |
| scf7180000051181 | 11777 | 0.6496 | 0 | 51.7696 | Eukaryota | 2323.0 | 0 | Chordata | 2323.0 | 0 | Carnivora | 2323.0 | 0 | Canidae | 2323.0 | 0 | Canis | 2323.0 | 0 | Canis lupus | 2323.0 | 0 |
| scf7180000041166 | 724713 | 0.4009 | 0 | 58.0236 | Eukaryota | 82729.0 | 0 | Chordata | 82729.0 | 0 | Carnivora | 82729.0 | 0 | Canidae | 82729.0 | 0 | Canis | 82729.0 | 0 | Canis lupus | 82729.0 | 0 |
| scf7180000042509 | 121276 | 0.3498 | 0 | 59.0971 | Eukaryota | 57053.0 | 0 | Chordata | 57053.0 | 0 | Carnivora | 57053.0 | 0 | Canidae | 57053.0 | 0 | Canis | 57053.0 | 0 | Canis lupus | 57053.0 | 0 |
| scf7180000046768 | 114359 | 0.5283 | 0 | 55.753 | Eukaryota | 18632.0 | 0 | Chordata | 18632.0 | 0 | Carnivora | 18632.0 | 0 | Canidae | 18632.0 | 0 | Canis | 18632.0 | 0 | Canis lupus | 18632.0 | 0 |
| scf7180000042790 | 284530 | 0.4184 | 0 | 60.2311 | Eukaryota | 57615.0 | 0 | Chordata | 57615.0 | 0 | Carnivora | 57615.0 | 0 | Canidae | 57615.0 | 0 | Canis | 57615.0 | 0 | Canis lupus | 57615.0 | 0 |
| scf7180000049047 | 57147 | 0.4096 | 0 | 57.5378 | Eukaryota | 35172.0 | 0 | Chordata | 35172.0 | 0 | Carnivora | 35172.0 | 0 | Otariidae | 18402.0 | 2 | Canis | 10438.0 | 3 | Canis lupus | 10438.0 | 3 |
| scf7180000049755 | 3147 | 0.4423 | 0 | 45.5195 | no-hit | 0.0 | 0 | no-hit | 0.0 | 0 | no-hit | 0.0 | 0 | no-hit | 0.0 | 0 | no-hit | 0.0 | 0 | no-hit | 0.0 | 0 |
| scf7180000043164 | 533383 | 0.4723 | 100 | 58.8003 | Eukaryota | 59415.0 | 0 | Chordata | 59415.0 | 0 | Carnivora | 59415.0 | 0 | Canidae | 59415.0 | 0 | Canis | 59415.0 | 0 | Canis lupus | 59415.0 | 0 |
| scf7180000052381 | 10215 | 0.4349 | 0 | 206.4676 | Eukaryota | 21233.0 | 0 | Chordata | 13070.0 | 1 | Strigeidida | 7658.0 | 6 | Schistosomatidae | 7658.0 | 6 | Schistosoma | 7658.0 | 6 | Schistosoma mansoni | 7658.0 | 6 |
| scf7180000046567 | 137822 | 0.3673 | 0 | 58.5425 | Eukaryota | 14235.0 | 1 | Chordata | 14235.0 | 1 | Primates | 5894.0 | 4 | Bovidae | 3796.0 | 6 | Homo | 3469.0 | 7 | Homo sapiens | 3469.0 | 7 |
| scf7180000049378 | 4311 | 0.4312 | 0 | 296.9177 | Eukaryota | 317.0 | 0 | Chordata | 165.0 | 1 | Kurtiformes | 165.0 | 1 | Apogonidae | 165.0 | 1 | Sphaeramia | 165.0 | 1 | Sphaeramia orbicularis | 165.0 | 1 |
| scf7180000041127 | 22048 | 0.4386 | 0 | 116.7665 | Eukaryota | 62375.0 | 0 | Chordata | 42366.0 | 2 | Gadiformes | 35194.0 | 5 | Gadidae | 35194.0 | 5 | Gadus | 35194.0 | 5 | Gadus morhua | 35194.0 | 5 |
| scf7180000044279 | 328672 | 0.5501 | 0 | 58.0386 | Eukaryota | 41660.0 | 0 | Chordata | 41660.0 | 0 | Carnivora | 41660.0 | 0 | Canidae | 41660.0 | 0 | Canis | 21049.0 | 1 | Canis lupus | 21049.0 | 1 |
| scf7180000043054 | 7728 | 0.4467 | 0 | 34.1603 | Eukaryota | 95308.0 | 0 | Chordata | 95308.0 | 0 | Carnivora | 95308.0 | 0 | Canidae | 95308.0 | 0 | Canis | 95308.0 | 0 | Canis lupus | 95308.0 | 0 |
| scf7180000055100 | 25959 | 0.442 | 0 | 204.298 | Eukaryota | 720.0 | 1 | Platyhelminthes | 720.0 | 1 | Polystomatidea | 720.0 | 1 | Polystomatidae | 720.0 | 1 | Protopolystoma | 720.0 | 1 | Protopolystoma xenopodis | 720.0 | 1 |
| scf7180000050315 | 3347 | 0.4443 | 0 | 582.4001 | no-hit | 0.0 | 0 | no-hit | 0.0 | 0 | no-hit | 0.0 | 0 | no-hit | 0.0 | 0 | no-hit | 0.0 | 0 | no-hit | 0.0 | 0 |
| scf7180000050575 | 25885 | 0.3861 | 0 | 58.6031 | Eukaryota | 41347.0 | 0 | Chordata | 41347.0 | 0 | Carnivora | 41347.0 | 0 | Canidae | 41347.0 | 0 | Canis | 41347.0 | 0 | Canis lupus | 41347.0 | 0 |
| scf7180000047993 | 72460 | 0.4161 | 0 | 61.7129 | Eukaryota | 24293.0 | 0 | Chordata | 24293.0 | 0 | Carnivora | 24293.0 | 0 | Canidae | 24293.0 | 0 | Canis | 24293.0 | 0 | Canis lupus | 24293.0 | 0 |
| scf7180000044987 | 15842 | 0.4471 | 0 | 45.9759 | Eukaryota | 6596.0 | 0 | Chordata | 6596.0 | 0 | Carnivora | 5568.0 | 1 | Phocidae | 3852.0 | 2 | Leptonychotes | 3852.0 | 3 | Leptonychotes weddellii | 3852.0 | 3 |
| scf7180000041602 | 455187 | 0.3971 | 0 | 59.1084 | Eukaryota | 62433.0 | 0 | Chordata | 62433.0 | 0 | Carnivora | 62433.0 | 0 | Canidae | 62433.0 | 0 | Canis | 62433.0 | 0 | Canis lupus | 62433.0 | 0 |
| scf7180000050129 | 10550 | 0.448 | 0 | 687.7525 | Eukaryota | 3053.0 | 0 | Platyhelminthes | 1933.0 | 2 | Polystomatidea | 1933.0 | 2 | Polystomatidae | 1933.0 | 2 | Protopolystoma | 1933.0 | 2 | Protopolystoma xenopodis | 1933.0 | 2 |
| scf7180000049669 | 30440 | 0.4657 | 0 | 61.9343 | Eukaryota | 15839.0 | 0 | Chordata | 15839.0 | 0 | Carnivora | 15839.0 | 0 | Canidae | 15839.0 | 0 | Canis | 15839.0 | 0 | Canis lupus | 15839.0 | 0 |
| scf7180000044850 | 32804 | 0.3979 | 0 | 65.701 | Eukaryota | 88586.0 | 0 | Chordata | 88586.0 | 0 | Carnivora | 88586.0 | 0 | Canidae | 88586.0 | 0 | Canis | 88586.0 | 0 | Canis lupus | 88586.0 | 0 |
| scf7180000043717 | 375522 | 0.337 | 0 | 58.6052 | Eukaryota | 93315.0 | 0 | Chordata | 93315.0 | 0 | Carnivora | 93315.0 | 0 | Canidae | 93315.0 | 0 | Canis | 93315.0 | 0 | Canis lupus | 93315.0 | 0 |
| scf7180000051297 | 21157 | 0.503 | 0 | 542.2859 | Eukaryota | 9468.0 | 0 | Chordata | 9468.0 | 0 | Carnivora | 9468.0 | 0 | Canidae | 9468.0 | 0 | Canis | 8225.0 | 1 | Canis lupus | 6915.0 | 2 |
| scf7180000053918 | 1887 | 0.4526 | 0 | 834.2972 | no-hit | 0.0 | 0 | no-hit | 0.0 | 0 | no-hit | 0.0 | 0 | no-hit | 0.0 | 0 | no-hit | 0.0 | 0 | no-hit | 0.0 | 0 |
| scf7180000048748 | 10526 | 0.4501 | 0 | 177.0798 | Eukaryota | 95621.0 | 0 | Chordata | 95621.0 | 0 | Carnivora | 95621.0 | 0 | Canidae | 95621.0 | 0 | Canis | 95621.0 | 0 | Canis lupus | 95621.0 | 0 |
| scf7180000051476 | 38322 | 0.506 | 0 | 275.4215 | Eukaryota | 11757.0 | 0 | Chordata | 11757.0 | 0 | Carnivora | 11757.0 | 0 | Canidae | 11757.0 | 0 | Canis | 8162.0 | 1 | Canis lupus | 8162.0 | 1 |
| scf7180000056445 | 450467 | 0.3662 | 0 | 59.4866 | Eukaryota | 37753.0 | 0 | Chordata | 37753.0 | 0 | Carnivora | 37753.0 | 0 | Canidae | 37753.0 | 0 | Canis | 37753.0 | 0 | Canis lupus | 37753.0 | 0 |
| scf7180000055316 | 36979 | 0.4393 | 0 | 124.0397 | Eukaryota | 58928.0 | 0 | Chordata | 58928.0 | 0 | Carnivora | 58928.0 | 0 | Canidae | 58928.0 | 0 | Canis | 58928.0 | 0 | Canis lupus | 58928.0 | 0 |
| scf7180000052563 | 13398 | 0.4349 | 0 | 289.4289 | Eukaryota | 1095.0 | 0 | Chordata | 595.0 | 1 | Tetraodontiformes | 595.0 | 2 | Tetraodontidae | 595.0 | 2 | Takifugu | 595.0 | 2 | Takifugu rubripes | 595.0 | 2 |
| scf7180000055709 | 60282 | 0.4425 | 0 | 392.1443 | Eukaryota | 4858.0 | 0 | Chordata | 4254.0 | 1 | Kurtiformes | 3758.0 | 2 | Apogonidae | 3758.0 | 2 | Sphaeramia | 3758.0 | 2 | Sphaeramia orbicularis | 3758.0 | 2 |
| scf7180000049557 | 33657 | 0.3618 | 0 | 57.669 | Eukaryota | 71108.0 | 0 | Chordata | 71108.0 | 0 | Carnivora | 71108.0 | 0 | Canidae | 71108.0 | 0 | Canis | 71108.0 | 0 | Canis lupus | 71108.0 | 0 |
| scf7180000056201 | 1473755 | 0.3842 | 0 | 60.5491 | Eukaryota | 88444.0 | 0 | Chordata | 88444.0 | 0 | Carnivora | 88444.0 | 0 | Canidae | 88444.0 | 0 | Canis | 88444.0 | 0 | Canis lupus | 88444.0 | 0 |
| scf7180000043467 | 7203 | 0.4039 | 0 | 52.8348 | Eukaryota | 14481.0 | 0 | Chordata | 14481.0 | 0 | Carnivora | 14481.0 | 0 | Canidae | 14481.0 | 0 | Canis | 14481.0 | 0 | Canis lupus | 14481.0 | 0 |
| scf7180000045962 | 134407 | 0.3429 | 0 | 59.516 | Eukaryota | 92572.0 | 0 | Chordata | 92572.0 | 0 | Carnivora | 92572.0 | 0 | Canidae | 92572.0 | 0 | Canis | 92572.0 | 0 | Canis lupus | 92572.0 | 0 |
| scf7180000043011 | 64718 | 0.3645 | 0 | 57.5256 | Eukaryota | 17883.0 | 0 | Chordata | 17883.0 | 0 | Carnivora | 17883.0 | 0 | Canidae | 17883.0 | 0 | Canis | 12807.0 | 1 | Canis lupus | 12807.0 | 1 |
| scf7180000044258 | 329510 | 0.3886 | 0 | 61.9533 | Eukaryota | 115128.0 | 0 | Chordata | 115128.0 | 0 | Carnivora | 115128.0 | 0 | Canidae | 115128.0 | 0 | Canis | 115128.0 | 0 | Canis lupus | 115128.0 | 0 |
| scf7180000051364 | 19704 | 0.4416 | 0 | 117.6092 | Eukaryota | 39313.0 | 0 | Chordata | 29003.0 | 1 | Gadiformes | 23188.0 | 4 | Gadidae | 23188.0 | 4 | Gadus | 23188.0 | 4 | Gadus morhua | 23188.0 | 4 |
| scf7180000047598 | 72879 | 0.4277 | 0 | 60.3298 | Eukaryota | 38853.0 | 0 | Chordata | 38853.0 | 0 | Carnivora | 38853.0 | 0 | Canidae | 18168.0 | 4 | Canis | 18168.0 | 5 | Canis lupus | 18168.0 | 5 |
| scf7180000054501 | 44213 | 0.3878 | 0 | 946.3706 | Eukaryota | 55588.0 | 0 | Chordata | 55588.0 | 0 | Carnivora | 55588.0 | 0 | Canidae | 55588.0 | 0 | Canis | 55588.0 | 0 | Canis lupus | 55588.0 | 0 |
| scf7180000043387 | 486692 | 0.3903 | 0 | 58.9318 | Eukaryota | 52250.0 | 0 | Chordata | 52250.0 | 0 | Carnivora | 52250.0 | 0 | Canidae | 52250.0 | 0 | Canis | 52250.0 | 0 | Canis lupus | 52250.0 | 0 |
| scf7180000051830 | 3386 | 0.4557 | 0 | 664.2245 | no-hit | 0.0 | 0 | no-hit | 0.0 | 0 | no-hit | 0.0 | 0 | no-hit | 0.0 | 0 | no-hit | 0.0 | 0 | no-hit | 0.0 | 0 |
| scf7180000049730 | 10105 | 0.4328 | 0 | 123.2205 | Eukaryota | 1285.0 | 0 | Platyhelminthes | 651.0 | 1 | Polystomatidea | 651.0 | 1 | Polystomatidae | 651.0 | 1 | Protopolystoma | 651.0 | 1 | Protopolystoma xenopodis | 651.0 | 1 |
| scf7180000045893 | 165470 | 0.4121 | 0 | 59.4623 | Eukaryota | 15629.0 | 0 | Chordata | 15629.0 | 0 | Apterygiformes | 13931.0 | 1 | Apterygidae | 13931.0 | 1 | Apteryx | 13931.0 | 1 | Apteryx mantelli | 13931.0 | 1 |
| scf7180000054238 | 56261 | 0.4456 | 0 | 524.4229 | Eukaryota | 15556.0 | 0 | Chordata | 13905.0 | 2 | Tetraodontiformes | 11496.0 | 3 | Tetraodontidae | 11496.0 | 3 | Takifugu | 11496.0 | 3 | Takifugu rubripes | 11496.0 | 3 |
| scf7180000047943 | 79517 | 0.4331 | 0 | 58.1455 | Eukaryota | 5897.0 | 0 | Chordata | 5300.0 | 1 | Primates | 2282.0 | 4 | Hominidae | 2282.0 | 5 | Homo | 2282.0 | 5 | Homo sapiens | 2282.0 | 5 |
| scf7180000053148 | 1958 | 0.4428 | 0 | 308.5936 | Eukaryota | 2612.0 | 0 | Chordata | 1282.0 | 2 | Polystomatidea | 997.0 | 4 | Polystomatidae | 997.0 | 4 | Protopolystoma | 997.0 | 4 | Protopolystoma xenopodis | 997.0 | 4 |
| scf7180000047114 | 24654 | 0.4369 | 0 | 164.9979 | no-hit | 0.0 | 0 | no-hit | 0.0 | 0 | no-hit | 0.0 | 0 | no-hit | 0.0 | 0 | no-hit | 0.0 | 0 | no-hit | 0.0 | 0 |
| scf7180000051531 | 2501 | 0.451 | 0 | 326.8072 | no-hit | 0.0 | 0 | no-hit | 0.0 | 0 | no-hit | 0.0 | 0 | no-hit | 0.0 | 0 | no-hit | 0.0 | 0 | no-hit | 0.0 | 0 |
| scf7180000043799 | 388227 | 0.517 | 0 | 60.1838 | Eukaryota | 43174.0 | 0 | Chordata | 43174.0 | 0 | Carnivora | 43174.0 | 0 | Canidae | 43174.0 | 0 | Canis | 43174.0 | 0 | Canis lupus | 43174.0 | 0 |
| scf7180000048010 | 2995 | 0.4337 | 0 | 94.2259 | no-hit | 0.0 | 0 | no-hit | 0.0 | 0 | no-hit | 0.0 | 0 | no-hit | 0.0 | 0 | no-hit | 0.0 | 0 | no-hit | 0.0 | 0 |
| scf7180000054347 | 83590 | 0.4418 | 0 | 250.9225 | Eukaryota | 28511.0 | 0 | Chordata | 24738.0 | 1 | Gonorynchiformes | 9409.0 | 4 | Chanidae | 9409.0 | 4 | Chanos | 9409.0 | 4 | Chanos chanos | 9409.0 | 4 |
| scf7180000040921 | 75906 | 0.4746 | 0 | 56.4944 | Eukaryota | 11559.0 | 1 | Chordata | 11559.0 | 1 | Primates | 6663.0 | 4 | Hominidae | 5477.0 | 5 | Homo | 5477.0 | 6 | Homo sapiens | 5477.0 | 6 |
| scf7180000041534 | 574787 | 0.4052 | 0 | 57.8652 | Eukaryota | 65816.0 | 0 | Chordata | 65816.0 | 0 | Carnivora | 65816.0 | 0 | Canidae | 65816.0 | 0 | Canis | 65816.0 | 0 | Canis lupus | 65816.0 | 0 |
| scf7180000051642 | 13322 | 0.4342 | 0 | 123.2432 | Eukaryota | 9059.0 | 0 | Platyhelminthes | 4787.0 | 2 | Poales | 2957.0 | 5 | Poaceae | 2957.0 | 5 | Oryza | 2957.0 | 6 | Oryza sativa | 2957.0 | 6 |
| scf7180000054389 | 32941 | 0.4 | 0 | 697.7881 | Eukaryota | 94052.0 | 0 | Chordata | 94052.0 | 0 | Carnivora | 94052.0 | 0 | Canidae | 94052.0 | 0 | Canis | 94052.0 | 0 | Canis lupus | 94052.0 | 0 |
| scf7180000044821 | 226390 | 0.4165 | 0 | 59.2844 | Eukaryota | 25415.0 | 0 | Chordata | 25415.0 | 0 | Carnivora | 20504.0 | 1 | Canidae | 20504.0 | 2 | Canis | 17868.0 | 3 | Canis lupus | 17868.0 | 3 |
| scf7180000055167 | 19822 | 0.4387 | 0 | 160.3422 | Eukaryota | 38743.0 | 0 | Chordata | 26580.0 | 1 | Cyclophyllidea | 12163.0 | 6 | Hymenolepididae | 12163.0 | 6 | Hymenolepis | 12163.0 | 6 | Hymenolepis microstoma | 12163.0 | 6 |
| scf7180000048168 | 8315 | 0.4423 | 0 | 135.1006 | Eukaryota | 3193.0 | 0 | Platyhelminthes | 2799.0 | 1 | Polystomatidea | 2799.0 | 1 | Polystomatidae | 2799.0 | 1 | Protopolystoma | 2799.0 | 1 | Protopolystoma xenopodis | 2799.0 | 1 |
| scf7180000055357 | 22771 | 0.6308 | 0 | 525.1042 | Eukaryota | 56621.0 | 0 | Chordata | 56621.0 | 0 | Primates | 48650.0 | 1 | Hominidae | 33408.0 | 2 | Homo | 33408.0 | 2 | Homo sapiens | 33408.0 | 2 |
| scf7180000056382 | 172337 | 0.3777 | 0 | 56.6574 | Eukaryota | 63552.0 | 0 | Chordata | 63552.0 | 0 | Carnivora | 63552.0 | 0 | Canidae | 45612.0 | 2 | Canis | 31750.0 | 4 | Canis lupus | 31750.0 | 4 |
| scf7180000046884 | 40174 | 0.4576 | 0 | 103.7303 | Eukaryota | 94878.0 | 0 | Chordata | 94878.0 | 0 | Carnivora | 94878.0 | 0 | Canidae | 94878.0 | 0 | Canis | 94878.0 | 0 | Canis lupus | 94878.0 | 0 |
| scf7180000056095 | 12497 | 0.4405 | 0 | 105.8613 | Eukaryota | 3515.0 | 0 | Platyhelminthes | 2068.0 | 2 | Polystomatidea | 2068.0 | 3 | Polystomatidae | 2068.0 | 3 | Protopolystoma | 2068.0 | 3 | Protopolystoma xenopodis | 2068.0 | 3 |
| scf7180000054594 | 6510 | 0.4458 | 0 | 397.9017 | Eukaryota | 7271.0 | 0 | Chordata | 6760.0 | 1 | Kurtiformes | 5375.0 | 2 | Apogonidae | 5375.0 | 2 | Sphaeramia | 5375.0 | 2 | Sphaeramia orbicularis | 5375.0 | 2 |
| scf7180000046490 | 97250 | 0.3937 | 0 | 62.3734 | Eukaryota | 56066.0 | 0 | Chordata | 56066.0 | 0 | Carnivora | 56066.0 | 0 | Canidae | 56066.0 | 0 | Canis | 56066.0 | 0 | Canis lupus | 56066.0 | 0 |
| scf7180000047835 | 16135 | 0.4404 | 0 | 106.2725 | Eukaryota | 3436.0 | 0 | Chordata | 2702.0 | 1 | Carnivora | 1783.0 | 3 | Canidae | 1783.0 | 3 | Canis | 1783.0 | 3 | Canis lupus | 1783.0 | 3 |
| scf7180000043148 | 332072 | 0.3956 | 0 | 56.1483 | Eukaryota | 67508.0 | 0 | Chordata | 67508.0 | 0 | Carnivora | 67508.0 | 0 | Canidae | 67508.0 | 0 | Canis | 67508.0 | 0 | Canis lupus | 67508.0 | 0 |
| scf7180000050605 | 23061 | 0.439 | 0 | 61.4803 | Eukaryota | 42310.0 | 0 | Platyhelminthes | 21867.0 | 1 | Strigeidida | 15654.0 | 5 | Schistosomatidae | 15654.0 | 5 | Schistosoma | 15654.0 | 5 | Schistosoma mansoni | 15654.0 | 5 |
| scf7180000044473 | 607810 | 0.4068 | 100 | 59.4034 | Eukaryota | 35192.0 | 0 | Chordata | 35192.0 | 0 | Carnivora | 35192.0 | 0 | Canidae | 35192.0 | 0 | Vulpes | 23233.0 | 1 | Vulpes vulpes | 23233.0 | 1 |
| scf7180000049548 | 4514 | 0.4548 | 0 | 352.943 | no-hit | 0.0 | 0 | no-hit | 0.0 | 0 | no-hit | 0.0 | 0 | no-hit | 0.0 | 0 | no-hit | 0.0 | 0 | no-hit | 0.0 | 0 |
| scf7180000041495 | 7895 | 0.4592 | 0 | 21.5063 | Eukaryota | 60570.0 | 0 | Chordata | 60570.0 | 0 | Carnivora | 60570.0 | 0 | Canidae | 60570.0 | 0 | Canis | 60570.0 | 0 | Canis lupus | 60570.0 | 0 |
| scf7180000045055 | 233506 | 0.3964 | 0 | 60.6694 | Eukaryota | 39418.0 | 0 | Chordata | 39418.0 | 0 | Carnivora | 39418.0 | 0 | Canidae | 39418.0 | 0 | Canis | 39418.0 | 0 | Canis lupus | 39418.0 | 0 |
| scf7180000045626 | 195830 | 0.4989 | 0 | 59.0424 | Eukaryota | 38903.0 | 0 | Chordata | 38903.0 | 0 | Carnivora | 38903.0 | 0 | Canidae | 38903.0 | 0 | Vulpes | 25992.0 | 1 | Vulpes vulpes | 25992.0 | 1 |
| scf7180000048831 | 12642 | 0.4492 | 0 | 405.1987 | Eukaryota | 4212.0 | 0 | Chordata | 1813.0 | 3 | Polystomatidea | 1152.0 | 4 | Polystomatidae | 1152.0 | 4 | Protopolystoma | 1152.0 | 4 | Protopolystoma xenopodis | 1152.0 | 4 |
| scf7180000040673 | 503008 | 0.503 | 0 | 57.4449 | Eukaryota | 67061.0 | 0 | Chordata | 67061.0 | 0 | Carnivora | 67061.0 | 0 | Canidae | 67061.0 | 0 | Canis | 67061.0 | 0 | Canis lupus | 67061.0 | 0 |
| scf7180000047016 | 62420 | 0.3793 | 0 | 68.1874 | Eukaryota | 82212.0 | 0 | Chordata | 82212.0 | 0 | Carnivora | 82212.0 | 0 | Canidae | 82212.0 | 0 | Canis | 82212.0 | 0 | Canis lupus | 82212.0 | 0 |
| scf7180000043234 | 78069 | 0.4134 | 0 | 617.2632 | Eukaryota | 94871.0 | 0 | Chordata | 94871.0 | 0 | Carnivora | 94871.0 | 0 | Canidae | 94871.0 | 0 | Canis | 94871.0 | 0 | Canis lupus | 94871.0 | 0 |
| scf7180000052889 | 11077 | 0.5108 | 0 | 365.3435 | Eukaryota | 34274.0 | 0 | Chordata | 34274.0 | 0 | Carnivora | 34274.0 | 0 | Canidae | 34274.0 | 0 | Canis | 34274.0 | 0 | Canis lupus | 34274.0 | 0 |
| scf7180000047339 | 13794 | 0.444 | 0 | 135.193 | Eukaryota | 2416.0 | 0 | Chordata | 1179.0 | 2 | Polystomatidea | 1042.0 | 4 | Polystomatidae | 1042.0 | 4 | Protopolystoma | 1042.0 | 5 | Protopolystoma xenopodis | 1042.0 | 5 |
| scf7180000049773 | 24461 | 0.4507 | 0 | 95.5725 | Eukaryota | 7480.0 | 0 | Chordata | 4007.0 | 2 | Forcipulatida | 2060.0 | 5 | Asteriidae | 2060.0 | 5 | Asterias | 2060.0 | 6 | Asterias rubens | 2060.0 | 6 |
| scf7180000045229 | 205285 | 0.4343 | 0 | 57.7883 | Eukaryota | 56201.0 | 0 | Chordata | 56201.0 | 0 | Carnivora | 56201.0 | 0 | Canidae | 56201.0 | 0 | Canis | 56201.0 | 0 | Canis lupus | 56201.0 | 0 |
| scf7180000044925 | 21514 | 0.4509 | 0 | 1706.6147 | Eukaryota | 2898.0 | 0 | Echinodermata | 933.0 | 3 | Forcipulatida | 933.0 | 3 | Asteriidae | 933.0 | 3 | Asterias | 933.0 | 3 | Asterias rubens | 933.0 | 3 |
| scf7180000051332 | 6349 | 0.4355 | 0 | 75.3112 | Eukaryota | 11067.0 | 0 | Chordata | 5947.0 | 3 | Strigeidida | 3422.0 | 7 | Schistosomatidae | 3422.0 | 7 | Schistosoma | 3422.0 | 7 | Schistosoma mansoni | 3422.0 | 7 |
| scf7180000048817 | 40534 | 0.4047 | 0 | 50.8829 | Eukaryota | 16218.0 | 0 | Chordata | 16218.0 | 0 | Carnivora | 16218.0 | 0 | Canidae | 16218.0 | 0 | Canis | 9901.0 | 1 | Canis lupus | 9901.0 | 1 |
| scf7180000042664 | 113254 | 0.4864 | 0 | 55.6617 | Eukaryota | 6281.0 | 0 | Chordata | 6281.0 | 0 | Carnivora | 2845.0 | 2 | Canidae | 2845.0 | 2 | Canis | 2845.0 | 3 | Canis lupus | 2845.0 | 3 |
| scf7180000048298 | 5621 | 0.4398 | 0 | 76.1766 | Eukaryota | 4968.0 | 0 | Platyhelminthes | 2640.0 | 1 | Polystomatidea | 2640.0 | 3 | Polystomatidae | 2640.0 | 3 | Protopolystoma | 2640.0 | 3 | Protopolystoma xenopodis | 2640.0 | 3 |
| scf7180000042594 | 641710 | 0.4269 | 0 | 63.0178 | Eukaryota | 92823.0 | 0 | Chordata | 92823.0 | 0 | Carnivora | 92823.0 | 0 | Canidae | 92823.0 | 0 | Canis | 92823.0 | 0 | Canis lupus | 92823.0 | 0 |
| scf7180000044621 | 215510 | 0.548 | 0 | 57.1143 | Eukaryota | 50622.0 | 0 | Chordata | 50622.0 | 0 | Carnivora | 50622.0 | 0 | Canidae | 44532.0 | 1 | Canis | 33738.0 | 3 | Canis lupus | 33738.0 | 3 |
| scf7180000048082 | 50334 | 0.4425 | 0 | 56.196 | Eukaryota | 9317.0 | 0 | Chordata | 9317.0 | 0 | Carnivora | 9317.0 | 0 | Canidae | 9317.0 | 0 | Canis | 9317.0 | 0 | Canis lupus | 9317.0 | 0 |
| scf7180000042631 | 186784 | 0.3408 | 0 | 59.8441 | Eukaryota | 93136.0 | 0 | Chordata | 93136.0 | 0 | Carnivora | 93136.0 | 0 | Canidae | 93136.0 | 0 | Canis | 93136.0 | 0 | Canis lupus | 93136.0 | 0 |
| scf7180000053920 | 9458 | 0.5022 | 0 | 217.3252 | Eukaryota | 8708.0 | 0 | Chordata | 8708.0 | 0 | Carnivora | 8708.0 | 0 | Canidae | 8708.0 | 0 | Canis | 7052.0 | 1 | Canis lupus | 5860.0 | 2 |
| scf7180000040812 | 1667194 | 0.3921 | 0 | 61.353 | Eukaryota | 94464.0 | 0 | Chordata | 94464.0 | 0 | Carnivora | 94464.0 | 0 | Canidae | 94464.0 | 0 | Canis | 94464.0 | 0 | Canis lupus | 94464.0 | 0 |
| scf7180000052642 | 4569 | 0.436 | 0 | 405.3911 | Eukaryota | 165.0 | 0 | Platyhelminthes | 165.0 | 0 | Polystomatidea | 165.0 | 0 | Polystomatidae | 165.0 | 0 | Protopolystoma | 165.0 | 0 | Protopolystoma xenopodis | 165.0 | 0 |
| scf7180000055028 | 18502 | 0.4105 | 0 | 194.1087 | Eukaryota | 81569.0 | 0 | Chordata | 81569.0 | 0 | Carnivora | 81569.0 | 0 | Canidae | 81569.0 | 0 | Canis | 81569.0 | 0 | Canis lupus | 81569.0 | 0 |
| scf7180000052615 | 11596 | 0.4411 | 0 | 407.3255 | Eukaryota | 4381.0 | 0 | Platyhelminthes | 2901.0 | 2 | Polystomatidea | 2901.0 | 2 | Polystomatidae | 2901.0 | 2 | Protopolystoma | 2901.0 | 2 | Protopolystoma xenopodis | 2901.0 | 2 |
| scf7180000043869 | 129795 | 0.3653 | 0 | 61.0904 | Eukaryota | 26797.0 | 0 | Chordata | 26797.0 | 0 | Carnivora | 26797.0 | 0 | Canidae | 26797.0 | 0 | Canis | 26797.0 | 0 | Canis lupus | 26797.0 | 0 |
| scf7180000043536 | 201624 | 0.4003 | 0 | 59.1351 | Eukaryota | 44294.0 | 0 | Chordata | 44294.0 | 0 | Carnivora | 44294.0 | 0 | Canidae | 44294.0 | 0 | Canis | 44294.0 | 0 | Canis lupus | 44294.0 | 0 |
| scf7180000040734 | 1817374 | 0.3623 | 0 | 59.683 | Eukaryota | 195632.0 | 0 | Chordata | 195632.0 | 0 | Carnivora | 195632.0 | 0 | Canidae | 195632.0 | 0 | Canis | 130946.0 | 1 | Canis lupus | 130946.0 | 1 |
| scf7180000046141 | 135741 | 0.3537 | 0 | 60.3439 | Eukaryota | 16612.0 | 0 | Chordata | 16612.0 | 0 | Carnivora | 16612.0 | 0 | Canidae | 16612.0 | 0 | Canis | 16612.0 | 0 | Canis lupus | 16612.0 | 0 |
| scf7180000047150 | 56713 | 0.4052 | 0 | 57.6111 | Eukaryota | 51886.0 | 0 | Chordata | 51886.0 | 0 | Carnivora | 51886.0 | 0 | Canidae | 51886.0 | 0 | Canis | 51886.0 | 0 | Canis lupus | 51886.0 | 0 |
| scf7180000047036 | 35061 | 0.4409 | 0 | 221.5056 | Eukaryota | 22103.0 | 0 | Chordata | 10601.0 | 2 | Cyclophyllidea | 10244.0 | 7 | Hymenolepididae | 10244.0 | 7 | Hymenolepis | 10244.0 | 8 | Hymenolepis microstoma | 10244.0 | 8 |
| scf7180000041884 | 466101 | 0.4402 | 0 | 58.4953 | Eukaryota | 193340.0 | 0 | Chordata | 193340.0 | 0 | Carnivora | 193340.0 | 0 | Canidae | 193340.0 | 0 | Canis | 193340.0 | 0 | Canis lupus | 193340.0 | 0 |
| scf7180000044454 | 234166 | 0.4285 | 0 | 59.9872 | Eukaryota | 41080.0 | 0 | Chordata | 41080.0 | 0 | Carnivora | 41080.0 | 0 | Canidae | 41080.0 | 0 | Canis | 41080.0 | 0 | Canis lupus | 41080.0 | 0 |
| scf7180000048430 | 2723 | 0.4389 | 0 | 90.8804 | Eukaryota | 185.0 | 0 | Chordata | 185.0 | 0 | Kurtiformes | 185.0 | 0 | Apogonidae | 185.0 | 0 | Sphaeramia | 185.0 | 0 | Sphaeramia orbicularis | 185.0 | 0 |
| scf7180000043854 | 405033 | 0.3737 | 0 | 60.34 | Eukaryota | 51091.0 | 0 | Chordata | 51091.0 | 0 | Carnivora | 51091.0 | 0 | Canidae | 51091.0 | 0 | Canis | 39283.0 | 1 | Canis lupus | 39283.0 | 1 |
| scf7180000044899 | 221328 | 0.4432 | 0 | 59.4605 | Eukaryota | 47082.0 | 0 | Chordata | 47082.0 | 0 | Carnivora | 47082.0 | 0 | Canidae | 47082.0 | 0 | Canis | 47082.0 | 0 | Canis lupus | 47082.0 | 0 |
| scf7180000044381 | 333570 | 0.4386 | 0 | 59.7011 | Eukaryota | 36813.0 | 0 | Chordata | 36813.0 | 0 | Carnivora | 36813.0 | 0 | Canidae | 36813.0 | 0 | Canis | 21300.0 | 1 | Canis lupus | 21300.0 | 1 |
| scf7180000054244 | 92518 | 0.4134 | 0 | 340.3819 | Eukaryota | 6528.0 | 0 | Chordata | 6528.0 | 0 | Carnivora | 5805.0 | 1 | Canidae | 5805.0 | 1 | Vulpes | 3225.0 | 2 | Vulpes vulpes | 3225.0 | 2 |
| scf7180000047583 | 11055 | 0.4469 | 0 | 412.9004 | Eukaryota | 4806.0 | 0 | Chordata | 2175.0 | 4 | Polystomatidea | 1479.0 | 7 | Polystomatidae | 1479.0 | 7 | Protopolystoma | 1479.0 | 7 | Protopolystoma xenopodis | 1479.0 | 7 |
| scf7180000053235 | 5859 | 0.4426 | 0 | 251.7795 | Eukaryota | 6803.0 | 0 | Chordata | 3769.0 | 2 | Polystomatidea | 2810.0 | 4 | Polystomatidae | 2810.0 | 4 | Protopolystoma | 2810.0 | 4 | Protopolystoma xenopodis | 2810.0 | 4 |
| scf7180000042861 | 146668 | 0.3597 | 0 | 59.9164 | Eukaryota | 12474.0 | 0 | Chordata | 12474.0 | 0 | Carnivora | 12474.0 | 0 | Canidae | 12474.0 | 0 | Canis | 12474.0 | 0 | Canis lupus | 12474.0 | 0 |
| scf7180000043398 | 75380 | 0.3546 | 0 | 60.6151 | Eukaryota | 44557.0 | 0 | Chordata | 44557.0 | 0 | Carnivora | 44557.0 | 0 | Canidae | 44557.0 | 0 | Canis | 44557.0 | 0 | Canis lupus | 44557.0 | 0 |
| scf7180000049246 | 3318 | 0.4439 | 0 | 184.1054 | Eukaryota | 20389.0 | 0 | Chordata | 15856.0 | 2 | Gadiformes | 10246.0 | 5 | Gadidae | 10246.0 | 5 | Gadus | 10246.0 | 5 | Gadus morhua | 10246.0 | 5 |
| scf7180000042685 | 561491 | 0.5291 | 0 | 56.4314 | Eukaryota | 45529.0 | 0 | Chordata | 45529.0 | 0 | Carnivora | 45529.0 | 0 | Canidae | 41935.0 | 1 | Vulpes | 24423.0 | 2 | Vulpes vulpes | 24423.0 | 2 |
| scf7180000049749 | 5840 | 0.449 | 0 | 492.9661 | Eukaryota | 309.0 | 0 | Platyhelminthes | 309.0 | 0 | Polystomatidea | 309.0 | 0 | Polystomatidae | 309.0 | 0 | Protopolystoma | 309.0 | 0 | Protopolystoma xenopodis | 309.0 | 0 |
| scf7180000046685 | 55598 | 0.4355 | 0 | 51.5891 | Eukaryota | 5776.0 | 0 | Chordata | 5776.0 | 0 | Carnivora | 3719.0 | 2 | Canidae | 2865.0 | 3 | Canis | 2865.0 | 4 | Canis lupus | 2865.0 | 4 |

| | | | | | | | | | | | | | | | | | | | | | | |
|---|---|---|---|---|---|---|---|---|---|---|---|---|---|---|---|---|---|---|---|---|---|---|
| scf7180000042127 | 107497 | 0.3461 | 0 | 57.9274 | Eukaryota | 10171.0 | 0 | Chordata | 10171.0 | 0 | Primates | 5189.0 | 3 | Hominidae | 5189.0 | 3 | Homo | 5189.0 | 5 | Homo sapiens | 5189.0 | 5 |
| scf7180000047627 | 10328 | 0.4396 | 0 | 110.1165 | Eukaryota | 4087.0 | 0 | Chordata | 2622.0 | 1 | Polystomatidea | 1465.0 | 4 | Polystomatidae | 1465.0 | 4 | Protopolystoma | 1465.0 | 4 | Protopolystoma xenopodis | 1465.0 | 4 |
| scf7180000045060 | 48270 | 0.4002 | 0 | 56.8307 | Eukaryota | 43344.0 | 0 | Chordata | 43344.0 | 0 | Carnivora | 43344.0 | 0 | Canidae | 43344.0 | 0 | Canis | 43344.0 | 0 | Canis lupus | 43344.0 | 0 |
| scf7180000051260 | 12302 | 0.4503 | 0 | 23.934 | Eukaryota | 32975.0 | 0 | Chordata | 32975.0 | 0 | Carnivora | 32975.0 | 0 | Canidae | 32975.0 | 0 | Canis | 32975.0 | 0 | Canis lupus | 32975.0 | 0 |
| scf7180000049602 | 6935 | 0.4287 | 0 | 124.3493 | Eukaryota | 15398.0 | 0 | Platyhelminthes | 8163.0 | 2 | Strigeidida | 8163.0 | 5 | Schistosomatidae | 8163.0 | 5 | Schistosoma | 8163.0 | 5 | Schistosoma mansoni | 8163.0 | 5 |
| scf7180000041841 | 140230 | 0.4143 | 0 | 58.9735 | Eukaryota | 75442.0 | 0 | Chordata | 75442.0 | 0 | Carnivora | 75442.0 | 0 | Canidae | 75442.0 | 0 | Canis | 75442.0 | 0 | Canis lupus | 75442.0 | 0 |
| scf7180000054783 | 23387 | 0.3559 | 0 | 461.0867 | Eukaryota | 8848.0 | 0 | Chordata | 8848.0 | 0 | Primates | 7640.0 | 1 | Hominidae | 6406.0 | 2 | Homo | 6406.0 | 2 | Homo sapiens | 6406.0 | 2 |
| scf7180000042552 | 683899 | 0.4682 | 0 | 59.1332 | Eukaryota | 90382.0 | 0 | Chordata | 90382.0 | 0 | Carnivora | 90382.0 | 0 | Canidae | 90382.0 | 0 | Canis | 90382.0 | 0 | Canis lupus | 90382.0 | 0 |
| scf7180000052903 | 7499 | 0.4366 | 0 | 111.4229 | Eukaryota | 20798.0 | 0 | Chordata | 12802.0 | 2 | Strigeidida | 7502.0 | 6 | Schistosomatidae | 7502.0 | 6 | Schistosoma | 7502.0 | 6 | Schistosoma mansoni | 7502.0 | 6 |
| scf7180000046044 | 138584 | 0.4063 | 0 | 116.3136 | Eukaryota | 64416.0 | 0 | Chordata | 64416.0 | 0 | Carnivora | 64416.0 | 0 | Canidae | 64416.0 | 0 | Canis | 64416.0 | 0 | Canis lupus | 64416.0 | 0 |
| scf7180000055096 | 66087 | 0.4444 | 0 | 148.3928 | Eukaryota | 9095.0 | 0 | Chordata | 9095.0 | 0 | Carnivora | 7683.0 | 2 | Canidae | 7683.0 | 2 | Canis | 4227.0 | 3 | Canis lupus | 4227.0 | 3 |
| scf7180000044933 | 137714 | 0.3817 | 0 | 57.6049 | Eukaryota | 75247.0 | 0 | Chordata | 75247.0 | 0 | Carnivora | 75247.0 | 0 | Canidae | 75247.0 | 0 | Canis | 75247.0 | 0 | Canis lupus | 75247.0 | 0 |
| scf7180000040768 | 304175 | 0.3502 | 0 | 58.1291 | Eukaryota | 15643.0 | 0 | Chordata | 15643.0 | 0 | Carnivora | 15643.0 | 0 | Otariidae | 10951.0 | 3 | Callorhinus | 4699.0 | 5 | Callorhinus ursinus | 4699.0 | 5 |
| scf7180000054569 | 23193 | 0.4455 | 0 | 182.8976 | Eukaryota | 47967.0 | 0 | Chordata | 39332.0 | 2 | Gadiformes | 27322.0 | 5 | Gadidae | 27322.0 | 5 | Gadus | 27322.0 | 5 | Gadus morhua | 27322.0 | 5 |
| scf7180000054742 | 13152 | 0.6179 | 0 | 379.4549 | Eukaryota | 60907.0 | 0 | Chordata | 60907.0 | 0 | Artiodactyla | 35666.0 | 3 | Bovidae | 35666.0 | 3 | Ovis | 22139.0 | 5 | Ovis aries | 22139.0 | 5 |
| scf7180000051478 | 20924 | 0.4414 | 0 | 80.4631 | Eukaryota | 1886.0 | 0 | Chordata | 1886.0 | 0 | Carnivora | 1886.0 | 0 | Canidae | 1886.0 | 0 | Canis | 1886.0 | 0 | Canis lupus | 1886.0 | 0 |
| scf7180000051068 | 9921 | 0.4375 | 0 | 155.1839 | Eukaryota | 355.0 | 0 | Platyhelminthes | 355.0 | 0 | Polystomatidea | 355.0 | 0 | Polystomatidae | 355.0 | 0 | Protopolystoma | 355.0 | 0 | Protopolystoma xenopodis | 355.0 | 0 |
| scf7180000046221 | 141438 | 0.3748 | 0 | 60.3612 | Eukaryota | 39622.0 | 0 | Chordata | 39622.0 | 0 | Carnivora | 39622.0 | 0 | Canidae | 39622.0 | 0 | Canis | 39622.0 | 0 | Canis lupus | 39622.0 | 0 |
| scf7180000056246 | 1064225 | 0.5042 | 0 | 58.3678 | Eukaryota | 116083.0 | 0 | Chordata | 116083.0 | 0 | Carnivora | 116083.0 | 0 | Canidae | 116083.0 | 0 | Canis | 62943.0 | 1 | Canis lupus | 62943.0 | 1 |
| scf7180000041904 | 837725 | 0.428 | 0 | 60.2916 | Eukaryota | 50531.0 | 0 | Chordata | 50531.0 | 0 | Carnivora | 50531.0 | 0 | Canidae | 50531.0 | 0 | Canis | 50531.0 | 0 | Canis lupus | 50531.0 | 0 |
| scf7180000047291 | 56583 | 0.4628 | 0 | 118.5198 | Eukaryota | 41466.0 | 0 | Chordata | 31402.0 | 2 | Gadiformes | 19842.0 | 5 | Gadidae | 19842.0 | 5 | Gadus | 19842.0 | 5 | Gadus morhua | 19842.0 | 5 |
| scf7180000051582 | 3491 | 0.4417 | 0 | 196.2257 | Eukaryota | 10655.0 | 0 | Chordata | 8805.0 | 2 | Gadiformes | 5450.0 | 5 | Gadidae | 5450.0 | 5 | Gadus | 5450.0 | 5 | Gadus morhua | 5450.0 | 5 |
| scf7180000042084 | 994032 | 0.3838 | 0 | 60.2012 | Eukaryota | 50586.0 | 0 | Chordata | 50586.0 | 0 | Carnivora | 50586.0 | 0 | Canidae | 50586.0 | 0 | Vulpes | 25774.0 | 1 | Vulpes vulpes | 25774.0 | 1 |
| scf7180000042759 | 8756 | 0.4366 | 0 | 143.8837 | no-hit | 0.0 | 0 | no-hit | 0.0 | 0 | no-hit | 0.0 | 0 | no-hit | 0.0 | 0 | no-hit | 0.0 | 0 | no-hit | 0.0 | 0 |
| scf7180000042367 | 1230491 | 0.4749 | 0 | 59.0159 | Eukaryota | 84979.0 | 0 | Chordata | 84979.0 | 0 | Carnivora | 84979.0 | 0 | Canidae | 84979.0 | 0 | Canis | 84979.0 | 0 | Canis lupus | 84979.0 | 0 |
| scf7180000053213 | 7209 | 0.4325 | 0 | 57.736 | Eukaryota | 3148.0 | 0 | Chordata | 3148.0 | 0 | Carnivora | 3148.0 | 0 | Canidae | 3148.0 | 0 | Canis | 3148.0 | 0 | Canis lupus | 3148.0 | 0 |
| scf7180000041369 | 1151725 | 0.4324 | 0 | 60.0294 | Eukaryota | 82961.0 | 0 | Chordata | 82961.0 | 0 | Carnivora | 82961.0 | 0 | Canidae | 82961.0 | 0 | Canis | 65084.0 | 1 | Canis lupus | 65084.0 | 1 |
| scf7180000041450 | 698361 | 0.4313 | 0 | 59.0186 | Eukaryota | 76961.0 | 0 | Chordata | 76961.0 | 0 | Carnivora | 76961.0 | 0 | Canidae | 76961.0 | 0 | Canis | 76961.0 | 0 | Canis lupus | 76961.0 | 0 |
| scf7180000054617 | 26160 | 0.3832 | 0 | 783.1856 | Eukaryota | 7003.0 | 0 | Chordata | 7003.0 | 0 | Carnivora | 5269.0 | 2 | Felidae | 4775.0 | 3 | Felis | 4021.0 | 4 | Felis catus | 4021.0 | 4 |
| scf7180000045843 | 25695 | 0.4517 | 0 | 157.2499 | Eukaryota | 25243.0 | 1 | Chordata | 25243.0 | 1 | Gadiformes | 10861.0 | 6 | Gadidae | 10861.0 | 6 | Gadus | 10861.0 | 6 | Gadus morhua | 10861.0 | 6 |
| scf7180000043729 | 83742 | 0.4099 | 0 | 58.1701 | Eukaryota | 35386.0 | 0 | Chordata | 35386.0 | 0 | Carnivora | 35386.0 | 0 | Canidae | 35386.0 | 0 | Canis | 35386.0 | 0 | Canis lupus | 35386.0 | 0 |
| scf7180000043824 | 62699 | 0.3757 | 0 | 69.9238 | Eukaryota | 89707.0 | 0 | Chordata | 89707.0 | 0 | Carnivora | 89707.0 | 0 | Canidae | 89707.0 | 0 | Canis | 89707.0 | 0 | Canis lupus | 89707.0 | 0 |
| scf7180000044749 | 261281 | 0.4953 | 0 | 60.4486 | Eukaryota | 61302.0 | 0 | Chordata | 61302.0 | 0 | Carnivora | 61302.0 | 0 | Ursidae | 35924.0 | 1 | Ursus | 31599.0 | 3 | Canis lupus | 21745.0 | 4 |
| scf7180000052261 | 34820 | 0.4435 | 0 | 52.4883 | Eukaryota | 25183.0 | 0 | Chordata | 14854.0 | 1 | Gonorynchiformes | 10108.0 | 4 | Chanidae | 10108.0 | 4 | Chanos | 10108.0 | 4 | Chanos chanos | 10108.0 | 4 |
| scf7180000040727 | 24569 | 0.4422 | 0 | 407.2943 | Eukaryota | 4081.0 | 0 | Chordata | 3650.0 | 1 | Tetraodontiformes | 2665.0 | 2 | Tetraodontidae | 2665.0 | 2 | Takifugu | 2665.0 | 2 | Takifugu rubripes | 2665.0 | 2 |
| scf7180000056490 | 1431757 | 0.382 | 0 | 61.7604 | Eukaryota | 92520.0 | 0 | Chordata | 92520.0 | 0 | Carnivora | 92520.0 | 0 | Canidae | 92520.0 | 0 | Canis | 92520.0 | 0 | Canis lupus | 92520.0 | 0 |
| scf7180000040629 | 1027077 | 0.3845 | 0 | 64.3072 | Eukaryota | 84323.0 | 0 | Chordata | 84323.0 | 0 | Carnivora | 84323.0 | 0 | Canidae | 84323.0 | 0 | Canis | 84323.0 | 0 | Canis lupus | 84323.0 | 0 |
| scf7180000041417 | 651192 | 0.3647 | 0 | 60.5532 | Eukaryota | 66714.0 | 0 | Chordata | 66714.0 | 0 | Carnivora | 66714.0 | 0 | Canidae | 66714.0 | 0 | Canis | 42895.0 | 1 | Canis lupus | 42895.0 | 1 |
| scf7180000051164 | 17336 | 0.4979 | 0 | 131.3549 | Eukaryota | 9224.0 | 0 | Chordata | 9224.0 | 0 | Carnivora | 9224.0 | 0 | Canidae | 9224.0 | 0 | Canis | 8601.0 | 1 | Canis lupus | 7453.0 | 2 |
| scf7180000041149 | 1320159 | 0.3773 | 0 | 59.4098 | Eukaryota | 22517.0 | 0 | Chordata | 22517.0 | 0 | Carnivora | 20179.0 | 1 | Canidae | 20179.0 | 1 | Canis | 20179.0 | 1 | Canis lupus | 20179.0 | 1 |
| scf7180000043632 | 415156 | 0.3867 | 0 | 59.739 | Eukaryota | 52600.0 | 0 | Chordata | 52600.0 | 0 | Carnivora | 52600.0 | 0 | Canidae | 52600.0 | 0 | Canis | 52600.0 | 0 | Canis lupus | 52600.0 | 0 |
| scf7180000043897 | 26972 | 0.4402 | 0 | 49.2226 | Eukaryota | 4653.0 | 0 | Chordata | 4653.0 | 0 | Primates | 2301.0 | 2 | Hominidae | 1741.0 | 3 | Canis | 1637.0 | 4 | Canis lupus | 1637.0 | 4 |
| scf7180000047246 | 81970 | 0.3846 | 0 | 60.3816 | Eukaryota | 53887.0 | 0 | Chordata | 53887.0 | 0 | Carnivora | 53887.0 | 0 | Canidae | 53887.0 | 0 | Canis | 53887.0 | 0 | Canis lupus | 53887.0 | 0 |
| scf7180000042864 | 623558 | 0.401 | 0 | 60.5665 | Eukaryota | 94184.0 | 0 | Chordata | 94184.0 | 0 | Carnivora | 94184.0 | 0 | Canidae | 94184.0 | 0 | Canis | 94184.0 | 0 | Canis lupus | 94184.0 | 0 |
| scf7180000041400 | 60948 | 0.4185 | 0 | 67.2576 | Eukaryota | 92530.0 | 0 | Chordata | 92530.0 | 0 | Carnivora | 92530.0 | 0 | Canidae | 92530.0 | 0 | Canis | 92530.0 | 0 | Canis lupus | 92530.0 | 0 |
| scf7180000043607 | 155232 | 0.3601 | 0 | 60.2938 | Eukaryota | 26737.0 | 0 | Chordata | 26737.0 | 0 | Carnivora | 26737.0 | 0 | Canidae | 8820.0 | 3 | Canis | 5922.0 | 7 | Canis lupus | 5922.0 | 7 |
| scf7180000049955 | 5314 | 0.453 | 0 | 311.4144 | Eukaryota | 1524.0 | 0 | Platyhelminthes | 701.0 | 2 | Polystomatidea | 701.0 | 2 | Polystomatidae | 701.0 | 2 | Protopolystoma | 701.0 | 2 | Protopolystoma xenopodis | 701.0 | 2 |
| scf7180000043572 | 33617 | 0.4889 | 0 | 55.2026 | Eukaryota | 3839.0 | 1 | Chordata | 3839.0 | 1 | Carnivora | 3110.0 | 3 | Felidae | 1611.0 | 4 | Canis | 1499.0 | 5 | Canis lupus | 1499.0 | 5 |
| scf7180000042946 | 556480 | 0.4313 | 0 | 59.4378 | Eukaryota | 58334.0 | 0 | Chordata | 58334.0 | 0 | Carnivora | 58334.0 | 0 | Canidae | 58334.0 | 0 | Canis | 58334.0 | 0 | Canis lupus | 58334.0 | 0 |
| scf7180000043951 | 258387 | 0.3872 | 0 | 58.5062 | Eukaryota | 65133.0 | 0 | Chordata | 65133.0 | 0 | Carnivora | 65133.0 | 0 | Canidae | 65133.0 | 0 | Canis | 65133.0 | 0 | Canis lupus | 65133.0 | 0 |
| scf7180000048867 | 52324 | 0.4188 | 0 | 60.6458 | Eukaryota | 15822.0 | 0 | Chordata | 15822.0 | 0 | Carnivora | 14350.0 | 1 | Felidae | 12863.0 | 2 | Felis | 12863.0 | 2 | Felis catus | 12863.0 | 2 |
| scf7180000055401 | 8387 | 0.4415 | 0 | 210.1415 | no-hit | 0.0 | 0 | no-hit | 0.0 | 0 | no-hit | 0.0 | 0 | no-hit | 0.0 | 0 | no-hit | 0.0 | 0 | no-hit | 0.0 | 0 |
| scf7180000054404 | 19504 | 0.4229 | 0 | 530.9693 | Eukaryota | 36749.0 | 0 | Chordata | 36749.0 | 0 | Carnivora | 36749.0 | 0 | Canidae | 36749.0 | 0 | Canis | 36749.0 | 0 | Canis lupus | 36749.0 | 0 |
| scf7180000046391 | 8888 | 0.3983 | 0 | 19.7184 | Eukaryota | 64888.0 | 0 | Chordata | 64888.0 | 0 | Carnivora | 64888.0 | 0 | Canidae | 64888.0 | 0 | Canis | 64888.0 | 0 | Canis lupus | 64888.0 | 0 |
| scf7180000050407 | 18328 | 0.4449 | 0 | 87.3593 | Eukaryota | 41725.0 | 0 | Platyhelminthes | 23098.0 | 2 | Cyclophyllidea | 12419.0 | 7 | Hymenolepididae | 12419.0 | 7 | Hymenolepis | 12419.0 | 7 | Hymenolepis microstoma | 12419.0 | 7 |
| scf7180000050783 | 5398 | 0.4298 | 0 | 90.7726 | Eukaryota | 6910.0 | 0 | Chordata | 6910.0 | 0 | Carnivora | 6910.0 | 0 | Canidae | 6910.0 | 0 | Canis | 6910.0 | 0 | Canis lupus | 6910.0 | 0 |
| scf7180000047031 | 29588 | 0.4316 | 0 | 98.5756 | Eukaryota | 25695.0 | 0 | Chordata | 25695.0 | 0 | Carnivora | 25695.0 | 0 | Canidae | 25695.0 | 0 | Canis | 25695.0 | 0 | Canis lupus | 25695.0 | 0 |
| scf7180000040962 | 536059 | 0.4734 | 0 | 58.2802 | Eukaryota | 89328.0 | 0 | Chordata | 89328.0 | 0 | Carnivora | 89328.0 | 0 | Canidae | 89328.0 | 0 | Canis | 89328.0 | 0 | Canis lupus | 89328.0 | 0 |
| scf7180000051772 | 20057 | 0.4523 | 0 | 61.0358 | Eukaryota | 3116.0 | 0 | Chordata | 3116.0 | 0 | Carnivora | 3116.0 | 0 | Canidae | 3116.0 | 0 | Canis | 2809.0 | 1 | Canis lupus | 2809.0 | 1 |
| scf7180000054562 | 14555 | 0.4131 | 0 | 497.4624 | Eukaryota | 33471.0 | 0 | Chordata | 33471.0 | 0 | Carnivora | 33471.0 | 0 | Canidae | 33471.0 | 0 | Canis | 33471.0 | 0 | Canis lupus | 33471.0 | 0 |
| scf7180000051497 | 16036 | 0.4133 | 0 | 60.0484 | Eukaryota | 18082.0 | 0 | Chordata | 18082.0 | 0 | Carnivora | 18082.0 | 0 | Canidae | 18082.0 | 0 | Canis | 18082.0 | 0 | Canis lupus | 18082.0 | 0 |
| scf7180000049254 | 7563 | 0.4416 | 0 | 400.2017 | no-hit | 0.0 | 0 | no-hit | 0.0 | 0 | no-hit | 0.0 | 0 | no-hit | 0.0 | 0 | no-hit | 0.0 | 0 | no-hit | 0.0 | 0 |
| scf7180000050934 | 11162 | 0.4561 | 0 | 1299.936 | Eukaryota | 2246.0 | 0 | Echinodermata | 957.0 | 3 | Forcipulatida | 957.0 | 3 | Asteriidae | 957.0 | 3 | Asterias | 957.0 | 3 | Asterias rubens | 957.0 | 3 |
| scf7180000046505 | 117531 | 0.4296 | 0 | 59.3207 | Eukaryota | 54474.0 | 0 | Chordata | 54474.0 | 0 | Carnivora | 54474.0 | 0 | Canidae | 54474.0 | 0 | Canis | 54474.0 | 0 | Canis lupus | 54474.0 | 0 |
| scf7180000044680 | 332365 | 0.3632 | 0 | 60.0479 | Eukaryota | 42768.0 | 0 | Chordata | 42768.0 | 0 | Carnivora | 42768.0 | 0 | Canidae | 42768.0 | 0 | Canis | 33466.0 | 1 | Canis lupus | 33466.0 | 1 |
| scf7180000043021 | 453356 | 0.4551 | 0 | 58.9313 | Eukaryota | 44765.0 | 0 | Chordata | 44765.0 | 0 | Carnivora | 44765.0 | 0 | Canidae | 41016.0 | 1 | Canis | 32670.0 | 2 | Canis lupus | 32670.0 | 2 |
| scf7180000056578 | 46311 | 0.3352 | 0 | 58.3778 | Eukaryota | 6076.0 | 0 | Chordata | 6076.0 | 0 | Carnivora | 6076.0 | 0 | Felidae | 4259.0 | 1 | Felis | 4259.0 | 1 | Felis catus | 4259.0 | 1 |
| scf7180000050108 | 9476 | 0.442 | 0 | 233.7184 | no-hit | 0.0 | 0 | no-hit | 0.0 | 0 | no-hit | 0.0 | 0 | no-hit | 0.0 | 0 | no-hit | 0.0 | 0 | no-hit | 0.0 | 0 |
| scf7180000043887 | 30711 | 0.3756 | 0 | 53.2959 | Eukaryota | 67075.0 | 0 | Chordata | 67075.0 | 0 | Carnivora | 67075.0 | 0 | Canidae | 67075.0 | 0 | Canis | 67075.0 | 0 | Canis lupus | 67075.0 | 0 |
| scf7180000051516 | 16062 | 0.4491 | 0 | 84.1148 | Eukaryota | 3802.0 | 0 | Platyhelminthes | 2305.0 | 3 | Cyclophyllidea | 2305.0 | 6 | Hymenolepididae | 2305.0 | 6 | Hymenolepis | 2305.0 | 6 | Hymenolepis microstoma | 2305.0 | 6 |
| scf7180000044490 | 314103 | 0.4973 | 0 | 57.9528 | Eukaryota | 34826.0 | 0 | Chordata | 34826.0 | 0 | Carnivora | 34826.0 | 0 | Canidae | 34826.0 | 0 | Canis | 31204.0 | 1 | Canis lupus | 31204.0 | 1 |
| scf7180000050822 | 9309 | 0.4331 | 0 | 421.481 | Eukaryota | 606.0 | 0 | Chordata | 606.0 | 0 | Tetraodontiformes | 606.0 | 0 | Tetraodontidae | 606.0 | 0 | Takifugu | 606.0 | 0 | Takifugu rubripes | 606.0 | 0 |
| scf7180000048880 | 32782 | 0.6044 | 0 | 59.6012 | Eukaryota | 48161.0 | 0 | Chordata | 48161.0 | 0 | Carnivora | 48161.0 | 0 | Canidae | 48161.0 | 0 | Canis | 38886.0 | 1 | Canis lupus | 38886.0 | 1 |
| scf7180000049842 | 16251 | 0.5016 | 0 | 776.9071 | Eukaryota | 9640.0 | 0 | Chordata | 9640.0 | 0 | Carnivora | 9640.0 | 0 | Canidae | 9640.0 | 0 | Canis | 8407.0 | 1 | Canis lupus | 7137.0 | 2 |
| scf7180000045353 | 220294 | 0.3706 | 0 | 58.7343 | Eukaryota | 59660.0 | 0 | Chordata | 59660.0 | 0 | Carnivora | 59660.0 | 0 | Canidae | 45307.0 | 2 | Canis | 39488.0 | 3 | Canis lupus | 39488.0 | 3 |
| scf7180000040743 | 3577 | 0.4543 | 0 | 472.4226 | Eukaryota | 2848.0 | 0 | Platyhelminthes | 1267.0 | 3 | Polystomatidea | 1267.0 | 3 | Polystomatidae | 1267.0 | 3 | Protopolystoma | 1267.0 | 3 | Protopolystoma xenopodis | 1267.0 | 3 |
| scf7180000043075 | 229347 | 0.4137 | 0 | 68.3707 | Eukaryota | 89257.0 | 0 | Chordata | 89257.0 | 0 | Carnivora | 89257.0 | 0 | Canidae | 89257.0 | 0 | Canis | 89257.0 | 0 | Canis lupus | 89257.0 | 0 |
| scf7180000043760 | 12312 | 0.3549 | 0 | 49.911 | Eukaryota | 4341.0 | 0 | Chordata | 4341.0 | 0 | Carnivora | 2501.0 | 2 | Canidae | 2501.0 | 2 | Canis | 2501.0 | 3 | Canis lupus | 2501.0 | 3 |
| scf7180000045027 | 206868 | 0.3453 | 0 | 58.9814 | Eukaryota | 82873.0 | 0 | Chordata | 82873.0 | 0 | Carnivora | 82873.0 | 0 | Canidae | 82873.0 | 0 | Canis | 82873.0 | 0 | Canis lupus | 82873.0 | 0 |
| scf7180000051782 | 11689 | 0.4464 | 0 | 297.6957 | Eukaryota | 4541.0 | 0 | Chordata | 2510.0 | 1 | Polystomatidea | 2031.0 | 2 | Polystomatidae | 2031.0 | 2 | Protopolystoma | 2031.0 | 3 | Protopolystoma xenopodis | 2031.0 | 3 |
| scf7180000048436 | 13171 | 0.3891 | 0 | 30.6825 | Eukaryota | 35226.0 | 0 | Chordata | 35226.0 | 0 | Carnivora | 35226.0 | 0 | Canidae | 35226.0 | 0 | Canis | 35226.0 | 0 | Canis lupus | 35226.0 | 0 |
| scf7180000048859 | 68824 | 0.4302 | 0 | 58.1846 | Eukaryota | 44219.0 | 0 | Chordata | 44219.0 | 0 | Carnivora | 44219.0 | 0 | Canidae | 23239.0 | 3 | Canis | 11705.0 | 7 | Canis lupus | 11705.0 | 7 |
| scf7180000048375 | 18261 | 0.4409 | 0 | 104.8225 | Eukaryota | 6898.0 | 0 | Chordata | 4282.0 | 2 | Kurtiformes | 4282.0 | 2 | Apogonidae | 4282.0 | 2 | Sphaeramia | 4282.0 | 2 | Sphaeramia orbicularis | 4282.0 | 2 |
| scf7180000045919 | 192270 | 0.4729 | 0 | 60.6241 | Eukaryota | 25979.0 | 0 | Chordata | 25979.0 | 0 | Carnivora | 13496.0 | 2 | Canidae | 13496.0 | 2 | Vulpes | 9180.0 | 3 | Vulpes vulpes | 9180.0 | 3 |
| scf7180000041738 | 13549 | 0.3833 | 0 | 240.4073 | Eukaryota | 90921.0 | 0 | Chordata | 90921.0 | 0 | Carnivora | 90921.0 | 0 | Canidae | 90921.0 | 0 | Canis | 90921.0 | 0 | Canis lupus | 90921.0 | 0 |
| scf7180000044834 | 184278 | 0.5332 | 0 | 56.3292 | Eukaryota | 53749.0 | 0 | Chordata | 53749.0 | 0 | Carnivora | 53749.0 | 0 | Canidae | 53749.0 | 0 | Canis | 53749.0 | 0 | Canis lupus | 53749.0 | 0 |
| scf7180000043198 | 5080 | 0.4346 | 0 | 62.8263 | Eukaryota | 182.0 | 0 | Platyhelminthes | 182.0 | 0 | Polystomatidea | 182.0 | 0 | Polystomatidae | 182.0 | 0 | Protopolystoma | 182.0 | 0 | Protopolystoma xenopodis | 182.0 | 0 |
| scf7180000041601 | 26890 | 0.3851 | 0 | 40.6488 | Eukaryota | 86456.0 | 0 | Chordata | 86456.0 | 0 | Carnivora | 86456.0 | 0 | Canidae | 86456.0 | 0 | Canis | 86456.0 | 0 | Canis lupus | 86456.0 | 0 |
| scf7180000043849 | 87125 | 0.4319 | 0 | 57.2017 | Eukaryota | 96304.0 | 0 | Chordata | 96304.0 | 0 | Carnivora | 93074.0 | 1 | Canidae | 79254.0 | 3 | Canis | 74387.0 | 5 | Canis lupus | 74387.0 | 5 |
| scf7180000055296 | 6459 | 0.4409 | 0 | 411.9413 | Eukaryota | 213.0 | 0 | Chordata | 213.0 | 0 | Tetraodontiformes | 213.0 | 0 | Tetraodontidae | 213.0 | 0 | Takifugu | 213.0 | 0 | Takifugu rubripes | 213.0 | 0 |
| scf7180000042988 | 304806 | 0.3765 | 0 | 62.8203 | Eukaryota | 96280.0 | 0 | Chordata | 96280.0 | 0 | Carnivora | 96280.0 | 0 | Canidae | 96280.0 | 0 | Canis | 96280.0 | 0 | Canis lupus | 96280.0 | 0 |
| scf7180000041789 | 189740 | 0.4391 | 0 | 59.5317 | Eukaryota | 43562.0 | 0 | Chordata | 43562.0 | 0 | Carnivora | 43562.0 | 0 | Canidae | 37600.0 | 1 | Vulpes | 23048.0 | 2 | Vulpes vulpes | 23048.0 | 2 |
| scf7180000046895 | 18963 | 0.3798 | 0 | 44.3165 | Eukaryota | 5347.0 | 0 | Chordata | 5347.0 | 0 | Carnivora | 4027.0 | 1 | Mustelidae | 2106.0 | 3 | Mustela | 2106.0 | 5 | Mustela putorius | 2106.0 | 5 |
| scf7180000041211 | 101395 | 0.3705 | 0 | 56.8633 | Eukaryota | 73335.0 | 0 | Chordata | 73335.0 | 0 | Carnivora | 73335.0 | 0 | Canidae | 73335.0 | 0 | Canis | 73335.0 | 0 | Canis lupus | 73335.0 | 0 |
| scf7180000042781 | 105498 | 0.3939 | 0 | 57.8608 | Eukaryota | 61845.0 | 0 | Chordata | 61845.0 | 0 | Carnivora | 61845.0 | 0 | Mustelidae | 24404.0 | 3 | Mustela | 12746.0 | 5 | Mustela putorius | 12746.0 | 5 |
| scf7180000048949 | 5951 | 0.4435 | 0 | 145.2822 | no-hit | 0.0 | 0 | no-hit | 0.0 | 0 | no-hit | 0.0 | 0 | no-hit | 0.0 | 0 | no-hit | 0.0 | 0 | no-hit | 0.0 | 0 |
| scf7180000049971 | 31882 | 0.4252 | 0 | 59.3412 | Eukaryota | 26837.0 | 0 | Chordata | 26837.0 | 0 | Carnivora | 26837.0 | 0 | Canidae | 26837.0 | 0 | Canis | 26837.0 | 0 | Canis lupus | 26837.0 | 0 |
| scf7180000043297 | 152456 | 0.4024 | 0 | 57.6647 | Eukaryota | 79550.0 | 0 | Chordata | 79550.0 | 0 | Carnivora | 79550.0 | 0 | Canidae | 79550.0 | 0 | Canis | 79550.0 | 0 | Canis lupus | 79550.0 | 0 |
| scf7180000045744 | 169264 | 0.3816 | 0 | 58.1472 | Eukaryota | 90972.0 | 0 | Chordata | 90972.0 | 0 | Carnivora | 90972.0 | 0 | Canidae | 90972.0 | 0 | Canis | 90972.0 | 0 | Canis lupus | 90972.0 | 0 |
| scf7180000053929 | 13044 | 0.5117 | 0 | 429.8165 | Eukaryota | 9694.0 | 0 | Chordata | 9694.0 | 0 | Carnivora | 9694.0 | 0 | Canidae | 9694.0 | 0 | Nyctereutes | 5300.0 | 1 | Nyctereutes procyonoides | 5300.0 | 1 |
| scf7180000055556 | 15201 | 0.4337 | 0 | 226.1608 | Eukaryota | 907.0 | 1 | Chordata | 907.0 | 1 | Gonorynchiformes | 422.0 | 3 | Chanidae | 422.0 | 3 | Chanos | 422.0 | 3 | Chanos chanos | 422.0 | 3 |
| scf7180000042086 | 784546 | 0.4592 | 0 | 58.4554 | Eukaryota | 77825.0 | 0 | Chordata | 77825.0 | 0 | Carnivora | 77825.0 | 0 | Canidae | 77825.0 | 0 | Canis | 70659.0 | 1 | Canis lupus | 70659.0 | 1 |
| scf7180000044851 | 25508 | 0.4363 | 0 | 160.6725 | Eukaryota | 1268.0 | 0 | Chordata | 828.0 | 1 | Tetraodontiformes | 828.0 | 1 | Tetraodontidae | 828.0 | 1 | Takifugu | 828.0 | 1 | Takifugu rubripes | 828.0 | 1 |
| scf7180000041579 | 1044605 | 0.3532 | 0 | 59.1664 | Eukaryota | 91704.0 | 0 | Chordata | 91704.0 | 0 | Carnivora | 91704.0 | 0 | Canidae | 91704.0 | 0 | Canis | 91704.0 | 0 | Canis lupus | 91704.0 | 0 |
| scf7180000047431 | 67323 | 0.4471 | 0 | 58.8054 | Eukaryota | 6506.0 | 0 | Chordata | 6506.0 | 0 | Carnivora | 6506.0 | 0 | Canidae | 6506.0 | 0 | Canis | 6506.0 | 0 | Canis lupus | 6506.0 | 0 |
| scf7180000056450 | 316671 | 0.3927 | 0 | 60.5393 | Eukaryota | 110865.0 | 0 | Chordata | 110865.0 | 0 | Carnivora | 110865.0 | 0 | Canidae | 110865.0 | 0 | Canis | 110865.0 | 0 | Canis lupus | 110865.0 | 0 |
| scf7180000056616 | 534773 | 0.3893 | 100 | 58.6479 | Eukaryota | 39243.0 | 0 | Chordata | 39243.0 | 0 | Carnivora | 39243.0 | 0 | Canidae | 39243.0 | 0 | Canis | 39243.0 | 0 | Canis lupus | 39243.0 | 0 |
| scf7180000046971 | 10176 | 0.4438 | 0 | 94.4109 | no-hit | 0.0 | 0 | no-hit | 0.0 | 0 | no-hit | 0.0 | 0 | no-hit | 0.0 | 0 | no-hit | 0.0 | 0 | no-hit | 0.0 | 0 |
| scf7180000046298 | 9052 | 0.4513 | 0 | 43.6811 | Eukaryota | 41785.0 | 0 | Chordata | 41785.0 | 0 | Carnivora | 41785.0 | 0 | Canidae | 41785.0 | 0 | Canis | 41785.0 | 0 | Canis lupus | 41785.0 | 0 |
| scf7180000041555 | 1008294 | 0.4349 | 0 | 58.4031 | Eukaryota | 53125.0 | 0 | Chordata | 53125.0 | 0 | Carnivora | 53125.0 | 0 | Canidae | 53125.0 | 0 | Canis | 53125.0 | 0 | Canis lupus | 53125.0 | 0 |
| scf7180000053605 | 5073 | 0.5105 | 0 | 336.8266 | Eukaryota | 27088.0 | 0 | Chordata | 27088.0 | 0 | Carnivora | 27088.0 | 0 | Canidae | 27088.0 | 0 | Canis | 27088.0 | 0 | Canis lupus | 27088.0 | 0 |
| scf7180000053762 | 2848 | 0.4417 | 0 | 225.0867 | no-hit | 0.0 | 0 | no-hit | 0.0 | 0 | no-hit | 0.0 | 0 | no-hit | 0.0 | 0 | no-hit | 0.0 | 0 | no-hit | 0.0 | 0 |
| scf7180000052965 | 3496 | 0.4502 | 0 | 433.5726 | no-hit | 0.0 | 0 | no-hit | 0.0 | 0 | no-hit | 0.0 | 0 | no-hit | 0.0 | 0 | no-hit | 0.0 | 0 | no-hit | 0.0 | 0 |
| scf7180000045760 | 52128 | 0.4084 | 0 | 57.1149 | Eukaryota | 63973.0 | 0 | Chordata | 63973.0 | 0 | Carnivora | 63973.0 | 0 | Canidae | 63973.0 | 0 | Canis | 63973.0 | 0 | Canis lupus | 63973.0 | 0 |
| scf7180000044257 | 321050 | 0.5115 | 0 | 57.5738 | Eukaryota | 101713.0 | 0 | Chordata | 101713.0 | 0 | Carnivora | 101713.0 | 0 | Canidae | 101713.0 | 0 | Canis | 71156.0 | 1 | Canis lupus | 71156.0 | 1 |
| scf7180000042432 | 121511 | 0.4308 | 0 | 58.8564 | Eukaryota | 89029.0 | 0 | Chordata | 89029.0 | 0 | Carnivora | 89029.0 | 0 | Canidae | 89029.0 | 0 | Canis | 89029.0 | 0 | Canis lupus | 89029.0 | 0 |
| scf7180000041871 | 869373 | 0.5122 | 0 | 68.9909 | Eukaryota | 119100.0 | 0 | Chordata | 119100.0 | 0 | Carnivora | 119100.0 | 0 | Canidae | 119100.0 | 0 | Canis | 119100.0 | 0 | Canis lupus | 119100.0 | 0 |
| scf7180000047012 | 25161 | 0.4364 | 0 | 111.9069 | Eukaryota | 1185.0 | 0 | Platyhelminthes | 664.0 | 2 | Polystomatidea | 664.0 | 2 | Polystomatidae | 664.0 | 2 | Protopolystoma | 664.0 | 2 | Protopolystoma xenopodis | 664.0 | 2 |
| scf7180000056597 | 767830 | 0.3908 | 100 | 59.1228 | Eukaryota | 118184.0 | 0 | Chordata | 118184.0 | 0 | Carnivora | 118184.0 | 0 | Canidae | 70900.0 | 2 | Canis | 58734.0 | 4 | Canis lupus | 58734.0 | 4 |
| scf7180000052510 | 4168 | 0.4407 | 0 | 271.9241 | Eukaryota | 693.0 | 0 | Chordata | 693.0 | 0 | Tetraodontiformes | 693.0 | 0 | Tetraodontidae | 693.0 | 0 | Takifugu | 693.0 | 0 | Takifugu rubripes | 693.0 | 0 |
| scf7180000051688 | 12286 | 0.4539 | 0 | 94.1029 | no-hit | 0.0 | 0 | no-hit | 0.0 | 0 | no-hit | 0.0 | 0 | no-hit | 0.0 | 0 | no-hit | 0.0 | 0 | no-hit | 0.0 | 0 |
| scf7180000045549 | 82302 | 0.4781 | 0 | 57.1217 | Eukaryota | 19494.0 | 0 | Chordata | 19494.0 | 0 | Carnivora | 19494.0 | 0 | Canidae | 19494.0 | 0 | Canis | 19494.0 | 0 | Canis lupus | 19494.0 | 0 |
| scf7180000047263 | 56879 | 0.5441 | 0 | 55.661 | Eukaryota | 8296.0 | 0 | Chordata | 8296.0 | 0 | Primates | 7656.0 | 1 | Hominidae | 6933.0 | 2 | Homo | 6933.0 | 2 | Homo sapiens | 6933.0 | 2 |
| scf7180000045376 | 102071 | 0.4051 | 0 | 58.0902 | Eukaryota | 91310.0 | 0 | Chordata | 91310.0 | 0 | Carnivora | 91310.0 | 0 | Canidae | 91310.0 | 0 | Canis | 91310.0 | 0 | Canis lupus | 91310.0 | 0 |
| scf7180000045090 | 9597 | 0.4337 | 0 | 24.6878 | Eukaryota | 44495.0 | 0 | Chordata | 44495.0 | 0 | Carnivora | 44495.0 | 0 | Canidae | 44495.0 | 0 | Canis | 44495.0 | 0 | Canis lupus | 44495.0 | 0 |
| scf7180000042481 | 14074 | 0.4695 | 0 | 96.0795 | Eukaryota | 24521.0 | 0 | Chordata | 24521.0 | 0 | Carnivora | 24521.0 | 0 | Canidae | 24521.0 | 0 | Canis | 24521.0 | 0 | Canis lupus | 24521.0 | 0 |
| scf7180000049505 | 628173 | 0.4557 | 0 | 60.2245 | Eukaryota | 127467.0 | 0 | Chordata | 127467.0 | 0 | Carnivora | 127467.0 | 0 | Canidae | 127467.0 | 0 | Canis | 127467.0 | 0 | Canis lupus | 127467.0 | 0 |
| scf7180000055953 | 30357 | 0.5866 | 0 | 733.9413 | Eukaryota | 40968.0 | 0 | Chordata | 40968.0 | 0 | Artiodactyla | 28808.0 | 1 | Bovidae | 28808.0 | 1 | Ovis | 16319.0 | 2 | Ovis aries | 16319.0 | 2 |
| scf7180000043408 | 209219 | 0.4415 | 0 | 60.9628 | Eukaryota | 88996.0 | 0 | Chordata | 88996.0 | 0 | Carnivora | 88996.0 | 0 | Canidae | 88996.0 | 0 | Canis | 88996.0 | 0 | Canis lupus | 88996.0 | 0 |
| scf7180000043620 | 658614 | 0.3435 | 0 | 59.6359 | Eukaryota | 87403.0 | 0 | Chordata | 87403.0 | 0 | Carnivora | 87403.0 | 0 | Canidae | 87403.0 | 0 | Canis | 87403.0 | 0 | Canis lupus | 87403.0 | 0 |
| scf7180000043288 | 119492 | 0.4209 | 0 | 61.4471 | Eukaryota | 49395.0 | 0 | Chordata | 49395.0 | 0 | Carnivora | 49395.0 | 0 | Canidae | 49395.0 | 0 | Canis | 28629.0 | 1 | Canis lupus | 28629.0 | 1 |
| scf7180000043290 | 504083 | 0.3776 | 0 | 60.214 | Eukaryota | 23780.0 | 0 | Chordata | 23780.0 | 0 | Carnivora | 23780.0 | 0 | Canidae | 23780.0 | 0 | Canis | 23780.0 | 0 | Canis lupus | 23780.0 | 0 |
| scf7180000045522 | 41192 | 0.3864 | 0 | 56.1662 | Eukaryota | 6712.0 | 1 | Chordata | 6712.0 | 1 | Carnivora | 4486.0 | 3 | Canidae | 3808.0 | 4 | Canis | 3808.0 | 5 | Canis lupus | 3808.0 | 5 |
| scf7180000050123 | 6366 | 0.4493 | 0 | 102.7219 | Eukaryota | 2180.0 | 0 | Chordata | 1198.0 | 1 | Polystomatidea | 982.0 | 2 | Polystomatidae | 982.0 | 2 | Protopolystoma | 982.0 | 2 | Protopolystoma xenopodis | 982.0 | 2 |
| scf7180000044683 | 669865 | 0.3974 | 100 | 60.3931 | Eukaryota | 54970.0 | 0 | Chordata | 54970.0 | 0 | Carnivora | 54970.0 | 0 | Canidae | 54970.0 | 0 | Canis | 54970.0 | 0 | Canis lupus | 54970.0 | 0 |
| scf7180000056226 | 1989577 | 0.4551 | 0 | 57.885 | Eukaryota | 69610.0 | 0 | Chordata | 69610.0 | 0 | Carnivora | 69610.0 | 0 | Canidae | 69610.0 | 0 | Canis | 69610.0 | 0 | Canis lupus | 69610.0 | 0 |
| scf7180000052295 | 3836 | 0.4466 | 0 | 928.943 | no-hit | 0.0 | 0 | no-hit | 0.0 | 0 | no-hit | 0.0 | 0 | no-hit | 0.0 | 0 | no-hit | 0.0 | 0 | no-hit | 0.0 | 0 |
| scf7180000041910 | 195069 | 0.3475 | 0 | 58.0628 | Eukaryota | 18985.0 | 0 | Chordata | 18985.0 | 0 | Carnivora | 18985.0 | 0 | Canidae | 18985.0 | 0 | Canis | 18985.0 | 0 | Canis lupus | 18985.0 | 0 |
| scf7180000040688 | 208891 | 0.4689 | 0 | 57.92 | Eukaryota | 54554.0 | 0 | Chordata | 54554.0 | 0 | Carnivora | 54554.0 | 0 | Canidae | 54554.0 | 0 | Canis | 54554.0 | 0 | Canis lupus | 54554.0 | 0 |
| scf7180000046405 | 95279 | 0.4358 | 0 | 221.8049 | Eukaryota | 42631.0 | 0 | Chordata | 24617.0 | 2 | Tetraodontiformes | 11638.0 | 7 | Tetraodontidae | 11638.0 | 7 | Takifugu | 11638.0 | 7 | Takifugu rubripes | 11638.0 | 7 |
| scf7180000042643 | 44470 | 0.4331 | 0 | 54.3579 | Eukaryota | 18022.0 | 0 | Chordata | 18022.0 | 0 | Carnivora | 18022.0 | 0 | Canidae | 13605.0 | 2 | Canis | 12125.0 | 4 | Canis lupus | 12125.0 | 4 |
| scf7180000044831 | 242507 | 0.4367 | 0 | 59.1622 | Eukaryota | 31512.0 | 0 | Chordata | 31512.0 | 0 | Carnivora | 31512.0 | 0 | Canidae | 29004.0 | 1 | Canis | 25594.0 | 2 | Canis lupus | 25594.0 | 2 |
| scf7180000047702 | 80166 | 0.3616 | 0 | 60.8298 | Eukaryota | 31804.0 | 0 | Chordata | 31804.0 | 0 | Carnivora | 31804.0 | 0 | Canidae | 31804.0 | 0 | Canis | 31804.0 | 0 | Canis lupus | 31804.0 | 0 |
| scf7180000041596 | 69449 | 0.3775 | 0 | 52.1736 | Eukaryota | 10024.0 | 0 | Chordata | 10024.0 | 0 | Primates | 6631.0 | 3 | Hominidae | 5644.0 | 4 | Pan | 3505.0 | 5 | Pan troglodytes | 3505.0 | 5 |
| scf7180000050919 | 9826 | 0.4998 | 0 | 189.9862 | Eukaryota | 8521.0 | 0 | Chordata | 8521.0 | 0 | Carnivora | 8521.0 | 0 | Canidae | 8521.0 | 0 | Canis | 7430.0 | 1 | Canis lupus | 6325.0 | 2 |
| scf7180000053537 | 6548 | 0.4487 | 0 | 80.4538 | Eukaryota | 300.0 | 0 | Chordata | 300.0 | 0 | Salmoniformes | 300.0 | 0 | Salmonidae | 300.0 | 0 | Coregonus | 300.0 | 0 | Coregonus sp. 'balchen' | 300.0 | 0 |
| scf7180000056443 | 330825 | 0.3835 | 0 | 58.8026 | Eukaryota | 94959.0 | 0 | Chordata | 94959.0 | 0 | Carnivora | 94959.0 | 0 | Canidae | 94959.0 | 0 | Canis | 94959.0 | 0 | Canis lupus | 94959.0 | 0 |
| scf7180000047655 | 76224 | 0.4423 | 0 | 58.0919 | Eukaryota | 32448.0 | 0 | Chordata | 32448.0 | 0 | Carnivora | 32448.0 | 0 | Canidae | 32448.0 | 0 | Canis | 22176.0 | 1 | Canis lupus | 22176.0 | 1 |
| scf7180000049531 | 3464 | 0.4538 | 0 | 433.8594 | Eukaryota | 3106.0 | 0 | Platyhelminthes | 1817.0 | 3 | Polystomatidea | 1817.0 | 3 | Polystomatidae | 1817.0 | 3 | Protopolystoma | 1817.0 | 3 | Protopolystoma xenopodis | 1817.0 | 3 |
| scf7180000045891 | 180399 | 0.4365 | 0 | 352.3168 | Eukaryota | 94499.0 | 0 | Chordata | 94499.0 | 0 | Carnivora | 94499.0 | 0 | Canidae | 94499.0 | 0 | Canis | 94499.0 | 0 | Canis lupus | 94499.0 | 0 |
| scf7180000051577 | 4245 | 0.437 | 0 | 152.5141 | Eukaryota | 1506.0 | 0 | Chordata | 1045.0 | 1 | Kurtiformes | 804.0 | 3 | Apogonidae | 804.0 | 3 | Sphaeramia | 804.0 | 3 | Sphaeramia orbicularis | 804.0 | 3 |
| scf7180000051300 | 25996 | 0.5606 | 0 | 56.6612 | Eukaryota | 6057.0 | 0 | Chordata | 6057.0 | 0 | Carnivora | 6057.0 | 0 | Canidae | 6057.0 | 0 | Canis | 6057.0 | 0 | Canis lupus | 6057.0 | 0 |
| scf7180000050612 | 8112 | 0.4514 | 0 | 1270.0383 | no-hit | 0.0 | 0 | no-hit | 0.0 | 0 | no-hit | 0.0 | 0 | no-hit | 0.0 | 0 | no-hit | 0.0 | 0 | no-hit | 0.0 | 0 |
| scf7180000042204 | 691292 | 0.3808 | 0 | 59.8207 | Eukaryota | 85931.0 | 0 | Chordata | 85931.0 | 0 | Carnivora | 85931.0 | 0 | Canidae | 76630.0 | 2 | Canis | 56373.0 | 3 | Canis lupus | 56373.0 | 3 |
| scf7180000047039 | 79516 | 0.3575 | 0 | 59.5698 | Eukaryota | 20669.0 | 0 | Chordata | 20669.0 | 0 | Primates | 11817.0 | 3 | Hominidae | 11817.0 | 4 | Homo | 8354.0 | 6 | Homo sapiens | 8354.0 | 6 |
| scf7180000045201 | 196148 | 0.6409 | 0 | 54.1217 | Eukaryota | 31618.0 | 0 | Chordata | 31618.0 | 0 | Carnivora | 31618.0 | 0 | Canidae | 31618.0 | 0 | Canis | 23580.0 | 1 | Canis lupus | 23580.0 | 1 |
| scf7180000054855 | 144787 | 0.4415 | 0 | 232.0632 | Eukaryota | 94369.0 | 0 | Chordata | 94369.0 | 0 | Carnivora | 94369.0 | 0 | Canidae | 94369.0 | 0 | Canis | 94369.0 | 0 | Canis lupus | 94369.0 | 0 |
| scf7180000045565 | 20253 | 0.4266 | 0 | 45.6952 | Eukaryota | 6629.0 | 0 | Chordata | 6629.0 | 0 | Artiodactyla | 4750.0 | 1 | Bovidae | 4168.0 | 2 | Bos | 2406.0 | 6 | Bos taurus | 1208.0 | 7 |
| scf7180000053073 | 4031 | 0.4294 | 0 | 139.1143 | no-hit | 0.0 | 0 | no-hit | 0.0 | 0 | no-hit | 0.0 | 0 | no-hit | 0.0 | 0 | no-hit | 0.0 | 0 | no-hit | 0.0 | 0 |
| scf7180000051251 | 6121 | 0.4458 | 0 | 248.4633 | Eukaryota | 2471.0 | 0 | Platyhelminthes | 927.0 | 3 | Polystomatidea | 927.0 | 3 | Polystomatidae | 927.0 | 3 | Protopolystoma | 927.0 | 3 | Protopolystoma xenopodis | 927.0 | 3 |
| scf7180000056329 | 634356 | 0.438 | 0 | 58.7568 | Eukaryota | 70966.0 | 0 | Chordata | 70966.0 | 0 | Carnivora | 70966.0 | 0 | Canidae | 70966.0 | 0 | Canis | 65004.0 | 1 | Canis lupus | 65004.0 | 1 |
| scf7180000041189 | 1311131 | 0.4033 | 0 | 58.6336 | Eukaryota | 79537.0 | 0 | Chordata | 79537.0 | 0 | Carnivora | 79537.0 | 0 | Canidae | 79537.0 | 0 | Canis | 57798.0 | 1 | Canis lupus | 57798.0 | 1 |
| scf7180000046770 | 17330 | 0.413 | 0 | 52.5842 | Eukaryota | 41258.0 | 0 | Chordata | 41258.0 | 0 | Carnivora | 41258.0 | 0 | Canidae | 38083.0 | 1 | Canis | 32376.0 | 2 | Canis lupus | 32376.0 | 2 |
| scf7180000045998 | 18075 | 0.4393 | 0 | 216.0349 | Eukaryota | 8254.0 | 0 | Chordata | 5604.0 | 1 | Salmoniformes | 2699.0 | 3 | Salmonidae | 2699.0 | 3 | Salmo | 2699.0 | 3 | Salmo trutta | 2699.0 | 3 |
| scf7180000046483 | 10858 | 0.5092 | 0 | 42.6827 | Eukaryota | 10073.0 | 0 | Chordata | 10073.0 | 0 | Carnivora | 8938.0 | 2 | Canidae | 3869.0 | 5 | Vulpes | 2793.0 | 8 | Vulpes vulpes | 2793.0 | 8 |
| scf7180000042882 | 581848 | 0.4104 | 0 | 60.2609 | Eukaryota | 88864.0 | 0 | Chordata | 88864.0 | 0 | Carnivora | 88864.0 | 0 | Canidae | 88864.0 | 0 | Canis | 70845.0 | 1 | Canis lupus | 70845.0 | 1 |
| scf7180000046235 | 168943 | 0.3367 | 0 | 60.3678 | Eukaryota | 25174.0 | 0 | Chordata | 25174.0 | 0 | Primates | 7855.0 | 3 | Hominidae | 7855.0 | 3 | Homo | 7855.0 | 4 | Homo sapiens | 7855.0 | 4 |
| scf7180000043740 | 252830 | 0.4155 | 0 | 58.9991 | Eukaryota | 35442.0 | 0 | Chordata | 35442.0 | 0 | Carnivora | 35442.0 | 0 | Canidae | 35442.0 | 0 | Canis | 35442.0 | 0 | Canis lupus | 35442.0 | 0 |
| scf7180000050462 | 3024 | 0.4517 | 0 | 724.4197 | no-hit | 0.0 | 0 | no-hit | 0.0 | 0 | no-hit | 0.0 | 0 | no-hit | 0.0 | 0 | no-hit | 0.0 | 0 | no-hit | 0.0 | 0 |
| scf7180000041374 | 58596 | 0.3786 | 0 | 54.4785 | Eukaryota | 16793.0 | 0 | Chordata | 16793.0 | 0 | Carnivora | 12648.0 | 2 | Canidae | 12648.0 | 2 | Canis | 12648.0 | 3 | Canis lupus | 12648.0 | 3 |
| scf7180000045819 | 147488 | 0.3971 | 0 | 59.0434 | Eukaryota | 65359.0 | 0 | Chordata | 65359.0 | 0 | Carnivora | 65359.0 | 0 | Canidae | 65359.0 | 0 | Canis | 65359.0 | 0 | Canis lupus | 65359.0 | 0 |
| scf7180000045636 | 110462 | 0.4145 | 0 | 59.3843 | Eukaryota | 4595.0 | 0 | Chordata | 4595.0 | 0 | Carnivora | 4007.0 | 1 | Canidae | 2556.0 | 2 | Canis | 2556.0 | 2 | Canis lupus | 2556.0 | 2 |
| scf7180000043889 | 411325 | 0.3867 | 0 | 61.2453 | Eukaryota | 28014.0 | 0 | Chordata | 28014.0 | 0 | Carnivora | 28014.0 | 0 | Canidae | 28014.0 | 0 | Canis | 28014.0 | 0 | Canis lupus | 28014.0 | 0 |
| scf7180000048538 | 3709 | 0.4492 | 0 | 44.3753 | no-hit | 0.0 | 0 | no-hit | 0.0 | 0 | no-hit | 0.0 | 0 | no-hit | 0.0 | 0 | no-hit | 0.0 | 0 | no-hit | 0.0 | 0 |
| scf7180000048387 | 49537 | 0.3773 | 0 | 59.7293 | Eukaryota | 18328.0 | 0 | Chordata | 18328.0 | 0 | Carnivora | 18328.0 | 0 | Phocidae | 5547.0 | 3 | Leptonychotes | 3678.0 | 8 | Leptonychotes weddellii | 3678.0 | 8 |
| scf7180000055608 | 12406 | 0.4389 | 0 | 218.8969 | Eukaryota | 1203.0 | 0 | Chordata | 957.0 | 1 | Kurtiformes | 957.0 | 1 | Apogonidae | 957.0 | 1 | Sphaeramia | 957.0 | 1 | Sphaeramia orbicularis | 957.0 | 1 |
| scf7180000043790 | 403863 | 0.5812 | 0 | 57.3231 | Eukaryota | 198895.0 | 0 | Chordata | 198895.0 | 0 | Carnivora | 198895.0 | 0 | Canidae | 198895.0 | 0 | Canis | 198895.0 | 0 | Canis lupus | 198895.0 | 0 |
| scf7180000053594 | 3042 | 0.4402 | 0 | 190.972 | no-hit | 0.0 | 0 | no-hit | 0.0 | 0 | no-hit | 0.0 | 0 | no-hit | 0.0 | 0 | no-hit | 0.0 | 0 | no-hit | 0.0 | 0 |

| | | | | | | | | | | | | | | | | | | | | | | |
|---|---|---|---|---|---|---|---|---|---|---|---|---|---|---|---|---|---|---|---|---|---|---|
| scf7180000051821 | 5865 | 0.4314 | 0 | 64.4444 | Eukaryota | 9801.0 | 0 | Chordata | 5941.0 | 3 | Gadiformes | 3997.0 | 7 | Gadidae | 3997.0 | 7 | Gadus | 3997.0 | 7 | Gadus morhua | 3997.0 | 8 |
| scf7180000042662 | 731380 | 0.4194 | 0 | 59.217 | Eukaryota | 39152.0 | 0 | Chordata | 39152.0 | 0 | Carnivora | 39152.0 | 0 | Canidae | 32876.0 | 1 | Canis | 21998.0 | 2 | Canis lupus | 21998.0 | 2 |
| scf7180000051640 | 16321 | 0.4481 | 0 | 41.7647 | Eukaryota | 9566.0 | 0 | Chordata | 9566.0 | 0 | Carnivora | 9566.0 | 0 | Canidae | 9566.0 | 0 | Canis | 5243.0 | 1 | Canis lupus | 5243.0 | 1 |
| scf7180000051480 | 12938 | 0.4437 | 0 | 106.1146 | Eukaryota | 8206.0 | 0 | Chordata | 7037.0 | 1 | Tetraodontiformes | 5583.0 | 2 | Tetraodontidae | 5583.0 | 2 | Takifugu | 5583.0 | 2 | Takifugu rubripes | 5583.0 | 2 |
| scf7180000046048 | 130630 | 0.4178 | 0 | 58.3025 | Eukaryota | 30084.0 | 0 | Chordata | 30084.0 | 0 | Carnivora | 30084.0 | 0 | Canidae | 30084.0 | 0 | Canis | 21776.0 | 1 | Canis lupus | 21776.0 | 1 |
| scf7180000054806 | 26664 | 0.4442 | 0 | 296.6245 | Eukaryota | 74169.0 | 0 | Chordata | 74169.0 | 0 | Carnivora | 74169.0 | 0 | Canidae | 74169.0 | 0 | Canis | 74169.0 | 0 | Canis lupus | 74169.0 | 0 |
| scf7180000054015 | 3603 | 0.4532 | 0 | 66.9142 | no-hit | 0.0 | 0 | no-hit | 0.0 | 0 | no-hit | 0.0 | 0 | no-hit | 0.0 | 0 | no-hit | 0.0 | 0 | no-hit | 0.0 | 0 |
| scf7180000042133 | 230519 | 0.3582 | 0 | 57.9516 | Eukaryota | 84639.0 | 0 | Chordata | 84639.0 | 0 | Carnivora | 84639.0 | 0 | Canidae | 84639.0 | 0 | Canis | 84639.0 | 0 | Canis lupus | 84639.0 | 0 |
| scf7180000055089 | 99784 | 0.4387 | 0 | 84.574 | Eukaryota | 34422.0 | 1 | Chordata | 30285.0 | 2 | Gadiformes | 15432.0 | 6 | Gadidae | 15432.0 | 6 | Gadus | 15432.0 | 6 | Gadus morhua | 15432.0 | 6 |
| scf7180000046442 | 126577 | 0.3451 | 0 | 59.3851 | Eukaryota | 43687.0 | 0 | Chordata | 43687.0 | 0 | Carnivora | 43687.0 | 0 | Canidae | 43687.0 | 0 | Canis | 43687.0 | 0 | Canis lupus | 43687.0 | 0 |
| scf7180000050268 | 18448 | 0.44 | 0 | 139.7219 | no-hit | 0.0 | 0 | no-hit | 0.0 | 0 | no-hit | 0.0 | 0 | no-hit | 0.0 | 0 | no-hit | 0.0 | 0 | no-hit | 0.0 | 0 |
| scf7180000042207 | 10102 | 0.464 | 0 | 32.5208 | Eukaryota | 19455.0 | 0 | Chordata | 19455.0 | 0 | Carnivora | 18946.0 | 1 | Canidae | 18667.0 | 2 | Canis | 18667.0 | 2 | Canis lupus | 18667.0 | 2 |
| scf7180000045448 | 25162 | 0.4687 | 0 | 160.6791 | Eukaryota | 62290.0 | 0 | Chordata | 62290.0 | 0 | Carnivora | 62290.0 | 0 | Canidae | 62290.0 | 0 | Canis | 62290.0 | 0 | Canis lupus | 62290.0 | 0 |
| scf7180000043507 | 18729 | 0.3609 | 0 | 54.8043 | Eukaryota | 2860.0 | 0 | Chordata | 2860.0 | 0 | Carnivora | 2860.0 | 0 | Canidae | 2860.0 | 0 | Canis | 2860.0 | 0 | Canis lupus | 2860.0 | 0 |
| scf7180000051454 | 16101 | 0.4002 | 0 | 58.5939 | Eukaryota | 3896.0 | 0 | Chordata | 3896.0 | 0 | Carnivora | 3896.0 | 0 | Canidae | 3896.0 | 0 | Canis | 3896.0 | 0 | Canis lupus | 3896.0 | 0 |
| scf7180000055564 | 8224 | 0.4335 | 0 | 133.9692 | Eukaryota | 211.0 | 0 | Platyhelminthes | 211.0 | 0 | Polystomatidea | 211.0 | 0 | Polystomatidae | 211.0 | 0 | Protopolystoma | 211.0 | 0 | Protopolystoma xenopodis | 211.0 | 0 |
| scf7180000054699 | 203659 | 0.3962 | 0 | 63.6896 | Eukaryota | 88827.0 | 0 | Chordata | 88827.0 | 0 | Carnivora | 88827.0 | 0 | Canidae | 88827.0 | 0 | Canis | 88827.0 | 0 | Canis lupus | 88827.0 | 0 |
| scf7180000053593 | 6859 | 0.4587 | 0 | 1262.1844 | no-hit | 0.0 | 0 | no-hit | 0.0 | 0 | no-hit | 0.0 | 0 | no-hit | 0.0 | 0 | no-hit | 0.0 | 0 | no-hit | 0.0 | 0 |
| scf7180000041081 | 688612 | 0.4945 | 0 | 58.8953 | Eukaryota | 27688.0 | 0 | Chordata | 27688.0 | 0 | Carnivora | 27688.0 | 0 | Canidae | 27688.0 | 0 | Canis | 27688.0 | 0 | Canis lupus | 27688.0 | 0 |
| scf7180000054668 | 26438 | 0.4469 | 0 | 165.0501 | Eukaryota | 8947.0 | 1 | Chordata | 8947.0 | 1 | Tetraodontiformes | 4656.0 | 2 | Tetraodontidae | 4656.0 | 2 | Takifugu | 4656.0 | 2 | Takifugu rubripes | 4656.0 | 2 |
| scf7180000041435 | 146086 | 0.3966 | 0 | 56.3895 | Eukaryota | 32110.0 | 0 | Chordata | 32110.0 | 0 | Carnivora | 32110.0 | 0 | Canidae | 30170.0 | 1 | Canis | 30170.0 | 1 | Canis lupus | 30170.0 | 1 |
| scf7180000054356 | 26387 | 0.3915 | 0 | 300.907 | Eukaryota | 95706.0 | 0 | Chordata | 95706.0 | 0 | Carnivora | 95706.0 | 0 | Canidae | 95706.0 | 0 | Canis | 95706.0 | 0 | Canis lupus | 95706.0 | 0 |
| scf7180000048255 | 2007 | 0.4534 | 0 | 227.0263 | no-hit | 0.0 | 0 | no-hit | 0.0 | 0 | no-hit | 0.0 | 0 | no-hit | 0.0 | 0 | no-hit | 0.0 | 0 | no-hit | 0.0 | 0 |
| scf7180000041713 | 649482 | 0.4699 | 0 | 58.2496 | Eukaryota | 57204.0 | 0 | Chordata | 57204.0 | 0 | Carnivora | 57204.0 | 0 | Canidae | 57204.0 | 0 | Vulpes | 28762.0 | 1 | Vulpes vulpes | 28762.0 | 1 |
| scf7180000054210 | 48372 | 0.4025 | 0 | 62.7252 | Eukaryota | 48085.0 | 0 | Chordata | 48085.0 | 0 | Carnivora | 48085.0 | 0 | Canidae | 48085.0 | 0 | Canis | 48085.0 | 0 | Canis lupus | 48085.0 | 0 |
| scf7180000055165 | 25419 | 0.4418 | 0 | 456.9094 | Eukaryota | 7891.0 | 0 | Chordata | 6295.0 | 1 | Synbranchiformes | 3018.0 | 4 | Mastacembelidae | 3018.0 | 4 | Mastacembelus | 3018.0 | 4 | Mastacembelus armatus | 3018.0 | 4 |
| scf7180000049206 | 6250 | 0.436 | 0 | 35.577 | Eukaryota | 1766.0 | 0 | Chordata | 1766.0 | 0 | Kurtiformes | 1766.0 | 0 | Apogonidae | 1766.0 | 0 | Sphaeramia | 1766.0 | 0 | Sphaeramia orbicularis | 1766.0 | 0 |
| scf7180000052559 | 2571 | 0.4461 | 0 | 169.6326 | Eukaryota | 8836.0 | 0 | Chordata | 7356.0 | 2 | Synbranchiformes | 2265.0 | 6 | Mastacembelidae | 2265.0 | 6 | Mastacembelus | 2265.0 | 6 | Mastacembelus armatus | 2265.0 | 6 |
| scf7180000051437 | 4143 | 0.4499 | 0 | 268.0063 | Eukaryota | 3726.0 | 0 | Platyhelminthes | 1399.0 | 3 | Polystomatidea | 1399.0 | 3 | Polystomatidae | 1399.0 | 3 | Protopolystoma | 1399.0 | 3 | Protopolystoma xenopodis | 1399.0 | 3 |
| scf7180000043689 | 471748 | 0.3901 | 0 | 62.5541 | Eukaryota | 33244.0 | 0 | Chordata | 33244.0 | 0 | Carnivora | 33244.0 | 0 | Canidae | 33244.0 | 0 | Canis | 33244.0 | 0 | Canis lupus | 33244.0 | 0 |
| scf7180000043603 | 405350 | 0.3738 | 0 | 60.1801 | Eukaryota | 15890.0 | 0 | Chordata | 15890.0 | 0 | Carnivora | 13961.0 | 1 | Canidae | 10608.0 | 2 | Canis | 10608.0 | 3 | Canis lupus | 10608.0 | 3 |
| scf7180000051932 | 5339 | 0.446 | 0 | 378.9952 | no-hit | 0.0 | 0 | no-hit | 0.0 | 0 | no-hit | 0.0 | 0 | no-hit | 0.0 | 0 | no-hit | 0.0 | 0 | no-hit | 0.0 | 0 |
| scf7180000041304 | 243206 | 0.3981 | 0 | 59.4453 | Eukaryota | 90091.0 | 0 | Chordata | 90091.0 | 0 | Carnivora | 90091.0 | 0 | Canidae | 90091.0 | 0 | Canis | 90091.0 | 0 | Canis lupus | 90091.0 | 0 |
| scf7180000048496 | 12518 | 0.3823 | 0 | 101.1936 | Eukaryota | 4811.0 | 0 | Chordata | 4811.0 | 0 | Carnivora | 4811.0 | 0 | Canidae | 4811.0 | 0 | Canis | 4811.0 | 0 | Canis lupus | 4811.0 | 0 |
| scf7180000047355 | 62094 | 0.6459 | 0 | 56.1932 | Eukaryota | 4640.0 | 0 | Chordata | 4640.0 | 0 | Carnivora | 4640.0 | 0 | Canidae | 4640.0 | 0 | Vulpes | 2784.0 | 1 | Vulpes vulpes | 2784.0 | 1 |
| scf7180000056404 | 445185 | 0.3592 | 0 | 60.3356 | Eukaryota | 96052.0 | 0 | Chordata | 96052.0 | 0 | Carnivora | 96052.0 | 0 | Canidae | 96052.0 | 0 | Canis | 85337.0 | 1 | Canis lupus | 85337.0 | 1 |
| scf7180000046485 | 119084 | 0.4443 | 0 | 56.9377 | Eukaryota | 24159.0 | 0 | Chordata | 24159.0 | 0 | Carnivora | 24159.0 | 0 | Canidae | 24159.0 | 0 | Canis | 21691.0 | 1 | Canis lupus | 21691.0 | 1 |
| scf7180000054320 | 21314 | 0.4573 | 0 | 396.8221 | Eukaryota | 11621.0 | 0 | Chordata | 11621.0 | 0 | Carnivora | 11621.0 | 0 | Canidae | 11621.0 | 0 | Canis | 8921.0 | 1 | Canis lupus | 7990.0 | 2 |
| scf7180000053294 | 11146 | 0.4472 | 0 | 57.5487 | Eukaryota | 25366.0 | 0 | Chordata | 25366.0 | 0 | Carnivora | 25366.0 | 0 | Canidae | 14001.0 | 4 | Canis | 9393.0 | 8 | Canis lupus | 9393.0 | 8 |
| scf7180000045540 | 79057 | 0.4382 | 0 | 129.4918 | Eukaryota | 63331.0 | 0 | Chordata | 44206.0 | 2 | Gadiformes | 37232.0 | 5 | Gadidae | 37232.0 | 5 | Gadus | 37232.0 | 5 | Gadus morhua | 37232.0 | 5 |
| scf7180000046224 | 128903 | 0.372 | 100 | 57.5689 | Eukaryota | 18565.0 | 0 | Chordata | 18565.0 | 0 | Primates | 6023.0 | 6 | Hominidae | 6023.0 | 6 | Homo | 6023.0 | 6 | Homo sapiens | 6023.0 | 6 |
| scf7180000054795 | 23751 | 0.4417 | 0 | 590.303 | Eukaryota | 40788.0 | 0 | Chordata | 31704.0 | 2 | Gadiformes | 26758.0 | 6 | Gadidae | 26758.0 | 6 | Gadus | 26758.0 | 6 | Gadus morhua | 26758.0 | 6 |
| scf7180000044865 | 100038 | 0.5509 | 0 | 56.2807 | Eukaryota | 60426.0 | 0 | Chordata | 60426.0 | 0 | Carnivora | 60426.0 | 0 | Canidae | 60426.0 | 0 | Canis | 55166.0 | 1 | Canis lupus | 55166.0 | 1 |
| scf7180000050903 | 15051 | 0.614 | 0 | 53.2158 | Eukaryota | 19296.0 | 0 | Chordata | 19296.0 | 0 | Carnivora | 16677.0 | 3 | Canidae | 16677.0 | 3 | Canis | 16677.0 | 4 | Canis lupus | 16677.0 | 4 |
| scf7180000042670 | 570618 | 0.3534 | 0 | 59.5696 | Eukaryota | 31058.0 | 0 | Chordata | 31058.0 | 0 | Carnivora | 31058.0 | 0 | Canidae | 31058.0 | 0 | Canis | 31058.0 | 0 | Canis lupus | 31058.0 | 0 |
| scf7180000041903 | 683199 | 0.3767 | 0 | 59.7542 | Eukaryota | 27335.0 | 0 | Chordata | 27335.0 | 0 | Carnivora | 27335.0 | 0 | Canidae | 27335.0 | 0 | Canis | 27335.0 | 0 | Canis lupus | 27335.0 | 0 |
| scf7180000048955 | 10376 | 0.4441 | 0 | 263.9422 | Eukaryota | 3197.0 | 0 | Platyhelminthes | 1838.0 | 2 | Polystomatidea | 1838.0 | 2 | Polystomatidae | 1838.0 | 2 | Protopolystoma | 1838.0 | 2 | Protopolystoma xenopodis | 1838.0 | 2 |
| scf7180000054796 | 11657 | 0.4366 | 0 | 235.4619 | Eukaryota | 4106.0 | 0 | Chordata | 1680.0 | 3 | Salmoniformes | 1386.0 | 4 | Salmonidae | 1386.0 | 4 | Salmo | 1386.0 | 4 | Salmo trutta | 1386.0 | 4 |
| scf7180000044990 | 233393 | 0.4893 | 0 | 59.2945 | Eukaryota | 46443.0 | 0 | Chordata | 46443.0 | 0 | Carnivora | 46443.0 | 0 | Canidae | 26957.0 | 3 | Canis | 20659.0 | 6 | Canis lupus | 20659.0 | 6 |
| scf7180000044305 | 13006 | 0.417 | 0 | 385.2951 | Eukaryota | 7102.0 | 0 | Chordata | 5260.0 | 3 | Primates | 2337.0 | 5 | Hominidae | 2337.0 | 5 | Homo | 2337.0 | 6 | Homo sapiens | 2337.0 | 6 |
| scf7180000040941 | 1462437 | 0.4595 | 0 | 70.0926 | Eukaryota | 209205.0 | 0 | Chordata | 209205.0 | 0 | Carnivora | 209205.0 | 0 | Canidae | 209205.0 | 0 | Canis | 209205.0 | 0 | Canis lupus | 209205.0 | 0 |
| scf7180000042624 | 331153 | 0.3508 | 0 | 59.1011 | Eukaryota | 59405.0 | 0 | Chordata | 59405.0 | 0 | Carnivora | 59405.0 | 0 | Canidae | 54887.0 | 1 | Canis | 49743.0 | 2 | Canis lupus | 49743.0 | 2 |
| scf7180000053909 | 2661 | 0.4551 | 0 | 347.2996 | Eukaryota | 2006.0 | 0 | Platyhelminthes | 1000.0 | 3 | Polystomatidea | 1000.0 | 3 | Polystomatidae | 1000.0 | 3 | Protopolystoma | 1000.0 | 3 | Protopolystoma xenopodis | 1000.0 | 3 |
| scf7180000048329 | 7211 | 0.4389 | 0 | 14.3623 | Eukaryota | 44677.0 | 0 | Chordata | 44677.0 | 0 | Carnivora | 44677.0 | 0 | Canidae | 44677.0 | 0 | Canis | 44677.0 | 0 | Canis lupus | 44677.0 | 0 |
| scf7180000048170 | 30135 | 0.4997 | 0 | 75.0993 | Eukaryota | 10340.0 | 0 | Chordata | 10340.0 | 0 | Carnivora | 10340.0 | 0 | Canidae | 10340.0 | 0 | Canis | 6441.0 | 1 | Canis lupus | 6441.0 | 1 |
| scf7180000048248 | 57683 | 0.444 | 0 | 58.7535 | Eukaryota | 11276.0 | 0 | Chordata | 10817.0 | 1 | Kurtiformes | 3095.0 | 6 | Apogonidae | 3095.0 | 6 | Sphaeramia | 3095.0 | 6 | Sphaeramia orbicularis | 3095.0 | 6 |
| scf7180000051191 | 8699 | 0.4525 | 0 | 554.7169 | Eukaryota | 3141.0 | 0 | Platyhelminthes | 1586.0 | 3 | Polystomatidea | 1586.0 | 3 | Polystomatidae | 1586.0 | 3 | Protopolystoma | 1586.0 | 3 | Protopolystoma xenopodis | 1586.0 | 3 |
| scf7180000045548 | 175110 | 0.4137 | 0 | 59.6231 | Eukaryota | 46277.0 | 0 | Chordata | 46277.0 | 0 | Carnivora | 46277.0 | 0 | Canidae | 46277.0 | 0 | Canis | 46277.0 | 0 | Canis lupus | 46277.0 | 0 |
| scf7180000045057 | 258000 | 0.4614 | 0 | 59.4003 | Eukaryota | 44765.0 | 0 | Chordata | 44765.0 | 0 | Carnivora | 44765.0 | 0 | Ursidae | 25188.0 | 3 | Ursus | 13014.0 | 4 | Ailuropoda melanoleuca | 12174.0 | 5 |
| scf7180000043973 | 446386 | 0.3515 | 0 | 59.9173 | Eukaryota | 62506.0 | 0 | Chordata | 62506.0 | 0 | Carnivora | 62506.0 | 0 | Canidae | 62506.0 | 0 | Canis | 62506.0 | 0 | Canis lupus | 62506.0 | 0 |
| scf7180000049427 | 13048 | 0.4407 | 0 | 74.4577 | Eukaryota | 78439.0 | 0 | Chordata | 52433.0 | 1 | Gadiformes | 52433.0 | 2 | Gadidae | 52433.0 | 2 | Gadus | 52433.0 | 2 | Gadus morhua | 52433.0 | 2 |
| scf7180000040913 | 620834 | 0.3805 | 0 | 59.4914 | Eukaryota | 48600.0 | 0 | Chordata | 48600.0 | 0 | Carnivora | 48600.0 | 0 | Canidae | 48600.0 | 0 | Canis | 48600.0 | 0 | Canis lupus | 48600.0 | 0 |
| scf7180000043940 | 363908 | 0.5041 | 0 | 55.5477 | Eukaryota | 22653.0 | 0 | Chordata | 22653.0 | 0 | Carnivora | 22653.0 | 0 | Canidae | 22653.0 | 0 | Canis | 22653.0 | 0 | Canis lupus | 22653.0 | 0 |
| scf7180000052298 | 8430 | 0.3731 | 0 | 30.0038 | Eukaryota | 39575.0 | 0 | Chordata | 39575.0 | 0 | Carnivora | 39575.0 | 0 | Canidae | 39575.0 | 0 | Canis | 39575.0 | 0 | Canis lupus | 39575.0 | 0 |
| scf7180000050085 | 2480 | 0.4472 | 0 | 525.0003 | no-hit | 0.0 | 0 | no-hit | 0.0 | 0 | no-hit | 0.0 | 0 | no-hit | 0.0 | 0 | no-hit | 0.0 | 0 | no-hit | 0.0 | 0 |
| scf7180000053061 | 8866 | 0.4478 | 0 | 1772.0124 | no-hit | 0.0 | 0 | no-hit | 0.0 | 0 | no-hit | 0.0 | 0 | no-hit | 0.0 | 0 | no-hit | 0.0 | 0 | no-hit | 0.0 | 0 |
| scf7180000043286 | 153199 | 0.3854 | 0 | 106.5737 | Eukaryota | 86189.0 | 0 | Chordata | 86189.0 | 0 | Carnivora | 86189.0 | 0 | Canidae | 86189.0 | 0 | Canis | 86189.0 | 0 | Canis lupus | 86189.0 | 0 |
| scf7180000044237 | 3348 | 0.4453 | 0 | 379.0182 | no-hit | 0.0 | 0 | no-hit | 0.0 | 0 | no-hit | 0.0 | 0 | no-hit | 0.0 | 0 | no-hit | 0.0 | 0 | no-hit | 0.0 | 0 |
| scf7180000040901 | 2218 | 0.4545 | 0 | 381.8212 | no-hit | 0.0 | 0 | no-hit | 0.0 | 0 | no-hit | 0.0 | 0 | no-hit | 0.0 | 0 | no-hit | 0.0 | 0 | no-hit | 0.0 | 0 |
| scf7180000052677 | 8860 | 0.5018 | 0 | 179.4946 | Eukaryota | 8256.0 | 0 | Chordata | 8256.0 | 0 | Carnivora | 8256.0 | 0 | Canidae | 8256.0 | 0 | Canis | 5929.0 | 1 | Canis lupus | 4666.0 | 2 |
| scf7180000050759 | 23710 | 0.508 | 0 | 734.7454 | Eukaryota | 9221.0 | 0 | Chordata | 9221.0 | 0 | Carnivora | 9221.0 | 0 | Canidae | 9221.0 | 0 | Canis | 7450.0 | 1 | Canis lupus | 6140.0 | 2 |
| scf7180000045714 | 50154 | 0.4447 | 0 | 55.5416 | Eukaryota | 9706.0 | 0 | Chordata | 9706.0 | 0 | Carnivora | 9706.0 | 0 | Canidae | 9706.0 | 0 | Canis | 9706.0 | 0 | Canis lupus | 9706.0 | 0 |
| scf7180000042398 | 138225 | 0.4654 | 0 | 59.8459 | Eukaryota | 20483.0 | 0 | Chordata | 20483.0 | 0 | Carnivora | 20483.0 | 0 | Canidae | 20483.0 | 0 | Canis | 16938.0 | 1 | Canis lupus | 16938.0 | 1 |
| scf7180000056504 | 277466 | 0.3676 | 0 | 57.3361 | Eukaryota | 89746.0 | 0 | Chordata | 89746.0 | 0 | Carnivora | 89746.0 | 0 | Canidae | 89746.0 | 0 | Canis | 89746.0 | 0 | Canis lupus | 89746.0 | 0 |
| scf7180000050292 | 22853 | 0.4402 | 0 | 418.338 | Eukaryota | 6334.0 | 0 | Chordata | 4538.0 | 1 | Kurtiformes | 4538.0 | 1 | Apogonidae | 4538.0 | 1 | Sphaeramia | 4538.0 | 1 | Sphaeramia orbicularis | 4538.0 | 1 |
| scf7180000056215 | 1328787 | 0.4226 | 0 | 60.2246 | Eukaryota | 70402.0 | 0 | Chordata | 70402.0 | 0 | Carnivora | 70402.0 | 0 | Canidae | 70402.0 | 0 | Canis | 58233.0 | 1 | Canis lupus | 58233.0 | 1 |
| scf7180000053243 | 4610 | 0.4462 | 0 | 760.8695 | no-hit | 0.0 | 0 | no-hit | 0.0 | 0 | no-hit | 0.0 | 0 | no-hit | 0.0 | 0 | no-hit | 0.0 | 0 | no-hit | 0.0 | 0 |
| scf7180000049259 | 5799 | 0.4468 | 0 | 651.5322 | Eukaryota | 1609.0 | 0 | Platyhelminthes | 751.0 | 2 | Polystomatidea | 751.0 | 2 | Polystomatidae | 751.0 | 2 | Protopolystoma | 751.0 | 2 | Protopolystoma xenopodis | 751.0 | 2 |
| scf7180000054253 | 16301 | 0.3948 | 0 | 96.3147 | Eukaryota | 5382.0 | 0 | Chordata | 5382.0 | 0 | Carnivora | 4438.0 | 1 | Canidae | 4438.0 | 1 | Canis | 4438.0 | 1 | Canis lupus | 4438.0 | 1 |
| scf7180000053885 | 3479 | 0.503 | 0 | 262.8359 | Eukaryota | 5758.0 | 0 | Chordata | 5758.0 | 0 | Carnivora | 5758.0 | 0 | Canidae | 5758.0 | 0 | Canis | 3292.0 | 1 | Nyctereutes procyonoides | 2466.0 | 2 |
| scf7180000042079 | 24464 | 0.4349 | 0 | 49.1366 | Eukaryota | 4323.0 | 0 | Chordata | 4323.0 | 0 | Carnivora | 2522.0 | 2 | Canidae | 2522.0 | 2 | Canis | 2522.0 | 3 | Canis lupus | 2522.0 | 3 |
| scf7180000041862 | 397452 | 0.3844 | 0 | 60.187 | Eukaryota | 91416.0 | 0 | Chordata | 91416.0 | 0 | Carnivora | 91416.0 | 0 | Canidae | 91416.0 | 0 | Canis | 91416.0 | 0 | Canis lupus | 91416.0 | 0 |
| scf7180000053234 | 3610 | 0.433 | 0 | 35.8971 | Eukaryota | 641.0 | 0 | Chordata | 641.0 | 0 | Kurtiformes | 641.0 | 0 | Apogonidae | 641.0 | 0 | Sphaeramia | 641.0 | 0 | Sphaeramia orbicularis | 641.0 | 0 |
| scf7180000045016 | 214413 | 0.405 | 0 | 60.7846 | Eukaryota | 67545.0 | 0 | Chordata | 67545.0 | 0 | Carnivora | 67545.0 | 0 | Canidae | 67545.0 | 0 | Canis | 67545.0 | 0 | Canis lupus | 67545.0 | 0 |
| scf7180000048342 | 74854 | 0.4645 | 0 | 59.515 | Eukaryota | 10448.0 | 0 | Chordata | 10448.0 | 0 | Carnivora | 6198.0 | 3 | Canidae | 6198.0 | 3 | Canis | 6198.0 | 4 | Canis lupus | 6198.0 | 4 |
| scf7180000049398 | 25460 | 0.4405 | 0 | 58.5235 | Eukaryota | 38747.0 | 0 | Chordata | 38747.0 | 0 | Carnivora | 38747.0 | 0 | Canidae | 38747.0 | 0 | Canis | 21857.0 | 1 | Canis lupus | 21857.0 | 1 |
| scf7180000047144 | 11491 | 0.4358 | 0 | 70.4724 | no-hit | 0.0 | 0 | no-hit | 0.0 | 0 | no-hit | 0.0 | 0 | no-hit | 0.0 | 0 | no-hit | 0.0 | 0 | no-hit | 0.0 | 0 |
| scf7180000046591 | 141162 | 0.3575 | 0 | 55.0651 | Eukaryota | 92187.0 | 0 | Chordata | 92187.0 | 0 | Carnivora | 92187.0 | 0 | Canidae | 92187.0 | 0 | Canis | 92187.0 | 0 | Canis lupus | 92187.0 | 0 |
| scf7180000049556 | 44998 | 0.3568 | 0 | 59.7024 | Eukaryota | 12369.0 | 0 | Chordata | 12369.0 | 0 | Carnivora | 12369.0 | 0 | Canidae | 12369.0 | 0 | Canis | 12369.0 | 0 | Canis lupus | 12369.0 | 0 |
| scf7180000055131 | 10895 | 0.4465 | 0 | 307.5684 | Eukaryota | 596.0 | 0 | Platyhelminthes | 596.0 | 0 | Polystomatidea | 596.0 | 0 | Polystomatidae | 596.0 | 0 | Protopolystoma | 596.0 | 0 | Protopolystoma xenopodis | 596.0 | 0 |
| scf7180000054759 | 15048 | 0.4464 | 0 | 233.8469 | Eukaryota | 257.0 | 0 | Platyhelminthes | 257.0 | 0 | Polystomatidea | 257.0 | 0 | Polystomatidae | 257.0 | 0 | Protopolystoma | 257.0 | 0 | Protopolystoma xenopodis | 257.0 | 0 |
| scf7180000047671 | 29247 | 0.3784 | 0 | 56.9113 | Eukaryota | 88989.0 | 0 | Chordata | 88989.0 | 0 | Carnivora | 88989.0 | 0 | Canidae | 88989.0 | 0 | Canis | 88989.0 | 0 | Canis lupus | 88989.0 | 0 |
| scf7180000051658 | 3300 | 0.4403 | 0 | 45.4199 | Eukaryota | 2560.0 | 0 | Chordata | 2207.0 | 1 | Syngnathiformes | 819.0 | 3 | Syngnathidae | 819.0 | 3 | Syngnathus | 819.0 | 3 | Syngnathus acus | 819.0 | 3 |
| scf7180000046877 | 83617 | 0.6028 | 0 | 55.5885 | Eukaryota | 5719.0 | 0 | Chordata | 5719.0 | 0 | Carnivora | 5719.0 | 0 | Canidae | 5719.0 | 0 | Canis | 5719.0 | 0 | Canis lupus | 5719.0 | 0 |
| scf7180000054661 | 66924 | 0.4412 | 0 | 228.6278 | Eukaryota | 56301.0 | 0 | Chordata | 48188.0 | 2 | Gadiformes | 39618.0 | 5 | Gadidae | 39618.0 | 5 | Gadus | 39618.0 | 5 | Gadus morhua | 39618.0 | 5 |
| scf7180000042679 | 531102 | 0.4259 | 0 | 61.1868 | Eukaryota | 33407.0 | 1 | Chordata | 33407.0 | 1 | Carnivora | 18471.0 | 3 | Canidae | 10788.0 | 5 | Canis | 10788.0 | 5 | Canis lupus | 10788.0 | 6 |
| scf7180000046361 | 44936 | 0.4126 | 0 | 51.8253 | Eukaryota | 5064.0 | 0 | Chordata | 5064.0 | 0 | Primates | 2591.0 | 3 | Hominidae | 2151.0 | 5 | Homo | 2151.0 | 6 | Homo sapiens | 2151.0 | 7 |
| scf7180000050398 | 8380 | 0.4425 | 0 | 97.6594 | Eukaryota | 5055.0 | 0 | Chordata | 4007.0 | 1 | Carnivora | 1921.0 | 3 | Canidae | 1921.0 | 3 | Canis | 1921.0 | 3 | Canis lupus | 1921.0 | 3 |
| scf7180000050244 | 7709 | 0.4401 | 0 | 183.9821 | Eukaryota | 40201.0 | 0 | Chordata | 27858.0 | 2 | Gadiformes | 18456.0 | 6 | Gadidae | 18456.0 | 6 | Gadus | 18456.0 | 6 | Gadus morhua | 18456.0 | 6 |
| scf7180000049642 | 9490 | 0.4373 | 0 | 279.5767 | Eukaryota | 523.0 | 0 | Platyhelminthes | 523.0 | 0 | Polystomatidea | 523.0 | 0 | Polystomatidae | 523.0 | 0 | Protopolystoma | 523.0 | 0 | Protopolystoma xenopodis | 523.0 | 0 |
| scf7180000043196 | 553361 | 0.3786 | 0 | 59.0462 | Eukaryota | 31103.0 | 0 | Chordata | 31103.0 | 0 | Carnivora | 31103.0 | 0 | Canidae | 19092.0 | 2 | Canis | 15345.0 | 4 | Canis lupus | 15345.0 | 4 |
| scf7180000042720 | 672590 | 0.4127 | 0 | 59.3833 | Eukaryota | 90109.0 | 0 | Chordata | 90109.0 | 0 | Carnivora | 90109.0 | 0 | Canidae | 90109.0 | 0 | Canis | 90109.0 | 0 | Canis lupus | 90109.0 | 0 |
| scf7180000040538 | 811394 | 0.3811 | 0 | 58.6313 | Eukaryota | 93949.0 | 0 | Chordata | 93949.0 | 0 | Carnivora | 93949.0 | 0 | Canidae | 93949.0 | 0 | Canis | 93949.0 | 0 | Canis lupus | 93949.0 | 0 |
| scf7180000041675 | 1008604 | 0.4175 | 0 | 59.1421 | Eukaryota | 71867.0 | 0 | Chordata | 71867.0 | 0 | Carnivora | 71867.0 | 0 | Canidae | 71867.0 | 0 | Canis | 71867.0 | 0 | Canis lupus | 71867.0 | 0 |
| scf7180000043176 | 257481 | 0.4247 | 0 | 59.3674 | Eukaryota | 63569.0 | 0 | Chordata | 63569.0 | 0 | Carnivora | 63569.0 | 0 | Canidae | 63569.0 | 0 | Canis | 63569.0 | 0 | Canis lupus | 63569.0 | 0 |
| scf7180000046351 | 27291 | 0.4391 | 0 | 183.5142 | Eukaryota | 6036.0 | 0 | Chordata | 4879.0 | 1 | Columbiformes | 1810.0 | 4 | Columbidae | 1810.0 | 4 | Streptopelia | 1810.0 | 4 | Streptopelia turtur | 1810.0 | 4 |
| scf7180000042653 | 437585 | 0.487 | 0 | 58.1675 | Eukaryota | 36827.0 | 0 | Chordata | 36827.0 | 0 | Carnivora | 36827.0 | 0 | Canidae | 36827.0 | 0 | Canis | 36827.0 | 0 | Canis lupus | 36827.0 | 0 |
| scf7180000043721 | 303438 | 0.4143 | 0 | 60.2466 | Eukaryota | 107438.0 | 0 | Chordata | 107438.0 | 0 | Carnivora | 107438.0 | 0 | Canidae | 101155.0 | 1 | Canis | 81599.0 | 2 | Canis lupus | 81599.0 | 2 |
| scf7180000048245 | 68149 | 0.4488 | 0 | 58.8812 | Eukaryota | 14711.0 | 1 | Chordata | 14711.0 | 1 | Carnivora | 13307.0 | 2 | Mustelidae | 7040.0 | 3 | Vulpes | 6267.0 | 4 | Vulpes vulpes | 6267.0 | 4 |
| scf7180000047000 | 37955 | 0.4988 | 0 | 200.4237 | Eukaryota | 10436.0 | 0 | Chordata | 10436.0 | 0 | Carnivora | 10436.0 | 0 | Canidae | 10436.0 | 0 | Canis | 5732.0 | 1 | Canis lupus | 5732.0 | 1 |
| scf7180000054194 | 159331 | 0.3625 | 0 | 55.7188 | Eukaryota | 90444.0 | 0 | Chordata | 90444.0 | 0 | Carnivora | 90444.0 | 0 | Canidae | 90444.0 | 0 | Canis | 90444.0 | 0 | Canis lupus | 90444.0 | 0 |
| scf7180000047687 | 90828 | 0.4439 | 0 | 58.3519 | Eukaryota | 83320.0 | 0 | Chordata | 83320.0 | 0 | Carnivora | 80383.0 | 1 | Canidae | 54634.0 | 3 | Canis | 50515.0 | 5 | Canis lupus | 50515.0 | 5 |
| scf7180000047307 | 83443 | 0.4661 | 0 | 58.2678 | Eukaryota | 22018.0 | 0 | Chordata | 22018.0 | 0 | Carnivora | 22018.0 | 0 | Otariidae | 19818.0 | 1 | Zalophus | 19818.0 | 1 | Zalophus californianus | 19818.0 | 1 |
| scf7180000040898 | 585190 | 0.4026 | 0 | 59.4254 | Eukaryota | 94830.0 | 0 | Chordata | 94830.0 | 0 | Carnivora | 94830.0 | 0 | Canidae | 94830.0 | 0 | Vulpes | 94830.0 | 0 | Vulpes vulpes | 94830.0 | 0 |
| scf7180000056417 | 501022 | 0.343 | 0 | 58.8926 | Eukaryota | 90630.0 | 0 | Chordata | 90630.0 | 0 | Carnivora | 90630.0 | 0 | Canidae | 90630.0 | 0 | Canis | 90630.0 | 0 | Canis lupus | 90630.0 | 0 |
| scf7180000045113 | 251966 | 0.389 | 0 | 58.384 | Eukaryota | 74528.0 | 0 | Chordata | 74528.0 | 0 | Carnivora | 74528.0 | 0 | Canidae | 34830.0 | 2 | Canis | 34830.0 | 2 | Canis lupus | 34830.0 | 2 |
| scf7180000051162 | 9861 | 0.4343 | 0 | 52.8564 | Eukaryota | 5195.0 | 0 | Chordata | 5195.0 | 0 | Kurtiformes | 3445.0 | 2 | Apogonidae | 3445.0 | 2 | Sphaeramia | 3445.0 | 2 | Sphaeramia orbicularis | 3445.0 | 3 |
| scf7180000045325 | 153848 | 0.4068 | 0 | 60.2628 | Eukaryota | 92358.0 | 0 | Chordata | 92358.0 | 0 | Carnivora | 92358.0 | 0 | Canidae | 92358.0 | 0 | Canis | 92358.0 | 0 | Canis lupus | 92358.0 | 0 |
| scf7180000054571 | 70406 | 0.4467 | 0 | 276.2437 | Eukaryota | 91044.0 | 0 | Chordata | 91044.0 | 0 | Carnivora | 91044.0 | 0 | Canidae | 91044.0 | 0 | Canis | 91044.0 | 0 | Canis lupus | 91044.0 | 0 |
| scf7180000043029 | 522171 | 0.522 | 0 | 57.6506 | Eukaryota | 48819.0 | 0 | Chordata | 48819.0 | 0 | Carnivora | 48819.0 | 0 | Otariidae | 28560.0 | 3 | Zalophus | 23797.0 | 4 | Zalophus californianus | 23797.0 | 4 |
| scf7180000043392 | 201709 | 0.3478 | 0 | 58.5793 | Eukaryota | 13666.0 | 0 | Chordata | 13666.0 | 0 | Primates | 11538.0 | 1 | Hominidae | 6637.0 | 2 | Homo | 5294.0 | 5 | Homo sapiens | 5294.0 | 6 |
| scf7180000049726 | 15946 | 0.4284 | 0 | 61.2114 | Eukaryota | 53364.0 | 0 | Chordata | 53364.0 | 0 | Carnivora | 53364.0 | 0 | Canidae | 50656.0 | 1 | Canis | 44960.0 | 2 | Canis lupus | 44960.0 | 2 |
| scf7180000045920 | 168241 | 0.3948 | 0 | 59.1728 | Eukaryota | 22914.0 | 0 | Chordata | 22914.0 | 0 | Carnivora | 22914.0 | 0 | Canidae | 22914.0 | 0 | Canis | 17968.0 | 1 | Canis lupus | 17968.0 | 1 |
| scf7180000051518 | 6476 | 0.4362 | 0 | 44.9033 | no-hit | 0.0 | 0 | no-hit | 0.0 | 0 | no-hit | 0.0 | 0 | no-hit | 0.0 | 0 | no-hit | 0.0 | 0 | no-hit | 0.0 | 0 |
| scf7180000052736 | 3670 | 0.4485 | 0 | 232.0336 | Eukaryota | 3265.0 | 0 | Chordata | 3265.0 | 0 | Kurtiformes | 2137.0 | 2 | Apogonidae | 2137.0 | 2 | Sphaeramia | 2137.0 | 2 | Sphaeramia orbicularis | 2137.0 | 2 |
| scf7180000046267 | 24642 | 0.4146 | 0 | 55.8462 | Eukaryota | 90979.0 | 0 | Chordata | 90979.0 | 0 | Carnivora | 90979.0 | 0 | Canidae | 90979.0 | 0 | Canis | 90979.0 | 0 | Canis lupus | 90979.0 | 0 |
| scf7180000056387 | 452530 | 0.4049 | 0 | 61.828 | Eukaryota | 48358.0 | 0 | Chordata | 48358.0 | 0 | Carnivora | 48358.0 | 0 | Canidae | 48358.0 | 0 | Canis | 48358.0 | 0 | Canis lupus | 48358.0 | 0 |
| scf7180000050737 | 27479 | 0.4136 | 0 | 62.5995 | Eukaryota | 44576.0 | 0 | Chordata | 44576.0 | 0 | Carnivora | 44576.0 | 0 | Canidae | 44576.0 | 0 | Canis | 44576.0 | 0 | Canis lupus | 44576.0 | 0 |
| scf7180000054369 | 35169 | 0.4057 | 0 | 249.8001 | Eukaryota | 8970.0 | 0 | Chordata | 8970.0 | 0 | Carnivora | 8970.0 | 0 | Canidae | 8970.0 | 0 | Canis | 8970.0 | 0 | Canis lupus | 8970.0 | 0 |
| scf7180000049813 | 6367 | 0.4556 | 0 | 680.9726 | no-hit | 0.0 | 0 | no-hit | 0.0 | 0 | no-hit | 0.0 | 0 | no-hit | 0.0 | 0 | no-hit | 0.0 | 0 | no-hit | 0.0 | 0 |
| scf7180000042886 | 558092 | 0.4391 | 0 | 59.4826 | Eukaryota | 30772.0 | 0 | Chordata | 30772.0 | 0 | Carnivora | 30772.0 | 0 | Canidae | 28317.0 | 1 | Canis | 23772.0 | 2 | Canis lupus | 23772.0 | 2 |
| scf7180000055271 | 25307 | 0.4426 | 0 | 156.4578 | Eukaryota | 3079.0 | 0 | Chordata | 1560.0 | 1 | Polystomatidea | 1519.0 | 3 | Polystomatidae | 1519.0 | 3 | Protopolystoma | 1519.0 | 4 | Protopolystoma xenopodis | 1519.0 | 4 |
| scf7180000041344 | 1166983 | 0.3892 | 0 | 60.9367 | Eukaryota | 98213.0 | 0 | Chordata | 98213.0 | 0 | Carnivora | 98213.0 | 0 | Canidae | 83158.0 | 2 | Canis | 62354.0 | 3 | Canis lupus | 62354.0 | 3 |
| scf7180000043564 | 186555 | 0.36 | 0 | 56.8349 | Eukaryota | 80698.0 | 0 | Chordata | 80698.0 | 0 | Carnivora | 80698.0 | 0 | Canidae | 80698.0 | 0 | Canis | 80698.0 | 0 | Canis lupus | 80698.0 | 0 |
| scf7180000055440 | 24133 | 0.4384 | 0 | 84.315 | Eukaryota | 3881.0 | 0 | Chordata | 3881.0 | 0 | Carnivora | 2394.0 | 1 | Canidae | 2394.0 | 1 | Canis | 2394.0 | 1 | Canis lupus | 2394.0 | 1 |
| scf7180000047223 | 80347 | 0.5123 | 0 | 62.1227 | Eukaryota | 18302.0 | 0 | Chordata | 18302.0 | 0 | Carnivora | 18302.0 | 0 | Canidae | 11816.0 | 1 | Canis | 11816.0 | 1 | Canis lupus | 11816.0 | 1 |
| scf7180000054777 | 25543 | 0.4434 | 0 | 414.782 | Eukaryota | 8581.0 | 0 | Chordata | 8581.0 | 0 | Anabantiformes | 3808.0 | 5 | Anabantidae | 3808.0 | 5 | Anabas | 3808.0 | 5 | Anabas testudineus | 3808.0 | 5 |
| scf7180000050520 | 5960 | 0.4549 | 0 | 599.3528 | Eukaryota | 487.0 | 0 | Chordata | 246.0 | 1 | Kurtiformes | 246.0 | 1 | Apogonidae | 246.0 | 1 | Sphaeramia | 246.0 | 1 | Sphaeramia orbicularis | 246.0 | 1 |
| scf7180000043422 | 473917 | 0.5653 | 0 | 58.8885 | Eukaryota | 95917.0 | 0 | Chordata | 95917.0 | 0 | Carnivora | 95917.0 | 0 | Canidae | 95917.0 | 0 | Canis | 95917.0 | 0 | Canis lupus | 95917.0 | 0 |
| scf7180000051171 | 4408 | 0.4503 | 0 | 248.7065 | Eukaryota | 1970.0 | 0 | Platyhelminthes | 766.0 | 3 | Polystomatidea | 766.0 | 3 | Polystomatidae | 766.0 | 3 | Protopolystoma | 766.0 | 3 | Protopolystoma xenopodis | 766.0 | 3 |
| scf7180000055072 | 13102 | 0.4356 | 0 | 116.8022 | Eukaryota | 22579.0 | 1 | Chordata | 18410.0 | 2 | Gadiformes | 10031.0 | 4 | Gadidae | 10031.0 | 4 | Gadus | 10031.0 | 4 | Gadus morhua | 10031.0 | 4 |
| scf7180000045649 | 185612 | 0.3455 | 0 | 59.6605 | Eukaryota | 37446.0 | 0 | Chordata | 37446.0 | 0 | Carnivora | 37446.0 | 0 | Canidae | 37446.0 | 0 | Canis | 37446.0 | 0 | Canis lupus | 37446.0 | 0 |
| scf7180000047609 | 82199 | 0.3659 | 0 | 58.0575 | Eukaryota | 22319.0 | 0 | Chordata | 22319.0 | 0 | Carnivora | 22319.0 | 0 | Canidae | 22319.0 | 0 | Canis | 22319.0 | 0 | Canis lupus | 22319.0 | 0 |
| scf7180000042628 | 60993 | 0.3918 | 0 | 347.5056 | Eukaryota | 89929.0 | 0 | Chordata | 89929.0 | 0 | Carnivora | 89929.0 | 0 | Canidae | 89929.0 | 0 | Canis | 89929.0 | 0 | Canis lupus | 89929.0 | 0 |
| scf7180000052101 | 2394 | 0.4457 | 0 | 434.5119 | no-hit | 0.0 | 0 | no-hit | 0.0 | 0 | no-hit | 0.0 | 0 | no-hit | 0.0 | 0 | no-hit | 0.0 | 0 | no-hit | 0.0 | 0 |
| scf7180000046829 | 99297 | 0.4418 | 0 | 87.7312 | Eukaryota | 71282.0 | 0 | Chordata | 71282.0 | 0 | Carnivora | 71282.0 | 0 | Canidae | 71282.0 | 0 | Canis | 71282.0 | 0 | Canis lupus | 71282.0 | 0 |
| scf7180000042693 | 52633 | 0.4367 | 0 | 52.0752 | Eukaryota | 69720.0 | 0 | Chordata | 69720.0 | 0 | Carnivora | 69720.0 | 0 | Canidae | 69720.0 | 0 | Canis | 69720.0 | 0 | Canis lupus | 69720.0 | 0 |
| scf7180000045710 | 98548 | 0.4738 | 0 | 58.4651 | Eukaryota | 67304.0 | 0 | Chordata | 67304.0 | 0 | Carnivora | 67304.0 | 0 | Canidae | 67304.0 | 0 | Canis | 67304.0 | 0 | Canis lupus | 67304.0 | 0 |
| scf7180000055680 | 14150 | 0.4394 | 0 | 217.8199 | Eukaryota | 8367.0 | 0 | Chordata | 7136.0 | 1 | Tetraodontiformes | 3314.0 | 4 | Tetraodontidae | 3314.0 | 4 | Takifugu | 3314.0 | 4 | Takifugu rubripes | 3314.0 | 4 |
| scf7180000046696 | 13107 | 0.4826 | 0 | 49.4646 | Eukaryota | 3433.0 | 0 | Chordata | 3433.0 | 0 | Carnivora | 2583.0 | 1 | Canidae | 2583.0 | 2 | Canis | 2583.0 | 2 | Canis lupus | 2583.0 | 2 |
| scf7180000041770 | 592725 | 0.3566 | 0 | 60.0485 | Eukaryota | 86998.0 | 0 | Chordata | 86998.0 | 0 | Carnivora | 86998.0 | 0 | Canidae | 86998.0 | 0 | Canis | 86998.0 | 0 | Canis lupus | 86998.0 | 0 |
| scf7180000050168 | 7129 | 0.4431 | 0 | 133.8139 | Eukaryota | 2388.0 | 0 | Chordata | 1944.0 | 1 | Kurtiformes | 1317.0 | 3 | Apogonidae | 1317.0 | 3 | Sphaeramia | 1317.0 | 3 | Sphaeramia orbicularis | 1317.0 | 3 |
| scf7180000056498 | 255104 | 0.4617 | 0 | 60.092 | Eukaryota | 34707.0 | 0 | Chordata | 34707.0 | 0 | Carnivora | 34707.0 | 0 | Canidae | 34707.0 | 0 | Canis | 27303.0 | 1 | Canis lupus | 27303.0 | 1 |
| scf7180000049617 | 22042 | 0.4511 | 0 | 1425.9101 | Eukaryota | 2474.0 | 0 | Platyhelminthes | 1563.0 | 1 | Polystomatidea | 1563.0 | 1 | Polystomatidae | 1563.0 | 1 | Protopolystoma | 1563.0 | 1 | Protopolystoma xenopodis | 1563.0 | 1 |
| scf7180000054147 | 185999 | 0.3705 | 0 | 81.7509 | Eukaryota | 51415.0 | 0 | Chordata | 43643.0 | 2 | Gadiformes | 35112.0 | 4 | Gadidae | 35112.0 | 4 | Gadus | 35112.0 | 4 | Gadus morhua | 35112.0 | 4 |
| scf7180000046162 | 7360 | 0.4346 | 0 | 103.6453 | no-hit | 0.0 | 0 | no-hit | 0.0 | 0 | no-hit | 0.0 | 0 | no-hit | 0.0 | 0 | no-hit | 0.0 | 0 | no-hit | 0.0 | 0 |
| scf7180000051137 | 30920 | 0.502 | 0 | 219.4053 | Eukaryota | 9623.0 | 0 | Chordata | 9623.0 | 0 | Carnivora | 9623.0 | 0 | Canidae | 9623.0 | 0 | Canis | 8941.0 | 1 | Canis lupus | 7673.0 | 2 |
| scf7180000051441 | 12826 | 0.4387 | 0 | 67.9963 | Eukaryota | 50297.0 | 0 | Chordata | 35783.0 | 2 | Gadiformes | 30895.0 | 5 | Gadidae | 30895.0 | 5 | Gadus | 30895.0 | 5 | Gadus morhua | 30895.0 | 5 |
| scf7180000042451 | 13394 | 0.3907 | 0 | 69.7561 | Eukaryota | 91261.0 | 0 | Chordata | 91261.0 | 0 | Carnivora | 91261.0 | 0 | Canidae | 91261.0 | 0 | Canis | 91261.0 | 0 | Canis lupus | 91261.0 | 0 |
| scf7180000047021 | 15060 | 0.4874 | 0 | 226.3862 | Eukaryota | 32299.0 | 0 | Chordata | 32299.0 | 0 | Carnivora | 32299.0 | 0 | Canidae | 32299.0 | 0 | Canis | 32299.0 | 0 | Canis lupus | 32299.0 | 0 |
| scf7180000041986 | 842902 | 0.4074 | 0 | 59.8735 | Eukaryota | 50626.0 | 0 | Chordata | 50626.0 | 0 | Carnivora | 50626.0 | 0 | Canidae | 50626.0 | 0 | Canis | 50626.0 | 0 | Canis lupus | 50626.0 | 0 |
| scf7180000056213 | 1808317 | 0.3584 | 0 | 60.3439 | Eukaryota | 59967.0 | 0 | Chordata | 59967.0 | 0 | Carnivora | 59967.0 | 0 | Canidae | 59967.0 | 0 | Canis | 59967.0 | 0 | Canis lupus | 59967.0 | 0 |
| scf7180000045974 | 9487 | 0.3788 | 0 | 30.0484 | Eukaryota | 2616.0 | 0 | Chordata | 2616.0 | 0 | Carnivora | 2616.0 | 0 | Canidae | 2079.0 | 1 | Canis | 2079.0 | 1 | Canis lupus | 2079.0 | 1 |
| scf7180000044325 | 319716 | 0.3987 | 0 | 57.9304 | Eukaryota | 60807.0 | 0 | Chordata | 60807.0 | 0 | Carnivora | 60807.0 | 0 | Canidae | 46744.0 | 2 | Canis | 31153.0 | 3 | Canis lupus | 31153.0 | 3 |
| scf7180000040604 | 863757 | 0.4507 | 0 | 59.9653 | Eukaryota | 45696.0 | 0 | Chordata | 45696.0 | 0 | Carnivora | 45696.0 | 0 | Canidae | 45696.0 | 0 | Canis | 40892.0 | 1 | Canis lupus | 40892.0 | 1 |
| scf7180000046022 | 154938 | 0.5634 | 0 | 59.298 | Eukaryota | 18211.0 | 0 | Chordata | 18211.0 | 0 | Carnivora | 18211.0 | 0 | Canidae | 18211.0 | 0 | Canis | 14827.0 | 1 | Canis lupus | 14827.0 | 1 |
| scf7180000047377 | 3365 | 0.4104 | 0 | 26.2538 | Eukaryota | 2580.0 | 0 | Chordata | 2580.0 | 0 | Carnivora | 2580.0 | 0 | Canidae | 2580.0 | 0 | Canis | 2580.0 | 0 | Canis lupus | 2580.0 | 0 |
| scf7180000044764 | 328366 | 0.389 | 0 | 58.0099 | Eukaryota | 89478.0 | 0 | Chordata | 89478.0 | 0 | Carnivora | 89478.0 | 0 | Canidae | 89478.0 | 0 | Canis | 89478.0 | 0 | Canis lupus | 89478.0 | 0 |
| scf7180000048144 | 67161 | 0.4041 | 0 | 58.5307 | Eukaryota | 42525.0 | 0 | Chordata | 42525.0 | 0 | Carnivora | 42525.0 | 0 | Canidae | 42525.0 | 0 | Canis | 42525.0 | 0 | Canis lupus | 42525.0 | 0 |
| scf7180000044217 | 235177 | 0.4238 | 0 | 61.2618 | Eukaryota | 77979.0 | 0 | Chordata | 77979.0 | 0 | Carnivora | 77979.0 | 0 | Canidae | 77979.0 | 0 | Canis | 77979.0 | 0 | Canis lupus | 77979.0 | 0 |
| scf7180000054109 | 4372 | 0.4536 | 0 | 573.051 | Eukaryota | 1988.0 | 0 | Echinodermata | 926.0 | 2 | Forcipulatida | 926.0 | 2 | Asteriidae | 926.0 | 2 | Asterias | 926.0 | 2 | Asterias rubens | 926.0 | 2 |
| scf7180000046563 | 61603 | 0.589 | 0 | 54.622 | Eukaryota | 24884.0 | 0 | Chordata | 24884.0 | 0 | Carnivora | 24884.0 | 0 | Canidae | 24884.0 | 0 | Canis | 24884.0 | 0 | Canis lupus | 24884.0 | 0 |
| scf7180000054036 | 6017 | 0.5056 | 0 | 68.244 | Eukaryota | 7165.0 | 0 | Chordata | 7165.0 | 0 | Carnivora | 7165.0 | 0 | Canidae | 7165.0 | 0 | Canis | 4673.0 | 1 | Canis lupus | 3597.0 | 2 |
| scf7180000049069 | 34751 | 0.5734 | 0 | 54.1762 | Eukaryota | 4679.0 | 0 | Chordata | 4679.0 | 0 | Carnivora | 3477.0 | 2 | Canidae | 3477.0 | 2 | Canis | 3052.0 | 3 | Canis lupus | 3052.0 | 3 |
| scf7180000043356 | 52435 | 0.3718 | 0 | 48.7236 | Eukaryota | 61182.0 | 0 | Chordata | 61182.0 | 0 | Carnivora | 61182.0 | 0 | Canidae | 39596.0 | 2 | Canis | 26442.0 | 3 | Canis lupus | 26442.0 | 3 |
| scf7180000046977 | 39929 | 0.4482 | 0 | 72.6778 | Eukaryota | 86461.0 | 0 | Chordata | 86461.0 | 0 | Carnivora | 86461.0 | 0 | Canidae | 86461.0 | 0 | Canis | 86461.0 | 0 | Canis lupus | 86461.0 | 0 |
| scf7180000052785 | 12665 | 0.3965 | 0 | 62.779 | Eukaryota | 22229.0 | 0 | Chordata | 22229.0 | 0 | Artiodactyla | 12680.0 | 1 | Canidae | 9549.0 | 3 | Canis | 6396.0 | 4 | Canis lupus | 6396.0 | 4 |
| scf7180000052513 | 4694 | 0.4474 | 0 | 464.4176 | Eukaryota | 2358.0 | 0 | Chordata | 1666.0 | 1 | Columbiformes | 846.0 | 3 | Columbidae | 846.0 | 3 | Streptopelia | 846.0 | 3 | Streptopelia turtur | 846.0 | 3 |

| | | | | | | | | | | | | | | | | | | | | | | |
|---|---|---|---|---|---|---|---|---|---|---|---|---|---|---|---|---|---|---|---|---|---|---|
| scf7180000055053 | 73010 | 0.4398 | 0 | 199.7167 | Eukaryota | 15707.0 | 0 | Chordata | 10088.0 | 3 | Gadiformes | 4499.0 | 8 | Gadidae | 4499.0 | 8 | Gadus | 4499.0 | 8 | Gadus morhua | 4499.0 | 8 |
| scf7180000051866 | 1586 | 0.4584 | 0 | 514.0372 | no-hit | 0.0 | 0 | no-hit | 0.0 | 0 | no-hit | 0.0 | 0 | no-hit | 0.0 | 0 | no-hit | 0.0 | 0 | no-hit | 0.0 | 0 |
| scf7180000051189 | 8506 | 0.4471 | 0 | 1017.2548 | Eukaryota | 2310.0 | 0 | Platyhelminthes | 1315.0 | 2 | Polystomatidea | 1315.0 | 2 | Polystomatidae | 1315.0 | 2 | Protopolystoma | 1315.0 | 2 | Protopolystoma xenopodis | 1315.0 | 2 |
| scf7180000044917 | 265775 | 0.3616 | 0 | 59.0417 | Eukaryota | 59491.0 | 0 | Chordata | 59491.0 | 0 | Carnivora | 59491.0 | 0 | Felidae | 37709.0 | 2 | Vulpes | 18718.0 | 4 | Vulpes vulpes | 18718.0 | 4 |
| scf7180000042522 | 60651 | 0.4928 | 0 | 52.7871 | Eukaryota | 21248.0 | 0 | Chordata | 21248.0 | 0 | Carnivora | 21248.0 | 0 | Canidae | 13749.0 | 1 | Vulpes | 11704.0 | 3 | Vulpes vulpes | 11704.0 | 4 |
| scf7180000046684 | 108204 | 0.3703 | 0 | 57.1809 | Eukaryota | 82636.0 | 0 | Chordata | 82636.0 | 0 | Carnivora | 82636.0 | 0 | Canidae | 82636.0 | 0 | Canis | 82636.0 | 0 | Canis lupus | 82636.0 | 0 |
| scf7180000042821 | 38963 | 0.3353 | 0 | 56.0 | Eukaryota | 24748.0 | 0 | Chordata | 24748.0 | 0 | Carnivora | 24748.0 | 0 | Canidae | 24748.0 | 0 | Canis | 24748.0 | 0 | Canis lupus | 24748.0 | 0 |
| scf7180000050010 | 10313 | 0.758 | 0 | 201.175 | Eukaryota | 2449.0 | 0 | Chordata | 2449.0 | 0 | Rodentia | 1221.0 | 4 | Caviidae | 723.0 | 6 | Cavia | 723.0 | 6 | Cavia porcellus | 723.0 | 6 |
| scf7180000046053 | 129171 | 0.6115 | 0 | 56.9697 | Eukaryota | 93237.0 | 0 | Chordata | 93237.0 | 0 | Carnivora | 93237.0 | 0 | Canidae | 93237.0 | 0 | Canis | 93237.0 | 0 | Canis lupus | 93237.0 | 0 |
| scf7180000043143 | 6113 | 0.5078 | 0 | 25.9877 | Eukaryota | 6220.0 | 0 | Chordata | 6220.0 | 0 | Carnivora | 6220.0 | 0 | Canidae | 6220.0 | 0 | Canis | 6220.0 | 0 | Canis lupus | 6220.0 | 0 |
| scf7180000055120 | 37987 | 0.4392 | 0 | 102.2116 | Eukaryota | 9420.0 | 0 | Chordata | 6150.0 | 2 | Kurtiformes | 4179.0 | 4 | Apogonidae | 4179.0 | 4 | Sphaeramia | 4179.0 | 4 | Sphaeramia orbicularis | 4179.0 | 4 |
| scf7180000040862 | 269922 | 0.4272 | 0 | 59.0923 | Eukaryota | 79039.0 | 0 | Chordata | 79039.0 | 0 | Carnivora | 79039.0 | 0 | Canidae | 79039.0 | 0 | Canis | 70749.0 | 1 | Canis lupus | 70749.0 | 1 |
| scf7180000054839 | 5418 | 0.4376 | 0 | 176.6726 | no-hit | 0.0 | 0 | no-hit | 0.0 | 0 | no-hit | 0.0 | 0 | no-hit | 0.0 | 0 | no-hit | 0.0 | 0 | no-hit | 0.0 | 0 |
| scf7180000043208 | 231971 | 0.3735 | 0 | 57.4828 | Eukaryota | 36133.0 | 0 | Chordata | 36133.0 | 0 | Carnivora | 36133.0 | 0 | Canidae | 36133.0 | 0 | Canis | 36133.0 | 0 | Canis lupus | 36133.0 | 0 |
| scf7180000040849 | 185446 | 0.3442 | 0 | 59.1014 | Eukaryota | 72016.0 | 0 | Chordata | 72016.0 | 0 | Carnivora | 72016.0 | 0 | Canidae | 72016.0 | 0 | Canis | 72016.0 | 0 | Canis lupus | 72016.0 | 0 |
| scf7180000054078 | 4029 | 0.455 | 0 | 72.5627 | no-hit | 0.0 | 0 | no-hit | 0.0 | 0 | no-hit | 0.0 | 0 | no-hit | 0.0 | 0 | no-hit | 0.0 | 0 | no-hit | 0.0 | 0 |
| scf7180000040902 | 4018 | 0.4515 | 0 | 477.2067 | no-hit | 0.0 | 0 | no-hit | 0.0 | 0 | no-hit | 0.0 | 0 | no-hit | 0.0 | 0 | no-hit | 0.0 | 0 | no-hit | 0.0 | 0 |
| scf7180000041225 | 244932 | 0.392 | 0 | 58.569 | Eukaryota | 102031.0 | 0 | Chordata | 102031.0 | 0 | Carnivora | 102031.0 | 0 | Canidae | 102031.0 | 0 | Canis | 102031.0 | 0 | Canis lupus | 102031.0 | 0 |
| scf7180000056155 | 5761 | 0.4444 | 0 | 390.2141 | Eukaryota | 1878.0 | 0 | Chordata | 966.0 | 2 | Tetraodontiformes | 966.0 | 2 | Tetraodontidae | 966.0 | 2 | Takifugu | 966.0 | 2 | Takifugu rubripes | 966.0 | 2 |
| scf7180000041595 | 23559 | 0.3828 | 0 | 54.8624 | Eukaryota | 4630.0 | 0 | Chordata | 4630.0 | 0 | Primates | 1537.0 | 4 | Hominidae | 1537.0 | 6 | Homo | 1537.0 | 6 | Homo sapiens | 1537.0 | 6 |
| scf7180000052050 | 4330 | 0.4524 | 0 | 286.8408 | no-hit | 0.0 | 0 | no-hit | 0.0 | 0 | no-hit | 0.0 | 0 | no-hit | 0.0 | 0 | no-hit | 0.0 | 0 | no-hit | 0.0 | 0 |
| scf7180000043844 | 748144 | 0.5303 | 0 | 58.4701 | Eukaryota | 72202.0 | 0 | Chordata | 72202.0 | 0 | Carnivora | 72202.0 | 0 | Canidae | 72202.0 | 0 | Canis | 72202.0 | 0 | Canis lupus | 72202.0 | 0 |
| scf7180000055457 | 28332 | 0.4394 | 0 | 185.9284 | no-hit | 0.0 | 0 | no-hit | 0.0 | 0 | no-hit | 0.0 | 0 | no-hit | 0.0 | 0 | no-hit | 0.0 | 0 | no-hit | 0.0 | 0 |
| scf7180000048959 | 44451 | 0.4488 | 0 | 149.7581 | Eukaryota | 1919.0 | 0 | Platyhelminthes | 1474.0 | 1 | Polystomatidea | 1474.0 | 2 | Polystomatidae | 1474.0 | 2 | Protopolystoma | 1474.0 | 2 | Protopolystoma xenopodis | 1474.0 | 2 |
| scf7180000045816 | 116552 | 0.4426 | 0 | 60.7221 | Eukaryota | 42092.0 | 0 | Chordata | 42092.0 | 0 | Carnivora | 42092.0 | 0 | Canidae | 42092.0 | 0 | Canis | 37030.0 | 1 | Canis lupus | 37030.0 | 1 |
| scf7180000046825 | 123165 | 0.5007 | 0 | 62.2387 | Eukaryota | 52566.0 | 0 | Chordata | 52566.0 | 0 | Carnivora | 52566.0 | 0 | Canidae | 52566.0 | 0 | Canis | 52566.0 | 0 | Canis lupus | 52566.0 | 0 |
| scf7180000044214 | 400685 | 0.3779 | 0 | 59.6449 | Eukaryota | 79660.0 | 0 | Chordata | 79660.0 | 0 | Carnivora | 79660.0 | 0 | Otariidae | 50562.0 | 2 | Zalophus | 50562.0 | 2 | Zalophus californianus | 50562.0 | 2 |
| scf7180000048749 | 63795 | 0.3961 | 0 | 61.3518 | Eukaryota | 8492.0 | 0 | Chordata | 8492.0 | 0 | Carnivora | 7732.0 | 1 | Canidae | 3579.0 | 4 | Canis | 2738.0 | 6 | Canis lupus | 2738.0 | 7 |
| scf7180000051602 | 12312 | 0.5082 | 0 | 564.9575 | Eukaryota | 9110.0 | 0 | Chordata | 9110.0 | 0 | Carnivora | 9110.0 | 0 | Canidae | 9110.0 | 0 | Canis | 7414.0 | 1 | Canis lupus | 6180.0 | 2 |
| scf7180000046848 | 100305 | 0.4393 | 0 | 60.0534 | Eukaryota | 23495.0 | 0 | Chordata | 23495.0 | 0 | Carnivora | 23495.0 | 0 | Canidae | 23495.0 | 0 | Canis | 23495.0 | 0 | Canis lupus | 23495.0 | 0 |
| scf7180000056428 | 559440 | 0.4103 | 0 | 60.0119 | Eukaryota | 150222.0 | 0 | Chordata | 150222.0 | 0 | Carnivora | 150222.0 | 0 | Canidae | 88596.0 | 3 | Canis | 70871.0 | 4 | Canis lupus | 70871.0 | 4 |
| scf7180000050037 | 15289 | 0.4075 | 0 | 1436.7645 | Eukaryota | 46911.0 | 0 | Chordata | 46911.0 | 0 | Carnivora | 46911.0 | 0 | Canidae | 46911.0 | 0 | Canis | 46911.0 | 0 | Canis lupus | 46911.0 | 0 |
| scf7180000045299 | 200890 | 0.4251 | 0 | 58.7158 | Eukaryota | 34063.0 | 0 | Chordata | 34063.0 | 0 | Primates | 19276.0 | 2 | Hominidae | 16308.0 | 3 | Homo | 16308.0 | 4 | Homo sapiens | 16308.0 | 4 |
| scf7180000055133 | 12279 | 0.432 | 0 | 239.7676 | no-hit | 0.0 | 0 | no-hit | 0.0 | 0 | no-hit | 0.0 | 0 | no-hit | 0.0 | 0 | no-hit | 0.0 | 0 | no-hit | 0.0 | 0 |
| scf7180000044227 | 58953 | 0.3903 | 0 | 58.3971 | Eukaryota | 17944.0 | 0 | Chordata | 17944.0 | 0 | Carnivora | 17944.0 | 0 | Canidae | 17944.0 | 0 | Canis | 17944.0 | 0 | Canis lupus | 17944.0 | 0 |
| scf7180000044610 | 163490 | 0.3697 | 0 | 57.4393 | Eukaryota | 72071.0 | 0 | Chordata | 72071.0 | 0 | Carnivora | 72071.0 | 0 | Canidae | 58863.0 | 2 | Canis | 44577.0 | 3 | Canis lupus | 44577.0 | 3 |
| scf7180000043796 | 420422 | 0.3709 | 0 | 61.1694 | Eukaryota | 87655.0 | 0 | Chordata | 87655.0 | 0 | Carnivora | 87655.0 | 0 | Canidae | 87655.0 | 0 | Canis | 87655.0 | 0 | Canis lupus | 87655.0 | 0 |
| scf7180000043962 | 363690 | 0.4424 | 0 | 59.0007 | Eukaryota | 64061.0 | 0 | Chordata | 64061.0 | 0 | Carnivora | 64061.0 | 0 | Canidae | 64061.0 | 0 | Canis | 64061.0 | 0 | Canis lupus | 64061.0 | 0 |
| scf7180000046467 | 93362 | 0.4335 | 0 | 54.7447 | Eukaryota | 17221.0 | 0 | Chordata | 17221.0 | 0 | Carnivora | 15882.0 | 1 | Canidae | 15882.0 | 1 | Canis | 14330.0 | 2 | Canis lupus | 14330.0 | 2 |
| scf7180000054267 | 1558311 | 0.3737 | 0 | 60.9889 | Eukaryota | 100861.0 | 0 | Chordata | 100861.0 | 0 | Carnivora | 100861.0 | 0 | Canidae | 100861.0 | 0 | Canis | 100861.0 | 0 | Canis lupus | 100861.0 | 0 |
| scf7180000051506 | 2105 | 0.429 | 0 | 119.1687 | Eukaryota | 3178.0 | 0 | Platyhelminthes | 1908.0 | 3 | Polystomatidea | 1908.0 | 3 | Polystomatidae | 1908.0 | 3 | Protopolystoma | 1908.0 | 3 | Protopolystoma xenopodis | 1908.0 | 3 |
| scf7180000050758 | 8727 | 0.4518 | 0 | 854.9743 | no-hit | 0.0 | 0 | no-hit | 0.0 | 0 | no-hit | 0.0 | 0 | no-hit | 0.0 | 0 | no-hit | 0.0 | 0 | no-hit | 0.0 | 0 |
| scf7180000044710 | 25577 | 0.4784 | 0 | 45.0755 | Eukaryota | 8022.0 | 0 | Chordata | 8022.0 | 0 | Carnivora | 8022.0 | 0 | Canidae | 5865.0 | 1 | Canis | 5510.0 | 2 | Canis lupus | 5510.0 | 2 |
| scf7180000054975 | 15279 | 0.4446 | 0 | 159.147 | no-hit | 0.0 | 0 | no-hit | 0.0 | 0 | no-hit | 0.0 | 0 | no-hit | 0.0 | 0 | no-hit | 0.0 | 0 | no-hit | 0.0 | 0 |
| scf7180000041465 | 1079791 | 0.3787 | 0 | 60.8328 | Eukaryota | 36147.0 | 0 | Chordata | 36147.0 | 0 | Carnivora | 36147.0 | 0 | Canidae | 36147.0 | 0 | Canis | 36147.0 | 0 | Canis lupus | 36147.0 | 0 |
| scf7180000047102 | 105983 | 0.353 | 0 | 58.8162 | Eukaryota | 36304.0 | 0 | Chordata | 36304.0 | 0 | Carnivora | 36304.0 | 0 | Canidae | 36304.0 | 0 | Canis | 36304.0 | 0 | Canis lupus | 36304.0 | 0 |
| scf7180000045728 | 48925 | 0.3513 | 0 | 54.6391 | Eukaryota | 16272.0 | 0 | Chordata | 16272.0 | 0 | Primates | 9654.0 | 2 | Cercopithecidae | 5241.0 | 3 | Macaca | 5241.0 | 3 | Macaca nemestrina | 5241.0 | 3 |
| scf7180000056463 | 377717 | 0.4466 | 0 | 59.9125 | Eukaryota | 35426.0 | 0 | Chordata | 35426.0 | 0 | Carnivora | 35426.0 | 0 | Canidae | 35426.0 | 0 | Canis | 35426.0 | 0 | Canis lupus | 35426.0 | 0 |
| scf7180000043692 | 40590 | 0.3847 | 0 | 59.6373 | Eukaryota | 13574.0 | 0 | Chordata | 13574.0 | 0 | Primates | 8077.0 | 3 | Hominidae | 6403.0 | 4 | Homo | 6403.0 | 4 | Homo sapiens | 6403.0 | 4 |
| scf7180000053878 | 7292 | 0.4457 | 0 | 318.9383 | no-hit | 0.0 | 0 | no-hit | 0.0 | 0 | no-hit | 0.0 | 0 | no-hit | 0.0 | 0 | no-hit | 0.0 | 0 | no-hit | 0.0 | 0 |
| scf7180000051439 | 13738 | 0.3544 | 0 | 59.6562 | Eukaryota | 5475.0 | 0 | Chordata | 5475.0 | 0 | Carnivora | 5475.0 | 0 | Canidae | 5475.0 | 0 | Canis | 5475.0 | 0 | Canis lupus | 5475.0 | 0 |
| scf7180000041717 | 313290 | 0.5331 | 0 | 55.1621 | Eukaryota | 30177.0 | 0 | Chordata | 30177.0 | 0 | Carnivora | 30177.0 | 0 | Canidae | 26427.0 | 1 | Canis | 22210.0 | 3 | Canis lupus | 22210.0 | 3 |
| scf7180000048456 | 28179 | 0.4367 | 0 | 122.1073 | Eukaryota | 12982.0 | 0 | Chordata | 12982.0 | 0 | Carnivora | 12982.0 | 0 | Canidae | 12982.0 | 0 | Canis | 12982.0 | 0 | Canis lupus | 12982.0 | 0 |
| scf7180000054606 | 44939 | 0.4473 | 0 | 910.265 | Eukaryota | 3946.0 | 0 | Chordata | 2344.0 | 1 | Polystomatidea | 1183.0 | 5 | Polystomatidae | 1183.0 | 5 | Protopolystoma | 1183.0 | 5 | Protopolystoma xenopodis | 1183.0 | 5 |
| scf7180000055341 | 6272 | 0.437 | 0 | 182.6339 | Eukaryota | 6155.0 | 0 | Platyhelminthes | 2860.0 | 2 | Polystomatidea | 2860.0 | 4 | Polystomatidae | 2860.0 | 4 | Protopolystoma | 2860.0 | 4 | Protopolystoma xenopodis | 2860.0 | 4 |
| scf7180000053435 | 2750 | 0.4455 | 0 | 463.8504 | Eukaryota | 431.0 | 0 | Platyhelminthes | 431.0 | 0 | Polystomatidea | 431.0 | 0 | Polystomatidae | 431.0 | 0 | Protopolystoma | 431.0 | 0 | Protopolystoma xenopodis | 431.0 | 0 |
| scf7180000041099 | 10810 | 0.3738 | 0 | 40.6335 | Eukaryota | 3958.0 | 0 | Chordata | 3958.0 | 0 | Carnivora | 3958.0 | 0 | Mustelidae | 2392.0 | 1 | Mustela | 1987.0 | 2 | Mustela putorius | 1987.0 | 2 |
| scf7180000042756 | 942246 | 0.4159 | 0 | 59.4707 | Eukaryota | 55940.0 | 0 | Chordata | 55940.0 | 0 | Carnivora | 55940.0 | 0 | Canidae | 55940.0 | 0 | Canis | 55940.0 | 0 | Canis lupus | 55940.0 | 0 |
| scf7180000047754 | 6555 | 0.4304 | 0 | 24.3148 | Eukaryota | 18098.0 | 0 | Chordata | 18098.0 | 0 | Carnivora | 18098.0 | 0 | Canidae | 18098.0 | 0 | Canis | 18098.0 | 0 | Canis lupus | 18098.0 | 0 |
| scf7180000050892 | 8361 | 0.4413 | 0 | 226.9321 | Eukaryota | 1712.0 | 0 | Platyhelminthes | 888.0 | 1 | Kurtiformes | 824.0 | 2 | Apogonidae | 824.0 | 2 | Sphaeramia | 824.0 | 2 | Sphaeramia orbicularis | 824.0 | 2 |
| scf7180000048433 | 1945 | 0.4488 | 0 | 30.8382 | Eukaryota | 7433.0 | 0 | Chordata | 7433.0 | 0 | Carnivora | 7433.0 | 0 | Canidae | 7433.0 | 0 | Canis | 7433.0 | 0 | Canis lupus | 7433.0 | 0 |
| scf7180000053818 | 5272 | 0.3619 | 0 | 30.9971 | Eukaryota | 11788.0 | 0 | Chordata | 11788.0 | 0 | Carnivora | 9931.0 | 3 | Canidae | 9931.0 | 5 | Canis | 9931.0 | 6 | Canis lupus | 9931.0 | 7 |
| scf7180000047108 | 81075 | 0.524 | 0 | 58.0841 | Eukaryota | 78316.0 | 0 | Chordata | 78316.0 | 0 | Carnivora | 78316.0 | 0 | Canidae | 78316.0 | 0 | Canis | 78316.0 | 0 | Canis lupus | 78316.0 | 0 |
| scf7180000051744 | 13374 | 0.4457 | 0 | 148.0678 | no-hit | 0.0 | 0 | no-hit | 0.0 | 0 | no-hit | 0.0 | 0 | no-hit | 0.0 | 0 | no-hit | 0.0 | 0 | no-hit | 0.0 | 0 |
| scf7180000046572 | 17542 | 0.442 | 0 | 398.5302 | no-hit | 0.0 | 0 | no-hit | 0.0 | 0 | no-hit | 0.0 | 0 | no-hit | 0.0 | 0 | no-hit | 0.0 | 0 | no-hit | 0.0 | 0 |
| scf7180000053597 | 7977 | 0.4531 | 0 | 60.4468 | no-hit | 0.0 | 0 | no-hit | 0.0 | 0 | no-hit | 0.0 | 0 | no-hit | 0.0 | 0 | no-hit | 0.0 | 0 | no-hit | 0.0 | 0 |
| scf7180000041483 | 51296 | 0.3638 | 0 | 51.1929 | Eukaryota | 88570.0 | 0 | Chordata | 88570.0 | 0 | Carnivora | 88570.0 | 0 | Canidae | 88570.0 | 0 | Canis | 88570.0 | 0 | Canis lupus | 88570.0 | 0 |
| scf7180000052996 | 5608 | 0.3738 | 0 | 61.3192 | Eukaryota | 45410.0 | 0 | Chordata | 45410.0 | 0 | Carnivora | 45410.0 | 0 | Canidae | 45410.0 | 0 | Canis | 45410.0 | 0 | Canis lupus | 45410.0 | 0 |
| scf7180000041128 | 1229895 | 0.3759 | 0 | 60.1156 | Eukaryota | 63045.0 | 0 | Chordata | 63045.0 | 0 | Carnivora | 63045.0 | 0 | Canidae | 63045.0 | 0 | Canis | 63045.0 | 0 | Canis lupus | 63045.0 | 0 |
| scf7180000049242 | 28142 | 0.4479 | 0 | 120.4879 | Eukaryota | 548.0 | 0 | Chordata | 548.0 | 0 | Kurtiformes | 548.0 | 0 | Apogonidae | 548.0 | 0 | Sphaeramia | 548.0 | 0 | Sphaeramia orbicularis | 548.0 | 0 |
| scf7180000055692 | 22015 | 0.4385 | 0 | 109.3353 | Eukaryota | 31380.0 | 0 | Chordata | 19562.0 | 2 | Strigeidida | 8573.0 | 6 | Schistosomatidae | 8573.0 | 6 | Schistosoma | 8573.0 | 6 | Schistosoma mansoni | 8573.0 | 6 |
| scf7180000052162 | 3490 | 0.4487 | 0 | 758.2484 | no-hit | 0.0 | 0 | no-hit | 0.0 | 0 | no-hit | 0.0 | 0 | no-hit | 0.0 | 0 | no-hit | 0.0 | 0 | no-hit | 0.0 | 0 |
| scf7180000041118 | 1007317 | 0.4383 | 0 | 60.4496 | Eukaryota | 78557.0 | 0 | Chordata | 78557.0 | 0 | Carnivora | 78557.0 | 0 | Canidae | 78557.0 | 0 | Canis | 70773.0 | 1 | Canis lupus | 70773.0 | 1 |
| scf7180000042961 | 272351 | 0.4288 | 0 | 60.7595 | Eukaryota | 99610.0 | 0 | Chordata | 99610.0 | 0 | Carnivora | 99610.0 | 0 | Canidae | 99610.0 | 0 | Canis | 75114.0 | 1 | Canis lupus | 75114.0 | 1 |
| scf7180000046232 | 122545 | 0.3775 | 0 | 59.8696 | Eukaryota | 30841.0 | 0 | Chordata | 30841.0 | 0 | Carnivora | 30841.0 | 0 | Canidae | 20033.0 | 2 | Canis | 20033.0 | 2 | Canis lupus | 20033.0 | 2 |
| scf7180000041971 | 315862 | 0.4629 | 0 | 58.4564 | Eukaryota | 34510.0 | 0 | Chordata | 34510.0 | 0 | Carnivora | 34510.0 | 0 | Canidae | 34510.0 | 0 | Canis | 24230.0 | 1 | Canis lupus | 24230.0 | 1 |
| scf7180000043976 | 24551 | 0.4496 | 0 | 73.9392 | Eukaryota | 42661.0 | 0 | Chordata | 42661.0 | 0 | Carnivora | 42661.0 | 0 | Canidae | 42661.0 | 0 | Canis | 42661.0 | 0 | Canis lupus | 42661.0 | 0 |
| scf7180000055590 | 15988 | 0.4356 | 0 | 157.0374 | Eukaryota | 20518.0 | 0 | Chordata | 12139.0 | 2 | Strigeidida | 7606.0 | 6 | Schistosomatidae | 7606.0 | 6 | Schistosoma | 7606.0 | 6 | Schistosoma mansoni | 7606.0 | 6 |
| scf7180000041297 | 401669 | 0.486 | 0 | 57.7677 | Eukaryota | 40803.0 | 0 | Chordata | 40803.0 | 0 | Carnivora | 40803.0 | 0 | Canidae | 40803.0 | 0 | Canis | 37381.0 | 1 | Canis lupus | 37381.0 | 1 |
| scf7180000056392 | 412759 | 0.3887 | 0 | 58.3626 | Eukaryota | 20367.0 | 0 | Chordata | 20367.0 | 0 | Carnivora | 20367.0 | 0 | Canidae | 20367.0 | 0 | Canis | 20367.0 | 0 | Canis lupus | 20367.0 | 0 |
| scf7180000041002 | 1479093 | 0.4314 | 0 | 60.0696 | Eukaryota | 79357.0 | 0 | Chordata | 79357.0 | 0 | Carnivora | 79357.0 | 0 | Canidae | 74245.0 | 1 | Vulpes | 57253.0 | 2 | Vulpes vulpes | 57253.0 | 2 |
| scf7180000051299 | 4175 | 0.4321 | 0 | 88.2871 | no-hit | 0.0 | 0 | no-hit | 0.0 | 0 | no-hit | 0.0 | 0 | no-hit | 0.0 | 0 | no-hit | 0.0 | 0 | no-hit | 0.0 | 0 |
| scf7180000056258 | 1049621 | 0.3561 | 0 | 60.4017 | Eukaryota | 78044.0 | 0 | Chordata | 78044.0 | 0 | Carnivora | 68265.0 | 1 | Canidae | 26298.0 | 6 | Canis | 26298.0 | 6 | Canis lupus | 26298.0 | 7 |
| scf7180000044358 | 247372 | 0.3833 | 0 | 74.1768 | Eukaryota | 19884.0 | 0 | Chordata | 19884.0 | 0 | Carnivora | 18029.0 | 1 | Canidae | 16130.0 | 2 | Canis | 16130.0 | 2 | Canis lupus | 16130.0 | 2 |
| scf7180000055944 | 32121 | 0.408 | 0 | 299.614 | Eukaryota | 95179.0 | 0 | Chordata | 95179.0 | 0 | Carnivora | 95179.0 | 0 | Canidae | 95179.0 | 0 | Canis | 95179.0 | 0 | Canis lupus | 95179.0 | 0 |
| scf7180000042506 | 657497 | 0.4664 | 0 | 59.0996 | Eukaryota | 30509.0 | 0 | Chordata | 30509.0 | 0 | Carnivora | 30509.0 | 0 | Canidae | 30509.0 | 0 | Canis | 30509.0 | 0 | Canis lupus | 30509.0 | 0 |
| scf7180000044976 | 203549 | 0.5238 | 0 | 58.4717 | Eukaryota | 67335.0 | 0 | Chordata | 67335.0 | 0 | Carnivora | 67335.0 | 0 | Canidae | 67335.0 | 0 | Canis | 67335.0 | 0 | Canis lupus | 67335.0 | 0 |
| scf7180000046697 | 120136 | 0.3868 | 0 | 58.9669 | Eukaryota | 85780.0 | 0 | Chordata | 85780.0 | 0 | Carnivora | 85780.0 | 0 | Canidae | 85780.0 | 0 | Canis | 85780.0 | 0 | Canis lupus | 85780.0 | 0 |
| scf7180000050100 | 2837 | 0.4491 | 0 | 225.0881 | no-hit | 0.0 | 0 | no-hit | 0.0 | 0 | no-hit | 0.0 | 0 | no-hit | 0.0 | 0 | no-hit | 0.0 | 0 | no-hit | 0.0 | 0 |
| scf7180000056459 | 544450 | 0.4315 | 0 | 60.0748 | Eukaryota | 77933.0 | 0 | Chordata | 77933.0 | 0 | Carnivora | 77933.0 | 0 | Canidae | 58522.0 | 2 | Canis | 50433.0 | 3 | Canis lupus | 50433.0 | 3 |
| scf7180000043231 | 16279 | 0.3757 | 0 | 53.8771 | Eukaryota | 6236.0 | 0 | Chordata | 6236.0 | 0 | Carnivora | 6236.0 | 0 | Canidae | 6236.0 | 0 | Canis | 6236.0 | 0 | Canis lupus | 6236.0 | 0 |
| scf7180000053156 | 7709 | 0.5058 | 0 | 93.5185 | Eukaryota | 8300.0 | 0 | Chordata | 8300.0 | 0 | Carnivora | 8300.0 | 0 | Canidae | 8300.0 | 0 | Canis | 7237.0 | 1 | Canis lupus | 6190.0 | 2 |
| scf7180000056436 | 504575 | 0.3863 | 0 | 59.4003 | Eukaryota | 31315.0 | 0 | Chordata | 31315.0 | 0 | Carnivora | 31315.0 | 0 | Canidae | 31315.0 | 0 | Canis | 31315.0 | 0 | Canis lupus | 31315.0 | 0 |
| scf7180000044844 | 281416 | 0.3713 | 0 | 60.1523 | Eukaryota | 28564.0 | 0 | Chordata | 28564.0 | 0 | Carnivora | 28564.0 | 0 | Canidae | 28564.0 | 0 | Canis | 28564.0 | 0 | Canis lupus | 28564.0 | 0 |
| scf7180000055425 | 23967 | 0.445 | 0 | 589.4181 | Eukaryota | 5354.0 | 0 | Chordata | 2664.0 | 3 | Salmoniformes | 2516.0 | 4 | Salmonidae | 2516.0 | 4 | Salmo | 2516.0 | 4 | Salmo trutta | 2516.0 | 4 |
| scf7180000050626 | 2711 | 0.4515 | 0 | 298.6604 | Eukaryota | 2592.0 | 0 | Platyhelminthes | 1450.0 | 2 | Polystomatidea | 1450.0 | 2 | Polystomatidae | 1450.0 | 2 | Protopolystoma | 1450.0 | 2 | Protopolystoma xenopodis | 1450.0 | 2 |
| scf7180000044830 | 1443574 | 0.3924 | 0 | 60.4001 | Eukaryota | 85154.0 | 0 | Chordata | 85154.0 | 0 | Carnivora | 85154.0 | 0 | Canidae | 85154.0 | 0 | Canis | 85154.0 | 0 | Canis lupus | 85154.0 | 0 |
| scf7180000052667 | 9418 | 0.4397 | 0 | 68.0523 | no-hit | 0.0 | 0 | no-hit | 0.0 | 0 | no-hit | 0.0 | 0 | no-hit | 0.0 | 0 | no-hit | 0.0 | 0 | no-hit | 0.0 | 0 |
| scf7180000049931 | 9339 | 0.4477 | 0 | 906.3606 | no-hit | 0.0 | 0 | no-hit | 0.0 | 0 | no-hit | 0.0 | 0 | no-hit | 0.0 | 0 | no-hit | 0.0 | 0 | no-hit | 0.0 | 0 |
| scf7180000040953 | 322817 | 0.3841 | 0 | 56.86 | Eukaryota | 94211.0 | 0 | Chordata | 94211.0 | 0 | Carnivora | 94211.0 | 0 | Canidae | 94211.0 | 0 | Canis | 94211.0 | 0 | Canis lupus | 94211.0 | 0 |
| scf7180000043825 | 32247 | 0.438 | 100 | 58.5849 | Eukaryota | 17756.0 | 0 | Chordata | 17142.0 | 1 | Carnivora | 12264.0 | 4 | Canidae | 12264.0 | 4 | Canis | 12264.0 | 6 | Canis lupus | 12264.0 | 6 |
| scf7180000051239 | 3551 | 0.4517 | 0 | 294.22 | no-hit | 0.0 | 0 | no-hit | 0.0 | 0 | no-hit | 0.0 | 0 | no-hit | 0.0 | 0 | no-hit | 0.0 | 0 | no-hit | 0.0 | 0 |
| scf7180000054014 | 3083 | 0.4538 | 0 | 596.8218 | Eukaryota | 2276.0 | 0 | Platyhelminthes | 1681.0 | 1 | Polystomatidea | 1681.0 | 1 | Polystomatidae | 1681.0 | 1 | Protopolystoma | 1681.0 | 1 | Protopolystoma xenopodis | 1681.0 | 1 |
| scf7180000043219 | 471118 | 0.5251 | 0 | 58.0232 | Eukaryota | 51353.0 | 0 | Chordata | 51353.0 | 0 | Carnivora | 51353.0 | 0 | Canidae | 51353.0 | 0 | Canis | 38257.0 | 1 | Canis lupus | 38257.0 | 1 |
| scf7180000055786 | 27256 | 0.4533 | 0 | 87.3901 | Eukaryota | 55372.0 | 0 | Chordata | 55372.0 | 0 | Carnivora | 55372.0 | 0 | Canidae | 55372.0 | 0 | Canis | 55372.0 | 0 | Canis lupus | 55372.0 | 0 |
| scf7180000045214 | 10943 | 0.4338 | 0 | 120.3238 | Eukaryota | 3252.0 | 0 | Echinodermata | 1693.0 | 2 | Forcipulatida | 1693.0 | 2 | Asteriidae | 1693.0 | 2 | Asterias | 1693.0 | 2 | Asterias rubens | 1693.0 | 2 |
| scf7180000047023 | 80448 | 0.5497 | 0 | 56.8612 | Eukaryota | 48600.0 | 0 | Chordata | 48600.0 | 0 | Carnivora | 48600.0 | 0 | Canidae | 48600.0 | 0 | Canis | 44125.0 | 1 | Canis lupus | 44125.0 | 1 |
| scf7180000040810 | 130585 | 0.38 | 0 | 56.0679 | Eukaryota | 90135.0 | 0 | Chordata | 90135.0 | 0 | Carnivora | 90135.0 | 0 | Canidae | 90135.0 | 0 | Canis | 90135.0 | 0 | Canis lupus | 90135.0 | 0 |
| scf7180000056099 | 40498 | 0.443 | 0 | 263.54 | Eukaryota | 96205.0 | 0 | Chordata | 96205.0 | 0 | Carnivora | 96205.0 | 0 | Canidae | 96205.0 | 0 | Canis | 96205.0 | 0 | Canis lupus | 96205.0 | 0 |
| scf7180000048902 | 41045 | 0.3575 | 0 | 59.7497 | Eukaryota | 14558.0 | 1 | Chordata | 14558.0 | 1 | Carnivora | 6807.0 | 3 | Canidae | 6807.0 | 3 | Canis | 6807.0 | 4 | Canis lupus | 6807.0 | 4 |
| scf7180000052263 | 3072 | 0.4938 | 0 | 715.176 | Eukaryota | 7408.0 | 0 | Chordata | 7408.0 | 0 | Carnivora | 7408.0 | 0 | Canidae | 7408.0 | 0 | Canis | 5860.0 | 1 | Canis lupus | 4742.0 | 2 |
| scf7180000046008 | 144026 | 0.5343 | 0 | 54.6007 | Eukaryota | 50940.0 | 0 | Chordata | 50940.0 | 0 | Carnivora | 50940.0 | 0 | Canidae | 50940.0 | 0 | Canis | 50940.0 | 0 | Canis lupus | 50940.0 | 0 |
| scf7180000044007 | 377779 | 0.3894 | 0 | 67.7459 | Eukaryota | 99199.0 | 0 | Chordata | 99199.0 | 0 | Carnivora | 99199.0 | 0 | Canidae | 99199.0 | 0 | Canis | 82322.0 | 1 | Canis lupus | 82322.0 | 1 |
| scf7180000053736 | 2738 | 0.4525 | 0 | 510.5994 | Eukaryota | 534.0 | 0 | Platyhelminthes | 534.0 | 0 | Polystomatidea | 534.0 | 0 | Polystomatidae | 534.0 | 0 | Protopolystoma | 534.0 | 0 | Protopolystoma xenopodis | 534.0 | 0 |
| scf7180000047369 | 3827 | 0.4591 | 0 | 214.3892 | Eukaryota | 135.0 | 0 | Chordata | 135.0 | 0 | Salmoniformes | 135.0 | 0 | Salmonidae | 135.0 | 0 | Salmo | 135.0 | 0 | Salmo trutta | 135.0 | 0 |
| scf7180000042179 | 949741 | 0.4171 | 0 | 60.0899 | Eukaryota | 95366.0 | 0 | Chordata | 95366.0 | 0 | Carnivora | 95366.0 | 0 | Canidae | 95366.0 | 0 | Canis | 95366.0 | 0 | Canis lupus | 95366.0 | 0 |
| scf7180000053184 | 2602 | 0.4523 | 0 | 187.313 | Eukaryota | 424.0 | 0 | Platyhelminthes | 424.0 | 0 | Polystomatidea | 424.0 | 0 | Polystomatidae | 424.0 | 0 | Protopolystoma | 424.0 | 0 | Protopolystoma xenopodis | 424.0 | 0 |
| scf7180000051224 | 5764 | 0.449 | 0 | 407.8208 | Eukaryota | 2913.0 | 0 | Platyhelminthes | 1483.0 | 3 | Polystomatidea | 1483.0 | 3 | Polystomatidae | 1483.0 | 3 | Protopolystoma | 1483.0 | 3 | Protopolystoma xenopodis | 1483.0 | 3 |
| scf7180000046915 | 38806 | 0.4464 | 0 | 205.4466 | Eukaryota | 189.0 | 0 | Chordata | 189.0 | 0 | Kurtiformes | 189.0 | 0 | Apogonidae | 189.0 | 0 | Sphaeramia | 189.0 | 0 | Sphaeramia orbicularis | 189.0 | 0 |
| scf7180000046661 | 117496 | 0.3429 | 0 | 60.0437 | Eukaryota | 45299.0 | 0 | Chordata | 45299.0 | 0 | Carnivora | 45299.0 | 0 | Canidae | 45299.0 | 0 | Canis | 45299.0 | 0 | Canis lupus | 45299.0 | 0 |
| scf7180000040805 | 21424 | 0.5045 | 0 | 81.1299 | Eukaryota | 8773.0 | 0 | Chordata | 8773.0 | 0 | Carnivora | 8773.0 | 0 | Canidae | 8773.0 | 0 | Nyctereutes | 4480.0 | 1 | Nyctereutes procyonoides | 4480.0 | 1 |
| scf7180000044294 | 48940 | 0.4014 | 0 | 57.8685 | Eukaryota | 86268.0 | 0 | Chordata | 86268.0 | 0 | Carnivora | 86268.0 | 0 | Canidae | 86268.0 | 0 | Canis | 86268.0 | 0 | Canis lupus | 86268.0 | 0 |
| scf7180000041792 | 835303 | 0.4725 | 0 | 167.614 | Eukaryota | 113001.0 | 0 | Chordata | 113001.0 | 0 | Carnivora | 113001.0 | 0 | Canidae | 113001.0 | 0 | Canis | 113001.0 | 0 | Canis lupus | 113001.0 | 0 |
| scf7180000041468 | 96956 | 0.369 | 0 | 57.5893 | Eukaryota | 74701.0 | 0 | Chordata | 74701.0 | 0 | Carnivora | 74701.0 | 0 | Canidae | 74701.0 | 0 | Canis | 74701.0 | 0 | Canis lupus | 74701.0 | 0 |
| scf7180000044446 | 354554 | 0.42 | 0 | 61.0135 | Eukaryota | 56466.0 | 0 | Chordata | 56466.0 | 0 | Carnivora | 56466.0 | 0 | Canidae | 22603.0 | 2 | Vulpes | 22603.0 | 3 | Vulpes vulpes | 22603.0 | 3 |
| scf7180000040719 | 324582 | 0.3534 | 0 | 54.3049 | Eukaryota | 26284.0 | 0 | Chordata | 26284.0 | 0 | Carnivora | 26284.0 | 0 | Canidae | 26284.0 | 0 | Canis | 26284.0 | 0 | Canis lupus | 26284.0 | 0 |
| scf7180000044731 | 229950 | 0.405 | 0 | 58.6874 | Eukaryota | 57615.0 | 0 | Chordata | 57615.0 | 0 | Carnivora | 57615.0 | 0 | Canidae | 57615.0 | 0 | Canis | 57615.0 | 0 | Canis lupus | 57615.0 | 0 |
| scf7180000046004 | 189916 | 0.3475 | 0 | 59.2758 | Eukaryota | 73070.0 | 0 | Chordata | 73070.0 | 0 | Carnivora | 73070.0 | 0 | Canidae | 73070.0 | 0 | Canis | 73070.0 | 0 | Canis lupus | 73070.0 | 0 |
| scf7180000046420 | 8741 | 0.5055 | 0 | 46.5898 | Eukaryota | 6953.0 | 0 | Chordata | 6953.0 | 0 | Carnivora | 6953.0 | 0 | Canidae | 6213.0 | 2 | Canis | 6213.0 | 2 | Canis lupus | 6213.0 | 2 |
| scf7180000052148 | 6315 | 0.5063 | 0 | 87.2956 | Eukaryota | 7786.0 | 0 | Chordata | 7786.0 | 0 | Carnivora | 7786.0 | 0 | Canidae | 7786.0 | 0 | Canis | 6719.0 | 1 | Canis lupus | 5672.0 | 2 |
| scf7180000043139 | 505162 | 0.5391 | 0 | 57.9412 | Eukaryota | 55909.0 | 0 | Chordata | 55909.0 | 0 | Carnivora | 55909.0 | 0 | Canidae | 55909.0 | 0 | Canis | 55909.0 | 0 | Canis lupus | 55909.0 | 0 |
| scf7180000055348 | 27982 | 0.4458 | 0 | 142.9981 | no-hit | 0.0 | 0 | no-hit | 0.0 | 0 | no-hit | 0.0 | 0 | no-hit | 0.0 | 0 | no-hit | 0.0 | 0 | no-hit | 0.0 | 0 |
| scf7180000040513 | 205562 | 0.3826 | 0 | 132.63 | Eukaryota | 32377.0 | 0 | Chordata | 32377.0 | 0 | Carnivora | 32377.0 | 0 | Canidae | 32377.0 | 0 | Canis | 32377.0 | 0 | Canis lupus | 32377.0 | 0 |
| scf7180000048325 | 18244 | 0.4455 | 0 | 209.0416 | Eukaryota | 2996.0 | 0 | Chordata | 2060.0 | 1 | Carnivora | 1747.0 | 2 | Canidae | 1747.0 | 2 | Canis | 1747.0 | 2 | Canis lupus | 1747.0 | 2 |
| scf7180000046878 | 9248 | 0.4385 | 0 | 196.2749 | no-hit | 0.0 | 0 | no-hit | 0.0 | 0 | no-hit | 0.0 | 0 | no-hit | 0.0 | 0 | no-hit | 0.0 | 0 | no-hit | 0.0 | 0 |
| scf7180000044971 | 247277 | 0.496 | 0 | 58.906 | Eukaryota | 34624.0 | 0 | Chordata | 34624.0 | 0 | Carnivora | 31126.0 | 1 | Canidae | 18661.0 | 4 | Canis | 13745.0 | 6 | Canis lupus | 13745.0 | 6 |
| scf7180000041405 | 693637 | 0.4009 | 0 | 59.8192 | Eukaryota | 91858.0 | 0 | Chordata | 91858.0 | 0 | Carnivora | 91858.0 | 0 | Canidae | 91858.0 | 0 | Canis | 91858.0 | 0 | Canis lupus | 91858.0 | 0 |
| scf7180000042178 | 49000 | 0.3718 | 0 | 52.7685 | Eukaryota | 33897.0 | 0 | Chordata | 33897.0 | 0 | Carnivora | 33897.0 | 0 | Canidae | 33897.0 | 0 | Canis | 33897.0 | 0 | Canis lupus | 33897.0 | 0 |
| scf7180000043052 | 377391 | 0.4423 | 0 | 57.963 | Eukaryota | 32115.0 | 0 | Chordata | 32115.0 | 0 | Carnivora | 32115.0 | 0 | Canidae | 32115.0 | 0 | Canis | 32115.0 | 0 | Canis lupus | 32115.0 | 0 |
| scf7180000047414 | 28377 | 0.4411 | 0 | 160.6544 | Eukaryota | 87522.0 | 0 | Chordata | 87522.0 | 0 | Carnivora | 87522.0 | 0 | Canidae | 87522.0 | 0 | Canis | 87522.0 | 0 | Canis lupus | 87522.0 | 0 |
| scf7180000043350 | 443312 | 0.4164 | 0 | 60.4569 | Eukaryota | 46471.0 | 0 | Chordata | 46471.0 | 0 | Carnivora | 46471.0 | 0 | Canidae | 23756.0 | 1 | Zalophus | 22715.0 | 2 | Zalophus californianus | 22715.0 | 2 |
| scf7180000056165 | 4694 | 0.4491 | 0 | 492.6624 | Eukaryota | 1206.0 | 0 | Echinodermata | 680.0 | 1 | Forcipulatida | 680.0 | 1 | Asteriidae | 680.0 | 1 | Asterias | 680.0 | 1 | Asterias rubens | 680.0 | 1 |
| scf7180000055495 | 4376 | 0.4463 | 0 | 75.2527 | no-hit | 0.0 | 0 | no-hit | 0.0 | 0 | no-hit | 0.0 | 0 | no-hit | 0.0 | 0 | no-hit | 0.0 | 0 | no-hit | 0.0 | 0 |
| scf7180000041000 | 1428012 | 0.3976 | 0 | 59.4805 | Eukaryota | 80208.0 | 0 | Chordata | 80208.0 | 0 | Carnivora | 80208.0 | 0 | Canidae | 80208.0 | 0 | Canis | 46308.0 | 1 | Canis lupus | 46308.0 | 1 |
| scf7180000053057 | 2888 | 0.4467 | 0 | 576.4351 | Eukaryota | 616.0 | 0 | Streptophyta | 482.0 | 1 | Fabales | 482.0 | 1 | Fabaceae | 482.0 | 1 | Cicer | 482.0 | 1 | Cicer arietinum | 482.0 | 1 |
| scf7180000041616 | 939702 | 0.4141 | 0 | 60.2576 | Eukaryota | 77572.0 | 0 | Chordata | 77572.0 | 0 | Carnivora | 77572.0 | 0 | Canidae | 77572.0 | 0 | Canis | 77572.0 | 0 | Canis lupus | 77572.0 | 0 |
| scf7180000055929 | 19195 | 0.4415 | 0 | 155.3784 | Eukaryota | 39712.0 | 0 | Chordata | 24049.0 | 2 | Gadiformes | 21784.0 | 5 | Gadidae | 21784.0 | 5 | Gadus | 21784.0 | 5 | Gadus morhua | 21784.0 | 5 |
| scf7180000044438 | 79347 | 0.4254 | 0 | 59.2801 | Eukaryota | 87964.0 | 0 | Chordata | 87964.0 | 0 | Carnivora | 87964.0 | 0 | Canidae | 87964.0 | 0 | Canis | 87964.0 | 0 | Canis lupus | 87964.0 | 0 |
| scf7180000045632 | 175309 | 0.4015 | 0 | 60.3139 | Eukaryota | 39202.0 | 0 | Chordata | 39202.0 | 0 | Carnivora | 32598.0 | 1 | Canidae | 29181.0 | 2 | Canis | 16964.0 | 3 | Canis lupus | 16964.0 | 3 |
| scf7180000053518 | 5201 | 0.4482 | 0 | 798.2153 | no-hit | 0.0 | 0 | no-hit | 0.0 | 0 | no-hit | 0.0 | 0 | no-hit | 0.0 | 0 | no-hit | 0.0 | 0 | no-hit | 0.0 | 0 |
| scf7180000045033 | 235441 | 0.4954 | 0 | 59.6319 | Eukaryota | 21022.0 | 0 | Chordata | 21022.0 | 0 | Carnivora | 21022.0 | 0 | Canidae | 21022.0 | 0 | Canis | 21022.0 | 0 | Canis lupus | 21022.0 | 0 |
| scf7180000041550 | 1018110 | 0.4891 | 0 | 58.39 | Eukaryota | 91984.0 | 0 | Chordata | 91984.0 | 0 | Carnivora | 91984.0 | 0 | Canidae | 79599.0 | 2 | Canis | 59451.0 | 3 | Canis lupus | 59451.0 | 3 |
| scf7180000045012 | 22802 | 0.453 | 0 | 179.7401 | Eukaryota | 18192.0 | 0 | Chordata | 11355.0 | 2 | Gadiformes | 5411.0 | 6 | Gadidae | 5411.0 | 6 | Gadus | 5411.0 | 6 | Gadus morhua | 5411.0 | 6 |
| scf7180000048308 | 18506 | 0.4483 | 0 | 172.0749 | Eukaryota | 8203.0 | 0 | Chordata | 8203.0 | 0 | Carnivora | 8203.0 | 0 | Canidae | 8203.0 | 0 | Canis | 8203.0 | 0 | Canis lupus | 8203.0 | 0 |
| scf7180000050568 | 29879 | 0.4506 | 0 | 60.0969 | Eukaryota | 14460.0 | 0 | Chordata | 14460.0 | 0 | Carnivora | 14460.0 | 0 | Canidae | 14460.0 | 0 | Canis | 14460.0 | 0 | Canis lupus | 14460.0 | 0 |
| scf7180000046307 | 92285 | 0.4421 | 0 | 97.7019 | Eukaryota | 6795.0 | 0 | Chordata | 6048.0 | 1 | Tetraodontiformes | 3540.0 | 4 | Tetraodontidae | 3540.0 | 4 | Takifugu | 3540.0 | 4 | Takifugu rubripes | 3540.0 | 4 |
| scf7180000045698 | 196257 | 0.3795 | 0 | 59.8056 | Eukaryota | 90568.0 | 0 | Chordata | 90568.0 | 0 | Carnivora | 90568.0 | 0 | Canidae | 90568.0 | 0 | Canis | 90568.0 | 0 | Canis lupus | 90568.0 | 0 |
| scf7180000048616 | 42745 | 0.3887 | 0 | 59.4462 | Eukaryota | 7242.0 | 0 | Chordata | 7242.0 | 0 | Primates | 6068.0 | 2 | Hominidae | 6068.0 | 2 | Homo | 4036.0 | 4 | Homo sapiens | 4036.0 | 4 |
| scf7180000043801 | 194159 | 0.4077 | 0 | 58.0193 | Eukaryota | 65767.0 | 0 | Chordata | 65767.0 | 0 | Carnivora | 65767.0 | 0 | Canidae | 65767.0 | 0 | Canis | 65767.0 | 0 | Canis lupus | 65767.0 | 0 |
| scf7180000052879 | 3015 | 0.4498 | 0 | 377.2082 | Eukaryota | 2416.0 | 0 | Platyhelminthes | 1358.0 | 2 | Polystomatidea | 1358.0 | 2 | Polystomatidae | 1358.0 | 2 | Protopolystoma | 1358.0 | 2 | Protopolystoma xenopodis | 1358.0 | 2 |
| scf7180000056338 | 597824 | 0.3812 | 0 | 61.1758 | Eukaryota | 83745.0 | 0 | Chordata | 83745.0 | 0 | Carnivora | 83745.0 | 0 | Canidae | 83745.0 | 0 | Canis | 48270.0 | 1 | Canis lupus | 48270.0 | 1 |
| scf7180000054421 | 26913 | 0.4051 | 0 | 258.9659 | Eukaryota | 42960.0 | 0 | Chordata | 42960.0 | 0 | Carnivora | 42960.0 | 0 | Canidae | 42960.0 | 0 | Canis | 42960.0 | 0 | Canis lupus | 42960.0 | 0 |
| scf7180000047714 | 67481 | 0.3985 | 0 | 53.0435 | Eukaryota | 38899.0 | 0 | Chordata | 38899.0 | 0 | Carnivora | 38899.0 | 0 | Canidae | 29992.0 | 4 | Canis | 28340.0 | 6 | Canis lupus | 28340.0 | 7 |
| scf7180000049000 | 37551 | 0.4461 | 0 | 57.9602 | Eukaryota | 3148.0 | 0 | Chordata | 1406.0 | 2 | Salmoniformes | 1252.0 | 4 | Salmonidae | 1252.0 | 4 | Coregonus | 915.0 | 6 | Coregonus sp. 'balchen' | 915.0 | 6 |
| scf7180000046191 | 128667 | 0.3898 | 0 | 57.9543 | Eukaryota | 11577.0 | 0 | Chordata | 11577.0 | 0 | Carnivora | 5347.0 | 2 | Bovidae | 4514.0 | 3 | Canis | 4405.0 | 5 | Canis lupus | 4405.0 | 5 |
| scf7180000046707 | 6242 | 0.4409 | 0 | 151.4325 | Eukaryota | 10368.0 | 0 | Chordata | 10368.0 | 0 | Tetraodontiformes | 3938.0 | 2 | Tetraodontidae | 3938.0 | 2 | Takifugu | 3938.0 | 2 | Sphaeramia orbicularis | 3846.0 | 3 |
| scf7180000040971 | 670654 | 0.452 | 0 | 59.2802 | Eukaryota | 72843.0 | 0 | Chordata | 72843.0 | 0 | Carnivora | 72843.0 | 0 | Canidae | 72843.0 | 0 | Canis | 43161.0 | 1 | Canis lupus | 43161.0 | 1 |
| scf7180000054248 | 8714 | 0.4536 | 0 | 156.298 | Eukaryota | 85768.0 | 0 | Chordata | 85768.0 | 0 | Carnivora | 85768.0 | 0 | Canidae | 42844.0 | 3 | Canis | 34229.0 | 5 | Canis lupus | 34229.0 | 6 |
| scf7180000054224 | 99977 | 0.4414 | 0 | 244.4387 | Eukaryota | 7068.0 | 0 | Chordata | 6462.0 | 1 | Salmoniformes | 2963.0 | 3 | Salmonidae | 2963.0 | 3 | Salmo | 2963.0 | 3 | Salmo trutta | 2963.0 | 3 |
| scf7180000049059 | 11762 | 0.4441 | 0 | 60.4638 | no-hit | 0.0 | 0 | no-hit | 0.0 | 0 | no-hit | 0.0 | 0 | no-hit | 0.0 | 0 | no-hit | 0.0 | 0 | no-hit | 0.0 | 0 |
| scf7180000050306 | 2897 | 0.457 | 0 | 84.5237 | no-hit | 0.0 | 0 | no-hit | 0.0 | 0 | no-hit | 0.0 | 0 | no-hit | 0.0 | 0 | no-hit | 0.0 | 0 | no-hit | 0.0 | 0 |
| scf7180000044106 | 933549 | 0.455 | 0 | 68.1356 | Eukaryota | 72275.0 | 0 | Chordata | 72275.0 | 0 | Carnivora | 72275.0 | 0 | Canidae | 72275.0 | 0 | Canis | 59378.0 | 2 | Canis lupus | 59378.0 | 2 |
| scf7180000056543 | 214149 | 0.5341 | 0 | 57.5587 | Eukaryota | 40091.0 | 0 | Chordata | 40091.0 | 0 | Carnivora | 40091.0 | 0 | Canidae | 40091.0 | 0 | Canis | 40091.0 | 0 | Canis lupus | 40091.0 | 0 |
| scf7180000045117 | 192778 | 0.4461 | 0 | 58.1704 | Eukaryota | 30072.0 | 0 | Chordata | 30072.0 | 0 | Carnivora | 30072.0 | 0 | Canidae | 30072.0 | 0 | Canis | 30072.0 | 0 | Canis lupus | 30072.0 | 0 |
| scf7180000053076 | 2583 | 0.5091 | 0 | 45.9386 | Eukaryota | 4283.0 | 0 | Chordata | 4283.0 | 0 | Carnivora | 4283.0 | 0 | Canidae | 4283.0 | 0 | Canis | 2906.0 | 1 | Canis lupus | 2579.0 | 2 |
| scf7180000048018 | 10317 | 0.4471 | 0 | 78.6052 | Eukaryota | 3743.0 | 0 | Chordata | 3336.0 | 1 | Kurtiformes | 2581.0 | 3 | Apogonidae | 2581.0 | 3 | Sphaeramia | 2581.0 | 3 | Sphaeramia orbicularis | 2581.0 | 3 |
| scf7180000048522 | 22837 | 0.4343 | 0 | 193.1153 | Eukaryota | 36923.0 | 0 | Chordata | 36923.0 | 0 | Carnivora | 36923.0 | 0 | Canidae | 36923.0 | 0 | Canis | 36923.0 | 0 | Canis lupus | 36923.0 | 0 |
| scf7180000056435 | 321671 | 0.3941 | 0 | 61.0115 | Eukaryota | 57697.0 | 0 | Chordata | 57697.0 | 0 | Carnivora | 57697.0 | 0 | Canidae | 57697.0 | 0 | Canis | 57697.0 | 0 | Canis lupus | 57697.0 | 0 |

| | | | | | | | | | | | | | | | | | | | | | | |
|---|---|---|---|---|---|---|---|---|---|---|---|---|---|---|---|---|---|---|---|---|---|---|
| scf7180000047382 | 70968 | 0.3361 | 0 | 59.6241 | Eukaryota | 8287.0 | 0 | Chordata | 8287.0 | 0 | Carnivora | 8287.0 | 0 | Canidae | 4194.0 | 2 | Canis | 3377.0 | 5 | Canis lupus | 3377.0 | 5 |
| scf7180000054027 | 2436 | 0.4433 | 0 | 90.6383 | no-hit | 0.0 | 0 | no-hit | 0.0 | 0 | no-hit | 0.0 | 0 | no-hit | 0.0 | 0 | no-hit | 0.0 | 0 | no-hit | 0.0 | 0 |
| scf7180000055374 | 54065 | 0.4379 | 0 | 140.9023 | Eukaryota | 8710.0 | 0 | Chordata | 8128.0 | 1 | Anabantiformes | 3199.0 | 4 | Anabantidae | 3199.0 | 4 | Anabas | 3199.0 | 4 | Anabas testudineus | 3199.0 | 4 |
| scf7180000048572 | 63382 | 0.3974 | 0 | 61.8522 | Eukaryota | 32198.0 | 0 | Chordata | 32198.0 | 0 | Carnivora | 32198.0 | 0 | Canidae | 32198.0 | 0 | Canis | 27368.0 | 1 | Canis lupus | 27368.0 | 1 |
| scf7180000045304 | 246847 | 0.4417 | 0 | 58.1832 | Eukaryota | 63456.0 | 0 | Chordata | 63456.0 | 0 | Carnivora | 63456.0 | 0 | Canidae | 63456.0 | 0 | Canis | 63456.0 | 0 | Canis lupus | 63456.0 | 0 |
| scf7180000056190 | 611343 | 0.3633 | 0 | 60.4629 | Eukaryota | 47919.0 | 0 | Chordata | 47919.0 | 0 | Carnivora | 47919.0 | 0 | Canidae | 47919.0 | 0 | Canis | 47919.0 | 0 | Canis lupus | 47919.0 | 0 |
| scf7180000054564 | 31987 | 0.3534 | 0 | 379.7188 | Eukaryota | 6881.0 | 0 | Chordata | 6881.0 | 0 | Carnivora | 4778.0 | 1 | Canidae | 4778.0 | 1 | Canis | 4778.0 | 1 | Canis lupus | 4778.0 | 1 |
| scf7180000051283 | 11298 | 0.3688 | 0 | 54.4712 | Eukaryota | 23587.0 | 0 | Chordata | 23587.0 | 0 | Carnivora | 23587.0 | 0 | Canidae | 23587.0 | 0 | Canis | 23587.0 | 0 | Canis lupus | 23587.0 | 0 |
| scf7180000042013 | 248618 | 0.4472 | 0 | 58.8776 | Eukaryota | 55141.0 | 0 | Chordata | 55141.0 | 0 | Carnivora | 55141.0 | 0 | Canidae | 55141.0 | 0 | Canis | 55141.0 | 0 | Canis lupus | 55141.0 | 0 |
| scf7180000054914 | 52551 | 0.4495 | 0 | 130.5571 | Eukaryota | 36739.0 | 0 | Chordata | 36739.0 | 0 | Carnivora | 36739.0 | 0 | Canidae | 36739.0 | 0 | Canis | 36739.0 | 0 | Canis lupus | 36739.0 | 0 |
| scf7180000052180 | 10142 | 0.4385 | 0 | 341.6035 | Eukaryota | 2092.0 | 0 | Platyhelminthes | 1547.0 | 2 | Polystomatidea | 1547.0 | 2 | Polystomatidae | 1547.0 | 2 | Protopolystoma | 1547.0 | 2 | Protopolystoma xenopodis | 1547.0 | 2 |
| scf7180000043504 | 369088 | 0.4345 | 0 | 57.0827 | Eukaryota | 19140.0 | 0 | Chordata | 19140.0 | 0 | Carnivora | 19140.0 | 0 | Canidae | 19140.0 | 0 | Canis | 11515.0 | 1 | Canis lupus | 11515.0 | 1 |
| scf7180000042920 | 62446 | 0.3881 | 0 | 58.1309 | Eukaryota | 77650.0 | 0 | Chordata | 77650.0 | 0 | Carnivora | 77650.0 | 0 | Canidae | 77650.0 | 0 | Canis | 77650.0 | 0 | Canis lupus | 77650.0 | 0 |
| scf7180000042157 | 741663 | 0.352 | 0 | 59.4051 | Eukaryota | 63874.0 | 0 | Chordata | 63874.0 | 0 | Carnivora | 63874.0 | 0 | Canidae | 63874.0 | 0 | Canis | 63874.0 | 0 | Canis lupus | 63874.0 | 0 |
| scf7180000053670 | 4430 | 0.4542 | 0 | 49.8702 | Eukaryota | 2703.0 | 0 | Chordata | 2556.0 | 1 | Salmoniformes | 1380.0 | 2 | Salmonidae | 1380.0 | 2 | Salmo | 1380.0 | 2 | Salmo trutta | 1380.0 | 2 |
| scf7180000048397 | 41371 | 0.4354 | 0 | 114.6296 | Eukaryota | 49575.0 | 0 | Chordata | 49575.0 | 0 | Carnivora | 49575.0 | 0 | Canidae | 49575.0 | 0 | Canis | 49575.0 | 0 | Canis lupus | 49575.0 | 0 |
| scf7180000054826 | 86137 | 0.4466 | 0 | 327.8033 | Eukaryota | 25767.0 | 0 | Chordata | 15297.0 | 2 | Strigeidida | 8274.0 | 7 | Schistosomatidae | 8274.0 | 7 | Schistosoma | 8274.0 | 7 | Schistosoma mansoni | 8274.0 | 7 |
| scf7180000056421 | 483222 | 0.4043 | 0 | 59.8824 | Eukaryota | 55336.0 | 0 | Chordata | 55336.0 | 0 | Carnivora | 55336.0 | 0 | Canidae | 55336.0 | 0 | Canis | 49712.0 | 1 | Canis lupus | 49712.0 | 1 |
| scf7180000041852 | 34764 | 0.403 | 0 | 59.0289 | Eukaryota | 48724.0 | 0 | Chordata | 48724.0 | 0 | Carnivora | 48724.0 | 0 | Canidae | 48724.0 | 0 | Canis | 48724.0 | 0 | Canis lupus | 48724.0 | 0 |
| scf7180000043447 | 37696 | 0.4232 | 0 | 270.1135 | Eukaryota | 15088.0 | 0 | Chordata | 15088.0 | 0 | Carnivora | 15088.0 | 0 | Canidae | 15088.0 | 0 | Canis | 15088.0 | 0 | Canis lupus | 15088.0 | 0 |
| scf7180000056529 | 181578 | 0.3924 | 0 | 60.1576 | Eukaryota | 40951.0 | 0 | Chordata | 40951.0 | 0 | Carnivora | 40951.0 | 0 | Canidae | 40951.0 | 0 | Canis | 32089.0 | 1 | Canis lupus | 32089.0 | 1 |
| scf7180000044520 | 60714 | 0.398 | 0 | 56.7312 | Eukaryota | 5533.0 | 0 | Chordata | 5533.0 | 0 | Carnivora | 5533.0 | 0 | Felidae | 3540.0 | 1 | Felis | 3540.0 | 1 | Felis catus | 3540.0 | 1 |
| scf7180000054174 | 19729 | 0.4454 | 0 | 199.1171 | no-hit | 0.0 | 0 | no-hit | 0.0 | 0 | no-hit | 0.0 | 0 | no-hit | 0.0 | 0 | no-hit | 0.0 | 0 | no-hit | 0.0 | 0 |
| scf7180000052019 | 9384 | 0.4358 | 0 | 163.0938 | Eukaryota | 365.0 | 0 | Platyhelminthes | 215.0 | 1 | Polystomatidea | 215.0 | 1 | Polystomatidae | 215.0 | 1 | Protopolystoma | 215.0 | 1 | Protopolystoma xenopodis | 215.0 | 1 |
| scf7180000041030 | 225614 | 0.3595 | 0 | 57.7588 | Eukaryota | 48254.0 | 0 | Chordata | 48254.0 | 0 | Carnivora | 48254.0 | 0 | Canidae | 48254.0 | 0 | Canis | 48254.0 | 0 | Canis lupus | 48254.0 | 0 |
| scf7180000044829 | 72360 | 0.3946 | 0 | 57.8424 | Eukaryota | 9294.0 | 0 | Chordata | 9294.0 | 0 | Carnivora | 9294.0 | 0 | Canidae | 9294.0 | 0 | Canis | 9294.0 | 0 | Canis lupus | 9294.0 | 0 |
| scf7180000051111 | 4669 | 0.4386 | 0 | 86.3864 | Eukaryota | 2922.0 | 0 | Chordata | 2096.0 | 1 | Columbiformes | 1746.0 | 2 | Columbidae | 1746.0 | 2 | Streptopelia | 1746.0 | 2 | Streptopelia turtur | 1746.0 | 2 |
| scf7180000041362 | 859206 | 0.3881 | 0 | 59.6306 | Eukaryota | 87538.0 | 0 | Chordata | 87538.0 | 0 | Carnivora | 87538.0 | 0 | Canidae | 87538.0 | 0 | Canis | 87538.0 | 0 | Canis lupus | 87538.0 | 0 |
| scf7180000053208 | 3612 | 0.441 | 0 | 631.6338 | no-hit | 0.0 | 0 | no-hit | 0.0 | 0 | no-hit | 0.0 | 0 | no-hit | 0.0 | 0 | no-hit | 0.0 | 0 | no-hit | 0.0 | 0 |
| scf7180000041547 | 997583 | 0.4527 | 0 | 59.5406 | Eukaryota | 87701.0 | 0 | Chordata | 87701.0 | 0 | Carnivora | 87701.0 | 0 | Canidae | 87701.0 | 0 | Canis | 87701.0 | 0 | Canis lupus | 87701.0 | 0 |
| scf7180000046631 | 196146 | 0.4318 | 0 | 59.2026 | Eukaryota | 12417.0 | 0 | Chordata | 12417.0 | 0 | Carnivora | 12417.0 | 0 | Canidae | 10568.0 | 2 | Canis | 9247.0 | 3 | Canis lupus | 9247.0 | 3 |
| scf7180000048934 | 17932 | 0.4317 | 0 | 71.737 | Eukaryota | 13190.0 | 0 | Chordata | 8075.0 | 3 | Gadiformes | 4477.0 | 7 | Gadidae | 4477.0 | 7 | Gadus | 4477.0 | 7 | Gadus morhua | 4477.0 | 7 |
| scf7180000049257 | 15368 | 0.4393 | 0 | 158.945 | Eukaryota | 5202.0 | 0 | Chordata | 4546.0 | 1 | Salmoniformes | 3242.0 | 3 | Salmonidae | 3242.0 | 3 | Salmo | 3242.0 | 3 | Salmo trutta | 3242.0 | 3 |
| scf7180000049962 | 19729 | 0.4518 | 0 | 767.4974 | Eukaryota | 2570.0 | 0 | Platyhelminthes | 1202.0 | 2 | Polystomatidea | 1202.0 | 2 | Polystomatidae | 1202.0 | 2 | Protopolystoma | 1202.0 | 2 | Protopolystoma xenopodis | 1202.0 | 2 |
| scf7180000047771 | 6228 | 0.4462 | 0 | 135.1646 | Eukaryota | 4396.0 | 0 | Platyhelminthes | 2530.0 | 1 | Polystomatidea | 2530.0 | 1 | Polystomatidae | 2530.0 | 1 | Protopolystoma | 2530.0 | 1 | Protopolystoma xenopodis | 2530.0 | 1 |
| scf7180000041223 | 1274035 | 0.3865 | 0 | 61.0641 | Eukaryota | 261169.0 | 0 | Chordata | 261169.0 | 0 | Carnivora | 261169.0 | 0 | Canidae | 261169.0 | 0 | Canis | 254153.0 | 1 | Canis lupus | 254153.0 | 1 |
| scf7180000045183 | 40579 | 0.4435 | 0 | 140.0133 | Eukaryota | 68980.0 | 0 | Chordata | 68980.0 | 0 | Carnivora | 68980.0 | 0 | Canidae | 68980.0 | 0 | Canis | 68980.0 | 0 | Canis lupus | 68980.0 | 0 |
| scf7180000045761 | 14732 | 0.3758 | 0 | 48.8571 | Eukaryota | 5423.0 | 0 | Chordata | 5423.0 | 0 | Primates | 1943.0 | 2 | Canidae | 1874.0 | 3 | Canis | 1874.0 | 5 | Canis lupus | 1874.0 | 5 |
| scf7180000051318 | 22151 | 0.4418 | 0 | 104.5876 | Eukaryota | 9615.0 | 0 | Chordata | 9095.0 | 1 | Gonorynchiformes | 4548.0 | 3 | Chanidae | 4548.0 | 3 | Chanos | 4548.0 | 3 | Chanos chanos | 4548.0 | 3 |
| scf7180000049884 | 9391 | 0.4426 | 0 | 118.1801 | Eukaryota | 12873.0 | 0 | Chordata | 7403.0 | 3 | Strigeidida | 4122.0 | 7 | Schistosomatidae | 4122.0 | 7 | Schistosoma | 4122.0 | 7 | Schistosoma mansoni | 4122.0 | 7 |
| scf7180000052778 | 11347 | 0.465 | 0 | 63.776 | Eukaryota | 27868.0 | 0 | Chordata | 27868.0 | 0 | Carnivora | 27868.0 | 0 | Canidae | 27868.0 | 0 | Canis | 27868.0 | 0 | Canis lupus | 27868.0 | 0 |
| scf7180000044872 | 3304 | 0.5079 | 0 | 83.3265 | Eukaryota | 7435.0 | 0 | Chordata | 7435.0 | 0 | Carnivora | 7435.0 | 0 | Canidae | 7435.0 | 0 | Canis | 5983.0 | 1 | Canis lupus | 4960.0 | 2 |
| scf7180000043884 | 431732 | 0.4562 | 0 | 58.1358 | Eukaryota | 39589.0 | 0 | Chordata | 39589.0 | 0 | Carnivora | 39589.0 | 0 | Canidae | 39589.0 | 0 | Canis | 39589.0 | 0 | Canis lupus | 39589.0 | 0 |
| scf7180000043719 | 124344 | 0.402 | 0 | 56.9609 | Eukaryota | 32653.0 | 0 | Chordata | 32653.0 | 0 | Carnivora | 32653.0 | 0 | Otariidae | 28150.0 | 2 | Zalophus | 20941.0 | 3 | Zalophus californianus | 20941.0 | 3 |
| scf7180000044916 | 15048 | 0.3591 | 0 | 48.1424 | Eukaryota | 22915.0 | 0 | Chordata | 22915.0 | 0 | Carnivora | 11409.0 | 4 | Canidae | 9032.0 | 6 | Vulpes | 9032.0 | 7 | Vulpes vulpes | 9032.0 | 7 |
| scf7180000054219 | 31170 | 0.3784 | 0 | 749.0722 | Eukaryota | 50351.0 | 0 | Chordata | 50351.0 | 0 | Carnivora | 50351.0 | 0 | Canidae | 50351.0 | 0 | Canis | 50351.0 | 0 | Canis lupus | 50351.0 | 0 |
| scf7180000052759 | 6263 | 0.4471 | 0 | 131.584 | Eukaryota | 294.0 | 0 | Chordata | 294.0 | 0 | Tetraodontiformes | 294.0 | 0 | Tetraodontidae | 294.0 | 0 | Takifugu | 294.0 | 0 | Takifugu rubripes | 294.0 | 0 |
| scf7180000052891 | 10674 | 0.441 | 0 | 141.055 | no-hit | 0.0 | 0 | no-hit | 0.0 | 0 | no-hit | 0.0 | 0 | no-hit | 0.0 | 0 | no-hit | 0.0 | 0 | no-hit | 0.0 | 0 |
| scf7180000041807 | 44249 | 0.3759 | 0 | 60.3572 | Eukaryota | 6235.0 | 0 | Chordata | 6235.0 | 0 | Carnivora | 6235.0 | 0 | Felidae | 4987.0 | 1 | Felis | 4333.0 | 2 | Felis catus | 4333.0 | 2 |
| scf7180000046147 | 3935 | 0.4442 | 0 | 438.0391 | no-hit | 0.0 | 0 | no-hit | 0.0 | 0 | no-hit | 0.0 | 0 | no-hit | 0.0 | 0 | no-hit | 0.0 | 0 | no-hit | 0.0 | 0 |
| scf7180000047586 | 45576 | 0.4451 | 0 | 257.7201 | Eukaryota | 45998.0 | 0 | Chordata | 35550.0 | 1 | Gadiformes | 28176.0 | 4 | Gadidae | 28176.0 | 4 | Gadus | 28176.0 | 4 | Gadus morhua | 28176.0 | 4 |
| scf7180000046115 | 126691 | 0.3881 | 0 | 60.2056 | Eukaryota | 13419.0 | 0 | Chordata | 13419.0 | 0 | Carnivora | 13419.0 | 0 | Canidae | 11231.0 | 1 | Canis | 6876.0 | 3 | Canis lupus | 6876.0 | 3 |
| scf7180000042537 | 622239 | 0.3618 | 0 | 60.264 | Eukaryota | 31967.0 | 0 | Chordata | 31967.0 | 0 | Carnivora | 31967.0 | 0 | Canidae | 24692.0 | 2 | Canis | 20834.0 | 3 | Canis lupus | 20834.0 | 3 |
| scf7180000041634 | 177676 | 0.3853 | 0 | 59.3449 | Eukaryota | 17258.0 | 0 | Chordata | 17258.0 | 0 | Carnivora | 9910.0 | 2 | Canidae | 7013.0 | 3 | Canis | 7013.0 | 4 | Canis lupus | 7013.0 | 4 |
| scf7180000051290 | 19254 | 0.5028 | 0 | 614.3721 | Eukaryota | 9313.0 | 0 | Chordata | 9313.0 | 0 | Carnivora | 9313.0 | 0 | Canidae | 9313.0 | 0 | Canis | 7537.0 | 1 | Canis lupus | 6251.0 | 2 |
| scf7180000044525 | 316052 | 0.3684 | 0 | 59.1631 | Eukaryota | 59595.0 | 0 | Chordata | 59595.0 | 0 | Carnivora | 59595.0 | 0 | Canidae | 59595.0 | 0 | Canis | 59595.0 | 0 | Canis lupus | 59595.0 | 0 |
| scf7180000041159 | 581493 | 0.4027 | 0 | 58.7343 | Eukaryota | 36708.0 | 0 | Chordata | 36708.0 | 0 | Carnivora | 36708.0 | 0 | Canidae | 36708.0 | 0 | Canis | 36708.0 | 0 | Canis lupus | 36708.0 | 0 |
| scf7180000055289 | 38237 | 0.4739 | 0 | 320.7078 | Eukaryota | 16179.0 | 0 | Chordata | 16179.0 | 0 | Carnivora | 16179.0 | 0 | Canidae | 16179.0 | 0 | Canis | 16179.0 | 0 | Canis lupus | 16179.0 | 0 |
| scf7180000053812 | 5576 | 0.4354 | 0 | 39.1307 | Eukaryota | 5704.0 | 0 | Platyhelminthes | 3345.0 | 2 | Cyclophyllidea | 2645.0 | 4 | Hymenolepididae | 2645.0 | 4 | Hymenolepis | 2645.0 | 5 | Hymenolepis microstoma | 2645.0 | 5 |
| scf7180000054111 | 9543 | 0.434 | 0 | 283.723 | Eukaryota | 154.0 | 0 | Platyhelminthes | 154.0 | 0 | Polystomatidea | 154.0 | 0 | Polystomatidae | 154.0 | 0 | Protopolystoma | 154.0 | 0 | Protopolystoma xenopodis | 154.0 | 0 |
| scf7180000055024 | 32715 | 0.4428 | 0 | 179.142 | Eukaryota | 12240.0 | 0 | Chordata | 8164.0 | 3 | Gadiformes | 4573.0 | 7 | Gadidae | 4573.0 | 7 | Gadus | 4573.0 | 7 | Gadus morhua | 4573.0 | 7 |
| scf7180000045768 | 180794 | 0.4994 | 0 | 57.4442 | Eukaryota | 78524.0 | 0 | Chordata | 78524.0 | 0 | Carnivora | 78524.0 | 0 | Canidae | 78524.0 | 0 | Canis | 62721.0 | 1 | Canis lupus | 62721.0 | 1 |
| scf7180000049534 | 4299 | 0.4382 | 0 | 85.7103 | Eukaryota | 2853.0 | 0 | Chordata | 2853.0 | 0 | Carnivora | 2853.0 | 0 | Canidae | 2853.0 | 0 | Canis | 2853.0 | 0 | Canis lupus | 2853.0 | 0 |
| scf7180000056471 | 310642 | 0.3991 | 0 | 59.0077 | Eukaryota | 12327.0 | 1 | Chordata | 12327.0 | 1 | Carnivora | 7903.0 | 3 | Canidae | 7903.0 | 3 | Canis | 7903.0 | 4 | Canis lupus | 7903.0 | 4 |
| scf7180000044679 | 50500 | 0.4347 | 0 | 67.8602 | Eukaryota | 76861.0 | 0 | Chordata | 76861.0 | 0 | Carnivora | 76861.0 | 0 | Canidae | 76861.0 | 0 | Canis | 76861.0 | 0 | Canis lupus | 76861.0 | 0 |
| scf7180000043115 | 474061 | 0.4286 | 0 | 59.9948 | Eukaryota | 59703.0 | 0 | Chordata | 59703.0 | 0 | Carnivora | 59703.0 | 0 | Canidae | 59703.0 | 0 | Canis | 59703.0 | 0 | Canis lupus | 59703.0 | 0 |
| scf7180000041608 | 1081863 | 0.3416 | 0 | 59.3903 | Eukaryota | 93481.0 | 0 | Chordata | 93481.0 | 0 | Carnivora | 93481.0 | 0 | Canidae | 93481.0 | 0 | Canis | 93481.0 | 0 | Canis lupus | 93481.0 | 0 |
| scf7180000046619 | 35426 | 0.3902 | 0 | 277.4723 | Eukaryota | 93838.0 | 0 | Chordata | 93838.0 | 0 | Carnivora | 93838.0 | 0 | Canidae | 93838.0 | 0 | Canis | 93838.0 | 0 | Canis lupus | 93838.0 | 0 |
| scf7180000049818 | 5195 | 0.4375 | 0 | 110.1231 | Eukaryota | 180.0 | 0 | Chordata | 180.0 | 0 | Blenniiformes | 180.0 | 0 | Gobiesocidae | 180.0 | 0 | Gouania | 180.0 | 0 | Gouania willdenowi | 180.0 | 0 |
| scf7180000055366 | 4062 | 0.4407 | 0 | 129.2121 | Eukaryota | 206.0 | 0 | Platyhelminthes | 206.0 | 0 | Polystomatidea | 206.0 | 0 | Polystomatidae | 206.0 | 0 | Protopolystoma | 206.0 | 0 | Protopolystoma xenopodis | 206.0 | 0 |
| scf7180000044377 | 302995 | 0.5535 | 0 | 56.0749 | Eukaryota | 22410.0 | 1 | Chordata | 22410.0 | 1 | Carnivora | 14049.0 | 3 | Canidae | 14049.0 | 3 | Canis | 14049.0 | 3 | Canis lupus | 14049.0 | 3 |
| scf7180000053027 | 11442 | 0.5006 | 0 | 149.9517 | Eukaryota | 8335.0 | 0 | Chordata | 8335.0 | 0 | Carnivora | 8335.0 | 0 | Canidae | 8335.0 | 0 | Canis | 6697.0 | 1 | Canis lupus | 5524.0 | 2 |
| scf7180000043899 | 7083 | 0.4477 | 0 | 651.7574 | Eukaryota | 3977.0 | 0 | Platyhelminthes | 2294.0 | 3 | Polystomatidea | 2294.0 | 3 | Polystomatidae | 2294.0 | 3 | Protopolystoma | 2294.0 | 3 | Protopolystoma xenopodis | 2294.0 | 3 |
| scf7180000044895 | 254398 | 0.4397 | 0 | 62.1127 | Eukaryota | 44865.0 | 0 | Chordata | 44865.0 | 0 | Carnivora | 44865.0 | 0 | Canidae | 44865.0 | 0 | Canis | 37597.0 | 1 | Canis lupus | 37597.0 | 1 |
| scf7180000046104 | 133940 | 0.5604 | 0 | 61.4409 | Eukaryota | 42968.0 | 0 | Chordata | 42968.0 | 0 | Carnivora | 42968.0 | 0 | Canidae | 42968.0 | 0 | Canis | 39913.0 | 1 | Canis lupus | 39913.0 | 1 |
| scf7180000046925 | 16501 | 0.4472 | 0 | 234.8601 | Eukaryota | 2953.0 | 0 | Chordata | 2006.0 | 1 | Carnivora | 1708.0 | 2 | Canidae | 1708.0 | 2 | Canis | 1708.0 | 2 | Canis lupus | 1708.0 | 2 |
| scf7180000046159 | 56883 | 0.3995 | 0 | 48.0404 | Eukaryota | 86888.0 | 0 | Chordata | 86888.0 | 0 | Carnivora | 86888.0 | 0 | Canidae | 86888.0 | 0 | Canis | 86888.0 | 0 | Canis lupus | 86888.0 | 0 |
| scf7180000044330 | 8291 | 0.3404 | 0 | 27.5252 | Eukaryota | 2562.0 | 0 | Chordata | 2562.0 | 0 | Carnivora | 2562.0 | 0 | Canidae | 2562.0 | 0 | Canis | 2562.0 | 0 | Canis lupus | 2562.0 | 0 |
| scf7180000046730 | 101912 | 0.4452 | 0 | 121.9548 | Eukaryota | 84546.0 | 0 | Platyhelminthes | 51433.0 | 1 | Strigeidida | 39064.0 | 5 | Schistosomatidae | 39064.0 | 5 | Schistosoma | 39064.0 | 5 | Schistosoma mansoni | 39064.0 | 5 |
| scf7180000050997 | 4011 | 0.5049 | 0 | 61.016 | Eukaryota | 6378.0 | 0 | Chordata | 6378.0 | 0 | Carnivora | 6378.0 | 0 | Canidae | 6378.0 | 0 | Canis | 3993.0 | 1 | Canis lupus | 3059.0 | 2 |
| scf7180000044913 | 236418 | 0.4289 | 0 | 60.6326 | Eukaryota | 25958.0 | 0 | Chordata | 25958.0 | 0 | Carnivora | 25958.0 | 0 | Canidae | 25958.0 | 0 | Canis | 20887.0 | 1 | Canis lupus | 20887.0 | 1 |
| scf7180000049861 | 2497 | 0.4377 | 0 | 188.051 | Eukaryota | 3448.0 | 0 | Platyhelminthes | 1730.0 | 1 | Polystomatidea | 1730.0 | 2 | Polystomatidae | 1730.0 | 2 | Protopolystoma | 1730.0 | 2 | Protopolystoma xenopodis | 1730.0 | 2 |
| scf7180000044644 | 258718 | 0.4701 | 0 | 55.6659 | Eukaryota | 33144.0 | 0 | Chordata | 33144.0 | 0 | Carnivora | 33144.0 | 0 | Canidae | 33144.0 | 0 | Canis | 33144.0 | 0 | Canis lupus | 33144.0 | 0 |
| scf7180000045352 | 232942 | 0.4698 | 0 | 58.5039 | Eukaryota | 42514.0 | 0 | Chordata | 42514.0 | 0 | Carnivora | 42514.0 | 0 | Canidae | 42514.0 | 0 | Canis | 42514.0 | 0 | Canis lupus | 42514.0 | 0 |
| scf7180000051229 | 13287 | 0.4387 | 0 | 120.1895 | Eukaryota | 51133.0 | 0 | Chordata | 31945.0 | 2 | Gadiformes | 29259.0 | 4 | Gadidae | 29259.0 | 4 | Gadus | 29259.0 | 4 | Gadus morhua | 29259.0 | 4 |
| scf7180000054191 | 1437846 | 0.4066 | 0 | 60.6298 | Eukaryota | 92619.0 | 0 | Chordata | 92619.0 | 0 | Carnivora | 92619.0 | 0 | Canidae | 92619.0 | 0 | Canis | 92619.0 | 0 | Canis lupus | 92619.0 | 0 |
| scf7180000040492 | 3036312 | 0.368 | 0 | 60.5728 | Eukaryota | 92994.0 | 0 | Chordata | 92994.0 | 0 | Carnivora | 92994.0 | 0 | Canidae | 92994.0 | 0 | Canis | 92994.0 | 0 | Canis lupus | 92994.0 | 0 |
| scf7180000041648 | 906436 | 0.4065 | 0 | 59.5528 | Eukaryota | 61714.0 | 0 | Chordata | 61714.0 | 0 | Carnivora | 61714.0 | 0 | Canidae | 61714.0 | 0 | Canis | 61714.0 | 0 | Canis lupus | 61714.0 | 0 |
| scf7180000050948 | 5024 | 0.4447 | 0 | 662.1212 | no-hit | 0.0 | 0 | no-hit | 0.0 | 0 | no-hit | 0.0 | 0 | no-hit | 0.0 | 0 | no-hit | 0.0 | 0 | no-hit | 0.0 | 0 |
| scf7180000044391 | 63035 | 0.4437 | 0 | 69.9722 | Eukaryota | 69273.0 | 0 | Chordata | 69273.0 | 0 | Carnivora | 69273.0 | 0 | Canidae | 69273.0 | 0 | Canis | 69273.0 | 0 | Canis lupus | 69273.0 | 0 |
| scf7180000052714 | 8575 | 0.5057 | 0 | 115.6794 | Eukaryota | 8518.0 | 0 | Chordata | 8518.0 | 0 | Carnivora | 8518.0 | 0 | Canidae | 8518.0 | 0 | Canis | 6925.0 | 1 | Canis lupus | 5740.0 | 2 |
| scf7180000042986 | 10297 | 0.4485 | 0 | 501.8995 | Eukaryota | 63416.0 | 0 | Chordata | 63416.0 | 0 | Carnivora | 63416.0 | 0 | Canidae | 63416.0 | 0 | Canis | 63416.0 | 0 | Canis lupus | 63416.0 | 0 |
| scf7180000041711 | 903311 | 0.425 | 0 | 61.6414 | Eukaryota | 33764.0 | 0 | Chordata | 33764.0 | 0 | Carnivora | 33764.0 | 0 | Canidae | 33764.0 | 0 | Canis | 21236.0 | 1 | Canis lupus | 21236.0 | 1 |
| scf7180000055840 | 11010 | 0.4416 | 0 | 144.0771 | Eukaryota | 50847.0 | 0 | Chordata | 36786.0 | 2 | Gadiformes | 29241.0 | 5 | Gadidae | 29241.0 | 5 | Gadus | 29241.0 | 5 | Gadus morhua | 29241.0 | 5 |
| scf7180000041736 | 532663 | 0.4167 | 0 | 58.9017 | Eukaryota | 88943.0 | 0 | Chordata | 88943.0 | 0 | Carnivora | 88943.0 | 0 | Canidae | 88943.0 | 0 | Canis | 79157.0 | 1 | Canis lupus | 79157.0 | 1 |
| scf7180000053460 | 9948 | 0.4502 | 0 | 67.3041 | Eukaryota | 161.0 | 0 | Platyhelminthes | 161.0 | 0 | Diphyllobothriidea | 161.0 | 0 | Diphyllobothriidae | 161.0 | 0 | Dibothriocephalus | 161.0 | 0 | Dibothriocephalus latus | 161.0 | 0 |
| scf7180000041600 | 898080 | 0.4667 | 0 | 60.0549 | Eukaryota | 71285.0 | 0 | Chordata | 71285.0 | 0 | Carnivora | 71285.0 | 0 | Canidae | 71285.0 | 0 | Canis | 71285.0 | 0 | Canis lupus | 71285.0 | 0 |
| scf7180000041653 | 584058 | 0.4105 | 0 | 58.2427 | Eukaryota | 52440.0 | 0 | Chordata | 52440.0 | 0 | Carnivora | 52440.0 | 0 | Canidae | 52440.0 | 0 | Canis | 52440.0 | 0 | Canis lupus | 52440.0 | 0 |
| scf7180000040875 | 555805 | 0.3827 | 0 | 60.0544 | Eukaryota | 65178.0 | 0 | Chordata | 65178.0 | 0 | Carnivora | 65178.0 | 0 | Canidae | 65178.0 | 0 | Canis | 65178.0 | 0 | Canis lupus | 65178.0 | 0 |
| scf7180000043032 | 535887 | 0.3685 | 0 | 60.7761 | Eukaryota | 39617.0 | 0 | Chordata | 39617.0 | 0 | Carnivora | 39617.0 | 0 | Canidae | 21498.0 | 2 | Vulpes | 12822.0 | 4 | Vulpes vulpes | 12822.0 | 4 |
| scf7180000041376 | 952570 | 0.4521 | 0 | 60.9903 | Eukaryota | 95294.0 | 0 | Chordata | 95294.0 | 0 | Carnivora | 95294.0 | 0 | Canidae | 95294.0 | 0 | Canis | 95294.0 | 0 | Canis lupus | 95294.0 | 0 |
| scf7180000049960 | 19472 | 0.4424 | 0 | 127.6568 | Eukaryota | 1909.0 | 0 | Chordata | 1422.0 | 1 | Tetraodontiformes | 1422.0 | 1 | Tetraodontidae | 1422.0 | 1 | Takifugu | 1422.0 | 1 | Takifugu rubripes | 1422.0 | 1 |
| scf7180000055899 | 11959 | 0.4338 | 0 | 77.8326 | Eukaryota | 983.0 | 0 | Platyhelminthes | 533.0 | 1 | Polystomatidea | 533.0 | 2 | Polystomatidae | 533.0 | 2 | Protopolystoma | 533.0 | 2 | Protopolystoma xenopodis | 533.0 | 2 |
| scf7180000049765 | 22194 | 0.3391 | 0 | 60.4291 | Eukaryota | 4278.0 | 0 | Chordata | 4278.0 | 0 | Carnivora | 4278.0 | 0 | Canidae | 3280.0 | 1 | Canis | 3280.0 | 1 | Canis lupus | 3280.0 | 1 |
| scf7180000049902 | 12699 | 0.4409 | 0 | 297.0715 | Eukaryota | 492.0 | 0 | Chordata | 492.0 | 0 | Kurtiformes | 492.0 | 0 | Apogonidae | 492.0 | 0 | Sphaeramia | 492.0 | 0 | Sphaeramia orbicularis | 492.0 | 0 |
| scf7180000044270 | 384313 | 0.4404 | 0 | 57.4593 | Eukaryota | 62785.0 | 0 | Chordata | 62785.0 | 0 | Carnivora | 62785.0 | 0 | Canidae | 62785.0 | 0 | Canis | 62785.0 | 0 | Canis lupus | 62785.0 | 0 |
| scf7180000049077 | 30965 | 0.4339 | 0 | 114.5707 | Eukaryota | 2087.0 | 0 | Chordata | 2087.0 | 0 | Kurtiformes | 1739.0 | 2 | Apogonidae | 1739.0 | 2 | Sphaeramia | 1739.0 | 2 | Sphaeramia orbicularis | 1739.0 | 2 |
| scf7180000055646 | 34472 | 0.4391 | 0 | 140.3412 | Eukaryota | 58946.0 | 0 | Chordata | 58946.0 | 0 | Carnivora | 58946.0 | 0 | Canidae | 58946.0 | 0 | Canis | 58946.0 | 0 | Canis lupus | 58946.0 | 0 |
| scf7180000041531 | 775425 | 0.558 | 0 | 56.707 | Eukaryota | 90641.0 | 0 | Chordata | 90641.0 | 0 | Carnivora | 90641.0 | 0 | Canidae | 90641.0 | 0 | Canis | 80471.0 | 1 | Canis lupus | 80471.0 | 1 |
| scf7180000044591 | 20249 | 0.5176 | 0 | 161.1118 | Eukaryota | 23353.0 | 0 | Chordata | 23353.0 | 0 | Carnivora | 23353.0 | 0 | Canidae | 23353.0 | 0 | Canis | 18759.0 | 1 | Canis lupus | 18759.0 | 1 |
| scf7180000050146 | 3425 | 0.4601 | 0 | 142.3032 | Eukaryota | 497.0 | 0 | Streptophyta | 325.0 | 1 | Fabales | 325.0 | 1 | Fabaceae | 325.0 | 1 | Cicer | 325.0 | 1 | Cicer arietinum | 325.0 | 1 |
| scf7180000041286 | 1107195 | 0.469 | 0 | 59.1679 | Eukaryota | 45148.0 | 0 | Chordata | 45148.0 | 0 | Carnivora | 45148.0 | 0 | Canidae | 45148.0 | 0 | Canis | 45148.0 | 0 | Canis lupus | 45148.0 | 0 |
| scf7180000056472 | 211333 | 0.3905 | 0 | 59.2251 | Eukaryota | 95121.0 | 0 | Chordata | 95121.0 | 0 | Carnivora | 95121.0 | 0 | Canidae | 95121.0 | 0 | Canis | 95121.0 | 0 | Canis lupus | 95121.0 | 0 |
| scf7180000044676 | 260105 | 0.4358 | 0 | 59.0471 | Eukaryota | 20631.0 | 0 | Chordata | 20631.0 | 0 | Carnivora | 12743.0 | 2 | Canidae | 12743.0 | 2 | Canis | 12743.0 | 3 | Canis lupus | 12743.0 | 3 |
| scf7180000042713 | 372242 | 0.3472 | 0 | 59.8011 | Eukaryota | 88831.0 | 0 | Chordata | 88831.0 | 0 | Carnivora | 88831.0 | 0 | Canidae | 88831.0 | 0 | Canis | 88831.0 | 0 | Canis lupus | 88831.0 | 0 |
| scf7180000044589 | 250845 | 0.3384 | 0 | 59.1693 | Eukaryota | 80572.0 | 0 | Chordata | 80572.0 | 0 | Carnivora | 80572.0 | 0 | Canidae | 80572.0 | 0 | Canis | 80572.0 | 0 | Canis lupus | 80572.0 | 0 |
| scf7180000051624 | 12456 | 0.4472 | 0 | 59.0275 | no-hit | 0.0 | 0 | no-hit | 0.0 | 0 | no-hit | 0.0 | 0 | no-hit | 0.0 | 0 | no-hit | 0.0 | 0 | no-hit | 0.0 | 0 |
| scf7180000051520 | 4510 | 0.4499 | 0 | 279.8642 | Eukaryota | 887.0 | 0 | Streptophyta | 528.0 | 2 | Fabales | 528.0 | 2 | Fabaceae | 528.0 | 2 | Cicer | 528.0 | 2 | Cicer arietinum | 528.0 | 2 |
| scf7180000042701 | 601604 | 0.4261 | 0 | 58.3486 | Eukaryota | 71143.0 | 0 | Chordata | 71143.0 | 0 | Carnivora | 71143.0 | 0 | Canidae | 30339.0 | 4 | Canis | 20001.0 | 7 | Canis lupus | 20001.0 | 8 |
| scf7180000042446 | 26874 | 0.4331 | 0 | 34.8971 | Eukaryota | 57107.0 | 0 | Chordata | 57107.0 | 0 | Carnivora | 57107.0 | 0 | Canidae | 57107.0 | 0 | Canis | 57107.0 | 0 | Canis lupus | 57107.0 | 0 |
| scf7180000049793 | 20614 | 0.5013 | 0 | 363.2941 | Eukaryota | 9173.0 | 0 | Chordata | 9173.0 | 0 | Carnivora | 9173.0 | 0 | Canidae | 9173.0 | 0 | Canis | 7420.0 | 1 | Canis lupus | 6132.0 | 2 |
| scf7180000054216 | 45598 | 0.4176 | 0 | 298.4392 | Eukaryota | 10298.0 | 0 | Chordata | 10298.0 | 0 | Carnivora | 10298.0 | 0 | Canidae | 10298.0 | 0 | Canis | 10298.0 | 0 | Canis lupus | 10298.0 | 0 |
| scf7180000040591 | 4888 | 0.4446 | 0 | 36.307 | Eukaryota | 14633.0 | 0 | Chordata | 14633.0 | 0 | Carnivora | 14633.0 | 0 | Canidae | 11516.0 | 2 | Canis | 7676.0 | 4 | Canis lupus | 7676.0 | 4 |
| scf7180000047161 | 360316 | 0.3483 | 0 | 59.1903 | Eukaryota | 14441.0 | 0 | Chordata | 14441.0 | 0 | Carnivora | 14441.0 | 0 | Canidae | 14441.0 | 0 | Canis | 14441.0 | 0 | Canis lupus | 14441.0 | 0 |
| scf7180000050447 | 14781 | 0.4358 | 0 | 138.2249 | Eukaryota | 23707.0 | 0 | Chordata | 20052.0 | 2 | Gadiformes | 6608.0 | 6 | Gadidae | 6608.0 | 6 | Gadus | 6608.0 | 7 | Gadus morhua | 6608.0 | 7 |
| scf7180000045002 | 29691 | 0.3651 | 0 | 58.225 | Eukaryota | 65307.0 | 0 | Chordata | 65307.0 | 0 | Carnivora | 65307.0 | 0 | Canidae | 65307.0 | 0 | Canis | 65307.0 | 0 | Canis lupus | 65307.0 | 0 |
| scf7180000055183 | 23521 | 0.4788 | 0 | 486.5715 | Eukaryota | 20062.0 | 0 | Chordata | 20062.0 | 0 | Carnivora | 20062.0 | 0 | Canidae | 20062.0 | 0 | Canis | 20062.0 | 0 | Canis lupus | 20062.0 | 0 |
| scf7180000048501 | 12289 | 0.4997 | 0 | 220.7375 | Eukaryota | 32993.0 | 0 | Chordata | 32993.0 | 0 | Carnivora | 32993.0 | 0 | Canidae | 32993.0 | 0 | Canis | 32993.0 | 0 | Canis lupus | 32993.0 | 0 |
| scf7180000049679 | 6767 | 0.4469 | 0 | 73.5048 | no-hit | 0.0 | 0 | no-hit | 0.0 | 0 | no-hit | 0.0 | 0 | no-hit | 0.0 | 0 | no-hit | 0.0 | 0 | no-hit | 0.0 | 0 |
| scf7180000041144 | 239264 | 0.4017 | 0 | 57.9466 | Eukaryota | 92130.0 | 0 | Chordata | 92130.0 | 0 | Carnivora | 92130.0 | 0 | Canidae | 92130.0 | 0 | Canis | 92130.0 | 0 | Canis lupus | 92130.0 | 0 |
| scf7180000044809 | 95060 | 0.3955 | 0 | 62.2391 | Eukaryota | 59835.0 | 0 | Chordata | 59835.0 | 0 | Carnivora | 59835.0 | 0 | Canidae | 59835.0 | 0 | Canis | 59835.0 | 0 | Canis lupus | 59835.0 | 0 |
| scf7180000053278 | 10705 | 0.4463 | 0 | 64.3192 | Eukaryota | 93589.0 | 0 | Chordata | 93589.0 | 0 | Carnivora | 93589.0 | 0 | Canidae | 93589.0 | 0 | Canis | 93589.0 | 0 | Canis lupus | 93589.0 | 0 |
| scf7180000055332 | 14536 | 0.4468 | 0 | 281.2077 | Eukaryota | 9226.0 | 0 | Chordata | 9226.0 | 0 | Carnivora | 9226.0 | 0 | Canidae | 9226.0 | 0 | Nyctereutes | 4968.0 | 1 | Nyctereutes procyonoides | 4968.0 | 1 |
| scf7180000041165 | 1370740 | 0.3526 | 0 | 59.9277 | Eukaryota | 67359.0 | 0 | Chordata | 67359.0 | 0 | Carnivora | 67359.0 | 0 | Canidae | 67359.0 | 0 | Canis | 67359.0 | 0 | Canis lupus | 67359.0 | 0 |
| scf7180000041029 | 1315863 | 0.4806 | 0 | 58.6861 | Eukaryota | 92833.0 | 0 | Chordata | 92833.0 | 0 | Carnivora | 92833.0 | 0 | Canidae | 92833.0 | 0 | Canis | 92833.0 | 0 | Canis lupus | 92833.0 | 0 |
| scf7180000041497 | 44485 | 0.3815 | 0 | 47.5775 | Eukaryota | 17026.0 | 0 | Chordata | 17026.0 | 0 | Carnivora | 17026.0 | 0 | Canidae | 17026.0 | 0 | Canis | 17026.0 | 0 | Canis lupus | 17026.0 | 0 |
| scf7180000043149 | 569959 | 0.3931 | 0 | 61.4597 | Eukaryota | 91830.0 | 0 | Chordata | 91830.0 | 0 | Carnivora | 91830.0 | 0 | Canidae | 91830.0 | 0 | Canis | 91830.0 | 0 | Canis lupus | 91830.0 | 0 |
| scf7180000049386 | 14198 | 0.4506 | 0 | 135.754 | Eukaryota | 2953.0 | 1 | Chordata | 2953.0 | 1 | Carnivora | 2953.0 | 1 | Canidae | 2953.0 | 1 | Canis | 2953.0 | 1 | Canis lupus | 2953.0 | 1 |
| scf7180000056216 | 1187181 | 0.3761 | 0 | 60.4851 | Eukaryota | 92092.0 | 0 | Chordata | 92092.0 | 0 | Carnivora | 92092.0 | 0 | Canidae | 92092.0 | 0 | Canis | 92092.0 | 0 | Canis lupus | 92092.0 | 0 |
| scf7180000040787 | 261218 | 0.3823 | 0 | 59.3778 | Eukaryota | 86659.0 | 0 | Chordata | 86659.0 | 0 | Carnivora | 86659.0 | 0 | Canidae | 86659.0 | 0 | Canis | 86659.0 | 0 | Canis lupus | 86659.0 | 0 |
| scf7180000050870 | 17804 | 0.5422 | 0 | 196.1208 | Eukaryota | 952.0 | 0 | Chordata | 952.0 | 0 | Carnivora | 952.0 | 0 | Canidae | 952.0 | 0 | Canis | 952.0 | 0 | Canis lupus | 952.0 | 0 |
| scf7180000049576 | 56589 | 0.5034 | 0 | 60.1561 | Eukaryota | 59121.0 | 0 | Chordata | 59121.0 | 0 | Carnivora | 59121.0 | 0 | Canidae | 59121.0 | 0 | Canis | 51141.0 | 1 | Canis lupus | 51141.0 | 1 |
| scf7180000054904 | 25817 | 0.4406 | 0 | 95.5974 | Eukaryota | 61103.0 | 0 | Chordata | 61103.0 | 0 | Carnivora | 61103.0 | 0 | Canidae | 61103.0 | 0 | Canis | 61103.0 | 0 | Canis lupus | 61103.0 | 0 |
| scf7180000048103 | 18452 | 0.4411 | 0 | 216.3877 | no-hit | 0.0 | 0 | no-hit | 0.0 | 0 | no-hit | 0.0 | 0 | no-hit | 0.0 | 0 | no-hit | 0.0 | 0 | no-hit | 0.0 | 0 |
| scf7180000042400 | 704491 | 0.3854 | 0 | 60.5837 | Eukaryota | 28671.0 | 0 | Chordata | 28671.0 | 0 | Carnivora | 28671.0 | 0 | Canidae | 28671.0 | 0 | Canis | 28671.0 | 0 | Canis lupus | 28671.0 | 0 |
| scf7180000055853 | 13002 | 0.5188 | 0 | 812.766 | Eukaryota | 16128.0 | 0 | Chordata | 16128.0 | 0 | Carnivora | 16128.0 | 0 | Canidae | 16128.0 | 0 | Canis | 16128.0 | 0 | Canis lupus | 16128.0 | 0 |
| scf7180000052825 | 19141 | 0.4408 | 0 | 116.7045 | Eukaryota | 7746.0 | 0 | Chordata | 4022.0 | 2 | Columbiformes | 2430.0 | 4 | Columbidae | 2430.0 | 4 | Streptopelia | 2430.0 | 5 | Streptopelia turtur | 2430.0 | 5 |
| scf7180000047237 | 13826 | 0.4398 | 0 | 110.6294 | Eukaryota | 5114.0 | 0 | Platyhelminthes | 3557.0 | 2 | Polystomatidea | 3557.0 | 2 | Polystomatidae | 3557.0 | 2 | Protopolystoma | 3557.0 | 2 | Protopolystoma xenopodis | 3557.0 | 2 |
| scf7180000056468 | 222398 | 0.415 | 0 | 70.2882 | Eukaryota | 56658.0 | 0 | Chordata | 56658.0 | 0 | Carnivora | 56658.0 | 0 | Canidae | 56658.0 | 0 | Canis | 51457.0 | 1 | Canis lupus | 51457.0 | 1 |
| scf7180000055050 | 18258 | 0.4402 | 0 | 220.4613 | Eukaryota | 3198.0 | 0 | Platyhelminthes | 1940.0 | 1 | Polystomatidea | 1940.0 | 2 | Polystomatidae | 1940.0 | 2 | Protopolystoma | 1940.0 | 2 | Protopolystoma xenopodis | 1940.0 | 2 |
| scf7180000049923 | 9971 | 0.4409 | 0 | 375.1252 | Eukaryota | 6317.0 | 0 | Platyhelminthes | 3843.0 | 2 | Polystomatidea | 3843.0 | 2 | Polystomatidae | 3843.0 | 2 | Protopolystoma | 3843.0 | 2 | Protopolystoma xenopodis | 3843.0 | 2 |
| scf7180000045945 | 130729 | 0.4113 | 0 | 57.5418 | Eukaryota | 28952.0 | 0 | Chordata | 28952.0 | 0 | Carnivora | 28952.0 | 0 | Canidae | 16182.0 | 3 | Canis | 12985.0 | 5 | Canis lupus | 12985.0 | 5 |
| scf7180000056082 | 4858 | 0.443 | 0 | 313.7293 | no-hit | 0.0 | 0 | no-hit | 0.0 | 0 | no-hit | 0.0 | 0 | no-hit | 0.0 | 0 | no-hit | 0.0 | 0 | no-hit | 0.0 | 0 |
| scf7180000052900 | 8678 | 0.4382 | 0 | 94.8006 | Eukaryota | 4634.0 | 0 | Chordata | 3206.0 | 1 | Salmoniformes | 3021.0 | 2 | Salmonidae | 3021.0 | 2 | Salmo | 3021.0 | 2 | Salmo trutta | 3021.0 | 2 |
| scf7180000043110 | 54596 | 0.4327 | 0 | 94.6815 | Eukaryota | 80637.0 | 0 | Chordata | 80637.0 | 0 | Carnivora | 80637.0 | 0 | Canidae | 80637.0 | 0 | Canis | 80637.0 | 0 | Canis lupus | 80637.0 | 0 |
| scf7180000046625 | 5118 | 0.4037 | 0 | 25.7015 | Eukaryota | 2908.0 | 0 | Chordata | 2908.0 | 0 | Carnivora | 2908.0 | 0 | Canidae | 2908.0 | 0 | Canis | 2908.0 | 0 | Canis lupus | 2908.0 | 0 |
| scf7180000043016 | 10441 | 0.4253 | 0 | 70.768 | Eukaryota | 49626.0 | 0 | Chordata | 49626.0 | 0 | Carnivora | 49626.0 | 0 | Canidae | 49626.0 | 0 | Canis | 49626.0 | 0 | Canis lupus | 49626.0 | 0 |
| scf7180000047682 | 10430 | 0.5335 | 0 | 79.6372 | Eukaryota | 30769.0 | 0 | Chordata | 30769.0 | 0 | Carnivora | 30769.0 | 0 | Canidae | 30769.0 | 0 | Canis | 30769.0 | 0 | Canis lupus | 30769.0 | 0 |
| scf7180000051222 | 11368 | 0.4298 | 0 | 88.6774 | Eukaryota | 6496.0 | 0 | Chordata | 5442.0 | 1 | Tetraodontiformes | 2395.0 | 3 | Tetraodontidae | 2395.0 | 3 | Takifugu | 2395.0 | 4 | Takifugu rubripes | 2395.0 | 4 |
| scf7180000043501 | 17584 | 0.417 | 0 | 56.1384 | Eukaryota | 5819.0 | 0 | Chordata | 5819.0 | 0 | Primates | 3932.0 | 2 | Hominidae | 3545.0 | 3 | Homo | 3545.0 | 3 | Homo sapiens | 3545.0 | 3 |
| scf7180000054283 | 29722 | 0.4362 | 0 | 161.8899 | Eukaryota | 5390.0 | 0 | Chordata | 2979.0 | 1 | Polystomatidea | 2411.0 | 2 | Polystomatidae | 2411.0 | 2 | Protopolystoma | 2411.0 | 2 | Protopolystoma xenopodis | 2411.0 | 2 |
| scf7180000050472 | 14103 | 0.4454 | 0 | 130.681 | Eukaryota | 283.0 | 0 | Chordata | 283.0 | 0 | Tetraodontiformes | 283.0 | 0 | Tetraodontidae | 283.0 | 0 | Takifugu | 283.0 | 0 | Takifugu rubripes | 283.0 | 0 |
| scf7180000043420 | 462785 | 0.3903 | 0 | 60.5625 | Eukaryota | 49663.0 | 0 | Chordata | 49663.0 | 0 | Carnivora | 49663.0 | 0 | Canidae | 49663.0 | 0 | Canis | 44444.0 | 1 | Canis lupus | 44444.0 | 1 |
| scf7180000044912 | 274673 | 0.4374 | 0 | 60.4955 | Eukaryota | 50664.0 | 0 | Chordata | 50664.0 | 0 | Carnivora | 50664.0 | 0 | Canidae | 50664.0 | 0 | Canis | 46776.0 | 1 | Canis lupus | 46776.0 | 1 |
| scf7180000043810 | 433851 | 0.4592 | 0 | 60.8954 | Eukaryota | 89998.0 | 0 | Chordata | 89998.0 | 0 | Carnivora | 89998.0 | 0 | Canidae | 89998.0 | 0 | Canis | 89998.0 | 0 | Canis lupus | 89998.0 | 0 |
| scf7180000045930 | 159263 | 0.3701 | 0 | 59.931 | Eukaryota | 45925.0 | 0 | Chordata | 45925.0 | 0 | Carnivora | 35563.0 | 1 | Ursidae | 20335.0 | 2 | Ursus | 20335.0 | 3 | Ursus arctos | 20335.0 | 3 |
| scf7180000045450 | 224187 | 0.4159 | 0 | 60.9725 | Eukaryota | 29164.0 | 0 | Chordata | 29164.0 | 0 | Carnivora | 29164.0 | 0 | Canidae | 29164.0 | 0 | Canis | 22671.0 | 1 | Canis lupus | 22671.0 | 1 |
| scf7180000049252 | 17110 | 0.4459 | 0 | 465.5104 | Eukaryota | 54312.0 | 0 | Chordata | 43311.0 | 1 | Gadiformes | 36538.0 | 3 | Gadidae | 36538.0 | 3 | Gadus | 36538.0 | 3 | Gadus morhua | 36538.0 | 3 |
| scf7180000045390 | 138364 | 0.4243 | 0 | 58.947 | Eukaryota | 72834.0 | 0 | Chordata | 72834.0 | 0 | Carnivora | 72834.0 | 0 | Canidae | 72834.0 | 0 | Canis | 72834.0 | 0 | Canis lupus | 72834.0 | 0 |
| scf7180000048466 | 9514 | 0.4386 | 0 | 53.9773 | Eukaryota | 5183.0 | 0 | Chordata | 4378.0 | 1 | Tetraodontiformes | 1833.0 | 5 | Tetraodontidae | 1833.0 | 5 | Takifugu | 1833.0 | 6 | Takifugu rubripes | 1833.0 | 6 |
| scf7180000052001 | 29726 | 0.3773 | 0 | 58.5372 | Eukaryota | 12524.0 | 0 | Chordata | 12524.0 | 0 | Primates | 5646.0 | 4 | Hominidae | 5646.0 | 4 | Homo | 5646.0 | 5 | Homo sapiens | 5646.0 | 6 |
| scf7180000052208 | 5049 | 0.4431 | 0 | 138.9736 | no-hit | 0.0 | 0 | no-hit | 0.0 | 0 | no-hit | 0.0 | 0 | no-hit | 0.0 | 0 | no-hit | 0.0 | 0 | no-hit | 0.0 | 0 |
| scf7180000041055 | 137162 | 0.3921 | 0 | 59.9957 | Eukaryota | 27012.0 | 0 | Chordata | 27012.0 | 0 | Carnivora | 27012.0 | 0 | Canidae | 27012.0 | 0 | Canis | 27012.0 | 0 | Canis lupus | 27012.0 | 0 |
| scf7180000041661 | 986971 | 0.3933 | 0 | 61.121 | Eukaryota | 86849.0 | 0 | Chordata | 86849.0 | 0 | Carnivora | 86849.0 | 0 | Canidae | 86849.0 | 0 | Canis | 55703.0 | 1 | Canis lupus | 55703.0 | 1 |
| scf7180000048379 | 58894 | 0.3532 | 0 | 68.2092 | Eukaryota | 57176.0 | 0 | Chordata | 57176.0 | 0 | Carnivora | 57176.0 | 0 | Canidae | 57176.0 | 0 | Canis | 57176.0 | 0 | Canis lupus | 57176.0 | 0 |
| scf7180000041734 | 888645 | 0.4163 | 0 | 59.8861 | Eukaryota | 67020.0 | 0 | Chordata | 67020.0 | 0 | Carnivora | 67020.0 | 0 | Canidae | 67020.0 | 0 | Canis | 67020.0 | 0 | Canis lupus | 67020.0 | 0 |
| scf7180000044521 | 46230 | 0.4226 | 0 | 56.5269 | Eukaryota | 8923.0 | 0 | Chordata | 8923.0 | 0 | Carnivora | 8923.0 | 0 | Canidae | 8923.0 | 0 | Canis | 8923.0 | 0 | Canis lupus | 8923.0 | 0 |
| scf7180000055187 | 30672 | 0.444 | 0 | 183.7824 | Eukaryota | 10381.0 | 0 | Chordata | 9970.0 | 2 | Anabantiformes | 3579.0 | 7 | Anabantidae | 3579.0 | 7 | Anabas | 3579.0 | 7 | Anabas testudineus | 3579.0 | 8 |

| | | | | | | | | | | | | | | | | | | | | | |
|---|---|---|---|---|---|---|---|---|---|---|---|---|---|---|---|---|---|---|---|---|---|
| scf7180000043116 | 119982 | 0.396 | 0 | 63.9289 | Eukaryota | 39025.0 | 0 | Chordata | 39025.0 | 0 | Carnivora | 39025.0 | 0 | Canidae | 39025.0 | 0 | Canis | 39025.0 | 0 | Canis lupus | 39025.0 | 0 |
| scf7180000050750 | 17046 | 0.5028 | 0 | 347.4153 | Eukaryota | 8937.0 | 0 | Chordata | 8937.0 | 0 | Carnivora | 8937.0 | 0 | Canidae | 8937.0 | 0 | Canis | 7750.0 | 1 | Canis lupus | 6525.0 | 2 |
| scf7180000050917 | 2667 | 0.4447 | 0 | 420.8175 | no-hit | 0.0 | 0 | no-hit | 0.0 | 0 | no-hit | 0.0 | 0 | no-hit | 0.0 | 0 | no-hit | 0.0 | 0 | no-hit | 0.0 | 0 |
| scf7180000054403 | 13034 | 0.4204 | 0 | 456.4995 | Eukaryota | 90533.0 | 0 | Chordata | 90533.0 | 0 | Carnivora | 90533.0 | 0 | Canidae | 90533.0 | 0 | Canis | 90533.0 | 0 | Canis lupus | 90533.0 | 0 |
| scf7180000045476 | 197194 | 0.3706 | 0 | 60.0598 | Eukaryota | 35391.0 | 0 | Chordata | 35391.0 | 0 | Carnivora | 35391.0 | 0 | Canidae | 35391.0 | 0 | Canis | 35391.0 | 0 | Canis lupus | 35391.0 | 0 |
| scf7180000042590 | 589429 | 0.5816 | 0 | 55.2583 | Eukaryota | 51351.0 | 0 | Chordata | 51351.0 | 0 | Carnivora | 51351.0 | 0 | Canidae | 51351.0 | 0 | Canis | 43491.0 | 1 | Canis lupus | 43491.0 | 1 |
| scf7180000042476 | 103989 | 0.3485 | 0 | 55.695 | Eukaryota | 91879.0 | 0 | Chordata | 91879.0 | 0 | Carnivora | 91879.0 | 0 | Canidae | 91879.0 | 0 | Canis | 91879.0 | 0 | Canis lupus | 91879.0 | 0 |
| scf7180000054824 | 26934 | 0.4364 | 0 | 171.9618 | Eukaryota | 55054.0 | 0 | Chordata | 39538.0 | 1 | Gadiformes | 35192.0 | 4 | Gadidae | 35192.0 | 4 | Gadus | 35192.0 | 4 | Gadus morhua | 35192.0 | 4 |
| scf7180000043521 | 446745 | 0.3817 | 0 | 60.3395 | Eukaryota | 69069.0 | 0 | Chordata | 69069.0 | 0 | Carnivora | 69069.0 | 0 | Canidae | 69069.0 | 0 | Vulpes | 34895.0 | 1 | Vulpes vulpes | 34895.0 | 1 |
| scf7180000047035 | 99085 | 0.4314 | 0 | 57.6033 | Eukaryota | 54926.0 | 0 | Chordata | 54926.0 | 0 | Primates | 32233.0 | 2 | Hominidae | 27056.0 | 4 | Homo | 27056.0 | 5 | Homo sapiens | 27056.0 | 5 |
| scf7180000054969 | 8776 | 0.4474 | 0 | 376.0203 | Eukaryota | 2542.0 | 0 | Chordata | 2542.0 | 0 | Carnivora | 2542.0 | 0 | Canidae | 2542.0 | 0 | Canis | 2542.0 | 0 | Canis lupus | 2542.0 | 0 |
| scf7180000045766 | 199490 | 0.3366 | 0 | 59.3679 | Eukaryota | 20936.0 | 0 | Chordata | 20936.0 | 0 | Carnivora | 20936.0 | 0 | Canidae | 20936.0 | 0 | Canis | 20936.0 | 0 | Canis lupus | 20936.0 | 0 |
| scf7180000041392 | 128247 | 0.446 | 0 | 57.8063 | Eukaryota | 46860.0 | 0 | Chordata | 46860.0 | 0 | Carnivora | 46860.0 | 0 | Canidae | 22349.0 | 4 | Canis | 12037.0 | 7 | Canis lupus | 12037.0 | 7 |
| scf7180000046528 | 94571 | 0.423 | 0 | 122.7197 | Eukaryota | 93610.0 | 0 | Chordata | 93610.0 | 0 | Carnivora | 93610.0 | 0 | Canidae | 93610.0 | 0 | Canis | 93610.0 | 0 | Canis lupus | 93610.0 | 0 |
| scf7180000051416 | 3593 | 0.4459 | 0 | 779.966 | no-hit | 0.0 | 0 | no-hit | 0.0 | 0 | no-hit | 0.0 | 0 | no-hit | 0.0 | 0 | no-hit | 0.0 | 0 | no-hit | 0.0 | 0 |
| scf7180000043492 | 128111 | 0.3955 | 0 | 58.9797 | Eukaryota | 31338.0 | 0 | Chordata | 31338.0 | 0 | Carnivora | 22943.0 | 1 | Canidae | 12149.0 | 5 | Canis | 9039.0 | 7 | Canis lupus | 9039.0 | 8 |
| scf7180000048592 | 90661 | 0.3262 | 0 | 52.3157 | Eukaryota | 10165.0 | 0 | Chordata | 10165.0 | 0 | Carnivora | 9027.0 | 1 | Canidae | 9027.0 | 1 | Canis | 9027.0 | 1 | Canis lupus | 9027.0 | 1 |
| scf7180000050950 | 4286 | 0.4489 | 0 | 974.3907 | no-hit | 0.0 | 0 | no-hit | 0.0 | 0 | no-hit | 0.0 | 0 | no-hit | 0.0 | 0 | no-hit | 0.0 | 0 | no-hit | 0.0 | 0 |
| scf7180000050174 | 4618 | 0.4426 | 0 | 86.6079 | no-hit | 0.0 | 0 | no-hit | 0.0 | 0 | no-hit | 0.0 | 0 | no-hit | 0.0 | 0 | no-hit | 0.0 | 0 | no-hit | 0.0 | 0 |
| scf7180000052224 | 9425 | 0.441 | 0 | 46.7564 | no-hit | 0.0 | 0 | no-hit | 0.0 | 0 | no-hit | 0.0 | 0 | no-hit | 0.0 | 0 | no-hit | 0.0 | 0 | no-hit | 0.0 | 0 |
| scf7180000045319 | 8231 | 0.434 | 0 | 59.3718 | Eukaryota | 745.0 | 0 | Platyhelminthes | 512.0 | 1 | Polystomatidea | 512.0 | 1 | Polystomatidae | 512.0 | 1 | Protopolystoma | 512.0 | 1 | Protopolystoma xenopodis | 512.0 | 1 |
| scf7180000043304 | 18750 | 0.4083 | 0 | 37.5685 | Eukaryota | 3789.0 | 0 | Chordata | 3789.0 | 0 | Carnivora | 3789.0 | 0 | Canidae | 3323.0 | 1 | Canis | 3323.0 | 1 | Canis lupus | 3323.0 | 1 |
| scf7180000040777 | 1746552 | 0.359 | 0 | 60.6886 | Eukaryota | 80322.0 | 0 | Chordata | 80322.0 | 0 | Carnivora | 80322.0 | 0 | Canidae | 27982.0 | 4 | Canis | 19345.0 | 8 | Canis lupus | 19345.0 | 8 |
| scf7180000043834 | 127046 | 0.3819 | 0 | 57.5877 | Eukaryota | 61809.0 | 0 | Chordata | 61809.0 | 0 | Carnivora | 61809.0 | 0 | Canidae | 61809.0 | 0 | Canis | 61809.0 | 0 | Canis lupus | 61809.0 | 0 |
| scf7180000055419 | 16409 | 0.5684 | 0 | 371.8918 | Eukaryota | 53778.0 | 0 | Chordata | 53778.0 | 0 | Primates | 26434.0 | 2 | Bovidae | 21469.0 | 3 | Homo | 21002.0 | 4 | Homo sapiens | 21002.0 | 4 |
| scf7180000044523 | 22021 | 0.3535 | 0 | 44.4903 | Eukaryota | 32556.0 | 0 | Chordata | 32556.0 | 0 | Carnivora | 32556.0 | 0 | Canidae | 32556.0 | 0 | Canis | 32556.0 | 0 | Canis lupus | 32556.0 | 0 |
| scf7180000053975 | 6413 | 0.4269 | 0 | 45.0198 | Eukaryota | 12979.0 | 0 | Chordata | 12979.0 | 0 | Carnivora | 12979.0 | 0 | Canidae | 12979.0 | 0 | Canis | 12979.0 | 0 | Canis lupus | 12979.0 | 0 |
| scf7180000048886 | 33360 | 0.4294 | 0 | 59.255 | Eukaryota | 6500.0 | 0 | Chordata | 6500.0 | 0 | Carnivora | 5862.0 | 1 | Canidae | 5862.0 | 1 | Canis | 5862.0 | 1 | Canis lupus | 5862.0 | 1 |
| scf7180000052628 | 9128 | 0.4533 | 0 | 98.665 | Eukaryota | 425.0 | 0 | Chordata | 425.0 | 0 | Tetraodontiformes | 425.0 | 0 | Tetraodontidae | 425.0 | 0 | Takifugu | 425.0 | 0 | Takifugu rubripes | 425.0 | 0 |
| scf7180000046557 | 8207 | 0.4379 | 0 | 55.3601 | Eukaryota | 3552.0 | 0 | Platyhelminthes | 2115.0 | 1 | Polystomatidea | 2115.0 | 3 | Polystomatidae | 2115.0 | 3 | Protopolystoma | 2115.0 | 3 | Protopolystoma xenopodis | 2115.0 | 3 |
| scf7180000051370 | 5716 | 0.4449 | 0 | 50.3729 | no-hit | 0.0 | 0 | no-hit | 0.0 | 0 | no-hit | 0.0 | 0 | no-hit | 0.0 | 0 | no-hit | 0.0 | 0 | no-hit | 0.0 | 0 |
| scf7180000043022 | 480487 | 0.3541 | 0 | 59.5937 | Eukaryota | 16915.0 | 0 | Chordata | 16915.0 | 0 | Carnivora | 16915.0 | 0 | Canidae | 16915.0 | 0 | Canis | 16915.0 | 0 | Canis lupus | 16915.0 | 0 |
| scf7180000045166 | 57871 | 0.4402 | 0 | 175.0067 | Eukaryota | 48342.0 | 0 | Chordata | 27512.0 | 2 | Cyclophyllidea | 11293.0 | 9 | Hymenolepididae | 11293.0 | 9 | Hymenolepis | 11293.0 | 9 | Hymenolepis microstoma | 11293.0 | 9 |
| scf7180000043030 | 29435 | 0.4317 | 0 | 142.9586 | Eukaryota | 32128.0 | 0 | Chordata | 32128.0 | 0 | Carnivora | 32128.0 | 0 | Canidae | 32128.0 | 0 | Canis | 32128.0 | 0 | Canis lupus | 32128.0 | 0 |
| scf7180000046478 | 102688 | 0.4418 | 0 | 80.1182 | Eukaryota | 92747.0 | 0 | Chordata | 92747.0 | 0 | Carnivora | 92747.0 | 0 | Canidae | 92747.0 | 0 | Canis | 92747.0 | 0 | Canis lupus | 92747.0 | 0 |
| scf7180000053667 | 2294 | 0.4568 | 0 | 677.7692 | no-hit | 0.0 | 0 | no-hit | 0.0 | 0 | no-hit | 0.0 | 0 | no-hit | 0.0 | 0 | no-hit | 0.0 | 0 | no-hit | 0.0 | 0 |
| scf7180000050930 | 1561 | 0.442 | 0 | 299.2257 | no-hit | 0.0 | 0 | no-hit | 0.0 | 0 | no-hit | 0.0 | 0 | no-hit | 0.0 | 0 | no-hit | 0.0 | 0 | no-hit | 0.0 | 0 |
| scf7180000051059 | 3849 | 0.4562 | 0 | 577.665 | Eukaryota | 313.0 | 0 | Chordata | 313.0 | 0 | Salmoniformes | 313.0 | 0 | Salmonidae | 313.0 | 0 | Coregonus | 313.0 | 0 | Coregonus sp. 'balchen' | 313.0 | 0 |
| scf7180000043629 | 4351 | 0.3765 | 0 | 28.438 | Eukaryota | 11119.0 | 0 | Chordata | 11119.0 | 0 | Carnivora | 11119.0 | 0 | Canidae | 11119.0 | 0 | Canis | 11119.0 | 0 | Canis lupus | 11119.0 | 0 |
| scf7180000046242 | 65958 | 0.37 | 0 | 58.2727 | Eukaryota | 4975.0 | 0 | Chordata | 4975.0 | 0 | Primates | 2896.0 | 1 | Canidae | 2079.0 | 2 | Homo | 2070.0 | 3 | Homo sapiens | 2070.0 | 3 |
| scf7180000045202 | 46508 | 0.389 | 0 | 55.8953 | Eukaryota | 7666.0 | 0 | Chordata | 7666.0 | 0 | Carnivora | 4562.0 | 1 | Hominidae | 3104.0 | 2 | Pan | 2082.0 | 4 | Pan troglodytes | 2082.0 | 5 |
| scf7180000052493 | 1845 | 0.4531 | 0 | 486.0929 | no-hit | 0.0 | 0 | no-hit | 0.0 | 0 | no-hit | 0.0 | 0 | no-hit | 0.0 | 0 | no-hit | 0.0 | 0 | no-hit | 0.0 | 0 |
| scf7180000045528 | 8011 | 0.4444 | 0 | 257.5219 | no-hit | 0.0 | 0 | no-hit | 0.0 | 0 | no-hit | 0.0 | 0 | no-hit | 0.0 | 0 | no-hit | 0.0 | 0 | no-hit | 0.0 | 0 |
| scf7180000042000 | 792032 | 0.4392 | 0 | 59.9386 | Eukaryota | 46721.0 | 0 | Chordata | 46721.0 | 0 | Carnivora | 46721.0 | 0 | Canidae | 26425.0 | 3 | Canis | 15935.0 | 5 | Canis lupus | 15935.0 | 6 |
| scf7180000043369 | 117584 | 0.3846 | 0 | 56.9326 | Eukaryota | 81610.0 | 0 | Chordata | 81610.0 | 0 | Carnivora | 81610.0 | 0 | Canidae | 81610.0 | 0 | Canis | 81610.0 | 0 | Canis lupus | 81610.0 | 0 |
| scf7180000052854 | 19471 | 0.4477 | 0 | 64.5567 | Eukaryota | 10073.0 | 0 | Chordata | 7354.0 | 1 | Salmoniformes | 2976.0 | 5 | Salmonidae | 2976.0 | 5 | Salmo | 2976.0 | 5 | Salmo trutta | 2976.0 | 5 |
| scf7180000052631 | 8794 | 0.4511 | 0 | 697.4162 | Eukaryota | 2723.0 | 0 | Chordata | 2244.0 | 1 | Carnivora | 1970.0 | 2 | Canidae | 1970.0 | 2 | Canis | 1970.0 | 2 | Canis lupus | 1970.0 | 2 |
| scf7180000047781 | 66967 | 0.5955 | 0 | 56.773 | Eukaryota | 30699.0 | 0 | Chordata | 30699.0 | 0 | Carnivora | 30699.0 | 0 | Canidae | 30699.0 | 0 | Canis | 30699.0 | 0 | Canis lupus | 30699.0 | 0 |
| scf7180000049340 | 41541 | 0.3816 | 0 | 62.209 | Eukaryota | 27601.0 | 0 | Chordata | 27601.0 | 0 | Carnivora | 27601.0 | 0 | Felidae | 13095.0 | 3 | Canis | 9044.0 | 5 | Canis lupus | 9044.0 | 5 |
| scf7180000045443 | 213546 | 0.4832 | 0 | 59.2103 | Eukaryota | 62165.0 | 0 | Chordata | 62165.0 | 0 | Carnivora | 62165.0 | 0 | Canidae | 62165.0 | 0 | Canis | 62165.0 | 0 | Canis lupus | 62165.0 | 0 |
| scf7180000056393 | 285317 | 0.386 | 0 | 58.5716 | Eukaryota | 95730.0 | 0 | Chordata | 95730.0 | 0 | Carnivora | 95730.0 | 0 | Canidae | 95730.0 | 0 | Canis | 95730.0 | 0 | Canis lupus | 95730.0 | 0 |
| scf7180000045507 | 67769 | 0.4376 | 0 | 171.4404 | Eukaryota | 29921.0 | 0 | Chordata | 29921.0 | 0 | Carnivora | 29921.0 | 0 | Canidae | 29921.0 | 0 | Canis | 29921.0 | 0 | Canis lupus | 29921.0 | 0 |
| scf7180000041182 | 197522 | 0.4179 | 0 | 61.0897 | Eukaryota | 10151.0 | 0 | Chordata | 10151.0 | 0 | Carnivora | 10151.0 | 0 | Canidae | 10151.0 | 0 | Canis | 10151.0 | 0 | Canis lupus | 10151.0 | 0 |
| scf7180000043793 | 14538 | 0.3555 | 0 | 33.1761 | Eukaryota | 15647.0 | 0 | Chordata | 15647.0 | 0 | Carnivora | 15647.0 | 0 | Canidae | 15647.0 | 0 | Canis | 9370.0 | 1 | Canis lupus | 9370.0 | 1 |
| scf7180000041995 | 786002 | 0.4892 | 0 | 58.854 | Eukaryota | 49392.0 | 0 | Chordata | 49392.0 | 0 | Carnivora | 49392.0 | 0 | Canidae | 49392.0 | 0 | Canis | 49392.0 | 0 | Canis lupus | 49392.0 | 0 |
| scf7180000046257 | 157634 | 0.3515 | 0 | 59.2826 | Eukaryota | 17307.0 | 0 | Chordata | 17307.0 | 0 | Artiodactyla | 4817.0 | 5 | Bovidae | 4817.0 | 5 | Canis | 4672.0 | 6 | Canis lupus | 4672.0 | 6 |
| scf7180000050118 | 27719 | 0.3847 | 0 | 60.7009 | Eukaryota | 5766.0 | 0 | Chordata | 5766.0 | 0 | Primates | 3879.0 | 2 | Atelidae | 1817.0 | 4 | Ateles | 1817.0 | 4 | Ateles belzebuth | 1817.0 | 4 |
| scf7180000044958 | 240450 | 0.5826 | 0 | 59.1305 | Eukaryota | 40290.0 | 0 | Chordata | 40290.0 | 0 | Carnivora | 40290.0 | 0 | Canidae | 40290.0 | 0 | Canis | 40290.0 | 0 | Canis lupus | 40290.0 | 0 |
| scf7180000046210 | 43042 | 0.3264 | 0 | 58.0899 | Eukaryota | 29275.0 | 0 | Chordata | 29275.0 | 0 | Carnivora | 29275.0 | 0 | Canidae | 29275.0 | 0 | Canis | 29275.0 | 0 | Canis lupus | 29275.0 | 0 |
| scf7180000042183 | 58730 | 0.4689 | 0 | 57.7105 | Eukaryota | 65007.0 | 0 | Chordata | 65007.0 | 0 | Carnivora | 65007.0 | 0 | Canidae | 65007.0 | 0 | Canis | 65007.0 | 0 | Canis lupus | 65007.0 | 0 |
| scf7180000051140 | 3780 | 0.4632 | 0 | 22.0635 | Eukaryota | 21610.0 | 0 | Chordata | 21610.0 | 0 | Carnivora | 21610.0 | 0 | Canidae | 21610.0 | 0 | Canis | 21610.0 | 0 | Canis lupus | 21610.0 | 0 |
| scf7180000054318 | 151266 | 0.4048 | 0 | 66.97 | Eukaryota | 89825.0 | 0 | Chordata | 89825.0 | 0 | Carnivora | 89825.0 | 0 | Canidae | 89825.0 | 0 | Canis | 89825.0 | 0 | Canis lupus | 89825.0 | 0 |
| scf7180000050808 | 6548 | 0.4386 | 0 | 506.7654 | no-hit | 0.0 | 0 | no-hit | 0.0 | 0 | no-hit | 0.0 | 0 | no-hit | 0.0 | 0 | no-hit | 0.0 | 0 | no-hit | 0.0 | 0 |
| scf7180000051629 | 11544 | 0.5036 | 0 | 1239.428 | Eukaryota | 8778.0 | 0 | Chordata | 8778.0 | 0 | Carnivora | 8778.0 | 0 | Canidae | 8778.0 | 0 | Canis | 7089.0 | 1 | Canis lupus | 5884.0 | 2 |
| scf7180000056524 | 141606 | 0.5672 | 0 | 89.1677 | Eukaryota | 21102.0 | 0 | Chordata | 21102.0 | 0 | Carnivora | 21102.0 | 0 | Canidae | 21102.0 | 0 | Canis | 14916.0 | 1 | Canis lupus | 14916.0 | 1 |
| scf7180000053813 | 2495 | 0.4481 | 0 | 341.8816 | no-hit | 0.0 | 0 | no-hit | 0.0 | 0 | no-hit | 0.0 | 0 | no-hit | 0.0 | 0 | no-hit | 0.0 | 0 | no-hit | 0.0 | 0 |
| scf7180000051022 | 23739 | 0.3786 | 0 | 42.3283 | Eukaryota | 7531.0 | 0 | Chordata | 7531.0 | 0 | Carnivora | 6578.0 | 1 | Canidae | 2811.0 | 3 | Canis | 2811.0 | 3 | Canis lupus | 2811.0 | 3 |
| scf7180000042312 | 26223 | 0.4158 | 0 | 57.7882 | Eukaryota | 6101.0 | 0 | Chordata | 6101.0 | 0 | Carnivora | 2362.0 | 3 | Bovidae | 1355.0 | 5 | Canis | 1292.0 | 7 | Canis lupus | 1292.0 | 7 |
| scf7180000046161 | 76151 | 0.4393 | 0 | 95.7463 | Eukaryota | 30502.0 | 0 | Chordata | 21840.0 | 1 | Gadiformes | 11076.0 | 6 | Gadidae | 11076.0 | 6 | Gadus | 11076.0 | 6 | Gadus morhua | 11076.0 | 6 |
| scf7180000050439 | 43598 | 0.3536 | 0 | 61.0645 | Eukaryota | 18475.0 | 0 | Chordata | 18475.0 | 0 | Carnivora | 18475.0 | 0 | Canidae | 18475.0 | 0 | Canis | 18475.0 | 0 | Canis lupus | 18475.0 | 0 |
| scf7180000041739 | 847857 | 0.4131 | 0 | 60.4053 | Eukaryota | 61489.0 | 0 | Chordata | 61489.0 | 0 | Carnivora | 61489.0 | 0 | Canidae | 61489.0 | 0 | Canis | 54045.0 | 1 | Canis lupus | 54045.0 | 1 |
| scf7180000053015 | 7106 | 0.4516 | 0 | 522.5416 | Eukaryota | 3408.0 | 0 | Chordata | 2538.0 | 1 | Gonorynchiformes | 1026.0 | 3 | Chanidae | 1026.0 | 3 | Chanos | 1026.0 | 3 | Chanos chanos | 1026.0 | 3 |
| scf7180000052999 | 4309 | 0.4539 | 0 | 469.0144 | no-hit | 0.0 | 0 | no-hit | 0.0 | 0 | no-hit | 0.0 | 0 | no-hit | 0.0 | 0 | no-hit | 0.0 | 0 | no-hit | 0.0 | 0 |
| scf7180000045772 | 159131 | 0.4163 | 0 | 58.7846 | Eukaryota | 19957.0 | 0 | Chordata | 19957.0 | 0 | Carnivora | 19957.0 | 0 | Canidae | 19957.0 | 0 | Canis | 19957.0 | 0 | Canis lupus | 19957.0 | 0 |
| scf7180000046324 | 137186 | 0.4461 | 0 | 70.6985 | Eukaryota | 82088.0 | 0 | Chordata | 82088.0 | 0 | Carnivora | 82088.0 | 0 | Canidae | 82088.0 | 0 | Canis | 82088.0 | 0 | Canis lupus | 82088.0 | 0 |
| scf7180000052138 | 14003 | 0.4562 | 0 | 57.8768 | Eukaryota | 9879.0 | 0 | Chordata | 9879.0 | 0 | Carnivora | 8671.0 | 1 | Canidae | 6508.0 | 4 | Canis | 3525.0 | 6 | Canis lupus | 3525.0 | 6 |
| scf7180000049343 | 5661 | 0.4524 | 0 | 88.5091 | no-hit | 0.0 | 0 | no-hit | 0.0 | 0 | no-hit | 0.0 | 0 | no-hit | 0.0 | 0 | no-hit | 0.0 | 0 | no-hit | 0.0 | 0 |
| scf7180000051657 | 7021 | 0.4338 | 0 | 81.3529 | Eukaryota | 9706.0 | 0 | Chordata | 9355.0 | 1 | Tetraodontiformes | 8819.0 | 2 | Tetraodontidae | 8819.0 | 2 | Takifugu | 8819.0 | 2 | Takifugu rubripes | 8819.0 | 2 |
| scf7180000050401 | 6492 | 0.4455 | 0 | 234.2042 | Eukaryota | 2933.0 | 0 | Chordata | 2396.0 | 1 | Salmoniformes | 2396.0 | 1 | Salmonidae | 2396.0 | 1 | Salmo | 2396.0 | 1 | Salmo trutta | 2396.0 | 1 |
| scf7180000054461 | 18017 | 0.6179 | 0 | 449.8679 | Eukaryota | 1723.0 | 0 | Chordata | 1723.0 | 0 | Carnivora | 1723.0 | 0 | Canidae | 1578.0 | 1 | Canis | 872.0 | 2 | Canis lupus | 872.0 | 2 |
| scf7180000049278 | 3403 | 0.4387 | 0 | 177.901 | Eukaryota | 530.0 | 0 | Streptophyta | 391.0 | 1 | Fabales | 391.0 | 1 | Fabaceae | 391.0 | 1 | Cicer | 391.0 | 1 | Cicer arietinum | 391.0 | 1 |
| scf7180000055865 | 17807 | 0.3791 | 0 | 113.9541 | Eukaryota | 8697.0 | 0 | Chordata | 8697.0 | 0 | Carnivora | 6086.0 | 2 | Canidae | 5755.0 | 4 | Canis | 5402.0 | 6 | Canis lupus | 5402.0 | 6 |
| scf7180000047024 | 90281 | 0.396 | 0 | 60.726 | Eukaryota | 66359.0 | 0 | Chordata | 66359.0 | 0 | Carnivora | 66359.0 | 0 | Canidae | 66359.0 | 0 | Canis | 59688.0 | 1 | Canis lupus | 59688.0 | 1 |
| scf7180000051080 | 4715 | 0.4492 | 0 | 222.9652 | no-hit | 0.0 | 0 | no-hit | 0.0 | 0 | no-hit | 0.0 | 0 | no-hit | 0.0 | 0 | no-hit | 0.0 | 0 | no-hit | 0.0 | 0 |
| scf7180000050576 | 22242 | 0.4438 | 0 | 975.3671 | Eukaryota | 5783.0 | 0 | Chordata | 3129.0 | 2 | Polystomatidea | 1693.0 | 3 | Polystomatidae | 1693.0 | 3 | Protopolystoma | 1693.0 | 3 | Protopolystoma xenopodis | 1693.0 | 3 |
| scf7180000055297 | 12544 | 0.4102 | 0 | 78.7977 | Eukaryota | 6457.0 | 0 | Chordata | 6457.0 | 0 | Carnivora | 6457.0 | 0 | Canidae | 6457.0 | 0 | Canis | 6457.0 | 0 | Canis lupus | 6457.0 | 0 |
| scf7180000050124 | 6454 | 0.3788 | 0 | 42.9799 | Eukaryota | 2139.0 | 0 | Chordata | 2139.0 | 0 | Carnivora | 2139.0 | 0 | Canidae | 2139.0 | 0 | Canis | 1933.0 | 1 | Canis lupus | 1933.0 | 1 |
| scf7180000043001 | 144035 | 0.3842 | 0 | 58.2855 | Eukaryota | 29111.0 | 0 | Chordata | 29111.0 | 0 | Carnivora | 29111.0 | 0 | Canidae | 29111.0 | 0 | Canis | 29111.0 | 0 | Canis lupus | 29111.0 | 0 |
| scf7180000041768 | 21409 | 0.449 | 0 | 85.8845 | Eukaryota | 8367.0 | 0 | Chordata | 8367.0 | 0 | Carnivora | 7456.0 | 1 | Canidae | 5550.0 | 4 | Canis | 4229.0 | 5 | Canis lupus | 4229.0 | 5 |
| scf7180000046151 | 15930 | 0.4041 | 0 | 57.4097 | Eukaryota | 69332.0 | 0 | Chordata | 69332.0 | 0 | Carnivora | 69332.0 | 0 | Canidae | 69332.0 | 0 | Canis | 69332.0 | 0 | Canis lupus | 69332.0 | 0 |
| scf7180000054520 | 155236 | 0.4393 | 0 | 171.2984 | Eukaryota | 40100.0 | 0 | Chordata | 36105.0 | 1 | Gadiformes | 23658.0 | 5 | Gadidae | 23658.0 | 5 | Gadus | 23658.0 | 5 | Gadus morhua | 23658.0 | 5 |
| scf7180000042545 | 718012 | 0.3666 | 0 | 61.0685 | Eukaryota | 66945.0 | 0 | Chordata | 66945.0 | 0 | Carnivora | 66945.0 | 0 | Canidae | 66945.0 | 0 | Canis | 66945.0 | 0 | Canis lupus | 66945.0 | 0 |
| scf7180000047964 | 101106 | 0.4296 | 0 | 61.1334 | Eukaryota | 62877.0 | 0 | Chordata | 62877.0 | 0 | Carnivora | 62877.0 | 0 | Otariidae | 41506.0 | 1 | Zalophus | 17724.0 | 4 | Zalophus californianus | 17724.0 | 4 |
| scf7180000051545 | 21659 | 0.4387 | 0 | 137.0353 | no-hit | 0.0 | 0 | no-hit | 0.0 | 0 | no-hit | 0.0 | 0 | no-hit | 0.0 | 0 | no-hit | 0.0 | 0 | no-hit | 0.0 | 0 |
| scf7180000041870 | 857482 | 0.4131 | 0 | 58.9974 | Eukaryota | 75162.0 | 0 | Chordata | 75162.0 | 0 | Carnivora | 75162.0 | 0 | Canidae | 38442.0 | 1 | Ursus | 36720.0 | 2 | Canis lupus | 30698.0 | 3 |
| scf7180000053491 | 2926 | 0.4545 | 0 | 325.0502 | Eukaryota | 2362.0 | 0 | Chordata | 1812.0 | 1 | Columbiformes | 891.0 | 3 | Columbidae | 891.0 | 3 | Streptopelia | 891.0 | 3 | Streptopelia turtur | 891.0 | 3 |
| scf7180000042724 | 593203 | 0.3882 | 0 | 59.9411 | Eukaryota | 88536.0 | 0 | Chordata | 88536.0 | 0 | Carnivora | 88536.0 | 0 | Canidae | 88536.0 | 0 | Canis | 88536.0 | 0 | Canis lupus | 88536.0 | 0 |
| scf7180000049673 | 12472 | 0.4377 | 0 | 59.9636 | Eukaryota | 10629.0 | 0 | Chordata | 8717.0 | 2 | Tetraodontiformes | 5584.0 | 3 | Tetraodontidae | 5584.0 | 3 | Takifugu | 5584.0 | 3 | Takifugu rubripes | 5584.0 | 3 |
| scf7180000056184 | 1153328 | 0.4126 | 0 | 63.38 | Eukaryota | 92978.0 | 0 | Chordata | 92978.0 | 0 | Carnivora | 92978.0 | 0 | Canidae | 92978.0 | 0 | Canis | 92978.0 | 0 | Canis lupus | 92978.0 | 0 |
| scf7180000049288 | 2693 | 0.4497 | 0 | 156.2468 | no-hit | 0.0 | 0 | no-hit | 0.0 | 0 | no-hit | 0.0 | 0 | no-hit | 0.0 | 0 | no-hit | 0.0 | 0 | no-hit | 0.0 | 0 |
| scf7180000056487 | 154103 | 0.4557 | 0 | 57.2232 | Eukaryota | 58948.0 | 0 | Chordata | 58948.0 | 0 | Carnivora | 58948.0 | 0 | Canidae | 58948.0 | 0 | Canis | 58948.0 | 0 | Canis lupus | 58948.0 | 0 |
| scf7180000055979 | 7562 | 0.4438 | 0 | 333.4796 | no-hit | 0.0 | 0 | no-hit | 0.0 | 0 | no-hit | 0.0 | 0 | no-hit | 0.0 | 0 | no-hit | 0.0 | 0 | no-hit | 0.0 | 0 |
| scf7180000047606 | 7577 | 0.4276 | 0 | 292.0541 | Eukaryota | 6313.0 | 1 | Chordata | 5576.0 | 2 | Carnivora | 3144.0 | 5 | Otariidae | 2182.0 | 6 | Callorhinus | 1498.0 | 7 | Callorhinus ursinus | 1498.0 | 7 |
| scf7180000045988 | 116740 | 0.4179 | 0 | 59.2166 | Eukaryota | 10676.0 | 0 | Chordata | 10676.0 | 0 | Carnivora | 10676.0 | 0 | Canidae | 10676.0 | 0 | Canis | 10676.0 | 0 | Canis lupus | 10676.0 | 0 |
| scf7180000045667 | 203500 | 0.3701 | 0 | 60.3501 | Eukaryota | 60858.0 | 0 | Chordata | 60858.0 | 0 | Carnivora | 60858.0 | 0 | Canidae | 60858.0 | 0 | Canis | 60858.0 | 0 | Canis lupus | 60858.0 | 0 |
| scf7180000045346 | 200566 | 0.3613 | 0 | 61.2492 | Eukaryota | 26077.0 | 0 | Chordata | 26077.0 | 0 | Carnivora | 26077.0 | 0 | Canidae | 26077.0 | 0 | Canis | 26077.0 | 0 | Canis lupus | 26077.0 | 0 |
| scf7180000049754 | 3748 | 0.4288 | 0 | 125.8044 | Eukaryota | 440.0 | 0 | Chordata | 440.0 | 0 | Salmoniformes | 440.0 | 0 | Salmonidae | 440.0 | 0 | Coregonus | 440.0 | 0 | Coregonus sp. 'balchen' | 440.0 | 0 |
| scf7180000054284 | 22197 | 0.4557 | 0 | 91.6455 | Eukaryota | 9263.0 | 0 | Chordata | 9263.0 | 0 | Carnivora | 9263.0 | 0 | Canidae | 9263.0 | 0 | Canis | 5546.0 | 1 | Canis lupus | 5546.0 | 1 |
| scf7180000054197 | 17151 | 0.4599 | 0 | 205.9775 | Eukaryota | 13290.0 | 0 | Chordata | 13290.0 | 0 | Carnivora | 13290.0 | 0 | Canidae | 13290.0 | 0 | Canis | 13290.0 | 0 | Canis lupus | 13290.0 | 0 |
| scf7180000040610 | 2242378 | 0.338 | 0 | 58.8202 | Eukaryota | 91733.0 | 0 | Chordata | 91733.0 | 0 | Carnivora | 91733.0 | 0 | Canidae | 91733.0 | 0 | Canis | 91733.0 | 0 | Canis lupus | 91733.0 | 0 |
| scf7180000049067 | 50162 | 0.4912 | 0 | 59.1952 | Eukaryota | 19843.0 | 0 | Chordata | 19843.0 | 0 | Carnivora | 19843.0 | 0 | Canidae | 9641.0 | 3 | Leptonychotes | 5076.0 | 5 | Leptonychotes weddellii | 5076.0 | 5 |
| scf7180000040664 | 43881 | 0.3714 | 0 | 56.6768 | Eukaryota | 50894.0 | 0 | Chordata | 50894.0 | 0 | Carnivora | 50894.0 | 0 | Canidae | 30099.0 | 2 | Canis | 25598.0 | 3 | Canis lupus | 25598.0 | 3 |
| scf7180000041025 | 45720 | 0.3643 | 0 | 56.5968 | Eukaryota | 36511.0 | 0 | Chordata | 36511.0 | 0 | Carnivora | 36511.0 | 0 | Canidae | 36511.0 | 0 | Canis | 36511.0 | 0 | Canis lupus | 36511.0 | 0 |
| scf7180000052984 | 4195 | 0.4505 | 0 | 625.1435 | Eukaryota | 858.0 | 0 | Platyhelminthes | 858.0 | 0 | Polystomatidea | 858.0 | 0 | Polystomatidae | 858.0 | 0 | Protopolystoma | 858.0 | 0 | Protopolystoma xenopodis | 858.0 | 0 |
| scf7180000044500 | 266531 | 0.4735 | 0 | 58.8054 | Eukaryota | 72544.0 | 0 | Chordata | 72544.0 | 0 | Carnivora | 72544.0 | 0 | Canidae | 72544.0 | 0 | Canis | 72544.0 | 0 | Canis lupus | 72544.0 | 0 |
| scf7180000053748 | 5779 | 0.4489 | 0 | 64.0301 | Eukaryota | 2155.0 | 0 | Chordata | 1201.0 | 1 | Polystomatidea | 954.0 | 3 | Polystomatidae | 954.0 | 3 | Protopolystoma | 954.0 | 3 | Protopolystoma xenopodis | 954.0 | 3 |
| scf7180000043017 | 477509 | 0.3703 | 0 | 60.1267 | Eukaryota | 85096.0 | 0 | Chordata | 85096.0 | 0 | Carnivora | 85096.0 | 0 | Canidae | 85096.0 | 0 | Canis | 85096.0 | 0 | Canis lupus | 85096.0 | 0 |
| scf7180000043289 | 396602 | 0.5138 | 0 | 57.7998 | Eukaryota | 67971.0 | 0 | Chordata | 67971.0 | 0 | Carnivora | 67971.0 | 0 | Canidae | 67971.0 | 0 | Canis | 41188.0 | 1 | Canis lupus | 41188.0 | 1 |
| scf7180000052716 | 2216 | 0.4472 | 0 | 416.6306 | no-hit | 0.0 | 0 | no-hit | 0.0 | 0 | no-hit | 0.0 | 0 | no-hit | 0.0 | 0 | no-hit | 0.0 | 0 | no-hit | 0.0 | 0 |
| scf7180000050053 | 19762 | 0.4419 | 0 | 69.9528 | Eukaryota | 3050.0 | 0 | Chordata | 3050.0 | 0 | Carnivora | 3050.0 | 0 | Canidae | 3050.0 | 0 | Canis | 3050.0 | 0 | Canis lupus | 3050.0 | 0 |
| scf7180000055335 | 32103 | 0.4478 | 0 | 562.6628 | Eukaryota | 11898.0 | 0 | Chordata | 11898.0 | 0 | Carnivora | 11898.0 | 0 | Canidae | 11898.0 | 0 | Canis | 10071.0 | 1 | Canis lupus | 9081.0 | 2 |
| scf7180000044504 | 29171 | 0.4864 | 0 | 565.4441 | Eukaryota | 90157.0 | 0 | Chordata | 90157.0 | 0 | Carnivora | 90157.0 | 0 | Canidae | 90157.0 | 0 | Canis | 90157.0 | 0 | Canis lupus | 90157.0 | 0 |
| scf7180000042998 | 97014 | 0.4278 | 0 | 58.9698 | Eukaryota | 25070.0 | 0 | Chordata | 25070.0 | 0 | Carnivora | 25070.0 | 0 | Canidae | 25070.0 | 0 | Canis | 25070.0 | 0 | Canis lupus | 25070.0 | 0 |
| scf7180000047553 | 14801 | 0.4404 | 0 | 111.4818 | Eukaryota | 1415.0 | 0 | Chordata | 1415.0 | 0 | Columbiformes | 1160.0 | 1 | Columbidae | 1160.0 | 1 | Streptopelia | 1160.0 | 1 | Streptopelia turtur | 1160.0 | 1 |
| scf7180000047485 | 34675 | 0.4407 | 0 | 88.2265 | Eukaryota | 943.0 | 0 | Chordata | 682.0 | 1 | Salmoniformes | 449.0 | 2 | Salmonidae | 449.0 | 2 | Coregonus | 449.0 | 2 | Coregonus sp. 'balchen' | 449.0 | 2 |
| scf7180000052866 | 2977 | 0.4501 | 0 | 433.8569 | no-hit | 0.0 | 0 | no-hit | 0.0 | 0 | no-hit | 0.0 | 0 | no-hit | 0.0 | 0 | no-hit | 0.0 | 0 | no-hit | 0.0 | 0 |
| scf7180000052689 | 4968 | 0.5225 | 0 | 56.7001 | Eukaryota | 815.0 | 1 | Chordata | 815.0 | 1 | Primates | 632.0 | 2 | Hominidae | 632.0 | 2 | Homo | 632.0 | 2 | Homo sapiens | 632.0 | 2 |
| scf7180000045278 | 58847 | 0.3516 | 0 | 58.3055 | Eukaryota | 40994.0 | 0 | Chordata | 40994.0 | 0 | Carnivora | 40994.0 | 0 | Canidae | 40994.0 | 0 | Canis | 40994.0 | 0 | Canis lupus | 40994.0 | 0 |
| scf7180000042985 | 183786 | 0.3926 | 0 | 58.8402 | Eukaryota | 88841.0 | 0 | Chordata | 88841.0 | 0 | Carnivora | 88841.0 | 0 | Canidae | 88841.0 | 0 | Canis | 88841.0 | 0 | Canis lupus | 88841.0 | 0 |
| scf7180000041747 | 303143 | 0.3891 | 0 | 61.5941 | Eukaryota | 17903.0 | 0 | Chordata | 17903.0 | 0 | Carnivora | 17903.0 | 0 | Canidae | 17903.0 | 0 | Canis | 17903.0 | 0 | Canis lupus | 17903.0 | 0 |
| scf7180000050329 | 7488 | 0.4439 | 0 | 70.6321 | no-hit | 0.0 | 0 | no-hit | 0.0 | 0 | no-hit | 0.0 | 0 | no-hit | 0.0 | 0 | no-hit | 0.0 | 0 | no-hit | 0.0 | 0 |
| scf7180000044047 | 74086 | 0.3727 | 0 | 56.1436 | Eukaryota | 89948.0 | 0 | Chordata | 89948.0 | 0 | Carnivora | 89948.0 | 0 | Canidae | 89948.0 | 0 | Canis | 89948.0 | 0 | Canis lupus | 89948.0 | 0 |
| scf7180000049915 | 12117 | 0.501 | 0 | 243.0806 | Eukaryota | 8251.0 | 0 | Chordata | 8251.0 | 0 | Carnivora | 8251.0 | 0 | Canidae | 8251.0 | 0 | Canis | 6569.0 | 1 | Canis lupus | 5457.0 | 2 |
| scf7180000047160 | 86942 | 0.4435 | 0 | 59.3842 | Eukaryota | 21384.0 | 0 | Chordata | 21384.0 | 0 | Carnivora | 21384.0 | 0 | Canidae | 21384.0 | 0 | Canis | 21384.0 | 0 | Canis lupus | 21384.0 | 0 |
| scf7180000050563 | 31864 | 0.4341 | 0 | 57.823 | Eukaryota | 7785.0 | 0 | Chordata | 7785.0 | 0 | Carnivora | 5136.0 | 1 | Canidae | 5136.0 | 1 | Canis | 5136.0 | 1 | Canis lupus | 5136.0 | 1 |
| scf7180000041527 | 47045 | 0.4324 | 0 | 285.0533 | Eukaryota | 88055.0 | 0 | Chordata | 88055.0 | 0 | Carnivora | 88055.0 | 0 | Canidae | 88055.0 | 0 | Canis | 88055.0 | 0 | Canis lupus | 88055.0 | 0 |
| scf7180000046481 | 113944 | 0.4523 | 0 | 58.5904 | Eukaryota | 73365.0 | 0 | Chordata | 73365.0 | 0 | Carnivora | 73365.0 | 0 | Canidae | 73365.0 | 0 | Canis | 73365.0 | 0 | Canis lupus | 73365.0 | 0 |
| scf7180000044002 | 136160 | 0.3719 | 0 | 59.8457 | Eukaryota | 30296.0 | 0 | Chordata | 30296.0 | 0 | Carnivora | 29300.0 | 1 | Canidae | 22738.0 | 2 | Vulpes | 15638.0 | 4 | Vulpes vulpes | 15638.0 | 4 |
| scf7180000050393 | 13169 | 0.4351 | 0 | 65.3163 | Eukaryota | 43620.0 | 0 | Chordata | 30673.0 | 2 | Gadiformes | 22401.0 | 5 | Gadidae | 22401.0 | 5 | Gadus | 22401.0 | 5 | Gadus morhua | 22401.0 | 5 |
| scf7180000044463 | 360264 | 0.3905 | 0 | 60.2776 | Eukaryota | 53447.0 | 0 | Chordata | 53447.0 | 0 | Carnivora | 53447.0 | 0 | Canidae | 53447.0 | 0 | Canis | 53447.0 | 0 | Canis lupus | 53447.0 | 0 |
| scf7180000041303 | 1246371 | 0.4033 | 0 | 58.6361 | Eukaryota | 88777.0 | 0 | Chordata | 88777.0 | 0 | Carnivora | 88777.0 | 0 | Canidae | 88777.0 | 0 | Canis | 88777.0 | 0 | Canis lupus | 88777.0 | 0 |
| scf7180000043241 | 169446 | 0.5905 | 0 | 53.7587 | Eukaryota | 52012.0 | 0 | Chordata | 52012.0 | 0 | Carnivora | 52012.0 | 0 | Canidae | 52012.0 | 0 | Canis | 47851.0 | 1 | Canis lupus | 47851.0 | 1 |
| scf7180000046396 | 100065 | 0.4584 | 0 | 58.6059 | Eukaryota | 40403.0 | 0 | Chordata | 40403.0 | 0 | Carnivora | 40403.0 | 0 | Canidae | 37254.0 | 1 | Canis | 18974.0 | 2 | Canis lupus | 18974.0 | 2 |
| scf7180000047994 | 53987 | 0.4215 | 0 | 61.5299 | Eukaryota | 10109.0 | 0 | Chordata | 10109.0 | 0 | Carnivora | 10109.0 | 0 | Canidae | 10109.0 | 0 | Canis | 10109.0 | 0 | Canis lupus | 10109.0 | 0 |
| scf7180000054638 | 237170 | 0.4839 | 0 | 57.8466 | Eukaryota | 44717.0 | 0 | Chordata | 44717.0 | 0 | Carnivora | 44717.0 | 0 | Canidae | 44717.0 | 0 | Canis | 36216.0 | 1 | Canis lupus | 36216.0 | 1 |
| scf7180000046700 | 52324 | 0.4363 | 0 | 113.6004 | Eukaryota | 76261.0 | 0 | Chordata | 46538.0 | 1 | Gadiformes | 37448.0 | 3 | Gadidae | 37448.0 | 3 | Gadus | 37448.0 | 3 | Gadus morhua | 37448.0 | 3 |
| scf7180000042034 | 808279 | 0.4041 | 0 | 60.2862 | Eukaryota | 111942.0 | 0 | Chordata | 111942.0 | 0 | Carnivora | 111942.0 | 0 | Canidae | 111942.0 | 0 | Canis | 111942.0 | 0 | Canis lupus | 111942.0 | 0 |
| scf7180000044465 | 263972 | 0.5487 | 0 | 58.7337 | Eukaryota | 41868.0 | 0 | Chordata | 41868.0 | 0 | Carnivora | 41868.0 | 0 | Canidae | 41868.0 | 0 | Canis | 33634.0 | 1 | Canis lupus | 33634.0 | 1 |
| scf7180000042229 | 317873 | 0.3559 | 0 | 72.7453 | Eukaryota | 96096.0 | 0 | Chordata | 96096.0 | 0 | Carnivora | 96096.0 | 0 | Canidae | 96096.0 | 0 | Canis | 96096.0 | 0 | Canis lupus | 96096.0 | 0 |
| scf7180000047537 | 76053 | 0.3799 | 0 | 58.1764 | Eukaryota | 85934.0 | 0 | Chordata | 85934.0 | 0 | Carnivora | 85934.0 | 0 | Canidae | 85934.0 | 0 | Canis | 85934.0 | 0 | Canis lupus | 85934.0 | 0 |
| scf7180000042517 | 647205 | 0.5504 | 0 | 55.4887 | Eukaryota | 15218.0 | 0 | Chordata | 15218.0 | 0 | Carnivora | 15218.0 | 0 | Canidae | 15218.0 | 0 | Canis | 13594.0 | 1 | Canis lupus | 13594.0 | 1 |
| scf7180000041265 | 1282193 | 0.3918 | 0 | 59.9682 | Eukaryota | 92499.0 | 0 | Chordata | 92499.0 | 0 | Carnivora | 92499.0 | 0 | Canidae | 92499.0 | 0 | Canis | 92499.0 | 0 | Canis lupus | 92499.0 | 0 |
| scf7180000043009 | 517911 | 0.4579 | 0 | 59.5853 | Eukaryota | 89973.0 | 0 | Chordata | 89973.0 | 0 | Carnivora | 89973.0 | 0 | Canidae | 89973.0 | 0 | Canis | 83782.0 | 1 | Canis lupus | 83782.0 | 1 |
| scf7180000042429 | 236146 | 0.4109 | 0 | 58.6737 | Eukaryota | 69190.0 | 0 | Chordata | 69190.0 | 0 | Carnivora | 69190.0 | 0 | Canidae | 69190.0 | 0 | Canis | 69190.0 | 0 | Canis lupus | 69190.0 | 0 |
| scf7180000054020 | 4275 | 0.4384 | 0 | 114.3133 | no-hit | 0.0 | 0 | no-hit | 0.0 | 0 | no-hit | 0.0 | 0 | no-hit | 0.0 | 0 | no-hit | 0.0 | 0 | no-hit | 0.0 | 0 |
| scf7180000044401 | 382280 | 0.3676 | 0 | 60.2239 | Eukaryota | 32483.0 | 0 | Chordata | 32483.0 | 0 | Carnivora | 32483.0 | 0 | Canidae | 32483.0 | 0 | Canis | 23272.0 | 1 | Canis lupus | 23272.0 | 1 |
| scf7180000040925 | 2312773 | 0.3838 | 100 | 58.9197 | Eukaryota | 90350.0 | 0 | Chordata | 90350.0 | 0 | Carnivora | 90350.0 | 0 | Canidae | 90350.0 | 0 | Canis | 90350.0 | 0 | Canis lupus | 90350.0 | 0 |
| scf7180000054663 | 19045 | 0.4396 | 0 | 527.4931 | Eukaryota | 2218.0 | 0 | Chordata | 2218.0 | 0 | Kurtiformes | 2044.0 | 1 | Apogonidae | 2044.0 | 1 | Sphaeramia | 2044.0 | 1 | Sphaeramia orbicularis | 2044.0 | 1 |
| scf7180000041498 | 238108 | 0.449 | 0 | 93.7322 | Eukaryota | 74579.0 | 0 | Chordata | 74579.0 | 0 | Carnivora | 74579.0 | 0 | Canidae | 74579.0 | 0 | Canis | 74579.0 | 0 | Canis lupus | 74579.0 | 0 |
| scf7180000048037 | 66379 | 0.5436 | 0 | 56.8447 | Eukaryota | 3375.0 | 0 | Chordata | 3375.0 | 0 | Carnivora | 2987.0 | 1 | Canidae | 2987.0 | 1 | Canis | 2987.0 | 1 | Canis lupus | 2987.0 | 1 |
| scf7180000052009 | 12856 | 0.448 | 0 | 89.1311 | no-hit | 0.0 | 0 | no-hit | 0.0 | 0 | no-hit | 0.0 | 0 | no-hit | 0.0 | 0 | no-hit | 0.0 | 0 | no-hit | 0.0 | 0 |
| scf7180000041131 | 948985 | 0.3719 | 0 | 58.8228 | Eukaryota | 74462.0 | 0 | Chordata | 74462.0 | 0 | Carnivora | 74462.0 | 0 | Canidae | 74462.0 | 0 | Canis | 74462.0 | 0 | Canis lupus | 74462.0 | 0 |
| scf7180000041686 | 610798 | 0.3383 | 0 | 58.87 | Eukaryota | 59987.0 | 0 | Chordata | 59987.0 | 0 | Carnivora | 59987.0 | 0 | Canidae | 59987.0 | 0 | Canis | 59987.0 | 0 | Canis lupus | 59987.0 | 0 |
| scf7180000046512 | 110281 | 0.4801 | 0 | 58.0176 | Eukaryota | 30027.0 | 0 | Chordata | 30027.0 | 0 | Carnivora | 30027.0 | 0 | Canidae | 25715.0 | 1 | Canis | 25715.0 | 1 | Canis lupus | 25715.0 | 1 |
| scf7180000055061 | 23419 | 0.4038 | 0 | 67.574 | Eukaryota | 92605.0 | 0 | Chordata | 92605.0 | 0 | Carnivora | 92605.0 | 0 | Canidae | 92605.0 | 0 | Canis | 92605.0 | 0 | Canis lupus | 92605.0 | 0 |
| scf7180000045181 | 91532 | 0.5128 | 0 | 54.7077 | Eukaryota | 25653.0 | 0 | Chordata | 25653.0 | 0 | Carnivora | 25653.0 | 0 | Canidae | 25653.0 | 0 | Canis | 25653.0 | 0 | Canis lupus | 25653.0 | 0 |
| scf7180000046264 | 98741 | 0.4146 | 0 | 58.8265 | Eukaryota | 23359.0 | 0 | Chordata | 23359.0 | 0 | Carnivora | 23359.0 | 0 | Canidae | 23359.0 | 0 | Canis | 23359.0 | 0 | Canis lupus | 23359.0 | 0 |
| scf7180000044733 | 276199 | 0.4041 | 0 | 59.6553 | Eukaryota | 21053.0 | 0 | Chordata | 21053.0 | 0 | Carnivora | 21053.0 | 0 | Canidae | 21053.0 | 0 | Canis | 13648.0 | 1 | Canis lupus | 13648.0 | 1 |
| scf7180000045106 | 255081 | 0.4278 | 0 | 57.7307 | Eukaryota | 34151.0 | 0 | Chordata | 34151.0 | 0 | Carnivora | 34151.0 | 0 | Canidae | 34151.0 | 0 | Vulpes | 18874.0 | 1 | Vulpes vulpes | 18874.0 | 1 |
| scf7180000056453 | 374163 | 0.4014 | 0 | 59.0378 | Eukaryota | 71436.0 | 0 | Chordata | 71436.0 | 0 | Carnivora | 71436.0 | 0 | Canidae | 71436.0 | 0 | Canis | 62791.0 | 1 | Canis lupus | 62791.0 | 1 |
| scf7180000045232 | 213743 | 0.5911 | 0 | 54.7821 | Eukaryota | 51120.0 | 0 | Chordata | 51120.0 | 0 | Carnivora | 51120.0 | 0 | Canidae | 51120.0 | 0 | Canis | 51120.0 | 0 | Canis lupus | 51120.0 | 0 |
| scf7180000051500 | 14505 | 0.4415 | 0 | 65.0321 | Eukaryota | 2390.0 | 0 | Chordata | 2390.0 | 0 | Salmoniformes | 2390.0 | 0 | Salmonidae | 2390.0 | 0 | Salmo | 2390.0 | 0 | Salmo trutta | 2390.0 | 0 |
| scf7180000053831 | 2387 | 0.4449 | 0 | 68.9518 | Eukaryota | 16233.0 | 0 | Chordata | 11363.0 | 2 | Gadiformes | 8505.0 | 5 | Gadidae | 8505.0 | 5 | Gadus | 8505.0 | 5 | Gadus morhua | 8505.0 | 5 |
| scf7180000046204 | 24500 | 0.4398 | 0 | 87.5873 | Eukaryota | 36690.0 | 1 | Chordata | 36690.0 | 1 | Gadiformes | 26431.0 | 4 | Gadidae | 26431.0 | 4 | Gadus | 26431.0 | 4 | Gadus morhua | 26431.0 | 4 |
| scf7180000043155 | 394770 | 0.4098 | 0 | 63.5266 | Eukaryota | 96162.0 | 0 | Chordata | 96162.0 | 0 | Carnivora | 96162.0 | 0 | Canidae | 96162.0 | 0 | Canis | 51930.0 | 1 | Canis lupus | 51930.0 | 1 |
| scf7180000050787 | 3228 | 0.4427 | 0 | 205.9253 | Eukaryota | 1112.0 | 0 | Chordata | 1112.0 | 0 | Salmoniformes | 1112.0 | 0 | Salmonidae | 1112.0 | 0 | Salmo | 1112.0 | 0 | Salmo trutta | 1112.0 | 0 |
| scf7180000041693 | 933483 | 0.3904 | 0 | 82.2541 | Eukaryota | 89038.0 | 0 | Chordata | 89038.0 | 0 | Carnivora | 89038.0 | 0 | Canidae | 89038.0 | 0 | Canis | 89038.0 | 0 | Canis lupus | 89038.0 | 0 |
| scf7180000050362 | 9723 | 0.4532 | 0 | 1297.9453 | no-hit | 0.0 | 0 | no-hit | 0.0 | 0 | no-hit | 0.0 | 0 | no-hit | 0.0 | 0 | no-hit | 0.0 | 0 | no-hit | 0.0 | 0 |

| | | | | | | | | | | | | | | | | | | | | | | |
|---|---|---|---|---|---|---|---|---|---|---|---|---|---|---|---|---|---|---|---|---|---|---|
| scf7180000048954 | 10993 | 0.4408 | 0 | 308.0694 | Eukaryota | 2812.0 | 0 | Platyhelminthes | 1704.0 | 2 | Polystomatidea | 1704.0 | 2 | Polystomatidae | 1704.0 | 2 | Protopolystoma | 1704.0 | 2 | Protopolystoma xenopodis | 1704.0 | 2 |
| scf7180000046621 | 3235 | 0.3981 | 0 | 27.3893 | Eukaryota | 50718.0 | 0 | Chordata | 50718.0 | 0 | Carnivora | 50718.0 | 0 | Canidae | 50718.0 | 0 | Canis | 50718.0 | 0 | Canis lupus | 50718.0 | 0 |
| scf7180000050590 | 12660 | 0.4472 | 0 | 173.7048 | Eukaryota | 3054.0 | 0 | Chordata | 3054.0 | 0 | Carnivora | 3054.0 | 0 | Canidae | 3054.0 | 0 | Canis | 3054.0 | 0 | Canis lupus | 3054.0 | 0 |
| scf7180000047621 | 73755 | 0.4445 | 0 | 161.0543 | Eukaryota | 5687.0 | 0 | Chordata | 3723.0 | 1 | Tetraodontiformes | 2018.0 | 4 | Tetraodontidae | 2018.0 | 4 | Takifugu | 2018.0 | 4 | Takifugu rubripes | 2018.0 | 4 |
| scf7180000052839 | 8377 | 0.4471 | 0 | 159.8522 | Eukaryota | 3395.0 | 0 | Chordata | 1910.0 | 1 | Polystomatidea | 1485.0 | 2 | Polystomatidae | 1485.0 | 2 | Protopolystoma | 1485.0 | 2 | Protopolystoma xenopodis | 1485.0 | 2 |
| scf7180000048093 | 5522 | 0.4506 | 0 | 615.8383 | Eukaryota | 204.0 | 0 | Streptophyta | 204.0 | 0 | Fabales | 204.0 | 0 | Fabaceae | 204.0 | 0 | Cicer | 204.0 | 0 | Cicer arietinum | 204.0 | 0 |
| scf7180000045718 | 165941 | 0.3745 | 0 | 57.8901 | Eukaryota | 84767.0 | 0 | Chordata | 84767.0 | 0 | Carnivora | 84767.0 | 0 | Canidae | 84767.0 | 0 | Canis | 84767.0 | 0 | Canis lupus | 84767.0 | 0 |
| scf7180000054811 | 17891 | 0.4318 | 0 | 120.2841 | Eukaryota | 32570.0 | 0 | Chordata | 32570.0 | 0 | Carnivora | 32570.0 | 0 | Canidae | 32570.0 | 0 | Canis | 32570.0 | 0 | Canis lupus | 32570.0 | 0 |
| scf7180000051055 | 10445 | 0.4364 | 0 | 907.1381 | no-hit | 0.0 | 0 | no-hit | 0.0 | 0 | no-hit | 0.0 | 0 | no-hit | 0.0 | 0 | no-hit | 0.0 | 0 | no-hit | 0.0 | 0 |
| scf7180000041960 | 1049720 | 0.3653 | 0 | 60.4012 | Eukaryota | 116239.0 | 0 | Chordata | 116239.0 | 0 | Carnivora | 116239.0 | 0 | Canidae | 116239.0 | 0 | Canis | 116239.0 | 0 | Canis lupus | 116239.0 | 0 |
| scf7180000041317 | 12867 | 0.3901 | 0 | 302.4046 | Eukaryota | 7801.0 | 1 | Chordata | 7801.0 | 1 | Primates | 4728.0 | 3 | Hominidae | 4728.0 | 3 | Homo | 4728.0 | 4 | Homo sapiens | 4728.0 | 4 |
| scf7180000051852 | 3991 | 0.4485 | 0 | 598.3514 | Eukaryota | 2625.0 | 0 | Chordata | 2318.0 | 1 | Columbiformes | 876.0 | 3 | Columbidae | 876.0 | 3 | Streptopelia | 876.0 | 3 | Streptopelia turtur | 876.0 | 3 |
| scf7180000041335 | 1157864 | 0.4137 | 0 | 60.6069 | Eukaryota | 55402.0 | 0 | Chordata | 55402.0 | 0 | Carnivora | 55402.0 | 0 | Canidae | 55402.0 | 0 | Canis | 55402.0 | 0 | Canis lupus | 55402.0 | 0 |
| scf7180000040983 | 221051 | 0.3465 | 0 | 58.771 | Eukaryota | 94896.0 | 0 | Chordata | 94896.0 | 0 | Carnivora | 94896.0 | 0 | Canidae | 94896.0 | 0 | Canis | 94896.0 | 0 | Canis lupus | 94896.0 | 0 |
| scf7180000045198 | 25756 | 0.4891 | 0 | 550.4324 | Eukaryota | 49814.0 | 0 | Chordata | 49814.0 | 0 | Carnivora | 49814.0 | 0 | Canidae | 49814.0 | 0 | Canis | 49814.0 | 0 | Canis lupus | 49814.0 | 0 |
| scf7180000051077 | 14526 | 0.503 | 0 | 429.6571 | Eukaryota | 8850.0 | 0 | Chordata | 8850.0 | 0 | Carnivora | 8850.0 | 0 | Canidae | 8850.0 | 0 | Canis | 7159.0 | 1 | Canis lupus | 5912.0 | 2 |
| scf7180000047617 | 88589 | 0.4085 | 0 | 61.2866 | Eukaryota | 30435.0 | 0 | Chordata | 30435.0 | 0 | Carnivora | 27912.0 | 1 | Felidae | 14337.0 | 3 | Mustela | 7369.0 | 6 | Mustela putorius | 7369.0 | 6 |
| scf7180000048484 | 26703 | 0.445 | 0 | 155.0413 | Eukaryota | 12111.0 | 0 | Chordata | 7160.0 | 3 | Gadiformes | 4556.0 | 7 | Gadidae | 4556.0 | 7 | Gadus | 4556.0 | 7 | Gadus morhua | 4556.0 | 7 |
| scf7180000045420 | 230194 | 0.4338 | 0 | 59.3153 | Eukaryota | 44276.0 | 0 | Chordata | 44276.0 | 0 | Carnivora | 44276.0 | 0 | Felidae | 22675.0 | 2 | Acinonyx | 13692.0 | 5 | Acinonyx jubatus | 13692.0 | 5 |
| scf7180000051655 | 11190 | 0.4389 | 0 | 92.8785 | Eukaryota | 544.0 | 0 | Chordata | 544.0 | 0 | Kurtiformes | 544.0 | 0 | Apogonidae | 544.0 | 0 | Sphaeramia | 544.0 | 0 | Sphaeramia orbicularis | 544.0 | 0 |
| scf7180000044832 | 384961 | 0.3977 | 0 | 59.9931 | Eukaryota | 87005.0 | 0 | Chordata | 87005.0 | 0 | Carnivora | 87005.0 | 0 | Canidae | 87005.0 | 0 | Vulpes | 46420.0 | 1 | Vulpes vulpes | 46420.0 | 1 |
| scf7180000053387 | 2300 | 0.4513 | 0 | 188.1489 | Eukaryota | 308.0 | 0 | Streptophyta | 308.0 | 0 | Fabales | 308.0 | 0 | Fabaceae | 308.0 | 0 | Cicer | 308.0 | 0 | Cicer arietinum | 308.0 | 0 |
| scf7180000044978 | 219423 | 0.4615 | 0 | 57.829 | Eukaryota | 15537.0 | 0 | Chordata | 15537.0 | 0 | Primates | 9265.0 | 3 | Hominidae | 7909.0 | 4 | Homo | 7909.0 | 5 | Homo sapiens | 7909.0 | 5 |
| scf7180000042484 | 665169 | 0.42 | 0 | 59.8617 | Eukaryota | 157904.0 | 0 | Chordata | 157904.0 | 0 | Carnivora | 157904.0 | 0 | Canidae | 139443.0 | 2 | Canis | 79527.0 | 3 | Canis lupus | 79527.0 | 3 |
| scf7180000045237 | 185434 | 0.3913 | 0 | 60.6257 | Eukaryota | 27802.0 | 0 | Chordata | 27802.0 | 0 | Carnivora | 27802.0 | 0 | Canidae | 13340.0 | 3 | Canis | 13340.0 | 4 | Canis lupus | 13340.0 | 4 |
| scf7180000041095 | 430920 | 0.3414 | 0 | 58.518 | Eukaryota | 38533.0 | 0 | Chordata | 38533.0 | 0 | Carnivora | 38533.0 | 0 | Canidae | 38533.0 | 0 | Canis | 38533.0 | 0 | Canis lupus | 38533.0 | 0 |
| scf7180000050594 | 15054 | 0.4396 | 0 | 91.8472 | Eukaryota | 17020.0 | 0 | Chordata | 13270.0 | 4 | Gadiformes | 10207.0 | 8 | Gadidae | 10207.0 | 8 | Gadus | 10207.0 | 8 | Gadus morhua | 10207.0 | 8 |
| scf7180000045618 | 184016 | 0.4261 | 0 | 57.839 | Eukaryota | 41608.0 | 0 | Chordata | 41608.0 | 0 | Carnivora | 41608.0 | 0 | Canidae | 21531.0 | 3 | Canis | 12952.0 | 4 | Canis lupus | 12952.0 | 5 |
| scf7180000043113 | 446239 | 0.3695 | 0 | 59.9169 | Eukaryota | 49167.0 | 0 | Chordata | 49167.0 | 0 | Carnivora | 42902.0 | 1 | Canidae | 42902.0 | 1 | Canis | 33122.0 | 2 | Canis lupus | 33122.0 | 2 |
| scf7180000055394 | 46616 | 0.3787 | 0 | 1042.5271 | Eukaryota | 53607.0 | 0 | Chordata | 53607.0 | 0 | Carnivora | 53607.0 | 0 | Canidae | 53607.0 | 0 | Canis | 53607.0 | 0 | Canis lupus | 53607.0 | 0 |
| scf7180000056538 | 60862 | 0.5645 | 0 | 56.9972 | Eukaryota | 76756.0 | 0 | Chordata | 76756.0 | 0 | Carnivora | 76756.0 | 0 | Canidae | 76756.0 | 0 | Canis | 76756.0 | 0 | Canis lupus | 76756.0 | 0 |
| scf7180000056295 | 1183736 | 0.5285 | 0 | 59.0095 | Eukaryota | 58691.0 | 0 | Chordata | 58691.0 | 0 | Carnivora | 58691.0 | 0 | Canidae | 58691.0 | 0 | Canis | 30684.0 | 1 | Canis lupus | 30684.0 | 1 |
| scf7180000042606 | 49834 | 0.5435 | 0 | 56.0419 | Eukaryota | 49410.0 | 0 | Chordata | 49410.0 | 0 | Carnivora | 49410.0 | 0 | Canidae | 49410.0 | 0 | Canis | 49410.0 | 0 | Canis lupus | 49410.0 | 0 |
| scf7180000050651 | 2249 | 0.4522 | 0 | 132.2495 | Eukaryota | 1262.0 | 0 | Platyhelminthes | 509.0 | 2 | Polystomatidea | 509.0 | 2 | Polystomatidae | 509.0 | 2 | Protopolystoma | 509.0 | 2 | Protopolystoma xenopodis | 509.0 | 2 |
| scf7180000047780 | 36340 | 0.4395 | 0 | 81.5813 | Eukaryota | 2866.0 | 0 | Chordata | 2866.0 | 0 | Carnivora | 2866.0 | 0 | Canidae | 2866.0 | 0 | Canis | 2866.0 | 0 | Canis lupus | 2866.0 | 0 |
| scf7180000045993 | 55370 | 0.3817 | 0 | 51.0708 | Eukaryota | 14318.0 | 0 | Chordata | 14318.0 | 0 | Carnivora | 14318.0 | 0 | Canidae | 14318.0 | 0 | Canis | 14318.0 | 0 | Canis lupus | 14318.0 | 0 |
| scf7180000041665 | 981347 | 0.3741 | 0 | 60.4377 | Eukaryota | 91594.0 | 0 | Chordata | 91594.0 | 0 | Carnivora | 91594.0 | 0 | Canidae | 91594.0 | 0 | Canis | 91594.0 | 0 | Canis lupus | 91594.0 | 0 |
| scf7180000049377 | 16835 | 0.4425 | 0 | 358.8875 | Eukaryota | 52768.0 | 0 | Chordata | 36124.0 | 1 | Chordata-undef | 9425.0 | 6 | Ambassidae | 9425.0 | 6 | Parambassis | 9425.0 | 6 | Parambassis ranga | 9425.0 | 6 |
| scf7180000044613 | 226269 | 0.4262 | 0 | 58.7601 | Eukaryota | 72870.0 | 0 | Chordata | 72870.0 | 0 | Carnivora | 72870.0 | 0 | Canidae | 72870.0 | 0 | Canis | 72870.0 | 0 | Canis lupus | 72870.0 | 0 |
| scf7180000053306 | 6241 | 0.5224 | 0 | 119.1641 | Eukaryota | 32404.0 | 0 | Chordata | 32404.0 | 0 | Carnivora | 32404.0 | 0 | Canidae | 32404.0 | 0 | Canis | 32404.0 | 0 | Canis lupus | 32404.0 | 0 |
| scf7180000043172 | 746637 | 0.5761 | 0 | 57.8582 | Eukaryota | 65675.0 | 0 | Chordata | 65675.0 | 0 | Carnivora | 65675.0 | 0 | Canidae | 65675.0 | 0 | Canis | 65675.0 | 0 | Canis lupus | 65675.0 | 0 |
| scf7180000054021 | 7341 | 0.4467 | 0 | 60.4375 | Eukaryota | 1030.0 | 1 | Chordata | 1030.0 | 1 | Columbiformes | 747.0 | 2 | Columbidae | 747.0 | 2 | Streptopelia | 747.0 | 2 | Streptopelia turtur | 747.0 | 2 |
| scf7180000051121 | 4282 | 0.4605 | 0 | 73.8966 | Eukaryota | 1993.0 | 0 | Chordata | 1780.0 | 1 | Columbiformes | 1310.0 | 3 | Columbidae | 1310.0 | 3 | Streptopelia | 1310.0 | 3 | Streptopelia turtur | 1310.0 | 3 |
| scf7180000051503 | 6669 | 0.4476 | 0 | 712.8194 | Eukaryota | 813.0 | 0 | Platyhelminthes | 813.0 | 0 | Polystomatidea | 813.0 | 0 | Polystomatidae | 813.0 | 0 | Protopolystoma | 813.0 | 0 | Protopolystoma xenopodis | 813.0 | 0 |
| scf7180000042909 | 30712 | 0.401 | 0 | 52.4639 | Eukaryota | 75984.0 | 0 | Chordata | 75984.0 | 0 | Carnivora | 75984.0 | 0 | Canidae | 75984.0 | 0 | Canis | 75984.0 | 0 | Canis lupus | 75984.0 | 0 |
| scf7180000043038 | 594604 | 0.3893 | 0 | 60.4064 | Eukaryota | 29067.0 | 0 | Chordata | 29067.0 | 0 | Carnivora | 29067.0 | 0 | Canidae | 29067.0 | 0 | Vulpes | 20412.0 | 1 | Vulpes vulpes | 20412.0 | 1 |
| scf7180000054198 | 15856 | 0.461 | 0 | 159.9202 | Eukaryota | 11484.0 | 0 | Chordata | 10671.0 | 1 | Carnivora | 10671.0 | 1 | Canidae | 10671.0 | 1 | Canis | 7984.0 | 2 | Canis lupus | 7093.0 | 3 |
| scf7180000056396 | 176833 | 0.4342 | 0 | 58.9391 | Eukaryota | 44934.0 | 0 | Chordata | 44934.0 | 0 | Carnivora | 44934.0 | 0 | Ursidae | 16336.0 | 3 | Ursus | 16336.0 | 5 | Ursus arctos | 16336.0 | 5 |
| scf7180000053522 | 3817 | 0.4483 | 0 | 475.0666 | no-hit | 0.0 | 0 | no-hit | 0.0 | 0 | no-hit | 0.0 | 0 | no-hit | 0.0 | 0 | no-hit | 0.0 | 0 | no-hit | 0.0 | 0 |
| scf7180000053644 | 3405 | 0.504 | 0 | 144.591 | Eukaryota | 5290.0 | 0 | Chordata | 5290.0 | 0 | Carnivora | 5290.0 | 0 | Canidae | 5290.0 | 0 | Canis | 4416.0 | 1 | Canis lupus | 4416.0 | 1 |
| scf7180000044577 | 315624 | 0.3873 | 0 | 58.5776 | Eukaryota | 89866.0 | 0 | Chordata | 89866.0 | 0 | Carnivora | 89866.0 | 0 | Canidae | 89866.0 | 0 | Canis | 89866.0 | 0 | Canis lupus | 89866.0 | 0 |
| scf7180000051100 | 6675 | 0.44 | 0 | 127.9064 | Eukaryota | 3725.0 | 0 | Platyhelminthes | 2419.0 | 2 | Polystomatidea | 2419.0 | 2 | Polystomatidae | 2419.0 | 2 | Protopolystoma | 2419.0 | 2 | Protopolystoma xenopodis | 2419.0 | 2 |
| scf7180000040502 | 2950737 | 0.4333 | 0 | 58.6593 | Eukaryota | 121520.0 | 0 | Chordata | 121520.0 | 0 | Carnivora | 121520.0 | 0 | Canidae | 121520.0 | 0 | Canis | 101658.0 | 1 | Canis lupus | 101658.0 | 1 |
| scf7180000051847 | 11894 | 0.5052 | 0 | 1096.7268 | Eukaryota | 8775.0 | 0 | Chordata | 8775.0 | 0 | Carnivora | 8775.0 | 0 | Canidae | 8775.0 | 0 | Canis | 7086.0 | 1 | Canis lupus | 5831.0 | 2 |
| scf7180000044949 | 23010 | 0.516 | 0 | 373.8952 | Eukaryota | 35077.0 | 0 | Chordata | 35077.0 | 0 | Carnivora | 35077.0 | 0 | Canidae | 35077.0 | 0 | Canis | 35077.0 | 0 | Canis lupus | 35077.0 | 0 |
| scf7180000051354 | 5964 | 0.4534 | 0 | 1255.5169 | Eukaryota | 263.0 | 0 | Chordata | 263.0 | 0 | Kurtiformes | 263.0 | 0 | Apogonidae | 263.0 | 0 | Sphaeramia | 263.0 | 0 | Sphaeramia orbicularis | 263.0 | 0 |
| scf7180000043606 | 496317 | 0.4868 | 100 | 59.2799 | Eukaryota | 59474.0 | 0 | Chordata | 59474.0 | 0 | Carnivora | 59474.0 | 0 | Canidae | 59474.0 | 0 | Canis | 43796.0 | 1 | Canis lupus | 43796.0 | 1 |
| scf7180000050227 | 3772 | 0.448 | 0 | 159.0489 | Eukaryota | 287.0 | 0 | Chordata | 287.0 | 0 | Salmoniformes | 287.0 | 0 | Salmonidae | 287.0 | 0 | Coregonus | 287.0 | 0 | Coregonus sp. 'balchen' | 287.0 | 0 |
| scf7180000052500 | 3056 | 0.4506 | 0 | 266.6042 | Eukaryota | 2184.0 | 0 | Platyhelminthes | 878.0 | 2 | Polystomatidea | 878.0 | 2 | Polystomatidae | 878.0 | 2 | Protopolystoma | 878.0 | 2 | Protopolystoma xenopodis | 878.0 | 2 |
| scf7180000042785 | 36370 | 0.4475 | 0 | 54.9699 | Eukaryota | 34063.0 | 0 | Chordata | 34063.0 | 0 | Carnivora | 34063.0 | 0 | Ursidae | 14093.0 | 3 | Ursus | 10626.0 | 4 | Ursus maritimus | 10626.0 | 4 |
| scf7180000042197 | 767983 | 0.4048 | 0 | 59.3264 | Eukaryota | 58886.0 | 0 | Chordata | 58886.0 | 0 | Carnivora | 58886.0 | 0 | Canidae | 58886.0 | 0 | Canis | 58886.0 | 0 | Canis lupus | 58886.0 | 0 |
| scf7180000056217 | 487266 | 0.395 | 0 | 59.8049 | Eukaryota | 83168.0 | 0 | Chordata | 83168.0 | 0 | Carnivora | 83168.0 | 0 | Canidae | 83168.0 | 0 | Canis | 83168.0 | 0 | Canis lupus | 83168.0 | 0 |
| scf7180000051898 | 2228 | 0.4461 | 0 | 616.8498 | no-hit | 0.0 | 0 | no-hit | 0.0 | 0 | no-hit | 0.0 | 0 | no-hit | 0.0 | 0 | no-hit | 0.0 | 0 | no-hit | 0.0 | 0 |
| scf7180000040648 | 1929304 | 0.4639 | 0 | 59.1741 | Eukaryota | 56667.0 | 0 | Chordata | 56667.0 | 0 | Carnivora | 56667.0 | 0 | Canidae | 56667.0 | 0 | Canis | 48903.0 | 1 | Canis lupus | 48903.0 | 1 |
| scf7180000044023 | 347767 | 0.3937 | 0 | 60.8241 | Eukaryota | 37822.0 | 0 | Chordata | 37822.0 | 0 | Carnivora | 37822.0 | 0 | Canidae | 37822.0 | 0 | Canis | 37822.0 | 0 | Canis lupus | 37822.0 | 0 |
| scf7180000040960 | 1412499 | 0.4884 | 0 | 59.0624 | Eukaryota | 92440.0 | 0 | Chordata | 92440.0 | 0 | Carnivora | 92440.0 | 0 | Canidae | 92440.0 | 0 | Canis | 92440.0 | 0 | Canis lupus | 92440.0 | 0 |
| scf7180000055236 | 25944 | 0.3765 | 0 | 808.6477 | Eukaryota | 16296.0 | 0 | Chordata | 16296.0 | 0 | Carnivora | 16296.0 | 0 | Canidae | 16296.0 | 0 | Canis | 16296.0 | 0 | Canis lupus | 16296.0 | 0 |
| scf7180000054368 | 16429 | 0.3723 | 0 | 222.1729 | Eukaryota | 27035.0 | 0 | Chordata | 27035.0 | 0 | Carnivora | 27035.0 | 0 | Canidae | 27035.0 | 0 | Canis | 27035.0 | 0 | Canis lupus | 27035.0 | 0 |
| scf7180000047500 | 83612 | 0.3869 | 0 | 58.9094 | Eukaryota | 25213.0 | 0 | Chordata | 25213.0 | 0 | Carnivora | 25213.0 | 0 | Canidae | 25213.0 | 0 | Canis | 25213.0 | 0 | Canis lupus | 25213.0 | 0 |
| scf7180000042929 | 549407 | 0.3818 | 0 | 60.1229 | Eukaryota | 80730.0 | 0 | Chordata | 80730.0 | 0 | Carnivora | 80730.0 | 0 | Canidae | 80730.0 | 0 | Canis | 80730.0 | 0 | Canis lupus | 80730.0 | 0 |
| scf7180000049543 | 10289 | 0.4358 | 0 | 69.4867 | Eukaryota | 4971.0 | 0 | Chordata | 4971.0 | 0 | Carnivora | 2562.0 | 2 | Canidae | 2562.0 | 2 | Canis | 2562.0 | 2 | Canis lupus | 2562.0 | 2 |
| scf7180000051627 | 26428 | 0.5124 | 0 | 693.6718 | Eukaryota | 32999.0 | 0 | Chordata | 32999.0 | 0 | Carnivora | 32999.0 | 0 | Canidae | 32999.0 | 0 | Canis | 32999.0 | 0 | Canis lupus | 32999.0 | 0 |
| scf7180000043855 | 20692 | 0.3464 | 0 | 44.5371 | Eukaryota | 8419.0 | 0 | Chordata | 8419.0 | 0 | Carnivora | 8419.0 | 0 | Canidae | 5889.0 | 1 | Canis | 5889.0 | 2 | Canis lupus | 5889.0 | 2 |
| scf7180000056362 | 851533 | 0.3611 | 0 | 60.708 | Eukaryota | 36572.0 | 0 | Chordata | 36572.0 | 0 | Carnivora | 36572.0 | 0 | Canidae | 36572.0 | 0 | Canis | 36572.0 | 0 | Canis lupus | 36572.0 | 0 |
| scf7180000042900 | 636778 | 0.4 | 0 | 59.9664 | Eukaryota | 47137.0 | 0 | Chordata | 47137.0 | 0 | Carnivora | 47137.0 | 0 | Canidae | 47137.0 | 0 | Canis | 47137.0 | 0 | Canis lupus | 47137.0 | 0 |
| scf7180000041652 | 1004178 | 0.4037 | 0 | 61.1524 | Eukaryota | 98265.0 | 0 | Chordata | 98265.0 | 0 | Carnivora | 98265.0 | 0 | Canidae | 98265.0 | 0 | Canis | 86799.0 | 1 | Canis lupus | 86799.0 | 1 |
| scf7180000054838 | 4836 | 0.4433 | 0 | 194.0335 | no-hit | 0.0 | 0 | no-hit | 0.0 | 0 | no-hit | 0.0 | 0 | no-hit | 0.0 | 0 | no-hit | 0.0 | 0 | no-hit | 0.0 | 0 |
| scf7180000054326 | 27725 | 0.4442 | 0 | 199.0515 | Eukaryota | 15363.0 | 0 | Platyhelminthes | 12421.0 | 1 | Strigeidida | 10113.0 | 5 | Schistosomatidae | 10113.0 | 5 | Schistosoma | 10113.0 | 5 | Schistosoma mansoni | 10113.0 | 5 |
| scf7180000046401 | 132633 | 0.5755 | 0 | 56.7805 | Eukaryota | 42246.0 | 0 | Chordata | 42246.0 | 0 | Carnivora | 42246.0 | 0 | Canidae | 42246.0 | 0 | Canis | 29983.0 | 1 | Canis lupus | 29983.0 | 1 |
| scf7180000050320 | 12701 | 0.4711 | 0 | 50.718 | Eukaryota | 16217.0 | 0 | Chordata | 16217.0 | 0 | Carnivora | 16217.0 | 0 | Canidae | 16217.0 | 0 | Canis | 10457.0 | 1 | Canis lupus | 10457.0 | 1 |
| scf7180000048138 | 73759 | 0.4425 | 0 | 66.516 | Eukaryota | 9721.0 | 0 | Chordata | 9721.0 | 0 | Carnivora | 9721.0 | 0 | Canidae | 9721.0 | 0 | Canis | 5804.0 | 1 | Canis lupus | 5804.0 | 1 |
| scf7180000052719 | 2578 | 0.4538 | 0 | 423.8889 | Eukaryota | 167.0 | 0 | Chordata | 167.0 | 0 | Salmoniformes | 167.0 | 0 | Salmonidae | 167.0 | 0 | Coregonus | 167.0 | 0 | Coregonus sp. 'balchen' | 167.0 | 0 |
| scf7180000050573 | 3467 | 0.4557 | 0 | 344.3929 | Eukaryota | 669.0 | 0 | Chordata | 669.0 | 0 | Kurtiformes | 669.0 | 0 | Apogonidae | 669.0 | 0 | Sphaeramia | 669.0 | 0 | Sphaeramia orbicularis | 669.0 | 0 |
| scf7180000040667 | 1753669 | 0.4222 | 0 | 59.8594 | Eukaryota | 96026.0 | 0 | Chordata | 96026.0 | 0 | Carnivora | 96026.0 | 0 | Canidae | 96026.0 | 0 | Canis | 96026.0 | 0 | Canis lupus | 96026.0 | 0 |
| scf7180000044220 | 285016 | 0.4013 | 0 | 60.4452 | Eukaryota | 42473.0 | 0 | Chordata | 42473.0 | 0 | Carnivora | 42473.0 | 0 | Canidae | 42473.0 | 0 | Canis | 42473.0 | 0 | Canis lupus | 42473.0 | 0 |
| scf7180000046041 | 165254 | 0.3618 | 0 | 60.3699 | Eukaryota | 51933.0 | 0 | Chordata | 51933.0 | 0 | Carnivora | 51933.0 | 0 | Canidae | 51933.0 | 0 | Canis | 51933.0 | 0 | Canis lupus | 51933.0 | 0 |
| scf7180000053637 | 6415 | 0.441 | 0 | 222.8586 | Eukaryota | 4486.0 | 0 | Chordata | 4486.0 | 0 | Kurtiformes | 4227.0 | 1 | Apogonidae | 4227.0 | 1 | Sphaeramia | 4227.0 | 1 | Sphaeramia orbicularis | 4227.0 | 1 |
| scf7180000044560 | 296066 | 0.4128 | 0 | 59.313 | Eukaryota | 32874.0 | 0 | Chordata | 32874.0 | 0 | Carnivora | 32874.0 | 0 | Canidae | 32874.0 | 0 | Canis | 24122.0 | 1 | Canis lupus | 24122.0 | 1 |
| scf7180000049613 | 25079 | 0.4441 | 0 | 332.0817 | Eukaryota | 3844.0 | 0 | Chordata | 2652.0 | 2 | Kurtiformes | 2652.0 | 2 | Apogonidae | 2652.0 | 2 | Sphaeramia | 2652.0 | 2 | Sphaeramia orbicularis | 2652.0 | 2 |
| scf7180000042071 | 348894 | 0.3765 | 0 | 59.6558 | Eukaryota | 85460.0 | 0 | Chordata | 85460.0 | 0 | Carnivora | 85460.0 | 0 | Canidae | 85460.0 | 0 | Canis | 85460.0 | 0 | Canis lupus | 85460.0 | 0 |
| scf7180000055659 | 89278 | 0.4367 | 0 | 255.0094 | Eukaryota | 7307.0 | 0 | Chordata | 7307.0 | 0 | Carnivora | 7307.0 | 0 | Canidae | 7307.0 | 0 | Canis | 7307.0 | 0 | Canis lupus | 7307.0 | 0 |
| scf7180000054972 | 29773 | 0.4393 | 0 | 298.281 | no-hit | 0.0 | 0 | no-hit | 0.0 | 0 | no-hit | 0.0 | 0 | no-hit | 0.0 | 0 | no-hit | 0.0 | 0 | no-hit | 0.0 | 0 |
| scf7180000055301 | 14457 | 0.4432 | 0 | 535.5195 | Eukaryota | 4236.0 | 0 | Chordata | 3625.0 | 1 | Kurtiformes | 2501.0 | 4 | Apogonidae | 2501.0 | 4 | Sphaeramia | 2501.0 | 4 | Sphaeramia orbicularis | 2501.0 | 4 |
| scf7180000044107 | 368802 | 0.3796 | 0 | 62.3225 | Eukaryota | 23888.0 | 0 | Chordata | 23888.0 | 0 | Carnivora | 23888.0 | 0 | Canidae | 9192.0 | 3 | Canis | 9192.0 | 4 | Canis lupus | 9192.0 | 4 |
| scf7180000042659 | 37636 | 0.367 | 0 | 47.2784 | Eukaryota | 33540.0 | 0 | Chordata | 33540.0 | 0 | Carnivora | 33540.0 | 0 | Canidae | 33540.0 | 0 | Canis | 33540.0 | 0 | Canis lupus | 33540.0 | 0 |
| scf7180000040873 | 591539 | 0.3591 | 0 | 58.6193 | Eukaryota | 40097.0 | 0 | Chordata | 40097.0 | 0 | Carnivora | 40097.0 | 0 | Canidae | 40097.0 | 0 | Canis | 40097.0 | 0 | Canis lupus | 40097.0 | 0 |
| scf7180000042843 | 429752 | 0.371 | 0 | 58.1396 | Eukaryota | 88910.0 | 0 | Chordata | 88910.0 | 0 | Carnivora | 88910.0 | 0 | Canidae | 88910.0 | 0 | Canis | 88910.0 | 0 | Canis lupus | 88910.0 | 0 |
| scf7180000040885 | 1581729 | 0.3835 | 0 | 60.9839 | Eukaryota | 95032.0 | 0 | Chordata | 95032.0 | 0 | Carnivora | 95032.0 | 0 | Canidae | 95032.0 | 0 | Canis | 95032.0 | 0 | Canis lupus | 95032.0 | 0 |
| scf7180000041058 | 1395776 | 0.4733 | 0 | 58.6856 | Eukaryota | 128452.0 | 0 | Chordata | 128452.0 | 0 | Carnivora | 128452.0 | 0 | Canidae | 128452.0 | 0 | Canis | 128452.0 | 0 | Canis lupus | 128452.0 | 0 |
| scf7180000049838 | 4908 | 0.4468 | 0 | 1073.0546 | no-hit | 0.0 | 0 | no-hit | 0.0 | 0 | no-hit | 0.0 | 0 | no-hit | 0.0 | 0 | no-hit | 0.0 | 0 | no-hit | 0.0 | 0 |
| scf7180000053255 | 7006 | 0.4448 | 0 | 431.5919 | no-hit | 0.0 | 0 | no-hit | 0.0 | 0 | no-hit | 0.0 | 0 | no-hit | 0.0 | 0 | no-hit | 0.0 | 0 | no-hit | 0.0 | 0 |
| scf7180000050536 | 7059 | 0.3904 | 0 | 28.1451 | Eukaryota | 3102.0 | 0 | Chordata | 3102.0 | 0 | Carnivora | 3102.0 | 0 | Canidae | 3102.0 | 0 | Canis | 3102.0 | 0 | Canis lupus | 3102.0 | 0 |
| scf7180000044550 | 262497 | 0.4264 | 0 | 60.6964 | Eukaryota | 61348.0 | 0 | Chordata | 61348.0 | 0 | Carnivora | 61348.0 | 0 | Canidae | 34419.0 | 2 | Canis | 26140.0 | 6 | Canis lupus | 26140.0 | 6 |
| scf7180000049147 | 43502 | 0.4812 | 0 | 55.0949 | Eukaryota | 5067.0 | 0 | Chordata | 5067.0 | 0 | Carnivora | 2403.0 | 2 | Canidae | 2403.0 | 2 | Canis | 2403.0 | 3 | Canis lupus | 2403.0 | 3 |
| scf7180000045165 | 9550 | 0.3695 | 0 | 34.3869 | Eukaryota | 6441.0 | 0 | Chordata | 6441.0 | 0 | Carnivora | 6441.0 | 0 | Canidae | 6441.0 | 0 | Canis | 6441.0 | 0 | Canis lupus | 6441.0 | 0 |
| scf7180000042639 | 515125 | 0.3684 | 0 | 59.8172 | Eukaryota | 49554.0 | 0 | Chordata | 49554.0 | 0 | Carnivora | 49554.0 | 0 | Canidae | 49554.0 | 0 | Vulpes | 28175.0 | 1 | Vulpes vulpes | 28175.0 | 1 |
| scf7180000048313 | 18640 | 0.5036 | 0 | 451.9054 | Eukaryota | 9575.0 | 0 | Chordata | 9575.0 | 0 | Carnivora | 9575.0 | 0 | Canidae | 9575.0 | 0 | Canis | 8368.0 | 1 | Canis lupus | 7106.0 | 2 |
| scf7180000056183 | 430041 | 0.4422 | 0 | 58.8681 | Eukaryota | 64187.0 | 0 | Chordata | 64187.0 | 0 | Carnivora | 64187.0 | 0 | Canidae | 45164.0 | 3 | Canis | 33962.0 | 5 | Canis lupus | 33962.0 | 5 |
| scf7180000043375 | 29029 | 0.4549 | 0 | 119.7376 | Eukaryota | 313.0 | 0 | Chordata | 313.0 | 0 | Kurtiformes | 313.0 | 0 | Apogonidae | 313.0 | 0 | Sphaeramia | 313.0 | 0 | Sphaeramia orbicularis | 313.0 | 0 |
| scf7180000040585 | 2114352 | 0.3636 | 0 | 61.1642 | Eukaryota | 99009.0 | 0 | Chordata | 99009.0 | 0 | Carnivora | 99009.0 | 0 | Canidae | 99009.0 | 0 | Canis | 89073.0 | 1 | Canis lupus | 89073.0 | 1 |
| scf7180000048963 | 27355 | 0.4465 | 0 | 87.9157 | Eukaryota | 56096.0 | 0 | Chordata | 56096.0 | 0 | Carnivora | 56096.0 | 0 | Canidae | 56096.0 | 0 | Canis | 56096.0 | 0 | Canis lupus | 56096.0 | 0 |
| scf7180000049969 | 6578 | 0.4953 | 0 | 105.2661 | Eukaryota | 8612.0 | 0 | Chordata | 8612.0 | 0 | Carnivora | 8612.0 | 0 | Canidae | 8612.0 | 0 | Canis | 6980.0 | 1 | Canis lupus | 5834.0 | 2 |
| scf7180000047770 | 20391 | 0.4348 | 0 | 111.0434 | Eukaryota | 8386.0 | 1 | Chordata | 5529.0 | 3 | Salmoniformes | 3242.0 | 6 | Salmonidae | 3242.0 | 6 | Salmo | 2828.0 | 7 | Salmo trutta | 2828.0 | 7 |
| scf7180000044524 | 564643 | 0.4448 | 0 | 59.4446 | Eukaryota | 60470.0 | 0 | Chordata | 60470.0 | 0 | Carnivora | 60470.0 | 0 | Canidae | 57531.0 | 1 | Canis | 36436.0 | 2 | Canis lupus | 36436.0 | 2 |
| scf7180000041298 | 353804 | 0.5116 | 0 | 58.516 | Eukaryota | 31065.0 | 0 | Chordata | 31065.0 | 0 | Carnivora | 31065.0 | 0 | Canidae | 31065.0 | 0 | Canis | 31065.0 | 0 | Canis lupus | 31065.0 | 0 |
| scf7180000045835 | 57212 | 0.4084 | 0 | 58.4481 | Eukaryota | 15631.0 | 0 | Chordata | 15631.0 | 0 | Carnivora | 15631.0 | 0 | Canidae | 5032.0 | 4 | Canis | 5032.0 | 5 | Canis lupus | 5032.0 | 5 |
| scf7180000054447 | 93884 | 0.444 | 0 | 387.1978 | Eukaryota | 50525.0 | 0 | Chordata | 47884.0 | 1 | Gadiformes | 31303.0 | 5 | Gadidae | 31303.0 | 5 | Gadus | 31303.0 | 5 | Gadus morhua | 31303.0 | 5 |
| scf7180000041883 | 272651 | 0.5632 | 0 | 56.4265 | Eukaryota | 44813.0 | 0 | Chordata | 44813.0 | 0 | Carnivora | 44813.0 | 0 | Canidae | 44813.0 | 0 | Canis | 39886.0 | 1 | Canis lupus | 39886.0 | 1 |
| scf7180000055032 | 14802 | 0.4447 | 0 | 161.0057 | Eukaryota | 11732.0 | 0 | Chordata | 11732.0 | 0 | Tetraodontiformes | 4062.0 | 3 | Tetraodontidae | 4062.0 | 3 | Takifugu | 4062.0 | 3 | Takifugu rubripes | 3302.0 | 4 |
| scf7180000043242 | 517021 | 0.4289 | 0 | 58.516 | Eukaryota | 42369.0 | 0 | Chordata | 42369.0 | 0 | Carnivora | 32357.0 | 1 | Canidae | 32357.0 | 1 | Canis | 23629.0 | 3 | Canis lupus | 23629.0 | 3 |
| scf7180000042167 | 76181 | 0.3748 | 0 | 58.9223 | Eukaryota | 32531.0 | 0 | Chordata | 32531.0 | 0 | Carnivora | 32531.0 | 0 | Canidae | 32531.0 | 0 | Canis | 32531.0 | 0 | Canis lupus | 32531.0 | 0 |
| scf7180000048367 | 23900 | 0.4374 | 0 | 136.56 | Eukaryota | 4666.0 | 0 | Platyhelminthes | 4272.0 | 1 | Polystomatidea | 4272.0 | 1 | Polystomatidae | 4272.0 | 1 | Protopolystoma | 4272.0 | 1 | Protopolystoma xenopodis | 4272.0 | 1 |
| scf7180000042461 | 4712 | 0.3971 | 0 | 30.125 | Eukaryota | 68119.0 | 0 | Chordata | 68119.0 | 0 | Carnivora | 68119.0 | 0 | Canidae | 68119.0 | 0 | Canis | 68119.0 | 0 | Canis lupus | 68119.0 | 0 |
| scf7180000051561 | 16700 | 0.4429 | 0 | 63.8002 | Eukaryota | 2204.0 | 0 | Chordata | 1971.0 | 1 | Tetraodontiformes | 1749.0 | 2 | Tetraodontidae | 1749.0 | 2 | Takifugu | 1749.0 | 2 | Takifugu rubripes | 1749.0 | 2 |
| scf7180000054309 | 18119 | 0.442 | 0 | 399.8416 | Eukaryota | 1990.0 | 0 | Platyhelminthes | 1289.0 | 1 | Polystomatidea | 1289.0 | 1 | Polystomatidae | 1289.0 | 1 | Protopolystoma | 1289.0 | 1 | Protopolystoma xenopodis | 1289.0 | 1 |
| scf7180000042048 | 576372 | 0.3631 | 0 | 59.2309 | Eukaryota | 85591.0 | 0 | Chordata | 85591.0 | 0 | Carnivora | 85591.0 | 0 | Canidae | 85591.0 | 0 | Canis | 85591.0 | 0 | Canis lupus | 85591.0 | 0 |
| scf7180000043271 | 34241 | 0.364 | 0 | 47.0111 | Eukaryota | 11987.0 | 0 | Chordata | 11987.0 | 0 | Carnivora | 11987.0 | 0 | Canidae | 11987.0 | 0 | Canis | 11987.0 | 0 | Canis lupus | 11987.0 | 0 |
| scf7180000050167 | 4379 | 0.4638 | 0 | 1120.3814 | Eukaryota | 2769.0 | 0 | Chordata | 2769.0 | 0 | Carnivora | 2769.0 | 0 | Canidae | 2769.0 | 0 | Canis | 2491.0 | 1 | Canis lupus | 2491.0 | 1 |
| scf7180000051853 | 6777 | 0.4526 | 0 | 1352.9839 | no-hit | 0.0 | 0 | no-hit | 0.0 | 0 | no-hit | 0.0 | 0 | no-hit | 0.0 | 0 | no-hit | 0.0 | 0 | no-hit | 0.0 | 0 |
| scf7180000052223 | 18075 | 0.4415 | 0 | 56.2474 | Eukaryota | 1782.0 | 0 | Chordata | 936.0 | 2 | Salmoniformes | 609.0 | 3 | Salmonidae | 609.0 | 3 | Coregonus | 609.0 | 3 | Coregonus sp. 'balchen' | 609.0 | 3 |
| scf7180000053037 | 5516 | 0.4365 | 0 | 209.6934 | Eukaryota | 2782.0 | 0 | Chordata | 2782.0 | 0 | Carnivora | 2782.0 | 0 | Canidae | 2782.0 | 0 | Canis | 2508.0 | 1 | Canis lupus | 2508.0 | 1 |
| scf7180000041470 | 8465 | 0.4412 | 0 | 46.5872 | Eukaryota | 8976.0 | 0 | Chordata | 8636.0 | 1 | Carnivora | 8636.0 | 1 | Canidae | 3012.0 | 6 | Vulpes | 2191.0 | 9 | Vulpes vulpes | 2191.0 | 9 |
| scf7180000049993 | 6511 | 0.4523 | 0 | 110.8454 | no-hit | 0.0 | 0 | no-hit | 0.0 | 0 | no-hit | 0.0 | 0 | no-hit | 0.0 | 0 | no-hit | 0.0 | 0 | no-hit | 0.0 | 0 |
| scf7180000041197 | 1183119 | 0.3847 | 0 | 61.3516 | Eukaryota | 80225.0 | 0 | Chordata | 80225.0 | 0 | Carnivora | 80225.0 | 0 | Canidae | 80225.0 | 0 | Canis | 59391.0 | 1 | Canis lupus | 59391.0 | 1 |
| scf7180000043963 | 390410 | 0.3855 | 0 | 60.7921 | Eukaryota | 81930.0 | 0 | Chordata | 81930.0 | 0 | Carnivora | 81930.0 | 0 | Canidae | 81930.0 | 0 | Canis | 54958.0 | 1 | Canis lupus | 54958.0 | 1 |
| scf7180000040529 | 280453 | 0.4132 | 0 | 56.1836 | Eukaryota | 93430.0 | 0 | Chordata | 93430.0 | 0 | Carnivora | 93430.0 | 0 | Canidae | 93430.0 | 0 | Canis | 93430.0 | 0 | Canis lupus | 93430.0 | 0 |
| scf7180000050846 | 3213 | 0.507 | 0 | 60.0939 | Eukaryota | 5925.0 | 0 | Chordata | 5925.0 | 0 | Carnivora | 5925.0 | 0 | Canidae | 5925.0 | 0 | Canis | 3866.0 | 1 | Canis lupus | 2806.0 | 2 |
| scf7180000045467 | 4175 | 0.4683 | 0 | 24.7752 | Eukaryota | 12238.0 | 0 | Chordata | 12238.0 | 0 | Carnivora | 12238.0 | 0 | Canidae | 12238.0 | 0 | Canis | 12238.0 | 0 | Canis lupus | 12238.0 | 0 |
| scf7180000043509 | 473671 | 0.3773 | 0 | 59.4426 | Eukaryota | 19919.0 | 0 | Chordata | 19919.0 | 0 | Carnivora | 19919.0 | 0 | Canidae | 19919.0 | 0 | Canis | 19919.0 | 0 | Canis lupus | 19919.0 | 0 |
| scf7180000051016 | 4881 | 0.4458 | 0 | 739.6834 | no-hit | 0.0 | 0 | no-hit | 0.0 | 0 | no-hit | 0.0 | 0 | no-hit | 0.0 | 0 | no-hit | 0.0 | 0 | no-hit | 0.0 | 0 |
| scf7180000049182 | 5088 | 0.5057 | 0 | 40.7869 | Eukaryota | 7547.0 | 0 | Chordata | 7547.0 | 0 | Carnivora | 7547.0 | 0 | Canidae | 7547.0 | 0 | Canis | 6055.0 | 1 | Canis lupus | 5010.0 | 2 |
| scf7180000052157 | 5024 | 0.4606 | 0 | 156.7057 | Eukaryota | 2818.0 | 0 | Chordata | 2818.0 | 0 | Carnivora | 2818.0 | 0 | Canidae | 2818.0 | 0 | Canis | 2818.0 | 0 | Canis lupus | 2818.0 | 0 |
| scf7180000056446 | 298823 | 0.4315 | 0 | 59.081 | Eukaryota | 79567.0 | 0 | Chordata | 79567.0 | 0 | Carnivora | 79567.0 | 0 | Canidae | 79567.0 | 0 | Canis | 58713.0 | 1 | Canis lupus | 58713.0 | 1 |
| scf7180000042593 | 716932 | 0.3438 | 0 | 58.707 | Eukaryota | 13404.0 | 0 | Chordata | 13404.0 | 0 | Carnivora | 10531.0 | 2 | Canidae | 10531.0 | 2 | Canis | 10531.0 | 2 | Canis lupus | 10531.0 | 2 |
| scf7180000055399 | 41740 | 0.4448 | 0 | 279.432 | Eukaryota | 48118.0 | 0 | Chordata | 39557.0 | 2 | Gadiformes | 27036.0 | 6 | Gadidae | 27036.0 | 6 | Gadus | 27036.0 | 6 | Gadus morhua | 27036.0 | 6 |
| scf7180000042650 | 493399 | 0.3957 | 0 | 60.0296 | Eukaryota | 87370.0 | 0 | Chordata | 87370.0 | 0 | Carnivora | 87370.0 | 0 | Canidae | 87370.0 | 0 | Canis | 87370.0 | 0 | Canis lupus | 87370.0 | 0 |
| scf7180000047999 | 10977 | 0.3616 | 0 | 87.4624 | Eukaryota | 6493.0 | 0 | Chordata | 6493.0 | 0 | Carnivora | 3481.0 | 1 | Canidae | 3481.0 | 3 | Canis | 3481.0 | 3 | Canis lupus | 3481.0 | 3 |
| scf7180000045859 | 13483 | 0.4033 | 0 | 37.5958 | Eukaryota | 76284.0 | 0 | Chordata | 76284.0 | 0 | Carnivora | 76284.0 | 0 | Canidae | 76284.0 | 0 | Canis | 76284.0 | 0 | Canis lupus | 76284.0 | 0 |
| scf7180000049675 | 1639 | 0.4606 | 0 | 124.7903 | no-hit | 0.0 | 0 | no-hit | 0.0 | 0 | no-hit | 0.0 | 0 | no-hit | 0.0 | 0 | no-hit | 0.0 | 0 | no-hit | 0.0 | 0 |
| scf7180000045662 | 48793 | 0.4328 | 0 | 45.262 | Eukaryota | 74490.0 | 0 | Chordata | 74490.0 | 0 | Carnivora | 74490.0 | 0 | Canidae | 74490.0 | 0 | Canis | 56738.0 | 1 | Canis lupus | 56738.0 | 1 |
| scf7180000041572 | 202562 | 0.3663 | 0 | 59.1188 | Eukaryota | 24974.0 | 0 | Chordata | 24974.0 | 0 | Carnivora | 24974.0 | 0 | Canidae | 24974.0 | 0 | Canis | 24974.0 | 0 | Canis lupus | 24974.0 | 0 |
| scf7180000044637 | 79094 | 0.4183 | 0 | 53.2336 | Eukaryota | 34524.0 | 0 | Chordata | 34524.0 | 0 | Carnivora | 34524.0 | 0 | Canidae | 28717.0 | 2 | Canis | 25644.0 | 3 | Canis lupus | 25644.0 | 4 |
| scf7180000041113 | 1523477 | 0.3953 | 100 | 64.6183 | Eukaryota | 93303.0 | 0 | Chordata | 93303.0 | 0 | Carnivora | 93303.0 | 0 | Canidae | 93303.0 | 0 | Canis | 93303.0 | 0 | Canis lupus | 93303.0 | 0 |
| scf7180000043066 | 577186 | 0.36 | 0 | 59.8915 | Eukaryota | 88402.0 | 0 | Chordata | 88402.0 | 0 | Carnivora | 88402.0 | 0 | Canidae | 88402.0 | 0 | Canis | 88402.0 | 0 | Canis lupus | 88402.0 | 0 |
| scf7180000044060 | 419493 | 0.4271 | 0 | 59.2848 | Eukaryota | 74904.0 | 0 | Chordata | 74904.0 | 0 | Carnivora | 74904.0 | 0 | Canidae | 74904.0 | 0 | Canis | 51188.0 | 1 | Canis lupus | 51188.0 | 1 |
| scf7180000044993 | 9894 | 0.4161 | 0 | 30.0747 | Eukaryota | 86037.0 | 0 | Chordata | 86037.0 | 0 | Carnivora | 86037.0 | 0 | Canidae | 86037.0 | 0 | Canis | 86037.0 | 0 | Canis lupus | 86037.0 | 0 |
| scf7180000048369 | 11211 | 0.4504 | 0 | 78.8592 | no-hit | 0.0 | 0 | no-hit | 0.0 | 0 | no-hit | 0.0 | 0 | no-hit | 0.0 | 0 | no-hit | 0.0 | 0 | no-hit | 0.0 | 0 |
| scf7180000056179 | 618217 | 0.4324 | 0 | 60.5283 | Eukaryota | 54892.0 | 0 | Chordata | 54892.0 | 0 | Carnivora | 54892.0 | 0 | Canidae | 54892.0 | 0 | Canis | 46509.0 | 1 | Canis lupus | 46509.0 | 1 |
| scf7180000045532 | 35176 | 0.3881 | 0 | 369.3659 | Eukaryota | 93941.0 | 0 | Chordata | 93941.0 | 0 | Carnivora | 93941.0 | 0 | Canidae | 93941.0 | 0 | Canis | 93941.0 | 0 | Canis lupus | 93941.0 | 0 |
| scf7180000055760 | 15215 | 0.4403 | 0 | 189.3443 | Eukaryota | 346.0 | 0 | Chordata | 346.0 | 0 | Tetraodontiformes | 346.0 | 0 | Tetraodontidae | 346.0 | 0 | Takifugu | 346.0 | 0 | Takifugu rubripes | 346.0 | 0 |
| scf7180000040609 | 2026413 | 0.4225 | 0 | 59.532 | Eukaryota | 52660.0 | 0 | Chordata | 52660.0 | 0 | Carnivora | 52660.0 | 0 | Canidae | 52660.0 | 0 | Canis | 35694.0 | 1 | Canis lupus | 35694.0 | 1 |
| scf7180000051566 | 11111 | 0.4454 | 0 | 791.4623 | Eukaryota | 1607.0 | 0 | Echinodermata | 881.0 | 2 | Forcipulatida | 881.0 | 2 | Asteriidae | 881.0 | 2 | Asterias | 881.0 | 2 | Asterias rubens | 881.0 | 2 |
| scf7180000043463 | 228209 | 0.4402 | 0 | 110.3505 | Eukaryota | 88403.0 | 0 | Platyhelminthes | 49005.0 | 1 | Strigeidida | 37182.0 | 4 | Schistosomatidae | 37182.0 | 4 | Schistosoma | 37182.0 | 4 | Schistosoma mansoni | 37182.0 | 4 |
| scf7180000045786 | 22410 | 0.4089 | 0 | 238.6725 | Eukaryota | 86151.0 | 0 | Chordata | 86151.0 | 0 | Carnivora | 86151.0 | 0 | Canidae | 86151.0 | 0 | Canis | 86151.0 | 0 | Canis lupus | 86151.0 | 0 |
| scf7180000042697 | 628140 | 0.3906 | 0 | 59.9168 | Eukaryota | 69687.0 | 0 | Chordata | 69687.0 | 0 | Carnivora | 69687.0 | 0 | Canidae | 69687.0 | 0 | Canis | 69687.0 | 0 | Canis lupus | 69687.0 | 0 |
| scf7180000041011 | 348687 | 0.459 | 0 | 66.0674 | Eukaryota | 82731.0 | 0 | Chordata | 82731.0 | 0 | Carnivora | 82731.0 | 0 | Canidae | 82731.0 | 0 | Canis | 82731.0 | 0 | Canis lupus | 82731.0 | 0 |
| scf7180000042610 | 507916 | 0.4827 | 0 | 55.7579 | Eukaryota | 86963.0 | 0 | Chordata | 86963.0 | 0 | Carnivora | 86963.0 | 0 | Canidae | 86963.0 | 0 | Canis | 86963.0 | 0 | Canis lupus | 86963.0 | 0 |
| scf7180000044587 | 350349 | 0.3513 | 0 | 60.495 | Eukaryota | 101045.0 | 0 | Chordata | 101045.0 | 0 | Carnivora | 101045.0 | 0 | Canidae | 101045.0 | 0 | Canis | 101045.0 | 0 | Canis lupus | 101045.0 | 0 |
| scf7180000053765 | 5083 | 0.5036 | 0 | 169.2722 | Eukaryota | 7154.0 | 0 | Chordata | 7154.0 | 0 | Carnivora | 7154.0 | 0 | Canidae | 7154.0 | 0 | Canis | 6633.0 | 1 | Canis lupus | 5668.0 | 2 |
| scf7180000051646 | 15394 | 0.4479 | 0 | 182.0123 | Eukaryota | 305.0 | 0 | Chordata | 305.0 | 0 | Kurtiformes | 305.0 | 0 | Apogonidae | 305.0 | 0 | Sphaeramia | 305.0 | 0 | Sphaeramia orbicularis | 305.0 | 0 |
| scf7180000043500 | 437428 | 0.3711 | 0 | 58.976 | Eukaryota | 63574.0 | 0 | Chordata | 63574.0 | 0 | Carnivora | 63574.0 | 0 | Canidae | 63574.0 | 0 | Canis | 54414.0 | 1 | Canis lupus | 54414.0 | 1 |
| scf7180000055545 | 31260 | 0.4402 | 0 | 247.1197 | Eukaryota | 53640.0 | 0 | Chordata | 38743.0 | 2 | Gadiformes | 32273.0 | 5 | Gadidae | 32273.0 | 5 | Gadus | 32273.0 | 5 | Gadus morhua | 32273.0 | 5 |
| scf7180000046668 | 99210 | 0.3968 | 0 | 59.768 | Eukaryota | 18759.0 | 1 | Chordata | 18759.0 | 1 | Carnivora | 15445.0 | 2 | Canidae | 15445.0 | 2 | Canis | 15445.0 | 2 | Canis lupus | 15445.0 | 2 |
| scf7180000041638 | 1004226 | 0.3841 | 0 | 60.3777 | Eukaryota | 75082.0 | 0 | Chordata | 75082.0 | 0 | Carnivora | 75082.0 | 0 | Canidae | 75082.0 | 0 | Canis | 75082.0 | 0 | Canis lupus | 75082.0 | 0 |
| scf7180000056049 | 451492 | 0.3709 | 0 | 536.5199 | Eukaryota | 88604.0 | 0 | Chordata | 88604.0 | 0 | Carnivora | 88604.0 | 0 | Canidae | 88604.0 | 0 | Canis | 88604.0 | 0 | Canis lupus | 88604.0 | 0 |
| scf7180000046383 | 129463 | 0.3572 | 0 | 60.9235 | Eukaryota | 69278.0 | 0 | Chordata | 69278.0 | 0 | Carnivora | 69278.0 | 0 | Canidae | 69278.0 | 0 | Canis | 69278.0 | 0 | Canis lupus | 69278.0 | 0 |
| scf7180000041204 | 119540 | 0.4718 | 0 | 58.3603 | Eukaryota | 60464.0 | 0 | Chordata | 60464.0 | 0 | Carnivora | 60464.0 | 0 | Canidae | 55839.0 | 1 | Canis | 50583.0 | 2 | Canis lupus | 50583.0 | 2 |
| scf7180000042957 | 586755 | 0.4216 | 0 | 60.8876 | Eukaryota | 90431.0 | 0 | Chordata | 90431.0 | 0 | Carnivora | 90431.0 | 0 | Canidae | 90431.0 | 0 | Canis | 90431.0 | 0 | Canis lupus | 90431.0 | 0 |

| | | | | | | | | | | | | | | | | | | | | | | |
|---|---|---|---|---|---|---|---|---|---|---|---|---|---|---|---|---|---|---|---|---|---|---|
| scf7180000053826 | 5592 | 0.4489 | 0 | 756.1539 | Eukaryota | 5330.0 | 0 | Chordata | 2885.0 | 2 | Tetraodontiformes | 2000.0 | 4 | Tetraodontidae | 2000.0 | 4 | Takifugu | 2000.0 | 4 | Takifugu rubripes | 2000.0 | 4 |
| scf7180000047756 | 5176 | 0.4484 | 0 | 103.7403 | no-hit | 0.0 | 0 | no-hit | 0.0 | 0 | no-hit | 0.0 | 0 | no-hit | 0.0 | 0 | no-hit | 0.0 | 0 | no-hit | 0.0 | 0 |
| scf7180000045469 | 226902 | 0.3883 | 0 | 59.2854 | Eukaryota | 30071.0 | 0 | Chordata | 30071.0 | 0 | Carnivora | 30071.0 | 0 | Canidae | 30071.0 | 0 | Canis | 30071.0 | 0 | Canis lupus | 30071.0 | 0 |
| scf7180000048759 | 43755 | 0.5073 | 0 | 57.6875 | Eukaryota | 79916.0 | 1 | Chordata | 79916.0 | 1 | Carnivora | 79916.0 | 1 | Canidae | 79916.0 | 1 | Canis | 79916.0 | 1 | Canis lupus | 79916.0 | 1 |
| scf7180000052039 | 2577 | 0.4517 | 0 | 160.4526 | Eukaryota | 900.0 | 0 | Chordata | 900.0 | 0 | Kurtiformes | 406.0 | 2 | Apogonidae | 406.0 | 2 | Sphaeramia | 406.0 | 2 | Sphaeramia orbicularis | 406.0 | 2 |
| scf7180000056327 | 243629 | 0.3748 | 0 | 58.1763 | Eukaryota | 23545.0 | 0 | Chordata | 23545.0 | 0 | Carnivora | 23545.0 | 0 | Canidae | 23545.0 | 0 | Canis | 23545.0 | 0 | Canis lupus | 23545.0 | 0 |
| scf7180000045506 | 197330 | 0.4405 | 0 | 59.4698 | Eukaryota | 44265.0 | 0 | Chordata | 44265.0 | 0 | Carnivora | 44265.0 | 0 | Canidae | 19994.0 | 3 | Ursus | 11753.0 | 4 | Canis lupus | 10002.0 | 5 |
| scf7180000042912 | 516835 | 0.4025 | 0 | 60.5799 | Eukaryota | 55524.0 | 0 | Chordata | 55524.0 | 0 | Carnivora | 55524.0 | 0 | Canidae | 55524.0 | 0 | Canis | 55524.0 | 0 | Canis lupus | 55524.0 | 0 |
| scf7180000049342 | 12226 | 0.444 | 0 | 159.1297 | no-hit | 0.0 | 0 | no-hit | 0.0 | 0 | no-hit | 0.0 | 0 | no-hit | 0.0 | 0 | no-hit | 0.0 | 0 | no-hit | 0.0 | 0 |
| scf7180000040583 | 725195 | 0.413 | 0 | 59.7164 | Eukaryota | 92533.0 | 0 | Chordata | 92533.0 | 0 | Carnivora | 92533.0 | 0 | Canidae | 92533.0 | 0 | Canis | 92533.0 | 0 | Canis lupus | 92533.0 | 0 |
| scf7180000054948 | 16585 | 0.4368 | 0 | 274.5429 | Eukaryota | 5109.0 | 0 | Chordata | 4185.0 | 2 | Salmoniformes | 2053.0 | 6 | Salmonidae | 2053.0 | 6 | Salmo | 1849.0 | 7 | Salmo trutta | 1849.0 | 7 |
| scf7180000056584 | 25669 | 0.3797 | 0 | 56.8558 | Eukaryota | 16545.0 | 0 | Chordata | 16545.0 | 0 | Carnivora | 16545.0 | 0 | Canidae | 16545.0 | 0 | Canis | 16545.0 | 0 | Canis lupus | 16545.0 | 0 |
| scf7180000056389 | 565494 | 0.4163 | 0 | 59.3619 | Eukaryota | 50546.0 | 0 | Chordata | 50546.0 | 0 | Carnivora | 50546.0 | 0 | Canidae | 50546.0 | 0 | Canis | 50546.0 | 0 | Canis lupus | 50546.0 | 0 |
| scf7180000048665 | 2768 | 0.4505 | 0 | 111.3541 | Eukaryota | 570.0 | 0 | Euglenozoa | 300.0 | 1 | Kinetoplastida | 300.0 | 1 | Trypanosomatidae | 300.0 | 1 | Leishmania | 300.0 | 1 | Leishmania infantum | 300.0 | 1 |
| scf7180000051694 | 4455 | 0.4352 | 0 | 21.9039 | Eukaryota | 26139.0 | 0 | Chordata | 13363.0 | 2 | Gadiformes | 8994.0 | 4 | Gadidae | 8994.0 | 5 | Gadus | 8994.0 | 5 | Gadus morhua | 8994.0 | 5 |
| scf7180000051462 | 9446 | 0.4439 | 0 | 72.7404 | Eukaryota | 882.0 | 0 | Platyhelminthes | 462.0 | 1 | Polystomatidea | 462.0 | 1 | Polystomatidae | 462.0 | 1 | Protopolystoma | 462.0 | 1 | Protopolystoma xenopodis | 462.0 | 1 |
| scf7180000043924 | 217934 | 0.4412 | 0 | 60.1101 | Eukaryota | 35681.0 | 0 | Chordata | 35681.0 | 0 | Carnivora | 35681.0 | 0 | Canidae | 35681.0 | 0 | Canis | 27372.0 | 1 | Canis lupus | 27372.0 | 1 |
| scf7180000052059 | 14250 | 0.5048 | 0 | 947.8692 | Eukaryota | 9157.0 | 0 | Chordata | 9157.0 | 0 | Carnivora | 9157.0 | 0 | Canidae | 9157.0 | 0 | Canis | 7409.0 | 1 | Canis lupus | 6126.0 | 2 |
| scf7180000053695 | 3204 | 0.4482 | 0 | 629.9972 | no-hit | 0.0 | 0 | no-hit | 0.0 | 0 | no-hit | 0.0 | 0 | no-hit | 0.0 | 0 | no-hit | 0.0 | 0 | no-hit | 0.0 | 0 |
| scf7180000052613 | 4753 | 0.4494 | 0 | 1053.4641 | Eukaryota | 2546.0 | 0 | Chordata | 2546.0 | 0 | Carnivora | 2546.0 | 0 | Canidae | 2546.0 | 0 | Canis | 2546.0 | 0 | Canis lupus | 2546.0 | 0 |
| scf7180000054576 | 64356 | 0.37 | 0 | 1352.1159 | Eukaryota | 10231.0 | 0 | Chordata | 10231.0 | 0 | Carnivora | 8856.0 | 1 | Canidae | 8856.0 | 1 | Canis | 8856.0 | 1 | Canis lupus | 8856.0 | 1 |
| scf7180000043411 | 416961 | 0.3935 | 0 | 59.2005 | Eukaryota | 33391.0 | 0 | Chordata | 33391.0 | 0 | Carnivora | 33391.0 | 0 | Canidae | 33391.0 | 0 | Canis | 33391.0 | 0 | Canis lupus | 33391.0 | 0 |
| scf7180000042531 | 611090 | 0.4109 | 0 | 59.83 | Eukaryota | 35881.0 | 0 | Chordata | 35881.0 | 0 | Carnivora | 35881.0 | 0 | Canidae | 35881.0 | 0 | Canis | 25152.0 | 1 | Canis lupus | 25152.0 | 1 |
| scf7180000054359 | 95399 | 0.4709 | 0 | 88.8569 | Eukaryota | 22669.0 | 0 | Chordata | 22669.0 | 0 | Carnivora | 22669.0 | 0 | Canidae | 22669.0 | 0 | Canis | 22669.0 | 0 | Canis lupus | 22669.0 | 0 |
| scf7180000046439 | 141216 | 0.3702 | 0 | 60.1041 | Eukaryota | 86435.0 | 0 | Chordata | 86435.0 | 0 | Carnivora | 86435.0 | 0 | Canidae | 86435.0 | 0 | Canis | 86435.0 | 0 | Canis lupus | 86435.0 | 0 |
| scf7180000055528 | 43378 | 0.4527 | 0 | 355.0568 | Eukaryota | 42344.0 | 0 | Chordata | 36390.0 | 2 | Gadiformes | 24674.0 | 5 | Gadidae | 24674.0 | 5 | Gadus | 24674.0 | 5 | Gadus morhua | 24674.0 | 5 |
| scf7180000044965 | 63690 | 0.3522 | 0 | 58.6461 | Eukaryota | 81190.0 | 0 | Chordata | 81190.0 | 0 | Carnivora | 81190.0 | 0 | Canidae | 81190.0 | 0 | Canis | 81190.0 | 0 | Canis lupus | 81190.0 | 0 |
| scf7180000041890 | 844794 | 0.4636 | 0 | 59.7104 | Eukaryota | 62697.0 | 0 | Chordata | 62697.0 | 0 | Carnivora | 62697.0 | 0 | Canidae | 62697.0 | 0 | Canis | 43843.0 | 1 | Canis lupus | 43843.0 | 1 |
| scf7180000048155 | 8658 | 0.441 | 0 | 124.684 | Eukaryota | 784.0 | 0 | Platyhelminthes | 407.0 | 1 | Polystomatidea | 407.0 | 1 | Polystomatidae | 407.0 | 1 | Protopolystoma | 407.0 | 1 | Protopolystoma xenopodis | 407.0 | 1 |
| scf7180000052711 | 4440 | 0.4538 | 0 | 281.6977 | Eukaryota | 1819.0 | 0 | Platyhelminthes | 760.0 | 2 | Polystomatidea | 760.0 | 2 | Polystomatidae | 760.0 | 2 | Protopolystoma | 760.0 | 2 | Protopolystoma xenopodis | 760.0 | 2 |
| scf7180000047973 | 32221 | 0.4501 | 0 | 1623.8609 | Eukaryota | 20325.0 | 0 | Chordata | 18915.0 | 2 | Gadiformes | 14140.0 | 7 | Gadidae | 14140.0 | 7 | Gadus | 14140.0 | 7 | Gadus morhua | 14140.0 | 7 |
| scf7180000053086 | 8847 | 0.4473 | 0 | 83.798 | no-hit | 0.0 | 0 | no-hit | 0.0 | 0 | no-hit | 0.0 | 0 | no-hit | 0.0 | 0 | no-hit | 0.0 | 0 | no-hit | 0.0 | 0 |
| scf7180000046897 | 107915 | 0.3869 | 0 | 59.2293 | Eukaryota | 7224.0 | 0 | Chordata | 7224.0 | 0 | Primates | 4589.0 | 1 | Canidae | 2635.0 | 2 | Canis | 2635.0 | 2 | Canis lupus | 2635.0 | 2 |
| scf7180000044049 | 74967 | 0.6203 | 0 | 69.8472 | Eukaryota | 38880.0 | 0 | Chordata | 38880.0 | 0 | Carnivora | 38880.0 | 0 | Canidae | 38880.0 | 0 | Canis | 34615.0 | 1 | Canis lupus | 34615.0 | 1 |
| scf7180000049916 | 4158 | 0.4483 | 0 | 81.8892 | Eukaryota | 12080.0 | 0 | Chordata | 8546.0 | 2 | Strigeidida | 2998.0 | 6 | Schistosomatidae | 2998.0 | 6 | Schistosoma | 2998.0 | 6 | Schistosoma mansoni | 2998.0 | 6 |
| scf7180000050381 | 14302 | 0.4518 | 0 | 1955.6378 | Eukaryota | 21374.0 | 1 | Chordata | 17375.0 | 2 | Tetraodontiformes | 7927.0 | 4 | Tetraodontidae | 7927.0 | 4 | Takifugu | 7927.0 | 4 | Takifugu rubripes | 7927.0 | 4 |
| scf7180000040659 | 101139 | 0.4669 | 0 | 55.1329 | Eukaryota | 10590.0 | 1 | Chordata | 10590.0 | 1 | Carnivora | 3998.0 | 4 | Canidae | 3998.0 | 4 | Canis | 3998.0 | 5 | Canis lupus | 3998.0 | 5 |
| scf7180000051557 | 2444 | 0.4574 | 0 | 303.9781 | Eukaryota | 450.0 | 0 | Streptophyta | 316.0 | 1 | Fabales | 316.0 | 1 | Fabaceae | 316.0 | 1 | Cicer | 316.0 | 1 | Cicer arietinum | 316.0 | 1 |
| scf7180000051450 | 9534 | 0.4514 | 0 | 963.74 | no-hit | 0.0 | 0 | no-hit | 0.0 | 0 | no-hit | 0.0 | 0 | no-hit | 0.0 | 0 | no-hit | 0.0 | 0 | no-hit | 0.0 | 0 |
| scf7180000046740 | 123063 | 0.384 | 0 | 61.0967 | Eukaryota | 26757.0 | 0 | Chordata | 26757.0 | 0 | Carnivora | 24507.0 | 1 | Canidae | 24507.0 | 1 | Canis | 24507.0 | 1 | Canis lupus | 24507.0 | 1 |
| scf7180000049653 | 6381 | 0.4292 | 0 | 124.359 | no-hit | 0.0 | 0 | no-hit | 0.0 | 0 | no-hit | 0.0 | 0 | no-hit | 0.0 | 0 | no-hit | 0.0 | 0 | no-hit | 0.0 | 0 |
| scf7180000048388 | 46241 | 0.4601 | 0 | 55.4105 | Eukaryota | 10380.0 | 0 | Chordata | 10380.0 | 0 | Carnivora | 10380.0 | 0 | Ursidae | 5373.0 | 4 | Ursus | 4414.0 | 5 | Ursus arctos | 3294.0 | 6 |
| scf7180000053825 | 6343 | 0.4588 | 0 | 126.0066 | Eukaryota | 374.0 | 0 | Platyhelminthes | 374.0 | 0 | Polystomatidea | 374.0 | 0 | Polystomatidae | 374.0 | 0 | Protopolystoma | 374.0 | 0 | Protopolystoma xenopodis | 374.0 | 0 |
| scf7180000053043 | 13121 | 0.5058 | 0 | 2352.2053 | Eukaryota | 9365.0 | 0 | Chordata | 9365.0 | 0 | Carnivora | 9365.0 | 0 | Canidae | 9365.0 | 0 | Canis | 7576.0 | 1 | Canis lupus | 6260.0 | 2 |
| scf7180000047398 | 119859 | 0.3702 | 0 | 59.9651 | Eukaryota | 74202.0 | 0 | Chordata | 74202.0 | 0 | Carnivora | 74202.0 | 0 | Canidae | 74202.0 | 0 | Canis | 65962.0 | 1 | Canis lupus | 65962.0 | 1 |
| scf7180000043278 | 313396 | 0.4827 | 0 | 59.8351 | Eukaryota | 43821.0 | 0 | Chordata | 43821.0 | 0 | Carnivora | 43821.0 | 0 | Canidae | 43821.0 | 0 | Canis | 29604.0 | 1 | Canis lupus | 29604.0 | 1 |
| scf7180000050477 | 14008 | 0.5116 | 0 | 615.7329 | Eukaryota | 21865.0 | 0 | Chordata | 21865.0 | 0 | Carnivora | 21865.0 | 0 | Canidae | 21865.0 | 0 | Canis | 21865.0 | 0 | Canis lupus | 21865.0 | 0 |
| scf7180000050259 | 33727 | 0.4412 | 0 | 136.4781 | Eukaryota | 3258.0 | 0 | Chordata | 2705.0 | 1 | Tetraodontiformes | 1827.0 | 5 | Tetraodontidae | 1827.0 | 5 | Takifugu | 1827.0 | 5 | Takifugu rubripes | 1827.0 | 5 |
| scf7180000042902 | 538235 | 0.4591 | 0 | 57.618 | Eukaryota | 56310.0 | 0 | Chordata | 56310.0 | 0 | Carnivora | 56310.0 | 0 | Canidae | 50440.0 | 1 | Canis | 50440.0 | 2 | Canis lupus | 50440.0 | 2 |
| scf7180000054346 | 34109 | 0.4629 | 0 | 139.0655 | Eukaryota | 46841.0 | 0 | Chordata | 46841.0 | 0 | Carnivora | 46841.0 | 0 | Canidae | 46841.0 | 0 | Canis | 46841.0 | 0 | Canis lupus | 46841.0 | 0 |
| scf7180000052037 | 4499 | 0.449 | 0 | 203.3911 | no-hit | 0.0 | 0 | no-hit | 0.0 | 0 | no-hit | 0.0 | 0 | no-hit | 0.0 | 0 | no-hit | 0.0 | 0 | no-hit | 0.0 | 0 |
| scf7180000052983 | 3588 | 0.444 | 0 | 606.4501 | no-hit | 0.0 | 0 | no-hit | 0.0 | 0 | no-hit | 0.0 | 0 | no-hit | 0.0 | 0 | no-hit | 0.0 | 0 | no-hit | 0.0 | 0 |
| scf7180000056353 | 686743 | 0.3417 | 0 | 58.7426 | Eukaryota | 41545.0 | 0 | Chordata | 41545.0 | 0 | Carnivora | 41545.0 | 0 | Canidae | 41545.0 | 0 | Vulpes | 20782.0 | 1 | Vulpes vulpes | 20782.0 | 1 |
| scf7180000044017 | 38422 | 0.3976 | 0 | 57.4725 | Eukaryota | 39667.0 | 0 | Chordata | 39667.0 | 0 | Carnivora | 39667.0 | 0 | Canidae | 39667.0 | 0 | Canis | 39667.0 | 0 | Canis lupus | 39667.0 | 0 |
| scf7180000055152 | 19523 | 0.4411 | 0 | 136.1203 | Eukaryota | 5643.0 | 0 | Chordata | 3888.0 | 2 | Salmoniformes | 3268.0 | 4 | Salmonidae | 3268.0 | 4 | Salmo | 3268.0 | 4 | Salmo trutta | 3268.0 | 4 |
| scf7180000046497 | 49666 | 0.3837 | 0 | 55.6001 | Eukaryota | 11404.0 | 0 | Chordata | 11404.0 | 0 | Carnivora | 11404.0 | 0 | Canidae | 11404.0 | 0 | Canis | 11404.0 | 0 | Canis lupus | 11404.0 | 0 |
| scf7180000045096 | 214007 | 0.5024 | 0 | 59.0274 | Eukaryota | 25921.0 | 0 | Chordata | 25921.0 | 0 | Carnivora | 25921.0 | 0 | Canidae | 25921.0 | 0 | Canis | 25921.0 | 0 | Canis lupus | 25921.0 | 0 |
| scf7180000052220 | 24258 | 0.3694 | 0 | 59.9377 | Eukaryota | 23517.0 | 1 | Chordata | 23517.0 | 1 | Primates | 12160.0 | 4 | Hominidae | 12160.0 | 4 | Homo | 12160.0 | 5 | Homo sapiens | 12160.0 | 5 |
| scf7180000055955 | 22391 | 0.4439 | 0 | 242.2162 | Eukaryota | 30176.0 | 0 | Chordata | 27233.0 | 2 | Gadiformes | 18871.0 | 6 | Gadidae | 18871.0 | 6 | Gadus | 18871.0 | 6 | Gadus morhua | 18871.0 | 6 |
| scf7180000051143 | 6708 | 0.4395 | 0 | 161.4531 | Eukaryota | 2880.0 | 0 | Platyhelminthes | 1910.0 | 2 | Polystomatidea | 1910.0 | 2 | Polystomatidae | 1910.0 | 2 | Protopolystoma | 1910.0 | 2 | Protopolystoma xenopodis | 1910.0 | 2 |
| scf7180000044472 | 369018 | 0.3481 | 0 | 59.3381 | Eukaryota | 44730.0 | 0 | Chordata | 44730.0 | 0 | Carnivora | 44730.0 | 0 | Canidae | 44730.0 | 0 | Canis | 31488.0 | 1 | Canis lupus | 31488.0 | 1 |
| scf7180000046089 | 96417 | 0.3891 | 0 | 56.5402 | Eukaryota | 85078.0 | 0 | Chordata | 85078.0 | 0 | Carnivora | 85078.0 | 0 | Canidae | 85078.0 | 0 | Canis | 85078.0 | 0 | Canis lupus | 85078.0 | 0 |
| scf7180000047887 | 59644 | 0.444 | 0 | 210.488 | Eukaryota | 58628.0 | 0 | Chordata | 58628.0 | 0 | Carnivora | 58628.0 | 0 | Canidae | 58628.0 | 0 | Canis | 58628.0 | 0 | Canis lupus | 58628.0 | 0 |
| scf7180000041039 | 1359457 | 0.3552 | 0 | 80.2434 | Eukaryota | 92106.0 | 0 | Chordata | 92106.0 | 0 | Carnivora | 92106.0 | 0 | Canidae | 92106.0 | 0 | Canis | 92106.0 | 0 | Canis lupus | 92106.0 | 0 |
| scf7180000046132 | 50617 | 0.4418 | 0 | 112.2237 | Eukaryota | 58288.0 | 0 | Chordata | 43402.0 | 1 | Gadiformes | 32752.0 | 5 | Gadidae | 32752.0 | 5 | Gadus | 32752.0 | 5 | Gadus morhua | 32752.0 | 5 |
| scf7180000049038 | 45839 | 0.3739 | 0 | 59.6346 | Eukaryota | 11807.0 | 0 | Chordata | 11807.0 | 0 | Carnivora | 10580.0 | 1 | Canidae | 9228.0 | 2 | Canis | 9228.0 | 2 | Canis lupus | 9228.0 | 2 |
| scf7180000042267 | 241285 | 0.3893 | 0 | 59.8787 | Eukaryota | 94679.0 | 0 | Chordata | 94679.0 | 0 | Carnivora | 94679.0 | 0 | Canidae | 94679.0 | 0 | Canis | 94679.0 | 0 | Canis lupus | 94679.0 | 0 |
| scf7180000046040 | 150965 | 0.4329 | 0 | 59.9786 | Eukaryota | 24346.0 | 0 | Chordata | 24346.0 | 0 | Carnivora | 24346.0 | 0 | Canidae | 13814.0 | 3 | Canis | 10362.0 | 6 | Canis lupus | 10362.0 | 7 |
| scf7180000046539 | 125877 | 0.4431 | 0 | 59.0414 | Eukaryota | 27048.0 | 0 | Chordata | 27048.0 | 0 | Carnivora | 27048.0 | 0 | Canidae | 27048.0 | 0 | Canis | 23856.0 | 1 | Canis lupus | 23856.0 | 1 |
| scf7180000043385 | 106817 | 0.445 | 0 | 56.4946 | Eukaryota | 11775.0 | 0 | Chordata | 11775.0 | 0 | Primates | 4633.0 | 3 | Hominidae | 4633.0 | 5 | Homo | 4633.0 | 5 | Homo sapiens | 4633.0 | 5 |
| scf7180000041840 | 536601 | 0.3371 | 0 | 58.1382 | Eukaryota | 56130.0 | 0 | Chordata | 56130.0 | 0 | Carnivora | 56130.0 | 0 | Canidae | 56130.0 | 0 | Canis | 56130.0 | 0 | Canis lupus | 56130.0 | 0 |
| scf7180000056106 | 26367 | 0.4302 | 0 | 142.5046 | Eukaryota | 54784.0 | 0 | Chordata | 54784.0 | 0 | Carnivora | 54784.0 | 0 | Canidae | 54784.0 | 0 | Canis | 54784.0 | 0 | Canis lupus | 54784.0 | 0 |
| scf7180000045282 | 13482 | 0.377 | 0 | 89.3546 | Eukaryota | 51036.0 | 0 | Chordata | 51036.0 | 0 | Carnivora | 51036.0 | 0 | Canidae | 51036.0 | 0 | Canis | 51036.0 | 0 | Canis lupus | 51036.0 | 0 |
| scf7180000053333 | 13279 | 0.4475 | 0 | 480.9986 | no-hit | 0.0 | 0 | no-hit | 0.0 | 0 | no-hit | 0.0 | 0 | no-hit | 0.0 | 0 | no-hit | 0.0 | 0 | no-hit | 0.0 | 0 |
| scf7180000043649 | 5854 | 0.3466 | 0 | 30.0269 | Eukaryota | 2968.0 | 0 | Chordata | 2968.0 | 0 | Carnivora | 2968.0 | 0 | Canidae | 2968.0 | 0 | Canis | 2968.0 | 0 | Canis lupus | 2968.0 | 0 |
| scf7180000047252 | 23991 | 0.382 | 0 | 50.7182 | Eukaryota | 47422.0 | 0 | Chordata | 47422.0 | 0 | Carnivora | 47422.0 | 0 | Canidae | 47422.0 | 0 | Canis | 47422.0 | 0 | Canis lupus | 47422.0 | 0 |
| scf7180000054313 | 783547 | 0.3714 | 0 | 60.2982 | Eukaryota | 178623.0 | 0 | Chordata | 178623.0 | 0 | Carnivora | 178623.0 | 0 | Canidae | 165192.0 | 1 | Canis | 165192.0 | 1 | Canis lupus | 165192.0 | 1 |
| scf7180000041663 | 859238 | 0.3781 | 0 | 60.215 | Eukaryota | 25297.0 | 1 | Chordata | 25297.0 | 1 | Carnivora | 15335.0 | 3 | Canidae | 15335.0 | 3 | Canis | 12841.0 | 4 | Canis lupus | 12841.0 | 4 |
| scf7180000055240 | 51278 | 0.4432 | 0 | 149.6515 | Eukaryota | 4493.0 | 0 | Platyhelminthes | 2258.0 | 2 | Polystomatidea | 2258.0 | 3 | Polystomatidae | 2258.0 | 3 | Protopolystoma | 2258.0 | 3 | Protopolystoma xenopodis | 2258.0 | 3 |
| scf7180000056177 | 177472 | 0.4013 | 0 | 59.6294 | Eukaryota | 29050.0 | 0 | Chordata | 29050.0 | 0 | Carnivora | 29050.0 | 0 | Canidae | 29050.0 | 0 | Vulpes | 15124.0 | 1 | Vulpes vulpes | 15124.0 | 1 |
| scf7180000055775 | 11434 | 0.4464 | 0 | 108.2497 | no-hit | 0.0 | 0 | no-hit | 0.0 | 0 | no-hit | 0.0 | 0 | no-hit | 0.0 | 0 | no-hit | 0.0 | 0 | no-hit | 0.0 | 0 |
| scf7180000045844 | 8999 | 0.3643 | 0 | 27.6585 | Eukaryota | 25096.0 | 0 | Chordata | 25096.0 | 0 | Carnivora | 25096.0 | 0 | Canidae | 25096.0 | 0 | Canis | 25096.0 | 0 | Canis lupus | 25096.0 | 0 |
| scf7180000050215 | 10566 | 0.4384 | 0 | 77.7625 | no-hit | 0.0 | 0 | no-hit | 0.0 | 0 | no-hit | 0.0 | 0 | no-hit | 0.0 | 0 | no-hit | 0.0 | 0 | no-hit | 0.0 | 0 |
| scf7180000050412 | 9105 | 0.4522 | 0 | 1780.9286 | Eukaryota | 3107.0 | 0 | Platyhelminthes | 1705.0 | 3 | Polystomatidea | 1705.0 | 3 | Polystomatidae | 1705.0 | 3 | Protopolystoma | 1705.0 | 3 | Protopolystoma xenopodis | 1705.0 | 3 |
| scf7180000043744 | 5899 | 0.4475 | 0 | 26.1585 | Eukaryota | 35210.0 | 0 | Chordata | 35210.0 | 0 | Carnivora | 35210.0 | 0 | Canidae | 35210.0 | 0 | Canis | 35210.0 | 0 | Canis lupus | 35210.0 | 0 |
| scf7180000043233 | 357430 | 0.4355 | 0 | 59.1084 | Eukaryota | 32020.0 | 0 | Chordata | 32020.0 | 0 | Carnivora | 32020.0 | 0 | Canidae | 32020.0 | 0 | Canis | 32020.0 | 0 | Canis lupus | 32020.0 | 0 |
| scf7180000055421 | 10620 | 0.4392 | 0 | 153.264 | Eukaryota | 765.0 | 0 | Streptophyta | 765.0 | 0 | Fabales | 765.0 | 0 | Fabaceae | 765.0 | 0 | Vigna | 765.0 | 0 | Vigna unguiculata | 765.0 | 0 |
| scf7180000044480 | 266876 | 0.394 | 0 | 60.9227 | Eukaryota | 39943.0 | 0 | Chordata | 39943.0 | 0 | Carnivora | 39943.0 | 0 | Canidae | 36090.0 | 1 | Canis | 36090.0 | 1 | Canis lupus | 36090.0 | 1 |
| scf7180000046712 | 97555 | 0.415 | 0 | 58.9588 | Eukaryota | 20667.0 | 0 | Chordata | 20667.0 | 0 | Carnivora | 20667.0 | 0 | Canidae | 14791.0 | 1 | Canis | 10558.0 | 2 | Canis lupus | 10558.0 | 2 |
| scf7180000040826 | 112650 | 0.3753 | 0 | 58.6617 | Eukaryota | 84535.0 | 0 | Chordata | 84535.0 | 0 | Carnivora | 84535.0 | 0 | Canidae | 84535.0 | 0 | Canis | 84535.0 | 0 | Canis lupus | 84535.0 | 0 |
| scf7180000053190 | 2638 | 0.4556 | 0 | 477.2801 | no-hit | 0.0 | 0 | no-hit | 0.0 | 0 | no-hit | 0.0 | 0 | no-hit | 0.0 | 0 | no-hit | 0.0 | 0 | no-hit | 0.0 | 0 |
| scf7180000053611 | 7532 | 0.4587 | 0 | 64.8299 | Eukaryota | 250.0 | 0 | Platyhelminthes | 250.0 | 0 | Polystomatidea | 250.0 | 0 | Polystomatidae | 250.0 | 0 | Protopolystoma | 250.0 | 0 | Protopolystoma xenopodis | 250.0 | 0 |
| scf7180000049887 | 19052 | 0.4491 | 0 | 569.9064 | Eukaryota | 27140.0 | 0 | Chordata | 23670.0 | 2 | Gadiformes | 18791.0 | 6 | Gadidae | 18791.0 | 6 | Gadus | 18791.0 | 6 | Gadus morhua | 18791.0 | 6 |
| scf7180000049972 | 41364 | 0.333 | 0 | 59.9211 | Eukaryota | 7143.0 | 0 | Chordata | 7143.0 | 0 | Primates | 5781.0 | 1 | Hominidae | 4538.0 | 3 | Homo | 4538.0 | 3 | Homo sapiens | 4538.0 | 3 |
| scf7180000048139 | 5479 | 0.4481 | 0 | 144.1434 | Eukaryota | 694.0 | 0 | Chordata | 492.0 | 1 | Tetraodontiformes | 492.0 | 1 | Tetraodontidae | 492.0 | 1 | Takifugu | 492.0 | 1 | Takifugu rubripes | 492.0 | 1 |
| scf7180000046802 | 55428 | 0.3984 | 0 | 55.3177 | Eukaryota | 24874.0 | 0 | Chordata | 24874.0 | 0 | Carnivora | 24874.0 | 0 | Canidae | 24874.0 | 0 | Canis | 24874.0 | 0 | Canis lupus | 24874.0 | 0 |
| scf7180000042772 | 1730 | 0.4445 | 0 | 26.1722 | Eukaryota | 3189.0 | 0 | Chordata | 3189.0 | 0 | Carnivora | 3189.0 | 0 | Canidae | 3189.0 | 0 | Canis | 3189.0 | 0 | Canis lupus | 3189.0 | 0 |
| scf7180000044155 | 415601 | 0.4255 | 0 | 60.6474 | Eukaryota | 36352.0 | 0 | Chordata | 36352.0 | 0 | Carnivora | 36352.0 | 0 | Canidae | 36352.0 | 0 | Canis | 36352.0 | 0 | Canis lupus | 36352.0 | 0 |
| scf7180000043461 | 105351 | 0.374 | 0 | 60.3646 | Eukaryota | 52658.0 | 0 | Chordata | 52658.0 | 0 | Carnivora | 52658.0 | 0 | Canidae | 52658.0 | 0 | Canis | 52658.0 | 0 | Canis lupus | 52658.0 | 0 |
| scf7180000047260 | 26568 | 0.4091 | 0 | 51.69 | Eukaryota | 5912.0 | 0 | Chordata | 5912.0 | 0 | Carnivora | 5202.0 | 1 | Canidae | 2305.0 | 3 | Canis | 2305.0 | 5 | Canis lupus | 2305.0 | 5 |
| scf7180000047192 | 92588 | 0.3961 | 0 | 60.8233 | Eukaryota | 44691.0 | 0 | Chordata | 44691.0 | 0 | Carnivora | 44691.0 | 0 | Canidae | 44691.0 | 0 | Canis | 28118.0 | 1 | Canis lupus | 28118.0 | 1 |
| scf7180000051647 | 9425 | 0.5091 | 0 | 146.8955 | Eukaryota | 8240.0 | 0 | Chordata | 8240.0 | 0 | Carnivora | 8240.0 | 0 | Canidae | 8240.0 | 0 | Canis | 6677.0 | 1 | Canis lupus | 5540.0 | 2 |
| scf7180000053581 | 4463 | 0.4508 | 0 | 342.254 | no-hit | 0.0 | 0 | no-hit | 0.0 | 0 | no-hit | 0.0 | 0 | no-hit | 0.0 | 0 | no-hit | 0.0 | 0 | no-hit | 0.0 | 0 |
| scf7180000052025 | 4836 | 0.4456 | 0 | 414.3525 | no-hit | 0.0 | 0 | no-hit | 0.0 | 0 | no-hit | 0.0 | 0 | no-hit | 0.0 | 0 | no-hit | 0.0 | 0 | no-hit | 0.0 | 0 |
| scf7180000053785 | 3769 | 0.4471 | 0 | 198.7364 | Eukaryota | 562.0 | 0 | Streptophyta | 417.0 | 1 | Fabales | 417.0 | 1 | Fabaceae | 417.0 | 1 | Cicer | 417.0 | 1 | Cicer arietinum | 417.0 | 1 |
| scf7180000041799 | 35505 | 0.405 | 0 | 65.1773 | Eukaryota | 37809.0 | 0 | Chordata | 37809.0 | 0 | Carnivora | 37809.0 | 0 | Canidae | 37809.0 | 0 | Canis | 37809.0 | 0 | Canis lupus | 37809.0 | 0 |
| scf7180000041927 | 86976 | 0.3679 | 0 | 52.1657 | Eukaryota | 83294.0 | 0 | Chordata | 83294.0 | 0 | Carnivora | 83294.0 | 0 | Canidae | 83294.0 | 0 | Canis | 83294.0 | 0 | Canis lupus | 83294.0 | 0 |
| scf7180000046079 | 140543 | 0.369 | 0 | 59.9827 | Eukaryota | 29165.0 | 0 | Chordata | 29165.0 | 0 | Carnivora | 29165.0 | 0 | Canidae | 29165.0 | 0 | Canis | 29165.0 | 0 | Canis lupus | 29165.0 | 0 |
| scf7180000056319 | 633391 | 0.4096 | 0 | 60.4843 | Eukaryota | 87865.0 | 0 | Chordata | 87865.0 | 0 | Carnivora | 87865.0 | 0 | Canidae | 87865.0 | 0 | Canis | 79037.0 | 1 | Canis lupus | 79037.0 | 1 |
| scf7180000051632 | 2768 | 0.4411 | 0 | 59.649 | Eukaryota | 474.0 | 0 | Platyhelminthes | 474.0 | 0 | Polystomatidea | 474.0 | 0 | Polystomatidae | 474.0 | 0 | Protopolystoma | 474.0 | 0 | Protopolystoma xenopodis | 474.0 | 0 |
| scf7180000053248 | 3677 | 0.4455 | 0 | 65.6383 | Eukaryota | 2314.0 | 0 | Chordata | 1550.0 | 1 | Salmoniformes | 1550.0 | 1 | Salmonidae | 1550.0 | 1 | Salmo | 1550.0 | 1 | Salmo trutta | 1550.0 | 1 |
| scf7180000053794 | 4587 | 0.4397 | 0 | 80.0453 | Eukaryota | 1917.0 | 0 | Chordata | 1630.0 | 1 | Columbiformes | 1630.0 | 1 | Columbidae | 1630.0 | 1 | Streptopelia | 1630.0 | 1 | Streptopelia turtur | 1630.0 | 1 |
| scf7180000055777 | 6654 | 0.4319 | 0 | 205.0576 | no-hit | 0.0 | 0 | no-hit | 0.0 | 0 | no-hit | 0.0 | 0 | no-hit | 0.0 | 0 | no-hit | 0.0 | 0 | no-hit | 0.0 | 0 |
| scf7180000055559 | 71355 | 0.4386 | 0 | 125.576 | Eukaryota | 11412.0 | 0 | Chordata | 11412.0 | 0 | Carnivora | 11412.0 | 0 | Otariidae | 4324.0 | 3 | Canis | 2596.0 | 5 | Canis lupus | 2596.0 | 5 |
| scf7180000040599 | 2516018 | 0.4021 | 0 | 60.7318 | Eukaryota | 104502.0 | 0 | Chordata | 104502.0 | 0 | Carnivora | 104502.0 | 0 | Canidae | 104502.0 | 0 | Canis | 82180.0 | 1 | Canis lupus | 82180.0 | 1 |
| scf7180000048711 | 24239 | 0.4496 | 0 | 2765.5905 | Eukaryota | 27085.0 | 1 | Chordata | 27085.0 | 1 | Gadiformes | 9116.0 | 4 | Gadidae | 9116.0 | 4 | Gadus | 9116.0 | 4 | Gadus morhua | 9116.0 | 4 |
| scf7180000056512 | 258855 | 0.5102 | 0 | 57.9671 | Eukaryota | 50784.0 | 0 | Chordata | 50784.0 | 0 | Carnivora | 50784.0 | 0 | Canidae | 50784.0 | 0 | Canis | 50784.0 | 0 | Canis lupus | 50784.0 | 0 |
| scf7180000050708 | 9757 | 0.4549 | 0 | 887.5869 | Eukaryota | 16635.0 | 0 | Chordata | 14965.0 | 1 | Salmoniformes | 4380.0 | 6 | Salmonidae | 4380.0 | 6 | Salmo | 4380.0 | 6 | Salmo trutta | 4380.0 | 6 |
| scf7180000041643 | 111831 | 0.3935 | 0 | 55.6486 | Eukaryota | 11940.0 | 0 | Chordata | 11940.0 | 0 | Carnivora | 11940.0 | 0 | Canidae | 11940.0 | 0 | Canis | 11940.0 | 0 | Canis lupus | 11940.0 | 0 |
| scf7180000052892 | 4789 | 0.4354 | 0 | 194.7424 | no-hit | 0.0 | 0 | no-hit | 0.0 | 0 | no-hit | 0.0 | 0 | no-hit | 0.0 | 0 | no-hit | 0.0 | 0 | no-hit | 0.0 | 0 |
| scf7180000051415 | 8829 | 0.5049 | 0 | 366.2917 | Eukaryota | 8468.0 | 0 | Chordata | 8468.0 | 0 | Carnivora | 8468.0 | 0 | Canidae | 8468.0 | 0 | Canis | 6830.0 | 1 | Canis lupus | 5669.0 | 2 |
| scf7180000043212 | 47571 | 0.4282 | 0 | 60.2941 | Eukaryota | 44252.0 | 0 | Chordata | 44252.0 | 0 | Carnivora | 44252.0 | 0 | Felidae | 17477.0 | 3 | Vulpes | 10355.0 | 6 | Vulpes vulpes | 10355.0 | 7 |
| scf7180000055832 | 6366 | 0.4384 | 0 | 32.8849 | Eukaryota | 2797.0 | 0 | Platyhelminthes | 1624.0 | 1 | Polystomatidea | 1624.0 | 3 | Polystomatidae | 1624.0 | 3 | Protopolystoma | 1624.0 | 3 | Protopolystoma xenopodis | 1624.0 | 3 |
| scf7180000047265 | 8670 | 0.4339 | 0 | 38.7476 | Eukaryota | 513.0 | 0 | Platyhelminthes | 289.0 | 1 | Polystomatidea | 289.0 | 1 | Polystomatidae | 289.0 | 1 | Protopolystoma | 289.0 | 1 | Protopolystoma xenopodis | 289.0 | 1 |
| scf7180000042066 | 849532 | 0.3637 | 0 | 60.4821 | Eukaryota | 35960.0 | 0 | Chordata | 35960.0 | 0 | Carnivora | 33538.0 | 1 | Canidae | 33538.0 | 1 | Canis | 30357.0 | 2 | Canis lupus | 30357.0 | 2 |
| scf7180000056297 | 1049316 | 0.4438 | 0 | 60.5163 | Eukaryota | 85695.0 | 0 | Chordata | 85695.0 | 0 | Carnivora | 85695.0 | 0 | Canidae | 85695.0 | 0 | Canis | 58442.0 | 1 | Canis lupus | 58442.0 | 1 |
| scf7180000046773 | 93288 | 0.5206 | 0 | 57.3069 | Eukaryota | 11636.0 | 1 | Chordata | 11636.0 | 1 | Primates | 5321.0 | 3 | Hominidae | 5321.0 | 3 | Homo | 5321.0 | 4 | Homo sapiens | 5321.0 | 5 |
| scf7180000052726 | 6516 | 0.4509 | 0 | 1262.7522 | no-hit | 0.0 | 0 | no-hit | 0.0 | 0 | no-hit | 0.0 | 0 | no-hit | 0.0 | 0 | no-hit | 0.0 | 0 | no-hit | 0.0 | 0 |
| scf7180000040627 | 26566 | 0.3844 | 0 | 49.5341 | Eukaryota | 2966.0 | 0 | Chordata | 2966.0 | 0 | Carnivora | 2966.0 | 0 | Canidae | 2966.0 | 0 | Canis | 2966.0 | 0 | Canis lupus | 2966.0 | 0 |
| scf7180000047276 | 35924 | 0.4391 | 0 | 174.5954 | no-hit | 0.0 | 0 | no-hit | 0.0 | 0 | no-hit | 0.0 | 0 | no-hit | 0.0 | 0 | no-hit | 0.0 | 0 | no-hit | 0.0 | 0 |
| scf7180000044745 | 4569 | 0.3861 | 0 | 26.056 | Eukaryota | 4758.0 | 0 | Chordata | 4758.0 | 0 | Carnivora | 4758.0 | 0 | Canidae | 4758.0 | 0 | Canis | 4758.0 | 0 | Canis lupus | 4758.0 | 0 |
| scf7180000042718 | 403857 | 0.3723 | 0 | 59.6394 | Eukaryota | 67717.0 | 0 | Chordata | 67717.0 | 0 | Carnivora | 67717.0 | 0 | Canidae | 67717.0 | 0 | Canis | 67717.0 | 0 | Canis lupus | 67717.0 | 0 |
| scf7180000056434 | 382677 | 0.3811 | 0 | 60.0322 | Eukaryota | 91682.0 | 0 | Chordata | 91682.0 | 0 | Carnivora | 91682.0 | 0 | Canidae | 91682.0 | 0 | Canis | 91682.0 | 0 | Canis lupus | 91682.0 | 0 |
| scf7180000052526 | 12139 | 0.4385 | 0 | 256.413 | Eukaryota | 4596.0 | 0 | Platyhelminthes | 3161.0 | 2 | Polystomatidea | 3161.0 | 2 | Polystomatidae | 3161.0 | 2 | Protopolystoma | 3161.0 | 2 | Protopolystoma xenopodis | 3161.0 | 2 |
| scf7180000046731 | 14978 | 0.4408 | 0 | 138.483 | Eukaryota | 86631.0 | 0 | Chordata | 86631.0 | 0 | Carnivora | 86631.0 | 0 | Canidae | 86631.0 | 0 | Canis | 86631.0 | 0 | Canis lupus | 86631.0 | 0 |
| scf7180000047502 | 4613 | 0.4821 | 0 | 74.3198 | Eukaryota | 4531.0 | 0 | Chordata | 4531.0 | 0 | Carnivora | 2031.0 | 2 | Cercopithecidae | 1039.0 | 4 | Macaca | 1039.0 | 6 | Macaca mulatta | 1039.0 | 6 |
| scf7180000048352 | 44429 | 0.4414 | 0 | 117.6132 | Eukaryota | 4621.0 | 0 | Chordata | 3930.0 | 1 | Tetraodontiformes | 1611.0 | 5 | Tetraodontidae | 1611.0 | 5 | Takifugu | 1611.0 | 6 | Sphaeramia orbicularis | 1122.0 | 7 |
| scf7180000042492 | 306265 | 0.3628 | 0 | 59.0821 | Eukaryota | 35943.0 | 0 | Chordata | 35943.0 | 0 | Primates | 26404.0 | 2 | Hominidae | 20189.0 | 4 | Homo | 20189.0 | 5 | Homo sapiens | 20189.0 | 5 |
| scf7180000054876 | 13186 | 0.5295 | 0 | 549.8205 | Eukaryota | 3588.0 | 0 | Chordata | 3588.0 | 0 | Carnivora | 3588.0 | 0 | Canidae | 3588.0 | 0 | Canis | 3588.0 | 0 | Canis lupus | 3588.0 | 0 |
| scf7180000053433 | 4742 | 0.438 | 0 | 190.3894 | Eukaryota | 6389.0 | 0 | Chordata | 5895.0 | 1 | Tetraodontiformes | 2821.0 | 4 | Tetraodontidae | 2821.0 | 4 | Takifugu | 2821.0 | 4 | Takifugu rubripes | 2821.0 | 4 |
| scf7180000042165 | 171455 | 0.4211 | 0 | 58.8406 | Eukaryota | 52263.0 | 0 | Chordata | 52263.0 | 0 | Carnivora | 52263.0 | 0 | Canidae | 52263.0 | 0 | Canis | 41201.0 | 1 | Canis lupus | 41201.0 | 1 |
| scf7180000052142 | 26475 | 0.4437 | 0 | 55.9899 | Eukaryota | 12948.0 | 1 | Chordata | 12948.0 | 1 | Primates | 11444.0 | 2 | Cercopithecidae | 6917.0 | 3 | Cercocebus | 5548.0 | 4 | Cercocebus atys | 5548.0 | 4 |
| scf7180000047505 | 9994 | 0.4418 | 0 | 98.0439 | Eukaryota | 568.0 | 0 | Platyhelminthes | 568.0 | 0 | Polystomatidea | 568.0 | 0 | Polystomatidae | 568.0 | 0 | Protopolystoma | 568.0 | 0 | Protopolystoma xenopodis | 568.0 | 0 |
| scf7180000052120 | 6397 | 0.4463 | 0 | 648.4094 | no-hit | 0.0 | 0 | no-hit | 0.0 | 0 | no-hit | 0.0 | 0 | no-hit | 0.0 | 0 | no-hit | 0.0 | 0 | no-hit | 0.0 | 0 |
| scf7180000040632 | 188628 | 0.3623 | 0 | 58.5231 | Eukaryota | 7880.0 | 0 | Chordata | 7880.0 | 0 | Carnivora | 5522.0 | 2 | Canidae | 5522.0 | 2 | Canis | 5522.0 | 2 | Canis lupus | 5522.0 | 2 |
| scf7180000050556 | 9404 | 0.4428 | 0 | 27.754 | Eukaryota | 7728.0 | 0 | Platyhelminthes | 4730.0 | 1 | Polystomatidea | 4730.0 | 1 | Polystomatidae | 4730.0 | 1 | Protopolystoma | 4730.0 | 1 | Protopolystoma xenopodis | 4730.0 | 1 |
| scf7180000040891 | 247166 | 0.3769 | 0 | 58.5745 | Eukaryota | 38942.0 | 0 | Chordata | 38942.0 | 0 | Carnivora | 38942.0 | 0 | Canidae | 38942.0 | 0 | Canis | 38942.0 | 0 | Canis lupus | 38942.0 | 0 |
| scf7180000042521 | 270351 | 0.4388 | 0 | 59.2074 | Eukaryota | 83921.0 | 0 | Chordata | 83921.0 | 0 | Carnivora | 83921.0 | 0 | Canidae | 83921.0 | 0 | Canis | 83921.0 | 0 | Canis lupus | 83921.0 | 0 |
| scf7180000050400 | 10519 | 0.4532 | 0 | 851.8249 | Eukaryota | 2094.0 | 0 | Echinodermata | 900.0 | 2 | Forcipulatida | 900.0 | 2 | Asteriidae | 900.0 | 2 | Asterias | 900.0 | 2 | Asterias rubens | 900.0 | 2 |
| scf7180000054263 | 17223 | 0.4421 | 0 | 342.2963 | Eukaryota | 1029.0 | 0 | Platyhelminthes | 639.0 | 1 | Polystomatidea | 639.0 | 1 | Polystomatidae | 639.0 | 1 | Protopolystoma | 639.0 | 1 | Protopolystoma xenopodis | 639.0 | 1 |
| scf7180000054130 | 1999 | 0.4562 | 0 | 262.1706 | no-hit | 0.0 | 0 | no-hit | 0.0 | 0 | no-hit | 0.0 | 0 | no-hit | 0.0 | 0 | no-hit | 0.0 | 0 | no-hit | 0.0 | 0 |
| scf7180000041545 | 1088546 | 0.4043 | 0 | 59.719 | Eukaryota | 88229.0 | 0 | Chordata | 88229.0 | 0 | Carnivora | 88229.0 | 0 | Canidae | 88229.0 | 0 | Canis | 88229.0 | 0 | Canis lupus | 88229.0 | 0 |
| scf7180000042378 | 965354 | 0.4732 | 0 | 60.0707 | Eukaryota | 97468.0 | 0 | Chordata | 97468.0 | 0 | Carnivora | 97468.0 | 0 | Canidae | 97468.0 | 0 | Vulpes | 66880.0 | 1 | Vulpes vulpes | 66880.0 | 1 |
| scf7180000052934 | 3159 | 0.4609 | 0 | 442.0164 | Eukaryota | 1607.0 | 0 | Chordata | 628.0 | 2 | Tetraodontiformes | 628.0 | 2 | Tetraodontidae | 628.0 | 2 | Takifugu | 628.0 | 2 | Takifugu rubripes | 628.0 | 2 |
| scf7180000047226 | 87434 | 0.428 | 0 | 61.7301 | Eukaryota | 105259.0 | 0 | Chordata | 105259.0 | 0 | Carnivora | 105259.0 | 0 | Canidae | 105259.0 | 0 | Canis | 64597.0 | 1 | Canis lupus | 64597.0 | 1 |
| scf7180000055729 | 21828 | 0.4104 | 0 | 83.008 | Eukaryota | 65908.0 | 0 | Chordata | 65908.0 | 0 | Carnivora | 65908.0 | 0 | Canidae | 65908.0 | 0 | Canis | 65908.0 | 0 | Canis lupus | 65908.0 | 0 |
| scf7180000051082 | 17489 | 0.4386 | 0 | 65.393 | Eukaryota | 3708.0 | 0 | Platyhelminthes | 1825.0 | 2 | Polystomatidea | 1825.0 | 2 | Polystomatidae | 1825.0 | 2 | Protopolystoma | 1825.0 | 2 | Protopolystoma xenopodis | 1825.0 | 2 |
| scf7180000055482 | 11465 | 0.4138 | 0 | 95.383 | Eukaryota | 6075.0 | 0 | Chordata | 6075.0 | 0 | Apterygiformes | 4446.0 | 2 | Apterygidae | 4446.0 | 2 | Apteryx | 4446.0 | 2 | Apteryx mantelli | 4446.0 | 2 |
| scf7180000041878 | 115904 | 0.4057 | 0 | 60.461 | Eukaryota | 87393.0 | 0 | Chordata | 87393.0 | 0 | Carnivora | 87393.0 | 0 | Canidae | 87393.0 | 0 | Canis | 87393.0 | 0 | Canis lupus | 87393.0 | 0 |
| scf7180000041519 | 217350 | 0.364 | 0 | 58.1585 | Eukaryota | 69289.0 | 0 | Chordata | 69289.0 | 0 | Carnivora | 69289.0 | 0 | Canidae | 69289.0 | 0 | Canis | 69289.0 | 0 | Canis lupus | 69289.0 | 0 |
| scf7180000042454 | 625119 | 0.3735 | 0 | 61.8423 | Eukaryota | 62516.0 | 0 | Chordata | 62516.0 | 0 | Carnivora | 62516.0 | 0 | Canidae | 62516.0 | 0 | Canis | 62516.0 | 0 | Canis lupus | 62516.0 | 0 |
| scf7180000042110 | 769145 | 0.386 | 0 | 58.8477 | Eukaryota | 91389.0 | 0 | Chordata | 91389.0 | 0 | Carnivora | 91389.0 | 0 | Canidae | 91389.0 | 0 | Canis | 91389.0 | 0 | Canis lupus | 91389.0 | 0 |
| scf7180000051541 | 6258 | 0.4506 | 0 | 642.5613 | no-hit | 0.0 | 0 | no-hit | 0.0 | 0 | no-hit | 0.0 | 0 | no-hit | 0.0 | 0 | no-hit | 0.0 | 0 | no-hit | 0.0 | 0 |
| scf7180000053084 | 3291 | 0.4369 | 0 | 249.1483 | Eukaryota | 292.0 | 0 | Chordata | 292.0 | 0 | Kurtiformes | 292.0 | 0 | Apogonidae | 292.0 | 0 | Sphaeramia | 292.0 | 0 | Sphaeramia orbicularis | 292.0 | 0 |
| scf7180000042371 | 739019 | 0.4183 | 0 | 60.4495 | Eukaryota | 60735.0 | 0 | Chordata | 60735.0 | 0 | Carnivora | 60735.0 | 0 | Canidae | 60735.0 | 0 | Canis | 52426.0 | 1 | Canis lupus | 52426.0 | 1 |
| scf7180000045395 | 170627 | 0.4288 | 0 | 61.5243 | Eukaryota | 16569.0 | 0 | Chordata | 16569.0 | 0 | Carnivora | 16569.0 | 0 | Canidae | 16569.0 | 0 | Canis | 16569.0 | 0 | Canis lupus | 16569.0 | 0 |
| scf7180000050165 | 7639 | 0.4269 | 0 | 98.0714 | Eukaryota | 196.0 | 0 | Platyhelminthes | 196.0 | 0 | Polystomatidea | 196.0 | 0 | Polystomatidae | 196.0 | 0 | Protopolystoma | 196.0 | 0 | Protopolystoma xenopodis | 196.0 | 0 |
| scf7180000052119 | 2012 | 0.4458 | 0 | 385.3477 | no-hit | 0.0 | 0 | no-hit | 0.0 | 0 | no-hit | 0.0 | 0 | no-hit | 0.0 | 0 | no-hit | 0.0 | 0 | no-hit | 0.0 | 0 |
| scf7180000045597 | 176459 | 0.3929 | 0 | 59.3339 | Eukaryota | 37539.0 | 0 | Chordata | 37539.0 | 0 | Carnivora | 37539.0 | 0 | Canidae | 14787.0 | 4 | Canis | 9878.0 | 6 | Canis lupus | 9878.0 | 7 |
| scf7180000053198 | 12081 | 0.44 | 0 | 76.6912 | Eukaryota | 3270.0 | 0 | Chordata | 2712.0 | 1 | Carnivora | 2712.0 | 1 | Canidae | 2712.0 | 1 | Canis | 2712.0 | 1 | Canis lupus | 2712.0 | 1 |
| scf7180000053801 | 4152 | 0.4627 | 0 | 115.6629 | Eukaryota | 27995.0 | 0 | Chordata | 27995.0 | 0 | Carnivora | 27995.0 | 0 | Canidae | 27995.0 | 0 | Canis | 27995.0 | 0 | Canis lupus | 27995.0 | 0 |
| scf7180000056571 | 98353 | 0.4627 | 0 | 57.6642 | Eukaryota | 29299.0 | 0 | Chordata | 29299.0 | 0 | Carnivora | 29299.0 | 0 | Canidae | 21275.0 | 2 | Vulpes | 13069.0 | 3 | Vulpes vulpes | 13069.0 | 4 |
| scf7180000048265 | 3425 | 0.4394 | 0 | 62.1864 | no-hit | 0.0 | 0 | no-hit | 0.0 | 0 | no-hit | 0.0 | 0 | no-hit | 0.0 | 0 | no-hit | 0.0 | 0 | no-hit | 0.0 | 0 |
| scf7180000047318 | 4734 | 0.4362 | 0 | 105.5384 | Eukaryota | 2921.0 | 0 | Platyhelminthes | 1936.0 | 1 | Polystomatidea | 1936.0 | 1 | Polystomatidae | 1936.0 | 1 | Protopolystoma | 1936.0 | 1 | Protopolystoma xenopodis | 1936.0 | 1 |
| scf7180000048966 | 16804 | 0.4542 | 0 | 1626.107 | no-hit | 0.0 | 0 | no-hit | 0.0 | 0 | no-hit | 0.0 | 0 | no-hit | 0.0 | 0 | no-hit | 0.0 | 0 | no-hit | 0.0 | 0 |
| scf7180000046542 | 68773 | 0.4072 | 0 | 57.9685 | Eukaryota | 89698.0 | 0 | Chordata | 89698.0 | 0 | Carnivora | 89698.0 | 0 | Canidae | 89698.0 | 0 | Canis | 89698.0 | 0 | Canis lupus | 89698.0 | 0 |
| scf7180000056050 | 49503 | 0.4633 | 0 | 527.8872 | Eukaryota | 90289.0 | 0 | Chordata | 90289.0 | 0 | Carnivora | 90289.0 | 0 | Canidae | 90289.0 | 0 | Canis | 90289.0 | 0 | Canis lupus | 90289.0 | 0 |
| scf7180000045175 | 231444 | 0.4016 | 0 | 59.7702 | Eukaryota | 80312.0 | 0 | Chordata | 80312.0 | 0 | Carnivora | 80312.0 | 0 | Canidae | 80312.0 | 0 | Canis | 80312.0 | 0 | Canis lupus | 80312.0 | 0 |

| | | | | | | | | | | | | | | | | | | | | | | |
|---|---|---|---|---|---|---|---|---|---|---|---|---|---|---|---|---|---|---|---|---|---|---|
| scf7180000054874 | 18450 | 0.4286 | 0 | 191.701 | Eukaryota | 715.0 | 0 | Chordata | 568.0 | 1 | Kurtiformes | 318.0 | 2 | Apogonidae | 318.0 | 2 | Sphaeramia | 318.0 | 2 | Sphaeramia orbicularis | 318.0 | 2 |
| scf7180000048217 | 13443 | 0.4486 | 0 | 72.0086 | Eukaryota | 2658.0 | 0 | Chordata | 2658.0 | 0 | Carnivora | 2658.0 | 0 | Canidae | 2658.0 | 0 | Canis | 2658.0 | 0 | Canis lupus | 2658.0 | 0 |
| scf7180000045095 | 189981 | 0.3527 | 0 | 57.7338 | Eukaryota | 90695.0 | 0 | Chordata | 90695.0 | 0 | Carnivora | 90695.0 | 0 | Canidae | 90695.0 | 0 | Canis | 90695.0 | 0 | Canis lupus | 90695.0 | 0 |
| scf7180000046443 | 55932 | 0.4433 | 0 | 136.2688 | Eukaryota | 2533.0 | 0 | Chordata | 2533.0 | 0 | Carnivora | 2533.0 | 0 | Canidae | 2533.0 | 0 | Canis | 2533.0 | 0 | Canis lupus | 2533.0 | 0 |
| scf7180000045846 | 147488 | 0.3796 | 0 | 60.9904 | Eukaryota | 19070.0 | 0 | Chordata | 19070.0 | 0 | Carnivora | 19070.0 | 0 | Canidae | 14128.0 | 1 | Canis | 14128.0 | 1 | Canis lupus | 14128.0 | 1 |
| scf7180000041605 | 51428 | 0.3915 | 0 | 56.9056 | Eukaryota | 9473.0 | 0 | Chordata | 9473.0 | 0 | Carnivora | 6248.0 | 2 | Canidae | 3287.0 | 4 | Canis | 3287.0 | 5 | Canis lupus | 3287.0 | 5 |
| scf7180000051161 | 27056 | 0.5055 | 0 | 798.6536 | Eukaryota | 9189.0 | 0 | Chordata | 9189.0 | 0 | Carnivora | 9189.0 | 0 | Canidae | 9189.0 | 0 | Canis | 7435.0 | 1 | Canis lupus | 6161.0 | 2 |
| scf7180000043275 | 493071 | 0.3989 | 0 | 61.2216 | Eukaryota | 69000.0 | 0 | Chordata | 69000.0 | 0 | Carnivora | 69000.0 | 0 | Canidae | 69000.0 | 0 | Canis | 69000.0 | 0 | Canis lupus | 69000.0 | 0 |
| scf7180000053352 | 13105 | 0.4652 | 0 | 81.6495 | Eukaryota | 9839.0 | 0 | Chordata | 9839.0 | 0 | Carnivora | 9839.0 | 0 | Canidae | 9839.0 | 0 | Canis | 5831.0 | 1 | Canis lupus | 5831.0 | 1 |
| scf7180000049052 | 16171 | 0.4473 | 0 | 1494.4311 | Eukaryota | 95909.0 | 0 | Chordata | 95909.0 | 0 | Carnivora | 95909.0 | 0 | Canidae | 95909.0 | 0 | Canis | 95909.0 | 0 | Canis lupus | 95909.0 | 0 |
| scf7180000041278 | 1152250 | 0.3695 | 0 | 59.3942 | Eukaryota | 78765.0 | 0 | Chordata | 78765.0 | 0 | Carnivora | 78765.0 | 0 | Canidae | 78765.0 | 0 | Canis | 78765.0 | 0 | Canis lupus | 78765.0 | 0 |
| scf7180000045956 | 170144 | 0.3946 | 0 | 60.2927 | Eukaryota | 92626.0 | 0 | Chordata | 92626.0 | 0 | Carnivora | 92626.0 | 0 | Canidae | 92626.0 | 0 | Canis | 92626.0 | 0 | Canis lupus | 92626.0 | 0 |
| scf7180000040672 | 665370 | 0.3404 | 0 | 58.7161 | Eukaryota | 73826.0 | 0 | Chordata | 73826.0 | 0 | Carnivora | 73826.0 | 0 | Canidae | 73826.0 | 0 | Canis | 73826.0 | 0 | Canis lupus | 73826.0 | 0 |
| scf7180000051788 | 2953 | 0.4646 | 0 | 569.5628 | no-hit | 0.0 | 0 | no-hit | 0.0 | 0 | no-hit | 0.0 | 0 | no-hit | 0.0 | 0 | no-hit | 0.0 | 0 | no-hit | 0.0 | 0 |
| scf7180000042563 | 20978 | 0.3547 | 0 | 48.0668 | Eukaryota | 11021.0 | 0 | Chordata | 11021.0 | 0 | Carnivora | 11021.0 | 0 | Canidae | 11021.0 | 0 | Canis | 9927.0 | 1 | Canis lupus | 9927.0 | 1 |
| scf7180000053047 | 16130 | 0.5038 | 0 | 150.8393 | Eukaryota | 64463.0 | 0 | Chordata | 64463.0 | 0 | Carnivora | 64463.0 | 0 | Canidae | 64463.0 | 0 | Canis | 64463.0 | 0 | Canis lupus | 64463.0 | 0 |
| scf7180000054513 | 151653 | 0.3786 | 0 | 446.6593 | Eukaryota | 38212.0 | 0 | Chordata | 38212.0 | 0 | Carnivora | 38212.0 | 0 | Canidae | 38212.0 | 0 | Canis | 38212.0 | 0 | Canis lupus | 38212.0 | 0 |
| scf7180000043706 | 48685 | 0.4098 | 0 | 55.8144 | Eukaryota | 49199.0 | 0 | Chordata | 49199.0 | 0 | Carnivora | 49199.0 | 0 | Canidae | 49199.0 | 0 | Canis | 49199.0 | 0 | Canis lupus | 49199.0 | 0 |
| scf7180000056514 | 150813 | 0.4804 | 0 | 58.4396 | Eukaryota | 93822.0 | 0 | Chordata | 93822.0 | 0 | Carnivora | 93822.0 | 0 | Canidae | 93822.0 | 0 | Canis | 93822.0 | 0 | Canis lupus | 93822.0 | 0 |
| scf7180000041607 | 943511 | 0.4391 | 0 | 58.8581 | Eukaryota | 101049.0 | 0 | Chordata | 101049.0 | 0 | Carnivora | 101049.0 | 0 | Canidae | 101049.0 | 0 | Canis | 101049.0 | 0 | Canis lupus | 101049.0 | 0 |
| scf7180000044938 | 280854 | 0.3645 | 0 | 56.9071 | Eukaryota | 86564.0 | 0 | Chordata | 86564.0 | 0 | Carnivora | 86564.0 | 0 | Canidae | 86564.0 | 0 | Canis | 86564.0 | 0 | Canis lupus | 86564.0 | 0 |
| scf7180000043328 | 14722 | 0.4893 | 0 | 28.3956 | Eukaryota | 24391.0 | 0 | Chordata | 24391.0 | 0 | Carnivora | 24391.0 | 0 | Canidae | 24391.0 | 0 | Canis | 24391.0 | 0 | Canis lupus | 24391.0 | 0 |
| scf7180000055203 | 13949 | 0.4774 | 0 | 79.9458 | Eukaryota | 9152.0 | 0 | Chordata | 9152.0 | 0 | Carnivora | 9152.0 | 0 | Canidae | 9152.0 | 0 | Canis | 5435.0 | 1 | Canis lupus | 5435.0 | 1 |
| scf7180000043454 | 440959 | 0.43 | 0 | 59.8451 | Eukaryota | 39316.0 | 0 | Chordata | 39316.0 | 0 | Carnivora | 36527.0 | 1 | Canidae | 22193.0 | 2 | Canis | 17209.0 | 3 | Canis lupus | 17209.0 | 3 |
| scf7180000052450 | 3373 | 0.4412 | 0 | 115.0653 | no-hit | 0.0 | 0 | no-hit | 0.0 | 0 | no-hit | 0.0 | 0 | no-hit | 0.0 | 0 | no-hit | 0.0 | 0 | no-hit | 0.0 | 0 |
| scf7180000049057 | 46422 | 0.3534 | 0 | 55.6036 | Eukaryota | 10948.0 | 0 | Chordata | 10948.0 | 0 | Carnivora | 10948.0 | 0 | Canidae | 10948.0 | 0 | Canis | 10948.0 | 0 | Canis lupus | 10948.0 | 0 |
| scf7180000053590 | 13336 | 0.4429 | 0 | 137.0359 | Eukaryota | 276.0 | 0 | Chordata | 276.0 | 0 | Salmoniformes | 276.0 | 0 | Salmonidae | 276.0 | 0 | Coregonus | 276.0 | 0 | Coregonus sp. 'balchen' | 276.0 | 0 |
| scf7180000055012 | 48780 | 0.4426 | 0 | 182.8594 | Eukaryota | 68739.0 | 0 | Chordata | 51994.0 | 1 | Gadiformes | 38571.0 | 5 | Gadidae | 38571.0 | 5 | Gadus | 38571.0 | 5 | Gadus morhua | 38571.0 | 5 |
| scf7180000050693 | 5548 | 0.4557 | 0 | 310.0056 | Eukaryota | 368.0 | 0 | Chordata | 368.0 | 0 | Kurtiformes | 368.0 | 0 | Apogonidae | 368.0 | 0 | Sphaeramia | 368.0 | 0 | Sphaeramia orbicularis | 368.0 | 0 |
| scf7180000047331 | 8960 | 0.4323 | 0 | 87.7074 | Eukaryota | 21432.0 | 0 | Platyhelminthes | 8749.0 | 2 | Strigeidida | 8749.0 | 4 | Schistosomatidae | 8749.0 | 4 | Schistosoma | 8749.0 | 4 | Schistosoma mansoni | 8749.0 | 5 |
| scf7180000046840 | 4603 | 0.3635 | 0 | 29.6794 | Eukaryota | 5207.0 | 0 | Chordata | 5207.0 | 0 | Carnivora | 5207.0 | 0 | Canidae | 5207.0 | 0 | Canis | 5207.0 | 0 | Canis lupus | 5207.0 | 0 |
| scf7180000041158 | 1296205 | 0.4746 | 0 | 59.0303 | Eukaryota | 60651.0 | 0 | Chordata | 60651.0 | 0 | Carnivora | 60651.0 | 0 | Canidae | 60651.0 | 0 | Canis | 45075.0 | 1 | Canis lupus | 45075.0 | 1 |
| scf7180000046272 | 137044 | 0.6329 | 0 | 56.7587 | Eukaryota | 123620.0 | 0 | Chordata | 123620.0 | 0 | Carnivora | 123620.0 | 0 | Canidae | 123620.0 | 0 | Canis | 110818.0 | 1 | Canis lupus | 110818.0 | 1 |
| scf7180000051567 | 2132 | 0.4381 | 0 | 256.2513 | no-hit | 0.0 | 0 | no-hit | 0.0 | 0 | no-hit | 0.0 | 0 | no-hit | 0.0 | 0 | no-hit | 0.0 | 0 | no-hit | 0.0 | 0 |
| scf7180000055681 | 13997 | 0.4354 | 0 | 222.6735 | Eukaryota | 363.0 | 0 | Chordata | 363.0 | 0 | Tetraodontiformes | 363.0 | 0 | Tetraodontidae | 363.0 | 0 | Takifugu | 363.0 | 0 | Takifugu rubripes | 363.0 | 0 |
| scf7180000050937 | 6667 | 0.4447 | 0 | 43.3436 | no-hit | 0.0 | 0 | no-hit | 0.0 | 0 | no-hit | 0.0 | 0 | no-hit | 0.0 | 0 | no-hit | 0.0 | 0 | no-hit | 0.0 | 0 |
| scf7180000045486 | 25904 | 0.4403 | 0 | 66.717 | Eukaryota | 91335.0 | 0 | Chordata | 91335.0 | 0 | Carnivora | 91335.0 | 0 | Canidae | 91335.0 | 0 | Canis | 91335.0 | 0 | Canis lupus | 91335.0 | 0 |
| scf7180000041576 | 952493 | 0.3852 | 0 | 59.6215 | Eukaryota | 77813.0 | 0 | Chordata | 77813.0 | 0 | Carnivora | 77813.0 | 0 | Canidae | 77813.0 | 0 | Canis | 69994.0 | 1 | Canis lupus | 69994.0 | 1 |
| scf7180000043183 | 699182 | 0.348 | 0 | 57.4219 | Eukaryota | 89344.0 | 0 | Chordata | 89344.0 | 0 | Carnivora | 89344.0 | 0 | Canidae | 89344.0 | 0 | Canis | 89344.0 | 0 | Canis lupus | 89344.0 | 0 |
| scf7180000050140 | 10562 | 0.4394 | 0 | 57.2199 | no-hit | 0.0 | 0 | no-hit | 0.0 | 0 | no-hit | 0.0 | 0 | no-hit | 0.0 | 0 | no-hit | 0.0 | 0 | no-hit | 0.0 | 0 |
| scf7180000050047 | 13035 | 0.4409 | 0 | 141.3582 | no-hit | 0.0 | 0 | no-hit | 0.0 | 0 | no-hit | 0.0 | 0 | no-hit | 0.0 | 0 | no-hit | 0.0 | 0 | no-hit | 0.0 | 0 |
| scf7180000042567 | 340322 | 0.3981 | 0 | 58.2432 | Eukaryota | 89087.0 | 0 | Chordata | 89087.0 | 0 | Carnivora | 89087.0 | 0 | Canidae | 89087.0 | 0 | Canis | 89087.0 | 0 | Canis lupus | 89087.0 | 0 |
| scf7180000043950 | 8840 | 0.4302 | 0 | 39.7392 | Eukaryota | 84539.0 | 0 | Chordata | 84539.0 | 0 | Carnivora | 84539.0 | 0 | Canidae | 84539.0 | 0 | Canis | 84539.0 | 0 | Canis lupus | 84539.0 | 0 |
| scf7180000044350 | 201867 | 0.396 | 0 | 60.4046 | Eukaryota | 89572.0 | 0 | Chordata | 89572.0 | 0 | Carnivora | 89572.0 | 0 | Canidae | 89572.0 | 0 | Canis | 89572.0 | 0 | Canis lupus | 89572.0 | 0 |
| scf7180000054517 | 327493 | 0.3668 | 0 | 63.1155 | Eukaryota | 43076.0 | 0 | Chordata | 43076.0 | 0 | Carnivora | 43076.0 | 0 | Canidae | 43076.0 | 0 | Canis | 43076.0 | 0 | Canis lupus | 43076.0 | 0 |
| scf7180000052653 | 2823 | 0.4552 | 0 | 190.0246 | no-hit | 0.0 | 0 | no-hit | 0.0 | 0 | no-hit | 0.0 | 0 | no-hit | 0.0 | 0 | no-hit | 0.0 | 0 | no-hit | 0.0 | 0 |
| scf7180000042067 | 15077 | 0.4102 | 0 | 104.712 | Eukaryota | 90107.0 | 0 | Chordata | 90107.0 | 0 | Carnivora | 90107.0 | 0 | Canidae | 90107.0 | 0 | Canis | 90107.0 | 0 | Canis lupus | 90107.0 | 0 |
| scf7180000046945 | 19798 | 0.4373 | 0 | 56.9646 | Eukaryota | 93837.0 | 0 | Chordata | 93837.0 | 0 | Carnivora | 93837.0 | 0 | Canidae | 93837.0 | 0 | Canis | 93837.0 | 0 | Canis lupus | 93837.0 | 0 |
| scf7180000042919 | 498154 | 0.4005 | 0 | 58.6373 | Eukaryota | 43670.0 | 0 | Chordata | 43670.0 | 0 | Carnivora | 39134.0 | 1 | Canidae | 39134.0 | 1 | Vulpes | 27254.0 | 2 | Vulpes vulpes | 27254.0 | 2 |
| scf7180000047209 | 117170 | 0.3787 | 0 | 61.4367 | Eukaryota | 28210.0 | 0 | Chordata | 28210.0 | 0 | Carnivora | 28210.0 | 0 | Canidae | 23168.0 | 1 | Canis | 23168.0 | 1 | Canis lupus | 23168.0 | 1 |
| scf7180000047767 | 173029 | 0.3475 | 0 | 57.0917 | Eukaryota | 61563.0 | 0 | Chordata | 61563.0 | 0 | Carnivora | 61563.0 | 0 | Canidae | 61563.0 | 0 | Canis | 61563.0 | 0 | Canis lupus | 61563.0 | 0 |
| scf7180000050166 | 5279 | 0.4476 | 0 | 886.0425 | Eukaryota | 592.0 | 0 | Chordata | 592.0 | 0 | Salmoniformes | 592.0 | 0 | Salmonidae | 592.0 | 0 | Coregonus | 592.0 | 0 | Coregonus sp. 'balchen' | 592.0 | 0 |
| scf7180000041681 | 893255 | 0.3538 | 0 | 60.3889 | Eukaryota | 56047.0 | 0 | Chordata | 56047.0 | 0 | Carnivora | 29777.0 | 1 | Hominidae | 19804.0 | 5 | Homo | 19804.0 | 7 | Homo sapiens | 19804.0 | 7 |
| scf7180000040815 | 1690912 | 0.3697 | 0 | 60.955 | Eukaryota | 42126.0 | 0 | Chordata | 42126.0 | 0 | Carnivora | 42126.0 | 0 | Canidae | 42126.0 | 0 | Canis | 29634.0 | 1 | Canis lupus | 29634.0 | 1 |
| scf7180000053580 | 5767 | 0.5048 | 0 | 96.5383 | Eukaryota | 7928.0 | 0 | Chordata | 7928.0 | 0 | Carnivora | 7928.0 | 0 | Canidae | 7928.0 | 0 | Canis | 6450.0 | 1 | Canis lupus | 5447.0 | 2 |
| scf7180000050233 | 17977 | 0.6622 | 0 | 366.8774 | Eukaryota | 35695.0 | 0 | Chordata | 35695.0 | 0 | Primates | 27523.0 | 1 | Cercopithecidae | 13783.0 | 2 | Macaca | 13783.0 | 3 | Macaca mulatta | 13783.0 | 3 |
| scf7180000050899 | 12555 | 0.4382 | 0 | 88.5218 | Eukaryota | 5216.0 | 1 | Chordata | 4860.0 | 2 | Salmoniformes | 3112.0 | 7 | Salmonidae | 3112.0 | 7 | Salmo | 2820.0 | 8 | Salmo trutta | 2820.0 | 8 |
| scf7180000048100 | 4989 | 0.4564 | 0 | 109.3761 | no-hit | 0.0 | 0 | no-hit | 0.0 | 0 | no-hit | 0.0 | 0 | no-hit | 0.0 | 0 | no-hit | 0.0 | 0 | no-hit | 0.0 | 0 |
| scf7180000052342 | 14174 | 0.5032 | 0 | 587.0694 | Eukaryota | 9020.0 | 0 | Chordata | 9020.0 | 0 | Carnivora | 9020.0 | 0 | Canidae | 9020.0 | 0 | Canis | 7343.0 | 1 | Canis lupus | 6125.0 | 2 |
| scf7180000044058 | 169268 | 0.4072 | 0 | 56.032 | Eukaryota | 21247.0 | 0 | Chordata | 21247.0 | 0 | Carnivora | 21247.0 | 0 | Canidae | 18160.0 | 1 | Canis | 10283.0 | 2 | Canis lupus | 10283.0 | 2 |
| scf7180000045785 | 169558 | 0.4275 | 0 | 58.7224 | Eukaryota | 35681.0 | 0 | Chordata | 35681.0 | 0 | Carnivora | 35681.0 | 0 | Canidae | 35681.0 | 0 | Canis | 35681.0 | 0 | Canis lupus | 35681.0 | 0 |
| scf7180000055995 | 70929 | 0.441 | 0 | 164.2298 | Eukaryota | 29370.0 | 0 | Chordata | 29370.0 | 0 | Gadiformes | 17949.0 | 5 | Gadidae | 17949.0 | 5 | Gadus | 17949.0 | 5 | Gadus morhua | 17949.0 | 5 |
| scf7180000044430 | 353238 | 0.37 | 0 | 59.545 | Eukaryota | 72363.0 | 0 | Chordata | 72363.0 | 0 | Carnivora | 72363.0 | 0 | Canidae | 65607.0 | 1 | Canis | 39355.0 | 2 | Canis lupus | 39355.0 | 2 |
| scf7180000046284 | 141496 | 0.3985 | 0 | 60.3668 | Eukaryota | 12169.0 | 0 | Chordata | 12169.0 | 0 | Primates | 9074.0 | 1 | Hominidae | 9074.0 | 1 | Homo | 9074.0 | 1 | Homo sapiens | 9074.0 | 1 |
| scf7180000046663 | 211794 | 0.3845 | 0 | 59.5017 | Eukaryota | 87395.0 | 0 | Chordata | 87395.0 | 0 | Carnivora | 87395.0 | 0 | Canidae | 87395.0 | 0 | Canis | 87395.0 | 0 | Canis lupus | 87395.0 | 0 |
| scf7180000049991 | 8840 | 0.5484 | 0 | 158.6035 | Eukaryota | 5607.0 | 0 | Chordata | 5607.0 | 0 | Artiodactyla | 4475.0 | 1 | Bovidae | 3907.0 | 2 | Bos | 2242.0 | 3 | Bos indicus x Bos taurus | 2242.0 | 3 |
| scf7180000044064 | 347526 | 0.4112 | 0 | 60.852 | Eukaryota | 80746.0 | 0 | Chordata | 80746.0 | 0 | Carnivora | 80746.0 | 0 | Canidae | 56928.0 | 1 | Vulpes | 34500.0 | 2 | Vulpes vulpes | 34500.0 | 2 |
| scf7180000056326 | 857183 | 0.3896 | 0 | 59.5402 | Eukaryota | 142880.0 | 0 | Chordata | 142880.0 | 0 | Carnivora | 142880.0 | 0 | Canidae | 142880.0 | 0 | Canis | 142880.0 | 0 | Canis lupus | 142880.0 | 0 |
| scf7180000054333 | 56412 | 0.4301 | 0 | 182.732 | Eukaryota | 89592.0 | 0 | Chordata | 89592.0 | 0 | Carnivora | 89592.0 | 0 | Canidae | 89592.0 | 0 | Canis | 89592.0 | 0 | Canis lupus | 89592.0 | 0 |
| scf7180000047037 | 18679 | 0.5639 | 0 | 52.1578 | Eukaryota | 3302.0 | 0 | Chordata | 3302.0 | 0 | Carnivora | 1881.0 | 2 | Canidae | 1881.0 | 2 | Canis | 1881.0 | 2 | Canis lupus | 1881.0 | 2 |
| scf7180000056269 | 1522610 | 0.3791 | 0 | 60.8518 | Eukaryota | 89230.0 | 0 | Chordata | 89230.0 | 0 | Carnivora | 89230.0 | 0 | Canidae | 89230.0 | 0 | Canis | 89230.0 | 0 | Canis lupus | 89230.0 | 0 |
| scf7180000048993 | 15758 | 0.4926 | 0 | 205.7899 | Eukaryota | 14377.0 | 0 | Chordata | 14377.0 | 0 | Carnivora | 14377.0 | 0 | Canidae | 14377.0 | 0 | Canis | 8890.0 | 1 | Canis lupus | 8890.0 | 1 |
| scf7180000045610 | 125545 | 0.3825 | 0 | 59.8028 | Eukaryota | 88865.0 | 0 | Chordata | 88865.0 | 0 | Carnivora | 88865.0 | 0 | Canidae | 88865.0 | 0 | Canis | 88865.0 | 0 | Canis lupus | 88865.0 | 0 |
| scf7180000041207 | 1475521 | 0.4849 | 0 | 59.8688 | Eukaryota | 86243.0 | 0 | Chordata | 86243.0 | 0 | Carnivora | 86243.0 | 0 | Canidae | 86243.0 | 0 | Canis | 86243.0 | 0 | Canis lupus | 86243.0 | 0 |
| scf7180000044969 | 472505 | 0.4203 | 0 | 59.0328 | Eukaryota | 43162.0 | 0 | Chordata | 43162.0 | 0 | Carnivora | 43162.0 | 0 | Canidae | 43162.0 | 0 | Canis | 43162.0 | 0 | Canis lupus | 43162.0 | 0 |
| scf7180000048720 | 5912 | 0.4438 | 0 | 102.7095 | no-hit | 0.0 | 0 | no-hit | 0.0 | 0 | no-hit | 0.0 | 0 | no-hit | 0.0 | 0 | no-hit | 0.0 | 0 | no-hit | 0.0 | 0 |
| scf7180000046409 | 25231 | 0.436 | 0 | 126.9376 | Eukaryota | 7676.0 | 0 | Chordata | 4989.0 | 1 | Polystomatidea | 2687.0 | 3 | Polystomatidae | 2687.0 | 3 | Protopolystoma | 2687.0 | 3 | Protopolystoma xenopodis | 2687.0 | 3 |
| scf7180000046882 | 35257 | 0.437 | 0 | 110.4471 | Eukaryota | 4690.0 | 1 | Platyhelminthes | 3065.0 | 2 | Polystomatidea | 3065.0 | 2 | Polystomatidae | 3065.0 | 2 | Protopolystoma | 3065.0 | 3 | Protopolystoma xenopodis | 3065.0 | 3 |
| scf7180000052209 | 7003 | 0.4494 | 0 | 703.198 | Eukaryota | 419.0 | 0 | Platyhelminthes | 254.0 | 1 | Polystomatidea | 254.0 | 1 | Polystomatidae | 254.0 | 1 | Protopolystoma | 254.0 | 1 | Protopolystoma xenopodis | 254.0 | 1 |
| scf7180000041493 | 500860 | 0.5288 | 0 | 59.9108 | Eukaryota | 103275.0 | 0 | Chordata | 103275.0 | 0 | Carnivora | 103275.0 | 0 | Canidae | 103275.0 | 0 | Canis | 103275.0 | 0 | Canis lupus | 103275.0 | 0 |
| scf7180000045220 | 88501 | 0.3694 | 0 | 56.6343 | Eukaryota | 9872.0 | 0 | Chordata | 9872.0 | 0 | Primates | 3596.0 | 2 | Hominidae | 3596.0 | 2 | Canis | 3558.0 | 4 | Canis lupus | 3558.0 | 4 |
| scf7180000041585 | 1018089 | 0.3725 | 0 | 59.7268 | Eukaryota | 89291.0 | 0 | Chordata | 89291.0 | 0 | Carnivora | 89291.0 | 0 | Canidae | 89291.0 | 0 | Canis | 89291.0 | 0 | Canis lupus | 89291.0 | 0 |
| scf7180000049907 | 33151 | 0.506 | 0 | 165.6907 | Eukaryota | 9026.0 | 0 | Chordata | 9026.0 | 0 | Carnivora | 9026.0 | 0 | Canidae | 9026.0 | 0 | Nyctereutes | 4807.0 | 1 | Nyctereutes procyonoides | 4807.0 | 1 |
| scf7180000046965 | 121419 | 0.362 | 0 | 59.5697 | Eukaryota | 14065.0 | 0 | Chordata | 14065.0 | 0 | Primates | 7340.0 | 2 | Hominidae | 5537.0 | 5 | Homo | 5537.0 | 7 | Homo sapiens | 5537.0 | 7 |
| scf7180000042665 | 22450 | 0.4503 | 0 | 51.9741 | Eukaryota | 36169.0 | 0 | Chordata | 36169.0 | 0 | Carnivora | 36169.0 | 0 | Canidae | 36169.0 | 0 | Canis | 36169.0 | 0 | Canis lupus | 36169.0 | 0 |
| scf7180000047620 | 29138 | 0.4376 | 0 | 92.6141 | Eukaryota | 15501.0 | 0 | Chordata | 9521.0 | 2 | Kurtiformes | 4150.0 | 6 | Apogonidae | 4150.0 | 6 | Sphaeramia | 4150.0 | 6 | Sphaeramia orbicularis | 4150.0 | 6 |
| scf7180000043073 | 261691 | 0.4519 | 0 | 55.6967 | Eukaryota | 24067.0 | 0 | Chordata | 24067.0 | 0 | Carnivora | 24067.0 | 0 | Canidae | 24067.0 | 0 | Canis | 24067.0 | 0 | Canis lupus | 24067.0 | 0 |
| scf7180000047885 | 47442 | 0.4336 | 0 | 282.7311 | Eukaryota | 3289.0 | 0 | Platyhelminthes | 2311.0 | 1 | Polystomatidea | 2311.0 | 2 | Polystomatidae | 2311.0 | 2 | Protopolystoma | 2311.0 | 2 | Protopolystoma xenopodis | 2311.0 | 2 |
| scf7180000046510 | 24686 | 0.4678 | 0 | 124.264 | Eukaryota | 11369.0 | 0 | Chordata | 11369.0 | 0 | Carnivora | 11369.0 | 0 | Canidae | 11369.0 | 0 | Canis | 8016.0 | 1 | Canis lupus | 8016.0 | 1 |
| scf7180000042353 | 723025 | 0.417 | 0 | 60.5092 | Eukaryota | 37361.0 | 0 | Chordata | 37361.0 | 0 | Carnivora | 37361.0 | 0 | Canidae | 37361.0 | 0 | Canis | 37361.0 | 0 | Canis lupus | 37361.0 | 0 |
| scf7180000042695 | 326203 | 0.3926 | 0 | 59.6258 | Eukaryota | 55150.0 | 0 | Chordata | 55150.0 | 0 | Carnivora | 55150.0 | 0 | Canidae | 55150.0 | 0 | Canis | 55150.0 | 0 | Canis lupus | 55150.0 | 0 |
| scf7180000052429 | 7369 | 0.4499 | 0 | 2298.2274 | Eukaryota | 375.0 | 0 | Chordata | 375.0 | 0 | Kurtiformes | 375.0 | 0 | Apogonidae | 375.0 | 0 | Sphaeramia | 375.0 | 0 | Sphaeramia orbicularis | 375.0 | 0 |
| scf7180000044908 | 265772 | 0.3737 | 0 | 60.8013 | Eukaryota | 38175.0 | 0 | Chordata | 38175.0 | 0 | Carnivora | 38175.0 | 0 | Canidae | 38175.0 | 0 | Canis | 38175.0 | 0 | Canis lupus | 38175.0 | 0 |
| scf7180000052095 | 3628 | 0.4551 | 0 | 775.7767 | Eukaryota | 3808.0 | 0 | Platyhelminthes | 2256.0 | 3 | Polystomatidea | 2256.0 | 3 | Polystomatidae | 2256.0 | 3 | Protopolystoma | 2256.0 | 3 | Protopolystoma xenopodis | 2256.0 | 3 |
| scf7180000041148 | 1194168 | 0.3951 | 0 | 60.8967 | Eukaryota | 124980.0 | 0 | Chordata | 124980.0 | 0 | Carnivora | 124980.0 | 0 | Canidae | 124980.0 | 0 | Canis | 97664.0 | 1 | Canis lupus | 97664.0 | 1 |
| scf7180000043892 | 330810 | 0.3994 | 0 | 59.9227 | Eukaryota | 28566.0 | 0 | Chordata | 28566.0 | 0 | Carnivora | 22752.0 | 1 | Canidae | 22752.0 | 1 | Canis | 22752.0 | 1 | Canis lupus | 22752.0 | 1 |
| scf7180000044551 | 25940 | 0.4444 | 0 | 183.542 | Eukaryota | 64874.0 | 0 | Chordata | 64874.0 | 0 | Carnivora | 64874.0 | 0 | Canidae | 64874.0 | 0 | Canis | 64874.0 | 0 | Canis lupus | 64874.0 | 0 |
| scf7180000049628 | 15869 | 0.4431 | 0 | 227.4304 | Eukaryota | 2019.0 | 0 | Chordata | 1804.0 | 1 | Kurtiformes | 1291.0 | 3 | Apogonidae | 1291.0 | 3 | Sphaeramia | 1291.0 | 3 | Sphaeramia orbicularis | 1291.0 | 3 |
| scf7180000047118 | 40732 | 0.4495 | 0 | 1571.7533 | Eukaryota | 25361.0 | 0 | Chordata | 22467.0 | 2 | Gadiformes | 15024.0 | 7 | Gadidae | 15024.0 | 7 | Gadus | 15024.0 | 7 | Gadus morhua | 15024.0 | 7 |
| scf7180000042830 | 10168 | 0.3906 | 0 | 51.6572 | Eukaryota | 60423.0 | 0 | Chordata | 60423.0 | 0 | Carnivora | 60423.0 | 0 | Canidae | 60423.0 | 0 | Canis | 60423.0 | 0 | Canis lupus | 60423.0 | 0 |
| scf7180000040890 | 9873 | 0.427 | 0 | 58.0503 | Eukaryota | 70777.0 | 0 | Chordata | 70777.0 | 0 | Carnivora | 70777.0 | 0 | Canidae | 70777.0 | 0 | Canis | 70777.0 | 0 | Canis lupus | 70777.0 | 0 |
| scf7180000047001 | 15009 | 0.4572 | 0 | 143.2421 | Eukaryota | 17362.0 | 0 | Chordata | 17362.0 | 0 | Carnivora | 17362.0 | 0 | Canidae | 17362.0 | 0 | Canis | 17362.0 | 0 | Canis lupus | 17362.0 | 0 |
| scf7180000046343 | 190972 | 0.3661 | 0 | 65.0075 | Eukaryota | 56275.0 | 0 | Chordata | 56275.0 | 0 | Carnivora | 56275.0 | 0 | Canidae | 56275.0 | 0 | Canis | 56275.0 | 0 | Canis lupus | 56275.0 | 0 |
| scf7180000048726 | 34737 | 0.444 | 0 | 127.8429 | Eukaryota | 3108.0 | 0 | Chordata | 2690.0 | 1 | Columbiformes | 2690.0 | 1 | Columbidae | 2690.0 | 1 | Streptopelia | 2690.0 | 1 | Streptopelia turtur | 2690.0 | 1 |
| scf7180000042445 | 746329 | 0.4205 | 0 | 60.5471 | Eukaryota | 92606.0 | 0 | Chordata | 92606.0 | 0 | Carnivora | 92606.0 | 0 | Canidae | 92606.0 | 0 | Canis | 80438.0 | 1 | Canis lupus | 80438.0 | 1 |
| scf7180000048182 | 63300 | 0.3709 | 0 | 58.9088 | Eukaryota | 46974.0 | 0 | Chordata | 46974.0 | 0 | Carnivora | 46974.0 | 0 | Canidae | 46974.0 | 0 | Canis | 46974.0 | 0 | Canis lupus | 46974.0 | 0 |
| scf7180000047191 | 22270 | 0.4383 | 0 | 94.4473 | Eukaryota | 991.0 | 0 | Platyhelminthes | 991.0 | 0 | Polystomatidea | 991.0 | 0 | Polystomatidae | 991.0 | 0 | Protopolystoma | 991.0 | 0 | Protopolystoma xenopodis | 991.0 | 0 |
| scf7180000044295 | 6201 | 0.4412 | 0 | 67.4569 | Eukaryota | 4504.0 | 0 | Platyhelminthes | 3312.0 | 1 | Polystomatidea | 3312.0 | 1 | Polystomatidae | 3312.0 | 1 | Protopolystoma | 3312.0 | 2 | Protopolystoma xenopodis | 3312.0 | 2 |
| scf7180000044522 | 24917 | 0.3498 | 0 | 51.7366 | Eukaryota | 89083.0 | 0 | Chordata | 89083.0 | 0 | Carnivora | 89083.0 | 0 | Canidae | 89083.0 | 0 | Canis | 89083.0 | 0 | Canis lupus | 89083.0 | 0 |
| scf7180000044221 | 362277 | 0.3943 | 0 | 59.3045 | Eukaryota | 62300.0 | 0 | Chordata | 62300.0 | 0 | Carnivora | 62300.0 | 0 | Canidae | 62300.0 | 0 | Canis | 57125.0 | 1 | Canis lupus | 57125.0 | 1 |
| scf7180000056493 | 364501 | 0.3841 | 0 | 60.5914 | Eukaryota | 42798.0 | 0 | Chordata | 42798.0 | 0 | Carnivora | 42798.0 | 0 | Canidae | 36072.0 | 1 | Canis | 24742.0 | 3 | Canis lupus | 24742.0 | 3 |
| scf7180000045634 | 210453 | 0.6363 | 0 | 56.0546 | Eukaryota | 58266.0 | 0 | Chordata | 58266.0 | 0 | Carnivora | 58266.0 | 0 | Canidae | 58266.0 | 0 | Canis | 58266.0 | 0 | Canis lupus | 58266.0 | 0 |
| scf7180000043471 | 472177 | 0.399 | 0 | 59.24 | Eukaryota | 90262.0 | 0 | Chordata | 90262.0 | 0 | Carnivora | 90262.0 | 0 | Canidae | 90262.0 | 0 | Canis | 90262.0 | 0 | Canis lupus | 90262.0 | 0 |
| scf7180000055861 | 18722 | 0.5189 | 0 | 398.3954 | Eukaryota | 15780.0 | 0 | Chordata | 15780.0 | 0 | Carnivora | 15780.0 | 0 | Canidae | 15780.0 | 0 | Canis | 15780.0 | 0 | Canis lupus | 15780.0 | 0 |
| scf7180000049152 | 5606 | 0.447 | 0 | 132.3371 | Eukaryota | 2835.0 | 0 | Chordata | 2256.0 | 1 | Carnivora | 2256.0 | 1 | Canidae | 2256.0 | 1 | Canis | 2256.0 | 1 | Canis lupus | 2256.0 | 1 |
| scf7180000044678 | 73343 | 0.3859 | 0 | 59.0522 | Eukaryota | 54662.0 | 0 | Chordata | 54662.0 | 0 | Carnivora | 54662.0 | 0 | Canidae | 54662.0 | 0 | Canis | 54662.0 | 0 | Canis lupus | 54662.0 | 0 |
| scf7180000041488 | 1022838 | 0.4913 | 0 | 58.6899 | Eukaryota | 70880.0 | 0 | Chordata | 70880.0 | 0 | Carnivora | 70880.0 | 0 | Canidae | 70880.0 | 0 | Canis | 56852.0 | 1 | Canis lupus | 56852.0 | 1 |
| scf7180000053620 | 2989 | 0.4483 | 0 | 392.4599 | no-hit | 0.0 | 0 | no-hit | 0.0 | 0 | no-hit | 0.0 | 0 | no-hit | 0.0 | 0 | no-hit | 0.0 | 0 | no-hit | 0.0 | 0 |
| scf7180000041260 | 1176948 | 0.4791 | 0 | 58.86 | Eukaryota | 85621.0 | 0 | Chordata | 85621.0 | 0 | Carnivora | 85621.0 | 0 | Canidae | 67982.0 | 2 | Canis | 38223.0 | 3 | Canis lupus | 38223.0 | 3 |
| scf7180000042462 | 538933 | 0.5196 | 0 | 58.9317 | Eukaryota | 255472.0 | 0 | Chordata | 255472.0 | 0 | Carnivora | 255472.0 | 0 | Canidae | 255472.0 | 0 | Canis | 238620.0 | 1 | Canis lupus | 238620.0 | 1 |
| scf7180000056427 | 156528 | 0.3895 | 0 | 57.8761 | Eukaryota | 22761.0 | 0 | Chordata | 22761.0 | 0 | Carnivora | 22761.0 | 0 | Canidae | 22761.0 | 0 | Canis | 11649.0 | 1 | Canis lupus | 11649.0 | 1 |
| scf7180000050624 | 59152 | 0.3955 | 0 | 58.8979 | Eukaryota | 13839.0 | 0 | Chordata | 13839.0 | 0 | Carnivora | 13839.0 | 0 | Canidae | 13839.0 | 0 | Canis | 13839.0 | 0 | Canis lupus | 13839.0 | 0 |
| scf7180000053654 | 5167 | 0.5038 | 0 | 61.6677 | Eukaryota | 7683.0 | 0 | Chordata | 7683.0 | 0 | Carnivora | 7683.0 | 0 | Canidae | 7683.0 | 0 | Canis | 5669.0 | 1 | Canis lupus | 4551.0 | 2 |
| scf7180000045999 | 46415 | 0.3956 | 0 | 56.2484 | Eukaryota | 52516.0 | 0 | Chordata | 52516.0 | 0 | Carnivora | 52516.0 | 0 | Canidae | 52516.0 | 0 | Canis | 52516.0 | 0 | Canis lupus | 52516.0 | 0 |
| scf7180000053366 | 5835 | 0.4488 | 0 | 727.9861 | Eukaryota | 3606.0 | 0 | Platyhelminthes | 1940.0 | 3 | Polystomatidea | 1940.0 | 3 | Polystomatidae | 1940.0 | 3 | Protopolystoma | 1940.0 | 3 | Protopolystoma xenopodis | 1940.0 | 3 |
| scf7180000055835 | 24010 | 0.4406 | 0 | 520.6799 | Eukaryota | 2701.0 | 0 | Platyhelminthes | 1261.0 | 2 | Polystomatidea | 1261.0 | 2 | Polystomatidae | 1261.0 | 2 | Protopolystoma | 1261.0 | 2 | Protopolystoma xenopodis | 1261.0 | 2 |
| scf7180000045948 | 133395 | 0.3715 | 0 | 59.8221 | Eukaryota | 11474.0 | 0 | Chordata | 10864.0 | 1 | Primates | 6798.0 | 4 | Hominidae | 5306.0 | 5 | Homo | 5306.0 | 5 | Homo sapiens | 5306.0 | 5 |
| scf7180000051579 | 6307 | 0.4279 | 0 | 103.9717 | Eukaryota | 2310.0 | 0 | Platyhelminthes | 1786.0 | 2 | Polystomatidea | 1786.0 | 2 | Polystomatidae | 1786.0 | 2 | Protopolystoma | 1786.0 | 2 | Protopolystoma xenopodis | 1786.0 | 2 |
| scf7180000053717 | 7759 | 0.4364 | 0 | 66.0858 | Eukaryota | 1768.0 | 0 | Chordata | 1001.0 | 1 | Polystomatidea | 767.0 | 3 | Polystomatidae | 767.0 | 3 | Protopolystoma | 767.0 | 4 | Protopolystoma xenopodis | 767.0 | 4 |
| scf7180000051665 | 4474 | 0.4443 | 0 | 61.9137 | Eukaryota | 14609.0 | 0 | Chordata | 14056.0 | 1 | Gadiformes | 6903.0 | 5 | Gadidae | 6903.0 | 5 | Gadus | 6903.0 | 5 | Gadus morhua | 6903.0 | 5 |
| scf7180000041868 | 853248 | 0.3919 | 0 | 59.6856 | Eukaryota | 56642.0 | 0 | Chordata | 56642.0 | 0 | Carnivora | 56642.0 | 0 | Canidae | 56642.0 | 0 | Vulpes | 28640.0 | 1 | Vulpes vulpes | 28640.0 | 1 |
| scf7180000040750 | 45694 | 0.4494 | 0 | 63.944 | Eukaryota | 84061.0 | 0 | Chordata | 84061.0 | 0 | Carnivora | 84061.0 | 0 | Canidae | 84061.0 | 0 | Canis | 84061.0 | 0 | Canis lupus | 84061.0 | 0 |
| scf7180000048390 | 11554 | 0.409 | 0 | 114.1072 | Eukaryota | 7678.0 | 0 | Chordata | 7678.0 | 0 | Carnivora | 7678.0 | 0 | Canidae | 6907.0 | 1 | Canis | 6907.0 | 1 | Canis lupus | 6907.0 | 1 |
| scf7180000040966 | 1476968 | 0.4887 | 0 | 59.279 | Eukaryota | 115770.0 | 0 | Chordata | 115770.0 | 0 | Carnivora | 115770.0 | 0 | Canidae | 115770.0 | 0 | Vulpes | 69498.0 | 1 | Vulpes vulpes | 69498.0 | 1 |
| scf7180000043747 | 473241 | 0.385 | 0 | 59.0163 | Eukaryota | 26346.0 | 0 | Chordata | 26346.0 | 0 | Carnivora | 26346.0 | 0 | Canidae | 21018.0 | 1 | Canis | 21018.0 | 2 | Canis lupus | 21018.0 | 2 |
| scf7180000040535 | 12848 | 0.3792 | 0 | 46.8769 | Eukaryota | 4451.0 | 1 | Chordata | 4451.0 | 1 | Artiodactyla | 2105.0 | 3 | Canidae | 1400.0 | 4 | Canis | 1400.0 | 5 | Canis lupus | 1400.0 | 5 |
| scf7180000043979 | 74862 | 0.35 | 0 | 58.3633 | Eukaryota | 12225.0 | 0 | Chordata | 12225.0 | 0 | Carnivora | 9378.0 | 1 | Canidae | 8585.0 | 2 | Canis | 5834.0 | 3 | Canis lupus | 5834.0 | 3 |
| scf7180000047527 | 9521 | 0.4287 | 0 | 119.1766 | no-hit | 0.0 | 0 | no-hit | 0.0 | 0 | no-hit | 0.0 | 0 | no-hit | 0.0 | 0 | no-hit | 0.0 | 0 | no-hit | 0.0 | 0 |
| scf7180000053476 | 4510 | 0.4455 | 0 | 200.3852 | no-hit | 0.0 | 0 | no-hit | 0.0 | 0 | no-hit | 0.0 | 0 | no-hit | 0.0 | 0 | no-hit | 0.0 | 0 | no-hit | 0.0 | 0 |
| scf7180000055518 | 41086 | 0.4422 | 0 | 106.9569 | Eukaryota | 77277.0 | 0 | Chordata | 77277.0 | 0 | Carnivora | 77277.0 | 0 | Canidae | 77277.0 | 0 | Canis | 77277.0 | 0 | Canis lupus | 77277.0 | 0 |
| scf7180000052882 | 6357 | 0.4515 | 0 | 678.1755 | Eukaryota | 4642.0 | 0 | Chordata | 4642.0 | 0 | Tetraodontiformes | 4642.0 | 0 | Tetraodontidae | 4642.0 | 0 | Takifugu | 4642.0 | 0 | Takifugu rubripes | 4642.0 | 0 |
| scf7180000053915 | 8886 | 0.507 | 0 | 59.4173 | Eukaryota | 8420.0 | 0 | Chordata | 8420.0 | 0 | Carnivora | 8420.0 | 0 | Canidae | 8420.0 | 0 | Canis | 6358.0 | 1 | Canis lupus | 5182.0 | 2 |
| scf7180000044201 | 1949 | 0.4433 | 0 | 325.7064 | Eukaryota | 5300.0 | 0 | Chordata | 5017.0 | 1 | Kurtiformes | 3625.0 | 2 | Apogonidae | 3625.0 | 2 | Sphaeramia | 3625.0 | 2 | Sphaeramia orbicularis | 3625.0 | 2 |
| scf7180000042487 | 20741 | 0.4308 | 0 | 56.909 | Eukaryota | 48322.0 | 0 | Chordata | 48322.0 | 0 | Carnivora | 48322.0 | 0 | Canidae | 48322.0 | 0 | Canis | 43481.0 | 1 | Canis lupus | 43481.0 | 1 |
| scf7180000053808 | 9274 | 0.5024 | 0 | 141.3045 | Eukaryota | 8801.0 | 0 | Chordata | 8801.0 | 0 | Carnivora | 8801.0 | 0 | Canidae | 8801.0 | 0 | Canis | 7077.0 | 1 | Canis lupus | 5866.0 | 2 |
| scf7180000048713 | 2647 | 0.4409 | 0 | 102.3865 | no-hit | 0.0 | 0 | no-hit | 0.0 | 0 | no-hit | 0.0 | 0 | no-hit | 0.0 | 0 | no-hit | 0.0 | 0 | no-hit | 0.0 | 0 |
| scf7180000052546 | 16370 | 0.461 | 0 | 82.3494 | Eukaryota | 36679.0 | 0 | Chordata | 36679.0 | 0 | Carnivora | 36679.0 | 0 | Canidae | 36679.0 | 0 | Canis | 36679.0 | 0 | Canis lupus | 36679.0 | 0 |
| scf7180000048289 | 30602 | 0.4381 | 0 | 109.48 | Eukaryota | 14000.0 | 0 | Chordata | 9846.0 | 1 | Carnivora | 7321.0 | 2 | Canidae | 7321.0 | 2 | Canis | 7321.0 | 2 | Canis lupus | 7321.0 | 2 |
| scf7180000045881 | 13962 | 0.4429 | 0 | 197.866 | Eukaryota | 87005.0 | 0 | Chordata | 87005.0 | 0 | Carnivora | 87005.0 | 0 | Canidae | 87005.0 | 0 | Canis | 87005.0 | 0 | Canis lupus | 87005.0 | 0 |
| scf7180000042947 | 202643 | 0.3427 | 0 | 57.8175 | Eukaryota | 70687.0 | 0 | Chordata | 70687.0 | 0 | Carnivora | 70687.0 | 0 | Canidae | 70687.0 | 0 | Canis | 70687.0 | 0 | Canis lupus | 70687.0 | 0 |
| scf7180000044646 | 15281 | 0.4179 | 0 | 47.817 | Eukaryota | 2897.0 | 0 | Chordata | 2897.0 | 0 | Carnivora | 2462.0 | 1 | Canidae | 2462.0 | 1 | Vulpes | 2192.0 | 2 | Vulpes vulpes | 2192.0 | 2 |
| scf7180000049817 | 38226 | 0.432 | 0 | 58.0974 | Eukaryota | 41830.0 | 0 | Chordata | 41830.0 | 0 | Carnivora | 41830.0 | 0 | Canidae | 41830.0 | 0 | Canis | 41830.0 | 0 | Canis lupus | 41830.0 | 0 |
| scf7180000047700 | 13789 | 0.4385 | 0 | 261.0389 | Eukaryota | 6648.0 | 1 | Chordata | 6648.0 | 1 | Apterygiformes | 2848.0 | 3 | Apterygidae | 2848.0 | 3 | Apteryx | 2848.0 | 3 | Apteryx mantelli | 2848.0 | 3 |
| scf7180000044538 | 268390 | 0.3529 | 0 | 60.5921 | Eukaryota | 90413.0 | 0 | Chordata | 90413.0 | 0 | Carnivora | 90413.0 | 0 | Canidae | 90413.0 | 0 | Canis | 90413.0 | 0 | Canis lupus | 90413.0 | 0 |
| scf7180000049736 | 43150 | 0.3457 | 0 | 58.7466 | Eukaryota | 14608.0 | 0 | Chordata | 14608.0 | 0 | Apterygiformes | 13043.0 | 1 | Apterygidae | 13043.0 | 1 | Apteryx | 13043.0 | 1 | Apteryx mantelli | 13043.0 | 1 |
| scf7180000052845 | 5793 | 0.4381 | 0 | 302.3299 | no-hit | 0.0 | 0 | no-hit | 0.0 | 0 | no-hit | 0.0 | 0 | no-hit | 0.0 | 0 | no-hit | 0.0 | 0 | no-hit | 0.0 | 0 |
| scf7180000045520 | 159088 | 0.3771 | 0 | 60.3445 | Eukaryota | 78102.0 | 0 | Chordata | 78102.0 | 0 | Carnivora | 78102.0 | 0 | Canidae | 78102.0 | 0 | Canis | 70430.0 | 1 | Canis lupus | 70430.0 | 1 |
| scf7180000041797 | 166141 | 0.4006 | 0 | 58.1039 | Eukaryota | 41397.0 | 0 | Chordata | 41397.0 | 0 | Carnivora | 41397.0 | 0 | Canidae | 41397.0 | 0 | Canis | 41397.0 | 0 | Canis lupus | 41397.0 | 0 |
| scf7180000050951 | 6258 | 0.4441 | 0 | 125.3479 | no-hit | 0.0 | 0 | no-hit | 0.0 | 0 | no-hit | 0.0 | 0 | no-hit | 0.0 | 0 | no-hit | 0.0 | 0 | no-hit | 0.0 | 0 |
| scf7180000049483 | 24860 | 0.442 | 0 | 39.017 | Eukaryota | 1053.0 | 0 | Platyhelminthes | 633.0 | 1 | Polystomatidea | 633.0 | 2 | Polystomatidae | 633.0 | 2 | Protopolystoma | 633.0 | 2 | Protopolystoma xenopodis | 633.0 | 2 |
| scf7180000054426 | 32353 | 0.3555 | 0 | 966.1155 | Eukaryota | 14129.0 | 0 | Chordata | 14129.0 | 0 | Carnivora | 14129.0 | 0 | Canidae | 9429.0 | 2 | Canis | 7595.0 | 4 | Canis lupus | 7595.0 | 4 |
| scf7180000053693 | 4719 | 0.4469 | 0 | 834.0596 | no-hit | 0.0 | 0 | no-hit | 0.0 | 0 | no-hit | 0.0 | 0 | no-hit | 0.0 | 0 | no-hit | 0.0 | 0 | no-hit | 0.0 | 0 |
| scf7180000050094 | 2744 | 0.4548 | 0 | 114.0825 | no-hit | 0.0 | 0 | no-hit | 0.0 | 0 | no-hit | 0.0 | 0 | no-hit | 0.0 | 0 | no-hit | 0.0 | 0 | no-hit | 0.0 | 0 |
| scf7180000041786 | 338175 | 0.4037 | 0 | 60.6221 | Eukaryota | 85495.0 | 0 | Chordata | 85495.0 | 0 | Carnivora | 85495.0 | 0 | Canidae | 85495.0 | 0 | Canis | 85495.0 | 0 | Canis lupus | 85495.0 | 0 |
| scf7180000044579 | 295665 | 0.3818 | 0 | 145.1219 | Eukaryota | 93483.0 | 0 | Chordata | 93483.0 | 0 | Carnivora | 93483.0 | 0 | Canidae | 93483.0 | 0 | Canis | 93483.0 | 0 | Canis lupus | 93483.0 | 0 |
| scf7180000047124 | 7142 | 0.4595 | 0 | 84.4379 | Eukaryota | 7907.0 | 0 | Chordata | 7907.0 | 0 | Carnivora | 7907.0 | 0 | Canidae | 7907.0 | 0 | Canis | 5911.0 | 1 | Canis lupus | 4876.0 | 2 |
| scf7180000047049 | 12955 | 0.5145 | 0 | 424.1724 | Eukaryota | 16119.0 | 0 | Chordata | 16119.0 | 0 | Carnivora | 16119.0 | 0 | Canidae | 16119.0 | 0 | Canis | 16119.0 | 0 | Canis lupus | 16119.0 | 0 |
| scf7180000043959 | 178540 | 0.4039 | 0 | 60.8198 | Eukaryota | 99708.0 | 1 | Chordata | 99708.0 | 1 | Carnivora | 75367.0 | 2 | Canidae | 67385.0 | 3 | Canis | 50545.0 | 4 | Canis lupus | 50545.0 | 4 |
| scf7180000045627 | 69365 | 0.4399 | 0 | 142.0461 | Eukaryota | 73950.0 | 0 | Chordata | 42403.0 | 1 | Gadiformes | 27887.0 | 6 | Gadidae | 27887.0 | 6 | Gadus | 27887.0 | 6 | Gadus morhua | 27887.0 | 6 |
| scf7180000042959 | 573533 | 0.3847 | 0 | 59.6573 | Eukaryota | 92824.0 | 0 | Chordata | 92824.0 | 0 | Carnivora | 92824.0 | 0 | Canidae | 92824.0 | 0 | Canis | 75309.0 | 1 | Canis lupus | 75309.0 | 1 |
| scf7180000045085 | 211035 | 0.5331 | 0 | 59.0837 | Eukaryota | 18940.0 | 0 | Chordata | 18940.0 | 0 | Carnivora | 18940.0 | 0 | Canidae | 18940.0 | 0 | Canis | 18940.0 | 0 | Canis lupus | 18940.0 | 0 |
| scf7180000047822 | 20168 | 0.4383 | 0 | 97.7753 | Eukaryota | 4430.0 | 0 | Platyhelminthes | 3249.0 | 2 | Polystomatidea | 3249.0 | 2 | Polystomatidae | 3249.0 | 2 | Protopolystoma | 3249.0 | 2 | Protopolystoma xenopodis | 3249.0 | 2 |
| scf7180000054853 | 40156 | 0.4483 | 0 | 281.8035 | Eukaryota | 84930.0 | 0 | Chordata | 84930.0 | 0 | Carnivora | 84930.0 | 0 | Canidae | 84930.0 | 0 | Canis | 84930.0 | 0 | Canis lupus | 84930.0 | 0 |
| scf7180000053292 | 6336 | 0.4591 | 0 | 57.6619 | no-hit | 0.0 | 0 | no-hit | 0.0 | 0 | no-hit | 0.0 | 0 | no-hit | 0.0 | 0 | no-hit | 0.0 | 0 | no-hit | 0.0 | 0 |
| scf7180000043757 | 526648 | 0.3835 | 0 | 58.2243 | Eukaryota | 17795.0 | 0 | Chordata | 17795.0 | 0 | Carnivora | 17795.0 | 0 | Canidae | 17795.0 | 0 | Canis | 17795.0 | 0 | Canis lupus | 17795.0 | 0 |
| scf7180000045010 | 248955 | 0.3857 | 0 | 60.8623 | Eukaryota | 16983.0 | 0 | Chordata | 16983.0 | 0 | Carnivora | 16983.0 | 0 | Canidae | 16983.0 | 0 | Canis | 16983.0 | 0 | Canis lupus | 16983.0 | 0 |
| scf7180000044977 | 292198 | 0.3937 | 0 | 122.342 | Eukaryota | 87871.0 | 0 | Chordata | 87871.0 | 0 | Carnivora | 87871.0 | 0 | Canidae | 87871.0 | 0 | Canis | 87871.0 | 0 | Canis lupus | 87871.0 | 0 |
| scf7180000055711 | 4608 | 0.4518 | 0 | 205.5259 | Eukaryota | 2341.0 | 0 | Chordata | 2006.0 | 1 | Carnivora | 1752.0 | 2 | Canidae | 1752.0 | 2 | Canis | 1322.0 | 3 | Canis lupus | 1322.0 | 3 |

| | | | | | | | | | | | | | | | | | | | | | | |
|---|---|---|---|---|---|---|---|---|---|---|---|---|---|---|---|---|---|---|---|---|---|---|
| scf7180000042493 | 731002 | 0.4515 | 0 | 58.5487 | Eukaryota | 78858.0 | 0 | Chordata | 78858.0 | 0 | Carnivora | 78858.0 | 0 | Canidae | 78858.0 | 0 | Canis | 41456.0 | 1 | Canis lupus | 41456.0 | 1 |
| scf7180000053539 | 5301 | 0.4493 | 0 | 430.5059 | no-hit | 0.0 | 0 | no-hit | 0.0 | 0 | no-hit | 0.0 | 0 | no-hit | 0.0 | 0 | no-hit | 0.0 | 0 | no-hit | 0.0 | 0 |
| scf7180000046920 | 131533 | 0.4077 | 0 | 59.7688 | Eukaryota | 93560.0 | 0 | Chordata | 93560.0 | 0 | Carnivora | 93560.0 | 0 | Canidae | 93560.0 | 0 | Canis | 93560.0 | 0 | Canis lupus | 93560.0 | 0 |
| scf7180000051572 | 18236 | 0.4394 | 0 | 234.0692 | Eukaryota | 30422.0 | 0 | Chordata | 30422.0 | 0 | Gadiformes | 13742.0 | 2 | Gadidae | 13742.0 | 2 | Gadus | 13742.0 | 2 | Gadus morhua | 13742.0 | 3 |
| scf7180000043360 | 37807 | 0.4426 | 0 | 189.8517 | Eukaryota | 42144.0 | 0 | Chordata | 42144.0 | 0 | Carnivora | 22283.0 | 1 | Canidae | 22283.0 | 1 | Canis | 22283.0 | 1 | Canis lupus | 22283.0 | 1 |
| scf7180000041777 | 348416 | 0.4177 | 0 | 58.8563 | Eukaryota | 62054.0 | 0 | Chordata | 62054.0 | 0 | Carnivora | 62054.0 | 0 | Canidae | 57365.0 | 1 | Canis | 43309.0 | 2 | Canis lupus | 43309.0 | 2 |
| scf7180000053821 | 5894 | 0.4428 | 0 | 69.0059 | Eukaryota | 5519.0 | 1 | Chordata | 4537.0 | 2 | Salmoniformes | 2842.0 | 5 | Salmonidae | 2842.0 | 5 | Salmo | 2612.0 | 6 | Salmo trutta | 2612.0 | 6 |
| scf7180000042108 | 735779 | 0.4117 | 0 | 60.2257 | Eukaryota | 52765.0 | 0 | Chordata | 52765.0 | 0 | Carnivora | 52765.0 | 0 | Canidae | 19894.0 | 4 | Canis | 11980.0 | 8 | Canis lupus | 11980.0 | 8 |
| scf7180000055070 | 26692 | 0.4393 | 0 | 95.4077 | Eukaryota | 8937.0 | 0 | Chordata | 6348.0 | 1 | Synbranchiformes | 2854.0 | 4 | Mastacembelidae | 2854.0 | 4 | Mastacembelus | 2854.0 | 4 | Mastacembelus armatus | 2854.0 | 4 |
| scf7180000047426 | 71307 | 0.4054 | 0 | 61.2164 | Eukaryota | 40483.0 | 0 | Chordata | 40483.0 | 0 | Carnivora | 40483.0 | 0 | Canidae | 22880.0 | 3 | Canis | 17563.0 | 7 | Canis lupus | 17563.0 | 8 |
| scf7180000041012 | 4594 | 0.429 | 0 | 77.7992 | Eukaryota | 2553.0 | 0 | Chordata | 2553.0 | 0 | Kurtiformes | 2553.0 | 0 | Apogonidae | 2553.0 | 0 | Sphaeramia | 2553.0 | 0 | Sphaeramia orbicularis | 2553.0 | 0 |
| scf7180000048837 | 384469 | 0.4431 | 0 | 61.0059 | Eukaryota | 91582.0 | 0 | Chordata | 91582.0 | 0 | Carnivora | 91582.0 | 0 | Canidae | 91582.0 | 0 | Canis | 91582.0 | 0 | Canis lupus | 91582.0 | 0 |
| scf7180000044055 | 345487 | 0.3967 | 0 | 60.1876 | Eukaryota | 42845.0 | 0 | Chordata | 42845.0 | 0 | Carnivora | 38835.0 | 1 | Canidae | 20694.0 | 4 | Canis | 16767.0 | 6 | Canis lupus | 16767.0 | 7 |
| scf7180000045845 | 151735 | 0.3762 | 0 | 57.0684 | Eukaryota | 26710.0 | 0 | Chordata | 26710.0 | 0 | Carnivora | 26710.0 | 0 | Canidae | 26710.0 | 0 | Canis | 23758.0 | 1 | Canis lupus | 23758.0 | 1 |
| scf7180000052596 | 2013 | 0.4501 | 0 | 478.6822 | no-hit | 0.0 | 0 | no-hit | 0.0 | 0 | no-hit | 0.0 | 0 | no-hit | 0.0 | 0 | no-hit | 0.0 | 0 | no-hit | 0.0 | 0 |
| scf7180000050764 | 6045 | 0.4518 | 0 | 122.9202 | no-hit | 0.0 | 0 | no-hit | 0.0 | 0 | no-hit | 0.0 | 0 | no-hit | 0.0 | 0 | no-hit | 0.0 | 0 | no-hit | 0.0 | 0 |
| scf7180000051774 | 3295 | 0.4473 | 0 | 624.7206 | Eukaryota | 285.0 | 0 | Platyhelminthes | 285.0 | 0 | Polystomatidea | 285.0 | 0 | Polystomatidae | 285.0 | 0 | Protopolystoma | 285.0 | 0 | Protopolystoma xenopodis | 285.0 | 0 |
| scf7180000047773 | 12229 | 0.4524 | 0 | 1502.7276 | Eukaryota | 4517.0 | 0 | Platyhelminthes | 2099.0 | 3 | Polystomatidea | 2099.0 | 3 | Polystomatidae | 2099.0 | 3 | Protopolystoma | 2099.0 | 3 | Protopolystoma xenopodis | 2099.0 | 3 |
| scf7180000042528 | 710282 | 0.5166 | 0 | 56.7665 | Eukaryota | 53150.0 | 0 | Chordata | 53150.0 | 0 | Carnivora | 53150.0 | 0 | Canidae | 53150.0 | 0 | Canis | 53150.0 | 0 | Canis lupus | 53150.0 | 0 |
| scf7180000056397 | 570117 | 0.3943 | 0 | 59.7751 | Eukaryota | 67446.0 | 0 | Chordata | 67446.0 | 0 | Carnivora | 36935.0 | 2 | Canidae | 36935.0 | 3 | Canis | 30743.0 | 6 | Canis lupus | 30743.0 | 6 |
| scf7180000041879 | 797221 | 0.5359 | 0 | 56.9196 | Eukaryota | 64248.0 | 0 | Chordata | 64248.0 | 0 | Carnivora | 64248.0 | 0 | Canidae | 64248.0 | 0 | Canis | 64248.0 | 0 | Canis lupus | 64248.0 | 0 |
| scf7180000047673 | 73511 | 0.4614 | 0 | 60.6323 | Eukaryota | 22437.0 | 0 | Chordata | 22437.0 | 0 | Carnivora | 22437.0 | 0 | Canidae | 13218.0 | 2 | Canis | 7902.0 | 6 | Canis lupus | 7902.0 | 6 |
| scf7180000055148 | 58389 | 0.4436 | 0 | 189.8568 | Eukaryota | 29489.0 | 0 | Chordata | 29489.0 | 0 | Gadiformes | 18930.0 | 5 | Gadidae | 18930.0 | 5 | Gadus | 18930.0 | 5 | Gadus morhua | 18930.0 | 5 |
| scf7180000053561 | 8857 | 0.4432 | 0 | 135.6912 | Eukaryota | 1942.0 | 0 | unresolved | 971.0 | 1 | Polystomatidea | 971.0 | 3 | Polystomatidae | 971.0 | 3 | Protopolystoma | 971.0 | 3 | Protopolystoma xenopodis | 971.0 | 3 |
| scf7180000050779 | 14516 | 0.475 | 0 | 63.4053 | Eukaryota | 56680.0 | 0 | Chordata | 56680.0 | 0 | Carnivora | 56680.0 | 0 | Canidae | 56680.0 | 0 | Canis | 56680.0 | 0 | Canis lupus | 56680.0 | 0 |
| scf7180000046739 | 572223 | 0.3967 | 0 | 60.4839 | Eukaryota | 63026.0 | 0 | Chordata | 63026.0 | 0 | Carnivora | 63026.0 | 0 | Canidae | 63026.0 | 0 | Canis | 63026.0 | 0 | Canis lupus | 63026.0 | 0 |
| scf7180000049048 | 5849 | 0.4464 | 0 | 145.6358 | Eukaryota | 2707.0 | 0 | Chordata | 2455.0 | 1 | Holocentriformes | 1760.0 | 2 | Holocentridae | 1760.0 | 2 | Myripristis | 1760.0 | 2 | Myripristis murdjan | 1760.0 | 2 |
| scf7180000047348 | 7804 | 0.4391 | 0 | 100.9495 | Eukaryota | 3191.0 | 0 | Chordata | 2723.0 | 1 | Carnivora | 2723.0 | 1 | Canidae | 2723.0 | 1 | Canis | 2723.0 | 1 | Canis lupus | 2723.0 | 1 |
| scf7180000046855 | 35861 | 0.445 | 0 | 591.3554 | Eukaryota | 7089.0 | 0 | Chordata | 7089.0 | 0 | Carnivora | 7089.0 | 0 | Canidae | 7089.0 | 0 | Nyctereutes | 4358.0 | 1 | Nyctereutes procyonoides | 4358.0 | 1 |
| scf7180000041178 | 1493500 | 0.3878 | 0 | 58.3675 | Eukaryota | 51368.0 | 0 | Chordata | 51368.0 | 0 | Carnivora | 51368.0 | 0 | Canidae | 51368.0 | 0 | Canis | 46531.0 | 1 | Canis lupus | 46531.0 | 1 |
| scf7180000054206 | 23142 | 0.5054 | 0 | 298.3422 | Eukaryota | 8640.0 | 0 | Chordata | 8640.0 | 0 | Carnivora | 8640.0 | 0 | Canidae | 8640.0 | 0 | Nyctereutes | 4356.0 | 1 | Nyctereutes procyonoides | 4356.0 | 1 |
| scf7180000053649 | 5626 | 0.4989 | 0 | 437.6898 | Eukaryota | 7598.0 | 0 | Chordata | 7598.0 | 0 | Carnivora | 7598.0 | 0 | Canidae | 7598.0 | 0 | Canis | 6581.0 | 1 | Canis lupus | 5586.0 | 2 |
| scf7180000042082 | 24282 | 0.423 | 0 | 55.774 | Eukaryota | 57574.0 | 0 | Chordata | 57574.0 | 0 | Carnivora | 57574.0 | 0 | Canidae | 57574.0 | 0 | Canis | 57574.0 | 0 | Canis lupus | 57574.0 | 0 |
| scf7180000045162 | 202284 | 0.4905 | 0 | 60.0517 | Eukaryota | 33300.0 | 0 | Chordata | 33300.0 | 0 | Carnivora | 33300.0 | 0 | Canidae | 33300.0 | 0 | Canis | 33300.0 | 0 | Canis lupus | 33300.0 | 0 |
| scf7180000043063 | 636952 | 0.4445 | 0 | 60.113 | Eukaryota | 137308.0 | 0 | Chordata | 137308.0 | 0 | Carnivora | 137308.0 | 0 | Canidae | 137308.0 | 0 | Canis | 111106.0 | 1 | Canis lupus | 111106.0 | 1 |
| scf7180000043325 | 30555 | 0.4716 | 0 | 54.6709 | Eukaryota | 5319.0 | 0 | Chordata | 5319.0 | 0 | Carnivora | 5319.0 | 0 | Canidae | 5319.0 | 0 | Canis | 5319.0 | 0 | Canis lupus | 5319.0 | 0 |
| scf7180000052284 | 7753 | 0.4512 | 0 | 708.6267 | Eukaryota | 2542.0 | 0 | Chordata | 2542.0 | 0 | Carnivora | 2542.0 | 0 | Canidae | 2542.0 | 0 | Canis | 2542.0 | 0 | Canis lupus | 2542.0 | 0 |
| scf7180000042765 | 5899 | 0.3616 | 0 | 35.7309 | Eukaryota | 3804.0 | 0 | Chordata | 3804.0 | 0 | Carnivora | 3804.0 | 0 | Canidae | 1194.0 | 3 | Callorhinus | 1119.0 | 4 | Callorhinus ursinus | 1119.0 | 4 |
| scf7180000052277 | 6080 | 0.4398 | 0 | 118.7673 | no-hit | 0.0 | 0 | no-hit | 0.0 | 0 | no-hit | 0.0 | 0 | no-hit | 0.0 | 0 | no-hit | 0.0 | 0 | no-hit | 0.0 | 0 |
| scf7180000046536 | 38151 | 0.4056 | 0 | 54.2215 | Eukaryota | 3552.0 | 0 | Chordata | 3552.0 | 0 | Carnivora | 3552.0 | 0 | Canidae | 3223.0 | 1 | Canis | 2860.0 | 2 | Canis lupus | 2860.0 | 2 |
| scf7180000040697 | 110617 | 0.4013 | 0 | 65.3738 | Eukaryota | 104163.0 | 0 | Chordata | 104163.0 | 0 | Carnivora | 104163.0 | 0 | Canidae | 104163.0 | 0 | Canis | 104163.0 | 0 | Canis lupus | 104163.0 | 0 |
| scf7180000041448 | 159776 | 0.4802 | 0 | 58.9137 | Eukaryota | 21302.0 | 1 | Chordata | 21302.0 | 1 | Carnivora | 13187.0 | 3 | Felidae | 5914.0 | 5 | Homo | 5775.0 | 7 | Homo sapiens | 5775.0 | 7 |
| scf7180000047935 | 15371 | 0.4453 | 0 | 795.0607 | no-hit | 0.0 | 0 | no-hit | 0.0 | 0 | no-hit | 0.0 | 0 | no-hit | 0.0 | 0 | no-hit | 0.0 | 0 | no-hit | 0.0 | 0 |
| scf7180000047957 | 68425 | 0.3738 | 0 | 57.6644 | Eukaryota | 15622.0 | 0 | Chordata | 15622.0 | 0 | Carnivora | 15622.0 | 0 | Canidae | 15622.0 | 0 | Canis | 15622.0 | 0 | Canis lupus | 15622.0 | 0 |
| scf7180000041416 | 1193085 | 0.4382 | 0 | 60.4316 | Eukaryota | 78452.0 | 0 | Chordata | 78452.0 | 0 | Carnivora | 78452.0 | 0 | Canidae | 78452.0 | 0 | Canis | 78452.0 | 0 | Canis lupus | 78452.0 | 0 |
| scf7180000052192 | 6223 | 0.4536 | 0 | 453.3221 | Eukaryota | 4594.0 | 0 | Chordata | 2115.0 | 3 | Polystomatidea | 1185.0 | 5 | Polystomatidae | 1185.0 | 5 | Protopolystoma | 1185.0 | 5 | Protopolystoma xenopodis | 1185.0 | 5 |
| scf7180000055693 | 14126 | 0.4414 | 0 | 227.4927 | Eukaryota | 55099.0 | 0 | Chordata | 39569.0 | 2 | Gadiformes | 32332.0 | 5 | Gadidae | 32332.0 | 5 | Gadus | 32332.0 | 5 | Gadus morhua | 32332.0 | 5 |
| scf7180000044213 | 197139 | 0.3875 | 0 | 58.8675 | Eukaryota | 46856.0 | 0 | Chordata | 46856.0 | 0 | Carnivora | 46856.0 | 0 | Canidae | 46856.0 | 0 | Canis | 46856.0 | 0 | Canis lupus | 46856.0 | 0 |
| scf7180000056531 | 191223 | 0.4237 | 0 | 58.8128 | Eukaryota | 40410.0 | 0 | Chordata | 40410.0 | 0 | Carnivora | 40410.0 | 0 | Canidae | 40410.0 | 0 | Canis | 40410.0 | 0 | Canis lupus | 40410.0 | 0 |
| scf7180000054400 | 11278 | 0.4148 | 0 | 51.8106 | Eukaryota | 50708.0 | 0 | Chordata | 50708.0 | 0 | Carnivora | 50708.0 | 0 | Canidae | 50708.0 | 0 | Canis | 50708.0 | 0 | Canis lupus | 50708.0 | 0 |
| scf7180000052535 | 11272 | 0.504 | 0 | 777.272 | Eukaryota | 9182.0 | 0 | Chordata | 9182.0 | 0 | Carnivora | 9182.0 | 0 | Canidae | 9182.0 | 0 | Canis | 7389.0 | 1 | Canis lupus | 6089.0 | 2 |
| scf7180000041275 | 262862 | 0.3658 | 0 | 59.4188 | Eukaryota | 22797.0 | 0 | Chordata | 22797.0 | 0 | Carnivora | 22797.0 | 0 | Canidae | 22797.0 | 0 | Canis | 22797.0 | 0 | Canis lupus | 22797.0 | 0 |
| scf7180000045852 | 12833 | 0.4222 | 0 | 225.8075 | Eukaryota | 28017.0 | 0 | Chordata | 28017.0 | 0 | Carnivora | 28017.0 | 0 | Canidae | 28017.0 | 0 | Canis | 28017.0 | 0 | Canis lupus | 28017.0 | 0 |
| scf7180000045344 | 196923 | 0.4091 | 0 | 58.6644 | Eukaryota | 7845.0 | 1 | Chordata | 7845.0 | 1 | Primates | 3204.0 | 4 | Bovidae | 2114.0 | 6 | Homo | 1901.0 | 7 | Homo sapiens | 1901.0 | 7 |
| scf7180000053188 | 3097 | 0.4462 | 0 | 192.0337 | Eukaryota | 1870.0 | 0 | Platyhelminthes | 702.0 | 2 | Polystomatidea | 702.0 | 2 | Polystomatidae | 702.0 | 2 | Protopolystoma | 702.0 | 2 | Protopolystoma xenopodis | 702.0 | 2 |
| scf7180000049605 | 17305 | 0.4402 | 0 | 570.4646 | Eukaryota | 2214.0 | 0 | Platyhelminthes | 1114.0 | 2 | Polystomatidea | 1114.0 | 2 | Polystomatidae | 1114.0 | 2 | Protopolystoma | 1114.0 | 2 | Protopolystoma xenopodis | 1114.0 | 2 |
| scf7180000045826 | 12207 | 0.4131 | 0 | 75.1322 | Eukaryota | 9170.0 | 0 | Chordata | 9170.0 | 0 | Carnivora | 9170.0 | 0 | Canidae | 9170.0 | 0 | Canis | 9170.0 | 0 | Canis lupus | 9170.0 | 0 |
| scf7180000043789 | 17671 | 0.4945 | 0 | 470.4717 | Eukaryota | 85227.0 | 0 | Chordata | 85227.0 | 0 | Carnivora | 85227.0 | 0 | Canidae | 85227.0 | 0 | Canis | 85227.0 | 0 | Canis lupus | 85227.0 | 0 |
| scf7180000042658 | 570628 | 0.3456 | 0 | 59.6953 | Eukaryota | 51182.0 | 0 | Chordata | 51182.0 | 0 | Carnivora | 42630.0 | 1 | Canidae | 42630.0 | 1 | Canis | 28341.0 | 2 | Canis lupus | 28341.0 | 2 |
| scf7180000053573 | 4549 | 0.5027 | 0 | 52.2267 | Eukaryota | 7708.0 | 0 | Chordata | 7708.0 | 0 | Carnivora | 7708.0 | 0 | Canidae | 7708.0 | 0 | Canis | 5685.0 | 1 | Canis lupus | 4595.0 | 2 |
| scf7180000040894 | 114904 | 0.3469 | 0 | 57.8416 | Eukaryota | 46881.0 | 0 | Chordata | 46881.0 | 0 | Carnivora | 46881.0 | 0 | Canidae | 46881.0 | 0 | Canis | 46881.0 | 0 | Canis lupus | 46881.0 | 0 |
| scf7180000042991 | 443537 | 0.4074 | 0 | 58.7445 | Eukaryota | 84630.0 | 0 | Chordata | 84630.0 | 0 | Carnivora | 84630.0 | 0 | Canidae | 84630.0 | 0 | Canis | 84630.0 | 0 | Canis lupus | 84630.0 | 0 |
| scf7180000044501 | 303753 | 0.3715 | 0 | 61.2871 | Eukaryota | 14473.0 | 0 | Chordata | 14473.0 | 0 | Carnivora | 12744.0 | 1 | Canidae | 12744.0 | 1 | Canis | 12744.0 | 1 | Canis lupus | 12744.0 | 1 |
| scf7180000049321 | 39474 | 0.4035 | 0 | 53.5776 | Eukaryota | 48689.0 | 0 | Chordata | 48689.0 | 0 | Carnivora | 48689.0 | 0 | Canidae | 48689.0 | 0 | Canis | 48689.0 | 0 | Canis lupus | 48689.0 | 0 |
| scf7180000053864 | 10031 | 0.3104 | 0 | 59.9116 | Eukaryota | 2772.0 | 0 | Chordata | 2772.0 | 0 | Carnivora | 1923.0 | 2 | Canidae | 1134.0 | 4 | Canis | 1134.0 | 5 | Canis lupus | 1134.0 | 5 |
| scf7180000040775 | 272943 | 0.3831 | 0 | 59.0937 | Eukaryota | 85597.0 | 0 | Chordata | 85597.0 | 0 | Carnivora | 85597.0 | 0 | Canidae | 85597.0 | 0 | Canis | 85597.0 | 0 | Canis lupus | 85597.0 | 0 |
| scf7180000043307 | 174000 | 0.3919 | 0 | 71.6908 | Eukaryota | 103477.0 | 0 | Chordata | 103477.0 | 0 | Carnivora | 103477.0 | 0 | Canidae | 103477.0 | 0 | Canis | 103477.0 | 0 | Canis lupus | 103477.0 | 0 |
| scf7180000047026 | 95022 | 0.5129 | 0 | 59.0188 | Eukaryota | 39358.0 | 0 | Chordata | 39358.0 | 0 | Carnivora | 39358.0 | 0 | Canidae | 36284.0 | 2 | Canis | 31533.0 | 3 | Canis lupus | 31533.0 | 3 |
| scf7180000045077 | 134676 | 0.3675 | 0 | 63.4614 | Eukaryota | 42756.0 | 0 | Chordata | 42756.0 | 0 | Carnivora | 42756.0 | 0 | Canidae | 39380.0 | 1 | Canis | 39380.0 | 1 | Canis lupus | 39380.0 | 1 |
| scf7180000052562 | 8805 | 0.4429 | 0 | 456.8222 | no-hit | 0.0 | 0 | no-hit | 0.0 | 0 | no-hit | 0.0 | 0 | no-hit | 0.0 | 0 | no-hit | 0.0 | 0 | no-hit | 0.0 | 0 |
| scf7180000040823 | 462257 | 0.5009 | 0 | 57.0903 | Eukaryota | 55739.0 | 0 | Chordata | 55739.0 | 0 | Carnivora | 55739.0 | 0 | Canidae | 55739.0 | 0 | Canis | 50527.0 | 1 | Canis lupus | 50527.0 | 1 |
| scf7180000045853 | 48145 | 0.4375 | 0 | 119.2155 | Eukaryota | 38596.0 | 0 | Chordata | 18803.0 | 2 | Cyclophyllidea | 12065.0 | 7 | Hymenolepididae | 12065.0 | 7 | Hymenolepis | 12065.0 | 7 | Hymenolepis microstoma | 12065.0 | 7 |
| scf7180000049621 | 3379 | 0.427 | 0 | 30.2651 | Eukaryota | 17325.0 | 0 | Chordata | 17325.0 | 0 | Carnivora | 17325.0 | 0 | Canidae | 17325.0 | 0 | Canis | 17325.0 | 0 | Canis lupus | 17325.0 | 0 |
| scf7180000041934 | 933974 | 0.4216 | 0 | 59.8901 | Eukaryota | 85543.0 | 0 | Chordata | 85543.0 | 0 | Carnivora | 85543.0 | 0 | Canidae | 85543.0 | 0 | Canis | 85543.0 | 0 | Canis lupus | 85543.0 | 0 |
| scf7180000042941 | 8552 | 0.3818 | 0 | 25.3463 | Eukaryota | 42167.0 | 0 | Chordata | 42167.0 | 0 | Carnivora | 42167.0 | 0 | Canidae | 42167.0 | 0 | Canis | 42167.0 | 0 | Canis lupus | 42167.0 | 0 |
| scf7180000053165 | 2862 | 0.4528 | 0 | 493.4777 | no-hit | 0.0 | 0 | no-hit | 0.0 | 0 | no-hit | 0.0 | 0 | no-hit | 0.0 | 0 | no-hit | 0.0 | 0 | no-hit | 0.0 | 0 |
| scf7180000051340 | 25546 | 0.5073 | 0 | 242.0101 | Eukaryota | 8665.0 | 0 | Chordata | 8665.0 | 0 | Carnivora | 8665.0 | 0 | Canidae | 8665.0 | 0 | Canis | 7541.0 | 1 | Canis lupus | 6375.0 | 2 |
| scf7180000041153 | 16748 | 0.3242 | 0 | 44.6152 | Eukaryota | 6543.0 | 0 | Chordata | 6543.0 | 0 | Primates | 2694.0 | 3 | Hominidae | 2694.0 | 3 | Homo | 2694.0 | 3 | Homo sapiens | 2694.0 | 3 |
| scf7180000041834 | 490612 | 0.3627 | 0 | 59.4668 | Eukaryota | 48077.0 | 0 | Chordata | 48077.0 | 0 | Carnivora | 42501.0 | 1 | Canidae | 39942.0 | 3 | Canis | 39942.0 | 3 | Canis lupus | 39942.0 | 3 |
| scf7180000053178 | 4149 | 0.4403 | 0 | 260.0298 | Eukaryota | 10574.0 | 0 | Chordata | 9053.0 | 2 | Gadiformes | 4454.0 | 6 | Gadidae | 4454.0 | 6 | Gadus | 4454.0 | 6 | Gadus morhua | 4454.0 | 6 |
| scf7180000042913 | 163291 | 0.3997 | 0 | 59.1953 | Eukaryota | 71156.0 | 0 | Chordata | 71156.0 | 0 | Carnivora | 71156.0 | 0 | Canidae | 71156.0 | 0 | Canis | 71156.0 | 0 | Canis lupus | 71156.0 | 0 |
| scf7180000044967 | 73361 | 0.4419 | 0 | 174.1337 | Eukaryota | 4924.0 | 0 | Chordata | 3019.0 | 1 | Polystomatidea | 1905.0 | 3 | Polystomatidae | 1905.0 | 3 | Protopolystoma | 1905.0 | 3 | Protopolystoma xenopodis | 1905.0 | 3 |
| scf7180000048383 | 63424 | 0.469 | 0 | 58.4243 | Eukaryota | 76554.0 | 0 | Chordata | 76554.0 | 0 | Carnivora | 76554.0 | 0 | Canidae | 71865.0 | 1 | Canis | 47547.0 | 2 | Canis lupus | 47547.0 | 2 |
| scf7180000042835 | 946713 | 0.4203 | 0 | 57.5602 | Eukaryota | 87430.0 | 0 | Chordata | 87430.0 | 0 | Carnivora | 87430.0 | 0 | Canidae | 87430.0 | 0 | Canis | 87430.0 | 0 | Canis lupus | 87430.0 | 0 |
| scf7180000055717 | 4435 | 0.4534 | 0 | 539.769 | Eukaryota | 1716.0 | 0 | Platyhelminthes | 1112.0 | 1 | Polystomatidea | 1112.0 | 1 | Polystomatidae | 1112.0 | 1 | Protopolystoma | 1112.0 | 1 | Protopolystoma xenopodis | 1112.0 | 1 |
| scf7180000043157 | 596706 | 0.4026 | 0 | 59.8729 | Eukaryota | 53384.0 | 0 | Chordata | 53384.0 | 0 | Carnivora | 53384.0 | 0 | Canidae | 53384.0 | 0 | Canis | 48335.0 | 1 | Canis lupus | 48335.0 | 1 |
| scf7180000048672 | 12722 | 0.3514 | 0 | 87.2882 | Eukaryota | 3877.0 | 0 | Chordata | 3877.0 | 0 | Carnivora | 2421.0 | 1 | Hominidae | 1456.0 | 3 | Homo | 1456.0 | 6 | Homo sapiens | 1456.0 | 6 |
| scf7180000042968 | 122349 | 0.381 | 0 | 56.7975 | Eukaryota | 51880.0 | 0 | Chordata | 51880.0 | 0 | Carnivora | 51880.0 | 0 | Canidae | 51880.0 | 0 | Canis | 51880.0 | 0 | Canis lupus | 51880.0 | 0 |
| scf7180000044168 | 610875 | 0.3668 | 0 | 60.2913 | Eukaryota | 87130.0 | 0 | Chordata | 87130.0 | 0 | Carnivora | 87130.0 | 0 | Canidae | 87130.0 | 0 | Canis | 87130.0 | 0 | Canis lupus | 87130.0 | 0 |
| scf7180000043473 | 469777 | 0.3657 | 0 | 60.2007 | Eukaryota | 40637.0 | 0 | Chordata | 40637.0 | 0 | Carnivora | 40637.0 | 0 | Canidae | 40637.0 | 0 | Canis | 40637.0 | 0 | Canis lupus | 40637.0 | 0 |
| scf7180000040801 | 537345 | 0.4082 | 0 | 59.8311 | Eukaryota | 94388.0 | 0 | Chordata | 94388.0 | 0 | Carnivora | 94388.0 | 0 | Canidae | 94388.0 | 0 | Canis | 94388.0 | 0 | Canis lupus | 94388.0 | 0 |
| scf7180000049682 | 14894 | 0.4085 | 0 | 46.9804 | Eukaryota | 3867.0 | 0 | Chordata | 3867.0 | 0 | Carnivora | 2944.0 | 1 | Canidae | 2944.0 | 1 | Canis | 2944.0 | 1 | Canis lupus | 2944.0 | 1 |
| scf7180000042956 | 864854 | 0.3925 | 0 | 60.317 | Eukaryota | 90979.0 | 0 | Chordata | 90979.0 | 0 | Carnivora | 90979.0 | 0 | Canidae | 90979.0 | 0 | Canis | 90979.0 | 0 | Canis lupus | 90979.0 | 0 |
| scf7180000040537 | 2396281 | 0.4231 | 0 | 59.6982 | Eukaryota | 92173.0 | 0 | Chordata | 92173.0 | 0 | Carnivora | 92173.0 | 0 | Canidae | 92173.0 | 0 | Canis | 92173.0 | 0 | Canis lupus | 92173.0 | 0 |
| scf7180000050247 | 4291 | 0.4421 | 0 | 68.2534 | Eukaryota | 1063.0 | 0 | Platyhelminthes | 883.0 | 1 | Polystomatidea | 883.0 | 1 | Polystomatidae | 883.0 | 1 | Protopolystoma | 883.0 | 1 | Protopolystoma xenopodis | 883.0 | 1 |
| scf7180000046991 | 105807 | 0.381 | 0 | 63.9705 | Eukaryota | 65246.0 | 0 | Chordata | 65246.0 | 0 | Carnivora | 65246.0 | 0 | Canidae | 65246.0 | 0 | Canis | 65246.0 | 0 | Canis lupus | 65246.0 | 0 |
| scf7180000053888 | 2281 | 0.4415 | 0 | 454.4403 | Eukaryota | 1382.0 | 0 | Platyhelminthes | 657.0 | 2 | Polystomatidea | 657.0 | 2 | Polystomatidae | 657.0 | 2 | Protopolystoma | 657.0 | 2 | Protopolystoma xenopodis | 657.0 | 2 |
| scf7180000050188 | 32215 | 0.3411 | 0 | 59.0872 | Eukaryota | 7708.0 | 0 | Chordata | 7708.0 | 0 | Carnivora | 6899.0 | 1 | Canidae | 6899.0 | 1 | Canis | 6899.0 | 1 | Canis lupus | 6899.0 | 1 |
| scf7180000044371 | 28200 | 0.4095 | 0 | 47.0422 | Eukaryota | 30985.0 | 0 | Chordata | 30985.0 | 0 | Carnivora | 30985.0 | 0 | Canidae | 30985.0 | 0 | Canis | 30985.0 | 0 | Canis lupus | 30985.0 | 0 |
| scf7180000041897 | 781114 | 0.3475 | 0 | 58.9962 | Eukaryota | 106563.0 | 0 | Chordata | 106563.0 | 0 | Carnivora | 106563.0 | 0 | Canidae | 106563.0 | 0 | Canis | 64129.0 | 1 | Canis lupus | 64129.0 | 1 |
| scf7180000053069 | 4067 | 0.5043 | 0 | 250.6206 | Eukaryota | 6060.0 | 0 | Chordata | 6060.0 | 0 | Carnivora | 6060.0 | 0 | Canidae | 6060.0 | 0 | Canis | 5146.0 | 1 | Canis lupus | 5146.0 | 1 |
| scf7180000041444 | 6477 | 0.388 | 0 | 33.9686 | Eukaryota | 61990.0 | 0 | Chordata | 61990.0 | 0 | Carnivora | 61990.0 | 0 | Canidae | 61990.0 | 0 | Canis | 61990.0 | 0 | Canis lupus | 61990.0 | 0 |
| scf7180000041880 | 909963 | 0.3792 | 0 | 60.79 | Eukaryota | 114243.0 | 0 | Chordata | 114243.0 | 0 | Carnivora | 114243.0 | 0 | Canidae | 114243.0 | 0 | Canis | 114243.0 | 0 | Canis lupus | 114243.0 | 0 |
| scf7180000052552 | 16308 | 0.5037 | 0 | 893.0513 | Eukaryota | 9180.0 | 0 | Chordata | 9180.0 | 0 | Carnivora | 9180.0 | 0 | Canidae | 9180.0 | 0 | Canis | 7436.0 | 1 | Canis lupus | 6201.0 | 2 |
| scf7180000052708 | 8146 | 0.4955 | 0 | 78.5829 | Eukaryota | 26232.0 | 0 | Chordata | 26232.0 | 0 | Carnivora | 26232.0 | 0 | Canidae | 26232.0 | 0 | Canis | 26232.0 | 0 | Canis lupus | 26232.0 | 0 |
| scf7180000044361 | 364222 | 0.425 | 0 | 60.2005 | Eukaryota | 51198.0 | 0 | Chordata | 51198.0 | 0 | Carnivora | 51198.0 | 0 | Canidae | 51198.0 | 0 | Canis | 51198.0 | 0 | Canis lupus | 51198.0 | 0 |
| scf7180000042682 | 34531 | 0.3438 | 0 | 59.9697 | Eukaryota | 44769.0 | 0 | Chordata | 44769.0 | 0 | Carnivora | 44769.0 | 0 | Canidae | 44769.0 | 0 | Canis | 44769.0 | 0 | Canis lupus | 44769.0 | 0 |
| scf7180000045205 | 217402 | 0.3693 | 0 | 61.1142 | Eukaryota | 98305.0 | 0 | Chordata | 98305.0 | 0 | Carnivora | 98305.0 | 0 | Canidae | 89841.0 | 1 | Canis | 59876.0 | 2 | Canis lupus | 59876.0 | 2 |
| scf7180000043403 | 125510 | 0.5224 | 0 | 56.8712 | Eukaryota | 151149.0 | 0 | Chordata | 151149.0 | 0 | Carnivora | 151149.0 | 0 | Canidae | 151149.0 | 0 | Canis | 139775.0 | 1 | Canis lupus | 139775.0 | 1 |
| scf7180000046182 | 8507 | 0.4358 | 0 | 95.2838 | no-hit | 0.0 | 0 | no-hit | 0.0 | 0 | no-hit | 0.0 | 0 | no-hit | 0.0 | 0 | no-hit | 0.0 | 0 | no-hit | 0.0 | 0 |
| scf7180000042696 | 618104 | 0.4476 | 0 | 59.1545 | Eukaryota | 90538.0 | 0 | Chordata | 90538.0 | 0 | Carnivora | 90538.0 | 0 | Canidae | 90538.0 | 0 | Canis | 90538.0 | 0 | Canis lupus | 90538.0 | 0 |
| scf7180000050645 | 5269 | 0.4511 | 0 | 224.3139 | Eukaryota | 2959.0 | 0 | Platyhelminthes | 1689.0 | 2 | Polystomatidea | 1689.0 | 2 | Polystomatidae | 1689.0 | 2 | Protopolystoma | 1689.0 | 2 | Protopolystoma xenopodis | 1689.0 | 2 |
| scf7180000042637 | 3204 | 0.5197 | 0 | 26.5984 | Eukaryota | 3588.0 | 0 | Chordata | 3588.0 | 0 | Carnivora | 2691.0 | 1 | Canidae | 2691.0 | 2 | Canis | 2691.0 | 2 | Canis lupus | 2691.0 | 2 |
| scf7180000045981 | 13363 | 0.5027 | 0 | 270.5874 | Eukaryota | 8769.0 | 0 | Chordata | 8769.0 | 0 | Carnivora | 8769.0 | 0 | Canidae | 8769.0 | 0 | Canis | 7176.0 | 1 | Canis lupus | 6025.0 | 2 |
| scf7180000056440 | 168151 | 0.3981 | 0 | 59.6741 | Eukaryota | 57496.0 | 0 | Chordata | 57496.0 | 0 | Carnivora | 57496.0 | 0 | Canidae | 57496.0 | 0 | Canis | 57496.0 | 0 | Canis lupus | 57496.0 | 0 |
| scf7180000042268 | 395473 | 0.3739 | 0 | 59.0863 | Eukaryota | 92172.0 | 0 | Chordata | 92172.0 | 0 | Carnivora | 92172.0 | 0 | Canidae | 92172.0 | 0 | Canis | 92172.0 | 0 | Canis lupus | 92172.0 | 0 |
| scf7180000043314 | 398194 | 0.379 | 0 | 59.7145 | Eukaryota | 41050.0 | 0 | Chordata | 41050.0 | 0 | Carnivora | 41050.0 | 0 | Canidae | 41050.0 | 0 | Canis | 23538.0 | 1 | Canis lupus | 23538.0 | 1 |
| scf7180000041269 | 1165602 | 0.3629 | 0 | 60.3405 | Eukaryota | 94424.0 | 0 | Chordata | 94424.0 | 0 | Carnivora | 94424.0 | 0 | Canidae | 94424.0 | 0 | Canis | 94424.0 | 0 | Canis lupus | 94424.0 | 0 |
| scf7180000045461 | 201422 | 0.3748 | 0 | 61.3177 | Eukaryota | 29759.0 | 0 | Chordata | 29759.0 | 0 | Carnivora | 29759.0 | 0 | Canidae | 10046.0 | 3 | Canis | 6720.0 | 7 | Canis lupus | 6720.0 | 7 |
| scf7180000053795 | 2451 | 0.4545 | 0 | 767.3299 | no-hit | 0.0 | 0 | no-hit | 0.0 | 0 | no-hit | 0.0 | 0 | no-hit | 0.0 | 0 | no-hit | 0.0 | 0 | no-hit | 0.0 | 0 |
| scf7180000045142 | 7127 | 0.4416 | 0 | 26.5622 | Eukaryota | 41099.0 | 0 | Chordata | 41099.0 | 0 | Carnivora | 41099.0 | 0 | Canidae | 41099.0 | 0 | Canis | 41099.0 | 0 | Canis lupus | 41099.0 | 0 |
| scf7180000040656 | 2002436 | 0.4316 | 0 | 59.9593 | Eukaryota | 124504.0 | 0 | Chordata | 124504.0 | 0 | Carnivora | 124504.0 | 0 | Canidae | 124504.0 | 0 | Canis | 71172.0 | 1 | Canis lupus | 71172.0 | 1 |
| scf7180000054033 | 2606 | 0.51 | 0 | 73.7417 | Eukaryota | 5360.0 | 0 | Chordata | 5360.0 | 0 | Carnivora | 5360.0 | 0 | Canidae | 5360.0 | 0 | Canis | 3276.0 | 1 | Canis lupus | 2373.0 | 2 |
| scf7180000042117 | 81322 | 0.393 | 0 | 52.3975 | Eukaryota | 58722.0 | 0 | Chordata | 58722.0 | 0 | Carnivora | 58722.0 | 0 | Canidae | 58722.0 | 0 | Canis | 58722.0 | 0 | Canis lupus | 58722.0 | 0 |
| scf7180000055868 | 16295 | 0.4057 | 0 | 64.3878 | Eukaryota | 80293.0 | 0 | Chordata | 80293.0 | 0 | Carnivora | 80293.0 | 0 | Canidae | 80293.0 | 0 | Canis | 80293.0 | 0 | Canis lupus | 80293.0 | 0 |
| scf7180000045518 | 219927 | 0.3438 | 0 | 59.799 | Eukaryota | 44460.0 | 0 | Chordata | 44460.0 | 0 | Carnivora | 44460.0 | 0 | Canidae | 44460.0 | 0 | Canis | 44460.0 | 0 | Canis lupus | 44460.0 | 0 |
| scf7180000040897 | 1549046 | 0.4391 | 0 | 58.9655 | Eukaryota | 52334.0 | 0 | Chordata | 52334.0 | 0 | Carnivora | 52334.0 | 0 | Canidae | 52334.0 | 0 | Canis | 47033.0 | 1 | Canis lupus | 47033.0 | 1 |
| scf7180000044171 | 325870 | 0.5378 | 0 | 56.1073 | Eukaryota | 91904.0 | 0 | Chordata | 91904.0 | 0 | Carnivora | 91904.0 | 0 | Canidae | 91904.0 | 0 | Canis | 91904.0 | 0 | Canis lupus | 91904.0 | 0 |
| scf7180000047083 | 116660 | 0.3662 | 0 | 61.248 | Eukaryota | 34336.0 | 0 | Chordata | 34336.0 | 0 | Carnivora | 34336.0 | 0 | Canidae | 15544.0 | 3 | Canis | 9558.0 | 5 | Canis lupus | 9558.0 | 5 |
| scf7180000046294 | 95141 | 0.3553 | 0 | 53.9562 | Eukaryota | 88654.0 | 0 | Chordata | 88654.0 | 0 | Carnivora | 88654.0 | 0 | Canidae | 88654.0 | 0 | Canis | 88654.0 | 0 | Canis lupus | 88654.0 | 0 |
| scf7180000040830 | 1737341 | 0.4826 | 0 | 59.4356 | Eukaryota | 93600.0 | 0 | Chordata | 93600.0 | 0 | Carnivora | 93600.0 | 0 | Canidae | 93600.0 | 0 | Canis | 93600.0 | 0 | Canis lupus | 93600.0 | 0 |
| scf7180000045559 | 172154 | 0.4205 | 0 | 61.2998 | Eukaryota | 68871.0 | 0 | Chordata | 68871.0 | 0 | Carnivora | 68871.0 | 0 | Canidae | 68871.0 | 0 | Canis | 68871.0 | 0 | Canis lupus | 68871.0 | 0 |
| scf7180000046629 | 5808 | 0.4146 | 0 | 29.8614 | Eukaryota | 14339.0 | 0 | Chordata | 14339.0 | 0 | Carnivora | 14339.0 | 0 | Canidae | 14339.0 | 0 | Canis | 12922.0 | 1 | Canis lupus | 12922.0 | 1 |
| scf7180000041979 | 862352 | 0.3567 | 0 | 59.5323 | Eukaryota | 34305.0 | 0 | Chordata | 34305.0 | 0 | Carnivora | 34305.0 | 0 | Canidae | 34305.0 | 0 | Canis | 30563.0 | 1 | Canis lupus | 30563.0 | 1 |
| scf7180000043372 | 276453 | 0.454 | 0 | 59.3746 | Eukaryota | 41139.0 | 0 | Chordata | 41139.0 | 0 | Primates | 20775.0 | 1 | Hominidae | 20775.0 | 1 | Canis | 20364.0 | 2 | Canis lupus | 20364.0 | 2 |
| scf7180000055647 | 15968 | 0.4409 | 0 | 174.7932 | no-hit | 0.0 | 0 | no-hit | 0.0 | 0 | no-hit | 0.0 | 0 | no-hit | 0.0 | 0 | no-hit | 0.0 | 0 | no-hit | 0.0 | 0 |
| scf7180000045327 | 205256 | 0.5426 | 0 | 56.1636 | Eukaryota | 13511.0 | 0 | Chordata | 13511.0 | 0 | Carnivora | 13511.0 | 0 | Canidae | 8643.0 | 1 | Canis | 5905.0 | 3 | Canis lupus | 5905.0 | 3 |
| scf7180000048793 | 12277 | 0.4382 | 0 | 111.2132 | Eukaryota | 24258.0 | 0 | Platyhelminthes | 14234.0 | 3 | Cyclophyllidea | 8408.0 | 7 | Hymenolepididae | 8408.0 | 7 | Hymenolepis | 8408.0 | 7 | Hymenolepis microstoma | 8408.0 | 7 |
| scf7180000055628 | 5125 | 0.4478 | 0 | 2021.8812 | no-hit | 0.0 | 0 | no-hit | 0.0 | 0 | no-hit | 0.0 | 0 | no-hit | 0.0 | 0 | no-hit | 0.0 | 0 | no-hit | 0.0 | 0 |
| scf7180000053039 | 4433 | 0.5062 | 0 | 28.9686 | Eukaryota | 6094.0 | 0 | Chordata | 6094.0 | 0 | Carnivora | 6094.0 | 0 | Canidae | 6094.0 | 0 | Canis | 4633.0 | 1 | Canis lupus | 3897.0 | 2 |
| scf7180000055618 | 25581 | 0.4356 | 0 | 155.7483 | no-hit | 0.0 | 0 | no-hit | 0.0 | 0 | no-hit | 0.0 | 0 | no-hit | 0.0 | 0 | no-hit | 0.0 | 0 | no-hit | 0.0 | 0 |
| scf7180000051084 | 6946 | 0.4171 | 0 | 27.8725 | Eukaryota | 3112.0 | 0 | Chordata | 3112.0 | 0 | Carnivora | 3112.0 | 0 | Canidae | 3112.0 | 0 | Canis | 3112.0 | 0 | Canis lupus | 3112.0 | 0 |
| scf7180000050660 | 9573 | 0.4987 | 0 | 67.7931 | Eukaryota | 8794.0 | 0 | Chordata | 8794.0 | 0 | Carnivora | 8794.0 | 0 | Canidae | 8794.0 | 0 | Canis | 7657.0 | 1 | Canis lupus | 6513.0 | 2 |
| scf7180000043132 | 322085 | 0.5181 | 0 | 57.4619 | Eukaryota | 31171.0 | 0 | Chordata | 31171.0 | 0 | Carnivora | 31171.0 | 0 | Canidae | 31171.0 | 0 | Canis | 21418.0 | 1 | Canis lupus | 21418.0 | 1 |
| scf7180000042490 | 661324 | 0.4118 | 0 | 60.4356 | Eukaryota | 46304.0 | 0 | Chordata | 46304.0 | 0 | Carnivora | 46304.0 | 0 | Canidae | 25224.0 | 3 | Canis | 20114.0 | 6 | Canis lupus | 20114.0 | 6 |
| scf7180000041101 | 1487004 | 0.3646 | 0 | 60.0405 | Eukaryota | 116253.0 | 0 | Chordata | 116253.0 | 0 | Carnivora | 116253.0 | 0 | Ursidae | 43068.0 | 3 | Canis | 29140.0 | 6 | Canis lupus | 29140.0 | 7 |
| scf7180000045163 | 209948 | 0.4461 | 0 | 59.1492 | Eukaryota | 11575.0 | 0 | Chordata | 11575.0 | 0 | Carnivora | 10391.0 | 1 | Canidae | 10391.0 | 1 | Canis | 10391.0 | 1 | Canis lupus | 10391.0 | 1 |
| scf7180000044211 | 326506 | 0.5899 | 0 | 57.9506 | Eukaryota | 77507.0 | 0 | Chordata | 77507.0 | 0 | Carnivora | 77507.0 | 0 | Canidae | 59210.0 | 2 | Canis | 53147.0 | 4 | Canis lupus | 53147.0 | 4 |
| scf7180000043349 | 532501 | 0.5382 | 0 | 57.2684 | Eukaryota | 32112.0 | 0 | Chordata | 32112.0 | 0 | Carnivora | 32112.0 | 0 | Canidae | 32112.0 | 0 | Canis | 16544.0 | 1 | Canis lupus | 16544.0 | 1 |
| scf7180000050647 | 48043 | 0.3556 | 0 | 63.8155 | Eukaryota | 23192.0 | 0 | Chordata | 23192.0 | 0 | Carnivora | 23192.0 | 0 | Otariidae | 12333.0 | 2 | Eumetopias | 12333.0 | 2 | Eumetopias jubatus | 12333.0 | 2 |
| scf7180000042260 | 1209704 | 0.3964 | 0 | 59.9803 | Eukaryota | 121928.0 | 0 | Chordata | 121928.0 | 0 | Carnivora | 121928.0 | 0 | Canidae | 121928.0 | 0 | Canis | 121928.0 | 0 | Canis lupus | 121928.0 | 0 |
| scf7180000050836 | 6635 | 0.4497 | 0 | 2279.7999 | no-hit | 0.0 | 0 | no-hit | 0.0 | 0 | no-hit | 0.0 | 0 | no-hit | 0.0 | 0 | no-hit | 0.0 | 0 | no-hit | 0.0 | 0 |
| scf7180000044594 | 47505 | 0.3661 | 0 | 56.067 | Eukaryota | 16778.0 | 0 | Chordata | 16778.0 | 0 | Artiodactyla | 9127.0 | 2 | Bovidae | 6233.0 | 3 | Homo | 4867.0 | 4 | Homo sapiens | 4867.0 | 4 |
| scf7180000044572 | 174734 | 0.4203 | 0 | 58.4528 | Eukaryota | 52248.0 | 0 | Chordata | 52248.0 | 0 | Carnivora | 52248.0 | 0 | Canidae | 52248.0 | 0 | Canis | 52248.0 | 0 | Canis lupus | 52248.0 | 0 |
| scf7180000051419 | 8887 | 0.4529 | 0 | 467.9063 | Eukaryota | 4332.0 | 0 | Chordata | 3305.0 | 1 | Salmoniformes | 2394.0 | 2 | Salmonidae | 2394.0 | 2 | Salmo | 2394.0 | 2 | Salmo trutta | 2394.0 | 2 |
| scf7180000045892 | 157814 | 0.3857 | 0 | 60.9611 | Eukaryota | 22846.0 | 0 | Chordata | 22846.0 | 0 | Carnivora | 22846.0 | 0 | Phocidae | 11449.0 | 3 | Neomonachus | 9155.0 | 5 | Neomonachus schauinslandi | 9155.0 | 5 |
| scf7180000044485 | 216021 | 0.3762 | 0 | 58.824 | Eukaryota | 50832.0 | 0 | Chordata | 50832.0 | 0 | Carnivora | 50832.0 | 0 | Canidae | 50832.0 | 0 | Canis | 50832.0 | 0 | Canis lupus | 50832.0 | 0 |
| scf7180000054604 | 2782 | 0.4335 | 0 | 131.4185 | no-hit | 0.0 | 0 | no-hit | 0.0 | 0 | no-hit | 0.0 | 0 | no-hit | 0.0 | 0 | no-hit | 0.0 | 0 | no-hit | 0.0 | 0 |
| scf7180000042225 | 185985 | 0.3968 | 0 | 58.905 | Eukaryota | 76285.0 | 0 | Chordata | 76285.0 | 0 | Carnivora | 71328.0 | 1 | Canidae | 71328.0 | 1 | Canis | 71328.0 | 1 | Canis lupus | 71328.0 | 1 |
| scf7180000054835 | 40819 | 0.441 | 0 | 317.1493 | Eukaryota | 6210.0 | 0 | Chordata | 5764.0 | 1 | Kurtiformes | 3415.0 | 3 | Apogonidae | 3415.0 | 3 | Sphaeramia | 3415.0 | 3 | Sphaeramia orbicularis | 3415.0 | 3 |
| scf7180000043519 | 369674 | 0.3592 | 0 | 66.5299 | Eukaryota | 92070.0 | 0 | Chordata | 92070.0 | 0 | Carnivora | 92070.0 | 0 | Canidae | 92070.0 | 0 | Canis | 92070.0 | 0 | Canis lupus | 92070.0 | 0 |
| scf7180000043718 | 118471 | 0.3926 | 0 | 57.7441 | Eukaryota | 41662.0 | 0 | Chordata | 41662.0 | 0 | Carnivora | 41662.0 | 0 | Canidae | 37852.0 | 1 | Canis | 23683.0 | 2 | Canis lupus | 23683.0 | 2 |
| scf7180000053747 | 4831 | 0.4972 | 0 | 80.3385 | Eukaryota | 7440.0 | 0 | Chordata | 7440.0 | 0 | Carnivora | 7440.0 | 0 | Canidae | 7440.0 | 0 | Canis | 5893.0 | 1 | Canis lupus | 4756.0 | 2 |
| scf7180000041727 | 8786 | 0.4552 | 0 | 49.0315 | Eukaryota | 46352.0 | 0 | Chordata | 46352.0 | 0 | Carnivora | 46352.0 | 0 | Canidae | 46352.0 | 0 | Canis | 46352.0 | 0 | Canis lupus | 46352.0 | 0 |
| scf7180000041524 | 144358 | 0.3764 | 0 | 55.8757 | Eukaryota | 91993.0 | 0 | Chordata | 91993.0 | 0 | Carnivora | 91993.0 | 0 | Canidae | 91993.0 | 0 | Canis | 91993.0 | 0 | Canis lupus | 91993.0 | 0 |
| scf7180000045908 | 186758 | 0.3796 | 0 | 58.4375 | Eukaryota | 71798.0 | 0 | Chordata | 71798.0 | 0 | Carnivora | 71798.0 | 0 | Canidae | 71798.0 | 0 | Canis | 71798.0 | 0 | Canis lupus | 71798.0 | 0 |
| scf7180000049747 | 4740 | 0.4498 | 0 | 283.1106 | no-hit | 0.0 | 0 | no-hit | 0.0 | 0 | no-hit | 0.0 | 0 | no-hit | 0.0 | 0 | no-hit | 0.0 | 0 | no-hit | 0.0 | 0 |
| scf7180000048362 | 63504 | 0.3908 | 0 | 58.6089 | Eukaryota | 49243.0 | 0 | Chordata | 49243.0 | 0 | Carnivora | 49243.0 | 0 | Canidae | 49243.0 | 0 | Canis | 47547.0 | 1 | Canis lupus | 47547.0 | 1 |
| scf7180000045087 | 203775 | 0.5332 | 0 | 57.5359 | Eukaryota | 15099.0 | 0 | Chordata | 15099.0 | 0 | Carnivora | 15099.0 | 0 | Canidae | 15099.0 | 0 | Canis | 15099.0 | 0 | Canis lupus | 15099.0 | 0 |
| scf7180000054274 | 22171 | 0.3608 | 0 | 65.0516 | Eukaryota | 92217.0 | 0 | Chordata | 92217.0 | 0 | Carnivora | 92217.0 | 0 | Canidae | 92217.0 | 0 | Canis | 92217.0 | 0 | Canis lupus | 92217.0 | 0 |
| scf7180000041061 | 15605 | 0.4281 | 0 | 51.3933 | Eukaryota | 6149.0 | 0 | Chordata | 6149.0 | 0 | Carnivora | 6149.0 | 0 | Canidae | 4846.0 | 1 | Canis | 4846.0 | 1 | Canis lupus | 4846.0 | 1 |
| scf7180000045860 | 171854 | 0.3344 | 0 | 60.1849 | Eukaryota | 19147.0 | 1 | Chordata | 19147.0 | 1 | Artiodactyla | 7565.0 | 3 | Bovidae | 7565.0 | 3 | unresolved | 7218.0 | 4 | unresolved | 7218.0 | 4 |
| scf7180000043823 | 448227 | 0.392 | 0 | 60.9647 | Eukaryota | 85860.0 | 0 | Chordata | 85860.0 | 0 | Carnivora | 85860.0 | 0 | Canidae | 85860.0 | 0 | Canis | 85860.0 | 0 | Canis lupus | 85860.0 | 0 |
| scf7180000046511 | 16159 | 0.3472 | 0 | 55.9278 | Eukaryota | 6002.0 | 0 | Chordata | 6002.0 | 0 | Carnivora | 5306.0 | 1 | Canidae | 5306.0 | 1 | Canis | 5306.0 | 2 | Canis lupus | 5306.0 | 2 |
| scf7180000041712 | 1040293 | 0.3541 | 0 | 59.7711 | Eukaryota | 88670.0 | 0 | Chordata | 88670.0 | 0 | Carnivora | 88670.0 | 0 | Canidae | 88670.0 | 0 | Canis | 88670.0 | 0 | Canis lupus | 88670.0 | 0 |
| scf7180000042317 | 635355 | 0.4759 | 0 | 59.8972 | Eukaryota | 65115.0 | 0 | Chordata | 65115.0 | 0 | Carnivora | 65115.0 | 0 | Canidae | 65115.0 | 0 | Canis | 65115.0 | 0 | Canis lupus | 65115.0 | 0 |
| scf7180000041193 | 112962 | 0.4416 | 0 | 53.8418 | Eukaryota | 109858.0 | 0 | Chordata | 109858.0 | 0 | Carnivora | 109858.0 | 0 | Canidae | 103268.0 | 1 | Canis | 103268.0 | 1 | Canis lupus | 103268.0 | 1 |
| scf7180000046131 | 120954 | 0.3797 | 0 | 60.2214 | Eukaryota | 39178.0 | 0 | Chordata | 39178.0 | 0 | Carnivora | 39178.0 | 0 | Otariidae | 15664.0 | 2 | Callorhinus | 15664.0 | 2 | Callorhinus ursinus | 15664.0 | 2 |

| | | | | | | | | | | | | | | | | | | | | | | |
|---|---|---|---|---|---|---|---|---|---|---|---|---|---|---|---|---|---|---|---|---|---|---|
| scf7180000045256 | 247779 | 0.426 | 0 | 59.5911 | Eukaryota | 50035.0 | 0 | Chordata | 50035.0 | 0 | Carnivora | 50035.0 | 0 | Canidae | 40051.0 | 1 | Canis | 35641.0 | 2 | Canis lupus | 35641.0 | 2 |
| scf7180000043203 | 503343 | 0.4067 | 0 | 59.9484 | Eukaryota | 59769.0 | 0 | Chordata | 59769.0 | 0 | Carnivora | 59769.0 | 0 | Canidae | 59769.0 | 0 | Canis | 59769.0 | 0 | Canis lupus | 59769.0 | 0 |
| scf7180000054767 | 15978 | 0.447 | 0 | 208.2086 | Eukaryota | 777.0 | 0 | Platyhelminthes | 777.0 | 0 | Polystomatidea | 507.0 | 1 | Polystomatidae | 507.0 | 1 | Protopolystoma | 507.0 | 1 | Protopolystoma xenopodis | 507.0 | 1 |
| scf7180000053927 | 5886 | 0.4514 | 0 | 104.3569 | Eukaryota | 3906.0 | 0 | Chordata | 3906.0 | 0 | Carnivora | 2563.0 | 1 | Canidae | 2563.0 | 1 | Canis | 2563.0 | 1 | Canis lupus | 2563.0 | 1 |
| scf7180000052140 | 14221 | 0.4411 | 0 | 60.4296 | no-hit | 0.0 | 0 | no-hit | 0.0 | 0 | no-hit | 0.0 | 0 | no-hit | 0.0 | 0 | no-hit | 0.0 | 0 | no-hit | 0.0 | 0 |
| scf7180000048692 | 20228 | 0.4482 | 0 | 1006.4601 | Eukaryota | 1470.0 | 0 | Platyhelminthes | 528.0 | 2 | Polystomatidea | 528.0 | 2 | Polystomatidae | 528.0 | 2 | Protopolystoma | 528.0 | 2 | Protopolystoma xenopodis | 528.0 | 2 |
| scf7180000047483 | 36885 | 0.4402 | 0 | 261.756 | Eukaryota | 14305.0 | 0 | Platyhelminthes | 8365.0 | 1 | Cyclophyllidea | 3971.0 | 6 | Hymenolepididae | 3971.0 | 6 | Hymenolepis | 3971.0 | 6 | Hymenolepis microstoma | 3971.0 | 6 |
| scf7180000047950 | 72287 | 0.4821 | 0 | 61.7814 | Eukaryota | 40193.0 | 0 | Chordata | 40193.0 | 0 | Carnivora | 40193.0 | 0 | Canidae | 30348.0 | 1 | Canis | 27380.0 | 2 | Canis lupus | 27380.0 | 2 |
| scf7180000055347 | 24009 | 0.347 | 0 | 118.7071 | Eukaryota | 11600.0 | 1 | Chordata | 11600.0 | 1 | Carnivora | 8107.0 | 3 | Canidae | 8107.0 | 3 | Vulpes | 4072.0 | 5 | Vulpes vulpes | 4072.0 | 5 |
| scf7180000042292 | 228746 | 0.4432 | 0 | 58.0902 | Eukaryota | 89891.0 | 0 | Chordata | 89891.0 | 0 | Carnivora | 89891.0 | 0 | Canidae | 89891.0 | 0 | Canis | 89891.0 | 0 | Canis lupus | 89891.0 | 0 |
| scf7180000040501 | 3639046 | 0.4274 | 0 | 59.3599 | Eukaryota | 103680.0 | 0 | Chordata | 103680.0 | 0 | Carnivora | 103680.0 | 0 | Canidae | 103680.0 | 0 | Canis | 86810.0 | 1 | Canis lupus | 86810.0 | 1 |
| scf7180000048818 | 53634 | 0.5013 | 0 | 317.6334 | Eukaryota | 9633.0 | 0 | Chordata | 9633.0 | 0 | Carnivora | 9633.0 | 0 | Canidae | 9633.0 | 0 | Canis | 8391.0 | 1 | Canis lupus | 7097.0 | 2 |
| scf7180000055981 | 12593 | 0.4413 | 0 | 70.8765 | Eukaryota | 1821.0 | 0 | Chordata | 1213.0 | 1 | Carnivora | 1213.0 | 1 | Canidae | 1213.0 | 1 | Canis | 1213.0 | 1 | Canis lupus | 1213.0 | 1 |
| scf7180000050065 | 5132 | 0.4452 | 0 | 302.8355 | no-hit | 0.0 | 0 | no-hit | 0.0 | 0 | no-hit | 0.0 | 0 | no-hit | 0.0 | 0 | no-hit | 0.0 | 0 | no-hit | 0.0 | 0 |
| scf7180000050172 | 10186 | 0.452 | 0 | 1254.4418 | Eukaryota | 4071.0 | 0 | Chordata | 1730.0 | 3 | Polystomatidea | 1167.0 | 4 | Polystomatidae | 1167.0 | 4 | Protopolystoma | 1167.0 | 4 | Protopolystoma xenopodis | 1167.0 | 4 |
| scf7180000040770 | 1370218 | 0.3697 | 0 | 60.7098 | Eukaryota | 92645.0 | 0 | Chordata | 92645.0 | 0 | Carnivora | 92645.0 | 0 | Canidae | 92645.0 | 0 | Canis | 92645.0 | 0 | Canis lupus | 92645.0 | 0 |
| scf7180000050801 | 5578 | 0.4516 | 0 | 1156.0761 | no-hit | 0.0 | 0 | no-hit | 0.0 | 0 | no-hit | 0.0 | 0 | no-hit | 0.0 | 0 | no-hit | 0.0 | 0 | no-hit | 0.0 | 0 |
| scf7180000045197 | 178109 | 0.4363 | 0 | 58.5812 | Eukaryota | 49117.0 | 0 | Chordata | 49117.0 | 0 | Carnivora | 49117.0 | 0 | Canidae | 49117.0 | 0 | Canis | 41804.0 | 1 | Canis lupus | 41804.0 | 1 |
| scf7180000047480 | 113966 | 0.3515 | 0 | 59.7367 | Eukaryota | 44516.0 | 0 | Chordata | 44516.0 | 0 | Carnivora | 44516.0 | 0 | Canidae | 44516.0 | 0 | Canis | 44516.0 | 0 | Canis lupus | 44516.0 | 0 |
| scf7180000053297 | 17302 | 0.4597 | 0 | 57.9633 | no-hit | 0.0 | 0 | no-hit | 0.0 | 0 | no-hit | 0.0 | 0 | no-hit | 0.0 | 0 | no-hit | 0.0 | 0 | no-hit | 0.0 | 0 |
| scf7180000047207 | 11365 | 0.4448 | 0 | 144.7627 | no-hit | 0.0 | 0 | no-hit | 0.0 | 0 | no-hit | 0.0 | 0 | no-hit | 0.0 | 0 | no-hit | 0.0 | 0 | no-hit | 0.0 | 0 |
| scf7180000056411 | 676868 | 0.4886 | 0 | 57.2113 | Eukaryota | 51519.0 | 0 | Chordata | 51519.0 | 0 | Carnivora | 51519.0 | 0 | Canidae | 51519.0 | 0 | Canis | 47378.0 | 1 | Canis lupus | 47378.0 | 1 |
| scf7180000053172 | 3842 | 0.4513 | 0 | 564.6974 | no-hit | 0.0 | 0 | no-hit | 0.0 | 0 | no-hit | 0.0 | 0 | no-hit | 0.0 | 0 | no-hit | 0.0 | 0 | no-hit | 0.0 | 0 |
| scf7180000046288 | 138256 | 0.4548 | 0 | 73.1308 | Eukaryota | 81590.0 | 0 | Platyhelminthes | 45583.0 | 1 | Strigeidida | 36227.0 | 5 | Schistosomatidae | 36227.0 | 5 | Schistosoma | 36227.0 | 5 | Schistosoma mansoni | 36227.0 | 5 |
| scf7180000051884 | 3608 | 0.4401 | 0 | 98.0866 | Eukaryota | 1102.0 | 0 | Platyhelminthes | 1102.0 | 0 | Polystomatidea | 1102.0 | 0 | Polystomatidae | 1102.0 | 0 | Protopolystoma | 1102.0 | 0 | Protopolystoma xenopodis | 1102.0 | 0 |
| scf7180000054245 | 18719 | 0.3902 | 0 | 79.0079 | Eukaryota | 6629.0 | 0 | Chordata | 6629.0 | 0 | Carnivora | 5904.0 | 1 | Canidae | 5904.0 | 1 | Vulpes | 3280.0 | 2 | Vulpes vulpes | 3280.0 | 2 |
| scf7180000043394 | 75934 | 0.3954 | 0 | 55.4335 | Eukaryota | 11667.0 | 0 | Chordata | 11667.0 | 0 | Carnivora | 11667.0 | 0 | Canidae | 11667.0 | 0 | Canis | 11667.0 | 0 | Canis lupus | 11667.0 | 0 |
| scf7180000040828 | 1678400 | 0.3447 | 0 | 59.2663 | Eukaryota | 93772.0 | 0 | Chordata | 93772.0 | 0 | Carnivora | 93772.0 | 0 | Canidae | 93772.0 | 0 | Canis | 93772.0 | 0 | Canis lupus | 93772.0 | 0 |
| scf7180000043419 | 445216 | 0.4195 | 0 | 59.6864 | Eukaryota | 43281.0 | 0 | Chordata | 43281.0 | 0 | Carnivora | 43281.0 | 0 | Canidae | 43281.0 | 0 | Canis | 29048.0 | 1 | Canis lupus | 29048.0 | 1 |
| scf7180000049464 | 3036 | 0.4499 | 0 | 664.3873 | no-hit | 0.0 | 0 | no-hit | 0.0 | 0 | no-hit | 0.0 | 0 | no-hit | 0.0 | 0 | no-hit | 0.0 | 0 | no-hit | 0.0 | 0 |
| scf7180000048559 | 44752 | 0.6079 | 0 | 53.0414 | Eukaryota | 12844.0 | 0 | Chordata | 12844.0 | 0 | Primates | 4331.0 | 5 | Hominidae | 4331.0 | 5 | Homo | 2886.0 | 7 | Homo sapiens | 2886.0 | 7 |
| scf7180000047576 | 3248 | 0.4329 | 0 | 241.3506 | Eukaryota | 15379.0 | 0 | Chordata | 10887.0 | 2 | Synbranchiformes | 2841.0 | 6 | Mastacembelidae | 2841.0 | 6 | Mastacembelus | 2841.0 | 7 | Mastacembelus armatus | 2841.0 | 8 |
| scf7180000056185 | 1641121 | 0.4784 | 0 | 58.9973 | Eukaryota | 109427.0 | 0 | Chordata | 109427.0 | 0 | Carnivora | 109427.0 | 0 | Canidae | 109427.0 | 0 | Canis | 91052.0 | 1 | Canis lupus | 91052.0 | 1 |
| scf7180000047497 | 29596 | 0.4376 | 0 | 196.7989 | Eukaryota | 25729.0 | 0 | Chordata | 25729.0 | 0 | Carnivora | 25729.0 | 0 | Canidae | 25729.0 | 0 | Canis | 25729.0 | 0 | Canis lupus | 25729.0 | 0 |
| scf7180000042927 | 54079 | 0.4075 | 0 | 58.0906 | Eukaryota | 37163.0 | 0 | Chordata | 37163.0 | 0 | Carnivora | 37163.0 | 0 | Canidae | 33912.0 | 1 | Canis | 29631.0 | 2 | Canis lupus | 29631.0 | 2 |
| scf7180000041699 | 915171 | 0.4001 | 0 | 59.7521 | Eukaryota | 98507.0 | 0 | Chordata | 98507.0 | 0 | Carnivora | 98507.0 | 0 | Canidae | 98507.0 | 0 | Canis | 68553.0 | 1 | Canis lupus | 68553.0 | 1 |
| scf7180000041130 | 1190753 | 0.4541 | 0 | 59.922 | Eukaryota | 66042.0 | 0 | Chordata | 66042.0 | 0 | Carnivora | 66042.0 | 0 | Canidae | 66042.0 | 0 | Canis | 66042.0 | 0 | Canis lupus | 66042.0 | 0 |
| scf7180000056247 | 662589 | 0.3601 | 0 | 59.5819 | Eukaryota | 54668.0 | 0 | Chordata | 54668.0 | 0 | Carnivora | 54668.0 | 0 | Canidae | 54668.0 | 0 | Canis | 54668.0 | 0 | Canis lupus | 54668.0 | 0 |
| scf7180000046448 | 128313 | 0.4183 | 0 | 60.8246 | Eukaryota | 16915.0 | 0 | Chordata | 16915.0 | 0 | Carnivora | 16915.0 | 0 | Canidae | 12715.0 | 1 | Canis | 8195.0 | 2 | Canis lupus | 8195.0 | 2 |
| scf7180000049300 | 11588 | 0.4416 | 0 | 91.4625 | no-hit | 0.0 | 0 | no-hit | 0.0 | 0 | no-hit | 0.0 | 0 | no-hit | 0.0 | 0 | no-hit | 0.0 | 0 | no-hit | 0.0 | 0 |
| scf7180000041354 | 257642 | 0.3659 | 0 | 59.7495 | Eukaryota | 47888.0 | 0 | Chordata | 47888.0 | 0 | Carnivora | 47888.0 | 0 | Canidae | 47888.0 | 0 | Canis | 47888.0 | 0 | Canis lupus | 47888.0 | 0 |
| scf7180000047556 | 21129 | 0.4349 | 0 | 68.6722 | Eukaryota | 80958.0 | 0 | Chordata | 80958.0 | 0 | Carnivora | 80958.0 | 0 | Canidae | 80958.0 | 0 | Canis | 80958.0 | 0 | Canis lupus | 80958.0 | 0 |
| scf7180000044048 | 83400 | 0.3876 | 0 | 55.4757 | Eukaryota | 8423.0 | 0 | Chordata | 8423.0 | 0 | Primates | 7028.0 | 2 | Cercopithecidae | 3536.0 | 3 | Homo | 2738.0 | 7 | Homo sapiens | 2738.0 | 8 |
| scf7180000053731 | 1864 | 0.4485 | 0 | 204.9356 | no-hit | 0.0 | 0 | no-hit | 0.0 | 0 | no-hit | 0.0 | 0 | no-hit | 0.0 | 0 | no-hit | 0.0 | 0 | no-hit | 0.0 | 0 |
| scf7180000051538 | 2905 | 0.4413 | 0 | 216.4154 | no-hit | 0.0 | 0 | no-hit | 0.0 | 0 | no-hit | 0.0 | 0 | no-hit | 0.0 | 0 | no-hit | 0.0 | 0 | no-hit | 0.0 | 0 |
| scf7180000052773 | 2675 | 0.5058 | 0 | 113.2313 | Eukaryota | 4637.0 | 0 | Chordata | 4637.0 | 0 | Carnivora | 4637.0 | 0 | Canidae | 4637.0 | 0 | Canis | 3940.0 | 2 | Canis lupus | 3940.0 | 2 |
| scf7180000053179 | 3074 | 0.4483 | 0 | 620.3245 | no-hit | 0.0 | 0 | no-hit | 0.0 | 0 | no-hit | 0.0 | 0 | no-hit | 0.0 | 0 | no-hit | 0.0 | 0 | no-hit | 0.0 | 0 |
| scf7180000043922 | 667395 | 0.4051 | 0 | 59.7636 | Eukaryota | 37372.0 | 0 | Chordata | 37372.0 | 0 | Carnivora | 37372.0 | 0 | Canidae | 12283.0 | 5 | Canis | 8198.0 | 7 | Canis lupus | 8198.0 | 7 |
| scf7180000046410 | 5952 | 0.4405 | 0 | 50.1025 | no-hit | 0.0 | 0 | no-hit | 0.0 | 0 | no-hit | 0.0 | 0 | no-hit | 0.0 | 0 | no-hit | 0.0 | 0 | no-hit | 0.0 | 0 |
| scf7180000051054 | 2743 | 0.4532 | 0 | 264.5818 | no-hit | 0.0 | 0 | no-hit | 0.0 | 0 | no-hit | 0.0 | 0 | no-hit | 0.0 | 0 | no-hit | 0.0 | 0 | no-hit | 0.0 | 0 |
| scf7180000056045 | 5617 | 0.438 | 0 | 346.0154 | no-hit | 0.0 | 0 | no-hit | 0.0 | 0 | no-hit | 0.0 | 0 | no-hit | 0.0 | 0 | no-hit | 0.0 | 0 | no-hit | 0.0 | 0 |
| scf7180000040678 | 22260 | 0.4766 | 0 | 149.8007 | Eukaryota | 76707.0 | 0 | Chordata | 76707.0 | 0 | Carnivora | 76707.0 | 0 | Canidae | 76707.0 | 0 | Canis | 76707.0 | 0 | Canis lupus | 76707.0 | 0 |
| scf7180000046506 | 86292 | 0.3778 | 0 | 58.8274 | Eukaryota | 38240.0 | 0 | Chordata | 38240.0 | 0 | Carnivora | 38240.0 | 0 | Canidae | 38240.0 | 0 | Canis | 38240.0 | 0 | Canis lupus | 38240.0 | 0 |
| scf7180000041833 | 965613 | 0.4016 | 0 | 60.7373 | Eukaryota | 51540.0 | 0 | Chordata | 51540.0 | 0 | Carnivora | 51540.0 | 0 | Canidae | 51540.0 | 0 | Canis | 51540.0 | 0 | Canis lupus | 51540.0 | 0 |
| scf7180000045223 | 255447 | 0.4521 | 0 | 58.3279 | Eukaryota | 43894.0 | 0 | Chordata | 43894.0 | 0 | Carnivora | 43894.0 | 0 | Canidae | 31813.0 | 4 | Canis | 21809.0 | 5 | Canis lupus | 21809.0 | 5 |
| scf7180000054627 | 30464 | 0.3913 | 0 | 756.7454 | Eukaryota | 69439.0 | 0 | Chordata | 69439.0 | 0 | Carnivora | 69439.0 | 0 | Canidae | 69439.0 | 0 | Canis | 69439.0 | 0 | Canis lupus | 69439.0 | 0 |
| scf7180000047694 | 84746 | 0.4224 | 0 | 59.6672 | Eukaryota | 40769.0 | 0 | Chordata | 40769.0 | 0 | Carnivora | 40769.0 | 0 | Canidae | 40769.0 | 0 | Canis | 24576.0 | 1 | Canis lupus | 24576.0 | 1 |
| scf7180000055814 | 15414 | 0.4516 | 0 | 1096.3038 | Eukaryota | 526.0 | 0 | Streptophyta | 526.0 | 0 | Fabales | 526.0 | 0 | Fabaceae | 526.0 | 0 | Cicer | 526.0 | 0 | Cicer arietinum | 526.0 | 0 |
| scf7180000052445 | 5541 | 0.4958 | 0 | 64.2679 | Eukaryota | 7581.0 | 0 | Chordata | 7581.0 | 0 | Carnivora | 7581.0 | 0 | Canidae | 7581.0 | 0 | Canis | 6475.0 | 1 | Canis lupus | 6475.0 | 1 |
| scf7180000041728 | 479642 | 0.3714 | 0 | 59.7236 | Eukaryota | 21056.0 | 0 | Chordata | 21056.0 | 0 | Carnivora | 13042.0 | 3 | Canidae | 13042.0 | 3 | Canis | 7769.0 | 4 | Canis lupus | 7769.0 | 4 |
| scf7180000055107 | 43158 | 0.4452 | 0 | 131.7309 | Eukaryota | 91068.0 | 0 | Chordata | 91068.0 | 0 | Carnivora | 91068.0 | 0 | Canidae | 91068.0 | 0 | Canis | 91068.0 | 0 | Canis lupus | 91068.0 | 0 |
| scf7180000044653 | 292012 | 0.4513 | 0 | 58.5787 | Eukaryota | 76941.0 | 0 | Chordata | 76941.0 | 0 | Carnivora | 76941.0 | 0 | Canidae | 76941.0 | 0 | Canis | 76941.0 | 0 | Canis lupus | 76941.0 | 0 |
| scf7180000050607 | 21154 | 0.4182 | 0 | 48.4172 | Eukaryota | 64410.0 | 0 | Chordata | 64410.0 | 0 | Carnivora | 64410.0 | 0 | Canidae | 64410.0 | 0 | Canis | 51504.0 | 1 | Canis lupus | 51504.0 | 1 |
| scf7180000051384 | 3118 | 0.4429 | 0 | 803.6046 | no-hit | 0.0 | 0 | no-hit | 0.0 | 0 | no-hit | 0.0 | 0 | no-hit | 0.0 | 0 | no-hit | 0.0 | 0 | no-hit | 0.0 | 0 |
| scf7180000053669 | 3636 | 0.5052 | 0 | 219.7826 | Eukaryota | 6145.0 | 0 | Chordata | 6145.0 | 0 | Carnivora | 6145.0 | 0 | Canidae | 6145.0 | 0 | Canis | 3717.0 | 1 | Canis lupus | 2588.0 | 2 |
| scf7180000042530 | 1136318 | 0.3422 | 0 | 59.5995 | Eukaryota | 117357.0 | 0 | Chordata | 117357.0 | 0 | Carnivora | 117357.0 | 0 | Canidae | 117357.0 | 0 | Canis | 117357.0 | 0 | Canis lupus | 117357.0 | 0 |
| scf7180000050643 | 38568 | 0.3675 | 0 | 60.8005 | Eukaryota | 4721.0 | 0 | Chordata | 4721.0 | 0 | Carnivora | 4083.0 | 1 | Canidae | 2861.0 | 3 | Canis | 2861.0 | 3 | Canis lupus | 2861.0 | 3 |
| scf7180000055701 | 20417 | 0.4508 | 0 | 165.993 | Eukaryota | 3448.0 | 0 | Chordata | 3448.0 | 0 | Carnivora | 3448.0 | 0 | Canidae | 3448.0 | 0 | Canis | 3448.0 | 0 | Canis lupus | 3448.0 | 0 |
| scf7180000043008 | 524844 | 0.3844 | 0 | 60.0008 | Eukaryota | 15979.0 | 0 | Chordata | 15979.0 | 0 | Carnivora | 15979.0 | 0 | Canidae | 15979.0 | 0 | Canis | 15979.0 | 0 | Canis lupus | 15979.0 | 0 |
| scf7180000051138 | 14063 | 0.5058 | 0 | 53.4573 | Eukaryota | 16990.0 | 0 | Chordata | 16990.0 | 0 | Carnivora | 16990.0 | 0 | Canidae | 16990.0 | 0 | Canis | 16990.0 | 0 | Canis lupus | 16990.0 | 0 |
| scf7180000044526 | 25193 | 0.4435 | 0 | 200.3047 | Eukaryota | 1070.0 | 0 | Chordata | 907.0 | 1 | Kurtiformes | 520.0 | 2 | Apogonidae | 520.0 | 2 | Sphaeramia | 520.0 | 2 | Sphaeramia orbicularis | 520.0 | 2 |
| scf7180000044673 | 15661 | 0.4329 | 0 | 104.1138 | Eukaryota | 88443.0 | 0 | Chordata | 88443.0 | 0 | Carnivora | 88443.0 | 0 | Canidae | 88443.0 | 0 | Canis | 88443.0 | 0 | Canis lupus | 88443.0 | 0 |
| scf7180000044206 | 123343 | 0.3556 | 0 | 55.844 | Eukaryota | 46556.0 | 0 | Chordata | 46556.0 | 0 | Carnivora | 46556.0 | 0 | Canidae | 46556.0 | 0 | Canis | 46556.0 | 0 | Canis lupus | 46556.0 | 0 |
| scf7180000050421 | 3847 | 0.4559 | 0 | 6.5274 | Eukaryota | 50175.0 | 0 | Chordata | 50175.0 | 0 | Carnivora | 50175.0 | 0 | Canidae | 50175.0 | 0 | Canis | 50175.0 | 0 | Canis lupus | 50175.0 | 0 |
| scf7180000041967 | 6875 | 0.4611 | 0 | 67.9216 | Eukaryota | 33939.0 | 0 | Chordata | 33939.0 | 0 | Carnivora | 33939.0 | 0 | Canidae | 33939.0 | 0 | Canis | 33939.0 | 0 | Canis lupus | 33939.0 | 0 |
| scf7180000048434 | 5470 | 0.4477 | 0 | 49.1179 | no-hit | 0.0 | 0 | no-hit | 0.0 | 0 | no-hit | 0.0 | 0 | no-hit | 0.0 | 0 | no-hit | 0.0 | 0 | no-hit | 0.0 | 0 |
| scf7180000042752 | 623614 | 0.3815 | 0 | 60.7553 | Eukaryota | 56338.0 | 0 | Chordata | 56338.0 | 0 | Carnivora | 36310.0 | 1 | Canidae | 31831.0 | 2 | Canis | 22271.0 | 3 | Canis lupus | 22271.0 | 3 |
| scf7180000056394 | 106787 | 0.4363 | 0 | 55.9025 | Eukaryota | 20585.0 | 0 | Chordata | 20585.0 | 0 | Carnivora | 20585.0 | 0 | Canidae | 20585.0 | 0 | Canis | 20585.0 | 0 | Canis lupus | 20585.0 | 0 |
| scf7180000050709 | 15369 | 0.4472 | 0 | 231.604 | Eukaryota | 3013.0 | 0 | Chordata | 2310.0 | 1 | Carnivora | 2310.0 | 1 | Canidae | 2310.0 | 1 | Canis | 2310.0 | 1 | Canis lupus | 2310.0 | 1 |
| scf7180000045511 | 175609 | 0.3987 | 0 | 62.3224 | Eukaryota | 22849.0 | 0 | Chordata | 22849.0 | 0 | Carnivora | 18745.0 | 1 | Canidae | 9756.0 | 5 | Canis | 9756.0 | 5 | Canis lupus | 9756.0 | 5 |
| scf7180000048982 | 13551 | 0.4389 | 0 | 128.5344 | Eukaryota | 23829.0 | 0 | Platyhelminthes | 11402.0 | 2 | Strigeidida | 11402.0 | 6 | Schistosomatidae | 11402.0 | 6 | Schistosoma | 11402.0 | 6 | Schistosoma mansoni | 11402.0 | 7 |
| scf7180000043863 | 413390 | 0.3959 | 0 | 58.631 | Eukaryota | 25811.0 | 0 | Chordata | 25811.0 | 0 | Carnivora | 25811.0 | 0 | Canidae | 25811.0 | 0 | Canis | 25811.0 | 0 | Canis lupus | 25811.0 | 0 |
| scf7180000051984 | 3643 | 0.4499 | 0 | 315.2902 | no-hit | 0.0 | 0 | no-hit | 0.0 | 0 | no-hit | 0.0 | 0 | no-hit | 0.0 | 0 | no-hit | 0.0 | 0 | no-hit | 0.0 | 0 |
| scf7180000043283 | 528848 | 0.4664 | 0 | 59.1312 | Eukaryota | 38092.0 | 0 | Chordata | 38092.0 | 0 | Carnivora | 38092.0 | 0 | Canidae | 38092.0 | 0 | Canis | 32575.0 | 1 | Canis lupus | 32575.0 | 1 |
| scf7180000044608 | 301648 | 0.4166 | 0 | 59.3433 | Eukaryota | 30082.0 | 0 | Chordata | 30082.0 | 0 | Carnivora | 30082.0 | 0 | Canidae | 30082.0 | 0 | Canis | 30082.0 | 0 | Canis lupus | 30082.0 | 0 |
| scf7180000050403 | 25060 | 0.4393 | 0 | 59.2551 | Eukaryota | 38564.0 | 1 | Platyhelminthes | 19531.0 | 2 | Strigeidida | 19531.0 | 5 | Schistosomatidae | 19531.0 | 5 | Schistosoma | 19531.0 | 5 | Schistosoma mansoni | 19531.0 | 5 |
| scf7180000052578 | 7563 | 0.5036 | 0 | 541.6477 | Eukaryota | 8880.0 | 0 | Chordata | 8880.0 | 0 | Carnivora | 8880.0 | 0 | Canidae | 8880.0 | 0 | Canis | 7204.0 | 1 | Canis lupus | 5969.0 | 2 |
| scf7180000042333 | 717611 | 0.4409 | 0 | 56.9575 | Eukaryota | 161815.0 | 0 | Chordata | 161815.0 | 0 | Carnivora | 161815.0 | 0 | Canidae | 161815.0 | 0 | Canis | 161815.0 | 0 | Canis lupus | 161815.0 | 0 |
| scf7180000055707 | 8860 | 0.4414 | 0 | 227.6614 | Eukaryota | 23422.0 | 0 | Chordata | 21423.0 | 1 | Gadiformes | 11676.0 | 7 | Gadidae | 11676.0 | 7 | Gadus | 11676.0 | 7 | Gadus morhua | 11676.0 | 7 |
| scf7180000044210 | 11883 | 0.4395 | 0 | 170.1392 | Eukaryota | 3901.0 | 0 | Platyhelminthes | 2351.0 | 3 | Polystomatidea | 2351.0 | 3 | Polystomatidae | 2351.0 | 3 | Protopolystoma | 2351.0 | 3 | Protopolystoma xenopodis | 2351.0 | 3 |
| scf7180000044898 | 76468 | 0.4302 | 0 | 64.7959 | Eukaryota | 93179.0 | 0 | Chordata | 93179.0 | 0 | Carnivora | 93179.0 | 0 | Canidae | 93179.0 | 0 | Canis | 93179.0 | 0 | Canis lupus | 93179.0 | 0 |
| scf7180000054906 | 27213 | 0.4439 | 0 | 303.296 | Eukaryota | 31887.0 | 1 | Chordata | 31887.0 | 1 | Gadiformes | 10717.0 | 6 | Gadidae | 10717.0 | 6 | Gadus | 10717.0 | 6 | Gadus morhua | 10717.0 | 6 |
| scf7180000049891 | 26256 | 0.4401 | 0 | 86.8632 | Eukaryota | 3370.0 | 1 | Viruses-undef | 3098.0 | 2 | Herpesvirales | 3098.0 | 3 | Herpesviridae | 3098.0 | 3 | Roseolovirus | 3098.0 | 3 | Human betaherpesvirus 7 | 3098.0 | 3 |
| scf7180000044746 | 249843 | 0.4477 | 0 | 59.9747 | Eukaryota | 30799.0 | 0 | Chordata | 30799.0 | 0 | Carnivora | 30799.0 | 0 | Canidae | 30799.0 | 0 | Canis | 19109.0 | 1 | Canis lupus | 19109.0 | 1 |
| scf7180000045138 | 12459 | 0.3631 | 0 | 46.2833 | Eukaryota | 3433.0 | 0 | Chordata | 3433.0 | 0 | Carnivora | 3433.0 | 0 | Canidae | 3017.0 | 1 | Canis | 3017.0 | 1 | Canis lupus | 3017.0 | 1 |
| scf7180000055536 | 27734 | 0.4392 | 0 | 118.1743 | no-hit | 0.0 | 0 | no-hit | 0.0 | 0 | no-hit | 0.0 | 0 | no-hit | 0.0 | 0 | no-hit | 0.0 | 0 | no-hit | 0.0 | 0 |
| scf7180000054062 | 3107 | 0.4442 | 0 | 54.7739 | Eukaryota | 176.0 | 0 | Platyhelminthes | 176.0 | 0 | Polystomatidea | 176.0 | 0 | Polystomatidae | 176.0 | 0 | Protopolystoma | 176.0 | 0 | Protopolystoma xenopodis | 176.0 | 0 |
| scf7180000047681 | 16329 | 0.4591 | 0 | 361.2997 | Eukaryota | 13003.0 | 0 | Chordata | 13003.0 | 0 | Carnivora | 13003.0 | 0 | Canidae | 13003.0 | 0 | Canis | 11152.0 | 1 | Canis lupus | 10150.0 | 2 |
| scf7180000047637 | 70482 | 0.4279 | 0 | 57.1149 | Eukaryota | 44871.0 | 0 | Chordata | 44871.0 | 0 | Carnivora | 44871.0 | 0 | Canidae | 44871.0 | 0 | Canis | 44871.0 | 0 | Canis lupus | 44871.0 | 0 |
| scf7180000047893 | 81334 | 0.3828 | 0 | 59.6124 | Eukaryota | 9473.0 | 0 | Chordata | 9473.0 | 0 | Primates | 3949.0 | 2 | Cervidae | 3236.0 | 3 | Odocoileus | 3236.0 | 3 | Odocoileus virginianus | 3236.0 | 3 |
| scf7180000051331 | 7703 | 0.4439 | 0 | 59.337 | Eukaryota | 4075.0 | 0 | Chordata | 2791.0 | 1 | Kurtiformes | 1299.0 | 4 | Apogonidae | 1299.0 | 4 | Sphaeramia | 1299.0 | 4 | Sphaeramia orbicularis | 1299.0 | 4 |
| scf7180000042654 | 172268 | 0.381 | 0 | 57.1914 | Eukaryota | 46879.0 | 0 | Chordata | 46879.0 | 0 | Carnivora | 46879.0 | 0 | Canidae | 46879.0 | 0 | Canis | 46879.0 | 0 | Canis lupus | 46879.0 | 0 |
| scf7180000050441 | 3245 | 0.4419 | 0 | 43.9043 | Eukaryota | 1815.0 | 0 | Chordata | 1515.0 | 1 | Columbiformes | 1515.0 | 1 | Columbidae | 1515.0 | 1 | Streptopelia | 1515.0 | 1 | Streptopelia turtur | 1515.0 | 1 |
| scf7180000041889 | 449547 | 0.4052 | 0 | 58.6463 | Eukaryota | 61799.0 | 0 | Chordata | 61799.0 | 0 | Carnivora | 61799.0 | 0 | Canidae | 61799.0 | 0 | Canis | 61799.0 | 0 | Canis lupus | 61799.0 | 0 |
| scf7180000046733 | 18398 | 0.439 | 0 | 80.2167 | Eukaryota | 5624.0 | 0 | Chordata | 2022.0 | 2 | Forcipulatida | 1962.0 | 3 | Asteriidae | 1962.0 | 3 | Asterias | 1962.0 | 4 | Asterias rubens | 1962.0 | 4 |
| scf7180000040563 | 222494 | 0.3578 | 0 | 58.8173 | Eukaryota | 84323.0 | 0 | Chordata | 84323.0 | 0 | Carnivora | 84323.0 | 0 | Canidae | 84323.0 | 0 | Canis | 84323.0 | 0 | Canis lupus | 84323.0 | 0 |
| scf7180000043189 | 94145 | 0.3866 | 0 | 56.9635 | Eukaryota | 86942.0 | 0 | Chordata | 86942.0 | 0 | Carnivora | 86942.0 | 0 | Canidae | 86942.0 | 0 | Canis | 86942.0 | 0 | Canis lupus | 86942.0 | 0 |
| scf7180000056195 | 2434891 | 0.4818 | 0 | 59.4266 | Eukaryota | 87085.0 | 0 | Chordata | 87085.0 | 0 | Carnivora | 87085.0 | 0 | Canidae | 87085.0 | 0 | Canis | 62230.0 | 1 | Canis lupus | 62230.0 | 1 |
| scf7180000048007 | 22210 | 0.4434 | 0 | 193.2127 | Eukaryota | 5367.0 | 0 | Chordata | 4857.0 | 1 | Tetraodontiformes | 3597.0 | 2 | Tetraodontidae | 3597.0 | 2 | Takifugu | 3597.0 | 2 | Takifugu rubripes | 3445.0 | 3 |
| scf7180000043911 | 403562 | 0.4021 | 0 | 58.1735 | Eukaryota | 61912.0 | 0 | Chordata | 61912.0 | 0 | Carnivora | 61912.0 | 0 | Canidae | 61912.0 | 0 | Canis | 48604.0 | 1 | Canis lupus | 48604.0 | 1 |
| scf7180000055126 | 24726 | 0.4376 | 0 | 161.5339 | Eukaryota | 4672.0 | 0 | Chordata | 2159.0 | 2 | Kurtiformes | 1724.0 | 3 | Apogonidae | 1724.0 | 3 | Sphaeramia | 1724.0 | 3 | Sphaeramia orbicularis | 1724.0 | 3 |
| scf7180000053525 | 5058 | 0.4541 | 0 | 270.3787 | Eukaryota | 200.0 | 0 | Streptophyta | 200.0 | 0 | Fabales | 200.0 | 0 | Fabaceae | 200.0 | 0 | Cicer | 200.0 | 0 | Cicer arietinum | 200.0 | 0 |
| scf7180000042753 | 44559 | 0.3889 | 0 | 58.5766 | Eukaryota | 14070.0 | 0 | Chordata | 14070.0 | 0 | Carnivora | 7059.0 | 1 | Suidae | 7011.0 | 4 | Sus | 7011.0 | 5 | Sus scrofa | 7011.0 | 5 |
| scf7180000040510 | 1025102 | 0.3981 | 0 | 59.8317 | Eukaryota | 60159.0 | 0 | Chordata | 60159.0 | 0 | Carnivora | 60159.0 | 0 | Canidae | 60159.0 | 0 | Canis | 60159.0 | 0 | Canis lupus | 60159.0 | 0 |
| scf7180000045805 | 38813 | 0.4192 | 0 | 124.7924 | Eukaryota | 33066.0 | 0 | Chordata | 33066.0 | 0 | Carnivora | 33066.0 | 0 | Canidae | 33066.0 | 0 | Canis | 33066.0 | 0 | Canis lupus | 33066.0 | 0 |
| scf7180000051949 | 16760 | 0.4008 | 0 | 61.7319 | Eukaryota | 6728.0 | 0 | Chordata | 6728.0 | 0 | Primates | 5800.0 | 1 | Cheirogaleidae | 3458.0 | 2 | Microcebus | 3458.0 | 2 | Microcebus murinus | 3458.0 | 2 |
| scf7180000045210 | 243792 | 0.3616 | 0 | 59.3798 | Eukaryota | 54576.0 | 0 | Chordata | 54576.0 | 0 | Carnivora | 54576.0 | 0 | Canidae | 54576.0 | 0 | Canis | 54576.0 | 0 | Canis lupus | 54576.0 | 0 |
| scf7180000042684 | 297987 | 0.3975 | 0 | 57.1452 | Eukaryota | 77557.0 | 0 | Chordata | 77557.0 | 0 | Carnivora | 77557.0 | 0 | Canidae | 77557.0 | 0 | Canis | 77557.0 | 0 | Canis lupus | 77557.0 | 0 |
| scf7180000052639 | 2352 | 0.4554 | 0 | 359.238 | no-hit | 0.0 | 0 | no-hit | 0.0 | 0 | no-hit | 0.0 | 0 | no-hit | 0.0 | 0 | no-hit | 0.0 | 0 | no-hit | 0.0 | 0 |
| scf7180000053853 | 6846 | 0.5061 | 0 | 116.1592 | Eukaryota | 8292.0 | 0 | Chordata | 8292.0 | 0 | Carnivora | 8292.0 | 0 | Canidae | 8292.0 | 0 | Canis | 6640.0 | 1 | Canis lupus | 5427.0 | 2 |
| scf7180000055249 | 12119 | 0.4401 | 0 | 125.9864 | Eukaryota | 3823.0 | 0 | Platyhelminthes | 1890.0 | 2 | Polystomatidea | 1890.0 | 3 | Polystomatidae | 1890.0 | 3 | Protopolystoma | 1890.0 | 3 | Protopolystoma xenopodis | 1890.0 | 3 |
| scf7180000043354 | 366272 | 0.3401 | 0 | 58.7326 | Eukaryota | 92495.0 | 0 | Chordata | 92495.0 | 0 | Carnivora | 92495.0 | 0 | Canidae | 92495.0 | 0 | Canis | 92495.0 | 0 | Canis lupus | 92495.0 | 0 |
| scf7180000040946 | 130630 | 0.4187 | 0 | 57.7687 | Eukaryota | 13109.0 | 0 | Chordata | 13109.0 | 0 | Carnivora | 13109.0 | 0 | Felidae | 11603.0 | 1 | Felis | 11603.0 | 1 | Felis catus | 11603.0 | 1 |
| scf7180000042636 | 82778 | 0.3851 | 0 | 59.0912 | Eukaryota | 17035.0 | 0 | Chordata | 17035.0 | 0 | Carnivora | 17035.0 | 0 | Canidae | 17035.0 | 0 | Canis | 13625.0 | 1 | Canis lupus | 13625.0 | 1 |
| scf7180000045733 | 105708 | 0.463 | 0 | 95.6333 | Eukaryota | 89734.0 | 0 | Chordata | 89734.0 | 0 | Carnivora | 89734.0 | 0 | Canidae | 89734.0 | 0 | Canis | 89734.0 | 0 | Canis lupus | 89734.0 | 0 |
| scf7180000052123 | 7515 | 0.4447 | 0 | 527.292 | no-hit | 0.0 | 0 | no-hit | 0.0 | 0 | no-hit | 0.0 | 0 | no-hit | 0.0 | 0 | no-hit | 0.0 | 0 | no-hit | 0.0 | 0 |
| scf7180000040482 | 1857237 | 0.4368 | 100 | 58.4793 | Eukaryota | 114333.0 | 0 | Chordata | 114333.0 | 0 | Carnivora | 114333.0 | 0 | Canidae | 114333.0 | 0 | Canis | 104739.0 | 1 | Canis lupus | 104739.0 | 1 |
| scf7180000054581 | 1622437 | 0.3807 | 0 | 60.8581 | Eukaryota | 93680.0 | 0 | Chordata | 93680.0 | 0 | Carnivora | 93680.0 | 0 | Canidae | 93680.0 | 0 | Canis | 83379.0 | 1 | Canis lupus | 83379.0 | 1 |
| scf7180000050747 | 11915 | 0.4429 | 0 | 89.2036 | no-hit | 0.0 | 0 | no-hit | 0.0 | 0 | no-hit | 0.0 | 0 | no-hit | 0.0 | 0 | no-hit | 0.0 | 0 | no-hit | 0.0 | 0 |
| scf7180000045898 | 125851 | 0.3992 | 0 | 58.3899 | Eukaryota | 86574.0 | 0 | Chordata | 86574.0 | 0 | Carnivora | 86574.0 | 0 | Canidae | 86574.0 | 0 | Canis | 86574.0 | 0 | Canis lupus | 86574.0 | 0 |
| scf7180000050741 | 19812 | 0.4426 | 0 | 110.1776 | Eukaryota | 4451.0 | 0 | Platyhelminthes | 3290.0 | 1 | Polystomatidea | 3290.0 | 2 | Polystomatidae | 3290.0 | 2 | Protopolystoma | 3290.0 | 2 | Protopolystoma xenopodis | 3290.0 | 2 |
| scf7180000046398 | 38088 | 0.4418 | 0 | 50.1812 | Eukaryota | 16402.0 | 0 | Chordata | 16402.0 | 0 | Carnivora | 16402.0 | 0 | Canidae | 6133.0 | 4 | Canis | 4188.0 | 7 | Canis lupus | 4188.0 | 8 |
| scf7180000053712 | 10879 | 0.5044 | 0 | 201.0177 | Eukaryota | 9073.0 | 0 | Chordata | 9073.0 | 0 | Carnivora | 9073.0 | 0 | Canidae | 9073.0 | 0 | Canis | 7347.0 | 1 | Canis lupus | 6114.0 | 2 |
| scf7180000042940 | 50198 | 0.4149 | 0 | 86.7359 | Eukaryota | 91641.0 | 0 | Chordata | 91641.0 | 0 | Carnivora | 91641.0 | 0 | Canidae | 91641.0 | 0 | Canis | 91641.0 | 0 | Canis lupus | 91641.0 | 0 |
| scf7180000044750 | 184238 | 0.3506 | 0 | 59.9028 | Eukaryota | 32064.0 | 0 | Chordata | 32064.0 | 0 | Carnivora | 32064.0 | 0 | Canidae | 32064.0 | 0 | Canis | 32064.0 | 0 | Canis lupus | 32064.0 | 0 |
| scf7180000046385 | 133850 | 0.4374 | 0 | 60.3653 | Eukaryota | 48009.0 | 0 | Chordata | 48009.0 | 0 | Carnivora | 48009.0 | 0 | Canidae | 48009.0 | 0 | Canis | 41071.0 | 1 | Canis lupus | 41071.0 | 1 |
| scf7180000047962 | 52482 | 0.4408 | 0 | 228.4085 | Eukaryota | 36167.0 | 0 | Chordata | 36167.0 | 0 | Carnivora | 36167.0 | 0 | Canidae | 36167.0 | 0 | Canis | 36167.0 | 0 | Canis lupus | 36167.0 | 0 |
| scf7180000053766 | 3023 | 0.4357 | 0 | 31.1177 | no-hit | 0.0 | 0 | no-hit | 0.0 | 0 | no-hit | 0.0 | 0 | no-hit | 0.0 | 0 | no-hit | 0.0 | 0 | no-hit | 0.0 | 0 |
| scf7180000045145 | 215422 | 0.3531 | 0 | 59.4031 | Eukaryota | 44426.0 | 0 | Chordata | 44426.0 | 0 | Carnivora | 44426.0 | 0 | Canidae | 44426.0 | 0 | Canis | 44426.0 | 0 | Canis lupus | 44426.0 | 0 |
| scf7180000046011 | 7691 | 0.3488 | 0 | 42.6648 | Eukaryota | 8818.0 | 0 | Chordata | 8818.0 | 0 | Carnivora | 8818.0 | 0 | Canidae | 8818.0 | 0 | Canis | 8818.0 | 0 | Canis lupus | 8818.0 | 0 |
| scf7180000047530 | 51674 | 0.4417 | 0 | 117.6858 | Eukaryota | 35350.0 | 0 | Chordata | 35350.0 | 0 | Carnivora | 35350.0 | 0 | Canidae | 35350.0 | 0 | Canis | 35350.0 | 0 | Canis lupus | 35350.0 | 0 |
| scf7180000041218 | 441067 | 0.4509 | 0 | 59.8555 | Eukaryota | 66931.0 | 0 | Chordata | 66931.0 | 0 | Carnivora | 66931.0 | 0 | Canidae | 66931.0 | 0 | Vulpes | 66931.0 | 0 | Vulpes vulpes | 66931.0 | 0 |
| scf7180000044826 | 15854 | 0.4232 | 0 | 98.743 | Eukaryota | 39707.0 | 0 | Chordata | 39707.0 | 0 | Carnivora | 39707.0 | 0 | Canidae | 39707.0 | 0 | Canis | 39707.0 | 0 | Canis lupus | 39707.0 | 0 |
| scf7180000052185 | 2889 | 0.4486 | 0 | 511.7891 | Eukaryota | 3474.0 | 0 | Chordata | 2617.0 | 1 | Columbiformes | 1013.0 | 3 | Columbidae | 1013.0 | 3 | Streptopelia | 1013.0 | 3 | Streptopelia turtur | 1013.0 | 3 |
| scf7180000056495 | 373500 | 0.4472 | 0 | 59.0329 | Eukaryota | 31903.0 | 0 | Chordata | 31903.0 | 0 | Carnivora | 31903.0 | 0 | Canidae | 28510.0 | 1 | Canis | 18951.0 | 2 | Canis lupus | 18951.0 | 2 |
| scf7180000042354 | 636592 | 0.4656 | 0 | 58.5559 | Eukaryota | 60328.0 | 0 | Chordata | 60328.0 | 0 | Carnivora | 60328.0 | 0 | Canidae | 60328.0 | 0 | Canis | 60328.0 | 0 | Canis lupus | 60328.0 | 0 |
| scf7180000048013 | 14513 | 0.449 | 0 | 621.3729 | Eukaryota | 2612.0 | 0 | Chordata | 2612.0 | 0 | Carnivora | 2331.0 | 1 | Canidae | 2331.0 | 1 | Canis | 2331.0 | 1 | Canis lupus | 2331.0 | 1 |
| scf7180000043413 | 74782 | 0.5124 | 0 | 55.3862 | Eukaryota | 28589.0 | 0 | Chordata | 28589.0 | 0 | Carnivora | 28589.0 | 0 | Canidae | 28589.0 | 0 | Canis | 16756.0 | 1 | Canis lupus | 16756.0 | 1 |
| scf7180000044432 | 344548 | 0.4068 | 0 | 59.7391 | Eukaryota | 77420.0 | 0 | Chordata | 77420.0 | 0 | Carnivora | 77420.0 | 0 | Canidae | 77420.0 | 0 | Vulpes | 38840.0 | 1 | Vulpes vulpes | 38840.0 | 1 |
| scf7180000045133 | 230494 | 0.3678 | 0 | 60.9344 | Eukaryota | 29351.0 | 1 | Chordata | 29351.0 | 1 | Carnivora | 29351.0 | 1 | Canidae | 26702.0 | 2 | Canis | 19642.0 | 3 | Canis lupus | 19642.0 | 3 |
| scf7180000042106 | 49853 | 0.3565 | 0 | 59.7839 | Eukaryota | 93506.0 | 0 | Chordata | 93506.0 | 0 | Carnivora | 93506.0 | 0 | Canidae | 93506.0 | 0 | Canis | 93506.0 | 0 | Canis lupus | 93506.0 | 0 |
| scf7180000040987 | 1488228 | 0.4006 | 0 | 59.9947 | Eukaryota | 73480.0 | 0 | Chordata | 73480.0 | 0 | Carnivora | 73480.0 | 0 | Canidae | 73480.0 | 0 | Canis | 66132.0 | 1 | Canis lupus | 66132.0 | 1 |
| scf7180000042275 | 195477 | 0.3766 | 100 | 59.2998 | Eukaryota | 66343.0 | 0 | Chordata | 66343.0 | 0 | Carnivora | 66343.0 | 0 | Canidae | 66343.0 | 0 | Canis | 66343.0 | 0 | Canis lupus | 66343.0 | 0 |
| scf7180000053474 | 9425 | 0.431 | 0 | 131.8622 | no-hit | 0.0 | 0 | no-hit | 0.0 | 0 | no-hit | 0.0 | 0 | no-hit | 0.0 | 0 | no-hit | 0.0 | 0 | no-hit | 0.0 | 0 |
| scf7180000042498 | 1147891 | 0.4166 | 0 | 60.1902 | Eukaryota | 115482.0 | 0 | Chordata | 115482.0 | 0 | Carnivora | 115482.0 | 0 | Canidae | 115482.0 | 0 | Canis | 115482.0 | 0 | Canis lupus | 115482.0 | 0 |
| scf7180000045941 | 135000 | 0.4569 | 0 | 56.9521 | Eukaryota | 17135.0 | 0 | Chordata | 17135.0 | 0 | Carnivora | 17135.0 | 0 | Canidae | 17135.0 | 0 | Canis | 17135.0 | 0 | Canis lupus | 17135.0 | 0 |
| scf7180000054955 | 74299 | 0.4446 | 0 | 177.9733 | Eukaryota | 5102.0 | 0 | Chordata | 3860.0 | 1 | Kurtiformes | 1822.0 | 5 | Apogonidae | 1822.0 | 5 | Sphaeramia | 1822.0 | 5 | Sphaeramia orbicularis | 1822.0 | 5 |
| scf7180000046006 | 130716 | 0.3303 | 0 | 58.8276 | Eukaryota | 15569.0 | 0 | Chordata | 15569.0 | 0 | Apterygiformes | 13879.0 | 1 | Apterygidae | 13879.0 | 1 | Apteryx | 13879.0 | 1 | Apteryx mantelli | 13879.0 | 1 |
| scf7180000045929 | 63961 | 0.3857 | 0 | 60.1979 | Eukaryota | 16700.0 | 0 | Chordata | 16700.0 | 0 | Carnivora | 16700.0 | 0 | Canidae | 16700.0 | 0 | Canis | 16700.0 | 0 | Canis lupus | 16700.0 | 0 |
| scf7180000055340 | 5533 | 0.4428 | 0 | 156.6575 | Eukaryota | 576.0 | 0 | Chordata | 576.0 | 0 | Kurtiformes | 341.0 | 1 | Apogonidae | 341.0 | 1 | Sphaeramia | 341.0 | 1 | Sphaeramia orbicularis | 341.0 | 1 |
| scf7180000054934 | 12781 | 0.4454 | 0 | 380.7579 | Eukaryota | 10488.0 | 0 | Chordata | 10488.0 | 0 | Tetraodontiformes | 3952.0 | 2 | Tetraodontidae | 3952.0 | 2 | Takifugu | 3952.0 | 2 | Sphaeramia orbicularis | 3869.0 | 3 |
| scf7180000042235 | 789797 | 0.3667 | 0 | 59.324 | Eukaryota | 52538.0 | 0 | Chordata | 52538.0 | 0 | Carnivora | 52538.0 | 0 | Canidae | 52538.0 | 0 | Canis | 52538.0 | 0 | Canis lupus | 52538.0 | 0 |
| scf7180000044667 | 262891 | 0.5003 | 0 | 59.5844 | Eukaryota | 46251.0 | 0 | Chordata | 46251.0 | 0 | Carnivora | 46251.0 | 0 | Canidae | 22133.0 | 3 | Canis | 17169.0 | 5 | Canis lupus | 17169.0 | 5 |
| scf7180000053159 | 8596 | 0.5054 | 0 | 155.3407 | Eukaryota | 8148.0 | 0 | Chordata | 8148.0 | 0 | Carnivora | 8148.0 | 0 | Canidae | 8148.0 | 0 | Canis | 6594.0 | 1 | Canis lupus | 5611.0 | 2 |
| scf7180000046449 | 25368 | 0.4271 | 0 | 109.5064 | Eukaryota | 37698.0 | 0 | Chordata | 37698.0 | 0 | Carnivora | 37698.0 | 0 | Canidae | 37698.0 | 0 | Canis | 37698.0 | 0 | Canis lupus | 37698.0 | 0 |
| scf7180000053632 | 4758 | 0.4344 | 0 | 38.9671 | no-hit | 0.0 | 0 | no-hit | 0.0 | 0 | no-hit | 0.0 | 0 | no-hit | 0.0 | 0 | no-hit | 0.0 | 0 | no-hit | 0.0 | 0 |
| scf7180000049215 | 11083 | 0.442 | 0 | 87.9407 | Eukaryota | 5199.0 | 0 | Chordata | 5199.0 | 0 | Kurtiformes | 2842.0 | 2 | Apogonidae | 2842.0 | 2 | Sphaeramia | 2842.0 | 2 | Sphaeramia orbicularis | 2842.0 | 2 |
| scf7180000042788 | 252138 | 0.3945 | 0 | 56.1242 | Eukaryota | 91579.0 | 0 | Chordata | 91579.0 | 0 | Carnivora | 91579.0 | 0 | Canidae | 91579.0 | 0 | Canis | 91579.0 | 0 | Canis lupus | 91579.0 | 0 |
| scf7180000041557 | 854545 | 0.3813 | 0 | 59.107 | Eukaryota | 88331.0 | 0 | Chordata | 88331.0 | 0 | Carnivora | 88331.0 | 0 | Canidae | 88331.0 | 0 | Canis | 88331.0 | 0 | Canis lupus | 88331.0 | 0 |
| scf7180000041425 | 30242 | 0.4187 | 0 | 65.8375 | Eukaryota | 79470.0 | 0 | Chordata | 79470.0 | 0 | Carnivora | 79470.0 | 0 | Canidae | 79470.0 | 0 | Canis | 79470.0 | 0 | Canis lupus | 79470.0 | 0 |
| scf7180000041132 | 1237217 | 0.4399 | 0 | 59.7463 | Eukaryota | 51572.0 | 0 | Chordata | 51572.0 | 0 | Carnivora | 51572.0 | 0 | Canidae | 51572.0 | 0 | Canis | 51572.0 | 0 | Canis lupus | 51572.0 | 0 |
| scf7180000048086 | 33557 | 0.3853 | 0 | 58.69 | Eukaryota | 11484.0 | 0 | Chordata | 11484.0 | 0 | Primates | 9622.0 | 1 | Hominidae | 9622.0 | 1 | Pan | 6401.0 | 2 | Pan troglodytes | 6401.0 | 2 |
| scf7180000053734 | 4745 | 0.4512 | 0 | 96.245 | Eukaryota | 1997.0 | 0 | Echinodermata | 1502.0 | 1 | Forcipulatida | 1502.0 | 2 | Asteriidae | 1502.0 | 2 | Asterias | 1502.0 | 2 | Asterias rubens | 1502.0 | 2 |
| scf7180000051098 | 6677 | 0.4486 | 0 | 1280.688 | Eukaryota | 150.0 | 0 | Platyhelminthes | 150.0 | 0 | Polystomatidea | 150.0 | 0 | Polystomatidae | 150.0 | 0 | Protopolystoma | 150.0 | 0 | Protopolystoma xenopodis | 150.0 | 0 |
| scf7180000044869 | 28698 | 0.448 | 0 | 121.6128 | Eukaryota | 57676.0 | 0 | Chordata | 57676.0 | 0 | Carnivora | 57676.0 | 0 | Canidae | 57676.0 | 0 | Canis | 57676.0 | 0 | Canis lupus | 57676.0 | 0 |
| scf7180000049878 | 50932 | 0.4723 | 0 | 58.1827 | Eukaryota | 28290.0 | 0 | Chordata | 28290.0 | 0 | Carnivora | 28290.0 | 0 | Canidae | 28290.0 | 0 | Vulpes | 18326.0 | 1 | Vulpes vulpes | 18326.0 | 1 |
| scf7180000053457 | 5282 | 0.5025 | 0 | 330.1497 | Eukaryota | 6607.0 | 0 | Chordata | 6607.0 | 0 | Carnivora | 6607.0 | 0 | Canidae | 6607.0 | 0 | Canis | 5171.0 | 2 | Canis lupus | 5171.0 | 2 |
| scf7180000052178 | 4094 | 0.4455 | 0 | 57.5225 | no-hit | 0.0 | 0 | no-hit | 0.0 | 0 | no-hit | 0.0 | 0 | no-hit | 0.0 | 0 | no-hit | 0.0 | 0 | no-hit | 0.0 | 0 |

| | | | | | | | | | | | | | | | | | | | | | | |
|---|---|---|---|---|---|---|---|---|---|---|---|---|---|---|---|---|---|---|---|---|---|---|
| scf7180000047630 | 16263 | 0.44 | 0 | 84.4992 | Eukaryota | 244.0 | 0 | Chordata | 244.0 | 0 | Kurtiformes | 244.0 | 0 | Apogonidae | 244.0 | 0 | Sphaeramia | 244.0 | 0 | Sphaeramia orbicularis | 244.0 | 0 |
| scf7180000052582 | 22103 | 0.5251 | 0 | 565.7619 | Eukaryota | 33151.0 | 0 | Chordata | 33151.0 | 0 | Carnivora | 33151.0 | 0 | Canidae | 33151.0 | 0 | Canis | 33151.0 | 0 | Canis lupus | 33151.0 | 0 |
| scf7180000046792 | 123377 | 0.58 | 0 | 54.5674 | Eukaryota | 16936.0 | 0 | Chordata | 16936.0 | 0 | Carnivora | 16936.0 | 0 | Canidae | 16936.0 | 0 | Canis | 14721.0 | 1 | Canis lupus | 14721.0 | 1 |
| scf7180000053166 | 3127 | 0.4535 | 0 | 197.9721 | Eukaryota | 243.0 | 0 | Streptophyta | 243.0 | 0 | Fabales | 243.0 | 0 | Fabaceae | 243.0 | 0 | Cicer | 243.0 | 0 | Cicer arietinum | 243.0 | 0 |
| scf7180000042455 | 718559 | 0.5079 | 0 | 59.5189 | Eukaryota | 67782.0 | 0 | Chordata | 67782.0 | 0 | Carnivora | 67782.0 | 0 | Canidae | 46340.0 | 2 | Canis | 46340.0 | 3 | Canis lupus | 46340.0 | 3 |
| scf7180000042816 | 98335 | 0.4044 | 0 | 83.7707 | Eukaryota | 106725.0 | 0 | Chordata | 106725.0 | 0 | Carnivora | 106725.0 | 0 | Canidae | 106725.0 | 0 | Canis | 106725.0 | 0 | Canis lupus | 106725.0 | 0 |
| scf7180000049178 | 6928 | 0.4463 | 0 | 84.6422 | no-hit | 0.0 | 0 | no-hit | 0.0 | 0 | no-hit | 0.0 | 0 | no-hit | 0.0 | 0 | no-hit | 0.0 | 0 | no-hit | 0.0 | 0 |
| scf7180000049232 | 25653 | 0.4401 | 0 | 65.4932 | no-hit | 0.0 | 0 | no-hit | 0.0 | 0 | no-hit | 0.0 | 0 | no-hit | 0.0 | 0 | no-hit | 0.0 | 0 | no-hit | 0.0 | 0 |
| scf7180000049583 | 35734 | 0.4429 | 0 | 72.2274 | Eukaryota | 1996.0 | 0 | Platyhelminthes | 1611.0 | 1 | Polystomatidea | 1611.0 | 1 | Polystomatidae | 1611.0 | 1 | Protopolystoma | 1611.0 | 2 | Protopolystoma xenopodis | 1611.0 | 2 |
| scf7180000043338 | 316762 | 0.3479 | 0 | 59.2198 | Eukaryota | 11158.0 | 0 | Chordata | 11158.0 | 0 | Carnivora | 11158.0 | 0 | Canidae | 11158.0 | 0 | Canis | 11158.0 | 0 | Canis lupus | 11158.0 | 0 |
| scf7180000041623 | 923084 | 0.4508 | 0 | 59.829 | Eukaryota | 106980.0 | 0 | Chordata | 106980.0 | 0 | Carnivora | 106980.0 | 0 | Canidae | 106980.0 | 0 | Canis | 91918.0 | 1 | Canis lupus | 91918.0 | 1 |
| scf7180000043624 | 370054 | 0.5043 | 0 | 57.5191 | Eukaryota | 64810.0 | 0 | Chordata | 64810.0 | 0 | Carnivora | 64810.0 | 0 | Canidae | 64810.0 | 0 | Canis | 64810.0 | 0 | Canis lupus | 64810.0 | 0 |
| scf7180000047534 | 18572 | 0.6357 | 0 | 545.1099 | Eukaryota | 34319.0 | 0 | Chordata | 34319.0 | 0 | Primates | 30889.0 | 1 | Hominidae | 13731.0 | 3 | Macaca | 13730.0 | 4 | Macaca mulatta | 13730.0 | 4 |
| scf7180000052731 | 9414 | 0.4409 | 0 | 213.7801 | Eukaryota | 42752.0 | 0 | Chordata | 31678.0 | 2 | Gadiformes | 23981.0 | 6 | Gadidae | 23981.0 | 6 | Gadus | 23981.0 | 6 | Gadus morhua | 23981.0 | 6 |
| scf7180000054973 | 24322 | 0.4415 | 0 | 302.4304 | Eukaryota | 39079.0 | 0 | Chordata | 39079.0 | 0 | Carnivora | 39079.0 | 0 | Canidae | 39079.0 | 0 | Canis | 39079.0 | 0 | Canis lupus | 39079.0 | 0 |
| scf7180000042211 | 10703 | 0.3745 | 0 | 49.5127 | Eukaryota | 4614.0 | 0 | Chordata | 4614.0 | 0 | Carnivora | 4614.0 | 0 | Canidae | 4614.0 | 0 | Canis | 4614.0 | 0 | Canis lupus | 4614.0 | 0 |
| scf7180000053678 | 8272 | 0.4543 | 0 | 75.813 | Eukaryota | 593.0 | 0 | Chordata | 593.0 | 0 | Salmoniformes | 593.0 | 0 | Salmonidae | 593.0 | 0 | Coregonus | 593.0 | 0 | Coregonus sp. 'balchen' | 593.0 | 0 |
| scf7180000041794 | 762577 | 0.4408 | 0 | 59.1522 | Eukaryota | 68373.0 | 0 | Chordata | 68373.0 | 0 | Carnivora | 68373.0 | 0 | Canidae | 68373.0 | 0 | Canis | 68373.0 | 0 | Canis lupus | 68373.0 | 0 |
| scf7180000050076 | 12716 | 0.4339 | 0 | 176.0605 | Eukaryota | 553.0 | 0 | Platyhelminthes | 553.0 | 0 | Polystomatidea | 553.0 | 0 | Polystomatidae | 553.0 | 0 | Protopolystoma | 553.0 | 0 | Protopolystoma xenopodis | 553.0 | 0 |
| scf7180000053443 | 3477 | 0.5116 | 0 | 46.7749 | Eukaryota | 6823.0 | 0 | Chordata | 6823.0 | 0 | Carnivora | 6823.0 | 0 | Canidae | 6823.0 | 0 | Canis | 4277.0 | 1 | Canis lupus | 3230.0 | 2 |
| scf7180000040957 | 19409 | 0.46 | 0 | 161.5084 | Eukaryota | 31101.0 | 0 | Chordata | 31101.0 | 0 | Carnivora | 31101.0 | 0 | Canidae | 31101.0 | 0 | Canis | 31101.0 | 0 | Canis lupus | 31101.0 | 0 |
| scf7180000054924 | 33469 | 0.4353 | 0 | 197.3774 | Eukaryota | 25316.0 | 0 | Chordata | 22017.0 | 1 | Gadiformes | 6730.0 | 5 | Gadidae | 6730.0 | 5 | Gadus | 6730.0 | 6 | Gadus morhua | 6730.0 | 6 |
| scf7180000044797 | 267254 | 0.3435 | 0 | 59.6141 | Eukaryota | 34654.0 | 0 | Chordata | 34654.0 | 0 | Primates | 16407.0 | 3 | Hominidae | 13293.0 | 4 | Homo | 11333.0 | 6 | Homo sapiens | 11333.0 | 6 |
| scf7180000051956 | 6967 | 0.4521 | 0 | 1629.0984 | no-hit | 0.0 | 0 | no-hit | 0.0 | 0 | no-hit | 0.0 | 0 | no-hit | 0.0 | 0 | no-hit | 0.0 | 0 | no-hit | 0.0 | 0 |
| scf7180000052972 | 4539 | 0.4085 | 0 | 57.358 | Eukaryota | 5930.0 | 0 | Chordata | 5930.0 | 0 | Carnivora | 5930.0 | 0 | Canidae | 5930.0 | 0 | Canis | 5930.0 | 0 | Canis lupus | 5930.0 | 0 |
| scf7180000045653 | 14012 | 0.4444 | 0 | 160.3984 | no-hit | 0.0 | 0 | no-hit | 0.0 | 0 | no-hit | 0.0 | 0 | no-hit | 0.0 | 0 | no-hit | 0.0 | 0 | no-hit | 0.0 | 0 |
| scf7180000044907 | 36811 | 0.353 | 0 | 54.3184 | Eukaryota | 9436.0 | 0 | Chordata | 9436.0 | 0 | Artiodactyla | 6220.0 | 2 | Bovidae | 6220.0 | 2 | Bos | 6220.0 | 2 | Bos mutus | 5337.0 | 3 |
| scf7180000040797 | 489299 | 0.3474 | 0 | 57.7533 | Eukaryota | 72475.0 | 0 | Chordata | 72475.0 | 0 | Carnivora | 72475.0 | 0 | Canidae | 72475.0 | 0 | Canis | 72475.0 | 0 | Canis lupus | 72475.0 | 0 |
| scf7180000052598 | 9357 | 0.5018 | 0 | 161.6872 | Eukaryota | 8235.0 | 0 | Chordata | 8235.0 | 0 | Carnivora | 8235.0 | 0 | Canidae | 8235.0 | 0 | Canis | 6164.0 | 1 | Canis lupus | 5055.0 | 2 |
| scf7180000045735 | 107285 | 0.3585 | 0 | 58.9502 | Eukaryota | 39592.0 | 0 | Chordata | 39592.0 | 0 | Carnivora | 39592.0 | 0 | Canidae | 39592.0 | 0 | Canis | 23736.0 | 1 | Canis lupus | 23736.0 | 1 |
| scf7180000056302 | 625228 | 0.3867 | 0 | 59.6034 | Eukaryota | 51264.0 | 0 | Chordata | 51264.0 | 0 | Carnivora | 51264.0 | 0 | Canidae | 51264.0 | 0 | Canis | 51264.0 | 0 | Canis lupus | 51264.0 | 0 |
| scf7180000046612 | 124458 | 0.3863 | 0 | 62.8048 | Eukaryota | 92616.0 | 0 | Chordata | 92616.0 | 0 | Carnivora | 92616.0 | 0 | Canidae | 92616.0 | 0 | Canis | 92616.0 | 0 | Canis lupus | 92616.0 | 0 |
| scf7180000043490 | 560980 | 0.484 | 0 | 56.5196 | Eukaryota | 25148.0 | 0 | Chordata | 25148.0 | 0 | Carnivora | 22126.0 | 2 | Canidae | 22126.0 | 2 | Canis | 20707.0 | 3 | Canis lupus | 20707.0 | 3 |
| scf7180000045049 | 78669 | 0.3762 | 0 | 60.656 | Eukaryota | 62685.0 | 0 | Chordata | 62685.0 | 0 | Carnivora | 62685.0 | 0 | Canidae | 62685.0 | 0 | Canis | 62685.0 | 0 | Canis lupus | 62685.0 | 0 |
| scf7180000048371 | 66966 | 0.4393 | 0 | 115.407 | Eukaryota | 6074.0 | 0 | Chordata | 3223.0 | 1 | Polystomatidea | 2851.0 | 3 | Polystomatidae | 2851.0 | 3 | Protopolystoma | 2851.0 | 3 | Protopolystoma xenopodis | 2851.0 | 3 |
| scf7180000043251 | 497678 | 0.3588 | 0 | 59.504 | Eukaryota | 91815.0 | 0 | Chordata | 91815.0 | 0 | Carnivora | 91815.0 | 0 | Canidae | 91815.0 | 0 | Canis | 91815.0 | 0 | Canis lupus | 91815.0 | 0 |
| scf7180000041625 | 16794 | 0.5269 | 0 | 50.2571 | Eukaryota | 3470.0 | 0 | Chordata | 3470.0 | 0 | Carnivora | 3470.0 | 0 | Canidae | 2928.0 | 1 | Canis | 2928.0 | 1 | Canis lupus | 2928.0 | 1 |
| scf7180000055170 | 41300 | 0.4405 | 0 | 261.6357 | Eukaryota | 50581.0 | 0 | Chordata | 29313.0 | 2 | Cyclophyllidea | 12004.0 | 9 | Hymenolepididae | 12004.0 | 9 | Hymenolepis | 12004.0 | 9 | Hymenolepis microstoma | 12004.0 | 9 |
| scf7180000049180 | 14010 | 0.4531 | 0 | 2032.1336 | no-hit | 0.0 | 0 | no-hit | 0.0 | 0 | no-hit | 0.0 | 0 | no-hit | 0.0 | 0 | no-hit | 0.0 | 0 | no-hit | 0.0 | 0 |
| scf7180000056267 | 1444840 | 0.3902 | 0 | 59.2854 | Eukaryota | 163034.0 | 0 | Chordata | 163034.0 | 0 | Carnivora | 163034.0 | 0 | Canidae | 163034.0 | 0 | Canis | 157994.0 | 1 | Canis lupus | 157994.0 | 1 |
| scf7180000053531 | 5924 | 0.446 | 0 | 200.3468 | no-hit | 0.0 | 0 | no-hit | 0.0 | 0 | no-hit | 0.0 | 0 | no-hit | 0.0 | 0 | no-hit | 0.0 | 0 | no-hit | 0.0 | 0 |
| scf7180000041532 | 231845 | 0.3899 | 100 | 60.894 | Eukaryota | 66119.0 | 0 | Chordata | 66119.0 | 0 | Carnivora | 66119.0 | 0 | Canidae | 66119.0 | 0 | Canis | 66119.0 | 0 | Canis lupus | 66119.0 | 0 |
| scf7180000046514 | 22144 | 0.4375 | 0 | 167.6775 | Eukaryota | 10058.0 | 0 | Chordata | 6595.0 | 2 | Tetraodontiformes | 3486.0 | 4 | Tetraodontidae | 3486.0 | 4 | Takifugu | 3486.0 | 4 | Takifugu rubripes | 3486.0 | 4 |
| scf7180000056429 | 374417 | 0.3715 | 0 | 59.8485 | Eukaryota | 60698.0 | 0 | Chordata | 60698.0 | 0 | Carnivora | 60698.0 | 0 | Canidae | 51064.0 | 2 | Canis | 29216.0 | 3 | Canis lupus | 29216.0 | 3 |
| scf7180000047399 | 4415 | 0.4924 | 0 | 48.5367 | Eukaryota | 5322.0 | 0 | Chordata | 5322.0 | 0 | Carnivora | 5322.0 | 0 | Canidae | 5322.0 | 0 | Canis | 4391.0 | 1 | Canis lupus | 4391.0 | 1 |
| scf7180000046590 | 22014 | 0.4459 | 0 | 170.3876 | Eukaryota | 11041.0 | 0 | Chordata | 10409.0 | 1 | Tetraodontiformes | 8052.0 | 3 | Tetraodontidae | 8052.0 | 3 | Takifugu | 8052.0 | 4 | Takifugu rubripes | 8052.0 | 4 |
| scf7180000041003 | 1576513 | 0.4426 | 0 | 60.5849 | Eukaryota | 100095.0 | 0 | Chordata | 100095.0 | 0 | Carnivora | 100095.0 | 0 | Canidae | 100095.0 | 0 | Canis | 100095.0 | 0 | Canis lupus | 100095.0 | 0 |
| scf7180000043171 | 514223 | 0.4107 | 0 | 60.1486 | Eukaryota | 64761.0 | 0 | Chordata | 64761.0 | 0 | Carnivora | 64761.0 | 0 | Canidae | 64761.0 | 0 | Canis | 49658.0 | 1 | Canis lupus | 49658.0 | 1 |
| scf7180000048758 | 6246 | 0.4456 | 0 | 283.9026 | Eukaryota | 2672.0 | 0 | Chordata | 2672.0 | 0 | Kurtiformes | 2168.0 | 1 | Apogonidae | 2168.0 | 1 | Sphaeramia | 2168.0 | 1 | Sphaeramia orbicularis | 2168.0 | 1 |
| scf7180000047542 | 23462 | 0.4439 | 0 | 201.6129 | Eukaryota | 17741.0 | 0 | Chordata | 13723.0 | 1 | Gadiformes | 8781.0 | 4 | Gadidae | 8781.0 | 4 | Gadus | 8781.0 | 5 | Gadus morhua | 8781.0 | 5 |
| scf7180000044194 | 357680 | 0.367 | 0 | 60.2496 | Eukaryota | 89830.0 | 0 | Chordata | 89830.0 | 0 | Carnivora | 89830.0 | 0 | Canidae | 89830.0 | 0 | Canis | 89830.0 | 0 | Canis lupus | 89830.0 | 0 |
| scf7180000042144 | 121786 | 0.4031 | 0 | 59.5611 | Eukaryota | 20385.0 | 0 | Chordata | 20385.0 | 0 | Carnivora | 8138.0 | 2 | Cheirogaleidae | 6867.0 | 4 | Microcebus | 6867.0 | 5 | Microcebus murinus | 6867.0 | 5 |
| scf7180000043838 | 176839 | 0.3781 | 0 | 59.3101 | Eukaryota | 85717.0 | 0 | Chordata | 85717.0 | 0 | Carnivora | 85717.0 | 0 | Canidae | 85717.0 | 0 | Canis | 85717.0 | 0 | Canis lupus | 85717.0 | 0 |
| scf7180000043994 | 330619 | 0.4084 | 0 | 58.5957 | Eukaryota | 65996.0 | 0 | Chordata | 65996.0 | 0 | Carnivora | 65996.0 | 0 | Canidae | 65996.0 | 0 | Canis | 65996.0 | 0 | Canis lupus | 65996.0 | 0 |
| scf7180000054115 | 3809 | 0.4563 | 0 | 644.2257 | Eukaryota | 822.0 | 0 | Echinodermata | 433.0 | 1 | Forcipulatida | 433.0 | 1 | Asteriidae | 433.0 | 1 | Asterias | 433.0 | 1 | Asterias rubens | 433.0 | 1 |
| scf7180000049913 | 15646 | 0.4391 | 0 | 230.805 | Eukaryota | 11729.0 | 0 | Chordata | 11729.0 | 0 | Tetraodontiformes | 8422.0 | 2 | Tetraodontidae | 8422.0 | 2 | Takifugu | 8422.0 | 2 | Takifugu rubripes | 8422.0 | 2 |
| scf7180000042539 | 606774 | 0.4001 | 0 | 60.2563 | Eukaryota | 87822.0 | 0 | Chordata | 87822.0 | 0 | Carnivora | 87822.0 | 0 | Canidae | 87822.0 | 0 | Canis | 87822.0 | 0 | Canis lupus | 87822.0 | 0 |
| scf7180000053233 | 4477 | 0.4458 | 0 | 1476.6514 | no-hit | 0.0 | 0 | no-hit | 0.0 | 0 | no-hit | 0.0 | 0 | no-hit | 0.0 | 0 | no-hit | 0.0 | 0 | no-hit | 0.0 | 0 |
| scf7180000046397 | 18543 | 0.5193 | 0 | 1753.079 | Eukaryota | 31739.0 | 0 | Chordata | 31739.0 | 0 | Carnivora | 31739.0 | 0 | Canidae | 31739.0 | 0 | Canis | 31739.0 | 0 | Canis lupus | 31739.0 | 0 |
| scf7180000042803 | 21490 | 0.4212 | 0 | 50.4573 | Eukaryota | 14925.0 | 0 | Chordata | 14925.0 | 0 | Primates | 9112.0 | 3 | Hominidae | 9112.0 | 3 | Pongo | 3690.0 | 5 | Pongo abelii | 3690.0 | 5 |
| scf7180000049889 | 35990 | 0.443 | 0 | 38.0783 | Eukaryota | 12626.0 | 0 | Chordata | 8454.0 | 2 | Synbranchiformes | 2976.0 | 5 | Mastacembelidae | 2976.0 | 5 | Mastacembelus | 2976.0 | 5 | Mastacembelus armatus | 2976.0 | 5 |
| scf7180000040522 | 2589086 | 0.4478 | 0 | 59.5864 | Eukaryota | 68104.0 | 0 | Chordata | 68104.0 | 0 | Carnivora | 68104.0 | 0 | Canidae | 68104.0 | 0 | Canis | 48156.0 | 1 | Canis lupus | 48156.0 | 1 |
| scf7180000040816 | 1549286 | 0.4304 | 0 | 60.4972 | Eukaryota | 65926.0 | 0 | Chordata | 65926.0 | 0 | Carnivora | 65926.0 | 0 | Canidae | 65926.0 | 0 | Canis | 39006.0 | 1 | Canis lupus | 39006.0 | 1 |
| scf7180000051719 | 4602 | 0.4535 | 0 | 1253.847 | Eukaryota | 15769.0 | 0 | Chordata | 13907.0 | 1 | Gadiformes | 7206.0 | 7 | Gadidae | 7206.0 | 7 | Gadus | 7206.0 | 7 | Gadus morhua | 7206.0 | 7 |
| scf7180000047669 | 111570 | 0.452 | 0 | 59.8896 | Eukaryota | 17472.0 | 0 | Chordata | 17472.0 | 0 | Carnivora | 17472.0 | 0 | Canidae | 17472.0 | 0 | Canis | 15741.0 | 1 | Canis lupus | 15741.0 | 1 |
| scf7180000050684 | 9396 | 0.4483 | 0 | 494.7632 | Eukaryota | 2217.0 | 0 | Platyhelminthes | 1015.0 | 3 | Polystomatidea | 1015.0 | 3 | Polystomatidae | 1015.0 | 3 | Protopolystoma | 1015.0 | 3 | Protopolystoma xenopodis | 1015.0 | 3 |
| scf7180000047486 | 98172 | 0.4099 | 0 | 57.1437 | Eukaryota | 6700.0 | 0 | Chordata | 6700.0 | 0 | Carnivora | 6700.0 | 0 | Canidae | 6700.0 | 0 | Canis | 6700.0 | 0 | Canis lupus | 6700.0 | 0 |
| scf7180000052044 | 31165 | 0.4399 | 0 | 227.2827 | Eukaryota | 98931.0 | 0 | Chordata | 55997.0 | 1 | Gadiformes | 55997.0 | 2 | Gadidae | 55997.0 | 2 | Gadus | 55997.0 | 2 | Gadus morhua | 55997.0 | 2 |
| scf7180000049291 | 45044 | 0.6423 | 0 | 53.5121 | Eukaryota | 10737.0 | 0 | Chordata | 10737.0 | 0 | Carnivora | 10737.0 | 0 | Canidae | 10286.0 | 1 | Canis | 6932.0 | 2 | Canis lupus | 6932.0 | 2 |
| scf7180000045539 | 266185 | 0.4399 | 0 | 59.5915 | Eukaryota | 15804.0 | 0 | Chordata | 15804.0 | 0 | Carnivora | 6186.0 | 3 | Canidae | 5039.0 | 4 | Homo | 4228.0 | 5 | Homo sapiens | 4228.0 | 5 |
| scf7180000041850 | 261038 | 0.3932 | 0 | 59.0719 | Eukaryota | 89940.0 | 0 | Chordata | 89940.0 | 0 | Carnivora | 89940.0 | 0 | Canidae | 89940.0 | 0 | Canis | 89940.0 | 0 | Canis lupus | 89940.0 | 0 |
| scf7180000056422 | 456786 | 0.3461 | 0 | 59.4988 | Eukaryota | 37618.0 | 0 | Chordata | 37618.0 | 0 | Carnivora | 37618.0 | 0 | Canidae | 37618.0 | 0 | Canis | 37618.0 | 0 | Canis lupus | 37618.0 | 0 |
| scf7180000052496 | 12693 | 0.5059 | 0 | 735.4495 | Eukaryota | 8835.0 | 0 | Chordata | 8835.0 | 0 | Carnivora | 8835.0 | 0 | Canidae | 8835.0 | 0 | Canis | 7184.0 | 1 | Canis lupus | 6005.0 | 2 |
| scf7180000056166 | 3049908 | 0.4208 | 0 | 58.926 | Eukaryota | 93702.0 | 0 | Chordata | 93702.0 | 0 | Carnivora | 93702.0 | 0 | Canidae | 93702.0 | 0 | Canis | 93702.0 | 0 | Canis lupus | 93702.0 | 0 |
| scf7180000045038 | 163631 | 0.3497 | 0 | 58.9968 | Eukaryota | 69669.0 | 0 | Chordata | 69669.0 | 0 | Carnivora | 69669.0 | 0 | Canidae | 69669.0 | 0 | Canis | 69669.0 | 0 | Canis lupus | 69669.0 | 0 |
| scf7180000047998 | 15657 | 0.5568 | 0 | 366.8264 | Eukaryota | 48746.0 | 0 | Chordata | 48746.0 | 0 | Primates | 24257.0 | 2 | Hominidae | 19315.0 | 3 | Ovis | 14745.0 | 4 | Ovis aries | 14745.0 | 4 |
| scf7180000055263 | 69231 | 0.4372 | 0 | 190.9171 | Eukaryota | 64127.0 | 0 | Chordata | 46164.0 | 2 | Gadiformes | 37068.0 | 5 | Gadidae | 37068.0 | 5 | Gadus | 37068.0 | 5 | Gadus morhua | 37068.0 | 5 |
| scf7180000043876 | 155358 | 0.3747 | 0 | 56.5499 | Eukaryota | 79962.0 | 0 | Chordata | 79962.0 | 0 | Carnivora | 79962.0 | 0 | Canidae | 79962.0 | 0 | Canis | 79962.0 | 0 | Canis lupus | 79962.0 | 0 |
| scf7180000047690 | 15801 | 0.4396 | 0 | 103.6113 | Eukaryota | 15246.0 | 0 | Chordata | 9658.0 | 2 | Gadiformes | 4630.0 | 6 | Gadidae | 4630.0 | 6 | Gadus | 4630.0 | 6 | Gadus morhua | 4630.0 | 6 |
| scf7180000041593 | 923828 | 0.4186 | 0 | 60.0879 | Eukaryota | 31130.0 | 0 | Chordata | 31130.0 | 0 | Primates | 22086.0 | 2 | Hominidae | 14020.0 | 4 | Homo | 14020.0 | 4 | Homo sapiens | 14020.0 | 4 |
| scf7180000053891 | 3977 | 0.5057 | 0 | 51.3304 | Eukaryota | 7200.0 | 0 | Chordata | 7200.0 | 0 | Carnivora | 7200.0 | 0 | Canidae | 7200.0 | 0 | Canis | 4718.0 | 1 | Canis lupus | 3699.0 | 2 |
| scf7180000054287 | 1553481 | 0.4408 | 0 | 63.6646 | Eukaryota | 126347.0 | 0 | Chordata | 126347.0 | 0 | Carnivora | 126347.0 | 0 | Canidae | 126347.0 | 0 | Canis | 126347.0 | 0 | Canis lupus | 126347.0 | 0 |
| scf7180000052807 | 5376 | 0.4622 | 0 | 71.2744 | Eukaryota | 3001.0 | 0 | Chordata | 3001.0 | 0 | Kurtiformes | 3001.0 | 0 | Apogonidae | 3001.0 | 0 | Sphaeramia | 3001.0 | 0 | Sphaeramia orbicularis | 3001.0 | 0 |
| scf7180000040899 | 26576 | 0.3687 | 0 | 53.6181 | Eukaryota | 96173.0 | 0 | Chordata | 96173.0 | 0 | Carnivora | 96173.0 | 0 | Canidae | 96173.0 | 0 | Canis | 96173.0 | 0 | Canis lupus | 96173.0 | 0 |
| scf7180000049693 | 38949 | 0.373 | 0 | 59.0633 | Eukaryota | 6125.0 | 1 | Chordata | 6125.0 | 1 | Carnivora | 4447.0 | 2 | Canidae | 2316.0 | 4 | Canis | 2316.0 | 4 | Canis lupus | 2316.0 | 4 |
| scf7180000041438 | 241090 | 0.3597 | 0 | 59.701 | Eukaryota | 45450.0 | 0 | Chordata | 45450.0 | 0 | Carnivora | 45450.0 | 0 | Canidae | 45450.0 | 0 | Canis | 34504.0 | 1 | Canis lupus | 34504.0 | 1 |
| scf7180000043832 | 372796 | 0.3843 | 0 | 62.2471 | Eukaryota | 91854.0 | 0 | Chordata | 91854.0 | 0 | Carnivora | 91854.0 | 0 | Canidae | 91854.0 | 0 | Canis | 91854.0 | 0 | Canis lupus | 91854.0 | 0 |
| scf7180000044517 | 353208 | 0.3458 | 0 | 59.3419 | Eukaryota | 51897.0 | 0 | Chordata | 51897.0 | 0 | Carnivora | 51897.0 | 0 | Canidae | 51897.0 | 0 | Canis | 51897.0 | 0 | Canis lupus | 51897.0 | 0 |
| scf7180000054344 | 553683 | 0.4509 | 0 | 58.8385 | Eukaryota | 71212.0 | 0 | Chordata | 71212.0 | 0 | Carnivora | 71212.0 | 0 | Canidae | 71212.0 | 0 | Canis | 59384.0 | 1 | Canis lupus | 59384.0 | 1 |
| scf7180000046421 | 28973 | 0.4411 | 0 | 132.4127 | no-hit | 0.0 | 0 | no-hit | 0.0 | 0 | no-hit | 0.0 | 0 | no-hit | 0.0 | 0 | no-hit | 0.0 | 0 | no-hit | 0.0 | 0 |
| scf7180000041542 | 1037511 | 0.3467 | 0 | 60.2695 | Eukaryota | 43222.0 | 0 | Chordata | 43222.0 | 0 | Carnivora | 43222.0 | 0 | Canidae | 43222.0 | 0 | Canis | 36217.0 | 1 | Canis lupus | 36217.0 | 1 |
| scf7180000050337 | 11524 | 0.4417 | 0 | 51.2578 | Eukaryota | 32006.0 | 0 | Chordata | 21777.0 | 2 | Gadiformes | 15209.0 | 5 | Gadidae | 15209.0 | 5 | Gadus | 15209.0 | 5 | Gadus morhua | 15209.0 | 5 |
| scf7180000055601 | 32503 | 0.4397 | 0 | 220.9567 | Eukaryota | 2511.0 | 0 | Streptophyta | 1066.0 | 2 | Fabales | 1066.0 | 4 | Fabaceae | 1066.0 | 4 | Vigna | 1066.0 | 4 | Vigna unguiculata | 1066.0 | 4 |
| scf7180000042170 | 278329 | 0.4677 | 0 | 57.7366 | Eukaryota | 35215.0 | 0 | Chordata | 35215.0 | 0 | Carnivora | 35215.0 | 0 | Canidae | 35215.0 | 0 | Canis | 35215.0 | 0 | Canis lupus | 35215.0 | 0 |
| scf7180000042854 | 98459 | 0.3759 | 0 | 66.4044 | Eukaryota | 12994.0 | 1 | Chordata | 12994.0 | 1 | Artiodactyla | 8750.0 | 3 | Suidae | 6008.0 | 4 | Sus | 6008.0 | 5 | Sus scrofa | 6008.0 | 5 |
| scf7180000050533 | 3708 | 0.4566 | 0 | 423.8367 | no-hit | 0.0 | 0 | no-hit | 0.0 | 0 | no-hit | 0.0 | 0 | no-hit | 0.0 | 0 | no-hit | 0.0 | 0 | no-hit | 0.0 | 0 |
| scf7180000052405 | 13633 | 0.4477 | 0 | 1326.1899 | no-hit | 0.0 | 0 | no-hit | 0.0 | 0 | no-hit | 0.0 | 0 | no-hit | 0.0 | 0 | no-hit | 0.0 | 0 | no-hit | 0.0 | 0 |
| scf7180000046673 | 42267 | 0.4401 | 0 | 75.7715 | Eukaryota | 4185.0 | 0 | Chordata | 2769.0 | 1 | Carnivora | 2259.0 | 2 | Canidae | 2259.0 | 2 | Canis | 2259.0 | 2 | Canis lupus | 2259.0 | 2 |
| scf7180000056103 | 19486 | 0.5176 | 0 | 646.1715 | Eukaryota | 16707.0 | 0 | Chordata | 16707.0 | 0 | Carnivora | 16707.0 | 0 | Canidae | 16707.0 | 0 | Canis | 16707.0 | 0 | Canis lupus | 16707.0 | 0 |
| scf7180000047075 | 91156 | 0.4333 | 0 | 73.1631 | Eukaryota | 60610.0 | 0 | Chordata | 60610.0 | 0 | Carnivora | 60610.0 | 0 | Canidae | 60610.0 | 0 | Canis | 60610.0 | 0 | Canis lupus | 60610.0 | 0 |
| scf7180000047662 | 63815 | 0.441 | 0 | 75.8349 | Eukaryota | 50139.0 | 0 | Chordata | 28253.0 | 2 | Cyclophyllidea | 12177.0 | 8 | Hymenolepididae | 12177.0 | 8 | Hymenolepis | 12177.0 | 8 | Hymenolepis microstoma | 12177.0 | 8 |
| scf7180000048729 | 8717 | 0.4409 | 0 | 339.5043 | Eukaryota | 366.0 | 0 | Platyhelminthes | 195.0 | 1 | Polystomatidea | 195.0 | 1 | Polystomatidae | 195.0 | 1 | Protopolystoma | 195.0 | 1 | Protopolystoma xenopodis | 195.0 | 1 |
| scf7180000042805 | 112331 | 0.4246 | 0 | 60.6768 | Eukaryota | 95000.0 | 0 | Chordata | 95000.0 | 0 | Carnivora | 95000.0 | 0 | Canidae | 95000.0 | 0 | Canis | 95000.0 | 0 | Canis lupus | 95000.0 | 0 |
| scf7180000051097 | 6495 | 0.4406 | 0 | 187.0405 | Eukaryota | 1347.0 | 0 | Chordata | 945.0 | 1 | Salmoniformes | 586.0 | 2 | Salmonidae | 586.0 | 2 | Protopolystoma | 402.0 | 3 | Protopolystoma xenopodis | 402.0 | 3 |
| scf7180000051438 | 8306 | 0.4467 | 0 | 416.1264 | no-hit | 0.0 | 0 | no-hit | 0.0 | 0 | no-hit | 0.0 | 0 | no-hit | 0.0 | 0 | no-hit | 0.0 | 0 | no-hit | 0.0 | 0 |
| scf7180000047509 | 55641 | 0.3732 | 0 | 59.1209 | Eukaryota | 86861.0 | 0 | Chordata | 86861.0 | 0 | Carnivora | 86861.0 | 0 | Canidae | 86861.0 | 0 | Canis | 86861.0 | 0 | Canis lupus | 86861.0 | 0 |
| scf7180000045527 | 152168 | 0.3518 | 0 | 58.9145 | Eukaryota | 27034.0 | 0 | Chordata | 27034.0 | 0 | Carnivora | 22486.0 | 1 | Canidae | 12908.0 | 3 | Canis | 12908.0 | 3 | Canis lupus | 12908.0 | 3 |
| scf7180000053074 | 2788 | 0.4523 | 0 | 753.2602 | no-hit | 0.0 | 0 | no-hit | 0.0 | 0 | no-hit | 0.0 | 0 | no-hit | 0.0 | 0 | no-hit | 0.0 | 0 | no-hit | 0.0 | 0 |
| scf7180000056300 | 743166 | 0.3807 | 0 | 60.6552 | Eukaryota | 88578.0 | 0 | Chordata | 88578.0 | 0 | Carnivora | 88578.0 | 0 | Canidae | 88578.0 | 0 | Canis | 79697.0 | 1 | Canis lupus | 79697.0 | 1 |
| scf7180000047092 | 29778 | 0.4495 | 0 | 102.9713 | Eukaryota | 8164.0 | 0 | Chordata | 5726.0 | 1 | Carnivora | 5726.0 | 1 | Canidae | 5726.0 | 1 | Canis | 5726.0 | 1 | Canis lupus | 5726.0 | 1 |
| scf7180000040596 | 557072 | 0.3614 | 0 | 60.6288 | Eukaryota | 90711.0 | 0 | Chordata | 90711.0 | 0 | Carnivora | 90711.0 | 0 | Canidae | 90711.0 | 0 | Canis | 90711.0 | 0 | Canis lupus | 90711.0 | 0 |
| scf7180000045990 | 12003 | 0.4424 | 0 | 88.4628 | Eukaryota | 54039.0 | 0 | Platyhelminthes | 31955.0 | 1 | Strigeidida | 25138.0 | 5 | Schistosomatidae | 25138.0 | 5 | Schistosoma | 25138.0 | 5 | Schistosoma mansoni | 25138.0 | 5 |
| scf7180000043644 | 144252 | 0.3494 | 0 | 58.3521 | Eukaryota | 11961.0 | 0 | Chordata | 11961.0 | 0 | Primates | 8801.0 | 2 | Hominidae | 7807.0 | 3 | Homo | 7807.0 | 3 | Homo sapiens | 7807.0 | 3 |
| scf7180000044053 | 376650 | 0.3445 | 0 | 60.2087 | Eukaryota | 34083.0 | 0 | Chordata | 34083.0 | 0 | Carnivora | 34083.0 | 0 | Canidae | 34083.0 | 0 | Canis | 34083.0 | 0 | Canis lupus | 34083.0 | 0 |
| scf7180000040917 | 7913 | 0.3817 | 0 | 32.4903 | Eukaryota | 3811.0 | 0 | Chordata | 3811.0 | 0 | Carnivora | 3811.0 | 0 | Canidae | 3811.0 | 0 | Canis | 2806.0 | 1 | Canis lupus | 2806.0 | 1 |
| scf7180000051746 | 4793 | 0.4459 | 0 | 53.9927 | no-hit | 0.0 | 0 | no-hit | 0.0 | 0 | no-hit | 0.0 | 0 | no-hit | 0.0 | 0 | no-hit | 0.0 | 0 | no-hit | 0.0 | 0 |
| scf7180000049963 | 4780 | 0.4349 | 0 | 149.2171 | Eukaryota | 494.0 | 0 | Platyhelminthes | 494.0 | 0 | Polystomatidea | 494.0 | 0 | Polystomatidae | 494.0 | 0 | Protopolystoma | 494.0 | 0 | Protopolystoma xenopodis | 494.0 | 0 |
| scf7180000045037 | 4618 | 0.4465 | 0 | 380.1279 | Eukaryota | 4724.0 | 0 | Platyhelminthes | 2630.0 | 1 | Polystomatidea | 2630.0 | 2 | Polystomatidae | 2630.0 | 2 | Protopolystoma | 2630.0 | 2 | Protopolystoma xenopodis | 2630.0 | 2 |
| scf7180000054808 | 46141 | 0.4452 | 0 | 404.223 | Eukaryota | 28341.0 | 0 | Platyhelminthes | 15337.0 | 2 | Cyclophyllidea | 12803.0 | 5 | Hymenolepididae | 12803.0 | 5 | Hymenolepis | 12803.0 | 5 | Hymenolepis microstoma | 12803.0 | 5 |
| scf7180000047905 | 46784 | 0.4174 | 0 | 55.8075 | Eukaryota | 88643.0 | 0 | Chordata | 88643.0 | 0 | Carnivora | 88643.0 | 0 | Canidae | 88643.0 | 0 | Canis | 88643.0 | 0 | Canis lupus | 88643.0 | 0 |
| scf7180000055015 | 51931 | 0.444 | 0 | 199.2711 | Eukaryota | 11274.0 | 0 | Chordata | 8228.0 | 1 | Chordata-undef | 4874.0 | 5 | Ambassidae | 4874.0 | 5 | Parambassis | 4874.0 | 5 | Parambassis ranga | 4874.0 | 5 |
| scf7180000049651 | 30758 | 0.4183 | 0 | 62.1231 | Eukaryota | 8462.0 | 1 | Chordata | 8462.0 | 1 | Carnivora | 3516.0 | 4 | Canidae | 3516.0 | 4 | Canis | 3516.0 | 5 | Canis lupus | 3516.0 | 5 |
| scf7180000051219 | 1839 | 0.4513 | 0 | 456.5489 | no-hit | 0.0 | 0 | no-hit | 0.0 | 0 | no-hit | 0.0 | 0 | no-hit | 0.0 | 0 | no-hit | 0.0 | 0 | no-hit | 0.0 | 0 |
| scf7180000041630 | 1008767 | 0.3815 | 0 | 59.8945 | Eukaryota | 91331.0 | 0 | Chordata | 91331.0 | 0 | Carnivora | 91331.0 | 0 | Canidae | 91331.0 | 0 | Canis | 91331.0 | 0 | Canis lupus | 91331.0 | 0 |
| scf7180000041805 | 26814 | 0.5284 | 0 | 795.1769 | Eukaryota | 32827.0 | 0 | Chordata | 32827.0 | 0 | Carnivora | 32827.0 | 0 | Canidae | 32827.0 | 0 | Canis | 32827.0 | 0 | Canis lupus | 32827.0 | 0 |
| scf7180000042700 | 60363 | 0.3839 | 0 | 57.6991 | Eukaryota | 92135.0 | 0 | Chordata | 92135.0 | 0 | Carnivora | 92135.0 | 0 | Canidae | 92135.0 | 0 | Canis | 92135.0 | 0 | Canis lupus | 92135.0 | 0 |
| scf7180000044335 | 16750 | 0.5297 | 0 | 1450.4318 | Eukaryota | 33196.0 | 0 | Chordata | 33196.0 | 0 | Carnivora | 33196.0 | 0 | Canidae | 33196.0 | 0 | Canis | 33196.0 | 0 | Canis lupus | 33196.0 | 0 |
| scf7180000055204 | 72111 | 0.4085 | 0 | 103.4976 | Eukaryota | 85891.0 | 0 | Chordata | 85891.0 | 0 | Carnivora | 85891.0 | 0 | Canidae | 85891.0 | 0 | Canis | 85891.0 | 0 | Canis lupus | 85891.0 | 0 |
| scf7180000051377 | 5683 | 0.4397 | 0 | 80.8402 | Eukaryota | 20387.0 | 0 | Chordata | 14380.0 | 1 | Strigeidida | 6007.0 | 5 | Schistosomatidae | 6007.0 | 5 | Schistosoma | 6007.0 | 5 | Schistosoma mansoni | 6007.0 | 6 |
| scf7180000045875 | 190159 | 0.439 | 0 | 148.2324 | Eukaryota | 94046.0 | 0 | Chordata | 94046.0 | 0 | Carnivora | 94046.0 | 0 | Canidae | 94046.0 | 0 | Canis | 94046.0 | 0 | Canis lupus | 94046.0 | 0 |
| scf7180000046601 | 110473 | 0.4125 | 0 | 59.3124 | Eukaryota | 94435.0 | 0 | Chordata | 94435.0 | 0 | Carnivora | 94435.0 | 0 | Canidae | 53451.0 | 2 | Canis | 42492.0 | 3 | Canis lupus | 42492.0 | 3 |
| scf7180000055197 | 19647 | 0.4399 | 0 | 158.5983 | Eukaryota | 40658.0 | 0 | Chordata | 30362.0 | 2 | Gadiformes | 22521.0 | 6 | Gadidae | 22521.0 | 6 | Gadus | 22521.0 | 6 | Gadus morhua | 22521.0 | 6 |
| scf7180000053290 | 8937 | 0.5182 | 0 | 366.8005 | Eukaryota | 13877.0 | 0 | Chordata | 13877.0 | 0 | Carnivora | 13877.0 | 0 | Canidae | 13877.0 | 0 | Canis | 13877.0 | 0 | Canis lupus | 13877.0 | 0 |
| scf7180000051564 | 13979 | 0.6783 | 0 | 54.3463 | Eukaryota | 11963.0 | 0 | Chordata | 11963.0 | 0 | Carnivora | 11963.0 | 0 | Canidae | 6396.0 | 4 | Canis | 4190.0 | 7 | Canis lupus | 4190.0 | 7 |
| scf7180000051235 | 7321 | 0.504 | 0 | 873.4951 | Eukaryota | 8959.0 | 0 | Chordata | 8959.0 | 0 | Carnivora | 8959.0 | 0 | Canidae | 8959.0 | 0 | Canis | 7316.0 | 1 | Canis lupus | 6103.0 | 2 |
| scf7180000048276 | 86372 | 0.3939 | 0 | 63.5617 | Eukaryota | 87898.0 | 0 | Chordata | 87898.0 | 0 | Carnivora | 87898.0 | 0 | Canidae | 87898.0 | 0 | Canis | 87898.0 | 0 | Canis lupus | 87898.0 | 0 |
| scf7180000055800 | 13493 | 0.4387 | 0 | 120.6784 | Eukaryota | 867.0 | 0 | Chordata | 867.0 | 0 | Columbiformes | 867.0 | 0 | Columbidae | 867.0 | 0 | Streptopelia | 867.0 | 0 | Streptopelia turtur | 867.0 | 0 |
| scf7180000046719 | 10404 | 0.4379 | 0 | 64.6293 | no-hit | 0.0 | 0 | no-hit | 0.0 | 0 | no-hit | 0.0 | 0 | no-hit | 0.0 | 0 | no-hit | 0.0 | 0 | no-hit | 0.0 | 0 |
| scf7180000044024 | 361853 | 0.4887 | 0 | 58.3916 | Eukaryota | 47891.0 | 0 | Chordata | 47891.0 | 0 | Carnivora | 47891.0 | 0 | Canidae | 47891.0 | 0 | Canis | 47891.0 | 0 | Canis lupus | 47891.0 | 0 |
| scf7180000051573 | 7901 | 0.524 | 0 | 79.8505 | Eukaryota | 10712.0 | 0 | Chordata | 10712.0 | 0 | Carnivora | 8602.0 | 1 | Canidae | 4808.0 | 3 | Canis | 3609.0 | 4 | Canis lupus | 3609.0 | 4 |
| scf7180000045433 | 224342 | 0.422 | 0 | 58.4552 | Eukaryota | 38080.0 | 0 | Chordata | 38080.0 | 0 | Carnivora | 38080.0 | 0 | Canidae | 34763.0 | 1 | Canis | 23403.0 | 2 | Canis lupus | 23403.0 | 2 |
| scf7180000049944 | 7470 | 0.4435 | 0 | 284.2194 | no-hit | 0.0 | 0 | no-hit | 0.0 | 0 | no-hit | 0.0 | 0 | no-hit | 0.0 | 0 | no-hit | 0.0 | 0 | no-hit | 0.0 | 0 |
| scf7180000049729 | 5461 | 0.4442 | 0 | 397.0457 | no-hit | 0.0 | 0 | no-hit | 0.0 | 0 | no-hit | 0.0 | 0 | no-hit | 0.0 | 0 | no-hit | 0.0 | 0 | no-hit | 0.0 | 0 |
| scf7180000056523 | 200382 | 0.4623 | 0 | 58.977 | Eukaryota | 64102.0 | 0 | Chordata | 64102.0 | 0 | Carnivora | 64102.0 | 0 | Canidae | 64102.0 | 0 | Canis | 64102.0 | 0 | Canis lupus | 64102.0 | 0 |
| scf7180000046691 | 136604 | 0.395 | 0 | 58.6131 | Eukaryota | 48095.0 | 0 | Chordata | 48095.0 | 0 | Carnivora | 48095.0 | 0 | Canidae | 48095.0 | 0 | Canis | 48095.0 | 0 | Canis lupus | 48095.0 | 0 |
| scf7180000044249 | 9289 | 0.3928 | 0 | 84.3348 | Eukaryota | 48809.0 | 0 | Chordata | 48809.0 | 0 | Carnivora | 48809.0 | 0 | Canidae | 48809.0 | 0 | Canis | 48809.0 | 0 | Canis lupus | 48809.0 | 0 |
| scf7180000043735 | 664853 | 0.4054 | 0 | 59.8129 | Eukaryota | 80293.0 | 0 | Chordata | 80293.0 | 0 | Carnivora | 80293.0 | 0 | Canidae | 80293.0 | 0 | Canis | 80293.0 | 0 | Canis lupus | 80293.0 | 0 |
| scf7180000040939 | 1964708 | 0.4056 | 100 | 67.6081 | Eukaryota | 92799.0 | 0 | Chordata | 92799.0 | 0 | Carnivora | 92799.0 | 0 | Canidae | 92799.0 | 0 | Canis | 84725.0 | 1 | Canis lupus | 84725.0 | 1 |
| scf7180000044729 | 313145 | 0.3731 | 0 | 60.1532 | Eukaryota | 72435.0 | 0 | Chordata | 72435.0 | 0 | Carnivora | 72435.0 | 0 | Canidae | 72435.0 | 0 | Canis | 72435.0 | 0 | Canis lupus | 72435.0 | 0 |
| scf7180000043780 | 390168 | 0.3677 | 0 | 60.0068 | Eukaryota | 91569.0 | 0 | Chordata | 91569.0 | 0 | Carnivora | 91569.0 | 0 | Canidae | 91569.0 | 0 | Canis | 91569.0 | 0 | Canis lupus | 91569.0 | 0 |
| scf7180000044066 | 340371 | 0.3662 | 0 | 59.0001 | Eukaryota | 93399.0 | 0 | Chordata | 93399.0 | 0 | Carnivora | 93399.0 | 0 | Canidae | 93399.0 | 0 | Canis | 93399.0 | 0 | Canis lupus | 93399.0 | 0 |
| scf7180000042551 | 293063 | 0.5064 | 0 | 59.1398 | Eukaryota | 55851.0 | 0 | Chordata | 55851.0 | 0 | Carnivora | 55851.0 | 0 | Canidae | 55851.0 | 0 | Canis | 50080.0 | 1 | Canis lupus | 50080.0 | 1 |
| scf7180000052558 | 13212 | 0.4197 | 0 | 25.7717 | Eukaryota | 89634.0 | 0 | Chordata | 89634.0 | 0 | Carnivora | 89634.0 | 0 | Canidae | 89634.0 | 0 | Canis | 89634.0 | 0 | Canis lupus | 89634.0 | 0 |
| scf7180000049137 | 7118 | 0.4441 | 0 | 833.2789 | no-hit | 0.0 | 0 | no-hit | 0.0 | 0 | no-hit | 0.0 | 0 | no-hit | 0.0 | 0 | no-hit | 0.0 | 0 | no-hit | 0.0 | 0 |
| scf7180000049859 | 5096 | 0.4429 | 0 | 74.2969 | Eukaryota | 14538.0 | 0 | Chordata | 14037.0 | 1 | Gadiformes | 8166.0 | 5 | Gadidae | 8166.0 | 5 | Gadus | 8166.0 | 5 | Gadus morhua | 8166.0 | 5 |
| scf7180000041342 | 29650 | 0.3785 | 0 | 100.3397 | Eukaryota | 74225.0 | 0 | Chordata | 74225.0 | 0 | Carnivora | 74225.0 | 0 | Canidae | 74225.0 | 0 | Canis | 74225.0 | 0 | Canis lupus | 74225.0 | 0 |
| scf7180000045389 | 169003 | 0.4605 | 0 | 59.0584 | Eukaryota | 29443.0 | 0 | Chordata | 29443.0 | 0 | Carnivora | 27888.0 | 1 | Canidae | 27888.0 | 1 | Canis | 18871.0 | 2 | Canis lupus | 18871.0 | 2 |
| scf7180000054311 | 20622 | 0.442 | 0 | 404.1379 | Eukaryota | 2775.0 | 1 | Platyhelminthes | 1500.0 | 2 | Polystomatidea | 1500.0 | 3 | Polystomatidae | 1500.0 | 3 | Protopolystoma | 1500.0 | 3 | Protopolystoma xenopodis | 1500.0 | 3 |
| scf7180000043150 | 251777 | 0.4047 | 0 | 59.3605 | Eukaryota | 103019.0 | 0 | Chordata | 103019.0 | 0 | Carnivora | 103019.0 | 0 | Canidae | 103019.0 | 0 | Canis | 76773.0 | 1 | Canis lupus | 76773.0 | 1 |
| scf7180000051167 | 4796 | 0.506 | 0 | 203.8989 | Eukaryota | 8690.0 | 0 | Chordata | 8690.0 | 0 | Carnivora | 8690.0 | 0 | Canidae | 8690.0 | 0 | Canis | 6998.0 | 1 | Canis lupus | 5782.0 | 2 |
| scf7180000051009 | 5913 | 0.4345 | 0 | 99.6776 | Eukaryota | 2214.0 | 0 | Chordata | 1216.0 | 1 | Polystomatidea | 998.0 | 2 | Polystomatidae | 998.0 | 2 | Protopolystoma | 998.0 | 2 | Protopolystoma xenopodis | 998.0 | 2 |
| scf7180000046474 | 5626 | 0.3841 | 0 | 30.9893 | Eukaryota | 15029.0 | 0 | Chordata | 15029.0 | 0 | Carnivora | 15029.0 | 0 | Canidae | 15029.0 | 0 | Canis | 15029.0 | 0 | Canis lupus | 15029.0 | 0 |
| scf7180000046339 | 5325 | 0.4515 | 0 | 28.0498 | Eukaryota | 9110.0 | 0 | Chordata | 9110.0 | 0 | Carnivora | 9110.0 | 0 | Canidae | 9110.0 | 0 | Canis | 8223.0 | 1 | Canis lupus | 8223.0 | 1 |
| scf7180000052695 | 5919 | 0.4393 | 0 | 79.5074 | Eukaryota | 916.0 | 0 | Chordata | 474.0 | 1 | Syngnathiformes | 474.0 | 1 | Syngnathidae | 474.0 | 1 | Syngnathus | 474.0 | 1 | Syngnathus acus | 474.0 | 1 |
| scf7180000043925 | 384616 | 0.5325 | 0 | 57.135 | Eukaryota | 79953.0 | 0 | Chordata | 79953.0 | 0 | Carnivora | 79953.0 | 0 | Canidae | 79953.0 | 0 | Vulpes | 79953.0 | 0 | Vulpes vulpes | 79953.0 | 0 |
| scf7180000052476 | 4473 | 0.4447 | 0 | 362.1411 | Eukaryota | 1894.0 | 0 | Echinodermata | 859.0 | 2 | Forcipulatida | 859.0 | 2 | Asteriidae | 859.0 | 2 | Asterias | 859.0 | 2 | Asterias rubens | 859.0 | 2 |
| scf7180000042525 | 642960 | 0.3942 | 0 | 59.7359 | Eukaryota | 80062.0 | 0 | Chordata | 80062.0 | 0 | Carnivora | 80062.0 | 0 | Canidae | 80062.0 | 0 | Vulpes | 48098.0 | 1 | Vulpes vulpes | 48098.0 | 1 |
| scf7180000040980 | 56107 | 0.4351 | 0 | 89.2724 | Eukaryota | 89523.0 | 0 | Chordata | 89523.0 | 0 | Carnivora | 89523.0 | 0 | Canidae | 89523.0 | 0 | Canis | 89523.0 | 0 | Canis lupus | 89523.0 | 0 |
| scf7180000049490 | 9164 | 0.5059 | 0 | 140.3598 | Eukaryota | 8136.0 | 0 | Chordata | 8136.0 | 0 | Carnivora | 8136.0 | 0 | Canidae | 8136.0 | 0 | Canis | 6046.0 | 1 | Canis lupus | 4942.0 | 2 |
| scf7180000052272 | 4347 | 0.4509 | 0 | 847.7288 | Eukaryota | 550.0 | 0 | Streptophyta | 550.0 | 0 | Fabales | 550.0 | 0 | Fabaceae | 550.0 | 0 | Cicer | 550.0 | 0 | Cicer arietinum | 550.0 | 0 |
| scf7180000048544 | 66860 | 0.4427 | 0 | 144.6981 | Eukaryota | 3979.0 | 0 | Chordata | 3428.0 | 1 | Kurtiformes | 2800.0 | 2 | Apogonidae | 2800.0 | 2 | Sphaeramia | 2800.0 | 2 | Sphaeramia orbicularis | 2800.0 | 2 |
| scf7180000047866 | 74398 | 0.4412 | 0 | 174.4714 | Eukaryota | 3619.0 | 0 | Chordata | 1623.0 | 2 | Kurtiformes | 1393.0 | 3 | Apogonidae | 1393.0 | 3 | Sphaeramia | 1393.0 | 3 | Sphaeramia orbicularis | 1393.0 | 3 |
| scf7180000049528 | 9015 | 0.4547 | 0 | 689.1375 | Eukaryota | 7287.0 | 0 | Chordata | 4484.0 | 4 | Gadiformes | 2017.0 | 8 | Gadidae | 2017.0 | 8 | Gadus | 2017.0 | 8 | Gadus morhua | 2017.0 | 8 |
| scf7180000054345 | 172429 | 0.5009 | 0 | 214.6905 | Eukaryota | 119409.0 | 0 | Chordata | 119409.0 | 0 | Carnivora | 119409.0 | 0 | Canidae | 119409.0 | 0 | Canis | 119409.0 | 0 | Canis lupus | 119409.0 | 0 |
| scf7180000052592 | 8961 | 0.4418 | 0 | 1578.049 | no-hit | 0.0 | 0 | no-hit | 0.0 | 0 | no-hit | 0.0 | 0 | no-hit | 0.0 | 0 | no-hit | 0.0 | 0 | no-hit | 0.0 | 0 |
| scf7180000046234 | 18711 | 0.4384 | 0 | 171.3578 | Eukaryota | 1056.0 | 0 | Platyhelminthes | 893.0 | 1 | Polystomatidea | 893.0 | 1 | Polystomatidae | 893.0 | 1 | Protopolystoma | 893.0 | 1 | Protopolystoma xenopodis | 893.0 | 1 |
| scf7180000045696 | 25848 | 0.4441 | 0 | 305.1783 | Eukaryota | 9534.0 | 0 | Chordata | 9534.0 | 0 | Carnivora | 7497.0 | 2 | Canidae | 7497.0 | 2 | Canis | 4331.0 | 3 | Canis lupus | 4331.0 | 3 |
| scf7180000040588 | 2154104 | 0.4537 | 0 | 59.9681 | Eukaryota | 103293.0 | 0 | Chordata | 103293.0 | 0 | Carnivora | 103293.0 | 0 | Canidae | 103293.0 | 0 | Canis | 63276.0 | 1 | Canis lupus | 63276.0 | 1 |

| | | | | | | | | | | | | | | | | | | | |
|---|---|---|---|---|---|---|---|---|---|---|---|---|---|---|---|---|---|---|---|
| scf7180000047239 | 84140 | 0.3415 | 0 | 62.636 | Eukaryota | 13520.0 | 0 | Chordata | 13520.0 | 0 | Artiodactyla | 5377.0 | 4 | Bovidae | 5377.0 | 5 | Homo | 3454.0 | 6 | Homo sapiens | 3454.0 | 6 |
| scf7180000053091 | 3612 | 0.4449 | 0 | 663.7648 | no-hit | 0.0 | 0 | no-hit | 0.0 | 0 | no-hit | 0.0 | 0 | no-hit | 0.0 | 0 | no-hit | 0.0 | 0 | no-hit | 0.0 | 0 |
| scf7180000056615 | 30237 | 0.4381 | 0 | 78.8262 | Eukaryota | 391.0 | 0 | Platyhelminthes | 198.0 | 1 | Polystomatidea | 198.0 | 1 | Polystomatidae | 198.0 | 1 | Protopolystoma | 198.0 | 1 | Protopolystoma xenopodis | 198.0 | 1 |
| scf7180000052137 | 11410 | 0.3776 | 0 | 58.3609 | Eukaryota | 3452.0 | 0 | Chordata | 3452.0 | 0 | Carnivora | 3452.0 | 0 | Canidae | 3452.0 | 0 | Canis | 3452.0 | 0 | Canis lupus | 3452.0 | 0 |
| scf7180000054430 | 18171 | 0.3943 | 0 | 417.1851 | Eukaryota | 27698.0 | 0 | Chordata | 27698.0 | 0 | Carnivora | 27698.0 | 0 | Canidae | 27698.0 | 0 | Canis | 27698.0 | 0 | Canis lupus | 27698.0 | 0 |
| scf7180000043317 | 477108 | 0.4098 | 0 | 58.5614 | Eukaryota | 52361.0 | 0 | Chordata | 52361.0 | 0 | Carnivora | 52361.0 | 0 | Canidae | 52361.0 | 0 | Canis | 47007.0 | 1 | Canis lupus | 47007.0 | 1 |
| scf7180000049453 | 28326 | 0.5043 | 0 | 2065.4425 | Eukaryota | 9373.0 | 0 | Chordata | 9373.0 | 0 | Carnivora | 9373.0 | 0 | Canidae | 9373.0 | 0 | Canis | 7582.0 | 1 | Canis lupus | 6269.0 | 2 |
| scf7180000048292 | 16467 | 0.4399 | 0 | 97.2637 | Eukaryota | 13414.0 | 0 | Chordata | 13168.0 | 1 | Tetraodontiformes | 9284.0 | 2 | Tetraodontidae | 9284.0 | 2 | Takifugu | 9284.0 | 2 | Takifugu rubripes | 8164.0 | 3 |
| scf7180000048806 | 4376 | 0.4541 | 0 | 171.883 | Eukaryota | 19349.0 | 0 | Chordata | 19349.0 | 0 | Carnivora | 19349.0 | 0 | Canidae | 19349.0 | 0 | Canis | 19349.0 | 0 | Canis lupus | 19349.0 | 0 |
| scf7180000046954 | 103549 | 0.483 | 0 | 58.8581 | Eukaryota | 53226.0 | 0 | Chordata | 53226.0 | 0 | Carnivora | 53226.0 | 0 | Canidae | 53226.0 | 0 | Canis | 41384.0 | 1 | Canis lupus | 41384.0 | 1 |
| scf7180000043559 | 436906 | 0.5376 | 0 | 58.5867 | Eukaryota | 91016.0 | 0 | Chordata | 91016.0 | 0 | Carnivora | 91016.0 | 0 | Canidae | 91016.0 | 0 | Canis | 91016.0 | 0 | Canis lupus | 91016.0 | 0 |
| scf7180000048363 | 14222 | 0.4377 | 0 | 273.2911 | Eukaryota | 3412.0 | 0 | Platyhelminthes | 3042.0 | 1 | Polystomatidea | 3042.0 | 1 | Polystomatidae | 3042.0 | 1 | Protopolystoma | 3042.0 | 1 | Protopolystoma xenopodis | 3042.0 | 1 |
| scf7180000056410 | 391002 | 0.3638 | 0 | 59.3201 | Eukaryota | 54755.0 | 0 | Chordata | 54755.0 | 0 | Carnivora | 54755.0 | 0 | Canidae | 54755.0 | 0 | Canis | 54755.0 | 0 | Canis lupus | 54755.0 | 0 |
| scf7180000043530 | 489079 | 0.3778 | 0 | 60.6048 | Eukaryota | 56794.0 | 0 | Chordata | 56794.0 | 0 | Carnivora | 56794.0 | 0 | Canidae | 56794.0 | 0 | Canis | 56794.0 | 0 | Canis lupus | 56794.0 | 0 |
| scf7180000041224 | 568189 | 0.3669 | 0 | 59.5587 | Eukaryota | 50547.0 | 0 | Chordata | 50547.0 | 0 | Carnivora | 50547.0 | 0 | Canidae | 50547.0 | 0 | Canis | 42857.0 | 1 | Canis lupus | 42857.0 | 1 |
| scf7180000056141 | 12457 | 0.4548 | 0 | 843.3323 | Eukaryota | 330.0 | 0 | Streptophyta | 180.0 | 1 | Fabales | 180.0 | 1 | Fabaceae | 180.0 | 1 | Cicer | 180.0 | 1 | Cicer arietinum | 180.0 | 1 |
| scf7180000045024 | 244704 | 0.409 | 0 | 59.3031 | Eukaryota | 89822.0 | 0 | Chordata | 89822.0 | 0 | Carnivora | 89822.0 | 0 | Canidae | 89822.0 | 0 | Canis | 89822.0 | 0 | Canis lupus | 89822.0 | 0 |
| scf7180000044090 | 177058 | 0.4573 | 0 | 59.2794 | Eukaryota | 17174.0 | 0 | Chordata | 17174.0 | 0 | Carnivora | 17174.0 | 0 | Canidae | 17174.0 | 0 | Canis | 13180.0 | 1 | Canis lupus | 13180.0 | 1 |
| scf7180000041044 | 569842 | 0.4017 | 0 | 68.976 | Eukaryota | 20901.0 | 0 | Chordata | 20901.0 | 0 | Carnivora | 15541.0 | 1 | Phocidae | 9696.0 | 3 | Neomonachus | 7767.0 | 4 | Neomonachus schauinslandi | 7767.0 | 4 |
| scf7180000053922 | 3514 | 0.4513 | 0 | 175.9841 | no-hit | 0.0 | 0 | no-hit | 0.0 | 0 | no-hit | 0.0 | 0 | no-hit | 0.0 | 0 | no-hit | 0.0 | 0 | no-hit | 0.0 | 0 |
| scf7180000047846 | 49690 | 0.4636 | 0 | 58.4666 | Eukaryota | 4720.0 | 0 | Chordata | 4720.0 | 0 | Carnivora | 4027.0 | 1 | Canidae | 4027.0 | 1 | Canis | 3581.0 | 2 | Canis lupus | 3581.0 | 2 |
| scf7180000051574 | 4171 | 0.4486 | 0 | 367.9611 | no-hit | 0.0 | 0 | no-hit | 0.0 | 0 | no-hit | 0.0 | 0 | no-hit | 0.0 | 0 | no-hit | 0.0 | 0 | no-hit | 0.0 | 0 |
| scf7180000043373 | 560261 | 0.3915 | 100 | 60.5741 | Eukaryota | 94281.0 | 0 | Chordata | 94281.0 | 0 | Carnivora | 94281.0 | 0 | Canidae | 94281.0 | 0 | Canis | 82892.0 | 1 | Canis lupus | 82892.0 | 1 |
| scf7180000046214 | 149940 | 0.3466 | 0 | 61.1557 | Eukaryota | 109552.0 | 0 | Chordata | 109552.0 | 0 | Carnivora | 109552.0 | 0 | Canidae | 109552.0 | 0 | Canis | 109552.0 | 0 | Canis lupus | 109552.0 | 0 |
| scf7180000056317 | 653856 | 0.5736 | 0 | 58.4843 | Eukaryota | 112649.0 | 0 | Chordata | 112649.0 | 0 | Carnivora | 112649.0 | 0 | Canidae | 112649.0 | 0 | Canis | 110451.0 | 1 | Canis lupus | 110451.0 | 1 |
| scf7180000046853 | 9361 | 0.4337 | 0 | 172.6957 | Eukaryota | 50793.0 | 0 | Chordata | 50793.0 | 0 | Carnivora | 50793.0 | 0 | Canidae | 50793.0 | 0 | Canis | 50793.0 | 0 | Canis lupus | 50793.0 | 0 |
| scf7180000046916 | 86614 | 0.3448 | 0 | 59.6407 | Eukaryota | 17110.0 | 0 | Chordata | 17110.0 | 0 | Carnivora | 17110.0 | 0 | Canidae | 17110.0 | 0 | Canis | 17110.0 | 0 | Canis lupus | 17110.0 | 0 |
| scf7180000049565 | 38767 | 0.3554 | 0 | 60.7569 | Eukaryota | 12470.0 | 0 | Chordata | 12470.0 | 0 | Carnivora | 12470.0 | 0 | Canidae | 12470.0 | 0 | Canis | 12470.0 | 0 | Canis lupus | 12470.0 | 0 |
| scf7180000050637 | 19222 | 0.5066 | 0 | 159.1063 | Eukaryota | 8770.0 | 0 | Chordata | 8770.0 | 0 | Carnivora | 8770.0 | 0 | Canidae | 8770.0 | 0 | Canis | 7064.0 | 1 | Canis lupus | 5833.0 | 2 |
| scf7180000051841 | 13670 | 0.4442 | 0 | 160.49 | Eukaryota | 3041.0 | 0 | Platyhelminthes | 1608.0 | 2 | Polystomatidea | 1608.0 | 2 | Polystomatidae | 1608.0 | 2 | Protopolystoma | 1608.0 | 2 | Protopolystoma xenopodis | 1608.0 | 2 |
| scf7180000042744 | 2797160 | 0.4647 | 100 | 59.3408 | Eukaryota | 68500.0 | 0 | Chordata | 68500.0 | 0 | Carnivora | 68500.0 | 0 | Canidae | 68500.0 | 0 | Canis | 45877.0 | 1 | Canis lupus | 45877.0 | 1 |
| scf7180000041594 | 2527 | 0.3862 | 0 | 21.9337 | Eukaryota | 23951.0 | 0 | Chordata | 23951.0 | 0 | Carnivora | 23951.0 | 0 | Canidae | 23951.0 | 0 | Canis | 23951.0 | 0 | Canis lupus | 23951.0 | 0 |
| scf7180000050246 | 10616 | 0.4424 | 0 | 119.189 | Eukaryota | 392.0 | 0 | Chordata | 392.0 | 0 | Kurtiformes | 392.0 | 0 | Apogonidae | 392.0 | 0 | Sphaeramia | 392.0 | 0 | Sphaeramia orbicularis | 392.0 | 0 |
| scf7180000046850 | 109224 | 0.4073 | 0 | 60.1944 | Eukaryota | 65236.0 | 0 | Chordata | 65236.0 | 0 | Carnivora | 65236.0 | 0 | Canidae | 42570.0 | 1 | Canis | 33944.0 | 3 | Canis lupus | 33944.0 | 3 |
| scf7180000053161 | 6234 | 0.4442 | 0 | 290.6686 | Eukaryota | 1195.0 | 0 | Platyhelminthes | 888.0 | 1 | Polystomatidea | 888.0 | 1 | Polystomatidae | 888.0 | 1 | Protopolystoma | 888.0 | 1 | Protopolystoma xenopodis | 888.0 | 1 |
| scf7180000043099 | 114193 | 0.4198 | 0 | 56.2306 | Eukaryota | 88995.0 | 0 | Chordata | 88995.0 | 0 | Carnivora | 88995.0 | 0 | Canidae | 88995.0 | 0 | Canis | 88995.0 | 0 | Canis lupus | 88995.0 | 0 |
| scf7180000056250 | 110910 | 0.4606 | 0 | 59.7603 | Eukaryota | 91417.0 | 0 | Chordata | 91417.0 | 0 | Carnivora | 91417.0 | 0 | Canidae | 91417.0 | 0 | Canis | 91417.0 | 0 | Canis lupus | 91417.0 | 0 |
| scf7180000051278 | 22961 | 0.3528 | 0 | 58.8808 | Eukaryota | 14048.0 | 0 | Chordata | 14048.0 | 0 | Carnivora | 14048.0 | 0 | Canidae | 14048.0 | 0 | Canis | 14048.0 | 0 | Canis lupus | 14048.0 | 0 |
| scf7180000046498 | 84595 | 0.3494 | 0 | 60.5796 | Eukaryota | 66151.0 | 0 | Chordata | 66151.0 | 0 | Carnivora | 66151.0 | 0 | Canidae | 66151.0 | 0 | Canis | 66151.0 | 0 | Canis lupus | 66151.0 | 0 |
| scf7180000049283 | 8636 | 0.448 | 0 | 64.7381 | no-hit | 0.0 | 0 | no-hit | 0.0 | 0 | no-hit | 0.0 | 0 | no-hit | 0.0 | 0 | no-hit | 0.0 | 0 | no-hit | 0.0 | 0 |
| scf7180000040545 | 862917 | 0.4062 | 0 | 59.9545 | Eukaryota | 17468.0 | 0 | Chordata | 17468.0 | 0 | Carnivora | 13683.0 | 2 | Canidae | 13683.0 | 2 | Canis | 13683.0 | 2 | Canis lupus | 13683.0 | 2 |
| scf7180000053337 | 7976 | 0.4349 | 0 | 200.5466 | no-hit | 0.0 | 0 | no-hit | 0.0 | 0 | no-hit | 0.0 | 0 | no-hit | 0.0 | 0 | no-hit | 0.0 | 0 | no-hit | 0.0 | 0 |
| scf7180000045245 | 115862 | 0.3834 | 0 | 59.5969 | Eukaryota | 38223.0 | 0 | Chordata | 38223.0 | 0 | Carnivora | 38223.0 | 0 | Canidae | 38223.0 | 0 | Canis | 38223.0 | 0 | Canis lupus | 38223.0 | 0 |
| scf7180000056132 | 10355 | 0.4459 | 0 | 302.8757 | Eukaryota | 2982.0 | 0 | Chordata | 2982.0 | 0 | Carnivora | 2982.0 | 0 | Canidae | 2982.0 | 0 | Canis | 2982.0 | 0 | Canis lupus | 2982.0 | 0 |
| scf7180000043556 | 92805 | 0.454 | 0 | 57.6739 | Eukaryota | 25691.0 | 0 | Chordata | 25691.0 | 0 | Carnivora | 25691.0 | 0 | Canidae | 23102.0 | 2 | Canis | 20459.0 | 3 | Canis lupus | 20459.0 | 3 |
| scf7180000040676 | 9249 | 0.4767 | 0 | 39.9513 | Eukaryota | 5457.0 | 0 | Chordata | 5457.0 | 0 | Carnivora | 5457.0 | 0 | Canidae | 5457.0 | 0 | Canis | 5457.0 | 0 | Canis lupus | 5457.0 | 0 |
| scf7180000045703 | 172508 | 0.3388 | 0 | 59.5834 | Eukaryota | 11022.0 | 0 | Chordata | 11022.0 | 0 | Carnivora | 5345.0 | 3 | Canidae | 5345.0 | 3 | Canis | 5345.0 | 3 | Canis lupus | 5345.0 | 3 |
| scf7180000056011 | 19137 | 0.6029 | 0 | 749.8689 | Eukaryota | 34404.0 | 0 | Chordata | 34404.0 | 0 | Primates | 30963.0 | 1 | Cercopithecidae | 13766.0 | 3 | Macaca | 13766.0 | 4 | Macaca mulatta | 13766.0 | 4 |
| scf7180000052616 | 4347 | 0.4536 | 0 | 905.1659 | Eukaryota | 2361.0 | 0 | Chordata | 2361.0 | 0 | Carnivora | 2098.0 | 1 | Canidae | 2098.0 | 1 | Canis | 2098.0 | 1 | Canis lupus | 2098.0 | 1 |
| scf7180000040907 | 80446 | 0.4135 | 0 | 57.5452 | Eukaryota | 38118.0 | 0 | Chordata | 38118.0 | 0 | Carnivora | 38118.0 | 0 | Canidae | 38118.0 | 0 | Canis | 38118.0 | 0 | Canis lupus | 38118.0 | 0 |
| scf7180000040592 | 2121543 | 0.3907 | 0 | 59.6655 | Eukaryota | 93510.0 | 0 | Chordata | 93510.0 | 0 | Carnivora | 93510.0 | 0 | Canidae | 93510.0 | 0 | Canis | 93510.0 | 0 | Canis lupus | 93510.0 | 0 |
| scf7180000044320 | 207938 | 0.4352 | 0 | 56.9296 | Eukaryota | 57003.0 | 0 | Chordata | 57003.0 | 0 | Carnivora | 57003.0 | 0 | Canidae | 57003.0 | 0 | Canis | 57003.0 | 0 | Canis lupus | 57003.0 | 0 |
| scf7180000041467 | 133845 | 0.4154 | 0 | 59.0903 | Eukaryota | 36867.0 | 0 | Chordata | 36867.0 | 0 | Carnivora | 36867.0 | 0 | Canidae | 36867.0 | 0 | Canis | 36867.0 | 0 | Canis lupus | 36867.0 | 0 |
| scf7180000044254 | 351296 | 0.3621 | 0 | 59.4023 | Eukaryota | 27916.0 | 0 | Chordata | 27916.0 | 0 | Carnivora | 20404.0 | 1 | Canidae | 20404.0 | 1 | Canis | 20404.0 | 1 | Canis lupus | 20404.0 | 1 |
| scf7180000041973 | 28884 | 0.3734 | 0 | 59.1175 | Eukaryota | 21422.0 | 0 | Chordata | 21422.0 | 0 | Carnivora | 21422.0 | 0 | Canidae | 21422.0 | 0 | Canis | 21422.0 | 0 | Canis lupus | 21422.0 | 0 |
| scf7180000046203 | 14032 | 0.4371 | 0 | 270.5566 | Eukaryota | 27223.0 | 0 | Chordata | 26425.0 | 1 | Gadiformes | 16998.0 | 6 | Gadidae | 16998.0 | 6 | Gadus | 16998.0 | 6 | Gadus morhua | 16998.0 | 6 |
| scf7180000044441 | 181563 | 0.3866 | 0 | 59.5298 | Eukaryota | 32049.0 | 0 | Chordata | 32049.0 | 0 | Carnivora | 32049.0 | 0 | Canidae | 27118.0 | 2 | Canis | 20568.0 | 3 | Canis lupus | 20568.0 | 3 |
| scf7180000040760 | 464748 | 0.3826 | 0 | 65.4534 | Eukaryota | 94043.0 | 0 | Chordata | 94043.0 | 0 | Carnivora | 94043.0 | 0 | Canidae | 94043.0 | 0 | Canis | 94043.0 | 0 | Canis lupus | 94043.0 | 0 |
| scf7180000056237 | 1178083 | 0.3597 | 0 | 59.1229 | Eukaryota | 140831.0 | 0 | Chordata | 140831.0 | 0 | Carnivora | 140831.0 | 0 | Canidae | 140831.0 | 0 | Canis | 140831.0 | 0 | Canis lupus | 140831.0 | 0 |
| scf7180000044614 | 59019 | 0.4381 | 0 | 164.2483 | Eukaryota | 43555.0 | 0 | Chordata | 43555.0 | 0 | Carnivora | 43555.0 | 0 | Canidae | 43555.0 | 0 | Canis | 43555.0 | 0 | Canis lupus | 43555.0 | 0 |
| scf7180000055415 | 20859 | 0.4467 | 0 | 154.3918 | Eukaryota | 516.0 | 0 | Chordata | 516.0 | 0 | Salmoniformes | 294.0 | 1 | Salmonidae | 294.0 | 1 | Coregonus | 294.0 | 1 | Coregonus sp. 'balchen' | 294.0 | 1 |
| scf7180000054258 | 20069 | 0.3636 | 0 | 104.7083 | Eukaryota | 3202.0 | 0 | Chordata | 3202.0 | 0 | Carnivora | 3202.0 | 0 | Canidae | 3202.0 | 0 | Canis | 3202.0 | 0 | Canis lupus | 3202.0 | 0 |
| scf7180000044876 | 2800 | 0.5104 | 0 | 526.6626 | Eukaryota | 6208.0 | 0 | Chordata | 6208.0 | 0 | Carnivora | 6208.0 | 0 | Canidae | 6208.0 | 0 | Canis | 4351.0 | 1 | Canis lupus | 3365.0 | 2 |
| scf7180000044322 | 408147 | 0.4141 | 0 | 59.8563 | Eukaryota | 47467.0 | 0 | Chordata | 47467.0 | 0 | Carnivora | 47467.0 | 0 | Canidae | 47467.0 | 0 | Canis | 43223.0 | 1 | Canis lupus | 43223.0 | 1 |
| scf7180000055172 | 21643 | 0.4399 | 0 | 158.628 | no-hit | 0.0 | 0 | no-hit | 0.0 | 0 | no-hit | 0.0 | 0 | no-hit | 0.0 | 0 | no-hit | 0.0 | 0 | no-hit | 0.0 | 0 |
| scf7180000052200 | 2521 | 0.4506 | 0 | 365.5026 | no-hit | 0.0 | 0 | no-hit | 0.0 | 0 | no-hit | 0.0 | 0 | no-hit | 0.0 | 0 | no-hit | 0.0 | 0 | no-hit | 0.0 | 0 |
| scf7180000049588 | 4476 | 0.4462 | 0 | 66.6823 | no-hit | 0.0 | 0 | no-hit | 0.0 | 0 | no-hit | 0.0 | 0 | no-hit | 0.0 | 0 | no-hit | 0.0 | 0 | no-hit | 0.0 | 0 |
| scf7180000045140 | 203152 | 0.4271 | 0 | 61.2547 | Eukaryota | 39271.0 | 0 | Chordata | 39271.0 | 0 | Carnivora | 39271.0 | 0 | Canidae | 37174.0 | 1 | Canis | 37174.0 | 1 | Canis lupus | 37174.0 | 1 |
| scf7180000044429 | 134314 | 0.3492 | 0 | 59.7703 | Eukaryota | 49827.0 | 0 | Chordata | 49827.0 | 0 | Carnivora | 49827.0 | 0 | Canidae | 49827.0 | 0 | Canis | 49827.0 | 0 | Canis lupus | 49827.0 | 0 |
| scf7180000043968 | 409635 | 0.3442 | 0 | 58.6241 | Eukaryota | 19164.0 | 0 | Chordata | 19164.0 | 0 | Carnivora | 19164.0 | 0 | Canidae | 19164.0 | 0 | Canis | 19164.0 | 0 | Canis lupus | 19164.0 | 0 |
| scf7180000042972 | 1467135 | 0.3699 | 0 | 59.4816 | Eukaryota | 141223.0 | 0 | Chordata | 141223.0 | 0 | Carnivora | 141223.0 | 0 | Canidae | 49552.0 | 3 | Canis | 32944.0 | 6 | Canis lupus | 32944.0 | 7 |
| scf7180000055156 | 23096 | 0.4473 | 0 | 238.09 | Eukaryota | 3042.0 | 0 | Chordata | 3042.0 | 0 | Carnivora | 3042.0 | 0 | Canidae | 3042.0 | 0 | Canis | 3042.0 | 0 | Canis lupus | 3042.0 | 0 |
| scf7180000053145 | 3344 | 0.4492 | 0 | 493.2781 | no-hit | 0.0 | 0 | no-hit | 0.0 | 0 | no-hit | 0.0 | 0 | no-hit | 0.0 | 0 | no-hit | 0.0 | 0 | no-hit | 0.0 | 0 |
| scf7180000041038 | 93577 | 0.3489 | 0 | 60.1068 | Eukaryota | 46109.0 | 0 | Chordata | 46109.0 | 0 | Carnivora | 46109.0 | 0 | Canidae | 46109.0 | 0 | Canis | 46109.0 | 0 | Canis lupus | 46109.0 | 0 |
| scf7180000044792 | 267660 | 0.3904 | 0 | 163.7727 | Eukaryota | 92849.0 | 0 | Chordata | 92849.0 | 0 | Carnivora | 92849.0 | 0 | Canidae | 92849.0 | 0 | Canis | 92849.0 | 0 | Canis lupus | 92849.0 | 0 |
| scf7180000044856 | 34251 | 0.3728 | 0 | 56.3574 | Eukaryota | 28683.0 | 0 | Chordata | 28683.0 | 0 | Carnivora | 28683.0 | 0 | Canidae | 28683.0 | 0 | Canis | 28683.0 | 0 | Canis lupus | 28683.0 | 0 |
| scf7180000044402 | 346697 | 0.3871 | 0 | 61.1724 | Eukaryota | 94115.0 | 0 | Chordata | 94115.0 | 0 | Carnivora | 94115.0 | 0 | Canidae | 94115.0 | 0 | Canis | 94115.0 | 0 | Canis lupus | 94115.0 | 0 |
| scf7180000050987 | 11082 | 0.4406 | 0 | 131.2596 | Eukaryota | 335.0 | 0 | Chordata | 335.0 | 0 | Salmoniformes | 335.0 | 0 | Salmonidae | 335.0 | 0 | Coregonus | 335.0 | 0 | Coregonus sp. 'balchen' | 335.0 | 0 |
| scf7180000048977 | 3849 | 0.4354 | 0 | 239.0571 | no-hit | 0.0 | 0 | no-hit | 0.0 | 0 | no-hit | 0.0 | 0 | no-hit | 0.0 | 0 | no-hit | 0.0 | 0 | no-hit | 0.0 | 0 |
| scf7180000055648 | 23814 | 0.4419 | 0 | 122.8643 | Eukaryota | 22045.0 | 0 | Platyhelminthes | 11676.0 | 2 | Cyclophyllidea | 11033.0 | 6 | Hymenolepididae | 11033.0 | 6 | Hymenolepis | 11033.0 | 6 | Hymenolepis microstoma | 11033.0 | 6 |
| scf7180000047275 | 105953 | 0.5671 | 0 | 56.8288 | Eukaryota | 38579.0 | 0 | Chordata | 38579.0 | 0 | Carnivora | 38579.0 | 0 | Canidae | 38579.0 | 0 | Vulpes | 19432.0 | 1 | Vulpes vulpes | 19432.0 | 1 |
| scf7180000050633 | 14899 | 0.4518 | 0 | 1710.5679 | Eukaryota | 2505.0 | 0 | Chordata | 1390.0 | 2 | Columbiformes | 859.0 | 3 | Columbidae | 859.0 | 3 | Streptopelia | 859.0 | 3 | Streptopelia turtur | 859.0 | 3 |
| scf7180000052696 | 15606 | 0.3743 | 0 | 59.7938 | Eukaryota | 4089.0 | 1 | Chordata | 4089.0 | 1 | Carnivora | 1725.0 | 3 | undef | 1508.0 | 4 | undef | 1508.0 | 4 | eukaryotic synthetic construct | 1508.0 | 4 |
| scf7180000043037 | 109624 | 0.3322 | 0 | 58.7439 | Eukaryota | 21742.0 | 0 | Chordata | 21742.0 | 0 | Carnivora | 21742.0 | 0 | Canidae | 21742.0 | 0 | Canis | 21742.0 | 0 | Canis lupus | 21742.0 | 0 |
| scf7180000047639 | 75378 | 0.4368 | 0 | 84.0173 | Eukaryota | 28331.0 | 0 | Platyhelminthes | 16603.0 | 2 | Cyclophyllidea | 11790.0 | 7 | Hymenolepididae | 11790.0 | 7 | Hymenolepis | 11790.0 | 7 | Hymenolepis microstoma | 11790.0 | 7 |
| scf7180000043709 | 22511 | 0.3556 | 0 | 28.032 | Eukaryota | 51961.0 | 0 | Chordata | 51961.0 | 0 | Carnivora | 51961.0 | 0 | Canidae | 51961.0 | 0 | Canis | 51961.0 | 0 | Canis lupus | 51961.0 | 0 |
| scf7180000043175 | 459882 | 0.4006 | 0 | 61.2606 | Eukaryota | 72830.0 | 0 | Chordata | 72830.0 | 0 | Carnivora | 72830.0 | 0 | Canidae | 72830.0 | 0 | Canis | 72830.0 | 0 | Canis lupus | 72830.0 | 0 |
| scf7180000051414 | 11832 | 0.5031 | 0 | 181.1431 | Eukaryota | 8727.0 | 0 | Chordata | 8727.0 | 0 | Carnivora | 8727.0 | 0 | Canidae | 8727.0 | 0 | Canis | 7018.0 | 1 | Canis lupus | 5781.0 | 2 |
| scf7180000040771 | 1723459 | 0.4261 | 0 | 58.5951 | Eukaryota | 92580.0 | 0 | Chordata | 92580.0 | 0 | Carnivora | 92580.0 | 0 | Canidae | 92580.0 | 0 | Canis | 92580.0 | 0 | Canis lupus | 92580.0 | 0 |
| scf7180000047867 | 76118 | 0.3795 | 0 | 59.3078 | Eukaryota | 24818.0 | 0 | Chordata | 24818.0 | 0 | Primates | 12332.0 | 4 | Cercopithecidae | 7220.0 | 5 | Homo | 5112.0 | 7 | Homo sapiens | 5112.0 | 8 |
| scf7180000041836 | 206283 | 0.3544 | 0 | 57.5889 | Eukaryota | 47179.0 | 0 | Chordata | 47179.0 | 0 | Carnivora | 47179.0 | 0 | Canidae | 47179.0 | 0 | Canis | 47179.0 | 0 | Canis lupus | 47179.0 | 0 |
| scf7180000046975 | 44235 | 0.3949 | 0 | 76.2771 | Eukaryota | 88720.0 | 0 | Chordata | 88720.0 | 0 | Carnivora | 88720.0 | 0 | Canidae | 88720.0 | 0 | Canis | 88720.0 | 0 | Canis lupus | 88720.0 | 0 |
| scf7180000055311 | 30191 | 0.4425 | 0 | 212.1064 | Eukaryota | 2799.0 | 0 | Chordata | 2497.0 | 1 | Carnivora | 2497.0 | 1 | Canidae | 2497.0 | 1 | Canis | 2497.0 | 1 | Canis lupus | 2497.0 | 1 |
| scf7180000044159 | 100229 | 0.3907 | 0 | 60.108 | Eukaryota | 45977.0 | 0 | Chordata | 45977.0 | 0 | Carnivora | 45977.0 | 0 | Canidae | 27136.0 | 1 | Suricata | 18841.0 | 2 | Suricata suricatta | 18841.0 | 2 |
| scf7180000052338 | 6003 | 0.4571 | 0 | 94.5853 | Eukaryota | 494.0 | 0 | Platyhelminthes | 250.0 | 1 | Polystomatidea | 250.0 | 1 | Polystomatidae | 250.0 | 1 | Protopolystoma | 250.0 | 1 | Protopolystoma xenopodis | 250.0 | 1 |
| scf7180000051674 | 13923 | 0.4403 | 0 | 200.8872 | Eukaryota | 38963.0 | 0 | Chordata | 38963.0 | 0 | Carnivora | 38963.0 | 0 | Canidae | 38963.0 | 0 | Canis | 38963.0 | 0 | Canis lupus | 38963.0 | 0 |
| scf7180000046175 | 56288 | 0.4395 | 0 | 126.4892 | Eukaryota | 9161.0 | 0 | Chordata | 8654.0 | 1 | Kurtiformes | 6689.0 | 3 | Apogonidae | 6689.0 | 3 | Sphaeramia | 6689.0 | 3 | Sphaeramia orbicularis | 6689.0 | 3 |
| scf7180000045056 | 7618 | 0.4366 | 0 | 41.7073 | Eukaryota | 3581.0 | 0 | Chordata | 3581.0 | 0 | Carnivora | 3581.0 | 0 | Canidae | 3581.0 | 0 | Canis | 3581.0 | 0 | Canis lupus | 3581.0 | 0 |
| scf7180000044732 | 45986 | 0.4452 | 0 | 137.4935 | Eukaryota | 2913.0 | 0 | Chordata | 1536.0 | 1 | Polystomatidea | 1377.0 | 4 | Polystomatidae | 1377.0 | 4 | Protopolystoma | 1377.0 | 4 | Protopolystoma xenopodis | 1377.0 | 4 |
| scf7180000053370 | 2780 | 0.4547 | 0 | 527.4563 | no-hit | 0.0 | 0 | no-hit | 0.0 | 0 | no-hit | 0.0 | 0 | no-hit | 0.0 | 0 | no-hit | 0.0 | 0 | no-hit | 0.0 | 0 |
| scf7180000049250 | 27565 | 0.4457 | 0 | 198.7814 | Eukaryota | 8956.0 | 0 | Chordata | 6119.0 | 1 | Polystomatidea | 2837.0 | 4 | Polystomatidae | 2837.0 | 4 | Protopolystoma | 2837.0 | 4 | Protopolystoma xenopodis | 2837.0 | 4 |
| scf7180000042162 | 75521 | 0.3709 | 0 | 57.9901 | Eukaryota | 90058.0 | 0 | Chordata | 90058.0 | 0 | Carnivora | 90058.0 | 0 | Canidae | 90058.0 | 0 | Canis | 90058.0 | 0 | Canis lupus | 90058.0 | 0 |
| scf7180000046180 | 40055 | 0.4437 | 0 | 220.599 | Eukaryota | 46492.0 | 0 | Chordata | 39863.0 | 2 | Gadiformes | 35839.0 | 4 | Gadidae | 35839.0 | 4 | Gadus | 35839.0 | 4 | Gadus morhua | 35839.0 | 4 |
| scf7180000054977 | 19868 | 0.4475 | 0 | 302.7909 | Eukaryota | 3664.0 | 0 | Chordata | 3664.0 | 0 | Primates | 2126.0 | 2 | Hominidae | 2126.0 | 2 | Homo | 2126.0 | 3 | Homo sapiens | 2126.0 | 3 |
| scf7180000056517 | 118410 | 0.376 | 0 | 57.1915 | Eukaryota | 71000.0 | 0 | Chordata | 71000.0 | 0 | Carnivora | 71000.0 | 0 | Canidae | 71000.0 | 0 | Canis | 71000.0 | 0 | Canis lupus | 71000.0 | 0 |
| scf7180000044497 | 261352 | 0.3711 | 0 | 63.0234 | Eukaryota | 93065.0 | 0 | Chordata | 93065.0 | 0 | Carnivora | 93065.0 | 0 | Canidae | 93065.0 | 0 | Canis | 93065.0 | 0 | Canis lupus | 93065.0 | 0 |
| scf7180000054476 | 55492 | 0.4027 | 0 | 2854.3723 | Eukaryota | 73368.0 | 0 | Chordata | 73368.0 | 0 | Carnivora | 73368.0 | 0 | Canidae | 73368.0 | 0 | Canis | 73368.0 | 0 | Canis lupus | 73368.0 | 0 |
| scf7180000050897 | 6137 | 0.4419 | 0 | 396.8144 | Eukaryota | 3460.0 | 0 | Platyhelminthes | 2436.0 | 2 | Polystomatidea | 2436.0 | 2 | Polystomatidae | 2436.0 | 2 | Protopolystoma | 2436.0 | 2 | Protopolystoma xenopodis | 2436.0 | 2 |
| scf7180000051570 | 11649 | 0.4569 | 0 | 59.6096 | Eukaryota | 4500.0 | 0 | Echinodermata | 2656.0 | 2 | Forcipulatida | 2656.0 | 4 | Asteriidae | 2656.0 | 4 | Asterias | 2656.0 | 4 | Asterias rubens | 2656.0 | 4 |
| scf7180000044400 | 346796 | 0.408 | 0 | 57.8316 | Eukaryota | 47095.0 | 0 | Chordata | 47095.0 | 0 | Carnivora | 47095.0 | 0 | Canidae | 47095.0 | 0 | Canis | 47095.0 | 0 | Canis lupus | 47095.0 | 0 |
| scf7180000046222 | 171674 | 0.3282 | 0 | 60.5845 | Eukaryota | 20106.0 | 0 | Chordata | 20106.0 | 0 | Carnivora | 20106.0 | 0 | Canidae | 20106.0 | 0 | Canis | 20106.0 | 0 | Canis lupus | 20106.0 | 0 |
| scf7180000040695 | 1735264 | 0.3642 | 0 | 59.7795 | Eukaryota | 70899.0 | 0 | Chordata | 70899.0 | 0 | Carnivora | 70899.0 | 0 | Canidae | 70899.0 | 0 | Canis | 70899.0 | 0 | Canis lupus | 70899.0 | 0 |
| scf7180000056254 | 671499 | 0.3704 | 0 | 60.2912 | Eukaryota | 89269.0 | 0 | Chordata | 89269.0 | 0 | Carnivora | 89269.0 | 0 | Canidae | 89269.0 | 0 | Canis | 89269.0 | 0 | Canis lupus | 89269.0 | 0 |
| scf7180000042730 | 20861 | 0.4765 | 0 | 237.4344 | Eukaryota | 53629.0 | 0 | Chordata | 53629.0 | 0 | Carnivora | 53629.0 | 0 | Canidae | 53629.0 | 0 | Canis | 53629.0 | 0 | Canis lupus | 53629.0 | 0 |
| scf7180000052491 | 9446 | 0.4387 | 0 | 45.7572 | Eukaryota | 10449.0 | 0 | Chordata | 6127.0 | 3 | Strigeidida | 2964.0 | 6 | Schistosomatidae | 2964.0 | 6 | Schistosoma | 2964.0 | 6 | Schistosoma mansoni | 2964.0 | 7 |
| scf7180000049151 | 1856 | 0.4623 | 0 | 428.9531 | no-hit | 0.0 | 0 | no-hit | 0.0 | 0 | no-hit | 0.0 | 0 | no-hit | 0.0 | 0 | no-hit | 0.0 | 0 | no-hit | 0.0 | 0 |
| scf7180000043875 | 341335 | 0.4346 | 0 | 58.8609 | Eukaryota | 92026.0 | 0 | Chordata | 92026.0 | 0 | Carnivora | 92026.0 | 0 | Canidae | 92026.0 | 0 | Canis | 92026.0 | 0 | Canis lupus | 92026.0 | 0 |
| scf7180000041541 | 940504 | 0.4711 | 0 | 58.8177 | Eukaryota | 63021.0 | 0 | Chordata | 63021.0 | 0 | Carnivora | 63021.0 | 0 | Canidae | 57516.0 | 1 | Canis | 38280.0 | 2 | Canis lupus | 38280.0 | 2 |
| scf7180000044926 | 249204 | 0.4869 | 0 | 59.3558 | Eukaryota | 49726.0 | 0 | Chordata | 49726.0 | 0 | Carnivora | 49726.0 | 0 | Canidae | 49726.0 | 0 | Vulpes | 49726.0 | 0 | Vulpes vulpes | 49726.0 | 0 |
| scf7180000052470 | 4993 | 0.4428 | 0 | 186.8639 | Eukaryota | 4355.0 | 0 | Platyhelminthes | 2258.0 | 1 | Polystomatidea | 2258.0 | 2 | Polystomatidae | 2258.0 | 2 | Protopolystoma | 2258.0 | 2 | Protopolystoma xenopodis | 2258.0 | 2 |
| scf7180000045958 | 16637 | 0.3642 | 0 | 46.1173 | Eukaryota | 93043.0 | 0 | Chordata | 93043.0 | 0 | Carnivora | 93043.0 | 0 | Canidae | 93043.0 | 0 | Canis | 93043.0 | 0 | Canis lupus | 93043.0 | 0 |
| scf7180000050254 | 30968 | 0.5009 | 0 | 252.3278 | Eukaryota | 9109.0 | 0 | Chordata | 9109.0 | 0 | Carnivora | 9109.0 | 0 | Canidae | 9109.0 | 0 | Canis | 7899.0 | 1 | Canis lupus | 6631.0 | 2 |
| scf7180000052420 | 9560 | 0.4589 | 0 | 57.8005 | no-hit | 0.0 | 0 | no-hit | 0.0 | 0 | no-hit | 0.0 | 0 | no-hit | 0.0 | 0 | no-hit | 0.0 | 0 | no-hit | 0.0 | 0 |
| scf7180000044705 | 32424 | 0.4428 | 0 | 155.0281 | Eukaryota | 4723.0 | 0 | Chordata | 2834.0 | 1 | Polystomatidea | 1889.0 | 4 | Polystomatidae | 1889.0 | 4 | Protopolystoma | 1889.0 | 4 | Protopolystoma xenopodis | 1889.0 | 4 |
| scf7180000046043 | 161656 | 0.3399 | 0 | 59.6523 | Eukaryota | 46067.0 | 0 | Chordata | 46067.0 | 0 | Carnivora | 46067.0 | 0 | Canidae | 46067.0 | 0 | Canis | 46067.0 | 0 | Canis lupus | 46067.0 | 0 |
| scf7180000050325 | 27729 | 0.5052 | 0 | 450.2694 | Eukaryota | 9693.0 | 0 | Chordata | 9693.0 | 0 | Carnivora | 9693.0 | 0 | Canidae | 9693.0 | 0 | Canis | 8456.0 | 1 | Canis lupus | 7156.0 | 2 |
| scf7180000043097 | 642893 | 0.3496 | 0 | 58.7685 | Eukaryota | 83771.0 | 0 | Chordata | 83771.0 | 0 | Carnivora | 83771.0 | 0 | Canidae | 83771.0 | 0 | Canis | 83771.0 | 0 | Canis lupus | 83771.0 | 0 |
| scf7180000042423 | 140361 | 0.3644 | 0 | 57.8803 | Eukaryota | 12149.0 | 0 | Chordata | 12149.0 | 0 | Carnivora | 7084.0 | 1 | Hominidae | 5065.0 | 2 | Homo | 5065.0 | 3 | Homo sapiens | 5065.0 | 3 |
| scf7180000046640 | 41993 | 0.4417 | 0 | 51.9748 | Eukaryota | 27559.0 | 0 | Chordata | 27559.0 | 0 | Carnivora | 27559.0 | 0 | Canidae | 27559.0 | 0 | Canis | 24033.0 | 1 | Canis lupus | 24033.0 | 1 |
| scf7180000040977 | 56584 | 0.4085 | 0 | 60.4373 | Eukaryota | 90887.0 | 0 | Chordata | 90887.0 | 0 | Carnivora | 90887.0 | 0 | Canidae | 90887.0 | 0 | Canis | 90887.0 | 0 | Canis lupus | 90887.0 | 0 |
| scf7180000052387 | 11410 | 0.5738 | 0 | 574.7602 | Eukaryota | 6006.0 | 0 | Chordata | 6006.0 | 0 | Carnivora | 6006.0 | 0 | Canidae | 6006.0 | 0 | Canis | 5317.0 | 1 | Canis lupus | 5317.0 | 1 |
| scf7180000041045 | 99110 | 0.5057 | 0 | 55.0052 | Eukaryota | 26901.0 | 0 | Chordata | 26901.0 | 0 | Carnivora | 26901.0 | 0 | Canidae | 20998.0 | 1 | Canis | 20998.0 | 1 | Canis lupus | 20998.0 | 1 |
| scf7180000047019 | 99777 | 0.3665 | 0 | 60.4586 | Eukaryota | 57474.0 | 0 | Chordata | 57474.0 | 0 | Carnivora | 57474.0 | 0 | Canidae | 44370.0 | 1 | Canis | 39177.0 | 2 | Canis lupus | 39177.0 | 2 |
| scf7180000056193 | 1265682 | 0.4441 | 0 | 89.1618 | Eukaryota | 105965.0 | 0 | Chordata | 105965.0 | 0 | Carnivora | 105965.0 | 0 | Canidae | 105965.0 | 0 | Canis | 105965.0 | 0 | Canis lupus | 105965.0 | 0 |
| scf7180000051382 | 6918 | 0.4441 | 0 | 266.1423 | Eukaryota | 2921.0 | 0 | Chordata | 2921.0 | 0 | Syngnathiformes | 2081.0 | 1 | Syngnathidae | 2081.0 | 1 | Syngnathus | 2081.0 | 1 | Syngnathus acus | 2081.0 | 1 |
| scf7180000040500 | 1634057 | 0.4081 | 0 | 60.1314 | Eukaryota | 87386.0 | 0 | Chordata | 87386.0 | 0 | Carnivora | 87386.0 | 0 | Phocidae | 58914.0 | 1 | Leptonychotes | 58914.0 | 1 | Leptonychotes weddellii | 58914.0 | 1 |
| scf7180000051317 | 18362 | 0.4412 | 0 | 61.71 | Eukaryota | 845.0 | 0 | Platyhelminthes | 845.0 | 0 | Polystomatidea | 845.0 | 0 | Polystomatidae | 845.0 | 0 | Protopolystoma | 845.0 | 0 | Protopolystoma xenopodis | 845.0 | 0 |
| scf7180000043046 | 13692 | 0.4543 | 0 | 1065.3031 | no-hit | 0.0 | 0 | no-hit | 0.0 | 0 | no-hit | 0.0 | 0 | no-hit | 0.0 | 0 | no-hit | 0.0 | 0 | no-hit | 0.0 | 0 |
| scf7180000044655 | 106928 | 0.4194 | 0 | 56.397 | Eukaryota | 10277.0 | 0 | Chordata | 10277.0 | 0 | Carnivora | 10277.0 | 0 | Canidae | 10277.0 | 0 | Canis | 10277.0 | 0 | Canis lupus | 10277.0 | 0 |
| scf7180000048076 | 73865 | 0.4553 | 0 | 180.5374 | Eukaryota | 26267.0 | 0 | Chordata | 26267.0 | 0 | Gadiformes | 16169.0 | 3 | Gadidae | 16169.0 | 3 | Gadus | 16169.0 | 3 | Gadus morhua | 16169.0 | 3 |
| scf7180000052518 | 11916 | 0.6044 | 0 | 335.2196 | Eukaryota | 65111.0 | 0 | Chordata | 65111.0 | 0 | Artiodactyla | 26466.0 | 3 | Bovidae | 26466.0 | 4 | Ovis | 26466.0 | 5 | Ovis aries | 26466.0 | 5 |
| scf7180000043257 | 265749 | 0.4281 | 0 | 87.7954 | Eukaryota | 91778.0 | 0 | Chordata | 91778.0 | 0 | Carnivora | 91778.0 | 0 | Canidae | 91778.0 | 0 | Canis | 91778.0 | 0 | Canis lupus | 91778.0 | 0 |
| scf7180000046593 | 22659 | 0.4476 | 0 | 326.3549 | Eukaryota | 5690.0 | 0 | Chordata | 5690.0 | 0 | Carnivora | 5690.0 | 0 | Canidae | 5690.0 | 0 | Canis | 5690.0 | 0 | Canis lupus | 5690.0 | 0 |
| scf7180000048054 | 9589 | 0.4359 | 0 | 104.4508 | Eukaryota | 3520.0 | 0 | Chordata | 2197.0 | 1 | Columbiformes | 1783.0 | 2 | Columbidae | 1783.0 | 2 | Streptopelia | 1783.0 | 2 | Streptopelia turtur | 1783.0 | 2 |
| scf7180000053129 | 5312 | 0.4443 | 0 | 662.6692 | no-hit | 0.0 | 0 | no-hit | 0.0 | 0 | no-hit | 0.0 | 0 | no-hit | 0.0 | 0 | no-hit | 0.0 | 0 | no-hit | 0.0 | 0 |
| scf7180000053700 | 4279 | 0.445 | 0 | 58.0458 | Eukaryota | 2997.0 | 0 | Chordata | 2997.0 | 0 | Carnivora | 2997.0 | 0 | Canidae | 2997.0 | 0 | Canis | 2997.0 | 0 | Canis lupus | 2997.0 | 0 |
| scf7180000049875 | 2459 | 0.4681 | 0 | 26.6673 | Eukaryota | 2832.0 | 0 | Chordata | 2832.0 | 0 | Carnivora | 2832.0 | 0 | Canidae | 2832.0 | 0 | Canis | 2832.0 | 0 | Canis lupus | 2832.0 | 0 |
| scf7180000049489 | 24528 | 0.4488 | 0 | 214.3552 | Eukaryota | 731.0 | 0 | Platyhelminthes | 731.0 | 0 | Polystomatidea | 731.0 | 0 | Polystomatidae | 731.0 | 0 | Protopolystoma | 731.0 | 0 | Protopolystoma xenopodis | 731.0 | 0 |
| scf7180000045513 | 188401 | 0.4004 | 0 | 59.1261 | Eukaryota | 46028.0 | 0 | Chordata | 46028.0 | 0 | Carnivora | 46028.0 | 0 | Canidae | 46028.0 | 0 | Canis | 32104.0 | 1 | Canis lupus | 32104.0 | 1 |
| scf7180000054836 | 44454 | 0.442 | 0 | 371.048 | Eukaryota | 19411.0 | 0 | Chordata | 17800.0 | 3 | Gadiformes | 10301.0 | 7 | Gadidae | 10301.0 | 7 | Gadus | 10301.0 | 7 | Gadus morhua | 10301.0 | 7 |
| scf7180000044376 | 186653 | 0.3601 | 0 | 58.7625 | Eukaryota | 49062.0 | 0 | Chordata | 49062.0 | 0 | Carnivora | 49062.0 | 0 | Canidae | 49062.0 | 0 | Canis | 49062.0 | 0 | Canis lupus | 49062.0 | 0 |
| scf7180000047152 | 80325 | 0.6674 | 0 | 53.7168 | Eukaryota | 56198.0 | 0 | Chordata | 56198.0 | 0 | Carnivora | 56198.0 | 0 | Canidae | 56198.0 | 0 | Canis | 56198.0 | 0 | Canis lupus | 56198.0 | 0 |
| scf7180000040746 | 4324 | 0.4526 | 0 | 688.4884 | no-hit | 0.0 | 0 | no-hit | 0.0 | 0 | no-hit | 0.0 | 0 | no-hit | 0.0 | 0 | no-hit | 0.0 | 0 | no-hit | 0.0 | 0 |
| scf7180000049775 | 31017 | 0.455 | 0 | 57.4389 | Eukaryota | 85940.0 | 0 | Chordata | 85940.0 | 0 | Carnivora | 85940.0 | 0 | Canidae | 85940.0 | 0 | Canis | 85940.0 | 0 | Canis lupus | 85940.0 | 0 |
| scf7180000045665 | 173350 | 0.484 | 0 | 59.8606 | Eukaryota | 45251.0 | 0 | Chordata | 45251.0 | 0 | Carnivora | 45251.0 | 0 | Canidae | 45251.0 | 0 | Canis | 36139.0 | 1 | Canis lupus | 36139.0 | 1 |
| scf7180000049805 | 2966 | 0.4346 | 0 | 111.9481 | no-hit | 0.0 | 0 | no-hit | 0.0 | 0 | no-hit | 0.0 | 0 | no-hit | 0.0 | 0 | no-hit | 0.0 | 0 | no-hit | 0.0 | 0 |
| scf7180000053639 | 3122 | 0.435 | 0 | 59.0116 | Eukaryota | 654.0 | 0 | Platyhelminthes | 654.0 | 0 | Polystomatidea | 654.0 | 0 | Polystomatidae | 654.0 | 0 | Protopolystoma | 654.0 | 0 | Protopolystoma xenopodis | 654.0 | 0 |
| scf7180000045491 | 157411 | 0.4712 | 0 | 58.4053 | Eukaryota | 57379.0 | 0 | Chordata | 57379.0 | 0 | Carnivora | 57379.0 | 0 | Canidae | 57379.0 | 0 | Canis | 45943.0 | 1 | Canis latrans | 34509.0 | 3 |
| scf7180000044992 | 76878 | 0.3432 | 0 | 54.5666 | Eukaryota | 73680.0 | 0 | Chordata | 73680.0 | 0 | Carnivora | 73680.0 | 0 | Canidae | 73680.0 | 0 | Canis | 73680.0 | 0 | Canis lupus | 73680.0 | 0 |
| scf7180000052252 | 3252 | 0.4499 | 0 | 795.5373 | no-hit | 0.0 | 0 | no-hit | 0.0 | 0 | no-hit | 0.0 | 0 | no-hit | 0.0 | 0 | no-hit | 0.0 | 0 | no-hit | 0.0 | 0 |
| scf7180000047921 | 8146 | 0.443 | 0 | 87.6551 | no-hit | 0.0 | 0 | no-hit | 0.0 | 0 | no-hit | 0.0 | 0 | no-hit | 0.0 | 0 | no-hit | 0.0 | 0 | no-hit | 0.0 | 0 |
| scf7180000044703 | 224369 | 0.39 | 0 | 60.407 | Eukaryota | 40318.0 | 0 | Chordata | 40318.0 | 0 | Carnivora | 40318.0 | 0 | Canidae | 40318.0 | 0 | Canis | 40318.0 | 0 | Canis lupus | 40318.0 | 0 |
| scf7180000049866 | 9488 | 0.5066 | 0 | 193.1888 | Eukaryota | 8029.0 | 0 | Chordata | 8029.0 | 0 | Carnivora | 8029.0 | 0 | Canidae | 8029.0 | 0 | Canis | 6402.0 | 1 | Canis lupus | 5250.0 | 2 |
| scf7180000045322 | 4706 | 0.438 | 0 | 57.0091 | Eukaryota | 1061.0 | 0 | Chordata | 743.0 | 1 | Kurtiformes | 388.0 | 2 | Apogonidae | 388.0 | 2 | Sphaeramia | 388.0 | 2 | Sphaeramia orbicularis | 388.0 | 2 |
| scf7180000054902 | 29304 | 0.4058 | 0 | 63.4339 | Eukaryota | 91237.0 | 0 | Chordata | 91237.0 | 0 | Carnivora | 91237.0 | 0 | Canidae | 91237.0 | 0 | Canis | 91237.0 | 0 | Canis lupus | 91237.0 | 0 |
| scf7180000052179 | 15038 | 0.4386 | 0 | 74.5112 | Eukaryota | 39318.0 | 0 | Chordata | 24063.0 | 2 | Cyclophyllidea | 7136.0 | 8 | Hymenolepididae | 7136.0 | 8 | Hymenolepis | 7136.0 | 8 | Hymenolepis microstoma | 7136.0 | 8 |
| scf7180000045153 | 170815 | 0.382 | 0 | 61.9799 | Eukaryota | 20229.0 | 0 | Chordata | 20229.0 | 0 | Carnivora | 20229.0 | 0 | Canidae | 20229.0 | 0 | Canis | 20229.0 | 0 | Canis lupus | 20229.0 | 0 |
| scf7180000048451 | 15115 | 0.4524 | 0 | 1249.3046 | Eukaryota | 2490.0 | 0 | Echinodermata | 1016.0 | 2 | Forcipulatida | 1016.0 | 2 | Asteriidae | 1016.0 | 2 | Asterias | 1016.0 | 2 | Asterias rubens | 1016.0 | 2 |
| scf7180000056602 | 543251 | 0.349 | 100 | 58.6319 | Eukaryota | 200638.0 | 0 | Chordata | 200638.0 | 0 | Carnivora | 200638.0 | 0 | Canidae | 200638.0 | 0 | Canis | 200638.0 | 0 | Canis lupus | 200638.0 | 0 |
| scf7180000055886 | 9002 | 0.4378 | 0 | 192.692 | Eukaryota | 20032.0 | 0 | Chordata | 16491.0 | 2 | Gadiformes | 5278.0 | 6 | Gadidae | 5278.0 | 6 | Gadus | 5278.0 | 7 | Gadus morhua | 5278.0 | 7 |
| scf7180000044178 | 335907 | 0.3373 | 0 | 59.0122 | Eukaryota | 51224.0 | 0 | Chordata | 51224.0 | 0 | Carnivora | 51224.0 | 0 | Canidae | 51224.0 | 0 | Canis | 51224.0 | 0 | Canis lupus | 51224.0 | 0 |
| scf7180000051366 | 4798 | 0.4441 | 0 | 100.6053 | Eukaryota | 28921.0 | 0 | Chordata | 20571.0 | 2 | Gadiformes | 15449.0 | 5 | Gadidae | 15449.0 | 5 | Gadus | 15449.0 | 5 | Gadus morhua | 15449.0 | 5 |
| scf7180000046900 | 64390 | 0.4159 | 0 | 59.8736 | Eukaryota | 27449.0 | 0 | Chordata | 27449.0 | 0 | Carnivora | 26309.0 | 1 | Canidae | 23093.0 | 3 | Canis | 23093.0 | 4 | Canis lupus | 23093.0 | 4 |
| scf7180000054445 | 24875 | 0.3902 | 0 | 63.6763 | Eukaryota | 11932.0 | 0 | Chordata | 11932.0 | 0 | Carnivora | 11932.0 | 0 | Canidae | 8045.0 | 1 | Canis | 8045.0 | 2 | Canis lupus | 8045.0 | 2 |
| scf7180000044041 | 346962 | 0.4794 | 0 | 59.9351 | Eukaryota | 82498.0 | 0 | Chordata | 82498.0 | 0 | Carnivora | 82498.0 | 0 | Canidae | 82498.0 | 0 | Canis | 58579.0 | 1 | Canis lupus | 58579.0 | 1 |
| scf7180000041936 | 458057 | 0.5314 | 0 | 57.2221 | Eukaryota | 41716.0 | 0 | Chordata | 41716.0 | 0 | Carnivora | 41716.0 | 0 | Canidae | 41716.0 | 0 | Canis | 31571.0 | 1 | Canis lupus | 31571.0 | 1 |
| scf7180000050082 | 20714 | 0.4382 | 0 | 58.9178 | Eukaryota | 17466.0 | 0 | Chordata | 11354.0 | 2 | Salmoniformes | 2832.0 | 7 | Salmonidae | 2832.0 | 7 | Salmo | 2832.0 | 7 | Salmo trutta | 2832.0 | 7 |
| scf7180000053420 | 7272 | 0.453 | 0 | 1228.3778 | no-hit | 0.0 | 0 | no-hit | 0.0 | 0 | no-hit | 0.0 | 0 | no-hit | 0.0 | 0 | no-hit | 0.0 | 0 | no-hit | 0.0 | 0 |

| | | | | | | | | | | | | | | | | | | | | | |
|---|---|---|---|---|---|---|---|---|---|---|---|---|---|---|---|---|---|---|---|---|---|
| scf7180000044030 | 83828 | 0.3977 | 0 | 58.7033 | Eukaryota | 42409.0 | 0 | Chordata | 42409.0 | 0 | Carnivora | 42409.0 | 0 | Canidae | 42409.0 | 0 | Canis | 42409.0 | 0 | Canis lupus | 42409.0 | 0 |
| scf7180000054929 | 20609 | 0.4749 | 0 | 106.3903 | Eukaryota | 9691.0 | 0 | Chordata | 9691.0 | 0 | Carnivora | 9691.0 | 0 | Canidae | 9691.0 | 0 | Canis | 5676.0 | 1 | Canis lupus | 5676.0 | 1 |
| scf7180000041944 | 789314 | 0.5169 | 0 | 59.1526 | Eukaryota | 39402.0 | 0 | Chordata | 39402.0 | 0 | Carnivora | 39402.0 | 0 | Canidae | 39402.0 | 0 | Canis | 39402.0 | 0 | Canis lupus | 39402.0 | 0 |
| scf7180000055837 | 6249 | 0.4378 | 0 | 494.2643 | Eukaryota | 1466.0 | 0 | Platyhelminthes | 1466.0 | 0 | Polystomatidea | 1466.0 | 0 | Polystomatidae | 1466.0 | 0 | Protopolystoma | 1466.0 | 0 | Protopolystoma xenopodis | 1466.0 | 0 |
| scf7180000050944 | 6039 | 0.4502 | 0 | 833.094 | Eukaryota | 483.0 | 0 | Platyhelminthes | 483.0 | 0 | Polystomatidea | 483.0 | 0 | Polystomatidae | 483.0 | 0 | Protopolystoma | 483.0 | 0 | Protopolystoma xenopodis | 483.0 | 0 |
| scf7180000044817 | 263969 | 0.3394 | 0 | 59.9577 | Eukaryota | 38047.0 | 0 | Chordata | 38047.0 | 0 | Carnivora | 38047.0 | 0 | Canidae | 38047.0 | 0 | Canis | 38047.0 | 0 | Canis lupus | 38047.0 | 0 |
| scf7180000041327 | 472502 | 0.4996 | 0 | 57.3256 | Eukaryota | 64627.0 | 0 | Chordata | 64627.0 | 0 | Carnivora | 64627.0 | 0 | Canidae | 64627.0 | 0 | Canis | 64627.0 | 0 | Canis lupus | 64627.0 | 0 |
| scf7180000052020 | 9748 | 0.4494 | 0 | 142.1482 | no-hit | 0.0 | 0 | no-hit | 0.0 | 0 | no-hit | 0.0 | 0 | no-hit | 0.0 | 0 | no-hit | 0.0 | 0 | no-hit | 0.0 | 0 |
| scf7180000043163 | 46656 | 0.3939 | 0 | 62.2045 | Eukaryota | 92939.0 | 0 | Chordata | 92939.0 | 0 | Carnivora | 92939.0 | 0 | Canidae | 92939.0 | 0 | Canis | 92939.0 | 0 | Canis lupus | 92939.0 | 0 |
| scf7180000045332 | 18961 | 0.4461 | 0 | 529.3085 | Eukaryota | 3436.0 | 0 | Chordata | 2069.0 | 1 | Salmoniformes | 2069.0 | 1 | Salmonidae | 2069.0 | 1 | Salmo | 2069.0 | 1 | Salmo trutta | 2069.0 | 1 |
| scf7180000053298 | 3189 | 0.4368 | 0 | 81.0274 | no-hit | 0.0 | 0 | no-hit | 0.0 | 0 | no-hit | 0.0 | 0 | no-hit | 0.0 | 0 | no-hit | 0.0 | 0 | no-hit | 0.0 | 0 |
| scf7180000049570 | 65286 | 0.4755 | 0 | 57.9099 | Eukaryota | 25064.0 | 0 | Chordata | 25064.0 | 0 | Carnivora | 22617.0 | 1 | Canidae | 15042.0 | 2 | Canis | 12742.0 | 3 | Canis lupus | 12742.0 | 3 |
| scf7180000047686 | 92530 | 0.4498 | 0 | 56.4743 | Eukaryota | 29635.0 | 0 | Chordata | 29635.0 | 0 | Carnivora | 29635.0 | 0 | Canidae | 29635.0 | 0 | Canis | 29635.0 | 0 | Canis lupus | 29635.0 | 0 |
| scf7180000042789 | 607860 | 0.411 | 0 | 59.4596 | Eukaryota | 86722.0 | 0 | Chordata | 86722.0 | 0 | Carnivora | 86722.0 | 0 | Canidae | 86722.0 | 0 | Canis | 86722.0 | 0 | Canis lupus | 86722.0 | 0 |
| scf7180000051862 | 15944 | 0.4403 | 0 | 227.1032 | no-hit | 0.0 | 0 | no-hit | 0.0 | 0 | no-hit | 0.0 | 0 | no-hit | 0.0 | 0 | no-hit | 0.0 | 0 | no-hit | 0.0 | 0 |
| scf7180000043774 | 451707 | 0.3693 | 0 | 61.1448 | Eukaryota | 94511.0 | 0 | Chordata | 94511.0 | 0 | Carnivora | 94511.0 | 0 | Canidae | 94511.0 | 0 | Canis | 94511.0 | 0 | Canis lupus | 94511.0 | 0 |
| scf7180000048186 | 66503 | 0.4126 | 0 | 60.2055 | Eukaryota | 12942.0 | 0 | Chordata | 12942.0 | 0 | Artiodactyla | 4969.0 | 4 | Suidae | 3166.0 | 6 | Sus | 3166.0 | 7 | Sus scrofa | 3166.0 | 7 |
| scf7180000045850 | 146971 | 0.3828 | 0 | 61.4563 | Eukaryota | 36691.0 | 0 | Chordata | 36691.0 | 0 | Carnivora | 36691.0 | 0 | Canidae | 36691.0 | 0 | Canis | 22035.0 | 1 | Canis lupus | 22035.0 | 1 |
| scf7180000051842 | 2468 | 0.4514 | 0 | 191.1569 | no-hit | 0.0 | 0 | no-hit | 0.0 | 0 | no-hit | 0.0 | 0 | no-hit | 0.0 | 0 | no-hit | 0.0 | 0 | no-hit | 0.0 | 0 |
| scf7180000041255 | 1168916 | 0.3845 | 0 | 59.7719 | Eukaryota | 87167.0 | 0 | Chordata | 87167.0 | 0 | Carnivora | 87167.0 | 0 | Canidae | 87167.0 | 0 | Canis | 79621.0 | 1 | Canis lupus | 79621.0 | 1 |
| scf7180000043661 | 170800 | 0.5239 | 0 | 57.5671 | Eukaryota | 47054.0 | 0 | Chordata | 47054.0 | 0 | Carnivora | 47054.0 | 0 | Canidae | 47054.0 | 0 | Canis | 47054.0 | 0 | Canis lupus | 47054.0 | 0 |
| scf7180000055875 | 10721 | 0.4357 | 0 | 85.5338 | Eukaryota | 1544.0 | 0 | Platyhelminthes | 1544.0 | 0 | Polystomatidea | 1544.0 | 0 | Polystomatidae | 1544.0 | 0 | Protopolystoma | 1544.0 | 0 | Protopolystoma xenopodis | 1544.0 | 0 |
| scf7180000049373 | 50825 | 0.5039 | 0 | 407.9087 | Eukaryota | 9378.0 | 0 | Chordata | 9378.0 | 0 | Carnivora | 9378.0 | 0 | Canidae | 9378.0 | 0 | Canis | 8132.0 | 1 | Canis lupus | 6835.0 | 2 |
| scf7180000042068 | 12633 | 0.4503 | 0 | 64.4829 | Eukaryota | 19685.0 | 0 | Chordata | 19685.0 | 0 | Carnivora | 19685.0 | 0 | Canidae | 19685.0 | 0 | Canis | 19685.0 | 0 | Canis lupus | 19685.0 | 0 |
| scf7180000056469 | 360524 | 0.3951 | 0 | 59.0843 | Eukaryota | 86586.0 | 0 | Chordata | 86586.0 | 0 | Carnivora | 86586.0 | 0 | Canidae | 86586.0 | 0 | Canis | 86586.0 | 0 | Canis lupus | 86586.0 | 0 |
| scf7180000043669 | 7537 | 0.4434 | 0 | 28.4317 | Eukaryota | 13939.0 | 0 | Chordata | 13939.0 | 0 | Carnivora | 13939.0 | 0 | Canidae | 11538.0 | 2 | Canis | 8670.0 | 3 | Canis lupus | 8670.0 | 3 |
| scf7180000052993 | 7326 | 0.4378 | 0 | 132.9683 | Eukaryota | 3378.0 | 0 | Platyhelminthes | 2289.0 | 3 | Polystomatidea | 2289.0 | 3 | Polystomatidae | 2289.0 | 3 | Protopolystoma | 2289.0 | 3 | Protopolystoma xenopodis | 2289.0 | 3 |
| scf7180000050225 | 2165 | 0.4536 | 0 | 54.0268 | Eukaryota | 2939.0 | 0 | Chordata | 2617.0 | 1 | Carnivora | 2617.0 | 1 | Canidae | 2617.0 | 1 | Canis | 2617.0 | 1 | Canis lupus | 2617.0 | 1 |
| scf7180000046927 | 114728 | 0.3969 | 0 | 60.5795 | Eukaryota | 34738.0 | 0 | Chordata | 34738.0 | 0 | Carnivora | 34738.0 | 0 | Canidae | 34738.0 | 0 | Canis | 29138.0 | 1 | Canis lupus | 29138.0 | 1 |
| scf7180000044324 | 54978 | 0.4125 | 0 | 56.2722 | Eukaryota | 38101.0 | 0 | Chordata | 38101.0 | 0 | Carnivora | 38101.0 | 0 | Canidae | 35894.0 | 1 | Canis | 28600.0 | 2 | Canis lupus | 28600.0 | 2 |
| scf7180000044133 | 423502 | 0.4315 | 0 | 58.1101 | Eukaryota | 24527.0 | 0 | Chordata | 24527.0 | 0 | Carnivora | 24527.0 | 0 | Canidae | 24527.0 | 0 | Canis | 24527.0 | 0 | Canis lupus | 24527.0 | 0 |
| scf7180000041326 | 12435 | 0.4715 | 0 | 114.5294 | Eukaryota | 4798.0 | 0 | Chordata | 4798.0 | 0 | Carnivora | 4798.0 | 0 | Canidae | 4798.0 | 0 | Canis | 4798.0 | 0 | Canis lupus | 4798.0 | 0 |
| scf7180000045708 | 202730 | 0.4276 | 0 | 59.3856 | Eukaryota | 20006.0 | 0 | Chordata | 20006.0 | 0 | Carnivora | 20006.0 | 0 | Canidae | 12370.0 | 2 | Canis | 8270.0 | 3 | Canis lupus | 8270.0 | 3 |
| scf7180000046453 | 7168 | 0.4375 | 0 | 131.0166 | Eukaryota | 664.0 | 0 | Platyhelminthes | 664.0 | 0 | Polystomatidea | 664.0 | 0 | Polystomatidae | 664.0 | 0 | Protopolystoma | 664.0 | 0 | Protopolystoma xenopodis | 664.0 | 0 |
| scf7180000048049 | 104151 | 0.382 | 0 | 60.839 | Eukaryota | 24774.0 | 0 | Chordata | 24774.0 | 0 | Carnivora | 24774.0 | 0 | Otariidae | 14022.0 | 2 | Zalophus | 9364.0 | 4 | Zalophus californianus | 9364.0 | 4 |
| scf7180000043128 | 207207 | 0.4866 | 0 | 58.6852 | Eukaryota | 70607.0 | 0 | Chordata | 70607.0 | 0 | Carnivora | 70607.0 | 0 | Canidae | 63988.0 | 1 | Vulpes | 35790.0 | 2 | Vulpes vulpes | 35790.0 | 2 |
| scf7180000043912 | 423286 | 0.3848 | 0 | 59.8733 | Eukaryota | 37764.0 | 0 | Chordata | 37764.0 | 0 | Carnivora | 37764.0 | 0 | Canidae | 37764.0 | 0 | Canis | 33719.0 | 1 | Canis lupus | 33719.0 | 1 |
| scf7180000046413 | 8836 | 0.4718 | 0 | 39.8455 | Eukaryota | 16302.0 | 0 | Chordata | 16302.0 | 0 | Carnivora | 16302.0 | 0 | Canidae | 6110.0 | 4 | Canis | 4176.0 | 7 | Canis lupus | 4176.0 | 8 |
| scf7180000047948 | 19615 | 0.343 | 0 | 50.7634 | Eukaryota | 3555.0 | 0 | Chordata | 2982.0 | 1 | Carnivora | 1683.0 | 3 | Canidae | 1683.0 | 3 | Canis | 1683.0 | 3 | Canis lupus | 1683.0 | 3 |
| scf7180000044452 | 98738 | 0.4876 | 0 | 56.2284 | Eukaryota | 12368.0 | 0 | Chordata | 12368.0 | 0 | Carnivora | 12368.0 | 0 | Canidae | 12368.0 | 0 | Canis | 12368.0 | 0 | Canis lupus | 12368.0 | 0 |
| scf7180000045971 | 115219 | 0.6614 | 0 | 51.3134 | Eukaryota | 33376.0 | 0 | Chordata | 33376.0 | 0 | Carnivora | 33376.0 | 0 | Canidae | 30191.0 | 2 | Canis | 19102.0 | 3 | Canis lupus | 19102.0 | 3 |
| scf7180000043235 | 467752 | 0.5689 | 0 | 56.4572 | Eukaryota | 56540.0 | 0 | Chordata | 56540.0 | 0 | Carnivora | 56540.0 | 0 | Canidae | 56540.0 | 0 | Canis | 56540.0 | 0 | Canis lupus | 56540.0 | 0 |
| scf7180000052871 | 5841 | 0.5074 | 0 | 159.6681 | Eukaryota | 8532.0 | 0 | Chordata | 8532.0 | 0 | Carnivora | 8532.0 | 0 | Canidae | 8532.0 | 0 | Canis | 6783.0 | 1 | Canis lupus | 5543.0 | 2 |
| scf7180000045393 | 16542 | 0.3244 | 0 | 28.803 | Eukaryota | 2612.0 | 1 | Chordata | 2612.0 | 1 | Carnivora | 1705.0 | 3 | Canidae | 1705.0 | 3 | Canis | 1705.0 | 3 | Canis lupus | 1705.0 | 3 |
| scf7180000052704 | 19368 | 0.673 | 0 | 54.239 | Eukaryota | 2556.0 | 0 | Chordata | 2556.0 | 0 | Carnivora | 2556.0 | 0 | Canidae | 2556.0 | 0 | Vulpes | 2097.0 | 1 | Vulpes vulpes | 2097.0 | 1 |
| scf7180000049145 | 26983 | 0.4462 | 0 | 89.1085 | undef | 339.0 | 0 | undef | 339.0 | 0 | undef | 339.0 | 0 | undef | 339.0 | 0 | undef | 339.0 | 0 | eukaryotic synthetic construct | 339.0 | 0 |
| scf7180000052952 | 7411 | 0.4821 | 0 | 57.8477 | Eukaryota | 15116.0 | 0 | Chordata | 15116.0 | 0 | Carnivora | 15116.0 | 0 | Canidae | 14635.0 | 1 | Canis | 14047.0 | 2 | Canis lupus | 14047.0 | 2 |
| scf7180000042041 | 824100 | 0.432 | 0 | 59.4586 | Eukaryota | 68378.0 | 0 | Chordata | 68378.0 | 0 | Carnivora | 68378.0 | 0 | Canidae | 68378.0 | 0 | Vulpes | 39442.0 | 1 | Vulpes vulpes | 39442.0 | 1 |
| scf7180000044615 | 230972 | 0.5045 | 0 | 58.7465 | Eukaryota | 40936.0 | 0 | Chordata | 40936.0 | 0 | Carnivora | 40936.0 | 0 | Canidae | 40936.0 | 0 | Canis | 34752.0 | 1 | Canis lupus | 34752.0 | 1 |
| scf7180000042195 | 162491 | 0.4877 | 0 | 61.3148 | Eukaryota | 25559.0 | 0 | Chordata | 25559.0 | 0 | Carnivora | 25559.0 | 0 | Canidae | 21826.0 | 2 | Canis | 19724.0 | 3 | Canis lupus | 19724.0 | 3 |
| scf7180000043650 | 420684 | 0.3637 | 0 | 58.2633 | Eukaryota | 90567.0 | 0 | Chordata | 90567.0 | 0 | Carnivora | 90567.0 | 0 | Canidae | 90567.0 | 0 | Canis | 90567.0 | 0 | Canis lupus | 90567.0 | 0 |
| scf7180000040838 | 53987 | 0.4346 | 0 | 171.2478 | Eukaryota | 3989.0 | 0 | Platyhelminthes | 2669.0 | 1 | Polystomatidea | 2669.0 | 2 | Polystomatidae | 2669.0 | 2 | Protopolystoma | 2669.0 | 3 | Protopolystoma xenopodis | 2669.0 | 3 |
| scf7180000051895 | 3703 | 0.4594 | 0 | 612.6683 | no-hit | 0.0 | 0 | no-hit | 0.0 | 0 | no-hit | 0.0 | 0 | no-hit | 0.0 | 0 | no-hit | 0.0 | 0 | no-hit | 0.0 | 0 |
| scf7180000042460 | 667804 | 0.3828 | 0 | 57.8756 | Eukaryota | 93131.0 | 0 | Chordata | 93131.0 | 0 | Carnivora | 93131.0 | 0 | Canidae | 93131.0 | 0 | Canis | 93131.0 | 0 | Canis lupus | 93131.0 | 0 |
| scf7180000043529 | 5807 | 0.7128 | 0 | 35.7817 | Eukaryota | 20195.0 | 0 | Chordata | 20195.0 | 0 | Carnivora | 20195.0 | 0 | Canidae | 11850.0 | 4 | Canis | 9113.0 | 6 | Canis lupus | 9113.0 | 6 |
| scf7180000046527 | 88490 | 0.6232 | 0 | 55.0681 | Eukaryota | 34610.0 | 0 | Chordata | 34610.0 | 0 | Carnivora | 34610.0 | 0 | Canidae | 34610.0 | 0 | Canis | 24152.0 | 1 | Canis lupus | 24152.0 | 1 |
| scf7180000046871 | 3413 | 0.4415 | 0 | 140.2493 | no-hit | 0.0 | 0 | no-hit | 0.0 | 0 | no-hit | 0.0 | 0 | no-hit | 0.0 | 0 | no-hit | 0.0 | 0 | no-hit | 0.0 | 0 |
| scf7180000040868 | 14758 | 0.4144 | 0 | 28.9699 | Eukaryota | 36021.0 | 0 | Chordata | 36021.0 | 0 | Carnivora | 36021.0 | 0 | Canidae | 36021.0 | 0 | Canis | 36021.0 | 0 | Canis lupus | 36021.0 | 0 |
| scf7180000045683 | 37239 | 0.4384 | 0 | 213.5885 | no-hit | 0.0 | 0 | no-hit | 0.0 | 0 | no-hit | 0.0 | 0 | no-hit | 0.0 | 0 | no-hit | 0.0 | 0 | no-hit | 0.0 | 0 |
| scf7180000046887 | 9076 | 0.4398 | 0 | 98.4132 | no-hit | 0.0 | 0 | no-hit | 0.0 | 0 | no-hit | 0.0 | 0 | no-hit | 0.0 | 0 | no-hit | 0.0 | 0 | no-hit | 0.0 | 0 |
| scf7180000056367 | 314312 | 0.4534 | 0 | 59.2182 | Eukaryota | 53896.0 | 0 | Chordata | 53896.0 | 0 | Carnivora | 53896.0 | 0 | Canidae | 22021.0 | 3 | Canis | 11039.0 | 6 | Canis lupus | 11039.0 | 7 |
| scf7180000045595 | 173306 | 0.3519 | 0 | 58.041 | Eukaryota | 89624.0 | 0 | Chordata | 89624.0 | 0 | Carnivora | 89624.0 | 0 | Canidae | 89624.0 | 0 | Canis | 89624.0 | 0 | Canis lupus | 89624.0 | 0 |
| scf7180000048606 | 28109 | 0.4394 | 0 | 89.1308 | Eukaryota | 544.0 | 0 | Platyhelminthes | 544.0 | 0 | Polystomatidea | 544.0 | 0 | Polystomatidae | 544.0 | 0 | Protopolystoma | 544.0 | 0 | Protopolystoma xenopodis | 544.0 | 0 |
| scf7180000053681 | 3699 | 0.4528 | 0 | 391.6284 | Eukaryota | 12893.0 | 0 | Chordata | 12311.0 | 1 | Gadiformes | 6460.0 | 5 | Gadidae | 6460.0 | 5 | Gadus | 6460.0 | 5 | Gadus morhua | 6460.0 | 5 |
| scf7180000052765 | 20574 | 0.4199 | 0 | 3978.2974 | Eukaryota | 182918.0 | 0 | Chordata | 182918.0 | 0 | Carnivora | 182918.0 | 0 | Canidae | 182918.0 | 0 | Canis | 86714.0 | 3 | Canis latrans | 86714.0 | 3 |
| scf7180000050345 | 4026 | 0.4414 | 0 | 50.4148 | no-hit | 0.0 | 0 | no-hit | 0.0 | 0 | no-hit | 0.0 | 0 | no-hit | 0.0 | 0 | no-hit | 0.0 | 0 | no-hit | 0.0 | 0 |
| scf7180000052116 | 4842 | 0.4502 | 0 | 949.6184 | no-hit | 0.0 | 0 | no-hit | 0.0 | 0 | no-hit | 0.0 | 0 | no-hit | 0.0 | 0 | no-hit | 0.0 | 0 | no-hit | 0.0 | 0 |
| scf7180000046336 | 191893 | 0.4042 | 0 | 103.095 | Eukaryota | 62916.0 | 0 | Chordata | 62916.0 | 0 | Carnivora | 62916.0 | 0 | Canidae | 62916.0 | 0 | Canis | 59767.0 | 1 | Canis lupus | 59767.0 | 1 |
| scf7180000041382 | 1094039 | 0.3714 | 0 | 59.8078 | Eukaryota | 87675.0 | 0 | Chordata | 87675.0 | 0 | Carnivora | 87675.0 | 0 | Canidae | 87675.0 | 0 | Canis | 87675.0 | 0 | Canis lupus | 87675.0 | 0 |
| scf7180000043491 | 329238 | 0.4332 | 0 | 58.3945 | Eukaryota | 55397.0 | 0 | Chordata | 55397.0 | 0 | Carnivora | 55397.0 | 0 | Canidae | 55397.0 | 0 | Canis | 55397.0 | 0 | Canis lupus | 55397.0 | 0 |
| scf7180000053257 | 4875 | 0.441 | 0 | 159.3602 | Eukaryota | 3723.0 | 0 | Platyhelminthes | 2607.0 | 1 | Polystomatidea | 2607.0 | 3 | Polystomatidae | 2607.0 | 3 | Protopolystoma | 2607.0 | 3 | Protopolystoma xenopodis | 2607.0 | 3 |
| scf7180000050625 | 13712 | 0.441 | 0 | 77.9307 | Eukaryota | 8807.0 | 0 | Chordata | 8267.0 | 1 | Salmoniformes | 7354.0 | 2 | Salmonidae | 7354.0 | 2 | Salmo | 5556.0 | 3 | Salmo trutta | 5556.0 | 3 |
| scf7180000054162 | 28284 | 0.4393 | 0 | 227.2031 | no-hit | 0.0 | 0 | no-hit | 0.0 | 0 | no-hit | 0.0 | 0 | no-hit | 0.0 | 0 | no-hit | 0.0 | 0 | no-hit | 0.0 | 0 |
| scf7180000043395 | 460160 | 0.4191 | 0 | 60.6484 | Eukaryota | 89663.0 | 0 | Chordata | 89663.0 | 0 | Carnivora | 89663.0 | 0 | Canidae | 89663.0 | 0 | Canis | 89663.0 | 0 | Canis lupus | 89663.0 | 0 |
| scf7180000055269 | 17773 | 0.4514 | 0 | 304.7346 | Eukaryota | 28655.0 | 0 | Chordata | 28655.0 | 0 | Carnivora | 28655.0 | 0 | Canidae | 28655.0 | 0 | Canis | 28655.0 | 0 | Canis lupus | 28655.0 | 0 |
| scf7180000056292 | 231729 | 0.3525 | 0 | 59.2043 | Eukaryota | 107200.0 | 0 | Chordata | 107200.0 | 0 | Carnivora | 107200.0 | 0 | Canidae | 107200.0 | 0 | Canis | 107200.0 | 0 | Canis lupus | 107200.0 | 0 |
| scf7180000042897 | 520730 | 0.4854 | 0 | 59.2001 | Eukaryota | 65395.0 | 0 | Chordata | 65395.0 | 0 | Carnivora | 65395.0 | 0 | Otariidae | 13925.0 | 6 | Canis | 7409.0 | 9 | Canis lupus | 7409.0 | 9 |
| scf7180000041533 | 1021741 | 0.3626 | 0 | 60.1227 | Eukaryota | 92004.0 | 0 | Chordata | 92004.0 | 0 | Carnivora | 92004.0 | 0 | Canidae | 92004.0 | 0 | Canis | 92004.0 | 0 | Canis lupus | 92004.0 | 0 |
| scf7180000050007 | 6657 | 0.4475 | 0 | 861.0609 | no-hit | 0.0 | 0 | no-hit | 0.0 | 0 | no-hit | 0.0 | 0 | no-hit | 0.0 | 0 | no-hit | 0.0 | 0 | no-hit | 0.0 | 0 |
| scf7180000046611 | 41421 | 0.4395 | 0 | 60.2281 | Eukaryota | 6749.0 | 0 | Chordata | 6749.0 | 0 | Carnivora | 6290.0 | 1 | Canidae | 6290.0 | 1 | Canis | 6290.0 | 1 | Canis lupus | 6290.0 | 1 |
| scf7180000049869 | 35531 | 0.4689 | 0 | 54.8581 | Eukaryota | 4428.0 | 0 | Chordata | 4428.0 | 0 | Carnivora | 4428.0 | 0 | Canidae | 4428.0 | 0 | Canis | 4428.0 | 0 | Canis lupus | 4428.0 | 0 |
| scf7180000053277 | 5238 | 0.3963 | 0 | 29.7825 | Eukaryota | 9721.0 | 0 | Chordata | 9721.0 | 0 | Carnivora | 9721.0 | 0 | Canidae | 9721.0 | 0 | Canis | 9721.0 | 0 | Canis lupus | 9721.0 | 0 |
| scf7180000047296 | 9159 | 0.4212 | 0 | 417.0619 | Eukaryota | 3084.0 | 0 | Platyhelminthes | 1666.0 | 2 | Polystomatidea | 1666.0 | 2 | Polystomatidae | 1666.0 | 2 | Protopolystoma | 1666.0 | 2 | Protopolystoma xenopodis | 1666.0 | 2 |
| scf7180000043160 | 43003 | 0.4295 | 0 | 59.9458 | Eukaryota | 107150.0 | 0 | Chordata | 107150.0 | 0 | Carnivora | 107150.0 | 0 | Canidae | 107150.0 | 0 | Canis | 107150.0 | 0 | Canis lupus | 107150.0 | 0 |
| scf7180000049660 | 47350 | 0.3868 | 0 | 58.5394 | Eukaryota | 73348.0 | 0 | Chordata | 73348.0 | 0 | Carnivora | 73348.0 | 0 | Canidae | 73348.0 | 0 | Canis | 73348.0 | 0 | Canis lupus | 73348.0 | 0 |
| scf7180000056248 | 1648975 | 0.421 | 100 | 59.1375 | Eukaryota | 90286.0 | 0 | Chordata | 90286.0 | 0 | Carnivora | 90286.0 | 0 | Canidae | 90286.0 | 0 | Canis | 90286.0 | 0 | Canis lupus | 90286.0 | 0 |
| scf7180000049115 | 6472 | 0.4611 | 0 | 229.3794 | Eukaryota | 46728.0 | 0 | Chordata | 46728.0 | 0 | Carnivora | 46728.0 | 0 | Canidae | 46728.0 | 0 | Canis | 46728.0 | 0 | Canis lupus | 46728.0 | 0 |
| scf7180000044274 | 306882 | 0.3755 | 0 | 60.0461 | Eukaryota | 21211.0 | 0 | Chordata | 21211.0 | 0 | Carnivora | 18885.0 | 1 | Canidae | 18885.0 | 1 | Canis | 18885.0 | 1 | Canis lupus | 18885.0 | 1 |
| scf7180000043053 | 945613 | 0.5306 | 100 | 57.5602 | Eukaryota | 32883.0 | 0 | Chordata | 32883.0 | 0 | Carnivora | 32883.0 | 0 | Canidae | 32883.0 | 0 | Canis | 24906.0 | 1 | Canis lupus | 24906.0 | 1 |
| scf7180000051967 | 26912 | 0.3637 | 0 | 59.6978 | Eukaryota | 5908.0 | 0 | Chordata | 5908.0 | 0 | Carnivora | 4850.0 | 2 | Felidae | 3236.0 | 3 | Felis | 3236.0 | 3 | Felis catus | 3236.0 | 3 |
| scf7180000047385 | 79964 | 0.387 | 0 | 59.6542 | Eukaryota | 15469.0 | 0 | Chordata | 15469.0 | 0 | Primates | 9106.0 | 2 | Hominidae | 9106.0 | 2 | Homo | 9106.0 | 3 | Homo sapiens | 9106.0 | 3 |
| scf7180000055668 | 19018 | 0.4374 | 0 | 163.057 | Eukaryota | 8218.0 | 0 | Chordata | 5676.0 | 2 | Salmoniformes | 3170.0 | 3 | Salmonidae | 3170.0 | 3 | Salmo | 3170.0 | 3 | Salmo trutta | 3170.0 | 3 |
| scf7180000043459 | 52278 | 0.4127 | 0 | 53.7853 | Eukaryota | 83552.0 | 0 | Chordata | 83552.0 | 0 | Carnivora | 83552.0 | 0 | Canidae | 83552.0 | 0 | Canis | 83552.0 | 0 | Canis lupus | 83552.0 | 0 |
| scf7180000043577 | 433964 | 0.3878 | 0 | 60.7213 | Eukaryota | 29966.0 | 0 | Chordata | 29966.0 | 0 | Carnivora | 29966.0 | 0 | Canidae | 29966.0 | 0 | Canis | 20978.0 | 1 | Canis lupus | 20978.0 | 1 |
| scf7180000046492 | 13515 | 0.407 | 0 | 173.6616 | Eukaryota | 30635.0 | 0 | Chordata | 30635.0 | 0 | Carnivora | 13060.0 | 2 | Canidae | 13060.0 | 4 | Canis | 13060.0 | 9 | Canis lupus | 13060.0 | 9 |
| scf7180000054133 | 241774 | 0.3858 | 0 | 81.138 | Eukaryota | 101431.0 | 0 | Chordata | 101431.0 | 0 | Carnivora | 101431.0 | 0 | Canidae | 101431.0 | 0 | Canis | 101431.0 | 0 | Canis lupus | 101431.0 | 0 |
| scf7180000052098 | 9332 | 0.663 | 0 | 53.9704 | Eukaryota | 35274.0 | 0 | Chordata | 35274.0 | 0 | Carnivora | 35274.0 | 0 | Canidae | 35274.0 | 0 | Canis | 19280.0 | 1 | Canis lupus | 19280.0 | 1 |
| scf7180000044052 | 283598 | 0.362 | 0 | 63.0189 | Eukaryota | 70474.0 | 0 | Chordata | 70474.0 | 0 | Carnivora | 70474.0 | 0 | Canidae | 70474.0 | 0 | Canis | 70474.0 | 0 | Canis lupus | 70474.0 | 0 |
| scf7180000043322 | 274284 | 0.3992 | 0 | 97.7691 | Eukaryota | 92541.0 | 0 | Chordata | 92541.0 | 0 | Carnivora | 92541.0 | 0 | Canidae | 92541.0 | 0 | Canis | 92541.0 | 0 | Canis lupus | 92541.0 | 0 |
| scf7180000047894 | 25212 | 0.4389 | 0 | 65.7254 | no-hit | 0.0 | 0 | no-hit | 0.0 | 0 | no-hit | 0.0 | 0 | no-hit | 0.0 | 0 | no-hit | 0.0 | 0 | no-hit | 0.0 | 0 |
| scf7180000044639 | 287469 | 0.3872 | 0 | 59.2999 | Eukaryota | 111045.0 | 0 | Chordata | 111045.0 | 0 | Carnivora | 97346.0 | 1 | Canidae | 97346.0 | 1 | Canis | 60713.0 | 2 | Canis lupus | 60713.0 | 2 |
| scf7180000041928 | 10190 | 0.3714 | 0 | 27.5029 | Eukaryota | 22276.0 | 0 | Chordata | 22276.0 | 0 | Carnivora | 22276.0 | 0 | Canidae | 22276.0 | 0 | Canis | 22276.0 | 0 | Canis lupus | 22276.0 | 0 |
| scf7180000055523 | 8240 | 0.4345 | 0 | 116.1507 | Eukaryota | 5600.0 | 1 | Chordata | 4209.0 | 2 | Tetraodontiformes | 3131.0 | 4 | Tetraodontidae | 3131.0 | 4 | Takifugu | 3131.0 | 4 | Takifugu rubripes | 3131.0 | 4 |
| scf7180000040840 | 400494 | 0.3865 | 0 | 59.2335 | Eukaryota | 76648.0 | 0 | Chordata | 76648.0 | 0 | Carnivora | 76648.0 | 0 | Canidae | 76648.0 | 0 | Canis | 76648.0 | 0 | Canis lupus | 76648.0 | 0 |
| scf7180000055535 | 23810 | 0.4414 | 0 | 269.0158 | Eukaryota | 1508.0 | 0 | Chordata | 1508.0 | 0 | Kurtiformes | 1508.0 | 0 | Apogonidae | 1508.0 | 0 | Sphaeramia | 1508.0 | 0 | Sphaeramia orbicularis | 1508.0 | 0 |
| scf7180000054128 | 2163 | 0.4549 | 0 | 495.0518 | Eukaryota | 509.0 | 0 | Streptophyta | 509.0 | 0 | Fabales | 509.0 | 0 | Fabaceae | 509.0 | 0 | Cicer | 509.0 | 0 | Cicer arietinum | 509.0 | 0 |
| scf7180000046325 | 148791 | 0.3897 | 0 | 58.8237 | Eukaryota | 95165.0 | 0 | Chordata | 95165.0 | 0 | Carnivora | 95165.0 | 0 | Canidae | 95165.0 | 0 | Canis | 95165.0 | 0 | Canis lupus | 95165.0 | 0 |
| scf7180000042003 | 107218 | 0.3514 | 0 | 58.811 | Eukaryota | 93508.0 | 0 | Chordata | 93508.0 | 0 | Carnivora | 93508.0 | 0 | Canidae | 93508.0 | 0 | Canis | 93508.0 | 0 | Canis lupus | 93508.0 | 0 |
| scf7180000053271 | 14560 | 0.3803 | 0 | 61.3818 | Eukaryota | 3166.0 | 0 | Chordata | 3166.0 | 0 | Carnivora | 762.0 | 5 | Canidae | 762.0 | 5 | Canis | 762.0 | 5 | Canis lupus | 762.0 | 5 |
| scf7180000054082 | 3300 | 0.4445 | 0 | 611.0392 | no-hit | 0.0 | 0 | no-hit | 0.0 | 0 | no-hit | 0.0 | 0 | no-hit | 0.0 | 0 | no-hit | 0.0 | 0 | no-hit | 0.0 | 0 |
| scf7180000044394 | 29375 | 0.5415 | 0 | 45.1384 | Eukaryota | 9703.0 | 0 | Chordata | 9703.0 | 0 | Carnivora | 9703.0 | 0 | Canidae | 5268.0 | 1 | Ailuropoda | 4435.0 | 2 | Ailuropoda melanoleuca | 4435.0 | 2 |
| scf7180000045273 | 32931 | 0.3956 | 0 | 41.6431 | Eukaryota | 74430.0 | 0 | Chordata | 74430.0 | 0 | Carnivora | 74430.0 | 0 | Canidae | 74430.0 | 0 | Canis | 74430.0 | 0 | Canis lupus | 74430.0 | 0 |
| scf7180000053746 | 3728 | 0.4464 | 0 | 66.0666 | no-hit | 0.0 | 0 | no-hit | 0.0 | 0 | no-hit | 0.0 | 0 | no-hit | 0.0 | 0 | no-hit | 0.0 | 0 | no-hit | 0.0 | 0 |
| scf7180000040852 | 1661666 | 0.3781 | 0 | 60.8488 | Eukaryota | 90794.0 | 0 | Chordata | 90794.0 | 0 | Carnivora | 90794.0 | 0 | Canidae | 90794.0 | 0 | Canis | 90794.0 | 0 | Canis lupus | 90794.0 | 0 |
| scf7180000041184 | 1265150 | 0.4152 | 0 | 60.5109 | Eukaryota | 56399.0 | 0 | Chordata | 56399.0 | 0 | Carnivora | 56399.0 | 0 | Canidae | 56399.0 | 0 | Canis | 56399.0 | 0 | Canis lupus | 56399.0 | 0 |
| scf7180000045644 | 176651 | 0.428 | 0 | 59.3688 | Eukaryota | 94452.0 | 0 | Chordata | 94452.0 | 0 | Carnivora | 94452.0 | 0 | Canidae | 94452.0 | 0 | Canis | 94452.0 | 0 | Canis lupus | 94452.0 | 0 |
| scf7180000041655 | 524962 | 0.3505 | 0 | 59.0767 | Eukaryota | 62963.0 | 0 | Chordata | 62963.0 | 0 | Carnivora | 62963.0 | 0 | Canidae | 62963.0 | 0 | Canis | 56344.0 | 1 | Canis lupus | 56344.0 | 1 |
| scf7180000053528 | 6974 | 0.5032 | 0 | 171.8804 | Eukaryota | 7932.0 | 0 | Chordata | 7932.0 | 0 | Carnivora | 7932.0 | 0 | Canidae | 7932.0 | 0 | Canis | 5863.0 | 1 | Canis lupus | 4704.0 | 2 |
| scf7180000049865 | 26296 | 0.5303 | 0 | 55.0499 | Eukaryota | 3628.0 | 0 | Chordata | 3628.0 | 0 | Carnivora | 3628.0 | 0 | Canidae | 2272.0 | 1 | Canis | 1518.0 | 2 | Canis lupus | 1518.0 | 3 |
| scf7180000045679 | 15557 | 0.3688 | 0 | 747.6354 | Eukaryota | 47223.0 | 0 | Chordata | 47223.0 | 0 | Carnivora | 47223.0 | 0 | Canidae | 47223.0 | 0 | Canis | 47223.0 | 0 | Canis lupus | 47223.0 | 0 |
| scf7180000050692 | 11990 | 0.4411 | 0 | 148.9363 | Eukaryota | 3299.0 | 0 | Chordata | 2868.0 | 1 | Carnivora | 2476.0 | 2 | Canidae | 2476.0 | 2 | Canis | 2476.0 | 2 | Canis lupus | 2476.0 | 2 |
| scf7180000050570 | 21251 | 0.5994 | 0 | 51.0493 | Eukaryota | 24483.0 | 0 | Chordata | 24483.0 | 0 | Carnivora | 24483.0 | 0 | Canidae | 24483.0 | 0 | Canis | 23415.0 | 1 | Canis lupus | 23415.0 | 1 |
| scf7180000048243 | 98286 | 0.4011 | 0 | 59.7422 | Eukaryota | 15377.0 | 0 | Chordata | 15377.0 | 0 | Carnivora | 15377.0 | 0 | Canidae | 15377.0 | 0 | Canis | 15377.0 | 0 | Canis lupus | 15377.0 | 0 |
| scf7180000045330 | 63751 | 0.4578 | 0 | 53.4163 | Eukaryota | 11777.0 | 0 | Chordata | 11777.0 | 0 | Carnivora | 11777.0 | 0 | Canidae | 5291.0 | 3 | Canis | 3689.0 | 4 | Canis lupus | 3689.0 | 4 |
| scf7180000043815 | 22566 | 0.4378 | 0 | 155.2936 | no-hit | 0.0 | 0 | no-hit | 0.0 | 0 | no-hit | 0.0 | 0 | no-hit | 0.0 | 0 | no-hit | 0.0 | 0 | no-hit | 0.0 | 0 |
| scf7180000043983 | 56259 | 0.3409 | 0 | 56.7254 | Eukaryota | 4276.0 | 0 | Chordata | 4276.0 | 0 | Carnivora | 2660.0 | 1 | Canidae | 2660.0 | 1 | Canis | 2660.0 | 1 | Canis lupus | 2660.0 | 1 |
| scf7180000050114 | 4227 | 0.4504 | 0 | 912.8561 | Eukaryota | 187.0 | 0 | Streptophyta | 187.0 | 0 | Fabales | 187.0 | 0 | Fabaceae | 187.0 | 0 | Cicer | 187.0 | 0 | Cicer arietinum | 187.0 | 0 |
| scf7180000056607 | 2234174 | 0.3809 | 0 | 60.0512 | Eukaryota | 40117.0 | 0 | Chordata | 40117.0 | 0 | Carnivora | 40117.0 | 0 | Canidae | 40117.0 | 0 | Canis | 31831.0 | 1 | Canis lupus | 31831.0 | 1 |
| scf7180000050144 | 18786 | 0.4457 | 0 | 751.2676 | Eukaryota | 6701.0 | 0 | Chordata | 4210.0 | 2 | Salmoniformes | 3268.0 | 3 | Salmonidae | 3268.0 | 3 | Salmo | 2894.0 | 4 | Salmo trutta | 2894.0 | 4 |
| scf7180000055498 | 13202 | 0.4458 | 0 | 177.0298 | Eukaryota | 11218.0 | 0 | Chordata | 11218.0 | 0 | Carnivora | 11218.0 | 0 | Otariidae | 4236.0 | 3 | Canis | 2570.0 | 5 | Canis lupus | 2570.0 | 5 |
| scf7180000053377 | 1669 | 0.4476 | 0 | 167.3721 | Eukaryota | 272.0 | 0 | Platyhelminthes | 272.0 | 0 | Polystomatidea | 272.0 | 0 | Polystomatidae | 272.0 | 0 | Protopolystoma | 272.0 | 0 | Protopolystoma xenopodis | 272.0 | 0 |
| scf7180000048136 | 5752 | 0.4404 | 0 | 174.4212 | Eukaryota | 4012.0 | 0 | Chordata | 3707.0 | 1 | Kurtiformes | 2488.0 | 3 | Apogonidae | 2488.0 | 3 | Sphaeramia | 2488.0 | 3 | Sphaeramia orbicularis | 2488.0 | 3 |
| scf7180000047623 | 60953 | 0.6336 | 0 | 53.4733 | Eukaryota | 3944.0 | 0 | Chordata | 3944.0 | 0 | Carnivora | 2741.0 | 1 | Canidae | 2741.0 | 1 | Canis | 2741.0 | 1 | Canis lupus | 2741.0 | 1 |
| scf7180000053126 | 2608 | 0.454 | 0 | 421.7068 | no-hit | 0.0 | 0 | no-hit | 0.0 | 0 | no-hit | 0.0 | 0 | no-hit | 0.0 | 0 | no-hit | 0.0 | 0 | no-hit | 0.0 | 0 |
| scf7180000052517 | 7203 | 0.5073 | 0 | 79.5502 | Eukaryota | 7618.0 | 0 | Chordata | 7618.0 | 0 | Carnivora | 7618.0 | 0 | Canidae | 7618.0 | 0 | Canis | 6634.0 | 1 | Canis lupus | 5617.0 | 2 |
| scf7180000052507 | 4041 | 0.4412 | 0 | 55.7012 | no-hit | 0.0 | 0 | no-hit | 0.0 | 0 | no-hit | 0.0 | 0 | no-hit | 0.0 | 0 | no-hit | 0.0 | 0 | no-hit | 0.0 | 0 |
| scf7180000047407 | 10136 | 0.5283 | 0 | 53.489 | Eukaryota | 5902.0 | 0 | Chordata | 5902.0 | 0 | Carnivora | 5902.0 | 0 | Canidae | 5902.0 | 0 | Canis | 5902.0 | 0 | Canis lupus | 5902.0 | 0 |
| scf7180000056499 | 89873 | 0.344 | 0 | 56.2489 | Eukaryota | 17339.0 | 0 | Chordata | 17339.0 | 0 | Carnivora | 17339.0 | 0 | Canidae | 17339.0 | 0 | Canis | 17339.0 | 0 | Canis lupus | 17339.0 | 0 |
| scf7180000051174 | 4552 | 0.4455 | 0 | 464.8424 | no-hit | 0.0 | 0 | no-hit | 0.0 | 0 | no-hit | 0.0 | 0 | no-hit | 0.0 | 0 | no-hit | 0.0 | 0 | no-hit | 0.0 | 0 |
| scf7180000042619 | 611084 | 0.4393 | 0 | 59.7014 | Eukaryota | 88959.0 | 0 | Chordata | 88959.0 | 0 | Carnivora | 88959.0 | 0 | Canidae | 60413.0 | 3 | Canis | 42686.0 | 5 | Canis lupus | 42686.0 | 5 |
| scf7180000041482 | 991506 | 0.3737 | 0 | 59.807 | Eukaryota | 90966.0 | 0 | Chordata | 90966.0 | 0 | Carnivora | 90966.0 | 0 | Canidae | 90966.0 | 0 | Canis | 90966.0 | 0 | Canis lupus | 90966.0 | 0 |
| scf7180000052133 | 3826 | 0.4482 | 0 | 517.3074 | no-hit | 0.0 | 0 | no-hit | 0.0 | 0 | no-hit | 0.0 | 0 | no-hit | 0.0 | 0 | no-hit | 0.0 | 0 | no-hit | 0.0 | 0 |
| scf7180000042289 | 633206 | 0.4869 | 0 | 59.5089 | Eukaryota | 82654.0 | 0 | Chordata | 82654.0 | 0 | Carnivora | 82654.0 | 0 | Canidae | 38034.0 | 3 | Canis | 28477.0 | 7 | Canis lupus | 28477.0 | 7 |
| scf7180000045025 | 280806 | 0.4394 | 0 | 60.4151 | Eukaryota | 91723.0 | 0 | Chordata | 91723.0 | 0 | Carnivora | 91723.0 | 0 | Canidae | 91723.0 | 0 | Canis | 91723.0 | 0 | Canis lupus | 91723.0 | 0 |
| scf7180000047992 | 66960 | 0.4501 | 0 | 57.3813 | Eukaryota | 55668.0 | 0 | Chordata | 55668.0 | 0 | Carnivora | 55668.0 | 0 | Canidae | 55668.0 | 0 | Canis | 55668.0 | 0 | Canis lupus | 55668.0 | 0 |
| scf7180000042159 | 769123 | 0.4388 | 0 | 58.7473 | Eukaryota | 44767.0 | 0 | Chordata | 44767.0 | 0 | Carnivora | 44767.0 | 0 | Canidae | 40061.0 | 1 | Canis | 31521.0 | 2 | Canis lupus | 31521.0 | 2 |
| scf7180000045472 | 128692 | 0.4101 | 0 | 58.8631 | Eukaryota | 46269.0 | 0 | Chordata | 46269.0 | 0 | Carnivora | 46269.0 | 0 | Canidae | 46269.0 | 0 | Canis | 38523.0 | 1 | Canis lupus | 38523.0 | 1 |
| scf7180000041121 | 1360806 | 0.412 | 0 | 59.7362 | Eukaryota | 97935.0 | 0 | Chordata | 97935.0 | 0 | Carnivora | 97935.0 | 0 | Canidae | 97935.0 | 0 | Canis | 97935.0 | 0 | Canis lupus | 97935.0 | 0 |
| scf7180000050876 | 21330 | 0.4779 | 0 | 57.7286 | Eukaryota | 2762.0 | 0 | Chordata | 2762.0 | 0 | Carnivora | 2762.0 | 0 | Canidae | 2762.0 | 0 | Canis | 2762.0 | 0 | Canis lupus | 2762.0 | 0 |
| scf7180000051169 | 2451 | 0.4561 | 0 | 141.9998 | Eukaryota | 966.0 | 0 | Platyhelminthes | 592.0 | 1 | Polystomatidea | 592.0 | 1 | Polystomatidae | 592.0 | 1 | Protopolystoma | 592.0 | 2 | Protopolystoma xenopodis | 592.0 | 2 |
| scf7180000049779 | 19286 | 0.4412 | 0 | 234.6226 | Eukaryota | 3607.0 | 0 | Platyhelminthes | 2907.0 | 1 | Polystomatidea | 2907.0 | 3 | Polystomatidae | 2907.0 | 3 | Protopolystoma | 2907.0 | 3 | Protopolystoma xenopodis | 2907.0 | 3 |
| scf7180000042716 | 660161 | 0.38 | 0 | 60.7642 | Eukaryota | 94946.0 | 0 | Chordata | 94946.0 | 0 | Carnivora | 94946.0 | 0 | Canidae | 89701.0 | 1 | Canis | 89701.0 | 1 | Canis lupus | 89701.0 | 1 |
| scf7180000040649 | 1937041 | 0.4201 | 0 | 58.5874 | Eukaryota | 93519.0 | 0 | Chordata | 93519.0 | 0 | Carnivora | 93519.0 | 0 | Canidae | 93519.0 | 0 | Canis | 93519.0 | 0 | Canis lupus | 93519.0 | 0 |
| scf7180000041526 | 1279841 | 0.3895 | 0 | 59.6181 | Eukaryota | 42351.0 | 0 | Chordata | 42351.0 | 0 | Carnivora | 42351.0 | 0 | Canidae | 42351.0 | 0 | Canis | 42351.0 | 0 | Canis lupus | 42351.0 | 0 |
| scf7180000056587 | 771799 | 0.3937 | 0 | 59.43 | Eukaryota | 58146.0 | 0 | Chordata | 58146.0 | 0 | Carnivora | 58146.0 | 0 | Canidae | 58146.0 | 0 | Canis | 58146.0 | 0 | Canis lupus | 58146.0 | 0 |
| scf7180000050457 | 7766 | 0.5049 | 0 | 1697.2473 | Eukaryota | 8005.0 | 0 | Chordata | 8005.0 | 0 | Carnivora | 8005.0 | 0 | Canidae | 8005.0 | 0 | Canis | 5307.0 | 1 | Canis lupus | 4227.0 | 2 |
| scf7180000055304 | 26507 | 0.4447 | 0 | 164.5616 | no-hit | 0.0 | 0 | no-hit | 0.0 | 0 | no-hit | 0.0 | 0 | no-hit | 0.0 | 0 | no-hit | 0.0 | 0 | no-hit | 0.0 | 0 |
| scf7180000049310 | 44723 | 0.4627 | 0 | 73.548 | Eukaryota | 14776.0 | 0 | Chordata | 14776.0 | 0 | Carnivora | 11771.0 | 1 | Canidae | 11771.0 | 1 | Canis | 10659.0 | 2 | Canis lupus | 10659.0 | 2 |
| scf7180000048092 | 2787 | 0.4377 | 0 | 107.6539 | Eukaryota | 470.0 | 0 | Platyhelminthes | 470.0 | 0 | Polystomatidea | 470.0 | 0 | Polystomatidae | 470.0 | 0 | Protopolystoma | 470.0 | 0 | Protopolystoma xenopodis | 470.0 | 0 |
| scf7180000056432 | 399529 | 0.4502 | 0 | 58.9248 | Eukaryota | 41648.0 | 0 | Chordata | 41648.0 | 0 | Carnivora | 41648.0 | 0 | Canidae | 41648.0 | 0 | Canis | 36231.0 | 1 | Canis lupus | 36231.0 | 1 |
| scf7180000052527 | 3602 | 0.4575 | 0 | 842.6382 | Eukaryota | 927.0 | 0 | Echinodermata | 623.0 | 2 | Forcipulatida | 623.0 | 2 | Asteriidae | 623.0 | 2 | Asterias | 623.0 | 2 | Asterias rubens | 623.0 | 2 |
| scf7180000055577 | 29519 | 0.4393 | 0 | 378.8054 | Eukaryota | 34708.0 | 0 | Chordata | 34708.0 | 0 | Gadiformes | 17581.0 | 2 | Gadidae | 17581.0 | 2 | Gadus | 17581.0 | 2 | Gadus morhua | 17581.0 | 3 |
| scf7180000047814 | 97788 | 0.397 | 0 | 61.8692 | Eukaryota | 85973.0 | 0 | Chordata | 85973.0 | 0 | Carnivora | 85973.0 | 0 | Canidae | 85973.0 | 0 | Canis | 85973.0 | 0 | Canis lupus | 85973.0 | 0 |
| scf7180000053481 | 2599 | 0.4436 | 0 | 522.3383 | Eukaryota | 154.0 | 0 | Chordata | 154.0 | 0 | Salmoniformes | 154.0 | 0 | Salmonidae | 154.0 | 0 | Salmo | 154.0 | 0 | Salmo trutta | 154.0 | 0 |
| scf7180000056228 | 635524 | 0.3421 | 0 | 58.7307 | Eukaryota | 94066.0 | 0 | Chordata | 94066.0 | 0 | Carnivora | 94066.0 | 0 | Canidae | 94066.0 | 0 | Canis | 94066.0 | 0 | Canis lupus | 94066.0 | 0 |
| scf7180000049062 | 9820 | 0.5164 | 0 | 453.7518 | Eukaryota | 15937.0 | 0 | Chordata | 15937.0 | 0 | Carnivora | 15937.0 | 0 | Canidae | 15937.0 | 0 | Canis | 15937.0 | 0 | Canis lupus | 15937.0 | 0 |
| scf7180000050386 | 7855 | 0.4511 | 0 | 89.5773 | no-hit | 0.0 | 0 | no-hit | 0.0 | 0 | no-hit | 0.0 | 0 | no-hit | 0.0 | 0 | no-hit | 0.0 | 0 | no-hit | 0.0 | 0 |
| scf7180000046588 | 14499 | 0.4419 | 0 | 162.3244 | Eukaryota | 1112.0 | 0 | Platyhelminthes | 562.0 | 1 | Polystomatidea | 562.0 | 2 | Polystomatidae | 562.0 | 2 | Protopolystoma | 562.0 | 2 | Protopolystoma xenopodis | 562.0 | 2 |
| scf7180000049203 | 30932 | 0.4316 | 0 | 62.2195 | Eukaryota | 17942.0 | 0 | Chordata | 17942.0 | 0 | Artiodactyla | 6879.0 | 2 | Ursidae | 6077.0 | 3 | Equus | 4986.0 | 4 | Camelus dromedarius | 4053.0 | 6 |
| scf7180000051443 | 7402 | 0.4572 | 0 | 1154.6866 | Eukaryota | 4788.0 | 0 | Chordata | 3163.0 | 1 | Salmoniformes | 2536.0 | 3 | Salmonidae | 2536.0 | 3 | Salmo | 2536.0 | 3 | Salmo trutta | 2536.0 | 3 |
| scf7180000041152 | 510845 | 0.3376 | 0 | 57.272 | Eukaryota | 92333.0 | 0 | Chordata | 92333.0 | 0 | Carnivora | 92333.0 | 0 | Canidae | 92333.0 | 0 | Canis | 92333.0 | 0 | Canis lupus | 92333.0 | 0 |
| scf7180000054634 | 25649 | 0.4369 | 0 | 360.4439 | Eukaryota | 4312.0 | 0 | Chordata | 2450.0 | 1 | Polystomatidea | 1862.0 | 3 | Polystomatidae | 1862.0 | 3 | Protopolystoma | 1862.0 | 3 | Protopolystoma xenopodis | 1862.0 | 3 |
| scf7180000045876 | 21101 | 0.4397 | 0 | 49.9015 | Eukaryota | 3875.0 | 0 | Platyhelminthes | 2050.0 | 1 | Polystomatidea | 2050.0 | 5 | Polystomatidae | 2050.0 | 5 | Protopolystoma | 2050.0 | 5 | Protopolystoma xenopodis | 2050.0 | 5 |
| scf7180000043345 | 485363 | 0.6197 | 0 | 55.7538 | Eukaryota | 57340.0 | 0 | Chordata | 57340.0 | 0 | Carnivora | 57340.0 | 0 | Canidae | 50872.0 | 1 | Canis | 45793.0 | 2 | Canis lupus | 45793.0 | 2 |
| scf7180000056263 | 253192 | 0.3808 | 0 | 58.2791 | Eukaryota | 44926.0 | 0 | Chordata | 44926.0 | 0 | Carnivora | 44926.0 | 0 | Canidae | 44926.0 | 0 | Canis | 44926.0 | 0 | Canis lupus | 44926.0 | 0 |
| scf7180000055932 | 13537 | 0.5426 | 0 | 69.4229 | Eukaryota | 31845.0 | 0 | Chordata | 31845.0 | 0 | Carnivora | 31845.0 | 0 | Canidae | 31845.0 | 0 | Canis | 31845.0 | 0 | Canis lupus | 31845.0 | 0 |
| scf7180000041958 | 778312 | 0.4069 | 0 | 60.1326 | Eukaryota | 68090.0 | 0 | Chordata | 68090.0 | 0 | Carnivora | 68090.0 | 0 | Canidae | 68090.0 | 0 | Canis | 46854.0 | 1 | Canis lupus | 46854.0 | 1 |
| scf7180000047656 | 28389 | 0.4443 | 0 | 99.2236 | Eukaryota | 29494.0 | 1 | Chordata | 29494.0 | 1 | Tetraodontiformes | 9433.0 | 5 | Tetraodontidae | 9433.0 | 5 | Takifugu | 9433.0 | 5 | Takifugu rubripes | 9433.0 | 5 |

| | | | | | | | | | | | | | | | | | | | | | | |
|---|---|---|---|---|---|---|---|---|---|---|---|---|---|---|---|---|---|---|---|---|---|---|
| scf7180000053106 | 2919 | 0.4519 | 0 | 484.51 | no-hit | 0.0 | 0 | no-hit | 0.0 | 0 | no-hit | 0.0 | 0 | no-hit | 0.0 | 0 | no-hit | 0.0 | 0 | no-hit | 0.0 | 0 |
| scf7180000045063 | 13283 | 0.5204 | 0 | 124.2221 | Eukaryota | 31973.0 | 0 | Chordata | 31973.0 | 0 | Carnivora | 31973.0 | 0 | Canidae | 31973.0 | 0 | Canis | 31973.0 | 0 | Canis lupus | 31973.0 | 0 |
| scf7180000040748 | 6660 | 0.4119 | 0 | 26.686 | Eukaryota | 3240.0 | 0 | Chordata | 3240.0 | 0 | Carnivora | 2233.0 | 2 | Canidae | 2233.0 | 2 | Canis | 2233.0 | 2 | Canis lupus | 2233.0 | 2 |
| scf7180000047864 | 71483 | 0.5084 | 0 | 60.1629 | Eukaryota | 41558.0 | 0 | Chordata | 41558.0 | 0 | Carnivora | 41558.0 | 0 | Canidae | 41558.0 | 0 | Vulpes | 25353.0 | 1 | Vulpes vulpes | 25353.0 | 1 |
| scf7180000056372 | 540172 | 0.4163 | 0 | 59.9893 | Eukaryota | 91440.0 | 0 | Chordata | 91440.0 | 0 | Carnivora | 91440.0 | 0 | Canidae | 91440.0 | 0 | Canis | 91440.0 | 0 | Canis lupus | 91440.0 | 0 |
| scf7180000050196 | 5986 | 0.446 | 0 | 162.4515 | no-hit | 0.0 | 0 | no-hit | 0.0 | 0 | no-hit | 0.0 | 0 | no-hit | 0.0 | 0 | no-hit | 0.0 | 0 | no-hit | 0.0 | 0 |
| scf7180000044756 | 421673 | 0.4042 | 0 | 59.1631 | Eukaryota | 64201.0 | 0 | Chordata | 64201.0 | 0 | Carnivora | 64201.0 | 0 | Canidae | 64201.0 | 0 | Canis | 64201.0 | 0 | Canis lupus | 64201.0 | 0 |
| scf7180000047853 | 26440 | 0.4456 | 0 | 594.5597 | Eukaryota | 5461.0 | 0 | Echinodermata | 2498.0 | 2 | Forcipulatida | 2498.0 | 2 | Asteriidae | 2498.0 | 2 | Asterias | 2498.0 | 2 | Asterias rubens | 2498.0 | 2 |
| scf7180000051242 | 22416 | 0.4994 | 0 | 405.5923 | Eukaryota | 9372.0 | 0 | Chordata | 9372.0 | 0 | Carnivora | 9372.0 | 0 | Canidae | 9372.0 | 0 | Canis | 7607.0 | 1 | Canis lupus | 6319.0 | 2 |
| scf7180000056334 | 921239 | 0.394 | 0 | 59.7139 | Eukaryota | 91244.0 | 0 | Chordata | 91244.0 | 0 | Carnivora | 91244.0 | 0 | Canidae | 91244.0 | 0 | Canis | 91244.0 | 0 | Canis lupus | 91244.0 | 0 |
| scf7180000045758 | 6360 | 0.4513 | 0 | 267.9714 | no-hit | 0.0 | 0 | no-hit | 0.0 | 0 | no-hit | 0.0 | 0 | no-hit | 0.0 | 0 | no-hit | 0.0 | 0 | no-hit | 0.0 | 0 |
| scf7180000050639 | 6896 | 0.4524 | 0 | 996.6659 | no-hit | 0.0 | 0 | no-hit | 0.0 | 0 | no-hit | 0.0 | 0 | no-hit | 0.0 | 0 | no-hit | 0.0 | 0 | no-hit | 0.0 | 0 |
| scf7180000047745 | 27516 | 0.4367 | 0 | 196.0903 | Eukaryota | 27932.0 | 0 | Chordata | 27932.0 | 0 | Carnivora | 27932.0 | 0 | Canidae | 27932.0 | 0 | Canis | 27932.0 | 0 | Canis lupus | 27932.0 | 0 |
| scf7180000051226 | 7696 | 0.4498 | 0 | 930.5938 | Eukaryota | 148.0 | 0 | Platyhelminthes | 148.0 | 0 | Polystomatidea | 148.0 | 0 | Polystomatidae | 148.0 | 0 | Protopolystoma | 148.0 | 0 | Protopolystoma xenopodis | 148.0 | 0 |
| scf7180000041172 | 1320327 | 0.4062 | 0 | 59.627 | Eukaryota | 93480.0 | 0 | Chordata | 93480.0 | 0 | Carnivora | 93480.0 | 0 | Canidae | 93480.0 | 0 | Canis | 93480.0 | 0 | Canis lupus | 93480.0 | 0 |
| scf7180000052135 | 12540 | 0.4537 | 0 | 2773.7703 | Eukaryota | 2510.0 | 0 | Chordata | 2510.0 | 0 | Carnivora | 2232.0 | 1 | Canidae | 2232.0 | 1 | Canis | 2232.0 | 1 | Canis lupus | 2232.0 | 1 |
| scf7180000042984 | 3153 | 0.4897 | 0 | 26.8636 | Eukaryota | 3148.0 | 0 | Chordata | 3148.0 | 0 | Carnivora | 3148.0 | 0 | Canidae | 3148.0 | 0 | Canis | 3148.0 | 0 | Canis lupus | 3148.0 | 0 |
| scf7180000041123 | 1401607 | 0.3727 | 0 | 61.2515 | Eukaryota | 76089.0 | 0 | Chordata | 76089.0 | 0 | Carnivora | 76089.0 | 0 | Canidae | 76089.0 | 0 | Canis | 76089.0 | 0 | Canis lupus | 76089.0 | 0 |
| scf7180000044038 | 345673 | 0.3659 | 0 | 59.2277 | Eukaryota | 91436.0 | 0 | Chordata | 91436.0 | 0 | Carnivora | 91436.0 | 0 | Canidae | 91436.0 | 0 | Canis | 91436.0 | 0 | Canis lupus | 91436.0 | 0 |
| scf7180000050102 | 28756 | 0.4374 | 0 | 66.9955 | Eukaryota | 37645.0 | 0 | Chordata | 37645.0 | 0 | Carnivora | 37645.0 | 0 | Canidae | 37645.0 | 0 | Canis | 37645.0 | 0 | Canis lupus | 37645.0 | 0 |
| scf7180000044406 | 199292 | 0.5286 | 0 | 58.7799 | Eukaryota | 62644.0 | 0 | Chordata | 62644.0 | 0 | Carnivora | 62644.0 | 0 | Canidae | 62644.0 | 0 | Canis | 62644.0 | 0 | Canis lupus | 62644.0 | 0 |
| scf7180000040842 | 456803 | 0.3528 | 0 | 59.6045 | Eukaryota | 72270.0 | 0 | Chordata | 72270.0 | 0 | Carnivora | 72270.0 | 0 | Canidae | 23547.0 | 3 | Odobenus | 20871.0 | 6 | Odobenus rosmarus | 20871.0 | 6 |
| scf7180000040906 | 1223622 | 0.3855 | 0 | 59.6527 | Eukaryota | 34544.0 | 0 | Chordata | 34544.0 | 0 | Carnivora | 34544.0 | 0 | Canidae | 34544.0 | 0 | Canis | 28134.0 | 1 | Canis lupus | 28134.0 | 1 |
| scf7180000046275 | 148163 | 0.3713 | 0 | 60.0126 | Eukaryota | 63329.0 | 0 | Chordata | 63329.0 | 0 | Carnivora | 63329.0 | 0 | Canidae | 63329.0 | 0 | Canis | 63329.0 | 0 | Canis lupus | 63329.0 | 0 |
| scf7180000047455 | 129559 | 0.5053 | 0 | 58.7353 | Eukaryota | 31653.0 | 0 | Chordata | 31653.0 | 0 | Carnivora | 31653.0 | 0 | Canidae | 31653.0 | 0 | Canis | 22927.0 | 1 | Canis lupus | 22927.0 | 1 |
| scf7180000042036 | 793850 | 0.4262 | 0 | 59.9639 | Eukaryota | 39514.0 | 0 | Chordata | 39514.0 | 0 | Carnivora | 39514.0 | 0 | Canidae | 39514.0 | 0 | Canis | 28915.0 | 1 | Canis lupus | 28915.0 | 1 |
| scf7180000046116 | 81098 | 0.3876 | 0 | 60.7308 | Eukaryota | 90906.0 | 0 | Chordata | 90906.0 | 0 | Carnivora | 90906.0 | 0 | Canidae | 90906.0 | 0 | Canis | 90906.0 | 0 | Canis lupus | 90906.0 | 0 |
| scf7180000044516 | 183301 | 0.3858 | 0 | 59.3544 | Eukaryota | 29103.0 | 0 | Chordata | 29103.0 | 0 | Carnivora | 29103.0 | 0 | Canidae | 29103.0 | 0 | Canis | 29103.0 | 0 | Canis lupus | 29103.0 | 0 |
| scf7180000056222 | 1732910 | 0.4358 | 0 | 59.2024 | Eukaryota | 57031.0 | 0 | Chordata | 57031.0 | 0 | Carnivora | 57031.0 | 0 | Canidae | 57031.0 | 0 | Canis | 57031.0 | 0 | Canis lupus | 57031.0 | 0 |
| scf7180000055552 | 13202 | 0.4409 | 0 | 212.4496 | Eukaryota | 1039.0 | 0 | Chordata | 1039.0 | 0 | Kurtiformes | 494.0 | 2 | Apogonidae | 494.0 | 2 | Sphaeramia | 494.0 | 2 | Sphaeramia orbicularis | 494.0 | 2 |
| scf7180000053342 | 3845 | 0.504 | 0 | 138.2979 | Eukaryota | 7027.0 | 0 | Chordata | 7027.0 | 0 | Carnivora | 7027.0 | 0 | Canidae | 7027.0 | 0 | Canis | 5272.0 | 1 | Canis lupus | 4354.0 | 2 |
| scf7180000052288 | 8522 | 0.4391 | 0 | 167.4531 | Eukaryota | 4338.0 | 0 | Chordata | 3807.0 | 1 | Tetraodontiformes | 2146.0 | 5 | Tetraodontidae | 2146.0 | 5 | Takifugu | 2146.0 | 5 | Takifugu rubripes | 2146.0 | 5 |
| scf7180000040651 | 214608 | 0.3694 | 0 | 61.928 | Eukaryota | 88097.0 | 0 | Chordata | 88097.0 | 0 | Carnivora | 88097.0 | 0 | Canidae | 88097.0 | 0 | Canis | 88097.0 | 0 | Canis lupus | 88097.0 | 0 |
| scf7180000052740 | 17840 | 0.4724 | 0 | 85.3366 | Eukaryota | 33283.0 | 0 | Chordata | 33283.0 | 0 | Carnivora | 33283.0 | 0 | Canidae | 33283.0 | 0 | Canis | 33283.0 | 0 | Canis lupus | 33283.0 | 0 |
| scf7180000042735 | 656015 | 0.3356 | 0 | 59.1584 | Eukaryota | 35121.0 | 0 | Chordata | 35121.0 | 0 | Carnivora | 35121.0 | 0 | Canidae | 35121.0 | 0 | Canis | 35121.0 | 0 | Canis lupus | 35121.0 | 0 |
| scf7180000052923 | 5209 | 0.5064 | 0 | 102.7334 | Eukaryota | 7518.0 | 0 | Chordata | 7518.0 | 0 | Carnivora | 7518.0 | 0 | Canidae | 7518.0 | 0 | Canis | 6938.0 | 1 | Canis lupus | 5853.0 | 2 |
| scf7180000050408 | 6707 | 0.4545 | 0 | 136.2003 | Eukaryota | 54322.0 | 0 | Chordata | 54322.0 | 0 | Carnivora | 54322.0 | 0 | Canidae | 54322.0 | 0 | Canis | 54322.0 | 0 | Canis lupus | 54322.0 | 0 |
| scf7180000043291 | 9741 | 0.4427 | 0 | 65.9561 | no-hit | 0.0 | 0 | no-hit | 0.0 | 0 | no-hit | 0.0 | 0 | no-hit | 0.0 | 0 | no-hit | 0.0 | 0 | no-hit | 0.0 | 0 |
| scf7180000047652 | 24762 | 0.4364 | 0 | 251.3276 | Eukaryota | 141.0 | 0 | Platyhelminthes | 141.0 | 0 | Polystomatidea | 141.0 | 0 | Polystomatidae | 141.0 | 0 | Protopolystoma | 141.0 | 0 | Protopolystoma xenopodis | 141.0 | 0 |
| scf7180000053869 | 2187 | 0.4417 | 0 | 87.5722 | no-hit | 0.0 | 0 | no-hit | 0.0 | 0 | no-hit | 0.0 | 0 | no-hit | 0.0 | 0 | no-hit | 0.0 | 0 | no-hit | 0.0 | 0 |
| scf7180000048595 | 21758 | 0.4371 | 0 | 535.7862 | Eukaryota | 300.0 | 0 | Chordata | 300.0 | 0 | Blenniiformes | 300.0 | 0 | Gobiesocidae | 300.0 | 0 | Gouania | 300.0 | 0 | Gouania willdenowi | 300.0 | 0 |
| scf7180000051702 | 13181 | 0.4396 | 0 | 62.8483 | Eukaryota | 52461.0 | 0 | Chordata | 39238.0 | 1 | Gadiformes | 28706.0 | 5 | Gadidae | 28706.0 | 5 | Gadus | 28706.0 | 5 | Gadus morhua | 28706.0 | 5 |
| scf7180000042687 | 617699 | 0.3713 | 0 | 58.9765 | Eukaryota | 106453.0 | 0 | Chordata | 106453.0 | 0 | Carnivora | 106453.0 | 0 | Canidae | 106453.0 | 0 | Canis | 106453.0 | 0 | Canis lupus | 106453.0 | 0 |
| scf7180000045989 | 12780 | 0.4418 | 0 | 117.1897 | Eukaryota | 3055.0 | 0 | Chordata | 3055.0 | 0 | Carnivora | 2726.0 | 1 | Canidae | 2726.0 | 1 | Canis | 2726.0 | 1 | Canis lupus | 2726.0 | 1 |
| scf7180000056128 | 4735 | 0.4437 | 0 | 148.0247 | Eukaryota | 4347.0 | 0 | Chordata | 4347.0 | 0 | Columbiformes | 2463.0 | 1 | Columbidae | 2463.0 | 1 | Streptopelia | 2463.0 | 1 | Streptopelia turtur | 2463.0 | 1 |
| scf7180000043829 | 35383 | 0.3992 | 0 | 57.4152 | Eukaryota | 42615.0 | 0 | Chordata | 42615.0 | 0 | Carnivora | 42615.0 | 0 | Canidae | 42615.0 | 0 | Canis | 42615.0 | 0 | Canis lupus | 42615.0 | 0 |
| scf7180000048532 | 13726 | 0.4372 | 0 | 383.2746 | Eukaryota | 2974.0 | 0 | Chordata | 2974.0 | 0 | Carnivora | 2974.0 | 0 | Canidae | 2974.0 | 0 | Canis | 2974.0 | 0 | Canis lupus | 2974.0 | 0 |
| scf7180000041281 | 1262525 | 0.3901 | 0 | 60.4621 | Eukaryota | 104979.0 | 0 | Chordata | 104979.0 | 0 | Carnivora | 104979.0 | 0 | Canidae | 104979.0 | 0 | Canis | 104979.0 | 0 | Canis lupus | 104979.0 | 0 |
| scf7180000055546 | 22035 | 0.4397 | 0 | 203.6042 | Eukaryota | 6556.0 | 0 | Platyhelminthes | 2487.0 | 3 | Polystomatidea | 2487.0 | 4 | Polystomatidae | 2487.0 | 4 | Protopolystoma | 2487.0 | 5 | Protopolystoma xenopodis | 2487.0 | 5 |
| scf7180000046709 | 120387 | 0.4366 | 0 | 60.4067 | Eukaryota | 16932.0 | 0 | Chordata | 16932.0 | 0 | Carnivora | 16932.0 | 0 | Canidae | 16932.0 | 0 | Canis | 14570.0 | 1 | Canis lupus | 14570.0 | 1 |
| scf7180000044251 | 7294 | 0.5457 | 0 | 46.8599 | Eukaryota | 3182.0 | 0 | Chordata | 3182.0 | 0 | Carnivora | 3182.0 | 0 | Canidae | 3182.0 | 0 | Canis | 3182.0 | 0 | Canis lupus | 3182.0 | 0 |
| scf7180000052862 | 3254 | 0.4582 | 0 | 440.2274 | no-hit | 0.0 | 0 | no-hit | 0.0 | 0 | no-hit | 0.0 | 0 | no-hit | 0.0 | 0 | no-hit | 0.0 | 0 | no-hit | 0.0 | 0 |
| scf7180000043256 | 10204 | 0.4255 | 0 | 64.6082 | Eukaryota | 57561.0 | 0 | Chordata | 57561.0 | 0 | Carnivora | 57561.0 | 0 | Canidae | 57561.0 | 0 | Canis | 57561.0 | 0 | Canis lupus | 57561.0 | 0 |
| scf7180000056244 | 751775 | 0.392 | 0 | 59.9298 | Eukaryota | 185151.0 | 0 | Chordata | 185151.0 | 0 | Carnivora | 185151.0 | 0 | Canidae | 89889.0 | 5 | Canis | 70556.0 | 8 | Canis lupus | 70556.0 | 8 |
| scf7180000045119 | 258098 | 0.3406 | 0 | 58.9019 | Eukaryota | 26573.0 | 0 | Chordata | 26573.0 | 0 | Carnivora | 26573.0 | 0 | Canidae | 26573.0 | 0 | Canis | 26573.0 | 0 | Canis lupus | 26573.0 | 0 |
| scf7180000052838 | 17065 | 0.5037 | 0 | 309.2729 | Eukaryota | 8733.0 | 0 | Chordata | 8733.0 | 0 | Carnivora | 8733.0 | 0 | Canidae | 8733.0 | 0 | Canis | 7543.0 | 1 | Canis lupus | 6303.0 | 2 |
| scf7180000053482 | 5856 | 0.4462 | 0 | 140.9292 | no-hit | 0.0 | 0 | no-hit | 0.0 | 0 | no-hit | 0.0 | 0 | no-hit | 0.0 | 0 | no-hit | 0.0 | 0 | no-hit | 0.0 | 0 |
| scf7180000045074 | 14806 | 0.3918 | 0 | 49.685 | Eukaryota | 16600.0 | 0 | Chordata | 16600.0 | 0 | Carnivora | 16600.0 | 0 | Canidae | 16600.0 | 0 | Canis | 14932.0 | 1 | Canis lupus | 14932.0 | 1 |
| scf7180000040562 | 2140043 | 0.5009 | 0 | 58.9489 | Eukaryota | 113353.0 | 0 | Chordata | 113353.0 | 0 | Carnivora | 113353.0 | 0 | Canidae | 113353.0 | 0 | Vulpes | 63264.0 | 1 | Vulpes vulpes | 63264.0 | 1 |
| scf7180000049201 | 33818 | 0.5335 | 0 | 56.4134 | Eukaryota | 12234.0 | 0 | Chordata | 12234.0 | 0 | Carnivora | 12234.0 | 0 | Canidae | 10301.0 | 1 | Canis | 8895.0 | 3 | Canis lupus | 8895.0 | 3 |
| scf7180000050721 | 8800 | 0.4351 | 0 | 52.428 | Eukaryota | 1083.0 | 0 | Chordata | 1083.0 | 0 | Kurtiformes | 595.0 | 1 | Apogonidae | 595.0 | 1 | Sphaeramia | 595.0 | 1 | Sphaeramia orbicularis | 595.0 | 1 |
| scf7180000042879 | 3616 | 0.4477 | 0 | 321.7103 | Eukaryota | 519.0 | 0 | Streptophyta | 519.0 | 0 | Fabales | 519.0 | 0 | Fabaceae | 519.0 | 0 | Cicer | 519.0 | 0 | Cicer arietinum | 519.0 | 0 |
| scf7180000043123 | 541704 | 0.4755 | 0 | 58.8885 | Eukaryota | 58968.0 | 0 | Chordata | 58968.0 | 0 | Carnivora | 58968.0 | 0 | Canidae | 54351.0 | 1 | Canis | 42800.0 | 2 | Canis lupus | 42800.0 | 2 |
| scf7180000041339 | 225801 | 0.3953 | 0 | 65.3838 | Eukaryota | 90944.0 | 0 | Chordata | 90944.0 | 0 | Carnivora | 90944.0 | 0 | Canidae | 90944.0 | 0 | Canis | 90944.0 | 0 | Canis lupus | 90944.0 | 0 |
| scf7180000043522 | 462482 | 0.3622 | 0 | 59.0991 | Eukaryota | 90073.0 | 0 | Chordata | 90073.0 | 0 | Carnivora | 90073.0 | 0 | Canidae | 90073.0 | 0 | Canis | 90073.0 | 0 | Canis lupus | 90073.0 | 0 |
| scf7180000053716 | 3606 | 0.4498 | 0 | 194.3336 | Eukaryota | 159.0 | 0 | Chordata | 159.0 | 0 | Tetraodontiformes | 159.0 | 0 | Tetraodontidae | 159.0 | 0 | Takifugu | 159.0 | 0 | Takifugu rubripes | 159.0 | 0 |
| scf7180000046602 | 121078 | 0.4765 | 0 | 61.3289 | Eukaryota | 9745.0 | 0 | Chordata | 9745.0 | 0 | Carnivora | 9745.0 | 0 | Canidae | 8808.0 | 1 | Canis | 8808.0 | 1 | Canis lupus | 8808.0 | 1 |
| scf7180000043568 | 380970 | 0.4842 | 0 | 59.8422 | Eukaryota | 41944.0 | 0 | Chordata | 41944.0 | 0 | Carnivora | 41944.0 | 0 | Canidae | 41944.0 | 0 | Canis | 33464.0 | 1 | Canis lupus | 33464.0 | 1 |
| scf7180000047987 | 61159 | 0.436 | 0 | 101.3829 | Eukaryota | 89435.0 | 0 | Chordata | 89435.0 | 0 | Carnivora | 89435.0 | 0 | Canidae | 89435.0 | 0 | Canis | 89435.0 | 0 | Canis lupus | 89435.0 | 0 |
| scf7180000047916 | 27489 | 0.4422 | 0 | 96.0825 | Eukaryota | 15654.0 | 0 | Chordata | 10281.0 | 2 | Gadiformes | 4599.0 | 7 | Gadidae | 4599.0 | 7 | Gadus | 4599.0 | 7 | Gadus morhua | 4599.0 | 7 |
| scf7180000050453 | 5077 | 0.4188 | 0 | 783.2936 | no-hit | 0.0 | 0 | no-hit | 0.0 | 0 | no-hit | 0.0 | 0 | no-hit | 0.0 | 0 | no-hit | 0.0 | 0 | no-hit | 0.0 | 0 |
| scf7180000045715 | 46142 | 0.3803 | 0 | 54.4156 | Eukaryota | 12557.0 | 0 | Chordata | 12557.0 | 0 | Carnivora | 12557.0 | 0 | Canidae | 9637.0 | 2 | Canis | 5522.0 | 4 | Canis lupus | 5522.0 | 4 |
| scf7180000049299 | 3540 | 0.437 | 0 | 138.6359 | Eukaryota | 1501.0 | 0 | Chordata | 1251.0 | 1 | Kurtiformes | 957.0 | 2 | Apogonidae | 957.0 | 2 | Sphaeramia | 957.0 | 2 | Sphaeramia orbicularis | 957.0 | 2 |
| scf7180000042613 | 363665 | 0.3971 | 0 | 60.0223 | Eukaryota | 112686.0 | 0 | Chordata | 112686.0 | 0 | Carnivora | 112686.0 | 0 | Canidae | 112686.0 | 0 | Canis | 101391.0 | 1 | Canis lupus | 101391.0 | 1 |
| scf7180000052973 | 11078 | 0.499 | 0 | 92.3025 | Eukaryota | 8393.0 | 0 | Chordata | 8393.0 | 0 | Carnivora | 8393.0 | 0 | Canidae | 8393.0 | 0 | Canis | 6852.0 | 1 | Canis lupus | 5735.0 | 2 |
| scf7180000051369 | 9172 | 0.4403 | 0 | 308.9876 | Eukaryota | 3038.0 | 0 | Platyhelminthes | 2273.0 | 1 | Polystomatidea | 2273.0 | 2 | Polystomatidae | 2273.0 | 2 | Protopolystoma | 2273.0 | 2 | Protopolystoma xenopodis | 2273.0 | 2 |
| scf7180000044836 | 88920 | 0.3733 | 0 | 55.9884 | Eukaryota | 36501.0 | 0 | Chordata | 36501.0 | 0 | Carnivora | 36501.0 | 0 | Canidae | 36501.0 | 0 | Canis | 36501.0 | 0 | Canis lupus | 36501.0 | 0 |
| scf7180000042570 | 280292 | 0.4077 | 0 | 58.2968 | Eukaryota | 17532.0 | 0 | Chordata | 17532.0 | 0 | Carnivora | 11637.0 | 2 | Canidae | 10604.0 | 3 | Canis | 10604.0 | 4 | Canis lupus | 10604.0 | 4 |
| scf7180000051726 | 6171 | 0.4515 | 0 | 708.315 | Eukaryota | 16643.0 | 0 | Chordata | 15369.0 | 2 | Gadiformes | 9834.0 | 7 | Gadidae | 9834.0 | 7 | Gadus | 9834.0 | 7 | Gadus morhua | 9834.0 | 7 |
| scf7180000048285 | 4495 | 0.3831 | 0 | 29.5531 | Eukaryota | 4020.0 | 0 | Chordata | 4020.0 | 0 | Carnivora | 4020.0 | 0 | Canidae | 1922.0 | 2 | Canis | 1922.0 | 5 | Canis lupus | 1922.0 | 5 |
| scf7180000044730 | 241815 | 0.4427 | 0 | 61.1571 | Eukaryota | 31496.0 | 0 | Chordata | 31496.0 | 0 | Carnivora | 31496.0 | 0 | Canidae | 31496.0 | 0 | Canis | 28269.0 | 1 | Canis lupus | 28269.0 | 1 |
| scf7180000048776 | 14476 | 0.4469 | 0 | 160.2814 | Eukaryota | 2318.0 | 0 | Chordata | 1336.0 | 1 | Polystomatidea | 982.0 | 3 | Polystomatidae | 982.0 | 3 | Protopolystoma | 982.0 | 3 | Protopolystoma xenopodis | 982.0 | 3 |
| scf7180000047603 | 78068 | 0.4012 | 0 | 62.0179 | Eukaryota | 7226.0 | 0 | Chordata | 7226.0 | 0 | Carnivora | 7226.0 | 0 | Canidae | 6516.0 | 1 | Canis | 6516.0 | 1 | Canis lupus | 6516.0 | 1 |
| scf7180000041079 | 625027 | 0.3796 | 0 | 60.3621 | Eukaryota | 85711.0 | 0 | Chordata | 85711.0 | 0 | Carnivora | 85711.0 | 0 | Canidae | 85711.0 | 0 | Canis | 68909.0 | 1 | Canis lupus | 68909.0 | 1 |
| scf7180000055334 | 26064 | 0.4383 | 0 | 582.111 | Eukaryota | 4553.0 | 0 | Chordata | 2473.0 | 1 | Polystomatidea | 2080.0 | 3 | Polystomatidae | 2080.0 | 3 | Protopolystoma | 2080.0 | 3 | Protopolystoma xenopodis | 2080.0 | 3 |
| scf7180000041984 | 789229 | 0.3947 | 0 | 59.6795 | Eukaryota | 25156.0 | 0 | Chordata | 25156.0 | 0 | Carnivora | 12255.0 | 2 | Hominidae | 7818.0 | 3 | Homo | 7818.0 | 5 | Homo sapiens | 7818.0 | 5 |
| scf7180000043651 | 146415 | 0.5297 | 0 | 55.5875 | Eukaryota | 48975.0 | 0 | Chordata | 48975.0 | 0 | Carnivora | 44199.0 | 1 | Canidae | 44199.0 | 1 | Canis | 39103.0 | 2 | Canis lupus | 39103.0 | 2 |
| scf7180000050797 | 32784 | 0.4413 | 0 | 157.8937 | Eukaryota | 94673.0 | 0 | Chordata | 94673.0 | 0 | Carnivora | 94673.0 | 0 | Canidae | 94673.0 | 0 | Canis | 94673.0 | 0 | Canis lupus | 94673.0 | 0 |
| scf7180000055166 | 24136 | 0.4426 | 0 | 153.3104 | no-hit | 0.0 | 0 | no-hit | 0.0 | 0 | no-hit | 0.0 | 0 | no-hit | 0.0 | 0 | no-hit | 0.0 | 0 | no-hit | 0.0 | 0 |
| scf7180000042958 | 316486 | 0.3989 | 0 | 60.9073 | Eukaryota | 81205.0 | 0 | Chordata | 81205.0 | 0 | Carnivora | 81205.0 | 0 | Canidae | 81205.0 | 0 | Canis | 81205.0 | 0 | Canis lupus | 81205.0 | 0 |
| scf7180000044260 | 6920 | 0.4205 | 0 | 43.2067 | Eukaryota | 27556.0 | 0 | Chordata | 27556.0 | 0 | Carnivora | 27556.0 | 0 | Canidae | 27556.0 | 0 | Canis | 27556.0 | 0 | Canis lupus | 27556.0 | 0 |
| scf7180000045072 | 74360 | 0.3798 | 0 | 56.8011 | Eukaryota | 24840.0 | 0 | Chordata | 24840.0 | 0 | Carnivora | 24840.0 | 0 | Canidae | 24840.0 | 0 | Canis | 24840.0 | 0 | Canis lupus | 24840.0 | 0 |
| scf7180000045206 | 210512 | 0.366 | 0 | 58.9487 | Eukaryota | 91612.0 | 0 | Chordata | 91612.0 | 0 | Carnivora | 91612.0 | 0 | Canidae | 91612.0 | 0 | Canis | 91612.0 | 0 | Canis lupus | 91612.0 | 0 |
| scf7180000043944 | 76026 | 0.4202 | 0 | 57.6374 | Eukaryota | 13541.0 | 0 | Chordata | 13541.0 | 0 | Carnivora | 13541.0 | 0 | Canidae | 13541.0 | 0 | Canis | 13541.0 | 0 | Canis lupus | 13541.0 | 0 |
| scf7180000041206 | 1374630 | 0.3549 | 0 | 58.5362 | Eukaryota | 27562.0 | 0 | Chordata | 27562.0 | 0 | Carnivora | 27562.0 | 0 | Canidae | 27562.0 | 0 | Canis | 27562.0 | 0 | Canis lupus | 27562.0 | 0 |
| scf7180000055583 | 15860 | 0.4561 | 0 | 80.4116 | Eukaryota | 9246.0 | 0 | Chordata | 9246.0 | 0 | Carnivora | 9246.0 | 0 | Canidae | 9246.0 | 0 | Canis | 5496.0 | 1 | Canis lupus | 5496.0 | 1 |
| scf7180000054045 | 4219 | 0.4416 | 0 | 66.2726 | Eukaryota | 3005.0 | 0 | Chordata | 3005.0 | 0 | Carnivora | 3005.0 | 0 | Canidae | 3005.0 | 0 | Canis | 3005.0 | 0 | Canis lupus | 3005.0 | 0 |
| scf7180000048347 | 48750 | 0.6326 | 0 | 54.0613 | Eukaryota | 24759.0 | 0 | Chordata | 24759.0 | 0 | Carnivora | 24759.0 | 0 | Felidae | 14020.0 | 1 | Felis | 9340.0 | 3 | Felis catus | 9340.0 | 3 |
| scf7180000055733 | 55746 | 0.4442 | 0 | 212.0287 | Eukaryota | 26348.0 | 0 | Platyhelminthes | 20366.0 | 1 | Strigeidida | 19719.0 | 2 | Schistosomatidae | 19719.0 | 2 | Schistosoma | 19719.0 | 3 | Schistosoma mansoni | 19719.0 | 3 |
| scf7180000050902 | 9743 | 0.4452 | 0 | 189.2543 | Eukaryota | 147.0 | 0 | Chordata | 147.0 | 0 | Kurtiformes | 147.0 | 0 | Apogonidae | 147.0 | 0 | Sphaeramia | 147.0 | 0 | Sphaeramia orbicularis | 147.0 | 0 |
| scf7180000056127 | 13712 | 0.4345 | 0 | 93.4542 | Eukaryota | 840.0 | 0 | Chordata | 840.0 | 0 | Carangiformes | 237.0 | 3 | Echeneidae | 237.0 | 3 | Echeneis | 237.0 | 3 | Echeneis naucrates | 237.0 | 3 |
| scf7180000044189 | 5466 | 0.4083 | 0 | 205.7798 | no-hit | 0.0 | 0 | no-hit | 0.0 | 0 | no-hit | 0.0 | 0 | no-hit | 0.0 | 0 | no-hit | 0.0 | 0 | no-hit | 0.0 | 0 |
| scf7180000048613 | 9608 | 0.445 | 0 | 117.8266 | Eukaryota | 1999.0 | 0 | Platyhelminthes | 1782.0 | 1 | Polystomatidea | 1782.0 | 1 | Polystomatidae | 1782.0 | 1 | Protopolystoma | 1782.0 | 1 | Protopolystoma xenopodis | 1782.0 | 1 |
| scf7180000048799 | 25722 | 0.4379 | 0 | 180.4338 | Eukaryota | 265.0 | 0 | Chordata | 265.0 | 0 | Kurtiformes | 265.0 | 0 | Apogonidae | 265.0 | 0 | Sphaeramia | 265.0 | 0 | Sphaeramia orbicularis | 265.0 | 0 |
| scf7180000056361 | 569139 | 0.4764 | 0 | 58.179 | Eukaryota | 66428.0 | 0 | Chordata | 66428.0 | 0 | Carnivora | 66428.0 | 0 | Canidae | 66428.0 | 0 | Canis | 66428.0 | 0 | Canis lupus | 66428.0 | 0 |
| scf7180000056315 | 525930 | 0.3978 | 0 | 60.2171 | Eukaryota | 42981.0 | 0 | Chordata | 42981.0 | 0 | Carnivora | 42981.0 | 0 | Canidae | 34518.0 | 2 | Vulpes | 21709.0 | 3 | Vulpes vulpes | 21709.0 | 3 |
| scf7180000049044 | 8690 | 0.4455 | 0 | 129.7646 | no-hit | 0.0 | 0 | no-hit | 0.0 | 0 | no-hit | 0.0 | 0 | no-hit | 0.0 | 0 | no-hit | 0.0 | 0 | no-hit | 0.0 | 0 |
| scf7180000048257 | 12687 | 0.4432 | 0 | 140.5284 | Eukaryota | 638.0 | 0 | Chordata | 377.0 | 1 | Kurtiformes | 377.0 | 1 | Apogonidae | 377.0 | 1 | Sphaeramia | 377.0 | 1 | Sphaeramia orbicularis | 377.0 | 1 |
| scf7180000051015 | 18880 | 0.5041 | 0 | 744.874 | Eukaryota | 9525.0 | 0 | Chordata | 9525.0 | 0 | Carnivora | 9525.0 | 0 | Canidae | 9525.0 | 0 | Canis | 7774.0 | 1 | Canis lupus | 6490.0 | 2 |
| scf7180000051900 | 28356 | 0.4103 | 0 | 57.791 | Eukaryota | 63242.0 | 0 | Chordata | 63242.0 | 0 | Carnivora | 63242.0 | 0 | Felidae | 29343.0 | 2 | Vulpes | 27792.0 | 3 | Vulpes vulpes | 27792.0 | 3 |
| scf7180000050849 | 4946 | 0.4515 | 0 | 583.0441 | Eukaryota | 870.0 | 0 | Streptophyta | 572.0 | 1 | Fabales | 572.0 | 1 | Fabaceae | 572.0 | 1 | Cicer | 572.0 | 1 | Cicer arietinum | 572.0 | 1 |
| scf7180000051234 | 13812 | 0.5372 | 0 | 47.279 | Eukaryota | 23591.0 | 0 | Chordata | 23591.0 | 0 | Carnivora | 23591.0 | 0 | Canidae | 20126.0 | 1 | Canis | 20126.0 | 2 | Canis lupus | 20126.0 | 2 |
| scf7180000043687 | 420606 | 0.4668 | 0 | 59.9984 | Eukaryota | 147970.0 | 0 | Chordata | 147970.0 | 0 | Carnivora | 147970.0 | 0 | Canidae | 147970.0 | 0 | Canis | 131850.0 | 1 | Canis lupus | 131850.0 | 1 |
| scf7180000043264 | 602067 | 0.3881 | 0 | 60.4793 | Eukaryota | 69972.0 | 0 | Chordata | 69972.0 | 0 | Carnivora | 69972.0 | 0 | Canidae | 47141.0 | 1 | Canis | 39314.0 | 4 | Canis lupus | 39314.0 | 5 |
| scf7180000046909 | 65669 | 0.4139 | 0 | 58.2993 | Eukaryota | 37156.0 | 0 | Chordata | 37156.0 | 0 | Carnivora | 37156.0 | 0 | Canidae | 37156.0 | 0 | Canis | 37156.0 | 0 | Canis lupus | 37156.0 | 0 |
| scf7180000052255 | 4661 | 0.4452 | 0 | 991.2123 | no-hit | 0.0 | 0 | no-hit | 0.0 | 0 | no-hit | 0.0 | 0 | no-hit | 0.0 | 0 | no-hit | 0.0 | 0 | no-hit | 0.0 | 0 |
| scf7180000055798 | 36426 | 0.4961 | 0 | 106.5989 | Eukaryota | 9463.0 | 0 | Chordata | 9463.0 | 0 | Carnivora | 9463.0 | 0 | Canidae | 9463.0 | 0 | Canis | 8820.0 | 1 | Canis lupus | 7602.0 | 2 |
| scf7180000047648 | 73647 | 0.4367 | 0 | 107.1495 | Eukaryota | 38359.0 | 0 | Chordata | 29636.0 | 1 | Gadiformes | 22670.0 | 6 | Gadidae | 22670.0 | 6 | Gadus | 22670.0 | 6 | Gadus morhua | 22670.0 | 6 |
| scf7180000053484 | 2973 | 0.4517 | 0 | 435.7098 | Eukaryota | 2518.0 | 0 | Platyhelminthes | 1463.0 | 2 | Polystomatidea | 1463.0 | 2 | Polystomatidae | 1463.0 | 2 | Protopolystoma | 1463.0 | 2 | Protopolystoma xenopodis | 1463.0 | 2 |
| scf7180000052447 | 2890 | 0.4408 | 0 | 441.11 | no-hit | 0.0 | 0 | no-hit | 0.0 | 0 | no-hit | 0.0 | 0 | no-hit | 0.0 | 0 | no-hit | 0.0 | 0 | no-hit | 0.0 | 0 |
| scf7180000047370 | 101583 | 0.451 | 0 | 59.6424 | Eukaryota | 7216.0 | 0 | Chordata | 7216.0 | 0 | Carnivora | 3063.0 | 3 | Muridae | 2700.0 | 4 | Rattus | 2700.0 | 5 | Rattus norvegicus | 2700.0 | 5 |
| scf7180000042499 | 27200 | 0.5084 | 0 | 58.3147 | Eukaryota | 8712.0 | 0 | Chordata | 8712.0 | 0 | Carnivora | 8712.0 | 0 | Canidae | 8712.0 | 0 | Canis | 8712.0 | 0 | Canis lupus | 8712.0 | 0 |
| scf7180000052254 | 4964 | 0.4519 | 0 | 495.4574 | no-hit | 0.0 | 0 | no-hit | 0.0 | 0 | no-hit | 0.0 | 0 | no-hit | 0.0 | 0 | no-hit | 0.0 | 0 | no-hit | 0.0 | 0 |
| scf7180000040712 | 452587 | 0.3729 | 0 | 59.4546 | Eukaryota | 78824.0 | 0 | Chordata | 78824.0 | 0 | Carnivora | 78824.0 | 0 | Canidae | 78824.0 | 0 | Canis | 78824.0 | 0 | Canis lupus | 78824.0 | 0 |
| scf7180000041556 | 428467 | 0.4235 | 0 | 58.2159 | Eukaryota | 36026.0 | 0 | Chordata | 36026.0 | 0 | Carnivora | 36026.0 | 0 | Canidae | 36026.0 | 0 | Canis | 36026.0 | 0 | Canis lupus | 36026.0 | 0 |
| scf7180000048160 | 53020 | 0.3989 | 0 | 58.5392 | Eukaryota | 15878.0 | 0 | Chordata | 15878.0 | 0 | Carnivora | 15878.0 | 0 | Canidae | 15878.0 | 0 | Canis | 15878.0 | 0 | Canis lupus | 15878.0 | 0 |
| scf7180000043442 | 332766 | 0.374 | 0 | 60.404 | Eukaryota | 30598.0 | 0 | Chordata | 30598.0 | 0 | Carnivora | 19495.0 | 3 | Canidae | 15716.0 | 4 | Canis | 11427.0 | 5 | Canis lupus | 11427.0 | 6 |
| scf7180000054912 | 32653 | 0.4449 | 0 | 133.4441 | Eukaryota | 1225.0 | 0 | Chordata | 890.0 | 1 | Kurtiformes | 460.0 | 3 | Apogonidae | 460.0 | 3 | Sphaeramia | 460.0 | 3 | Sphaeramia orbicularis | 460.0 | 3 |
| scf7180000044902 | 103273 | 0.3633 | 0 | 59.2157 | Eukaryota | 60278.0 | 0 | Chordata | 60278.0 | 0 | Carnivora | 60278.0 | 0 | Canidae | 60278.0 | 0 | Canis | 60278.0 | 0 | Canis lupus | 60278.0 | 0 |
| scf7180000049161 | 27556 | 0.4398 | 0 | 95.7262 | no-hit | 0.0 | 0 | no-hit | 0.0 | 0 | no-hit | 0.0 | 0 | no-hit | 0.0 | 0 | no-hit | 0.0 | 0 | no-hit | 0.0 | 0 |
| scf7180000053059 | 2835 | 0.4589 | 0 | 279.0345 | Eukaryota | 1818.0 | 0 | Echinodermata | 915.0 | 2 | Forcipulatida | 915.0 | 2 | Asteriidae | 915.0 | 2 | Asterias | 915.0 | 2 | Asterias rubens | 915.0 | 2 |
| scf7180000049249 | 11505 | 0.4505 | 0 | 81.6051 | no-hit | 0.0 | 0 | no-hit | 0.0 | 0 | no-hit | 0.0 | 0 | no-hit | 0.0 | 0 | no-hit | 0.0 | 0 | no-hit | 0.0 | 0 |
| scf7180000051758 | 4492 | 0.4495 | 0 | 458.9731 | no-hit | 0.0 | 0 | no-hit | 0.0 | 0 | no-hit | 0.0 | 0 | no-hit | 0.0 | 0 | no-hit | 0.0 | 0 | no-hit | 0.0 | 0 |
| scf7180000055811 | 26440 | 0.4367 | 0 | 203.0173 | Eukaryota | 4978.0 | 0 | Chordata | 4512.0 | 1 | Tetraodontiformes | 2353.0 | 2 | Tetraodontidae | 2353.0 | 2 | Takifugu | 2353.0 | 2 | Canis lupus | 2159.0 | 3 |
| scf7180000053138 | 10235 | 0.5024 | 0 | 285.6678 | Eukaryota | 8264.0 | 0 | Chordata | 8264.0 | 0 | Carnivora | 8264.0 | 0 | Canidae | 8264.0 | 0 | Canis | 6684.0 | 1 | Canis lupus | 5541.0 | 2 |
| scf7180000052877 | 3338 | 0.453 | 0 | 274.2501 | Eukaryota | 18645.0 | 0 | Chordata | 12817.0 | 2 | Gadiformes | 10472.0 | 5 | Gadidae | 10472.0 | 5 | Gadus | 10472.0 | 5 | Gadus morhua | 10472.0 | 5 |
| scf7180000046552 | 43844 | 0.4221 | 0 | 55.2824 | Eukaryota | 21463.0 | 0 | Chordata | 21463.0 | 0 | Carnivora | 21463.0 | 0 | Canidae | 21463.0 | 0 | Canis | 21463.0 | 0 | Canis lupus | 21463.0 | 0 |
| scf7180000054424 | 33485 | 0.386 | 0 | 1034.7988 | Eukaryota | 22579.0 | 0 | Chordata | 22579.0 | 0 | Carnivora | 22579.0 | 0 | Canidae | 22579.0 | 0 | Canis | 22579.0 | 0 | Canis lupus | 22579.0 | 0 |
| scf7180000040856 | 49461 | 0.4307 | 0 | 56.2409 | Eukaryota | 45828.0 | 0 | Chordata | 45828.0 | 0 | Carnivora | 45828.0 | 0 | Canidae | 45828.0 | 0 | Canis | 45828.0 | 0 | Canis lupus | 45828.0 | 0 |
| scf7180000052575 | 5714 | 0.4543 | 0 | 407.2908 | no-hit | 0.0 | 0 | no-hit | 0.0 | 0 | no-hit | 0.0 | 0 | no-hit | 0.0 | 0 | no-hit | 0.0 | 0 | no-hit | 0.0 | 0 |
| scf7180000046020 | 82966 | 0.4995 | 0 | 75.869 | Eukaryota | 34506.0 | 0 | Chordata | 34506.0 | 0 | Carnivora | 14446.0 | 2 | Cricetidae | 10085.0 | 3 | Cricetulus | 10085.0 | 4 | Cricetulus griseus | 10085.0 | 4 |
| scf7180000042356 | 337045 | 0.4129 | 100 | 59.444 | Eukaryota | 48028.0 | 0 | Chordata | 48028.0 | 0 | Carnivora | 48028.0 | 0 | Canidae | 48028.0 | 0 | Canis | 39096.0 | 1 | Canis lupus | 39096.0 | 1 |
| scf7180000043548 | 435537 | 0.4236 | 0 | 59.5508 | Eukaryota | 56552.0 | 0 | Chordata | 56552.0 | 0 | Carnivora | 56552.0 | 0 | Canidae | 56552.0 | 0 | Canis | 51301.0 | 1 | Canis lupus | 51301.0 | 1 |
| scf7180000043079 | 145069 | 0.3909 | 0 | 60.297 | Eukaryota | 67493.0 | 0 | Chordata | 67493.0 | 0 | Carnivora | 67493.0 | 0 | Canidae | 67493.0 | 0 | Canis | 67493.0 | 0 | Canis lupus | 67493.0 | 0 |
| scf7180000041314 | 42682 | 0.5539 | 0 | 57.7263 | Eukaryota | 11869.0 | 0 | Chordata | 11869.0 | 0 | Carnivora | 6428.0 | 2 | Felidae | 4809.0 | 4 | Homo | 4375.0 | 6 | Homo sapiens | 4375.0 | 6 |
| scf7180000048826 | 60753 | 0.3945 | 0 | 58.6051 | Eukaryota | 35358.0 | 0 | Chordata | 35358.0 | 0 | Carnivora | 35358.0 | 0 | Canidae | 35358.0 | 0 | Canis | 35358.0 | 0 | Canis lupus | 35358.0 | 0 |
| scf7180000051459 | 4358 | 0.455 | 0 | 747.2162 | no-hit | 0.0 | 0 | no-hit | 0.0 | 0 | no-hit | 0.0 | 0 | no-hit | 0.0 | 0 | no-hit | 0.0 | 0 | no-hit | 0.0 | 0 |
| scf7180000041307 | 1264181 | 0.3984 | 0 | 60.6711 | Eukaryota | 131784.0 | 0 | Chordata | 131784.0 | 0 | Carnivora | 131784.0 | 0 | Canidae | 131784.0 | 0 | Canis | 131784.0 | 0 | Canis lupus | 131784.0 | 0 |
| scf7180000043268 | 521279 | 0.3502 | 0 | 59.6504 | Eukaryota | 92885.0 | 0 | Chordata | 92885.0 | 0 | Carnivora | 92885.0 | 0 | Canidae | 92885.0 | 0 | Canis | 92885.0 | 0 | Canis lupus | 92885.0 | 0 |
| scf7180000049309 | 14912 | 0.4476 | 0 | 1656.0454 | Eukaryota | 24950.0 | 0 | Chordata | 24950.0 | 0 | Gadiformes | 15689.0 | 4 | Gadidae | 15689.0 | 4 | Gadus | 15689.0 | 4 | Gadus morhua | 15689.0 | 4 |
| scf7180000043985 | 3221 | 0.4508 | 0 | 186.0143 | Eukaryota | 1158.0 | 0 | Chordata | 720.0 | 1 | Fabales | 438.0 | 3 | Fabaceae | 438.0 | 3 | Cicer | 438.0 | 3 | Cicer arietinum | 438.0 | 3 |
| scf7180000045281 | 44355 | 0.4158 | 0 | 61.7941 | Eukaryota | 92354.0 | 0 | Chordata | 92354.0 | 0 | Carnivora | 92354.0 | 0 | Canidae | 92354.0 | 0 | Canis | 92354.0 | 0 | Canis lupus | 92354.0 | 0 |
| scf7180000050552 | 22265 | 0.4119 | 0 | 110.2347 | Eukaryota | 9539.0 | 0 | Chordata | 9539.0 | 0 | Primates | 5308.0 | 2 | Hominidae | 3744.0 | 3 | Canis | 2432.0 | 5 | Canis lupus | 2432.0 | 5 |
| scf7180000046119 | 137386 | 0.3921 | 0 | 58.6791 | Eukaryota | 72964.0 | 0 | Chordata | 72964.0 | 0 | Carnivora | 72964.0 | 0 | Canidae | 47467.0 | 4 | Canis | 42148.0 | 5 | Canis lupus | 42148.0 | 5 |
| scf7180000045219 | 34433 | 0.3963 | 0 | 71.9901 | Eukaryota | 81036.0 | 0 | Chordata | 81036.0 | 0 | Carnivora | 81036.0 | 0 | Canidae | 81036.0 | 0 | Canis | 81036.0 | 0 | Canis lupus | 81036.0 | 0 |
| scf7180000051747 | 18972 | 0.4423 | 0 | 52.4016 | Eukaryota | 4570.0 | 0 | Chordata | 3591.0 | 1 | Kurtiformes | 3354.0 | 2 | Apogonidae | 3354.0 | 2 | Sphaeramia | 3354.0 | 2 | Sphaeramia orbicularis | 3354.0 | 2 |
| scf7180000041200 | 799014 | 0.3423 | 0 | 58.8625 | Eukaryota | 89749.0 | 0 | Chordata | 89749.0 | 0 | Carnivora | 89749.0 | 0 | Canidae | 89749.0 | 0 | Canis | 89749.0 | 0 | Canis lupus | 89749.0 | 0 |
| scf7180000055346 | 32257 | 0.348 | 0 | 154.4472 | Eukaryota | 13634.0 | 0 | Chordata | 13634.0 | 0 | Carnivora | 13634.0 | 0 | Canidae | 13634.0 | 0 | Canis | 9525.0 | 1 | Canis lupus | 9525.0 | 1 |
| scf7180000047151 | 6307 | 0.4441 | 0 | 42.4964 | Eukaryota | 3087.0 | 0 | Chordata | 3087.0 | 0 | Carnivora | 3087.0 | 0 | Canidae | 3087.0 | 0 | Canis | 3087.0 | 0 | Canis lupus | 3087.0 | 0 |
| scf7180000052229 | 11730 | 0.4419 | 0 | 231.1903 | no-hit | 0.0 | 0 | no-hit | 0.0 | 0 | no-hit | 0.0 | 0 | no-hit | 0.0 | 0 | no-hit | 0.0 | 0 | no-hit | 0.0 | 0 |
| scf7180000052109 | 7522 | 0.4543 | 0 | 735.889 | no-hit | 0.0 | 0 | no-hit | 0.0 | 0 | no-hit | 0.0 | 0 | no-hit | 0.0 | 0 | no-hit | 0.0 | 0 | no-hit | 0.0 | 0 |
| scf7180000050751 | 21775 | 0.5334 | 0 | 361.3594 | Eukaryota | 23265.0 | 0 | Chordata | 23265.0 | 0 | Artiodactyla | 16344.0 | 1 | Bovidae | 16344.0 | 1 | Ovis | 9330.0 | 3 | Ovis aries | 9330.0 | 3 |
| scf7180000050867 | 6807 | 0.4534 | 0 | 965.5246 | Eukaryota | 1336.0 | 0 | Echinodermata | 512.0 | 2 | Forcipulatida | 512.0 | 2 | Asteriidae | 512.0 | 2 | Asterias | 512.0 | 2 | Asterias rubens | 512.0 | 2 |
| scf7180000044307 | 315330 | 0.5295 | 0 | 56.6211 | Eukaryota | 201259.0 | 0 | Chordata | 201259.0 | 0 | Carnivora | 201259.0 | 0 | Canidae | 193331.0 | 1 | Canis | 157663.0 | 2 | Canis lupus | 157663.0 | 2 |
| scf7180000042681 | 25718 | 0.3297 | 0 | 55.9785 | Eukaryota | 18049.0 | 0 | Chordata | 18049.0 | 0 | Carnivora | 18049.0 | 0 | Canidae | 18049.0 | 0 | Canis | 18049.0 | 0 | Canis lupus | 18049.0 | 0 |
| scf7180000055071 | 25522 | 0.441 | 0 | 140.4833 | Eukaryota | 83030.0 | 0 | Chordata | 83030.0 | 0 | Carnivora | 83030.0 | 0 | Canidae | 83030.0 | 0 | Canis | 83030.0 | 0 | Canis lupus | 83030.0 | 0 |
| scf7180000056069 | 24123 | 0.4715 | 0 | 1008.9583 | Eukaryota | 56630.0 | 0 | Chordata | 56630.0 | 0 | Carnivora | 56630.0 | 0 | Canidae | 56630.0 | 0 | Canis | 56630.0 | 0 | Canis lupus | 56630.0 | 0 |
| scf7180000045333 | 51180 | 0.4383 | 0 | 55.7018 | Eukaryota | 17214.0 | 0 | Chordata | 17214.0 | 0 | Carnivora | 14599.0 | 2 | Canidae | 14599.0 | 2 | Canis | 14599.0 | 2 | Canis lupus | 14599.0 | 2 |
| scf7180000054103 | 7264 | 0.4489 | 0 | 164.8035 | Eukaryota | 342.0 | 0 | Chordata | 342.0 | 0 | Salmoniformes | 342.0 | 0 | Salmonidae | 342.0 | 0 | Coregonus | 342.0 | 0 | Coregonus sp. 'balchen' | 342.0 | 0 |
| scf7180000056223 | 408789 | 0.4087 | 0 | 58.0051 | Eukaryota | 85587.0 | 0 | Chordata | 85587.0 | 0 | Carnivora | 85587.0 | 0 | Canidae | 85587.0 | 0 | Canis | 85587.0 | 0 | Canis lupus | 85587.0 | 0 |
| scf7180000040624 | 475116 | 0.3799 | 0 | 146.9911 | Eukaryota | 95324.0 | 0 | Chordata | 95324.0 | 0 | Carnivora | 95324.0 | 0 | Canidae | 95324.0 | 0 | Canis | 95324.0 | 0 | Canis lupus | 95324.0 | 0 |
| scf7180000046866 | 109041 | 0.4096 | 0 | 58.9435 | Eukaryota | 16652.0 | 0 | Chordata | 16652.0 | 0 | Primates | 6528.0 | 3 | Hominidae | 6528.0 | 3 | Canis | 5589.0 | 5 | Canis lupus | 5589.0 | 5 |
| scf7180000046782 | 62301 | 0.4441 | 0 | 143.387 | Eukaryota | 4701.0 | 0 | Chordata | 2889.0 | 2 | unresolved | 1496.0 | 6 | unresolved | 1496.0 | 6 | unresolved | 1496.0 | 6 | unresolved | 1496.0 | 6 |
| scf7180000053056 | 3632 | 0.4433 | 0 | 249.1845 | no-hit | 0.0 | 0 | no-hit | 0.0 | 0 | no-hit | 0.0 | 0 | no-hit | 0.0 | 0 | no-hit | 0.0 | 0 | no-hit | 0.0 | 0 |
| scf7180000045551 | 11292 | 0.4327 | 0 | 33.4951 | no-hit | 0.0 | 0 | no-hit | 0.0 | 0 | no-hit | 0.0 | 0 | no-hit | 0.0 | 0 | no-hit | 0.0 | 0 | no-hit | 0.0 | 0 |
| scf7180000053002 | 4024 | 0.4476 | 0 | 475.4928 | Eukaryota | 2685.0 | 0 | Platyhelminthes | 1542.0 | 2 | Polystomatidea | 1542.0 | 2 | Polystomatidae | 1542.0 | 2 | Protopolystoma | 1542.0 | 2 | Protopolystoma xenopodis | 1542.0 | 2 |
| scf7180000042119 | 857616 | 0.4073 | 0 | 60.6036 | Eukaryota | 39302.0 | 0 | Chordata | 39302.0 | 0 | Carnivora | 39302.0 | 0 | Otariidae | 22811.0 | 2 | Zalophus | 15196.0 | 5 | Zalophus californianus | 15196.0 | 5 |
| scf7180000055290 | 76416 | 0.4394 | 0 | 72.6227 | Eukaryota | 95795.0 | 0 | Chordata | 95795.0 | 0 | Carnivora | 95795.0 | 0 | Canidae | 95795.0 | 0 | Canis | 95795.0 | 0 | Canis lupus | 95795.0 | 0 |

| | | | | | | | | | | | | | | | | | | | | | | |
|---|---|---|---|---|---|---|---|---|---|---|---|---|---|---|---|---|---|---|---|---|---|---|
| scf7180000048604 | 48248 | 0.3769 | 0 | 62.3228 | Eukaryota | 39149.0 | 0 | Chordata | 39149.0 | 0 | Carnivora | 39149.0 | 0 | Felidae | 20168.0 | 3 | Canis | 11248.0 | 5 | Canis lupus | 11248.0 | 5 |
| scf7180000046174 | 159122 | 0.4859 | 0 | 59.8893 | Eukaryota | 26240.0 | 0 | Chordata | 26240.0 | 0 | Carnivora | 26240.0 | 0 | Canidae | 22009.0 | 3 | Canis | 17308.0 | 4 | Canis lupus | 17308.0 | 4 |
| scf7180000043340 | 21501 | 0.4033 | 0 | 43.1276 | Eukaryota | 75413.0 | 0 | Chordata | 75413.0 | 0 | Carnivora | 75413.0 | 0 | Canidae | 75413.0 | 0 | Canis | 75413.0 | 0 | Canis lupus | 75413.0 | 0 |
| scf7180000054995 | 27974 | 0.4621 | 0 | 449.7816 | Eukaryota | 15328.0 | 0 | Chordata | 15328.0 | 0 | Carnivora | 13664.0 | 1 | Canidae | 13664.0 | 1 | Canis | 8856.0 | 2 | Canis lupus | 8856.0 | 2 |
| scf7180000047038 | 130212 | 0.3833 | 0 | 64.8447 | Eukaryota | 90882.0 | 0 | Chordata | 90882.0 | 0 | Carnivora | 90882.0 | 0 | Canidae | 90882.0 | 0 | Canis | 90882.0 | 0 | Canis lupus | 90882.0 | 0 |
| scf7180000051481 | 3722 | 0.4436 | 0 | 250.9195 | no-hit | 0.0 | 0 | no-hit | 0.0 | 0 | no-hit | 0.0 | 0 | no-hit | 0.0 | 0 | no-hit | 0.0 | 0 | no-hit | 0.0 | 0 |
| scf7180000056205 | 1385707 | 0.3688 | 0 | 58.6824 | Eukaryota | 46642.0 | 0 | Chordata | 46642.0 | 0 | Carnivora | 46642.0 | 0 | Canidae | 46642.0 | 0 | Canis | 35628.0 | 1 | Canis lupus | 35628.0 | 1 |
| scf7180000042060 | 818964 | 0.4046 | 0 | 58.8506 | Eukaryota | 131087.0 | 0 | Chordata | 131087.0 | 0 | Carnivora | 131087.0 | 0 | Canidae | 131087.0 | 0 | Canis | 131087.0 | 0 | Canis lupus | 131087.0 | 0 |
| scf7180000051678 | 3835 | 0.4558 | 0 | 34.266 | Eukaryota | 3109.0 | 0 | Chordata | 2124.0 | 1 | Tetraodontiformes | 1674.0 | 3 | Tetraodontidae | 1674.0 | 3 | Takifugu | 1674.0 | 3 | Takifugu rubripes | 1674.0 | 3 |
| scf7180000044696 | 265602 | 0.47 | 0 | 59.1928 | Eukaryota | 59155.0 | 0 | Chordata | 59155.0 | 0 | Carnivora | 59155.0 | 0 | Canidae | 59155.0 | 0 | Vulpes | 29745.0 | 1 | Vulpes vulpes | 29745.0 | 1 |
| scf7180000043846 | 338850 | 0.5125 | 0 | 59.8705 | Eukaryota | 74928.0 | 0 | Chordata | 74928.0 | 0 | Carnivora | 74928.0 | 0 | Canidae | 74928.0 | 0 | Canis | 51233.0 | 1 | Canis lupus | 51233.0 | 1 |
| scf7180000050232 | 10921 | 0.414 | 0 | 57.4953 | Eukaryota | 20177.0 | 0 | Chordata | 20177.0 | 0 | Carnivora | 7659.0 | 2 | Hominidae | 5884.0 | 4 | Homo | 5884.0 | 4 | Homo sapiens | 5884.0 | 4 |
| scf7180000040959 | 184050 | 0.4348 | 0 | 59.8247 | Eukaryota | 36912.0 | 0 | Chordata | 36912.0 | 0 | Carnivora | 36912.0 | 0 | Canidae | 16334.0 | 3 | Canis | 11179.0 | 5 | Canis lupus | 11179.0 | 5 |
| scf7180000042549 | 155699 | 0.4279 | 0 | 82.3661 | Eukaryota | 43928.0 | 0 | Chordata | 43928.0 | 0 | Carnivora | 43928.0 | 0 | Canidae | 43928.0 | 0 | Canis | 43928.0 | 0 | Canis lupus | 43928.0 | 0 |
| scf7180000046125 | 137545 | 0.4219 | 0 | 57.7807 | Eukaryota | 24156.0 | 0 | Chordata | 24156.0 | 0 | Carnivora | 24156.0 | 0 | Canidae | 24156.0 | 0 | Canis | 24156.0 | 0 | Canis lupus | 24156.0 | 0 |
| scf7180000052974 | 8355 | 0.4369 | 0 | 64.1435 | Eukaryota | 28916.0 | 0 | Chordata | 28916.0 | 0 | Carnivora | 28916.0 | 0 | Otariidae | 25427.0 | 1 | Callorhinus | 11265.0 | 3 | Callorhinus ursinus | 11265.0 | 3 |
| scf7180000044657 | 56864 | 0.4376 | 0 | 163.0023 | Eukaryota | 19299.0 | 0 | Chordata | 17677.0 | 1 | Tetraodontiformes | 10565.0 | 2 | Tetraodontidae | 10565.0 | 2 | Takifugu | 10565.0 | 2 | Takifugu rubripes | 10565.0 | 2 |
| scf7180000041810 | 932277 | 0.436 | 0 | 58.6285 | Eukaryota | 53490.0 | 0 | Chordata | 53490.0 | 0 | Carnivora | 53490.0 | 0 | Canidae | 53490.0 | 0 | Canis | 37461.0 | 1 | Canis lupus | 37461.0 | 1 |
| scf7180000041793 | 982374 | 0.463 | 0 | 57.8831 | Eukaryota | 59015.0 | 0 | Chordata | 59015.0 | 0 | Carnivora | 59015.0 | 0 | Canidae | 59015.0 | 0 | Canis | 33242.0 | 1 | Canis lupus | 33242.0 | 1 |
| scf7180000050258 | 26690 | 0.4568 | 0 | 66.5803 | Eukaryota | 1298.0 | 0 | Platyhelminthes | 651.0 | 1 | Polystomatidea | 651.0 | 1 | Polystomatidae | 651.0 | 1 | Protopolystoma | 651.0 | 1 | Protopolystoma xenopodis | 651.0 | 1 |
| scf7180000055456 | 17686 | 0.5176 | 0 | 340.9862 | Eukaryota | 15866.0 | 0 | Chordata | 15866.0 | 0 | Carnivora | 15866.0 | 0 | Canidae | 15866.0 | 0 | Canis | 15866.0 | 0 | Canis lupus | 15866.0 | 0 |
| scf7180000049468 | 35303 | 0.404 | 0 | 58.7578 | Eukaryota | 9860.0 | 0 | Chordata | 9860.0 | 0 | Primates | 7710.0 | 2 | Hominidae | 7710.0 | 2 | Pan | 4075.0 | 3 | Pan troglodytes | 4075.0 | 3 |
| scf7180000054187 | 212263 | 0.3953 | 0 | 171.9197 | Eukaryota | 87514.0 | 0 | Chordata | 87514.0 | 0 | Carnivora | 87514.0 | 0 | Canidae | 87514.0 | 0 | Canis | 87514.0 | 0 | Canis lupus | 87514.0 | 0 |
| scf7180000045468 | 20977 | 0.4953 | 0 | 185.6086 | Eukaryota | 9231.0 | 0 | Chordata | 9231.0 | 0 | Carnivora | 9231.0 | 0 | Canidae | 9231.0 | 0 | Canis | 8588.0 | 1 | Canis lupus | 7357.0 | 2 |
| scf7180000051778 | 4478 | 0.4395 | 0 | 309.9604 | no-hit | 0.0 | 0 | no-hit | 0.0 | 0 | no-hit | 0.0 | 0 | no-hit | 0.0 | 0 | no-hit | 0.0 | 0 | no-hit | 0.0 | 0 |
| scf7180000051508 | 22780 | 0.4438 | 0 | 112.0262 | Eukaryota | 3125.0 | 0 | Chordata | 2673.0 | 1 | Tetraodontiformes | 2673.0 | 2 | Tetraodontidae | 2673.0 | 2 | Takifugu | 2673.0 | 2 | Takifugu rubripes | 2673.0 | 2 |
| scf7180000044530 | 7639 | 0.4165 | 0 | 111.7823 | Eukaryota | 67921.0 | 0 | Chordata | 67921.0 | 0 | Carnivora | 67921.0 | 0 | Canidae | 67921.0 | 0 | Canis | 67921.0 | 0 | Canis lupus | 67921.0 | 0 |
| scf7180000050704 | 14189 | 0.4396 | 0 | 38.3752 | Eukaryota | 1933.0 | 0 | Platyhelminthes | 1933.0 | 0 | Polystomatidea | 1933.0 | 0 | Polystomatidae | 1933.0 | 0 | Protopolystoma | 1933.0 | 0 | Protopolystoma xenopodis | 1933.0 | 0 |
| scf7180000042754 | 553393 | 0.4108 | 0 | 58.7849 | Eukaryota | 32801.0 | 0 | Chordata | 32801.0 | 0 | Carnivora | 32801.0 | 0 | Canidae | 32801.0 | 0 | Canis | 32801.0 | 0 | Canis lupus | 32801.0 | 0 |
| scf7180000040766 | 1795544 | 0.384 | 0 | 59.631 | Eukaryota | 94567.0 | 0 | Chordata | 94567.0 | 0 | Carnivora | 94567.0 | 0 | Canidae | 94567.0 | 0 | Canis | 94567.0 | 0 | Canis lupus | 94567.0 | 0 |
| scf7180000052486 | 7664 | 0.5034 | 0 | 70.5719 | Eukaryota | 8157.0 | 0 | Chordata | 8157.0 | 0 | Carnivora | 8157.0 | 0 | Canidae | 8157.0 | 0 | Canis | 6682.0 | 1 | Canis lupus | 5639.0 | 2 |
| scf7180000044018 | 369583 | 0.442 | 0 | 59.5695 | Eukaryota | 38517.0 | 0 | Chordata | 38517.0 | 0 | Carnivora | 38517.0 | 0 | Canidae | 38517.0 | 0 | Canis | 38517.0 | 0 | Canis lupus | 38517.0 | 0 |
| scf7180000041119 | 1524345 | 0.5114 | 0 | 58.5281 | Eukaryota | 65244.0 | 0 | Chordata | 65244.0 | 0 | Carnivora | 65244.0 | 0 | Canidae | 65244.0 | 0 | Canis | 52040.0 | 1 | Canis lupus | 52040.0 | 1 |
| scf7180000045690 | 164963 | 0.4594 | 0 | 58.9849 | Eukaryota | 21015.0 | 0 | Chordata | 21015.0 | 0 | Carnivora | 21015.0 | 0 | Canidae | 8651.0 | 3 | Canis | 7672.0 | 6 | Canis lupus | 7672.0 | 7 |
| scf7180000051122 | 7623 | 0.4482 | 0 | 700.3322 | no-hit | 0.0 | 0 | no-hit | 0.0 | 0 | no-hit | 0.0 | 0 | no-hit | 0.0 | 0 | no-hit | 0.0 | 0 | no-hit | 0.0 | 0 |
| scf7180000044498 | 329107 | 0.3962 | 0 | 60.3596 | Eukaryota | 85730.0 | 0 | Chordata | 85730.0 | 0 | Carnivora | 85730.0 | 0 | Canidae | 85730.0 | 0 | Canis | 85730.0 | 0 | Canis lupus | 85730.0 | 0 |
| scf7180000052021 | 4087 | 0.439 | 0 | 221.093 | Eukaryota | 4563.0 | 0 | Chordata | 1883.0 | 3 | Polystomatidea | 1627.0 | 4 | Polystomatidae | 1627.0 | 4 | Protopolystoma | 1627.0 | 4 | Protopolystoma xenopodis | 1627.0 | 4 |
| scf7180000056281 | 833015 | 0.4867 | 0 | 58.93 | Eukaryota | 81846.0 | 0 | Chordata | 81846.0 | 0 | Carnivora | 81846.0 | 0 | Canidae | 81846.0 | 0 | Canis | 63297.0 | 1 | Canis lupus | 63297.0 | 1 |
| scf7180000052644 | 2287 | 0.4609 | 0 | 223.5235 | Eukaryota | 999.0 | 0 | Platyhelminthes | 398.0 | 2 | Polystomatidea | 398.0 | 2 | Polystomatidae | 398.0 | 2 | Protopolystoma | 398.0 | 2 | Protopolystoma xenopodis | 398.0 | 2 |
| scf7180000051795 | 21070 | 0.4436 | 0 | 61.263 | Eukaryota | 16288.0 | 0 | Platyhelminthes | 11450.0 | 1 | Cyclophyllidea | 10067.0 | 4 | Hymenolepididae | 10067.0 | 4 | Hymenolepis | 10067.0 | 4 | Hymenolepis microstoma | 10067.0 | 4 |
| scf7180000045231 | 253589 | 0.3872 | 0 | 59.0718 | Eukaryota | 109353.0 | 0 | Chordata | 109353.0 | 0 | Carnivora | 109353.0 | 0 | Canidae | 109353.0 | 0 | Canis | 78361.0 | 1 | Canis lupus | 78361.0 | 1 |
| scf7180000050125 | 15856 | 0.4518 | 0 | 1412.605 | Eukaryota | 3824.0 | 0 | Platyhelminthes | 1960.0 | 3 | Polystomatidea | 1960.0 | 4 | Polystomatidae | 1960.0 | 4 | Protopolystoma | 1960.0 | 4 | Protopolystoma xenopodis | 1960.0 | 4 |
| scf7180000055429 | 56931 | 0.442 | 0 | 277.1024 | Eukaryota | 81907.0 | 0 | Platyhelminthes | 54479.0 | 1 | Strigeidida | 42290.0 | 5 | Schistosomatidae | 42290.0 | 5 | Schistosoma | 42290.0 | 5 | Schistosoma mansoni | 42290.0 | 5 |
| scf7180000051701 | 7303 | 0.4546 | 0 | 324.4975 | Eukaryota | 3421.0 | 0 | Platyhelminthes | 1561.0 | 3 | Polystomatidea | 1561.0 | 4 | Polystomatidae | 1561.0 | 4 | Protopolystoma | 1561.0 | 4 | Protopolystoma xenopodis | 1561.0 | 4 |
| scf7180000047513 | 33850 | 0.4393 | 0 | 123.4556 | Eukaryota | 3361.0 | 0 | Chordata | 2806.0 | 1 | Kurtiformes | 1374.0 | 3 | Apogonidae | 1374.0 | 3 | Sphaeramia | 1374.0 | 3 | Sphaeramia orbicularis | 1374.0 | 4 |
| scf7180000050791 | 5884 | 0.4262 | 0 | 28.149 | Eukaryota | 3097.0 | 0 | Chordata | 3097.0 | 0 | Carnivora | 3097.0 | 0 | Canidae | 3097.0 | 0 | Canis | 3097.0 | 0 | Canis lupus | 3097.0 | 0 |
| scf7180000051781 | 5448 | 0.4396 | 0 | 358.2774 | Eukaryota | 1632.0 | 0 | Platyhelminthes | 1632.0 | 0 | Polystomatidea | 1632.0 | 0 | Polystomatidae | 1632.0 | 0 | Protopolystoma | 1632.0 | 0 | Protopolystoma xenopodis | 1632.0 | 0 |
| scf7180000052464 | 3624 | 0.449 | 0 | 274.5429 | Eukaryota | 1949.0 | 0 | Platyhelminthes | 1125.0 | 2 | Polystomatidea | 1125.0 | 2 | Polystomatidae | 1125.0 | 2 | Protopolystoma | 1125.0 | 2 | Protopolystoma xenopodis | 1125.0 | 2 |
| scf7180000047193 | 35371 | 0.4459 | 0 | 55.544 | Eukaryota | 4114.0 | 0 | Chordata | 2082.0 | 1 | Polystomatidea | 2032.0 | 2 | Polystomatidae | 2032.0 | 2 | Protopolystoma | 2032.0 | 2 | Protopolystoma xenopodis | 2032.0 | 2 |
| scf7180000044911 | 287786 | 0.3954 | 0 | 59.4113 | Eukaryota | 30995.0 | 0 | Chordata | 30995.0 | 0 | Carnivora | 30995.0 | 0 | Felidae | 19126.0 | 1 | Canis | 11869.0 | 4 | Canis lupus | 11869.0 | 4 |
| scf7180000044040 | 128520 | 0.4078 | 0 | 57.6187 | Eukaryota | 70940.0 | 0 | Chordata | 70940.0 | 0 | Carnivora | 70940.0 | 0 | Canidae | 70940.0 | 0 | Canis | 70940.0 | 0 | Canis lupus | 70940.0 | 0 |
| scf7180000047735 | 82298 | 0.3791 | 0 | 57.5276 | Eukaryota | 34688.0 | 0 | Chordata | 34688.0 | 0 | Carnivora | 32891.0 | 1 | Canidae | 31522.0 | 2 | Canis | 24705.0 | 3 | Canis lupus | 24705.0 | 3 |
| scf7180000048121 | 2481 | 0.3853 | 0 | 29.0005 | Eukaryota | 3343.0 | 0 | Chordata | 3343.0 | 0 | Carnivora | 3343.0 | 0 | Canidae | 3343.0 | 0 | Canis | 3343.0 | 0 | Canis lupus | 3343.0 | 0 |
| scf7180000048885 | 14780 | 0.4427 | 0 | 132.9348 | Eukaryota | 81824.0 | 0 | Chordata | 55011.0 | 1 | Gadiformes | 55011.0 | 2 | Gadidae | 55011.0 | 2 | Gadus | 55011.0 | 2 | Gadus morhua | 55011.0 | 2 |
| scf7180000052313 | 2706 | 0.4557 | 0 | 2207.718 | no-hit | 0.0 | 0 | no-hit | 0.0 | 0 | no-hit | 0.0 | 0 | no-hit | 0.0 | 0 | no-hit | 0.0 | 0 | no-hit | 0.0 | 0 |
| scf7180000045478 | 175941 | 0.4894 | 0 | 58.8763 | Eukaryota | 43594.0 | 0 | Chordata | 43594.0 | 0 | Carnivora | 43594.0 | 0 | Canidae | 37248.0 | 1 | Canis | 29420.0 | 2 | Canis lupus | 29420.0 | 2 |
| scf7180000053759 | 3518 | 0.446 | 0 | 101.6685 | Eukaryota | 1017.0 | 0 | Platyhelminthes | 688.0 | 1 | Polystomatidea | 532.0 | 2 | Polystomatidae | 532.0 | 2 | Protopolystoma | 532.0 | 2 | Protopolystoma xenopodis | 532.0 | 2 |
| scf7180000044314 | 343726 | 0.381 | 0 | 59.6242 | Eukaryota | 39651.0 | 0 | Chordata | 39651.0 | 0 | Carnivora | 39651.0 | 0 | Canidae | 39651.0 | 0 | Canis | 39651.0 | 0 | Canis lupus | 39651.0 | 0 |
| scf7180000044132 | 387252 | 0.354 | 0 | 61.0755 | Eukaryota | 91384.0 | 0 | Chordata | 91384.0 | 0 | Carnivora | 91384.0 | 0 | Canidae | 91384.0 | 0 | Canis | 91384.0 | 0 | Canis lupus | 91384.0 | 0 |
| scf7180000042116 | 237180 | 0.4012 | 0 | 58.9369 | Eukaryota | 50258.0 | 0 | Chordata | 50258.0 | 0 | Carnivora | 50258.0 | 0 | Canidae | 50258.0 | 0 | Canis | 50258.0 | 0 | Canis lupus | 50258.0 | 0 |
| scf7180000042558 | 593461 | 0.3848 | 0 | 61.1921 | Eukaryota | 63273.0 | 0 | Chordata | 63273.0 | 0 | Carnivora | 63273.0 | 0 | Canidae | 63273.0 | 0 | Canis | 63273.0 | 0 | Canis lupus | 63273.0 | 0 |
| scf7180000046105 | 16739 | 0.442 | 0 | 162.7703 | Eukaryota | 4639.0 | 0 | Platyhelminthes | 3025.0 | 1 | Polystomatidea | 3025.0 | 3 | Polystomatidae | 3025.0 | 3 | Protopolystoma | 3025.0 | 3 | Protopolystoma xenopodis | 3025.0 | 3 |
| scf7180000048901 | 13735 | 0.4831 | 0 | 283.1517 | Eukaryota | 9618.0 | 0 | Chordata | 9618.0 | 0 | Carnivora | 9618.0 | 0 | Canidae | 9618.0 | 0 | Canis | 5606.0 | 1 | Canis lupus | 5606.0 | 1 |
| scf7180000042221 | 540362 | 0.3767 | 0 | 59.3716 | Eukaryota | 40976.0 | 0 | Chordata | 40976.0 | 0 | Carnivora | 40976.0 | 0 | Canidae | 40976.0 | 0 | Canis | 40976.0 | 0 | Canis lupus | 40976.0 | 0 |
| scf7180000050326 | 6043 | 0.4387 | 0 | 50.4676 | Eukaryota | 16672.0 | 1 | Chordata | 12507.0 | 3 | Gadiformes | 7216.0 | 6 | Gadidae | 7216.0 | 6 | Gadus | 7216.0 | 6 | Gadus morhua | 7216.0 | 6 |
| scf7180000043381 | 84790 | 0.4 | 0 | 57.5509 | Eukaryota | 58848.0 | 0 | Chordata | 58848.0 | 0 | Carnivora | 58848.0 | 0 | Canidae | 58848.0 | 0 | Canis | 58848.0 | 0 | Canis lupus | 58848.0 | 0 |
| scf7180000056605 | 880445 | 0.4755 | 0 | 57.2833 | Eukaryota | 63918.0 | 0 | Chordata | 63918.0 | 0 | Carnivora | 63918.0 | 0 | Canidae | 63918.0 | 0 | Canis | 44277.0 | 1 | Canis lupus | 44277.0 | 1 |
| scf7180000043177 | 237669 | 0.4098 | 0 | 60.3983 | Eukaryota | 90999.0 | 0 | Chordata | 90999.0 | 0 | Carnivora | 90999.0 | 0 | Canidae | 90999.0 | 0 | Canis | 90999.0 | 0 | Canis lupus | 90999.0 | 0 |
| scf7180000055699 | 38395 | 0.4418 | 0 | 66.0457 | Eukaryota | 2020.0 | 0 | Chordata | 1521.0 | 1 | Kurtiformes | 1521.0 | 1 | Apogonidae | 1521.0 | 1 | Sphaeramia | 1521.0 | 1 | Sphaeramia orbicularis | 1521.0 | 1 |
| scf7180000041540 | 37174 | 0.4721 | 0 | 57.2985 | Eukaryota | 17461.0 | 0 | Chordata | 17461.0 | 0 | Carnivora | 17461.0 | 0 | Canidae | 9043.0 | 2 | Canis | 6738.0 | 5 | Canis lupus | 6738.0 | 5 |
| scf7180000054743 | 14103 | 0.7266 | 0 | 326.458 | Eukaryota | 39932.0 | 0 | Chordata | 39932.0 | 0 | Primates | 28742.0 | 1 | Cercopithecidae | 18581.0 | 3 | Macaca | 18581.0 | 3 | Macaca mulatta | 18581.0 | 3 |
| scf7180000048678 | 7838 | 0.4481 | 0 | 31.8716 | Eukaryota | 89467.0 | 0 | Chordata | 89467.0 | 0 | Carnivora | 89467.0 | 0 | Canidae | 89467.0 | 0 | Canis | 89467.0 | 0 | Canis lupus | 89467.0 | 0 |
| scf7180000047130 | 24078 | 0.4449 | 0 | 198.811 | Eukaryota | 9129.0 | 0 | Chordata | 6802.0 | 1 | Carnivora | 3160.0 | 3 | Hymenolepididae | 2327.0 | 5 | Hymenolepis | 2327.0 | 5 | Hymenolepis microstoma | 2327.0 | 6 |
| scf7180000041290 | 1610241 | 0.4703 | 0 | 57.9068 | Eukaryota | 155653.0 | 0 | Chordata | 155653.0 | 0 | Carnivora | 155653.0 | 0 | Canidae | 155653.0 | 0 | Canis | 113603.0 | 1 | Canis lupus | 113603.0 | 1 |
| scf7180000044268 | 288569 | 0.4311 | 0 | 57.315 | Eukaryota | 79708.0 | 0 | Chordata | 79708.0 | 0 | Carnivora | 79708.0 | 0 | Canidae | 74616.0 | 1 | Canis | 53975.0 | 2 | Canis lupus | 53975.0 | 2 |
| scf7180000048134 | 61498 | 0.5445 | 0 | 56.6955 | Eukaryota | 4311.0 | 0 | Chordata | 4311.0 | 0 | Primates | 1863.0 | 2 | Hominidae | 1863.0 | 2 | Homo | 1863.0 | 3 | Homo sapiens | 1863.0 | 3 |
| scf7180000050285 | 8182 | 0.4495 | 0 | 57.9941 | no-hit | 0.0 | 0 | no-hit | 0.0 | 0 | no-hit | 0.0 | 0 | no-hit | 0.0 | 0 | no-hit | 0.0 | 0 | no-hit | 0.0 | 0 |
| scf7180000042951 | 39012 | 0.4231 | 0 | 52.2877 | Eukaryota | 23645.0 | 0 | Chordata | 23645.0 | 0 | Carnivora | 23645.0 | 0 | Canidae | 18147.0 | 1 | Canis | 18147.0 | 2 | Canis lupus | 18147.0 | 2 |
| scf7180000049690 | 14933 | 0.4448 | 0 | 74.0505 | no-hit | 0.0 | 0 | no-hit | 0.0 | 0 | no-hit | 0.0 | 0 | no-hit | 0.0 | 0 | no-hit | 0.0 | 0 | no-hit | 0.0 | 0 |
| scf7180000048530 | 23681 | 0.4379 | 0 | 315.7318 | Eukaryota | 92926.0 | 0 | Chordata | 54191.0 | 1 | Gadiformes | 54191.0 | 2 | Gadidae | 54191.0 | 2 | Gadus | 54191.0 | 2 | Gadus morhua | 54191.0 | 2 |
| scf7180000048259 | 80266 | 0.3733 | 0 | 61.43 | Eukaryota | 14356.0 | 0 | Chordata | 14356.0 | 0 | Carnivora | 6715.0 | 3 | Canidae | 6715.0 | 3 | Canis | 6715.0 | 4 | Canis lupus | 6715.0 | 4 |
| scf7180000054070 | 11013 | 0.4422 | 0 | 166.591 | Eukaryota | 3492.0 | 0 | Platyhelminthes | 1788.0 | 1 | Kurtiformes | 1704.0 | 2 | Apogonidae | 1704.0 | 2 | Sphaeramia | 1704.0 | 2 | Sphaeramia orbicularis | 1704.0 | 2 |
| scf7180000051460 | 5983 | 0.4499 | 0 | 1042.503 | no-hit | 0.0 | 0 | no-hit | 0.0 | 0 | no-hit | 0.0 | 0 | no-hit | 0.0 | 0 | no-hit | 0.0 | 0 | no-hit | 0.0 | 0 |
| scf7180000045382 | 189738 | 0.4861 | 0 | 58.6614 | Eukaryota | 86972.0 | 0 | Chordata | 86972.0 | 0 | Carnivora | 86972.0 | 0 | Canidae | 86972.0 | 0 | Canis | 76134.0 | 1 | Canis lupus | 76134.0 | 1 |
| scf7180000053117 | 3700 | 0.4432 | 0 | 388.2523 | Eukaryota | 267.0 | 0 | Platyhelminthes | 267.0 | 0 | Polystomatidea | 267.0 | 0 | Polystomatidae | 267.0 | 0 | Protopolystoma | 267.0 | 0 | Protopolystoma xenopodis | 267.0 | 0 |
| scf7180000044224 | 378911 | 0.3615 | 0 | 59.0291 | Eukaryota | 29200.0 | 0 | Chordata | 29200.0 | 0 | Carnivora | 29200.0 | 0 | Canidae | 29200.0 | 0 | Canis | 29200.0 | 0 | Canis lupus | 29200.0 | 0 |
| scf7180000045635 | 175824 | 0.5166 | 0 | 54.8152 | Eukaryota | 27305.0 | 0 | Chordata | 27305.0 | 0 | Carnivora | 27305.0 | 0 | Canidae | 27305.0 | 0 | Canis | 27305.0 | 0 | Canis lupus | 27305.0 | 0 |
| scf7180000041702 | 97519 | 0.361 | 0 | 54.4323 | Eukaryota | 84767.0 | 0 | Chordata | 84767.0 | 0 | Carnivora | 84767.0 | 0 | Canidae | 84767.0 | 0 | Canis | 84767.0 | 0 | Canis lupus | 84767.0 | 0 |
| scf7180000047619 | 101641 | 0.4236 | 0 | 59.5674 | Eukaryota | 26179.0 | 0 | Chordata | 26179.0 | 0 | Carnivora | 26179.0 | 0 | Canidae | 26179.0 | 0 | Canis | 26179.0 | 0 | Canis lupus | 26179.0 | 0 |
| scf7180000051072 | 4270 | 0.4253 | 0 | 31.5674 | Eukaryota | 8825.0 | 0 | Chordata | 8825.0 | 0 | Carnivora | 8825.0 | 0 | Canidae | 8825.0 | 0 | Canis | 8825.0 | 0 | Canis lupus | 8825.0 | 0 |
| scf7180000047469 | 55563 | 0.4429 | 0 | 56.3002 | Eukaryota | 7244.0 | 0 | Chordata | 6283.0 | 1 | Kurtiformes | 3680.0 | 3 | Apogonidae | 3680.0 | 3 | Sphaeramia | 3680.0 | 4 | Sphaeramia orbicularis | 3680.0 | 4 |
| scf7180000046074 | 8133 | 0.4402 | 0 | 54.3601 | Eukaryota | 3137.0 | 0 | Chordata | 2564.0 | 1 | Carnivora | 2050.0 | 2 | Canidae | 2050.0 | 2 | Canis | 2050.0 | 2 | Canis lupus | 2050.0 | 2 |
| scf7180000042836 | 524406 | 0.3848 | 0 | 57.3116 | Eukaryota | 127743.0 | 0 | Chordata | 127743.0 | 0 | Carnivora | 119004.0 | 1 | Canidae | 84335.0 | 2 | Canis | 84335.0 | 2 | Canis lupus | 84335.0 | 2 |
| scf7180000049037 | 49282 | 0.4351 | 0 | 59.3994 | Eukaryota | 7034.0 | 0 | Chordata | 7034.0 | 0 | Primates | 3884.0 | 2 | Hominidae | 3884.0 | 3 | Homo | 3884.0 | 3 | Homo sapiens | 3884.0 | 3 |
| scf7180000055763 | 17075 | 0.5195 | 0 | 411.373 | Eukaryota | 15936.0 | 0 | Chordata | 15936.0 | 0 | Carnivora | 15936.0 | 0 | Canidae | 15936.0 | 0 | Canis | 15936.0 | 0 | Canis lupus | 15936.0 | 0 |
| scf7180000049518 | 60010 | 0.4729 | 0 | 58.5089 | Eukaryota | 30507.0 | 0 | Chordata | 30507.0 | 0 | Carnivora | 30507.0 | 0 | Canidae | 19894.0 | 2 | Canis | 14701.0 | 4 | Canis lupus | 14701.0 | 4 |
| scf7180000050991 | 17846 | 0.4683 | 0 | 69.3779 | Eukaryota | 7097.0 | 0 | Chordata | 7097.0 | 0 | Carnivora | 7097.0 | 0 | Canidae | 7097.0 | 0 | Canis | 5826.0 | 1 | Canis lupus | 5061.0 | 2 |
| scf7180000046974 | 128072 | 0.3665 | 0 | 63.042 | Eukaryota | 89158.0 | 0 | Chordata | 89158.0 | 0 | Carnivora | 89158.0 | 0 | Canidae | 89158.0 | 0 | Canis | 89158.0 | 0 | Canis lupus | 89158.0 | 0 |
| scf7180000055185 | 28752 | 0.4477 | 0 | 181.5607 | Eukaryota | 42311.0 | 0 | Chordata | 25736.0 | 1 | Strigeidida | 16575.0 | 6 | Schistosomatidae | 16575.0 | 6 | Schistosoma | 16575.0 | 6 | Schistosoma mansoni | 16575.0 | 6 |
| scf7180000056478 | 110353 | 0.4071 | 0 | 60.7811 | Eukaryota | 51835.0 | 0 | Chordata | 51835.0 | 0 | Carnivora | 51835.0 | 0 | Canidae | 51835.0 | 0 | Canis | 51835.0 | 0 | Canis lupus | 51835.0 | 0 |
| scf7180000041478 | 1051101 | 0.4201 | 0 | 75.0969 | Eukaryota | 89600.0 | 0 | Chordata | 89600.0 | 0 | Carnivora | 89600.0 | 0 | Canidae | 89600.0 | 0 | Canis | 89600.0 | 0 | Canis lupus | 89600.0 | 0 |
| scf7180000053353 | 1950 | 0.4215 | 0 | 436.8993 | no-hit | 0.0 | 0 | no-hit | 0.0 | 0 | no-hit | 0.0 | 0 | no-hit | 0.0 | 0 | no-hit | 0.0 | 0 | no-hit | 0.0 | 0 |
| scf7180000046642 | 108039 | 0.414 | 0 | 59.9369 | Eukaryota | 26093.0 | 1 | Chordata | 26093.0 | 1 | Carnivora | 14238.0 | 3 | Canidae | 9779.0 | 4 | Canis | 9779.0 | 6 | Canis lupus | 9779.0 | 6 |
| scf7180000053185 | 11778 | 0.4306 | 0 | 71.7047 | Eukaryota | 45855.0 | 0 | Chordata | 45855.0 | 0 | Carnivora | 45855.0 | 0 | Canidae | 45855.0 | 0 | Canis | 45855.0 | 0 | Canis lupus | 45855.0 | 0 |
| scf7180000056594 | 27337 | 0.4499 | 100 | 59.2386 | Eukaryota | 88448.0 | 0 | Chordata | 88448.0 | 0 | Carnivora | 88448.0 | 0 | Canidae | 88448.0 | 0 | Canis | 88448.0 | 0 | Canis lupus | 88448.0 | 0 |
| scf7180000042844 | 567809 | 0.4328 | 0 | 60.0632 | Eukaryota | 53690.0 | 0 | Chordata | 53690.0 | 0 | Carnivora | 53690.0 | 0 | Canidae | 53690.0 | 0 | Canis | 53690.0 | 0 | Canis lupus | 53690.0 | 0 |
| scf7180000043658 | 1610 | 0.3559 | 0 | 20.8498 | Eukaryota | 2977.0 | 0 | Chordata | 2977.0 | 0 | Carnivora | 2977.0 | 0 | Canidae | 2977.0 | 0 | Canis | 2977.0 | 0 | Canis lupus | 2977.0 | 0 |
| scf7180000050311 | 7045 | 0.4372 | 0 | 471.7721 | Eukaryota | 1231.0 | 0 | Platyhelminthes | 807.0 | 1 | Polystomatidea | 625.0 | 2 | Polystomatidae | 625.0 | 2 | Protopolystoma | 625.0 | 2 | Protopolystoma xenopodis | 625.0 | 2 |
| scf7180000042580 | 778376 | 0.3814 | 0 | 61.395 | Eukaryota | 49445.0 | 0 | Chordata | 49445.0 | 0 | Carnivora | 49445.0 | 0 | Canidae | 49445.0 | 0 | Canis | 44484.0 | 1 | Canis lupus | 44484.0 | 1 |
| scf7180000042811 | 28285 | 0.421 | 0 | 55.5395 | Eukaryota | 11023.0 | 0 | Chordata | 11023.0 | 0 | Carnivora | 8947.0 | 1 | Canidae | 8947.0 | 1 | Canis | 8947.0 | 1 | Canis lupus | 8947.0 | 1 |
| scf7180000054848 | 38203 | 0.443 | 0 | 242.8931 | Eukaryota | 9867.0 | 0 | Chordata | 9867.0 | 0 | Carnivora | 8214.0 | 1 | Canidae | 8214.0 | 1 | Canis | 5462.0 | 2 | Canis lupus | 5462.0 | 2 |
| scf7180000047421 | 84921 | 0.5278 | 0 | 57.505 | Eukaryota | 5770.0 | 0 | Chordata | 5770.0 | 0 | Carnivora | 5770.0 | 0 | Canidae | 4408.0 | 1 | Canis | 4408.0 | 2 | Canis lupus | 4408.0 | 2 |
| scf7180000042388 | 1409807 | 0.384 | 0 | 60.7535 | Eukaryota | 62942.0 | 0 | Chordata | 62942.0 | 0 | Carnivora | 62942.0 | 0 | Canidae | 62942.0 | 0 | Canis | 62942.0 | 0 | Canis lupus | 62942.0 | 0 |
| scf7180000041603 | 1865164 | 0.4235 | 0 | 58.6596 | Eukaryota | 64950.0 | 0 | Chordata | 64950.0 | 0 | Carnivora | 64950.0 | 0 | Canidae | 64950.0 | 0 | Canis | 64950.0 | 0 | Canis lupus | 64950.0 | 0 |
| scf7180000048232 | 50362 | 0.4446 | 0 | 62.5589 | Eukaryota | 89336.0 | 0 | Chordata | 89336.0 | 0 | Carnivora | 89336.0 | 0 | Canidae | 89336.0 | 0 | Canis | 89336.0 | 0 | Canis lupus | 89336.0 | 0 |
| scf7180000044590 | 276045 | 0.4487 | 0 | 58.2614 | Eukaryota | 44158.0 | 0 | Chordata | 44158.0 | 0 | Carnivora | 44158.0 | 0 | Canidae | 44158.0 | 0 | Canis | 35799.0 | 1 | Canis lupus | 35799.0 | 1 |
| scf7180000046005 | 135396 | 0.4832 | 0 | 57.9886 | Eukaryota | 17925.0 | 0 | Chordata | 17925.0 | 0 | Carnivora | 17925.0 | 0 | Ursidae | 6882.0 | 3 | Ailuropoda | 3923.0 | 7 | Ailuropoda melanoleuca | 3923.0 | 7 |
| scf7180000046438 | 4938 | 0.405 | 0 | 28.6408 | Eukaryota | 10316.0 | 0 | Chordata | 10316.0 | 0 | Carnivora | 10316.0 | 0 | Canidae | 10316.0 | 0 | Canis | 10316.0 | 0 | Canis lupus | 10316.0 | 0 |
| scf7180000043443 | 481082 | 0.3742 | 0 | 59.6495 | Eukaryota | 89291.0 | 0 | Chordata | 89291.0 | 0 | Carnivora | 89291.0 | 0 | Canidae | 89291.0 | 0 | Canis | 89291.0 | 0 | Canis lupus | 89291.0 | 0 |
| scf7180000045572 | 190230 | 0.3794 | 0 | 59.721 | Eukaryota | 13669.0 | 0 | Chordata | 13669.0 | 0 | Carnivora | 10196.0 | 1 | Canidae | 7910.0 | 4 | Canis | 6332.0 | 5 | Canis lupus | 6332.0 | 6 |
| scf7180000042008 | 556265 | 0.3725 | 0 | 59.697 | Eukaryota | 91635.0 | 0 | Chordata | 91635.0 | 0 | Carnivora | 91635.0 | 0 | Canidae | 91635.0 | 0 | Canis | 91635.0 | 0 | Canis lupus | 91635.0 | 0 |
| scf7180000043598 | 96077 | 0.413 | 0 | 59.5896 | Eukaryota | 8901.0 | 0 | Chordata | 8901.0 | 0 | Carnivora | 8901.0 | 0 | Canidae | 8901.0 | 0 | Canis | 8901.0 | 0 | Canis lupus | 8901.0 | 0 |
| scf7180000049728 | 2965 | 0.4486 | 0 | 42.4774 | no-hit | 0.0 | 0 | no-hit | 0.0 | 0 | no-hit | 0.0 | 0 | no-hit | 0.0 | 0 | no-hit | 0.0 | 0 | no-hit | 0.0 | 0 |
| scf7180000045354 | 156945 | 0.4769 | 0 | 58.3151 | Eukaryota | 61860.0 | 1 | Chordata | 61860.0 | 1 | Carnivora | 58026.0 | 2 | Canidae | 58026.0 | 2 | Canis | 58026.0 | 2 | Canis lupus | 58026.0 | 2 |
| scf7180000047056 | 91994 | 0.4104 | 0 | 50.627 | Eukaryota | 17856.0 | 0 | Chordata | 17856.0 | 0 | Carnivora | 17856.0 | 0 | Canidae | 17856.0 | 0 | Canis | 17856.0 | 0 | Canis lupus | 17856.0 | 0 |
| scf7180000040631 | 145575 | 0.3596 | 0 | 60.2751 | Eukaryota | 38408.0 | 0 | Chordata | 38408.0 | 0 | Carnivora | 38408.0 | 0 | Otariidae | 15111.0 | 3 | Canis | 11970.0 | 4 | Canis lupus | 11970.0 | 4 |
| scf7180000045977 | 103692 | 0.3869 | 0 | 57.2486 | Eukaryota | 20314.0 | 0 | Chordata | 20314.0 | 0 | Carnivora | 20314.0 | 0 | Canidae | 14264.0 | 1 | Canis | 14264.0 | 2 | Canis lupus | 14264.0 | 2 |
| scf7180000044344 | 351431 | 0.481 | 0 | 58.1677 | Eukaryota | 30681.0 | 0 | Chordata | 30681.0 | 0 | Carnivora | 30681.0 | 0 | Canidae | 30681.0 | 0 | Canis | 30681.0 | 0 | Canis lupus | 30681.0 | 0 |
| scf7180000040998 | 954761 | 0.3877 | 0 | 60.1627 | Eukaryota | 87834.0 | 0 | Chordata | 87834.0 | 0 | Carnivora | 87834.0 | 0 | Canidae | 87834.0 | 0 | Canis | 60904.0 | 1 | Canis lupus | 60904.0 | 1 |
| scf7180000048135 | 91526 | 0.4216 | 0 | 62.0749 | Eukaryota | 10290.0 | 0 | Chordata | 10290.0 | 0 | Carnivora | 5703.0 | 1 | Hominidae | 4587.0 | 2 | Homo | 4587.0 | 2 | Homo sapiens | 4587.0 | 2 |
| scf7180000051653 | 9520 | 0.3833 | 0 | 39.839 | Eukaryota | 9037.0 | 1 | Chordata | 9037.0 | 1 | Primates | 4413.0 | 3 | Hominidae | 3432.0 | 5 | Homo | 3432.0 | 6 | Homo sapiens | 3432.0 | 6 |
| scf7180000046524 | 117620 | 0.4226 | 0 | 59.1099 | Eukaryota | 57288.0 | 0 | Chordata | 57288.0 | 0 | Carnivora | 57288.0 | 0 | Canidae | 57288.0 | 0 | Canis | 57288.0 | 0 | Canis lupus | 57288.0 | 0 |
| scf7180000043468 | 354786 | 0.3689 | 0 | 61.1283 | Eukaryota | 72118.0 | 0 | Chordata | 72118.0 | 0 | Carnivora | 72118.0 | 0 | Canidae | 72118.0 | 0 | Canis | 72118.0 | 0 | Canis lupus | 72118.0 | 0 |
| scf7180000049664 | 16611 | 0.4377 | 0 | 82.1729 | Eukaryota | 52674.0 | 0 | Chordata | 34228.0 | 1 | Gadiformes | 16269.0 | 7 | Gadidae | 16269.0 | 7 | Gadus | 16269.0 | 7 | Gadus morhua | 16269.0 | 7 |
| scf7180000048230 | 7782 | 0.4481 | 0 | 97.8911 | Eukaryota | 9109.0 | 0 | Chordata | 6716.0 | 3 | Gadiformes | 3496.0 | 6 | Gadidae | 3496.0 | 6 | Gadus | 3496.0 | 6 | Gadus morhua | 3496.0 | 6 |
| scf7180000041954 | 2342 | 0.4065 | 0 | 23.6107 | Eukaryota | 3820.0 | 0 | Chordata | 3820.0 | 0 | Carnivora | 3820.0 | 0 | Canidae | 3820.0 | 0 | Canis | 3820.0 | 0 | Canis lupus | 3820.0 | 0 |
| scf7180000046067 | 172039 | 0.3691 | 0 | 60.5308 | Eukaryota | 28483.0 | 0 | Chordata | 28483.0 | 0 | Carnivora | 28483.0 | 0 | Canidae | 28483.0 | 0 | Canis | 28483.0 | 0 | Canis lupus | 28483.0 | 0 |
| scf7180000048870 | 3973 | 0.4372 | 0 | 66.3808 | Eukaryota | 2516.0 | 0 | Chordata | 2516.0 | 0 | Tetraodontiformes | 2345.0 | 1 | Tetraodontidae | 2345.0 | 1 | Takifugu | 2345.0 | 1 | Takifugu rubripes | 2345.0 | 1 |
| scf7180000046269 | 15033 | 0.4255 | 0 | 143.1606 | Eukaryota | 36522.0 | 0 | Chordata | 36522.0 | 0 | Carnivora | 36522.0 | 0 | Canidae | 36522.0 | 0 | Canis | 36522.0 | 0 | Canis lupus | 36522.0 | 0 |
| scf7180000053781 | 4228 | 0.5071 | 0 | 49.5448 | Eukaryota | 6517.0 | 0 | Chordata | 6517.0 | 0 | Carnivora | 6517.0 | 0 | Canidae | 6517.0 | 0 | Canis | 4345.0 | 1 | Canis lupus | 3397.0 | 2 |
| scf7180000041970 | 796859 | 0.3615 | 0 | 60.385 | Eukaryota | 93119.0 | 0 | Chordata | 93119.0 | 0 | Carnivora | 93119.0 | 0 | Canidae | 93119.0 | 0 | Canis | 93119.0 | 0 | Canis lupus | 93119.0 | 0 |
| scf7180000044529 | 65904 | 0.3665 | 0 | 55.294 | Eukaryota | 21090.0 | 0 | Chordata | 21090.0 | 0 | Carnivora | 21090.0 | 0 | Canidae | 21090.0 | 0 | Canis | 21090.0 | 0 | Canis lupus | 21090.0 | 0 |
| scf7180000054169 | 38014 | 0.4418 | 0 | 324.7043 | Eukaryota | 16107.0 | 0 | Chordata | 9913.0 | 2 | Strigeidida | 4342.0 | 6 | Schistosomatidae | 4342.0 | 6 | Schistosoma | 4342.0 | 6 | Schistosoma mansoni | 4342.0 | 6 |
| scf7180000053396 | 7366 | 0.4336 | 0 | 255.8375 | no-hit | 0.0 | 0 | no-hit | 0.0 | 0 | no-hit | 0.0 | 0 | no-hit | 0.0 | 0 | no-hit | 0.0 | 0 | no-hit | 0.0 | 0 |
| scf7180000045497 | 7633 | 0.4707 | 0 | 39.7844 | Eukaryota | 44555.0 | 0 | Chordata | 44555.0 | 0 | Carnivora | 44555.0 | 0 | Canidae | 44555.0 | 0 | Canis | 44555.0 | 0 | Canis lupus | 44555.0 | 0 |
| scf7180000056347 | 762001 | 0.3505 | 0 | 59.1112 | Eukaryota | 30686.0 | 0 | Chordata | 30686.0 | 0 | Carnivora | 30686.0 | 0 | Canidae | 30686.0 | 0 | Canis | 30686.0 | 0 | Canis lupus | 30686.0 | 0 |
| scf7180000041539 | 32312 | 0.4812 | 0 | 77.823 | Eukaryota | 51311.0 | 0 | Chordata | 51311.0 | 0 | Carnivora | 51311.0 | 0 | Canidae | 51311.0 | 0 | Canis | 51311.0 | 0 | Canis lupus | 51311.0 | 0 |
| scf7180000041947 | 458369 | 0.3889 | 0 | 59.3177 | Eukaryota | 92454.0 | 0 | Chordata | 92454.0 | 0 | Carnivora | 92454.0 | 0 | Canidae | 92454.0 | 0 | Canis | 92454.0 | 0 | Canis lupus | 92454.0 | 0 |
| scf7180000056261 | 963236 | 0.4207 | 0 | 60.4174 | Eukaryota | 55446.0 | 0 | Chordata | 55446.0 | 0 | Carnivora | 44088.0 | 1 | Canidae | 35758.0 | 2 | Canis | 27990.0 | 3 | Canis lupus | 27990.0 | 3 |
| scf7180000046197 | 131983 | 0.3423 | 0 | 61.8817 | Eukaryota | 31264.0 | 0 | Chordata | 31264.0 | 0 | Chiroptera | 20766.0 | 1 | Pteropodidae | 20766.0 | 1 | Rousettus | 20766.0 | 3 | Rousettus aegyptiacus | 20766.0 | 3 |
| scf7180000047462 | 4361 | 0.416 | 0 | 24.5838 | Eukaryota | 15824.0 | 0 | Chordata | 15824.0 | 0 | Carnivora | 15824.0 | 0 | Canidae | 15824.0 | 0 | Canis | 15824.0 | 0 | Canis lupus | 15824.0 | 0 |
| scf7180000042689 | 353564 | 0.4545 | 0 | 84.0654 | Eukaryota | 94399.0 | 0 | Chordata | 94399.0 | 0 | Carnivora | 94399.0 | 0 | Canidae | 94399.0 | 0 | Canis | 84829.0 | 1 | Canis lupus | 84829.0 | 1 |
| scf7180000049030 | 55388 | 0.3782 | 0 | 59.8584 | Eukaryota | 14469.0 | 1 | Chordata | 14469.0 | 1 | Carnivora | 12603.0 | 2 | Canidae | 12603.0 | 2 | Canis | 12603.0 | 2 | Canis lupus | 12603.0 | 2 |
| scf7180000041573 | 350386 | 0.422 | 0 | 93.9013 | Eukaryota | 92197.0 | 0 | Chordata | 67645.0 | 1 | Carnivora | 38309.0 | 3 | Canidae | 38309.0 | 3 | Canis | 38309.0 | 3 | Canis lupus | 38309.0 | 3 |
| scf7180000049060 | 5009 | 0.4566 | 0 | 888.2686 | no-hit | 0.0 | 0 | no-hit | 0.0 | 0 | no-hit | 0.0 | 0 | no-hit | 0.0 | 0 | no-hit | 0.0 | 0 | no-hit | 0.0 | 0 |
| scf7180000051608 | 46683 | 0.4015 | 0 | 61.6027 | Eukaryota | 6865.0 | 1 | Chordata | 6865.0 | 1 | Primates | 3061.0 | 3 | Hominidae | 3061.0 | 3 | Homo | 3061.0 | 4 | Homo sapiens | 3061.0 | 4 |
| scf7180000049920 | 9806 | 0.4597 | 0 | 68.8227 | Eukaryota | 11486.0 | 0 | Chordata | 11486.0 | 0 | Carnivora | 11486.0 | 0 | Canidae | 11486.0 | 0 | Canis | 5931.0 | 1 | Canis lupus | 5931.0 | 1 |
| scf7180000049802 | 18965 | 0.443 | 0 | 350.1314 | Eukaryota | 2774.0 | 0 | Platyhelminthes | 2294.0 | 1 | Polystomatidea | 2294.0 | 2 | Polystomatidae | 2294.0 | 2 | Protopolystoma | 2294.0 | 2 | Protopolystoma xenopodis | 2294.0 | 2 |
| scf7180000042326 | 531193 | 0.39 | 0 | 59.628 | Eukaryota | 65114.0 | 0 | Chordata | 65114.0 | 0 | Carnivora | 65114.0 | 0 | Canidae | 65114.0 | 0 | Canis | 65114.0 | 0 | Canis lupus | 65114.0 | 0 |
| scf7180000044450 | 20288 | 0.4388 | 0 | 124.3116 | Eukaryota | 6254.0 | 0 | Chordata | 2277.0 | 2 | Polystomatidea | 2145.0 | 4 | Polystomatidae | 2145.0 | 4 | Protopolystoma | 2145.0 | 5 | Protopolystoma xenopodis | 2145.0 | 5 |
| scf7180000042527 | 617119 | 0.3992 | 0 | 60.2033 | Eukaryota | 89131.0 | 0 | Chordata | 89131.0 | 0 | Carnivora | 89131.0 | 0 | Canidae | 89131.0 | 0 | Canis | 89131.0 | 0 | Canis lupus | 89131.0 | 0 |
| scf7180000049796 | 30625 | 0.4015 | 0 | 58.2166 | Eukaryota | 16119.0 | 0 | Chordata | 16119.0 | 0 | Carnivora | 16119.0 | 0 | Canidae | 16119.0 | 0 | Canis | 16119.0 | 0 | Canis lupus | 16119.0 | 0 |
| scf7180000043213 | 347677 | 0.377 | 0 | 59.3425 | Eukaryota | 69350.0 | 0 | Chordata | 69350.0 | 0 | Carnivora | 69350.0 | 0 | Canidae | 69350.0 | 0 | Canis | 69350.0 | 0 | Canis lupus | 69350.0 | 0 |
| scf7180000048127 | 81666 | 0.3815 | 100 | 60.2771 | Eukaryota | 20341.0 | 0 | Chordata | 20341.0 | 0 | Carnivora | 20341.0 | 0 | Canidae | 20341.0 | 0 | Canis | 20341.0 | 0 | Canis lupus | 20341.0 | 0 |
| scf7180000052390 | 4777 | 0.4503 | 0 | 940.8866 | no-hit | 0.0 | 0 | no-hit | 0.0 | 0 | no-hit | 0.0 | 0 | no-hit | 0.0 | 0 | no-hit | 0.0 | 0 | no-hit | 0.0 | 0 |
| scf7180000041668 | 234079 | 0.3908 | 0 | 58.0279 | Eukaryota | 58795.0 | 0 | Chordata | 58795.0 | 0 | Carnivora | 58795.0 | 0 | Canidae | 58795.0 | 0 | Canis | 58795.0 | 0 | Canis lupus | 58795.0 | 0 |
| scf7180000041294 | 628613 | 0.3975 | 0 | 58.9699 | Eukaryota | 39081.0 | 0 | Chordata | 39081.0 | 0 | Primates | 24393.0 | 2 | Hominidae | 19451.0 | 3 | Homo | 19451.0 | 4 | Homo sapiens | 19451.0 | 4 |
| scf7180000040559 | 2302324 | 0.4048 | 0 | 60.2314 | Eukaryota | 97971.0 | 0 | Chordata | 97971.0 | 0 | Carnivora | 97971.0 | 0 | Canidae | 97971.0 | 0 | Canis | 97971.0 | 0 | Canis lupus | 97971.0 | 0 |
| scf7180000053439 | 2223 | 0.439 | 0 | 432.5261 | no-hit | 0.0 | 0 | no-hit | 0.0 | 0 | no-hit | 0.0 | 0 | no-hit | 0.0 | 0 | no-hit | 0.0 | 0 | no-hit | 0.0 | 0 |
| scf7180000044565 | 295987 | 0.3914 | 0 | 59.2356 | Eukaryota | 23591.0 | 0 | Chordata | 23591.0 | 0 | Carnivora | 23591.0 | 0 | Canidae | 23591.0 | 0 | Canis | 14125.0 | 1 | Canis lupus | 14125.0 | 1 |
| scf7180000044474 | 110116 | 0.3592 | 0 | 58.4815 | Eukaryota | 57424.0 | 0 | Chordata | 57424.0 | 0 | Carnivora | 52657.0 | 2 | Canidae | 52657.0 | 2 | Canis | 52657.0 | 2 | Canis lupus | 52657.0 | 2 |
| scf7180000044364 | 78553 | 0.4502 | 0 | 77.4214 | Eukaryota | 81949.0 | 0 | Chordata | 81949.0 | 0 | Carnivora | 81949.0 | 0 | Canidae | 81949.0 | 0 | Canis | 81949.0 | 0 | Canis lupus | 81949.0 | 0 |
| scf7180000042641 | 582229 | 0.4253 | 0 | 58.4071 | Eukaryota | 82075.0 | 0 | Chordata | 82075.0 | 0 | Carnivora | 82075.0 | 0 | Canidae | 82075.0 | 0 | Canis | 43040.0 | 1 | Canis lupus | 43040.0 | 1 |
| scf7180000045652 | 156584 | 0.3928 | 0 | 76.036 | Eukaryota | 19033.0 | 0 | Chordata | 19033.0 | 0 | Carnivora | 17453.0 | 1 | Canidae | 10772.0 | 3 | Canis | 10772.0 | 4 | Canis lupus | 10772.0 | 4 |
| scf7180000043798 | 734133 | 0.3641 | 100 | 59.7048 | Eukaryota | 94128.0 | 0 | Chordata | 94128.0 | 0 | Carnivora | 94128.0 | 0 | Canidae | 94128.0 | 0 | Canis | 94128.0 | 0 | Canis lupus | 94128.0 | 0 |
| scf7180000046873 | 25888 | 0.4514 | 0 | 195.6711 | no-hit | 0.0 | 0 | no-hit | 0.0 | 0 | no-hit | 0.0 | 0 | no-hit | 0.0 | 0 | no-hit | 0.0 | 0 | no-hit | 0.0 | 0 |
| scf7180000048492 | 89095 | 0.4092 | 0 | 60.2624 | Eukaryota | 21754.0 | 0 | Chordata | 21754.0 | 0 | Carnivora | 18480.0 | 2 | Otariidae | 4975.0 | 6 | Ursus | 4621.0 | 7 | Canis lupus | 4292.0 | 8 |
| scf7180000045315 | 255670 | 0.5514 | 0 | 56.6089 | Eukaryota | 28093.0 | 0 | Chordata | 28093.0 | 0 | Carnivora | 28093.0 | 0 | Canidae | 28093.0 | 0 | Canis | 25327.0 | 1 | Canis lupus | 25327.0 | 1 |

| | | | | | | | | | | | | | | | | | | | | | |
|---|---|---|---|---|---|---|---|---|---|---|---|---|---|---|---|---|---|---|---|---|---|
| scf7180000050395 | 7660 | 0.4402 | 0 | 170.3647 | Eukaryota | 1451.0 | 0 | Platyhelminthes | 1046.0 | 1 | Polystomatidea | 1046.0 | 1 | Polystomatidae | 1046.0 | 1 | Protopolystoma | 1046.0 | 1 | Protopolystoma xenopodis | 1046.0 | 1 |
| scf7180000054669 | 20277 | 0.45 | 0 | 172.221 | Eukaryota | 10149.0 | 1 | Chordata | 10149.0 | 1 | Tetraodontiformes | 5867.0 | 2 | Tetraodontidae | 5867.0 | 2 | Takifugu | 5867.0 | 2 | Takifugu rubripes | 5867.0 | 2 |
| scf7180000056220 | 488444 | 0.4013 | 0 | 60.1276 | Eukaryota | 40360.0 | 0 | Chordata | 40360.0 | 0 | Carnivora | 40360.0 | 0 | Canidae | 40360.0 | 0 | Canis | 40360.0 | 0 | Canis lupus | 40360.0 | 0 |
| scf7180000042950 | 107136 | 0.4019 | 0 | 57.4692 | Eukaryota | 115029.0 | 0 | Chordata | 115029.0 | 0 | Carnivora | 115029.0 | 0 | Canidae | 115029.0 | 0 | Canis | 115029.0 | 0 | Canis lupus | 115029.0 | 0 |
| scf7180000048814 | 43029 | 0.4351 | 0 | 62.2349 | Eukaryota | 25267.0 | 0 | Chordata | 25267.0 | 0 | Carnivora | 25267.0 | 0 | Canidae | 13793.0 | 2 | Vulpes | 7827.0 | 5 | Vulpes vulpes | 7827.0 | 6 |
| scf7180000041262 | 1122530 | 0.4512 | 0 | 60.1862 | Eukaryota | 92362.0 | 0 | Chordata | 92362.0 | 0 | Carnivora | 92362.0 | 0 | Canidae | 92362.0 | 0 | Canis | 92362.0 | 0 | Canis lupus | 92362.0 | 0 |
| scf7180000040511 | 1361923 | 0.3629 | 0 | 58.3759 | Eukaryota | 94565.0 | 0 | Chordata | 94565.0 | 0 | Carnivora | 94565.0 | 0 | Canidae | 94565.0 | 0 | Canis | 94565.0 | 0 | Canis lupus | 94565.0 | 0 |
| scf7180000053839 | 2260 | 0.4451 | 0 | 297.2933 | no-hit | 0.0 | 0 | no-hit | 0.0 | 0 | no-hit | 0.0 | 0 | no-hit | 0.0 | 0 | no-hit | 0.0 | 0 | no-hit | 0.0 | 0 |
| scf7180000055714 | 19694 | 0.436 | 0 | 128.6688 | Eukaryota | 11294.0 | 0 | Chordata | 10232.0 | 1 | Tetraodontiformes | 9435.0 | 2 | Tetraodontidae | 9435.0 | 2 | Takifugu | 9435.0 | 2 | Takifugu rubripes | 9435.0 | 2 |
| scf7180000042431 | 680870 | 0.391 | 0 | 60.2481 | Eukaryota | 74244.0 | 0 | Chordata | 74244.0 | 0 | Carnivora | 51300.0 | 1 | Canidae | 38634.0 | 2 | Canis | 38634.0 | 2 | Canis lupus | 38634.0 | 2 |
| scf7180000054443 | 26675 | 0.4497 | 0 | 372.241 | Eukaryota | 9543.0 | 0 | Chordata | 8101.0 | 1 | Carnivora | 4820.0 | 3 | Canidae | 4820.0 | 3 | Canis | 4197.0 | 4 | Canis lupus | 4197.0 | 4 |
| scf7180000048858 | 17062 | 0.4529 | 0 | 1611.8585 | no-hit | 0.0 | 0 | no-hit | 0.0 | 0 | no-hit | 0.0 | 0 | no-hit | 0.0 | 0 | no-hit | 0.0 | 0 | no-hit | 0.0 | 0 |
| scf7180000048286 | 64989 | 0.4458 | 0 | 59.3618 | Eukaryota | 13286.0 | 0 | Chordata | 13286.0 | 0 | Carnivora | 12615.0 | 1 | Canidae | 12615.0 | 1 | Canis | 11024.0 | 2 | Canis lupus | 11024.0 | 2 |
| scf7180000045673 | 152399 | 0.3555 | 0 | 59.0335 | Eukaryota | 11650.0 | 0 | Chordata | 11650.0 | 0 | Carnivora | 11650.0 | 0 | Canidae | 11650.0 | 0 | Canis | 11650.0 | 0 | Canis lupus | 11650.0 | 0 |
| scf7180000040623 | 1635950 | 0.3786 | 0 | 60.5626 | Eukaryota | 92103.0 | 0 | Chordata | 92103.0 | 0 | Carnivora | 92103.0 | 0 | Canidae | 92103.0 | 0 | Canis | 92103.0 | 0 | Canis lupus | 92103.0 | 0 |
| scf7180000043342 | 863464 | 0.4041 | 0 | 59.8 | Eukaryota | 89070.0 | 0 | Chordata | 89070.0 | 0 | Carnivora | 89070.0 | 0 | Canidae | 65113.0 | 2 | Canis | 35598.0 | 3 | Canis lupus | 35598.0 | 3 |
| scf7180000043245 | 373599 | 0.3612 | 0 | 59.4313 | Eukaryota | 26318.0 | 0 | Chordata | 26318.0 | 0 | Carnivora | 26318.0 | 0 | Canidae | 26318.0 | 0 | Canis | 26318.0 | 0 | Canis lupus | 26318.0 | 0 |
| scf7180000041345 | 543794 | 0.338 | 0 | 58.7157 | Eukaryota | 16750.0 | 0 | Chordata | 16750.0 | 0 | Carnivora | 16750.0 | 0 | Canidae | 16750.0 | 0 | Canis | 16750.0 | 0 | Canis lupus | 16750.0 | 0 |
| scf7180000041726 | 9652 | 0.4628 | 0 | 27.7418 | Eukaryota | 4233.0 | 1 | Chordata | 4233.0 | 1 | Primates | 1305.0 | 4 | Hominidae | 1305.0 | 4 | Homo | 1305.0 | 5 | Homo sapiens | 1305.0 | 5 |
| scf7180000043886 | 422484 | 0.3936 | 0 | 60.7597 | Eukaryota | 27993.0 | 0 | Chordata | 27993.0 | 0 | Carnivora | 27993.0 | 0 | Canidae | 27993.0 | 0 | Canis | 25821.0 | 1 | Canis lupus | 25821.0 | 1 |
| scf7180000046139 | 144643 | 0.3931 | 0 | 61.098 | Eukaryota | 25313.0 | 0 | Chordata | 25313.0 | 0 | Carnivora | 25313.0 | 0 | Canidae | 22812.0 | 1 | Canis | 15243.0 | 2 | Canis lupus | 15243.0 | 2 |
| scf7180000042390 | 67361 | 0.4267 | 0 | 54.2467 | Eukaryota | 7103.0 | 0 | Chordata | 7103.0 | 0 | Carnivora | 7103.0 | 0 | Canidae | 7103.0 | 0 | Canis | 7103.0 | 0 | Canis lupus | 7103.0 | 0 |
| scf7180000041645 | 634814 | 0.3601 | 0 | 59.849 | Eukaryota | 87272.0 | 0 | Chordata | 87272.0 | 0 | Carnivora | 87272.0 | 0 | Canidae | 87272.0 | 0 | Canis | 87272.0 | 0 | Canis lupus | 87272.0 | 0 |
| scf7180000054171 | 59619 | 0.4435 | 0 | 359.5429 | Eukaryota | 89322.0 | 0 | Chordata | 89322.0 | 0 | Carnivora | 89322.0 | 0 | Canidae | 89322.0 | 0 | Canis | 89322.0 | 0 | Canis lupus | 89322.0 | 0 |
| scf7180000042125 | 147281 | 0.3754 | 0 | 59.1617 | Eukaryota | 93844.0 | 0 | Chordata | 93844.0 | 0 | Carnivora | 93844.0 | 0 | Canidae | 93844.0 | 0 | Canis | 93844.0 | 0 | Canis lupus | 93844.0 | 0 |
| scf7180000052524 | 7643 | 0.4469 | 0 | 661.4399 | no-hit | 0.0 | 0 | no-hit | 0.0 | 0 | no-hit | 0.0 | 0 | no-hit | 0.0 | 0 | no-hit | 0.0 | 0 | no-hit | 0.0 | 0 |
| scf7180000046831 | 35618 | 0.4423 | 0 | 199.4155 | Eukaryota | 92046.0 | 0 | Chordata | 92046.0 | 0 | Carnivora | 92046.0 | 0 | Canidae | 92046.0 | 0 | Canis | 92046.0 | 0 | Canis lupus | 92046.0 | 0 |
| scf7180000051492 | 11736 | 0.4524 | 0 | 1365.3574 | Eukaryota | 1044.0 | 0 | Chordata | 825.0 | 1 | Tetraodontiformes | 825.0 | 1 | Tetraodontidae | 825.0 | 1 | Takifugu | 825.0 | 1 | Takifugu rubripes | 825.0 | 1 |
| scf7180000044188 | 123881 | 0.4332 | 0 | 58.9676 | Eukaryota | 37409.0 | 0 | Chordata | 37409.0 | 0 | Carnivora | 37409.0 | 0 | Canidae | 37409.0 | 0 | Canis | 37409.0 | 0 | Canis lupus | 37409.0 | 0 |
| scf7180000053472 | 3384 | 0.4453 | 0 | 240.4171 | no-hit | 0.0 | 0 | no-hit | 0.0 | 0 | no-hit | 0.0 | 0 | no-hit | 0.0 | 0 | no-hit | 0.0 | 0 | no-hit | 0.0 | 0 |
| scf7180000050471 | 12732 | 0.4355 | 0 | 368.0086 | no-hit | 0.0 | 0 | no-hit | 0.0 | 0 | no-hit | 0.0 | 0 | no-hit | 0.0 | 0 | no-hit | 0.0 | 0 | no-hit | 0.0 | 0 |
| scf7180000049966 | 16958 | 0.4462 | 0 | 71.3382 | Eukaryota | 235.0 | 0 | Platyhelminthes | 235.0 | 0 | Polystomatidea | 235.0 | 0 | Polystomatidae | 235.0 | 0 | Protopolystoma | 235.0 | 0 | Protopolystoma xenopodis | 235.0 | 0 |
| scf7180000056405 | 512188 | 0.4322 | 0 | 59.7487 | Eukaryota | 66199.0 | 0 | Chordata | 66199.0 | 0 | Carnivora | 66199.0 | 0 | Canidae | 66199.0 | 0 | Canis | 45560.0 | 1 | Canis lupus | 45560.0 | 1 |
| scf7180000041295 | 30144 | 0.3577 | 0 | 56.6508 | Eukaryota | 25211.0 | 0 | Chordata | 25211.0 | 0 | Carnivora | 25211.0 | 0 | Canidae | 25211.0 | 0 | Canis | 25211.0 | 0 | Canis lupus | 25211.0 | 0 |
| scf7180000046046 | 140119 | 0.4579 | 0 | 119.6878 | Eukaryota | 65206.0 | 0 | Chordata | 65206.0 | 0 | Carnivora | 65206.0 | 0 | Canidae | 65206.0 | 0 | Canis | 65206.0 | 0 | Canis lupus | 65206.0 | 0 |
| scf7180000047705 | 72236 | 0.4007 | 0 | 59.6852 | Eukaryota | 14715.0 | 0 | Chordata | 14715.0 | 0 | Carnivora | 14715.0 | 0 | Canidae | 14715.0 | 0 | Canis | 14715.0 | 0 | Canis lupus | 14715.0 | 0 |
| scf7180000041337 | 35574 | 0.3859 | 0 | 86.9295 | Eukaryota | 65930.0 | 0 | Chordata | 65930.0 | 0 | Carnivora | 65930.0 | 0 | Canidae | 65930.0 | 0 | Canis | 65930.0 | 0 | Canis lupus | 65930.0 | 0 |
| scf7180000044927 | 258340 | 0.4229 | 0 | 61.0743 | Eukaryota | 47733.0 | 0 | Chordata | 47733.0 | 0 | Carnivora | 47733.0 | 0 | Canidae | 47733.0 | 0 | Canis | 47733.0 | 0 | Canis lupus | 47733.0 | 0 |
| scf7180000051495 | 2062 | 0.4433 | 0 | 265.96 | no-hit | 0.0 | 0 | no-hit | 0.0 | 0 | no-hit | 0.0 | 0 | no-hit | 0.0 | 0 | no-hit | 0.0 | 0 | no-hit | 0.0 | 0 |
| scf7180000052700 | 10487 | 0.5051 | 0 | 134.1908 | Eukaryota | 7510.0 | 0 | Chordata | 7510.0 | 0 | Carnivora | 7510.0 | 0 | Canidae | 7510.0 | 0 | Canis | 6053.0 | 1 | Canis lupus | 5034.0 | 2 |
| scf7180000043218 | 591992 | 0.368 | 0 | 65.2714 | Eukaryota | 101394.0 | 0 | Chordata | 101394.0 | 0 | Carnivora | 101394.0 | 0 | Canidae | 97104.0 | 1 | Canis | 92274.0 | 2 | Canis lupus | 92274.0 | 3 |
| scf7180000042480 | 679689 | 0.4885 | 0 | 58.4583 | Eukaryota | 71689.0 | 0 | Chordata | 71689.0 | 0 | Carnivora | 71689.0 | 0 | Canidae | 71689.0 | 0 | Canis | 65146.0 | 1 | Canis lupus | 65146.0 | 1 |
| scf7180000051132 | 27365 | 0.5792 | 0 | 53.7695 | Eukaryota | 15540.0 | 0 | Chordata | 15540.0 | 0 | Carnivora | 15540.0 | 0 | Canidae | 15540.0 | 0 | Canis | 15540.0 | 0 | Canis lupus | 15540.0 | 0 |
| scf7180000055949 | 2736 | 0.458 | 0 | 1244.4526 | Eukaryota | 12116.0 | 0 | Chordata | 10226.0 | 2 | Gadiformes | 4938.0 | 6 | Gadidae | 4938.0 | 6 | Gadus | 4938.0 | 6 | Gadus morhua | 4938.0 | 6 |
| scf7180000056560 | 100672 | 0.477 | 0 | 58.6325 | Eukaryota | 22137.0 | 0 | Chordata | 22137.0 | 0 | Rodentia | 10434.0 | 2 | Muridae | 10434.0 | 2 | Mus | 8337.0 | 4 | Mus musculus | 8337.0 | 4 |
| scf7180000050592 | 4452 | 0.4483 | 0 | 245.7225 | Eukaryota | 13590.0 | 0 | Chordata | 12366.0 | 2 | Gadiformes | 7115.0 | 7 | Gadidae | 7115.0 | 7 | Gadus | 7115.0 | 7 | Gadus morhua | 7115.0 | 7 |
| scf7180000040825 | 1611086 | 0.484 | 0 | 59.4096 | Eukaryota | 68153.0 | 0 | Chordata | 68153.0 | 0 | Carnivora | 68153.0 | 0 | Canidae | 68153.0 | 0 | Canis | 41729.0 | 1 | Canis lupus | 41729.0 | 1 |
| scf7180000040652 | 12184 | 0.3506 | 0 | 28.448 | Eukaryota | 2963.0 | 0 | Chordata | 2963.0 | 0 | Carnivora | 1812.0 | 1 | Canidae | 1812.0 | 1 | Canis | 1812.0 | 1 | Canis lupus | 1812.0 | 1 |
| scf7180000052401 | 20418 | 0.433 | 0 | 176.3589 | no-hit | 0.0 | 0 | no-hit | 0.0 | 0 | no-hit | 0.0 | 0 | no-hit | 0.0 | 0 | no-hit | 0.0 | 0 | no-hit | 0.0 | 0 |
| scf7180000054471 | 16238 | 0.4191 | 0 | 354.759 | Eukaryota | 33878.0 | 0 | Chordata | 33878.0 | 0 | Carnivora | 33878.0 | 0 | Canidae | 33878.0 | 0 | Canis | 33878.0 | 0 | Canis lupus | 33878.0 | 0 |
| scf7180000049375 | 22564 | 0.4509 | 0 | 2226.9028 | Eukaryota | 971.0 | 0 | Platyhelminthes | 765.0 | 1 | Polystomatidea | 765.0 | 1 | Polystomatidae | 765.0 | 1 | Protopolystoma | 765.0 | 1 | Protopolystoma xenopodis | 765.0 | 1 |
| scf7180000044784 | 273028 | 0.418 | 0 | 60.7185 | Eukaryota | 33402.0 | 0 | Chordata | 33402.0 | 0 | Carnivora | 33402.0 | 0 | Canidae | 33402.0 | 0 | Canis | 33402.0 | 0 | Canis lupus | 33402.0 | 0 |
| scf7180000051342 | 22314 | 0.4364 | 0 | 415.7319 | Eukaryota | 20172.0 | 0 | Chordata | 20172.0 | 0 | Carnivora | 20172.0 | 0 | Canidae | 20172.0 | 0 | Canis | 20172.0 | 0 | Canis lupus | 20172.0 | 0 |
| scf7180000041212 | 298314 | 0.3773 | 0 | 60.8854 | Eukaryota | 49930.0 | 0 | Chordata | 49930.0 | 0 | Carnivora | 49930.0 | 0 | Canidae | 49930.0 | 0 | Canis | 49930.0 | 0 | Canis lupus | 49930.0 | 0 |
| scf7180000042856 | 22712 | 0.3694 | 0 | 42.8274 | Eukaryota | 3154.0 | 0 | Chordata | 3154.0 | 0 | Carnivora | 3154.0 | 0 | Canidae | 2286.0 | 1 | Canis | 2286.0 | 1 | Canis lupus | 2286.0 | 1 |
| scf7180000051644 | 19535 | 0.384 | 0 | 59.9227 | Eukaryota | 7419.0 | 0 | Chordata | 7419.0 | 0 | Carnivora | 5414.0 | 4 | Canidae | 3959.0 | 6 | Canis | 3959.0 | 6 | Canis lupus | 3959.0 | 6 |
| scf7180000042466 | 446404 | 0.3964 | 0 | 60.4331 | Eukaryota | 128893.0 | 0 | Chordata | 128893.0 | 0 | Carnivora | 128893.0 | 0 | Canidae | 128893.0 | 0 | Canis | 128893.0 | 0 | Canis lupus | 128893.0 | 0 |
| scf7180000048525 | 50090 | 0.4004 | 0 | 61.0466 | Eukaryota | 22905.0 | 0 | Chordata | 22905.0 | 0 | Carnivora | 22905.0 | 0 | Canidae | 22905.0 | 0 | Canis | 22905.0 | 0 | Canis lupus | 22905.0 | 0 |
| scf7180000056236 | 958375 | 0.4377 | 0 | 61.414 | Eukaryota | 78610.0 | 0 | Chordata | 78610.0 | 0 | Carnivora | 78610.0 | 0 | Canidae | 52482.0 | 3 | Canis | 52482.0 | 3 | Canis lupus | 52482.0 | 3 |
| scf7180000047451 | 76411 | 0.3988 | 0 | 60.9926 | Eukaryota | 16125.0 | 1 | Chordata | 16125.0 | 1 | Carnivora | 8269.0 | 3 | Canidae | 8269.0 | 3 | Canis | 8269.0 | 4 | Canis lupus | 8269.0 | 4 |
| scf7180000052980 | 13578 | 0.5217 | 0 | 1196.4221 | Eukaryota | 34360.0 | 0 | Chordata | 34360.0 | 0 | Carnivora | 34360.0 | 0 | Canidae | 34360.0 | 0 | Canis | 34360.0 | 0 | Canis lupus | 34360.0 | 0 |
| scf7180000052678 | 7316 | 0.4569 | 0 | 54.1172 | Eukaryota | 2960.0 | 0 | Chordata | 2960.0 | 0 | Carnivora | 2391.0 | 1 | Canidae | 2391.0 | 1 | Canis | 2391.0 | 1 | Canis lupus | 2391.0 | 1 |
| scf7180000051872 | 12101 | 0.5047 | 0 | 844.1011 | Eukaryota | 9223.0 | 0 | Chordata | 9223.0 | 0 | Carnivora | 9223.0 | 0 | Canidae | 9223.0 | 0 | Canis | 7458.0 | 1 | Canis lupus | 6170.0 | 2 |
| scf7180000041514 | 423261 | 0.3867 | 0 | 59.1148 | Eukaryota | 47518.0 | 0 | Chordata | 47518.0 | 0 | Carnivora | 47518.0 | 0 | Canidae | 29779.0 | 1 | Canis | 24820.0 | 2 | Canis lupus | 24820.0 | 2 |
| scf7180000044790 | 408312 | 0.4009 | 0 | 59.0071 | Eukaryota | 53843.0 | 0 | Chordata | 53843.0 | 0 | Carnivora | 53843.0 | 0 | Canidae | 53843.0 | 0 | Canis | 53843.0 | 0 | Canis lupus | 53843.0 | 0 |
| scf7180000045136 | 215400 | 0.3823 | 0 | 58.2597 | Eukaryota | 18250.0 | 0 | Chordata | 18250.0 | 0 | Carnivora | 18250.0 | 0 | Canidae | 10518.0 | 1 | Canis | 10518.0 | 1 | Canis lupus | 10518.0 | 1 |
| scf7180000041882 | 865213 | 0.4498 | 0 | 60.4502 | Eukaryota | 58898.0 | 0 | Chordata | 58898.0 | 0 | Carnivora | 58898.0 | 0 | Canidae | 39575.0 | 2 | Canis | 33408.0 | 3 | Canis lupus | 33408.0 | 3 |
| scf7180000049909 | 13084 | 0.4463 | 0 | 149.2017 | no-hit | 0.0 | 0 | no-hit | 0.0 | 0 | no-hit | 0.0 | 0 | no-hit | 0.0 | 0 | no-hit | 0.0 | 0 | no-hit | 0.0 | 0 |
| scf7180000052231 | 10911 | 0.4609 | 0 | 50.8344 | no-hit | 0.0 | 0 | no-hit | 0.0 | 0 | no-hit | 0.0 | 0 | no-hit | 0.0 | 0 | no-hit | 0.0 | 0 | no-hit | 0.0 | 0 |
| scf7180000051740 | 10626 | 0.5044 | 0 | 159.4588 | Eukaryota | 8166.0 | 0 | Chordata | 8166.0 | 0 | Carnivora | 8166.0 | 0 | Canidae | 8166.0 | 0 | Canis | 6497.0 | 1 | Canis lupus | 5275.0 | 2 |
| scf7180000054028 | 6689 | 0.4451 | 0 | 64.6363 | no-hit | 0.0 | 0 | no-hit | 0.0 | 0 | no-hit | 0.0 | 0 | no-hit | 0.0 | 0 | no-hit | 0.0 | 0 | no-hit | 0.0 | 0 |
| scf7180000040786 | 38705 | 0.5096 | 0 | 790.3588 | Eukaryota | 8907.0 | 0 | Chordata | 8907.0 | 0 | Carnivora | 8907.0 | 0 | Canidae | 8907.0 | 0 | Nyctereutes | 4500.0 | 1 | Nyctereutes procyonoides | 4500.0 | 1 |
| scf7180000050878 | 21054 | 0.4417 | 0 | 60.123 | Eukaryota | 41089.0 | 1 | Chordata | 36786.0 | 2 | Gadiformes | 22357.0 | 6 | Gadidae | 22357.0 | 6 | Gadus | 22357.0 | 6 | Gadus morhua | 22357.0 | 6 |
| scf7180000042078 | 1019643 | 0.4434 | 0 | 60.3439 | Eukaryota | 41598.0 | 0 | Chordata | 41598.0 | 0 | Carnivora | 41598.0 | 0 | Canidae | 41598.0 | 0 | Canis | 37221.0 | 1 | Canis lupus | 37221.0 | 1 |
| scf7180000053343 | 2960 | 0.4382 | 0 | 326.1516 | no-hit | 0.0 | 0 | no-hit | 0.0 | 0 | no-hit | 0.0 | 0 | no-hit | 0.0 | 0 | no-hit | 0.0 | 0 | no-hit | 0.0 | 0 |
| scf7180000053082 | 8558 | 0.4401 | 0 | 276.0597 | Eukaryota | 2193.0 | 0 | Platyhelminthes | 1864.0 | 1 | Polystomatidea | 1864.0 | 1 | Polystomatidae | 1864.0 | 1 | Protopolystoma | 1864.0 | 1 | Protopolystoma xenopodis | 1864.0 | 1 |
| scf7180000040579 | 2286898 | 0.3782 | 0 | 60.5482 | Eukaryota | 95914.0 | 0 | Chordata | 95914.0 | 0 | Carnivora | 95914.0 | 0 | Canidae | 95914.0 | 0 | Canis | 95914.0 | 0 | Canis lupus | 95914.0 | 0 |
| scf7180000048745 | 56273 | 0.4485 | 0 | 68.3388 | Eukaryota | 51011.0 | 0 | Chordata | 51011.0 | 0 | Carnivora | 51011.0 | 0 | Canidae | 51011.0 | 0 | Canis | 51011.0 | 0 | Canis lupus | 51011.0 | 0 |
| scf7180000044840 | 239014 | 0.4277 | 0 | 57.9294 | Eukaryota | 44297.0 | 0 | Chordata | 44297.0 | 0 | Carnivora | 44297.0 | 0 | Canidae | 44297.0 | 0 | Canis | 44297.0 | 0 | Canis lupus | 44297.0 | 0 |
| scf7180000054352 | 22628 | 0.3957 | 0 | 1181.5033 | Eukaryota | 55964.0 | 0 | Chordata | 55964.0 | 0 | Carnivora | 55964.0 | 0 | Canidae | 55964.0 | 0 | Canis | 55964.0 | 0 | Canis lupus | 55964.0 | 0 |
| scf7180000049045 | 65432 | 0.3582 | 0 | 59.1647 | Eukaryota | 10273.0 | 1 | Chordata | 10273.0 | 1 | Carnivora | 3795.0 | 3 | Bovidae | 3697.0 | 4 | Homo | 2781.0 | 5 | Homo sapiens | 2781.0 | 5 |
| scf7180000056447 | 303340 | 0.5013 | 0 | 58.2059 | Eukaryota | 20698.0 | 0 | Chordata | 20698.0 | 0 | Carnivora | 20698.0 | 0 | Canidae | 20698.0 | 0 | Vulpes | 12438.0 | 1 | Vulpes vulpes | 12438.0 | 1 |
| scf7180000041751 | 895565 | 0.3648 | 0 | 61.2199 | Eukaryota | 34416.0 | 0 | Chordata | 34416.0 | 0 | Carnivora | 34416.0 | 0 | Canidae | 34416.0 | 0 | Canis | 34416.0 | 0 | Canis lupus | 34416.0 | 0 |
| scf7180000040621 | 2023125 | 0.3839 | 0 | 60.5142 | Eukaryota | 91099.0 | 0 | Chordata | 91099.0 | 0 | Carnivora | 91099.0 | 0 | Canidae | 91099.0 | 0 | Canis | 79278.0 | 1 | Canis lupus | 79278.0 | 1 |
| scf7180000047272 | 28520 | 0.3844 | 0 | 58.3301 | Eukaryota | 18462.0 | 0 | Chordata | 18462.0 | 0 | Carnivora | 18462.0 | 0 | Canidae | 18462.0 | 0 | Canis | 18462.0 | 0 | Canis lupus | 18462.0 | 0 |
| scf7180000046752 | 44615 | 0.4403 | 0 | 304.6006 | Eukaryota | 14500.0 | 0 | Chordata | 13667.0 | 1 | Tetraodontiformes | 12841.0 | 2 | Tetraodontidae | 12841.0 | 2 | Takifugu | 12841.0 | 2 | Takifugu rubripes | 12841.0 | 2 |
| scf7180000043456 | 52223 | 0.3817 | 0 | 103.1669 | Eukaryota | 60882.0 | 0 | Chordata | 60882.0 | 0 | Carnivora | 60882.0 | 0 | Canidae | 60882.0 | 0 | Canis | 60882.0 | 0 | Canis lupus | 60882.0 | 0 |
| scf7180000049527 | 5066 | 0.4568 | 0 | 263.454 | Eukaryota | 955.0 | 0 | Platyhelminthes | 955.0 | 0 | Polystomatidea | 955.0 | 0 | Polystomatidae | 955.0 | 0 | Protopolystoma | 955.0 | 0 | Protopolystoma xenopodis | 955.0 | 0 |
| scf7180000054458 | 178123 | 0.5888 | 0 | 111.4607 | Eukaryota | 48622.0 | 0 | Chordata | 48622.0 | 0 | Gadiformes | 26169.0 | 2 | Gadidae | 26169.0 | 2 | Gadus | 26169.0 | 3 | Gadus morhua | 26169.0 | 3 |
| scf7180000049750 | 28965 | 0.3757 | 0 | 59.1948 | Eukaryota | 37049.0 | 0 | Chordata | 37049.0 | 0 | Carnivora | 37049.0 | 0 | Canidae | 37049.0 | 0 | Canis | 37049.0 | 0 | Canis lupus | 37049.0 | 0 |
| scf7180000045034 | 219233 | 0.6025 | 0 | 55.7254 | Eukaryota | 57069.0 | 0 | Chordata | 57069.0 | 0 | Carnivora | 57069.0 | 0 | Canidae | 57069.0 | 0 | Canis | 42606.0 | 1 | Canis lupus | 42606.0 | 1 |
| scf7180000052163 | 7571 | 0.4418 | 0 | 402.5285 | Eukaryota | 4957.0 | 0 | Chordata | 4184.0 | 1 | Tetraodontiformes | 1873.0 | 4 | Tetraodontidae | 1873.0 | 4 | Takifugu | 1873.0 | 4 | Takifugu rubripes | 1873.0 | 4 |
| scf7180000044877 | 299457 | 0.3594 | 0 | 60.152 | Eukaryota | 39359.0 | 0 | Chordata | 39359.0 | 0 | Carnivora | 19645.0 | 2 | Canidae | 19645.0 | 2 | Canis | 16422.0 | 3 | Canis lupus | 16422.0 | 3 |
| scf7180000041138 | 1274570 | 0.3999 | 0 | 60.0374 | Eukaryota | 188088.0 | 0 | Chordata | 188088.0 | 0 | Carnivora | 188088.0 | 0 | Canidae | 101956.0 | 4 | Canis | 101956.0 | 6 | Canis lupus | 101956.0 | 6 |
| scf7180000051327 | 12625 | 0.4418 | 0 | 52.6207 | Eukaryota | 92536.0 | 0 | Chordata | 92536.0 | 0 | Carnivora | 92536.0 | 0 | Canidae | 92536.0 | 0 | Canis | 92536.0 | 0 | Canis lupus | 92536.0 | 0 |
| scf7180000042649 | 183376 | 0.3814 | 0 | 59.6629 | Eukaryota | 55685.0 | 0 | Chordata | 55685.0 | 0 | Carnivora | 55685.0 | 0 | Canidae | 55685.0 | 0 | Canis | 55685.0 | 0 | Canis lupus | 55685.0 | 0 |
| scf7180000045357 | 199282 | 0.3833 | 0 | 62.77 | Eukaryota | 124829.0 | 0 | Chordata | 124829.0 | 0 | Carnivora | 124829.0 | 0 | Canidae | 124829.0 | 0 | Canis | 124829.0 | 0 | Canis lupus | 124829.0 | 0 |
| scf7180000041364 | 1061807 | 0.3479 | 0 | 58.8519 | Eukaryota | 86507.0 | 0 | Chordata | 86507.0 | 0 | Carnivora | 86507.0 | 0 | Canidae | 86507.0 | 0 | Canis | 86507.0 | 0 | Canis lupus | 86507.0 | 0 |
| scf7180000055392 | 23742 | 0.3966 | 0 | 316.2193 | Eukaryota | 40353.0 | 0 | Chordata | 40353.0 | 0 | Carnivora | 40353.0 | 0 | Canidae | 40353.0 | 0 | Canis | 40353.0 | 0 | Canis lupus | 40353.0 | 0 |
| scf7180000044528 | 249425 | 0.5095 | 0 | 60.1503 | Eukaryota | 42031.0 | 0 | Chordata | 42031.0 | 0 | Carnivora | 33753.0 | 1 | Canidae | 33753.0 | 1 | Canis | 23479.0 | 2 | Canis lupus | 23479.0 | 2 |
| scf7180000049766 | 35695 | 0.5544 | 0 | 57.6569 | Eukaryota | 3321.0 | 0 | Chordata | 3321.0 | 0 | Rodentia | 722.0 | 7 | Muridae | 722.0 | 7 | Mus | 722.0 | 8 | Mus musculus | 722.0 | 8 |
| scf7180000042832 | 143824 | 0.3895 | 0 | 56.8883 | Eukaryota | 80718.0 | 0 | Chordata | 80718.0 | 0 | Carnivora | 80718.0 | 0 | Canidae | 80718.0 | 0 | Canis | 80718.0 | 0 | Canis lupus | 80718.0 | 0 |
| scf7180000051208 | 2164 | 0.4506 | 0 | 424.177 | no-hit | 0.0 | 0 | no-hit | 0.0 | 0 | no-hit | 0.0 | 0 | no-hit | 0.0 | 0 | no-hit | 0.0 | 0 | no-hit | 0.0 | 0 |
| scf7180000045421 | 195820 | 0.3894 | 0 | 59.0866 | Eukaryota | 83211.0 | 0 | Chordata | 83211.0 | 0 | Carnivora | 83211.0 | 0 | Canidae | 83211.0 | 0 | Canis | 83211.0 | 0 | Canis lupus | 83211.0 | 0 |
| scf7180000047221 | 2683 | 0.4394 | 0 | 218.2223 | no-hit | 0.0 | 0 | no-hit | 0.0 | 0 | no-hit | 0.0 | 0 | no-hit | 0.0 | 0 | no-hit | 0.0 | 0 | no-hit | 0.0 | 0 |
| scf7180000048671 | 33738 | 0.4419 | 0 | 125.6856 | Eukaryota | 109767.0 | 0 | Chordata | 62622.0 | 1 | Gadiformes | 62622.0 | 2 | Gadidae | 62622.0 | 2 | Gadus | 62622.0 | 2 | Gadus morhua | 62622.0 | 2 |
| scf7180000049228 | 4571 | 0.4634 | 0 | 109.8711 | Eukaryota | 726.0 | 0 | Chordata | 726.0 | 0 | Syngnathiformes | 726.0 | 0 | Syngnathidae | 726.0 | 0 | Syngnathus | 726.0 | 0 | Syngnathus acus | 726.0 | 0 |
| scf7180000047663 | 107035 | 0.3605 | 0 | 61.4971 | Eukaryota | 11003.0 | 0 | Chordata | 11003.0 | 0 | Artiodactyla | 7523.0 | 1 | Delphinidae | 3799.0 | 3 | Sus | 3724.0 | 5 | Sus scrofa | 3724.0 | 5 |
| scf7180000041929 | 37077 | 0.3744 | 0 | 59.396 | Eukaryota | 35671.0 | 0 | Chordata | 35671.0 | 0 | Carnivora | 35671.0 | 0 | Canidae | 35671.0 | 0 | Canis | 35671.0 | 0 | Canis lupus | 35671.0 | 0 |
| scf7180000050069 | 1988 | 0.4396 | 0 | 106.4826 | no-hit | 0.0 | 0 | no-hit | 0.0 | 0 | no-hit | 0.0 | 0 | no-hit | 0.0 | 0 | no-hit | 0.0 | 0 | no-hit | 0.0 | 0 |
| scf7180000043441 | 161977 | 0.5183 | 0 | 55.3797 | Eukaryota | 5742.0 | 0 | Chordata | 5742.0 | 0 | Carnivora | 5742.0 | 0 | Canidae | 5742.0 | 0 | Canis | 5742.0 | 0 | Canis lupus | 5742.0 | 0 |
| scf7180000047741 | 20186 | 0.3941 | 0 | 43.6296 | Eukaryota | 45162.0 | 0 | Chordata | 45162.0 | 0 | Carnivora | 45162.0 | 0 | Canidae | 45162.0 | 0 | Canis | 45162.0 | 0 | Canis lupus | 45162.0 | 0 |
| scf7180000054997 | 33213 | 0.4897 | 0 | 429.6421 | Eukaryota | 15373.0 | 0 | Chordata | 15373.0 | 0 | Carnivora | 15373.0 | 0 | Canidae | 15373.0 | 0 | Canis | 9379.0 | 1 | Canis lupus | 9379.0 | 1 |
| scf7180000049353 | 5516 | 0.4578 | 0 | 118.8106 | Eukaryota | 809.0 | 0 | Chordata | 809.0 | 0 | Syngnathiformes | 809.0 | 0 | Syngnathidae | 809.0 | 0 | Syngnathus | 809.0 | 0 | Syngnathus acus | 809.0 | 0 |
| scf7180000054166 | 369805 | 0.3934 | 0 | 60.9474 | Eukaryota | 95221.0 | 0 | Chordata | 95221.0 | 0 | Carnivora | 95221.0 | 0 | Canidae | 95221.0 | 0 | Canis | 95221.0 | 0 | Canis lupus | 95221.0 | 0 |
| scf7180000054442 | 33922 | 0.4466 | 0 | 593.2229 | Eukaryota | 78809.0 | 0 | Chordata | 44208.0 | 1 | Gadiformes | 25804.0 | 5 | Gadidae | 25804.0 | 6 | Gadus | 25804.0 | 6 | Gadus morhua | 25804.0 | 6 |
| scf7180000040477 | 2597830 | 0.4482 | 0 | 63.8116 | Eukaryota | 97797.0 | 0 | Chordata | 97797.0 | 0 | Carnivora | 97797.0 | 0 | Canidae | 97797.0 | 0 | Canis | 87969.0 | 1 | Canis lupus | 87969.0 | 1 |
| scf7180000051035 | 5802 | 0.4483 | 0 | 67.0503 | no-hit | 0.0 | 0 | no-hit | 0.0 | 0 | no-hit | 0.0 | 0 | no-hit | 0.0 | 0 | no-hit | 0.0 | 0 | no-hit | 0.0 | 0 |
| scf7180000046395 | 155118 | 0.3438 | 0 | 58.6739 | Eukaryota | 12279.0 | 0 | Chordata | 12279.0 | 0 | Carnivora | 12279.0 | 0 | Canidae | 9617.0 | 1 | Canis | 5999.0 | 2 | Canis lupus | 5999.0 | 2 |
| scf7180000046365 | 24887 | 0.4371 | 0 | 219.7586 | Eukaryota | 292.0 | 0 | Chordata | 292.0 | 0 | Kurtiformes | 292.0 | 0 | Apogonidae | 292.0 | 0 | Sphaeramia | 292.0 | 0 | Sphaeramia orbicularis | 292.0 | 0 |
| scf7180000042256 | 635274 | 0.3938 | 0 | 59.9503 | Eukaryota | 154592.0 | 0 | Chordata | 154592.0 | 0 | Carnivora | 154592.0 | 0 | Canidae | 154592.0 | 0 | Canis | 154592.0 | 0 | Canis lupus | 154592.0 | 0 |
| scf7180000055265 | 10576 | 0.4393 | 0 | 100.8419 | Eukaryota | 43827.0 | 0 | Chordata | 29615.0 | 2 | Gadiformes | 24530.0 | 5 | Gadidae | 24530.0 | 5 | Gadus | 24530.0 | 5 | Gadus morhua | 24530.0 | 5 |
| scf7180000046648 | 138732 | 0.4372 | 0 | 61.0621 | Eukaryota | 27113.0 | 0 | Chordata | 27113.0 | 0 | Carnivora | 27113.0 | 0 | Canidae | 18145.0 | 1 | Canis | 14054.0 | 2 | Canis lupus | 14054.0 | 2 |
| scf7180000043558 | 386435 | 0.442 | 0 | 60.2218 | Eukaryota | 74746.0 | 0 | Chordata | 74746.0 | 0 | Carnivora | 74746.0 | 0 | Canidae | 74746.0 | 0 | Canis | 74746.0 | 0 | Canis lupus | 74746.0 | 0 |
| scf7180000056527 | 249148 | 0.5381 | 0 | 58.9827 | Eukaryota | 53655.0 | 0 | Chordata | 53655.0 | 0 | Carnivora | 53655.0 | 0 | Canidae | 53655.0 | 0 | Canis | 53655.0 | 0 | Canis lupus | 53655.0 | 0 |
| scf7180000047688 | 22748 | 0.4579 | 0 | 277.4797 | Eukaryota | 11427.0 | 0 | Chordata | 11427.0 | 0 | Carnivora | 11427.0 | 0 | Canidae | 11427.0 | 0 | Canis | 8797.0 | 1 | Canis lupus | 7866.0 | 2 |
| scf7180000055004 | 22424 | 0.4389 | 0 | 197.438 | Eukaryota | 427.0 | 0 | Chordata | 279.0 | 1 | Kurtiformes | 279.0 | 1 | Apogonidae | 279.0 | 1 | Sphaeramia | 279.0 | 1 | Sphaeramia orbicularis | 279.0 | 1 |
| scf7180000049549 | 5145 | 0.4416 | 0 | 145.8131 | Eukaryota | 795.0 | 0 | Platyhelminthes | 795.0 | 0 | Polystomatidea | 795.0 | 0 | Polystomatidae | 795.0 | 0 | Protopolystoma | 795.0 | 0 | Protopolystoma xenopodis | 795.0 | 0 |
| scf7180000050933 | 17309 | 0.4324 | 0 | 800.0775 | Eukaryota | 45767.0 | 0 | Chordata | 45767.0 | 0 | Carnivora | 45767.0 | 0 | Canidae | 45767.0 | 0 | Canis | 45767.0 | 0 | Canis lupus | 45767.0 | 0 |
| scf7180000042652 | 565531 | 0.3852 | 0 | 60.8925 | Eukaryota | 26449.0 | 0 | Chordata | 26449.0 | 0 | Carnivora | 26449.0 | 0 | Canidae | 26449.0 | 0 | Canis | 22814.0 | 1 | Canis lupus | 22814.0 | 1 |
| scf7180000055507 | 28787 | 0.4104 | 0 | 365.2245 | Eukaryota | 87872.0 | 0 | Chordata | 87872.0 | 0 | Carnivora | 87872.0 | 0 | Canidae | 87872.0 | 0 | Canis | 87872.0 | 0 | Canis lupus | 87872.0 | 0 |
| scf7180000044690 | 55843 | 0.4912 | 0 | 176.5813 | Eukaryota | 81470.0 | 0 | Chordata | 81470.0 | 0 | Carnivora | 81470.0 | 0 | Canidae | 81470.0 | 0 | Canis | 81470.0 | 0 | Canis lupus | 81470.0 | 0 |
| scf7180000049209 | 56077 | 0.4633 | 0 | 60.6682 | Eukaryota | 47240.0 | 0 | Chordata | 47240.0 | 0 | Carnivora | 47240.0 | 0 | Canidae | 47240.0 | 0 | Canis | 43018.0 | 1 | Canis lupus | 43018.0 | 1 |
| scf7180000052958 | 8974 | 0.5021 | 0 | 57.4138 | Eukaryota | 7904.0 | 0 | Chordata | 7904.0 | 0 | Carnivora | 7904.0 | 0 | Canidae | 7904.0 | 0 | Canis | 6228.0 | 1 | Canis lupus | 5010.0 | 2 |
| scf7180000047963 | 68987 | 0.4395 | 0 | 61.3231 | Eukaryota | 26797.0 | 0 | Chordata | 26797.0 | 0 | Carnivora | 26797.0 | 0 | Canidae | 26797.0 | 0 | Canis | 20906.0 | 1 | Canis lupus | 20906.0 | 1 |
| scf7180000055136 | 19123 | 0.4508 | 0 | 168.8362 | Eukaryota | 2820.0 | 0 | Chordata | 2820.0 | 0 | Carnivora | 2820.0 | 0 | Canidae | 2820.0 | 0 | Canis | 2820.0 | 0 | Canis lupus | 2820.0 | 0 |
| scf7180000052522 | 1484 | 0.4535 | 0 | 249.7847 | no-hit | 0.0 | 0 | no-hit | 0.0 | 0 | no-hit | 0.0 | 0 | no-hit | 0.0 | 0 | no-hit | 0.0 | 0 | no-hit | 0.0 | 0 |
| scf7180000050608 | 10637 | 0.4522 | 0 | 102.3307 | Eukaryota | 368.0 | 0 | Platyhelminthes | 368.0 | 0 | Polystomatidea | 368.0 | 0 | Polystomatidae | 368.0 | 0 | Protopolystoma | 368.0 | 0 | Protopolystoma xenopodis | 368.0 | 0 |
| scf7180000041654 | 297371 | 0.3998 | 0 | 59.6298 | Eukaryota | 43489.0 | 0 | Chordata | 43489.0 | 0 | Carnivora | 43489.0 | 0 | Canidae | 43489.0 | 0 | Canis | 28283.0 | 1 | Canis lupus | 28283.0 | 1 |
| scf7180000048306 | 81062 | 0.4439 | 0 | 75.3911 | Eukaryota | 7025.0 | 0 | Chordata | 2621.0 | 2 | Carnivora | 2621.0 | 2 | Canidae | 2621.0 | 2 | Canis | 2621.0 | 2 | Canis lupus | 2621.0 | 2 |
| scf7180000054163 | 199861 | 0.4122 | 0 | 102.1493 | Eukaryota | 103629.0 | 0 | Chordata | 103629.0 | 0 | Carnivora | 103629.0 | 0 | Canidae | 103629.0 | 0 | Canis | 103629.0 | 0 | Canis lupus | 103629.0 | 0 |
| scf7180000044597 | 262525 | 0.5377 | 0 | 59.9698 | Eukaryota | 21237.0 | 0 | Chordata | 21237.0 | 0 | Carnivora | 21237.0 | 0 | Canidae | 17013.0 | 1 | Canis | 15131.0 | 4 | Canis lupus | 15131.0 | 4 |
| scf7180000041083 | 155322 | 0.3517 | 0 | 59.4295 | Eukaryota | 76983.0 | 0 | Chordata | 76983.0 | 0 | Carnivora | 76983.0 | 0 | Canidae | 76983.0 | 0 | Canis | 76983.0 | 0 | Canis lupus | 76983.0 | 0 |
| scf7180000040935 | 1541987 | 0.4323 | 0 | 58.994 | Eukaryota | 92115.0 | 0 | Chordata | 92115.0 | 0 | Carnivora | 92115.0 | 0 | Canidae | 92115.0 | 0 | Canis | 92115.0 | 0 | Canis lupus | 92115.0 | 0 |
| scf7180000046605 | 55537 | 0.4389 | 0 | 174.3752 | Eukaryota | 48791.0 | 0 | Chordata | 48791.0 | 0 | Carnivora | 48791.0 | 0 | Canidae | 48791.0 | 0 | Canis | 48791.0 | 0 | Canis lupus | 48791.0 | 0 |
| scf7180000052415 | 10482 | 0.508 | 0 | 304.0469 | Eukaryota | 8948.0 | 0 | Chordata | 8948.0 | 0 | Carnivora | 8948.0 | 0 | Canidae | 8948.0 | 0 | Canis | 6656.0 | 1 | Canis lupus | 5379.0 | 2 |
| scf7180000042970 | 36168 | 0.4412 | 0 | 146.6065 | Eukaryota | 82862.0 | 0 | Chordata | 82862.0 | 0 | Carnivora | 82862.0 | 0 | Canidae | 82862.0 | 0 | Canis | 82862.0 | 0 | Canis lupus | 82862.0 | 0 |
| scf7180000042859 | 212644 | 0.3822 | 0 | 64.2499 | Eukaryota | 70377.0 | 0 | Chordata | 70377.0 | 0 | Carnivora | 70377.0 | 0 | Canidae | 70377.0 | 0 | Canis | 70377.0 | 0 | Canis lupus | 70377.0 | 0 |
| scf7180000043061 | 5198 | 0.4198 | 0 | 137.7684 | Eukaryota | 3434.0 | 0 | Chordata | 3434.0 | 0 | Carnivora | 3434.0 | 0 | Canidae | 3434.0 | 0 | Canis | 3434.0 | 0 | Canis lupus | 3434.0 | 0 |
| scf7180000041939 | 98544 | 0.4273 | 0 | 107.698 | Eukaryota | 87379.0 | 0 | Chordata | 87379.0 | 0 | Carnivora | 87379.0 | 0 | Canidae | 87379.0 | 0 | Canis | 87379.0 | 0 | Canis lupus | 87379.0 | 0 |
| scf7180000043415 | 526506 | 0.3952 | 0 | 59.5576 | Eukaryota | 89531.0 | 0 | Chordata | 89531.0 | 0 | Carnivora | 89531.0 | 0 | Canidae | 89531.0 | 0 | Canis | 89531.0 | 0 | Canis lupus | 89531.0 | 0 |
| scf7180000040871 | 633296 | 0.339 | 0 | 59.2463 | Eukaryota | 66477.0 | 0 | Chordata | 66477.0 | 0 | Carnivora | 66477.0 | 0 | Canidae | 66477.0 | 0 | Canis | 66477.0 | 0 | Canis lupus | 66477.0 | 0 |
| scf7180000056238 | 1252623 | 0.3689 | 0 | 59.6168 | Eukaryota | 95941.0 | 0 | Chordata | 95941.0 | 0 | Carnivora | 95941.0 | 0 | Canidae | 95941.0 | 0 | Canis | 95941.0 | 0 | Canis lupus | 95941.0 | 0 |
| scf7180000044396 | 312320 | 0.4253 | 0 | 59.3272 | Eukaryota | 28327.0 | 0 | Chordata | 28327.0 | 0 | Carnivora | 28327.0 | 0 | Canidae | 21577.0 | 1 | Canis | 16993.0 | 2 | Canis lupus | 16993.0 | 2 |
| scf7180000049732 | 8903 | 0.4531 | 0 | 964.7766 | Eukaryota | 419.0 | 0 | Streptophyta | 419.0 | 0 | Fabales | 419.0 | 0 | Fabaceae | 419.0 | 0 | Cicer | 419.0 | 0 | Cicer arietinum | 419.0 | 0 |
| scf7180000045062 | 59148 | 0.3777 | 0 | 53.4752 | Eukaryota | 70279.0 | 0 | Chordata | 70279.0 | 0 | Carnivora | 70279.0 | 0 | Canidae | 70279.0 | 0 | Canis | 70279.0 | 0 | Canis lupus | 70279.0 | 0 |
| scf7180000043282 | 244189 | 0.5214 | 0 | 59.4543 | Eukaryota | 33962.0 | 0 | Chordata | 33962.0 | 0 | Carnivora | 33962.0 | 0 | Canidae | 33962.0 | 0 | Canis | 28351.0 | 1 | Canis lupus | 28351.0 | 1 |
| scf7180000045716 | 161019 | 0.4138 | 0 | 57.0764 | Eukaryota | 28856.0 | 0 | Chordata | 28856.0 | 0 | Primates | 24238.0 | 2 | Hominidae | 21025.0 | 3 | Homo | 19691.0 | 5 | Homo sapiens | 19691.0 | 5 |
| scf7180000048823 | 363655 | 0.5188 | 0 | 56.5194 | Eukaryota | 43685.0 | 0 | Chordata | 43685.0 | 0 | Carnivora | 43685.0 | 0 | Canidae | 43685.0 | 0 | Canis | 40589.0 | 1 | Canis lupus | 40589.0 | 1 |
| scf7180000046148 | 106817 | 0.6188 | 0 | 54.229 | Eukaryota | 75281.0 | 0 | Chordata | 75281.0 | 0 | Carnivora | 75281.0 | 0 | Canidae | 75281.0 | 0 | Canis | 72832.0 | 1 | Canis lupus | 72832.0 | 1 |
| scf7180000044668 | 282677 | 0.3996 | 0 | 58.245 | Eukaryota | 89080.0 | 0 | Chordata | 89080.0 | 0 | Carnivora | 89080.0 | 0 | Canidae | 89080.0 | 0 | Canis | 89080.0 | 0 | Canis lupus | 89080.0 | 0 |
| scf7180000045064 | 191430 | 0.3371 | 0 | 56.4608 | Eukaryota | 75306.0 | 0 | Chordata | 75306.0 | 0 | Carnivora | 75306.0 | 0 | Canidae | 75306.0 | 0 | Canis | 75306.0 | 0 | Canis lupus | 75306.0 | 0 |
| scf7180000048623 | 51482 | 0.4854 | 0 | 60.2603 | Eukaryota | 4054.0 | 0 | Chordata | 4054.0 | 0 | Primates | 2187.0 | 2 | Canidae | 1525.0 | 3 | Canis | 1525.0 | 4 | Canis lupus | 1525.0 | 4 |
| scf7180000046143 | 49874 | 0.3993 | 0 | 56.096 | Eukaryota | 27147.0 | 0 | Chordata | 27147.0 | 0 | Carnivora | 25500.0 | 1 | Canidae | 24159.0 | 2 | Canis | 24159.0 | 2 | Canis lupus | 24159.0 | 2 |
| scf7180000040725 | 202081 | 0.3529 | 0 | 61.6894 | Eukaryota | 94807.0 | 0 | Chordata | 94807.0 | 0 | Carnivora | 94807.0 | 0 | Canidae | 94807.0 | 0 | Canis | 94807.0 | 0 | Canis lupus | 94807.0 | 0 |
| scf7180000055428 | 26405 | 0.4423 | 0 | 191.7611 | Eukaryota | 11914.0 | 0 | Chordata | 8777.0 | 2 | Synbranchiformes | 2835.0 | 7 | Mastacembelidae | 2835.0 | 7 | Mastacembelus | 2835.0 | 7 | Mastacembelus armatus | 2835.0 | 7 |
| scf7180000044413 | 349284 | 0.4458 | 0 | 59.8845 | Eukaryota | 51349.0 | 0 | Chordata | 51349.0 | 0 | Carnivora | 51349.0 | 0 | Canidae | 51349.0 | 0 | Canis | 35077.0 | 1 | Canis lupus | 35077.0 | 1 |
| scf7180000047818 | 70556 | 0.4114 | 0 | 97.6168 | Eukaryota | 58325.0 | 0 | Chordata | 58325.0 | 0 | Carnivora | 58325.0 | 0 | Canidae | 58325.0 | 0 | Canis | 58325.0 | 0 | Canis lupus | 58325.0 | 0 |
| scf7180000056233 | 1562192 | 0.4035 | 0 | 60.1139 | Eukaryota | 58085.0 | 0 | Chordata | 58085.0 | 0 | Carnivora | 58085.0 | 0 | Canidae | 58085.0 | 0 | Canis | 58085.0 | 0 | Canis lupus | 58085.0 | 0 |
| scf7180000043112 | 50626 | 0.3702 | 0 | 66.3241 | Eukaryota | 74522.0 | 0 | Chordata | 74522.0 | 0 | Carnivora | 74522.0 | 0 | Canidae | 74522.0 | 0 | Canis | 74522.0 | 0 | Canis lupus | 74522.0 | 0 |
| scf7180000042871 | 235530 | 0.395 | 0 | 59.4387 | Eukaryota | 90771.0 | 0 | Chordata | 90771.0 | 0 | Carnivora | 90771.0 | 0 | Canidae | 90771.0 | 0 | Canis | 90771.0 | 0 | Canis lupus | 90771.0 | 0 |
| scf7180000051751 | 3846 | 0.5023 | 0 | 46.482 | Eukaryota | 6369.0 | 0 | Chordata | 6369.0 | 0 | Carnivora | 6369.0 | 0 | Canidae | 6369.0 | 0 | Canis | 5329.0 | 1 | Canis lupus | 5329.0 | 1 |
| scf7180000048979 | 9809 | 0.4392 | 0 | 108.0638 | Eukaryota | 20736.0 | 0 | Chordata | 18915.0 | 1 | Gadiformes | 10989.0 | 6 | Gadidae | 10989.0 | 6 | Gadus | 10989.0 | 6 | Gadus morhua | 10989.0 | 6 |
| scf7180000049049 | 71314 | 0.423 | 0 | 58.1252 | Eukaryota | 18788.0 | 0 | Chordata | 18788.0 | 0 | Primates | 9836.0 | 2 | Hominidae | 9836.0 | 2 | Homo | 6514.0 | 5 | Homo sapiens | 6514.0 | 5 |
| scf7180000052540 | 6000 | 0.5222 | 0 | 194.4913 | Eukaryota | 47080.0 | 0 | Chordata | 47080.0 | 0 | Carnivora | 47080.0 | 0 | Canidae | 47080.0 | 0 | Canis | 47080.0 | 0 | Canis lupus | 47080.0 | 0 |
| scf7180000051670 | 5021 | 0.4517 | 0 | 462.5983 | no-hit | 0.0 | 0 | no-hit | 0.0 | 0 | no-hit | 0.0 | 0 | no-hit | 0.0 | 0 | no-hit | 0.0 | 0 | no-hit | 0.0 | 0 |
| scf7180000043986 | 400655 | 0.3837 | 0 | 59.7974 | Eukaryota | 34631.0 | 0 | Chordata | 34631.0 | 0 | Carnivora | 34631.0 | 0 | Canidae | 17674.0 | 2 | Canis | 17674.0 | 2 | Canis lupus | 17674.0 | 2 |
| scf7180000050077 | 4709 | 0.4392 | 0 | 79.5464 | Eukaryota | 16173.0 | 0 | Chordata | 12963.0 | 3 | Gadiformes | 7336.0 | 8 | Gadidae | 7336.0 | 8 | Gadus | 7336.0 | 8 | Gadus morhua | 7336.0 | 8 |

| | | | | | | | | | | | | | | | | | | | | |
|---|---|---|---|---|---|---|---|---|---|---|---|---|---|---|---|---|---|---|---|---|
| scf7180000046816 | 101778 | 0.5239 | 0 | 58.531 | Eukaryota | 22062.0 | 0 | Chordata | 22062.0 | 0 | Carnivora | 22062.0 | 0 | Canidae | 22062.0 | 0 | Canis | 16347.0 | 1 | Canis lupus | 16347.0 | 1 |
| scf7180000047801 | 76925 | 0.3606 | 0 | 59.7461 | Eukaryota | 55652.0 | 0 | Chordata | 55652.0 | 0 | Carnivora | 55652.0 | 0 | Canidae | 55652.0 | 0 | Canis | 55652.0 | 0 | Canis lupus | 55652.0 | 0 |
| scf7180000048181 | 73606 | 0.3775 | 0 | 58.4524 | Eukaryota | 9410.0 | 0 | Chordata | 9410.0 | 0 | Carnivora | 6840.0 | 1 | Canidae | 5033.0 | 2 | Canis | 5033.0 | 3 | Canis lupus | 5033.0 | 3 |
| scf7180000048389 | 64996 | 0.3737 | 0 | 59.8959 | Eukaryota | 60696.0 | 0 | Chordata | 60696.0 | 0 | Carnivora | 60696.0 | 0 | Canidae | 60696.0 | 0 | Canis | 60696.0 | 0 | Canis lupus | 60696.0 | 0 |
| scf7180000052709 | 11013 | 0.5079 | 0 | 268.1807 | Eukaryota | 8904.0 | 0 | Chordata | 8904.0 | 0 | Carnivora | 8904.0 | 0 | Canidae | 8904.0 | 0 | Canis | 7183.0 | 1 | Canis lupus | 5929.0 | 2 |
| scf7180000048802 | 10558 | 0.4338 | 0 | 165.6863 | Eukaryota | 4054.0 | 0 | Platyhelminthes | 2764.0 | 1 | Polystomatidea | 2764.0 | 2 | Polystomatidae | 2764.0 | 2 | Protopolystoma | 2764.0 | 3 | Protopolystoma xenopodis | 2764.0 | 3 |
| scf7180000052520 | 6523 | 0.4552 | 0 | 1448.7915 | no-hit | 0.0 | 0 | no-hit | 0.0 | 0 | no-hit | 0.0 | 0 | no-hit | 0.0 | 0 | no-hit | 0.0 | 0 | no-hit | 0.0 | 0 |
| scf7180000046484 | 7455 | 0.4482 | 0 | 95.3784 | no-hit | 0.0 | 0 | no-hit | 0.0 | 0 | no-hit | 0.0 | 0 | no-hit | 0.0 | 0 | no-hit | 0.0 | 0 | no-hit | 0.0 | 0 |
| scf7180000051474 | 8704 | 0.4485 | 0 | 427.4433 | Eukaryota | 1601.0 | 0 | Platyhelminthes | 834.0 | 1 | Polystomatidea | 834.0 | 2 | Polystomatidae | 834.0 | 2 | Protopolystoma | 834.0 | 2 | Protopolystoma xenopodis | 834.0 | 2 |
| scf7180000045134 | 236273 | 0.3706 | 0 | 59.9335 | Eukaryota | 27312.0 | 0 | Chordata | 27312.0 | 0 | Carnivora | 27312.0 | 0 | Canidae | 27312.0 | 0 | Canis | 27312.0 | 0 | Canis lupus | 27312.0 | 0 |
| scf7180000054335 | 23931 | 0.4297 | 0 | 88.9141 | Eukaryota | 46282.0 | 0 | Chordata | 46282.0 | 0 | Carnivora | 46282.0 | 0 | Canidae | 46282.0 | 0 | Canis | 46282.0 | 0 | Canis lupus | 46282.0 | 0 |
| scf7180000040687 | 363653 | 0.3945 | 0 | 58.5097 | Eukaryota | 35229.0 | 0 | Chordata | 35229.0 | 0 | Carnivora | 35229.0 | 0 | Canidae | 35229.0 | 0 | Canis | 35229.0 | 0 | Canis lupus | 35229.0 | 0 |
| scf7180000054549 | 7977 | 0.4437 | 0 | 89.2675 | Eukaryota | 2041.0 | 0 | Streptophyta | 1062.0 | 1 | Fabales | 1062.0 | 2 | Fabaceae | 1062.0 | 2 | Vigna | 1062.0 | 2 | Vigna unguiculata | 1062.0 | 2 |
| scf7180000048154 | 7288 | 0.4391 | 0 | 132.9297 | no-hit | 0.0 | 0 | no-hit | 0.0 | 0 | no-hit | 0.0 | 0 | no-hit | 0.0 | 0 | no-hit | 0.0 | 0 | no-hit | 0.0 | 0 |
| scf7180000048579 | 40674 | 0.3853 | 0 | 62.2203 | Eukaryota | 7672.0 | 0 | Chordata | 7672.0 | 0 | Carnivora | 4339.0 | 2 | Hominidae | 2583.0 | 4 | Homo | 2583.0 | 4 | Homo sapiens | 2583.0 | 4 |
| scf7180000043388 | 122193 | 0.3501 | 0 | 60.1176 | Eukaryota | 12187.0 | 0 | Chordata | 12187.0 | 0 | Carnivora | 12187.0 | 0 | Canidae | 8621.0 | 1 | Canis | 8621.0 | 1 | Canis lupus | 8621.0 | 1 |
| scf7180000045553 | 5620 | 0.4559 | 0 | 75.7271 | Eukaryota | 165.0 | 0 | Platyhelminthes | 165.0 | 0 | Diphyllobothriidea | 165.0 | 0 | Diphyllobothriidae | 165.0 | 0 | Dibothriocephalus | 165.0 | 0 | Dibothriocephalus latus | 165.0 | 0 |
| scf7180000045435 | 190464 | 0.4195 | 0 | 59.1606 | Eukaryota | 21968.0 | 0 | Chordata | 21968.0 | 0 | Carnivora | 21968.0 | 0 | Canidae | 20926.0 | 1 | Canis | 19810.0 | 2 | Canis lupus | 19810.0 | 2 |
| scf7180000045608 | 195311 | 0.5466 | 0 | 58.5757 | Eukaryota | 30549.0 | 0 | Chordata | 30549.0 | 0 | Carnivora | 30549.0 | 0 | Canidae | 30549.0 | 0 | Canis | 26861.0 | 1 | Canis lupus | 26861.0 | 1 |
| scf7180000048808 | 10041 | 0.437 | 0 | 69.7906 | Eukaryota | 531.0 | 0 | Platyhelminthes | 531.0 | 0 | Polystomatidea | 531.0 | 0 | Polystomatidae | 531.0 | 0 | Protopolystoma | 531.0 | 0 | Protopolystoma xenopodis | 531.0 | 0 |
| scf7180000056274 | 1178759 | 0.3653 | 0 | 60.5674 | Eukaryota | 94544.0 | 0 | Chordata | 94544.0 | 0 | Carnivora | 94544.0 | 0 | Canidae | 94544.0 | 0 | Canis | 94544.0 | 0 | Canis lupus | 94544.0 | 0 |
| scf7180000054504 | 17022 | 0.4183 | 0 | 482.3863 | Eukaryota | 92219.0 | 0 | Chordata | 92219.0 | 0 | Carnivora | 92219.0 | 0 | Canidae | 92219.0 | 0 | Canis | 92219.0 | 0 | Canis lupus | 92219.0 | 0 |
| scf7180000056164 | 2514 | 0.4348 | 0 | 267.1013 | no-hit | 0.0 | 0 | no-hit | 0.0 | 0 | no-hit | 0.0 | 0 | no-hit | 0.0 | 0 | no-hit | 0.0 | 0 | no-hit | 0.0 | 0 |
| scf7180000040837 | 431508 | 0.366 | 0 | 60.0263 | Eukaryota | 62213.0 | 0 | Chordata | 62213.0 | 0 | Carnivora | 62213.0 | 0 | Canidae | 62213.0 | 0 | Canis | 62213.0 | 0 | Canis lupus | 62213.0 | 0 |
| scf7180000049871 | 42198 | 0.4689 | 100 | 53.0443 | Eukaryota | 21930.0 | 0 | Chordata | 21930.0 | 0 | Carnivora | 21930.0 | 0 | Canidae | 21930.0 | 0 | Canis | 13158.0 | 1 | Canis lupus | 13158.0 | 1 |
| scf7180000045091 | 191108 | 0.3777 | 0 | 59.7407 | Eukaryota | 57297.0 | 0 | Chordata | 57297.0 | 0 | Carnivora | 57297.0 | 0 | Canidae | 57297.0 | 0 | Canis | 57297.0 | 0 | Canis lupus | 57297.0 | 0 |
| scf7180000044616 | 557755 | 0.4235 | 0 | 60.7073 | Eukaryota | 70627.0 | 0 | Chordata | 70627.0 | 0 | Carnivora | 70627.0 | 0 | Canidae | 66169.0 | 1 | Canis | 59629.0 | 2 | Canis lupus | 59629.0 | 2 |
| scf7180000055023 | 36341 | 0.4422 | 0 | 170.2433 | Eukaryota | 28302.0 | 0 | Chordata | 27459.0 | 1 | Gadiformes | 16782.0 | 5 | Gadidae | 16782.0 | 5 | Gadus | 16782.0 | 5 | Gadus morhua | 16782.0 | 5 |
| scf7180000045098 | 35252 | 0.4729 | 0 | 114.8511 | Eukaryota | 29360.0 | 0 | Chordata | 29360.0 | 0 | Carnivora | 29360.0 | 0 | Canidae | 29360.0 | 0 | Canis | 29360.0 | 0 | Canis lupus | 29360.0 | 0 |
| scf7180000046529 | 130854 | 0.3725 | 0 | 61.7266 | Eukaryota | 21241.0 | 0 | Chordata | 21241.0 | 0 | Primates | 10413.0 | 3 | Hominidae | 10413.0 | 3 | Homo | 10413.0 | 3 | Homo sapiens | 10413.0 | 3 |
| scf7180000040952 | 1511787 | 0.3686 | 0 | 59.4884 | Eukaryota | 92345.0 | 0 | Chordata | 92345.0 | 0 | Carnivora | 92345.0 | 0 | Canidae | 92345.0 | 0 | Canis | 92345.0 | 0 | Canis lupus | 92345.0 | 0 |
| scf7180000051692 | 6632 | 0.4448 | 0 | 544.4881 | Eukaryota | 2648.0 | 0 | Platyhelminthes | 1492.0 | 2 | Polystomatidea | 1492.0 | 2 | Polystomatidae | 1492.0 | 2 | Protopolystoma | 1492.0 | 2 | Protopolystoma xenopodis | 1492.0 | 2 |
| scf7180000043216 | 636449 | 0.356 | 0 | 59.0311 | Eukaryota | 67329.0 | 0 | Chordata | 67329.0 | 0 | Carnivora | 67329.0 | 0 | Canidae | 67329.0 | 0 | Canis | 67329.0 | 0 | Canis lupus | 67329.0 | 0 |
| scf7180000047443 | 30557 | 0.5044 | 0 | 49.5962 | Eukaryota | 3208.0 | 0 | Chordata | 3208.0 | 0 | Carnivora | 2093.0 | 2 | Canidae | 2093.0 | 2 | Canis | 2093.0 | 2 | Canis lupus | 2093.0 | 2 |
| scf7180000055683 | 5508 | 0.4336 | 0 | 181.888 | no-hit | 0.0 | 0 | no-hit | 0.0 | 0 | no-hit | 0.0 | 0 | no-hit | 0.0 | 0 | no-hit | 0.0 | 0 | no-hit | 0.0 | 0 |
| scf7180000051375 | 3824 | 0.4417 | 0 | 123.4062 | Eukaryota | 4285.0 | 0 | Chordata | 2295.0 | 2 | Salmoniformes | 2295.0 | 2 | Salmonidae | 2295.0 | 2 | Salmo | 2295.0 | 2 | Salmo trutta | 2295.0 | 2 |
| scf7180000047495 | 29370 | 0.4133 | 0 | 53.7181 | Eukaryota | 87592.0 | 0 | Chordata | 87592.0 | 0 | Carnivora | 87592.0 | 0 | Canidae | 87592.0 | 0 | Canis | 87592.0 | 0 | Canis lupus | 87592.0 | 0 |
| scf7180000043679 | 251806 | 0.4414 | 0 | 60.1231 | Eukaryota | 70156.0 | 0 | Chordata | 70156.0 | 0 | Carnivora | 70156.0 | 0 | Canidae | 70156.0 | 0 | Canis | 43839.0 | 1 | Canis lupus | 43839.0 | 1 |
| scf7180000047977 | 29327 | 0.4425 | 0 | 141.1381 | no-hit | 0.0 | 0 | no-hit | 0.0 | 0 | no-hit | 0.0 | 0 | no-hit | 0.0 | 0 | no-hit | 0.0 | 0 | no-hit | 0.0 | 0 |
| scf7180000043156 | 159563 | 0.3965 | 0 | 60.6672 | Eukaryota | 61916.0 | 0 | Chordata | 61916.0 | 0 | Carnivora | 61916.0 | 0 | Canidae | 61916.0 | 0 | Canis | 61916.0 | 0 | Canis lupus | 61916.0 | 0 |
| scf7180000042917 | 1951 | 0.4131 | 0 | 7.4691 | Eukaryota | 12251.0 | 0 | Chordata | 12251.0 | 0 | Carnivora | 12251.0 | 0 | Canidae | 12251.0 | 0 | Canis | 12251.0 | 0 | Canis lupus | 12251.0 | 0 |
| scf7180000052104 | 9931 | 0.4511 | 0 | 1751.7747 | Eukaryota | 3525.0 | 0 | Platyhelminthes | 1762.0 | 3 | Polystomatidea | 1762.0 | 3 | Polystomatidae | 1762.0 | 3 | Protopolystoma | 1762.0 | 3 | Protopolystoma xenopodis | 1762.0 | 3 |
| scf7180000046913 | 3937 | 0.3686 | 0 | 27.0331 | Eukaryota | 3316.0 | 0 | Chordata | 3316.0 | 0 | Carnivora | 3316.0 | 0 | Canidae | 3316.0 | 0 | Canis | 3316.0 | 0 | Canis lupus | 3316.0 | 0 |
| scf7180000040993 | 1878141 | 0.3807 | 0 | 60.194 | Eukaryota | 88745.0 | 0 | Chordata | 88745.0 | 0 | Carnivora | 88745.0 | 0 | Canidae | 88745.0 | 0 | Canis | 88745.0 | 0 | Canis lupus | 88745.0 | 0 |
| scf7180000044859 | 22696 | 0.55 | 0 | 282.7589 | Eukaryota | 1474.0 | 0 | Chordata | 1474.0 | 0 | Carnivora | 1474.0 | 0 | Canidae | 1474.0 | 0 | Canis | 1474.0 | 0 | Canis lupus | 1474.0 | 0 |
| scf7180000046195 | 116888 | 0.3856 | 0 | 88.137 | Eukaryota | 78575.0 | 0 | Chordata | 78575.0 | 0 | Carnivora | 78575.0 | 0 | Canidae | 78575.0 | 0 | Canis | 78575.0 | 0 | Canis lupus | 78575.0 | 0 |
| scf7180000043971 | 257412 | 0.3534 | 0 | 59.0108 | Eukaryota | 11691.0 | 0 | Chordata | 11691.0 | 0 | Primates | 4884.0 | 2 | Hominidae | 4884.0 | 3 | Homo | 4884.0 | 4 | Homo sapiens | 4884.0 | 4 |
| scf7180000056352 | 560306 | 0.3892 | 0 | 60.8575 | Eukaryota | 61603.0 | 0 | Chordata | 61603.0 | 0 | Carnivora | 61603.0 | 0 | Canidae | 61603.0 | 0 | Canis | 61603.0 | 0 | Canis lupus | 61603.0 | 0 |
| scf7180000047353 | 26867 | 0.4423 | 0 | 85.3769 | Eukaryota | 37120.0 | 0 | Chordata | 28074.0 | 1 | Gadiformes | 23021.0 | 4 | Gadidae | 23021.0 | 4 | Gadus | 23021.0 | 4 | Gadus morhua | 23021.0 | 4 |
| scf7180000051126 | 2474 | 0.4555 | 0 | 159.1916 | no-hit | 0.0 | 0 | no-hit | 0.0 | 0 | no-hit | 0.0 | 0 | no-hit | 0.0 | 0 | no-hit | 0.0 | 0 | no-hit | 0.0 | 0 |
| scf7180000042895 | 5916 | 0.5112 | 0 | 27.6009 | Eukaryota | 2369.0 | 0 | Chordata | 2369.0 | 0 | Carnivora | 2369.0 | 0 | Canidae | 2369.0 | 0 | Canis | 2369.0 | 0 | Canis lupus | 2369.0 | 0 |
| scf7180000042305 | 270944 | 0.4236 | 0 | 58.8843 | Eukaryota | 58123.0 | 0 | Chordata | 58123.0 | 0 | Carnivora | 58123.0 | 0 | Canidae | 49667.0 | 1 | Canis | 26000.0 | 2 | Canis lupus | 26000.0 | 2 |
| scf7180000056283 | 1160990 | 0.3893 | 0 | 61.245 | Eukaryota | 92199.0 | 0 | Chordata | 92199.0 | 0 | Carnivora | 92199.0 | 0 | Canidae | 92199.0 | 0 | Canis | 92199.0 | 0 | Canis lupus | 92199.0 | 0 |
| scf7180000042274 | 723370 | 0.5891 | 0 | 56.6843 | Eukaryota | 60012.0 | 0 | Chordata | 60012.0 | 0 | Carnivora | 60012.0 | 0 | Phocidae | 31140.0 | 1 | Leptonychotes | 31140.0 | 2 | Leptonychotes weddellii | 31140.0 | 2 |
| scf7180000045594 | 51036 | 0.3986 | 0 | 56.3586 | Eukaryota | 3938.0 | 1 | Chordata | 3938.0 | 1 | Primates | 3938.0 | 1 | Hominidae | 2717.0 | 2 | Homo | 1773.0 | 3 | Homo sapiens | 1773.0 | 3 |
| scf7180000045456 | 203842 | 0.4963 | 0 | 56.8768 | Eukaryota | 31185.0 | 0 | Chordata | 31185.0 | 0 | Carnivora | 27189.0 | 2 | Canidae | 27189.0 | 2 | Canis | 27189.0 | 2 | Canis lupus | 27189.0 | 2 |
| scf7180000042164 | 289605 | 0.45 | 0 | 61.6677 | Eukaryota | 85027.0 | 0 | Chordata | 85027.0 | 0 | Carnivora | 85027.0 | 0 | Canidae | 85027.0 | 0 | Canis | 85027.0 | 0 | Canis lupus | 85027.0 | 0 |
| scf7180000051840 | 9372 | 0.4489 | 0 | 1269.1458 | Eukaryota | 1566.0 | 0 | Echinodermata | 592.0 | 2 | Forcipulatida | 592.0 | 2 | Asteriidae | 592.0 | 2 | Asterias | 592.0 | 2 | Asterias rubens | 592.0 | 2 |
| scf7180000046025 | 134613 | 0.3779 | 0 | 58.901 | Eukaryota | 44811.0 | 0 | Chordata | 44811.0 | 0 | Carnivora | 44811.0 | 0 | Canidae | 44811.0 | 0 | Canis | 38253.0 | 1 | Canis lupus | 38253.0 | 1 |
| scf7180000046876 | 16945 | 0.395 | 0 | 47.0681 | Eukaryota | 53248.0 | 0 | Chordata | 53248.0 | 0 | Carnivora | 53248.0 | 0 | Canidae | 53248.0 | 0 | Canis | 53248.0 | 0 | Canis lupus | 53248.0 | 0 |
| scf7180000046309 | 18648 | 0.4447 | 0 | 52.4748 | Eukaryota | 4942.0 | 0 | Platyhelminthes | 3104.0 | 1 | Polystomatidea | 3104.0 | 2 | Polystomatidae | 3104.0 | 2 | Protopolystoma | 3104.0 | 2 | Protopolystoma xenopodis | 3104.0 | 2 |
| scf7180000048698 | 78478 | 0.4587 | 0 | 60.4339 | Eukaryota | 24897.0 | 0 | Chordata | 24897.0 | 0 | Carnivora | 24897.0 | 0 | Canidae | 24897.0 | 0 | Canis | 17923.0 | 1 | Canis lupus | 17923.0 | 1 |
| scf7180000047997 | 19445 | 0.434 | 0 | 70.8323 | Eukaryota | 3574.0 | 0 | Chordata | 3134.0 | 1 | Kurtiformes | 2651.0 | 2 | Apogonidae | 2651.0 | 2 | Sphaeramia | 2651.0 | 2 | Sphaeramia orbicularis | 2651.0 | 2 |
| scf7180000044491 | 290307 | 0.4474 | 0 | 59.1204 | Eukaryota | 59980.0 | 0 | Chordata | 59980.0 | 0 | Carnivora | 59980.0 | 0 | Canidae | 59980.0 | 0 | Canis | 59980.0 | 0 | Canis lupus | 59980.0 | 0 |
| scf7180000043126 | 967817 | 0.4259 | 0 | 59.7928 | Eukaryota | 94978.0 | 0 | Chordata | 94978.0 | 0 | Carnivora | 94978.0 | 0 | Canidae | 94978.0 | 0 | Canis | 94978.0 | 0 | Canis lupus | 94978.0 | 0 |
| scf7180000050812 | 4258 | 0.4436 | 0 | 63.4045 | Eukaryota | 2588.0 | 0 | Chordata | 2588.0 | 0 | Carnivora | 2588.0 | 0 | Canidae | 2588.0 | 0 | Canis | 2588.0 | 0 | Canis lupus | 2588.0 | 0 |
| scf7180000042949 | 214324 | 0.3732 | 0 | 58.2335 | Eukaryota | 11374.0 | 0 | Chordata | 11374.0 | 0 | Carnivora | 11374.0 | 0 | Canidae | 11374.0 | 0 | Canis | 11374.0 | 0 | Canis lupus | 11374.0 | 0 |
| scf7180000043486 | 172686 | 0.4027 | 0 | 59.0036 | Eukaryota | 22607.0 | 0 | Chordata | 22607.0 | 0 | Carnivora | 11340.0 | 2 | Hominidae | 8869.0 | 3 | Homo | 8869.0 | 4 | Homo sapiens | 8869.0 | 4 |
| scf7180000056021 | 9936 | 0.441 | 0 | 649.0653 | no-hit | 0.0 | 0 | no-hit | 0.0 | 0 | no-hit | 0.0 | 0 | no-hit | 0.0 | 0 | no-hit | 0.0 | 0 | no-hit | 0.0 | 0 |
| scf7180000055117 | 26649 | 0.4528 | 0 | 112.1958 | Eukaryota | 2748.0 | 0 | Chordata | 2748.0 | 0 | Carnivora | 2748.0 | 0 | Canidae | 2748.0 | 0 | Canis | 2748.0 | 0 | Canis lupus | 2748.0 | 0 |
| scf7180000048141 | 14007 | 0.4123 | 0 | 49.2588 | Eukaryota | 40710.0 | 0 | Chordata | 40710.0 | 0 | Carnivora | 40710.0 | 0 | Canidae | 40710.0 | 0 | Canis | 40710.0 | 0 | Canis lupus | 40710.0 | 0 |
| scf7180000047408 | 40769 | 0.4398 | 0 | 99.2325 | Eukaryota | 660.0 | 0 | Platyhelminthes | 429.0 | 1 | Polystomatidea | 429.0 | 1 | Polystomatidae | 429.0 | 1 | Protopolystoma | 429.0 | 1 | Protopolystoma xenopodis | 429.0 | 1 |
| scf7180000045263 | 219555 | 0.5172 | 0 | 56.0407 | Eukaryota | 162724.0 | 0 | Chordata | 162724.0 | 0 | Carnivora | 162724.0 | 0 | Canidae | 162724.0 | 0 | Canis | 162724.0 | 0 | Canis lupus | 162724.0 | 0 |
| scf7180000041815 | 904403 | 0.3895 | 0 | 60.9301 | Eukaryota | 67777.0 | 0 | Chordata | 67777.0 | 0 | Carnivora | 67777.0 | 0 | Canidae | 67777.0 | 0 | Canis | 55922.0 | 1 | Canis lupus | 55922.0 | 1 |
| scf7180000052577 | 9611 | 0.4449 | 0 | 395.4848 | no-hit | 0.0 | 0 | no-hit | 0.0 | 0 | no-hit | 0.0 | 0 | no-hit | 0.0 | 0 | no-hit | 0.0 | 0 | no-hit | 0.0 | 0 |
| scf7180000040938 | 29452 | 0.4316 | 0 | 57.1911 | Eukaryota | 7915.0 | 0 | Chordata | 7915.0 | 0 | Artiodactyla | 4600.0 | 1 | Delphinidae | 2258.0 | 5 | Vulpes | 1615.0 | 8 | Vulpes vulpes | 1615.0 | 8 |
| scf7180000042131 | 788991 | 0.463 | 0 | 61.018 | Eukaryota | 94957.0 | 0 | Chordata | 94957.0 | 0 | Carnivora | 94957.0 | 0 | Canidae | 47572.0 | 2 | Ailuropoda | 39244.0 | 3 | Ailuropoda melanoleuca | 39244.0 | 3 |
| scf7180000056341 | 719089 | 0.3581 | 0 | 59.5237 | Eukaryota | 41914.0 | 0 | Chordata | 41914.0 | 0 | Carnivora | 36300.0 | 1 | Canidae | 36300.0 | 1 | Canis | 36300.0 | 1 | Canis lupus | 36300.0 | 1 |
| scf7180000054419 | 12299 | 0.4094 | 0 | 300.9571 | Eukaryota | 54935.0 | 0 | Chordata | 54935.0 | 0 | Carnivora | 54935.0 | 0 | Canidae | 54935.0 | 0 | Canis | 54935.0 | 0 | Canis lupus | 54935.0 | 0 |
| scf7180000042750 | 546638 | 0.4176 | 0 | 59.8941 | Eukaryota | 77312.0 | 0 | Chordata | 77312.0 | 0 | Carnivora | 77312.0 | 0 | Canidae | 77312.0 | 0 | Canis | 77312.0 | 0 | Canis lupus | 77312.0 | 0 |
| scf7180000042705 | 562458 | 0.3742 | 0 | 59.302 | Eukaryota | 96663.0 | 0 | Chordata | 96663.0 | 0 | Carnivora | 96663.0 | 0 | Canidae | 96663.0 | 0 | Canis | 96663.0 | 0 | Canis lupus | 96663.0 | 0 |
| scf7180000051062 | 16985 | 0.4331 | 0 | 284.9376 | Eukaryota | 1706.0 | 0 | Chordata | 1706.0 | 0 | Carnivora | 1706.0 | 0 | Canidae | 1706.0 | 0 | Canis | 1706.0 | 0 | Canis lupus | 1706.0 | 0 |
| scf7180000041529 | 1026347 | 0.4472 | 0 | 59.55 | Eukaryota | 63754.0 | 0 | Chordata | 63754.0 | 0 | Carnivora | 63754.0 | 0 | Canidae | 63754.0 | 0 | Canis | 63754.0 | 0 | Canis lupus | 63754.0 | 0 |
| scf7180000046657 | 30186 | 0.4418 | 0 | 215.2751 | Eukaryota | 7938.0 | 0 | Chordata | 3693.0 | 2 | Polystomatidea | 3378.0 | 3 | Polystomatidae | 3378.0 | 3 | Protopolystoma | 3378.0 | 3 | Protopolystoma xenopodis | 3378.0 | 3 |
| scf7180000050213 | 16665 | 0.4365 | 0 | 75.2548 | Eukaryota | 8159.0 | 0 | Chordata | 8159.0 | 0 | Kurtiformes | 3456.0 | 3 | Apogonidae | 3456.0 | 3 | Sphaeramia | 3456.0 | 4 | Sphaeramia orbicularis | 3456.0 | 4 |
| scf7180000045523 | 198075 | 0.4295 | 0 | 58.4113 | Eukaryota | 68447.0 | 0 | Chordata | 68447.0 | 0 | Carnivora | 68447.0 | 0 | Canidae | 61365.0 | 1 | Canis | 61365.0 | 1 | Canis lupus | 61365.0 | 1 |
| scf7180000049036 | 7402 | 0.4406 | 0 | 135.2416 | no-hit | 0.0 | 0 | no-hit | 0.0 | 0 | no-hit | 0.0 | 0 | no-hit | 0.0 | 0 | no-hit | 0.0 | 0 | no-hit | 0.0 | 0 |
| scf7180000042898 | 13374 | 0.3732 | 0 | 176.5334 | Eukaryota | 5205.0 | 0 | Chordata | 5205.0 | 0 | Carnivora | 4713.0 | 1 | Felidae | 3522.0 | 3 | Felis | 3050.0 | 4 | Felis catus | 3050.0 | 4 |
| scf7180000056378 | 387139 | 0.4212 | 0 | 56.2622 | Eukaryota | 89207.0 | 0 | Chordata | 89207.0 | 0 | Carnivora | 89207.0 | 0 | Canidae | 89207.0 | 0 | Canis | 66575.0 | 1 | Canis lupus | 66575.0 | 1 |
| scf7180000041552 | 892278 | 0.396 | 0 | 62.5409 | Eukaryota | 149652.0 | 0 | Chordata | 149652.0 | 0 | Carnivora | 149652.0 | 0 | Canidae | 149652.0 | 0 | Canis | 149652.0 | 0 | Canis lupus | 149652.0 | 0 |
| scf7180000047360 | 99796 | 0.4372 | 0 | 62.2462 | Eukaryota | 30872.0 | 0 | Chordata | 30872.0 | 0 | Carnivora | 30872.0 | 0 | Canidae | 30872.0 | 0 | Canis | 30872.0 | 0 | Canis lupus | 30872.0 | 0 |
| scf7180000055409 | 24248 | 0.4271 | 0 | 103.1836 | Eukaryota | 78944.0 | 0 | Chordata | 78944.0 | 0 | Carnivora | 78944.0 | 0 | Canidae | 56500.0 | 1 | Canis | 48075.0 | 3 | Canis lupus | 48075.0 | 4 |
| scf7180000041496 | 48693 | 0.4582 | 0 | 61.2429 | Eukaryota | 83857.0 | 0 | Chordata | 83857.0 | 0 | Carnivora | 83857.0 | 0 | Canidae | 83857.0 | 0 | Canis | 83857.0 | 0 | Canis lupus | 83857.0 | 0 |
| scf7180000046150 | 27060 | 0.4691 | 0 | 53.5966 | Eukaryota | 13560.0 | 0 | Chordata | 13560.0 | 0 | Carnivora | 13560.0 | 0 | Canidae | 12839.0 | 1 | Canis | 11717.0 | 2 | Canis lupus | 11717.0 | 2 |
| scf7180000040992 | 461675 | 0.4692 | 100 | 59.156 | Eukaryota | 69415.0 | 0 | Chordata | 69415.0 | 0 | Carnivora | 69415.0 | 0 | Canidae | 69415.0 | 0 | Canis | 69415.0 | 0 | Canis lupus | 69415.0 | 0 |
| scf7180000053603 | 6169 | 0.5545 | 0 | 141.6667 | Eukaryota | 30425.0 | 0 | Chordata | 30425.0 | 0 | Carnivora | 30425.0 | 0 | Canidae | 30425.0 | 0 | Canis | 30425.0 | 0 | Canis lupus | 30425.0 | 0 |
| scf7180000043766 | 527814 | 0.3587 | 0 | 59.7845 | Eukaryota | 96097.0 | 0 | Chordata | 96097.0 | 0 | Carnivora | 96097.0 | 0 | Canidae | 96097.0 | 0 | Canis | 85622.0 | 1 | Canis lupus | 85622.0 | 1 |
| scf7180000052012 | 3308 | 0.4589 | 0 | 294.0432 | Eukaryota | 322.0 | 0 | Streptophyta | 322.0 | 0 | Fabales | 322.0 | 0 | Fabaceae | 322.0 | 0 | Cicer | 322.0 | 0 | Cicer arietinum | 322.0 | 0 |
| scf7180000056496 | 86279 | 0.4197 | 0 | 58.0053 | Eukaryota | 32897.0 | 0 | Chordata | 32897.0 | 0 | Carnivora | 32897.0 | 0 | Canidae | 32897.0 | 0 | Canis | 24221.0 | 1 | Canis lupus | 24221.0 | 1 |
| scf7180000055216 | 31528 | 0.4428 | 0 | 159.4297 | Eukaryota | 15118.0 | 0 | Chordata | 15118.0 | 0 | Carnivora | 15118.0 | 0 | Canidae | 8797.0 | 3 | Canis | 8797.0 | 3 | Canis lupus | 8797.0 | 4 |
| scf7180000052074 | 13842 | 0.4293 | 0 | 599.0165 | Eukaryota | 62630.0 | 0 | Chordata | 62630.0 | 0 | Carnivora | 62630.0 | 0 | Canidae | 62630.0 | 0 | Canis | 62630.0 | 0 | Canis lupus | 62630.0 | 0 |
| scf7180000043301 | 415718 | 0.4883 | 0 | 56.9973 | Eukaryota | 30778.0 | 0 | Chordata | 30778.0 | 0 | Carnivora | 30778.0 | 0 | Canidae | 30778.0 | 0 | Canis | 30778.0 | 0 | Canis lupus | 30778.0 | 0 |
| scf7180000046370 | 9055 | 0.3687 | 0 | 36.4679 | Eukaryota | 7024.0 | 0 | Chordata | 7024.0 | 0 | Carnivora | 7024.0 | 0 | Felidae | 5588.0 | 1 | Felis | 5588.0 | 1 | Felis catus | 5588.0 | 1 |
| scf7180000049282 | 12610 | 0.445 | 0 | 250.7977 | Eukaryota | 2443.0 | 0 | Platyhelminthes | 1269.0 | 2 | Polystomatidea | 1269.0 | 2 | Polystomatidae | 1269.0 | 2 | Protopolystoma | 1269.0 | 2 | Protopolystoma xenopodis | 1269.0 | 2 |
| scf7180000042513 | 66450 | 0.3789 | 0 | 57.62 | Eukaryota | 11259.0 | 0 | Chordata | 11259.0 | 0 | Carnivora | 11259.0 | 0 | Canidae | 11259.0 | 0 | Canis | 11259.0 | 0 | Canis lupus | 11259.0 | 0 |
| scf7180000047233 | 73874 | 0.4893 | 0 | 58.2667 | Eukaryota | 14353.0 | 0 | Chordata | 14353.0 | 0 | Carnivora | 14353.0 | 0 | Canidae | 9845.0 | 1 | Vulpes | 6514.0 | 2 | Vulpes vulpes | 6514.0 | 2 |
| scf7180000051129 | 19167 | 0.6199 | 0 | 52.2622 | Eukaryota | 15758.0 | 0 | Chordata | 15758.0 | 0 | Carnivora | 15758.0 | 0 | Canidae | 15758.0 | 0 | Canis | 10879.0 | 1 | Canis lupus | 10879.0 | 1 |
| scf7180000041389 | 1032429 | 0.4182 | 0 | 59.3168 | Eukaryota | 68483.0 | 0 | Chordata | 68483.0 | 0 | Carnivora | 68483.0 | 0 | Canidae | 68483.0 | 0 | Canis | 68483.0 | 0 | Canis lupus | 68483.0 | 0 |
| scf7180000052754 | 7136 | 0.4434 | 0 | 57.3056 | no-hit | 0.0 | 0 | no-hit | 0.0 | 0 | no-hit | 0.0 | 0 | no-hit | 0.0 | 0 | no-hit | 0.0 | 0 | no-hit | 0.0 | 0 |
| scf7180000040843 | 1659923 | 0.3879 | 0 | 61.2772 | Eukaryota | 96953.0 | 0 | Chordata | 96953.0 | 0 | Carnivora | 96953.0 | 0 | Canidae | 96953.0 | 0 | Canis | 86982.0 | 1 | Canis lupus | 86982.0 | 1 |
| scf7180000055757 | 45753 | 0.5019 | 0 | 441.6705 | Eukaryota | 9173.0 | 0 | Chordata | 9173.0 | 0 | Carnivora | 9173.0 | 0 | Canidae | 9173.0 | 0 | Canis | 7331.0 | 1 | Canis lupus | 6021.0 | 2 |
| scf7180000053089 | 3099 | 0.455 | 0 | 363.667 | no-hit | 0.0 | 0 | no-hit | 0.0 | 0 | no-hit | 0.0 | 0 | no-hit | 0.0 | 0 | no-hit | 0.0 | 0 | no-hit | 0.0 | 0 |
| scf7180000051684 | 3711 | 0.4573 | 0 | 864.9742 | no-hit | 0.0 | 0 | no-hit | 0.0 | 0 | no-hit | 0.0 | 0 | no-hit | 0.0 | 0 | no-hit | 0.0 | 0 | no-hit | 0.0 | 0 |
| scf7180000044352 | 144049 | 0.5281 | 0 | 57.5798 | Eukaryota | 54791.0 | 0 | Chordata | 54791.0 | 0 | Carnivora | 54791.0 | 0 | Otariidae | 26249.0 | 3 | Zalophus | 15753.0 | 6 | Zalophus californianus | 15753.0 | 6 |
| scf7180000043121 | 8901 | 0.4365 | 0 | 153.9645 | Eukaryota | 452.0 | 0 | Chordata | 309.0 | 1 | Kurtiformes | 309.0 | 1 | Apogonidae | 309.0 | 1 | Sphaeramia | 309.0 | 1 | Sphaeramia orbicularis | 309.0 | 1 |
| scf7180000056363 | 484586 | 0.3835 | 0 | 60.8101 | Eukaryota | 52910.0 | 0 | Chordata | 52910.0 | 0 | Carnivora | 52910.0 | 0 | Canidae | 52910.0 | 0 | Canis | 45630.0 | 1 | Canis lupus | 45630.0 | 1 |
| scf7180000055616 | 18641 | 0.4416 | 0 | 144.1033 | Eukaryota | 2661.0 | 0 | Chordata | 2222.0 | 1 | Kurtiformes | 1430.0 | 3 | Apogonidae | 1430.0 | 3 | Sphaeramia | 1430.0 | 3 | Sphaeramia orbicularis | 1430.0 | 3 |
| scf7180000046708 | 111198 | 0.4124 | 0 | 161.3018 | Eukaryota | 96407.0 | 0 | Chordata | 96407.0 | 0 | Carnivora | 96407.0 | 0 | Canidae | 96407.0 | 0 | Canis | 96407.0 | 0 | Canis lupus | 96407.0 | 0 |
| scf7180000054841 | 11831 | 0.4326 | 0 | 163.8817 | no-hit | 0.0 | 0 | no-hit | 0.0 | 0 | no-hit | 0.0 | 0 | no-hit | 0.0 | 0 | no-hit | 0.0 | 0 | no-hit | 0.0 | 0 |
| scf7180000045889 | 12520 | 0.4315 | 0 | 171.0746 | Eukaryota | 94324.0 | 0 | Chordata | 94324.0 | 0 | Carnivora | 94324.0 | 0 | Canidae | 94324.0 | 0 | Canis | 94324.0 | 0 | Canis lupus | 94324.0 | 0 |
| scf7180000040575 | 2324916 | 0.3878 | 0 | 60.7085 | Eukaryota | 90047.0 | 0 | Chordata | 90047.0 | 0 | Carnivora | 90047.0 | 0 | Canidae | 90047.0 | 0 | Canis | 90047.0 | 0 | Canis lupus | 90047.0 | 0 |
| scf7180000041423 | 291165 | 0.3917 | 0 | 64.7285 | Eukaryota | 93847.0 | 0 | Chordata | 93847.0 | 0 | Carnivora | 93847.0 | 0 | Canidae | 93847.0 | 0 | Canis | 93847.0 | 0 | Canis lupus | 93847.0 | 0 |
| scf7180000043770 | 384193 | 0.4796 | 0 | 60.2359 | Eukaryota | 66695.0 | 0 | Chordata | 66695.0 | 0 | Carnivora | 66695.0 | 0 | Canidae | 66695.0 | 0 | Canis | 66695.0 | 0 | Canis lupus | 66695.0 | 0 |
| scf7180000051134 | 13252 | 0.4319 | 0 | 117.3062 | Eukaryota | 6139.0 | 0 | Chordata | 5767.0 | 1 | Synbranchiformes | 2632.0 | 3 | Mastacembelidae | 2632.0 | 3 | Mastacembelus | 2632.0 | 3 | Mastacembelus armatus | 2632.0 | 3 |
| scf7180000043244 | 106574 | 0.3285 | 0 | 54.9741 | Eukaryota | 14302.0 | 0 | Chordata | 14302.0 | 0 | Carnivora | 14302.0 | 0 | Canidae | 14302.0 | 0 | Canis | 14302.0 | 0 | Canis lupus | 14302.0 | 0 |
| scf7180000048063 | 3996 | 0.4572 | 0 | 29.955 | Eukaryota | 9338.0 | 0 | Chordata | 9338.0 | 0 | Carnivora | 3691.0 | 3 | Equidae | 3004.0 | 5 | Equus | 3004.0 | 5 | Canis lupus | 2653.0 | 6 |
| scf7180000053263 | 5666 | 0.4352 | 0 | 99.6369 | Eukaryota | 13016.0 | 0 | Chordata | 10359.0 | 2 | Synbranchiformes | 4016.0 | 6 | Mastacembelidae | 4016.0 | 6 | Mastacembelus | 4016.0 | 7 | Mastacembelus armatus | 4016.0 | 8 |
| scf7180000043190 | 7108 | 0.4532 | 0 | 36.0051 | Eukaryota | 3326.0 | 1 | Chordata | 3326.0 | 1 | Primates | 1470.0 | 3 | Hominidae | 1470.0 | 3 | Homo | 1470.0 | 3 | Homo sapiens | 1470.0 | 3 |
| scf7180000044275 | 106513 | 0.3913 | 0 | 57.4621 | Eukaryota | 9159.0 | 0 | Chordata | 9159.0 | 0 | Carnivora | 9159.0 | 0 | Canidae | 9159.0 | 0 | Canis | 9159.0 | 0 | Canis lupus | 9159.0 | 0 |
| scf7180000045675 | 164356 | 0.4368 | 0 | 60.881 | Eukaryota | 76462.0 | 0 | Chordata | 76462.0 | 0 | Carnivora | 76462.0 | 0 | Canidae | 76462.0 | 0 | Canis | 53532.0 | 1 | Canis lupus | 53532.0 | 1 |
| scf7180000043064 | 651194 | 0.3421 | 0 | 59.0199 | Eukaryota | 37680.0 | 0 | Chordata | 37680.0 | 0 | Carnivora | 37680.0 | 0 | Canidae | 37680.0 | 0 | Canis | 37680.0 | 0 | Canis lupus | 37680.0 | 0 |
| scf7180000043005 | 479887 | 0.447 | 0 | 57.7853 | Eukaryota | 54938.0 | 0 | Chordata | 54938.0 | 0 | Carnivora | 39345.0 | 1 | Canidae | 39345.0 | 2 | Canis | 21720.0 | 3 | Canis lupus | 21720.0 | 3 |
| scf7180000045726 | 14213 | 0.4124 | 0 | 54.6141 | Eukaryota | 20524.0 | 0 | Chordata | 20524.0 | 0 | Carnivora | 20524.0 | 0 | Canidae | 20524.0 | 0 | Canis | 20524.0 | 0 | Canis lupus | 20524.0 | 0 |
| scf7180000047871 | 60600 | 0.3417 | 0 | 63.4357 | Eukaryota | 11193.0 | 0 | Chordata | 11193.0 | 0 | Carnivora | 11193.0 | 0 | Canidae | 6737.0 | 1 | Canis | 4509.0 | 3 | Canis lupus | 4509.0 | 3 |
| scf7180000056482 | 302310 | 0.3664 | 0 | 58.7042 | Eukaryota | 74576.0 | 0 | Chordata | 74576.0 | 0 | Carnivora | 74576.0 | 0 | Canidae | 74576.0 | 0 | Canis | 74576.0 | 0 | Canis lupus | 74576.0 | 0 |
| scf7180000041447 | 1051050 | 0.3928 | 0 | 65.2797 | Eukaryota | 63382.0 | 0 | Chordata | 63382.0 | 0 | Carnivora | 63382.0 | 0 | Canidae | 63382.0 | 0 | Canis | 47700.0 | 1 | Canis lupus | 47700.0 | 1 |
| scf7180000056002 | 9923 | 0.5085 | 0 | 302.2677 | Eukaryota | 3523.0 | 0 | Chordata | 3523.0 | 0 | Carnivora | 3523.0 | 0 | Canidae | 2979.0 | 1 | Canis | 2979.0 | 1 | Canis lupus | 2979.0 | 1 |
| scf7180000046061 | 70013 | 0.4078 | 0 | 55.6856 | Eukaryota | 50284.0 | 0 | Chordata | 50284.0 | 0 | Carnivora | 50284.0 | 0 | Canidae | 50284.0 | 0 | Canis | 50284.0 | 0 | Canis lupus | 50284.0 | 0 |
| scf7180000051662 | 2788 | 0.401 | 0 | 18.7603 | Eukaryota | 39231.0 | 0 | Chordata | 39231.0 | 0 | Carnivora | 39231.0 | 0 | Canidae | 39231.0 | 0 | Canis | 39231.0 | 0 | Canis lupus | 39231.0 | 0 |
| scf7180000051089 | 4936 | 0.4473 | 0 | 425.2002 | Eukaryota | 546.0 | 0 | Streptophyta | 546.0 | 0 | Fabales | 546.0 | 0 | Fabaceae | 546.0 | 0 | Cicer | 546.0 | 0 | Cicer arietinum | 546.0 | 0 |
| scf7180000049502 | 23599 | 0.4867 | 0 | 59.3234 | Eukaryota | 4568.0 | 0 | Chordata | 4568.0 | 0 | Carnivora | 4568.0 | 0 | Felidae | 3138.0 | 1 | Panthera | 2078.0 | 2 | Canis lupus | 1430.0 | 3 |
| scf7180000047844 | 3295 | 0.4807 | 0 | 23.8895 | Eukaryota | 4454.0 | 0 | Chordata | 3988.0 | 1 | Primates | 1340.0 | 4 | Hominidae | 1340.0 | 4 | Homo | 1340.0 | 5 | Homo sapiens | 1340.0 | 5 |
| scf7180000041390 | 1157490 | 0.366 | 0 | 60.5376 | Eukaryota | 85870.0 | 0 | Chordata | 85870.0 | 0 | Carnivora | 85870.0 | 0 | Canidae | 85870.0 | 0 | Canis | 85870.0 | 0 | Canis lupus | 85870.0 | 0 |
| scf7180000056224 | 1538502 | 0.5473 | 0 | 58.5395 | Eukaryota | 94716.0 | 0 | Chordata | 94716.0 | 0 | Carnivora | 94716.0 | 0 | Canidae | 94716.0 | 0 | Canis | 94716.0 | 0 | Canis lupus | 94716.0 | 0 |
| scf7180000054065 | 7941 | 0.5047 | 0 | 245.7505 | Eukaryota | 8468.0 | 0 | Chordata | 8468.0 | 0 | Carnivora | 8468.0 | 0 | Canidae | 8468.0 | 0 | Canis | 6894.0 | 1 | Canis lupus | 5764.0 | 2 |
| scf7180000052283 | 6388 | 0.4497 | 0 | 1693.2297 | no-hit | 0.0 | 0 | no-hit | 0.0 | 0 | no-hit | 0.0 | 0 | no-hit | 0.0 | 0 | no-hit | 0.0 | 0 | no-hit | 0.0 | 0 |
| scf7180000042849 | 108048 | 0.363 | 0 | 59.2576 | Eukaryota | 76781.0 | 0 | Chordata | 76781.0 | 0 | Carnivora | 76781.0 | 0 | Canidae | 76781.0 | 0 | Canis | 76781.0 | 0 | Canis lupus | 76781.0 | 0 |
| scf7180000040747 | 217664 | 0.3815 | 0 | 59.554 | Eukaryota | 60130.0 | 0 | Chordata | 60130.0 | 0 | Carnivora | 60130.0 | 0 | Canidae | 60130.0 | 0 | Canis | 44834.0 | 1 | Canis lupus | 44834.0 | 1 |
| scf7180000041434 | 373694 | 0.4398 | 0 | 59.7181 | Eukaryota | 38729.0 | 0 | Chordata | 38729.0 | 0 | Carnivora | 38729.0 | 0 | Canidae | 38729.0 | 0 | Canis | 38729.0 | 0 | Canis lupus | 38729.0 | 0 |
| scf7180000054804 | 62103 | 0.4968 | 0 | 261.7013 | Eukaryota | 15186.0 | 0 | Chordata | 15186.0 | 0 | Carnivora | 15186.0 | 0 | Canidae | 15186.0 | 0 | Canis | 9313.0 | 1 | Canis lupus | 9313.0 | 1 |
| scf7180000051807 | 4671 | 0.4457 | 0 | 271.332 | Eukaryota | 1419.0 | 0 | Echinodermata | 749.0 | 2 | Forcipulatida | 749.0 | 2 | Asteriidae | 749.0 | 2 | Asterias | 749.0 | 2 | Asterias rubens | 749.0 | 2 |
| scf7180000048670 | 7393 | 0.4376 | 0 | 117.7017 | no-hit | 0.0 | 0 | no-hit | 0.0 | 0 | no-hit | 0.0 | 0 | no-hit | 0.0 | 0 | no-hit | 0.0 | 0 | no-hit | 0.0 | 0 |
| scf7180000048036 | 12832 | 0.3886 | 0 | 35.8428 | Eukaryota | 6596.0 | 0 | Chordata | 6596.0 | 0 | Carnivora | 6596.0 | 0 | Canidae | 6596.0 | 0 | Canis | 6596.0 | 0 | Canis lupus | 6596.0 | 0 |
| scf7180000056580 | 64683 | 0.4371 | 0 | 81.7529 | Eukaryota | 40493.0 | 0 | Chordata | 27745.0 | 2 | Gadiformes | 19503.0 | 4 | Gadidae | 19503.0 | 4 | Gadus | 19503.0 | 4 | Gadus morhua | 19503.0 | 4 |
| scf7180000041020 | 87893 | 0.3825 | 0 | 57.4983 | Eukaryota | 17717.0 | 0 | Chordata | 17717.0 | 0 | Carnivora | 17717.0 | 0 | Canidae | 17717.0 | 0 | Canis | 17717.0 | 0 | Canis lupus | 17717.0 | 0 |
| scf7180000046701 | 114355 | 0.3807 | 0 | 60.0785 | Eukaryota | 27136.0 | 0 | Chordata | 27136.0 | 0 | Carnivora | 25257.0 | 1 | Canidae | 15706.0 | 2 | Canis | 15706.0 | 2 | Canis lupus | 15706.0 | 2 |
| scf7180000046098 | 140722 | 0.4837 | 0 | 57.9158 | Eukaryota | 18724.0 | 0 | Chordata | 18724.0 | 0 | Carnivora | 18724.0 | 0 | Canidae | 18724.0 | 0 | Canis | 18724.0 | 0 | Canis lupus | 18724.0 | 0 |
| scf7180000054908 | 92252 | 0.4413 | 0 | 81.6658 | Eukaryota | 66027.0 | 0 | Chordata | 46046.0 | 2 | Gadiformes | 38778.0 | 5 | Gadidae | 38778.0 | 5 | Gadus | 38778.0 | 5 | Gadus morhua | 38778.0 | 5 |
| scf7180000056550 | 126963 | 0.4016 | 0 | 57.858 | Eukaryota | 54078.0 | 0 | Chordata | 54078.0 | 0 | Carnivora | 54078.0 | 0 | Canidae | 51439.0 | 1 | Canis | 26959.0 | 2 | Canis lupus | 26959.0 | 2 |
| scf7180000042448 | 34622 | 0.4384 | 0 | 64.8891 | Eukaryota | 88828.0 | 0 | Chordata | 88828.0 | 0 | Carnivora | 88828.0 | 0 | Canidae | 88828.0 | 0 | Canis | 88828.0 | 0 | Canis lupus | 88828.0 | 0 |
| scf7180000045094 | 212424 | 0.517 | 0 | 58.235 | Eukaryota | 66823.0 | 0 | Chordata | 66823.0 | 0 | Carnivora | 66823.0 | 0 | Canidae | 66823.0 | 0 | Canis | 60019.0 | 1 | Canis lupus | 60019.0 | 1 |
| scf7180000049862 | 3164 | 0.5054 | 0 | 163.2531 | Eukaryota | 4424.0 | 0 | Chordata | 4424.0 | 0 | Carnivora | 4424.0 | 0 | Canidae | 4424.0 | 0 | Canis | 3249.0 | 2 | Canis lupus | 3249.0 | 2 |
| scf7180000049686 | 9348 | 0.4427 | 0 | 219.3258 | Eukaryota | 3574.0 | 0 | Platyhelminthes | 2075.0 | 2 | Polystomatidea | 2075.0 | 2 | Polystomatidae | 2075.0 | 2 | Protopolystoma | 2075.0 | 2 | Protopolystoma xenopodis | 2075.0 | 2 |
| scf7180000047248 | 52469 | 0.3768 | 0 | 79.8126 | Eukaryota | 74962.0 | 0 | Chordata | 74962.0 | 0 | Carnivora | 74962.0 | 0 | Canidae | 74962.0 | 0 | Canis | 74962.0 | 0 | Canis lupus | 74962.0 | 0 |
| scf7180000045290 | 3220 | 0.4339 | 0 | 80.409 | no-hit | 0.0 | 0 | no-hit | 0.0 | 0 | no-hit | 0.0 | 0 | no-hit | 0.0 | 0 | no-hit | 0.0 | 0 | no-hit | 0.0 | 0 |
| scf7180000046761 | 126852 | 0.5368 | 0 | 56.565 | Eukaryota | 16045.0 | 0 | Chordata | 16045.0 | 0 | Carnivora | 16045.0 | 0 | Canidae | 16045.0 | 0 | Canis | 11233.0 | 1 | Canis lupus | 11233.0 | 1 |
| scf7180000053718 | 2030 | 0.4611 | 0 | 439.2941 | Eukaryota | 574.0 | 0 | Platyhelminthes | 574.0 | 0 | Polystomatidea | 574.0 | 0 | Polystomatidae | 574.0 | 0 | Protopolystoma | 574.0 | 0 | Protopolystoma xenopodis | 574.0 | 0 |
| scf7180000044804 | 15319 | 0.5197 | 0 | 1071.295 | Eukaryota | 33137.0 | 0 | Chordata | 33137.0 | 0 | Carnivora | 33137.0 | 0 | Canidae | 33137.0 | 0 | Canis | 33137.0 | 0 | Canis lupus | 33137.0 | 0 |
| scf7180000044848 | 275905 | 0.4395 | 0 | 58.9762 | Eukaryota | 26306.0 | 0 | Chordata | 26306.0 | 0 | Carnivora | 26306.0 | 0 | Canidae | 26306.0 | 0 | Canis | 26306.0 | 0 | Canis lupus | 26306.0 | 0 |
| scf7180000049874 | 35502 | 0.3536 | 0 | 59.0735 | Eukaryota | 7038.0 | 0 | Chordata | 7038.0 | 0 | Carnivora | 5024.0 | 1 | Canidae | 5024.0 | 1 | Canis | 5024.0 | 1 | Canis lupus | 5024.0 | 1 |
| scf7180000056567 | 122732 | 0.4367 | 0 | 58.4871 | Eukaryota | 21491.0 | 0 | Chordata | 21491.0 | 0 | Carnivora | 21491.0 | 0 | Phocidae | 8833.0 | 3 | Neomonachus | 7097.0 | 4 | Neomonachus schauinslandi | 7097.0 | 4 |
| scf7180000052783 | 8241 | 0.3415 | 0 | 61.1312 | Eukaryota | 3240.0 | 0 | Chordata | 3240.0 | 0 | Carnivora | 3240.0 | 0 | Canidae | 3240.0 | 0 | Canis | 3240.0 | 0 | Canis lupus | 3240.0 | 0 |
| scf7180000045287 | 5701 | 0.4345 | 0 | 32.6357 | no-hit | 0.0 | 0 | no-hit | 0.0 | 0 | no-hit | 0.0 | 0 | no-hit | 0.0 | 0 | no-hit | 0.0 | 0 | no-hit | 0.0 | 0 |
| scf7180000046254 | 110831 | 0.3917 | 0 | 58.0936 | Eukaryota | 46504.0 | 0 | Chordata | 46504.0 | 0 | Carnivora | 46504.0 | 0 | Canidae | 46504.0 | 0 | Canis | 46504.0 | 0 | Canis lupus | 46504.0 | 0 |
| scf7180000049031 | 53371 | 0.3544 | 0 | 60.2849 | Eukaryota | 80521.0 | 0 | Chordata | 80521.0 | 0 | Carnivora | 80521.0 | 0 | Canidae | 58387.0 | 2 | Canis | 50014.0 | 3 | Canis lupus | 50014.0 | 3 |
| scf7180000056424 | 685699 | 0.4171 | 0 | 58.7006 | Eukaryota | 98950.0 | 0 | Chordata | 98950.0 | 0 | Carnivora | 98950.0 | 0 | Canidae | 98950.0 | 0 | Canis | 98950.0 | 0 | Canis lupus | 98950.0 | 0 |
| scf7180000044006 | 398626 | 0.3876 | 0 | 59.0779 | Eukaryota | 50163.0 | 0 | Chordata | 50163.0 | 0 | Carnivora | 50163.0 | 0 | Canidae | 43729.0 | 1 | Canis | 43729.0 | 1 | Canis lupus | 43729.0 | 1 |
| scf7180000041261 | 65619 | 0.3607 | 0 | 57.6656 | Eukaryota | 106786.0 | 0 | Chordata | 106786.0 | 0 | Carnivora | 106786.0 | 0 | Canidae | 106786.0 | 0 | Canis | 106786.0 | 0 | Canis lupus | 106786.0 | 0 |

| | | | | | | | | | | | | | | | | | | | |
|---|---|---|---|---|---|---|---|---|---|---|---|---|---|---|---|---|---|---|---|
| scf7180000047079 | 92073 | 0.416 | 0 | 59.8324 | Eukaryota | 12414.0 | 0 | Chordata | 12414.0 | 0 | Carnivora | 12414.0 | 0 | Otariidae | 4500.0 | 2 | Canis | 3007.0 | 5 | Canis lupus | 3007.0 | 5 |
| scf7180000052541 | 12295 | 0.51 | 0 | 115.3996 | Eukaryota | 8490.0 | 0 | Chordata | 8490.0 | 0 | Carnivora | 8490.0 | 0 | Canidae | 8490.0 | 0 | Canis | 6893.0 | 1 | Canis lupus | 5780.0 | 2 |
| scf7180000047544 | 79596 | 0.5502 | 0 | 55.8876 | Eukaryota | 22298.0 | 0 | Chordata | 22298.0 | 0 | Carnivora | 22298.0 | 0 | Canidae | 18671.0 | 2 | Canis | 13160.0 | 5 | Canis lupus | 13160.0 | 5 |
| scf7180000055441 | 56786 | 0.4469 | 0 | 80.0253 | Eukaryota | 19865.0 | 0 | Chordata | 19865.0 | 0 | Carnivora | 19865.0 | 0 | Canidae | 19865.0 | 0 | Canis | 19865.0 | 0 | Canis lupus | 19865.0 | 0 |
| scf7180000046466 | 34685 | 0.5177 | 0 | 55.2959 | Eukaryota | 35239.0 | 0 | Chordata | 35239.0 | 0 | Carnivora | 35239.0 | 0 | Canidae | 35239.0 | 0 | Canis | 35239.0 | 0 | Canis lupus | 35239.0 | 0 |
| scf7180000052691 | 8490 | 0.4578 | 0 | 61.1773 | Eukaryota | 4377.0 | 0 | Chordata | 4377.0 | 0 | Carnivora | 4377.0 | 0 | Canidae | 4377.0 | 0 | Canis | 4377.0 | 0 | Canis lupus | 4377.0 | 0 |
| scf7180000049674 | 13340 | 0.443 | 0 | 167.0057 | Eukaryota | 54922.0 | 0 | Chordata | 37914.0 | 2 | Gadiformes | 29941.0 | 5 | Gadidae | 29941.0 | 5 | Gadus | 29941.0 | 5 | Gadus morhua | 29941.0 | 5 |
| scf7180000043611 | 118684 | 0.3955 | 0 | 59.0086 | Eukaryota | 39481.0 | 0 | Chordata | 39481.0 | 0 | Carnivora | 39481.0 | 0 | Canidae | 39481.0 | 0 | Canis | 39481.0 | 0 | Canis lupus | 39481.0 | 0 |
| scf7180000049764 | 54237 | 0.3347 | 0 | 57.0026 | Eukaryota | 3558.0 | 0 | Chordata | 3558.0 | 0 | Carnivora | 2933.0 | 1 | Canidae | 2933.0 | 1 | Canis | 2933.0 | 1 | Canis lupus | 2933.0 | 1 |
| scf7180000054180 | 14565 | 0.4376 | 0 | 185.9491 | Eukaryota | 5427.0 | 0 | Chordata | 4225.0 | 1 | Salmoniformes | 2303.0 | 3 | Salmonidae | 2303.0 | 3 | Salmo | 2303.0 | 3 | Salmo trutta | 2303.0 | 3 |
| scf7180000052959 | 5302 | 0.4536 | 0 | 1030.0991 | no-hit | 0.0 | 0 | no-hit | 0.0 | 0 | no-hit | 0.0 | 0 | no-hit | 0.0 | 0 | no-hit | 0.0 | 0 | no-hit | 0.0 | 0 |
| scf7180000052920 | 2933 | 0.4576 | 0 | 233.7349 | no-hit | 0.0 | 0 | no-hit | 0.0 | 0 | no-hit | 0.0 | 0 | no-hit | 0.0 | 0 | no-hit | 0.0 | 0 | no-hit | 0.0 | 0 |
| scf7180000051854 | 9400 | 0.4473 | 0 | 2189.3746 | Eukaryota | 222.0 | 0 | Streptophyta | 222.0 | 0 | Fabales | 222.0 | 0 | Fabaceae | 222.0 | 0 | Cicer | 222.0 | 0 | Cicer arietinum | 222.0 | 0 |
| scf7180000049450 | 6482 | 0.4408 | 0 | 73.6455 | no-hit | 0.0 | 0 | no-hit | 0.0 | 0 | no-hit | 0.0 | 0 | no-hit | 0.0 | 0 | no-hit | 0.0 | 0 | no-hit | 0.0 | 0 |
| scf7180000044315 | 183557 | 0.3637 | 0 | 59.2149 | Eukaryota | 16998.0 | 0 | Chordata | 16998.0 | 0 | Carnivora | 16998.0 | 0 | Canidae | 16998.0 | 0 | Canis | 16998.0 | 0 | Canis lupus | 16998.0 | 0 |
| scf7180000044240 | 340684 | 0.363 | 0 | 58.1867 | Eukaryota | 36297.0 | 0 | Chordata | 36297.0 | 0 | Carnivora | 36297.0 | 0 | Canidae | 36297.0 | 0 | Canis | 36297.0 | 0 | Canis lupus | 36297.0 | 0 |
| scf7180000054946 | 26901 | 0.4388 | 0 | 168.5986 | Eukaryota | 8010.0 | 0 | Chordata | 7208.0 | 1 | Kurtiformes | 5270.0 | 5 | Apogonidae | 5270.0 | 5 | Sphaeramia | 5270.0 | 5 | Sphaeramia orbicularis | 5270.0 | 5 |
| scf7180000054688 | 47204 | 0.4423 | 0 | 172.242 | Eukaryota | 53663.0 | 0 | Chordata | 53663.0 | 0 | Carnivora | 53663.0 | 0 | Canidae | 53663.0 | 0 | Canis | 53663.0 | 0 | Canis lupus | 53663.0 | 0 |
| scf7180000047855 | 12223 | 0.4366 | 0 | 119.6773 | Eukaryota | 2744.0 | 0 | Chordata | 2744.0 | 0 | Carnivora | 2339.0 | 1 | Canidae | 2339.0 | 1 | Canis | 2339.0 | 1 | Canis lupus | 2339.0 | 1 |
| scf7180000050252 | 22327 | 0.442 | 0 | 59.498 | Eukaryota | 2487.0 | 0 | Platyhelminthes | 1952.0 | 1 | Polystomatidea | 1952.0 | 2 | Polystomatidae | 1952.0 | 2 | Protopolystoma | 1952.0 | 2 | Protopolystoma xenopodis | 1952.0 | 2 |
| scf7180000048066 | 8719 | 0.5852 | 0 | 224.3892 | Eukaryota | 35025.0 | 0 | Chordata | 35025.0 | 0 | Artiodactyla | 24563.0 | 1 | Bovidae | 24563.0 | 2 | Ovis | 13892.0 | 3 | Ovis aries | 13892.0 | 3 |
| scf7180000049532 | 34263 | 0.4243 | 0 | 57.4047 | Eukaryota | 88491.0 | 0 | Chordata | 88491.0 | 0 | Carnivora | 88491.0 | 0 | Canidae | 53124.0 | 2 | Canis | 42518.0 | 4 | Canis lupus | 42518.0 | 4 |
| scf7180000048426 | 13122 | 0.4463 | 0 | 154.114 | Eukaryota | 5253.0 | 0 | Chordata | 5253.0 | 0 | Salmoniformes | 4275.0 | 2 | Salmonidae | 4275.0 | 2 | Salmo | 4275.0 | 2 | Salmo trutta | 4275.0 | 2 |
| scf7180000054866 | 17481 | 0.4103 | 0 | 1169.7083 | Eukaryota | 49042.0 | 0 | Chordata | 49042.0 | 0 | Carnivora | 49042.0 | 0 | Canidae | 49042.0 | 0 | Canis | 49042.0 | 0 | Canis lupus | 49042.0 | 0 |
| scf7180000040660 | 360706 | 0.3717 | 0 | 60.1695 | Eukaryota | 90313.0 | 0 | Chordata | 90313.0 | 0 | Carnivora | 90313.0 | 0 | Canidae | 90313.0 | 0 | Canis | 90313.0 | 0 | Canis lupus | 90313.0 | 0 |
| scf7180000047317 | 22616 | 0.5007 | 0 | 67.7469 | Eukaryota | 10865.0 | 0 | Chordata | 10865.0 | 0 | Carnivora | 10865.0 | 0 | Canidae | 10865.0 | 0 | Canis | 7676.0 | 1 | Canis lupus | 7676.0 | 1 |
| scf7180000049656 | 17101 | 0.4511 | 0 | 84.2692 | Eukaryota | 2789.0 | 0 | Chordata | 2789.0 | 0 | Carnivora | 2789.0 | 0 | Canidae | 2789.0 | 0 | Canis | 2789.0 | 0 | Canis lupus | 2789.0 | 0 |
| scf7180000056259 | 733353 | 0.3701 | 0 | 59.5385 | Eukaryota | 90212.0 | 0 | Chordata | 90212.0 | 0 | Carnivora | 90212.0 | 0 | Canidae | 90212.0 | 0 | Canis | 90212.0 | 0 | Canis lupus | 90212.0 | 0 |
| scf7180000054840 | 19132 | 0.5317 | 0 | 86.3268 | Eukaryota | 5236.0 | 0 | Chordata | 4611.0 | 1 | Carnivora | 3580.0 | 2 | Canidae | 2189.0 | 4 | Canis | 1880.0 | 5 | Canis lupus | 1880.0 | 5 |
| scf7180000047597 | 48512 | 0.4394 | 0 | 232.336 | Eukaryota | 94247.0 | 0 | Chordata | 94247.0 | 0 | Carnivora | 94247.0 | 0 | Canidae | 94247.0 | 0 | Canis | 94247.0 | 0 | Canis lupus | 94247.0 | 0 |
| scf7180000056500 | 209307 | 0.5065 | 0 | 57.8714 | Eukaryota | 26568.0 | 0 | Chordata | 26568.0 | 0 | Carnivora | 26568.0 | 0 | Canidae | 14751.0 | 1 | Canis | 11009.0 | 4 | Canis lupus | 11009.0 | 4 |
| scf7180000042205 | 9991 | 0.4552 | 0 | 1421.9139 | Eukaryota | 27949.0 | 0 | Chordata | 27949.0 | 0 | Carnivora | 27949.0 | 0 | Canidae | 27949.0 | 0 | Canis | 27949.0 | 0 | Canis lupus | 27949.0 | 0 |
| scf7180000042109 | 13762 | 0.4275 | 0 | 33.4112 | Eukaryota | 6964.0 | 0 | Chordata | 6964.0 | 0 | Primates | 3954.0 | 3 | Hominidae | 3954.0 | 3 | Homo | 2554.0 | 4 | Homo sapiens | 2554.0 | 4 |
| scf7180000051907 | 7069 | 0.4477 | 0 | 894.0139 | no-hit | 0.0 | 0 | no-hit | 0.0 | 0 | no-hit | 0.0 | 0 | no-hit | 0.0 | 0 | no-hit | 0.0 | 0 | no-hit | 0.0 | 0 |
| scf7180000049939 | 14585 | 0.4549 | 0 | 93.525 | no-hit | 0.0 | 0 | no-hit | 0.0 | 0 | no-hit | 0.0 | 0 | no-hit | 0.0 | 0 | no-hit | 0.0 | 0 | no-hit | 0.0 | 0 |
| scf7180000042572 | 575779 | 0.3776 | 0 | 57.9998 | Eukaryota | 93515.0 | 0 | Chordata | 93515.0 | 0 | Carnivora | 93515.0 | 0 | Canidae | 93515.0 | 0 | Canis | 93515.0 | 0 | Canis lupus | 93515.0 | 0 |
| scf7180000043981 | 16661 | 0.3662 | 0 | 43.4426 | Eukaryota | 30652.0 | 0 | Chordata | 30652.0 | 0 | Carnivora | 30652.0 | 0 | Canidae | 30652.0 | 0 | Canis | 30652.0 | 0 | Canis lupus | 30652.0 | 0 |
| scf7180000042379 | 143127 | 0.3702 | 0 | 59.4043 | Eukaryota | 71094.0 | 0 | Chordata | 71094.0 | 0 | Carnivora | 71094.0 | 0 | Canidae | 71094.0 | 0 | Canis | 71094.0 | 0 | Canis lupus | 71094.0 | 0 |
| scf7180000053601 | 6800 | 0.3646 | 0 | 63.4855 | Eukaryota | 2387.0 | 0 | Chordata | 2387.0 | 0 | Carnivora | 2387.0 | 0 | Canidae | 2387.0 | 0 | Canis | 2387.0 | 0 | Canis lupus | 2387.0 | 0 |
| scf7180000049614 | 30958 | 0.5071 | 0 | 579.8 | Eukaryota | 9177.0 | 0 | Chordata | 9177.0 | 0 | Carnivora | 9177.0 | 0 | Canidae | 9177.0 | 0 | Canis | 6863.0 | 1 | Canis lupus | 5575.0 | 2 |
| scf7180000046905 | 94275 | 0.3684 | 0 | 57.6772 | Eukaryota | 12209.0 | 0 | Chordata | 12209.0 | 0 | Carnivora | 12209.0 | 0 | Canidae | 12209.0 | 0 | Canis | 12209.0 | 0 | Canis lupus | 12209.0 | 0 |
| scf7180000042007 | 867777 | 0.3551 | 0 | 59.2048 | Eukaryota | 42483.0 | 0 | Chordata | 42483.0 | 0 | Carnivora | 42483.0 | 0 | Canidae | 42483.0 | 0 | Canis | 42483.0 | 0 | Canis lupus | 42483.0 | 0 |
| scf7180000042831 | 520872 | 0.3579 | 0 | 58.3356 | Eukaryota | 86691.0 | 0 | Chordata | 86691.0 | 0 | Carnivora | 86691.0 | 0 | Canidae | 86691.0 | 0 | Canis | 74270.0 | 1 | Canis lupus | 74270.0 | 1 |
| scf7180000053207 | 3361 | 0.4335 | 0 | 238.5783 | no-hit | 0.0 | 0 | no-hit | 0.0 | 0 | no-hit | 0.0 | 0 | no-hit | 0.0 | 0 | no-hit | 0.0 | 0 | no-hit | 0.0 | 0 |
| scf7180000049105 | 13240 | 0.4451 | 0 | 103.2814 | Eukaryota | 3180.0 | 0 | Chordata | 1721.0 | 1 | Polystomatidea | 1459.0 | 3 | Polystomatidae | 1459.0 | 3 | Protopolystoma | 1459.0 | 3 | Protopolystoma xenopodis | 1459.0 | 3 |
| scf7180000040819 | 379477 | 0.3742 | 0 | 60.4696 | Eukaryota | 85895.0 | 0 | Chordata | 85895.0 | 0 | Carnivora | 85895.0 | 0 | Canidae | 85895.0 | 0 | Canis | 85895.0 | 0 | Canis lupus | 85895.0 | 0 |
| scf7180000043873 | 39024 | 0.4445 | 0 | 174.799 | Eukaryota | 65341.0 | 0 | Chordata | 65341.0 | 0 | Carnivora | 65341.0 | 0 | Canidae | 65341.0 | 0 | Canis | 65341.0 | 0 | Canis lupus | 65341.0 | 0 |
| scf7180000043285 | 427171 | 0.5189 | 0 | 58.454 | Eukaryota | 75309.0 | 0 | Chordata | 75309.0 | 0 | Carnivora | 75309.0 | 0 | Canidae | 75309.0 | 0 | Canis | 67774.0 | 1 | Canis lupus | 67774.0 | 1 |
| scf7180000047920 | 10689 | 0.4402 | 0 | 94.3105 | Eukaryota | 670.0 | 0 | Chordata | 670.0 | 0 | Tetraodontiformes | 670.0 | 0 | Tetraodontidae | 670.0 | 0 | Takifugu | 670.0 | 0 | Takifugu rubripes | 670.0 | 0 |
| scf7180000042450 | 20181 | 0.3829 | 0 | 38.5924 | Eukaryota | 15760.0 | 0 | Chordata | 15760.0 | 0 | Carnivora | 15760.0 | 0 | Canidae | 15760.0 | 0 | Canis | 15760.0 | 0 | Canis lupus | 15760.0 | 0 |
| scf7180000042866 | 15915 | 0.4294 | 0 | 51.2261 | Eukaryota | 4688.0 | 0 | Chordata | 4688.0 | 0 | Carnivora | 1594.0 | 4 | Canidae | 1594.0 | 6 | Canis | 1594.0 | 6 | Canis lupus | 1594.0 | 6 |
| scf7180000044139 | 102134 | 0.5481 | 0 | 53.5015 | Eukaryota | 38423.0 | 0 | Chordata | 38423.0 | 0 | Carnivora | 38423.0 | 0 | Canidae | 38423.0 | 0 | Canis | 36635.0 | 1 | Canis lupus | 36635.0 | 1 |
| scf7180000048874 | 26780 | 0.4476 | 0 | 1176.1358 | Eukaryota | 916.0 | 0 | Chordata | 916.0 | 0 | Tetraodontiformes | 429.0 | 2 | Tetraodontidae | 429.0 | 2 | Takifugu | 429.0 | 2 | Takifugu rubripes | 429.0 | 2 |
| scf7180000051177 | 5576 | 0.4467 | 0 | 1197.9782 | Eukaryota | 241.0 | 0 | Chordata | 241.0 | 0 | Blenniiformes | 241.0 | 0 | Gobiesocidae | 241.0 | 0 | Gouania | 241.0 | 0 | Gouania willdenowi | 241.0 | 0 |
| scf7180000046186 | 140163 | 0.3765 | 0 | 59.943 | Eukaryota | 29631.0 | 0 | Chordata | 29631.0 | 0 | Carnivora | 29631.0 | 0 | Canidae | 19807.0 | 1 | Canis | 16953.0 | 4 | Canis lupus | 16953.0 | 4 |
| scf7180000047989 | 66290 | 0.3883 | 0 | 57.865 | Eukaryota | 41082.0 | 0 | Chordata | 41082.0 | 0 | Carnivora | 41082.0 | 0 | Canidae | 19506.0 | 4 | Canis | 14715.0 | 6 | Canis lupus | 14715.0 | 7 |
| scf7180000043645 | 110330 | 0.4269 | 0 | 59.3751 | Eukaryota | 33651.0 | 0 | Chordata | 33651.0 | 0 | Carnivora | 33651.0 | 0 | Canidae | 20461.0 | 4 | Canis | 16606.0 | 5 | Canis lupus | 16606.0 | 5 |
| scf7180000044404 | 313975 | 0.4308 | 0 | 59.3618 | Eukaryota | 51575.0 | 0 | Chordata | 51575.0 | 0 | Carnivora | 51575.0 | 0 | Canidae | 51575.0 | 0 | Canis | 51575.0 | 0 | Canis lupus | 51575.0 | 0 |
| scf7180000050318 | 4832 | 0.4489 | 0 | 246.7547 | Eukaryota | 1977.0 | 0 | Platyhelminthes | 1036.0 | 2 | Polystomatidea | 1036.0 | 2 | Polystomatidae | 1036.0 | 2 | Protopolystoma | 1036.0 | 2 | Protopolystoma xenopodis | 1036.0 | 2 |
| scf7180000046705 | 29850 | 0.4392 | 0 | 183.5125 | Eukaryota | 93077.0 | 0 | Chordata | 93077.0 | 0 | Carnivora | 93077.0 | 0 | Canidae | 93077.0 | 0 | Canis | 93077.0 | 0 | Canis lupus | 93077.0 | 0 |
| scf7180000041922 | 650213 | 0.3523 | 0 | 59.4904 | Eukaryota | 70435.0 | 0 | Chordata | 70435.0 | 0 | Carnivora | 70435.0 | 0 | Canidae | 70435.0 | 0 | Canis | 70435.0 | 0 | Canis lupus | 70435.0 | 0 |
| scf7180000040755 | 39517 | 0.3631 | 0 | 60.5948 | Eukaryota | 57618.0 | 0 | Chordata | 57618.0 | 0 | Carnivora | 57618.0 | 0 | Canidae | 39732.0 | 2 | Canis | 29722.0 | 4 | Canis lupus | 29722.0 | 5 |
| scf7180000046610 | 158651 | 0.3601 | 0 | 58.5655 | Eukaryota | 64255.0 | 0 | Chordata | 64255.0 | 0 | Carnivora | 64255.0 | 0 | Canidae | 64255.0 | 0 | Canis | 64255.0 | 0 | Canis lupus | 64255.0 | 0 |
| scf7180000056540 | 197709 | 0.382 | 0 | 60.9076 | Eukaryota | 33087.0 | 0 | Chordata | 33087.0 | 0 | Carnivora | 33087.0 | 0 | Canidae | 33087.0 | 0 | Canis | 29362.0 | 1 | Canis lupus | 29362.0 | 1 |
| scf7180000056388 | 748108 | 0.445 | 0 | 59.4801 | Eukaryota | 133519.0 | 0 | Chordata | 133519.0 | 0 | Carnivora | 133519.0 | 0 | Canidae | 133519.0 | 0 | Canis | 93340.0 | 1 | Canis lupus | 93340.0 | 1 |
| scf7180000051491 | 3819 | 0.4318 | 0 | 121.0467 | no-hit | 0.0 | 0 | no-hit | 0.0 | 0 | no-hit | 0.0 | 0 | no-hit | 0.0 | 0 | no-hit | 0.0 | 0 | no-hit | 0.0 | 0 |
| scf7180000056452 | 396776 | 0.4113 | 0 | 59.4694 | Eukaryota | 39244.0 | 0 | Chordata | 39244.0 | 0 | Carnivora | 39244.0 | 0 | Ursidae | 17360.0 | 3 | Ailuropoda | 17360.0 | 4 | Ailuropoda melanoleuca | 17360.0 | 4 |
| scf7180000041041 | 1430224 | 0.375 | 0 | 59.7422 | Eukaryota | 43044.0 | 0 | Chordata | 43044.0 | 0 | Carnivora | 43044.0 | 0 | Canidae | 43044.0 | 0 | Canis | 38872.0 | 1 | Canis lupus | 38872.0 | 1 |
| scf7180000047783 | 21281 | 0.4408 | 0 | 83.0017 | Eukaryota | 9945.0 | 0 | Chordata | 5606.0 | 2 | Strigeidida | 2680.0 | 7 | Schistosomatidae | 2680.0 | 7 | Schistosoma | 2680.0 | 7 | Schistosoma mansoni | 2680.0 | 7 |
| scf7180000044581 | 275203 | 0.3922 | 0 | 58.3305 | Eukaryota | 45065.0 | 0 | Chordata | 45065.0 | 0 | Carnivora | 45065.0 | 0 | Canidae | 45065.0 | 0 | Canis | 45065.0 | 0 | Canis lupus | 45065.0 | 0 |
| scf7180000048357 | 5599 | 0.4522 | 0 | 747.4238 | no-hit | 0.0 | 0 | no-hit | 0.0 | 0 | no-hit | 0.0 | 0 | no-hit | 0.0 | 0 | no-hit | 0.0 | 0 | no-hit | 0.0 | 0 |
| scf7180000055325 | 27831 | 0.4447 | 0 | 144.218 | Eukaryota | 5054.0 | 0 | Chordata | 2959.0 | 2 | Carnivora | 1557.0 | 4 | Canidae | 1557.0 | 4 | Canis | 1557.0 | 5 | Canis lupus | 1557.0 | 5 |
| scf7180000048843 | 7842 | 0.4597 | 0 | 772.1427 | Eukaryota | 400.0 | 0 | Chordata | 400.0 | 0 | Kurtiformes | 250.0 | 1 | Apogonidae | 250.0 | 1 | Sphaeramia | 250.0 | 1 | Sphaeramia orbicularis | 250.0 | 1 |
| scf7180000056547 | 186306 | 0.4153 | 0 | 59.0214 | Eukaryota | 37859.0 | 0 | Chordata | 37859.0 | 0 | Carnivora | 37859.0 | 0 | Canidae | 21008.0 | 3 | Canis | 15126.0 | 4 | Canis lupus | 15126.0 | 4 |
| scf7180000050874 | 21816 | 0.3887 | 0 | 60.4205 | Eukaryota | 3482.0 | 0 | Chordata | 3482.0 | 0 | Carnivora | 2670.0 | 2 | Ursidae | 1605.0 | 3 | Ursus | 1276.0 | 5 | Ursus maritimus | 952.0 | 6 |
| scf7180000044846 | 283759 | 0.4229 | 0 | 60.6048 | Eukaryota | 54577.0 | 0 | Chordata | 54577.0 | 0 | Carnivora | 54577.0 | 0 | Canidae | 49952.0 | 1 | Canis | 28456.0 | 2 | Canis lupus | 28456.0 | 2 |
| scf7180000046083 | 10215 | 0.5122 | 0 | 33.5239 | Eukaryota | 5873.0 | 0 | Chordata | 5873.0 | 0 | Carnivora | 5873.0 | 0 | Canidae | 5344.0 | 1 | Canis | 3569.0 | 2 | Canis lupus | 3569.0 | 2 |
| scf7180000054064 | 4829 | 0.4446 | 0 | 313.1076 | Eukaryota | 405.0 | 0 | Chordata | 405.0 | 0 | Columbiformes | 405.0 | 0 | Columbidae | 405.0 | 0 | Streptopelia | 405.0 | 0 | Streptopelia turtur | 405.0 | 0 |
| scf7180000045873 | 113435 | 0.4476 | 0 | 103.8888 | Eukaryota | 90572.0 | 0 | Chordata | 90572.0 | 0 | Carnivora | 90572.0 | 0 | Canidae | 90572.0 | 0 | Canis | 90572.0 | 0 | Canis lupus | 90572.0 | 0 |
| scf7180000044437 | 34632 | 0.3707 | 0 | 57.7661 | Eukaryota | 39162.0 | 0 | Chordata | 39162.0 | 0 | Carnivora | 39162.0 | 0 | Canidae | 39162.0 | 0 | Canis | 39162.0 | 0 | Canis lupus | 39162.0 | 0 |
| scf7180000047668 | 6549 | 0.4349 | 0 | 88.4876 | Eukaryota | 945.0 | 0 | Chordata | 945.0 | 0 | Kurtiformes | 945.0 | 0 | Apogonidae | 945.0 | 0 | Sphaeramia | 945.0 | 0 | Sphaeramia orbicularis | 945.0 | 0 |
| scf7180000055116 | 12164 | 0.4514 | 0 | 186.5851 | Eukaryota | 2700.0 | 0 | Chordata | 2700.0 | 0 | Carnivora | 2700.0 | 0 | Canidae | 2700.0 | 0 | Canis | 2700.0 | 0 | Canis lupus | 2700.0 | 0 |
| scf7180000049957 | 1548 | 0.4522 | 0 | 689.376 | no-hit | 0.0 | 0 | no-hit | 0.0 | 0 | no-hit | 0.0 | 0 | no-hit | 0.0 | 0 | no-hit | 0.0 | 0 | no-hit | 0.0 | 0 |
| scf7180000047713 | 20935 | 0.445 | 0 | 283.2691 | Eukaryota | 17352.0 | 0 | Chordata | 17352.0 | 0 | Tetraodontiformes | 17352.0 | 0 | Tetraodontidae | 17352.0 | 0 | Takifugu | 17352.0 | 0 | Takifugu rubripes | 15909.0 | 1 |
| scf7180000043436 | 122358 | 0.4281 | 0 | 58.2533 | Eukaryota | 84507.0 | 0 | Chordata | 84507.0 | 0 | Carnivora | 84507.0 | 0 | Canidae | 50489.0 | 2 | Canis | 40256.0 | 5 | Canis lupus | 40256.0 | 5 |
| scf7180000044199 | 342514 | 0.3758 | 0 | 58.1391 | Eukaryota | 22798.0 | 0 | Chordata | 22798.0 | 0 | Carnivora | 22798.0 | 0 | Canidae | 22798.0 | 0 | Canis | 22798.0 | 0 | Canis lupus | 22798.0 | 0 |
| scf7180000053206 | 2286 | 0.444 | 0 | 134.4346 | Eukaryota | 340.0 | 0 | Platyhelminthes | 340.0 | 0 | Polystomatidea | 193.0 | 1 | Polystomatidae | 193.0 | 1 | Protopolystoma | 193.0 | 1 | Protopolystoma xenopodis | 193.0 | 1 |
| scf7180000054518 | 50750 | 0.4528 | 0 | 238.4033 | Eukaryota | 57217.0 | 0 | Chordata | 57217.0 | 0 | Carnivora | 57217.0 | 0 | Canidae | 57217.0 | 0 | Canis | 57217.0 | 0 | Canis lupus | 57217.0 | 0 |
| scf7180000049436 | 18055 | 0.4448 | 0 | 129.3812 | Eukaryota | 1935.0 | 0 | Chordata | 979.0 | 1 | Polystomatidea | 956.0 | 4 | Polystomatidae | 956.0 | 4 | Protopolystoma | 956.0 | 4 | Protopolystoma xenopodis | 956.0 | 4 |
| scf7180000046334 | 57364 | 0.3653 | 0 | 57.4903 | Eukaryota | 68690.0 | 0 | Chordata | 68690.0 | 0 | Carnivora | 68690.0 | 0 | Canidae | 68690.0 | 0 | Canis | 68690.0 | 0 | Canis lupus | 68690.0 | 0 |
| scf7180000051819 | 9616 | 0.452 | 0 | 2336.7798 | Eukaryota | 498.0 | 0 | Streptophyta | 498.0 | 0 | Fabales | 498.0 | 0 | Fabaceae | 498.0 | 0 | Cicer | 498.0 | 0 | Cicer arietinum | 498.0 | 0 |
| scf7180000048610 | 53025 | 0.3607 | 0 | 61.9223 | Eukaryota | 17126.0 | 0 | Chordata | 17126.0 | 0 | Carnivora | 13083.0 | 1 | Canidae | 13083.0 | 1 | Canis | 11463.0 | 3 | Canis lupus | 11463.0 | 3 |
| scf7180000055531 | 50783 | 0.4537 | 0 | 78.8243 | Eukaryota | 4495.0 | 0 | Platyhelminthes | 2886.0 | 1 | Polystomatidea | 2241.0 | 4 | Polystomatidae | 2241.0 | 4 | Protopolystoma | 2241.0 | 4 | Protopolystoma xenopodis | 2241.0 | 4 |
| scf7180000055658 | 15427 | 0.4411 | 0 | 241.537 | no-hit | 0.0 | 0 | no-hit | 0.0 | 0 | no-hit | 0.0 | 0 | no-hit | 0.0 | 0 | no-hit | 0.0 | 0 | no-hit | 0.0 | 0 |
| scf7180000045795 | 143780 | 0.4957 | 0 | 58.1011 | Eukaryota | 24949.0 | 0 | Chordata | 24949.0 | 0 | Carnivora | 24949.0 | 0 | Canidae | 24949.0 | 0 | Canis | 24949.0 | 0 | Canis lupus | 24949.0 | 0 |
| scf7180000048227 | 6072 | 0.4448 | 0 | 56.6334 | no-hit | 0.0 | 0 | no-hit | 0.0 | 0 | no-hit | 0.0 | 0 | no-hit | 0.0 | 0 | no-hit | 0.0 | 0 | no-hit | 0.0 | 0 |
| scf7180000043674 | 497293 | 0.4083 | 0 | 61.3766 | Eukaryota | 184378.0 | 0 | Chordata | 184378.0 | 0 | Carnivora | 184378.0 | 0 | Canidae | 170504.0 | 1 | Canis | 170504.0 | 1 | Canis lupus | 170504.0 | 1 |
| scf7180000042186 | 152945 | 0.388 | 0 | 55.9849 | Eukaryota | 75269.0 | 0 | Chordata | 75269.0 | 0 | Carnivora | 75269.0 | 0 | Canidae | 75269.0 | 0 | Canis | 75269.0 | 0 | Canis lupus | 75269.0 | 0 |
| scf7180000041719 | 874278 | 0.5175 | 0 | 56.9236 | Eukaryota | 84703.0 | 0 | Chordata | 84703.0 | 0 | Carnivora | 84703.0 | 0 | Canidae | 84703.0 | 0 | Canis | 69591.0 | 1 | Canis lupus | 69591.0 | 1 |
| scf7180000044184 | 359595 | 0.3972 | 0 | 59.6248 | Eukaryota | 191910.0 | 0 | Chordata | 191910.0 | 0 | Carnivora | 191910.0 | 0 | Canidae | 180238.0 | 1 | Canis | 180238.0 | 1 | Canis lupus | 180238.0 | 1 |
| scf7180000053095 | 2252 | 0.4516 | 0 | 737.3806 | no-hit | 0.0 | 0 | no-hit | 0.0 | 0 | no-hit | 0.0 | 0 | no-hit | 0.0 | 0 | no-hit | 0.0 | 0 | no-hit | 0.0 | 0 |
| scf7180000047222 | 89253 | 0.4324 | 0 | 58.5055 | Eukaryota | 18526.0 | 0 | Chordata | 18526.0 | 0 | Carnivora | 18526.0 | 0 | Canidae | 18526.0 | 0 | Canis | 16405.0 | 1 | Canis lupus | 16405.0 | 1 |
| scf7180000048539 | 30647 | 0.4989 | 0 | 62.6155 | Eukaryota | 7596.0 | 0 | Chordata | 7596.0 | 0 | Carnivora | 7596.0 | 0 | Canidae | 3680.0 | 2 | Canis | 3680.0 | 2 | Canis lupus | 3680.0 | 3 |
| scf7180000052717 | 3685 | 0.4505 | 0 | 400.3191 | no-hit | 0.0 | 0 | no-hit | 0.0 | 0 | no-hit | 0.0 | 0 | no-hit | 0.0 | 0 | no-hit | 0.0 | 0 | no-hit | 0.0 | 0 |
| scf7180000048702 | 10940 | 0.4545 | 0 | 157.2725 | Eukaryota | 8627.0 | 0 | Chordata | 8627.0 | 0 | Carnivora | 8627.0 | 0 | Canidae | 8627.0 | 0 | Canis | 8627.0 | 0 | Canis lupus | 8627.0 | 0 |
| scf7180000043359 | 515272 | 0.4159 | 0 | 58.8484 | Eukaryota | 19972.0 | 0 | Chordata | 19972.0 | 0 | Carnivora | 19972.0 | 0 | Canidae | 19972.0 | 0 | Canis | 19972.0 | 0 | Canis lupus | 19972.0 | 0 |
| scf7180000041398 | 620898 | 0.4058 | 0 | 59.8581 | Eukaryota | 58189.0 | 0 | Chordata | 58189.0 | 0 | Carnivora | 58189.0 | 0 | Canidae | 58189.0 | 0 | Canis | 52264.0 | 1 | Canis lupus | 52264.0 | 1 |
| scf7180000051123 | 7253 | 0.4508 | 0 | 1144.7368 | no-hit | 0.0 | 0 | no-hit | 0.0 | 0 | no-hit | 0.0 | 0 | no-hit | 0.0 | 0 | no-hit | 0.0 | 0 | no-hit | 0.0 | 0 |
| scf7180000043745 | 442798 | 0.3901 | 0 | 60.0602 | Eukaryota | 61208.0 | 0 | Chordata | 61208.0 | 0 | Carnivora | 61208.0 | 0 | Canidae | 53297.0 | 2 | Canis | 44407.0 | 3 | Canis lupus | 44407.0 | 3 |
| scf7180000056403 | 517808 | 0.4388 | 0 | 59.8319 | Eukaryota | 53746.0 | 0 | Chordata | 53746.0 | 0 | Carnivora | 53746.0 | 0 | Canidae | 53746.0 | 0 | Vulpes | 32412.0 | 1 | Vulpes vulpes | 32412.0 | 1 |
| scf7180000043246 | 9942 | 0.4398 | 0 | 76.5055 | no-hit | 0.0 | 0 | no-hit | 0.0 | 0 | no-hit | 0.0 | 0 | no-hit | 0.0 | 0 | no-hit | 0.0 | 0 | no-hit | 0.0 | 0 |
| scf7180000043127 | 79396 | 0.3326 | 0 | 51.8365 | Eukaryota | 7031.0 | 1 | Chordata | 7031.0 | 1 | Carnivora | 4276.0 | 3 | Canidae | 2188.0 | 4 | Canis | 2188.0 | 5 | Canis lupus | 2188.0 | 5 |
| scf7180000043673 | 397758 | 0.3876 | 0 | 60.9552 | Eukaryota | 89206.0 | 0 | Chordata | 89206.0 | 0 | Carnivora | 89206.0 | 0 | Canidae | 89206.0 | 0 | Canis | 89206.0 | 0 | Canis lupus | 89206.0 | 0 |
| scf7180000050924 | 25924 | 0.5041 | 0 | 175.9413 | Eukaryota | 9351.0 | 0 | Chordata | 9351.0 | 0 | Carnivora | 9351.0 | 0 | Canidae | 9351.0 | 0 | Canis | 7621.0 | 1 | Canis lupus | 6369.0 | 2 |
| scf7180000040618 | 442936 | 0.4006 | 0 | 62.2869 | Eukaryota | 110490.0 | 0 | Chordata | 110490.0 | 0 | Carnivora | 110490.0 | 0 | Canidae | 110490.0 | 0 | Canis | 110490.0 | 0 | Canis lupus | 110490.0 | 0 |
| scf7180000052724 | 5382 | 0.455 | 0 | 1028.3736 | Eukaryota | 2967.0 | 0 | Chordata | 2313.0 | 1 | Carnivora | 2313.0 | 1 | Canidae | 2313.0 | 1 | Canis | 2313.0 | 1 | Canis lupus | 2313.0 | 1 |
| scf7180000042428 | 89237 | 0.3926 | 0 | 58.1486 | Eukaryota | 15177.0 | 0 | Chordata | 15177.0 | 0 | Carnivora | 15177.0 | 0 | Canidae | 15177.0 | 0 | Canis | 15177.0 | 0 | Canis lupus | 15177.0 | 0 |
| scf7180000041867 | 838816 | 0.4156 | 0 | 59.1519 | Eukaryota | 44890.0 | 0 | Chordata | 44890.0 | 0 | Carnivora | 44890.0 | 0 | Canidae | 44890.0 | 0 | Canis | 38526.0 | 1 | Canis lupus | 38526.0 | 1 |
| scf7180000044546 | 11734 | 0.4413 | 0 | 71.7074 | no-hit | 0.0 | 0 | no-hit | 0.0 | 0 | no-hit | 0.0 | 0 | no-hit | 0.0 | 0 | no-hit | 0.0 | 0 | no-hit | 0.0 | 0 |
| scf7180000048689 | 50202 | 0.4435 | 0 | 73.348 | Eukaryota | 7213.0 | 0 | Platyhelminthes | 4204.0 | 1 | Polystomatidea | 4204.0 | 2 | Polystomatidae | 4204.0 | 2 | Protopolystoma | 4204.0 | 2 | Protopolystoma xenopodis | 4204.0 | 2 |
| scf7180000041548 | 79435 | 0.4196 | 0 | 176.5607 | Eukaryota | 91312.0 | 0 | Chordata | 91312.0 | 0 | Carnivora | 91312.0 | 0 | Canidae | 91312.0 | 0 | Canis | 91312.0 | 0 | Canis lupus | 91312.0 | 0 |
| scf7180000056553 | 126829 | 0.4639 | 0 | 56.3003 | Eukaryota | 41824.0 | 0 | Chordata | 41824.0 | 0 | Carnivora | 41824.0 | 0 | Canidae | 41824.0 | 0 | Canis | 14423.0 | 4 | Canis lupus | 10805.0 | 5 |
| scf7180000043201 | 34670 | 0.3858 | 0 | 50.6175 | Eukaryota | 13199.0 | 0 | Chordata | 13199.0 | 0 | Carnivora | 13199.0 | 0 | Canidae | 6760.0 | 2 | Canis | 6760.0 | 2 | Canis lupus | 6760.0 | 2 |
| scf7180000043311 | 430152 | 0.4077 | 0 | 59.1412 | Eukaryota | 38608.0 | 0 | Chordata | 38608.0 | 0 | Carnivora | 38608.0 | 0 | Canidae | 23154.0 | 2 | Canis | 15219.0 | 3 | Canis lupus | 15219.0 | 4 |
| scf7180000050819 | 21376 | 0.4466 | 0 | 118.4286 | Eukaryota | 92518.0 | 0 | Chordata | 92518.0 | 0 | Carnivora | 92518.0 | 0 | Canidae | 92518.0 | 0 | Canis | 92518.0 | 0 | Canis lupus | 92518.0 | 0 |
| scf7180000053173 | 3805 | 0.4489 | 0 | 768.1389 | no-hit | 0.0 | 0 | no-hit | 0.0 | 0 | no-hit | 0.0 | 0 | no-hit | 0.0 | 0 | no-hit | 0.0 | 0 | no-hit | 0.0 | 0 |
| scf7180000049618 | 52430 | 0.4407 | 0 | 52.0312 | Eukaryota | 66130.0 | 0 | Chordata | 45780.0 | 2 | Gadiformes | 38480.0 | 5 | Gadidae | 38480.0 | 5 | Gadus | 38480.0 | 5 | Gadus morhua | 38480.0 | 5 |
| scf7180000042209 | 49256 | 0.5196 | 0 | 50.2173 | Eukaryota | 58737.0 | 0 | Chordata | 58737.0 | 0 | Carnivora | 58737.0 | 0 | Canidae | 58737.0 | 0 | Canis | 58737.0 | 0 | Canis lupus | 58737.0 | 0 |
| scf7180000046776 | 10412 | 0.4366 | 0 | 147.9159 | Eukaryota | 2787.0 | 0 | Platyhelminthes | 1347.0 | 2 | Polystomatidea | 1347.0 | 2 | Polystomatidae | 1347.0 | 2 | Protopolystoma | 1347.0 | 2 | Protopolystoma xenopodis | 1347.0 | 2 |
| scf7180000042632 | 622050 | 0.3764 | 0 | 59.8591 | Eukaryota | 21670.0 | 0 | Chordata | 21670.0 | 0 | Carnivora | 21670.0 | 0 | Canidae | 21670.0 | 0 | Canis | 21670.0 | 0 | Canis lupus | 21670.0 | 0 |
| scf7180000042967 | 551433 | 0.3899 | 0 | 61.2936 | Eukaryota | 94377.0 | 0 | Chordata | 94377.0 | 0 | Carnivora | 94377.0 | 0 | Canidae | 94377.0 | 0 | Canis | 84879.0 | 1 | Canis lupus | 84879.0 | 1 |
| scf7180000041196 | 256110 | 0.4017 | 0 | 57.7918 | Eukaryota | 20294.0 | 0 | Chordata | 20294.0 | 0 | Carnivora | 20294.0 | 0 | Canidae | 20294.0 | 0 | Canis | 20294.0 | 0 | Canis lupus | 20294.0 | 0 |
| scf7180000045150 | 3811 | 0.3322 | 0 | 29.3949 | Eukaryota | 5023.0 | 0 | Chordata | 5023.0 | 0 | Carnivora | 3230.0 | 1 | Felidae | 2044.0 | 3 | Felis | 2044.0 | 3 | Felis catus | 2044.0 | 3 |
| scf7180000053997 | 5174 | 0.3384 | 0 | 27.8877 | Eukaryota | 2781.0 | 0 | Chordata | 2781.0 | 0 | Carnivora | 2781.0 | 0 | Canidae | 2781.0 | 0 | Canis | 2781.0 | 0 | Canis lupus | 2781.0 | 0 |
| scf7180000056574 | 15997 | 0.4555 | 0 | 128.0431 | Eukaryota | 2808.0 | 0 | Platyhelminthes | 1887.0 | 1 | Polystomatidea | 1887.0 | 1 | Polystomatidae | 1887.0 | 1 | Protopolystoma | 1887.0 | 1 | Protopolystoma xenopodis | 1887.0 | 1 |
| scf7180000048972 | 54159 | 0.4549 | 0 | 56.2775 | Eukaryota | 10523.0 | 0 | Chordata | 10523.0 | 0 | Carnivora | 10523.0 | 0 | Felidae | 4479.0 | 2 | Felis | 4479.0 | 2 | Felis catus | 4479.0 | 2 |
| scf7180000051408 | 18483 | 0.505 | 0 | 387.5605 | Eukaryota | 9273.0 | 0 | Chordata | 9273.0 | 0 | Carnivora | 9273.0 | 0 | Canidae | 9273.0 | 0 | Canis | 8063.0 | 1 | Canis lupus | 6797.0 | 2 |
| scf7180000050731 | 27375 | 0.439 | 0 | 278.2503 | no-hit | 0.0 | 0 | no-hit | 0.0 | 0 | no-hit | 0.0 | 0 | no-hit | 0.0 | 0 | no-hit | 0.0 | 0 | no-hit | 0.0 | 0 |
| scf7180000054273 | 11961 | 0.3305 | 0 | 50.0269 | Eukaryota | 3638.0 | 1 | Chordata | 3638.0 | 1 | Primates | 2097.0 | 3 | Hominidae | 2097.0 | 4 | Homo | 1814.0 | 5 | Homo sapiens | 1814.0 | 5 |
| scf7180000042579 | 555223 | 0.3608 | 0 | 58.9291 | Eukaryota | 102886.0 | 0 | Chordata | 102886.0 | 0 | Carnivora | 102886.0 | 0 | Canidae | 102886.0 | 0 | Canis | 102886.0 | 0 | Canis lupus | 102886.0 | 0 |
| scf7180000044957 | 113893 | 0.4617 | 0 | 60.2457 | Eukaryota | 26157.0 | 0 | Chordata | 26157.0 | 0 | Carnivora | 26157.0 | 0 | Canidae | 26157.0 | 0 | Canis | 19944.0 | 1 | Canis lupus | 19944.0 | 1 |
| scf7180000044334 | 299183 | 0.5636 | 0 | 58.0423 | Eukaryota | 42407.0 | 0 | Chordata | 42407.0 | 0 | Carnivora | 42407.0 | 0 | Canidae | 42407.0 | 0 | Canis | 42407.0 | 0 | Canis lupus | 42407.0 | 0 |
| scf7180000051067 | 5276 | 0.4447 | 0 | 112.1808 | no-hit | 0.0 | 0 | no-hit | 0.0 | 0 | no-hit | 0.0 | 0 | no-hit | 0.0 | 0 | no-hit | 0.0 | 0 | no-hit | 0.0 | 0 |
| scf7180000053214 | 4902 | 0.5059 | 0 | 300.5884 | Eukaryota | 8059.0 | 0 | Chordata | 8059.0 | 0 | Carnivora | 8059.0 | 0 | Canidae | 8059.0 | 0 | Canis | 6421.0 | 1 | Canis lupus | 5218.0 | 2 |
| scf7180000042626 | 563390 | 0.3922 | 0 | 59.6249 | Eukaryota | 90952.0 | 0 | Chordata | 90952.0 | 0 | Carnivora | 90952.0 | 0 | Canidae | 90952.0 | 0 | Canis | 90952.0 | 0 | Canis lupus | 90952.0 | 0 |
| scf7180000046154 | 137391 | 0.3944 | 0 | 60.5643 | Eukaryota | 15318.0 | 0 | Chordata | 15318.0 | 0 | Carnivora | 15318.0 | 0 | Canidae | 11705.0 | 1 | Vulpes | 6611.0 | 3 | Vulpes vulpes | 6611.0 | 3 |
| scf7180000054522 | 14620 | 0.3993 | 0 | 252.2264 | Eukaryota | 82351.0 | 0 | Chordata | 82351.0 | 0 | Carnivora | 82351.0 | 0 | Canidae | 82351.0 | 0 | Canis | 82351.0 | 0 | Canis lupus | 82351.0 | 0 |
| scf7180000056595 | 1572625 | 0.4407 | 100 | 59.3351 | Eukaryota | 126297.0 | 0 | Chordata | 126297.0 | 0 | Carnivora | 126297.0 | 0 | Canidae | 126297.0 | 0 | Canis | 88958.0 | 1 | Canis lupus | 88958.0 | 1 |
| scf7180000043250 | 504070 | 0.4388 | 100 | 60.906 | Eukaryota | 48794.0 | 0 | Chordata | 48794.0 | 0 | Carnivora | 48794.0 | 0 | Canidae | 48794.0 | 0 | Canis | 30640.0 | 1 | Canis lupus | 30640.0 | 1 |
| scf7180000043093 | 4868 | 0.4634 | 0 | 231.7533 | Eukaryota | 7978.0 | 0 | Chordata | 7978.0 | 0 | Carnivora | 7978.0 | 0 | Canidae | 7978.0 | 0 | Canis | 7978.0 | 0 | Canis lupus | 7978.0 | 0 |
| scf7180000040857 | 12358 | 0.4385 | 0 | 33.6275 | no-hit | 0.0 | 0 | no-hit | 0.0 | 0 | no-hit | 0.0 | 0 | no-hit | 0.0 | 0 | no-hit | 0.0 | 0 | no-hit | 0.0 | 0 |
| scf7180000045579 | 31121 | 0.3995 | 0 | 56.3314 | Eukaryota | 4989.0 | 0 | Chordata | 4989.0 | 0 | Carnivora | 3157.0 | 1 | Canidae | 3157.0 | 1 | Vulpes | 3157.0 | 1 | Vulpes vulpes | 3157.0 | 1 |
| scf7180000041991 | 1423289 | 0.3962 | 100 | 60.3626 | Eukaryota | 98374.0 | 0 | Chordata | 98374.0 | 0 | Carnivora | 98374.0 | 0 | Canidae | 98374.0 | 0 | Canis | 86239.0 | 1 | Canis lupus | 86239.0 | 1 |
| scf7180000040988 | 155355 | 0.3571 | 0 | 57.4647 | Eukaryota | 56745.0 | 0 | Chordata | 56745.0 | 0 | Carnivora | 56745.0 | 0 | Canidae | 56745.0 | 0 | Canis | 56745.0 | 0 | Canis lupus | 56745.0 | 0 |
| scf7180000050778 | 2197 | 0.4529 | 0 | 183.7634 | Eukaryota | 1715.0 | 0 | Platyhelminthes | 846.0 | 2 | Polystomatidea | 846.0 | 2 | Polystomatidae | 846.0 | 2 | Protopolystoma | 846.0 | 2 | Protopolystoma xenopodis | 846.0 | 2 |
| scf7180000045338 | 2683 | 0.4898 | 0 | 41.3168 | no-hit | 0.0 | 0 | no-hit | 0.0 | 0 | no-hit | 0.0 | 0 | no-hit | 0.0 | 0 | no-hit | 0.0 | 0 | no-hit | 0.0 | 0 |
| scf7180000045487 | 77473 | 0.4164 | 0 | 60.5092 | Eukaryota | 83515.0 | 0 | Chordata | 83515.0 | 0 | Carnivora | 83515.0 | 0 | Canidae | 83515.0 | 0 | Canis | 83515.0 | 0 | Canis lupus | 83515.0 | 0 |
| scf7180000044020 | 172740 | 0.4419 | 0 | 62.6209 | Eukaryota | 32607.0 | 0 | Chordata | 32607.0 | 0 | Carnivora | 32607.0 | 0 | Felidae | 11502.0 | 2 | Zalophus | 10809.0 | 3 | Zalophus californianus | 10809.0 | 3 |
| scf7180000042608 | 636975 | 0.3623 | 0 | 59.1602 | Eukaryota | 91453.0 | 0 | Chordata | 91453.0 | 0 | Carnivora | 91453.0 | 0 | Canidae | 91453.0 | 0 | Canis | 91453.0 | 0 | Canis lupus | 91453.0 | 0 |
| scf7180000044984 | 12031 | 0.4289 | 0 | 60.2583 | Eukaryota | 61831.0 | 0 | Chordata | 61831.0 | 0 | Carnivora | 61831.0 | 0 | Canidae | 61831.0 | 0 | Canis | 61831.0 | 0 | Canis lupus | 61831.0 | 0 |
| scf7180000046562 | 146290 | 0.4052 | 0 | 60.1922 | Eukaryota | 50451.0 | 0 | Chordata | 50451.0 | 0 | Carnivora | 50451.0 | 0 | Canidae | 27511.0 | 1 | Callorhinus | 22940.0 | 2 | Callorhinus ursinus | 22940.0 | 2 |
| scf7180000048675 | 49416 | 0.3537 | 0 | 60.0072 | Eukaryota | 7224.0 | 0 | Chordata | 7224.0 | 0 | Carnivora | 3954.0 | 2 | Canidae | 3954.0 | 3 | Canis | 3954.0 | 3 | Canis lupus | 3954.0 | 3 |
| scf7180000049495 | 43946 | 0.447 | 0 | 77.6935 | Eukaryota | 3154.0 | 0 | Chordata | 3154.0 | 0 | Carnivora | 3154.0 | 0 | Canidae | 3154.0 | 0 | Canis | 3154.0 | 0 | Canis lupus | 3154.0 | 0 |
| scf7180000054516 | 12831 | 0.4376 | 0 | 275.8035 | Eukaryota | 662.0 | 0 | Chordata | 662.0 | 0 | Salmoniformes | 384.0 | 1 | Salmonidae | 384.0 | 1 | Sphaeramia | 278.0 | 2 | Sphaeramia orbicularis | 278.0 | 2 |
| scf7180000042335 | 235116 | 0.3845 | 0 | 60.25 | Eukaryota | 83095.0 | 0 | Chordata | 83095.0 | 0 | Carnivora | 83095.0 | 0 | Canidae | 83095.0 | 0 | Canis | 83095.0 | 0 | Canis lupus | 83095.0 | 0 |
| scf7180000056230 | 203465 | 0.3636 | 0 | 58.6333 | Eukaryota | 95668.0 | 0 | Chordata | 95668.0 | 0 | Carnivora | 95668.0 | 0 | Canidae | 95668.0 | 0 | Canis | 95668.0 | 0 | Canis lupus | 95668.0 | 0 |
| scf7180000043423 | 467163 | 0.4517 | 0 | 58.8183 | Eukaryota | 87969.0 | 0 | Chordata | 87969.0 | 0 | Carnivora | 87969.0 | 0 | Canidae | 87969.0 | 0 | Canis | 75893.0 | 1 | Canis lupus | 75893.0 | 1 |
| scf7180000056189 | 1827431 | 0.394 | 0 | 59.5119 | Eukaryota | 121714.0 | 0 | Chordata | 121714.0 | 0 | Carnivora | 121714.0 | 0 | Canidae | 121714.0 | 0 | Canis | 121714.0 | 0 | Canis lupus | 121714.0 | 0 |
| scf7180000055830 | 6512 | 0.4387 | 0 | 101.7775 | Eukaryota | 1448.0 | 0 | Platyhelminthes | 919.0 | 1 | Polystomatidea | 919.0 | 1 | Polystomatidae | 919.0 | 1 | Protopolystoma | 919.0 | 1 | Protopolystoma xenopodis | 919.0 | 1 |
| scf7180000050145 | 21487 | 0.4526 | 0 | 57.9123 | Eukaryota | 26293.0 | 0 | Chordata | 26293.0 | 0 | Carnivora | 26293.0 | 0 | Canidae | 26293.0 | 0 | Canis | 26293.0 | 0 | Canis lupus | 26293.0 | 0 |
| scf7180000054595 | 12215 | 0.4397 | 0 | 202.8273 | Eukaryota | 84806.0 | 0 | Chordata | 84806.0 | 0 | Carnivora | 84806.0 | 0 | Canidae | 84806.0 | 0 | Canis | 84806.0 | 0 | Canis lupus | 84806.0 | 0 |
| scf7180000041135 | 1342933 | 0.4437 | 0 | 59.5107 | Eukaryota | 94436.0 | 0 | Chordata | 94436.0 | 0 | Carnivora | 94436.0 | 0 | Canidae | 94436.0 | 0 | Canis | 94436.0 | 0 | Canis lupus | 94436.0 | 0 |
| scf7180000045071 | 2913 | 0.4195 | 0 | 28.2729 | Eukaryota | 31079.0 | 0 | Chordata | 31079.0 | 0 | Carnivora | 31079.0 | 0 | Canidae | 31079.0 | 0 | Canis | 31079.0 | 0 | Canis lupus | 31079.0 | 0 |
| scf7180000053139 | 2648 | 0.4468 | 0 | 390.8675 | no-hit | 0.0 | 0 | no-hit | 0.0 | 0 | no-hit | 0.0 | 0 | no-hit | 0.0 | 0 | no-hit | 0.0 | 0 | no-hit | 0.0 | 0 |
| scf7180000043738 | 104742 | 0.3526 | 0 | 58.9405 | Eukaryota | 39249.0 | 0 | Chordata | 39249.0 | 0 | Carnivora | 39249.0 | 0 | Canidae | 20179.0 | 3 | Canis | 16132.0 | 5 | Canis lupus | 16132.0 | 5 |
| scf7180000046911 | 28973 | 0.4089 | 0 | 84.0704 | Eukaryota | 10491.0 | 0 | Chordata | 10491.0 | 0 | Carnivora | 10491.0 | 0 | Canidae | 10491.0 | 0 | Canis | 10491.0 | 0 | Canis lupus | 10491.0 | 0 |
| scf7180000046523 | 16964 | 0.4157 | 0 | 54.5163 | Eukaryota | 20896.0 | 0 | Chordata | 20896.0 | 0 | Carnivora | 20896.0 | 0 | Canidae | 20896.0 | 0 | Canis | 20896.0 | 0 | Canis lupus | 20896.0 | 0 |
| scf7180000041049 | 1376086 | 0.4272 | 0 | 60.4527 | Eukaryota | 47534.0 | 0 | Chordata | 47534.0 | 0 | Carnivora | 47534.0 | 0 | Canidae | 47534.0 | 0 | Canis | 38921.0 | 1 | Canis lupus | 38921.0 | 1 |

| | | | | | | | | | | | | | | | | | | | | | | |
|---|---|---|---|---|---|---|---|---|---|---|---|---|---|---|---|---|---|---|---|---|---|---|
| scf7180000043474 | 610031 | 0.4046 | 0 | 60.5452 | Eukaryota | 62055.0 | 0 | Chordata | 62055.0 | 0 | Carnivora | 62055.0 | 0 | Canidae | 18717.0 | 5 | Canis | 12820.0 | 7 | Canis lupus | 12820.0 | 7 |
| scf7180000052660 | 2559 | 0.4525 | 0 | 586.9395 | no-hit | 0.0 | 0 | no-hit | 0.0 | 0 | no-hit | 0.0 | 0 | no-hit | 0.0 | 0 | no-hit | 0.0 | 0 | no-hit | 0.0 | 0 |
| scf7180000055592 | 28552 | 0.4336 | 0 | 162.0211 | Eukaryota | 2566.0 | 0 | Platyhelminthes | 1877.0 | 1 | Polystomatidea | 1877.0 | 1 | Polystomatidae | 1877.0 | 1 | Protopolystoma | 1877.0 | 1 | Protopolystoma xenopodis | 1877.0 | 1 |
| scf7180000041428 | 596277 | 0.3947 | 0 | 58.9247 | Eukaryota | 70812.0 | 0 | Chordata | 70812.0 | 0 | Carnivora | 70812.0 | 0 | Canidae | 70812.0 | 0 | Canis | 42513.0 | 1 | Canis lupus | 42513.0 | 1 |
| scf7180000049009 | 29464 | 0.4465 | 0 | 65.8494 | Eukaryota | 1968.0 | 0 | Platyhelminthes | 1241.0 | 1 | Polystomatidea | 1028.0 | 2 | Polystomatidae | 1028.0 | 2 | Protopolystoma | 1028.0 | 2 | Protopolystoma xenopodis | 1028.0 | 2 |
| scf7180000046902 | 47155 | 0.4408 | 0 | 137.9865 | Eukaryota | 4710.0 | 0 | Chordata | 4710.0 | 0 | Carnivora | 2369.0 | 2 | Canidae | 2369.0 | 2 | Canis | 2369.0 | 2 | Canis lupus | 2369.0 | 2 |
| scf7180000042504 | 747245 | 0.3689 | 0 | 61.2672 | Eukaryota | 44461.0 | 0 | Chordata | 44461.0 | 0 | Carnivora | 44461.0 | 0 | Canidae | 44461.0 | 0 | Canis | 40014.0 | 1 | Canis lupus | 40014.0 | 1 |
| scf7180000046382 | 136610 | 0.4255 | 0 | 58.3752 | Eukaryota | 10464.0 | 0 | Chordata | 10464.0 | 0 | Sirenia | 3492.0 | 5 | Trichechidae | 3492.0 | 6 | Trichechus | 3492.0 | 6 | Trichechus manatus | 3492.0 | 6 |
| scf7180000045309 | 182894 | 0.4872 | 0 | 60.9 | Eukaryota | 34732.0 | 0 | Chordata | 34732.0 | 0 | Carnivora | 34732.0 | 0 | Canidae | 26345.0 | 2 | Canis | 22904.0 | 4 | Canis lupus | 22904.0 | 4 |
| scf7180000047939 | 3406 | 0.3714 | 0 | 31.0948 | Eukaryota | 7485.0 | 0 | Chordata | 7485.0 | 0 | Carnivora | 7485.0 | 0 | Felidae | 5311.0 | 1 | Felis | 5311.0 | 1 | Felis catus | 5311.0 | 1 |
| scf7180000056521 | 97891 | 0.4016 | 0 | 55.4124 | Eukaryota | 86545.0 | 0 | Chordata | 86545.0 | 0 | Carnivora | 86545.0 | 0 | Canidae | 86545.0 | 0 | Canis | 86545.0 | 0 | Canis lupus | 86545.0 | 0 |
| scf7180000041853 | 428568 | 0.3849 | 0 | 60.1798 | Eukaryota | 83917.0 | 0 | Chordata | 83917.0 | 0 | Carnivora | 83917.0 | 0 | Canidae | 83917.0 | 0 | Canis | 83917.0 | 0 | Canis lupus | 83917.0 | 0 |
| scf7180000047612 | 2757 | 0.4324 | 0 | 84.3325 | no-hit | 0.0 | 0 | no-hit | 0.0 | 0 | no-hit | 0.0 | 0 | no-hit | 0.0 | 0 | no-hit | 0.0 | 0 | no-hit | 0.0 | 0 |
| scf7180000052955 | 14942 | 0.5023 | 0 | 433.6909 | Eukaryota | 9043.0 | 0 | Chordata | 9043.0 | 0 | Carnivora | 9043.0 | 0 | Canidae | 9043.0 | 0 | Canis | 7851.0 | 1 | Canis lupus | 6611.0 | 2 |
| scf7180000048366 | 8972 | 0.5085 | 0 | 96.8132 | Eukaryota | 32367.0 | 0 | Chordata | 32367.0 | 0 | Carnivora | 32367.0 | 0 | Canidae | 32367.0 | 0 | Canis | 32367.0 | 0 | Canis lupus | 32367.0 | 0 |
| scf7180000041716 | 1525134 | 0.4122 | 0 | 59.8976 | Eukaryota | 136000.0 | 0 | Chordata | 136000.0 | 0 | Carnivora | 136000.0 | 0 | Canidae | 136000.0 | 0 | Canis | 136000.0 | 0 | Canis lupus | 136000.0 | 0 |
| scf7180000052665 | 3986 | 0.5068 | 0 | 39.7247 | Eukaryota | 5473.0 | 0 | Chordata | 5473.0 | 0 | Carnivora | 5473.0 | 0 | Canidae | 5473.0 | 0 | Canis | 4479.0 | 2 | Canis lupus | 4479.0 | 2 |
| scf7180000047587 | 20326 | 0.4477 | 0 | 111.949 | Eukaryota | 224.0 | 0 | Chordata | 224.0 | 0 | Blenniiformes | 224.0 | 0 | Gobiesocidae | 224.0 | 0 | Gouania | 224.0 | 0 | Gouania willdenowi | 224.0 | 0 |
| scf7180000055920 | 6230 | 0.4364 | 0 | 112.9839 | Eukaryota | 749.0 | 0 | Platyhelminthes | 438.0 | 1 | Polystomatidea | 438.0 | 1 | Polystomatidae | 438.0 | 1 | Protopolystoma | 438.0 | 1 | Protopolystoma xenopodis | 438.0 | 1 |
| scf7180000041103 | 1359447 | 0.354 | 0 | 60.4359 | Eukaryota | 64183.0 | 0 | Chordata | 64183.0 | 0 | Carnivora | 64183.0 | 0 | Canidae | 50252.0 | 2 | Canis | 50252.0 | 3 | Canis lupus | 50252.0 | 3 |
| scf7180000042557 | 63568 | 0.4581 | 0 | 57.3546 | Eukaryota | 46764.0 | 0 | Chordata | 46764.0 | 0 | Carnivora | 46764.0 | 0 | Canidae | 46764.0 | 0 | Canis | 46764.0 | 0 | Canis lupus | 46764.0 | 0 |
| scf7180000041776 | 932254 | 0.3785 | 0 | 60.9315 | Eukaryota | 30467.0 | 0 | Chordata | 30467.0 | 0 | Carnivora | 30467.0 | 0 | Canidae | 30467.0 | 0 | Canis | 30467.0 | 0 | Canis lupus | 30467.0 | 0 |
| scf7180000043380 | 490415 | 0.3818 | 0 | 60.267 | Eukaryota | 122817.0 | 0 | Chordata | 122817.0 | 0 | Carnivora | 122817.0 | 0 | Canidae | 122817.0 | 0 | Canis | 122817.0 | 0 | Canis lupus | 122817.0 | 0 |
| scf7180000046058 | 50220 | 0.4384 | 0 | 232.6391 | Eukaryota | 4298.0 | 0 | Chordata | 3947.0 | 1 | Synbranchiformes | 2726.0 | 4 | Mastacembelidae | 2726.0 | 4 | Mastacembelus | 2726.0 | 4 | Mastacembelus armatus | 2726.0 | 4 |
| scf7180000049002 | 3655 | 0.4014 | 0 | 67.9201 | Eukaryota | 28043.0 | 0 | Chordata | 28043.0 | 0 | Carnivora | 28043.0 | 0 | Canidae | 28043.0 | 0 | Canis | 28043.0 | 0 | Canis lupus | 28043.0 | 0 |
| scf7180000051475 | 13489 | 0.5062 | 0 | 288.5374 | Eukaryota | 8171.0 | 0 | Chordata | 8171.0 | 0 | Carnivora | 8171.0 | 0 | Canidae | 8171.0 | 0 | Canis | 6467.0 | 1 | Canis lupus | 5260.0 | 2 |
| scf7180000042960 | 110877 | 0.3746 | 0 | 59.1432 | Eukaryota | 66009.0 | 0 | Chordata | 66009.0 | 0 | Carnivora | 66009.0 | 0 | Canidae | 66009.0 | 0 | Canis | 66009.0 | 0 | Canis lupus | 66009.0 | 0 |
| scf7180000042038 | 6774 | 0.3571 | 0 | 52.5315 | Eukaryota | 6407.0 | 1 | Chordata | 6407.0 | 1 | Primates | 2734.0 | 4 | Hominidae | 2250.0 | 5 | Homo | 2250.0 | 6 | Homo sapiens | 2250.0 | 7 |
| scf7180000056110 | 20022 | 0.4485 | 0 | 186.3648 | Eukaryota | 3381.0 | 0 | Chordata | 2856.0 | 1 | Carnivora | 2856.0 | 1 | Canidae | 2856.0 | 1 | Canis | 2856.0 | 1 | Canis lupus | 2856.0 | 1 |
| scf7180000042782 | 47382 | 0.6521 | 0 | 55.1388 | Eukaryota | 27226.0 | 0 | Chordata | 27226.0 | 0 | Carnivora | 27226.0 | 0 | Canidae | 27226.0 | 0 | Canis | 23899.0 | 1 | Canis lupus | 23899.0 | 1 |
| scf7180000048813 | 8182 | 0.445 | 0 | 124.4797 | Eukaryota | 2822.0 | 0 | Chordata | 2197.0 | 1 | Carnivora | 2197.0 | 1 | Canidae | 2197.0 | 1 | Canis | 2197.0 | 1 | Canis lupus | 2197.0 | 1 |
| scf7180000042858 | 4561 | 0.4036 | 0 | 20.6531 | Eukaryota | 25563.0 | 0 | Chordata | 25563.0 | 0 | Carnivora | 25563.0 | 0 | Canidae | 25563.0 | 0 | Canis | 25563.0 | 0 | Canis lupus | 25563.0 | 0 |
| scf7180000044760 | 248304 | 0.3956 | 0 | 60.0598 | Eukaryota | 53054.0 | 0 | Chordata | 53054.0 | 0 | Carnivora | 53054.0 | 0 | Canidae | 50730.0 | 1 | Canis | 35524.0 | 2 | Canis lupus | 35524.0 | 2 |
| scf7180000042212 | 765991 | 0.4122 | 0 | 60.3779 | Eukaryota | 62053.0 | 0 | Chordata | 62053.0 | 0 | Carnivora | 62053.0 | 0 | Canidae | 62053.0 | 0 | Canis | 41028.0 | 1 | Canis lupus | 41028.0 | 1 |
| scf7180000049702 | 47682 | 0.3358 | 0 | 59.4061 | Eukaryota | 7733.0 | 0 | Chordata | 7733.0 | 0 | Carnivora | 6761.0 | 1 | Felidae | 4003.0 | 2 | Felis | 4003.0 | 2 | Felis catus | 4003.0 | 2 |
| scf7180000049129 | 22202 | 0.4511 | 0 | 1656.1911 | Eukaryota | 3511.0 | 0 | Platyhelminthes | 2102.0 | 2 | Polystomatidea | 2102.0 | 2 | Polystomatidae | 2102.0 | 2 | Protopolystoma | 2102.0 | 2 | Protopolystoma xenopodis | 2102.0 | 2 |
| scf7180000042304 | 259993 | 0.4267 | 0 | 57.0103 | Eukaryota | 17114.0 | 0 | Chordata | 17114.0 | 0 | Carnivora | 17114.0 | 0 | Canidae | 17114.0 | 0 | Canis | 17114.0 | 0 | Canis lupus | 17114.0 | 0 |
| scf7180000049694 | 3392 | 0.4531 | 0 | 344.1828 | Eukaryota | 924.0 | 0 | Echinodermata | 394.0 | 2 | Forcipulatida | 394.0 | 2 | Asteriidae | 394.0 | 2 | Asterias | 394.0 | 2 | Asterias rubens | 394.0 | 2 |
| scf7180000054566 | 30599 | 0.439 | 0 | 158.1641 | Eukaryota | 411.0 | 0 | Chordata | 411.0 | 0 | Salmoniformes | 255.0 | 1 | Salmonidae | 255.0 | 1 | Coregonus | 255.0 | 1 | Coregonus sp. 'balchen' | 255.0 | 1 |
| scf7180000048734 | 36672 | 0.5016 | 0 | 554.8011 | Eukaryota | 9399.0 | 0 | Chordata | 9399.0 | 0 | Carnivora | 9399.0 | 0 | Canidae | 9399.0 | 0 | Canis | 8185.0 | 1 | Canis lupus | 6911.0 | 2 |
| scf7180000056200 | 927795 | 0.343 | 0 | 60.1594 | Eukaryota | 53850.0 | 0 | Chordata | 53850.0 | 0 | Carnivora | 53850.0 | 0 | Felidae | 21728.0 | 3 | Panthera | 21728.0 | 5 | Panthera pardus | 21728.0 | 5 |
| scf7180000041489 | 778775 | 0.3519 | 0 | 59.9598 | Eukaryota | 93758.0 | 0 | Chordata | 93758.0 | 0 | Carnivora | 93758.0 | 0 | Canidae | 93758.0 | 0 | Canis | 93758.0 | 0 | Canis lupus | 93758.0 | 0 |
| scf7180000050591 | 32476 | 0.5002 | 0 | 374.7625 | Eukaryota | 9198.0 | 0 | Chordata | 9198.0 | 0 | Carnivora | 9198.0 | 0 | Canidae | 9198.0 | 0 | Canis | 7969.0 | 1 | Canis lupus | 6729.0 | 2 |
| scf7180000049330 | 31000 | 0.499 | 0 | 52.5976 | Eukaryota | 5337.0 | 1 | Chordata | 5337.0 | 1 | Carnivora | 2866.0 | 3 | Canidae | 2866.0 | 5 | Canis | 2866.0 | 5 | Canis lupus | 2866.0 | 5 |
| scf7180000054362 | 24586 | 0.5315 | 0 | 206.3719 | Eukaryota | 7919.0 | 0 | Chordata | 7919.0 | 0 | Carnivora | 7919.0 | 0 | Canidae | 7919.0 | 0 | Canis | 7919.0 | 0 | Canis lupus | 7919.0 | 0 |
| scf7180000042741 | 579328 | 0.4577 | 0 | 59.6945 | Eukaryota | 62991.0 | 0 | Chordata | 62991.0 | 0 | Carnivora | 62991.0 | 0 | Canidae | 57489.0 | 1 | Canis | 51444.0 | 2 | Canis lupus | 51444.0 | 2 |
| scf7180000048646 | 59472 | 0.4416 | 0 | 65.7918 | Eukaryota | 25372.0 | 1 | Chordata | 25372.0 | 1 | Gadiformes | 10813.0 | 5 | Gadidae | 10813.0 | 5 | Gadus | 10813.0 | 5 | Gadus morhua | 10813.0 | 5 |
| scf7180000044624 | 126208 | 0.3494 | 0 | 57.501 | Eukaryota | 14030.0 | 0 | Chordata | 14030.0 | 0 | Carnivora | 14030.0 | 0 | Canidae | 14030.0 | 0 | Canis | 14030.0 | 0 | Canis lupus | 14030.0 | 0 |
| scf7180000041226 | 1258633 | 0.3764 | 0 | 59.7867 | Eukaryota | 64416.0 | 0 | Chordata | 64416.0 | 0 | Carnivora | 64416.0 | 0 | Canidae | 64416.0 | 0 | Canis | 64416.0 | 0 | Canis lupus | 64416.0 | 0 |
| scf7180000048957 | 38484 | 0.4031 | 0 | 302.2725 | Eukaryota | 61907.0 | 0 | Chordata | 61907.0 | 0 | Carnivora | 61907.0 | 0 | Canidae | 61907.0 | 0 | Canis | 61907.0 | 0 | Canis lupus | 61907.0 | 0 |
| scf7180000041222 | 463029 | 0.3853 | 0 | 58.8921 | Eukaryota | 58214.0 | 0 | Chordata | 58214.0 | 0 | Carnivora | 58214.0 | 0 | Otariidae | 28160.0 | 3 | Zalophus | 16848.0 | 6 | Zalophus californianus | 16848.0 | 6 |
| scf7180000045643 | 7037 | 0.3351 | 0 | 32.0419 | Eukaryota | 3402.0 | 0 | Chordata | 3402.0 | 0 | Carnivora | 1636.0 | 3 | Canidae | 1636.0 | 3 | Canis | 1636.0 | 3 | Canis lupus | 1636.0 | 3 |
| scf7180000049501 | 53748 | 0.4422 | 0 | 58.7751 | Eukaryota | 90968.0 | 0 | Chordata | 90968.0 | 0 | Carnivora | 90968.0 | 0 | Canidae | 90968.0 | 0 | Canis | 90968.0 | 0 | Canis lupus | 90968.0 | 0 |
| scf7180000048970 | 29387 | 0.4408 | 0 | 86.7998 | no-hit | 0.0 | 0 | no-hit | 0.0 | 0 | no-hit | 0.0 | 0 | no-hit | 0.0 | 0 | no-hit | 0.0 | 0 | no-hit | 0.0 | 0 |
| scf7180000054441 | 18182 | 0.4434 | 0 | 317.3867 | Eukaryota | 248.0 | 0 | Chordata | 248.0 | 0 | Kurtiformes | 248.0 | 0 | Apogonidae | 248.0 | 0 | Sphaeramia | 248.0 | 0 | Sphaeramia orbicularis | 248.0 | 0 |
| scf7180000052402 | 10036 | 0.4394 | 0 | 58.728 | Eukaryota | 5221.0 | 0 | Chordata | 4139.0 | 1 | Tetraodontiformes | 2703.0 | 4 | Tetraodontidae | 2703.0 | 4 | Takifugu | 2703.0 | 4 | Takifugu rubripes | 2703.0 | 4 |
| scf7180000048304 | 5103 | 0.4331 | 0 | 122.2561 | Eukaryota | 5980.0 | 0 | Chordata | 5980.0 | 0 | Tetraodontiformes | 5980.0 | 0 | Tetraodontidae | 5980.0 | 0 | Takifugu | 5980.0 | 0 | Takifugu rubripes | 5387.0 | 1 |
| scf7180000049864 | 3796 | 0.4497 | 0 | 509.9478 | no-hit | 0.0 | 0 | no-hit | 0.0 | 0 | no-hit | 0.0 | 0 | no-hit | 0.0 | 0 | no-hit | 0.0 | 0 | no-hit | 0.0 | 0 |
| scf7180000052437 | 33152 | 0.4967 | 0 | 369.2636 | Eukaryota | 9294.0 | 0 | Chordata | 9294.0 | 0 | Carnivora | 9294.0 | 0 | Canidae | 9294.0 | 0 | Canis | 7495.0 | 1 | Canis lupus | 6195.0 | 2 |
| scf7180000045590 | 4313 | 0.3573 | 0 | 31.9164 | Eukaryota | 3338.0 | 0 | Chordata | 3338.0 | 0 | Carnivora | 3338.0 | 0 | Canidae | 3338.0 | 0 | Canis | 3338.0 | 0 | Canis lupus | 3338.0 | 0 |
| scf7180000044988 | 426460 | 0.4677 | 0 | 59.9262 | Eukaryota | 119406.0 | 0 | Chordata | 119406.0 | 0 | Carnivora | 119406.0 | 0 | Canidae | 94263.0 | 1 | Canis | 66241.0 | 2 | Canis lupus | 66241.0 | 2 |
| scf7180000050931 | 17300 | 0.5032 | 0 | 626.2379 | Eukaryota | 9165.0 | 0 | Chordata | 9165.0 | 0 | Carnivora | 9165.0 | 0 | Canidae | 9165.0 | 0 | Canis | 7452.0 | 1 | Canis lupus | 6212.0 | 2 |
| scf7180000050465 | 18785 | 0.4392 | 0 | 66.3532 | Eukaryota | 12177.0 | 0 | Platyhelminthes | 6724.0 | 2 | Cyclophyllidea | 5158.0 | 6 | Hymenolepididae | 5158.0 | 6 | Hymenolepis | 5158.0 | 7 | Hymenolepis microstoma | 5158.0 | 7 |
| scf7180000055522 | 7026 | 0.4328 | 0 | 99.0539 | no-hit | 0.0 | 0 | no-hit | 0.0 | 0 | no-hit | 0.0 | 0 | no-hit | 0.0 | 0 | no-hit | 0.0 | 0 | no-hit | 0.0 | 0 |
| scf7180000042868 | 154891 | 0.3392 | 0 | 57.4084 | Eukaryota | 49953.0 | 0 | Chordata | 49953.0 | 0 | Carnivora | 49953.0 | 0 | Canidae | 49953.0 | 0 | Canis | 49953.0 | 0 | Canis lupus | 49953.0 | 0 |
| scf7180000049260 | 10564 | 0.441 | 0 | 145.9354 | Eukaryota | 630.0 | 0 | Platyhelminthes | 630.0 | 0 | Polystomatidea | 472.0 | 1 | Polystomatidae | 472.0 | 1 | Protopolystoma | 472.0 | 1 | Protopolystoma xenopodis | 472.0 | 1 |
| scf7180000040679 | 340808 | 0.377 | 0 | 59.2623 | Eukaryota | 92357.0 | 0 | Chordata | 92357.0 | 0 | Carnivora | 92357.0 | 0 | Canidae | 92357.0 | 0 | Canis | 92357.0 | 0 | Canis lupus | 92357.0 | 0 |
| scf7180000052383 | 11919 | 0.6822 | 0 | 48.1946 | Eukaryota | 7912.0 | 0 | Chordata | 7912.0 | 0 | Carnivora | 7912.0 | 0 | Canidae | 5763.0 | 1 | Canis | 3812.0 | 3 | Canis lupus | 3812.0 | 3 |
| scf7180000056366 | 167655 | 0.3964 | 0 | 57.4131 | Eukaryota | 61121.0 | 0 | Chordata | 61121.0 | 0 | Carnivora | 61121.0 | 0 | Canidae | 61121.0 | 0 | Canis | 61121.0 | 0 | Canis lupus | 61121.0 | 0 |
| scf7180000043181 | 515270 | 0.3448 | 0 | 59.0305 | Eukaryota | 91057.0 | 0 | Chordata | 91057.0 | 0 | Carnivora | 91057.0 | 0 | Canidae | 91057.0 | 0 | Canis | 91057.0 | 0 | Canis lupus | 91057.0 | 0 |
| scf7180000043726 | 51932 | 0.4393 | 0 | 58.2123 | Eukaryota | 24492.0 | 0 | Chordata | 24492.0 | 0 | Carnivora | 24492.0 | 0 | Otariidae | 8525.0 | 4 | Canis | 7279.0 | 6 | Canis lupus | 7279.0 | 7 |
| scf7180000050048 | 3525 | 0.5067 | 0 | 50.4195 | Eukaryota | 6613.0 | 0 | Chordata | 6613.0 | 0 | Carnivora | 6613.0 | 0 | Canidae | 6613.0 | 0 | Canis | 4223.0 | 1 | Canis lupus | 3228.0 | 2 |
| scf7180000045471 | 205068 | 0.4409 | 0 | 59.7085 | Eukaryota | 73692.0 | 0 | Chordata | 73692.0 | 0 | Carnivora | 73692.0 | 0 | Canidae | 65690.0 | 1 | Canis | 53662.0 | 2 | Canis lupus | 53662.0 | 2 |
| scf7180000045670 | 171578 | 0.4718 | 0 | 59.1192 | Eukaryota | 34058.0 | 0 | Chordata | 34058.0 | 0 | Carnivora | 34058.0 | 0 | Canidae | 34058.0 | 0 | Canis | 34058.0 | 0 | Canis lupus | 34058.0 | 0 |
| scf7180000048631 | 39374 | 0.4413 | 0 | 99.9758 | Eukaryota | 180.0 | 0 | Chordata | 180.0 | 0 | Gadiformes | 180.0 | 0 | Gadidae | 180.0 | 0 | Gadus | 180.0 | 0 | Gadus morhua | 180.0 | 0 |
| scf7180000047753 | 38478 | 0.5793 | 0 | 55.8132 | Eukaryota | 8451.0 | 0 | Chordata | 8451.0 | 0 | Carnivora | 6339.0 | 1 | Canidae | 6339.0 | 1 | Canis | 5622.0 | 2 | Canis lupus | 5622.0 | 2 |
| scf7180000042806 | 588431 | 0.401 | 100 | 58.9847 | Eukaryota | 62001.0 | 0 | Chordata | 62001.0 | 0 | Carnivora | 62001.0 | 0 | Canidae | 62001.0 | 0 | Canis | 52420.0 | 1 | Canis lupus | 52420.0 | 1 |
| scf7180000051263 | 14992 | 0.4606 | 0 | 58.9535 | Eukaryota | 13702.0 | 0 | Chordata | 13702.0 | 0 | Carnivora | 13702.0 | 0 | Canidae | 13702.0 | 0 | Canis | 13702.0 | 0 | Canis lupus | 13702.0 | 0 |
| scf7180000055565 | 22198 | 0.4396 | 0 | 201.3589 | Eukaryota | 97737.0 | 0 | Chordata | 57622.0 | 1 | Gadiformes | 57622.0 | 2 | Gadidae | 57622.0 | 2 | Gadus | 57622.0 | 2 | Gadus morhua | 57622.0 | 2 |
| scf7180000048070 | 4064 | 0.4456 | 0 | 98.9904 | no-hit | 0.0 | 0 | no-hit | 0.0 | 0 | no-hit | 0.0 | 0 | no-hit | 0.0 | 0 | no-hit | 0.0 | 0 | no-hit | 0.0 | 0 |
| scf7180000043668 | 233902 | 0.4971 | 0 | 58.464 | Eukaryota | 51175.0 | 0 | Chordata | 51175.0 | 0 | Carnivora | 51175.0 | 0 | Canidae | 51175.0 | 0 | Canis | 41842.0 | 1 | Canis lupus | 41842.0 | 1 |
| scf7180000056415 | 444437 | 0.539 | 0 | 57.0315 | Eukaryota | 61430.0 | 0 | Chordata | 61430.0 | 0 | Carnivora | 61430.0 | 0 | Canidae | 61430.0 | 0 | Canis | 50154.0 | 1 | Canis lupus | 50154.0 | 1 |
| scf7180000046077 | 126461 | 0.3502 | 0 | 59.2904 | Eukaryota | 87898.0 | 0 | Chordata | 87898.0 | 0 | Carnivora | 87898.0 | 0 | Canidae | 87898.0 | 0 | Canis | 87898.0 | 0 | Canis lupus | 87898.0 | 0 |
| scf7180000041522 | 1352757 | 0.457 | 0 | 58.7931 | Eukaryota | 103231.0 | 0 | Chordata | 103231.0 | 0 | Carnivora | 103231.0 | 0 | Canidae | 103231.0 | 0 | Canis | 92559.0 | 1 | Canis lupus | 92559.0 | 1 |
| scf7180000046624 | 108276 | 0.443 | 0 | 54.3611 | Eukaryota | 39419.0 | 0 | Chordata | 39419.0 | 0 | Carnivora | 39419.0 | 0 | Canidae | 39419.0 | 0 | Canis | 32198.0 | 1 | Canis lupus | 32198.0 | 1 |
| scf7180000045717 | 28221 | 0.4376 | 0 | 137.4072 | Eukaryota | 19004.0 | 0 | Chordata | 10204.0 | 1 | Cyclophyllidea | 8050.0 | 4 | Hymenolepididae | 8050.0 | 4 | Hymenolepis | 8050.0 | 5 | Hymenolepis microstoma | 8050.0 | 5 |
| scf7180000044791 | 98438 | 0.4194 | 0 | 56.68 | Eukaryota | 24706.0 | 0 | Chordata | 24706.0 | 0 | Carnivora | 24706.0 | 0 | Canidae | 10374.0 | 4 | Canis | 6905.0 | 8 | Canis lupus | 6905.0 | 8 |
| scf7180000050222 | 3300 | 0.4424 | 0 | 196.228 | no-hit | 0.0 | 0 | no-hit | 0.0 | 0 | no-hit | 0.0 | 0 | no-hit | 0.0 | 0 | no-hit | 0.0 | 0 | no-hit | 0.0 | 0 |
| scf7180000040931 | 430752 | 0.4124 | 0 | 60.3431 | Eukaryota | 24101.0 | 0 | Chordata | 24101.0 | 0 | Carnivora | 24101.0 | 0 | Canidae | 19837.0 | 2 | Canis | 12385.0 | 4 | Canis lupus | 12385.0 | 4 |
| scf7180000056379 | 604054 | 0.377 | 0 | 60.4873 | Eukaryota | 79431.0 | 0 | Chordata | 79431.0 | 0 | Carnivora | 79431.0 | 0 | Canidae | 79431.0 | 0 | Canis | 79431.0 | 0 | Canis lupus | 79431.0 | 0 |
| scf7180000042168 | 13914 | 0.3893 | 0 | 41.8361 | Eukaryota | 28596.0 | 0 | Chordata | 28596.0 | 0 | Carnivora | 28596.0 | 0 | Canidae | 24613.0 | 1 | Canis | 23196.0 | 2 | Canis lupus | 23196.0 | 2 |
| scf7180000047347 | 82959 | 0.4654 | 0 | 59.5566 | Eukaryota | 22726.0 | 0 | Chordata | 22726.0 | 0 | Carnivora | 22726.0 | 0 | Canidae | 15118.0 | 3 | Canis | 12536.0 | 5 | Canis lupus | 12536.0 | 6 |
| scf7180000046960 | 104388 | 0.4406 | 0 | 160.1933 | Eukaryota | 17283.0 | 0 | Chordata | 13096.0 | 1 | Gadiformes | 4590.0 | 6 | Gadidae | 4590.0 | 6 | Gadus | 4590.0 | 6 | Gadus morhua | 4590.0 | 6 |
| scf7180000051660 | 4211 | 0.4519 | 0 | 211.2147 | no-hit | 0.0 | 0 | no-hit | 0.0 | 0 | no-hit | 0.0 | 0 | no-hit | 0.0 | 0 | no-hit | 0.0 | 0 | no-hit | 0.0 | 0 |
| scf7180000052042 | 2946 | 0.4525 | 0 | 380.6549 | no-hit | 0.0 | 0 | no-hit | 0.0 | 0 | no-hit | 0.0 | 0 | no-hit | 0.0 | 0 | no-hit | 0.0 | 0 | no-hit | 0.0 | 0 |
| scf7180000055009 | 14769 | 0.3756 | 0 | 521.3434 | Eukaryota | 30403.0 | 0 | Chordata | 30403.0 | 0 | Carnivora | 30403.0 | 0 | Canidae | 30403.0 | 0 | Canis | 30403.0 | 0 | Canis lupus | 30403.0 | 0 |
| scf7180000052196 | 3367 | 0.4464 | 0 | 446.3971 | Eukaryota | 303.0 | 0 | Chordata | 303.0 | 0 | Kurtiformes | 303.0 | 0 | Apogonidae | 303.0 | 0 | Sphaeramia | 303.0 | 0 | Sphaeramia orbicularis | 303.0 | 0 |
| scf7180000056255 | 1573959 | 0.4914 | 0 | 59.3527 | Eukaryota | 82697.0 | 0 | Chordata | 82697.0 | 0 | Carnivora | 82697.0 | 0 | Canidae | 82697.0 | 0 | Canis | 76514.0 | 1 | Canis lupus | 76514.0 | 1 |
| scf7180000056386 | 566650 | 0.4179 | 0 | 60.3858 | Eukaryota | 97394.0 | 0 | Chordata | 97394.0 | 0 | Carnivora | 97394.0 | 0 | Canidae | 97394.0 | 0 | Canis | 87815.0 | 1 | Canis lupus | 87815.0 | 1 |
| scf7180000044665 | 54727 | 0.5423 | 0 | 54.2765 | Eukaryota | 29397.0 | 0 | Chordata | 29397.0 | 0 | Carnivora | 29397.0 | 0 | Canidae | 29397.0 | 0 | Canis | 22039.0 | 1 | Canis lupus | 22039.0 | 1 |
| scf7180000045622 | 234798 | 0.3993 | 0 | 59.9903 | Eukaryota | 12542.0 | 0 | Chordata | 12542.0 | 0 | Carnivora | 10807.0 | 1 | Canidae | 9456.0 | 2 | Canis | 9456.0 | 2 | Canis lupus | 9456.0 | 2 |
| scf7180000042435 | 754876 | 0.4047 | 0 | 59.7185 | Eukaryota | 67844.0 | 0 | Chordata | 67844.0 | 0 | Carnivora | 67844.0 | 0 | Canidae | 67844.0 | 0 | Canis | 53197.0 | 1 | Canis lupus | 53197.0 | 1 |
| scf7180000044071 | 362583 | 0.38 | 0 | 59.4733 | Eukaryota | 50398.0 | 0 | Chordata | 50398.0 | 0 | Carnivora | 50398.0 | 0 | Canidae | 50398.0 | 0 | Canis | 50398.0 | 0 | Canis lupus | 50398.0 | 0 |
| scf7180000047591 | 53890 | 0.3972 | 0 | 61.771 | Eukaryota | 15926.0 | 0 | Chordata | 15926.0 | 0 | Carnivora | 15926.0 | 0 | Canidae | 13232.0 | 1 | Canis | 7183.0 | 2 | Canis lupus | 7183.0 | 2 |
| scf7180000048508 | 19258 | 0.4461 | 0 | 208.2366 | no-hit | 0.0 | 0 | no-hit | 0.0 | 0 | no-hit | 0.0 | 0 | no-hit | 0.0 | 0 | no-hit | 0.0 | 0 | no-hit | 0.0 | 0 |
| scf7180000040553 | 219907 | 0.3679 | 0 | 61.6871 | Eukaryota | 90361.0 | 0 | Chordata | 90361.0 | 0 | Carnivora | 90361.0 | 0 | Canidae | 90361.0 | 0 | Canis | 90361.0 | 0 | Canis lupus | 90361.0 | 0 |
| scf7180000044160 | 410165 | 0.3711 | 0 | 60.7051 | Eukaryota | 91591.0 | 0 | Chordata | 91591.0 | 0 | Carnivora | 91591.0 | 0 | Canidae | 91591.0 | 0 | Canis | 91591.0 | 0 | Canis lupus | 91591.0 | 0 |
| scf7180000054252 | 43139 | 0.369 | 0 | 318.9916 | Eukaryota | 22158.0 | 0 | Chordata | 22158.0 | 0 | Carnivora | 22158.0 | 0 | Canidae | 22158.0 | 0 | Canis | 22158.0 | 0 | Canis lupus | 22158.0 | 0 |
| scf7180000044033 | 158563 | 0.3642 | 0 | 59.3078 | Eukaryota | 72239.0 | 0 | Chordata | 72239.0 | 0 | Carnivora | 72239.0 | 0 | Canidae | 72239.0 | 0 | Canis | 72239.0 | 0 | Canis lupus | 72239.0 | 0 |
| scf7180000048378 | 5807 | 0.5096 | 0 | 30.8972 | Eukaryota | 2053.0 | 0 | Chordata | 2053.0 | 0 | Carnivora | 2053.0 | 0 | Canidae | 2053.0 | 0 | Canis | 2053.0 | 0 | Canis lupus | 2053.0 | 0 |
| scf7180000041830 | 842532 | 0.448 | 0 | 59.1772 | Eukaryota | 85703.0 | 0 | Chordata | 85703.0 | 0 | Carnivora | 85703.0 | 0 | Canidae | 85703.0 | 0 | Canis | 85703.0 | 0 | Canis lupus | 85703.0 | 0 |
| scf7180000041347 | 253284 | 0.4055 | 0 | 57.6178 | Eukaryota | 91574.0 | 0 | Chordata | 91574.0 | 0 | Carnivora | 91574.0 | 0 | Canidae | 91574.0 | 0 | Canis | 91574.0 | 0 | Canis lupus | 91574.0 | 0 |
| scf7180000043826 | 44484 | 0.4972 | 0 | 45.5575 | Eukaryota | 3582.0 | 1 | Chordata | 3582.0 | 1 | Carnivora | 1087.0 | 6 | Canidae | 1087.0 | 6 | Homo | 780.0 | 7 | Homo sapiens | 780.0 | 7 |
| scf7180000044935 | 241903 | 0.4097 | 0 | 60.1646 | Eukaryota | 61447.0 | 0 | Chordata | 61447.0 | 0 | Carnivora | 61447.0 | 0 | Canidae | 56239.0 | 1 | Canis | 48778.0 | 2 | Canis lupus | 48778.0 | 2 |
| scf7180000048130 | 58828 | 0.391 | 0 | 60.0074 | Eukaryota | 5738.0 | 0 | Chordata | 5738.0 | 0 | Carnivora | 3783.0 | 1 | Canidae | 3783.0 | 2 | Canis | 3783.0 | 2 | Canis lupus | 3783.0 | 2 |
| scf7180000041563 | 1016801 | 0.4157 | 0 | 60.1562 | Eukaryota | 91410.0 | 0 | Chordata | 91410.0 | 0 | Carnivora | 91410.0 | 0 | Canidae | 91410.0 | 0 | Canis | 91410.0 | 0 | Canis lupus | 91410.0 | 0 |
| scf7180000047277 | 56623 | 0.4446 | 0 | 109.1736 | Eukaryota | 5989.0 | 0 | Chordata | 3181.0 | 2 | Forcipulatida | 1834.0 | 4 | Asteriidae | 1834.0 | 4 | Asterias | 1834.0 | 5 | Asterias rubens | 1834.0 | 5 |
| scf7180000044893 | 243107 | 0.4193 | 0 | 74.167 | Eukaryota | 20593.0 | 0 | Chordata | 20593.0 | 0 | Carnivora | 20593.0 | 0 | Canidae | 20593.0 | 0 | Canis | 20593.0 | 0 | Canis lupus | 20593.0 | 0 |
| scf7180000041180 | 1294226 | 0.3946 | 0 | 61.6961 | Eukaryota | 69336.0 | 0 | Chordata | 69336.0 | 0 | Carnivora | 69336.0 | 0 | Canidae | 69336.0 | 0 | Canis | 52510.0 | 1 | Canis lupus | 52510.0 | 1 |
| scf7180000045693 | 59968 | 0.445 | 0 | 191.7767 | Eukaryota | 16121.0 | 0 | Chordata | 15638.0 | 1 | Tetraodontiformes | 13493.0 | 3 | Tetraodontidae | 13493.0 | 3 | Takifugu | 13493.0 | 3 | Takifugu rubripes | 13493.0 | 3 |
| scf7180000047271 | 13393 | 0.4683 | 0 | 117.9073 | Eukaryota | 9717.0 | 0 | Chordata | 9717.0 | 0 | Carnivora | 5586.0 | 2 | Canidae | 3956.0 | 4 | Canis | 3956.0 | 5 | Canis lupus | 3956.0 | 5 |
| scf7180000046377 | 126178 | 0.3688 | 0 | 59.345 | Eukaryota | 16423.0 | 0 | Chordata | 16423.0 | 0 | Carnivora | 16423.0 | 0 | Canidae | 16423.0 | 0 | Canis | 16423.0 | 0 | Canis lupus | 16423.0 | 0 |
| scf7180000041236 | 660617 | 0.3821 | 0 | 60.058 | Eukaryota | 92083.0 | 0 | Chordata | 92083.0 | 0 | Carnivora | 92083.0 | 0 | Canidae | 92083.0 | 0 | Canis | 92083.0 | 0 | Canis lupus | 92083.0 | 0 |
| scf7180000044057 | 392323 | 0.579 | 0 | 55.348 | Eukaryota | 113294.0 | 0 | Chordata | 113294.0 | 0 | Carnivora | 113294.0 | 0 | Canidae | 113294.0 | 0 | Canis | 113294.0 | 0 | Canis lupus | 113294.0 | 0 |
| scf7180000048043 | 68756 | 0.367 | 0 | 63.9361 | Eukaryota | 18551.0 | 0 | Chordata | 18551.0 | 0 | Primates | 9395.0 | 2 | Hominidae | 9395.0 | 2 | Canis | 7975.0 | 3 | Canis lupus | 7975.0 | 3 |
| scf7180000041365 | 82584 | 0.4511 | 0 | 60.0524 | Eukaryota | 90663.0 | 0 | Chordata | 90663.0 | 0 | Carnivora | 90663.0 | 0 | Canidae | 90663.0 | 0 | Canis | 90663.0 | 0 | Canis lupus | 90663.0 | 0 |
| scf7180000040706 | 55961 | 0.493 | 0 | 49.6642 | Eukaryota | 40801.0 | 0 | Chordata | 40801.0 | 0 | Carnivora | 40801.0 | 0 | Canidae | 38077.0 | 1 | Vulpes | 21459.0 | 2 | Vulpes vulpes | 21459.0 | 2 |
| scf7180000050517 | 26875 | 0.4489 | 0 | 56.6525 | Eukaryota | 12040.0 | 0 | Chordata | 12040.0 | 0 | Carnivora | 12040.0 | 0 | Canidae | 12040.0 | 0 | Canis | 10939.0 | 1 | Canis lupus | 10939.0 | 1 |
| scf7180000056279 | 842798 | 0.4449 | 0 | 59.0952 | Eukaryota | 47823.0 | 0 | Chordata | 47823.0 | 0 | Carnivora | 47823.0 | 0 | Canidae | 47823.0 | 0 | Canis | 47823.0 | 0 | Canis lupus | 47823.0 | 0 |
| scf7180000045697 | 175273 | 0.3824 | 100 | 60.9668 | Eukaryota | 49135.0 | 0 | Chordata | 49135.0 | 0 | Carnivora | 49135.0 | 0 | Canidae | 49135.0 | 0 | Canis | 49135.0 | 0 | Canis lupus | 49135.0 | 0 |
| scf7180000041724 | 913398 | 0.3826 | 0 | 61.5009 | Eukaryota | 99276.0 | 0 | Chordata | 99276.0 | 0 | Carnivora | 99276.0 | 0 | Canidae | 99276.0 | 0 | Canis | 99276.0 | 0 | Canis lupus | 99276.0 | 0 |
| scf7180000052908 | 7396 | 0.4475 | 0 | 287.8371 | Eukaryota | 733.0 | 0 | Chordata | 733.0 | 0 | Kurtiformes | 579.0 | 1 | Apogonidae | 579.0 | 1 | Sphaeramia | 579.0 | 1 | Sphaeramia orbicularis | 579.0 | 1 |
| scf7180000048085 | 84984 | 0.4733 | 0 | 56.233 | Eukaryota | 15128.0 | 0 | Chordata | 15128.0 | 0 | Carnivora | 10537.0 | 2 | Canidae | 7297.0 | 4 | Vulpes | 5764.0 | 5 | Vulpes vulpes | 5764.0 | 5 |
| scf7180000051625 | 16355 | 0.3506 | 0 | 56.9043 | Eukaryota | 4101.0 | 0 | Chordata | 4101.0 | 0 | Carnivora | 1290.0 | 5 | Canidae | 1290.0 | 5 | Canis | 1290.0 | 6 | Canis lupus | 1290.0 | 6 |
| scf7180000055526 | 13179 | 0.4406 | 0 | 114.6093 | no-hit | 0.0 | 0 | no-hit | 0.0 | 0 | no-hit | 0.0 | 0 | no-hit | 0.0 | 0 | no-hit | 0.0 | 0 | no-hit | 0.0 | 0 |
| scf7180000047518 | 58177 | 0.3835 | 0 | 58.7869 | Eukaryota | 44641.0 | 0 | Chordata | 44641.0 | 0 | Carnivora | 44641.0 | 0 | Canidae | 44641.0 | 0 | Canis | 44641.0 | 0 | Canis lupus | 44641.0 | 0 |
| scf7180000042776 | 677841 | 0.3724 | 0 | 60.5529 | Eukaryota | 86667.0 | 0 | Chordata | 86667.0 | 0 | Carnivora | 86667.0 | 0 | Canidae | 86667.0 | 0 | Canis | 81549.0 | 1 | Canis lupus | 81549.0 | 1 |
| scf7180000043318 | 225666 | 0.5338 | 0 | 57.0525 | Eukaryota | 25920.0 | 0 | Chordata | 25920.0 | 0 | Carnivora | 25920.0 | 0 | Canidae | 19017.0 | 1 | Canis | 16891.0 | 2 | Canis lupus | 16891.0 | 2 |
| scf7180000045899 | 25186 | 0.3963 | 0 | 32.4317 | Eukaryota | 8214.0 | 0 | Chordata | 8214.0 | 0 | Carnivora | 8214.0 | 0 | Felidae | 5151.0 | 1 | Felis | 5151.0 | 2 | Felis catus | 5151.0 | 2 |
| scf7180000055805 | 32967 | 0.447 | 0 | 297.3322 | Eukaryota | 9700.0 | 0 | Chordata | 9700.0 | 0 | Carnivora | 9700.0 | 0 | Canidae | 9700.0 | 0 | Canis | 9700.0 | 0 | Canis lupus | 9700.0 | 0 |
| scf7180000043146 | 93849 | 0.4232 | 0 | 52.1412 | Eukaryota | 25120.0 | 0 | Chordata | 25120.0 | 0 | Carnivora | 25120.0 | 0 | Canidae | 25120.0 | 0 | Canis | 22813.0 | 1 | Canis lupus | 22813.0 | 1 |
| scf7180000042540 | 9329 | 0.4331 | 0 | 29.4136 | Eukaryota | 25028.0 | 0 | Chordata | 25028.0 | 0 | Carnivora | 25028.0 | 0 | Canidae | 25028.0 | 0 | Canis | 25028.0 | 0 | Canis lupus | 25028.0 | 0 |
| scf7180000041887 | 1037138 | 0.3937 | 0 | 60.6141 | Eukaryota | 81042.0 | 0 | Chordata | 81042.0 | 0 | Carnivora | 81042.0 | 0 | Canidae | 81042.0 | 0 | Canis | 81042.0 | 0 | Canis lupus | 81042.0 | 0 |
| scf7180000045067 | 244608 | 0.38 | 0 | 60.9466 | Eukaryota | 26463.0 | 0 | Chordata | 26463.0 | 0 | Carnivora | 26463.0 | 0 | Canidae | 26463.0 | 0 | Canis | 26463.0 | 0 | Canis lupus | 26463.0 | 0 |
| scf7180000041232 | 1233892 | 0.3608 | 0 | 61.6794 | Eukaryota | 92673.0 | 0 | Chordata | 92673.0 | 0 | Carnivora | 92673.0 | 0 | Canidae | 92673.0 | 0 | Canis | 92673.0 | 0 | Canis lupus | 92673.0 | 0 |
| scf7180000054444 | 28106 | 0.44 | 0 | 142.7838 | Eukaryota | 94999.0 | 0 | Chordata | 94999.0 | 0 | Carnivora | 94999.0 | 0 | Canidae | 94999.0 | 0 | Canis | 94999.0 | 0 | Canis lupus | 94999.0 | 0 |
| scf7180000055279 | 15536 | 0.3838 | 0 | 122.4294 | Eukaryota | 4690.0 | 0 | Chordata | 3879.0 | 1 | Carnivora | 1884.0 | 3 | Canidae | 1884.0 | 3 | Canis | 1884.0 | 3 | Canis lupus | 1884.0 | 3 |
| scf7180000050115 | 15970 | 0.4773 | 0 | 59.4493 | Eukaryota | 3052.0 | 0 | Chordata | 3052.0 | 0 | Carnivora | 2285.0 | 1 | Canidae | 2285.0 | 1 | Canis | 1996.0 | 2 | Canis lupus | 1996.0 | 2 |
| scf7180000053036 | 2414 | 0.4495 | 0 | 1049.3339 | Eukaryota | 1728.0 | 0 | Platyhelminthes | 664.0 | 2 | Polystomatidea | 664.0 | 2 | Polystomatidae | 664.0 | 2 | Protopolystoma | 664.0 | 2 | Protopolystoma xenopodis | 664.0 | 2 |
| scf7180000046698 | 13171 | 0.4234 | 0 | 1538.3307 | Eukaryota | 8035.0 | 0 | Chordata | 8035.0 | 0 | Carnivora | 8035.0 | 0 | Canidae | 8035.0 | 0 | Canis | 8035.0 | 0 | Canis lupus | 8035.0 | 0 |
| scf7180000049327 | 3610 | 0.4515 | 0 | 400.0581 | no-hit | 0.0 | 0 | no-hit | 0.0 | 0 | no-hit | 0.0 | 0 | no-hit | 0.0 | 0 | no-hit | 0.0 | 0 | no-hit | 0.0 | 0 |
| scf7180000050110 | 30628 | 0.3474 | 0 | 60.3647 | Eukaryota | 16421.0 | 0 | Chordata | 16421.0 | 0 | Carnivora | 16421.0 | 0 | Phocidae | 8091.0 | 2 | Canis | 6712.0 | 3 | Canis lupus | 6712.0 | 3 |
| scf7180000043998 | 381372 | 0.4169 | 0 | 58.6925 | Eukaryota | 44344.0 | 0 | Chordata | 44344.0 | 0 | Carnivora | 44344.0 | 0 | Canidae | 44344.0 | 0 | Canis | 39206.0 | 1 | Canis lupus | 39206.0 | 1 |
| scf7180000047283 | 13843 | 0.4485 | 0 | 209.3135 | Eukaryota | 2912.0 | 0 | Chordata | 2912.0 | 0 | Carnivora | 2614.0 | 1 | Canidae | 2614.0 | 1 | Canis | 2614.0 | 1 | Canis lupus | 2614.0 | 1 |
| scf7180000049498 | 7721 | 0.4382 | 0 | 98.7656 | Eukaryota | 902.0 | 0 | Chordata | 902.0 | 0 | Salmoniformes | 405.0 | 2 | Salmonidae | 405.0 | 2 | Coregonus | 405.0 | 2 | Coregonus sp. 'balchen' | 405.0 | 2 |
| scf7180000047946 | 17209 | 0.4481 | 0 | 80.4515 | Eukaryota | 2833.0 | 0 | Chordata | 2833.0 | 0 | Carnivora | 2833.0 | 0 | Canidae | 2833.0 | 0 | Canis | 2833.0 | 0 | Canis lupus | 2833.0 | 0 |
| scf7180000052443 | 4262 | 0.4789 | 0 | 40.9987 | Eukaryota | 27404.0 | 0 | Chordata | 27404.0 | 0 | Carnivora | 27404.0 | 0 | Canidae | 27404.0 | 0 | Canis | 27404.0 | 0 | Canis lupus | 27404.0 | 0 |
| scf7180000041167 | 32215 | 0.4048 | 0 | 55.7027 | Eukaryota | 24540.0 | 0 | Chordata | 24540.0 | 0 | Carnivora | 24540.0 | 0 | Canidae | 24540.0 | 0 | Canis | 24540.0 | 0 | Canis lupus | 24540.0 | 0 |
| scf7180000047022 | 107112 | 0.3609 | 0 | 60.503 | Eukaryota | 64874.0 | 0 | Chordata | 64874.0 | 0 | Carnivora | 64874.0 | 0 | Canidae | 64874.0 | 0 | Canis | 64874.0 | 0 | Canis lupus | 64874.0 | 0 |
| scf7180000046296 | 146070 | 0.6262 | 0 | 55.5495 | Eukaryota | 33482.0 | 0 | Chordata | 33482.0 | 0 | Carnivora | 18598.0 | 1 | Monodontidae | 7460.0 | 6 | Canis | 6102.0 | 8 | Canis lupus | 6102.0 | 8 |
| scf7180000049496 | 2692 | 0.4435 | 0 | 760.4513 | Eukaryota | 1533.0 | 0 | Echinodermata | 627.0 | 2 | Forcipulatida | 627.0 | 2 | Asteriidae | 627.0 | 2 | Asterias | 627.0 | 2 | Asterias rubens | 627.0 | 2 |
| scf7180000043424 | 535020 | 0.4018 | 0 | 61.7686 | Eukaryota | 66305.0 | 0 | Chordata | 66305.0 | 0 | Carnivora | 66305.0 | 0 | Canidae | 66305.0 | 0 | Canis | 56818.0 | 1 | Canis lupus | 56818.0 | 1 |
| scf7180000053684 | 4825 | 0.447 | 0 | 57.9221 | Eukaryota | 2775.0 | 1 | Chordata | 2775.0 | 1 | Carnivora | 1692.0 | 2 | Canidae | 1692.0 | 2 | Canis | 1692.0 | 2 | Canis lupus | 1692.0 | 2 |
| scf7180000052167 | 7159 | 0.4347 | 0 | 33.4587 | Eukaryota | 374.0 | 0 | Chordata | 374.0 | 0 | Kurtiformes | 374.0 | 0 | Apogonidae | 374.0 | 0 | Sphaeramia | 374.0 | 0 | Sphaeramia orbicularis | 374.0 | 0 |
| scf7180000042978 | 9673 | 0.334 | 0 | 29.0401 | Eukaryota | 5258.0 | 0 | Chordata | 4905.0 | 1 | Primates | 3183.0 | 3 | Hominidae | 3183.0 | 3 | Pan | 2337.0 | 4 | Pan troglodytes | 2337.0 | 4 |
| scf7180000051058 | 16849 | 0.5204 | 0 | 222.156 | Eukaryota | 3816.0 | 0 | Chordata | 3816.0 | 0 | Carnivora | 3816.0 | 0 | Canidae | 3555.0 | 1 | Canis | 3139.0 | 2 | Canis lupus | 3139.0 | 2 |
| scf7180000052538 | 8892 | 0.5053 | 0 | 636.2787 | Eukaryota | 8799.0 | 0 | Chordata | 8799.0 | 0 | Carnivora | 8799.0 | 0 | Canidae | 8799.0 | 0 | Canis | 7128.0 | 1 | Canis lupus | 5928.0 | 2 |
| scf7180000042180 | 834021 | 0.4588 | 0 | 58.6303 | Eukaryota | 68874.0 | 0 | Chordata | 68874.0 | 0 | Carnivora | 68874.0 | 0 | Canidae | 68874.0 | 0 | Canis | 59880.0 | 1 | Canis lupus | 59880.0 | 1 |
| scf7180000054584 | 21146 | 0.5428 | 0 | 298.5813 | Eukaryota | 43541.0 | 0 | Chordata | 43541.0 | 0 | Carnivora | 43541.0 | 0 | Canidae | 43541.0 | 0 | Canis | 43541.0 | 0 | Canis lupus | 43541.0 | 0 |
| scf7180000043197 | 431256 | 0.4301 | 0 | 59.7407 | Eukaryota | 51031.0 | 0 | Chordata | 51031.0 | 0 | Carnivora | 51031.0 | 0 | Canidae | 51031.0 | 0 | Canis | 51031.0 | 0 | Canis lupus | 51031.0 | 0 |
| scf7180000053314 | 6426 | 0.5736 | 0 | 191.0331 | Eukaryota | 13733.0 | 0 | Chordata | 13733.0 | 0 | Artiodactyla | 10975.0 | 1 | Bovidae | 9597.0 | 2 | Ovis | 5484.0 | 3 | Ovis aries | 5484.0 | 3 |
| scf7180000047420 | 2792 | 0.3857 | 0 | 26.0032 | Eukaryota | 3105.0 | 0 | Chordata | 3105.0 | 0 | Carnivora | 3105.0 | 0 | Canidae | 3105.0 | 0 | Canis | 3105.0 | 0 | Canis lupus | 3105.0 | 0 |
| scf7180000045228 | 21149 | 0.4456 | 0 | 705.8824 | Eukaryota | 4860.0 | 0 | Chordata | 2594.0 | 1 | Polystomatidea | 2266.0 | 2 | Polystomatidae | 2266.0 | 2 | Protopolystoma | 2266.0 | 2 | Protopolystoma xenopodis | 2266.0 | 2 |
| scf7180000046305 | 13813 | 0.4455 | 0 | 43.9411 | Eukaryota | 7051.0 | 0 | Chordata | 7051.0 | 0 | Carnivora | 7051.0 | 0 | Canidae | 7051.0 | 0 | Canis | 7051.0 | 0 | Canis lupus | 7051.0 | 0 |
| scf7180000045495 | 11605 | 0.3777 | 0 | 38.758 | Eukaryota | 4090.0 | 0 | Chordata | 4090.0 | 0 | Carnivora | 3032.0 | 1 | Canidae | 3032.0 | 1 | Canis | 3032.0 | 1 | Canis lupus | 3032.0 | 1 |
| scf7180000041753 | 7034 | 0.4562 | 0 | 32.1666 | Eukaryota | 6834.0 | 0 | Chordata | 6834.0 | 0 | Primates | 3027.0 | 2 | Hominidae | 3027.0 | 3 | Homo | 3027.0 | 3 | Homo sapiens | 3027.0 | 3 |
| scf7180000055108 | 107944 | 0.3726 | 0 | 842.2432 | Eukaryota | 65921.0 | 0 | Chordata | 65921.0 | 0 | Carnivora | 65921.0 | 0 | Canidae | 65921.0 | 0 | Canis | 65921.0 | 0 | Canis lupus | 65921.0 | 0 |
| scf7180000047974 | 24525 | 0.4435 | 0 | 163.2185 | Eukaryota | 7604.0 | 1 | Chordata | 5410.0 | 3 | Salmoniformes | 3120.0 | 6 | Salmonidae | 3120.0 | 6 | Salmo | 2796.0 | 7 | Salmo trutta | 2796.0 | 7 |

| | | | | | | | | | | | | | | | | | | | | | | |
|---|---|---|---|---|---|---|---|---|---|---|---|---|---|---|---|---|---|---|---|---|---|---|
| scf7180000044539 | 173918 | 0.3478 | 0 | 59.2118 | Eukaryota | 79538.0 | 0 | Chordata | 79538.0 | 0 | Carnivora | 79538.0 | 0 | Canidae | 79538.0 | 0 | Canis | 79538.0 | 0 | Canis lupus | 79538.0 | 0 |
| scf7180000045943 | 17218 | 0.4131 | 0 | 170.4384 | Eukaryota | 74777.0 | 0 | Chordata | 74777.0 | 0 | Carnivora | 74777.0 | 0 | Canidae | 74777.0 | 0 | Canis | 74777.0 | 0 | Canis lupus | 74777.0 | 0 |
| scf7180000051418 | 9775 | 0.4444 | 0 | 95.9546 | no-hit | 0.0 | 0 | no-hit | 0.0 | 0 | no-hit | 0.0 | 0 | no-hit | 0.0 | 0 | no-hit | 0.0 | 0 | no-hit | 0.0 | 0 |
| scf7180000045084 | 208017 | 0.3377 | 0 | 58.852 | Eukaryota | 13517.0 | 0 | Chordata | 13517.0 | 0 | Carnivora | 11814.0 | 1 | Canidae | 7900.0 | 4 | Canis | 7900.0 | 4 | Canis lupus | 7900.0 | 4 |
| scf7180000050138 | 3328 | 0.4468 | 0 | 430.1651 | no-hit | 0.0 | 0 | no-hit | 0.0 | 0 | no-hit | 0.0 | 0 | no-hit | 0.0 | 0 | no-hit | 0.0 | 0 | no-hit | 0.0 | 0 |
| scf7180000044416 | 183020 | 0.3658 | 0 | 57.841 | Eukaryota | 78021.0 | 0 | Chordata | 78021.0 | 0 | Carnivora | 78021.0 | 0 | Canidae | 78021.0 | 0 | Canis | 78021.0 | 0 | Canis lupus | 78021.0 | 0 |
| scf7180000040752 | 707493 | 0.3535 | 0 | 58.8593 | Eukaryota | 81100.0 | 0 | Chordata | 81100.0 | 0 | Carnivora | 81100.0 | 0 | Canidae | 81100.0 | 0 | Canis | 81100.0 | 0 | Canis lupus | 81100.0 | 0 |
| scf7180000044384 | 22251 | 0.4394 | 0 | 484.5685 | Eukaryota | 1377.0 | 0 | Platyhelminthes | 1377.0 | 0 | Polystomatidea | 1377.0 | 0 | Polystomatidae | 1377.0 | 0 | Protopolystoma | 1377.0 | 0 | Protopolystoma xenopodis | 1377.0 | 0 |
| scf7180000040945 | 747794 | 0.3941 | 0 | 59.2992 | Eukaryota | 290957.0 | 0 | Chordata | 290957.0 | 0 | Carnivora | 290957.0 | 0 | Canidae | 214216.0 | 3 | Vulpes | 152960.0 | 4 | Vulpes vulpes | 152960.0 | 4 |
| scf7180000042845 | 322049 | 0.4342 | 0 | 60.2386 | Eukaryota | 59964.0 | 0 | Chordata | 59964.0 | 0 | Carnivora | 59964.0 | 0 | Canidae | 59964.0 | 0 | Canis | 59964.0 | 0 | Canis lupus | 59964.0 | 0 |
| scf7180000041688 | 866696 | 0.481 | 0 | 59.5171 | Eukaryota | 88833.0 | 0 | Chordata | 88833.0 | 0 | Carnivora | 88833.0 | 0 | Canidae | 88833.0 | 0 | Canis | 57310.0 | 1 | Canis lupus | 57310.0 | 1 |
| scf7180000051187 | 9546 | 0.4515 | 0 | 1233.0779 | Eukaryota | 326.0 | 0 | Platyhelminthes | 326.0 | 0 | Polystomatidea | 326.0 | 0 | Polystomatidae | 326.0 | 0 | Protopolystoma | 326.0 | 0 | Protopolystoma xenopodis | 326.0 | 0 |
| scf7180000048602 | 49075 | 0.3437 | 0 | 60.7078 | Eukaryota | 3383.0 | 0 | Chordata | 3383.0 | 0 | Carnivora | 1741.0 | 2 | Canidae | 1741.0 | 2 | Canis | 1741.0 | 2 | Canis lupus | 1741.0 | 2 |
| scf7180000053638 | 3154 | 0.4372 | 0 | 206.8378 | Eukaryota | 670.0 | 0 | Echinodermata | 440.0 | 1 | Forcipulatida | 440.0 | 1 | Asteriidae | 440.0 | 1 | Asterias | 440.0 | 1 | Asterias rubens | 440.0 | 1 |
| scf7180000041876 | 868766 | 0.4382 | 0 | 58.8611 | Eukaryota | 90528.0 | 0 | Chordata | 90528.0 | 0 | Carnivora | 90528.0 | 0 | Canidae | 90528.0 | 0 | Vulpes | 47049.0 | 1 | Vulpes vulpes | 47049.0 | 1 |
| scf7180000043162 | 372331 | 0.357 | 0 | 59.1414 | Eukaryota | 18127.0 | 0 | Chordata | 18127.0 | 0 | Carnivora | 18127.0 | 0 | Canidae | 18127.0 | 0 | Canis | 18127.0 | 0 | Canis lupus | 18127.0 | 0 |
| scf7180000052672 | 6787 | 0.4461 | 0 | 1333.3345 | no-hit | 0.0 | 0 | no-hit | 0.0 | 0 | no-hit | 0.0 | 0 | no-hit | 0.0 | 0 | no-hit | 0.0 | 0 | no-hit | 0.0 | 0 |
| scf7180000042112 | 47549 | 0.4008 | 0 | 72.4795 | Eukaryota | 93383.0 | 0 | Chordata | 93383.0 | 0 | Carnivora | 93383.0 | 0 | Canidae | 93383.0 | 0 | Canis | 93383.0 | 0 | Canis lupus | 93383.0 | 0 |
| scf7180000045254 | 111600 | 0.5171 | 0 | 55.9131 | Eukaryota | 18880.0 | 0 | Chordata | 18880.0 | 0 | Carnivora | 18880.0 | 0 | Canidae | 18880.0 | 0 | Canis | 18880.0 | 0 | Canis lupus | 18880.0 | 0 |
| scf7180000047918 | 17974 | 0.4503 | 0 | 232.9931 | Eukaryota | 3277.0 | 0 | Chordata | 3277.0 | 0 | Carnivora | 3277.0 | 0 | Canidae | 3277.0 | 0 | Canis | 3277.0 | 0 | Canis lupus | 3277.0 | 0 |
| scf7180000043514 | 451849 | 0.6491 | 0 | 54.5615 | Eukaryota | 36591.0 | 0 | Chordata | 36591.0 | 0 | Carnivora | 36591.0 | 0 | Canidae | 36591.0 | 0 | Canis | 27220.0 | 1 | Canis lupus | 27220.0 | 1 |
| scf7180000041662 | 938984 | 0.4294 | 0 | 59.2807 | Eukaryota | 25909.0 | 0 | Chordata | 25909.0 | 0 | Carnivora | 25909.0 | 0 | Canidae | 10660.0 | 4 | Vulpes | 8004.0 | 6 | Vulpes vulpes | 8004.0 | 6 |
| scf7180000045847 | 177456 | 0.381 | 0 | 67.5818 | Eukaryota | 11062.0 | 0 | Chordata | 11062.0 | 0 | Carnivora | 11062.0 | 0 | Canidae | 11062.0 | 0 | Canis | 11062.0 | 0 | Canis lupus | 11062.0 | 0 |
| scf7180000043896 | 262998 | 0.3442 | 0 | 59.4998 | Eukaryota | 18296.0 | 0 | Chordata | 18296.0 | 0 | Carnivora | 18296.0 | 0 | Canidae | 18296.0 | 0 | Canis | 18296.0 | 0 | Canis lupus | 18296.0 | 0 |
| scf7180000054643 | 30739 | 0.4306 | 0 | 122.7907 | Eukaryota | 49687.0 | 0 | Chordata | 49687.0 | 0 | Carnivora | 49687.0 | 0 | Canidae | 49687.0 | 0 | Canis | 49687.0 | 0 | Canis lupus | 49687.0 | 0 |
| scf7180000056351 | 94773 | 0.3418 | 0 | 58.876 | Eukaryota | 9061.0 | 0 | Chordata | 9061.0 | 0 | Carnivora | 5385.0 | 1 | Hominidae | 3676.0 | 2 | Felis | 3037.0 | 3 | Felis catus | 3037.0 | 3 |
| scf7180000055082 | 25951 | 0.4061 | 0 | 139.1035 | Eukaryota | 96445.0 | 0 | Chordata | 96445.0 | 0 | Carnivora | 96445.0 | 0 | Canidae | 96445.0 | 0 | Canis | 96445.0 | 0 | Canis lupus | 96445.0 | 0 |
| scf7180000047532 | 74723 | 0.4061 | 0 | 58.0522 | Eukaryota | 24479.0 | 0 | Chordata | 24479.0 | 0 | Carnivora | 24479.0 | 0 | Canidae | 22833.0 | 1 | Canis | 18485.0 | 2 | Canis lupus | 18485.0 | 2 |
| scf7180000052687 | 4829 | 0.5086 | 0 | 895.3245 | Eukaryota | 5086.0 | 0 | Chordata | 5086.0 | 0 | Carnivora | 5086.0 | 0 | Canidae | 5086.0 | 0 | Canis | 4295.0 | 2 | Canis lupus | 4295.0 | 2 |
| scf7180000048640 | 11807 | 0.395 | 0 | 65.3575 | Eukaryota | 85302.0 | 0 | Chordata | 85302.0 | 0 | Carnivora | 85302.0 | 0 | Canidae | 85302.0 | 0 | Canis | 85302.0 | 0 | Canis lupus | 85302.0 | 0 |
| scf7180000049781 | 26528 | 0.4424 | 0 | 338.456 | Eukaryota | 7177.0 | 0 | Chordata | 6652.0 | 1 | Kurtiformes | 5278.0 | 2 | Apogonidae | 5278.0 | 2 | Sphaeramia | 5278.0 | 2 | Sphaeramia orbicularis | 5278.0 | 2 |
| scf7180000053494 | 3872 | 0.4419 | 0 | 211.0575 | no-hit | 0.0 | 0 | no-hit | 0.0 | 0 | no-hit | 0.0 | 0 | no-hit | 0.0 | 0 | no-hit | 0.0 | 0 | no-hit | 0.0 | 0 |
| scf7180000045840 | 17132 | 0.4421 | 0 | 197.6621 | Eukaryota | 84393.0 | 0 | Chordata | 84393.0 | 0 | Carnivora | 84393.0 | 0 | Canidae | 84393.0 | 0 | Canis | 84393.0 | 0 | Canis lupus | 84393.0 | 0 |
| scf7180000054265 | 37719 | 0.4406 | 0 | 318.0235 | Eukaryota | 55842.0 | 0 | Chordata | 41224.0 | 2 | Gadiformes | 34187.0 | 5 | Gadidae | 34187.0 | 5 | Gadus | 34187.0 | 5 | Gadus morhua | 34187.0 | 5 |
| scf7180000043946 | 393905 | 0.3494 | 0 | 58.3032 | Eukaryota | 87646.0 | 0 | Chordata | 87646.0 | 0 | Carnivora | 87646.0 | 0 | Canidae | 87646.0 | 0 | Canis | 87646.0 | 0 | Canis lupus | 87646.0 | 0 |
| scf7180000047967 | 57813 | 0.4305 | 0 | 62.364 | Eukaryota | 9405.0 | 0 | Chordata | 9405.0 | 0 | Carnivora | 6383.0 | 2 | Canidae | 5181.0 | 4 | Canis | 5181.0 | 5 | Canis lupus | 5181.0 | 5 |
| scf7180000047463 | 42647 | 0.4091 | 0 | 57.8866 | Eukaryota | 4382.0 | 0 | Chordata | 4382.0 | 0 | Carnivora | 3573.0 | 1 | Ursidae | 1979.0 | 2 | Ursus | 1979.0 | 2 | Ursus arctos | 1979.0 | 2 |
| scf7180000047386 | 18984 | 0.4455 | 0 | 347.0601 | Eukaryota | 105654.0 | 0 | Chordata | 55633.0 | 1 | Gadiformes | 55633.0 | 2 | Gadidae | 55633.0 | 2 | Gadus | 55633.0 | 2 | Gadus morhua | 55633.0 | 2 |
| scf7180000043274 | 294658 | 0.365 | 0 | 58.5724 | Eukaryota | 51628.0 | 0 | Chordata | 51628.0 | 0 | Carnivora | 51628.0 | 0 | Canidae | 51628.0 | 0 | Canis | 51628.0 | 0 | Canis lupus | 51628.0 | 0 |
| scf7180000052568 | 13741 | 0.5182 | 0 | 86.5721 | Eukaryota | 55131.0 | 0 | Chordata | 55131.0 | 0 | Carnivora | 55131.0 | 0 | Canidae | 55131.0 | 0 | Canis | 55131.0 | 0 | Canis lupus | 55131.0 | 0 |
| scf7180000042783 | 673762 | 0.3438 | 0 | 60.2811 | Eukaryota | 42520.0 | 0 | Chordata | 42520.0 | 0 | Carnivora | 42520.0 | 0 | Canidae | 42520.0 | 0 | Canis | 42520.0 | 0 | Canis lupus | 42520.0 | 0 |
| scf7180000045855 | 21511 | 0.4414 | 0 | 200.6888 | no-hit | 0.0 | 0 | no-hit | 0.0 | 0 | no-hit | 0.0 | 0 | no-hit | 0.0 | 0 | no-hit | 0.0 | 0 | no-hit | 0.0 | 0 |
| scf7180000048879 | 45920 | 0.4448 | 0 | 61.1444 | Eukaryota | 25527.0 | 1 | Chordata | 25527.0 | 1 | Gadiformes | 10756.0 | 5 | Gadidae | 10756.0 | 5 | Gadus | 10756.0 | 5 | Gadus morhua | 10756.0 | 5 |
| scf7180000042501 | 676437 | 0.381 | 0 | 59.0823 | Eukaryota | 124454.0 | 0 | Chordata | 124454.0 | 0 | Carnivora | 124454.0 | 0 | Canidae | 124454.0 | 0 | Canis | 110295.0 | 1 | Canis lupus | 110295.0 | 1 |
| scf7180000045430 | 59237 | 0.3848 | 0 | 56.9508 | Eukaryota | 23181.0 | 0 | Chordata | 23181.0 | 0 | Carnivora | 23181.0 | 0 | Canidae | 14763.0 | 1 | Canis | 14763.0 | 1 | Canis lupus | 14763.0 | 1 |
| scf7180000042721 | 173638 | 0.3865 | 0 | 58.7221 | Eukaryota | 25951.0 | 0 | Chordata | 25951.0 | 0 | Carnivora | 25951.0 | 0 | Canidae | 25951.0 | 0 | Canis | 25951.0 | 0 | Canis lupus | 25951.0 | 0 |
| scf7180000055694 | 8172 | 0.4422 | 0 | 127.6412 | Eukaryota | 231.0 | 0 | Chordata | 231.0 | 0 | Salmoniformes | 231.0 | 0 | Salmonidae | 231.0 | 0 | Coregonus | 231.0 | 0 | Coregonus sp. 'balchen' | 231.0 | 0 |
| scf7180000051730 | 3879 | 0.5063 | 0 | 234.5249 | Eukaryota | 5434.0 | 0 | Chordata | 5434.0 | 0 | Carnivora | 5434.0 | 0 | Canidae | 5434.0 | 0 | Canis | 3603.0 | 1 | Canis lupus | 2685.0 | 2 |
| scf7180000041784 | 269511 | 0.4306 | 0 | 57.7274 | Eukaryota | 65256.0 | 0 | Chordata | 65256.0 | 0 | Carnivora | 65256.0 | 0 | Canidae | 65256.0 | 0 | Canis | 65256.0 | 0 | Canis lupus | 65256.0 | 0 |
| scf7180000045380 | 27899 | 0.3826 | 0 | 139.9771 | Eukaryota | 4257.0 | 0 | Chordata | 4257.0 | 0 | Carnivora | 2546.0 | 2 | Canidae | 1864.0 | 4 | Canis | 1864.0 | 4 | Canis lupus | 1864.0 | 4 |
| scf7180000055645 | 17125 | 0.516 | 0 | 235.1763 | Eukaryota | 15614.0 | 0 | Chordata | 15614.0 | 0 | Carnivora | 15614.0 | 0 | Canidae | 15614.0 | 0 | Canis | 15614.0 | 0 | Canis lupus | 15614.0 | 0 |
| scf7180000049795 | 39822 | 0.4399 | 0 | 91.0652 | Eukaryota | 1190.0 | 0 | Streptophyta | 634.0 | 1 | Fabales | 634.0 | 1 | Fabaceae | 634.0 | 1 | Cicer | 634.0 | 1 | Cicer arietinum | 634.0 | 1 |
| scf7180000043714 | 199592 | 0.5026 | 0 | 56.8539 | Eukaryota | 35934.0 | 0 | Chordata | 35934.0 | 0 | Carnivora | 35934.0 | 0 | Canidae | 35934.0 | 0 | Canis | 26383.0 | 1 | Canis lupus | 26383.0 | 1 |
| scf7180000055317 | 28365 | 0.4373 | 0 | 149.5393 | Eukaryota | 1931.0 | 0 | Chordata | 1219.0 | 1 | Kurtiformes | 1219.0 | 1 | Apogonidae | 1219.0 | 1 | Sphaeramia | 1219.0 | 1 | Sphaeramia orbicularis | 1219.0 | 1 |
| scf7180000043528 | 457756 | 0.3629 | 0 | 59.6902 | Eukaryota | 32331.0 | 0 | Chordata | 32331.0 | 0 | Primates | 23541.0 | 2 | Hominidae | 23541.0 | 2 | Homo | 23541.0 | 2 | Homo sapiens | 23541.0 | 2 |
| scf7180000046094 | 163650 | 0.4423 | 100 | 59.5622 | Eukaryota | 37408.0 | 0 | Chordata | 37408.0 | 0 | Carnivora | 37408.0 | 0 | Canidae | 37408.0 | 0 | Canis | 37408.0 | 0 | Canis lupus | 37408.0 | 0 |
| scf7180000054034 | 8070 | 0.4404 | 0 | 56.574 | no-hit | 0.0 | 0 | no-hit | 0.0 | 0 | no-hit | 0.0 | 0 | no-hit | 0.0 | 0 | no-hit | 0.0 | 0 | no-hit | 0.0 | 0 |
| scf7180000052930 | 5169 | 0.5045 | 0 | 247.0284 | Eukaryota | 8226.0 | 0 | Chordata | 8226.0 | 0 | Carnivora | 8226.0 | 0 | Canidae | 8226.0 | 0 | Canis | 6545.0 | 1 | Canis lupus | 5347.0 | 2 |
| scf7180000048898 | 28610 | 0.4481 | 0 | 57.2871 | Eukaryota | 39736.0 | 0 | Chordata | 31061.0 | 2 | Gadiformes | 21462.0 | 6 | Gadidae | 21462.0 | 6 | Gadus | 21462.0 | 6 | Gadus morhua | 21462.0 | 6 |
| scf7180000050858 | 13386 | 0.3638 | 0 | 52.1175 | Eukaryota | 58320.0 | 0 | Chordata | 58320.0 | 0 | Carnivora | 58320.0 | 0 | Canidae | 58320.0 | 0 | Canis | 58320.0 | 0 | Canis lupus | 58320.0 | 0 |
| scf7180000053830 | 3935 | 0.4394 | 0 | 1920.7858 | no-hit | 0.0 | 0 | no-hit | 0.0 | 0 | no-hit | 0.0 | 0 | no-hit | 0.0 | 0 | no-hit | 0.0 | 0 | no-hit | 0.0 | 0 |
| scf7180000048855 | 2257 | 0.4448 | 0 | 93.215 | no-hit | 0.0 | 0 | no-hit | 0.0 | 0 | no-hit | 0.0 | 0 | no-hit | 0.0 | 0 | no-hit | 0.0 | 0 | no-hit | 0.0 | 0 |
| scf7180000043279 | 349936 | 0.4585 | 0 | 190.2411 | Eukaryota | 61247.0 | 0 | Chordata | 61247.0 | 0 | Carnivora | 61247.0 | 0 | Canidae | 61247.0 | 0 | Canis | 61247.0 | 0 | Canis lupus | 61247.0 | 0 |
| scf7180000054270 | 207597 | 0.401 | 0 | 196.7864 | Eukaryota | 91775.0 | 0 | Chordata | 91775.0 | 0 | Carnivora | 91775.0 | 0 | Canidae | 91775.0 | 0 | Canis | 91775.0 | 0 | Canis lupus | 91775.0 | 0 |
| scf7180000042629 | 612592 | 0.355 | 0 | 60.0724 | Eukaryota | 41241.0 | 0 | Chordata | 41241.0 | 0 | Primates | 18187.0 | 2 | Hominidae | 18187.0 | 3 | Homo | 18187.0 | 4 | Homo sapiens | 18187.0 | 4 |
| scf7180000052566 | 5510 | 0.4503 | 0 | 752.5542 | Eukaryota | 2762.0 | 0 | Platyhelminthes | 1431.0 | 2 | Polystomatidea | 1431.0 | 2 | Polystomatidae | 1431.0 | 2 | Protopolystoma | 1431.0 | 2 | Protopolystoma xenopodis | 1431.0 | 2 |
| scf7180000041064 | 1535 | 0.4476 | 0 | 269.118 | no-hit | 0.0 | 0 | no-hit | 0.0 | 0 | no-hit | 0.0 | 0 | no-hit | 0.0 | 0 | no-hit | 0.0 | 0 | no-hit | 0.0 | 0 |
| scf7180000052567 | 14238 | 0.3836 | 0 | 29.9092 | Eukaryota | 70750.0 | 0 | Chordata | 70750.0 | 0 | Carnivora | 70750.0 | 0 | Canidae | 70750.0 | 0 | Canis | 70750.0 | 0 | Canis lupus | 70750.0 | 0 |
| scf7180000053732 | 4049 | 0.4539 | 0 | 761.181 | Eukaryota | 278.0 | 0 | Euglenozoa | 278.0 | 0 | Kinetoplastida | 278.0 | 0 | Trypanosomatidae | 278.0 | 0 | Leishmania | 278.0 | 0 | Leishmania infantum | 278.0 | 0 |
| scf7180000052863 | 7329 | 0.4377 | 0 | 52.9383 | Eukaryota | 1263.0 | 0 | Chordata | 998.0 | 1 | Salmoniformes | 998.0 | 1 | Salmonidae | 998.0 | 1 | Coregonus | 667.0 | 2 | Coregonus sp. 'balchen' | 667.0 | 2 |
| scf7180000052532 | 6552 | 0.4947 | 0 | 57.3298 | Eukaryota | 7470.0 | 0 | Chordata | 7470.0 | 0 | Carnivora | 7470.0 | 0 | Canidae | 7470.0 | 0 | Canis | 5297.0 | 1 | Canis lupus | 4160.0 | 2 |
| scf7180000048491 | 54362 | 0.3879 | 0 | 59.7579 | Eukaryota | 19438.0 | 0 | Chordata | 19438.0 | 0 | Carnivora | 19438.0 | 0 | Mustelidae | 9460.0 | 3 | Mustela | 9460.0 | 3 | Mustela putorius | 9460.0 | 3 |
| scf7180000041551 | 590231 | 0.3923 | 0 | 59.668 | Eukaryota | 56408.0 | 1 | Chordata | 56408.0 | 1 | Carnivora | 34918.0 | 2 | Canidae | 34918.0 | 2 | Canis | 34918.0 | 2 | Canis lupus | 34918.0 | 2 |
| scf7180000047070 | 20650 | 0.4668 | 0 | 107.9936 | Eukaryota | 95254.0 | 0 | Chordata | 95254.0 | 0 | Carnivora | 95254.0 | 0 | Canidae | 95254.0 | 0 | Canis | 95254.0 | 0 | Canis lupus | 95254.0 | 0 |
| scf7180000044386 | 316630 | 0.4269 | 0 | 59.5412 | Eukaryota | 36460.0 | 0 | Chordata | 36460.0 | 0 | Carnivora | 36460.0 | 0 | Canidae | 36460.0 | 0 | Canis | 36460.0 | 0 | Canis lupus | 36460.0 | 0 |
| scf7180000050397 | 4097 | 0.453 | 0 | 60.3068 | Eukaryota | 2771.0 | 0 | Chordata | 2443.0 | 1 | Tetraodontiformes | 1392.0 | 5 | Tetraodontidae | 1392.0 | 5 | Takifugu | 1392.0 | 5 | Takifugu rubripes | 1392.0 | 5 |
| scf7180000044564 | 281603 | 0.4047 | 0 | 58.663 | Eukaryota | 19955.0 | 0 | Chordata | 19955.0 | 0 | Carnivora | 19955.0 | 0 | Canidae | 11517.0 | 2 | Canis | 11517.0 | 2 | Canis lupus | 11517.0 | 2 |
| scf7180000049645 | 12229 | 0.4367 | 0 | 253.9195 | Eukaryota | 3115.0 | 0 | Platyhelminthes | 2121.0 | 2 | Polystomatidea | 2121.0 | 2 | Polystomatidae | 2121.0 | 2 | Protopolystoma | 2121.0 | 2 | Protopolystoma xenopodis | 2121.0 | 2 |
| scf7180000054847 | 9836 | 0.5187 | 0 | 622.1713 | Eukaryota | 16408.0 | 0 | Chordata | 16408.0 | 0 | Carnivora | 16408.0 | 0 | Canidae | 16408.0 | 0 | Canis | 16408.0 | 0 | Canis lupus | 16408.0 | 0 |
| scf7180000052869 | 4741 | 0.4444 | 0 | 123.6049 | Eukaryota | 259.0 | 0 | Chordata | 259.0 | 0 | Salmoniformes | 259.0 | 0 | Salmonidae | 259.0 | 0 | Coregonus | 259.0 | 0 | Coregonus sp. 'balchen' | 259.0 | 0 |
| scf7180000043557 | 398211 | 0.4621 | 0 | 57.6966 | Eukaryota | 87651.0 | 0 | Chordata | 87651.0 | 0 | Carnivora | 69411.0 | 1 | Canidae | 69411.0 | 1 | Canis | 48774.0 | 2 | Canis lupus | 48774.0 | 2 |
| scf7180000050097 | 1767 | 0.446 | 0 | 609.237 | no-hit | 0.0 | 0 | no-hit | 0.0 | 0 | no-hit | 0.0 | 0 | no-hit | 0.0 | 0 | no-hit | 0.0 | 0 | no-hit | 0.0 | 0 |
| scf7180000041615 | 382165 | 0.3384 | 0 | 58.1939 | Eukaryota | 57588.0 | 0 | Chordata | 57588.0 | 0 | Carnivora | 57588.0 | 0 | Canidae | 57588.0 | 0 | Canis | 57588.0 | 0 | Canis lupus | 57588.0 | 0 |
| scf7180000051087 | 11065 | 0.5706 | 0 | 271.7885 | Eukaryota | 29668.0 | 0 | Chordata | 29668.0 | 0 | Primates | 16698.0 | 2 | Hominidae | 16698.0 | 2 | Ovis | 9213.0 | 3 | Ovis aries | 9213.0 | 3 |
| scf7180000050543 | 15844 | 0.5194 | 0 | 703.1174 | Eukaryota | 16115.0 | 0 | Chordata | 16115.0 | 0 | Carnivora | 16115.0 | 0 | Canidae | 16115.0 | 0 | Canis | 16115.0 | 0 | Canis lupus | 16115.0 | 0 |
| scf7180000044424 | 3703 | 0.3927 | 0 | 3.3731 | Eukaryota | 53654.0 | 0 | Chordata | 53654.0 | 0 | Carnivora | 53654.0 | 0 | Canidae | 53654.0 | 0 | Canis | 53654.0 | 0 | Canis lupus | 53654.0 | 0 |
| scf7180000041140 | 1402215 | 0.368 | 0 | 59.4728 | Eukaryota | 203068.0 | 0 | Chordata | 203068.0 | 0 | Carnivora | 203068.0 | 0 | Canidae | 203068.0 | 0 | Canis | 203068.0 | 0 | Canis lupus | 203068.0 | 0 |
| scf7180000049357 | 5382 | 0.4526 | 0 | 92.2754 | no-hit | 0.0 | 0 | no-hit | 0.0 | 0 | no-hit | 0.0 | 0 | no-hit | 0.0 | 0 | no-hit | 0.0 | 0 | no-hit | 0.0 | 0 |
| scf7180000042464 | 402864 | 0.3922 | 0 | 59.3097 | Eukaryota | 92102.0 | 0 | Chordata | 92102.0 | 0 | Carnivora | 92102.0 | 0 | Canidae | 92102.0 | 0 | Canis | 92102.0 | 0 | Canis lupus | 92102.0 | 0 |
| scf7180000056039 | 14124 | 0.627 | 0 | 290.8937 | Eukaryota | 66912.0 | 0 | Chordata | 66912.0 | 0 | Artiodactyla | 27682.0 | 3 | Bovidae | 27682.0 | 3 | Ovis | 27682.0 | 4 | Ovis aries | 27682.0 | 4 |
| scf7180000053565 | 2942 | 0.4483 | 0 | 453.5683 | no-hit | 0.0 | 0 | no-hit | 0.0 | 0 | no-hit | 0.0 | 0 | no-hit | 0.0 | 0 | no-hit | 0.0 | 0 | no-hit | 0.0 | 0 |
| scf7180000045081 | 97521 | 0.3727 | 0 | 62.7137 | Eukaryota | 83448.0 | 0 | Chordata | 83448.0 | 0 | Carnivora | 83448.0 | 0 | Canidae | 83448.0 | 0 | Canis | 83448.0 | 0 | Canis lupus | 83448.0 | 0 |
| scf7180000043779 | 356617 | 0.3694 | 0 | 57.5736 | Eukaryota | 55404.0 | 0 | Chordata | 55404.0 | 0 | Carnivora | 55404.0 | 0 | Canidae | 55404.0 | 0 | Canis | 55404.0 | 0 | Canis lupus | 55404.0 | 0 |
| scf7180000044244 | 268824 | 0.5005 | 0 | 58.4674 | Eukaryota | 39463.0 | 0 | Chordata | 39463.0 | 0 | Carnivora | 39463.0 | 0 | Canidae | 39463.0 | 0 | Canis | 34785.0 | 1 | Canis lupus | 34785.0 | 1 |
| scf7180000048376 | 18053 | 0.4437 | 0 | 201.9273 | Eukaryota | 14551.0 | 0 | Chordata | 9054.0 | 3 | Gadiformes | 4279.0 | 7 | Gadidae | 4279.0 | 7 | Gadus | 4279.0 | 7 | Gadus morhua | 4279.0 | 7 |
| scf7180000046823 | 13838 | 0.3992 | 0 | 81.1888 | Eukaryota | 26911.0 | 0 | Chordata | 26911.0 | 0 | Carnivora | 22375.0 | 2 | Canidae | 22375.0 | 3 | Canis | 22375.0 | 3 | Canis lupus | 22375.0 | 3 |
| scf7180000044233 | 78033 | 0.3476 | 0 | 58.4326 | Eukaryota | 19337.0 | 0 | Chordata | 19337.0 | 0 | Carnivora | 19337.0 | 0 | Canidae | 19337.0 | 0 | Canis | 19337.0 | 0 | Canis lupus | 19337.0 | 0 |
| scf7180000047922 | 3103 | 0.4457 | 0 | 46.794 | Eukaryota | 2798.0 | 0 | Chordata | 2027.0 | 1 | Anabantiformes | 1099.0 | 2 | Osphronemidae | 1099.0 | 2 | Betta | 1099.0 | 2 | Betta splendens | 1099.0 | 2 |
| scf7180000044904 | 144302 | 0.4365 | 0 | 162.3022 | Eukaryota | 60834.0 | 0 | Chordata | 48553.0 | 1 | Gadiformes | 23619.0 | 3 | Gadidae | 23619.0 | 3 | Gadus | 23619.0 | 3 | Gadus morhua | 23619.0 | 3 |
| scf7180000043582 | 546250 | 0.3884 | 0 | 59.1461 | Eukaryota | 108331.0 | 0 | Chordata | 108331.0 | 0 | Carnivora | 108331.0 | 0 | Canidae | 108331.0 | 0 | Canis | 108331.0 | 0 | Canis lupus | 108331.0 | 0 |
| scf7180000040487 | 587478 | 0.4809 | 0 | 57.6239 | Eukaryota | 91974.0 | 0 | Chordata | 91974.0 | 0 | Carnivora | 91974.0 | 0 | Canidae | 91974.0 | 0 | Canis | 91974.0 | 0 | Canis lupus | 91974.0 | 0 |
| scf7180000044874 | 258049 | 0.4733 | 0 | 58.6539 | Eukaryota | 28539.0 | 0 | Chordata | 28539.0 | 0 | Carnivora | 28539.0 | 0 | Canidae | 23602.0 | 1 | Canis | 20909.0 | 2 | Canis lupus | 20909.0 | 2 |
| scf7180000049689 | 3504 | 0.4458 | 0 | 90.9352 | no-hit | 0.0 | 0 | no-hit | 0.0 | 0 | no-hit | 0.0 | 0 | no-hit | 0.0 | 0 | no-hit | 0.0 | 0 | no-hit | 0.0 | 0 |
| scf7180000041918 | 829161 | 0.3829 | 0 | 60.6099 | Eukaryota | 41311.0 | 0 | Chordata | 41311.0 | 0 | Carnivora | 41311.0 | 0 | Canidae | 41311.0 | 0 | Canis | 41311.0 | 0 | Canis lupus | 41311.0 | 0 |
| scf7180000055915 | 25851 | 0.442 | 0 | 611.7403 | Eukaryota | 3560.0 | 0 | Chordata | 2100.0 | 1 | Kurtiformes | 1839.0 | 2 | Apogonidae | 1839.0 | 2 | Sphaeramia | 1839.0 | 2 | Sphaeramia orbicularis | 1839.0 | 2 |
| scf7180000051254 | 5921 | 0.4302 | 0 | 185.6597 | Eukaryota | 2189.0 | 0 | Platyhelminthes | 1115.0 | 2 | Polystomatidea | 1115.0 | 2 | Polystomatidae | 1115.0 | 2 | Protopolystoma | 1115.0 | 2 | Protopolystoma xenopodis | 1115.0 | 2 |
| scf7180000046881 | 20100 | 0.3977 | 0 | 59.6365 | Eukaryota | 68754.0 | 0 | Chordata | 68754.0 | 0 | Carnivora | 68754.0 | 0 | Canidae | 68754.0 | 0 | Canis | 68754.0 | 0 | Canis lupus | 68754.0 | 0 |
| scf7180000047774 | 15154 | 0.4326 | 0 | 61.8091 | Eukaryota | 63986.0 | 0 | Chordata | 63986.0 | 0 | Carnivora | 63986.0 | 0 | Canidae | 63986.0 | 0 | Canis | 63986.0 | 0 | Canis lupus | 63986.0 | 0 |
| scf7180000041732 | 40836 | 0.3242 | 0 | 51.8958 | Eukaryota | 14325.0 | 0 | Chordata | 14325.0 | 0 | Carnivora | 14325.0 | 0 | Canidae | 14325.0 | 0 | Canis | 14325.0 | 0 | Canis lupus | 14325.0 | 0 |
| scf7180000056423 | 463010 | 0.3722 | 0 | 58.6436 | Eukaryota | 20039.0 | 0 | Chordata | 20039.0 | 0 | Carnivora | 14263.0 | 2 | Canidae | 14263.0 | 2 | Canis | 12165.0 | 3 | Canis lupus | 12165.0 | 3 |
| scf7180000047854 | 10731 | 0.3991 | 0 | 36.8466 | Eukaryota | 8533.0 | 0 | Chordata | 8533.0 | 0 | Carnivora | 8533.0 | 0 | Canidae | 8533.0 | 0 | Canis | 8533.0 | 0 | Canis lupus | 8533.0 | 0 |
| scf7180000042231 | 93633 | 0.3968 | 0 | 58.4311 | Eukaryota | 89661.0 | 0 | Chordata | 89661.0 | 0 | Carnivora | 89661.0 | 0 | Canidae | 89661.0 | 0 | Canis | 89661.0 | 0 | Canis lupus | 89661.0 | 0 |
| scf7180000056350 | 531319 | 0.4487 | 0 | 58.401 | Eukaryota | 107257.0 | 0 | Chordata | 107257.0 | 0 | Carnivora | 107257.0 | 0 | Canidae | 107257.0 | 0 | Canis | 107257.0 | 0 | Canis lupus | 107257.0 | 0 |
| scf7180000040713 | 1887852 | 0.3963 | 100 | 60.3939 | Eukaryota | 74152.0 | 0 | Chordata | 74152.0 | 0 | Carnivora | 74152.0 | 0 | Canidae | 74152.0 | 0 | Canis | 59292.0 | 1 | Canis lupus | 59292.0 | 1 |
| scf7180000053602 | 3558 | 0.4514 | 0 | 296.2741 | no-hit | 0.0 | 0 | no-hit | 0.0 | 0 | no-hit | 0.0 | 0 | no-hit | 0.0 | 0 | no-hit | 0.0 | 0 | no-hit | 0.0 | 0 |
| scf7180000049958 | 15761 | 0.4397 | 0 | 36.7518 | Eukaryota | 84373.0 | 0 | Chordata | 84373.0 | 0 | Carnivora | 84373.0 | 0 | Canidae | 84373.0 | 0 | Canis | 84373.0 | 0 | Canis lupus | 84373.0 | 0 |
| scf7180000054484 | 26558 | 0.3875 | 0 | 817.926 | Eukaryota | 26170.0 | 0 | Chordata | 26170.0 | 0 | Carnivora | 26170.0 | 0 | Canidae | 26170.0 | 0 | Canis | 26170.0 | 0 | Canis lupus | 26170.0 | 0 |
| scf7180000050460 | 10489 | 0.478 | 0 | 55.4598 | Eukaryota | 6282.0 | 0 | Chordata | 6282.0 | 0 | Carnivora | 4628.0 | 1 | Canidae | 3418.0 | 2 | Canis | 2620.0 | 3 | Canis lupus | 2620.0 | 3 |
| scf7180000047491 | 60166 | 0.4485 | 0 | 99.3842 | Eukaryota | 93469.0 | 0 | Chordata | 93469.0 | 0 | Carnivora | 93469.0 | 0 | Canidae | 93469.0 | 0 | Canis | 93469.0 | 0 | Canis lupus | 93469.0 | 0 |
| scf7180000053886 | 4209 | 0.4524 | 0 | 461.7958 | Eukaryota | 2709.0 | 0 | Platyhelminthes | 1526.0 | 2 | Polystomatidea | 1526.0 | 2 | Polystomatidae | 1526.0 | 2 | Protopolystoma | 1526.0 | 2 | Protopolystoma xenopodis | 1526.0 | 2 |
| scf7180000052303 | 7661 | 0.4567 | 0 | 1323.8306 | Eukaryota | 1207.0 | 0 | Echinodermata | 998.0 | 1 | Forcipulatida | 998.0 | 1 | Asteriidae | 998.0 | 1 | Asterias | 998.0 | 1 | Asterias rubens | 998.0 | 1 |
| scf7180000054583 | 22074 | 0.5014 | 0 | 256.9113 | Eukaryota | 43311.0 | 0 | Chordata | 43311.0 | 0 | Carnivora | 43311.0 | 0 | Canidae | 43311.0 | 0 | Canis | 43311.0 | 0 | Canis lupus | 43311.0 | 0 |
| scf7180000044661 | 267961 | 0.3427 | 0 | 59.6204 | Eukaryota | 28294.0 | 0 | Chordata | 28294.0 | 0 | Carnivora | 28294.0 | 0 | Canidae | 28294.0 | 0 | Canis | 28294.0 | 0 | Canis lupus | 28294.0 | 0 |
| scf7180000041175 | 20676 | 0.3797 | 0 | 49.3981 | Eukaryota | 83209.0 | 0 | Chordata | 83209.0 | 0 | Carnivora | 83209.0 | 0 | Canidae | 83209.0 | 0 | Canis | 83209.0 | 0 | Canis lupus | 83209.0 | 0 |
| scf7180000056414 | 428861 | 0.3617 | 0 | 59.1833 | Eukaryota | 40726.0 | 0 | Chordata | 40726.0 | 0 | Primates | 29462.0 | 1 | Hominidae | 20456.0 | 2 | Homo | 20456.0 | 3 | Homo sapiens | 20456.0 | 3 |
| scf7180000051020 | 4816 | 0.4448 | 0 | 134.2424 | no-hit | 0.0 | 0 | no-hit | 0.0 | 0 | no-hit | 0.0 | 0 | no-hit | 0.0 | 0 | no-hit | 0.0 | 0 | no-hit | 0.0 | 0 |
| scf7180000042261 | 746649 | 0.4572 | 0 | 60.6469 | Eukaryota | 97995.0 | 0 | Chordata | 97995.0 | 0 | Carnivora | 97995.0 | 0 | Canidae | 97995.0 | 0 | Canis | 97995.0 | 0 | Canis lupus | 97995.0 | 0 |
| scf7180000041316 | 1160055 | 0.3813 | 0 | 59.6791 | Eukaryota | 83560.0 | 0 | Chordata | 83560.0 | 0 | Carnivora | 83560.0 | 0 | Canidae | 83560.0 | 0 | Canis | 74361.0 | 1 | Canis lupus | 74361.0 | 1 |
| scf7180000047884 | 16658 | 0.4286 | 0 | 77.075 | Eukaryota | 40894.0 | 0 | Chordata | 40894.0 | 0 | Carnivora | 40894.0 | 0 | Canidae | 40894.0 | 0 | Canis | 40894.0 | 0 | Canis lupus | 40894.0 | 0 |
| scf7180000047067 | 12731 | 0.4445 | 0 | 81.8917 | Eukaryota | 3757.0 | 0 | Chordata | 3757.0 | 0 | Carnivora | 3326.0 | 1 | Canidae | 3326.0 | 1 | Canis | 3326.0 | 1 | Canis lupus | 3326.0 | 1 |
| scf7180000053111 | 5307 | 0.5039 | 0 | 112.1629 | Eukaryota | 8529.0 | 0 | Chordata | 8529.0 | 0 | Carnivora | 8529.0 | 0 | Canidae | 8529.0 | 0 | Canis | 6912.0 | 1 | Canis lupus | 5783.0 | 2 |
| scf7180000044548 | 95708 | 0.3719 | 0 | 56.6872 | Eukaryota | 56512.0 | 0 | Chordata | 56512.0 | 0 | Carnivora | 56512.0 | 0 | Canidae | 56512.0 | 0 | Canis | 56512.0 | 0 | Canis lupus | 56512.0 | 0 |
| scf7180000040912 | 701885 | 0.4246 | 0 | 61.2389 | Eukaryota | 65425.0 | 0 | Chordata | 65425.0 | 0 | Carnivora | 65425.0 | 0 | Canidae | 65425.0 | 0 | Vulpes | 37120.0 | 1 | Vulpes vulpes | 37120.0 | 1 |
| scf7180000046245 | 5206 | 0.4335 | 0 | 161.6241 | Eukaryota | 7606.0 | 0 | Chordata | 4562.0 | 4 | Gadiformes | 3467.0 | 8 | Gadidae | 3467.0 | 8 | Gadus | 3467.0 | 8 | Gadus morhua | 3467.0 | 8 |
| scf7180000047206 | 67730 | 0.3715 | 0 | 58.5404 | Eukaryota | 84754.0 | 0 | Chordata | 84754.0 | 0 | Carnivora | 84754.0 | 0 | Canidae | 84754.0 | 0 | Canis | 84754.0 | 0 | Canis lupus | 84754.0 | 0 |
| scf7180000046852 | 103879 | 0.3456 | 0 | 60.1466 | Eukaryota | 9884.0 | 0 | Chordata | 9884.0 | 0 | Primates | 3604.0 | 3 | Hominidae | 3604.0 | 4 | Homo | 3604.0 | 5 | Homo sapiens | 3604.0 | 5 |
| scf7180000044606 | 267138 | 0.3946 | 0 | 61.4771 | Eukaryota | 18357.0 | 0 | Chordata | 18357.0 | 0 | Carnivora | 18357.0 | 0 | Canidae | 18357.0 | 0 | Canis | 18357.0 | 0 | Canis lupus | 18357.0 | 0 |
| scf7180000043047 | 250766 | 0.4347 | 0 | 59.8692 | Eukaryota | 38356.0 | 0 | Chordata | 38356.0 | 0 | Carnivora | 38356.0 | 0 | Canidae | 38356.0 | 0 | Canis | 38356.0 | 0 | Canis lupus | 38356.0 | 0 |
| scf7180000047143 | 110418 | 0.4635 | 0 | 58.6585 | Eukaryota | 31582.0 | 0 | Chordata | 31582.0 | 0 | Carnivora | 16008.0 | 1 | Hominidae | 15574.0 | 2 | Homo | 15574.0 | 2 | Homo sapiens | 15574.0 | 2 |
| scf7180000046231 | 80879 | 0.4548 | 0 | 157.1719 | Eukaryota | 4889.0 | 0 | Platyhelminthes | 2596.0 | 1 | Polystomatidea | 2596.0 | 3 | Polystomatidae | 2596.0 | 3 | Protopolystoma | 2596.0 | 4 | Protopolystoma xenopodis | 2596.0 | 4 |
| scf7180000048783 | 54745 | 0.4246 | 0 | 54.7981 | Eukaryota | 21853.0 | 0 | Chordata | 21853.0 | 0 | Primates | 13613.0 | 2 | Hominidae | 11690.0 | 3 | Pan | 5864.0 | 4 | Pan troglodytes | 5864.0 | 4 |
| scf7180000043743 | 43598 | 0.4401 | 0 | 50.6344 | Eukaryota | 44381.0 | 0 | Chordata | 44381.0 | 0 | Carnivora | 44381.0 | 0 | Canidae | 44381.0 | 0 | Canis | 44381.0 | 0 | Canis lupus | 44381.0 | 0 |
| scf7180000056477 | 314414 | 0.4521 | 0 | 59.6791 | Eukaryota | 43302.0 | 0 | Chordata | 43302.0 | 0 | Carnivora | 36642.0 | 1 | Otariidae | 18450.0 | 2 | Zalophus | 18450.0 | 3 | Zalophus californianus | 18450.0 | 3 |
| scf7180000052674 | 4000 | 0.4505 | 0 | 186.251 | no-hit | 0.0 | 0 | no-hit | 0.0 | 0 | no-hit | 0.0 | 0 | no-hit | 0.0 | 0 | no-hit | 0.0 | 0 | no-hit | 0.0 | 0 |
| scf7180000046525 | 34938 | 0.4546 | 0 | 55.1971 | Eukaryota | 9805.0 | 0 | Chordata | 9805.0 | 0 | Carnivora | 8173.0 | 1 | Felidae | 5134.0 | 4 | Felis | 5134.0 | 4 | Felis catus | 5134.0 | 4 |
| scf7180000040821 | 1555573 | 0.4508 | 0 | 59.4667 | Eukaryota | 95849.0 | 0 | Chordata | 95849.0 | 0 | Carnivora | 95849.0 | 0 | Canidae | 95849.0 | 0 | Canis | 95849.0 | 0 | Canis lupus | 95849.0 | 0 |
| scf7180000050489 | 9274 | 0.4382 | 0 | 82.942 | no-hit | 0.0 | 0 | no-hit | 0.0 | 0 | no-hit | 0.0 | 0 | no-hit | 0.0 | 0 | no-hit | 0.0 | 0 | no-hit | 0.0 | 0 |
| scf7180000040972 | 1502628 | 0.3466 | 0 | 58.5847 | Eukaryota | 95813.0 | 0 | Chordata | 95813.0 | 0 | Carnivora | 95813.0 | 0 | Canidae | 95813.0 | 0 | Canis | 95813.0 | 0 | Canis lupus | 95813.0 | 0 |
| scf7180000055324 | 28575 | 0.4405 | 0 | 137.7448 | Eukaryota | 1801.0 | 0 | Platyhelminthes | 1263.0 | 1 | Polystomatidea | 1263.0 | 2 | Polystomatidae | 1263.0 | 2 | Protopolystoma | 1263.0 | 2 | Protopolystoma xenopodis | 1263.0 | 2 |
| scf7180000048564 | 7780 | 0.4506 | 0 | 1257.0904 | no-hit | 0.0 | 0 | no-hit | 0.0 | 0 | no-hit | 0.0 | 0 | no-hit | 0.0 | 0 | no-hit | 0.0 | 0 | no-hit | 0.0 | 0 |
| scf7180000056573 | 74880 | 0.5013 | 0 | 58.5595 | Eukaryota | 35217.0 | 0 | Chordata | 35217.0 | 0 | Carnivora | 35217.0 | 0 | Canidae | 31635.0 | 1 | Canis | 30098.0 | 2 | Canis lupus | 30098.0 | 3 |
| scf7180000046082 | 86650 | 0.5391 | 0 | 59.269 | Eukaryota | 19192.0 | 0 | Chordata | 19192.0 | 0 | Carnivora | 19192.0 | 0 | Canidae | 14180.0 | 2 | Canis | 10883.0 | 3 | Canis lupus | 10883.0 | 3 |
| scf7180000046604 | 4306 | 0.4285 | 0 | 26.3783 | Eukaryota | 4761.0 | 0 | Chordata | 4761.0 | 0 | Carnivora | 4761.0 | 0 | Canidae | 4761.0 | 0 | Canis | 4761.0 | 0 | Canis lupus | 4761.0 | 0 |
| scf7180000048642 | 11502 | 0.4391 | 0 | 116.761 | no-hit | 0.0 | 0 | no-hit | 0.0 | 0 | no-hit | 0.0 | 0 | no-hit | 0.0 | 0 | no-hit | 0.0 | 0 | no-hit | 0.0 | 0 |
| scf7180000053775 | 7878 | 0.4494 | 0 | 967.1469 | Eukaryota | 268.0 | 0 | Chordata | 268.0 | 0 | Kurtiformes | 268.0 | 0 | Apogonidae | 268.0 | 0 | Sphaeramia | 268.0 | 0 | Sphaeramia orbicularis | 268.0 | 0 |
| scf7180000053607 | 5780 | 0.5036 | 0 | 87.6998 | Eukaryota | 7216.0 | 0 | Chordata | 7216.0 | 0 | Carnivora | 7216.0 | 0 | Canidae | 7216.0 | 0 | Canis | 5279.0 | 1 | Canis lupus | 4232.0 | 2 |
| scf7180000049313 | 40690 | 0.3689 | 0 | 61.5596 | Eukaryota | 22388.0 | 0 | Chordata | 22388.0 | 0 | Carnivora | 19918.0 | 1 | Ursidae | 8372.0 | 4 | Ursus | 8372.0 | 4 | Canis lupus | 6875.0 | 5 |
| scf7180000044985 | 48556 | 0.4436 | 0 | 146.2855 | Eukaryota | 89617.0 | 0 | Chordata | 89617.0 | 0 | Carnivora | 89617.0 | 0 | Canidae | 89617.0 | 0 | Canis | 89617.0 | 0 | Canis lupus | 89617.0 | 0 |
| scf7180000045740 | 176852 | 0.4051 | 0 | 132.3043 | Eukaryota | 89872.0 | 0 | Chordata | 89872.0 | 0 | Carnivora | 89872.0 | 0 | Canidae | 89872.0 | 0 | Canis | 89872.0 | 0 | Canis lupus | 89872.0 | 0 |
| scf7180000047005 | 70371 | 0.343 | 0 | 54.391 | Eukaryota | 22600.0 | 0 | Chordata | 22600.0 | 0 | Carnivora | 22600.0 | 0 | Canidae | 22600.0 | 0 | Canis | 22600.0 | 0 | Canis lupus | 22600.0 | 0 |
| scf7180000056168 | 1496352 | 0.4372 | 0 | 59.7209 | Eukaryota | 89671.0 | 0 | Chordata | 89671.0 | 0 | Carnivora | 89671.0 | 0 | Canidae | 89671.0 | 0 | Canis | 89671.0 | 0 | Canis lupus | 89671.0 | 0 |
| scf7180000056488 | 370892 | 0.3904 | 0 | 59.2575 | Eukaryota | 43911.0 | 0 | Chordata | 43911.0 | 0 | Carnivora | 24873.0 | 3 | Canidae | 24873.0 | 4 | Canis | 24873.0 | 4 | Canis lupus | 24873.0 | 4 |
| scf7180000049341 | 24696 | 0.5023 | 0 | 715.7734 | Eukaryota | 9542.0 | 0 | Chordata | 9542.0 | 0 | Carnivora | 9542.0 | 0 | Canidae | 9542.0 | 0 | Canis | 7755.0 | 1 | Canis lupus | 6459.0 | 2 |
| scf7180000054927 | 5614 | 0.4875 | 0 | 149.5303 | Eukaryota | 8143.0 | 0 | Chordata | 8143.0 | 0 | Carnivora | 8143.0 | 0 | Canidae | 8143.0 | 0 | Canis | 8143.0 | 0 | Canis lupus | 8143.0 | 0 |
| scf7180000054960 | 25456 | 0.4396 | 0 | 299.6964 | Eukaryota | 94981.0 | 0 | Chordata | 94981.0 | 0 | Carnivora | 94981.0 | 0 | Canidae | 94981.0 | 0 | Canis | 94981.0 | 0 | Canis lupus | 94981.0 | 0 |
| scf7180000042873 | 299452 | 0.4578 | 0 | 58.2478 | Eukaryota | 52115.0 | 0 | Chordata | 52115.0 | 0 | Carnivora | 52115.0 | 0 | Canidae | 32148.0 | 4 | Vulpes | 21542.0 | 6 | Vulpes vulpes | 21542.0 | 6 |
| scf7180000050388 | 16727 | 0.4467 | 0 | 966.5962 | Eukaryota | 4349.0 | 0 | Platyhelminthes | 2868.0 | 3 | Polystomatidea | 2868.0 | 3 | Polystomatidae | 2868.0 | 3 | Protopolystoma | 2868.0 | 3 | Protopolystoma xenopodis | 2868.0 | 3 |
| scf7180000054586 | 18714 | 0.462 | 0 | 121.9537 | Eukaryota | 29897.0 | 0 | Chordata | 29897.0 | 0 | Carnivora | 29897.0 | 0 | Canidae | 29897.0 | 0 | Canis | 29897.0 | 0 | Canis lupus | 29897.0 | 0 |
| scf7180000056203 | 1049655 | 0.3707 | 0 | 59.6323 | Eukaryota | 44607.0 | 0 | Chordata | 44607.0 | 0 | Carnivora | 44607.0 | 0 | Canidae | 44607.0 | 0 | Canis | 44607.0 | 0 | Canis lupus | 44607.0 | 0 |
| scf7180000046317 | 94750 | 0.4038 | 0 | 60.501 | Eukaryota | 60456.0 | 0 | Chordata | 60456.0 | 0 | Carnivora | 60456.0 | 0 | Canidae | 60456.0 | 0 | Canis | 60456.0 | 0 | Canis lupus | 60456.0 | 0 |
| scf7180000042604 | 616083 | 0.3636 | 0 | 59.0649 | Eukaryota | 31108.0 | 0 | Chordata | 31108.0 | 0 | Carnivora | 31108.0 | 0 | Canidae | 31108.0 | 0 | Canis | 28775.0 | 1 | Canis lupus | 28775.0 | 1 |
| scf7180000051618 | 12157 | 0.4528 | 0 | 128.3003 | Eukaryota | 3554.0 | 0 | Chordata | 3128.0 | 1 | Tetraodontiformes | 3128.0 | 1 | Tetraodontidae | 3128.0 | 1 | Takifugu | 3128.0 | 1 | Takifugu rubripes | 3128.0 | 1 |
| scf7180000045438 | 104277 | 0.4368 | 0 | 96.4042 | Eukaryota | 78097.0 | 0 | Chordata | 78097.0 | 0 | Carnivora | 78097.0 | 0 | Canidae | 78097.0 | 0 | Canis | 78097.0 | 0 | Canis lupus | 78097.0 | 0 |
| scf7180000050555 | 5888 | 0.4496 | 0 | 170.1272 | Eukaryota | 22167.0 | 0 | Chordata | 17884.0 | 2 | Gadiformes | 12432.0 | 6 | Gadidae | 12432.0 | 6 | Gadus | 12432.0 | 6 | Gadus morhua | 12432.0 | 6 |
| scf7180000055019 | 29821 | 0.4434 | 0 | 159.4851 | Eukaryota | 34891.0 | 0 | Chordata | 34891.0 | 0 | Gadiformes | 19861.0 | 6 | Gadidae | 19861.0 | 6 | Gadus | 19861.0 | 6 | Gadus morhua | 19861.0 | 6 |
| scf7180000045770 | 184954 | 0.3536 | 0 | 60.3089 | Eukaryota | 19748.0 | 0 | Chordata | 19748.0 | 0 | Carnivora | 19748.0 | 0 | Canidae | 19748.0 | 0 | Canis | 19748.0 | 0 | Canis lupus | 19748.0 | 0 |
| scf7180000041221 | 1279804 | 0.3818 | 0 | 59.5973 | Eukaryota | 49198.0 | 0 | Chordata | 49198.0 | 0 | Carnivora | 49198.0 | 0 | Canidae | 49198.0 | 0 | Canis | 49198.0 | 0 | Canis lupus | 49198.0 | 0 |
| scf7180000043438 | 456787 | 0.4479 | 0 | 59.8125 | Eukaryota | 89490.0 | 0 | Chordata | 89490.0 | 0 | Carnivora | 89490.0 | 0 | Canidae | 89490.0 | 0 | Canis | 89490.0 | 0 | Canis lupus | 89490.0 | 0 |

| | | | | | | | | | | | | | | | | | | | | | | |
|---|---|---|---|---|---|---|---|---|---|---|---|---|---|---|---|---|---|---|---|---|---|---|
| scf7180000048324 | 32934 | 0.4418 | 0 | 97.2203 | Eukaryota | 17023.0 | 0 | Chordata | 12965.0 | 1 | Tetraodontiformes | 8359.0 | 5 | Tetraodontidae | 8359.0 | 5 | Takifugu | 8359.0 | 5 | Takifugu rubripes | 7651.0 | 6 |
| scf7180000054597 | 45136 | 0.4425 | 0 | 434.4769 | Eukaryota | 93372.0 | 0 | Chordata | 93372.0 | 0 | Carnivora | 93372.0 | 0 | Canidae | 93372.0 | 0 | Canis | 93372.0 | 0 | Canis lupus | 93372.0 | 0 |
| scf7180000047153 | 4389 | 0.4361 | 0 | 267.0955 | no-hit | 0.0 | 0 | no-hit | 0.0 | 0 | no-hit | 0.0 | 0 | no-hit | 0.0 | 0 | no-hit | 0.0 | 0 | no-hit | 0.0 | 0 |
| scf7180000045691 | 141919 | 0.358 | 0 | 59.3012 | Eukaryota | 63748.0 | 0 | Chordata | 63748.0 | 0 | Carnivora | 63748.0 | 0 | Canidae | 63748.0 | 0 | Canis | 63748.0 | 0 | Canis lupus | 63748.0 | 0 |
| scf7180000055328 | 31449 | 0.4463 | 0 | 291.7761 | Eukaryota | 2960.0 | 0 | Chordata | 2960.0 | 0 | Carnivora | 2446.0 | 1 | Canidae | 2446.0 | 1 | Canis | 2446.0 | 1 | Canis lupus | 2446.0 | 1 |
| scf7180000045241 | 17308 | 0.5061 | 0 | 397.2429 | Eukaryota | 9296.0 | 0 | Chordata | 9296.0 | 0 | Carnivora | 9296.0 | 0 | Canidae | 9296.0 | 0 | Canis | 7530.0 | 1 | Canis lupus | 6222.0 | 2 |
| scf7180000052080 | 15951 | 0.5001 | 0 | 290.5776 | Eukaryota | 9133.0 | 0 | Chordata | 9133.0 | 0 | Carnivora | 9133.0 | 0 | Canidae | 9133.0 | 0 | Canis | 7350.0 | 1 | Canis lupus | 6050.0 | 2 |
| scf7180000042409 | 611732 | 0.4413 | 0 | 67.2599 | Eukaryota | 107246.0 | 0 | Chordata | 107246.0 | 0 | Carnivora | 107246.0 | 0 | Canidae | 107246.0 | 0 | Canis | 107246.0 | 0 | Canis lupus | 107246.0 | 0 |
| scf7180000053007 | 6755 | 0.4511 | 0 | 1526.3298 | no-hit | 0.0 | 0 | no-hit | 0.0 | 0 | no-hit | 0.0 | 0 | no-hit | 0.0 | 0 | no-hit | 0.0 | 0 | no-hit | 0.0 | 0 |
| scf7180000054552 | 22721 | 0.4388 | 0 | 195.1549 | Eukaryota | 1779.0 | 0 | Chordata | 1439.0 | 1 | Tetraodontiformes | 656.0 | 3 | Tetraodontidae | 656.0 | 3 | Takifugu | 656.0 | 3 | Takifugu rubripes | 656.0 | 3 |
| scf7180000043769 | 390822 | 0.3596 | 0 | 61.5323 | Eukaryota | 58392.0 | 0 | Chordata | 58392.0 | 0 | Carnivora | 58392.0 | 0 | Canidae | 58392.0 | 0 | Canis | 45168.0 | 1 | Canis lupus | 45168.0 | 1 |
| scf7180000049485 | 34021 | 0.5005 | 0 | 271.6567 | Eukaryota | 8731.0 | 0 | Chordata | 8731.0 | 0 | Carnivora | 8731.0 | 0 | Canidae | 8731.0 | 0 | Canis | 7043.0 | 2 | Canis lupus | 5871.0 | 3 |
| scf7180000041420 | 1135146 | 0.3589 | 0 | 59.4702 | Eukaryota | 90963.0 | 0 | Chordata | 90963.0 | 0 | Carnivora | 90963.0 | 0 | Canidae | 90963.0 | 0 | Canis | 90963.0 | 0 | Canis lupus | 90963.0 | 0 |
| scf7180000044527 | 335536 | 0.3629 | 0 | 61.4619 | Eukaryota | 90907.0 | 0 | Chordata | 90907.0 | 0 | Carnivora | 90907.0 | 0 | Canidae | 90907.0 | 0 | Canis | 90907.0 | 0 | Canis lupus | 90907.0 | 0 |
| scf7180000041356 | 1195471 | 0.4008 | 0 | 60.2327 | Eukaryota | 93572.0 | 0 | Chordata | 93572.0 | 0 | Carnivora | 93572.0 | 0 | Canidae | 93572.0 | 0 | Canis | 93572.0 | 0 | Canis lupus | 93572.0 | 0 |
| scf7180000053704 | 3972 | 0.4587 | 0 | 352.1656 | Eukaryota | 2125.0 | 0 | Echinodermata | 809.0 | 3 | Forcipulatida | 809.0 | 3 | Asteriidae | 809.0 | 3 | Asterias | 809.0 | 3 | Asterias rubens | 809.0 | 3 |
| scf7180000045661 | 30107 | 0.3824 | 0 | 52.1352 | Eukaryota | 54399.0 | 0 | Chordata | 54399.0 | 0 | Carnivora | 54399.0 | 0 | Canidae | 54399.0 | 0 | Canis | 54399.0 | 0 | Canis lupus | 54399.0 | 0 |
| scf7180000041227 | 15906 | 0.399 | 0 | 52.5066 | Eukaryota | 9060.0 | 0 | Chordata | 9060.0 | 0 | Carnivora | 9060.0 | 0 | Canidae | 9060.0 | 0 | Canis | 9060.0 | 0 | Canis lupus | 9060.0 | 0 |
| scf7180000045933 | 4851 | 0.4424 | 0 | 78.3175 | Eukaryota | 494.0 | 0 | Platyhelminthes | 494.0 | 0 | Polystomatidea | 494.0 | 0 | Polystomatidae | 494.0 | 0 | Protopolystoma | 494.0 | 0 | Protopolystoma xenopodis | 494.0 | 0 |
| scf7180000043012 | 517119 | 0.4507 | 0 | 58.8609 | Eukaryota | 128300.0 | 0 | Chordata | 128300.0 | 0 | Carnivora | 128300.0 | 0 | Canidae | 128300.0 | 0 | Canis | 128300.0 | 0 | Canis lupus | 128300.0 | 0 |
| scf7180000054603 | 19330 | 0.4353 | 0 | 126.1916 | Eukaryota | 2601.0 | 0 | Platyhelminthes | 2181.0 | 1 | Polystomatidea | 2181.0 | 1 | Polystomatidae | 2181.0 | 1 | Protopolystoma | 2181.0 | 1 | Protopolystoma xenopodis | 2181.0 | 1 |
| scf7180000055908 | 17637 | 0.4451 | 0 | 138.5262 | Eukaryota | 1831.0 | 0 | Chordata | 1193.0 | 1 | Salmoniformes | 806.0 | 3 | Salmonidae | 806.0 | 3 | Salmo | 806.0 | 3 | Salmo trutta | 806.0 | 3 |
| scf7180000040518 | 2741928 | 0.3909 | 0 | 60.1696 | Eukaryota | 103730.0 | 0 | Chordata | 103730.0 | 0 | Carnivora | 103730.0 | 0 | Canidae | 103730.0 | 0 | Canis | 103730.0 | 0 | Canis lupus | 103730.0 | 0 |
| scf7180000053408 | 2856 | 0.451 | 0 | 432.4211 | no-hit | 0.0 | 0 | no-hit | 0.0 | 0 | no-hit | 0.0 | 0 | no-hit | 0.0 | 0 | no-hit | 0.0 | 0 | no-hit | 0.0 | 0 |
| scf7180000049845 | 8144 | 0.4498 | 0 | 934.0082 | Eukaryota | 274.0 | 0 | Chordata | 274.0 | 0 | Kurtiformes | 274.0 | 0 | Apogonidae | 274.0 | 0 | Sphaeramia | 274.0 | 0 | Sphaeramia orbicularis | 274.0 | 0 |
| scf7180000051517 | 3540 | 0.4446 | 0 | 849.7514 | no-hit | 0.0 | 0 | no-hit | 0.0 | 0 | no-hit | 0.0 | 0 | no-hit | 0.0 | 0 | no-hit | 0.0 | 0 | no-hit | 0.0 | 0 |
| scf7180000047251 | 8787 | 0.4501 | 0 | 617.1404 | Eukaryota | 7645.0 | 0 | Chordata | 6922.0 | 1 | Kurtiformes | 5101.0 | 3 | Apogonidae | 5101.0 | 3 | Sphaeramia | 5101.0 | 3 | Sphaeramia orbicularis | 5101.0 | 3 |
| scf7180000051053 | 8608 | 0.4372 | 0 | 68.6972 | Eukaryota | 1782.0 | 0 | Platyhelminthes | 1782.0 | 0 | Polystomatidea | 1782.0 | 0 | Polystomatidae | 1782.0 | 0 | Protopolystoma | 1782.0 | 0 | Protopolystoma xenopodis | 1782.0 | 0 |
| scf7180000041893 | 9823 | 0.3925 | 0 | 49.0475 | Eukaryota | 6618.0 | 0 | Chordata | 6618.0 | 0 | Carnivora | 6618.0 | 0 | Canidae | 6618.0 | 0 | Canis | 6618.0 | 0 | Canis lupus | 6618.0 | 0 |
| scf7180000041351 | 1092026 | 0.4207 | 0 | 63.2212 | Eukaryota | 90525.0 | 0 | Chordata | 90525.0 | 0 | Carnivora | 90525.0 | 0 | Canidae | 90525.0 | 0 | Canis | 90525.0 | 0 | Canis lupus | 90525.0 | 0 |
| scf7180000056412 | 250246 | 0.4676 | 0 | 56.627 | Eukaryota | 87771.0 | 0 | Chordata | 87771.0 | 0 | Carnivora | 87771.0 | 0 | Canidae | 63502.0 | 2 | Canis | 51858.0 | 3 | Canis lupus | 51858.0 | 3 |
| scf7180000049898 | 64020 | 0.4098 | 0 | 59.5957 | Eukaryota | 12081.0 | 0 | Chordata | 12081.0 | 0 | Artiodactyla | 4325.0 | 3 | Canidae | 2687.0 | 6 | Canis | 2687.0 | 6 | Canis lupus | 2687.0 | 6 |
| scf7180000041300 | 427810 | 0.3675 | 0 | 60.6875 | Eukaryota | 93269.0 | 0 | Chordata | 93269.0 | 0 | Carnivora | 93269.0 | 0 | Canidae | 93269.0 | 0 | Canis | 93269.0 | 0 | Canis lupus | 93269.0 | 0 |
| scf7180000044630 | 258266 | 0.3839 | 0 | 60.2893 | Eukaryota | 53863.0 | 0 | Chordata | 53863.0 | 0 | Carnivora | 53863.0 | 0 | Canidae | 53863.0 | 0 | Canis | 37855.0 | 1 | Canis lupus | 37855.0 | 1 |
| scf7180000045437 | 192020 | 0.3466 | 0 | 58.7618 | Eukaryota | 58974.0 | 0 | Chordata | 58974.0 | 0 | Carnivora | 58974.0 | 0 | Canidae | 53330.0 | 1 | Canis | 53330.0 | 1 | Canis lupus | 53330.0 | 1 |
| scf7180000049900 | 19193 | 0.4491 | 0 | 2215.4526 | Eukaryota | 3189.0 | 0 | Platyhelminthes | 1738.0 | 2 | Polystomatidea | 1738.0 | 2 | Polystomatidae | 1738.0 | 2 | Protopolystoma | 1738.0 | 2 | Protopolystoma xenopodis | 1738.0 | 2 |
| scf7180000045755 | 52154 | 0.4426 | 0 | 102.9366 | Eukaryota | 15062.0 | 0 | Chordata | 13837.0 | 1 | Kurtiformes | 5199.0 | 4 | Apogonidae | 5199.0 | 4 | Sphaeramia | 5199.0 | 4 | Sphaeramia orbicularis | 5199.0 | 4 |
| scf7180000043541 | 256025 | 0.387 | 0 | 59.2487 | Eukaryota | 65111.0 | 0 | Chordata | 65111.0 | 0 | Carnivora | 65111.0 | 0 | Canidae | 65111.0 | 0 | Canis | 65111.0 | 0 | Canis lupus | 65111.0 | 0 |
| scf7180000048809 | 3117 | 0.4405 | 0 | 36.6492 | Eukaryota | 2798.0 | 0 | Chordata | 2798.0 | 0 | Carnivora | 2798.0 | 0 | Canidae | 2798.0 | 0 | Canis | 2798.0 | 0 | Canis lupus | 2798.0 | 0 |
| scf7180000049358 | 6539 | 0.4418 | 0 | 97.736 | no-hit | 0.0 | 0 | no-hit | 0.0 | 0 | no-hit | 0.0 | 0 | no-hit | 0.0 | 0 | no-hit | 0.0 | 0 | no-hit | 0.0 | 0 |
| scf7180000043762 | 21060 | 0.3663 | 0 | 44.7783 | Eukaryota | 63846.0 | 0 | Chordata | 63846.0 | 0 | Carnivora | 63846.0 | 0 | Canidae | 63846.0 | 0 | Canis | 63846.0 | 0 | Canis lupus | 63846.0 | 0 |
| scf7180000045946 | 59203 | 0.3645 | 0 | 56.4517 | Eukaryota | 14326.0 | 0 | Chordata | 14326.0 | 0 | Carnivora | 14326.0 | 0 | Canidae | 14326.0 | 0 | Canis | 14326.0 | 0 | Canis lupus | 14326.0 | 0 |
| scf7180000052533 | 3515 | 0.4481 | 0 | 812.8944 | no-hit | 0.0 | 0 | no-hit | 0.0 | 0 | no-hit | 0.0 | 0 | no-hit | 0.0 | 0 | no-hit | 0.0 | 0 | no-hit | 0.0 | 0 |
| scf7180000040696 | 1679403 | 0.4155 | 0 | 59.5406 | Eukaryota | 87838.0 | 0 | Chordata | 87838.0 | 0 | Carnivora | 87838.0 | 0 | Canidae | 87838.0 | 0 | Canis | 87838.0 | 0 | Canis lupus | 87838.0 | 0 |
| scf7180000046033 | 9248 | 0.4365 | 0 | 148.9894 | Eukaryota | 333.0 | 0 | Chordata | 333.0 | 0 | Salmoniformes | 333.0 | 0 | Salmonidae | 333.0 | 0 | Coregonus | 333.0 | 0 | Coregonus sp. 'balchen' | 333.0 | 0 |
| scf7180000045749 | 116421 | 0.3734 | 0 | 59.9321 | Eukaryota | 67138.0 | 0 | Chordata | 67138.0 | 0 | Carnivora | 67138.0 | 0 | Otariidae | 25634.0 | 3 | Zalophus | 25634.0 | 3 | Zalophus californianus | 25634.0 | 3 |
| scf7180000049454 | 19136 | 0.4369 | 0 | 53.3667 | no-hit | 0.0 | 0 | no-hit | 0.0 | 0 | no-hit | 0.0 | 0 | no-hit | 0.0 | 0 | no-hit | 0.0 | 0 | no-hit | 0.0 | 0 |
| scf7180000056171 | 540430 | 0.4501 | 0 | 59.0194 | Eukaryota | 94570.0 | 0 | Chordata | 94570.0 | 0 | Carnivora | 94570.0 | 0 | Canidae | 94570.0 | 0 | Canis | 94570.0 | 0 | Canis lupus | 94570.0 | 0 |
| scf7180000053999 | 6186 | 0.4381 | 0 | 34.4518 | no-hit | 0.0 | 0 | no-hit | 0.0 | 0 | no-hit | 0.0 | 0 | no-hit | 0.0 | 0 | no-hit | 0.0 | 0 | no-hit | 0.0 | 0 |
| scf7180000053995 | 1809 | 0.4334 | 0 | 313.2254 | no-hit | 0.0 | 0 | no-hit | 0.0 | 0 | no-hit | 0.0 | 0 | no-hit | 0.0 | 0 | no-hit | 0.0 | 0 | no-hit | 0.0 | 0 |
| scf7180000055342 | 8514 | 0.4294 | 0 | 79.4805 | Eukaryota | 3385.0 | 0 | Chordata | 2788.0 | 2 | Salmoniformes | 2595.0 | 3 | Salmonidae | 2595.0 | 3 | Salmo | 2349.0 | 4 | Salmo trutta | 2349.0 | 4 |
| scf7180000047624 | 14758 | 0.4223 | 0 | 47.2458 | Eukaryota | 16644.0 | 0 | Chordata | 16644.0 | 0 | Carnivora | 16644.0 | 0 | Felidae | 9667.0 | 2 | Felis | 9667.0 | 2 | Felis catus | 9667.0 | 2 |
| scf7180000046177 | 153960 | 0.4256 | 0 | 59.5217 | Eukaryota | 80669.0 | 0 | Chordata | 80669.0 | 0 | Carnivora | 80669.0 | 0 | Canidae | 80669.0 | 0 | Canis | 80669.0 | 0 | Canis lupus | 80669.0 | 0 |
| scf7180000055140 | 24514 | 0.4404 | 0 | 308.3574 | Eukaryota | 2911.0 | 0 | Platyhelminthes | 1734.0 | 2 | Polystomatidea | 1734.0 | 2 | Polystomatidae | 1734.0 | 2 | Protopolystoma | 1734.0 | 2 | Protopolystoma xenopodis | 1734.0 | 2 |
| scf7180000043475 | 425007 | 0.4804 | 0 | 59.5127 | Eukaryota | 78514.0 | 0 | Chordata | 78514.0 | 0 | Carnivora | 78514.0 | 0 | Canidae | 78514.0 | 0 | Canis | 78514.0 | 0 | Canis lupus | 78514.0 | 0 |
| scf7180000048840 | 55247 | 0.4535 | 0 | 59.2458 | Eukaryota | 15959.0 | 0 | Chordata | 15959.0 | 0 | Carnivora | 15959.0 | 0 | Canidae | 6687.0 | 3 | Canis | 4993.0 | 4 | Canis lupus | 4993.0 | 4 |
| scf7180000049363 | 6907 | 0.451 | 0 | 629.2966 | no-hit | 0.0 | 0 | no-hit | 0.0 | 0 | no-hit | 0.0 | 0 | no-hit | 0.0 | 0 | no-hit | 0.0 | 0 | no-hit | 0.0 | 0 |
| scf7180000042401 | 377010 | 0.3627 | 0 | 59.6339 | Eukaryota | 87738.0 | 0 | Chordata | 87738.0 | 0 | Carnivora | 87738.0 | 0 | Canidae | 87738.0 | 0 | Canis | 87738.0 | 0 | Canis lupus | 87738.0 | 0 |
| scf7180000050375 | 18089 | 0.451 | 0 | 2540.8118 | Eukaryota | 2318.0 | 0 | Echinodermata | 832.0 | 3 | Forcipulatida | 832.0 | 3 | Asteriidae | 832.0 | 3 | Asterias | 832.0 | 3 | Asterias rubens | 832.0 | 3 |
| scf7180000044643 | 118173 | 0.3727 | 0 | 58.2024 | Eukaryota | 83644.0 | 0 | Chordata | 83644.0 | 0 | Carnivora | 83644.0 | 0 | Canidae | 83644.0 | 0 | Canis | 83644.0 | 0 | Canis lupus | 83644.0 | 0 |
| scf7180000052901 | 5117 | 0.4522 | 0 | 618.3038 | no-hit | 0.0 | 0 | no-hit | 0.0 | 0 | no-hit | 0.0 | 0 | no-hit | 0.0 | 0 | no-hit | 0.0 | 0 | no-hit | 0.0 | 0 |
| scf7180000050596 | 5900 | 0.4449 | 0 | 286.0634 | no-hit | 0.0 | 0 | no-hit | 0.0 | 0 | no-hit | 0.0 | 0 | no-hit | 0.0 | 0 | no-hit | 0.0 | 0 | no-hit | 0.0 | 0 |
| scf7180000052408 | 2874 | 0.4398 | 0 | 286.7468 | no-hit | 0.0 | 0 | no-hit | 0.0 | 0 | no-hit | 0.0 | 0 | no-hit | 0.0 | 0 | no-hit | 0.0 | 0 | no-hit | 0.0 | 0 |
| scf7180000054399 | 27937 | 0.4364 | 0 | 169.0621 | Eukaryota | 5408.0 | 0 | Platyhelminthes | 2245.0 | 2 | Polystomatidea | 2245.0 | 3 | Polystomatidae | 2245.0 | 3 | Protopolystoma | 2245.0 | 3 | Protopolystoma xenopodis | 2245.0 | 3 |
| scf7180000050414 | 8658 | 0.4453 | 0 | 199.1158 | Eukaryota | 1941.0 | 0 | Echinodermata | 859.0 | 2 | Forcipulatida | 859.0 | 2 | Asteriidae | 859.0 | 2 | Asterias | 859.0 | 2 | Asterias rubens | 859.0 | 2 |
| scf7180000042308 | 142793 | 0.3863 | 0 | 60.6768 | Eukaryota | 94045.0 | 0 | Chordata | 94045.0 | 0 | Carnivora | 94045.0 | 0 | Canidae | 94045.0 | 0 | Canis | 94045.0 | 0 | Canis lupus | 94045.0 | 0 |
| scf7180000049051 | 4931 | 0.5216 | 0 | 224.0165 | Eukaryota | 12482.0 | 0 | Chordata | 12482.0 | 0 | Carnivora | 12482.0 | 0 | Canidae | 12482.0 | 0 | Canis | 12482.0 | 0 | Canis lupus | 12482.0 | 0 |
| scf7180000050005 | 17610 | 0.4497 | 0 | 88.2392 | no-hit | 0.0 | 0 | no-hit | 0.0 | 0 | no-hit | 0.0 | 0 | no-hit | 0.0 | 0 | no-hit | 0.0 | 0 | no-hit | 0.0 | 0 |
| scf7180000045083 | 239375 | 0.452 | 0 | 59.6091 | Eukaryota | 52966.0 | 0 | Chordata | 52966.0 | 0 | Carnivora | 52966.0 | 0 | Canidae | 52966.0 | 0 | Canis | 37175.0 | 1 | Canis lupus | 37175.0 | 1 |
| scf7180000044061 | 223877 | 0.3501 | 0 | 58.3127 | Eukaryota | 76411.0 | 0 | Chordata | 76411.0 | 0 | Carnivora | 76411.0 | 0 | Canidae | 76411.0 | 0 | Canis | 76411.0 | 0 | Canis lupus | 76411.0 | 0 |
| scf7180000043389 | 467914 | 0.4565 | 0 | 59.5758 | Eukaryota | 30891.0 | 0 | Chordata | 30891.0 | 0 | Carnivora | 30891.0 | 0 | Canidae | 30891.0 | 0 | Canis | 27729.0 | 1 | Canis lupus | 27729.0 | 1 |
| scf7180000053442 | 2170 | 0.4548 | 0 | 514.3802 | Eukaryota | 172.0 | 0 | Streptophyta | 172.0 | 0 | Fabales | 172.0 | 0 | Fabaceae | 172.0 | 0 | Cicer | 172.0 | 0 | Cicer arietinum | 172.0 | 0 |
| scf7180000052174 | 10461 | 0.3441 | 0 | 59.1653 | Eukaryota | 9103.0 | 0 | Chordata | 9103.0 | 0 | Carnivora | 9103.0 | 0 | Canidae | 9103.0 | 0 | Canis | 9103.0 | 0 | Canis lupus | 9103.0 | 0 |
| scf7180000044972 | 252848 | 0.3763 | 0 | 57.7553 | Eukaryota | 49074.0 | 0 | Chordata | 49074.0 | 0 | Carnivora | 49074.0 | 0 | Canidae | 49074.0 | 0 | Canis | 49074.0 | 0 | Canis lupus | 49074.0 | 0 |
| scf7180000046103 | 215685 | 0.4124 | 0 | 57.9119 | Eukaryota | 40440.0 | 0 | Chordata | 40440.0 | 0 | Artiodactyla | 19292.0 | 2 | Canidae | 14349.0 | 4 | Canis | 11209.0 | 6 | Canis lupus | 11209.0 | 6 |
| scf7180000045342 | 38595 | 0.5188 | 0 | 966.5216 | Eukaryota | 16365.0 | 0 | Chordata | 16365.0 | 0 | Carnivora | 16365.0 | 0 | Canidae | 16365.0 | 0 | Canis | 16365.0 | 0 | Canis lupus | 16365.0 | 0 |
| scf7180000047786 | 94415 | 0.3798 | 0 | 62.1875 | Eukaryota | 62664.0 | 0 | Chordata | 62664.0 | 0 | Carnivora | 62664.0 | 0 | Canidae | 62664.0 | 0 | Canis | 62664.0 | 0 | Canis lupus | 62664.0 | 0 |
| scf7180000047748 | 53110 | 0.4416 | 0 | 66.2525 | Eukaryota | 26617.0 | 0 | Chordata | 20306.0 | 1 | Tetraodontiformes | 7050.0 | 7 | Tetraodontidae | 7050.0 | 7 | Takifugu | 7050.0 | 7 | Takifugu rubripes | 7050.0 | 7 |
| scf7180000049652 | 26954 | 0.373 | 0 | 61.9845 | Eukaryota | 7504.0 | 1 | Chordata | 7504.0 | 1 | Artiodactyla | 2747.0 | 4 | Bovidae | 2747.0 | 4 | Homo | 2720.0 | 5 | Homo sapiens | 2720.0 | 5 |
| scf7180000043050 | 503025 | 0.3862 | 0 | 60.1855 | Eukaryota | 51116.0 | 0 | Chordata | 51116.0 | 0 | Carnivora | 48353.0 | 1 | Canidae | 48353.0 | 1 | Canis | 39939.0 | 2 | Canis lupus | 39939.0 | 2 |
| scf7180000042977 | 52369 | 0.4076 | 0 | 54.7253 | Eukaryota | 20231.0 | 0 | Chordata | 20231.0 | 0 | Carnivora | 20231.0 | 0 | Canidae | 13124.0 | 1 | Canis | 7950.0 | 4 | Canis lupus | 7950.0 | 4 |
| scf7180000046886 | 12477 | 0.4423 | 0 | 149.1086 | Eukaryota | 8934.0 | 0 | Chordata | 8934.0 | 0 | Tetraodontiformes | 4314.0 | 2 | Tetraodontidae | 4314.0 | 2 | Takifugu | 4314.0 | 2 | Takifugu rubripes | 4314.0 | 2 |
| scf7180000049298 | 3282 | 0.45 | 0 | 206.9375 | no-hit | 0.0 | 0 | no-hit | 0.0 | 0 | no-hit | 0.0 | 0 | no-hit | 0.0 | 0 | no-hit | 0.0 | 0 | no-hit | 0.0 | 0 |
| scf7180000041321 | 219135 | 0.4209 | 0 | 59.1447 | Eukaryota | 91181.0 | 0 | Chordata | 91181.0 | 0 | Carnivora | 91181.0 | 0 | Canidae | 91181.0 | 0 | Canis | 91181.0 | 0 | Canis lupus | 91181.0 | 0 |
| scf7180000041782 | 703059 | 0.3761 | 0 | 59.9314 | Eukaryota | 46025.0 | 0 | Chordata | 46025.0 | 0 | Carnivora | 46025.0 | 0 | Canidae | 46025.0 | 0 | Canis | 46025.0 | 0 | Canis lupus | 46025.0 | 0 |
| scf7180000042827 | 55829 | 0.6175 | 0 | 55.7057 | Eukaryota | 25402.0 | 0 | Chordata | 25402.0 | 0 | Carnivora | 25402.0 | 0 | Canidae | 25402.0 | 0 | Canis | 14993.0 | 1 | Canis lupus | 14993.0 | 1 |
| scf7180000040608 | 553254 | 0.4617 | 0 | 59.207 | Eukaryota | 43658.0 | 0 | Chordata | 43658.0 | 0 | Carnivora | 43658.0 | 0 | Canidae | 43658.0 | 0 | Canis | 43658.0 | 0 | Canis lupus | 43658.0 | 0 |
| scf7180000051915 | 4758 | 0.4563 | 0 | 434.1068 | Eukaryota | 9616.0 | 0 | Chordata | 9616.0 | 0 | Tetraodontiformes | 2658.0 | 5 | Tetraodontidae | 2658.0 | 5 | Takifugu | 2658.0 | 5 | Gouania willdenowi | 2145.0 | 6 |
| scf7180000056055 | 22222 | 0.4364 | 0 | 168.9742 | Eukaryota | 8420.0 | 0 | Chordata | 8420.0 | 0 | Carnivora | 6594.0 | 1 | Canidae | 6594.0 | 1 | Canis | 6594.0 | 1 | Canis lupus | 6594.0 | 1 |
| scf7180000052485 | 3378 | 0.4464 | 0 | 164.7739 | Eukaryota | 819.0 | 0 | Platyhelminthes | 660.0 | 1 | Polystomatidea | 501.0 | 2 | Polystomatidae | 501.0 | 2 | Protopolystoma | 501.0 | 2 | Protopolystoma xenopodis | 501.0 | 2 |
| scf7180000043167 | 459134 | 0.3801 | 0 | 60.0888 | Eukaryota | 49565.0 | 0 | Chordata | 49565.0 | 0 | Carnivora | 49565.0 | 0 | Canidae | 49565.0 | 0 | Canis | 49565.0 | 0 | Canis lupus | 49565.0 | 0 |
| scf7180000049191 | 38083 | 0.4079 | 0 | 61.035 | Eukaryota | 6769.0 | 0 | Chordata | 6769.0 | 0 | Artiodactyla | 2455.0 | 4 | Bovidae | 2455.0 | 4 | Homo | 1380.0 | 5 | Homo sapiens | 1380.0 | 5 |
| scf7180000055951 | 13528 | 0.7572 | 0 | 247.5129 | Eukaryota | 2425.0 | 0 | Chordata | 2425.0 | 0 | Rodentia | 984.0 | 4 | Caviidae | 723.0 | 6 | Cavia | 723.0 | 6 | Cavia porcellus | 723.0 | 6 |
| scf7180000042824 | 547426 | 0.344 | 0 | 59.2897 | Eukaryota | 41770.0 | 0 | Chordata | 41770.0 | 0 | Carnivora | 27893.0 | 2 | Canidae | 14236.0 | 4 | Canis | 14236.0 | 4 | Canis lupus | 14236.0 | 4 |
| scf7180000056333 | 641842 | 0.3857 | 0 | 59.0461 | Eukaryota | 115533.0 | 0 | Chordata | 115533.0 | 0 | Carnivora | 115533.0 | 0 | Canidae | 115533.0 | 0 | Canis | 115533.0 | 0 | Canis lupus | 115533.0 | 0 |
| scf7180000043626 | 60838 | 0.415 | 0 | 52.8537 | Eukaryota | 8902.0 | 1 | Chordata | 8902.0 | 1 | Carnivora | 3118.0 | 5 | Bovidae | 1841.0 | 7 | Canis | 1760.0 | 8 | Canis lupus | 1760.0 | 8 |
| scf7180000056485 | 237769 | 0.5502 | 0 | 56.5594 | Eukaryota | 37489.0 | 0 | Chordata | 37489.0 | 0 | Carnivora | 37489.0 | 0 | Canidae | 35627.0 | 1 | Canis | 27149.0 | 2 | Canis lupus | 27149.0 | 2 |
| scf7180000056206 | 1441838 | 0.4392 | 0 | 59.5057 | Eukaryota | 59936.0 | 0 | Chordata | 59936.0 | 0 | Carnivora | 59936.0 | 0 | Canidae | 59936.0 | 0 | Canis | 59936.0 | 0 | Canis lupus | 59936.0 | 0 |
| scf7180000050855 | 4959 | 0.4473 | 0 | 270.1356 | Eukaryota | 1600.0 | 0 | Echinodermata | 693.0 | 2 | Forcipulatida | 693.0 | 2 | Asteriidae | 693.0 | 2 | Asterias | 693.0 | 2 | Asterias rubens | 693.0 | 2 |
| scf7180000047562 | 7112 | 0.4447 | 0 | 54.4987 | Eukaryota | 473.0 | 0 | Chordata | 473.0 | 0 | Columbiformes | 473.0 | 0 | Columbidae | 473.0 | 0 | Streptopelia | 473.0 | 0 | Streptopelia turtur | 473.0 | 0 |
| scf7180000045014 | 15972 | 0.3547 | 0 | 29.0 | Eukaryota | 19116.0 | 0 | Chordata | 19116.0 | 0 | Carnivora | 19116.0 | 0 | Canidae | 19116.0 | 0 | Canis | 19116.0 | 0 | Canis lupus | 19116.0 | 0 |
| scf7180000054861 | 21981 | 0.3781 | 0 | 131.5474 | Eukaryota | 33791.0 | 0 | Chordata | 33791.0 | 0 | Carnivora | 33791.0 | 0 | Canidae | 33791.0 | 0 | Canis | 33791.0 | 0 | Canis lupus | 33791.0 | 0 |
| scf7180000056052 | 944965 | 0.4408 | 0 | 82.5746 | Eukaryota | 51825.0 | 0 | Chordata | 51825.0 | 0 | Carnivora | 51825.0 | 0 | Canidae | 51825.0 | 0 | Canis | 46137.0 | 1 | Canis lupus | 46137.0 | 1 |
| scf7180000043619 | 396761 | 0.4767 | 0 | 59.6806 | Eukaryota | 90964.0 | 0 | Chordata | 90964.0 | 0 | Carnivora | 90964.0 | 0 | Canidae | 90964.0 | 0 | Canis | 66118.0 | 1 | Canis lupus | 66118.0 | 1 |
| scf7180000054379 | 35427 | 0.4354 | 0 | 213.4769 | Eukaryota | 1159.0 | 0 | Chordata | 942.0 | 1 | Carnivora | 503.0 | 3 | Canidae | 503.0 | 3 | Canis | 503.0 | 3 | Canis lupus | 503.0 | 3 |
| scf7180000053683 | 3537 | 0.4558 | 0 | 276.7345 | no-hit | 0.0 | 0 | no-hit | 0.0 | 0 | no-hit | 0.0 | 0 | no-hit | 0.0 | 0 | no-hit | 0.0 | 0 | no-hit | 0.0 | 0 |
| scf7180000051820 | 1924 | 0.4569 | 0 | 711.0687 | no-hit | 0.0 | 0 | no-hit | 0.0 | 0 | no-hit | 0.0 | 0 | no-hit | 0.0 | 0 | no-hit | 0.0 | 0 | no-hit | 0.0 | 0 |
| scf7180000041950 | 11526 | 0.4545 | 0 | 241.1746 | Eukaryota | 30136.0 | 0 | Chordata | 30136.0 | 0 | Carnivora | 25323.0 | 1 | Canidae | 9199.0 | 3 | Canis | 9199.0 | 5 | Canis lupus | 9199.0 | 6 |
| scf7180000041424 | 1376071 | 0.4035 | 0 | 59.3312 | Eukaryota | 91809.0 | 0 | Chordata | 91809.0 | 0 | Carnivora | 91809.0 | 0 | Canidae | 91809.0 | 0 | Canis | 91809.0 | 0 | Canis lupus | 91809.0 | 0 |
| scf7180000040751 | 2644 | 0.455 | 0 | 620.9703 | no-hit | 0.0 | 0 | no-hit | 0.0 | 0 | no-hit | 0.0 | 0 | no-hit | 0.0 | 0 | no-hit | 0.0 | 0 | no-hit | 0.0 | 0 |
| scf7180000047430 | 51376 | 0.6791 | 0 | 58.0218 | Eukaryota | 20422.0 | 0 | Chordata | 20422.0 | 0 | Carnivora | 20422.0 | 0 | Felidae | 7904.0 | 3 | Canis | 4886.0 | 8 | Canis lupus | 4886.0 | 8 |
| scf7180000044889 | 28424 | 0.4372 | 0 | 214.1827 | Eukaryota | 1265.0 | 0 | Platyhelminthes | 967.0 | 1 | Polystomatidea | 809.0 | 2 | Polystomatidae | 809.0 | 2 | Protopolystoma | 809.0 | 2 | Protopolystoma xenopodis | 809.0 | 2 |
| scf7180000052664 | 4059 | 0.458 | 0 | 885.0553 | no-hit | 0.0 | 0 | no-hit | 0.0 | 0 | no-hit | 0.0 | 0 | no-hit | 0.0 | 0 | no-hit | 0.0 | 0 | no-hit | 0.0 | 0 |
| scf7180000042369 | 188118 | 0.3724 | 0 | 58.0427 | Eukaryota | 109487.0 | 0 | Chordata | 109487.0 | 0 | Carnivora | 109487.0 | 0 | Canidae | 109487.0 | 0 | Canis | 109487.0 | 0 | Canis lupus | 109487.0 | 0 |
| scf7180000040527 | 2486956 | 0.3637 | 0 | 59.4535 | Eukaryota | 109362.0 | 0 | Chordata | 109362.0 | 0 | Carnivora | 109362.0 | 0 | Canidae | 109362.0 | 0 | Canis | 109362.0 | 0 | Canis lupus | 109362.0 | 0 |
| scf7180000045452 | 118028 | 0.4386 | 0 | 61.4288 | Eukaryota | 99609.0 | 0 | Chordata | 99609.0 | 0 | Carnivora | 99609.0 | 0 | Canidae | 99609.0 | 0 | Canis | 99609.0 | 0 | Canis lupus | 99609.0 | 0 |
| scf7180000052846 | 5747 | 0.4495 | 0 | 1675.2524 | Eukaryota | 2787.0 | 0 | Platyhelminthes | 1293.0 | 2 | Polystomatidea | 1293.0 | 2 | Polystomatidae | 1293.0 | 2 | Protopolystoma | 1293.0 | 2 | Protopolystoma xenopodis | 1293.0 | 2 |
| scf7180000054758 | 6661 | 0.4364 | 0 | 155.1698 | Eukaryota | 1087.0 | 0 | Chordata | 787.0 | 1 | Kurtiformes | 429.0 | 3 | Apogonidae | 429.0 | 3 | Sphaeramia | 429.0 | 3 | Sphaeramia orbicularis | 429.0 | 3 |
| scf7180000056549 | 166673 | 0.5183 | 0 | 57.8146 | Eukaryota | 22953.0 | 0 | Chordata | 22953.0 | 0 | Carnivora | 22953.0 | 0 | Canidae | 17894.0 | 2 | Canis | 12626.0 | 4 | Canis lupus | 12626.0 | 4 |
| scf7180000053487 | 8309 | 0.4353 | 0 | 61.2259 | no-hit | 0.0 | 0 | no-hit | 0.0 | 0 | no-hit | 0.0 | 0 | no-hit | 0.0 | 0 | no-hit | 0.0 | 0 | no-hit | 0.0 | 0 |
| scf7180000044448 | 283490 | 0.4044 | 0 | 60.627 | Eukaryota | 94469.0 | 0 | Chordata | 94469.0 | 0 | Carnivora | 94469.0 | 0 | Canidae | 94469.0 | 0 | Canis | 94469.0 | 0 | Canis lupus | 94469.0 | 0 |
| scf7180000046402 | 133432 | 0.33 | 0 | 58.5643 | Eukaryota | 23156.0 | 0 | Chordata | 23156.0 | 0 | Carnivora | 23156.0 | 0 | Canidae | 23156.0 | 0 | Canis | 23156.0 | 0 | Canis lupus | 23156.0 | 0 |
| scf7180000053702 | 9179 | 0.4385 | 0 | 31.2976 | Eukaryota | 3367.0 | 0 | Chordata | 2865.0 | 1 | Columbiformes | 2359.0 | 3 | Columbidae | 2359.0 | 3 | Streptopelia | 2359.0 | 3 | Streptopelia turtur | 2359.0 | 3 |
| scf7180000048945 | 25009 | 0.5415 | 0 | 56.0152 | Eukaryota | 21556.0 | 0 | Chordata | 21556.0 | 0 | Carnivora | 21556.0 | 0 | Canidae | 21556.0 | 0 | Canis | 21556.0 | 0 | Canis lupus | 21556.0 | 0 |
| scf7180000053245 | 5456 | 0.4542 | 0 | 1213.6909 | Eukaryota | 1950.0 | 0 | Chordata | 1300.0 | 1 | Gonorynchiformes | 1085.0 | 2 | Chanidae | 1085.0 | 2 | Chanos | 1085.0 | 2 | Chanos chanos | 1085.0 | 2 |
| scf7180000056336 | 882382 | 0.4006 | 0 | 59.7502 | Eukaryota | 87672.0 | 0 | Chordata | 87672.0 | 0 | Carnivora | 87672.0 | 0 | Canidae | 51535.0 | 3 | Canis | 41066.0 | 5 | Canis lupus | 41066.0 | 5 |
| scf7180000049325 | 35419 | 0.4432 | 0 | 86.7883 | Eukaryota | 2576.0 | 0 | Chordata | 2341.0 | 1 | Columbiformes | 1247.0 | 3 | Columbidae | 1247.0 | 3 | Streptopelia | 1247.0 | 3 | Streptopelia turtur | 1247.0 | 3 |
| scf7180000047139 | 9725 | 0.4252 | 0 | 35.3335 | Eukaryota | 17637.0 | 0 | Chordata | 17637.0 | 0 | Carnivora | 17637.0 | 0 | Canidae | 15873.0 | 1 | Canis | 13878.0 | 2 | Canis lupus | 13878.0 | 2 |
| scf7180000051595 | 16255 | 0.4301 | 0 | 55.0692 | Eukaryota | 3175.0 | 0 | Chordata | 3175.0 | 0 | Carnivora | 3175.0 | 0 | Canidae | 1142.0 | 4 | Canis | 754.0 | 7 | Canis lupus | 754.0 | 7 |
| scf7180000048838 | 46398 | 0.4443 | 0 | 240.8775 | Eukaryota | 4542.0 | 0 | Platyhelminthes | 2459.0 | 3 | Polystomatidea | 2459.0 | 3 | Polystomatidae | 2459.0 | 3 | Protopolystoma | 2459.0 | 3 | Protopolystoma xenopodis | 2459.0 | 3 |
| scf7180000051925 | 4040 | 0.447 | 0 | 113.008 | no-hit | 0.0 | 0 | no-hit | 0.0 | 0 | no-hit | 0.0 | 0 | no-hit | 0.0 | 0 | no-hit | 0.0 | 0 | no-hit | 0.0 | 0 |
| scf7180000045923 | 170914 | 0.3808 | 0 | 60.9635 | Eukaryota | 19725.0 | 0 | Chordata | 19725.0 | 0 | Carnivora | 14457.0 | 2 | Canidae | 14457.0 | 2 | Canis | 8500.0 | 3 | Canis lupus | 8500.0 | 3 |
| scf7180000043481 | 476869 | 0.3416 | 0 | 59.9591 | Eukaryota | 62619.0 | 0 | Chordata | 62619.0 | 0 | Carnivora | 62619.0 | 0 | Canidae | 62619.0 | 0 | Canis | 50067.0 | 1 | Canis lupus | 50067.0 | 1 |
| scf7180000042444 | 8766 | 0.4099 | 0 | 41.6034 | Eukaryota | 13430.0 | 0 | Chordata | 13430.0 | 0 | Carnivora | 13430.0 | 0 | Canidae | 13430.0 | 0 | Canis | 13430.0 | 0 | Canis lupus | 13430.0 | 0 |
| scf7180000054225 | 19851 | 0.4009 | 0 | 563.5435 | Eukaryota | 53856.0 | 0 | Chordata | 53856.0 | 0 | Carnivora | 53856.0 | 0 | Canidae | 53856.0 | 0 | Canis | 53856.0 | 0 | Canis lupus | 53856.0 | 0 |
| scf7180000044553 | 71195 | 0.3984 | 0 | 52.9263 | Eukaryota | 40849.0 | 0 | Chordata | 40849.0 | 0 | Carnivora | 40849.0 | 0 | Canidae | 37541.0 | 1 | Canis | 33029.0 | 2 | Canis lupus | 33029.0 | 2 |
| scf7180000051092 | 16436 | 0.5194 | 0 | 492.3734 | Eukaryota | 16077.0 | 0 | Chordata | 16077.0 | 0 | Carnivora | 16077.0 | 0 | Canidae | 16077.0 | 0 | Canis | 16077.0 | 0 | Canis lupus | 16077.0 | 0 |
| scf7180000040506 | 2926779 | 0.3699 | 0 | 59.6737 | Eukaryota | 73949.0 | 0 | Chordata | 73949.0 | 0 | Carnivora | 73949.0 | 0 | Canidae | 73949.0 | 0 | Canis | 57373.0 | 1 | Canis lupus | 57373.0 | 1 |
| scf7180000043939 | 410171 | 0.4067 | 0 | 61.5283 | Eukaryota | 90310.0 | 0 | Chordata | 90310.0 | 0 | Carnivora | 90310.0 | 0 | Canidae | 90310.0 | 0 | Canis | 90310.0 | 0 | Canis lupus | 90310.0 | 0 |
| scf7180000044222 | 331744 | 0.5213 | 0 | 57.8416 | Eukaryota | 33597.0 | 0 | Chordata | 33597.0 | 0 | Carnivora | 33597.0 | 0 | Canidae | 33597.0 | 0 | Canis | 30019.0 | 1 | Canis lupus | 30019.0 | 1 |
| scf7180000046598 | 84880 | 0.4056 | 0 | 57.7918 | Eukaryota | 10766.0 | 0 | Chordata | 10766.0 | 0 | Primates | 9217.0 | 2 | Hominidae | 9217.0 | 2 | Homo | 6589.0 | 3 | Homo sapiens | 6589.0 | 3 |
| scf7180000044067 | 357042 | 0.4491 | 0 | 58.6172 | Eukaryota | 101804.0 | 0 | Chordata | 101804.0 | 0 | Carnivora | 101804.0 | 0 | Canidae | 98428.0 | 1 | Canis | 88629.0 | 2 | Canis lupus | 88629.0 | 2 |
| scf7180000049016 | 24529 | 0.4385 | 0 | 94.1936 | Eukaryota | 89293.0 | 0 | Chordata | 89293.0 | 0 | Carnivora | 89293.0 | 0 | Canidae | 89293.0 | 0 | Canis | 89293.0 | 0 | Canis lupus | 89293.0 | 0 |
| scf7180000041191 | 165528 | 0.3758 | 0 | 57.2029 | Eukaryota | 72781.0 | 0 | Chordata | 72781.0 | 0 | Carnivora | 72781.0 | 0 | Canidae | 72781.0 | 0 | Canis | 72781.0 | 0 | Canis lupus | 72781.0 | 0 |
| scf7180000053894 | 3717 | 0.4385 | 0 | 183.1047 | no-hit | 0.0 | 0 | no-hit | 0.0 | 0 | no-hit | 0.0 | 0 | no-hit | 0.0 | 0 | no-hit | 0.0 | 0 | no-hit | 0.0 | 0 |
| scf7180000043590 | 448494 | 0.3715 | 0 | 60.0885 | Eukaryota | 19815.0 | 1 | Chordata | 19815.0 | 1 | Carnivora | 11785.0 | 3 | Canidae | 9321.0 | 4 | Canis | 9321.0 | 5 | Canis lupus | 9321.0 | 5 |
| scf7180000041476 | 458052 | 0.4075 | 200 | 58.7933 | Eukaryota | 65775.0 | 0 | Chordata | 65775.0 | 0 | Carnivora | 65775.0 | 0 | Canidae | 65775.0 | 0 | Canis | 35791.0 | 1 | Canis lupus | 35791.0 | 1 |
| scf7180000047122 | 8033 | 0.4775 | 0 | 178.6185 | Eukaryota | 36504.0 | 0 | Chordata | 36504.0 | 0 | Carnivora | 36504.0 | 0 | Canidae | 36504.0 | 0 | Canis | 36504.0 | 0 | Canis lupus | 36504.0 | 0 |
| scf7180000045305 | 224885 | 0.3895 | 0 | 60.7208 | Eukaryota | 73335.0 | 0 | Chordata | 73335.0 | 0 | Carnivora | 73335.0 | 0 | Canidae | 73335.0 | 0 | Canis | 73335.0 | 0 | Canis lupus | 73335.0 | 0 |
| scf7180000042839 | 493521 | 0.4256 | 0 | 59.7068 | Eukaryota | 97279.0 | 0 | Chordata | 97279.0 | 0 | Carnivora | 97279.0 | 0 | Canidae | 97279.0 | 0 | Canis | 97279.0 | 0 | Canis lupus | 97279.0 | 0 |
| scf7180000047220 | 18177 | 0.4338 | 0 | 54.0564 | Eukaryota | 63871.0 | 0 | Chordata | 63871.0 | 0 | Carnivora | 63871.0 | 0 | Canidae | 63871.0 | 0 | Canis | 63871.0 | 0 | Canis lupus | 63871.0 | 0 |
| scf7180000041399 | 748491 | 0.3812 | 0 | 59.5413 | Eukaryota | 47676.0 | 0 | Chordata | 47676.0 | 0 | Carnivora | 47676.0 | 0 | Canidae | 47676.0 | 0 | Canis | 47676.0 | 0 | Canis lupus | 47676.0 | 0 |
| scf7180000055092 | 44521 | 0.4396 | 0 | 188.6889 | Eukaryota | 5153.0 | 0 | Chordata | 3839.0 | 1 | Kurtiformes | 1652.0 | 6 | Apogonidae | 1652.0 | 6 | Sphaeramia | 1652.0 | 6 | Sphaeramia orbicularis | 1652.0 | 6 |
| scf7180000049366 | 30053 | 0.5051 | 0 | 373.079 | Eukaryota | 8355.0 | 0 | Chordata | 8355.0 | 0 | Carnivora | 8355.0 | 0 | Canidae | 8355.0 | 0 | Canis | 6635.0 | 1 | Canis lupus | 5425.0 | 2 |
| scf7180000054277 | 19088 | 0.4323 | 0 | 90.9495 | Eukaryota | 58876.0 | 0 | Chordata | 58876.0 | 0 | Carnivora | 58876.0 | 0 | Canidae | 42988.0 | 1 | Canis | 36810.0 | 3 | Canis lupus | 36810.0 | 4 |
| scf7180000048040 | 101569 | 0.4585 | 100 | 58.4015 | Eukaryota | 39430.0 | 0 | Chordata | 39430.0 | 0 | Carnivora | 39430.0 | 0 | Canidae | 39430.0 | 0 | Canis | 31544.0 | 1 | Canis lupus | 31544.0 | 1 |
| scf7180000054858 | 18485 | 0.4437 | 0 | 257.7052 | Eukaryota | 3313.0 | 0 | Chordata | 2270.0 | 1 | Tetraodontiformes | 1776.0 | 2 | Tetraodontidae | 1776.0 | 2 | Takifugu | 1776.0 | 2 | Takifugu rubripes | 1776.0 | 2 |
| scf7180000041201 | 18339 | 0.4499 | 0 | 2332.0224 | Eukaryota | 11200.0 | 1 | Chordata | 11200.0 | 1 | Artiodactyla | 3353.0 | 7 | Bovidae | 3353.0 | 7 | Mus | 2176.0 | 8 | Mus musculus | 2176.0 | 8 |
| scf7180000043182 | 513462 | 0.4475 | 0 | 59.2854 | Eukaryota | 53613.0 | 0 | Chordata | 53613.0 | 0 | Carnivora | 53613.0 | 0 | Canidae | 53613.0 | 0 | Canis | 31806.0 | 1 | Canis lupus | 31806.0 | 1 |
| scf7180000045995 | 144840 | 0.394 | 0 | 60.0474 | Eukaryota | 26185.0 | 0 | Chordata | 26185.0 | 0 | Carnivora | 26185.0 | 0 | Canidae | 16844.0 | 3 | Canis | 16844.0 | 4 | Canis lupus | 16844.0 | 4 |
| scf7180000051750 | 2176 | 0.4513 | 0 | 912.6764 | no-hit | 0.0 | 0 | no-hit | 0.0 | 0 | no-hit | 0.0 | 0 | no-hit | 0.0 | 0 | no-hit | 0.0 | 0 | no-hit | 0.0 | 0 |
| scf7180000053535 | 1997 | 0.4517 | 0 | 243.1481 | no-hit | 0.0 | 0 | no-hit | 0.0 | 0 | no-hit | 0.0 | 0 | no-hit | 0.0 | 0 | no-hit | 0.0 | 0 | no-hit | 0.0 | 0 |
| scf7180000043837 | 310685 | 0.4138 | 0 | 59.8195 | Eukaryota | 56807.0 | 0 | Chordata | 56807.0 | 0 | Carnivora | 56807.0 | 0 | Canidae | 56807.0 | 0 | Vulpes | 40040.0 | 1 | Vulpes vulpes | 40040.0 | 1 |
| scf7180000048923 | 12302 | 0.4419 | 0 | 73.8116 | Eukaryota | 1235.0 | 0 | Platyhelminthes | 681.0 | 1 | Polystomatidea | 681.0 | 2 | Polystomatidae | 681.0 | 2 | Protopolystoma | 681.0 | 2 | Protopolystoma xenopodis | 681.0 | 2 |
| scf7180000048008 | 19952 | 0.4421 | 0 | 143.0599 | Eukaryota | 4643.0 | 0 | Chordata | 4334.0 | 1 | Tetraodontiformes | 1791.0 | 5 | Tetraodontidae | 1791.0 | 5 | Takifugu | 1791.0 | 6 | Sphaeramia orbicularis | 1270.0 | 7 |
| scf7180000056461 | 307957 | 0.4631 | 0 | 60.1914 | Eukaryota | 97444.0 | 0 | Chordata | 97444.0 | 0 | Carnivora | 97444.0 | 0 | Canidae | 97444.0 | 0 | Canis | 97444.0 | 0 | Canis lupus | 97444.0 | 0 |
| scf7180000046329 | 6714 | 0.4611 | 0 | 273.1436 | Eukaryota | 1135.0 | 0 | Chordata | 723.0 | 1 | Kurtiformes | 564.0 | 2 | Apogonidae | 564.0 | 2 | Sphaeramia | 564.0 | 2 | Sphaeramia orbicularis | 564.0 | 2 |
| scf7180000041422 | 1064167 | 0.3988 | 0 | 61.3677 | Eukaryota | 88580.0 | 0 | Chordata | 88580.0 | 0 | Carnivora | 88580.0 | 0 | Canidae | 88580.0 | 0 | Canis | 88580.0 | 0 | Canis lupus | 88580.0 | 0 |
| scf7180000048177 | 7924 | 0.4464 | 0 | 924.4898 | no-hit | 0.0 | 0 | no-hit | 0.0 | 0 | no-hit | 0.0 | 0 | no-hit | 0.0 | 0 | no-hit | 0.0 | 0 | no-hit | 0.0 | 0 |
| scf7180000055525 | 6375 | 0.4387 | 0 | 125.0679 | no-hit | 0.0 | 0 | no-hit | 0.0 | 0 | no-hit | 0.0 | 0 | no-hit | 0.0 | 0 | no-hit | 0.0 | 0 | no-hit | 0.0 | 0 |
| scf7180000044242 | 5860 | 0.4012 | 0 | 52.0436 | Eukaryota | 61997.0 | 0 | Chordata | 61997.0 | 0 | Carnivora | 61997.0 | 0 | Canidae | 61997.0 | 0 | Canis | 61997.0 | 0 | Canis lupus | 61997.0 | 0 |
| scf7180000045557 | 10051 | 0.4383 | 0 | 83.4427 | Eukaryota | 2077.0 | 0 | Chordata | 1829.0 | 1 | Tetraodontiformes | 1018.0 | 3 | Tetraodontidae | 1018.0 | 3 | Takifugu | 1018.0 | 3 | Takifugu rubripes | 1018.0 | 3 |
| scf7180000042255 | 296977 | 0.4742 | 0 | 56.3722 | Eukaryota | 70146.0 | 0 | Chordata | 70146.0 | 0 | Carnivora | 70146.0 | 0 | Canidae | 55333.0 | 2 | Canis | 55333.0 | 3 | Canis lupus | 55333.0 | 4 |
| scf7180000049295 | 59453 | 0.3981 | 0 | 60.8888 | Eukaryota | 19101.0 | 0 | Chordata | 19101.0 | 0 | Carnivora | 17499.0 | 1 | Felidae | 11740.0 | 3 | Felis | 5100.0 | 5 | Felis catus | 5100.0 | 5 |
| scf7180000052514 | 22523 | 0.4396 | 0 | 78.8868 | Eukaryota | 16827.0 | 0 | Chordata | 14095.0 | 2 | Synbranchiformes | 3885.0 | 6 | Mastacembelidae | 3885.0 | 6 | Mastacembelus | 3885.0 | 6 | Mastacembelus armatus | 3885.0 | 6 |
| scf7180000046435 | 133727 | 0.5494 | 0 | 55.4053 | Eukaryota | 10094.0 | 0 | Chordata | 10094.0 | 0 | Carnivora | 8954.0 | 1 | Canidae | 4140.0 | 5 | Canis | 2657.0 | 7 | Canis lupus | 2657.0 | 7 |
| scf7180000046090 | 103874 | 0.3787 | 0 | 60.683 | Eukaryota | 10725.0 | 0 | Chordata | 10725.0 | 0 | Carnivora | 8533.0 | 1 | Felidae | 7657.0 | 2 | Felis | 7657.0 | 2 | Felis catus | 7657.0 | 2 |
| scf7180000041129 | 157999 | 0.4457 | 0 | 58.808 | Eukaryota | 31105.0 | 0 | Chordata | 31105.0 | 0 | Carnivora | 31105.0 | 0 | Canidae | 31105.0 | 0 | Canis | 31105.0 | 0 | Canis lupus | 31105.0 | 0 |

| | | | | | | | | | | | | | | | | | | | | | |
|---|---|---|---|---|---|---|---|---|---|---|---|---|---|---|---|---|---|---|---|---|---|
| scf7180000050093 | 27065 | 0.3924 | 0 | 56.2788 | Eukaryota | 89743.0 | 0 | Chordata | 89743.0 | 0 | Carnivora | 89743.0 | 0 | Canidae | 89743.0 | 0 | Canis | 89743.0 | 0 | Canis lupus | 89743.0 | 0 |
| scf7180000044727 | 270611 | 0.5204 | 0 | 53.9416 | Eukaryota | 27244.0 | 0 | Chordata | 27244.0 | 0 | Carnivora | 27244.0 | 0 | Canidae | 27244.0 | 0 | Canis | 25753.0 | 1 | Canis lupus | 25753.0 | 1 |
| scf7180000048537 | 11810 | 0.4448 | 0 | 83.6069 | Eukaryota | 1123.0 | 0 | Chordata | 1123.0 | 0 | Tetraodontiformes | 1123.0 | 0 | Tetraodontidae | 1123.0 | 0 | Takifugu | 1123.0 | 0 | Takifugu bimaculatus | 1123.0 | 0 |
| scf7180000042389 | 30998 | 0.438 | 0 | 51.8917 | Eukaryota | 83651.0 | 0 | Chordata | 83651.0 | 0 | Carnivora | 83651.0 | 0 | Canidae | 83651.0 | 0 | Canis | 83651.0 | 0 | Canis lupus | 83651.0 | 0 |
| scf7180000042230 | 634606 | 0.4522 | 0 | 59.6302 | Eukaryota | 54575.0 | 0 | Chordata | 54575.0 | 0 | Carnivora | 54575.0 | 0 | Canidae | 54575.0 | 0 | Canis | 34416.0 | 1 | Canis lupus | 34416.0 | 1 |
| scf7180000048673 | 5248 | 0.4369 | 0 | 70.8311 | no-hit | 0.0 | 0 | no-hit | 0.0 | 0 | no-hit | 0.0 | 0 | no-hit | 0.0 | 0 | no-hit | 0.0 | 0 | no-hit | 0.0 | 0 |
| scf7180000047375 | 55490 | 0.4413 | 0 | 54.4727 | Eukaryota | 45544.0 | 0 | Chordata | 45544.0 | 0 | Carnivora | 45544.0 | 0 | Canidae | 45544.0 | 0 | Canis | 45544.0 | 0 | Canis lupus | 45544.0 | 0 |
| scf7180000042310 | 146516 | 0.4018 | 0 | 55.273 | Eukaryota | 81824.0 | 0 | Chordata | 81824.0 | 0 | Carnivora | 81824.0 | 0 | Canidae | 81824.0 | 0 | Canis | 81824.0 | 0 | Canis lupus | 81824.0 | 0 |
| scf7180000050207 | 4367 | 0.4431 | 0 | 197.5917 | no-hit | 0.0 | 0 | no-hit | 0.0 | 0 | no-hit | 0.0 | 0 | no-hit | 0.0 | 0 | no-hit | 0.0 | 0 | no-hit | 0.0 | 0 |
| scf7180000054971 | 14630 | 0.4481 | 0 | 797.8359 | no-hit | 0.0 | 0 | no-hit | 0.0 | 0 | no-hit | 0.0 | 0 | no-hit | 0.0 | 0 | no-hit | 0.0 | 0 | no-hit | 0.0 | 0 |
| scf7180000041673 | 966741 | 0.3744 | 0 | 61.1229 | Eukaryota | 137596.0 | 0 | Chordata | 137596.0 | 0 | Carnivora | 137596.0 | 0 | Canidae | 137596.0 | 0 | Canis | 137596.0 | 0 | Canis lupus | 137596.0 | 0 |
| scf7180000049700 | 4727 | 0.4519 | 0 | 898.5696 | Eukaryota | 355.0 | 0 | Platyhelminthes | 355.0 | 0 | Polystomatidea | 355.0 | 0 | Polystomatidae | 355.0 | 0 | Protopolystoma | 355.0 | 0 | Protopolystoma xenopodis | 355.0 | 0 |
| scf7180000054551 | 12117 | 0.4259 | 0 | 87.3836 | Eukaryota | 73668.0 | 0 | Chordata | 73668.0 | 0 | Carnivora | 73668.0 | 0 | Canidae | 73668.0 | 0 | Canis | 73668.0 | 0 | Canis lupus | 73668.0 | 0 |
| scf7180000041855 | 241370 | 0.4143 | 0 | 58.49 | Eukaryota | 23595.0 | 0 | Chordata | 23595.0 | 0 | Carnivora | 23595.0 | 0 | Canidae | 23595.0 | 0 | Canis | 15000.0 | 1 | Canis lupus | 15000.0 | 1 |
| scf7180000056456 | 434192 | 0.4273 | 0 | 59.9678 | Eukaryota | 87849.0 | 0 | Chordata | 87849.0 | 0 | Carnivora | 87849.0 | 0 | Canidae | 87849.0 | 0 | Canis | 52731.0 | 1 | Canis lupus | 52731.0 | 1 |
| scf7180000053446 | 4281 | 0.4536 | 0 | 412.179 | no-hit | 0.0 | 0 | no-hit | 0.0 | 0 | no-hit | 0.0 | 0 | no-hit | 0.0 | 0 | no-hit | 0.0 | 0 | no-hit | 0.0 | 0 |
| scf7180000049622 | 7746 | 0.4454 | 0 | 96.6159 | Eukaryota | 19507.0 | 1 | Chordata | 19507.0 | 1 | Gadiformes | 5616.0 | 5 | Gadidae | 5616.0 | 5 | Gadus | 5616.0 | 5 | Gadus morhua | 5616.0 | 5 |
| scf7180000047314 | 27254 | 0.3845 | 0 | 46.5459 | Eukaryota | 16376.0 | 0 | Chordata | 16376.0 | 0 | Carnivora | 14554.0 | 2 | Canidae | 14204.0 | 3 | Canis | 14204.0 | 3 | Canis lupus | 14204.0 | 3 |
| scf7180000040772 | 1069460 | 0.4019 | 0 | 59.3322 | Eukaryota | 68128.0 | 0 | Chordata | 68128.0 | 0 | Carnivora | 68128.0 | 0 | Canidae | 57018.0 | 1 | Canis | 57018.0 | 1 | Canis lupus | 57018.0 | 2 |
| scf7180000054297 | 623187 | 0.356 | 0 | 61.4584 | Eukaryota | 26132.0 | 0 | Chordata | 26132.0 | 0 | Carnivora | 26132.0 | 0 | Canidae | 26132.0 | 0 | Canis | 26132.0 | 0 | Canis lupus | 26132.0 | 0 |
| scf7180000052692 | 16694 | 0.5616 | 0 | 53.9718 | Eukaryota | 3033.0 | 0 | Chordata | 3033.0 | 0 | Carnivora | 3033.0 | 0 | Canidae | 3033.0 | 0 | Canis | 3033.0 | 0 | Canis lupus | 3033.0 | 0 |
| scf7180000047041 | 13457 | 0.5131 | 0 | 428.4982 | Eukaryota | 37447.0 | 0 | Chordata | 37447.0 | 0 | Carnivora | 37447.0 | 0 | Canidae | 37447.0 | 0 | Canis | 37447.0 | 0 | Canis lupus | 37447.0 | 0 |
| scf7180000051166 | 4107 | 0.457 | 0 | 434.1719 | no-hit | 0.0 | 0 | no-hit | 0.0 | 0 | no-hit | 0.0 | 0 | no-hit | 0.0 | 0 | no-hit | 0.0 | 0 | no-hit | 0.0 | 0 |
| scf7180000042770 | 80426 | 0.346 | 0 | 57.4171 | Eukaryota | 20034.0 | 0 | Chordata | 20034.0 | 0 | Carnivora | 20034.0 | 0 | Canidae | 20034.0 | 0 | Canis | 20034.0 | 0 | Canis lupus | 20034.0 | 0 |
| scf7180000055054 | 32031 | 0.4357 | 0 | 276.9655 | Eukaryota | 9650.0 | 0 | Chordata | 7820.0 | 1 | Kurtiformes | 3260.0 | 4 | Apogonidae | 3260.0 | 4 | Sphaeramia | 3260.0 | 4 | Sphaeramia orbicularis | 3260.0 | 4 |
| scf7180000056142 | 2328 | 0.4424 | 0 | 131.0032 | no-hit | 0.0 | 0 | no-hit | 0.0 | 0 | no-hit | 0.0 | 0 | no-hit | 0.0 | 0 | no-hit | 0.0 | 0 | no-hit | 0.0 | 0 |
| scf7180000043797 | 372116 | 0.3829 | 0 | 60.1657 | Eukaryota | 40534.0 | 0 | Chordata | 40534.0 | 0 | Carnivora | 40534.0 | 0 | Canidae | 40534.0 | 0 | Canis | 40534.0 | 0 | Canis lupus | 40534.0 | 0 |
| scf7180000044365 | 29301 | 0.407 | 0 | 122.7246 | Eukaryota | 19912.0 | 0 | Chordata | 19912.0 | 0 | Carnivora | 12101.0 | 1 | Canidae | 8227.0 | 3 | Canis | 6254.0 | 4 | Canis lupus | 6254.0 | 7 |
| scf7180000056536 | 304186 | 0.4408 | 0 | 60.0424 | Eukaryota | 29535.0 | 0 | Chordata | 29535.0 | 0 | Carnivora | 29535.0 | 0 | Canidae | 29535.0 | 0 | Canis | 17611.0 | 1 | Canis lupus | 17611.0 | 1 |
| scf7180000040491 | 3023915 | 0.422 | 0 | 59.8465 | Eukaryota | 95612.0 | 0 | Chordata | 95612.0 | 0 | Carnivora | 95612.0 | 0 | Canidae | 95612.0 | 0 | Canis | 95612.0 | 0 | Canis lupus | 95612.0 | 0 |
| scf7180000053568 | 3033 | 0.4451 | 0 | 629.7396 | no-hit | 0.0 | 0 | no-hit | 0.0 | 0 | no-hit | 0.0 | 0 | no-hit | 0.0 | 0 | no-hit | 0.0 | 0 | no-hit | 0.0 | 0 |
| scf7180000046007 | 176449 | 0.4364 | 0 | 59.1006 | Eukaryota | 49641.0 | 0 | Chordata | 49641.0 | 0 | Carnivora | 49641.0 | 0 | Canidae | 49641.0 | 0 | Canis | 49641.0 | 0 | Canis lupus | 49641.0 | 0 |
| scf7180000051942 | 15581 | 0.4458 | 0 | 155.1187 | Eukaryota | 19744.0 | 0 | Chordata | 11533.0 | 3 | Gadiformes | 8036.0 | 7 | Gadidae | 8036.0 | 7 | Gadus | 8036.0 | 7 | Gadus morhua | 8036.0 | 7 |
| scf7180000046919 | 59807 | 0.4166 | 0 | 55.118 | Eukaryota | 23721.0 | 0 | Chordata | 23721.0 | 0 | Carnivora | 21327.0 | 1 | Canidae | 21327.0 | 1 | Canis | 21327.0 | 1 | Canis lupus | 21327.0 | 1 |
| scf7180000040581 | 2090911 | 0.5228 | 0 | 59.0314 | Eukaryota | 74952.0 | 0 | Chordata | 74952.0 | 0 | Carnivora | 74952.0 | 0 | Canidae | 74952.0 | 0 | Canis | 58824.0 | 1 | Canis lupus | 58824.0 | 1 |
| scf7180000046937 | 7359 | 0.4297 | 0 | 54.8795 | Eukaryota | 1004.0 | 0 | Chordata | 1004.0 | 0 | Salmoniformes | 1004.0 | 0 | Salmonidae | 1004.0 | 0 | Coregonus | 697.0 | 1 | Coregonus sp. 'balchen' | 697.0 | 1 |
| scf7180000040616 | 605804 | 0.4705 | 0 | 58.6945 | Eukaryota | 89921.0 | 0 | Chordata | 89921.0 | 0 | Carnivora | 89921.0 | 0 | Canidae | 89921.0 | 0 | Canis | 89921.0 | 0 | Canis lupus | 89921.0 | 0 |
| scf7180000045842 | 164536 | 0.4364 | 0 | 102.7518 | Eukaryota | 91020.0 | 0 | Chordata | 91020.0 | 0 | Carnivora | 91020.0 | 0 | Canidae | 91020.0 | 0 | Canis | 91020.0 | 0 | Canis lupus | 91020.0 | 0 |
| scf7180000045326 | 161173 | 0.44 | 0 | 58.6896 | Eukaryota | 48164.0 | 0 | Chordata | 48164.0 | 0 | Carnivora | 48164.0 | 0 | Canidae | 48164.0 | 0 | Canis | 37736.0 | 1 | Canis lupus | 37736.0 | 1 |
| scf7180000043455 | 179518 | 0.4333 | 0 | 59.2172 | Eukaryota | 59579.0 | 0 | Chordata | 59579.0 | 0 | Carnivora | 59579.0 | 0 | Canidae | 59579.0 | 0 | Canis | 59579.0 | 0 | Canis lupus | 59579.0 | 0 |
| scf7180000049716 | 19948 | 0.4366 | 0 | 76.5339 | Eukaryota | 2920.0 | 0 | Chordata | 2020.0 | 1 | Carnivora | 2020.0 | 1 | Canidae | 2020.0 | 1 | Canis | 2020.0 | 1 | Canis lupus | 2020.0 | 1 |
| scf7180000044225 | 6101 | 0.3514 | 0 | 25.4631 | Eukaryota | 40592.0 | 0 | Chordata | 40592.0 | 0 | Carnivora | 40592.0 | 0 | Canidae | 40592.0 | 0 | Canis | 40592.0 | 0 | Canis lupus | 40592.0 | 0 |
| scf7180000053035 | 3884 | 0.714 | 0 | 45.1228 | Eukaryota | 2507.0 | 0 | Chordata | 2507.0 | 0 | Carnivora | 2507.0 | 0 | Canidae | 1661.0 | 2 | Canis | 1446.0 | 3 | Canis lupus | 1446.0 | 3 |
| scf7180000051924 | 25762 | 0.3974 | 0 | 57.4536 | Eukaryota | 30519.0 | 0 | Chordata | 30519.0 | 0 | Carnivora | 30519.0 | 0 | Canidae | 30519.0 | 0 | Canis | 30519.0 | 0 | Canis lupus | 30519.0 | 0 |
| scf7180000042415 | 278436 | 0.5538 | 0 | 57.7116 | Eukaryota | 55154.0 | 0 | Chordata | 55154.0 | 0 | Carnivora | 55154.0 | 0 | Canidae | 55154.0 | 0 | Vulpes | 33190.0 | 1 | Vulpes vulpes | 33190.0 | 1 |
| scf7180000042171 | 154693 | 0.3966 | 0 | 59.1733 | Eukaryota | 110617.0 | 0 | Chordata | 110617.0 | 0 | Carnivora | 110617.0 | 0 | Canidae | 110617.0 | 0 | Canis | 110617.0 | 0 | Canis lupus | 110617.0 | 0 |
| scf7180000054370 | 15016 | 0.3917 | 0 | 103.3611 | Eukaryota | 8717.0 | 0 | Chordata | 8717.0 | 0 | Carnivora | 7216.0 | 1 | Canidae | 7216.0 | 1 | Canis | 7216.0 | 1 | Canis lupus | 7216.0 | 2 |
| scf7180000044605 | 42390 | 0.3952 | 0 | 62.9275 | Eukaryota | 94403.0 | 0 | Chordata | 94403.0 | 0 | Carnivora | 94403.0 | 0 | Canidae | 94403.0 | 0 | Canis | 94403.0 | 0 | Canis lupus | 94403.0 | 0 |
| scf7180000053480 | 2577 | 0.5029 | 0 | 25.2443 | Eukaryota | 3818.0 | 0 | Chordata | 3818.0 | 0 | Carnivora | 3818.0 | 0 | Canidae | 3818.0 | 0 | Canis | 3066.0 | 2 | Canis lupus | 3066.0 | 2 |
| scf7180000046790 | 4778 | 0.3198 | 0 | 30.4441 | Eukaryota | 7404.0 | 0 | Chordata | 7404.0 | 0 | Carnivora | 6357.0 | 2 | Canidae | 6357.0 | 2 | Canis | 6357.0 | 2 | Canis lupus | 6357.0 | 2 |
| scf7180000042544 | 219917 | 0.3688 | 0 | 57.6973 | Eukaryota | 89421.0 | 0 | Chordata | 89421.0 | 0 | Carnivora | 89421.0 | 0 | Canidae | 89421.0 | 0 | Canis | 89421.0 | 0 | Canis lupus | 89421.0 | 0 |
| scf7180000041700 | 33738 | 0.5031 | 0 | 85.7637 | Eukaryota | 34386.0 | 0 | Chordata | 34386.0 | 0 | Carnivora | 34386.0 | 0 | Canidae | 34386.0 | 0 | Canis | 34386.0 | 0 | Canis lupus | 34386.0 | 0 |
| scf7180000041156 | 417531 | 0.4139 | 0 | 59.4546 | Eukaryota | 35412.0 | 0 | Chordata | 35412.0 | 0 | Carnivora | 35412.0 | 0 | Canidae | 35412.0 | 0 | Canis | 35412.0 | 0 | Canis lupus | 35412.0 | 0 |
| scf7180000051458 | 3082 | 0.4536 | 0 | 387.3297 | Eukaryota | 1649.0 | 0 | Echinodermata | 593.0 | 2 | Forcipulatida | 593.0 | 2 | Asteriidae | 593.0 | 2 | Asterias | 593.0 | 2 | Asterias rubens | 593.0 | 2 |
| scf7180000056032 | 2370 | 0.4405 | 0 | 215.1465 | no-hit | 0.0 | 0 | no-hit | 0.0 | 0 | no-hit | 0.0 | 0 | no-hit | 0.0 | 0 | no-hit | 0.0 | 0 | no-hit | 0.0 | 0 |
| scf7180000049850 | 4321 | 0.4406 | 0 | 156.727 | no-hit | 0.0 | 0 | no-hit | 0.0 | 0 | no-hit | 0.0 | 0 | no-hit | 0.0 | 0 | no-hit | 0.0 | 0 | no-hit | 0.0 | 0 |
| scf7180000047755 | 94298 | 0.4456 | 0 | 59.1753 | Eukaryota | 40222.0 | 0 | Chordata | 40222.0 | 0 | Carnivora | 40222.0 | 0 | Canidae | 19868.0 | 2 | Canis | 15070.0 | 4 | Canis lupus | 15070.0 | 4 |
| scf7180000054779 | 108266 | 0.4434 | 0 | 198.9526 | Eukaryota | 87939.0 | 0 | Chordata | 87939.0 | 0 | Carnivora | 87939.0 | 0 | Canidae | 87939.0 | 0 | Canis | 87939.0 | 0 | Canis lupus | 87939.0 | 0 |
| scf7180000054435 | 567895 | 0.3934 | 0 | 64.5957 | Eukaryota | 88391.0 | 0 | Chordata | 88391.0 | 0 | Carnivora | 88391.0 | 0 | Canidae | 88391.0 | 0 | Canis | 88391.0 | 0 | Canis lupus | 88391.0 | 0 |
| scf7180000047607 | 2919 | 0.4515 | 0 | 138.9345 | no-hit | 0.0 | 0 | no-hit | 0.0 | 0 | no-hit | 0.0 | 0 | no-hit | 0.0 | 0 | no-hit | 0.0 | 0 | no-hit | 0.0 | 0 |
| scf7180000048812 | 50447 | 0.441 | 0 | 140.9052 | Eukaryota | 139.0 | 0 | Chordata | 139.0 | 0 | Blenniiformes | 139.0 | 0 | Gobiesocidae | 139.0 | 0 | Gouania | 139.0 | 0 | Gouania willdenowi | 139.0 | 0 |
| scf7180000049687 | 35825 | 0.4756 | 0 | 57.1208 | Eukaryota | 14595.0 | 0 | Chordata | 14595.0 | 0 | Carnivora | 14595.0 | 0 | Canidae | 14595.0 | 0 | Canis | 11239.0 | 1 | Canis lupus | 11239.0 | 1 |
| scf7180000046693 | 118910 | 0.4342 | 0 | 59.5355 | Eukaryota | 19392.0 | 0 | Chordata | 19392.0 | 0 | Carnivora | 19392.0 | 0 | Canidae | 19392.0 | 0 | Canis | 11649.0 | 1 | Canis lupus | 11649.0 | 1 |
| scf7180000041134 | 136955 | 0.3414 | 0 | 55.9121 | Eukaryota | 89001.0 | 0 | Chordata | 89001.0 | 0 | Carnivora | 89001.0 | 0 | Canidae | 89001.0 | 0 | Canis | 89001.0 | 0 | Canis lupus | 89001.0 | 0 |
| scf7180000049742 | 8454 | 0.4403 | 0 | 79.1058 | no-hit | 0.0 | 0 | no-hit | 0.0 | 0 | no-hit | 0.0 | 0 | no-hit | 0.0 | 0 | no-hit | 0.0 | 0 | no-hit | 0.0 | 0 |
| scf7180000044135 | 128001 | 0.4825 | 0 | 58.3301 | Eukaryota | 6657.0 | 0 | Chordata | 6657.0 | 0 | Carnivora | 3363.0 | 2 | Canidae | 3363.0 | 3 | Canis | 2414.0 | 4 | Canis lupus | 2414.0 | 4 |
| scf7180000048625 | 53073 | 0.3995 | 0 | 60.6945 | Eukaryota | 33512.0 | 0 | Chordata | 33512.0 | 0 | Carnivora | 33512.0 | 0 | Canidae | 33512.0 | 0 | Canis | 33512.0 | 0 | Canis lupus | 33512.0 | 0 |
| scf7180000049541 | 54548 | 0.3759 | 0 | 59.8977 | Eukaryota | 10176.0 | 0 | Chordata | 10176.0 | 0 | Carnivora | 8390.0 | 1 | Canidae | 5755.0 | 4 | Canis | 4683.0 | 5 | Canis lupus | 4683.0 | 5 |
| scf7180000049281 | 14716 | 0.4437 | 0 | 142.7167 | Eukaryota | 8570.0 | 0 | Chordata | 6786.0 | 1 | Synbranchiformes | 2787.0 | 4 | Mastacembelidae | 2787.0 | 4 | Mastacembelus | 2787.0 | 4 | Mastacembelus armatus | 2787.0 | 4 |
| scf7180000046166 | 24605 | 0.4673 | 0 | 525.0629 | Eukaryota | 87208.0 | 0 | Chordata | 87208.0 | 0 | Carnivora | 87208.0 | 0 | Canidae | 87208.0 | 0 | Canis | 87208.0 | 0 | Canis lupus | 87208.0 | 0 |
| scf7180000045519 | 201210 | 0.3753 | 0 | 60.1078 | Eukaryota | 111956.0 | 0 | Chordata | 111956.0 | 0 | Carnivora | 111956.0 | 0 | Canidae | 111956.0 | 0 | Canis | 78239.0 | 1 | Canis lupus | 78239.0 | 1 |
| scf7180000047720 | 3598 | 0.4433 | 0 | 66.6456 | Eukaryota | 2891.0 | 0 | Chordata | 2891.0 | 0 | Carnivora | 2891.0 | 0 | Canidae | 2891.0 | 0 | Canis | 2891.0 | 0 | Canis lupus | 2891.0 | 0 |
| scf7180000044886 | 156577 | 0.3394 | 0 | 57.5264 | Eukaryota | 18660.0 | 0 | Chordata | 18660.0 | 0 | Carnivora | 18660.0 | 0 | Odobenidae | 5664.0 | 4 | Odobenus | 5664.0 | 5 | Odobenus rosmarus | 5664.0 | 5 |
| scf7180000046253 | 31596 | 0.3565 | 0 | 45.1239 | Eukaryota | 80063.0 | 0 | Chordata | 80063.0 | 0 | Carnivora | 80063.0 | 0 | Canidae | 80063.0 | 0 | Canis | 80063.0 | 0 | Canis lupus | 80063.0 | 0 |
| scf7180000048566 | 44162 | 0.4241 | 0 | 57.639 | Eukaryota | 18155.0 | 0 | Chordata | 18155.0 | 0 | Carnivora | 18155.0 | 0 | Canidae | 9890.0 | 1 | Canis | 7810.0 | 3 | Canis lupus | 7810.0 | 3 |
| scf7180000056272 | 387567 | 0.3837 | 0 | 59.2966 | Eukaryota | 45371.0 | 0 | Chordata | 45371.0 | 0 | Carnivora | 45371.0 | 0 | Canidae | 45371.0 | 0 | Canis | 39404.0 | 1 | Canis lupus | 39404.0 | 1 |
| scf7180000054536 | 34228 | 0.4452 | 0 | 293.3801 | no-hit | 0.0 | 0 | no-hit | 0.0 | 0 | no-hit | 0.0 | 0 | no-hit | 0.0 | 0 | no-hit | 0.0 | 0 | no-hit | 0.0 | 0 |
| scf7180000055212 | 7158 | 0.4579 | 0 | 162.4999 | Eukaryota | 8318.0 | 0 | Chordata | 8318.0 | 0 | Carnivora | 8318.0 | 0 | Canidae | 8318.0 | 0 | Canis | 5248.0 | 1 | Canis lupus | 5248.0 | 1 |
| scf7180000041510 | 962960 | 0.3804 | 0 | 60.5091 | Eukaryota | 94943.0 | 0 | Chordata | 94943.0 | 0 | Carnivora | 94943.0 | 0 | Canidae | 94943.0 | 0 | Canis | 94943.0 | 0 | Canis lupus | 94943.0 | 0 |
| scf7180000047446 | 22214 | 0.4452 | 0 | 179.5837 | Eukaryota | 7943.0 | 0 | Chordata | 7943.0 | 0 | Anabantiformes | 2891.0 | 5 | Anabantidae | 2891.0 | 5 | Anabas | 2891.0 | 5 | Anabas testudineus | 2891.0 | 5 |
| scf7180000042087 | 645838 | 0.3716 | 0 | 60.6283 | Eukaryota | 33495.0 | 0 | Chordata | 33495.0 | 0 | Carnivora | 33495.0 | 0 | Canidae | 33495.0 | 0 | Canis | 33495.0 | 0 | Canis lupus | 33495.0 | 0 |
| scf7180000041111 | 20021 | 0.3849 | 0 | 51.1182 | Eukaryota | 53331.0 | 0 | Chordata | 53331.0 | 0 | Carnivora | 53331.0 | 0 | Canidae | 53331.0 | 0 | Canis | 53331.0 | 0 | Canis lupus | 53331.0 | 0 |
| scf7180000040756 | 270358 | 0.3887 | 0 | 59.5741 | Eukaryota | 87868.0 | 0 | Chordata | 87868.0 | 0 | Carnivora | 87868.0 | 0 | Canidae | 87868.0 | 0 | Canis | 87868.0 | 0 | Canis lupus | 87868.0 | 0 |
| scf7180000045275 | 18467 | 0.4082 | 0 | 55.1239 | Eukaryota | 91993.0 | 0 | Chordata | 91993.0 | 0 | Carnivora | 91993.0 | 0 | Canidae | 91993.0 | 0 | Canis | 91993.0 | 0 | Canis lupus | 91993.0 | 0 |
| scf7180000050566 | 12592 | 0.4485 | 0 | 773.1628 | Eukaryota | 2432.0 | 0 | Chordata | 2156.0 | 1 | Carnivora | 2156.0 | 1 | Canidae | 2156.0 | 1 | Canis | 2156.0 | 1 | Canis lupus | 2156.0 | 1 |
| scf7180000040534 | 493210 | 0.4619 | 0 | 58.9676 | Eukaryota | 73041.0 | 0 | Chordata | 73041.0 | 0 | Carnivora | 73041.0 | 0 | Canidae | 73041.0 | 0 | Canis | 44389.0 | 1 | Canis lupus | 44389.0 | 1 |
| scf7180000049982 | 40664 | 0.4606 | 0 | 44.8394 | Eukaryota | 74579.0 | 0 | Chordata | 74579.0 | 0 | Carnivora | 74579.0 | 0 | Canidae | 74579.0 | 0 | Canis | 74579.0 | 0 | Canis lupus | 74579.0 | 0 |
| scf7180000044134 | 209154 | 0.3862 | 0 | 58.7125 | Eukaryota | 17650.0 | 0 | Chordata | 17650.0 | 0 | Primates | 10048.0 | 2 | Hominidae | 8633.0 | 3 | Homo | 8633.0 | 5 | Homo sapiens | 8633.0 | 5 |
| scf7180000051757 | 3144 | 0.4504 | 0 | 1345.3287 | no-hit | 0.0 | 0 | no-hit | 0.0 | 0 | no-hit | 0.0 | 0 | no-hit | 0.0 | 0 | no-hit | 0.0 | 0 | no-hit | 0.0 | 0 |
| scf7180000053665 | 2797 | 0.5066 | 0 | 43.3542 | Eukaryota | 6910.0 | 0 | Chordata | 6910.0 | 0 | Carnivora | 6910.0 | 0 | Canidae | 6910.0 | 0 | Canis | 5209.0 | 1 | Canis lupus | 4785.0 | 2 |
| scf7180000052071 | 7204 | 0.4359 | 0 | 55.1573 | Eukaryota | 32704.0 | 0 | Chordata | 32704.0 | 0 | Carnivora | 32704.0 | 0 | Canidae | 32704.0 | 0 | Canis | 32704.0 | 0 | Canis lupus | 32704.0 | 0 |
| scf7180000042983 | 412275 | 0.3751 | 0 | 58.1023 | Eukaryota | 91118.0 | 0 | Chordata | 91118.0 | 0 | Carnivora | 91118.0 | 0 | Canidae | 91118.0 | 0 | Canis | 91118.0 | 0 | Canis lupus | 91118.0 | 0 |
| scf7180000044502 | 341539 | 0.3905 | 0 | 60.6536 | Eukaryota | 31919.0 | 0 | Chordata | 31919.0 | 0 | Carnivora | 31919.0 | 0 | Canidae | 22665.0 | 2 | Canis | 22665.0 | 3 | Canis lupus | 22665.0 | 3 |
| scf7180000042404 | 724180 | 0.4255 | 0 | 60.6531 | Eukaryota | 70995.0 | 0 | Chordata | 70995.0 | 0 | Carnivora | 70995.0 | 0 | Canidae | 63364.0 | 1 | Canis | 37678.0 | 2 | Canis lupus | 37678.0 | 2 |
| scf7180000042410 | 639753 | 0.4129 | 0 | 59.498 | Eukaryota | 41675.0 | 0 | Chordata | 41675.0 | 0 | Carnivora | 41675.0 | 0 | Canidae | 41675.0 | 0 | Canis | 41675.0 | 0 | Canis lupus | 41675.0 | 0 |
| scf7180000045218 | 201754 | 0.5345 | 0 | 58.3109 | Eukaryota | 76420.0 | 0 | Chordata | 76420.0 | 0 | Carnivora | 76420.0 | 0 | Canidae | 76420.0 | 0 | Canis | 65960.0 | 1 | Canis lupus | 65960.0 | 1 |
| scf7180000054788 | 6944 | 0.4435 | 0 | 128.5164 | Eukaryota | 532.0 | 0 | Chordata | 532.0 | 0 | Columbiformes | 532.0 | 0 | Columbidae | 532.0 | 0 | Streptopelia | 532.0 | 0 | Streptopelia turtur | 532.0 | 0 |
| scf7180000045419 | 170879 | 0.4414 | 0 | 55.5807 | Eukaryota | 24505.0 | 0 | Chordata | 24505.0 | 0 | Carnivora | 24505.0 | 0 | Canidae | 24505.0 | 0 | Canis | 24505.0 | 0 | Canis lupus | 24505.0 | 0 |
| scf7180000040653 | 692479 | 0.3713 | 0 | 59.6214 | Eukaryota | 90754.0 | 0 | Chordata | 90754.0 | 0 | Carnivora | 90754.0 | 0 | Canidae | 90754.0 | 0 | Canis | 90754.0 | 0 | Canis lupus | 90754.0 | 0 |
| scf7180000042503 | 159646 | 0.4535 | 0 | 59.4947 | Eukaryota | 88376.0 | 0 | Chordata | 88376.0 | 0 | Carnivora | 88376.0 | 0 | Canidae | 88376.0 | 0 | Canis | 88376.0 | 0 | Canis lupus | 88376.0 | 0 |
| scf7180000053332 | 13771 | 0.5045 | 0 | 2623.3153 | Eukaryota | 9200.0 | 0 | Chordata | 9200.0 | 0 | Carnivora | 9200.0 | 0 | Canidae | 9200.0 | 0 | Canis | 7389.0 | 1 | Canis lupus | 6099.0 | 2 |
| scf7180000042301 | 169609 | 0.3454 | 0 | 59.5102 | Eukaryota | 92402.0 | 0 | Chordata | 92402.0 | 0 | Carnivora | 92402.0 | 0 | Canidae | 92402.0 | 0 | Canis | 92402.0 | 0 | Canis lupus | 92402.0 | 0 |
| scf7180000052648 | 5090 | 0.455 | 0 | 488.0238 | no-hit | 0.0 | 0 | no-hit | 0.0 | 0 | no-hit | 0.0 | 0 | no-hit | 0.0 | 0 | no-hit | 0.0 | 0 | no-hit | 0.0 | 0 |
| scf7180000042512 | 792041 | 0.4596 | 0 | 59.1548 | Eukaryota | 91925.0 | 0 | Chordata | 91925.0 | 0 | Carnivora | 91925.0 | 0 | Canidae | 91925.0 | 0 | Canis | 91925.0 | 0 | Canis lupus | 91925.0 | 0 |
| scf7180000044740 | 112834 | 0.394 | 0 | 59.3824 | Eukaryota | 89122.0 | 0 | Chordata | 89122.0 | 0 | Carnivora | 89122.0 | 0 | Canidae | 89122.0 | 0 | Canis | 89122.0 | 0 | Canis lupus | 89122.0 | 0 |
| scf7180000041086 | 1380635 | 0.3445 | 0 | 59.4262 | Eukaryota | 93013.0 | 0 | Chordata | 93013.0 | 0 | Carnivora | 93013.0 | 0 | Canidae | 93013.0 | 0 | Canis | 93013.0 | 0 | Canis lupus | 93013.0 | 0 |
| scf7180000040630 | 4153 | 0.53 | 0 | 25.909 | Eukaryota | 722.0 | 0 | Chordata | 722.0 | 0 | Primates | 722.0 | 0 | Hominidae | 722.0 | 0 | Homo | 722.0 | 0 | Homo sapiens | 722.0 | 0 |
| scf7180000047464 | 13046 | 0.408 | 0 | 52.648 | Eukaryota | 10733.0 | 0 | Chordata | 10733.0 | 0 | Carnivora | 10733.0 | 0 | Canidae | 10733.0 | 0 | Canis | 10733.0 | 0 | Canis lupus | 10733.0 | 0 |
| scf7180000049500 | 43729 | 0.4031 | 0 | 61.2059 | Eukaryota | 11134.0 | 0 | Chordata | 11134.0 | 0 | Carnivora | 11134.0 | 0 | Canidae | 11134.0 | 0 | Canis | 9865.0 | 1 | Canis lupus | 9865.0 | 1 |
| scf7180000046868 | 95322 | 0.4899 | 0 | 58.874 | Eukaryota | 48187.0 | 0 | Chordata | 48187.0 | 0 | Carnivora | 48187.0 | 0 | Canidae | 43258.0 | 1 | Canis | 31086.0 | 2 | Canis lupus | 31086.0 | 2 |
| scf7180000044681 | 37306 | 0.3979 | 0 | 49.974 | Eukaryota | 38364.0 | 0 | Chordata | 38364.0 | 0 | Carnivora | 38364.0 | 0 | Canidae | 38364.0 | 0 | Canis | 38364.0 | 0 | Canis lupus | 38364.0 | 0 |
| scf7180000056048 | 10579 | 0.5034 | 0 | 746.0575 | Eukaryota | 16143.0 | 0 | Chordata | 16143.0 | 0 | Carnivora | 16143.0 | 0 | Canidae | 16143.0 | 0 | Canis | 16143.0 | 0 | Canis lupus | 16143.0 | 0 |
| scf7180000044625 | 4452 | 0.327 | 0 | 29.9262 | Eukaryota | 5080.0 | 0 | Chordata | 5080.0 | 0 | Carnivora | 4599.0 | 1 | Felidae | 3454.0 | 2 | Felis | 2728.0 | 3 | Felis catus | 2728.0 | 3 |
| scf7180000042411 | 385792 | 0.3837 | 0 | 59.7178 | Eukaryota | 102146.0 | 0 | Chordata | 102146.0 | 0 | Carnivora | 102146.0 | 0 | Canidae | 102146.0 | 0 | Canis | 102146.0 | 0 | Canis lupus | 102146.0 | 0 |
| scf7180000048268 | 54814 | 0.4722 | 0 | 59.4812 | Eukaryota | 15889.0 | 0 | Chordata | 15889.0 | 0 | Carnivora | 15889.0 | 0 | Ursidae | 8154.0 | 2 | Canis | 4219.0 | 4 | Canis lupus | 4219.0 | 4 |
| scf7180000056194 | 1474043 | 0.3937 | 0 | 61.1 | Eukaryota | 94308.0 | 0 | Chordata | 94308.0 | 0 | Carnivora | 94308.0 | 0 | Canidae | 94308.0 | 0 | Canis | 94308.0 | 0 | Canis lupus | 94308.0 | 0 |
| scf7180000043117 | 443556 | 0.3553 | 0 | 59.92 | Eukaryota | 17968.0 | 0 | Chordata | 17968.0 | 0 | Carnivora | 17968.0 | 0 | Canidae | 12959.0 | 2 | Canis | 11208.0 | 3 | Canis lupus | 11208.0 | 3 |
| scf7180000040762 | 357476 | 0.3931 | 0 | 59.3053 | Eukaryota | 70870.0 | 0 | Chordata | 70870.0 | 0 | Carnivora | 70870.0 | 0 | Canidae | 70870.0 | 0 | Canis | 62672.0 | 1 | Canis lupus | 62672.0 | 1 |
| scf7180000041740 | 148338 | 0.4435 | 0 | 57.1333 | Eukaryota | 13411.0 | 0 | Chordata | 13411.0 | 0 | Carnivora | 13411.0 | 0 | Canidae | 13411.0 | 0 | Canis | 10900.0 | 2 | Canis lupus | 7174.0 | 3 |
| scf7180000045459 | 22184 | 0.4425 | 0 | 748.0446 | Eukaryota | 74273.0 | 0 | Chordata | 74273.0 | 0 | Carnivora | 74273.0 | 0 | Canidae | 74273.0 | 0 | Canis | 74273.0 | 0 | Canis lupus | 74273.0 | 0 |
| scf7180000044584 | 211322 | 0.4471 | 0 | 59.8932 | Eukaryota | 39646.0 | 0 | Chordata | 39646.0 | 0 | Carnivora | 39646.0 | 0 | Canidae | 39646.0 | 0 | Canis | 23606.0 | 1 | Canis lupus | 23606.0 | 1 |
| scf7180000043988 | 374925 | 0.3558 | 0 | 59.048 | Eukaryota | 149070.0 | 0 | Chordata | 149070.0 | 0 | Carnivora | 149070.0 | 0 | Canidae | 149070.0 | 0 | Canis | 149070.0 | 0 | Canis lupus | 149070.0 | 0 |
| scf7180000040476 | 2847344 | 0.4085 | 0 | 60.8716 | Eukaryota | 106464.0 | 0 | Chordata | 106464.0 | 0 | Carnivora | 106464.0 | 0 | Canidae | 106464.0 | 0 | Canis | 106464.0 | 0 | Canis lupus | 106464.0 | 0 |
| scf7180000046993 | 4030 | 0.4635 | 0 | 27.4746 | Eukaryota | 2361.0 | 0 | Chordata | 2361.0 | 0 | Carnivora | 2361.0 | 0 | Canidae | 2361.0 | 0 | Canis | 1943.0 | 1 | Canis lupus | 1943.0 | 1 |
| scf7180000047078 | 29613 | 0.4201 | 0 | 57.4733 | Eukaryota | 12161.0 | 0 | Chordata | 12161.0 | 0 | Carnivora | 5808.0 | 3 | Canidae | 5808.0 | 3 | Canis | 5808.0 | 3 | Canis lupus | 5808.0 | 3 |
| scf7180000041925 | 419868 | 0.4219 | 0 | 89.5084 | Eukaryota | 116692.0 | 0 | Chordata | 116692.0 | 0 | Carnivora | 116692.0 | 0 | Canidae | 116692.0 | 0 | Canis | 116692.0 | 0 | Canis lupus | 116692.0 | 0 |
| scf7180000047166 | 26210 | 0.4381 | 0 | 145.2545 | Eukaryota | 1742.0 | 0 | Platyhelminthes | 936.0 | 2 | Polystomatidea | 936.0 | 2 | Polystomatidae | 936.0 | 2 | Protopolystoma | 936.0 | 2 | Protopolystoma xenopodis | 936.0 | 2 |
| scf7180000047082 | 81283 | 0.4401 | 0 | 82.0498 | Eukaryota | 12839.0 | 0 | Chordata | 12233.0 | 1 | Kurtiformes | 5642.0 | 3 | Apogonidae | 5642.0 | 3 | Sphaeramia | 5642.0 | 3 | Sphaeramia orbicularis | 5642.0 | 3 |
| scf7180000042707 | 560558 | 0.4492 | 0 | 59.7 | Eukaryota | 67662.0 | 0 | Chordata | 67662.0 | 0 | Carnivora | 67662.0 | 0 | Canidae | 67662.0 | 0 | Canis | 67662.0 | 0 | Canis lupus | 67662.0 | 0 |
| scf7180000042738 | 612178 | 0.3752 | 0 | 60.119 | Eukaryota | 85394.0 | 0 | Chordata | 85394.0 | 0 | Carnivora | 85394.0 | 0 | Canidae | 85394.0 | 0 | Canis | 85394.0 | 0 | Canis lupus | 85394.0 | 0 |
| scf7180000054764 | 8496 | 0.4409 | 0 | 75.3456 | Eukaryota | 1410.0 | 0 | Platyhelminthes | 750.0 | 1 | Polystomatidea | 750.0 | 1 | Polystomatidae | 750.0 | 1 | Protopolystoma | 750.0 | 1 | Protopolystoma xenopodis | 750.0 | 1 |
| scf7180000043836 | 16172 | 0.3997 | 0 | 48.272 | Eukaryota | 10000.0 | 0 | Chordata | 10000.0 | 0 | Apterygiformes | 7997.0 | 2 | Apterygidae | 7997.0 | 2 | Apteryx | 7997.0 | 2 | Apteryx mantelli | 7997.0 | 2 |
| scf7180000045942 | 162502 | 0.3456 | 0 | 59.8478 | Eukaryota | 19838.0 | 0 | Chordata | 19838.0 | 0 | Rodentia | 7188.0 | 4 | Muridae | 7188.0 | 4 | Mus | 7188.0 | 5 | Mus musculus | 7188.0 | 5 |
| scf7180000049676 | 4971 | 0.4506 | 0 | 412.3914 | Eukaryota | 182.0 | 0 | Streptophyta | 182.0 | 0 | Fabales | 182.0 | 0 | Fabaceae | 182.0 | 0 | Cicer | 182.0 | 0 | Cicer arietinum | 182.0 | 0 |
| scf7180000045868 | 151122 | 0.4679 | 0 | 59.2508 | Eukaryota | 23219.0 | 0 | Chordata | 23219.0 | 0 | Carnivora | 23219.0 | 0 | Canidae | 12483.0 | 2 | Vulpes | 10584.0 | 4 | Vulpes vulpes | 10584.0 | 5 |
| scf7180000046777 | 121905 | 0.3408 | 0 | 58.8818 | Eukaryota | 18997.0 | 0 | Chordata | 18997.0 | 0 | Carnivora | 18997.0 | 0 | Canidae | 18997.0 | 0 | Canis | 18997.0 | 0 | Canis lupus | 18997.0 | 0 |
| scf7180000042810 | 552494 | 0.4268 | 0 | 59.3739 | Eukaryota | 50886.0 | 0 | Chordata | 50886.0 | 0 | Carnivora | 50886.0 | 0 | Canidae | 50886.0 | 0 | Canis | 30598.0 | 1 | Canis lupus | 30598.0 | 1 |
| scf7180000046508 | 125461 | 0.4221 | 0 | 175.8812 | Eukaryota | 20180.0 | 0 | Chordata | 16263.0 | 1 | Carnivora | 7788.0 | 4 | Gadidae | 4241.0 | 7 | Gadus | 4241.0 | 8 | Gadus morhua | 4241.0 | 8 |
| scf7180000040615 | 575919 | 0.3823 | 0 | 59.493 | Eukaryota | 73829.0 | 0 | Chordata | 73829.0 | 0 | Carnivora | 73829.0 | 0 | Canidae | 73829.0 | 0 | Canis | 73829.0 | 0 | Canis lupus | 73829.0 | 0 |
| scf7180000043281 | 424340 | 0.3547 | 0 | 60.0272 | Eukaryota | 83133.0 | 0 | Chordata | 83133.0 | 0 | Carnivora | 83133.0 | 0 | Canidae | 83133.0 | 0 | Canis | 83133.0 | 0 | Canis lupus | 83133.0 | 0 |
| scf7180000046223 | 3344 | 0.5072 | 0 | 57.4674 | Eukaryota | 7089.0 | 0 | Chordata | 7089.0 | 0 | Carnivora | 7089.0 | 0 | Canidae | 7089.0 | 0 | Canis | 4586.0 | 1 | Canis lupus | 3582.0 | 2 |
| scf7180000040860 | 928600 | 0.4345 | 0 | 60.3254 | Eukaryota | 58317.0 | 0 | Chordata | 58317.0 | 0 | Carnivora | 58317.0 | 0 | Canidae | 58317.0 | 0 | Canis | 58317.0 | 0 | Canis lupus | 58317.0 | 0 |
| scf7180000055867 | 28949 | 0.4623 | 0 | 59.5825 | Eukaryota | 90950.0 | 0 | Chordata | 90950.0 | 0 | Carnivora | 90950.0 | 0 | Canidae | 90950.0 | 0 | Canis | 90950.0 | 0 | Canis lupus | 90950.0 | 0 |
| scf7180000051154 | 2494 | 0.4503 | 0 | 568.2748 | no-hit | 0.0 | 0 | no-hit | 0.0 | 0 | no-hit | 0.0 | 0 | no-hit | 0.0 | 0 | no-hit | 0.0 | 0 | no-hit | 0.0 | 0 |
| scf7180000048967 | 57454 | 0.3747 | 0 | 60.3419 | Eukaryota | 12977.0 | 0 | Chordata | 12977.0 | 0 | Carnivora | 12977.0 | 0 | Canidae | 12977.0 | 0 | Canis | 12977.0 | 0 | Canis lupus | 12977.0 | 0 |
| scf7180000053317 | 3886 | 0.4241 | 0 | 25.8812 | Eukaryota | 8032.0 | 0 | Chordata | 8032.0 | 0 | Carnivora | 8032.0 | 0 | Canidae | 8032.0 | 0 | Canis | 8032.0 | 0 | Canis lupus | 8032.0 | 0 |
| scf7180000050742 | 25003 | 0.4467 | 0 | 60.4102 | Eukaryota | 14203.0 | 0 | Chordata | 14203.0 | 0 | Carnivora | 14203.0 | 0 | Canidae | 12969.0 | 1 | Canis | 12969.0 | 1 | Canis lupus | 12969.0 | 1 |
| scf7180000049411 | 11138 | 0.4404 | 0 | 64.3302 | Eukaryota | 21443.0 | 0 | Chordata | 19372.0 | 1 | Gadiformes | 14940.0 | 6 | Gadidae | 14940.0 | 6 | Gadus | 14940.0 | 6 | Gadus morhua | 14940.0 | 6 |
| scf7180000050827 | 3833 | 0.4388 | 0 | 65.6146 | no-hit | 0.0 | 0 | no-hit | 0.0 | 0 | no-hit | 0.0 | 0 | no-hit | 0.0 | 0 | no-hit | 0.0 | 0 | no-hit | 0.0 | 0 |
| scf7180000051455 | 14139 | 0.5048 | 0 | 400.0076 | Eukaryota | 9209.0 | 0 | Chordata | 9209.0 | 0 | Carnivora | 9209.0 | 0 | Canidae | 9209.0 | 0 | Canis | 7959.0 | 1 | Canis lupus | 6649.0 | 2 |
| scf7180000054001 | 6859 | 0.5007 | 0 | 311.8898 | Eukaryota | 8902.0 | 0 | Chordata | 8902.0 | 0 | Carnivora | 8902.0 | 0 | Canidae | 8902.0 | 0 | Canis | 7179.0 | 1 | Canis lupus | 5931.0 | 2 |
| scf7180000053012 | 3459 | 0.4516 | 0 | 419.2903 | Eukaryota | 1839.0 | 0 | Platyhelminthes | 745.0 | 2 | Polystomatidea | 745.0 | 2 | Polystomatidae | 745.0 | 2 | Protopolystoma | 745.0 | 2 | Protopolystoma xenopodis | 745.0 | 2 |
| scf7180000053501 | 14876 | 0.4372 | 0 | 196.4492 | Eukaryota | 3570.0 | 0 | Platyhelminthes | 2071.0 | 3 | Polystomatidea | 2071.0 | 3 | Polystomatidae | 2071.0 | 3 | Protopolystoma | 2071.0 | 3 | Protopolystoma xenopodis | 2071.0 | 3 |
| scf7180000041975 | 102429 | 0.3986 | 0 | 81.6711 | Eukaryota | 84752.0 | 0 | Chordata | 84752.0 | 0 | Carnivora | 84752.0 | 0 | Canidae | 84752.0 | 0 | Canis | 84752.0 | 0 | Canis lupus | 84752.0 | 0 |
| scf7180000045053 | 274899 | 0.3456 | 0 | 57.5729 | Eukaryota | 39678.0 | 0 | Chordata | 39678.0 | 0 | Carnivora | 39678.0 | 0 | Canidae | 39678.0 | 0 | Canis | 39678.0 | 0 | Canis lupus | 39678.0 | 0 |
| scf7180000040735 | 6815 | 0.4386 | 0 | 321.5974 | Eukaryota | 1529.0 | 0 | Platyhelminthes | 875.0 | 1 | Polystomatidea | 875.0 | 1 | Polystomatidae | 875.0 | 1 | Protopolystoma | 875.0 | 1 | Protopolystoma xenopodis | 875.0 | 1 |
| scf7180000046023 | 170662 | 0.3638 | 0 | 57.6381 | Eukaryota | 67105.0 | 0 | Chordata | 67105.0 | 0 | Carnivora | 67105.0 | 0 | Canidae | 67105.0 | 0 | Canis | 67105.0 | 0 | Canis lupus | 67105.0 | 0 |
| scf7180000044945 | 4031 | 0.3746 | 0 | 22.2389 | Eukaryota | 2972.0 | 0 | Chordata | 2972.0 | 0 | Carnivora | 2972.0 | 0 | Canidae | 2972.0 | 0 | Canis | 2972.0 | 0 | Canis lupus | 2972.0 | 0 |
| scf7180000048662 | 5156 | 0.4517 | 0 | 594.0702 | Eukaryota | 3699.0 | 0 | Platyhelminthes | 1951.0 | 3 | Polystomatidea | 1951.0 | 3 | Polystomatidae | 1951.0 | 3 | Protopolystoma | 1951.0 | 3 | Protopolystoma xenopodis | 1951.0 | 3 |
| scf7180000055753 | 22270 | 0.4455 | 0 | 384.9053 | Eukaryota | 3141.0 | 0 | Chordata | 3141.0 | 0 | Carnivora | 3141.0 | 0 | Canidae | 3141.0 | 0 | Canis | 3141.0 | 0 | Canis lupus | 3141.0 | 0 |
| scf7180000041065 | 1451460 | 0.374 | 0 | 60.997 | Eukaryota | 56501.0 | 0 | Chordata | 56501.0 | 0 | Carnivora | 56501.0 | 0 | Canidae | 42584.0 | 2 | Canis | 42584.0 | 2 | Canis lupus | 42584.0 | 2 |
| scf7180000042966 | 34087 | 0.362 | 0 | 56.598 | Eukaryota | 86513.0 | 0 | Chordata | 86513.0 | 0 | Carnivora | 86513.0 | 0 | Canidae | 86513.0 | 0 | Canis | 86513.0 | 0 | Canis lupus | 86513.0 | 0 |
| scf7180000041404 | 2597 | 0.4655 | 0 | 22.8956 | Eukaryota | 4530.0 | 0 | Chordata | 4530.0 | 0 | Carnivora | 4530.0 | 0 | Canidae | 4530.0 | 0 | Canis | 2718.0 | 1 | Canis lupus | 2718.0 | 1 |
| scf7180000054336 | 30805 | 0.452 | 0 | 93.1311 | Eukaryota | 15923.0 | 0 | Chordata | 15923.0 | 0 | Carnivora | 14617.0 | 2 | Canidae | 14617.0 | 3 | Vulpes | 14617.0 | 3 | Vulpes vulpes | 14617.0 | 3 |
| scf7180000042979 | 24219 | 0.4185 | 0 | 56.5799 | Eukaryota | 61325.0 | 0 | Chordata | 61325.0 | 0 | Carnivora | 61325.0 | 0 | Canidae | 61325.0 | 0 | Canis | 61325.0 | 0 | Canis lupus | 61325.0 | 0 |
| scf7180000041284 | 480879 | 0.4193 | 0 | 59.2269 | Eukaryota | 66195.0 | 0 | Chordata | 66195.0 | 0 | Carnivora | 66195.0 | 0 | Canidae | 63005.0 | 1 | Canis | 56996.0 | 2 | Canis lupus | 56996.0 | 2 |
| scf7180000041240 | 1360829 | 0.4181 | 0 | 59.4115 | Eukaryota | 90827.0 | 0 | Chordata | 90827.0 | 0 | Carnivora | 90827.0 | 0 | Canidae | 90827.0 | 0 | Canis | 90827.0 | 0 | Canis lupus | 90827.0 | 0 |
| scf7180000049655 | 18786 | 0.4355 | 0 | 234.5439 | Eukaryota | 328.0 | 0 | Chordata | 328.0 | 0 | Salmoniformes | 328.0 | 0 | Salmonidae | 328.0 | 0 | Salmo | 185.0 | 1 | Salmo trutta | 185.0 | 1 |
| scf7180000045568 | 29292 | 0.4383 | 0 | 136.5318 | Eukaryota | 4911.0 | 0 | Chordata | 4911.0 | 0 | Carnivora | 2198.0 | 2 | Canidae | 2198.0 | 2 | Canis | 2198.0 | 2 | Canis lupus | 2198.0 | 3 |
| scf7180000042486 | 812401 | 0.4521 | 0 | 59.5242 | Eukaryota | 70201.0 | 0 | Chordata | 70201.0 | 0 | Carnivora | 70201.0 | 0 | Canidae | 70201.0 | 0 | Canis | 63660.0 | 1 | Canis lupus | 63660.0 | 1 |
| scf7180000046997 | 85528 | 0.4534 | 0 | 139.0752 | Eukaryota | 3158.0 | 0 | Chordata | 3158.0 | 0 | Carnivora | 3158.0 | 0 | Canidae | 3158.0 | 0 | Canis | 2840.0 | 1 | Canis lupus | 2840.0 | 1 |
| scf7180000046093 | 134723 | 0.3363 | 0 | 60.0775 | Eukaryota | 10977.0 | 0 | Chordata | 10977.0 | 0 | Carnivora | 10977.0 | 0 | Canidae | 10977.0 | 0 | Canis | 10977.0 | 0 | Canis lupus | 10977.0 | 0 |
| scf7180000055871 | 10006 | 0.4327 | 0 | 201.9511 | no-hit | 0.0 | 0 | no-hit | 0.0 | 0 | no-hit | 0.0 | 0 | no-hit | 0.0 | 0 | no-hit | 0.0 | 0 | no-hit | 0.0 | 0 |
| scf7180000043653 | 88881 | 0.4369 | 0 | 54.3339 | Eukaryota | 17700.0 | 0 | Chordata | 17700.0 | 0 | Carnivora | 12206.0 | 2 | Canidae | 9448.0 | 3 | Canis | 7521.0 | 5 | Canis lupus | 7521.0 | 5 |
| scf7180000043007 | 32356 | 0.3856 | 0 | 159.5601 | Eukaryota | 91162.0 | 0 | Chordata | 91162.0 | 0 | Carnivora | 91162.0 | 0 | Canidae | 91162.0 | 0 | Canis | 91162.0 | 0 | Canis lupus | 91162.0 | 0 |
| scf7180000052490 | 7064 | 0.5093 | 0 | 81.4655 | Eukaryota | 7979.0 | 0 | Chordata | 7979.0 | 0 | Carnivora | 7979.0 | 0 | Canidae | 7979.0 | 0 | Canis | 6407.0 | 1 | Canis lupus | 5287.0 | 2 |
| scf7180000052456 | 7950 | 0.4367 | 0 | 49.6214 | Eukaryota | 285.0 | 0 | Chordata | 285.0 | 0 | Salmoniformes | 285.0 | 0 | Salmonidae | 285.0 | 0 | Coregonus | 285.0 | 0 | Coregonus sp. 'balchen' | 285.0 | 0 |
| scf7180000040620 | 2067040 | 0.3829 | 0 | 60.7351 | Eukaryota | 88883.0 | 0 | Chordata | 88883.0 | 0 | Carnivora | 88883.0 | 0 | Canidae | 88883.0 | 0 | Canis | 88883.0 | 0 | Canis lupus | 88883.0 | 0 |

| | | | | | | | | | | | | | | | | | | | |
|---|---|---|---|---|---|---|---|---|---|---|---|---|---|---|---|---|---|---|---|
| scf7180000053634 | 3797 | 0.4475 | 0 | 268.5104 | no-hit | 0.0 | 0 | no-hit | 0.0 | 0 | no-hit | 0.0 | 0 | no-hit | 0.0 | 0 | no-hit | 0.0 | 0 | no-hit | 0.0 | 0 |
| scf7180000049286 | 8024 | 0.4518 | 0 | 660.1281 | Eukaryota | 3273.0 | 0 | Platyhelminthes | 1707.0 | 3 | Polystomatidea | 1707.0 | 3 | Polystomatidae | 1707.0 | 3 | Protopolystoma | 1707.0 | 3 | Protopolystoma xenopodis | 1707.0 | 3 |
| scf7180000043660 | 506179 | 0.3646 | 0 | 59.4861 | Eukaryota | 48511.0 | 0 | Chordata | 48511.0 | 0 | Carnivora | 48511.0 | 0 | Canidae | 48511.0 | 0 | Canis | 48511.0 | 0 | Canis lupus | 48511.0 | 0 |
| scf7180000042672 | 514167 | 0.427 | 0 | 58.9063 | Eukaryota | 89802.0 | 0 | Chordata | 89802.0 | 0 | Carnivora | 89802.0 | 0 | Canidae | 89802.0 | 0 | Canis | 53596.0 | 1 | Canis lupus | 53596.0 | 1 |
| scf7180000045809 | 19584 | 0.3917 | 0 | 84.806 | Eukaryota | 81872.0 | 0 | Chordata | 81872.0 | 0 | Carnivora | 81872.0 | 0 | Canidae | 81872.0 | 0 | Canis | 81872.0 | 0 | Canis lupus | 81872.0 | 0 |
| scf7180000055914 | 6003 | 0.4408 | 0 | 465.2323 | Eukaryota | 270.0 | 0 | Chordata | 270.0 | 0 | Tetraodontiformes | 270.0 | 0 | Tetraodontidae | 270.0 | 0 | Takifugu | 270.0 | 0 | Takifugu rubripes | 270.0 | 0 |
| scf7180000040597 | 1597170 | 0.4912 | 0 | 59.0155 | Eukaryota | 102516.0 | 0 | Chordata | 102516.0 | 0 | Carnivora | 102516.0 | 0 | Canidae | 102516.0 | 0 | Canis | 102516.0 | 0 | Canis lupus | 102516.0 | 0 |
| scf7180000046418 | 15247 | 0.4384 | 0 | 240.0535 | no-hit | 0.0 | 0 | no-hit | 0.0 | 0 | no-hit | 0.0 | 0 | no-hit | 0.0 | 0 | no-hit | 0.0 | 0 | no-hit | 0.0 | 0 |
| scf7180000043284 | 494267 | 0.4916 | 0 | 58.8059 | Eukaryota | 14768.0 | 0 | Chordata | 14768.0 | 0 | Carnivora | 10260.0 | 2 | Canidae | 10260.0 | 2 | Canis | 10260.0 | 2 | Canis lupus | 10260.0 | 2 |
| scf7180000041891 | 882920 | 0.3823 | 0 | 60.8128 | Eukaryota | 84254.0 | 0 | Chordata | 84254.0 | 0 | Carnivora | 84254.0 | 0 | Canidae | 78715.0 | 1 | Canis | 60991.0 | 2 | Canis lupus | 60991.0 | 2 |
| scf7180000043306 | 189364 | 0.4284 | 0 | 58.3506 | Eukaryota | 17923.0 | 0 | Chordata | 17923.0 | 0 | Primates | 8689.0 | 2 | Hominidae | 6729.0 | 5 | Canis | 3086.0 | 8 | Canis lupus | 3086.0 | 8 |
| scf7180000045900 | 142215 | 0.4694 | 0 | 57.768 | Eukaryota | 26068.0 | 0 | Chordata | 26068.0 | 0 | Carnivora | 26068.0 | 0 | Canidae | 26068.0 | 0 | Canis | 16752.0 | 1 | Canis lupus | 16752.0 | 1 |
| scf7180000056331 | 577834 | 0.4643 | 0 | 60.0506 | Eukaryota | 92695.0 | 0 | Chordata | 92695.0 | 0 | Carnivora | 92695.0 | 0 | Canidae | 92695.0 | 0 | Canis | 52972.0 | 1 | Canis lupus | 52972.0 | 1 |
| scf7180000056018 | 5542 | 0.4894 | 0 | 37.7509 | Eukaryota | 7222.0 | 0 | Chordata | 7222.0 | 0 | Carnivora | 7222.0 | 0 | Canidae | 7222.0 | 0 | Canis | 5752.0 | 1 | Canis lupus | 4713.0 | 2 |
| scf7180000053469 | 2152 | 0.5107 | 0 | 64.4499 | Eukaryota | 6240.0 | 0 | Chordata | 6240.0 | 0 | Carnivora | 6240.0 | 0 | Canidae | 6240.0 | 0 | Canis | 4303.0 | 1 | Canis lupus | 3301.0 | 2 |
| scf7180000053406 | 8313 | 0.4409 | 0 | 77.912 | Eukaryota | 619.0 | 0 | Streptophyta | 619.0 | 0 | Fabales | 619.0 | 0 | Fabaceae | 619.0 | 0 | Cicer | 619.0 | 0 | Cicer arietinum | 619.0 | 0 |
| scf7180000045155 | 221340 | 0.5838 | 0 | 56.1484 | Eukaryota | 70392.0 | 0 | Chordata | 70392.0 | 0 | Carnivora | 70392.0 | 0 | Canidae | 70392.0 | 0 | Vulpes | 35285.0 | 1 | Vulpes vulpes | 35285.0 | 1 |
| scf7180000048649 | 7954 | 0.4443 | 0 | 75.0163 | Eukaryota | 3081.0 | 0 | Chordata | 3081.0 | 0 | Carnivora | 2353.0 | 1 | Canidae | 2353.0 | 1 | Canis | 2353.0 | 1 | Canis lupus | 2353.0 | 1 |
| scf7180000051180 | 18245 | 0.5051 | 0 | 419.7219 | Eukaryota | 9429.0 | 0 | Chordata | 9429.0 | 0 | Carnivora | 9429.0 | 0 | Canidae | 9429.0 | 0 | Canis | 8177.0 | 1 | Canis lupus | 6870.0 | 2 |
| scf7180000040900 | 1740908 | 0.3559 | 0 | 58.9268 | Eukaryota | 93377.0 | 0 | Chordata | 93377.0 | 0 | Carnivora | 93377.0 | 0 | Canidae | 93377.0 | 0 | Canis | 93377.0 | 0 | Canis lupus | 93377.0 | 0 |
| scf7180000049611 | 7683 | 0.4423 | 0 | 46.2824 | Eukaryota | 21578.0 | 0 | Chordata | 15257.0 | 2 | Gadiformes | 11492.0 | 5 | Gadidae | 11492.0 | 5 | Gadus | 11492.0 | 5 | Gadus morhua | 11492.0 | 5 |
| scf7180000047510 | 18266 | 0.4234 | 0 | 44.7824 | Eukaryota | 21585.0 | 1 | Chordata | 19468.0 | 2 | Gadiformes | 11441.0 | 5 | Gadidae | 11441.0 | 5 | Gadus | 11441.0 | 5 | Gadus morhua | 11441.0 | 5 |
| scf7180000045221 | 220314 | 0.3925 | 0 | 57.2927 | Eukaryota | 90708.0 | 0 | Chordata | 90708.0 | 0 | Carnivora | 90708.0 | 0 | Canidae | 90708.0 | 0 | Canis | 90708.0 | 0 | Canis lupus | 90708.0 | 0 |
| scf7180000041641 | 950938 | 0.376 | 0 | 59.3111 | Eukaryota | 68480.0 | 0 | Chordata | 68480.0 | 0 | Carnivora | 68480.0 | 0 | Canidae | 53297.0 | 2 | Canis | 47267.0 | 3 | Canis lupus | 47267.0 | 4 |
| scf7180000053410 | 1518 | 0.4592 | 0 | 85.6251 | no-hit | 0.0 | 0 | no-hit | 0.0 | 0 | no-hit | 0.0 | 0 | no-hit | 0.0 | 0 | no-hit | 0.0 | 0 | no-hit | 0.0 | 0 |
| scf7180000051267 | 7718 | 0.4438 | 0 | 75.9953 | Eukaryota | 12479.0 | 0 | Chordata | 7664.0 | 3 | Gadiformes | 4314.0 | 7 | Gadidae | 4314.0 | 7 | Gadus | 4314.0 | 7 | Gadus morhua | 4314.0 | 7 |
| scf7180000042964 | 491093 | 0.3544 | 0 | 60.441 | Eukaryota | 37212.0 | 0 | Chordata | 37212.0 | 0 | Carnivora | 37212.0 | 0 | Canidae | 37212.0 | 0 | Canis | 33326.0 | 1 | Canis lupus | 33326.0 | 1 |
| scf7180000053900 | 3743 | 0.5052 | 0 | 443.6498 | Eukaryota | 7061.0 | 0 | Chordata | 7061.0 | 0 | Carnivora | 7061.0 | 0 | Canidae | 7061.0 | 0 | Canis | 5349.0 | 1 | Canis lupus | 4456.0 | 2 |
| scf7180000042142 | 473813 | 0.3527 | 0 | 60.2129 | Eukaryota | 79738.0 | 0 | Chordata | 79738.0 | 0 | Carnivora | 79738.0 | 0 | Canidae | 79738.0 | 0 | Canis | 79738.0 | 0 | Canis lupus | 79738.0 | 0 |
| scf7180000040675 | 1966658 | 0.3547 | 0 | 60.1352 | Eukaryota | 101419.0 | 0 | Chordata | 101419.0 | 0 | Carnivora | 101419.0 | 0 | Canidae | 101419.0 | 0 | Canis | 74827.0 | 1 | Canis lupus | 74827.0 | 1 |
| scf7180000050101 | 23720 | 0.4428 | 0 | 59.0317 | Eukaryota | 26771.0 | 0 | Chordata | 26771.0 | 0 | Carnivora | 26771.0 | 0 | Canidae | 16533.0 | 4 | Canis | 14294.0 | 5 | Canis lupus | 14294.0 | 6 |
| scf7180000042033 | 405172 | 0.4 | 0 | 59.3498 | Eukaryota | 90195.0 | 0 | Chordata | 90195.0 | 0 | Carnivora | 90195.0 | 0 | Canidae | 90195.0 | 0 | Canis | 90195.0 | 0 | Canis lupus | 90195.0 | 0 |
| scf7180000043065 | 39120 | 0.3579 | 0 | 66.9004 | Eukaryota | 65363.0 | 0 | Chordata | 65363.0 | 0 | Carnivora | 65363.0 | 0 | Canidae | 65363.0 | 0 | Canis | 65363.0 | 0 | Canis lupus | 65363.0 | 0 |
| scf7180000055880 | 7518 | 0.4705 | 0 | 202.5706 | Eukaryota | 17764.0 | 0 | Chordata | 17764.0 | 0 | Carnivora | 17764.0 | 0 | Canidae | 17764.0 | 0 | Canis | 17764.0 | 0 | Canis lupus | 17764.0 | 0 |
| scf7180000054544 | 22002 | 0.4361 | 0 | 56.4326 | Eukaryota | 88883.0 | 0 | Chordata | 88883.0 | 0 | Carnivora | 88883.0 | 0 | Canidae | 88883.0 | 0 | Canis | 88883.0 | 0 | Canis lupus | 88883.0 | 0 |
| scf7180000045110 | 271911 | 0.4821 | 0 | 58.3495 | Eukaryota | 119404.0 | 0 | Chordata | 119404.0 | 0 | Carnivora | 119404.0 | 0 | Canidae | 119404.0 | 0 | Canis | 119404.0 | 0 | Canis lupus | 119404.0 | 0 |
| scf7180000051829 | 3518 | 0.4417 | 0 | 232.0733 | no-hit | 0.0 | 0 | no-hit | 0.0 | 0 | no-hit | 0.0 | 0 | no-hit | 0.0 | 0 | no-hit | 0.0 | 0 | no-hit | 0.0 | 0 |
| scf7180000049600 | 4151 | 0.4423 | 0 | 480.0963 | no-hit | 0.0 | 0 | no-hit | 0.0 | 0 | no-hit | 0.0 | 0 | no-hit | 0.0 | 0 | no-hit | 0.0 | 0 | no-hit | 0.0 | 0 |
| scf7180000040888 | 1725053 | 0.3529 | 0 | 59.5632 | Eukaryota | 81720.0 | 0 | Chordata | 81720.0 | 0 | Carnivora | 81720.0 | 0 | Canidae | 81720.0 | 0 | Canis | 81720.0 | 0 | Canis lupus | 81720.0 | 0 |
| scf7180000042550 | 471624 | 0.3881 | 0 | 59.9453 | Eukaryota | 93988.0 | 0 | Chordata | 93988.0 | 0 | Carnivora | 93988.0 | 0 | Canidae | 93988.0 | 0 | Canis | 93988.0 | 0 | Canis lupus | 93988.0 | 0 |
| scf7180000053171 | 2183 | 0.4475 | 0 | 163.3068 | Eukaryota | 3150.0 | 0 | Chordata | 1639.0 | 3 | Polystomatidea | 664.0 | 6 | Polystomatidae | 664.0 | 6 | Protopolystoma | 664.0 | 6 | Protopolystoma xenopodis | 664.0 | 6 |
| scf7180000046315 | 121455 | 0.4973 | 0 | 55.4887 | Eukaryota | 46119.0 | 0 | Chordata | 46119.0 | 0 | Carnivora | 46119.0 | 0 | Canidae | 21513.0 | 2 | Zalophus | 18720.0 | 3 | Zalophus californianus | 18720.0 | 4 |
| scf7180000048777 | 10298 | 0.4494 | 0 | 168.2017 | no-hit | 0.0 | 0 | no-hit | 0.0 | 0 | no-hit | 0.0 | 0 | no-hit | 0.0 | 0 | no-hit | 0.0 | 0 | no-hit | 0.0 | 0 |
| scf7180000051436 | 10816 | 0.4311 | 0 | 38.5419 | Eukaryota | 4540.0 | 0 | Chordata | 4540.0 | 0 | Carnivora | 4540.0 | 0 | Canidae | 4540.0 | 0 | Canis | 4540.0 | 0 | Canis lupus | 4540.0 | 0 |
| scf7180000047917 | 46646 | 0.4554 | 0 | 214.8893 | no-hit | 0.0 | 0 | no-hit | 0.0 | 0 | no-hit | 0.0 | 0 | no-hit | 0.0 | 0 | no-hit | 0.0 | 0 | no-hit | 0.0 | 0 |
| scf7180000048848 | 7941 | 0.4474 | 0 | 1289.965 | Eukaryota | 2861.0 | 0 | Chordata | 2861.0 | 0 | Salmoniformes | 1280.0 | 2 | Salmonidae | 1280.0 | 2 | Salmo | 1280.0 | 2 | Salmo trutta | 1280.0 | 2 |
| scf7180000056593 | 186125 | 0.4756 | 0 | 57.912 | Eukaryota | 33463.0 | 0 | Chordata | 33463.0 | 0 | Carnivora | 33463.0 | 0 | Canidae | 33463.0 | 0 | Canis | 27714.0 | 1 | Canis lupus | 27714.0 | 1 |
| scf7180000048854 | 39526 | 0.4431 | 0 | 67.2172 | Eukaryota | 37790.0 | 1 | Chordata | 32832.0 | 2 | Gadiformes | 22505.0 | 5 | Gadidae | 22505.0 | 5 | Gadus | 22505.0 | 5 | Gadus morhua | 22505.0 | 5 |
| scf7180000056291 | 780822 | 0.4471 | 0 | 81.4655 | Eukaryota | 83700.0 | 0 | Chordata | 83700.0 | 0 | Carnivora | 83700.0 | 0 | Canidae | 83700.0 | 0 | Canis | 83700.0 | 0 | Canis lupus | 83700.0 | 0 |
| scf7180000040776 | 1935776 | 0.385 | 0 | 60.1931 | Eukaryota | 139649.0 | 0 | Chordata | 139649.0 | 0 | Carnivora | 139649.0 | 0 | Canidae | 139649.0 | 0 | Canis | 139649.0 | 0 | Canis lupus | 139649.0 | 0 |
| scf7180000052956 | 2788 | 0.4408 | 0 | 87.5949 | Eukaryota | 1088.0 | 0 | Echinodermata | 592.0 | 2 | Forcipulatida | 592.0 | 2 | Asteriidae | 592.0 | 2 | Asterias | 592.0 | 2 | Asterias rubens | 592.0 | 2 |
| scf7180000046206 | 22504 | 0.4587 | 0 | 49.953 | Eukaryota | 44302.0 | 0 | Chordata | 44302.0 | 0 | Carnivora | 44302.0 | 0 | Canidae | 44302.0 | 0 | Canis | 44302.0 | 0 | Canis lupus | 44302.0 | 0 |
| scf7180000048725 | 12889 | 0.5311 | 0 | 45.9909 | Eukaryota | 8779.0 | 0 | Chordata | 8779.0 | 0 | Carnivora | 8212.0 | 1 | Canidae | 7443.0 | 2 | Canis | 5946.0 | 3 | Canis lupus | 5946.0 | 3 |
| scf7180000056566 | 171974 | 0.412 | 0 | 61.2112 | Eukaryota | 37661.0 | 0 | Chordata | 37661.0 | 0 | Carnivora | 37661.0 | 0 | Canidae | 37661.0 | 0 | Canis | 37661.0 | 0 | Canis lupus | 37661.0 | 0 |
| scf7180000048732 | 2538 | 0.448 | 0 | 252.2208 | no-hit | 0.0 | 0 | no-hit | 0.0 | 0 | no-hit | 0.0 | 0 | no-hit | 0.0 | 0 | no-hit | 0.0 | 0 | no-hit | 0.0 | 0 |
| scf7180000053612 | 5043 | 0.4436 | 0 | 92.1315 | Eukaryota | 1909.0 | 0 | Echinodermata | 1607.0 | 1 | Forcipulatida | 1607.0 | 1 | Asteriidae | 1607.0 | 1 | Asterias | 1607.0 | 1 | Asterias rubens | 1607.0 | 1 |
| scf7180000047256 | 22926 | 0.4423 | 0 | 108.8044 | Eukaryota | 2404.0 | 1 | Chordata | 1330.0 | 2 | Polystomatidea | 1074.0 | 4 | Polystomatidae | 1074.0 | 4 | Protopolystoma | 1074.0 | 4 | Protopolystoma xenopodis | 1074.0 | 4 |
| scf7180000050396 | 12183 | 0.4585 | 0 | 575.3182 | Eukaryota | 44799.0 | 0 | Chordata | 44799.0 | 0 | Carnivora | 44799.0 | 0 | Canidae | 44799.0 | 0 | Canis | 44799.0 | 0 | Canis lupus | 44799.0 | 0 |
| scf7180000053322 | 10617 | 0.4435 | 0 | 126.0159 | Eukaryota | 566.0 | 0 | Platyhelminthes | 566.0 | 0 | Polystomatidea | 566.0 | 0 | Polystomatidae | 566.0 | 0 | Protopolystoma | 566.0 | 0 | Protopolystoma xenopodis | 566.0 | 0 |
| scf7180000044508 | 92122 | 0.4975 | 0 | 368.4637 | Eukaryota | 80693.0 | 0 | Chordata | 80693.0 | 0 | Carnivora | 80693.0 | 0 | Canidae | 80693.0 | 0 | Canis | 80693.0 | 0 | Canis lupus | 80693.0 | 0 |
| scf7180000047940 | 97625 | 0.4941 | 0 | 54.232 | Eukaryota | 20518.0 | 0 | Chordata | 20518.0 | 0 | Carnivora | 20518.0 | 0 | Canidae | 20518.0 | 0 | Canis | 17973.0 | 1 | Canis lupus | 17973.0 | 1 |
| scf7180000045426 | 8323 | 0.4427 | 0 | 141.7708 | Eukaryota | 2093.0 | 0 | Platyhelminthes | 1782.0 | 1 | Polystomatidea | 1782.0 | 1 | Polystomatidae | 1782.0 | 1 | Protopolystoma | 1782.0 | 1 | Protopolystoma xenopodis | 1782.0 | 1 |
| scf7180000054387 | 18881 | 0.4069 | 0 | 206.7852 | Eukaryota | 88528.0 | 0 | Chordata | 88528.0 | 0 | Carnivora | 88528.0 | 0 | Canidae | 88528.0 | 0 | Canis | 88528.0 | 0 | Canis lupus | 88528.0 | 0 |
| scf7180000047699 | 4590 | 0.4429 | 0 | 67.0104 | no-hit | 0.0 | 0 | no-hit | 0.0 | 0 | no-hit | 0.0 | 0 | no-hit | 0.0 | 0 | no-hit | 0.0 | 0 | no-hit | 0.0 | 0 |
| scf7180000043074 | 527383 | 0.4304 | 0 | 106.1404 | Eukaryota | 84683.0 | 0 | Chordata | 84683.0 | 0 | Carnivora | 84683.0 | 0 | Canidae | 84683.0 | 0 | Canis | 84683.0 | 0 | Canis lupus | 84683.0 | 0 |
| scf7180000049486 | 16041 | 0.4383 | 0 | 59.5268 | Eukaryota | 5233.0 | 0 | Platyhelminthes | 3272.0 | 1 | Polystomatidea | 3272.0 | 2 | Polystomatidae | 3272.0 | 2 | Protopolystoma | 3272.0 | 2 | Protopolystoma xenopodis | 3272.0 | 2 |
| scf7180000053538 | 2841 | 0.4555 | 0 | 205.9487 | Eukaryota | 774.0 | 0 | Platyhelminthes | 418.0 | 1 | Polystomatidea | 418.0 | 1 | Polystomatidae | 418.0 | 1 | Protopolystoma | 418.0 | 1 | Protopolystoma xenopodis | 418.0 | 1 |
| scf7180000051610 | 15178 | 0.3582 | 0 | 50.9446 | Eukaryota | 55695.0 | 0 | Chordata | 55695.0 | 0 | Carnivora | 55695.0 | 0 | Canidae | 55695.0 | 0 | Canis | 55695.0 | 0 | Canis lupus | 55695.0 | 0 |
| scf7180000042542 | 297644 | 0.3915 | 0 | 60.271 | Eukaryota | 44741.0 | 0 | Chordata | 44741.0 | 0 | Carnivora | 44741.0 | 0 | Canidae | 44741.0 | 0 | Canis | 30171.0 | 1 | Canis lupus | 30171.0 | 1 |
| scf7180000051921 | 15092 | 0.4421 | 0 | 67.3218 | Eukaryota | 11045.0 | 0 | Chordata | 10208.0 | 1 | Kurtiformes | 6156.0 | 4 | Apogonidae | 6156.0 | 4 | Sphaeramia | 6156.0 | 4 | Sphaeramia orbicularis | 6156.0 | 4 |
| scf7180000043204 | 284732 | 0.4256 | 0 | 60.1327 | Eukaryota | 28458.0 | 0 | Chordata | 28458.0 | 0 | Carnivora | 28458.0 | 0 | Canidae | 25628.0 | 1 | Canis | 25628.0 | 1 | Canis lupus | 25628.0 | 1 |
| scf7180000048534 | 2487 | 0.431 | 0 | 68.115 | no-hit | 0.0 | 0 | no-hit | 0.0 | 0 | no-hit | 0.0 | 0 | no-hit | 0.0 | 0 | no-hit | 0.0 | 0 | no-hit | 0.0 | 0 |
| scf7180000040995 | 1314766 | 0.4771 | 0 | 59.4226 | Eukaryota | 80392.0 | 0 | Chordata | 80392.0 | 0 | Carnivora | 80392.0 | 0 | Canidae | 80392.0 | 0 | Canis | 56176.0 | 1 | Canis lupus | 56176.0 | 1 |
| scf7180000053887 | 9867 | 0.5052 | 0 | 90.3919 | Eukaryota | 7089.0 | 0 | Chordata | 7089.0 | 0 | Carnivora | 7089.0 | 0 | Canidae | 7089.0 | 0 | Canis | 5335.0 | 1 | Canis lupus | 4440.0 | 2 |
| scf7180000043941 | 112165 | 0.4476 | 0 | 58.9432 | Eukaryota | 73967.0 | 0 | Chordata | 73967.0 | 0 | Carnivora | 73967.0 | 0 | Canidae | 73967.0 | 0 | Canis | 73967.0 | 0 | Canis lupus | 73967.0 | 0 |
| scf7180000047187 | 43938 | 0.4346 | 0 | 117.1982 | Eukaryota | 14941.0 | 0 | Chordata | 8020.0 | 3 | Kinetoplastida | 3245.0 | 6 | Trypanosomatidae | 3245.0 | 6 | Leishmania | 3245.0 | 7 | Leishmania braziliensis | 3245.0 | 7 |
| scf7180000056565 | 78079 | 0.4036 | 0 | 56.4652 | Eukaryota | 14408.0 | 0 | Chordata | 14408.0 | 0 | Carnivora | 14408.0 | 0 | Canidae | 14408.0 | 0 | Canis | 14408.0 | 0 | Canis lupus | 14408.0 | 0 |
| scf7180000053400 | 2732 | 0.4447 | 0 | 998.4866 | no-hit | 0.0 | 0 | no-hit | 0.0 | 0 | no-hit | 0.0 | 0 | no-hit | 0.0 | 0 | no-hit | 0.0 | 0 | no-hit | 0.0 | 0 |
| scf7180000042905 | 497188 | 0.5388 | 0 | 57.4922 | Eukaryota | 63974.0 | 0 | Chordata | 63974.0 | 0 | Carnivora | 63974.0 | 0 | Canidae | 63974.0 | 0 | Canis | 51256.0 | 1 | Canis lupus | 51256.0 | 1 |
| scf7180000043970 | 164367 | 0.4095 | 0 | 57.8452 | Eukaryota | 92223.0 | 0 | Chordata | 92223.0 | 0 | Carnivora | 92223.0 | 0 | Canidae | 92223.0 | 0 | Canis | 92223.0 | 0 | Canis lupus | 92223.0 | 0 |
| scf7180000042023 | 65599 | 0.3907 | 0 | 56.5864 | Eukaryota | 32955.0 | 0 | Chordata | 32955.0 | 0 | Carnivora | 32955.0 | 0 | Canidae | 32955.0 | 0 | Canis | 32955.0 | 0 | Canis lupus | 32955.0 | 0 |
| scf7180000041052 | 23760 | 0.4194 | 0 | 60.6902 | Eukaryota | 73320.0 | 0 | Chordata | 73320.0 | 0 | Carnivora | 73320.0 | 0 | Canidae | 73320.0 | 0 | Canis | 73320.0 | 0 | Canis lupus | 73320.0 | 0 |
| scf7180000053191 | 2624 | 0.4535 | 0 | 153.5128 | Eukaryota | 1974.0 | 0 | Platyhelminthes | 914.0 | 2 | Polystomatidea | 914.0 | 2 | Polystomatidae | 914.0 | 2 | Protopolystoma | 914.0 | 2 | Protopolystoma xenopodis | 914.0 | 2 |
| scf7180000054498 | 61286 | 0.439 | 0 | 237.9055 | Eukaryota | 78018.0 | 0 | Chordata | 78018.0 | 0 | Carnivora | 78018.0 | 0 | Canidae | 78018.0 | 0 | Canis | 78018.0 | 0 | Canis lupus | 78018.0 | 0 |
| scf7180000042815 | 680348 | 0.3487 | 0 | 60.3699 | Eukaryota | 54706.0 | 0 | Chordata | 54706.0 | 0 | Carnivora | 54706.0 | 0 | Canidae | 54706.0 | 0 | Canis | 54706.0 | 0 | Canis lupus | 54706.0 | 0 |
| scf7180000056316 | 766366 | 0.4231 | 0 | 58.5999 | Eukaryota | 27366.0 | 0 | Chordata | 27366.0 | 0 | Carnivora | 27366.0 | 0 | Canidae | 27366.0 | 0 | Canis | 27366.0 | 0 | Canis lupus | 27366.0 | 0 |
| scf7180000043489 | 492587 | 0.4394 | 0 | 58.6771 | Eukaryota | 14546.0 | 0 | Chordata | 14546.0 | 0 | Carnivora | 14546.0 | 0 | Canidae | 14546.0 | 0 | Canis | 13221.0 | 1 | Canis lupus | 13221.0 | 1 |
| scf7180000047297 | 59847 | 0.6097 | 0 | 55.7877 | Eukaryota | 5888.0 | 0 | Chordata | 5888.0 | 0 | Carnivora | 5888.0 | 0 | Canidae | 3907.0 | 1 | Canis | 3907.0 | 1 | Canis lupus | 3907.0 | 2 |
| scf7180000046059 | 13880 | 0.4137 | 0 | 47.0661 | Eukaryota | 83910.0 | 0 | Chordata | 83910.0 | 0 | Carnivora | 83910.0 | 0 | Canidae | 83910.0 | 0 | Canis | 83910.0 | 0 | Canis lupus | 83910.0 | 0 |
| scf7180000041363 | 1205948 | 0.3842 | 0 | 60.4338 | Eukaryota | 74988.0 | 0 | Chordata | 74988.0 | 0 | Carnivora | 74988.0 | 0 | Canidae | 74988.0 | 0 | Canis | 68581.0 | 1 | Canis lupus | 68581.0 | 1 |
| scf7180000052855 | 5624 | 0.4422 | 0 | 84.3025 | Eukaryota | 4601.0 | 0 | Chordata | 3309.0 | 1 | Kurtiformes | 2718.0 | 3 | Apogonidae | 2718.0 | 3 | Sphaeramia | 2718.0 | 3 | Sphaeramia orbicularis | 2718.0 | 3 |
| scf7180000041825 | 888221 | 0.4134 | 0 | 59.7434 | Eukaryota | 88456.0 | 0 | Chordata | 88456.0 | 0 | Carnivora | 88456.0 | 0 | Canidae | 88456.0 | 0 | Canis | 88456.0 | 0 | Canis lupus | 88456.0 | 0 |
| scf7180000043910 | 363146 | 0.3557 | 0 | 59.1303 | Eukaryota | 59746.0 | 0 | Chordata | 59746.0 | 0 | Carnivora | 59746.0 | 0 | Canidae | 23928.0 | 3 | Canis | 16351.0 | 6 | Canis lupus | 16351.0 | 7 |
| scf7180000046651 | 115623 | 0.3549 | 0 | 58.5374 | Eukaryota | 7372.0 | 0 | Chordata | 7372.0 | 0 | Artiodactyla | 3212.0 | 2 | Bovidae | 2024.0 | 4 | Felis | 1661.0 | 5 | Felis catus | 1661.0 | 5 |
| scf7180000046599 | 124162 | 0.4346 | 0 | 59.9209 | Eukaryota | 20474.0 | 0 | Chordata | 20474.0 | 0 | Carnivora | 16375.0 | 1 | Canidae | 12086.0 | 3 | Canis | 12086.0 | 3 | Canis lupus | 12086.0 | 4 |
| scf7180000046487 | 115448 | 0.3645 | 0 | 59.6235 | Eukaryota | 17217.0 | 0 | Chordata | 17217.0 | 0 | Carnivora | 14987.0 | 1 | Canidae | 13079.0 | 2 | Canis | 11016.0 | 3 | Canis lupus | 11016.0 | 3 |
| scf7180000052265 | 3817 | 0.4449 | 0 | 460.1707 | no-hit | 0.0 | 0 | no-hit | 0.0 | 0 | no-hit | 0.0 | 0 | no-hit | 0.0 | 0 | no-hit | 0.0 | 0 | no-hit | 0.0 | 0 |
| scf7180000048126 | 73747 | 0.3491 | 0 | 61.3328 | Eukaryota | 4974.0 | 1 | Chordata | 4974.0 | 1 | Carnivora | 2155.0 | 3 | Felidae | 1739.0 | 4 | undef | 1522.0 | 6 | eukaryotic synthetic construct | 1522.0 | 6 |
| scf7180000043287 | 121676 | 0.6403 | 0 | 52.3052 | Eukaryota | 16715.0 | 0 | Chordata | 16715.0 | 0 | Carnivora | 16715.0 | 0 | Canidae | 13830.0 | 2 | Canis | 13830.0 | 2 | Canis lupus | 13830.0 | 2 |
| scf7180000053110 | 11904 | 0.5056 | 0 | 897.986 | Eukaryota | 8924.0 | 0 | Chordata | 8924.0 | 0 | Carnivora | 8924.0 | 0 | Canidae | 8924.0 | 0 | Canis | 6620.0 | 1 | Canis lupus | 5354.0 | 2 |
| scf7180000042222 | 431055 | 0.4078 | 0 | 88.4109 | Eukaryota | 138184.0 | 0 | Chordata | 138184.0 | 0 | Carnivora | 138184.0 | 0 | Canidae | 138184.0 | 0 | Canis | 138184.0 | 0 | Canis lupus | 138184.0 | 0 |
| scf7180000044094 | 390309 | 0.3775 | 0 | 60.7151 | Eukaryota | 67107.0 | 0 | Chordata | 67107.0 | 0 | Carnivora | 67107.0 | 0 | Canidae | 67107.0 | 0 | Canis | 67107.0 | 0 | Canis lupus | 67107.0 | 0 |
| scf7180000047954 | 23535 | 0.5044 | 0 | 1307.3812 | Eukaryota | 9246.0 | 0 | Chordata | 9246.0 | 0 | Carnivora | 9246.0 | 0 | Canidae | 9246.0 | 0 | Canis | 7465.0 | 1 | Canis lupus | 6181.0 | 2 |
| scf7180000042742 | 25790 | 0.4055 | 0 | 57.8756 | Eukaryota | 85565.0 | 0 | Chordata | 85565.0 | 0 | Carnivora | 85565.0 | 0 | Canidae | 85565.0 | 0 | Canis | 85565.0 | 0 | Canis lupus | 85565.0 | 0 |
| scf7180000050768 | 13918 | 0.4348 | 0 | 89.36 | Eukaryota | 19221.0 | 0 | Chordata | 13980.0 | 2 | Gadiformes | 5947.0 | 6 | Gadidae | 5947.0 | 6 | Gadus | 5947.0 | 7 | Gadus morhua | 5947.0 | 8 |
| scf7180000042597 | 356062 | 0.3607 | 0 | 58.5568 | Eukaryota | 86415.0 | 0 | Chordata | 86415.0 | 0 | Carnivora | 86415.0 | 0 | Canidae | 86415.0 | 0 | Canis | 86415.0 | 0 | Canis lupus | 86415.0 | 0 |
| scf7180000042252 | 322280 | 0.3418 | 0 | 57.4102 | Eukaryota | 87256.0 | 0 | Chordata | 87256.0 | 0 | Carnivora | 87256.0 | 0 | Canidae | 87256.0 | 0 | Canis | 87256.0 | 0 | Canis lupus | 87256.0 | 0 |
| scf7180000053471 | 2300 | 0.507 | 0 | 24.7773 | Eukaryota | 14968.0 | 0 | Chordata | 14968.0 | 0 | Carnivora | 14968.0 | 0 | Canidae | 14968.0 | 0 | Canis | 14968.0 | 0 | Canis lupus | 14968.0 | 0 |
| scf7180000041757 | 930864 | 0.3936 | 0 | 60.0744 | Eukaryota | 94584.0 | 0 | Chordata | 94584.0 | 0 | Carnivora | 94584.0 | 0 | Canidae | 94584.0 | 0 | Canis | 94584.0 | 0 | Canis lupus | 94584.0 | 0 |
| scf7180000041885 | 133452 | 0.3789 | 0 | 62.2632 | Eukaryota | 91642.0 | 0 | Chordata | 91642.0 | 0 | Carnivora | 91642.0 | 0 | Canidae | 91642.0 | 0 | Canis | 91642.0 | 0 | Canis lupus | 91642.0 | 0 |
| scf7180000056358 | 518259 | 0.41 | 0 | 59.5295 | Eukaryota | 47571.0 | 0 | Chordata | 47571.0 | 0 | Carnivora | 47571.0 | 0 | Canidae | 47571.0 | 0 | Canis | 34853.0 | 1 | Canis lupus | 34853.0 | 1 |
| scf7180000051449 | 5477 | 0.4517 | 0 | 518.488 | Eukaryota | 2601.0 | 0 | Platyhelminthes | 1363.0 | 2 | Polystomatidea | 1363.0 | 2 | Polystomatidae | 1363.0 | 2 | Protopolystoma | 1363.0 | 2 | Protopolystoma xenopodis | 1363.0 | 2 |
| scf7180000056473 | 293869 | 0.5145 | 0 | 56.1241 | Eukaryota | 24213.0 | 0 | Chordata | 24213.0 | 0 | Carnivora | 24213.0 | 0 | Canidae | 24213.0 | 0 | Canis | 24213.0 | 0 | Canis lupus | 24213.0 | 0 |
| scf7180000047378 | 102297 | 0.3659 | 0 | 59.8519 | Eukaryota | 31146.0 | 0 | Chordata | 31146.0 | 0 | Carnivora | 31146.0 | 0 | Canidae | 31146.0 | 0 | Canis | 31146.0 | 0 | Canis lupus | 31146.0 | 0 |
| scf7180000048269 | 54654 | 0.4098 | 0 | 104.8152 | Eukaryota | 82351.0 | 0 | Chordata | 82351.0 | 0 | Carnivora | 82351.0 | 0 | Canidae | 82351.0 | 0 | Canis | 82351.0 | 0 | Canis lupus | 82351.0 | 0 |
| scf7180000048619 | 46923 | 0.4107 | 0 | 61.027 | Eukaryota | 16119.0 | 0 | Chordata | 16119.0 | 0 | Carnivora | 16119.0 | 0 | Canidae | 16119.0 | 0 | Canis | 11724.0 | 1 | Canis lupus | 11724.0 | 1 |
| scf7180000047995 | 72614 | 0.569 | 0 | 57.2763 | Eukaryota | 19801.0 | 0 | Chordata | 19801.0 | 0 | Carnivora | 19801.0 | 0 | Canidae | 19801.0 | 0 | Canis | 16489.0 | 1 | Canis lupus | 16489.0 | 1 |
| scf7180000042263 | 808632 | 0.3984 | 100 | 59.0298 | Eukaryota | 78007.0 | 0 | Chordata | 78007.0 | 0 | Carnivora | 78007.0 | 0 | Canidae | 78007.0 | 0 | Canis | 78007.0 | 0 | Canis lupus | 78007.0 | 0 |
| scf7180000051780 | 5371 | 0.4526 | 0 | 1070.3288 | Eukaryota | 1772.0 | 0 | Platyhelminthes | 889.0 | 2 | Polystomatidea | 889.0 | 2 | Polystomatidae | 889.0 | 2 | Protopolystoma | 889.0 | 2 | Protopolystoma xenopodis | 889.0 | 2 |
| scf7180000050615 | 9324 | 0.4402 | 0 | 470.3544 | Eukaryota | 2838.0 | 0 | Platyhelminthes | 1846.0 | 1 | Polystomatidea | 1846.0 | 2 | Polystomatidae | 1846.0 | 2 | Protopolystoma | 1846.0 | 2 | Protopolystoma xenopodis | 1846.0 | 2 |
| scf7180000041773 | 406515 | 0.3673 | 0 | 58.6848 | Eukaryota | 80231.0 | 0 | Chordata | 80231.0 | 0 | Carnivora | 80231.0 | 0 | Canidae | 80231.0 | 0 | Canis | 80231.0 | 0 | Canis lupus | 80231.0 | 0 |
| scf7180000044433 | 334139 | 0.4283 | 0 | 60.0157 | Eukaryota | 91798.0 | 0 | Chordata | 91798.0 | 0 | Carnivora | 91798.0 | 0 | Canidae | 91798.0 | 0 | Canis | 91798.0 | 0 | Canis lupus | 91798.0 | 0 |
| scf7180000040594 | 2203702 | 0.3827 | 0 | 59.7589 | Eukaryota | 169572.0 | 0 | Chordata | 169572.0 | 0 | Carnivora | 169572.0 | 0 | Otariidae | 84597.0 | 3 | Zalophus | 69024.0 | 5 | Zalophus californianus | 69024.0 | 5 |
| scf7180000044979 | 237560 | 0.4124 | 0 | 59.0294 | Eukaryota | 58844.0 | 0 | Chordata | 58844.0 | 0 | Carnivora | 58844.0 | 0 | Canidae | 58844.0 | 0 | Canis | 49553.0 | 1 | Canis lupus | 49553.0 | 1 |
| scf7180000043265 | 137337 | 0.335 | 0 | 57.6911 | Eukaryota | 9936.0 | 0 | Chordata | 9936.0 | 0 | Carnivora | 8864.0 | 1 | Canidae | 5197.0 | 4 | Canis | 5197.0 | 4 | Canis lupus | 5197.0 | 4 |
| scf7180000042908 | 45695 | 0.4244 | 0 | 112.7801 | Eukaryota | 5554.0 | 0 | Chordata | 5554.0 | 0 | Primates | 3087.0 | 2 | Hominidae | 2412.0 | 3 | Homo | 2412.0 | 5 | Homo sapiens | 2412.0 | 5 |
| scf7180000052933 | 5081 | 0.4414 | 0 | 374.2714 | Eukaryota | 909.0 | 0 | Chordata | 418.0 | 2 | Kurtiformes | 418.0 | 2 | Apogonidae | 418.0 | 2 | Sphaeramia | 418.0 | 2 | Sphaeramia orbicularis | 418.0 | 2 |
| scf7180000046165 | 65962 | 0.501 | 0 | 54.3479 | Eukaryota | 26931.0 | 0 | Chordata | 26931.0 | 0 | Carnivora | 26931.0 | 0 | Canidae | 26931.0 | 0 | Canis | 26931.0 | 0 | Canis lupus | 26931.0 | 0 |
| scf7180000054636 | 32138 | 0.4432 | 0 | 328.2157 | Eukaryota | 440.0 | 0 | Platyhelminthes | 440.0 | 0 | Polystomatidea | 440.0 | 0 | Polystomatidae | 440.0 | 0 | Protopolystoma | 440.0 | 0 | Protopolystoma xenopodis | 440.0 | 0 |
| scf7180000046138 | 22401 | 0.445 | 0 | 624.141 | Eukaryota | 9059.0 | 0 | Chordata | 9059.0 | 0 | Carnivora | 9059.0 | 0 | Canidae | 9059.0 | 0 | Nyctereutes | 4968.0 | 1 | Nyctereutes procyonoides | 4968.0 | 1 |
| scf7180000047344 | 81411 | 0.4388 | 0 | 137.4462 | Eukaryota | 3654.0 | 0 | Chordata | 2261.0 | 1 | Carnivora | 1544.0 | 2 | Canidae | 1544.0 | 2 | Canis | 1544.0 | 2 | Canis lupus | 1544.0 | 2 |
| scf7180000046311 | 269601 | 0.3728 | 0 | 59.7206 | Eukaryota | 73344.0 | 0 | Chordata | 73344.0 | 0 | Carnivora | 73344.0 | 0 | Canidae | 73344.0 | 0 | Canis | 73344.0 | 0 | Canis lupus | 73344.0 | 0 |
| scf7180000051899 | 6186 | 0.4531 | 0 | 574.8269 | Eukaryota | 1475.0 | 0 | Platyhelminthes | 835.0 | 1 | Polystomatidea | 835.0 | 1 | Polystomatidae | 835.0 | 1 | Protopolystoma | 835.0 | 1 | Protopolystoma xenopodis | 835.0 | 1 |
| scf7180000045114 | 234127 | 0.4643 | 0 | 60.2467 | Eukaryota | 61694.0 | 0 | Chordata | 61694.0 | 0 | Carnivora | 61694.0 | 0 | Canidae | 61694.0 | 0 | Canis | 61694.0 | 0 | Canis lupus | 61694.0 | 0 |
| scf7180000054888 | 15087 | 0.4389 | 0 | 95.5875 | Eukaryota | 1557.0 | 0 | Platyhelminthes | 1557.0 | 0 | Polystomatidea | 1557.0 | 0 | Polystomatidae | 1557.0 | 0 | Protopolystoma | 1557.0 | 0 | Protopolystoma xenopodis | 1557.0 | 0 |
| scf7180000043019 | 552020 | 0.4561 | 0 | 57.2098 | Eukaryota | 14243.0 | 0 | Chordata | 14243.0 | 0 | Carnivora | 14243.0 | 0 | Canidae | 14243.0 | 0 | Canis | 12610.0 | 1 | Canis lupus | 12610.0 | 1 |
| scf7180000055491 | 28061 | 0.4361 | 0 | 260.9738 | Eukaryota | 2106.0 | 0 | Platyhelminthes | 1491.0 | 1 | Polystomatidea | 1491.0 | 2 | Polystomatidae | 1491.0 | 2 | Protopolystoma | 1491.0 | 2 | Protopolystoma xenopodis | 1491.0 | 2 |
| scf7180000055137 | 12465 | 0.4489 | 0 | 180.8687 | Eukaryota | 2712.0 | 0 | Chordata | 2712.0 | 0 | Carnivora | 2712.0 | 0 | Canidae | 2712.0 | 0 | Canis | 2712.0 | 0 | Canis lupus | 2712.0 | 0 |
| scf7180000042294 | 742946 | 0.4684 | 0 | 56.9901 | Eukaryota | 22593.0 | 0 | Chordata | 22593.0 | 0 | Carnivora | 22593.0 | 0 | Canidae | 22593.0 | 0 | Canis | 22593.0 | 0 | Canis lupus | 22593.0 | 0 |
| scf7180000042031 | 15725 | 0.4028 | 0 | 58.7621 | Eukaryota | 38532.0 | 0 | Chordata | 38532.0 | 0 | Carnivora | 38532.0 | 0 | Canidae | 38532.0 | 0 | Canis | 38532.0 | 0 | Canis lupus | 38532.0 | 0 |
| scf7180000046941 | 3285 | 0.4384 | 0 | 44.0826 | Eukaryota | 3459.0 | 0 | Chordata | 2852.0 | 1 | Tetraodontiformes | 1607.0 | 4 | Tetraodontidae | 1607.0 | 4 | Takifugu | 1607.0 | 4 | Takifugu rubripes | 1607.0 | 4 |
| scf7180000042226 | 802290 | 0.3544 | 0 | 59.6451 | Eukaryota | 80864.0 | 0 | Chordata | 80864.0 | 0 | Carnivora | 80864.0 | 0 | Canidae | 80864.0 | 0 | Canis | 80864.0 | 0 | Canis lupus | 80864.0 | 0 |
| scf7180000041028 | 1354546 | 0.39 | 0 | 60.6274 | Eukaryota | 179261.0 | 0 | Chordata | 179261.0 | 0 | Carnivora | 179261.0 | 0 | Canidae | 179261.0 | 0 | Canis | 179261.0 | 0 | Canis lupus | 179261.0 | 0 |
| scf7180000043702 | 23270 | 0.319 | 0 | 49.4045 | Eukaryota | 14031.0 | 0 | Chordata | 14031.0 | 0 | Carnivora | 14031.0 | 0 | Canidae | 8518.0 | 2 | Canis | 5670.0 | 3 | Canis lupus | 5670.0 | 3 |
| scf7180000044853 | 302349 | 0.4006 | 0 | 58.756 | Eukaryota | 61283.0 | 0 | Chordata | 61283.0 | 0 | Carnivora | 61283.0 | 0 | Canidae | 61283.0 | 0 | Canis | 61283.0 | 0 | Canis lupus | 61283.0 | 0 |
| scf7180000050686 | 24515 | 0.4106 | 0 | 58.3752 | Eukaryota | 58890.0 | 0 | Chordata | 58890.0 | 0 | Carnivora | 58890.0 | 0 | Canidae | 58890.0 | 0 | Canis | 58890.0 | 0 | Canis lupus | 58890.0 | 0 |
| scf7180000052382 | 8343 | 0.4719 | 0 | 44.3833 | Eukaryota | 9130.0 | 0 | Chordata | 9130.0 | 0 | Carnivora | 9130.0 | 0 | Canidae | 8496.0 | 1 | Vulpes | 4851.0 | 2 | Vulpes vulpes | 4851.0 | 2 |
| scf7180000045222 | 79913 | 0.3395 | 0 | 59.5229 | Eukaryota | 52529.0 | 0 | Chordata | 52529.0 | 0 | Carnivora | 52529.0 | 0 | Canidae | 52529.0 | 0 | Canis | 52529.0 | 0 | Canis lupus | 52529.0 | 0 |
| scf7180000040714 | 1913508 | 0.4055 | 0 | 59.1967 | Eukaryota | 93584.0 | 0 | Chordata | 93584.0 | 0 | Carnivora | 93584.0 | 0 | Canidae | 93584.0 | 0 | Canis | 93584.0 | 0 | Canis lupus | 93584.0 | 0 |
| scf7180000050113 | 37871 | 0.4495 | 0 | 59.9441 | Eukaryota | 3749.0 | 0 | Chordata | 3749.0 | 0 | Carnivora | 2086.0 | 2 | Canidae | 2086.0 | 3 | Canis | 2086.0 | 3 | Canis lupus | 2086.0 | 3 |
| scf7180000048703 | 55445 | 0.4409 | 0 | 55.6222 | Eukaryota | 49080.0 | 0 | Chordata | 49080.0 | 0 | Carnivora | 49080.0 | 0 | Canidae | 49080.0 | 0 | Canis | 49080.0 | 0 | Canis lupus | 49080.0 | 0 |
| scf7180000041752 | 996593 | 0.3953 | 0 | 59.4626 | Eukaryota | 88864.0 | 0 | Chordata | 88864.0 | 0 | Carnivora | 88864.0 | 0 | Canidae | 88864.0 | 0 | Canis | 88864.0 | 0 | Canis lupus | 88864.0 | 0 |
| scf7180000042514 | 250854 | 0.3716 | 0 | 59.7055 | Eukaryota | 86778.0 | 0 | Chordata | 86778.0 | 0 | Carnivora | 86778.0 | 0 | Canidae | 86778.0 | 0 | Canis | 86778.0 | 0 | Canis lupus | 86778.0 | 0 |
| scf7180000045687 | 67303 | 0.4387 | 0 | 142.1972 | Eukaryota | 59773.0 | 0 | Chordata | 42738.0 | 2 | Gadiformes | 34395.0 | 5 | Gadidae | 34395.0 | 5 | Gadus | 34395.0 | 5 | Gadus morhua | 34395.0 | 5 |
| scf7180000041485 | 335826 | 0.4115 | 0 | 60.0113 | Eukaryota | 36421.0 | 0 | Chordata | 36421.0 | 0 | Carnivora | 36421.0 | 0 | Canidae | 36421.0 | 0 | Canis | 36421.0 | 0 | Canis lupus | 36421.0 | 0 |
| scf7180000053217 | 2673 | 0.4523 | 0 | 366.3788 | Eukaryota | 644.0 | 0 | Streptophyta | 505.0 | 1 | Fabales | 505.0 | 1 | Fabaceae | 505.0 | 1 | Cicer | 505.0 | 1 | Cicer arietinum | 505.0 | 1 |
| scf7180000052600 | 5509 | 0.4576 | 0 | 900.017 | Eukaryota | 1943.0 | 0 | Chordata | 1943.0 | 0 | Carnivora | 1943.0 | 0 | Canidae | 1943.0 | 0 | Canis | 1943.0 | 0 | Canis lupus | 1943.0 | 0 |
| scf7180000045005 | 290344 | 0.3809 | 0 | 60.0494 | Eukaryota | 65769.0 | 0 | Chordata | 65769.0 | 0 | Carnivora | 65769.0 | 0 | Canidae | 65769.0 | 0 | Canis | 58282.0 | 1 | Canis lupus | 58282.0 | 1 |
| scf7180000045157 | 50858 | 0.3789 | 0 | 58.009 | Eukaryota | 72264.0 | 0 | Chordata | 72264.0 | 0 | Carnivora | 72264.0 | 0 | Canidae | 72264.0 | 0 | Canis | 65058.0 | 1 | Canis lupus | 65058.0 | 1 |
| scf7180000049506 | 22321 | 0.446 | 0 | 647.9394 | Eukaryota | 30183.0 | 0 | Chordata | 27013.0 | 1 | Gadiformes | 18485.0 | 6 | Gadidae | 18485.0 | 6 | Gadus | 18485.0 | 6 | Gadus morhua | 18485.0 | 6 |
| scf7180000052607 | 1946 | 0.4445 | 0 | 510.0059 | no-hit | 0.0 | 0 | no-hit | 0.0 | 0 | no-hit | 0.0 | 0 | no-hit | 0.0 | 0 | no-hit | 0.0 | 0 | no-hit | 0.0 | 0 |
| scf7180000046626 | 110646 | 0.3681 | 0 | 58.5541 | Eukaryota | 48489.0 | 0 | Chordata | 48489.0 | 0 | Carnivora | 48489.0 | 0 | Canidae | 48489.0 | 0 | Canis | 48489.0 | 0 | Canis lupus | 48489.0 | 0 |
| scf7180000047032 | 4306 | 0.3516 | 0 | 29.6416 | Eukaryota | 2974.0 | 0 | Chordata | 2974.0 | 0 | Carnivora | 2974.0 | 0 | Canidae | 2178.0 | 2 | Vulpes | 1165.0 | 4 | Vulpes vulpes | 1165.0 | 4 |
| scf7180000049025 | 7853 | 0.4471 | 0 | 101.38 | no-hit | 0.0 | 0 | no-hit | 0.0 | 0 | no-hit | 0.0 | 0 | no-hit | 0.0 | 0 | no-hit | 0.0 | 0 | no-hit | 0.0 | 0 |
| scf7180000048173 | 105302 | 0.3892 | 0 | 61.024 | Eukaryota | 85811.0 | 0 | Chordata | 85811.0 | 0 | Carnivora | 85811.0 | 0 | Canidae | 85811.0 | 0 | Canis | 85811.0 | 0 | Canis lupus | 85811.0 | 0 |
| scf7180000044879 | 263687 | 0.4856 | 0 | 58.7079 | Eukaryota | 38399.0 | 0 | Chordata | 38399.0 | 0 | Carnivora | 38399.0 | 0 | Canidae | 38399.0 | 0 | Canis | 30302.0 | 1 | Canis lupus | 30302.0 | 1 |
| scf7180000044765 | 162716 | 0.3557 | 0 | 59.2118 | Eukaryota | 56116.0 | 0 | Chordata | 56116.0 | 0 | Carnivora | 56116.0 | 0 | Canidae | 47881.0 | 1 | Canis | 31202.0 | 2 | Canis lupus | 31202.0 | 2 |
| scf7180000044428 | 469070 | 0.4733 | 0 | 58.6465 | Eukaryota | 60314.0 | 0 | Chordata | 60314.0 | 0 | Carnivora | 54785.0 | 1 | Canidae | 43994.0 | 3 | Canis | 25256.0 | 4 | Canis lupus | 25256.0 | 4 |
| scf7180000052348 | 13009 | 0.4379 | 0 | 55.3581 | Eukaryota | 494.0 | 0 | Platyhelminthes | 494.0 | 0 | Polystomatidea | 494.0 | 0 | Polystomatidae | 494.0 | 0 | Protopolystoma | 494.0 | 0 | Protopolystoma xenopodis | 494.0 | 0 |
| scf7180000048228 | 20144 | 0.4236 | 0 | 49.8243 | Eukaryota | 19747.0 | 0 | Chordata | 19747.0 | 0 | Carnivora | 19747.0 | 0 | Canidae | 19747.0 | 0 | Canis | 19747.0 | 0 | Canis lupus | 19747.0 | 0 |
| scf7180000048441 | 89422 | 0.4244 | 0 | 61.4839 | Eukaryota | 16270.0 | 0 | Chordata | 16270.0 | 0 | Carnivora | 16270.0 | 0 | Canidae | 16270.0 | 0 | Canis | 16270.0 | 0 | Canis lupus | 16270.0 | 0 |
| scf7180000050116 | 11292 | 0.4506 | 0 | 2030.0207 | Eukaryota | 2255.0 | 0 | Platyhelminthes | 925.0 | 2 | Polystomatidea | 925.0 | 2 | Polystomatidae | 925.0 | 2 | Protopolystoma | 925.0 | 2 | Protopolystoma xenopodis | 925.0 | 2 |
| scf7180000048487 | 26439 | 0.4504 | 0 | 2515.5834 | Eukaryota | 3612.0 | 0 | Platyhelminthes | 2217.0 | 2 | Polystomatidea | 2217.0 | 2 | Polystomatidae | 2217.0 | 2 | Protopolystoma | 2217.0 | 2 | Protopolystoma xenopodis | 2217.0 | 2 |
| scf7180000045713 | 9130 | 0.3887 | 0 | 37.0946 | Eukaryota | 12120.0 | 0 | Chordata | 12120.0 | 0 | Carnivora | 12120.0 | 0 | Phocidae | 4944.0 | 3 | Leptonychotes | 4944.0 | 5 | Leptonychotes weddellii | 4944.0 | 6 |
| scf7180000051539 | 2782 | 0.4375 | 0 | 189.0369 | no-hit | 0.0 | 0 | no-hit | 0.0 | 0 | no-hit | 0.0 | 0 | no-hit | 0.0 | 0 | no-hit | 0.0 | 0 | no-hit | 0.0 | 0 |
| scf7180000056031 | 8764 | 0.439 | 0 | 281.9226 | no-hit | 0.0 | 0 | no-hit | 0.0 | 0 | no-hit | 0.0 | 0 | no-hit | 0.0 | 0 | no-hit | 0.0 | 0 | no-hit | 0.0 | 0 |
| scf7180000045224 | 213259 | 0.3573 | 0 | 59.3278 | Eukaryota | 90221.0 | 0 | Chordata | 90221.0 | 0 | Carnivora | 90221.0 | 0 | Canidae | 90221.0 | 0 | Canis | 90221.0 | 0 | Canis lupus | 90221.0 | 0 |
| scf7180000043302 | 76946 | 0.4639 | 0 | 56.4156 | Eukaryota | 81729.0 | 0 | Chordata | 81729.0 | 0 | Carnivora | 81729.0 | 0 | Canidae | 81729.0 | 0 | Canis | 81729.0 | 0 | Canis lupus | 81729.0 | 0 |

| | | | | | | | | | | | | | | | | | | | | | | |
|---|---|---|---|---|---|---|---|---|---|---|---|---|---|---|---|---|---|---|---|---|---|---|
| scf7180000047969 | 18607 | 0.4526 | 0 | 98.9645 | Eukaryota | 2395.0 | 0 | Platyhelminthes | 1491.0 | 1 | Polystomatidea | 1491.0 | 1 | Polystomatidae | 1491.0 | 1 | Protopolystoma | 1491.0 | 1 | Protopolystoma xenopodis | 1491.0 | 1 |
| scf7180000045625 | 176069 | 0.4148 | 0 | 61.1162 | Eukaryota | 30923.0 | 0 | Chordata | 30923.0 | 0 | Carnivora | 29135.0 | 1 | Canidae | 16847.0 | 4 | Canis | 12640.0 | 6 | Canis lupus | 12640.0 | 7 |
| scf7180000054212 | 17369 | 0.4782 | 0 | 253.4612 | Eukaryota | 53873.0 | 0 | Chordata | 53873.0 | 0 | Carnivora | 53873.0 | 0 | Canidae | 53873.0 | 0 | Canis | 53873.0 | 0 | Canis lupus | 53873.0 | 0 |
| scf7180000046577 | 147611 | 0.3407 | 0 | 59.7813 | Eukaryota | 32899.0 | 0 | Chordata | 32899.0 | 0 | Carnivora | 32899.0 | 0 | Canidae | 32899.0 | 0 | Canis | 32899.0 | 0 | Canis lupus | 32899.0 | 0 |
| scf7180000046144 | 8119 | 0.4244 | 0 | 21.7463 | Eukaryota | 82092.0 | 0 | Chordata | 82092.0 | 0 | Carnivora | 82092.0 | 0 | Canidae | 82092.0 | 0 | Canis | 82092.0 | 0 | Canis lupus | 82092.0 | 0 |
| scf7180000048017 | 22621 | 0.4345 | 0 | 231.7759 | Eukaryota | 14366.0 | 0 | Chordata | 8842.0 | 2 | Gadiformes | 4420.0 | 6 | Gadidae | 4420.0 | 6 | Gadus | 4420.0 | 6 | Gadus morhua | 4420.0 | 6 |
| scf7180000045536 | 180246 | 0.5405 | 0 | 58.1248 | Eukaryota | 82538.0 | 0 | Chordata | 82538.0 | 0 | Carnivora | 82538.0 | 0 | Canidae | 82538.0 | 0 | Canis | 82538.0 | 0 | Canis lupus | 82538.0 | 0 |
| scf7180000054376 | 19206 | 0.4252 | 0 | 161.3931 | Eukaryota | 3076.0 | 0 | Chordata | 3076.0 | 0 | Carnivora | 3076.0 | 0 | Canidae | 3076.0 | 0 | Canis | 3076.0 | 0 | Canis lupus | 3076.0 | 0 |
| scf7180000049899 | 12097 | 0.4391 | 0 | 71.5454 | no-hit | 0.0 | 0 | no-hit | 0.0 | 0 | no-hit | 0.0 | 0 | no-hit | 0.0 | 0 | no-hit | 0.0 | 0 | no-hit | 0.0 | 0 |
| scf7180000050426 | 3099 | 0.4434 | 0 | 83.1287 | no-hit | 0.0 | 0 | no-hit | 0.0 | 0 | no-hit | 0.0 | 0 | no-hit | 0.0 | 0 | no-hit | 0.0 | 0 | no-hit | 0.0 | 0 |
| scf7180000042823 | 456632 | 0.3631 | 0 | 59.9619 | Eukaryota | 31222.0 | 0 | Chordata | 31222.0 | 0 | Carnivora | 31222.0 | 0 | Canidae | 10908.0 | 3 | Mustela | 8688.0 | 5 | Mustela putorius | 8688.0 | 5 |
| scf7180000041411 | 72036 | 0.53 | 0 | 56.6775 | Eukaryota | 15621.0 | 0 | Chordata | 15621.0 | 0 | Carnivora | 15621.0 | 0 | Canidae | 15621.0 | 0 | Canis | 15621.0 | 0 | Canis lupus | 15621.0 | 0 |
| scf7180000043980 | 332043 | 0.4129 | 0 | 61.733 | Eukaryota | 63734.0 | 0 | Chordata | 63734.0 | 0 | Carnivora | 63734.0 | 0 | Canidae | 63734.0 | 0 | Canis | 63734.0 | 0 | Canis lupus | 63734.0 | 0 |
| scf7180000056569 | 120026 | 0.5154 | 0 | 58.4898 | Eukaryota | 120590.0 | 0 | Chordata | 120590.0 | 0 | Carnivora | 120590.0 | 0 | Mustelidae | 77133.0 | 1 | Mustela | 44064.0 | 3 | Mustela putorius | 44064.0 | 3 |
| scf7180000055021 | 40632 | 0.4425 | 0 | 195.8135 | Eukaryota | 75932.0 | 0 | Chordata | 75932.0 | 0 | Carnivora | 75932.0 | 0 | Canidae | 75932.0 | 0 | Canis | 75932.0 | 0 | Canis lupus | 75932.0 | 0 |
| scf7180000043495 | 474295 | 0.4639 | 0 | 57.7818 | Eukaryota | 36655.0 | 0 | Chordata | 36655.0 | 0 | Carnivora | 36655.0 | 0 | Canidae | 36655.0 | 0 | Canis | 36655.0 | 0 | Canis lupus | 36655.0 | 0 |
| scf7180000047923 | 3542 | 0.4376 | 0 | 163.3893 | Eukaryota | 4452.0 | 1 | Chordata | 2248.0 | 3 | Polystomatidea | 1553.0 | 5 | Polystomatidae | 1553.0 | 5 | Protopolystoma | 1553.0 | 5 | Protopolystoma xenopodis | 1553.0 | 5 |
| scf7180000051268 | 3091 | 0.4478 | 0 | 388.4328 | no-hit | 0.0 | 0 | no-hit | 0.0 | 0 | no-hit | 0.0 | 0 | no-hit | 0.0 | 0 | no-hit | 0.0 | 0 | no-hit | 0.0 | 0 |
| scf7180000049005 | 3965 | 0.4499 | 0 | 1161.4976 | no-hit | 0.0 | 0 | no-hit | 0.0 | 0 | no-hit | 0.0 | 0 | no-hit | 0.0 | 0 | no-hit | 0.0 | 0 | no-hit | 0.0 | 0 |
| scf7180000043095 | 714621 | 0.5208 | 0 | 58.009 | Eukaryota | 103296.0 | 0 | Chordata | 103296.0 | 0 | Carnivora | 103296.0 | 0 | Canidae | 103296.0 | 0 | Canis | 94437.0 | 1 | Canis lupus | 94437.0 | 1 |
| scf7180000043485 | 271729 | 0.4279 | 0 | 59.487 | Eukaryota | 80573.0 | 0 | Chordata | 80573.0 | 0 | Carnivora | 80573.0 | 0 | Canidae | 39114.0 | 4 | Canis | 26148.0 | 7 | Canis lupus | 26148.0 | 7 |
| scf7180000046030 | 139072 | 0.3914 | 0 | 58.4673 | Eukaryota | 23622.0 | 0 | Chordata | 23622.0 | 0 | Carnivora | 23622.0 | 0 | Canidae | 23622.0 | 0 | Canis | 18994.0 | 1 | Canis lupus | 18994.0 | 1 |
| scf7180000041254 | 1499652 | 0.3569 | 0 | 60.7393 | Eukaryota | 86825.0 | 0 | Chordata | 86825.0 | 0 | Carnivora | 86825.0 | 0 | Canidae | 86825.0 | 0 | Canis | 86825.0 | 0 | Canis lupus | 86825.0 | 0 |
| scf7180000044629 | 19854 | 0.4443 | 0 | 252.9701 | Eukaryota | 1338.0 | 0 | Platyhelminthes | 1338.0 | 0 | Polystomatidea | 1338.0 | 0 | Polystomatidae | 1338.0 | 0 | Protopolystoma | 1338.0 | 0 | Protopolystoma xenopodis | 1338.0 | 0 |
| scf7180000056419 | 712392 | 0.4945 | 0 | 58.9371 | Eukaryota | 65653.0 | 0 | Chordata | 65653.0 | 0 | Carnivora | 65653.0 | 0 | Canidae | 65653.0 | 0 | Canis | 65653.0 | 0 | Canis lupus | 65653.0 | 0 |
| scf7180000050498 | 32974 | 0.4427 | 0 | 79.2357 | Eukaryota | 8847.0 | 0 | Chordata | 3806.0 | 2 | Polystomatidea | 3059.0 | 3 | Polystomatidae | 3059.0 | 3 | Protopolystoma | 3059.0 | 3 | Protopolystoma xenopodis | 3059.0 | 3 |
| scf7180000055994 | 13330 | 0.4298 | 0 | 170.0592 | Eukaryota | 940.0 | 0 | Chordata | 940.0 | 0 | Kurtiformes | 940.0 | 0 | Apogonidae | 940.0 | 0 | Sphaeramia | 940.0 | 0 | Sphaeramia orbicularis | 940.0 | 0 |
| scf7180000044316 | 286679 | 0.5263 | 0 | 58.9795 | Eukaryota | 63350.0 | 0 | Chordata | 63350.0 | 0 | Carnivora | 63350.0 | 0 | Canidae | 63350.0 | 0 | Canis | 58055.0 | 1 | Canis lupus | 58055.0 | 1 |
| scf7180000051146 | 10347 | 0.4551 | 0 | 52.021 | Eukaryota | 2795.0 | 0 | Chordata | 2795.0 | 0 | Salmoniformes | 2795.0 | 0 | Salmonidae | 2795.0 | 0 | Salmo | 2795.0 | 0 | Salmo trutta | 2795.0 | 0 |
| scf7180000053140 | 3262 | 0.4577 | 0 | 530.607 | no-hit | 0.0 | 0 | no-hit | 0.0 | 0 | no-hit | 0.0 | 0 | no-hit | 0.0 | 0 | no-hit | 0.0 | 0 | no-hit | 0.0 | 0 |
| scf7180000046803 | 106298 | 0.3647 | 0 | 61.1769 | Eukaryota | 90788.0 | 0 | Chordata | 90788.0 | 0 | Carnivora | 90788.0 | 0 | Canidae | 90788.0 | 0 | Canis | 90788.0 | 0 | Canis lupus | 90788.0 | 0 |
| scf7180000043267 | 3211 | 0.4394 | 0 | 44.7744 | Eukaryota | 2186.0 | 0 | Platyhelminthes | 1316.0 | 1 | Polystomatidea | 1316.0 | 3 | Polystomatidae | 1316.0 | 3 | Protopolystoma | 1316.0 | 3 | Protopolystoma xenopodis | 1316.0 | 3 |
| scf7180000050509 | 4263 | 0.441 | 0 | 211.4157 | Eukaryota | 1648.0 | 0 | Platyhelminthes | 999.0 | 1 | Polystomatidea | 862.0 | 2 | Polystomatidae | 862.0 | 2 | Protopolystoma | 862.0 | 2 | Protopolystoma xenopodis | 862.0 | 2 |
| scf7180000044319 | 289591 | 0.5262 | 0 | 57.0341 | Eukaryota | 36274.0 | 0 | Chordata | 36274.0 | 0 | Carnivora | 36274.0 | 0 | Canidae | 36274.0 | 0 | Canis | 36274.0 | 0 | Canis lupus | 36274.0 | 0 |
| scf7180000048042 | 18890 | 0.4392 | 0 | 93.06 | Eukaryota | 4979.0 | 0 | Chordata | 4552.0 | 1 | Syngnathiformes | 1701.0 | 4 | Syngnathidae | 1701.0 | 4 | Syngnathus | 1701.0 | 4 | Syngnathus acus | 1701.0 | 4 |
| scf7180000048911 | 9340 | 0.4331 | 0 | 124.3279 | Eukaryota | 3987.0 | 0 | Platyhelminthes | 2110.0 | 1 | Kurtiformes | 1620.0 | 4 | Apogonidae | 1620.0 | 4 | Sphaeramia | 1620.0 | 4 | Sphaeramia orbicularis | 1620.0 | 4 |
| scf7180000052483 | 5794 | 0.5003 | 0 | 84.8605 | Eukaryota | 7984.0 | 0 | Chordata | 7984.0 | 0 | Carnivora | 7984.0 | 0 | Canidae | 7984.0 | 0 | Canis | 6393.0 | 1 | Canis lupus | 5254.0 | 2 |
| scf7180000042589 | 111888 | 0.4655 | 0 | 57.8355 | Eukaryota | 26219.0 | 0 | Chordata | 26219.0 | 0 | Carnivora | 26219.0 | 0 | Canidae | 26219.0 | 0 | Canis | 21995.0 | 1 | Canis lupus | 21995.0 | 1 |
| scf7180000042474 | 637696 | 0.4136 | 0 | 59.9748 | Eukaryota | 64213.0 | 0 | Chordata | 64213.0 | 0 | Carnivora | 64213.0 | 0 | Canidae | 60235.0 | 1 | Canis | 60235.0 | 1 | Canis lupus | 60235.0 | 1 |
| scf7180000046799 | 118987 | 0.3401 | 0 | 59.3143 | Eukaryota | 9303.0 | 0 | Chordata | 9303.0 | 0 | Carnivora | 6632.0 | 2 | Canidae | 4050.0 | 4 | Canis | 2125.0 | 6 | Canis lupus | 2125.0 | 6 |
| scf7180000051207 | 6551 | 0.4425 | 0 | 133.0484 | Eukaryota | 893.0 | 0 | Platyhelminthes | 893.0 | 0 | Polystomatidea | 893.0 | 0 | Polystomatidae | 893.0 | 0 | Protopolystoma | 893.0 | 0 | Protopolystoma xenopodis | 893.0 | 0 |
| scf7180000043258 | 478588 | 0.4154 | 0 | 59.778 | Eukaryota | 79851.0 | 0 | Chordata | 79851.0 | 0 | Carnivora | 79851.0 | 0 | Canidae | 79851.0 | 0 | Canis | 79851.0 | 0 | Canis lupus | 79851.0 | 0 |
| scf7180000042676 | 420160 | 0.3908 | 0 | 61.1332 | Eukaryota | 96521.0 | 0 | Chordata | 96521.0 | 0 | Carnivora | 96521.0 | 0 | Canidae | 96521.0 | 0 | Canis | 96521.0 | 0 | Canis lupus | 96521.0 | 0 |
| scf7180000047751 | 570367 | 0.5193 | 0 | 57.572 | Eukaryota | 33277.0 | 0 | Chordata | 33277.0 | 0 | Carnivora | 33277.0 | 0 | Canidae | 33277.0 | 0 | Canis | 21577.0 | 1 | Canis lupus | 21577.0 | 1 |
| scf7180000055381 | 36181 | 0.4416 | 0 | 223.9661 | Eukaryota | 28228.0 | 0 | Chordata | 28228.0 | 0 | Gadiformes | 18433.0 | 5 | Gadidae | 18433.0 | 5 | Gadus | 18433.0 | 5 | Gadus morhua | 18433.0 | 5 |
| scf7180000052145 | 10254 | 0.4479 | 0 | 472.841 | Eukaryota | 2345.0 | 0 | Platyhelminthes | 1528.0 | 2 | Polystomatidea | 1528.0 | 2 | Polystomatidae | 1528.0 | 2 | Protopolystoma | 1528.0 | 2 | Protopolystoma xenopodis | 1528.0 | 2 |
| scf7180000046052 | 190649 | 0.3491 | 0 | 59.284 | Eukaryota | 14271.0 | 0 | Chordata | 14271.0 | 0 | Carnivora | 11794.0 | 1 | Canidae | 10410.0 | 2 | Canis | 10410.0 | 3 | Canis lupus | 10410.0 | 3 |
| scf7180000056172 | 2542562 | 0.4503 | 0 | 58.5052 | Eukaryota | 77045.0 | 0 | Chordata | 77045.0 | 0 | Carnivora | 77045.0 | 0 | Canidae | 66056.0 | 2 | Canis | 49196.0 | 3 | Canis lupus | 49196.0 | 3 |
| scf7180000043578 | 7743 | 0.3504 | 0 | 29.9941 | Eukaryota | 6659.0 | 0 | Chordata | 6659.0 | 0 | Carnivora | 6038.0 | 1 | Felidae | 4663.0 | 2 | Felis | 4663.0 | 2 | Felis catus | 4663.0 | 2 |
| scf7180000041048 | 226784 | 0.3687 | 0 | 60.2881 | Eukaryota | 89545.0 | 0 | Chordata | 89545.0 | 0 | Carnivora | 89545.0 | 0 | Canidae | 89545.0 | 0 | Canis | 89545.0 | 0 | Canis lupus | 89545.0 | 0 |
| scf7180000042290 | 833595 | 0.3942 | 0 | 59.781 | Eukaryota | 36417.0 | 0 | Chordata | 36417.0 | 0 | Carnivora | 36417.0 | 0 | Canidae | 36417.0 | 0 | Canis | 36417.0 | 0 | Canis lupus | 36417.0 | 0 |
| scf7180000047788 | 9471 | 0.448 | 0 | 73.249 | Eukaryota | 1658.0 | 0 | Chordata | 1658.0 | 0 | Carnivora | 1658.0 | 0 | Canidae | 1658.0 | 0 | Canis | 1658.0 | 0 | Canis lupus | 1658.0 | 0 |
| scf7180000052608 | 5058 | 0.4476 | 0 | 638.3521 | no-hit | 0.0 | 0 | no-hit | 0.0 | 0 | no-hit | 0.0 | 0 | no-hit | 0.0 | 0 | no-hit | 0.0 | 0 | no-hit | 0.0 | 0 |
| scf7180000045422 | 233720 | 0.3768 | 0 | 62.3406 | Eukaryota | 96841.0 | 0 | Chordata | 96841.0 | 0 | Carnivora | 96841.0 | 0 | Canidae | 96841.0 | 0 | Canis | 96841.0 | 0 | Canis lupus | 96841.0 | 0 |
| scf7180000055667 | 19582 | 0.4388 | 0 | 336.7472 | Eukaryota | 346.0 | 0 | Chordata | 346.0 | 0 | Kurtiformes | 346.0 | 0 | Apogonidae | 346.0 | 0 | Sphaeramia | 346.0 | 0 | Sphaeramia orbicularis | 346.0 | 0 |
| scf7180000045537 | 180395 | 0.4942 | 0 | 57.9448 | Eukaryota | 17297.0 | 0 | Chordata | 17297.0 | 0 | Carnivora | 17297.0 | 0 | Canidae | 17297.0 | 0 | Canis | 16168.0 | 1 | Canis lupus | 16168.0 | 1 |
| scf7180000044942 | 177519 | 0.3884 | 0 | 129.6451 | Eukaryota | 55768.0 | 0 | Chordata | 55768.0 | 0 | Carnivora | 55768.0 | 0 | Canidae | 55768.0 | 0 | Canis | 55768.0 | 0 | Canis lupus | 55768.0 | 0 |
| scf7180000044806 | 4399 | 0.5619 | 0 | 33.5994 | Eukaryota | 3314.0 | 0 | Chordata | 3314.0 | 0 | Carnivora | 3314.0 | 0 | Canidae | 2546.0 | 2 | Canis | 2546.0 | 2 | Canis lupus | 2546.0 | 2 |
| scf7180000043229 | 432897 | 0.3523 | 0 | 57.9664 | Eukaryota | 9308.0 | 0 | Chordata | 9308.0 | 0 | Carnivora | 9308.0 | 0 | Canidae | 9308.0 | 0 | Canis | 9308.0 | 0 | Canis lupus | 9308.0 | 0 |
| scf7180000044113 | 36372 | 0.3768 | 0 | 51.3181 | Eukaryota | 11157.0 | 0 | Chordata | 11157.0 | 0 | Primates | 5851.0 | 3 | Hominidae | 5202.0 | 5 | Homo | 5202.0 | 6 | Homo sapiens | 5202.0 | 6 |
| scf7180000053925 | 6366 | 0.4505 | 0 | 665.1582 | Eukaryota | 658.0 | 0 | Platyhelminthes | 658.0 | 0 | Polystomatidea | 658.0 | 0 | Polystomatidae | 658.0 | 0 | Protopolystoma | 658.0 | 0 | Protopolystoma xenopodis | 658.0 | 0 |
| scf7180000049893 | 6969 | 0.4464 | 0 | 308.4044 | Eukaryota | 20688.0 | 0 | Chordata | 15446.0 | 2 | Gadiformes | 10811.0 | 6 | Gadidae | 10811.0 | 6 | Gadus | 10811.0 | 6 | Gadus morhua | 10811.0 | 6 |
| scf7180000045066 | 185840 | 0.3567 | 0 | 60.3826 | Eukaryota | 82512.0 | 0 | Chordata | 82512.0 | 0 | Carnivora | 82512.0 | 0 | Canidae | 82512.0 | 0 | Canis | 82512.0 | 0 | Canis lupus | 82512.0 | 0 |
| scf7180000053239 | 4066 | 0.4469 | 0 | 176.6679 | no-hit | 0.0 | 0 | no-hit | 0.0 | 0 | no-hit | 0.0 | 0 | no-hit | 0.0 | 0 | no-hit | 0.0 | 0 | no-hit | 0.0 | 0 |
| scf7180000042223 | 74209 | 0.3834 | 0 | 54.4708 | Eukaryota | 87676.0 | 0 | Chordata | 87676.0 | 0 | Carnivora | 87676.0 | 0 | Canidae | 87676.0 | 0 | Canis | 87676.0 | 0 | Canis lupus | 87676.0 | 0 |
| scf7180000044342 | 16974 | 0.4618 | 0 | 220.9339 | Eukaryota | 41769.0 | 0 | Chordata | 41769.0 | 0 | Carnivora | 41769.0 | 0 | Canidae | 41769.0 | 0 | Canis | 41769.0 | 0 | Canis lupus | 41769.0 | 0 |
| scf7180000052523 | 11697 | 0.6561 | 0 | 53.4474 | Eukaryota | 6976.0 | 0 | Chordata | 6976.0 | 0 | Carnivora | 6976.0 | 0 | Canidae | 5300.0 | 2 | Canis | 3631.0 | 4 | Canis lupus | 3631.0 | 4 |
| scf7180000051012 | 17979 | 0.4449 | 0 | 133.7853 | Eukaryota | 11809.0 | 0 | Chordata | 7434.0 | 3 | Gadiformes | 4492.0 | 7 | Gadidae | 4492.0 | 7 | Gadus | 4492.0 | 7 | Gadus morhua | 4492.0 | 7 |
| scf7180000048282 | 62867 | 0.3337 | 0 | 60.6701 | Eukaryota | 14115.0 | 0 | Chordata | 14115.0 | 0 | Carnivora | 14115.0 | 0 | Canidae | 14115.0 | 0 | Vulpes | 8478.0 | 1 | Vulpes vulpes | 8478.0 | 1 |
| scf7180000052519 | 2988 | 0.4508 | 0 | 239.2734 | no-hit | 0.0 | 0 | no-hit | 0.0 | 0 | no-hit | 0.0 | 0 | no-hit | 0.0 | 0 | no-hit | 0.0 | 0 | no-hit | 0.0 | 0 |
| scf7180000042829 | 15935 | 0.3906 | 0 | 45.283 | Eukaryota | 77430.0 | 0 | Chordata | 77430.0 | 0 | Carnivora | 77430.0 | 0 | Canidae | 77430.0 | 0 | Canis | 77430.0 | 0 | Canis lupus | 77430.0 | 0 |
| scf7180000046970 | 20744 | 0.4246 | 0 | 73.8731 | Eukaryota | 12196.0 | 0 | Chordata | 12196.0 | 0 | Carnivora | 12196.0 | 0 | Canidae | 12196.0 | 0 | Canis | 12196.0 | 0 | Canis lupus | 12196.0 | 0 |
| scf7180000055454 | 14496 | 0.4405 | 0 | 241.2707 | Eukaryota | 3497.0 | 0 | Chordata | 3497.0 | 0 | Kurtiformes | 3497.0 | 0 | Apogonidae | 3497.0 | 0 | Sphaeramia | 3497.0 | 0 | Sphaeramia orbicularis | 3497.0 | 0 |
| scf7180000048470 | 64260 | 0.413 | 0 | 60.5211 | Eukaryota | 34414.0 | 0 | Chordata | 34414.0 | 0 | Carnivora | 34414.0 | 0 | Canidae | 26407.0 | 1 | Canis | 15108.0 | 2 | Canis lupus | 15108.0 | 2 |
| scf7180000056307 | 979580 | 0.3974 | 0 | 61.2603 | Eukaryota | 54491.0 | 0 | Chordata | 54491.0 | 0 | Carnivora | 54491.0 | 0 | Canidae | 54491.0 | 0 | Canis | 45992.0 | 1 | Canis lupus | 45992.0 | 1 |
| scf7180000040904 | 41346 | 0.3913 | 0 | 54.8147 | Eukaryota | 20768.0 | 0 | Chordata | 20768.0 | 0 | Carnivora | 20768.0 | 0 | Canidae | 20768.0 | 0 | Canis | 20768.0 | 0 | Canis lupus | 20768.0 | 0 |
| scf7180000044190 | 32370 | 0.3828 | 0 | 56.7463 | Eukaryota | 5757.0 | 1 | Chordata | 5757.0 | 1 | Artiodactyla | 2839.0 | 3 | Hominidae | 2520.0 | 4 | Homo | 2520.0 | 5 | Homo sapiens | 2520.0 | 5 |
| scf7180000042943 | 757449 | 0.4383 | 100 | 58.9306 | Eukaryota | 183611.0 | 0 | Chordata | 183611.0 | 0 | Carnivora | 183611.0 | 0 | Canidae | 183611.0 | 0 | Canis | 154364.0 | 1 | Canis lupus | 154364.0 | 1 |
| scf7180000055058 | 11858 | 0.4412 | 0 | 68.5876 | Eukaryota | 33598.0 | 0 | Chordata | 33598.0 | 0 | Carnivora | 33598.0 | 0 | Canidae | 33598.0 | 0 | Canis | 33598.0 | 0 | Canis lupus | 33598.0 | 0 |
| scf7180000043082 | 568749 | 0.4361 | 0 | 60.435 | Eukaryota | 113987.0 | 0 | Chordata | 113987.0 | 0 | Carnivora | 113987.0 | 0 | Canidae | 113987.0 | 0 | Canis | 86315.0 | 1 | Canis lupus | 86315.0 | 1 |
| scf7180000046096 | 125361 | 0.3971 | 0 | 58.2735 | Eukaryota | 24793.0 | 0 | Chordata | 24793.0 | 0 | Carnivora | 21671.0 | 1 | Canidae | 21671.0 | 1 | Canis | 20487.0 | 2 | Canis lupus | 20487.0 | 2 |
| scf7180000053003 | 4592 | 0.4425 | 0 | 254.2679 | no-hit | 0.0 | 0 | no-hit | 0.0 | 0 | no-hit | 0.0 | 0 | no-hit | 0.0 | 0 | no-hit | 0.0 | 0 | no-hit | 0.0 | 0 |
| scf7180000041586 | 1002416 | 0.4121 | 0 | 58.7532 | Eukaryota | 87856.0 | 0 | Chordata | 87856.0 | 0 | Carnivora | 87856.0 | 0 | Canidae | 87856.0 | 0 | Canis | 87856.0 | 0 | Canis lupus | 87856.0 | 0 |
| scf7180000042051 | 704718 | 0.4267 | 0 | 60.4579 | Eukaryota | 148360.0 | 0 | Chordata | 148360.0 | 0 | Carnivora | 148360.0 | 0 | Canidae | 148360.0 | 0 | Canis | 128840.0 | 1 | Canis lupus | 128840.0 | 1 |
| scf7180000054057 | 5811 | 0.4443 | 0 | 254.307 | Eukaryota | 4428.0 | 0 | Chordata | 3631.0 | 1 | Kurtiformes | 1281.0 | 4 | Apogonidae | 1281.0 | 4 | Sphaeramia | 1281.0 | 4 | Sphaeramia orbicularis | 1281.0 | 4 |
| scf7180000047428 | 1380238 | 0.4059 | 0 | 59.9709 | Eukaryota | 59736.0 | 0 | Chordata | 59736.0 | 0 | Carnivora | 59736.0 | 0 | Canidae | 55791.0 | 1 | Canis | 50707.0 | 2 | Canis lupus | 50707.0 | 2 |
| scf7180000041859 | 793351 | 0.4747 | 0 | 57.9972 | Eukaryota | 45388.0 | 0 | Chordata | 45388.0 | 0 | Carnivora | 45388.0 | 0 | Otariidae | 17811.0 | 3 | Canis | 10336.0 | 6 | Canis lupus | 10336.0 | 6 |
| scf7180000043464 | 491205 | 0.3456 | 0 | 58.8133 | Eukaryota | 135923.0 | 0 | Chordata | 135923.0 | 0 | Carnivora | 135923.0 | 0 | Canidae | 135923.0 | 0 | Canis | 135923.0 | 0 | Canis lupus | 135923.0 | 0 |
| scf7180000048822 | 39569 | 0.623 | 0 | 54.4486 | Eukaryota | 46044.0 | 0 | Chordata | 46044.0 | 0 | Carnivora | 46044.0 | 0 | Canidae | 46044.0 | 0 | Canis | 46044.0 | 0 | Canis lupus | 46044.0 | 0 |
| scf7180000053008 | 2346 | 0.4501 | 0 | 164.9378 | no-hit | 0.0 | 0 | no-hit | 0.0 | 0 | no-hit | 0.0 | 0 | no-hit | 0.0 | 0 | no-hit | 0.0 | 0 | no-hit | 0.0 | 0 |
| scf7180000056559 | 68928 | 0.3479 | 0 | 55.2352 | Eukaryota | 5758.0 | 0 | Chordata | 5758.0 | 0 | Carnivora | 5229.0 | 1 | Canidae | 3516.0 | 2 | Canis | 1974.0 | 3 | Canis lupus | 1974.0 | 3 |
| scf7180000053455 | 7178 | 0.4383 | 0 | 195.4898 | no-hit | 0.0 | 0 | no-hit | 0.0 | 0 | no-hit | 0.0 | 0 | no-hit | 0.0 | 0 | no-hit | 0.0 | 0 | no-hit | 0.0 | 0 |
| scf7180000041077 | 13322 | 0.4402 | 0 | 43.3241 | Eukaryota | 4096.0 | 0 | Chordata | 4096.0 | 0 | Carnivora | 2009.0 | 2 | Hominidae | 1531.0 | 4 | Homo | 1531.0 | 7 | Homo sapiens | 1531.0 | 7 |
| scf7180000041370 | 1345614 | 0.3977 | 0 | 60.6388 | Eukaryota | 87170.0 | 0 | Chordata | 87170.0 | 0 | Carnivora | 87170.0 | 0 | Canidae | 87170.0 | 0 | Canis | 87170.0 | 0 | Canis lupus | 87170.0 | 0 |
| scf7180000050823 | 1596 | 0.4536 | 0 | 469.4848 | no-hit | 0.0 | 0 | no-hit | 0.0 | 0 | no-hit | 0.0 | 0 | no-hit | 0.0 | 0 | no-hit | 0.0 | 0 | no-hit | 0.0 | 0 |
| scf7180000052739 | 5488 | 0.428 | 0 | 35.3011 | no-hit | 0.0 | 0 | no-hit | 0.0 | 0 | no-hit | 0.0 | 0 | no-hit | 0.0 | 0 | no-hit | 0.0 | 0 | no-hit | 0.0 | 0 |
| scf7180000053275 | 7104 | 0.4524 | 0 | 56.4282 | Eukaryota | 2681.0 | 0 | Chordata | 2681.0 | 0 | Tetraodontiformes | 2187.0 | 1 | Tetraodontidae | 2187.0 | 1 | Takifugu | 2187.0 | 1 | Takifugu rubripes | 2187.0 | 1 |
| scf7180000044059 | 418982 | 0.3433 | 0 | 59.0006 | Eukaryota | 94057.0 | 0 | Chordata | 94057.0 | 0 | Carnivora | 94057.0 | 0 | Canidae | 94057.0 | 0 | Canis | 94057.0 | 0 | Canis lupus | 94057.0 | 0 |
| scf7180000046874 | 101323 | 0.3577 | 0 | 57.1335 | Eukaryota | 14558.0 | 0 | Chordata | 14558.0 | 0 | Carnivora | 8787.0 | 2 | Canidae | 8787.0 | 2 | Canis | 4889.0 | 3 | Canis lupus | 4889.0 | 3 |
| scf7180000053174 | 2648 | 0.4464 | 0 | 202.8285 | no-hit | 0.0 | 0 | no-hit | 0.0 | 0 | no-hit | 0.0 | 0 | no-hit | 0.0 | 0 | no-hit | 0.0 | 0 | no-hit | 0.0 | 0 |
| scf7180000042383 | 17059 | 0.4473 | 0 | 136.3021 | Eukaryota | 38452.0 | 0 | Chordata | 22523.0 | 2 | Cyclophyllidea | 8582.0 | 8 | Hymenolepididae | 8582.0 | 8 | Hymenolepis | 8582.0 | 8 | Hymenolepis microstoma | 8582.0 | 8 |
| scf7180000040536 | 2222441 | 0.4001 | 0 | 59.917 | Eukaryota | 120888.0 | 0 | Chordata | 120888.0 | 0 | Carnivora | 120888.0 | 0 | Canidae | 120888.0 | 0 | Canis | 120888.0 | 0 | Canis lupus | 120888.0 | 0 |
| scf7180000050429 | 2797 | 0.4462 | 0 | 373.8599 | no-hit | 0.0 | 0 | no-hit | 0.0 | 0 | no-hit | 0.0 | 0 | no-hit | 0.0 | 0 | no-hit | 0.0 | 0 | no-hit | 0.0 | 0 |
| scf7180000048435 | 36303 | 0.4397 | 0 | 366.3036 | Eukaryota | 55471.0 | 0 | Chordata | 27259.0 | 2 | Gadiformes | 15794.0 | 6 | Gadidae | 15794.0 | 6 | Gadus | 15794.0 | 6 | Gadus morhua | 15794.0 | 6 |
| scf7180000050600 | 5070 | 0.4314 | 0 | 41.9326 | Eukaryota | 29053.0 | 0 | Chordata | 20015.0 | 2 | Gadiformes | 16782.0 | 5 | Gadidae | 16782.0 | 5 | Gadus | 16782.0 | 5 | Gadus morhua | 16782.0 | 5 |
| scf7180000055144 | 28302 | 0.4295 | 0 | 466.7043 | Eukaryota | 252.0 | 0 | Chordata | 252.0 | 0 | Salmoniformes | 252.0 | 0 | Salmonidae | 252.0 | 0 | Salmo | 252.0 | 0 | Salmo trutta | 252.0 | 0 |
| scf7180000048750 | 49182 | 0.3399 | 0 | 58.2213 | Eukaryota | 5712.0 | 1 | Chordata | 5712.0 | 1 | Carnivora | 2738.0 | 3 | Canidae | 2738.0 | 3 | Canis | 2738.0 | 4 | Canis lupus | 2738.0 | 4 |
| scf7180000056033 | 25938 | 0.3391 | 0 | 309.7124 | Eukaryota | 5161.0 | 0 | Chordata | 5161.0 | 0 | Apterygiformes | 2689.0 | 1 | Apterygidae | 2689.0 | 1 | Apteryx | 2689.0 | 1 | Apteryx mantelli | 2689.0 | 1 |
| scf7180000045935 | 141036 | 0.4628 | 0 | 55.6826 | Eukaryota | 10024.0 | 0 | Chordata | 10024.0 | 0 | Carnivora | 10024.0 | 0 | Canidae | 7172.0 | 2 | Canis | 5786.0 | 3 | Canis lupus | 5786.0 | 3 |
| scf7180000046451 | 14514 | 0.4174 | 0 | 29.6004 | Eukaryota | 4306.0 | 0 | Chordata | 4306.0 | 0 | Carnivora | 2665.0 | 2 | Canidae | 2665.0 | 2 | Canis | 2665.0 | 2 | Canis lupus | 2665.0 | 2 |
| scf7180000043913 | 371883 | 0.4007 | 0 | 60.5432 | Eukaryota | 61867.0 | 0 | Chordata | 61867.0 | 0 | Carnivora | 61867.0 | 0 | Canidae | 61867.0 | 0 | Vulpes | 47019.0 | 1 | Vulpes vulpes | 47019.0 | 1 |
| scf7180000051298 | 10868 | 0.4357 | 0 | 47.6356 | no-hit | 0.0 | 0 | no-hit | 0.0 | 0 | no-hit | 0.0 | 0 | no-hit | 0.0 | 0 | no-hit | 0.0 | 0 | no-hit | 0.0 | 0 |
| scf7180000050418 | 13008 | 0.4476 | 0 | 882.4467 | Eukaryota | 5069.0 | 0 | Chordata | 3074.0 | 2 | Polystomatidea | 1458.0 | 4 | Polystomatidae | 1458.0 | 4 | Protopolystoma | 1458.0 | 4 | Protopolystoma xenopodis | 1458.0 | 4 |
| scf7180000044174 | 391773 | 0.4964 | 0 | 58.0592 | Eukaryota | 40673.0 | 0 | Chordata | 40673.0 | 0 | Carnivora | 40673.0 | 0 | Canidae | 30013.0 | 1 | Canis | 25737.0 | 3 | Canis lupus | 25737.0 | 3 |
| scf7180000044772 | 27494 | 0.4709 | 0 | 57.5681 | Eukaryota | 37214.0 | 0 | Chordata | 37214.0 | 0 | Carnivora | 37214.0 | 0 | Canidae | 37214.0 | 0 | Canis | 37214.0 | 0 | Canis lupus | 37214.0 | 0 |
| scf7180000041791 | 864985 | 0.4485 | 0 | 59.9882 | Eukaryota | 157965.0 | 0 | Chordata | 157965.0 | 0 | Carnivora | 157965.0 | 0 | Canidae | 157965.0 | 0 | Canis | 157965.0 | 0 | Canis lupus | 157965.0 | 0 |
| scf7180000052867 | 2417 | 0.4402 | 0 | 61.6901 | no-hit | 0.0 | 0 | no-hit | 0.0 | 0 | no-hit | 0.0 | 0 | no-hit | 0.0 | 0 | no-hit | 0.0 | 0 | no-hit | 0.0 | 0 |
| scf7180000049840 | 6191 | 0.4432 | 0 | 280.6454 | no-hit | 0.0 | 0 | no-hit | 0.0 | 0 | no-hit | 0.0 | 0 | no-hit | 0.0 | 0 | no-hit | 0.0 | 0 | no-hit | 0.0 | 0 |
| scf7180000054671 | 17935 | 0.4109 | 0 | 405.2422 | Eukaryota | 95190.0 | 0 | Chordata | 95190.0 | 0 | Carnivora | 95190.0 | 0 | Canidae | 95190.0 | 0 | Canis | 95190.0 | 0 | Canis lupus | 95190.0 | 0 |
| scf7180000042278 | 449142 | 0.5869 | 0 | 55.2994 | Eukaryota | 67219.0 | 0 | Chordata | 67219.0 | 0 | Carnivora | 67219.0 | 0 | Canidae | 67219.0 | 0 | Canis | 67219.0 | 0 | Canis lupus | 67219.0 | 0 |
| scf7180000045669 | 192831 | 0.338 | 0 | 59.1867 | Eukaryota | 41851.0 | 0 | Chordata | 41851.0 | 0 | Carnivora | 41851.0 | 0 | Canidae | 41851.0 | 0 | Canis | 41851.0 | 0 | Canis lupus | 41851.0 | 0 |
| scf7180000047140 | 6086 | 0.4512 | 0 | 96.1601 | Eukaryota | 37189.0 | 0 | Chordata | 37189.0 | 0 | Carnivora | 37189.0 | 0 | Canidae | 37189.0 | 0 | Canis | 37189.0 | 0 | Canis lupus | 37189.0 | 0 |
| scf7180000047545 | 22795 | 0.4419 | 0 | 76.2218 | Eukaryota | 446.0 | 0 | Platyhelminthes | 446.0 | 0 | Polystomatidea | 446.0 | 0 | Polystomatidae | 446.0 | 0 | Protopolystoma | 446.0 | 0 | Protopolystoma xenopodis | 446.0 | 0 |
| scf7180000049099 | 5451 | 0.43 | 0 | 36.0181 | no-hit | 0.0 | 0 | no-hit | 0.0 | 0 | no-hit | 0.0 | 0 | no-hit | 0.0 | 0 | no-hit | 0.0 | 0 | no-hit | 0.0 | 0 |
| scf7180000044192 | 449481 | 0.3517 | 0 | 58.5301 | Eukaryota | 35550.0 | 0 | Chordata | 35550.0 | 0 | Carnivora | 35550.0 | 0 | Canidae | 35550.0 | 0 | Canis | 35550.0 | 0 | Canis lupus | 35550.0 | 0 |
| scf7180000040685 | 1940076 | 0.4269 | 0 | 59.9878 | Eukaryota | 81136.0 | 0 | Chordata | 81136.0 | 0 | Carnivora | 81136.0 | 0 | Canidae | 81136.0 | 0 | Canis | 64450.0 | 1 | Canis lupus | 64450.0 | 1 |
| scf7180000042128 | 189746 | 0.4765 | 0 | 57.8636 | Eukaryota | 62900.0 | 0 | Chordata | 62900.0 | 0 | Carnivora | 62900.0 | 0 | Canidae | 62900.0 | 0 | Vulpes | 62900.0 | 0 | Vulpes vulpes | 62900.0 | 0 |
| scf7180000047882 | 16066 | 0.5184 | 0 | 838.2904 | Eukaryota | 4021.0 | 0 | Chordata | 4021.0 | 0 | Carnivora | 4021.0 | 0 | Canidae | 4021.0 | 0 | Canis | 4021.0 | 0 | Canis lupus | 4021.0 | 0 |
| scf7180000041491 | 433926 | 0.4082 | 0 | 58.228 | Eukaryota | 37108.0 | 0 | Chordata | 37108.0 | 0 | Carnivora | 37108.0 | 0 | Canidae | 37108.0 | 0 | Canis | 37108.0 | 0 | Canis lupus | 37108.0 | 0 |
| scf7180000044356 | 130538 | 0.4371 | 0 | 159.9117 | Eukaryota | 67258.0 | 0 | Chordata | 50922.0 | 1 | Gadiformes | 43340.0 | 4 | Gadidae | 43340.0 | 4 | Gadus | 43340.0 | 4 | Gadus morhua | 43340.0 | 4 |
| scf7180000043497 | 314635 | 0.5339 | 0 | 56.2036 | Eukaryota | 36486.0 | 0 | Chordata | 36486.0 | 0 | Carnivora | 36486.0 | 0 | Canidae | 36486.0 | 0 | Canis | 36486.0 | 0 | Canis lupus | 36486.0 | 0 |
| scf7180000042611 | 631623 | 0.3574 | 0 | 59.4438 | Eukaryota | 57207.0 | 0 | Chordata | 57207.0 | 0 | Carnivora | 57207.0 | 0 | Canidae | 57207.0 | 0 | Canis | 57207.0 | 0 | Canis lupus | 57207.0 | 0 |
| scf7180000043852 | 335639 | 0.374 | 0 | 60.8953 | Eukaryota | 33767.0 | 0 | Chordata | 33767.0 | 0 | Carnivora | 33767.0 | 0 | Canidae | 33767.0 | 0 | Canis | 31116.0 | 1 | Canis lupus | 31116.0 | 1 |
| scf7180000046348 | 119659 | 0.3981 | 0 | 58.8605 | Eukaryota | 68783.0 | 0 | Chordata | 68783.0 | 0 | Carnivora | 68783.0 | 0 | Canidae | 24389.0 | 4 | Canis | 16243.0 | 7 | Canis lupus | 16243.0 | 8 |
| scf7180000054618 | 49287 | 0.4338 | 0 | 312.3017 | Eukaryota | 4154.0 | 0 | Chordata | 3197.0 | 1 | Polystomatidea | 957.0 | 4 | Polystomatidae | 957.0 | 4 | Protopolystoma | 957.0 | 5 | Protopolystoma xenopodis | 957.0 | 5 |
| scf7180000046106 | 157085 | 0.3614 | 0 | 59.2371 | Eukaryota | 38483.0 | 0 | Chordata | 38483.0 | 0 | Carnivora | 38483.0 | 0 | Canidae | 38483.0 | 0 | Canis | 38483.0 | 0 | Canis lupus | 38483.0 | 0 |
| scf7180000044451 | 353098 | 0.3629 | 0 | 60.9515 | Eukaryota | 89690.0 | 0 | Chordata | 89690.0 | 0 | Carnivora | 89690.0 | 0 | Canidae | 89690.0 | 0 | Canis | 89690.0 | 0 | Canis lupus | 89690.0 | 0 |
| scf7180000053441 | 7779 | 0.4492 | 0 | 381.5293 | Eukaryota | 1733.0 | 0 | Chordata | 1057.0 | 1 | Tetraodontiformes | 1057.0 | 2 | Tetraodontidae | 1057.0 | 2 | Takifugu | 1057.0 | 2 | Takifugu rubripes | 1057.0 | 2 |
| scf7180000050389 | 30748 | 0.3533 | 0 | 59.7271 | Eukaryota | 6159.0 | 0 | Chordata | 6159.0 | 0 | Carnivora | 3622.0 | 1 | Canidae | 3622.0 | 1 | Canis | 3622.0 | 1 | Canis lupus | 3622.0 | 1 |
| scf7180000043786 | 376918 | 0.3764 | 0 | 60.8411 | Eukaryota | 43400.0 | 0 | Chordata | 43400.0 | 0 | Carnivora | 43400.0 | 0 | Canidae | 36042.0 | 1 | Canis | 32036.0 | 2 | Canis lupus | 32036.0 | 3 |
| scf7180000043180 | 637992 | 0.4181 | 0 | 57.285 | Eukaryota | 26027.0 | 0 | Chordata | 26027.0 | 0 | Carnivora | 26027.0 | 0 | Canidae | 26027.0 | 0 | Canis | 23142.0 | 1 | Canis lupus | 23142.0 | 1 |
| scf7180000049213 | 28274 | 0.4388 | 0 | 63.4347 | Eukaryota | 29602.0 | 1 | Chordata | 21467.0 | 2 | Gadiformes | 17116.0 | 5 | Gadidae | 17116.0 | 5 | Gadus | 17116.0 | 5 | Gadus morhua | 17116.0 | 5 |
| scf7180000047476 | 29264 | 0.4391 | 0 | 51.3473 | Eukaryota | 14454.0 | 0 | Chordata | 13443.0 | 1 | Tetraodontiformes | 13443.0 | 1 | Tetraodontidae | 13443.0 | 1 | Takifugu | 13443.0 | 1 | Takifugu rubripes | 13443.0 | 1 |
| scf7180000042468 | 395580 | 0.3639 | 0 | 59.0043 | Eukaryota | 22519.0 | 0 | Chordata | 22519.0 | 0 | Primates | 12255.0 | 1 | Hominidae | 12255.0 | 1 | Homo | 12255.0 | 1 | Homo sapiens | 12255.0 | 1 |
| scf7180000050835 | 2374073 | 0.4027 | 0 | 59.7095 | Eukaryota | 94194.0 | 0 | Chordata | 94194.0 | 0 | Carnivora | 94194.0 | 0 | Canidae | 94194.0 | 0 | Canis | 94194.0 | 0 | Canis lupus | 94194.0 | 0 |
| scf7180000053186 | 4285 | 0.4975 | 0 | 201.4033 | Eukaryota | 8193.0 | 0 | Chordata | 8193.0 | 0 | Carnivora | 8193.0 | 0 | Canidae | 8193.0 | 0 | Canis | 6451.0 | 1 | Canis lupus | 5273.0 | 2 |
| scf7180000052847 | 4082 | 0.5061 | 0 | 69.3893 | Eukaryota | 4708.0 | 0 | Chordata | 4708.0 | 0 | Carnivora | 4708.0 | 0 | Canidae | 4708.0 | 0 | Canis | 2727.0 | 1 | Nyctereutes procyonoides | 1981.0 | 2 |
| scf7180000050499 | 5186 | 0.4553 | 0 | 887.4809 | Eukaryota | 719.0 | 0 | Platyhelminthes | 719.0 | 0 | Polystomatidea | 719.0 | 0 | Polystomatidae | 719.0 | 0 | Protopolystoma | 719.0 | 0 | Protopolystoma xenopodis | 719.0 | 0 |
| scf7180000051432 | 4347 | 0.444 | 0 | 375.9902 | no-hit | 0.0 | 0 | no-hit | 0.0 | 0 | no-hit | 0.0 | 0 | no-hit | 0.0 | 0 | no-hit | 0.0 | 0 | no-hit | 0.0 | 0 |
| scf7180000048090 | 8989 | 0.43 | 0 | 25.0839 | Eukaryota | 18515.0 | 0 | Chordata | 18515.0 | 0 | Carnivora | 18515.0 | 0 | Canidae | 18515.0 | 0 | Canis | 17140.0 | 1 | Canis lupus | 17140.0 | 1 |
| scf7180000051274 | 38087 | 0.4162 | 0 | 50.8691 | Eukaryota | 18155.0 | 0 | Chordata | 18155.0 | 0 | Carnivora | 18155.0 | 0 | Canidae | 14945.0 | 1 | Canis | 11230.0 | 2 | Canis lupus | 11230.0 | 2 |
| scf7180000051691 | 8231 | 0.4424 | 0 | 78.1666 | Eukaryota | 9021.0 | 1 | Chordata | 8485.0 | 2 | Salmoniformes | 4389.0 | 5 | Salmonidae | 4389.0 | 5 | Salmo | 3962.0 | 6 | Salmo trutta | 3962.0 | 7 |
| scf7180000044883 | 73990 | 0.3793 | 0 | 57.0007 | Eukaryota | 56500.0 | 0 | Chordata | 56500.0 | 0 | Carnivora | 56500.0 | 0 | Canidae | 56500.0 | 0 | Canis | 56500.0 | 0 | Canis lupus | 56500.0 | 0 |
| scf7180000052837 | 8998 | 0.4344 | 0 | 34.9782 | Eukaryota | 63886.0 | 0 | Chordata | 44882.0 | 1 | Gadiformes | 44882.0 | 2 | Gadidae | 44882.0 | 2 | Gadus | 44882.0 | 2 | Gadus morhua | 44882.0 | 2 |
| scf7180000045441 | 9855 | 0.4368 | 0 | 251.4517 | Eukaryota | 9591.0 | 0 | Chordata | 7523.0 | 2 | Tetraodontiformes | 6841.0 | 3 | Tetraodontidae | 6841.0 | 3 | Takifugu | 6841.0 | 3 | Takifugu rubripes | 6841.0 | 3 |
| scf7180000055238 | 15148 | 0.6402 | 0 | 259.1358 | Eukaryota | 35334.0 | 0 | Chordata | 35334.0 | 0 | Primates | 27557.0 | 1 | Hominidae | 27557.0 | 1 | Homo | 27557.0 | 1 | Homo sapiens | 27557.0 | 1 |
| scf7180000047883 | 43012 | 0.4381 | 0 | 262.2054 | Eukaryota | 94020.0 | 0 | Chordata | 94020.0 | 0 | Carnivora | 94020.0 | 0 | Canidae | 94020.0 | 0 | Canis | 94020.0 | 0 | Canis lupus | 94020.0 | 0 |
| scf7180000056516 | 260427 | 0.5365 | 0 | 57.917 | Eukaryota | 116550.0 | 0 | Chordata | 116550.0 | 0 | Carnivora | 116550.0 | 0 | Canidae | 116550.0 | 0 | Canis | 91665.0 | 1 | Canis lupus | 91665.0 | 1 |
| scf7180000050644 | 32067 | 0.4359 | 0 | 58.4389 | Eukaryota | 2807.0 | 0 | Chordata | 1762.0 | 1 | Carnivora | 1433.0 | 2 | Canidae | 1433.0 | 2 | Canis | 1433.0 | 2 | Canis lupus | 1433.0 | 2 |
| scf7180000040484 | 430658 | 0.3678 | 0 | 59.8058 | Eukaryota | 83709.0 | 0 | Chordata | 83709.0 | 0 | Carnivora | 83709.0 | 0 | Canidae | 83709.0 | 0 | Canis | 83709.0 | 0 | Canis lupus | 83709.0 | 0 |
| scf7180000042362 | 15412 | 0.49 | 0 | 641.073 | Eukaryota | 33536.0 | 0 | Chordata | 33536.0 | 0 | Carnivora | 33536.0 | 0 | Canidae | 33536.0 | 0 | Canis | 33536.0 | 0 | Canis lupus | 33536.0 | 0 |
| scf7180000056268 | 145659 | 0.3726 | 0 | 58.8469 | Eukaryota | 49849.0 | 0 | Chordata | 49849.0 | 0 | Carnivora | 49849.0 | 0 | Canidae | 49849.0 | 0 | Canis | 49849.0 | 0 | Canis lupus | 49849.0 | 0 |
| scf7180000055572 | 8923 | 0.4412 | 0 | 220.134 | Eukaryota | 3103.0 | 0 | Platyhelminthes | 1971.0 | 3 | Polystomatidea | 1971.0 | 3 | Polystomatidae | 1971.0 | 3 | Protopolystoma | 1971.0 | 3 | Protopolystoma xenopodis | 1971.0 | 3 |
| scf7180000045368 | 218089 | 0.4399 | 0 | 60.5357 | Eukaryota | 58121.0 | 0 | Chordata | 58121.0 | 0 | Carnivora | 58121.0 | 0 | Canidae | 58121.0 | 0 | Canis | 58121.0 | 0 | Canis lupus | 58121.0 | 0 |
| scf7180000041914 | 6243 | 0.3561 | 0 | 34.6582 | Eukaryota | 7836.0 | 0 | Chordata | 7836.0 | 0 | Apterygiformes | 4870.0 | 2 | Apterygidae | 4870.0 | 2 | Apteryx | 4870.0 | 2 | Apteryx mantelli | 4870.0 | 2 |
| scf7180000051487 | 9412 | 0.4561 | 0 | 1469.6428 | Eukaryota | 1269.0 | 0 | Echinodermata | 501.0 | 2 | Forcipulatida | 501.0 | 2 | Asteriidae | 501.0 | 2 | Asterias | 501.0 | 2 | Asterias rubens | 501.0 | 2 |
| scf7180000051798 | 14773 | 0.4508 | 0 | 1000.5438 | Eukaryota | 18734.0 | 0 | Chordata | 15933.0 | 3 | Gadiformes | 11970.0 | 6 | Gadidae | 11970.0 | 6 | Gadus | 11970.0 | 6 | Gadus morhua | 11970.0 | 6 |
| scf7180000046875 | 113555 | 0.3462 | 0 | 61.2133 | Eukaryota | 12247.0 | 0 | Chordata | 12247.0 | 0 | Carnivora | 12247.0 | 0 | Canidae | 12247.0 | 0 | Canis | 12247.0 | 0 | Canis lupus | 12247.0 | 0 |
| scf7180000056384 | 532066 | 0.5012 | 0 | 58.0405 | Eukaryota | 41951.0 | 0 | Chordata | 41951.0 | 0 | Carnivora | 41951.0 | 0 | Canidae | 24293.0 | 2 | Canis | 19715.0 | 5 | Canis lupus | 19715.0 | 5 |
| scf7180000053429 | 2264 | 0.4488 | 0 | 654.1655 | Eukaryota | 4075.0 | 0 | Platyhelminthes | 2061.0 | 1 | Polystomatidea | 2061.0 | 2 | Polystomatidae | 2061.0 | 2 | Protopolystoma | 2061.0 | 2 | Protopolystoma xenopodis | 2061.0 | 2 |
| scf7180000049383 | 39095 | 0.3915 | 0 | 57.2153 | Eukaryota | 6402.0 | 1 | Chordata | 6402.0 | 1 | Artiodactyla | 2220.0 | 3 | Bovidae | 2220.0 | 3 | Canis | 2137.0 | 5 | Canis lupus | 2137.0 | 6 |
| scf7180000052944 | 2135 | 0.5082 | 0 | 20.8418 | Eukaryota | 4196.0 | 0 | Chordata | 4196.0 | 0 | Carnivora | 4196.0 | 0 | Canidae | 4196.0 | 0 | Canis | 2605.0 | 2 | Canis lupus | 2605.0 | 2 |
| scf7180000042406 | 40888 | 0.4379 | 0 | 122.8348 | Eukaryota | 92643.0 | 0 | Chordata | 92643.0 | 0 | Carnivora | 92643.0 | 0 | Canidae | 92643.0 | 0 | Canis | 92643.0 | 0 | Canis lupus | 92643.0 | 0 |
| scf7180000046019 | 8145 | 0.4996 | 0 | 35.3986 | Eukaryota | 5982.0 | 0 | Chordata | 5982.0 | 0 | Carnivora | 5982.0 | 0 | Canidae | 5982.0 | 0 | Canis | 5982.0 | 0 | Canis lupus | 5982.0 | 0 |
| scf7180000046996 | 4548 | 0.4398 | 0 | 222.6271 | Eukaryota | 3044.0 | 1 | Chordata | 2026.0 | 2 | Gonorynchiformes | 1044.0 | 4 | Chanidae | 1044.0 | 4 | Chanos | 1044.0 | 4 | Chanos chanos | 1044.0 | 4 |
| scf7180000052067 | 6952 | 0.443 | 0 | 1183.6326 | no-hit | 0.0 | 0 | no-hit | 0.0 | 0 | no-hit | 0.0 | 0 | no-hit | 0.0 | 0 | no-hit | 0.0 | 0 | no-hit | 0.0 | 0 |
| scf7180000041215 | 1317936 | 0.3869 | 100 | 60.2415 | Eukaryota | 72865.0 | 0 | Chordata | 72865.0 | 0 | Carnivora | 72865.0 | 0 | Canidae | 72865.0 | 0 | Canis | 43721.0 | 1 | Canis lupus | 43721.0 | 1 |
| scf7180000046149 | 133394 | 0.334 | 0 | 59.5719 | Eukaryota | 39908.0 | 0 | Chordata | 39908.0 | 0 | Carnivora | 39908.0 | 0 | Canidae | 39908.0 | 0 | Canis | 39908.0 | 0 | Canis lupus | 39908.0 | 0 |

| | | | | | | | | | | | | | | | | | | | | | | |
|---|---|---|---|---|---|---|---|---|---|---|---|---|---|---|---|---|---|---|---|---|---|---|
| scf7180000056282 | 801207 | 0.4475 | 0 | 58.5742 | Eukaryota | 71677.0 | 0 | Chordata | 71677.0 | 0 | Carnivora | 71677.0 | 0 | Canidae | 71677.0 | 0 | Canis | 71677.0 | 0 | Canis lupus | 71677.0 | 0 |
| scf7180000042787 | 154726 | 0.3822 | 0 | 58.0919 | Eukaryota | 91345.0 | 0 | Chordata | 91345.0 | 0 | Carnivora | 91345.0 | 0 | Canidae | 91345.0 | 0 | Canis | 91345.0 | 0 | Canis lupus | 91345.0 | 0 |
| scf7180000040817 | 2454003 | 0.446 | 0 | 59.5045 | Eukaryota | 112865.0 | 0 | Chordata | 112865.0 | 0 | Carnivora | 112865.0 | 0 | Canidae | 112865.0 | 0 | Canis | 112865.0 | 0 | Canis lupus | 112865.0 | 0 |
| scf7180000051546 | 7462 | 0.4623 | 0 | 957.7536 | Eukaryota | 1931.0 | 0 | Platyhelminthes | 718.0 | 2 | Polystomatidea | 718.0 | 2 | Polystomatidae | 718.0 | 2 | Protopolystoma | 718.0 | 2 | Protopolystoma xenopodis | 718.0 | 2 |
| scf7180000043262 | 350667 | 0.5145 | 0 | 58.0537 | Eukaryota | 54448.0 | 0 | Chordata | 54448.0 | 0 | Carnivora | 54448.0 | 0 | Canidae | 54448.0 | 0 | Canis | 54448.0 | 0 | Canis lupus | 54448.0 | 0 |
| scf7180000049706 | 42415 | 0.4276 | 0 | 59.2185 | Eukaryota | 52429.0 | 0 | Chordata | 52429.0 | 0 | Carnivora | 52429.0 | 0 | Canidae | 52429.0 | 0 | Canis | 52429.0 | 0 | Canis lupus | 52429.0 | 0 |
| scf7180000053029 | 7074 | 0.3527 | 0 | 64.5579 | Eukaryota | 3836.0 | 0 | Chordata | 3836.0 | 0 | Carnivora | 3836.0 | 0 | Canidae | 3836.0 | 0 | Canis | 2982.0 | 1 | Canis lupus | 2982.0 | 1 |
| scf7180000047461 | 69112 | 0.429 | 0 | 59.8209 | Eukaryota | 79873.0 | 0 | Chordata | 79873.0 | 0 | Carnivora | 79873.0 | 0 | Canidae | 79873.0 | 0 | Canis | 79873.0 | 0 | Canis lupus | 79873.0 | 0 |
| scf7180000040602 | 444645 | 0.396 | 0 | 59.0359 | Eukaryota | 74406.0 | 0 | Chordata | 74406.0 | 0 | Carnivora | 74406.0 | 0 | Canidae | 74406.0 | 0 | Canis | 74406.0 | 0 | Canis lupus | 74406.0 | 0 |
| scf7180000047602 | 17497 | 0.4424 | 0 | 180.3248 | Eukaryota | 2579.0 | 0 | Chordata | 2579.0 | 0 | Carnivora | 2305.0 | 1 | Canidae | 2305.0 | 1 | Canis | 2305.0 | 1 | Canis lupus | 2305.0 | 1 |
| scf7180000046463 | 118986 | 0.413 | 0 | 59.6713 | Eukaryota | 26850.0 | 0 | Chordata | 26850.0 | 0 | Carnivora | 26850.0 | 0 | Canidae | 26850.0 | 0 | Canis | 23952.0 | 1 | Canis lupus | 23952.0 | 1 |
| scf7180000050349 | 8324 | 0.4402 | 0 | 248.3075 | no-hit | 0.0 | 0 | no-hit | 0.0 | 0 | no-hit | 0.0 | 0 | no-hit | 0.0 | 0 | no-hit | 0.0 | 0 | no-hit | 0.0 | 0 |
| scf7180000045692 | 12564 | 0.5176 | 0 | 484.0564 | Eukaryota | 16085.0 | 0 | Chordata | 16085.0 | 0 | Carnivora | 16085.0 | 0 | Canidae | 16085.0 | 0 | Canis | 16085.0 | 0 | Canis lupus | 16085.0 | 0 |
| scf7180000043031 | 473669 | 0.4015 | 0 | 58.2051 | Eukaryota | 73615.0 | 0 | Chordata | 73615.0 | 0 | Carnivora | 73615.0 | 0 | Canidae | 73615.0 | 0 | Canis | 73615.0 | 0 | Canis lupus | 73615.0 | 0 |
| scf7180000051869 | 6114 | 0.4436 | 0 | 627.4131 | Eukaryota | 3720.0 | 0 | Platyhelminthes | 2225.0 | 2 | Polystomatidea | 2225.0 | 2 | Polystomatidae | 2225.0 | 2 | Protopolystoma | 2225.0 | 2 | Protopolystoma xenopodis | 2225.0 | 2 |
| scf7180000056563 | 666082 | 0.3646 | 0 | 68.1486 | Eukaryota | 88968.0 | 0 | Chordata | 88968.0 | 0 | Carnivora | 88968.0 | 0 | Canidae | 88968.0 | 0 | Canis | 88968.0 | 0 | Canis lupus | 88968.0 | 0 |
| scf7180000053630 | 4935 | 0.4991 | 0 | 92.6186 | Eukaryota | 8301.0 | 0 | Chordata | 8301.0 | 0 | Carnivora | 8301.0 | 0 | Canidae | 8301.0 | 0 | Canis | 6750.0 | 1 | Canis lupus | 6240.0 | 2 |
| scf7180000044092 | 349082 | 0.3918 | 0 | 60.2647 | Eukaryota | 41148.0 | 0 | Chordata | 41148.0 | 0 | Carnivora | 41148.0 | 0 | Canidae | 41148.0 | 0 | Canis | 26688.0 | 1 | Canis lupus | 26688.0 | 1 |
| scf7180000045917 | 141249 | 0.5904 | 0 | 55.1657 | Eukaryota | 69900.0 | 0 | Chordata | 69900.0 | 0 | Carnivora | 69900.0 | 0 | Canidae | 69900.0 | 0 | Canis | 57833.0 | 1 | Canis lupus | 57833.0 | 1 |
| scf7180000050190 | 30407 | 0.5256 | 0 | 57.4487 | Eukaryota | 43397.0 | 0 | Chordata | 43397.0 | 0 | Carnivora | 43397.0 | 0 | Canidae | 43397.0 | 0 | Canis | 40752.0 | 1 | Canis lupus | 40752.0 | 1 |
| scf7180000050893 | 4167 | 0.4454 | 0 | 452.9304 | Eukaryota | 6105.0 | 0 | Chordata | 5374.0 | 1 | Tetraodontiformes | 3491.0 | 3 | Tetraodontidae | 3491.0 | 3 | Takifugu | 3491.0 | 3 | Takifugu rubripes | 3491.0 | 3 |
| scf7180000044043 | 11895 | 0.5154 | 0 | 41.5649 | Eukaryota | 8769.0 | 0 | Chordata | 8769.0 | 0 | Carnivora | 5126.0 | 1 | Hominidae | 3643.0 | 2 | Pan | 2606.0 | 5 | Pan troglodytes | 2606.0 | 5 |
| scf7180000044218 | 144971 | 0.409 | 0 | 60.7147 | Eukaryota | 84767.0 | 0 | Chordata | 84767.0 | 0 | Carnivora | 84767.0 | 0 | Canidae | 84767.0 | 0 | Canis | 84767.0 | 0 | Canis lupus | 84767.0 | 0 |
| scf7180000048921 | 11971 | 0.441 | 0 | 63.0601 | Eukaryota | 463.0 | 0 | Platyhelminthes | 268.0 | 1 | Polystomatidea | 268.0 | 1 | Polystomatidae | 268.0 | 1 | Protopolystoma | 268.0 | 1 | Protopolystoma xenopodis | 268.0 | 1 |
| scf7180000050697 | 3541 | 0.4485 | 0 | 318.8607 | Eukaryota | 3813.0 | 0 | Platyhelminthes | 1953.0 | 2 | Polystomatidea | 1953.0 | 3 | Polystomatidae | 1953.0 | 3 | Protopolystoma | 1953.0 | 3 | Protopolystoma xenopodis | 1953.0 | 3 |
| scf7180000051637 | 29515 | 0.5154 | 0 | 1345.6352 | Eukaryota | 33881.0 | 0 | Chordata | 33881.0 | 0 | Carnivora | 33881.0 | 0 | Canidae | 33881.0 | 0 | Canis | 33881.0 | 0 | Canis lupus | 33881.0 | 0 |
| scf7180000043758 | 21545 | 0.4381 | 0 | 113.828 | Eukaryota | 12944.0 | 0 | Chordata | 6420.0 | 3 | Strigeidida | 3919.0 | 7 | Schistosomatidae | 3919.0 | 7 | Schistosoma | 3919.0 | 7 | Schistosoma mansoni | 3919.0 | 7 |
| scf7180000054165 | 27809 | 0.3805 | 0 | 95.0845 | Eukaryota | 92288.0 | 0 | Chordata | 92288.0 | 0 | Carnivora | 92288.0 | 0 | Canidae | 92288.0 | 0 | Canis | 92288.0 | 0 | Canis lupus | 92288.0 | 0 |
| scf7180000044499 | 10141 | 0.4279 | 0 | 41.6205 | Eukaryota | 4303.0 | 0 | Chordata | 4303.0 | 0 | Carnivora | 2595.0 | 1 | Canidae | 2595.0 | 1 | Canis | 2595.0 | 1 | Canis lupus | 2595.0 | 1 |
| scf7180000044290 | 123423 | 0.3764 | 0 | 58.4918 | Eukaryota | 91193.0 | 0 | Chordata | 91193.0 | 0 | Carnivora | 91193.0 | 0 | Canidae | 91193.0 | 0 | Canis | 91193.0 | 0 | Canis lupus | 91193.0 | 0 |
| scf7180000041715 | 54984 | 0.361 | 0 | 57.5699 | Eukaryota | 50710.0 | 0 | Chordata | 50710.0 | 0 | Carnivora | 50710.0 | 0 | Canidae | 50710.0 | 0 | Canis | 50710.0 | 0 | Canis lupus | 50710.0 | 0 |
| scf7180000052473 | 3744 | 0.4287 | 0 | 126.0083 | no-hit | 0.0 | 0 | no-hit | 0.0 | 0 | no-hit | 0.0 | 0 | no-hit | 0.0 | 0 | no-hit | 0.0 | 0 | no-hit | 0.0 | 0 |
| scf7180000041733 | 225910 | 0.3955 | 0 | 71.7261 | Eukaryota | 49857.0 | 0 | Chordata | 49857.0 | 0 | Carnivora | 49857.0 | 0 | Canidae | 49857.0 | 0 | Canis | 49857.0 | 0 | Canis lupus | 49857.0 | 0 |
| scf7180000042524 | 218795 | 0.3753 | 0 | 58.3785 | Eukaryota | 26373.0 | 0 | Chordata | 26373.0 | 0 | Carnivora | 26373.0 | 0 | Canidae | 26373.0 | 0 | Canis | 26373.0 | 0 | Canis lupus | 26373.0 | 0 |
| scf7180000049446 | 7411 | 0.3902 | 0 | 12.7634 | Eukaryota | 79357.0 | 0 | Chordata | 79357.0 | 0 | Carnivora | 79357.0 | 0 | Canidae | 79357.0 | 0 | Canis | 79357.0 | 0 | Canis lupus | 79357.0 | 0 |
| scf7180000043111 | 1773394 | 0.3621 | 0 | 60.6156 | Eukaryota | 86033.0 | 0 | Chordata | 86033.0 | 0 | Carnivora | 86033.0 | 0 | Canidae | 86033.0 | 0 | Canis | 86033.0 | 0 | Canis lupus | 86033.0 | 0 |
| scf7180000056494 | 154483 | 0.4367 | 0 | 58.9118 | Eukaryota | 55798.0 | 0 | Chordata | 55798.0 | 0 | Carnivora | 55798.0 | 0 | Canidae | 43686.0 | 1 | Canis | 43686.0 | 1 | Canis lupus | 43686.0 | 1 |
| scf7180000052369 | 9162 | 0.4998 | 0 | 196.1576 | Eukaryota | 9030.0 | 0 | Chordata | 9030.0 | 0 | Carnivora | 9030.0 | 0 | Canidae | 9030.0 | 0 | Canis | 7916.0 | 1 | Canis lupus | 6788.0 | 2 |
| scf7180000044230 | 43989 | 0.4749 | 0 | 54.9168 | Eukaryota | 34236.0 | 0 | Chordata | 34236.0 | 0 | Carnivora | 34236.0 | 0 | Canidae | 34236.0 | 0 | Canis | 34236.0 | 0 | Canis lupus | 34236.0 | 0 |
| scf7180000042176 | 337206 | 0.3738 | 0 | 60.1473 | Eukaryota | 77283.0 | 0 | Chordata | 77283.0 | 0 | Carnivora | 77283.0 | 0 | Canidae | 77283.0 | 0 | Canis | 77283.0 | 0 | Canis lupus | 77283.0 | 0 |
| scf7180000043033 | 468825 | 0.5655 | 0 | 56.8686 | Eukaryota | 29580.0 | 0 | Chordata | 29580.0 | 0 | Carnivora | 29580.0 | 0 | Canidae | 29580.0 | 0 | Canis | 22006.0 | 1 | Canis lupus | 22006.0 | 1 |
| scf7180000044403 | 299367 | 0.4972 | 0 | 57.7645 | Eukaryota | 22185.0 | 0 | Chordata | 22185.0 | 0 | Carnivora | 22185.0 | 0 | Canidae | 22185.0 | 0 | Canis | 22185.0 | 0 | Canis lupus | 22185.0 | 0 |
| scf7180000046179 | 27527 | 0.35 | 0 | 52.5125 | Eukaryota | 6230.0 | 0 | Chordata | 6230.0 | 0 | Carnivora | 5236.0 | 1 | Canidae | 5236.0 | 1 | Canis | 5236.0 | 1 | Canis lupus | 5236.0 | 1 |
| scf7180000041872 | 244303 | 0.3862 | 0 | 69.2871 | Eukaryota | 93878.0 | 0 | Chordata | 93878.0 | 0 | Carnivora | 93878.0 | 0 | Canidae | 93878.0 | 0 | Canis | 93878.0 | 0 | Canis lupus | 93878.0 | 0 |
| scf7180000056454 | 455092 | 0.3957 | 0 | 60.5742 | Eukaryota | 28991.0 | 0 | Chordata | 28991.0 | 0 | Carnivora | 28991.0 | 0 | Canidae | 23491.0 | 1 | Canis | 23491.0 | 1 | Canis lupus | 23491.0 | 1 |
| scf7180000041707 | 82269 | 0.3421 | 0 | 54.7518 | Eukaryota | 88423.0 | 0 | Chordata | 88423.0 | 0 | Carnivora | 88423.0 | 0 | Canidae | 88423.0 | 0 | Canis | 88423.0 | 0 | Canis lupus | 88423.0 | 0 |
| scf7180000042982 | 562735 | 0.3781 | 0 | 59.7919 | Eukaryota | 99994.0 | 0 | Chordata | 99994.0 | 0 | Carnivora | 99994.0 | 0 | Canidae | 99994.0 | 0 | Canis | 99994.0 | 0 | Canis lupus | 99994.0 | 0 |
| scf7180000042749 | 384111 | 0.4014 | 0 | 59.4331 | Eukaryota | 37285.0 | 0 | Chordata | 37285.0 | 0 | Carnivora | 37285.0 | 0 | Canidae | 37285.0 | 0 | Canis | 37285.0 | 0 | Canis lupus | 37285.0 | 0 |
| scf7180000053613 | 4919 | 0.4515 | 0 | 57.8732 | no-hit | 0.0 | 0 | no-hit | 0.0 | 0 | no-hit | 0.0 | 0 | no-hit | 0.0 | 0 | no-hit | 0.0 | 0 | no-hit | 0.0 | 0 |
| scf7180000054312 | 24240 | 0.4401 | 0 | 461.9675 | Eukaryota | 202.0 | 0 | Platyhelminthes | 202.0 | 0 | Polystomatidea | 202.0 | 0 | Polystomatidae | 202.0 | 0 | Protopolystoma | 202.0 | 0 | Protopolystoma xenopodis | 202.0 | 0 |
| scf7180000048231 | 65016 | 0.6082 | 0 | 58.1484 | Eukaryota | 23617.0 | 0 | Chordata | 23617.0 | 0 | Carnivora | 23617.0 | 0 | Canidae | 23617.0 | 0 | Vulpes | 12075.0 | 1 | Vulpes vulpes | 12075.0 | 1 |
| scf7180000054215 | 7291 | 0.4485 | 0 | 603.9513 | Eukaryota | 710.0 | 0 | Chordata | 710.0 | 0 | Salmoniformes | 710.0 | 0 | Salmonidae | 710.0 | 0 | Coregonus | 710.0 | 0 | Coregonus sp. 'balchen' | 710.0 | 0 |
| scf7180000049418 | 56954 | 0.4422 | 0 | 181.2877 | Eukaryota | 92988.0 | 0 | Chordata | 92988.0 | 0 | Carnivora | 92988.0 | 0 | Canidae | 92988.0 | 0 | Canis | 92988.0 | 0 | Canis lupus | 92988.0 | 0 |
| scf7180000051761 | 10844 | 0.5057 | 0 | 155.8092 | Eukaryota | 8468.0 | 0 | Chordata | 8468.0 | 0 | Carnivora | 8468.0 | 0 | Canidae | 8468.0 | 0 | Canis | 7398.0 | 1 | Canis lupus | 6283.0 | 2 |
| scf7180000052937 | 17441 | 0.4845 | 0 | 49.3774 | Eukaryota | 4046.0 | 0 | Chordata | 4046.0 | 0 | Carnivora | 4046.0 | 0 | Canidae | 4046.0 | 0 | Canis | 4046.0 | 0 | Canis lupus | 4046.0 | 0 |
| scf7180000053510 | 2688 | 0.4501 | 0 | 256.3941 | Eukaryota | 254.0 | 0 | Platyhelminthes | 254.0 | 0 | Polystomatidea | 254.0 | 0 | Polystomatidae | 254.0 | 0 | Protopolystoma | 254.0 | 0 | Protopolystoma xenopodis | 254.0 | 0 |
| scf7180000047020 | 124061 | 0.3959 | 0 | 58.2172 | Eukaryota | 89372.0 | 0 | Chordata | 89372.0 | 0 | Carnivora | 89372.0 | 0 | Canidae | 89372.0 | 0 | Canis | 89372.0 | 0 | Canis lupus | 89372.0 | 0 |
| scf7180000046190 | 132464 | 0.4901 | 0 | 58.398 | Eukaryota | 30482.0 | 0 | Chordata | 30482.0 | 0 | Carnivora | 30482.0 | 0 | Canidae | 25969.0 | 3 | Canis | 23372.0 | 4 | Canis lupus | 23372.0 | 4 |
| scf7180000041412 | 365340 | 0.3766 | 0 | 59.1855 | Eukaryota | 31131.0 | 0 | Chordata | 31131.0 | 0 | Carnivora | 31131.0 | 0 | Canidae | 31131.0 | 0 | Canis | 24811.0 | 1 | Canis lupus | 24811.0 | 1 |
| scf7180000043594 | 79269 | 0.4358 | 0 | 104.4874 | Eukaryota | 33184.0 | 0 | Chordata | 21249.0 | 2 | Gadiformes | 10735.0 | 6 | Gadidae | 10735.0 | 6 | Gadus | 10735.0 | 6 | Gadus morhua | 10735.0 | 6 |
| scf7180000051709 | 6988 | 0.437 | 0 | 446.3886 | Eukaryota | 2047.0 | 0 | Platyhelminthes | 1085.0 | 2 | Polystomatidea | 1085.0 | 2 | Polystomatidae | 1085.0 | 2 | Protopolystoma | 1085.0 | 2 | Protopolystoma xenopodis | 1085.0 | 2 |
| scf7180000053394 | 6544 | 0.4456 | 0 | 687.0089 | Eukaryota | 1997.0 | 0 | Platyhelminthes | 819.0 | 2 | Polystomatidea | 819.0 | 2 | Polystomatidae | 819.0 | 2 | Protopolystoma | 819.0 | 2 | Protopolystoma xenopodis | 819.0 | 2 |
| scf7180000053658 | 3307 | 0.4518 | 0 | 128.1418 | no-hit | 0.0 | 0 | no-hit | 0.0 | 0 | no-hit | 0.0 | 0 | no-hit | 0.0 | 0 | no-hit | 0.0 | 0 | no-hit | 0.0 | 0 |
| scf7180000050063 | 7745 | 0.4505 | 0 | 1195.7093 | Eukaryota | 749.0 | 0 | Platyhelminthes | 749.0 | 0 | Polystomatidea | 749.0 | 0 | Polystomatidae | 749.0 | 0 | Protopolystoma | 749.0 | 0 | Protopolystoma xenopodis | 749.0 | 0 |
| scf7180000050436 | 5422 | 0.4386 | 0 | 30.8115 | Eukaryota | 3002.0 | 0 | Chordata | 3002.0 | 0 | Carnivora | 3002.0 | 0 | Canidae | 3002.0 | 0 | Canis | 3002.0 | 0 | Canis lupus | 3002.0 | 0 |
| scf7180000047839 | 9223 | 0.4392 | 0 | 449.3084 | no-hit | 0.0 | 0 | no-hit | 0.0 | 0 | no-hit | 0.0 | 0 | no-hit | 0.0 | 0 | no-hit | 0.0 | 0 | no-hit | 0.0 | 0 |
| scf7180000048650 | 6917 | 0.4473 | 0 | 259.9547 | no-hit | 0.0 | 0 | no-hit | 0.0 | 0 | no-hit | 0.0 | 0 | no-hit | 0.0 | 0 | no-hit | 0.0 | 0 | no-hit | 0.0 | 0 |
| scf7180000042833 | 575060 | 0.3613 | 0 | 61.2783 | Eukaryota | 94486.0 | 0 | Chordata | 94486.0 | 0 | Carnivora | 94486.0 | 0 | Canidae | 94486.0 | 0 | Canis | 94486.0 | 0 | Canis lupus | 94486.0 | 0 |
| scf7180000046266 | 142962 | 0.6336 | 0 | 54.1807 | Eukaryota | 30672.0 | 0 | Chordata | 30672.0 | 0 | Carnivora | 30672.0 | 0 | Canidae | 30672.0 | 0 | Canis | 30672.0 | 0 | Canis lupus | 30672.0 | 0 |
| scf7180000053633 | 3169 | 0.508 | 0 | 51.6084 | Eukaryota | 5665.0 | 0 | Chordata | 5665.0 | 0 | Carnivora | 5665.0 | 0 | Canidae | 5665.0 | 0 | Canis | 3714.0 | 1 | Canis lupus | 2980.0 | 2 |
| scf7180000045739 | 3034 | 0.3368 | 0 | 25.9446 | Eukaryota | 2963.0 | 0 | Chordata | 2963.0 | 0 | Carnivora | 2963.0 | 0 | Canidae | 2963.0 | 0 | Canis | 2963.0 | 0 | Canis lupus | 2963.0 | 0 |
| scf7180000049774 | 12775 | 0.4557 | 0 | 89.5741 | Eukaryota | 11441.0 | 0 | Chordata | 11441.0 | 0 | Carnivora | 9625.0 | 1 | Canidae | 9625.0 | 1 | Nyctereutes | 4967.0 | 2 | Nyctereutes procyonoides | 4967.0 | 2 |
| scf7180000054261 | 23291 | 0.395 | 0 | 598.5539 | Eukaryota | 91613.0 | 0 | Chordata | 91613.0 | 0 | Carnivora | 91613.0 | 0 | Canidae | 91613.0 | 0 | Canis | 91613.0 | 0 | Canis lupus | 91613.0 | 0 |
| scf7180000041490 | 149455 | 0.3492 | 0 | 57.852 | Eukaryota | 29843.0 | 0 | Chordata | 29843.0 | 0 | Carnivora | 29843.0 | 0 | Canidae | 26683.0 | 1 | Canis | 26683.0 | 1 | Canis lupus | 26683.0 | 1 |
| scf7180000051406 | 15383 | 0.5067 | 0 | 196.4449 | Eukaryota | 8987.0 | 0 | Chordata | 8987.0 | 0 | Carnivora | 8987.0 | 0 | Canidae | 8987.0 | 0 | Canis | 7327.0 | 1 | Canis lupus | 6139.0 | 2 |
| scf7180000053055 | 9887 | 0.4518 | 0 | 763.2833 | Eukaryota | 1308.0 | 0 | Echinodermata | 669.0 | 2 | Forcipulatida | 669.0 | 2 | Asteriidae | 669.0 | 2 | Asterias | 669.0 | 2 | Asterias rubens | 669.0 | 2 |
| scf7180000047088 | 27352 | 0.4501 | 0 | 128.387 | Eukaryota | 1857.0 | 0 | Chordata | 1857.0 | 0 | Kurtiformes | 1857.0 | 0 | Apogonidae | 1857.0 | 0 | Sphaeramia | 1857.0 | 0 | Sphaeramia orbicularis | 1857.0 | 0 |
| scf7180000056368 | 740931 | 0.4314 | 0 | 59.849 | Eukaryota | 49436.0 | 0 | Chordata | 49436.0 | 0 | Carnivora | 49436.0 | 0 | Canidae | 49436.0 | 0 | Canis | 49436.0 | 0 | Canis lupus | 49436.0 | 0 |
| scf7180000043616 | 11477 | 0.4516 | 0 | 79.8316 | no-hit | 0.0 | 0 | no-hit | 0.0 | 0 | no-hit | 0.0 | 0 | no-hit | 0.0 | 0 | no-hit | 0.0 | 0 | no-hit | 0.0 | 0 |
| scf7180000054645 | 19342 | 0.4413 | 0 | 119.1309 | no-hit | 0.0 | 0 | no-hit | 0.0 | 0 | no-hit | 0.0 | 0 | no-hit | 0.0 | 0 | no-hit | 0.0 | 0 | no-hit | 0.0 | 0 |
| scf7180000042975 | 206144 | 0.5557 | 0 | 58.1875 | Eukaryota | 41425.0 | 0 | Chordata | 41425.0 | 0 | Carnivora | 41425.0 | 0 | Canidae | 41425.0 | 0 | Canis | 25684.0 | 1 | Canis lupus | 25684.0 | 1 |
| scf7180000042394 | 74905 | 0.4669 | 0 | 60.3273 | Eukaryota | 31847.0 | 0 | Chordata | 31847.0 | 0 | Carnivora | 31847.0 | 0 | Felidae | 17843.0 | 2 | Panthera | 6852.0 | 6 | Panthera pardus | 6852.0 | 6 |
| scf7180000051533 | 5904 | 0.4565 | 0 | 462.4114 | no-hit | 0.0 | 0 | no-hit | 0.0 | 0 | no-hit | 0.0 | 0 | no-hit | 0.0 | 0 | no-hit | 0.0 | 0 | no-hit | 0.0 | 0 |
| scf7180000042712 | 1101059 | 0.3654 | 0 | 59.8995 | Eukaryota | 36469.0 | 0 | Chordata | 36469.0 | 0 | Carnivora | 36469.0 | 0 | Canidae | 36469.0 | 0 | Canis | 36469.0 | 0 | Canis lupus | 36469.0 | 0 |
| scf7180000051944 | 5475 | 0.4544 | 0 | 116.2782 | Eukaryota | 4240.0 | 0 | Chordata | 3600.0 | 1 | Carnivora | 1494.0 | 3 | Canidae | 1494.0 | 3 | Canis | 1494.0 | 3 | Canis lupus | 1494.0 | 3 |
| scf7180000044010 | 684186 | 0.4614 | 0 | 58.6893 | Eukaryota | 105085.0 | 0 | Chordata | 105085.0 | 0 | Carnivora | 105085.0 | 0 | Canidae | 105085.0 | 0 | Canis | 105085.0 | 0 | Canis lupus | 105085.0 | 0 |
| scf7180000047112 | 23916 | 0.4409 | 0 | 159.383 | no-hit | 0.0 | 0 | no-hit | 0.0 | 0 | no-hit | 0.0 | 0 | no-hit | 0.0 | 0 | no-hit | 0.0 | 0 | no-hit | 0.0 | 0 |
| scf7180000050104 | 17418 | 0.4467 | 0 | 826.2497 | Eukaryota | 3846.0 | 0 | Platyhelminthes | 2297.0 | 2 | Polystomatidea | 2297.0 | 2 | Polystomatidae | 2297.0 | 2 | Protopolystoma | 2297.0 | 2 | Protopolystoma xenopodis | 2297.0 | 2 |
| scf7180000043239 | 558454 | 0.3816 | 0 | 67.1763 | Eukaryota | 90779.0 | 0 | Chordata | 90779.0 | 0 | Carnivora | 90779.0 | 0 | Canidae | 90779.0 | 0 | Canis | 90779.0 | 0 | Canis lupus | 90779.0 | 0 |
| scf7180000049510 | 4612 | 0.4441 | 0 | 94.3939 | no-hit | 0.0 | 0 | no-hit | 0.0 | 0 | no-hit | 0.0 | 0 | no-hit | 0.0 | 0 | no-hit | 0.0 | 0 | no-hit | 0.0 | 0 |
| scf7180000043187 | 521548 | 0.4432 | 0 | 57.6665 | Eukaryota | 33863.0 | 0 | Chordata | 33863.0 | 0 | Carnivora | 33863.0 | 0 | Canidae | 33863.0 | 0 | Canis | 33863.0 | 0 | Canis lupus | 33863.0 | 0 |
| scf7180000051428 | 7671 | 0.4402 | 0 | 112.7577 | no-hit | 0.0 | 0 | no-hit | 0.0 | 0 | no-hit | 0.0 | 0 | no-hit | 0.0 | 0 | no-hit | 0.0 | 0 | no-hit | 0.0 | 0 |
| scf7180000041459 | 1102285 | 0.3899 | 0 | 57.8632 | Eukaryota | 62327.0 | 0 | Chordata | 62327.0 | 0 | Carnivora | 62327.0 | 0 | Canidae | 62327.0 | 0 | Canis | 62327.0 | 0 | Canis lupus | 62327.0 | 0 |
| scf7180000047850 | 26605 | 0.4477 | 0 | 203.1764 | Eukaryota | 993.0 | 0 | Platyhelminthes | 993.0 | 0 | Polystomatidea | 993.0 | 0 | Polystomatidae | 993.0 | 0 | Protopolystoma | 993.0 | 0 | Protopolystoma xenopodis | 993.0 | 0 |
| scf7180000042152 | 401593 | 0.4518 | 0 | 59.0111 | Eukaryota | 23314.0 | 0 | Chordata | 23314.0 | 0 | Carnivora | 23314.0 | 0 | Canidae | 18470.0 | 1 | Canis | 11936.0 | 2 | Canis lupus | 11936.0 | 2 |
| scf7180000047224 | 91905 | 0.4065 | 0 | 59.7237 | Eukaryota | 55932.0 | 0 | Chordata | 55932.0 | 0 | Carnivora | 55932.0 | 0 | Canidae | 55932.0 | 0 | Canis | 44006.0 | 1 | Canis lupus | 44006.0 | 1 |
| scf7180000051411 | 6549 | 0.4544 | 0 | 690.9744 | no-hit | 0.0 | 0 | no-hit | 0.0 | 0 | no-hit | 0.0 | 0 | no-hit | 0.0 | 0 | no-hit | 0.0 | 0 | no-hit | 0.0 | 0 |
| scf7180000050711 | 25898 | 0.4606 | 0 | 56.323 | Eukaryota | 14424.0 | 0 | Chordata | 14424.0 | 0 | Carnivora | 14424.0 | 0 | Canidae | 14424.0 | 0 | Canis | 8421.0 | 1 | Canis lupus | 8421.0 | 1 |
| scf7180000051037 | 13413 | 0.4515 | 0 | 1195.7166 | Eukaryota | 5202.0 | 0 | Chordata | 3655.0 | 2 | Tetraodontiformes | 2117.0 | 5 | Tetraodontidae | 2117.0 | 5 | Takifugu | 2117.0 | 5 | Takifugu rubripes | 2117.0 | 5 |
| scf7180000045177 | 148697 | 0.3765 | 0 | 58.8434 | Eukaryota | 17719.0 | 0 | Chordata | 17719.0 | 0 | Carnivora | 17719.0 | 0 | Felidae | 15894.0 | 1 | Panthera | 15894.0 | 1 | Panthera pardus | 15894.0 | 1 |
| scf7180000054343 | 632313 | 0.3399 | 0 | 59.7211 | Eukaryota | 89992.0 | 0 | Chordata | 89992.0 | 0 | Carnivora | 89992.0 | 0 | Canidae | 89992.0 | 0 | Canis | 89992.0 | 0 | Canis lupus | 89992.0 | 0 |
| scf7180000052311 | 4917 | 0.5034 | 0 | 343.0411 | Eukaryota | 8323.0 | 0 | Chordata | 8323.0 | 0 | Carnivora | 8323.0 | 0 | Canidae | 8323.0 | 0 | Canis | 5607.0 | 1 | Canis lupus | 4367.0 | 2 |
| scf7180000050434 | 28330 | 0.3555 | 0 | 55.0472 | Eukaryota | 11513.0 | 0 | Chordata | 11513.0 | 0 | Primates | 8318.0 | 2 | Hominidae | 4614.0 | 4 | Homo | 4614.0 | 4 | Homo sapiens | 4614.0 | 4 |
| scf7180000054110 | 5824 | 0.345 | 0 | 28.9081 | Eukaryota | 2874.0 | 0 | Chordata | 2874.0 | 0 | Carnivora | 1705.0 | 2 | Canidae | 1705.0 | 2 | Canis | 1705.0 | 2 | Canis lupus | 1705.0 | 2 |
| scf7180000050938 | 11629 | 0.4368 | 0 | 30.1413 | Eukaryota | 2078.0 | 0 | Chordata | 2078.0 | 0 | Kurtiformes | 1301.0 | 2 | Apogonidae | 1301.0 | 2 | Sphaeramia | 1301.0 | 2 | Sphaeramia orbicularis | 1301.0 | 2 |
| scf7180000045262 | 171110 | 0.4867 | 0 | 57.6316 | Eukaryota | 13983.0 | 0 | Chordata | 13983.0 | 0 | Carnivora | 12854.0 | 1 | Canidae | 9335.0 | 2 | Canis | 9335.0 | 2 | Canis lupus | 9335.0 | 2 |
| scf7180000041813 | 121758 | 0.3338 | 0 | 57.8779 | Eukaryota | 11661.0 | 1 | Chordata | 11661.0 | 1 | Carnivora | 6601.0 | 3 | undef | 3592.0 | 5 | undef | 3592.0 | 7 | eukaryotic synthetic construct | 3592.0 | 7 |
| scf7180000048526 | 7558 | 0.4407 | 0 | 48.1324 | Eukaryota | 749.0 | 0 | Platyhelminthes | 749.0 | 0 | Polystomatidea | 749.0 | 0 | Polystomatidae | 749.0 | 0 | Protopolystoma | 749.0 | 0 | Protopolystoma xenopodis | 749.0 | 0 |
| scf7180000045170 | 23402 | 0.4414 | 0 | 108.6151 | Eukaryota | 9909.0 | 0 | Chordata | 5571.0 | 2 | Kurtiformes | 4102.0 | 4 | Apogonidae | 4102.0 | 4 | Sphaeramia | 4102.0 | 4 | Sphaeramia orbicularis | 4102.0 | 4 |
| scf7180000045020 | 261751 | 0.3814 | 0 | 60.2292 | Eukaryota | 46507.0 | 0 | Chordata | 46507.0 | 0 | Carnivora | 46507.0 | 0 | Canidae | 46507.0 | 0 | Canis | 46507.0 | 0 | Canis lupus | 46507.0 | 0 |
| scf7180000047142 | 19590 | 0.4391 | 0 | 178.4732 | Eukaryota | 93198.0 | 0 | Chordata | 93198.0 | 0 | Carnivora | 93198.0 | 0 | Canidae | 93198.0 | 0 | Canis | 93198.0 | 0 | Canis lupus | 93198.0 | 0 |
| scf7180000055292 | 18163 | 0.4485 | 0 | 205.6686 | Eukaryota | 4081.0 | 0 | Chordata | 4081.0 | 0 | Carnivora | 2688.0 | 1 | Canidae | 2688.0 | 1 | Canis | 2688.0 | 1 | Canis lupus | 2688.0 | 1 |
| scf7180000041818 | 8402 | 0.4175 | 0 | 28.1526 | Eukaryota | 8852.0 | 0 | Chordata | 8852.0 | 0 | Carnivora | 8852.0 | 0 | Canidae | 8852.0 | 0 | Canis | 8852.0 | 0 | Canis lupus | 8852.0 | 0 |
| scf7180000041051 | 1079690 | 0.3911 | 0 | 58.7475 | Eukaryota | 92230.0 | 0 | Chordata | 92230.0 | 0 | Carnivora | 92230.0 | 0 | Canidae | 92230.0 | 0 | Canis | 92230.0 | 0 | Canis lupus | 92230.0 | 0 |
| scf7180000044788 | 222977 | 0.4886 | 0 | 59.3294 | Eukaryota | 48640.0 | 0 | Chordata | 48640.0 | 0 | Carnivora | 48640.0 | 0 | Canidae | 48640.0 | 0 | Canis | 27792.0 | 1 | Canis lupus | 27792.0 | 1 |
| scf7180000046153 | 15883 | 0.4085 | 0 | 551.8348 | Eukaryota | 87579.0 | 0 | Chordata | 87579.0 | 0 | Carnivora | 87579.0 | 0 | Canidae | 87579.0 | 0 | Canis | 87579.0 | 0 | Canis lupus | 87579.0 | 0 |
| scf7180000043933 | 374163 | 0.4072 | 0 | 60.2021 | Eukaryota | 49842.0 | 0 | Chordata | 49842.0 | 0 | Carnivora | 49842.0 | 0 | Canidae | 49842.0 | 0 | Canis | 43640.0 | 1 | Canis lupus | 43640.0 | 1 |
| scf7180000049597 | 12916 | 0.4395 | 0 | 59.1514 | Eukaryota | 733.0 | 0 | Platyhelminthes | 533.0 | 1 | Polystomatidea | 533.0 | 1 | Polystomatidae | 533.0 | 1 | Protopolystoma | 533.0 | 1 | Protopolystoma xenopodis | 533.0 | 1 |
| scf7180000056425 | 150434 | 0.4548 | 0 | 58.3066 | Eukaryota | 9552.0 | 0 | Chordata | 9552.0 | 0 | Carnivora | 5102.0 | 2 | Canidae | 3396.0 | 3 | Canis | 3396.0 | 4 | Canis lupus | 3396.0 | 4 |
| scf7180000050769 | 25362 | 0.5749 | 0 | 56.3436 | Eukaryota | 21508.0 | 0 | Chordata | 21508.0 | 0 | Carnivora | 21508.0 | 0 | Canidae | 16704.0 | 4 | Canis | 14914.0 | 7 | Canis lupus | 14914.0 | 7 |
| scf7180000055543 | 9649 | 0.4351 | 0 | 261.081 | Eukaryota | 39770.0 | 0 | Chordata | 27566.0 | 2 | Gadiformes | 22769.0 | 5 | Gadidae | 22769.0 | 5 | Gadus | 22769.0 | 5 | Gadus morhua | 22769.0 | 5 |
| scf7180000051755 | 2225 | 0.4467 | 0 | 145.1958 | no-hit | 0.0 | 0 | no-hit | 0.0 | 0 | no-hit | 0.0 | 0 | no-hit | 0.0 | 0 | no-hit | 0.0 | 0 | no-hit | 0.0 | 0 |
| scf7180000040869 | 269779 | 0.3724 | 100 | 60.2905 | Eukaryota | 86351.0 | 0 | Chordata | 86351.0 | 0 | Carnivora | 86351.0 | 0 | Canidae | 86351.0 | 0 | Canis | 86351.0 | 0 | Canis lupus | 86351.0 | 0 |
| scf7180000051939 | 2067 | 0.4514 | 0 | 373.8981 | no-hit | 0.0 | 0 | no-hit | 0.0 | 0 | no-hit | 0.0 | 0 | no-hit | 0.0 | 0 | no-hit | 0.0 | 0 | no-hit | 0.0 | 0 |
| scf7180000042973 | 63593 | 0.5343 | 0 | 57.0149 | Eukaryota | 59676.0 | 0 | Chordata | 59676.0 | 0 | Carnivora | 59676.0 | 0 | Canidae | 59676.0 | 0 | Canis | 59676.0 | 0 | Canis lupus | 59676.0 | 0 |
| scf7180000053650 | 2605 | 0.4683 | 0 | 134.9038 | no-hit | 0.0 | 0 | no-hit | 0.0 | 0 | no-hit | 0.0 | 0 | no-hit | 0.0 | 0 | no-hit | 0.0 | 0 | no-hit | 0.0 | 0 |
| scf7180000048377 | 3254 | 0.4459 | 0 | 293.5249 | no-hit | 0.0 | 0 | no-hit | 0.0 | 0 | no-hit | 0.0 | 0 | no-hit | 0.0 | 0 | no-hit | 0.0 | 0 | no-hit | 0.0 | 0 |
| scf7180000045701 | 179028 | 0.3747 | 0 | 61.7725 | Eukaryota | 41483.0 | 0 | Chordata | 41483.0 | 0 | Carnivora | 41483.0 | 0 | Canidae | 34252.0 | 2 | Canis | 25620.0 | 3 | Canis lupus | 25620.0 | 3 |
| scf7180000056051 | 26290 | 0.4251 | 0 | 219.3195 | Eukaryota | 6598.0 | 0 | Chordata | 6598.0 | 0 | Carnivora | 6598.0 | 0 | Canidae | 6598.0 | 0 | Canis | 6598.0 | 0 | Canis lupus | 6598.0 | 0 |
| scf7180000042380 | 575120 | 0.3913 | 0 | 59.3019 | Eukaryota | 37771.0 | 0 | Chordata | 37771.0 | 0 | Carnivora | 37771.0 | 0 | Canidae | 37771.0 | 0 | Canis | 37771.0 | 0 | Canis lupus | 37771.0 | 0 |
| scf7180000045001 | 263474 | 0.3655 | 0 | 60.5785 | Eukaryota | 18536.0 | 0 | Chordata | 18536.0 | 0 | Carnivora | 8652.0 | 2 | Hominidae | 8303.0 | 4 | Homo | 8303.0 | 5 | Homo sapiens | 8303.0 | 5 |
| scf7180000053506 | 2577 | 0.4505 | 0 | 36.3518 | Eukaryota | 2209.0 | 0 | Chordata | 2209.0 | 0 | Carnivora | 2209.0 | 0 | Canidae | 2209.0 | 0 | Canis | 2209.0 | 0 | Canis lupus | 2209.0 | 0 |
| scf7180000053464 | 5705 | 0.4503 | 0 | 40.4353 | no-hit | 0.0 | 0 | no-hit | 0.0 | 0 | no-hit | 0.0 | 0 | no-hit | 0.0 | 0 | no-hit | 0.0 | 0 | no-hit | 0.0 | 0 |
| scf7180000054012 | 2827 | 0.4627 | 0 | 55.2284 | no-hit | 0.0 | 0 | no-hit | 0.0 | 0 | no-hit | 0.0 | 0 | no-hit | 0.0 | 0 | no-hit | 0.0 | 0 | no-hit | 0.0 | 0 |
| scf7180000043843 | 394192 | 0.3618 | 0 | 59.6946 | Eukaryota | 100467.0 | 0 | Chordata | 100467.0 | 0 | Chiroptera | 58899.0 | 1 | Pteropodidae | 58899.0 | 1 | Rousettus | 41615.0 | 3 | Rousettus aegyptiacus | 41615.0 | 4 |
| scf7180000040806 | 1687423 | 0.4015 | 0 | 65.2963 | Eukaryota | 94776.0 | 0 | Chordata | 94776.0 | 0 | Carnivora | 94776.0 | 0 | Canidae | 94776.0 | 0 | Canis | 94776.0 | 0 | Canis lupus | 94776.0 | 0 |
| scf7180000046488 | 78431 | 0.3918 | 0 | 59.5312 | Eukaryota | 11126.0 | 0 | Chordata | 11126.0 | 0 | Carnivora | 9308.0 | 1 | Canidae | 6448.0 | 3 | Canis | 5229.0 | 5 | Canis lupus | 5229.0 | 5 |
| scf7180000054349 | 35159 | 0.442 | 0 | 261.2209 | Eukaryota | 527.0 | 0 | Chordata | 527.0 | 0 | Tetraodontiformes | 318.0 | 1 | Tetraodontidae | 318.0 | 1 | Takifugu | 318.0 | 1 | Takifugu rubripes | 318.0 | 1 |
| scf7180000042264 | 485424 | 0.4088 | 0 | 59.217 | Eukaryota | 63266.0 | 0 | Chordata | 63266.0 | 0 | Carnivora | 63266.0 | 0 | Canidae | 63266.0 | 0 | Vulpes | 58056.0 | 1 | Vulpes vulpes | 58056.0 | 1 |
| scf7180000052979 | 4057 | 0.4501 | 0 | 1066.6565 | no-hit | 0.0 | 0 | no-hit | 0.0 | 0 | no-hit | 0.0 | 0 | no-hit | 0.0 | 0 | no-hit | 0.0 | 0 | no-hit | 0.0 | 0 |
| scf7180000050700 | 9250 | 0.4373 | 0 | 93.7207 | no-hit | 0.0 | 0 | no-hit | 0.0 | 0 | no-hit | 0.0 | 0 | no-hit | 0.0 | 0 | no-hit | 0.0 | 0 | no-hit | 0.0 | 0 |
| scf7180000044568 | 296546 | 0.4483 | 0 | 78.7465 | Eukaryota | 23550.0 | 0 | Chordata | 23550.0 | 0 | Carnivora | 23550.0 | 0 | Canidae | 17565.0 | 1 | Canis | 14375.0 | 2 | Canis lupus | 14375.0 | 3 |
| scf7180000042146 | 325792 | 0.3399 | 0 | 57.7434 | Eukaryota | 91960.0 | 0 | Chordata | 91960.0 | 0 | Carnivora | 91960.0 | 0 | Canidae | 91960.0 | 0 | Canis | 91960.0 | 0 | Canis lupus | 91960.0 | 0 |
| scf7180000055829 | 43771 | 0.4007 | 0 | 91.8708 | Eukaryota | 94940.0 | 0 | Chordata | 94940.0 | 0 | Carnivora | 94940.0 | 0 | Canidae | 94940.0 | 0 | Canis | 94940.0 | 0 | Canis lupus | 94940.0 | 0 |
| scf7180000053049 | 4149 | 0.4391 | 0 | 694.8868 | no-hit | 0.0 | 0 | no-hit | 0.0 | 0 | no-hit | 0.0 | 0 | no-hit | 0.0 | 0 | no-hit | 0.0 | 0 | no-hit | 0.0 | 0 |
| scf7180000056075 | 251029 | 0.3951 | 0 | 69.5951 | Eukaryota | 92338.0 | 0 | Chordata | 92338.0 | 0 | Carnivora | 92338.0 | 0 | Canidae | 92338.0 | 0 | Canis | 92338.0 | 0 | Canis lupus | 92338.0 | 0 |
| scf7180000050099 | 31454 | 0.4384 | 0 | 57.402 | Eukaryota | 429.0 | 0 | Chordata | 429.0 | 0 | Salmoniformes | 429.0 | 0 | Salmonidae | 429.0 | 0 | Coregonus | 429.0 | 0 | Coregonus sp. 'balchen' | 429.0 | 0 |
| scf7180000046227 | 32993 | 0.4328 | 0 | 105.5605 | Eukaryota | 9203.0 | 0 | Chordata | 4811.0 | 2 | Forcipulatida | 3874.0 | 4 | Asteriidae | 3874.0 | 4 | Asterias | 3874.0 | 4 | Asterias rubens | 3874.0 | 4 |
| scf7180000043292 | 550313 | 0.3752 | 0 | 59.2839 | Eukaryota | 62734.0 | 0 | Chordata | 62734.0 | 0 | Carnivora | 62734.0 | 0 | Canidae | 62734.0 | 0 | Canis | 62734.0 | 0 | Canis lupus | 62734.0 | 0 |
| scf7180000053914 | 6546 | 0.3961 | 0 | 96.4587 | Eukaryota | 1236.0 | 0 | Chordata | 1082.0 | 1 | Artiodactyla | 791.0 | 2 | Bovidae | 791.0 | 2 | Ovis | 791.0 | 2 | Ovis canadensis | 791.0 | 2 |
| scf7180000046372 | 151480 | 0.3676 | 0 | 60.7986 | Eukaryota | 43725.0 | 0 | Chordata | 43725.0 | 0 | Carnivora | 43725.0 | 0 | Canidae | 43725.0 | 0 | Canis | 43725.0 | 0 | Canis lupus | 43725.0 | 0 |
| scf7180000042686 | 612837 | 0.4702 | 0 | 58.498 | Eukaryota | 84876.0 | 0 | Chordata | 84876.0 | 0 | Carnivora | 84876.0 | 0 | Canidae | 84876.0 | 0 | Canis | 63177.0 | 1 | Canis lupus | 63177.0 | 1 |
| scf7180000050282 | 11676 | 0.4435 | 0 | 44.8406 | Eukaryota | 55152.0 | 0 | Chordata | 55152.0 | 0 | Carnivora | 55152.0 | 0 | Canidae | 55152.0 | 0 | Canis | 55152.0 | 0 | Canis lupus | 55152.0 | 0 |
| scf7180000052264 | 4723 | 0.4461 | 0 | 85.4262 | no-hit | 0.0 | 0 | no-hit | 0.0 | 0 | no-hit | 0.0 | 0 | no-hit | 0.0 | 0 | no-hit | 0.0 | 0 | no-hit | 0.0 | 0 |
| scf7180000049198 | 37949 | 0.54 | 0 | 59.022 | Eukaryota | 40797.0 | 0 | Chordata | 40797.0 | 0 | Carnivora | 40797.0 | 0 | Canidae | 34440.0 | 1 | Canis | 25986.0 | 2 | Canis lupus | 25986.0 | 2 |
| scf7180000042857 | 24680 | 0.3916 | 0 | 59.8986 | Eukaryota | 29706.0 | 0 | Chordata | 29706.0 | 0 | Carnivora | 29706.0 | 0 | Canidae | 29706.0 | 0 | Canis | 29706.0 | 0 | Canis lupus | 29706.0 | 0 |
| scf7180000051115 | 12663 | 0.4374 | 0 | 119.2589 | Eukaryota | 4150.0 | 0 | Chordata | 2542.0 | 1 | Polystomatidea | 1608.0 | 4 | Polystomatidae | 1608.0 | 4 | Protopolystoma | 1608.0 | 4 | Protopolystoma xenopodis | 1608.0 | 4 |
| scf7180000042469 | 30581 | 0.3849 | 0 | 57.9643 | Eukaryota | 85187.0 | 0 | Chordata | 85187.0 | 0 | Carnivora | 85187.0 | 0 | Canidae | 85187.0 | 0 | Canis | 85187.0 | 0 | Canis lupus | 85187.0 | 0 |
| scf7180000047264 | 31961 | 0.4214 | 0 | 53.4546 | Eukaryota | 93405.0 | 0 | Chordata | 93405.0 | 0 | Carnivora | 93405.0 | 0 | Canidae | 93405.0 | 0 | Canis | 93405.0 | 0 | Canis lupus | 93405.0 | 0 |
| scf7180000046063 | 22580 | 0.4332 | 0 | 51.3752 | Eukaryota | 87637.0 | 0 | Chordata | 87637.0 | 0 | Carnivora | 87637.0 | 0 | Canidae | 87637.0 | 0 | Canis | 87637.0 | 0 | Canis lupus | 87637.0 | 0 |
| scf7180000054008 | 3544 | 0.5996 | 0 | 51.067 | no-hit | 0.0 | 0 | no-hit | 0.0 | 0 | no-hit | 0.0 | 0 | no-hit | 0.0 | 0 | no-hit | 0.0 | 0 | no-hit | 0.0 | 0 |
| scf7180000047831 | 27501 | 0.5022 | 0 | 327.4843 | Eukaryota | 9832.0 | 0 | Chordata | 9832.0 | 0 | Carnivora | 9832.0 | 0 | Canidae | 9832.0 | 0 | Canis | 9143.0 | 1 | Canis lupus | 7855.0 | 2 |
| scf7180000042987 | 633217 | 0.3518 | 0 | 59.3977 | Eukaryota | 91107.0 | 0 | Chordata | 91107.0 | 0 | Carnivora | 91107.0 | 0 | Canidae | 91107.0 | 0 | Canis | 91107.0 | 0 | Canis lupus | 91107.0 | 0 |
| scf7180000040780 | 51892 | 0.4823 | 0 | 151.569 | Eukaryota | 95043.0 | 0 | Chordata | 95043.0 | 0 | Carnivora | 95043.0 | 0 | Canidae | 95043.0 | 0 | Canis | 95043.0 | 0 | Canis lupus | 95043.0 | 0 |
| scf7180000051388 | 3229 | 0.4487 | 0 | 391.0523 | no-hit | 0.0 | 0 | no-hit | 0.0 | 0 | no-hit | 0.0 | 0 | no-hit | 0.0 | 0 | no-hit | 0.0 | 0 | no-hit | 0.0 | 0 |
| scf7180000044640 | 180896 | 0.3665 | 0 | 59.9781 | Eukaryota | 26367.0 | 0 | Chordata | 26367.0 | 0 | Carnivora | 26367.0 | 0 | Canidae | 21985.0 | 2 | Canis | 16350.0 | 3 | Canis lupus | 16350.0 | 3 |
| scf7180000051005 | 29850 | 0.4747 | 0 | 57.5487 | Eukaryota | 35140.0 | 0 | Chordata | 35140.0 | 0 | Carnivora | 35140.0 | 0 | Canidae | 35140.0 | 0 | Canis | 35140.0 | 0 | Canis lupus | 35140.0 | 0 |
| scf7180000042107 | 805237 | 0.4386 | 0 | 56.5127 | Eukaryota | 34436.0 | 0 | Chordata | 34436.0 | 0 | Carnivora | 34436.0 | 0 | Canidae | 34436.0 | 0 | Canis | 34436.0 | 0 | Canis lupus | 34436.0 | 0 |
| scf7180000043713 | 396994 | 0.5601 | 0 | 57.9492 | Eukaryota | 38163.0 | 0 | Chordata | 38163.0 | 0 | Carnivora | 38163.0 | 0 | Canidae | 38163.0 | 0 | Vulpes | 25050.0 | 1 | Vulpes vulpes | 25050.0 | 1 |
| scf7180000042786 | 8524 | 0.3932 | 0 | 30.6956 | Eukaryota | 6491.0 | 0 | Chordata | 6491.0 | 0 | Carnivora | 6491.0 | 0 | Canidae | 5778.0 | 1 | Canis | 5778.0 | 1 | Canis lupus | 5778.0 | 1 |
| scf7180000042412 | 206513 | 0.4425 | 0 | 58.2713 | Eukaryota | 19914.0 | 0 | Chordata | 19914.0 | 0 | Carnivora | 19914.0 | 0 | Canidae | 19914.0 | 0 | Canis | 19914.0 | 0 | Canis lupus | 19914.0 | 0 |
| scf7180000045267 | 22134 | 0.3571 | 0 | 52.4457 | Eukaryota | 6357.0 | 0 | Chordata | 6357.0 | 0 | Primates | 2096.0 | 3 | Hominidae | 2096.0 | 3 | Homo | 2096.0 | 4 | Homo sapiens | 2096.0 | 4 |
| scf7180000044001 | 411781 | 0.353 | 0 | 60.4681 | Eukaryota | 73171.0 | 0 | Chordata | 73171.0 | 0 | Carnivora | 73171.0 | 0 | Canidae | 68705.0 | 1 | Canis | 55156.0 | 2 | Canis lupus | 55156.0 | 2 |
| scf7180000042302 | 39520 | 0.3787 | 0 | 29.7289 | Eukaryota | 90640.0 | 0 | Chordata | 90640.0 | 0 | Carnivora | 90640.0 | 0 | Canidae | 90640.0 | 0 | Canis | 90640.0 | 0 | Canis lupus | 90640.0 | 0 |
| scf7180000045526 | 2580 | 0.3798 | 0 | 48.3118 | Eukaryota | 6337.0 | 0 | Chordata | 6337.0 | 0 | Carnivora | 6337.0 | 0 | Canidae | 6337.0 | 0 | Canis | 6337.0 | 0 | Canis lupus | 6337.0 | 0 |
| scf7180000045686 | 169694 | 0.5344 | 0 | 57.0131 | Eukaryota | 16411.0 | 0 | Chordata | 16411.0 | 0 | Carnivora | 16411.0 | 0 | Canidae | 16411.0 | 0 | Canis | 16411.0 | 0 | Canis lupus | 16411.0 | 0 |
| scf7180000054658 | 50405 | 0.3908 | 0 | 1159.5253 | Eukaryota | 89482.0 | 0 | Chordata | 89482.0 | 0 | Carnivora | 89482.0 | 0 | Canidae | 89482.0 | 0 | Canis | 89482.0 | 0 | Canis lupus | 89482.0 | 0 |
| scf7180000047117 | 105818 | 0.56 | 0 | 55.9435 | Eukaryota | 7873.0 | 0 | Chordata | 7873.0 | 0 | Carnivora | 7873.0 | 0 | Canidae | 2572.0 | 4 | Canis | 1720.0 | 7 | Canis lupus | 1720.0 | 8 |

| | | | | | | | | | | | | | | | | | | | | | | |
|---|---|---|---|---|---|---|---|---|---|---|---|---|---|---|---|---|---|---|---|---|---|---|
| scf7180000045040 | 268627 | 0.3791 | 0 | 61.3073 | Eukaryota | 33481.0 | 0 | Chordata | 33481.0 | 0 | Primates | 20897.0 | 2 | Hominidae | 20897.0 | 2 | Homo | 20897.0 | 3 | Homo sapiens | 20897.0 | 3 |
| scf7180000051030 | 13931 | 0.4787 | 0 | 56.5228 | Eukaryota | 19093.0 | 0 | Chordata | 19093.0 | 0 | Carnivora | 19093.0 | 0 | Canidae | 19093.0 | 0 | Canis | 19093.0 | 0 | Canis lupus | 19093.0 | 0 |
| scf7180000046747 | 23025 | 0.4344 | 0 | 106.6244 | Eukaryota | 12284.0 | 0 | Chordata | 11842.0 | 1 | Tetraodontiformes | 7600.0 | 3 | Tetraodontidae | 7600.0 | 3 | Takifugu | 7600.0 | 3 | Takifugu rubripes | 7600.0 | 3 |
| scf7180000055752 | 26327 | 0.4477 | 0 | 205.3812 | Eukaryota | 41968.0 | 0 | Chordata | 26922.0 | 2 | Gadiformes | 7845.0 | 8 | Gadidae | 7845.0 | 8 | Gadus | 7845.0 | 8 | Gadus morhua | 7845.0 | 8 |
| scf7180000045867 | 3001 | 0.4848 | 0 | 47.1271 | Eukaryota | 18861.0 | 0 | Chordata | 18861.0 | 0 | Carnivora | 18861.0 | 0 | Canidae | 18861.0 | 0 | Canis | 18861.0 | 0 | Canis lupus | 18861.0 | 0 |
| scf7180000050702 | 16774 | 0.3235 | 0 | 60.1291 | Eukaryota | 4156.0 | 0 | Chordata | 4156.0 | 0 | Carnivora | 4156.0 | 0 | Canidae | 4156.0 | 0 | Canis | 4156.0 | 0 | Canis lupus | 4156.0 | 0 |
| scf7180000052656 | 16632 | 0.5022 | 0 | 589.84 | Eukaryota | 9570.0 | 0 | Chordata | 9570.0 | 0 | Carnivora | 9570.0 | 0 | Canidae | 9570.0 | 0 | Canis | 8894.0 | 1 | Canis lupus | 7632.0 | 2 |
| scf7180000051955 | 17669 | 0.5188 | 0 | 367.9856 | Eukaryota | 15925.0 | 0 | Chordata | 15925.0 | 0 | Carnivora | 15925.0 | 0 | Canidae | 15925.0 | 0 | Canis | 15925.0 | 0 | Canis lupus | 15925.0 | 0 |
| scf7180000054462 | 530230 | 0.4173 | 0 | 60.7966 | Eukaryota | 17503.0 | 0 | Chordata | 17503.0 | 0 | Carnivora | 17503.0 | 0 | Canidae | 17503.0 | 0 | Canis | 17503.0 | 0 | Canis lupus | 17503.0 | 0 |
| scf7180000048571 | 11779 | 0.4431 | 0 | 130.5619 | no-hit | 0.0 | 0 | no-hit | 0.0 | 0 | no-hit | 0.0 | 0 | no-hit | 0.0 | 0 | no-hit | 0.0 | 0 | no-hit | 0.0 | 0 |
| scf7180000053355 | 9602 | 0.5042 | 0 | 140.3123 | Eukaryota | 8705.0 | 0 | Chordata | 8705.0 | 0 | Carnivora | 8705.0 | 0 | Canidae | 8705.0 | 0 | Canis | 7568.0 | 1 | Canis lupus | 6370.0 | 2 |
| scf7180000047367 | 81897 | 0.4468 | 0 | 167.8372 | Eukaryota | 1000.0 | 0 | Chordata | 1000.0 | 0 | Kurtiformes | 824.0 | 1 | Apogonidae | 824.0 | 1 | Sphaeramia | 824.0 | 1 | Sphaeramia orbicularis | 824.0 | 1 |
| scf7180000043236 | 552253 | 0.3725 | 0 | 59.9575 | Eukaryota | 95176.0 | 0 | Chordata | 95176.0 | 0 | Carnivora | 95176.0 | 0 | Canidae | 95176.0 | 0 | Canis | 95176.0 | 0 | Canis lupus | 95176.0 | 0 |
| scf7180000052832 | 6130 | 0.4444 | 0 | 205.5425 | Eukaryota | 246.0 | 0 | Streptophyta | 246.0 | 0 | Fabales | 246.0 | 0 | Fabaceae | 246.0 | 0 | Cicer | 246.0 | 0 | Cicer arietinum | 246.0 | 0 |
| scf7180000049608 | 31108 | 0.4442 | 0 | 79.4841 | Eukaryota | 14556.0 | 0 | Chordata | 7795.0 | 3 | Gadiformes | 4819.0 | 8 | Gadidae | 4819.0 | 8 | Gadus | 4819.0 | 8 | Gadus morhua | 4819.0 | 8 |
| scf7180000055088 | 3933 | 0.4422 | 0 | 109.2677 | no-hit | 0.0 | 0 | no-hit | 0.0 | 0 | no-hit | 0.0 | 0 | no-hit | 0.0 | 0 | no-hit | 0.0 | 0 | no-hit | 0.0 | 0 |
| scf7180000041549 | 1100033 | 0.386 | 0 | 59.8197 | Eukaryota | 53734.0 | 0 | Chordata | 53734.0 | 0 | Carnivora | 24268.0 | 2 | Canidae | 24268.0 | 2 | Canis | 24268.0 | 2 | Canis lupus | 24268.0 | 2 |
| scf7180000044034 | 167440 | 0.3833 | 0 | 57.9383 | Eukaryota | 38422.0 | 0 | Chordata | 38422.0 | 0 | Carnivora | 38422.0 | 0 | Canidae | 34772.0 | 1 | Vulpes | 34772.0 | 1 | Vulpes vulpes | 34772.0 | 1 |
| scf7180000043867 | 382746 | 0.3743 | 0 | 61.1296 | Eukaryota | 49114.0 | 0 | Chordata | 49114.0 | 0 | Carnivora | 49114.0 | 0 | Canidae | 49114.0 | 0 | Canis | 49114.0 | 0 | Canis lupus | 49114.0 | 0 |
| scf7180000043237 | 283459 | 0.3476 | 0 | 59.4044 | Eukaryota | 48957.0 | 0 | Chordata | 48957.0 | 0 | Carnivora | 48957.0 | 0 | Canidae | 48957.0 | 0 | Canis | 48957.0 | 0 | Canis lupus | 48957.0 | 0 |
| scf7180000043784 | 390391 | 0.389 | 0 | 61.0043 | Eukaryota | 82099.0 | 0 | Chordata | 82099.0 | 0 | Carnivora | 82099.0 | 0 | Canidae | 82099.0 | 0 | Canis | 73260.0 | 1 | Canis lupus | 73260.0 | 1 |
| scf7180000044459 | 17516 | 0.4325 | 0 | 153.1655 | Eukaryota | 41223.0 | 0 | Chordata | 41223.0 | 0 | Carnivora | 41223.0 | 0 | Canidae | 41223.0 | 0 | Canis | 41223.0 | 0 | Canis lupus | 41223.0 | 0 |
| scf7180000052987 | 5313 | 0.4734 | 0 | 224.195 | Eukaryota | 6710.0 | 0 | Chordata | 6710.0 | 0 | Carnivora | 6710.0 | 0 | Canidae | 6710.0 | 0 | Canis | 4302.0 | 1 | Canis lupus | 3336.0 | 2 |
| scf7180000051490 | 3604 | 0.4351 | 0 | 222.0228 | no-hit | 0.0 | 0 | no-hit | 0.0 | 0 | no-hit | 0.0 | 0 | no-hit | 0.0 | 0 | no-hit | 0.0 | 0 | no-hit | 0.0 | 0 |
| scf7180000040690 | 999213 | 0.3839 | 0 | 59.4181 | Eukaryota | 93466.0 | 0 | Chordata | 93466.0 | 0 | Carnivora | 93466.0 | 0 | Canidae | 93466.0 | 0 | Canis | 93466.0 | 0 | Canis lupus | 93466.0 | 0 |
| scf7180000055571 | 46491 | 0.4394 | 0 | 453.533 | Eukaryota | 22827.0 | 0 | Chordata | 20155.0 | 2 | Gadiformes | 12887.0 | 6 | Gadidae | 12887.0 | 6 | Gadus | 12887.0 | 6 | Gadus morhua | 12887.0 | 6 |
| scf7180000051793 | 5543 | 0.4541 | 0 | 437.8122 | Eukaryota | 444.0 | 0 | Streptophyta | 444.0 | 0 | Fabales | 444.0 | 0 | Fabaceae | 444.0 | 0 | Cicer | 444.0 | 0 | Cicer arietinum | 444.0 | 0 |
| scf7180000042965 | 270923 | 0.3722 | 0 | 59.2087 | Eukaryota | 17473.0 | 0 | Chordata | 17473.0 | 0 | Carnivora | 17473.0 | 0 | Canidae | 17473.0 | 0 | Canis | 17473.0 | 0 | Canis lupus | 17473.0 | 0 |
| scf7180000056270 | 847018 | 0.4169 | 0 | 59.1171 | Eukaryota | 82527.0 | 0 | Chordata | 82527.0 | 0 | Carnivora | 82527.0 | 0 | Canidae | 82527.0 | 0 | Canis | 74663.0 | 1 | Canis lupus | 74663.0 | 1 |
| scf7180000045682 | 118853 | 0.3948 | 0 | 60.6653 | Eukaryota | 58390.0 | 0 | Chordata | 58390.0 | 0 | Carnivora | 58390.0 | 0 | Canidae | 58390.0 | 0 | Canis | 52175.0 | 1 | Canis lupus | 52175.0 | 1 |
| scf7180000045200 | 82339 | 0.4404 | 0 | 102.66 | Eukaryota | 9627.0 | 0 | Chordata | 7055.0 | 1 | Tetraodontiformes | 3046.0 | 6 | Tetraodontidae | 3046.0 | 6 | Takifugu | 3046.0 | 6 | Takifugu rubripes | 3046.0 | 6 |
| scf7180000046842 | 94216 | 0.4632 | 0 | 58.7614 | Eukaryota | 24861.0 | 1 | Chordata | 24861.0 | 1 | Carnivora | 16019.0 | 3 | Canidae | 16019.0 | 3 | Canis | 16019.0 | 4 | Canis lupus | 16019.0 | 4 |
| scf7180000041938 | 838069 | 0.3631 | 0 | 60.8534 | Eukaryota | 90682.0 | 0 | Chordata | 90682.0 | 0 | Carnivora | 90682.0 | 0 | Canidae | 90682.0 | 0 | Canis | 90682.0 | 0 | Canis lupus | 90682.0 | 0 |
| scf7180000040948 | 674817 | 0.4281 | 0 | 59.1029 | Eukaryota | 56801.0 | 0 | Chordata | 56801.0 | 0 | Carnivora | 56801.0 | 0 | Canidae | 56801.0 | 0 | Canis | 49966.0 | 1 | Canis lupus | 49966.0 | 1 |
| scf7180000056121 | 2731 | 0.4405 | 0 | 442.2086 | Eukaryota | 11064.0 | 0 | Chordata | 11064.0 | 0 | Gadiformes | 4748.0 | 5 | Gadidae | 4748.0 | 5 | Gadus | 4748.0 | 5 | Gadus morhua | 4748.0 | 5 |
| scf7180000044475 | 323615 | 0.5117 | 0 | 58.9247 | Eukaryota | 29621.0 | 0 | Chordata | 29621.0 | 0 | Carnivora | 29621.0 | 0 | Canidae | 29621.0 | 0 | Canis | 26716.0 | 1 | Canis lupus | 26716.0 | 1 |
| scf7180000052528 | 9454 | 0.4479 | 0 | 1287.1901 | no-hit | 0.0 | 0 | no-hit | 0.0 | 0 | no-hit | 0.0 | 0 | no-hit | 0.0 | 0 | no-hit | 0.0 | 0 | no-hit | 0.0 | 0 |
| scf7180000056210 | 824513 | 0.5027 | 0 | 58.8491 | Eukaryota | 71617.0 | 0 | Chordata | 71617.0 | 0 | Carnivora | 71617.0 | 0 | Canidae | 71617.0 | 0 | Canis | 56418.0 | 1 | Canis lupus | 56418.0 | 1 |
| scf7180000046632 | 102828 | 0.5973 | 0 | 55.671 | Eukaryota | 19057.0 | 0 | Chordata | 19057.0 | 0 | Carnivora | 19057.0 | 0 | Canidae | 17429.0 | 1 | Canis | 14113.0 | 2 | Canis lupus | 14113.0 | 2 |
| scf7180000052794 | 3619 | 0.4518 | 0 | 1193.2351 | no-hit | 0.0 | 0 | no-hit | 0.0 | 0 | no-hit | 0.0 | 0 | no-hit | 0.0 | 0 | no-hit | 0.0 | 0 | no-hit | 0.0 | 0 |
| scf7180000044103 | 316076 | 0.4379 | 0 | 59.2434 | Eukaryota | 43252.0 | 0 | Chordata | 43252.0 | 0 | Carnivora | 43252.0 | 0 | Canidae | 43252.0 | 0 | Canis | 36418.0 | 1 | Canis lupus | 36418.0 | 1 |
| scf7180000042419 | 684768 | 0.4727 | 0 | 59.0849 | Eukaryota | 44761.0 | 0 | Chordata | 44761.0 | 0 | Carnivora | 44761.0 | 0 | Canidae | 44761.0 | 0 | Canis | 44761.0 | 0 | Canis lupus | 44761.0 | 0 |
| scf7180000047028 | 56799 | 0.4372 | 0 | 171.4056 | Eukaryota | 16505.0 | 0 | Chordata | 10555.0 | 2 | Gadiformes | 4482.0 | 7 | Gadidae | 4482.0 | 7 | Gadus | 4482.0 | 7 | Gadus morhua | 4482.0 | 7 |
| scf7180000040753 | 927513 | 0.4791 | 0 | 58.2744 | Eukaryota | 81026.0 | 0 | Chordata | 81026.0 | 0 | Carnivora | 81026.0 | 0 | Canidae | 81026.0 | 0 | Canis | 81026.0 | 0 | Canis lupus | 81026.0 | 0 |
| scf7180000040783 | 1563330 | 0.4379 | 0 | 57.9183 | Eukaryota | 80235.0 | 0 | Chordata | 80235.0 | 0 | Carnivora | 80235.0 | 0 | Canidae | 80235.0 | 0 | Canis | 80235.0 | 0 | Canis lupus | 80235.0 | 0 |
| scf7180000048041 | 12885 | 0.4502 | 0 | 1310.7838 | Eukaryota | 3541.0 | 0 | Platyhelminthes | 1956.0 | 3 | Polystomatidea | 1956.0 | 3 | Polystomatidae | 1956.0 | 3 | Protopolystoma | 1956.0 | 3 | Protopolystoma xenopodis | 1956.0 | 3 |
| scf7180000041170 | 1219475 | 0.3467 | 0 | 59.0806 | Eukaryota | 91560.0 | 0 | Chordata | 91560.0 | 0 | Carnivora | 91560.0 | 0 | Canidae | 91560.0 | 0 | Canis | 91560.0 | 0 | Canis lupus | 91560.0 | 0 |
| scf7180000051947 | 17519 | 0.5301 | 0 | 398.8083 | Eukaryota | 4426.0 | 0 | Chordata | 4426.0 | 0 | Carnivora | 4426.0 | 0 | Canidae | 4426.0 | 0 | Canis | 3725.0 | 1 | Canis lupus | 3725.0 | 1 |
| scf7180000045583 | 219616 | 0.426 | 0 | 59.4791 | Eukaryota | 42654.0 | 0 | Chordata | 42654.0 | 0 | Carnivora | 42654.0 | 0 | Canidae | 42654.0 | 0 | Canis | 31510.0 | 1 | Canis lupus | 31510.0 | 1 |
| scf7180000051387 | 9779 | 0.4371 | 0 | 324.88 | Eukaryota | 3535.0 | 0 | Platyhelminthes | 2461.0 | 2 | Polystomatidea | 2461.0 | 2 | Polystomatidae | 2461.0 | 2 | Protopolystoma | 2461.0 | 2 | Protopolystoma xenopodis | 2461.0 | 2 |
| scf7180000056314 | 734867 | 0.3532 | 0 | 60.0125 | Eukaryota | 68528.0 | 0 | Chordata | 68528.0 | 0 | Carnivora | 68528.0 | 0 | Canidae | 68528.0 | 0 | Canis | 68528.0 | 0 | Canis lupus | 68528.0 | 0 |
| scf7180000040963 | 245419 | 0.4474 | 0 | 59.5735 | Eukaryota | 136623.0 | 0 | Chordata | 136623.0 | 0 | Carnivora | 136623.0 | 0 | Canidae | 136623.0 | 0 | Canis | 117234.0 | 1 | Canis lupus | 117234.0 | 1 |
| scf7180000043545 | 209117 | 0.4098 | 0 | 59.518 | Eukaryota | 41949.0 | 0 | Chordata | 41949.0 | 0 | Carnivora | 41949.0 | 0 | Canidae | 21529.0 | 1 | Ailuropoda | 20420.0 | 2 | Ailuropoda melanoleuca | 20420.0 | 2 |
| scf7180000042259 | 728073 | 0.3946 | 0 | 59.3758 | Eukaryota | 73380.0 | 0 | Chordata | 73380.0 | 0 | Carnivora | 73380.0 | 0 | Canidae | 73380.0 | 0 | Canis | 73380.0 | 0 | Canis lupus | 73380.0 | 0 |
| scf7180000042101 | 86301 | 0.3875 | 0 | 61.4732 | Eukaryota | 91180.0 | 0 | Chordata | 91180.0 | 0 | Carnivora | 91180.0 | 0 | Canidae | 91180.0 | 0 | Canis | 91180.0 | 0 | Canis lupus | 91180.0 | 0 |
| scf7180000043952 | 353166 | 0.4561 | 0 | 59.4562 | Eukaryota | 51649.0 | 0 | Chordata | 51649.0 | 0 | Carnivora | 51649.0 | 0 | Canidae | 51649.0 | 0 | Canis | 33947.0 | 1 | Canis lupus | 33947.0 | 1 |
| scf7180000050169 | 5143 | 0.4527 | 0 | 1408.2029 | Eukaryota | 5275.0 | 0 | Chordata | 5275.0 | 0 | Tetraodontiformes | 5275.0 | 0 | Tetraodontidae | 5275.0 | 0 | Takifugu | 5275.0 | 0 | Takifugu rubripes | 5275.0 | 0 |
| scf7180000046293 | 14510 | 0.4992 | 0 | 420.0969 | Eukaryota | 33484.0 | 0 | Chordata | 33484.0 | 0 | Carnivora | 33484.0 | 0 | Canidae | 33484.0 | 0 | Canis | 33484.0 | 0 | Canis lupus | 33484.0 | 0 |
| scf7180000045832 | 40840 | 0.3291 | 0 | 55.3154 | Eukaryota | 44779.0 | 0 | Chordata | 44779.0 | 0 | Carnivora | 44779.0 | 0 | Canidae | 44779.0 | 0 | Canis | 44779.0 | 0 | Canis lupus | 44779.0 | 0 |
| scf7180000055386 | 12853 | 0.4492 | 0 | 186.0175 | Eukaryota | 640.0 | 0 | Chordata | 337.0 | 1 | Kurtiformes | 337.0 | 1 | Apogonidae | 337.0 | 1 | Sphaeramia | 337.0 | 1 | Sphaeramia orbicularis | 337.0 | 1 |
| scf7180000047849 | 59750 | 0.3316 | 0 | 58.953 | Eukaryota | 9799.0 | 0 | Chordata | 9799.0 | 0 | Carnivora | 9799.0 | 0 | Canidae | 9799.0 | 0 | Canis | 9799.0 | 0 | Canis lupus | 9799.0 | 0 |
| scf7180000044091 | 435303 | 0.3891 | 0 | 61.0716 | Eukaryota | 36863.0 | 0 | Chordata | 36863.0 | 0 | Carnivora | 36863.0 | 0 | Canidae | 36863.0 | 0 | Canis | 31104.0 | 1 | Canis lupus | 31104.0 | 1 |
| scf7180000053721 | 2252 | 0.4387 | 0 | 430.1339 | no-hit | 0.0 | 0 | no-hit | 0.0 | 0 | no-hit | 0.0 | 0 | no-hit | 0.0 | 0 | no-hit | 0.0 | 0 | no-hit | 0.0 | 0 |
| scf7180000043534 | 452975 | 0.4625 | 0 | 58.8378 | Eukaryota | 68829.0 | 0 | Chordata | 68829.0 | 0 | Carnivora | 68829.0 | 0 | Canidae | 68829.0 | 0 | Canis | 60317.0 | 1 | Canis lupus | 60317.0 | 1 |
| scf7180000041998 | 826049 | 0.4741 | 0 | 59.7056 | Eukaryota | 49776.0 | 0 | Chordata | 49776.0 | 0 | Carnivora | 49776.0 | 0 | Canidae | 49776.0 | 0 | Canis | 36285.0 | 1 | Canis lupus | 36285.0 | 1 |
| scf7180000053782 | 5584 | 0.4473 | 0 | 445.7585 | Eukaryota | 1951.0 | 0 | Platyhelminthes | 1099.0 | 1 | Polystomatidea | 1099.0 | 1 | Polystomatidae | 1099.0 | 1 | Protopolystoma | 1099.0 | 1 | Protopolystoma xenopodis | 1099.0 | 1 |
| scf7180000049320 | 3788 | 0.4461 | 0 | 333.3099 | Eukaryota | 1271.0 | 0 | Echinodermata | 821.0 | 1 | Forcipulatida | 821.0 | 1 | Asteriidae | 821.0 | 1 | Asterias | 821.0 | 1 | Asterias rubens | 821.0 | 1 |
| scf7180000047284 | 116756 | 0.3334 | 0 | 59.4715 | Eukaryota | 37725.0 | 0 | Chordata | 37725.0 | 0 | Carnivora | 37725.0 | 0 | Canidae | 37725.0 | 0 | Canis | 37725.0 | 0 | Canis lupus | 37725.0 | 0 |
| scf7180000042694 | 11934 | 0.3757 | 0 | 29.48 | Eukaryota | 37795.0 | 0 | Chordata | 37795.0 | 0 | Carnivora | 37795.0 | 0 | Canidae | 37795.0 | 0 | Canis | 37795.0 | 0 | Canis lupus | 37795.0 | 0 |
| scf7180000046202 | 18849 | 0.3732 | 0 | 55.2423 | Eukaryota | 62699.0 | 0 | Chordata | 62699.0 | 0 | Carnivora | 62699.0 | 0 | Canidae | 62699.0 | 0 | Canis | 62699.0 | 0 | Canis lupus | 62699.0 | 0 |
| scf7180000041407 | 19903 | 0.4354 | 0 | 214.2226 | Eukaryota | 3323.0 | 0 | Chordata | 3323.0 | 0 | Carnivora | 2857.0 | 1 | Canidae | 2857.0 | 1 | Canis | 2522.0 | 2 | Canis lupus | 2522.0 | 2 |
| scf7180000056199 | 2159397 | 0.394 | 0 | 60.7015 | Eukaryota | 117410.0 | 0 | Chordata | 117410.0 | 0 | Carnivora | 117410.0 | 0 | Canidae | 83307.0 | 1 | Vulpes | 83307.0 | 1 | Vulpes vulpes | 83307.0 | 1 |
| scf7180000046381 | 134433 | 0.3789 | 0 | 60.0658 | Eukaryota | 15344.0 | 0 | Chordata | 15344.0 | 0 | Carnivora | 15344.0 | 0 | Canidae | 15344.0 | 0 | Canis | 15344.0 | 0 | Canis lupus | 15344.0 | 0 |
| scf7180000050376 | 5205 | 0.4423 | 0 | 178.928 | Eukaryota | 7278.0 | 1 | Chordata | 3540.0 | 3 | Strigeidida | 2464.0 | 8 | Schistosomatidae | 2464.0 | 8 | Schistosoma | 2464.0 | 8 | Schistosoma mansoni | 2464.0 | 8 |
| scf7180000056451 | 369484 | 0.3625 | 0 | 60.8039 | Eukaryota | 33344.0 | 0 | Chordata | 33344.0 | 0 | Carnivora | 33344.0 | 0 | Canidae | 33344.0 | 0 | Canis | 33344.0 | 0 | Canis lupus | 33344.0 | 0 |
| scf7180000041181 | 843012 | 0.3831 | 0 | 57.3794 | Eukaryota | 52868.0 | 0 | Chordata | 52868.0 | 0 | Carnivora | 52868.0 | 0 | Canidae | 52868.0 | 0 | Canis | 52868.0 | 0 | Canis lupus | 52868.0 | 0 |
| scf7180000054074 | 2599 | 0.4575 | 0 | 68.6839 | no-hit | 0.0 | 0 | no-hit | 0.0 | 0 | no-hit | 0.0 | 0 | no-hit | 0.0 | 0 | no-hit | 0.0 | 0 | no-hit | 0.0 | 0 |
| scf7180000042295 | 298490 | 0.4278 | 0 | 59.2981 | Eukaryota | 64274.0 | 0 | Chordata | 64274.0 | 0 | Carnivora | 35661.0 | 2 | Canidae | 20893.0 | 5 | Canis | 11282.0 | 7 | Canis lupus | 11282.0 | 7 |
| scf7180000054607 | 9286 | 0.4213 | 0 | 134.6066 | Eukaryota | 35396.0 | 0 | Chordata | 35396.0 | 0 | Carnivora | 35396.0 | 0 | Canidae | 35396.0 | 0 | Canis | 35396.0 | 0 | Canis lupus | 35396.0 | 0 |
| scf7180000049319 | 5587 | 0.4428 | 0 | 104.0499 | no-hit | 0.0 | 0 | no-hit | 0.0 | 0 | no-hit | 0.0 | 0 | no-hit | 0.0 | 0 | no-hit | 0.0 | 0 | no-hit | 0.0 | 0 |
| scf7180000053350 | 4427 | 0.4488 | 0 | 622.7422 | Eukaryota | 1586.0 | 0 | Platyhelminthes | 1369.0 | 1 | Polystomatidea | 1369.0 | 1 | Polystomatidae | 1369.0 | 1 | Protopolystoma | 1369.0 | 1 | Protopolystoma xenopodis | 1369.0 | 1 |
| scf7180000041964 | 96432 | 0.3959 | 0 | 56.9591 | Eukaryota | 66315.0 | 0 | Chordata | 66315.0 | 0 | Carnivora | 66315.0 | 0 | Canidae | 66315.0 | 0 | Canis | 66315.0 | 0 | Canis lupus | 66315.0 | 0 |
| scf7180000046400 | 132164 | 0.3524 | 0 | 61.0498 | Eukaryota | 9195.0 | 0 | Chordata | 8674.0 | 1 | Primates | 5819.0 | 4 | Hominidae | 5819.0 | 4 | Homo | 5819.0 | 4 | Homo sapiens | 5819.0 | 4 |
| scf7180000045129 | 106570 | 0.3407 | 0 | 57.2751 | Eukaryota | 6098.0 | 0 | Chordata | 6098.0 | 0 | Primates | 3664.0 | 2 | Hominidae | 3664.0 | 3 | Homo | 3664.0 | 4 | Homo sapiens | 3664.0 | 4 |
| scf7180000046934 | 18249 | 0.4425 | 0 | 64.2239 | Eukaryota | 1030.0 | 0 | Chordata | 564.0 | 1 | Polystomatidea | 466.0 | 2 | Polystomatidae | 466.0 | 2 | Protopolystoma | 466.0 | 2 | Protopolystoma xenopodis | 466.0 | 2 |
| scf7180000050087 | 34855 | 0.4479 | 0 | 54.9291 | Eukaryota | 71934.0 | 0 | Chordata | 71934.0 | 0 | Carnivora | 71934.0 | 0 | Canidae | 71934.0 | 0 | Canis | 71934.0 | 0 | Canis lupus | 71934.0 | 0 |
| scf7180000054556 | 5667 | 0.4339 | 0 | 382.8653 | Eukaryota | 1271.0 | 0 | Chordata | 1271.0 | 0 | Salmoniformes | 1271.0 | 0 | Salmonidae | 1271.0 | 0 | Salmo | 1271.0 | 0 | Salmo trutta | 1271.0 | 0 |
| scf7180000048011 | 46365 | 0.38 | 0 | 57.9815 | Eukaryota | 51599.0 | 0 | Chordata | 51599.0 | 0 | Carnivora | 51599.0 | 0 | Canidae | 51599.0 | 0 | Canis | 51599.0 | 0 | Canis lupus | 51599.0 | 0 |
| scf7180000040808 | 1767961 | 0.4102 | 0 | 60.1738 | Eukaryota | 85998.0 | 0 | Chordata | 85998.0 | 0 | Carnivora | 85998.0 | 0 | Canidae | 85998.0 | 0 | Canis | 85998.0 | 0 | Canis lupus | 85998.0 | 0 |
| scf7180000046414 | 29019 | 0.3728 | 0 | 380.7815 | Eukaryota | 25410.0 | 0 | Chordata | 25410.0 | 0 | Carnivora | 25410.0 | 0 | Canidae | 17921.0 | 1 | Canis | 15754.0 | 3 | Canis lupus | 15754.0 | 3 |
| scf7180000048003 | 73454 | 0.4436 | 0 | 125.9376 | Eukaryota | 66971.0 | 0 | Chordata | 66971.0 | 0 | Carnivora | 66971.0 | 0 | Canidae | 66971.0 | 0 | Canis | 66971.0 | 0 | Canis lupus | 66971.0 | 0 |
| scf7180000043828 | 344034 | 0.3808 | 0 | 58.5228 | Eukaryota | 25529.0 | 0 | Chordata | 25529.0 | 0 | Carnivora | 25529.0 | 0 | Canidae | 25529.0 | 0 | Canis | 25529.0 | 0 | Canis lupus | 25529.0 | 0 |
| scf7180000053646 | 2666 | 0.4542 | 0 | 242.9417 | Eukaryota | 1719.0 | 0 | Chordata | 1377.0 | 1 | Columbiformes | 907.0 | 2 | Columbidae | 907.0 | 2 | Streptopelia | 907.0 | 2 | Streptopelia turtur | 907.0 | 2 |
| scf7180000050185 | 13477 | 0.4498 | 0 | 1630.2869 | Eukaryota | 3305.0 | 0 | Platyhelminthes | 1622.0 | 3 | Polystomatidea | 1622.0 | 3 | Polystomatidae | 1622.0 | 3 | Protopolystoma | 1622.0 | 3 | Protopolystoma xenopodis | 1622.0 | 3 |
| scf7180000043641 | 222422 | 0.3694 | 0 | 60.1426 | Eukaryota | 84003.0 | 0 | Chordata | 84003.0 | 0 | Carnivora | 84003.0 | 0 | Canidae | 84003.0 | 0 | Canis | 61492.0 | 1 | Canis lupus | 61492.0 | 1 |
| scf7180000052788 | 3740 | 0.4489 | 0 | 636.3513 | Eukaryota | 3717.0 | 0 | Platyhelminthes | 2172.0 | 3 | Polystomatidea | 2172.0 | 3 | Polystomatidae | 2172.0 | 3 | Protopolystoma | 2172.0 | 3 | Protopolystoma xenopodis | 2172.0 | 3 |
| scf7180000046894 | 66221 | 0.4952 | 0 | 60.1043 | Eukaryota | 26087.0 | 0 | Chordata | 26087.0 | 0 | Carnivora | 26087.0 | 0 | Felidae | 8193.0 | 4 | Odobenus | 6762.0 | 5 | Odobenus rosmarus | 6762.0 | 6 |
| scf7180000045261 | 3472 | 0.4683 | 0 | 24.6203 | Eukaryota | 2728.0 | 0 | Chordata | 2728.0 | 0 | Carnivora | 2728.0 | 0 | Canidae | 2147.0 | 2 | Canis | 2147.0 | 2 | Canis lupus | 2147.0 | 2 |
| scf7180000045152 | 41614 | 0.343 | 0 | 58.6513 | Eukaryota | 78156.0 | 0 | Chordata | 78156.0 | 0 | Carnivora | 78156.0 | 0 | Canidae | 78156.0 | 0 | Canis | 78156.0 | 0 | Canis lupus | 78156.0 | 0 |
| scf7180000041164 | 440909 | 0.4466 | 0 | 59.2649 | Eukaryota | 109148.0 | 0 | Chordata | 109148.0 | 0 | Carnivora | 109148.0 | 0 | Canidae | 109148.0 | 0 | Canis | 109148.0 | 0 | Canis lupus | 109148.0 | 0 |
| scf7180000045727 | 7267 | 0.5829 | 0 | 25.0299 | Eukaryota | 3275.0 | 0 | Chordata | 3275.0 | 0 | Carnivora | 3275.0 | 0 | Canidae | 3275.0 | 0 | Canis | 3275.0 | 0 | Canis lupus | 3275.0 | 0 |
| scf7180000041779 | 821768 | 0.429 | 0 | 61.3135 | Eukaryota | 89466.0 | 0 | Chordata | 89466.0 | 0 | Carnivora | 89466.0 | 0 | Canidae | 89466.0 | 0 | Canis | 89466.0 | 0 | Canis lupus | 89466.0 | 0 |
| scf7180000048810 | 1370 | 0.446 | 0 | 144.8294 | no-hit | 0.0 | 0 | no-hit | 0.0 | 0 | no-hit | 0.0 | 0 | no-hit | 0.0 | 0 | no-hit | 0.0 | 0 | no-hit | 0.0 | 0 |
| scf7180000045585 | 161983 | 0.3976 | 0 | 59.4749 | Eukaryota | 45802.0 | 0 | Chordata | 45802.0 | 0 | Carnivora | 45802.0 | 0 | Canidae | 45802.0 | 0 | Canis | 45802.0 | 0 | Canis lupus | 45802.0 | 0 |
| scf7180000050885 | 8565 | 0.6076 | 0 | 248.2021 | Eukaryota | 30264.0 | 0 | Chordata | 30264.0 | 0 | Primates | 13457.0 | 2 | Hominidae | 13457.0 | 2 | Homo | 10753.0 | 4 | Homo sapiens | 10753.0 | 4 |
| scf7180000056486 | 249467 | 0.4083 | 0 | 57.9085 | Eukaryota | 50830.0 | 0 | Chordata | 50830.0 | 0 | Carnivora | 50830.0 | 0 | Canidae | 50830.0 | 0 | Canis | 50830.0 | 0 | Canis lupus | 50830.0 | 0 |
| scf7180000048129 | 60272 | 0.666 | 0 | 52.8773 | Eukaryota | 22687.0 | 0 | Chordata | 22687.0 | 0 | Carnivora | 22687.0 | 0 | Canidae | 22687.0 | 0 | Canis | 17576.0 | 1 | Canis lupus | 17576.0 | 1 |
| scf7180000051537 | 5518 | 0.4523 | 0 | 1132.4057 | no-hit | 0.0 | 0 | no-hit | 0.0 | 0 | no-hit | 0.0 | 0 | no-hit | 0.0 | 0 | no-hit | 0.0 | 0 | no-hit | 0.0 | 0 |
| scf7180000046780 | 11457 | 0.4448 | 0 | 120.706 | no-hit | 0.0 | 0 | no-hit | 0.0 | 0 | no-hit | 0.0 | 0 | no-hit | 0.0 | 0 | no-hit | 0.0 | 0 | no-hit | 0.0 | 0 |
| scf7180000045684 | 19204 | 0.3937 | 0 | 39.6479 | Eukaryota | 34748.0 | 0 | Chordata | 34748.0 | 0 | Carnivora | 34748.0 | 0 | Canidae | 34748.0 | 0 | Canis | 34748.0 | 0 | Canis lupus | 34748.0 | 0 |
| scf7180000048274 | 74642 | 0.3571 | 0 | 58.8159 | Eukaryota | 24006.0 | 0 | Chordata | 24006.0 | 0 | Carnivora | 24006.0 | 0 | Canidae | 22286.0 | 1 | Canis | 18187.0 | 2 | Canis lupus | 18187.0 | 2 |
| scf7180000046959 | 56620 | 0.4394 | 0 | 169.9919 | Eukaryota | 7120.0 | 1 | Chordata | 4828.0 | 3 | Tetraodontiformes | 2617.0 | 4 | Tetraodontidae | 2617.0 | 4 | Takifugu | 2617.0 | 4 | Takifugu rubripes | 2617.0 | 4 |
| scf7180000052525 | 7594 | 0.4481 | 0 | 504.605 | no-hit | 0.0 | 0 | no-hit | 0.0 | 0 | no-hit | 0.0 | 0 | no-hit | 0.0 | 0 | no-hit | 0.0 | 0 | no-hit | 0.0 | 0 |
| scf7180000053882 | 5760 | 0.4415 | 0 | 51.9307 | Eukaryota | 9743.0 | 0 | Chordata | 5482.0 | 2 | Strigeidida | 2545.0 | 6 | Schistosomatidae | 2545.0 | 6 | Schistosoma | 2545.0 | 6 | Schistosoma mansoni | 2545.0 | 6 |
| scf7180000052395 | 9201 | 0.4561 | 0 | 53.3955 | no-hit | 0.0 | 0 | no-hit | 0.0 | 0 | no-hit | 0.0 | 0 | no-hit | 0.0 | 0 | no-hit | 0.0 | 0 | no-hit | 0.0 | 0 |
| scf7180000055787 | 6590 | 0.4533 | 0 | 1706.0271 | Eukaryota | 2461.0 | 0 | Platyhelminthes | 1395.0 | 2 | Polystomatidea | 1395.0 | 2 | Polystomatidae | 1395.0 | 2 | Protopolystoma | 1395.0 | 2 | Protopolystoma xenopodis | 1395.0 | 2 |
| scf7180000047596 | 8082 | 0.4317 | 0 | 229.7111 | no-hit | 0.0 | 0 | no-hit | 0.0 | 0 | no-hit | 0.0 | 0 | no-hit | 0.0 | 0 | no-hit | 0.0 | 0 | no-hit | 0.0 | 0 |
| scf7180000042508 | 30706 | 0.4118 | 0 | 48.453 | Eukaryota | 86839.0 | 0 | Chordata | 86839.0 | 0 | Carnivora | 86839.0 | 0 | Canidae | 86839.0 | 0 | Canis | 86839.0 | 0 | Canis lupus | 86839.0 | 0 |
| scf7180000051816 | 4416 | 0.4506 | 0 | 514.6318 | no-hit | 0.0 | 0 | no-hit | 0.0 | 0 | no-hit | 0.0 | 0 | no-hit | 0.0 | 0 | no-hit | 0.0 | 0 | no-hit | 0.0 | 0 |
| scf7180000046725 | 142933 | 0.4309 | 0 | 56.573 | Eukaryota | 12979.0 | 0 | Chordata | 12979.0 | 0 | Carnivora | 12979.0 | 0 | Canidae | 12979.0 | 0 | Canis | 12979.0 | 0 | Canis lupus | 12979.0 | 0 |
| scf7180000054964 | 31273 | 0.4429 | 0 | 1506.0284 | Eukaryota | 1334.0 | 0 | Platyhelminthes | 773.0 | 1 | Polystomatidea | 773.0 | 1 | Polystomatidae | 773.0 | 1 | Protopolystoma | 773.0 | 1 | Protopolystoma xenopodis | 773.0 | 1 |
| scf7180000046964 | 110792 | 0.4969 | 0 | 59.9989 | Eukaryota | 32835.0 | 0 | Chordata | 32835.0 | 0 | Carnivora | 32835.0 | 0 | Canidae | 32835.0 | 0 | Vulpes | 17602.0 | 1 | Vulpes vulpes | 17602.0 | 1 |
| scf7180000046333 | 28412 | 0.4449 | 0 | 269.3676 | Eukaryota | 747.0 | 0 | Platyhelminthes | 381.0 | 1 | Polystomatidea | 381.0 | 1 | Polystomatidae | 381.0 | 1 | Protopolystoma | 381.0 | 1 | Protopolystoma xenopodis | 381.0 | 1 |
| scf7180000053231 | 3160 | 0.4538 | 0 | 180.4233 | no-hit | 0.0 | 0 | no-hit | 0.0 | 0 | no-hit | 0.0 | 0 | no-hit | 0.0 | 0 | no-hit | 0.0 | 0 | no-hit | 0.0 | 0 |
| scf7180000054278 | 13511 | 0.4505 | 0 | 463.2635 | no-hit | 0.0 | 0 | no-hit | 0.0 | 0 | no-hit | 0.0 | 0 | no-hit | 0.0 | 0 | no-hit | 0.0 | 0 | no-hit | 0.0 | 0 |
| scf7180000040686 | 32215 | 0.3843 | 0 | 58.4249 | Eukaryota | 87246.0 | 0 | Chordata | 87246.0 | 0 | Carnivora | 87246.0 | 0 | Canidae | 87246.0 | 0 | Canis | 87246.0 | 0 | Canis lupus | 87246.0 | 0 |
| scf7180000043929 | 5003 | 0.4169 | 0 | 156.1932 | Eukaryota | 44809.0 | 0 | Chordata | 44809.0 | 0 | Carnivora | 44809.0 | 0 | Canidae | 44809.0 | 0 | Canis | 44809.0 | 0 | Canis lupus | 44809.0 | 0 |
| scf7180000046535 | 125634 | 0.365 | 0 | 59.9158 | Eukaryota | 45509.0 | 0 | Chordata | 45509.0 | 0 | Carnivora | 45509.0 | 0 | Otariidae | 13543.0 | 4 | Canis | 9947.0 | 7 | Canis lupus | 9947.0 | 8 |
| scf7180000047886 | 98240 | 0.348 | 0 | 62.0508 | Eukaryota | 56908.0 | 0 | Chordata | 56908.0 | 0 | Carnivora | 56908.0 | 0 | Felidae | 33457.0 | 1 | Vulpes | 23451.0 | 2 | Vulpes vulpes | 23451.0 | 3 |
| scf7180000041940 | 857239 | 0.3945 | 0 | 61.3887 | Eukaryota | 23300.0 | 1 | Chordata | 23300.0 | 1 | Primates | 9946.0 | 3 | Canidae | 7896.0 | 4 | Homo | 7691.0 | 6 | Homo sapiens | 7691.0 | 6 |
| scf7180000049919 | 42957 | 0.4093 | 0 | 54.4127 | Eukaryota | 24327.0 | 0 | Chordata | 24327.0 | 0 | Carnivora | 24327.0 | 0 | Canidae | 24327.0 | 0 | Canis | 24327.0 | 0 | Canis lupus | 24327.0 | 0 |
| scf7180000041778 | 918750 | 0.3876 | 0 | 61.2605 | Eukaryota | 39065.0 | 0 | Chordata | 39065.0 | 0 | Carnivora | 39065.0 | 0 | Canidae | 39065.0 | 0 | Canis | 34957.0 | 1 | Canis lupus | 34957.0 | 1 |
| scf7180000051185 | 2563 | 0.4503 | 0 | 554.245 | no-hit | 0.0 | 0 | no-hit | 0.0 | 0 | no-hit | 0.0 | 0 | no-hit | 0.0 | 0 | no-hit | 0.0 | 0 | no-hit | 0.0 | 0 |
| scf7180000046869 | 10065 | 0.4126 | 0 | 62.3851 | Eukaryota | 85602.0 | 0 | Chordata | 85602.0 | 0 | Carnivora | 85602.0 | 0 | Canidae | 85602.0 | 0 | Canis | 85602.0 | 0 | Canis lupus | 85602.0 | 0 |
| scf7180000045825 | 128556 | 0.3511 | 0 | 59.8635 | Eukaryota | 48571.0 | 0 | Chordata | 48571.0 | 0 | Carnivora | 39283.0 | 1 | Canidae | 10110.0 | 6 | Canis | 10110.0 | 7 | Canis lupus | 10110.0 | 8 |
| scf7180000046921 | 19158 | 0.4445 | 0 | 170.2664 | Eukaryota | 8938.0 | 0 | Chordata | 6865.0 | 1 | Kurtiformes | 3980.0 | 2 | Apogonidae | 3980.0 | 2 | Sphaeramia | 3980.0 | 2 | Sphaeramia orbicularis | 3980.0 | 2 |
| scf7180000043434 | 473960 | 0.4047 | 0 | 60.6052 | Eukaryota | 40621.0 | 0 | Chordata | 40621.0 | 0 | Carnivora | 40621.0 | 0 | Canidae | 40621.0 | 0 | Canis | 40621.0 | 0 | Canis lupus | 40621.0 | 0 |
| scf7180000049610 | 16689 | 0.4456 | 0 | 359.0179 | Eukaryota | 395.0 | 0 | Platyhelminthes | 206.0 | 1 | Polystomatidea | 206.0 | 1 | Polystomatidae | 206.0 | 1 | Protopolystoma | 206.0 | 1 | Protopolystoma xenopodis | 206.0 | 1 |
| scf7180000043418 | 195177 | 0.3664 | 0 | 57.0361 | Eukaryota | 79635.0 | 0 | Chordata | 79635.0 | 0 | Carnivora | 79635.0 | 0 | Canidae | 79635.0 | 0 | Canis | 79635.0 | 0 | Canis lupus | 79635.0 | 0 |
| scf7180000049537 | 26415 | 0.4452 | 0 | 924.2935 | no-hit | 0.0 | 0 | no-hit | 0.0 | 0 | no-hit | 0.0 | 0 | no-hit | 0.0 | 0 | no-hit | 0.0 | 0 | no-hit | 0.0 | 0 |
| scf7180000048393 | 59154 | 0.4496 | 0 | 240.7745 | Eukaryota | 14258.0 | 0 | Chordata | 14258.0 | 0 | Carnivora | 14258.0 | 0 | Canidae | 14258.0 | 0 | Canis | 14258.0 | 0 | Canis lupus | 14258.0 | 0 |
| scf7180000051559 | 2985 | 0.4506 | 0 | 807.6783 | Eukaryota | 883.0 | 0 | Chordata | 883.0 | 0 | Tetraodontiformes | 883.0 | 0 | Tetraodontidae | 883.0 | 0 | Takifugu | 883.0 | 0 | Takifugu rubripes | 883.0 | 0 |
| scf7180000056395 | 696965 | 0.4099 | 0 | 60.1425 | Eukaryota | 90366.0 | 0 | Chordata | 90366.0 | 0 | Carnivora | 90366.0 | 0 | Canidae | 90366.0 | 0 | Canis | 90366.0 | 0 | Canis lupus | 90366.0 | 0 |
| scf7180000054393 | 24014 | 0.4123 | 0 | 796.1984 | Eukaryota | 87629.0 | 0 | Chordata | 87629.0 | 0 | Carnivora | 87629.0 | 0 | Canidae | 87629.0 | 0 | Canis | 87629.0 | 0 | Canis lupus | 87629.0 | 0 |
| scf7180000056176 | 767415 | 0.3779 | 0 | 60.7557 | Eukaryota | 90780.0 | 0 | Chordata | 90780.0 | 0 | Carnivora | 90780.0 | 0 | Canidae | 90780.0 | 0 | Canis | 90780.0 | 0 | Canis lupus | 90780.0 | 0 |
| scf7180000050531 | 25780 | 0.4527 | 0 | 152.9513 | Eukaryota | 13533.0 | 0 | Chordata | 13533.0 | 0 | Carnivora | 13533.0 | 0 | Canidae | 13145.0 | 1 | Canis | 13145.0 | 1 | Canis lupus | 13145.0 | 1 |
| scf7180000049392 | 44029 | 0.501 | 0 | 295.9124 | Eukaryota | 10236.0 | 0 | Chordata | 10236.0 | 0 | Carnivora | 10236.0 | 0 | Canidae | 10236.0 | 0 | Canis | 9015.0 | 1 | Canis lupus | 7727.0 | 2 |
| scf7180000054093 | 3271 | 0.4439 | 0 | 153.0998 | Eukaryota | 505.0 | 0 | Platyhelminthes | 505.0 | 0 | Polystomatidea | 505.0 | 0 | Polystomatidae | 505.0 | 0 | Protopolystoma | 505.0 | 0 | Protopolystoma xenopodis | 505.0 | 0 |
| scf7180000048137 | 59686 | 0.488 | 0 | 57.608 | Eukaryota | 42899.0 | 0 | Chordata | 42899.0 | 0 | Carnivora | 42899.0 | 0 | Canidae | 34303.0 | 3 | Canis | 28901.0 | 4 | Canis lupus | 28901.0 | 4 |
| scf7180000056375 | 1065257 | 0.3969 | 0 | 60.3548 | Eukaryota | 57895.0 | 0 | Chordata | 57895.0 | 0 | Carnivora | 57895.0 | 0 | Canidae | 57895.0 | 0 | Canis | 52059.0 | 1 | Canis lupus | 52059.0 | 1 |
| scf7180000053282 | 5242 | 0.4508 | 0 | 68.2812 | Eukaryota | 1563.0 | 1 | Chordata | 1339.0 | 2 | Salmoniformes | 1161.0 | 3 | Salmonidae | 1161.0 | 3 | Coregonus | 1161.0 | 3 | Coregonus sp. 'balchen' | 1161.0 | 3 |
| scf7180000053051 | 2495 | 0.4521 | 0 | 171.6187 | no-hit | 0.0 | 0 | no-hit | 0.0 | 0 | no-hit | 0.0 | 0 | no-hit | 0.0 | 0 | no-hit | 0.0 | 0 | no-hit | 0.0 | 0 |
| scf7180000044598 | 48828 | 0.3577 | 0 | 54.8436 | Eukaryota | 40325.0 | 0 | Chordata | 40325.0 | 0 | Carnivora | 40325.0 | 0 | Canidae | 40325.0 | 0 | Canis | 40325.0 | 0 | Canis lupus | 40325.0 | 0 |
| scf7180000044595 | 258422 | 0.3867 | 0 | 57.8825 | Eukaryota | 27703.0 | 0 | Chordata | 27703.0 | 0 | Carnivora | 27703.0 | 0 | Otariidae | 6999.0 | 5 | Canis | 6900.0 | 7 | Canis lupus | 6900.0 | 8 |
| scf7180000046127 | 91721 | 0.4463 | 0 | 95.8756 | Eukaryota | 5848.0 | 0 | Platyhelminthes | 3597.0 | 1 | Polystomatidea | 3597.0 | 2 | Polystomatidae | 3597.0 | 2 | Protopolystoma | 3597.0 | 2 | Protopolystoma xenopodis | 3597.0 | 2 |
| scf7180000056325 | 256734 | 0.3386 | 0 | 60.0323 | Eukaryota | 90198.0 | 0 | Chordata | 90198.0 | 0 | Carnivora | 90198.0 | 0 | Canidae | 90198.0 | 0 | Canis | 90198.0 | 0 | Canis lupus | 90198.0 | 0 |
| scf7180000053389 | 2673 | 0.4471 | 0 | 483.8809 | Eukaryota | 2917.0 | 0 | Platyhelminthes | 1506.0 | 3 | Polystomatidea | 1506.0 | 3 | Polystomatidae | 1506.0 | 3 | Protopolystoma | 1506.0 | 3 | Protopolystoma xenopodis | 1506.0 | 3 |
| scf7180000042137 | 374749 | 0.3743 | 0 | 59.3976 | Eukaryota | 87675.0 | 0 | Chordata | 87675.0 | 0 | Carnivora | 87675.0 | 0 | Canidae | 87675.0 | 0 | Canis | 87675.0 | 0 | Canis lupus | 87675.0 | 0 |
| scf7180000042801 | 599650 | 0.4388 | 0 | 58.619 | Eukaryota | 86289.0 | 0 | Chordata | 86289.0 | 0 | Carnivora | 86289.0 | 0 | Canidae | 86289.0 | 0 | Canis | 86289.0 | 0 | Canis lupus | 86289.0 | 0 |
| scf7180000040737 | 234307 | 0.4168 | 0 | 58.3679 | Eukaryota | 22257.0 | 0 | Chordata | 22257.0 | 0 | Carnivora | 22257.0 | 0 | Canidae | 22257.0 | 0 | Canis | 15149.0 | 1 | Canis lupus | 15149.0 | 1 |
| scf7180000043512 | 457805 | 0.3815 | 0 | 59.0148 | Eukaryota | 106290.0 | 0 | Chordata | 106290.0 | 0 | Carnivora | 106290.0 | 0 | Canidae | 106290.0 | 0 | Canis | 63857.0 | 1 | Canis lupus | 63857.0 | 1 |
| scf7180000041380 | 867563 | 0.4544 | 0 | 59.5261 | Eukaryota | 148551.0 | 0 | Chordata | 148551.0 | 0 | Carnivora | 148551.0 | 0 | Canidae | 148551.0 | 0 | Canis | 148551.0 | 0 | Canis lupus | 148551.0 | 0 |
| scf7180000053176 | 2984 | 0.4544 | 0 | 268.11 | no-hit | 0.0 | 0 | no-hit | 0.0 | 0 | no-hit | 0.0 | 0 | no-hit | 0.0 | 0 | no-hit | 0.0 | 0 | no-hit | 0.0 | 0 |
| scf7180000045424 | 210494 | 0.36 | 0 | 60.0676 | Eukaryota | 11595.0 | 0 | Chordata | 11595.0 | 0 | Primates | 7740.0 | 2 | Hominidae | 7740.0 | 2 | Homo | 7740.0 | 2 | Homo sapiens | 7740.0 | 2 |
| scf7180000044542 | 42110 | 0.3862 | 0 | 53.1652 | Eukaryota | 82176.0 | 0 | Chordata | 82176.0 | 0 | Carnivora | 82176.0 | 0 | Canidae | 82176.0 | 0 | Canis | 82176.0 | 0 | Canis lupus | 82176.0 | 0 |
| scf7180000043989 | 256751 | 0.3807 | 0 | 60.0942 | Eukaryota | 16115.0 | 0 | Chordata | 16115.0 | 0 | Carnivora | 14006.0 | 1 | Canidae | 14006.0 | 1 | Canis | 14006.0 | 1 | Canis lupus | 14006.0 | 1 |
| scf7180000041480 | 461761 | 0.3581 | 0 | 59.7735 | Eukaryota | 93745.0 | 0 | Chordata | 93745.0 | 0 | Carnivora | 93745.0 | 0 | Canidae | 93745.0 | 0 | Canis | 93745.0 | 0 | Canis lupus | 93745.0 | 0 |
| scf7180000041431 | 339771 | 0.3913 | 0 | 59.5164 | Eukaryota | 49655.0 | 0 | Chordata | 49655.0 | 0 | Carnivora | 49655.0 | 0 | Canidae | 49655.0 | 0 | Canis | 49655.0 | 0 | Canis lupus | 49655.0 | 0 |
| scf7180000044762 | 13744 | 0.4023 | 0 | 50.0499 | Eukaryota | 35347.0 | 0 | Chordata | 35347.0 | 0 | Carnivora | 35347.0 | 0 | Canidae | 35347.0 | 0 | Canis | 35347.0 | 0 | Canis lupus | 35347.0 | 0 |
| scf7180000054528 | 28713 | 0.436 | 0 | 280.5683 | Eukaryota | 150.0 | 0 | Chordata | 150.0 | 0 | Osteoglossiformes | 150.0 | 0 | Osteoglossidae | 150.0 | 0 | Scleropages | 150.0 | 0 | Scleropages formosus | 150.0 | 0 |
| scf7180000045003 | 298547 | 0.3643 | 0 | 59.3448 | Eukaryota | 14558.0 | 0 | Chordata | 14558.0 | 0 | Carnivora | 9079.0 | 2 | Canidae | 8330.0 | 3 | Homo | 4546.0 | 4 | Homo sapiens | 4546.0 | 4 |
| scf7180000049821 | 17402 | 0.4445 | 0 | 86.8864 | Eukaryota | 627.0 | 0 | Chordata | 627.0 | 0 | Tetraodontiformes | 381.0 | 1 | Tetraodontidae | 381.0 | 1 | Takifugu | 381.0 | 1 | Takifugu rubripes | 381.0 | 1 |
| scf7180000054164 | 29039 | 0.4056 | 0 | 52.7488 | Eukaryota | 47303.0 | 0 | Chordata | 47303.0 | 0 | Carnivora | 47303.0 | 0 | Canidae | 47303.0 | 0 | Canis | 47303.0 | 0 | Canis lupus | 47303.0 | 0 |
| scf7180000044702 | 126086 | 0.3546 | 0 | 58.7031 | Eukaryota | 45644.0 | 0 | Chordata | 45644.0 | 0 | Carnivora | 45644.0 | 0 | Canidae | 45644.0 | 0 | Canis | 45644.0 | 0 | Canis lupus | 45644.0 | 0 |
| scf7180000042206 | 760103 | 0.358 | 0 | 60.7943 | Eukaryota | 63043.0 | 0 | Chordata | 63043.0 | 0 | Carnivora | 63043.0 | 0 | Canidae | 63043.0 | 0 | Canis | 44050.0 | 1 | Canis lupus | 44050.0 | 1 |
| scf7180000043252 | 447780 | 0.3717 | 0 | 59.6157 | Eukaryota | 31187.0 | 0 | Chordata | 31187.0 | 0 | Carnivora | 31187.0 | 0 | Otariidae | 17978.0 | 2 | Zalophus | 14980.0 | 3 | Zalophus californianus | 14980.0 | 3 |
| scf7180000044532 | 98668 | 0.3947 | 0 | 57.521 | Eukaryota | 33818.0 | 0 | Chordata | 33818.0 | 0 | Carnivora | 33818.0 | 0 | Canidae | 33818.0 | 0 | Canis | 33818.0 | 0 | Canis lupus | 33818.0 | 0 |
| scf7180000047809 | 98434 | 0.5005 | 0 | 179.5082 | Eukaryota | 8969.0 | 0 | Chordata | 8969.0 | 0 | Carnivora | 8969.0 | 0 | Canidae | 8969.0 | 0 | Canis | 7849.0 | 1 | Canis lupus | 6714.0 | 2 |

| | | | | | | | | | | | | | | | | | | | |
|---|---|---|---|---|---|---|---|---|---|---|---|---|---|---|---|---|---|---|---|
| scf7180000048156 | 6431 | 0.4419 | 0 | 98.568 | Eukaryota | 257.0 | 0 | Chordata | 257.0 | 0 | Kurtiformes | 257.0 | 0 | Apogonidae | 257.0 | 0 | Sphaeramia | 257.0 | 0 | Sphaeramia orbicularis | 257.0 | 0 |
| scf7180000043174 | 548078 | 0.3404 | 0 | 59.1005 | Eukaryota | 94213.0 | 0 | Chordata | 94213.0 | 0 | Carnivora | 94213.0 | 0 | Canidae | 94213.0 | 0 | Canis | 94213.0 | 0 | Canis lupus | 94213.0 | 0 |
| scf7180000041611 | 969936 | 0.392 | 0 | 60.9794 | Eukaryota | 89703.0 | 0 | Chordata | 89703.0 | 0 | Carnivora | 89703.0 | 0 | Canidae | 89703.0 | 0 | Canis | 78711.0 | 1 | Canis lupus | 78711.0 | 1 |
| scf7180000056588 | 341018 | 0.4629 | 0 | 57.5025 | Eukaryota | 43300.0 | 0 | Chordata | 43300.0 | 0 | Carnivora | 43300.0 | 0 | Canidae | 39685.0 | 1 | Canis | 30653.0 | 2 | Canis lupus | 30653.0 | 2 |
| scf7180000056391 | 189915 | 0.4065 | 0 | 63.1421 | Eukaryota | 85238.0 | 0 | Chordata | 85238.0 | 0 | Carnivora | 85238.0 | 0 | Canidae | 85238.0 | 0 | Canis | 85238.0 | 0 | Canis lupus | 85238.0 | 0 |
| scf7180000043571 | 9609 | 0.4435 | 0 | 42.8551 | Eukaryota | 2235.0 | 0 | Chordata | 2235.0 | 0 | Carnivora | 1329.0 | 1 | Canidae | 1329.0 | 1 | Canis | 1329.0 | 2 | Canis lupus | 1329.0 | 2 |
| scf7180000054778 | 33994 | 0.4449 | 0 | 367.3204 | Eukaryota | 4280.0 | 0 | Chordata | 3102.0 | 1 | Tetraodontiformes | 3102.0 | 1 | Tetraodontidae | 3102.0 | 1 | Takifugu | 3102.0 | 1 | Takifugu rubripes | 3102.0 | 1 |
| scf7180000056552 | 185647 | 0.4052 | 0 | 60.6234 | Eukaryota | 33234.0 | 0 | Chordata | 33234.0 | 0 | Carnivora | 33234.0 | 0 | Canidae | 33234.0 | 0 | Canis | 33234.0 | 0 | Canis lupus | 33234.0 | 0 |
| scf7180000042284 | 724752 | 0.5053 | 0 | 58.1293 | Eukaryota | 96551.0 | 0 | Chordata | 96551.0 | 0 | Carnivora | 96551.0 | 0 | Canidae | 96551.0 | 0 | Canis | 89311.0 | 1 | Canis lupus | 89311.0 | 1 |
| scf7180000048580 | 52403 | 0.4822 | 0 | 58.2353 | Eukaryota | 52515.0 | 0 | Chordata | 52515.0 | 0 | Carnivora | 52515.0 | 0 | Canidae | 52515.0 | 0 | Canis | 52515.0 | 0 | Canis lupus | 52515.0 | 0 |
| scf7180000046411 | 117415 | 0.3494 | 0 | 59.2186 | Eukaryota | 89706.0 | 0 | Chordata | 89706.0 | 0 | Carnivora | 89706.0 | 0 | Canidae | 89706.0 | 0 | Canis | 89706.0 | 0 | Canis lupus | 89706.0 | 0 |
| scf7180000052286 | 3911 | 0.4505 | 0 | 572.932 | Eukaryota | 3635.0 | 0 | Chordata | 1637.0 | 3 | Kurtiformes | 1238.0 | 4 | Apogonidae | 1238.0 | 4 | Sphaeramia | 1238.0 | 4 | Sphaeramia orbicularis | 1238.0 | 4 |
| scf7180000046415 | 6065 | 0.402 | 0 | 41.6392 | Eukaryota | 73432.0 | 0 | Chordata | 73432.0 | 0 | Carnivora | 73432.0 | 0 | Canidae | 73432.0 | 0 | Canis | 73432.0 | 0 | Canis lupus | 73432.0 | 0 |
| scf7180000054963 | 27282 | 0.4439 | 0 | 423.7484 | Eukaryota | 48776.0 | 0 | Chordata | 31275.0 | 2 | Gadiformes | 21224.0 | 6 | Gadidae | 21224.0 | 6 | Gadus | 21224.0 | 6 | Gadus morhua | 21224.0 | 6 |
| scf7180000040970 | 132490 | 0.4525 | 0 | 67.3839 | Eukaryota | 92626.0 | 0 | Chordata | 92626.0 | 0 | Carnivora | 92626.0 | 0 | Canidae | 92626.0 | 0 | Canis | 92626.0 | 0 | Canis lupus | 92626.0 | 0 |
| scf7180000053987 | 4328 | 0.4108 | 0 | 42.4478 | Eukaryota | 500.0 | 0 | Chordata | 500.0 | 0 | Primates | 500.0 | 0 | Hominidae | 500.0 | 0 | Pongo | 500.0 | 0 | Pongo abelii | 500.0 | 0 |
| scf7180000042633 | 568848 | 0.4951 | 0 | 59.2568 | Eukaryota | 47895.0 | 0 | Chordata | 47895.0 | 0 | Carnivora | 47895.0 | 0 | Canidae | 47895.0 | 0 | Canis | 38009.0 | 1 | Canis lupus | 38009.0 | 1 |
| scf7180000045481 | 14791 | 0.4399 | 0 | 202.3458 | Eukaryota | 500.0 | 0 | Platyhelminthes | 326.0 | 1 | Polystomatidea | 326.0 | 1 | Polystomatidae | 326.0 | 1 | Protopolystoma | 326.0 | 1 | Protopolystoma xenopodis | 326.0 | 1 |
| scf7180000040704 | 1824304 | 0.4425 | 0 | 59.0618 | Eukaryota | 111073.0 | 0 | Chordata | 111073.0 | 0 | Carnivora | 111073.0 | 0 | Canidae | 111073.0 | 0 | Canis | 111073.0 | 0 | Canis lupus | 111073.0 | 0 |
| scf7180000044875 | 14629 | 0.4395 | 0 | 59.2423 | Eukaryota | 2751.0 | 0 | Chordata | 2751.0 | 0 | Carnivora | 2751.0 | 0 | Canidae | 2751.0 | 0 | Canis | 2199.0 | 1 | Canis lupus | 2199.0 | 1 |
| scf7180000045324 | 215615 | 0.3976 | 0 | 60.561 | Eukaryota | 36012.0 | 0 | Chordata | 36012.0 | 0 | Carnivora | 36012.0 | 0 | Canidae | 36012.0 | 0 | Canis | 36012.0 | 0 | Canis lupus | 36012.0 | 0 |
| scf7180000047425 | 102249 | 0.4633 | 0 | 57.3236 | Eukaryota | 27173.0 | 1 | Chordata | 27173.0 | 1 | Gadiformes | 7073.0 | 6 | Gadidae | 7073.0 | 6 | Gadus | 7073.0 | 6 | Gadus morhua | 7073.0 | 6 |
| scf7180000056036 | 5933 | 0.4379 | 0 | 104.0784 | Eukaryota | 1249.0 | 0 | Platyhelminthes | 925.0 | 1 | Polystomatidea | 925.0 | 1 | Polystomatidae | 925.0 | 1 | Protopolystoma | 925.0 | 1 | Protopolystoma xenopodis | 925.0 | 1 |
| scf7180000055881 | 11262 | 0.4536 | 0 | 237.6898 | Eukaryota | 17601.0 | 0 | Chordata | 17601.0 | 0 | Carnivora | 17601.0 | 0 | Canidae | 17601.0 | 0 | Canis | 17601.0 | 0 | Canis lupus | 17601.0 | 0 |
| scf7180000043723 | 40823 | 0.4065 | 0 | 59.4053 | Eukaryota | 22479.0 | 0 | Chordata | 22479.0 | 0 | Carnivora | 22479.0 | 0 | Canidae | 22479.0 | 0 | Canis | 22479.0 | 0 | Canis lupus | 22479.0 | 0 |
| scf7180000048047 | 36279 | 0.4418 | 0 | 215.7079 | Eukaryota | 5287.0 | 0 | Platyhelminthes | 2839.0 | 3 | Polystomatidea | 2839.0 | 3 | Polystomatidae | 2839.0 | 3 | Protopolystoma | 2839.0 | 3 | Protopolystoma xenopodis | 2839.0 | 3 |
| scf7180000055329 | 25825 | 0.4437 | 0 | 535.2581 | Eukaryota | 4394.0 | 1 | Platyhelminthes | 1992.0 | 4 | Strigeidida | 1644.0 | 7 | Schistosomatidae | 1644.0 | 7 | Schistosoma | 1644.0 | 7 | Schistosoma mansoni | 1644.0 | 7 |
| scf7180000053786 | 4268 | 0.5021 | 0 | 35.3928 | Eukaryota | 4173.0 | 0 | Chordata | 4173.0 | 0 | Carnivora | 4173.0 | 0 | Canidae | 4173.0 | 0 | Canis | 3453.0 | 1 | Canis lupus | 3453.0 | 1 |
| scf7180000046111 | 171186 | 0.3655 | 0 | 60.2335 | Eukaryota | 22621.0 | 0 | Chordata | 22621.0 | 0 | Primates | 15071.0 | 1 | Hominidae | 15071.0 | 1 | Homo | 15071.0 | 2 | Homo sapiens | 15071.0 | 2 |
| scf7180000055051 | 14700 | 0.4378 | 0 | 194.8537 | Eukaryota | 748.0 | 0 | Platyhelminthes | 748.0 | 0 | Polystomatidea | 748.0 | 0 | Polystomatidae | 748.0 | 0 | Protopolystoma | 748.0 | 0 | Protopolystoma xenopodis | 748.0 | 0 |
| scf7180000048429 | 7934 | 0.4363 | 0 | 90.03 | no-hit | 0.0 | 0 | no-hit | 0.0 | 0 | no-hit | 0.0 | 0 | no-hit | 0.0 | 0 | no-hit | 0.0 | 0 | no-hit | 0.0 | 0 |
| scf7180000045949 | 2407 | 0.4491 | 0 | 211.515 | no-hit | 0.0 | 0 | no-hit | 0.0 | 0 | no-hit | 0.0 | 0 | no-hit | 0.0 | 0 | no-hit | 0.0 | 0 | no-hit | 0.0 | 0 |
| scf7180000054769 | 34624 | 0.4432 | 0 | 342.9754 | Eukaryota | 63626.0 | 0 | Chordata | 63626.0 | 0 | Carnivora | 63626.0 | 0 | Canidae | 63626.0 | 0 | Canis | 63626.0 | 0 | Canis lupus | 63626.0 | 0 |
| scf7180000050084 | 7437 | 0.4351 | 0 | 117.8801 | Eukaryota | 967.0 | 0 | Chordata | 967.0 | 0 | Kurtiformes | 833.0 | 1 | Apogonidae | 833.0 | 1 | Sphaeramia | 833.0 | 1 | Sphaeramia orbicularis | 833.0 | 1 |
| scf7180000041217 | 301933 | 0.3403 | 0 | 58.0415 | Eukaryota | 62196.0 | 0 | Chordata | 62196.0 | 0 | Carnivora | 62196.0 | 0 | Canidae | 62196.0 | 0 | Canis | 62196.0 | 0 | Canis lupus | 62196.0 | 0 |
| scf7180000055268 | 14286 | 0.432 | 0 | 264.2435 | Eukaryota | 90706.0 | 0 | Chordata | 90706.0 | 0 | Carnivora | 90706.0 | 0 | Canidae | 90706.0 | 0 | Canis | 90706.0 | 0 | Canis lupus | 90706.0 | 0 |
| scf7180000041926 | 470162 | 0.3551 | 100 | 58.974 | Eukaryota | 55126.0 | 0 | Chordata | 55126.0 | 0 | Carnivora | 55126.0 | 0 | Canidae | 55126.0 | 0 | Vulpes | 39550.0 | 1 | Vulpes vulpes | 39550.0 | 1 |
| scf7180000046462 | 58332 | 0.3474 | 0 | 56.7861 | Eukaryota | 90689.0 | 0 | Chordata | 90689.0 | 0 | Carnivora | 90689.0 | 0 | Canidae | 90689.0 | 0 | Canis | 90689.0 | 0 | Canis lupus | 90689.0 | 0 |
| scf7180000055488 | 15316 | 0.4414 | 0 | 211.1171 | Eukaryota | 41578.0 | 0 | Chordata | 34827.0 | 2 | Gadiformes | 27292.0 | 5 | Gadidae | 27292.0 | 5 | Gadus | 27292.0 | 5 | Gadus morhua | 27292.0 | 5 |
| scf7180000052003 | 2438 | 0.4537 | 0 | 718.9616 | Eukaryota | 8655.0 | 0 | Chordata | 7527.0 | 2 | Gadiformes | 3797.0 | 6 | Gadidae | 3797.0 | 6 | Gadus | 3797.0 | 6 | Gadus morhua | 3797.0 | 6 |
| scf7180000054480 | 138406 | 0.3866 | 0 | 154.5006 | Eukaryota | 93180.0 | 0 | Chordata | 93180.0 | 0 | Carnivora | 93180.0 | 0 | Canidae | 93180.0 | 0 | Canis | 93180.0 | 0 | Canis lupus | 93180.0 | 0 |
| scf7180000051946 | 28141 | 0.3522 | 0 | 55.3522 | Eukaryota | 56078.0 | 0 | Chordata | 56078.0 | 0 | Carnivora | 56078.0 | 0 | Canidae | 56078.0 | 0 | Canis | 56078.0 | 0 | Canis lupus | 56078.0 | 0 |
| scf7180000054139 | 1290428 | 0.3677 | 0 | 60.7843 | Eukaryota | 52606.0 | 0 | Chordata | 52606.0 | 0 | Carnivora | 52606.0 | 0 | Canidae | 52606.0 | 0 | Canis | 35501.0 | 1 | Canis lupus | 35501.0 | 1 |
| scf7180000050879 | 19957 | 0.4114 | 0 | 58.9947 | Eukaryota | 18884.0 | 0 | Chordata | 18884.0 | 0 | Carnivora | 18884.0 | 0 | Canidae | 18884.0 | 0 | Canis | 18884.0 | 0 | Canis lupus | 18884.0 | 0 |
| scf7180000042762 | 337213 | 0.4993 | 0 | 57.5976 | Eukaryota | 47916.0 | 0 | Chordata | 47916.0 | 0 | Carnivora | 47916.0 | 0 | Canidae | 44324.0 | 1 | Canis | 44324.0 | 1 | Canis lupus | 44324.0 | 1 |
| scf7180000046450 | 88973 | 0.4417 | 0 | 56.4186 | Eukaryota | 36783.0 | 0 | Chordata | 36783.0 | 0 | Carnivora | 36783.0 | 0 | Canidae | 36783.0 | 0 | Canis | 36783.0 | 0 | Canis lupus | 36783.0 | 0 |
| scf7180000049607 | 24823 | 0.4499 | 0 | 826.5169 | Eukaryota | 12489.0 | 0 | Platyhelminthes | 8995.0 | 2 | Cyclophyllidea | 7504.0 | 5 | Hymenolepididae | 7504.0 | 5 | Hymenolepis | 7504.0 | 5 | Hymenolepis microstoma | 7504.0 | 5 |
| scf7180000044372 | 206693 | 0.501 | 0 | 58.3515 | Eukaryota | 28230.0 | 0 | Chordata | 28230.0 | 0 | Carnivora | 28230.0 | 0 | Canidae | 28230.0 | 0 | Vulpes | 16874.0 | 1 | Vulpes vulpes | 16874.0 | 1 |
| scf7180000055961 | 14026 | 0.4465 | 0 | 122.3911 | Eukaryota | 1855.0 | 0 | Chordata | 1217.0 | 1 | Salmoniformes | 806.0 | 3 | Salmonidae | 806.0 | 3 | Salmo | 806.0 | 3 | Salmo trutta | 806.0 | 3 |
| scf7180000042241 | 233598 | 0.3873 | 0 | 60.5814 | Eukaryota | 91491.0 | 0 | Chordata | 91491.0 | 0 | Carnivora | 91491.0 | 0 | Canidae | 91491.0 | 0 | Canis | 91491.0 | 0 | Canis lupus | 91491.0 | 0 |
| scf7180000042088 | 344531 | 0.392 | 0 | 62.222 | Eukaryota | 25880.0 | 0 | Chordata | 25880.0 | 0 | Carnivora | 25880.0 | 0 | Canidae | 25880.0 | 0 | Canis | 25880.0 | 0 | Canis lupus | 25880.0 | 0 |
| scf7180000046243 | 157431 | 0.3964 | 0 | 60.6177 | Eukaryota | 16274.0 | 0 | Chordata | 16274.0 | 0 | Carnivora | 16274.0 | 0 | Canidae | 13155.0 | 1 | Canis | 9881.0 | 3 | Canis lupus | 9881.0 | 3 |
| scf7180000055177 | 22225 | 0.3968 | 0 | 235.7593 | Eukaryota | 80031.0 | 0 | Chordata | 80031.0 | 0 | Carnivora | 80031.0 | 0 | Canidae | 80031.0 | 0 | Canis | 80031.0 | 0 | Canis lupus | 80031.0 | 0 |
| scf7180000041843 | 924102 | 0.4593 | 0 | 58.6635 | Eukaryota | 88620.0 | 0 | Chordata | 88620.0 | 0 | Carnivora | 88620.0 | 0 | Canidae | 88620.0 | 0 | Canis | 52932.0 | 1 | Canis lupus | 52932.0 | 1 |
| scf7180000055977 | 4548 | 0.4529 | 0 | 1399.015 | Eukaryota | 4791.0 | 0 | Platyhelminthes | 2008.0 | 3 | Polystomatidea | 2008.0 | 4 | Polystomatidae | 2008.0 | 4 | Protopolystoma | 2008.0 | 4 | Protopolystoma xenopodis | 2008.0 | 4 |
| scf7180000051333 | 27624 | 0.5008 | 0 | 99.998 | Eukaryota | 9108.0 | 0 | Chordata | 9108.0 | 0 | Carnivora | 9108.0 | 0 | Canidae | 9108.0 | 0 | Canis | 5214.0 | 1 | Canis lupus | 5214.0 | 1 |
| scf7180000040549 | 677064 | 0.3976 | 0 | 59.4088 | Eukaryota | 88460.0 | 0 | Chordata | 88460.0 | 0 | Carnivora | 88460.0 | 0 | Canidae | 88460.0 | 0 | Canis | 88460.0 | 0 | Canis lupus | 88460.0 | 0 |
| scf7180000041372 | 1214031 | 0.3701 | 0 | 59.9574 | Eukaryota | 47402.0 | 0 | Chordata | 47402.0 | 0 | Carnivora | 47402.0 | 0 | Canidae | 47402.0 | 0 | Canis | 42362.0 | 1 | Canis lupus | 42362.0 | 1 |
| scf7180000052919 | 3205 | 0.449 | 0 | 459.5426 | Eukaryota | 908.0 | 0 | Chordata | 603.0 | 1 | Tetraodontiformes | 603.0 | 1 | Tetraodontidae | 603.0 | 1 | Takifugu | 603.0 | 1 | Takifugu rubripes | 603.0 | 1 |
| scf7180000041320 | 65987 | 0.4069 | 0 | 58.4048 | Eukaryota | 13795.0 | 0 | Chordata | 13795.0 | 0 | Carnivora | 6201.0 | 3 | Canidae | 6201.0 | 7 | Canis | 6201.0 | 8 | Canis lupus | 6201.0 | 8 |
| scf7180000054649 | 30015 | 0.4171 | 0 | 124.2787 | Eukaryota | 30477.0 | 0 | Chordata | 30477.0 | 0 | Carnivora | 30477.0 | 0 | Canidae | 30477.0 | 0 | Canis | 30477.0 | 0 | Canis lupus | 30477.0 | 0 |
| scf7180000042495 | 48243 | 0.5439 | 0 | 52.7551 | Eukaryota | 26315.0 | 0 | Chordata | 26315.0 | 0 | Carnivora | 26315.0 | 0 | Canidae | 26315.0 | 0 | Canis | 26315.0 | 0 | Canis lupus | 26315.0 | 0 |
| scf7180000051879 | 4065 | 0.4517 | 0 | 498.5431 | Eukaryota | 2612.0 | 0 | Platyhelminthes | 1147.0 | 3 | Polystomatidea | 1147.0 | 3 | Polystomatidae | 1147.0 | 3 | Protopolystoma | 1147.0 | 3 | Protopolystoma xenopodis | 1147.0 | 3 |
| scf7180000049306 | 60668 | 0.4155 | 0 | 55.2729 | Eukaryota | 10730.0 | 0 | Chordata | 10730.0 | 0 | Carnivora | 10730.0 | 0 | Canidae | 6058.0 | 3 | Canis | 4868.0 | 6 | Canis lupus | 4868.0 | 6 |
| scf7180000051957 | 4719 | 0.4524 | 0 | 75.4902 | no-hit | 0.0 | 0 | no-hit | 0.0 | 0 | no-hit | 0.0 | 0 | no-hit | 0.0 | 0 | no-hit | 0.0 | 0 | no-hit | 0.0 | 0 |
| scf7180000042215 | 794566 | 0.3746 | 0 | 59.8597 | Eukaryota | 115671.0 | 0 | Chordata | 115671.0 | 0 | Carnivora | 115671.0 | 0 | Canidae | 110239.0 | 1 | Canis | 102420.0 | 2 | Canis lupus | 102420.0 | 2 |
| scf7180000044956 | 46451 | 0.4567 | 0 | 54.2385 | Eukaryota | 5847.0 | 0 | Chordata | 5847.0 | 0 | Carnivora | 5847.0 | 0 | Canidae | 5847.0 | 0 | Canis | 4792.0 | 1 | Canis lupus | 4792.0 | 1 |
| scf7180000043573 | 460550 | 0.3608 | 0 | 60.0261 | Eukaryota | 53800.0 | 0 | Chordata | 53800.0 | 0 | Carnivora | 53800.0 | 0 | Canidae | 53800.0 | 0 | Canis | 53800.0 | 0 | Canis lupus | 53800.0 | 0 |
| scf7180000050585 | 3936 | 0.455 | 0 | 1375.1358 | Eukaryota | 542.0 | 0 | Platyhelminthes | 542.0 | 0 | Polystomatidea | 542.0 | 0 | Polystomatidae | 542.0 | 0 | Protopolystoma | 542.0 | 0 | Protopolystoma xenopodis | 542.0 | 0 |
| scf7180000050845 | 331888 | 0.5612 | 0 | 55.6184 | Eukaryota | 48420.0 | 0 | Chordata | 48420.0 | 0 | Carnivora | 48420.0 | 0 | Canidae | 48420.0 | 0 | Canis | 39387.0 | 1 | Canis lupus | 39387.0 | 1 |
| scf7180000052075 | 2383 | 0.4612 | 0 | 419.9196 | no-hit | 0.0 | 0 | no-hit | 0.0 | 0 | no-hit | 0.0 | 0 | no-hit | 0.0 | 0 | no-hit | 0.0 | 0 | no-hit | 0.0 | 0 |
| scf7180000047706 | 8547 | 0.4352 | 0 | 119.4378 | no-hit | 0.0 | 0 | no-hit | 0.0 | 0 | no-hit | 0.0 | 0 | no-hit | 0.0 | 0 | no-hit | 0.0 | 0 | no-hit | 0.0 | 0 |
| scf7180000043807 | 365660 | 0.4037 | 0 | 61.1675 | Eukaryota | 31331.0 | 0 | Chordata | 31331.0 | 0 | Carnivora | 31331.0 | 0 | Canidae | 31331.0 | 0 | Canis | 31331.0 | 0 | Canis lupus | 31331.0 | 0 |
| scf7180000041472 | 1073587 | 0.5002 | 0 | 58.2642 | Eukaryota | 151535.0 | 0 | Chordata | 151535.0 | 0 | Carnivora | 151535.0 | 0 | Canidae | 101339.0 | 1 | Canis | 101339.0 | 1 | Canis lupus | 101339.0 | 1 |
| scf7180000042813 | 345598 | 0.4451 | 0 | 57.705 | Eukaryota | 53776.0 | 0 | Chordata | 53776.0 | 0 | Carnivora | 53776.0 | 0 | Canidae | 53776.0 | 0 | Canis | 53776.0 | 0 | Canis lupus | 53776.0 | 0 |
| scf7180000056448 | 350100 | 0.4074 | 0 | 59.2953 | Eukaryota | 34644.0 | 0 | Chordata | 34644.0 | 0 | Carnivora | 34644.0 | 0 | Canidae | 25544.0 | 2 | Canis | 19160.0 | 4 | Canis lupus | 19160.0 | 4 |
| scf7180000055632 | 5102 | 0.45 | 0 | 463.8225 | no-hit | 0.0 | 0 | no-hit | 0.0 | 0 | no-hit | 0.0 | 0 | no-hit | 0.0 | 0 | no-hit | 0.0 | 0 | no-hit | 0.0 | 0 |
| scf7180000056030 | 17519 | 0.5027 | 0 | 836.4882 | Eukaryota | 8844.0 | 0 | Chordata | 8844.0 | 0 | Carnivora | 8844.0 | 0 | Canidae | 8844.0 | 0 | Canis | 7161.0 | 1 | Canis lupus | 5944.0 | 2 |
| scf7180000046426 | 124358 | 0.5623 | 0 | 54.7961 | Eukaryota | 90627.0 | 0 | Chordata | 90627.0 | 0 | Carnivora | 90627.0 | 0 | Canidae | 90627.0 | 0 | Canis | 83298.0 | 1 | Canis lupus | 83298.0 | 1 |
| scf7180000050677 | 24586 | 0.442 | 0 | 58.9528 | Eukaryota | 20757.0 | 0 | Chordata | 20757.0 | 0 | Carnivora | 20757.0 | 0 | Otariidae | 8956.0 | 4 | Callorhinus | 6771.0 | 6 | Callorhinus ursinus | 6771.0 | 6 |
| scf7180000042251 | 1581341 | 0.4175 | 0 | 58.9599 | Eukaryota | 89592.0 | 0 | Chordata | 89592.0 | 0 | Carnivora | 89592.0 | 0 | Canidae | 89592.0 | 0 | Canis | 89592.0 | 0 | Canis lupus | 89592.0 | 0 |
| scf7180000050075 | 6332 | 0.4272 | 0 | 107.8968 | no-hit | 0.0 | 0 | no-hit | 0.0 | 0 | no-hit | 0.0 | 0 | no-hit | 0.0 | 0 | no-hit | 0.0 | 0 | no-hit | 0.0 | 0 |
| scf7180000041187 | 1239005 | 0.3843 | 0 | 59.0331 | Eukaryota | 94652.0 | 0 | Chordata | 94652.0 | 0 | Carnivora | 94652.0 | 0 | Canidae | 94652.0 | 0 | Canis | 84434.0 | 1 | Canis lupus | 84434.0 | 1 |
| scf7180000044890 | 46291 | 0.3869 | 0 | 54.5187 | Eukaryota | 18031.0 | 0 | Chordata | 18031.0 | 0 | Carnivora | 18031.0 | 0 | Canidae | 18031.0 | 0 | Canis | 16235.0 | 1 | Canis lupus | 16235.0 | 1 |
| scf7180000041102 | 312706 | 0.3658 | 0 | 58.6532 | Eukaryota | 90760.0 | 0 | Chordata | 90760.0 | 0 | Carnivora | 90760.0 | 0 | Canidae | 90760.0 | 0 | Canis | 90760.0 | 0 | Canis lupus | 90760.0 | 0 |
| scf7180000056080 | 25424 | 0.3591 | 0 | 411.611 | Eukaryota | 6917.0 | 0 | Chordata | 6917.0 | 0 | Carnivora | 4822.0 | 1 | Canidae | 4822.0 | 1 | Canis | 4822.0 | 1 | Canis lupus | 4822.0 | 1 |
| scf7180000052301 | 4798 | 0.4535 | 0 | 321.1507 | Eukaryota | 605.0 | 0 | Platyhelminthes | 605.0 | 0 | Polystomatidea | 605.0 | 0 | Polystomatidae | 605.0 | 0 | Protopolystoma | 605.0 | 0 | Protopolystoma xenopodis | 605.0 | 0 |
| scf7180000052772 | 22412 | 0.3709 | 0 | 59.7891 | Eukaryota | 7612.0 | 0 | Chordata | 7612.0 | 0 | Carnivora | 6055.0 | 2 | Canidae | 6055.0 | 2 | Canis | 6055.0 | 2 | Canis lupus | 6055.0 | 2 |
| scf7180000040707 | 94349 | 0.4781 | 0 | 55.6393 | Eukaryota | 34016.0 | 0 | Chordata | 34016.0 | 0 | Carnivora | 34016.0 | 0 | Canidae | 25301.0 | 1 | Canis | 25301.0 | 1 | Canis lupus | 25301.0 | 1 |
| scf7180000055149 | 38832 | 0.444 | 0 | 110.2804 | Eukaryota | 4583.0 | 0 | Chordata | 3504.0 | 1 | Kurtiformes | 1630.0 | 3 | Apogonidae | 1630.0 | 3 | Sphaeramia | 1630.0 | 4 | Sphaeramia orbicularis | 1630.0 | 4 |
| scf7180000055985 | 16295 | 0.4252 | 0 | 280.5707 | Eukaryota | 70662.0 | 0 | Chordata | 70662.0 | 0 | Carnivora | 70662.0 | 0 | Canidae | 70662.0 | 0 | Canis | 70662.0 | 0 | Canis lupus | 70662.0 | 0 |
| scf7180000046581 | 15615 | 0.3708 | 0 | 47.0359 | Eukaryota | 50767.0 | 0 | Chordata | 50767.0 | 0 | Carnivora | 50767.0 | 0 | Canidae | 50767.0 | 0 | Canis | 50767.0 | 0 | Canis lupus | 50767.0 | 0 |
| scf7180000041781 | 197740 | 0.3893 | 0 | 60.4355 | Eukaryota | 21263.0 | 0 | Chordata | 21263.0 | 0 | Carnivora | 21263.0 | 0 | Canidae | 12147.0 | 3 | Canis | 7341.0 | 4 | Canis lupus | 7341.0 | 5 |
| scf7180000042158 | 855451 | 0.4007 | 0 | 59.3795 | Eukaryota | 109274.0 | 0 | Chordata | 109274.0 | 0 | Carnivora | 109274.0 | 0 | Canidae | 109274.0 | 0 | Canis | 109274.0 | 0 | Canis lupus | 109274.0 | 0 |
| scf7180000043147 | 426880 | 0.5058 | 0 | 58.7664 | Eukaryota | 43839.0 | 0 | Chordata | 43839.0 | 0 | Carnivora | 43839.0 | 0 | Canidae | 43839.0 | 0 | Canis | 26629.0 | 1 | Canis lupus | 26629.0 | 1 |
| scf7180000055199 | 35062 | 0.443 | 0 | 431.0941 | Eukaryota | 5950.0 | 0 | Chordata | 4765.0 | 1 | Kurtiformes | 3061.0 | 2 | Apogonidae | 3061.0 | 2 | Sphaeramia | 3061.0 | 2 | Sphaeramia orbicularis | 3061.0 | 2 |
| scf7180000046054 | 15751 | 0.4434 | 0 | 36.2547 | Eukaryota | 3452.0 | 0 | Chordata | 3452.0 | 0 | Carnivora | 2019.0 | 1 | Canidae | 1693.0 | 2 | Canis | 1356.0 | 4 | Canis lupus | 1356.0 | 4 |
| scf7180000056598 | 885929 | 0.466 | 0 | 58.8221 | Eukaryota | 47260.0 | 0 | Chordata | 47260.0 | 0 | Carnivora | 47260.0 | 0 | Canidae | 43272.0 | 1 | Canis | 35375.0 | 2 | Canis lupus | 35375.0 | 2 |
| scf7180000040691 | 1928741 | 0.3763 | 0 | 59.4606 | Eukaryota | 95073.0 | 0 | Chordata | 95073.0 | 0 | Carnivora | 95073.0 | 0 | Canidae | 95073.0 | 0 | Canis | 95073.0 | 0 | Canis lupus | 95073.0 | 0 |
| scf7180000050309 | 24375 | 0.5193 | 0 | 53.1058 | Eukaryota | 5134.0 | 1 | Chordata | 5134.0 | 1 | Carnivora | 3408.0 | 3 | Canidae | 3071.0 | 4 | Vulpes | 1663.0 | 6 | Vulpes vulpes | 1663.0 | 6 |
| scf7180000050553 | 9406 | 0.4392 | 0 | 137.5232 | no-hit | 0.0 | 0 | no-hit | 0.0 | 0 | no-hit | 0.0 | 0 | no-hit | 0.0 | 0 | no-hit | 0.0 | 0 | no-hit | 0.0 | 0 |
| scf7180000051309 | 7750 | 0.4511 | 0 | 553.7775 | no-hit | 0.0 | 0 | no-hit | 0.0 | 0 | no-hit | 0.0 | 0 | no-hit | 0.0 | 0 | no-hit | 0.0 | 0 | no-hit | 0.0 | 0 |
| scf7180000051286 | 15951 | 0.4444 | 0 | 83.6724 | Eukaryota | 206.0 | 0 | Chordata | 206.0 | 0 | Salmoniformes | 206.0 | 0 | Salmonidae | 206.0 | 0 | Coregonus | 206.0 | 0 | Coregonus sp. 'balchen' | 206.0 | 0 |
| scf7180000047287 | 45745 | 0.4012 | 0 | 55.8236 | Eukaryota | 22889.0 | 0 | Chordata | 22889.0 | 0 | Carnivora | 22889.0 | 0 | Canidae | 22889.0 | 0 | Canis | 22889.0 | 0 | Canis lupus | 22889.0 | 0 |
| scf7180000048191 | 16452 | 0.4399 | 0 | 100.9191 | Eukaryota | 4162.0 | 0 | Chordata | 4162.0 | 0 | Carnivora | 3530.0 | 1 | Canidae | 3530.0 | 1 | Canis | 3530.0 | 1 | Canis lupus | 3530.0 | 1 |
| scf7180000043902 | 366341 | 0.4852 | 0 | 59.8648 | Eukaryota | 33666.0 | 0 | Chordata | 33666.0 | 0 | Carnivora | 33666.0 | 0 | Canidae | 33666.0 | 0 | Canis | 26084.0 | 1 | Canis lupus | 26084.0 | 1 |
| scf7180000042599 | 106054 | 0.3907 | 0 | 65.7121 | Eukaryota | 89435.0 | 0 | Chordata | 89435.0 | 0 | Carnivora | 89435.0 | 0 | Canidae | 89435.0 | 0 | Canis | 89435.0 | 0 | Canis lupus | 89435.0 | 0 |
| scf7180000056251 | 1034442 | 0.48 | 0 | 59.2963 | Eukaryota | 128115.0 | 0 | Chordata | 128115.0 | 0 | Carnivora | 128115.0 | 0 | Canidae | 128115.0 | 0 | Canis | 116144.0 | 1 | Canis lupus | 116144.0 | 1 |
| scf7180000050361 | 5354 | 0.4466 | 0 | 437.35 | Eukaryota | 570.0 | 0 | Streptophyta | 570.0 | 0 | Fabales | 570.0 | 0 | Fabaceae | 570.0 | 0 | Cicer | 570.0 | 0 | Cicer arietinum | 570.0 | 0 |
| scf7180000043547 | 490437 | 0.4075 | 0 | 56.5658 | Eukaryota | 18371.0 | 0 | Chordata | 18371.0 | 0 | Carnivora | 18371.0 | 0 | Canidae | 9516.0 | 2 | Canis | 9516.0 | 4 | Canis lupus | 9516.0 | 4 |
| scf7180000040480 | 2050206 | 0.38 | 0 | 58.9182 | Eukaryota | 117885.0 | 0 | Chordata | 117885.0 | 0 | Carnivora | 117885.0 | 0 | Canidae | 117885.0 | 0 | Canis | 117885.0 | 0 | Canis lupus | 117885.0 | 0 |
| scf7180000056416 | 452273 | 0.3861 | 0 | 59.0525 | Eukaryota | 123940.0 | 0 | Chordata | 123940.0 | 0 | Carnivora | 123940.0 | 0 | Canidae | 123940.0 | 0 | Canis | 123940.0 | 0 | Canis lupus | 123940.0 | 0 |
| scf7180000042149 | 52787 | 0.3725 | 0 | 59.1945 | Eukaryota | 72731.0 | 0 | Chordata | 72731.0 | 0 | Carnivora | 72731.0 | 0 | Canidae | 72731.0 | 0 | Canis | 72731.0 | 0 | Canis lupus | 72731.0 | 0 |
| scf7180000052215 | 7626 | 0.4491 | 0 | 1908.9307 | Eukaryota | 6096.0 | 0 | Chordata | 3981.0 | 3 | Tetraodontiformes | 3981.0 | 3 | Tetraodontidae | 3981.0 | 3 | Takifugu | 3981.0 | 3 | Takifugu rubripes | 3981.0 | 3 |
| scf7180000045711 | 5566 | 0.4896 | 0 | 30.9961 | Eukaryota | 3935.0 | 0 | Chordata | 3935.0 | 0 | Carnivora | 3935.0 | 0 | Canidae | 3935.0 | 0 | Canis | 3458.0 | 1 | Canis lupus | 3458.0 | 1 |
| scf7180000041809 | 179073 | 0.3631 | 0 | 57.6112 | Eukaryota | 46560.0 | 0 | Chordata | 46560.0 | 0 | Carnivora | 46560.0 | 0 | Canidae | 46560.0 | 0 | Canis | 46560.0 | 0 | Canis lupus | 46560.0 | 0 |
| scf7180000052633 | 5594 | 0.4487 | 0 | 779.0724 | Eukaryota | 2628.0 | 0 | Platyhelminthes | 1219.0 | 2 | Polystomatidea | 1219.0 | 2 | Polystomatidae | 1219.0 | 2 | Protopolystoma | 1219.0 | 2 | Protopolystoma xenopodis | 1219.0 | 2 |
| scf7180000052324 | 3968 | 0.4609 | 0 | 674.8749 | Eukaryota | 139.0 | 0 | Platyhelminthes | 139.0 | 0 | Polystomatidea | 139.0 | 0 | Polystomatidae | 139.0 | 0 | Protopolystoma | 139.0 | 0 | Protopolystoma xenopodis | 139.0 | 0 |
| scf7180000040958 | 1493306 | 0.4171 | 0 | 60.3648 | Eukaryota | 43472.0 | 0 | Chordata | 43472.0 | 0 | Carnivora | 43472.0 | 0 | Canidae | 43472.0 | 0 | Vulpes | 34856.0 | 1 | Vulpes vulpes | 34856.0 | 1 |
| scf7180000050746 | 25953 | 0.4556 | 0 | 57.853 | Eukaryota | 5588.0 | 0 | Platyhelminthes | 4602.0 | 1 | Polystomatidea | 4602.0 | 2 | Polystomatidae | 4602.0 | 2 | Protopolystoma | 4602.0 | 2 | Protopolystoma xenopodis | 4602.0 | 2 |
| scf7180000047095 | 83703 | 0.6328 | 0 | 55.7506 | Eukaryota | 78393.0 | 0 | Chordata | 78393.0 | 0 | Carnivora | 78393.0 | 0 | Canidae | 78393.0 | 0 | Canis | 55651.0 | 1 | Canis lupus | 55651.0 | 1 |
| scf7180000042072 | 293872 | 0.4003 | 0 | 58.9659 | Eukaryota | 69494.0 | 0 | Chordata | 69494.0 | 0 | Carnivora | 69494.0 | 0 | Canidae | 69494.0 | 0 | Canis | 69494.0 | 0 | Canis lupus | 69494.0 | 0 |
| scf7180000056070 | 128153 | 0.4305 | 0 | 664.181 | Eukaryota | 34023.0 | 0 | Chordata | 34023.0 | 0 | Carnivora | 34023.0 | 0 | Canidae | 34023.0 | 0 | Canis | 34023.0 | 0 | Canis lupus | 34023.0 | 0 |
| scf7180000042337 | 747453 | 0.42 | 0 | 70.2803 | Eukaryota | 73760.0 | 0 | Chordata | 73760.0 | 0 | Carnivora | 73760.0 | 0 | Canidae | 73760.0 | 0 | Vulpes | 73760.0 | 0 | Vulpes vulpes | 73760.0 | 0 |
| scf7180000042497 | 82700 | 0.3725 | 0 | 60.8319 | Eukaryota | 73068.0 | 0 | Chordata | 73068.0 | 0 | Carnivora | 73068.0 | 0 | Canidae | 73068.0 | 0 | Canis | 73068.0 | 0 | Canis lupus | 73068.0 | 0 |
| scf7180000041477 | 996671 | 0.3909 | 0 | 58.954 | Eukaryota | 136436.0 | 0 | Chordata | 136436.0 | 0 | Carnivora | 136436.0 | 0 | Canidae | 136436.0 | 0 | Canis | 136436.0 | 0 | Canis lupus | 136436.0 | 0 |
| scf7180000042585 | 116904 | 0.4536 | 0 | 58.4447 | Eukaryota | 39352.0 | 0 | Chordata | 39352.0 | 0 | Carnivora | 39352.0 | 0 | Canidae | 39352.0 | 0 | Canis | 39352.0 | 0 | Canis lupus | 39352.0 | 0 |
| scf7180000046574 | 89151 | 0.3524 | 0 | 59.4058 | Eukaryota | 26199.0 | 0 | Chordata | 26199.0 | 0 | Carnivora | 26199.0 | 0 | Canidae | 26199.0 | 0 | Canis | 26199.0 | 0 | Canis lupus | 26199.0 | 0 |
| scf7180000046999 | 28610 | 0.4919 | 0 | 168.2958 | Eukaryota | 9258.0 | 0 | Chordata | 9258.0 | 0 | Carnivora | 9258.0 | 0 | Canidae | 9258.0 | 0 | Canis | 7430.0 | 2 | Canis lupus | 6159.0 | 3 |
| scf7180000041082 | 1294974 | 0.4753 | 0 | 59.7713 | Eukaryota | 130521.0 | 0 | Chordata | 130521.0 | 0 | Carnivora | 117656.0 | 1 | Canidae | 117656.0 | 2 | Canis | 117656.0 | 2 | Canis lupus | 117656.0 | 2 |
| scf7180000046156 | 144463 | 0.4211 | 0 | 59.5061 | Eukaryota | 20047.0 | 0 | Chordata | 20047.0 | 0 | Carnivora | 20047.0 | 0 | Canidae | 20047.0 | 0 | Vulpes | 16066.0 | 1 | Vulpes vulpes | 16066.0 | 1 |
| scf7180000053756 | 2968 | 0.4444 | 0 | 446.6579 | Eukaryota | 2069.0 | 0 | Platyhelminthes | 1045.0 | 3 | Polystomatidea | 1045.0 | 3 | Polystomatidae | 1045.0 | 3 | Protopolystoma | 1045.0 | 3 | Protopolystoma xenopodis | 1045.0 | 3 |
| scf7180000045616 | 165807 | 0.4531 | 0 | 60.546 | Eukaryota | 14458.0 | 0 | Chordata | 14458.0 | 0 | Carnivora | 8354.0 | 2 | Canidae | 8354.0 | 2 | Canis | 7685.0 | 4 | Canis lupus | 7685.0 | 4 |
| scf7180000052049 | 5455 | 0.4367 | 0 | 65.615 | Eukaryota | 185.0 | 0 | Chordata | 185.0 | 0 | Salmoniformes | 185.0 | 0 | Salmonidae | 185.0 | 0 | Salmo | 185.0 | 0 | Salmo trutta | 185.0 | 0 |
| scf7180000049243 | 7580 | 0.4427 | 0 | 96.8265 | Eukaryota | 29451.0 | 0 | Chordata | 25479.0 | 2 | Gadiformes | 17176.0 | 5 | Gadidae | 17176.0 | 5 | Gadus | 17176.0 | 5 | Gadus morhua | 17176.0 | 5 |
| scf7180000043818 | 231171 | 0.3673 | 0 | 57.3514 | Eukaryota | 81296.0 | 0 | Chordata | 81296.0 | 0 | Carnivora | 81296.0 | 0 | Canidae | 81296.0 | 0 | Canis | 81296.0 | 0 | Canis lupus | 81296.0 | 0 |
| scf7180000044141 | 303800 | 0.5652 | 0 | 57.2608 | Eukaryota | 36379.0 | 0 | Chordata | 36379.0 | 0 | Carnivora | 36379.0 | 0 | Canidae | 36379.0 | 0 | Canis | 26889.0 | 1 | Canis lupus | 26889.0 | 1 |
| scf7180000045951 | 35595 | 0.4305 | 0 | 54.7704 | Eukaryota | 16164.0 | 0 | Chordata | 16164.0 | 0 | Carnivora | 16164.0 | 0 | Ursidae | 7829.0 | 3 | Ursus | 7829.0 | 4 | Ursus arctos | 6528.0 | 5 |
| scf7180000048735 | 12727 | 0.4433 | 0 | 186.3052 | Eukaryota | 16452.0 | 1 | Chordata | 14704.0 | 3 | Gadiformes | 8977.0 | 8 | Gadidae | 8977.0 | 8 | Gadus | 8977.0 | 8 | Gadus morhua | 8977.0 | 8 |
| scf7180000044672 | 230051 | 0.343 | 0 | 59.3494 | Eukaryota | 37500.0 | 0 | Chordata | 37500.0 | 0 | Carnivora | 37500.0 | 0 | Canidae | 37500.0 | 0 | Canis | 37500.0 | 0 | Canis lupus | 37500.0 | 0 |
| scf7180000046812 | 14252 | 0.4478 | 0 | 135.5317 | Eukaryota | 6980.0 | 0 | Chordata | 6980.0 | 0 | Tetraodontiformes | 5281.0 | 2 | Tetraodontidae | 5281.0 | 2 | Takifugu | 5281.0 | 2 | Takifugu rubripes | 4603.0 | 3 |
| scf7180000047178 | 35779 | 0.3894 | 0 | 53.4487 | Eukaryota | 13671.0 | 0 | Chordata | 13671.0 | 0 | Carnivora | 13671.0 | 0 | Canidae | 13671.0 | 0 | Canis | 13671.0 | 0 | Canis lupus | 13671.0 | 0 |
| scf7180000045316 | 281616 | 0.3482 | 100 | 58.8711 | Eukaryota | 32214.0 | 0 | Chordata | 32214.0 | 0 | Carnivora | 32214.0 | 0 | Canidae | 32214.0 | 0 | Canis | 32214.0 | 0 | Canis lupus | 32214.0 | 0 |
| scf7180000041774 | 163361 | 0.4015 | 0 | 57.4719 | Eukaryota | 57314.0 | 0 | Chordata | 57314.0 | 0 | Carnivora | 57314.0 | 0 | Canidae | 57314.0 | 0 | Canis | 57314.0 | 0 | Canis lupus | 57314.0 | 0 |
| scf7180000045323 | 194673 | 0.4236 | 0 | 59.8183 | Eukaryota | 38815.0 | 0 | Chordata | 38815.0 | 0 | Carnivora | 38815.0 | 0 | Canidae | 35513.0 | 1 | Canis | 30940.0 | 2 | Canis lupus | 30940.0 | 2 |
| scf7180000055519 | 19312 | 0.4214 | 0 | 83.09 | Eukaryota | 18510.0 | 0 | Chordata | 18510.0 | 0 | Carnivora | 18510.0 | 0 | Canidae | 18510.0 | 0 | Vulpes | 16194.0 | 1 | Vulpes vulpes | 16194.0 | 1 |
| scf7180000043453 | 81578 | 0.404 | 0 | 58.7371 | Eukaryota | 10558.0 | 0 | Chordata | 10558.0 | 0 | Carnivora | 10558.0 | 0 | Canidae | 10558.0 | 0 | Canis | 10558.0 | 0 | Canis lupus | 10558.0 | 0 |
| scf7180000045970 | 30734 | 0.429 | 0 | 46.3133 | Eukaryota | 66438.0 | 0 | Chordata | 66438.0 | 0 | Carnivora | 66438.0 | 0 | Canidae | 66438.0 | 0 | Canis | 66438.0 | 0 | Canis lupus | 66438.0 | 0 |
| scf7180000045188 | 283821 | 0.429 | 0 | 58.7157 | Eukaryota | 25700.0 | 0 | Chordata | 25700.0 | 0 | Carnivora | 25700.0 | 0 | Canidae | 25700.0 | 0 | Canis | 25700.0 | 0 | Canis lupus | 25700.0 | 0 |
| scf7180000045294 | 228027 | 0.5285 | 0 | 55.6772 | Eukaryota | 60853.0 | 0 | Chordata | 60853.0 | 0 | Carnivora | 60853.0 | 0 | Canidae | 26749.0 | 3 | Canis | 14763.0 | 6 | Canis lupus | 14763.0 | 6 |
| scf7180000052812 | 11605 | 0.384 | 0 | 61.024 | Eukaryota | 97426.0 | 0 | Chordata | 97426.0 | 0 | Carnivora | 97426.0 | 0 | Canidae | 97426.0 | 0 | Canis | 97426.0 | 0 | Canis lupus | 97426.0 | 0 |
| scf7180000042677 | 507486 | 0.3498 | 0 | 60.9668 | Eukaryota | 91571.0 | 0 | Chordata | 91571.0 | 0 | Carnivora | 91571.0 | 0 | Canidae | 91571.0 | 0 | Canis | 91571.0 | 0 | Canis lupus | 91571.0 | 0 |
| scf7180000052643 | 7971 | 0.4514 | 0 | 50.8618 | Eukaryota | 3121.0 | 0 | Chordata | 3121.0 | 0 | Carnivora | 3121.0 | 0 | Canidae | 3121.0 | 0 | Canis | 3121.0 | 0 | Canis lupus | 3121.0 | 0 |
| scf7180000048030 | 6208 | 0.4546 | 0 | 838.2683 | Eukaryota | 2762.0 | 0 | Chordata | 2287.0 | 1 | Carnivora | 2287.0 | 1 | Canidae | 2287.0 | 1 | Canis | 2287.0 | 1 | Canis lupus | 2287.0 | 1 |
| scf7180000042586 | 206402 | 0.343 | 0 | 58.3644 | Eukaryota | 91565.0 | 0 | Chordata | 91565.0 | 0 | Carnivora | 91565.0 | 0 | Canidae | 91565.0 | 0 | Canis | 91565.0 | 0 | Canis lupus | 91565.0 | 0 |
| scf7180000044648 | 288115 | 0.4942 | 0 | 60.2656 | Eukaryota | 55946.0 | 0 | Chordata | 55946.0 | 0 | Carnivora | 55946.0 | 0 | Canidae | 55946.0 | 0 | Canis | 34334.0 | 1 | Canis lupus | 34334.0 | 1 |
| scf7180000044291 | 545328 | 0.4047 | 0 | 59.5813 | Eukaryota | 94579.0 | 0 | Chordata | 94579.0 | 0 | Carnivora | 94579.0 | 0 | Canidae | 94579.0 | 0 | Canis | 94579.0 | 0 | Canis lupus | 94579.0 | 0 |
| scf7180000047096 | 6691 | 0.4446 | 0 | 99.7548 | no-hit | 0.0 | 0 | no-hit | 0.0 | 0 | no-hit | 0.0 | 0 | no-hit | 0.0 | 0 | no-hit | 0.0 | 0 | no-hit | 0.0 | 0 |
| scf7180000050567 | 8313 | 0.4509 | 0 | 61.713 | no-hit | 0.0 | 0 | no-hit | 0.0 | 0 | no-hit | 0.0 | 0 | no-hit | 0.0 | 0 | no-hit | 0.0 | 0 | no-hit | 0.0 | 0 |
| scf7180000056278 | 436223 | 0.5927 | 0 | 56.669 | Eukaryota | 48858.0 | 0 | Chordata | 48858.0 | 0 | Carnivora | 48858.0 | 0 | Canidae | 45748.0 | 1 | Canis | 23850.0 | 2 | Canis lupus | 23850.0 | 2 |
| scf7180000050569 | 2588 | 0.4552 | 0 | 96.4254 | no-hit | 0.0 | 0 | no-hit | 0.0 | 0 | no-hit | 0.0 | 0 | no-hit | 0.0 | 0 | no-hit | 0.0 | 0 | no-hit | 0.0 | 0 |
| scf7180000041504 | 1046243 | 0.3827 | 0 | 59.923 | Eukaryota | 92052.0 | 0 | Chordata | 92052.0 | 0 | Carnivora | 92052.0 | 0 | Canidae | 92052.0 | 0 | Canis | 92052.0 | 0 | Canis lupus | 92052.0 | 0 |
| scf7180000045048 | 211762 | 0.4059 | 0 | 59.4812 | Eukaryota | 83142.0 | 0 | Chordata | 83142.0 | 0 | Carnivora | 83142.0 | 0 | Canidae | 83142.0 | 0 | Canis | 83142.0 | 0 | Canis lupus | 83142.0 | 0 |
| scf7180000042200 | 754780 | 0.4283 | 0 | 59.3218 | Eukaryota | 77033.0 | 0 | Chordata | 77033.0 | 0 | Carnivora | 77033.0 | 0 | Canidae | 77033.0 | 0 | Canis | 77033.0 | 0 | Canis lupus | 77033.0 | 0 |
| scf7180000048161 | 12876 | 0.4471 | 0 | 112.4459 | Eukaryota | 315.0 | 0 | Chordata | 315.0 | 0 | Kurtiformes | 315.0 | 0 | Apogonidae | 315.0 | 0 | Sphaeramia | 315.0 | 0 | Sphaeramia orbicularis | 315.0 | 0 |
| scf7180000044808 | 29962 | 0.4274 | 0 | 235.2879 | Eukaryota | 2685.0 | 0 | Chordata | 2685.0 | 0 | Carnivora | 2685.0 | 0 | Canidae | 2685.0 | 0 | Canis | 2685.0 | 0 | Canis lupus | 2685.0 | 0 |
| scf7180000045167 | 121592 | 0.3397 | 0 | 59.4271 | Eukaryota | 86392.0 | 0 | Chordata | 86392.0 | 0 | Carnivora | 86392.0 | 0 | Canidae | 86392.0 | 0 | Canis | 86392.0 | 0 | Canis lupus | 86392.0 | 0 |
| scf7180000043733 | 561879 | 0.5047 | 0 | 57.9656 | Eukaryota | 31921.0 | 0 | Chordata | 31921.0 | 0 | Carnivora | 31921.0 | 0 | Canidae | 31921.0 | 0 | Canis | 25035.0 | 1 | Canis lupus | 25035.0 | 1 |
| scf7180000045189 | 7305 | 0.4424 | 0 | 166.3187 | Eukaryota | 16041.0 | 1 | Chordata | 11312.0 | 3 | Gadiformes | 5639.0 | 6 | Gadidae | 5639.0 | 6 | Gadus | 5639.0 | 6 | Gadus morhua | 5639.0 | 6 |
| scf7180000041846 | 793496 | 0.3662 | 0 | 59.9718 | Eukaryota | 72461.0 | 0 | Chordata | 72461.0 | 0 | Carnivora | 72461.0 | 0 | Canidae | 72461.0 | 0 | Canis | 72461.0 | 0 | Canis lupus | 72461.0 | 0 |
| scf7180000045109 | 202627 | 0.3773 | 0 | 58.3396 | Eukaryota | 24019.0 | 0 | Chordata | 24019.0 | 0 | Carnivora | 24019.0 | 0 | Canidae | 24019.0 | 0 | Canis | 24019.0 | 0 | Canis lupus | 24019.0 | 0 |
| scf7180000052809 | 20890 | 0.5118 | 0 | 184.9304 | Eukaryota | 33313.0 | 0 | Chordata | 33313.0 | 0 | Carnivora | 33313.0 | 0 | Canidae | 33313.0 | 0 | Canis | 33313.0 | 0 | Canis lupus | 33313.0 | 0 |
| scf7180000043452 | 38613 | 0.3979 | 0 | 55.6079 | Eukaryota | 5204.0 | 0 | Chordata | 5204.0 | 0 | Carnivora | 3679.0 | 1 | Canidae | 3679.0 | 2 | Canis | 3679.0 | 2 | Canis lupus | 3679.0 | 2 |
| scf7180000043524 | 384933 | 0.5226 | 0 | 59.3066 | Eukaryota | 46555.0 | 0 | Chordata | 46555.0 | 0 | Carnivora | 46555.0 | 0 | Canidae | 43537.0 | 1 | Canis | 40120.0 | 2 | Canis lupus | 40120.0 | 2 |
| scf7180000051501 | 5293 | 0.4485 | 0 | 606.9377 | Eukaryota | 145.0 | 0 | Chordata | 145.0 | 0 | Salmoniformes | 145.0 | 0 | Salmonidae | 145.0 | 0 | Salmo | 145.0 | 0 | Salmo trutta | 145.0 | 0 |
| scf7180000051951 | 4395 | 0.4491 | 0 | 2071.205 | no-hit | 0.0 | 0 | no-hit | 0.0 | 0 | no-hit | 0.0 | 0 | no-hit | 0.0 | 0 | no-hit | 0.0 | 0 | no-hit | 0.0 | 0 |
| scf7180000041419 | 29416 | 0.3912 | 0 | 186.5606 | Eukaryota | 4067.0 | 0 | Chordata | 4067.0 | 0 | Carnivora | 2859.0 | 2 | Canidae | 2088.0 | 3 | Canis | 2088.0 | 3 | Canis lupus | 2088.0 | 4 |
| scf7180000042798 | 466973 | 0.3965 | 100 | 60.732 | Eukaryota | 14613.0 | 0 | Chordata | 14613.0 | 0 | Carnivora | 14613.0 | 0 | Canidae | 14613.0 | 0 | Canis | 13207.0 | 1 | Canis lupus | 13207.0 | 1 |

| | | | | | | | | | | | | | | | | | | | | | |
|---|---|---|---|---|---|---|---|---|---|---|---|---|---|---|---|---|---|---|---|---|---|
| scf7180000052550 | 16930 | 0.498 | 0 | 5962.6734 | Eukaryota | 8541.0 | 0 | Chordata | 8541.0 | 0 | Carnivora | 8541.0 | 0 | Canidae | 8541.0 | 0 | Canis | 6371.0 | 1 | Canis lupus | 5199.0 | 2 |
| scf7180000047901 | 10483 | 0.3456 | 0 | 39.1829 | Eukaryota | 3659.0 | 0 | Chordata | 3659.0 | 0 | Carnivora | 3659.0 | 0 | Felidae | 1897.0 | 1 | Felis | 1897.0 | 1 | Felis catus | 1897.0 | 1 |
| scf7180000043856 | 437180 | 0.3729 | 0 | 59.3493 | Eukaryota | 67801.0 | 0 | Chordata | 67801.0 | 0 | Carnivora | 67801.0 | 0 | Canidae | 67801.0 | 0 | Canis | 62984.0 | 1 | Canis lupus | 62984.0 | 1 |
| scf7180000055797 | 28778 | 0.4582 | 0 | 294.6238 | Eukaryota | 17120.0 | 0 | Chordata | 17120.0 | 0 | Carnivora | 10796.0 | 2 | Canidae | 10796.0 | 2 | Canis | 8428.0 | 3 | Canis lupus | 8428.0 | 4 |
| scf7180000044752 | 196382 | 0.4865 | 0 | 59.5184 | Eukaryota | 77566.0 | 0 | Chordata | 77566.0 | 0 | Carnivora | 77566.0 | 0 | Canidae | 77566.0 | 0 | Canis | 46934.0 | 1 | Canis lupus | 46934.0 | 1 |
| scf7180000056359 | 662324 | 0.4972 | 0 | 58.3301 | Eukaryota | 60730.0 | 0 | Chordata | 60730.0 | 0 | Carnivora | 60730.0 | 0 | Canidae | 60730.0 | 0 | Canis | 60730.0 | 0 | Canis lupus | 60730.0 | 0 |
| scf7180000050586 | 5527 | 0.4511 | 0 | 532.583 | Eukaryota | 1883.0 | 0 | Echinodermata | 867.0 | 2 | Forcipulatida | 867.0 | 2 | Asteriidae | 867.0 | 2 | Asterias | 867.0 | 2 | Asterias rubens | 867.0 | 2 |
| scf7180000050851 | 2675 | 0.4535 | 0 | 666.9363 | Eukaryota | 413.0 | 0 | Chordata | 215.0 | 1 | Kurtiformes | 215.0 | 1 | Apogonidae | 215.0 | 1 | Sphaeramia | 215.0 | 1 | Sphaeramia orbicularis | 215.0 | 1 |
| scf7180000050051 | 4369 | 0.4557 | 0 | 596.5538 | no-hit | 0.0 | 0 | no-hit | 0.0 | 0 | no-hit | 0.0 | 0 | no-hit | 0.0 | 0 | no-hit | 0.0 | 0 | no-hit | 0.0 | 0 |
| scf7180000056285 | 725237 | 0.4517 | 0 | 59.2055 | Eukaryota | 104007.0 | 0 | Chordata | 104007.0 | 0 | Carnivora | 104007.0 | 0 | Canidae | 57973.0 | 2 | Canis | 46414.0 | 4 | Canis lupus | 46414.0 | 4 |
| scf7180000056027 | 1666 | 0.4496 | 0 | 263.9666 | no-hit | 0.0 | 0 | no-hit | 0.0 | 0 | no-hit | 0.0 | 0 | no-hit | 0.0 | 0 | no-hit | 0.0 | 0 | no-hit | 0.0 | 0 |
| scf7180000046603 | 27738 | 0.443 | 0 | 205.2718 | Eukaryota | 91239.0 | 0 | Chordata | 91239.0 | 0 | Carnivora | 91239.0 | 0 | Canidae | 91239.0 | 0 | Canis | 91239.0 | 0 | Canis lupus | 91239.0 | 0 |
| scf7180000040514 | 202847 | 0.4723 | 0 | 57.7279 | Eukaryota | 29267.0 | 0 | Chordata | 29267.0 | 0 | Carnivora | 21132.0 | 2 | Felidae | 14281.0 | 3 | Canis | 6851.0 | 5 | Canis lupus | 6851.0 | 5 |
| scf7180000053828 | 3833 | 0.4503 | 0 | 465.7176 | no-hit | 0.0 | 0 | no-hit | 0.0 | 0 | no-hit | 0.0 | 0 | no-hit | 0.0 | 0 | no-hit | 0.0 | 0 | no-hit | 0.0 | 0 |
| scf7180000053123 | 8519 | 0.4171 | 0 | 61.5188 | Eukaryota | 2579.0 | 0 | Chordata | 2579.0 | 0 | Carnivora | 2579.0 | 0 | Canidae | 2579.0 | 0 | Canis | 2579.0 | 0 | Canis lupus | 2579.0 | 0 |
| scf7180000055487 | 21158 | 0.4403 | 0 | 533.854 | Eukaryota | 7270.0 | 0 | Chordata | 7270.0 | 0 | Carnivora | 7270.0 | 0 | Canidae | 7270.0 | 0 | Canis | 7270.0 | 0 | Canis lupus | 7270.0 | 0 |
| scf7180000052988 | 3785 | 0.4486 | 0 | 225.3619 | no-hit | 0.0 | 0 | no-hit | 0.0 | 0 | no-hit | 0.0 | 0 | no-hit | 0.0 | 0 | no-hit | 0.0 | 0 | no-hit | 0.0 | 0 |
| scf7180000051199 | 16937 | 0.438 | 0 | 43.8101 | Eukaryota | 13150.0 | 0 | Chordata | 13150.0 | 0 | Carnivora | 7190.0 | 3 | Canidae | 7190.0 | 3 | Canis | 7190.0 | 4 | Canis lupus | 7190.0 | 4 |
| scf7180000041208 | 205029 | 0.4377 | 0 | 58.3307 | Eukaryota | 60175.0 | 0 | Chordata | 60175.0 | 0 | Carnivora | 60175.0 | 0 | Canidae | 60175.0 | 0 | Canis | 60175.0 | 0 | Canis lupus | 60175.0 | 0 |
| scf7180000054519 | 12909 | 0.4464 | 0 | 153.2099 | Eukaryota | 2792.0 | 0 | Chordata | 2792.0 | 0 | Carnivora | 2792.0 | 0 | Canidae | 2792.0 | 0 | Canis | 2792.0 | 0 | Canis lupus | 2792.0 | 0 |
| scf7180000047101 | 105221 | 0.4757 | 0 | 59.4598 | Eukaryota | 4922.0 | 0 | Chordata | 4922.0 | 0 | Carnivora | 4487.0 | 1 | Canidae | 4487.0 | 1 | Canis | 4487.0 | 1 | Canis lupus | 4487.0 | 1 |
| scf7180000040683 | 1977176 | 0.3604 | 0 | 59.7886 | Eukaryota | 45821.0 | 0 | Chordata | 45821.0 | 0 | Carnivora | 45821.0 | 0 | Canidae | 41370.0 | 1 | Canis | 36599.0 | 2 | Canis lupus | 36599.0 | 2 |
| scf7180000044635 | 134825 | 0.362 | 0 | 61.2808 | Eukaryota | 63752.0 | 0 | Chordata | 63752.0 | 0 | Carnivora | 63752.0 | 0 | Canidae | 63752.0 | 0 | Canis | 63752.0 | 0 | Canis lupus | 63752.0 | 0 |
| scf7180000056537 | 291005 | 0.4296 | 0 | 58.3844 | Eukaryota | 91529.0 | 0 | Chordata | 91529.0 | 0 | Carnivora | 91529.0 | 0 | Canidae | 91529.0 | 0 | Canis | 91529.0 | 0 | Canis lupus | 91529.0 | 0 |
| scf7180000043819 | 150191 | 0.4294 | 0 | 58.1913 | Eukaryota | 33122.0 | 0 | Chordata | 33122.0 | 0 | Carnivora | 33122.0 | 0 | Canidae | 33122.0 | 0 | Canis | 33122.0 | 0 | Canis lupus | 33122.0 | 0 |
| scf7180000044100 | 358687 | 0.4222 | 0 | 59.9089 | Eukaryota | 40894.0 | 0 | Chordata | 40894.0 | 0 | Carnivora | 40894.0 | 0 | Canidae | 26379.0 | 3 | Canis | 18811.0 | 6 | Canis lupus | 18811.0 | 6 |
| scf7180000055180 | 6923 | 0.3987 | 0 | 353.6416 | Eukaryota | 6684.0 | 0 | Chordata | 6684.0 | 0 | Carnivora | 6684.0 | 0 | Canidae | 4699.0 | 1 | Canis | 4699.0 | 1 | Canis lupus | 4699.0 | 1 |
| scf7180000053896 | 13156 | 0.4419 | 0 | 58.8635 | no-hit | 0.0 | 0 | no-hit | 0.0 | 0 | no-hit | 0.0 | 0 | no-hit | 0.0 | 0 | no-hit | 0.0 | 0 | no-hit | 0.0 | 0 |
| scf7180000041760 | 960528 | 0.4401 | 0 | 59.4978 | Eukaryota | 56720.0 | 0 | Chordata | 56720.0 | 0 | Carnivora | 56720.0 | 0 | Canidae | 56720.0 | 0 | Canis | 56720.0 | 0 | Canis lupus | 56720.0 | 0 |
| scf7180000047364 | 100944 | 0.3503 | 0 | 57.2668 | Eukaryota | 46059.0 | 0 | Chordata | 46059.0 | 0 | Carnivora | 46059.0 | 0 | Canidae | 46059.0 | 0 | Canis | 46059.0 | 0 | Canis lupus | 46059.0 | 0 |
| scf7180000049117 | 36501 | 0.6879 | 0 | 54.9045 | Eukaryota | 8869.0 | 0 | Chordata | 8869.0 | 0 | Carnivora | 8869.0 | 0 | Canidae | 8869.0 | 0 | Canis | 7169.0 | 1 | Canis lupus | 7169.0 | 1 |
| scf7180000041660 | 1014113 | 0.3943 | 0 | 60.4743 | Eukaryota | 99985.0 | 0 | Chordata | 99985.0 | 0 | Carnivora | 99985.0 | 0 | Canidae | 74542.0 | 2 | Canis | 63908.0 | 3 | Canis lupus | 63908.0 | 3 |
| scf7180000043561 | 48269 | 0.5343 | 0 | 56.5084 | Eukaryota | 14685.0 | 0 | Chordata | 14685.0 | 0 | Carnivora | 14685.0 | 0 | Canidae | 13203.0 | 1 | Canis | 12599.0 | 2 | Canis lupus | 12599.0 | 2 |
| scf7180000042187 | 642012 | 0.4158 | 0 | 62.2193 | Eukaryota | 70017.0 | 0 | Chordata | 70017.0 | 0 | Carnivora | 70017.0 | 0 | Canidae | 49398.0 | 2 | Canis | 37922.0 | 5 | Canis lupus | 37922.0 | 5 |
| scf7180000041900 | 819572 | 0.4352 | 0 | 59.5667 | Eukaryota | 59340.0 | 0 | Chordata | 59340.0 | 0 | Carnivora | 59340.0 | 0 | Canidae | 59340.0 | 0 | Canis | 59340.0 | 0 | Canis lupus | 59340.0 | 0 |
| scf7180000051312 | 22092 | 0.5078 | 0 | 700.7758 | Eukaryota | 8974.0 | 0 | Chordata | 8974.0 | 0 | Carnivora | 8974.0 | 0 | Canidae | 8974.0 | 0 | Canis | 7315.0 | 1 | Canis lupus | 6121.0 | 2 |
| scf7180000045604 | 220429 | 0.4439 | 0 | 60.8752 | Eukaryota | 58985.0 | 0 | Chordata | 58985.0 | 0 | Carnivora | 58985.0 | 0 | Canidae | 54117.0 | 1 | Canis | 48085.0 | 2 | Canis lupus | 48085.0 | 2 |
| scf7180000048164 | 34526 | 0.4429 | 0 | 1143.0381 | Eukaryota | 17884.0 | 0 | Platyhelminthes | 12556.0 | 2 | Cyclophyllidea | 11793.0 | 5 | Hymenolepididae | 11793.0 | 5 | Hymenolepis | 11793.0 | 5 | Hymenolepis microstoma | 11793.0 | 5 |
| scf7180000041242 | 24892 | 0.3979 | 0 | 102.9237 | Eukaryota | 5998.0 | 0 | Chordata | 5998.0 | 0 | Carnivora | 5998.0 | 0 | Felidae | 1934.0 | 5 | Felis | 1934.0 | 7 | Felis catus | 1934.0 | 7 |
| scf7180000045428 | 6192 | 0.4436 | 0 | 29.0602 | Eukaryota | 2700.0 | 0 | Chordata | 2700.0 | 0 | Carnivora | 2700.0 | 0 | Canidae | 2700.0 | 0 | Canis | 1890.0 | 1 | Canis lupus | 1890.0 | 1 |
| scf7180000050206 | 11085 | 0.4355 | 0 | 55.6822 | no-hit | 0.0 | 0 | no-hit | 0.0 | 0 | no-hit | 0.0 | 0 | no-hit | 0.0 | 0 | no-hit | 0.0 | 0 | no-hit | 0.0 | 0 |
| scf7180000041856 | 25950 | 0.3671 | 0 | 52.4798 | Eukaryota | 16481.0 | 0 | Chordata | 16481.0 | 0 | Carnivora | 16481.0 | 0 | Canidae | 16481.0 | 0 | Canis | 16481.0 | 0 | Canis lupus | 16481.0 | 0 |
| scf7180000049544 | 36690 | 0.4442 | 0 | 315.7546 | Eukaryota | 6246.0 | 0 | Chordata | 3332.0 | 2 | Polystomatidea | 2027.0 | 4 | Polystomatidae | 2027.0 | 4 | Protopolystoma | 2027.0 | 4 | Protopolystoma xenopodis | 2027.0 | 4 |
| scf7180000044794 | 245600 | 0.4236 | 0 | 60.2771 | Eukaryota | 61369.0 | 0 | Chordata | 61369.0 | 0 | Carnivora | 61369.0 | 0 | Canidae | 61369.0 | 0 | Canis | 52756.0 | 1 | Canis lupus | 52756.0 | 1 |
| scf7180000043084 | 593462 | 0.4639 | 0 | 59.0014 | Eukaryota | 46424.0 | 0 | Chordata | 46424.0 | 0 | Carnivora | 46424.0 | 0 | Canidae | 23798.0 | 1 | Vulpes | 23798.0 | 1 | Vulpes vulpes | 23798.0 | 1 |
| scf7180000043820 | 24909 | 0.4281 | 0 | 55.2319 | Eukaryota | 5350.0 | 0 | Chordata | 5350.0 | 0 | Primates | 2067.0 | 3 | Hominidae | 2067.0 | 3 | Homo | 2067.0 | 4 | Homo sapiens | 2067.0 | 4 |
| scf7180000046279 | 150405 | 0.4778 | 0 | 55.8637 | Eukaryota | 23848.0 | 0 | Chordata | 23848.0 | 0 | Carnivora | 23848.0 | 0 | Canidae | 21633.0 | 1 | Canis | 21633.0 | 1 | Canis lupus | 21633.0 | 1 |
| scf7180000042575 | 285158 | 0.5126 | 0 | 55.6239 | Eukaryota | 53959.0 | 0 | Chordata | 53959.0 | 0 | Carnivora | 53959.0 | 0 | Canidae | 53959.0 | 0 | Canis | 40443.0 | 1 | Canis lupus | 40443.0 | 1 |
| scf7180000048739 | 53885 | 0.3953 | 0 | 59.9851 | Eukaryota | 15019.0 | 0 | Chordata | 15019.0 | 0 | Carnivora | 15019.0 | 0 | Canidae | 15019.0 | 0 | Canis | 13988.0 | 1 | Canis lupus | 13988.0 | 1 |
| scf7180000041619 | 61330 | 0.379 | 0 | 55.9513 | Eukaryota | 15263.0 | 0 | Chordata | 15263.0 | 0 | Carnivora | 15263.0 | 0 | Canidae | 15263.0 | 0 | Canis | 15263.0 | 0 | Canis lupus | 15263.0 | 0 |
| scf7180000048358 | 24591 | 0.4464 | 0 | 355.6926 | Eukaryota | 817.0 | 0 | Platyhelminthes | 502.0 | 1 | Polystomatidea | 346.0 | 2 | Polystomatidae | 346.0 | 2 | Protopolystoma | 346.0 | 2 | Protopolystoma xenopodis | 346.0 | 2 |
| scf7180000054930 | 17578 | 0.4636 | 0 | 292.4455 | Eukaryota | 9402.0 | 0 | Chordata | 9402.0 | 0 | Carnivora | 9402.0 | 0 | Canidae | 9402.0 | 0 | Canis | 5656.0 | 1 | Canis lupus | 5656.0 | 1 |
| scf7180000043926 | 212236 | 0.4248 | 0 | 59.0718 | Eukaryota | 83873.0 | 0 | Chordata | 83873.0 | 0 | Carnivora | 83873.0 | 0 | Canidae | 83873.0 | 0 | Canis | 83873.0 | 0 | Canis lupus | 83873.0 | 0 |
| scf7180000045355 | 20831 | 0.4373 | 0 | 72.3716 | Eukaryota | 51263.0 | 0 | Chordata | 51263.0 | 0 | Carnivora | 51263.0 | 0 | Canidae | 51263.0 | 0 | Canis | 51263.0 | 0 | Canis lupus | 51263.0 | 0 |
| scf7180000042288 | 433921 | 0.4034 | 0 | 59.6631 | Eukaryota | 50347.0 | 0 | Chordata | 50347.0 | 0 | Carnivora | 50347.0 | 0 | Ursidae | 22240.0 | 2 | Ursus | 22240.0 | 3 | Canis lupus | 13802.0 | 4 |
| scf7180000052502 | 16711 | 0.3849 | 0 | 66.4754 | Eukaryota | 11227.0 | 0 | Chordata | 11227.0 | 0 | Carnivora | 9119.0 | 2 | Canidae | 9119.0 | 2 | Canis | 9119.0 | 3 | Canis lupus | 9119.0 | 3 |
| scf7180000045982 | 12302 | 0.4298 | 0 | 108.7133 | Eukaryota | 39169.0 | 0 | Chordata | 39169.0 | 0 | Carnivora | 39169.0 | 0 | Canidae | 39169.0 | 0 | Canis | 39169.0 | 0 | Canis lupus | 39169.0 | 0 |
| scf7180000051401 | 5340 | 0.4408 | 0 | 168.9711 | Eukaryota | 661.0 | 0 | Chordata | 466.0 | 1 | Salmoniformes | 466.0 | 1 | Salmonidae | 466.0 | 1 | Coregonus | 466.0 | 1 | Coregonus sp. 'balchen' | 466.0 | 1 |
| scf7180000047805 | 58903 | 0.4234 | 0 | 57.1523 | Eukaryota | 8321.0 | 0 | Chordata | 8321.0 | 0 | Carnivora | 6564.0 | 1 | Canidae | 6564.0 | 1 | Canis | 6564.0 | 1 | Canis lupus | 6564.0 | 1 |
| scf7180000042475 | 242598 | 0.4059 | 0 | 57.5638 | Eukaryota | 39794.0 | 0 | Chordata | 39794.0 | 0 | Carnivora | 39794.0 | 0 | Canidae | 39794.0 | 0 | Canis | 39794.0 | 0 | Canis lupus | 39794.0 | 0 |
| scf7180000052668 | 4954 | 0.4386 | 0 | 61.4301 | no-hit | 0.0 | 0 | no-hit | 0.0 | 0 | no-hit | 0.0 | 0 | no-hit | 0.0 | 0 | no-hit | 0.0 | 0 | no-hit | 0.0 | 0 |
| scf7180000040589 | 2218728 | 0.4059 | 0 | 59.6301 | Eukaryota | 75332.0 | 0 | Chordata | 75332.0 | 0 | Carnivora | 75332.0 | 0 | Canidae | 75332.0 | 0 | Canis | 75332.0 | 0 | Canis lupus | 75332.0 | 0 |
| scf7180000044464 | 324262 | 0.3676 | 0 | 58.9965 | Eukaryota | 30274.0 | 0 | Chordata | 30274.0 | 0 | Carnivora | 30274.0 | 0 | Canidae | 30274.0 | 0 | Canis | 30274.0 | 0 | Canis lupus | 30274.0 | 0 |
| scf7180000052287 | 13367 | 0.4465 | 0 | 75.8442 | Eukaryota | 1962.0 | 1 | Chordata | 1017.0 | 3 | Kurtiformes | 1017.0 | 3 | Apogonidae | 1017.0 | 3 | Sphaeramia | 1017.0 | 3 | Sphaeramia orbicularis | 1017.0 | 3 |
| scf7180000054530 | 14239 | 0.3407 | 0 | 120.5694 | Eukaryota | 4604.0 | 0 | Chordata | 4604.0 | 0 | Carnivora | 3088.0 | 1 | Felidae | 2360.0 | 2 | Acinonyx | 2360.0 | 2 | Acinonyx jubatus | 2360.0 | 2 |
| scf7180000051423 | 7426 | 0.5048 | 0 | 156.4952 | Eukaryota | 8757.0 | 0 | Chordata | 8757.0 | 0 | Carnivora | 8757.0 | 0 | Canidae | 8757.0 | 0 | Canis | 6548.0 | 1 | Canis lupus | 5418.0 | 2 |
| scf7180000044408 | 133647 | 0.3377 | 0 | 57.4089 | Eukaryota | 77996.0 | 0 | Chordata | 77996.0 | 0 | Carnivora | 77996.0 | 0 | Canidae | 77996.0 | 0 | Canis | 77996.0 | 0 | Canis lupus | 77996.0 | 0 |
| scf7180000041861 | 499159 | 0.4702 | 100 | 57.7934 | Eukaryota | 62153.0 | 0 | Chordata | 62153.0 | 0 | Carnivora | 62153.0 | 0 | Canidae | 62153.0 | 0 | Canis | 62153.0 | 0 | Canis lupus | 62153.0 | 0 |
| scf7180000045485 | 153313 | 0.5357 | 0 | 65.0643 | Eukaryota | 58429.0 | 0 | Chordata | 58429.0 | 0 | Carnivora | 58429.0 | 0 | Canidae | 58429.0 | 0 | Canis | 56013.0 | 1 | Canis lupus | 56013.0 | 1 |
| scf7180000040932 | 1510502 | 0.4174 | 0 | 60.2922 | Eukaryota | 146101.0 | 0 | Chordata | 146101.0 | 0 | Carnivora | 146101.0 | 0 | Canidae | 146101.0 | 0 | Canis | 146101.0 | 0 | Canis lupus | 146101.0 | 0 |
| scf7180000055104 | 20672 | 0.4055 | 0 | 158.4571 | Eukaryota | 82288.0 | 0 | Chordata | 82288.0 | 0 | Carnivora | 82288.0 | 0 | Canidae | 66720.0 | 1 | Canis | 48664.0 | 3 | Canis lupus | 48664.0 | 3 |
| scf7180000041575 | 909007 | 0.4144 | 0 | 59.5475 | Eukaryota | 98803.0 | 0 | Chordata | 98803.0 | 0 | Carnivora | 98803.0 | 0 | Canidae | 98803.0 | 0 | Canis | 87639.0 | 1 | Canis lupus | 87639.0 | 1 |
| scf7180000055225 | 42140 | 0.4441 | 0 | 128.556 | Eukaryota | 42647.0 | 0 | Chordata | 30520.0 | 2 | Gadiformes | 15137.0 | 7 | Gadidae | 15137.0 | 7 | Gadus | 15137.0 | 7 | Gadus morhua | 15137.0 | 7 |
| scf7180000055390 | 37793 | 0.4632 | 0 | 446.7593 | Eukaryota | 35025.0 | 0 | Chordata | 35025.0 | 0 | Carnivora | 35025.0 | 0 | Canidae | 35025.0 | 0 | Canis | 35025.0 | 0 | Canis lupus | 35025.0 | 0 |
| scf7180000046633 | 26082 | 0.4293 | 0 | 50.3286 | Eukaryota | 17008.0 | 0 | Chordata | 17008.0 | 0 | Carnivora | 17008.0 | 0 | Canidae | 15989.0 | 1 | Canis | 15989.0 | 1 | Canis lupus | 15989.0 | 1 |
| scf7180000046788 | 33311 | 0.4416 | 0 | 62.1153 | Eukaryota | 59666.0 | 0 | Chordata | 38251.0 | 1 | Gadiformes | 18931.0 | 6 | Gadidae | 18931.0 | 7 | Gadus | 18931.0 | 7 | Gadus morhua | 18931.0 | 7 |
| scf7180000040565 | 176535 | 0.4667 | 0 | 57.2749 | Eukaryota | 55698.0 | 0 | Chordata | 55698.0 | 0 | Carnivora | 55698.0 | 0 | Canidae | 55698.0 | 0 | Canis | 38910.0 | 1 | Canis lupus | 38910.0 | 1 |
| scf7180000055285 | 17391 | 0.5557 | 0 | 454.1059 | Eukaryota | 35613.0 | 0 | Chordata | 35613.0 | 0 | Artiodactyla | 25015.0 | 1 | Bovidae | 25015.0 | 2 | Ovis | 14182.0 | 3 | Ovis aries | 14182.0 | 3 |
| scf7180000053627 | 4431 | 0.4536 | 0 | 377.0841 | Eukaryota | 1880.0 | 0 | Platyhelminthes | 841.0 | 3 | Polystomatidea | 841.0 | 3 | Polystomatidae | 841.0 | 3 | Protopolystoma | 841.0 | 3 | Protopolystoma xenopodis | 841.0 | 3 |
| scf7180000044955 | 100556 | 0.4087 | 0 | 57.5848 | Eukaryota | 62769.0 | 0 | Chordata | 62769.0 | 0 | Carnivora | 62769.0 | 0 | Canidae | 62769.0 | 0 | Canis | 62769.0 | 0 | Canis lupus | 62769.0 | 0 |
| scf7180000049873 | 34103 | 0.5207 | 0 | 859.693 | Eukaryota | 21919.0 | 0 | Chordata | 21919.0 | 0 | Carnivora | 21919.0 | 0 | Canidae | 21919.0 | 0 | Canis | 21919.0 | 0 | Canis lupus | 21919.0 | 0 |
| scf7180000055435 | 25719 | 0.4366 | 0 | 175.1987 | Eukaryota | 24413.0 | 0 | Chordata | 20549.0 | 2 | Gadiformes | 6756.0 | 6 | Gadidae | 6756.0 | 6 | Gadus | 6756.0 | 7 | Gadus morhua | 6756.0 | 7 |
| scf7180000049734 | 7639 | 0.4972 | 0 | 292.1363 | Eukaryota | 24625.0 | 0 | Chordata | 24625.0 | 0 | Carnivora | 24625.0 | 0 | Canidae | 24625.0 | 0 | Canis | 24625.0 | 0 | Canis lupus | 24625.0 | 0 |
| scf7180000054642 | 71338 | 0.5341 | 0 | 103.9497 | Eukaryota | 41602.0 | 0 | Chordata | 41602.0 | 0 | Carnivora | 41602.0 | 0 | Canidae | 41602.0 | 0 | Canis | 41602.0 | 0 | Canis lupus | 41602.0 | 0 |
| scf7180000040984 | 1451431 | 0.4507 | 0 | 66.5656 | Eukaryota | 127228.0 | 0 | Chordata | 127228.0 | 0 | Carnivora | 127228.0 | 0 | Canidae | 127228.0 | 0 | Canis | 118323.0 | 1 | Canis lupus | 118323.0 | 1 |
| scf7180000045924 | 155398 | 0.3899 | 0 | 60.2335 | Eukaryota | 18209.0 | 1 | Chordata | 18209.0 | 1 | Carnivora | 10416.0 | 4 | Canidae | 10416.0 | 4 | Vulpes | 8552.0 | 7 | Vulpes vulpes | 8552.0 | 7 |
| scf7180000049807 | 5461 | 0.4333 | 0 | 196.462 | no-hit | 0.0 | 0 | no-hit | 0.0 | 0 | no-hit | 0.0 | 0 | no-hit | 0.0 | 0 | no-hit | 0.0 | 0 | no-hit | 0.0 | 0 |
| scf7180000047313 | 72086 | 0.5103 | 0 | 55.7285 | Eukaryota | 26358.0 | 0 | Chordata | 26358.0 | 0 | Carnivora | 26358.0 | 0 | Canidae | 26358.0 | 0 | Canis | 26358.0 | 0 | Canis lupus | 26358.0 | 0 |
| scf7180000051210 | 2421 | 0.4424 | 0 | 497.542 | no-hit | 0.0 | 0 | no-hit | 0.0 | 0 | no-hit | 0.0 | 0 | no-hit | 0.0 | 0 | no-hit | 0.0 | 0 | no-hit | 0.0 | 0 |
| scf7180000046992 | 102105 | 0.4432 | 0 | 58.7157 | Eukaryota | 75030.0 | 0 | Chordata | 39496.0 | 1 | Strigeidida | 35534.0 | 3 | Schistosomatidae | 35534.0 | 3 | Schistosoma | 35534.0 | 3 | Schistosoma mansoni | 35534.0 | 3 |
| scf7180000056402 | 451913 | 0.3682 | 0 | 58.4074 | Eukaryota | 14255.0 | 0 | Chordata | 14255.0 | 0 | Carnivora | 14255.0 | 0 | Canidae | 14255.0 | 0 | Canis | 14255.0 | 0 | Canis lupus | 14255.0 | 0 |
| scf7180000052803 | 2340 | 0.4474 | 0 | 79.2886 | Eukaryota | 311.0 | 0 | Chordata | 311.0 | 0 | Salmoniformes | 311.0 | 0 | Salmonidae | 311.0 | 0 | Coregonus | 311.0 | 0 | Coregonus sp. 'balchen' | 311.0 | 0 |
| scf7180000042240 | 740354 | 0.5147 | 0 | 57.5735 | Eukaryota | 78925.0 | 0 | Chordata | 78925.0 | 0 | Carnivora | 78925.0 | 0 | Canidae | 78925.0 | 0 | Canis | 70448.0 | 1 | Canis lupus | 70448.0 | 1 |
| scf7180000048863 | 34682 | 0.4857 | 0 | 56.4936 | Eukaryota | 51175.0 | 0 | Chordata | 51175.0 | 0 | Carnivora | 51175.0 | 0 | Canidae | 51175.0 | 0 | Canis | 51175.0 | 0 | Canis lupus | 51175.0 | 0 |
| scf7180000042098 | 486654 | 0.4568 | 0 | 58.6067 | Eukaryota | 35120.0 | 0 | Chordata | 35120.0 | 0 | Carnivora | 35120.0 | 0 | Canidae | 35120.0 | 0 | Canis | 35120.0 | 0 | Canis lupus | 35120.0 | 0 |
| scf7180000044763 | 322116 | 0.3642 | 0 | 59.5826 | Eukaryota | 79206.0 | 0 | Chordata | 79206.0 | 0 | Carnivora | 79206.0 | 0 | Canidae | 79206.0 | 0 | Canis | 79206.0 | 0 | Canis lupus | 79206.0 | 0 |
| scf7180000056092 | 33611 | 0.4514 | 0 | 276.4957 | Eukaryota | 18002.0 | 0 | Chordata | 18002.0 | 0 | Carnivora | 18002.0 | 0 | Canidae | 18002.0 | 0 | Canis | 18002.0 | 0 | Canis lupus | 18002.0 | 0 |
| scf7180000052907 | 3579 | 0.4476 | 0 | 623.7525 | no-hit | 0.0 | 0 | no-hit | 0.0 | 0 | no-hit | 0.0 | 0 | no-hit | 0.0 | 0 | no-hit | 0.0 | 0 | no-hit | 0.0 | 0 |
| scf7180000056167 | 2815536 | 0.3661 | 0 | 59.6933 | Eukaryota | 94624.0 | 0 | Chordata | 94624.0 | 0 | Carnivora | 94624.0 | 0 | Canidae | 94624.0 | 0 | Canis | 94624.0 | 0 | Canis lupus | 94624.0 | 0 |
| scf7180000042175 | 788421 | 0.3592 | 0 | 61.2805 | Eukaryota | 119207.0 | 0 | Chordata | 119207.0 | 0 | Carnivora | 119207.0 | 0 | Canidae | 119207.0 | 0 | Canis | 83405.0 | 1 | Canis lupus | 83405.0 | 1 |
| scf7180000044901 | 4554 | 0.4238 | 0 | 25.1704 | Eukaryota | 3022.0 | 0 | Chordata | 3022.0 | 0 | Carnivora | 3022.0 | 0 | Canidae | 3022.0 | 0 | Canis | 2722.0 | 1 | Canis lupus | 2722.0 | 1 |
| scf7180000041263 | 11389 | 0.4299 | 0 | 159.9145 | Eukaryota | 85542.0 | 0 | Chordata | 85542.0 | 0 | Carnivora | 85542.0 | 0 | Canidae | 85542.0 | 0 | Canis | 85542.0 | 0 | Canis lupus | 85542.0 | 0 |
| scf7180000046766 | 119302 | 0.4022 | 0 | 59.9538 | Eukaryota | 75759.0 | 0 | Chordata | 75759.0 | 0 | Carnivora | 75759.0 | 0 | Canidae | 75759.0 | 0 | Canis | 53416.0 | 1 | Canis lupus | 53416.0 | 1 |
| scf7180000045118 | 214162 | 0.4162 | 0 | 59.0734 | Eukaryota | 72266.0 | 0 | Chordata | 72266.0 | 0 | Carnivora | 72266.0 | 0 | Canidae | 72266.0 | 0 | Canis | 65106.0 | 1 | Canis lupus | 65106.0 | 1 |
| scf7180000046185 | 151641 | 0.4573 | 0 | 59.0206 | Eukaryota | 17746.0 | 0 | Chordata | 17746.0 | 0 | Primates | 17746.0 | 0 | Hominidae | 14298.0 | 1 | Homo | 14298.0 | 1 | Homo sapiens | 14298.0 | 1 |
| scf7180000045890 | 163922 | 0.3526 | 0 | 58.8023 | Eukaryota | 20839.0 | 0 | Chordata | 20839.0 | 0 | Carnivora | 17646.0 | 1 | Canidae | 17646.0 | 1 | Canis | 17646.0 | 1 | Canis lupus | 17646.0 | 1 |
| scf7180000054367 | 186115 | 0.4021 | 0 | 61.1666 | Eukaryota | 73934.0 | 0 | Chordata | 73934.0 | 0 | Carnivora | 73934.0 | 0 | Canidae | 73934.0 | 0 | Canis | 73934.0 | 0 | Canis lupus | 73934.0 | 0 |
| scf7180000041765 | 12334 | 0.4972 | 0 | 40.9505 | Eukaryota | 5355.0 | 0 | Chordata | 5355.0 | 0 | Carnivora | 5355.0 | 0 | Canidae | 3549.0 | 1 | Canis | 3549.0 | 2 | Canis lupus | 3549.0 | 2 |
| scf7180000052637 | 2829 | 0.4521 | 0 | 245.4088 | no-hit | 0.0 | 0 | no-hit | 0.0 | 0 | no-hit | 0.0 | 0 | no-hit | 0.0 | 0 | no-hit | 0.0 | 0 | no-hit | 0.0 | 0 |
| scf7180000045848 | 181074 | 0.3491 | 0 | 59.4204 | Eukaryota | 39561.0 | 0 | Chordata | 39561.0 | 0 | Carnivora | 39561.0 | 0 | Canidae | 39561.0 | 0 | Canis | 39561.0 | 0 | Canis lupus | 39561.0 | 0 |
| scf7180000044541 | 230583 | 0.4767 | 0 | 58.1934 | Eukaryota | 37621.0 | 0 | Chordata | 37621.0 | 0 | Carnivora | 35225.0 | 1 | Canidae | 35225.0 | 1 | Canis | 20920.0 | 2 | Canis lupus | 20920.0 | 2 |
| scf7180000043270 | 469784 | 0.4863 | 0 | 59.4398 | Eukaryota | 52191.0 | 0 | Chordata | 52191.0 | 0 | Carnivora | 52191.0 | 0 | Canidae | 52191.0 | 0 | Canis | 52191.0 | 0 | Canis lupus | 52191.0 | 0 |
| scf7180000042080 | 711847 | 0.4772 | 0 | 59.4812 | Eukaryota | 81481.0 | 0 | Chordata | 81481.0 | 0 | Carnivora | 81481.0 | 0 | Canidae | 81481.0 | 0 | Canis | 81481.0 | 0 | Canis lupus | 81481.0 | 0 |
| scf7180000040929 | 893939 | 0.3739 | 0 | 58.9044 | Eukaryota | 93834.0 | 0 | Chordata | 93834.0 | 0 | Carnivora | 93834.0 | 0 | Canidae | 93834.0 | 0 | Canis | 93834.0 | 0 | Canis lupus | 93834.0 | 0 |
| scf7180000045741 | 35130 | 0.4375 | 0 | 260.7958 | Eukaryota | 2577.0 | 0 | Platyhelminthes | 1376.0 | 1 | Polystomatidea | 1376.0 | 1 | Polystomatidae | 1376.0 | 1 | Protopolystoma | 1376.0 | 1 | Protopolystoma xenopodis | 1376.0 | 1 |
| scf7180000043816 | 50815 | 0.3706 | 0 | 54.0894 | Eukaryota | 71235.0 | 0 | Chordata | 71235.0 | 0 | Carnivora | 71235.0 | 0 | Canidae | 71235.0 | 0 | Canis | 71235.0 | 0 | Canis lupus | 71235.0 | 0 |
| scf7180000043931 | 265782 | 0.446 | 0 | 58.7713 | Eukaryota | 50098.0 | 0 | Chordata | 50098.0 | 0 | Carnivora | 50098.0 | 0 | Canidae | 50098.0 | 0 | Canis | 50098.0 | 0 | Canis lupus | 50098.0 | 0 |
| scf7180000042089 | 789426 | 0.411 | 0 | 59.5751 | Eukaryota | 71282.0 | 0 | Chordata | 71282.0 | 0 | Carnivora | 71282.0 | 0 | Canidae | 71282.0 | 0 | Canis | 61767.0 | 1 | Canis lupus | 61767.0 | 1 |
| scf7180000053080 | 3950 | 0.4461 | 0 | 622.6127 | Eukaryota | 3497.0 | 0 | Platyhelminthes | 2025.0 | 2 | Polystomatidea | 2025.0 | 2 | Polystomatidae | 2025.0 | 2 | Protopolystoma | 2025.0 | 2 | Protopolystoma xenopodis | 2025.0 | 2 |
| scf7180000045017 | 287287 | 0.4476 | 0 | 59.9022 | Eukaryota | 32001.0 | 0 | Chordata | 32001.0 | 0 | Carnivora | 32001.0 | 0 | Felidae | 12570.0 | 3 | Canis | 9976.0 | 5 | Canis lupus | 9976.0 | 5 |
| scf7180000041328 | 1317305 | 0.448 | 0 | 59.4905 | Eukaryota | 93104.0 | 0 | Chordata | 93104.0 | 0 | Carnivora | 93104.0 | 0 | Canidae | 93104.0 | 0 | Canis | 79892.0 | 1 | Canis lupus | 79892.0 | 1 |
| scf7180000053016 | 5156 | 0.4474 | 0 | 112.8738 | Eukaryota | 610.0 | 0 | Eukaryota-undef | 316.0 | 1 | Craspedida | 316.0 | 1 | Salpingoecidae | 316.0 | 1 | Monosiga | 316.0 | 1 | Monosiga brevicollis | 316.0 | 1 |
| scf7180000049961 | 25674 | 0.4378 | 0 | 161.4428 | Eukaryota | 765.0 | 0 | Platyhelminthes | 558.0 | 1 | Polystomatidea | 558.0 | 1 | Polystomatidae | 558.0 | 1 | Protopolystoma | 558.0 | 1 | Protopolystoma xenopodis | 558.0 | 1 |
| scf7180000043217 | 469049 | 0.4115 | 0 | 60.5637 | Eukaryota | 84920.0 | 0 | Chordata | 84920.0 | 0 | Carnivora | 84920.0 | 0 | Canidae | 84920.0 | 0 | Vulpes | 84920.0 | 0 | Vulpes vulpes | 84920.0 | 0 |
| scf7180000041093 | 1205663 | 0.5447 | 0 | 58.6197 | Eukaryota | 72808.0 | 0 | Chordata | 72808.0 | 0 | Carnivora | 72808.0 | 0 | Canidae | 72808.0 | 0 | Canis | 72808.0 | 0 | Canis lupus | 72808.0 | 0 |
| scf7180000041451 | 29381 | 0.3854 | 0 | 30.0241 | Eukaryota | 32601.0 | 0 | Chordata | 32601.0 | 0 | Carnivora | 32601.0 | 0 | Canidae | 32601.0 | 0 | Canis | 32601.0 | 0 | Canis lupus | 32601.0 | 0 |
| scf7180000046428 | 25757 | 0.4455 | 0 | 360.8672 | Eukaryota | 462.0 | 0 | Platyhelminthes | 462.0 | 0 | Polystomatidea | 462.0 | 0 | Polystomatidae | 462.0 | 0 | Protopolystoma | 462.0 | 0 | Protopolystoma xenopodis | 462.0 | 0 |
| scf7180000056620 | 28181 | 0.4559 | 0 | 93.183 | no-hit | 0.0 | 0 | no-hit | 0.0 | 0 | no-hit | 0.0 | 0 | no-hit | 0.0 | 0 | no-hit | 0.0 | 0 | no-hit | 0.0 | 0 |
| scf7180000052494 | 14003 | 0.4345 | 0 | 58.8338 | Eukaryota | 2434.0 | 0 | Chordata | 2434.0 | 0 | Carnivora | 1788.0 | 2 | Canidae | 1571.0 | 3 | Canis | 1571.0 | 3 | Canis lupus | 1571.0 | 3 |
| scf7180000047045 | 35047 | 0.4342 | 0 | 96.2592 | Eukaryota | 8750.0 | 0 | Chordata | 7784.0 | 1 | Kurtiformes | 3042.0 | 4 | Apogonidae | 3042.0 | 4 | Sphaeramia | 3042.0 | 5 | Sphaeramia orbicularis | 3042.0 | 5 |
| scf7180000042584 | 188188 | 0.3915 | 0 | 60.8924 | Eukaryota | 118998.0 | 0 | Chordata | 118998.0 | 0 | Carnivora | 118998.0 | 0 | Canidae | 118998.0 | 0 | Vulpes | 100520.0 | 1 | Vulpes vulpes | 100520.0 | 1 |
| scf7180000043410 | 119579 | 0.3934 | 0 | 71.8402 | Eukaryota | 46604.0 | 0 | Chordata | 46604.0 | 0 | Carnivora | 46604.0 | 0 | Canidae | 46604.0 | 0 | Canis | 46604.0 | 0 | Canis lupus | 46604.0 | 0 |
| scf7180000049276 | 49939 | 0.5056 | 0 | 422.2505 | Eukaryota | 9740.0 | 0 | Chordata | 9740.0 | 0 | Carnivora | 9740.0 | 0 | Canidae | 9740.0 | 0 | Canis | 9041.0 | 1 | Canis lupus | 7745.0 | 2 |
| scf7180000052894 | 4416 | 0.4484 | 0 | 176.5428 | Eukaryota | 1942.0 | 0 | Chordata | 1942.0 | 0 | Salmoniformes | 848.0 | 2 | Salmonidae | 848.0 | 2 | Takifugu | 798.0 | 3 | Takifugu rubripes | 798.0 | 3 |
| scf7180000053739 | 2797 | 0.5109 | 0 | 572.1633 | Eukaryota | 5426.0 | 0 | Chordata | 5426.0 | 0 | Carnivora | 5426.0 | 0 | Canidae | 5426.0 | 0 | Canis | 3261.0 | 1 | Canis lupus | 2460.0 | 2 |
| scf7180000053229 | 4175 | 0.4505 | 0 | 272.2472 | no-hit | 0.0 | 0 | no-hit | 0.0 | 0 | no-hit | 0.0 | 0 | no-hit | 0.0 | 0 | no-hit | 0.0 | 0 | no-hit | 0.0 | 0 |
| scf7180000050974 | 9729 | 0.4414 | 0 | 187.7747 | Eukaryota | 600.0 | 0 | Chordata | 600.0 | 0 | Kurtiformes | 387.0 | 1 | Apogonidae | 387.0 | 1 | Sphaeramia | 387.0 | 1 | Sphaeramia orbicularis | 387.0 | 1 |
| scf7180000041494 | 1123630 | 0.4559 | 0 | 58.7456 | Eukaryota | 93204.0 | 0 | Chordata | 93204.0 | 0 | Carnivora | 93204.0 | 0 | Canidae | 93204.0 | 0 | Canis | 93204.0 | 0 | Canis lupus | 93204.0 | 0 |
| scf7180000042834 | 3275 | 0.4495 | 0 | 26.7369 | Eukaryota | 3491.0 | 0 | Chordata | 3491.0 | 0 | Carnivora | 3491.0 | 0 | Canidae | 3491.0 | 0 | Canis | 3491.0 | 0 | Canis lupus | 3491.0 | 0 |
| scf7180000056277 | 1201109 | 0.4481 | 0 | 58.9674 | Eukaryota | 87986.0 | 0 | Chordata | 87986.0 | 0 | Carnivora | 87986.0 | 0 | Canidae | 87986.0 | 0 | Canis | 87986.0 | 0 | Canis lupus | 87986.0 | 0 |
| scf7180000053319 | 9241 | 0.4349 | 0 | 49.7148 | Eukaryota | 35212.0 | 0 | Chordata | 35212.0 | 0 | Carnivora | 35212.0 | 0 | Canidae | 35212.0 | 0 | Canis | 35212.0 | 0 | Canis lupus | 35212.0 | 0 |
| scf7180000053155 | 7019 | 0.4508 | 0 | 1670.3037 | Eukaryota | 16576.0 | 0 | Chordata | 13940.0 | 2 | Gadiformes | 9157.0 | 7 | Gadidae | 9157.0 | 7 | Gadus | 9157.0 | 7 | Gadus morhua | 9157.0 | 7 |
| scf7180000041015 | 1477000 | 0.4266 | 0 | 60.269 | Eukaryota | 76212.0 | 0 | Chordata | 76212.0 | 0 | Carnivora | 76212.0 | 0 | Canidae | 33791.0 | 3 | Canis | 25567.0 | 6 | Canis lupus | 25567.0 | 7 |
| scf7180000041143 | 1219693 | 0.3953 | 0 | 60.5639 | Eukaryota | 95551.0 | 0 | Chordata | 95551.0 | 0 | Carnivora | 95551.0 | 0 | Canidae | 95551.0 | 0 | Canis | 85023.0 | 1 | Canis lupus | 85023.0 | 1 |
| scf7180000056525 | 348657 | 0.3731 | 0 | 59.5269 | Eukaryota | 33097.0 | 0 | Chordata | 33097.0 | 0 | Carnivora | 33097.0 | 0 | Canidae | 28893.0 | 1 | Canis | 28893.0 | 1 | Canis lupus | 28893.0 | 1 |
| scf7180000049289 | 3641 | 0.4864 | 0 | 30.5887 | Eukaryota | 33158.0 | 0 | Chordata | 33158.0 | 0 | Carnivora | 33158.0 | 0 | Canidae | 33158.0 | 0 | Canis | 33158.0 | 0 | Canis lupus | 33158.0 | 0 |
| scf7180000042045 | 197652 | 0.3967 | 0 | 58.922 | Eukaryota | 13026.0 | 0 | Chordata | 13026.0 | 0 | Primates | 11908.0 | 1 | Hominidae | 11908.0 | 1 | Homo | 8195.0 | 4 | Homo sapiens | 8195.0 | 4 |
| scf7180000049339 | 11382 | 0.452 | 0 | 1952.9953 | no-hit | 0.0 | 0 | no-hit | 0.0 | 0 | no-hit | 0.0 | 0 | no-hit | 0.0 | 0 | no-hit | 0.0 | 0 | no-hit | 0.0 | 0 |
| scf7180000045439 | 225503 | 0.333 | 0 | 59.5978 | Eukaryota | 69900.0 | 0 | Chordata | 69900.0 | 0 | Carnivora | 69900.0 | 0 | Canidae | 69900.0 | 0 | Canis | 69900.0 | 0 | Canis lupus | 69900.0 | 0 |
| scf7180000044712 | 261574 | 0.4335 | 0 | 61.2031 | Eukaryota | 62040.0 | 0 | Chordata | 62040.0 | 0 | Carnivora | 62040.0 | 0 | Canidae | 62040.0 | 0 | Canis | 62040.0 | 0 | Canis lupus | 62040.0 | 0 |
| scf7180000054842 | 3263 | 0.4364 | 0 | 45.4963 | no-hit | 0.0 | 0 | no-hit | 0.0 | 0 | no-hit | 0.0 | 0 | no-hit | 0.0 | 0 | no-hit | 0.0 | 0 | no-hit | 0.0 | 0 |
| scf7180000055661 | 6932 | 0.4365 | 0 | 398.9421 | Eukaryota | 174.0 | 0 | Chordata | 174.0 | 0 | Kurtiformes | 174.0 | 0 | Apogonidae | 174.0 | 0 | Sphaeramia | 174.0 | 0 | Sphaeramia orbicularis | 174.0 | 0 |
| scf7180000048348 | 4838 | 0.4477 | 0 | 147.8585 | Eukaryota | 3061.0 | 0 | Chordata | 2726.0 | 1 | Carnivora | 2726.0 | 1 | Canidae | 2726.0 | 1 | Canis | 2726.0 | 1 | Canis lupus | 2726.0 | 1 |
| scf7180000050275 | 17993 | 0.4382 | 0 | 44.2326 | Eukaryota | 635.0 | 0 | Chordata | 396.0 | 1 | Polystomatidea | 239.0 | 2 | Polystomatidae | 239.0 | 2 | Protopolystoma | 239.0 | 2 | Protopolystoma xenopodis | 239.0 | 2 |
| scf7180000045363 | 5277 | 0.4395 | 0 | 321.8577 | Eukaryota | 272.0 | 0 | Platyhelminthes | 272.0 | 0 | Polystomatidea | 272.0 | 0 | Polystomatidae | 272.0 | 0 | Protopolystoma | 272.0 | 0 | Protopolystoma xenopodis | 272.0 | 0 |
| scf7180000050995 | 5067 | 0.4541 | 0 | 470.6469 | no-hit | 0.0 | 0 | no-hit | 0.0 | 0 | no-hit | 0.0 | 0 | no-hit | 0.0 | 0 | no-hit | 0.0 | 0 | no-hit | 0.0 | 0 |
| scf7180000054550 | 17676 | 0.4423 | 0 | 112.3622 | Eukaryota | 774.0 | 0 | Platyhelminthes | 774.0 | 0 | Polystomatidea | 774.0 | 0 | Polystomatidae | 774.0 | 0 | Protopolystoma | 774.0 | 0 | Protopolystoma xenopodis | 774.0 | 0 |
| scf7180000051728 | 3279 | 0.4288 | 0 | 217.0467 | Eukaryota | 759.0 | 0 | Echinodermata | 416.0 | 2 | Forcipulatida | 416.0 | 2 | Asteriidae | 416.0 | 2 | Asterias | 416.0 | 2 | Asterias rubens | 416.0 | 2 |
| scf7180000053302 | 7683 | 0.5059 | 0 | 66.0621 | Eukaryota | 8385.0 | 0 | Chordata | 8385.0 | 0 | Carnivora | 8385.0 | 0 | Canidae | 8385.0 | 0 | Canis | 6253.0 | 1 | Canis lupus | 5066.0 | 2 |
| scf7180000052183 | 12799 | 0.4421 | 0 | 395.3793 | Eukaryota | 1801.0 | 0 | Platyhelminthes | 1235.0 | 1 | Polystomatidea | 1235.0 | 1 | Polystomatidae | 1235.0 | 1 | Protopolystoma | 1235.0 | 1 | Protopolystoma xenopodis | 1235.0 | 1 |
| scf7180000043409 | 473633 | 0.3546 | 0 | 60.7995 | Eukaryota | 78193.0 | 0 | Chordata | 78193.0 | 0 | Carnivora | 78193.0 | 0 | Canidae | 78193.0 | 0 | Canis | 78193.0 | 0 | Canis lupus | 78193.0 | 0 |
| scf7180000048800 | 27639 | 0.4424 | 0 | 101.3936 | Eukaryota | 33422.0 | 1 | Chordata | 29183.0 | 2 | Gadiformes | 15579.0 | 6 | Gadidae | 15579.0 | 6 | Gadus | 15579.0 | 6 | Gadus morhua | 15579.0 | 6 |
| scf7180000043215 | 60834 | 0.3969 | 0 | 58.0159 | Eukaryota | 15856.0 | 0 | Chordata | 15856.0 | 0 | Carnivora | 15856.0 | 0 | Canidae | 15856.0 | 0 | Canis | 15856.0 | 0 | Canis lupus | 15856.0 | 0 |
| scf7180000045695 | 8654 | 0.3966 | 0 | 58.2739 | Eukaryota | 22359.0 | 0 | Chordata | 22359.0 | 0 | Carnivora | 22359.0 | 0 | Canidae | 22359.0 | 0 | Canis | 22359.0 | 0 | Canis lupus | 22359.0 | 0 |
| scf7180000041087 | 1302006 | 0.3567 | 0 | 60.5263 | Eukaryota | 92580.0 | 0 | Chordata | 92580.0 | 0 | Carnivora | 92580.0 | 0 | Canidae | 92580.0 | 0 | Canis | 92580.0 | 0 | Canis lupus | 92580.0 | 0 |
| scf7180000042239 | 536334 | 0.384 | 0 | 59.7944 | Eukaryota | 49224.0 | 0 | Chordata | 49224.0 | 0 | Carnivora | 49224.0 | 0 | Canidae | 49224.0 | 0 | Canis | 49224.0 | 0 | Canis lupus | 49224.0 | 0 |
| scf7180000046658 | 19798 | 0.4358 | 0 | 175.7175 | no-hit | 0.0 | 0 | no-hit | 0.0 | 0 | no-hit | 0.0 | 0 | no-hit | 0.0 | 0 | no-hit | 0.0 | 0 | no-hit | 0.0 | 0 |
| scf7180000044981 | 241263 | 0.3401 | 0 | 59.6021 | Eukaryota | 12098.0 | 0 | Chordata | 12098.0 | 0 | Primates | 10660.0 | 2 | Hominidae | 7647.0 | 3 | Homo | 7647.0 | 3 | Homo sapiens | 7647.0 | 3 |
| scf7180000051855 | 6284 | 0.4453 | 0 | 130.5438 | Eukaryota | 1026.0 | 0 | Chordata | 809.0 | 1 | Syngnathiformes | 809.0 | 1 | Syngnathidae | 809.0 | 1 | Syngnathus | 809.0 | 1 | Syngnathus acus | 809.0 | 1 |
| scf7180000045092 | 16148 | 0.4473 | 0 | 287.5886 | Eukaryota | 1327.0 | 0 | Chordata | 1327.0 | 0 | Primates | 544.0 | 2 | Hominidae | 544.0 | 2 | Homo | 544.0 | 3 | Homo sapiens | 544.0 | 3 |
| scf7180000042800 | 55972 | 0.5189 | 0 | 53.0041 | Eukaryota | 21426.0 | 0 | Chordata | 21426.0 | 0 | Carnivora | 21426.0 | 0 | Canidae | 21426.0 | 0 | Canis | 21426.0 | 0 | Canis lupus | 21426.0 | 0 |
| scf7180000053676 | 3195 | 0.4532 | 0 | 229.9203 | Eukaryota | 1635.0 | 0 | Platyhelminthes | 777.0 | 2 | Polystomatidea | 777.0 | 2 | Polystomatidae | 777.0 | 2 | Protopolystoma | 777.0 | 2 | Protopolystoma xenopodis | 777.0 | 2 |
| scf7180000040701 | 1771949 | 0.4149 | 0 | 60.2464 | Eukaryota | 107640.0 | 0 | Chordata | 107640.0 | 0 | Carnivora | 107640.0 | 0 | Canidae | 107640.0 | 0 | Canis | 65408.0 | 1 | Canis lupus | 65408.0 | 1 |
| scf7180000055178 | 30746 | 0.3802 | 0 | 443.046 | Eukaryota | 57071.0 | 0 | Chordata | 57071.0 | 0 | Carnivora | 57071.0 | 0 | Canidae | 57071.0 | 0 | Canis | 57071.0 | 0 | Canis lupus | 57071.0 | 0 |
| scf7180000042341 | 105623 | 0.376 | 0 | 59.2594 | Eukaryota | 93101.0 | 0 | Chordata | 93101.0 | 0 | Carnivora | 93101.0 | 0 | Canidae | 93101.0 | 0 | Canis | 93101.0 | 0 | Canis lupus | 93101.0 | 0 |
| scf7180000046494 | 21137 | 0.3407 | 0 | 50.9615 | Eukaryota | 9027.0 | 0 | Chordata | 9027.0 | 0 | Carnivora | 8301.0 | 1 | Felidae | 6701.0 | 2 | Felis | 6701.0 | 2 | Felis catus | 6701.0 | 2 |
| scf7180000054785 | 23735 | 0.4755 | 0 | 235.8901 | Eukaryota | 17594.0 | 0 | Chordata | 17594.0 | 0 | Carnivora | 17594.0 | 0 | Canidae | 17594.0 | 0 | Canis | 17594.0 | 0 | Canis lupus | 17594.0 | 0 |
| scf7180000054243 | 30960 | 0.4251 | 0 | 910.5024 | Eukaryota | 80339.0 | 0 | Chordata | 80339.0 | 0 | Carnivora | 80339.0 | 0 | Canidae | 80339.0 | 0 | Canis | 80339.0 | 0 | Canis lupus | 80339.0 | 0 |

| | | | | | | | | | | | | | | | | | | | | | | |
|---|---|---|---|---|---|---|---|---|---|---|---|---|---|---|---|---|---|---|---|---|---|---|
| scf7180000048611 | 70894 | 0.6343 | 0 | 54.5531 | Eukaryota | 12154.0 | 0 | Chordata | 12154.0 | 0 | Carnivora | 12154.0 | 0 | Canidae | 12154.0 | 0 | Canis | 10525.0 | 1 | Canis lupus | 10525.0 | 1 |
| scf7180000042808 | 182374 | 0.3782 | 0 | 60.724 | Eukaryota | 81626.0 | 0 | Chordata | 81626.0 | 0 | Carnivora | 81626.0 | 0 | Canidae | 81626.0 | 0 | Canis | 81626.0 | 0 | Canis lupus | 81626.0 | 0 |
| scf7180000056001 | 2168 | 0.4405 | 0 | 910.643 | no-hit | 0.0 | 0 | no-hit | 0.0 | 0 | no-hit | 0.0 | 0 | no-hit | 0.0 | 0 | no-hit | 0.0 | 0 | no-hit | 0.0 | 0 |
| scf7180000050058 | 24083 | 0.4432 | 0 | 346.8583 | Eukaryota | 3325.0 | 0 | Platyhelminthes | 2139.0 | 1 | Polystomatidea | 2139.0 | 3 | Polystomatidae | 2139.0 | 3 | Protopolystoma | 2139.0 | 3 | Protopolystoma xenopodis | 2139.0 | 3 |
| scf7180000052202 | 10549 | 0.4115 | 0 | 65.332 | Eukaryota | 4577.0 | 0 | Chordata | 4577.0 | 0 | Carnivora | 3313.0 | 1 | Canidae | 1996.0 | 2 | Canis | 1750.0 | 3 | Canis lupus | 1750.0 | 4 |
| scf7180000048681 | 34891 | 0.4756 | 0 | 59.1533 | Eukaryota | 45285.0 | 0 | Chordata | 45285.0 | 0 | Carnivora | 45285.0 | 0 | Canidae | 31858.0 | 2 | Canis | 16576.0 | 3 | Canis lupus | 16576.0 | 3 |
| scf7180000042945 | 378731 | 0.3737 | 0 | 60.0308 | Eukaryota | 91952.0 | 0 | Chordata | 91952.0 | 0 | Carnivora | 91952.0 | 0 | Canidae | 91952.0 | 0 | Canis | 91952.0 | 0 | Canis lupus | 91952.0 | 0 |
| scf7180000050841 | 4871 | 0.4432 | 0 | 188.4991 | Eukaryota | 26746.0 | 0 | Chordata | 20838.0 | 2 | Gadiformes | 13509.0 | 5 | Gadidae | 13509.0 | 5 | Gadus | 13509.0 | 5 | Gadus morhua | 13509.0 | 5 |
| scf7180000042511 | 423120 | 0.473 | 0 | 57.6791 | Eukaryota | 50264.0 | 0 | Chordata | 50264.0 | 0 | Carnivora | 50264.0 | 0 | Canidae | 41026.0 | 2 | Canis | 34712.0 | 3 | Canis lupus | 34712.0 | 3 |
| scf7180000055842 | 18023 | 0.4392 | 0 | 148.5753 | Eukaryota | 23732.0 | 0 | Chordata | 21087.0 | 2 | Gadiformes | 9584.0 | 6 | Gadidae | 9584.0 | 6 | Gadus | 9584.0 | 6 | Gadus morhua | 9584.0 | 6 |
| scf7180000048167 | 6871 | 0.4459 | 0 | 882.1318 | Eukaryota | 2072.0 | 0 | Platyhelminthes | 1018.0 | 2 | Polystomatidea | 1018.0 | 3 | Polystomatidae | 1018.0 | 3 | Protopolystoma | 1018.0 | 3 | Protopolystoma xenopodis | 1018.0 | 3 |
| scf7180000056260 | 898828 | 0.5829 | 0 | 54.8592 | Eukaryota | 36815.0 | 0 | Chordata | 36815.0 | 0 | Carnivora | 36815.0 | 0 | Canidae | 36815.0 | 0 | Canis | 36815.0 | 0 | Canis lupus | 36815.0 | 0 |
| scf7180000056532 | 195616 | 0.3796 | 0 | 68.5424 | Eukaryota | 56968.0 | 0 | Chordata | 56968.0 | 0 | Carnivora | 56968.0 | 0 | Canidae | 56968.0 | 0 | Canis | 56968.0 | 0 | Canis lupus | 56968.0 | 0 |
| scf7180000047539 | 49539 | 0.4474 | 0 | 61.2449 | Eukaryota | 7990.0 | 0 | Chordata | 7990.0 | 0 | Carnivora | 7990.0 | 0 | Canidae | 7990.0 | 0 | Canis | 7990.0 | 0 | Canis lupus | 7990.0 | 0 |
| scf7180000044892 | 270510 | 0.396 | 0 | 60.2534 | Eukaryota | 58672.0 | 0 | Chordata | 58672.0 | 0 | Carnivora | 58672.0 | 0 | Canidae | 58672.0 | 0 | Canis | 47060.0 | 1 | Canis lupus | 47060.0 | 1 |
| scf7180000044868 | 9353 | 0.3911 | 0 | 37.323 | Eukaryota | 5607.0 | 0 | Chordata | 5607.0 | 0 | Carnivora | 5607.0 | 0 | Canidae | 4399.0 | 1 | Canis | 4399.0 | 1 | Canis lupus | 4399.0 | 1 |
| scf7180000045191 | 32147 | 0.3864 | 0 | 44.824 | Eukaryota | 85787.0 | 0 | Chordata | 85787.0 | 0 | Carnivora | 85787.0 | 0 | Canidae | 85787.0 | 0 | Canis | 85787.0 | 0 | Canis lupus | 85787.0 | 0 |
| scf7180000041110 | 105706 | 0.3556 | 0 | 57.113 | Eukaryota | 5951.0 | 0 | Chordata | 5951.0 | 0 | Carnivora | 5223.0 | 1 | Canidae | 5223.0 | 1 | Canis | 5223.0 | 1 | Canis lupus | 5223.0 | 1 |
| scf7180000053846 | 4334 | 0.4522 | 0 | 309.962 | Eukaryota | 695.0 | 0 | Platyhelminthes | 441.0 | 1 | Polystomatidea | 441.0 | 1 | Polystomatidae | 441.0 | 1 | Protopolystoma | 441.0 | 1 | Protopolystoma xenopodis | 441.0 | 1 |
| scf7180000042910 | 144518 | 0.3378 | 0 | 58.0862 | Eukaryota | 75191.0 | 0 | Chordata | 75191.0 | 0 | Carnivora | 75191.0 | 0 | Canidae | 75191.0 | 0 | Canis | 75191.0 | 0 | Canis lupus | 75191.0 | 0 |
| scf7180000043750 | 59761 | 0.3703 | 0 | 58.2128 | Eukaryota | 59455.0 | 0 | Chordata | 59455.0 | 0 | Carnivora | 59455.0 | 0 | Canidae | 59455.0 | 0 | Canis | 59455.0 | 0 | Canis lupus | 59455.0 | 0 |
| scf7180000047393 | 100985 | 0.4183 | 0 | 60.6346 | Eukaryota | 7133.0 | 0 | Chordata | 7133.0 | 0 | Primates | 4725.0 | 2 | Hominidae | 4725.0 | 2 | Homo | 4725.0 | 3 | Homo sapiens | 4725.0 | 3 |
| scf7180000040779 | 1652456 | 0.3991 | 0 | 59.8187 | Eukaryota | 99942.0 | 0 | Chordata | 99942.0 | 0 | Carnivora | 99942.0 | 0 | Canidae | 99942.0 | 0 | Canis | 90924.0 | 1 | Canis lupus | 90924.0 | 1 |
| scf7180000050926 | 31204 | 0.3524 | 0 | 63.37 | Eukaryota | 72688.0 | 0 | Chordata | 72688.0 | 0 | Carnivora | 72688.0 | 0 | Canidae | 72688.0 | 0 | Canis | 72688.0 | 0 | Canis lupus | 72688.0 | 0 |
| scf7180000045521 | 11334 | 0.3972 | 0 | 40.3161 | Eukaryota | 90397.0 | 0 | Chordata | 90397.0 | 0 | Carnivora | 90397.0 | 0 | Canidae | 90397.0 | 0 | Canis | 90397.0 | 0 | Canis lupus | 90397.0 | 0 |
| scf7180000048464 | 17658 | 0.4391 | 0 | 278.0787 | Eukaryota | 3114.0 | 0 | Platyhelminthes | 2340.0 | 1 | Polystomatidea | 2340.0 | 1 | Polystomatidae | 2340.0 | 1 | Protopolystoma | 2340.0 | 1 | Protopolystoma xenopodis | 2340.0 | 1 |
| scf7180000052624 | 3923 | 0.4507 | 0 | 307.5123 | no-hit | 0.0 | 0 | no-hit | 0.0 | 0 | no-hit | 0.0 | 0 | no-hit | 0.0 | 0 | no-hit | 0.0 | 0 | no-hit | 0.0 | 0 |
| scf7180000051990 | 16371 | 0.5044 | 0 | 446.214 | Eukaryota | 9183.0 | 0 | Chordata | 9183.0 | 0 | Carnivora | 9183.0 | 0 | Canidae | 9183.0 | 0 | Canis | 7400.0 | 1 | Canis lupus | 6093.0 | 2 |
| scf7180000043698 | 2805 | 0.3932 | 0 | 17.7189 | Eukaryota | 16667.0 | 0 | Chordata | 16667.0 | 0 | Carnivora | 16667.0 | 0 | Canidae | 16667.0 | 0 | Canis | 16667.0 | 0 | Canis lupus | 16667.0 | 0 |
| scf7180000046906 | 7008 | 0.4362 | 0 | 140.5058 | no-hit | 0.0 | 0 | no-hit | 0.0 | 0 | no-hit | 0.0 | 0 | no-hit | 0.0 | 0 | no-hit | 0.0 | 0 | no-hit | 0.0 | 0 |
| scf7180000042963 | 520921 | 0.3965 | 0 | 60.8933 | Eukaryota | 61384.0 | 0 | Chordata | 61384.0 | 0 | Carnivora | 61384.0 | 0 | Canidae | 61384.0 | 0 | Canis | 61384.0 | 0 | Canis lupus | 61384.0 | 0 |
| scf7180000052280 | 8018 | 0.4509 | 0 | 144.113 | Eukaryota | 4112.0 | 0 | Chordata | 4112.0 | 0 | Carnivora | 2009.0 | 2 | Canidae | 2009.0 | 2 | Canis | 2009.0 | 2 | Canis lupus | 2009.0 | 2 |
| scf7180000044117 | 6995 | 0.4179 | 0 | 139.3905 | Eukaryota | 1738.0 | 0 | Chordata | 1738.0 | 0 | Primates | 1738.0 | 0 | Hominidae | 1738.0 | 0 | Homo | 1738.0 | 0 | Homo sapiens | 1738.0 | 0 |
| scf7180000047721 | 68154 | 0.4903 | 0 | 61.5495 | Eukaryota | 41037.0 | 0 | Chordata | 41037.0 | 0 | Carnivora | 41037.0 | 0 | Canidae | 18923.0 | 4 | Canis | 18923.0 | 6 | Canis lupus | 18923.0 | 6 |
| scf7180000049061 | 26480 | 0.4414 | 0 | 176.3842 | Eukaryota | 8839.0 | 0 | Chordata | 7664.0 | 1 | Tetraodontiformes | 5397.0 | 5 | Tetraodontidae | 5397.0 | 5 | Takifugu | 5397.0 | 5 | Takifugu rubripes | 5397.0 | 5 |
| scf7180000047349 | 82560 | 0.4242 | 0 | 58.3993 | Eukaryota | 6615.0 | 0 | Chordata | 6615.0 | 0 | Carnivora | 4889.0 | 2 | Canidae | 4889.0 | 2 | Canis | 4312.0 | 3 | Canis lupus | 4312.0 | 3 |
| scf7180000050142 | 9448 | 0.4388 | 0 | 510.2454 | Eukaryota | 3238.0 | 0 | Chordata | 3238.0 | 0 | Carnivora | 3238.0 | 0 | Canidae | 3238.0 | 0 | Canis | 3238.0 | 0 | Canis lupus | 3238.0 | 0 |
| scf7180000054921 | 9970 | 0.4103 | 0 | 414.9655 | Eukaryota | 6997.0 | 0 | Chordata | 6997.0 | 0 | Carnivora | 6997.0 | 0 | Canidae | 6997.0 | 0 | Canis | 6997.0 | 0 | Canis lupus | 6997.0 | 0 |
| scf7180000045379 | 173872 | 0.3755 | 0 | 61.1933 | Eukaryota | 79887.0 | 0 | Chordata | 79887.0 | 0 | Carnivora | 79887.0 | 0 | Canidae | 79887.0 | 0 | Canis | 79887.0 | 0 | Canis lupus | 79887.0 | 0 |
| scf7180000042740 | 600385 | 0.372 | 0 | 59.7201 | Eukaryota | 58349.0 | 0 | Chordata | 58349.0 | 0 | Carnivora | 58349.0 | 0 | Canidae | 58349.0 | 0 | Canis | 58349.0 | 0 | Canis lupus | 58349.0 | 0 |
| scf7180000044493 | 289997 | 0.484 | 0 | 59.4393 | Eukaryota | 40184.0 | 0 | Chordata | 40184.0 | 0 | Carnivora | 40184.0 | 0 | Canidae | 33299.0 | 1 | Canis | 25667.0 | 2 | Canis lupus | 25667.0 | 3 |
| scf7180000045059 | 228786 | 0.3625 | 0 | 59.0747 | Eukaryota | 21041.0 | 0 | Chordata | 21041.0 | 0 | Artiodactyla | 12893.0 | 1 | Suidae | 12893.0 | 1 | Sus | 12893.0 | 1 | Sus scrofa | 12893.0 | 1 |
| scf7180000053419 | 13445 | 0.5029 | 0 | 251.0538 | Eukaryota | 9016.0 | 0 | Chordata | 9016.0 | 0 | Carnivora | 9016.0 | 0 | Canidae | 9016.0 | 0 | Canis | 7234.0 | 1 | Canis lupus | 5972.0 | 2 |
| scf7180000043768 | 387648 | 0.4387 | 0 | 59.451 | Eukaryota | 60147.0 | 0 | Chordata | 60147.0 | 0 | Carnivora | 60147.0 | 0 | Canidae | 60147.0 | 0 | Canis | 48650.0 | 1 | Canis lupus | 48650.0 | 1 |
| scf7180000051547 | 9563 | 0.5007 | 0 | 541.47 | Eukaryota | 8771.0 | 0 | Chordata | 8771.0 | 0 | Carnivora | 8771.0 | 0 | Canidae | 8771.0 | 0 | Canis | 7096.0 | 1 | Canis lupus | 5863.0 | 2 |
| scf7180000041429 | 1020562 | 0.4683 | 0 | 59.378 | Eukaryota | 90574.0 | 0 | Chordata | 90574.0 | 0 | Carnivora | 90574.0 | 0 | Canidae | 90574.0 | 0 | Canis | 90574.0 | 0 | Canis lupus | 90574.0 | 0 |
| scf7180000056508 | 195262 | 0.4501 | 0 | 58.3312 | Eukaryota | 22613.0 | 0 | Chordata | 22613.0 | 0 | Carnivora | 22613.0 | 0 | Canidae | 22613.0 | 0 | Canis | 22613.0 | 0 | Canis lupus | 22613.0 | 0 |
| scf7180000051790 | 5051 | 0.4486 | 0 | 470.6695 | no-hit | 0.0 | 0 | no-hit | 0.0 | 0 | no-hit | 0.0 | 0 | no-hit | 0.0 | 0 | no-hit | 0.0 | 0 | no-hit | 0.0 | 0 |
| scf7180000052622 | 1354 | 0.4461 | 0 | 128.0911 | no-hit | 0.0 | 0 | no-hit | 0.0 | 0 | no-hit | 0.0 | 0 | no-hit | 0.0 | 0 | no-hit | 0.0 | 0 | no-hit | 0.0 | 0 |
| scf7180000050208 | 29940 | 0.4735 | 0 | 59.0749 | Eukaryota | 4771.0 | 0 | Chordata | 4771.0 | 0 | Carnivora | 2706.0 | 1 | Canidae | 2706.0 | 1 | Canis | 2238.0 | 2 | Canis lupus | 2238.0 | 2 |
| scf7180000044569 | 337596 | 0.4728 | 0 | 59.4019 | Eukaryota | 44867.0 | 0 | Chordata | 44867.0 | 0 | Carnivora | 44867.0 | 0 | Canidae | 44867.0 | 0 | Canis | 44867.0 | 0 | Canis lupus | 44867.0 | 0 |
| scf7180000053844 | 7972 | 0.4454 | 0 | 57.1063 | Eukaryota | 2492.0 | 0 | Platyhelminthes | 1270.0 | 1 | Polystomatidea | 1270.0 | 2 | Polystomatidae | 1270.0 | 2 | Protopolystoma | 1270.0 | 2 | Protopolystoma xenopodis | 1270.0 | 2 |
| scf7180000048868 | 7050 | 0.4485 | 0 | 107.0216 | Eukaryota | 372.0 | 0 | Platyhelminthes | 372.0 | 0 | Polystomatidea | 372.0 | 0 | Polystomatidae | 372.0 | 0 | Protopolystoma | 372.0 | 0 | Protopolystoma xenopodis | 372.0 | 0 |
| scf7180000052370 | 16691 | 0.4411 | 0 | 60.6541 | Eukaryota | 174.0 | 0 | Platyhelminthes | 174.0 | 0 | Polystomatidea | 174.0 | 0 | Polystomatidae | 174.0 | 0 | Protopolystoma | 174.0 | 0 | Protopolystoma xenopodis | 174.0 | 0 |
| scf7180000042315 | 380000 | 0.3913 | 0 | 59.4708 | Eukaryota | 32677.0 | 0 | Chordata | 32677.0 | 0 | Carnivora | 29687.0 | 1 | Canidae | 15864.0 | 3 | Canis | 15864.0 | 5 | Canis lupus | 15864.0 | 5 |
| scf7180000055975 | 5957 | 0.4531 | 0 | 1440.8747 | Eukaryota | 2558.0 | 0 | Chordata | 2558.0 | 0 | Carnivora | 2558.0 | 0 | Canidae | 2558.0 | 0 | Canis | 2558.0 | 0 | Canis lupus | 2558.0 | 0 |
| scf7180000053170 | 5149 | 0.4488 | 0 | 827.5023 | no-hit | 0.0 | 0 | no-hit | 0.0 | 0 | no-hit | 0.0 | 0 | no-hit | 0.0 | 0 | no-hit | 0.0 | 0 | no-hit | 0.0 | 0 |
| scf7180000051615 | 20732 | 0.4538 | 0 | 58.3018 | Eukaryota | 43245.0 | 0 | Chordata | 29155.0 | 2 | Gadiformes | 20284.0 | 5 | Gadidae | 20284.0 | 5 | Gadus | 20284.0 | 5 | Gadus morhua | 20284.0 | 5 |
| scf7180000053604 | 12743 | 0.4393 | 0 | 38.2475 | no-hit | 0.0 | 0 | no-hit | 0.0 | 0 | no-hit | 0.0 | 0 | no-hit | 0.0 | 0 | no-hit | 0.0 | 0 | no-hit | 0.0 | 0 |
| scf7180000042322 | 706555 | 0.3444 | 0 | 58.9063 | Eukaryota | 58159.0 | 0 | Chordata | 58159.0 | 0 | Carnivora | 58159.0 | 0 | Canidae | 58159.0 | 0 | Canis | 58159.0 | 0 | Canis lupus | 58159.0 | 0 |
| scf7180000049881 | 22521 | 0.452 | 0 | 1238.6381 | Eukaryota | 3267.0 | 0 | Chordata | 994.0 | 3 | Polystomatidea | 942.0 | 5 | Polystomatidae | 942.0 | 5 | Protopolystoma | 942.0 | 5 | Protopolystoma xenopodis | 942.0 | 5 |
| scf7180000054342 | 16418 | 0.4428 | 0 | 1089.1894 | no-hit | 0.0 | 0 | no-hit | 0.0 | 0 | no-hit | 0.0 | 0 | no-hit | 0.0 | 0 | no-hit | 0.0 | 0 | no-hit | 0.0 | 0 |
| scf7180000053094 | 1961 | 0.4656 | 0 | 200.5045 | Eukaryota | 2041.0 | 0 | Chordata | 1460.0 | 1 | Columbiformes | 872.0 | 2 | Columbidae | 872.0 | 2 | Streptopelia | 872.0 | 2 | Streptopelia turtur | 872.0 | 2 |
| scf7180000045440 | 22034 | 0.4448 | 0 | 639.0776 | Eukaryota | 22041.0 | 0 | Chordata | 12016.0 | 2 | Strigeidida | 7845.0 | 6 | Schistosomatidae | 7845.0 | 6 | Schistosoma | 7845.0 | 6 | Schistosoma mansoni | 7845.0 | 6 |
| scf7180000050042 | 16547 | 0.4493 | 0 | 113.2979 | Eukaryota | 5947.0 | 1 | Chordata | 4852.0 | 2 | Tetraodontiformes | 3654.0 | 4 | Tetraodontidae | 3654.0 | 4 | Takifugu | 3654.0 | 4 | Takifugu rubripes | 3654.0 | 4 |
| scf7180000047170 | 4010 | 0.3753 | 0 | 25.7947 | Eukaryota | 2335.0 | 0 | Chordata | 2335.0 | 0 | Carnivora | 2335.0 | 0 | Canidae | 2335.0 | 0 | Canis | 2335.0 | 0 | Canis lupus | 2335.0 | 0 |
| scf7180000046674 | 85371 | 0.6822 | 0 | 51.5644 | Eukaryota | 27298.0 | 0 | Chordata | 27298.0 | 0 | Carnivora | 27298.0 | 0 | Canidae | 27298.0 | 0 | Canis | 25870.0 | 1 | Canis lupus | 25870.0 | 1 |
| scf7180000042061 | 130969 | 0.3835 | 0 | 57.7771 | Eukaryota | 15407.0 | 0 | Chordata | 15407.0 | 0 | Carnivora | 15407.0 | 0 | Canidae | 15407.0 | 0 | Canis | 15407.0 | 0 | Canis lupus | 15407.0 | 0 |
| scf7180000051399 | 6219 | 0.4366 | 0 | 156.9928 | no-hit | 0.0 | 0 | no-hit | 0.0 | 0 | no-hit | 0.0 | 0 | no-hit | 0.0 | 0 | no-hit | 0.0 | 0 | no-hit | 0.0 | 0 |
| scf7180000044081 | 6854 | 0.4669 | 0 | 27.4683 | Eukaryota | 3356.0 | 0 | Chordata | 3356.0 | 0 | Carnivora | 3356.0 | 0 | Canidae | 2737.0 | 3 | Canis | 2101.0 | 4 | Canis lupus | 2101.0 | 4 |
| scf7180000043402 | 8347 | 0.4541 | 0 | 77.9376 | no-hit | 0.0 | 0 | no-hit | 0.0 | 0 | no-hit | 0.0 | 0 | no-hit | 0.0 | 0 | no-hit | 0.0 | 0 | no-hit | 0.0 | 0 |
| scf7180000050734 | 14665 | 0.4826 | 0 | 58.9195 | Eukaryota | 2550.0 | 0 | Chordata | 2550.0 | 0 | Carnivora | 2550.0 | 0 | Felidae | 1281.0 | 1 | Felis | 1281.0 | 1 | Felis catus | 1281.0 | 1 |
| scf7180000054272 | 19372 | 0.3411 | 0 | 78.1178 | Eukaryota | 3855.0 | 1 | Chordata | 3855.0 | 1 | Primates | 1781.0 | 3 | Hominidae | 1781.0 | 4 | Homo | 1496.0 | 6 | Homo sapiens | 1496.0 | 6 |
| scf7180000055439 | 23367 | 0.4485 | 0 | 541.6336 | no-hit | 0.0 | 0 | no-hit | 0.0 | 0 | no-hit | 0.0 | 0 | no-hit | 0.0 | 0 | no-hit | 0.0 | 0 | no-hit | 0.0 | 0 |
| scf7180000045829 | 20041 | 0.4211 | 0 | 116.2807 | Eukaryota | 38614.0 | 0 | Chordata | 38614.0 | 0 | Carnivora | 38614.0 | 0 | Canidae | 38614.0 | 0 | Canis | 38614.0 | 0 | Canis lupus | 38614.0 | 0 |
| scf7180000041379 | 79702 | 0.4827 | 0 | 54.2379 | Eukaryota | 10287.0 | 1 | Chordata | 10287.0 | 1 | Carnivora | 8391.0 | 2 | Canidae | 8391.0 | 2 | Canis | 7382.0 | 3 | Canis lupus | 7382.0 | 3 |
| scf7180000041500 | 994028 | 0.4935 | 0 | 59.8386 | Eukaryota | 66734.0 | 0 | Chordata | 66734.0 | 0 | Carnivora | 66734.0 | 0 | Canidae | 57310.0 | 1 | Canis | 43760.0 | 2 | Canis lupus | 43760.0 | 2 |
| scf7180000049596 | 50630 | 0.5009 | 0 | 324.5555 | Eukaryota | 9303.0 | 0 | Chordata | 9303.0 | 0 | Carnivora | 9303.0 | 0 | Canidae | 9303.0 | 0 | Canis | 8057.0 | 1 | Canis lupus | 6750.0 | 2 |
| scf7180000046170 | 16481 | 0.5047 | 0 | 334.7025 | Eukaryota | 9145.0 | 0 | Chordata | 9145.0 | 0 | Carnivora | 9145.0 | 0 | Canidae | 9145.0 | 0 | Canis | 7424.0 | 1 | Canis lupus | 6150.0 | 2 |
| scf7180000043018 | 551727 | 0.4044 | 0 | 59.3065 | Eukaryota | 33726.0 | 0 | Chordata | 33726.0 | 0 | Carnivora | 22716.0 | 1 | Canidae | 22716.0 | 2 | Canis | 18900.0 | 3 | Canis lupus | 18900.0 | 3 |
| scf7180000042242 | 694072 | 0.4778 | 0 | 58.3906 | Eukaryota | 80700.0 | 0 | Chordata | 80700.0 | 0 | Carnivora | 80700.0 | 0 | Canidae | 80700.0 | 0 | Canis | 80700.0 | 0 | Canis lupus | 80700.0 | 0 |
| scf7180000056596 | 1397664 | 0.3892 | 100 | 60.0506 | Eukaryota | 92589.0 | 0 | Chordata | 92589.0 | 0 | Carnivora | 92589.0 | 0 | Canidae | 92589.0 | 0 | Canis | 92589.0 | 0 | Canis lupus | 92589.0 | 0 |
| scf7180000041920 | 98860 | 0.3804 | 0 | 56.9382 | Eukaryota | 80867.0 | 0 | Chordata | 80867.0 | 0 | Carnivora | 80867.0 | 0 | Canidae | 80867.0 | 0 | Canis | 80867.0 | 0 | Canis lupus | 80867.0 | 0 |
| scf7180000053042 | 10903 | 0.5068 | 0 | 184.4056 | Eukaryota | 8508.0 | 0 | Chordata | 8508.0 | 0 | Carnivora | 8508.0 | 0 | Canidae | 8508.0 | 0 | Canis | 6369.0 | 1 | Canis lupus | 5172.0 | 2 |
| scf7180000050990 | 32751 | 0.4284 | 0 | 58.2235 | Eukaryota | 81450.0 | 0 | Chordata | 81450.0 | 0 | Carnivora | 81450.0 | 0 | Canidae | 81450.0 | 0 | Canis | 81450.0 | 0 | Canis lupus | 81450.0 | 0 |
| scf7180000044858 | 139334 | 0.4244 | 0 | 58.0034 | Eukaryota | 17107.0 | 0 | Chordata | 17107.0 | 0 | Carnivora | 10802.0 | 1 | Canidae | 10802.0 | 1 | Canis | 10802.0 | 1 | Canis lupus | 10802.0 | 1 |
| scf7180000042181 | 45921 | 0.369 | 0 | 56.2188 | Eukaryota | 91061.0 | 0 | Chordata | 91061.0 | 0 | Carnivora | 91061.0 | 0 | Canidae | 91061.0 | 0 | Canis | 91061.0 | 0 | Canis lupus | 91061.0 | 0 |
| scf7180000052289 | 13747 | 0.5047 | 0 | 60.4784 | Eukaryota | 8052.0 | 0 | Chordata | 8052.0 | 0 | Carnivora | 8052.0 | 0 | Canidae | 8052.0 | 0 | Canis | 6493.0 | 1 | Canis lupus | 5343.0 | 2 |
| scf7180000044250 | 347604 | 0.5334 | 0 | 58.539 | Eukaryota | 38849.0 | 0 | Chordata | 38849.0 | 0 | Carnivora | 38849.0 | 0 | Canidae | 38849.0 | 0 | Canis | 38849.0 | 0 | Canis lupus | 38849.0 | 0 |
| scf7180000043140 | 439344 | 0.4841 | 0 | 58.9024 | Eukaryota | 65167.0 | 0 | Chordata | 65167.0 | 0 | Carnivora | 65167.0 | 0 | Canidae | 65167.0 | 0 | Canis | 48275.0 | 1 | Canis lupus | 48275.0 | 1 |
| scf7180000056592 | 715858 | 0.3636 | 0 | 57.6668 | Eukaryota | 13108.0 | 0 | Chordata | 13108.0 | 0 | Carnivora | 11538.0 | 1 | Canidae | 11538.0 | 1 | Canis | 11538.0 | 1 | Canis lupus | 11538.0 | 1 |
| scf7180000043546 | 6662 | 0.3756 | 0 | 26.9905 | Eukaryota | 5451.0 | 0 | Chordata | 5451.0 | 0 | Carnivora | 5451.0 | 0 | Canidae | 4702.0 | 1 | Canis | 4702.0 | 3 | Canis lupus | 4702.0 | 3 |
| scf7180000042799 | 577514 | 0.4175 | 0 | 60.1292 | Eukaryota | 27981.0 | 0 | Chordata | 27981.0 | 0 | Carnivora | 27981.0 | 0 | Canidae | 24376.0 | 1 | Canis | 24376.0 | 1 | Canis lupus | 24376.0 | 1 |
| scf7180000053077 | 4121 | 0.4506 | 0 | 64.9124 | no-hit | 0.0 | 0 | no-hit | 0.0 | 0 | no-hit | 0.0 | 0 | no-hit | 0.0 | 0 | no-hit | 0.0 | 0 | no-hit | 0.0 | 0 |
| scf7180000045036 | 309780 | 0.3583 | 100 | 60.0258 | Eukaryota | 40915.0 | 0 | Chordata | 40915.0 | 0 | Carnivora | 40915.0 | 0 | Canidae | 40915.0 | 0 | Canis | 40915.0 | 0 | Canis lupus | 40915.0 | 0 |
| scf7180000050440 | 45960 | 0.3948 | 0 | 59.0162 | Eukaryota | 7288.0 | 0 | Chordata | 7288.0 | 0 | Carnivora | 7288.0 | 0 | Canidae | 7288.0 | 0 | Canis | 7288.0 | 0 | Canis lupus | 7288.0 | 0 |
| scf7180000044317 | 339988 | 0.3894 | 0 | 60.9563 | Eukaryota | 88476.0 | 0 | Chordata | 88476.0 | 0 | Carnivora | 88476.0 | 0 | Canidae | 88476.0 | 0 | Canis | 88476.0 | 0 | Canis lupus | 88476.0 | 0 |
| scf7180000055035 | 100218 | 0.4379 | 0 | 300.5095 | Eukaryota | 26604.0 | 0 | Chordata | 22213.0 | 2 | Gadiformes | 14191.0 | 7 | Gadidae | 14191.0 | 7 | Gadus | 14191.0 | 7 | Gadus morhua | 14191.0 | 7 |
| scf7180000046985 | 80937 | 0.3851 | 0 | 61.6731 | Eukaryota | 37323.0 | 1 | Chordata | 37323.0 | 1 | Primates | 26583.0 | 3 | Hominidae | 26583.0 | 3 | Homo | 26583.0 | 4 | Homo sapiens | 26583.0 | 4 |
| scf7180000041034 | 99615 | 0.409 | 0 | 59.158 | Eukaryota | 58632.0 | 0 | Chordata | 58632.0 | 0 | Carnivora | 58632.0 | 0 | Canidae | 58632.0 | 0 | Canis | 35068.0 | 1 | Canis lupus | 35068.0 | 1 |
| scf7180000045542 | 57019 | 0.3798 | 0 | 59.7298 | Eukaryota | 55657.0 | 0 | Chordata | 55657.0 | 0 | Carnivora | 55657.0 | 0 | Canidae | 55657.0 | 0 | Canis | 55657.0 | 0 | Canis lupus | 55657.0 | 0 |
| scf7180000047572 | 9135 | 0.4397 | 0 | 48.4686 | Eukaryota | 263.0 | 0 | Platyhelminthes | 263.0 | 0 | Polystomatidea | 263.0 | 0 | Polystomatidae | 263.0 | 0 | Protopolystoma | 263.0 | 0 | Protopolystoma xenopodis | 263.0 | 0 |
| scf7180000053547 | 2892 | 0.4402 | 0 | 414.8012 | no-hit | 0.0 | 0 | no-hit | 0.0 | 0 | no-hit | 0.0 | 0 | no-hit | 0.0 | 0 | no-hit | 0.0 | 0 | no-hit | 0.0 | 0 |
| scf7180000049696 | 19760 | 0.5246 | 0 | 284.304 | Eukaryota | 35227.0 | 0 | Chordata | 35227.0 | 0 | Carnivora | 35227.0 | 0 | Canidae | 35227.0 | 0 | Canis | 35227.0 | 0 | Canis lupus | 35227.0 | 0 |
| scf7180000042784 | 628826 | 0.4127 | 0 | 60.1026 | Eukaryota | 93439.0 | 0 | Chordata | 93439.0 | 0 | Carnivora | 93439.0 | 0 | Canidae | 93439.0 | 0 | Canis | 93439.0 | 0 | Canis lupus | 93439.0 | 0 |
| scf7180000052461 | 3495 | 0.4378 | 0 | 201.883 | no-hit | 0.0 | 0 | no-hit | 0.0 | 0 | no-hit | 0.0 | 0 | no-hit | 0.0 | 0 | no-hit | 0.0 | 0 | no-hit | 0.0 | 0 |
| scf7180000054691 | 19263 | 0.4419 | 0 | 418.8911 | Eukaryota | 3478.0 | 0 | Platyhelminthes | 2117.0 | 2 | Polystomatidea | 2117.0 | 2 | Polystomatidae | 2117.0 | 2 | Protopolystoma | 2117.0 | 2 | Protopolystoma xenopodis | 2117.0 | 2 |
| scf7180000045239 | 218862 | 0.4465 | 0 | 60.2047 | Eukaryota | 20830.0 | 0 | Chordata | 20830.0 | 0 | Primates | 14923.0 | 2 | Hominidae | 14923.0 | 2 | Homo | 14923.0 | 3 | Homo sapiens | 14923.0 | 3 |
| scf7180000052565 | 6156 | 0.7034 | 0 | 46.5188 | no-hit | 0.0 | 0 | no-hit | 0.0 | 0 | no-hit | 0.0 | 0 | no-hit | 0.0 | 0 | no-hit | 0.0 | 0 | no-hit | 0.0 | 0 |
| scf7180000054807 | 18234 | 0.4409 | 0 | 205.5243 | Eukaryota | 94172.0 | 0 | Chordata | 94172.0 | 0 | Carnivora | 94172.0 | 0 | Canidae | 94172.0 | 0 | Canis | 94172.0 | 0 | Canis lupus | 94172.0 | 0 |
| scf7180000046908 | 50362 | 0.4373 | 0 | 135.6712 | Eukaryota | 7254.0 | 0 | Chordata | 3868.0 | 1 | Kurtiformes | 3467.0 | 2 | Apogonidae | 3467.0 | 2 | Sphaeramia | 3467.0 | 2 | Sphaeramia orbicularis | 3467.0 | 2 |
| scf7180000053892 | 2517 | 0.4374 | 0 | 821.8526 | no-hit | 0.0 | 0 | no-hit | 0.0 | 0 | no-hit | 0.0 | 0 | no-hit | 0.0 | 0 | no-hit | 0.0 | 0 | no-hit | 0.0 | 0 |
| scf7180000046056 | 22326 | 0.4352 | 0 | 231.0038 | Eukaryota | 19227.0 | 0 | Chordata | 16248.0 | 3 | Gadiformes | 10218.0 | 6 | Gadidae | 10218.0 | 6 | Gadus | 10218.0 | 7 | Gadus morhua | 10218.0 | 7 |
| scf7180000046126 | 123363 | 0.4305 | 0 | 58.2576 | Eukaryota | 22739.0 | 0 | Chordata | 22739.0 | 0 | Carnivora | 22739.0 | 0 | Canidae | 19199.0 | 1 | Canis | 14901.0 | 2 | Canis lupus | 14901.0 | 2 |
| scf7180000041709 | 836542 | 0.4502 | 0 | 58.8723 | Eukaryota | 24620.0 | 0 | Chordata | 24620.0 | 0 | Carnivora | 24620.0 | 0 | Canidae | 24620.0 | 0 | Canis | 21840.0 | 1 | Canis lupus | 21840.0 | 1 |
| scf7180000055118 | 72514 | 0.4384 | 0 | 203.1175 | Eukaryota | 3933.0 | 0 | Chordata | 3393.0 | 1 | Kurtiformes | 1022.0 | 4 | Apogonidae | 1022.0 | 4 | Sphaeramia | 1022.0 | 4 | Sphaeramia orbicularis | 1022.0 | 4 |
| scf7180000047106 | 29345 | 0.493 | 0 | 455.397 | Eukaryota | 8834.0 | 0 | Chordata | 8834.0 | 0 | Carnivora | 8834.0 | 0 | Canidae | 8834.0 | 0 | Canis | 7129.0 | 1 | Canis lupus | 5900.0 | 2 |
| scf7180000046578 | 19861 | 0.4084 | 0 | 344.6929 | Eukaryota | 90729.0 | 0 | Chordata | 90729.0 | 0 | Carnivora | 90729.0 | 0 | Canidae | 90729.0 | 0 | Canis | 90729.0 | 0 | Canis lupus | 90729.0 | 0 |
| scf7180000051127 | 22732 | 0.5347 | 0 | 48.1527 | Eukaryota | 21930.0 | 0 | Chordata | 21930.0 | 0 | Carnivora | 21930.0 | 0 | Canidae | 21930.0 | 0 | Canis | 13158.0 | 1 | Canis lupus | 13158.0 | 1 |
| scf7180000052234 | 15012 | 0.4423 | 0 | 68.4866 | Eukaryota | 772.0 | 0 | Chordata | 424.0 | 1 | Kurtiformes | 424.0 | 1 | Apogonidae | 424.0 | 1 | Sphaeramia | 424.0 | 1 | Sphaeramia orbicularis | 424.0 | 1 |
| scf7180000049345 | 10500 | 0.4537 | 0 | 938.9037 | Eukaryota | 3703.0 | 0 | Platyhelminthes | 2270.0 | 2 | Polystomatidea | 2270.0 | 2 | Polystomatidae | 2270.0 | 2 | Protopolystoma | 2270.0 | 2 | Protopolystoma xenopodis | 2270.0 | 2 |
| scf7180000043551 | 198854 | 0.4713 | 0 | 57.22 | Eukaryota | 41252.0 | 0 | Chordata | 41252.0 | 0 | Carnivora | 41252.0 | 0 | Canidae | 37763.0 | 1 | Canis | 22724.0 | 2 | Canis lupus | 22724.0 | 2 |
| scf7180000046864 | 4337 | 0.439 | 0 | 70.0495 | Eukaryota | 233.0 | 0 | Chordata | 233.0 | 0 | Kurtiformes | 233.0 | 0 | Apogonidae | 233.0 | 0 | Sphaeramia | 233.0 | 0 | Sphaeramia orbicularis | 233.0 | 0 |
| scf7180000046814 | 19796 | 0.3644 | 0 | 57.9233 | Eukaryota | 3747.0 | 0 | Chordata | 3747.0 | 0 | Carnivora | 2117.0 | 1 | Canidae | 2117.0 | 1 | Canis | 2117.0 | 1 | Canis lupus | 2117.0 | 1 |
| scf7180000050469 | 6648 | 0.4498 | 0 | 909.9802 | Eukaryota | 1251.0 | 0 | Echinodermata | 556.0 | 2 | Forcipulatida | 556.0 | 2 | Asteriidae | 556.0 | 2 | Asterias | 556.0 | 2 | Asterias rubens | 556.0 | 2 |
| scf7180000041368 | 1137702 | 0.3835 | 0 | 60.7672 | Eukaryota | 89125.0 | 0 | Chordata | 89125.0 | 0 | Carnivora | 89125.0 | 0 | Canidae | 89125.0 | 0 | Canis | 79261.0 | 1 | Canis lupus | 79261.0 | 1 |
| scf7180000051696 | 6535 | 0.4459 | 0 | 46.5087 | Eukaryota | 444.0 | 0 | Chordata | 303.0 | 1 | Tetraodontiformes | 303.0 | 1 | Tetraodontidae | 303.0 | 1 | Takifugu | 303.0 | 1 | Takifugu rubripes | 303.0 | 1 |
| scf7180000041353 | 1168781 | 0.3793 | 0 | 60.4466 | Eukaryota | 222838.0 | 0 | Chordata | 222838.0 | 0 | Carnivora | 222838.0 | 0 | Canidae | 222838.0 | 0 | Canis | 222838.0 | 0 | Canis lupus | 222838.0 | 0 |
| scf7180000041338 | 868667 | 0.4165 | 0 | 60.202 | Eukaryota | 95043.0 | 0 | Chordata | 95043.0 | 0 | Carnivora | 95043.0 | 0 | Canidae | 95043.0 | 0 | Canis | 95043.0 | 0 | Canis lupus | 95043.0 | 0 |
| scf7180000045601 | 140723 | 0.4041 | 0 | 60.8019 | Eukaryota | 25374.0 | 0 | Chordata | 25374.0 | 0 | Carnivora | 25374.0 | 0 | Canidae | 25374.0 | 0 | Canis | 17556.0 | 1 | Canis lupus | 17556.0 | 1 |
| scf7180000050717 | 41794 | 0.5049 | 0 | 59.0773 | Eukaryota | 24476.0 | 0 | Chordata | 24476.0 | 0 | Carnivora | 24476.0 | 0 | Canidae | 19074.0 | 1 | Canis | 19074.0 | 1 | Canis lupus | 19074.0 | 1 |
| scf7180000053763 | 2273 | 0.4417 | 0 | 301.0309 | Eukaryota | 1606.0 | 0 | Platyhelminthes | 1471.0 | 1 | Strigeidida | 1471.0 | 1 | Schistosomatidae | 1471.0 | 1 | Schistosoma | 1471.0 | 1 | Schistosoma mansoni | 1471.0 | 1 |
| scf7180000046062 | 10593 | 0.4594 | 0 | 114.068 | Eukaryota | 88448.0 | 0 | Chordata | 88448.0 | 0 | Carnivora | 88448.0 | 0 | Canidae | 88448.0 | 0 | Canis | 88448.0 | 0 | Canis lupus | 88448.0 | 0 |
| scf7180000043194 | 46387 | 0.5863 | 0 | 57.4323 | Eukaryota | 14250.0 | 0 | Chordata | 14250.0 | 0 | Carnivora | 14250.0 | 0 | Canidae | 11717.0 | 1 | Canis | 11717.0 | 2 | Canis lupus | 11717.0 | 2 |
| scf7180000048507 | 39930 | 0.4193 | 0 | 55.4649 | Eukaryota | 96333.0 | 0 | Chordata | 96333.0 | 0 | Carnivora | 96333.0 | 0 | Canidae | 96333.0 | 0 | Canis | 87932.0 | 1 | Canis lupus | 87932.0 | 1 |
| scf7180000051421 | 11102 | 0.4427 | 0 | 35.4512 | Eukaryota | 187.0 | 0 | Chordata | 187.0 | 0 | Kurtiformes | 187.0 | 0 | Apogonidae | 187.0 | 0 | Sphaeramia | 187.0 | 0 | Sphaeramia orbicularis | 187.0 | 0 |
| scf7180000049551 | 5855 | 0.5023 | 0 | 170.2417 | Eukaryota | 7671.0 | 0 | Chordata | 7671.0 | 0 | Carnivora | 7671.0 | 0 | Canidae | 7671.0 | 0 | Canis | 6427.0 | 1 | Canis lupus | 5510.0 | 2 |
| scf7180000051335 | 3828 | 0.4501 | 0 | 241.6261 | Eukaryota | 1926.0 | 0 | Platyhelminthes | 798.0 | 3 | Polystomatidea | 798.0 | 3 | Polystomatidae | 798.0 | 3 | Protopolystoma | 798.0 | 3 | Protopolystoma xenopodis | 798.0 | 3 |
| scf7180000053592 | 2904 | 0.438 | 0 | 35.1706 | Eukaryota | 963.0 | 0 | Chordata | 963.0 | 0 | Kurtiformes | 963.0 | 0 | Apogonidae | 963.0 | 0 | Sphaeramia | 963.0 | 0 | Sphaeramia orbicularis | 963.0 | 0 |
| scf7180000045172 | 115385 | 0.3961 | 0 | 59.6493 | Eukaryota | 228210.0 | 0 | Chordata | 228210.0 | 0 | Carnivora | 228210.0 | 0 | Canidae | 208177.0 | 1 | Canis | 185078.0 | 2 | Canis lupus | 185078.0 | 2 |
| scf7180000056357 | 526266 | 0.4781 | 0 | 59.9121 | Eukaryota | 69698.0 | 0 | Chordata | 69698.0 | 0 | Carnivora | 69698.0 | 0 | Canidae | 69698.0 | 0 | Canis | 40893.0 | 1 | Canis lupus | 40893.0 | 1 |
| scf7180000054158 | 725687 | 0.3545 | 0 | 59.2694 | Eukaryota | 33018.0 | 0 | Chordata | 33018.0 | 0 | Carnivora | 33018.0 | 0 | Canidae | 33018.0 | 0 | Canis | 33018.0 | 0 | Canis lupus | 33018.0 | 0 |
| scf7180000050871 | 5853 | 0.4511 | 0 | 259.151 | no-hit | 0.0 | 0 | no-hit | 0.0 | 0 | no-hit | 0.0 | 0 | no-hit | 0.0 | 0 | no-hit | 0.0 | 0 | no-hit | 0.0 | 0 |
| scf7180000049663 | 43498 | 0.3963 | 0 | 60.2358 | Eukaryota | 7443.0 | 0 | Chordata | 7443.0 | 0 | Carnivora | 7443.0 | 0 | Canidae | 7443.0 | 0 | Canis | 7443.0 | 0 | Canis lupus | 7443.0 | 0 |
| scf7180000044418 | 12605 | 0.526 | 0 | 242.4745 | Eukaryota | 3438.0 | 0 | Chordata | 3438.0 | 0 | Carnivora | 3438.0 | 0 | Canidae | 3438.0 | 0 | Canis | 2749.0 | 1 | Canis lupus | 2749.0 | 1 |
| scf7180000048075 | 14717 | 0.3571 | 0 | 143.706 | Eukaryota | 13033.0 | 0 | Chordata | 13033.0 | 0 | Carnivora | 7865.0 | 2 | Canidae | 7865.0 | 3 | Vulpes | 4494.0 | 5 | Vulpes vulpes | 4494.0 | 5 |
| scf7180000045320 | 190897 | 0.4141 | 0 | 59.6248 | Eukaryota | 49429.0 | 0 | Chordata | 49429.0 | 0 | Carnivora | 49429.0 | 0 | Canidae | 49429.0 | 0 | Canis | 34499.0 | 1 | Canis lupus | 34499.0 | 1 |
| scf7180000050143 | 2177 | 0.4355 | 0 | 199.377 | no-hit | 0.0 | 0 | no-hit | 0.0 | 0 | no-hit | 0.0 | 0 | no-hit | 0.0 | 0 | no-hit | 0.0 | 0 | no-hit | 0.0 | 0 |
| scf7180000043567 | 477195 | 0.4508 | 0 | 57.8828 | Eukaryota | 57754.0 | 0 | Chordata | 57754.0 | 0 | Carnivora | 57754.0 | 0 | Canidae | 57754.0 | 0 | Canis | 57754.0 | 0 | Canis lupus | 57754.0 | 0 |
| scf7180000045806 | 15627 | 0.4845 | 0 | 31.014 | Eukaryota | 23418.0 | 0 | Chordata | 23418.0 | 0 | Carnivora | 23418.0 | 0 | Canidae | 23418.0 | 0 | Canis | 23418.0 | 0 | Canis lupus | 23418.0 | 0 |
| scf7180000056078 | 41171 | 0.3753 | 0 | 53.5972 | Eukaryota | 42848.0 | 0 | Chordata | 42848.0 | 0 | Carnivora | 42848.0 | 0 | Canidae | 42848.0 | 0 | Canis | 42848.0 | 0 | Canis lupus | 42848.0 | 0 |
| scf7180000046167 | 174373 | 0.3681 | 0 | 58.5141 | Eukaryota | 32748.0 | 0 | Chordata | 32748.0 | 0 | Carnivora | 32748.0 | 0 | Canidae | 32748.0 | 0 | Canis | 26912.0 | 1 | Canis lupus | 26912.0 | 1 |
| scf7180000045947 | 180729 | 0.3736 | 0 | 59.8312 | Eukaryota | 15265.0 | 0 | Chordata | 15265.0 | 0 | Carnivora | 15265.0 | 0 | Canidae | 15265.0 | 0 | Canis | 15265.0 | 0 | Canis lupus | 15265.0 | 0 |
| scf7180000040647 | 8498 | 0.4088 | 0 | 55.1792 | Eukaryota | 2966.0 | 0 | Chordata | 2966.0 | 0 | Carnivora | 2966.0 | 0 | Canidae | 2966.0 | 0 | Canis | 2966.0 | 0 | Canis lupus | 2966.0 | 0 |
| scf7180000041698 | 79482 | 0.5795 | 0 | 51.8246 | Eukaryota | 26138.0 | 0 | Chordata | 26138.0 | 0 | Carnivora | 26138.0 | 0 | Canidae | 26138.0 | 0 | Canis | 22891.0 | 1 | Canis lupus | 22891.0 | 1 |
| scf7180000041324 | 2213752 | 0.4392 | 0 | 60.0044 | Eukaryota | 64935.0 | 0 | Chordata | 64935.0 | 0 | Carnivora | 64935.0 | 0 | Canidae | 64935.0 | 0 | Canis | 58021.0 | 1 | Canis lupus | 58021.0 | 1 |
| scf7180000045446 | 197858 | 0.4362 | 0 | 58.0545 | Eukaryota | 70412.0 | 0 | Chordata | 70412.0 | 0 | Carnivora | 70412.0 | 0 | Canidae | 57536.0 | 1 | Canis | 49126.0 | 2 | Canis lupus | 49126.0 | 2 |
| scf7180000043700 | 546327 | 0.4057 | 0 | 58.9124 | Eukaryota | 89720.0 | 0 | Chordata | 89720.0 | 0 | Carnivora | 89720.0 | 0 | Canidae | 89720.0 | 0 | Canis | 89720.0 | 0 | Canis lupus | 89720.0 | 0 |
| scf7180000051368 | 11951 | 0.459 | 0 | 77.5036 | Eukaryota | 55672.0 | 0 | Chordata | 34257.0 | 2 | Cyclophyllidea | 9932.0 | 8 | Hymenolepididae | 9932.0 | 8 | Hymenolepis | 9932.0 | 8 | Hymenolepis microstoma | 9932.0 | 8 |
| scf7180000043731 | 503095 | 0.4159 | 0 | 59.1243 | Eukaryota | 58962.0 | 0 | Chordata | 58962.0 | 0 | Carnivora | 58962.0 | 0 | Canidae | 58962.0 | 0 | Canis | 58962.0 | 0 | Canis lupus | 58962.0 | 0 |
| scf7180000045888 | 138700 | 0.4642 | 0 | 79.2089 | Eukaryota | 26236.0 | 0 | Chordata | 26236.0 | 0 | Carnivora | 17521.0 | 1 | Canidae | 17521.0 | 1 | Canis | 11683.0 | 2 | Canis lupus | 11683.0 | 2 |
| scf7180000055513 | 13314 | 0.4327 | 0 | 94.4497 | no-hit | 0.0 | 0 | no-hit | 0.0 | 0 | no-hit | 0.0 | 0 | no-hit | 0.0 | 0 | no-hit | 0.0 | 0 | no-hit | 0.0 | 0 |
| scf7180000053742 | 5449 | 0.4438 | 0 | 94.7702 | no-hit | 0.0 | 0 | no-hit | 0.0 | 0 | no-hit | 0.0 | 0 | no-hit | 0.0 | 0 | no-hit | 0.0 | 0 | no-hit | 0.0 | 0 |
| scf7180000044753 | 116362 | 0.3769 | 0 | 56.134 | Eukaryota | 73671.0 | 0 | Chordata | 73671.0 | 0 | Carnivora | 73671.0 | 0 | Canidae | 73671.0 | 0 | Canis | 73671.0 | 0 | Canis lupus | 73671.0 | 0 |
| scf7180000050019 | 5453 | 0.4432 | 0 | 17.2508 | Eukaryota | 44871.0 | 0 | Chordata | 44871.0 | 0 | Carnivora | 44871.0 | 0 | Canidae | 44871.0 | 0 | Canis | 44871.0 | 0 | Canis lupus | 44871.0 | 0 |
| scf7180000052193 | 4985 | 0.4506 | 0 | 1297.2011 | Eukaryota | 17065.0 | 0 | Chordata | 14373.0 | 1 | Gadiformes | 7224.0 | 5 | Gadidae | 7224.0 | 5 | Gadus | 7224.0 | 5 | Gadus morhua | 7224.0 | 5 |
| scf7180000044469 | 54646 | 0.4318 | 0 | 68.7759 | Eukaryota | 26559.0 | 0 | Chordata | 26559.0 | 0 | Carnivora | 26559.0 | 0 | Canidae | 26559.0 | 0 | Canis | 26559.0 | 0 | Canis lupus | 26559.0 | 0 |
| scf7180000052940 | 4036 | 0.4492 | 0 | 485.1364 | no-hit | 0.0 | 0 | no-hit | 0.0 | 0 | no-hit | 0.0 | 0 | no-hit | 0.0 | 0 | no-hit | 0.0 | 0 | no-hit | 0.0 | 0 |
| scf7180000052331 | 17345 | 0.4461 | 0 | 60.8125 | Eukaryota | 2458.0 | 0 | Chordata | 1723.0 | 1 | Columbiformes | 1212.0 | 4 | Columbidae | 1212.0 | 4 | Streptopelia | 1212.0 | 4 | Streptopelia turtur | 1212.0 | 4 |
| scf7180000049719 | 13492 | 0.4458 | 0 | 108.9899 | Eukaryota | 4954.0 | 0 | Chordata | 4110.0 | 1 | Tetraodontiformes | 2599.0 | 4 | Tetraodontidae | 2599.0 | 4 | Takifugu | 2599.0 | 4 | Takifugu rubripes | 2599.0 | 4 |
| scf7180000054135 | 104146 | 0.3909 | 0 | 65.3081 | Eukaryota | 94430.0 | 0 | Chordata | 94430.0 | 0 | Carnivora | 94430.0 | 0 | Canidae | 94430.0 | 0 | Canis | 94430.0 | 0 | Canis lupus | 94430.0 | 0 |
| scf7180000055540 | 25142 | 0.4395 | 0 | 426.8623 | Eukaryota | 26109.0 | 0 | Chordata | 18130.0 | 2 | Gadiformes | 14126.0 | 5 | Gadidae | 14126.0 | 5 | Gadus | 14126.0 | 5 | Gadus morhua | 14126.0 | 5 |
| scf7180000042343 | 652823 | 0.4131 | 0 | 61.0233 | Eukaryota | 63770.0 | 0 | Chordata | 63770.0 | 0 | Carnivora | 63770.0 | 0 | Canidae | 63770.0 | 0 | Canis | 63770.0 | 0 | Canis lupus | 63770.0 | 0 |
| scf7180000056266 | 172015 | 0.3581 | 0 | 57.575 | Eukaryota | 93139.0 | 0 | Chordata | 93139.0 | 0 | Carnivora | 93139.0 | 0 | Canidae | 93139.0 | 0 | Canis | 93139.0 | 0 | Canis lupus | 93139.0 | 0 |
| scf7180000041146 | 171875 | 0.3762 | 0 | 57.9792 | Eukaryota | 68012.0 | 0 | Chordata | 68012.0 | 0 | Carnivora | 68012.0 | 0 | Canidae | 68012.0 | 0 | Canis | 68012.0 | 0 | Canis lupus | 68012.0 | 0 |
| scf7180000046724 | 49935 | 0.384 | 0 | 55.7227 | Eukaryota | 3588.0 | 0 | Chordata | 3588.0 | 0 | Carnivora | 3081.0 | 1 | Canidae | 3081.0 | 1 | Canis | 2746.0 | 2 | Canis lupus | 2746.0 | 2 |
| scf7180000042253 | 320272 | 0.3917 | 0 | 58.8739 | Eukaryota | 93244.0 | 0 | Chordata | 93244.0 | 0 | Carnivora | 93244.0 | 0 | Canidae | 93244.0 | 0 | Canis | 93244.0 | 0 | Canis lupus | 93244.0 | 0 |
| scf7180000046896 | 26351 | 0.4552 | 0 | 160.0947 | Eukaryota | 18353.0 | 0 | Chordata | 18353.0 | 0 | Carnivora | 18353.0 | 0 | Canidae | 12340.0 | 2 | Canis | 9346.0 | 6 | Canis lupus | 9346.0 | 6 |

| | | | | | | | | | | | | | | | | | | | | | | |
|---|---|---|---|---|---|---|---|---|---|---|---|---|---|---|---|---|---|---|---|---|---|---|
| scf7180000054472 | 18259 | 0.4172 | 0 | 624.2296 | Eukaryota | 73839.0 | 0 | Chordata | 73839.0 | 0 | Carnivora | 73839.0 | 0 | Canidae | 73839.0 | 0 | Canis | 73839.0 | 0 | Canis lupus | 73839.0 | 0 |
| scf7180000043767 | 368510 | 0.4255 | 0 | 59.416 | Eukaryota | 58146.0 | 0 | Chordata | 58146.0 | 0 | Carnivora | 58146.0 | 0 | Canidae | 58146.0 | 0 | Vulpes | 40232.0 | 1 | Vulpes vulpes | 40232.0 | 1 |
| scf7180000044578 | 69867 | 0.3796 | 0 | 138.1986 | Eukaryota | 92026.0 | 0 | Chordata | 92026.0 | 0 | Carnivora | 92026.0 | 0 | Canidae | 92026.0 | 0 | Canis | 92026.0 | 0 | Canis lupus | 92026.0 | 0 |
| scf7180000048438 | 33053 | 0.427 | 0 | 54.1488 | Eukaryota | 7599.0 | 1 | Chordata | 7599.0 | 1 | Carnivora | 3687.0 | 4 | Felidae | 2801.0 | 5 | Rousettus | 1345.0 | 8 | Rousettus aegyptiacus | 1345.0 | 8 |
| scf7180000051110 | 4869 | 0.4444 | 0 | 215.3519 | no-hit | 0.0 | 0 | no-hit | 0.0 | 0 | no-hit | 0.0 | 0 | no-hit | 0.0 | 0 | no-hit | 0.0 | 0 | no-hit | 0.0 | 0 |
| scf7180000048952 | 8929 | 0.4506 | 0 | 2285.9054 | Eukaryota | 1880.0 | 0 | Platyhelminthes | 1264.0 | 2 | Polystomatidea | 1264.0 | 2 | Polystomatidae | 1264.0 | 2 | Protopolystoma | 1264.0 | 2 | Protopolystoma xenopodis | 1264.0 | 2 |
| scf7180000042282 | 767668 | 0.4084 | 0 | 60.6631 | Eukaryota | 64137.0 | 0 | Chordata | 64137.0 | 0 | Carnivora | 64137.0 | 0 | Canidae | 64137.0 | 0 | Canis | 64137.0 | 0 | Canis lupus | 64137.0 | 0 |
| scf7180000041795 | 853256 | 0.4708 | 0 | 58.7849 | Eukaryota | 116880.0 | 0 | Chordata | 116880.0 | 0 | Carnivora | 116880.0 | 0 | Canidae | 116880.0 | 0 | Canis | 91530.0 | 1 | Canis lupus | 91530.0 | 1 |
| scf7180000041004 | 871162 | 0.3529 | 0 | 59.8169 | Eukaryota | 73513.0 | 0 | Chordata | 73513.0 | 0 | Carnivora | 73513.0 | 0 | Canidae | 73513.0 | 0 | Canis | 73513.0 | 0 | Canis lupus | 73513.0 | 0 |
| scf7180000046012 | 30546 | 0.4314 | 0 | 211.6901 | Eukaryota | 55011.0 | 0 | Chordata | 55011.0 | 0 | Carnivora | 55011.0 | 0 | Canidae | 55011.0 | 0 | Canis | 55011.0 | 0 | Canis lupus | 55011.0 | 0 |
| scf7180000040665 | 1906426 | 0.389 | 0 | 59.2122 | Eukaryota | 32260.0 | 0 | Chordata | 32260.0 | 0 | Carnivora | 32260.0 | 0 | Canidae | 32260.0 | 0 | Vulpes | 22351.0 | 1 | Vulpes vulpes | 22351.0 | 1 |
| scf7180000051882 | 20577 | 0.5069 | 0 | 260.1042 | Eukaryota | 8671.0 | 0 | Chordata | 8671.0 | 0 | Carnivora | 8671.0 | 0 | Canidae | 8671.0 | 0 | Canis | 7027.0 | 1 | Canis lupus | 5868.0 | 2 |
| scf7180000048253 | 70940 | 0.4737 | 0 | 58.5472 | Eukaryota | 87404.0 | 0 | Chordata | 87404.0 | 0 | Carnivora | 87404.0 | 0 | Canidae | 87404.0 | 0 | Canis | 87404.0 | 0 | Canis lupus | 87404.0 | 0 |
| scf7180000051247 | 3005 | 0.4519 | 0 | 292.4746 | no-hit | 0.0 | 0 | no-hit | 0.0 | 0 | no-hit | 0.0 | 0 | no-hit | 0.0 | 0 | no-hit | 0.0 | 0 | no-hit | 0.0 | 0 |
| scf7180000044726 | 103394 | 0.4323 | 0 | 60.2588 | Eukaryota | 51495.0 | 0 | Chordata | 51495.0 | 0 | Carnivora | 51495.0 | 0 | Canidae | 51495.0 | 0 | Vulpes | 37611.0 | 1 | Vulpes vulpes | 37611.0 | 1 |
| scf7180000053621 | 6123 | 0.5012 | 0 | 88.248 | Eukaryota | 8454.0 | 0 | Chordata | 8454.0 | 0 | Carnivora | 8454.0 | 0 | Canidae | 8454.0 | 0 | Canis | 7395.0 | 1 | Canis lupus | 6319.0 | 2 |
| scf7180000049271 | 44784 | 0.5184 | 0 | 55.0604 | Eukaryota | 7105.0 | 0 | Chordata | 7105.0 | 0 | Carnivora | 7105.0 | 0 | Canidae | 4559.0 | 2 | Canis | 2348.0 | 3 | Canis lupus | 2348.0 | 4 |
| scf7180000052377 | 6491 | 0.4434 | 0 | 117.5411 | no-hit | 0.0 | 0 | no-hit | 0.0 | 0 | no-hit | 0.0 | 0 | no-hit | 0.0 | 0 | no-hit | 0.0 | 0 | no-hit | 0.0 | 0 |
| scf7180000046568 | 89764 | 0.3678 | 0 | 58.8871 | Eukaryota | 24167.0 | 0 | Chordata | 24167.0 | 0 | Carnivora | 24167.0 | 0 | Canidae | 24167.0 | 0 | Canis | 24167.0 | 0 | Canis lupus | 24167.0 | 0 |
| scf7180000041109 | 19535 | 0.4469 | 0 | 179.9014 | Eukaryota | 45940.0 | 0 | Chordata | 45940.0 | 0 | Carnivora | 45940.0 | 0 | Canidae | 45940.0 | 0 | Canis | 45940.0 | 0 | Canis lupus | 45940.0 | 0 |
| scf7180000053211 | 12742 | 0.5052 | 0 | 205.0682 | Eukaryota | 8935.0 | 0 | Chordata | 8935.0 | 0 | Carnivora | 8935.0 | 0 | Canidae | 8935.0 | 0 | Canis | 7202.0 | 1 | Canis lupus | 5956.0 | 2 |
| scf7180000048178 | 7904 | 0.4398 | 0 | 133.7469 | Eukaryota | 20856.0 | 0 | Chordata | 17839.0 | 1 | Gadiformes | 11578.0 | 5 | Gadidae | 11578.0 | 5 | Gadus | 11578.0 | 5 | Gadus morhua | 11578.0 | 5 |
| scf7180000041270 | 405243 | 0.4001 | 100 | 59.4754 | Eukaryota | 80279.0 | 0 | Chordata | 80279.0 | 0 | Carnivora | 80279.0 | 0 | Canidae | 80279.0 | 0 | Canis | 80279.0 | 0 | Canis lupus | 80279.0 | 0 |
| scf7180000044281 | 244542 | 0.3619 | 0 | 60.1054 | Eukaryota | 49840.0 | 0 | Chordata | 49840.0 | 0 | Carnivora | 49840.0 | 0 | Canidae | 49840.0 | 0 | Canis | 49840.0 | 0 | Canis lupus | 49840.0 | 0 |
| scf7180000042526 | 629103 | 0.4891 | 0 | 59.2803 | Eukaryota | 66782.0 | 0 | Chordata | 66782.0 | 0 | Carnivora | 66782.0 | 0 | Canidae | 66782.0 | 0 | Canis | 66782.0 | 0 | Canis lupus | 66782.0 | 0 |
| scf7180000053479 | 4295 | 0.4482 | 0 | 55.5576 | Eukaryota | 3145.0 | 0 | Chordata | 3145.0 | 0 | Carnivora | 3145.0 | 0 | Canidae | 3145.0 | 0 | Canis | 3145.0 | 0 | Canis lupus | 3145.0 | 0 |
| scf7180000045815 | 139451 | 0.4156 | 0 | 61.3761 | Eukaryota | 117549.0 | 0 | Chordata | 117549.0 | 0 | Carnivora | 117549.0 | 0 | Canidae | 117549.0 | 0 | Canis | 117549.0 | 0 | Canis lupus | 117549.0 | 0 |
| scf7180000049445 | 6338 | 0.4446 | 0 | 72.301 | Eukaryota | 2297.0 | 0 | Chordata | 2062.0 | 1 | Tetraodontiformes | 1059.0 | 3 | Tetraodontidae | 1059.0 | 3 | Takifugu | 1059.0 | 3 | Takifugu rubripes | 1059.0 | 3 |
| scf7180000043724 | 280010 | 0.3854 | 100 | 60.046 | Eukaryota | 93475.0 | 0 | Chordata | 93475.0 | 0 | Carnivora | 93475.0 | 0 | Canidae | 93475.0 | 0 | Canis | 93475.0 | 0 | Canis lupus | 93475.0 | 0 |
| scf7180000049644 | 8307 | 0.446 | 0 | 248.565 | Eukaryota | 1281.0 | 0 | Chordata | 652.0 | 1 | Tetraodontiformes | 652.0 | 1 | Tetraodontidae | 652.0 | 1 | Takifugu | 652.0 | 1 | Takifugu rubripes | 652.0 | 1 |
| scf7180000043588 | 466446 | 0.3773 | 0 | 58.2222 | Eukaryota | 93192.0 | 0 | Chordata | 93192.0 | 0 | Carnivora | 93192.0 | 0 | Canidae | 93192.0 | 0 | Canis | 93192.0 | 0 | Canis lupus | 93192.0 | 0 |
| scf7180000042914 | 132932 | 0.3832 | 0 | 62.1505 | Eukaryota | 18103.0 | 0 | Chordata | 18103.0 | 0 | Carnivora | 7689.0 | 2 | Hominidae | 7281.0 | 4 | Homo | 7281.0 | 5 | Homo sapiens | 7281.0 | 5 |
| scf7180000042795 | 552975 | 0.4128 | 0 | 60.8707 | Eukaryota | 49891.0 | 0 | Chordata | 49891.0 | 0 | Carnivora | 49891.0 | 0 | Canidae | 42585.0 | 1 | Canis | 23219.0 | 2 | Canis lupus | 23219.0 | 2 |
| scf7180000050322 | 36011 | 0.3999 | 0 | 59.5717 | Eukaryota | 47266.0 | 0 | Chordata | 47266.0 | 0 | Carnivora | 47266.0 | 0 | Canidae | 47266.0 | 0 | Canis | 42822.0 | 1 | Canis lupus | 42822.0 | 1 |
| scf7180000046241 | 101038 | 0.5469 | 0 | 57.5342 | Eukaryota | 84561.0 | 0 | Chordata | 84561.0 | 0 | Carnivora | 81316.0 | 1 | Canidae | 43471.0 | 4 | Canis | 28948.0 | 7 | Canis lupus | 28948.0 | 7 |
| scf7180000046060 | 27252 | 0.4378 | 0 | 211.2529 | Eukaryota | 37130.0 | 0 | Chordata | 35157.0 | 1 | Gadiformes | 18348.0 | 5 | Gadidae | 18348.0 | 5 | Gadus | 18348.0 | 5 | Gadus morhua | 18348.0 | 5 |
| scf7180000050193 | 24663 | 0.3504 | 0 | 60.8051 | Eukaryota | 6996.0 | 0 | Chordata | 6996.0 | 0 | Carnivora | 5164.0 | 1 | Canidae | 5164.0 | 1 | Canis | 5164.0 | 1 | Canis lupus | 5164.0 | 1 |
| scf7180000048162 | 37501 | 0.4509 | 0 | 1769.293 | Eukaryota | 5873.0 | 0 | Platyhelminthes | 3258.0 | 2 | Polystomatidea | 3258.0 | 3 | Polystomatidae | 3258.0 | 3 | Protopolystoma | 3258.0 | 3 | Protopolystoma xenopodis | 3258.0 | 3 |
| scf7180000054428 | 14316 | 0.4139 | 0 | 343.2856 | Eukaryota | 92224.0 | 0 | Chordata | 92224.0 | 0 | Carnivora | 92224.0 | 0 | Canidae | 92224.0 | 0 | Canis | 92224.0 | 0 | Canis lupus | 92224.0 | 0 |
| scf7180000055833 | 7911 | 0.4419 | 0 | 121.935 | Eukaryota | 642.0 | 0 | Streptophyta | 374.0 | 1 | Fabales | 374.0 | 1 | Fabaceae | 374.0 | 1 | Vigna | 374.0 | 1 | Vigna unguiculata | 374.0 | 1 |
| scf7180000051376 | 19976 | 0.4508 | 0 | 131.8415 | Eukaryota | 95738.0 | 0 | Chordata | 95738.0 | 0 | Carnivora | 95738.0 | 0 | Canidae | 95738.0 | 0 | Canis | 95738.0 | 0 | Canis lupus | 95738.0 | 0 |
| scf7180000052994 | 6004 | 0.4414 | 0 | 218.3508 | no-hit | 0.0 | 0 | no-hit | 0.0 | 0 | no-hit | 0.0 | 0 | no-hit | 0.0 | 0 | no-hit | 0.0 | 0 | no-hit | 0.0 | 0 |
| scf7180000052547 | 5853 | 0.5052 | 0 | 91.0518 | Eukaryota | 6572.0 | 0 | Chordata | 6572.0 | 0 | Carnivora | 6572.0 | 0 | Canidae | 6572.0 | 0 | Canis | 4612.0 | 1 | Canis lupus | 3584.0 | 2 |
| scf7180000047660 | 16440 | 0.4501 | 0 | 64.1681 | Eukaryota | 763.0 | 0 | Platyhelminthes | 448.0 | 1 | Polystomatidea | 448.0 | 1 | Polystomatidae | 448.0 | 1 | Protopolystoma | 448.0 | 1 | Protopolystoma xenopodis | 448.0 | 1 |
| scf7180000044065 | 91080 | 0.3837 | 0 | 55.7349 | Eukaryota | 11789.0 | 0 | Chordata | 11789.0 | 0 | Carnivora | 7907.0 | 2 | Otariidae | 3520.0 | 4 | Zalophus | 3520.0 | 7 | Zalophus californianus | 3520.0 | 7 |
| scf7180000047365 | 33653 | 0.3678 | 0 | 47.6624 | Eukaryota | 92676.0 | 0 | Chordata | 92676.0 | 0 | Carnivora | 92676.0 | 0 | Canidae | 92676.0 | 0 | Canis | 92676.0 | 0 | Canis lupus | 92676.0 | 0 |
| scf7180000046379 | 147917 | 0.3957 | 0 | 58.9452 | Eukaryota | 62994.0 | 0 | Chordata | 62994.0 | 0 | Carnivora | 62994.0 | 0 | Canidae | 62994.0 | 0 | Canis | 62994.0 | 0 | Canis lupus | 62994.0 | 0 |
| scf7180000052713 | 3222 | 0.4438 | 0 | 198.394 | no-hit | 0.0 | 0 | no-hit | 0.0 | 0 | no-hit | 0.0 | 0 | no-hit | 0.0 | 0 | no-hit | 0.0 | 0 | no-hit | 0.0 | 0 |
| scf7180000048699 | 39839 | 0.4983 | 0 | 52.5185 | Eukaryota | 40091.0 | 0 | Chordata | 40091.0 | 0 | Carnivora | 40091.0 | 0 | Canidae | 40091.0 | 0 | Canis | 40091.0 | 0 | Canis lupus | 40091.0 | 0 |
| scf7180000048340 | 57152 | 0.4625 | 0 | 134.8237 | Eukaryota | 20485.0 | 0 | Chordata | 20485.0 | 0 | Gadiformes | 11014.0 | 3 | Gadidae | 11014.0 | 3 | Gadus | 11014.0 | 4 | Gadus morhua | 11014.0 | 5 |
| scf7180000045277 | 208853 | 0.3598 | 0 | 58.9983 | Eukaryota | 24586.0 | 0 | Chordata | 24586.0 | 0 | Carnivora | 14914.0 | 2 | Canidae | 14914.0 | 2 | Canis | 14914.0 | 2 | Canis lupus | 14914.0 | 2 |
| scf7180000041072 | 3436584 | 0.4064 | 0 | 60.2278 | Eukaryota | 148807.0 | 0 | Chordata | 148807.0 | 0 | Carnivora | 148807.0 | 0 | Canidae | 148807.0 | 0 | Canis | 148807.0 | 0 | Canis lupus | 148807.0 | 0 |
| scf7180000048189 | 21270 | 0.4449 | 0 | 95.3521 | Eukaryota | 988.0 | 0 | Platyhelminthes | 514.0 | 1 | Polystomatidea | 514.0 | 2 | Polystomatidae | 514.0 | 2 | Protopolystoma | 514.0 | 2 | Protopolystoma xenopodis | 514.0 | 2 |
| scf7180000041642 | 38145 | 0.3599 | 0 | 53.8309 | Eukaryota | 12590.0 | 0 | Chordata | 12590.0 | 0 | Carnivora | 11302.0 | 1 | Canidae | 11302.0 | 1 | Canis | 11302.0 | 1 | Canis lupus | 11302.0 | 1 |
| scf7180000054422 | 19201 | 0.4418 | 0 | 296.5518 | Eukaryota | 79667.0 | 0 | Chordata | 79667.0 | 0 | Carnivora | 79667.0 | 0 | Canidae | 79667.0 | 0 | Canis | 79667.0 | 0 | Canis lupus | 79667.0 | 0 |
| scf7180000043794 | 33092 | 0.3571 | 0 | 45.8204 | Eukaryota | 6246.0 | 0 | Chordata | 6246.0 | 0 | Carnivora | 2564.0 | 2 | Canidae | 2564.0 | 2 | Canis | 2564.0 | 3 | Canis lupus | 2564.0 | 3 |
| scf7180000052413 | 11225 | 0.4422 | 0 | 1545.1849 | Eukaryota | 4705.0 | 0 | Chordata | 3224.0 | 1 | Salmoniformes | 1737.0 | 3 | Salmonidae | 1737.0 | 3 | Protopolystoma | 1481.0 | 4 | Protopolystoma xenopodis | 1481.0 | 4 |
| scf7180000046507 | 56976 | 0.3593 | 0 | 58.9825 | Eukaryota | 19186.0 | 0 | Chordata | 19186.0 | 0 | Carnivora | 19186.0 | 0 | Canidae | 19186.0 | 0 | Canis | 19186.0 | 0 | Canis lupus | 19186.0 | 0 |
| scf7180000043871 | 348489 | 0.4639 | 0 | 59.9861 | Eukaryota | 34998.0 | 0 | Chordata | 34998.0 | 0 | Carnivora | 31740.0 | 1 | Canidae | 31740.0 | 1 | Canis | 31740.0 | 1 | Canis lupus | 31740.0 | 1 |
| scf7180000051737 | 5368 | 0.4493 | 0 | 257.2367 | no-hit | 0.0 | 0 | no-hit | 0.0 | 0 | no-hit | 0.0 | 0 | no-hit | 0.0 | 0 | no-hit | 0.0 | 0 | no-hit | 0.0 | 0 |
| scf7180000051743 | 6688 | 0.4978 | 0 | 65.3074 | Eukaryota | 33949.0 | 0 | Chordata | 33949.0 | 0 | Carnivora | 33949.0 | 0 | Canidae | 33949.0 | 0 | Canis | 33949.0 | 0 | Canis lupus | 33949.0 | 0 |
| scf7180000052171 | 12559 | 0.4461 | 0 | 62.7184 | Eukaryota | 3855.0 | 0 | Chordata | 3855.0 | 0 | Tetraodontiformes | 3855.0 | 0 | Tetraodontidae | 3855.0 | 0 | Takifugu | 3855.0 | 0 | Takifugu rubripes | 3855.0 | 0 |
| scf7180000052943 | 5782 | 0.4568 | 0 | 463.0323 | Eukaryota | 15900.0 | 0 | Chordata | 13834.0 | 2 | Gadiformes | 8936.0 | 7 | Gadidae | 8936.0 | 7 | Gadus | 8936.0 | 7 | Gadus morhua | 8936.0 | 7 |
| scf7180000043232 | 13417 | 0.4397 | 0 | 79.8866 | Eukaryota | 1653.0 | 0 | Chordata | 1653.0 | 0 | Salmoniformes | 1653.0 | 0 | Salmonidae | 1653.0 | 0 | Coregonus | 1653.0 | 0 | Coregonus sp. 'balchen' | 1653.0 | 0 |
| scf7180000052997 | 4231 | 0.5079 | 0 | 43.1134 | Eukaryota | 5771.0 | 0 | Chordata | 5771.0 | 0 | Carnivora | 5771.0 | 0 | Canidae | 5771.0 | 0 | Canis | 3884.0 | 1 | Canis lupus | 2769.0 | 2 |
| scf7180000041068 | 1322570 | 0.4273 | 0 | 59.5338 | Eukaryota | 68170.0 | 0 | Chordata | 68170.0 | 0 | Carnivora | 68170.0 | 0 | Canidae | 68170.0 | 0 | Canis | 62005.0 | 1 | Canis lupus | 62005.0 | 1 |
| scf7180000040916 | 34257 | 0.3936 | 0 | 47.8589 | Eukaryota | 72731.0 | 0 | Chordata | 72731.0 | 0 | Carnivora | 72731.0 | 0 | Canidae | 72731.0 | 0 | Canis | 72731.0 | 0 | Canis lupus | 72731.0 | 0 |
| scf7180000046898 | 118360 | 0.4277 | 0 | 60.5981 | Eukaryota | 42916.0 | 0 | Chordata | 42916.0 | 0 | Carnivora | 42916.0 | 0 | Canidae | 16229.0 | 3 | Canis | 10722.0 | 6 | Canis lupus | 10722.0 | 7 |
| scf7180000041725 | 317591 | 0.3777 | 0 | 59.7511 | Eukaryota | 57053.0 | 0 | Chordata | 57053.0 | 0 | Carnivora | 57053.0 | 0 | Canidae | 57053.0 | 0 | Canis | 50242.0 | 1 | Canis lupus | 50242.0 | 1 |
| scf7180000042364 | 670454 | 0.4127 | 0 | 60.5991 | Eukaryota | 49163.0 | 0 | Chordata | 49163.0 | 0 | Carnivora | 49163.0 | 0 | Canidae | 49163.0 | 0 | Canis | 44260.0 | 1 | Canis lupus | 44260.0 | 1 |
| scf7180000043617 | 164741 | 0.4553 | 0 | 59.0889 | Eukaryota | 40094.0 | 0 | Chordata | 40094.0 | 0 | Carnivora | 40094.0 | 0 | Canidae | 40094.0 | 0 | Canis | 40094.0 | 0 | Canis lupus | 40094.0 | 0 |
| scf7180000053990 | 3231 | 0.5831 | 0 | 64.9941 | Eukaryota | 7390.0 | 0 | Chordata | 7390.0 | 0 | Primates | 3956.0 | 1 | Hominidae | 3956.0 | 3 | Homo | 3956.0 | 5 | Homo sapiens | 3956.0 | 5 |
| scf7180000042767 | 636149 | 0.3426 | 0 | 58.774 | Eukaryota | 63715.0 | 0 | Chordata | 63715.0 | 0 | Carnivora | 63715.0 | 0 | Canidae | 63715.0 | 0 | Canis | 63715.0 | 0 | Canis lupus | 63715.0 | 0 |
| scf7180000044931 | 467140 | 0.3668 | 0 | 59.3172 | Eukaryota | 21269.0 | 0 | Chordata | 21269.0 | 0 | Carnivora | 21269.0 | 0 | Canidae | 21269.0 | 0 | Canis | 21269.0 | 0 | Canis lupus | 21269.0 | 0 |
| scf7180000042298 | 730254 | 0.4064 | 0 | 61.3361 | Eukaryota | 94741.0 | 0 | Chordata | 94741.0 | 0 | Carnivora | 94741.0 | 0 | Canidae | 94741.0 | 0 | Canis | 94741.0 | 0 | Canis lupus | 94741.0 | 0 |
| scf7180000044510 | 341677 | 0.4851 | 0 | 58.4308 | Eukaryota | 52618.0 | 0 | Chordata | 52618.0 | 0 | Carnivora | 52618.0 | 0 | Canidae | 17267.0 | 5 | Canis | 11344.0 | 7 | Canis lupus | 11344.0 | 8 |
| scf7180000053250 | 3723 | 0.4432 | 0 | 72.9701 | Eukaryota | 465.0 | 0 | Platyhelminthes | 465.0 | 0 | Polystomatidea | 465.0 | 0 | Polystomatidae | 465.0 | 0 | Protopolystoma | 465.0 | 0 | Protopolystoma xenopodis | 465.0 | 0 |
| scf7180000053725 | 5870 | 0.5053 | 0 | 257.403 | Eukaryota | 7207.0 | 0 | Chordata | 7207.0 | 0 | Carnivora | 7207.0 | 0 | Canidae | 7207.0 | 0 | Canis | 5202.0 | 1 | Canis lupus | 4095.0 | 2 |
| scf7180000044601 | 31465 | 0.3728 | 0 | 55.1531 | Eukaryota | 16516.0 | 0 | Chordata | 16516.0 | 0 | Carnivora | 16516.0 | 0 | Felidae | 7077.0 | 3 | Canis | 3811.0 | 7 | Canis lupus | 3811.0 | 7 |
| scf7180000047843 | 65098 | 0.443 | 0 | 61.4629 | Eukaryota | 69753.0 | 0 | Chordata | 33972.0 | 2 | Strigeidida | 20079.0 | 8 | Schistosomatidae | 20079.0 | 8 | Schistosoma | 20079.0 | 8 | Schistosoma mansoni | 20079.0 | 8 |
| scf7180000045075 | 147851 | 0.3852 | 0 | 57.1578 | Eukaryota | 99478.0 | 0 | Chordata | 99478.0 | 0 | Carnivora | 99478.0 | 0 | Canidae | 99478.0 | 0 | Canis | 99478.0 | 0 | Canis lupus | 99478.0 | 0 |
| scf7180000042797 | 130835 | 0.3802 | 0 | 58.9839 | Eukaryota | 62785.0 | 0 | Chordata | 62785.0 | 0 | Carnivora | 62785.0 | 0 | Canidae | 62785.0 | 0 | Canis | 62785.0 | 0 | Canis lupus | 62785.0 | 0 |
| scf7180000044313 | 7646 | 0.4766 | 0 | 41.027 | Eukaryota | 11351.0 | 0 | Chordata | 11351.0 | 0 | Carnivora | 6905.0 | 2 | Canidae | 6905.0 | 2 | Canis | 6905.0 | 4 | Canis lupus | 6905.0 | 5 |
| scf7180000054664 | 247662 | 0.4397 | 0 | 194.2499 | Eukaryota | 58711.0 | 0 | Chordata | 41946.0 | 1 | Gadiformes | 37484.0 | 3 | Gadidae | 37484.0 | 3 | Gadus | 37484.0 | 3 | Gadus morhua | 37484.0 | 3 |
| scf7180000052875 | 3465 | 0.4401 | 0 | 34.6957 | no-hit | 0.0 | 0 | no-hit | 0.0 | 0 | no-hit | 0.0 | 0 | no-hit | 0.0 | 0 | no-hit | 0.0 | 0 | no-hit | 0.0 | 0 |
| scf7180000044736 | 332821 | 0.3924 | 0 | 59.102 | Eukaryota | 89351.0 | 0 | Chordata | 89351.0 | 0 | Carnivora | 89351.0 | 0 | Canidae | 89351.0 | 0 | Canis | 89351.0 | 0 | Canis lupus | 89351.0 | 0 |
| scf7180000049166 | 31897 | 0.5807 | 0 | 57.2656 | Eukaryota | 21244.0 | 0 | Chordata | 21244.0 | 0 | Carnivora | 21244.0 | 0 | Canidae | 21244.0 | 0 | Canis | 14833.0 | 1 | Canis lupus | 14833.0 | 1 |
| scf7180000043523 | 4639 | 0.3678 | 0 | 32.6745 | Eukaryota | 3326.0 | 0 | Chordata | 3326.0 | 0 | Carnivora | 3326.0 | 0 | Canidae | 3326.0 | 0 | Canis | 3326.0 | 0 | Canis lupus | 3326.0 | 0 |
| scf7180000054061 | 5972 | 0.4287 | 0 | 109.4336 | Eukaryota | 3160.0 | 0 | Platyhelminthes | 1691.0 | 2 | Polystomatidea | 1691.0 | 4 | Polystomatidae | 1691.0 | 4 | Protopolystoma | 1691.0 | 4 | Protopolystoma xenopodis | 1691.0 | 4 |
| scf7180000055783 | 30912 | 0.4364 | 0 | 487.6267 | Eukaryota | 1846.0 | 0 | Platyhelminthes | 1846.0 | 0 | Polystomatidea | 1846.0 | 0 | Polystomatidae | 1846.0 | 0 | Protopolystoma | 1846.0 | 0 | Protopolystoma xenopodis | 1846.0 | 0 |
| scf7180000055143 | 12214 | 0.4448 | 0 | 239.4777 | Eukaryota | 12208.0 | 0 | Chordata | 8395.0 | 1 | Polystomatidea | 3813.0 | 3 | Polystomatidae | 3813.0 | 3 | Protopolystoma | 3813.0 | 3 | Protopolystoma xenopodis | 3813.0 | 3 |
| scf7180000052537 | 9785 | 0.4458 | 0 | 660.4644 | no-hit | 0.0 | 0 | no-hit | 0.0 | 0 | no-hit | 0.0 | 0 | no-hit | 0.0 | 0 | no-hit | 0.0 | 0 | no-hit | 0.0 | 0 |
| scf7180000052448 | 16049 | 0.4325 | 0 | 159.7319 | Eukaryota | 6856.0 | 0 | Chordata | 5944.0 | 1 | Columbiformes | 2217.0 | 6 | Columbidae | 2217.0 | 6 | Streptopelia | 2217.0 | 6 | Streptopelia turtur | 2217.0 | 6 |
| scf7180000045790 | 23247 | 0.3961 | 0 | 49.1232 | Eukaryota | 7845.0 | 0 | Chordata | 7845.0 | 0 | Carnivora | 3097.0 | 3 | Bovidae | 2551.0 | 4 | Canis | 2095.0 | 5 | Canis lupus | 2095.0 | 5 |
| scf7180000043026 | 532397 | 0.3777 | 0 | 61.5037 | Eukaryota | 35782.0 | 0 | Chordata | 35782.0 | 0 | Carnivora | 35782.0 | 0 | Canidae | 35782.0 | 0 | Canis | 35782.0 | 0 | Canis lupus | 35782.0 | 0 |
| scf7180000048645 | 17624 | 0.4443 | 0 | 625.2969 | Eukaryota | 29304.0 | 0 | Chordata | 28531.0 | 1 | Gadiformes | 19105.0 | 6 | Gadidae | 19105.0 | 6 | Gadus | 19105.0 | 6 | Gadus morhua | 19105.0 | 6 |
| scf7180000044918 | 271584 | 0.385 | 0 | 59.0019 | Eukaryota | 88732.0 | 0 | Chordata | 88732.0 | 0 | Carnivora | 88732.0 | 0 | Canidae | 88732.0 | 0 | Canis | 88732.0 | 0 | Canis lupus | 88732.0 | 0 |
| scf7180000055629 | 16775 | 0.4488 | 0 | 1574.664 | no-hit | 0.0 | 0 | no-hit | 0.0 | 0 | no-hit | 0.0 | 0 | no-hit | 0.0 | 0 | no-hit | 0.0 | 0 | no-hit | 0.0 | 0 |
| scf7180000042556 | 584300 | 0.3962 | 0 | 58.7677 | Eukaryota | 93229.0 | 0 | Chordata | 93229.0 | 0 | Carnivora | 93229.0 | 0 | Canidae | 93229.0 | 0 | Canis | 93229.0 | 0 | Canis lupus | 93229.0 | 0 |
| scf7180000045550 | 131457 | 0.4228 | 0 | 59.066 | Eukaryota | 13770.0 | 0 | Chordata | 13770.0 | 0 | Carnivora | 13770.0 | 0 | Canidae | 13770.0 | 0 | Canis | 13770.0 | 0 | Canis lupus | 13770.0 | 0 |
| scf7180000050535 | 7231 | 0.4528 | 0 | 825.6551 | Eukaryota | 8054.0 | 0 | Chordata | 6962.0 | 1 | Tetraodontiformes | 6496.0 | 3 | Tetraodontidae | 6496.0 | 3 | Takifugu | 6496.0 | 3 | Takifugu rubripes | 6496.0 | 3 |
| scf7180000051978 | 30927 | 0.3689 | 0 | 57.7196 | Eukaryota | 16194.0 | 0 | Chordata | 16194.0 | 0 | Carnivora | 16194.0 | 0 | Canidae | 16194.0 | 0 | Canis | 16194.0 | 0 | Canis lupus | 16194.0 | 0 |
| scf7180000056355 | 52353 | 0.4248 | 0 | 58.701 | Eukaryota | 73428.0 | 0 | Chordata | 73428.0 | 0 | Carnivora | 73428.0 | 0 | Canidae | 73428.0 | 0 | Canis | 73428.0 | 0 | Canis lupus | 73428.0 | 0 |
| scf7180000054239 | 16073 | 0.4851 | 0 | 196.0094 | Eukaryota | 9330.0 | 0 | Chordata | 9330.0 | 0 | Carnivora | 9330.0 | 0 | Canidae | 9330.0 | 0 | Canis | 7286.0 | 1 | Canis lupus | 7286.0 | 1 |
| scf7180000056618 | 41355 | 0.4472 | 0 | 135.0359 | Eukaryota | 13876.0 | 0 | Chordata | 8969.0 | 2 | Tetraodontiformes | 5605.0 | 4 | Tetraodontidae | 5605.0 | 4 | Takifugu | 5605.0 | 4 | Takifugu rubripes | 5605.0 | 4 |
| scf7180000042699 | 116238 | 0.3323 | 0 | 57.3289 | Eukaryota | 20940.0 | 0 | Chordata | 20940.0 | 0 | Carnivora | 20940.0 | 0 | Canidae | 20940.0 | 0 | Canis | 20940.0 | 0 | Canis lupus | 20940.0 | 0 |
| scf7180000045797 | 19089 | 0.413 | 0 | 43.8536 | Eukaryota | 7058.0 | 0 | Chordata | 7058.0 | 0 | Carnivora | 7058.0 | 0 | Canidae | 6598.0 | 1 | Canis | 3805.0 | 2 | Canis lupus | 3805.0 | 2 |
| scf7180000043040 | 533927 | 0.4071 | 0 | 61.2761 | Eukaryota | 36210.0 | 1 | Chordata | 36210.0 | 1 | Primates | 28444.0 | 2 | Hominidae | 28444.0 | 2 | Homo | 25012.0 | 4 | Homo sapiens | 25012.0 | 4 |
| scf7180000043431 | 432508 | 0.4171 | 0 | 59.8737 | Eukaryota | 40726.0 | 0 | Chordata | 40726.0 | 0 | Carnivora | 40726.0 | 0 | Canidae | 40726.0 | 0 | Canis | 32042.0 | 1 | Canis lupus | 32042.0 | 1 |
| scf7180000053451 | 3081 | 0.4489 | 0 | 466.2556 | no-hit | 0.0 | 0 | no-hit | 0.0 | 0 | no-hit | 0.0 | 0 | no-hit | 0.0 | 0 | no-hit | 0.0 | 0 | no-hit | 0.0 | 0 |
| scf7180000047506 | 118604 | 0.4045 | 0 | 59.1757 | Eukaryota | 40635.0 | 0 | Chordata | 40635.0 | 0 | Carnivora | 40635.0 | 0 | Canidae | 40635.0 | 0 | Canis | 25531.0 | 1 | Canis lupus | 25531.0 | 1 |
| scf7180000054453 | 16540 | 0.4425 | 0 | 389.9494 | Eukaryota | 769.0 | 0 | Streptophyta | 769.0 | 0 | Fabales | 769.0 | 0 | Fabaceae | 769.0 | 0 | Vigna | 769.0 | 0 | Vigna unguiculata | 769.0 | 0 |
| scf7180000055252 | 20241 | 0.4419 | 0 | 182.5678 | Eukaryota | 8262.0 | 0 | Chordata | 6110.0 | 1 | Strigeidida | 2152.0 | 5 | Schistosomatidae | 2152.0 | 5 | Schistosoma | 2152.0 | 5 | Schistosoma mansoni | 2152.0 | 5 |
| scf7180000048693 | 11666 | 0.4834 | 0 | 81.0142 | Eukaryota | 18379.0 | 0 | Chordata | 18379.0 | 0 | Carnivora | 18379.0 | 0 | Canidae | 18379.0 | 0 | Canis | 18379.0 | 0 | Canis lupus | 18379.0 | 0 |
| scf7180000051918 | 2622 | 0.4375 | 0 | 62.9208 | no-hit | 0.0 | 0 | no-hit | 0.0 | 0 | no-hit | 0.0 | 0 | no-hit | 0.0 | 0 | no-hit | 0.0 | 0 | no-hit | 0.0 | 0 |
| scf7180000043579 | 14218 | 0.3474 | 0 | 39.5781 | Eukaryota | 21905.0 | 0 | Chordata | 21905.0 | 0 | Carnivora | 21905.0 | 0 | Otariidae | 12343.0 | 3 | Zalophus | 6631.0 | 4 | Zalophus californianus | 6631.0 | 5 |
| scf7180000052204 | 7090 | 0.4511 | 0 | 1733.8446 | no-hit | 0.0 | 0 | no-hit | 0.0 | 0 | no-hit | 0.0 | 0 | no-hit | 0.0 | 0 | no-hit | 0.0 | 0 | no-hit | 0.0 | 0 |
| scf7180000045227 | 35232 | 0.4395 | 0 | 163.216 | Eukaryota | 66831.0 | 0 | Chordata | 46902.0 | 2 | Gadiformes | 38557.0 | 5 | Gadidae | 38557.0 | 5 | Gadus | 38557.0 | 5 | Gadus morhua | 38557.0 | 5 |
| scf7180000056275 | 1102137 | 0.3799 | 0 | 60.3548 | Eukaryota | 91758.0 | 0 | Chordata | 91758.0 | 0 | Carnivora | 91758.0 | 0 | Canidae | 91758.0 | 0 | Canis | 91758.0 | 0 | Canis lupus | 91758.0 | 0 |
| scf7180000050799 | 5788 | 0.4352 | 0 | 156.949 | no-hit | 0.0 | 0 | no-hit | 0.0 | 0 | no-hit | 0.0 | 0 | no-hit | 0.0 | 0 | no-hit | 0.0 | 0 | no-hit | 0.0 | 0 |
| scf7180000048925 | 8774 | 0.4363 | 0 | 69.7108 | Eukaryota | 3433.0 | 0 | Chordata | 2369.0 | 2 | Columbiformes | 2162.0 | 3 | Columbidae | 2162.0 | 3 | Streptopelia | 2162.0 | 3 | Streptopelia turtur | 2162.0 | 3 |
| scf7180000047613 | 78314 | 0.3674 | 0 | 61.7716 | Eukaryota | 119068.0 | 0 | Chordata | 119068.0 | 0 | Carnivora | 119068.0 | 0 | Canidae | 85290.0 | 1 | Canis | 74703.0 | 2 | Canis lupus | 74703.0 | 2 |
| scf7180000056489 | 369049 | 0.5546 | 0 | 56.4175 | Eukaryota | 57758.0 | 0 | Chordata | 57758.0 | 0 | Carnivora | 57758.0 | 0 | Canidae | 57758.0 | 0 | Canis | 42616.0 | 1 | Canis lupus | 42616.0 | 1 |
| scf7180000052659 | 4078 | 0.4492 | 0 | 752.2904 | no-hit | 0.0 | 0 | no-hit | 0.0 | 0 | no-hit | 0.0 | 0 | no-hit | 0.0 | 0 | no-hit | 0.0 | 0 | no-hit | 0.0 | 0 |
| scf7180000050296 | 9908 | 0.4502 | 0 | 280.4277 | no-hit | 0.0 | 0 | no-hit | 0.0 | 0 | no-hit | 0.0 | 0 | no-hit | 0.0 | 0 | no-hit | 0.0 | 0 | no-hit | 0.0 | 0 |
| scf7180000043469 | 489805 | 0.4016 | 0 | 60.0397 | Eukaryota | 48957.0 | 0 | Chordata | 48957.0 | 0 | Carnivora | 48957.0 | 0 | Canidae | 25320.0 | 3 | Canis | 20055.0 | 4 | Canis lupus | 20055.0 | 4 |
| scf7180000044148 | 214304 | 0.4569 | 0 | 57.3015 | Eukaryota | 59153.0 | 0 | Chordata | 59153.0 | 0 | Carnivora | 59153.0 | 0 | Canidae | 59153.0 | 0 | Canis | 56139.0 | 1 | Canis lupus | 56139.0 | 1 |
| scf7180000048400 | 25641 | 0.4396 | 0 | 323.4585 | Eukaryota | 28693.0 | 0 | Chordata | 25837.0 | 1 | Gadiformes | 18397.0 | 6 | Gadidae | 18397.0 | 6 | Gadus | 18397.0 | 6 | Gadus morhua | 18397.0 | 6 |
| scf7180000054825 | 24328 | 0.4567 | 0 | 251.0139 | Eukaryota | 93482.0 | 0 | Chordata | 93482.0 | 0 | Carnivora | 93482.0 | 0 | Canidae | 93482.0 | 0 | Canis | 93482.0 | 0 | Canis lupus | 93482.0 | 0 |
| scf7180000043202 | 151875 | 0.3451 | 0 | 59.3331 | Eukaryota | 35501.0 | 0 | Chordata | 35501.0 | 0 | Carnivora | 35501.0 | 0 | Canidae | 35501.0 | 0 | Canis | 35501.0 | 0 | Canis lupus | 35501.0 | 0 |
| scf7180000052325 | 7176 | 0.4569 | 0 | 2296.8517 | Eukaryota | 3464.0 | 0 | Platyhelminthes | 1788.0 | 3 | Polystomatidea | 1788.0 | 3 | Polystomatidae | 1788.0 | 3 | Protopolystoma | 1788.0 | 3 | Protopolystoma xenopodis | 1788.0 | 3 |
| scf7180000045865 | 189151 | 0.4427 | 0 | 58.3441 | Eukaryota | 26292.0 | 0 | Chordata | 26292.0 | 0 | Carnivora | 26292.0 | 0 | Canidae | 26292.0 | 0 | Vulpes | 17412.0 | 1 | Vulpes vulpes | 17412.0 | 1 |
| scf7180000052873 | 2353 | 0.445 | 0 | 416.5921 | no-hit | 0.0 | 0 | no-hit | 0.0 | 0 | no-hit | 0.0 | 0 | no-hit | 0.0 | 0 | no-hit | 0.0 | 0 | no-hit | 0.0 | 0 |
| scf7180000045779 | 88391 | 0.4564 | 0 | 59.5854 | Eukaryota | 46035.0 | 0 | Chordata | 46035.0 | 0 | Carnivora | 46035.0 | 0 | Canidae | 46035.0 | 0 | Canis | 46035.0 | 0 | Canis lupus | 46035.0 | 0 |
| scf7180000055824 | 6013 | 0.4417 | 0 | 128.9189 | Eukaryota | 20379.0 | 0 | Chordata | 14409.0 | 2 | Gadiformes | 8692.0 | 7 | Gadidae | 8692.0 | 7 | Gadus | 8692.0 | 7 | Gadus morhua | 8692.0 | 7 |
| scf7180000044533 | 64355 | 0.4498 | 0 | 58.4889 | Eukaryota | 20668.0 | 0 | Chordata | 20668.0 | 0 | Carnivora | 18660.0 | 1 | Canidae | 4535.0 | 6 | Panthera | 4012.0 | 8 | Panthera pardus | 4012.0 | 8 |
| scf7180000047775 | 20378 | 0.4366 | 0 | 203.3999 | Eukaryota | 751.0 | 0 | Chordata | 466.0 | 1 | Kurtiformes | 466.0 | 1 | Apogonidae | 466.0 | 1 | Sphaeramia | 466.0 | 1 | Sphaeramia orbicularis | 466.0 | 1 |
| scf7180000054612 | 27293 | 0.4373 | 0 | 425.1127 | Eukaryota | 3637.0 | 0 | Chordata | 2591.0 | 2 | Kurtiformes | 1280.0 | 5 | Apogonidae | 1280.0 | 5 | Sphaeramia | 1280.0 | 5 | Sphaeramia orbicularis | 1280.0 | 5 |
| scf7180000045702 | 217247 | 0.3867 | 0 | 60.4049 | Eukaryota | 10458.0 | 0 | Chordata | 10458.0 | 0 | Carnivora | 5790.0 | 1 | Canidae | 5790.0 | 1 | Canis | 5790.0 | 2 | Canis lupus | 5790.0 | 2 |
| scf7180000042021 | 410751 | 0.384 | 0 | 57.8921 | Eukaryota | 86511.0 | 0 | Chordata | 86511.0 | 0 | Carnivora | 86511.0 | 0 | Canidae | 86511.0 | 0 | Canis | 86511.0 | 0 | Canis lupus | 86511.0 | 0 |
| scf7180000044658 | 189428 | 0.4376 | 0 | 58.7286 | Eukaryota | 16877.0 | 0 | Chordata | 16877.0 | 0 | Carnivora | 15423.0 | 1 | Canidae | 14213.0 | 2 | Canis | 14213.0 | 2 | Canis lupus | 14213.0 | 2 |
| scf7180000046085 | 154954 | 0.3798 | 0 | 59.7067 | Eukaryota | 24437.0 | 0 | Chordata | 24437.0 | 0 | Carnivora | 22477.0 | 1 | Canidae | 13611.0 | 3 | Vulpes | 9161.0 | 6 | Vulpes vulpes | 9161.0 | 7 |
| scf7180000046655 | 101647 | 0.3489 | 0 | 61.0692 | Eukaryota | 21759.0 | 0 | Chordata | 21759.0 | 0 | Carnivora | 21759.0 | 0 | Canidae | 10748.0 | 2 | Canis | 10748.0 | 3 | Canis lupus | 10748.0 | 3 |
| scf7180000056116 | 21247 | 0.4461 | 0 | 562.5973 | Eukaryota | 8314.0 | 0 | Chordata | 8314.0 | 0 | Carnivora | 8314.0 | 0 | Canidae | 8314.0 | 0 | Canis | 5780.0 | 1 | Canis lupus | 5780.0 | 1 |
| scf7180000046460 | 98834 | 0.346 | 0 | 56.9479 | Eukaryota | 17861.0 | 0 | Chordata | 17861.0 | 0 | Carnivora | 14441.0 | 1 | Felidae | 12839.0 | 3 | Felis | 12839.0 | 3 | Felis catus | 12839.0 | 3 |
| scf7180000053758 | 5643 | 0.5237 | 0 | 163.0109 | Eukaryota | 15961.0 | 0 | Chordata | 15961.0 | 0 | Carnivora | 15961.0 | 0 | Canidae | 15961.0 | 0 | Canis | 15961.0 | 0 | Canis lupus | 15961.0 | 0 |
| scf7180000041916 | 57485 | 0.4071 | 0 | 141.084 | Eukaryota | 29167.0 | 0 | Chordata | 29167.0 | 0 | Carnivora | 29167.0 | 0 | Canidae | 29167.0 | 0 | Canis | 29167.0 | 0 | Canis lupus | 29167.0 | 0 |
| scf7180000048251 | 22411 | 0.4454 | 0 | 207.9271 | Eukaryota | 16126.0 | 0 | Chordata | 9384.0 | 2 | Gadiformes | 4732.0 | 5 | Gadidae | 4732.0 | 5 | Gadus | 4732.0 | 5 | Gadus morhua | 4732.0 | 5 |
| scf7180000045161 | 6285 | 0.3933 | 0 | 22.2334 | Eukaryota | 55111.0 | 0 | Chordata | 55111.0 | 0 | Carnivora | 55111.0 | 0 | Canidae | 55111.0 | 0 | Canis | 55111.0 | 0 | Canis lupus | 55111.0 | 0 |
| scf7180000049148 | 33771 | 0.4789 | 0 | 57.1507 | Eukaryota | 3673.0 | 0 | Chordata | 3673.0 | 0 | Carnivora | 2700.0 | 1 | Canidae | 2291.0 | 2 | Canis | 2291.0 | 3 | Canis lupus | 2291.0 | 3 |
| scf7180000044051 | 90867 | 0.3904 | 0 | 58.2954 | Eukaryota | 88911.0 | 0 | Chordata | 88911.0 | 0 | Carnivora | 88911.0 | 0 | Canidae | 88911.0 | 0 | Canis | 88911.0 | 0 | Canis lupus | 88911.0 | 0 |
| scf7180000043186 | 51205 | 0.3661 | 0 | 54.6862 | Eukaryota | 87659.0 | 0 | Chordata | 87659.0 | 0 | Carnivora | 87659.0 | 0 | Canidae | 87659.0 | 0 | Canis | 87659.0 | 0 | Canis lupus | 87659.0 | 0 |
| scf7180000046804 | 49322 | 0.3776 | 0 | 54.0411 | Eukaryota | 70500.0 | 0 | Chordata | 70500.0 | 0 | Carnivora | 70500.0 | 0 | Canidae | 70500.0 | 0 | Canis | 70500.0 | 0 | Canis lupus | 70500.0 | 0 |
| scf7180000056231 | 1415294 | 0.4093 | 0 | 59.7011 | Eukaryota | 92167.0 | 0 | Chordata | 92167.0 | 0 | Carnivora | 92167.0 | 0 | Canidae | 92167.0 | 0 | Canis | 92167.0 | 0 | Canis lupus | 92167.0 | 0 |
| scf7180000046208 | 13076 | 0.4308 | 0 | 97.4325 | Eukaryota | 40862.0 | 0 | Chordata | 40862.0 | 0 | Carnivora | 40862.0 | 0 | Canidae | 40862.0 | 0 | Canis | 40862.0 | 0 | Canis lupus | 40862.0 | 0 |
| scf7180000050068 | 2862 | 0.4364 | 0 | 106.333 | no-hit | 0.0 | 0 | no-hit | 0.0 | 0 | no-hit | 0.0 | 0 | no-hit | 0.0 | 0 | no-hit | 0.0 | 0 | no-hit | 0.0 | 0 |
| scf7180000047634 | 40168 | 0.4436 | 0 | 475.4155 | Eukaryota | 94434.0 | 0 | Chordata | 94434.0 | 0 | Carnivora | 94434.0 | 0 | Canidae | 94434.0 | 0 | Canis | 94434.0 | 0 | Canis lupus | 94434.0 | 0 |
| scf7180000043179 | 487081 | 0.4461 | 0 | 59.7229 | Eukaryota | 12914.0 | 0 | Chordata | 12914.0 | 0 | Carnivora | 11320.0 | 1 | Canidae | 11320.0 | 1 | Canis | 11320.0 | 1 | Canis lupus | 11320.0 | 1 |
| scf7180000044137 | 233032 | 0.4896 | 0 | 57.024 | Eukaryota | 40998.0 | 0 | Chordata | 40998.0 | 0 | Carnivora | 40998.0 | 0 | Otariidae | 15730.0 | 4 | Canis | 8362.0 | 7 | Canis lupus | 8362.0 | 7 |
| scf7180000049474 | 34622 | 0.4564 | 0 | 59.4401 | Eukaryota | 17318.0 | 0 | Platyhelminthes | 15338.0 | 1 | Cyclophyllidea | 14198.0 | 3 | Hymenolepididae | 14198.0 | 3 | Hymenolepis | 14198.0 | 3 | Hymenolepis microstoma | 14198.0 | 3 |
| scf7180000044862 | 32572 | 0.3886 | 0 | 52.4399 | Eukaryota | 49868.0 | 0 | Chordata | 49868.0 | 0 | Carnivora | 49868.0 | 0 | Canidae | 49868.0 | 0 | Canis | 49868.0 | 0 | Canis lupus | 49868.0 | 0 |
| scf7180000044559 | 167181 | 0.3976 | 0 | 58.2475 | Eukaryota | 55891.0 | 0 | Chordata | 55891.0 | 0 | Carnivora | 55891.0 | 0 | Canidae | 55891.0 | 0 | Canis | 55891.0 | 0 | Canis lupus | 55891.0 | 0 |
| scf7180000044215 | 299561 | 0.4239 | 0 | 60.0742 | Eukaryota | 28091.0 | 0 | Chordata | 28091.0 | 0 | Carnivora | 28091.0 | 0 | Canidae | 28091.0 | 0 | Canis | 28091.0 | 0 | Canis lupus | 28091.0 | 0 |
| scf7180000043656 | 353956 | 0.4992 | 0 | 56.8864 | Eukaryota | 71805.0 | 0 | Chordata | 71805.0 | 0 | Carnivora | 71805.0 | 0 | Canidae | 71805.0 | 0 | Vulpes | 35995.0 | 1 | Vulpes vulpes | 35995.0 | 1 |
| scf7180000041393 | 262359 | 0.3523 | 0 | 59.2713 | Eukaryota | 45507.0 | 0 | Chordata | 45507.0 | 0 | Carnivora | 45507.0 | 0 | Canidae | 45507.0 | 0 | Canis | 45507.0 | 0 | Canis lupus | 45507.0 | 0 |
| scf7180000051571 | 3466 | 0.4455 | 0 | 1385.8898 | no-hit | 0.0 | 0 | no-hit | 0.0 | 0 | no-hit | 0.0 | 0 | no-hit | 0.0 | 0 | no-hit | 0.0 | 0 | no-hit | 0.0 | 0 |
| scf7180000056564 | 99325 | 0.3629 | 0 | 57.8728 | Eukaryota | 20790.0 | 0 | Chordata | 20790.0 | 0 | Carnivora | 12912.0 | 2 | Canidae | 12912.0 | 2 | Canis | 12912.0 | 2 | Canis lupus | 12912.0 | 2 |
| scf7180000049338 | 8310 | 0.4439 | 0 | 675.8779 | no-hit | 0.0 | 0 | no-hit | 0.0 | 0 | no-hit | 0.0 | 0 | no-hit | 0.0 | 0 | no-hit | 0.0 | 0 | no-hit | 0.0 | 0 |
| scf7180000046283 | 108684 | 0.4866 | 0 | 57.4826 | Eukaryota | 36529.0 | 0 | Chordata | 36529.0 | 0 | Carnivora | 36529.0 | 0 | Canidae | 36529.0 | 0 | Canis | 29067.0 | 1 | Canis lupus | 29067.0 | 1 |
| scf7180000041520 | 1442887 | 0.4454 | 0 | 60.0394 | Eukaryota | 97389.0 | 0 | Chordata | 97389.0 | 0 | Carnivora | 97389.0 | 0 | Canidae | 97389.0 | 0 | Canis | 97389.0 | 0 | Canis lupus | 97389.0 | 0 |
| scf7180000052905 | 2322 | 0.4543 | 0 | 778.3566 | no-hit | 0.0 | 0 | no-hit | 0.0 | 0 | no-hit | 0.0 | 0 | no-hit | 0.0 | 0 | no-hit | 0.0 | 0 | no-hit | 0.0 | 0 |
| scf7180000041046 | 2012842 | 0.4841 | 0 | 59.4504 | Eukaryota | 96348.0 | 0 | Chordata | 96348.0 | 0 | Carnivora | 96348.0 | 0 | Canidae | 96348.0 | 0 | Canis | 84643.0 | 1 | Canis lupus | 84643.0 | 1 |
| scf7180000041229 | 1296024 | 0.4203 | 0 | 58.3511 | Eukaryota | 81060.0 | 0 | Chordata | 81060.0 | 0 | Carnivora | 81060.0 | 0 | Canidae | 81060.0 | 0 | Canis | 81060.0 | 0 | Canis lupus | 81060.0 | 0 |
| scf7180000052149 | 8911 | 0.4514 | 0 | 190.3087 | Eukaryota | 1930.0 | 0 | Platyhelminthes | 922.0 | 2 | Polystomatidea | 922.0 | 2 | Polystomatidae | 922.0 | 2 | Protopolystoma | 922.0 | 2 | Protopolystoma xenopodis | 922.0 | 2 |
| scf7180000040605 | 2109365 | 0.3675 | 0 | 59.9415 | Eukaryota | 92342.0 | 0 | Chordata | 92342.0 | 0 | Carnivora | 92342.0 | 0 | Canidae | 92342.0 | 0 | Canis | 92342.0 | 0 | Canis lupus | 92342.0 | 0 |
| scf7180000044183 | 323293 | 0.337 | 0 | 58.6888 | Eukaryota | 11202.0 | 0 | Chordata | 11202.0 | 0 | Primates | 8111.0 | 2 | Hominidae | 8111.0 | 3 | Homo | 8111.0 | 3 | Homo sapiens | 8111.0 | 3 |
| scf7180000055957 | 6103 | 0.4445 | 0 | 126.5914 | no-hit | 0.0 | 0 | no-hit | 0.0 | 0 | no-hit | 0.0 | 0 | no-hit | 0.0 | 0 | no-hit | 0.0 | 0 | no-hit | 0.0 | 0 |
| scf7180000041817 | 22734 | 0.3618 | 0 | 59.6233 | Eukaryota | 27619.0 | 0 | Chordata | 27619.0 | 0 | Carnivora | 27619.0 | 0 | Canidae | 27619.0 | 0 | Canis | 27619.0 | 0 | Canis lupus | 27619.0 | 0 |

| | | | | | | | | | | | | | | | | | | | | | | |
|---|---|---|---|---|---|---|---|---|---|---|---|---|---|---|---|---|---|---|---|---|---|---|
| scf7180000053859 | 2977 | 0.4454 | 0 | 532.8481 | no-hit | 0.0 | 0 | no-hit | 0.0 | 0 | no-hit | 0.0 | 0 | no-hit | 0.0 | 0 | no-hit | 0.0 | 0 | no-hit | 0.0 | 0 |
| scf7180000054119 | 6907 | 0.4451 | 0 | 1513.273 | Eukaryota | 220.0 | 0 | Platyhelminthes | 220.0 | 0 | Polystomatidea | 220.0 | 0 | Polystomatidae | 220.0 | 0 | Protopolystoma | 220.0 | 0 | Protopolystoma xenopodis | 220.0 | 0 |
| scf7180000047281 | 35064 | 0.4348 | 0 | 80.9767 | Eukaryota | 57606.0 | 0 | Chordata | 41024.0 | 2 | Gadiformes | 33830.0 | 5 | Gadidae | 33830.0 | 5 | Gadus | 33830.0 | 5 | Gadus morhua | 33830.0 | 5 |
| scf7180000046819 | 105865 | 0.4899 | 0 | 60.9662 | Eukaryota | 18720.0 | 0 | Chordata | 18720.0 | 0 | Carnivora | 18720.0 | 0 | Canidae | 18720.0 | 0 | Canis | 14452.0 | 1 | Canis lupus | 14452.0 | 1 |
| scf7180000044618 | 302641 | 0.4406 | 0 | 57.9203 | Eukaryota | 68137.0 | 0 | Chordata | 68137.0 | 0 | Carnivora | 68137.0 | 0 | Canidae | 57302.0 | 1 | Canis | 35763.0 | 2 | Canis lupus | 35763.0 | 3 |
| scf7180000046769 | 88043 | 0.392 | 0 | 59.4781 | Eukaryota | 8945.0 | 0 | Chordata | 8945.0 | 0 | Carnivora | 5284.0 | 2 | Canidae | 5284.0 | 2 | Canis | 5284.0 | 3 | Canis lupus | 5284.0 | 3 |
| scf7180000048936 | 9958 | 0.518 | 0 | 296.9067 | Eukaryota | 15672.0 | 0 | Chordata | 15672.0 | 0 | Carnivora | 15672.0 | 0 | Canidae | 15672.0 | 0 | Canis | 15672.0 | 0 | Canis lupus | 15672.0 | 0 |
| scf7180000040638 | 2076666 | 0.3436 | 0 | 58.9823 | Eukaryota | 93190.0 | 0 | Chordata | 93190.0 | 0 | Carnivora | 93190.0 | 0 | Canidae | 93190.0 | 0 | Canis | 93190.0 | 0 | Canis lupus | 93190.0 | 0 |
| scf7180000048498 | 10973 | 0.396 | 0 | 41.9982 | Eukaryota | 38234.0 | 0 | Chordata | 38234.0 | 0 | Carnivora | 38234.0 | 0 | Canidae | 38234.0 | 0 | Canis | 38234.0 | 0 | Canis lupus | 38234.0 | 0 |
| scf7180000048569 | 69072 | 0.4702 | 0 | 57.744 | Eukaryota | 15073.0 | 0 | Chordata | 15073.0 | 0 | Carnivora | 15073.0 | 0 | Canidae | 15073.0 | 0 | Canis | 12009.0 | 1 | Canis lupus | 12009.0 | 1 |
| scf7180000046775 | 24559 | 0.4581 | 0 | 85.8934 | Eukaryota | 7071.0 | 0 | Chordata | 5866.0 | 1 | Tetraodontiformes | 4352.0 | 2 | Tetraodontidae | 4352.0 | 2 | Takifugu | 4352.0 | 2 | Takifugu rubripes | 4352.0 | 2 |
| scf7180000056591 | 136130 | 0.3734 | 0 | 60.4372 | Eukaryota | 68061.0 | 0 | Chordata | 68061.0 | 0 | Carnivora | 68061.0 | 0 | Canidae | 68061.0 | 0 | Canis | 68061.0 | 0 | Canis lupus | 68061.0 | 0 |
| scf7180000042761 | 509300 | 0.4488 | 0 | 59.927 | Eukaryota | 84838.0 | 0 | Chordata | 84838.0 | 0 | Carnivora | 84838.0 | 0 | Canidae | 53938.0 | 1 | Canis | 37598.0 | 3 | Canis lupus | 37598.0 | 3 |
| scf7180000044682 | 20928 | 0.4233 | 0 | 51.8372 | Eukaryota | 48952.0 | 0 | Chordata | 48952.0 | 0 | Carnivora | 48952.0 | 0 | Canidae | 48952.0 | 0 | Canis | 48952.0 | 0 | Canis lupus | 48952.0 | 0 |
| scf7180000046531 | 32337 | 0.4384 | 0 | 109.7097 | Eukaryota | 34342.0 | 0 | Chordata | 17238.0 | 2 | Gadiformes | 10510.0 | 6 | Gadidae | 10510.0 | 6 | Gadus | 10510.0 | 6 | Gadus morhua | 10510.0 | 6 |
| scf7180000041050 | 1273310 | 0.4061 | 0 | 61.0331 | Eukaryota | 56383.0 | 0 | Chordata | 56383.0 | 0 | Carnivora | 56383.0 | 0 | Canidae | 56383.0 | 0 | Canis | 32419.0 | 1 | Canis lupus | 32419.0 | 1 |
| scf7180000050973 | 6224 | 0.4393 | 0 | 258.9637 | Eukaryota | 505.0 | 0 | Chordata | 505.0 | 0 | Kurtiformes | 505.0 | 0 | Apogonidae | 505.0 | 0 | Sphaeramia | 505.0 | 0 | Sphaeramia orbicularis | 505.0 | 0 |
| scf7180000045894 | 164179 | 0.3564 | 0 | 59.2135 | Eukaryota | 92348.0 | 0 | Chordata | 92348.0 | 0 | Carnivora | 92348.0 | 0 | Canidae | 92348.0 | 0 | Canis | 92348.0 | 0 | Canis lupus | 92348.0 | 0 |
| scf7180000046717 | 128090 | 0.3748 | 0 | 60.1545 | Eukaryota | 9592.0 | 1 | Chordata | 9592.0 | 1 | Primates | 3505.0 | 5 | Hominidae | 3505.0 | 5 | Homo | 3505.0 | 6 | Homo sapiens | 3505.0 | 6 |
| scf7180000044382 | 262604 | 0.4025 | 0 | 57.8471 | Eukaryota | 39919.0 | 0 | Chordata | 39919.0 | 0 | Carnivora | 39919.0 | 0 | Canidae | 39919.0 | 0 | Canis | 24035.0 | 1 | Canis lupus | 24035.0 | 1 |
| scf7180000050621 | 13481 | 0.4469 | 0 | 2302.9322 | Eukaryota | 3200.0 | 0 | Platyhelminthes | 1853.0 | 2 | Polystomatidea | 1853.0 | 2 | Polystomatidae | 1853.0 | 2 | Protopolystoma | 1853.0 | 2 | Protopolystoma xenopodis | 1853.0 | 2 |
| scf7180000051008 | 8509 | 0.5019 | 0 | 160.4917 | Eukaryota | 8953.0 | 0 | Chordata | 8953.0 | 0 | Carnivora | 8953.0 | 0 | Canidae | 8953.0 | 0 | Canis | 7218.0 | 1 | Canis lupus | 6012.0 | 2 |
| scf7180000047305 | 72702 | 0.4305 | 0 | 59.3635 | Eukaryota | 74745.0 | 0 | Chordata | 74745.0 | 0 | Carnivora | 74745.0 | 0 | Canidae | 71291.0 | 1 | Canis | 66642.0 | 2 | Canis lupus | 66642.0 | 2 |
| scf7180000056530 | 194355 | 0.4031 | 0 | 57.122 | Eukaryota | 23587.0 | 0 | Chordata | 23587.0 | 0 | Carnivora | 23587.0 | 0 | Canidae | 23587.0 | 0 | Canis | 23587.0 | 0 | Canis lupus | 23587.0 | 0 |
| scf7180000047924 | 26983 | 0.4419 | 0 | 205.5474 | no-hit | 0.0 | 0 | no-hit | 0.0 | 0 | no-hit | 0.0 | 0 | no-hit | 0.0 | 0 | no-hit | 0.0 | 0 | no-hit | 0.0 | 0 |
| scf7180000042596 | 12722 | 0.4171 | 0 | 47.4537 | Eukaryota | 6473.0 | 0 | Chordata | 6473.0 | 0 | Carnivora | 6473.0 | 0 | Canidae | 6473.0 | 0 | Canis | 6473.0 | 0 | Canis lupus | 6473.0 | 0 |
| scf7180000045488 | 187271 | 0.4449 | 0 | 58.4692 | Eukaryota | 90015.0 | 0 | Chordata | 90015.0 | 0 | Carnivora | 90015.0 | 0 | Canidae | 90015.0 | 0 | Canis | 90015.0 | 0 | Canis lupus | 90015.0 | 0 |
| scf7180000044852 | 249230 | 0.3826 | 0 | 59.5783 | Eukaryota | 54605.0 | 0 | Chordata | 54605.0 | 0 | Carnivora | 54605.0 | 0 | Canidae | 54605.0 | 0 | Canis | 48651.0 | 1 | Canis lupus | 48651.0 | 1 |
| scf7180000043688 | 591889 | 0.3615 | 100 | 60.2173 | Eukaryota | 90964.0 | 0 | Chordata | 90964.0 | 0 | Carnivora | 90964.0 | 0 | Canidae | 90964.0 | 0 | Canis | 90964.0 | 0 | Canis lupus | 90964.0 | 0 |
| scf7180000044409 | 29816 | 0.5229 | 0 | 437.6168 | Eukaryota | 4389.0 | 0 | Chordata | 4389.0 | 0 | Carnivora | 4389.0 | 0 | Canidae | 4389.0 | 0 | Canis | 3666.0 | 1 | Canis lupus | 3666.0 | 1 |
| scf7180000042363 | 184673 | 0.3634 | 0 | 59.0205 | Eukaryota | 41321.0 | 0 | Chordata | 41321.0 | 0 | Carnivora | 41321.0 | 0 | Canidae | 41321.0 | 0 | Canis | 41321.0 | 0 | Canis lupus | 41321.0 | 0 |
| scf7180000042094 | 802641 | 0.3388 | 0 | 59.3261 | Eukaryota | 31132.0 | 0 | Chordata | 31132.0 | 0 | Carnivora | 31132.0 | 0 | Canidae | 31132.0 | 0 | Canis | 31132.0 | 0 | Canis lupus | 31132.0 | 0 |
| scf7180000042774 | 365727 | 0.4134 | 0 | 59.3046 | Eukaryota | 53695.0 | 0 | Chordata | 53695.0 | 0 | Carnivora | 53695.0 | 0 | Canidae | 53695.0 | 0 | Canis | 53695.0 | 0 | Canis lupus | 53695.0 | 0 |
| scf7180000045672 | 32685 | 0.4427 | 0 | 116.3836 | no-hit | 0.0 | 0 | no-hit | 0.0 | 0 | no-hit | 0.0 | 0 | no-hit | 0.0 | 0 | no-hit | 0.0 | 0 | no-hit | 0.0 | 0 |
| scf7180000052421 | 5019 | 0.4463 | 0 | 1581.0097 | Eukaryota | 396.0 | 0 | Chordata | 396.0 | 0 | Kurtiformes | 396.0 | 0 | Apogonidae | 396.0 | 0 | Sphaeramia | 396.0 | 0 | Sphaeramia orbicularis | 396.0 | 0 |
| scf7180000049499 | 3946 | 0.448 | 0 | 2936.0583 | no-hit | 0.0 | 0 | no-hit | 0.0 | 0 | no-hit | 0.0 | 0 | no-hit | 0.0 | 0 | no-hit | 0.0 | 0 | no-hit | 0.0 | 0 |
| scf7180000044964 | 211417 | 0.4118 | 0 | 59.5087 | Eukaryota | 75587.0 | 0 | Chordata | 75587.0 | 0 | Carnivora | 75587.0 | 0 | Canidae | 75587.0 | 0 | Canis | 66940.0 | 1 | Canis lupus | 66940.0 | 1 |
| scf7180000045818 | 201298 | 0.3747 | 0 | 59.7602 | Eukaryota | 41880.0 | 0 | Chordata | 41880.0 | 0 | Carnivora | 41880.0 | 0 | Canidae | 41880.0 | 0 | Canis | 41880.0 | 0 | Canis lupus | 41880.0 | 0 |
| scf7180000055074 | 35111 | 0.3863 | 0 | 633.9141 | Eukaryota | 65673.0 | 0 | Chordata | 65673.0 | 0 | Carnivora | 65673.0 | 0 | Canidae | 65673.0 | 0 | Canis | 65673.0 | 0 | Canis lupus | 65673.0 | 0 |
| scf7180000051831 | 5807 | 0.4491 | 0 | 1519.6337 | Eukaryota | 17093.0 | 0 | Chordata | 14333.0 | 2 | Gadiformes | 8496.0 | 6 | Gadidae | 8496.0 | 6 | Gadus | 8496.0 | 6 | Gadus morhua | 8496.0 | 6 |
| scf7180000040543 | 447334 | 0.3622 | 0 | 58.7773 | Eukaryota | 21494.0 | 0 | Chordata | 21494.0 | 0 | Carnivora | 21494.0 | 0 | Canidae | 21494.0 | 0 | Canis | 21494.0 | 0 | Canis lupus | 21494.0 | 0 |
| scf7180000050346 | 22556 | 0.3535 | 0 | 61.1321 | Eukaryota | 5883.0 | 0 | Chordata | 5883.0 | 0 | Primates | 2950.0 | 1 | Hominidae | 2512.0 | 5 | Homo | 2512.0 | 6 | Homo sapiens | 2512.0 | 6 |
| scf7180000043691 | 419789 | 0.3538 | 0 | 59.7153 | Eukaryota | 54268.0 | 0 | Chordata | 54268.0 | 0 | Carnivora | 54268.0 | 0 | Canidae | 54268.0 | 0 | Canis | 54268.0 | 0 | Canis lupus | 54268.0 | 0 |
| scf7180000053838 | 3860 | 0.4383 | 0 | 77.2227 | no-hit | 0.0 | 0 | no-hit | 0.0 | 0 | no-hit | 0.0 | 0 | no-hit | 0.0 | 0 | no-hit | 0.0 | 0 | no-hit | 0.0 | 0 |
| scf7180000043002 | 531856 | 0.4355 | 0 | 60.8921 | Eukaryota | 42160.0 | 0 | Chordata | 42160.0 | 0 | Carnivora | 42160.0 | 0 | Canidae | 34646.0 | 1 | Canis | 31074.0 | 3 | Canis lupus | 31074.0 | 3 |
| scf7180000042709 | 558650 | 0.3982 | 0 | 58.6251 | Eukaryota | 92114.0 | 0 | Chordata | 92114.0 | 0 | Carnivora | 92114.0 | 0 | Canidae | 92114.0 | 0 | Canis | 92114.0 | 0 | Canis lupus | 92114.0 | 0 |
