## Supplementary material for "High-quality carnivore genomes from roadkill samples enable species delimitation in aardwolf and bat-eared fox": Table S8

| Contigs name | length | GC | N | bam0 | superkingdom.t. | superkingdom.s. | superkingdom.c. | phylum.t. | phylum.s. | phylum.c. | order.t. | order.s. | order.c. | family.t. | family.s. | family.c. | genus.t. | genus.s. | genus.c | species.t. | species.s. | species.c. |
|---|---|---|---|---|---|---|---|---|---|---|---|---|---|---|---|---|---|---|---|---|---|---|
| scf7180000019764 | 243696 | 0.549 | 0 | 26.627 | Eukaryota | 86243.0 | 0 | Chordata | 86243.0 | 0 | Carnivora | 63847.0 | 2 | Felidae | 40403.0 | 5 | Myotis | 14178.0 | 7 | yotis lucifugu | 14178.0 | 7 |
| scf7180000018798 | 577059 | 0.3806 | 0 | 51.8235 | Eukaryota | 36159.0 | 0 | Chordata | 36159.0 | 0 | Carnivora | 36159.0 | 0 | Felidae | 36159.0 | 0 | Felis | 32511.0 | 1 | Felis catus | 32511.0 | 1 |
| scf7180000018868 | 2190210 | 0.4599 | 0 | 52.2105 | Eukaryota | 70011.0 | 0 | Chordata | 70011.0 | 0 | Carnivora | 70011.0 | 0 | Felidae | 70011.0 | 0 | Panthera | 35446.0 | 3 | Panthera tigri | 20908.0 | 4 |
| scf7180000020080 | 191413 | 0.3863 | 0 | 50.2408 | Eukaryota | 22926.0 | 0 | Chordata | 22926.0 | 0 | Carnivora | 22926.0 | 0 | Felidae | 22926.0 | 0 | Felis | 20606.0 | 1 | Felis catus | 20606.0 | 1 |
| scf7180000018202 | 1533268 | 0.4004 | 0 | 51.1355 | Eukaryota | 87671.0 | 0 | Chordata | 87671.0 | 0 | Chiroptera | 77540.0 | 1 | Pteropodidae | 64576.0 | 2 | Pteropus | 64576.0 | 2 | ropus vampy | 64576.0 | 2 |
| scf7180000019289 | 1200804 | 0.4881 | 0 | 51.7095 | Eukaryota | 48033.0 | 0 | Chordata | 48033.0 | 0 | Carnivora | 48033.0 | 0 | Felidae | 48033.0 | 0 | Felis | 48033.0 | 0 | Felis catus | 48033.0 | 0 |
| scf7180000018426 | 747129 | 0.3934 | 0 | 50.2791 | Eukaryota | 47238.0 | 0 | Chordata | 47238.0 | 0 | Carnivora | 47238.0 | 0 | Felidae | 47238.0 | 0 | Felis | 47238.0 | 0 | Felis catus | 47238.0 | 0 |
| scf7180000018222 | 1146792 | 0.4271 | 0 | 51.3568 | Eukaryota | 47154.0 | 0 | Chordata | 47154.0 | 0 | Carnivora | 47154.0 | 0 | Felidae | 47154.0 | 0 | Felis | 47154.0 | 0 | Felis catus | 47154.0 | 0 |
| scf7180000022685 | 19858 | 0.3658 | 0 | 24.0854 | Eukaryota | 16434.0 | 0 | Chordata | 16434.0 | 0 | Carnivora | 16434.0 | 0 | Felidae | 16434.0 | 0 | Felis | 13135.0 | 1 | Felis catus | 13135.0 | 1 |
| scf7180000023524 | 16013 | 0.4635 | 0 | 264.7461 | no-hit | 0.0 | 0 | no-hit | 0.0 | 0 | no-hit | 0.0 | 0 | no-hit | 0.0 | 0 | no-hit | 0.0 | 0 | no-hit | 0.0 | 0 |
| scf7180000018077 | 1220306 | 0.4075 | 0 | 51.7414 | Eukaryota | 45867.0 | 0 | Chordata | 45867.0 | 0 | Carnivora | 45867.0 | 0 | Felidae | 45867.0 | 0 | Felis | 45867.0 | 0 | Felis catus | 45867.0 | 0 |
| scf7180000019404 | 26794 | 0.3691 | 0 | 40.6337 | Eukaryota | 45503.0 | 0 | Chordata | 45503.0 | 0 | Carnivora | 45503.0 | 0 | Felidae | 45503.0 | 0 | Felis | 45503.0 | 0 | Felis catus | 45503.0 | 0 |
| scf7180000024076 | 270767 | 0.5343 | 0 | 48.3743 | Eukaryota | 39635.0 | 0 | Chordata | 39635.0 | 0 | Carnivora | 39635.0 | 0 | Felidae | 35683.0 | 1 | Felis | 12831.0 | 3 | Felis catus | 12831.0 | 4 |
| scf7180000017716 | 1827474 | 0.4385 | 0 | 54.8365 | Eukaryota | 46722.0 | 0 | Chordata | 46722.0 | 0 | Carnivora | 46722.0 | 0 | Felidae | 41994.0 | 1 | Felis | 41994.0 | 1 | Felis catus | 41994.0 | 1 |
| scf7180000021069 | 77262 | 0.5219 | 0 | 25.7213 | Eukaryota | 25577.0 | 0 | Chordata | 25577.0 | 0 | Carnivora | 25577.0 | 0 | Felidae | 23335.0 | 1 | Felis | 10880.0 | 3 | Felis catus | 10880.0 | 3 |
| scf7180000020487 | 86335 | 0.3782 | 0 | 24.8687 | Eukaryota | 41644.0 | 0 | Chordata | 41644.0 | 0 | Carnivora | 41644.0 | 0 | Felidae | 41644.0 | 0 | Felis | 41644.0 | 0 | Felis catus | 41644.0 | 0 |
| scf7180000019981 | 190780 | 0.502 | 0 | 48.4149 | Eukaryota | 30442.0 | 0 | Chordata | 30442.0 | 0 | Carnivora | 30442.0 | 0 | Felidae | 30442.0 | 0 | Felis | 27317.0 | 1 | Felis catus | 27317.0 | 1 |
| scf7180000017734 | 2171113 | 0.4033 | 0 | 149.4 | Eukaryota | 47744.0 | 0 | Chordata | 47744.0 | 0 | Carnivora | 47744.0 | 0 | Felidae | 47744.0 | 0 | Felis | 47744.0 | 0 | Felis catus | 47744.0 | 0 |
| scf7180000024217 | 40114 | 0.4147 | 0 | 25.8054 | Eukaryota | 13864.0 | 0 | Chordata | 13864.0 | 0 | Carnivora | 9364.0 | 1 | Felidae | 5014.0 | 2 | Homo | 4500.0 | 5 | Homo sapiens | 4500.0 | 5 |
| scf7180000023617 | 36846 | 0.4314 | 0 | 219.0538 | Eukaryota | 25965.0 | 0 | Chordata | 25965.0 | 0 | Carnivora | 25965.0 | 0 | Felidae | 25965.0 | 0 | Felis | 23340.0 | 1 | Felis catus | 23340.0 | 1 |
| scf7180000018291 | 49098 | 0.4353 | 0 | 23.0116 | Eukaryota | 8791.0 | 0 | Chordata | 8791.0 | 0 | Carnivora | 8791.0 | 0 | Felidae | 8791.0 | 0 | Felis | 8791.0 | 0 | Felis catus | 8791.0 | 0 |
| scf7180000023401 | 21449 | 0.5084 | 0 | 444.7426 | no-hit | 0.0 | 0 | no-hit | 0.0 | 0 | no-hit | 0.0 | 0 | no-hit | 0.0 | 0 | no-hit | 0.0 | 0 | no-hit | 0.0 | 0 |
| scf7180000022724 | 753140 | 0.381 | 0 | 49.9012 | Eukaryota | 77751.0 | 0 | Chordata | 77751.0 | 0 | Carnivora | 77751.0 | 0 | Felidae | 77751.0 | 0 | Felis | 31723.0 | 3 | Felis catus | 31723.0 | 3 |
| scf7180000022010 | 17094 | 0.6245 | 0 | 43.4068 | Eukaryota | 3077.0 | 0 | Chordata | 3077.0 | 0 | Carnivora | 3077.0 | 0 | Felidae | 3077.0 | 0 | Neofelis | 1052.0 | 3 | eofelis nebulo | 1052.0 | 3 |
| scf7180000019399 | 459695 | 0.525 | 0 | 50.6346 | Eukaryota | 15148.0 | 0 | Chordata | 15148.0 | 0 | Carnivora | 15148.0 | 0 | Felidae | 15148.0 | 0 | Felis | 15148.0 | 0 | Felis catus | 15148.0 | 0 |
| scf7180000021504 | 34952 | 0.4437 | 0 | 24.1287 | Eukaryota | 3031.0 | 0 | Chordata | 3031.0 | 0 | Carnivora | 2029.0 | 2 | Felidae | 1775.0 | 3 | Felis | 1529.0 | 4 | Felis catus | 1529.0 | 4 |
| scf7180000017553 | 3454165 | 0.4388 | 100 | 50.8685 | Eukaryota | 59366.0 | 0 | Chordata | 59366.0 | 0 | Carnivora | 59366.0 | 0 | Otariidae | 31099.0 | 2 | Zalophus | 19799.0 | 3 | phus californi | 19799.0 | 3 |
| scf7180000020837 | 493940 | 0.4851 | 0 | 50.8566 | Eukaryota | 45361.0 | 0 | Chordata | 45361.0 | 0 | Carnivora | 45361.0 | 0 | Felidae | 45361.0 | 0 | Felis | 45361.0 | 0 | Felis catus | 45361.0 | 0 |
| scf7180000023964 | 424273 | 0.4021 | 0 | 50.0544 | Eukaryota | 31409.0 | 0 | Chordata | 31409.0 | 0 | Carnivora | 31409.0 | 0 | Felidae | 31409.0 | 0 | Felis | 28212.0 | 1 | Felis catus | 28212.0 | 1 |
| scf7180000017801 | 157278 | 0.4087 | 0 | 50.0343 | Eukaryota | 24293.0 | 0 | Chordata | 24293.0 | 0 | Carnivora | 24293.0 | 0 | Ursidae | 11785.0 | 3 | Ursus | 9428.0 | 4 | Enhydra lutris | 6972.0 | 5 |
| scf7180000018883 | 1766658 | 0.355 | 0 | 49.6766 | Eukaryota | 45895.0 | 0 | Chordata | 45895.0 | 0 | Carnivora | 45895.0 | 0 | Felidae | 45895.0 | 0 | Felis | 45895.0 | 0 | Felis catus | 45895.0 | 0 |
| scf7180000017950 | 6746 | 0.4363 | 0 | 31.0747 | Eukaryota | 6061.0 | 0 | Chordata | 6061.0 | 0 | Carnivora | 6061.0 | 0 | Felidae | 6061.0 | 0 | Panthera | 2630.0 | 3 | anthera pardu | 2630.0 | 3 |
| scf7180000019486 | 36037 | 0.4013 | 0 | 26.2249 | Eukaryota | 18183.0 | 0 | Chordata | 18183.0 | 0 | Carnivora | 18183.0 | 0 | Felidae | 18183.0 | 0 | Felis | 18183.0 | 0 | Felis catus | 18183.0 | 0 |
| scf7180000022089 | 13796 | 0.3723 | 0 | 23.1244 | Eukaryota | 22813.0 | 0 | Chordata | 22813.0 | 0 | Carnivora | 22813.0 | 0 | Felidae | 22813.0 | 0 | Felis | 20478.0 | 1 | Felis catus | 20478.0 | 1 |
| scf7180000021905 | 22830 | 0.378 | 0 | 139.4408 | Eukaryota | 7224.0 | 0 | Chordata | 7224.0 | 0 | Artiodactyla | 7224.0 | 0 | Bovidae | 7224.0 | 0 | Ovis | 7224.0 | 0 | vis canadens | 7224.0 | 0 |
| scf7180000020293 | 95889 | 0.5046 | 0 | 51.0563 | Eukaryota | 11552.0 | 0 | Chordata | 11552.0 | 0 | Carnivora | 11552.0 | 0 | Felidae | 3802.0 | 4 | Panthera | 2782.0 | 7 | anthera pardu | 2782.0 | 8 |
| scf7180000021729 | 18284 | 0.6844 | 0 | 37.1841 | Eukaryota | 4293.0 | 0 | Chordata | 4293.0 | 0 | Carnivora | 2328.0 | 2 | Felidae | 2328.0 | 2 | Panthera | 2328.0 | 3 | anthera pardu | 2328.0 | 3 |
| scf7180000018056 | 1248890 | 0.4313 | 0 | 52.3533 | Eukaryota | 110664.0 | 0 | Chordata | 110664.0 | 0 | Carnivora | 110664.0 | 0 | Herpestidae | 88648.0 | 1 | Suricata | 88648.0 | 1 | ricata suricat | 88648.0 | 1 |
| scf7180000018589 | 8497 | 0.6368 | 0 | 23.2147 | Eukaryota | 2810.0 | 0 | Chordata | 2810.0 | 0 | Carnivora | 2810.0 | 0 | Felidae | 2810.0 | 0 | Acinonyx | 1686.0 | 2 | cinonyx jubat | 1686.0 | 2 |
| scf7180000023405 | 49458 | 0.4252 | 0 | 447.1669 | Eukaryota | 257405.0 | 0 | Chordata | 257405.0 | 0 | Carnivora | 257405.0 | 0 | Hyaenidae | 257405.0 | 0 | Proteles | 155760.0 | 1 | roteles cristat | 155760.0 | 1 |
| scf7180000024056 | 403309 | 0.4882 | 0 | 50.0853 | Eukaryota | 49237.0 | 0 | Chordata | 49237.0 | 0 | Carnivora | 49237.0 | 0 | Felidae | 30317.0 | 1 | Suricata | 18920.0 | 4 | ricata suricat | 18920.0 | 4 |
| scf7180000022064 | 494461 | 0.6029 | 0 | 56.5757 | Eukaryota | 37657.0 | 0 | Chordata | 37657.0 | 0 | Carnivora | 37657.0 | 0 | Herpestidae | 14379.0 | 3 | Suricata | 14379.0 | 3 | ricata suricat | 14379.0 | 3 |
| scf7180000020286 | 179851 | 0.5497 | 0 | 48.9308 | Eukaryota | 29598.0 | 0 | Chordata | 29598.0 | 0 | Carnivora | 29598.0 | 0 | Felidae | 17720.0 | 3 | Panthera | 7754.0 | 7 | anthera pardu | 7754.0 | 7 |
| scf7180000017567 | 3007162 | 0.4136 | 0 | 51.7504 | Eukaryota | 68649.0 | 0 | Chordata | 68649.0 | 0 | Carnivora | 68649.0 | 0 | Otariidae | 48204.0 | 1 | Zalophus | 34316.0 | 2 | phus californi | 34316.0 | 2 |
| scf7180000021066 | 89970 | 0.4061 | 0 | 27.0949 | Eukaryota | 25058.0 | 0 | Chordata | 25058.0 | 0 | Carnivora | 25058.0 | 0 | Felidae | 25058.0 | 0 | Felis | 22535.0 | 1 | Felis catus | 22535.0 | 1 |
| scf7180000017564 | 2626601 | 0.3964 | 0 | 51.5511 | Eukaryota | 46831.0 | 0 | Chordata | 46831.0 | 0 | Carnivora | 46831.0 | 0 | Felidae | 46831.0 | 0 | Felis | 42367.0 | 1 | Felis catus | 42367.0 | 1 |
| scf7180000018292 | 831567 | 0.3757 | 0 | 50.0144 | Eukaryota | 47588.0 | 0 | Chordata | 47588.0 | 0 | Carnivora | 47588.0 | 0 | Felidae | 33293.0 | 1 | Panthera | 19016.0 | 2 | ricata suricat | 14295.0 | 3 |
| scf7180000022488 | 25035 | 0.4088 | 0 | 641.2338 | no-hit | 0.0 | 0 | no-hit | 0.0 | 0 | no-hit | 0.0 | 0 | no-hit | 0.0 | 0 | no-hit | 0.0 | 0 | no-hit | 0.0 | 0 |
| scf7180000018584 | 633984 | 0.5653 | 0 | 60.3949 | Eukaryota | 24751.0 | 0 | Chordata | 24751.0 | 0 | Carnivora | 24751.0 | 0 | Felidae | 24751.0 | 0 | Felis | 22244.0 | 1 | Felis catus | 22244.0 | 1 |
| scf7180000020950 | 121823 | 0.3504 | 0 | 25.1808 | Eukaryota | 35319.0 | 0 | Chordata | 35319.0 | 0 | Carnivora | 35319.0 | 0 | Felidae | 35319.0 | 0 | Felis | 35319.0 | 0 | Felis catus | 35319.0 | 0 |
| scf7180000024095 | 99731 | 0.3798 | 0 | 45.0903 | Eukaryota | 27732.0 | 0 | Chordata | 27732.0 | 0 | Carnivora | 27732.0 | 0 | Felidae | 10855.0 | 4 | Enhydra | 5718.0 | 8 | Enhydra lutris | 5718.0 | 8 |
| scf7180000022926 | 56260 | 0.3883 | 0 | 389.4935 | Eukaryota | 22508.0 | 0 | Chordata | 22508.0 | 0 | Carnivora | 22508.0 | 0 | Felidae | 22508.0 | 0 | Felis | 20232.0 | 1 | Felis catus | 20232.0 | 1 |
| scf7180000018820 | 710686 | 0.4333 | 0 | 52.2269 | Eukaryota | 44836.0 | 0 | Chordata | 44836.0 | 0 | Carnivora | 44836.0 | 0 | Felidae | 44836.0 | 0 | Felis | 44836.0 | 0 | Felis catus | 44836.0 | 0 |
| scf7180000024169 | 201225 | 0.583 | 0 | 49.4164 | Eukaryota | 15963.0 | 0 | Chordata | 15963.0 | 0 | Carnivora | 15963.0 | 0 | Felidae | 15963.0 | 0 | Lynx | 6291.0 | 2 | ynx canadens | 6291.0 | 2 |
| scf7180000018857 | 615695 | 0.399 | 0 | 50.8342 | Eukaryota | 46871.0 | 0 | Chordata | 46871.0 | 0 | Carnivora | 46871.0 | 0 | Felidae | 46871.0 | 0 | Felis | 46871.0 | 0 | Felis catus | 46871.0 | 0 |
| scf7180000017887 | 564532 | 0.4542 | 0 | 54.4798 | Eukaryota | 47046.0 | 0 | Chordata | 47046.0 | 0 | Carnivora | 47046.0 | 0 | Felidae | 47046.0 | 0 | Felis | 41866.0 | 1 | Felis catus | 41866.0 | 1 |
| scf7180000020307 | 195061 | 0.4904 | 0 | 50.7788 | Eukaryota | 8314.0 | 0 | Chordata | 8314.0 | 0 | Primates | 4470.0 | 3 | Hominidae | 4470.0 | 3 | Homo | 3640.0 | 6 | Homo sapiens | 3640.0 | 6 |
| scf7180000020748 | 27449 | 0.4235 | 0 | 236.79 | Eukaryota | 10456.0 | 0 | Chordata | 10456.0 | 0 | Carnivora | 10456.0 | 0 | Felidae | 10456.0 | 0 | Felis | 9408.0 | 1 | Felis catus | 9408.0 | 1 |
| scf7180000021684 | 19970 | 0.3918 | 0 | 26.0572 | Eukaryota | 3333.0 | 0 | Chordata | 3333.0 | 0 | Carnivora | 2111.0 | 2 | Felidae | 2111.0 | 2 | Felis | 1602.0 | 3 | Felis catus | 1602.0 | 3 |
| scf7180000020317 | 124926 | 0.5093 | 0 | 53.8853 | Eukaryota | 9791.0 | 0 | Chordata | 9791.0 | 0 | Carnivora | 9791.0 | 0 | Otariidae | 2888.0 | 4 | Suricata | 2025.0 | 7 | ricata suricat | 2025.0 | 7 |
| scf7180000023341 | 32699 | 0.3971 | 0 | 298.9623 | Eukaryota | 6641.0 | 0 | Chordata | 6641.0 | 0 | Carnivora | 6641.0 | 0 | Felidae | 6641.0 | 0 | Felis | 6641.0 | 0 | Felis catus | 6641.0 | 0 |
| scf7180000022500 | 12180 | 0.4664 | 0 | 26.355 | Eukaryota | 1999.0 | 0 | Chordata | 1999.0 | 0 | Carnivora | 1999.0 | 0 | Felidae | 1795.0 | 1 | Felis | 1373.0 | 2 | Felis catus | 1373.0 | 2 |
| scf7180000023646 | 31873 | 0.4908 | 0 | 153.0658 | no-hit | 0.0 | 0 | no-hit | 0.0 | 0 | no-hit | 0.0 | 0 | no-hit | 0.0 | 0 | no-hit | 0.0 | 0 | no-hit | 0.0 | 0 |
| scf7180000018442 | 324316 | 0.4153 | 0 | 47.0553 | Eukaryota | 32568.0 | 0 | Chordata | 32568.0 | 0 | Carnivora | 32568.0 | 0 | Felidae | 32568.0 | 0 | Felis | 29269.0 | 1 | Felis catus | 29269.0 | 1 |
| scf7180000021325 | 15783 | 0.4609 | 0 | 108.8593 | Eukaryota | 1999.0 | 0 | Chordata | 1999.0 | 0 | Carnivora | 1999.0 | 0 | Felidae | 1999.0 | 0 | Felis | 1786.0 | 1 | Felis catus | 1786.0 | 1 |
| scf7180000021446 | 38346 | 0.5913 | 0 | 53.303 | Eukaryota | 4919.0 | 0 | Chordata | 4919.0 | 0 | Carnivora | 4438.0 | 1 | Felidae | 1501.0 | 5 | Eumetopias | 1443.0 | 7 | netopias juba | 1443.0 | 7 |
| scf7180000017488 | 2918418 | 0.4262 | 0 | 50.2882 | Eukaryota | 46542.0 | 0 | Chordata | 46542.0 | 0 | Carnivora | 46542.0 | 0 | Felidae | 46542.0 | 0 | Felis | 46542.0 | 0 | Felis catus | 46542.0 | 0 |
| scf7180000022444 | 8275 | 0.3768 | 0 | 46.359 | Eukaryota | 2838.0 | 0 | Chordata | 2838.0 | 0 | Artiodactyla | 2168.0 | 1 | Suidae | 2168.0 | 2 | Sus | 2168.0 | 3 | Sus scrofa | 2168.0 | 3 |
| scf7180000023790 | 288043 | 0.4508 | 0 | 51.168 | Eukaryota | 23615.0 | 0 | Chordata | 23615.0 | 0 | Carnivora | 23615.0 | 0 | Felidae | 21040.0 | 1 | Panthera | 10309.0 | 3 | cinonyx jubat | 6405.0 | 4 |
| scf7180000021443 | 17030 | 0.6402 | 0 | 100.8082 | Eukaryota | 2278.0 | 0 | Chordata | 2278.0 | 0 | Carnivora | 2278.0 | 0 | Felidae | 2061.0 | 1 | Felis | 1818.0 | 2 | Felis catus | 1818.0 | 2 |
| scf7180000020030 | 143999 | 0.3678 | 0 | 26.8141 | Eukaryota | 29525.0 | 0 | Chordata | 29525.0 | 0 | Carnivora | 29525.0 | 0 | Felidae | 29525.0 | 0 | Felis | 26592.0 | 1 | Felis catus | 26592.0 | 1 |
| scf7180000019700 | 282420 | 0.3532 | 0 | 25.5411 | Eukaryota | 13650.0 | 0 | Chordata | 13650.0 | 0 | Carnivora | 6797.0 | 2 | Felidae | 6797.0 | 2 | Felis | 5685.0 | 4 | Felis catus | 5685.0 | 4 |
| scf7180000018297 | 697474 | 0.4384 | 0 | 49.2584 | Eukaryota | 25341.0 | 0 | Chordata | 25341.0 | 0 | Carnivora | 16561.0 | 2 | Felidae | 16561.0 | 2 | Panthera | 7093.0 | 6 | Mus musculus | 5583.0 | 7 |
| scf7180000019929 | 41564 | 0.414 | 0 | 41.7716 | Eukaryota | 17263.0 | 0 | Chordata | 17263.0 | 0 | Carnivora | 17263.0 | 0 | Felidae | 17263.0 | 0 | Felis | 15532.0 | 1 | Felis catus | 15532.0 | 1 |
| scf7180000018896 | 527229 | 0.3771 | 0 | 49.0611 | Eukaryota | 31042.0 | 0 | Chordata | 31042.0 | 0 | Carnivora | 31042.0 | 0 | Felidae | 31042.0 | 0 | Felis | 27850.0 | 1 | Felis catus | 27850.0 | 1 |
| scf7180000020326 | 76052 | 0.4843 | 0 | 25.6879 | Eukaryota | 4194.0 | 0 | Chordata | 4194.0 | 0 | Carnivora | 3916.0 | 1 | Felidae | 3302.0 | 2 | Lynx | 1756.0 | 4 | ynx canadens | 1756.0 | 4 |
| scf7180000018979 | 466745 | 0.4018 | 0 | 51.2276 | Eukaryota | 44735.0 | 0 | Chordata | 44735.0 | 0 | Carnivora | 44735.0 | 0 | Felidae | 44735.0 | 0 | Felis | 44735.0 | 0 | Felis catus | 44735.0 | 0 |
| scf7180000017679 | 2648614 | 0.4787 | 0 | 51.6572 | Eukaryota | 56062.0 | 0 | Chordata | 56062.0 | 0 | Carnivora | 56062.0 | 0 | Felidae | 56062.0 | 0 | Panthera | 25255.0 | 2 | anthera pardu | 25255.0 | 2 |
| scf7180000021860 | 68107 | 0.3768 | 0 | 50.4065 | Eukaryota | 45811.0 | 0 | Chordata | 45811.0 | 0 | Carnivora | 45811.0 | 0 | Felidae | 45811.0 | 0 | Felis | 41321.0 | 1 | Felis catus | 41321.0 | 1 |
| scf7180000019127 | 103679 | 0.3736 | 0 | 47.6642 | Eukaryota | 24996.0 | 0 | Chordata | 24996.0 | 0 | Carnivora | 24996.0 | 0 | Felidae | 24996.0 | 0 | Felis | 22477.0 | 1 | Felis catus | 22477.0 | 1 |
| scf7180000020731 | 126174 | 0.4113 | 0 | 26.07 | Eukaryota | 16383.0 | 0 | Chordata | 16383.0 | 0 | Carnivora | 16383.0 | 0 | Ursidae | 4597.0 | 4 | Enhydra | 3085.0 | 7 | Enhydra lutris | 3085.0 | 8 |
| scf7180000021171 | 57804 | 0.3913 | 100 | 23.3069 | Eukaryota | 35121.0 | 0 | Chordata | 35121.0 | 0 | Carnivora | 35121.0 | 0 | Felidae | 35121.0 | 0 | Felis | 35121.0 | 0 | Felis catus | 35121.0 | 0 |
| scf7180000022458 | 36566 | 0.3755 | 100 | 25.3972 | Eukaryota | 30152.0 | 0 | Chordata | 30152.0 | 0 | Carnivora | 30152.0 | 0 | Felidae | 30152.0 | 0 | Felis | 27193.0 | 1 | Felis catus | 27193.0 | 1 |
| scf7180000024208 | 60126 | 0.469 | 0 | 39.354 | Eukaryota | 28534.0 | 0 | Chordata | 28534.0 | 0 | Carnivora | 28534.0 | 0 | Felidae | 28534.0 | 0 | Felis | 25680.0 | 1 | Felis catus | 25680.0 | 1 |
| scf7180000020295 | 129833 | 0.3967 | 0 | 25.789 | Eukaryota | 40911.0 | 0 | Chordata | 40911.0 | 0 | Carnivora | 40911.0 | 0 | Otariidae | 11619.0 | 4 | Zalophus | 11619.0 | 4 | phus californi | 11619.0 | 4 |
| scf7180000020213 | 181269 | 0.4538 | 0 | 25.6701 | Eukaryota | 70658.0 | 0 | Chordata | 70658.0 | 0 | Carnivora | 70658.0 | 0 | Otariidae | 32275.0 | 2 | Zalophus | 19365.0 | 6 | phus californi | 19365.0 | 6 |
| scf7180000023980 | 564333 | 0.4101 | 0 | 49.4376 | Eukaryota | 77180.0 | 0 | Chordata | 77180.0 | 0 | Carnivora | 77180.0 | 0 | Felidae | 77180.0 | 0 | Felis | 30944.0 | 2 | Felis catus | 30944.0 | 3 |
| scf7180000019219 | 96989 | 0.4 | 0 | 46.8639 | Eukaryota | 19985.0 | 0 | Chordata | 19985.0 | 0 | Carnivora | 19985.0 | 0 | Felidae | 16822.0 | 1 | Felis | 11387.0 | 5 | Felis catus | 11387.0 | 5 |
| scf7180000024164 | 162693 | 0.3931 | 0 | 26.3603 | Eukaryota | 44423.0 | 0 | Chordata | 44423.0 | 0 | Carnivora | 44423.0 | 0 | Felidae | 44423.0 | 0 | Felis | 44423.0 | 0 | Felis catus | 44423.0 | 0 |
| scf7180000022321 | 12749 | 0.3629 | 0 | 20.607 | Eukaryota | 38815.0 | 0 | Chordata | 38815.0 | 0 | Carnivora | 38815.0 | 0 | Felidae | 38815.0 | 0 | Felis | 38815.0 | 0 | Felis catus | 38815.0 | 0 |
| scf7180000018786 | 14867 | 0.3721 | 0 | 20.5679 | Eukaryota | 42283.0 | 0 | Chordata | 42283.0 | 0 | Carnivora | 42283.0 | 0 | Felidae | 42283.0 | 0 | Felis | 42283.0 | 0 | Felis catus | 42283.0 | 0 |
| scf7180000021799 | 43634 | 0.4228 | 0 | 53.7462 | Eukaryota | 37191.0 | 0 | Chordata | 37191.0 | 0 | Carnivora | 37191.0 | 0 | Felidae | 28743.0 | 2 | Felis | 18015.0 | 3 | Felis catus | 18015.0 | 3 |
| scf7180000021686 | 46452 | 0.4554 | 100 | 46.0453 | Eukaryota | 13718.0 | 0 | Chordata | 13718.0 | 0 | Carnivora | 13718.0 | 0 | Felidae | 13718.0 | 0 | Felis | 12340.0 | 1 | Felis catus | 12340.0 | 1 |
| scf7180000021851 | 39131 | 0.4153 | 0 | 47.6608 | Eukaryota | 15941.0 | 0 | Chordata | 15941.0 | 0 | Carnivora | 15941.0 | 0 | Felidae | 15941.0 | 0 | Felis | 14319.0 | 1 | Felis catus | 14319.0 | 1 |
| scf7180000022762 | 5910 | 0.3937 | 0 | 26.8017 | Eukaryota | 16253.0 | 0 | Chordata | 16253.0 | 0 | Carnivora | 16253.0 | 0 | Felidae | 11474.0 | 2 | Panthera | 4846.0 | 6 | Panthera tigri | 3246.0 | 8 |
| scf7180000021173 | 67799 | 0.3986 | 0 | 24.9673 | Eukaryota | 10760.0 | 0 | Chordata | 10760.0 | 0 | Carnivora | 10760.0 | 0 | Felidae | 10760.0 | 0 | Felis | 9672.0 | 1 | Felis catus | 9672.0 | 1 |
| scf7180000019226 | 356145 | 0.475 | 0 | 49.4477 | Eukaryota | 36737.0 | 0 | Chordata | 36737.0 | 0 | Carnivora | 36737.0 | 0 | Felidae | 36737.0 | 0 | Felis | 36737.0 | 0 | Felis catus | 36737.0 | 0 |
| scf7180000018509 | 11053 | 0.4241 | 0 | 37.428 | Eukaryota | 4720.0 | 1 | Chordata | 4720.0 | 1 | Primates | 2409.0 | 6 | Hominidae | 2409.0 | 6 | Homo | 2409.0 | 7 | Homo sapiens | 2409.0 | 7 |
| scf7180000023154 | 94934 | 0.4879 | 0 | 356.9305 | Eukaryota | 37454.0 | 0 | Chordata | 37454.0 | 0 | Carnivora | 37454.0 | 0 | Felidae | 25884.0 | 2 | Acinonyx | 11154.0 | 5 | cinonyx jubat | 11154.0 | 5 |
| scf7180000020167 | 143775 | 0.3707 | 0 | 25.4125 | Eukaryota | 45638.0 | 0 | Chordata | 45638.0 | 0 | Carnivora | 45638.0 | 0 | Felidae | 45638.0 | 0 | Felis | 45638.0 | 0 | Felis catus | 45638.0 | 0 |
| scf7180000020815 | 77625 | 0.394 | 0 | 24.7481 | Eukaryota | 32206.0 | 0 | Chordata | 32206.0 | 0 | Carnivora | 32206.0 | 0 | Felidae | 32206.0 | 0 | Felis | 32206.0 | 0 | Felis catus | 32206.0 | 0 |
| scf7180000024015 | 519630 | 0.4481 | 0 | 50.5294 | Eukaryota | 28424.0 | 0 | Chordata | 28424.0 | 0 | Carnivora | 28424.0 | 0 | Felidae | 28424.0 | 0 | Lynx | 11382.0 | 3 | ynx canadens | 11382.0 | 3 |
| scf7180000023979 | 868964 | 0.4264 | 0 | 49.7009 | Eukaryota | 27099.0 | 0 | Chordata | 27099.0 | 0 | Carnivora | 27099.0 | 0 | Felidae | 27099.0 | 0 | Felis | 21659.0 | 1 | Felis catus | 21659.0 | 1 |
| scf7180000022027 | 13782 | 0.4307 | 0 | 166.4797 | Eukaryota | 2216.0 | 0 | Chordata | 2216.0 | 0 | Carnivora | 2216.0 | 0 | Felidae | 2216.0 | 0 | Felis | 1567.0 | 2 | Felis catus | 1567.0 | 2 |
| scf7180000019391 | 278999 | 0.4232 | 0 | 50.121 | Eukaryota | 33587.0 | 0 | Chordata | 33587.0 | 0 | Carnivora | 33587.0 | 0 | Felidae | 33587.0 | 0 | Felis | 20191.0 | 3 | Felis catus | 20191.0 | 3 |
| scf7180000021733 | 61142 | 0.3896 | 0 | 25.5646 | Eukaryota | 42721.0 | 0 | Chordata | 42721.0 | 0 | Carnivora | 42721.0 | 0 | Felidae | 42721.0 | 0 | Felis | 38473.0 | 1 | Felis catus | 38473.0 | 1 |
| scf7180000023472 | 17012 | 0.4778 | 0 | 119.5587 | no-hit | 0.0 | 0 | no-hit | 0.0 | 0 | no-hit | 0.0 | 0 | no-hit | 0.0 | 0 | no-hit | 0.0 | 0 | no-hit | 0.0 | 0 |
| scf7180000021972 | 31023 | 0.3955 | 0 | 25.7446 | Eukaryota | 22881.0 | 0 | Chordata | 22881.0 | 0 | Carnivora | 22881.0 | 0 | Felidae | 22881.0 | 0 | Felis | 20585.0 | 1 | Felis catus | 20585.0 | 1 |
| scf7180000019042 | 16784 | 0.5807 | 0 | 32.6551 | Eukaryota | 7106.0 | 1 | Chordata | 7106.0 | 1 | Carnivora | 4579.0 | 2 | Felidae | 2313.0 | 5 | Homo | 1740.0 | 7 | Homo sapiens | 1740.0 | 7 |
| scf7180000019589 | 282655 | 0.5226 | 0 | 52.5781 | Eukaryota | 27623.0 | 0 | Chordata | 27623.0 | 0 | Carnivora | 27623.0 | 0 | Felidae | 27623.0 | 0 | Felis | 22043.0 | 1 | Felis catus | 22043.0 | 1 |
| scf7180000021690 | 40861 | 0.3794 | 0 | 24.946 | Eukaryota | 12781.0 | 0 | Chordata | 12781.0 | 0 | Carnivora | 12781.0 | 0 | Felidae | 12781.0 | 0 | Lynx | 5100.0 | 3 | ynx canadens | 5100.0 | 4 |
| scf7180000022669 | 9649 | 0.396 | 0 | 22.0077 | Eukaryota | 3596.0 | 0 | Chordata | 3596.0 | 0 | Carnivora | 3596.0 | 0 | Felidae | 3596.0 | 0 | Felis | 3259.0 | 1 | Felis catus | 3259.0 | 1 |
| scf7180000021097 | 90879 | 0.5595 | 0 | 46.33 | Eukaryota | 14438.0 | 0 | Chordata | 14438.0 | 0 | Primates | 4701.0 | 6 | Hominidae | 4701.0 | 6 | Homo | 4701.0 | 6 | Homo sapiens | 4701.0 | 6 |
| scf7180000023333 | 28284 | 0.6988 | 0 | 226.6362 | no-hit | 0.0 | 0 | no-hit | 0.0 | 0 | no-hit | 0.0 | 0 | no-hit | 0.0 | 0 | no-hit | 0.0 | 0 | no-hit | 0.0 | 0 |
| scf7180000020131 | 205951 | 0.4842 | 0 | 54.6258 | Eukaryota | 37342.0 | 0 | Chordata | 37342.0 | 0 | Carnivora | 37342.0 | 0 | Felidae | 37342.0 | 0 | Felis | 37342.0 | 0 | Felis catus | 37342.0 | 0 |
| scf7180000017802 | 4531 | 0.4083 | 0 | 17.1243 | Eukaryota | 16627.0 | 0 | Chordata | 16627.0 | 0 | Carnivora | 16627.0 | 0 | Felidae | 16627.0 | 0 | Felis | 14983.0 | 1 | Felis catus | 14983.0 | 1 |
| scf7180000019540 | 17359 | 0.3932 | 0 | 310.9078 | Eukaryota | 8155.0 | 0 | Chordata | 8155.0 | 0 | Carnivora | 8155.0 | 0 | Felidae | 8155.0 | 0 | Felis | 8155.0 | 0 | Felis catus | 8155.0 | 0 |
| scf7180000019238 | 432620 | 0.3878 | 0 | 51.5375 | Eukaryota | 36402.0 | 0 | Chordata | 36402.0 | 0 | Carnivora | 36402.0 | 0 | Felidae | 36402.0 | 0 | Lynx | 32778.0 | 1 | ynx canadens | 32778.0 | 1 |
| scf7180000021148 | 98883 | 0.4933 | 0 | 28.1855 | Eukaryota | 21948.0 | 0 | Chordata | 21948.0 | 0 | Carnivora | 21948.0 | 0 | Felidae | 21948.0 | 0 | Felis | 13176.0 | 2 | Felis catus | 13176.0 | 3 |
| scf7180000018835 | 535201 | 0.3878 | 0 | 51.3609 | Eukaryota | 46848.0 | 0 | Chordata | 46848.0 | 0 | Carnivora | 46848.0 | 0 | Felidae | 46848.0 | 0 | Felis | 46848.0 | 0 | Felis catus | 46848.0 | 0 |
| scf7180000017609 | 559565 | 0.4227 | 0 | 50.0026 | Eukaryota | 45253.0 | 0 | Chordata | 45253.0 | 0 | Carnivora | 45253.0 | 0 | Felidae | 45253.0 | 0 | Felis | 45253.0 | 0 | Felis catus | 45253.0 | 0 |
| scf7180000021579 | 64327 | 0.3885 | 0 | 24.3103 | Eukaryota | 34248.0 | 0 | Chordata | 34248.0 | 0 | Carnivora | 34248.0 | 0 | Felidae | 34248.0 | 0 | Felis | 34248.0 | 0 | Felis catus | 34248.0 | 0 |
| scf7180000021505 | 22591 | 0.4316 | 0 | 75.4181 | Eukaryota | 2218.0 | 0 | Chordata | 2218.0 | 0 | Carnivora | 2218.0 | 0 | Felidae | 2218.0 | 0 | Felis | 1358.0 | 2 | Felis catus | 1358.0 | 2 |
| scf7180000020412 | 288925 | 0.4018 | 0 | 25.5592 | Eukaryota | 35187.0 | 0 | Chordata | 35187.0 | 0 | Primates | 25458.0 | 3 | Hominidae | 22524.0 | 4 | Pan | 11284.0 | 5 | an troglodyte | 11284.0 | 5 |
| scf7180000021376 | 48471 | 0.3962 | 0 | 25.0607 | Eukaryota | 23765.0 | 0 | Chordata | 23765.0 | 0 | Carnivora | 23765.0 | 0 | Felidae | 23765.0 | 0 | Felis | 21367.0 | 1 | Felis catus | 21367.0 | 1 |
| scf7180000024103 | 222960 | 0.4353 | 0 | 25.8775 | Eukaryota | 9372.0 | 0 | Chordata | 9372.0 | 0 | Carnivora | 6417.0 | 1 | Felidae | 6417.0 | 1 | Felis | 6417.0 | 2 | Felis catus | 6417.0 | 2 |
| scf7180000017633 | 2030613 | 0.354 | 0 | 49.6539 | Eukaryota | 47360.0 | 0 | Chordata | 47360.0 | 0 | Carnivora | 47360.0 | 0 | Felidae | 47360.0 | 0 | Felis | 47360.0 | 0 | Felis catus | 47360.0 | 0 |
| scf7180000023282 | 160462 | 0.4231 | 0 | 193.5061 | Eukaryota | 24366.0 | 0 | Chordata | 24366.0 | 0 | Carnivora | 24366.0 | 0 | Felidae | 24366.0 | 0 | Felis | 20226.0 | 1 | Felis catus | 20226.0 | 1 |
| scf7180000023612 | 32879 | 0.4249 | 0 | 413.2004 | Eukaryota | 256605.0 | 0 | Chordata | 256605.0 | 0 | Carnivora | 256605.0 | 0 | Hyaenidae | 256605.0 | 0 | Proteles | 155086.0 | 1 | roteles cristat | 155086.0 | 1 |
| scf7180000023672 | 26493 | 0.5572 | 0 | 231.5131 | Eukaryota | 3702.0 | 0 | Chordata | 3702.0 | 0 | Carnivora | 3702.0 | 0 | Felidae | 3702.0 | 0 | unresolved | 1104.0 | 4 | unresolved | 1104.0 | 4 |
| scf7180000021740 | 22993 | 0.5103 | 0 | 45.7094 | Eukaryota | 18785.0 | 0 | Chordata | 18785.0 | 0 | Carnivora | 18785.0 | 0 | Felidae | 18785.0 | 0 | Felis | 18785.0 | 0 | Felis catus | 18785.0 | 0 |
| scf7180000017947 | 1458176 | 0.4748 | 0 | 52.7043 | Eukaryota | 65405.0 | 0 | Chordata | 65405.0 | 0 | Carnivora | 53172.0 | 1 | Otariidae | 13890.0 | 7 | Zalophus | 7304.0 | 9 | phus californi | 7304.0 | 9 |
| scf7180000018116 | 443227 | 0.38 | 0 | 48.3902 | Eukaryota | 46490.0 | 0 | Chordata | 46490.0 | 0 | Carnivora | 46490.0 | 0 | Felidae | 46490.0 | 0 | Felis | 46490.0 | 0 | Felis catus | 46490.0 | 0 |
| scf7180000018409 | 1439750 | 0.347 | 0 | 48.3084 | Eukaryota | 46843.0 | 0 | Chordata | 46843.0 | 0 | Carnivora | 46843.0 | 0 | Felidae | 46843.0 | 0 | Felis | 46843.0 | 0 | Felis catus | 46843.0 | 0 |
| scf7180000019874 | 254373 | 0.4898 | 0 | 60.4681 | Eukaryota | 40372.0 | 0 | Chordata | 40372.0 | 0 | Carnivora | 40372.0 | 0 | Felidae | 40372.0 | 0 | Felis | 40372.0 | 0 | Felis catus | 40372.0 | 0 |
| scf7180000020456 | 200900 | 0.4391 | 0 | 51.4508 | Eukaryota | 37449.0 | 0 | Chordata | 37449.0 | 0 | Carnivora | 37449.0 | 0 | Felidae | 37449.0 | 0 | Felis | 37449.0 | 0 | Felis catus | 37449.0 | 0 |
| scf7180000019014 | 45606 | 0.4311 | 0 | 42.2237 | Eukaryota | 45638.0 | 0 | Chordata | 45638.0 | 0 | Carnivora | 45638.0 | 0 | Felidae | 45638.0 | 0 | Felis | 45638.0 | 0 | Felis catus | 45638.0 | 0 |
| scf7180000021143 | 85587 | 0.5049 | 0 | 51.0859 | Eukaryota | 40704.0 | 0 | Chordata | 40704.0 | 0 | Carnivora | 40704.0 | 0 | Herpestidae | 29511.0 | 1 | Suricata | 29511.0 | 1 | ricata suricat | 29511.0 | 1 |
| scf7180000018788 | 990746 | 0.4361 | 0 | 52.2592 | Eukaryota | 45141.0 | 0 | Chordata | 45141.0 | 0 | Carnivora | 45141.0 | 0 | Felidae | 45141.0 | 0 | Felis | 45141.0 | 0 | Felis catus | 45141.0 | 0 |
| scf7180000020567 | 121068 | 0.4427 | 0 | 47.9773 | Eukaryota | 34626.0 | 0 | Chordata | 34626.0 | 0 | Carnivora | 34626.0 | 0 | Felidae | 34626.0 | 0 | Felis | 34626.0 | 0 | Felis catus | 34626.0 | 0 |
| scf7180000022055 | 45088 | 0.3624 | 0 | 24.6026 | Eukaryota | 10523.0 | 0 | Chordata | 10523.0 | 0 | Carnivora | 9509.0 | 1 | Felidae | 8509.0 | 2 | Felis | 8509.0 | 2 | Felis catus | 8509.0 | 2 |
| scf7180000022398 | 6922 | 0.4522 | 0 | 54.7948 | Eukaryota | 2043.0 | 0 | Chordata | 2043.0 | 0 | Carnivora | 2043.0 | 0 | Felidae | 2043.0 | 0 | Felis | 1633.0 | 1 | Felis catus | 1633.0 | 1 |
| scf7180000018485 | 684302 | 0.358 | 0 | 51.9701 | Eukaryota | 46683.0 | 0 | Chordata | 46683.0 | 0 | Carnivora | 46683.0 | 0 | Felidae | 46683.0 | 0 | Felis | 46683.0 | 0 | Felis catus | 46683.0 | 0 |
| scf7180000020867 | 191725 | 0.3589 | 0 | 24.8459 | Eukaryota | 45695.0 | 0 | Chordata | 45695.0 | 0 | Carnivora | 45695.0 | 0 | Felidae | 45695.0 | 0 | Felis | 45695.0 | 0 | Felis catus | 45695.0 | 0 |
| scf7180000017869 | 1399245 | 0.4172 | 0 | 50.9823 | Eukaryota | 63277.0 | 0 | Chordata | 63277.0 | 0 | Carnivora | 63277.0 | 0 | Felidae | 35966.0 | 2 | Felis | 30160.0 | 3 | Felis catus | 30160.0 | 3 |
| scf7180000017644 | 326107 | 0.4285 | 0 | 49.6858 | Eukaryota | 45521.0 | 0 | Chordata | 45521.0 | 0 | Carnivora | 45521.0 | 0 | Felidae | 45521.0 | 0 | Felis | 45521.0 | 0 | Felis catus | 45521.0 | 0 |
| scf7180000018632 | 24872 | 0.3868 | 0 | 26.0662 | Eukaryota | 29226.0 | 0 | Chordata | 29226.0 | 0 | Carnivora | 29226.0 | 0 | Felidae | 29226.0 | 0 | Felis | 29226.0 | 0 | Felis catus | 29226.0 | 0 |
| scf7180000018755 | 3538416 | 0.369 | 100 | 47.6326 | Eukaryota | 47476.0 | 0 | Chordata | 47476.0 | 0 | Carnivora | 47476.0 | 0 | Felidae | 47476.0 | 0 | Felis | 47476.0 | 0 | Felis catus | 47476.0 | 0 |
| scf7180000018065 | 1150586 | 0.4496 | 0 | 52.1986 | Eukaryota | 39668.0 | 0 | Chordata | 39668.0 | 0 | Carnivora | 39668.0 | 0 | Felidae | 35625.0 | 1 | Felis | 18378.0 | 4 | Felis catus | 18378.0 | 5 |
| scf7180000022596 | 13360 | 0.4793 | 0 | 26.4911 | Eukaryota | 5801.0 | 0 | Chordata | 5801.0 | 0 | Primates | 3248.0 | 3 | Hominidae | 3248.0 | 3 | Homo | 2451.0 | 4 | Homo sapiens | 2451.0 | 4 |
| scf7180000021040 | 81077 | 0.4294 | 0 | 108.8076 | Eukaryota | 47722.0 | 0 | Chordata | 47722.0 | 0 | Carnivora | 47722.0 | 0 | Felidae | 47722.0 | 0 | Felis | 47722.0 | 0 | Felis catus | 47722.0 | 0 |
| scf7180000022189 | 22380 | 0.3839 | 0 | 283.7979 | Eukaryota | 10083.0 | 0 | Chordata | 10083.0 | 0 | pterygiforme | 9019.0 | 1 | Apterygidae | 9019.0 | 1 | Apteryx | 9019.0 | 1 | pteryx mante | 9019.0 | 1 |
| scf7180000019205 | 397960 | 0.3925 | 0 | 50.3888 | Eukaryota | 43024.0 | 0 | Chordata | 43024.0 | 0 | Carnivora | 43024.0 | 0 | Felidae | 43024.0 | 0 | Acinonyx | 16377.0 | 2 | cinonyx jubat | 16377.0 | 2 |
| scf7180000017589 | 905364 | 0.38 | 0 | 49.4284 | Eukaryota | 47785.0 | 0 | Chordata | 47785.0 | 0 | Carnivora | 47785.0 | 0 | Felidae | 47785.0 | 0 | Felis | 47785.0 | 0 | Felis catus | 47785.0 | 0 |
| scf7180000021843 | 43478 | 0.383 | 0 | 26.285 | Eukaryota | 5803.0 | 0 | Chordata | 5247.0 | 1 | Carnivora | 5247.0 | 1 | Felidae | 5247.0 | 1 | Felis | 4080.0 | 3 | Felis catus | 4080.0 | 3 |
| scf7180000018088 | 307265 | 0.3718 | 0 | 49.8909 | Eukaryota | 44553.0 | 0 | Chordata | 44553.0 | 0 | Carnivora | 44553.0 | 0 | Felidae | 44553.0 | 0 | Felis | 40115.0 | 1 | Felis catus | 40115.0 | 1 |
| scf7180000017702 | 1816899 | 0.3949 | 0 | 50.9003 | Eukaryota | 63437.0 | 0 | Chordata | 63437.0 | 0 | Carnivora | 63437.0 | 0 | Felidae | 37929.0 | 1 | Suricata | 25508.0 | 3 | ricata suricat | 25508.0 | 3 |

| | | | | | | | | | | | | | | | | | | | | | | |
|---|---|---|---|---|---|---|---|---|---|---|---|---|---|---|---|---|---|---|---|---|---|---|
| scf7180000020757 | 95670 | 0.3578 | 0 | 47.4221 | Eukaryota | 12593.0 | 0 | Chordata | 12593.0 | 0 | Carnivora | 12593.0 | 0 | Felidae | 12593.0 | 0 | Felis | 11337.0 | 1 | Felis catus | 11337.0 | 1 |
| scf7180000022339 | 22602 | 0.4707 | 0 | 26.5324 | Eukaryota | 7001.0 | 0 | Chordata | 7001.0 | 0 | Carnivora | 7001.0 | 0 | Felidae | 7001.0 | 0 | Felis | 7001.0 | 0 | Felis catus | 7001.0 | 0 |
| scf7180000019614 | 163139 | 0.4579 | 0 | 86.8506 | Eukaryota | 2580.0 | 0 | Chordata | 2580.0 | 0 | Carnivora | 2580.0 | 0 | Felidae | 2580.0 | 0 | Felis | 2341.0 | 1 | Felis catus | 2341.0 | 1 |
| scf7180000023941 | 616232 | 0.3822 | 0 | 49.4469 | Eukaryota | 77537.0 | 0 | Chordata | 77537.0 | 0 | Carnivora | 77537.0 | 0 | Felidae | 62234.0 | 2 | Acinonyx | 15698.0 | 6 | :inonyx jubat | 15698.0 | 6 |
| scf7180000019158 | 610481 | 0.3874 | 0 | 50.6785 | Eukaryota | 47177.0 | 0 | Chordata | 47177.0 | 0 | Carnivora | 47177.0 | 0 | Felidae | 47177.0 | 0 | Felis | 47177.0 | 0 | Felis catus | 47177.0 | 0 |
| scf7180000021697 | 49515 | 0.4065 | 0 | 24.3224 | Eukaryota | 42428.0 | 0 | Chordata | 42428.0 | 0 | Carnivora | 42428.0 | 0 | Felidae | 42428.0 | 0 | Felis | 42428.0 | 0 | Felis catus | 42428.0 | 0 |
| scf7180000023828 | 2085795 | 0.4054 | 0 | 51.341 | Eukaryota | 52144.0 | 0 | Chordata | 52144.0 | 0 | Carnivora | 52144.0 | 0 | Felidae | 52144.0 | 0 | Panthera | 30195.0 | 1 | Panthera tigri | 30195.0 | 1 |
| scf7180000019140 | 7600 | 0.4036 | 0 | 23.1214 | Eukaryota | 38307.0 | 0 | Chordata | 38307.0 | 0 | Carnivora | 38307.0 | 0 | Felidae | 38307.0 | 0 | Felis | 38307.0 | 0 | Felis catus | 38307.0 | 0 |
| scf7180000017467 | 2537210 | 0.3743 | 0 | 49.5808 | Eukaryota | 61891.0 | 0 | Chordata | 61891.0 | 0 | Carnivora | 61891.0 | 0 | Felidae | 35111.0 | 1 | Suricata | 26780.0 | 3 | ıricata suricat | 26780.0 | 3 |
| scf7180000021018 | 43325 | 0.4762 | 0 | 53.9124 | Eukaryota | 6829.0 | 0 | Chordata | 6829.0 | 0 | Carnivora | 5144.0 | 1 | Felidae | 3955.0 | 2 | Apteryx | 1685.0 | 4 | pteryx mante | 1685.0 | 4 |
| scf7180000020605 | 89527 | 0.4155 | 0 | 48.3013 | Eukaryota | 12936.0 | 0 | Chordata | 12936.0 | 0 | Carnivora | 12936.0 | 0 | Felidae | 10377.0 | 1 | Felis | 10377.0 | 1 | Felis catus | 10377.0 | 1 |
| scf7180000018021 | 925919 | 0.352 | 0 | 49.7989 | Eukaryota | 47635.0 | 0 | Chordata | 47635.0 | 0 | Carnivora | 47635.0 | 0 | Felidae | 47635.0 | 0 | Felis | 47635.0 | 0 | Felis catus | 47635.0 | 0 |
| scf7180000017530 | 134959 | 0.4555 | 0 | 43.9679 | Eukaryota | 5849.0 | 0 | Chordata | 5849.0 | 0 | Carnivora | 5849.0 | 0 | Felidae | 2972.0 | 2 | Felis | 2347.0 | 3 | Felis catus | 2347.0 | 4 |
| scf7180000018909 | 423202 | 0.3577 | 0 | 48.257 | Eukaryota | 41373.0 | 0 | Chordata | 41373.0 | 0 | Carnivora | 41373.0 | 0 | Felidae | 41373.0 | 0 | Felis | 41373.0 | 0 | Felis catus | 41373.0 | 0 |
| scf7180000022295 | 9598 | 0.5206 | 0 | 75.1543 | no-hit | 0.0 | 0 | no-hit | 0.0 | 0 | no-hit | 0.0 | 0 | no-hit | 0.0 | 0 | no-hit | 0.0 | 0 | no-hit | 0.0 | 0 |
| scf7180000023871 | 1805027 | 0.4112 | 0 | 52.4478 | Eukaryota | 60390.0 | 0 | Chordata | 60390.0 | 0 | Carnivora | 60390.0 | 0 | Felidae | 60390.0 | 0 | Felis | 55675.0 | 1 | Felis catus | 55675.0 | 1 |
| scf7180000024247 | 4903089 | 0.4449 | 100 | 51.3268 | Eukaryota | 65448.0 | 1 | Chordata | 65448.0 | 1 | Carnivora | 39441.0 | 2 | Felidae | 23627.0 | 5 | Homo | 18066.0 | 7 | Homo sapiens | 18066.0 | 7 |
| scf7180000020045 | 31822 | 0.5043 | 0 | 341.4967 | Eukaryota | 5925.0 | 0 | Chordata | 5925.0 | 0 | Carnivora | 5925.0 | 0 | Phocidae | 2416.0 | 2 | .eptonychote | 1812.0 | 4 | ınychotes wed | 1812.0 | 4 |
| scf7180000022410 | 14010 | 0.6187 | 0 | 53.1033 | Eukaryota | 2880.0 | 0 | Chordata | 2880.0 | 0 | Primates | 1380.0 | 3 | Hominidae | 1380.0 | 4 | Homo | 1380.0 | 4 | Homo sapiens | 1380.0 | 4 |
| scf7180000021932 | 50222 | 0.632 | 0 | 43.3177 | Eukaryota | 2338.0 | 0 | Chordata | 2338.0 | 0 | Carnivora | 2338.0 | 0 | Felidae | 2092.0 | 1 | Felis | 1386.0 | 3 | Felis catus | 1386.0 | 3 |
| scf7180000022309 | 30289 | 0.4833 | 0 | 26.4175 | Eukaryota | 3075.0 | 0 | Chordata | 3075.0 | 0 | Carnivora | 1677.0 | 2 | Felidae | 1396.0 | 3 | Felis | 1396.0 | 3 | Felis catus | 1396.0 | 3 |
| scf7180000023789 | 1204302 | 0.3591 | 0 | 49.7919 | Eukaryota | 45452.0 | 0 | Chordata | 45452.0 | 0 | Carnivora | 45452.0 | 0 | Felidae | 45452.0 | 0 | Felis | 45452.0 | 0 | Felis catus | 45452.0 | 0 |
| scf7180000023682 | 81091 | 0.4402 | 0 | 250.18 | Eukaryota | 22446.0 | 0 | Chordata | 22446.0 | 0 | Carnivora | 22446.0 | 0 | Felidae | 22446.0 | 0 | Felis | 20250.0 | 1 | Felis catus | 20250.0 | 1 |
| scf7180000019029 | 478818 | 0.405 | 0 | 51.3615 | Eukaryota | 26062.0 | 0 | Chordata | 26062.0 | 0 | Carnivora | 26062.0 | 0 | Felidae | 26062.0 | 0 | Felis | 23406.0 | 1 | Felis catus | 23406.0 | 1 |
| scf7180000021147 | 51749 | 0.5904 | 0 | 27.1415 | Eukaryota | 13551.0 | 0 | Chordata | 13551.0 | 0 | Primates | 7223.0 | 1 | Hominidae | 7223.0 | 1 | Homo | 7223.0 | 3 | Homo sapiens | 7223.0 | 4 |
| scf7180000022587 | 5130 | 0.6413 | 0 | 46.9907 | Eukaryota | 1765.0 | 0 | Chordata | 1765.0 | 0 | Carnivora | 1765.0 | 0 | Felidae | 1554.0 | 1 | Felis | 1374.0 | 2 | Felis catus | 1374.0 | 2 |
| scf7180000024014 | 546622 | 0.3859 | 0 | 49.5061 | Eukaryota | 41149.0 | 0 | Chordata | 41149.0 | 0 | Carnivora | 41149.0 | 0 | Felidae | 41149.0 | 0 | Felis | 41149.0 | 0 | Felis catus | 41149.0 | 0 |
| scf7180000018330 | 6996 | 0.3836 | 0 | 35.8015 | Eukaryota | 31896.0 | 0 | Chordata | 31896.0 | 0 | Carnivora | 31896.0 | 0 | Felidae | 31896.0 | 0 | Felis | 28621.0 | 1 | Felis catus | 28621.0 | 1 |
| scf7180000021585 | 22492 | 0.4611 | 0 | 48.3066 | Eukaryota | 3374.0 | 0 | Chordata | 3374.0 | 0 | Carnivora | 2774.0 | 1 | Felidae | 1563.0 | 3 | Ursus | 915.0 | 5 | :inonyx jubat | 646.0 | 7 |
| scf7180000023719 | 20262 | 0.5621 | 0 | 60.2803 | Eukaryota | 2760.0 | 0 | Chordata | 2760.0 | 0 | Carnivora | 2760.0 | 0 | Felidae | 2488.0 | 1 | Felis | 1366.0 | 2 | Felis catus | 1366.0 | 2 |
| scf7180000020276 | 147945 | 0.4515 | 0 | 50.9858 | Eukaryota | 31090.0 | 0 | Chordata | 31090.0 | 0 | Carnivora | 31090.0 | 0 | Felidae | 18514.0 | 1 | Suricata | 12576.0 | 3 | ıricata suricat | 12576.0 | 3 |
| scf7180000024032 | 457909 | 0.3776 | 0 | 50.1927 | Eukaryota | 52986.0 | 0 | Chordata | 52986.0 | 0 | Carnivora | 52986.0 | 0 | Felidae | 47739.0 | 1 | Acinonyx | 31770.0 | 2 | :inonyx jubat | 31770.0 | 2 |
| scf7180000020038 | 26310 | 0.5271 | 0 | 143.2775 | Eukaryota | 12702.0 | 0 | Chordata | 12702.0 | 0 | Carnivora | 12702.0 | 0 | Felidae | 12702.0 | 0 | Felis | 12702.0 | 0 | Felis catus | 12702.0 | 0 |
| scf7180000021352 | 54820 | 0.5941 | 0 | 46.6552 | Eukaryota | 3310.0 | 1 | Chordata | 3310.0 | 1 | Primates | 1298.0 | 3 | Hominidae | 1298.0 | 4 | Homo | 1298.0 | 5 | Homo sapiens | 1298.0 | 5 |
| scf7180000022317 | 30644 | 0.3883 | 0 | 26.1114 | Eukaryota | 10003.0 | 0 | Chordata | 10003.0 | 0 | Carnivora | 5680.0 | 2 | Felidae | 5680.0 | 3 | Felis | 4760.0 | 5 | Felis catus | 4760.0 | 5 |
| scf7180000017834 | 1591862 | 0.3823 | 0 | 49.5783 | Eukaryota | 40178.0 | 0 | Chordata | 40178.0 | 0 | Carnivora | 40178.0 | 0 | Felidae | 40178.0 | 0 | Felis | 36207.0 | 1 | Felis catus | 36207.0 | 1 |
| scf7180000024279 | 1139041 | 0.4538 | 100 | 50.3311 | Eukaryota | 38744.0 | 0 | Chordata | 38744.0 | 0 | Carnivora | 31422.0 | 1 | Felidae | 27222.0 | 2 | Felis | 17551.0 | 5 | Felis catus | 17551.0 | 5 |
| scf7180000021189 | 158736 | 0.3977 | 0 | 51.3561 | Eukaryota | 14061.0 | 0 | Chordata | 14061.0 | 0 | Carnivora | 8526.0 | 1 | Felidae | 6385.0 | 2 | Homo | 5535.0 | 3 | Homo sapiens | 5535.0 | 3 |
| scf7180000020063 | 177436 | 0.436 | 0 | 54.1775 | Eukaryota | 29165.0 | 0 | Chordata | 29165.0 | 0 | Carnivora | 29165.0 | 0 | Felidae | 29165.0 | 0 | Felis | 29165.0 | 0 | Felis catus | 29165.0 | 0 |
| scf7180000019766 | 188734 | 0.4508 | 0 | 26.2068 | Eukaryota | 56803.0 | 0 | Chordata | 56803.0 | 0 | Carnivora | 56803.0 | 0 | Felidae | 45862.0 | 2 | Felis | 16800.0 | 6 | Felis catus | 16800.0 | 6 |
| scf7180000021978 | 833325 | 0.3607 | 0 | 48.1858 | Eukaryota | 79009.0 | 0 | Chordata | 79009.0 | 0 | Perissodactyla | 43746.0 | 2 | Equidae | 43746.0 | 2 | Equus | 43746.0 | 2 | Equus caballu | 43746.0 | 2 |
| scf7180000023193 | 26309 | 0.4202 | 0 | 180.0185 | Eukaryota | 12155.0 | 0 | Chordata | 12155.0 | 0 | Carnivora | 12155.0 | 0 | Felidae | 12155.0 | 0 | Felis | 10958.0 | 1 | Felis catus | 10958.0 | 1 |
| scf7180000021583 | 63329 | 0.4266 | 0 | 28.6035 | Eukaryota | 45623.0 | 0 | Chordata | 45623.0 | 0 | Carnivora | 45623.0 | 0 | Felidae | 45623.0 | 0 | Felis | 45623.0 | 0 | Felis catus | 45623.0 | 0 |
| scf7180000019351 | 402163 | 0.3717 | 0 | 50.649 | Eukaryota | 47873.0 | 0 | Chordata | 47873.0 | 0 | Carnivora | 47873.0 | 0 | Felidae | 47873.0 | 0 | Felis | 47873.0 | 0 | Felis catus | 47873.0 | 0 |
| scf7180000023869 | 1418468 | 0.3515 | 0 | 49.0297 | Eukaryota | 44645.0 | 0 | Chordata | 44645.0 | 0 | Carnivora | 44645.0 | 0 | Felidae | 44645.0 | 0 | Felis | 44645.0 | 0 | Felis catus | 44645.0 | 0 |
| scf7180000018178 | 1007144 | 0.4145 | 0 | 51.5457 | Eukaryota | 51756.0 | 0 | Chordata | 51756.0 | 0 | Carnivora | 46583.0 | 1 | Herpestidae | 28161.0 | 2 | Suricata | 28161.0 | 3 | ıricata suricat | 28161.0 | 3 |
| scf7180000022316 | 10964 | 0.4048 | 0 | 17.9799 | Eukaryota | 17470.0 | 0 | Chordata | 17470.0 | 0 | Carnivora | 17470.0 | 0 | Felidae | 17470.0 | 0 | Felis | 17470.0 | 0 | Felis catus | 17470.0 | 0 |
| scf7180000018335 | 915963 | 0.3607 | 0 | 49.5363 | Eukaryota | 51299.0 | 0 | Chordata | 51299.0 | 0 | Carnivora | 37942.0 | 2 | Felidae | 32401.0 | 3 | Panthera | 14529.0 | 6 | anthera pardı | 9688.0 | 7 |
| scf7180000021991 | 17875 | 0.5604 | 0 | 238.7119 | Eukaryota | 3628.0 | 0 | Chordata | 3628.0 | 0 | Carnivora | 3628.0 | 0 | Felidae | 3628.0 | 0 | Panthera | 1474.0 | 4 | anthera pardı | 1110.0 | 5 |
| scf7180000023957 | 1143563 | 0.4201 | 0 | 50.9764 | Eukaryota | 47431.0 | 0 | Chordata | 47431.0 | 0 | Carnivora | 47431.0 | 0 | Felidae | 47431.0 | 0 | Felis | 47431.0 | 0 | Felis catus | 47431.0 | 0 |
| scf7180000020497 | 117319 | 0.3731 | 0 | 25.5389 | Eukaryota | 46345.0 | 0 | Chordata | 46345.0 | 0 | Carnivora | 46345.0 | 0 | Felidae | 46345.0 | 0 | Felis | 46345.0 | 0 | Felis catus | 46345.0 | 0 |
| scf7180000020471 | 20202 | 0.4385 | 0 | 35.3016 | Eukaryota | 45182.0 | 0 | Chordata | 45182.0 | 0 | Carnivora | 45182.0 | 0 | Felidae | 45182.0 | 0 | Felis | 45182.0 | 0 | Felis catus | 45182.0 | 0 |
| scf7180000024188 | 62815 | 0.5152 | 0 | 50.321 | Eukaryota | 8188.0 | 1 | Chordata | 8188.0 | 1 | Artiodactyla | 2984.0 | 4 | Bovidae | 2984.0 | 5 | Homo | 1837.0 | 7 | Homo sapiens | 1837.0 | 7 |
| scf7180000020998 | 107798 | 0.3597 | 0 | 23.2486 | Eukaryota | 30532.0 | 0 | Chordata | 30532.0 | 0 | Carnivora | 30532.0 | 0 | Felidae | 30532.0 | 0 | Felis | 30532.0 | 0 | Felis catus | 30532.0 | 0 |
| scf7180000023878 | 1294451 | 0.3848 | 0 | 50.0066 | Eukaryota | 82753.0 | 0 | Chordata | 82753.0 | 0 | Carnivora | 82753.0 | 0 | Felidae | 82753.0 | 0 | Panthera | 57988.0 | 2 | anthera pardı | 33124.0 | 3 |
| scf7180000023186 | 45055 | 0.3912 | 0 | 457.1076 | Eukaryota | 45834.0 | 0 | Chordata | 45834.0 | 0 | Carnivora | 45834.0 | 0 | Felidae | 45834.0 | 0 | Felis | 45834.0 | 0 | Felis catus | 45834.0 | 0 |
| scf7180000018674 | 1001290 | 0.5112 | 0 | 54.3116 | Eukaryota | 38767.0 | 0 | Chordata | 38767.0 | 0 | Carnivora | 38767.0 | 0 | Felidae | 38767.0 | 0 | Felis | 20819.0 | 1 | Felis catus | 20819.0 | 1 |
| scf7180000020969 | 66899 | 0.3633 | 0 | 24.2469 | Eukaryota | 20287.0 | 0 | Chordata | 20287.0 | 0 | Carnivora | 20287.0 | 0 | Felidae | 20287.0 | 0 | Felis | 18281.0 | 1 | Felis catus | 18281.0 | 1 |
| scf7180000020734 | 67512 | 0.6631 | 0 | 52.4711 | Eukaryota | 5524.0 | 0 | Chordata | 5524.0 | 0 | Carnivora | 3000.0 | 2 | Ursidae | 1564.0 | 4 | Ailuropoda | 1564.0 | 5 | ›poda melanc | 1564.0 | 5 |
| scf7180000019556 | 29813 | 0.4628 | 0 | 252.6318 | Eukaryota | 16638.0 | 0 | Chordata | 16638.0 | 0 | Carnivora | 16638.0 | 0 | Felidae | 16638.0 | 0 | Felis | 14957.0 | 1 | Felis catus | 14957.0 | 1 |
| scf7180000017569 | 2460404 | 0.389 | 0 | 50.8581 | Eukaryota | 47786.0 | 0 | Chordata | 47786.0 | 0 | Carnivora | 47786.0 | 0 | Felidae | 47786.0 | 0 | Felis | 47786.0 | 0 | Felis catus | 47786.0 | 0 |
| scf7180000019946 | 470430 | 0.5389 | 0 | 49.9114 | Eukaryota | 16723.0 | 0 | Chordata | 16723.0 | 0 | Carnivora | 16723.0 | 0 | Felidae | 16723.0 | 0 | Felis | 16723.0 | 0 | Felis catus | 16723.0 | 0 |
| scf7180000022663 | 21240 | 0.3462 | 0 | 23.6952 | Eukaryota | 7492.0 | 0 | Chordata | 7492.0 | 0 | Carnivora | 7492.0 | 0 | Felidae | 7492.0 | 0 | Felis | 4490.0 | 1 | Felis catus | 4490.0 | 1 |
| scf7180000020785 | 62416 | 0.5356 | 0 | 50.2818 | Eukaryota | 9322.0 | 0 | Chordata | 9322.0 | 0 | Carnivora | 9322.0 | 0 | Felidae | 6585.0 | 2 | Lynx | 1890.0 | 7 | ynx canadens | 1890.0 | 8 |
| scf7180000022241 | 20332 | 0.3917 | 0 | 31.6493 | Eukaryota | 34390.0 | 0 | Chordata | 34390.0 | 0 | Carnivora | 34390.0 | 0 | Felidae | 34390.0 | 0 | Felis | 34390.0 | 0 | Felis catus | 34390.0 | 0 |
| scf7180000017953 | 1617104 | 0.3962 | 0 | 50.5772 | Eukaryota | 51055.0 | 0 | Chordata | 51055.0 | 0 | Carnivora | 42078.0 | 1 | Felidae | 42078.0 | 1 | Felis | 42078.0 | 1 | Felis catus | 42078.0 | 1 |
| scf7180000021155 | 44387 | 0.4331 | 0 | 26.0255 | Eukaryota | 11585.0 | 0 | Chordata | 11585.0 | 0 | Carnivora | 5761.0 | 2 | Felidae | 5761.0 | 2 | Felis | 5761.0 | 3 | Felis catus | 5761.0 | 3 |
| scf7180000022481 | 35953 | 0.4316 | 0 | 371.886 | Eukaryota | 47065.0 | 0 | Chordata | 47065.0 | 0 | Carnivora | 47065.0 | 0 | Felidae | 47065.0 | 0 | Felis | 47065.0 | 0 | Felis catus | 47065.0 | 0 |
| scf7180000019529 | 336502 | 0.4849 | 0 | 51.9642 | Eukaryota | 21121.0 | 0 | Chordata | 21121.0 | 0 | Carnivora | 21121.0 | 0 | Felidae | 11805.0 | 1 | Felis | 9838.0 | 2 | Felis catus | 9838.0 | 2 |
| scf7180000021467 | 47903 | 0.3833 | 0 | 24.4898 | Eukaryota | 42098.0 | 0 | Chordata | 42098.0 | 0 | Carnivora | 42098.0 | 0 | Felidae | 42098.0 | 0 | Felis | 37911.0 | 1 | Felis catus | 37911.0 | 1 |
| scf7180000017743 | 144661 | 0.3793 | 0 | 53.4991 | Eukaryota | 49588.0 | 0 | Chordata | 49588.0 | 0 | Carnivora | 49588.0 | 0 | Felidae | 49588.0 | 0 | Felis | 44601.0 | 1 | Felis catus | 44601.0 | 1 |
| scf7180000020563 | 703959 | 0.4224 | 0 | 49.0875 | Eukaryota | 46346.0 | 0 | Chordata | 46346.0 | 0 | Carnivora | 46346.0 | 0 | Felidae | 46346.0 | 0 | Lynx | 27986.0 | 1 | ynx canadens | 27986.0 | 1 |
| scf7180000017500 | 3537076 | 0.3806 | 0 | 50.5448 | Eukaryota | 49002.0 | 0 | Chordata | 49002.0 | 0 | Carnivora | 49002.0 | 0 | Felidae | 49002.0 | 0 | Panthera | 34230.0 | 2 | anthera pardı | 29310.0 | 3 |
| scf7180000017608 | 51545 | 0.3736 | 0 | 22.6682 | Eukaryota | 45678.0 | 0 | Chordata | 45678.0 | 0 | Carnivora | 45678.0 | 0 | Felidae | 45678.0 | 0 | Felis | 45678.0 | 0 | Felis catus | 45678.0 | 0 |
| scf7180000023932 | 928190 | 0.6257 | 0 | 50.6331 | Eukaryota | 34862.0 | 0 | Chordata | 34862.0 | 0 | Carnivora | 32427.0 | 1 | Felidae | 15544.0 | 3 | Acinonyx | 15544.0 | 3 | :inonyx jubat | 15544.0 | 3 |
| scf7180000019624 | 288939 | 0.4357 | 0 | 52.4552 | Eukaryota | 19971.0 | 0 | Chordata | 19971.0 | 0 | Carnivora | 19971.0 | 0 | Felidae | 19971.0 | 0 | Felis | 17961.0 | 1 | Felis catus | 17961.0 | 1 |
| scf7180000020669 | 103366 | 0.5378 | 0 | 49.5457 | Eukaryota | 9265.0 | 1 | Chordata | 9265.0 | 1 | Carnivora | 8345.0 | 2 | Felidae | 8345.0 | 2 | Felis | 7150.0 | 3 | Felis catus | 7150.0 | 3 |
| scf7180000018093 | 400926 | 0.4516 | 0 | 52.1849 | Eukaryota | 58823.0 | 0 | Chordata | 58823.0 | 0 | Carnivora | 58823.0 | 0 | Felidae | 41977.0 | 3 | Panthera | 13270.0 | 6 | anthera pardı | 13270.0 | 6 |
| scf7180000018650 | 619964 | 0.4009 | 0 | 50.5578 | Eukaryota | 63887.0 | 0 | Chordata | 63887.0 | 0 | Carnivora | 63887.0 | 0 | Felidae | 52333.0 | 1 | Lynx | 29625.0 | 4 | ynx canadens | 29625.0 | 4 |
| scf7180000019893 | 176759 | 0.5672 | 0 | 50.1956 | Eukaryota | 13500.0 | 0 | Chordata | 13500.0 | 0 | Carnivora | 13500.0 | 0 | Felidae | 13500.0 | 0 | Felis | 10800.0 | 1 | Felis catus | 10800.0 | 1 |
| scf7180000020768 | 95493 | 0.4753 | 0 | 27.0621 | Eukaryota | 49118.0 | 0 | Chordata | 49118.0 | 0 | Carnivora | 49118.0 | 0 | Herpestidae | 29428.0 | 1 | Suricata | 29428.0 | 2 | ıricata suricat | 29428.0 | 2 |
| scf7180000018028 | 1132451 | 0.5161 | 0 | 52.4253 | Eukaryota | 182148.0 | 0 | Chordata | 182148.0 | 0 | Carnivora | 182148.0 | 0 | Felidae | 114954.0 | 2 | Panthera | 76756.0 | 3 | anthera pardı | 76756.0 | 3 |
| scf7180000018878 | 346506 | 0.3727 | 0 | 48.854 | Eukaryota | 42168.0 | 0 | Chordata | 42168.0 | 0 | Carnivora | 42168.0 | 0 | Felidae | 42168.0 | 0 | Felis | 42168.0 | 0 | Felis catus | 42168.0 | 0 |
| scf7180000021386 | 17152 | 0.5209 | 0 | 49.3927 | no-hit | 0.0 | 0 | no-hit | 0.0 | 0 | no-hit | 0.0 | 0 | no-hit | 0.0 | 0 | no-hit | 0.0 | 0 | no-hit | 0.0 | 0 |
| scf7180000017864 | 1476014 | 0.3623 | 0 | 49.7984 | Eukaryota | 47951.0 | 0 | Chordata | 47951.0 | 0 | Carnivora | 47951.0 | 0 | Felidae | 41788.0 | 1 | Felis | 37204.0 | 2 | Felis catus | 37204.0 | 2 |
| scf7180000020532 | 132224 | 0.4857 | 0 | 26.0368 | Eukaryota | 13742.0 | 0 | Chordata | 13742.0 | 0 | Rodentia | 3126.0 | 4 | Muridae | 3126.0 | 4 | Rousettus | 3083.0 | 6 | ɛettus aegypti | 3083.0 | 6 |
| scf7180000018648 | 432020 | 0.3912 | 0 | 55.8115 | Eukaryota | 46722.0 | 0 | Chordata | 46722.0 | 0 | Carnivora | 46722.0 | 0 | Felidae | 46722.0 | 0 | Felis | 46722.0 | 0 | Felis catus | 46722.0 | 0 |
| scf7180000021144 | 79320 | 0.4397 | 0 | 50.8957 | Eukaryota | 15927.0 | 0 | Chordata | 15927.0 | 0 | Carnivora | 15927.0 | 0 | Felidae | 15927.0 | 0 | Felis | 9093.0 | 1 | Felis catus | 9093.0 | 1 |
| scf7180000023393 | 31028 | 0.5148 | 0 | 203.291 | Eukaryota | 4562.0 | 0 | Chordata | 4562.0 | 0 | Carnivora | 4562.0 | 0 | Felidae | 4562.0 | 0 | Felis | 4290.0 | 1 | Felis catus | 4290.0 | 1 |
| scf7180000020957 | 64367 | 0.3917 | 0 | 25.1142 | Eukaryota | 43653.0 | 0 | Chordata | 43653.0 | 0 | Carnivora | 43653.0 | 0 | Felidae | 43653.0 | 0 | Felis | 43653.0 | 0 | Felis catus | 43653.0 | 0 |
| scf7180000022301 | 17967 | 0.4996 | 0 | 49.0738 | Eukaryota | 11843.0 | 0 | Chordata | 11843.0 | 0 | Carnivora | 11843.0 | 0 | Felidae | 11843.0 | 0 | Felis | 11843.0 | 0 | Felis catus | 11843.0 | 0 |
| scf7180000018984 | 403956 | 0.4215 | 0 | 50.9832 | Eukaryota | 38709.0 | 0 | Chordata | 38709.0 | 0 | Carnivora | 38709.0 | 0 | Felidae | 38709.0 | 0 | Felis | 38709.0 | 0 | Felis catus | 38709.0 | 0 |
| scf7180000019928 | 164527 | 0.3705 | 0 | 24.7938 | Eukaryota | 45874.0 | 0 | Chordata | 45874.0 | 0 | Carnivora | 45874.0 | 0 | Felidae | 45874.0 | 0 | Felis | 45874.0 | 0 | Felis catus | 45874.0 | 0 |
| scf7180000018981 | 670599 | 0.3987 | 0 | 49.5501 | Eukaryota | 46183.0 | 0 | Chordata | 46183.0 | 0 | Carnivora | 46183.0 | 0 | Herpestidae | 27750.0 | 1 | Suricata | 27750.0 | 3 | ıricata suricat | 27750.0 | 3 |
| scf7180000018623 | 599175 | 0.4025 | 0 | 49.9476 | Eukaryota | 33328.0 | 0 | Chordata | 33328.0 | 0 | Carnivora | 33328.0 | 0 | Felidae | 33328.0 | 0 | Felis | 29350.0 | 1 | Felis catus | 29350.0 | 1 |
| scf7180000018600 | 1194294 | 0.5322 | 100 | 49.9799 | Eukaryota | 47486.0 | 0 | Chordata | 47486.0 | 0 | Carnivora | 47486.0 | 0 | Canidae | 21953.0 | 3 | Canis | 12899.0 | 6 | Canis lupus | 12899.0 | 6 |
| scf7180000022552 | 17998 | 0.3652 | 0 | 25.7219 | Eukaryota | 43693.0 | 0 | Chordata | 43693.0 | 0 | Carnivora | 43693.0 | 0 | Felidae | 25873.0 | 2 | Panthera | 12497.0 | 4 | ıricata suricat | 10118.0 | 5 |
| scf7180000021496 | 36888 | 0.38 | 0 | 29.1021 | Eukaryota | 48747.0 | 0 | Chordata | 48747.0 | 0 | Carnivora | 48747.0 | 0 | Felidae | 48747.0 | 0 | Felis | 43930.0 | 1 | Felis catus | 43930.0 | 1 |
| scf7180000019634 | 328923 | 0.4511 | 0 | 52.4448 | Eukaryota | 50410.0 | 0 | Chordata | 50410.0 | 0 | Carnivora | 50410.0 | 0 | Canidae | 18797.0 | 4 | Vulpes | 14054.0 | 5 | Vulpes vulpes | 14054.0 | 5 |
| scf7180000022281 | 17864 | 0.4016 | 0 | 24.2385 | Eukaryota | 33550.0 | 0 | Chordata | 33550.0 | 0 | Carnivora | 33550.0 | 0 | Felidae | 33550.0 | 0 | Felis | 33550.0 | 0 | Felis catus | 33550.0 | 0 |
| scf7180000021331 | 76041 | 0.6671 | 0 | 47.9414 | Eukaryota | 15843.0 | 0 | Chordata | 15843.0 | 0 | Carnivora | 15843.0 | 0 | Felidae | 14191.0 | 1 | Acinonyx | 12589.0 | 2 | :inonyx jubat | 12589.0 | 2 |
| scf7180000018709 | 600526 | 0.4029 | 0 | 49.2752 | Eukaryota | 105192.0 | 0 | Chordata | 105192.0 | 0 | Carnivora | 105192.0 | 0 | Felidae | 105192.0 | 0 | Lynx | 36700.0 | 3 | ynx canadens | 36700.0 | 3 |
| scf7180000022279 | 14519 | 0.4721 | 0 | 46.9203 | Eukaryota | 1922.0 | 0 | Chordata | 1922.0 | 0 | Carnivora | 1735.0 | 1 | Felidae | 1735.0 | 1 | Felis | 1735.0 | 1 | Felis catus | 1735.0 | 1 |
| scf7180000023060 | 41136 | 0.3831 | 0 | 356.8752 | Eukaryota | 7148.0 | 0 | Chordata | 7148.0 | 0 | Carnivora | 7148.0 | 0 | Felidae | 7148.0 | 0 | Felis | 7148.0 | 0 | Felis catus | 7148.0 | 0 |
| scf7180000020953 | 97541 | 0.3959 | 0 | 26.0562 | Eukaryota | 18360.0 | 0 | Chordata | 18360.0 | 0 | Carnivora | 18360.0 | 0 | Felidae | 18360.0 | 0 | Felis | 16520.0 | 1 | Felis catus | 16520.0 | 1 |
| scf7180000023624 | 19589 | 0.5976 | 0 | 115.6815 | Eukaryota | 2486.0 | 0 | Chordata | 2486.0 | 0 | Carnivora | 2486.0 | 0 | Felidae | 2486.0 | 0 | Felis | 1983.0 | 1 | Felis catus | 1983.0 | 1 |
| scf7180000021100 | 79348 | 0.6637 | 0 | 49.5506 | Eukaryota | 17343.0 | 0 | Chordata | 17343.0 | 0 | Carnivora | 17343.0 | 0 | Felidae | 6798.0 | 3 | Suricata | 5847.0 | 6 | ıricata suricat | 5847.0 | 6 |
| scf7180000022227 | 12617 | 0.3583 | 0 | 23.3236 | Eukaryota | 39003.0 | 0 | Chordata | 39003.0 | 0 | Carnivora | 39003.0 | 0 | Felidae | 39003.0 | 0 | Felis | 34980.0 | 1 | Felis catus | 34980.0 | 1 |
| scf7180000019702 | 284347 | 0.423 | 0 | 49.4551 | Eukaryota | 15991.0 | 0 | Chordata | 15991.0 | 0 | Carnivora | 15991.0 | 0 | Felidae | 15991.0 | 0 | Felis | 11835.0 | 2 | Felis catus | 11835.0 | 2 |
| scf7180000022574 | 19246 | 0.5275 | 0 | 52.9818 | Eukaryota | 6799.0 | 0 | Chordata | 6799.0 | 0 | Carnivora | 6799.0 | 0 | Felidae | 4578.0 | 3 | Lynx | 4578.0 | 3 | ynx canadens | 4578.0 | 3 |
| scf7180000020414 | 138303 | 0.3845 | 0 | 24.3984 | Eukaryota | 25830.0 | 0 | Chordata | 25830.0 | 0 | Carnivora | 25830.0 | 0 | Felidae | 25830.0 | 0 | Felis | 23272.0 | 1 | Felis catus | 23272.0 | 1 |
| scf7180000018089 | 7463 | 0.4104 | 0 | 21.5229 | Eukaryota | 5728.0 | 0 | Chordata | 5728.0 | 0 | Carnivora | 5728.0 | 0 | Felidae | 5728.0 | 0 | Felis | 4578.0 | 2 | Felis catus | 4578.0 | 2 |
| scf7180000022003 | 20837 | 0.4452 | 0 | 25.0502 | Eukaryota | 13104.0 | 0 | Chordata | 13104.0 | 0 | Primates | 8790.0 | 2 | Hominidae | 8790.0 | 3 | Homo | 5250.0 | 5 | Homo sapiens | 5250.0 | 5 |
| scf7180000020952 | 67783 | 0.5307 | 0 | 48.2923 | Eukaryota | 28194.0 | 0 | Chordata | 28194.0 | 0 | Carnivora | 28194.0 | 0 | Felidae | 28194.0 | 0 | Lynx | 11196.0 | 3 | ynx canadens | 11196.0 | 3 |
| scf7180000020027 | 165652 | 0.425 | 0 | 46.6534 | Eukaryota | 38176.0 | 0 | Chordata | 38176.0 | 0 | Carnivora | 38176.0 | 0 | Felidae | 38176.0 | 0 | Felis | 38176.0 | 0 | Felis catus | 38176.0 | 0 |
| scf7180000022348 | 27720 | 0.442 | 0 | 54.986 | Eukaryota | 14171.0 | 0 | Chordata | 14171.0 | 0 | Carnivora | 14171.0 | 0 | Felidae | 14171.0 | 0 | Felis | 12712.0 | 1 | Felis catus | 12712.0 | 1 |
| scf7180000023943 | 883197 | 0.3731 | 0 | 50.0739 | Eukaryota | 45324.0 | 0 | Chordata | 45324.0 | 0 | Carnivora | 45324.0 | 0 | Felidae | 45324.0 | 0 | Felis | 40821.0 | 1 | Felis catus | 40821.0 | 1 |
| scf7180000023905 | 1021069 | 0.5031 | 0 | 52.9214 | Eukaryota | 24407.0 | 0 | Chordata | 24407.0 | 0 | Carnivora | 14248.0 | 2 | Felidae | 7475.0 | 4 | Homo | 7353.0 | 6 | Homo sapiens | 7353.0 | 6 |
| scf7180000020174 | 1164711 | 0.4277 | 0 | 51.388 | Eukaryota | 44970.0 | 0 | Chordata | 44970.0 | 0 | Carnivora | 44970.0 | 0 | Felidae | 44970.0 | 0 | Felis | 40484.0 | 1 | Felis catus | 40484.0 | 1 |
| scf7180000021506 | 47254 | 0.4134 | 0 | 24.6114 | Eukaryota | 11568.0 | 0 | Chordata | 11568.0 | 0 | Carnivora | 11568.0 | 0 | Felidae | 11568.0 | 0 | Felis | 10432.0 | 1 | Felis catus | 10432.0 | 1 |
| scf7180000019340 | 292500 | 0.4209 | 0 | 50.5567 | Eukaryota | 9505.0 | 1 | Chordata | 9505.0 | 1 | Chiroptera | 4210.0 | 4 | Cercopithecidae | 1962.0 | 8 | undef | 1230.0 | 9 | tic synthetic c | 1230.0 | 9 |
| scf7180000024058 | 286728 | 0.4877 | 0 | 50.5766 | Eukaryota | 24093.0 | 0 | Chordata | 24093.0 | 0 | Carnivora | 24093.0 | 0 | Felidae | 24093.0 | 0 | Felis | 21644.0 | 1 | Felis catus | 21644.0 | 1 |
| scf7180000021473 | 31045 | 0.4105 | 0 | 45.4854 | Eukaryota | 16389.0 | 0 | Chordata | 16389.0 | 0 | Carnivora | 16389.0 | 0 | Felidae | 16389.0 | 0 | Felis | 16389.0 | 0 | Felis catus | 16389.0 | 0 |
| scf7180000017922 | 1993201 | 0.3784 | 0 | 62.0798 | Eukaryota | 47488.0 | 0 | Chordata | 47488.0 | 0 | Carnivora | 47488.0 | 0 | Felidae | 47488.0 | 0 | Felis | 47488.0 | 0 | Felis catus | 47488.0 | 0 |
| scf7180000019278 | 311002 | 0.3544 | 0 | 47.8442 | Eukaryota | 35820.0 | 0 | Chordata | 35820.0 | 0 | Carnivora | 35820.0 | 0 | Felidae | 35820.0 | 0 | Felis | 35820.0 | 0 | Felis catus | 35820.0 | 0 |
| scf7180000019583 | 385347 | 0.3938 | 0 | 51.8753 | Eukaryota | 46718.0 | 0 | Chordata | 46718.0 | 0 | Carnivora | 46718.0 | 0 | Felidae | 46718.0 | 0 | Felis | 46718.0 | 0 | Felis catus | 46718.0 | 0 |
| scf7180000019069 | 720692 | 0.3595 | 0 | 49.6398 | Eukaryota | 41100.0 | 0 | Chordata | 41100.0 | 0 | Carnivora | 41100.0 | 0 | Felidae | 41100.0 | 0 | Felis | 41100.0 | 0 | Felis catus | 41100.0 | 0 |
| scf7180000021835 | 40836 | 0.4171 | 0 | 25.0491 | Eukaryota | 13532.0 | 0 | Chordata | 13532.0 | 0 | Carnivora | 13532.0 | 0 | Felidae | 13532.0 | 0 | Felis | 12189.0 | 1 | Felis catus | 12189.0 | 1 |
| scf7180000023437 | 11577 | 0.5222 | 0 | 131.3165 | no-hit | 0.0 | 0 | no-hit | 0.0 | 0 | no-hit | 0.0 | 0 | no-hit | 0.0 | 0 | no-hit | 0.0 | 0 | no-hit | 0.0 | 0 |
| scf7180000022770 | 6448 | 0.3766 | 0 | 23.8938 | Eukaryota | 8029.0 | 0 | Chordata | 8029.0 | 0 | Carnivora | 8029.0 | 0 | Felidae | 8029.0 | 0 | Felis | 8029.0 | 0 | Felis catus | 8029.0 | 0 |
| scf7180000019628 | 326318 | 0.5053 | 0 | 62.3608 | Eukaryota | 14295.0 | 0 | Chordata | 14295.0 | 0 | Carnivora | 14295.0 | 0 | Felidae | 14295.0 | 0 | Felis | 14295.0 | 0 | Felis catus | 14295.0 | 0 |
| scf7180000021559 | 66474 | 0.3572 | 0 | 24.8972 | Eukaryota | 15363.0 | 0 | Chordata | 15363.0 | 0 | Carnivora | 15363.0 | 0 | Felidae | 15363.0 | 0 | Felis | 12257.0 | 1 | Felis catus | 12257.0 | 1 |
| scf7180000020120 | 153279 | 0.3785 | 0 | 24.2178 | Eukaryota | 37883.0 | 0 | Chordata | 37883.0 | 0 | Carnivora | 37883.0 | 0 | Felidae | 37883.0 | 0 | Felis | 37883.0 | 0 | Felis catus | 37883.0 | 0 |
| scf7180000021297 | 44480 | 0.3897 | 0 | 24.7219 | Eukaryota | 44395.0 | 0 | Chordata | 44395.0 | 0 | Carnivora | 44395.0 | 0 | Felidae | 44395.0 | 0 | Felis | 44395.0 | 0 | Felis catus | 44395.0 | 0 |
| scf7180000017539 | 1259026 | 0.3616 | 0 | 49.1649 | Eukaryota | 46264.0 | 0 | Chordata | 46264.0 | 0 | Carnivora | 46264.0 | 0 | Felidae | 46264.0 | 0 | Felis | 46264.0 | 0 | Felis catus | 46264.0 | 0 |
| scf7180000020243 | 142752 | 0.4252 | 0 | 25.2158 | Eukaryota | 14697.0 | 0 | Chordata | 14697.0 | 0 | Carnivora | 14697.0 | 0 | Felidae | 14697.0 | 0 | Felis | 14697.0 | 0 | Felis catus | 14697.0 | 0 |
| scf7180000021033 | 53784 | 0.4193 | 0 | 27.0395 | Eukaryota | 11485.0 | 0 | Chordata | 11485.0 | 0 | Carnivora | 11485.0 | 0 | Felidae | 11485.0 | 0 | Felis | 9190.0 | 1 | Felis catus | 9190.0 | 1 |
| scf7180000021941 | 21502 | 0.3655 | 0 | 48.1251 | Eukaryota | 17697.0 | 0 | Chordata | 17697.0 | 0 | Carnivora | 17697.0 | 0 | Felidae | 17697.0 | 0 | Felis | 17697.0 | 0 | Felis catus | 17697.0 | 0 |
| scf7180000023267 | 37141 | 0.4138 | 0 | 485.5319 | Eukaryota | 4585.0 | 0 | Chordata | 4585.0 | 0 | Carnivora | 4585.0 | 0 | Felidae | 4585.0 | 0 | Felis | 3694.0 | 1 | Felis catus | 3694.0 | 1 |
| scf7180000022478 | 9969 | 0.3832 | 0 | 25.646 | Eukaryota | 29543.0 | 0 | Chordata | 29543.0 | 0 | Carnivora | 26881.0 | 1 | Felidae | 14497.0 | 3 | Felis | 6237.0 | 6 | Felis catus | 6237.0 | 6 |
| scf7180000018919 | 63690 | 0.6122 | 0 | 49.6335 | Eukaryota | 6107.0 | 0 | Chordata | 6107.0 | 0 | Carnivora | 6107.0 | 0 | Felidae | 4277.0 | 1 | Suricata | 1830.0 | 5 | ıricata suricat | 1830.0 | 5 |
| scf7180000020397 | 137076 | 0.4659 | 0 | 48.1846 | Eukaryota | 39732.0 | 0 | Chordata | 39732.0 | 0 | Carnivora | 39732.0 | 0 | Felidae | 39732.0 | 0 | Felis | 35730.0 | 1 | Felis catus | 35730.0 | 1 |
| scf7180000021420 | 46948 | 0.365 | 0 | 24.387 | Eukaryota | 21156.0 | 0 | Chordata | 21156.0 | 0 | Carnivora | 21156.0 | 0 | Felidae | 21156.0 | 0 | Felis | 19039.0 | 1 | Felis catus | 19039.0 | 1 |
| scf7180000023191 | 35287 | 0.4174 | 0 | 196.959 | Eukaryota | 27132.0 | 0 | Chordata | 27132.0 | 0 | Carnivora | 27132.0 | 0 | Felidae | 27132.0 | 0 | Felis | 24374.0 | 1 | Felis catus | 24374.0 | 1 |
| scf7180000021619 | 36491 | 0.4572 | 0 | 24.0471 | Eukaryota | 40492.0 | 0 | Chordata | 40492.0 | 0 | Carnivora | 13017.0 | 4 | Otariidae | 8562.0 | 6 | Pteropus | 7936.0 | 7 | nachus schau | 4455.0 | 9 |
| scf7180000020130 | 170169 | 0.3532 | 0 | 47.9151 | Eukaryota | 31820.0 | 0 | Chordata | 31820.0 | 0 | Carnivora | 31820.0 | 0 | Felidae | 31820.0 | 0 | Felis | 28628.0 | 1 | Felis catus | 28628.0 | 1 |
| scf7180000024189 | 117547 | 0.3708 | 0 | 25.1936 | Eukaryota | 17528.0 | 0 | Chordata | 17528.0 | 0 | Carnivora | 17528.0 | 0 | Felidae | 17528.0 | 0 | Felis | 15762.0 | 1 | Felis catus | 15762.0 | 1 |
| scf7180000023690 | 21237 | 0.4074 | 0 | 196.4 | Eukaryota | 6041.0 | 0 | Chordata | 6041.0 | 0 | Carnivora | 6041.0 | 0 | Felidae | 5246.0 | 1 | Felis | 5246.0 | 1 | Felis catus | 5246.0 | 1 |
| scf7180000023698 | 1241007 | 0.4514 | 0 | 79.2593 | Eukaryota | 64587.0 | 0 | Chordata | 64587.0 | 0 | Carnivora | 64587.0 | 0 | Felidae | 64587.0 | 0 | Panthera | 45252.0 | 1 | anthera pardı | 25848.0 | 2 |
| scf7180000020580 | 128314 | 0.4686 | 0 | 25.4134 | Eukaryota | 14373.0 | 0 | Chordata | 14373.0 | 0 | Carnivora | 14373.0 | 0 | Felidae | 14373.0 | 0 | Felis | 12877.0 | 1 | Felis catus | 12877.0 | 1 |
| scf7180000018036 | 273511 | 0.4876 | 0 | 77.1676 | Eukaryota | 41995.0 | 0 | Chordata | 41995.0 | 0 | Carnivora | 41995.0 | 0 | Felidae | 41995.0 | 0 | Lynx | 24616.0 | 2 | ynx canadens | 24616.0 | 3 |
| scf7180000018927 | 175987 | 0.4189 | 0 | 51.9572 | Eukaryota | 8464.0 | 1 | Chordata | 8464.0 | 1 | Carnivora | 6509.0 | 2 | Felidae | 6509.0 | 3 | Felis | 6509.0 | 3 | Felis catus | 6509.0 | 3 |
| scf7180000019318 | 329398 | 0.3713 | 0 | 50.5178 | Eukaryota | 47600.0 | 0 | Chordata | 47600.0 | 0 | Carnivora | 47600.0 | 0 | Felidae | 47600.0 | 0 | Felis | 47600.0 | 0 | Felis catus | 47600.0 | 0 |
| scf7180000022859 | 38693 | 0.4388 | 0 | 170.7528 | Eukaryota | 5566.0 | 0 | Chordata | 5566.0 | 0 | Artiodactyla | 5566.0 | 0 | Bovidae | 5566.0 | 0 | Ovis | 5566.0 | 0 | Ovis canadens | 5566.0 | 0 |
| scf7180000018298 | 2783327 | 0.4479 | 0 | 51.4652 | Eukaryota | 45890.0 | 0 | Chordata | 45890.0 | 0 | Carnivora | 45890.0 | 0 | Felidae | 45890.0 | 0 | Acinonyx | 26094.0 | 3 | :inonyx jubat | 26094.0 | 3 |
| scf7180000017517 | 393032 | 0.3761 | 0 | 49.7555 | Eukaryota | 45047.0 | 0 | Chordata | 45047.0 | 0 | Carnivora | 45047.0 | 0 | Felidae | 45047.0 | 0 | Felis | 45047.0 | 0 | Felis catus | 45047.0 | 0 |
| scf7180000020435 | 98459 | 0.3786 | 0 | 24.8116 | Eukaryota | 25735.0 | 0 | Chordata | 25735.0 | 0 | Primates | 16674.0 | 2 | Hominidae | 16674.0 | 3 | Homo | 10014.0 | 4 | Homo sapiens | 10014.0 | 4 |
| scf7180000019332 | 387799 | 0.4202 | 0 | 50.8191 | Eukaryota | 26729.0 | 0 | Chordata | 26729.0 | 0 | Carnivora | 26729.0 | 0 | Felidae | 26729.0 | 0 | Acinonyx | 8097.0 | 4 | :inonyx jubat | 8097.0 | 4 |
| scf7180000018101 | 25304 | 0.4317 | 0 | 24.7898 | Eukaryota | 69432.0 | 0 | Chordata | 69432.0 | 0 | Carnivora | 69432.0 | 0 | Felidae | 69432.0 | 0 | Felis | 41700.0 | 1 | Felis catus | 41700.0 | 1 |

| | | | | | | | | | | | | | | | | | | | | | | |
|---|---|---|---|---|---|---|---|---|---|---|---|---|---|---|---|---|---|---|---|---|---|---|
| scf7180000018044 | 1330552 | 0.4113 | 0 | 50.6845 | Eukaryota | 46198.0 | 0 | Chordata | 46198.0 | 0 | Carnivora | 46198.0 | 0 | Felidae | 46198.0 | 0 | Felis | 46198.0 | 0 | Felis catus | 46198.0 | 0 |
| scf7180000018151 | 942215 | 0.3707 | 0 | 50.4636 | Eukaryota | 51840.0 | 0 | Chordata | 51840.0 | 0 | Carnivora | 51840.0 | 0 | Felidae | 51840.0 | 0 | Felis | 46106.0 | 1 | Felis catus | 46106.0 | 1 |
| scf7180000023751 | 38706 | 0.3779 | 0 | 417.4957 | Eukaryota | 7262.0 | 0 | Chordata | 7262.0 | 0 | Carnivora | 7262.0 | 0 | Felidae | 7262.0 | 0 | Felis | 7262.0 | 0 | Felis catus | 7262.0 | 0 |
| scf7180000019862 | 307836 | 0.4261 | 0 | 51.3151 | Eukaryota | 16614.0 | 0 | Chordata | 16614.0 | 0 | Carnivora | 16614.0 | 0 | Felidae | 16614.0 | 0 | Felis | 14922.0 | 1 | Felis catus | 14922.0 | 1 |
| scf7180000019193 | 418308 | 0.477 | 0 | 55.2864 | Eukaryota | 25323.0 | 0 | Chordata | 25323.0 | 0 | Carnivora | 25323.0 | 0 | Felidae | 25323.0 | 0 | Felis | 11028.0 | 3 | Felis catus | 11028.0 | 3 |
| scf7180000018948 | 614785 | 0.3785 | 0 | 50.6202 | Eukaryota | 44905.0 | 0 | Chordata | 44905.0 | 0 | Carnivora | 44905.0 | 0 | Felidae | 44905.0 | 0 | Felis | 44905.0 | 0 | Felis catus | 44905.0 | 0 |
| scf7180000018071 | 24260 | 0.375 | 0 | 29.3865 | Eukaryota | 45781.0 | 0 | Chordata | 45781.0 | 0 | Carnivora | 45781.0 | 0 | Felidae | 45781.0 | 0 | Felis | 45781.0 | 0 | Felis catus | 45781.0 | 0 |
| scf7180000023861 | 388900 | 0.4868 | 0 | 50.1925 | Eukaryota | 41778.0 | 0 | Chordata | 41778.0 | 0 | Carnivora | 41778.0 | 0 | Felidae | 41778.0 | 0 | Felis | 37624.0 | 1 | Felis catus | 37624.0 | 1 |
| scf7180000018855 | 57358 | 0.3938 | 0 | 24.66 | Eukaryota | 15104.0 | 0 | Chordata | 15104.0 | 0 | Carnivora | 15104.0 | 0 | Felidae | 15104.0 | 0 | Felis | 10550.0 | 1 | Felis catus | 10550.0 | 1 |
| scf7180000021241 | 32837 | 0.6355 | 0 | 51.5352 | Eukaryota | 9718.0 | 0 | Chordata | 9718.0 | 0 | Carnivora | 9718.0 | 0 | Felidae | 9718.0 | 0 | Lynx | 4864.0 | 1 | ynx canadens | 4864.0 | 1 |
| scf7180000021391 | 8633 | 0.5568 | 0 | 39.4768 | Eukaryota | 3364.0 | 0 | Chordata | 3364.0 | 0 | Carnivora | 3364.0 | 0 | Felidae | 3364.0 | 0 | Felis | 3064.0 | 1 | Felis catus | 3064.0 | 1 |
| scf7180000020569 | 109553 | 0.49 | 0 | 45.5897 | Eukaryota | 21560.0 | 0 | Chordata | 21560.0 | 0 | Carnivora | 21560.0 | 0 | Felidae | 21560.0 | 0 | Lynx | 17288.0 | 1 | ynx canadens | 17288.0 | 1 |
| scf7180000018846 | 504009 | 0.4661 | 0 | 48.9004 | Eukaryota | 47404.0 | 0 | Chordata | 47404.0 | 0 | Carnivora | 47404.0 | 0 | Otariidae | 37202.0 | 1 | Eumetopias | 18556.0 | 3 | netopias juba | 18556.0 | 3 |
| scf7180000018004 | 1237767 | 0.5336 | 0 | 50.5521 | Eukaryota | 23037.0 | 0 | Chordata | 23037.0 | 0 | Carnivora | 23037.0 | 0 | Felidae | 19295.0 | 1 | Panthera | 11539.0 | 5 | anthera pardu | 7597.0 | 7 |
| scf7180000020750 | 63692 | 0.4524 | 0 | 23.216 | Eukaryota | 35570.0 | 0 | Chordata | 35570.0 | 0 | Carnivora | 35570.0 | 0 | Felidae | 35570.0 | 0 | Lynx | 19465.0 | 1 | ynx canadens | 19465.0 | 1 |
| scf7180000020486 | 58460 | 0.3681 | 0 | 26.8094 | Eukaryota | 45582.0 | 0 | Chordata | 45582.0 | 0 | Carnivora | 45582.0 | 0 | Felidae | 45582.0 | 0 | Felis | 45582.0 | 0 | Felis catus | 45582.0 | 0 |
| scf7180000018068 | 263063 | 0.426 | 0 | 49.8112 | Eukaryota | 45315.0 | 0 | Chordata | 45315.0 | 0 | Carnivora | 45315.0 | 0 | Felidae | 45315.0 | 0 | Felis | 45315.0 | 0 | Felis catus | 45315.0 | 0 |
| scf7180000018170 | 881460 | 0.4842 | 0 | 50.3565 | Eukaryota | 32152.0 | 0 | Chordata | 32152.0 | 0 | Carnivora | 32152.0 | 0 | Felidae | 32152.0 | 0 | Felis | 28879.0 | 1 | Felis catus | 28879.0 | 1 |
| scf7180000018809 | 328649 | 0.4258 | 0 | 78.1981 | Eukaryota | 45290.0 | 0 | Chordata | 45290.0 | 0 | Carnivora | 45290.0 | 0 | Felidae | 45290.0 | 0 | Felis | 45290.0 | 0 | Felis catus | 45290.0 | 0 |
| scf7180000021984 | 22676 | 0.419 | 0 | 27.8593 | Eukaryota | 5134.0 | 0 | Chordata | 5134.0 | 0 | Carnivora | 5134.0 | 0 | Felidae | 5134.0 | 0 | Felis | 4716.0 | 1 | Felis catus | 4716.0 | 1 |
| scf7180000019026 | 441933 | 0.526 | 0 | 52.672 | Eukaryota | 11089.0 | 0 | Chordata | 11089.0 | 0 | Carnivora | 11089.0 | 0 | Felidae | 6351.0 | 2 | Suricata | 3853.0 | 6 | ıricata suricat | 3853.0 | 6 |
| scf7180000018227 | 949157 | 0.3751 | 0 | 49.7074 | Eukaryota | 47227.0 | 0 | Chordata | 47227.0 | 0 | Carnivora | 47227.0 | 0 | Felidae | 47227.0 | 0 | Felis | 47227.0 | 0 | Felis catus | 47227.0 | 0 |
| scf7180000020374 | 90070 | 0.6237 | 0 | 47.9607 | Eukaryota | 12515.0 | 0 | Chordata | 12515.0 | 0 | Carnivora | 4524.0 | 2 | Delphinidae | 3729.0 | 4 | Homo | 3471.0 | 5 | Homo sapien | 3471.0 | 5 |
| scf7180000021816 | 51592 | 0.4578 | 0 | 25.4108 | Eukaryota | 66840.0 | 0 | Chordata | 66840.0 | 0 | Carnivora | 56370.0 | 1 | Felidae | 51160.0 | 2 | Acinonyx | 27456.0 | 5 | cinonyx jubat | 27456.0 | 5 |
| scf7180000019224 | 348427 | 0.4221 | 0 | 50.8047 | Eukaryota | 77556.0 | 0 | Chordata | 77556.0 | 0 | Carnivora | 77556.0 | 0 | Felidae | 77556.0 | 0 | Panthera | 77556.0 | 0 | anthera pardu | 69811.0 | 1 |
| scf7180000020303 | 273767 | 0.4099 | 0 | 30.0029 | Eukaryota | 37106.0 | 0 | Chordata | 37106.0 | 0 | Carnivora | 37106.0 | 0 | Felidae | 37106.0 | 0 | Felis | 37106.0 | 0 | Felis catus | 37106.0 | 0 |
| scf7180000018685 | 643493 | 0.3774 | 0 | 51.0513 | Eukaryota | 40665.0 | 0 | Chordata | 40665.0 | 0 | Carnivora | 40665.0 | 0 | Felidae | 24779.0 | 2 | Panthera | 24779.0 | 2 | anthera pardu | 20683.0 | 3 |
| scf7180000019362 | 364588 | 0.4304 | 0 | 51.6241 | Eukaryota | 17355.0 | 0 | Chordata | 17355.0 | 0 | Carnivora | 17355.0 | 0 | Felidae | 15644.0 | 1 | Felis | 13882.0 | 2 | Felis catus | 13882.0 | 2 |
| scf7180000021644 | 52492 | 0.3648 | 0 | 24.8384 | Eukaryota | 14382.0 | 0 | Chordata | 14382.0 | 0 | Carnivora | 14382.0 | 0 | Felidae | 14382.0 | 0 | Felis | 14382.0 | 0 | Felis catus | 14382.0 | 0 |
| scf7180000023831 | 1258344 | 0.4892 | 0 | 51.207 | Eukaryota | 23838.0 | 1 | Chordata | 23838.0 | 1 | Artiodactyla | 19709.0 | 3 | Bovidae | 19709.0 | 3 | Ovis | 19709.0 | 3 | Ovis canadens | 19709.0 | 3 |
| scf7180000019321 | 614852 | 0.6126 | 0 | 52.4772 | Eukaryota | 30602.0 | 0 | Chordata | 30602.0 | 0 | Carnivora | 30602.0 | 0 | Felidae | 30602.0 | 0 | Acinonyx | 21476.0 | 1 | cinonyx jubat | 21476.0 | 1 |
| scf7180000018018 | 17350 | 0.527 | 0 | 584.7211 | Eukaryota | 1818.0 | 0 | Chordata | 1818.0 | 0 | Carnivora | 1818.0 | 0 | Felidae | 1818.0 | 0 | Felis | 1818.0 | 0 | Felis catus | 1818.0 | 0 |
| scf7180000021146 | 43792 | 0.4384 | 0 | 24.4377 | Eukaryota | 16034.0 | 0 | Chordata | 16034.0 | 0 | Carnivora | 16034.0 | 0 | Felidae | 16034.0 | 0 | Felis | 16034.0 | 0 | Felis catus | 16034.0 | 0 |
| scf7180000018561 | 592305 | 0.417 | 0 | 50.501 | Eukaryota | 36215.0 | 0 | Chordata | 36215.0 | 0 | Carnivora | 36215.0 | 0 | Felidae | 19972.0 | 2 | Acinonyx | 10027.0 | 4 | cinonyx jubat | 10027.0 | 5 |
| scf7180000020162 | 105710 | 0.53 | 0 | 48.3569 | Eukaryota | 12708.0 | 0 | Chordata | 12708.0 | 0 | Carnivora | 5469.0 | 2 | Hominidae | 4784.0 | 4 | Homo | 3576.0 | 7 | Homo sapien | 3576.0 | 7 |
| scf7180000018527 | 667898 | 0.3935 | 0 | 51.0469 | Eukaryota | 103352.0 | 0 | Chordata | 103352.0 | 0 | Carnivora | 103352.0 | 0 | Felidae | 61956.0 | 1 | Felis | 61956.0 | 1 | Felis catus | 61956.0 | 1 |
| scf7180000023456 | 123997 | 0.3576 | 0 | 630.1741 | Eukaryota | 43560.0 | 0 | Chordata | 43560.0 | 0 | Carnivora | 43560.0 | 0 | Felidae | 43560.0 | 0 | Felis | 39188.0 | 1 | Felis catus | 39188.0 | 1 |
| scf7180000020061 | 706707 | 0.3944 | 100 | 52.2431 | Eukaryota | 47130.0 | 0 | Chordata | 47130.0 | 0 | Carnivora | 47130.0 | 0 | Felidae | 47130.0 | 0 | Felis | 47130.0 | 0 | Felis catus | 47130.0 | 0 |
| scf7180000023819 | 2405975 | 0.4344 | 0 | 50.8595 | Eukaryota | 87691.0 | 0 | Chordata | 87691.0 | 0 | Carnivora | 87691.0 | 0 | Felidae | 87691.0 | 0 | Panthera | 26439.0 | 3 | anthera pardu | 26439.0 | 3 |
| scf7180000023786 | 1494169 | 0.3541 | 0 | 48.5629 | Eukaryota | 48152.0 | 0 | Chordata | 48152.0 | 0 | Carnivora | 48152.0 | 0 | Felidae | 48152.0 | 0 | Felis | 48152.0 | 0 | Felis catus | 48152.0 | 0 |
| scf7180000022823 | 62200 | 0.4421 | 0 | 88.7548 | Eukaryota | 42292.0 | 0 | Chordata | 42292.0 | 0 | Carnivora | 42292.0 | 0 | Felidae | 42292.0 | 0 | Felis | 42292.0 | 0 | Felis catus | 42292.0 | 0 |
| scf7180000018204 | 955527 | 0.4526 | 0 | 50.6004 | Eukaryota | 53607.0 | 0 | Chordata | 53607.0 | 0 | Carnivora | 53607.0 | 0 | Felidae | 33082.0 | 1 | Felis | 33082.0 | 3 | Felis catus | 33082.0 | 3 |
| scf7180000024081 | 142864 | 0.3809 | 0 | 48.8883 | Eukaryota | 25619.0 | 0 | Chordata | 25619.0 | 0 | Carnivora | 25619.0 | 0 | Felidae | 25619.0 | 0 | Felis | 23018.0 | 1 | Felis catus | 23018.0 | 1 |
| scf7180000022423 | 10249 | 0.4993 | 0 | 26.5633 | Eukaryota | 9458.0 | 0 | Chordata | 9458.0 | 0 | Primates | 5574.0 | 3 | Hominidae | 5574.0 | 3 | Homo | 4185.0 | 4 | Homo sapien | 4185.0 | 4 |
| scf7180000022660 | 6721 | 0.6341 | 0 | 42.6611 | Eukaryota | 2447.0 | 0 | Chordata | 2447.0 | 0 | Carnivora | 2447.0 | 0 | Felidae | 2447.0 | 0 | Felis | 1475.0 | 2 | Felis catus | 1475.0 | 3 |
| scf7180000024150 | 102728 | 0.5096 | 0 | 48.6138 | Eukaryota | 8516.0 | 0 | Chordata | 8516.0 | 0 | Carnivora | 8516.0 | 0 | Felidae | 8516.0 | 0 | Felis | 8516.0 | 0 | Felis catus | 8516.0 | 0 |
| scf7180000017718 | 1711656 | 0.4263 | 0 | 51.81 | Eukaryota | 46130.0 | 0 | Chordata | 46130.0 | 0 | Carnivora | 46130.0 | 0 | Felidae | 46130.0 | 0 | Felis | 46130.0 | 0 | Felis catus | 46130.0 | 0 |
| scf7180000022183 | 21220 | 0.4366 | 0 | 26.3359 | Eukaryota | 3850.0 | 0 | Chordata | 3850.0 | 0 | Carnivora | 1495.0 | 3 | Hominidae | 1350.0 | 4 | Felis | 1001.0 | 6 | Felis catus | 1001.0 | 6 |
| scf7180000023680 | 82468 | 0.3856 | 0 | 396.5678 | Eukaryota | 12367.0 | 0 | Chordata | 12367.0 | 0 | Carnivora | 12367.0 | 0 | Felidae | 12367.0 | 0 | Felis | 7458.0 | 1 | Felis catus | 7458.0 | 1 |
| scf7180000022235 | 13579 | 0.3589 | 0 | 20.5595 | Eukaryota | 28429.0 | 0 | Chordata | 28429.0 | 0 | Carnivora | 28429.0 | 0 | Felidae | 28429.0 | 0 | Felis | 25627.0 | 1 | Felis catus | 25627.0 | 1 |
| scf7180000018158 | 1278198 | 0.4058 | 0 | 51.9172 | Eukaryota | 55658.0 | 0 | Chordata | 55658.0 | 0 | Carnivora | 55658.0 | 0 | Felidae | 55658.0 | 0 | Acinonyx | 44504.0 | 2 | cinonyx jubat | 44504.0 | 2 |
| scf7180000023885 | 1041764 | 0.4559 | 0 | 52.0323 | Eukaryota | 58930.0 | 0 | Chordata | 58930.0 | 0 | Carnivora | 58930.0 | 0 | Felidae | 34530.0 | 1 | Felis | 34530.0 | 1 | Felis catus | 34530.0 | 1 |
| scf7180000020660 | 90524 | 0.4844 | 0 | 26.6023 | Eukaryota | 24876.0 | 0 | Chordata | 24876.0 | 0 | Carnivora | 24876.0 | 0 | Felidae | 24876.0 | 0 | Felis | 22368.0 | 1 | Felis catus | 22368.0 | 1 |
| scf7180000022840 | 24385 | 0.4574 | 0 | 479.9629 | no-hit | 0.0 | 0 | no-hit | 0.0 | 0 | no-hit | 0.0 | 0 | no-hit | 0.0 | 0 | no-hit | 0.0 | 0 | no-hit | 0.0 | 0 |
| scf7180000024168 | 154611 | 0.3725 | 0 | 25.4795 | Eukaryota | 20639.0 | 0 | Chordata | 20639.0 | 0 | Carnivora | 20639.0 | 0 | Felidae | 20639.0 | 0 | Felis | 16455.0 | 1 | Felis catus | 16455.0 | 1 |
| scf7180000022659 | 5478 | 0.4356 | 0 | 204.8471 | Eukaryota | 2156.0 | 0 | Chordata | 2156.0 | 0 | Carnivora | 2156.0 | 0 | Felidae | 2156.0 | 0 | Felis | 1317.0 | 2 | Felis catus | 1317.0 | 3 |
| scf7180000019386 | 291264 | 0.4258 | 0 | 49.4516 | Eukaryota | 9085.0 | 0 | Chordata | 9085.0 | 0 | Carnivora | 9085.0 | 0 | Felidae | 9085.0 | 0 | Felis | 8202.0 | 1 | Felis catus | 8202.0 | 1 |
| scf7180000020882 | 74089 | 0.34 | 0 | 49.1143 | Eukaryota | 5711.0 | 1 | Chordata | 5711.0 | 1 | Carnivora | 1865.0 | 4 | Bovidae | 1836.0 | 5 | Felis | 1519.0 | 6 | Felis catus | 1519.0 | 6 |
| scf7180000019092 | 61183 | 0.4277 | 0 | 55.2432 | Eukaryota | 32570.0 | 0 | Chordata | 32570.0 | 0 | Carnivora | 32570.0 | 0 | Felidae | 32570.0 | 0 | Felis | 29340.0 | 1 | Felis catus | 29340.0 | 1 |
| scf7180000022448 | 19407 | 0.405 | 0 | 24.575 | Eukaryota | 44885.0 | 0 | Chordata | 44885.0 | 0 | Carnivora | 44885.0 | 0 | Felidae | 44885.0 | 0 | Felis | 44885.0 | 0 | Felis catus | 44885.0 | 0 |
| scf7180000022341 | 20369 | 0.4038 | 0 | 23.6999 | Eukaryota | 40288.0 | 0 | Chordata | 40288.0 | 0 | Carnivora | 40288.0 | 0 | Felidae | 40288.0 | 0 | Felis | 40288.0 | 0 | Felis catus | 40288.0 | 0 |
| scf7180000019878 | 155019 | 0.4631 | 0 | 52.1948 | Eukaryota | 19464.0 | 0 | Chordata | 19464.0 | 0 | Carnivora | 19464.0 | 0 | Felidae | 19464.0 | 0 | Felis | 17454.0 | 1 | Felis catus | 17454.0 | 1 |
| scf7180000024159 | 114245 | 0.441 | 0 | 26.3777 | Eukaryota | 31697.0 | 0 | Chordata | 31697.0 | 0 | Carnivora | 28945.0 | 1 | Felidae | 18184.0 | 5 | Panthera | 7229.0 | 8 | ıricata suricat | 3788.0 | 9 |
| scf7180000020011 | 208976 | 0.5337 | 0 | 53.5532 | Eukaryota | 33977.0 | 0 | Chordata | 33977.0 | 0 | Carnivora | 33977.0 | 0 | Felidae | 10659.0 | 5 | Panthera | 3565.0 | 9 | anthera pardu | 3565.0 | 9 |
| scf7180000018035 | 44205 | 0.4561 | 0 | 37.2604 | Eukaryota | 7283.0 | 1 | Chordata | 7283.0 | 1 | Carnivora | 6214.0 | 3 | Felidae | 6214.0 | 3 | Felis | 6214.0 | 3 | Felis catus | 6214.0 | 3 |
| scf7180000020733 | 168470 | 0.4331 | 0 | 54.0007 | Eukaryota | 37721.0 | 0 | Chordata | 37721.0 | 0 | Carnivora | 37721.0 | 0 | Felidae | 31651.0 | 2 | Puma | 11876.0 | 6 | Puma concolo | 11876.0 | 7 |
| scf7180000022521 | 20637 | 0.4272 | 0 | 24.3396 | Eukaryota | 3525.0 | 0 | Chordata | 3183.0 | 1 | Primates | 2493.0 | 2 | Hominidae | 2493.0 | 2 | Homo | 2084.0 | 3 | Homo sapien | 2084.0 | 3 |
| scf7180000021212 | 52848 | 0.3872 | 0 | 24.9356 | Eukaryota | 20939.0 | 0 | Chordata | 20939.0 | 0 | Carnivora | 20939.0 | 0 | Felidae | 12809.0 | 2 | Panthera | 4274.0 | 6 | ynx canadens | 4248.0 | 7 |
| scf7180000020223 | 107149 | 0.4029 | 0 | 44.8456 | Eukaryota | 25123.0 | 0 | Chordata | 25123.0 | 0 | Carnivora | 25123.0 | 0 | Felidae | 25123.0 | 0 | Felis | 22565.0 | 1 | Felis catus | 22565.0 | 1 |
| scf7180000022409 | 14100 | 0.37 | 0 | 25.6274 | Eukaryota | 6454.0 | 0 | Chordata | 6454.0 | 0 | Carnivora | 4406.0 | 1 | Felidae | 4406.0 | 1 | Felis | 3851.0 | 2 | Felis catus | 3851.0 | 2 |
| scf7180000021899 | 25482 | 0.4641 | 0 | 51.0933 | Eukaryota | 28891.0 | 0 | Chordata | 28891.0 | 0 | Carnivora | 28891.0 | 0 | Felidae | 28891.0 | 0 | Felis | 28891.0 | 0 | Felis catus | 28891.0 | 0 |
| scf7180000018064 | 2135801 | 0.4205 | 0 | 51.6553 | Eukaryota | 59298.0 | 0 | Chordata | 59298.0 | 0 | Carnivora | 44780.0 | 1 | Felidae | 44780.0 | 2 | Felis | 26238.0 | 3 | Felis catus | 26238.0 | 3 |
| scf7180000021058 | 178893 | 0.6064 | 0 | 49.0683 | Eukaryota | 18613.0 | 0 | Chordata | 18613.0 | 0 | Carnivora | 14077.0 | 1 | Felidae | 7099.0 | 4 | Homo | 4536.0 | 7 | Homo sapien | 4536.0 | 8 |
| scf7180000022714 | 25554 | 0.4025 | 0 | 25.8306 | Eukaryota | 23282.0 | 0 | Chordata | 23282.0 | 0 | Carnivora | 23282.0 | 0 | Felidae | 23282.0 | 0 | Felis | 16324.0 | 1 | Felis catus | 16324.0 | 1 |
| scf7180000022371 | 7494 | 0.4029 | 0 | 28.0556 | Eukaryota | 10559.0 | 0 | Chordata | 10559.0 | 0 | Carnivora | 9517.0 | 1 | Felidae | 9517.0 | 2 | Felis | 9517.0 | 2 | Felis catus | 9517.0 | 2 |
| scf7180000020320 | 219670 | 0.5769 | 0 | 48.8972 | Eukaryota | 10557.0 | 0 | Chordata | 10557.0 | 0 | Carnivora | 10557.0 | 0 | Felidae | 6139.0 | 4 | Felis | 2830.0 | 7 | Felis catus | 2830.0 | 8 |
| scf7180000018746 | 672661 | 0.394 | 0 | 51.3784 | Eukaryota | 57804.0 | 0 | Chordata | 57804.0 | 0 | Carnivora | 57804.0 | 0 | Felidae | 57804.0 | 0 | Felis | 57804.0 | 0 | Felis catus | 57804.0 | 0 |
| scf7180000019186 | 667083 | 0.3905 | 0 | 49.71 | Eukaryota | 45560.0 | 0 | Chordata | 45560.0 | 0 | Carnivora | 45560.0 | 0 | Felidae | 45560.0 | 0 | Felis | 45560.0 | 0 | Felis catus | 45560.0 | 0 |
| scf7180000019891 | 166370 | 0.3831 | 0 | 25.594 | Eukaryota | 42903.0 | 0 | Chordata | 42903.0 | 0 | Carnivora | 42903.0 | 0 | Felidae | 37903.0 | 1 | Felis | 37903.0 | 1 | Felis catus | 37903.0 | 1 |
| scf7180000021101 | 2388 | 0.3882 | 0 | 24.4077 | Eukaryota | 2198.0 | 0 | Chordata | 2198.0 | 0 | Carnivora | 2198.0 | 0 | Felidae | 1959.0 | 1 | Felis | 1519.0 | 3 | Felis catus | 1519.0 | 3 |
| scf7180000021771 | 26872 | 0.3749 | 0 | 24.2689 | Eukaryota | 44266.0 | 0 | Chordata | 44266.0 | 0 | Carnivora | 44266.0 | 0 | Felidae | 44266.0 | 0 | Felis | 44266.0 | 0 | Felis catus | 44266.0 | 0 |
| scf7180000023663 | 64175 | 0.3853 | 0 | 294.4416 | Eukaryota | 7127.0 | 0 | Chordata | 7127.0 | 0 | Carnivora | 7127.0 | 0 | Felidae | 7127.0 | 0 | Felis | 7127.0 | 0 | Felis catus | 7127.0 | 0 |
| scf7180000023040 | 23186 | 0.3996 | 0 | 314.8848 | Eukaryota | 10163.0 | 0 | Chordata | 10163.0 | 0 | .pterygiforme | 9093.0 | 1 | Apterygidae | 9093.0 | 1 | Apteryx | 9093.0 | 1 | pteryx mante | 9093.0 | 1 |
| scf7180000018564 | 724430 | 0.3431 | 0 | 48.1411 | Eukaryota | 45211.0 | 0 | Chordata | 45211.0 | 0 | Carnivora | 45211.0 | 0 | Felidae | 45211.0 | 0 | Felis | 45211.0 | 0 | Felis catus | 45211.0 | 0 |
| scf7180000022726 | 5425 | 0.3504 | 0 | 23.6092 | Eukaryota | 16371.0 | 0 | Chordata | 16371.0 | 0 | Carnivora | 16371.0 | 0 | Felidae | 16371.0 | 0 | Felis | 13117.0 | 1 | Felis catus | 13117.0 | 1 |
| scf7180000019353 | 296864 | 0.5372 | 0 | 54.8674 | Eukaryota | 39228.0 | 0 | Chordata | 39228.0 | 0 | Carnivora | 39228.0 | 0 | Felidae | 39228.0 | 0 | Felis | 35348.0 | 1 | Felis catus | 35348.0 | 1 |
| scf7180000018387 | 878651 | 0.4086 | 0 | 48.2331 | Eukaryota | 48048.0 | 0 | Chordata | 48048.0 | 0 | Carnivora | 48048.0 | 0 | Felidae | 48048.0 | 0 | Felis | 48048.0 | 0 | Felis catus | 48048.0 | 0 |
| scf7180000020635 | 94935 | 0.5379 | 0 | 50.1629 | Eukaryota | 8268.0 | 0 | Chordata | 8268.0 | 0 | Carnivora | 8268.0 | 0 | Felidae | 8268.0 | 0 | Felis | 6623.0 | 1 | Felis catus | 6623.0 | 1 |
| scf7180000024017 | 354819 | 0.5024 | 0 | 52.613 | Eukaryota | 34857.0 | 0 | Chordata | 34857.0 | 0 | Carnivora | 26946.0 | 1 | Felidae | 17304.0 | 4 | Panthera | 6913.0 | 7 | ıricata suricat | 6820.0 | 8 |
| scf7180000020794 | 102697 | 0.568 | 0 | 49.1273 | Eukaryota | 5945.0 | 1 | Chordata | 5945.0 | 1 | Primates | 2811.0 | 5 | Hominidae | 1419.0 | 6 | Homo | 1419.0 | 8 | Homo sapien | 1419.8 | 8 |
| scf7180000019046 | 73754 | 0.4515 | 0 | 47.8072 | Eukaryota | 6296.0 | 0 | Chordata | 6296.0 | 0 | Carnivora | 6296.0 | 0 | Felidae | 5727.0 | 1 | Acinonyx | 1900.0 | 5 | cinonyx jubat | 1900.0 | 5 |
| scf7180000021333 | 64510 | 0.3639 | 0 | 24.7574 | Eukaryota | 40951.0 | 0 | Chordata | 40951.0 | 0 | Carnivora | 40951.0 | 0 | Felidae | 40951.0 | 0 | Felis | 36816.0 | 1 | Felis catus | 36816.0 | 1 |
| scf7180000019068 | 106454 | 0.3823 | 0 | 52.3165 | Eukaryota | 38797.0 | 0 | Chordata | 38797.0 | 0 | Carnivora | 38797.0 | 0 | Felidae | 38797.0 | 0 | Felis | 38797.0 | 0 | Felis catus | 38797.0 | 0 |
| scf7180000021747 | 28774 | 0.391 | 0 | 24.6646 | Eukaryota | 21501.0 | 0 | Chordata | 21501.0 | 0 | Carnivora | 21501.0 | 0 | Felidae | 21501.0 | 0 | Felis | 19384.0 | 1 | Felis catus | 19384.0 | 1 |
| scf7180000020115 | 163963 | 0.4379 | 0 | 26.7342 | Eukaryota | 37680.0 | 0 | Chordata | 37680.0 | 0 | Carnivora | 37680.0 | 0 | Felidae | 37680.0 | 0 | Felis | 37680.0 | 0 | Felis catus | 37680.0 | 0 |
| scf7180000018234 | 288630 | 0.443 | 0 | 51.6059 | Eukaryota | 45333.0 | 0 | Chordata | 45333.0 | 0 | Carnivora | 45333.0 | 0 | Felidae | 45333.0 | 0 | Felis | 45333.0 | 0 | Felis catus | 45333.0 | 0 |
| scf7180000019742 | 248255 | 0.3871 | 0 | 50.2043 | Eukaryota | 36326.0 | 0 | Chordata | 36326.0 | 0 | Carnivora | 36326.0 | 0 | Felidae | 36326.0 | 0 | Felis | 24350.0 | 2 | Felis catus | 24350.0 | 2 |
| scf7180000018817 | 47906 | 0.3888 | 0 | 26.2526 | Eukaryota | 30839.0 | 0 | Chordata | 30839.0 | 0 | Carnivora | 30839.0 | 0 | Felidae | 30839.0 | 0 | Felis | 30839.0 | 0 | Felis catus | 30839.0 | 0 |
| scf7180000023960 | 1446515 | 0.4479 | 0 | 52.5921 | Eukaryota | 42277.0 | 0 | Chordata | 42277.0 | 0 | Carnivora | 42277.0 | 0 | Felidae | 42277.0 | 0 | Felis | 42277.0 | 0 | Felis catus | 42277.0 | 0 |
| scf7180000021870 | 26839 | 0.4289 | 0 | 24.6123 | Eukaryota | 5027.0 | 0 | Chordata | 5027.0 | 0 | Carnivora | 4021.0 | 1 | Felidae | 4021.0 | 1 | Felis | 4021.0 | 2 | Felis catus | 4021.0 | 2 |
| scf7180000020730 | 76684 | 0.5458 | 0 | 46.1177 | Eukaryota | 30771.0 | 0 | Chordata | 30771.0 | 0 | Carnivora | 30771.0 | 0 | Felidae | 15242.0 | 2 | Eumetopias | 12155.0 | 4 | netopias juba | 12155.0 | 4 |
| scf7180000020127 | 172160 | 0.4622 | 0 | 50.403 | Eukaryota | 46402.0 | 0 | Chordata | 46402.0 | 0 | Carnivora | 46402.0 | 0 | Otariidae | 17938.0 | 5 | Zalophus | 8980.0 | 8 | phus californ | 8980.0 | 8 |
| scf7180000021430 | 47057 | 0.6212 | 0 | 47.6013 | Eukaryota | 8460.0 | 0 | Chordata | 8460.0 | 0 | Carnivora | 8460.0 | 0 | Felidae | 7588.0 | 1 | Panthera | 2577.0 | 4 | ynx canadens | 2512.0 | 5 |
| scf7180000023856 | 1075279 | 0.4468 | 0 | 49.9878 | Eukaryota | 39825.0 | 1 | Chordata | 39825.0 | 1 | Carnivora | 35080.2 | 2 | Felidae | 35080.0 | 2 | Felis | 35080.0 | 2 | Felis catus | 35080.2 | 2 |
| scf7180000021844 | 24272 | 0.5231 | 0 | 44.5754 | Eukaryota | 3206.0 | 1 | Chordata | 3206.0 | 1 | Carnivora | 2823.0 | 2 | Felidae | 2823.0 | 2 | Felis | 2823.0 | 2 | Felis catus | 2823.0 | 2 |
| scf7180000022212 | 17677 | 0.3614 | 0 | 21.8207 | Eukaryota | 38640.0 | 0 | Chordata | 38640.0 | 0 | Carnivora | 38640.0 | 0 | Felidae | 38640.0 | 0 | Felis | 34734.0 | 1 | Felis catus | 34734.0 | 1 |
| scf7180000022361 | 24802 | 0.3753 | 0 | 332.0037 | Eukaryota | 6530.0 | 0 | Chordata | 6530.0 | 0 | Carnivora | 6530.0 | 0 | Felidae | 6530.0 | 0 | Felis | 6530.0 | 0 | Felis catus | 6530.0 | 0 |
| scf7180000018543 | 780209 | 0.3954 | 0 | 50.0925 | Eukaryota | 46205.0 | 0 | Chordata | 46205.0 | 0 | Carnivora | 46205.0 | 0 | Felidae | 46205.0 | 0 | Felis | 46205.0 | 0 | Felis catus | 46205.0 | 0 |
| scf7180000018312 | 948950 | 0.5211 | 0 | 57.9553 | Eukaryota | 44410.0 | 0 | Chordata | 44410.0 | 0 | Carnivora | 44410.0 | 0 | Felidae | 22257.0 | 1 | Suricata | 22153.0 | 2 | ıricata suricat | 22153.0 | 2 |
| scf7180000018263 | 56685 | 0.391 | 0 | 24.8851 | Eukaryota | 18827.0 | 0 | Chordata | 18827.0 | 0 | Carnivora | 18827.0 | 0 | Felidae | 18827.0 | 0 | Felis | 16924.0 | 1 | Felis catus | 16924.0 | 1 |
| scf7180000022846 | 38963 | 0.415 | 0 | 145.025 | Eukaryota | 17056.0 | 0 | Chordata | 17056.0 | 0 | Carnivora | 17056.0 | 0 | Felidae | 17056.0 | 0 | Felis | 13684.0 | 1 | Felis catus | 13684.0 | 1 |
| scf7180000022401 | 15728 | 0.3843 | 0 | 25.28 | Eukaryota | 6748.0 | 0 | Chordata | 6748.0 | 0 | Carnivora | 3103.0 | 2 | Felidae | 2655.0 | 3 | Homo | 2254.0 | 4 | Homo sapien | 2254.0 | 4 |
| scf7180000018736 | 719089 | 0.3966 | 0 | 51.0994 | Eukaryota | 47015.0 | 0 | Chordata | 47015.0 | 0 | Carnivora | 47015.0 | 0 | Felidae | 47015.0 | 0 | Felis | 47015.0 | 0 | Felis catus | 47015.0 | 0 |
| scf7180000022274 | 13315 | 0.3976 | 0 | 46.3953 | Eukaryota | 4543.0 | 0 | Chordata | 4543.0 | 0 | Carnivora | 3541.0 | 1 | Canidae | 2642.0 | 2 | Canis | 2642.0 | 2 | Canis lupus | 2642.0 | 2 |
| scf7180000021720 | 27501 | 0.4277 | 0 | 146.729 | Eukaryota | 2197.0 | 0 | Chordata | 2197.0 | 0 | Carnivora | 2197.0 | 0 | Felidae | 2197.0 | 0 | Felis | 1548.0 | 1 | Felis catus | 1548.0 | 1 |
| scf7180000018132 | 1124916 | 0.3844 | 0 | 49.9933 | Eukaryota | 46712.0 | 0 | Chordata | 46712.0 | 0 | Carnivora | 46712.0 | 0 | Felidae | 46712.0 | 0 | Felis | 46712.0 | 0 | Felis catus | 46712.0 | 0 |
| scf7180000022399 | 10733 | 0.3916 | 0 | 25.2501 | Eukaryota | 3523.0 | 0 | Chordata | 3523.0 | 0 | Primates | 2010.0 | 3 | Hominidae | 2010.0 | 3 | Homo | 1503.0 | 5 | Homo sapien | 1503.0 | 5 |
| scf7180000019569 | 226609 | 0.5166 | 0 | 52.3175 | Eukaryota | 31946.0 | 0 | Chordata | 31946.0 | 0 | Carnivora | 18964.0 | 1 | Felidae | 13156.0 | 3 | agenorhynchu | 6532.0 | 5 | rhynchus obli | 6532.0 | 5 |
| scf7180000023112 | 37319 | 0.4428 | 0 | 179.9841 | Eukaryota | 7435.0 | 0 | Chordata | 7435.0 | 0 | Carnivora | 7435.0 | 0 | Felidae | 7435.0 | 0 | Felis | 7435.0 | 0 | Felis catus | 7435.0 | 0 |
| scf7180000022102 | 17454 | 0.3789 | 0 | 22.9716 | Eukaryota | 7347.0 | 0 | Chordata | 7347.0 | 0 | Carnivora | 5889.0 | 1 | Felidae | 5889.0 | 1 | Felis | 5889.0 | 1 | Felis catus | 5889.0 | 1 |
| scf7180000023232 | 26360 | 0.4388 | 0 | 237.9438 | Eukaryota | 4357.0 | 0 | Chordata | 4357.0 | 0 | Carnivora | 4357.0 | 0 | Felidae | 2708.0 | 1 | Felis | 2138.0 | 2 | Felis catus | 2138.0 | 2 |
| scf7180000021185 | 60292 | 0.3765 | 0 | 23.9464 | Eukaryota | 39388.0 | 0 | Chordata | 39388.0 | 0 | Carnivora | 39388.0 | 0 | Felidae | 39388.0 | 0 | Felis | 39388.0 | 0 | Felis catus | 39388.0 | 0 |
| scf7180000023652 | 42207 | 0.3931 | 0 | 280.1737 | Eukaryota | 11829.0 | 0 | Chordata | 11829.0 | 0 | Carnivora | 11829.0 | 0 | Felidae | 11829.0 | 0 | Felis | 11829.0 | 0 | Felis catus | 11829.0 | 0 |
| scf7180000022232 | 16733 | 0.5478 | 0 | 39.6324 | Eukaryota | 2774.0 | 0 | Chordata | 2774.0 | 0 | Carnivora | 2774.0 | 0 | Felidae | 2774.0 | 0 | Felis | 2216.0 | 1 | Felis catus | 2216.0 | 1 |
| scf7180000021553 | 40024 | 0.5431 | 0 | 47.9857 | Eukaryota | 8693.0 | 0 | Chordata | 8693.0 | 0 | Carnivora | 5953.0 | 2 | Felidae | 5164.0 | 3 | Homo | 1836.0 | 6 | Homo sapien | 1836.0 | 6 |
| scf7180000023494 | 17850 | 0.4899 | 0 | 85.8406 | no-hit | 0.0 | 0 | no-hit | 0.0 | 0 | no-hit | 0.0 | 0 | no-hit | 0.0 | 0 | no-hit | 0.0 | 0 | no-hit | 0.0 | 0 |
| scf7180000018350 | 863866 | 0.3941 | 0 | 52.3022 | Eukaryota | 65510.0 | 0 | Chordata | 65510.0 | 0 | Rodentia | 39144.0 | 1 | Bathyergidae | 39144.0 | 2 | leterocephalu | 39144.0 | 2 | rocephalus gl | 39144.0 | 2 |
| scf7180000024139 | 194261 | 0.4178 | 0 | 26.1982 | Eukaryota | 48141.0 | 0 | Chordata | 48141.0 | 0 | Carnivora | 48141.0 | 0 | Felidae | 30684.0 | 2 | Lynx | 15356.0 | 5 | ynx canadens | 15356.0 | 6 |
| scf7180000022777 | 20921 | 0.4292 | 0 | 26.5999 | Eukaryota | 22501.0 | 0 | Chordata | 22501.0 | 0 | Carnivora | 22501.0 | 0 | Felidae | 20262.0 | 1 | Lynx | 15449.0 | 3 | ynx canadens | 15449.0 | 3 |
| scf7180000023956 | 852521 | 0.4365 | 0 | 52.3489 | Eukaryota | 47917.0 | 0 | Chordata | 47917.0 | 0 | Carnivora | 47917.0 | 0 | Felidae | 47917.0 | 0 | Felis | 47917.0 | 0 | Felis catus | 47917.0 | 0 |
| scf7180000024251 | 1428010 | 0.3614 | 0 | 50.1744 | Eukaryota | 47246.0 | 0 | Chordata | 47246.0 | 0 | Carnivora | 47246.0 | 0 | Felidae | 47246.0 | 0 | Felis | 47246.0 | 0 | Felis catus | 47246.0 | 0 |
| scf7180000021498 | 12263 | 0.4844 | 0 | 23.6437 | Eukaryota | 5609.0 | 0 | Chordata | 5609.0 | 0 | Carnivora | 5609.0 | 0 | Felidae | 5609.0 | 0 | Felis | 4923.0 | 1 | Felis catus | 4923.0 | 1 |
| scf7180000017593 | 4860826 | 0.4303 | 0 | 51.3708 | Eukaryota | 68986.0 | 0 | Chordata | 68986.0 | 0 | Carnivora | 68986.0 | 0 | Mustelidae | 20853.0 | 4 | Mustela | 13848.0 | 7 | ıustela putori | 13848.0 | 7 |
| scf7180000022084 | 15270 | 0.4993 | 0 | 27.4889 | Eukaryota | 3646.0 | 0 | Chordata | 3646.0 | 0 | Carnivora | 3646.0 | 0 | Felidae | 3646.0 | 0 | Felis | 3289.0 | 1 | Felis catus | 3289.0 | 1 |
| scf7180000017558 | 2665743 | 0.4194 | 0 | 51.9503 | Eukaryota | 69707.0 | 0 | Chordata | 69707.0 | 0 | Carnivora | 69707.0 | 0 | Ursidae | 53023.0 | 1 | Ursus | 32773.0 | 2 | Ursus arctos | 26824.0 | 3 |
| scf7180000017871 | 575418 | 0.3725 | 0 | 47.9284 | Eukaryota | 47456.0 | 0 | Chordata | 47456.0 | 0 | Carnivora | 47456.0 | 0 | Felidae | 47456.0 | 0 | Felis | 47456.0 | 0 | Felis catus | 47456.0 | 0 |
| scf7180000019846 | 27237 | 0.4059 | 0 | 41.0769 | Eukaryota | 21211.0 | 0 | Chordata | 21211.0 | 0 | Carnivora | 21211.0 | 0 | Felidae | 21211.0 | 0 | Felis | 14886.0 | 1 | Felis catus | 14886.1 | 1 |
| scf7180000020590 | 30007 | 0.4662 | 0 | 43.4669 | Eukaryota | 3045.0 | 0 | Chordata | 3045.0 | 0 | Primates | 1495.0 | 3 | Equidae | 632.0 | 8 | Equus | 632.0 | 8 | unresolved | 316.0 | 9 |
| scf7180000018141 | 996707 | 0.4394 | 0 | 50.0418 | Eukaryota | 45898.0 | 0 | Chordata | 45898.0 | 0 | Carnivora | 45898.0 | 0 | Felidae | 45898.0 | 0 | Felis | 45898.0 | 0 | Felis catus | 45898.0 | 0 |
| scf7180000021688 | 23578 | 0.3683 | 0 | 24.0118 | Eukaryota | 38934.0 | 0 | Chordata | 38934.0 | 0 | Carnivora | 38934.0 | 0 | Felidae | 38934.0 | 0 | Felis | 38934.0 | 0 | Felis catus | 38934.0 | 0 |
| scf7180000019377 | 566883 | 0.4322 | 100 | 53.1432 | Eukaryota | 32058.0 | 0 | Chordata | 32058.0 | 0 | Carnivora | 32058.0 | 0 | Felidae | 29457.0 | 1 | Felis | 11766.0 | 5 | Felis catus | 11766.0 | 6 |
| scf7180000022847 | 30120 | 0.4101 | 0 | 264.3668 | Eukaryota | 11946.0 | 0 | Chordata | 11946.0 | 0 | Carnivora | 11946.0 | 0 | Felidae | 11946.0 | 0 | Felis | 10771.0 | 1 | Felis catus | 10771.0 | 1 |
| scf7180000024198 | 61663 | 0.3512 | 0 | 22.9303 | Eukaryota | 22516.0 | 0 | Chordata | 22516.0 | 0 | Carnivora | 22516.0 | 0 | Felidae | 22516.0 | 0 | Felis | 20259.0 | 1 | Felis catus | 20259.0 | 1 |
| scf7180000020481 | 95946 | 0.4338 | 0 | 25.495 | Eukaryota | 5521.0 | 0 | Chordata | 5521.0 | 0 | Primates | 2479.0 | 2 | Hominidae | 2479.0 | 2 | Felis | 1822.0 | 3 | Felis catus | 1822.0 | 3 |
| scf7180000022427 | 29331 | 0.3598 | 0 | 23.0444 | Eukaryota | 15561.0 | 0 | Chordata | 15561.0 | 0 | Carnivora | 15561.0 | 0 | Felidae | 15561.0 | 0 | Felis | 15561.0 | 0 | Felis catus | 15561.0 | 0 |
| scf7180000022378 | 22283 | 0.4311 | 0 | 274.2317 | Eukaryota | 2343.0 | 0 | Chordata | 2343.0 | 0 | Carnivora | 2343.0 | 0 | Felidae | 2343.0 | 0 | Felis | 1702.0 | 2 | Felis catus | 1702.0 | 2 |
| scf7180000023966 | 830641 | 0.3957 | 0 | 50.6041 | Eukaryota | 44688.0 | 0 | Chordata | 44688.0 | 0 | Carnivora | 44688.0 | 0 | Felidae | 44688.0 | 0 | Felis | 40248.0 | 1 | Felis catus | 40248.0 | 1 |
| scf7180000018592 | 623261 | 0.4413 | 0 | 50.1633 | Eukaryota | 44903.0 | 0 | Chordata | 44903.0 | 0 | Carnivora | 44903.0 | 0 | Felidae | 44903.0 | 0 | Felis | 44903.0 | 0 | Felis catus | 44903.0 | 0 |
| scf7180000020924 | 49020 | 0.4069 | 0 | 64.8229 | Eukaryota | 8004.0 | 0 | Chordata | 8004.0 | 0 | Primates | 4270.0 | 2 | Hominidae | 4270.0 | 2 | Felis | 2997.0 | 3 | Felis catus | 2997.0 | 3 |
| scf7180000023036 | 27592 | 0.5566 | 0 | 116.1287 | Eukaryota | 6343.0 | 0 | Chordata | 6343.0 | 0 | Carnivora | 6343.0 | 0 | Felidae | 6113.0 | 1 | Felis | 2479.0 | 3 | Felis catus | 2479.0 | 3 |
| scf7180000022735 | 10928 | 0.3944 | 0 | 25.864 | Eukaryota | 6277.0 | 0 | Chordata | 6277.0 | 0 | Carnivora | 6277.0 | 0 | Felidae | 6277.0 | 0 | Felis | 6277.0 | 0 | Felis catus | 6277.0 | 0 |
| scf7180000019013 | 1090757 | 0.4581 | 0 | 50.8272 | Eukaryota | 69249.0 | 0 | Chordata | 69249.0 | 0 | Carnivora | 69249.0 | 0 | Felidae | 35176.0 | 4 | Acinonyx | 14054.0 | 8 | cinonyx jubat | 14054.0 | 8 |
| scf7180000024092 | 229210 | 0.5199 | 0 | 52.22 | Eukaryota | 40991.0 | 0 | Chordata | 40991.0 | 0 | Carnivora | 40991.0 | 0 | Felidae | 40991.0 | 0 | Felis | 40991.0 | 0 | Felis catus | 40991.0 | 0 |
| scf7180000024211 | 73246 | 0.3852 | 0 | 25.1532 | Eukaryota | 32042.0 | 0 | Chordata | 32042.0 | 0 | Carnivora | 32042.0 | 0 | Felidae | 32042.0 | 0 | Felis | 28871.0 | 1 | Felis catus | 28871.0 | 1 |
| scf7180000020654 | 75086 | 0.6277 | 0 | 52.8938 | Eukaryota | 16578.0 | 0 | Chordata | 16578.0 | 0 | Carnivora | 16578.0 | 0 | Felidae | 16578.0 | 0 | Panthera | 11619.0 | 2 | anthera pardu | 11619.0 | 2 |
| scf7180000022612 | 7196 | 0.3883 | 0 | 20.7222 | Eukaryota | 2477.0 | 0 | Chordata | 2477.0 | 0 | Carnivora | 2477.0 | 0 | Felidae | 2477.0 | 0 | Felis | 2222.0 | 1 | Felis catus | 2222.0 | 1 |
| scf7180000021798 | 35456 | 0.5241 | 0 | 49.7249 | Eukaryota | 4634.0 | 0 | Chordata | 4634.0 | 0 | Carnivora | 4634.0 | 0 | Felidae | 2238.0 | 2 | Felis | 2238.0 | 2 | Felis catus | 2238.0 | 2 |
| scf7180000020232 | 6730 | 0.4955 | 0 | 26.6436 | Eukaryota | 12620.0 | 0 | Chordata | 12620.0 | 0 | Carnivora | 12620.0 | 0 | Felidae | 5742.0 | 2 | .eptonychote | 2830.0 | 5 | onychotes we | 2830.0 | 5 |
| scf7180000024136 | 124118 | 0.3934 | 0 | 26.478 | Eukaryota | 44766.0 | 0 | Chordata | 44766.0 | 0 | Carnivora | 44766.0 | 0 | Felidae | 44766.0 | 0 | Felis | 44766.0 | 0 | Felis catus | 44766.0 | 0 |

| | | | | | | | | | | | | | | | | | | | | | | |
|---|---|---|---|---|---|---|---|---|---|---|---|---|---|---|---|---|---|---|---|---|---|---|
| scf7180000019683 | 280855 | 0.3733 | 0 | 49.4439 | Eukaryota | 46561.0 | 0 | Chordata | 46561.0 | 0 | Carnivora | 46561.0 | 0 | Felidae | 46561.0 | 0 | Felis | 46561.0 | 0 | Felis catus | 46561.0 | 0 |
| scf7180000023037 | 42261 | 0.3767 | 0 | 354.681 | Eukaryota | 10025.0 | 0 | Chordata | 10025.0 | 0 | pterygiforme | 8966.0 | 1 | Apterygidae | 8966.0 | 1 | Apteryx | 8966.0 | 1 | pteryx mante | 8966.0 | 1 |
| scf7180000020648 | 73221 | 0.4055 | 0 | 28.9002 | Eukaryota | 29060.0 | 0 | Chordata | 29060.0 | 0 | Carnivora | 26360.0 | 1 | Felidae | 14888.0 | 4 | Panthera | 5900.0 | 7 | anthera pardu | 5900.0 | 8 |
| scf7180000020985 | 91110 | 0.3918 | 0 | 52.3612 | Eukaryota | 10932.0 | 0 | Chordata | 10932.0 | 0 | Carnivora | 10030.0 | 1 | Felidae | 5500.0 | 2 | Canis | 4530.0 | 4 | Canis lupus | 4530.0 | 5 |
| scf7180000018941 | 823967 | 0.3523 | 0 | 47.9069 | Eukaryota | 47208.0 | 0 | Chordata | 47208.0 | 0 | Carnivora | 47208.0 | 0 | Felidae | 47208.0 | 0 | Felis | 47208.0 | 0 | Felis catus | 47208.0 | 0 |
| scf7180000018199 | 2113116 | 0.4283 | 0 | 51.908 | Eukaryota | 46710.0 | 0 | Chordata | 46710.0 | 0 | Carnivora | 46710.0 | 0 | Felidae | 46710.0 | 0 | Felis | 46710.0 | 0 | Felis catus | 46710.0 | 0 |
| scf7180000019114 | 451927 | 0.4015 | 0 | 46.8886 | Eukaryota | 46279.0 | 0 | Chordata | 46279.0 | 0 | Carnivora | 46279.0 | 0 | Felidae | 46279.0 | 0 | Felis | 46279.0 | 0 | Felis catus | 46279.0 | 0 |
| scf7180000021091 | 16830 | 0.4926 | 0 | 365.3267 | Eukaryota | 6039.0 | 0 | Chordata | 6039.0 | 0 | Carnivora | 6039.0 | 0 | Phocidae | 2464.0 | 2 | .eptonychote | 1848.0 | 4 | nychotes wed | 1848.0 | 4 |
| scf7180000020098 | 57460 | 0.4981 | 0 | 56.852 | Eukaryota | 23826.0 | 0 | Chordata | 23826.0 | 0 | Carnivora | 23826.0 | 0 | Felidae | 23826.0 | 0 | Felis | 23826.0 | 0 | Felis catus | 23826.0 | 0 |
| scf7180000022695 | 17935 | 0.4146 | 0 | 24.7221 | Eukaryota | 8376.0 | 0 | Chordata | 8376.0 | 0 | Primates | 4644.0 | 2 | Hominidae | 4644.0 | 2 | Homo | 3495.0 | 3 | Homo sapiens | 3495.0 | 3 |
| scf7180000019293 | 302324 | 0.4136 | 0 | 50.349 | Eukaryota | 96600.0 | 0 | Chordata | 96600.0 | 0 | Carnivora | 96600.0 | 0 | Felidae | 96600.0 | 0 | Acinonyx | 45577.0 | 2 | cinonyx jubat | 45577.0 | 2 |
| scf7180000018062 | 1214253 | 0.349 | 0 | 48.2807 | Eukaryota | 46452.0 | 0 | Chordata | 46452.0 | 0 | Carnivora | 46452.0 | 0 | Felidae | 46452.0 | 0 | Felis | 46452.0 | 0 | Felis catus | 46452.0 | 0 |
| scf7180000021859 | 16843 | 0.3723 | 0 | 48.4841 | Eukaryota | 3994.0 | 0 | Chordata | 3994.0 | 0 | Carnivora | 2517.0 | 2 | Felidae | 1156.0 | 4 | Neofelis | 867.0 | 7 | eofelis nebulo | 867.0 | 7 |
| scf7180000018289 | 961982 | 0.3807 | 0 | 49.9722 | Eukaryota | 47126.0 | 0 | Chordata | 47126.0 | 0 | Carnivora | 47126.0 | 0 | Felidae | 47126.0 | 0 | Felis | 47126.0 | 0 | Felis catus | 47126.0 | 0 |
| scf7180000018494 | 1818085 | 0.4605 | 100 | 51.8237 | Eukaryota | 43344.0 | 0 | Chordata | 43344.0 | 0 | Carnivora | 43344.0 | 0 | Felidae | 43344.0 | 0 | Felis | 43344.0 | 0 | Felis catus | 43344.0 | 0 |
| scf7180000018338 | 858454 | 0.3456 | 0 | 49.0206 | Eukaryota | 46070.0 | 0 | Chordata | 46070.0 | 0 | Carnivora | 46070.0 | 0 | Felidae | 46070.0 | 0 | Felis | 46070.0 | 0 | Felis catus | 46070.0 | 0 |
| scf7180000018458 | 24324 | 0.426 | 0 | 28.2815 | Eukaryota | 6115.0 | 0 | Chordata | 6115.0 | 0 | Carnivora | 6115.0 | 0 | Felidae | 3045.0 | 2 | Suricata | 2440.0 | 4 | ıricata suricat | 2440.0 | 4 |
| scf7180000023599 | 50528 | 0.4282 | 0 | 522.4636 | Eukaryota | 29775.0 | 0 | Chordata | 29775.0 | 0 | Carnivora | 29775.0 | 0 | Felidae | 29775.0 | 0 | Felis | 29775.0 | 0 | Felis catus | 29775.0 | 0 |
| scf7180000020250 | 179941 | 0.4591 | 0 | 48.9627 | Eukaryota | 28262.0 | 0 | Chordata | 28262.0 | 0 | Carnivora | 28262.0 | 0 | Felidae | 28262.0 | 0 | Felis | 24033.0 | 1 | Felis catus | 24033.0 | 1 |
| scf7180000021628 | 19102 | 0.7505 | 100 | 62.4018 | no-hit | 0.0 | 0 | no-hit | 0.0 | 0 | no-hit | 0.0 | 0 | no-hit | 0.0 | 0 | no-hit | 0.0 | 0 | no-hit | 0.0 | 0 |
| scf7180000018464 | 605653 | 0.4218 | 0 | 51.1456 | Eukaryota | 46450.0 | 0 | Chordata | 46450.0 | 0 | Carnivora | 46450.0 | 0 | Felidae | 46450.0 | 0 | Felis | 46450.0 | 0 | Felis catus | 46450.0 | 0 |
| scf7180000022988 | 33551 | 0.5538 | 0 | 110.2787 | Eukaryota | 6470.0 | 0 | Chordata | 6470.0 | 0 | Carnivora | 6470.0 | 0 | Felidae | 6220.0 | 1 | Felis | 2548.0 | 3 | Felis catus | 2548.0 | 3 |
| scf7180000021673 | 23972 | 0.3971 | 0 | 26.6215 | Eukaryota | 33090.0 | 0 | Chordata | 33090.0 | 0 | Carnivora | 33090.0 | 0 | Felidae | 33090.0 | 0 | Felis | 33090.0 | 0 | Felis catus | 33090.0 | 0 |
| scf7180000022491 | 5128 | 0.6673 | 0 | 111.0793 | Eukaryota | 374.0 | 0 | Chordata | 374.0 | 0 | Carnivora | 374.0 | 0 | Felidae | 374.0 | 0 | Panthera | 374.0 | 0 | anthera pardu | 374.0 | 0 |
| scf7180000021191 | 44164 | 0.3726 | 0 | 24.7671 | Eukaryota | 4324.0 | 0 | Chordata | 4324.0 | 0 | Carnivora | 2102.0 | 2 | Hominidae | 1560.0 | 4 | Homo | 1560.0 | 4 | Homo sapiens | 1560.0 | 4 |
| scf7180000020626 | 11716 | 0.4007 | 0 | 16.5195 | Eukaryota | 34854.0 | 0 | Chordata | 34854.0 | 0 | Carnivora | 34854.0 | 0 | Felidae | 34854.0 | 0 | Felis | 34854.0 | 0 | Felis catus | 34854.0 | 0 |
| scf7180000022201 | 9744 | 0.3659 | 0 | 40.5764 | Eukaryota | 44387.0 | 0 | Chordata | 44387.0 | 0 | Carnivora | 44387.0 | 0 | Felidae | 44387.0 | 0 | Felis | 39984.0 | 1 | Felis catus | 39984.0 | 1 |
| scf7180000019315 | 25044 | 0.3848 | 0 | 19.8539 | Eukaryota | 33351.0 | 0 | Chordata | 33351.0 | 0 | Carnivora | 33351.0 | 0 | Herpestidae | 14935.0 | 2 | Suricata | 14935.0 | 4 | ıricata suricat | 14935.0 | 5 |
| scf7180000018854 | 547899 | 0.4589 | 0 | 50.8007 | Eukaryota | 47240.0 | 0 | Chordata | 47240.0 | 0 | Carnivora | 47240.0 | 0 | Felidae | 47240.0 | 0 | Felis | 47240.0 | 0 | Felis catus | 47240.0 | 0 |
| scf7180000019863 | 83383 | 0.4132 | 0 | 26.831 | Eukaryota | 46752.0 | 0 | Chordata | 46752.0 | 0 | Carnivora | 46752.0 | 0 | Felidae | 46752.0 | 0 | Felis | 46752.0 | 0 | Felis catus | 46752.0 | 0 |
| scf7180000020245 | 22767 | 0.3719 | 0 | 20.3557 | Eukaryota | 39723.0 | 0 | Chordata | 39723.0 | 0 | Carnivora | 39723.0 | 0 | Felidae | 39723.0 | 0 | Felis | 39723.0 | 0 | Felis catus | 39723.0 | 0 |
| scf7180000017712 | 1803121 | 0.3756 | 0 | 50.3525 | Eukaryota | 47031.0 | 0 | Chordata | 47031.0 | 0 | Carnivora | 47031.0 | 0 | Felidae | 47031.0 | 0 | Felis | 47031.0 | 0 | Felis catus | 47031.0 | 0 |
| scf7180000020630 | 115344 | 0.3817 | 0 | 25.3325 | Eukaryota | 45102.0 | 0 | Chordata | 45102.0 | 0 | Carnivora | 45102.0 | 0 | Felidae | 45102.0 | 0 | Felis | 45102.0 | 0 | Felis catus | 45102.0 | 0 |
| scf7180000018261 | 946472 | 0.365 | 0 | 50.5049 | Eukaryota | 61700.0 | 0 | Chordata | 61700.0 | 0 | Carnivora | 61700.0 | 0 | Felidae | 33555.0 | 1 | Acinonyx | 33555.0 | 1 | cinonyx jubat | 33555.0 | 1 |
| scf7180000019155 | 561616 | 0.4193 | 0 | 54.4388 | Eukaryota | 31059.0 | 0 | Chordata | 31059.0 | 0 | Carnivora | 31059.0 | 0 | Felidae | 31059.0 | 0 | Felis | 24825.0 | 1 | Felis catus | 24825.0 | 1 |
| scf7180000019297 | 346974 | 0.4274 | 0 | 51.4858 | Eukaryota | 26873.0 | 0 | Chordata | 26873.0 | 0 | Carnivora | 26873.0 | 0 | Felidae | 21726.0 | 1 | Panthera | 8104.0 | 4 | anthera pardu | 6536.0 | 5 |
| scf7180000022384 | 12555 | 0.4951 | 0 | 34.7048 | Eukaryota | 2994.0 | 0 | Chordata | 2994.0 | 0 | Carnivora | 2994.0 | 0 | Felidae | 2994.0 | 0 | Felis | 1794.0 | 2 | Felis catus | 1794.0 | 2 |
| scf7180000020575 | 71122 | 0.4191 | 0 | 51.064 | Eukaryota | 6480.0 | 0 | Chordata | 6480.0 | 0 | Carnivora | 6480.0 | 0 | Felidae | 4842.0 | 1 | Felis | 3618.0 | 2 | Felis catus | 3618.0 | 2 |
| scf7180000019741 | 96979 | 0.3874 | 0 | 47.5387 | Eukaryota | 20450.0 | 0 | Chordata | 20450.0 | 0 | Chiroptera | 20450.0 | 0 | Pteropodidae | 20450.0 | 0 | Rousettus | 20450.0 | 0 | settus aegypti | 20450.0 | 0 |
| scf7180000022681 | 15693 | 0.4763 | 0 | 81.2058 | Eukaryota | 11954.0 | 0 | Chordata | 11954.0 | 0 | Carnivora | 11954.0 | 0 | Felidae | 11954.0 | 0 | Felis | 10783.0 | 1 | Felis catus | 10783.0 | 1 |
| scf7180000020053 | 16539 | 0.5233 | 0 | 193.3707 | Eukaryota | 12181.0 | 0 | Chordata | 12181.0 | 0 | Carnivora | 12181.0 | 0 | Felidae | 12181.0 | 0 | Felis | 10973.0 | 1 | Felis catus | 10973.0 | 1 |
| scf7180000019059 | 227816 | 0.367 | 0 | 51.9664 | Eukaryota | 44687.0 | 0 | Chordata | 44687.0 | 0 | Carnivora | 44687.0 | 0 | Felidae | 44687.0 | 0 | Felis | 44687.0 | 0 | Felis catus | 44687.0 | 0 |
| scf7180000019757 | 54070 | 0.3693 | 0 | 46.9064 | Eukaryota | 37680.0 | 0 | Chordata | 37680.0 | 0 | Carnivora | 37680.0 | 0 | Felidae | 37680.0 | 0 | Felis | 37680.0 | 0 | Felis catus | 37680.0 | 0 |
| scf7180000022839 | 1315485 | 0.3959 | 0 | 52.8649 | Eukaryota | 46820.0 | 0 | Chordata | 46820.0 | 0 | Carnivora | 46820.0 | 0 | Felidae | 46820.0 | 0 | Felis | 46820.0 | 0 | Felis catus | 46820.0 | 0 |
| scf7180000021092 | 69723 | 0.363 | 0 | 25.1746 | Eukaryota | 15448.0 | 0 | Chordata | 15448.0 | 0 | Primates | 10172.0 | 1 | Hominidae | 10172.0 | 1 | Pan | 6017.0 | 2 | an troglodyte | 6017.0 | 2 |
| scf7180000023039 | 28427 | 0.3723 | 0 | 229.9668 | Eukaryota | 10148.0 | 0 | Chordata | 10148.0 | 0 | pterygiforme | 9076.0 | 1 | Apterygidae | 9076.0 | 1 | Apteryx | 9076.0 | 1 | pteryx mante | 9076.0 | 1 |
| scf7180000021281 | 59546 | 0.3694 | 0 | 24.1134 | Eukaryota | 23689.0 | 0 | Chordata | 23689.0 | 0 | Carnivora | 23689.0 | 0 | Felidae | 23689.0 | 0 | Felis | 21269.0 | 1 | Felis catus | 21269.0 | 1 |
| scf7180000019652 | 742445 | 0.3964 | 200 | 51.6665 | Eukaryota | 98794.0 | 0 | Chordata | 98794.0 | 0 | Carnivora | 98794.0 | 0 | Felidae | 98794.0 | 0 | Panthera | 79040.0 | 1 | anthera pardu | 79040.0 | 1 |
| scf7180000023803 | 1476319 | 0.4745 | 0 | 52.8584 | Eukaryota | 60260.0 | 0 | Chordata | 60260.0 | 0 | Carnivora | 60260.0 | 0 | Felidae | 44377.0 | 2 | Acinonyx | 38007.0 | 3 | cinonyx jubat | 38007.0 | 3 |
| scf7180000020416 | 140770 | 0.3514 | 0 | 49.7702 | Eukaryota | 12924.0 | 0 | Chordata | 12924.0 | 0 | Primates | 6837.0 | 2 | Hominidae | 6837.0 | 3 | Homo | 3909.0 | 4 | Homo sapiens | 3909.0 | 4 |
| scf7180000017643 | 2159564 | 0.4589 | 0 | 52.8911 | Eukaryota | 48534.0 | 0 | Chordata | 48534.0 | 0 | Carnivora | 48534.0 | 0 | Felidae | 48534.0 | 0 | Felis | 48534.0 | 0 | Felis catus | 48534.0 | 0 |
| scf7180000019783 | 198359 | 0.5579 | 0 | 45.0778 | Eukaryota | 5123.0 | 0 | Chordata | 5123.0 | 0 | Primates | 3133.0 | 2 | Hominidae | 3133.0 | 3 | Homo | 2484.0 | 5 | Homo sapiens | 2484.0 | 5 |
| scf7180000018590 | 736127 | 0.5606 | 0 | 54.2761 | Eukaryota | 24151.0 | 0 | Chordata | 24151.0 | 0 | Carnivora | 8291.0 | 5 | Felidae | 6010.0 | 6 | Homo | 5380.0 | 9 | Homo sapiens | 5380.0 | 9 |
| scf7180000019531 | 92041 | 0.398 | 0 | 48.9033 | Eukaryota | 9884.0 | 0 | Chordata | 9884.0 | 0 | Carnivora | 9884.0 | 0 | Felidae | 7556.0 | 1 | Felis | 7556.0 | 1 | Felis catus | 7556.0 | 1 |
| scf7180000023986 | 577970 | 0.4623 | 0 | 54.1565 | Eukaryota | 35524.0 | 0 | Chordata | 35524.0 | 0 | Carnivora | 27124.0 | 1 | Herpestidae | 20856.0 | 2 | Suricata | 20856.0 | 2 | ıricata suricat | 20856.0 | 2 |
| scf7180000022973 | 35201 | 0.5296 | 0 | 190.5006 | Eukaryota | 16413.0 | 0 | Chordata | 16413.0 | 0 | Carnivora | 16413.0 | 0 | Felidae | 16413.0 | 0 | Felis | 16413.0 | 0 | Felis catus | 16413.0 | 0 |
| scf7180000023797 | 1731768 | 0.496 | 0 | 52.5237 | Eukaryota | 44062.0 | 0 | Chordata | 44062.0 | 0 | Carnivora | 44062.0 | 0 | Felidae | 44062.0 | 0 | Felis | 17645.0 | 3 | Felis catus | 17645.0 | 3 |
| scf7180000020463 | 145267 | 0.5288 | 0 | 50.4188 | Eukaryota | 19463.0 | 1 | Chordata | 19463.0 | 1 | Primates | 12085.0 | 3 | undef | 6268.0 | 6 | undef | 6268.0 | 7 | tic synthetic c | 6268.0 | 7 |
| scf7180000017747 | 3028908 | 0.4086 | 0 | 51.3425 | Eukaryota | 76027.0 | 0 | Chordata | 76027.0 | 0 | Carnivora | 76027.0 | 0 | Felidae | 76027.0 | 0 | Felis | 31795.0 | 2 | Felis catus | 31795.0 | 2 |
| scf7180000017820 | 307514 | 0.4571 | 100 | 52.9615 | Eukaryota | 45125.0 | 0 | Chordata | 45125.0 | 0 | Carnivora | 45125.0 | 0 | Felidae | 45125.0 | 0 | Felis | 45125.0 | 0 | Felis catus | 45125.0 | 0 |
| scf7180000020401 | 146143 | 0.4099 | 0 | 52.1623 | Eukaryota | 26593.0 | 0 | Chordata | 26593.0 | 0 | Artiodactyla | 8458.0 | 3 | Bovidae | 8458.0 | 4 | Crocidura | 5866.0 | 5 | Bos mutus | 4362.0 | 7 |
| scf7180000023900 | 786667 | 0.4564 | 0 | 52.67 | Eukaryota | 45597.0 | 0 | Chordata | 45597.0 | 0 | Carnivora | 45597.0 | 0 | Felidae | 45597.0 | 0 | Felis | 36457.0 | 1 | Felis catus | 36457.0 | 2 |
| scf7180000018686 | 620127 | 0.4756 | 0 | 53.6508 | Eukaryota | 33036.0 | 0 | Chordata | 33036.0 | 0 | Carnivora | 25919.0 | 1 | Felidae | 14311.0 | 4 | unresolved | 5804.0 | 7 | Canis lupus | 5804.0 | 8 |
| scf7180000020422 | 102316 | 0.3722 | 0 | 23.817 | Eukaryota | 32853.0 | 0 | Chordata | 32853.0 | 0 | Carnivora | 32853.0 | 0 | Felidae | 32853.0 | 0 | Felis | 32853.0 | 0 | Felis catus | 32853.0 | 0 |
| scf7180000021371 | 4346 | 0.4257 | 0 | 19.6187 | Eukaryota | 9964.0 | 0 | Chordata | 9964.0 | 0 | Carnivora | 9964.0 | 0 | Felidae | 9964.0 | 0 | Felis | 7965.0 | 1 | Felis catus | 7965.0 | 1 |
| scf7180000020705 | 109126 | 0.3807 | 0 | 23.95 | Eukaryota | 11279.0 | 0 | Chordata | 11279.0 | 0 | Carnivora | 11279.0 | 0 | Felidae | 11279.0 | 0 | Felis | 9059.0 | 1 | Felis catus | 9059.0 | 1 |
| scf7180000022379 | 30263 | 0.5371 | 0 | 54.6713 | Eukaryota | 2744.0 | 0 | Chordata | 2744.0 | 0 | Carnivora | 2744.0 | 0 | Felidae | 1014.0 | 4 | Zalophus | 512.0 | 7 | phus californi | 512.0 | 7 |
| scf7180000021678 | 44782 | 0.5032 | 0 | 49.2734 | Eukaryota | 16429.0 | 0 | Chordata | 16429.0 | 0 | Carnivora | 16429.0 | 0 | Canidae | 9135.0 | 3 | Canis | 7626.0 | 5 | Canis lupus | 7626.0 | 5 |
| scf7180000019853 | 188035 | 0.4371 | 0 | 51.508 | Eukaryota | 45020.0 | 0 | Chordata | 45020.0 | 0 | Carnivora | 45020.0 | 0 | Felidae | 45020.0 | 0 | Lynx | 22499.0 | 2 | ynx canadens | 22499.0 | 2 |
| scf7180000020857 | 57833 | 0.4331 | 0 | 25.4396 | Eukaryota | 5964.0 | 0 | Chordata | 5964.0 | 0 | Carnivora | 5964.0 | 0 | Felidae | 5964.0 | 0 | Felis | 5964.0 | 0 | Felis catus | 5964.0 | 0 |
| scf7180000018224 | 279742 | 0.4617 | 0 | 51.3557 | Eukaryota | 43664.0 | 0 | Chordata | 43664.0 | 0 | Carnivora | 43664.0 | 0 | Felidae | 30856.0 | 4 | Panthera | 13334.0 | 7 | Panthera tigri | 6728.0 | 9 |
| scf7180000022934 | 26777 | 0.3849 | 0 | 203.4363 | Eukaryota | 12513.0 | 0 | Chordata | 12513.0 | 0 | Carnivora | 12513.0 | 0 | Felidae | 12513.0 | 0 | Felis | 10030.0 | 1 | Felis catus | 10030.0 | 1 |
| scf7180000018266 | 1956408 | 0.4037 | 100 | 50.6945 | Eukaryota | 46186.0 | 0 | Chordata | 46186.0 | 0 | Carnivora | 46186.0 | 0 | Felidae | 46186.0 | 0 | Felis | 46186.0 | 0 | Felis catus | 46186.0 | 0 |
| scf7180000019852 | 177554 | 0.4112 | 0 | 47.6263 | Eukaryota | 38698.0 | 0 | Chordata | 38698.0 | 0 | Carnivora | 38698.0 | 0 | Felidae | 38698.0 | 0 | Felis | 38698.0 | 0 | Felis catus | 38698.0 | 0 |
| scf7180000019358 | 3574 | 0.399 | 0 | 19.5283 | Eukaryota | 6956.0 | 0 | Chordata | 6956.0 | 0 | Carnivora | 6956.0 | 0 | Felidae | 6956.0 | 0 | Felis | 6248.0 | 1 | Felis catus | 6248.0 | 1 |
| scf7180000022474 | 12728 | 0.368 | 0 | 47.5044 | Eukaryota | 3980.0 | 1 | Chordata | 3980.0 | 1 | Primates | 2476.0 | 3 | Hominidae | 2476.0 | 4 | Homo | 2476.0 | 4 | Homo sapiens | 2476.0 | 4 |
| scf7180000018138 | 1077100 | 0.4522 | 0 | 52.8768 | Eukaryota | 21717.0 | 0 | Chordata | 21717.0 | 0 | Carnivora | 21717.0 | 0 | Felidae | 21717.0 | 0 | Felis | 19528.0 | 1 | Felis catus | 19528.0 | 1 |
| scf7180000017806 | 1691299 | 0.3639 | 0 | 49.969 | Eukaryota | 47351.0 | 0 | Chordata | 47351.0 | 0 | Carnivora | 47351.0 | 0 | Felidae | 47351.0 | 0 | Felis | 47351.0 | 0 | Felis catus | 47351.0 | 0 |
| scf7180000019497 | 278427 | 0.3874 | 0 | 51.7009 | Eukaryota | 17396.0 | 0 | Chordata | 17396.0 | 0 | Carnivora | 17396.0 | 0 | Felidae | 17396.0 | 0 | Felis | 15671.0 | 1 | Felis catus | 15671.0 | 1 |
| scf7180000023906 | 1974075 | 0.3863 | 0 | 50.8275 | Eukaryota | 54459.0 | 0 | Chordata | 54459.0 | 0 | Carnivora | 37565.0 | 1 | Felidae | 37565.0 | 2 | Felis | 33073.0 | 3 | Felis catus | 33073.0 | 3 |
| scf7180000018959 | 553087 | 0.3593 | 0 | 49.6778 | Eukaryota | 47531.0 | 0 | Chordata | 47531.0 | 0 | Carnivora | 47531.0 | 0 | Felidae | 47531.0 | 0 | Felis | 47531.0 | 0 | Felis catus | 47531.0 | 0 |
| scf7180000021291 | 57964 | 0.3656 | 0 | 26.0781 | Eukaryota | 6885.0 | 0 | Chordata | 6885.0 | 0 | Primates | 3268.0 | 2 | Hominidae | 3268.0 | 3 | Homo | 2484.0 | 5 | Homo sapiens | 2484.0 | 5 |
| scf7180000022147 | 9969 | 0.5213 | 0 | 37.5628 | no-hit | 0.0 | 0 | no-hit | 0.0 | 0 | no-hit | 0.0 | 0 | no-hit | 0.0 | 0 | no-hit | 0.0 | 0 | no-hit | 0.0 | 0 |
| scf7180000023771 | 370805 | 0.3659 | 0 | 49.2708 | Eukaryota | 46628.0 | 0 | Chordata | 46628.0 | 0 | Carnivora | 46628.0 | 0 | Felidae | 46628.0 | 0 | Felis | 46628.0 | 0 | Felis catus | 46628.0 | 0 |
| scf7180000024278 | 862044 | 0.3617 | 0 | 49.4032 | Eukaryota | 69047.0 | 0 | Chordata | 69047.0 | 0 | Carnivora | 69047.0 | 0 | Felidae | 24107.0 | 3 | Suricata | 18684.0 | 5 | ıricata suricat | 18684.0 | 5 |
| scf7180000021026 | 30562 | 0.4091 | 0 | 221.6801 | Eukaryota | 36917.0 | 0 | Chordata | 36917.0 | 0 | Carnivora | 36917.0 | 0 | Felidae | 36917.0 | 0 | Felis | 36917.0 | 0 | Felis catus | 36917.0 | 0 |
| scf7180000018910 | 69269 | 0.4436 | 0 | 78.059 | Eukaryota | 9384.0 | 0 | Chordata | 9384.0 | 0 | Carnivora | 9384.0 | 0 | Felidae | 8903.0 | 1 | Felis | 7615.0 | 2 | Felis catus | 7615.0 | 2 |
| scf7180000020325 | 85594 | 0.4314 | 0 | 51.6643 | Eukaryota | 28282.0 | 0 | Chordata | 28282.0 | 0 | Carnivora | 28282.0 | 0 | Felidae | 28282.0 | 0 | Felis | 11198.0 | 2 | Felis catus | 11198.0 | 2 |
| scf7180000024192 | 77016 | 0.3749 | 0 | 22.5063 | Eukaryota | 38631.0 | 0 | Chordata | 38631.0 | 0 | Carnivora | 38631.0 | 0 | Felidae | 38631.0 | 0 | Felis | 38631.0 | 0 | Felis catus | 38631.0 | 0 |
| scf7180000018577 | 86777 | 0.3783 | 0 | 48.5043 | Eukaryota | 45429.0 | 0 | Chordata | 45429.0 | 0 | Carnivora | 45429.0 | 0 | Felidae | 45429.0 | 0 | Felis | 45429.0 | 0 | Felis catus | 45429.0 | 0 |
| scf7180000020824 | 106755 | 0.4164 | 0 | 26.5557 | Eukaryota | 13265.0 | 0 | Chordata | 13265.0 | 0 | Carnivora | 13265.0 | 0 | Felidae | 13265.0 | 0 | Felis | 10630.0 | 1 | Felis catus | 10630.0 | 1 |
| scf7180000019965 | 212907 | 0.5206 | 0 | 51.4072 | Eukaryota | 45927.0 | 0 | Chordata | 45927.0 | 0 | Carnivora | 45927.0 | 0 | Felidae | 45927.0 | 0 | Felis | 45927.0 | 0 | Felis catus | 45927.0 | 0 |
| scf7180000020197 | 183898 | 0.5652 | 0 | 52.9924 | Eukaryota | 37181.0 | 0 | Chordata | 37181.0 | 0 | Carnivora | 37181.0 | 0 | Felidae | 32446.0 | 1 | Felis | 28891.0 | 2 | Felis catus | 28891.0 | 2 |
| scf7180000020033 | 133258 | 0.3828 | 0 | 49.5717 | Eukaryota | 28018.0 | 0 | Chordata | 28018.0 | 0 | Carnivora | 28018.0 | 0 | Felidae | 28018.0 | 0 | Felis | 25271.0 | 1 | Felis catus | 25271.0 | 1 |
| scf7180000022490 | 12855 | 0.3924 | 0 | 23.1806 | Eukaryota | 23071.0 | 0 | Chordata | 23071.0 | 0 | Carnivora | 23071.0 | 0 | Felidae | 23071.0 | 0 | Felis | 23071.0 | 0 | Felis catus | 23071.0 | 0 |
| scf7180000020638 | 71064 | 0.4795 | 0 | 25.2255 | Eukaryota | 13606.0 | 0 | Chordata | 13606.0 | 0 | Carnivora | 10578.0 | 1 | Felidae | 10578.0 | 1 | Acinonyx | 4156.0 | 4 | cinonyx jubat | 4156.0 | 4 |
| scf7180000017458 | 3748321 | 0.3683 | 0 | 50.3636 | Eukaryota | 82196.0 | 0 | Chordata | 82196.0 | 0 | Carnivora | 82196.0 | 0 | Felidae | 82196.0 | 0 | Felis | 32942.0 | 2 | Felis catus | 32942.0 | 2 |
| scf7180000018033 | 1374974 | 0.3817 | 0 | 49.8298 | Eukaryota | 51669.0 | 0 | Chordata | 51669.0 | 0 | Carnivora | 51669.0 | 0 | Odobenidae | 25933.0 | 1 | Odobenus | 25933.0 | 2 | obenus rosma | 25933.0 | 2 |
| scf7180000023887 | 824885 | 0.3612 | 0 | 49.2204 | Eukaryota | 35425.0 | 0 | Chordata | 35425.0 | 0 | Carnivora | 35425.0 | 0 | Felidae | 35425.0 | 0 | Felis | 32555.0 | 1 | Felis catus | 32555.0 | 1 |
| scf7180000021478 | 39147 | 0.5877 | 0 | 46.9079 | Eukaryota | 2441.0 | 0 | Chordata | 2441.0 | 0 | Carnivora | 2441.0 | 0 | Felidae | 2441.0 | 0 | Felis | 2202.0 | 1 | Felis catus | 2202.0 | 1 |
| scf7180000022817 | 49991 | 0.4387 | 0 | 469.6461 | no-hit | 0.0 | 0 | no-hit | 0.0 | 0 | no-hit | 0.0 | 0 | no-hit | 0.0 | 0 | no-hit | 0.0 | 0 | no-hit | 0.0 | 0 |
| scf7180000022689 | 6996 | 0.4182 | 0 | 24.4278 | Eukaryota | 10567.0 | 0 | Chordata | 10567.0 | 0 | Carnivora | 10567.0 | 0 | Felidae | 10567.0 | 0 | Felis | 10567.0 | 0 | Felis catus | 10567.0 | 0 |
| scf7180000020613 | 107450 | 0.444 | 0 | 25.0411 | Eukaryota | 7380.0 | 0 | Chordata | 7380.0 | 0 | Carnivora | 7380.0 | 0 | Ursidae | 5912.0 | 1 | Ursus | 5912.0 | 1 | Ursus arctos | 5912.0 | 1 |
| scf7180000017464 | 281370 | 0.4097 | 0 | 47.7415 | Eukaryota | 46166.0 | 0 | Chordata | 46166.0 | 0 | Carnivora | 46166.0 | 0 | Felidae | 46166.0 | 0 | Felis | 46166.0 | 0 | Felis catus | 46166.0 | 0 |
| scf7180000020364 | 109540 | 0.4564 | 0 | 26.5143 | Eukaryota | 16038.0 | 0 | Chordata | 16038.0 | 0 | Artiodactyla | 5740.0 | 4 | Suidae | 5740.0 | 4 | Sus | 5740.0 | 5 | Sus scrofa | 5740.0 | 5 |
| scf7180000021582 | 71121 | 0.4038 | 0 | 25.2627 | Eukaryota | 9816.0 | 0 | Chordata | 9816.0 | 0 | Carnivora | 5085.0 | 2 | Felidae | 5085.0 | 2 | Felis | 5085.0 | 3 | Felis catus | 5085.0 | 3 |
| scf7180000022627 | 6348 | 0.5235 | 0 | 23.9193 | no-hit | 0.0 | 0 | no-hit | 0.0 | 0 | no-hit | 0.0 | 0 | no-hit | 0.0 | 0 | no-hit | 0.0 | 0 | no-hit | 0.0 | 0 |
| scf7180000022550 | 20455 | 0.4089 | 0 | 24.2597 | Eukaryota | 21714.0 | 0 | Chordata | 21714.0 | 0 | Carnivora | 21714.0 | 0 | Felidae | 21714.0 | 0 | Felis | 19531.0 | 1 | Felis catus | 19531.0 | 1 |
| scf7180000019608 | 144424 | 0.4386 | 0 | 46.6902 | Eukaryota | 10701.0 | 0 | Chordata | 10701.0 | 0 | Carnivora | 10701.0 | 0 | Felidae | 6469.0 | 3 | Panthera | 2197.0 | 6 | Felis catus | 2119.0 | 7 |
| scf7180000023413 | 78274 | 0.3957 | 0 | 308.951 | Eukaryota | 6748.0 | 0 | Chordata | 6748.0 | 0 | Carnivora | 6748.0 | 0 | Felidae | 6748.0 | 0 | Felis | 6107.0 | 1 | Felis catus | 6107.0 | 1 |
| scf7180000017583 | 37362 | 0.3596 | 0 | 40.2625 | Eukaryota | 4285.0 | 0 | Chordata | 4285.0 | 0 | Carnivora | 2544.0 | 2 | Felidae | 1464.0 | 3 | Homo | 1460.0 | 5 | Homo sapiens | 1460.0 | 5 |
| scf7180000018902 | 1041954 | 0.3736 | 0 | 50.6425 | Eukaryota | 50320.0 | 0 | Chordata | 50320.0 | 0 | Carnivora | 50320.0 | 0 | Felidae | 19504.0 | 2 | Panthera | 19504.0 | 2 | anthera pardu | 19504.0 | 2 |
| scf7180000020922 | 80103 | 0.3984 | 0 | 25.1922 | Eukaryota | 30634.0 | 0 | Chordata | 30634.0 | 0 | Carnivora | 19897.0 | 1 | Felidae | 11732.0 | 3 | Canis | 8165.0 | 5 | Canis lupus | 8165.0 | 6 |
| scf7180000022577 | 12993 | 0.5226 | 0 | 23.7725 | Eukaryota | 8402.0 | 0 | Chordata | 8402.0 | 0 | Carnivora | 8402.0 | 0 | Felidae | 8402.0 | 0 | Panthera | 7812.0 | 1 | anthera pardu | 7812.0 | 1 |
| scf7180000017595 | 278633 | 0.3553 | 0 | 48.2784 | Eukaryota | 45546.0 | 0 | Chordata | 45546.0 | 0 | Carnivora | 45546.0 | 0 | Felidae | 45546.0 | 0 | Felis | 45546.0 | 0 | Felis catus | 45546.0 | 0 |
| scf7180000023166 | 30658 | 0.3899 | 0 | 173.8164 | Eukaryota | 10039.0 | 0 | Chordata | 10039.0 | 0 | pterygiforme | 8975.0 | 1 | Apterygidae | 8975.0 | 1 | Apteryx | 8975.0 | 1 | pteryx mante | 8975.0 | 1 |
| scf7180000022583 | 6092 | 0.6906 | 0 | 154.5639 | Eukaryota | 337.0 | 0 | Chordata | 337.0 | 0 | Carnivora | 337.0 | 0 | Felidae | 337.0 | 0 | Panthera | 337.0 | 0 | anthera pardu | 337.0 | 0 |
| scf7180000019579 | 236421 | 0.4355 | 0 | 50.3074 | Eukaryota | 28843.0 | 0 | Chordata | 28843.0 | 0 | Carnivora | 28843.0 | 0 | Felidae | 28843.0 | 0 | Felis | 25908.0 | 1 | Felis catus | 25908.0 | 1 |
| scf7180000023241 | 25959 | 0.379 | 0 | 287.0687 | Eukaryota | 10088.0 | 0 | Chordata | 10088.0 | 0 | pterygiforme | 9022.0 | 1 | Apterygidae | 9022.0 | 1 | Apteryx | 9022.0 | 1 | pteryx mante | 9022.0 | 1 |
| scf7180000020490 | 12259 | 0.4324 | 0 | 28.0829 | Eukaryota | 21056.0 | 0 | Chordata | 21056.0 | 0 | Carnivora | 21056.0 | 0 | Felidae | 21056.0 | 0 | Felis | 21056.0 | 0 | Felis catus | 21056.0 | 0 |
| scf7180000019490 | 220249 | 0.4699 | 0 | 52.7607 | Eukaryota | 29692.0 | 0 | Chordata | 29692.0 | 0 | Carnivora | 29692.0 | 0 | Felidae | 29692.0 | 0 | Felis | 23743.0 | 1 | Felis catus | 23743.0 | 1 |
| scf7180000021448 | 64649 | 0.4658 | 0 | 25.5005 | Eukaryota | 11434.0 | 0 | Chordata | 11434.0 | 0 | Primates | 4834.0 | 2 | Hominidae | 4834.0 | 4 | Homo | 3860.0 | 5 | Homo sapiens | 3860.0 | 5 |
| scf7180000017912 | 362338 | 0.3806 | 0 | 49.6838 | Eukaryota | 20752.0 | 0 | Chordata | 20752.0 | 0 | Carnivora | 20752.0 | 0 | Felidae | 20752.0 | 0 | Felis | 20752.0 | 0 | Felis catus | 20752.0 | 0 |
| scf7180000022536 | 22736 | 0.364 | 0 | 24.7269 | Eukaryota | 12864.0 | 0 | Chordata | 12864.0 | 0 | Carnivora | 12864.0 | 0 | Felidae | 12864.0 | 0 | Felis | 11600.0 | 1 | Felis catus | 11600.0 | 1 |
| scf7180000023213 | 95069 | 0.5176 | 0 | 486.5986 | Eukaryota | 15311.0 | 0 | Chordata | 15311.0 | 0 | Carnivora | 15311.0 | 0 | Felidae | 15311.0 | 0 | Felis | 15311.0 | 0 | Felis catus | 15311.0 | 0 |
| scf7180000021248 | 65915 | 0.3577 | 0 | 25.572 | Eukaryota | 16652.0 | 0 | Chordata | 16652.0 | 0 | Primates | 13260.0 | 2 | Hominidae | 13260.0 | 2 | Homo | 8048.0 | 3 | Homo sapiens | 8048.0 | 3 |
| scf7180000023868 | 1248250 | 0.435 | 0 | 51.8372 | Eukaryota | 42670.0 | 0 | Chordata | 42670.0 | 0 | Carnivora | 42670.0 | 0 | Felidae | 42670.0 | 0 | Felis | 42670.0 | 0 | Felis catus | 42670.0 | 0 |
| scf7180000017937 | 4044 | 0.4881 | 0 | 22.5526 | Eukaryota | 2224.0 | 0 | Chordata | 2224.0 | 0 | Carnivora | 2224.0 | 0 | Felidae | 2224.0 | 0 | Felis | 1776.0 | 1 | Felis catus | 1776.0 | 1 |
| scf7180000018428 | 313692 | 0.3848 | 0 | 51.115 | Eukaryota | 46940.0 | 0 | Chordata | 46940.0 | 0 | Carnivora | 46940.0 | 0 | Felidae | 46940.0 | 0 | Felis | 46940.0 | 0 | Felis catus | 46940.0 | 0 |
| scf7180000024118 | 338819 | 0.3993 | 0 | 51.2038 | Eukaryota | 62966.0 | 0 | Chordata | 62966.0 | 0 | Carnivora | 62966.0 | 0 | Felidae | 62966.0 | 0 | Felis | 38254.0 | 1 | Felis catus | 38254.0 | 1 |
| scf7180000022254 | 15412 | 0.5511 | 0 | 53.307 | Eukaryota | 2384.0 | 0 | Chordata | 2384.0 | 0 | Carnivora | 1311.0 | 1 | Felidae | 1311.0 | 2 | Felis | 1311.0 | 2 | Felis catus | 1311.0 | 2 |
| scf7180000017528 | 6766 | 0.4005 | 0 | 22.4964 | Eukaryota | 33556.0 | 0 | Chordata | 33556.0 | 0 | Carnivora | 33556.0 | 0 | Felidae | 33556.0 | 0 | Felis | 33556.0 | 0 | Felis catus | 33556.0 | 0 |
| scf7180000018339 | 708864 | 0.4854 | 0 | 51.416 | Eukaryota | 32676.0 | 0 | Chordata | 32676.0 | 0 | Carnivora | 32676.0 | 0 | Felidae | 22781.0 | 1 | Suricata | 9895.0 | 5 | ıricata suricat | 9895.0 | 6 |
| scf7180000018988 | 2074181 | 0.3815 | 100 | 50.8666 | Eukaryota | 46866.0 | 0 | Chordata | 46866.0 | 0 | Carnivora | 46866.0 | 0 | Felidae | 46866.0 | 0 | Felis | 46866.0 | 0 | Felis catus | 46866.0 | 0 |
| scf7180000022273 | 6409 | 0.3862 | 0 | 27.1672 | Eukaryota | 17494.0 | 0 | Chordata | 17494.0 | 0 | Carnivora | 17494.0 | 0 | Felidae | 17494.0 | 0 | Felis | 15761.0 | 1 | Felis catus | 15761.0 | 1 |
| scf7180000018867 | 532726 | 0.419 | 0 | 50.6246 | Eukaryota | 54343.0 | 0 | Chordata | 54343.0 | 0 | Carnivora | 50195.0 | 1 | Felidae | 46014.0 | 2 | Panthera | 14939.0 | 5 | ynx canadens | 10951.0 | 6 |
| scf7180000022885 | 32818 | 0.3968 | 0 | 240.5251 | Eukaryota | 45640.0 | 0 | Chordata | 45640.0 | 0 | Carnivora | 45640.0 | 0 | Felidae | 45640.0 | 0 | Felis | 41139.0 | 1 | Felis catus | 41139.0 | 1 |
| scf7180000018505 | 1548840 | 0.4059 | 0 | 51.4525 | Eukaryota | 54295.0 | 0 | Chordata | 54295.0 | 0 | Carnivora | 54295.0 | 0 | Felidae | 48721.0 | 1 | Lynx | 16245.0 | 4 | ynx canadens | 16245.0 | 5 |
| scf7180000022732 | 21250 | 0.4544 | 0 | 25.2156 | Eukaryota | 3086.0 | 0 | Chordata | 3086.0 | 0 | Artiodactyla | 1579.0 | 3 | Bovidae | 786.0 | 5 | Suricata | 715.0 | 6 | ıricata suricat | 715.0 | 6 |
| scf7180000017735 | 1860029 | 0.3849 | 0 | 51.331 | Eukaryota | 49299.0 | 0 | Chordata | 49299.0 | 0 | Carnivora | 49299.0 | 0 | Felidae | 42898.0 | 1 | Felis | 42898.0 | 1 | Felis catus | 42898.0 | 1 |
| scf7180000018087 | 325200 | 0.3661 | 0 | 45.9582 | Eukaryota | 46806.0 | 0 | Chordata | 46806.0 | 0 | Carnivora | 46806.0 | 0 | Felidae | 37025.0 | 2 | Felis | 37025.0 | 2 | Felis catus | 37025.0 | 2 |
| scf7180000021939 | 14957 | 0.6375 | 0 | 51.131 | Eukaryota | 12190.0 | 0 | Chordata | 12190.0 | 0 | Carnivora | 11041.0 | 1 | Felidae | 11041.0 | 3 | Acinonyx | 5916.0 | 4 | cinonyx jubat | 5916.0 | 4 |
| scf7180000018437 | 893995 | 0.4197 | 0 | 52.5993 | Eukaryota | 44585.0 | 0 | Chordata | 44585.0 | 0 | Carnivora | 44585.0 | 0 | Felidae | 44585.0 | 0 | Felis | 40121.0 | 1 | Felis catus | 40121.0 | 1 |
| scf7180000018220 | 1010002 | 0.4458 | 0 | 50.4257 | Eukaryota | 18411.0 | 0 | Chordata | 18411.0 | 0 | Carnivora | 18411.0 | 0 | Felidae | 18411.0 | 0 | Felis | 14737.0 | 1 | Felis catus | 14737.0 | 1 |
| scf7180000021412 | 61161 | 0.4909 | 0 | 52.2949 | Eukaryota | 37547.0 | 0 | Chordata | 37547.0 | 0 | Carnivora | 37547.0 | 0 | Felidae | 30508.0 | 2 | Acinonyx | 11489.0 | 4 | cinonyx jubat | 11489.0 | 4 |
| scf7180000019352 | 772584 | 0.416 | 100 | 50.77 | Eukaryota | 48612.0 | 0 | Chordata | 48612.0 | 0 | Carnivora | 48612.0 | 0 | Felidae | 48612.0 | 0 | Felis | 41973.0 | 1 | Felis catus | 41973.0 | 1 |
| scf7180000022852 | 882149 | 0.4944 | 0 | 66.2106 | Eukaryota | 44178.0 | 0 | Chordata | 44178.0 | 0 | Carnivora | 44178.0 | 0 | Felidae | 44178.0 | 0 | Felis | 44178.0 | 0 | Felis catus | 44178.0 | 0 |
| scf7180000023247 | 23491 | 0.3904 | 0 | 269.5856 | Eukaryota | 4074.0 | 0 | Chordata | 4074.0 | 0 | Primates | 2577.0 | 2 | Hominidae | 2200.0 | 3 | Homo | 2200.0 | 4 | Homo sapiens | 2200.0 | 4 |
| scf7180000020847 | 77900 | 0.3873 | 0 | 25.5091 | Eukaryota | 8650.0 | 0 | Chordata | 8650.0 | 0 | Carnivora | 4729.0 | 3 | Felidae | 2708.0 | 5 | Vulpes | 2021.0 | 8 | Vulpes vulpes | 2021.0 | 8 |
| scf7180000020666 | 153422 | 0.3624 | 0 | 26.1101 | Eukaryota | 45310.0 | 0 | Chordata | 45310.0 | 0 | Carnivora | 45310.0 | 0 | Felidae | 45310.0 | 0 | Felis | 45310.0 | 0 | Felis catus | 45310.0 | 0 |
| scf7180000024241 | 355738 | 0.4341 | 0 | 51.1646 | Eukaryota | 47179.0 | 0 | Chordata | 47179.0 | 0 | Carnivora | 47179.0 | 0 | Felidae | 47179.0 | 0 | Felis | 47179.0 | 0 | Felis catus | 47179.0 | 0 |
| scf7180000018051 | 24792 | 0.3777 | 0 | 43.0015 | Eukaryota | 20639.0 | 0 | Chordata | 20639.0 | 0 | Carnivora | 20639.0 | 0 | Felidae | 20639.0 | 0 | Felis | 18583.0 | 1 | Felis catus | 18583.0 | 1 |
| scf7180000021389 | 38546 | 0.3484 | 0 | 23.9946 | Eukaryota | 10618.0 | 0 | Chordata | 10618.0 | 0 | Carnivora | 4861.0 | 2 | Felidae | 4861.0 | 2 | Felis | 4013.0 | 4 | Felis catus | 4013.0 | 4 |
| scf7180000022210 | 15085 | 0.4325 | 0 | 30.7054 | Eukaryota | 17494.0 | 0 | Chordata | 17494.0 | 0 | Carnivora | 17494.0 | 0 | Felidae | 17494.0 | 0 | Felis | 15741.0 | 1 | Felis catus | 15741.0 | 1 |
| scf7180000019341 | 317194 | 0.38 | 0 | 50.6166 | Eukaryota | 45999.0 | 0 | Chordata | 45999.0 | 0 | Carnivora | 45999.0 | 0 | Felidae | 45999.0 | 0 | Felis | 45999.0 | 0 | Felis catus | 45999.0 | 0 |
| scf7180000017884 | 1462275 | 0.3632 | 0 | 49.8107 | Eukaryota | 47009.0 | 0 | Chordata | 47009.0 | 0 | Carnivora | 47009.0 | 0 | Felidae | 47009.0 | 0 | Felis | 47009.0 | 0 | Felis catus | 47009.0 | 0 |
| scf7180000019093 | 42133 | 0.3676 | 0 | 24.325 | Eukaryota | 42369.0 | 0 | Chordata | 42369.0 | 0 | Carnivora | 42369.0 | 0 | Felidae | 42369.0 | 0 | Felis | 42369.0 | 0 | Felis catus | 42369.0 | 0 |
| scf7180000024260 | 470885 | 0.5218 | 0 | 50.6904 | Eukaryota | 90188.0 | 0 | Chordata | 90188.0 | 0 | Carnivora | 90188.0 | 0 | Felidae | 62952.0 | 1 | Panthera | 53648.0 | 2 | anthera pardu | 53648.0 | 2 |
| scf7180000017623 | 1797590 | 0.4732 | 0 | 52.7816 | Eukaryota | 81403.0 | 0 | Chordata | 81403.0 | 0 | Carnivora | 81403.0 | 0 | Felidae | 48612.0 | 1 | Lynx | 48612.0 | 1 | ynx canadens | 48612.0 | 1 |

| | | | | | | | | | | | | | | | | | | | | |
|---|---|---|---|---|---|---|---|---|---|---|---|---|---|---|---|---|---|---|---|---|
| scf7180000020285 | 178506 | 0.3785 | 0 | 23.7506 | Eukaryota | 45382.0 | 0 | Chordata | 45382.0 | 0 | Carnivora | 45382.0 | 0 | Felidae | 45382.0 | 0 | Felis | 45382.0 | 0 | Felis catus | 45382.0 | 0 |
| scf7180000023146 | 42185 | 0.4342 | 0 | 138.7475 | Eukaryota | 5743.0 | 0 | Chordata | 5743.0 | 0 | Carnivora | 5743.0 | 0 | Felidae | 5743.0 | 0 | Felis | 5292.0 | 1 | Felis catus | 5292.0 | 1 |
| scf7180000018264 | 91481 | 0.367 | 0 | 47.0315 | Eukaryota | 29066.0 | 0 | Chordata | 29066.0 | 0 | Carnivora | 29066.0 | 0 | Felidae | 29066.0 | 0 | Felis | 29066.0 | 0 | Felis catus | 29066.0 | 0 |
| scf7180000018221 | 1467801 | 0.354 | 0 | 49.0166 | Eukaryota | 47076.0 | 0 | Chordata | 47076.0 | 0 | Carnivora | 47076.0 | 0 | Felidae | 47076.0 | 0 | Felis | 47076.0 | 0 | Felis catus | 47076.0 | 0 |
| scf7180000019768 | 164746 | 0.4188 | 0 | 47.3141 | Eukaryota | 36414.0 | 0 | Chordata | 36414.0 | 0 | Carnivora | 36414.0 | 0 | Felidae | 36414.0 | 0 | Felis | 36414.0 | 0 | Felis catus | 36414.0 | 0 |
| scf7180000018010 | 23084 | 0.5269 | 0 | 475.3203 | Eukaryota | 2192.0 | 0 | Chordata | 2192.0 | 0 | Carnivora | 2192.0 | 0 | Felidae | 2192.0 | 0 | Felis | 1818.0 | 2 | Felis catus | 1818.0 | 2 |
| scf7180000018664 | 268016 | 0.5365 | 0 | 47.8823 | Eukaryota | 15786.0 | 0 | Chordata | 15786.0 | 0 | Carnivora | 15786.0 | 0 | Felidae | 15786.0 | 0 | Panthera | 15786.0 | 0 | anthera pardu | 15786.0 | 0 |
| scf7180000023363 | 19861 | 0.5086 | 0 | 385.5266 | no-hit | 0.0 | 0 | no-hit | 0.0 | 0 | no-hit | 0.0 | 0 | no-hit | 0.0 | 0 | no-hit | 0.0 | 0 | no-hit | 0.0 | 0 |
| scf7180000023910 | 604122 | 0.4005 | 0 | 49.2008 | Eukaryota | 45335.0 | 0 | Chordata | 45335.0 | 0 | Carnivora | 45335.0 | 0 | Felidae | 45335.0 | 0 | Felis | 45335.0 | 0 | Felis catus | 45335.0 | 0 |
| scf7180000023469 | 37519 | 0.6727 | 0 | 287.1789 | Eukaryota | 2579.0 | 0 | Chordata | 2579.0 | 0 | Rodentia | 1046.0 | 3 | Caviidae | 783.0 | 4 | Cavia | 783.0 | 4 | Cavia porcellu | 783.0 | 4 |
| scf7180000018894 | 515371 | 0.4287 | 0 | 49.5584 | Eukaryota | 57187.0 | 1 | Chordata | 57187.0 | 1 | Carnivora | 39180.0 | 3 | Felidae | 23577.0 | 4 | Canis | 15603.0 | 5 | Canis lupus | 15603.0 | 5 |
| scf7180000020332 | 23091 | 0.4752 | 0 | 441.333 | Eukaryota | 24577.0 | 0 | Chordata | 24577.0 | 0 | Carnivora | 24577.0 | 0 | Felidae | 24577.0 | 0 | Felis | 22085.0 | 1 | Felis catus | 22085.0 | 1 |
| scf7180000020864 | 12452 | 0.4083 | 0 | 42.0876 | Eukaryota | 4855.0 | 0 | Chordata | 4855.0 | 0 | Carnivora | 3973.0 | 2 | Felidae | 3973.0 | 2 | Felis | 2320.0 | 4 | Felis catus | 2320.0 | 4 |
| scf7180000023471 | 18767 | 0.4816 | 0 | 116.0139 | no-hit | 0.0 | 0 | no-hit | 0.0 | 0 | no-hit | 0.0 | 0 | no-hit | 0.0 | 0 | no-hit | 0.0 | 0 | no-hit | 0.0 | 0 |
| scf7180000018519 | 196835 | 0.4337 | 0 | 52.2122 | Eukaryota | 35811.0 | 0 | Chordata | 35811.0 | 0 | Carnivora | 35811.0 | 0 | Felidae | 35811.0 | 0 | Felis | 32281.0 | 1 | Felis catus | 32281.0 | 1 |
| scf7180000021877 | 34298 | 0.367 | 0 | 25.0182 | Eukaryota | 14010.0 | 0 | Chordata | 14010.0 | 0 | Carnivora | 14010.0 | 0 | Felidae | 14010.0 | 0 | Felis | 12510.0 | 1 | Felis catus | 12510.0 | 1 |
| scf7180000021210 | 48714 | 0.3956 | 0 | 28.9951 | Eukaryota | 45476.0 | 0 | Chordata | 45476.0 | 0 | Carnivora | 45476.0 | 0 | Felidae | 45476.0 | 0 | Felis | 45476.0 | 0 | Felis catus | 45476.0 | 0 |
| scf7180000023812 | 1576473 | 0.4192 | 0 | 51.533 | Eukaryota | 81041.0 | 0 | Chordata | 81041.0 | 0 | Carnivora | 81041.0 | 0 | Felidae | 32485.0 | 5 | Lynx | 16248.0 | 7 | ynx canadens | 16248.0 | 7 |
| scf7180000019214 | 408506 | 0.4387 | 0 | 51.3411 | Eukaryota | 33270.0 | 0 | Chordata | 33270.0 | 0 | Carnivora | 33270.0 | 0 | Felidae | 29386.0 | 1 | Felis | 8768.0 | 6 | Felis catus | 8768.0 | 7 |
| scf7180000019978 | 209951 | 0.5255 | 0 | 50.9421 | Eukaryota | 11165.0 | 0 | Chordata | 11165.0 | 0 | Carnivora | 11165.0 | 0 | Felidae | 8916.0 | 1 | Felis | 6623.0 | 2 | Felis catus | 6623.0 | 2 |
| scf7180000023990 | 580676 | 0.4049 | 0 | 50.098 | Eukaryota | 45913.0 | 0 | Chordata | 45913.0 | 0 | Carnivora | 45913.0 | 0 | Felidae | 45913.0 | 0 | Felis | 45913.0 | 0 | Felis catus | 45913.0 | 0 |
| scf7180000017523 | 2591125 | 0.3738 | 0 | 50.0569 | Eukaryota | 47677.0 | 0 | Chordata | 47677.0 | 0 | Carnivora | 47677.0 | 0 | Felidae | 47677.0 | 0 | Felis | 47677.0 | 0 | Felis catus | 47677.0 | 0 |
| scf7180000021344 | 53547 | 0.5388 | 0 | 47.8013 | Eukaryota | 5891.0 | 1 | Chordata | 5891.0 | 1 | Primates | 3074.0 | 4 | Hominidae | 3074.0 | 4 | Homo | 1574.0 | 5 | Homo sapiens | 1574.0 | 5 |
| scf7180000018389 | 792480 | 0.3512 | 0 | 49.4813 | Eukaryota | 47547.0 | 0 | Chordata | 47547.0 | 0 | Carnivora | 47547.0 | 0 | Felidae | 47547.0 | 0 | Felis | 47547.0 | 0 | Felis catus | 47547.0 | 0 |
| scf7180000020594 | 48988 | 0.4212 | 0 | 46.6621 | Eukaryota | 37317.0 | 0 | Chordata | 37317.0 | 0 | Carnivora | 37317.0 | 0 | Felidae | 37317.0 | 0 | Acinonyx | 18785.0 | 2 | cinonyx jubat | 18785.0 | 3 |
| scf7180000019538 | 331113 | 0.4055 | 0 | 52.2971 | Eukaryota | 14702.0 | 0 | Chordata | 14702.0 | 0 | Carnivora | 14702.0 | 0 | Felidae | 14702.0 | 0 | Felis | 12053.0 | 2 | Felis catus | 12053.0 | 2 |
| scf7180000020400 | 97171 | 0.3807 | 0 | 25.1659 | Eukaryota | 19981.0 | 0 | Chordata | 19981.0 | 0 | Carnivora | 19981.0 | 0 | Felidae | 19981.0 | 0 | Felis | 18344.0 | 1 | Felis catus | 18344.0 | 1 |
| scf7180000018273 | 1114061 | 0.5144 | 0 | 52.2148 | Eukaryota | 46900.0 | 0 | Chordata | 46900.0 | 0 | Carnivora | 46900.0 | 0 | Felidae | 46900.0 | 0 | Felis | 46900.0 | 0 | Felis catus | 46900.0 | 0 |
| scf7180000022615 | 7649 | 0.6139 | 0 | 199.3665 | Eukaryota | 2027.0 | 0 | Chordata | 2027.0 | 0 | Carnivora | 2027.0 | 0 | Felidae | 1594.0 | 2 | Felis | 621.0 | 4 | Felis catus | 621.0 | 6 |
| scf7180000019899 | 193632 | 0.4 | 0 | 47.055 | Eukaryota | 46597.0 | 0 | Chordata | 46597.0 | 0 | Carnivora | 46597.0 | 0 | Felidae | 46597.0 | 0 | Felis | 46597.0 | 0 | Felis catus | 46597.0 | 0 |
| scf7180000023418 | 31826 | 0.5099 | 0 | 101.6476 | Eukaryota | 6998.0 | 0 | Chordata | 6998.0 | 0 | Carnivora | 6998.0 | 0 | Felidae | 6998.0 | 0 | Felis | 6560.0 | 1 | Felis catus | 6560.0 | 1 |
| scf7180000019547 | 313849 | 0.5384 | 0 | 46.8089 | Eukaryota | 18993.0 | 0 | Chordata | 18993.0 | 0 | Carnivora | 18993.0 | 0 | Felidae | 13511.0 | 2 | Felis | 13511.0 | 2 | Felis catus | 13511.0 | 2 |
| scf7180000022020 | 21536 | 0.4102 | 0 | 24.513 | Eukaryota | 24220.0 | 0 | Chordata | 24220.0 | 0 | Carnivora | 24220.0 | 0 | Felidae | 24220.0 | 0 | Felis | 24220.0 | 0 | Felis catus | 24220.0 | 0 |
| scf7180000023860 | 2033022 | 0.4297 | 0 | 51.5953 | Eukaryota | 50053.0 | 0 | Chordata | 50053.0 | 0 | Carnivora | 50053.0 | 0 | Felidae | 50053.0 | 0 | Lynx | 24580.0 | 2 | ynx canadens | 24580.0 | 2 |
| scf7180000020736 | 83163 | 0.383 | 0 | 25.478 | Eukaryota | 23182.0 | 0 | Chordata | 23182.0 | 0 | Carnivora | 23182.0 | 0 | Felidae | 18669.0 | 1 | Panthera | 9322.0 | 5 | anthera pardu | 6998.0 | 6 |
| scf7180000020122 | 200302 | 0.4439 | 0 | 48.5803 | Eukaryota | 50379.0 | 0 | Chordata | 50379.0 | 0 | Carnivora | 50379.0 | 0 | Felidae | 50379.0 | 0 | Panthera | 25210.0 | 2 | anthera pardu | 25210.0 | 2 |
| scf7180000023924 | 778945 | 0.3733 | 0 | 50.0798 | Eukaryota | 47676.0 | 0 | Chordata | 47676.0 | 0 | Carnivora | 47676.0 | 0 | Felidae | 47676.0 | 0 | Felis | 42883.0 | 1 | Felis catus | 42883.0 | 1 |
| scf7180000022960 | 21150 | 0.3949 | 0 | 210.5774 | Eukaryota | 7002.0 | 0 | Chordata | 7002.0 | 0 | Carnivora | 7002.0 | 0 | Felidae | 7002.0 | 0 | Felis | 7002.0 | 0 | Felis catus | 7002.0 | 0 |
| scf7180000019845 | 236468 | 0.3551 | 0 | 49.4083 | Eukaryota | 30028.0 | 0 | Chordata | 30028.0 | 0 | Carnivora | 30028.0 | 0 | Felidae | 30028.0 | 0 | Felis | 27003.0 | 1 | Felis catus | 27003.0 | 1 |
| scf7180000022262 | 21022 | 0.3623 | 0 | 23.5936 | Eukaryota | 7975.0 | 0 | Chordata | 7975.0 | 0 | Carnivora | 7975.0 | 0 | Felidae | 5548.0 | 1 | Felis | 5548.0 | 1 | Felis catus | 5548.0 | 1 |
| scf7180000022858 | 676473 | 0.4886 | 0 | 58.0012 | Eukaryota | 53694.0 | 0 | Chordata | 53694.0 | 0 | Carnivora | 53694.0 | 0 | Ursidae | 53694.0 | 0 | Ursus | 32250.0 | 1 | Ursus arctos | 26870.0 | 2 |
| scf7180000018877 | 647093 | 0.3951 | 0 | 51.5282 | Eukaryota | 47139.0 | 0 | Chordata | 47139.0 | 0 | Carnivora | 47139.0 | 0 | Felidae | 47139.0 | 0 | Felis | 47139.0 | 0 | Felis catus | 47139.0 | 0 |
| scf7180000018801 | 1308801 | 0.3965 | 100 | 50.7598 | Eukaryota | 38491.0 | 0 | Chordata | 38491.0 | 0 | Carnivora | 38491.0 | 0 | Felidae | 38491.0 | 0 | Felis | 34668.0 | 1 | Felis catus | 34668.0 | 1 |
| scf7180000020483 | 97496 | 0.5358 | 0 | 49.3174 | Eukaryota | 6200.0 | 0 | Chordata | 6200.0 | 0 | Carnivora | 6200.0 | 0 | Herpestidae | 2729.0 | 2 | Suricata | 2729.0 | 2 | uricata suricat | 2729.0 | 2 |
| scf7180000023756 | 3351750 | 0.3797 | 0 | 49.6322 | Eukaryota | 46780.0 | 0 | Chordata | 46780.0 | 0 | Carnivora | 46780.0 | 0 | Felidae | 46780.0 | 0 | Felis | 41797.0 | 1 | Felis catus | 41797.0 | 1 |
| scf7180000018753 | 877537 | 0.4565 | 0 | 50.9006 | Eukaryota | 44881.0 | 0 | Chordata | 44881.0 | 0 | Carnivora | 44881.0 | 0 | Felidae | 41872.0 | 1 | Acinonyx | 13348.0 | 4 | cinonyx jubat | 13348.0 | 5 |
| scf7180000023053 | 232420 | 0.5421 | 0 | 57.5834 | Eukaryota | 9270.0 | 0 | Chordata | 9270.0 | 0 | pterygiforme | 8326.0 | 1 | Apterygidae | 8326.0 | 1 | Apteryx | 8326.0 | 1 | pteryx mante | 8326.0 | 1 |
| scf7180000023578 | 26447 | 0.4629 | 0 | 301.0497 | no-hit | 0.0 | 0 | no-hit | 0.0 | 0 | no-hit | 0.0 | 0 | no-hit | 0.0 | 0 | no-hit | 0.0 | 0 | no-hit | 0.0 | 0 |
| scf7180000018915 | 593894 | 0.3753 | 0 | 51.6718 | Eukaryota | 74935.0 | 0 | Chordata | 74935.0 | 0 | Carnivora | 74935.0 | 0 | Herpestidae | 44240.0 | 1 | Suricata | 44240.0 | 4 | uricata suricat | 44240.0 | 4 |
| scf7180000017655 | 2838666 | 0.3819 | 0 | 49.947 | Eukaryota | 43983.0 | 0 | Chordata | 43983.0 | 0 | Carnivora | 43983.0 | 0 | Felidae | 43983.0 | 0 | Felis | 39419.0 | 1 | Felis catus | 39419.0 | 1 |
| scf7180000018469 | 164078 | 0.3743 | 0 | 50.976 | Eukaryota | 46403.0 | 0 | Chordata | 46403.0 | 0 | Carnivora | 46403.0 | 0 | Felidae | 46403.0 | 0 | Felis | 46403.0 | 0 | Felis catus | 46403.0 | 0 |
| scf7180000019576 | 187313 | 0.3523 | 0 | 46.3936 | Eukaryota | 45861.0 | 0 | Chordata | 45861.0 | 0 | Carnivora | 45861.0 | 0 | Felidae | 45861.0 | 0 | Felis | 45861.0 | 0 | Felis catus | 45861.0 | 0 |
| scf7180000018393 | 1117674 | 0.424 | 0 | 52.2664 | Eukaryota | 45972.0 | 0 | Chordata | 45972.0 | 0 | Carnivora | 45972.0 | 0 | Felidae | 45972.0 | 0 | Felis | 45972.0 | 0 | Felis catus | 45972.0 | 0 |
| scf7180000020354 | 254233 | 0.3885 | 0 | 51.5705 | Eukaryota | 46991.0 | 0 | Chordata | 46991.0 | 0 | Carnivora | 46991.0 | 0 | Felidae | 46991.0 | 0 | Felis | 46991.0 | 0 | Felis catus | 46991.0 | 0 |
| scf7180000020028 | 6736 | 0.5338 | 0 | 162.8242 | Eukaryota | 12009.0 | 0 | Chordata | 12009.0 | 0 | Carnivora | 12009.0 | 0 | Felidae | 12009.0 | 0 | Felis | 12009.0 | 0 | Felis catus | 12009.0 | 0 |
| scf7180000019037 | 912986 | 0.4138 | 0 | 52.8229 | Eukaryota | 83196.0 | 0 | Chordata | 83196.0 | 0 | Carnivora | 83196.0 | 0 | Felidae | 33730.0 | 3 | Suricata | 26079.0 | 5 | uricata suricat | 26079.0 | 5 |
| scf7180000018079 | 972422 | 0.4548 | 0 | 52.0316 | Eukaryota | 56981.0 | 0 | Chordata | 56981.0 | 0 | Carnivora | 56981.0 | 0 | Felidae | 56981.0 | 0 | Felis | 36689.0 | 1 | Felis catus | 36689.0 | 1 |
| scf7180000019543 | 254073 | 0.3995 | 0 | 51.3492 | Eukaryota | 88423.0 | 0 | Chordata | 88423.0 | 0 | Carnivora | 88423.0 | 0 | Felidae | 77825.0 | 1 | Panthera | 29128.0 | 4 | Felis catus | 29097.0 | 5 |
| scf7180000024085 | 297857 | 0.5721 | 0 | 48.1642 | Eukaryota | 29134.0 | 0 | Chordata | 29134.0 | 0 | Carnivora | 29134.0 | 0 | Felidae | 23272.0 | 1 | Felis | 23272.0 | 1 | Felis catus | 23272.0 | 1 |
| scf7180000018960 | 508464 | 0.3732 | 0 | 50.778 | Eukaryota | 47400.0 | 0 | Chordata | 47400.0 | 0 | Carnivora | 47400.0 | 0 | Felidae | 47400.0 | 0 | Felis | 37039.0 | 2 | Felis catus | 37039.0 | 2 |
| scf7180000020906 | 71227 | 0.3645 | 0 | 23.9064 | Eukaryota | 29900.0 | 0 | Chordata | 29900.0 | 0 | Carnivora | 29900.0 | 0 | Felidae | 29900.0 | 0 | Felis | 26863.0 | 1 | Felis catus | 26863.0 | 1 |
| scf7180000019879 | 27566 | 0.4241 | 0 | 24.7533 | Eukaryota | 5709.0 | 0 | Chordata | 5709.0 | 0 | Carnivora | 5709.0 | 0 | Felidae | 5709.0 | 0 | Felis | 5709.0 | 0 | Felis catus | 5709.0 | 0 |
| scf7180000022622 | 6530 | 0.6513 | 0 | 43.909 | no-hit | 0.0 | 0 | no-hit | 0.0 | 0 | no-hit | 0.0 | 0 | no-hit | 0.0 | 0 | no-hit | 0.0 | 0 | no-hit | 0.0 | 0 |
| scf7180000018844 | 558216 | 0.3469 | 0 | 47.915 | Eukaryota | 45526.0 | 0 | Chordata | 45526.0 | 0 | Carnivora | 45526.0 | 0 | Felidae | 45526.0 | 0 | Felis | 45526.0 | 0 | Felis catus | 45526.0 | 0 |
| scf7180000022184 | 28985 | 0.4225 | 0 | 25.3432 | Eukaryota | 5618.0 | 0 | Chordata | 5618.0 | 0 | Carnivora | 3778.0 | 1 | Delphinidae | 1840.0 | 3 | Globicephala | 1840.0 | 5 | obicephala me | 1840.0 | 5 |
| scf7180000021237 | 77919 | 0.4576 | 0 | 24.1322 | Eukaryota | 11399.0 | 0 | Chordata | 11399.0 | 0 | Primates | 8254.0 | 1 | Hominidae | 8254.0 | 3 | Homo | 5476.0 | 4 | Homo sapiens | 5476.0 | 4 |
| scf7180000024129 | 193955 | 0.4418 | 0 | 51.1928 | Eukaryota | 11989.0 | 0 | Chordata | 11989.0 | 0 | Carnivora | 11989.0 | 0 | Felidae | 11989.0 | 0 | Felis | 9785.0 | 1 | Felis catus | 9785.0 | 2 |
| scf7180000022505 | 7492 | 0.5039 | 0 | 56.7374 | Eukaryota | 2817.0 | 0 | Chordata | 2817.0 | 0 | Carnivora | 2817.0 | 0 | Felidae | 2817.0 | 0 | Felis | 1678.0 | 1 | Felis catus | 1678.0 | 1 |
| scf7180000021813 | 13600 | 0.5212 | 0 | 98.0617 | no-hit | 0.0 | 0 | no-hit | 0.0 | 0 | no-hit | 0.0 | 0 | no-hit | 0.0 | 0 | no-hit | 0.0 | 0 | no-hit | 0.0 | 0 |
| scf7180000017842 | 595214 | 0.3902 | 0 | 52.2166 | Eukaryota | 45430.0 | 0 | Chordata | 45430.0 | 0 | Carnivora | 45430.0 | 0 | Felidae | 45430.0 | 0 | Felis | 45430.0 | 0 | Felis catus | 45430.0 | 0 |
| scf7180000019390 | 381817 | 0.3589 | 0 | 50.7836 | Eukaryota | 38538.0 | 0 | Chordata | 38538.0 | 0 | Primates | 25147.0 | 2 | Hominidae | 25147.0 | 3 | Homo | 25147.0 | 3 | Homo sapiens | 25147.0 | 3 |
| scf7180000022005 | 18406 | 0.4628 | 0 | 352.945 | no-hit | 0.0 | 0 | no-hit | 0.0 | 0 | no-hit | 0.0 | 0 | no-hit | 0.0 | 0 | no-hit | 0.0 | 0 | no-hit | 0.0 | 0 |
| scf7180000017873 | 46603 | 0.4134 | 0 | 24.5782 | Eukaryota | 5743.0 | 1 | Chordata | 5743.0 | 1 | Carnivora | 3832.0 | 3 | Felidae | 3832.0 | 3 | Felis | 3832.0 | 3 | Felis catus | 3832.0 | 3 |
| scf7180000018496 | 187054 | 0.4415 | 0 | 51.7332 | Eukaryota | 13047.0 | 0 | Chordata | 13047.0 | 0 | Carnivora | 13047.0 | 0 | Felidae | 13047.0 | 0 | Felis | 13047.0 | 0 | Felis catus | 13047.0 | 0 |
| scf7180000018542 | 663958 | 0.4573 | 0 | 52.995 | Eukaryota | 38708.0 | 0 | Chordata | 38708.0 | 0 | Carnivora | 38708.0 | 0 | Felidae | 32271.0 | 2 | Panthera | 15088.0 | 5 | Panthera tigri | 9118.0 | 6 |
| scf7180000019032 | 454904 | 0.4844 | 0 | 47.9003 | Eukaryota | 29955.0 | 0 | Chordata | 29955.0 | 0 | Carnivora | 29955.0 | 0 | Felidae | 24307.0 | 1 | Felis | 24307.0 | 1 | Felis catus | 24307.0 | 1 |
| scf7180000019073 | 4165772 | 0.411 | 100 | 50.3504 | Eukaryota | 47409.0 | 0 | Chordata | 47409.0 | 0 | Carnivora | 47409.0 | 0 | Felidae | 47409.0 | 0 | Felis | 47409.0 | 0 | Felis catus | 47409.0 | 0 |
| scf7180000022417 | 13330 | 0.3672 | 0 | 23.515 | Eukaryota | 4632.0 | 0 | Chordata | 4632.0 | 0 | Carnivora | 4632.0 | 0 | Felidae | 3260.0 | 1 | Felis | 3260.0 | 1 | Felis catus | 3260.0 | 1 |
| scf7180000017627 | 2131931 | 0.438 | 0 | 51.5598 | Eukaryota | 47145.0 | 0 | Chordata | 47145.0 | 0 | Carnivora | 47145.0 | 0 | Felidae | 47145.0 | 0 | Felis | 47145.0 | 0 | Felis catus | 47145.0 | 0 |
| scf7180000022436 | 20578 | 0.6061 | 0 | 43.8394 | Eukaryota | 5509.0 | 0 | Chordata | 5509.0 | 0 | Carnivora | 5509.0 | 0 | Felidae | 4241.0 | 2 | Panthera | 1766.0 | 4 | anthera pardu | 1766.0 | 4 |
| scf7180000017749 | 79981 | 0.403 | 0 | 45.0088 | Eukaryota | 35952.0 | 0 | Chordata | 35952.0 | 0 | Carnivora | 35952.0 | 0 | Felidae | 35952.0 | 0 | Felis | 35952.0 | 0 | Felis catus | 35952.0 | 0 |
| scf7180000019273 | 341516 | 0.4394 | 0 | 49.1159 | Eukaryota | 52204.0 | 0 | Chordata | 52204.0 | 0 | Carnivora | 52204.0 | 0 | Felidae | 23516.0 | 4 | Felis | 6394.0 | 9 | Felis catus | 6394.0 | 9 |
| scf7180000019516 | 150658 | 0.3915 | 0 | 25.5985 | Eukaryota | 27042.0 | 0 | Chordata | 27042.0 | 0 | Carnivora | 27042.0 | 0 | Felidae | 27042.0 | 0 | Felis | 18911.0 | 2 | Felis catus | 18911.0 | 2 |
| scf7180000021763 | 45615 | 0.3926 | 0 | 25.0319 | Eukaryota | 19942.0 | 0 | Chordata | 19942.0 | 0 | Carnivora | 19942.0 | 0 | Felidae | 6304.0 | 5 | Acinonyx | 2117.0 | 9 | cinonyx jubat | 2117.0 | 9 |
| scf7180000018807 | 743748 | 0.3926 | 0 | 50.3538 | Eukaryota | 29837.0 | 0 | Chordata | 29837.0 | 0 | Carnivora | 29837.0 | 0 | Felidae | 29837.0 | 0 | Felis | 26839.0 | 1 | Felis catus | 26839.0 | 1 |
| scf7180000022692 | 5586 | 0.6459 | 0 | 50.8419 | Eukaryota | 2084.0 | 0 | Chordata | 2084.0 | 0 | Carnivora | 2084.0 | 0 | Felidae | 2084.0 | 0 | Felis | 2084.0 | 0 | Felis catus | 2084.0 | 0 |
| scf7180000022794 | 13418 | 0.3679 | 0 | 27.2158 | Eukaryota | 13384.0 | 0 | Chordata | 13384.0 | 0 | Carnivora | 11527.0 | 1 | Felidae | 6979.0 | 2 | Felis | 5051.0 | 3 | Felis catus | 5051.0 | 3 |
| scf7180000023903 | 1039718 | 0.4591 | 0 | 51.7102 | Eukaryota | 76254.0 | 0 | Chordata | 76254.0 | 0 | Carnivora | 76254.0 | 0 | Felidae | 62436.0 | 1 | Acinonyx | 31264.0 | 2 | cinonyx jubat | 31264.0 | 2 |
| scf7180000018843 | 569523 | 0.4178 | 0 | 50.845 | Eukaryota | 31575.0 | 0 | Chordata | 31575.0 | 0 | Carnivora | 31575.0 | 0 | Felidae | 16150.0 | 1 | Suricata | 15425.0 | 2 | uricata suricat | 15425.0 | 3 |
| scf7180000022144 | 74527 | 0.4372 | 0 | 24.6561 | Eukaryota | 8331.0 | 0 | Chordata | 8331.0 | 0 | Carnivora | 4583.0 | 2 | Felidae | 3875.0 | 3 | Felis | 3875.0 | 4 | Felis catus | 3875.0 | 4 |
| scf7180000022701 | 12545 | 0.4007 | 0 | 24.1085 | Eukaryota | 31067.0 | 0 | Chordata | 31067.0 | 0 | Carnivora | 31067.0 | 0 | Felidae | 31067.0 | 0 | Felis | 27944.0 | 1 | Felis catus | 27944.0 | 1 |
| scf7180000021789 | 29444 | 0.4071 | 0 | 26.7721 | Eukaryota | 8462.0 | 0 | Chordata | 8462.0 | 0 | Artiodactyla | 8462.0 | 0 | Bovidae | 7538.0 | 1 | Ovis | 7538.0 | 1 | Ovis canadens | 7538.0 | 1 |
| scf7180000019983 | 550183 | 0.4753 | 0 | 49.7104 | Eukaryota | 40474.0 | 0 | Chordata | 40474.0 | 0 | Carnivora | 40474.0 | 0 | Felidae | 40474.0 | 0 | Lynx | 20960.0 | 1 | ynx canadens | 20960.0 | 2 |
| scf7180000018593 | 975131 | 0.5185 | 100 | 51.5285 | Eukaryota | 23078.0 | 0 | Chordata | 23078.0 | 0 | Carnivora | 23078.0 | 0 | Otariidae | 16100.0 | 1 | Zalophus | 9200.0 | 2 | phus californi | 9200.0 | 2 |
| scf7180000019554 | 317087 | 0.3682 | 0 | 49.1385 | Eukaryota | 45266.0 | 0 | Chordata | 45266.0 | 0 | Carnivora | 45266.0 | 0 | Felidae | 45266.0 | 0 | Felis | 45266.0 | 0 | Felis catus | 45266.0 | 0 |
| scf7180000018837 | 844977 | 0.4026 | 0 | 51.6015 | Eukaryota | 47551.0 | 0 | Chordata | 47551.0 | 0 | Carnivora | 47551.0 | 0 | Felidae | 47551.0 | 0 | Felis | 47551.0 | 0 | Felis catus | 47551.0 | 0 |
| scf7180000020361 | 212915 | 0.5281 | 0 | 48.5933 | Eukaryota | 20257.0 | 0 | Chordata | 20257.0 | 0 | Carnivora | 20257.0 | 0 | Felidae | 20257.0 | 0 | Felis | 20257.0 | 0 | Felis catus | 20257.0 | 0 |
| scf7180000023676 | 157620 | 0.493 | 0 | 534.3911 | Eukaryota | 15167.0 | 0 | Chordata | 15167.0 | 0 | Carnivora | 15167.0 | 0 | Felidae | 15167.0 | 0 | Felis | 15167.0 | 0 | Felis catus | 15167.0 | 0 |
| scf7180000023879 | 804016 | 0.3622 | 0 | 49.9664 | Eukaryota | 46972.0 | 0 | Chordata | 46972.0 | 0 | Carnivora | 46972.0 | 0 | Felidae | 46972.0 | 0 | Felis | 46972.0 | 0 | Felis catus | 46972.0 | 0 |
| scf7180000019593 | 266329 | 0.3737 | 0 | 48.3636 | Eukaryota | 42162.0 | 0 | Chordata | 42162.0 | 0 | Carnivora | 42162.0 | 0 | Felidae | 42162.0 | 0 | Felis | 42162.0 | 0 | Felis catus | 42162.0 | 0 |
| scf7180000020584 | 108472 | 0.3732 | 0 | 25.2558 | Eukaryota | 41910.0 | 0 | Chordata | 41910.0 | 0 | Carnivora | 41910.0 | 0 | Felidae | 41910.0 | 0 | Felis | 41910.0 | 0 | Felis catus | 41910.0 | 0 |
| scf7180000018544 | 59906 | 0.4155 | 0 | 52.1035 | Eukaryota | 37218.0 | 0 | Chordata | 37218.0 | 0 | Carnivora | 37218.0 | 0 | Felidae | 37218.0 | 0 | Felis | 37218.0 | 0 | Felis catus | 37218.0 | 0 |
| scf7180000022719 | 16928 | 0.528 | 0 | 26.0685 | Eukaryota | 12643.0 | 0 | Chordata | 12643.0 | 0 | Carnivora | 12643.0 | 0 | Felidae | 3952.0 | 4 | Neomonachus | 2435.0 | 8 | nachus schau | 2435.0 | 8 |
| scf7180000023870 | 1066766 | 0.4322 | 0 | 52.0515 | Eukaryota | 45587.0 | 0 | Chordata | 45587.0 | 0 | Carnivora | 45587.0 | 0 | Felidae | 45587.0 | 0 | Felis | 45587.0 | 0 | Felis catus | 45587.0 | 0 |
| scf7180000019950 | 1476330 | 0.4673 | 0 | 52.7309 | Eukaryota | 40186.0 | 0 | Chordata | 40186.0 | 0 | Carnivora | 40186.0 | 0 | Felidae | 22741.0 | 3 | Felis | 11266.0 | 7 | Felis catus | 11266.0 | 7 |
| scf7180000021090 | 20058 | 0.519 | 0 | 81.1541 | no-hit | 0.0 | 0 | no-hit | 0.0 | 0 | no-hit | 0.0 | 0 | no-hit | 0.0 | 0 | no-hit | 0.0 | 0 | no-hit | 0.0 | 0 |
| scf7180000023653 | 40733 | 0.3882 | 0 | 427.6471 | Eukaryota | 6143.0 | 0 | Chordata | 6143.0 | 0 | Carnivora | 6143.0 | 0 | Felidae | 6143.0 | 0 | Felis | 6143.0 | 0 | Felis catus | 6143.0 | 0 |
| scf7180000020135 | 129953 | 0.443 | 0 | 25.9376 | Eukaryota | 21976.0 | 0 | Chordata | 21976.0 | 0 | Carnivora | 21976.0 | 0 | Felidae | 21976.0 | 0 | Felis | 19765.0 | 1 | Felis catus | 19765.0 | 1 |
| scf7180000023947 | 963517 | 0.4129 | 0 | 50.8306 | Eukaryota | 44648.0 | 0 | Chordata | 44648.0 | 0 | Carnivora | 44648.0 | 0 | Felidae | 27602.0 | 2 | Panthera | 13802.0 | 4 | anthera pardu | 9220.0 | 5 |
| scf7180000020645 | 115437 | 0.4611 | 0 | 50.7839 | Eukaryota | 28001.0 | 0 | Chordata | 28001.0 | 0 | Carnivora | 28001.0 | 0 | Felidae | 28001.0 | 0 | Felis | 13559.0 | 3 | Felis catus | 13559.0 | 3 |
| scf7180000017968 | 1608973 | 0.3719 | 0 | 50.0785 | Eukaryota | 47495.0 | 0 | Chordata | 47495.0 | 0 | Carnivora | 47495.0 | 0 | Felidae | 47495.0 | 0 | Felis | 47495.0 | 0 | Felis catus | 47495.0 | 0 |
| scf7180000024061 | 298264 | 0.5876 | 0 | 50.9073 | Eukaryota | 25654.0 | 0 | Chordata | 25654.0 | 0 | Carnivora | 25654.0 | 0 | Otariidae | 25654.0 | 0 | Callorhinus | 12845.0 | 2 | llorhinus ursir | 12845.0 | 2 |
| scf7180000020407 | 115467 | 0.3848 | 0 | 24.4117 | Eukaryota | 42803.0 | 0 | Chordata | 42803.0 | 0 | Carnivora | 42803.0 | 0 | Felidae | 42803.0 | 0 | Felis | 42803.0 | 0 | Felis catus | 42803.0 | 0 |
| scf7180000019261 | 415573 | 0.4975 | 0 | 52.6805 | Eukaryota | 39778.0 | 0 | Chordata | 39778.0 | 0 | Carnivora | 39778.0 | 0 | Felidae | 14079.0 | 5 | Suricata | 9469.0 | 8 | uricata suricat | 9469.0 | 8 |
| scf7180000022099 | 72647 | 0.403 | 0 | 25.6655 | Eukaryota | 43579.0 | 0 | Chordata | 43579.0 | 0 | Carnivora | 43579.0 | 0 | Felidae | 43579.0 | 0 | Felis | 43579.0 | 0 | Felis catus | 43579.0 | 0 |
| scf7180000023804 | 2378379 | 0.417 | 0 | 51.9745 | Eukaryota | 64400.0 | 0 | Chordata | 64400.0 | 0 | Carnivora | 64400.0 | 0 | Felidae | 39076.0 | 3 | Panthera | 13071.0 | 8 | uricata suricat | 6647.0 | 9 |
| scf7180000024179 | 118879 | 0.4603 | 0 | 51.6505 | Eukaryota | 14495.0 | 0 | Chordata | 14495.0 | 0 | Carnivora | 14495.0 | 0 | Felidae | 14495.0 | 0 | Felis | 10912.0 | 1 | Felis catus | 10912.0 | 1 |
| scf7180000018694 | 1955875 | 0.4163 | 0 | 51.1722 | Eukaryota | 47642.0 | 0 | Chordata | 47642.0 | 0 | Carnivora | 47642.0 | 0 | Felidae | 47642.0 | 0 | Felis | 47642.0 | 0 | Felis catus | 47642.0 | 0 |
| scf7180000017651 | 2138670 | 0.3895 | 0 | 50.837 | Eukaryota | 63965.0 | 0 | Chordata | 63965.0 | 0 | Carnivora | 38781.0 | 2 | Ursidae | 21164.0 | 5 | Ailuropoda | 21164.0 | 5 | opoda melanc | 21164.0 | 5 |
| scf7180000024183 | 59073 | 0.4005 | 0 | 25.4125 | Eukaryota | 16995.0 | 0 | Chordata | 16995.0 | 0 | Primates | 14701.0 | 1 | Hominidae | 7422.0 | 3 | Chlorocebus | 4434.0 | 5 | rocebus saba | 4434.0 | 5 |
| scf7180000021875 | 91102 | 0.3672 | 0 | 26.1238 | Eukaryota | 45579.0 | 0 | Chordata | 45579.0 | 0 | Carnivora | 45579.0 | 0 | Felidae | 45579.0 | 0 | Felis | 45579.0 | 0 | Felis catus | 45579.0 | 0 |
| scf7180000017661 | 3170902 | 0.4056 | 0 | 50.3343 | Eukaryota | 55700.0 | 0 | Chordata | 55700.0 | 0 | Carnivora | 55700.0 | 0 | Felidae | 55700.0 | 0 | Felis | 31613.0 | 2 | Felis catus | 31613.0 | 2 |
| scf7180000021776 | 12245 | 0.5205 | 0 | 313.1057 | no-hit | 0.0 | 0 | no-hit | 0.0 | 0 | no-hit | 0.0 | 0 | no-hit | 0.0 | 0 | no-hit | 0.0 | 0 | no-hit | 0.0 | 0 |
| scf7180000022122 | 28038 | 0.3749 | 0 | 23.3347 | Eukaryota | 33184.0 | 0 | Chordata | 33184.0 | 0 | Carnivora | 33184.0 | 0 | Felidae | 33184.0 | 0 | Felis | 29887.0 | 1 | Felis catus | 29887.0 | 1 |
| scf7180000020683 | 98483 | 0.3863 | 0 | 25.2415 | Eukaryota | 28367.0 | 0 | Chordata | 28367.0 | 0 | Carnivora | 28367.0 | 0 | Felidae | 28367.0 | 0 | Felis | 25491.0 | 1 | Felis catus | 25491.0 | 1 |
| scf7180000018872 | 511825 | 0.3694 | 0 | 50.0006 | Eukaryota | 45868.0 | 0 | Chordata | 45868.0 | 0 | Carnivora | 45868.0 | 0 | Felidae | 45868.0 | 0 | Felis | 45868.0 | 0 | Felis catus | 45868.0 | 0 |
| scf7180000021689 | 1020707 | 0.5041 | 0 | 51.8631 | Eukaryota | 112804.0 | 0 | Chordata | 112804.0 | 0 | Carnivora | 112804.0 | 0 | Felidae | 61632.0 | 2 | Felis | 24728.0 | 5 | Felis catus | 24728.0 | 5 |
| scf7180000020862 | 84717 | 0.4014 | 0 | 51.3461 | Eukaryota | 12031.0 | 0 | Chordata | 12031.0 | 0 | Carnivora | 12031.0 | 0 | Felidae | 12031.0 | 0 | Felis | 10850.0 | 1 | Felis catus | 10850.0 | 1 |
| scf7180000021413 | 56225 | 0.4551 | 0 | 27.4371 | Eukaryota | 13755.0 | 0 | Chordata | 13755.0 | 0 | Carnivora | 13755.0 | 0 | Felidae | 13755.0 | 0 | Felis | 12377.0 | 1 | Felis catus | 12377.0 | 1 |
| scf7180000021556 | 117017 | 0.5058 | 100 | 47.8453 | Eukaryota | 28347.0 | 0 | Chordata | 28347.0 | 0 | Carnivora | 20672.0 | 2 | Felidae | 9385.0 | 7 | Panthera | 3168.0 | 9 | anthera pardu | 3168.0 | 9 |
| scf7180000020946 | 11492 | 0.3804 | 0 | 33.8661 | Eukaryota | 2480.0 | 0 | Chordata | 2480.0 | 0 | Carnivora | 2480.0 | 0 | Felidae | 2480.0 | 0 | Felis | 2201.0 | 1 | Felis catus | 2201.0 | 1 |
| scf7180000019243 | 12477 | 0.4526 | 0 | 26.0153 | Eukaryota | 12461.0 | 0 | Chordata | 12461.0 | 0 | Carnivora | 12461.0 | 0 | Felidae | 12461.0 | 0 | Felis | 11227.0 | 1 | Felis catus | 11227.0 | 1 |
| scf7180000021279 | 61710 | 0.3857 | 0 | 25.3633 | Eukaryota | 42574.0 | 0 | Chordata | 42574.0 | 0 | Carnivora | 42574.0 | 0 | Felidae | 42574.0 | 0 | Felis | 42574.0 | 0 | Felis catus | 42574.0 | 0 |
| scf7180000023681 | 102215 | 0.4074 | 0 | 775.1664 | Eukaryota | 30362.0 | 0 | Chordata | 30362.0 | 0 | Carnivora | 30362.0 | 0 | Felidae | 30362.0 | 0 | Felis | 30362.0 | 0 | Felis catus | 30362.0 | 0 |
| scf7180000021001 | 74854 | 0.3834 | 0 | 25.9503 | Eukaryota | 11746.0 | 0 | Chordata | 11746.0 | 0 | Carnivora | 9962.0 | 1 | Felidae | 9962.0 | 1 | Felis | 8975.0 | 2 | Felis catus | 8975.0 | 2 |
| scf7180000020873 | 79073 | 0.4364 | 0 | 26.8427 | Eukaryota | 46420.0 | 0 | Chordata | 46420.0 | 0 | Carnivora | 46420.0 | 0 | Felidae | 46420.0 | 0 | Felis | 46420.0 | 0 | Felis catus | 46420.0 | 0 |
| scf7180000017640 | 1935070 | 0.4349 | 0 | 52.309 | Eukaryota | 95805.0 | 0 | Chordata | 95805.0 | 0 | Carnivora | 95805.0 | 0 | Felidae | 95805.0 | 0 | Panthera | 47750.0 | 2 | anthera pardu | 47750.0 | 2 |
| scf7180000020173 | 135815 | 0.3796 | 0 | 50.6366 | Eukaryota | 29887.0 | 0 | Chordata | 29887.0 | 0 | Carnivora | 29887.0 | 0 | Felidae | 29887.0 | 0 | Felis | 29887.0 | 0 | Felis catus | 29887.0 | 0 |
| scf7180000022330 | 13924 | 0.456 | 0 | 138.6528 | Eukaryota | 5552.0 | 0 | Chordata | 5552.0 | 0 | Carnivora | 5552.0 | 0 | Felidae | 5552.0 | 0 | Felis | 5552.0 | 0 | Felis catus | 5552.0 | 0 |
| scf7180000017642 | 537239 | 0.4237 | 0 | 50.8797 | Eukaryota | 43128.0 | 0 | Chordata | 43128.0 | 0 | Carnivora | 43128.0 | 0 | Felidae | 43128.0 | 0 | Felis | 43128.0 | 0 | Felis catus | 43128.0 | 0 |
| scf7180000020425 | 16545 | 0.445 | 0 | 57.2407 | Eukaryota | 3401.0 | 0 | Chordata | 3401.0 | 0 | Carnivora | 3401.0 | 0 | Felidae | 2577.0 | 1 | Felis | 1941.0 | 2 | Felis catus | 1941.0 | 2 |
| scf7180000023899 | 1664876 | 0.3818 | 0 | 49.1667 | Eukaryota | 45776.0 | 0 | Chordata | 45776.0 | 0 | Carnivora | 45776.0 | 0 | Felidae | 45776.0 | 0 | Felis | 45776.0 | 0 | Felis catus | 45776.0 | 0 |
| scf7180000022106 | 19416 | 0.3706 | 0 | 206.7066 | Eukaryota | 12382.0 | 0 | Chordata | 12382.0 | 0 | Carnivora | 12382.0 | 0 | Felidae | 12382.0 | 0 | Felis | 11159.0 | 1 | Felis catus | 11159.0 | 1 |
| scf7180000017858 | 19985 | 0.4007 | 0 | 32.4155 | Eukaryota | 19058.0 | 0 | Chordata | 19058.0 | 0 | Carnivora | 19058.0 | 0 | Felidae | 19058.0 | 0 | Felis | 17141.0 | 1 | Felis catus | 17141.0 | 1 |
| scf7180000018558 | 8113 | 0.4307 | 0 | 23.0322 | Eukaryota | 5818.0 | 0 | Chordata | 5818.0 | 0 | Carnivora | 5818.0 | 0 | Felidae | 5818.0 | 0 | Felis | 5818.0 | 0 | Felis catus | 5818.0 | 0 |
| scf7180000021828 | 61947 | 0.4807 | 0 | 54.1763 | Eukaryota | 8382.0 | 0 | Chordata | 8382.0 | 0 | Carnivora | 8382.0 | 0 | Felidae | 8382.0 | 0 | Felis | 6716.0 | 1 | Felis catus | 6716.0 | 1 |
| scf7180000020843 | 90122 | 0.4144 | 0 | 23.2731 | Eukaryota | 7155.0 | 0 | Chordata | 7155.0 | 0 | Primates | 3238.0 | 2 | Hominidae | 3238.0 | 2 | Felis | 2579.0 | 3 | Felis catus | 2579.0 | 3 |
| scf7180000017748 | 1810933 | 0.3747 | 0 | 49.8723 | Eukaryota | 54150.0 | 0 | Chordata | 54150.0 | 0 | Carnivora | 54150.0 | 0 | Felidae | 54150.0 | 0 | Panthera | 54150.0 | 0 | anthera pardu | 54150.0 | 0 |
| scf7180000021007 | 71840 | 0.3882 | 0 | 25.1149 | Eukaryota | 28740.0 | 0 | Chordata | 28740.0 | 0 | Carnivora | 28740.0 | 0 | Felidae | 28740.0 | 0 | Felis | 25796.0 | 1 | Felis catus | 25796.0 | 1 |
| scf7180000020517 | 24094 | 0.6199 | 0 | 28.1354 | Eukaryota | 6406.0 | 0 | Chordata | 6406.0 | 0 | Carnivora | 6406.0 | 0 | Felidae | 6406.0 | 0 | Acinonyx | 3847.0 | 3 | cinonyx jubat | 3847.0 | 3 |
| scf7180000019400 | 74338 | 0.4154 | 0 | 26.3568 | Eukaryota | 46065.0 | 0 | Chordata | 46065.0 | 0 | Carnivora | 46065.0 | 0 | Herpestidae | 23235.0 | 1 | Suricata | 23235.0 | 1 | uricata suricat | 23235.0 | 1 |
| scf7180000021963 | 43014 | 0.3915 | 0 | 45.3664 | Eukaryota | 35689.0 | 0 | Chordata | 35689.0 | 0 | Carnivora | 35689.0 | 0 | Felidae | 35689.0 | 0 | Felis | 32110.0 | 1 | Felis catus | 32110.0 | 1 |
| scf7180000021606 | 52294 | 0.3939 | 0 | 25.3205 | Eukaryota | 31891.0 | 0 | Chordata | 31891.0 | 0 | Carnivora | 31891.0 | 0 | Felidae | 31891.0 | 0 | Felis | 28622.0 | 1 | Felis catus | 28622.0 | 1 |
| scf7180000019555 | 269515 | 0.619 | 0 | 49.7084 | Eukaryota | 22347.0 | 0 | Chordata | 22347.0 | 0 | Carnivora | 22347.0 | 0 | Felidae | 22347.0 | 0 | Panthera | 13453.0 | 3 | anthera pardu | 13453.0 | 3 |
| scf7180000024178 | 45697 | 0.4535 | 0 | 46.4968 | Eukaryota | 22827.0 | 0 | Chordata | 22827.0 | 0 | Carnivora | 22827.0 | 0 | Canidae | 20400.0 | 1 | Canis | 18100.0 | 2 | Canis lupus | 18100.0 | 2 |
| scf7180000018508 | 507884 | 0.3653 | 0 | 47.773 | Eukaryota | 39715.0 | 0 | Chordata | 39715.0 | 0 | Carnivora | 39715.0 | 0 | Felidae | 39715.0 | 0 | Felis | 39715.0 | 0 | Felis catus | 39715.0 | 0 |
| scf7180000023271 | 21197 | 0.3723 | 0 | 149.4417 | Eukaryota | 44355.0 | 0 | Chordata | 44355.0 | 0 | Carnivora | 44355.0 | 0 | Felidae | 44355.0 | 0 | Felis | 44355.0 | 0 | Felis catus | 44355.0 | 0 |
| scf7180000019183 | 530494 | 0.3642 | 0 | 48.3579 | Eukaryota | 20769.0 | 0 | Chordata | 20769.0 | 0 | Carnivora | 20769.0 | 0 | Felidae | 20769.0 | 0 | Felis | 18682.0 | 1 | Felis catus | 18682.0 | 1 |
| scf7180000021196 | 78902 | 0.4343 | 0 | 25.5458 | Eukaryota | 20931.0 | 0 | Chordata | 20931.0 | 0 | Carnivora | 20931.0 | 0 | Felidae | 15030.0 | 2 | Felis | 4316.0 | 5 | Felis catus | 4316.0 | 6 |

| | | | | | | | | | | | | | | | | | | | | | |
|---|---|---|---|---|---|---|---|---|---|---|---|---|---|---|---|---|---|---|---|---|---|
| scf7180000019087 | 387618 | 0.4094 | 0 | 49.8895 | Eukaryota | 42512.0 | 0 | Chordata | 42512.0 | 0 | Carnivora | 42512.0 | 0 | Felidae | 42512.0 | 0 | Felis | 42512.0 | 0 | Felis catus | 42512.0 | 0 |
| scf7180000023840 | 403674 | 0.4376 | 0 | 52.1779 | Eukaryota | 57613.0 | 0 | Chordata | 57613.0 | 0 | Carnivora | 57613.0 | 0 | Felidae | 57613.0 | 0 | Felis | 57613.0 | 0 | Felis catus | 57613.0 | 0 |
| scf7180000019977 | 152649 | 0.3702 | 0 | 28.7927 | Eukaryota | 47182.0 | 0 | Chordata | 47182.0 | 0 | Carnivora | 47182.0 | 0 | Felidae | 47182.0 | 0 | Felis | 47182.0 | 0 | Felis catus | 47182.0 | 0 |
| scf7180000018243 | 72719 | 0.3932 | 0 | 24.9304 | Eukaryota | 46504.0 | 0 | Chordata | 46504.0 | 0 | Carnivora | 46504.0 | 0 | Felidae | 46504.0 | 0 | Felis | 46504.0 | 0 | Felis catus | 46504.0 | 0 |
| scf7180000023197 | 64876 | 0.3871 | 0 | 300.7498 | Eukaryota | 46128.0 | 0 | Chordata | 46128.0 | 0 | Carnivora | 46128.0 | 0 | Felidae | 46128.0 | 0 | Felis | 46128.0 | 0 | Felis catus | 46128.0 | 0 |
| scf7180000017529 | 2083815 | 0.3755 | 0 | 50.7341 | Eukaryota | 47101.0 | 0 | Chordata | 47101.0 | 0 | Carnivora | 47101.0 | 0 | Felidae | 47101.0 | 0 | Felis | 47101.0 | 0 | Felis catus | 47101.0 | 0 |
| scf7180000019370 | 665068 | 0.4116 | 0 | 49.6901 | Eukaryota | 46456.0 | 0 | Chordata | 46456.0 | 0 | Carnivora | 46456.0 | 0 | Felidae | 46456.0 | 0 | Felis | 46456.0 | 0 | Felis catus | 46456.0 | 0 |
| scf7180000021580 | 35579 | 0.394 | 0 | 24.2252 | Eukaryota | 14308.0 | 0 | Chordata | 14308.0 | 0 | Carnivora | 14308.0 | 0 | Felidae | 14308.0 | 0 | Felis | 12847.0 | 1 | Felis catus | 12847.0 | 1 |
| scf7180000020727 | 97139 | 0.3869 | 0 | 29.5209 | Eukaryota | 49365.0 | 0 | Chordata | 49365.0 | 0 | Carnivora | 49365.0 | 0 | Felidae | 49365.0 | 0 | Felis | 44399.0 | 1 | Felis catus | 44399.0 | 1 |
| scf7180000021318 | 5545 | 0.4575 | 0 | 23.7678 | Eukaryota | 6960.0 | 0 | Chordata | 6960.0 | 0 | Carnivora | 6211.0 | 1 | Felidae | 6053.0 | 2 | Felis | 4184.0 | 5 | Felis catus | 4184.0 | 5 |
| scf7180000019357 | 24883 | 0.55 | 0 | 266.2114 | Eukaryota | 7162.0 | 0 | Chordata | 7162.0 | 0 | Carnivora | 7162.0 | 0 | Felidae | 6239.0 | 2 | Felis | 2589.0 | 4 | Felis catus | 2589.0 | 4 |
| scf7180000022333 | 22775 | 0.4177 | 0 | 22.8534 | Eukaryota | 10652.0 | 0 | Chordata | 10652.0 | 0 | Artiodactyla | 9143.0 | 1 | Bovidae | 9143.0 | 1 | Ovis | 9143.0 | 1 | vis canadens | 9143.0 | 1 |
| scf7180000019610 | 3453220 | 0.4023 | 0 | 50.4107 | Eukaryota | 47137.0 | 0 | Chordata | 47137.0 | 0 | Carnivora | 47137.0 | 0 | Felidae | 37253.0 | 1 | Felis | 37253.0 | 1 | Felis catus | 37253.0 | 1 |
| scf7180000019389 | 57324 | 0.5264 | 0 | 317.5726 | Eukaryota | 12423.0 | 0 | Chordata | 12423.0 | 0 | Carnivora | 12423.0 | 0 | Felidae | 12423.0 | 0 | Felis | 12423.0 | 0 | Felis catus | 12423.0 | 0 |
| scf7180000018971 | 643537 | 0.4603 | 0 | 51.5234 | Eukaryota | 36493.0 | 0 | Chordata | 36493.0 | 0 | Carnivora | 36493.0 | 0 | Felidae | 30240.0 | 1 | Felis | 30240.0 | 1 | Felis catus | 30240.0 | 1 |
| scf7180000018513 | 713164 | 0.3873 | 0 | 49.6397 | Eukaryota | 47576.0 | 0 | Chordata | 47576.0 | 0 | Carnivora | 47576.0 | 0 | Felidae | 47576.0 | 0 | Felis | 47576.0 | 0 | Felis catus | 47576.0 | 0 |
| scf7180000017507 | 3869470 | 0.3754 | 0 | 50.8143 | Eukaryota | 61000.0 | 0 | Chordata | 61000.0 | 0 | Carnivora | 61000.0 | 0 | Felidae | 61000.0 | 0 | Felis | 61000.0 | 0 | Felis catus | 61000.0 | 0 |
| scf7180000022222 | 17172 | 0.6438 | 0 | 50.5818 | Eukaryota | 3202.0 | 0 | Chordata | 3202.0 | 0 | Carnivora | 3202.0 | 0 | Felidae | 2950.0 | 1 | Felis | 2232.0 | 3 | Felis catus | 2232.0 | 3 |
| scf7180000017698 | 3049275 | 0.3609 | 0 | 50.1103 | Eukaryota | 47904.0 | 0 | Chordata | 47904.0 | 0 | Carnivora | 47904.0 | 0 | Felidae | 47904.0 | 0 | Felis | 47904.0 | 0 | Felis catus | 47904.0 | 0 |
| scf7180000020470 | 123433 | 0.421 | 0 | 53.4172 | Eukaryota | 18228.0 | 0 | Chordata | 18228.0 | 0 | Primates | 9141.0 | 1 | Hominidae | 9141.0 | 1 | Felis | 9087.0 | 2 | Felis catus | 9087.0 | 2 |
| scf7180000021636 | 59198 | 0.4444 | 0 | 24.7025 | Eukaryota | 6828.0 | 0 | Chordata | 6828.0 | 0 | Primates | 4483.0 | 3 | Hominidae | 4483.0 | 3 | Pan | 2157.0 | 5 | an troglodyte | 2157.0 | 5 |
| scf7180000018017 | 3672147 | 0.3628 | 0 | 49.6029 | Eukaryota | 46821.0 | 0 | Chordata | 46821.0 | 0 | Carnivora | 46821.0 | 0 | Felidae | 46821.0 | 0 | Felis | 46821.0 | 0 | Felis catus | 46821.0 | 0 |
| scf7180000021610 | 38495 | 0.3638 | 0 | 24.0682 | Eukaryota | 32973.0 | 0 | Chordata | 32973.0 | 0 | Carnivora | 32973.0 | 0 | Felidae | 32973.0 | 0 | Felis | 32973.0 | 0 | Felis catus | 32973.0 | 0 |
| scf7180000020861 | 67953 | 0.4054 | 0 | 25.9342 | Eukaryota | 20750.0 | 0 | Chordata | 20750.0 | 0 | Carnivora | 8495.0 | 2 | Felidae | 8495.0 | 2 | Felis | 7216.0 | 3 | Felis catus | 7216.0 | 3 |
| scf7180000018097 | 78301 | 0.3864 | 0 | 49.5794 | Eukaryota | 38114.0 | 0 | Chordata | 38114.0 | 0 | Carnivora | 38114.0 | 0 | Felidae | 24799.0 | 2 | Acinonyx | 12435.0 | 5 | cinonyx jubat | 12435.0 | 5 |
| scf7180000022136 | 31420 | 0.3862 | 0 | 24.1038 | Eukaryota | 36803.0 | 0 | Chordata | 36803.0 | 0 | Carnivora | 36803.0 | 0 | Felidae | 36803.0 | 0 | Felis | 33105.0 | 1 | Felis catus | 33105.0 | 1 |
| scf7180000021263 | 73890 | 0.403 | 0 | 27.3992 | Eukaryota | 51235.0 | 0 | Chordata | 51235.0 | 0 | Carnivora | 51235.0 | 0 | Felidae | 30216.0 | 2 | Lynx | 16318.0 | 5 | ynx canadens | 16318.0 | 5 |
| scf7180000021622 | 8145 | 0.6268 | 0 | 33.3187 | Eukaryota | 4225.0 | 0 | Chordata | 4225.0 | 0 | Crocodylia | 843.0 | 7 | unresolved | 427.0 | 9 | unresolved | 427.0 | 9 | unresolved | 427.0 | 9 |
| scf7180000023262 | 17343 | 0.4263 | 0 | 384.6817 | Eukaryota | 131834.0 | 0 | Chordata | 131834.0 | 0 | Carnivora | 131834.0 | 0 | Hyaenidae | 131834.0 | 0 | Proteles | 79231.0 | 1 | roteles cristat | 79231.0 | 1 |
| scf7180000017758 | 1654680 | 0.3978 | 0 | 49.7283 | Eukaryota | 45647.0 | 0 | Chordata | 45647.0 | 0 | Carnivora | 45647.0 | 0 | Felidae | 45647.0 | 0 | Felis | 45647.0 | 0 | Felis catus | 45647.0 | 0 |
| scf7180000018897 | 529764 | 0.4926 | 0 | 50.1207 | Eukaryota | 10012.0 | 0 | Chordata | 10012.0 | 0 | Carnivora | 10012.0 | 0 | Felidae | 10012.0 | 0 | Felis | 9016.0 | 1 | Felis catus | 9016.0 | 1 |
| scf7180000024141 | 178149 | 0.4229 | 0 | 49.8575 | Eukaryota | 6586.0 | 0 | Chordata | 6586.0 | 0 | Carnivora | 6586.0 | 0 | Felidae | 6586.0 | 0 | Felis | 5939.0 | 1 | Felis catus | 5939.0 | 1 |
| scf7180000020979 | 8557 | 0.596 | 0 | 27.7809 | no-hit | 0.0 | 0 | no-hit | 0.0 | 0 | no-hit | 0.0 | 0 | no-hit | 0.0 | 0 | no-hit | 0.0 | 0 | no-hit | 0.0 | 0 |
| scf7180000023999 | 590764 | 0.3555 | 0 | 51.2036 | Eukaryota | 46090.0 | 0 | Chordata | 46090.0 | 0 | Carnivora | 46090.0 | 0 | Felidae | 46090.0 | 0 | Felis | 46090.0 | 0 | Felis catus | 46090.0 | 0 |
| scf7180000019023 | 484568 | 0.364 | 0 | 49.599 | Eukaryota | 46506.0 | 0 | Chordata | 46506.0 | 0 | Carnivora | 46506.0 | 0 | Felidae | 46506.0 | 0 | Felis | 41885.0 | 1 | Felis catus | 41885.0 | 1 |
| scf7180000021540 | 71123 | 0.4343 | 0 | 26.8731 | Eukaryota | 44492.0 | 0 | Chordata | 44492.0 | 0 | Carnivora | 44492.0 | 0 | Felidae | 38870.0 | 1 | Felis | 34600.0 | 2 | Felis catus | 34600.0 | 2 |
| scf7180000019713 | 200972 | 0.4478 | 0 | 26.9452 | Eukaryota | 12235.0 | 0 | Chordata | 12235.0 | 0 | Carnivora | 10979.0 | 1 | Felidae | 8536.0 | 3 | Felis | 8536.0 | 3 | Felis catus | 8536.0 | 3 |
| scf7180000021205 | 56464 | 0.4143 | 0 | 38.4632 | Eukaryota | 36661.0 | 0 | Chordata | 36661.0 | 0 | Carnivora | 36661.0 | 0 | Felidae | 36661.0 | 0 | Felis | 36661.0 | 0 | Felis catus | 36661.0 | 0 |
| scf7180000018357 | 850857 | 0.3978 | 0 | 51.2139 | Eukaryota | 74058.0 | 0 | Chordata | 74058.0 | 0 | Carnivora | 74058.0 | 0 | Felidae | 24260.0 | 4 | Suricata | 15441.0 | 8 | ricata suricat | 15441.0 | 8 |
| scf7180000021134 | 76183 | 0.3787 | 0 | 51.5458 | Eukaryota | 16378.0 | 0 | Chordata | 16378.0 | 0 | Carnivora | 16378.0 | 0 | Felidae | 11800.0 | 1 | Panthera | 7000.0 | 4 | anthera pardu | 4752.0 | 5 |
| scf7180000023207 | 25945 | 0.5383 | 0 | 288.0202 | Eukaryota | 22118.0 | 0 | Chordata | 22118.0 | 0 | Carnivora | 22118.0 | 0 | Felidae | 22118.0 | 0 | Felis | 19885.0 | 1 | Felis catus | 19885.0 | 1 |
| scf7180000017660 | 1978249 | 0.5011 | 0 | 52.0661 | Eukaryota | 40741.0 | 0 | Chordata | 40741.0 | 0 | Carnivora | 40741.0 | 0 | Canidae | 20135.0 | 3 | Canis | 12040.0 | 4 | Canis lupus | 12040.0 | 4 |
| scf7180000018520 | 959425 | 0.4165 | 0 | 52.4295 | Eukaryota | 46939.0 | 0 | Chordata | 46939.0 | 0 | Carnivora | 46939.0 | 0 | Felidae | 46939.0 | 0 | Felis | 42322.0 | 1 | Felis catus | 42322.0 | 1 |
| scf7180000018531 | 726640 | 0.4335 | 0 | 51.2255 | Eukaryota | 40109.0 | 0 | Chordata | 40109.0 | 0 | Carnivora | 40109.0 | 0 | Felidae | 40109.0 | 0 | Felis | 35383.0 | 1 | Felis catus | 35383.0 | 1 |
| scf7180000019265 | 325138 | 0.4594 | 0 | 55.3013 | Eukaryota | 36051.0 | 0 | Chordata | 36051.0 | 0 | Carnivora | 26183.0 | 1 | Felidae | 26183.0 | 2 | Lynx | 7854.0 | 5 | ynx canadens | 7854.0 | 6 |
| scf7180000021357 | 39509 | 0.3836 | 0 | 23.6165 | Eukaryota | 33381.0 | 0 | Chordata | 33381.0 | 0 | Carnivora | 33381.0 | 0 | Felidae | 33381.0 | 0 | Felis | 24109.0 | 3 | Felis catus | 24109.0 | 3 |
| scf7180000024224 | 33977 | 0.4159 | 0 | 26.0558 | Eukaryota | 5367.0 | 0 | Chordata | 5367.0 | 0 | Primates | 2075.0 | 3 | Hominidae | 2075.0 | 3 | Felis | 1950.0 | 4 | Felis catus | 1950.0 | 4 |
| scf7180000017448 | 272749 | 0.4174 | 0 | 49.9647 | Eukaryota | 44902.0 | 0 | Chordata | 44902.0 | 0 | Carnivora | 44902.0 | 0 | Felidae | 35358.0 | 1 | Felis | 30973.0 | 2 | Felis catus | 30973.0 | 2 |
| scf7180000022865 | 150351 | 0.4569 | 0 | 574.2728 | Eukaryota | 31868.0 | 0 | Chordata | 31868.0 | 0 | Carnivora | 31868.0 | 0 | Felidae | 31868.0 | 0 | Felis | 31868.0 | 0 | Felis catus | 31868.0 | 0 |
| scf7180000024006 | 573033 | 0.5117 | 0 | 51.2609 | Eukaryota | 44011.0 | 0 | Chordata | 44011.0 | 0 | Carnivora | 44011.0 | 0 | Felidae | 44011.0 | 0 | Felis | 44011.0 | 0 | Felis catus | 44011.0 | 0 |
| scf7180000022509 | 14405 | 0.3741 | 0 | 22.6192 | Eukaryota | 26748.0 | 0 | Chordata | 26748.0 | 0 | Carnivora | 26748.0 | 0 | Felidae | 26748.0 | 0 | Felis | 20359.0 | 1 | Felis catus | 20359.0 | 1 |
| scf7180000020118 | 130749 | 0.4567 | 0 | 24.7414 | Eukaryota | 28462.0 | 0 | Chordata | 28462.0 | 0 | Carnivora | 28462.0 | 0 | Felidae | 8932.0 | 5 | Panthera | 8932.0 | 5 | anthera pardu | 8932.0 | 5 |
| scf7180000018396 | 745820 | 0.4923 | 0 | 52.8328 | Eukaryota | 46996.0 | 0 | Chordata | 46996.0 | 0 | Carnivora | 46996.0 | 0 | Felidae | 46996.0 | 0 | Felis | 46996.0 | 0 | Felis catus | 46996.0 | 0 |
| scf7180000020610 | 6235 | 0.4223 | 0 | 56.6247 | Eukaryota | 2126.0 | 0 | Chordata | 2126.0 | 0 | Carnivora | 2126.0 | 0 | Felidae | 2126.0 | 0 | Felis | 1503.0 | 1 | Felis catus | 1503.0 | 1 |
| scf7180000019571 | 433114 | 0.5661 | 0 | 49.0558 | Eukaryota | 10796.0 | 0 | Chordata | 10796.0 | 0 | Primates | 5246.0 | 4 | Hominidae | 4627.0 | 6 | Homo | 4627.0 | 6 | Homo sapien | 4627.0 | 6 |
| scf7180000018468 | 105833 | 0.3702 | 0 | 50.5703 | Eukaryota | 36950.0 | 0 | Chordata | 36950.0 | 0 | Carnivora | 36950.0 | 0 | Felidae | 36950.0 | 0 | Felis | 36950.0 | 0 | Felis catus | 36950.0 | 0 |
| scf7180000020810 | 72562 | 0.5378 | 0 | 50.0268 | Eukaryota | 5077.0 | 0 | Chordata | 5077.0 | 0 | Carnivora | 5077.0 | 0 | Felidae | 5077.0 | 0 | Felis | 4552.0 | 1 | Felis catus | 4552.0 | 1 |
| scf7180000018103 | 1465102 | 0.3595 | 0 | 49.6709 | Eukaryota | 50997.0 | 0 | Chordata | 50997.0 | 0 | Carnivora | 50997.0 | 0 | Felidae | 50997.0 | 0 | Felis | 50997.0 | 0 | Felis catus | 50997.0 | 0 |
| scf7180000018244 | 675589 | 0.3559 | 0 | 49.4548 | Eukaryota | 46678.0 | 0 | Chordata | 46678.0 | 0 | Carnivora | 46678.0 | 0 | Felidae | 46678.0 | 0 | Felis | 46678.0 | 0 | Felis catus | 46678.0 | 0 |
| scf7180000019567 | 284252 | 0.4371 | 0 | 47.6162 | Eukaryota | 38686.0 | 0 | Chordata | 38686.0 | 0 | Carnivora | 38686.0 | 0 | Felidae | 38686.0 | 0 | Panthera | 20410.0 | 1 | Panthera tigri | 20410.0 | 1 |
| scf7180000022426 | 21104 | 0.3636 | 0 | 25.4428 | Eukaryota | 8893.0 | 0 | Chordata | 8893.0 | 0 | Primates | 4481.0 | 3 | Hominidae | 4481.0 | 3 | Homo | 4481.0 | 3 | Homo sapien | 4481.0 | 3 |
| scf7180000020905 | 65534 | 0.3823 | 0 | 25.3808 | Eukaryota | 19195.0 | 0 | Chordata | 19195.0 | 0 | Carnivora | 19195.0 | 0 | Felidae | 19195.0 | 0 | Felis | 19195.0 | 0 | Felis catus | 19195.0 | 0 |
| scf7180000017803 | 1610790 | 0.3464 | 0 | 48.4916 | Eukaryota | 44981.0 | 0 | Chordata | 44981.0 | 0 | Carnivora | 44981.0 | 0 | Felidae | 44981.0 | 0 | Felis | 44981.0 | 0 | Felis catus | 44981.0 | 0 |
| scf7180000023300 | 44126 | 0.4041 | 0 | 130.8663 | Eukaryota | 12954.0 | 0 | Chordata | 12954.0 | 0 | Carnivora | 12954.0 | 0 | Felidae | 12954.0 | 0 | Felis | 10392.0 | 1 | Felis catus | 10392.0 | 1 |
| scf7180000022260 | 78423 | 0.4269 | 0 | 1014.5293 | no-hit | 0.0 | 0 | no-hit | 0.0 | 0 | no-hit | 0.0 | 0 | no-hit | 0.0 | 0 | no-hit | 0.0 | 0 | no-hit | 0.0 | 0 |
| scf7180000019443 | 276729 | 0.423 | 0 | 24.8433 | Eukaryota | 21968.0 | 0 | Chordata | 21968.0 | 0 | Carnivora | 21968.0 | 0 | Felidae | 21968.0 | 0 | Felis | 19759.0 | 1 | Felis catus | 19759.0 | 1 |
| scf7180000023507 | 45530 | 0.4862 | 0 | 631.9142 | no-hit | 0.0 | 0 | no-hit | 0.0 | 0 | no-hit | 0.0 | 0 | no-hit | 0.0 | 0 | no-hit | 0.0 | 0 | no-hit | 0.0 | 0 |
| scf7180000023749 | 18789 | 0.4314 | 0 | 493.7233 | no-hit | 0.0 | 0 | no-hit | 0.0 | 0 | no-hit | 0.0 | 0 | no-hit | 0.0 | 0 | no-hit | 0.0 | 0 | no-hit | 0.0 | 0 |
| scf7180000018203 | 506035 | 0.372 | 0 | 49.1067 | Eukaryota | 47259.0 | 0 | Chordata | 47259.0 | 0 | Carnivora | 47259.0 | 0 | Felidae | 47259.0 | 0 | Felis | 47259.0 | 0 | Felis catus | 47259.0 | 0 |
| scf7180000020986 | 17014 | 0.5219 | 0 | 259.6 | no-hit | 0.0 | 0 | no-hit | 0.0 | 0 | no-hit | 0.0 | 0 | no-hit | 0.0 | 0 | no-hit | 0.0 | 0 | no-hit | 0.0 | 0 |
| scf7180000021934 | 9191 | 0.3751 | 0 | 24.4777 | Eukaryota | 4128.0 | 0 | Chordata | 4128.0 | 0 | Carnivora | 1812.0 | 3 | Hominidae | 1397.0 | 4 | Felis | 1112.0 | 5 | Felis catus | 1112.0 | 5 |
| scf7180000019368 | 426715 | 0.3682 | 0 | 50.6441 | Eukaryota | 47054.0 | 0 | Chordata | 47054.0 | 0 | Carnivora | 47054.0 | 0 | Felidae | 47054.0 | 0 | Felis | 47054.0 | 0 | Felis catus | 47054.0 | 0 |
| scf7180000021049 | 111566 | 0.3926 | 0 | 49.6665 | Eukaryota | 17779.0 | 0 | Chordata | 17779.0 | 0 | Carnivora | 17779.0 | 0 | Felidae | 17779.0 | 0 | Felis | 15806.0 | 1 | Felis catus | 15806.0 | 1 |
| scf7180000021668 | 14785 | 0.5213 | 0 | 231.4689 | no-hit | 0.0 | 0 | no-hit | 0.0 | 0 | no-hit | 0.0 | 0 | no-hit | 0.0 | 0 | no-hit | 0.0 | 0 | no-hit | 0.0 | 0 |
| scf7180000019033 | 100703 | 0.4595 | 0 | 41.6395 | Eukaryota | 17173.0 | 0 | Chordata | 17173.0 | 0 | Carnivora | 16297.0 | 1 | Felidae | 16297.0 | 1 | Lynx | 7131.0 | 4 | ynx canadens | 7131.0 | 4 |
| scf7180000021010 | 31540 | 0.3962 | 0 | 25.2676 | Eukaryota | 45788.0 | 0 | Chordata | 45788.0 | 0 | Carnivora | 45788.0 | 0 | Felidae | 45788.0 | 0 | Felis | 45788.0 | 0 | Felis catus | 45788.0 | 0 |
| scf7180000020264 | 25652 | 0.6981 | 0 | 140.7537 | no-hit | 0.0 | 0 | no-hit | 0.0 | 0 | no-hit | 0.0 | 0 | no-hit | 0.0 | 0 | no-hit | 0.0 | 0 | no-hit | 0.0 | 0 |
| scf7180000020787 | 83430 | 0.4135 | 0 | 25.5003 | Eukaryota | 15744.0 | 0 | Chordata | 15744.0 | 0 | Artiodactyla | 9721.0 | 1 | Suidae | 5027.0 | 5 | Sus | 5027.0 | 7 | Sus scrofa | 5027.0 | 7 |
| scf7180000018410 | 758205 | 0.3486 | 0 | 47.4498 | Eukaryota | 44981.0 | 0 | Chordata | 44981.0 | 0 | Carnivora | 44981.0 | 0 | Felidae | 44981.0 | 0 | Felis | 40685.0 | 1 | Felis catus | 40685.0 | 1 |
| scf7180000024170 | 137713 | 0.3857 | 0 | 25.9695 | Eukaryota | 18037.0 | 0 | Chordata | 18037.0 | 0 | Carnivora | 18037.0 | 0 | Felidae | 18037.0 | 0 | Felis | 14432.0 | 1 | Felis catus | 14432.0 | 1 |
| scf7180000020696 | 109158 | 0.4862 | 0 | 49.7033 | Eukaryota | 9044.0 | 0 | Chordata | 9044.0 | 0 | Carnivora | 9044.0 | 0 | Felidae | 9044.0 | 0 | Felis | 8211.0 | 1 | Felis catus | 8211.0 | 1 |
| scf7180000018281 | 6641 | 0.435 | 0 | 22.7894 | Eukaryota | 2548.0 | 0 | Chordata | 2548.0 | 0 | Carnivora | 1988.0 | 2 | Felidae | 1988.0 | 2 | Felis | 1242.0 | 3 | Felis catus | 1242.0 | 3 |
| scf7180000019895 | 221108 | 0.5787 | 0 | 51.466 | Eukaryota | 29237.0 | 0 | Chordata | 29237.0 | 0 | Carnivora | 26527.0 | 1 | Felidae | 14985.0 | 4 | Panthera | 5936.0 | 7 | anthera pardu | 5936.0 | 8 |
| scf7180000020564 | 114859 | 0.3702 | 0 | 24.2996 | Eukaryota | 33655.0 | 0 | Chordata | 33655.0 | 0 | Carnivora | 33655.0 | 0 | Felidae | 33655.0 | 0 | Felis | 33655.0 | 0 | Felis catus | 33655.0 | 0 |
| scf7180000018662 | 819201 | 0.3702 | 0 | 48.5136 | Eukaryota | 47421.0 | 0 | Chordata | 47421.0 | 0 | Carnivora | 47421.0 | 0 | Felidae | 47421.0 | 0 | Felis | 47421.0 | 0 | Felis catus | 47421.0 | 0 |
| scf7180000019623 | 276857 | 0.4015 | 0 | 53.4803 | Eukaryota | 45747.0 | 0 | Chordata | 45747.0 | 0 | Carnivora | 45747.0 | 0 | Felidae | 45747.0 | 0 | Felis | 41167.0 | 1 | Felis catus | 41167.0 | 1 |
| scf7180000019189 | 415117 | 0.4013 | 0 | 50.6898 | Eukaryota | 20588.0 | 0 | Chordata | 20588.0 | 0 | Carnivora | 20588.0 | 0 | Felidae | 20588.0 | 0 | Felis | 18569.0 | 1 | Felis catus | 18569.0 | 1 |
| scf7180000021558 | 1975545 | 0.4181 | 0 | 51.296 | Eukaryota | 92035.0 | 0 | Chordata | 92035.0 | 0 | Carnivora | 92035.0 | 0 | Felidae | 92035.0 | 0 | Panthera | 36952.0 | 2 | anthera pardu | 36952.0 | 2 |
| scf7180000024096 | 169420 | 0.3748 | 0 | 49.3563 | Eukaryota | 16426.0 | 0 | Chordata | 16426.0 | 0 | Carnivora | 16426.0 | 0 | Felidae | 16426.0 | 0 | Felis | 13163.0 | 1 | Felis catus | 13163.0 | 1 |
| scf7180000021485 | 26580 | 0.4452 | 0 | 26.3932 | Eukaryota | 7593.0 | 0 | Chordata | 7593.0 | 0 | Primates | 4125.0 | 5 | Hominidae | 4125.0 | 5 | Homo | 2070.0 | 6 | Homo sapien | 2070.0 | 6 |
| scf7180000019731 | 193056 | 0.5259 | 0 | 46.8801 | Eukaryota | 39981.0 | 0 | Chordata | 39981.0 | 0 | Carnivora | 21684.0 | 2 | Felidae | 13594.0 | 4 | Felis | 8498.0 | 6 | Felis catus | 8498.0 | 7 |
| scf7180000018682 | 618833 | 0.4254 | 0 | 50.7874 | Eukaryota | 44190.0 | 0 | Chordata | 44190.0 | 0 | Carnivora | 44190.0 | 0 | Felidae | 44190.0 | 0 | Felis | 39820.0 | 1 | Felis catus | 39820.0 | 1 |
| scf7180000018718 | 618654 | 0.432 | 0 | 52.9498 | Eukaryota | 29701.0 | 0 | Chordata | 29701.0 | 0 | Carnivora | 29701.0 | 0 | Felidae | 26360.0 | 1 | Panthera | 11464.0 | 3 | Felis catus | 9658.0 | 4 |
| scf7180000022974 | 59936 | 0.5524 | 0 | 182.6745 | Eukaryota | 13715.0 | 0 | Chordata | 13715.0 | 0 | Carnivora | 13715.0 | 0 | Felidae | 13715.0 | 0 | Felis | 13715.0 | 0 | Felis catus | 13715.0 | 0 |
| scf7180000020496 | 20970 | 0.5159 | 0 | 234.8627 | Eukaryota | 12236.0 | 0 | Chordata | 12236.0 | 0 | Carnivora | 12236.0 | 0 | Felidae | 12236.0 | 0 | Felis | 11024.0 | 1 | Felis catus | 11024.0 | 1 |
| scf7180000020242 | 141273 | 0.4674 | 0 | 52.7911 | Eukaryota | 25584.0 | 0 | Chordata | 25584.0 | 0 | Carnivora | 25584.0 | 0 | Felidae | 9101.0 | 4 | Canis | 3800.0 | 8 | Canis lupus | 3800.0 | 8 |
| scf7180000018784 | 218878 | 0.3471 | 0 | 43.3695 | Eukaryota | 46546.0 | 0 | Chordata | 46546.0 | 0 | Carnivora | 46546.0 | 0 | Felidae | 46546.0 | 0 | Felis | 46546.0 | 0 | Felis catus | 46546.0 | 0 |
| scf7180000019181 | 419114 | 0.41 | 0 | 50.6558 | Eukaryota | 54503.0 | 0 | Chordata | 54503.0 | 0 | Carnivora | 54503.0 | 0 | Felidae | 54503.0 | 0 | Panthera | 22240.0 | 3 | anthera pardu | 22240.0 | 3 |
| scf7180000020318 | 186781 | 0.491 | 0 | 51.5974 | Eukaryota | 17300.0 | 0 | Chordata | 17300.0 | 0 | Carnivora | 17300.0 | 0 | Herpestidae | 9210.0 | 1 | Suricata | 9210.0 | 5 | ricata suricat | 9210.0 | 5 |
| scf7180000017928 | 2136956 | 0.479 | 0 | 51.7455 | Eukaryota | 51126.0 | 0 | Chordata | 51126.0 | 0 | Carnivora | 51126.0 | 0 | Felidae | 51126.0 | 0 | Panthera | 51126.0 | 0 | anthera pardu | 35776.0 | 1 |
| scf7180000020624 | 81946 | 0.4504 | 0 | 71.4122 | Eukaryota | 35505.0 | 0 | Chordata | 35505.0 | 0 | Carnivora | 35505.0 | 0 | Felidae | 32400.0 | 1 | Felis | 25762.0 | 3 | Felis catus | 25762.0 | 3 |
| scf7180000021468 | 38181 | 0.3898 | 0 | 24.6745 | Eukaryota | 41949.0 | 0 | Chordata | 41949.0 | 0 | Carnivora | 41949.0 | 0 | Felidae | 41949.0 | 0 | Felis | 41949.0 | 0 | Felis catus | 41949.0 | 0 |
| scf7180000019871 | 194627 | 0.4243 | 0 | 51.3231 | Eukaryota | 23747.0 | 0 | Chordata | 23747.0 | 0 | Carnivora | 23747.0 | 0 | Felidae | 23747.0 | 0 | Felis | 21349.0 | 1 | Felis catus | 21349.0 | 1 |
| scf7180000017697 | 2115090 | 0.3754 | 0 | 50.7904 | Eukaryota | 52708.0 | 0 | Chordata | 52708.0 | 0 | Carnivora | 52708.0 | 0 | Felidae | 32060.0 | 1 | Acinonyx | 32060.0 | 1 | cinonyx jubat | 32060.0 | 1 |
| scf7180000018419 | 57743 | 0.4072 | 0 | 26.1843 | Eukaryota | 36064.0 | 0 | Chordata | 36064.0 | 0 | Carnivora | 36064.0 | 0 | Felidae | 24399.0 | 2 | Felis | 14270.0 | 5 | Felis catus | 14270.0 | 6 |
| scf7180000022899 | 634865 | 0.3579 | 0 | 50.1043 | Eukaryota | 47526.0 | 0 | Chordata | 47526.0 | 0 | Carnivora | 47526.0 | 0 | Felidae | 47526.0 | 0 | Felis | 47526.0 | 0 | Felis catus | 47526.0 | 0 |
| scf7180000022605 | 13442 | 0.3427 | 0 | 23.1994 | Eukaryota | 4577.0 | 0 | Chordata | 4577.0 | 0 | Primates | 2033.0 | 2 | Hominidae | 2033.0 | 2 | Homo | 2033.0 | 2 | Homo sapien | 2033.0 | 2 |
| scf7180000019621 | 296364 | 0.3922 | 100 | 52.5425 | Eukaryota | 47453.0 | 0 | Chordata | 47453.0 | 0 | Carnivora | 47453.0 | 0 | Felidae | 47453.0 | 0 | Felis | 47453.0 | 0 | Felis catus | 47453.0 | 0 |
| scf7180000024142 | 149189 | 0.4216 | 0 | 25.6142 | Eukaryota | 43943.0 | 0 | Chordata | 43943.0 | 0 | Carnivora | 43943.0 | 0 | Felidae | 43943.0 | 0 | Felis | 39584.0 | 1 | Felis catus | 39584.0 | 1 |
| scf7180000021380 | 40613 | 0.5951 | 0 | 49.2063 | Eukaryota | 3954.0 | 0 | Chordata | 3954.0 | 0 | Carnivora | 3954.0 | 0 | Felidae | 1233.0 | 3 | Zalophus | 1089.0 | 6 | phus californi | 1089.0 | 6 |
| scf7180000018537 | 989013 | 0.4006 | 0 | 50.8014 | Eukaryota | 46082.0 | 0 | Chordata | 46082.0 | 0 | Carnivora | 46082.0 | 0 | Felidae | 46082.0 | 0 | Felis | 46082.0 | 0 | Felis catus | 46082.0 | 0 |
| scf7180000022323 | 19673 | 0.3684 | 0 | 24.2553 | Eukaryota | 6299.0 | 0 | Chordata | 6299.0 | 0 | Primates | 3410.0 | 3 | Hominidae | 3410.0 | 4 | Homo | 2276.0 | 6 | Homo sapien | 2276.0 | 6 |
| scf7180000024181 | 73478 | 0.5262 | 0 | 43.4968 | Eukaryota | 6703.0 | 0 | Chordata | 6703.0 | 0 | Passeriformes | 2916.0 | 3 | Corvidae | 2916.0 | 3 | Aphelocoma | 2916.0 | 3 | Sus scrofa | 2062.0 | 5 |
| scf7180000020859 | 123785 | 0.4092 | 0 | 26.3702 | Eukaryota | 16844.0 | 0 | Chordata | 16844.0 | 0 | Carnivora | 9190.0 | 2 | Felidae | 7474.0 | 3 | Equus | 5868.0 | 5 | quus caballu | 5868.0 | 6 |
| scf7180000020430 | 113672 | 0.4331 | 0 | 25.2566 | Eukaryota | 15805.0 | 0 | Chordata | 15805.0 | 0 | Primates | 12271.0 | 3 | Hominidae | 11469.0 | 4 | Homo | 7645.0 | 5 | Homo sapien | 7645.0 | 5 |
| scf7180000024122 | 156882 | 0.3838 | 0 | 48.9577 | Eukaryota | 47480.0 | 0 | Chordata | 47480.0 | 0 | Carnivora | 47480.0 | 0 | Felidae | 47480.0 | 0 | Felis | 47480.0 | 0 | Felis catus | 47480.0 | 0 |
| scf7180000019231 | 77277 | 0.428 | 0 | 24.7811 | Eukaryota | 4585.0 | 0 | Chordata | 4585.0 | 0 | Primates | 3494.0 | 2 | Hominidae | 3494.0 | 2 | Homo | 3019.0 | 3 | Homo sapien | 3019.0 | 3 |
| scf7180000019000 | 754797 | 0.3446 | 0 | 48.1098 | Eukaryota | 45742.0 | 0 | Chordata | 45742.0 | 0 | Carnivora | 45742.0 | 0 | Felidae | 45742.0 | 0 | Felis | 45742.0 | 0 | Felis catus | 45742.0 | 0 |
| scf7180000020300 | 19321 | 0.5165 | 0 | 28.3352 | Eukaryota | 2693.0 | 0 | Chordata | 2693.0 | 0 | Carnivora | 2693.0 | 0 | Felidae | 2693.0 | 0 | Felis | 1935.0 | 1 | Felis catus | 1935.0 | 1 |
| scf7180000023951 | 533997 | 0.3793 | 0 | 251.0183 | Eukaryota | 45273.0 | 0 | Chordata | 45273.0 | 0 | Carnivora | 45273.0 | 0 | Felidae | 45273.0 | 0 | Felis | 45273.0 | 0 | Felis catus | 45273.0 | 0 |
| scf7180000023965 | 417229 | 0.5768 | 0 | 48.6578 | Eukaryota | 13954.0 | 0 | Chordata | 13954.0 | 0 | Carnivora | 9175.0 | 2 | Felidae | 9175.0 | 2 | Felis | 4017.0 | 5 | Felis catus | 4017.0 | 5 |
| scf7180000020951 | 57637 | 0.396 | 0 | 26.941 | Eukaryota | 10344.0 | 0 | Chordata | 10344.0 | 0 | Carnivora | 10344.0 | 0 | Otariidae | 4811.0 | 3 | Zalophus | 2904.0 | 5 | phus californi | 2904.0 | 6 |
| scf7180000019771 | 239047 | 0.5364 | 0 | 50.1264 | Eukaryota | 16031.0 | 0 | Chordata | 16031.0 | 0 | Carnivora | 16031.0 | 0 | Felidae | 16031.0 | 0 | Felis | 6410.0 | 2 | Felis catus | 6410.0 | 2 |
| scf7180000021284 | 81700 | 0.3928 | 0 | 26.0463 | Eukaryota | 29245.0 | 0 | Chordata | 29245.0 | 0 | Carnivora | 29245.0 | 0 | Felidae | 29245.0 | 0 | Felis | 26247.0 | 1 | Felis catus | 26247.0 | 1 |
| scf7180000019721 | 194426 | 0.631 | 0 | 53.3359 | Eukaryota | 33250.0 | 0 | Chordata | 33250.0 | 0 | Carnivora | 33250.0 | 0 | Felidae | 33250.0 | 0 | Felis | 20082.0 | 2 | Felis catus | 20082.0 | 2 |
| scf7180000019085 | 31682 | 0.5474 | 0 | 141.6574 | Eukaryota | 6405.0 | 0 | Chordata | 6405.0 | 0 | Carnivora | 6405.0 | 0 | Felidae | 6166.0 | 1 | Felis | 1836.0 | 6 | Felis catus | 1836.0 | 6 |
| scf7180000020917 | 63121 | 0.4153 | 0 | 50.8651 | Eukaryota | 30276.0 | 0 | Chordata | 30276.0 | 0 | Carnivora | 30276.0 | 0 | Felidae | 20353.0 | 2 | Panthera | 11265.0 | 4 | anthera pardu | 11265.0 | 4 |
| scf7180000018940 | 118593 | 0.3503 | 0 | 46.9542 | Eukaryota | 34284.0 | 0 | Chordata | 34284.0 | 0 | Carnivora | 34284.0 | 0 | Felidae | 34284.0 | 0 | Felis | 34284.0 | 0 | Felis catus | 34284.0 | 0 |
| scf7180000019553 | 324178 | 0.3947 | 0 | 52.0302 | Eukaryota | 47015.0 | 0 | Chordata | 47015.0 | 0 | Carnivora | 47015.0 | 0 | Felidae | 47015.0 | 0 | Felis | 47015.0 | 0 | Felis catus | 47015.0 | 0 |
| scf7180000019074 | 353751 | 0.3759 | 0 | 50.6093 | Eukaryota | 26532.0 | 0 | Chordata | 26532.0 | 0 | Carnivora | 14000.0 | 2 | Hominidae | 7314.0 | 5 | Homo | 7314.0 | 7 | Homo sapien | 7314.0 | 7 |
| scf7180000018521 | 226515 | 0.3856 | 0 | 51.1182 | Eukaryota | 46015.0 | 0 | Chordata | 46015.0 | 0 | Carnivora | 46015.0 | 0 | Felidae | 46015.0 | 0 | Felis | 46015.0 | 0 | Felis catus | 46015.0 | 0 |
| scf7180000019545 | 341517 | 0.6327 | 100 | 51.1683 | Eukaryota | 24290.0 | 0 | Chordata | 24290.0 | 0 | Carnivora | 24290.0 | 0 | Felidae | 19545.0 | 2 | Acinonyx | 11975.0 | 5 | cinonyx jubat | 11975.0 | 5 |
| scf7180000020870 | 85314 | 0.4127 | 0 | 24.898 | Eukaryota | 30928.0 | 0 | Chordata | 30928.0 | 0 | Carnivora | 30928.0 | 0 | Felidae | 24638.0 | 1 | Felis | 18354.0 | 2 | Felis catus | 18354.0 | 2 |
| scf7180000019259 | 381048 | 0.4764 | 0 | 51.0527 | Eukaryota | 25583.0 | 0 | Chordata | 25583.0 | 0 | Carnivora | 13593.0 | 3 | Delphinidae | 7378.0 | 5 | Odobenus | 4634.0 | 8 | obenus rosma | 4634.0 | 8 |
| scf7180000018724 | 218854 | 0.3674 | 0 | 49.3566 | Eukaryota | 48445.0 | 0 | Chordata | 48445.0 | 0 | Carnivora | 48445.0 | 0 | Felidae | 48445.0 | 0 | Felis | 43652.0 | 1 | Felis catus | 43652.0 | 1 |
| scf7180000019164 | 1210790 | 0.3959 | 0 | 50.34 | Eukaryota | 69720.0 | 0 | Chordata | 69720.0 | 0 | Carnivora | 69720.0 | 0 | Felidae | 69720.0 | 0 | Felis | 27880.0 | 2 | Felis catus | 27880.0 | 2 |
| scf7180000019433 | 276887 | 0.5504 | 0 | 46.3371 | Eukaryota | 18953.0 | 0 | Chordata | 18953.0 | 0 | Carnivora | 18953.0 | 0 | Felidae | 18953.0 | 0 | Felis | 17065.0 | 1 | Felis catus | 17065.0 | 1 |
| scf7180000018573 | 1423404 | 0.4753 | 0 | 50.4864 | Eukaryota | 45220.0 | 0 | Chordata | 45220.0 | 0 | Carnivora | 45220.0 | 0 | Felidae | 45220.0 | 0 | Felis | 45220.0 | 0 | Felis catus | 45220.0 | 0 |
| scf7180000024285 | 79142 | 0.6455 | 0 | 51.4293 | Eukaryota | 12387.0 | 0 | Chordata | 12387.0 | 0 | Carnivora | 12387.0 | 0 | Felidae | 9826.0 | 1 | Acinonyx | 3678.0 | 5 | cinonyx jubat | 3678.0 | 5 |
| scf7180000017872 | 1407510 | 0.4212 | 0 | 54.2532 | Eukaryota | 53473.0 | 0 | Chordata | 53473.0 | 0 | Carnivora | 53473.0 | 0 | Felidae | 53473.0 | 0 | Panthera | 48150.0 | 1 | anthera pardu | 48150.0 | 1 |
| scf7180000022324 | 12245 | 0.5232 | 0 | 398.0572 | no-hit | 0.0 | 0 | no-hit | 0.0 | 0 | no-hit | 0.0 | 0 | no-hit | 0.0 | 0 | no-hit | 0.0 | 0 | no-hit | 0.0 | 0 |
| scf7180000018210 | 15588 | 0.3489 | 0 | 23.5542 | Eukaryota | 9467.0 | 0 | Chordata | 9467.0 | 0 | Carnivora | 9467.0 | 0 | Felidae | 9467.0 | 0 | Felis | 8526.0 | 1 | Felis catus | 8526.0 | 1 |
| scf7180000022575 | 30900 | 0.4861 | 0 | 1218.8436 | Eukaryota | 2323.0 | 0 | Chordata | 2323.0 | 0 | Carnivora | 1555.0 | 1 | Felidae | 1555.0 | 3 | Felis | 1555.0 | 3 | Felis catus | 1555.0 | 3 |
| scf7180000019485 | 292906 | 0.3469 | 0 | 47.5051 | Eukaryota | 47505.0 | 0 | Chordata | 47505.0 | 0 | Carnivora | 47505.0 | 0 | Felidae | 47505.0 | 0 | Felis | 47505.0 | 0 | Felis catus | 47505.0 | 0 |
| scf7180000023554 | 39140 | 0.5 | 0 | 5792.696 | Eukaryota | 17951.0 | 0 | Chordata | 17951.0 | 0 | Carnivora | 17951.0 | 0 | Felidae | 14462.0 | 1 | Acinonyx | 3554.0 | 5 | cinonyx jubat | 3554.0 | 5 |
| scf7180000018885 | 492110 | 0.5405 | 0 | 50.3705 | Eukaryota | 34657.0 | 0 | Chordata | 34657.0 | 0 | Perissodactyla | 19872.0 | 2 | Equidae | 19872.0 | 3 | Equus | 19872.0 | 3 | quus caballu | 19872.0 | 3 |
| scf7180000020771 | 132424 | 0.4354 | 0 | 50.1297 | Eukaryota | 14827.0 | 1 | Chordata | 14827.0 | 1 | Primates | 7483.0 | 3 | Hominidae | 6229.0 | 5 | Homo | 6229.0 | 6 | Homo sapien | 6229.0 | 6 |
| scf7180000022270 | 10170 | 0.5218 | 0 | 266.966 | no-hit | 0.0 | 0 | no-hit | 0.0 | 0 | no-hit | 0.0 | 0 | no-hit | 0.0 | 0 | no-hit | 0.0 | 0 | no-hit | 0.0 | 0 |
| scf7180000023662 | 100919 | 0.3953 | 0 | 373.5486 | Eukaryota | 46120.0 | 0 | Chordata | 46120.0 | 0 | Carnivora | 46120.0 | 0 | Felidae | 46120.0 | 0 | Felis | 46120.0 | 0 | Felis catus | 46120.0 | 0 |
| scf7180000019364 | 453051 | 0.6465 | 0 | 50.4081 | Eukaryota | 42000.0 | 0 | Chordata | 42000.0 | 0 | Carnivora | 42000.0 | 0 | Felidae | 42000.0 | 0 | Acinonyx | 18144.0 | 3 | cinonyx jubat | 18144.0 | 3 |
| scf7180000024281 | 219707 | 0.4979 | 0 | 27.012 | Eukaryota | 15780.0 | 0 | Chordata | 15780.0 | 0 | Carnivora | 15780.0 | 0 | Felidae | 13793.0 | 2 | Felis | 6500.0 | 4 | Felis catus | 6500.0 | 4 |
| scf7180000019154 | 611139 | 0.4096 | 0 | 51.817 | Eukaryota | 73321.0 | 0 | Chordata | 73321.0 | 0 | Carnivora | 73321.0 | 0 | Ursidae | 41165.0 | 1 | Ursus | 34576.0 | 3 | rsus maritimu | 20754.0 | 4 |
| scf7180000024120 | 260260 | 0.4353 | 0 | 50.7818 | Eukaryota | 13469.0 | 0 | Chordata | 13469.0 | 0 | Carnivora | 13469.0 | 0 | Felidae | 13469.0 | 0 | Felis | 13469.0 | 0 | Felis catus | 13469.0 | 0 |
| scf7180000019159 | 378972 | 0.4454 | 0 | 51.8176 | Eukaryota | 47875.0 | 0 | Chordata | 47875.0 | 0 | Carnivora | 47875.0 | 0 | Felidae | 33567.0 | 1 | Lynx | 19549.0 | 3 | ynx canadens | 19549.0 | 4 |
| scf7180000023020 | 25749 | 0.4023 | 0 | 248.0259 | Eukaryota | 6555.0 | 0 | Chordata | 6555.0 | 0 | Carnivora | 6555.0 | 0 | Felidae | 6555.0 | 0 | Felis | 6555.0 | 0 | Felis catus | 6555.0 | 0 |
| scf7180000018963 | 626132 | 0.4531 | 0 | 49.8325 | Eukaryota | 46129.0 | 0 | Chordata | 46129.0 | 0 | Carnivora | 46129.0 | 0 | Felidae | 46129.0 | 0 | Felis | 41558.0 | 1 | Felis catus | 41558.0 | 1 |
| scf7180000017969 | 54737 | 0.3938 | 0 | 25.0698 | Eukaryota | 46385.0 | 0 | Chordata | 46385.0 | 0 | Carnivora | 46385.0 | 0 | Felidae | 46385.0 | 0 | Panthera | 23325.0 | 1 | anthera pardu | 23325.0 | 1 |
| scf7180000021274 | 59533 | 0.4119 | 0 | 26.3532 | Eukaryota | 42174.0 | 0 | Chordata | 42174.0 | 0 | Carnivora | 42174.0 | 0 | Felidae | 42174.0 | 0 | Felis | 38020.0 | 1 | Felis catus | 38020.0 | 1 |
| scf7180000019600 | 140129 | 0.4394 | 0 | 48.5496 | Eukaryota | 51893.0 | 0 | Chordata | 51893.0 | 0 | Carnivora | 39947.0 | 2 | Felidae | 35677.0 | 3 | Lynx | 11972.0 | 6 | ynx canadens | 11972.0 | 7 |

| | | | | | | | | | | | | | | | | | | | | | | |
|---|---|---|---|---|---|---|---|---|---|---|---|---|---|---|---|---|---|---|---|---|---|---|
| scf7180000024280 | 329869 | 0.4667 | 0 | 52.2089 | Eukaryota | 51425.0 | 0 | Chordata | 51425.0 | 0 | Carnivora | 51425.0 | 0 | Felidae | 45838.0 | 1 | Felis | 20332.0 | 3 | Felis catus | 20332.0 | 3 |
| scf7180000024112 | 1235015 | 0.3986 | 0 | 50.6317 | Eukaryota | 51788.0 | 0 | Chordata | 51788.0 | 0 | Carnivora | 51788.0 | 0 | Felidae | 51788.0 | 0 | Felis | 51788.0 | 0 | Felis catus | 51788.0 | 0 |
| scf7180000022265 | 39823 | 0.3523 | 0 | 24.375 | Eukaryota | 9114.0 | 0 | Chordata | 9114.0 | 0 | Carnivora | 4580.0 | 2 | Felidae | 3630.0 | 3 | Felis | 3630.0 | 3 | Felis catus | 3630.0 | 3 |
| scf7180000017794 | 795365 | 0.3863 | 0 | 50.9956 | Eukaryota | 45411.0 | 0 | Chordata | 45411.0 | 0 | Carnivora | 45411.0 | 0 | Felidae | 45411.0 | 0 | Felis | 45411.0 | 0 | Felis catus | 45411.0 | 0 |
| scf7180000018802 | 333877 | 0.4997 | 0 | 50.0596 | Eukaryota | 26796.0 | 0 | Chordata | 26796.0 | 0 | Carnivora | 26796.0 | 0 | Felidae | 26796.0 | 0 | Panthera | 21440.0 | 1 | unresolved | 10720.0 | 2 |
| scf7180000017886 | 1254791 | 0.4562 | 0 | 51.6605 | Eukaryota | 55850.0 | 0 | Chordata | 55850.0 | 0 | Carnivora | 55850.0 | 0 | Herpestidae | 29120.0 | 1 | Suricata | 29120.0 | 1 | ricata suricat | 29120.0 | 1 |
| scf7180000018344 | 487342 | 0.3891 | 0 | 48.2723 | Eukaryota | 47376.0 | 0 | Chordata | 47376.0 | 0 | Carnivora | 47376.0 | 0 | Felidae | 47376.0 | 0 | Felis | 47376.0 | 0 | Felis catus | 47376.0 | 0 |
| scf7180000018692 | 700308 | 0.3472 | 0 | 48.5183 | Eukaryota | 53611.0 | 0 | Chordata | 53611.0 | 0 | Carnivora | 53611.0 | 0 | Mustelidae | 11785.0 | 4 | Canis | 10614.0 | 7 | Canis lupus | 10614.0 | 7 |
| scf7180000024144 | 55911 | 0.4663 | 0 | 39.6272 | Eukaryota | 18449.0 | 0 | Chordata | 18449.0 | 0 | Carnivora | 18449.0 | 0 | Otariidae | 6884.0 | 3 | Suricata | 4994.0 | 6 | ricata suricat | 4994.0 | 7 |
| scf7180000023996 | 529448 | 0.469 | 0 | 53.0547 | Eukaryota | 22319.0 | 0 | Chordata | 22319.0 | 0 | Carnivora | 22319.0 | 0 | Felidae | 20141.0 | 1 | Acinonyx | 5736.0 | 5 | cinonyx jubat | 5736.0 | 6 |
| scf7180000021784 | 18460 | 0.6178 | 0 | 33.6789 | Eukaryota | 2440.0 | 0 | Chordata | 2440.0 | 0 | Carnivora | 2440.0 | 0 | Felidae | 2186.0 | 1 | Neofelis | 977.0 | 3 | eofelis nebulo | 977.0 | 4 |
| scf7180000021825 | 52242 | 0.4016 | 0 | 25.168 | Eukaryota | 45730.0 | 0 | Chordata | 45730.0 | 0 | Carnivora | 45730.0 | 0 | Felidae | 45730.0 | 0 | Felis | 45730.0 | 0 | Felis catus | 45730.0 | 0 |
| scf7180000021474 | 55909 | 0.581 | 0 | 52.2516 | Eukaryota | 23352.0 | 0 | Chordata | 23352.0 | 0 | Carnivora | 23352.0 | 0 | Felidae | 18777.0 | 2 | Panthera | 9383.0 | 6 | anthera pardu | 7045.0 | 7 |
| scf7180000020200 | 197767 | 0.3958 | 0 | 24.5786 | Eukaryota | 17600.0 | 0 | Chordata | 17600.0 | 0 | Carnivora | 17600.0 | 0 | Felidae | 17600.0 | 0 | Felis | 14005.0 | 1 | Felis catus | 14005.0 | 1 |
| scf7180000022015 | 33016 | 0.3887 | 0 | 27.8787 | Eukaryota | 45380.0 | 0 | Chordata | 45380.0 | 0 | Carnivora | 45380.0 | 0 | Felidae | 45380.0 | 0 | Felis | 45380.0 | 0 | Felis catus | 45380.0 | 0 |
| scf7180000018515 | 1403116 | 0.5085 | 0 | 50.6747 | Eukaryota | 40508.0 | 0 | Chordata | 40508.0 | 0 | Carnivora | 25851.0 | 1 | Felidae | 22259.0 | 3 | Felis | 10865.0 | 8 | Felis catus | 10865.0 | 8 |
| scf7180000019137 | 391147 | 0.344 | 0 | 48.3707 | Eukaryota | 33682.0 | 0 | Chordata | 33682.0 | 0 | Carnivora | 33682.0 | 0 | Felidae | 33682.0 | 0 | Felis | 30505.0 | 1 | Felis catus | 30505.0 | 1 |
| scf7180000023427 | 39403 | 0.526 | 0 | 165.487 | Eukaryota | 3725.0 | 0 | Chordata | 3725.0 | 0 | Carnivora | 3725.0 | 0 | Felidae | 3481.0 | 1 | Felis | 2840.0 | 2 | Felis catus | 2840.0 | 2 |
| scf7180000019834 | 142247 | 0.3917 | 0 | 46.0241 | Eukaryota | 21682.0 | 0 | Chordata | 21682.0 | 0 | Carnivora | 21682.0 | 0 | Felidae | 21682.0 | 0 | Felis | 21682.0 | 0 | Felis catus | 21682.0 | 0 |
| scf7180000024007 | 557715 | 0.3908 | 0 | 50.5072 | Eukaryota | 75394.0 | 0 | Chordata | 75394.0 | 0 | Carnivora | 75394.0 | 0 | Felidae | 61472.0 | 2 | Felis | 30924.0 | 3 | Felis catus | 30924.0 | 3 |
| scf7180000022391 | 12055 | 0.368 | 0 | 43.4737 | Eukaryota | 7999.0 | 0 | Chordata | 7999.0 | 0 | Carnivora | 7999.0 | 0 | Felidae | 7999.0 | 0 | Felis | 7999.0 | 0 | Felis catus | 7999.0 | 0 |
| scf7180000020745 | 79030 | 0.3892 | 0 | 24.9345 | Eukaryota | 48198.0 | 0 | Chordata | 48198.0 | 0 | Carnivora | 48198.0 | 0 | Felidae | 48198.0 | 0 | Felis | 37434.0 | 2 | Felis catus | 37434.0 | 2 |
| scf7180000020826 | 176839 | 0.5673 | 0 | 51.0385 | Eukaryota | 17871.0 | 0 | Chordata | 17871.0 | 0 | Carnivora | 17871.0 | 0 | Felidae | 17871.0 | 0 | Felis | 14310.0 | 1 | Felis catus | 14310.0 | 1 |
| scf7180000024238 | 67024 | 0.4126 | 0 | 56.6067 | Eukaryota | 41018.0 | 0 | Chordata | 41018.0 | 0 | Carnivora | 41018.0 | 0 | Felidae | 41018.0 | 0 | Felis | 36896.0 | 1 | Felis catus | 36896.0 | 1 |
| scf7180000021791 | 29816 | 0.3671 | 0 | 23.4291 | Eukaryota | 19202.0 | 0 | Chordata | 19202.0 | 0 | Carnivora | 19202.0 | 0 | Felidae | 19202.0 | 0 | Felis | 15334.0 | 1 | Felis catus | 15334.0 | 1 |
| scf7180000022518 | 22075 | 0.3701 | 0 | 25.6298 | Eukaryota | 7845.0 | 0 | Chordata | 7845.0 | 0 | Carnivora | 6568.0 | 1 | Felidae | 6568.0 | 1 | Felis | 3949.0 | 5 | Felis catus | 3949.0 | 6 |
| scf7180000019101 | 33691 | 0.4531 | 0 | 30.1337 | Eukaryota | 21039.0 | 0 | Chordata | 21039.0 | 0 | Carnivora | 21039.0 | 0 | Felidae | 21039.0 | 0 | Felis | 18935.0 | 1 | Felis catus | 18935.0 | 1 |
| scf7180000020018 | 2363582 | 0.3874 | 100 | 51.705 | Eukaryota | 50290.0 | 0 | Chordata | 50290.0 | 0 | Carnivora | 50290.0 | 0 | Felidae | 50290.0 | 0 | Panthera | 50290.0 | 0 | anthera pardu | 50290.0 | 0 |
| scf7180000022192 | 35111 | 0.4881 | 0 | 2887.2817 | Eukaryota | 2234.0 | 0 | Chordata | 2234.0 | 0 | Carnivora | 1758.0 | 1 | Felidae | 1552.0 | 3 | Felis | 1552.0 | 3 | Felis catus | 1552.0 | 3 |
| scf7180000018466 | 663100 | 0.3452 | 0 | 48.4307 | Eukaryota | 36082.0 | 0 | Chordata | 36082.0 | 0 | Carnivora | 36082.0 | 0 | Felidae | 36082.0 | 0 | Felis | 36082.0 | 0 | Felis catus | 36082.0 | 0 |
| scf7180000018319 | 450535 | 0.4249 | 0 | 52.2431 | Eukaryota | 49948.0 | 0 | Chordata | 49948.0 | 0 | Carnivora | 49948.0 | 0 | Felidae | 49948.0 | 0 | Felis | 29319.0 | 3 | Felis catus | 29319.0 | 3 |
| scf7180000022228 | 27991 | 0.3996 | 0 | 24.6383 | Eukaryota | 35714.0 | 0 | Chordata | 35714.0 | 0 | Carnivora | 35714.0 | 0 | Felidae | 35714.0 | 0 | Felis | 35714.0 | 0 | Felis catus | 35714.0 | 0 |
| scf7180000022093 | 42711 | 0.406 | 0 | 2388.8541 | no-hit | 0.0 | 0 | no-hit | 0.0 | 0 | no-hit | 0.0 | 0 | no-hit | 0.0 | 0 | no-hit | 0.0 | 0 | no-hit | 0.0 | 0 |
| scf7180000018167 | 72818 | 0.3785 | 0 | 50.8535 | Eukaryota | 45136.0 | 0 | Chordata | 45136.0 | 0 | Carnivora | 45136.0 | 0 | Felidae | 45136.0 | 0 | Felis | 40613.0 | 1 | Felis catus | 40613.0 | 1 |
| scf7180000020874 | 97576 | 0.4291 | 0 | 336.3112 | Eukaryota | 2200.0 | 0 | Chordata | 2200.0 | 0 | Carnivora | 2200.0 | 0 | Felidae | 2200.0 | 0 | Felis | 1770.0 | 2 | Felis catus | 1770.0 | 2 |
| scf7180000021998 | 52550 | 0.4131 | 0 | 23.5464 | Eukaryota | 4987.0 | 0 | Chordata | 4987.0 | 0 | Carnivora | 2085.0 | 2 | Felidae | 2085.0 | 2 | Pan | 1346.0 | 5 | an troglodyte | 1346.0 | 6 |
| scf7180000018072 | 47608 | 0.4029 | 0 | 44.3263 | Eukaryota | 9939.0 | 0 | Chordata | 9939.0 | 0 | Carnivora | 9939.0 | 0 | Felidae | 9939.0 | 0 | Felis | 9166.0 | 1 | Felis catus | 9166.0 | 1 |
| scf7180000018096 | 1122530 | 0.3535 | 0 | 48.5179 | Eukaryota | 44741.0 | 0 | Chordata | 44741.0 | 0 | Carnivora | 44741.0 | 0 | Felidae | 44741.0 | 0 | Felis | 40323.0 | 1 | Felis catus | 40323.0 | 1 |
| scf7180000020740 | 61609 | 0.5722 | 0 | 44.4566 | Eukaryota | 9157.0 | 0 | Chordata | 9157.0 | 0 | Carnivora | 9157.0 | 0 | Felidae | 6315.0 | 1 | Felis | 6315.0 | 1 | Felis catus | 6315.0 | 1 |
| scf7180000017548 | 1310174 | 0.4079 | 0 | 49.2888 | Eukaryota | 47667.0 | 0 | Chordata | 47667.0 | 0 | Carnivora | 47667.0 | 0 | Felidae | 47667.0 | 0 | Felis | 47667.0 | 0 | Felis catus | 47667.0 | 0 |
| scf7180000021562 | 71003 | 0.4196 | 0 | 25.3796 | Eukaryota | 43947.0 | 0 | Chordata | 43947.0 | 0 | Carnivora | 43947.0 | 0 | Felidae | 43947.0 | 0 | Felis | 43947.0 | 0 | Felis catus | 43947.0 | 0 |
| scf7180000021047 | 86841 | 0.3763 | 0 | 24.7794 | Eukaryota | 40777.0 | 0 | Chordata | 40777.0 | 0 | Carnivora | 40777.0 | 0 | Felidae | 40777.0 | 0 | Felis | 40777.0 | 0 | Felis catus | 40777.0 | 0 |
| scf7180000024131 | 233156 | 0.382 | 0 | 25.533 | Eukaryota | 45979.0 | 0 | Chordata | 45979.0 | 0 | Carnivora | 45979.0 | 0 | Felidae | 45979.0 | 0 | Felis | 45979.0 | 0 | Felis catus | 45979.0 | 0 |
| scf7180000017700 | 285853 | 0.3816 | 0 | 50.6327 | Eukaryota | 23158.0 | 0 | Chordata | 23158.0 | 0 | Carnivora | 23158.0 | 0 | Felidae | 23158.0 | 0 | Felis | 20825.0 | 1 | Felis catus | 20825.0 | 1 |
| scf7180000019947 | 159300 | 0.6071 | 0 | 53.0247 | Eukaryota | 7722.0 | 0 | Chordata | 7722.0 | 0 | Carnivora | 7722.0 | 0 | Felidae | 6990.0 | 1 | Acinonyx | 5422.0 | 3 | cinonyx jubat | 5422.0 | 3 |
| scf7180000019699 | 73894 | 0.3748 | 0 | 22.4809 | Eukaryota | 18618.0 | 0 | Chordata | 18618.0 | 0 | Carnivora | 18618.0 | 0 | Felidae | 18618.0 | 0 | Felis | 16562.0 | 1 | Felis catus | 16562.0 | 1 |
| scf7180000022675 | 8861 | 0.6428 | 0 | 51.2395 | no-hit | 0.0 | 0 | no-hit | 0.0 | 0 | no-hit | 0.0 | 0 | no-hit | 0.0 | 0 | no-hit | 0.0 | 0 | no-hit | 0.0 | 0 |
| scf7180000018862 | 2481386 | 0.526 | 0 | 51.8127 | Eukaryota | 48289.0 | 0 | Chordata | 48289.0 | 0 | Carnivora | 48289.0 | 0 | Felidae | 38261.0 | 2 | Acinonyx | 20309.0 | 3 | cinonyx jubat | 20309.0 | 3 |
| scf7180000018921 | 394523 | 0.4086 | 0 | 50.7954 | Eukaryota | 47155.0 | 0 | Chordata | 47155.0 | 0 | Carnivora | 47155.0 | 0 | Felidae | 47155.0 | 0 | Felis | 47155.0 | 0 | Felis catus | 47155.0 | 0 |
| scf7180000023774 | 3273761 | 0.3953 | 0 | 51.1254 | Eukaryota | 56959.0 | 1 | Chordata | 56959.0 | 1 | Carnivora | 39037.0 | 2 | Felidae | 39037.0 | 2 | Panthera | 27895.0 | 4 | anthera pardu | 22316.0 | 5 |
| scf7180000021723 | 45321 | 0.4106 | 0 | 26.1226 | Eukaryota | 45624.0 | 0 | Chordata | 45624.0 | 0 | Carnivora | 45624.0 | 0 | Felidae | 45624.0 | 0 | Felis | 45624.0 | 0 | Felis catus | 45624.0 | 0 |
| scf7180000018608 | 678871 | 0.3747 | 0 | 52.9059 | Eukaryota | 45616.0 | 0 | Chordata | 45616.0 | 0 | Carnivora | 45616.0 | 0 | Felidae | 45616.0 | 0 | Felis | 45616.0 | 0 | Felis catus | 45616.0 | 0 |
| scf7180000020269 | 114323 | 0.642 | 0 | 50.9649 | Eukaryota | 15040.0 | 0 | Chordata | 15040.0 | 0 | Carnivora | 15040.0 | 0 | Felidae | 15040.0 | 0 | Felis | 15040.0 | 0 | Felis catus | 15040.0 | 0 |
| scf7180000021398 | 12014 | 0.3617 | 0 | 25.8745 | Eukaryota | 45984.0 | 0 | Chordata | 45984.0 | 0 | Carnivora | 45984.0 | 0 | Felidae | 45984.0 | 0 | Felis | 45984.0 | 0 | Felis catus | 45984.0 | 0 |
| scf7180000022648 | 5563 | 0.3786 | 0 | 32.3664 | Eukaryota | 2034.0 | 0 | Chordata | 2034.0 | 0 | Carnivora | 2034.0 | 0 | Felidae | 2034.0 | 0 | Felis | 1219.0 | 3 | Felis catus | 1219.0 | 3 |
| scf7180000022069 | 15561 | 0.6556 | 0 | 54.1246 | Eukaryota | 2616.0 | 0 | Chordata | 2616.0 | 0 | Carnivora | 2616.0 | 0 | Felidae | 2616.0 | 0 | Felis | 1553.0 | 1 | Felis catus | 1553.0 | 1 |
| scf7180000020301 | 117915 | 0.4135 | 0 | 24.585 | Eukaryota | 19821.0 | 0 | Chordata | 19821.0 | 0 | Primates | 9978.0 | 2 | Hominidae | 9978.0 | 2 | Pan | 6024.0 | 3 | an troglodyte | 6024.0 | 3 |
| scf7180000020573 | 113139 | 0.4681 | 0 | 50.6647 | Eukaryota | 17368.0 | 0 | Chordata | 17368.0 | 0 | Carnivora | 17368.0 | 0 | Felidae | 17368.0 | 0 | Felis | 13897.0 | 1 | Felis catus | 13897.0 | 1 |
| scf7180000017769 | 3870749 | 0.3924 | 0 | 50.3558 | Eukaryota | 60085.0 | 0 | Chordata | 60085.0 | 0 | Carnivora | 54583.0 | 1 | Felidae | 26183.0 | 3 | Canis | 21919.0 | 5 | Canis lupus | 21919.0 | 6 |
| scf7180000022598 | 15033 | 0.4019 | 0 | 23.9545 | Eukaryota | 14310.0 | 0 | Chordata | 14310.0 | 0 | Carnivora | 14310.0 | 0 | Felidae | 14310.0 | 0 | Felis | 12854.0 | 1 | Felis catus | 12854.0 | 1 |
| scf7180000019264 | 315510 | 0.4264 | 0 | 53.0078 | Eukaryota | 31411.0 | 0 | Chordata | 31411.0 | 0 | Carnivora | 31411.0 | 0 | Felidae | 24590.0 | 2 | Acinonyx | 8818.0 | 6 | cinonyx jubat | 8818.0 | 6 |
| scf7180000022258 | 25793 | 0.3517 | 0 | 22.4727 | Eukaryota | 37617.0 | 0 | Chordata | 37617.0 | 0 | Carnivora | 37617.0 | 0 | Felidae | 37617.0 | 0 | Felis | 33823.0 | 1 | Felis catus | 33823.0 | 1 |
| scf7180000019319 | 31175 | 0.4293 | 0 | 140.5911 | Eukaryota | 5209.0 | 0 | Chordata | 5209.0 | 0 | Carnivora | 4285.0 | 1 | Felidae | 2893.0 | 2 | Felis | 2893.0 | 2 | Felis catus | 2893.0 | 2 |
| scf7180000017849 | 1469037 | 0.4133 | 0 | 51.283 | Eukaryota | 49658.0 | 0 | Chordata | 49658.0 | 0 | Carnivora | 49658.0 | 0 | Herpestidae | 44847.0 | 1 | Suricata | 44847.0 | 1 | ricata suricat | 44847.0 | 1 |
| scf7180000023735 | 218230 | 0.4668 | 0 | 53.0169 | Eukaryota | 36129.0 | 0 | Chordata | 36129.0 | 0 | Carnivora | 36129.0 | 0 | Felidae | 36129.0 | 0 | Felis | 36129.0 | 0 | Felis catus | 36129.0 | 0 |
| scf7180000018953 | 142559 | 0.4006 | 0 | 49.4451 | Eukaryota | 45974.0 | 0 | Chordata | 45974.0 | 0 | Carnivora | 45974.0 | 0 | Felidae | 45974.0 | 0 | Felis | 45974.0 | 0 | Felis catus | 45974.0 | 0 |
| scf7180000017492 | 3095458 | 0.3775 | 0 | 50.5366 | Eukaryota | 48005.0 | 0 | Chordata | 48005.0 | 0 | Carnivora | 48005.0 | 0 | Felidae | 48005.0 | 0 | Felis | 48005.0 | 0 | Felis catus | 48005.0 | 0 |
| scf7180000020312 | 20416 | 0.4994 | 0 | 27.0276 | Eukaryota | 3286.0 | 0 | Chordata | 3286.0 | 0 | Carnivora | 3286.0 | 0 | Felidae | 3016.0 | 1 | Felis | 1647.0 | 5 | Felis catus | 1647.0 | 5 |
| scf7180000020869 | 138850 | 0.3627 | 0 | 25.9725 | Eukaryota | 46186.0 | 0 | Chordata | 46186.0 | 0 | Carnivora | 46186.0 | 0 | Felidae | 46186.0 | 0 | Felis | 46186.0 | 0 | Felis catus | 46186.0 | 0 |
| scf7180000024138 | 199102 | 0.4632 | 0 | 51.5133 | Eukaryota | 16290.0 | 0 | Chordata | 16290.0 | 0 | Carnivora | 16290.0 | 0 | Felidae | 14871.0 | 1 | Panthera | 7635.0 | 5 | anthera pardu | 6240.0 | 6 |
| scf7180000023923 | 854839 | 0.3406 | 0 | 48.0328 | Eukaryota | 46638.0 | 0 | Chordata | 46638.0 | 0 | Carnivora | 46638.0 | 0 | Felidae | 46638.0 | 0 | Felis | 46638.0 | 0 | Felis catus | 46638.0 | 0 |
| scf7180000017998 | 924545 | 0.3512 | 0 | 48.0246 | Eukaryota | 45309.0 | 0 | Chordata | 45309.0 | 0 | Carnivora | 45309.0 | 0 | Felidae | 45309.0 | 0 | Felis | 45309.0 | 0 | Felis catus | 45309.0 | 0 |
| scf7180000018702 | 695464 | 0.4823 | 0 | 51.102 | Eukaryota | 17646.0 | 0 | Chordata | 17646.0 | 0 | Carnivora | 17646.0 | 0 | Felidae | 17646.0 | 0 | Felis | 15882.0 | 1 | Felis catus | 15882.0 | 1 |
| scf7180000018889 | 525983 | 0.3449 | 0 | 48.5393 | Eukaryota | 46309.0 | 0 | Chordata | 46309.0 | 0 | Carnivora | 46309.0 | 0 | Felidae | 46309.0 | 0 | Felis | 46309.0 | 0 | Felis catus | 46309.0 | 0 |
| scf7180000024107 | 112472 | 0.4374 | 0 | 49.7615 | Eukaryota | 10225.0 | 0 | Chordata | 10225.0 | 0 | Carnivora | 10225.0 | 0 | Felidae | 10225.0 | 0 | Felis | 9198.0 | 1 | Felis catus | 9198.0 | 1 |
| scf7180000019327 | 297545 | 0.5118 | 0 | 51.3236 | Eukaryota | 14230.0 | 0 | Chordata | 14230.0 | 0 | Carnivora | 14230.0 | 0 | Felidae | 14230.0 | 0 | Felis | 9090.0 | 2 | Felis catus | 9090.0 | 2 |
| scf7180000018287 | 1852828 | 0.426 | 0 | 51.0482 | Eukaryota | 44631.0 | 0 | Chordata | 44631.0 | 0 | Carnivora | 44631.0 | 0 | Felidae | 44631.0 | 0 | Felis | 44631.0 | 0 | Felis catus | 44631.0 | 0 |
| scf7180000020134 | 153314 | 0.4726 | 0 | 52.0108 | Eukaryota | 75080.0 | 0 | Chordata | 75080.0 | 0 | Carnivora | 75080.0 | 0 | Canidae | 35583.0 | 2 | Canis | 28512.0 | 5 | Canis lupus | 28512.0 | 5 |
| scf7180000017923 | 55475 | 0.4327 | 0 | 56.8809 | Eukaryota | 14091.0 | 0 | Chordata | 14091.0 | 0 | Carnivora | 12700.0 | 1 | Felidae | 8358.0 | 3 | Panthera | 4173.0 | 6 | ricata suricat | 2908.0 | 7 |
| scf7180000021769 | 15300 | 0.3793 | 0 | 24.3938 | Eukaryota | 3368.0 | 1 | Chordata | 3368.0 | 1 | Primates | 3368.0 | 1 | Hominidae | 3368.0 | 1 | Homo | 2989.0 | 2 | Homo sapiens | 2989.0 | 2 |
| scf7180000022420 | 14341 | 0.6512 | 0 | 49.5305 | Eukaryota | 2593.0 | 0 | Chordata | 2593.0 | 0 | Carnivora | 2593.0 | 0 | Felidae | 2593.0 | 0 | Felis | 1820.0 | 1 | Felis catus | 1820.0 | 1 |
| scf7180000022060 | 57123 | 0.5296 | 0 | 51.2884 | Eukaryota | 3762.0 | 0 | Chordata | 3762.0 | 0 | Primates | 1425.0 | 3 | Hominidae | 1425.0 | 3 | Homo | 1425.0 | 4 | Homo sapiens | 1425.0 | 4 |
| scf7180000018355 | 16300 | 0.4025 | 0 | 22.2823 | Eukaryota | 6803.0 | 0 | Chordata | 6803.0 | 0 | Carnivora | 6803.0 | 0 | Felidae | 6803.0 | 0 | Felis | 6803.0 | 0 | Felis catus | 6803.0 | 0 |
| scf7180000021377 | 39490 | 0.4758 | 0 | 46.6572 | Eukaryota | 11285.0 | 0 | Chordata | 11285.0 | 0 | Carnivora | 11285.0 | 0 | Felidae | 10112.0 | 1 | Lynx | 4524.0 | 3 | ynx canadens | 4524.0 | 3 |
| scf7180000019612 | 273264 | 0.5134 | 0 | 51.1516 | Eukaryota | 20820.0 | 1 | Chordata | 20820.0 | 1 | Artiodactyla | 11312.0 | 4 | Monodontidae | 6792.0 | 5 | Monodon | 4529.0 | 7 | odon monoc | 4529.0 | 7 |
| scf7180000019992 | 190400 | 0.4187 | 0 | 25.4991 | Eukaryota | 24911.0 | 0 | Chordata | 24911.0 | 0 | Carnivora | 24911.0 | 0 | Felidae | 21695.0 | 1 | Felis | 19248.0 | 2 | Felis catus | 19248.0 | 2 |
| scf7180000019328 | 469708 | 0.459 | 0 | 52.8095 | Eukaryota | 32989.0 | 0 | Chordata | 32989.0 | 0 | Carnivora | 32989.0 | 0 | Felidae | 32989.0 | 0 | Felis | 32989.0 | 0 | Felis catus | 32989.0 | 0 |
| scf7180000019536 | 105553 | 0.3743 | 0 | 25.327 | Eukaryota | 23604.0 | 0 | Chordata | 23604.0 | 0 | Carnivora | 23604.0 | 0 | Felidae | 23604.0 | 0 | Felis | 18841.0 | 1 | Felis catus | 18841.0 | 1 |
| scf7180000023285 | 76960 | 0.4175 | 0 | 74.656 | Eukaryota | 45507.0 | 0 | Chordata | 45507.0 | 0 | Carnivora | 45507.0 | 0 | Felidae | 45507.0 | 0 | Felis | 45507.0 | 0 | Felis catus | 45507.0 | 0 |
| scf7180000017874 | 10624 | 0.4133 | 0 | 38.0424 | Eukaryota | 43349.0 | 0 | Chordata | 43349.0 | 0 | Carnivora | 43349.0 | 0 | Felidae | 43349.0 | 0 | Felis | 43349.0 | 0 | Felis catus | 43349.0 | 0 |
| scf7180000022485 | 13754 | 0.5523 | 0 | 47.9105 | Eukaryota | 2501.0 | 0 | Chordata | 2501.0 | 0 | Carnivora | 2501.0 | 0 | Felidae | 2251.0 | 1 | Felis | 2251.0 | 1 | Felis catus | 2251.0 | 1 |
| scf7180000021665 | 54152 | 0.4977 | 0 | 48.5538 | Eukaryota | 9176.0 | 0 | Chordata | 9176.0 | 0 | Carnivora | 9176.0 | 0 | Felidae | 8503.0 | 1 | Felis | 8503.0 | 1 | Felis catus | 8503.0 | 1 |
| scf7180000020744 | 51009 | 0.4039 | 0 | 49.0176 | Eukaryota | 7570.0 | 0 | Chordata | 7570.0 | 0 | Carnivora | 7570.0 | 0 | Felidae | 7570.0 | 0 | Felis | 7570.0 | 0 | Felis catus | 7570.0 | 0 |
| scf7180000018474 | 93663 | 0.3857 | 0 | 50.823 | Eukaryota | 36997.0 | 0 | Chordata | 36997.0 | 0 | Carnivora | 36997.0 | 0 | Felidae | 36997.0 | 0 | Felis | 33490.0 | 1 | Felis catus | 33490.0 | 1 |
| scf7180000020700 | 109414 | 0.3866 | 0 | 49.3992 | Eukaryota | 46787.0 | 0 | Chordata | 46787.0 | 0 | Carnivora | 46787.0 | 0 | Felidae | 46787.0 | 0 | Felis | 42131.0 | 1 | Felis catus | 42131.0 | 1 |
| scf7180000018187 | 1154125 | 0.416 | 0 | 50.8435 | Eukaryota | 46260.0 | 0 | Chordata | 46260.0 | 0 | Carnivora | 46260.0 | 0 | Felidae | 46260.0 | 0 | Felis | 46260.0 | 0 | Felis catus | 46260.0 | 0 |
| scf7180000018949 | 497661 | 0.3477 | 0 | 47.982 | Eukaryota | 42332.0 | 0 | Chordata | 42332.0 | 0 | Carnivora | 42332.0 | 0 | Felidae | 42332.0 | 0 | Felis | 38106.0 | 1 | Felis catus | 38106.0 | 1 |
| scf7180000023894 | 236171 | 0.3905 | 0 | 47.735 | Eukaryota | 46840.0 | 0 | Chordata | 46840.0 | 0 | Carnivora | 46840.0 | 0 | Felidae | 46840.0 | 0 | Felis | 46840.0 | 0 | Felis catus | 46840.0 | 0 |
| scf7180000024132 | 212983 | 0.551 | 0 | 51.7339 | Eukaryota | 15408.0 | 0 | Chordata | 15408.0 | 0 | Carnivora | 15408.0 | 0 | Felidae | 15408.0 | 0 | Felis | 15408.0 | 0 | Felis catus | 15408.0 | 0 |
| scf7180000018421 | 119964 | 0.3598 | 0 | 45.5274 | Eukaryota | 44567.0 | 0 | Chordata | 44567.0 | 0 | Carnivora | 44567.0 | 0 | Felidae | 44567.0 | 0 | Felis | 44567.0 | 0 | Felis catus | 44567.0 | 0 |
| scf7180000021050 | 81294 | 0.5988 | 0 | 63.6721 | Eukaryota | 4894.0 | 0 | Chordata | 4894.0 | 0 | Primates | 3454.0 | 1 | Hominidae | 1824.0 | 3 | Homo | 1824.0 | 3 | Homo sapiens | 1824.0 | 3 |
| scf7180000022667 | 9288 | 0.3925 | 0 | 23.7158 | Eukaryota | 4631.0 | 0 | Chordata | 4631.0 | 0 | Carnivora | 4631.0 | 0 | Felidae | 4631.0 | 0 | Felis | 4631.0 | 0 | Felis catus | 4631.0 | 0 |
| scf7180000019004 | 361868 | 0.4662 | 0 | 52.7047 | Eukaryota | 28119.0 | 0 | Chordata | 28119.0 | 0 | Carnivora | 28119.0 | 0 | Felidae | 28119.0 | 0 | Felis | 25304.0 | 1 | Felis catus | 25304.0 | 1 |
| scf7180000020898 | 62909 | 0.3762 | 0 | 23.1568 | Eukaryota | 32884.0 | 0 | Chordata | 32884.0 | 0 | Carnivora | 32884.0 | 0 | Felidae | 32884.0 | 0 | Felis | 32884.0 | 0 | Felis catus | 32884.0 | 0 |
| scf7180000017980 | 1349300 | 0.4491 | 0 | 51.6342 | Eukaryota | 59277.0 | 0 | Chordata | 59277.0 | 0 | Carnivora | 59277.0 | 0 | Felidae | 30749.0 | 3 | Canis | 11604.0 | 7 | Canis lupus | 11604.0 | 7 |
| scf7180000018006 | 1293091 | 0.3995 | 0 | 51.9601 | Eukaryota | 59841.0 | 0 | Chordata | 59841.0 | 0 | Carnivora | 59841.0 | 0 | Felidae | 59841.0 | 0 | Felis | 49054.0 | 1 | Felis catus | 49054.0 | 1 |
| scf7180000017959 | 407856 | 0.454 | 0 | 51.3268 | Eukaryota | 46055.0 | 0 | Chordata | 46055.0 | 0 | Carnivora | 46055.0 | 0 | Felidae | 46055.0 | 0 | Felis | 41508.0 | 1 | Felis catus | 41508.0 | 1 |
| scf7180000021856 | 30325 | 0.4649 | 0 | 160.7691 | Eukaryota | 2297.0 | 0 | Chordata | 2297.0 | 0 | Carnivora | 2297.0 | 0 | Felidae | 2297.0 | 0 | Felis | 1859.0 | 1 | Felis catus | 1859.0 | 1 |
| scf7180000021741 | 9914 | 0.5231 | 0 | 35.7366 | no-hit | 0.0 | 0 | no-hit | 0.0 | 0 | no-hit | 0.0 | 0 | no-hit | 0.0 | 0 | no-hit | 0.0 | 0 | no-hit | 0.0 | 0 |
| scf7180000019675 | 250484 | 0.4151 | 0 | 53.8511 | Eukaryota | 46406.0 | 0 | Chordata | 46406.0 | 0 | Carnivora | 46406.0 | 0 | Felidae | 46406.0 | 0 | Felis | 46406.0 | 0 | Felis catus | 46406.0 | 0 |
| scf7180000021403 | 34142 | 0.4909 | 0 | 243.7313 | Eukaryota | 3282.0 | 0 | Chordata | 3282.0 | 0 | Carnivora | 2349.0 | 1 | Felidae | 1874.0 | 2 | Homo | 933.0 | 4 | Homo sapiens | 933.0 | 4 |
| scf7180000018310 | 23436 | 0.4178 | 0 | 27.5789 | Eukaryota | 16023.0 | 0 | Chordata | 16023.0 | 0 | Carnivora | 16023.0 | 0 | Felidae | 16023.0 | 0 | Felis | 12816.0 | 1 | Felis catus | 12816.0 | 1 |
| scf7180000024046 | 341923 | 0.5534 | 0 | 50.7954 | Eukaryota | 60072.0 | 0 | Chordata | 60072.0 | 0 | Carnivora | 60072.0 | 0 | Felidae | 60072.0 | 0 | Lynx | 30327.0 | 1 | ynx canadens | 30327.0 | 1 |
| scf7180000018349 | 752942 | 0.3892 | 0 | 51.3846 | Eukaryota | 47081.0 | 0 | Chordata | 47081.0 | 0 | Carnivora | 47081.0 | 0 | Felidae | 47081.0 | 0 | Felis | 47081.0 | 0 | Felis catus | 47081.0 | 0 |
| scf7180000019241 | 37976 | 0.3962 | 0 | 41.029 | Eukaryota | 18073.0 | 0 | Chordata | 18073.0 | 0 | Carnivora | 18073.0 | 0 | Felidae | 18073.0 | 0 | Felis | 18073.0 | 0 | Felis catus | 18073.0 | 0 |
| scf7180000020253 | 143348 | 0.541 | 0 | 49.2127 | Eukaryota | 7649.0 | 1 | Chordata | 7649.0 | 1 | Carnivora | 5344.0 | 2 | Felidae | 5344.0 | 3 | Felis | 2698.0 | 4 | Felis catus | 2698.0 | 4 |
| scf7180000018288 | 1090570 | 0.4159 | 0 | 57.0791 | Eukaryota | 118910.0 | 0 | Chordata | 118910.0 | 0 | Carnivora | 118910.0 | 0 | Felidae | 118910.0 | 0 | Lynx | 56593.0 | 3 | ynx canadens | 56593.0 | 3 |
| scf7180000023094 | 12823 | 0.5113 | 0 | 351.6745 | Eukaryota | 21685.0 | 0 | Chordata | 21685.0 | 0 | Carnivora | 21685.0 | 0 | Felidae | 21685.0 | 0 | Felis | 19481.0 | 1 | Felis catus | 19481.0 | 1 |
| scf7180000017814 | 1649496 | 0.3694 | 0 | 49.8573 | Eukaryota | 45320.0 | 0 | Chordata | 45320.0 | 0 | Carnivora | 45320.0 | 0 | Felidae | 45320.0 | 0 | Felis | 45320.0 | 0 | Felis catus | 45320.0 | 0 |
| scf7180000021985 | 21944 | 0.3889 | 0 | 25.1287 | Eukaryota | 15910.0 | 0 | Chordata | 15910.0 | 0 | Carnivora | 15910.0 | 0 | Felidae | 15910.0 | 0 | Felis | 14288.0 | 1 | Felis catus | 14288.0 | 1 |
| scf7180000018073 | 1805534 | 0.3942 | 0 | 51.0092 | Eukaryota | 30272.0 | 0 | Chordata | 30272.0 | 0 | Carnivora | 30272.0 | 0 | Felidae | 30272.0 | 0 | Felis | 30272.0 | 0 | Felis catus | 30272.0 | 0 |
| scf7180000021837 | 43312 | 0.3626 | 0 | 23.6743 | Eukaryota | 34091.0 | 0 | Chordata | 34091.0 | 0 | Carnivora | 34091.0 | 0 | Felidae | 34091.0 | 0 | Felis | 34091.0 | 0 | Felis catus | 34091.0 | 0 |
| scf7180000017856 | 2781099 | 0.383 | 0 | 49.8669 | Eukaryota | 52722.0 | 0 | Chordata | 52722.0 | 0 | Carnivora | 52722.0 | 0 | Felidae | 47407.0 | 1 | Acinonyx | 31548.0 | 3 | cinonyx jubat | 31548.0 | 3 |
| scf7180000023837 | 1614864 | 0.531 | 0 | 50.1612 | Eukaryota | 28517.0 | 0 | Chordata | 28517.0 | 0 | Carnivora | 28517.0 | 0 | Felidae | 28517.0 | 0 | Felis | 25645.0 | 1 | Felis catus | 25645.0 | 1 |
| scf7180000018219 | 961526 | 0.4443 | 0 | 52.0272 | Eukaryota | 32351.0 | 0 | Chordata | 32351.0 | 0 | Carnivora | 32351.0 | 0 | Felidae | 25964.0 | 2 | Acinonyx | 9794.0 | 5 | cinonyx jubat | 9794.0 | 5 |
| scf7180000022897 | 69478 | 0.3553 | 0 | 504.86 | Eukaryota | 12692.0 | 0 | Chordata | 12692.0 | 0 | Carnivora | 12692.0 | 0 | Felidae | 12692.0 | 0 | Felis | 7650.0 | 1 | Felis catus | 7650.0 | 1 |
| scf7180000022806 | 3868 | 0.3976 | 0 | 25.1144 | Eukaryota | 3863.0 | 0 | Chordata | 3863.0 | 0 | Carnivora | 3004.0 | 1 | Felidae | 3004.0 | 1 | Felis | 2627.0 | 2 | Felis catus | 2627.0 | 2 |
| scf7180000018467 | 78606 | 0.3979 | 0 | 50.0955 | Eukaryota | 44942.0 | 0 | Chordata | 44942.0 | 0 | Carnivora | 44942.0 | 0 | Felidae | 44942.0 | 0 | Felis | 40513.0 | 1 | Felis catus | 40513.0 | 1 |
| scf7180000024199 | 88021 | 0.4723 | 0 | 53.3173 | Eukaryota | 24424.0 | 0 | Chordata | 24424.0 | 0 | Carnivora | 24424.0 | 0 | Felidae | 24424.0 | 0 | Felis | 19116.0 | 1 | Felis catus | 19116.0 | 1 |
| scf7180000021568 | 54263 | 0.4279 | 0 | 25.4713 | Eukaryota | 47410.0 | 0 | Chordata | 47410.0 | 0 | Carnivora | 42704.0 | 1 | Felidae | 32932.0 | 2 | Lynx | 14151.0 | 4 | ynx canadens | 14151.0 | 4 |
| scf7180000022400 | 30108 | 0.5664 | 0 | 52.7333 | Eukaryota | 22273.0 | 0 | Chordata | 22273.0 | 0 | Carnivora | 22273.0 | 0 | Felidae | 14744.0 | 3 | Lynx | 5124.0 | 6 | ynx canadens | 5124.0 | 6 |
| scf7180000020849 | 137618 | 0.3641 | 0 | 49.698 | Eukaryota | 45373.0 | 0 | Chordata | 45373.0 | 0 | Carnivora | 45373.0 | 0 | Felidae | 45373.0 | 0 | Felis | 45373.0 | 0 | Felis catus | 45373.0 | 0 |
| scf7180000019179 | 183189 | 0.3723 | 100 | 25.0324 | Eukaryota | 46221.0 | 0 | Chordata | 46221.0 | 0 | Carnivora | 46221.0 | 0 | Felidae | 46221.0 | 0 | Felis | 46221.0 | 0 | Felis catus | 46221.0 | 0 |
| scf7180000019110 | 514191 | 0.3714 | 0 | 50.6904 | Eukaryota | 47247.0 | 0 | Chordata | 47247.0 | 0 | Carnivora | 47247.0 | 0 | Felidae | 47247.0 | 0 | Felis | 42584.0 | 1 | Felis catus | 42584.0 | 1 |
| scf7180000022807 | 3747 | 0.4406 | 0 | 24.7035 | Eukaryota | 15649.0 | 0 | Chordata | 15649.0 | 0 | Carnivora | 15649.0 | 0 | Felidae | 15649.0 | 0 | Felis | 15649.0 | 0 | Felis catus | 15649.0 | 0 |
| scf7180000022307 | 13247 | 0.6469 | 0 | 54.9475 | Eukaryota | 2760.0 | 0 | Chordata | 2760.0 | 0 | Artiodactyla | 2492.0 | 1 | Physeteridae | 1683.0 | 2 | Physeter | 1683.0 | 3 | yseter catod | 1683.0 | 4 |
| scf7180000023581 | 20997 | 0.4756 | 0 | 412.4777 | no-hit | 0.0 | 0 | no-hit | 0.0 | 0 | no-hit | 0.0 | 0 | no-hit | 0.0 | 0 | no-hit | 0.0 | 0 | no-hit | 0.0 | 0 |
| scf7180000021477 | 53578 | 0.4139 | 0 | 25.3073 | Eukaryota | 10974.0 | 0 | Chordata | 10974.0 | 0 | Artiodactyla | 5591.0 | 2 | Delphinidae | 3553.0 | 5 | Pseudorca | 3553.0 | 5 | udorca crassic | 3553.0 | 5 |
| scf7180000021086 | 61700 | 0.4148 | 0 | 25.3374 | Eukaryota | 9502.0 | 0 | Chordata | 9502.0 | 0 | Artiodactyla | 3639.0 | 3 | Felidae | 3539.0 | 4 | Felis | 2814.0 | 5 | Felis catus | 2814.0 | 5 |
| scf7180000023284 | 48515 | 0.3897 | 0 | 146.2588 | Eukaryota | 14705.0 | 0 | Chordata | 14705.0 | 0 | Carnivora | 14705.0 | 0 | Felidae | 14705.0 | 0 | Felis | 11768.0 | 1 | Felis catus | 11768.0 | 1 |
| scf7180000020026 | 189353 | 0.4336 | 0 | 50.0783 | Eukaryota | 30892.0 | 0 | Chordata | 30892.0 | 0 | Primates | 17243.0 | 2 | Hominidae | 10178.0 | 4 | Homo | 10178.0 | 5 | Homo sapiens | 10178.0 | 5 |
| scf7180000022098 | 4483 | 0.5206 | 0 | 36.6581 | no-hit | 0.0 | 0 | no-hit | 0.0 | 0 | no-hit | 0.0 | 0 | no-hit | 0.0 | 0 | no-hit | 0.0 | 0 | no-hit | 0.0 | 0 |
| scf7180000023849 | 323123 | 0.5371 | 0 | 52.2043 | Eukaryota | 21203.0 | 0 | Chordata | 21203.0 | 0 | Carnivora | 16329.0 | 1 | Felidae | 10591.0 | 4 | Felis | 6473.0 | 6 | anthera pardu | 4118.0 | 7 |
| scf7180000019837 | 339650 | 0.4595 | 0 | 25.7972 | Eukaryota | 32163.0 | 0 | Chordata | 32163.0 | 0 | Carnivora | 32163.0 | 0 | Felidae | 16777.0 | 3 | Felis | 11535.0 | 7 | Felis catus | 11535.0 | 7 |
| scf7180000018345 | 551425 | 0.4885 | 0 | 49.0508 | Eukaryota | 37078.0 | 0 | Chordata | 37078.0 | 0 | Carnivora | 37078.0 | 0 | Felidae | 30666.0 | 1 | Acinonyx | 11541.0 | 4 | cinonyx jubat | 11541.0 | 4 |
| scf7180000019194 | 4932 | 0.382 | 0 | 24.1124 | Eukaryota | 11802.0 | 0 | Chordata | 11802.0 | 0 | Carnivora | 11802.0 | 0 | Felidae | 11802.0 | 0 | Felis | 9441.0 | 1 | Felis catus | 9441.0 | 1 |
| scf7180000017767 | 2319926 | 0.3728 | 0 | 50.1521 | Eukaryota | 56026.0 | 0 | Chordata | 56026.0 | 0 | Carnivora | 56026.0 | 0 | Felidae | 33619.0 | 3 | Suricata | 12355.0 | 7 | ricata suricat | 12355.0 | 8 |
| scf7180000021832 | 41640 | 0.3619 | 0 | 24.827 | Eukaryota | 59677.0 | 0 | Chordata | 59677.0 | 0 | Carnivora | 59677.0 | 0 | Felidae | 43045.0 | 2 | Felis | 22023.0 | 4 | Felis catus | 22023.0 | 4 |
| scf7180000018001 | 2528617 | 0.396 | 0 | 50.7416 | Eukaryota | 70978.0 | 0 | Chordata | 70978.0 | 0 | Carnivora | 70978.0 | 0 | Felidae | 44210.0 | 2 | Felis | 15124.0 | 5 | Felis catus | 15124.0 | 5 |
| scf7180000024246 | 1036456 | 0.4824 | 0 | 51.0694 | Eukaryota | 46206.0 | 0 | Chordata | 46206.0 | 0 | Carnivora | 46206.0 | 0 | Felidae | 37568.0 | 1 | Panthera | 13737.0 | 5 | ynx canadens | 9702.0 | 6 |
| scf7180000021266 | 77519 | 0.4048 | 0 | 24.584 | Eukaryota | 30002.0 | 0 | Chordata | 30002.0 | 0 | Carnivora | 30002.0 | 0 | Canidae | 30002.0 | 0 | Canis | 30002.0 | 0 | Canis lupus | 30002.0 | 0 |
| scf7180000020533 | 180228 | 0.3993 | 0 | 51.2516 | Eukaryota | 15332.0 | 0 | Chordata | 15332.0 | 0 | pterygiforme | 13651.0 | 1 | Apterygidae | 13651.0 | 1 | Apteryx | 13651.0 | 1 | pteryx mante | 13651.0 | 1 |
| scf7180000020109 | 30928 | 0.4109 | 0 | 286.5167 | Eukaryota | 8913.0 | 0 | Chordata | 8913.0 | 0 | Carnivora | 8913.0 | 0 | Felidae | 8913.0 | 0 | Felis | 7032.0 | 1 | Felis catus | 7032.0 | 1 |
| scf7180000018655 | 7488 | 0.472 | 0 | 25.1199 | Eukaryota | 11120.0 | 0 | Chordata | 11120.0 | 0 | Carnivora | 11120.0 | 0 | Felidae | 11120.0 | 0 | Felis | 11120.0 | 0 | Felis catus | 11120.0 | 0 |
| scf7180000020475 | 97613 | 0.5582 | 0 | 47.06 | Eukaryota | 27293.0 | 1 | Chordata | 27293.0 | 1 | Primates | 11659.0 | 3 | Hominidae | 11659.0 | 5 | Homo | 8881.0 | 7 | Homo sapiens | 8881.0 | 7 |
| scf7180000020277 | 181373 | 0.4111 | 100 | 47.069 | Eukaryota | 26020.0 | 0 | Chordata | 26020.0 | 0 | Carnivora | 26020.0 | 0 | Felidae | 26020.0 | 0 | Felis | 23366.0 | 1 | Felis catus | 23366.0 | 1 |
| scf7180000018211 | 1031184 | 0.4723 | 0 | 51.5961 | Eukaryota | 28690.0 | 0 | Chordata | 28690.0 | 0 | Carnivora | 28690.0 | 0 | Felidae | 28690.0 | 0 | Felis | 25794.0 | 1 | Felis catus | 25794.0 | 1 |
| scf7180000022285 | 42274 | 0.3694 | 0 | 22.8359 | Eukaryota | 25083.0 | 0 | Chordata | 25083.0 | 0 | Carnivora | 25083.0 | 0 | Felidae | 25083.0 | 0 | Felis | 25083.0 | 0 | Felis catus | 25083.0 | 0 |
| scf7180000019626 | 4125 | 0.3862 | 0 | 27.1797 | Eukaryota | 2096.0 | 1 | Chordata | 2096.0 | 1 | Carnivora | 1285.0 | 2 | Felidae | 1033.0 | 3 | Felis | 1033.0 | 4 | Felis catus | 1033.0 | 4 |

| | | | | | | | | | | | | | | | | | | | | | | |
|---|---|---|---|---|---|---|---|---|---|---|---|---|---|---|---|---|---|---|---|---|---|---|
| scf7180000017710 | 1676063 | 0.4476 | 0 | 50.5605 | Eukaryota | 80288.0 | 0 | Chordata | 80288.0 | 0 | Carnivora | 80288.0 | 0 | Felidae | 80288.0 | 0 | Acinonyx | 38087.0 | 2 | cinonyx jubat | 38087.0 | 2 |
| scf7180000019349 | 298093 | 0.4915 | 0 | 50.9997 | Eukaryota | 18215.0 | 0 | Chordata | 18215.0 | 0 | Carnivora | 18215.0 | 0 | Felidae | 18215.0 | 0 | Felis | 12785.0 | 3 | Felis catus | 12785.0 | 3 |
| scf7180000021577 | 26473 | 0.4315 | 0 | 187.4576 | Eukaryota | 2288.0 | 0 | Chordata | 2288.0 | 0 | Carnivora | 2288.0 | 0 | Felidae | 2288.0 | 0 | Felis | 1643.0 | 2 | Felis catus | 1643.0 | 2 |
| scf7180000022785 | 8763 | 0.3822 | 0 | 22.0864 | Eukaryota | 15806.0 | 0 | Chordata | 15806.0 | 0 | Carnivora | 15806.0 | 0 | Felidae | 15806.0 | 0 | Felis | 14252.0 | 1 | Felis catus | 14252.0 | 1 |
| scf7180000017446 | 4096778 | 0.3861 | 0 | 51.0061 | Eukaryota | 65915.0 | 0 | Chordata | 65915.0 | 0 | Carnivora | 65915.0 | 0 | Felidae | 65915.0 | 0 | Panthera | 29752.0 | 3 | anthera pardu | 29752.0 | 3 |
| scf7180000018936 | 13874 | 0.3685 | 0 | 24.8139 | Eukaryota | 11169.0 | 0 | Chordata | 11169.0 | 0 | Carnivora | 11169.0 | 0 | Felidae | 11169.0 | 0 | Felis | 9903.0 | 1 | Felis catus | 9903.0 | 1 |
| scf7180000020907 | 76687 | 0.5755 | 0 | 50.3615 | Eukaryota | 13099.0 | 0 | Chordata | 13099.0 | 0 | Carnivora | 13099.0 | 0 | Felidae | 13099.0 | 0 | Felis | 11828.0 | 1 | Felis catus | 11828.0 | 1 |
| scf7180000020995 | 77921 | 0.403 | 0 | 57.9111 | Eukaryota | 38828.0 | 0 | Chordata | 38828.0 | 0 | Carnivora | 38828.0 | 0 | Felidae | 38828.0 | 0 | Felis | 34979.0 | 1 | Felis catus | 34979.0 | 1 |
| scf7180000020279 | 58503 | 0.4151 | 0 | 62.6432 | Eukaryota | 13551.0 | 0 | Chordata | 13551.0 | 0 | Carnivora | 13551.0 | 0 | Felidae | 13551.0 | 0 | Felis | 12226.0 | 1 | Felis catus | 12226.0 | 1 |
| scf7180000020468 | 124034 | 0.5682 | 0 | 26.9698 | Eukaryota | 31286.0 | 0 | Chordata | 31286.0 | 0 | Carnivora | 31286.0 | 0 | Felidae | 18918.0 | 1 | Felis | 12676.0 | 2 | Felis catus | 12676.0 | 2 |
| scf7180000017762 | 30631 | 0.3691 | 0 | 47.4253 | Eukaryota | 21703.0 | 0 | Chordata | 21703.0 | 0 | Carnivora | 21703.0 | 0 | Felidae | 21703.0 | 0 | Felis | 19555.0 | 1 | Felis catus | 19555.0 | 1 |
| scf7180000021803 | 45458 | 0.4648 | 0 | 26.9613 | Eukaryota | 13748.0 | 0 | Chordata | 13748.0 | 0 | Primates | 10025.0 | 3 | Hominidae | 7149.0 | 4 | Homo | 7149.0 | 5 | Homo sapiens | 7149.0 | 5 |
| scf7180000024171 | 93421 | 0.3836 | 0 | 51.1348 | Eukaryota | 10650.0 | 0 | Chordata | 10650.0 | 0 | Carnivora | 10650.0 | 0 | Felidae | 10650.0 | 0 | Felis | 4272.0 | 2 | Felis catus | 4272.0 | 2 |
| scf7180000022655 | 17473 | 0.5146 | 0 | 28.2937 | Eukaryota | 17033.0 | 0 | Chordata | 17033.0 | 0 | Carnivora | 12400.0 | 3 | Felidae | 9099.0 | 5 | Felis | 2534.0 | 8 | Felis catus | 2534.0 | 8 |
| scf7180000019985 | 155710 | 0.3687 | 0 | 48.5036 | Eukaryota | 44696.0 | 0 | Chordata | 44696.0 | 0 | Carnivora | 44696.0 | 0 | Felidae | 44696.0 | 0 | Felis | 44696.0 | 0 | Felis catus | 44696.0 | 0 |
| scf7180000018432 | 181123 | 0.3823 | 0 | 49.5061 | Eukaryota | 29700.0 | 0 | Chordata | 29700.0 | 0 | Carnivora | 29700.0 | 0 | Felidae | 29700.0 | 0 | Acinonyx | 14820.0 | 2 | cinonyx jubat | 14820.0 | 2 |
| scf7180000022225 | 255492 | 0.5611 | 0 | 52.0012 | Eukaryota | 20811.0 | 0 | Chordata | 20811.0 | 0 | Carnivora | 20811.0 | 0 | Felidae | 20811.0 | 0 | Lynx | 14378.0 | 1 | ynx canadens | 14378.0 | 1 |
| scf7180000017585 | 1991287 | 0.4321 | 0 | 51.7769 | Eukaryota | 51902.0 | 0 | Chordata | 51902.0 | 0 | Carnivora | 51902.0 | 0 | Felidae | 26818.0 | 4 | Panthera | 10697.0 | 7 | ynx canadens | 5398.0 | 9 |
| scf7180000018848 | 3083677 | 0.3593 | 0 | 48.9879 | Eukaryota | 47003.0 | 0 | Chordata | 47003.0 | 0 | Carnivora | 47003.0 | 0 | Phocidae | 18679.0 | 2 | Zalophus | 13978.0 | 4 | phus californi | 13978.0 | 4 |
| scf7180000018732 | 677648 | 0.4369 | 0 | 51.3009 | Eukaryota | 22977.0 | 0 | Chordata | 22977.0 | 0 | Carnivora | 22977.0 | 0 | Felidae | 22977.0 | 0 | Felis | 22977.0 | 0 | Felis catus | 22977.0 | 0 |
| scf7180000020128 | 89581 | 0.4285 | 0 | 48.9449 | Eukaryota | 13632.0 | 0 | Chordata | 13632.0 | 0 | Carnivora | 13632.0 | 0 | Felidae | 13632.0 | 0 | Felis | 10934.0 | 1 | Felis catus | 10934.0 | 1 |
| scf7180000017745 | 2106719 | 0.4187 | 0 | 51.3462 | Eukaryota | 46622.0 | 0 | Chordata | 46622.0 | 0 | Carnivora | 46622.0 | 0 | Felidae | 46622.0 | 0 | Felis | 46622.0 | 0 | Felis catus | 46622.0 | 0 |
| scf7180000019687 | 238985 | 0.3737 | 0 | 25.4348 | Eukaryota | 51541.0 | 0 | Chordata | 51541.0 | 0 | Carnivora | 51541.0 | 0 | Felidae | 26881.0 | 4 | Panthera | 10849.0 | 8 | anthera pardu | 5432.0 | 9 |
| scf7180000024257 | 3204198 | 0.4122 | 0 | 52.5559 | Eukaryota | 44703.0 | 0 | Chordata | 44703.0 | 0 | Carnivora | 44703.0 | 0 | Felidae | 44703.0 | 0 | Felis | 35721.0 | 2 | Felis catus | 35721.0 | 2 |
| scf7180000019530 | 374484 | 0.3751 | 0 | 50.5864 | Eukaryota | 46379.0 | 0 | Chordata | 46379.0 | 0 | Carnivora | 46379.0 | 0 | Felidae | 46379.0 | 0 | Felis | 46379.0 | 0 | Felis catus | 46379.0 | 0 |
| scf7180000020793 | 119361 | 0.3798 | 0 | 27.979 | Eukaryota | 22970.0 | 0 | Chordata | 22970.0 | 0 | Carnivora | 22970.0 | 0 | Felidae | 22970.0 | 0 | Felis | 16095.0 | 1 | Felis catus | 16095.0 | 1 |
| scf7180000018408 | 14525 | 0.4927 | 0 | 39.9403 | Eukaryota | 13957.0 | 0 | Chordata | 13957.0 | 0 | Carnivora | 13957.0 | 0 | Felidae | 13957.0 | 0 | Felis | 9856.0 | 1 | Felis catus | 9856.0 | 1 |
| scf7180000021584 | 39995 | 0.4487 | 0 | 47.2964 | Eukaryota | 39220.0 | 0 | Chordata | 39220.0 | 0 | Carnivora | 39220.0 | 0 | Felidae | 30444.0 | 1 | Panthera | 12827.0 | 4 | ricata suricat | 8776.0 | 5 |
| scf7180000022251 | 10392 | 0.3685 | 0 | 23.054 | Eukaryota | 2428.0 | 0 | Chordata | 2428.0 | 0 | Carnivora | 2428.0 | 0 | Felidae | 2163.0 | 1 | Felis | 1935.0 | 2 | Felis catus | 1935.0 | 2 |
| scf7180000020927 | 95757 | 0.3927 | 0 | 49.2684 | Eukaryota | 7044.0 | 0 | Chordata | 7044.0 | 0 | Carnivora | 4673.0 | 1 | Felidae | 4673.0 | 1 | Felis | 4673.0 | 1 | Felis catus | 4673.0 | 1 |
| scf7180000020631 | 124985 | 0.4662 | 0 | 53.6985 | Eukaryota | 17796.0 | 0 | Chordata | 17796.0 | 0 | Carnivora | 10639.0 | 2 | Felidae | 7161.0 | 4 | Equus | 5393.0 | 7 | Equus caballu | 5393.0 | 7 |
| scf7180000019826 | 275249 | 0.3867 | 0 | 52.6162 | Eukaryota | 48521.0 | 0 | Chordata | 48521.0 | 0 | Carnivora | 48521.0 | 0 | Felidae | 48521.0 | 0 | Panthera | 22834.0 | 2 | ynx canadens | 22420.0 | 3 |
| scf7180000021457 | 46423 | 0.3792 | 0 | 25.6331 | Eukaryota | 29462.0 | 0 | Chordata | 29462.0 | 0 | Carnivora | 29462.0 | 0 | Felidae | 29462.0 | 0 | Felis | 26453.0 | 1 | Felis catus | 26453.0 | 1 |
| scf7180000020259 | 120408 | 0.3946 | 0 | 49.9761 | Eukaryota | 6794.0 | 0 | Chordata | 6794.0 | 0 | Primates | 3349.0 | 2 | Hominidae | 3349.0 | 3 | Homo | 3349.0 | 3 | Homo sapiens | 3349.0 | 3 |
| scf7180000018085 | 1257586 | 0.424 | 0 | 51.7992 | Eukaryota | 44063.0 | 0 | Chordata | 44063.0 | 0 | Carnivora | 35436.0 | 1 | Felidae | 31007.0 | 3 | Acinonyx | 16792.0 | 4 | cinonyx jubat | 16792.0 | 4 |
| scf7180000020543 | 77622 | 0.4254 | 0 | 44.6453 | Eukaryota | 13852.0 | 0 | Chordata | 13852.0 | 0 | Carnivora | 13852.0 | 0 | Felidae | 13852.0 | 0 | Leopardus | 5476.0 | 5 | Felis catus | 1533.0 | 9 |
| scf7180000020531 | 107026 | 0.4165 | 0 | 25.4606 | Eukaryota | 9566.0 | 0 | Chordata | 9566.0 | 0 | Carnivora | 7988.0 | 1 | Felidae | 7988.0 | 1 | Felis | 7988.0 | 1 | Felis catus | 7988.0 | 1 |
| scf7180000019044 | 890077 | 0.4893 | 0 | 49.4462 | Eukaryota | 32630.0 | 0 | Chordata | 32630.0 | 0 | Artiodactyla | 25294.0 | 2 | Delphinidae | 19314.0 | 3 | agenorhynchu | 6574.0 | 6 | rhynchus obli | 6574.0 | 6 |
| scf7180000023781 | 2235489 | 0.3999 | 0 | 51.5537 | Eukaryota | 52240.0 | 0 | Chordata | 52240.0 | 0 | Carnivora | 52240.0 | 0 | Felidae | 29640.0 | 3 | Felis | 18430.0 | 5 | Felis catus | 18430.0 | 5 |
| scf7180000023994 | 642721 | 0.3542 | 0 | 49.4449 | Eukaryota | 22196.0 | 0 | Chordata | 22196.0 | 0 | Carnivora | 22196.0 | 0 | Herpestidae | 13620.0 | 1 | Suricata | 13620.0 | 2 | ricata suricat | 13620.0 | 2 |
| scf7180000022579 | 17324 | 0.5249 | 0 | 49.6452 | Eukaryota | 2180.0 | 0 | Chordata | 2180.0 | 0 | Carnivora | 2180.0 | 0 | Felidae | 1954.0 | 1 | Felis | 1954.0 | 1 | Felis catus | 1954.0 | 1 |
| scf7180000017990 | 1208474 | 0.4573 | 0 | 47.5951 | Eukaryota | 36236.0 | 0 | Chordata | 36236.0 | 0 | Carnivora | 36236.0 | 0 | Felidae | 28774.0 | 1 | Acinonyx | 7750.0 | 4 | cinonyx jubat | 7750.0 | 5 |
| scf7180000019671 | 242963 | 0.5235 | 0 | 51.3747 | Eukaryota | 17074.0 | 0 | Chordata | 17074.0 | 0 | Carnivora | 17074.0 | 0 | Felidae | 10298.0 | 1 | Suricata | 6776.0 | 2 | ricata suricat | 6776.0 | 2 |
| scf7180000020372 | 977988 | 0.368 | 0 | 49.9982 | Eukaryota | 47438.0 | 0 | Chordata | 47438.0 | 0 | Carnivora | 47438.0 | 0 | Felidae | 47438.0 | 0 | Felis | 47438.0 | 0 | Felis catus | 47438.0 | 0 |
| scf7180000022068 | 10385 | 0.5198 | 0 | 148.2838 | no-hit | 0.0 | 0 | no-hit | 0.0 | 0 | no-hit | 0.0 | 0 | no-hit | 0.0 | 0 | no-hit | 0.0 | 0 | no-hit | 0.0 | 0 |
| scf7180000020386 | 18785 | 0.4259 | 0 | 422.4251 | Eukaryota | 197378.0 | 0 | Chordata | 197378.0 | 0 | Carnivora | 197378.0 | 0 | Hyaenidae | 197378.0 | 0 | Proteles | 121338.0 | 1 | roteles cristat | 121338.0 | 1 |
| scf7180000021958 | 21920 | 0.4296 | 0 | 26.6522 | Eukaryota | 41683.0 | 0 | Chordata | 41683.0 | 0 | Carnivora | 41683.0 | 0 | Felidae | 41683.0 | 0 | Felis | 41683.0 | 0 | Felis catus | 41683.0 | 0 |
| scf7180000018512 | 953672 | 0.4515 | 0 | 53.0123 | Eukaryota | 41450.0 | 0 | Chordata | 41450.0 | 0 | Carnivora | 41450.0 | 0 | Felidae | 41450.0 | 0 | Felis | 37331.0 | 1 | Felis catus | 37331.0 | 1 |
| scf7180000020291 | 122846 | 0.4471 | 0 | 25.6561 | Eukaryota | 37065.0 | 0 | Chordata | 37065.0 | 0 | Carnivora | 37065.0 | 0 | Ursidae | 18525.0 | 3 | Ursus | 9675.0 | 4 | Ursus arctos | 9675.0 | 4 |
| scf7180000018476 | 29584 | 0.382 | 0 | 61.4811 | Eukaryota | 13647.0 | 0 | Chordata | 13647.0 | 0 | Carnivora | 13647.0 | 0 | Felidae | 10904.0 | 2 | Panthera | 6753.0 | 4 | anthera pardu | 4085.0 | 5 |
| scf7180000022637 | 15234 | 0.3736 | 0 | 26.1323 | Eukaryota | 5710.0 | 0 | Chordata | 5710.0 | 0 | Primates | 4134.0 | 2 | Hominidae | 4134.0 | 2 | Homo | 3759.0 | 3 | Homo sapiens | 3759.0 | 3 |
| scf7180000020072 | 33283 | 0.4596 | 0 | 247.3745 | Eukaryota | 23845.0 | 0 | Chordata | 23845.0 | 0 | Carnivora | 23845.0 | 0 | Felidae | 23845.0 | 0 | Felis | 21442.0 | 1 | Felis catus | 21442.0 | 1 |
| scf7180000021920 | 5669 | 0.5209 | 0 | 84.9148 | no-hit | 0.0 | 0 | no-hit | 0.0 | 0 | no-hit | 0.0 | 0 | no-hit | 0.0 | 0 | no-hit | 0.0 | 0 | no-hit | 0.0 | 0 |
| scf7180000023909 | 639121 | 0.3754 | 0 | 49.0217 | Eukaryota | 46498.0 | 0 | Chordata | 46498.0 | 0 | Carnivora | 46498.0 | 0 | Felidae | 46498.0 | 0 | Felis | 46498.0 | 0 | Felis catus | 46498.0 | 0 |
| scf7180000021509 | 54580 | 0.4529 | 0 | 25.9174 | Eukaryota | 8229.0 | 0 | Chordata | 8229.0 | 0 | Carnivora | 8229.0 | 0 | Felidae | 8229.0 | 0 | Felis | 6608.0 | 1 | Felis catus | 6608.0 | 1 |
| scf7180000018045 | 1039060 | 0.3954 | 0 | 50.6737 | Eukaryota | 46530.0 | 0 | Chordata | 46530.0 | 0 | Carnivora | 46530.0 | 0 | Felidae | 46530.0 | 0 | Felis | 41340.0 | 1 | Felis catus | 41340.0 | 1 |
| scf7180000021595 | 35016 | 0.4826 | 0 | 49.0845 | Eukaryota | 3903.0 | 1 | Chordata | 3903.0 | 1 | Carnivora | 1563.0 | 3 | Felidae | 1563.0 | 4 | undef | 1473.0 | 8 | tic synthetic c | 1473.0 | 8 |
| scf7180000020013 | 190116 | 0.4184 | 0 | 50.3434 | Eukaryota | 6769.0 | 0 | Chordata | 6769.0 | 0 | Primates | 3203.0 | 3 | Hominidae | 3203.0 | 3 | Homo | 2510.0 | 4 | Homo sapiens | 2510.0 | 4 |
| scf7180000020582 | 60308 | 0.5106 | 0 | 48.2658 | Eukaryota | 4061.0 | 0 | Chordata | 4061.0 | 0 | Carnivora | 4061.0 | 0 | Felidae | 3700.0 | 1 | Felis | 2151.0 | 3 | Felis catus | 2151.0 | 3 |
| scf7180000023526 | 17498 | 0.4628 | 0 | 239.7461 | no-hit | 0.0 | 0 | no-hit | 0.0 | 0 | no-hit | 0.0 | 0 | no-hit | 0.0 | 0 | no-hit | 0.0 | 0 | no-hit | 0.0 | 0 |
| scf7180000017509 | 204610 | 0.3852 | 0 | 50.2736 | Eukaryota | 45081.0 | 0 | Chordata | 45081.0 | 0 | Carnivora | 45081.0 | 0 | Felidae | 45081.0 | 0 | Felis | 45081.0 | 0 | Felis catus | 45081.0 | 0 |
| scf7180000018252 | 26356 | 0.3592 | 0 | 31.3456 | Eukaryota | 19338.0 | 0 | Chordata | 19338.0 | 0 | Carnivora | 19338.0 | 0 | Felidae | 19338.0 | 0 | Felis | 17328.0 | 1 | Felis catus | 17328.0 | 1 |
| scf7180000021035 | 41462 | 0.5335 | 0 | 134.6186 | Eukaryota | 19785.0 | 0 | Chordata | 19785.0 | 0 | Carnivora | 19785.0 | 0 | Felidae | 19785.0 | 0 | Felis | 19785.0 | 0 | Felis catus | 19785.0 | 0 |
| scf7180000019094 | 407503 | 0.4066 | 0 | 50.0663 | Eukaryota | 29604.0 | 0 | Chordata | 29604.0 | 0 | Carnivora | 29604.0 | 0 | Felidae | 29604.0 | 0 | Felis | 26543.0 | 1 | Felis catus | 26543.0 | 1 |
| scf7180000022440 | 11190 | 0.4371 | 0 | 21.5205 | Eukaryota | 3392.0 | 0 | Chordata | 3392.0 | 0 | Carnivora | 2153.0 | 2 | Felidae | 1894.0 | 3 | Felis | 831.0 | 5 | Felis catus | 831.0 | 5 |
| scf7180000018360 | 834847 | 0.4674 | 0 | 51.406 | Eukaryota | 53730.0 | 0 | Chordata | 53730.0 | 0 | Carnivora | 53730.0 | 0 | Felidae | 48398.0 | 1 | Acinonyx | 26870.0 | 2 | cinonyx jubat | 26870.0 | 2 |
| scf7180000022123 | 36503 | 0.4245 | 100 | 27.1443 | Eukaryota | 11769.0 | 0 | Chordata | 11769.0 | 0 | Carnivora | 11769.0 | 0 | Felidae | 11769.0 | 0 | Felis | 9424.0 | 1 | Felis catus | 9424.0 | 1 |
| scf7180000021124 | 57110 | 0.3889 | 0 | 26.4204 | Eukaryota | 17150.0 | 0 | Chordata | 17150.0 | 0 | Carnivora | 17150.0 | 0 | Felidae | 17150.0 | 0 | Felis | 15438.0 | 1 | Felis catus | 15438.0 | 1 |
| scf7180000018568 | 706542 | 0.4486 | 0 | 53.0472 | Eukaryota | 43951.0 | 0 | Chordata | 43951.0 | 0 | Carnivora | 35801.0 | 1 | Felidae | 27798.0 | 2 | Panthera | 8846.0 | 5 | anthera pardu | 8846.0 | 5 |
| scf7180000019888 | 529518 | 0.6161 | 0 | 51.0019 | Eukaryota | 22699.0 | 0 | Chordata | 22699.0 | 0 | Carnivora | 22699.0 | 0 | Felidae | 17690.0 | 1 | Felis | 6319.0 | 4 | Felis catus | 6319.0 | 4 |
| scf7180000021138 | 53045 | 0.473 | 0 | 47.5701 | Eukaryota | 5686.0 | 0 | Chordata | 5686.0 | 0 | Carnivora | 5686.0 | 0 | Felidae | 5686.0 | 0 | Felis | 5686.0 | 0 | Felis catus | 5686.0 | 0 |
| scf7180000022557 | 18123 | 0.3549 | 0 | 70.3174 | Eukaryota | 2559.0 | 0 | Chordata | 2559.0 | 0 | Carnivora | 2154.0 | 1 | Felidae | 1906.0 | 2 | Felis | 1678.0 | 3 | Felis catus | 1678.0 | 3 |
| scf7180000017835 | 979283 | 0.3493 | 0 | 48.736 | Eukaryota | 47732.0 | 0 | Chordata | 47732.0 | 0 | Carnivora | 47732.0 | 0 | Felidae | 47732.0 | 0 | Felis | 47732.0 | 0 | Felis catus | 47732.0 | 0 |
| scf7180000021902 | 28111 | 0.3583 | 0 | 25.0614 | Eukaryota | 14206.0 | 0 | Chordata | 14206.0 | 0 | Carnivora | 14206.0 | 0 | Phocidae | 4365.0 | 4 | Neomonachus | 2906.0 | 7 | nachus schau | 2906.0 | 7 |
| scf7180000020872 | 62000 | 0.3775 | 0 | 25.5341 | Eukaryota | 43111.0 | 0 | Chordata | 43111.0 | 0 | Carnivora | 43111.0 | 0 | Felidae | 43111.0 | 0 | Felis | 43111.0 | 0 | Felis catus | 43111.0 | 0 |
| scf7180000020353 | 107589 | 0.5598 | 0 | 49.133 | Eukaryota | 14369.0 | 0 | Chordata | 14369.0 | 0 | Carnivora | 7212.0 | 1 | Felidae | 7212.0 | 1 | Eptesicus | 5764.0 | 5 | ptesicus fuscu | 5764.0 | 5 |
| scf7180000023280 | 45746 | 0.5195 | 0 | 219.5614 | Eukaryota | 4591.0 | 0 | Chordata | 4591.0 | 0 | Carnivora | 4591.0 | 0 | Felidae | 4591.0 | 0 | Felis | 4319.0 | 1 | Felis catus | 4319.0 | 1 |
| scf7180000022280 | 23952 | 0.3701 | 0 | 50.5493 | Eukaryota | 6362.0 | 0 | Chordata | 6362.0 | 0 | Primates | 2784.0 | 2 | Hominidae | 2784.0 | 3 | Homo | 2784.0 | 4 | Homo sapiens | 2784.0 | 4 |
| scf7180000017819 | 1482206 | 0.4405 | 0 | 52.7437 | Eukaryota | 46764.0 | 0 | Chordata | 46764.0 | 0 | Carnivora | 46764.0 | 0 | Felidae | 46764.0 | 0 | Felis | 32823.0 | 1 | Felis catus | 32823.0 | 1 |
| scf7180000021742 | 29878 | 0.3858 | 0 | 23.7656 | Eukaryota | 34652.0 | 0 | Chordata | 34652.0 | 0 | Carnivora | 34652.0 | 0 | Felidae | 34652.0 | 0 | Felis | 34652.0 | 0 | Felis catus | 34652.0 | 0 |
| scf7180000018991 | 1080828 | 0.4362 | 0 | 52.3652 | Eukaryota | 47159.0 | 0 | Chordata | 47159.0 | 0 | Carnivora | 47159.0 | 0 | Felidae | 47159.0 | 0 | Felis | 47159.0 | 0 | Felis catus | 47159.0 | 0 |
| scf7180000020848 | 71365 | 0.4048 | 0 | 51.1866 | Eukaryota | 23203.0 | 0 | Chordata | 23203.0 | 0 | Perissodactyla | 13039.0 | 1 | Equidae | 13039.0 | 1 | Equus | 13039.0 | 4 | uus przewals | 8652.0 | 6 |
| scf7180000017497 | 3865597 | 0.3561 | 0 | 49.6901 | Eukaryota | 47038.0 | 0 | Chordata | 47038.0 | 0 | Carnivora | 47038.0 | 0 | Felidae | 47038.0 | 0 | Felis | 47038.0 | 0 | Felis catus | 47038.0 | 0 |
| scf7180000020585 | 154909 | 0.4958 | 0 | 26.8764 | Eukaryota | 35893.0 | 0 | Chordata | 35893.0 | 0 | Carnivora | 35893.0 | 0 | Felidae | 35893.0 | 0 | Felis | 35893.0 | 0 | Felis catus | 35893.0 | 0 |
| scf7180000017961 | 271390 | 0.3681 | 0 | 48.8114 | Eukaryota | 45928.0 | 0 | Chordata | 45928.0 | 0 | Carnivora | 45928.0 | 0 | Felidae | 45928.0 | 0 | Felis | 45928.0 | 0 | Felis catus | 45928.0 | 0 |
| scf7180000019930 | 185701 | 0.4002 | 0 | 25.8665 | Eukaryota | 45654.0 | 0 | Chordata | 45654.0 | 0 | Carnivora | 45654.0 | 0 | Felidae | 45654.0 | 0 | Felis | 41146.0 | 1 | Felis catus | 41146.0 | 1 |
| scf7180000023908 | 177707 | 0.3545 | 0 | 49.1911 | Eukaryota | 47470.0 | 0 | Chordata | 47470.0 | 0 | Carnivora | 47470.0 | 0 | Felidae | 47470.0 | 0 | Felis | 47470.0 | 0 | Felis catus | 47470.0 | 0 |
| scf7180000017985 | 1255180 | 0.4438 | 0 | 51.6684 | Eukaryota | 45895.0 | 0 | Chordata | 45895.0 | 0 | Carnivora | 45895.0 | 0 | Felidae | 40428.0 | 1 | Felis | 40428.0 | 1 | Felis catus | 40428.0 | 1 |
| scf7180000019263 | 21207 | 0.4382 | 0 | 33.3311 | Eukaryota | 23284.0 | 0 | Chordata | 23284.0 | 0 | Carnivora | 23284.0 | 0 | Felidae | 23284.0 | 0 | Felis | 23284.0 | 0 | Felis catus | 23284.0 | 0 |
| scf7180000023601 | 39997 | 0.4316 | 0 | 444.5569 | no-hit | 0.0 | 0 | no-hit | 0.0 | 0 | no-hit | 0.0 | 0 | no-hit | 0.0 | 0 | no-hit | 0.0 | 0 | no-hit | 0.0 | 0 |
| scf7180000023865 | 1023733 | 0.3827 | 0 | 50.5321 | Eukaryota | 47058.0 | 0 | Chordata | 47058.0 | 0 | Carnivora | 47058.0 | 0 | Felidae | 47058.0 | 0 | Felis | 47058.0 | 0 | Felis catus | 47058.0 | 0 |
| scf7180000020619 | 29119 | 0.3929 | 0 | 50.2023 | Eukaryota | 7881.0 | 0 | Chordata | 7881.0 | 0 | Carnivora | 7881.0 | 0 | Felidae | 7881.0 | 0 | Felis | 7105.0 | 1 | Felis catus | 7105.0 | 1 |
| scf7180000022336 | 22125 | 0.4367 | 0 | 26.5507 | Eukaryota | 23099.0 | 0 | Chordata | 23099.0 | 0 | Carnivora | 23099.0 | 0 | Canidae | 8554.0 | 3 | Lynx | 7614.0 | 5 | ynx canadens | 7614.0 | 5 |
| scf7180000021224 | 13294 | 0.4047 | 0 | 20.0694 | Eukaryota | 21292.0 | 0 | Chordata | 21292.0 | 0 | Carnivora | 21292.0 | 0 | Felidae | 21292.0 | 0 | Felis | 21292.0 | 0 | Felis catus | 21292.0 | 0 |
| scf7180000019416 | 275403 | 0.5047 | 0 | 49.5011 | Eukaryota | 48123.0 | 0 | Chordata | 48123.0 | 0 | Carnivora | 48123.0 | 0 | Felidae | 48123.0 | 0 | Panthera | 32886.0 | 2 | anthera pardu | 32886.0 | 2 |
| scf7180000023136 | 28527 | 0.3903 | 0 | 401.5203 | Eukaryota | 6229.0 | 0 | Chordata | 6229.0 | 0 | Carnivora | 6229.0 | 0 | Felidae | 6229.0 | 0 | Felis | 6229.0 | 0 | Felis catus | 6229.0 | 0 |
| scf7180000020424 | 111020 | 0.4145 | 0 | 47.259 | Eukaryota | 45998.0 | 0 | Chordata | 45998.0 | 0 | Carnivora | 45998.0 | 0 | Felidae | 45998.0 | 0 | Felis | 45998.0 | 0 | Felis catus | 45998.0 | 0 |
| scf7180000020278 | 653795 | 0.3967 | 0 | 50.542 | Eukaryota | 33063.0 | 0 | Chordata | 33063.0 | 0 | Carnivora | 33063.0 | 0 | Felidae | 33063.0 | 0 | Felis | 33063.0 | 0 | Felis catus | 33063.0 | 0 |
| scf7180000021638 | 34551 | 0.5671 | 0 | 50.5868 | Eukaryota | 2998.0 | 0 | Chordata | 2998.0 | 0 | Carnivora | 2685.0 | 1 | Canidae | 1776.0 | 3 | Canis | 1480.0 | 4 | Canis lupus | 1480.0 | 5 |
| scf7180000020896 | 157402 | 0.4694 | 0 | 47.6725 | Eukaryota | 22275.0 | 0 | Chordata | 22275.0 | 0 | Carnivora | 22275.0 | 0 | Felidae | 22275.0 | 0 | Felis | 20060.0 | 1 | Felis catus | 20060.0 | 1 |
| scf7180000019115 | 483306 | 0.3437 | 0 | 47.4589 | Eukaryota | 45962.0 | 0 | Chordata | 45962.0 | 0 | Carnivora | 45962.0 | 0 | Felidae | 45962.0 | 0 | Felis | 45962.0 | 0 | Felis catus | 45962.0 | 0 |
| scf7180000022382 | 14817 | 0.4821 | 0 | 24.3956 | Eukaryota | 7061.0 | 0 | Chordata | 7061.0 | 0 | Primates | 3604.0 | 2 | Hominidae | 3604.0 | 4 | Homo | 1808.0 | 5 | Homo sapiens | 1808.0 | 5 |
| scf7180000019728 | 232415 | 0.3879 | 0 | 24.6502 | Eukaryota | 16587.0 | 0 | Chordata | 16587.0 | 0 | Carnivora | 16587.0 | 0 | Felidae | 14490.0 | 1 | Felis | 10253.0 | 3 | Felis catus | 10253.0 | 3 |
| scf7180000022288 | 18090 | 0.4397 | 0 | 27.0345 | Eukaryota | 3873.0 | 0 | Chordata | 3873.0 | 0 | Carnivora | 2400.0 | 1 | Canidae | 1956.0 | 2 | Homo | 1473.0 | 3 | Homo sapiens | 1473.0 | 3 |
| scf7180000023824 | 1802253 | 0.3934 | 0 | 50.8121 | Eukaryota | 47110.0 | 0 | Chordata | 47110.0 | 0 | Carnivora | 47110.0 | 0 | Felidae | 47110.0 | 0 | Felis | 47110.0 | 0 | Felis catus | 47110.0 | 0 |
| scf7180000019769 | 187023 | 0.4245 | 0 | 52.2228 | Eukaryota | 53627.0 | 0 | Chordata | 53627.0 | 0 | Carnivora | 53627.0 | 0 | Felidae | 43257.0 | 2 | Lynx | 16284.0 | 5 | ynx canadens | 16284.0 | 5 |
| scf7180000021887 | 33546 | 0.5129 | 0 | 50.1472 | Eukaryota | 4888.0 | 0 | Chordata | 4888.0 | 0 | Carnivora | 4888.0 | 0 | Felidae | 4553.0 | 1 | Panthera | 1411.0 | 5 | ynx canadens | 998.0 | 6 |
| scf7180000017863 | 30223 | 0.3819 | 0 | 50.3789 | Eukaryota | 8244.0 | 0 | Chordata | 8244.0 | 0 | Carnivora | 8244.0 | 0 | Felidae | 8244.0 | 0 | Panthera | 7422.0 | 1 | anthera pardu | 6576.0 | 2 |
| scf7180000017431 | 2964028 | 0.3804 | 0 | 52.1472 | Eukaryota | 62015.0 | 0 | Chordata | 62015.0 | 0 | Carnivora | 62015.0 | 0 | Ursidae | 23487.0 | 3 | Ursus | 23487.0 | 4 | Ursus arctos | 17756.0 | 5 |
| scf7180000024261 | 347269 | 0.3864 | 100 | 48.2303 | Eukaryota | 45463.0 | 0 | Chordata | 45463.0 | 0 | Carnivora | 45463.0 | 0 | Felidae | 45463.0 | 0 | Felis | 45463.0 | 0 | Felis catus | 45463.0 | 0 |
| scf7180000018759 | 28065 | 0.409 | 0 | 40.7557 | Eukaryota | 46233.0 | 0 | Chordata | 46233.0 | 0 | Carnivora | 46233.0 | 0 | Felidae | 46233.0 | 0 | Felis | 46233.0 | 0 | Felis catus | 46233.0 | 0 |
| scf7180000020915 | 79657 | 0.3768 | 0 | 25.9904 | Eukaryota | 47232.0 | 0 | Chordata | 47232.0 | 0 | Carnivora | 47232.0 | 0 | Felidae | 47232.0 | 0 | Felis | 47232.0 | 0 | Felis catus | 47232.0 | 0 |
| scf7180000018117 | 207292 | 0.3637 | 0 | 47.8895 | Eukaryota | 44285.0 | 0 | Chordata | 44285.0 | 0 | Carnivora | 44285.0 | 0 | Felidae | 44285.0 | 0 | Felis | 44285.0 | 0 | Felis catus | 44285.0 | 0 |
| scf7180000024047 | 395093 | 0.5269 | 0 | 52.3327 | Eukaryota | 33918.0 | 0 | Chordata | 33918.0 | 0 | Carnivora | 33918.0 | 0 | Felidae | 33918.0 | 0 | Felis | 33918.0 | 0 | Felis catus | 33918.0 | 0 |
| scf7180000021600 | 33952 | 0.3652 | 0 | 23.5929 | Eukaryota | 40938.0 | 0 | Chordata | 40938.0 | 0 | Carnivora | 40938.0 | 0 | Felidae | 40938.0 | 0 | Felis | 40938.0 | 0 | Felis catus | 40938.0 | 0 |
| scf7180000020491 | 99872 | 0.6472 | 0 | 51.5578 | Eukaryota | 94201.0 | 0 | Chordata | 94201.0 | 0 | Carnivora | 60187.0 | 3 | Felidae | 33079.0 | 4 | Canis | 20048.0 | 7 | Canis lupus | 20048.0 | 7 |
| scf7180000020339 | 113550 | 0.398 | 0 | 51.2562 | Eukaryota | 42538.0 | 0 | Chordata | 42538.0 | 0 | Carnivora | 42538.0 | 0 | Felidae | 42538.0 | 0 | Felis | 42538.0 | 0 | Felis catus | 42538.0 | 0 |
| scf7180000018069 | 252601 | 0.4011 | 0 | 49.801 | Eukaryota | 46852.0 | 0 | Chordata | 46852.0 | 0 | Carnivora | 46852.0 | 0 | Felidae | 46852.0 | 0 | Felis | 46852.0 | 0 | Felis catus | 46852.0 | 0 |
| scf7180000023802 | 2799608 | 0.4073 | 0 | 51.1958 | Eukaryota | 51291.0 | 0 | Chordata | 51291.0 | 0 | Carnivora | 51291.0 | 0 | Felidae | 39951.0 | 1 | Felis | 28477.0 | 2 | Felis catus | 28477.0 | 2 |
| scf7180000018704 | 698215 | 0.4031 | 0 | 51.7751 | Eukaryota | 48554.0 | 0 | Chordata | 48554.0 | 0 | Carnivora | 48554.0 | 0 | Felidae | 48554.0 | 0 | Panthera | 38892.0 | 1 | anthera pardu | 34027.0 | 2 |
| scf7180000023757 | 4322580 | 0.362 | 0 | 49.3905 | Eukaryota | 46966.0 | 0 | Chordata | 46966.0 | 0 | Carnivora | 46966.0 | 0 | Felidae | 46966.0 | 0 | Felis | 32617.0 | 1 | Felis catus | 32617.0 | 1 |
| scf7180000021534 | 39123 | 0.4147 | 0 | 24.4726 | Eukaryota | 30312.0 | 0 | Chordata | 30312.0 | 0 | Carnivora | 30312.0 | 0 | Felidae | 30312.0 | 0 | Felis | 27215.0 | 1 | Felis catus | 27215.0 | 1 |
| scf7180000020586 | 25655 | 0.4606 | 0 | 1399.5096 | Eukaryota | 22916.0 | 0 | Chordata | 22916.0 | 0 | Carnivora | 22916.0 | 0 | Felidae | 22916.0 | 0 | Felis | 22916.0 | 0 | Felis catus | 22916.0 | 0 |
| scf7180000022921 | 31195 | 0.5779 | 0 | 239.4483 | Eukaryota | 3645.0 | 0 | Chordata | 3645.0 | 0 | Carnivora | 3645.0 | 0 | Felidae | 2427.0 | 1 | Felis | 1861.0 | 2 | Felis catus | 1861.0 | 2 |
| scf7180000020670 | 78298 | 0.5704 | 0 | 45.6633 | Eukaryota | 18074.0 | 0 | Chordata | 18074.0 | 0 | Carnivora | 18074.0 | 0 | Felidae | 14246.0 | 1 | Panthera | 10718.0 | 2 | anthera pardu | 10718.0 | 2 |
| scf7180000020309 | 124210 | 0.3871 | 0 | 46.4199 | Eukaryota | 27959.0 | 0 | Chordata | 27959.0 | 0 | Carnivora | 27959.0 | 0 | Felidae | 27959.0 | 0 | Panthera | 11270.0 | 2 | anthera pardu | 8466.0 | 3 |
| scf7180000020406 | 111484 | 0.393 | 0 | 25.4204 | Eukaryota | 80275.0 | 0 | Chordata | 80275.0 | 0 | Carnivora | 80275.0 | 0 | Felidae | 80275.0 | 0 | Lynx | 40050.0 | 2 | ynx canadens | 40050.0 | 2 |
| scf7180000018436 | 92461 | 0.4016 | 0 | 50.8948 | Eukaryota | 33819.0 | 0 | Chordata | 33819.0 | 0 | Carnivora | 33819.0 | 0 | Felidae | 33819.0 | 0 | Felis | 30465.0 | 1 | Felis catus | 30465.0 | 1 |
| scf7180000023106 | 15170 | 0.5013 | 0 | 259.766 | Eukaryota | 5984.0 | 0 | Chordata | 5984.0 | 0 | Carnivora | 5984.0 | 0 | Phocidae | 2440.0 | 2 | Leptonychote | 1830.0 | 4 | nychotes wed | 1830.0 | 4 |
| scf7180000018037 | 1849665 | 0.3988 | 0 | 51.2777 | Eukaryota | 48611.0 | 0 | Chordata | 48611.0 | 0 | Carnivora | 48611.0 | 0 | Felidae | 48611.0 | 0 | Felis | 29041.0 | 2 | Felis catus | 29041.0 | 2 |
| scf7180000021879 | 44024 | 0.3562 | 0 | 24.7587 | Eukaryota | 26412.0 | 0 | Chordata | 26412.0 | 0 | Carnivora | 26412.0 | 0 | Felidae | 26412.0 | 0 | Felis | 23758.0 | 1 | Felis catus | 23758.0 | 1 |
| scf7180000022322 | 14545 | 0.3704 | 0 | 26.8512 | Eukaryota | 22042.0 | 0 | Chordata | 22042.0 | 0 | Carnivora | 22042.0 | 0 | Felidae | 22042.0 | 0 | Felis | 19814.0 | 1 | Felis catus | 19814.0 | 1 |
| scf7180000019053 | 824068 | 0.5054 | 0 | 52.4233 | Eukaryota | 33658.0 | 0 | Chordata | 33658.0 | 0 | Carnivora | 33658.0 | 0 | Felidae | 21240.0 | 1 | Canis | 12418.0 | 4 | Canis lupus | 12418.0 | 4 |
| scf7180000020708 | 85345 | 0.4357 | 0 | 25.2978 | Eukaryota | 44674.0 | 0 | Chordata | 44674.0 | 0 | Carnivora | 44674.0 | 0 | Felidae | 44674.0 | 0 | Felis | 44674.0 | 0 | Felis catus | 44674.0 | 0 |
| scf7180000022169 | 2716 | 0.4091 | 0 | 433.758 | no-hit | 0.0 | 0 | no-hit | 0.0 | 0 | no-hit | 0.0 | 0 | no-hit | 0.0 | 0 | no-hit | 0.0 | 0 | no-hit | 0.0 | 0 |
| scf7180000019371 | 183302 | 0.4017 | 0 | 46.9011 | Eukaryota | 38011.0 | 0 | Chordata | 38011.0 | 0 | Carnivora | 38011.0 | 0 | Felidae | 38011.0 | 0 | Felis | 34230.0 | 1 | Felis catus | 34230.0 | 1 |
| scf7180000017579 | 3833756 | 0.4528 | 100 | 50.5951 | Eukaryota | 46719.0 | 0 | Chordata | 46719.0 | 0 | Carnivora | 46719.0 | 0 | Felidae | 46719.0 | 0 | Felis | 46719.0 | 0 | Felis catus | 46719.0 | 0 |
| scf7180000020844 | 63754 | 0.3998 | 0 | 23.7718 | Eukaryota | 44661.0 | 0 | Chordata | 44661.0 | 0 | Carnivora | 44661.0 | 0 | Felidae | 44661.0 | 0 | Felis | 44661.0 | 0 | Felis catus | 44661.0 | 0 |
| scf7180000023364 | 28127 | 0.5109 | 0 | 435.9161 | no-hit | 0.0 | 0 | no-hit | 0.0 | 0 | no-hit | 0.0 | 0 | no-hit | 0.0 | 0 | no-hit | 0.0 | 0 | no-hit | 0.0 | 0 |
| scf7180000021109 | 103186 | 0.4145 | 0 | 27.4323 | Eukaryota | 69600.0 | 0 | Chordata | 69600.0 | 0 | Carnivora | 69600.0 | 0 | Felidae | 55498.0 | 1 | Acinonyx | 41628.0 | 2 | cinonyx jubat | 41628.0 | 2 |
| scf7180000019761 | 225827 | 0.5041 | 0 | 51.0273 | Eukaryota | 6472.0 | 1 | Chordata | 6472.0 | 1 | Artiodactyla | 3037.0 | 4 | Hominidae | 2157.0 | 5 | Homo | 2157.0 | 6 | Homo sapiens | 2157.0 | 6 |
| scf7180000019751 | 17531 | 0.6624 | 0 | 276.8832 | Eukaryota | 2542.0 | 0 | Chordata | 2542.0 | 0 | Carnivora | 2299.0 | 1 | Felidae | 2299.0 | 1 | Felis | 1673.0 | 3 | Felis catus | 1673.0 | 3 |
| scf7180000021987 | 43407 | 0.3816 | 0 | 27.0008 | Eukaryota | 8775.0 | 0 | Chordata | 8775.0 | 0 | Carnivora | 8775.0 | 0 | Felidae | 7888.0 | 1 | Felis | 7888.0 | 1 | Felis catus | 7888.0 | 1 |
| scf7180000020719 | 171825 | 0.4362 | 100 | 25.448 | Eukaryota | 13169.0 | 0 | Chordata | 13169.0 | 0 | Carnivora | 6240.0 | 3 | Felidae | 6240.0 | 3 | Felis | 6240.0 | 3 | Felis catus | 6240.0 | 3 |
| scf7180000020270 | 160974 | 0.5096 | 0 | 50.4907 | Eukaryota | 19509.0 | 0 | Chordata | 19509.0 | 0 | Carnivora | 17874.0 | 1 | Felidae | 13238.0 | 2 | Suricata | 4636.0 | 5 | ricata suricat | 4636.0 | 6 |
| scf7180000019160 | 391604 | 0.3481 | 0 | 47.3334 | Eukaryota | 32221.0 | 0 | Chordata | 32221.0 | 0 | Carnivora | 32221.0 | 0 | Felidae | 32221.0 | 0 | Felis | 29018.0 | 1 | Felis catus | 29018.0 | 1 |
| scf7180000023298 | 27022 | 0.3838 | 0 | 227.8784 | Eukaryota | 25357.0 | 0 | Chordata | 25357.0 | 0 | Carnivora | 25357.0 | 0 | Felidae | 25357.0 | 0 | Felis | 22808.0 | 1 | Felis catus | 22808.0 | 1 |
| scf7180000021829 | 30660 | 0.3776 | 0 | 23.6963 | Eukaryota | 18124.0 | 0 | Chordata | 18124.0 | 0 | Carnivora | 18124.0 | 0 | Felidae | 18124.0 | 0 | Felis | 14499.0 | 1 | Felis catus | 14499.0 | 1 |
| scf7180000020082 | 245348 | 0.3877 | 0 | 49.6708 | Eukaryota | 47464.0 | 0 | Chordata | 47464.0 | 0 | Carnivora | 47464.0 | 0 | Felidae | 47464.0 | 0 | Felis | 47464.0 | 0 | Felis catus | 47464.0 | 0 |
| scf7180000017909 | 1597114 | 0.4539 | 0 | 52.8941 | Eukaryota | 64282.0 | 0 | Chordata | 64282.0 | 0 | Carnivora | 64282.0 | 0 | Felidae | 52552.0 | 1 | Felis | 21970.0 | 4 | Felis catus | 21970.0 | 5 |
| scf7180000021378 | 38855 | 0.3449 | 0 | 47.3815 | Eukaryota | 5734.0 | 0 | Chordata | 5734.0 | 0 | Rodentia | 1980.0 | 4 | Muridae | 1980.0 | 4 | Mus | 1980.0 | 4 | Mus musculus | 1980.0 | 4 |
| scf7180000020220 | 9810 | 0.5083 | 0 | 24.0811 | Eukaryota | 16721.0 | 1 | Chordata | 16721.0 | 1 | Carnivora | 14284.0 | 3 | Felidae | 14284.0 | 3 | Lynx | 11160.0 | 6 | ynx canadens | 11160.0 | 6 |
| scf7180000024145 | 206188 | 0.4624 | 0 | 51.1042 | Eukaryota | 19170.0 | 0 | Chordata | 19170.0 | 0 | Carnivora | 19170.0 | 0 | Felidae | 19170.0 | 0 | Lynx | 19170.0 | 0 | ynx canadens | 19170.0 | 0 |
| scf7180000018091 | 1223426 | 0.3895 | 0 | 48.9874 | Eukaryota | 50687.0 | 0 | Chordata | 50687.0 | 0 | Carnivora | 50687.0 | 0 | Felidae | 50687.0 | 0 | Felis | 37595.0 | 2 | Felis catus | 37595.0 | 2 |
| scf7180000021850 | 7659 | 0.5526 | 0 | 262.0549 | Eukaryota | 2012.0 | 0 | Chordata | 2012.0 | 0 | Carnivora | 2012.0 | 0 | Felidae | 2012.0 | 0 | Felis | 1279.0 | 2 | Felis catus | 1279.0 | 2 |
| scf7180000023245 | 29303 | 0.3888 | 0 | 295.8091 | Eukaryota | 6888.0 | 0 | Chordata | 6888.0 | 0 | Carnivora | 6888.0 | 0 | Felidae | 6888.0 | 0 | Felis | 6888.0 | 0 | Felis catus | 6888.0 | 0 |
| scf7180000020314 | 132447 | 0.4213 | 0 | 49.7332 | Eukaryota | 7198.0 | 0 | Chordata | 7198.0 | 0 | Carnivora | 6442.0 | 1 | Felidae | 6442.0 | 1 | Felis | 5730.0 | 2 | Felis catus | 5730.0 | 2 |
| scf7180000017580 | 2101507 | 0.4087 | 0 | 51.4937 | Eukaryota | 45824.0 | 0 | Chordata | 45824.0 | 0 | Carnivora | 45824.0 | 0 | Felidae | 45824.0 | 0 | Felis | 45824.0 | 0 | Felis catus | 45824.0 | 0 |
| scf7180000021444 | 61799 | 0.5462 | 0 | 25.3168 | Eukaryota | 32558.0 | 0 | Chordata | 32558.0 | 0 | Artiodactyla | 20205.0 | 1 | Suidae | 20205.0 | 1 | Sus | 20205.0 | 2 | Sus scrofa | 20205.0 | 2 |

| | | | | | | | | | | | | | | | | | | | | | | |
|---|---|---|---|---|---|---|---|---|---|---|---|---|---|---|---|---|---|---|---|---|---|---|
| scf7180000018379 | 915405 | 0.3649 | 0 | 50.4103 | Eukaryota | 50463.0 | 0 | Chordata | 50463.0 | 0 | Carnivora | 50463.0 | 0 | Felidae | 50463.0 | 0 | Felis | 50463.0 | 0 | Felis catus | 50463.0 | 0 |
| scf7180000018614 | 28318 | 0.3729 | 0 | 47.1694 | Eukaryota | 43510.0 | 0 | Chordata | 43510.0 | 0 | Carnivora | 43510.0 | 0 | Felidae | 43510.0 | 0 | Felis | 39212.0 | 1 | Felis catus | 39212.0 | 1 |
| scf7180000021806 | 16853 | 0.4038 | 0 | 22.5278 | Eukaryota | 14668.0 | 0 | Chordata | 14668.0 | 0 | Carnivora | 14668.0 | 0 | Felidae | 14668.0 | 0 | Felis | 11747.0 | 1 | Felis catus | 11747.0 | 1 |
| scf7180000022044 | 23032 | 0.3829 | 0 | 26.5529 | Eukaryota | 35899.0 | 0 | Chordata | 35899.0 | 0 | Carnivora | 35899.0 | 0 | Felidae | 35899.0 | 0 | Felis | 35899.0 | 0 | Felis catus | 35899.0 | 0 |
| scf7180000020199 | 103293 | 0.4902 | 0 | 49.5591 | Eukaryota | 14984.0 | 0 | Chordata | 14984.0 | 0 | Carnivora | 14984.0 | 0 | Felidae | 14984.0 | 0 | Felis | 13318.0 | 1 | Felis catus | 13318.0 | 1 |
| scf7180000022712 | 12491 | 0.4058 | 0 | 25.2728 | Eukaryota | 9492.0 | 0 | Chordata | 9492.0 | 0 | Primates | 5411.0 | 3 | Hominidae | 5411.0 | 3 | Homo | 3606.0 | 4 | Homo sapiens | 3606.0 | 4 |
| scf7180000021257 | 45532 | 0.3961 | 0 | 25.9907 | Eukaryota | 45792.0 | 0 | Chordata | 45792.0 | 0 | Carnivora | 45792.0 | 0 | Felidae | 45792.0 | 0 | Felis | 41267.0 | 1 | Felis catus | 41267.0 | 1 |
| scf7180000022234 | 17856 | 0.3869 | 0 | 24.7483 | Eukaryota | 19800.0 | 0 | Chordata | 19800.0 | 0 | Carnivora | 19800.0 | 0 | Felidae | 19800.0 | 0 | Felis | 17801.0 | 1 | Felis catus | 17801.0 | 1 |
| scf7180000018721 | 599896 | 0.4645 | 100 | 50.343 | Eukaryota | 27542.0 | 0 | Chordata | 27542.0 | 0 | Carnivora | 27542.0 | 0 | Felidae | 16790.0 | 3 | Felis | 5628.0 | 6 | Felis catus | 5628.0 | 7 |
| scf7180000018470 | 4214 | 0.4089 | 0 | 27.7509 | Eukaryota | 7500.0 | 0 | Chordata | 7500.0 | 0 | Carnivora | 7500.0 | 0 | Felidae | 7500.0 | 0 | Felis | 6559.0 | 1 | Felis catus | 6559.0 | 1 |
| scf7180000018643 | 1122161 | 0.4314 | 0 | 50.6486 | Eukaryota | 47892.0 | 0 | Chordata | 47892.0 | 0 | Carnivora | 47892.0 | 0 | Felidae | 47892.0 | 0 | Felis | 42965.0 | 1 | Felis catus | 42965.0 | 1 |
| scf7180000023858 | 713747 | 0.4325 | 0 | 52.4963 | Eukaryota | 31836.0 | 0 | Chordata | 31836.0 | 0 | Carnivora | 31836.0 | 0 | Felidae | 31836.0 | 0 | Panthera | 19161.0 | 2 | anthera pardu | 15960.0 | 3 |
| scf7180000020394 | 67385 | 0.3998 | 0 | 44.4162 | Eukaryota | 46002.0 | 0 | Chordata | 46002.0 | 0 | Carnivora | 46002.0 | 0 | Felidae | 46002.0 | 0 | Felis | 41468.0 | 1 | Felis catus | 41468.0 | 1 |
| scf7180000021797 | 16457 | 0.5557 | 0 | 52.7494 | Eukaryota | 2785.0 | 0 | Chordata | 2785.0 | 0 | Carnivora | 1477.0 | 2 | Felidae | 1477.0 | 2 | Felis | 1477.0 | 3 | Felis catus | 1477.0 | 3 |
| scf7180000023716 | 13105 | 0.4069 | 0 | 263.0401 | Eukaryota | 14520.0 | 0 | Chordata | 14520.0 | 0 | Carnivora | 14520.0 | 0 | Felidae | 14520.0 | 0 | Felis | 11628.0 | 1 | Felis catus | 11628.0 | 1 |
| scf7180000021898 | 33997 | 0.3862 | 0 | 24.8975 | Eukaryota | 44132.0 | 0 | Chordata | 44132.0 | 0 | Carnivora | 44132.0 | 0 | Felidae | 44132.0 | 0 | Felis | 39790.0 | 1 | Felis catus | 39790.0 | 1 |
| scf7180000024244 | 200278 | 0.3991 | 0 | 49.5636 | Eukaryota | 39573.0 | 0 | Chordata | 39573.0 | 0 | Carnivora | 39573.0 | 0 | Felidae | 39573.0 | 0 | Felis | 35626.0 | 1 | Felis catus | 35626.0 | 1 |
| scf7180000023369 | 69284 | 0.4273 | 0 | 98.1823 | Eukaryota | 2306.0 | 0 | Chordata | 2306.0 | 0 | Carnivora | 2306.0 | 0 | Felidae | 2306.0 | 0 | Felis | 1634.0 | 2 | Felis catus | 1634.0 | 2 |
| scf7180000019901 | 160512 | 0.4804 | 0 | 50.3646 | Eukaryota | 15315.0 | 0 | Chordata | 15315.0 | 0 | Carnivora | 13222.0 | 1 | Felidae | 8382.0 | 5 | Mustela | 2542.0 | 8 | ustela putori | 2542.0 | 8 |
| scf7180000018257 | 136363 | 0.4419 | 0 | 52.5276 | Eukaryota | 46262.0 | 0 | Chordata | 46262.0 | 0 | Carnivora | 46262.0 | 0 | Felidae | 46262.0 | 0 | Felis | 46262.0 | 0 | Felis catus | 46262.0 | 0 |
| scf7180000020461 | 940276 | 0.4372 | 0 | 49.3165 | Eukaryota | 37534.0 | 0 | Chordata | 37534.0 | 0 | Carnivora | 37534.0 | 0 | Felidae | 37534.0 | 0 | Felis | 33823.0 | 1 | Felis catus | 33823.0 | 1 |
| scf7180000020217 | 12165 | 0.3577 | 0 | 21.2809 | Eukaryota | 24001.0 | 0 | Chordata | 24001.0 | 0 | Carnivora | 24001.0 | 0 | Felidae | 24001.0 | 0 | Felis | 24001.0 | 0 | Felis catus | 24001.0 | 0 |
| scf7180000020601 | 17374 | 0.4254 | 0 | 419.5729 | Eukaryota | 201644.0 | 0 | Chordata | 201644.0 | 0 | Carnivora | 201644.0 | 0 | Hyaenidae | 201644.0 | 0 | Proteles | 123739.0 | 1 | roteles cristat | 123739.0 | 1 |
| scf7180000017704 | 10217545 | 0.4197 | 0 | 51.0186 | Eukaryota | 70003.0 | 0 | Chordata | 70003.0 | 0 | Carnivora | 70003.0 | 0 | Felidae | 70003.0 | 0 | Felis | 56568.0 | 2 | Felis catus | 56568.0 | 2 |
| scf7180000019388 | 148182 | 0.4372 | 0 | 44.0797 | Eukaryota | 13865.0 | 0 | Chordata | 13865.0 | 0 | Carnivora | 13865.0 | 0 | Felidae | 13865.0 | 0 | Felis | 12489.0 | 1 | Felis catus | 12489.0 | 1 |
| scf7180000020155 | 209400 | 0.5904 | 0 | 48.2773 | Eukaryota | 31840.0 | 0 | Chordata | 31840.0 | 0 | Carnivora | 31840.0 | 0 | Felidae | 31840.0 | 0 | Lynx | 31840.0 | 0 | ynx canadens | 31840.0 | 0 |
| scf7180000020037 | 114410 | 0.4321 | 0 | 48.4557 | Eukaryota | 22086.0 | 0 | Chordata | 22086.0 | 0 | Carnivora | 22086.0 | 0 | Felidae | 22086.0 | 0 | Felis | 15272.0 | 2 | Felis catus | 15272.0 | 2 |
| scf7180000017484 | 155211 | 0.4275 | 0 | 50.9574 | Eukaryota | 47223.0 | 0 | Chordata | 47223.0 | 0 | Carnivora | 47223.0 | 0 | Felidae | 47223.0 | 0 | Felis | 47223.0 | 0 | Felis catus | 47223.0 | 0 |
| scf7180000021203 | 84747 | 0.6242 | 0 | 52.3238 | Eukaryota | 5067.0 | 0 | Chordata | 5067.0 | 0 | Carnivora | 5067.0 | 0 | Otariidae | 2970.0 | 2 | Zalophus | 2491.0 | 3 | phus californi | 2491.0 | 3 |
| scf7180000017709 | 23120 | 0.4334 | 0 | 34.6196 | Eukaryota | 27739.0 | 0 | Chordata | 27739.0 | 0 | Carnivora | 27739.0 | 0 | Felidae | 27739.0 | 0 | Felis | 24917.0 | 1 | Felis catus | 24917.0 | 1 |
| scf7180000022815 | 14309 | 0.6423 | 0 | 28.2021 | Eukaryota | 20452.0 | 0 | Chordata | 20452.0 | 0 | Primates | 16430.0 | 2 | Hominidae | 16430.0 | 2 | Homo | 13975.0 | 3 | Homo sapiens | 13975.0 | 3 |
| scf7180000019510 | 135008 | 0.3623 | 0 | 43.4726 | Eukaryota | 45823.0 | 0 | Chordata | 45823.0 | 0 | Carnivora | 45823.0 | 0 | Felidae | 45823.0 | 0 | Felis | 45823.0 | 0 | Felis catus | 45823.0 | 0 |
| scf7180000020444 | 184212 | 0.3787 | 0 | 24.8569 | Eukaryota | 45298.0 | 0 | Chordata | 45298.0 | 0 | Carnivora | 45298.0 | 0 | Felidae | 45298.0 | 0 | Felis | 45298.0 | 0 | Felis catus | 45298.0 | 0 |
| scf7180000017611 | 2936649 | 0.3605 | 0 | 49.1853 | Eukaryota | 46555.0 | 0 | Chordata | 46555.0 | 0 | Carnivora | 46555.0 | 0 | Felidae | 46555.0 | 0 | Felis | 46555.0 | 0 | Felis catus | 46555.0 | 0 |
| scf7180000023855 | 1949502 | 0.4273 | 0 | 51.2599 | Eukaryota | 30192.0 | 0 | Chordata | 30192.0 | 0 | Carnivora | 30192.0 | 0 | Felidae | 30192.0 | 0 | Lynx | 30192.0 | 0 | ynx canadens | 30192.0 | 0 |
| scf7180000021296 | 72234 | 0.3994 | 0 | 25.3462 | Eukaryota | 32690.0 | 0 | Chordata | 32690.0 | 0 | Carnivora | 32690.0 | 0 | Felidae | 22850.0 | 1 | Panthera | 13180.0 | 3 | ricata suricat | 9840.0 | 4 |
| scf7180000018812 | 498261 | 0.3673 | 0 | 49.5798 | Eukaryota | 37372.0 | 0 | Chordata | 37372.0 | 0 | Carnivora | 37372.0 | 0 | Felidae | 37372.0 | 0 | Felis | 37372.0 | 0 | Felis catus | 37372.0 | 0 |
| scf7180000019134 | 1607420 | 0.4496 | 0 | 51.7865 | Eukaryota | 88188.0 | 0 | Chordata | 88188.0 | 0 | Carnivora | 53504.0 | 1 | Felidae | 35148.0 | 2 | Felis | 35148.0 | 2 | Felis catus | 35148.0 | 2 |
| scf7180000022982 | 36268 | 0.5325 | 0 | 691.8771 | Eukaryota | 12122.0 | 0 | Chordata | 12122.0 | 0 | Carnivora | 12122.0 | 0 | Felidae | 12122.0 | 0 | Felis | 12122.0 | 0 | Felis catus | 12122.0 | 0 |
| scf7180000023816 | 1717503 | 0.4293 | 0 | 51.2159 | Eukaryota | 52490.0 | 0 | Chordata | 52490.0 | 0 | Carnivora | 52490.0 | 0 | Felidae | 52490.0 | 0 | Felis | 52490.0 | 0 | Felis catus | 52490.0 | 0 |
| scf7180000018415 | 2939148 | 0.3977 | 100 | 50.6448 | Eukaryota | 96071.0 | 0 | Chordata | 96071.0 | 0 | Carnivora | 96071.0 | 0 | Felidae | 41727.0 | 4 | Panthera | 25084.0 | 7 | anthera pardu | 16740.0 | 8 |
| scf7180000017545 | 2701704 | 0.4095 | 0 | 51.6024 | Eukaryota | 45307.0 | 0 | Chordata | 45307.0 | 0 | Carnivora | 45307.0 | 0 | Felidae | 45307.0 | 0 | Felis | 45307.0 | 0 | Felis catus | 45307.0 | 0 |
| scf7180000024190 | 102932 | 0.3711 | 0 | 23.6792 | Eukaryota | 42317.0 | 0 | Chordata | 42317.0 | 0 | Carnivora | 42317.0 | 0 | Felidae | 42317.0 | 0 | Felis | 42317.0 | 0 | Felis catus | 42317.0 | 0 |
| scf7180000018882 | 475052 | 0.4403 | 0 | 50.7467 | Eukaryota | 46512.0 | 0 | Chordata | 46512.0 | 0 | Carnivora | 46512.0 | 0 | Felidae | 46512.0 | 0 | Felis | 46512.0 | 0 | Felis catus | 46512.0 | 0 |
| scf7180000022970 | 32175 | 0.536 | 0 | 285.4122 | Eukaryota | 2452.0 | 0 | Chordata | 2452.0 | 0 | Carnivora | 2452.0 | 0 | Felidae | 2224.0 | 1 | Felis | 1010.0 | 5 | Felis catus | 1010.0 | 5 |
| scf7180000018557 | 105785 | 0.4653 | 0 | 48.2372 | Eukaryota | 31743.0 | 0 | Chordata | 31743.0 | 0 | Carnivora | 31743.0 | 0 | Felidae | 31743.0 | 0 | Felis | 28522.0 | 1 | Felis catus | 28522.0 | 1 |
| scf7180000021526 | 55430 | 0.3542 | 0 | 22.9481 | Eukaryota | 42901.0 | 0 | Chordata | 42901.0 | 0 | Carnivora | 42901.0 | 0 | Felidae | 42901.0 | 0 | Felis | 42901.0 | 0 | Felis catus | 42901.0 | 0 |
| scf7180000023013 | 50297 | 0.4094 | 0 | 463.9914 | Eukaryota | 6583.0 | 0 | Chordata | 6583.0 | 0 | Carnivora | 6583.0 | 0 | Felidae | 6583.0 | 0 | Felis | 6583.0 | 0 | Felis catus | 6583.0 | 0 |
| scf7180000019506 | 573961 | 0.456 | 0 | 53.167 | Eukaryota | 23530.0 | 0 | Chordata | 23530.0 | 0 | Carnivora | 23530.0 | 0 | Felidae | 23530.0 | 0 | Felis | 9605.0 | 3 | Felis catus | 9605.0 | 3 |
| scf7180000021261 | 59968 | 0.3839 | 0 | 31.711 | Eukaryota | 47360.0 | 0 | Chordata | 47360.0 | 0 | Carnivora | 47360.0 | 0 | Felidae | 47360.0 | 0 | Felis | 47360.0 | 0 | Felis catus | 47360.0 | 0 |
| scf7180000018676 | 512683 | 0.3749 | 0 | 48.8363 | Eukaryota | 39760.0 | 0 | Chordata | 39760.0 | 0 | Carnivora | 39760.0 | 0 | Felidae | 33826.0 | 1 | Felis | 33826.0 | 1 | Felis catus | 33826.0 | 1 |
| scf7180000017792 | 79055 | 0.601 | 0 | 47.5788 | Eukaryota | 14947.0 | 0 | Chordata | 14947.0 | 0 | Carnivora | 14947.0 | 0 | Felidae | 14947.0 | 0 | Felis | 14947.0 | 0 | Felis catus | 14947.0 | 0 |
| scf7180000018560 | 32835 | 0.3947 | 0 | 26.7133 | Eukaryota | 31370.0 | 0 | Chordata | 31370.0 | 0 | Carnivora | 31370.0 | 0 | Felidae | 31370.0 | 0 | Felis | 28273.0 | 1 | Felis catus | 28273.0 | 1 |
| scf7180000019827 | 271815 | 0.4104 | 100 | 45.3882 | Eukaryota | 27600.0 | 0 | Chordata | 27600.0 | 0 | Carnivora | 27600.0 | 0 | Felidae | 27600.0 | 0 | Felis | 21933.0 | 2 | Felis catus | 21933.0 | 2 |
| scf7180000019083 | 605897 | 0.4103 | 0 | 49.1956 | Eukaryota | 47239.0 | 0 | Chordata | 47239.0 | 0 | Carnivora | 47239.0 | 0 | Felidae | 47239.0 | 0 | Felis | 47239.0 | 0 | Felis catus | 47239.0 | 0 |
| scf7180000019773 | 22111 | 0.385 | 0 | 24.8885 | Eukaryota | 24842.0 | 0 | Chordata | 24842.0 | 0 | Carnivora | 24842.0 | 0 | Felidae | 24842.0 | 0 | Felis | 22954.0 | 1 | Felis catus | 22954.0 | 1 |
| scf7180000022672 | 17697 | 0.3848 | 0 | 22.8799 | Eukaryota | 40461.0 | 0 | Chordata | 40461.0 | 0 | Carnivora | 40461.0 | 0 | Felidae | 40461.0 | 0 | Felis | 40461.0 | 0 | Felis catus | 40461.0 | 0 |
| scf7180000021176 | 46168 | 0.3618 | 0 | 23.223 | Eukaryota | 37282.0 | 0 | Chordata | 37282.0 | 0 | Carnivora | 37282.0 | 0 | Felidae | 37282.0 | 0 | Felis | 37282.0 | 0 | Felis catus | 37282.0 | 0 |
| scf7180000021561 | 35109 | 0.3575 | 0 | 25.0994 | Eukaryota | 59952.0 | 0 | Chordata | 59952.0 | 0 | Carnivora | 59952.0 | 0 | Felidae | 59952.0 | 0 | Panthera | 47992.0 | 1 | anthera pardu | 47992.0 | 1 |
| scf7180000017888 | 1607486 | 0.3535 | 0 | 49.4216 | Eukaryota | 46249.0 | 0 | Chordata | 46249.0 | 0 | Carnivora | 46249.0 | 0 | Felidae | 46249.0 | 0 | Felis | 46249.0 | 0 | Felis catus | 46249.0 | 0 |
| scf7180000021618 | 40040 | 0.3689 | 0 | 27.7167 | Eukaryota | 19531.0 | 0 | Chordata | 19531.0 | 0 | Carnivora | 19531.0 | 0 | Felidae | 19531.0 | 0 | Felis | 17508.0 | 1 | Felis catus | 17508.0 | 1 |
| scf7180000018228 | 640058 | 0.4106 | 0 | 52.9227 | Eukaryota | 31869.0 | 0 | Chordata | 31869.0 | 0 | Carnivora | 29383.0 | 1 | Felidae | 20627.0 | 5 | Felis | 14055.0 | 8 | Felis catus | 14055.0 | 8 |
| scf7180000018745 | 602406 | 0.401 | 0 | 51.1828 | Eukaryota | 43705.0 | 0 | Chordata | 43705.0 | 0 | Carnivora | 43705.0 | 0 | Felidae | 24745.0 | 2 | Acinonyx | 19792.0 | 4 | inonyx jubat | 19792.0 | 4 |
| scf7180000023008 | 83426 | 0.5052 | 0 | 389.5472 | Eukaryota | 21794.0 | 0 | Chordata | 21794.0 | 0 | Carnivora | 21794.0 | 0 | Felidae | 21794.0 | 0 | Felis | 21794.0 | 0 | Felis catus | 21794.0 | 0 |
| scf7180000022248 | 19756 | 0.6249 | 0 | 51.4638 | Eukaryota | 3027.0 | 0 | Chordata | 3027.0 | 0 | Carnivora | 3027.0 | 0 | Felidae | 2768.0 | 1 | Felis | 2268.0 | 2 | Felis catus | 2268.0 | 2 |
| scf7180000018214 | 74988 | 0.3662 | 0 | 44.0814 | Eukaryota | 46034.0 | 0 | Chordata | 46034.0 | 0 | Carnivora | 46034.0 | 0 | Felidae | 46034.0 | 0 | Felis | 46034.0 | 0 | Felis catus | 46034.0 | 0 |
| scf7180000017944 | 351046 | 0.3867 | 0 | 49.3696 | Eukaryota | 32480.0 | 0 | Chordata | 32480.0 | 0 | Carnivora | 24227.0 | 2 | Felidae | 18080.0 | 4 | Panthera | 7992.0 | 8 | inonyx jubat | 4043.0 | 9 |
| scf7180000022597 | 14804 | 0.6558 | 0 | 44.6461 | Eukaryota | 4354.0 | 0 | Chordata | 4354.0 | 0 | Primates | 1940.0 | 4 | Hominidae | 1577.0 | 5 | Homo | 1577.0 | 5 | Homo sapiens | 1577.0 | 5 |
| scf7180000020675 | 54004 | 0.6077 | 0 | 49.0306 | Eukaryota | 42380.0 | 0 | Chordata | 42380.0 | 0 | Carnivora | 42380.0 | 0 | Otariidae | 20603.0 | 2 | Callorhinus | 12343.0 | 6 | llorhinus ursir | 12343.0 | 6 |
| scf7180000022564 | 29814 | 0.3864 | 0 | 332.4161 | Eukaryota | 9977.0 | 0 | Chordata | 9977.0 | 0 | pterygiforme | 8924.0 | 1 | Apterygidae | 8924.0 | 1 | Apteryx | 8924.0 | 1 | pteryx mante | 8924.0 | 1 |
| scf7180000019131 | 385424 | 0.3956 | 0 | 51.9477 | Eukaryota | 73149.0 | 0 | Chordata | 73149.0 | 0 | Carnivora | 73149.0 | 0 | Felidae | 73149.0 | 0 | Acinonyx | 69957.0 | 1 | inonyx jubat | 69957.0 | 1 |
| scf7180000024020 | 452214 | 0.4821 | 0 | 50.85 | Eukaryota | 46499.0 | 0 | Chordata | 46499.0 | 0 | Carnivora | 46499.0 | 0 | Felidae | 46499.0 | 0 | Felis | 46499.0 | 0 | Felis catus | 46499.0 | 0 |
| scf7180000017927 | 740082 | 0.3676 | 0 | 47.7013 | Eukaryota | 45481.0 | 0 | Chordata | 45481.0 | 0 | Carnivora | 45481.0 | 0 | Felidae | 45481.0 | 0 | Felis | 45481.0 | 0 | Felis catus | 45481.0 | 0 |
| scf7180000017527 | 1328143 | 0.3738 | 0 | 50.1709 | Eukaryota | 47115.0 | 0 | Chordata | 47115.0 | 0 | Carnivora | 47115.0 | 0 | Felidae | 47115.0 | 0 | Felis | 47115.0 | 0 | Felis catus | 47115.0 | 0 |
| scf7180000018368 | 1702409 | 0.3753 | 0 | 50.3966 | Eukaryota | 51414.0 | 0 | Chordata | 51414.0 | 0 | Carnivora | 51414.0 | 0 | Felidae | 51414.0 | 0 | Felis | 51414.0 | 0 | Felis catus | 51414.0 | 0 |
| scf7180000020737 | 48775 | 0.4305 | 0 | 167.7881 | Eukaryota | 2349.0 | 0 | Chordata | 2349.0 | 0 | Carnivora | 2349.0 | 0 | Felidae | 2349.0 | 0 | Felis | 1690.0 | 2 | Felis catus | 1690.0 | 2 |
| scf7180000019359 | 8160 | 0.4211 | 0 | 23.7488 | Eukaryota | 2336.0 | 0 | Chordata | 2336.0 | 0 | Carnivora | 2336.0 | 0 | Felidae | 1836.0 | 2 | Felis | 925.0 | 4 | Felis catus | 925.0 | 6 |
| scf7180000017427 | 2184763 | 0.4154 | 0 | 51.2836 | Eukaryota | 46736.0 | 0 | Chordata | 46736.0 | 0 | Carnivora | 46736.0 | 0 | Felidae | 46736.0 | 0 | Felis | 46736.0 | 0 | Felis catus | 46736.0 | 0 |
| scf7180000019281 | 50752 | 0.526 | 0 | 201.4036 | Eukaryota | 12347.0 | 0 | Chordata | 12347.0 | 0 | Carnivora | 12347.0 | 0 | Felidae | 12347.0 | 0 | Felis | 12347.0 | 0 | Felis catus | 12347.0 | 0 |
| scf7180000017943 | 1176852 | 0.409 | 100 | 51.6735 | Eukaryota | 27800.0 | 0 | Chordata | 27800.0 | 0 | Carnivora | 27800.0 | 0 | Felidae | 22815.0 | 1 | Felis | 20281.0 | 2 | Felis catus | 20281.0 | 2 |
| scf7180000024268 | 3129060 | 0.3704 | 0 | 49.6096 | Eukaryota | 75091.0 | 0 | Chordata | 75091.0 | 0 | Carnivora | 75091.0 | 0 | Felidae | 60247.0 | 1 | Acinonyx | 30083.0 | 3 | inonyx jubat | 30083.0 | 3 |
| scf7180000021167 | 86582 | 0.4088 | 0 | 26.4628 | Eukaryota | 43228.0 | 0 | Chordata | 43228.0 | 0 | Carnivora | 43228.0 | 0 | Felidae | 43228.0 | 0 | Felis | 38869.0 | 1 | Felis catus | 38869.0 | 1 |
| scf7180000022219 | 14291 | 0.6403 | 0 | 52.5613 | Eukaryota | 2980.0 | 0 | Chordata | 2980.0 | 0 | Carnivora | 2980.0 | 0 | Felidae | 2980.0 | 0 | Felis | 2980.0 | 0 | Felis catus | 2980.0 | 0 |
| scf7180000022520 | 13816 | 0.423 | 0 | 24.1478 | Eukaryota | 10707.0 | 0 | Chordata | 10707.0 | 0 | Carnivora | 10707.0 | 0 | Felidae | 10707.0 | 0 | Felis | 10707.0 | 0 | Felis catus | 10707.0 | 0 |
| scf7180000019987 | 192384 | 0.4483 | 0 | 51.1991 | Eukaryota | 10223.0 | 0 | Chordata | 10223.0 | 0 | Primates | 4415.0 | 2 | Hominidae | 4415.0 | 4 | Homo | 4415.0 | 5 | Homo sapiens | 4415.0 | 5 |
| scf7180000019233 | 48012 | 0.4343 | 0 | 71.1119 | Eukaryota | 3279.0 | 0 | Chordata | 3279.0 | 0 | Perissodactyla | 1691.0 | 2 | Equidae | 1691.0 | 2 | Equus | 1691.0 | 3 | quus caballu | 1404.0 | 4 |
| scf7180000023773 | 2589175 | 0.3996 | 0 | 50.8897 | Eukaryota | 48390.0 | 0 | Chordata | 48390.0 | 0 | Carnivora | 48390.0 | 0 | Felidae | 48390.0 | 0 | Panthera | 48390.0 | 0 | anthera pardu | 48390.0 | 0 |
| scf7180000022043 | 20344 | 0.5447 | 0 | 50.4265 | Eukaryota | 13237.0 | 0 | Chordata | 13237.0 | 0 | Carnivora | 13237.0 | 0 | Felidae | 10728.0 | 2 | Panthera | 5205.0 | 4 | anthera pardu | 5205.0 | 4 |
| scf7180000018911 | 527977 | 0.4203 | 0 | 51.1587 | Eukaryota | 26099.0 | 0 | Chordata | 26099.0 | 0 | Carnivora | 26099.0 | 0 | Felidae | 26099.0 | 0 | Felis | 20030.0 | 1 | Felis catus | 20030.0 | 1 |
| scf7180000019099 | 17819 | 0.3866 | 0 | 31.1472 | Eukaryota | 43956.0 | 0 | Chordata | 43956.0 | 0 | Carnivora | 43956.0 | 0 | Felidae | 43956.0 | 0 | Felis | 39621.0 | 1 | Felis catus | 39621.0 | 1 |
| scf7180000020387 | 111007 | 0.4822 | 0 | 48.7405 | Eukaryota | 9854.0 | 0 | Chordata | 9854.0 | 0 | Carnivora | 9854.0 | 0 | Canidae | 5256.0 | 3 | Vulpes | 5256.0 | 4 | Vulpes vulpes | 5256.0 | 4 |
| scf7180000021230 | 58955 | 0.5375 | 0 | 49.1426 | Eukaryota | 4897.0 | 0 | Chordata | 4897.0 | 0 | Rodentia | 1308.0 | 5 | Muridae | 1308.0 | 6 | Mus | 1308.0 | 7 | Mus musculus | 1308.0 | 7 |
| scf7180000018958 | 518931 | 0.3794 | 0 | 25.5752 | Eukaryota | 46161.0 | 0 | Chordata | 46161.0 | 0 | Carnivora | 29531.0 | 3 | Felidae | 18533.0 | 5 | Vulpes | 7158.0 | 8 | Vulpes vulpes | 7158.0 | 8 |
| scf7180000022650 | 12043 | 0.4191 | 0 | 25.4731 | Eukaryota | 15642.0 | 0 | Chordata | 15642.0 | 0 | Carnivora | 15642.0 | 0 | Felidae | 15642.0 | 0 | Panthera | 9330.0 | 1 | anthera pardu | 9330.0 | 1 |
| scf7180000022014 | 35055 | 0.3638 | 0 | 24.7356 | Eukaryota | 39046.0 | 0 | Chordata | 39046.0 | 0 | Carnivora | 39046.0 | 0 | Felidae | 39046.0 | 0 | Felis | 35166.0 | 1 | Felis catus | 35166.0 | 1 |
| scf7180000024055 | 93288 | 0.3697 | 0 | 21.6767 | Eukaryota | 22537.0 | 0 | Chordata | 22537.0 | 0 | Carnivora | 22537.0 | 0 | Felidae | 22537.0 | 0 | Felis | 17760.0 | 1 | Felis catus | 17760.0 | 1 |
| scf7180000018453 | 683256 | 0.5313 | 0 | 50.3015 | Eukaryota | 30354.0 | 0 | Chordata | 30354.0 | 0 | Artiodactyla | 23834.0 | 1 | Bovidae | 23834.0 | 1 | Bos | 14869.0 | 5 | Bos mutus | 5981.0 | 8 |
| scf7180000021381 | 49044 | 0.4605 | 0 | 26.5468 | Eukaryota | 86734.0 | 0 | Chordata | 86734.0 | 0 | Carnivora | 86734.0 | 0 | Felidae | 53462.0 | 1 | Lynx | 35592.0 | 2 | ynx canadens | 35592.0 | 2 |
| scf7180000022326 | 29188 | 0.4343 | 0 | 33.693 | Eukaryota | 24956.0 | 0 | Chordata | 24956.0 | 0 | Carnivora | 24956.0 | 0 | Felidae | 24956.0 | 0 | Felis | 22409.0 | 1 | Felis catus | 22409.0 | 1 |
| scf7180000022137 | 22852 | 0.3614 | 0 | 23.3883 | Eukaryota | 7894.0 | 0 | Chordata | 7894.0 | 0 | Carnivora | 7894.0 | 0 | Felidae | 7894.0 | 0 | Felis | 7894.0 | 0 | Felis catus | 7894.0 | 0 |
| scf7180000019693 | 215240 | 0.4657 | 0 | 50.6976 | Eukaryota | 40672.0 | 0 | Chordata | 40672.0 | 0 | Carnivora | 40672.0 | 0 | Felidae | 24312.0 | 1 | Lynx | 24312.0 | 1 | ynx canadens | 24312.0 | 1 |
| scf7180000019162 | 398439 | 0.3528 | 0 | 48.4103 | Eukaryota | 24904.0 | 0 | Chordata | 24904.0 | 0 | Carnivora | 24904.0 | 0 | Felidae | 24904.0 | 0 | Felis | 19704.0 | 1 | Felis catus | 19704.0 | 1 |
| scf7180000019880 | 237798 | 0.3576 | 0 | 49.3869 | Eukaryota | 10784.0 | 0 | Chordata | 10784.0 | 0 | Carnivora | 9467.0 | 1 | Felidae | 9467.0 | 1 | Felis | 7388.0 | 2 | Felis catus | 7388.0 | 2 |
| scf7180000019427 | 389641 | 0.5197 | 0 | 50.5496 | Eukaryota | 19099.0 | 0 | Chordata | 19099.0 | 0 | Carnivora | 19099.0 | 0 | Felidae | 19099.0 | 0 | Felis | 15299.0 | 1 | Felis catus | 15299.0 | 1 |
| scf7180000022862 | 68230 | 0.6539 | 0 | 74.0932 | Eukaryota | 11778.0 | 0 | Chordata | 11778.0 | 0 | Carnivora | 11778.0 | 0 | Felidae | 7380.0 | 2 | Felis | 7380.0 | 2 | Felis catus | 7380.0 | 2 |
| scf7180000021812 | 65993 | 0.3597 | 100 | 23.6143 | Eukaryota | 25192.0 | 0 | Chordata | 25192.0 | 0 | Carnivora | 25192.0 | 0 | Felidae | 25192.0 | 0 | Felis | 22615.0 | 1 | Felis catus | 22615.0 | 1 |
| scf7180000018461 | 70992 | 0.3981 | 0 | 44.5197 | Eukaryota | 47186.0 | 0 | Chordata | 47186.0 | 0 | Carnivora | 47186.0 | 0 | Felidae | 47186.0 | 0 | Felis | 47186.0 | 0 | Felis catus | 47186.0 | 0 |
| scf7180000020222 | 93155 | 0.5134 | 0 | 47.9332 | Eukaryota | 19494.0 | 0 | Chordata | 19494.0 | 0 | Carnivora | 19494.0 | 0 | Felidae | 19494.0 | 0 | Felis | 17575.0 | 1 | Felis catus | 17575.0 | 1 |
| scf7180000019174 | 341332 | 0.3823 | 0 | 49.675 | Eukaryota | 62869.0 | 0 | Chordata | 62869.0 | 0 | Carnivora | 62869.0 | 0 | Felidae | 23488.0 | 5 | Suricata | 7890.0 | 9 | ricata suricat | 7890.0 | 9 |
| scf7180000023695 | 951942 | 0.442 | 0 | 53.5604 | Eukaryota | 39998.0 | 0 | Chordata | 39998.0 | 0 | Carnivora | 39998.0 | 0 | Felidae | 23730.0 | 1 | Suricata | 16268.0 | 2 | ricata suricat | 16268.0 | 2 |
| scf7180000020479 | 86491 | 0.6181 | 0 | 46.6602 | Eukaryota | 9217.0 | 0 | Chordata | 9217.0 | 0 | Carnivora | 9217.0 | 0 | Felidae | 7805.0 | 2 | Felis | 5810.0 | 3 | Felis catus | 5810.0 | 3 |
| scf7180000019270 | 246267 | 0.3654 | 0 | 49.8734 | Eukaryota | 36343.0 | 0 | Chordata | 36343.0 | 0 | Carnivora | 36343.0 | 0 | Otariidae | 34344.0 | 1 | Zalophus | 34344.0 | 1 | phus californi | 34344.0 | 1 |
| scf7180000019222 | 4043811 | 0.3627 | 100 | 48.9667 | Eukaryota | 50941.0 | 0 | Chordata | 50941.0 | 0 | Carnivora | 50941.0 | 0 | Felidae | 42293.0 | 1 | Felis | 42293.0 | 1 | Felis catus | 42293.0 | 1 |
| scf7180000023475 | 21039 | 0.4832 | 0 | 142.4235 | no-hit | 0.0 | 0 | no-hit | 0.0 | 0 | no-hit | 0.0 | 0 | no-hit | 0.0 | 0 | no-hit | 0.0 | 0 | no-hit | 0.0 | 0 |
| scf7180000020087 | 148678 | 0.5782 | 0 | 50.6554 | Eukaryota | 6020.0 | 1 | Chordata | 6020.0 | 1 | Carnivora | 2506.0 | 4 | Felidae | 2506.0 | 4 | Felis | 1390.0 | 7 | Felis catus | 1390.0 | 7 |
| scf7180000021706 | 31544 | 0.3593 | 0 | 22.0985 | Eukaryota | 40536.0 | 0 | Chordata | 40536.0 | 0 | Carnivora | 40536.0 | 0 | Felidae | 40536.0 | 0 | Felis | 40536.0 | 0 | Felis catus | 40536.0 | 0 |
| scf7180000020168 | 229955 | 0.3971 | 0 | 24.1841 | Eukaryota | 9444.0 | 0 | Chordata | 9444.0 | 0 | Carnivora | 9444.0 | 0 | Felidae | 9444.0 | 0 | Felis | 9444.0 | 0 | Felis catus | 9444.0 | 0 |
| scf7180000021072 | 69784 | 0.3979 | 100 | 25.2864 | Eukaryota | 30319.0 | 0 | Chordata | 30319.0 | 0 | Carnivora | 30319.0 | 0 | Felidae | 30319.0 | 0 | Felis | 27271.0 | 1 | Felis catus | 27271.0 | 1 |
| scf7180000018215 | 1064613 | 0.3751 | 0 | 50.7873 | Eukaryota | 39201.0 | 0 | Chordata | 39201.0 | 0 | Carnivora | 39201.0 | 0 | Felidae | 35232.0 | 1 | Felis | 31328.0 | 2 | Felis catus | 31328.0 | 2 |
| scf7180000024218 | 42187 | 0.4432 | 0 | 24.6371 | Eukaryota | 15577.0 | 0 | Chordata | 15577.0 | 0 | Primates | 11980.0 | 3 | Hominidae | 9390.0 | 5 | Homo | 9390.0 | 5 | Homo sapiens | 9390.0 | 5 |
| scf7180000019599 | 342201 | 0.5036 | 0 | 51.8061 | Eukaryota | 31377.0 | 0 | Chordata | 31377.0 | 0 | Carnivora | 31377.0 | 0 | Felidae | 31377.0 | 0 | Puma | 11992.0 | 3 | Puma concolo | 11992.0 | 3 |
| scf7180000022853 | 28558 | 0.4885 | 0 | 476.8555 | no-hit | 0.0 | 0 | no-hit | 0.0 | 0 | no-hit | 0.0 | 0 | no-hit | 0.0 | 0 | no-hit | 0.0 | 0 | no-hit | 0.0 | 0 |
| scf7180000017675 | 1801277 | 0.4436 | 0 | 51.9488 | Eukaryota | 49665.0 | 0 | Chordata | 49665.0 | 0 | Carnivora | 49665.0 | 0 | Felidae | 49665.0 | 0 | Felis | 49665.0 | 0 | Felis catus | 49665.0 | 0 |
| scf7180000023466 | 23268 | 0.4874 | 0 | 274.9455 | Eukaryota | 9292.0 | 0 | Chordata | 9292.0 | 0 | Carnivora | 9292.0 | 0 | Felidae | 9292.0 | 0 | Felis | 9292.0 | 0 | Felis catus | 9292.0 | 0 |
| scf7180000022180 | 37234 | 0.4424 | 0 | 33.9666 | Eukaryota | 6092.0 | 0 | Chordata | 6092.0 | 0 | Carnivora | 6092.0 | 0 | Felidae | 6092.0 | 0 | Felis | 2823.0 | 2 | Felis catus | 2823.0 | 2 |
| scf7180000018731 | 1197692 | 0.4391 | 0 | 50.8724 | Eukaryota | 46098.0 | 0 | Chordata | 46098.0 | 0 | Carnivora | 46098.0 | 0 | Felidae | 46098.0 | 0 | Felis | 46098.0 | 0 | Felis catus | 46098.0 | 0 |
| scf7180000021280 | 51374 | 0.3839 | 0 | 25.2536 | Eukaryota | 32496.0 | 0 | Chordata | 32496.0 | 0 | Carnivora | 32496.0 | 0 | Felidae | 32496.0 | 0 | Felis | 29223.0 | 1 | Felis catus | 29223.0 | 1 |
| scf7180000023934 | 128975 | 0.3591 | 0 | 50.3985 | Eukaryota | 31469.0 | 0 | Chordata | 31469.0 | 0 | Carnivora | 31469.0 | 0 | Felidae | 31469.0 | 0 | Felis | 28264.0 | 1 | Felis catus | 28264.0 | 1 |
| scf7180000021929 | 13854 | 0.5213 | 0 | 219.137 | no-hit | 0.0 | 0 | no-hit | 0.0 | 0 | no-hit | 0.0 | 0 | no-hit | 0.0 | 0 | no-hit | 0.0 | 0 | no-hit | 0.0 | 0 |
| scf7180000018739 | 1442463 | 0.3547 | 0 | 49.8507 | Eukaryota | 62941.0 | 0 | Chordata | 62941.0 | 0 | Artiodactyla | 47609.0 | 1 | Monodontidae | 24118.0 | 5 | elphinapteru | 24118.0 | 5 | hinapterus le | 24118.0 | 5 |
| scf7180000020933 | 54828 | 0.3719 | 0 | 25.9739 | Eukaryota | 39246.0 | 0 | Chordata | 39246.0 | 0 | Carnivora | 39246.0 | 0 | Felidae | 39246.0 | 0 | Felis | 39246.0 | 0 | Felis catus | 39246.0 | 0 |
| scf7180000023731 | 7652 | 0.6373 | 0 | 92.8276 | Eukaryota | 2454.0 | 0 | Chordata | 2454.0 | 0 | Carnivora | 2217.0 | 1 | Felidae | 1993.0 | 2 | Felis | 1366.0 | 4 | Felis catus | 1366.0 | 4 |
| scf7180000022141 | 23459 | 0.4571 | 0 | 46.0042 | Eukaryota | 6228.0 | 0 | Chordata | 6228.0 | 0 | Carnivora | 6228.0 | 0 | Felidae | 6228.0 | 0 | Felis | 5607.0 | 1 | Felis catus | 5607.0 | 1 |
| scf7180000020044 | 86546 | 0.6519 | 0 | 46.8479 | Eukaryota | 9851.0 | 0 | Chordata | 9851.0 | 0 | Carnivora | 9851.0 | 0 | Felidae | 5887.0 | 1 | Suricata | 3964.0 | 5 | ricata suricat | 3964.0 | 5 |
| scf7180000020021 | 146532 | 0.3594 | 0 | 47.9237 | Eukaryota | 46027.0 | 0 | Chordata | 46027.0 | 0 | Carnivora | 46027.0 | 0 | Felidae | 46027.0 | 0 | Felis | 46027.0 | 0 | Felis catus | 46027.0 | 0 |
| scf7180000020194 | 298018 | 0.5497 | 0 | 48.2154 | Eukaryota | 15169.0 | 0 | Chordata | 15169.0 | 0 | Primates | 7934.0 | 2 | Hominidae | 4186.0 | 4 | Felis | 3597.0 | 7 | Felis catus | 3597.0 | 7 |
| scf7180000018778 | 2680074 | 0.4307 | 0 | 51.0792 | Eukaryota | 112720.0 | 0 | Chordata | 112720.0 | 0 | Carnivora | 112720.0 | 0 | Felidae | 51505.0 | 2 | Zalophus | 49599.0 | 3 | phus californi | 49599.0 | 3 |
| scf7180000017905 | 1352522 | 0.4334 | 0 | 51.4843 | Eukaryota | 55343.0 | 0 | Chordata | 55343.0 | 0 | Carnivora | 55343.0 | 0 | Felidae | 55343.0 | 0 | Felis | 41307.0 | 2 | Felis catus | 41307.0 | 2 |
| scf7180000020729 | 92364 | 0.5098 | 0 | 49.7693 | Eukaryota | 46704.0 | 0 | Chordata | 46704.0 | 0 | Carnivora | 46704.0 | 0 | Felidae | 46704.0 | 0 | Felis | 46704.0 | 0 | Felis catus | 46704.0 | 0 |
| scf7180000019605 | 246706 | 0.529 | 0 | 47.8675 | Eukaryota | 16118.0 | 0 | Chordata | 16118.0 | 0 | Carnivora | 14546.0 | 1 | Felidae | 14546.0 | 1 | Felis | 14546.0 | 1 | Felis catus | 14546.0 | 1 |
| scf7180000017526 | 1290611 | 0.3571 | 0 | 48.2714 | Eukaryota | 46828.0 | 0 | Chordata | 46828.0 | 0 | Carnivora | 46828.0 | 0 | Felidae | 46828.0 | 0 | Felis | 46828.0 | 0 | Felis catus | 46828.0 | 0 |
| scf7180000017844 | 3105628 | 0.3615 | 100 | 49.2861 | Eukaryota | 47028.0 | 0 | Chordata | 47028.0 | 0 | Carnivora | 47028.0 | 0 | Felidae | 42649.0 | 1 | Felis | 42649.0 | 1 | Felis catus | 42649.0 | 1 |
| scf7180000019931 | 20987 | 0.6606 | 0 | 38.6335 | Eukaryota | 11209.0 | 0 | Chordata | 11209.0 | 0 | Carnivora | 11209.0 | 0 | Felidae | 11209.0 | 0 | Panthera | 4604.0 | 3 | anthera pardu | 4604.0 | 3 |
| scf7180000023366 | 27552 | 0.6965 | 0 | 113.1083 | no-hit | 0.0 | 0 | no-hit | 0.0 | 0 | no-hit | 0.0 | 0 | no-hit | 0.0 | 0 | no-hit | 0.0 | 0 | no-hit | 0.0 | 0 |
| scf7180000019818 | 196706 | 0.4781 | 0 | 48.7166 | Eukaryota | 13558.0 | 0 | Chordata | 13558.0 | 0 | Artiodactyla | 5147.0 | 2 | Bovidae | 4005.0 | 4 | Homo | 3858.0 | 5 | Homo sapiens | 3858.0 | 5 |
| scf7180000021544 | 51778 | 0.3682 | 0 | 28.2544 | Eukaryota | 42672.0 | 0 | Chordata | 42672.0 | 0 | Carnivora | 42672.0 | 0 | Felidae | 42672.0 | 0 | Felis | 42672.0 | 0 | Felis catus | 42672.0 | 0 |
| scf7180000017524 | 1787209 | 0.3409 | 0 | 47.1832 | Eukaryota | 31996.0 | 0 | Chordata | 31996.0 | 0 | Carnivora | 31996.0 | 0 | Felidae | 31996.0 | 0 | Felis | 28749.0 | 1 | Felis catus | 28749.0 | 1 |
| scf7180000019035 | 379930 | 0.3902 | 0 | 51.2068 | Eukaryota | 13124.0 | 0 | Chordata | 13124.0 | 0 | Carnivora | 13124.0 | 0 | Felidae | 13124.0 | 0 | Felis | 11773.0 | 1 | Felis catus | 11773.0 | 1 |
| scf7180000023817 | 1874823 | 0.3768 | 0 | 51.4388 | Eukaryota | 296732.0 | 0 | Chordata | 296732.0 | 0 | Carnivora | 201038.0 | 1 | Felidae | 172069.0 | 2 | Puma | 143405.0 | 3 | Puma concolo | 143405.0 | 3 |
| scf7180000022825 | 342911 | 0.3987 | 0 | 53.0243 | Eukaryota | 46515.0 | 0 | Chordata | 46515.0 | 0 | Carnivora | 46515.0 | 0 | Felidae | 46515.0 | 0 | Felis | 46515.0 | 0 | Felis catus | 46515.0 | 0 |
| scf7180000018604 | 23225 | 0.4262 | 0 | 50.0269 | Eukaryota | 20173.0 | 0 | Chordata | 20173.0 | 0 | Carnivora | 20173.0 | 0 | Felidae | 20173.0 | 0 | Felis | 18141.0 | 1 | Felis catus | 18141.0 | 1 |
| scf7180000017941 | 1434136 | 0.3965 | 0 | 51.0176 | Eukaryota | 45834.0 | 0 | Chordata | 45834.0 | 0 | Carnivora | 45834.0 | 0 | Felidae | 45834.0 | 0 | Felis | 45834.0 | 0 | Felis catus | 45834.0 | 0 |
| scf7180000020450 | 42189 | 0.5105 | 0 | 49.0929 | Eukaryota | 10853.0 | 0 | Chordata | 10853.0 | 0 | Carnivora | 10156.0 | 1 | Felidae | 9465.0 | 2 | Panthera | 4755.0 | 3 | anthera pardu | 4755.0 | 3 |
| scf7180000024186 | 85719 | 0.6041 | 0 | 50.7011 | Eukaryota | 15770.0 | 0 | Chordata | 15770.0 | 0 | Carnivora | 15770.0 | 0 | Felidae | 15770.0 | 0 | Lynx | 7900.0 | 1 | ynx canadens | 7900.0 | 1 |
| scf7180000023967 | 739870 | 0.4193 | 0 | 49.799 | Eukaryota | 101292.0 | 0 | Chordata | 101292.0 | 0 | Carnivora | 101292.0 | 0 | Felidae | 101292.0 | 0 | Panthera | 60900.0 | 1 | anthera pardu | 60900.0 | 1 |
| scf7180000017764 | 1908829 | 0.4325 | 0 | 52.5095 | Eukaryota | 44195.0 | 0 | Chordata | 44195.0 | 0 | Carnivora | 44195.0 | 0 | Felidae | 31324.0 | 1 | Suricata | 12871.0 | 5 | ricata suricat | 12871.0 | 5 |
| scf7180000017920 | 1466746 | 0.3618 | 0 | 49.1718 | Eukaryota | 46639.0 | 0 | Chordata | 46639.0 | 0 | Carnivora | 46639.0 | 0 | Felidae | 46639.0 | 0 | Felis | 41900.0 | 1 | Felis catus | 41900.0 | 1 |
| scf7180000019496 | 334496 | 0.4258 | 0 | 49.7816 | Eukaryota | 24511.0 | 0 | Chordata | 24511.0 | 0 | Carnivora | 24511.0 | 0 | Felidae | 24511.0 | 0 | Felis | 10392.0 | 4 | Felis catus | 10392.0 | 5 |
| scf7180000021161 | 55977 | 0.3701 | 0 | 26.0509 | Eukaryota | 45372.0 | 0 | Chordata | 45372.0 | 0 | Carnivora | 45372.0 | 0 | Felidae | 45372.0 | 0 | Felis | 45372.0 | 0 | Felis catus | 45372.0 | 0 |

| | | | | | | | | | | | | | | | | | | | | | | |
|---|---|---|---|---|---|---|---|---|---|---|---|---|---|---|---|---|---|---|---|---|---|---|
| scf7180000018570 | 670164 | 0.3596 | 0 | 49.0624 | Eukaryota | 30555.0 | 0 | Chordata | 30555.0 | 0 | Artiodactyla | 18113.0 | 1 | Bovidae | 18113.0 | 3 | Bos | 9647.0 | 7 | ıricata suricat | 4148.0 | 9 |
| scf7180000020136 | 268853 | 0.4672 | 0 | 25.3984 | Eukaryota | 36362.0 | 0 | Chordata | 36362.0 | 0 | Carnivora | 36362.0 | 0 | Felidae | 36362.0 | 0 | Felis | 32735.0 | 1 | Felis catus | 32735.0 | 1 |
| scf7180000017983 | 1919985 | 0.4036 | 0 | 51.2566 | Eukaryota | 47823.0 | 0 | Chordata | 47823.0 | 0 | Carnivora | 47823.0 | 0 | Felidae | 47823.0 | 0 | Felis | 47823.0 | 0 | Felis catus | 47823.0 | 0 |
| scf7180000020308 | 110522 | 0.4091 | 0 | 50.5163 | Eukaryota | 49332.0 | 0 | Chordata | 49332.0 | 0 | Carnivora | 49332.0 | 0 | Phocidae | 14838.0 | 3 | Neomonachus | 14838.0 | 4 | nachus schau | 14838.0 | 4 |
| scf7180000022042 | 20847 | 0.4713 | 0 | 46.1824 | Eukaryota | 2265.0 | 0 | Chordata | 2265.0 | 0 | Carnivora | 2265.0 | 0 | Felidae | 2032.0 | 1 | Felis | 1582.0 | 2 | Felis catus | 1582.0 | 2 |
| scf7180000021855 | 29858 | 0.4951 | 0 | 27.9178 | Eukaryota | 5672.0 | 0 | Chordata | 5127.0 | 1 | Primates | 3507.0 | 3 | Hominidae | 3507.0 | 4 | Homo | 2712.0 | 5 | Homo sapiens | 2712.0 | 5 |
| scf7180000017932 | 83313 | 0.4075 | 0 | 46.2076 | Eukaryota | 43848.0 | 0 | Chordata | 43848.0 | 0 | Carnivora | 43848.0 | 0 | Felidae | 43848.0 | 0 | Felis | 43848.0 | 0 | Felis catus | 43848.0 | 0 |
| scf7180000018772 | 493052 | 0.476 | 0 | 50.1089 | Eukaryota | 46351.0 | 0 | Chordata | 46351.0 | 0 | Carnivora | 46351.0 | 0 | Felidae | 46351.0 | 0 | Felis | 46351.0 | 0 | Felis catus | 46351.0 | 0 |
| scf7180000023375 | 14833 | 0.504 | 0 | 173.5968 | Eukaryota | 7090.0 | 0 | Chordata | 7090.0 | 0 | Carnivora | 7090.0 | 0 | Felidae | 7090.0 | 0 | Felis | 7090.0 | 0 | Felis catus | 7090.0 | 0 |
| scf7180000022375 | 33992 | 0.3748 | 0 | 18.7159 | Eukaryota | 19993.0 | 0 | Chordata | 19993.0 | 0 | Carnivora | 19993.0 | 0 | Felidae | 19993.0 | 0 | Felis | 17978.0 | 1 | Felis catus | 17978.0 | 1 |
| scf7180000019382 | 224201 | 0.6125 | 0 | 50.6776 | Eukaryota | 28129.0 | 0 | Chordata | 28129.0 | 0 | Carnivora | 28129.0 | 0 | Felidae | 28129.0 | 0 | Felis | 25587.0 | 1 | Felis catus | 25587.0 | 1 |
| scf7180000021744 | 64818 | 0.4206 | 0 | 26.5432 | Eukaryota | 45851.0 | 0 | Chordata | 45851.0 | 0 | Carnivora | 45851.0 | 0 | Felidae | 45851.0 | 0 | Felis | 45851.0 | 0 | Felis catus | 45851.0 | 0 |
| scf7180000021110 | 91170 | 0.3993 | 0 | 27.3259 | Eukaryota | 36942.0 | 0 | Chordata | 36942.0 | 0 | Carnivora | 36942.0 | 0 | Felidae | 36942.0 | 0 | Felis | 36942.0 | 0 | Felis catus | 36942.0 | 0 |
| scf7180000018701 | 32302 | 0.3828 | 0 | 24.9206 | Eukaryota | 24515.0 | 0 | Chordata | 24515.0 | 0 | Carnivora | 24515.0 | 0 | Felidae | 24515.0 | 0 | Felis | 22053.0 | 1 | Felis catus | 22053.0 | 1 |
| scf7180000024062 | 482459 | 0.4403 | 0 | 50.4837 | Eukaryota | 25371.0 | 0 | Chordata | 25371.0 | 0 | Carnivora | 25371.0 | 0 | Felidae | 25371.0 | 0 | Felis | 20515.0 | 1 | Felis catus | 20515.0 | 1 |
| scf7180000019512 | 170004 | 0.401 | 0 | 24.6356 | Eukaryota | 16889.0 | 0 | Chordata | 16889.0 | 0 | Carnivora | 16889.0 | 0 | Felidae | 14412.0 | 1 | Felis | 9201.0 | 4 | Felis catus | 9201.0 | 4 |
| scf7180000024263 | 86804 | 0.61 | 0 | 50.9556 | Eukaryota | 3666.0 | 0 | Chordata | 3666.0 | 0 | Carnivora | 3066.0 | 1 | Felidae | 1935.0 | 2 | Felis | 1137.0 | 4 | Felis catus | 1137.0 | 5 |
| scf7180000021590 | 34711 | 0.3837 | 0 | 25.5119 | Eukaryota | 5204.0 | 0 | Chordata | 5204.0 | 0 | Carnivora | 1853.0 | 2 | Hominidae | 1810.0 | 3 | Sus | 1541.0 | 5 | Sus scrofa | 1541.0 | 5 |
| scf7180000019984 | 173611 | 0.4287 | 0 | 26.6941 | Eukaryota | 13881.0 | 0 | Chordata | 13881.0 | 0 | Primates | 7662.0 | 3 | Hominidae | 7662.0 | 3 | Homo | 7662.0 | 4 | Homo sapiens | 7662.0 | 4 |
| scf7180000019395 | 301825 | 0.6084 | 0 | 51.0136 | Eukaryota | 21836.0 | 0 | Chordata | 21836.0 | 0 | Carnivora | 19893.0 | 1 | Phocidae | 7123.0 | 5 | Neomonachus | 4748.0 | 8 | nachus schau | 4748.0 | 8 |
| scf7180000023665 | 64633 | 0.3861 | 0 | 270.9974 | Eukaryota | 7156.0 | 0 | Chordata | 7156.0 | 0 | Carnivora | 7156.0 | 0 | Felidae | 7156.0 | 0 | Felis | 6455.0 | 1 | Felis catus | 6455.0 | 1 |
| scf7180000022383 | 11809 | 0.49 | 0 | 26.1875 | Eukaryota | 14492.0 | 0 | Chordata | 14492.0 | 0 | Primates | 9442.0 | 3 | Hominidae | 9442.0 | 4 | Homo | 5009.0 | 6 | Homo sapiens | 5009.0 | 6 |
| scf7180000019269 | 278369 | 0.3604 | 0 | 48.6501 | Eukaryota | 48280.0 | 0 | Chordata | 48280.0 | 0 | Carnivora | 48280.0 | 0 | Felidae | 48280.0 | 0 | Felis | 48280.0 | 0 | Felis catus | 48280.0 | 0 |
| scf7180000022774 | 7313 | 0.3497 | 0 | 22.6244 | Eukaryota | 2260.0 | 0 | Chordata | 2260.0 | 0 | Carnivora | 1886.0 | 1 | Felidae | 1886.0 | 1 | Felis | 1456.0 | 2 | Felis catus | 1456.0 | 2 |
| scf7180000022980 | 40435 | 0.5686 | 0 | 95.7609 | Eukaryota | 6267.0 | 0 | Chordata | 6267.0 | 0 | Carnivora | 6267.0 | 0 | Felidae | 3596.0 | 3 | Ursus | 894.0 | 9 | Ursus arctos | 894.0 | 9 |
| scf7180000018722 | 691428 | 0.3998 | 0 | 51.0187 | Eukaryota | 47220.0 | 0 | Chordata | 47220.0 | 0 | Carnivora | 47220.0 | 0 | Felidae | 47220.0 | 0 | Felis | 47220.0 | 0 | Felis catus | 47220.0 | 0 |
| scf7180000024193 | 122244 | 0.4119 | 0 | 25.0293 | Eukaryota | 7746.0 | 0 | Chordata | 7746.0 | 0 | Carnivora | 6807.0 | 1 | Felidae | 5979.0 | 2 | Felis | 5979.0 | 2 | Felis catus | 5979.0 | 2 |
| scf7180000022765 | 4959 | 0.4543 | 0 | 26.8509 | Eukaryota | 5267.0 | 0 | Chordata | 5267.0 | 0 | Carnivora | 5267.0 | 0 | Felidae | 5267.0 | 0 | Felis | 3693.0 | 1 | Felis catus | 3693.0 | 1 |
| scf7180000020634 | 81664 | 0.3925 | 0 | 24.7258 | Eukaryota | 41968.0 | 0 | Chordata | 41968.0 | 0 | Carnivora | 41968.0 | 0 | Felidae | 41968.0 | 0 | Felis | 37805.0 | 1 | Felis catus | 37805.0 | 1 |
| scf7180000018381 | 1495243 | 0.4176 | 0 | 51.9126 | Eukaryota | 46769.0 | 0 | Chordata | 46769.0 | 0 | Carnivora | 46769.0 | 0 | Felidae | 46769.0 | 0 | Felis | 46769.0 | 0 | Felis catus | 46769.0 | 0 |
| scf7180000020304 | 57712 | 0.4035 | 0 | 42.9985 | Eukaryota | 31411.0 | 0 | Chordata | 31411.0 | 0 | Carnivora | 31411.0 | 0 | Felidae | 31411.0 | 0 | Felis | 28321.0 | 1 | Felis catus | 28321.0 | 1 |
| scf7180000019150 | 150708 | 0.3588 | 0 | 132.4541 | Eukaryota | 46598.0 | 0 | Chordata | 46598.0 | 0 | Carnivora | 46598.0 | 0 | Felidae | 46598.0 | 0 | Felis | 46598.0 | 0 | Felis catus | 46598.0 | 0 |
| scf7180000017823 | 896812 | 0.3747 | 0 | 49.6078 | Eukaryota | 55599.0 | 0 | Chordata | 55599.0 | 0 | Carnivora | 55599.0 | 0 | Felidae | 55599.0 | 0 | Felis | 55599.0 | 0 | Felis catus | 55599.0 | 0 |
| scf7180000018642 | 627185 | 0.3757 | 0 | 50.4801 | Eukaryota | 25866.0 | 0 | Chordata | 25866.0 | 0 | Carnivora | 25866.0 | 0 | Felidae | 25866.0 | 0 | Felis | 23241.0 | 1 | Felis catus | 23241.0 | 1 |
| scf7180000019760 | 50706 | 0.3967 | 0 | 51.3703 | Eukaryota | 45953.0 | 0 | Chordata | 45953.0 | 0 | Carnivora | 45953.0 | 0 | Felidae | 45953.0 | 0 | Felis | 45953.0 | 0 | Felis catus | 45953.0 | 0 |
| scf7180000022143 | 15829 | 0.4103 | 0 | 49.7136 | Eukaryota | 2880.0 | 0 | Chordata | 2880.0 | 0 | Carnivora | 2880.0 | 0 | Felidae | 2880.0 | 0 | Felis | 2318.0 | 1 | Felis catus | 2318.0 | 1 |
| scf7180000023233 | 16133 | 0.4305 | 0 | 134.6254 | Eukaryota | 4232.0 | 0 | Chordata | 4232.0 | 0 | Carnivora | 3897.0 | 1 | Felidae | 2271.0 | 2 | Felis | 2271.0 | 2 | Felis catus | 2271.0 | 2 |
| scf7180000018886 | 460565 | 0.3886 | 0 | 49.4081 | Eukaryota | 43449.0 | 0 | Chordata | 43449.0 | 0 | Carnivora | 43449.0 | 0 | Felidae | 43449.0 | 0 | Felis | 39164.0 | 1 | Felis catus | 39164.0 | 1 |
| scf7180000022496 | 11360 | 0.3704 | 0 | 39.9351 | Eukaryota | 9012.0 | 0 | Chordata | 9012.0 | 0 | Carnivora | 9012.0 | 0 | Felidae | 9012.0 | 0 | Felis | 7109.0 | 1 | Felis catus | 7109.0 | 1 |
| scf7180000018658 | 563904 | 0.3911 | 0 | 50.2784 | Eukaryota | 24845.0 | 0 | Chordata | 24845.0 | 0 | Carnivora | 24845.0 | 0 | Otariidae | 14954.0 | 1 | Zalophus | 9964.0 | 3 | phus californi | 9964.0 | 3 |
| scf7180000021745 | 58131 | 0.3898 | 0 | 25.1559 | Eukaryota | 42086.0 | 0 | Chordata | 42086.0 | 0 | Carnivora | 42086.0 | 0 | Felidae | 25763.0 | 3 | Lynx | 10242.0 | 6 | ynx canadens | 10242.0 | 6 |
| scf7180000018954 | 134124 | 0.5357 | 0 | 43.9861 | Eukaryota | 12921.0 | 0 | Chordata | 12921.0 | 0 | Carnivora | 8349.0 | 1 | Felidae | 8349.0 | 1 | Felis | 8349.0 | 1 | Felis catus | 8349.0 | 1 |
| scf7180000024001 | 1441014 | 0.4477 | 0 | 51.4714 | Eukaryota | 56727.0 | 0 | Chordata | 56727.0 | 0 | Carnivora | 56727.0 | 0 | Felidae | 39856.0 | 1 | Panthera | 17132.0 | 5 | Felis catus | 11420.0 | 6 |
| scf7180000022121 | 11302 | 0.4207 | 0 | 24.3512 | Eukaryota | 7106.0 | 0 | Chordata | 7106.0 | 0 | Carnivora | 4998.0 | 1 | Felidae | 4998.0 | 1 | Felis | 4998.0 | 1 | Felis catus | 4998.0 | 1 |
| scf7180000023982 | 544153 | 0.3981 | 0 | 50.3749 | Eukaryota | 47483.0 | 0 | Chordata | 47483.0 | 0 | Carnivora | 47483.0 | 0 | Mustelidae | 20396.0 | 3 | Mustela | 20396.0 | 4 | ıustela putori | 20396.0 | 4 |
| scf7180000017648 | 1400214 | 0.4009 | 100 | 50.109 | Eukaryota | 51215.0 | 0 | Chordata | 51215.0 | 0 | Carnivora | 51215.0 | 0 | Felidae | 51215.0 | 0 | Felis | 19688.0 | 3 | Felis catus | 19688.0 | 3 |
| scf7180000023431 | 20253 | 0.4709 | 0 | 210.4209 | Eukaryota | 11836.0 | 0 | Chordata | 11836.0 | 0 | Carnivora | 11836.0 | 0 | Felidae | 11836.0 | 0 | Felis | 11836.0 | 0 | Felis catus | 11836.0 | 0 |
| scf7180000018149 | 1075338 | 0.3793 | 0 | 71.9442 | Eukaryota | 45989.0 | 0 | Chordata | 45989.0 | 0 | Carnivora | 45989.0 | 0 | Felidae | 45989.0 | 0 | Felis | 45989.0 | 0 | Felis catus | 45989.0 | 0 |
| scf7180000024037 | 285833 | 0.4441 | 0 | 26.8371 | Eukaryota | 24422.0 | 0 | Chordata | 24422.0 | 0 | Carnivora | 24422.0 | 0 | Felidae | 14802.0 | 2 | Acinonyx | 7449.0 | 3 | :inonyx jubat | 7449.0 | 3 |
| scf7180000021857 | 32142 | 0.3579 | 0 | 23.1581 | Eukaryota | 6852.0 | 0 | Chordata | 6852.0 | 0 | Carnivora | 4372.0 | 2 | Felidae | 4372.0 | 2 | Felis | 3836.0 | 3 | Felis catus | 3836.0 | 3 |
| scf7180000020671 | 115688 | 0.3544 | 0 | 24.947 | Eukaryota | 42163.0 | 0 | Chordata | 42163.0 | 0 | Carnivora | 42163.0 | 0 | Felidae | 42163.0 | 0 | Felis | 38026.0 | 1 | Felis catus | 38026.0 | 1 |
| scf7180000018391 | 946927 | 0.4205 | 0 | 50.6908 | Eukaryota | 82982.0 | 0 | Chordata | 82982.0 | 0 | Carnivora | 82982.0 | 0 | Felidae | 64537.0 | 2 | Felis | 24434.0 | 4 | Felis catus | 24434.0 | 4 |
| scf7180000023105 | 41590 | 0.5152 | 0 | 169.9423 | Eukaryota | 15407.0 | 0 | Chordata | 15407.0 | 0 | Carnivora | 15407.0 | 0 | Felidae | 15407.0 | 0 | Felis | 15407.0 | 0 | Felis catus | 15407.0 | 0 |
| scf7180000018156 | 922249 | 0.3624 | 0 | 48.033 | Eukaryota | 46499.0 | 0 | Chordata | 46499.0 | 0 | Carnivora | 46499.0 | 0 | Felidae | 46499.0 | 0 | Felis | 46499.0 | 0 | Felis catus | 46499.0 | 0 |
| scf7180000018192 | 945861 | 0.3507 | 0 | 48.9927 | Eukaryota | 40040.0 | 0 | Chordata | 40040.0 | 0 | Carnivora | 40040.0 | 0 | Felidae | 40040.0 | 0 | Panthera | 40040.0 | 0 | anthera pardı | 40040.0 | 0 |
| scf7180000022515 | 14035 | 0.471 | 0 | 24.2031 | Eukaryota | 2789.0 | 0 | Chordata | 2789.0 | 0 | Carnivora | 2789.0 | 0 | Felidae | 2789.0 | 0 | Felis | 1962.0 | 2 | Felis catus | 1962.0 | 2 |
| scf7180000021988 | 32571 | 0.354 | 0 | 24.5552 | Eukaryota | 18082.0 | 0 | Chordata | 18082.0 | 0 | Carnivora | 18082.0 | 0 | Felidae | 18082.0 | 0 | Felis | 14526.0 | 1 | Felis catus | 14526.0 | 1 |
| scf7180000022278 | 23757 | 0.432 | 0 | 24.5757 | Eukaryota | 6926.0 | 0 | Chordata | 6926.0 | 0 | Carnivora | 6926.0 | 0 | Felidae | 6926.0 | 0 | Felis | 6926.0 | 0 | Felis catus | 6926.0 | 0 |
| scf7180000018026 | 59481 | 0.481 | 0 | 124.6807 | Eukaryota | 15408.0 | 0 | Chordata | 15408.0 | 0 | Carnivora | 15408.0 | 0 | Felidae | 15408.0 | 0 | Felis | 13847.0 | 1 | Felis catus | 13847.0 | 1 |
| scf7180000023351 | 50621 | 0.4168 | 0 | 424.1838 | Eukaryota | 6039.0 | 0 | Chordata | 6039.0 | 0 | Carnivora | 6039.0 | 0 | Phocidae | 2464.0 | 2 | .eptonychote | 1848.0 | 4 | ınychotes wec | 1848.0 | 4 |
| scf7180000018587 | 1771019 | 0.4869 | 0 | 52.7278 | Eukaryota | 51871.0 | 0 | Chordata | 51871.0 | 0 | Carnivora | 51871.0 | 0 | Felidae | 51871.0 | 0 | Felis | 39079.0 | 1 | Felis catus | 39079.0 | 1 |
| scf7180000022431 | 6845 | 0.5005 | 0 | 3399.9679 | no-hit | 0.0 | 0 | no-hit | 0.0 | 0 | no-hit | 0.0 | 0 | no-hit | 0.0 | 0 | no-hit | 0.0 | 0 | no-hit | 0.0 | 0 |
| scf7180000019663 | 348992 | 0.3535 | 0 | 49.4644 | Eukaryota | 48005.0 | 0 | Chordata | 48005.0 | 0 | Carnivora | 48005.0 | 0 | Felidae | 48005.0 | 0 | Felis | 48005.0 | 0 | Felis catus | 48005.0 | 0 |
| scf7180000023702 | 78816 | 0.405 | 0 | 408.1839 | Eukaryota | 11808.0 | 0 | Chordata | 11808.0 | 0 | Carnivora | 11808.0 | 0 | Felidae | 11808.0 | 0 | Felis | 11808.0 | 0 | Felis catus | 11808.0 | 0 |
| scf7180000018826 | 475956 | 0.3536 | 0 | 45.5567 | Eukaryota | 45449.0 | 0 | Chordata | 45449.0 | 0 | Carnivora | 45449.0 | 0 | Felidae | 45449.0 | 0 | Felis | 45449.0 | 0 | Felis catus | 45449.0 | 0 |
| scf7180000021121 | 57758 | 0.4108 | 0 | 27.1502 | Eukaryota | 9330.0 | 0 | Chordata | 9330.0 | 0 | Primates | 9330.0 | 0 | Cercopithecidae | 9330.0 | 0 | Macaca | 9330.0 | 0 | Aacaca mulatt | 9330.0 | 0 |
| scf7180000024191 | 69311 | 0.604 | 0 | 43.8104 | Eukaryota | 15703.0 | 0 | Chordata | 15703.0 | 0 | Carnivora | 15703.0 | 0 | Felidae | 13913.0 | 1 | Panthera | 6256.0 | 3 | anthera pardı | 6256.0 | 3 |
| scf7180000018839 | 595646 | 0.3475 | 0 | 47.7742 | Eukaryota | 45598.0 | 0 | Chordata | 45598.0 | 0 | Carnivora | 45598.0 | 0 | Felidae | 45598.0 | 0 | Felis | 45598.0 | 0 | Felis catus | 45598.0 | 0 |
| scf7180000018239 | 1038251 | 0.4918 | 0 | 50.7075 | Eukaryota | 46916.0 | 0 | Chordata | 46916.0 | 0 | Carnivora | 46916.0 | 0 | Felidae | 46916.0 | 0 | Felis | 46916.0 | 0 | Felis catus | 46916.0 | 0 |
| scf7180000017859 | 173528 | 0.3952 | 0 | 25.3269 | Eukaryota | 44450.0 | 0 | Chordata | 44450.0 | 0 | Carnivora | 28081.0 | 2 | Canidae | 17825.0 | 6 | Canis | 9646.0 | 7 | Canis lupus | 9646.0 | 7 |
| scf7180000022595 | 8678 | 0.3932 | 0 | 26.0246 | Eukaryota | 9976.0 | 0 | Chordata | 9490.0 | 1 | Primates | 4580.0 | 4 | Hominidae | 4580.0 | 4 | Homo | 3060.0 | 5 | Homo sapiens | 3060.0 | 5 |
| scf7180000022614 | 15239 | 0.3949 | 0 | 22.8917 | Eukaryota | 30024.0 | 0 | Chordata | 30024.0 | 0 | Carnivora | 30024.0 | 0 | Felidae | 30024.0 | 0 | Felis | 27010.0 | 1 | Felis catus | 27010.0 | 1 |
| scf7180000023829 | 481551 | 0.4494 | 0 | 52.0535 | Eukaryota | 41955.0 | 0 | Chordata | 41955.0 | 0 | Carnivora | 41955.0 | 0 | Felidae | 41955.0 | 0 | Panthera | 25237.0 | 2 | anthera pardı | 21035.0 | 3 |
| scf7180000018126 | 580801 | 0.3725 | 0 | 46.4022 | Eukaryota | 46553.0 | 0 | Chordata | 46553.0 | 0 | Carnivora | 46553.0 | 0 | Felidae | 46553.0 | 0 | Felis | 46553.0 | 0 | Felis catus | 46553.0 | 0 |
| scf7180000021995 | 16848 | 0.3299 | 0 | 25.1199 | Eukaryota | 18332.0 | 0 | Chordata | 18332.0 | 0 | Primates | 10082.0 | 1 | Hominidae | 10082.0 | 1 | Ceratotheriun | 8250.0 | 2 | ıtotherium sir | 8250.0 | 2 |
| scf7180000021993 | 24578 | 0.3841 | 0 | 24.7618 | Eukaryota | 16049.0 | 0 | Chordata | 16049.0 | 0 | Carnivora | 16049.0 | 0 | Felidae | 16049.0 | 0 | Felis | 14412.0 | 1 | Felis catus | 14412.0 | 1 |
| scf7180000018488 | 1551580 | 0.402 | 0 | 50.9887 | Eukaryota | 44464.0 | 0 | Chordata | 44464.0 | 0 | Carnivora | 44464.0 | 0 | Felidae | 44464.0 | 0 | Panthera | 39897.0 | 1 | anthera pardı | 39897.0 | 1 |
| scf7180000021550 | 15866 | 0.4152 | 0 | 27.4078 | Eukaryota | 4668.0 | 0 | Chordata | 4668.0 | 0 | Carnivora | 3760.0 | 1 | Felidae | 2734.0 | 2 | Acinonyx | 1161.0 | 6 | :inonyx jubat | 1161.0 | 6 |
| scf7180000021200 | 60352 | 0.4323 | 0 | 25.3251 | Eukaryota | 33762.0 | 0 | Chordata | 33762.0 | 0 | Carnivora | 33762.0 | 0 | Felidae | 33762.0 | 0 | Felis | 30428.0 | 1 | Felis catus | 30428.0 | 1 |
| scf7180000018969 | 83684 | 0.3808 | 0 | 46.5469 | Eukaryota | 15617.0 | 0 | Chordata | 15617.0 | 0 | Carnivora | 15617.0 | 0 | Felidae | 15617.0 | 0 | Felis | 15617.0 | 0 | Felis catus | 15617.0 | 0 |
| scf7180000022849 | 27081 | 0.4526 | 0 | 322.7306 | Eukaryota | 12325.0 | 0 | Chordata | 12325.0 | 0 | Carnivora | 12325.0 | 0 | Felidae | 12325.0 | 0 | Felis | 11109.0 | 1 | Felis catus | 11109.0 | 1 |
| scf7180000022151 | 22654 | 0.3874 | 0 | 25.4719 | Eukaryota | 10385.0 | 0 | Chordata | 10385.0 | 0 | Primates | 7089.0 | 3 | Hominidae | 7089.0 | 4 | Homo | 4263.0 | 5 | Homo sapiens | 4263.0 | 5 |
| scf7180000021102 | 79117 | 0.4239 | 0 | 24.4378 | Eukaryota | 45879.0 | 0 | Chordata | 45879.0 | 0 | Carnivora | 45879.0 | 0 | Felidae | 45879.0 | 0 | Felis | 45879.0 | 0 | Felis catus | 45879.0 | 0 |
| scf7180000020909 | 106922 | 0.3929 | 0 | 25.5872 | Eukaryota | 43960.0 | 0 | Chordata | 43960.0 | 0 | Carnivora | 43960.0 | 0 | Felidae | 43960.0 | 0 | Felis | 43960.0 | 0 | Felis catus | 43960.0 | 0 |
| scf7180000017831 | 2567473 | 0.4606 | 0 | 53.359 | Eukaryota | 51461.0 | 0 | Chordata | 51461.0 | 0 | Carnivora | 51461.0 | 0 | Felidae | 46199.0 | 1 | Panthera | 30761.0 | 4 | anthera pardı | 25625.0 | 5 |
| scf7180000018552 | 86300 | 0.4214 | 0 | 46.124 | Eukaryota | 41078.0 | 0 | Chordata | 41078.0 | 0 | Carnivora | 41078.0 | 0 | Felidae | 41078.0 | 0 | Felis | 37009.0 | 1 | Felis catus | 37009.0 | 1 |
| scf7180000021145 | 89160 | 0.3868 | 0 | 24.6692 | Eukaryota | 42961.0 | 0 | Chordata | 42961.0 | 0 | Carnivora | 42961.0 | 0 | Felidae | 42961.0 | 0 | Felis | 42961.0 | 0 | Felis catus | 42961.0 | 0 |
| scf7180000020538 | 79872 | 0.3891 | 0 | 24.5152 | Eukaryota | 16254.0 | 0 | Chordata | 16254.0 | 0 | Carnivora | 14220.0 | 1 | Felidae | 14220.0 | 1 | Felis | 12679.0 | 2 | Felis catus | 12679.0 | 2 |
| scf7180000022710 | 10504 | 0.4048 | 0 | 23.5975 | Eukaryota | 5376.0 | 0 | Chordata | 5376.0 | 0 | Primates | 2972.0 | 3 | Hominidae | 2247.0 | 4 | Homo | 2247.0 | 5 | Homo sapiens | 2247.0 | 5 |
| scf7180000023888 | 1046732 | 0.4032 | 0 | 50.1333 | Eukaryota | 39295.0 | 0 | Chordata | 39295.0 | 0 | Carnivora | 39295.0 | 0 | Felidae | 39295.0 | 0 | Felis | 35427.0 | 1 | Felis catus | 35427.0 | 1 |
| scf7180000022944 | 58912 | 0.4929 | 0 | 370.5169 | Eukaryota | 7381.0 | 0 | Chordata | 7381.0 | 0 | Carnivora | 7381.0 | 0 | Otariidae | 5263.0 | 1 | Eumetopias | 2980.0 | 2 | netopias juba | 2980.0 | 2 |
| scf7180000020294 | 199178 | 0.3695 | 0 | 24.5878 | Eukaryota | 44050.0 | 0 | Chordata | 44050.0 | 0 | Carnivora | 44050.0 | 0 | Felidae | 44050.0 | 0 | Felis | 44050.0 | 0 | Felis catus | 44050.0 | 0 |
| scf7180000017890 | 753233 | 0.3653 | 0 | 50.0071 | Eukaryota | 47845.0 | 0 | Chordata | 47845.0 | 0 | Carnivora | 47845.0 | 0 | Felidae | 47845.0 | 0 | Felis | 47845.0 | 0 | Felis catus | 47845.0 | 0 |
| scf7180000021353 | 81750 | 0.3624 | 0 | 25.2704 | Eukaryota | 39842.0 | 0 | Chordata | 39842.0 | 0 | Carnivora | 39842.0 | 0 | Felidae | 39842.0 | 0 | Felis | 39842.0 | 0 | Felis catus | 39842.0 | 0 |
| scf7180000021252 | 37693 | 0.6066 | 0 | 53.9966 | Eukaryota | 2712.0 | 0 | Chordata | 2712.0 | 0 | Carnivora | 2712.0 | 0 | Felidae | 2013.0 | 2 | Felis | 1504.0 | 3 | Felis catus | 1504.0 | 3 |
| scf7180000018983 | 546463 | 0.3893 | 0 | 50.107 | Eukaryota | 57648.0 | 0 | Chordata | 57648.0 | 0 | Carnivora | 57648.0 | 0 | Felidae | 36733.0 | 1 | Mustela | 20915.0 | 5 | ıustela putori | 20915.0 | 5 |
| scf7180000020305 | 137697 | 0.4256 | 0 | 25.4917 | Eukaryota | 11268.0 | 0 | Chordata | 11268.0 | 0 | Primates | 7448.0 | 2 | Hominidae | 7448.0 | 2 | Pan | 5015.0 | 3 | an troglodyte | 5015.0 | 3 |
| scf7180000024160 | 149097 | 0.3746 | 0 | 24.2871 | Eukaryota | 44995.0 | 0 | Chordata | 44995.0 | 0 | Carnivora | 44995.0 | 0 | Felidae | 44995.0 | 0 | Felis | 44995.0 | 0 | Felis catus | 44995.0 | 0 |
| scf7180000019360 | 470177 | 0.3767 | 0 | 48.9231 | Eukaryota | 44582.0 | 0 | Chordata | 44582.0 | 0 | Carnivora | 44582.0 | 0 | Felidae | 44582.0 | 0 | Felis | 44582.0 | 0 | Felis catus | 44582.0 | 0 |
| scf7180000021738 | 62646 | 0.4525 | 0 | 26.7919 | Eukaryota | 31790.0 | 0 | Chordata | 31790.0 | 0 | Carnivora | 31790.0 | 0 | Felidae | 31790.0 | 0 | Felis | 28578.0 | 1 | Felis catus | 28578.0 | 1 |
| scf7180000020967 | 28043 | 0.4319 | 0 | 88.7594 | Eukaryota | 2370.0 | 0 | Chordata | 2370.0 | 0 | Carnivora | 2370.0 | 0 | Felidae | 2370.0 | 0 | Felis | 1698.0 | 2 | Felis catus | 1698.0 | 2 |
| scf7180000018302 | 35925 | 0.4813 | 0 | 44.2911 | Eukaryota | 31777.0 | 0 | Chordata | 31777.0 | 0 | Carnivora | 31777.0 | 0 | Felidae | 31777.0 | 0 | Felis | 31777.0 | 0 | Felis catus | 31777.0 | 0 |
| scf7180000019789 | 210996 | 0.4273 | 0 | 52.1162 | Eukaryota | 18187.0 | 0 | Chordata | 18187.0 | 0 | Carnivora | 18187.0 | 0 | Felidae | 14501.0 | 1 | Felis | 10893.0 | 3 | Felis catus | 10893.0 | 3 |
| scf7180000023365 | 23863 | 0.6991 | 0 | 183.1 | no-hit | 0.0 | 0 | no-hit | 0.0 | 0 | no-hit | 0.0 | 0 | no-hit | 0.0 | 0 | no-hit | 0.0 | 0 | no-hit | 0.0 | 0 |
| scf7180000017657 | 2262031 | 0.4428 | 100 | 51.7189 | Eukaryota | 47510.0 | 0 | Chordata | 47510.0 | 0 | Carnivora | 47510.0 | 0 | Felidae | 47510.0 | 0 | Felis | 42607.0 | 1 | Felis catus | 42607.0 | 1 |
| scf7180000020233 | 25908 | 0.3866 | 0 | 326.7483 | Eukaryota | 3716.0 | 0 | Chordata | 3716.0 | 0 | Primates | 2547.0 | 2 | Hominidae | 2170.0 | 4 | Homo | 2170.0 | 5 | Homo sapiens | 2170.0 | 5 |
| scf7180000020327 | 101666 | 0.4476 | 0 | 46.9087 | Eukaryota | 46949.0 | 0 | Chordata | 46949.0 | 0 | Carnivora | 46949.0 | 0 | Felidae | 37412.0 | 2 | Lynx | 18720.0 | 4 | ynx canadens | 18720.0 | 4 |
| scf7180000018194 | 1215156 | 0.4137 | 0 | 51.2162 | Eukaryota | 45096.0 | 0 | Chordata | 45096.0 | 0 | Carnivora | 45096.0 | 0 | Felidae | 45096.0 | 0 | Felis | 39760.0 | 1 | Felis catus | 39760.0 | 1 |
| scf7180000019714 | 185724 | 0.4112 | 0 | 50.2959 | Eukaryota | 40540.0 | 0 | Chordata | 40540.0 | 0 | Carnivora | 40540.0 | 0 | Mustelidae | 40540.0 | 0 | Enhydra | 40540.0 | 0 | Enhydra lutris | 40540.0 | 0 |
| scf7180000020833 | 58696 | 0.5489 | 0 | 48.7712 | Eukaryota | 5396.0 | 0 | Chordata | 5396.0 | 0 | Carnivora | 5396.0 | 0 | Felidae | 3076.0 | 1 | Suricata | 2320.0 | 2 | ıricata suricat | 2320.0 | 2 |
| scf7180000017521 | 39518 | 0.3837 | 0 | 24.5067 | Eukaryota | 39947.0 | 0 | Chordata | 39947.0 | 0 | Carnivora | 39947.0 | 0 | Felidae | 39947.0 | 0 | Felis | 39947.0 | 0 | Felis catus | 39947.0 | 0 |
| scf7180000024203 | 71762 | 0.3446 | 0 | 23.5247 | Eukaryota | 37753.0 | 0 | Chordata | 37753.0 | 0 | Carnivora | 37753.0 | 0 | Felidae | 23085.0 | 2 | Panthera | 19273.0 | 3 | anthera pardı | 15374.0 | 5 |
| scf7180000020852 | 94623 | 0.3783 | 0 | 26.8297 | Eukaryota | 47373.0 | 0 | Chordata | 47373.0 | 0 | Carnivora | 47373.0 | 0 | Felidae | 47373.0 | 0 | Felis | 47373.0 | 0 | Felis catus | 47373.0 | 0 |
| scf7180000021507 | 50871 | 0.6181 | 0 | 44.0422 | Eukaryota | 4225.0 | 0 | Chordata | 4225.0 | 0 | Crocodylia | 843.0 | 7 | unresolved | 427.0 | 9 | unresolved | 427.0 | 9 | unresolved | 427.0 | 9 |
| scf7180000019232 | 300397 | 0.4404 | 0 | 50.0515 | Eukaryota | 13270.0 | 0 | Chordata | 13270.0 | 0 | Carnivora | 13270.0 | 0 | Felidae | 13270.0 | 0 | Felis | 10601.0 | 1 | Felis catus | 10601.0 | 1 |
| scf7180000021587 | 5239 | 0.6064 | 0 | 145.4796 | no-hit | 0.0 | 0 | no-hit | 0.0 | 0 | no-hit | 0.0 | 0 | no-hit | 0.0 | 0 | no-hit | 0.0 | 0 | no-hit | 0.0 | 0 |
| scf7180000018804 | 15245 | 0.5863 | 0 | 36.2601 | Eukaryota | 11778.0 | 0 | Chordata | 11778.0 | 0 | Carnivora | 11778.0 | 0 | Felidae | 7380.0 | 2 | Felis | 7380.0 | 2 | Felis catus | 7380.0 | 2 |
| scf7180000021013 | 57575 | 0.354 | 0 | 25.6245 | Eukaryota | 21273.0 | 0 | Chordata | 21273.0 | 0 | Carnivora | 21273.0 | 0 | Felidae | 21273.0 | 0 | Felis | 19132.0 | 1 | Felis catus | 19132.0 | 1 |
| scf7180000020776 | 52636 | 0.4409 | 0 | 70.4273 | Eukaryota | 31274.0 | 0 | Chordata | 31274.0 | 0 | Carnivora | 31274.0 | 0 | Felidae | 31274.0 | 0 | Felis | 28171.0 | 1 | Felis catus | 28171.0 | 1 |
| scf7180000024114 | 190551 | 0.604 | 0 | 51.4059 | Eukaryota | 10556.0 | 0 | Chordata | 10556.0 | 0 | Carnivora | 10556.0 | 0 | Felidae | 6493.0 | 1 | Suricata | 4063.0 | 5 | ıricata suricat | 4063.0 | 6 |
| scf7180000020621 | 93708 | 0.3888 | 0 | 48.7108 | Eukaryota | 51568.0 | 0 | Chordata | 51568.0 | 0 | Primates | 31783.0 | 1 | Hominidae | 27020.0 | 4 | Pan | 27020.0 | 4 | an troglodyte | 27020.0 | 4 |
| scf7180000017498 | 2981175 | 0.4047 | 0 | 51.4459 | Eukaryota | 49930.0 | 0 | Chordata | 49930.0 | 0 | Carnivora | 49930.0 | 0 | Felidae | 49930.0 | 0 | Felis | 49930.0 | 0 | Felis catus | 49930.0 | 0 |
| scf7180000020549 | 69841 | 0.5066 | 0 | 48.6925 | Eukaryota | 44693.0 | 0 | Chordata | 44693.0 | 0 | Carnivora | 44693.0 | 0 | Felidae | 44693.0 | 0 | Felis | 44693.0 | 0 | Felis catus | 44693.0 | 0 |
| scf7180000017632 | 176535 | 0.3505 | 0 | 47.1497 | Eukaryota | 27804.0 | 0 | Chordata | 27804.0 | 0 | Carnivora | 27804.0 | 0 | Felidae | 27804.0 | 0 | Felis | 25061.0 | 1 | Felis catus | 25061.0 | 1 |
| scf7180000017626 | 651192 | 0.3491 | 0 | 46.5929 | Eukaryota | 46411.0 | 0 | Chordata | 46411.0 | 0 | Carnivora | 46411.0 | 0 | Felidae | 46411.0 | 0 | Felis | 46411.0 | 0 | Felis catus | 46411.0 | 0 |
| scf7180000022665 | 8232 | 0.6279 | 0 | 126.1238 | Eukaryota | 398.0 | 0 | Chordata | 398.0 | 0 | Carnivora | 398.0 | 0 | Felidae | 398.0 | 0 | Panthera | 398.0 | 0 | anthera pardı | 398.0 | 0 |
| scf7180000017951 | 151939 | 0.3923 | 0 | 44.3735 | Eukaryota | 41292.0 | 0 | Chordata | 41292.0 | 0 | Carnivora | 41292.0 | 0 | Felidae | 41292.0 | 0 | Felis | 41292.0 | 0 | Felis catus | 41292.0 | 0 |
| scf7180000018711 | 446567 | 0.4917 | 0 | 51.3342 | Eukaryota | 46956.0 | 1 | Chordata | 46956.0 | 1 | Carnivora | 36074.0 | 2 | Felidae | 21137.0 | 5 | Homo | 10882.0 | 8 | Homo sapiens | 10882.0 | 8 |
| scf7180000021494 | 38480 | 0.3541 | 0 | 23.2897 | Eukaryota | 19099.0 | 0 | Chordata | 19099.0 | 0 | Carnivora | 19099.0 | 0 | Felidae | 19099.0 | 0 | Felis | 19099.0 | 0 | Felis catus | 19099.0 | 0 |
| scf7180000021096 | 70940 | 0.4461 | 0 | 26.0301 | Eukaryota | 7085.0 | 0 | Chordata | 7085.0 | 0 | Carnivora | 3963.0 | 1 | Herpestidae | 3394.0 | 2 | Suricata | 3394.0 | 3 | ıricata suricat | 3394.0 | 3 |
| scf7180000023793 | 3008591 | 0.455 | 0 | 52.4452 | Eukaryota | 47675.0 | 0 | Chordata | 47675.0 | 0 | Carnivora | 47675.0 | 0 | Felidae | 47675.0 | 0 | Felis | 47675.0 | 0 | Felis catus | 47675.0 | 0 |
| scf7180000018787 | 575637 | 0.5442 | 0 | 51.3214 | Eukaryota | 42364.0 | 0 | Chordata | 42364.0 | 0 | Carnivora | 42364.0 | 0 | Herpestidae | 33432.0 | 1 | Suricata | 33432.0 | 1 | ıricata suricat | 33432.0 | 1 |
| scf7180000021912 | 32008 | 0.5615 | 0 | 54.2618 | Eukaryota | 3664.0 | 0 | Chordata | 3664.0 | 0 | Carnivora | 2665.0 | 1 | Felidae | 1074.0 | 5 | Homo | 999.0 | 6 | Homo sapiens | 999.0 | 6 |
| scf7180000019283 | 341706 | 0.4815 | 0 | 53.3833 | Eukaryota | 14198.0 | 0 | Chordata | 14198.0 | 0 | Carnivora | 8714.0 | 1 | Hominidae | 5484.0 | 2 | Homo | 5484.0 | 2 | Homo sapiens | 5484.0 | 2 |
| scf7180000023321 | 20516 | 0.421 | 0 | 618.1639 | Eukaryota | 13477.0 | 0 | Chordata | 13477.0 | 0 | Carnivora | 13477.0 | 0 | Felidae | 13477.0 | 0 | Felis | 12145.0 | 1 | Felis catus | 12145.0 | 1 |
| scf7180000022127 | 21171 | 0.3412 | 0 | 24.2751 | Eukaryota | 8255.0 | 0 | Chordata | 8255.0 | 0 | Carnivora | 8255.0 | 0 | Felidae | 5748.0 | 2 | Felis | 5748.0 | 2 | Felis catus | 5748.0 | 2 |
| scf7180000020623 | 93521 | 0.3677 | 0 | 48.8968 | Eukaryota | 20080.0 | 0 | Chordata | 20080.0 | 0 | Carnivora | 20080.0 | 0 | Felidae | 20080.0 | 0 | Felis | 20080.0 | 0 | Felis catus | 20080.0 | 0 |
| scf7180000018500 | 19863 | 0.391 | 0 | 35.8667 | Eukaryota | 47654.0 | 0 | Chordata | 47654.0 | 0 | Carnivora | 47654.0 | 0 | Felidae | 47654.0 | 0 | Felis | 47654.0 | 0 | Felis catus | 47654.0 | 0 |
| scf7180000019812 | 259796 | 0.3495 | 0 | 48.397 | Eukaryota | 43318.0 | 0 | Chordata | 43318.0 | 0 | Carnivora | 43318.0 | 0 | Felidae | 43318.0 | 0 | Felis | 39022.0 | 1 | Felis catus | 39022.0 | 1 |
| scf7180000021682 | 48840 | 0.3706 | 0 | 49.4184 | Eukaryota | 5187.0 | 0 | Chordata | 5187.0 | 0 | Carnivora | 4043.0 | 1 | Felidae | 3097.0 | 2 | Felis | 3097.0 | 2 | Felis catus | 3097.0 | 2 |
| scf7180000017600 | 2378078 | 0.3956 | 0 | 50.8981 | Eukaryota | 46494.0 | 0 | Chordata | 46494.0 | 0 | Carnivora | 46494.0 | 0 | Felidae | 46494.0 | 0 | Felis | 46494.0 | 0 | Felis catus | 46494.0 | 0 |
| scf7180000018591 | 620299 | 0.5571 | 0 | 50.4128 | Eukaryota | 40835.0 | 0 | Chordata | 40835.0 | 0 | Carnivora | 40835.0 | 0 | Felidae | 40835.0 | 0 | Panthera | 20425.0 | 1 | anthera pardı | 20425.0 | 1 |
| scf7180000022198 | 5468 | 0.5216 | 0 | 118.4302 | no-hit | 0.0 | 0 | no-hit | 0.0 | 0 | no-hit | 0.0 | 0 | no-hit | 0.0 | 0 | no-hit | 0.0 | 0 | no-hit | 0.0 | 0 |
| scf7180000020954 | 92322 | 0.3607 | 0 | 25.103 | Eukaryota | 35567.0 | 0 | Chordata | 35567.0 | 0 | Carnivora | 35567.0 | 0 | Felidae | 35567.0 | 0 | Felis | 35567.0 | 0 | Felis catus | 35567.0 | 0 |
| scf7180000020477 | 216918 | 0.3944 | 0 | 25.4459 | Eukaryota | 126431.0 | 0 | Chordata | 126431.0 | 0 | Carnivora | 126431.0 | 0 | Felidae | 63921.0 | 2 | Ursus | 24774.0 | 8 | ynx canadens | 12839.0 | 9 |
| scf7180000020520 | 132329 | 0.3746 | 0 | 24.7817 | Eukaryota | 38782.0 | 0 | Chordata | 38782.0 | 0 | Carnivora | 38782.0 | 0 | Felidae | 38782.0 | 0 | Felis | 34946.0 | 1 | Felis catus | 34946.0 | 1 |
| scf7180000019406 | 27219 | 0.4273 | 0 | 26.1191 | Eukaryota | 13492.0 | 0 | Chordata | 13492.0 | 0 | Primates | 5749.0 | 3 | Hominidae | 5749.0 | 3 | Mus | 5452.0 | 4 | Mus musculus | 5452.0 | 4 |
| scf7180000018631 | 655193 | 0.4654 | 0 | 50.7403 | Eukaryota | 59655.0 | 0 | Chordata | 59655.0 | 0 | Carnivora | 59655.0 | 0 | Felidae | 59655.0 | 0 | Acinonyx | 59655.0 | 0 | :inonyx jubat | 59655.0 | 0 |
| scf7180000018254 | 914731 | 0.3846 | 0 | 49.6203 | Eukaryota | 36042.0 | 0 | Chordata | 36042.0 | 0 | Carnivora | 36042.0 | 0 | Felidae | 20906.0 | 1 | Felis | 20906.0 | 1 | Felis catus | 20906.0 | 1 |
| scf7180000017997 | 1218459 | 0.3836 | 0 | 48.3641 | Eukaryota | 47022.0 | 0 | Chordata | 47022.0 | 0 | Carnivora | 47022.0 | 0 | Felidae | 47022.0 | 0 | Felis | 47022.0 | 0 | Felis catus | 47022.0 | 0 |
| scf7180000019952 | 321633 | 0.4537 | 0 | 49.1692 | Eukaryota | 38792.0 | 0 | Chordata | 38792.0 | 0 | Carnivora | 38792.0 | 0 | Felidae | 38792.0 | 0 | Felis | 38792.0 | 0 | Felis catus | 38792.0 | 0 |
| scf7180000023147 | 38597 | 0.5832 | 0 | 182.0753 | Eukaryota | 4020.0 | 0 | Chordata | 4020.0 | 0 | Carnivora | 4020.0 | 0 | Felidae | 2534.0 | 2 | Felis | 2242.0 | 3 | Felis catus | 2242.0 | 3 |
| scf7180000021805 | 27896 | 0.6026 | 0 | 53.6853 | Eukaryota | 4069.0 | 0 | Chordata | 4069.0 | 0 | Carnivora | 4069.0 | 0 | Felidae | 3081.0 | 3 | Panthera | 1133.0 | 7 | anthera pardı | 902.0 | 8 |
| scf7180000017799 | 305539 | 0.4019 | 0 | 53.1184 | Eukaryota | 47846.0 | 0 | Chordata | 47846.0 | 0 | Carnivora | 47846.0 | 0 | Felidae | 47846.0 | 0 | Felis | 47846.0 | 0 | Felis catus | 47846.0 | 0 |
| scf7180000022428 | 21428 | 0.4246 | 0 | 25.1967 | Eukaryota | 7308.0 | 0 | Chordata | 7308.0 | 0 | Carnivora | 7308.0 | 0 | Mustelidae | 2007.0 | 4 | Felis | 1724.0 | 7 | Felis catus | 1724.0 | 7 |
| scf7180000022763 | 9356 | 0.5147 | 0 | 51.822 | Eukaryota | 3152.0 | 0 | Chordata | 3152.0 | 0 | Carnivora | 3152.0 | 0 | Felidae | 2909.0 | 1 | Felis | 2633.0 | 2 | Felis catus | 2633.0 | 2 |
| scf7180000021113 | 17995 | 0.4054 | 0 | 23.2231 | Eukaryota | 10377.0 | 0 | Chordata | 10377.0 | 0 | Carnivora | 10377.0 | 0 | Felidae | 10377.0 | 0 | Felis | 10377.0 | 0 | Felis catus | 10377.0 | 0 |
| scf7180000021521 | 53535 | 0.4612 | 0 | 25.9748 | Eukaryota | 46217.0 | 0 | Chordata | 46217.0 | 0 | Carnivora | 46217.0 | 0 | Felidae | 40296.0 | 1 | Acinonyx | 31104.0 | 2 | :inonyx jubat | 31104.0 | 2 |
| scf7180000021796 | 55783 | 0.6335 | 0 | 46.9811 | Eukaryota | 9287.0 | 0 | Chordata | 9287.0 | 0 | Carnivora | 9287.0 | 0 | Felidae | 4804.0 | 3 | Acinonyx | 4804.0 | 5 | :inonyx jubat | 4804.0 | 5 |

| | | | | | | | | | | | | | | | | | | | | | | |
|---|---|---|---|---|---|---|---|---|---|---|---|---|---|---|---|---|---|---|---|---|---|---|
| scf7180000019753 | 16470 | 0.6054 | 0 | 173.1729 | Eukaryota | 2388.0 | 0 | Chordata | 2388.0 | 0 | Carnivora | 1945.0 | 1 | Felidae | 1710.0 | 3 | Felis | 1094.0 | 5 | Felis catus | 1094.0 | 5 |
| scf7180000020049 | 143669 | 0.6273 | 0 | 43.7891 | Eukaryota | 15378.0 | 0 | Chordata | 15378.0 | 0 | Carnivora | 13484.0 | 1 | Felidae | 13484.0 | 2 | Panthera | 4212.0 | 6 | anthera pardu | 4212.0 | 6 |
| scf7180000018683 | 558840 | 0.4258 | 0 | 49.4288 | Eukaryota | 48277.0 | 0 | Chordata | 48277.0 | 0 | Carnivora | 44217.0 | 1 | Herpestidae | 44217.0 | 1 | Suricata | 44217.0 | 1 | ricata suricat | 44217.0 | 1 |
| scf7180000021884 | 31134 | 0.3644 | 0 | 25.5216 | Eukaryota | 7026.0 | 0 | Chordata | 7026.0 | 0 | Carnivora | 2526.0 | 2 | Hominidae | 2155.0 | 4 | Homo | 2155.0 | 4 | Homo sapiens | 2155.0 | 4 |
| scf7180000018822 | 600165 | 0.354 | 0 | 48.374 | Eukaryota | 44156.0 | 0 | Chordata | 44156.0 | 0 | Carnivora | 44156.0 | 0 | Felidae | 44156.0 | 0 | Felis | 39792.0 | 1 | Felis catus | 39792.0 | 1 |
| scf7180000021153 | 83096 | 0.6069 | 0 | 46.1809 | Eukaryota | 11948.0 | 0 | Chordata | 11948.0 | 0 | Carnivora | 11948.0 | 0 | Felidae | 5091.0 | 4 | Mustela | 2299.0 | 8 | ustela putori | 2299.0 | 8 |
| scf7180000022632 | 16129 | 0.4223 | 0 | 25.0351 | Eukaryota | 5632.0 | 0 | Chordata | 5632.0 | 0 | Carnivora | 3531.0 | 1 | Felidae | 3531.0 | 1 | Felis | 3531.0 | 2 | Felis catus | 3531.0 | 2 |
| scf7180000021004 | 100110 | 0.4678 | 0 | 28.1651 | Eukaryota | 44280.0 | 0 | Chordata | 44280.0 | 0 | Carnivora | 44280.0 | 0 | Felidae | 44280.0 | 0 | Felis | 35334.0 | 2 | Felis catus | 35334.0 | 2 |
| scf7180000019552 | 331620 | 0.5104 | 0 | 52.8516 | Eukaryota | 26761.0 | 0 | Chordata | 26761.0 | 0 | Carnivora | 26761.0 | 0 | Felidae | 26761.0 | 0 | Acinonyx | 16146.0 | 2 | cinonyx jubat | 16146.0 | 2 |
| scf7180000017472 | 3328613 | 0.3784 | 0 | 50.2979 | Eukaryota | 47266.0 | 0 | Chordata | 47266.0 | 0 | Carnivora | 47266.0 | 0 | Felidae | 47266.0 | 0 | Felis | 47266.0 | 0 | Felis catus | 47266.0 | 0 |
| scf7180000019602 | 13043 | 0.3866 | 0 | 23.596 | Eukaryota | 11406.0 | 0 | Chordata | 11406.0 | 0 | Carnivora | 11406.0 | 0 | Felidae | 4948.0 | 2 | Suricata | 4138.0 | 4 | ricata suricat | 4138.0 | 4 |
| scf7180000017830 | 1614564 | 0.3687 | 0 | 49.3908 | Eukaryota | 45877.0 | 0 | Chordata | 45877.0 | 0 | Carnivora | 45877.0 | 0 | Felidae | 45877.0 | 0 | Felis | 45877.0 | 0 | Felis catus | 45877.0 | 0 |
| scf7180000020749 | 48964 | 0.4511 | 0 | 55.2451 | Eukaryota | 14550.0 | 0 | Chordata | 14550.0 | 0 | Carnivora | 14550.0 | 0 | Felidae | 14550.0 | 0 | Felis | 13111.0 | 1 | Felis catus | 13111.0 | 1 |
| scf7180000019198 | 430938 | 0.3664 | 0 | 49.2455 | Eukaryota | 30172.0 | 0 | Chordata | 30172.0 | 0 | Carnivora | 30172.0 | 0 | Canidae | 21406.0 | 1 | Canis | 21406.0 | 1 | Canis lupus | 21406.0 | 1 |
| scf7180000022647 | 11960 | 0.3855 | 0 | 23.6736 | Eukaryota | 11156.0 | 0 | Chordata | 11156.0 | 0 | Carnivora | 8636.0 | 1 | Phocidae | 4330.0 | 4 | Neomonachus | 2167.0 | 7 | nachus schau | 2167.0 | 7 |
| scf7180000020482 | 23350 | 0.3771 | 0 | 24.9776 | Eukaryota | 18475.0 | 0 | Chordata | 18475.0 | 0 | Carnivora | 18475.0 | 0 | Felidae | 18475.0 | 0 | Felis | 16611.0 | 1 | Felis catus | 16611.0 | 1 |
| scf7180000018303 | 918646 | 0.4166 | 0 | 51.3548 | Eukaryota | 42650.0 | 0 | Chordata | 42650.0 | 0 | Carnivora | 42650.0 | 0 | Felidae | 42650.0 | 0 | Felis | 42650.0 | 0 | Felis catus | 42650.0 | 0 |
| scf7180000023544 | 19281 | 0.4958 | 0 | 246.5987 | Eukaryota | 3699.0 | 0 | Chordata | 3699.0 | 0 | Carnivora | 3699.0 | 0 | Felidae | 3699.0 | 0 | unresolved | 1104.0 | 4 | unresolved | 1104.0 | 4 |
| scf7180000017776 | 22419 | 0.6686 | 0 | 1419.1319 | Eukaryota | 636.0 | 0 | Chordata | 495.0 | 1 | Carnivora | 495.0 | 1 | Felidae | 356.0 | 2 | Felis | 356.0 | 2 | Felis catus | 356.0 | 2 |
| scf7180000022782 | 3727 | 0.3944 | 0 | 24.6036 | Eukaryota | 1651.0 | 0 | Chordata | 1651.0 | 0 | Carnivora | 1651.0 | 0 | Felidae | 1325.0 | 1 | Felis | 1151.0 | 3 | Felis catus | 1151.0 | 3 |
| scf7180000021437 | 76085 | 0.5025 | 0 | 49.2555 | Eukaryota | 12628.0 | 0 | Chordata | 12628.0 | 0 | Primates | 12628.0 | 0 | Hominidae | 7611.0 | 2 | Homo | 3845.0 | 5 | Homo sapiens | 3845.0 | 5 |
| scf7180000018549 | 524754 | 0.3951 | 0 | 68.0297 | Eukaryota | 45054.0 | 0 | Chordata | 45054.0 | 0 | Carnivora | 45054.0 | 0 | Felidae | 45054.0 | 0 | Felis | 45054.0 | 0 | Felis catus | 45054.0 | 0 |
| scf7180000023176 | 12425 | 0.5129 | 0 | 213.2061 | Eukaryota | 2854.0 | 0 | Chordata | 2854.0 | 0 | Carnivora | 2854.0 | 0 | Felidae | 2600.0 | 1 | Felis | 988.0 | 3 | Felis catus | 988.0 | 3 |
| scf7180000023627 | 47818 | 0.4018 | 0 | 375.86 | Eukaryota | 12161.0 | 0 | Chordata | 12161.0 | 0 | Carnivora | 12161.0 | 0 | Felidae | 12161.0 | 0 | Felis | 12161.0 | 0 | Felis catus | 12161.0 | 0 |
| scf7180000019287 | 434904 | 0.4307 | 0 | 52.3595 | Eukaryota | 40157.0 | 0 | Chordata | 40157.0 | 0 | Carnivora | 28281.0 | 1 | Phocidae | 14690.0 | 3 | eptonychote | 14690.0 | 4 | nychotes we | 14690.0 | 4 |
| scf7180000017667 | 1856774 | 0.446 | 0 | 52.7789 | Eukaryota | 48120.0 | 0 | Chordata | 48120.0 | 0 | Carnivora | 43237.0 | 1 | Mustelidae | 14630.0 | 6 | Mustela | 14630.0 | 6 | ustela putori | 14630.0 | 6 |
| scf7180000021178 | 68121 | 0.3918 | 0 | 25.6704 | Eukaryota | 70959.0 | 0 | Chordata | 70959.0 | 0 | erissodactyla | 48834.0 | 2 | Equidae | 48834.0 | 2 | Equus | 48834.0 | 2 | Equus caballu | 48834.0 | 2 |
| scf7180000022120 | 34418 | 0.4193 | 0 | 26.0527 | Eukaryota | 20777.0 | 0 | Chordata | 20777.0 | 0 | Carnivora | 20777.0 | 0 | Felidae | 20777.0 | 0 | Felis | 20777.0 | 0 | Felis catus | 20777.0 | 0 |
| scf7180000021641 | 30300 | 0.4367 | 0 | 25.8402 | Eukaryota | 5616.0 | 0 | Chordata | 5616.0 | 0 | Carnivora | 3352.0 | 2 | Felidae | 3352.0 | 2 | Felis | 2770.0 | 4 | Felis catus | 2770.0 | 4 |
| scf7180000021589 | 41879 | 0.3761 | 0 | 24.6439 | Eukaryota | 37303.0 | 0 | Chordata | 37303.0 | 0 | Carnivora | 37303.0 | 0 | Felidae | 37303.0 | 0 | Felis | 33609.0 | 1 | Felis catus | 33609.0 | 1 |
| scf7180000022631 | 9535 | 0.5227 | 0 | 155.273 | Eukaryota | 2303.0 | 0 | Chordata | 2303.0 | 0 | Carnivora | 2303.0 | 0 | Felidae | 1822.0 | 2 | Felis | 986.0 | 4 | Felis catus | 986.0 | 4 |
| scf7180000019407 | 309355 | 0.6212 | 0 | 48.8758 | Eukaryota | 14613.0 | 0 | Chordata | 14613.0 | 0 | Carnivora | 14613.0 | 0 | Felidae | 9548.0 | 4 | Felis | 3711.0 | 8 | Felis catus | 3711.0 | 8 |
| scf7180000022729 | 23235 | 0.4007 | 0 | 27.94 | Eukaryota | 38886.0 | 0 | Chordata | 38886.0 | 0 | Carnivora | 38886.0 | 0 | Felidae | 38886.0 | 0 | Felis | 35054.0 | 1 | Felis catus | 35054.0 | 1 |
| scf7180000020281 | 86046 | 0.5791 | 0 | 50.7601 | Eukaryota | 8160.0 | 0 | Chordata | 8160.0 | 0 | Carnivora | 8160.0 | 0 | Felidae | 8160.0 | 0 | Felis | 7363.0 | 1 | Felis catus | 7363.0 | 1 |
| scf7180000024005 | 422974 | 0.4354 | 0 | 52.7444 | Eukaryota | 51928.0 | 0 | Chordata | 51928.0 | 0 | Carnivora | 51928.0 | 0 | Felidae | 51928.0 | 0 | Lynx | 31236.0 | 1 | ynx canadens | 31236.0 | 1 |
| scf7180000020672 | 100237 | 0.4037 | 0 | 48.2232 | Eukaryota | 28559.0 | 0 | Chordata | 28559.0 | 0 | Carnivora | 28559.0 | 0 | Felidae | 28559.0 | 0 | Felis | 25663.0 | 1 | Felis catus | 25663.0 | 1 |
| scf7180000024283 | 1846948 | 0.3955 | 100 | 52.0112 | Eukaryota | 35899.0 | 0 | Chordata | 35899.0 | 0 | Carnivora | 35899.0 | 0 | Felidae | 25156.0 | 1 | Acinonyx | 17960.0 | 2 | cinonyx jubat | 17960.0 | 2 |
| scf7180000021707 | 35228 | 0.3809 | 0 | 46.274 | Eukaryota | 35559.0 | 0 | Chordata | 35559.0 | 0 | Carnivora | 35559.0 | 0 | Felidae | 35559.0 | 0 | Felis | 35559.0 | 0 | Felis catus | 35559.0 | 0 |
| scf7180000019379 | 415179 | 0.4195 | 0 | 50.5976 | Eukaryota | 35858.0 | 0 | Chordata | 35858.0 | 0 | Carnivora | 35858.0 | 0 | Felidae | 35858.0 | 0 | Felis | 35858.0 | 0 | Felis catus | 35858.0 | 0 |
| scf7180000023611 | 22210 | 0.4252 | 0 | 408.6044 | Eukaryota | 237156.0 | 0 | Chordata | 237156.0 | 0 | Carnivora | 237156.0 | 0 | Hyaenidae | 237156.0 | 0 | Proteles | 142792.0 | 2 | roteles cristat | 142792.0 | 2 |
| scf7180000021080 | 50634 | 0.4468 | 0 | 25.2292 | Eukaryota | 7858.0 | 0 | Chordata | 7858.0 | 0 | Primates | 3232.0 | 4 | Hominidae | 3232.0 | 5 | Homo | 3232.0 | 6 | Homo sapiens | 3232.0 | 6 |
| scf7180000022599 | 23431 | 0.6695 | 0 | 49.7086 | Eukaryota | 326.0 | 0 | Chordata | 326.0 | 0 | Carnivora | 326.0 | 0 | Felidae | 326.0 | 0 | Acinonyx | 176.0 | 1 | cinonyx jubat | 176.0 | 1 |
| scf7180000018773 | 446739 | 0.528 | 0 | 49.3392 | Eukaryota | 6722.0 | 1 | Chordata | 6722.0 | 1 | Carnivora | 5098.0 | 2 | Felidae | 5098.0 | 2 | Felis | 5098.0 | 2 | Felis catus | 5098.0 | 2 |
| scf7180000024042 | 434726 | 0.4191 | 0 | 48.4663 | Eukaryota | 44743.0 | 0 | Chordata | 44743.0 | 0 | Carnivora | 44743.0 | 0 | Felidae | 44743.0 | 0 | Felis | 44743.0 | 0 | Felis catus | 44743.0 | 0 |
| scf7180000017733 | 1675958 | 0.3714 | 0 | 50.8054 | Eukaryota | 45846.0 | 0 | Chordata | 45846.0 | 0 | Carnivora | 45846.0 | 0 | Felidae | 45846.0 | 0 | Felis | 45846.0 | 0 | Felis catus | 45846.0 | 0 |
| scf7180000021842 | 21945 | 0.3677 | 0 | 24.417 | Eukaryota | 38117.0 | 0 | Chordata | 38117.0 | 0 | Carnivora | 38117.0 | 0 | Felidae | 38117.0 | 0 | Felis | 38117.0 | 0 | Felis catus | 38117.0 | 0 |
| scf7180000021055 | 82563 | 0.5314 | 0 | 53.8023 | Eukaryota | 3098.0 | 0 | Chordata | 3098.0 | 0 | Carnivora | 2423.0 | 1 | Bovidae | 675.0 | 5 | Odobenus | 614.0 | 7 | obenus rosma | 614.0 | 7 |
| scf7180000019745 | 185291 | 0.4396 | 0 | 52.1606 | Eukaryota | 28962.0 | 0 | Chordata | 28962.0 | 0 | Carnivora | 28962.0 | 0 | Felidae | 28962.0 | 0 | Felis | 26020.0 | 1 | Felis catus | 26020.0 | 1 |
| scf7180000022808 | 9977 | 0.3637 | 0 | 22.1028 | Eukaryota | 25469.0 | 0 | Chordata | 25469.0 | 0 | Carnivora | 25469.0 | 0 | Felidae | 25469.0 | 0 | Felis | 22972.0 | 1 | Felis catus | 22972.0 | 1 |
| scf7180000018606 | 329104 | 0.5101 | 0 | 49.9252 | Eukaryota | 19294.0 | 0 | Chordata | 19294.0 | 0 | Carnivora | 19294.0 | 0 | Felidae | 19294.0 | 0 | Felis | 17449.0 | 1 | Felis catus | 17449.0 | 1 |
| scf7180000020493 | 183735 | 0.3743 | 0 | 24.5086 | Eukaryota | 42686.0 | 0 | Chordata | 42686.0 | 0 | Carnivora | 42686.0 | 0 | Felidae | 42686.0 | 0 | Felis | 42686.0 | 0 | Felis catus | 42686.0 | 0 |
| scf7180000020714 | 107473 | 0.3992 | 0 | 24.6569 | Eukaryota | 45699.0 | 0 | Chordata | 45699.0 | 0 | Carnivora | 45699.0 | 0 | Felidae | 45699.0 | 0 | Felis | 45699.0 | 0 | Felis catus | 45699.0 | 0 |
| scf7180000019218 | 67820 | 0.398 | 0 | 36.7305 | Eukaryota | 46057.0 | 0 | Chordata | 46057.0 | 0 | Carnivora | 46057.0 | 0 | Felidae | 46057.0 | 0 | Felis | 46057.0 | 0 | Felis catus | 46057.0 | 0 |
| scf7180000018533 | 229163 | 0.445 | 0 | 50.6393 | Eukaryota | 20305.0 | 0 | Chordata | 20305.0 | 0 | Carnivora | 20305.0 | 0 | Felidae | 14235.0 | 2 | Suricata | 4396.0 | 5 | ricata suricat | 4396.0 | 6 |
| scf7180000023488 | 97112 | 0.3875 | 0 | 369.1585 | Eukaryota | 23029.0 | 0 | Chordata | 23029.0 | 0 | Carnivora | 23029.0 | 0 | Felidae | 23029.0 | 0 | Felis | 20733.0 | 1 | Felis catus | 20733.0 | 1 |
| scf7180000020695 | 121137 | 0.3814 | 0 | 49.0011 | Eukaryota | 7300.0 | 0 | Chordata | 7300.0 | 0 | Carnivora | 5156.0 | 2 | Felidae | 5156.0 | 2 | Felis | 3873.0 | 3 | Felis catus | 3873.0 | 3 |
| scf7180000022684 | 17278 | 0.4004 | 0 | 25.8054 | Eukaryota | 18728.0 | 0 | Chordata | 18728.0 | 0 | Carnivora | 18728.0 | 0 | Felidae | 18728.0 | 0 | Felis | 16870.0 | 1 | Felis catus | 16870.0 | 1 |
| scf7180000019872 | 156909 | 0.5142 | 0 | 47.338 | Eukaryota | 9430.0 | 0 | Chordata | 9430.0 | 0 | Carnivora | 9430.0 | 0 | Felidae | 9430.0 | 0 | Felis | 4720.0 | 1 | Felis catus | 4720.0 | 1 |
| scf7180000020723 | 136584 | 0.5177 | 0 | 48.0103 | Eukaryota | 11610.0 | 0 | Chordata | 11610.0 | 0 | Carnivora | 11610.0 | 0 | Felidae | 11610.0 | 0 | Felis | 10389.0 | 1 | Felis catus | 10389.0 | 1 |
| scf7180000019435 | 323293 | 0.4683 | 0 | 50.4398 | Eukaryota | 45467.0 | 0 | Chordata | 45467.0 | 0 | Carnivora | 45467.0 | 0 | Felidae | 45467.0 | 0 | Felis | 45467.0 | 0 | Felis catus | 45467.0 | 0 |
| scf7180000022021 | 18834 | 0.4546 | 0 | 22.2214 | Eukaryota | 8045.0 | 0 | Chordata | 8045.0 | 0 | Carnivora | 8045.0 | 0 | Felidae | 8045.0 | 0 | Felis | 7164.0 | 1 | Felis catus | 7164.0 | 1 |
| scf7180000022494 | 29232 | 0.4019 | 0 | 24.5829 | Eukaryota | 5281.0 | 0 | Chordata | 5281.0 | 0 | Carnivora | 5281.0 | 0 | Felidae | 5281.0 | 0 | Felis | 5281.0 | 0 | Felis catus | 5281.0 | 0 |
| scf7180000024106 | 240719 | 0.6035 | 0 | 51.0779 | Eukaryota | 14767.0 | 0 | Chordata | 14767.0 | 0 | Carnivora | 13392.0 | 1 | Felidae | 10491.0 | 3 | Lynx | 7497.0 | 4 | ynx canadens | 7497.0 | 4 |
| scf7180000020446 | 64063 | 0.384 | 0 | 25.3068 | Eukaryota | 38170.0 | 0 | Chordata | 38170.0 | 0 | Carnivora | 38170.0 | 0 | Felidae | 38170.0 | 0 | Felis | 38170.0 | 0 | Felis catus | 38170.0 | 0 |
| scf7180000020928 | 14695 | 0.5219 | 0 | 56.5058 | no-hit | 0.0 | 0 | no-hit | 0.0 | 0 | no-hit | 0.0 | 0 | no-hit | 0.0 | 0 | no-hit | 0.0 | 0 | no-hit | 0.0 | 0 |
| scf7180000020170 | 111067 | 0.3843 | 0 | 27.1214 | Eukaryota | 40048.0 | 0 | Chordata | 40048.0 | 0 | Carnivora | 40048.0 | 0 | Felidae | 40048.0 | 0 | Felis | 36048.0 | 1 | Felis catus | 36048.0 | 1 |
| scf7180000022700 | 12173 | 0.481 | 0 | 24.337 | Eukaryota | 4548.0 | 0 | Chordata | 4548.0 | 0 | Artiodactyla | 2542.0 | 3 | Hominidae | 1284.0 | 6 | Homo | 1284.0 | 8 | Homo sapiens | 1284.0 | 8 |
| scf7180000020899 | 97893 | 0.4189 | 0 | 25.9524 | Eukaryota | 26205.0 | 0 | Chordata | 26205.0 | 0 | Carnivora | 26205.0 | 0 | Felidae | 26205.0 | 0 | Felis | 20944.0 | 1 | Felis catus | 20944.0 | 1 |
| scf7180000017965 | 524944 | 0.3538 | 0 | 49.6143 | Eukaryota | 47029.0 | 0 | Chordata | 47029.0 | 0 | Carnivora | 47029.0 | 0 | Felidae | 47029.0 | 0 | Felis | 47029.0 | 0 | Felis catus | 47029.0 | 0 |
| scf7180000022168 | 18763 | 0.4046 | 0 | 27.6147 | Eukaryota | 10251.0 | 0 | Chordata | 10251.0 | 0 | Primates | 5887.0 | 2 | Hominidae | 5887.0 | 2 | Homo | 4400.0 | 4 | Homo sapiens | 4400.0 | 4 |
| scf7180000023317 | 187843 | 0.4488 | 0 | 151.3221 | Eukaryota | 44848.0 | 0 | Chordata | 44848.0 | 0 | Carnivora | 44848.0 | 0 | Felidae | 42866.0 | 1 | Acinonyx | 17292.0 | 3 | cinonyx jubat | 17292.0 | 3 |
| scf7180000021287 | 78249 | 0.4041 | 0 | 25.0141 | Eukaryota | 41301.0 | 0 | Chordata | 41301.0 | 0 | Carnivora | 41301.0 | 0 | Felidae | 41301.0 | 0 | Felis | 33769.0 | 1 | Felis catus | 33769.0 | 1 |
| scf7180000024182 | 104149 | 0.3846 | 0 | 23.9562 | Eukaryota | 44172.0 | 0 | Chordata | 44172.0 | 0 | Carnivora | 44172.0 | 0 | Felidae | 44172.0 | 0 | Felis | 39810.0 | 1 | Felis catus | 39810.0 | 1 |
| scf7180000021709 | 33070 | 0.4068 | 0 | 28.1886 | Eukaryota | 45919.0 | 0 | Chordata | 45919.0 | 0 | Carnivora | 45919.0 | 0 | Felidae | 45919.0 | 0 | Felis | 45919.0 | 0 | Felis catus | 45919.0 | 0 |
| scf7180000023893 | 792137 | 0.3902 | 0 | 49.9752 | Eukaryota | 46430.0 | 0 | Chordata | 46430.0 | 0 | Carnivora | 46430.0 | 0 | Felidae | 46430.0 | 0 | Felis | 46430.0 | 0 | Felis catus | 46430.0 | 0 |
| scf7180000018304 | 1247196 | 0.3877 | 0 | 51.2102 | Eukaryota | 47211.0 | 0 | Chordata | 47211.0 | 0 | Carnivora | 47211.0 | 0 | Felidae | 47211.0 | 0 | Felis | 32747.0 | 3 | Felis catus | 32747.0 | 3 |
| scf7180000020894 | 69511 | 0.6368 | 0 | 43.9015 | Eukaryota | 4189.0 | 0 | Chordata | 4189.0 | 0 | Carnivora | 4189.0 | 0 | Felidae | 4189.0 | 0 | Felis | 3340.0 | 1 | Felis catus | 3340.0 | 1 |
| scf7180000017493 | 1733809 | 0.3651 | 0 | 50.1903 | Eukaryota | 46541.0 | 0 | Chordata | 46541.0 | 0 | Carnivora | 46541.0 | 0 | Felidae | 46541.0 | 0 | Felis | 46541.0 | 0 | Felis catus | 46541.0 | 0 |
| scf7180000020578 | 69355 | 0.4102 | 0 | 46.9319 | Eukaryota | 8956.0 | 0 | Chordata | 8956.0 | 0 | Primates | 4063.0 | 2 | Hominidae | 4063.0 | 4 | Homo | 4063.0 | 7 | Homo sapiens | 4063.0 | 7 |
| scf7180000017515 | 21837 | 0.3509 | 0 | 25.1385 | Eukaryota | 7587.0 | 0 | Chordata | 7587.0 | 0 | pterygiforme | 6053.0 | 2 | Apterygidae | 6053.0 | 2 | Apteryx | 6053.0 | 2 | pteryx mante | 6053.0 | 2 |
| scf7180000023929 | 1170892 | 0.4509 | 0 | 52.4307 | Eukaryota | 47275.0 | 0 | Chordata | 47275.0 | 0 | Carnivora | 47275.0 | 0 | Felidae | 38087.0 | 2 | Panthera | 9604.0 | 6 | anthera pardu | 9604.0 | 6 |
| scf7180000023464 | 51682 | 0.4197 | 0 | 294.1501 | Eukaryota | 9392.0 | 0 | Chordata | 9392.0 | 0 | Carnivora | 9392.0 | 0 | Felidae | 9392.0 | 0 | Felis | 7465.0 | 1 | Felis catus | 7465.0 | 1 |
| scf7180000020323 | 2305895 | 0.4476 | 0 | 51.9079 | Eukaryota | 144860.0 | 0 | Chordata | 144860.0 | 0 | Carnivora | 144860.0 | 0 | Felidae | 144860.0 | 0 | Lynx | 144860.0 | 0 | ynx canadens | 144860.0 | 0 |
| scf7180000019672 | 235906 | 0.3566 | 0 | 49.303 | Eukaryota | 45883.0 | 0 | Chordata | 45883.0 | 0 | Carnivora | 45883.0 | 0 | Felidae | 45883.0 | 0 | Felis | 45883.0 | 0 | Felis catus | 45883.0 | 0 |
| scf7180000022702 | 4867 | 0.5194 | 0 | 70.5319 | no-hit | 0.0 | 0 | no-hit | 0.0 | 0 | no-hit | 0.0 | 0 | no-hit | 0.0 | 0 | no-hit | 0.0 | 0 | no-hit | 0.0 | 0 |
| scf7180000020527 | 142398 | 0.4196 | 0 | 26.2613 | Eukaryota | 11209.0 | 0 | Chordata | 11209.0 | 0 | pterygiforme | 7681.0 | 3 | Apterygidae | 7681.0 | 3 | Apteryx | 7681.0 | 3 | pteryx mante | 7681.0 | 3 |
| scf7180000019698 | 452480 | 0.4874 | 0 | 50.0582 | Eukaryota | 16731.0 | 0 | Chordata | 16731.0 | 0 | Carnivora | 16731.0 | 0 | Felidae | 13136.0 | 2 | Felis | 7116.0 | 4 | Felis catus | 7116.0 | 4 |
| scf7180000018198 | 993662 | 0.5152 | 0 | 52.2708 | Eukaryota | 35390.0 | 0 | Chordata | 35390.0 | 0 | Carnivora | 32619.0 | 1 | Felidae | 32619.0 | 1 | Lynx | 14084.0 | 4 | ynx canadens | 14084.0 | 4 |
| scf7180000018571 | 627307 | 0.4819 | 0 | 50.7861 | Eukaryota | 43135.0 | 0 | Chordata | 43135.0 | 0 | Carnivora | 43135.0 | 0 | Felidae | 43135.0 | 0 | Panthera | 17321.0 | 3 | anthera pardu | 13005.0 | 4 |
| scf7180000022784 | 12328 | 0.3625 | 0 | 24.3221 | Eukaryota | 21369.0 | 0 | Chordata | 21369.0 | 0 | Carnivora | 21369.0 | 0 | Felidae | 21369.0 | 0 | Felis | 19258.0 | 1 | Felis catus | 19258.0 | 1 |
| scf7180000017666 | 1531446 | 0.4476 | 0 | 60.7663 | Eukaryota | 45516.0 | 0 | Chordata | 45516.0 | 0 | Carnivora | 45516.0 | 0 | Felidae | 22761.0 | 1 | Suricata | 22755.0 | 2 | ricata suricat | 22755.0 | 2 |
| scf7180000023985 | 542211 | 0.4954 | 0 | 51.4708 | Eukaryota | 25745.0 | 0 | Chordata | 25745.0 | 0 | Carnivora | 25745.0 | 0 | Felidae | 6280.0 | 5 | Panthera | 6280.0 | 7 | ricata suricat | 6004.0 | 8 |
| scf7180000020012 | 185772 | 0.4775 | 0 | 51.2888 | Eukaryota | 9547.0 | 0 | Chordata | 9547.0 | 0 | Carnivora | 9547.0 | 0 | Felidae | 4494.0 | 2 | Acinonyx | 2672.0 | 4 | cinonyx jubat | 2672.0 | 6 |
| scf7180000023712 | 38314 | 0.5468 | 0 | 210.0761 | Eukaryota | 12577.0 | 0 | Chordata | 12577.0 | 0 | Carnivora | 12577.0 | 0 | Felidae | 12577.0 | 0 | Felis | 12577.0 | 0 | Felis catus | 12577.0 | 0 |
| scf7180000022392 | 7392 | 0.63 | 0 | 58.5763 | Eukaryota | 3203.0 | 0 | Chordata | 3203.0 | 0 | Carnivora | 1681.0 | 2 | Hominidae | 588.0 | 8 | Homo | 588.0 | 8 | Homo sapiens | 588.0 | 8 |
| scf7180000021950 | 32096 | 0.4505 | 0 | 27.3553 | Eukaryota | 6460.0 | 0 | Chordata | 6460.0 | 0 | Primates | 4909.0 | 3 | Hominidae | 4909.0 | 3 | Pan | 3371.0 | 4 | an troglodyte | 3371.0 | 4 |
| scf7180000020244 | 166162 | 0.4418 | 0 | 24.2095 | Eukaryota | 21198.0 | 0 | Chordata | 21198.0 | 0 | Carnivora | 21198.0 | 0 | Canidae | 7407.0 | 4 | Canis | 5539.0 | 8 | Canis lupus | 5539.0 | 8 |
| scf7180000022131 | 26308 | 0.4516 | 0 | 23.7591 | Eukaryota | 3602.0 | 0 | Chordata | 3602.0 | 0 | Carnivora | 2074.0 | 3 | Felidae | 1491.0 | 4 | Felis | 1491.0 | 4 | Felis catus | 1491.0 | 4 |
| scf7180000021983 | 29019 | 0.3846 | 0 | 23.2495 | Eukaryota | 32972.0 | 0 | Chordata | 32972.0 | 0 | Carnivora | 32972.0 | 0 | Felidae | 32972.0 | 0 | Felis | 32972.0 | 0 | Felis catus | 32972.0 | 0 |
| scf7180000020653 | 75985 | 0.4169 | 0 | 1019.437 | Eukaryota | 13555.0 | 0 | Chordata | 13555.0 | 0 | Carnivora | 13555.0 | 0 | Felidae | 13555.0 | 0 | Felis | 13555.0 | 0 | Felis catus | 13555.0 | 0 |
| scf7180000018888 | 474220 | 0.5292 | 0 | 51.962 | Eukaryota | 19603.0 | 0 | Chordata | 19603.0 | 0 | Carnivora | 14421.0 | 1 | Felidae | 14421.0 | 1 | Felis | 12631.0 | 2 | Felis catus | 12631.0 | 2 |
| scf7180000023889 | 226671 | 0.503 | 0 | 51.5279 | Eukaryota | 18774.0 | 1 | Chordata | 18774.0 | 1 | Carnivora | 11832.0 | 2 | Hominidae | 6942.0 | 3 | Homo | 6942.0 | 5 | Homo sapiens | 6942.0 | 5 |
| scf7180000018413 | 1008811 | 0.3936 | 100 | 51.364 | Eukaryota | 48441.0 | 0 | Chordata | 48441.0 | 0 | Carnivora | 48441.0 | 0 | Felidae | 41809.0 | 1 | Felis | 41809.0 | 1 | Felis catus | 41809.0 | 1 |
| scf7180000022294 | 22132 | 0.4303 | 0 | 27.2916 | Eukaryota | 29634.0 | 0 | Chordata | 29634.0 | 0 | Carnivora | 19594.0 | 2 | Felidae | 16960.0 | 5 | Felis | 9996.0 | 6 | Felis catus | 9996.0 | 6 |
| scf7180000019544 | 277661 | 0.6162 | 0 | 49.6499 | Eukaryota | 13170.0 | 0 | Chordata | 13170.0 | 0 | Carnivora | 13170.0 | 0 | Felidae | 13170.0 | 0 | Felis | 9289.0 | 1 | Felis catus | 9289.0 | 1 |
| scf7180000022034 | 22849 | 0.4557 | 0 | 27.1826 | Eukaryota | 4192.0 | 0 | Chordata | 4192.0 | 0 | Primates | 1416.0 | 6 | Hominidae | 1416.0 | 6 | Homo | 1416.0 | 6 | Homo sapiens | 1416.0 | 6 |
| scf7180000022654 | 14048 | 0.3921 | 0 | 48.4494 | Eukaryota | 2739.0 | 1 | Chordata | 2739.0 | 1 | Primates | 1841.0 | 2 | Hominidae | 1841.0 | 3 | Homo | 1841.0 | 4 | Homo sapiens | 1841.0 | 4 |
| scf7180000018707 | 492488 | 0.4767 | 0 | 53.3353 | Eukaryota | 52361.0 | 0 | Chordata | 52361.0 | 0 | Carnivora | 52361.0 | 0 | Felidae | 52361.0 | 0 | Felis | 26310.0 | 2 | Felis catus | 26310.0 | 2 |
| scf7180000019976 | 228696 | 0.5667 | 0 | 48.1806 | Eukaryota | 13144.0 | 0 | Chordata | 13144.0 | 0 | Carnivora | 13144.0 | 0 | Felidae | 13144.0 | 0 | Felis | 13144.0 | 0 | Felis catus | 13144.0 | 0 |
| scf7180000021915 | 21713 | 0.4261 | 0 | 198.6144 | Eukaryota | 12999.0 | 0 | Chordata | 12999.0 | 0 | Carnivora | 12999.0 | 0 | Felidae | 12999.0 | 0 | Felis | 11719.0 | 1 | Felis catus | 11719.0 | 1 |
| scf7180000020509 | 119860 | 0.5258 | 0 | 52.0434 | Eukaryota | 26432.0 | 0 | Chordata | 26432.0 | 0 | Carnivora | 26432.0 | 0 | Felidae | 23076.0 | 1 | Panthera | 7669.0 | 5 | cinonyx jubat | 5128.0 | 6 |
| scf7180000019271 | 386992 | 0.3846 | 0 | 50.7661 | Eukaryota | 77178.0 | 0 | Chordata | 77178.0 | 0 | Carnivora | 77178.0 | 0 | Felidae | 61594.0 | 1 | Panthera | 30822.0 | 2 | anthera pardu | 30822.0 | 2 |
| scf7180000020676 | 125666 | 0.391 | 0 | 50.8472 | Eukaryota | 46511.0 | 0 | Chordata | 46511.0 | 0 | Carnivora | 46511.0 | 0 | Felidae | 46511.0 | 0 | Felis | 46511.0 | 0 | Felis catus | 46511.0 | 0 |
| scf7180000020728 | 94497 | 0.3676 | 0 | 23.3571 | Eukaryota | 45183.0 | 0 | Chordata | 45183.0 | 0 | Carnivora | 45183.0 | 0 | Felidae | 45183.0 | 0 | Felis | 45183.0 | 0 | Felis catus | 45183.0 | 0 |
| scf7180000020235 | 133899 | 0.4407 | 0 | 48.1496 | Eukaryota | 12647.0 | 1 | Chordata | 12647.0 | 1 | Carnivora | 8019.0 | 2 | Hominidae | 4628.0 | 3 | Suricata | 3879.0 | 6 | ricata suricat | 3879.0 | 7 |
| scf7180000018250 | 1212801 | 0.4133 | 0 | 52.851 | Eukaryota | 90957.0 | 0 | Chordata | 90957.0 | 0 | Carnivora | 90957.0 | 0 | Felidae | 81551.0 | 1 | Felis | 46370.0 | 3 | Felis catus | 46370.0 | 3 |
| scf7180000023591 | 10399 | 0.7211 | 0 | 479.6195 | Eukaryota | 2544.0 | 0 | Chordata | 2544.0 | 0 | Carnivora | 2544.0 | 0 | Felidae | 2314.0 | 1 | Panthera | 1164.0 | 3 | anthera pardu | 1164.0 | 3 |
| scf7180000020205 | 129294 | 0.4428 | 0 | 49.8922 | Eukaryota | 18707.0 | 1 | Chordata | 18707.0 | 1 | Primates | 9116.0 | 3 | Hominidae | 7182.0 | 4 | Homo | 7182.0 | 5 | Homo sapiens | 7182.0 | 5 |
| scf7180000020031 | 153270 | 0.5288 | 0 | 48.8091 | Eukaryota | 10360.0 | 0 | Chordata | 10360.0 | 0 | Carnivora | 5897.0 | 1 | Felidae | 5897.0 | 2 | Felis | 5897.0 | 3 | Felis catus | 5897.0 | 3 |
| scf7180000020888 | 23481 | 0.3812 | 0 | 216.7353 | Eukaryota | 6451.0 | 0 | Chordata | 6451.0 | 0 | Carnivora | 6451.0 | 0 | Felidae | 6451.0 | 0 | Felis | 5804.0 | 1 | Felis catus | 5804.0 | 1 |
| scf7180000017592 | 603623 | 0.3812 | 0 | 51.5491 | Eukaryota | 46635.0 | 0 | Chordata | 46635.0 | 0 | Carnivora | 46635.0 | 0 | Felidae | 46635.0 | 0 | Felis | 46635.0 | 0 | Felis catus | 46635.0 | 0 |
| scf7180000017847 | 4851842 | 0.403 | 0 | 50.6371 | Eukaryota | 49611.0 | 0 | Chordata | 49611.0 | 0 | Carnivora | 49611.0 | 0 | Felidae | 49611.0 | 0 | Felis | 42593.0 | 1 | Felis catus | 42593.0 | 1 |
| scf7180000020663 | 62207 | 0.4354 | 0 | 211.9961 | Eukaryota | 22544.0 | 0 | Chordata | 22544.0 | 0 | Carnivora | 22544.0 | 0 | Felidae | 22544.0 | 0 | Felis | 20322.0 | 1 | Felis catus | 20322.0 | 1 |
| scf7180000017531 | 3501263 | 0.3985 | 0 | 50.68 | Eukaryota | 56308.0 | 0 | Chordata | 56308.0 | 0 | Carnivora | 56308.0 | 0 | Felidae | 28779.0 | 2 | Panthera | 17304.0 | 6 | anthera pardu | 17304.0 | 6 |
| scf7180000020775 | 22740 | 0.5207 | 0 | 180.0123 | no-hit | 0.0 | 0 | no-hit | 0.0 | 0 | no-hit | 0.0 | 0 | no-hit | 0.0 | 0 | no-hit | 0.0 | 0 | no-hit | 0.0 | 0 |
| scf7180000023505 | 13686 | 0.4326 | 0 | 381.0383 | Eukaryota | 2294.0 | 0 | Chordata | 2294.0 | 0 | Carnivora | 2294.0 | 0 | Felidae | 2294.0 | 0 | Felis | 1622.0 | 2 | Felis catus | 1622.0 | 2 |
| scf7180000021821 | 40069 | 0.4816 | 0 | 24.7998 | Eukaryota | 20964.0 | 0 | Chordata | 20964.0 | 0 | Chiroptera | 13150.0 | 2 | Pteropodidae | 13150.0 | 2 | Pteropus | 13150.0 | 3 | ropus vampy | 7089.0 | 4 |
| scf7180000024233 | 116822 | 0.458 | 0 | 51.5049 | Eukaryota | 9183.0 | 0 | Chordata | 9183.0 | 0 | Carnivora | 9183.0 | 0 | Felidae | 9183.0 | 0 | Felis | 9183.0 | 0 | Felis catus | 9183.0 | 0 |
| scf7180000019002 | 581696 | 0.4919 | 100 | 47.7963 | Eukaryota | 29004.0 | 0 | Chordata | 29004.0 | 0 | Carnivora | 29004.0 | 0 | Felidae | 20286.0 | 3 | Felis | 5844.0 | 7 | Felis catus | 5844.0 | 7 |
| scf7180000020651 | 128176 | 0.4585 | 0 | 52.1106 | Eukaryota | 46673.0 | 0 | Chordata | 46673.0 | 0 | Carnivora | 46673.0 | 0 | Felidae | 46673.0 | 0 | Felis | 46673.0 | 0 | Felis catus | 46673.0 | 0 |
| scf7180000021453 | 29867 | 0.5977 | 0 | 48.0833 | Eukaryota | 15090.0 | 0 | Chordata | 15090.0 | 0 | Carnivora | 15090.0 | 0 | Felidae | 10389.0 | 1 | Felis | 9186.0 | 2 | Felis catus | 9186.0 | 2 |
| scf7180000023468 | 21640 | 0.6692 | 0 | 223.794 | Eukaryota | 2553.0 | 0 | Chordata | 2553.0 | 0 | Rodentia | 1040.0 | 3 | Cercopithecidae | 1000.0 | 4 | Macaca | 1000.0 | 4 | Cavia porcellu | 783.0 | 5 |
| scf7180000018451 | 436218 | 0.3637 | 0 | 47.9374 | Eukaryota | 46567.0 | 0 | Chordata | 46567.0 | 0 | Carnivora | 46567.0 | 0 | Felidae | 46567.0 | 0 | Felis | 41924.0 | 1 | Felis catus | 41924.0 | 1 |
| scf7180000022812 | 14474 | 0.7229 | 0 | 318.6821 | Eukaryota | 2451.0 | 0 | Chordata | 2451.0 | 0 | Carnivora | 2451.0 | 0 | Felidae | 1789.0 | 1 | Panthera | 1132.0 | 2 | anthera pardu | 1132.0 | 2 |
| scf7180000017684 | 2912771 | 0.4745 | 0 | 53.5633 | Eukaryota | 107198.0 | 0 | Chordata | 107198.0 | 0 | Carnivora | 107198.0 | 0 | Felidae | 96681.0 | 1 | Panthera | 42977.0 | 5 | anthera pardu | 32205.0 | 6 |
| scf7180000019419 | 286407 | 0.5144 | 0 | 51.37 | Eukaryota | 29106.0 | 0 | Chordata | 29106.0 | 0 | Carnivora | 29106.0 | 0 | Felidae | 25768.0 | 1 | Acinonyx | 20055.0 | 3 | cinonyx jubat | 20055.0 | 3 |
| scf7180000022709 | 11412 | 0.4409 | 0 | 2162.8245 | Eukaryota | 6141.0 | 0 | Chordata | 6141.0 | 0 | Artiodactyla | 6141.0 | 0 | Bovidae | 6141.0 | 0 | Ovis | 6141.0 | 0 | vis canadensi | 6141.0 | 0 |
| scf7180000021149 | 60045 | 0.4401 | 0 | 24.5086 | Eukaryota | 12234.0 | 0 | Chordata | 12234.0 | 0 | Carnivora | 12234.0 | 0 | Felidae | 8809.0 | 1 | Felis | 8809.0 | 1 | Felis catus | 8809.0 | 1 |
| scf7180000022983 | 20974 | 0.5095 | 0 | 140.4046 | Eukaryota | 16434.0 | 0 | Chordata | 16434.0 | 0 | Carnivora | 16434.0 | 0 | Felidae | 16434.0 | 0 | Felis | 16434.0 | 0 | Felis catus | 16434.0 | 0 |
| scf7180000022289 | 18320 | 0.3777 | 0 | 25.101 | Eukaryota | 22650.0 | 0 | Chordata | 22650.0 | 0 | Carnivora | 22650.0 | 0 | Felidae | 22650.0 | 0 | Felis | 20594.0 | 1 | Felis catus | 20594.0 | 1 |
| scf7180000021994 | 22792 | 0.4108 | 0 | 25.7683 | Eukaryota | 45925.0 | 0 | Chordata | 45925.0 | 0 | Carnivora | 45925.0 | 0 | Felidae | 45925.0 | 0 | Felis | 45925.0 | 0 | Felis catus | 45925.0 | 0 |
| scf7180000024044 | 235935 | 0.5116 | 0 | 49.4014 | Eukaryota | 30512.0 | 0 | Chordata | 30512.0 | 0 | Carnivora | 30512.0 | 0 | Felidae | 30512.0 | 0 | Felis | 27557.0 | 1 | Felis catus | 27557.0 | 1 |
| scf7180000019963 | 186402 | 0.4913 | 0 | 25.6709 | Eukaryota | 37468.0 | 0 | Chordata | 37468.0 | 0 | Carnivora | 37468.0 | 0 | Felidae | 34853.0 | 1 | Felis | 15472.0 | 3 | Felis catus | 15472.0 | 3 |
| scf7180000021317 | 18149 | 0.3974 | 0 | 396.6992 | Eukaryota | 6437.0 | 0 | Chordata | 6437.0 | 0 | Carnivora | 6437.0 | 0 | Felidae | 6437.0 | 0 | Felis | 6437.0 | 0 | Felis catus | 6437.0 | 0 |
| scf7180000020204 | 62716 | 0.4 | 0 | 48.8337 | Eukaryota | 13339.0 | 0 | Chordata | 13339.0 | 0 | Carnivora | 13339.0 | 0 | Felidae | 13339.0 | 0 | Felis | 12009.0 | 1 | Felis catus | 12009.0 | 1 |
| scf7180000024147 | 134698 | 0.4396 | 0 | 24.8695 | Eukaryota | 10355.0 | 0 | Chordata | 10355.0 | 0 | Artiodactyla | 4128.0 | 4 | Canidae | 3139.0 | 5 | Canis | 2102.0 | 7 | Canis lupus | 2102.0 | 7 |
| scf7180000017537 | 2725435 | 0.4861 | 0 | 50.1083 | Eukaryota | 38476.0 | 0 | Chordata | 38476.0 | 0 | Carnivora | 38476.0 | 0 | Canidae | 30544.0 | 1 | Canis | 30544.0 | 2 | Canis lupus | 30544.0 | 2 |
| scf7180000022742 | 13771 | 0.4101 | 0 | 312.2336 | no-hit | 0.0 | 0 | no-hit | 0.0 | 0 | no-hit | 0.0 | 0 | no-hit | 0.0 | 0 | no-hit | 0.0 | 0 | no-hit | 0.0 | 0 |
| scf7180000021392 | 52665 | 0.5624 | 0 | 50.8012 | Eukaryota | 6943.0 | 0 | Chordata | 6943.0 | 0 | Carnivora | 3250.0 | 2 | Hominidae | 2629.0 | 4 | Pan | 2102.0 | 6 | Canis lupus | 2073.0 | 7 |
| scf7180000023671 | 31093 | 0.6082 | 0 | 325.0505 | Eukaryota | 2987.0 | 0 | Chordata | 2987.0 | 0 | Carnivora | 2987.0 | 0 | Felidae | 2432.0 | 2 | Panthera | 1651.0 | 3 | anthera pardu | 1324.0 | 4 |
| scf7180000018440 | 961050 | 0.4248 | 100 | 52.1297 | Eukaryota | 73739.0 | 0 | Chordata | 73739.0 | 0 | Carnivora | 73739.0 | 0 | Phocidae | 29364.0 | 3 | Callorhinus | 22002.0 | 5 | llorhinus ursi | 22002.0 | 5 |
| scf7180000019934 | 168184 | 0.4737 | 0 | 27.7762 | Eukaryota | 35655.0 | 0 | Chordata | 35655.0 | 0 | Carnivora | 35655.0 | 0 | Felidae | 32079.0 | 1 | Acinonyx | 28520.0 | 2 | cinonyx jubat | 28520.0 | 2 |
| scf7180000022415 | 361617 | 0.475 | 0 | 47.2649 | Eukaryota | 41401.0 | 0 | Chordata | 41401.0 | 0 | Carnivora | 41401.0 | 0 | Felidae | 37281.0 | 1 | Acinonyx | 20744.0 | 2 | cinonyx jubat | 20744.0 | 2 |
| scf7180000022866 | 24129 | 0.489 | 0 | 122.3639 | Eukaryota | 4345.0 | 0 | Chordata | 4345.0 | 0 | Artiodactyla | 2867.0 | 1 | Bovidae | 2867.0 | 2 | Ovis | 2867.0 | 2 | vis canadensi | 2144.0 | 3 |
| scf7180000019907 | 184294 | 0.3709 | 0 | 25.7812 | Eukaryota | 46905.0 | 0 | Chordata | 46905.0 | 0 | Carnivora | 34712.0 | 1 | Felidae | 34712.0 | 1 | Felis | 34712.0 | 1 | Felis catus | 34712.0 | 1 |
| scf7180000018605 | 345149 | 0.3503 | 0 | 45.8855 | Eukaryota | 45179.0 | 0 | Chordata | 45179.0 | 0 | Carnivora | 45179.0 | 0 | Felidae | 45179.0 | 0 | Felis | 45179.0 | 0 | Felis catus | 45179.0 | 0 |
| scf7180000019345 | 392470 | 0.3848 | 0 | 51.6514 | Eukaryota | 31227.0 | 0 | Chordata | 31227.0 | 0 | Carnivora | 31227.0 | 0 | Canidae | 11756.0 | 3 | Mustela | 9354.0 | 4 | ustela putori | 9354.0 | 5 |
| scf7180000017652 | 1275946 | 0.3542 | 0 | 48.1929 | Eukaryota | 26880.0 | 0 | Chordata | 26880.0 | 0 | Carnivora | 26880.0 | 0 | Felidae | 26880.0 | 0 | Felis | 24143.0 | 1 | Felis catus | 24143.0 | 1 |
| scf7180000021029 | 73149 | 0.3613 | 0 | 26.0375 | Eukaryota | 32565.0 | 0 | Chordata | 32565.0 | 0 | Carnivora | 32565.0 | 0 | Felidae | 32565.0 | 0 | Felis | 32565.0 | 0 | Felis catus | 32565.0 | 0 |

| | | | | | | | | | | | | | | | | | | | | | | |
|---|---|---|---|---|---|---|---|---|---|---|---|---|---|---|---|---|---|---|---|---|---|---|
| scf7180000023392 | 37401 | 0.5228 | 0 | 187.2557 | Eukaryota | 4612.0 | 0 | Chordata | 4612.0 | 0 | Carnivora | 4612.0 | 0 | Felidae | 4612.0 | 0 | Felis | 4340.0 | 1 | Felis catus | 4340.0 | 1 |
| scf7180000020697 | 102869 | 0.3707 | 0 | 23.5313 | Eukaryota | 42765.0 | 0 | Chordata | 42765.0 | 0 | Carnivora | 42765.0 | 0 | Felidae | 42765.0 | 0 | Felis | 38526.0 | 1 | Felis catus | 38526.0 | 1 |
| scf7180000018043 | 320537 | 0.3755 | 0 | 47.5136 | Eukaryota | 47296.0 | 0 | Chordata | 47296.0 | 0 | Carnivora | 47296.0 | 0 | Felidae | 47296.0 | 0 | Felis | 47296.0 | 0 | Felis catus | 47296.0 | 0 |
| scf7180000023407 | 29459 | 0.4262 | 0 | 436.4912 | Eukaryota | 256970.0 | 0 | Chordata | 256970.0 | 0 | Carnivora | 256970.0 | 0 | Hyaenidae | 256970.0 | 0 | Proteles | 155516.0 | 1 | roteles cristat | 155516.0 | 1 |
| scf7180000019566 | 264565 | 0.394 | 0 | 50.325 | Eukaryota | 26243.0 | 0 | Chordata | 26243.0 | 0 | Carnivora | 21975.0 | 1 | Felidae | 21975.0 | 1 | Panthera | 7476.0 | 4 | anthera pardu | 7476.0 | 4 |
| scf7180000024148 | 172681 | 0.4034 | 0 | 25.4203 | Eukaryota | 44036.0 | 0 | Chordata | 44036.0 | 0 | Carnivora | 44036.0 | 0 | Felidae | 31678.0 | 2 | Panthera | 10594.0 | 5 | anthera pardu | 10594.0 | 6 |
| scf7180000023974 | 484271 | 0.5219 | 0 | 53.072 | Eukaryota | 27143.0 | 0 | Chordata | 27143.0 | 0 | Carnivora | 27143.0 | 0 | Felidae | 27143.0 | 0 | Felis | 24441.0 | 1 | Felis catus | 24441.0 | 1 |
| scf7180000024174 | 72119 | 0.383 | 0 | 24.2181 | Eukaryota | 32148.0 | 0 | Chordata | 32148.0 | 0 | Carnivora | 32148.0 | 0 | Felidae | 32148.0 | 0 | Felis | 32148.0 | 0 | Felis catus | 32148.0 | 0 |
| scf7180000020881 | 79110 | 0.5051 | 0 | 49.0677 | Eukaryota | 4135.0 | 0 | Chordata | 4135.0 | 0 | Carnivora | 3741.0 | 1 | Canidae | 2557.0 | 2 | Canis | 2557.0 | 2 | Canis lupus | 2557.0 | 2 |
| scf7180000023917 | 1381266 | 0.3902 | 0 | 51.9365 | Eukaryota | 48674.0 | 0 | Chordata | 48674.0 | 0 | Carnivora | 48674.0 | 0 | Felidae | 48674.0 | 0 | Felis | 39066.0 | 2 | Felis catus | 39066.0 | 2 |
| scf7180000018567 | 1924453 | 0.4017 | 0 | 49.9099 | Eukaryota | 67267.0 | 0 | Chordata | 67267.0 | 0 | Carnivora | 67267.0 | 0 | Ursidae | 44185.0 | 1 | Ursus | 44185.0 | 1 | Ursus arctos | 44185.0 | 1 |
| scf7180000019833 | 315101 | 0.3676 | 0 | 24.4865 | Eukaryota | 37077.0 | 0 | Chordata | 37077.0 | 0 | Carnivora | 37077.0 | 0 | Felidae | 37077.0 | 0 | Felis | 37077.0 | 0 | Felis catus | 37077.0 | 0 |
| scf7180000018530 | 64863 | 0.4521 | 0 | 486.9128 | Eukaryota | 42605.0 | 0 | Chordata | 42605.0 | 0 | Carnivora | 42605.0 | 0 | Felidae | 42605.0 | 0 | Felis | 38376.0 | 1 | Felis catus | 38376.0 | 1 |
| scf7180000024200 | 76937 | 0.6324 | 0 | 50.3941 | Eukaryota | 5419.0 | 0 | Chordata | 5419.0 | 0 | Carnivora | 4875.0 | 1 | Felidae | 3755.0 | 2 | Acinonyx | 3261.0 | 3 | inonyx jubat | 3261.0 | 3 |
| scf7180000022109 | 35495 | 0.3817 | 0 | 24.7183 | Eukaryota | 41807.0 | 0 | Chordata | 41807.0 | 0 | Carnivora | 41807.0 | 0 | Felidae | 41807.0 | 0 | Felis | 37683.0 | 1 | Felis catus | 37683.0 | 1 |
| scf7180000019489 | 297099 | 0.3585 | 0 | 49.4126 | Eukaryota | 45929.0 | 0 | Chordata | 45929.0 | 0 | Carnivora | 45929.0 | 0 | Felidae | 45929.0 | 0 | Felis | 45929.0 | 0 | Felis catus | 45929.0 | 0 |
| scf7180000021136 | 29425 | 0.5821 | 0 | 42.4593 | Eukaryota | 2850.0 | 0 | Chordata | 2850.0 | 0 | Carnivora | 2128.0 | 1 | Felidae | 1865.0 | 2 | Felis | 1865.0 | 2 | Felis catus | 1865.0 | 2 |
| scf7180000021785 | 31190 | 0.6893 | 0 | 49.6264 | Eukaryota | 7995.0 | 1 | Chordata | 7995.0 | 1 | Carnivora | 4307.0 | 2 | Hominidae | 3688.0 | 3 | Homo | 3688.0 | 6 | Homo sapiens | 3688.0 | 6 |
| scf7180000023748 | 12154 | 0.432 | 0 | 421.5333 | no-hit | 0.0 | 0 | no-hit | 0.0 | 0 | no-hit | 0.0 | 0 | no-hit | 0.0 | 0 | no-hit | 0.0 | 0 | no-hit | 0.0 | 0 |
| scf7180000021190 | 14885 | 0.5234 | 0 | 69.951 | no-hit | 0.0 | 0 | no-hit | 0.0 | 0 | no-hit | 0.0 | 0 | no-hit | 0.0 | 0 | no-hit | 0.0 | 0 | no-hit | 0.0 | 0 |
| scf7180000020735 | 79064 | 0.3863 | 0 | 25.6688 | Eukaryota | 23758.0 | 0 | Chordata | 23758.0 | 0 | Primates | 9923.0 | 3 | Hominidae | 9923.0 | 3 | Rousettus | 6692.0 | 4 | ettus aegypti | 6692.0 | 4 |
| scf7180000024039 | 909529 | 0.4498 | 0 | 52.0609 | Eukaryota | 46754.0 | 0 | Chordata | 46754.0 | 0 | Carnivora | 46754.0 | 0 | Felidae | 46754.0 | 0 | Felis | 46754.0 | 0 | Felis catus | 46754.0 | 0 |
| scf7180000024033 | 311040 | 0.443 | 0 | 51.2842 | Eukaryota | 41089.0 | 0 | Chordata | 41089.0 | 0 | Carnivora | 41089.0 | 0 | Felidae | 28885.0 | 2 | Suricata | 8348.0 | 6 | ricata suricat | 8348.0 | 7 |
| scf7180000019670 | 17308 | 0.423 | 0 | 26.6544 | Eukaryota | 13901.0 | 0 | Chordata | 13901.0 | 0 | Carnivora | 13901.0 | 0 | Felidae | 13901.0 | 0 | Felis | 11112.0 | 1 | Felis catus | 11112.0 | 1 |
| scf7180000018343 | 773145 | 0.4077 | 0 | 50.7501 | Eukaryota | 46712.0 | 0 | Chordata | 46712.0 | 0 | Carnivora | 46712.0 | 0 | Felidae | 46712.0 | 0 | Felis | 46712.0 | 0 | Felis catus | 46712.0 | 0 |
| scf7180000017741 | 2339019 | 0.4056 | 0 | 51.6882 | Eukaryota | 85825.0 | 0 | Chordata | 85825.0 | 0 | Carnivora | 85825.0 | 0 | Felidae | 59752.0 | 1 | Acinonyx | 51168.0 | 2 | inonyx jubat | 51168.0 | 2 |
| scf7180000023850 | 798385 | 0.4384 | 0 | 52.8414 | Eukaryota | 50800.0 | 0 | Chordata | 50800.0 | 0 | Carnivora | 50800.0 | 0 | Felidae | 50800.0 | 0 | Felis | 50800.0 | 0 | Felis catus | 50800.0 | 0 |
| scf7180000022957 | 38131 | 0.4252 | 0 | 200.9783 | Eukaryota | 6705.0 | 0 | Chordata | 6705.0 | 0 | Carnivora | 6705.0 | 0 | Felidae | 6705.0 | 0 | Felis | 6705.0 | 0 | Felis catus | 6705.0 | 0 |
| scf7180000018032 | 3344260 | 0.4347 | 0 | 52.2631 | Eukaryota | 60878.0 | 0 | Chordata | 60878.0 | 0 | Carnivora | 60878.0 | 0 | Felidae | 43317.0 | 2 | Felis | 24888.0 | 3 | Felis catus | 24888.0 | 3 |
| scf7180000023189 | 35846 | 0.3911 | 0 | 167.768 | Eukaryota | 6814.0 | 0 | Chordata | 6814.0 | 0 | Carnivora | 6814.0 | 0 | Felidae | 6814.0 | 0 | Felis | 6110.0 | 1 | Felis catus | 6110.0 | 1 |
| scf7180000022572 | 19079 | 0.6362 | 0 | 37.7687 | Eukaryota | 4114.0 | 0 | Chordata | 4114.0 | 0 | Passeriformes | 3120.0 | 1 | Corvidae | 3120.0 | 1 | Aphelocoma | 3120.0 | 1 | locoma califo | 1560.0 | 4 |
| scf7180000018438 | 575981 | 0.471 | 0 | 51.237 | Eukaryota | 33624.0 | 0 | Chordata | 33624.0 | 0 | Carnivora | 33624.0 | 0 | Felidae | 33624.0 | 0 | Felis | 13586.0 | 2 | Felis catus | 13586.0 | 2 |
| scf7180000022715 | 20717 | 0.3683 | 0 | 24.7175 | Eukaryota | 17662.0 | 0 | Chordata | 17662.0 | 0 | Carnivora | 17662.0 | 0 | Felidae | 17662.0 | 0 | Felis | 17662.0 | 0 | Felis catus | 17662.0 | 0 |
| scf7180000021117 | 85441 | 0.3463 | 0 | 49.5655 | Eukaryota | 36382.0 | 0 | Chordata | 36382.0 | 0 | Carnivora | 36382.0 | 0 | Felidae | 36382.0 | 0 | Felis | 36382.0 | 0 | Felis catus | 36382.0 | 0 |
| scf7180000024035 | 453454 | 0.3978 | 0 | 49.1461 | Eukaryota | 47721.0 | 0 | Chordata | 47721.0 | 0 | Carnivora | 47721.0 | 0 | Felidae | 47721.0 | 0 | Felis | 42901.0 | 1 | Felis catus | 42901.0 | 1 |
| scf7180000022009 | 48827 | 0.3818 | 0 | 28.6821 | Eukaryota | 15023.0 | 0 | Chordata | 15023.0 | 0 | Carnivora | 15023.0 | 0 | Felidae | 15023.0 | 0 | Felis | 12056.0 | 1 | Felis catus | 12056.0 | 1 |
| scf7180000022624 | 12094 | 0.4187 | 0 | 22.163 | Eukaryota | 2826.0 | 0 | Chordata | 2826.0 | 0 | Carnivora | 1172.0 | 2 | Felidae | 1172.0 | 2 | Felis | 1172.0 | 3 | Felis catus | 1172.0 | 3 |
| scf7180000020465 | 99568 | 0.3943 | 0 | 24.9951 | Eukaryota | 5777.0 | 0 | Chordata | 5777.0 | 0 | Primates | 2254.0 | 2 | Hominidae | 2254.0 | 4 | Sus | 1576.0 | 5 | Sus scrofa | 1576.0 | 5 |
| scf7180000017613 | 437271 | 0.419 | 0 | 53.0835 | Eukaryota | 44958.0 | 0 | Chordata | 44958.0 | 0 | Carnivora | 44958.0 | 0 | Felidae | 44958.0 | 0 | Acinonyx | 25208.0 | 1 | inonyx jubat | 25208.0 | 1 |
| scf7180000019923 | 698796 | 0.4827 | 0 | 50.2743 | Eukaryota | 41477.0 | 0 | Chordata | 41477.0 | 0 | Carnivora | 41477.0 | 0 | Felidae | 32973.0 | 1 | Acinonyx | 12417.0 | 4 | inonyx jubat | 12417.0 | 5 |
| scf7180000024166 | 121572 | 0.5106 | 0 | 52.6022 | Eukaryota | 38075.0 | 0 | Chordata | 38075.0 | 0 | Carnivora | 38075.0 | 0 | Felidae | 38075.0 | 0 | Felis | 19115.0 | 1 | Felis catus | 19115.0 | 1 |
| scf7180000019970 | 25608 | 0.362 | 0 | 133.6502 | Eukaryota | 7309.0 | 0 | Chordata | 7309.0 | 0 | Artiodactyla | 7309.0 | 0 | Bovidae | 7309.0 | 0 | Ovis | 7309.0 | 0 | vis canadens | 7309.0 | 0 |
| scf7180000017903 | 1262683 | 0.436 | 0 | 53.8611 | Eukaryota | 47453.0 | 0 | Chordata | 47453.0 | 0 | Carnivora | 47453.0 | 0 | Felidae | 47453.0 | 0 | Felis | 47453.0 | 0 | Felis catus | 47453.0 | 0 |
| scf7180000023857 | 1477263 | 0.385 | 0 | 50.7378 | Eukaryota | 65599.0 | 0 | Chordata | 65599.0 | 0 | Carnivora | 37315.0 | 1 | Felidae | 37315.0 | 1 | Felis | 31100.0 | 2 | Felis catus | 31100.0 | 2 |
| scf7180000017956 | 207859 | 0.4555 | 0 | 50.1915 | Eukaryota | 33457.0 | 0 | Chordata | 33457.0 | 0 | Carnivora | 33457.0 | 0 | Felidae | 33457.0 | 0 | Felis | 30046.0 | 1 | Felis catus | 30046.0 | 1 |
| scf7180000020841 | 72601 | 0.4086 | 0 | 49.5311 | Eukaryota | 44503.0 | 0 | Chordata | 44503.0 | 0 | Carnivora | 44503.0 | 0 | Felidae | 44503.0 | 0 | Felis | 44503.0 | 0 | Felis catus | 44503.0 | 0 |
| scf7180000021126 | 39606 | 0.4518 | 0 | 24.4148 | Eukaryota | 8202.0 | 0 | Chordata | 8202.0 | 0 | Primates | 4577.0 | 3 | Hominidae | 3461.0 | 4 | Pan | 2306.0 | 6 | an troglodyte | 2306.0 | 6 |
| scf7180000024050 | 434689 | 0.4329 | 0 | 52.2385 | Eukaryota | 83595.0 | 0 | Chordata | 83595.0 | 0 | Carnivora | 83595.0 | 0 | Felidae | 83595.0 | 0 | Panthera | 83595.0 | 0 | anthera pardu | 83595.0 | 0 |
| scf7180000022471 | 19748 | 0.4363 | 0 | 25.1407 | Eukaryota | 9759.0 | 0 | Chordata | 9759.0 | 0 | Carnivora | 9759.0 | 0 | Felidae | 9759.0 | 0 | Felis | 8772.0 | 1 | Felis catus | 8772.0 | 1 |
| scf7180000023770 | 3071863 | 0.4424 | 0 | 51.761 | Eukaryota | 48499.0 | 0 | Chordata | 48499.0 | 0 | Carnivora | 48499.0 | 0 | Felidae | 48499.0 | 0 | Felis | 28108.0 | 3 | Felis catus | 28108.0 | 3 |
| scf7180000019743 | 355319 | 0.397 | 0 | 50.4605 | Eukaryota | 47058.0 | 0 | Chordata | 47058.0 | 0 | Carnivora | 47058.0 | 0 | Felidae | 47058.0 | 0 | Felis | 47058.0 | 0 | Felis catus | 47058.0 | 0 |
| scf7180000023657 | 34463 | 0.3956 | 0 | 304.9064 | Eukaryota | 6746.0 | 0 | Chordata | 6746.0 | 0 | Carnivora | 6746.0 | 0 | Felidae | 6746.0 | 0 | Felis | 6746.0 | 0 | Felis catus | 6746.0 | 0 |
| scf7180000021931 | 18971 | 0.4277 | 0 | 76.8088 | Eukaryota | 2263.0 | 0 | Chordata | 2263.0 | 0 | Carnivora | 2263.0 | 0 | Felidae | 2263.0 | 0 | Felis | 1622.0 | 1 | Felis catus | 1622.0 | 1 |
| scf7180000017810 | 1401507 | 0.4489 | 0 | 51.2512 | Eukaryota | 47441.0 | 0 | Chordata | 47441.0 | 0 | Carnivora | 47441.0 | 0 | Felidae | 47441.0 | 0 | Felis | 47441.0 | 0 | Felis catus | 47441.0 | 0 |
| scf7180000022407 | 28427 | 0.37 | 0 | 23.4943 | Eukaryota | 3575.0 | 0 | Chordata | 3575.0 | 0 | Carnivora | 1780.0 | 2 | Felidae | 1515.0 | 3 | Sus | 1444.0 | 4 | Sus scrofa | 1444.0 | 4 |
| scf7180000020298 | 28798 | 0.6981 | 0 | 141.2293 | no-hit | 0.0 | 0 | no-hit | 0.0 | 0 | no-hit | 0.0 | 0 | no-hit | 0.0 | 0 | no-hit | 0.0 | 0 | no-hit | 0.0 | 0 |
| scf7180000017753 | 113522 | 0.3702 | 0 | 47.9256 | Eukaryota | 40596.0 | 0 | Chordata | 40596.0 | 0 | Carnivora | 40596.0 | 0 | Felidae | 40596.0 | 0 | Felis | 40596.0 | 0 | Felis catus | 40596.0 | 0 |
| scf7180000022086 | 28505 | 0.6396 | 0 | 52.8553 | Eukaryota | 6991.0 | 0 | Chordata | 6991.0 | 0 | Carnivora | 6991.0 | 0 | Felidae | 4072.0 | 3 | Acinonyx | 1848.0 | 6 | inonyx jubat | 1848.0 | 6 |
| scf7180000018781 | 572718 | 0.3525 | 0 | 48.405 | Eukaryota | 32794.0 | 0 | Chordata | 32794.0 | 0 | Carnivora | 32794.0 | 0 | Felidae | 32794.0 | 0 | Felis | 29442.0 | 1 | Felis catus | 29442.0 | 1 |
| scf7180000022017 | 35025 | 0.3758 | 0 | 24.5342 | Eukaryota | 35849.0 | 0 | Chordata | 35849.0 | 0 | Carnivora | 35849.0 | 0 | Felidae | 35849.0 | 0 | Felis | 35849.0 | 0 | Felis catus | 35849.0 | 0 |
| scf7180000018697 | 646366 | 0.5991 | 0 | 57.1766 | Eukaryota | 21561.0 | 0 | Chordata | 21561.0 | 0 | Carnivora | 21561.0 | 0 | Felidae | 9624.0 | 3 | Zalophus | 5886.0 | 4 | phus californi | 5886.0 | 4 |
| scf7180000022696 | 6424 | 0.398 | 0 | 23.3859 | Eukaryota | 13780.0 | 0 | Chordata | 13780.0 | 0 | Carnivora | 13780.0 | 0 | Felidae | 13780.0 | 0 | Felis | 12409.0 | 1 | Felis catus | 12409.0 | 1 |
| scf7180000020124 | 223407 | 0.4283 | 0 | 25.9764 | Eukaryota | 22465.0 | 0 | Chordata | 22465.0 | 0 | Carnivora | 22465.0 | 0 | Felidae | 22465.0 | 0 | Felis | 20182.0 | 1 | Felis catus | 20182.0 | 1 |
| scf7180000018346 | 262713 | 0.4232 | 0 | 51.3727 | Eukaryota | 62740.0 | 0 | Chordata | 62740.0 | 0 | Carnivora | 62740.0 | 0 | Felidae | 56381.0 | 1 | Panthera | 25068.0 | 4 | anthera pardu | 18828.0 | 5 |
| scf7180000019727 | 286601 | 0.3824 | 0 | 50.9254 | Eukaryota | 44655.0 | 0 | Chordata | 44655.0 | 0 | Carnivora | 44655.0 | 0 | Felidae | 44655.0 | 0 | Felis | 40204.0 | 1 | Felis catus | 40204.0 | 1 |
| scf7180000019945 | 43089 | 0.5776 | 0 | 109.9576 | Eukaryota | 2738.0 | 0 | Chordata | 2738.0 | 0 | Carnivora | 2738.0 | 0 | Felidae | 2738.0 | 0 | Felis | 1640.0 | 1 | Felis catus | 1640.0 | 1 |
| scf7180000023577 | 17194 | 0.4626 | 0 | 336.8504 | no-hit | 0.0 | 0 | no-hit | 0.0 | 0 | no-hit | 0.0 | 0 | no-hit | 0.0 | 0 | no-hit | 0.0 | 0 | no-hit | 0.0 | 0 |
| scf7180000021861 | 45632 | 0.5723 | 0 | 47.9678 | Eukaryota | 4941.0 | 0 | Chordata | 4941.0 | 0 | Carnivora | 4941.0 | 0 | Felidae | 4941.0 | 0 | Felis | 4941.0 | 0 | Felis catus | 4941.0 | 0 |
| scf7180000018832 | 49524 | 0.3656 | 0 | 48.9207 | Eukaryota | 45307.0 | 0 | Chordata | 45307.0 | 0 | Carnivora | 45307.0 | 0 | Felidae | 45307.0 | 0 | Felis | 45307.0 | 0 | Felis catus | 45307.0 | 0 |
| scf7180000020480 | 173292 | 0.3894 | 0 | 50.7878 | Eukaryota | 43586.0 | 0 | Chordata | 43586.0 | 0 | Carnivora | 43586.0 | 0 | Felidae | 43586.0 | 0 | Felis | 43586.0 | 0 | Felis catus | 43586.0 | 0 |
| scf7180000022623 | 34150 | 0.5888 | 0 | 45.7091 | Eukaryota | 20841.0 | 0 | Chordata | 20841.0 | 0 | Carnivora | 20841.0 | 0 | Felidae | 18425.0 | 1 | Acinonyx | 8216.0 | 3 | inonyx jubat | 8216.0 | 3 |
| scf7180000018218 | 1562134 | 0.5138 | 0 | 51.8869 | Eukaryota | 56960.0 | 0 | Chordata | 56960.0 | 0 | Carnivora | 56960.0 | 0 | Felidae | 35673.0 | 2 | Ursus | 15717.0 | 5 | Ursus arctos | 15717.0 | 6 |
| scf7180000020077 | 78745 | 0.5051 | 0 | 46.6047 | Eukaryota | 7670.0 | 0 | Chordata | 7670.0 | 0 | Carnivora | 7670.0 | 0 | Felidae | 7670.0 | 0 | Felis | 5279.0 | 1 | Felis catus | 5279.0 | 1 |
| scf7180000022722 | 12612 | 0.4319 | 0 | 26.3451 | Eukaryota | 2917.0 | 0 | Chordata | 2584.0 | 1 | Carnivora | 1818.0 | 3 | Felidae | 1563.0 | 4 | Felis | 1563.0 | 4 | Felis catus | 1563.0 | 4 |
| scf7180000024161 | 132745 | 0.3776 | 0 | 29.0883 | Eukaryota | 48154.0 | 0 | Chordata | 48154.0 | 0 | Carnivora | 48154.0 | 0 | Felidae | 48154.0 | 0 | Felis | 48154.0 | 0 | Felis catus | 48154.0 | 0 |
| scf7180000023455 | 30861 | 0.3369 | 0 | 279.7165 | Eukaryota | 3478.0 | 1 | Chordata | 3478.0 | 1 | Carnivora | 2434.0 | 2 | Felidae | 1740.0 | 4 | Felis | 1472.0 | 5 | Felis catus | 1472.0 | 5 |
| scf7180000021778 | 26511 | 0.383 | 0 | 30.6909 | Eukaryota | 46105.0 | 0 | Chordata | 46105.0 | 0 | Carnivora | 46105.0 | 0 | Felidae | 46105.0 | 0 | Felis | 46105.0 | 0 | Felis catus | 46105.0 | 0 |
| scf7180000022422 | 16298 | 0.4119 | 0 | 27.0382 | Eukaryota | 10152.0 | 0 | Chordata | 10152.0 | 0 | Carnivora | 10152.0 | 0 | Felidae | 10152.0 | 0 | Felis | 10152.0 | 0 | Felis catus | 10152.0 | 0 |
| scf7180000019038 | 56046 | 0.4423 | 0 | 43.0755 | Eukaryota | 43498.0 | 0 | Chordata | 43498.0 | 0 | Carnivora | 43498.0 | 0 | Felidae | 43498.0 | 0 | Felis | 43498.0 | 0 | Felis catus | 43498.0 | 0 |
| scf7180000019842 | 220664 | 0.5408 | 0 | 52.1001 | Eukaryota | 21608.0 | 0 | Chordata | 21608.0 | 0 | Carnivora | 21608.0 | 0 | Mustelidae | 6114.0 | 4 | Mustela | 4584.0 | 6 | ustela putori | 4584.0 | 6 |
| scf7180000022286 | 17212 | 0.4287 | 0 | 26.3789 | Eukaryota | 23535.0 | 0 | Chordata | 23535.0 | 0 | Carnivora | 23535.0 | 0 | Felidae | 9803.0 | 4 | Panthera | 4917.0 | 8 | poda melanc | 2660.0 | 9 |
| scf7180000022079 | 35896 | 0.4243 | 0 | 44.4269 | Eukaryota | 12112.0 | 0 | Chordata | 12112.0 | 0 | Carnivora | 12112.0 | 0 | Felidae | 5674.0 | 2 | Suricata | 5335.0 | 3 | ricata suricat | 5335.0 | 3 |
| scf7180000023754 | 27342 | 0.5522 | 0 | 126.7538 | Eukaryota | 2172.0 | 0 | Chordata | 2172.0 | 0 | Carnivora | 2172.0 | 0 | Felidae | 1931.0 | 1 | Felis | 1931.0 | 1 | Felis catus | 1931.0 | 1 |
| scf7180000019402 | 326830 | 0.4184 | 0 | 52.7739 | Eukaryota | 21708.0 | 0 | Chordata | 21708.0 | 0 | Carnivora | 21708.0 | 0 | Felidae | 21708.0 | 0 | Panthera | 13032.0 | 1 | anthera pardu | 13032.0 | 1 |
| scf7180000021005 | 121505 | 0.3597 | 0 | 24.2761 | Eukaryota | 39223.0 | 0 | Chordata | 39223.0 | 0 | Carnivora | 39223.0 | 0 | Felidae | 39223.0 | 0 | Felis | 35346.0 | 1 | Felis catus | 35346.0 | 1 |
| scf7180000017461 | 1429517 | 0.3601 | 0 | 48.2722 | Eukaryota | 45804.0 | 0 | Chordata | 45804.0 | 0 | Carnivora | 45804.0 | 0 | Felidae | 45804.0 | 0 | Felis | 45804.0 | 0 | Felis catus | 45804.0 | 0 |
| scf7180000021750 | 45798 | 0.4543 | 0 | 25.5803 | Eukaryota | 3903.0 | 0 | Chordata | 3903.0 | 0 | Carnivora | 1328.0 | 3 | Felidae | 1328.0 | 3 | Homo | 1264.0 | 6 | Homo sapiens | 1264.0 | 6 |
| scf7180000021555 | 62053 | 0.581 | 0 | 45.6143 | Eukaryota | 4134.0 | 0 | Chordata | 4134.0 | 0 | Carnivora | 4134.0 | 0 | Felidae | 2926.0 | 1 | Panthera | 1770.0 | 3 | anthera pardu | 1483.0 | 4 |
| scf7180000019129 | 370911 | 0.4564 | 0 | 50.1779 | Eukaryota | 33120.0 | 0 | Chordata | 33120.0 | 0 | Carnivora | 33120.0 | 0 | Herpestidae | 33120.0 | 0 | Suricata | 33120.0 | 0 | ricata suricat | 33120.0 | 0 |
| scf7180000017989 | 368598 | 0.3915 | 0 | 48.7262 | Eukaryota | 34033.0 | 0 | Chordata | 34033.0 | 0 | Carnivora | 34033.0 | 0 | Felidae | 34033.0 | 0 | Felis | 34033.0 | 0 | Felis catus | 34033.0 | 0 |
| scf7180000017840 | 1681422 | 0.425 | 0 | 50.5328 | Eukaryota | 112164.0 | 0 | Chordata | 112164.0 | 0 | Primates | 72509.0 | 1 | Hominidae | 72509.0 | 2 | Pan | 42095.0 | 4 | an troglodyte | 42095.0 | 4 |
| scf7180000020284 | 127992 | 0.4588 | 0 | 51.2545 | Eukaryota | 33633.0 | 0 | Chordata | 33633.0 | 0 | Carnivora | 33633.0 | 0 | Felidae | 33633.0 | 0 | Felis | 30192.0 | 1 | Felis catus | 30192.0 | 1 |
| scf7180000019677 | 151240 | 0.472 | 0 | 46.7919 | Eukaryota | 28439.0 | 0 | Chordata | 28439.0 | 0 | Carnivora | 17252.0 | 1 | Otariidae | 8657.0 | 4 | Zalophus | 5781.0 | 6 | phus californi | 5781.0 | 7 |
| scf7180000020451 | 100750 | 0.3939 | 0 | 24.4967 | Eukaryota | 41995.0 | 0 | Chordata | 41995.0 | 0 | Carnivora | 41995.0 | 0 | Felidae | 41995.0 | 0 | Felis | 41995.0 | 0 | Felis catus | 41995.0 | 0 |
| scf7180000021065 | 1213065 | 0.3623 | 0 | 50.3618 | Eukaryota | 47960.0 | 0 | Chordata | 47960.0 | 0 | Carnivora | 47960.0 | 0 | Felidae | 47960.0 | 0 | Felis | 47960.0 | 0 | Felis catus | 47960.0 | 0 |
| scf7180000018657 | 592603 | 0.3952 | 0 | 48.3743 | Eukaryota | 36758.0 | 0 | Chordata | 36758.0 | 0 | Carnivora | 36758.0 | 0 | Herpestidae | 14916.0 | 2 | Suricata | 14916.0 | 2 | ricata suricat | 14916.0 | 2 |
| scf7180000019815 | 225303 | 0.4523 | 0 | 51.4074 | Eukaryota | 13940.0 | 1 | Chordata | 13940.0 | 1 | Primates | 9879.0 | 2 | Hominidae | 5143.0 | 4 | Homo | 5143.0 | 4 | Homo sapiens | 5143.0 | 4 |
| scf7180000024029 | 332163 | 0.4181 | 0 | 54.1317 | Eukaryota | 43787.0 | 0 | Chordata | 43787.0 | 0 | Carnivora | 43787.0 | 0 | Felidae | 43787.0 | 0 | Felis | 39415.0 | 1 | Felis catus | 39415.0 | 1 |
| scf7180000017787 | 1862097 | 0.5249 | 100 | 52.8463 | Eukaryota | 45124.0 | 0 | Chordata | 45124.0 | 0 | Primates | 23554.0 | 3 | Cercopithecidae | 12968.0 | 4 | Pan | 10586.0 | 8 | an troglodyte | 10586.0 | 8 |
| scf7180000021681 | 33551 | 0.4351 | 0 | 25.9108 | Eukaryota | 4134.0 | 0 | Chordata | 4134.0 | 0 | Primates | 2356.0 | 3 | Hominidae | 2356.0 | 3 | Pan | 2017.0 | 5 | an troglodyte | 2017.0 | 5 |
| scf7180000022792 | 16487 | 0.5025 | 0 | 26.1128 | Eukaryota | 3413.0 | 0 | Chordata | 3413.0 | 0 | Carnivora | 2265.0 | 2 | Felidae | 2265.0 | 2 | Felis | 1095.0 | 4 | Felis catus | 1095.0 | 4 |
| scf7180000023872 | 771280 | 0.4419 | 0 | 50.9556 | Eukaryota | 40649.0 | 0 | Chordata | 40649.0 | 0 | Carnivora | 40649.0 | 0 | Felidae | 40649.0 | 0 | Felis | 36512.0 | 1 | Felis catus | 36512.0 | 1 |
| scf7180000018849 | 544389 | 0.4437 | 0 | 51.6499 | Eukaryota | 42846.0 | 0 | Chordata | 42846.0 | 0 | Carnivora | 42846.0 | 0 | Felidae | 39158.0 | 1 | Panthera | 20751.0 | 5 | Panthera tigri | 11742.0 | 6 |
| scf7180000017817 | 1464461 | 0.4351 | 0 | 51.2804 | Eukaryota | 120692.0 | 0 | Chordata | 120692.0 | 0 | Carnivora | 120692.0 | 0 | Felidae | 120692.0 | 0 | Acinonyx | 108666.0 | 1 | inonyx jubat | 108666.0 | 1 |
| scf7180000024094 | 575381 | 0.4383 | 0 | 48.7006 | Eukaryota | 33681.0 | 0 | Chordata | 33681.0 | 0 | Carnivora | 33681.0 | 0 | Felidae | 33681.0 | 0 | Felis | 16657.0 | 2 | Felis catus | 16657.0 | 2 |
| scf7180000022492 | 9032 | 0.3499 | 0 | 25.032 | Eukaryota | 3835.0 | 0 | Chordata | 3835.0 | 0 | Artiodactyla | 2032.0 | 2 | Hominidae | 1063.0 | 6 | Pan | 708.0 | 8 | an troglodyte | 708.0 | 8 |
| scf7180000018027 | 1283669 | 0.3644 | 0 | 49.1702 | Eukaryota | 57667.0 | 0 | Chordata | 57667.0 | 0 | Carnivora | 23729.0 | 2 | Pteropodidae | 22628.0 | 5 | Rousettus | 22628.0 | 5 | ettus aegypti | 22628.0 | 5 |
| scf7180000019869 | 205147 | 0.4453 | 0 | 51.6742 | Eukaryota | 24120.0 | 0 | Chordata | 24120.0 | 0 | Carnivora | 24120.0 | 0 | Felidae | 24120.0 | 0 | Felis | 14472.0 | 1 | Felis catus | 14472.0 | 1 |
| scf7180000021911 | 52001 | 0.457 | 0 | 153.3382 | Eukaryota | 4921.0 | 0 | Chordata | 4921.0 | 0 | Carnivora | 4627.0 | 1 | Felidae | 3610.0 | 3 | Felis | 3138.0 | 5 | Felis catus | 3138.0 | 5 |
| scf7180000020056 | 3177583 | 0.3912 | 0 | 50.9401 | Eukaryota | 50051.0 | 0 | Chordata | 50051.0 | 0 | Carnivora | 50051.0 | 0 | Felidae | 45340.0 | 1 | Felis | 38237.0 | 2 | Felis catus | 38237.0 | 2 |
| scf7180000017759 | 1848458 | 0.38 | 0 | 50.3969 | Eukaryota | 48095.0 | 0 | Chordata | 48095.0 | 0 | Carnivora | 48095.0 | 0 | Felidae | 48095.0 | 0 | Felis | 48095.0 | 0 | Felis catus | 48095.0 | 0 |
| scf7180000022465 | 54816 | 0.4372 | 0 | 54.4561 | Eukaryota | 15525.0 | 0 | Chordata | 15525.0 | 0 | Carnivora | 8655.0 | 2 | Felidae | 6503.0 | 3 | Acinonyx | 6503.0 | 5 | inonyx jubat | 6503.0 | 5 |
| scf7180000019712 | 116549 | 0.3903 | 0 | 25.7852 | Eukaryota | 44942.0 | 0 | Chordata | 44942.0 | 0 | Carnivora | 44942.0 | 0 | Felidae | 44942.0 | 0 | Felis | 44942.0 | 0 | Felis catus | 44942.0 | 0 |
| scf7180000018620 | 2245126 | 0.5181 | 0 | 52.3835 | Eukaryota | 31450.0 | 0 | Chordata | 31450.0 | 0 | Carnivora | 31450.0 | 0 | Felidae | 31450.0 | 0 | Acinonyx | 10703.0 | 3 | inonyx jubat | 10703.0 | 3 |
| scf7180000023359 | 32224 | 0.3877 | 0 | 228.9142 | Eukaryota | 44081.0 | 0 | Chordata | 44081.0 | 0 | Carnivora | 44081.0 | 0 | Felidae | 44081.0 | 0 | Felis | 44081.0 | 0 | Felis catus | 44081.0 | 0 |
| scf7180000022071 | 19245 | 0.4326 | 0 | 25.2384 | Eukaryota | 4359.0 | 0 | Chordata | 4359.0 | 0 | Primates | 2023.0 | 2 | Hominidae | 2023.0 | 2 | Homo | 1205.0 | 5 | Homo sapiens | 1205.0 | 5 |
| scf7180000022938 | 43832 | 0.3832 | 0 | 337.1636 | Eukaryota | 7482.0 | 0 | Chordata | 7482.0 | 0 | Carnivora | 7482.0 | 0 | Felidae | 7482.0 | 0 | Felis | 7482.0 | 0 | Felis catus | 7482.0 | 0 |
| scf7180000020838 | 42666 | 0.6109 | 0 | 44.9679 | Eukaryota | 25285.0 | 0 | Chordata | 25285.0 | 0 | Carnivora | 25285.0 | 0 | Felidae | 25285.0 | 0 | Felis | 22723.0 | 1 | Felis catus | 22723.0 | 1 |
| scf7180000021339 | 74442 | 0.3875 | 0 | 49.154 | Eukaryota | 23668.0 | 0 | Chordata | 23668.0 | 0 | Carnivora | 23668.0 | 0 | Felidae | 19052.0 | 2 | Panthera | 9503.0 | 6 | anthera pardu | 7124.0 | 7 |
| scf7180000019440 | 336562 | 0.3685 | 0 | 50.7485 | Eukaryota | 44891.0 | 0 | Chordata | 44891.0 | 0 | Carnivora | 44891.0 | 0 | Felidae | 44891.0 | 0 | Felis | 44891.0 | 0 | Felis catus | 44891.0 | 0 |
| scf7180000019631 | 262216 | 0.4229 | 0 | 25.6845 | Eukaryota | 44696.0 | 0 | Chordata | 44696.0 | 0 | Carnivora | 44696.0 | 0 | Felidae | 44696.0 | 0 | Felis | 40269.0 | 1 | Felis catus | 40269.0 | 1 |
| scf7180000019141 | 11436 | 0.4122 | 0 | 27.6368 | Eukaryota | 33739.0 | 0 | Chordata | 33739.0 | 0 | Carnivora | 14973.0 | 3 | Felidae | 11493.0 | 6 | Equus | 6655.0 | 8 | anthera pardu | 3904.0 | 9 |
| scf7180000018761 | 13637 | 0.4153 | 0 | 37.16 | Eukaryota | 28217.0 | 0 | Chordata | 28217.0 | 0 | Carnivora | 28217.0 | 0 | Felidae | 14715.0 | 4 | Lynx | 2953.0 | 9 | ynx canadens | 2953.0 | 9 |
| scf7180000022990 | 29348 | 0.5515 | 0 | 169.1149 | Eukaryota | 6580.0 | 0 | Chordata | 6580.0 | 0 | Carnivora | 6580.0 | 0 | Felidae | 6334.0 | 1 | Felis | 2578.0 | 3 | Felis catus | 2578.0 | 3 |
| scf7180000022138 | 14296 | 0.4312 | 0 | 258.7589 | Eukaryota | 2340.0 | 0 | Chordata | 2340.0 | 0 | Carnivora | 2340.0 | 0 | Felidae | 2340.0 | 0 | Felis | 1890.0 | 1 | Felis catus | 1890.0 | 1 |
| scf7180000021727 | 80613 | 0.4732 | 0 | 27.6916 | Eukaryota | 28617.0 | 0 | Chordata | 28617.0 | 0 | Carnivora | 26144.0 | 1 | Felidae | 10992.0 | 6 | Canis | 5658.0 | 8 | Canis lupus | 5658.0 | 8 |
| scf7180000019528 | 274880 | 0.4342 | 0 | 50.9337 | Eukaryota | 15708.0 | 0 | Chordata | 15708.0 | 0 | Carnivora | 15708.0 | 0 | Felidae | 15708.0 | 0 | Acinonyx | 8556.0 | 2 | inonyx jubat | 8556.0 | 2 |
| scf7180000017807 | 156927 | 0.3904 | 0 | 47.8239 | Eukaryota | 22746.0 | 0 | Chordata | 22746.0 | 0 | Carnivora | 22746.0 | 0 | Felidae | 22746.0 | 0 | Felis | 20467.0 | 1 | Felis catus | 20467.0 | 1 |
| scf7180000023580 | 29976 | 0.4757 | 0 | 482.6303 | no-hit | 0.0 | 0 | no-hit | 0.0 | 0 | no-hit | 0.0 | 0 | no-hit | 0.0 | 0 | no-hit | 0.0 | 0 | no-hit | 0.0 | 0 |
| scf7180000018477 | 716763 | 0.4295 | 0 | 48.2425 | Eukaryota | 34922.0 | 0 | Chordata | 34922.0 | 0 | Carnivora | 34922.0 | 0 | Felidae | 34922.0 | 0 | Panthera | 28006.0 | 1 | anthera pardu | 17590.0 | 2 |
| scf7180000021882 | 32113 | 0.3817 | 0 | 24.1655 | Eukaryota | 38554.0 | 0 | Chordata | 38554.0 | 0 | Carnivora | 38554.0 | 0 | Felidae | 38554.0 | 0 | Felis | 34722.0 | 1 | Felis catus | 34722.0 | 1 |
| scf7180000019635 | 330969 | 0.5041 | 0 | 52.9948 | Eukaryota | 21017.0 | 1 | Chordata | 21017.0 | 1 | Carnivora | 16826.0 | 3 | Felidae | 9931.0 | 5 | Zalophus | 4588.0 | 8 | phus californi | 4588.0 | 8 |
| scf7180000019680 | 140681 | 0.387 | 0 | 48.8262 | Eukaryota | 15946.0 | 0 | Chordata | 15946.0 | 0 | Carnivora | 15946.0 | 0 | Felidae | 15946.0 | 0 | Felis | 14361.0 | 1 | Felis catus | 14361.0 | 1 |
| scf7180000023458 | 203311 | 0.3885 | 0 | 539.48 | Eukaryota | 48198.0 | 0 | Chordata | 48198.0 | 0 | Carnivora | 48198.0 | 0 | Felidae | 48198.0 | 0 | Felis | 48198.0 | 0 | Felis catus | 48198.0 | 0 |
| scf7180000018268 | 299400 | 0.3788 | 0 | 50.1339 | Eukaryota | 107278.0 | 0 | Chordata | 107278.0 | 0 | Carnivora | 107278.0 | 0 | Felidae | 107278.0 | 0 | Panthera | 98506.0 | 1 | anthera pardu | 98506.0 | 1 |
| scf7180000022489 | 14374 | 0.3732 | 0 | 23.5375 | Eukaryota | 20671.0 | 0 | Chordata | 20671.0 | 0 | Carnivora | 20671.0 | 0 | Felidae | 20671.0 | 0 | Felis | 18595.0 | 1 | Felis catus | 18595.0 | 1 |
| scf7180000021675 | 40817 | 0.6124 | 0 | 48.4116 | Eukaryota | 9673.0 | 0 | Chordata | 9673.0 | 0 | Carnivora | 9673.0 | 0 | Felidae | 7300.0 | 2 | Panthera | 3084.0 | 5 | ynx canadens | 1751.0 | 6 |
| scf7180000023768 | 2771805 | 0.4345 | 0 | 50.9152 | Eukaryota | 52860.0 | 0 | Chordata | 52860.0 | 0 | Carnivora | 52860.0 | 0 | Felidae | 29961.0 | 2 | Panthera | 19561.0 | 5 | anthera pardu | 12726.0 | 6 |
| scf7180000020526 | 30059 | 0.4388 | 0 | 353.531 | Eukaryota | 45969.0 | 0 | Chordata | 45969.0 | 0 | Carnivora | 45969.0 | 0 | Felidae | 45969.0 | 0 | Felis | 45969.0 | 0 | Felis catus | 45969.0 | 0 |
| scf7180000021961 | 16604 | 0.4181 | 0 | 46.9237 | Eukaryota | 26106.0 | 1 | Chordata | 26106.0 | 1 | Primates | 20020.0 | 3 | Hominidae | 20020.0 | 3 | Homo | 11460.0 | 4 | Homo sapiens | 11460.0 | 4 |
| scf7180000018003 | 1243113 | 0.4253 | 0 | 51.3948 | Eukaryota | 44851.0 | 0 | Chordata | 44851.0 | 0 | Carnivora | 44851.0 | 0 | Felidae | 38393.0 | 1 | Felis | 34136.0 | 2 | Felis catus | 34136.0 | 2 |
| scf7180000021827 | 29047 | 0.5425 | 0 | 50.9744 | Eukaryota | 3514.0 | 1 | Chordata | 3514.0 | 1 | Rodentia | 1908.0 | 3 | Muridae | 1908.0 | 3 | Mus | 1908.0 | 4 | Mus musculus | 1908.0 | 4 |
| scf7180000022161 | 26493 | 0.4874 | 0 | 768.098 | Eukaryota | 17850.0 | 0 | Chordata | 17850.0 | 0 | Carnivora | 17850.0 | 0 | Felidae | 14338.0 | 1 | Acinonyx | 3526.0 | 5 | inonyx jubat | 3526.0 | 5 |
| scf7180000019225 | 416028 | 0.3465 | 0 | 48.4869 | Eukaryota | 58753.0 | 0 | Chordata | 58753.0 | 0 | Carnivora | 58753.0 | 0 | Felidae | 30434.0 | 3 | Panthera | 12159.0 | 7 | unresolved | 6115.0 | 9 |
| scf7180000020331 | 220781 | 0.5992 | 0 | 51.716 | Eukaryota | 21788.0 | 0 | Chordata | 21788.0 | 0 | Carnivora | 21788.0 | 0 | Felidae | 21788.0 | 0 | Lynx | 13212.0 | 2 | ynx canadens | 13212.0 | 2 |
| scf7180000019882 | 172136 | 0.4422 | 0 | 51.6386 | Eukaryota | 23013.0 | 0 | Chordata | 23013.0 | 0 | Carnivora | 23013.0 | 0 | Felidae | 23013.0 | 0 | Panthera | 16086.0 | 1 | anthera pardu | 16086.0 | 1 |
| scf7180000019132 | 39966 | 0.4674 | 0 | 39.7323 | Eukaryota | 6382.0 | 0 | Chordata | 6382.0 | 0 | Carnivora | 4039.0 | 2 | Felidae | 3281.0 | 3 | Homo | 1630.0 | 5 | Homo sapiens | 1630.0 | 5 |
| scf7180000020743 | 48376 | 0.3707 | 0 | 22.007 | Eukaryota | 45389.0 | 0 | Chordata | 45389.0 | 0 | Carnivora | 45389.0 | 0 | Felidae | 45389.0 | 0 | Felis | 45389.0 | 0 | Felis catus | 45389.0 | 0 |
| scf7180000020596 | 94008 | 0.397 | 0 | 24.6805 | Eukaryota | 11025.0 | 0 | Chordata | 11025.0 | 0 | Carnivora | 9375.0 | 1 | Felidae | 8342.0 | 2 | Panthera | 4154.0 | 5 | Panthera tigri | 2088.0 | 6 |
| scf7180000020273 | 75941 | 0.5656 | 0 | 48.5797 | Eukaryota | 9839.0 | 0 | Chordata | 9839.0 | 0 | Carnivora | 7957.0 | 1 | Otariidae | 2924.0 | 5 | Zalophus | 2924.0 | 7 | phus californi | 2924.0 | 7 |
| scf7180000023766 | 3160036 | 0.3984 | 0 | 50.873 | Eukaryota | 49469.0 | 0 | Chordata | 49469.0 | 0 | Carnivora | 49469.0 | 0 | Felidae | 37535.0 | 1 | Felis | 37535.0 | 1 | Felis catus | 37535.0 | 1 |
| scf7180000021415 | 45045 | 0.3969 | 0 | 47.8896 | Eukaryota | 16453.0 | 0 | Chordata | 16453.0 | 0 | Carnivora | 16453.0 | 0 | Felidae | 16453.0 | 0 | Felis | 13297.0 | 1 | Felis catus | 13297.0 | 1 |
| scf7180000019228 | 513945 | 0.425 | 0 | 53.3799 | Eukaryota | 51837.0 | 0 | Chordata | 51837.0 | 0 | Carnivora | 51837.0 | 0 | Ursidae | 25941.0 | 2 | Canis | 20902.0 | 3 | Canis lupus | 20902.0 | 4 |
| scf7180000020365 | 113171 | 0.3575 | 0 | 24.7692 | Eukaryota | 16874.0 | 0 | Chordata | 16874.0 | 0 | Carnivora | 8539.0 | 2 | Felidae | 8539.0 | 2 | Felis | 7032.0 | 4 | Felis catus | 7032.0 | 4 |
| scf7180000023221 | 21798 | 0.5445 | 0 | 154.8362 | Eukaryota | 3340.0 | 0 | Chordata | 3340.0 | 0 | Carnivora | 3340.0 | 0 | Felidae | 3340.0 | 0 | Acinonyx | 1836.0 | 1 | inonyx jubat | 1836.0 | 1 |
| scf7180000018860 | 803017 | 0.3852 | 0 | 49.5361 | Eukaryota | 37742.0 | 0 | Chordata | 37742.0 | 0 | Carnivora | 37742.0 | 0 | Felidae | 30296.0 | 1 | Acinonyx | 11420.0 | 3 | inonyx jubat | 11420.0 | 3 |
| scf7180000020079 | 226342 | 0.6257 | 0 | 48.9862 | Eukaryota | 15033.0 | 0 | Chordata | 15033.0 | 0 | Carnivora | 8536.0 | 2 | Otariidae | 3497.0 | 5 | Acinonyx | 3474.0 | 7 | inonyx jubat | 3474.0 | 7 |
| scf7180000017599 | 49635 | 0.3941 | 0 | 27.1556 | Eukaryota | 24241.0 | 0 | Chordata | 24241.0 | 0 | Carnivora | 24241.0 | 0 | Felidae | 24241.0 | 0 | Felis | 21799.0 | 1 | Felis catus | 21799.0 | 1 |
| scf7180000018325 | 1318877 | 0.5026 | 0 | 50.7826 | Eukaryota | 39804.0 | 0 | Chordata | 39804.0 | 0 | Carnivora | 39804.0 | 0 | Felidae | 39804.0 | 0 | Felis | 39804.0 | 0 | Felis catus | 39804.0 | 0 |

| | | | | | | | | | | | | | | | | | | | | | | |
|---|---|---|---|---|---|---|---|---|---|---|---|---|---|---|---|---|---|---|---|---|---|---|
| scf7180000022764 | 3177 | 0.5212 | 0 | 34.9005 | no-hit | 0.0 | 0 | no-hit | 0.0 | 0 | no-hit | 0.0 | 0 | no-hit | 0.0 | 0 | no-hit | 0.0 | 0 | no-hit | 0.0 | 0 |
| scf7180000020644 | 136156 | 0.3666 | 0 | 49.1946 | Eukaryota | 18916.0 | 0 | Chordata | 18916.0 | 0 | Carnivora | 15806.0 | 1 | Felidae | 12283.0 | 3 | Panthera | 4131.0 | 6 | Felis catus | 4096.0 | 7 |
| scf7180000024034 | 749567 | 0.3834 | 100 | 49.7452 | Eukaryota | 47042.0 | 0 | Chordata | 47042.0 | 0 | Carnivora | 47042.0 | 0 | Felidae | 47042.0 | 0 | Felis | 47042.0 | 0 | Felis catus | 47042.0 | 0 |
| scf7180000019334 | 80413 | 0.4071 | 0 | 49.8392 | Eukaryota | 47527.0 | 0 | Chordata | 47527.0 | 0 | Carnivora | 47527.0 | 0 | Felidae | 47527.0 | 0 | Felis | 47527.0 | 0 | Felis catus | 47527.0 | 0 |
| scf7180000019919 | 237786 | 0.463 | 0 | 51.4888 | Eukaryota | 49054.0 | 0 | Chordata | 49054.0 | 0 | Carnivora | 49054.0 | 0 | Felidae | 42627.0 | 1 | Lynx | 12500.0 | 4 | ynx canadens | 12500.0 | 5 |
| scf7180000018277 | 862965 | 0.3587 | 0 | 48.8257 | Eukaryota | 45980.0 | 0 | Chordata | 45980.0 | 0 | Carnivora | 45980.0 | 0 | Felidae | 45980.0 | 0 | Felis | 45980.0 | 0 | Felis catus | 45980.0 | 0 |
| scf7180000020210 | 168387 | 0.4353 | 0 | 50.3153 | Eukaryota | 32239.0 | 0 | Chordata | 32239.0 | 0 | Carnivora | 32239.0 | 0 | Felidae | 32239.0 | 0 | Felis | 28964.0 | 1 | Felis catus | 28964.0 | 1 |
| scf7180000022271 | 13399 | 0.4267 | 0 | 23.9388 | Eukaryota | 4176.0 | 0 | Chordata | 4176.0 | 0 | Carnivora | 3549.0 | 1 | Felidae | 2390.0 | 2 | Felis | 2390.0 | 2 | Felis catus | 2390.0 | 2 |
| scf7180000022223 | 18845 | 0.3863 | 0 | 24.1784 | Eukaryota | 23548.0 | 0 | Chordata | 23548.0 | 0 | Carnivora | 23548.0 | 0 | Felidae | 23548.0 | 0 | Felis | 21147.0 | 1 | Felis catus | 21147.0 | 1 |
| scf7180000024273 | 1110040 | 0.4904 | 0 | 52.0061 | Eukaryota | 40667.0 | 0 | Chordata | 40667.0 | 0 | Carnivora | 27412.0 | 1 | Otariidae | 7226.0 | 6 | Zalophus | 7226.0 | 7 | phus californi | 7226.0 | 7 |
| scf7180000022687 | 13609 | 0.5205 | 0 | 43.7926 | Eukaryota | 2400.0 | 0 | Chordata | 2400.0 | 0 | Carnivora | 2400.0 | 0 | Felidae | 2150.0 | 1 | Felis | 1457.0 | 3 | Felis catus | 1233.0 | 5 |
| scf7180000023021 | 65535 | 0.3876 | 0 | 233.4106 | Eukaryota | 7010.0 | 0 | Chordata | 7010.0 | 0 | Carnivora | 7010.0 | 0 | Felidae | 7010.0 | 0 | Felis | 7010.0 | 0 | Felis catus | 7010.0 | 0 |
| scf7180000020440 | 132711 | 0.4158 | 0 | 26.4867 | Eukaryota | 47246.0 | 0 | Chordata | 47246.0 | 0 | Carnivora | 47246.0 | 0 | Felidae | 39068.0 | 1 | Felis | 18548.0 | 4 | Felis catus | 18548.0 | 5 |
| scf7180000017902 | 1434109 | 0.3495 | 0 | 48.4408 | Eukaryota | 47152.0 | 0 | Chordata | 47152.0 | 0 | Carnivora | 47152.0 | 0 | Felidae | 47152.0 | 0 | Felis | 47152.0 | 0 | Felis catus | 47152.0 | 0 |
| scf7180000019572 | 392573 | 0.3745 | 0 | 48.3438 | Eukaryota | 44041.0 | 0 | Chordata | 44041.0 | 0 | Carnivora | 44041.0 | 0 | Felidae | 44041.0 | 0 | Felis | 39588.0 | 1 | Felis catus | 39588.0 | 1 |
| scf7180000020686 | 90861 | 0.4239 | 0 | 50.9998 | Eukaryota | 28166.0 | 0 | Chordata | 28166.0 | 0 | Carnivora | 28166.0 | 0 | Felidae | 28166.0 | 0 | Felis | 25298.0 | 1 | Felis catus | 25298.0 | 1 |
| scf7180000021295 | 49739 | 0.5071 | 0 | 50.9012 | Eukaryota | 42240.0 | 0 | Chordata | 42240.0 | 0 | Carnivora | 42240.0 | 0 | Felidae | 42240.0 | 0 | Felis | 38047.0 | 1 | Felis catus | 38047.0 | 1 |
| scf7180000017919 | 1422044 | 0.3767 | 0 | 51.3582 | Eukaryota | 47922.0 | 0 | Chordata | 47922.0 | 0 | Carnivora | 47922.0 | 0 | Felidae | 47922.0 | 0 | Felis | 47922.0 | 0 | Felis catus | 47922.0 | 0 |
| scf7180000020759 | 8506 | 0.5225 | 0 | 286.7051 | no-hit | 0.0 | 0 | no-hit | 0.0 | 0 | no-hit | 0.0 | 0 | no-hit | 0.0 | 0 | no-hit | 0.0 | 0 | no-hit | 0.0 | 0 |
| scf7180000020640 | 28970 | 0.602 | 0 | 46.9749 | Eukaryota | 2743.0 | 0 | Chordata | 2743.0 | 0 | Carnivora | 2743.0 | 0 | Felidae | 2491.0 | 1 | Felis | 1279.0 | 4 | Felis catus | 1279.0 | 4 |
| scf7180000018166 | 1060051 | 0.4832 | 0 | 51.8407 | Eukaryota | 55213.0 | 0 | Chordata | 55213.0 | 0 | Carnivora | 40379.0 | 1 | Herpestidae | 20748.0 | 4 | Suricata | 20748.0 | 7 | ıricata suricat | 20748.0 | 7 |
| scf7180000017851 | 1432925 | 0.407 | 0 | 51.3701 | Eukaryota | 47163.0 | 0 | Chordata | 47163.0 | 0 | Carnivora | 47163.0 | 0 | Felidae | 47163.0 | 0 | Felis | 47163.0 | 0 | Felis catus | 47163.0 | 0 |
| scf7180000019786 | 285474 | 0.3906 | 0 | 51.2483 | Eukaryota | 44896.0 | 0 | Chordata | 44896.0 | 0 | Carnivora | 44896.0 | 0 | Felidae | 44896.0 | 0 | Felis | 39793.0 | 1 | Felis catus | 39793.0 | 1 |
| scf7180000020746 | 76909 | 0.3878 | 0 | 25.4323 | Eukaryota | 47049.0 | 0 | Chordata | 47049.0 | 0 | Carnivora | 47049.0 | 0 | Felidae | 47049.0 | 0 | Felis | 42602.0 | 1 | Felis catus | 42602.0 | 1 |
| scf7180000020505 | 99957 | 0.6038 | 0 | 48.1886 | Eukaryota | 3770.0 | 0 | Chordata | 3770.0 | 0 | Carnivora | 2179.0 | 1 | Felidae | 2179.0 | 1 | Homo | 1591.0 | 3 | Homo sapien: | 1591.0 | 4 |
| scf7180000018076 | 1033474 | 0.4281 | 0 | 51.0781 | Eukaryota | 34357.0 | 0 | Chordata | 34357.0 | 0 | Carnivora | 34357.0 | 0 | Felidae | 34357.0 | 0 | Felis | 34357.0 | 0 | Felis catus | 34357.0 | 0 |
| scf7180000024053 | 614177 | 0.4222 | 0 | 51.0268 | Eukaryota | 47934.0 | 0 | Chordata | 47934.0 | 0 | Carnivora | 43974.0 | 1 | Felidae | 33982.0 | 2 | Lynx | 19708.0 | 3 | ynx canadens | 19708.0 | 3 |
| scf7180000020158 | 102907 | 0.3741 | 0 | 25.799 | Eukaryota | 66112.0 | 1 | Chordata | 66112.0 | 1 | Carnivora | 47694.0 | 2 | Felidae | 24241.0 | 3 | Homo | 18418.0 | 6 | Homo sapien: | 18418.0 | 6 |
| scf7180000022311 | 24458 | 0.3539 | 0 | 24.0738 | Eukaryota | 4095.0 | 0 | Chordata | 4095.0 | 0 | Carnivora | 3502.0 | 1 | Felidae | 1935.0 | 3 | Felis | 1935.0 | 3 | Felis catus | 1935.0 | 3 |
| scf7180000019005 | 1036290 | 0.3958 | 0 | 50.028 | Eukaryota | 47344.0 | 0 | Chordata | 47344.0 | 0 | Carnivora | 47344.0 | 0 | Felidae | 47344.0 | 0 | Felis | 47344.0 | 0 | Felis catus | 47344.0 | 0 |
| scf7180000019409 | 195425 | 0.3985 | 0 | 49.2936 | Eukaryota | 25045.0 | 0 | Chordata | 25045.0 | 0 | Carnivora | 25045.0 | 0 | Felidae | 25045.0 | 0 | Felis | 22581.0 | 1 | Felis catus | 22581.0 | 1 |
| scf7180000024196 | 64599 | 0.5873 | 0 | 49.2301 | Eukaryota | 9063.0 | 0 | Chordata | 9063.0 | 0 | Carnivora | 9063.0 | 0 | Felidae | 7623.0 | 2 | Acinonyx | 2600.0 | 5 | :inonyx jubat | 2600.0 | 6 |
| scf7180000017949 | 387953 | 0.3707 | 0 | 47.1348 | Eukaryota | 45291.0 | 0 | Chordata | 45291.0 | 0 | Carnivora | 45291.0 | 0 | Felidae | 45291.0 | 0 | Felis | 45291.0 | 0 | Felis catus | 45291.0 | 0 |
| scf7180000023490 | 50554 | 0.4329 | 0 | 216.3483 | Eukaryota | 28200.0 | 0 | Chordata | 28200.0 | 0 | Carnivora | 28200.0 | 0 | Felidae | 28200.0 | 0 | Felis | 28200.0 | 0 | Felis catus | 28200.0 | 0 |
| scf7180000021694 | 70665 | 0.3971 | 0 | 40.2251 | Eukaryota | 31785.0 | 0 | Chordata | 31785.0 | 0 | Carnivora | 31785.0 | 0 | Felidae | 31785.0 | 0 | Felis | 31785.0 | 0 | Felis catus | 31785.0 | 0 |
| scf7180000023864 | 374038 | 0.3996 | 0 | 50.435 | Eukaryota | 44562.0 | 0 | Chordata | 44562.0 | 0 | Carnivora | 44562.0 | 0 | Felidae | 44562.0 | 0 | Felis | 44562.0 | 0 | Felis catus | 44562.0 | 0 |
| scf7180000021710 | 44240 | 0.4158 | 0 | 25.3931 | Eukaryota | 50922.0 | 0 | Chordata | 50922.0 | 0 | Carnivora | 31354.0 | 2 | Canidae | 10132.0 | 8 | Canis | 10132.0 | 8 | Canis lupus | 10132.0 | 8 |
| scf7180000021781 | 66831 | 0.5203 | 0 | 45.338 | Eukaryota | 6813.0 | 0 | Chordata | 6813.0 | 0 | Carnivora | 6813.0 | 0 | Felidae | 6813.0 | 0 | Felis | 4068.0 | 1 | Felis catus | 4068.0 | 1 |
| scf7180000020478 | 189280 | 0.5602 | 0 | 47.7545 | Eukaryota | 13921.0 | 0 | Chordata | 13921.0 | 0 | Carnivora | 10233.0 | 1 | Felidae | 7679.0 | 2 | Felis | 7679.0 | 2 | Felis catus | 7679.0 | 2 |
| scf7180000022297 | 12019 | 0.5589 | 0 | 26.0697 | Eukaryota | 2960.0 | 0 | Chordata | 2960.0 | 0 | Carnivora | 2960.0 | 0 | Felidae | 2960.0 | 0 | Felis | 1751.0 | 1 | Felis catus | 1751.0 | 1 |
| scf7180000022090 | 14531 | 0.4971 | 0 | 26.8685 | Eukaryota | 2920.0 | 0 | Chordata | 2920.0 | 0 | Carnivora | 2920.0 | 0 | Felidae | 2044.0 | 1 | Suricata | 876.0 | 4 | ıricata suricat | 876.0 | 4 |
| scf7180000018548 | 1467901 | 0.4253 | 0 | 94.4967 | Eukaryota | 61854.0 | 0 | Chordata | 61854.0 | 0 | Carnivora | 61854.0 | 0 | Otariidae | 29900.0 | 2 | Callorhinus | 29900.0 | 3 | llorhinus ursir | 29900.0 | 3 |
| scf7180000021839 | 48317 | 0.382 | 0 | 23.86 | Eukaryota | 41308.0 | 0 | Chordata | 41308.0 | 0 | Carnivora | 41308.0 | 0 | Felidae | 41308.0 | 0 | Felis | 37221.0 | 1 | Felis catus | 37221.0 | 1 |
| scf7180000022940 | 49635 | 0.3958 | 0 | 432.1964 | Eukaryota | 7061.0 | 0 | Chordata | 7061.0 | 0 | Carnivora | 7061.0 | 0 | Felidae | 4982.0 | 1 | Felis | 4982.0 | 1 | Felis catus | 4982.0 | 1 |
| scf7180000018884 | 572555 | 0.4292 | 0 | 50.6653 | Eukaryota | 28585.0 | 0 | Chordata | 28585.0 | 0 | Carnivora | 28585.0 | 0 | Felidae | 28585.0 | 0 | Felis | 25713.0 | 1 | Felis catus | 25713.0 | 1 |
| scf7180000017942 | 2016347 | 0.4109 | 0 | 50.8726 | Eukaryota | 47962.0 | 0 | Chordata | 47962.0 | 0 | Carnivora | 47962.0 | 0 | Felidae | 47962.0 | 0 | Felis | 47962.0 | 0 | Felis catus | 47962.0 | 0 |
| scf7180000020280 | 178392 | 0.4116 | 0 | 51.2511 | Eukaryota | 43116.0 | 0 | Chordata | 43116.0 | 0 | Carnivora | 43116.0 | 0 | Felidae | 43116.0 | 0 | Panthera | 13058.0 | 3 | anthera pardı | 13058.0 | 3 |
| scf7180000019204 | 382455 | 0.3959 | 0 | 51.2496 | Eukaryota | 45672.0 | 0 | Chordata | 45672.0 | 0 | Carnivora | 45672.0 | 0 | Felidae | 45672.0 | 0 | Felis | 45672.0 | 0 | Felis catus | 45672.0 | 0 |
| scf7180000022933 | 32739 | 0.3986 | 0 | 197.9396 | Eukaryota | 6949.0 | 0 | Chordata | 6949.0 | 0 | Carnivora | 6949.0 | 0 | Felidae | 6949.0 | 0 | Felis | 6273.0 | 1 | Felis catus | 6273.0 | 1 |
| scf7180000018944 | 34632 | 0.5723 | 0 | 40.6921 | Eukaryota | 10792.0 | 0 | Chordata | 10792.0 | 0 | Carnivora | 10792.0 | 0 | Felidae | 10792.0 | 0 | Felis | 8627.0 | 1 | Felis catus | 8627.0 | 1 |
| scf7180000020156 | 142812 | 0.4159 | 0 | 50.4051 | Eukaryota | 17396.0 | 0 | Chordata | 17396.0 | 0 | Carnivora | 17396.0 | 0 | Felidae | 17396.0 | 0 | Panthera | 7088.0 | 4 | unresolved | 3544.0 | 5 |
| scf7180000022484 | 13740 | 0.4435 | 0 | 26.596 | Eukaryota | 8441.0 | 0 | Chordata | 8441.0 | 0 | Primates | 5187.0 | 3 | Hominidae | 5187.0 | 3 | Homo | 3436.0 | 4 | Homo sapien: | 3436.0 | 4 |
| scf7180000024019 | 550367 | 0.3998 | 0 | 49.0842 | Eukaryota | 185705.0 | 0 | Chordata | 185705.0 | 0 | Carnivora | 185705.0 | 0 | Felidae | 145851.0 | 1 | Acinonyx | 91143.0 | 2 | :inonyx jubat | 91143.0 | 2 |
| scf7180000020034 | 145228 | 0.3778 | 0 | 25.1505 | Eukaryota | 39817.0 | 0 | Chordata | 39817.0 | 0 | Carnivora | 39817.0 | 0 | Felidae | 39817.0 | 0 | Felis | 39817.0 | 0 | Felis catus | 39817.0 | 0 |
| scf7180000018951 | 42001 | 0.4531 | 0 | 26.4138 | Eukaryota | 11197.0 | 0 | Chordata | 11197.0 | 0 | Carnivora | 11197.0 | 0 | Felidae | 11197.0 | 0 | Felis | 10325.0 | 1 | Felis catus | 10325.0 | 1 |
| scf7180000020783 | 58656 | 0.3831 | 0 | 26.4519 | Eukaryota | 24181.0 | 0 | Chordata | 24181.0 | 0 | Carnivora | 24181.0 | 0 | Felidae | 24181.0 | 0 | Felis | 21752.0 | 1 | Felis catus | 21752.0 | 1 |
| scf7180000022331 | 15254 | 0.4308 | 0 | 46.931 | Eukaryota | 2517.0 | 0 | Chordata | 2517.0 | 0 | Carnivora | 2517.0 | 0 | Felidae | 2267.0 | 1 | Felis | 1775.0 | 2 | Felis catus | 1775.0 | 2 |
| scf7180000022698 | 13989 | 0.5384 | 0 | 47.203 | Eukaryota | 2130.0 | 1 | Chordata | 2130.0 | 1 | Carnivora | 982.0 | 3 | Felidae | 982.0 | 3 | Felis | 667.0 | 6 | Felis catus | 667.0 | 6 |
| scf7180000022613 | 6998 | 0.393 | 0 | 25.7117 | Eukaryota | 8536.0 | 0 | Chordata | 8101.0 | 1 | Primates | 3620.0 | 4 | Hominidae | 3620.0 | 4 | Sus | 3332.0 | 5 | Sus scrofa | 3332.0 | 5 |
| scf7180000021962 | 25335 | 0.3981 | 0 | 26.9868 | Eukaryota | 10189.0 | 0 | Chordata | 10189.0 | 0 | Carnivora | 10189.0 | 0 | Felidae | 10189.0 | 0 | Felis | 8172.0 | 1 | Felis catus | 8172.0 | 1 |
| scf7180000020324 | 139855 | 0.5171 | 0 | 49.9976 | Eukaryota | 21146.0 | 0 | Chordata | 21146.0 | 0 | Carnivora | 21146.0 | 0 | Felidae | 9928.0 | 4 | Ursus | 4079.0 | 8 | Felis catus | 3317.0 | 9 |
| scf7180000019638 | 312567 | 0.3841 | 0 | 24.1414 | Eukaryota | 24257.0 | 0 | Chordata | 24257.0 | 0 | Carnivora | 24257.0 | 0 | Felidae | 24257.0 | 0 | Felis | 24257.0 | 0 | Felis catus | 24257.0 | 0 |
| scf7180000021364 | 50293 | 0.4119 | 0 | 25.9859 | Eukaryota | 23022.0 | 0 | Chordata | 23022.0 | 0 | Carnivora | 23022.0 | 0 | Felidae | 23022.0 | 0 | Felis | 20532.0 | 1 | Felis catus | 20532.0 | 1 |
| scf7180000018946 | 456613 | 0.5014 | 0 | 49.5254 | Eukaryota | 19503.0 | 0 | Chordata | 19503.0 | 0 | Carnivora | 19503.0 | 0 | Felidae | 19503.0 | 0 | Felis | 7781.0 | 3 | Felis catus | 7781.0 | 4 |
| scf7180000017680 | 1901712 | 0.4336 | 0 | 50.3429 | Eukaryota | 35706.0 | 0 | Chordata | 35706.0 | 0 | Carnivora | 35706.0 | 0 | Felidae | 35706.0 | 0 | Felis | 32114.0 | 1 | Felis catus | 32114.0 | 1 |
| scf7180000018743 | 118055 | 0.5155 | 0 | 48.9439 | Eukaryota | 11767.0 | 0 | Chordata | 11767.0 | 0 | Carnivora | 11767.0 | 0 | Felidae | 10152.0 | 1 | Felis | 5894.0 | 3 | Felis catus | 5894.0 | 4 |
| scf7180000021435 | 85539 | 0.5528 | 0 | 46.7282 | Eukaryota | 9596.0 | 0 | Chordata | 9596.0 | 0 | Carnivora | 9596.0 | 0 | Felidae | 9596.0 | 0 | Felis | 9596.0 | 0 | Felis catus | 9596.0 | 0 |
| scf7180000018994 | 964839 | 0.3482 | 0 | 48.8764 | Eukaryota | 95300.0 | 0 | Chordata | 95300.0 | 0 | Carnivora | 95300.0 | 0 | Felidae | 77740.0 | 1 | Panthera | 45066.0 | 3 | anthera pardı | 45066.0 | 3 |
| scf7180000022656 | 15662 | 0.4947 | 0 | 51.8161 | Eukaryota | 2858.0 | 0 | Chordata | 2858.0 | 0 | Carnivora | 2858.0 | 0 | Felidae | 2858.0 | 0 | Felis | 2571.0 | 1 | Felis catus | 2571.0 | 1 |
| scf7180000021053 | 67532 | 0.5415 | 0 | 25.4233 | Eukaryota | 21302.0 | 0 | Chordata | 21302.0 | 0 | Carnivora | 8717.0 | 2 | Hominidae | 7741.0 | 3 | Pan | 5164.0 | 5 | an troglodyte | 5164.0 | 5 |
| scf7180000019793 | 112640 | 0.3791 | 0 | 45.9992 | Eukaryota | 27658.0 | 0 | Chordata | 27658.0 | 0 | Carnivora | 27658.0 | 0 | Felidae | 24908.0 | 1 | Felis | 24908.0 | 1 | Felis catus | 24908.0 | 1 |
| scf7180000021253 | 54438 | 0.4063 | 0 | 24.6733 | Eukaryota | 8181.0 | 0 | Chordata | 8181.0 | 0 | Carnivora | 5202.0 | 1 | Felidae | 5202.0 | 1 | Felis | 4622.0 | 2 | Felis catus | 4622.0 | 2 |
| scf7180000020910 | 104603 | 0.3573 | 0 | 24.697 | Eukaryota | 43209.0 | 0 | Chordata | 43209.0 | 0 | Carnivora | 43209.0 | 0 | Felidae | 43209.0 | 0 | Felis | 43209.0 | 0 | Felis catus | 43209.0 | 0 |
| scf7180000019890 | 263984 | 0.458 | 0 | 24.8262 | Eukaryota | 43641.0 | 0 | Chordata | 43641.0 | 0 | Carnivora | 23886.0 | 2 | Hominidae | 12850.0 | 5 | Homo | 12850.0 | 7 | Homo sapien: | 12850.0 | 7 |
| scf7180000017926 | 1452623 | 0.4492 | 0 | 52.7427 | Eukaryota | 117815.0 | 0 | Chordata | 117815.0 | 0 | Carnivora | 112618.0 | 1 | Felidae | 93932.0 | 2 | Panthera | 47893.0 | 6 | anthera pardı | 40546.0 | 8 |
| scf7180000019746 | 187306 | 0.5795 | 0 | 51.6774 | Eukaryota | 19952.0 | 0 | Chordata | 19952.0 | 0 | Carnivora | 19952.0 | 0 | Felidae | 18642.0 | 1 | Lynx | 6326.0 | 5 | ynx canadens | 6326.0 | 5 |
| scf7180000020145 | 46716 | 0.5721 | 0 | 164.9213 | Eukaryota | 3808.0 | 0 | Chordata | 3808.0 | 0 | Carnivora | 2980.0 | 1 | Felidae | 1950.0 | 2 | Acinonyx | 1116.0 | 4 | :inonyx jubat | 1116.0 | 4 |
| scf7180000019938 | 11048 | 0.5511 | 0 | 154.6297 | Eukaryota | 2989.0 | 0 | Chordata | 2989.0 | 0 | Carnivora | 2710.0 | 1 | Felidae | 2259.0 | 2 | unresolved | 849.0 | 5 | unresolved | 849.0 | 5 |
| scf7180000018945 | 8373 | 0.4001 | 0 | 33.2243 | Eukaryota | 41557.0 | 0 | Chordata | 41557.0 | 0 | Carnivora | 41557.0 | 0 | Felidae | 41557.0 | 0 | Felis | 41557.0 | 0 | Felis catus | 41557.0 | 0 |
| scf7180000019067 | 11600 | 0.3722 | 0 | 20.0131 | Eukaryota | 21209.0 | 0 | Chordata | 21209.0 | 0 | Carnivora | 21209.0 | 0 | Felidae | 21209.0 | 0 | Felis | 18815.0 | 1 | Felis catus | 18815.0 | 1 |
| scf7180000017934 | 1079165 | 0.378 | 0 | 51.9157 | Eukaryota | 98836.0 | 0 | Chordata | 98836.0 | 0 | Carnivora | 98836.0 | 0 | Felidae | 98836.0 | 0 | Neofelis | 52337.0 | 1 | eofelis nebulo | 52337.0 | 1 |
| scf7180000024124 | 145501 | 0.4899 | 0 | 50.0085 | Eukaryota | 20512.0 | 0 | Chordata | 20512.0 | 0 | Carnivora | 20512.0 | 0 | Felidae | 20512.0 | 0 | Panthera | 12356.0 | 1 | anthera pardı | 10300.0 | 2 |
| scf7180000021883 | 888435 | 0.4244 | 100 | 50.8955 | Eukaryota | 36336.0 | 0 | Chordata | 36336.0 | 0 | Carnivora | 36336.0 | 0 | Felidae | 36336.0 | 0 | Felis | 24307.0 | 4 | Felis catus | 24307.0 | 4 |
| scf7180000021406 | 119680 | 0.372 | 0 | 23.877 | Eukaryota | 43241.0 | 0 | Chordata | 43241.0 | 0 | Carnivora | 43241.0 | 0 | Felidae | 43241.0 | 0 | Felis | 43241.0 | 0 | Felis catus | 43241.0 | 0 |
| scf7180000019644 | 275885 | 0.586 | 0 | 51.3318 | Eukaryota | 8135.0 | 0 | Chordata | 8135.0 | 0 | Carnivora | 3908.0 | 3 | Felidae | 3908.0 | 3 | Homo | 2706.0 | 4 | Homo sapien: | 2706.0 | 4 |
| scf7180000021557 | 41318 | 0.5885 | 0 | 53.9704 | Eukaryota | 2628.0 | 0 | Chordata | 2628.0 | 0 | Carnivora | 2628.0 | 0 | Felidae | 2628.0 | 0 | Felis | 1602.0 | 1 | Felis catus | 1602.0 | 1 |
| scf7180000022130 | 15618 | 0.5409 | 0 | 46.3602 | Eukaryota | 2717.0 | 0 | Chordata | 2717.0 | 0 | Carnivora | 2329.0 | 1 | Felidae | 2070.0 | 2 | Leopardus | 1040.0 | 3 | Felis catus | 1030.0 | 6 |
| scf7180000020535 | 5760 | 0.4962 | 0 | 22.988 | Eukaryota | 2589.0 | 0 | Chordata | 2589.0 | 0 | Carnivora | 2293.0 | 1 | Felidae | 2293.0 | 1 | Felis | 1531.0 | 3 | Felis catus | 1279.0 | 4 |
| scf7180000022081 | 39632 | 0.3751 | 0 | 23.5682 | Eukaryota | 7454.0 | 0 | Chordata | 7454.0 | 0 | Carnivora | 5816.0 | 3 | Felidae | 4611.0 | 5 | Felis | 2436.0 | 6 | Felis catus | 2436.0 | 6 |
| scf7180000018342 | 8106 | 0.5263 | 0 | 22.8078 | Eukaryota | 7988.0 | 0 | Chordata | 7988.0 | 0 | Carnivora | 7988.0 | 0 | Felidae | 7988.0 | 0 | Acinonyx | 4576.0 | 2 | :inonyx jubat | 4576.0 | 2 |
| scf7180000020851 | 117179 | 0.3843 | 0 | 24.7689 | Eukaryota | 43773.0 | 0 | Chordata | 43773.0 | 0 | Carnivora | 43773.0 | 0 | Felidae | 43773.0 | 0 | Felis | 43773.0 | 0 | Felis catus | 43773.0 | 0 |
| scf7180000021549 | 47994 | 0.4542 | 0 | 46.3213 | Eukaryota | 28764.0 | 0 | Chordata | 28764.0 | 0 | Carnivora | 28764.0 | 0 | Felidae | 28764.0 | 0 | Felis | 28764.0 | 0 | Felis catus | 28764.0 | 0 |
| scf7180000018499 | 735401 | 0.3913 | 0 | 50.6255 | Eukaryota | 45846.0 | 0 | Chordata | 45846.0 | 0 | Carnivora | 45846.0 | 0 | Felidae | 45846.0 | 0 | Felis | 45846.0 | 0 | Felis catus | 45846.0 | 0 |
| scf7180000019494 | 413899 | 0.356 | 0 | 48.4723 | Eukaryota | 30671.0 | 0 | Chordata | 30671.0 | 0 | Carnivora | 30671.0 | 0 | Felidae | 30671.0 | 0 | Felis | 27535.0 | 1 | Felis catus | 27535.0 | 1 |
| scf7180000023415 | 20970 | 0.4261 | 0 | 411.681 | Eukaryota | 171327.0 | 0 | Chordata | 171327.0 | 0 | Carnivora | 171327.0 | 0 | Hyaenidae | 171327.0 | 0 | Proteles | 101016.0 | 1 | roteles cristat | 101016.0 | 1 |
| scf7180000018594 | 742512 | 0.3612 | 0 | 49.8161 | Eukaryota | 60285.0 | 0 | Chordata | 60285.0 | 0 | Carnivora | 60285.0 | 0 | Felidae | 42214.0 | 1 | Panthera | 24060.0 | 2 | anthera pardı | 24060.0 | 2 |
| scf7180000022593 | 6392 | 0.395 | 0 | 22.0703 | Eukaryota | 2935.0 | 0 | Chordata | 2935.0 | 0 | Carnivora | 2318.0 | 1 | Felidae | 2318.0 | 1 | Felis | 1284.0 | 3 | Felis catus | 1284.0 | 4 |
| scf7180000021603 | 32534 | 0.3831 | 0 | 25.6329 | Eukaryota | 16714.0 | 0 | Chordata | 16714.0 | 0 | Carnivora | 16714.0 | 0 | Felidae | 16714.0 | 0 | Felis | 16714.0 | 0 | Felis catus | 16714.0 | 0 |
| scf7180000020830 | 77920 | 0.4147 | 0 | 25.944 | Eukaryota | 20306.0 | 0 | Chordata | 20306.0 | 0 | Primates | 16328.0 | 3 | Hominidae | 14941.0 | 4 | Pan | 7780.0 | 5 | an troglodyte | 7780.0 | 5 |
| scf7180000023589 | 9012 | 0.7254 | 0 | 361.4552 | Eukaryota | 2485.0 | 0 | Chordata | 2485.0 | 0 | Carnivora | 2485.0 | 0 | Felidae | 2255.0 | 1 | Panthera | 1164.0 | 3 | anthera pardı | 1164.0 | 3 |
| scf7180000022771 | 4174 | 0.4329 | 0 | 24.8149 | Eukaryota | 3005.0 | 0 | Chordata | 3005.0 | 0 | Carnivora | 3005.0 | 0 | Felidae | 3005.0 | 0 | Felis | 1809.0 | 1 | Felis catus | 1809.0 | 1 |
| scf7180000021054 | 65642 | 0.588 | 0 | 51.5302 | Eukaryota | 17920.0 | 0 | Chordata | 17920.0 | 0 | Carnivora | 17920.0 | 0 | Herpestidae | 11598.0 | 1 | Suricata | 11598.0 | 2 | ıricata suricat | 11598.0 | 2 |
| scf7180000018670 | 2582018 | 0.4085 | 0 | 51.148 | Eukaryota | 47773.0 | 0 | Chordata | 47773.0 | 0 | Carnivora | 47773.0 | 0 | Felidae | 47773.0 | 0 | Felis | 47773.0 | 0 | Felis catus | 47773.0 | 0 |
| scf7180000020436 | 17211 | 0.4267 | 0 | 397.8923 | Eukaryota | 136991.0 | 0 | Chordata | 136991.0 | 0 | Carnivora | 136991.0 | 0 | Hyaenidae | 136991.0 | 0 | Proteles | 85759.0 | 1 | roteles cristat | 85759.0 | 1 |
| scf7180000020521 | 115731 | 0.3667 | 0 | 24.5303 | Eukaryota | 43438.0 | 0 | Chordata | 43438.0 | 0 | Carnivora | 43438.0 | 0 | Felidae | 43438.0 | 0 | Felis | 43438.0 | 0 | Felis catus | 43438.0 | 0 |
| scf7180000018539 | 737895 | 0.4723 | 0 | 51.5664 | Eukaryota | 38452.0 | 0 | Chordata | 38452.0 | 0 | Carnivora | 38452.0 | 0 | Felidae | 31349.0 | 1 | Panthera | 9063.0 | 4 | Felis catus | 8766.0 | 5 |
| scf7180000022551 | 18502 | 0.4178 | 0 | 25.1684 | Eukaryota | 8580.0 | 0 | Chordata | 8580.0 | 0 | Carnivora | 7652.0 | 1 | Felidae | 4629.0 | 2 | Felis | 4629.0 | 2 | Felis catus | 4629.0 | 2 |
| scf7180000020187 | 44180 | 0.4194 | 0 | 262.7991 | Eukaryota | 22241.0 | 0 | Chordata | 22241.0 | 0 | Carnivora | 22241.0 | 0 | Felidae | 22241.0 | 0 | Felis | 20074.0 | 1 | Felis catus | 20074.0 | 1 |
| scf7180000018689 | 506294 | 0.4694 | 0 | 49.5672 | Eukaryota | 53306.0 | 0 | Chordata | 53306.0 | 0 | Carnivora | 53306.0 | 0 | Felidae | 53306.0 | 0 | Panthera | 22251.0 | 2 | ynx canadens | 18164.0 | 3 |
| scf7180000021909 | 26375 | 0.4596 | 0 | 51.3152 | Eukaryota | 14884.0 | 0 | Chordata | 14884.0 | 0 | Primates | 5640.0 | 5 | Hominidae | 5640.0 | 5 | Homo | 5640.0 | 5 | Homo sapien: | 5640.0 | 5 |
| scf7180000020201 | 186516 | 0.448 | 0 | 48.9657 | Eukaryota | 46121.0 | 0 | Chordata | 46121.0 | 0 | Carnivora | 46121.0 | 0 | Felidae | 46121.0 | 0 | Felis | 46121.0 | 0 | Felis catus | 46121.0 | 0 |
| scf7180000018395 | 618115 | 0.3949 | 0 | 49.9797 | Eukaryota | 45613.0 | 0 | Chordata | 45613.0 | 0 | Carnivora | 45613.0 | 0 | Felidae | 45613.0 | 0 | Felis | 45613.0 | 0 | Felis catus | 45613.0 | 0 |
| scf7180000022609 | 6715 | 0.4144 | 0 | 21.6906 | Eukaryota | 3787.0 | 0 | Chordata | 3787.0 | 0 | Carnivora | 3787.0 | 0 | Felidae | 1811.0 | 2 | Canis | 1647.0 | 3 | Canis lupus | 1647.0 | 3 |
| scf7180000024108 | 581204 | 0.3553 | 0 | 48.7205 | Eukaryota | 45031.0 | 0 | Chordata | 45031.0 | 0 | Carnivora | 45031.0 | 0 | Felidae | 45031.0 | 0 | Felis | 45031.0 | 0 | Felis catus | 45031.0 | 0 |
| scf7180000020113 | 12167 | 0.651 | 0 | 31.9059 | Eukaryota | 5157.0 | 0 | Chordata | 5157.0 | 0 | Carnivora | 5157.0 | 0 | Canidae | 2597.0 | 1 | Acinonyx | 2560.0 | 2 | :inonyx jubat | 2560.0 | 2 |
| scf7180000024137 | 165117 | 0.3617 | 0 | 23.3457 | Eukaryota | 41853.0 | 0 | Chordata | 41853.0 | 0 | Carnivora | 41853.0 | 0 | Felidae | 41853.0 | 0 | Felis | 37712.0 | 1 | Felis catus | 37712.0 | 1 |
| scf7180000017650 | 1785184 | 0.3823 | 0 | 49.4688 | Eukaryota | 47238.0 | 0 | Chordata | 47238.0 | 0 | Carnivora | 47238.0 | 0 | Felidae | 47238.0 | 0 | Felis | 47238.0 | 0 | Felis catus | 47238.0 | 0 |
| scf7180000020562 | 100154 | 0.4279 | 0 | 50.4686 | Eukaryota | 11415.0 | 0 | Chordata | 11415.0 | 0 | Carnivora | 11415.0 | 0 | Felidae | 11415.0 | 0 | Felis | 11415.0 | 0 | Felis catus | 11415.0 | 0 |
| scf7180000019749 | 21174 | 0.4301 | 0 | 38.6839 | Eukaryota | 13862.0 | 0 | Chordata | 13862.0 | 0 | Carnivora | 13862.0 | 0 | Felidae | 13862.0 | 0 | Felis | 12467.0 | 1 | Felis catus | 12467.0 | 1 |
| scf7180000018190 | 1225721 | 0.4315 | 0 | 50.0559 | Eukaryota | 45429.0 | 0 | Chordata | 45429.0 | 0 | Carnivora | 45429.0 | 0 | Felidae | 45429.0 | 0 | Felis | 45429.0 | 0 | Felis catus | 45429.0 | 0 |
| scf7180000018782 | 444694 | 0.554 | 0 | 54.4109 | Eukaryota | 32223.0 | 0 | Chordata | 32223.0 | 0 | Carnivora | 29142.0 | 1 | Felidae | 22840.0 | 3 | Panthera | 13484.0 | 6 | anthera pardı | 9958.0 | 7 |
| scf7180000021518 | 39178 | 0.3898 | 0 | 24.2077 | Eukaryota | 23927.0 | 0 | Chordata | 23927.0 | 0 | Carnivora | 23927.0 | 0 | Felidae | 23927.0 | 0 | Felis | 21491.0 | 1 | Felis catus | 21491.0 | 1 |
| scf7180000023989 | 1230196 | 0.3957 | 0 | 51.9267 | Eukaryota | 47248.0 | 0 | Chordata | 47248.0 | 0 | Carnivora | 47248.0 | 0 | Felidae | 47248.0 | 0 | Felis | 47248.0 | 0 | Felis catus | 47248.0 | 0 |
| scf7180000019548 | 55604 | 0.4225 | 0 | 46.0129 | Eukaryota | 16393.0 | 0 | Chordata | 16393.0 | 0 | Carnivora | 16393.0 | 0 | Felidae | 16393.0 | 0 | Felis | 14773.0 | 1 | Felis catus | 14773.0 | 1 |
| scf7180000021947 | 19096 | 0.5455 | 0 | 27.9562 | Eukaryota | 5760.0 | 0 | Chordata | 5760.0 | 0 | Carnivora | 5178.0 | 1 | Felidae | 2759.0 | 4 | Panthera | 1387.0 | 6 | Canis lupus | 1094.0 | 7 |
| scf7180000019838 | 33172 | 0.4185 | 0 | 31.6385 | Eukaryota | 9575.0 | 0 | Chordata | 9575.0 | 0 | Carnivora | 9575.0 | 0 | Felidae | 9575.0 | 0 | Felis | 9575.0 | 0 | Felis catus | 9575.0 | 0 |
| scf7180000020659 | 47509 | 0.4312 | 0 | 31.4054 | Eukaryota | 46890.0 | 0 | Chordata | 46890.0 | 0 | Carnivora | 46890.0 | 0 | Felidae | 46890.0 | 0 | Felis | 46890.0 | 0 | Felis catus | 46890.0 | 0 |
| scf7180000018279 | 1204745 | 0.4158 | 0 | 50.9401 | Eukaryota | 29720.0 | 0 | Chordata | 29720.0 | 0 | Carnivora | 29720.0 | 0 | Felidae | 29720.0 | 0 | Panthera | 11900.0 | 3 | anthera pardı | 8934.0 | 4 |
| scf7180000020139 | 158003 | 0.3961 | 0 | 45.5903 | Eukaryota | 38328.0 | 0 | Chordata | 38328.0 | 0 | Carnivora | 38328.0 | 0 | Felidae | 18483.0 | 3 | Zalophus | 8906.0 | 6 | phus californi | 8906.0 | 7 |
| scf7180000021112 | 45236 | 0.5218 | 0 | 354.1618 | no-hit | 0.0 | 0 | no-hit | 0.0 | 0 | no-hit | 0.0 | 0 | no-hit | 0.0 | 0 | no-hit | 0.0 | 0 | no-hit | 0.0 | 0 |
| scf7180000020688 | 124876 | 0.5933 | 0 | 51.1639 | Eukaryota | 8174.0 | 0 | Chordata | 8174.0 | 0 | Carnivora | 8174.0 | 0 | Felidae | 7368.0 | 1 | Acinonyx | 2538.0 | 3 | :inonyx jubat | 2538.0 | 3 |
| scf7180000023553 | 17125 | 0.7236 | 0 | 486.2724 | Eukaryota | 2556.0 | 0 | Chordata | 2556.0 | 0 | Carnivora | 2556.0 | 0 | Felidae | 2556.0 | 0 | Panthera | 1176.0 | 2 | anthera pardı | 1176.0 | 2 |
| scf7180000021631 | 9801 | 0.4415 | 0 | 40.8202 | Eukaryota | 12662.0 | 0 | Chordata | 12662.0 | 0 | Carnivora | 12662.0 | 0 | Felidae | 12662.0 | 0 | Felis | 11422.0 | 1 | Felis catus | 11422.0 | 1 |
| scf7180000022424 | 12881 | 0.6143 | 0 | 44.3174 | Eukaryota | 2707.0 | 1 | Chordata | 2707.0 | 1 | Artiodactyla | 828.0 | 6 | Suidae | 828.0 | 6 | Sus | 828.0 | 7 | Sus scrofa | 828.0 | 7 |
| scf7180000021758 | 47957 | 0.3796 | 0 | 25.5718 | Eukaryota | 14995.0 | 0 | Chordata | 14995.0 | 0 | Primates | 11071.0 | 3 | Hominidae | 11071.0 | 3 | Homo | 6789.0 | 5 | Homo sapien: | 6789.0 | 5 |
| scf7180000021464 | 42599 | 0.4069 | 0 | 23.4053 | Eukaryota | 7360.0 | 0 | Chordata | 7360.0 | 0 | Carnivora | 7360.0 | 0 | Felidae | 3998.0 | 2 | Felis | 3009.0 | 3 | Felis catus | 3009.0 | 3 |
| scf7180000024059 | 273369 | 0.514 | 0 | 51.9228 | Eukaryota | 39076.0 | 0 | Chordata | 39076.0 | 0 | Carnivora | 39076.0 | 0 | Felidae | 39076.0 | 0 | Felis | 39076.0 | 0 | Felis catus | 39076.0 | 0 |
| scf7180000019461 | 21101 | 0.5692 | 0 | 28.3553 | Eukaryota | 7618.0 | 0 | Chordata | 7618.0 | 0 | Primates | 6001.0 | 1 | Hominidae | 5365.0 | 2 | Homo | 5365.0 | 3 | Homo sapien: | 5365.0 | 3 |
| scf7180000021904 | 27357 | 0.3776 | 0 | 23.6668 | Eukaryota | 4311.0 | 0 | Chordata | 4311.0 | 0 | Primates | 2200.0 | 2 | Hominidae | 2200.0 | 2 | Felis | 1512.0 | 3 | Felis catus | 1512.0 | 3 |
| scf7180000018966 | 465931 | 0.4081 | 0 | 49.6401 | Eukaryota | 40512.0 | 0 | Chordata | 40512.0 | 0 | Carnivora | 40512.0 | 0 | Felidae | 40512.0 | 0 | Felis | 40512.0 | 0 | Felis catus | 40512.0 | 0 |
| scf7180000018361 | 1950810 | 0.4323 | 0 | 50.2528 | Eukaryota | 69380.0 | 0 | Chordata | 69380.0 | 0 | Carnivora | 49276.0 | 1 | Felidae | 49276.0 | 2 | Panthera | 25799.0 | 4 | anthera pardı | 25799.0 | 4 |
| scf7180000022744 | 22996 | 0.4563 | 0 | 4766.8288 | Eukaryota | 4170.0 | 0 | Chordata | 4170.0 | 0 | Carnivora | 4170.0 | 0 | Felidae | 4170.0 | 0 | Felis | 4170.0 | 0 | Felis catus | 4170.0 | 0 |
| scf7180000018183 | 36589 | 0.4223 | 0 | 24.8924 | Eukaryota | 6492.0 | 0 | Chordata | 6492.0 | 0 | Carnivora | 6492.0 | 0 | Felidae | 6492.0 | 0 | Felis | 5845.0 | 1 | Felis catus | 5845.0 | 1 |
| scf7180000018534 | 119063 | 0.3421 | 0 | 42.5065 | Eukaryota | 25629.0 | 0 | Chordata | 25629.0 | 0 | Carnivora | 25629.0 | 0 | Felidae | 25629.0 | 0 | Felis | 23039.0 | 1 | Felis catus | 23039.0 | 1 |
| scf7180000020287 | 110422 | 0.3622 | 0 | 48.4922 | Eukaryota | 29485.0 | 0 | Chordata | 29485.0 | 0 | Carnivora | 29485.0 | 0 | Felidae | 29485.0 | 0 | Felis | 26574.0 | 1 | Felis catus | 26574.0 | 1 |
| scf7180000023015 | 37442 | 0.5678 | 0 | 229.4548 | Eukaryota | 6171.0 | 0 | Chordata | 6171.0 | 0 | Carnivora | 6171.0 | 0 | Felidae | 3763.0 | 2 | Felis | 2103.0 | 4 | Felis catus | 2103.0 | 4 |
| scf7180000021685 | 44388 | 0.405 | 0 | 25.2471 | Eukaryota | 12309.0 | 0 | Chordata | 12309.0 | 0 | Carnivora | 12309.0 | 0 | Felidae | 8889.0 | 1 | Acinonyx | 3891.0 | 3 | :inonyx jubat | 3891.0 | 3 |
| scf7180000020209 | 250947 | 0.3954 | 100 | 24.7599 | Eukaryota | 92342.0 | 0 | Chordata | 92342.0 | 0 | Carnivora | 92342.0 | 0 | Felidae | 82294.0 | 1 | Panthera | 38504.0 | 4 | anthera pardı | 38504.0 | 4 |
| scf7180000019182 | 380134 | 0.3519 | 0 | 48.0708 | Eukaryota | 44208.0 | 0 | Chordata | 44208.0 | 0 | Carnivora | 44208.0 | 0 | Felidae | 44208.0 | 0 | Felis | 39655.0 | 1 | Felis catus | 39655.0 | 1 |
| scf7180000022040 | 30836 | 0.4266 | 0 | 420.6353 | Eukaryota | 243750.0 | 0 | Chordata | 243750.0 | 0 | Carnivora | 243750.0 | 0 | Hyaenidae | 243750.0 | 0 | Proteles | 146965.0 | 1 | roteles cristat | 146965.0 | 1 |
| scf7180000021563 | 7709 | 0.6797 | 0 | 152.3862 | no-hit | 0.0 | 0 | no-hit | 0.0 | 0 | no-hit | 0.0 | 0 | no-hit | 0.0 | 0 | no-hit | 0.0 | 0 | no-hit | 0.0 | 0 |
| scf7180000017866 | 160671 | 0.4781 | 0 | 42.3478 | Eukaryota | 46007.0 | 0 | Chordata | 46007.0 | 0 | Carnivora | 46007.0 | 0 | Felidae | 46007.0 | 0 | Felis | 46007.0 | 0 | Felis catus | 46007.0 | 0 |
| scf7180000018861 | 553495 | 0.4392 | 0 | 50.0266 | Eukaryota | 41762.0 | 0 | Chordata | 41762.0 | 0 | Carnivora | 41762.0 | 0 | Felidae | 41762.0 | 0 | Acinonyx | 26419.0 | 2 | :inonyx jubat | 26419.0 | 2 |
| scf7180000019045 | 351587 | 0.3467 | 0 | 47.9899 | Eukaryota | 35665.0 | 0 | Chordata | 35665.0 | 0 | Carnivora | 35665.0 | 0 | Felidae | 35665.0 | 0 | Felis | 35665.0 | 0 | Felis catus | 35665.0 | 0 |
| scf7180000020751 | 105970 | 0.3792 | 0 | 24.1542 | Eukaryota | 37378.0 | 0 | Chordata | 37378.0 | 0 | Carnivora | 37378.0 | 0 | Felidae | 37378.0 | 0 | Felis | 33642.0 | 1 | Felis catus | 33642.0 | 1 |
| scf7180000018181 | 825888 | 0.5777 | 0 | 49.4892 | Eukaryota | 33630.0 | 0 | Chordata | 33630.0 | 0 | Carnivora | 33630.0 | 0 | Felidae | 13884.0 | 5 | Suricata | 4217.0 | 9 | ıricata suricat | 4217.0 | 9 |
| scf7180000020311 | 4298737 | 0.3915 | 0 | 51.2127 | Eukaryota | 48186.0 | 0 | Chordata | 48186.0 | 0 | Carnivora | 48186.0 | 0 | Felidae | 48186.0 | 0 | Felis | 48186.0 | 0 | Felis catus | 48186.0 | 0 |
| scf7180000018460 | 1362652 | 0.4392 | 0 | 51.2497 | Eukaryota | 32974.0 | 0 | Chordata | 32974.0 | 0 | Carnivora | 32974.0 | 0 | Felidae | 32974.0 | 0 | Felis | 29638.0 | 1 | Felis catus | 29638.0 | 1 |
| scf7180000020984 | 84931 | 0.377 | 0 | 45.5386 | Eukaryota | 46479.0 | 0 | Chordata | 46479.0 | 0 | Carnivora | 46479.0 | 0 | Felidae | 46479.0 | 0 | Felis | 46479.0 | 0 | Felis catus | 46479.0 | 0 |

| | | | | | | | | | | | | | | | | | | | | | | |
|---|---|---|---|---|---|---|---|---|---|---|---|---|---|---|---|---|---|---|---|---|---|---|
| scf7180000023585 | 30838 | 0.433 | 0 | 199.9849 | Eukaryota | 7666.0 | 0 | Chordata | 7666.0 | 0 | Carnivora | 7666.0 | 0 | Felidae | 7666.0 | 0 | Felis | 6881.0 | 1 | Felis catus | 6881.0 | 1 |
| scf7180000018246 | 1740360 | 0.3994 | 0 | 50.9023 | Eukaryota | 29414.0 | 0 | Chordata | 29414.0 | 0 | Carnivora | 29414.0 | 0 | Felidae | 29414.0 | 0 | Felis | 29414.0 | 0 | Felis catus | 29414.0 | 0 |
| scf7180000018740 | 1686496 | 0.544 | 100 | 51.3964 | Eukaryota | 56476.0 | 0 | Chordata | 56476.0 | 0 | Carnivora | 56476.0 | 0 | Mustelidae | 43560.0 | 1 | Mustela | 43560.0 | 1 | ustela putori | 43560.0 | 1 |
| scf7180000022708 | 9565 | 0.4817 | 0 | 25.0343 | Eukaryota | 2373.0 | 0 | Chordata | 2373.0 | 0 | Carnivora | 1303.0 | 2 | Felidae | 1303.0 | 2 | Felis | 813.0 | 4 | Felis catus | 813.0 | 4 |
| scf7180000022163 | 17839 | 0.4359 | 0 | 26.3744 | Eukaryota | 9740.0 | 0 | Chordata | 9740.0 | 0 | Primates | 5802.0 | 5 | Hominidae | 5802.0 | 5 | Homo | 3486.0 | 6 | Homo sapiens | 3486.0 | 6 |
| scf7180000023602 | 59293 | 0.4322 | 0 | 462.9537 | no-hit | 0.0 | 0 | no-hit | 0.0 | 0 | no-hit | 0.0 | 0 | no-hit | 0.0 | 0 | no-hit | 0.0 | 0 | no-hit | 0.0 | 0 |
| scf7180000022989 | 33516 | 0.5536 | 0 | 154.7448 | Eukaryota | 3850.0 | 0 | Chordata | 3850.0 | 0 | Carnivora | 3850.0 | 0 | Felidae | 3598.0 | 1 | Acinonyx | 1926.0 | 3 | cinonyx jubat | 1926.0 | 3 |
| scf7180000021334 | 67492 | 0.4058 | 0 | 29.2686 | Eukaryota | 76136.0 | 0 | Chordata | 76136.0 | 0 | Carnivora | 76136.0 | 0 | Felidae | 56267.0 | 1 | Lynx | 38648.0 | 2 | ynx canadens | 38648.0 | 2 |
| scf7180000020617 | 122561 | 0.3972 | 0 | 24.7021 | Eukaryota | 44674.0 | 0 | Chordata | 44674.0 | 0 | Carnivora | 44674.0 | 0 | Felidae | 44674.0 | 0 | Felis | 44674.0 | 0 | Felis catus | 44674.0 | 0 |
| scf7180000020172 | 157732 | 0.3707 | 0 | 24.2104 | Eukaryota | 10198.0 | 0 | Chordata | 10198.0 | 0 | Carnivora | 5742.0 | 2 | Felidae | 5742.0 | 2 | Felis | 5742.0 | 3 | Felis catus | 5742.0 | 3 |
| scf7180000022179 | 32736 | 0.3858 | 0 | 27.1872 | Eukaryota | 44064.0 | 0 | Chordata | 44064.0 | 0 | Carnivora | 44064.0 | 0 | Felidae | 44064.0 | 0 | Felis | 44064.0 | 0 | Felis catus | 44064.0 | 0 |
| scf7180000020722 | 38717 | 0.5207 | 0 | 324.5648 | no-hit | 0.0 | 0 | no-hit | 0.0 | 0 | no-hit | 0.0 | 0 | no-hit | 0.0 | 0 | no-hit | 0.0 | 0 | no-hit | 0.0 | 0 |
| scf7180000024057 | 135727 | 0.3685 | 0 | 24.7977 | Eukaryota | 31820.0 | 0 | Chordata | 31820.0 | 0 | Carnivora | 21872.0 | 1 | Felidae | 21872.0 | 1 | Felis | 21872.0 | 1 | Felis catus | 21872.0 | 1 |
| scf7180000021630 | 61321 | 0.611 | 0 | 62.2971 | Eukaryota | 5870.0 | 0 | Chordata | 5870.0 | 0 | Carnivora | 5870.0 | 0 | Felidae | 4186.0 | 1 | Felis | 4186.0 | 1 | Felis catus | 4186.0 | 1 |
| scf7180000020219 | 154954 | 0.3534 | 0 | 47.8299 | Eukaryota | 42535.0 | 0 | Chordata | 42535.0 | 0 | Carnivora | 42535.0 | 0 | Felidae | 42535.0 | 0 | Felis | 42535.0 | 0 | Felis catus | 42535.0 | 0 |
| scf7180000020963 | 11486 | 0.4701 | 0 | 25.1462 | Eukaryota | 7782.0 | 0 | Chordata | 7782.0 | 0 | Carnivora | 7782.0 | 0 | Felidae | 4638.0 | 2 | Suricata | 2672.0 | 5 | ricata suricat | 2672.0 | 5 |
| scf7180000020698 | 106020 | 0.6121 | 0 | 46.781 | Eukaryota | 4690.0 | 0 | Chordata | 4690.0 | 0 | Carnivora | 2439.0 | 2 | Otariidae | 1132.0 | 5 | Zalophus | 1132.0 | 6 | phus californi | 1132.0 | 6 |
| scf7180000018934 | 16458 | 0.4367 | 0 | 36.3471 | Eukaryota | 30364.0 | 0 | Chordata | 30364.0 | 0 | Carnivora | 30364.0 | 0 | Felidae | 30364.0 | 0 | Felis | 27222.0 | 1 | Felis catus | 27222.0 | 1 |
| scf7180000020560 | 161084 | 0.5814 | 0 | 51.1883 | Eukaryota | 46167.0 | 0 | Chordata | 46167.0 | 0 | Carnivora | 46167.0 | 0 | Felidae | 46167.0 | 0 | Felis | 46167.0 | 0 | Felis catus | 46167.0 | 0 |
| scf7180000024143 | 122799 | 0.3709 | 0 | 24.0022 | Eukaryota | 25959.0 | 0 | Chordata | 25959.0 | 0 | Carnivora | 25959.0 | 0 | Felidae | 25959.0 | 0 | Felis | 23290.0 | 1 | Felis catus | 23290.0 | 1 |
| scf7180000018375 | 747266 | 0.3701 | 0 | 50.384 | Eukaryota | 46765.0 | 0 | Chordata | 46765.0 | 0 | Carnivora | 46765.0 | 0 | Felidae | 46765.0 | 0 | Felis | 46765.0 | 0 | Felis catus | 46765.0 | 0 |
| scf7180000020091 | 156691 | 0.4551 | 0 | 50.3852 | Eukaryota | 45928.0 | 0 | Chordata | 45928.0 | 0 | Carnivora | 45928.0 | 0 | Felidae | 30104.0 | 1 | Suricata | 15824.0 | 3 | ricata suricat | 15824.0 | 3 |
| scf7180000018930 | 582326 | 0.3596 | 0 | 49.1727 | Eukaryota | 46395.0 | 0 | Chordata | 46395.0 | 0 | Carnivora | 46395.0 | 0 | Felidae | 46395.0 | 0 | Felis | 46395.0 | 0 | Felis catus | 46395.0 | 0 |
| scf7180000019696 | 795062 | 0.4839 | 100 | 49.6084 | Eukaryota | 24410.0 | 0 | Chordata | 24410.0 | 0 | Carnivora | 24410.0 | 0 | Felidae | 12006.0 | 2 | Canis | 6892.0 | 4 | Canis lupus | 6892.0 | 4 |
| scf7180000020940 | 91155 | 0.5462 | 0 | 53.0454 | Eukaryota | 21930.0 | 0 | Chordata | 21930.0 | 0 | Carnivora | 21930.0 | 0 | Felidae | 21930.0 | 0 | Felis | 6606.0 | 4 | Felis catus | 6606.0 | 4 |
| scf7180000022869 | 19787 | 0.5094 | 0 | 479.1842 | no-hit | 0.0 | 0 | no-hit | 0.0 | 0 | no-hit | 0.0 | 0 | no-hit | 0.0 | 0 | no-hit | 0.0 | 0 | no-hit | 0.0 | 0 |
| scf7180000020973 | 76447 | 0.3584 | 0 | 23.3845 | Eukaryota | 39390.0 | 0 | Chordata | 39390.0 | 0 | Carnivora | 39390.0 | 0 | Felidae | 39390.0 | 0 | Felis | 35478.0 | 1 | Felis catus | 35478.0 | 1 |
| scf7180000019413 | 1290287 | 0.4521 | 0 | 51.2102 | Eukaryota | 76574.0 | 0 | Chordata | 76574.0 | 0 | Carnivora | 76574.0 | 0 | Felidae | 76574.0 | 0 | Felis | 52588.0 | 1 | Felis catus | 52588.0 | 1 |
| scf7180000024210 | 45484 | 0.4194 | 0 | 26.8887 | Eukaryota | 4948.0 | 0 | Chordata | 4948.0 | 0 | Carnivora | 3014.0 | 3 | Felidae | 2196.0 | 4 | Felis | 1288.0 | 6 | Felis catus | 1288.0 | 6 |
| scf7180000021358 | 10973 | 0.5367 | 0 | 240.0427 | Eukaryota | 3675.0 | 0 | Chordata | 3675.0 | 0 | Carnivora | 3675.0 | 0 | Felidae | 3675.0 | 0 | unresolved | 1104.0 | 4 | unresolved | 1104.0 | 4 |
| scf7180000021951 | 17678 | 0.3814 | 0 | 26.2996 | Eukaryota | 11984.0 | 0 | Chordata | 11984.0 | 0 | Carnivora | 11984.0 | 0 | Felidae | 11984.0 | 0 | Felis | 11984.0 | 0 | Felis catus | 11984.0 | 0 |
| scf7180000019816 | 255397 | 0.35 | 0 | 48.0469 | Eukaryota | 63683.0 | 0 | Chordata | 63683.0 | 0 | Carnivora | 63683.0 | 0 | Felidae | 34464.0 | 2 | Canis | 22986.0 | 4 | Canis lupus | 22986.0 | 4 |
| scf7180000021471 | 122963 | 0.4337 | 0 | 27.4845 | Eukaryota | 38667.0 | 0 | Chordata | 38667.0 | 0 | Carnivora | 38667.0 | 0 | Felidae | 38667.0 | 0 | Felis | 38667.0 | 0 | Felis catus | 38667.0 | 0 |
| scf7180000024116 | 141522 | 0.5232 | 0 | 52.9124 | Eukaryota | 26994.0 | 0 | Chordata | 26994.0 | 0 | Carnivora | 26994.0 | 0 | Felidae | 26994.0 | 0 | Lynx | 9321.0 | 3 | ynx canadens | 9321.0 | 3 |
| scf7180000024237 | 10553 | 0.641 | 0 | 47.187 | Eukaryota | 2144.0 | 0 | Chordata | 2144.0 | 0 | Carnivora | 2144.0 | 0 | Felidae | 2144.0 | 0 | Felis | 1714.0 | 1 | Felis catus | 1714.0 | 1 |
| scf7180000021737 | 48149 | 0.3986 | 0 | 25.2514 | Eukaryota | 7433.0 | 0 | Chordata | 7433.0 | 0 | Primates | 3310.0 | 2 | Hominidae | 3310.0 | 2 | Felis | 3072.0 | 3 | Felis catus | 3072.0 | 3 |
| scf7180000023992 | 505073 | 0.4448 | 0 | 51.7052 | Eukaryota | 44891.0 | 0 | Chordata | 44891.0 | 0 | Carnivora | 44891.0 | 0 | Felidae | 44891.0 | 0 | Felis | 44891.0 | 0 | Felis catus | 44891.0 | 0 |
| scf7180000020384 | 106927 | 0.4029 | 0 | 28.1657 | Eukaryota | 37596.0 | 0 | Chordata | 37596.0 | 0 | Carnivora | 37596.0 | 0 | Felidae | 37596.0 | 0 | Felis | 37596.0 | 0 | Felis catus | 37596.0 | 0 |
| scf7180000020720 | 22167 | 0.6785 | 0 | 33.2776 | Eukaryota | 3240.0 | 0 | Chordata | 3240.0 | 0 | Carnivora | 3240.0 | 0 | Felidae | 2914.0 | 1 | Felis | 960.0 | 5 | Felis catus | 960.0 | 6 |
| scf7180000018511 | 1182474 | 0.3754 | 0 | 49.1032 | Eukaryota | 44806.0 | 0 | Chordata | 44806.0 | 0 | Carnivora | 44806.0 | 0 | Felidae | 44806.0 | 0 | Felis | 44806.0 | 0 | Felis catus | 44806.0 | 0 |
| scf7180000024030 | 761759 | 0.4075 | 0 | 50.9341 | Eukaryota | 44473.0 | 0 | Chordata | 44473.0 | 0 | Carnivora | 44473.0 | 0 | Felidae | 44473.0 | 0 | Felis | 44473.0 | 0 | Felis catus | 44473.0 | 0 |
| scf7180000019462 | 14522 | 0.3772 | 0 | 23.7152 | Eukaryota | 3547.0 | 1 | Chordata | 3547.0 | 1 | Carnivora | 2664.0 | 2 | Felidae | 2664.0 | 2 | Felis | 1809.0 | 3 | Felis catus | 1809.0 | 3 |
| scf7180000019246 | 48972 | 0.5385 | 0 | 125.381 | Eukaryota | 12702.0 | 0 | Chordata | 12702.0 | 0 | Carnivora | 12702.0 | 0 | Felidae | 12702.0 | 0 | Felis | 12702.0 | 0 | Felis catus | 12702.0 | 0 |
| scf7180000024234 | 28424 | 0.4061 | 0 | 50.4416 | Eukaryota | 4943.0 | 1 | Chordata | 4943.0 | 1 | Carnivora | 3698.0 | 2 | Felidae | 3698.0 | 2 | Felis | 3195.0 | 3 | Felis catus | 3195.0 | 3 |
| scf7180000021944 | 26726 | 0.3752 | 0 | 50.169 | Eukaryota | 9295.0 | 0 | Chordata | 9295.0 | 0 | Primates | 9295.0 | 0 | Cercopithecidae | 6284.0 | 1 | Papio | 5495.0 | 2 | Papio anubis | 5495.0 | 2 |
| scf7180000018427 | 656472 | 0.3606 | 0 | 49.9994 | Eukaryota | 46479.0 | 0 | Chordata | 46479.0 | 0 | Carnivora | 46479.0 | 0 | Felidae | 46479.0 | 0 | Felis | 46479.0 | 0 | Felis catus | 46479.0 | 0 |
| scf7180000019146 | 467506 | 0.3984 | 0 | 50.3779 | Eukaryota | 38020.0 | 0 | Chordata | 38020.0 | 0 | Carnivora | 15290.0 | 2 | Felidae | 15290.0 | 3 | Sus | 8339.0 | 6 | Sus scrofa | 8339.0 | 6 |
| scf7180000017829 | 1102276 | 0.3718 | 0 | 51.3103 | Eukaryota | 58950.0 | 0 | Chordata | 58950.0 | 0 | Carnivora | 58950.0 | 0 | Felidae | 58950.0 | 0 | Acinonyx | 58950.0 | 0 | cinonyx jubat | 58950.0 | 0 |
| scf7180000021765 | 48372 | 0.3865 | 0 | 343.7905 | Eukaryota | 6143.0 | 0 | Chordata | 6143.0 | 0 | Carnivora | 6143.0 | 0 | Felidae | 6143.0 | 0 | Felis | 6143.0 | 0 | Felis catus | 6143.0 | 0 |
| scf7180000018800 | 596704 | 0.391 | 0 | 50.1992 | Eukaryota | 42926.0 | 0 | Chordata | 42926.0 | 0 | Carnivora | 42926.0 | 0 | Felidae | 42926.0 | 0 | Acinonyx | 12946.0 | 3 | cinonyx jubat | 12946.0 | 4 |
| scf7180000023968 | 453099 | 0.4008 | 0 | 51.5072 | Eukaryota | 61267.0 | 0 | Chordata | 61267.0 | 0 | Carnivora | 61267.0 | 0 | Herpestidae | 54856.0 | 1 | Suricata | 54856.0 | 2 | ricata suricat | 54856.0 | 2 |
| scf7180000020839 | 55437 | 0.418 | 0 | 25.644 | Eukaryota | 18894.0 | 0 | Chordata | 18894.0 | 0 | Carnivora | 18894.0 | 0 | Felidae | 18894.0 | 0 | Felis | 16988.0 | 1 | Felis catus | 16988.0 | 1 |
| scf7180000017581 | 3223824 | 0.4329 | 0 | 50.3864 | Eukaryota | 56834.0 | 0 | Chordata | 56834.0 | 0 | Carnivora | 56834.0 | 0 | Felidae | 32063.0 | 1 | Felis | 27529.0 | 2 | Felis catus | 27529.0 | 2 |
| scf7180000021594 | 31667 | 0.3859 | 0 | 23.1381 | Eukaryota | 39600.0 | 0 | Chordata | 39600.0 | 0 | Carnivora | 39600.0 | 0 | Felidae | 39600.0 | 0 | Felis | 39600.0 | 0 | Felis catus | 39600.0 | 0 |
| scf7180000019828 | 179375 | 0.5389 | 0 | 49.8789 | Eukaryota | 7270.0 | 0 | Chordata | 7270.0 | 0 | Carnivora | 7270.0 | 0 | Felidae | 7270.0 | 0 | Felis | 6232.0 | 1 | Felis catus | 6232.0 | 1 |
| scf7180000018100 | 259138 | 0.3743 | 0 | 52.0862 | Eukaryota | 44479.0 | 0 | Chordata | 44479.0 | 0 | Carnivora | 44479.0 | 0 | Felidae | 44479.0 | 0 | Felis | 44479.0 | 0 | Felis catus | 44479.0 | 0 |
| scf7180000022032 | 32660 | 0.3425 | 0 | 23.7873 | Eukaryota | 11016.0 | 0 | Chordata | 11016.0 | 0 | Primates | 3859.0 | 5 | Hominidae | 3859.0 | 5 | Sus | 2540.0 | 6 | Sus scrofa | 2540.0 | 6 |
| scf7180000019803 | 57229 | 0.4441 | 0 | 23.0073 | Eukaryota | 35570.0 | 0 | Chordata | 35570.0 | 0 | Carnivora | 35570.0 | 0 | Felidae | 35570.0 | 0 | Lynx | 19465.0 | 1 | ynx canadens | 19465.0 | 1 |
| scf7180000021476 | 38793 | 0.4419 | 0 | 25.137 | Eukaryota | 7418.0 | 0 | Chordata | 7418.0 | 0 | Carnivora | 6625.0 | 1 | Felidae | 6625.0 | 1 | Felis | 6625.0 | 1 | Felis catus | 6625.0 | 1 |
| scf7180000023836 | 586347 | 0.3981 | 0 | 49.7808 | Eukaryota | 42004.0 | 0 | Chordata | 42004.0 | 0 | Carnivora | 42004.0 | 0 | Felidae | 42004.0 | 0 | Felis | 37878.0 | 1 | Felis catus | 37878.0 | 1 |
| scf7180000019337 | 393278 | 0.3952 | 0 | 50.1385 | Eukaryota | 45996.0 | 0 | Chordata | 45996.0 | 0 | Carnivora | 45996.0 | 0 | Felidae | 45996.0 | 0 | Felis | 45996.0 | 0 | Felis catus | 45996.0 | 0 |
| scf7180000022112 | 16904 | 0.4058 | 0 | 23.902 | Eukaryota | 45983.0 | 0 | Chordata | 45983.0 | 0 | Carnivora | 45983.0 | 0 | Felidae | 45983.0 | 0 | Felis | 45983.0 | 0 | Felis catus | 45983.0 | 0 |
| scf7180000022037 | 19852 | 0.4082 | 0 | 49.5443 | Eukaryota | 3589.0 | 1 | Chordata | 3589.0 | 1 | Primates | 2171.0 | 3 | Hominidae | 2171.0 | 3 | Homo | 1892.0 | 5 | Homo sapiens | 1892.0 | 5 |
| scf7180000019330 | 381449 | 0.3491 | 0 | 48.1039 | Eukaryota | 45048.0 | 0 | Chordata | 45048.0 | 0 | Carnivora | 45048.0 | 0 | Felidae | 45048.0 | 0 | Felis | 45048.0 | 0 | Felis catus | 45048.0 | 0 |
| scf7180000022811 | 12766 | 0.3867 | 0 | 22.305 | Eukaryota | 45290.0 | 0 | Chordata | 45290.0 | 0 | Carnivora | 45290.0 | 0 | Felidae | 45290.0 | 0 | Felis | 40813.0 | 1 | Felis catus | 40813.0 | 1 |
| scf7180000021071 | 58469 | 0.4773 | 0 | 48.9606 | Eukaryota | 15979.0 | 1 | Chordata | 15979.0 | 1 | Primates | 11380.0 | 3 | Hominidae | 11380.0 | 4 | Homo | 9080.0 | 6 | Homo sapiens | 9080.0 | 6 |
| scf7180000020178 | 8656 | 0.3923 | 0 | 27.4632 | Eukaryota | 7286.0 | 0 | Chordata | 7286.0 | 0 | Carnivora | 7286.0 | 0 | Felidae | 4437.0 | 2 | Enhydra | 2124.0 | 6 | Enhydra lutris | 2124.0 | 7 |
| scf7180000020530 | 111801 | 0.424 | 0 | 52.2053 | Eukaryota | 14885.0 | 0 | Chordata | 14885.0 | 0 | Carnivora | 14885.0 | 0 | Felidae | 12025.0 | 1 | Felis | 10569.0 | 2 | Felis catus | 10569.0 | 2 |
| scf7180000022643 | 11421 | 0.6344 | 0 | 55.4653 | Eukaryota | 2883.0 | 0 | Chordata | 2883.0 | 0 | Carnivora | 2883.0 | 0 | Felidae | 2883.0 | 0 | Felis | 1165.0 | 5 | Felis catus | 1165.0 | 6 |
| scf7180000024204 | 77557 | 0.3674 | 0 | 23.8492 | Eukaryota | 43349.0 | 0 | Chordata | 43349.0 | 0 | Carnivora | 43349.0 | 0 | Felidae | 43349.0 | 0 | Felis | 39077.0 | 1 | Felis catus | 39077.0 | 1 |
| scf7180000019661 | 232238 | 0.4317 | 0 | 51.197 | Eukaryota | 23650.0 | 0 | Chordata | 23650.0 | 0 | Carnivora | 16443.0 | 2 | Felidae | 10577.0 | 3 | Acinonyx | 8202.0 | 5 | cinonyx jubat | 8202.0 | 5 |
| scf7180000021031 | 1189686 | 0.3898 | 0 | 50.7613 | Eukaryota | 43950.0 | 0 | Chordata | 43950.0 | 0 | Carnivora | 43950.0 | 0 | Felidae | 43950.0 | 0 | Felis | 43950.0 | 0 | Felis catus | 43950.0 | 0 |
| scf7180000021361 | 75349 | 0.5513 | 0 | 48.7676 | Eukaryota | 5141.0 | 0 | Chordata | 5141.0 | 0 | Carnivora | 5141.0 | 0 | Felidae | 2758.0 | 1 | Felis | 2447.0 | 2 | Felis catus | 2447.0 | 2 |
| scf7180000023955 | 566287 | 0.4797 | 0 | 50.9422 | Eukaryota | 26270.0 | 0 | Chordata | 26270.0 | 0 | Carnivora | 26270.0 | 0 | Felidae | 26270.0 | 0 | Felis | 23618.0 | 1 | Felis catus | 23618.0 | 1 |
| scf7180000022476 | 4966 | 0.6979 | 0 | 167.2575 | Eukaryota | 272.0 | 0 | Chordata | 272.0 | 0 | Carnivora | 272.0 | 0 | Felidae | 272.0 | 0 | Panthera | 272.0 | 0 | anthera pardu | 272.0 | 0 |
| scf7180000021082 | 75172 | 0.3836 | 0 | 25.1952 | Eukaryota | 41327.0 | 0 | Chordata | 41327.0 | 0 | Carnivora | 41327.0 | 0 | Felidae | 41327.0 | 0 | Felis | 37260.0 | 1 | Felis catus | 37260.0 | 1 |
| scf7180000022243 | 20105 | 0.4184 | 0 | 25.2237 | Eukaryota | 28950.0 | 0 | Chordata | 28950.0 | 0 | Carnivora | 28950.0 | 0 | Felidae | 28950.0 | 0 | Panthera | 11554.0 | 3 | cinonyx jubat | 5822.0 | 4 |
| scf7180000021923 | 9152 | 0.5577 | 0 | 138.4465 | Eukaryota | 3166.0 | 0 | Chordata | 3166.0 | 0 | Carnivora | 2881.0 | 1 | Felidae | 2424.0 | 2 | Panthera | 1268.0 | 4 | anthera pardu | 1268.0 | 4 |
| scf7180000018622 | 182446 | 0.3806 | 0 | 45.4031 | Eukaryota | 46538.0 | 0 | Chordata | 46538.0 | 0 | Carnivora | 46538.0 | 0 | Felidae | 46538.0 | 0 | Felis | 46538.0 | 0 | Felis catus | 46538.0 | 0 |
| scf7180000019734 | 252891 | 0.4335 | 0 | 50.6942 | Eukaryota | 30300.0 | 0 | Chordata | 30300.0 | 0 | Carnivora | 30300.0 | 0 | Felidae | 30300.0 | 0 | Felis | 18132.0 | 1 | Felis catus | 18132.0 | 1 |
| scf7180000021304 | 76315 | 0.3947 | 0 | 25.8329 | Eukaryota | 45725.0 | 0 | Chordata | 45725.0 | 0 | Carnivora | 45725.0 | 0 | Felidae | 45725.0 | 0 | Felis | 30729.0 | 1 | Felis catus | 30729.0 | 2 |
| scf7180000024127 | 195994 | 0.4149 | 0 | 51.6458 | Eukaryota | 42192.0 | 0 | Chordata | 42192.0 | 0 | Carnivora | 42192.0 | 0 | Felidae | 42192.0 | 0 | Felis | 38036.0 | 1 | Felis catus | 38036.0 | 1 |
| scf7180000017502 | 12650 | 0.3683 | 0 | 21.453 | Eukaryota | 18995.0 | 0 | Chordata | 18995.0 | 0 | Carnivora | 18995.0 | 0 | Felidae | 18995.0 | 0 | Felis | 18995.0 | 0 | Felis catus | 18995.0 | 0 |
| scf7180000021916 | 9754 | 0.5245 | 0 | 80.6383 | no-hit | 0.0 | 0 | no-hit | 0.0 | 0 | no-hit | 0.0 | 0 | no-hit | 0.0 | 0 | no-hit | 0.0 | 0 | no-hit | 0.0 | 0 |
| scf7180000022946 | 27286 | 0.4111 | 0 | 367.1323 | Eukaryota | 5925.0 | 0 | Chordata | 5925.0 | 0 | Carnivora | 5925.0 | 0 | Phocidae | 2416.0 | 2 | eptonychote | 1812.0 | 4 | nychotes wei | 1812.0 | 4 |
| scf7180000020788 | 39831 | 0.3826 | 0 | 23.379 | Eukaryota | 24480.0 | 0 | Chordata | 24480.0 | 0 | Carnivora | 24480.0 | 0 | Felidae | 24480.0 | 0 | Felis | 21999.0 | 1 | Felis catus | 21999.0 | 1 |
| scf7180000022693 | 6007 | 0.4262 | 0 | 23.6084 | Eukaryota | 3173.0 | 0 | Chordata | 3173.0 | 0 | Carnivora | 3173.0 | 0 | Felidae | 1868.0 | 1 | Canis | 1305.0 | 2 | Canis lupus | 1305.0 | 2 |
| scf7180000023897 | 1299389 | 0.3569 | 0 | 49.5558 | Eukaryota | 48009.0 | 0 | Chordata | 48009.0 | 0 | Carnivora | 48009.0 | 0 | Felidae | 48009.0 | 0 | Felis | 43253.0 | 1 | Felis catus | 43253.0 | 1 |
| scf7180000019411 | 159800 | 0.42 | 0 | 47.6812 | Eukaryota | 26485.0 | 0 | Chordata | 26485.0 | 0 | Carnivora | 26485.0 | 0 | Felidae | 26485.0 | 0 | Felis | 23812.0 | 1 | Felis catus | 23812.0 | 1 |
| scf7180000023761 | 482238 | 0.3859 | 0 | 51.3786 | Eukaryota | 37613.0 | 0 | Chordata | 37613.0 | 0 | Carnivora | 37613.0 | 0 | Felidae | 37613.0 | 0 | Felis | 29475.0 | 1 | Felis catus | 29475.0 | 1 |
| scf7180000020182 | 12990 | 0.382 | 0 | 23.0731 | Eukaryota | 11823.0 | 0 | Chordata | 11823.0 | 0 | Carnivora | 11823.0 | 0 | Felidae | 11823.0 | 0 | Felis | 10620.0 | 1 | Felis catus | 10620.0 | 1 |
| scf7180000021362 | 70626 | 0.4126 | 0 | 48.8925 | Eukaryota | 32815.0 | 0 | Chordata | 32815.0 | 0 | Carnivora | 32815.0 | 0 | Felidae | 32815.0 | 0 | Felis | 29537.0 | 1 | Felis catus | 29537.0 | 1 |
| scf7180000023938 | 722477 | 0.4507 | 0 | 51.0251 | Eukaryota | 24970.0 | 0 | Chordata | 24970.0 | 0 | Carnivora | 24970.0 | 0 | Felidae | 24970.0 | 0 | Felis | 13289.0 | 2 | Felis catus | 13289.0 | 3 |
| scf7180000020964 | 77646 | 0.3891 | 0 | 25.1415 | Eukaryota | 45725.0 | 0 | Chordata | 45725.0 | 0 | Carnivora | 45725.0 | 0 | Felidae | 45725.0 | 0 | Felis | 41198.0 | 1 | Felis catus | 41198.0 | 1 |
| scf7180000022542 | 12013 | 0.4172 | 0 | 26.8759 | Eukaryota | 44524.0 | 0 | Chordata | 44524.0 | 0 | Carnivora | 44524.0 | 0 | Felidae | 44524.0 | 0 | Felis | 44524.0 | 0 | Felis catus | 44524.0 | 0 |
| scf7180000017756 | 669509 | 0.3816 | 0 | 49.9195 | Eukaryota | 46380.0 | 0 | Chordata | 46380.0 | 0 | Carnivora | 46380.0 | 0 | Felidae | 46380.0 | 0 | Felis | 46380.0 | 0 | Felis catus | 46380.0 | 0 |
| scf7180000021125 | 49496 | 0.3757 | 0 | 24.0242 | Eukaryota | 36298.0 | 0 | Chordata | 36298.0 | 0 | Carnivora | 36298.0 | 0 | Felidae | 11033.0 | 5 | Canis | 7184.0 | 8 | Canis lupus | 7184.0 | 8 |
| scf7180000021393 | 63203 | 0.376 | 0 | 26.6399 | Eukaryota | 38716.0 | 0 | Chordata | 38716.0 | 0 | Carnivora | 38716.0 | 0 | Felidae | 38716.0 | 0 | Felis | 34570.0 | 1 | Felis catus | 34570.0 | 1 |
| scf7180000018497 | 656911 | 0.4098 | 0 | 49.6781 | Eukaryota | 44943.0 | 0 | Chordata | 44943.0 | 0 | Carnivora | 44943.0 | 0 | Felidae | 44943.0 | 0 | Felis | 40501.0 | 1 | Felis catus | 40501.0 | 1 |
| scf7180000019526 | 244755 | 0.3681 | 0 | 49.1005 | Eukaryota | 17448.0 | 0 | Chordata | 17448.0 | 0 | Carnivora | 17448.0 | 0 | Felidae | 17448.0 | 0 | Felis | 17448.0 | 0 | Felis catus | 17448.0 | 0 |
| scf7180000022083 | 30492 | 0.4149 | 0 | 23.9892 | Eukaryota | 26524.0 | 0 | Chordata | 26524.0 | 0 | Carnivora | 26524.0 | 0 | Felidae | 26524.0 | 0 | Felis | 24040.0 | 1 | Felis catus | 24040.0 | 1 |
| scf7180000020850 | 71195 | 0.3921 | 0 | 47.8916 | Eukaryota | 42889.0 | 0 | Chordata | 42889.0 | 0 | Carnivora | 42889.0 | 0 | Felidae | 42889.0 | 0 | Felis | 38580.0 | 1 | Felis catus | 38580.0 | 1 |
| scf7180000017962 | 631759 | 0.3792 | 0 | 49.0372 | Eukaryota | 40805.0 | 0 | Chordata | 40805.0 | 0 | Carnivora | 40805.0 | 0 | Felidae | 40805.0 | 0 | Felis | 36799.0 | 1 | Felis catus | 36799.0 | 1 |
| scf7180000022727 | 2701 | 0.5268 | 0 | 352.9195 | Eukaryota | 694.0 | 0 | Chordata | 694.0 | 0 | Carnivora | 694.0 | 0 | Felidae | 694.0 | 0 | unresolved | 233.0 | 2 | unresolved | 233.0 | 2 |
| scf7180000018899 | 513732 | 0.3907 | 0 | 48.5304 | Eukaryota | 47071.0 | 0 | Chordata | 47071.0 | 0 | Carnivora | 40646.0 | 1 | Felidae | 40646.0 | 1 | Felis | 40646.0 | 1 | Felis catus | 40646.0 | 1 |
| scf7180000018703 | 2960008 | 0.4408 | 0 | 52.0788 | Eukaryota | 46367.0 | 0 | Chordata | 46367.0 | 0 | Carnivora | 46367.0 | 0 | Felidae | 41585.0 | 1 | Felis | 32367.0 | 2 | Felis catus | 32367.0 | 2 |
| scf7180000021209 | 36011 | 0.3903 | 0 | 23.5119 | Eukaryota | 41549.0 | 0 | Chordata | 41549.0 | 0 | Carnivora | 41549.0 | 0 | Felidae | 41549.0 | 0 | Felis | 41549.0 | 0 | Felis catus | 41549.0 | 0 |
| scf7180000024162 | 389387 | 0.5369 | 0 | 54.2551 | Eukaryota | 52219.0 | 0 | Chordata | 52219.0 | 0 | Carnivora | 52219.0 | 0 | Felidae | 52219.0 | 0 | Acinonyx | 20920.0 | 2 | cinonyx jubat | 20920.0 | 2 |
| scf7180000020095 | 159747 | 0.4495 | 0 | 25.0562 | Eukaryota | 10252.0 | 0 | Chordata | 10252.0 | 0 | Carnivora | 8294.0 | 1 | Felidae | 8294.0 | 1 | Felis | 8294.0 | 1 | Felis catus | 8294.0 | 1 |
| scf7180000017795 | 2481395 | 0.3631 | 0 | 50.8185 | Eukaryota | 108581.0 | 0 | Chordata | 108581.0 | 0 | Carnivora | 108581.0 | 0 | Felidae | 88373.0 | 1 | Lynx | 33342.0 | 3 | ynx canadens | 33342.0 | 3 |
| scf7180000021441 | 58005 | 0.5201 | 0 | 49.9987 | Eukaryota | 11497.0 | 1 | Chordata | 11497.0 | 1 | Primates | 2656.0 | 7 | Hominidae | 2656.0 | 7 | Homo | 2656.0 | 7 | Homo sapiens | 2656.0 | 7 |
| scf7180000020717 | 85605 | 0.4099 | 0 | 25.8729 | Eukaryota | 20609.0 | 0 | Chordata | 20609.0 | 0 | Carnivora | 20609.0 | 0 | Felidae | 20609.0 | 0 | Felis | 18477.0 | 1 | Felis catus | 18477.0 | 1 |
| scf7180000021734 | 22913 | 0.4572 | 0 | 26.6098 | Eukaryota | 16758.0 | 0 | Chordata | 16758.0 | 0 | Carnivora | 16758.0 | 0 | Felidae | 16758.0 | 0 | Felis | 15073.0 | 1 | Felis catus | 15073.0 | 1 |
| scf7180000021880 | 15754 | 0.5872 | 0 | 194.994 | Eukaryota | 2515.0 | 0 | Chordata | 2515.0 | 0 | Carnivora | 2278.0 | 1 | Felidae | 2278.0 | 1 | Felis | 1412.0 | 3 | Felis catus | 1412.0 | 3 |
| scf7180000023901 | 1324049 | 0.3887 | 0 | 50.4091 | Eukaryota | 46562.0 | 0 | Chordata | 46562.0 | 0 | Carnivora | 46562.0 | 0 | Felidae | 46562.0 | 0 | Felis | 46562.0 | 0 | Felis catus | 46562.0 | 0 |
| scf7180000022931 | 64766 | 0.4366 | 0 | 449.363 | Eukaryota | 7397.0 | 0 | Chordata | 7397.0 | 0 | Carnivora | 7397.0 | 0 | Otariidae | 5252.0 | 1 | Eumetopias | 2972.0 | 2 | netopias juba | 2972.0 | 2 |
| scf7180000023760 | 489525 | 0.3942 | 0 | 52.6045 | Eukaryota | 44776.0 | 0 | Chordata | 44776.0 | 0 | Carnivora | 44776.0 | 0 | Felidae | 44776.0 | 0 | Felis | 44776.0 | 0 | Felis catus | 44776.0 | 0 |
| scf7180000023187 | 43129 | 0.3921 | 0 | 418.494 | Eukaryota | 40883.0 | 0 | Chordata | 40883.0 | 0 | Carnivora | 40883.0 | 0 | Felidae | 40883.0 | 0 | Felis | 40883.0 | 0 | Felis catus | 40883.0 | 0 |
| scf7180000020668 | 35358 | 0.438 | 0 | 55.221 | Eukaryota | 33190.0 | 0 | Chordata | 33190.0 | 0 | Carnivora | 33190.0 | 0 | Felidae | 33190.0 | 0 | Felis | 33190.0 | 0 | Felis catus | 33190.0 | 0 |
| scf7180000019020 | 502830 | 0.4697 | 0 | 51.2524 | Eukaryota | 40582.0 | 0 | Chordata | 40582.0 | 0 | Carnivora | 40582.0 | 0 | Felidae | 40582.0 | 0 | Felis | 36569.0 | 1 | Felis catus | 36569.0 | 1 |
| scf7180000017696 | 722989 | 0.3758 | 0 | 51.141 | Eukaryota | 45519.0 | 0 | Chordata | 45519.0 | 0 | Carnivora | 45519.0 | 0 | Felidae | 45519.0 | 0 | Felis | 45519.0 | 0 | Felis catus | 45519.0 | 0 |
| scf7180000020188 | 39251 | 0.3888 | 0 | 24.2762 | Eukaryota | 40683.0 | 0 | Chordata | 40683.0 | 0 | Carnivora | 40683.0 | 0 | Felidae | 40683.0 | 0 | Felis | 40683.0 | 0 | Felis catus | 40683.0 | 0 |
| scf7180000018225 | 1047910 | 0.4472 | 0 | 50.5352 | Eukaryota | 27048.0 | 0 | Chordata | 27048.0 | 0 | Carnivora | 24102.0 | 1 | Ursidae | 11260.0 | 3 | Ursus | 11260.0 | 4 | Ursus arctos | 11260.0 | 4 |
| scf7180000024070 | 544785 | 0.3865 | 0 | 48.6172 | Eukaryota | 42642.0 | 0 | Chordata | 42642.0 | 0 | Carnivora | 42642.0 | 0 | Felidae | 42642.0 | 0 | Felis | 42642.0 | 0 | Felis catus | 42642.0 | 0 |
| scf7180000021493 | 53698 | 0.3865 | 0 | 23.5062 | Eukaryota | 42810.0 | 0 | Chordata | 42810.0 | 0 | Carnivora | 42810.0 | 0 | Felidae | 42810.0 | 0 | Felis | 42810.0 | 0 | Felis catus | 42810.0 | 0 |
| scf7180000020807 | 87889 | 0.6483 | 0 | 49.6661 | Eukaryota | 5450.0 | 1 | Chordata | 5450.0 | 1 | Rodentia | 2410.0 | 3 | Felidae | 1585.0 | 5 | Mus | 1492.0 | 7 | Mus musculus | 1492.0 | 7 |
| scf7180000019758 | 152152 | 0.5662 | 0 | 50.9766 | Eukaryota | 32298.0 | 0 | Chordata | 32298.0 | 0 | Carnivora | 26092.0 | 1 | Hyaenidae | 21196.0 | 4 | Hyaena | 6039.0 | 8 | Hyaena hyaen | 6039.0 | 8 |
| scf7180000022107 | 14547 | 0.4246 | 0 | 23.1896 | Eukaryota | 34228.0 | 0 | Chordata | 34228.0 | 0 | Carnivora | 34228.0 | 0 | Felidae | 23788.0 | 1 | Suricata | 10440.0 | 4 | ricata suricat | 10440.0 | 4 |
| scf7180000020883 | 105935 | 0.3955 | 0 | 51.7073 | Eukaryota | 8943.0 | 0 | Chordata | 8943.0 | 0 | Carnivora | 3881.0 | 2 | Canidae | 2644.0 | 4 | Canis | 2644.0 | 6 | Canis lupus | 2644.0 | 6 |
| scf7180000020025 | 157590 | 0.4588 | 0 | 24.7639 | Eukaryota | 17382.0 | 0 | Chordata | 17382.0 | 0 | Carnivora | 17382.0 | 0 | Felidae | 12721.0 | 2 | Panthera | 5771.0 | 4 | anthera pardu | 3950.0 | 5 |
| scf7180000023692 | 344804 | 0.4008 | 0 | 48.6171 | Eukaryota | 20484.0 | 0 | Chordata | 20484.0 | 0 | Carnivora | 20484.0 | 0 | Felidae | 20484.0 | 0 | Felis | 18423.0 | 1 | Felis catus | 18423.0 | 1 |
| scf7180000017971 | 1255286 | 0.3926 | 0 | 50.4958 | Eukaryota | 45510.0 | 0 | Chordata | 45510.0 | 0 | Carnivora | 45510.0 | 0 | Felidae | 45510.0 | 0 | Felis | 41020.0 | 1 | Felis catus | 41020.0 | 1 |
| scf7180000017688 | 1814974 | 0.3874 | 100 | 50.3787 | Eukaryota | 46894.0 | 0 | Chordata | 46894.0 | 0 | Carnivora | 46894.0 | 0 | Felidae | 46894.0 | 0 | Felis | 46894.0 | 0 | Felis catus | 46894.0 | 0 |
| scf7180000021528 | 58390 | 0.4059 | 0 | 25.8009 | Eukaryota | 43953.0 | 0 | Chordata | 43953.0 | 0 | Carnivora | 43953.0 | 0 | Felidae | 43953.0 | 0 | Felis | 43953.0 | 0 | Felis catus | 43953.0 | 0 |
| scf7180000018207 | 11314 | 0.4014 | 0 | 26.666 | Eukaryota | 44539.0 | 0 | Chordata | 44539.0 | 0 | Carnivora | 44539.0 | 0 | Felidae | 44539.0 | 0 | Felis | 44539.0 | 0 | Felis catus | 44539.0 | 0 |
| scf7180000021451 | 123322 | 0.3872 | 0 | 51.8066 | Eukaryota | 27000.0 | 0 | Chordata | 27000.0 | 0 | Carnivora | 27000.0 | 0 | Felidae | 27000.0 | 0 | Acinonyx | 27000.0 | 0 | cinonyx jubat | 27000.0 | 0 |
| scf7180000022959 | 23447 | 0.4075 | 0 | 315.3222 | Eukaryota | 3974.0 | 0 | Chordata | 3974.0 | 0 | Primates | 2566.0 | 2 | Hominidae | 2192.0 | 4 | Homo | 2192.0 | 4 | Homo sapiens | 2192.0 | 4 |
| scf7180000018430 | 808758 | 0.4023 | 0 | 52.8513 | Eukaryota | 36637.0 | 0 | Chordata | 36637.0 | 0 | Carnivora | 36637.0 | 0 | Felidae | 36637.0 | 0 | Felis | 36637.0 | 0 | Felis catus | 36637.0 | 0 |
| scf7180000021949 | 21957 | 0.495 | 0 | 1749.9462 | no-hit | 0.0 | 0 | no-hit | 0.0 | 0 | no-hit | 0.0 | 0 | no-hit | 0.0 | 0 | no-hit | 0.0 | 0 | no-hit | 0.0 | 0 |
| scf7180000017561 | 2635277 | 0.3613 | 0 | 50.2641 | Eukaryota | 153200.0 | 0 | Chordata | 153200.0 | 0 | Primates | 109757.0 | 1 | Hominidae | 109757.0 | 1 | Pan | 66189.0 | 3 | an troglodyte | 66189.0 | 3 |
| scf7180000021907 | 16674 | 0.409 | 0 | 24.3653 | Eukaryota | 9889.0 | 0 | Chordata | 9889.0 | 0 | Carnivora | 9889.0 | 0 | Felidae | 9889.0 | 0 | Felis | 8904.0 | 1 | Felis catus | 8904.0 | 1 |
| scf7180000019348 | 1430763 | 0.4322 | 100 | 52.4274 | Eukaryota | 47051.0 | 0 | Chordata | 47051.0 | 0 | Carnivora | 47051.0 | 0 | Felidae | 47051.0 | 0 | Felis | 47051.0 | 0 | Felis catus | 47051.0 | 0 |
| scf7180000019383 | 9650 | 0.6062 | 0 | 125.826 | Eukaryota | 2610.0 | 0 | Chordata | 2610.0 | 0 | Carnivora | 2610.0 | 0 | Felidae | 2343.0 | 1 | Felis | 1817.0 | 2 | Felis catus | 1817.0 | 2 |
| scf7180000019744 | 57850 | 0.3733 | 0 | 42.3975 | Eukaryota | 45481.0 | 0 | Chordata | 45481.0 | 0 | Carnivora | 45481.0 | 0 | Felidae | 45481.0 | 0 | Felis | 45481.0 | 0 | Felis catus | 45481.0 | 0 |
| scf7180000021087 | 66701 | 0.4043 | 0 | 49.0163 | Eukaryota | 12274.0 | 0 | Chordata | 12274.0 | 0 | Carnivora | 10278.0 | 1 | Felidae | 10278.0 | 1 | Felis | 4321.0 | 5 | Felis catus | 4321.0 | 6 |
| scf7180000021221 | 12725 | 0.5213 | 0 | 76.1455 | no-hit | 0.0 | 0 | no-hit | 0.0 | 0 | no-hit | 0.0 | 0 | no-hit | 0.0 | 0 | no-hit | 0.0 | 0 | no-hit | 0.0 | 0 |
| scf7180000019701 | 3684883 | 0.452 | 0 | 51.2267 | Eukaryota | 45628.0 | 0 | Chordata | 45628.0 | 0 | Carnivora | 45628.0 | 0 | Felidae | 45628.0 | 0 | Felis | 18484.0 | 2 | Felis catus | 18484.0 | 2 |
| scf7180000020119 | 133613 | 0.3447 | 0 | 49.8484 | Eukaryota | 23896.0 | 0 | Chordata | 23896.0 | 0 | Carnivora | 23896.0 | 0 | Felidae | 23896.0 | 0 | Felis | 21491.0 | 1 | Felis catus | 21491.0 | 1 |
| scf7180000023255 | 59055 | 0.3946 | 0 | 335.0736 | Eukaryota | 45780.0 | 0 | Chordata | 45780.0 | 0 | Carnivora | 45780.0 | 0 | Felidae | 45780.0 | 0 | Felis | 45780.0 | 0 | Felis catus | 45780.0 | 0 |
| scf7180000022630 | 54064 | 0.4069 | 0 | 3841.1759 | no-hit | 0.0 | 0 | no-hit | 0.0 | 0 | no-hit | 0.0 | 0 | no-hit | 0.0 | 0 | no-hit | 0.0 | 0 | no-hit | 0.0 | 0 |
| scf7180000021175 | 14365 | 0.5201 | 0 | 96.8022 | no-hit | 0.0 | 0 | no-hit | 0.0 | 0 | no-hit | 0.0 | 0 | no-hit | 0.0 | 0 | no-hit | 0.0 | 0 | no-hit | 0.0 | 0 |
| scf7180000020755 | 75018 | 0.4651 | 0 | 27.2742 | Eukaryota | 42447.0 | 0 | Chordata | 42447.0 | 0 | Carnivora | 42447.0 | 0 | Felidae | 42447.0 | 0 | Felis | 38212.0 | 1 | Felis catus | 38212.0 | 1 |
| scf7180000023970 | 2189719 | 0.3908 | 0 | 50.4761 | Eukaryota | 47856.0 | 0 | Chordata | 47856.0 | 0 | Carnivora | 47856.0 | 0 | Otariidae | 38120.0 | 1 | Callorhinus | 38120.0 | 1 | llorhinus ursir | 38120.0 | 1 |
| scf7180000022096 | 23096 | 0.6094 | 0 | 50.0813 | Eukaryota | 5864.0 | 0 | Chordata | 5864.0 | 0 | Carnivora | 5864.0 | 0 | Felidae | 4716.0 | 1 | Felis | 3787.0 | 2 | Felis catus | 3787.0 | 2 |
| scf7180000021834 | 23576 | 0.4018 | 0 | 22.2449 | Eukaryota | 35042.0 | 0 | Chordata | 35042.0 | 0 | Carnivora | 35042.0 | 0 | Felidae | 35042.0 | 0 | Felis | 31553.0 | 1 | Felis catus | 31553.0 | 1 |
| scf7180000021410 | 63699 | 0.3743 | 0 | 24.4762 | Eukaryota | 15949.0 | 0 | Chordata | 15949.0 | 0 | Carnivora | 15949.0 | 0 | Felidae | 15949.0 | 0 | Felis | 14024.0 | 1 | Felis catus | 14024.0 | 1 |
| scf7180000019799 | 97477 | 0.3903 | 0 | 44.3188 | Eukaryota | 31799.0 | 0 | Chordata | 31799.0 | 0 | Carnivora | 31799.0 | 0 | Felidae | 31799.0 | 0 | Felis | 31799.0 | 0 | Felis catus | 31799.0 | 0 |
| scf7180000021283 | 52122 | 0.4471 | 0 | 48.3152 | Eukaryota | 2594.0 | 0 | Chordata | 2594.0 | 0 | Carnivora | 2594.0 | 0 | Felidae | 2331.0 | 1 | Felis | 1822.0 | 2 | Felis catus | 1822.0 | 2 |
| scf7180000017730 | 213979 | 0.398 | 0 | 50.7188 | Eukaryota | 45821.0 | 0 | Chordata | 45821.0 | 0 | Carnivora | 45821.0 | 0 | Felidae | 45821.0 | 0 | Felis | 45821.0 | 0 | Felis catus | 45821.0 | 0 |

| | | | | | | | | | | | | | | | | | | | | | | |
|---|---|---|---|---|---|---|---|---|---|---|---|---|---|---|---|---|---|---|---|---|---|---|
| scf7180000017671 | 149601 | 0.368 | 0 | 47.2756 | Eukaryota | 42552.0 | 0 | Chordata | 42552.0 | 0 | Carnivora | 42552.0 | 0 | Felidae | 42552.0 | 0 | Felis | 42552.0 | 0 | Felis catus | 42552.0 | 0 |
| scf7180000017616 | 1062770 | 0.4161 | 0 | 50.9138 | Eukaryota | 46025.0 | 0 | Chordata | 46025.0 | 0 | Carnivora | 46025.0 | 0 | Felidae | 46025.0 | 0 | Felis | 46025.0 | 0 | Felis catus | 46025.0 | 0 |
| scf7180000019914 | 314926 | 0.4345 | 0 | 50.1064 | Eukaryota | 25682.0 | 0 | Chordata | 25682.0 | 0 | Carnivora | 25682.0 | 0 | Felidae | 25682.0 | 0 | Felis | 23109.0 | 1 | Felis catus | 23109.0 | 1 |
| scf7180000017654 | 1055723 | 0.3795 | 0 | 51.5669 | Eukaryota | 47063.0 | 0 | Chordata | 47063.0 | 0 | Carnivora | 47063.0 | 0 | Felidae | 47063.0 | 0 | Felis | 47063.0 | 0 | Felis catus | 47063.0 | 0 |
| scf7180000021321 | 54373 | 0.4413 | 0 | 96.1531 | Eukaryota | 8666.0 | 0 | Chordata | 8666.0 | 0 | Carnivora | 8666.0 | 0 | Felidae | 8666.0 | 0 | Felis | 8666.0 | 0 | Felis catus | 8666.0 | 0 |
| scf7180000018108 | 1933161 | 0.4185 | 0 | 51.9797 | Eukaryota | 47173.0 | 0 | Chordata | 47173.0 | 0 | Carnivora | 47173.0 | 0 | Felidae | 47173.0 | 0 | Felis | 47173.0 | 0 | Felis catus | 47173.0 | 0 |
| scf7180000023461 | 57693 | 0.4297 | 0 | 285.5829 | Eukaryota | 9238.0 | 0 | Chordata | 9238.0 | 0 | Carnivora | 9238.0 | 0 | Felidae | 9238.0 | 0 | Felis | 7346.0 | 1 | Felis catus | 7346.0 | 1 |
| scf7180000021372 | 44341 | 0.3622 | 0 | 50.4305 | Eukaryota | 34088.0 | 0 | Chordata | 34088.0 | 0 | Carnivora | 34088.0 | 0 | Felidae | 34088.0 | 0 | Felis | 30673.0 | 1 | Felis catus | 30673.0 | 1 |
| scf7180000020420 | 148203 | 0.4083 | 0 | 25.6819 | Eukaryota | 38848.0 | 0 | Chordata | 38848.0 | 0 | Primates | 31047.0 | 2 | Hominidae | 28328.0 | 3 | Homo | 19910.0 | 4 | Homo sapiens | 19910.0 | 4 |
| scf7180000020738 | 89293 | 0.3718 | 0 | 47.2305 | Eukaryota | 30952.0 | 0 | Chordata | 30952.0 | 0 | Carnivora | 30952.0 | 0 | Felidae | 30952.0 | 0 | Felis | 27890.0 | 1 | Felis catus | 27890.0 | 1 |
| scf7180000019195 | 473331 | 0.4588 | 0 | 51.2552 | Eukaryota | 50200.0 | 0 | Chordata | 50200.0 | 0 | Carnivora | 50200.0 | 0 | Herpestidae | 50200.0 | 0 | Suricata | 50200.0 | 0 | uricata suricat | 50200.0 | 0 |
| scf7180000020240 | 173861 | 0.4405 | 100 | 50.3837 | Eukaryota | 35384.0 | 0 | Chordata | 35384.0 | 0 | Carnivora | 35384.0 | 0 | Felidae | 32035.0 | 1 | Acinonyx | 16825.0 | 3 | cinonyx jubat | 16825.0 | 4 |
| scf7180000018182 | 1222320 | 0.361 | 0 | 48.9936 | Eukaryota | 46489.0 | 0 | Chordata | 46489.0 | 0 | Carnivora | 46489.0 | 0 | Felidae | 46489.0 | 0 | Felis | 46489.0 | 0 | Felis catus | 46489.0 | 0 |
| scf7180000022676 | 3705 | 0.6704 | 0 | 528.2612 | Eukaryota | 1345.0 | 0 | Chordata | 1211.0 | 1 | Carnivora | 1211.0 | 1 | Otariidae | 804.0 | 4 | unresolved | 402.0 | 5 | unresolved | 402.0 | 5 |
| scf7180000021786 | 71359 | 0.5975 | 0 | 51.2482 | Eukaryota | 10928.0 | 0 | Chordata | 10928.0 | 0 | Carnivora | 10928.0 | 0 | Felidae | 10928.0 | 0 | Panthera | 10187.0 | 1 | anthera pardu | 7941.0 | 2 |
| scf7180000020003 | 32407 | 0.408 | 0 | 24.6833 | Eukaryota | 22764.0 | 0 | Chordata | 22764.0 | 0 | Carnivora | 22764.0 | 0 | Felidae | 22764.0 | 0 | Felis | 22764.0 | 0 | Felis catus | 22764.0 | 0 |
| scf7180000022743 | 39558 | 0.4759 | 0 | 23.8563 | Eukaryota | 3649.0 | 0 | Chordata | 3649.0 | 0 | Primates | 1636.0 | 3 | Hominidae | 1636.0 | 4 | unresolved | 818.0 | 5 | unresolved | 818.0 | 5 |
| scf7180000022644 | 22462 | 0.4187 | 0 | 24.8908 | Eukaryota | 9395.0 | 0 | Chordata | 9395.0 | 0 | Carnivora | 5527.0 | 1 | Felidae | 4662.0 | 5 | Panthera | 1818.0 | 8 | coileus virgini | 1040.0 | 9 |
| scf7180000023355 | 127122 | 0.3896 | 0 | 53.0859 | Eukaryota | 24824.0 | 0 | Chordata | 24824.0 | 0 | Carnivora | 24824.0 | 0 | Felidae | 24824.0 | 0 | Felis | 22327.0 | 1 | Felis catus | 22327.0 | 1 |
| scf7180000022503 | 9954 | 0.5214 | 0 | 82.4866 | no-hit | 0.0 | 0 | no-hit | 0.0 | 0 | no-hit | 0.0 | 0 | no-hit | 0.0 | 0 | no-hit | 0.0 | 0 | no-hit | 0.0 | 0 |
| scf7180000019509 | 328112 | 0.4324 | 0 | 51.5316 | Eukaryota | 54215.0 | 0 | Chordata | 54215.0 | 0 | Carnivora | 54215.0 | 0 | Felidae | 54215.0 | 0 | Felis | 35476.0 | 2 | Felis catus | 35476.0 | 2 |
| scf7180000018111 | 145957 | 0.3945 | 0 | 48.8179 | Eukaryota | 29675.0 | 0 | Chordata | 29675.0 | 0 | Carnivora | 29675.0 | 0 | Felidae | 29675.0 | 0 | Felis | 29675.0 | 0 | Felis catus | 29675.0 | 0 |
| scf7180000021354 | 59362 | 0.3772 | 0 | 24.653 | Eukaryota | 45052.0 | 0 | Chordata | 45052.0 | 0 | Carnivora | 45052.0 | 0 | Felidae | 45052.0 | 0 | Felis | 45052.0 | 0 | Felis catus | 45052.0 | 0 |
| scf7180000020432 | 119154 | 0.3791 | 0 | 25.6483 | Eukaryota | 81325.0 | 0 | Chordata | 81325.0 | 0 | Carnivora | 45747.0 | 2 | Felidae | 45747.0 | 3 | Homo | 15380.0 | 7 | Homo sapiens | 15380.0 | 8 |
| scf7180000018162 | 790392 | 0.4252 | 0 | 51.0149 | Eukaryota | 47415.0 | 0 | Chordata | 47415.0 | 0 | Carnivora | 47415.0 | 0 | Felidae | 47415.0 | 0 | Felis | 42717.0 | 1 | Felis catus | 42717.0 | 1 |
| scf7180000024130 | 175965 | 0.4038 | 0 | 50.605 | Eukaryota | 27845.0 | 0 | Chordata | 27845.0 | 0 | Carnivora | 27845.0 | 0 | Felidae | 18704.0 | 3 | Felis | 5700.0 | 6 | Felis catus | 5700.0 | 6 |
| scf7180000019956 | 233514 | 0.4428 | 0 | 24.7435 | Eukaryota | 30577.0 | 0 | Chordata | 30577.0 | 0 | Carnivora | 30577.0 | 0 | Felidae | 30577.0 | 0 | Panthera | 24426.0 | 1 | anthera pardu | 24426.0 | 1 |
| scf7180000020721 | 133655 | 0.6178 | 0 | 47.119 | Eukaryota | 9207.0 | 0 | Chordata | 9207.0 | 0 | Carnivora | 9207.0 | 0 | Felidae | 9207.0 | 0 | Felis | 7385.0 | 1 | Felis catus | 7385.0 | 1 |
| scf7180000021831 | 30743 | 0.4027 | 0 | 24.7232 | Eukaryota | 26432.0 | 0 | Chordata | 26432.0 | 0 | Carnivora | 26432.0 | 0 | Felidae | 26432.0 | 0 | Felis | 21120.0 | 1 | Felis catus | 21120.0 | 1 |
| scf7180000017444 | 4671033 | 0.3726 | 0 | 50.3825 | Eukaryota | 95029.0 | 0 | Chordata | 95029.0 | 0 | Carnivora | 67574.0 | 1 | Felidae | 39695.0 | 5 | Acinonyx | 11064.0 | 9 | cinonyx jubat | 11064.0 | 9 |
| scf7180000021270 | 44603 | 0.351 | 0 | 24.5067 | Eukaryota | 7785.0 | 0 | Chordata | 7785.0 | 0 | Primates | 3076.0 | 2 | Hominidae | 3076.0 | 4 | Sus | 2389.0 | 5 | Sus scrofa | 2389.0 | 5 |
| scf7180000018282 | 848681 | 0.4057 | 0 | 50.4724 | Eukaryota | 35597.0 | 0 | Chordata | 35597.0 | 0 | Carnivora | 35597.0 | 0 | Felidae | 35597.0 | 0 | Felis | 35597.0 | 0 | Felis catus | 35597.0 | 0 |
| scf7180000022877 | 21912 | 0.3855 | 0 | 315.2732 | Eukaryota | 6143.0 | 0 | Chordata | 6143.0 | 0 | Carnivora | 6143.0 | 0 | Felidae | 6143.0 | 0 | Felis | 6143.0 | 0 | Felis catus | 6143.0 | 0 |
| scf7180000021228 | 80438 | 0.4138 | 0 | 49.2068 | Eukaryota | 5597.0 | 0 | Chordata | 5597.0 | 0 | Carnivora | 4898.0 | 1 | Felidae | 2990.0 | 2 | Panthera | 2541.0 | 3 | Canis lupus | 1908.0 | 4 |
| scf7180000020942 | 72295 | 0.3753 | 0 | 23.2725 | Eukaryota | 11535.0 | 0 | Chordata | 11535.0 | 0 | Carnivora | 11535.0 | 0 | Felidae | 11535.0 | 0 | Felis | 9181.0 | 1 | Felis catus | 9181.0 | 1 |
| scf7180000018681 | 691348 | 0.4293 | 0 | 50.839 | Eukaryota | 39665.0 | 0 | Chordata | 39665.0 | 0 | Carnivora | 22041.0 | 1 | Felidae | 22041.0 | 1 | Felis | 18829.0 | 2 | Felis catus | 18829.0 | 2 |
| scf7180000020358 | 160752 | 0.4433 | 0 | 26.2829 | Eukaryota | 14097.0 | 0 | Chordata | 14097.0 | 0 | Primates | 8974.0 | 2 | Hominidae | 8974.0 | 2 | Homo | 7345.0 | 3 | Homo sapiens | 7345.0 | 3 |
| scf7180000020778 | 112531 | 0.5729 | 0 | 48.6731 | Eukaryota | 8599.0 | 0 | Chordata | 8599.0 | 0 | Carnivora | 7552.0 | 2 | Felidae | 3984.0 | 4 | Acinonyx | 3984.0 | 4 | cinonyx jubat | 3984.0 | 4 |
| scf7180000024125 | 281633 | 0.4899 | 0 | 49.9954 | Eukaryota | 11979.0 | 0 | Chordata | 11979.0 | 0 | Carnivora | 11979.0 | 0 | Felidae | 11979.0 | 0 | Felis | 8664.0 | 2 | Felis catus | 8664.0 | 2 |
| scf7180000019866 | 272939 | 0.4414 | 0 | 49.3219 | Eukaryota | 20420.0 | 0 | Chordata | 20420.0 | 0 | Carnivora | 20420.0 | 0 | Felidae | 18336.0 | 1 | Felis | 16337.0 | 2 | Felis catus | 16337.0 | 2 |
| scf7180000017477 | 1050227 | 0.3697 | 0 | 48.4456 | Eukaryota | 45818.0 | 0 | Chordata | 45818.0 | 0 | Carnivora | 45818.0 | 0 | Felidae | 45818.0 | 0 | Felis | 45818.0 | 0 | Felis catus | 45818.0 | 0 |
| scf7180000023991 | 765627 | 0.3815 | 0 | 50.8557 | Eukaryota | 47994.0 | 0 | Chordata | 47994.0 | 0 | Carnivora | 47994.0 | 0 | Felidae | 47994.0 | 0 | Felis | 47994.0 | 0 | Felis catus | 47994.0 | 0 |
| scf7180000018454 | 905495 | 0.4049 | 0 | 50.7214 | Eukaryota | 45665.0 | 0 | Chordata | 45665.0 | 0 | Carnivora | 45665.0 | 0 | Felidae | 45665.0 | 0 | Felis | 45665.0 | 0 | Felis catus | 45665.0 | 0 |
| scf7180000022238 | 11690 | 0.3582 | 0 | 23.924 | Eukaryota | 10536.0 | 0 | Chordata | 10536.0 | 0 | Primates | 6204.0 | 3 | Hominidae | 5393.0 | 7 | Homo | 3594.0 | 8 | Homo sapiens | 3594.0 | 8 |
| scf7180000017705 | 2393025 | 0.3989 | 0 | 51.0656 | Eukaryota | 46973.0 | 0 | Chordata | 46973.0 | 0 | Carnivora | 46973.0 | 0 | Felidae | 46973.0 | 0 | Felis | 46973.0 | 0 | Felis catus | 46973.0 | 0 |
| scf7180000022267 | 20647 | 0.4068 | 0 | 2024.6295 | Eukaryota | 13207.0 | 0 | Chordata | 13207.0 | 0 | Carnivora | 13207.0 | 0 | Felidae | 13207.0 | 0 | Felis | 13207.0 | 0 | Felis catus | 13207.0 | 0 |
| scf7180000019491 | 255942 | 0.4182 | 0 | 50.3365 | Eukaryota | 41635.0 | 0 | Chordata | 41635.0 | 0 | Carnivora | 41635.0 | 0 | Felidae | 41635.0 | 0 | Felis | 37579.0 | 1 | Felis catus | 37579.0 | 1 |
| scf7180000018212 | 793225 | 0.4255 | 0 | 51.877 | Eukaryota | 46833.0 | 0 | Chordata | 46833.0 | 0 | Carnivora | 46833.0 | 0 | Felidae | 46833.0 | 0 | Felis | 46833.0 | 0 | Felis catus | 46833.0 | 0 |
| scf7180000020313 | 160663 | 0.4253 | 0 | 25.0373 | Eukaryota | 25820.0 | 0 | Chordata | 25820.0 | 0 | Carnivora | 25820.0 | 0 | Felidae | 25820.0 | 0 | Felis | 23182.0 | 1 | Felis catus | 23182.0 | 1 |
| scf7180000019229 | 21901 | 0.3875 | 0 | 25.3362 | Eukaryota | 3250.0 | 0 | Chordata | 3250.0 | 0 | Carnivora | 2135.0 | 1 | Felidae | 1531.0 | 2 | Felis | 1531.0 | 3 | Felis catus | 1531.0 | 3 |
| scf7180000017496 | 4979459 | 0.3746 | 0 | 50.802 | Eukaryota | 53147.0 | 0 | Chordata | 53147.0 | 0 | Carnivora | 53147.0 | 0 | Felidae | 42767.0 | 1 | Felis | 32229.0 | 2 | Felis catus | 32229.0 | 2 |
| scf7180000022543 | 12117 | 0.631 | 0 | 52.6777 | Eukaryota | 4105.0 | 0 | Chordata | 4105.0 | 0 | Carnivora | 4105.0 | 0 | Felidae | 3563.0 | 1 | Panthera | 1271.0 | 4 | Felis catus | 1195.0 | 5 |
| scf7180000021452 | 68468 | 0.459 | 0 | 50.8203 | Eukaryota | 9025.0 | 1 | Chordata | 9025.0 | 1 | Artiodactyla | 4521.0 | 5 | Bovidae | 3721.0 | 6 | Bos | 1622.0 | 8 | Bos mutus | 1622.0 | 8 |
| scf7180000021794 | 59486 | 0.3545 | 0 | 46.736 | Eukaryota | 37171.0 | 0 | Chordata | 37171.0 | 0 | Carnivora | 37171.0 | 0 | Felidae | 37171.0 | 0 | Felis | 37171.0 | 0 | Felis catus | 37171.0 | 0 |
| scf7180000018892 | 550893 | 0.4164 | 0 | 50.8106 | Eukaryota | 44127.0 | 0 | Chordata | 44127.0 | 0 | Carnivora | 44127.0 | 0 | Felidae | 44127.0 | 0 | Felis | 44127.0 | 0 | Felis catus | 44127.0 | 0 |
| scf7180000019849 | 239538 | 0.3924 | 0 | 25.6013 | Eukaryota | 45080.0 | 0 | Chordata | 45080.0 | 0 | Carnivora | 45080.0 | 0 | Felidae | 45080.0 | 0 | Felis | 45080.0 | 0 | Felis catus | 45080.0 | 0 |
| scf7180000019534 | 104994 | 0.419 | 0 | 25.1542 | Eukaryota | 40002.0 | 0 | Chordata | 40002.0 | 0 | Carnivora | 40002.0 | 0 | Felidae | 40002.0 | 0 | Felis | 36011.0 | 1 | Felis catus | 36011.0 | 1 |
| scf7180000017565 | 544062 | 0.399 | 0 | 51.8784 | Eukaryota | 46699.0 | 0 | Chordata | 46699.0 | 0 | Carnivora | 46699.0 | 0 | Felidae | 46699.0 | 0 | Felis | 46699.0 | 0 | Felis catus | 46699.0 | 0 |
| scf7180000020637 | 104585 | 0.3942 | 0 | 23.854 | Eukaryota | 40471.0 | 0 | Chordata | 40471.0 | 0 | Carnivora | 40471.0 | 0 | Felidae | 40471.0 | 0 | Felis | 36478.0 | 1 | Felis catus | 36478.0 | 1 |
| scf7180000019302 | 326740 | 0.478 | 0 | 50.4315 | Eukaryota | 17252.0 | 0 | Chordata | 17252.0 | 0 | Carnivora | 17252.0 | 0 | Otariidae | 11444.0 | 1 | Callorhinus | 6584.0 | 4 | llorhinus ursir | 6584.0 | 4 |
| scf7180000019932 | 919247 | 0.5692 | 0 | 51.7848 | Eukaryota | 55124.0 | 0 | Chordata | 55124.0 | 0 | Carnivora | 55124.0 | 0 | Felidae | 55124.0 | 0 | Acinonyx | 28045.0 | 1 | cinonyx jubat | 28045.0 | 2 |
| scf7180000022149 | 37728 | 0.3972 | 0 | 25.2365 | Eukaryota | 44141.0 | 0 | Chordata | 44141.0 | 0 | Carnivora | 44141.0 | 0 | Felidae | 44141.0 | 0 | Felis | 44141.0 | 0 | Felis catus | 44141.0 | 0 |
| scf7180000018452 | 518134 | 0.3724 | 0 | 50.223 | Eukaryota | 47652.0 | 0 | Chordata | 47652.0 | 0 | Carnivora | 47652.0 | 0 | Felidae | 47652.0 | 0 | Felis | 47652.0 | 0 | Felis catus | 47652.0 | 0 |
| scf7180000024071 | 614396 | 0.4696 | 0 | 52.5695 | Eukaryota | 25841.0 | 0 | Chordata | 25841.0 | 0 | Carnivora | 25841.0 | 0 | Felidae | 7730.0 | 5 | Panthera | 5476.0 | 7 | opoda melanc | 4947.0 | 8 |
| scf7180000020246 | 118895 | 0.3866 | 0 | 49.4588 | Eukaryota | 37797.0 | 0 | Chordata | 37797.0 | 0 | Carnivora | 37797.0 | 0 | Felidae | 37797.0 | 0 | Felis | 37797.0 | 0 | Felis catus | 37797.0 | 0 |
| scf7180000021989 | 32049 | 0.3775 | 0 | 24.959 | Eukaryota | 8597.0 | 0 | Chordata | 8597.0 | 0 | Carnivora | 8597.0 | 0 | Felidae | 7509.0 | 1 | Felis | 6940.0 | 2 | Felis catus | 6940.0 | 2 |
| scf7180000022780 | 47095 | 0.4617 | 0 | 26.6337 | Eukaryota | 20728.0 | 0 | Chordata | 20728.0 | 0 | Carnivora | 10528.0 | 3 | Felidae | 5158.0 | 5 | Myotis | 4788.0 | 7 | Homo sapiens | 3026.0 | 8 |
| scf7180000019314 | 348798 | 0.5009 | 0 | 49.9907 | Eukaryota | 43117.0 | 0 | Chordata | 43117.0 | 0 | Carnivora | 43117.0 | 0 | Felidae | 43117.0 | 0 | Felis | 38841.0 | 1 | Felis catus | 38841.0 | 1 |
| scf7180000024221 | 40956 | 0.4426 | 0 | 29.6016 | Eukaryota | 27999.0 | 0 | Chordata | 27999.0 | 0 | Carnivora | 27999.0 | 0 | Felidae | 17181.0 | 2 | Ursus | 8143.0 | 5 | rsus maritimu | 5502.0 | 7 |
| scf7180000022453 | 18568 | 0.4243 | 0 | 23.4794 | Eukaryota | 15366.0 | 0 | Chordata | 15366.0 | 0 | Carnivora | 15366.0 | 0 | Felidae | 15366.0 | 0 | Felis | 13847.0 | 1 | Felis catus | 13847.0 | 1 |
| scf7180000020206 | 124219 | 0.3854 | 0 | 46.8091 | Eukaryota | 44880.0 | 0 | Chordata | 44880.0 | 0 | Carnivora | 44880.0 | 0 | Felidae | 44880.0 | 0 | Felis | 44880.0 | 0 | Felis catus | 44880.0 | 0 |
| scf7180000018107 | 861596 | 0.4402 | 0 | 52.6309 | Eukaryota | 68783.0 | 0 | Chordata | 68783.0 | 0 | Carnivora | 68783.0 | 0 | Felidae | 48083.0 | 2 | Zalophus | 17940.0 | 5 | phus californi | 17940.0 | 5 |
| scf7180000023654 | 73623 | 0.4018 | 0 | 280.5812 | Eukaryota | 6973.0 | 0 | Chordata | 6973.0 | 0 | Carnivora | 6973.0 | 0 | Felidae | 6973.0 | 0 | Felis | 6973.0 | 0 | Felis catus | 6973.0 | 0 |
| scf7180000019794 | 212888 | 0.5773 | 0 | 53.0486 | Eukaryota | 17585.0 | 0 | Chordata | 17585.0 | 0 | Carnivora | 17585.0 | 0 | Felidae | 15712.0 | 1 | Panthera | 8722.0 | 3 | anthera pardu | 6984.0 | 4 |
| scf7180000022443 | 26903 | 0.4844 | 0 | 22.2125 | Eukaryota | 2981.0 | 0 | Chordata | 2981.0 | 0 | Carnivora | 2981.0 | 0 | Felidae | 2072.0 | 1 | Panthera | 1188.0 | 4 | uricata suricat | 909.0 | 5 |
| scf7180000024254 | 198532 | 0.4089 | 0 | 48.6893 | Eukaryota | 15726.0 | 0 | Chordata | 15726.0 | 0 | Carnivora | 15726.0 | 0 | Felidae | 11298.0 | 1 | Lynx | 9312.0 | 2 | ynx canadens | 9312.0 | 2 |
| scf7180000024013 | 427211 | 0.4288 | 0 | 25.285 | Eukaryota | 34139.0 | 0 | Chordata | 34139.0 | 0 | Primates | 22357.0 | 2 | Hominidae | 22357.0 | 2 | Homo | 16514.0 | 3 | Homo sapiens | 16514.0 | 3 |
| scf7180000017832 | 874167 | 0.3854 | 0 | 50.4853 | Eukaryota | 47099.0 | 0 | Chordata | 47099.0 | 0 | Carnivora | 47099.0 | 0 | Felidae | 47099.0 | 0 | Felis | 47099.0 | 0 | Felis catus | 47099.0 | 0 |
| scf7180000019722 | 290314 | 0.3859 | 0 | 51.1684 | Eukaryota | 17440.0 | 0 | Chordata | 17440.0 | 0 | Carnivora | 17440.0 | 0 | Felidae | 17440.0 | 0 | Felis | 17440.0 | 0 | Felis catus | 17440.0 | 0 |
| scf7180000018351 | 908661 | 0.4241 | 0 | 50.7726 | Eukaryota | 94313.0 | 0 | Chordata | 94313.0 | 0 | Carnivora | 94313.0 | 0 | Felidae | 55473.0 | 1 | Felis | 46095.0 | 2 | Felis catus | 46095.0 | 2 |
| scf7180000021337 | 48902 | 0.6078 | 0 | 46.2634 | Eukaryota | 5552.0 | 0 | Chordata | 5552.0 | 0 | Primates | 2380.0 | 6 | Hominidae | 2380.0 | 6 | Homo | 2380.0 | 6 | Homo sapiens | 2380.0 | 6 |
| scf7180000024040 | 264315 | 0.4905 | 0 | 49.8304 | Eukaryota | 8666.0 | 0 | Chordata | 8666.0 | 0 | Carnivora | 8666.0 | 0 | Felidae | 5728.0 | 3 | Panthera | 2144.0 | 7 | ynx canadens | 1986.0 | 8 |
| scf7180000020712 | 49748 | 0.5421 | 0 | 56.8271 | Eukaryota | 41597.0 | 0 | Chordata | 41597.0 | 0 | Carnivora | 17654.0 | 3 | Felidae | 13233.0 | 5 | Lynx | 9945.0 | 6 | ynx canadens | 9945.0 | 6 |
| scf7180000021170 | 58735 | 0.4122 | 0 | 23.888 | Eukaryota | 5679.0 | 0 | Chordata | 5679.0 | 0 | Carnivora | 4912.0 | 1 | Felidae | 4912.0 | 1 | Felis | 4912.0 | 1 | Felis catus | 4912.0 | 1 |
| scf7180000019800 | 12581 | 0.3485 | 0 | 26.8232 | Eukaryota | 8462.0 | 1 | Chordata | 8462.0 | 1 | Artiodactyla | 4753.0 | 4 | Bovidae | 4753.0 | 5 | Bos | 2446.0 | 7 | Bos mutus | 2446.0 | 7 |
| scf7180000022292 | 30075 | 0.6263 | 0 | 50.7009 | Eukaryota | 2431.0 | 0 | Chordata | 2431.0 | 0 | Carnivora | 2431.0 | 0 | Felidae | 2174.0 | 1 | Felis | 1915.0 | 2 | Felis catus | 1915.0 | 2 |
| scf7180000020542 | 142666 | 0.3939 | 0 | 26.1799 | Eukaryota | 34393.0 | 0 | Chordata | 34393.0 | 0 | Carnivora | 34393.0 | 0 | Felidae | 34393.0 | 0 | Felis | 34393.0 | 0 | Felis catus | 34393.0 | 0 |
| scf7180000018660 | 1295354 | 0.4437 | 0 | 52.3978 | Eukaryota | 47639.0 | 0 | Chordata | 47639.0 | 0 | Carnivora | 47639.0 | 0 | Felidae | 47639.0 | 0 | Felis | 47639.0 | 0 | Felis catus | 47639.0 | 0 |
| scf7180000022103 | 41942 | 0.4448 | 0 | 42.0914 | Eukaryota | 26325.0 | 0 | Chordata | 26325.0 | 0 | Carnivora | 26325.0 | 0 | Felidae | 26325.0 | 0 | Felis | 23663.0 | 1 | Felis catus | 23663.0 | 1 |
| scf7180000019637 | 205764 | 0.38 | 0 | 49.4167 | Eukaryota | 42446.0 | 0 | Chordata | 42446.0 | 0 | Carnivora | 42446.0 | 0 | Felidae | 42446.0 | 0 | Felis | 38148.0 | 1 | Felis catus | 38148.0 | 1 |
| scf7180000020932 | 959002 | 0.4681 | 200 | 48.7411 | Eukaryota | 79029.0 | 0 | Chordata | 79029.0 | 0 | Carnivora | 79029.0 | 0 | Herpestidae | 40584.0 | 1 | Suricata | 40584.0 | 2 | uricata suricat | 40584.0 | 2 |
| scf7180000020895 | 82918 | 0.6594 | 0 | 48.6216 | Eukaryota | 8406.0 | 0 | Chordata | 8406.0 | 0 | Carnivora | 8406.0 | 0 | Felidae | 6267.0 | 1 | Felis | 2897.0 | 4 | Felis catus | 2897.0 | 4 |
| scf7180000022553 | 5554 | 0.5126 | 0 | 25.6677 | Eukaryota | 2509.0 | 0 | Chordata | 2509.0 | 0 | Carnivora | 2509.0 | 0 | Felidae | 2509.0 | 0 | Felis | 1256.0 | 1 | Felis catus | 1256.0 | 1 |
| scf7180000018873 | 531028 | 0.3568 | 0 | 49.7094 | Eukaryota | 45043.0 | 0 | Chordata | 45043.0 | 0 | Carnivora | 45043.0 | 0 | Felidae | 45043.0 | 0 | Felis | 45043.0 | 0 | Felis catus | 45043.0 | 0 |
| scf7180000019493 | 273006 | 0.5513 | 0 | 48.2491 | Eukaryota | 11409.0 | 0 | Chordata | 11409.0 | 0 | Carnivora | 11409.0 | 0 | Felidae | 11409.0 | 0 | Felis | 11409.0 | 0 | Felis catus | 11409.0 | 0 |
| scf7180000021192 | 54390 | 0.4732 | 0 | 48.3232 | Eukaryota | 8286.0 | 1 | Chordata | 8286.0 | 1 | Carnivora | 4226.0 | 3 | Hominidae | 3212.0 | 4 | Canis | 1849.0 | 7 | Canis lupus | 1849.0 | 7 |
| scf7180000020370 | 193478 | 0.5821 | 0 | 48.8597 | Eukaryota | 9858.0 | 0 | Chordata | 9858.0 | 0 | Carnivora | 9858.0 | 0 | Felidae | 7590.0 | 1 | Panthera | 4785.0 | 4 | anthera pardu | 3828.0 | 5 |
| scf7180000024253 | 2006224 | 0.4544 | 100 | 52.902 | Eukaryota | 75200.0 | 0 | Chordata | 75200.0 | 0 | Carnivora | 75200.0 | 0 | Felidae | 75200.0 | 0 | Acinonyx | 45044.0 | 1 | cinonyx jubat | 45044.0 | 1 |
| scf7180000017602 | 573124 | 0.3796 | 0 | 49.4393 | Eukaryota | 46368.0 | 0 | Chordata | 46368.0 | 0 | Carnivora | 46368.0 | 0 | Felidae | 46368.0 | 0 | Felis | 46368.0 | 0 | Felis catus | 46368.0 | 0 |
| scf7180000018811 | 494959 | 0.4329 | 0 | 50.0983 | Eukaryota | 41296.0 | 0 | Chordata | 41296.0 | 0 | Carnivora | 41296.0 | 0 | Felidae | 36865.0 | 1 | Panthera | 17576.0 | 3 | anthera pardu | 17576.0 | 3 |
| scf7180000019690 | 112441 | 0.4164 | 0 | 48.5624 | Eukaryota | 46035.0 | 0 | Chordata | 46035.0 | 0 | Carnivora | 46035.0 | 0 | Felidae | 46035.0 | 0 | Felis | 41449.0 | 1 | Felis catus | 41449.0 | 1 |
| scf7180000023981 | 585380 | 0.4414 | 0 | 52.3544 | Eukaryota | 45094.0 | 0 | Chordata | 45094.0 | 0 | Carnivora | 45094.0 | 0 | Felidae | 45094.0 | 0 | Acinonyx | 18036.0 | 3 | cinonyx jubat | 18036.0 | 4 |
| scf7180000022381 | 6498 | 0.4432 | 0 | 408.5913 | Eukaryota | 3345.0 | 0 | Chordata | 3345.0 | 0 | Artiodactyla | 2608.0 | 1 | Bovidae | 2608.0 | 2 | Ovis | 2608.0 | 3 | vis canadens | 2608.0 | 3 |
| scf7180000021503 | 27418 | 0.4619 | 0 | 25.6407 | Eukaryota | 15249.0 | 0 | Chordata | 15249.0 | 0 | Artiodactyla | 10760.0 | 1 | Delphinidae | 8290.0 | 2 | Tursiops | 6150.0 | 4 | rsiops truncat | 6150.0 | 4 |
| scf7180000018405 | 208464 | 0.3559 | 0 | 48.5641 | Eukaryota | 41771.0 | 0 | Chordata | 41771.0 | 0 | Carnivora | 41771.0 | 0 | Felidae | 41771.0 | 0 | Felis | 41771.0 | 0 | Felis catus | 41771.0 | 0 |
| scf7180000017470 | 3376522 | 0.3921 | 0 | 50.5168 | Eukaryota | 86241.0 | 0 | Chordata | 86241.0 | 0 | Carnivora | 86241.0 | 0 | Felidae | 86241.0 | 0 | Acinonyx | 33864.0 | 2 | cinonyx jubat | 33864.0 | 2 |
| scf7180000022370 | 22977 | 0.4546 | 0 | 24.5806 | Eukaryota | 3377.0 | 0 | Chordata | 3377.0 | 0 | Carnivora | 2048.0 | 2 | Felidae | 1793.0 | 3 | Felis | 1282.0 | 5 | Felis catus | 1282.0 | 5 |
| scf7180000019667 | 14310 | 0.413 | 0 | 24.3519 | Eukaryota | 11795.0 | 0 | Chordata | 11795.0 | 0 | Carnivora | 11795.0 | 0 | Felidae | 11795.0 | 0 | Felis | 10613.0 | 1 | Felis catus | 10613.0 | 1 |
| scf7180000022733 | 7152 | 0.4999 | 0 | 48.1879 | Eukaryota | 20740.0 | 0 | Chordata | 20740.0 | 0 | Carnivora | 20740.0 | 0 | Felidae | 20740.0 | 0 | Felis | 20740.0 | 0 | Felis catus | 20740.0 | 0 |
| scf7180000019808 | 272007 | 0.5902 | 0 | 48.3585 | Eukaryota | 12208.0 | 0 | Chordata | 12208.0 | 0 | Carnivora | 12208.0 | 0 | Felidae | 12208.0 | 0 | Felis | 10981.0 | 1 | Felis catus | 10981.0 | 1 |
| scf7180000022346 | 28016 | 0.3719 | 0 | 24.6472 | Eukaryota | 16598.0 | 0 | Chordata | 16598.0 | 0 | Primates | 8068.0 | 2 | Hominidae | 8068.0 | 2 | Homo | 5368.0 | 3 | Homo sapiens | 5368.0 | 3 |
| scf7180000017973 | 228403 | 0.4385 | 0 | 49.2577 | Eukaryota | 29410.0 | 0 | Chordata | 29410.0 | 0 | Primates | 11820.0 | 2 | Bovidae | 10058.0 | 3 | Felis | 7532.0 | 4 | Felis catus | 7532.0 | 4 |
| scf7180000017441 | 337939 | 0.5503 | 0 | 49.7277 | Eukaryota | 13752.0 | 0 | Chordata | 13752.0 | 0 | Carnivora | 13752.0 | 0 | Felidae | 13752.0 | 0 | Felis | 13752.0 | 0 | Felis catus | 13752.0 | 0 |
| scf7180000017701 | 1311539 | 0.3678 | 0 | 47.8155 | Eukaryota | 47063.0 | 0 | Chordata | 47063.0 | 0 | Carnivora | 47063.0 | 0 | Felidae | 47063.0 | 0 | Felis | 47063.0 | 0 | Felis catus | 47063.0 | 0 |
| scf7180000020568 | 152856 | 0.4854 | 0 | 48.0702 | Eukaryota | 18990.0 | 0 | Chordata | 18990.0 | 0 | Carnivora | 18990.0 | 0 | Felidae | 18990.0 | 0 | Felis | 18990.0 | 0 | Felis catus | 18990.0 | 0 |
| scf7180000019860 | 46762 | 0.3855 | 0 | 304.0073 | Eukaryota | 10099.0 | 0 | Chordata | 10099.0 | 0 | pterygiforme | 9033.0 | 1 | Apterygidae | 9033.0 | 1 | Apteryx | 9033.0 | 1 | pteryx mante | 9033.0 | 1 |
| scf7180000019955 | 182148 | 0.4678 | 0 | 611.5891 | Eukaryota | 17884.0 | 0 | Chordata | 17884.0 | 0 | Carnivora | 17884.0 | 0 | Felidae | 17884.0 | 0 | Felis | 14461.0 | 1 | Felis catus | 14461.0 | 1 |
| scf7180000021024 | 62800 | 0.5739 | 0 | 24.8817 | Eukaryota | 43373.0 | 0 | Chordata | 43373.0 | 0 | Carnivora | 43373.0 | 0 | Herpestidae | 43373.0 | 0 | Suricata | 43373.0 | 0 | uricata suricat | 43373.0 | 0 |
| scf7180000022468 | 31609 | 0.4923 | 0 | 24.4216 | Eukaryota | 5523.0 | 0 | Chordata | 5523.0 | 0 | Carnivora | 3397.0 | 2 | Felidae | 3397.0 | 3 | Panthera | 1139.0 | 7 | Felis catus | 1119.0 | 8 |
| scf7180000020981 | 67142 | 0.4646 | 0 | 49.6675 | Eukaryota | 22545.0 | 0 | Chordata | 22545.0 | 0 | Carnivora | 22545.0 | 0 | Felidae | 9916.0 | 2 | Panthera | 9916.0 | 2 | phus californi | 7768.0 | 3 |
| scf7180000020504 | 34045 | 0.4438 | 0 | 49.8666 | Eukaryota | 33632.0 | 0 | Chordata | 33632.0 | 0 | Carnivora | 33632.0 | 0 | Felidae | 33632.0 | 0 | Felis | 30130.0 | 1 | Felis catus | 30130.0 | 1 |
| scf7180000020367 | 219568 | 0.3997 | 0 | 25.6325 | Eukaryota | 43379.0 | 0 | Chordata | 43379.0 | 0 | Carnivora | 15392.0 | 2 | Felidae | 15392.0 | 2 | Sus | 13386.0 | 4 | Sus scrofa | 13386.0 | 4 |
| scf7180000020588 | 110286 | 0.3667 | 0 | 25.5338 | Eukaryota | 24814.0 | 0 | Chordata | 24814.0 | 0 | Carnivora | 24814.0 | 0 | Felidae | 24814.0 | 0 | Felis | 22313.0 | 1 | Felis catus | 22313.0 | 1 |
| scf7180000020524 | 94692 | 0.5691 | 0 | 48.2634 | Eukaryota | 10484.0 | 0 | Chordata | 10484.0 | 0 | Carnivora | 10484.0 | 0 | Felidae | 5529.0 | 3 | Canis | 2917.0 | 6 | Canis lupus | 2917.0 | 6 |
| scf7180000020256 | 145574 | 0.5352 | 0 | 48.8537 | Eukaryota | 14731.0 | 0 | Chordata | 14731.0 | 0 | Carnivora | 14731.0 | 0 | Felidae | 14731.0 | 0 | Felis | 8886.0 | 3 | Felis catus | 8886.0 | 3 |
| scf7180000018054 | 984171 | 0.4804 | 0 | 60.2262 | Eukaryota | 47916.0 | 0 | Chordata | 47916.0 | 0 | Carnivora | 47916.0 | 0 | Felidae | 47916.0 | 0 | Felis | 47916.0 | 0 | Felis catus | 47916.0 | 0 |
| scf7180000022221 | 19628 | 0.65 | 0 | 48.2057 | no-hit | 0.0 | 0 | no-hit | 0.0 | 0 | no-hit | 0.0 | 0 | no-hit | 0.0 | 0 | no-hit | 0.0 | 0 | no-hit | 0.0 | 0 |
| scf7180000023549 | 20095 | 0.3666 | 0 | 265.787 | Eukaryota | 9951.0 | 0 | Chordata | 9951.0 | 0 | pterygiforme | 8898.0 | 1 | Apterygidae | 8898.0 | 1 | Apteryx | 8898.0 | 1 | pteryx mante | 8898.0 | 1 |
| scf7180000024018 | 190547 | 0.4097 | 0 | 47.474 | Eukaryota | 44703.0 | 0 | Chordata | 44703.0 | 0 | Carnivora | 44703.0 | 0 | Felidae | 44703.0 | 0 | Felis | 44703.0 | 0 | Felis catus | 44703.0 | 0 |
| scf7180000022216 | 67310 | 0.4714 | 0 | 126.3795 | Eukaryota | 5078.0 | 0 | Chordata | 5078.0 | 0 | Carnivora | 5078.0 | 0 | Felidae | 5078.0 | 0 | Felis | 5078.0 | 0 | Felis catus | 5078.0 | 0 |
| scf7180000021569 | 68113 | 0.3901 | 0 | 25.7897 | Eukaryota | 12836.0 | 0 | Chordata | 12836.0 | 0 | Carnivora | 12836.0 | 0 | Felidae | 12836.0 | 0 | Acinonyx | 8990.0 | 1 | cinonyx jubat | 8990.0 | 1 |
| scf7180000018305 | 481614 | 0.3979 | 0 | 65.0602 | Eukaryota | 41832.0 | 0 | Chordata | 41832.0 | 0 | Carnivora | 41832.0 | 0 | Felidae | 41832.0 | 0 | Felis | 37706.0 | 1 | Felis catus | 37706.0 | 1 |
| scf7180000022405 | 11067 | 0.3694 | 0 | 24.1303 | Eukaryota | 4951.0 | 0 | Chordata | 4951.0 | 0 | Carnivora | 4951.0 | 0 | Felidae | 4951.0 | 0 | Felis | 4428.0 | 1 | Felis catus | 4428.0 | 1 |
| scf7180000023618 | 40625 | 0.4348 | 0 | 323.44 | Eukaryota | 5309.0 | 0 | Chordata | 5309.0 | 0 | Carnivora | 4396.0 | 1 | Felidae | 3049.0 | 2 | Felis | 3049.0 | 2 | Felis catus | 3049.0 | 2 |
| scf7180000019173 | 387937 | 0.4336 | 0 | 49.5946 | Eukaryota | 45389.0 | 0 | Chordata | 45389.0 | 0 | Carnivora | 45389.0 | 0 | Felidae | 45389.0 | 0 | Felis | 45389.0 | 0 | Felis catus | 45389.0 | 0 |
| scf7180000018316 | 1025551 | 0.3495 | 0 | 48.1421 | Eukaryota | 45891.0 | 0 | Chordata | 45891.0 | 0 | Carnivora | 45891.0 | 0 | Felidae | 45891.0 | 0 | Felis | 45891.0 | 0 | Felis catus | 45891.0 | 0 |
| scf7180000019108 | 606061 | 0.5365 | 0 | 51.0434 | Eukaryota | 20147.0 | 1 | Chordata | 20147.0 | 1 | Primates | 11365.0 | 2 | Hominidae | 11365.0 | 2 | Homo | 11365.0 | 2 | Homo sapiens | 11365.0 | 2 |
| scf7180000023292 | 21251 | 0.4516 | 0 | 340.5692 | Eukaryota | 12325.0 | 0 | Chordata | 12325.0 | 0 | Carnivora | 12325.0 | 0 | Felidae | 12325.0 | 0 | Felis | 11109.0 | 1 | Felis catus | 11109.0 | 1 |
| scf7180000017885 | 489262 | 0.3979 | 0 | 50.3822 | Eukaryota | 65267.0 | 0 | Chordata | 65267.0 | 0 | Carnivora | 43827.0 | 1 | Felidae | 43827.0 | 1 | Felis | 43827.0 | 1 | Felis catus | 43827.0 | 1 |
| scf7180000018716 | 672649 | 0.4375 | 0 | 49.9808 | Eukaryota | 45314.0 | 0 | Chordata | 45314.0 | 0 | Carnivora | 45314.0 | 0 | Felidae | 45314.0 | 0 | Felis | 45314.0 | 0 | Felis catus | 45314.0 | 0 |
| scf7180000021259 | 12042 | 0.3881 | 0 | 17.0075 | Eukaryota | 42796.0 | 0 | Chordata | 42796.0 | 0 | Carnivora | 42796.0 | 0 | Felidae | 42796.0 | 0 | Felis | 42796.0 | 0 | Felis catus | 42796.0 | 0 |
| scf7180000023119 | 19516 | 0.5228 | 0 | 357.1437 | Eukaryota | 22236.0 | 0 | Chordata | 22236.0 | 0 | Carnivora | 22236.0 | 0 | Felidae | 22236.0 | 0 | Felis | 19992.0 | 1 | Felis catus | 19992.0 | 1 |
| scf7180000023952 | 616150 | 0.4505 | 0 | 51.6266 | Eukaryota | 45757.0 | 0 | Chordata | 45757.0 | 0 | Carnivora | 45757.0 | 0 | Felidae | 37605.0 | 1 | Panthera | 20534.0 | 4 | Panthera tigri | 11580.0 | 5 |
| scf7180000020903 | 75817 | 0.3733 | 0 | 51.3507 | Eukaryota | 45450.0 | 0 | Chordata | 45450.0 | 0 | Carnivora | 45450.0 | 0 | Felidae | 45450.0 | 0 | Felis | 45450.0 | 0 | Felis catus | 45450.0 | 0 |
| scf7180000023918 | 1422634 | 0.3962 | 0 | 51.0971 | Eukaryota | 62743.0 | 0 | Chordata | 62743.0 | 0 | Carnivora | 62743.0 | 0 | Felidae | 44692.0 | 1 | Lynx | 44692.0 | 1 | ynx canadens | 44692.0 | 1 |
| scf7180000020405 | 105202 | 0.3725 | 0 | 24.8963 | Eukaryota | 43797.0 | 0 | Chordata | 43797.0 | 0 | Carnivora | 43797.0 | 0 | Felidae | 43797.0 | 0 | Felis | 43797.0 | 0 | Felis catus | 43797.0 | 0 |
| scf7180000020878 | 13575 | 0.6494 | 0 | 83.0734 | Eukaryota | 1248.0 | 0 | Chordata | 1248.0 | 0 | Carnivora | 1248.0 | 0 | Felidae | 1248.0 | 0 | Felis | 1248.0 | 0 | Felis catus | 1248.0 | 0 |
| scf7180000018372 | 200718 | 0.3841 | 0 | 47.8236 | Eukaryota | 44604.0 | 0 | Chordata | 44604.0 | 0 | Carnivora | 44604.0 | 0 | Felidae | 44604.0 | 0 | Lynx | 31171.0 | 1 | ynx canadens | 31171.0 | 2 |
| scf7180000019459 | 270614 | 0.4004 | 0 | 51.2931 | Eukaryota | 50176.0 | 0 | Chordata | 50176.0 | 0 | Carnivora | 50176.0 | 0 | Felidae | 39684.0 | 2 | Felis | 28012.0 | 4 | Felis catus | 28012.0 | 4 |
| scf7180000020681 | 112600 | 0.4214 | 0 | 50.5446 | Eukaryota | 38986.0 | 0 | Chordata | 38986.0 | 0 | Carnivora | 38986.0 | 0 | Felidae | 38986.0 | 0 | Acinonyx | 15714.0 | 2 | cinonyx jubat | 15714.0 | 2 |
| scf7180000019802 | 90176 | 0.4194 | 0 | 49.3397 | Eukaryota | 45036.0 | 0 | Chordata | 45036.0 | 0 | Carnivora | 45036.0 | 0 | Felidae | 45036.0 | 0 | Felis | 40589.0 | 1 | Felis catus | 40589.0 | 1 |
| scf7180000021093 | 60202 | 0.3498 | 0 | 50.0179 | Eukaryota | 3841.0 | 0 | Chordata | 3841.0 | 0 | Carnivora | 2206.0 | 2 | Felidae | 1240.0 | 3 | Canis | 966.0 | 6 | Canis lupus | 966.0 | 6 |
| scf7180000022664 | 14195 | 0.3691 | 0 | 22.8231 | Eukaryota | 31813.0 | 0 | Chordata | 31813.0 | 0 | Carnivora | 31813.0 | 0 | Felidae | 31813.0 | 0 | Felis | 31813.0 | 0 | Felis catus | 31813.0 | 0 |
| scf7180000021691 | 27056 | 0.6252 | 0 | 46.2639 | Eukaryota | 2595.0 | 0 | Chordata | 2595.0 | 0 | Carnivora | 2595.0 | 0 | Felidae | 2345.0 | 1 | Felis | 1623.0 | 2 | Felis catus | 1623.0 | 2 |
| scf7180000023667 | 29463 | 0.4062 | 0 | 234.9934 | Eukaryota | 5925.0 | 0 | Chordata | 5925.0 | 0 | Carnivora | 5925.0 | 0 | Phocidae | 2416.0 | 2 | eptonychote | 1812.0 | 4 | nychotes wed | 1812.0 | 4 |
| scf7180000021964 | 24576 | 0.3615 | 0 | 24.3358 | Eukaryota | 28629.0 | 0 | Chordata | 28629.0 | 0 | Carnivora | 28629.0 | 0 | Felidae | 28629.0 | 0 | Felis | 25751.0 | 1 | Felis catus | 25751.0 | 1 |
| scf7180000018823 | 768193 | 0.3499 | 0 | 48.3573 | Eukaryota | 25247.0 | 0 | Chordata | 25247.0 | 0 | Carnivora | 25247.0 | 0 | Felidae | 25247.0 | 0 | Felis | 25247.0 | 0 | Felis catus | 25247.0 | 0 |
| scf7180000022740 | 12075 | 0.4385 | 0 | 22.864 | Eukaryota | 3205.0 | 0 | Chordata | 3205.0 | 0 | Carnivora | 2471.0 | 1 | Felidae | 2105.0 | 2 | Felis | 1831.0 | 4 | Felis catus | 1831.0 | 4 |
| scf7180000023772 | 2643422 | 0.3986 | 0 | 50.6326 | Eukaryota | 92268.0 | 0 | Chordata | 92268.0 | 0 | Carnivora | 92268.0 | 0 | Felidae | 92268.0 | 0 | Panthera | 64533.0 | 1 | anthera pardu | 64533.0 | 1 |

| | | | | | | | | | | | | | | | | | | | | | |
|---|---|---|---|---|---|---|---|---|---|---|---|---|---|---|---|---|---|---|---|---|---|
| scf7180000020536 | 50400 | 0.4065 | 0 | 42.5705 | Eukaryota | 10472.0 | 0 | Chordata | 10472.0 | 0 | Carnivora | 10472.0 | 0 | Felidae | 6606.0 | 2 | Acinonyx | 2236.0 | 6 | cinonyx jubat | 2236.0 | 7 |
| scf7180000019574 | 261184 | 0.4214 | 0 | 52.9948 | Eukaryota | 22238.0 | 0 | Chordata | 22238.0 | 0 | Carnivora | 22238.0 | 0 | Felidae | 18341.0 | 1 | Acinonyx | 8662.0 | 3 | cinonyx jubat | 8662.0 | 3 |
| scf7180000019415 | 250949 | 0.3683 | 0 | 24.7597 | Eukaryota | 37460.0 | 0 | Chordata | 37460.0 | 0 | Carnivora | 37460.0 | 0 | Felidae | 37460.0 | 0 | Felis | 33705.0 | 1 | Felis catus | 33705.0 | 1 |
| scf7180000018363 | 772695 | 0.4165 | 0 | 50.4965 | Eukaryota | 27338.0 | 0 | Chordata | 27338.0 | 0 | Carnivora | 27338.0 | 0 | Felidae | 22276.0 | 1 | Felis | 22276.0 | 1 | Felis catus | 22276.0 | 1 |
| scf7180000019926 | 202158 | 0.4171 | 0 | 25.9644 | Eukaryota | 37073.0 | 0 | Chordata | 37073.0 | 0 | Carnivora | 31789.0 | 1 | Felidae | 31789.0 | 1 | Panthera | 17665.0 | 2 | anthera pard | 17665.0 | 2 |
| scf7180000023486 | 25711 | 0.4575 | 0 | 487.3801 | no-hit | 0.0 | 0 | no-hit | 0.0 | 0 | no-hit | 0.0 | 0 | no-hit | 0.0 | 0 | no-hit | 0.0 | 0 | no-hit | 0.0 | 0 |
| scf7180000021388 | 106609 | 0.3755 | 0 | 25.0518 | Eukaryota | 18210.0 | 0 | Chordata | 18210.0 | 0 | Carnivora | 18210.0 | 0 | Felidae | 18210.0 | 0 | Felis | 16381.0 | 1 | Felis catus | 16381.0 | 1 |
| scf7180000022802 | 13087 | 0.365 | 0 | 24.4055 | Eukaryota | 18025.0 | 0 | Chordata | 18025.0 | 0 | Carnivora | 18025.0 | 0 | Felidae | 18025.0 | 0 | Felis | 16547.0 | 1 | Felis catus | 16547.0 | 1 |
| scf7180000018005 | 1178466 | 0.3811 | 0 | 51.8606 | Eukaryota | 46224.0 | 0 | Chordata | 46224.0 | 0 | Carnivora | 46224.0 | 0 | Felidae | 46224.0 | 0 | Felis | 46224.0 | 0 | Felis catus | 46224.0 | 0 |
| scf7180000020357 | 107657 | 0.5877 | 0 | 45.8834 | Eukaryota | 6675.0 | 0 | Chordata | 6675.0 | 0 | Primates | 4503.0 | 3 | Hominidae | 3508.0 | 4 | Homo | 3033.0 | 5 | Homo sapiens | 3033.0 | 6 |
| scf7180000023785 | 622531 | 0.3857 | 0 | 48.9414 | Eukaryota | 49315.0 | 0 | Chordata | 49315.0 | 0 | Carnivora | 49315.0 | 0 | Felidae | 49315.0 | 0 | Felis | 41608.0 | 1 | Felis catus | 41608.0 | 1 |
| scf7180000018599 | 1165030 | 0.4263 | 0 | 50.754 | Eukaryota | 60266.0 | 0 | Chordata | 60266.0 | 0 | Carnivora | 60266.0 | 0 | Felidae | 60266.0 | 0 | Panthera | 50068.0 | 1 | Panthera tigri | 25152.0 | 2 |
| scf7180000022469 | 21757 | 0.5329 | 0 | 49.6428 | Eukaryota | 16022.0 | 0 | Chordata | 16022.0 | 0 | Carnivora | 16022.0 | 0 | Felidae | 16022.0 | 0 | Felis | 14317.0 | 1 | Felis catus | 14317.0 | 1 |
| scf7180000020464 | 126606 | 0.4817 | 0 | 49.2762 | Eukaryota | 8044.0 | 0 | Chordata | 8044.0 | 0 | Carnivora | 5477.0 | 3 | Felidae | 3300.0 | 6 | unresolved | 1079.0 | 9 | unresolved | 1079.0 | 9 |
| scf7180000021730 | 40282 | 0.6322 | 0 | 46.7782 | Eukaryota | 8952.0 | 0 | Chordata | 8952.0 | 0 | Carnivora | 4934.0 | 2 | Felidae | 3774.0 | 3 | Mus | 2424.0 | 4 | Mus musculus | 2424.0 | 4 |
| scf7180000019723 | 222765 | 0.4167 | 0 | 25.9857 | Eukaryota | 27110.0 | 0 | Chordata | 27110.0 | 0 | Carnivora | 27110.0 | 0 | Felidae | 27110.0 | 0 | Felis | 24343.0 | 1 | Felis catus | 24343.0 | 1 |
| scf7180000021731 | 38221 | 0.5295 | 0 | 35.8188 | Eukaryota | 38371.0 | 0 | Chordata | 38371.0 | 0 | Carnivora | 38371.0 | 0 | Felidae | 38371.0 | 0 | Felis | 38371.0 | 0 | Felis catus | 38371.0 | 0 |
| scf7180000018693 | 618663 | 0.3851 | 0 | 50.8722 | Eukaryota | 46256.0 | 0 | Chordata | 46256.0 | 0 | Carnivora | 46256.0 | 0 | Felidae | 46256.0 | 0 | Felis | 46256.0 | 0 | Felis catus | 46256.0 | 0 |
| scf7180000021440 | 39491 | 0.6826 | 0 | 45.882 | Eukaryota | 8972.0 | 0 | Chordata | 8972.0 | 0 | Carnivora | 8972.0 | 0 | Felidae | 7339.0 | 1 | Panthera | 2852.0 | 4 | anthera pard | 2852.0 | 4 |
| scf7180000020255 | 26661 | 0.3932 | 0 | 28.7545 | Eukaryota | 9144.0 | 0 | Chordata | 9144.0 | 0 | Primates | 3814.0 | 3 | Felidae | 3080.0 | 4 | Felis | 3080.0 | 4 | Felis catus | 3080.0 | 4 |
| scf7180000018610 | 707050 | 0.3844 | 0 | 51.6412 | Eukaryota | 63576.0 | 0 | Chordata | 63576.0 | 0 | Carnivora | 43371.0 | 3 | Felidae | 33729.0 | 5 | Panthera | 13572.0 | 7 | Panthera tigri | 13572.0 | 7 |
| scf7180000017766 | 12591 | 0.3394 | 0 | 23.1654 | Eukaryota | 9764.0 | 0 | Chordata | 9764.0 | 0 | Carnivora | 9764.0 | 0 | Felidae | 9764.0 | 0 | Felis | 9764.0 | 0 | Felis catus | 9764.0 | 0 |
| scf7180000017514 | 2915107 | 0.4034 | 0 | 51.3853 | Eukaryota | 129506.0 | 0 | Chordata | 129506.0 | 0 | Carnivora | 129506.0 | 0 | Canidae | 83039.0 | 1 | Canis | 83039.0 | 2 | Canis lupus | 83039.0 | 2 |
| scf7180000020203 | 102895 | 0.4189 | 0 | 48.85 | Eukaryota | 15872.0 | 0 | Chordata | 15872.0 | 0 | Carnivora | 15872.0 | 0 | Herpestidae | 8348.0 | 1 | Suricata | 8348.0 | 1 | ricata suricat | 8348.0 | 1 |
| scf7180000019396 | 331042 | 0.4225 | 0 | 51.5693 | Eukaryota | 31214.0 | 0 | Chordata | 31214.0 | 0 | Carnivora | 28765.0 | 1 | Felidae | 17144.0 | 4 | Panthera | 8215.0 | 7 | ricata suricat | 5960.0 | 8 |
| scf7180000019381 | 462577 | 0.614 | 0 | 51.382 | Eukaryota | 29901.0 | 0 | Chordata | 29901.0 | 0 | Carnivora | 19090.0 | 2 | Felidae | 19090.0 | 3 | Felis | 14428.0 | 4 | Felis catus | 14428.0 | 4 |
| scf7180000019682 | 1778000 | 0.3933 | 100 | 50.3625 | Eukaryota | 46159.0 | 0 | Chordata | 46159.0 | 0 | Carnivora | 46159.0 | 0 | Felidae | 46159.0 | 0 | Felis | 46159.0 | 0 | Felis catus | 46159.0 | 0 |
| scf7180000024101 | 275448 | 0.4147 | 0 | 25.7895 | Eukaryota | 50423.0 | 0 | Chordata | 50423.0 | 0 | Primates | 29079.0 | 2 | Hominidae | 29079.0 | 3 | Homo | 11668.0 | 5 | Homo sapiens | 11668.0 | 5 |
| scf7180000018498 | 2224695 | 0.4337 | 0 | 50.5512 | Eukaryota | 48788.0 | 0 | Chordata | 48788.0 | 0 | Carnivora | 48788.0 | 0 | Felidae | 37600.0 | 1 | Lynx | 23710.0 | 2 | ynx canadens | 23710.0 | 2 |
| scf7180000021014 | 87525 | 0.3636 | 0 | 25.1993 | Eukaryota | 24092.0 | 0 | Chordata | 24092.0 | 0 | Carnivora | 24092.0 | 0 | Felidae | 24092.0 | 0 | Felis | 24092.0 | 0 | Felis catus | 24092.0 | 0 |
| scf7180000017751 | 28553 | 0.439 | 0 | 30.3466 | Eukaryota | 5811.0 | 0 | Chordata | 5811.0 | 0 | Artiodactyla | 1811.0 | 3 | Bovidae | 1811.0 | 3 | Homo | 1746.0 | 4 | Homo sapiens | 1746.0 | 4 |
| scf7180000024214 | 56193 | 0.413 | 0 | 24.9796 | Eukaryota | 17537.0 | 0 | Chordata | 17537.0 | 0 | Carnivora | 17537.0 | 0 | Felidae | 17537.0 | 0 | Felis | 15765.0 | 1 | Felis catus | 15765.0 | 1 |
| scf7180000019843 | 241759 | 0.3999 | 0 | 50.2582 | Eukaryota | 45383.0 | 0 | Chordata | 45383.0 | 0 | Carnivora | 45383.0 | 0 | Felidae | 45383.0 | 0 | Felis | 45383.0 | 0 | Felis catus | 45383.0 | 0 |
| scf7180000019064 | 852846 | 0.4821 | 0 | 53.6638 | Eukaryota | 54177.0 | 0 | Chordata | 54177.0 | 0 | Carnivora | 54177.0 | 0 | Felidae | 27630.0 | 2 | Suricata | 16251.0 | 5 | ricata suricat | 16251.0 | 7 |
| scf7180000022404 | 108457 | 0.3708 | 0 | 41.9109 | Eukaryota | 40564.0 | 0 | Chordata | 40564.0 | 0 | Carnivora | 40564.0 | 0 | Felidae | 40564.0 | 0 | Felis | 36521.0 | 1 | Felis catus | 36521.0 | 1 |
| scf7180000024008 | 615099 | 0.4561 | 0 | 50.5889 | Eukaryota | 39734.0 | 0 | Chordata | 39734.0 | 0 | Carnivora | 39734.0 | 0 | Felidae | 39734.0 | 0 | Felis | 36452.0 | 1 | Felis catus | 36452.0 | 1 |
| scf7180000020866 | 101498 | 0.399 | 0 | 24.6551 | Eukaryota | 28411.0 | 0 | Chordata | 28411.0 | 0 | Carnivora | 28411.0 | 0 | Felidae | 28411.0 | 0 | Felis | 16635.0 | 2 | Felis catus | 16635.0 | 2 |
| scf7180000024223 | 19440 | 0.394 | 0 | 24.3996 | Eukaryota | 15248.0 | 0 | Chordata | 15248.0 | 0 | Carnivora | 15248.0 | 0 | Felidae | 15248.0 | 0 | Felis | 10420.0 | 1 | Felis catus | 10420.0 | 1 |
| scf7180000021936 | 19218 | 0.41 | 0 | 25.8985 | Eukaryota | 10928.0 | 0 | Chordata | 10928.0 | 0 | Carnivora | 10928.0 | 0 | Felidae | 8561.0 | 1 | Felis | 8561.0 | 1 | Felis catus | 8561.0 | 1 |
| scf7180000020752 | 60303 | 0.4382 | 0 | 24.769 | Eukaryota | 26050.0 | 0 | Chordata | 26050.0 | 0 | Carnivora | 26050.0 | 0 | Felidae | 21074.0 | 2 | Panthera | 10596.0 | 4 | unresolved | 5298.0 | 5 |
| scf7180000024109 | 241835 | 0.4058 | 0 | 27.6849 | Eukaryota | 45977.0 | 0 | Chordata | 45977.0 | 0 | Carnivora | 45977.0 | 0 | Felidae | 45977.0 | 0 | Felis | 41445.0 | 1 | Felis catus | 41445.0 | 1 |
| scf7180000017972 | 315287 | 0.4755 | 0 | 49.7217 | Eukaryota | 11669.0 | 0 | Chordata | 11669.0 | 0 | Carnivora | 8509.0 | 2 | Felidae | 8509.0 | 2 | Felis | 6384.0 | 3 | Felis catus | 6384.0 | 3 |
| scf7180000023128 | 30759 | 0.4433 | 0 | 213.8313 | Eukaryota | 12074.0 | 0 | Chordata | 12074.0 | 0 | Carnivora | 12074.0 | 0 | Felidae | 12074.0 | 0 | Felis | 10875.0 | 1 | Felis catus | 10875.0 | 1 |
| scf7180000022628 | 7330 | 0.434 | 0 | 23.6355 | Eukaryota | 20559.0 | 0 | Chordata | 20559.0 | 0 | Carnivora | 20559.0 | 0 | Felidae | 20559.0 | 0 | Felis | 16421.0 | 1 | Felis catus | 16421.0 | 1 |
| scf7180000019429 | 349795 | 0.4297 | 0 | 53.1147 | Eukaryota | 38973.0 | 0 | Chordata | 38973.0 | 0 | Carnivora | 38973.0 | 0 | Felidae | 38973.0 | 0 | Felis | 14221.0 | 2 | Felis catus | 14221.0 | 2 |
| scf7180000018161 | 8909 | 0.3922 | 0 | 23.3774 | Eukaryota | 2593.0 | 0 | Chordata | 2593.0 | 0 | Carnivora | 2593.0 | 0 | Felidae | 2326.0 | 1 | Felis | 1800.0 | 2 | Felis catus | 1800.0 | 2 |
| scf7180000022376 | 9157 | 0.5278 | 0 | 479.9262 | no-hit | 0.0 | 0 | no-hit | 0.0 | 0 | no-hit | 0.0 | 0 | no-hit | 0.0 | 0 | no-hit | 0.0 | 0 | no-hit | 0.0 | 0 |
| scf7180000022797 | 15937 | 0.3755 | 0 | 24.7078 | Eukaryota | 4783.0 | 0 | Chordata | 4783.0 | 0 | Carnivora | 4783.0 | 0 | Felidae | 4783.0 | 0 | Felis | 3363.0 | 1 | Felis catus | 3363.0 | 1 |
| scf7180000017812 | 1725402 | 0.4199 | 0 | 51.5673 | Eukaryota | 118846.0 | 0 | Chordata | 118846.0 | 0 | Carnivora | 118846.0 | 0 | Felidae | 118846.0 | 0 | Lynx | 71394.0 | 1 | ynx canadens | 71394.0 | 1 |
| scf7180000019047 | 703024 | 0.3664 | 100 | 50.0618 | Eukaryota | 44023.0 | 0 | Chordata | 44023.0 | 0 | Carnivora | 44023.0 | 0 | Felidae | 44023.0 | 0 | Felis | 44023.0 | 0 | Felis catus | 44023.0 | 0 |
| scf7180000020216 | 220523 | 0.3865 | 0 | 50.9213 | Eukaryota | 21674.0 | 1 | Chordata | 21674.0 | 1 | Primates | 7804.0 | 4 | Delphinidae | 5753.0 | 5 | Homo | 5419.0 | 6 | Homo sapiens | 5419.0 | 6 |
| scf7180000021194 | 5737 | 0.522 | 0 | 195.4913 | no-hit | 0.0 | 0 | no-hit | 0.0 | 0 | no-hit | 0.0 | 0 | no-hit | 0.0 | 0 | no-hit | 0.0 | 0 | no-hit | 0.0 | 0 |
| scf7180000018362 | 831463 | 0.3673 | 0 | 49.1154 | Eukaryota | 47282.0 | 0 | Chordata | 47282.0 | 0 | Carnivora | 47282.0 | 0 | Felidae | 47282.0 | 0 | Felis | 47282.0 | 0 | Felis catus | 47282.0 | 0 |
| scf7180000024063 | 371925 | 0.4541 | 0 | 49.7939 | Eukaryota | 22355.0 | 0 | Chordata | 22355.0 | 0 | Carnivora | 22355.0 | 0 | Felidae | 22355.0 | 0 | Panthera | 7403.0 | 4 | cinonyx jubat | 5252.0 | 5 |
| scf7180000021129 | 7576 | 0.479 | 0 | 23.505 | Eukaryota | 1988.0 | 0 | Chordata | 1988.0 | 0 | Carnivora | 1988.0 | 0 | Felidae | 1988.0 | 0 | Felis | 1988.0 | 0 | Felis catus | 1988.0 | 0 |
| scf7180000022004 | 17563 | 0.3628 | 0 | 25.5135 | Eukaryota | 9488.0 | 0 | Chordata | 9488.0 | 0 | Carnivora | 9488.0 | 0 | Felidae | 9488.0 | 0 | Felis | 8394.0 | 1 | Felis catus | 8394.0 | 1 |
| scf7180000021135 | 64204 | 0.3668 | 0 | 47.6769 | Eukaryota | 41033.0 | 0 | Chordata | 41033.0 | 0 | Carnivora | 41033.0 | 0 | Felidae | 41033.0 | 0 | Felis | 36973.0 | 1 | Felis catus | 36973.0 | 1 |
| scf7180000022447 | 8598 | 0.5295 | 0 | 123.8637 | Eukaryota | 12002.0 | 0 | Chordata | 12002.0 | 0 | Carnivora | 12002.0 | 0 | Felidae | 12002.0 | 0 | Felis | 12002.0 | 0 | Felis catus | 12002.0 | 0 |
| scf7180000019639 | 277807 | 0.3495 | 0 | 49.2626 | Eukaryota | 39766.0 | 0 | Chordata | 39766.0 | 0 | Carnivora | 39766.0 | 0 | Felidae | 39766.0 | 0 | Felis | 35823.0 | 1 | Felis catus | 35823.0 | 1 |
| scf7180000023643 | 44510 | 0.4533 | 0 | 251.0643 | Eukaryota | 2636.0 | 0 | Chordata | 2636.0 | 0 | Carnivora | 2221.0 | 1 | Hyaenidae | 774.0 | 4 | Crocuta | 774.0 | 5 | rocuta crocut | 774.0 | 5 |
| scf7180000018708 | 694983 | 0.4923 | 0 | 54.7921 | Eukaryota | 53532.0 | 0 | Chordata | 53532.0 | 0 | Carnivora | 53532.0 | 0 | Felidae | 41853.0 | 1 | Felis | 22405.0 | 3 | Felis catus | 22405.0 | 4 |
| scf7180000023294 | 78662 | 0.4465 | 0 | 435.6262 | Eukaryota | 13305.0 | 0 | Chordata | 13305.0 | 0 | Carnivora | 13305.0 | 0 | Felidae | 13305.0 | 0 | Felis | 10671.0 | 1 | Felis catus | 10671.0 | 1 |
| scf7180000019236 | 480826 | 0.3489 | 0 | 48.7326 | Eukaryota | 47616.0 | 0 | Chordata | 47616.0 | 0 | Carnivora | 47616.0 | 0 | Felidae | 47616.0 | 0 | Felis | 47616.0 | 0 | Felis catus | 47616.0 | 0 |
| scf7180000018866 | 538928 | 0.3589 | 0 | 49.5977 | Eukaryota | 46858.0 | 0 | Chordata | 46858.0 | 0 | Carnivora | 46858.0 | 0 | Felidae | 46858.0 | 0 | Felis | 46858.0 | 0 | Felis catus | 46858.0 | 0 |
| scf7180000020296 | 108273 | 0.475 | 0 | 46.595 | Eukaryota | 26518.0 | 0 | Chordata | 26518.0 | 0 | Primates | 10730.0 | 3 | Hominidae | 7263.0 | 5 | Homo | 7263.0 | 6 | Homo sapiens | 7263.0 | 7 |
| scf7180000018142 | 2484635 | 0.4512 | 100 | 52.0011 | Eukaryota | 47753.0 | 0 | Chordata | 47753.0 | 0 | Carnivora | 47753.0 | 0 | Felidae | 47753.0 | 0 | Felis | 38035.0 | 1 | Felis catus | 38035.0 | 1 |
| scf7180000019217 | 401490 | 0.3786 | 0 | 50.7388 | Eukaryota | 45279.0 | 0 | Chordata | 45279.0 | 0 | Carnivora | 45279.0 | 0 | Felidae | 45279.0 | 0 | Felis | 40821.0 | 1 | Felis catus | 40821.0 | 1 |
| scf7180000017456 | 1334876 | 0.3706 | 0 | 50.1142 | Eukaryota | 46924.0 | 0 | Chordata | 46924.0 | 0 | Carnivora | 46924.0 | 0 | Felidae | 41958.0 | 1 | Felis | 37435.0 | 2 | Felis catus | 37435.0 | 2 |
| scf7180000020550 | 66207 | 0.4195 | 0 | 826.899 | Eukaryota | 13569.0 | 0 | Chordata | 13569.0 | 0 | Carnivora | 13569.0 | 0 | Felidae | 13569.0 | 0 | Felis | 12226.0 | 1 | Felis catus | 12226.0 | 1 |
| scf7180000021432 | 48829 | 0.5354 | 0 | 49.4776 | Eukaryota | 7647.0 | 0 | Chordata | 7647.0 | 0 | Artiodactyla | 1966.0 | 5 | Bovidae | 1966.0 | 5 | Felis | 1456.0 | 7 | Felis catus | 1456.0 | 7 |
| scf7180000020426 | 121486 | 0.3945 | 0 | 25.5176 | Eukaryota | 18084.0 | 0 | Chordata | 18084.0 | 0 | Carnivora | 18084.0 | 0 | Felidae | 18084.0 | 0 | Felis | 10898.0 | 2 | Felis catus | 10898.0 | 2 |
| scf7180000021571 | 45665 | 0.533 | 0 | 48.8734 | Eukaryota | 3507.0 | 0 | Chordata | 3507.0 | 0 | Carnivora | 3507.0 | 0 | Felidae | 3196.0 | 1 | Felis | 2403.0 | 2 | Felis catus | 2403.0 | 2 |
| scf7180000018417 | 185479 | 0.3855 | 0 | 52.8723 | Eukaryota | 61473.0 | 0 | Chordata | 61473.0 | 0 | Carnivora | 61473.0 | 0 | Felidae | 51256.0 | 1 | Panthera | 45989.0 | 3 | Panthera tigri | 24818.0 | 4 |
| scf7180000022430 | 14303 | 0.4316 | 0 | 26.4144 | Eukaryota | 4349.0 | 0 | Chordata | 4349.0 | 0 | Artiodactyla | 3306.0 | 1 | Bovidae | 3306.0 | 2 | Ovis | 3306.0 | 2 | Ovis canadensi | 3306.0 | 2 |
| scf7180000021640 | 42658 | 0.4735 | 0 | 27.6431 | Eukaryota | 3580.0 | 0 | Chordata | 3580.0 | 0 | Carnivora | 1743.0 | 2 | Felidae | 1743.0 | 2 | Felis | 1427.0 | 3 | Felis catus | 1427.0 | 3 |
| scf7180000019864 | 245848 | 0.5462 | 0 | 52.5628 | Eukaryota | 14005.0 | 0 | Chordata | 14005.0 | 0 | Carnivora | 14005.0 | 0 | Otariidae | 4056.0 | 4 | Suricata | 2699.0 | 8 | ricata suricat | 2699.0 | 8 |
| scf7180000020467 | 127511 | 0.382 | 0 | 26.28 | Eukaryota | 38865.0 | 0 | Chordata | 38865.0 | 0 | Carnivora | 38865.0 | 0 | Felidae | 38865.0 | 0 | Felis | 38865.0 | 0 | Felis catus | 38865.0 | 0 |
| scf7180000019625 | 203748 | 0.442 | 0 | 50.9175 | Eukaryota | 16111.0 | 0 | Chordata | 16111.0 | 0 | Primates | 7911.0 | 2 | Hominidae | 7911.0 | 2 | Homo | 7911.0 | 3 | Homo sapiens | 7911.0 | 3 |
| scf7180000022638 | 10372 | 0.5168 | 0 | 44.1567 | Eukaryota | 6770.0 | 0 | Chordata | 6770.0 | 0 | Carnivora | 6770.0 | 0 | Felidae | 4205.0 | 1 | Canis | 2565.0 | 4 | Canis lupus | 2565.0 | 4 |
| scf7180000021458 | 80449 | 0.6058 | 0 | 47.045 | Eukaryota | 10174.0 | 0 | Chordata | 10174.0 | 0 | Rodentia | 2594.0 | 7 | Muridae | 2594.0 | 7 | Mus | 2594.0 | 7 | Mus musculus | 2594.0 | 7 |
| scf7180000019732 | 354911 | 0.4742 | 0 | 52.167 | Eukaryota | 8345.0 | 0 | Chordata | 8345.0 | 0 | Carnivora | 8345.0 | 0 | Felidae | 8345.0 | 0 | Felis | 8345.0 | 0 | Felis catus | 8345.0 | 0 |
| scf7180000022437 | 6152 | 0.6702 | 0 | 370.059 | Eukaryota | 606.0 | 0 | Chordata | 471.0 | 1 | Carnivora | 471.0 | 1 | Felidae | 334.0 | 2 | Felis | 334.0 | 2 | Felis catus | 334.0 | 2 |
| scf7180000020970 | 28007 | 0.3897 | 0 | 25.15 | Eukaryota | 42871.0 | 0 | Chordata | 42871.0 | 0 | Carnivora | 42871.0 | 0 | Felidae | 42871.0 | 0 | Felis | 38610.0 | 1 | Felis catus | 38610.0 | 1 |
| scf7180000019403 | 797743 | 0.4182 | 0 | 79.5231 | Eukaryota | 46998.0 | 0 | Chordata | 46998.0 | 0 | Carnivora | 46998.0 | 0 | Felidae | 46998.0 | 0 | Felis | 46998.0 | 0 | Felis catus | 46998.0 | 0 |
| scf7180000023898 | 1486229 | 0.4277 | 0 | 49.1958 | Eukaryota | 57221.0 | 0 | Chordata | 57221.0 | 0 | Carnivora | 57221.0 | 0 | Felidae | 57221.0 | 0 | Lynx | 40616.0 | 2 | ynx canadens | 40616.0 | 2 |
| scf7180000020741 | 88214 | 0.363 | 0 | 25.2747 | Eukaryota | 26956.0 | 0 | Chordata | 26956.0 | 0 | Carnivora | 26956.0 | 0 | Felidae | 26956.0 | 0 | Felis | 23238.0 | 1 | Felis catus | 23238.0 | 1 |
| scf7180000020297 | 99649 | 0.4495 | 0 | 48.4543 | Eukaryota | 39289.0 | 0 | Chordata | 39289.0 | 0 | Carnivora | 39289.0 | 0 | Felidae | 39289.0 | 0 | Felis | 39289.0 | 0 | Felis catus | 39289.0 | 0 |
| scf7180000020337 | 150445 | 0.3772 | 0 | 25.3831 | Eukaryota | 76995.0 | 0 | Chordata | 76995.0 | 0 | Carnivora | 76995.0 | 0 | Felidae | 76995.0 | 0 | Felis | 34157.0 | 3 | Felis catus | 34157.0 | 4 |
| scf7180000022527 | 7744 | 0.4595 | 0 | 24.8488 | Eukaryota | 4873.0 | 0 | Chordata | 4873.0 | 0 | Carnivora | 4873.0 | 0 | Felidae | 4873.0 | 0 | Felis | 4322.0 | 1 | Felis catus | 4322.0 | 1 |
| scf7180000020473 | 125001 | 0.3826 | 0 | 27.4868 | Eukaryota | 46236.0 | 0 | Chordata | 46236.0 | 0 | Carnivora | 46236.0 | 0 | Felidae | 46236.0 | 0 | Felis | 46236.0 | 0 | Felis catus | 46236.0 | 0 |
| scf7180000018168 | 12547 | 0.3611 | 0 | 26.955 | Eukaryota | 8332.0 | 0 | Chordata | 8332.0 | 0 | Carnivora | 7401.0 | 1 | Felidae | 7401.0 | 1 | Felis | 7401.0 | 1 | Felis catus | 7401.0 | 1 |
| scf7180000021401 | 36104 | 0.3789 | 0 | 23.3965 | Eukaryota | 42683.0 | 0 | Chordata | 42683.0 | 0 | Carnivora | 42683.0 | 0 | Felidae | 42683.0 | 0 | Felis | 42683.0 | 0 | Felis catus | 42683.0 | 0 |
| scf7180000019525 | 240300 | 0.4821 | 0 | 49.2852 | Eukaryota | 19861.0 | 0 | Chordata | 19861.0 | 0 | Carnivora | 19861.0 | 0 | Felidae | 19861.0 | 0 | Felis | 17870.0 | 1 | Felis catus | 17870.0 | 1 |
| scf7180000021472 | 39271 | 0.4273 | 0 | 26.2553 | Eukaryota | 45383.0 | 0 | Chordata | 45383.0 | 0 | Carnivora | 45383.0 | 0 | Felidae | 45383.0 | 0 | Felis | 45383.0 | 0 | Felis catus | 45383.0 | 0 |
| scf7180000019740 | 543064 | 0.4642 | 0 | 49.8457 | Eukaryota | 44023.0 | 0 | Chordata | 44023.0 | 0 | Carnivora | 44023.0 | 0 | Felidae | 23907.0 | 1 | Suricata | 20116.0 | 2 | ricata suricat | 20116.0 | 3 |
| scf7180000020577 | 115133 | 0.4341 | 0 | 24.7039 | Eukaryota | 7467.0 | 0 | Chordata | 6999.0 | 1 | Primates | 3138.0 | 4 | Hominidae | 3138.0 | 5 | Homo | 3138.0 | 6 | Homo sapiens | 3138.0 | 6 |
| scf7180000018677 | 117054 | 0.375 | 0 | 23.7128 | Eukaryota | 34923.0 | 0 | Chordata | 34923.0 | 0 | Carnivora | 34923.0 | 0 | Felidae | 34923.0 | 0 | Felis | 34923.0 | 0 | Felis catus | 34923.0 | 0 |
| scf7180000017442 | 1342318 | 0.3642 | 0 | 49.804 | Eukaryota | 45484.0 | 0 | Chordata | 45484.0 | 0 | Carnivora | 45484.0 | 0 | Felidae | 45484.0 | 0 | Felis | 40922.0 | 1 | Felis catus | 40922.0 | 1 |
| scf7180000018306 | 525638 | 0.3727 | 0 | 49.6602 | Eukaryota | 46694.0 | 0 | Chordata | 46694.0 | 0 | Carnivora | 46694.0 | 0 | Felidae | 46694.0 | 0 | Felis | 46694.0 | 0 | Felis catus | 46694.0 | 0 |
| scf7180000021028 | 103129 | 0.3663 | 0 | 24.348 | Eukaryota | 31559.0 | 0 | Chordata | 31559.0 | 0 | Carnivora | 31559.0 | 0 | Felidae | 31559.0 | 0 | Felis | 31559.0 | 0 | Felis catus | 31559.0 | 0 |
| scf7180000021025 | 97515 | 0.3768 | 0 | 23.8322 | Eukaryota | 37475.0 | 0 | Chordata | 37475.0 | 0 | Carnivora | 37475.0 | 0 | Felidae | 37475.0 | 0 | Felis | 33824.0 | 1 | Felis catus | 33824.0 | 1 |
| scf7180000020875 | 75436 | 0.43 | 0 | 263.5129 | Eukaryota | 2155.0 | 0 | Chordata | 2155.0 | 0 | Carnivora | 2155.0 | 0 | Felidae | 2155.0 | 0 | Felis | 1729.0 | 2 | Felis catus | 1729.0 | 2 |
| scf7180000024267 | 1731863 | 0.4357 | 100 | 49.4478 | Eukaryota | 39051.0 | 0 | Chordata | 39051.0 | 0 | Carnivora | 39051.0 | 0 | Felidae | 39051.0 | 0 | Felis | 39051.0 | 0 | Felis catus | 39051.0 | 0 |
| scf7180000020608 | 124068 | 0.4351 | 0 | 24.2733 | Eukaryota | 40359.0 | 0 | Chordata | 40359.0 | 0 | Primates | 24037.0 | 3 | Hominidae | 22273.0 | 4 | Homo | 14176.0 | 5 | Homo sapiens | 14176.0 | 5 |
| scf7180000022349 | 14109 | 0.3867 | 0 | 25.2251 | Eukaryota | 38494.0 | 0 | Chordata | 38494.0 | 0 | Carnivora | 38494.0 | 0 | Felidae | 38494.0 | 0 | Felis | 38494.0 | 0 | Felis catus | 38494.0 | 0 |
| scf7180000020157 | 120963 | 0.3738 | 0 | 49.2742 | Eukaryota | 9009.0 | 0 | Chordata | 9009.0 | 0 | Carnivora | 6644.0 | 2 | Felidae | 6644.0 | 2 | Felis | 6644.0 | 2 | Felis catus | 6644.0 | 2 |
| scf7180000020448 | 144134 | 0.442 | 0 | 26.0163 | Eukaryota | 53371.0 | 0 | Chordata | 53371.0 | 0 | Primates | 32268.0 | 2 | Hominidae | 27154.0 | 3 | Homo | 27154.0 | 3 | Homo sapiens | 27154.0 | 5 |
| scf7180000019178 | 769009 | 0.3775 | 0 | 50.3252 | Eukaryota | 47724.0 | 0 | Chordata | 47724.0 | 0 | Carnivora | 47724.0 | 0 | Felidae | 47724.0 | 0 | Felis | 47724.0 | 0 | Felis catus | 47724.0 | 0 |
| scf7180000023454 | 81809 | 0.3482 | 0 | 439.5058 | Eukaryota | 5904.0 | 0 | Chordata | 5904.0 | 0 | Carnivora | 5074.0 | 1 | Felidae | 5074.0 | 1 | Felis | 5074.0 | 1 | Felis catus | 5074.0 | 1 |
| scf7180000021960 | 35563 | 0.4507 | 0 | 25.8403 | Eukaryota | 8971.0 | 0 | Chordata | 8971.0 | 0 | Carnivora | 5331.0 | 2 | Felidae | 5331.0 | 2 | Felis | 5331.0 | 2 | Felis catus | 5331.0 | 2 |
| scf7180000017510 | 3245181 | 0.4619 | 0 | 51.3882 | Eukaryota | 53415.0 | 0 | Chordata | 53415.0 | 0 | Carnivora | 53415.0 | 0 | Felidae | 23025.0 | 4 | Felis | 18423.0 | 6 | Felis catus | 18423.0 | 6 |
| scf7180000020667 | 44445 | 0.4325 | 0 | 30.5244 | Eukaryota | 22943.0 | 0 | Chordata | 22943.0 | 0 | Carnivora | 22943.0 | 0 | Felidae | 22943.0 | 0 | Felis | 18469.0 | 1 | Felis catus | 18469.0 | 1 |
| scf7180000021422 | 65401 | 0.4166 | 0 | 25.3965 | Eukaryota | 33536.0 | 0 | Chordata | 33536.0 | 0 | Carnivora | 33536.0 | 0 | Felidae | 33536.0 | 0 | Felis | 30529.0 | 1 | Felis catus | 30529.0 | 1 |
| scf7180000017960 | 1285793 | 0.4033 | 0 | 50.1853 | Eukaryota | 47694.0 | 0 | Chordata | 47694.0 | 0 | Carnivora | 47694.0 | 0 | Felidae | 47694.0 | 0 | Felis | 47694.0 | 0 | Felis catus | 47694.0 | 0 |
| scf7180000021057 | 66087 | 0.4421 | 0 | 25.1197 | Eukaryota | 5031.0 | 0 | Chordata | 5031.0 | 0 | Primates | 2245.0 | 3 | Hominidae | 2245.0 | 4 | Homo | 1526.0 | 7 | Homo sapiens | 1526.0 | 7 |
| scf7180000018092 | 1206822 | 0.3887 | 0 | 51.9404 | Eukaryota | 47477.0 | 0 | Chordata | 47477.0 | 0 | Carnivora | 47477.0 | 0 | Felidae | 47477.0 | 0 | Felis | 47477.0 | 0 | Felis catus | 47477.0 | 0 |
| scf7180000024184 | 89871 | 0.4425 | 0 | 51.4583 | Eukaryota | 6492.0 | 1 | Chordata | 6492.0 | 1 | Carnivora | 3802.0 | 2 | Felidae | 3802.0 | 2 | Felis | 3802.0 | 2 | Felis catus | 3802.0 | 2 |
| scf7180000022097 | 25311 | 0.3694 | 0 | 24.8221 | Eukaryota | 4241.0 | 0 | Chordata | 4241.0 | 0 | Carnivora | 4241.0 | 0 | Felidae | 3499.0 | 1 | Felis | 3499.0 | 1 | Felis catus | 3499.0 | 1 |
| scf7180000022778 | 9105 | 0.3807 | 0 | 24.1381 | Eukaryota | 3066.0 | 0 | Chordata | 3066.0 | 0 | Carnivora | 3066.0 | 0 | Felidae | 1836.0 | 1 | Felis | 1545.0 | 2 | Felis catus | 1545.0 | 2 |
| scf7180000018153 | 1079582 | 0.5023 | 0 | 51.2526 | Eukaryota | 29280.0 | 0 | Chordata | 29280.0 | 0 | Carnivora | 29280.0 | 0 | Canidae | 29280.0 | 0 | Canis | 29280.0 | 0 | Canis lupus | 29280.0 | 0 |
| scf7180000018406 | 847059 | 0.4084 | 0 | 50.9296 | Eukaryota | 47037.0 | 0 | Chordata | 47037.0 | 0 | Carnivora | 47037.0 | 0 | Felidae | 47037.0 | 0 | Felis | 47037.0 | 0 | Felis catus | 47037.0 | 0 |
| scf7180000020175 | 15576 | 0.37 | 0 | 24.2794 | Eukaryota | 14242.0 | 0 | Chordata | 14242.0 | 0 | Carnivora | 14242.0 | 0 | Felidae | 14242.0 | 0 | Felis | 7551.0 | 3 | Felis catus | 7551.0 | 3 |
| scf7180000017525 | 3579852 | 0.4182 | 0 | 52.3542 | Eukaryota | 62853.0 | 0 | Chordata | 62853.0 | 0 | Carnivora | 56986.0 | 1 | Canidae | 18438.0 | 5 | Canis | 18438.0 | 6 | Canis lupus | 18438.0 | 6 |
| scf7180000019408 | 288870 | 0.3974 | 0 | 48.829 | Eukaryota | 45423.0 | 0 | Chordata | 45423.0 | 0 | Carnivora | 45423.0 | 0 | Felidae | 45423.0 | 0 | Felis | 45423.0 | 0 | Felis catus | 45423.0 | 0 |
| scf7180000017455 | 2531333 | 0.3588 | 0 | 50.8071 | Eukaryota | 47261.0 | 0 | Chordata | 47261.0 | 0 | Carnivora | 47261.0 | 0 | Felidae | 47261.0 | 0 | Felis | 47261.0 | 0 | Felis catus | 47261.0 | 0 |
| scf7180000018208 | 983898 | 0.478 | 0 | 53.1076 | Eukaryota | 52740.0 | 0 | Chordata | 52740.0 | 0 | Carnivora | 52740.0 | 0 | Felidae | 43059.0 | 2 | Panthera | 22832.0 | 5 | Panthera tigri | 11421.0 | 6 |
| scf7180000019475 | 300315 | 0.3818 | 0 | 49.5769 | Eukaryota | 21962.0 | 0 | Chordata | 21962.0 | 0 | Carnivora | 20144.0 | 1 | Felidae | 15079.0 | 3 | Panthera | 4413.0 | 8 | Canis lupus | 2839.0 | 9 |
| scf7180000018439 | 861307 | 0.3875 | 0 | 52.3338 | Eukaryota | 63770.0 | 0 | Chordata | 63770.0 | 0 | Carnivora | 63770.0 | 0 | Felidae | 43771.0 | 3 | Felis | 31144.0 | 5 | Felis catus | 31144.0 | 5 |
| scf7180000024206 | 63738 | 0.3508 | 0 | 23.2419 | Eukaryota | 22126.0 | 0 | Chordata | 22126.0 | 0 | Carnivora | 22126.0 | 0 | Felidae | 17554.0 | 2 | Felis | 17554.0 | 2 | Felis catus | 17554.0 | 2 |
| scf7180000022411 | 16612 | 0.4698 | 0 | 25.4693 | Eukaryota | 13909.0 | 0 | Chordata | 13909.0 | 0 | Carnivora | 13909.0 | 0 | Felidae | 13909.0 | 0 | Felis | 12540.0 | 1 | Felis catus | 12540.0 | 1 |
| scf7180000024104 | 344857 | 0.3968 | 0 | 50.505 | Eukaryota | 14383.0 | 0 | Chordata | 14383.0 | 0 | Carnivora | 7310.0 | 2 | Felidae | 7310.0 | 3 | Felis | 3967.0 | 7 | Felis catus | 3967.0 | 7 |
| scf7180000021841 | 35332 | 0.563 | 0 | 48.4599 | Eukaryota | 9521.0 | 0 | Chordata | 9521.0 | 0 | Squamata | 2983.0 | 4 | Bovidae | 2345.0 | 6 | Homo | 1310.0 | 9 | Homo sapiens | 1310.0 | 9 |
| scf7180000021735 | 27399 | 0.4205 | 0 | 22.8175 | Eukaryota | 7775.0 | 0 | Chordata | 7775.0 | 0 | Carnivora | 7775.0 | 0 | Felidae | 7775.0 | 0 | Felis | 7775.0 | 0 | Felis catus | 7775.0 | 0 |
| scf7180000024242 | 346216 | 0.4941 | 0 | 49.7402 | Eukaryota | 30641.0 | 0 | Chordata | 30641.0 | 0 | Carnivora | 21319.0 | 1 | Felidae | 17893.0 | 4 | Physeter | 4968.0 | 7 | yseter catod | 4968.0 | 7 |
| scf7180000022148 | 7926 | 0.5276 | 0 | 343.4078 | Eukaryota | 145.0 | 0 | latyhelminthe | 145.0 | 0 | olystomatide | 145.0 | 0 | Polystomatidae | 145.0 | 0 | rotopolystom | 145.0 | 0 | olystoma xen | 145.0 | 0 |
| scf7180000020677 | 21979 | 0.4526 | 0 | 27.9263 | Eukaryota | 15678.0 | 0 | Chordata | 15678.0 | 0 | Carnivora | 15678.0 | 0 | Felidae | 12747.0 | 1 | Felis | 12747.0 | 1 | Felis catus | 12747.0 | 1 |
| scf7180000023395 | 9290 | 0.5236 | 0 | 204.5115 | no-hit | 0.0 | 0 | no-hit | 0.0 | 0 | no-hit | 0.0 | 0 | no-hit | 0.0 | 0 | no-hit | 0.0 | 0 | no-hit | 0.0 | 0 |
| scf7180000021823 | 30383 | 0.4877 | 0 | 48.7777 | Eukaryota | 46374.0 | 0 | Chordata | 46374.0 | 0 | Carnivora | 46374.0 | 0 | Felidae | 46374.0 | 0 | Felis | 41762.0 | 1 | Felis catus | 41762.0 | 1 |
| scf7180000022231 | 9417 | 0.5211 | 0 | 130.871 | no-hit | 0.0 | 0 | no-hit | 0.0 | 0 | no-hit | 0.0 | 0 | no-hit | 0.0 | 0 | no-hit | 0.0 | 0 | no-hit | 0.0 | 0 |
| scf7180000022705 | 14030 | 0.4091 | 0 | 26.2668 | Eukaryota | 11132.0 | 0 | Chordata | 11132.0 | 0 | Carnivora | 11132.0 | 0 | Felidae | 11132.0 | 0 | Felis | 10095.0 | 1 | Felis catus | 10095.0 | 1 |
| scf7180000019831 | 79332 | 0.3847 | 0 | 38.8926 | Eukaryota | 44887.0 | 0 | Chordata | 44887.0 | 0 | Carnivora | 44887.0 | 0 | Felidae | 44887.0 | 0 | Felis | 44887.0 | 0 | Felis catus | 44887.0 | 0 |
| scf7180000023353 | 33196 | 0.3772 | 0 | 232.7574 | Eukaryota | 6664.0 | 0 | Chordata | 6664.0 | 0 | Carnivora | 6664.0 | 0 | Felidae | 6664.0 | 0 | Felis | 6664.0 | 0 | Felis catus | 6664.0 | 0 |
| scf7180000017555 | 916244 | 0.4511 | 0 | 51.2492 | Eukaryota | 44272.0 | 0 | Chordata | 44272.0 | 0 | Carnivora | 44272.0 | 0 | Felidae | 44272.0 | 0 | Felis | 30732.0 | 3 | Felis catus | 30732.0 | 3 |
| scf7180000020126 | 158142 | 0.4931 | 0 | 46.224 | Eukaryota | 31611.0 | 0 | Chordata | 31611.0 | 0 | Carnivora | 31611.0 | 0 | Felidae | 12764.0 | 3 | Ursus | 11061.0 | 4 | Ursus arctos | 11061.0 | 4 |
| scf7180000018157 | 937915 | 0.5449 | 0 | 51.0129 | Eukaryota | 22031.0 | 0 | Chordata | 22031.0 | 0 | Carnivora | 22031.0 | 0 | Felidae | 22031.0 | 0 | Lynx | 8836.0 | 3 | ynx canadens | 8836.0 | 3 |
| scf7180000020770 | 88542 | 0.5343 | 0 | 45.3907 | Eukaryota | 3062.0 | 0 | Chordata | 3062.0 | 0 | Carnivora | 3062.0 | 0 | Felidae | 3062.0 | 0 | Felis | 3062.0 | 0 | Felis catus | 3062.0 | 0 |
| scf7180000022013 | 39561 | 0.36 | 0 | 25.7438 | Eukaryota | 9917.0 | 0 | Chordata | 9917.0 | 0 | Carnivora | 5351.0 | 2 | Felidae | 5351.0 | 3 | Felis | 5351.0 | 4 | Felis catus | 5351.0 | 4 |
| scf7180000021436 | 1134220 | 0.5368 | 0 | 50.8162 | Eukaryota | 24721.0 | 0 | Chordata | 24721.0 | 0 | Carnivora | 24721.0 | 0 | Felidae | 22069.0 | 1 | Felis | 22069.0 | 1 | Felis catus | 22069.0 | 1 |
| scf7180000019587 | 212785 | 0.3853 | 0 | 45.6041 | Eukaryota | 44503.0 | 0 | Chordata | 44503.0 | 0 | Carnivora | 44503.0 | 0 | Felidae | 44503.0 | 0 | Felis | 44503.0 | 0 | Felis catus | 44503.0 | 0 |
| scf7180000021419 | 56329 | 0.3676 | 0 | 25.1412 | Eukaryota | 14418.0 | 0 | Chordata | 14418.0 | 0 | Carnivora | 14418.0 | 0 | Felidae | 14418.0 | 0 | Felis | 11558.0 | 1 | Felis catus | 11558.0 | 1 |
| scf7180000022237 | 12893 | 0.5221 | 0 | 90.1648 | no-hit | 0.0 | 0 | no-hit | 0.0 | 0 | no-hit | 0.0 | 0 | no-hit | 0.0 | 0 | no-hit | 0.0 | 0 | no-hit | 0.0 | 0 |
| scf7180000021800 | 22343 | 0.5036 | 0 | 49.022 | Eukaryota | 2536.0 | 0 | Chordata | 2536.0 | 0 | Carnivora | 2536.0 | 0 | Felidae | 2266.0 | 1 | Felis | 1498.0 | 2 | Felis catus | 1498.0 | 2 |
| scf7180000020048 | 20866 | 0.5624 | 0 | 99.4923 | Eukaryota | 6295.0 | 0 | Chordata | 6295.0 | 0 | Carnivora | 6295.0 | 0 | Felidae | 6041.0 | 1 | Felis | 2453.0 | 3 | Felis catus | 2453.0 | 3 |
| scf7180000022394 | 29445 | 0.3764 | 0 | 24.8716 | Eukaryota | 27031.0 | 0 | Chordata | 27031.0 | 0 | Carnivora | 27031.0 | 0 | Felidae | 27031.0 | 0 | Felis | 24257.0 | 1 | Felis catus | 24257.0 | 1 |
| scf7180000023884 | 870547 | 0.5575 | 0 | 51.953 | Eukaryota | 66117.0 | 0 | Chordata | 66117.0 | 0 | Carnivora | 66117.0 | 0 | Felidae | 53054.0 | 2 | Panthera | 13400.0 | 5 | anthera pard | 13400.0 | 5 |
| scf7180000021151 | 44829 | 0.4699 | 0 | 50.3259 | Eukaryota | 5274.0 | 1 | Chordata | 5274.0 | 1 | Carnivora | 2832.0 | 3 | Herpestidae | 2000.0 | 4 | Suricata | 2000.0 | 4 | ricata suricat | 2000.0 | 4 |
| scf7180000022602 | 7275 | 0.4047 | 0 | 25.3281 | Eukaryota | 2593.0 | 0 | Chordata | 2593.0 | 0 | Primates | 1830.0 | 1 | Hominidae | 1830.0 | 1 | Homo | 1228.0 | 3 | Homo sapiens | 1228.0 | 3 |
| scf7180000018972 | 392505 | 0.4906 | 0 | 51.283 | Eukaryota | 31001.0 | 0 | Chordata | 31001.0 | 0 | Carnivora | 31001.0 | 0 | Felidae | 31001.0 | 0 | Felis | 27907.0 | 1 | Felis catus | 27907.0 | 1 |
| scf7180000021752 | 26839 | 0.3845 | 0 | 24.7599 | Eukaryota | 35124.0 | 0 | Chordata | 35124.0 | 0 | Carnivora | 35124.0 | 0 | Felidae | 35124.0 | 0 | Felis | 26807.0 | 3 | Felis catus | 26807.0 | 3 |

| | | | | | | | | | | | | | | | | | | | |
|---|---|---|---|---|---|---|---|---|---|---|---|---|---|---|---|---|---|---|---|
| scf7180000021059 | 62698 | 0.3837 | 0 | 32.5349 | Eukaryota | 47556.0 | 0 | Chordata | 47556.0 | 0 | Carnivora | 47556.0 | 0 | Felidae | 47556.0 | 0 | Felis | 47556.0 | 0 | Felis catus | 47556.0 | 0 |
| scf7180000018965 | 195987 | 0.4124 | 0 | 81.9074 | Eukaryota | 14431.0 | 0 | Chordata | 14431.0 | 0 | Carnivora | 14431.0 | 0 | Herpestidae | 8099.0 | 1 | Suricata | 8099.0 | 4 | ricata suricat | 8099.0 | 5 |
| scf7180000017546 | 222418 | 0.3995 | 0 | 50.2059 | Eukaryota | 49724.0 | 0 | Chordata | 49724.0 | 0 | Carnivora | 49724.0 | 0 | Felidae | 49724.0 | 0 | Felis | 49724.0 | 0 | Felis catus | 49724.0 | 0 |
| scf7180000022233 | 28278 | 0.3881 | 0 | 24.9965 | Eukaryota | 42515.0 | 0 | Chordata | 42515.0 | 0 | Carnivora | 42515.0 | 0 | Felidae | 42515.0 | 0 | Felis | 38317.0 | 1 | Felis catus | 38317.0 | 1 |
| scf7180000018929 | 730041 | 0.4504 | 0 | 51.9667 | Eukaryota | 45114.0 | 0 | Chordata | 45114.0 | 0 | Carnivora | 45114.0 | 0 | Felidae | 45114.0 | 0 | Felis | 45114.0 | 0 | Felis catus | 45114.0 | 0 |
| scf7180000017815 | 1754006 | 0.3653 | 0 | 50.1573 | Eukaryota | 47568.0 | 0 | Chordata | 47568.0 | 0 | Carnivora | 47568.0 | 0 | Felidae | 47568.0 | 0 | Felis | 47568.0 | 0 | Felis catus | 47568.0 | 0 |
| scf7180000021698 | 31355 | 0.5132 | 0 | 44.0846 | Eukaryota | 3647.0 | 0 | Chordata | 3647.0 | 0 | Carnivora | 3647.0 | 0 | Felidae | 3647.0 | 0 | Felis | 3647.0 | 0 | Felis catus | 3647.0 | 0 |
| scf7180000021365 | 120176 | 0.3942 | 0 | 25.6289 | Eukaryota | 28237.0 | 0 | Chordata | 28237.0 | 0 | Carnivora | 28237.0 | 0 | Felidae | 28237.0 | 0 | Felis | 25374.0 | 1 | Felis catus | 25374.0 | 1 |
| scf7180000017443 | 3946964 | 0.3739 | 0 | 50.258 | Eukaryota | 47744.0 | 0 | Chordata | 47744.0 | 0 | Carnivora | 47744.0 | 0 | Felidae | 47744.0 | 0 | Felis | 47744.0 | 0 | Felis catus | 47744.0 | 0 |
| scf7180000024250 | 1915783 | 0.4947 | 0 | 52.6848 | Eukaryota | 37938.0 | 0 | Chordata | 37938.0 | 0 | Carnivora | 37938.0 | 0 | Felidae | 18919.0 | 5 | Felis | 7624.0 | 7 | Felis catus | 7624.0 | 8 |
| scf7180000018603 | 1008479 | 0.4192 | 0 | 50.3881 | Eukaryota | 47628.0 | 0 | Chordata | 47628.0 | 0 | Carnivora | 47628.0 | 0 | Felidae | 47628.0 | 0 | Felis | 47628.0 | 0 | Felis catus | 47628.0 | 0 |
| scf7180000017805 | 1707726 | 0.3822 | 0 | 50.7305 | Eukaryota | 42813.0 | 0 | Chordata | 42813.0 | 0 | Carnivora | 42813.0 | 0 | Felidae | 18606.0 | 3 | Ursus | 11874.0 | 5 | Felis catus | 9303.0 | 6 |
| scf7180000019260 | 355420 | 0.4763 | 0 | 50.8169 | Eukaryota | 52217.0 | 0 | Chordata | 52217.0 | 0 | Carnivora | 52217.0 | 0 | Felidae | 37776.0 | 2 | Felis | 27260.0 | 4 | Felis catus | 27260.0 | 4 |
| scf7180000023330 | 34215 | 0.428 | 0 | 400.267 | Eukaryota | 4015.0 | 0 | Chordata | 4015.0 | 0 | Carnivora | 2662.0 | 1 | Felidae | 2192.0 | 2 | Panthera | 1474.0 | 4 | Homo sapien | 1353.0 | 5 |
| scf7180000020884 | 14831 | 0.4911 | 0 | 101.7707 | Eukaryota | 3847.0 | 0 | Chordata | 3847.0 | 0 | Carnivora | 3847.0 | 0 | Felidae | 3551.0 | 1 | Felis | 3225.0 | 2 | Felis catus | 3225.0 | 2 |
| scf7180000021808 | 19719 | 0.453 | 0 | 45.7554 | Eukaryota | 24772.0 | 0 | Chordata | 24772.0 | 0 | Carnivora | 24772.0 | 0 | Felidae | 24772.0 | 0 | Felis | 22321.0 | 1 | Felis catus | 22321.0 | 1 |
| scf7180000017706 | 1707051 | 0.46 | 0 | 51.1733 | Eukaryota | 47490.0 | 0 | Chordata | 47490.0 | 0 | Carnivora | 47490.0 | 0 | Felidae | 47490.0 | 0 | Felis | 47490.0 | 0 | Felis catus | 47490.0 | 0 |
| scf7180000019615 | 311522 | 0.4213 | 0 | 50.2725 | Eukaryota | 46291.0 | 0 | Chordata | 46291.0 | 0 | Carnivora | 46291.0 | 0 | Felidae | 46291.0 | 0 | Felis | 46291.0 | 0 | Felis catus | 46291.0 | 0 |
| scf7180000022639 | 12355 | 0.4214 | 0 | 29.3872 | Eukaryota | 2482.0 | 0 | Chordata | 2482.0 | 0 | Carnivora | 1702.0 | 2 | Felidae | 1702.0 | 2 | Felis | 1487.0 | 3 | Felis catus | 1487.0 | 3 |
| scf7180000023916 | 668371 | 0.4156 | 0 | 49.9048 | Eukaryota | 122625.0 | 0 | Chordata | 122625.0 | 0 | Carnivora | 122625.0 | 0 | Felidae | 122625.0 | 0 | Panthera | 49097.0 | 2 | ynx canadens | 49018.0 | 3 |
| scf7180000019586 | 141675 | 0.392 | 0 | 32.5218 | Eukaryota | 44107.0 | 0 | Chordata | 44107.0 | 0 | Carnivora | 44107.0 | 0 | Felidae | 44107.0 | 0 | Felis | 44107.0 | 0 | Felis catus | 44107.0 | 0 |
| scf7180000021724 | 6040 | 0.5237 | 0 | 30.2815 | no-hit | 0.0 | 0 | no-hit | 0.0 | 0 | no-hit | 0.0 | 0 | no-hit | 0.0 | 0 | no-hit | 0.0 | 0 | no-hit | 0.0 | 0 |
| scf7180000021106 | 54371 | 0.4421 | 0 | 53.2152 | Eukaryota | 51616.0 | 0 | Chordata | 51616.0 | 0 | Carnivora | 51616.0 | 0 | Felidae | 17216.0 | 4 | Panthera | 17216.0 | 4 | Panthera tigri | 17216.0 | 4 |
| scf7180000021367 | 52272 | 0.3593 | 0 | 24.5689 | Eukaryota | 14007.0 | 0 | Chordata | 14007.0 | 0 | Carnivora | 14007.0 | 0 | Felidae | 14007.0 | 0 | Felis | 12608.0 | 1 | Felis catus | 12608.0 | 1 |
| scf7180000018630 | 38738 | 0.3912 | 0 | 38.1935 | Eukaryota | 43235.0 | 0 | Chordata | 43235.0 | 0 | Carnivora | 43235.0 | 0 | Felidae | 43235.0 | 0 | Felis | 38972.0 | 1 | Felis catus | 38972.0 | 1 |
| scf7180000018705 | 678461 | 0.411 | 0 | 48.7205 | Eukaryota | 47148.0 | 0 | Chordata | 47148.0 | 0 | Carnivora | 47148.0 | 0 | Felidae | 47148.0 | 0 | Felis | 42475.0 | 1 | Felis catus | 42475.0 | 1 |
| scf7180000023954 | 841252 | 0.4207 | 0 | 51.2489 | Eukaryota | 47601.0 | 0 | Chordata | 47601.0 | 0 | Carnivora | 47601.0 | 0 | Felidae | 47601.0 | 0 | Felis | 47601.0 | 0 | Felis catus | 47601.0 | 0 |
| scf7180000022617 | 13134 | 0.3563 | 0 | 22.4809 | Eukaryota | 12184.0 | 0 | Chordata | 12184.0 | 0 | Carnivora | 12184.0 | 0 | Felidae | 12184.0 | 0 | Felis | 12184.0 | 0 | Felis catus | 12184.0 | 0 |
| scf7180000021067 | 91267 | 0.4419 | 0 | 48.7187 | Eukaryota | 4308.0 | 0 | Chordata | 4308.0 | 0 | Carnivora | 2559.0 | 2 | Felidae | 1599.0 | 4 | Felis | 1286.0 | 7 | Felis catus | 1286.0 | 7 |
| scf7180000020024 | 181915 | 0.5195 | 0 | 50.3953 | Eukaryota | 7112.0 | 0 | Chordata | 7112.0 | 0 | Carnivora | 7112.0 | 0 | Felidae | 7112.0 | 0 | Felis | 6360.0 | 1 | Felis catus | 6360.0 | 1 |
| scf7180000022332 | 13026 | 0.5193 | 0 | 242.2626 | no-hit | 0.0 | 0 | no-hit | 0.0 | 0 | no-hit | 0.0 | 0 | no-hit | 0.0 | 0 | no-hit | 0.0 | 0 | no-hit | 0.0 | 0 |
| scf7180000020428 | 97624 | 0.5752 | 0 | 51.6397 | Eukaryota | 10199.0 | 0 | Chordata | 10199.0 | 0 | Primates | 5890.0 | 2 | Hominidae | 5890.0 | 3 | Homo | 5890.0 | 3 | Homo sapien | 5890.0 | 3 |
| scf7180000023496 | 24055 | 0.4751 | 0 | 408.2804 | no-hit | 0.0 | 0 | no-hit | 0.0 | 0 | no-hit | 0.0 | 0 | no-hit | 0.0 | 0 | no-hit | 0.0 | 0 | no-hit | 0.0 | 0 |
| scf7180000020070 | 4109 | 0.6179 | 0 | 104.715 | no-hit | 0.0 | 0 | no-hit | 0.0 | 0 | no-hit | 0.0 | 0 | no-hit | 0.0 | 0 | no-hit | 0.0 | 0 | no-hit | 0.0 | 0 |
| scf7180000019892 | 1806304 | 0.3887 | 100 | 50.0433 | Eukaryota | 57753.0 | 0 | Chordata | 57753.0 | 0 | Carnivora | 57753.0 | 0 | Canidae | 27786.0 | 2 | Canis | 27786.0 | 3 | Canis lupus | 27786.0 | 3 |
| scf7180000018392 | 773924 | 0.4547 | 0 | 50.8345 | Eukaryota | 56122.0 | 0 | Chordata | 56122.0 | 0 | Artiodactyla | 27020.0 | 2 | Delphinidae | 10818.0 | 6 | Orcinus | 10818.0 | 6 | Orcinus orca | 10818.0 | 6 |
| scf7180000022134 | 76625 | 0.3729 | 0 | 23.729 | Eukaryota | 5714.0 | 0 | Chordata | 5714.0 | 0 | Carnivora | 2920.0 | 1 | Hominidae | 2794.0 | 2 | Felis | 1845.0 | 4 | Felis catus | 1845.0 | 4 |
| scf7180000019640 | 275554 | 0.4676 | 0 | 28.0318 | Eukaryota | 45337.0 | 0 | Chordata | 45337.0 | 0 | Carnivora | 45337.0 | 0 | Felidae | 45337.0 | 0 | Felis | 40869.0 | 1 | Felis catus | 40869.0 | 1 |
| scf7180000020237 | 105882 | 0.5225 | 0 | 48.2795 | Eukaryota | 5431.0 | 0 | Chordata | 5431.0 | 0 | Artiodactyla | 1259.0 | 7 | Bovidae | 1259.0 | 8 | Homo | 756.0 | 9 | Homo sapien | 756.0 | 9 |
| scf7180000019242 | 37497 | 0.486 | 0 | 66.5656 | Eukaryota | 10093.0 | 0 | Chordata | 10093.0 | 0 | Carnivora | 10093.0 | 0 | Felidae | 10093.0 | 0 | Felis | 8088.0 | 2 | Felis catus | 8088.0 | 2 |
| scf7180000019584 | 245318 | 0.3522 | 0 | 50.6126 | Eukaryota | 47831.0 | 0 | Chordata | 47831.0 | 0 | Carnivora | 47831.0 | 0 | Felidae | 47831.0 | 0 | Felis | 47831.0 | 0 | Felis catus | 47831.0 | 0 |
| scf7180000021896 | 44052 | 0.3969 | 0 | 23.3805 | Eukaryota | 40914.0 | 0 | Chordata | 40914.0 | 0 | Carnivora | 40914.0 | 0 | Felidae | 40914.0 | 0 | Felis | 36867.0 | 1 | Felis catus | 36867.0 | 1 |
| scf7180000019303 | 6633 | 0.4172 | 0 | 26.884 | Eukaryota | 22326.0 | 0 | Chordata | 22326.0 | 0 | Carnivora | 22326.0 | 0 | Felidae | 22326.0 | 0 | Felis | 20080.0 | 1 | Felis catus | 20080.0 | 1 |
| scf7180000024202 | 51306 | 0.3709 | 0 | 23.8737 | Eukaryota | 18628.0 | 0 | Chordata | 18628.0 | 0 | Carnivora | 18628.0 | 0 | Felidae | 18628.0 | 0 | Felis | 14917.0 | 1 | Felis catus | 14917.0 | 1 |
| scf7180000020036 | 258923 | 0.3894 | 0 | 49.8387 | Eukaryota | 16859.0 | 0 | Chordata | 16859.0 | 0 | Carnivora | 16859.0 | 0 | Felidae | 16859.0 | 0 | Felis | 13494.0 | 1 | Felis catus | 13494.0 | 1 |
| scf7180000018370 | 740023 | 0.466 | 0 | 51.6295 | Eukaryota | 28175.0 | 0 | Chordata | 28175.0 | 0 | Carnivora | 28175.0 | 0 | Felidae | 28175.0 | 0 | Felis | 25329.0 | 1 | Felis catus | 25329.0 | 1 |
| scf7180000021531 | 55691 | 0.3584 | 0 | 23.3916 | Eukaryota | 29610.0 | 0 | Chordata | 29610.0 | 0 | Carnivora | 29610.0 | 0 | Felidae | 29610.0 | 0 | Felis | 26670.0 | 1 | Felis catus | 26670.0 | 1 |
| scf7180000020443 | 123826 | 0.4457 | 0 | 49.604 | Eukaryota | 20872.0 | 0 | Chordata | 20872.0 | 0 | Carnivora | 14663.0 | 1 | Felidae | 14663.0 | 1 | Panthera | 6300.0 | 3 | anthera pardu | 6300.0 | 3 |
| scf7180000024128 | 171912 | 0.3717 | 0 | 24.6429 | Eukaryota | 38228.0 | 0 | Chordata | 38228.0 | 0 | Carnivora | 38228.0 | 0 | Felidae | 38228.0 | 0 | Felis | 38228.0 | 0 | Felis catus | 38228.0 | 0 |
| scf7180000020252 | 140114 | 0.3946 | 0 | 25.5465 | Eukaryota | 43305.0 | 0 | Chordata | 43305.0 | 0 | Carnivora | 43305.0 | 0 | Felidae | 43305.0 | 0 | Felis | 39015.0 | 1 | Felis catus | 39015.0 | 1 |
| scf7180000019620 | 417364 | 0.4274 | 0 | 48.9653 | Eukaryota | 43446.0 | 0 | Chordata | 43446.0 | 0 | Carnivora | 43446.0 | 0 | Felidae | 43446.0 | 0 | Felis | 39145.0 | 1 | Felis catus | 39145.0 | 1 |
| scf7180000023848 | 1458471 | 0.4618 | 0 | 51.5797 | Eukaryota | 86181.0 | 0 | Chordata | 86181.0 | 0 | Carnivora | 86181.0 | 0 | Ursidae | 32370.0 | 3 | Ursus | 32370.0 | 3 | ricata suricat | 28255.0 | 4 |
| scf7180000022250 | 24336 | 0.3875 | 0 | 24.5606 | Eukaryota | 3739.0 | 0 | Chordata | 3739.0 | 0 | Carnivora | 2676.0 | 2 | Felidae | 1680.0 | 3 | Felis | 1680.0 | 5 | Felis catus | 1680.0 | 5 |
| scf7180000017628 | 28757 | 0.5301 | 0 | 46.4235 | Eukaryota | 27074.0 | 0 | Chordata | 27074.0 | 0 | Carnivora | 27074.0 | 0 | Felidae | 27074.0 | 0 | Felis | 24327.0 | 1 | Felis catus | 24327.0 | 1 |
| scf7180000019432 | 280407 | 0.4525 | 0 | 50.6845 | Eukaryota | 22738.0 | 1 | Chordata | 22738.0 | 1 | Primates | 12241.0 | 2 | Hominidae | 12241.0 | 3 | Homo | 12241.0 | 4 | Homo sapien | 12241.0 | 4 |
| scf7180000021788 | 31042 | 0.3649 | 0 | 24.19 | Eukaryota | 27444.0 | 0 | Chordata | 27444.0 | 0 | Carnivora | 27444.0 | 0 | Felidae | 27444.0 | 0 | Felis | 24721.0 | 1 | Felis catus | 24721.0 | 1 |
| scf7180000020539 | 107836 | 0.4251 | 0 | 50.1233 | Eukaryota | 43215.0 | 0 | Chordata | 43215.0 | 0 | Carnivora | 43215.0 | 0 | Felidae | 38107.0 | 1 | Felis | 38107.0 | 1 | Felis catus | 38107.0 | 1 |
| scf7180000020947 | 64321 | 0.4153 | 100 | 25.364 | Eukaryota | 39138.0 | 0 | Chordata | 39138.0 | 0 | Carnivora | 39138.0 | 0 | Felidae | 39138.0 | 0 | Felis | 39138.0 | 0 | Felis catus | 39138.0 | 0 |
| scf7180000018084 | 1101675 | 0.3856 | 0 | 50.3513 | Eukaryota | 46870.0 | 0 | Chordata | 46870.0 | 0 | Carnivora | 46870.0 | 0 | Felidae | 46870.0 | 0 | Felis | 46870.0 | 0 | Felis catus | 46870.0 | 0 |
| scf7180000019889 | 241565 | 0.3694 | 0 | 49.9528 | Eukaryota | 42330.0 | 0 | Chordata | 42330.0 | 0 | Carnivora | 42330.0 | 0 | Felidae | 42330.0 | 0 | Felis | 42330.0 | 0 | Felis catus | 42330.0 | 0 |
| scf7180000022699 | 6921 | 0.6807 | 0 | 22.7078 | Eukaryota | 4136.0 | 0 | Chordata | 4136.0 | 0 | Carnivora | 3727.0 | 1 | Felidae | 3727.0 | 1 | Lynx | 2055.0 | 4 | ynx canadens | 2055.0 | 5 |
| scf7180000022576 | 11127 | 0.5189 | 0 | 150.8739 | no-hit | 0.0 | 0 | no-hit | 0.0 | 0 | no-hit | 0.0 | 0 | no-hit | 0.0 | 0 | no-hit | 0.0 | 0 | no-hit | 0.0 | 0 |
| scf7180000018011 | 793645 | 0.3754 | 0 | 51.0393 | Eukaryota | 47463.0 | 0 | Chordata | 47463.0 | 0 | Carnivora | 47463.0 | 0 | Felidae | 47463.0 | 0 | Felis | 47463.0 | 0 | Felis catus | 47463.0 | 0 |
| scf7180000018231 | 974409 | 0.4162 | 0 | 49.9084 | Eukaryota | 37380.0 | 0 | Chordata | 37380.0 | 0 | Carnivora | 37380.0 | 0 | Ursidae | 29749.0 | 1 | Ursus | 29749.0 | 1 | Ursus arctos | 18580.0 | 2 |
| scf7180000019163 | 31472 | 0.508 | 0 | 874.4503 | Eukaryota | 2477.0 | 0 | Chordata | 2477.0 | 0 | Carnivora | 2477.0 | 0 | Felidae | 2223.0 | 1 | Felis | 1492.0 | 2 | Felis catus | 1492.0 | 2 |
| scf7180000023984 | 906794 | 0.4523 | 0 | 51.8114 | Eukaryota | 48550.0 | 0 | Chordata | 48550.0 | 0 | Carnivora | 48550.0 | 0 | Felidae | 29562.0 | 1 | Panthera | 19960.0 | 2 | Panthera tigri | 19960.0 | 2 |
| scf7180000023809 | 768264 | 0.3665 | 0 | 50.8424 | Eukaryota | 47425.0 | 0 | Chordata | 47425.0 | 0 | Carnivora | 47425.0 | 0 | Felidae | 47425.0 | 0 | Felis | 47425.0 | 0 | Felis catus | 47425.0 | 0 |
| scf7180000019116 | 272943 | 0.4594 | 0 | 50.4487 | Eukaryota | 45736.0 | 0 | Chordata | 45736.0 | 0 | Carnivora | 45736.0 | 0 | Felidae | 45736.0 | 0 | Felis | 45736.0 | 0 | Felis catus | 45736.0 | 0 |
| scf7180000022618 | 12548 | 0.4401 | 0 | 26.3908 | Eukaryota | 6140.0 | 0 | Chordata | 6140.0 | 0 | Primates | 5021.0 | 2 | Hominidae | 2936.0 | 3 | Homo | 2936.0 | 5 | Homo sapien | 2936.0 | 6 |
| scf7180000018206 | 177032 | 0.4273 | 0 | 47.4039 | Eukaryota | 39917.0 | 0 | Chordata | 39917.0 | 0 | Carnivora | 39917.0 | 0 | Felidae | 39917.0 | 0 | Felis | 35957.0 | 1 | Felis catus | 35957.0 | 1 |
| scf7180000023807 | 1436972 | 0.417 | 0 | 50.6029 | Eukaryota | 46721.0 | 0 | Chordata | 46721.0 | 0 | Carnivora | 46721.0 | 0 | Felidae | 46721.0 | 0 | Felis | 46721.0 | 0 | Felis catus | 46721.0 | 0 |
| scf7180000019030 | 170695 | 0.4859 | 0 | 52.0941 | Eukaryota | 19353.0 | 0 | Chordata | 19353.0 | 0 | Carnivora | 19353.0 | 0 | Felidae | 19353.0 | 0 | Felis | 17419.0 | 1 | Felis catus | 17419.0 | 1 |
| scf7180000020039 | 161650 | 0.4084 | 0 | 51.2885 | Eukaryota | 42453.0 | 0 | Chordata | 42453.0 | 0 | Carnivora | 42453.0 | 0 | Felidae | 42453.0 | 0 | Felis | 42453.0 | 0 | Felis catus | 42453.0 | 0 |
| scf7180000018824 | 21524 | 0.3612 | 0 | 41.5217 | Eukaryota | 15487.0 | 0 | Chordata | 15487.0 | 0 | Carnivora | 15487.0 | 0 | Felidae | 15487.0 | 0 | Felis | 12415.0 | 1 | Felis catus | 12415.0 | 1 |
| scf7180000022758 | 8144 | 0.3692 | 0 | 52.6813 | Eukaryota | 6645.0 | 0 | Chordata | 6645.0 | 0 | Carnivora | 6645.0 | 0 | Canidae | 5874.0 | 1 | Canis | 5874.0 | 1 | Canis lupus | 5874.0 | 1 |
| scf7180000018735 | 131969 | 0.3581 | 0 | 46.1972 | Eukaryota | 49602.0 | 0 | Chordata | 49602.0 | 0 | Carnivora | 49602.0 | 0 | Felidae | 25146.0 | 4 | Panthera | 10066.0 | 7 | obenus rosma | 9748.0 | 8 |
| scf7180000018205 | 928201 | 0.4858 | 0 | 54.5513 | Eukaryota | 62274.0 | 0 | Chordata | 62274.0 | 0 | Carnivora | 62274.0 | 0 | Felidae | 39006.0 | 1 | Suricata | 23268.0 | 4 | ricata suricat | 23268.0 | 5 |
| scf7180000020798 | 71090 | 0.393 | 0 | 27.3312 | Eukaryota | 46930.0 | 0 | Chordata | 46930.0 | 0 | Carnivora | 46930.0 | 0 | Felidae | 46930.0 | 0 | Felis | 46930.0 | 0 | Felis catus | 46930.0 | 0 |
| scf7180000020379 | 104391 | 0.3673 | 0 | 24.8455 | Eukaryota | 21538.0 | 0 | Chordata | 21538.0 | 0 | Carnivora | 21538.0 | 0 | Felidae | 21538.0 | 0 | Felis | 17197.0 | 1 | Felis catus | 17197.0 | 1 |
| scf7180000019446 | 88358 | 0.3969 | 0 | 46.2163 | Eukaryota | 46614.0 | 0 | Chordata | 46614.0 | 0 | Carnivora | 46614.0 | 0 | Felidae | 46614.0 | 0 | Felis | 46614.0 | 0 | Felis catus | 46614.0 | 0 |
| scf7180000023205 | 46173 | 0.3887 | 0 | 354.8435 | Eukaryota | 12272.0 | 0 | Chordata | 12272.0 | 0 | Carnivora | 12272.0 | 0 | Felidae | 12272.0 | 0 | Felis | 11054.0 | 1 | Felis catus | 11054.0 | 1 |
| scf7180000021743 | 49526 | 0.5192 | 0 | 46.8973 | Eukaryota | 42247.0 | 0 | Chordata | 42247.0 | 0 | Carnivora | 42247.0 | 0 | Felidae | 42247.0 | 0 | Felis | 38088.0 | 1 | Felis catus | 38088.0 | 1 |
| scf7180000023439 | 30930 | 0.4543 | 0 | 194.625 | Eukaryota | 3000.0 | 0 | Chordata | 3000.0 | 0 | Carnivora | 3000.0 | 0 | Felidae | 2227.0 | 1 | Lynx | 1215.0 | 4 | ynx canadens | 1215.0 | 4 |
| scf7180000020319 | 114892 | 0.5037 | 0 | 52.7137 | Eukaryota | 20120.0 | 0 | Chordata | 20120.0 | 0 | Carnivora | 20120.0 | 0 | Felidae | 20120.0 | 0 | Felis | 18119.0 | 1 | Felis catus | 18119.0 | 1 |
| scf7180000017468 | 3719737 | 0.3537 | 0 | 48.9514 | Eukaryota | 53725.0 | 0 | Chordata | 53725.0 | 0 | Rodentia | 24559.0 | 4 | Caviidae | 24559.0 | 4 | Cavia | 24559.0 | 4 | Cavia porcellu | 24559.0 | 4 |
| scf7180000024201 | 74387 | 0.4214 | 0 | 25.8104 | Eukaryota | 11223.0 | 0 | Chordata | 11223.0 | 0 | Carnivora | 11223.0 | 0 | Felidae | 11223.0 | 0 | Felis | 11223.0 | 0 | Felis catus | 11223.0 | 0 |
| scf7180000017821 | 51178 | 0.3937 | 0 | 41.7009 | Eukaryota | 45589.0 | 0 | Chordata | 45589.0 | 0 | Carnivora | 45589.0 | 0 | Felidae | 45589.0 | 0 | Felis | 45589.0 | 0 | Felis catus | 45589.0 | 0 |
| scf7180000021817 | 55168 | 0.4041 | 0 | 24.2414 | Eukaryota | 33081.0 | 0 | Chordata | 33081.0 | 0 | Carnivora | 33081.0 | 0 | Felidae | 33081.0 | 0 | Felis | 29732.0 | 1 | Felis catus | 29732.0 | 1 |
| scf7180000023586 | 59463 | 0.3977 | 0 | 361.9329 | Eukaryota | 43088.0 | 0 | Chordata | 43088.0 | 0 | Carnivora | 43088.0 | 0 | Felidae | 43088.0 | 0 | Felis | 43088.0 | 0 | Felis catus | 43088.0 | 0 |
| scf7180000020711 | 76001 | 0.3852 | 0 | 25.5016 | Eukaryota | 13638.0 | 0 | Chordata | 13638.0 | 0 | Carnivora | 13638.0 | 0 | Felidae | 12949.0 | 1 | Felis | 8694.0 | 5 | Felis catus | 8694.0 | 5 |
| scf7180000018163 | 1130187 | 0.4101 | 0 | 50.9209 | Eukaryota | 46053.0 | 0 | Chordata | 46053.0 | 0 | Carnivora | 46053.0 | 0 | Felidae | 46053.0 | 0 | Felis | 41493.0 | 1 | Felis catus | 41493.0 | 1 |
| scf7180000019412 | 389338 | 0.3563 | 0 | 49.9323 | Eukaryota | 47775.0 | 0 | Chordata | 47775.0 | 0 | Carnivora | 47775.0 | 0 | Felidae | 47775.0 | 0 | Felis | 47775.0 | 0 | Felis catus | 47775.0 | 0 |
| scf7180000020452 | 127155 | 0.3652 | 0 | 23.6169 | Eukaryota | 24000.0 | 0 | Chordata | 24000.0 | 0 | Carnivora | 24000.0 | 0 | Felidae | 24000.0 | 0 | Felis | 21674.0 | 1 | Felis catus | 21674.0 | 1 |
| scf7180000018374 | 905956 | 0.3856 | 0 | 51.2149 | Eukaryota | 37481.0 | 0 | Chordata | 37481.0 | 0 | Carnivora | 37481.0 | 0 | Felidae | 37481.0 | 0 | Felis | 37481.0 | 0 | Felis catus | 37481.0 | 0 |
| scf7180000018275 | 3382179 | 0.3898 | 0 | 51.2757 | Eukaryota | 56040.0 | 0 | Chordata | 56040.0 | 0 | Carnivora | 56040.0 | 0 | Felidae | 56040.0 | 0 | Felis | 56040.0 | 0 | Felis catus | 56040.0 | 0 |
| scf7180000020429 | 31866 | 0.405 | 0 | 38.803 | Eukaryota | 12974.0 | 0 | Chordata | 12974.0 | 0 | Carnivora | 5537.0 | 2 | Felidae | 5537.0 | 2 | Felis | 4471.0 | 4 | Felis catus | 4471.0 | 4 |
| scf7180000018575 | 86120 | 0.4774 | 0 | 45.6283 | Eukaryota | 21620.0 | 0 | Chordata | 21620.0 | 0 | Carnivora | 21620.0 | 0 | Felidae | 21620.0 | 0 | Felis | 17329.0 | 1 | Felis catus | 17329.0 | 1 |
| scf7180000019057 | 60834 | 0.465 | 0 | 48.0194 | Eukaryota | 28158.0 | 0 | Chordata | 28158.0 | 0 | Carnivora | 28158.0 | 0 | Felidae | 28158.0 | 0 | Panthera | 16950.0 | 1 | anthera pardu | 14120.0 | 2 |
| scf7180000023933 | 608505 | 0.5819 | 0 | 49.8791 | Eukaryota | 56009.0 | 0 | Chordata | 56009.0 | 0 | Carnivora | 56009.0 | 0 | Otariidae | 43840.0 | 2 | Zalophus | 43840.0 | 2 | phus californi | 43840.0 | 2 |
| scf7180000017620 | 2192130 | 0.5058 | 0 | 53.111 | Eukaryota | 47334.0 | 0 | Chordata | 47334.0 | 0 | Carnivora | 47334.0 | 0 | Felidae | 47334.0 | 0 | Felis | 47334.0 | 0 | Felis catus | 47334.0 | 0 |
| scf7180000018856 | 337104 | 0.3762 | 0 | 46.7343 | Eukaryota | 45315.0 | 0 | Chordata | 45315.0 | 0 | Carnivora | 45315.0 | 0 | Felidae | 45315.0 | 0 | Felis | 45315.0 | 0 | Felis catus | 45315.0 | 0 |
| scf7180000018039 | 3306226 | 0.4143 | 0 | 50.805 | Eukaryota | 90356.0 | 0 | Chordata | 90356.0 | 0 | Carnivora | 90356.0 | 0 | Felidae | 80869.0 | 1 | Panthera | 71864.0 | 2 | anthera pardu | 71864.0 | 2 |
| scf7180000022798 | 17659 | 0.3846 | 0 | 42.3334 | Eukaryota | 41617.0 | 0 | Chordata | 41617.0 | 0 | Carnivora | 41617.0 | 0 | Felidae | 41617.0 | 0 | Felis | 41617.0 | 0 | Felis catus | 41617.0 | 0 |
| scf7180000021748 | 36633 | 0.4295 | 0 | 25.197 | Eukaryota | 5199.0 | 0 | Chordata | 5199.0 | 0 | Carnivora | 4641.0 | 1 | Felidae | 4641.0 | 1 | Felis | 4131.0 | 2 | Felis catus | 4131.0 | 2 |
| scf7180000020055 | 14704 | 0.4169 | 0 | 23.5732 | Eukaryota | 36578.0 | 0 | Chordata | 36578.0 | 0 | Carnivora | 36578.0 | 0 | Felidae | 36578.0 | 0 | Felis | 36578.0 | 0 | Felis catus | 36578.0 | 0 |
| scf7180000021658 | 57171 | 0.3704 | 0 | 48.7128 | Eukaryota | 46760.0 | 0 | Chordata | 46760.0 | 0 | Carnivora | 46760.0 | 0 | Felidae | 46760.0 | 0 | Felis | 46760.0 | 0 | Felis catus | 46760.0 | 0 |
| scf7180000021179 | 43403 | 0.6381 | 0 | 50.6365 | Eukaryota | 8744.0 | 1 | Chordata | 8744.0 | 1 | Carnivora | 8256.0 | 2 | Felidae | 4221.0 | 3 | Suricata | 4035.0 | 4 | ricata suricat | 4035.0 | 4 |
| scf7180000019061 | 454885 | 0.4445 | 0 | 56.9436 | Eukaryota | 42034.0 | 0 | Chordata | 42034.0 | 0 | Carnivora | 42034.0 | 0 | Felidae | 33863.0 | 2 | Panthera | 12630.0 | 6 | cinonyx jubat | 8468.0 | 7 |
| scf7180000021542 | 60492 | 0.4188 | 0 | 26.2832 | Eukaryota | 11061.0 | 0 | Chordata | 11061.0 | 0 | Primates | 7115.0 | 3 | Hominidae | 7115.0 | 4 | Pan | 4269.0 | 5 | an troglodyte | 4269.0 | 5 |
| scf7180000022095 | 21632 | 0.5906 | 0 | 42.939 | Eukaryota | 7281.0 | 0 | Chordata | 7281.0 | 0 | Carnivora | 7281.0 | 0 | Felidae | 6027.0 | 1 | Lynx | 1656.0 | 5 | ynx canadens | 1656.0 | 5 |
| scf7180000022290 | 24534 | 0.4 | 0 | 26.0571 | Eukaryota | 12348.0 | 0 | Chordata | 12348.0 | 0 | Carnivora | 12348.0 | 0 | Felidae | 12348.0 | 0 | Felis | 11110.0 | 1 | Felis catus | 11110.0 | 1 |
| scf7180000021642 | 37954 | 0.3888 | 0 | 27.1273 | Eukaryota | 5436.0 | 0 | Chordata | 5436.0 | 0 | Carnivora | 5436.0 | 0 | Felidae | 5436.0 | 0 | Felis | 4351.0 | 1 | Felis catus | 4351.0 | 1 |
| scf7180000018359 | 853138 | 0.5126 | 0 | 49.1527 | Eukaryota | 19828.0 | 1 | Chordata | 19828.0 | 1 | Carnivora | 17840.0 | 2 | Felidae | 17840.0 | 2 | Panthera | 7613.0 | 4 | anthera pardu | 7613.0 | 4 |
| scf7180000020159 | 169863 | 0.5026 | 0 | 49.2225 | Eukaryota | 9776.0 | 0 | Chordata | 9776.0 | 0 | Carnivora | 9776.0 | 0 | Felidae | 9776.0 | 0 | Felis | 7834.0 | 1 | Felis catus | 7834.0 | 1 |
| scf7180000021459 | 45997 | 0.5351 | 0 | 49.1771 | Eukaryota | 10404.0 | 1 | Chordata | 10404.0 | 1 | Artiodactyla | 3096.0 | 7 | Bovidae | 3096.0 | 7 | Homo | 2520.0 | 8 | Homo sapien | 2520.0 | 8 |
| scf7180000018471 | 1189228 | 0.404 | 100 | 50.6551 | Eukaryota | 47511.0 | 0 | Chordata | 47511.0 | 0 | Carnivora | 47511.0 | 0 | Felidae | 47511.0 | 0 | Felis | 47511.0 | 0 | Felis catus | 47511.0 | 0 |
| scf7180000022166 | 27464 | 0.3632 | 0 | 24.1513 | Eukaryota | 20211.0 | 0 | Chordata | 20211.0 | 0 | Carnivora | 20211.0 | 0 | Felidae | 20211.0 | 0 | Felis | 18168.0 | 1 | Felis catus | 18168.0 | 1 |
| scf7180000020360 | 92533 | 0.4271 | 0 | 26.4193 | Eukaryota | 17089.0 | 0 | Chordata | 17089.0 | 0 | Carnivora | 11678.0 | 1 | Felidae | 11678.0 | 1 | Acinonyx | 5291.0 | 4 | cinonyx jubat | 5291.0 | 4 |
| scf7180000022429 | 43817 | 0.3815 | 0 | 27.7507 | Eukaryota | 35659.0 | 0 | Chordata | 35659.0 | 0 | Carnivora | 35659.0 | 0 | Felidae | 35659.0 | 0 | Felis | 35659.0 | 0 | Felis catus | 35659.0 | 0 |
| scf7180000020593 | 98606 | 0.4544 | 0 | 49.2454 | Eukaryota | 26615.0 | 0 | Chordata | 26615.0 | 0 | Carnivora | 26615.0 | 0 | Felidae | 26615.0 | 0 | Felis | 20191.0 | 2 | Felis catus | 20191.0 | 2 |
| scf7180000019666 | 345496 | 0.5536 | 0 | 50.8185 | Eukaryota | 18122.0 | 0 | Chordata | 18122.0 | 0 | Carnivora | 18122.0 | 0 | Herpestidae | 11082.0 | 1 | Suricata | 11082.0 | 1 | ricata suricat | 11082.0 | 1 |
| scf7180000024172 | 189421 | 0.3819 | 0 | 25.0965 | Eukaryota | 13068.0 | 0 | Chordata | 13068.0 | 0 | Carnivora | 13068.0 | 0 | Felidae | 13068.0 | 0 | Felis | 13068.0 | 0 | Felis catus | 13068.0 | 0 |
| scf7180000021567 | 82121 | 0.3919 | 0 | 24.5204 | Eukaryota | 33136.0 | 0 | Chordata | 33136.0 | 0 | Carnivora | 33136.0 | 0 | Felidae | 33136.0 | 0 | Felis | 33136.0 | 0 | Felis catus | 33136.0 | 0 |
| scf7180000021383 | 32132 | 0.5044 | 0 | 41.4857 | Eukaryota | 2700.0 | 0 | Chordata | 2700.0 | 0 | Carnivora | 2700.0 | 0 | Felidae | 2700.0 | 0 | Felis | 2167.0 | 1 | Felis catus | 2167.0 | 1 |
| scf7180000018990 | 516094 | 0.4878 | 0 | 49.7796 | Eukaryota | 45408.0 | 0 | Chordata | 45408.0 | 0 | Carnivora | 45408.0 | 0 | Felidae | 45408.0 | 0 | Panthera | 22810.0 | 2 | anthera pardu | 22810.0 | 2 |
| scf7180000022019 | 19835 | 0.4688 | 0 | 30.4583 | Eukaryota | 45752.0 | 0 | Chordata | 45752.0 | 0 | Carnivora | 45752.0 | 0 | Felidae | 45752.0 | 0 | Felis | 45752.0 | 0 | Felis catus | 45752.0 | 0 |
| scf7180000024152 | 195828 | 0.3928 | 0 | 25.4034 | Eukaryota | 46037.0 | 0 | Chordata | 46037.0 | 0 | Carnivora | 46037.0 | 0 | Felidae | 46037.0 | 0 | Felis | 25528.0 | 2 | Felis catus | 25528.0 | 2 |
| scf7180000021023 | 69504 | 0.3955 | 0 | 25.6777 | Eukaryota | 21914.0 | 0 | Chordata | 21914.0 | 0 | Carnivora | 21914.0 | 0 | Felidae | 21914.0 | 0 | Felis | 21914.0 | 0 | Felis catus | 21914.0 | 0 |
| scf7180000020485 | 10440 | 0.3852 | 0 | 16.6895 | Eukaryota | 15932.0 | 0 | Chordata | 15932.0 | 0 | Carnivora | 15932.0 | 0 | Felidae | 15932.0 | 0 | Felis | 14358.0 | 1 | Felis catus | 14358.0 | 1 |
| scf7180000023324 | 50525 | 0.4241 | 0 | 417.7449 | Eukaryota | 6943.0 | 0 | Chordata | 6943.0 | 0 | Carnivora | 6943.0 | 0 | Felidae | 6943.0 | 0 | Felis | 6943.0 | 0 | Felis catus | 6943.0 | 0 |
| scf7180000024021 | 357809 | 0.559 | 0 | 48.0214 | Eukaryota | 11777.0 | 0 | Chordata | 11777.0 | 0 | Carnivora | 11777.0 | 0 | Felidae | 11777.0 | 0 | Panthera | 10476.0 | 1 | anthera pardu | 10476.0 | 1 |
| scf7180000018933 | 537067 | 0.3794 | 0 | 52.7043 | Eukaryota | 47957.0 | 0 | Chordata | 47957.0 | 0 | Carnivora | 47957.0 | 0 | Felidae | 47957.0 | 0 | Felis | 47957.0 | 0 | Felis catus | 47957.0 | 0 |
| scf7180000020877 | 99374 | 0.4302 | 0 | 28.4522 | Eukaryota | 13429.0 | 0 | Chordata | 13429.0 | 0 | Carnivora | 13429.0 | 0 | Felidae | 6727.0 | 2 | Felis | 6727.0 | 2 | Felis catus | 6727.0 | 2 |
| scf7180000017591 | 2066658 | 0.5154 | 0 | 50.475 | Eukaryota | 38164.0 | 0 | Chordata | 38164.0 | 0 | Carnivora | 38164.0 | 0 | Felidae | 38164.0 | 0 | Acinonyx | 30656.0 | 2 | cinonyx jubat | 30656.0 | 2 |
| scf7180000018416 | 300422 | 0.4862 | 0 | 51.8297 | Eukaryota | 58843.0 | 0 | Chordata | 58843.0 | 0 | Carnivora | 58843.0 | 0 | Felidae | 47027.0 | 1 | Panthera | 18003.0 | 5 | anthera pardu | 18003.0 | 5 |
| scf7180000019367 | 239328 | 0.5421 | 0 | 50.8799 | Eukaryota | 26057.0 | 0 | Chordata | 26057.0 | 0 | Carnivora | 26057.0 | 0 | Felidae | 11956.0 | 2 | Suricata | 10064.0 | 3 | ricata suricat | 10064.0 | 3 |
| scf7180000022486 | 14621 | 0.4718 | 0 | 236.342 | Eukaryota | 24462.0 | 0 | Chordata | 24462.0 | 0 | Carnivora | 24462.0 | 0 | Felidae | 24462.0 | 0 | Felis | 24462.0 | 0 | Felis catus | 24462.0 | 0 |
| scf7180000020410 | 150233 | 0.5296 | 0 | 50.7365 | Eukaryota | 22843.0 | 0 | Chordata | 22843.0 | 0 | Carnivora | 22843.0 | 0 | Felidae | 20154.0 | 1 | Panthera | 5346.0 | 5 | ynx canadens | 5327.0 | 6 |
| scf7180000019276 | 236388 | 0.4824 | 100 | 47.2657 | Eukaryota | 34780.0 | 0 | Chordata | 34780.0 | 0 | Carnivora | 34780.0 | 0 | Felidae | 34780.0 | 0 | Felis | 34780.0 | 0 | Felis catus | 34780.0 | 0 |
| scf7180000018803 | 648408 | 0.3952 | 100 | 52.1093 | Eukaryota | 47291.0 | 0 | Chordata | 47291.0 | 0 | Carnivora | 47291.0 | 0 | Felidae | 47291.0 | 0 | Felis | 47291.0 | 0 | Felis catus | 47291.0 | 0 |
| scf7180000023508 | 18970 | 0.4858 | 0 | 621.5024 | no-hit | 0.0 | 0 | no-hit | 0.0 | 0 | no-hit | 0.0 | 0 | no-hit | 0.0 | 0 | no-hit | 0.0 | 0 | no-hit | 0.0 | 0 |
| scf7180000018364 | 1025892 | 0.4284 | 0 | 51.3613 | Eukaryota | 48066.0 | 0 | Chordata | 48066.0 | 0 | Carnivora | 48066.0 | 0 | Felidae | 48066.0 | 0 | Felis | 48066.0 | 0 | Felis catus | 48066.0 | 0 |
| scf7180000022224 | 5377 | 0.5429 | 0 | 50.7008 | no-hit | 0.0 | 0 | no-hit | 0.0 | 0 | no-hit | 0.0 | 0 | no-hit | 0.0 | 0 | no-hit | 0.0 | 0 | no-hit | 0.0 | 0 |
| scf7180000018134 | 326488 | 0.3878 | 0 | 48.9718 | Eukaryota | 45308.0 | 0 | Chordata | 45308.0 | 0 | Carnivora | 45308.0 | 0 | Felidae | 45308.0 | 0 | Felis | 45308.0 | 0 | Felis catus | 45308.0 | 0 |
| scf7180000020655 | 6445 | 0.5341 | 0 | 21.812 | Eukaryota | 4825.0 | 0 | Chordata | 4825.0 | 0 | Carnivora | 4825.0 | 0 | Felidae | 4825.0 | 0 | Felis | 4825.0 | 0 | Felis catus | 4825.0 | 0 |
| scf7180000019951 | 200951 | 0.4175 | 0 | 49.5318 | Eukaryota | 19888.0 | 0 | Chordata | 19888.0 | 0 | Artiodactyla | 11952.0 | 1 | Rhinocerotidae | 7936.0 | 2 | Ceratotherium | 7936.0 | 2 | totherium sir | 7936.0 | 2 |
| scf7180000021511 | 62361 | 0.4678 | 0 | 24.5608 | Eukaryota | 19498.0 | 0 | Chordata | 19498.0 | 0 | Carnivora | 19498.0 | 0 | Canidae | 15870.0 | 1 | Canis | 15870.0 | 1 | Canis lupus | 15870.0 | 1 |
| scf7180000020597 | 20898 | 0.4256 | 0 | 399.4087 | Eukaryota | 257074.0 | 0 | Chordata | 257074.0 | 0 | Carnivora | 257074.0 | 0 | Hyaenidae | 257074.0 | 0 | Proteles | 155539.0 | 1 | roteles cristat | 155539.0 | 1 |
| scf7180000020754 | 102184 | 0.4372 | 0 | 26.1778 | Eukaryota | 11976.0 | 0 | Chordata | 11976.0 | 0 | Primates | 4945.0 | 3 | Hominidae | 4202.0 | 4 | Felis | 2682.0 | 6 | Felis catus | 2682.0 | 6 |
| scf7180000020138 | 28102 | 0.3804 | 0 | 188.8023 | Eukaryota | 47659.0 | 0 | Chordata | 47659.0 | 0 | Carnivora | 47659.0 | 0 | Felidae | 47659.0 | 0 | Felis | 47659.0 | 0 | Felis catus | 47659.0 | 0 |
| scf7180000021256 | 66598 | 0.45 | 0 | 26.9932 | Eukaryota | 15190.0 | 0 | Chordata | 15190.0 | 0 | Artiodactyla | 10347.0 | 2 | Bovidae | 2690.0 | 6 | Manis | 2560.0 | 8 | Manis javanica | 2560.0 | 8 |
| scf7180000017658 | 1896571 | 0.3946 | 0 | 50.2117 | Eukaryota | 54534.0 | 0 | Chordata | 54534.0 | 0 | Carnivora | 54534.0 | 0 | Felidae | 54534.0 | 0 | Panthera | 21840.0 | 3 | anthera pardu | 16395.0 | 4 |
| scf7180000020338 | 158542 | 0.3843 | 0 | 25.5596 | Eukaryota | 46699.0 | 0 | Chordata | 46699.0 | 0 | Carnivora | 46699.0 | 0 | Felidae | 46699.0 | 0 | Felis | 46699.0 | 0 | Felis catus | 46699.0 | 0 |
| scf7180000020646 | 181599 | 0.4628 | 0 | 50.8312 | Eukaryota | 24653.0 | 0 | Chordata | 24653.0 | 0 | Carnivora | 24653.0 | 0 | Felidae | 24653.0 | 0 | Felis | 22176.0 | 1 | Felis catus | 22176.0 | 1 |
| scf7180000021906 | 21398 | 0.4423 | 0 | 3244.9742 | Eukaryota | 5836.0 | 0 | Chordata | 5836.0 | 0 | Artiodactyla | 5836.0 | 0 | Bovidae | 5836.0 | 0 | Ovis | 5836.0 | 0 | Ovis canadens | 5836.0 | 0 |
| scf7180000022522 | 8961 | 0.4406 | 0 | 55.5287 | Eukaryota | 15626.0 | 0 | Chordata | 15626.0 | 0 | Carnivora | 15626.0 | 0 | Otariidae | 5600.0 | 4 | Puma | 2820.0 | 7 | Puma concolo | 2820.0 | 7 |
| scf7180000020726 | 58721 | 0.6351 | 0 | 52.2719 | Eukaryota | 8378.0 | 0 | Chordata | 8378.0 | 0 | Carnivora | 6801.0 | 2 | Felidae | 3331.0 | 4 | Ursus | 2727.0 | 6 | Ursus arctos | 2727.0 | 6 |
| scf7180000023023 | 77143 | 0.3856 | 0 | 500.1549 | Eukaryota | 7072.0 | 0 | Chordata | 7072.0 | 0 | Carnivora | 7072.0 | 0 | Felidae | 7072.0 | 0 | Felis | 7072.0 | 0 | Felis catus | 7072.0 | 0 |
| scf7180000018997 | 635981 | 0.3787 | 0 | 48.6977 | Eukaryota | 35024.0 | 0 | Chordata | 35024.0 | 0 | Carnivora | 35024.0 | 0 | Felidae | 35024.0 | 0 | Felis | 35024.0 | 0 | Felis catus | 35024.0 | 0 |

| | | | | | | | | | | | | | | | | | | | | | | |
|---|---|---|---|---|---|---|---|---|---|---|---|---|---|---|---|---|---|---|---|---|---|---|
| scf7180000020959 | 66108 | 0.4612 | 0 | 24.1908 | Eukaryota | 39496.0 | 0 | Chordata | 39496.0 | 0 | Carnivora | 39496.0 | 0 | Felidae | 22284.0 | 2 | Lynx | 13777.0 | 4 | ynx canadens | 13777.0 | 4 |
| scf7180000024163 | 106656 | 0.6338 | 0 | 53.9683 | Eukaryota | 23580.0 | 0 | Chordata | 23580.0 | 0 | Carnivora | 23580.0 | 0 | Felidae | 9446.0 | 4 | Ursus | 6974.0 | 6 | Ursus arctos | 6974.0 | 6 |
| scf7180000023673 | 41077 | 0.5216 | 0 | 325.9664 | Eukaryota | 15306.0 | 0 | Chordata | 15306.0 | 0 | Carnivora | 15306.0 | 0 | Felidae | 15306.0 | 0 | Felis | 15306.0 | 0 | Felis catus | 15306.0 | 0 |
| scf7180000021539 | 20518 | 0.3835 | 0 | 47.8819 | Eukaryota | 14333.0 | 0 | Chordata | 14333.0 | 0 | Carnivora | 14333.0 | 0 | Felidae | 14333.0 | 0 | Felis | 11506.0 | 1 | Felis catus | 11506.0 | 1 |
| scf7180000023919 | 614340 | 0.4763 | 0 | 52.7089 | Eukaryota | 36803.0 | 0 | Chordata | 36803.0 | 0 | Carnivora | 36803.0 | 0 | Felidae | 26103.0 | 1 | Suricata | 10700.0 | 4 | ıricata suricat | 10700.0 | 4 |
| scf7180000022563 | 6970 | 0.5626 | 0 | 45.5047 | Eukaryota | 2366.0 | 0 | Chordata | 2366.0 | 0 | Carnivora | 2366.0 | 0 | Felidae | 2366.0 | 0 | Felis | 1898.0 | 2 | Felis catus | 1898.0 | 2 |
| scf7180000018932 | 36720 | 0.3686 | 0 | 20.7023 | Eukaryota | 26362.0 | 0 | Chordata | 26362.0 | 0 | Carnivora | 26362.0 | 0 | Felidae | 26362.0 | 0 | Felis | 23699.0 | 1 | Felis catus | 23699.0 | 1 |
| scf7180000019277 | 90091 | 0.45 | 0 | 95.9223 | Eukaryota | 7767.0 | 0 | Chordata | 7767.0 | 0 | Carnivora | 6993.0 | 1 | Felidae | 6274.0 | 2 | Felis | 2733.0 | 5 | Felis catus | 2733.0 | 5 |
| scf7180000019791 | 692759 | 0.4403 | 100 | 53.0984 | Eukaryota | 46112.0 | 0 | Chordata | 46112.0 | 0 | Carnivora | 46112.0 | 0 | Felidae | 46112.0 | 0 | Felis | 46112.0 | 0 | Felis catus | 46112.0 | 0 |
| scf7180000017782 | 660796 | 0.4007 | 0 | 49.8936 | Eukaryota | 47748.0 | 0 | Chordata | 47748.0 | 0 | Carnivora | 47748.0 | 0 | Felidae | 47748.0 | 0 | Felis | 47748.0 | 0 | Felis catus | 47748.0 | 0 |
| scf7180000018898 | 86946 | 0.4391 | 0 | 26.399 | Eukaryota | 74661.0 | 0 | Chordata | 74661.0 | 0 | Carnivora | 66664.0 | 1 | Felidae | 39789.0 | 3 | Suricata | 18010.0 | 7 | ıricata suricat | 18010.0 | 7 |
| scf7180000023854 | 561301 | 0.4185 | 0 | 50.6989 | Eukaryota | 34126.0 | 0 | Chordata | 34126.0 | 0 | Carnivora | 34126.0 | 0 | Felidae | 34126.0 | 0 | Felis | 34126.0 | 0 | Felis catus | 34126.0 | 0 |
| scf7180000022313 | 31039 | 0.3589 | 0 | 24.4316 | Eukaryota | 5528.0 | 0 | Chordata | 5528.0 | 0 | Carnivora | 5528.0 | 0 | Felidae | 4490.0 | 1 | Felis | 3921.0 | 2 | Felis catus | 3921.0 | 2 |
| scf7180000021649 | 28318 | 0.3616 | 0 | 24.5155 | Eukaryota | 4370.0 | 0 | Chordata | 4370.0 | 0 | Primates | 1510.0 | 3 | Hominidae | 1510.0 | 4 | Sus | 1297.0 | 6 | Sus scrofa | 1297.0 | 7 |
| scf7180000020445 | 15429 | 0.4551 | 0 | 25.778 | Eukaryota | 2511.0 | 0 | Chordata | 2511.0 | 0 | Carnivora | 2511.0 | 0 | Felidae | 2257.0 | 1 | Felis | 2011.0 | 2 | Felis catus | 2011.0 | 2 |
| scf7180000019694 | 351361 | 0.4536 | 0 | 51.149 | Eukaryota | 34864.0 | 0 | Chordata | 34864.0 | 0 | Carnivora | 22523.0 | 2 | Otariidae | 7989.0 | 5 | Ursus | 7214.0 | 7 | Homo sapiens | 6336.0 | 8 |
| scf7180000017508 | 55028 | 0.3852 | 0 | 38.1626 | Eukaryota | 45291.0 | 0 | Chordata | 45291.0 | 0 | Carnivora | 45291.0 | 0 | Felidae | 45291.0 | 0 | Felis | 45291.0 | 0 | Felis catus | 45291.0 | 0 |
| scf7180000020088 | 114932 | 0.6292 | 0 | 49.7882 | Eukaryota | 17252.0 | 0 | Chordata | 17252.0 | 0 | Carnivora | 17252.0 | 0 | Felidae | 17252.0 | 0 | Felis | 17252.0 | 0 | Felis catus | 17252.0 | 0 |
| scf7180000021608 | 25005 | 0.4366 | 0 | 25.7841 | Eukaryota | 20223.0 | 0 | Chordata | 20223.0 | 0 | Carnivora | 20223.0 | 0 | Felidae | 15620.0 | 3 | Panthera | 6588.0 | 5 | Felis catus | 4562.0 | 6 |
| scf7180000019798 | 178557 | 0.6137 | 0 | 52.5483 | Eukaryota | 48035.0 | 0 | Chordata | 48035.0 | 0 | Carnivora | 48035.0 | 0 | Felidae | 48035.0 | 0 | Panthera | 23915.0 | 3 | anthera pardu | 23915.0 | 3 |
| scf7180000021311 | 53879 | 0.349 | 0 | 49.8012 | Eukaryota | 17044.0 | 0 | Chordata | 17044.0 | 0 | Carnivora | 17044.0 | 0 | Felidae | 17044.0 | 0 | Felis | 15474.0 | 1 | Felis catus | 15474.0 | 1 |
| scf7180000022287 | 21913 | 0.3907 | 0 | 43.4727 | Eukaryota | 34586.0 | 0 | Chordata | 34586.0 | 0 | Carnivora | 34586.0 | 0 | Felidae | 34586.0 | 0 | Felis | 34586.0 | 0 | Felis catus | 34586.0 | 0 |
| scf7180000021529 | 55171 | 0.473 | 0 | 26.5818 | Eukaryota | 9768.0 | 0 | Chordata | 9768.0 | 0 | Carnivora | 9768.0 | 0 | Felidae | 3518.0 | 4 | Suricata | 1788.0 | 9 | ıricata suricat | 1788.0 | 9 |
| scf7180000021802 | 38505 | 0.4429 | 0 | 49.1202 | Eukaryota | 45640.0 | 0 | Chordata | 45640.0 | 0 | Carnivora | 45640.0 | 0 | Felidae | 45640.0 | 0 | Felis | 45640.0 | 0 | Felis catus | 45640.0 | 0 |
| scf7180000017566 | 2363606 | 0.3704 | 0 | 49.7522 | Eukaryota | 46624.0 | 0 | Chordata | 46624.0 | 0 | Carnivora | 46624.0 | 0 | Felidae | 46624.0 | 0 | Felis | 46624.0 | 0 | Felis catus | 46624.0 | 0 |
| scf7180000018688 | 599042 | 0.5825 | 0 | 51.0879 | Eukaryota | 10690.0 | 0 | Chordata | 10690.0 | 0 | Carnivora | 9618.0 | 1 | Felidae | 4313.0 | 4 | Zalophus | 2106.0 | 8 | phus californi | 2106.0 | 8 |
| scf7180000024236 | 19945 | 0.3721 | 0 | 23.8873 | Eukaryota | 29429.0 | 0 | Chordata | 29429.0 | 0 | Carnivora | 26958.0 | 1 | Felidae | 17408.0 | 4 | Panthera | 6801.0 | 7 | :inonyx jubat | 3736.0 | 9 |
| scf7180000021925 | 46082 | 0.3747 | 0 | 23.9753 | Eukaryota | 35359.0 | 0 | Chordata | 35359.0 | 0 | Carnivora | 35359.0 | 0 | Felidae | 35359.0 | 0 | Felis | 31853.0 | 1 | Felis catus | 31853.0 | 1 |
| scf7180000021530 | 28977 | 0.3889 | 0 | 295.5327 | Eukaryota | 12301.0 | 0 | Chordata | 12301.0 | 0 | Carnivora | 12301.0 | 0 | Felidae | 12301.0 | 0 | Felis | 8638.0 | 1 | Felis catus | 8638.0 | 1 |
| scf7180000019118 | 13428 | 0.3925 | 0 | 22.1177 | Eukaryota | 45366.0 | 0 | Chordata | 45366.0 | 0 | Carnivora | 45366.0 | 0 | Felidae | 45366.0 | 0 | Felis | 45366.0 | 0 | Felis catus | 45366.0 | 0 |
| scf7180000022272 | 12503 | 0.5374 | 0 | 47.7951 | Eukaryota | 3267.0 | 0 | Chordata | 3267.0 | 0 | Carnivora | 3267.0 | 0 | Felidae | 1855.0 | 1 | Felis | 1071.0 | 5 | Felis catus | 1071.0 | 5 |
| scf7180000017746 | 1696197 | 0.3678 | 0 | 49.9698 | Eukaryota | 47492.0 | 0 | Chordata | 47492.0 | 0 | Carnivora | 47492.0 | 0 | Felidae | 47492.0 | 0 | Felis | 47492.0 | 0 | Felis catus | 47492.0 | 0 |
| scf7180000020476 | 34062 | 0.4202 | 0 | 32.3849 | Eukaryota | 11105.0 | 0 | Chordata | 11105.0 | 0 | Artiodactyla | 9143.0 | 1 | Bovidae | 9143.0 | 1 | Ovis | 9143.0 | 1 | )vis canadens | 9143.0 | 1 |
| scf7180000021217 | 44446 | 0.4854 | 0 | 27.5863 | Eukaryota | 14861.0 | 0 | Chordata | 14861.0 | 0 | Carnivora | 14861.0 | 0 | Phocidae | 7716.0 | 2 | .eptonychote | 6430.0 | 3 | ›nychotes wed | 6430.0 | 3 |
| scf7180000021749 | 82013 | 0.6186 | 0 | 44.3312 | Eukaryota | 9186.0 | 0 | Chordata | 9186.0 | 0 | Carnivora | 9186.0 | 0 | Felidae | 4951.0 | 5 | Panthera | 2057.0 | 8 | Panthera tigri | 1120.0 | 9 |
| scf7180000023843 | 1256476 | 0.395 | 0 | 51.8643 | Eukaryota | 54960.0 | 0 | Chordata | 54960.0 | 0 | Chiroptera | 54960.0 | 0 | Vespertilionidae | 54960.0 | 0 | Myotis | 54960.0 | 0 | Myotis brandt | 54960.0 | 0 |
| scf7180000018478 | 681526 | 0.472 | 0 | 52.4365 | Eukaryota | 42982.0 | 0 | Chordata | 42982.0 | 0 | Carnivora | 42982.0 | 0 | Felidae | 42982.0 | 0 | Panthera | 26634.0 | 1 | anthera pardu | 26634.0 | 1 |
| scf7180000020989 | 65279 | 0.5402 | 0 | 52.175 | Eukaryota | 9171.0 | 0 | Chordata | 9171.0 | 0 | Carnivora | 9171.0 | 0 | Felidae | 7027.0 | 2 | Felis | 1834.0 | 6 | Felis catus | 1834.0 | 7 |
| scf7180000017503 | 28853 | 0.3713 | 0 | 20.7927 | Eukaryota | 41824.0 | 0 | Chordata | 41824.0 | 0 | Carnivora | 41824.0 | 0 | Felidae | 41824.0 | 0 | Felis | 41824.0 | 0 | Felis catus | 41824.0 | 0 |
| scf7180000019055 | 377995 | 0.5514 | 0 | 51.8458 | Eukaryota | 13849.0 | 0 | Chordata | 13849.0 | 0 | Carnivora | 13849.0 | 0 | Canidae | 7893.0 | 2 | Canis | 7893.0 | 4 | Canis lupus | 7893.0 | 4 |
| scf7180000022419 | 15735 | 0.4099 | 0 | 1652.8362 | no-hit | 0.0 | 0 | no-hit | 0.0 | 0 | no-hit | 0.0 | 0 | no-hit | 0.0 | 0 | no-hit | 0.0 | 0 | no-hit | 0.0 | 0 |
| scf7180000023131 | 20537 | 0.4844 | 0 | 229.1431 | Eukaryota | 11883.0 | 0 | Chordata | 11883.0 | 0 | Carnivora | 11883.0 | 0 | Felidae | 11883.0 | 0 | Felis | 10706.0 | 1 | Felis catus | 10706.0 | 1 |
| scf7180000019060 | 383979 | 0.4107 | 0 | 52.0913 | Eukaryota | 47321.0 | 0 | Chordata | 47321.0 | 0 | Carnivora | 47321.0 | 0 | Felidae | 47321.0 | 0 | Felis | 47321.0 | 0 | Felis catus | 47321.0 | 0 |
| scf7180000023744 | 23854 | 0.4411 | 0 | 204.9369 | Eukaryota | 3002.0 | 0 | Chordata | 3002.0 | 0 | Artiodactyla | 2865.0 | 1 | Bovidae | 2865.0 | 1 | Ovis | 2865.0 | 1 | )vis canadens | 2142.0 | 2 |
| scf7180000022308 | 65351 | 0.3514 | 0 | 23.5052 | Eukaryota | 20885.0 | 0 | Chordata | 20885.0 | 0 | Carnivora | 20885.0 | 0 | Felidae | 20885.0 | 0 | Felis | 20885.0 | 0 | Felis catus | 20885.0 | 0 |
| scf7180000018353 | 829742 | 0.5102 | 100 | 51.6745 | Eukaryota | 49948.0 | 0 | Chordata | 49948.0 | 0 | Carnivora | 49948.0 | 0 | Felidae | 30965.0 | 1 | Crocuta | 18983.0 | 3 | rocuta crocut | 18983.0 | 3 |
| scf7180000024276 | 562587 | 0.3808 | 0 | 50.2295 | Eukaryota | 45400.0 | 0 | Chordata | 45400.0 | 0 | Carnivora | 45400.0 | 0 | Felidae | 45400.0 | 0 | Felis | 40882.0 | 1 | Felis catus | 40882.0 | 1 |
| scf7180000023780 | 167603 | 0.3746 | 0 | 48.6952 | Eukaryota | 44741.0 | 0 | Chordata | 44741.0 | 0 | Carnivora | 44741.0 | 0 | Felidae | 44741.0 | 0 | Felis | 44741.0 | 0 | Felis catus | 44741.0 | 0 |
| scf7180000020043 | 169038 | 0.4861 | 0 | 51.4203 | Eukaryota | 19715.0 | 0 | Chordata | 19715.0 | 0 | Carnivora | 19715.0 | 0 | Felidae | 19715.0 | 0 | Felis | 19715.0 | 0 | Felis catus | 19715.0 | 0 |
| scf7180000017469 | 3383909 | 0.3873 | 0 | 52.2621 | Eukaryota | 46615.0 | 0 | Chordata | 46615.0 | 0 | Carnivora | 46615.0 | 0 | Felidae | 46615.0 | 0 | Felis | 28167.0 | 1 | Felis catus | 28167.0 | 1 |
| scf7180000022246 | 2069553 | 0.4218 | 0 | 51.8168 | Eukaryota | 48283.0 | 0 | Chordata | 48283.0 | 0 | Carnivora | 48283.0 | 0 | Felidae | 28391.0 | 1 | Felis | 23670.0 | 2 | Felis catus | 23670.0 | 2 |
| scf7180000017544 | 2650683 | 0.3657 | 0 | 49.6757 | Eukaryota | 57137.0 | 1 | Chordata | 57137.0 | 1 | Primates | 32651.0 | 2 | Hominidae | 25753.0 | 4 | Homo | 20393.0 | 5 | Homo sapiens | 20393.0 | 5 |
| scf7180000022566 | 10990 | 0.4261 | 0 | 25.3015 | Eukaryota | 45945.0 | 0 | Chordata | 45945.0 | 0 | Carnivora | 45945.0 | 0 | Felidae | 45945.0 | 0 | Felis | 45945.0 | 0 | Felis catus | 45945.0 | 0 |
| scf7180000018102 | 1138300 | 0.3944 | 0 | 57.7367 | Eukaryota | 46683.0 | 0 | Chordata | 46683.0 | 0 | Carnivora | 46683.0 | 0 | Felidae | 46683.0 | 0 | Felis | 42086.0 | 1 | Felis catus | 42086.0 | 1 |
| scf7180000019288 | 431999 | 0.4163 | 100 | 49.4465 | Eukaryota | 45764.0 | 0 | Chordata | 45764.0 | 0 | Carnivora | 45764.0 | 0 | Felidae | 45764.0 | 0 | Felis | 45764.0 | 0 | Felis catus | 45764.0 | 0 |
| scf7180000019449 | 269079 | 0.4467 | 0 | 49.1552 | Eukaryota | 31990.0 | 0 | Chordata | 31990.0 | 0 | Carnivora | 31990.0 | 0 | Felidae | 31990.0 | 0 | Felis | 28734.0 | 1 | Felis catus | 28734.0 | 1 |
| scf7180000021276 | 61873 | 0.3823 | 0 | 25.5205 | Eukaryota | 42728.0 | 0 | Chordata | 42728.0 | 0 | Carnivora | 42728.0 | 0 | Felidae | 42728.0 | 0 | Felis | 42728.0 | 0 | Felis catus | 42728.0 | 0 |
| scf7180000022783 | 16875 | 0.3517 | 0 | 24.1557 | Eukaryota | 22675.0 | 0 | Chordata | 22675.0 | 0 | Carnivora | 22675.0 | 0 | Felidae | 22675.0 | 0 | Felis | 22675.0 | 0 | Felis catus | 22675.0 | 0 |
| scf7180000019078 | 46849 | 0.5344 | 0 | 155.4353 | Eukaryota | 2893.0 | 0 | Chordata | 2893.0 | 0 | Carnivora | 2893.0 | 0 | Felidae | 1900.0 | 1 | Felis | 1676.0 | 3 | Felis catus | 1676.0 | 3 |
| scf7180000020189 | 21094 | 0.3966 | 0 | 442.09 | Eukaryota | 6531.0 | 0 | Chordata | 6531.0 | 0 | Carnivora | 6531.0 | 0 | Felidae | 6531.0 | 0 | Felis | 6531.0 | 0 | Felis catus | 6531.0 | 0 |
| scf7180000024119 | 150685 | 0.4089 | 0 | 49.3483 | Eukaryota | 46266.0 | 0 | Chordata | 46266.0 | 0 | Carnivora | 46266.0 | 0 | Felidae | 46266.0 | 0 | Felis | 46266.0 | 0 | Felis catus | 46266.0 | 0 |
| scf7180000021062 | 64360 | 0.4072 | 0 | 26.2264 | Eukaryota | 14750.0 | 0 | Chordata | 14750.0 | 0 | Primates | 8592.0 | 5 | Hominidae | 8592.0 | 5 | Homo | 5259.0 | 6 | Homo sapiens | 5259.0 | 6 |
| scf7180000022900 | 38305 | 0.4769 | 0 | 151.2831 | Eukaryota | 3919.0 | 0 | Chordata | 3919.0 | 0 | Carnivora | 2533.0 | 1 | Felidae | 2085.0 | 2 | Homo | 1386.0 | 4 | Homo sapiens | 1386.0 | 5 |
| scf7180000017432 | 1131093 | 0.422 | 0 | 51.8607 | Eukaryota | 77940.0 | 0 | Chordata | 77940.0 | 0 | Carnivora | 51528.0 | 2 | Herpestidae | 29142.0 | 4 | Suricata | 29142.0 | 4 | ıricata suricat | 29142.0 | 4 |
| scf7180000022725 | 4653 | 0.4083 | 0 | 667.5988 | no-hit | 0.0 | 0 | no-hit | 0.0 | 0 | no-hit | 0.0 | 0 | no-hit | 0.0 | 0 | no-hit | 0.0 | 0 | no-hit | 0.0 | 0 |
| scf7180000022472 | 8286 | 0.521 | 0 | 164.2365 | no-hit | 0.0 | 0 | no-hit | 0.0 | 0 | no-hit | 0.0 | 0 | no-hit | 0.0 | 0 | no-hit | 0.0 | 0 | no-hit | 0.0 | 0 |
| scf7180000020474 | 110685 | 0.4179 | 0 | 51.5711 | Eukaryota | 164190.0 | 0 | Chordata | 164190.0 | 0 | Carnivora | 164190.0 | 0 | Felidae | 164190.0 | 0 | Felis | 164190.0 | 0 | Felis catus | 164190.0 | 0 |
| scf7180000018579 | 291096 | 0.3539 | 0 | 48.976 | Eukaryota | 46341.0 | 0 | Chordata | 46341.0 | 0 | Carnivora | 46341.0 | 0 | Felidae | 46341.0 | 0 | Felis | 46341.0 | 0 | Felis catus | 46341.0 | 0 |
| scf7180000022997 | 40894 | 0.4734 | 0 | 228.9867 | Eukaryota | 24489.0 | 0 | Chordata | 24489.0 | 0 | Carnivora | 24489.0 | 0 | Felidae | 24489.0 | 0 | Felis | 21979.0 | 1 | Felis catus | 21979.0 | 1 |
| scf7180000019894 | 162658 | 0.3873 | 0 | 49.2361 | Eukaryota | 11285.0 | 0 | Chordata | 11285.0 | 0 | Carnivora | 11285.0 | 0 | Felidae | 11285.0 | 0 | Felis | 10169.0 | 1 | Felis catus | 10169.0 | 1 |
| scf7180000018586 | 654052 | 0.4062 | 0 | 49.9933 | Eukaryota | 52706.0 | 0 | Chordata | 52706.0 | 0 | Artiodactyla | 30723.0 | 2 | Monodontidae | 30723.0 | 2 | )elphinapteru | 30723.0 | 2 | ›hinapterus le | 30723.0 | 3 |
| scf7180000023733 | 28753 | 0.4014 | 0 | 312.8426 | Eukaryota | 3842.0 | 0 | Chordata | 3842.0 | 0 | Primates | 2471.0 | 1 | Hominidae | 2103.0 | 3 | Homo | 2103.0 | 3 | Homo sapiens | 2103.0 | 3 |
| scf7180000018986 | 605972 | 0.5231 | 0 | 52.7743 | Eukaryota | 43789.0 | 0 | Chordata | 43789.0 | 0 | Carnivora | 43789.0 | 0 | Felidae | 13800.0 | 4 | Canis | 8246.0 | 8 | Canis lupus | 8246.0 | 8 |
| scf7180000020589 | 173966 | 0.5491 | 0 | 48.0573 | Eukaryota | 14868.0 | 0 | Chordata | 14868.0 | 0 | Carnivora | 14868.0 | 0 | Felidae | 14868.0 | 0 | Panthera | 7544.0 | 4 | anthera pardu | 4506.0 | 5 |
| scf7180000019105 | 1407994 | 0.4249 | 0 | 50.8008 | Eukaryota | 48683.0 | 0 | Chordata | 48683.0 | 0 | Carnivora | 48683.0 | 0 | Felidae | 48683.0 | 0 | Felis | 43949.0 | 1 | Felis catus | 43949.0 | 1 |
| scf7180000022028 | 24958 | 0.4391 | 0 | 24.9748 | Eukaryota | 2999.0 | 0 | Chordata | 2999.0 | 0 | Carnivora | 2341.0 | 1 | Felidae | 2063.0 | 2 | Felis | 1557.0 | 3 | Felis catus | 1557.0 | 3 |
| scf7180000021502 | 42491 | 0.3697 | 0 | 25.4918 | Eukaryota | 11898.0 | 0 | Chordata | 11898.0 | 0 | Carnivora | 10483.0 | 1 | Felidae | 10483.0 | 1 | Felis | 9319.0 | 2 | Felis catus | 9319.0 | 2 |
| scf7180000020949 | 64155 | 0.4125 | 0 | 25.2667 | Eukaryota | 45535.0 | 0 | Chordata | 45535.0 | 0 | Carnivora | 45535.0 | 0 | Felidae | 45535.0 | 0 | Felis | 45535.0 | 0 | Felis catus | 45535.0 | 0 |
| scf7180000021160 | 44770 | 0.4641 | 0 | 24.0145 | Eukaryota | 35782.0 | 0 | Chordata | 35782.0 | 0 | Artiodactyla | 13351.0 | 3 | Hominidae | 12378.0 | 5 | Homo | 12378.0 | 5 | Homo sapiens | 12378.0 | 6 |
| scf7180000017452 | 1936113 | 0.3653 | 0 | 48.8213 | Eukaryota | 46786.0 | 0 | Chordata | 46786.0 | 0 | Carnivora | 46786.0 | 0 | Felidae | 46786.0 | 0 | Felis | 46786.0 | 0 | Felis catus | 46786.0 | 0 |
| scf7180000022464 | 6922 | 0.4546 | 0 | 24.6353 | Eukaryota | 5388.0 | 0 | Chordata | 5388.0 | 0 | Primates | 2931.0 | 2 | Hominidae | 2931.0 | 3 | Homo | 2931.0 | 4 | Homo sapiens | 2931.0 | 4 |
| scf7180000024026 | 375629 | 0.6226 | 0 | 50.4777 | Eukaryota | 13109.0 | 0 | Chordata | 13109.0 | 0 | Carnivora | 11836.0 | 1 | Felidae | 10816.0 | 2 | Felis | 7536.0 | 4 | Felis catus | 7536.0 | 4 |
| scf7180000019940 | 203709 | 0.4453 | 0 | 53.5437 | Eukaryota | 47868.0 | 0 | Chordata | 47868.0 | 0 | Carnivora | 47868.0 | 0 | Felidae | 47868.0 | 0 | Felis | 47868.0 | 0 | Felis catus | 47868.0 | 0 |
| scf7180000019157 | 1194934 | 0.4119 | 0 | 51.1013 | Eukaryota | 69228.0 | 0 | Chordata | 69228.0 | 0 | Carnivora | 69228.0 | 0 | Felidae | 53185.0 | 1 | Panthera | 33256.0 | 2 | anthera pardu | 26596.0 | 3 |
| scf7180000024099 | 250250 | 0.6589 | 0 | 49.0237 | Eukaryota | 19525.0 | 0 | Chordata | 19525.0 | 0 | Carnivora | 19525.0 | 0 | Felidae | 19525.0 | 0 | Acinonyx | 7869.0 | 2 | :inonyx jubat | 7869.0 | 3 |
| scf7180000018908 | 128418 | 0.4108 | 0 | 48.2995 | Eukaryota | 14188.0 | 0 | Chordata | 14188.0 | 0 | Carnivora | 14188.0 | 0 | Felidae | 14188.0 | 0 | Felis | 9906.0 | 2 | Felis catus | 9906.0 | 2 |
| scf7180000020836 | 207239 | 0.3653 | 0 | 47.3375 | Eukaryota | 27277.0 | 0 | Chordata | 27277.0 | 0 | Carnivora | 27277.0 | 0 | Felidae | 27277.0 | 0 | Felis | 24482.0 | 1 | Felis catus | 24482.0 | 1 |
| scf7180000022191 | 25974 | 0.4285 | 0 | 26.9041 | Eukaryota | 34684.0 | 0 | Chordata | 34684.0 | 0 | Carnivora | 34684.0 | 0 | Felidae | 34684.0 | 0 | Felis | 31214.0 | 1 | Felis catus | 31214.0 | 1 |
| scf7180000021811 | 34542 | 0.4457 | 0 | 27.2274 | Eukaryota | 14067.0 | 0 | Chordata | 14067.0 | 0 | Primates | 12037.0 | 1 | Hominidae | 8729.0 | 2 | Homo | 7046.0 | 4 | Homo sapiens | 7046.0 | 5 |
| scf7180000024134 | 280507 | 0.4617 | 0 | 25.7638 | Eukaryota | 23358.0 | 0 | Chordata | 23358.0 | 0 | Carnivora | 23358.0 | 0 | Felidae | 14670.0 | 3 | Lynx | 7254.0 | 6 | ynx canadens | 7254.0 | 7 |
| scf7180000018022 | 1171045 | 0.4087 | 0 | 50.957 | Eukaryota | 55178.0 | 0 | Chordata | 55178.0 | 0 | Carnivora | 55178.0 | 0 | Felidae | 55178.0 | 0 | Acinonyx | 33030.0 | 1 | :inonyx jubat | 33030.0 | 1 |
| scf7180000022633 | 12857 | 0.4191 | 0 | 25.256 | Eukaryota | 43847.0 | 0 | Chordata | 43847.0 | 0 | Carnivora | 43847.0 | 0 | Felidae | 43847.0 | 0 | Felis | 43847.0 | 0 | Felis catus | 43847.0 | 0 |
| scf7180000020093 | 189368 | 0.4654 | 0 | 53.1699 | Eukaryota | 10435.0 | 0 | Chordata | 10435.0 | 0 | Carnivora | 8584.0 | 1 | Felidae | 8584.0 | 1 | Panthera | 5090.0 | 3 | anthera pardu | 5090.0 | 3 |
| scf7180000022125 | 23026 | 0.3902 | 0 | 35.5349 | Eukaryota | 47626.0 | 0 | Chordata | 47626.0 | 0 | Carnivora | 47626.0 | 0 | Felidae | 47626.0 | 0 | Felis | 47626.0 | 0 | Felis catus | 47626.0 | 0 |
| scf7180000018309 | 1008852 | 0.3722 | 0 | 51.5014 | Eukaryota | 32336.0 | 0 | Chordata | 32336.0 | 0 | Carnivora | 32336.0 | 0 | Felidae | 32336.0 | 0 | Felis | 29150.0 | 1 | Felis catus | 29150.0 | 1 |
| scf7180000023460 | 71676 | 0.4262 | 0 | 243.2189 | Eukaryota | 8893.0 | 0 | Chordata | 8893.0 | 0 | Carnivora | 8893.0 | 0 | Felidae | 8152.0 | 1 | Felis | 6277.0 | 2 | Felis catus | 6277.0 | 2 |
| scf7180000022209 | 12439 | 0.4529 | 0 | 11.9879 | Eukaryota | 7924.0 | 0 | Chordata | 7924.0 | 0 | Carnivora | 7924.0 | 0 | Felidae | 5515.0 | 1 | Panthera | 2381.0 | 4 | anthera pardu | 2381.0 | 4 |
| scf7180000022146 | 21489 | 0.3482 | 0 | 24.3996 | Eukaryota | 22376.0 | 0 | Chordata | 22376.0 | 0 | Carnivora | 22376.0 | 0 | Felidae | 22376.0 | 0 | Felis | 20121.0 | 1 | Felis catus | 20121.0 | 1 |
| scf7180000020409 | 117230 | 0.4481 | 0 | 25.686 | Eukaryota | 24630.0 | 0 | Chordata | 24630.0 | 0 | Carnivora | 24630.0 | 0 | Felidae | 15186.0 | 1 | Acinonyx | 15186.0 | 2 | :inonyx jubat | 15186.0 | 2 |
| scf7180000022677 | 5596 | 0.3815 | 0 | 23.7944 | Eukaryota | 2056.0 | 0 | Chordata | 2056.0 | 0 | Carnivora | 2056.0 | 0 | Felidae | 1832.0 | 1 | Felis | 1236.0 | 3 | Felis catus | 1236.0 | 4 |
| scf7180000017811 | 2144107 | 0.4156 | 0 | 51.1038 | Eukaryota | 62888.0 | 0 | Chordata | 62888.0 | 0 | Carnivora | 62888.0 | 0 | Felidae | 62888.0 | 0 | Panthera | 37730.0 | 2 | anthera pardu | 25168.0 | 3 |
| scf7180000022449 | 19934 | 0.3791 | 0 | 50.2158 | Eukaryota | 33061.0 | 0 | Chordata | 33061.0 | 0 | Carnivora | 33061.0 | 0 | Felidae | 33061.0 | 0 | Felis | 33061.0 | 0 | Felis catus | 33061.0 | 0 |
| scf7180000018816 | 55334 | 0.3454 | 0 | 39.0743 | Eukaryota | 15683.0 | 0 | Chordata | 15683.0 | 0 | Carnivora | 9722.0 | 2 | Felidae | 9722.0 | 2 | Felis | 9722.0 | 2 | Felis catus | 9722.0 | 2 |
| scf7180000017695 | 649732 | 0.3851 | 100 | 49.6955 | Eukaryota | 47743.0 | 0 | Chordata | 47743.0 | 0 | Carnivora | 47743.0 | 0 | Felidae | 47743.0 | 0 | Felis | 47743.0 | 0 | Felis catus | 47743.0 | 0 |
| scf7180000022734 | 29527 | 0.4294 | 0 | 220.8841 | Eukaryota | 2150.0 | 0 | Chordata | 2150.0 | 0 | Carnivora | 2150.0 | 0 | Felidae | 2150.0 | 0 | Felis | 1310.0 | 2 | Felis catus | 1310.0 | 3 |
| scf7180000018484 | 166887 | 0.5356 | 0 | 50.818 | Eukaryota | 17313.0 | 0 | Chordata | 17313.0 | 0 | Carnivora | 9381.0 | 2 | Hominidae | 5919.0 | 4 | Homo | 5919.0 | 5 | Homo sapiens | 5919.0 | 5 |
| scf7180000018891 | 577127 | 0.4332 | 0 | 51.3718 | Eukaryota | 29416.0 | 0 | Chordata | 29416.0 | 0 | Carnivora | 29416.0 | 0 | Felidae | 29416.0 | 0 | Felis | 26429.0 | 1 | Felis catus | 26429.0 | 1 |
| scf7180000022892 | 42131 | 0.5641 | 0 | 364.8926 | Eukaryota | 14940.0 | 0 | Chordata | 14940.0 | 0 | Carnivora | 10721.0 | 3 | Felidae | 6047.0 | 6 | Neomonachus | 1655.0 | 9 | nachus schau | 1655.0 | 9 |
| scf7180000023034 | 24573 | 0.56 | 0 | 146.1838 | Eukaryota | 6553.0 | 0 | Chordata | 6553.0 | 0 | Carnivora | 6553.0 | 0 | Felidae | 6314.0 | 1 | Felis | 2548.0 | 3 | Felis catus | 2548.0 | 3 |
| scf7180000019375 | 374095 | 0.5707 | 0 | 51.7462 | Eukaryota | 23567.0 | 0 | Chordata | 23567.0 | 0 | Carnivora | 23567.0 | 0 | Felidae | 23567.0 | 0 | Felis | 21175.0 | 1 | Felis catus | 21175.0 | 1 |
| scf7180000023880 | 1494787 | 0.4519 | 0 | 50.8176 | Eukaryota | 100566.0 | 0 | Chordata | 100566.0 | 0 | Carnivora | 100566.0 | 0 | Felidae | 100566.0 | 0 | Felis | 40324.0 | 2 | Felis catus | 40324.0 | 2 |
| scf7180000022188 | 24387 | 0.458 | 0 | 144.4317 | Eukaryota | 3726.0 | 0 | Chordata | 3726.0 | 0 | Carnivora | 3726.0 | 0 | Felidae | 3267.0 | 1 | unresolved | 1089.0 | 4 | unresolved | 1089.0 | 4 |
| scf7180000020960 | 120753 | 0.4627 | 0 | 28.4107 | Eukaryota | 21847.0 | 0 | Chordata | 21847.0 | 0 | Carnivora | 21847.0 | 0 | Felidae | 21847.0 | 0 | Acinonyx | 10914.0 | 2 | :inonyx jubat | 10914.0 | 2 |
| scf7180000023907 | 109761 | 0.3674 | 0 | 45.9307 | Eukaryota | 48013.0 | 0 | Chordata | 48013.0 | 0 | Carnivora | 48013.0 | 0 | Felidae | 48013.0 | 0 | Felis | 48013.0 | 0 | Felis catus | 48013.0 | 0 |
| scf7180000021098 | 49395 | 0.5204 | 0 | 49.4418 | Eukaryota | 20854.0 | 0 | Chordata | 20854.0 | 0 | Carnivora | 20854.0 | 0 | Felidae | 20854.0 | 0 | Felis | 18715.0 | 1 | Felis catus | 18715.0 | 1 |
| scf7180000022282 | 9615 | 0.5096 | 0 | 25.8694 | Eukaryota | 4088.0 | 0 | Chordata | 4088.0 | 0 | Carnivora | 4088.0 | 0 | Felidae | 4088.0 | 0 | Felis | 1914.0 | 2 | Felis catus | 1914.0 | 2 |
| scf7180000021289 | 84668 | 0.5142 | 0 | 50.4857 | Eukaryota | 17554.0 | 0 | Chordata | 17554.0 | 0 | Carnivora | 17554.0 | 0 | Felidae | 17554.0 | 0 | Panthera | 14056.0 | 1 | anthera pardu | 14056.0 | 1 |
| scf7180000022479 | 17151 | 0.413 | 0 | 26.2563 | Eukaryota | 18635.0 | 0 | Chordata | 18635.0 | 0 | Carnivora | 18635.0 | 0 | Felidae | 18635.0 | 0 | Felis | 18635.0 | 0 | Felis catus | 18635.0 | 0 |
| scf7180000023379 | 25341 | 0.4383 | 0 | 174.6085 | Eukaryota | 3153.0 | 0 | Chordata | 3153.0 | 0 | Artiodactyla | 2871.0 | 1 | Bovidae | 2871.0 | 1 | Ovis | 2871.0 | 1 | )vis canadens | 2148.0 | 2 |
| scf7180000017621 | 1926039 | 0.4857 | 0 | 51.9162 | Eukaryota | 89815.0 | 0 | Chordata | 89815.0 | 0 | Primates | 80411.0 | 1 | Cercopithecidae | 80411.0 | 1 | Rhinopithecus | 26801.0 | 6 | ›pithecus roxe | 17876.0 | 8 |
| scf7180000019730 | 220691 | 0.3767 | 0 | 50.1418 | Eukaryota | 45982.0 | 0 | Chordata | 45982.0 | 0 | Carnivora | 45982.0 | 0 | Felidae | 45982.0 | 0 | Felis | 45982.0 | 0 | Felis catus | 45982.0 | 0 |
| scf7180000022748 | 8921 | 0.4603 | 0 | 25.7041 | Eukaryota | 11166.0 | 0 | Chordata | 11166.0 | 0 | Carnivora | 11166.0 | 0 | Otariidae | 8784.0 | 2 | Zalophus | 8784.0 | 2 | phus californi | 8784.0 | 2 |
| scf7180000019308 | 387299 | 0.3585 | 0 | 49.3155 | Eukaryota | 32863.0 | 0 | Chordata | 32863.0 | 0 | Carnivora | 32863.0 | 0 | Felidae | 32863.0 | 0 | Felis | 29952.0 | 1 | Felis catus | 29952.0 | 1 |
| scf7180000022795 | 12724 | 0.4029 | 0 | 28.1751 | Eukaryota | 32353.0 | 0 | Chordata | 32353.0 | 0 | Carnivora | 28399.0 | 1 | Otariidae | 16353.0 | 3 | Callorhinus | 9807.0 | 4 | llorhinus ursir | 9807.0 | 4 |
| scf7180000019697 | 89933 | 0.3517 | 0 | 45.4262 | Eukaryota | 14753.0 | 0 | Chordata | 14753.0 | 0 | Carnivora | 14753.0 | 0 | Felidae | 14753.0 | 0 | Felis | 14753.0 | 0 | Felis catus | 14753.0 | 0 |
| scf7180000020351 | 56886 | 0.4214 | 0 | 50.2338 | Eukaryota | 45322.0 | 0 | Chordata | 45322.0 | 0 | Carnivora | 45322.0 | 0 | Felidae | 45322.0 | 0 | Felis | 45322.0 | 0 | Felis catus | 45322.0 | 0 |
| scf7180000018328 | 927780 | 0.4468 | 0 | 50.2748 | Eukaryota | 43161.0 | 0 | Chordata | 43161.0 | 0 | Carnivora | 43161.0 | 0 | Felidae | 34597.0 | 2 | Acinonyx | 13480.0 | 4 | :inonyx jubat | 13480.0 | 4 |
| scf7180000021623 | 43246 | 0.3692 | 0 | 24.6089 | Eukaryota | 9209.0 | 0 | Chordata | 9209.0 | 0 | Carnivora | 9209.0 | 0 | Felidae | 9209.0 | 0 | Felis | 7480.0 | 1 | Felis catus | 7480.0 | 1 |
| scf7180000023832 | 1238985 | 0.4293 | 0 | 50.9597 | Eukaryota | 29223.0 | 0 | Chordata | 29223.0 | 0 | Carnivora | 29223.0 | 0 | Felidae | 29223.0 | 0 | Felis | 29223.0 | 0 | Felis catus | 29223.0 | 0 |
| scf7180000018423 | 644761 | 0.5131 | 0 | 50.9269 | Eukaryota | 67653.0 | 0 | Chordata | 67653.0 | 0 | Carnivora | 67653.0 | 0 | Felidae | 38893.0 | 2 | Felis | 31076.0 | 3 | Felis catus | 31076.0 | 3 |
| scf7180000018140 | 2156393 | 0.3672 | 0 | 50.4904 | Eukaryota | 77656.0 | 0 | Chordata | 77656.0 | 0 | Carnivora | 65970.0 | 1 | Phocidae | 65970.0 | 1 | .eptonychote | 65970.0 | 1 | ›nychotes wed | 65970.0 | 1 |
| scf7180000022059 | 25023 | 0.3936 | 0 | 24.1978 | Eukaryota | 45200.0 | 0 | Chordata | 45200.0 | 0 | Carnivora | 45200.0 | 0 | Felidae | 45200.0 | 0 | Felis | 45200.0 | 0 | Felis catus | 45200.0 | 0 |
| scf7180000018401 | 381706 | 0.4646 | 0 | 52.9387 | Eukaryota | 35026.0 | 0 | Chordata | 35026.0 | 0 | Carnivora | 35026.0 | 0 | Felidae | 23726.0 | 2 | Felis | 10560.0 | 4 | Felis catus | 10560.0 | 4 |
| scf7180000022157 | 72903 | 0.3997 | 0 | 75.8756 | Eukaryota | 31567.0 | 0 | Chordata | 31567.0 | 0 | Carnivora | 31567.0 | 0 | Felidae | 31567.0 | 0 | Felis | 31567.0 | 0 | Felis catus | 31567.0 | 0 |
| scf7180000018240 | 1014438 | 0.4319 | 0 | 51.3245 | Eukaryota | 49645.0 | 0 | Chordata | 49645.0 | 0 | Carnivora | 49645.0 | 0 | Felidae | 49645.0 | 0 | Felis | 44686.0 | 1 | Felis catus | 44686.0 | 1 |
| scf7180000018831 | 755913 | 0.4689 | 0 | 52.5228 | Eukaryota | 19059.0 | 1 | Chordata | 19059.0 | 1 | Primates | 17223.0 | 2 | Cercopithecidae | 9172.0 | 4 | Homo | 6368.0 | 6 | Homo sapiens | 6368.0 | 8 |
| scf7180000022075 | 24379 | 0.4302 | 0 | 25.5324 | Eukaryota | 20197.0 | 0 | Chordata | 20197.0 | 0 | Carnivora | 20197.0 | 0 | Felidae | 20197.0 | 0 | Panthera | 11525.0 | 2 | anthera pardu | 11525.0 | 2 |
| scf7180000023120 | 21722 | 0.4027 | 0 | 341.9579 | Eukaryota | 5445.0 | 0 | Chordata | 5445.0 | 0 | Carnivora | 5445.0 | 0 | Felidae | 5445.0 | 0 | Felis | 5445.0 | 0 | Felis catus | 5445.0 | 0 |
| scf7180000024003 | 1957412 | 0.3841 | 0 | 51.1261 | Eukaryota | 51635.0 | 0 | Chordata | 51635.0 | 0 | Carnivora | 51635.0 | 0 | Felidae | 51635.0 | 0 | Felis | 51635.0 | 0 | Felis catus | 51635.0 | 0 |
| scf7180000018529 | 6173193 | 0.374 | 0 | 50.4901 | Eukaryota | 95850.0 | 0 | Chordata | 95850.0 | 0 | Carnivora | 95850.0 | 0 | Felidae | 95850.0 | 0 | Acinonyx | 48120.0 | 1 | :inonyx jubat | 48120.0 | 1 |
| scf7180000019897 | 265512 | 0.4361 | 0 | 51.2277 | Eukaryota | 14186.0 | 1 | Chordata | 14186.0 | 1 | Primates | 10431.0 | 2 | Cercopithecidae | 10431.0 | 2 | Cercocebus | 5964.0 | 4 | ercocebus aty | 5964.0 | 4 |
| scf7180000022303 | 4060 | 0.5232 | 0 | 86.1363 | no-hit | 0.0 | 0 | no-hit | 0.0 | 0 | no-hit | 0.0 | 0 | no-hit | 0.0 | 0 | no-hit | 0.0 | 0 | no-hit | 0.0 | 0 |
| scf7180000024025 | 359094 | 0.4383 | 0 | 51.9177 | Eukaryota | 45553.0 | 0 | Chordata | 45553.0 | 0 | Carnivora | 45553.0 | 0 | Felidae | 45553.0 | 0 | Felis | 45553.0 | 0 | Felis catus | 45553.0 | 0 |
| scf7180000021251 | 38630 | 0.4609 | 0 | 84.1638 | Eukaryota | 2445.0 | 0 | Chordata | 2445.0 | 0 | Carnivora | 2445.0 | 0 | Felidae | 2199.0 | 1 | Felis | 2199.0 | 1 | Felis catus | 2199.0 | 1 |
| scf7180000019645 | 169337 | 0.403 | 0 | 30.6753 | Eukaryota | 40441.0 | 0 | Chordata | 40441.0 | 0 | Carnivora | 40441.0 | 0 | Felidae | 40441.0 | 0 | Felis | 36372.0 | 1 | Felis catus | 36372.0 | 1 |
| scf7180000021792 | 18791 | 0.6574 | 0 | 50.5571 | Eukaryota | 6312.0 | 0 | Chordata | 6312.0 | 0 | Primates | 4960.0 | 1 | Hominidae | 4960.0 | 1 | Homo | 4960.0 | 1 | Homo sapiens | 4960.0 | 1 |
| scf7180000023065 | 58816 | 0.3888 | 0 | 475.747 | Eukaryota | 8155.0 | 0 | Chordata | 8155.0 | 0 | Carnivora | 8155.0 | 0 | Felidae | 8155.0 | 0 | Felis | 8155.0 | 0 | Felis catus | 8155.0 | 0 |
| scf7180000018757 | 3388795 | 0.436 | 100 | 51.4932 | Eukaryota | 46937.0 | 0 | Chordata | 46937.0 | 0 | Carnivora | 46937.0 | 0 | Felidae | 46937.0 | 0 | Felis | 41978.0 | 1 | Felis catus | 41978.0 | 1 |
| scf7180000021840 | 52663 | 0.4326 | 0 | 24.7483 | Eukaryota | 21044.0 | 0 | Chordata | 21044.0 | 0 | Carnivora | 21044.0 | 0 | Felidae | 21044.0 | 0 | Felis | 18920.0 | 1 | Felis catus | 18920.0 | 1 |
| scf7180000020000 | 169962 | 0.4709 | 0 | 51.7465 | Eukaryota | 17920.0 | 0 | Chordata | 17920.0 | 0 | Carnivora | 17920.0 | 0 | Felidae | 17920.0 | 0 | Felis | 17920.0 | 0 | Felis catus | 17920.0 | 0 |
| scf7180000020806 | 113300 | 0.4105 | 0 | 36.0444 | Eukaryota | 42946.0 | 0 | Chordata | 42946.0 | 0 | Carnivora | 42946.0 | 0 | Felidae | 42946.0 | 0 | Felis | 38702.0 | 1 | Felis catus | 38702.0 | 1 |
| scf7180000017868 | 594040 | 0.3852 | 0 | 50.1438 | Eukaryota | 45865.0 | 0 | Chordata | 45865.0 | 0 | Carnivora | 45865.0 | 0 | Felidae | 45865.0 | 0 | Felis | 41333.0 | 1 | Felis catus | 41333.0 | 1 |
| scf7180000021236 | 58656 | 0.4244 | 0 | 24.6248 | Eukaryota | 45180.0 | 0 | Chordata | 45180.0 | 0 | Carnivora | 45180.0 | 0 | Felidae | 45180.0 | 0 | Felis | 45180.0 | 0 | Felis catus | 45180.0 | 0 |
| scf7180000020434 | 78237 | 0.3803 | 0 | 38.4515 | Eukaryota | 38325.0 | 0 | Chordata | 38325.0 | 0 | Carnivora | 38325.0 | 0 | Felidae | 38325.0 | 0 | Felis | 34432.0 | 1 | Felis catus | 34432.0 | 1 |
| scf7180000022939 | 62525 | 0.3812 | 0 | 504.4806 | Eukaryota | 7145.0 | 0 | Chordata | 7145.0 | 0 | Carnivora | 7145.0 | 0 | Felidae | 7145.0 | 0 | Felis | 7145.0 | 0 | Felis catus | 7145.0 | 0 |
| scf7180000023814 | 2095574 | 0.4561 | 0 | 51.4395 | Eukaryota | 46601.0 | 0 | Chordata | 46601.0 | 0 | Carnivora | 46601.0 | 0 | Felidae | 46601.0 | 0 | Felis | 46601.0 | 0 | Felis catus | 46601.0 | 0 |
| scf7180000024121 | 224830 | 0.632 | 0 | 48.5225 | Eukaryota | 6147.0 | 0 | Chordata | 6147.0 | 0 | Carnivora | 5751.0 | 1 | Felidae | 4779.0 | 2 | Lynx | 2769.0 | 5 | ynx canadens | 2769.0 | 5 |
| scf7180000019775 | 122207 | 0.4995 | 0 | 52.5243 | Eukaryota | 72705.0 | 0 | Chordata | 72705.0 | 0 | Carnivora | 72705.0 | 0 | Canidae | 72705.0 | 0 | Canis | 72705.0 | 0 | Canis lupus | 72705.0 | 0 |
| scf7180000022194 | 21828 | 0.4102 | 0 | 24.8257 | Eukaryota | 12838.0 | 0 | Chordata | 12838.0 | 0 | Carnivora | 6032.0 | 2 | Felidae | 6032.0 | 2 | Apteryx | 5651.0 | 3 | pteryx mante | 5651.0 | 3 |

| | | | | | | | | | | | | | | | | | | | | | | |
|---|---|---|---|---|---|---|---|---|---|---|---|---|---|---|---|---|---|---|---|---|---|---|
| scf7180000018790 | 577407 | 0.3722 | 0 | 50.4425 | Eukaryota | 47043.0 | 0 | Chordata | 47043.0 | 0 | Carnivora | 47043.0 | 0 | Felidae | 47043.0 | 0 | Felis | 47043.0 | 0 | Felis catus | 47043.0 | 0 |
| scf7180000019453 | 330260 | 0.5823 | 0 | 50.4339 | Eukaryota | 46817.0 | 0 | Chordata | 46817.0 | 0 | Carnivora | 46817.0 | 0 | Felidae | 42006.0 | 1 | Felis | 23365.0 | 3 | Felis catus | 23365.0 | 3 |
| scf7180000021159 | 57837 | 0.5085 | 0 | 50.1589 | Eukaryota | 12677.0 | 0 | Chordata | 12677.0 | 0 | Carnivora | 12677.0 | 0 | Felidae | 10134.0 | 2 | Panthera | 5450.0 | 4 | anthera pardu | 4908.0 | 6 |
| scf7180000022717 | 8660 | 0.5939 | 0 | 51.6143 | Eukaryota | 2111.0 | 0 | Chordata | 2111.0 | 0 | Carnivora | 2111.0 | 0 | Felidae | 2111.0 | 0 | Felis | 1488.0 | 3 | Felis catus | 1488.0 | 3 |
| scf7180000022253 | 22105 | 0.396 | 0 | 25.4831 | Eukaryota | 37891.0 | 0 | Chordata | 37891.0 | 0 | Carnivora | 37891.0 | 0 | Felidae | 37891.0 | 0 | Felis | 37891.0 | 0 | Felis catus | 37891.0 | 0 |
| scf7180000017978 | 24490 | 0.4327 | 0 | 25.7452 | Eukaryota | 5391.0 | 0 | Chordata | 5391.0 | 0 | Carnivora | 5391.0 | 0 | Felidae | 5391.0 | 0 | Felis | 5391.0 | 0 | Felis catus | 5391.0 | 0 |
| scf7180000023018 | 50569 | 0.5301 | 0 | 320.9109 | Eukaryota | 3020.0 | 0 | Chordata | 3020.0 | 0 | Carnivora | 3020.0 | 0 | Felidae | 2269.0 | 2 | Felis | 1332.0 | 4 | Felis catus | 1332.0 | 4 |
| scf7180000021677 | 55635 | 0.3953 | 100 | 26.2627 | Eukaryota | 33674.0 | 0 | Chordata | 33674.0 | 0 | Carnivora | 33674.0 | 0 | Felidae | 33674.0 | 0 | Felis | 33674.0 | 0 | Felis catus | 33674.0 | 0 |
| scf7180000021233 | 49861 | 0.3508 | 0 | 48.9847 | Eukaryota | 8109.0 | 0 | Chordata | 8109.0 | 0 | Carnivora | 8109.0 | 0 | Felidae | 8109.0 | 0 | Felis | 7311.0 | 1 | Felis catus | 7311.0 | 1 |
| scf7180000022338 | 20792 | 0.3799 | 0 | 24.176 | Eukaryota | 18137.0 | 0 | Chordata | 18137.0 | 0 | Carnivora | 18137.0 | 0 | Felidae | 18137.0 | 0 | Felis | 16290.0 | 1 | Felis catus | 16290.0 | 1 |
| scf7180000023782 | 3194906 | 0.3812 | 0 | 50.7208 | Eukaryota | 63493.0 | 0 | Chordata | 63493.0 | 0 | Carnivora | 63493.0 | 0 | Felidae | 63493.0 | 0 | Acinonyx | 23612.0 | 3 | :inonyx jubat | 23612.0 | 3 |
| scf7180000021525 | 42898 | 0.4117 | 0 | 24.6989 | Eukaryota | 11271.0 | 0 | Chordata | 11271.0 | 0 | Carnivora | 5678.0 | 2 | Felidae | 5678.0 | 2 | Felis | 5678.0 | 2 | Felis catus | 5678.0 | 2 |
| scf7180000020009 | 274494 | 0.4206 | 0 | 26.7325 | Eukaryota | 42467.0 | 0 | Chordata | 42467.0 | 0 | Carnivora | 42467.0 | 0 | Felidae | 42467.0 | 0 | Felis | 42467.0 | 0 | Felis catus | 42467.0 | 0 |
| scf7180000020725 | 71274 | 0.6563 | 0 | 46.8246 | Eukaryota | 9920.0 | 0 | Chordata | 9920.0 | 0 | Carnivora | 9920.0 | 0 | Felidae | 7625.0 | 2 | Panthera | 2218.0 | 6 | anthera pardu | 2218.0 | 6 |
| scf7180000017618 | 25720 | 0.3967 | 0 | 29.7872 | Eukaryota | 46154.0 | 0 | Chordata | 46154.0 | 0 | Carnivora | 46154.0 | 0 | Felidae | 46154.0 | 0 | Felis | 46154.0 | 0 | Felis catus | 46154.0 | 0 |
| scf7180000021615 | 39660 | 0.4105 | 0 | 25.2022 | Eukaryota | 18559.0 | 0 | Chordata | 18559.0 | 0 | Carnivora | 18559.0 | 0 | Felidae | 18559.0 | 0 | Felis | 16666.0 | 1 | Felis catus | 16666.0 | 1 |
| scf7180000018139 | 1374267 | 0.4473 | 0 | 53.0042 | Eukaryota | 43161.0 | 1 | Chordata | 43161.0 | 1 | Carnivora | 37236.0 | 2 | Felidae | 37236.0 | 2 | Felis | 37236.0 | 2 | Felis catus | 37236.0 | 2 |
| scf7180000019138 | 380136 | 0.3556 | 0 | 48.8893 | Eukaryota | 46581.0 | 0 | Chordata | 46581.0 | 0 | Carnivora | 46581.0 | 0 | Felidae | 46581.0 | 0 | Felis | 41905.0 | 1 | Felis catus | 41905.0 | 1 |
| scf7180000022117 | 10521 | 0.4272 | 0 | 24.3945 | Eukaryota | 19888.0 | 0 | Chordata | 19888.0 | 0 | Carnivora | 19888.0 | 0 | Otariidae | 10944.0 | 1 | Eumetopias | 5547.0 | 5 | netopias juba | 5547.0 | 5 |
| scf7180000020821 | 106397 | 0.4899 | 0 | 48.7169 | Eukaryota | 8973.0 | 1 | Chordata | 8973.0 | 1 | Artiodactyla | 3310.0 | 6 | Bovidae | 3310.0 | 7 | unresolved | 1362.0 | 9 | unresolved | 1362.0 | 9 |
| scf7180000018023 | 947398 | 0.3392 | 0 | 47.8604 | Eukaryota | 46616.0 | 0 | Chordata | 46616.0 | 0 | Carnivora | 46616.0 | 0 | Felidae | 46616.0 | 0 | Felis | 46616.0 | 0 | Felis catus | 46616.0 | 0 |
| scf7180000021924 | 47601 | 0.3728 | 0 | 24.9338 | Eukaryota | 36744.0 | 0 | Chordata | 36744.0 | 0 | Carnivora | 36744.0 | 0 | Felidae | 36744.0 | 0 | Felis | 33094.0 | 1 | Felis catus | 33094.0 | 1 |
| scf7180000024195 | 82502 | 0.4182 | 0 | 24.9231 | Eukaryota | 45707.0 | 0 | Chordata | 45707.0 | 0 | Carnivora | 45707.0 | 0 | Felidae | 45707.0 | 0 | Felis | 45707.0 | 0 | Felis catus | 45707.0 | 0 |
| scf7180000022775 | 12642 | 0.3723 | 0 | 21.5351 | Eukaryota | 3390.0 | 0 | Chordata | 3390.0 | 0 | Carnivora | 2287.0 | 1 | Felidae | 2024.0 | 2 | Felis | 1533.0 | 4 | Felis catus | 1533.0 | 4 |
| scf7180000019689 | 363126 | 0.4509 | 0 | 51.7652 | Eukaryota | 29821.0 | 0 | Chordata | 29821.0 | 0 | Carnivora | 29821.0 | 0 | Felidae | 29821.0 | 0 | Felis | 26801.0 | 1 | Felis catus | 26801.0 | 1 |
| scf7180000021048 | 85534 | 0.471 | 0 | 50.0975 | Eukaryota | 8622.0 | 1 | Chordata | 8622.0 | 1 | Artiodactyla | 4142.0 | 3 | Bovidae | 4142.0 | 3 | Homo | 3896.0 | 5 | Homo sapiens | 3896.0 | 5 |
| scf7180000021708 | 84361 | 0.4343 | 0 | 25.634 | Eukaryota | 10265.0 | 0 | Chordata | 10265.0 | 0 | Carnivora | 10265.0 | 0 | Felidae | 10265.0 | 0 | Felis | 6199.0 | 1 | Felis catus | 6199.0 | 1 |
| scf7180000018112 | 2009006 | 0.3697 | 0 | 48.9748 | Eukaryota | 46448.0 | 0 | Chordata | 46448.0 | 0 | Carnivora | 46448.0 | 0 | Felidae | 46448.0 | 0 | Felis | 46448.0 | 0 | Felis catus | 46448.0 | 0 |
| scf7180000022992 | 42401 | 0.5156 | 0 | 307.292 | Eukaryota | 15829.0 | 0 | Chordata | 15829.0 | 0 | Carnivora | 15829.0 | 0 | Felidae | 15829.0 | 0 | Felis | 15829.0 | 0 | Felis catus | 15829.0 | 0 |
| scf7180000020014 | 26832 | 0.4821 | 0 | 294.544 | Eukaryota | 7011.0 | 0 | Chordata | 7011.0 | 0 | Carnivora | 7011.0 | 0 | Felidae | 7011.0 | 0 | Felis | 7011.0 | 0 | Felis catus | 7011.0 | 0 |
| scf7180000018876 | 1055977 | 0.4663 | 0 | 52.0605 | Eukaryota | 47610.0 | 0 | Chordata | 47610.0 | 0 | Carnivora | 47610.0 | 0 | Felidae | 33661.0 | 1 | Panthera | 19281.0 | 3 | anthera pardu | 14483.0 | 4 |
| scf7180000021766 | 18087 | 0.4264 | 0 | 384.8406 | Eukaryota | 188513.0 | 0 | Chordata | 188513.0 | 0 | Carnivora | 188513.0 | 0 | Hyaenidae | 188513.0 | 0 | Proteles | 111576.0 | 2 | roteles cristat | 111576.0 | 2 |
| scf7180000022512 | 7826 | 0.4392 | 0 | 25.2526 | Eukaryota | 2351.0 | 0 | Chordata | 2351.0 | 0 | Carnivora | 2351.0 | 0 | Felidae | 2351.0 | 0 | Felis | 2125.0 | 1 | Felis catus | 2125.0 | 1 |
| scf7180000018834 | 445801 | 0.6483 | 0 | 50.6949 | Eukaryota | 63246.0 | 0 | Chordata | 63246.0 | 0 | Carnivora | 63246.0 | 0 | Felidae | 58368.0 | 1 | Acinonyx | 18970.0 | 5 | :inonyx jubat | 18970.0 | 5 |
| scf7180000022094 | 15357 | 0.3602 | 0 | 21.7811 | Eukaryota | 13390.0 | 0 | Chordata | 13390.0 | 0 | Carnivora | 13390.0 | 0 | Felidae | 13390.0 | 0 | Felis | 12067.0 | 1 | Felis catus | 12067.0 | 1 |
| scf7180000019438 | 154475 | 0.3574 | 0 | 45.937 | Eukaryota | 46144.0 | 0 | Chordata | 46144.0 | 0 | Carnivora | 46144.0 | 0 | Felidae | 46144.0 | 0 | Felis | 46144.0 | 0 | Felis catus | 46144.0 | 0 |
| scf7180000021019 | 90536 | 0.4046 | 0 | 26.022 | Eukaryota | 38541.0 | 0 | Chordata | 38541.0 | 0 | Carnivora | 38541.0 | 0 | Felidae | 34574.0 | 1 | Felis | 14793.0 | 4 | Felis catus | 14793.0 | 4 |
| scf7180000021188 | 54303 | 0.3945 | 0 | 25.5072 | Eukaryota | 24434.0 | 0 | Chordata | 24434.0 | 0 | Primates | 11597.0 | 2 | Hominidae | 9700.0 | 4 | Homo | 9700.0 | 4 | Homo sapiens | 9700.0 | 4 |
| scf7180000024252 | 2187837 | 0.3988 | 0 | 50.5201 | Eukaryota | 43001.0 | 0 | Chordata | 43001.0 | 0 | Carnivora | 43001.0 | 0 | Felidae | 43001.0 | 0 | Felis | 38181.0 | 1 | Felis catus | 38181.0 | 1 |
| scf7180000020897 | 301726 | 0.4925 | 0 | 49.4349 | Eukaryota | 8380.0 | 0 | Chordata | 8380.0 | 0 | Carnivora | 7589.0 | 1 | Felidae | 7589.0 | 1 | unresolved | 2529.0 | 5 | unresolved | 2529.0 | 5 |
| scf7180000018491 | 7650 | 0.4637 | 0 | 34.2452 | Eukaryota | 3479.0 | 1 | Chordata | 3479.0 | 1 | Artiodactyla | 1211.0 | 3 | Bovidae | 1211.0 | 4 | Homo | 1120.0 | 6 | Homo sapiens | 1120.0 | 6 |
| scf7180000021454 | 28927 | 0.4347 | 0 | 26.5405 | Eukaryota | 5017.0 | 0 | Chordata | 5017.0 | 0 | Primates | 2410.0 | 2 | Hominidae | 2410.0 | 2 | Felis | 2254.0 | 3 | Felis catus | 2254.0 | 3 |
| scf7180000019166 | 93386 | 0.3677 | 0 | 44.3985 | Eukaryota | 29163.0 | 0 | Chordata | 29163.0 | 0 | Carnivora | 17961.0 | 1 | Ursidae | 12009.0 | 2 | Ursus | 12009.0 | 2 | Ursus arctos | 9015.0 | 4 |
| scf7180000023521 | 17585 | 0.538 | 0 | 217.2022 | Eukaryota | 3693.0 | 0 | Chordata | 3693.0 | 0 | Carnivora | 3693.0 | 0 | Felidae | 3693.0 | 0 | unresolved | 1104.0 | 4 | unresolved | 1104.0 | 4 |
| scf7180000020767 | 79753 | 0.386 | 0 | 24.9521 | Eukaryota | 33009.0 | 0 | Chordata | 33009.0 | 0 | Carnivora | 33009.0 | 0 | Felidae | 33009.0 | 0 | Felis | 33009.0 | 0 | Felis catus | 33009.0 | 0 |
| scf7180000020999 | 36731 | 0.4276 | 0 | 28.5797 | Eukaryota | 12968.0 | 0 | Chordata | 12968.0 | 0 | Carnivora | 12968.0 | 0 | Felidae | 12968.0 | 0 | Felis | 10341.0 | 1 | Felis catus | 10341.0 | 1 |
| scf7180000024153 | 142969 | 0.4601 | 0 | 26.1772 | Eukaryota | 9053.0 | 0 | Chordata | 9053.0 | 0 | Carnivora | 9053.0 | 0 | Felidae | 9053.0 | 0 | Felis | 8114.0 | 1 | Felis catus | 8114.0 | 1 |
| scf7180000020020 | 78150 | 0.4085 | 0 | 50.4416 | Eukaryota | 13528.0 | 0 | Chordata | 13528.0 | 0 | Carnivora | 13528.0 | 0 | Otariidae | 6784.0 | 3 | Eumetopias | 2760.0 | 7 | netopias juba | 2760.0 | 7 |
| scf7180000022678 | 502846 | 0.3772 | 0 | 51.5956 | Eukaryota | 47543.0 | 0 | Chordata | 47543.0 | 0 | Carnivora | 47543.0 | 0 | Felidae | 47543.0 | 0 | Felis | 47543.0 | 0 | Felis catus | 47543.0 | 0 |
| scf7180000021115 | 83550 | 0.3439 | 0 | 49.5942 | Eukaryota | 8954.0 | 0 | Chordata | 8954.0 | 0 | Carnivora | 4488.0 | 2 | Felidae | 3780.0 | 3 | Felis | 3780.0 | 4 | Felis catus | 3780.0 | 4 |
| scf7180000023649 | 26108 | 0.3869 | 0 | 303.5557 | Eukaryota | 8965.0 | 0 | Chordata | 8965.0 | 0 | .pterygiforme | 6316.0 | 3 | Apterygidae | 6316.0 | 3 | Apteryx | 6316.0 | 3 | pteryx mante | 6316.0 | 3 |
| scf7180000024146 | 120792 | 0.3652 | 0 | 25.5207 | Eukaryota | 41872.0 | 0 | Chordata | 41872.0 | 0 | Carnivora | 41872.0 | 0 | Felidae | 41872.0 | 0 | Felis | 41872.0 | 0 | Felis catus | 41872.0 | 0 |
| scf7180000022634 | 8262 | 0.4096 | 0 | 25.0583 | Eukaryota | 10376.0 | 0 | Chordata | 10376.0 | 0 | Carnivora | 10376.0 | 0 | Felidae | 10376.0 | 0 | Felis | 9369.0 | 1 | Felis catus | 9369.0 | 1 |
| scf7180000018061 | 1255987 | 0.3783 | 0 | 50.3717 | Eukaryota | 45782.0 | 0 | Chordata | 45782.0 | 0 | Carnivora | 45782.0 | 0 | Felidae | 45782.0 | 0 | Felis | 45782.0 | 0 | Felis catus | 45782.0 | 0 |
| scf7180000018272 | 836348 | 0.4751 | 0 | 50.7251 | Eukaryota | 40170.0 | 0 | Chordata | 40170.0 | 0 | Carnivora | 40170.0 | 0 | Felidae | 40170.0 | 0 | Felis | 36173.0 | 1 | Felis catus | 36173.0 | 1 |
| scf7180000019840 | 25279 | 0.3611 | 0 | 23.4779 | Eukaryota | 16893.0 | 0 | Chordata | 16893.0 | 0 | Carnivora | 16893.0 | 0 | Felidae | 16893.0 | 0 | Felis | 15177.0 | 1 | Felis catus | 15177.0 | 1 |
| scf7180000021838 | 28898 | 0.3662 | 0 | 48.0341 | Eukaryota | 8347.0 | 0 | Chordata | 8347.0 | 0 | Primates | 4807.0 | 3 | Cercopithecidae | 2739.0 | 4 | Hipposideros | 2130.0 | 6 | osideros arm | 2130.0 | 6 |
| scf7180000018525 | 749003 | 0.5031 | 0 | 52.0358 | Eukaryota | 71950.0 | 0 | Chordata | 71950.0 | 0 | Carnivora | 71950.0 | 0 | Felidae | 71950.0 | 0 | Panthera | 28931.0 | 3 | anthera pardu | 21717.0 | 4 |
| scf7180000017506 | 152580 | 0.4231 | 0 | 49.8247 | Eukaryota | 47001.0 | 0 | Chordata | 47001.0 | 0 | Carnivora | 47001.0 | 0 | Felidae | 47001.0 | 0 | Felis | 47001.0 | 0 | Felis catus | 47001.0 | 0 |
| scf7180000018922 | 460500 | 0.3898 | 0 | 49.2536 | Eukaryota | 60574.0 | 0 | Chordata | 60574.0 | 0 | Carnivora | 60574.0 | 0 | Felidae | 60574.0 | 0 | Felis | 22360.0 | 2 | Felis catus | 22360.0 | 2 |
| scf7180000021783 | 39840 | 0.4102 | 0 | 31.3285 | Eukaryota | 6896.0 | 0 | Chordata | 6896.0 | 0 | Carnivora | 4965.0 | 2 | Felidae | 4163.0 | 3 | Felis | 2209.0 | 5 | Felis catus | 2209.0 | 5 |
| scf7180000018271 | 335375 | 0.3671 | 0 | 50.1083 | Eukaryota | 45368.0 | 0 | Chordata | 45368.0 | 0 | Carnivora | 45368.0 | 0 | Felidae | 45368.0 | 0 | Felis | 45368.0 | 0 | Felis catus | 45368.0 | 0 |
| scf7180000019237 | 232314 | 0.4863 | 0 | 38.5521 | Eukaryota | 32073.0 | 0 | Chordata | 32073.0 | 0 | Carnivora | 32073.0 | 0 | Felidae | 32073.0 | 0 | Felis | 28826.0 | 1 | Felis catus | 28826.0 | 1 |
| scf7180000023799 | 1870979 | 0.3646 | 0 | 50.2442 | Eukaryota | 43849.0 | 0 | Chordata | 43849.0 | 0 | Carnivora | 43849.0 | 0 | Felidae | 43849.0 | 0 | Felis | 22425.0 | 2 | Felis catus | 22425.0 | 2 |
| scf7180000024080 | 375937 | 0.5622 | 0 | 47.2081 | Eukaryota | 25652.0 | 0 | Chordata | 25652.0 | 0 | Carnivora | 25652.0 | 0 | Ursidae | 13022.0 | 2 | Ailuropoda | 7617.0 | 3 | ›poda melanc | 7617.0 | 4 |
| scf7180000020472 | 113313 | 0.3765 | 0 | 48.2062 | Eukaryota | 17614.0 | 0 | Chordata | 17614.0 | 0 | Carnivora | 17614.0 | 0 | Felidae | 17614.0 | 0 | Felis | 16230.0 | 1 | Felis catus | 16230.0 | 1 |
| scf7180000017921 | 33647 | 0.4053 | 0 | 22.8615 | Eukaryota | 25889.0 | 0 | Chordata | 25889.0 | 0 | Carnivora | 25889.0 | 0 | Felidae | 25889.0 | 0 | Felis | 23327.0 | 1 | Felis catus | 23327.0 | 1 |
| scf7180000022178 | 51504 | 0.3805 | 0 | 24.0414 | Eukaryota | 43435.0 | 0 | Chordata | 43435.0 | 0 | Carnivora | 43435.0 | 0 | Felidae | 43435.0 | 0 | Felis | 43435.0 | 0 | Felis catus | 43435.0 | 0 |
| scf7180000018441 | 723741 | 0.4467 | 0 | 52.422 | Eukaryota | 47070.0 | 0 | Chordata | 47070.0 | 0 | Carnivora | 47070.0 | 0 | Felidae | 47070.0 | 0 | Felis | 47070.0 | 0 | Felis catus | 47070.0 | 0 |
| scf7180000018637 | 381035 | 0.4092 | 0 | 406.5168 | Eukaryota | 44796.0 | 0 | Chordata | 44796.0 | 0 | Carnivora | 44796.0 | 0 | Felidae | 44796.0 | 0 | Felis | 44796.0 | 0 | Felis catus | 44796.0 | 0 |
| scf7180000022530 | 15722 | 0.3816 | 0 | 24.1248 | Eukaryota | 6460.0 | 0 | Chordata | 6460.0 | 0 | Carnivora | 5567.0 | 1 | Felidae | 4315.0 | 2 | Felis | 4315.0 | 2 | Felis catus | 4315.0 | 2 |
| scf7180000022033 | 22059 | 0.5095 | 0 | 31.4883 | Eukaryota | 4042.0 | 0 | Chordata | 4042.0 | 0 | Carnivora | 4042.0 | 0 | Felidae | 4042.0 | 0 | Felis | 3657.0 | 1 | Felis catus | 3657.0 | 1 |
| scf7180000021450 | 59019 | 0.3629 | 0 | 25.1716 | Eukaryota | 34180.0 | 0 | Chordata | 34180.0 | 0 | Carnivora | 34180.0 | 0 | Felidae | 34180.0 | 0 | Felis | 30789.0 | 1 | Felis catus | 30789.0 | 1 |
| scf7180000017793 | 620952 | 0.4172 | 0 | 52.3338 | Eukaryota | 35965.0 | 0 | Chordata | 35965.0 | 0 | Carnivora | 35965.0 | 0 | Felidae | 35965.0 | 0 | Felis | 35965.0 | 0 | Felis catus | 35965.0 | 0 |
| scf7180000021368 | 191591 | 0.4348 | 0 | 23.9842 | Eukaryota | 23843.0 | 0 | Chordata | 23843.0 | 0 | Carnivora | 23843.0 | 0 | Felidae | 23843.0 | 0 | Felis | 19002.0 | 1 | Felis catus | 19002.0 | 1 |
| scf7180000022470 | 23090 | 0.5219 | 0 | 25.8806 | Eukaryota | 5045.0 | 0 | Chordata | 5045.0 | 0 | Primates | 2212.0 | 3 | Hominidae | 2212.0 | 3 | Pan | 1399.0 | 5 | an troglodyte | 1399.0 | 5 |
| scf7180000018150 | 670309 | 0.393 | 0 | 50.3035 | Eukaryota | 53614.0 | 0 | Chordata | 53614.0 | 0 | Carnivora | 53614.0 | 0 | Felidae | 37336.0 | 1 | Felis | 32674.0 | 2 | Felis catus | 32674.0 | 2 |
| scf7180000019502 | 301912 | 0.4213 | 0 | 55.515 | Eukaryota | 37982.0 | 0 | Chordata | 37982.0 | 0 | Carnivora | 37982.0 | 0 | Felidae | 33507.0 | 1 | Felis | 33507.0 | 1 | Felis catus | 33507.0 | 1 |
| scf7180000017925 | 2160431 | 0.42 | 0 | 51.6195 | Eukaryota | 64752.0 | 0 | Chordata | 64752.0 | 0 | Carnivora | 64752.0 | 0 | Felidae | 64752.0 | 0 | Felis | 51936.0 | 1 | Felis catus | 51936.0 | 1 |
| scf7180000024051 | 530999 | 0.446 | 0 | 52.8137 | Eukaryota | 107544.0 | 0 | Chordata | 107544.0 | 0 | Carnivora | 107544.0 | 0 | Felidae | 107544.0 | 0 | Felis | 107544.0 | 0 | Felis catus | 107544.0 | 0 |
| scf7180000020192 | 230859 | 0.5973 | 0 | 53.0021 | Eukaryota | 14554.0 | 0 | Chordata | 14554.0 | 0 | Carnivora | 14554.0 | 0 | Mustelidae | 9898.0 | 2 | Mustela | 8448.0 | 4 | lustela putori | 8448.0 | 4 |
| scf7180000024117 | 166369 | 0.3877 | 0 | 50.6073 | Eukaryota | 28701.0 | 0 | Chordata | 28701.0 | 0 | Carnivora | 28701.0 | 0 | Phocidae | 24264.0 | 2 | Neomonachus | 12236.0 | 3 | nachus schau | 12236.0 | 3 |
| scf7180000021515 | 65401 | 0.4177 | 0 | 27.8298 | Eukaryota | 30021.0 | 0 | Chordata | 30021.0 | 0 | Carnivora | 30021.0 | 0 | Felidae | 21963.0 | 1 | Vulpes | 8058.0 | 4 | Vulpes vulpes | 8058.0 | 4 |
| scf7180000022766 | 9594 | 0.3797 | 0 | 24.8297 | Eukaryota | 8300.0 | 0 | Chordata | 8300.0 | 0 | Primates | 4352.0 | 2 | Hominidae | 4352.0 | 2 | Sus | 2524.0 | 3 | Sus scrofa | 2524.0 | 3 |
| scf7180000022217 | 20160 | 0.5657 | 0 | 48.2411 | Eukaryota | 2640.0 | 0 | Chordata | 2640.0 | 0 | Carnivora | 2640.0 | 0 | Felidae | 2640.0 | 0 | Felis | 2640.0 | 0 | Felis catus | 2640.0 | 0 |
| scf7180000019747 | 161376 | 0.4009 | 0 | 24.6982 | Eukaryota | 14921.0 | 0 | Chordata | 14921.0 | 0 | Carnivora | 14921.0 | 0 | Felidae | 14921.0 | 0 | Felis | 11965.0 | 1 | Felis catus | 11965.0 | 1 |
| scf7180000017612 | 2207930 | 0.3726 | 0 | 49.3898 | Eukaryota | 47285.0 | 0 | Chordata | 47285.0 | 0 | Carnivora | 47285.0 | 0 | Felidae | 47285.0 | 0 | Felis | 42583.0 | 1 | Felis catus | 42583.0 | 1 |
| scf7180000023576 | 24539 | 0.4807 | 0 | 241.4709 | Eukaryota | 4020.0 | 0 | Chordata | 4020.0 | 0 | Carnivora | 2682.0 | 1 | Felidae | 2223.0 | 2 | Panthera | 1864.0 | 3 | anthera pardu | 1500.0 | 4 |
| scf7180000017631 | 2541636 | 0.3512 | 0 | 48.5712 | Eukaryota | 47523.0 | 0 | Chordata | 47523.0 | 0 | Carnivora | 47523.0 | 0 | Felidae | 47523.0 | 0 | Felis | 47523.0 | 0 | Felis catus | 47523.0 | 0 |
| scf7180000024284 | 110481 | 0.3758 | 0 | 25.7811 | Eukaryota | 42680.0 | 0 | Chordata | 42680.0 | 0 | Carnivora | 42680.0 | 0 | Felidae | 42680.0 | 0 | Felis | 42680.0 | 0 | Felis catus | 42680.0 | 0 |
| scf7180000021499 | 35960 | 0.4262 | 0 | 21.9554 | Eukaryota | 7446.0 | 0 | Chordata | 7446.0 | 0 | Carnivora | 4234.0 | 2 | Felidae | 3012.0 | 3 | Felis | 3012.0 | 4 | Felis catus | 3012.0 | 4 |
| scf7180000022483 | 8638 | 0.5883 | 0 | 52.5194 | no-hit | 0.0 | 0 | no-hit | 0.0 | 0 | no-hit | 0.0 | 0 | no-hit | 0.0 | 0 | no-hit | 0.0 | 0 | no-hit | 0.0 | 0 |
| scf7180000019385 | 25124 | 0.5383 | 0 | 33.7299 | Eukaryota | 4881.0 | 0 | Chordata | 4881.0 | 0 | Carnivora | 4881.0 | 0 | Herpestidae | 3391.0 | 1 | Suricata | 3391.0 | 1 | ıricata suricat | 3391.0 | 1 |
| scf7180000023493 | 50198 | 0.4564 | 0 | 75.8816 | Eukaryota | 2338.0 | 0 | Chordata | 2338.0 | 0 | Carnivora | 2338.0 | 0 | Felidae | 1912.0 | 1 | Felis | 1912.0 | 1 | Felis catus | 1912.0 | 1 |
| scf7180000021312 | 48013 | 0.3679 | 0 | 24.9415 | Eukaryota | 23882.0 | 0 | Chordata | 23882.0 | 0 | Carnivora | 23882.0 | 0 | Felidae | 23882.0 | 0 | Felis | 21451.0 | 1 | Felis catus | 21451.0 | 1 |
| scf7180000023902 | 1048583 | 0.4242 | 0 | 51.1751 | Eukaryota | 79792.0 | 0 | Chordata | 79792.0 | 0 | Carnivora | 79792.0 | 0 | Felidae | 54937.0 | 1 | Lynx | 47094.0 | 2 | ynx canadens | 47094.0 | 2 |
| scf7180000020165 | 148580 | 0.5958 | 0 | 47.6904 | Eukaryota | 8278.0 | 0 | Chordata | 8278.0 | 0 | Carnivora | 8278.0 | 0 | Felidae | 6168.0 | 1 | Felis | 3845.0 | 4 | Felis catus | 3845.0 | 4 |
| scf7180000019724 | 195366 | 0.3782 | 0 | 22.736 | Eukaryota | 43033.0 | 0 | Chordata | 43033.0 | 0 | Carnivora | 43033.0 | 0 | Felidae | 43033.0 | 0 | Felis | 43033.0 | 0 | Felis catus | 43033.0 | 0 |
| scf7180000019267 | 362862 | 0.4331 | 0 | 50.4339 | Eukaryota | 26925.0 | 1 | Chordata | 26925.0 | 1 | Artiodactyla | 12914.0 | 3 | Hominidae | 10673.0 | 4 | Homo | 10673.0 | 5 | Homo sapiens | 10673.0 | 5 |
| scf7180000022263 | 21907 | 0.473 | 0 | 31.4187 | Eukaryota | 7129.0 | 0 | Chordata | 7129.0 | 0 | Carnivora | 6390.0 | 1 | Felidae | 4232.0 | 2 | Panthera | 2850.0 | 5 | rsus maritimu | 1426.0 | 6 |
| scf7180000019595 | 45740 | 0.402 | 0 | 44.2346 | Eukaryota | 24562.0 | 0 | Chordata | 24562.0 | 0 | Carnivora | 24562.0 | 0 | Felidae | 24562.0 | 0 | Felis | 22098.0 | 1 | Felis catus | 22098.0 | 1 |
| scf7180000022493 | 13699 | 0.3685 | 0 | 24.4444 | Eukaryota | 27547.0 | 0 | Chordata | 27547.0 | 0 | Carnivora | 27547.0 | 0 | Felidae | 27547.0 | 0 | Felis | 24738.0 | 1 | Felis catus | 24738.0 | 1 |
| scf7180000021979 | 30798 | 0.37 | 0 | 27.0497 | Eukaryota | 5537.0 | 0 | Chordata | 5537.0 | 0 | Carnivora | 4622.0 | 1 | Felidae | 4622.0 | 1 | Felis | 4622.0 | 1 | Felis catus | 4622.0 | 1 |
| scf7180000021660 | 56390 | 0.4794 | 0 | 28.5155 | Eukaryota | 46667.0 | 0 | Chordata | 46667.0 | 0 | Carnivora | 46667.0 | 0 | Felidae | 46667.0 | 0 | Felis | 46667.0 | 0 | Felis catus | 46667.0 | 0 |
| scf7180000021310 | 38744 | 0.3929 | 0 | 26.8616 | Eukaryota | 28698.0 | 0 | Chordata | 28698.0 | 0 | Carnivora | 28698.0 | 0 | Felidae | 28698.0 | 0 | Felis | 25798.0 | 1 | Felis catus | 25798.0 | 1 |
| scf7180000018756 | 573752 | 0.4537 | 0 | 52.0454 | Eukaryota | 41506.0 | 0 | Chordata | 41506.0 | 0 | Carnivora | 41506.0 | 0 | Felidae | 41506.0 | 0 | Felis | 37378.0 | 1 | Felis catus | 37378.0 | 1 |
| scf7180000021286 | 59656 | 0.3835 | 0 | 25.774 | Eukaryota | 21574.0 | 0 | Chordata | 21574.0 | 0 | Primates | 18766.0 | 1 | Hominidae | 18766.0 | 1 | Pan | 10842.0 | 2 | an troglodyte | 10842.0 | 2 |
| scf7180000023806 | 1543240 | 0.4338 | 0 | 51.4323 | Eukaryota | 47568.0 | 0 | Chordata | 47568.0 | 0 | Carnivora | 47568.0 | 0 | Felidae | 47568.0 | 0 | Felis | 47568.0 | 0 | Felis catus | 47568.0 | 0 |
| scf7180000019527 | 247203 | 0.3786 | 0 | 50.1066 | Eukaryota | 30192.0 | 0 | Chordata | 30192.0 | 0 | Carnivora | 30192.0 | 0 | Felidae | 30192.0 | 0 | Lynx | 30192.0 | 0 | ynx canadens | 30192.0 | 0 |
| scf7180000022397 | 19700 | 0.4237 | 0 | 430.2305 | Eukaryota | 236485.0 | 0 | Chordata | 236485.0 | 0 | Carnivora | 236485.0 | 0 | Hyaenidae | 236485.0 | 0 | Proteles | 142875.0 | 1 | roteles cristat | 142875.0 | 1 |
| scf7180000022789 | 12003 | 0.3899 | 0 | 20.5268 | Eukaryota | 43968.0 | 0 | Chordata | 43968.0 | 0 | Carnivora | 43968.0 | 0 | Felidae | 43968.0 | 0 | Felis | 39646.0 | 1 | Felis catus | 39646.0 | 1 |
| scf7180000021132 | 14501 | 0.4468 | 0 | 25.4393 | Eukaryota | 5899.0 | 0 | Chordata | 5899.0 | 0 | Carnivora | 3339.0 | 2 | Felidae | 3339.0 | 2 | Felis | 2375.0 | 3 | Felis catus | 2375.0 | 3 |
| scf7180000022718 | 4845 | 0.5234 | 0 | 21.5059 | no-hit | 0.0 | 0 | no-hit | 0.0 | 0 | no-hit | 0.0 | 0 | no-hit | 0.0 | 0 | no-hit | 0.0 | 0 | no-hit | 0.0 | 0 |
| scf7180000022373 | 20848 | 0.5868 | 0 | 28.7112 | Eukaryota | 2529.0 | 0 | Chordata | 2529.0 | 0 | Carnivora | 2529.0 | 0 | Felidae | 2281.0 | 1 | Felis | 2038.0 | 2 | Felis catus | 2038.0 | 2 |
| scf7180000019365 | 75303 | 0.3973 | 0 | 40.7023 | Eukaryota | 32566.0 | 0 | Chordata | 32566.0 | 0 | Carnivora | 32566.0 | 0 | Felidae | 30170.0 | 1 | Felis | 30170.0 | 1 | Felis catus | 30170.0 | 1 |
| scf7180000019460 | 187652 | 0.4013 | 0 | 45.3635 | Eukaryota | 32610.0 | 0 | Chordata | 32610.0 | 0 | Carnivora | 32610.0 | 0 | Felidae | 32610.0 | 0 | Felis | 29367.0 | 1 | Felis catus | 29367.0 | 1 |
| scf7180000018818 | 20200 | 0.4313 | 0 | 31.2986 | Eukaryota | 15029.0 | 0 | Chordata | 15029.0 | 0 | Carnivora | 15029.0 | 0 | Felidae | 15029.0 | 0 | Felis | 13536.0 | 1 | Felis catus | 13536.0 | 1 |
| scf7180000024229 | 314496 | 0.3758 | 0 | 50.3913 | Eukaryota | 43070.0 | 0 | Chordata | 43070.0 | 0 | Carnivora | 43070.0 | 0 | Felidae | 43070.0 | 0 | Felis | 43070.0 | 0 | Felis catus | 43070.0 | 0 |
| scf7180000018159 | 1229482 | 0.3957 | 0 | 51.9135 | Eukaryota | 71933.0 | 0 | Chordata | 71933.0 | 0 | Carnivora | 71933.0 | 0 | Felidae | 43655.0 | 1 | Ursus | 28278.0 | 3 | Ursus arctos | 28278.0 | 4 |
| scf7180000021407 | 39245 | 0.3825 | 0 | 25.4434 | Eukaryota | 5462.0 | 0 | Chordata | 5462.0 | 0 | Carnivora | 2932.0 | 2 | Felidae | 2451.0 | 3 | Felis | 1523.0 | 4 | Felis catus | 1523.0 | 4 |
| scf7180000023086 | 29904 | 0.5718 | 0 | 197.1188 | Eukaryota | 3139.0 | 0 | Chordata | 3139.0 | 0 | Carnivora | 3139.0 | 0 | Felidae | 2588.0 | 1 | Felis | 1521.0 | 3 | Felis catus | 1521.0 | 3 |
| scf7180000022356 | 14077 | 0.5028 | 0 | 27.4802 | Eukaryota | 22869.0 | 0 | Chordata | 22869.0 | 0 | Carnivora | 18835.0 | 1 | Otariidae | 7377.0 | 3 | Zalophus | 7377.0 | 3 | phus californi | 7377.0 | 3 |
| scf7180000021970 | 13939 | 0.5533 | 0 | 41.9746 | Eukaryota | 10320.0 | 0 | Chordata | 10320.0 | 0 | Carnivora | 7249.0 | 3 | Felidae | 4159.0 | 6 | Acinonyx | 2054.0 | 8 | :inonyx jubat | 2054.0 | 8 |
| scf7180000021073 | 86969 | 0.4059 | 0 | 27.4051 | Eukaryota | 42631.0 | 0 | Chordata | 42631.0 | 0 | Carnivora | 42631.0 | 0 | Felidae | 42631.0 | 0 | Felis | 38400.0 | 1 | Felis catus | 38400.0 | 1 |
| scf7180000018931 | 497893 | 0.4787 | 0 | 53.0956 | Eukaryota | 64868.0 | 0 | Chordata | 64868.0 | 0 | Carnivora | 58214.0 | 1 | Canidae | 34712.0 | 2 | Canis | 34712.0 | 2 | Canis lupus | 34712.0 | 2 |
| scf7180000021163 | 79567 | 0.4206 | 0 | 25.1939 | Eukaryota | 30379.0 | 0 | Chordata | 30379.0 | 0 | Carnivora | 30379.0 | 0 | Felidae | 30379.0 | 0 | Acinonyx | 9138.0 | 3 | :inonyx jubat | 9138.0 | 3 |
| scf7180000021399 | 138553 | 0.3948 | 0 | 26.4506 | Eukaryota | 11521.0 | 0 | Chordata | 11521.0 | 0 | Carnivora | 6424.0 | 3 | Felidae | 6424.0 | 3 | Felis | 6424.0 | 3 | Felis catus | 6424.0 | 3 |
| scf7180000023822 | 4262793 | 0.3657 | 100 | 50.5173 | Eukaryota | 90960.0 | 0 | Chordata | 90960.0 | 0 | Carnivora | 82014.0 | 1 | Felidae | 82014.0 | 1 | Acinonyx | 53154.0 | 2 | :inonyx jubat | 53154.0 | 2 |
| scf7180000021232 | 22488 | 0.4627 | 0 | 27.3273 | Eukaryota | 22003.0 | 0 | Chordata | 22003.0 | 0 | Carnivora | 22003.0 | 0 | Felidae | 22003.0 | 0 | Felis | 19810.0 | 1 | Felis catus | 19810.0 | 1 |
| scf7180000018906 | 12109 | 0.3662 | 0 | 41.6549 | Eukaryota | 2924.0 | 0 | Chordata | 2924.0 | 0 | Carnivora | 2005.0 | 1 | Felidae | 1192.0 | 2 | Felis | 1192.0 | 3 | Felis catus | 1192.0 | 3 |
| scf7180000021959 | 31754 | 0.3898 | 0 | 49.2555 | Eukaryota | 7381.0 | 0 | Chordata | 7381.0 | 0 | Perissodactyla | 3971.0 | 1 | Felidae | 3410.0 | 2 | Felis | 3410.0 | 2 | Felis catus | 3410.0 | 3 |
| scf7180000020860 | 56505 | 0.4119 | 0 | 46.775 | Eukaryota | 18625.0 | 0 | Chordata | 18625.0 | 0 | Carnivora | 18625.0 | 0 | Felidae | 18625.0 | 0 | Felis | 14923.0 | 1 | Felis catus | 14923.0 | 1 |
| scf7180000020010 | 142593 | 0.3896 | 0 | 50.3298 | Eukaryota | 10032.0 | 1 | Chordata | 10032.0 | 1 | Carnivora | 5871.0 | 3 | Felidae | 5871.0 | 3 | Felis | 4729.0 | 4 | Felis catus | 4729.0 | 4 |
| scf7180000022160 | 30686 | 0.4439 | 0 | 24.6037 | Eukaryota | 4390.0 | 0 | Chordata | 4390.0 | 0 | Carnivora | 2754.0 | 1 | Felidae | 1686.0 | 2 | Felis | 1686.0 | 3 | Felis catus | 1686.0 | 4 |
| scf7180000020611 | 84403 | 0.4161 | 0 | 51.8245 | Eukaryota | 46889.0 | 0 | Chordata | 46889.0 | 0 | Carnivora | 46889.0 | 0 | Felidae | 46889.0 | 0 | Felis | 46889.0 | 0 | Felis catus | 46889.0 | 0 |
| scf7180000017703 | 859746 | 0.3982 | 0 | 50.3867 | Eukaryota | 45993.0 | 0 | Chordata | 45993.0 | 0 | Carnivora | 45993.0 | 0 | Felidae | 40619.0 | 1 | Panthera | 17970.0 | 3 | anthera pardu | 13536.0 | 4 |
| scf7180000020581 | 104657 | 0.3841 | 0 | 49.1367 | Eukaryota | 34846.0 | 0 | Chordata | 34846.0 | 0 | Carnivora | 34846.0 | 0 | Felidae | 34846.0 | 0 | Panthera | 24420.0 | 1 | anthera pardu | 13980.0 | 2 |
| scf7180000017779 | 1486638 | 0.4967 | 0 | 50.3967 | Eukaryota | 29857.0 | 0 | Chordata | 29857.0 | 0 | Carnivora | 29857.0 | 0 | Felidae | 29857.0 | 0 | Felis | 26826.0 | 1 | Felis catus | 26826.0 | 1 |
| scf7180000022252 | 45081 | 0.387 | 0 | 24.3072 | Eukaryota | 27187.0 | 0 | Chordata | 27187.0 | 0 | Carnivora | 27187.0 | 0 | Felidae | 27187.0 | 0 | Felis | 24389.0 | 1 | Felis catus | 24389.0 | 1 |
| scf7180000018999 | 1100977 | 0.3842 | 0 | 50.4617 | Eukaryota | 47401.0 | 0 | Chordata | 47401.0 | 0 | Carnivora | 47401.0 | 0 | Felidae | 47401.0 | 0 | Felis | 47401.0 | 0 | Felis catus | 47401.0 | 0 |
| scf7180000021275 | 38947 | 0.6227 | 0 | 46.793 | Eukaryota | 4361.0 | 0 | Chordata | 4361.0 | 0 | Primates | 1529.0 | 6 | Hominidae | 1529.0 | 6 | Homo | 1529.0 | 6 | Homo sapiens | 1529.0 | 6 |
| scf7180000023332 | 16902 | 0.7026 | 0 | 196.5759 | no-hit | 0.0 | 0 | no-hit | 0.0 | 0 | no-hit | 0.0 | 0 | no-hit | 0.0 | 0 | no-hit | 0.0 | 0 | no-hit | 0.0 | 0 |
| scf7180000019570 | 337930 | 0.4506 | 0 | 51.0411 | Eukaryota | 28950.0 | 0 | Chordata | 28950.0 | 0 | Carnivora | 28950.0 | 0 | Felidae | 28950.0 | 0 | Felis | 11825.0 | 3 | Felis catus | 11825.0 | 3 |
| scf7180000020408 | 181863 | 0.3704 | 0 | 24.7411 | Eukaryota | 19554.0 | 0 | Chordata | 19554.0 | 0 | Carnivora | 19554.0 | 0 | Felidae | 19554.0 | 0 | Felis | 15662.0 | 1 | Felis catus | 15662.0 | 1 |
| scf7180000021679 | 24402 | 0.4676 | 0 | 48.8463 | Eukaryota | 2693.0 | 0 | Chordata | 2693.0 | 0 | Carnivora | 2693.0 | 0 | Felidae | 2693.0 | 0 | Felis | 2159.0 | 1 | Felis catus | 2159.0 | 1 |
| scf7180000020625 | 120139 | 0.3771 | 0 | 48.8046 | Eukaryota | 31153.0 | 0 | Chordata | 31153.0 | 0 | Carnivora | 27895.0 | 1 | Mustelidae | 16843.0 | 3 | Enhydra | 16843.0 | 5 | Enhydra lutris | 16843.0 | 5 |
| scf7180000022319 | 92883 | 0.4662 | 0 | 602.8859 | Eukaryota | 40920.0 | 0 | Chordata | 40920.0 | 0 | Carnivora | 40920.0 | 0 | Felidae | 40920.0 | 0 | Felis | 40920.0 | 0 | Felis catus | 40920.0 | 0 |
| scf7180000021416 | 115324 | 0.3634 | 0 | 24.7564 | Eukaryota | 45670.0 | 0 | Chordata | 45670.0 | 0 | Carnivora | 45670.0 | 0 | Felidae | 45670.0 | 0 | Felis | 45670.0 | 0 | Felis catus | 45670.0 | 0 |
| scf7180000019223 | 381293 | 0.5355 | 0 | 52.124 | Eukaryota | 15343.0 | 0 | Chordata | 15343.0 | 0 | Carnivora | 15343.0 | 0 | Felidae | 12488.0 | 1 | Felis | 5445.0 | 4 | Felis catus | 5445.0 | 4 |
| scf7180000017577 | 381777 | 0.402 | 0 | 48.2658 | Eukaryota | 72015.0 | 0 | Chordata | 72015.0 | 0 | Carnivora | 72015.0 | 0 | Felidae | 67115.0 | 1 | Panthera | 31320.0 | 4 | anthera pardu | 23973.0 | 5 |
| scf7180000018918 | 495931 | 0.4504 | 0 | 51.0756 | Eukaryota | 57261.0 | 0 | Chordata | 57261.0 | 0 | Carnivora | 57261.0 | 0 | Felidae | 29166.0 | 2 | Suricata | 22292.0 | 5 | ıricata suricat | 22292.0 | 6 |
| scf7180000019175 | 424089 | 0.5578 | 0 | 52.2002 | Eukaryota | 32628.0 | 0 | Chordata | 32628.0 | 0 | Carnivora | 32628.0 | 0 | Felidae | 17910.0 | 2 | Felis | 17910.0 | 2 | Felis catus | 17910.0 | 2 |
| scf7180000021624 | 59248 | 0.3961 | 0 | 24.7297 | Eukaryota | 15221.0 | 0 | Chordata | 15221.0 | 0 | Carnivora | 15221.0 | 0 | Felidae | 15221.0 | 0 | Felis | 13725.0 | 1 | Felis catus | 13725.0 | 1 |
| scf7180000022532 | 6166 | 0.6635 | 0 | 884.4633 | Eukaryota | 1418.0 | 0 | Chordata | 1277.0 | 1 | Carnivora | 1277.0 | 1 | Otariidae | 670.0 | 4 | Eumetopias | 402.0 | 5 | netopias juba | 402.0 | 5 |
| scf7180000019111 | 20103 | 0.5014 | 0 | 47.6761 | Eukaryota | 22689.0 | 0 | Chordata | 22689.0 | 0 | Carnivora | 22689.0 | 0 | Felidae | 22689.0 | 0 | Felis | 11325.0 | 2 | Felis catus | 11325.0 | 2 |
| scf7180000022276 | 30057 | 0.3642 | 0 | 23.5541 | Eukaryota | 41239.0 | 0 | Chordata | 41239.0 | 0 | Carnivora | 41239.0 | 0 | Felidae | 41239.0 | 0 | Felis | 41239.0 | 0 | Felis catus | 41239.0 | 0 |
| scf7180000023835 | 550085 | 0.4067 | 0 | 50.3111 | Eukaryota | 45344.0 | 0 | Chordata | 45344.0 | 0 | Carnivora | 45344.0 | 0 | Felidae | 45344.0 | 0 | Felis | 45344.0 | 0 | Felis catus | 45344.0 | 0 |
| scf7180000022025 | 32094 | 0.4742 | 0 | 25.0821 | Eukaryota | 13293.0 | 0 | Chordata | 13293.0 | 0 | Carnivora | 13293.0 | 0 | Felidae | 13293.0 | 0 | Felis | 13293.0 | 0 | Felis catus | 13293.0 | 0 |
| scf7180000020935 | 81395 | 0.3766 | 0 | 24.7391 | Eukaryota | 27601.0 | 0 | Chordata | 27601.0 | 0 | Carnivora | 17734.0 | 2 | Canidae | 9306.0 | 5 | Canis | 9306.0 | 5 | Canis lupus | 9306.0 | 5 |

| | | | | | | | | | | | | | | | | | | | | | | |
|---|---|---|---|---|---|---|---|---|---|---|---|---|---|---|---|---|---|---|---|---|---|---|
| scf7180000022061 | 20940 | 0.4933 | 0 | 45.67 | Eukaryota | 3262.0 | 0 | Chordata | 3262.0 | 0 | Carnivora | 3262.0 | 0 | Felidae | 2970.0 | 1 | Felis | 2970.0 | 1 | Felis catus | 2970.0 | 1 |
| scf7180000024115 | 99327 | 0.3828 | 0 | 34.7087 | Eukaryota | 45241.0 | 0 | Chordata | 45241.0 | 0 | Carnivora | 45241.0 | 0 | Felidae | 45241.0 | 0 | Felis | 45241.0 | 0 | Felis catus | 45241.0 | 0 |
| scf7180000021427 | 197940 | 0.5332 | 0 | 49.3106 | Eukaryota | 10381.0 | 0 | Chordata | 10381.0 | 0 | Carnivora | 10381.0 | 0 | Felidae | 10381.0 | 0 | Felis | 9357.0 | 1 | Felis catus | 9357.0 | 1 |
| scf7180000020330 | 102709 | 0.4716 | 0 | 49.7963 | Eukaryota | 10337.0 | 0 | Chordata | 10337.0 | 0 | Carnivora | 10337.0 | 0 | Felidae | 10337.0 | 0 | Felis | 6886.0 | 1 | Felis catus | 6886.0 | 1 |
| scf7180000023452 | 25929 | 0.3536 | 0 | 227.6862 | Eukaryota | 12265.0 | 0 | Chordata | 12265.0 | 0 | Carnivora | 12265.0 | 0 | Felidae | 12265.0 | 0 | Felis | 9823.0 | 1 | Felis catus | 9823.0 | 1 |
| scf7180000021633 | 52596 | 0.3909 | 0 | 25.968 | Eukaryota | 23474.0 | 0 | Chordata | 23474.0 | 0 | Carnivora | 23474.0 | 0 | Felidae | 23474.0 | 0 | Felis | 21073.0 | 1 | Felis catus | 21073.0 | 1 |
| scf7180000019056 | 47500 | 0.4193 | 0 | 43.4814 | Eukaryota | 55659.0 | 0 | Chordata | 55659.0 | 0 | Carnivora | 55659.0 | 0 | Felidae | 55659.0 | 0 | Panthera | 27148.0 | 2 | anthera pardu | 27148.0 | 2 |
| scf7180000022197 | 14647 | 0.4011 | 0 | 24.8393 | Eukaryota | 20051.0 | 0 | Chordata | 20051.0 | 0 | Carnivora | 20051.0 | 0 | Felidae | 20051.0 | 0 | Felis | 16030.0 | 1 | Felis catus | 16030.0 | 1 |
| scf7180000021130 | 29072 | 0.5216 | 0 | 278.5408 | no-hit | 0.0 | 0 | no-hit | 0.0 | 0 | no-hit | 0.0 | 0 | no-hit | 0.0 | 0 | no-hit | 0.0 | 0 | no-hit | 0.0 | 0 |
| scf7180000019581 | 58172 | 0.3833 | 0 | 390.1042 | Eukaryota | 7365.0 | 0 | Chordata | 7365.0 | 0 | Carnivora | 4201.0 | 2 | Felidae | 4201.0 | 2 | Felis | 3574.0 | 3 | Felis catus | 3574.0 | 3 |
| scf7180000020713 | 119632 | 0.3689 | 0 | 24.6934 | Eukaryota | 40458.0 | 0 | Chordata | 40458.0 | 0 | Carnivora | 40458.0 | 0 | Felidae | 40458.0 | 0 | Felis | 36400.0 | 1 | Felis catus | 36400.0 | 1 |
| scf7180000021215 | 38457 | 0.4384 | 0 | 26.4088 | Eukaryota | 8997.0 | 0 | Chordata | 8997.0 | 0 | Artiodactyla | 4943.0 | 2 | Delphinidae | 3864.0 | 3 | Felis | 2947.0 | 4 | Felis catus | 2947.0 | 4 |
| scf7180000018404 | 788404 | 0.3605 | 0 | 50.4381 | Eukaryota | 47791.0 | 0 | Chordata | 47791.0 | 0 | Carnivora | 47791.0 | 0 | Felidae | 47791.0 | 0 | Felis | 47791.0 | 0 | Felis catus | 47791.0 | 0 |
| scf7180000022257 | 11516 | 0.3594 | 0 | 24.8335 | Eukaryota | 2892.0 | 0 | Chordata | 2892.0 | 0 | Carnivora | 2260.0 | 1 | Felidae | 1986.0 | 2 | Felis | 1492.0 | 3 | Felis catus | 1492.0 | 3 |
| scf7180000023394 | 18284 | 0.5229 | 0 | 536.9501 | no-hit | 0.0 | 0 | no-hit | 0.0 | 0 | no-hit | 0.0 | 0 | no-hit | 0.0 | 0 | no-hit | 0.0 | 0 | no-hit | 0.0 | 0 |
| scf7180000022904 | 65324 | 0.3994 | 0 | 358.0439 | Eukaryota | 46314.0 | 0 | Chordata | 46314.0 | 0 | Carnivora | 46314.0 | 0 | Felidae | 46314.0 | 0 | Felis | 46314.0 | 0 | Felis catus | 46314.0 | 0 |
| scf7180000018517 | 3622513 | 0.369 | 100 | 49.5474 | Eukaryota | 45868.0 | 0 | Chordata | 45868.0 | 0 | Carnivora | 45868.0 | 0 | Felidae | 45868.0 | 0 | Felis | 45868.0 | 0 | Felis catus | 45868.0 | 0 |
| scf7180000023274 | 36121 | 0.5599 | 0 | 180.2801 | Eukaryota | 3740.0 | 0 | Chordata | 3740.0 | 0 | Carnivora | 3740.0 | 0 | Felidae | 3496.0 | 1 | Acinonyx | 1858.0 | 3 | cinonyx jubat | 1858.0 | 3 |
| scf7180000019207 | 350933 | 0.4499 | 0 | 50.7882 | Eukaryota | 15615.0 | 0 | Chordata | 15615.0 | 0 | Carnivora | 14767.0 | 1 | Felidae | 14767.0 | 1 | Lynx | 6397.0 | 3 | ynx canadens | 6397.0 | 3 |
| scf7180000020469 | 91809 | 0.6433 | 0 | 72.9776 | Eukaryota | 6473.0 | 0 | Chordata | 6473.0 | 0 | Carnivora | 6473.0 | 0 | Felidae | 6473.0 | 0 | Acinonyx | 3869.0 | 3 | cinonyx jubat | 3869.0 | 3 |
| scf7180000019861 | 62665 | 0.3705 | 0 | 23.9598 | Eukaryota | 26218.0 | 0 | Chordata | 26218.0 | 0 | Carnivora | 26218.0 | 0 | Felidae | 26218.0 | 0 | Felis | 21050.0 | 1 | Felis catus | 21050.0 | 1 |
| scf7180000023579 | 38298 | 0.4753 | 0 | 429.149 | no-hit | 0.0 | 0 | no-hit | 0.0 | 0 | no-hit | 0.0 | 0 | no-hit | 0.0 | 0 | no-hit | 0.0 | 0 | no-hit | 0.0 | 0 |
| scf7180000018479 | 738758 | 0.3728 | 0 | 48.791 | Eukaryota | 46190.0 | 0 | Chordata | 46190.0 | 0 | Carnivora | 46190.0 | 0 | Felidae | 46190.0 | 0 | Felis | 46190.0 | 0 | Felis catus | 46190.0 | 0 |
| scf7180000021885 | 34845 | 0.4238 | 0 | 24.5864 | Eukaryota | 44749.0 | 0 | Chordata | 44749.0 | 0 | Carnivora | 44749.0 | 0 | Felidae | 44749.0 | 0 | Felis | 44749.0 | 0 | Felis catus | 44749.0 | 0 |
| scf7180000020262 | 68118 | 0.4015 | 0 | 46.7323 | Eukaryota | 34261.0 | 0 | Chordata | 34261.0 | 0 | Carnivora | 34261.0 | 0 | Felidae | 34261.0 | 0 | Felis | 30761.0 | 1 | Felis catus | 30761.0 | 1 |
| scf7180000018652 | 630687 | 0.415 | 0 | 50.6632 | Eukaryota | 47889.0 | 0 | Chordata | 47889.0 | 0 | Carnivora | 47889.0 | 0 | Felidae | 47889.0 | 0 | Felis | 47889.0 | 0 | Felis catus | 47889.0 | 0 |
| scf7180000018938 | 607822 | 0.6136 | 0 | 49.4706 | Eukaryota | 12638.0 | 1 | Chordata | 12638.0 | 1 | Carnivora | 9966.0 | 2 | Herpestidae | 6811.0 | 4 | Suricata | 6811.0 | 4 | ricata suricat | 6811.0 | 4 |
| scf7180000020627 | 78756 | 0.4962 | 0 | 50.3536 | Eukaryota | 15921.0 | 0 | Chordata | 15921.0 | 0 | Carnivora | 15921.0 | 0 | Felidae | 15921.0 | 0 | Felis | 14336.0 | 1 | Felis catus | 14336.0 | 1 |
| scf7180000018402 | 94483 | 0.352 | 0 | 46.4462 | Eukaryota | 30592.0 | 0 | Chordata | 30592.0 | 0 | Carnivora | 30592.0 | 0 | Felidae | 30592.0 | 0 | Felis | 30592.0 | 0 | Felis catus | 30592.0 | 0 |
| scf7180000018717 | 313071 | 0.3681 | 0 | 49.5941 | Eukaryota | 51349.0 | 0 | Chordata | 51349.0 | 0 | Carnivora | 51349.0 | 0 | Herpestidae | 36589.0 | 1 | Suricata | 36589.0 | 1 | ricata suricat | 36589.0 | 1 |
| scf7180000019759 | 262402 | 0.4354 | 0 | 51.6406 | Eukaryota | 49472.0 | 1 | Chordata | 49472.0 | 1 | Carnivora | 21354.0 | 3 | Felidae | 21354.0 | 3 | Panthera | 14835.0 | 6 | anthera pardu | 14835.0 | 6 |
| scf7180000021429 | 52448 | 0.5024 | 0 | 41.5783 | Eukaryota | 12235.0 | 0 | Chordata | 12235.0 | 0 | Carnivora | 12235.0 | 0 | Felidae | 12235.0 | 0 | Felis | 6301.0 | 2 | Felis catus | 6301.0 | 2 |
| scf7180000018172 | 1079429 | 0.3882 | 0 | 50.3938 | Eukaryota | 44447.0 | 0 | Chordata | 44447.0 | 0 | Carnivora | 44447.0 | 0 | Felidae | 44447.0 | 0 | Felis | 44447.0 | 0 | Felis catus | 44447.0 | 0 |
| scf7180000020506 | 139943 | 0.395 | 0 | 24.8863 | Eukaryota | 46317.0 | 0 | Chordata | 46317.0 | 0 | Carnivora | 46317.0 | 0 | Felidae | 46317.0 | 0 | Felis | 46317.0 | 0 | Felis catus | 46317.0 | 0 |
| scf7180000019107 | 255155 | 0.5279 | 0 | 51.3715 | Eukaryota | 28455.0 | 0 | Chordata | 28455.0 | 0 | Carnivora | 28455.0 | 0 | Felidae | 28455.0 | 0 | Felis | 25548.0 | 1 | Felis catus | 25548.0 | 1 |
| scf7180000021260 | 33306 | 0.6479 | 0 | 41.8143 | Eukaryota | 4961.0 | 0 | Chordata | 4961.0 | 0 | Carnivora | 3547.0 | 1 | Felidae | 2615.0 | 3 | Panthera | 1080.0 | 8 | anthera pardu | 1080.0 | 8 |
| scf7180000018850 | 996357 | 0.394 | 0 | 50.8801 | Eukaryota | 66389.0 | 0 | Chordata | 66389.0 | 0 | Carnivora | 66389.0 | 0 | Felidae | 66389.0 | 0 | Felis | 39894.0 | 2 | Felis catus | 39894.0 | 2 |
| scf7180000019908 | 188673 | 0.3575 | 0 | 24.9461 | Eukaryota | 17372.0 | 0 | Chordata | 17372.0 | 0 | Carnivora | 17372.0 | 0 | Felidae | 10252.0 | 3 | Felis | 5091.0 | 5 | Felis catus | 5091.0 | 5 |
| scf7180000018719 | 388394 | 0.605 | 0 | 50.7415 | Eukaryota | 33535.0 | 0 | Chordata | 33535.0 | 0 | Artiodactyla | 17496.0 | 1 | Monodontidae | 8051.0 | 6 | Zalophus | 5720.0 | 8 | phus californi | 5720.0 | 8 |
| scf7180000024064 | 343485 | 0.5808 | 0 | 49.8608 | Eukaryota | 14779.0 | 0 | Chordata | 14779.0 | 0 | Carnivora | 14779.0 | 0 | Felidae | 7761.0 | 3 | Ursus | 4106.0 | 6 | Ursus arctos | 2748.0 | 8 |
| scf7180000019147 | 124153 | 0.4175 | 0 | 51.728 | Eukaryota | 42956.0 | 0 | Chordata | 42956.0 | 0 | Carnivora | 42956.0 | 0 | Felidae | 42956.0 | 0 | Felis | 38719.0 | 1 | Felis catus | 38719.0 | 1 |
| scf7180000019654 | 183966 | 0.3682 | 0 | 25.024 | Eukaryota | 45455.0 | 0 | Chordata | 45455.0 | 0 | Carnivora | 45455.0 | 0 | Felidae | 45455.0 | 0 | Felis | 45455.0 | 0 | Felis catus | 45455.0 | 0 |
| scf7180000020081 | 216989 | 0.3991 | 0 | 51.3258 | Eukaryota | 24421.0 | 0 | Chordata | 24421.0 | 0 | Carnivora | 24421.0 | 0 | Felidae | 18936.0 | 1 | Felis | 8010.0 | 5 | Felis catus | 8010.0 | 5 |
| scf7180000017614 | 530811 | 0.3503 | 0 | 49.2963 | Eukaryota | 46830.0 | 0 | Chordata | 46830.0 | 0 | Carnivora | 46830.0 | 0 | Felidae | 46830.0 | 0 | Felis | 46830.0 | 0 | Felis catus | 46830.0 | 0 |
| scf7180000019212 | 380736 | 0.4655 | 0 | 49.9228 | Eukaryota | 17841.0 | 0 | Chordata | 17841.0 | 0 | Carnivora | 17841.0 | 0 | Felidae | 17841.0 | 0 | Felis | 12512.0 | 1 | Felis catus | 12512.0 | 1 |
| scf7180000017800 | 1597938 | 0.3692 | 0 | 50.0945 | Eukaryota | 45346.0 | 0 | Chordata | 45346.0 | 0 | Carnivora | 45346.0 | 0 | Felidae | 45346.0 | 0 | Felis | 40463.0 | 1 | Felis catus | 40463.0 | 1 |
| scf7180000022296 | 41658 | 0.3541 | 0 | 49.2315 | Eukaryota | 39392.0 | 0 | Chordata | 39392.0 | 0 | Carnivora | 39392.0 | 0 | Felidae | 39392.0 | 0 | Felis | 35464.0 | 1 | Felis catus | 35464.0 | 1 |
| scf7180000021400 | 111864 | 0.3895 | 0 | 25.8518 | Eukaryota | 8944.0 | 0 | Chordata | 8944.0 | 0 | Carnivora | 6245.0 | 1 | Felidae | 4772.0 | 2 | Felis | 4772.0 | 2 | Felis catus | 4772.0 | 2 |
| scf7180000022116 | 35566 | 0.3821 | 0 | 25.6224 | Eukaryota | 41965.0 | 0 | Chordata | 41965.0 | 0 | Carnivora | 41965.0 | 0 | Felidae | 41965.0 | 0 | Felis | 37830.0 | 1 | Felis catus | 37830.0 | 1 |
| scf7180000022600 | 8492 | 0.4531 | 0 | 25.3414 | Eukaryota | 4088.0 | 0 | Chordata | 4088.0 | 0 | Primates | 1672.0 | 2 | Hominidae | 1672.0 | 2 | Sus | 1286.0 | 5 | Sus scrofa | 1286.0 | 5 |
| scf7180000018094 | 1039016 | 0.3934 | 0 | 50.9578 | Eukaryota | 45621.0 | 0 | Chordata | 45621.0 | 0 | Carnivora | 45621.0 | 0 | Felidae | 45621.0 | 0 | Felis | 45621.0 | 0 | Felis catus | 45621.0 | 0 |
| scf7180000021746 | 23778 | 0.3674 | 0 | 23.1437 | Eukaryota | 8868.0 | 0 | Chordata | 8868.0 | 0 | Carnivora | 8868.0 | 0 | Felidae | 8868.0 | 0 | Felis | 8868.0 | 0 | Felis catus | 8868.0 | 0 |
| scf7180000017848 | 221627 | 0.4179 | 0 | 49.9049 | Eukaryota | 55577.0 | 0 | Chordata | 55577.0 | 0 | Carnivora | 55577.0 | 0 | Felidae | 55577.0 | 0 | Felis | 16676.0 | 3 | Felis catus | 16676.0 | 4 |
| scf7180000021764 | 29861 | 0.3712 | 0 | 24.5583 | Eukaryota | 39315.0 | 0 | Chordata | 39315.0 | 0 | Carnivora | 39315.0 | 0 | Felidae | 39315.0 | 0 | Felis | 35422.0 | 1 | Felis catus | 35422.0 | 1 |
| scf7180000022269 | 19863 | 0.4127 | 0 | 23.9049 | Eukaryota | 14577.0 | 0 | Chordata | 14577.0 | 0 | Carnivora | 14577.0 | 0 | Felidae | 14577.0 | 0 | Felis | 11671.0 | 1 | Felis catus | 11671.0 | 1 |
| scf7180000020236 | 137724 | 0.5277 | 0 | 52.517 | Eukaryota | 9065.0 | 0 | Chordata | 9065.0 | 0 | Primates | 4936.0 | 2 | Felidae | 2248.0 | 4 | Homo | 2158.0 | 6 | Homo sapiens | 2158.0 | 6 |
| scf7180000019347 | 378207 | 0.381 | 0 | 49.4867 | Eukaryota | 46919.0 | 0 | Chordata | 46919.0 | 0 | Carnivora | 46919.0 | 0 | Felidae | 46919.0 | 0 | Felis | 46919.0 | 0 | Felis catus | 46919.0 | 0 |
| scf7180000022995 | 41968 | 0.4733 | 0 | 228.8146 | Eukaryota | 24065.0 | 0 | Chordata | 24065.0 | 0 | Carnivora | 24065.0 | 0 | Felidae | 24065.0 | 0 | Felis | 24065.0 | 0 | Felis catus | 24065.0 | 0 |
| scf7180000021847 | 31085 | 0.4685 | 0 | 24.1088 | Eukaryota | 9267.0 | 0 | Chordata | 9267.0 | 0 | Primates | 5149.0 | 3 | Hominidae | 4284.0 | 4 | Homo | 2568.0 | 5 | Homo sapiens | 2568.0 | 5 |
| scf7180000021084 | 88161 | 0.3689 | 0 | 25.4218 | Eukaryota | 21274.0 | 0 | Chordata | 21274.0 | 0 | Carnivora | 21274.0 | 0 | Felidae | 21274.0 | 0 | Felis | 19213.0 | 1 | Felis catus | 19213.0 | 1 |
| scf7180000022507 | 20902 | 0.6677 | 0 | 618.6838 | Eukaryota | 624.0 | 0 | Chordata | 483.0 | 1 | Carnivora | 483.0 | 1 | Felidae | 344.0 | 2 | Felis | 344.0 | 2 | Felis catus | 344.0 | 2 |
| scf7180000018726 | 1214664 | 0.4332 | 100 | 51.1704 | Eukaryota | 44772.0 | 0 | Chordata | 44772.0 | 0 | Carnivora | 44772.0 | 0 | Felidae | 28087.0 | 2 | Panthera | 9386.0 | 6 | Canis lupus | 8296.0 | 8 |
| scf7180000017453 | 4779948 | 0.3618 | 0 | 49.3074 | Eukaryota | 47856.0 | 0 | Chordata | 47856.0 | 0 | Carnivora | 47856.0 | 0 | Felidae | 47856.0 | 0 | Felis | 47856.0 | 0 | Felis catus | 47856.0 | 0 |
| scf7180000019541 | 4933 | 0.3673 | 0 | 24.4086 | Eukaryota | 26548.0 | 0 | Chordata | 26548.0 | 0 | Carnivora | 26548.0 | 0 | Felidae | 26548.0 | 0 | Felis | 23846.0 | 1 | Felis catus | 23846.0 | 1 |
| scf7180000018323 | 1021858 | 0.3746 | 0 | 49.7994 | Eukaryota | 46114.0 | 0 | Chordata | 46114.0 | 0 | Carnivora | 46114.0 | 0 | Felidae | 46114.0 | 0 | Felis | 46114.0 | 0 | Felis catus | 46114.0 | 0 |
| scf7180000022150 | 35424 | 0.4322 | 0 | 75.6172 | Eukaryota | 40973.0 | 0 | Chordata | 40973.0 | 0 | Carnivora | 40973.0 | 0 | Felidae | 40973.0 | 0 | Felis | 36919.0 | 1 | Felis catus | 36919.0 | 1 |
| scf7180000019884 | 29992 | 0.4039 | 0 | 315.3708 | Eukaryota | 12715.0 | 0 | Chordata | 12715.0 | 0 | Carnivora | 12715.0 | 0 | Felidae | 12715.0 | 0 | Felis | 11449.0 | 1 | Felis catus | 11449.0 | 1 |
| scf7180000018935 | 10150 | 0.3712 | 0 | 30.8823 | Eukaryota | 8670.0 | 0 | Chordata | 8670.0 | 0 | Carnivora | 8670.0 | 0 | Felidae | 8670.0 | 0 | Felis | 8670.0 | 0 | Felis catus | 8670.0 | 0 |
| scf7180000017911 | 1379959 | 0.3841 | 0 | 50.9914 | Eukaryota | 48059.0 | 0 | Chordata | 48059.0 | 0 | Carnivora | 48059.0 | 0 | Felidae | 42253.0 | 1 | Felis | 42253.0 | 1 | Felis catus | 42253.0 | 1 |
| scf7180000018887 | 604509 | 0.6124 | 0 | 48.8813 | Eukaryota | 29142.0 | 0 | Chordata | 29142.0 | 0 | Carnivora | 29142.0 | 0 | Felidae | 23682.0 | 2 | Felis | 7548.0 | 5 | Felis catus | 7548.0 | 5 |
| scf7180000018762 | 655322 | 0.3945 | 0 | 52.106 | Eukaryota | 47653.0 | 0 | Chordata | 47653.0 | 0 | Carnivora | 47653.0 | 0 | Felidae | 47653.0 | 0 | Felis | 47653.0 | 0 | Felis catus | 47653.0 | 0 |
| scf7180000023813 | 1461991 | 0.5051 | 0 | 51.6596 | Eukaryota | 54759.0 | 0 | Chordata | 54759.0 | 0 | Carnivora | 54759.0 | 0 | Felidae | 41169.0 | 1 | Felis | 26657.0 | 2 | Felis catus | 26657.0 | 2 |
| scf7180000017604 | 10409 | 0.3515 | 0 | 26.4284 | Eukaryota | 7219.0 | 0 | Chordata | 7219.0 | 0 | Carnivora | 7219.0 | 0 | Felidae | 7219.0 | 0 | Felis | 6415.0 | 1 | Felis catus | 6415.0 | 1 |
| scf7180000021222 | 33872 | 0.3722 | 0 | 24.9602 | Eukaryota | 26398.0 | 0 | Chordata | 26398.0 | 0 | Carnivora | 26398.0 | 0 | Felidae | 26398.0 | 0 | Felis | 23707.0 | 1 | Felis catus | 23707.0 | 1 |
| scf7180000022054 | 18524 | 0.6307 | 0 | 62.0747 | Eukaryota | 2409.0 | 0 | Chordata | 2409.0 | 0 | Carnivora | 2207.0 | 1 | Felidae | 2207.0 | 1 | Felis | 1012.0 | 4 | Felis catus | 1012.0 | 4 |
| scf7180000018602 | 597659 | 0.3966 | 0 | 49.8681 | Eukaryota | 43552.0 | 0 | Chordata | 43552.0 | 0 | Carnivora | 43552.0 | 0 | Felidae | 43552.0 | 0 | Felis | 39225.0 | 1 | Felis catus | 39225.0 | 1 |
| scf7180000020375 | 119740 | 0.3826 | 0 | 50.3097 | Eukaryota | 34049.0 | 0 | Chordata | 34049.0 | 0 | Carnivora | 34049.0 | 0 | Felidae | 34049.0 | 0 | Felis | 34049.0 | 0 | Felis catus | 34049.0 | 0 |
| scf7180000024177 | 100551 | 0.3712 | 0 | 23.5532 | Eukaryota | 40267.0 | 0 | Chordata | 40267.0 | 0 | Carnivora | 40267.0 | 0 | Felidae | 40267.0 | 0 | Felis | 40267.0 | 0 | Felis catus | 40267.0 | 0 |
| scf7180000022558 | 15642 | 0.3963 | 0 | 333.0738 | Eukaryota | 3510.0 | 0 | Chordata | 3510.0 | 0 | Carnivora | 3510.0 | 0 | Canidae | 2034.0 | 2 | Canis | 2034.0 | 2 | Canis lupus | 2034.0 | 2 |
| scf7180000019619 | 479334 | 0.4684 | 0 | 50.4447 | Eukaryota | 35305.0 | 0 | Chordata | 35305.0 | 0 | Carnivora | 35305.0 | 0 | Herpestidae | 17770.0 | 1 | Suricata | 17770.0 | 1 | ricata suricat | 17770.0 | 1 |
| scf7180000022791 | 4530 | 0.3786 | 0 | 24.5858 | Eukaryota | 3432.0 | 0 | Chordata | 3432.0 | 0 | Carnivora | 3432.0 | 0 | Felidae | 2788.0 | 1 | Felis | 2788.0 | 1 | Felis catus | 2788.0 | 1 |
| scf7180000019165 | 473871 | 0.3441 | 0 | 48.8846 | Eukaryota | 44866.0 | 0 | Chordata | 44866.0 | 0 | Carnivora | 44866.0 | 0 | Felidae | 44866.0 | 0 | Felis | 40430.0 | 1 | Felis catus | 40430.0 | 1 |
| scf7180000018874 | 419476 | 0.4249 | 0 | 49.8067 | Eukaryota | 43376.0 | 0 | Chordata | 43376.0 | 0 | Carnivora | 43376.0 | 0 | Felidae | 43376.0 | 0 | Felis | 39143.0 | 1 | Felis catus | 39143.0 | 1 |
| scf7180000023297 | 30077 | 0.3973 | 0 | 135.3142 | Eukaryota | 25637.0 | 0 | Chordata | 25637.0 | 0 | Carnivora | 25637.0 | 0 | Felidae | 25637.0 | 0 | Felis | 23040.0 | 1 | Felis catus | 23040.0 | 1 |
| scf7180000021243 | 111202 | 0.4491 | 0 | 356.6255 | Eukaryota | 43246.0 | 0 | Chordata | 43246.0 | 0 | Carnivora | 43246.0 | 0 | Felidae | 43246.0 | 0 | Felis | 43246.0 | 0 | Felis catus | 43246.0 | 0 |
| scf7180000018341 | 781996 | 0.3871 | 0 | 50.6055 | Eukaryota | 65885.0 | 0 | Chordata | 65885.0 | 0 | Carnivora | 65885.0 | 0 | Felidae | 65885.0 | 0 | Acinonyx | 33060.0 | 1 | cinonyx jubat | 33060.0 | 1 |
| scf7180000020449 | 94921 | 0.4113 | 0 | 23.6778 | Eukaryota | 5431.0 | 0 | Chordata | 5431.0 | 0 | Carnivora | 1924.0 | 3 | Felidae | 1924.0 | 3 | Equus | 1689.0 | 5 | Equus caballu | 1689.0 | 5 |
| scf7180000023826 | 1342284 | 0.4342 | 0 | 51.6969 | Eukaryota | 53890.0 | 0 | Chordata | 53890.0 | 0 | Carnivora | 53890.0 | 0 | Felidae | 53890.0 | 0 | Panthera | 53890.0 | 0 | anthera pardu | 53890.0 | 0 |
| scf7180000022088 | 17055 | 0.3411 | 0 | 48.0918 | Eukaryota | 12476.0 | 0 | Chordata | 12476.0 | 0 | Carnivora | 12476.0 | 0 | Felidae | 12476.0 | 0 | Felis | 11299.0 | 1 | Felis catus | 11299.0 | 1 |
| scf7180000024048 | 452265 | 0.3891 | 0 | 50.9734 | Eukaryota | 45569.0 | 0 | Chordata | 45569.0 | 0 | Carnivora | 45569.0 | 0 | Felidae | 34211.0 | 1 | Felis | 34211.0 | 1 | Felis catus | 34211.0 | 1 |
| scf7180000018489 | 726887 | 0.4405 | 0 | 48.3181 | Eukaryota | 44624.0 | 0 | Chordata | 44624.0 | 0 | Carnivora | 44624.0 | 0 | Felidae | 44624.0 | 0 | Felis | 40121.0 | 1 | Felis catus | 40121.0 | 1 |
| scf7180000020514 | 109360 | 0.5643 | 0 | 51.7852 | Eukaryota | 28147.0 | 0 | Chordata | 28147.0 | 0 | Carnivora | 26113.0 | 1 | Felidae | 12663.0 | 4 | Felis | 3459.0 | 9 | Felis catus | 3459.0 | 9 |
| scf7180000017619 | 2002061 | 0.3687 | 0 | 49.8907 | Eukaryota | 46321.0 | 0 | Chordata | 46321.0 | 0 | Carnivora | 46321.0 | 0 | Felidae | 46321.0 | 0 | Felis | 46321.0 | 0 | Felis catus | 46321.0 | 0 |
| scf7180000021775 | 48557 | 0.4101 | 0 | 23.793 | Eukaryota | 4796.0 | 0 | Chordata | 4796.0 | 0 | Primates | 1850.0 | 3 | Hominidae | 1850.0 | 3 | Homo | 1534.0 | 4 | Homo sapiens | 1534.0 | 4 |
| scf7180000023225 | 167563 | 0.4199 | 0 | 245.5785 | Eukaryota | 45283.0 | 0 | Chordata | 45283.0 | 0 | Carnivora | 45283.0 | 0 | Felidae | 45283.0 | 0 | Felis | 45283.0 | 0 | Felis catus | 45283.0 | 0 |
| scf7180000023886 | 402705 | 0.3788 | 0 | 50.7206 | Eukaryota | 13112.0 | 0 | Chordata | 13112.0 | 0 | Carnivora | 13112.0 | 0 | Felidae | 13112.0 | 0 | Felis | 10496.0 | 1 | Felis catus | 10496.0 | 1 |
| scf7180000022421 | 30992 | 0.3971 | 0 | 24.9278 | Eukaryota | 8667.0 | 0 | Chordata | 8667.0 | 0 | Primates | 4156.0 | 2 | Hominidae | 4156.0 | 3 | Pan | 3326.0 | 4 | an troglodyte | 3326.0 | 4 |
| scf7180000018939 | 609251 | 0.3755 | 0 | 51.221 | Eukaryota | 73144.0 | 0 | Chordata | 73144.0 | 0 | Primates | 57452.0 | 1 | Hominidae | 57452.0 | 2 | Pan | 34343.0 | 3 | an troglodyte | 27488.0 | 4 |
| scf7180000020100 | 169327 | 0.4264 | 0 | 25.3538 | Eukaryota | 41535.0 | 0 | Chordata | 41535.0 | 0 | Carnivora | 20327.0 | 3 | Felidae | 16245.0 | 4 | Homo | 8282.0 | 7 | Homo sapiens | 8282.0 | 8 |
| scf7180000022159 | 15571 | 0.4262 | 0 | 49.4009 | Eukaryota | 19105.0 | 0 | Chordata | 19105.0 | 0 | Carnivora | 19105.0 | 0 | Felidae | 19105.0 | 0 | Felis | 15288.0 | 1 | Felis catus | 15288.0 | 1 |
| scf7180000019266 | 21120 | 0.44 | 0 | 66.925 | Eukaryota | 10416.0 | 0 | Chordata | 10416.0 | 0 | Carnivora | 10416.0 | 0 | Felidae | 10416.0 | 0 | Felis | 8373.0 | 1 | Felis catus | 8373.0 | 1 |
| scf7180000020395 | 156294 | 0.592 | 0 | 51.5386 | Eukaryota | 27800.0 | 0 | Chordata | 27800.0 | 0 | Carnivora | 27800.0 | 0 | Felidae | 27800.0 | 0 | Acinonyx | 13900.0 | 2 | cinonyx jubat | 13900.0 | 2 |
| scf7180000023251 | 32966 | 0.465 | 0 | 454.4831 | Eukaryota | 13566.0 | 0 | Chordata | 13566.0 | 0 | Carnivora | 13566.0 | 0 | Felidae | 13566.0 | 0 | Felis | 13566.0 | 0 | Felis catus | 13566.0 | 0 |
| scf7180000017578 | 18894 | 0.4062 | 0 | 27.1678 | Eukaryota | 10191.0 | 0 | Chordata | 10191.0 | 0 | Carnivora | 10191.0 | 0 | Felidae | 10191.0 | 0 | Felis | 10191.0 | 0 | Felis catus | 10191.0 | 0 |
| scf7180000017637 | 1167139 | 0.3847 | 0 | 45.1286 | Eukaryota | 58659.0 | 0 | Chordata | 58659.0 | 0 | Carnivora | 52090.0 | 1 | Canidae | 28146.0 | 2 | Canis | 28146.0 | 2 | Canis lupus | 28146.0 | 2 |
| scf7180000020114 | 17883 | 0.4801 | 0 | 161.7181 | Eukaryota | 12910.0 | 0 | Chordata | 12910.0 | 0 | Artiodactyla | 6455.0 | 3 | Delphinidae | 5164.0 | 4 | Globicephala | 3873.0 | 6 | bicephala me | 3873.0 | 6 |
| scf7180000020169 | 103169 | 0.3912 | 0 | 49.783 | Eukaryota | 44837.0 | 0 | Chordata | 44837.0 | 0 | Carnivora | 44837.0 | 0 | Felidae | 44837.0 | 0 | Felis | 44837.0 | 0 | Felis catus | 44837.0 | 0 |
| scf7180000021482 | 15012 | 0.5228 | 0 | 233.7309 | no-hit | 0.0 | 0 | no-hit | 0.0 | 0 | no-hit | 0.0 | 0 | no-hit | 0.0 | 0 | no-hit | 0.0 | 0 | no-hit | 0.0 | 0 |
| scf7180000021385 | 41717 | 0.4255 | 0 | 55.9401 | Eukaryota | 29590.0 | 0 | Chordata | 29590.0 | 0 | erissodactyla | 29590.0 | 0 | Equidae | 29590.0 | 0 | Equus | 29590.0 | 0 | Equus caballu | 29590.0 | 0 |
| scf7180000020292 | 9276 | 0.5497 | 0 | 26.2012 | Eukaryota | 272.0 | 0 | Chordata | 272.0 | 0 | Carnivora | 272.0 | 0 | Felidae | 272.0 | 0 | Felis | 137.0 | 1 | Felis catus | 137.0 | 1 |
| scf7180000020908 | 85625 | 0.3849 | 0 | 24.7078 | Eukaryota | 43325.0 | 0 | Chordata | 43325.0 | 0 | Carnivora | 43325.0 | 0 | Felidae | 43325.0 | 0 | Felis | 43325.0 | 0 | Felis catus | 43325.0 | 0 |
| scf7180000019573 | 235514 | 0.4008 | 0 | 25.0294 | Eukaryota | 32434.0 | 0 | Chordata | 32434.0 | 0 | Carnivora | 32434.0 | 0 | Felidae | 32434.0 | 0 | Felis | 29241.0 | 1 | Felis catus | 29241.0 | 1 |
| scf7180000020929 | 87400 | 0.3559 | 0 | 48.463 | Eukaryota | 20400.0 | 0 | Chordata | 20400.0 | 0 | Carnivora | 20400.0 | 0 | Felidae | 20400.0 | 0 | Felis | 20400.0 | 0 | Felis catus | 20400.0 | 0 |
| scf7180000018265 | 32043 | 0.3857 | 0 | 24.4925 | Eukaryota | 13716.0 | 0 | Chordata | 13716.0 | 0 | Carnivora | 13716.0 | 0 | Felidae | 10976.0 | 2 | Panthera | 5504.0 | 5 | Panthera tigri | 2755.0 | 6 |
| scf7180000017883 | 2188608 | 0.3694 | 0 | 49.6549 | Eukaryota | 46842.0 | 0 | Chordata | 46842.0 | 0 | Carnivora | 46842.0 | 0 | Felidae | 46842.0 | 0 | Felis | 46842.0 | 0 | Felis catus | 46842.0 | 0 |
| scf7180000020350 | 202220 | 0.4239 | 0 | 26.1187 | Eukaryota | 10978.0 | 0 | Chordata | 10978.0 | 0 | Carnivora | 6783.0 | 2 | Felidae | 6783.0 | 2 | Felis | 6783.0 | 3 | Felis catus | 6783.0 | 3 |
| scf7180000022377 | 20994 | 0.408 | 0 | 1235.0869 | no-hit | 0.0 | 0 | no-hit | 0.0 | 0 | no-hit | 0.0 | 0 | no-hit | 0.0 | 0 | no-hit | 0.0 | 0 | no-hit | 0.0 | 0 |
| scf7180000021382 | 58944 | 0.3675 | 0 | 24.5948 | Eukaryota | 34186.0 | 0 | Chordata | 34186.0 | 0 | Carnivora | 34186.0 | 0 | Felidae | 34186.0 | 0 | Felis | 30764.0 | 1 | Felis catus | 30764.0 | 1 |
| scf7180000019417 | 345454 | 0.3473 | 0 | 46.8305 | Eukaryota | 40970.0 | 0 | Chordata | 40970.0 | 0 | Carnivora | 40970.0 | 0 | Felidae | 40970.0 | 0 | Felis | 36837.0 | 1 | Felis catus | 36837.0 | 1 |
| scf7180000019954 | 748813 | 0.4227 | 0 | 53.3042 | Eukaryota | 32675.0 | 0 | Chordata | 32675.0 | 0 | Carnivora | 32675.0 | 0 | Felidae | 20136.0 | 2 | Acinonyx | 20136.0 | 3 | cinonyx jubat | 20136.0 | 3 |
| scf7180000021081 | 51733 | 0.368 | 0 | 48.224 | Eukaryota | 8743.0 | 0 | Chordata | 8743.0 | 0 | Carnivora | 8743.0 | 0 | Felidae | 8743.0 | 0 | Felis | 7910.0 | 1 | Felis catus | 7910.0 | 1 |
| scf7180000017804 | 1547856 | 0.4519 | 0 | 51.3913 | Eukaryota | 46409.0 | 0 | Chordata | 46409.0 | 0 | Carnivora | 46409.0 | 0 | Felidae | 46409.0 | 0 | Felis | 46409.0 | 0 | Felis catus | 46409.0 | 0 |
| scf7180000022293 | 9133 | 0.6671 | 0 | 31.1641 | Eukaryota | 4136.0 | 0 | Chordata | 4136.0 | 0 | Carnivora | 3727.0 | 1 | Felidae | 3727.0 | 1 | Lynx | 2055.0 | 4 | ynx canadens | 2055.0 | 5 |
| scf7180000018661 | 62367 | 0.3754 | 0 | 51.1963 | Eukaryota | 15189.0 | 0 | Chordata | 15189.0 | 0 | Carnivora | 15189.0 | 0 | Felidae | 15189.0 | 0 | Felis | 13829.0 | 1 | Felis catus | 13829.0 | 1 |
| scf7180000023859 | 1273684 | 0.3882 | 0 | 51.0213 | Eukaryota | 45116.0 | 0 | Chordata | 45116.0 | 0 | Primates | 26684.0 | 1 | Hominidae | 26684.0 | 1 | Homo | 26684.0 | 2 | Homo sapiens | 26684.0 | 3 |
| scf7180000021751 | 21219 | 0.5256 | 0 | 92.4018 | Eukaryota | 7065.0 | 0 | Chordata | 7065.0 | 0 | Carnivora | 7065.0 | 0 | Felidae | 7065.0 | 0 | Felis | 6455.0 | 1 | Felis catus | 6455.0 | 1 |
| scf7180000019312 | 302683 | 0.4839 | 0 | 45.485 | Eukaryota | 8589.0 | 0 | Chordata | 8589.0 | 0 | Carnivora | 5704.0 | 1 | Felidae | 5704.0 | 1 | Felis | 5704.0 | 1 | Felis catus | 5704.0 | 1 |
| scf7180000017740 | 17739 | 0.3941 | 0 | 30.7366 | Eukaryota | 27194.0 | 0 | Chordata | 27194.0 | 0 | Carnivora | 27194.0 | 0 | Felidae | 27194.0 | 0 | Felis | 27194.0 | 0 | Felis catus | 27194.0 | 0 |
| scf7180000019474 | 242814 | 0.4058 | 0 | 26.2818 | Eukaryota | 29347.0 | 0 | Chordata | 29347.0 | 0 | Carnivora | 29347.0 | 0 | Felidae | 29347.0 | 0 | Felis | 23424.0 | 1 | Felis catus | 23424.0 | 1 |
| scf7180000018481 | 746746 | 0.4262 | 0 | 54.7803 | Eukaryota | 25443.0 | 0 | Chordata | 25443.0 | 0 | Carnivora | 25443.0 | 0 | Felidae | 25443.0 | 0 | Felis | 22876.0 | 1 | Felis catus | 22876.0 | 1 |
| scf7180000021254 | 45829 | 0.5818 | 100 | 52.0199 | Eukaryota | 11844.0 | 0 | Chordata | 11844.0 | 0 | Carnivora | 11844.0 | 0 | Felidae | 3810.0 | 4 | eptonychote | 2364.0 | 7 | nychotes we | 2364.0 | 7 |
| scf7180000022245 | 17809 | 0.573 | 0 | 46.4448 | Eukaryota | 6346.0 | 0 | Chordata | 6346.0 | 0 | Primates | 3972.0 | 2 | Hominidae | 3972.0 | 2 | Homo | 3972.0 | 2 | Homo sapiens | 3972.0 | 2 |
| scf7180000020508 | 50898 | 0.3905 | 0 | 31.3371 | Eukaryota | 23633.0 | 0 | Chordata | 23633.0 | 0 | Carnivora | 23633.0 | 0 | Felidae | 23633.0 | 0 | Felis | 18882.0 | 1 | Felis catus | 18882.0 | 1 |
| scf7180000021581 | 36595 | 0.5978 | 0 | 49.2424 | Eukaryota | 26236.0 | 0 | Chordata | 26236.0 | 0 | Carnivora | 26236.0 | 0 | Felidae | 15100.0 | 1 | Suricata | 11136.0 | 2 | ricata suricat | 11136.0 | 2 |
| scf7180000021488 | 50520 | 0.6461 | 0 | 54.3372 | Eukaryota | 14969.0 | 0 | Chordata | 14969.0 | 0 | Carnivora | 14969.0 | 0 | Felidae | 11263.0 | 1 | Felis | 8436.0 | 3 | Felis catus | 8436.0 | 3 |
| scf7180000018617 | 1092272 | 0.3919 | 0 | 51.4999 | Eukaryota | 37886.0 | 0 | Chordata | 37886.0 | 0 | Carnivora | 37886.0 | 0 | Felidae | 37886.0 | 0 | Felis | 34340.0 | 1 | Felis catus | 34340.0 | 1 |
| scf7180000018904 | 651371 | 0.3789 | 0 | 50.1073 | Eukaryota | 45877.0 | 0 | Chordata | 45877.0 | 0 | Carnivora | 45877.0 | 0 | Felidae | 45877.0 | 0 | Felis | 45877.0 | 0 | Felis catus | 45877.0 | 0 |
| scf7180000018177 | 1034266 | 0.3859 | 0 | 50.3572 | Eukaryota | 41543.0 | 0 | Chordata | 41543.0 | 0 | Carnivora | 36580.0 | 1 | Canidae | 25146.0 | 3 | Canis | 25146.0 | 3 | Canis lupus | 25146.0 | 3 |
| scf7180000020829 | 84895 | 0.4503 | 0 | 28.0499 | Eukaryota | 41220.0 | 0 | Chordata | 41220.0 | 0 | Carnivora | 41220.0 | 0 | Felidae | 41220.0 | 0 | Felis | 41220.0 | 0 | Felis catus | 41220.0 | 0 |
| scf7180000020636 | 107046 | 0.3622 | 0 | 48.8135 | Eukaryota | 6138.0 | 0 | Chordata | 6138.0 | 0 | Carnivora | 4674.0 | 1 | Felidae | 4674.0 | 1 | Felis | 4674.0 | 2 | Felis catus | 4674.0 | 2 |
| scf7180000022045 | 23007 | 0.3721 | 0 | 24.7368 | Eukaryota | 33185.0 | 0 | Chordata | 33185.0 | 0 | Carnivora | 33185.0 | 0 | Felidae | 33185.0 | 0 | Felis | 29888.0 | 1 | Felis catus | 29888.0 | 1 |
| scf7180000017534 | 2909658 | 0.3861 | 0 | 50.9328 | Eukaryota | 47653.0 | 0 | Chordata | 47653.0 | 0 | Carnivora | 47653.0 | 0 | Felidae | 47653.0 | 0 | Felis | 42862.0 | 1 | Felis catus | 42862.0 | 1 |
| scf7180000022646 | 11321 | 0.4574 | 0 | 25.4143 | Eukaryota | 6193.0 | 0 | Chordata | 6193.0 | 0 | Carnivora | 2701.0 | 2 | Felidae | 2701.0 | 3 | Pan | 1176.0 | 7 | an troglodyte | 1176.0 | 8 |
| scf7180000017463 | 548243 | 0.4151 | 0 | 48.4331 | Eukaryota | 65534.0 | 0 | Chordata | 65534.0 | 0 | Carnivora | 65534.0 | 0 | Felidae | 60051.0 | 1 | Acinonyx | 33591.0 | 2 | cinonyx jubat | 33591.0 | 2 |
| scf7180000017838 | 1528949 | 0.3903 | 0 | 50.6472 | Eukaryota | 47162.0 | 0 | Chordata | 47162.0 | 0 | Carnivora | 47162.0 | 0 | Felidae | 47162.0 | 0 | Felis | 47162.0 | 0 | Felis catus | 47162.0 | 0 |
| scf7180000018457 | 1083190 | 0.4044 | 0 | 49.9897 | Eukaryota | 51168.0 | 0 | Chordata | 51168.0 | 0 | Carnivora | 51168.0 | 0 | Felidae | 41734.0 | 1 | Felis | 23168.0 | 3 | Felis catus | 23168.0 | 3 |
| scf7180000022781 | 20130 | 0.4082 | 0 | 44.1819 | Eukaryota | 16946.0 | 0 | Chordata | 16946.0 | 0 | Carnivora | 16946.0 | 0 | Felidae | 16946.0 | 0 | Felis | 15226.0 | 1 | Felis catus | 15226.0 | 1 |
| scf7180000021463 | 89328 | 0.3903 | 0 | 25.9527 | Eukaryota | 43795.0 | 0 | Chordata | 43795.0 | 0 | Carnivora | 43795.0 | 0 | Felidae | 43795.0 | 0 | Felis | 39446.0 | 1 | Felis catus | 39446.0 | 1 |
| scf7180000022154 | 22083 | 0.3689 | 0 | 23.1605 | Eukaryota | 6185.0 | 0 | Chordata | 6185.0 | 0 | Carnivora | 6185.0 | 0 | Felidae | 3759.0 | 1 | Felis | 3143.0 | 2 | Felis catus | 3143.0 | 2 |
| scf7180000021002 | 51466 | 0.4261 | 0 | 50.2198 | Eukaryota | 9661.0 | 0 | Chordata | 9661.0 | 0 | Carnivora | 9661.0 | 0 | Felidae | 9661.0 | 0 | Felis | 9661.0 | 0 | Felis catus | 9661.0 | 0 |
| scf7180000020393 | 129149 | 0.3612 | 0 | 47.937 | Eukaryota | 18369.0 | 0 | Chordata | 18369.0 | 0 | Carnivora | 18369.0 | 0 | Felidae | 18369.0 | 0 | Felis | 16509.0 | 1 | Felis catus | 16509.0 | 1 |
| scf7180000017670 | 1867955 | 0.4102 | 0 | 51.0449 | Eukaryota | 103160.0 | 0 | Chordata | 103160.0 | 0 | Carnivora | 103160.0 | 0 | Felidae | 75381.0 | 1 | Panthera | 22084.0 | 6 | anthera pardu | 22084.0 | 7 |
| scf7180000021330 | 78811 | 0.5619 | 0 | 46.2391 | Eukaryota | 19065.0 | 0 | Chordata | 19065.0 | 0 | Carnivora | 15285.0 | 1 | Otariidae | 13048.0 | 2 | Eumetopias | 13048.0 | 2 | netopias juba | 13048.0 | 2 |
| scf7180000019422 | 400585 | 0.4186 | 100 | 49.5526 | Eukaryota | 49500.0 | 0 | Chordata | 49500.0 | 0 | Carnivora | 49500.0 | 0 | Felidae | 49500.0 | 0 | Lynx | 45911.0 | 1 | ynx canadens | 45911.0 | 1 |
| scf7180000020546 | 140947 | 0.4449 | 0 | 26.5519 | Eukaryota | 32642.0 | 0 | Chordata | 32642.0 | 0 | Carnivora | 32642.0 | 0 | Felidae | 26632.0 | 1 | Lynx | 13320.0 | 2 | ynx canadens | 13320.0 | 2 |
| scf7180000020016 | 4515 | 0.6315 | 0 | 127.9969 | no-hit | 0.0 | 0 | no-hit | 0.0 | 0 | no-hit | 0.0 | 0 | no-hit | 0.0 | 0 | no-hit | 0.0 | 0 | no-hit | 0.0 | 0 |
| scf7180000021008 | 87184 | 0.3948 | 0 | 25.7209 | Eukaryota | 34605.0 | 0 | Chordata | 34605.0 | 0 | Carnivora | 34605.0 | 0 | Felidae | 24080.0 | 1 | Felis | 13611.0 | 4 | Felis catus | 13611.0 | 4 |
| scf7180000022635 | 11107 | 0.4128 | 0 | 49.2343 | Eukaryota | 5616.0 | 0 | Chordata | 5616.0 | 0 | Carnivora | 5616.0 | 0 | Felidae | 5616.0 | 0 | Felis | 5616.0 | 0 | Felis catus | 5616.0 | 0 |
| scf7180000019239 | 231480 | 0.424 | 0 | 49.2097 | Eukaryota | 45382.0 | 0 | Chordata | 45382.0 | 0 | Carnivora | 45382.0 | 0 | Felidae | 45382.0 | 0 | Felis | 45382.0 | 0 | Felis catus | 45382.0 | 0 |

| | | | | | | | | | | | | | | | | | | | | | | |
|---|---|---|---|---|---|---|---|---|---|---|---|---|---|---|---|---|---|---|---|---|---|---|
| scf7180000022713 | 12730 | 0.4174 | 0 | 19.4226 | Eukaryota | 3350.0 | 0 | Chordata | 3350.0 | 0 | Carnivora | 3350.0 | 0 | Felidae | 2082.0 | 2 | Felis | 1223.0 | 4 | Felis catus | 1223.0 | 4 |
| scf7180000019772 | 247759 | 0.3525 | 0 | 47.1496 | Eukaryota | 45650.0 | 0 | Chordata | 45650.0 | 0 | Carnivora | 45650.0 | 0 | Felidae | 45650.0 | 0 | Felis | 45650.0 | 0 | Felis catus | 45650.0 | 0 |
| scf7180000017867 | 383378 | 0.3929 | 0 | 51.8444 | Eukaryota | 26464.0 | 0 | Chordata | 26464.0 | 0 | Carnivora | 26464.0 | 0 | Felidae | 26464.0 | 0 | Felis | 19983.0 | 2 | Felis catus | 19983.0 | 2 |
| scf7180000018947 | 489986 | 0.4188 | 0 | 49.741 | Eukaryota | 28487.0 | 1 | Chordata | 28487.0 | 1 | Carnivora | 11182.0 | 3 | Felidae | 11182.0 | 3 | Homo | 8872.0 | 5 | Homo sapiens | 8872.0 | 5 |
| scf7180000021070 | 131870 | 0.5532 | 0 | 49.6608 | Eukaryota | 27766.0 | 0 | Chordata | 27766.0 | 0 | Carnivora | 27766.0 | 0 | Felidae | 26390.0 | 1 | Lynx | 11696.0 | 3 | ynx canadens | 11696.0 | 3 |
| scf7180000021971 | 21175 | 0.389 | 0 | 26.7484 | Eukaryota | 10052.0 | 1 | Chordata | 10052.0 | 1 | Primates | 6621.0 | 4 | Hominidae | 3963.0 | 5 | Homo | 3963.0 | 5 | Homo sapiens | 3963.0 | 5 |
| scf7180000019618 | 26037 | 0.4199 | 0 | 42.3522 | Eukaryota | 7805.0 | 0 | Chordata | 7805.0 | 0 | Carnivora | 7805.0 | 0 | Felidae | 3589.0 | 2 | Panthera | 1623.0 | 6 | phus californi | 1466.0 | 7 |
| scf7180000019855 | 274965 | 0.4164 | 0 | 49.7951 | Eukaryota | 14485.0 | 0 | Chordata | 14485.0 | 0 | Carnivora | 13203.0 | 1 | Canidae | 6828.0 | 2 | Felis | 6375.0 | 3 | Felis catus | 6375.0 | 3 |
| scf7180000022746 | 2153 | 0.3683 | 0 | 25.5774 | Eukaryota | 21289.0 | 0 | Chordata | 21289.0 | 0 | Carnivora | 21289.0 | 0 | Felidae | 21289.0 | 0 | Felis | 21289.0 | 0 | Felis catus | 21289.0 | 0 |
| scf7180000023784 | 4388236 | 0.4054 | 0 | 51.667 | Eukaryota | 53238.0 | 0 | Chordata | 53238.0 | 0 | Carnivora | 53238.0 | 0 | Felidae | 53238.0 | 0 | Acinonyx | 42600.0 | 1 | cinonyx jubat | 42600.0 | 1 |
| scf7180000019953 | 224186 | 0.3689 | 0 | 50.1208 | Eukaryota | 35207.0 | 0 | Chordata | 35207.0 | 0 | Carnivora | 35207.0 | 0 | Felidae | 35207.0 | 0 | Felis | 35207.0 | 0 | Felis catus | 35207.0 | 0 |
| scf7180000017625 | 1905473 | 0.3975 | 0 | 49.9627 | Eukaryota | 45785.0 | 0 | Chordata | 45785.0 | 0 | Carnivora | 45785.0 | 0 | Felidae | 45785.0 | 0 | Felis | 45785.0 | 0 | Felis catus | 45785.0 | 0 |
| scf7180000017914 | 76266 | 0.4354 | 100 | 26.3969 | Eukaryota | 21283.0 | 0 | Chordata | 21283.0 | 0 | Carnivora | 21283.0 | 0 | Otariidae | 6057.0 | 5 | Acinonyx | 2477.0 | 9 | cinonyx jubat | 2477.0 | 9 |
| scf7180000018793 | 518386 | 0.4015 | 0 | 25.4718 | Eukaryota | 44539.0 | 0 | Chordata | 44539.0 | 0 | Carnivora | 37582.0 | 1 | Felidae | 37582.0 | 1 | Felis | 33434.0 | 2 | Felis catus | 33434.0 | 2 |
| scf7180000024219 | 37145 | 0.3784 | 0 | 24.2067 | Eukaryota | 23731.0 | 0 | Chordata | 23731.0 | 0 | Carnivora | 23731.0 | 0 | Felidae | 23731.0 | 0 | Felis | 23731.0 | 0 | Felis catus | 23731.0 | 0 |
| scf7180000019646 | 259040 | 0.6005 | 0 | 52.3357 | Eukaryota | 26626.0 | 0 | Chordata | 26626.0 | 0 | Carnivora | 26626.0 | 0 | Otariidae | 7941.0 | 5 | Lynx | 3061.0 | 9 | ynx canadens | 3061.0 | 9 |
| scf7180000022132 | 20632 | 0.3579 | 0 | 24.0336 | Eukaryota | 32492.0 | 0 | Chordata | 32492.0 | 0 | Carnivora | 32492.0 | 0 | Felidae | 32492.0 | 0 | Felis | 32492.0 | 0 | Felis catus | 32492.0 | 0 |
| scf7180000018095 | 7010373 | 0.3552 | 100 | 48.8711 | Eukaryota | 70634.0 | 1 | Chordata | 70634.0 | 1 | Primates | 35204.0 | 4 | Hominidae | 27723.0 | 5 | Homo | 27723.0 | 5 | Homo sapiens | 27723.0 | 6 |
| scf7180000020404 | 97329 | 0.4453 | 0 | 50.4535 | Eukaryota | 4025.0 | 0 | Chordata | 4025.0 | 0 | Carnivora | 1208.0 | 3 | Bovidae | 1191.0 | 4 | Mus | 1153.0 | 6 | Mus musculus | 1153.0 | 6 |
| scf7180000022668 | 20175 | 0.445 | 0 | 24.5841 | Eukaryota | 10327.0 | 0 | Chordata | 10327.0 | 0 | Primates | 3370.0 | 4 | Hominidae | 3370.0 | 6 | Homo | 3370.0 | 6 | Homo sapiens | 3370.0 | 7 |
| scf7180000018229 | 1452490 | 0.4389 | 0 | 52.4431 | Eukaryota | 48079.0 | 0 | Chordata | 48079.0 | 0 | Carnivora | 48079.0 | 0 | Felidae | 48079.0 | 0 | Felis | 48079.0 | 0 | Felis catus | 48079.0 | 0 |
| scf7180000019323 | 382232 | 0.3944 | 0 | 59.5724 | Eukaryota | 46080.0 | 0 | Chordata | 46080.0 | 0 | Carnivora | 46080.0 | 0 | Felidae | 46080.0 | 0 | Felis | 46080.0 | 0 | Felis catus | 46080.0 | 0 |
| scf7180000023383 | 28148 | 0.4392 | 0 | 88.2964 | Eukaryota | 4715.0 | 0 | Chordata | 4715.0 | 0 | Carnivora | 2669.0 | 1 | Bovidae | 2046.0 | 2 | Ovis | 2046.0 | 2 | Ovis canadens | 2046.0 | 2 |
| scf7180000019307 | 96669 | 0.4175 | 0 | 47.0216 | Eukaryota | 47425.0 | 0 | Chordata | 47425.0 | 0 | Carnivora | 47425.0 | 0 | Felidae | 47425.0 | 0 | Felis | 47425.0 | 0 | Felis catus | 47425.0 | 0 |
| scf7180000020274 | 138052 | 0.5305 | 0 | 48.683 | Eukaryota | 28605.0 | 0 | Chordata | 28605.0 | 0 | Carnivora | 28605.0 | 0 | Felidae | 28605.0 | 0 | Felis | 25827.0 | 1 | Felis catus | 25827.0 | 1 |
| scf7180000019444 | 267853 | 0.3665 | 0 | 49.4915 | Eukaryota | 35615.0 | 0 | Chordata | 35615.0 | 0 | Carnivora | 35615.0 | 0 | Felidae | 35615.0 | 0 | Felis | 32067.0 | 1 | Felis catus | 32067.0 | 1 |
| scf7180000020347 | 22167 | 0.4281 | 0 | 415.6606 | Eukaryota | 169670.0 | 0 | Chordata | 169670.0 | 0 | Carnivora | 169670.0 | 0 | Hyaenidae | 169670.0 | 0 | Proteles | 100404.0 | 1 | roteles cristat | 100404.0 | 1 |
| scf7180000019903 | 462461 | 0.4707 | 0 | 51.464 | Eukaryota | 33949.0 | 0 | Chordata | 33949.0 | 0 | Carnivora | 33949.0 | 0 | Felidae | 23704.0 | 1 | Panthera | 23704.0 | 1 | anthera pardu | 20322.0 | 2 |
| scf7180000020099 | 220584 | 0.3988 | 0 | 50.5504 | Eukaryota | 35197.0 | 0 | Chordata | 35197.0 | 0 | Carnivora | 35197.0 | 0 | Felidae | 35197.0 | 0 | Felis | 31697.0 | 1 | Felis catus | 31697.0 | 1 |
| scf7180000023370 | 36759 | 0.4122 | 0 | 190.2973 | Eukaryota | 11787.0 | 0 | Chordata | 11787.0 | 0 | Carnivora | 11787.0 | 0 | Felidae | 11787.0 | 0 | Felis | 11787.0 | 0 | Felis catus | 11787.0 | 0 |
| scf7180000019479 | 351241 | 0.5409 | 0 | 49.3954 | Eukaryota | 12242.0 | 0 | Chordata | 12242.0 | 0 | Carnivora | 12242.0 | 0 | Felidae | 12242.0 | 0 | Acinonyx | 9624.0 | 2 | cinonyx jubat | 9624.0 | 2 |
| scf7180000021032 | 70343 | 0.4726 | 0 | 25.0829 | Eukaryota | 6591.0 | 0 | Chordata | 6591.0 | 0 | Primates | 3155.0 | 5 | Hominidae | 3155.0 | 5 | Homo | 3155.0 | 5 | Homo sapiens | 3155.0 | 5 |
| scf7180000020572 | 83707 | 0.4201 | 0 | 51.9656 | Eukaryota | 10442.0 | 0 | Chordata | 10442.0 | 0 | Carnivora | 9409.0 | 1 | Felidae | 5418.0 | 2 | Felis | 5418.0 | 3 | Felis catus | 5418.0 | 3 |
| scf7180000022418 | 38098 | 0.6188 | 0 | 48.9466 | Eukaryota | 3163.0 | 0 | Chordata | 3163.0 | 0 | Carnivora | 3163.0 | 0 | Felidae | 2482.0 | 1 | Felis | 1155.0 | 3 | Felis catus | 1155.0 | 3 |
| scf7180000023334 | 23740 | 0.6926 | 0 | 89.2564 | no-hit | 0.0 | 0 | no-hit | 0.0 | 0 | no-hit | 0.0 | 0 | no-hit | 0.0 | 0 | no-hit | 0.0 | 0 | no-hit | 0.0 | 0 |
| scf7180000021893 | 61182 | 0.3887 | 0 | 24.0127 | Eukaryota | 43225.0 | 0 | Chordata | 43225.0 | 0 | Carnivora | 43225.0 | 0 | Felidae | 43225.0 | 0 | Felis | 38953.0 | 1 | Felis catus | 38953.0 | 1 |
| scf7180000020363 | 170313 | 0.3714 | 0 | 23.6136 | Eukaryota | 42376.0 | 0 | Chordata | 42376.0 | 0 | Carnivora | 42376.0 | 0 | Felidae | 42376.0 | 0 | Felis | 42376.0 | 0 | Felis catus | 42376.0 | 0 |
| scf7180000020247 | 162254 | 0.5999 | 0 | 49.5928 | Eukaryota | 5123.0 | 0 | Chordata | 5123.0 | 0 | Carnivora | 5123.0 | 0 | Felidae | 3805.0 | 2 | Felis | 1553.0 | 5 | Felis catus | 1553.0 | 6 |
| scf7180000017882 | 448379 | 0.3926 | 0 | 49.8365 | Eukaryota | 44079.0 | 0 | Chordata | 44079.0 | 0 | Carnivora | 44079.0 | 0 | Felidae | 44079.0 | 0 | Felis | 39709.0 | 1 | Felis catus | 39709.0 | 1 |
| scf7180000024149 | 166274 | 0.3622 | 0 | 51.475 | Eukaryota | 31767.0 | 0 | Chordata | 31767.0 | 0 | Carnivora | 31767.0 | 0 | Felidae | 31767.0 | 0 | Felis | 28465.0 | 1 | Felis catus | 28465.0 | 1 |
| scf7180000024031 | 472256 | 0.4761 | 0 | 49.5287 | Eukaryota | 39900.0 | 0 | Chordata | 39900.0 | 0 | Carnivora | 39900.0 | 0 | Felidae | 39900.0 | 0 | Felis | 39900.0 | 0 | Felis catus | 39900.0 | 0 |
| scf7180000020019 | 145964 | 0.3599 | 0 | 45.4388 | Eukaryota | 39019.0 | 0 | Chordata | 39019.0 | 0 | Carnivora | 39019.0 | 0 | Felidae | 39019.0 | 0 | Felis | 39019.0 | 0 | Felis catus | 39019.0 | 0 |
| scf7180000020628 | 149246 | 0.4779 | 0 | 52.5786 | Eukaryota | 23555.0 | 0 | Chordata | 23555.0 | 0 | Carnivora | 23555.0 | 0 | Felidae | 21643.0 | 1 | Acinonyx | 6276.0 | 4 | cinonyx jubat | 6276.0 | 4 |
| scf7180000020196 | 192114 | 0.5668 | 0 | 50.5838 | Eukaryota | 42180.0 | 0 | Chordata | 42180.0 | 0 | Carnivora | 42180.0 | 0 | Felidae | 42180.0 | 0 | Acinonyx | 42180.0 | 0 | cinonyx jubat | 42180.0 | 0 |
| scf7180000018274 | 1046152 | 0.4405 | 0 | 49.2074 | Eukaryota | 18729.0 | 0 | Chordata | 18729.0 | 0 | Carnivora | 18729.0 | 0 | Felidae | 15217.0 | 2 | Acinonyx | 8518.0 | 4 | cinonyx jubat | 8518.0 | 4 |
| scf7180000017677 | 1360988 | 0.3628 | 0 | 48.4344 | Eukaryota | 46657.0 | 0 | Chordata | 46657.0 | 0 | Carnivora | 46657.0 | 0 | Felidae | 46657.0 | 0 | Felis | 46657.0 | 0 | Felis catus | 46657.0 | 0 |
| scf7180000018358 | 905308 | 0.4283 | 0 | 52.714 | Eukaryota | 45676.0 | 0 | Chordata | 45676.0 | 0 | Carnivora | 45676.0 | 0 | Felidae | 39892.0 | 1 | Felis | 39892.0 | 1 | Felis catus | 39892.0 | 1 |
| scf7180000019009 | 516553 | 0.427 | 0 | 52.872 | Eukaryota | 26851.0 | 0 | Chordata | 26851.0 | 0 | Carnivora | 12788.0 | 3 | Felidae | 10703.0 | 5 | Panthera | 3603.0 | 9 | anthera pardu | 3603.0 | 9 |
| scf7180000020732 | 74351 | 0.4005 | 0 | 29.3298 | Eukaryota | 45790.0 | 0 | Chordata | 45790.0 | 0 | Carnivora | 45790.0 | 0 | Felidae | 45790.0 | 0 | Felis | 45790.0 | 0 | Felis catus | 45790.0 | 0 |
| scf7180000019483 | 184467 | 0.3925 | 0 | 27.4174 | Eukaryota | 46504.0 | 0 | Chordata | 46504.0 | 0 | Carnivora | 46504.0 | 0 | Felidae | 46504.0 | 0 | Felis | 46504.0 | 0 | Felis catus | 46504.0 | 0 |
| scf7180000023853 | 1726163 | 0.4264 | 0 | 51.6469 | Eukaryota | 83361.0 | 0 | Chordata | 83361.0 | 0 | Carnivora | 83361.0 | 0 | Felidae | 83361.0 | 0 | Acinonyx | 75187.0 | 1 | cinonyx jubat | 75187.0 | 1 |
| scf7180000018114 | 571319 | 0.3835 | 0 | 50.1608 | Eukaryota | 31227.0 | 0 | Chordata | 31227.0 | 0 | Carnivora | 31227.0 | 0 | Felidae | 31227.0 | 0 | Felis | 31227.0 | 0 | Felis catus | 31227.0 | 0 |
| scf7180000021938 | 13189 | 0.5157 | 0 | 48.762 | Eukaryota | 11599.0 | 0 | Chordata | 11599.0 | 0 | Carnivora | 9555.0 | 1 | Ursidae | 7034.0 | 3 | Ursus | 4795.0 | 4 | rsus maritimu | 3552.0 | 5 |
| scf7180000021449 | 86862 | 0.3471 | 0 | 24.2206 | Eukaryota | 47240.0 | 0 | Chordata | 47240.0 | 0 | Carnivora | 47240.0 | 0 | Felidae | 47240.0 | 0 | Felis | 47240.0 | 0 | Felis catus | 47240.0 | 0 |
| scf7180000020974 | 57711 | 0.4114 | 0 | 30.1636 | Eukaryota | 43729.0 | 0 | Chordata | 43729.0 | 0 | Carnivora | 43729.0 | 0 | Felidae | 43729.0 | 0 | Felis | 43729.0 | 0 | Felis catus | 43729.0 | 0 |
| scf7180000021779 | 5765 | 0.5197 | 0 | 53.5779 | no-hit | 0.0 | 0 | no-hit | 0.0 | 0 | no-hit | 0.0 | 0 | no-hit | 0.0 | 0 | no-hit | 0.0 | 0 | no-hit | 0.0 | 0 |
| scf7180000018188 | 178970 | 0.3749 | 0 | 47.1589 | Eukaryota | 46344.0 | 0 | Chordata | 46344.0 | 0 | Carnivora | 46344.0 | 0 | Felidae | 46344.0 | 0 | Felis | 46344.0 | 0 | Felis catus | 46344.0 | 0 |
| scf7180000024194 | 75140 | 0.6313 | 0 | 50.061 | Eukaryota | 4163.0 | 0 | Chordata | 4163.0 | 0 | Carnivora | 4163.0 | 0 | Felidae | 3206.0 | 2 | Lynx | 1036.0 | 5 | ynx canadens | 1036.0 | 5 |
| scf7180000018048 | 90402 | 0.4093 | 0 | 27.1899 | Eukaryota | 44795.0 | 0 | Chordata | 44795.0 | 0 | Carnivora | 44795.0 | 0 | Felidae | 44795.0 | 0 | Felis | 44795.0 | 0 | Felis catus | 44795.0 | 0 |
| scf7180000018598 | 153207 | 0.3565 | 0 | 25.8964 | Eukaryota | 23058.0 | 0 | Chordata | 23058.0 | 0 | Carnivora | 23058.0 | 0 | Felidae | 23058.0 | 0 | Felis | 20732.0 | 1 | Felis catus | 20732.0 | 1 |
| scf7180000018047 | 2620629 | 0.384 | 0 | 50.8923 | Eukaryota | 82943.0 | 0 | Chordata | 82943.0 | 0 | Carnivora | 82943.0 | 0 | Felidae | 75377.0 | 1 | Felis | 27079.0 | 3 | Felis catus | 27079.0 | 3 |
| scf7180000022452 | 11224 | 0.3784 | 0 | 26.2624 | Eukaryota | 3963.0 | 0 | Chordata | 3963.0 | 0 | Primates | 3963.0 | 0 | Hominidae | 3658.0 | 1 | Homo | 2852.0 | 3 | Homo sapiens | 2852.0 | 3 |
| scf7180000020937 | 122316 | 0.6301 | 0 | 53.3709 | Eukaryota | 42655.0 | 0 | Chordata | 42655.0 | 0 | Carnivora | 42655.0 | 0 | Felidae | 21625.0 | 2 | Felis | 17344.0 | 3 | Felis catus | 17344.0 | 3 |
| scf7180000024072 | 550378 | 0.3541 | 0 | 49.0825 | Eukaryota | 34362.0 | 0 | Chordata | 34362.0 | 0 | Carnivora | 34362.0 | 0 | Felidae | 34362.0 | 0 | Felis | 34362.0 | 0 | Felis catus | 34362.0 | 0 |
| scf7180000020060 | 134898 | 0.5928 | 0 | 52.0954 | Eukaryota | 37985.0 | 0 | Chordata | 37985.0 | 0 | Carnivora | 37985.0 | 0 | Felidae | 37985.0 | 0 | Lynx | 11760.0 | 3 | ynx canadens | 11760.0 | 4 |
| scf7180000022706 | 19380 | 0.396 | 0 | 25.1712 | Eukaryota | 4421.0 | 0 | Chordata | 4421.0 | 0 | Primates | 2162.0 | 2 | Hominidae | 2162.0 | 2 | Suricata | 1773.0 | 3 | ricata suricat | 1773.0 | 3 |
| scf7180000018147 | 1004116 | 0.4065 | 0 | 51.498 | Eukaryota | 47780.0 | 0 | Chordata | 47780.0 | 0 | Carnivora | 47780.0 | 0 | Felidae | 47780.0 | 0 | Panthera | 38224.0 | 1 | anthera pardu | 38224.0 | 1 |
| scf7180000022590 | 3954 | 0.3493 | 0 | 19.0286 | Eukaryota | 2780.0 | 0 | Chordata | 2780.0 | 0 | Carnivora | 2780.0 | 0 | Felidae | 2536.0 | 1 | Felis | 2058.0 | 2 | Felis catus | 2058.0 | 2 |
| scf7180000022105 | 9916 | 0.3843 | 0 | 24.6474 | Eukaryota | 13097.0 | 0 | Chordata | 13097.0 | 0 | Carnivora | 13097.0 | 0 | Felidae | 13097.0 | 0 | Felis | 10493.0 | 1 | Felis catus | 10493.0 | 1 |
| scf7180000019632 | 209173 | 0.4873 | 0 | 51.7321 | Eukaryota | 30169.0 | 0 | Chordata | 30169.0 | 0 | Carnivora | 30169.0 | 0 | Felidae | 30169.0 | 0 | Felis | 27112.0 | 1 | Felis catus | 27112.0 | 1 |
| scf7180000020267 | 152267 | 0.6037 | 0 | 47.9327 | Eukaryota | 4296.0 | 0 | Chordata | 4296.0 | 0 | Carnivora | 4296.0 | 0 | Felidae | 2533.0 | 2 | Felis | 2230.0 | 3 | Felis catus | 2230.0 | 3 |
| scf7180000021316 | 11704 | 0.5866 | 0 | 140.1265 | Eukaryota | 2469.0 | 0 | Chordata | 2469.0 | 0 | Carnivora | 2228.0 | 1 | Felidae | 2228.0 | 1 | Felis | 1645.0 | 3 | Felis catus | 1645.0 | 3 |
| scf7180000019575 | 333999 | 0.387 | 0 | 48.4377 | Eukaryota | 45598.0 | 0 | Chordata | 45598.0 | 0 | Carnivora | 45598.0 | 0 | Felidae | 45598.0 | 0 | Felis | 36512.0 | 1 | Felis catus | 36512.0 | 1 |
| scf7180000018123 | 1211755 | 0.3503 | 0 | 49.5342 | Eukaryota | 47113.0 | 0 | Chordata | 47113.0 | 0 | Carnivora | 47113.0 | 0 | Felidae | 47113.0 | 0 | Felis | 47113.0 | 0 | Felis catus | 47113.0 | 0 |
| scf7180000019797 | 168980 | 0.4218 | 0 | 51.4701 | Eukaryota | 13755.0 | 0 | Chordata | 13755.0 | 0 | Carnivora | 11834.0 | 1 | Felidae | 10103.0 | 2 | Felis | 6545.0 | 3 | Felis catus | 6545.0 | 3 |
| scf7180000020359 | 144311 | 0.6468 | 0 | 52.7551 | Eukaryota | 49593.0 | 0 | Chordata | 49593.0 | 0 | Carnivora | 49593.0 | 0 | Felidae | 49593.0 | 0 | Acinonyx | 23613.0 | 2 | cinonyx jubat | 23613.0 | 2 |
| scf7180000018852 | 457483 | 0.4018 | 0 | 52.0434 | Eukaryota | 46291.0 | 0 | Chordata | 46291.0 | 0 | Carnivora | 46291.0 | 0 | Felidae | 46291.0 | 0 | Felis | 41718.0 | 1 | Felis catus | 41718.0 | 1 |
| scf7180000020103 | 177564 | 0.6264 | 0 | 49.5394 | Eukaryota | 13438.0 | 0 | Chordata | 13438.0 | 0 | Carnivora | 13438.0 | 0 | Felidae | 10914.0 | 2 | Lynx | 7159.0 | 3 | ynx canadens | 7159.0 | 3 |
| scf7180000018748 | 628276 | 0.3499 | 0 | 49.8116 | Eukaryota | 43178.0 | 0 | Chordata | 43178.0 | 0 | Carnivora | 43178.0 | 0 | Felidae | 43178.0 | 0 | Felis | 43178.0 | 0 | Felis catus | 43178.0 | 0 |
| scf7180000017892 | 1471908 | 0.396 | 0 | 49.2645 | Eukaryota | 45722.0 | 0 | Chordata | 45722.0 | 0 | Carnivora | 45722.0 | 0 | Felidae | 45722.0 | 0 | Felis | 41188.0 | 1 | Felis catus | 41188.0 | 1 |
| scf7180000019398 | 243282 | 0.5067 | 0 | 63.1678 | Eukaryota | 31876.0 | 0 | Chordata | 31876.0 | 0 | Carnivora | 31876.0 | 0 | Felidae | 31876.0 | 0 | Felis | 28614.0 | 1 | Felis catus | 28614.0 | 1 |
| scf7180000020519 | 10559 | 0.5288 | 0 | 165.131 | Eukaryota | 12293.0 | 0 | Chordata | 12293.0 | 0 | Carnivora | 12293.0 | 0 | Felidae | 12293.0 | 0 | Felis | 11077.0 | 1 | Felis catus | 11077.0 | 1 |
| scf7180000019071 | 4236353 | 0.4263 | 0 | 51.7691 | Eukaryota | 111444.0 | 0 | Chordata | 111444.0 | 0 | Carnivora | 111444.0 | 0 | Herpestidae | 71281.0 | 1 | Suricata | 71281.0 | 3 | ricata suricat | 71281.0 | 3 |
| scf7180000019658 | 371044 | 0.3929 | 100 | 51.8708 | Eukaryota | 40198.0 | 0 | Chordata | 40198.0 | 0 | Carnivora | 40198.0 | 0 | Felidae | 40198.0 | 0 | Felis | 40198.0 | 0 | Felis catus | 40198.0 | 0 |
| scf7180000018493 | 685674 | 0.3815 | 0 | 66.3103 | Eukaryota | 43597.0 | 0 | Chordata | 43597.0 | 0 | Carnivora | 43597.0 | 0 | Felidae | 43597.0 | 0 | Felis | 43597.0 | 0 | Felis catus | 43597.0 | 0 |
| scf7180000018173 | 28354 | 0.3882 | 0 | 54.5334 | Eukaryota | 20350.0 | 0 | Chordata | 20350.0 | 0 | Carnivora | 20350.0 | 0 | Felidae | 20350.0 | 0 | Felis | 18346.0 | 1 | Felis catus | 18346.0 | 1 |
| scf7180000020306 | 218615 | 0.4326 | 100 | 24.991 | Eukaryota | 45587.0 | 0 | Chordata | 45587.0 | 0 | Carnivora | 45587.0 | 0 | Felidae | 45587.0 | 0 | Felis | 45587.0 | 0 | Felis catus | 45587.0 | 0 |
| scf7180000021355 | 78802 | 0.3647 | 0 | 24.3251 | Eukaryota | 45817.0 | 0 | Chordata | 45817.0 | 0 | Carnivora | 45817.0 | 0 | Felidae | 45817.0 | 0 | Felis | 45817.0 | 0 | Felis catus | 45817.0 | 0 |
| scf7180000017945 | 7025731 | 0.3659 | 0 | 50.3756 | Eukaryota | 126409.0 | 0 | Chordata | 126409.0 | 0 | Carnivora | 126409.0 | 0 | Otariidae | 87534.0 | 2 | Zalophus | 75024.0 | 4 | phus californi | 75024.0 | 4 |
| scf7180000021917 | 5881 | 0.6742 | 0 | 25.2295 | Eukaryota | 2275.0 | 0 | Chordata | 2275.0 | 0 | Carnivora | 2097.0 | 1 | Felidae | 2097.0 | 1 | Felis | 890.0 | 4 | Felis catus | 890.0 | 4 |
| scf7180000022335 | 20285 | 0.3745 | 0 | 27.0618 | Eukaryota | 5247.0 | 0 | Chordata | 5247.0 | 0 | Primates | 2705.0 | 3 | Hominidae | 2705.0 | 4 | Pan | 1625.0 | 5 | an troglodyte | 1625.0 | 5 |
| scf7180000018518 | 762291 | 0.4251 | 0 | 51.7419 | Eukaryota | 41981.0 | 0 | Chordata | 41981.0 | 0 | Carnivora | 41981.0 | 0 | Felidae | 41981.0 | 0 | Felis | 41981.0 | 0 | Felis catus | 41981.0 | 0 |
| scf7180000021438 | 56707 | 0.5411 | 0 | 49.1922 | Eukaryota | 8202.0 | 1 | Chordata | 8202.0 | 1 | Artiodactyla | 3038.0 | 6 | undef | 2266.0 | 7 | undef | 2266.0 | 8 | tic synthetic c | 2266.0 | 8 |
| scf7180000020966 | 46167 | 0.5655 | 0 | 159.1628 | Eukaryota | 4315.0 | 0 | Chordata | 4315.0 | 0 | Carnivora | 3127.0 | 1 | Felidae | 2062.0 | 2 | Acinonyx | 1490.0 | 4 | cinonyx jubat | 1490.0 | 4 |
| scf7180000020961 | 19099 | 0.3956 | 0 | 329.0136 | Eukaryota | 6240.0 | 0 | Chordata | 6240.0 | 0 | Carnivora | 6240.0 | 0 | Felidae | 6240.0 | 0 | Felis | 5599.0 | 1 | Felis catus | 5599.0 | 1 |
| scf7180000022213 | 10041 | 0.6709 | 0 | 2206.7699 | Eukaryota | 622.0 | 0 | Chordata | 481.0 | 1 | Carnivora | 481.0 | 1 | Felidae | 342.0 | 2 | Felis | 342.0 | 2 | Felis catus | 342.0 | 2 |
| scf7180000019616 | 217790 | 0.3604 | 0 | 24.6861 | Eukaryota | 24015.0 | 0 | Chordata | 24015.0 | 0 | Carnivora | 24015.0 | 0 | Felidae | 24015.0 | 0 | Felis | 21660.0 | 1 | Felis catus | 21660.0 | 1 |
| scf7180000019326 | 348462 | 0.432 | 0 | 49.7172 | Eukaryota | 45448.0 | 0 | Chordata | 45448.0 | 0 | Carnivora | 45448.0 | 0 | Felidae | 45448.0 | 0 | Felis | 40875.0 | 1 | Felis catus | 40875.0 | 1 |
| scf7180000024090 | 330894 | 0.6104 | 0 | 53.6908 | Eukaryota | 32026.0 | 0 | Chordata | 32026.0 | 0 | Carnivora | 32026.0 | 0 | Felidae | 32026.0 | 0 | Lynx | 19230.0 | 1 | ynx canadens | 19230.0 | 1 |
| scf7180000018217 | 36014 | 0.3566 | 0 | 36.0759 | Eukaryota | 3434.0 | 1 | Chordata | 3434.0 | 1 | Primates | 1115.0 | 4 | Hominidae | 1115.0 | 5 | Homo | 1115.0 | 6 | Homo sapiens | 1115.0 | 6 |
| scf7180000017518 | 229314 | 0.3484 | 0 | 47.1133 | Eukaryota | 45779.0 | 0 | Chordata | 45779.0 | 0 | Carnivora | 45779.0 | 0 | Felidae | 45779.0 | 0 | Felis | 45779.0 | 0 | Felis catus | 45779.0 | 0 |
| scf7180000024176 | 107292 | 0.6301 | 0 | 48.6427 | Eukaryota | 9492.0 | 0 | Chordata | 9492.0 | 0 | Carnivora | 9492.0 | 0 | Felidae | 8466.0 | 1 | Panthera | 1968.0 | 5 | anthera pardu | 1968.0 | 5 |
| scf7180000019546 | 271983 | 0.5114 | 0 | 53.5237 | Eukaryota | 19445.0 | 0 | Chordata | 19445.0 | 0 | Carnivora | 19445.0 | 0 | Herpestidae | 15632.0 | 2 | Suricata | 15632.0 | 2 | ricata suricat | 15632.0 | 2 |
| scf7180000017669 | 2066307 | 0.3876 | 0 | 50.9115 | Eukaryota | 104401.0 | 0 | Chordata | 104401.0 | 0 | Carnivora | 61286.0 | 2 | Felidae | 40008.0 | 4 | Acinonyx | 40008.0 | 4 | cinonyx jubat | 40008.0 | 4 |
| scf7180000018284 | 926778 | 0.3605 | 0 | 49.0456 | Eukaryota | 48090.0 | 0 | Chordata | 48090.0 | 0 | Carnivora | 31940.0 | 1 | Felidae | 31940.0 | 2 | Felis | 31940.0 | 2 | Felis catus | 31940.0 | 2 |
| scf7180000023268 | 34738 | 0.4117 | 0 | 574.2066 | Eukaryota | 1635.0 | 0 | Chordata | 1635.0 | 0 | Carnivora | 1635.0 | 0 | Felidae | 1635.0 | 0 | Felis | 1474.0 | 1 | Felis catus | 1474.0 | 1 |
| scf7180000023569 | 29474 | 0.4872 | 0 | 758.9458 | no-hit | 0.0 | 0 | no-hit | 0.0 | 0 | no-hit | 0.0 | 0 | no-hit | 0.0 | 0 | no-hit | 0.0 | 0 | no-hit | 0.0 | 0 |
| scf7180000019192 | 370975 | 0.4662 | 0 | 53.9347 | Eukaryota | 20291.0 | 0 | Chordata | 20291.0 | 0 | Carnivora | 20291.0 | 0 | Felidae | 12698.0 | 4 | Panthera | 4286.0 | 6 | anthera pardu | 4286.0 | 6 |
| scf7180000018607 | 572598 | 0.462 | 0 | 48.1662 | Eukaryota | 45754.0 | 0 | Chordata | 45754.0 | 0 | Carnivora | 45754.0 | 0 | Felidae | 45754.0 | 0 | Felis | 41436.0 | 1 | Felis catus | 41436.0 | 1 |
| scf7180000018540 | 45990 | 0.4626 | 0 | 41.3572 | Eukaryota | 46582.0 | 0 | Chordata | 46582.0 | 0 | Carnivora | 46582.0 | 0 | Felidae | 46582.0 | 0 | Felis | 46582.0 | 0 | Felis catus | 46582.0 | 0 |
| scf7180000020402 | 127419 | 0.3711 | 0 | 50.6429 | Eukaryota | 13260.0 | 0 | Chordata | 13260.0 | 0 | Carnivora | 13260.0 | 0 | Felidae | 13260.0 | 0 | Felis | 9687.0 | 2 | Felis catus | 9687.0 | 2 |
| scf7180000020355 | 110104 | 0.4248 | 0 | 50.0922 | Eukaryota | 9318.0 | 1 | Chordata | 9318.0 | 1 | Artiodactyla | 2690.0 | 4 | Bovidae | 2690.0 | 4 | Homo | 2369.0 | 6 | Homo sapiens | 2369.0 | 6 |
| scf7180000018390 | 261976 | 0.3821 | 0 | 50.0736 | Eukaryota | 46254.0 | 0 | Chordata | 46254.0 | 0 | Carnivora | 46254.0 | 0 | Felidae | 46254.0 | 0 | Felis | 46254.0 | 0 | Felis catus | 46254.0 | 0 |
| scf7180000024259 | 166839 | 0.4984 | 100 | 45.4839 | Eukaryota | 7031.0 | 0 | Chordata | 7031.0 | 0 | Carnivora | 7031.0 | 0 | Felidae | 7031.0 | 0 | Felis | 6356.0 | 1 | Felis catus | 6356.0 | 1 |
| scf7180000018136 | 1197402 | 0.4143 | 0 | 50.8965 | Eukaryota | 32296.0 | 0 | Chordata | 32296.0 | 0 | Carnivora | 32296.0 | 0 | Felidae | 21633.0 | 2 | Felis | 18901.0 | 3 | Felis catus | 18901.0 | 3 |
| scf7180000019184 | 14090 | 0.3693 | 0 | 23.1513 | Eukaryota | 2602.0 | 0 | Chordata | 2602.0 | 0 | Carnivora | 1914.0 | 1 | Felidae | 1662.0 | 2 | Felis | 938.0 | 3 | Felis catus | 938.0 | 3 |
| scf7180000017689 | 644539 | 0.3924 | 0 | 50.125 | Eukaryota | 53953.0 | 0 | Chordata | 53953.0 | 0 | Carnivora | 53953.0 | 0 | Felidae | 53953.0 | 0 | Felis | 35966.0 | 3 | Felis catus | 35966.0 | 3 |
| scf7180000019935 | 3109 | 0.6874 | 0 | 104.8818 | Eukaryota | 350.0 | 0 | Chordata | 350.0 | 0 | Carnivora | 350.0 | 0 | Felidae | 350.0 | 0 | Panthera | 350.0 | 0 | anthera pardu | 350.0 | 0 |
| scf7180000018070 | 786075 | 0.4824 | 0 | 53.4433 | Eukaryota | 42639.0 | 0 | Chordata | 42639.0 | 0 | Carnivora | 42639.0 | 0 | Phocidae | 23781.0 | 2 | eptonychote | 14277.0 | 3 | nychotes wec | 14277.0 | 3 |
| scf7180000021535 | 31464 | 0.4113 | 0 | 24.1724 | Eukaryota | 11721.0 | 0 | Chordata | 11721.0 | 0 | Carnivora | 11721.0 | 0 | Felidae | 11721.0 | 0 | Felis | 10507.0 | 1 | Felis catus | 10507.0 | 1 |
| scf7180000021127 | 58893 | 0.5314 | 0 | 50.6274 | Eukaryota | 19993.0 | 0 | Chordata | 19993.0 | 0 | Carnivora | 19993.0 | 0 | Felidae | 19993.0 | 0 | Felis | 16064.0 | 1 | Felis catus | 16064.0 | 1 |
| scf7180000019431 | 35124 | 0.4019 | 0 | 633.2057 | Eukaryota | 24799.0 | 0 | Chordata | 24799.0 | 0 | Carnivora | 24799.0 | 0 | Felidae | 24799.0 | 0 | Felis | 22294.0 | 1 | Felis catus | 22294.0 | 1 |
| scf7180000023322 | 35207 | 0.3947 | 0 | 319.4856 | Eukaryota | 12574.0 | 0 | Chordata | 12574.0 | 0 | Carnivora | 12574.0 | 0 | Felidae | 12574.0 | 0 | Felis | 11323.0 | 1 | Felis catus | 11323.0 | 1 |
| scf7180000019941 | 241288 | 0.4542 | 0 | 26.6234 | Eukaryota | 28326.0 | 0 | Chordata | 28326.0 | 0 | Carnivora | 28326.0 | 0 | Felidae | 28326.0 | 0 | Felis | 28326.0 | 0 | Felis catus | 28326.0 | 0 |
| scf7180000021060 | 89464 | 0.3449 | 0 | 43.1289 | Eukaryota | 42749.0 | 0 | Chordata | 42749.0 | 0 | Carnivora | 42749.0 | 0 | Felidae | 42749.0 | 0 | Felis | 42749.0 | 0 | Felis catus | 42749.0 | 0 |
| scf7180000022129 | 14500 | 0.4192 | 0 | 23.4525 | Eukaryota | 14163.0 | 0 | Chordata | 14163.0 | 0 | Carnivora | 14163.0 | 0 | Felidae | 14163.0 | 0 | Felis | 9918.0 | 1 | Felis catus | 9918.0 | 1 |
| scf7180000017540 | 2561783 | 0.3776 | 0 | 50.3307 | Eukaryota | 53417.0 | 0 | Chordata | 53417.0 | 0 | Carnivora | 53417.0 | 0 | Felidae | 53417.0 | 0 | Felis | 53417.0 | 0 | Felis catus | 53417.0 | 0 |
| scf7180000019999 | 217428 | 0.428 | 0 | 53.4082 | Eukaryota | 16147.0 | 0 | Chordata | 16147.0 | 0 | Carnivora | 14645.0 | 1 | Felidae | 13114.0 | 2 | Panthera | 5006.0 | 5 | anthera pardu | 3358.0 | 6 |
| scf7180000020996 | 70275 | 0.6084 | 0 | 49.5347 | Eukaryota | 15485.0 | 0 | Chordata | 15485.0 | 0 | Carnivora | 15485.0 | 0 | Felidae | 12860.0 | 2 | Felis | 7796.0 | 3 | Felis catus | 7796.0 | 3 |
| scf7180000020620 | 109620 | 0.3909 | 0 | 48.0116 | Eukaryota | 16900.0 | 0 | Chordata | 16900.0 | 0 | Carnivora | 16900.0 | 0 | Felidae | 8965.0 | 4 | Felis | 3568.0 | 7 | Felis catus | 3568.0 | 7 |
| scf7180000021293 | 36505 | 0.3979 | 0 | 24.5633 | Eukaryota | 24838.0 | 0 | Chordata | 24838.0 | 0 | Carnivora | 24838.0 | 0 | Felidae | 24838.0 | 0 | Felis | 22298.0 | 1 | Felis catus | 22298.0 | 1 |
| scf7180000017948 | 1294079 | 0.3876 | 0 | 51.0084 | Eukaryota | 48575.0 | 0 | Chordata | 48575.0 | 0 | Carnivora | 48575.0 | 0 | Felidae | 42913.0 | 1 | Felis | 42913.0 | 1 | Felis catus | 42913.0 | 1 |
| scf7180000018734 | 448204 | 0.4703 | 0 | 49.6169 | Eukaryota | 52196.0 | 0 | Chordata | 52196.0 | 0 | Carnivora | 39398.0 | 1 | Herpestidae | 26457.0 | 2 | Suricata | 26457.0 | 3 | ricata suricat | 26457.0 | 3 |
| scf7180000018680 | 611662 | 0.5033 | 0 | 50.5631 | Eukaryota | 31804.0 | 0 | Chordata | 31804.0 | 0 | Carnivora | 31804.0 | 0 | Felidae | 31804.0 | 0 | Felis | 29824.0 | 1 | Felis catus | 29824.0 | 1 |
| scf7180000021351 | 46989 | 0.4421 | 0 | 26.5779 | Eukaryota | 26147.0 | 0 | Chordata | 26147.0 | 0 | Carnivora | 26147.0 | 0 | Felidae | 26147.0 | 0 | Felis | 20844.0 | 1 | Felis catus | 20844.0 | 1 |
| scf7180000020948 | 15844 | 0.5167 | 0 | 31.5229 | Eukaryota | 44175.0 | 0 | Chordata | 44175.0 | 0 | Carnivora | 44175.0 | 0 | Felidae | 44175.0 | 0 | Felis | 44175.0 | 0 | Felis catus | 44175.0 | 0 |
| scf7180000019006 | 1059316 | 0.5521 | 0 | 50.1555 | Eukaryota | 30702.0 | 0 | Chordata | 30702.0 | 0 | Carnivora | 30702.0 | 0 | Herpestidae | 17574.0 | 1 | Suricata | 17574.0 | 2 | ricata suricat | 17574.0 | 2 |
| scf7180000020976 | 86502 | 0.5413 | 0 | 46.557 | Eukaryota | 5084.0 | 0 | Chordata | 5084.0 | 0 | Carnivora | 2769.0 | 3 | Herpestidae | 2188.0 | 5 | Suricata | 2188.0 | 5 | ricata suricat | 2188.0 | 5 |
| scf7180000023800 | 320273 | 0.3497 | 0 | 48.2857 | Eukaryota | 45744.0 | 0 | Chordata | 45744.0 | 0 | Carnivora | 45744.0 | 0 | Felidae | 45744.0 | 0 | Felis | 45744.0 | 0 | Felis catus | 45744.0 | 0 |
| scf7180000018425 | 753261 | 0.5053 | 0 | 50.882 | Eukaryota | 64881.0 | 0 | Chordata | 64881.0 | 0 | Carnivora | 64881.0 | 0 | Felidae | 64881.0 | 0 | Felis | 25716.0 | 2 | Felis catus | 25716.0 | 2 |
| scf7180000023007 | 65599 | 0.517 | 0 | 193.8091 | Eukaryota | 12472.0 | 0 | Chordata | 12472.0 | 0 | Carnivora | 12472.0 | 0 | Felidae | 12472.0 | 0 | Felis | 12472.0 | 0 | Felis catus | 12472.0 | 0 |
| scf7180000021370 | 49767 | 0.4502 | 0 | 24.3011 | Eukaryota | 14327.0 | 0 | Chordata | 14327.0 | 0 | Carnivora | 14327.0 | 0 | Felidae | 14327.0 | 0 | Felis | 4783.0 | 3 | Felis catus | 4783.0 | 3 |
| scf7180000022185 | 18093 | 0.363 | 0 | 24.4994 | Eukaryota | 10518.0 | 0 | Chordata | 10518.0 | 0 | Carnivora | 10518.0 | 0 | Felidae | 10518.0 | 0 | Felis | 9485.0 | 1 | Felis catus | 9485.0 | 1 |
| scf7180000019517 | 313808 | 0.3691 | 0 | 25.121 | Eukaryota | 47308.0 | 0 | Chordata | 47308.0 | 0 | Carnivora | 47308.0 | 0 | Felidae | 47308.0 | 0 | Felis | 42632.0 | 1 | Felis catus | 42632.0 | 1 |
| scf7180000023445 | 31101 | 0.4063 | 0 | 367.4419 | Eukaryota | 23886.0 | 0 | Chordata | 23886.0 | 0 | Carnivora | 23886.0 | 0 | Felidae | 23886.0 | 0 | Felis | 21492.0 | 1 | Felis catus | 21492.0 | 1 |
| scf7180000022360 | 18207 | 0.3719 | 0 | 23.2517 | Eukaryota | 22526.0 | 0 | Chordata | 22526.0 | 0 | Carnivora | 22526.0 | 0 | Felidae | 22526.0 | 0 | Felis | 20296.0 | 1 | Felis catus | 20296.0 | 1 |
| scf7180000024227 | 154008 | 0.4087 | 0 | 45.5476 | Eukaryota | 44162.0 | 0 | Chordata | 44162.0 | 0 | Carnivora | 44162.0 | 0 | Felidae | 44162.0 | 0 | Felis | 44162.0 | 0 | Felis catus | 44162.0 | 0 |
| scf7180000020808 | 86203 | 0.4028 | 0 | 24.9599 | Eukaryota | 16904.0 | 0 | Chordata | 16904.0 | 0 | Carnivora | 16904.0 | 0 | Felidae | 16904.0 | 0 | Felis | 15219.0 | 1 | Felis catus | 15219.0 | 1 |
| scf7180000018377 | 996269 | 0.3921 | 0 | 51.1002 | Eukaryota | 45084.0 | 0 | Chordata | 45084.0 | 0 | Carnivora | 45084.0 | 0 | Felidae | 45084.0 | 0 | Felis | 40332.0 | 1 | Felis catus | 40332.0 | 1 |
| scf7180000021756 | 42521 | 0.6355 | 0 | 56.3462 | Eukaryota | 8587.0 | 0 | Chordata | 8587.0 | 0 | Carnivora | 8587.0 | 0 | Otariidae | 5371.0 | 2 | Zalophus | 5371.0 | 2 | phus californi | 5371.0 | 2 |
| scf7180000020699 | 95322 | 0.356 | 0 | 45.9841 | Eukaryota | 37451.0 | 0 | Chordata | 37451.0 | 0 | Carnivora | 37451.0 | 0 | Felidae | 37451.0 | 0 | Felis | 33742.0 | 1 | Felis catus | 33742.0 | 1 |
| scf7180000024069 | 283303 | 0.4165 | 0 | 51.0308 | Eukaryota | 20749.0 | 0 | Chordata | 20749.0 | 0 | Carnivora | 20749.0 | 0 | Felidae | 9605.0 | 3 | Panthera | 4723.0 | 6 | Ursus arctos | 4175.0 | 7 |
| scf7180000018053 | 1352074 | 0.4386 | 0 | 52.1971 | Eukaryota | 42150.0 | 0 | Chordata | 42150.0 | 0 | Carnivora | 42150.0 | 0 | Herpestidae | 42150.0 | 0 | Suricata | 42150.0 | 0 | ricata suricat | 42150.0 | 0 |
| scf7180000018690 | 757717 | 0.3511 | 0 | 48.8991 | Eukaryota | 46673.0 | 0 | Chordata | 46673.0 | 0 | Carnivora | 46673.0 | 0 | Felidae | 46673.0 | 0 | Felis | 46673.0 | 0 | Felis catus | 46673.0 | 0 |
| scf7180000018865 | 190757 | 0.3626 | 0 | 49.8226 | Eukaryota | 37045.0 | 0 | Chordata | 37045.0 | 0 | Carnivora | 37045.0 | 0 | Felidae | 37045.0 | 0 | Felis | 37045.0 | 0 | Felis catus | 37045.0 | 0 |
| scf7180000020551 | 20768 | 0.5333 | 0 | 340.0524 | Eukaryota | 2338.0 | 0 | Chordata | 2338.0 | 0 | Carnivora | 2338.0 | 0 | Felidae | 1861.0 | 1 | Felis | 1700.0 | 2 | Felis catus | 1700.0 | 2 |
| scf7180000018336 | 893745 | 0.5871 | 0 | 50.5178 | Eukaryota | 27826.0 | 0 | Chordata | 27826.0 | 0 | Carnivora | 27826.0 | 0 | Felidae | 11978.0 | 5 | Canis | 5318.0 | 8 | Canis lupus | 5318.0 | 8 |
| scf7180000022023 | 45340 | 0.4049 | 0 | 26.6412 | Eukaryota | 45091.0 | 0 | Chordata | 45091.0 | 0 | Carnivora | 45091.0 | 0 | Felidae | 45091.0 | 0 | Felis | 45091.0 | 0 | Felis catus | 45091.0 | 0 |
| scf7180000020334 | 100070 | 0.4159 | 0 | 25.9666 | Eukaryota | 45036.0 | 0 | Chordata | 45036.0 | 0 | Carnivora | 45036.0 | 0 | Felidae | 45036.0 | 0 | Felis | 45036.0 | 0 | Felis catus | 45036.0 | 0 |

| | | | | | | | | | | | | | | | | | | | | | |
|---|---|---|---|---|---|---|---|---|---|---|---|---|---|---|---|---|---|---|---|---|---|
| scf7180000020271 | 167416 | 0.3661 | 0 | 46.5247 | Eukaryota | 44597.0 | 0 | Chordata | 44597.0 | 0 | Carnivora | 44597.0 | 0 | Felidae | 44597.0 | 0 | Felis | 44597.0 | 0 | Felis catus | 44597.0 | 0 |
| scf7180000023397 | 21336 | 0.4369 | 0 | 198.4757 | Eukaryota | 4816.0 | 0 | Chordata | 4816.0 | 0 | Artiodactyla | 4673.0 | 1 | Bovidae | 4673.0 | 1 | Ovis | 4673.0 | 1 | vis canadens | 3874.0 | 2 |
| scf7180000017957 | 59760 | 0.398 | 0 | 50.6425 | Eukaryota | 40254.0 | 0 | Chordata | 40254.0 | 0 | Carnivora | 40254.0 | 0 | Felidae | 40254.0 | 0 | Felis | 40254.0 | 0 | Felis catus | 40254.0 | 0 |
| scf7180000023881 | 917936 | 0.4384 | 0 | 51.0176 | Eukaryota | 37513.0 | 0 | Chordata | 37513.0 | 0 | Carnivora | 37513.0 | 0 | Felidae | 37513.0 | 0 | Felis | 33786.0 | 1 | Felis catus | 33786.0 | 1 |
| scf7180000022836 | 77561 | 0.5098 | 0 | 49.4765 | Eukaryota | 21265.0 | 0 | Chordata | 21265.0 | 0 | Carnivora | 21265.0 | 0 | Felidae | 21265.0 | 0 | Felis | 19141.0 | 1 | Felis catus | 19141.0 | 1 |
| scf7180000020820 | 80971 | 0.5097 | 0 | 46.9618 | Eukaryota | 29265.0 | 0 | Chordata | 29265.0 | 0 | Carnivora | 29265.0 | 0 | Felidae | 29265.0 | 0 | Felis | 29265.0 | 0 | Felis catus | 29265.0 | 0 |
| scf7180000023758 | 4770918 | 0.3796 | 0 | 50.296 | Eukaryota | 71096.0 | 0 | Chordata | 71096.0 | 0 | Carnivora | 71096.0 | 0 | Felidae | 71096.0 | 0 | Panthera | 26139.0 | 3 | Felis catus | 25683.0 | 4 |
| scf7180000019754 | 1390165 | 0.3626 | 0 | 48.4977 | Eukaryota | 71968.0 | 0 | Chordata | 71968.0 | 0 | Carnivora | 71968.0 | 0 | Felidae | 46619.0 | 2 | Acinonyx | 19923.0 | 4 | cinonyx jubat | 19923.0 | 4 |
| scf7180000020774 | 123849 | 0.3837 | 0 | 29.8088 | Eukaryota | 46912.0 | 0 | Chordata | 46912.0 | 0 | Carnivora | 46912.0 | 0 | Felidae | 46912.0 | 0 | Felis | 46912.0 | 0 | Felis catus | 46912.0 | 0 |
| scf7180000023440 | 22620 | 0.4578 | 0 | 220.964 | Eukaryota | 2991.0 | 0 | Chordata | 2991.0 | 0 | Carnivora | 2761.0 | 1 | Felidae | 1981.0 | 2 | Lynx | 974.0 | 5 | ynx canadens | 974.0 | 5 |
| scf7180000017541 | 1127594 | 0.358 | 0 | 49.9032 | Eukaryota | 56243.0 | 0 | Chordata | 56243.0 | 0 | Carnivora | 56243.0 | 0 | Felidae | 51107.0 | 1 | Panthera | 22775.0 | 4 | anthera pardu | 17070.0 | 5 |
| scf7180000019058 | 776194 | 0.3528 | 0 | 48.6384 | Eukaryota | 46241.0 | 0 | Chordata | 46241.0 | 0 | Carnivora | 46241.0 | 0 | Felidae | 46241.0 | 0 | Felis | 46241.0 | 0 | Felis catus | 46241.0 | 0 |
| scf7180000017845 | 1623375 | 0.3519 | 0 | 49.1409 | Eukaryota | 44317.0 | 0 | Chordata | 44317.0 | 0 | Carnivora | 44317.0 | 0 | Felidae | 34290.0 | 2 | Felis | 34290.0 | 2 | Felis catus | 34290.0 | 2 |
| scf7180000022787 | 20616 | 0.3722 | 0 | 23.7401 | Eukaryota | 14564.0 | 0 | Chordata | 14564.0 | 0 | Carnivora | 14564.0 | 0 | Felidae | 14564.0 | 0 | Felis | 13119.0 | 1 | Felis catus | 13119.0 | 1 |
| scf7180000019325 | 137063 | 0.4112 | 0 | 51.816 | Eukaryota | 18945.0 | 0 | Chordata | 18945.0 | 0 | Carnivora | 18945.0 | 0 | Felidae | 10183.0 | 3 | Panthera | 4141.0 | 7 | anthera pardu | 2078.0 | 9 |
| scf7180000019711 | 204058 | 0.5759 | 0 | 48.1779 | Eukaryota | 25460.0 | 0 | Chordata | 25460.0 | 0 | Primates | 20115.0 | 2 | Hominidae | 15980.0 | 3 | Homo | 15980.0 | 4 | Homo sapiens | 15980.0 | 4 |
| scf7180000022872 | 24872 | 0.437 | 0 | 684.4832 | no-hit | 0.0 | 0 | no-hit | 0.0 | 0 | no-hit | 0.0 | 0 | no-hit | 0.0 | 0 | no-hit | 0.0 | 0 | no-hit | 0.0 | 0 |
| scf7180000024140 | 116788 | 0.4814 | 0 | 50.4391 | Eukaryota | 18913.0 | 0 | Chordata | 18913.0 | 0 | Carnivora | 18913.0 | 0 | Felidae | 18913.0 | 0 | Felis | 17021.0 | 1 | Felis catus | 17021.0 | 1 |
| scf7180000020413 | 149958 | 0.6313 | 0 | 51.7761 | Eukaryota | 18259.0 | 0 | Chordata | 18259.0 | 0 | Carnivora | 16587.0 | 1 | Canidae | 7596.0 | 5 | Canis | 7596.0 | 6 | Canis lupus | 7596.0 | 6 |
| scf7180000019821 | 270124 | 0.4334 | 0 | 51.2217 | Eukaryota | 45624.0 | 0 | Chordata | 45624.0 | 0 | Carnivora | 45624.0 | 0 | Felidae | 45624.0 | 0 | Felis | 45624.0 | 0 | Felis catus | 45624.0 | 0 |
| scf7180000022501 | 23253 | 0.518 | 0 | 38.8228 | Eukaryota | 11621.0 | 0 | Chordata | 11621.0 | 0 | Carnivora | 11621.0 | 0 | Felidae | 11621.0 | 0 | Felis | 10426.0 | 1 | Felis catus | 10426.0 | 1 |
| scf7180000024207 | 30297 | 0.6472 | 0 | 48.2831 | Eukaryota | 4912.0 | 0 | Chordata | 4912.0 | 0 | Carnivora | 3226.0 | 2 | Felidae | 2767.0 | 3 | Panthera | 1831.0 | 6 | anthera pardu | 1831.0 | 6 |
| scf7180000018236 | 940230 | 0.4438 | 0 | 52.2387 | Eukaryota | 86428.0 | 0 | Chordata | 86428.0 | 0 | Carnivora | 86428.0 | 0 | Felidae | 59234.0 | 1 | Lynx | 33806.0 | 3 | ynx canadens | 33806.0 | 3 |
| scf7180000023958 | 746501 | 0.4063 | 0 | 50.5273 | Eukaryota | 81942.0 | 0 | Chordata | 81942.0 | 0 | Carnivora | 81942.0 | 0 | Felidae | 81942.0 | 0 | Acinonyx | 40917.0 | 2 | cinonyx jubat | 40917.0 | 2 |
| scf7180000023140 | 51310 | 0.3799 | 0 | 288.36 | Eukaryota | 24376.0 | 0 | Chordata | 24376.0 | 0 | Carnivora | 24376.0 | 0 | Felidae | 24376.0 | 0 | Felis | 21899.0 | 1 | Felis catus | 21899.0 | 1 |
| scf7180000019156 | 442672 | 0.3584 | 0 | 50.4321 | Eukaryota | 46658.0 | 0 | Chordata | 46658.0 | 0 | Carnivora | 46658.0 | 0 | Felidae | 46658.0 | 0 | Felis | 46658.0 | 0 | Felis catus | 46658.0 | 0 |
| scf7180000017901 | 309518 | 0.3646 | 0 | 48.7193 | Eukaryota | 46265.0 | 0 | Chordata | 46265.0 | 0 | Carnivora | 46265.0 | 0 | Felidae | 46265.0 | 0 | Felis | 46265.0 | 0 | Felis catus | 46265.0 | 0 |
| scf7180000020333 | 18672 | 0.5446 | 0 | 47.6252 | Eukaryota | 3431.0 | 0 | Chordata | 3431.0 | 0 | Carnivora | 3431.0 | 0 | Felidae | 2758.0 | 1 | Acinonyx | 1002.0 | 4 | cinonyx jubat | 1002.0 | 5 |
| scf7180000022072 | 26233 | 0.4018 | 0 | 24.6148 | Eukaryota | 15836.0 | 0 | Chordata | 15836.0 | 0 | Carnivora | 15836.0 | 0 | Felidae | 11731.0 | 2 | Panthera | 4883.0 | 5 | cinonyx jubat | 2742.0 | 6 |
| scf7180000019424 | 255452 | 0.5175 | 0 | 49.9074 | Eukaryota | 26361.0 | 0 | Chordata | 26361.0 | 0 | Carnivora | 26361.0 | 0 | Felidae | 26361.0 | 0 | Felis | 23688.0 | 1 | Felis catus | 23688.0 | 1 |
| scf7180000017861 | 197468 | 0.4533 | 0 | 49.4009 | Eukaryota | 52034.0 | 0 | Chordata | 52034.0 | 0 | Carnivora | 52034.0 | 0 | Felidae | 48419.0 | 1 | Panthera | 37611.0 | 2 | Panthera tigri | 37611.0 | 2 |
| scf7180000019128 | 558794 | 0.5038 | 0 | 49.492 | Eukaryota | 36691.0 | 0 | Chordata | 36691.0 | 0 | Carnivora | 36691.0 | 0 | Felidae | 36691.0 | 0 | Felis | 33080.0 | 1 | Felis catus | 33080.0 | 1 |
| scf7180000018913 | 563334 | 0.4918 | 0 | 50.2892 | Eukaryota | 25507.0 | 0 | Chordata | 25507.0 | 0 | Carnivora | 25507.0 | 0 | Felidae | 23231.0 | 1 | Acinonyx | 10316.0 | 4 | cinonyx jubat | 10316.0 | 4 |
| scf7180000019221 | 18598 | 0.4117 | 0 | 30.5745 | Eukaryota | 18554.0 | 0 | Chordata | 18554.0 | 0 | Carnivora | 18554.0 | 0 | Felidae | 12980.0 | 1 | Acinonyx | 11118.0 | 2 | cinonyx jubat | 11118.0 | 2 |
| scf7180000017954 | 192878 | 0.3773 | 0 | 47.3166 | Eukaryota | 22902.0 | 0 | Chordata | 22902.0 | 0 | Carnivora | 22902.0 | 0 | Felidae | 22902.0 | 0 | Felis | 20593.0 | 1 | Felis catus | 20593.0 | 1 |
| scf7180000023302 | 67197 | 0.4282 | 0 | 397.5362 | Eukaryota | 13435.0 | 0 | Chordata | 13435.0 | 0 | Carnivora | 13435.0 | 0 | Felidae | 13435.0 | 0 | Felis | 12105.0 | 1 | Felis catus | 12105.0 | 1 |
| scf7180000020047 | 285622 | 0.4194 | 0 | 51.0177 | Eukaryota | 27550.0 | 0 | Chordata | 27550.0 | 0 | Carnivora | 27550.0 | 0 | Herpestidae | 15895.0 | 1 | Suricata | 15895.0 | 2 | ricata suricat | 15895.0 | 3 |
| scf7180000018465 | 445476 | 0.3577 | 0 | 49.486 | Eukaryota | 31325.0 | 0 | Chordata | 31325.0 | 0 | Carnivora | 31325.0 | 0 | Felidae | 28213.0 | 1 | Lynx | 12568.0 | 5 | ynx canadens | 12568.0 | 5 |
| scf7180000017445 | 9847 | 0.3608 | 0 | 22.5706 | Eukaryota | 2281.0 | 0 | Chordata | 2281.0 | 0 | Carnivora | 2281.0 | 0 | Felidae | 2057.0 | 1 | Felis | 1123.0 | 2 | Felis catus | 1123.0 | 2 |
| scf7180000019142 | 1053297 | 0.5032 | 0 | 50.407 | Eukaryota | 35159.0 | 0 | Chordata | 35159.0 | 0 | Carnivora | 35159.0 | 0 | Felidae | 31570.0 | 1 | Panthera | 21039.0 | 2 | anthera pardu | 17528.0 | 3 |
| scf7180000021692 | 4389 | 0.5256 | 0 | 49.2852 | no-hit | 0.0 | 0 | no-hit | 0.0 | 0 | no-hit | 0.0 | 0 | no-hit | 0.0 | 0 | no-hit | 0.0 | 0 | no-hit | 0.0 | 0 |
| scf7180000023382 | 32766 | 0.4334 | 0 | 185.317 | Eukaryota | 5278.0 | 0 | Chordata | 5278.0 | 0 | Carnivora | 4382.0 | 1 | Felidae | 2967.0 | 2 | Felis | 2967.0 | 2 | Felis catus | 2967.0 | 2 |
| scf7180000020321 | 134729 | 0.3635 | 0 | 25.6785 | Eukaryota | 15664.0 | 0 | Chordata | 15664.0 | 0 | Carnivora | 15664.0 | 0 | Felidae | 15664.0 | 0 | Felis | 14110.0 | 1 | Felis catus | 14110.0 | 1 |
| scf7180000020512 | 86550 | 0.5469 | 0 | 51.1337 | Eukaryota | 15699.0 | 0 | Chordata | 15699.0 | 0 | Carnivora | 15699.0 | 0 | Felidae | 15699.0 | 0 | Felis | 15699.0 | 0 | Felis catus | 15699.0 | 0 |
| scf7180000018923 | 591672 | 0.4102 | 0 | 50.7563 | Eukaryota | 43213.0 | 0 | Chordata | 43213.0 | 0 | Carnivora | 43213.0 | 0 | Felidae | 43213.0 | 0 | Felis | 38948.0 | 1 | Felis catus | 38948.0 | 1 |
| scf7180000018663 | 416847 | 0.3887 | 0 | 49.6435 | Eukaryota | 45488.0 | 0 | Chordata | 45488.0 | 0 | Carnivora | 45488.0 | 0 | Felidae | 45488.0 | 0 | Felis | 45488.0 | 0 | Felis catus | 45488.0 | 0 |
| scf7180000019822 | 9629 | 0.6455 | 0 | 22.8888 | Eukaryota | 4669.0 | 0 | Chordata | 4669.0 | 0 | Carnivora | 4669.0 | 0 | Herpestidae | 3185.0 | 1 | Suricata | 3185.0 | 2 | ricata suricat | 3185.0 | 3 |
| scf7180000019306 | 47531 | 0.3573 | 0 | 279.2707 | Eukaryota | 12515.0 | 0 | Chordata | 12515.0 | 0 | Carnivora | 12515.0 | 0 | Felidae | 12515.0 | 0 | Felis | 7545.0 | 1 | Felis catus | 7545.0 | 1 |
| scf7180000023412 | 35639 | 0.4524 | 0 | 338.214 | Eukaryota | 5984.0 | 0 | Chordata | 5984.0 | 0 | Carnivora | 5984.0 | 0 | Phocidae | 2440.0 | 2 | .eptonychote | 1830.0 | 4 | nychotes wed | 1830.0 | 4 |
| scf7180000018332 | 131460 | 0.4978 | 0 | 50.2657 | Eukaryota | 28574.0 | 0 | Chordata | 28574.0 | 0 | Perissodactyla | 17442.0 | 2 | Equidae | 14533.0 | 4 | Equus | 14533.0 | 4 | quus caballu | 5872.0 | 6 |
| scf7180000022239 | 15907 | 0.4384 | 0 | 26.8004 | Eukaryota | 13233.0 | 0 | Chordata | 13233.0 | 0 | Primates | 5690.0 | 3 | Hominidae | 5690.0 | 4 | Equus | 4116.0 | 5 | Equus asinus | 4116.0 | 5 |
| scf7180000022039 | 30496 | 0.5422 | 0 | 50.4636 | Eukaryota | 9033.0 | 0 | Chordata | 9033.0 | 0 | Carnivora | 7577.0 | 1 | Felidae | 4425.0 | 3 | Ursus | 2374.0 | 7 | cinonyx jubat | 2127.0 | 8 |
| scf7180000020687 | 133083 | 0.3664 | 0 | 24.9221 | Eukaryota | 26519.0 | 0 | Chordata | 26519.0 | 0 | Carnivora | 26519.0 | 0 | Felidae | 16371.0 | 2 | Felis | 7764.0 | 6 | Felis catus | 7764.0 | 6 |
| scf7180000021011 | 95679 | 0.4452 | 0 | 52.0557 | Eukaryota | 22005.0 | 0 | Chordata | 22005.0 | 0 | Carnivora | 22005.0 | 0 | Felidae | 22005.0 | 0 | Felis | 17531.0 | 1 | Felis catus | 17531.0 | 1 |
| scf7180000017722 | 389426 | 0.4217 | 0 | 50.97 | Eukaryota | 46254.0 | 0 | Chordata | 46254.0 | 0 | Carnivora | 46254.0 | 0 | Felidae | 46254.0 | 0 | Felis | 46254.0 | 0 | Felis catus | 46254.0 | 0 |
| scf7180000019647 | 15087 | 0.406 | 0 | 24.771 | Eukaryota | 26456.0 | 0 | Chordata | 26456.0 | 0 | Carnivora | 26456.0 | 0 | Felidae | 26456.0 | 0 | Felis | 23772.0 | 1 | Felis catus | 23772.0 | 1 |
| scf7180000021085 | 49059 | 0.3805 | 0 | 25.6507 | Eukaryota | 11268.0 | 0 | Chordata | 11268.0 | 0 | Carnivora | 7831.0 | 1 | Felidae | 7831.0 | 1 | Lynx | 4928.0 | 3 | ynx canadens | 4928.0 | 4 |
| scf7180000017917 | 2069791 | 0.3735 | 100 | 50.7563 | Eukaryota | 48632.0 | 0 | Chordata | 48632.0 | 0 | Carnivora | 41919.0 | 1 | Felidae | 41919.0 | 1 | Felis | 41919.0 | 1 | Felis catus | 41919.0 | 1 |
| scf7180000021501 | 66800 | 0.418 | 0 | 416.6827 | Eukaryota | 27002.0 | 0 | Chordata | 27002.0 | 0 | Carnivora | 27002.0 | 0 | Felidae | 27002.0 | 0 | Felis | 13435.0 | 2 | Felis catus | 13435.0 | 2 |
| scf7180000018634 | 18974 | 0.4029 | 0 | 35.3558 | Eukaryota | 29002.0 | 0 | Chordata | 29002.0 | 0 | Carnivora | 29002.0 | 0 | Felidae | 29002.0 | 0 | Felis | 26290.0 | 1 | Felis catus | 26290.0 | 1 |
| scf7180000022930 | 62049 | 0.4582 | 0 | 679.9163 | Eukaryota | 15600.0 | 0 | Chordata | 15600.0 | 0 | Carnivora | 14093.0 | 1 | Felidae | 9351.0 | 4 | Felis | 4691.0 | 7 | Felis catus | 4691.0 | 7 |
| scf7180000018750 | 513651 | 0.4436 | 0 | 51.6915 | Eukaryota | 45885.0 | 0 | Chordata | 45885.0 | 0 | Carnivora | 45885.0 | 0 | Felidae | 45885.0 | 0 | Felis | 45885.0 | 0 | Felis catus | 45885.0 | 0 |
| scf7180000020438 | 36499 | 0.5469 | 0 | 59.2179 | Eukaryota | 9641.0 | 0 | Chordata | 9641.0 | 0 | Carnivora | 9641.0 | 0 | Felidae | 8833.0 | 1 | Felis | 4446.0 | 3 | Felis catus | 4446.0 | 3 |
| scf7180000021094 | 76773 | 0.4624 | 0 | 26.135 | Eukaryota | 50478.0 | 0 | Chordata | 50478.0 | 0 | Carnivora | 21676.0 | 2 | Equidae | 14583.0 | 5 | Equus | 14583.0 | 6 | Equus asinus | 9704.0 | 7 |
| scf7180000022087 | 27455 | 0.38 | 0 | 24.8309 | Eukaryota | 17919.0 | 0 | Chordata | 17919.0 | 0 | Carnivora | 17919.0 | 0 | Felidae | 17919.0 | 0 | Felis | 16109.0 | 1 | Felis catus | 16109.0 | 1 |
| scf7180000020283 | 194849 | 0.3503 | 0 | 46.8513 | Eukaryota | 23032.0 | 0 | Chordata | 23032.0 | 0 | Carnivora | 23032.0 | 0 | Felidae | 23032.0 | 0 | Felis | 18514.0 | 1 | Felis catus | 18514.0 | 1 |
| scf7180000019452 | 472896 | 0.4237 | 0 | 50.0277 | Eukaryota | 16622.0 | 0 | Chordata | 16622.0 | 0 | Carnivora | 16622.0 | 0 | Felidae | 16622.0 | 0 | Acinonyx | 8395.0 | 1 | cinonyx jubat | 8395.0 | 2 |
| scf7180000022569 | 14871 | 0.6688 | 0 | 772.0495 | Eukaryota | 610.0 | 0 | Chordata | 469.0 | 1 | Carnivora | 469.0 | 1 | Felidae | 330.0 | 2 | Felis | 330.0 | 2 | Felis catus | 330.0 | 2 |
| scf7180000023634 | 30136 | 0.5526 | 0 | 161.9088 | Eukaryota | 6341.0 | 0 | Chordata | 6341.0 | 0 | Carnivora | 6341.0 | 0 | Felidae | 6113.0 | 1 | Felis | 2437.0 | 3 | Felis catus | 2437.0 | 3 |
| scf7180000020348 | 22005 | 0.4727 | 0 | 414.5817 | Eukaryota | 7132.0 | 0 | Chordata | 7132.0 | 0 | Carnivora | 7132.0 | 0 | Felidae | 7132.0 | 0 | Felis | 7132.0 | 0 | Felis catus | 7132.0 | 0 |
| scf7180000022155 | 32349 | 0.3732 | 0 | 24.4636 | Eukaryota | 7918.0 | 0 | Chordata | 7918.0 | 0 | Carnivora | 7918.0 | 0 | Felidae | 7918.0 | 0 | Felis | 7147.0 | 1 | Felis catus | 7147.0 | 1 |
| scf7180000019286 | 296024 | 0.498 | 0 | 49.7839 | Eukaryota | 24741.0 | 0 | Chordata | 24741.0 | 0 | Carnivora | 24741.0 | 0 | Felidae | 14313.0 | 2 | Felis | 7512.0 | 7 | Felis catus | 7512.0 | 7 |
| scf7180000017557 | 3055474 | 0.444 | 100 | 50.793 | Eukaryota | 46719.0 | 0 | Chordata | 46719.0 | 0 | Carnivora | 46719.0 | 0 | Felidae | 46719.0 | 0 | Felis | 46719.0 | 0 | Felis catus | 46719.0 | 0 |
| scf7180000022457 | 13655 | 0.3706 | 0 | 23.4447 | Eukaryota | 5858.0 | 0 | Chordata | 5858.0 | 0 | Carnivora | 5858.0 | 0 | Felidae | 4750.0 | 1 | Felis | 4168.0 | 2 | Felis catus | 4168.0 | 2 |
| scf7180000021240 | 60213 | 0.3618 | 0 | 23.873 | Eukaryota | 39294.0 | 0 | Chordata | 39294.0 | 0 | Carnivora | 39294.0 | 0 | Felidae | 39294.0 | 0 | Felis | 39294.0 | 0 | Felis catus | 39294.0 | 0 |
| scf7180000021184 | 49338 | 0.3945 | 0 | 30.2687 | Eukaryota | 44085.0 | 0 | Chordata | 44085.0 | 0 | Carnivora | 44085.0 | 0 | Felidae | 44085.0 | 0 | Felis | 39736.0 | 1 | Felis catus | 39736.0 | 1 |
| scf7180000019050 | 48158 | 0.4278 | 0 | 43.432 | Eukaryota | 31093.0 | 0 | Chordata | 31093.0 | 0 | Carnivora | 31093.0 | 0 | Felidae | 31093.0 | 0 | Felis | 27888.0 | 1 | Felis catus | 27888.0 | 1 |
| scf7180000023866 | 945366 | 0.4118 | 0 | 52.0018 | Eukaryota | 34692.0 | 0 | Chordata | 34692.0 | 0 | Carnivora | 34692.0 | 0 | Felidae | 21376.0 | 3 | Felis | 9676.0 | 6 | Felis catus | 9676.0 | 6 |
| scf7180000018450 | 748718 | 0.4068 | 0 | 52.9358 | Eukaryota | 47069.0 | 0 | Chordata | 47069.0 | 0 | Carnivora | 47069.0 | 0 | Felidae | 47069.0 | 0 | Felis | 47069.0 | 0 | Felis catus | 47069.0 | 0 |
| scf7180000019960 | 34327 | 0.5515 | 0 | 183.9123 | Eukaryota | 3642.0 | 0 | Chordata | 3642.0 | 0 | Carnivora | 3642.0 | 0 | Felidae | 3392.0 | 1 | Acinonyx | 1814.0 | 4 | cinonyx jubat | 1814.0 | 4 |
| scf7180000020266 | 111094 | 0.6024 | 0 | 49.247 | Eukaryota | 35187.0 | 0 | Chordata | 35187.0 | 0 | Carnivora | 35187.0 | 0 | Felidae | 15469.0 | 3 | Felis | 7739.0 | 7 | Felis catus | 7739.0 | 7 |
| scf7180000023026 | 38540 | 0.4032 | 0 | 366.7248 | Eukaryota | 7073.0 | 0 | Chordata | 7073.0 | 0 | Carnivora | 7073.0 | 0 | Felidae | 7073.0 | 0 | Felis | 7073.0 | 0 | Felis catus | 7073.0 | 0 |
| scf7180000017895 | 1380266 | 0.3828 | 0 | 51.456 | Eukaryota | 47277.0 | 0 | Chordata | 47277.0 | 0 | Carnivora | 47277.0 | 0 | Felidae | 28233.0 | 1 | Panthera | 23475.0 | 2 | anthera pardu | 23475.0 | 2 |
| scf7180000019710 | 172371 | 0.4632 | 0 | 51.014 | Eukaryota | 10080.0 | 0 | Chordata | 10080.0 | 0 | Carnivora | 10080.0 | 0 | Felidae | 10080.0 | 0 | Felis | 9138.0 | 1 | Felis catus | 9138.0 | 1 |
| scf7180000020574 | 118923 | 0.4424 | 0 | 24.7741 | Eukaryota | 24458.0 | 0 | Chordata | 24458.0 | 0 | Carnivora | 24458.0 | 0 | Felidae | 24458.0 | 0 | Felis | 22011.0 | 1 | Felis catus | 22011.0 | 1 |
| scf7180000017665 | 518499 | 0.5048 | 0 | 55.6528 | Eukaryota | 38809.0 | 0 | Chordata | 38809.0 | 0 | Carnivora | 38809.0 | 0 | Felidae | 27198.0 | 2 | Felis | 11604.0 | 4 | Felis catus | 11604.0 | 4 |
| scf7180000017535 | 5482578 | 0.3785 | 0 | 50.626 | Eukaryota | 70916.0 | 0 | Chordata | 70916.0 | 0 | Carnivora | 70916.0 | 0 | Felidae | 28956.0 | 4 | Suricata | 15100.0 | 5 | ricata suricat | 15100.0 | 6 |
| scf7180000019613 | 264530 | 0.3554 | 0 | 48.6078 | Eukaryota | 45407.0 | 0 | Chordata | 45407.0 | 0 | Carnivora | 45407.0 | 0 | Felidae | 45407.0 | 0 | Felis | 45407.0 | 0 | Felis catus | 45407.0 | 0 |
| scf7180000018758 | 717433 | 0.344 | 0 | 47.8114 | Eukaryota | 46805.0 | 0 | Chordata | 46805.0 | 0 | Carnivora | 46805.0 | 0 | Felidae | 46805.0 | 0 | Felis | 46805.0 | 0 | Felis catus | 46805.0 | 0 |
| scf7180000023890 | 1798775 | 0.4573 | 0 | 51.0898 | Eukaryota | 115442.0 | 0 | Chordata | 115442.0 | 0 | Carnivora | 115442.0 | 0 | Felidae | 76535.0 | 2 | Felis | 29456.0 | 6 | Felis catus | 29456.0 | 6 |
| scf7180000020958 | 112237 | 0.5223 | 0 | 26.7846 | Eukaryota | 12817.0 | 0 | Chordata | 12817.0 | 0 | Carnivora | 7890.0 | 2 | Herpestidae | 7890.0 | 2 | Suricata | 7890.0 | 3 | ricata suricat | 7890.0 | 3 |
| scf7180000021262 | 39459 | 0.3892 | 0 | 24.697 | Eukaryota | 40543.0 | 0 | Chordata | 40543.0 | 0 | Primates | 31868.0 | 1 | Hominidae | 31868.0 | 2 | Homo | 23918.0 | 3 | Homo sapiens | 23918.0 | 3 |
| scf7180000018799 | 586029 | 0.4568 | 0 | 50.6464 | Eukaryota | 46999.0 | 0 | Chordata | 46999.0 | 0 | Carnivora | 46999.0 | 0 | Felidae | 46999.0 | 0 | Felis | 46999.0 | 0 | Felis catus | 46999.0 | 0 |
| scf7180000018104 | 827919 | 0.3715 | 0 | 51.1839 | Eukaryota | 47622.0 | 0 | Chordata | 47622.0 | 0 | Carnivora | 47622.0 | 0 | Felidae | 47622.0 | 0 | Felis | 47622.0 | 0 | Felis catus | 47622.0 | 0 |
| scf7180000018841 | 738800 | 0.3673 | 100 | 49.9471 | Eukaryota | 46552.0 | 0 | Chordata | 46552.0 | 0 | Carnivora | 46552.0 | 0 | Felidae | 46552.0 | 0 | Felis | 46552.0 | 0 | Felis catus | 46552.0 | 0 |
| scf7180000019980 | 276514 | 0.4617 | 0 | 49.6607 | Eukaryota | 28547.0 | 0 | Chordata | 28547.0 | 0 | Carnivora | 23183.0 | 1 | Otariidae | 10160.0 | 4 | Zalophus | 7059.0 | 7 | phus californi | 7059.0 | 8 |
| scf7180000020171 | 231198 | 0.4851 | 0 | 25.3178 | Eukaryota | 37755.0 | 0 | Chordata | 37755.0 | 0 | Carnivora | 37755.0 | 0 | Felidae | 19449.0 | 3 | Panthera | 7721.0 | 8 | ricata suricat | 3978.0 | 9 |
| scf7180000022757 | 4483 | 0.6246 | 0 | 42.9242 | no-hit | 0.0 | 0 | no-hit | 0.0 | 0 | no-hit | 0.0 | 0 | no-hit | 0.0 | 0 | no-hit | 0.0 | 0 | no-hit | 0.0 | 0 |
| scf7180000020107 | 139718 | 0.4949 | 0 | 27.0635 | Eukaryota | 14404.0 | 0 | Chordata | 14404.0 | 0 | Primates | 8278.0 | 1 | Hominidae | 8278.0 | 2 | Sus | 4842.0 | 3 | Sus scrofa | 4842.0 | 3 |
| scf7180000023047 | 25837 | 0.4379 | 0 | 199.7737 | Eukaryota | 23728.0 | 0 | Chordata | 23728.0 | 0 | Carnivora | 23728.0 | 0 | Felidae | 23728.0 | 0 | Felis | 21332.0 | 1 | Felis catus | 21332.0 | 1 |
| scf7180000020616 | 99384 | 0.4275 | 0 | 49.2297 | Eukaryota | 21090.0 | 0 | Chordata | 21090.0 | 0 | Carnivora | 21090.0 | 0 | Felidae | 21090.0 | 0 | Felis | 18912.0 | 1 | Felis catus | 18912.0 | 1 |
| scf7180000023210 | 28877 | 0.4797 | 0 | 168.0979 | Eukaryota | 12072.0 | 0 | Chordata | 12072.0 | 0 | Carnivora | 12072.0 | 0 | Felidae | 12072.0 | 0 | Felis | 12072.0 | 0 | Felis catus | 12072.0 | 0 |
| scf7180000017774 | 38678 | 0.3894 | 0 | 47.9087 | Eukaryota | 26590.0 | 0 | Chordata | 26590.0 | 0 | Carnivora | 26590.0 | 0 | Felidae | 26590.0 | 0 | Felis | 23884.0 | 1 | Felis catus | 23884.0 | 1 |
| scf7180000019974 | 220602 | 0.4226 | 0 | 50.5676 | Eukaryota | 52010.0 | 0 | Chordata | 52010.0 | 0 | Carnivora | 52010.0 | 0 | Felidae | 52010.0 | 0 | Panthera | 26015.0 | 1 | anthera pardu | 26015.0 | 1 |
| scf7180000019868 | 271416 | 0.4009 | 0 | 51.7169 | Eukaryota | 18845.0 | 0 | Chordata | 18845.0 | 0 | Carnivora | 18845.0 | 0 | Felidae | 18845.0 | 0 | Felis | 16928.0 | 1 | Felis catus | 16928.0 | 1 |
| scf7180000018482 | 119648 | 0.5049 | 0 | 43.1726 | Eukaryota | 33975.0 | 0 | Chordata | 33975.0 | 0 | Carnivora | 33975.0 | 0 | Felidae | 33975.0 | 0 | Felis | 30519.0 | 1 | Felis catus | 30519.0 | 1 |
| scf7180000021328 | 34072 | 0.4303 | 0 | 263.7176 | Eukaryota | 16643.0 | 0 | Chordata | 16643.0 | 0 | Carnivora | 16643.0 | 0 | Felidae | 16643.0 | 0 | Felis | 14991.0 | 1 | Felis catus | 14991.0 | 1 |
| scf7180000021342 | 33493 | 0.6466 | 0 | 50.5739 | Eukaryota | 17552.0 | 0 | Chordata | 17552.0 | 0 | Carnivora | 16083.0 | 1 | Felidae | 12533.0 | 2 | Panthera | 5371.0 | 6 | anthera pardu | 3588.0 | 7 |
| scf7180000022050 | 22547 | 0.3827 | 0 | 17.4424 | Eukaryota | 36201.0 | 0 | Chordata | 36201.0 | 0 | Carnivora | 36201.0 | 0 | Felidae | 36201.0 | 0 | Felis | 36201.0 | 0 | Felis catus | 36201.0 | 0 |
| scf7180000017754 | 1825325 | 0.3563 | 0 | 50.0072 | Eukaryota | 59042.0 | 0 | Chordata | 59042.0 | 0 | Carnivora | 59042.0 | 0 | Felidae | 59042.0 | 0 | Felis | 42236.0 | 2 | Felis catus | 42236.0 | 3 |
| scf7180000017634 | 3173366 | 0.3556 | 0 | 49.1416 | Eukaryota | 47934.0 | 0 | Chordata | 47934.0 | 0 | Carnivora | 47934.0 | 0 | Felidae | 47934.0 | 0 | Felis | 47934.0 | 0 | Felis catus | 47934.0 | 0 |
| scf7180000019927 | 167604 | 0.5648 | 0 | 48.0009 | Eukaryota | 24569.0 | 0 | Chordata | 24569.0 | 0 | Carnivora | 24569.0 | 0 | Felidae | 24569.0 | 0 | Felis | 22081.0 | 1 | Felis catus | 22081.0 | 1 |
| scf7180000020046 | 235463 | 0.4426 | 0 | 53.1598 | Eukaryota | 10688.0 | 0 | Chordata | 10688.0 | 0 | Carnivora | 10688.0 | 0 | Felidae | 10688.0 | 0 | Felis | 10688.0 | 0 | Felis catus | 10688.0 | 0 |
| scf7180000017672 | 27765 | 0.5023 | 0 | 69.5515 | Eukaryota | 4654.0 | 0 | Chordata | 4654.0 | 0 | Carnivora | 2803.0 | 2 | Felidae | 2006.0 | 3 | Acinonyx | 1334.0 | 5 | cinonyx jubat | 1334.0 | 5 |
| scf7180000024049 | 493415 | 0.5241 | 0 | 52.6643 | Eukaryota | 45653.0 | 0 | Chordata | 45653.0 | 0 | Carnivora | 45653.0 | 0 | Felidae | 45653.0 | 0 | Panthera | 22979.0 | 2 | anthera pardu | 22979.0 | 2 |
| scf7180000019805 | 45297 | 0.5225 | 0 | 160.6544 | Eukaryota | 4859.0 | 0 | Chordata | 4859.0 | 0 | Carnivora | 3803.0 | 1 | Felidae | 2880.0 | 2 | Felis | 1416.0 | 4 | Felis catus | 1416.0 | 4 |
| scf7180000019309 | 278591 | 0.367 | 0 | 49.0315 | Eukaryota | 45533.0 | 0 | Chordata | 45533.0 | 0 | Carnivora | 45533.0 | 0 | Felidae | 45533.0 | 0 | Felis | 45533.0 | 0 | Felis catus | 45533.0 | 0 |
| scf7180000018580 | 791974 | 0.388 | 0 | 52.3433 | Eukaryota | 46842.0 | 0 | Chordata | 46842.0 | 0 | Carnivora | 46842.0 | 0 | Felidae | 46842.0 | 0 | Felis | 46842.0 | 0 | Felis catus | 46842.0 | 0 |
| scf7180000020352 | 76660 | 0.3965 | 0 | 29.9331 | Eukaryota | 20220.0 | 0 | Chordata | 20220.0 | 0 | Carnivora | 20220.0 | 0 | Felidae | 20220.0 | 0 | Felis | 13073.0 | 1 | Felis catus | 13073.0 | 1 |
| scf7180000021687 | 37392 | 0.3676 | 0 | 23.3948 | Eukaryota | 33442.0 | 0 | Chordata | 33442.0 | 0 | Carnivora | 33442.0 | 0 | Felidae | 33442.0 | 0 | Felis | 30304.0 | 1 | Felis catus | 30304.0 | 1 |
| scf7180000017808 | 2308432 | 0.4385 | 0 | 51.5158 | Eukaryota | 30906.0 | 0 | Chordata | 30906.0 | 0 | Carnivora | 30906.0 | 0 | Felidae | 30906.0 | 0 | Lynx | 15460.0 | 2 | ynx canadens | 15460.0 | 2 |
| scf7180000019678 | 271684 | 0.5087 | 0 | 50.0102 | Eukaryota | 17495.0 | 0 | Chordata | 17495.0 | 0 | Carnivora | 17495.0 | 0 | Felidae | 7078.0 | 3 | Suricata | 6003.0 | 4 | ricata suricat | 6003.0 | 4 |
| scf7180000024083 | 321569 | 0.425 | 0 | 96.6529 | Eukaryota | 33518.0 | 0 | Chordata | 33518.0 | 0 | Carnivora | 33518.0 | 0 | Felidae | 19860.0 | 1 | Suricata | 13658.0 | 2 | ricata suricat | 13658.0 | 2 |
| scf7180000024262 | 63066 | 0.3859 | 0 | 25.4106 | Eukaryota | 39342.0 | 0 | Chordata | 39342.0 | 0 | Carnivora | 39342.0 | 0 | Felidae | 39342.0 | 0 | Felis | 35426.0 | 1 | Felis catus | 35426.0 | 1 |
| scf7180000020612 | 101594 | 0.522 | 0 | 51.8106 | Eukaryota | 7716.0 | 0 | Chordata | 7716.0 | 0 | Carnivora | 7716.0 | 0 | Felidae | 7716.0 | 0 | Felis | 6170.0 | 1 | Felis catus | 6170.0 | 1 |
| scf7180000019916 | 35875 | 0.5234 | 0 | 41.2794 | Eukaryota | 18451.0 | 0 | Chordata | 18451.0 | 0 | Carnivora | 18451.0 | 0 | Felidae | 18451.0 | 0 | Felis | 8161.0 | 4 | Felis catus | 8161.0 | 4 |
| scf7180000017559 | 11379 | 0.4541 | 0 | 33.974 | Eukaryota | 4853.0 | 0 | Chordata | 4853.0 | 0 | Primates | 2525.0 | 2 | Cercopithecidae | 2525.0 | 2 | Macaca | 2028.0 | 5 | Macaca mulatt | 2028.0 | 5 |
| scf7180000019200 | 413294 | 0.4088 | 0 | 50.1583 | Eukaryota | 47145.0 | 0 | Chordata | 47145.0 | 0 | Carnivora | 47145.0 | 0 | Felidae | 47145.0 | 0 | Felis | 47145.0 | 0 | Felis catus | 47145.0 | 0 |
| scf7180000017827 | 403799 | 0.4195 | 0 | 49.4147 | Eukaryota | 47646.0 | 0 | Chordata | 47646.0 | 0 | Carnivora | 47646.0 | 0 | Felidae | 47646.0 | 0 | Felis | 42929.0 | 1 | Felis catus | 42929.0 | 1 |
| scf7180000020856 | 91996 | 0.3681 | 0 | 24.7575 | Eukaryota | 39664.0 | 0 | Chordata | 39664.0 | 0 | Carnivora | 39664.0 | 0 | Felidae | 39664.0 | 0 | Felis | 39664.0 | 0 | Felis catus | 39664.0 | 0 |
| scf7180000018725 | 577434 | 0.5052 | 0 | 52.0922 | Eukaryota | 79712.0 | 0 | Chordata | 79712.0 | 0 | Carnivora | 79712.0 | 0 | Ursidae | 79712.0 | 0 | Ursus | 63866.0 | 1 | Ursus arctos | 55888.0 | 2 |
| scf7180000017974 | 321557 | 0.4318 | 0 | 50.7822 | Eukaryota | 31983.0 | 0 | Chordata | 31983.0 | 0 | Carnivora | 31983.0 | 0 | Felidae | 31983.0 | 0 | Felis | 28825.0 | 1 | Felis catus | 28825.0 | 1 |
| scf7180000018262 | 532538 | 0.4742 | 0 | 48.6375 | Eukaryota | 18266.0 | 0 | Chordata | 18266.0 | 0 | Carnivora | 18266.0 | 0 | Felidae | 8364.0 | 3 | Felis | 4949.0 | 4 | Felis catus | 4949.0 | 4 |
| scf7180000023714 | 25949 | 0.4379 | 0 | 255.5996 | Eukaryota | 24049.0 | 0 | Chordata | 24049.0 | 0 | Carnivora | 24049.0 | 0 | Felidae | 24049.0 | 0 | Felis | 21613.0 | 1 | Felis catus | 21613.0 | 1 |
| scf7180000022716 | 16248 | 0.6857 | 0 | 51.5771 | Eukaryota | 95855.0 | 0 | Chordata | 95855.0 | 0 | Carnivora | 95855.0 | 0 | Felidae | 95855.0 | 0 | Felis | 48220.0 | 1 | Felis catus | 48220.0 | 1 |
| scf7180000017822 | 593308 | 0.398 | 0 | 50.9371 | Eukaryota | 46321.0 | 0 | Chordata | 46321.0 | 0 | Carnivora | 46321.0 | 0 | Felidae | 46321.0 | 0 | Felis | 41746.0 | 1 | Felis catus | 41746.0 | 1 |
| scf7180000021119 | 54294 | 0.4019 | 0 | 182.783 | Eukaryota | 42958.0 | 0 | Chordata | 42958.0 | 0 | Carnivora | 42958.0 | 0 | Felidae | 42958.0 | 0 | Felis | 42958.0 | 0 | Felis catus | 42958.0 | 0 |
| scf7180000018975 | 468870 | 0.4848 | 0 | 51.144 | Eukaryota | 47490.0 | 0 | Chordata | 47490.0 | 0 | Carnivora | 47490.0 | 0 | Felidae | 43141.0 | 1 | Lynx | 13947.0 | 4 | ynx canadens | 13947.0 | 5 |
| scf7180000020911 | 97228 | 0.5147 | 0 | 49.2383 | Eukaryota | 4455.0 | 1 | Chordata | 4455.0 | 1 | Primates | 1575.0 | 6 | Hominidae | 1575.0 | 6 | Homo | 1575.0 | 7 | Homo sapiens | 1575.0 | 7 |
| scf7180000020058 | 182072 | 0.4096 | 0 | 24.9901 | Eukaryota | 30320.0 | 0 | Chordata | 30320.0 | 0 | Carnivora | 30320.0 | 0 | Felidae | 30320.0 | 0 | Felis | 27223.0 | 1 | Felis catus | 27223.0 | 1 |
| scf7180000020335 | 12937 | 0.4207 | 0 | 28.5912 | Eukaryota | 7482.0 | 0 | Chordata | 7482.0 | 0 | Primates | 3289.0 | 3 | Cercopithecidae | 2332.0 | 6 | Felis | 1642.0 | 8 | Felis catus | 1642.0 | 8 |
| scf7180000017935 | 537787 | 0.3601 | 0 | 47.9352 | Eukaryota | 46286.0 | 0 | Chordata | 46286.0 | 0 | Carnivora | 46286.0 | 0 | Felidae | 46286.0 | 0 | Felis | 46286.0 | 0 | Felis catus | 46286.0 | 0 |
| scf7180000022662 | 6325 | 0.4555 | 0 | 26.3149 | Eukaryota | 4901.0 | 0 | Chordata | 4901.0 | 0 | Primates | 2272.0 | 2 | Hominidae | 1542.0 | 4 | Sus | 1406.0 | 6 | Sus scrofa | 1406.0 | 6 |
| scf7180000022284 | 14993 | 0.4087 | 0 | 1228.6462 | no-hit | 0.0 | 0 | no-hit | 0.0 | 0 | no-hit | 0.0 | 0 | no-hit | 0.0 | 0 | no-hit | 0.0 | 0 | no-hit | 0.0 | 0 |
| scf7180000019958 | 325868 | 0.3773 | 0 | 25.8899 | Eukaryota | 45123.0 | 0 | Chordata | 45123.0 | 0 | Carnivora | 45123.0 | 0 | Felidae | 45123.0 | 0 | Felis | 40637.0 | 1 | Felis catus | 40637.0 | 1 |
| scf7180000019562 | 269650 | 0.4793 | 0 | 52.1584 | Eukaryota | 43530.0 | 0 | Chordata | 43530.0 | 0 | Carnivora | 43530.0 | 0 | Felidae | 43530.0 | 0 | Felis | 39195.0 | 1 | Felis catus | 39195.0 | 1 |
| scf7180000017875 | 1382035 | 0.3977 | 0 | 50.9691 | Eukaryota | 49375.0 | 0 | Chordata | 49375.0 | 0 | Carnivora | 49375.0 | 0 | Felidae | 34867.0 | 1 | Felis | 34867.0 | 1 | Felis catus | 34867.0 | 1 |
| scf7180000023398 | 29631 | 0.4361 | 0 | 174.8877 | Eukaryota | 7329.0 | 0 | Chordata | 7329.0 | 0 | Artiodactyla | 7329.0 | 0 | Bovidae | 7329.0 | 0 | Ovis | 6601.0 | 1 | vis canadens | 6601.0 | 1 |
| scf7180000024187 | 101845 | 0.4035 | 0 | 24.2212 | Eukaryota | 26038.0 | 0 | Chordata | 26038.0 | 0 | Carnivora | 26038.0 | 0 | Felidae | 26038.0 | 0 | Felis | 23454.0 | 1 | Felis catus | 23454.0 | 1 |
| scf7180000017571 | 2480510 | 0.3971 | 0 | 50.4454 | Eukaryota | 52869.0 | 0 | Chordata | 52869.0 | 0 | Carnivora | 47663.0 | 1 | Felidae | 33121.0 | 3 | Felis | 33121.0 | 3 | Felis catus | 33121.0 | 3 |
| scf7180000020694 | 111499 | 0.4499 | 0 | 25.7788 | Eukaryota | 14603.0 | 0 | Chordata | 14603.0 | 0 | Primates | 6538.0 | 3 | Hominidae | 6538.0 | 5 | Homo | 6538.0 | 7 | Homo sapiens | 6538.0 | 7 |
| scf7180000022018 | 22771 | 0.3586 | 0 | 25.5203 | Eukaryota | 34090.0 | 0 | Chordata | 34090.0 | 0 | Carnivora | 34090.0 | 0 | Felidae | 34090.0 | 0 | Lynx | 17050.0 | 2 | ynx canadens | 17050.0 | 2 |
| scf7180000023779 | 2648402 | 0.4481 | 0 | 51.9179 | Eukaryota | 57166.0 | 0 | Chordata | 57166.0 | 0 | Carnivora | 57166.0 | 0 | Felidae | 57166.0 | 0 | Felis | 28600.0 | 2 | Felis catus | 28600.0 | 2 |
| scf7180000018955 | 202930 | 0.5458 | 0 | 52.135 | Eukaryota | 14870.0 | 0 | Chordata | 14870.0 | 0 | Carnivora | 14870.0 | 0 | Otariidae | 7248.0 | 3 | Eumetopias | 4362.0 | 5 | netopias juba | 4362.0 | 5 |
| scf7180000018615 | 693937 | 0.358 | 0 | 48.225 | Eukaryota | 45948.0 | 0 | Chordata | 45948.0 | 0 | Carnivora | 45948.0 | 0 | Felidae | 45948.0 | 0 | Felis | 45948.0 | 0 | Felis catus | 45948.0 | 0 |
| scf7180000021672 | 52151 | 0.4308 | 0 | 24.6401 | Eukaryota | 10696.0 | 0 | Chordata | 9973.0 | 1 | Primates | 4354.0 | 4 | Hominidae | 4354.0 | 4 | Homo | 4354.0 | 4 | Homo sapiens | 4354.0 | 4 |
| scf7180000021235 | 56110 | 0.6048 | 0 | 47.1881 | Eukaryota | 23944.0 | 0 | Chordata | 23944.0 | 0 | Carnivora | 23944.0 | 0 | Felidae | 21412.0 | 1 | Felis | 9508.0 | 4 | Felis catus | 9508.0 | 5 |
| scf7180000018656 | 486146 | 0.4164 | 0 | 51.9438 | Eukaryota | 35656.0 | 0 | Chordata | 35656.0 | 0 | Carnivora | 35656.0 | 0 | Felidae | 35656.0 | 0 | Felis | 35656.0 | 0 | Felis catus | 35656.0 | 0 |
| scf7180000023678 | 40786 | 0.5411 | 0 | 186.7 | Eukaryota | 6549.0 | 0 | Chordata | 6549.0 | 0 | Carnivora | 6549.0 | 0 | Felidae | 6294.0 | 1 | Felis | 2334.0 | 4 | Felis catus | 2334.0 | 4 |

| | | | | | | | | | | | | | | | | | | | | | | |
|---|---|---|---|---|---|---|---|---|---|---|---|---|---|---|---|---|---|---|---|---|---|---|
| scf7180000019011 | 89676 | 0.3879 | 0 | 45.7068 | Eukaryota | 18462.0 | 1 | Chordata | 18462.0 | 1 | Primates | 6794.0 | 5 | Hominidae | 6794.0 | 5 | Homo | 6794.0 | 7 | Homo sapien | 6794.0 | 7 |
| scf7180000021490 | 27511 | 0.4805 | 0 | 53.0782 | Eukaryota | 20392.0 | 0 | Chordata | 20392.0 | 0 | Carnivora | 20392.0 | 0 | Felidae | 20392.0 | 0 | Felis | 16354.0 | 1 | Felis catus | 16354.0 | 1 |
| scf7180000023845 | 1302927 | 0.5451 | 0 | 50.1464 | Eukaryota | 31304.0 | 0 | Chordata | 31304.0 | 0 | Carnivora | 31304.0 | 0 | Felidae | 31304.0 | 0 | Lynx | 28179.0 | 1 | ynx canadens | 28179.0 | 1 |
| scf7180000022873 | 34628 | 0.4377 | 0 | 788.5096 | no-hit | 0.0 | 0 | no-hit | 0.0 | 0 | no-hit | 0.0 | 0 | no-hit | 0.0 | 0 | no-hit | 0.0 | 0 | no-hit | 0.0 | 0 |
| scf7180000019048 | 28079 | 0.3586 | 0 | 34.6887 | Eukaryota | 12081.0 | 0 | Chordata | 12081.0 | 0 | Carnivora | 12081.0 | 0 | Felidae | 12081.0 | 0 | Felis | 12081.0 | 0 | Felis catus | 12081.0 | 0 |
| scf7180000021366 | 87064 | 0.3857 | 0 | 24.0905 | Eukaryota | 25840.0 | 0 | Chordata | 25840.0 | 0 | Carnivora | 25840.0 | 0 | Felidae | 25840.0 | 0 | Felis | 23265.0 | 1 | Felis catus | 23265.0 | 1 |
| scf7180000022182 | 2761 | 0.5194 | 0 | 24.8304 | no-hit | 0.0 | 0 | no-hit | 0.0 | 0 | no-hit | 0.0 | 0 | no-hit | 0.0 | 0 | no-hit | 0.0 | 0 | no-hit | 0.0 | 0 |
| scf7180000022358 | 26868 | 0.5566 | 0 | 245.9036 | Eukaryota | 3493.0 | 0 | Chordata | 3493.0 | 0 | Carnivora | 3493.0 | 0 | Felidae | 3052.0 | 2 | Acinonyx | 1892.0 | 3 | cinonyx jubat | 1892.0 | 3 |
| scf7180000024270 | 3316815 | 0.4047 | 200 | 50.1676 | Eukaryota | 46741.0 | 0 | Chordata | 46741.0 | 0 | Carnivora | 46741.0 | 0 | Felidae | 46741.0 | 0 | Felis | 42109.0 | 1 | Felis catus | 42109.0 | 1 |
| scf7180000024084 | 265445 | 0.3922 | 0 | 24.9287 | Eukaryota | 41853.0 | 0 | Chordata | 41853.0 | 0 | Carnivora | 41853.0 | 0 | Felidae | 41853.0 | 0 | Felis | 41853.0 | 0 | Felis catus | 41853.0 | 0 |
| scf7180000021815 | 15285 | 0.568 | 0 | 55.7091 | Eukaryota | 2310.0 | 1 | Chordata | 2310.0 | 1 | Artiodactyla | 1071.0 | 3 | Bovidae | 838.0 | 5 | Bos | 547.0 | 7 | Bos mutus | 547.0 | 7 |
| scf7180000019681 | 17563 | 0.3811 | 0 | 222.7517 | Eukaryota | 17775.0 | 0 | Chordata | 17775.0 | 0 | Carnivora | 17775.0 | 0 | Felidae | 17775.0 | 0 | Felis | 14228.0 | 1 | Felis catus | 14228.0 | 1 |
| scf7180000019809 | 217035 | 0.4362 | 0 | 51.5019 | Eukaryota | 8123.0 | 0 | Chordata | 8123.0 | 0 | Primates | 3264.0 | 2 | Hominidae | 3264.0 | 2 | Homo | 3264.0 | 3 | Homo sapien | 3264.0 | 3 |
| scf7180000023659 | 46972 | 0.4467 | 0 | 252.9138 | Eukaryota | 23040.0 | 0 | Chordata | 23040.0 | 0 | Carnivora | 23040.0 | 0 | Felidae | 23040.0 | 0 | Felis | 20698.0 | 1 | Felis catus | 20698.0 | 1 |
| scf7180000021833 | 55138 | 0.4017 | 0 | 24.304 | Eukaryota | 40890.0 | 0 | Chordata | 40890.0 | 0 | Carnivora | 40890.0 | 0 | Felidae | 40890.0 | 0 | Felis | 36847.0 | 1 | Felis catus | 36847.0 | 1 |
| scf7180000022953 | 43661 | 0.3935 | 0 | 293.8012 | Eukaryota | 46026.0 | 0 | Chordata | 46026.0 | 0 | Carnivora | 46026.0 | 0 | Felidae | 46026.0 | 0 | Felis | 46026.0 | 0 | Felis catus | 46026.0 | 0 |
| scf7180000021586 | 28961 | 0.4661 | 0 | 25.9453 | Eukaryota | 20823.0 | 0 | Chordata | 20823.0 | 0 | Carnivora | 20823.0 | 0 | Felidae | 13269.0 | 1 | Suricata | 7554.0 | 4 | ricata suricat | 7554.0 | 5 |
| scf7180000022861 | 162338 | 0.6239 | 0 | 48.9367 | Eukaryota | 33792.0 | 0 | Chordata | 33792.0 | 0 | Carnivora | 33792.0 | 0 | Felidae | 33792.0 | 0 | Felis | 30403.0 | 1 | Felis catus | 30403.0 | 1 |
| scf7180000019737 | 16668 | 0.406 | 0 | 24.5077 | Eukaryota | 29277.0 | 0 | Chordata | 29277.0 | 0 | Carnivora | 29277.0 | 0 | Felidae | 29277.0 | 0 | Felis | 26307.0 | 1 | Felis catus | 26307.0 | 1 |
| scf7180000020633 | 127779 | 0.4062 | 0 | 52.9175 | Eukaryota | 20437.0 | 0 | Chordata | 20437.0 | 0 | Carnivora | 12249.0 | 1 | Felidae | 12249.0 | 1 | Homo | 8188.0 | 3 | Homo sapien | 8188.0 | 3 |
| scf7180000018492 | 685198 | 0.4668 | 0 | 53.2765 | Eukaryota | 37436.0 | 0 | Chordata | 37436.0 | 0 | Carnivora | 34246.0 | 1 | Otariidae | 9680.0 | 6 | Canis | 6537.0 | 8 | Canis lupus | 6537.0 | 8 |
| scf7180000021323 | 73656 | 0.4949 | 0 | 51.2175 | Eukaryota | 46666.0 | 0 | Chordata | 46666.0 | 0 | Carnivora | 46666.0 | 0 | Felidae | 46666.0 | 0 | Felis | 46666.0 | 0 | Felis catus | 46666.0 | 0 |
| scf7180000019905 | 228466 | 0.3976 | 0 | 49.9466 | Eukaryota | 46223.0 | 0 | Chordata | 46223.0 | 0 | Carnivora | 46223.0 | 0 | Felidae | 46223.0 | 0 | Felis | 41681.0 | 1 | Felis catus | 41681.0 | 1 |
| scf7180000018403 | 12161 | 0.4467 | 0 | 26.3148 | Eukaryota | 5381.0 | 0 | Chordata | 5381.0 | 0 | Carnivora | 5381.0 | 0 | Felidae | 2335.0 | 3 | Acinonyx | 1656.0 | 4 | cinonyx jubat | 1656.0 | 4 |
| scf7180000022388 | 10033 | 0.4002 | 0 | 24.6066 | Eukaryota | 7242.0 | 0 | Chordata | 7242.0 | 0 | Carnivora | 3560.0 | 2 | Felidae | 3560.0 | 2 | Felis | 3560.0 | 2 | Felis catus | 3560.0 | 2 |
| scf7180000024113 | 282271 | 0.3959 | 0 | 50.9245 | Eukaryota | 17364.0 | 0 | Chordata | 17364.0 | 0 | Carnivora | 8957.0 | 1 | Equidae | 8407.0 | 2 | Equus | 8407.0 | 2 | ustela putori | 5247.0 | 4 |
| scf7180000022657 | 11790 | 0.455 | 0 | 25.8833 | Eukaryota | 15943.0 | 0 | Chordata | 15943.0 | 0 | Carnivora | 14565.0 | 1 | Felidae | 8677.0 | 4 | Felis | 1753.0 | 9 | Felis catus | 1753.0 | 9 |
| scf7180000022524 | 33134 | 0.3803 | 0 | 24.9409 | Eukaryota | 29464.0 | 0 | Chordata | 29464.0 | 0 | Carnivora | 29464.0 | 0 | Felidae | 29464.0 | 0 | Felis | 26560.0 | 1 | Felis catus | 26560.0 | 1 |
| scf7180000020828 | 81363 | 0.5604 | 0 | 47.9782 | Eukaryota | 7211.0 | 0 | Chordata | 7211.0 | 0 | Carnivora | 7211.0 | 0 | Felidae | 2902.0 | 3 | Suricata | 2235.0 | 7 | ricata suricat | 2235.0 | 7 |
| scf7180000017641 | 1831588 | 0.5177 | 0 | 52.2557 | Eukaryota | 76340.0 | 0 | Chordata | 76340.0 | 0 | Carnivora | 76340.0 | 0 | Felidae | 62051.0 | 2 | Panthera | 22296.0 | 5 | anthera pardu | 22296.0 | 5 |
| scf7180000019750 | 7703 | 0.4236 | 0 | 50.3959 | Eukaryota | 2293.0 | 0 | Chordata | 1736.0 | 1 | Carnivora | 1736.0 | 1 | Herpestidae | 772.0 | 3 | Suricata | 772.0 | 4 | ricata suricat | 772.0 | 4 |
| scf7180000020110 | 43532 | 0.5314 | 0 | 231.9675 | Eukaryota | 4082.0 | 0 | Chordata | 4082.0 | 0 | Carnivora | 4082.0 | 0 | Felidae | 3366.0 | 2 | Acinonyx | 2056.0 | 3 | cinonyx jubat | 2056.0 | 3 |
| scf7180000019498 | 126026 | 0.3803 | 0 | 49.8358 | Eukaryota | 39829.0 | 0 | Chordata | 39829.0 | 0 | Carnivora | 39829.0 | 0 | Felidae | 39829.0 | 0 | Felis | 35869.0 | 1 | Felis catus | 35869.0 | 1 |
| scf7180000017663 | 1849782 | 0.4394 | 0 | 49.2405 | Eukaryota | 47628.0 | 0 | Chordata | 47628.0 | 0 | Carnivora | 47628.0 | 0 | Felidae | 47628.0 | 0 | Felis | 47628.0 | 0 | Felis catus | 47628.0 | 0 |
| scf7180000017685 | 1852497 | 0.3929 | 0 | 48.259 | Eukaryota | 46708.0 | 0 | Chordata | 46708.0 | 0 | Carnivora | 46708.0 | 0 | Felidae | 46708.0 | 0 | Felis | 46708.0 | 0 | Felis catus | 46708.0 | 0 |
| scf7180000018583 | 1011602 | 0.5247 | 0 | 50.4913 | Eukaryota | 18394.0 | 0 | Chordata | 18394.0 | 0 | Carnivora | 18394.0 | 0 | Felidae | 18394.0 | 0 | Panthera | 7296.0 | 3 | Felis catus | 5559.0 | 4 |
| scf7180000018766 | 477846 | 0.5191 | 0 | 48.5906 | Eukaryota | 57882.0 | 0 | Chordata | 57882.0 | 0 | Carnivora | 57882.0 | 0 | Felidae | 37122.0 | 4 | Panthera | 12350.0 | 8 | ricata suricat | 6383.0 | 9 |
| scf7180000019629 | 181178 | 0.3565 | 0 | 48.6742 | Eukaryota | 44417.0 | 0 | Chordata | 44417.0 | 0 | Carnivora | 44417.0 | 0 | Felidae | 44417.0 | 0 | Felis | 44417.0 | 0 | Felis catus | 44417.0 | 0 |
| scf7180000018546 | 31553 | 0.395 | 0 | 23.2923 | Eukaryota | 26200.0 | 0 | Chordata | 26200.0 | 0 | Carnivora | 26200.0 | 0 | Felidae | 26200.0 | 0 | Felis | 23599.0 | 1 | Felis catus | 23599.0 | 1 |
| scf7180000018858 | 460862 | 0.5499 | 0 | 50.3322 | Eukaryota | 31097.0 | 0 | Chordata | 31097.0 | 0 | Carnivora | 28162.0 | 1 | Felidae | 22122.0 | 3 | Panthera | 9567.0 | 6 | anthera pardu | 6427.0 | 7 |
| scf7180000022561 | 3819 | 0.5177 | 0 | 51.6834 | no-hit | 0.0 | 0 | no-hit | 0.0 | 0 | no-hit | 0.0 | 0 | no-hit | 0.0 | 0 | no-hit | 0.0 | 0 | no-hit | 0.0 | 0 |
| scf7180000020772 | 112490 | 0.6033 | 0 | 46.9362 | Eukaryota | 26040.0 | 0 | Chordata | 26040.0 | 0 | Carnivora | 26040.0 | 0 | Felidae | 22184.0 | 1 | Lynx | 14656.0 | 2 | ynx canadens | 14656.0 | 2 |
| scf7180000021790 | 32275 | 0.356 | 0 | 23.7988 | Eukaryota | 4156.0 | 0 | Chordata | 4156.0 | 0 | Artiodactyla | 3331.0 | 1 | Delphinidae | 2469.0 | 3 | agenorhynchu | 1260.0 | 4 | rhynchus obli | 1260.0 | 4 |
| scf7180000017967 | 1403286 | 0.3745 | 0 | 50.3536 | Eukaryota | 100944.0 | 0 | Chordata | 100944.0 | 0 | Carnivora | 100944.0 | 0 | Felidae | 100944.0 | 0 | Acinonyx | 40524.0 | 3 | cinonyx jubat | 40524.0 | 3 |
| scf7180000022275 | 16234 | 0.3905 | 0 | 25.3306 | Eukaryota | 5386.0 | 0 | Chordata | 5386.0 | 0 | Primates | 2002.0 | 3 | Hominidae | 2002.0 | 3 | Sus | 1757.0 | 6 | Sus scrofa | 1757.0 | 6 |
| scf7180000018678 | 17985 | 0.5067 | 0 | 79.6505 | Eukaryota | 12153.0 | 0 | Chordata | 12153.0 | 0 | Carnivora | 12153.0 | 0 | Felidae | 12153.0 | 0 | Felis | 10928.0 | 1 | Felis catus | 10928.0 | 1 |
| scf7180000018322 | 678772 | 0.381 | 0 | 50.5091 | Eukaryota | 46588.0 | 0 | Chordata | 46588.0 | 0 | Carnivora | 46588.0 | 0 | Felidae | 46588.0 | 0 | Felis | 46588.0 | 0 | Felis catus | 46588.0 | 0 |
| scf7180000018255 | 955597 | 0.3571 | 0 | 49.6927 | Eukaryota | 46136.0 | 0 | Chordata | 46136.0 | 0 | Carnivora | 46136.0 | 0 | Felidae | 46136.0 | 0 | Felis | 46136.0 | 0 | Felis catus | 46136.0 | 0 |
| scf7180000017780 | 252420 | 0.372 | 0 | 46.906 | Eukaryota | 45249.0 | 0 | Chordata | 45249.0 | 0 | Carnivora | 45249.0 | 0 | Felidae | 45249.0 | 0 | Felis | 45249.0 | 0 | Felis catus | 45249.0 | 0 |
| scf7180000019971 | 38545 | 0.3859 | 0 | 566.0523 | Eukaryota | 12432.0 | 0 | Chordata | 12432.0 | 0 | Carnivora | 12432.0 | 0 | Felidae | 12432.0 | 0 | Felis | 11205.0 | 1 | Felis catus | 11205.0 | 1 |
| scf7180000019627 | 14238 | 0.474 | 0 | 27.9256 | Eukaryota | 2998.0 | 0 | Chordata | 2998.0 | 0 | Carnivora | 2465.0 | 2 | Felidae | 2465.0 | 2 | Lynx | 994.0 | 4 | ynx canadens | 994.0 | 4 |
| scf7180000017899 | 733524 | 0.3732 | 0 | 50.6442 | Eukaryota | 45572.0 | 0 | Chordata | 45572.0 | 0 | Carnivora | 39880.0 | 1 | Felidae | 39880.0 | 1 | Felis | 39880.0 | 1 | Felis catus | 39880.0 | 1 |
| scf7180000023642 | 26015 | 0.4576 | 0 | 180.6259 | Eukaryota | 3060.0 | 0 | Chordata | 3060.0 | 0 | Carnivora | 2830.0 | 1 | Felidae | 2019.0 | 2 | Lynx | 996.0 | 5 | ynx canadens | 996.0 | 5 |
| scf7180000022508 | 11330 | 0.3649 | 0 | 24.3238 | Eukaryota | 42526.0 | 0 | Chordata | 42526.0 | 0 | Carnivora | 42526.0 | 0 | Felidae | 42526.0 | 0 | Felis | 38228.0 | 1 | Felis catus | 38228.0 | 1 |
| scf7180000018063 | 17325 | 0.4196 | 0 | 27.014 | Eukaryota | 2548.0 | 0 | Chordata | 2548.0 | 0 | Carnivora | 2071.0 | 1 | Felidae | 1838.0 | 2 | Felis | 1838.0 | 2 | Felis catus | 1838.0 | 2 |
| scf7180000019245 | 30597 | 0.3989 | 0 | 389.4621 | Eukaryota | 6698.0 | 0 | Chordata | 6698.0 | 0 | Carnivora | 6698.0 | 0 | Felidae | 6698.0 | 0 | Felis | 6698.0 | 0 | Felis catus | 6698.0 | 0 |
| scf7180000022357 | 33161 | 0.3643 | 0 | 24.9356 | Eukaryota | 10211.0 | 0 | Chordata | 10211.0 | 0 | Carnivora | 10211.0 | 0 | Felidae | 10211.0 | 0 | Felis | 9152.0 | 1 | Felis catus | 9152.0 | 1 |
| scf7180000019653 | 152037 | 0.3836 | 0 | 24.2391 | Eukaryota | 40344.0 | 0 | Chordata | 40344.0 | 0 | Carnivora | 40344.0 | 0 | Felidae | 40344.0 | 0 | Felis | 40344.0 | 0 | Felis catus | 40344.0 | 0 |
| scf7180000023852 | 1305713 | 0.4226 | 0 | 52.0626 | Eukaryota | 46993.0 | 0 | Chordata | 46993.0 | 0 | Carnivora | 46993.0 | 0 | Felidae | 46993.0 | 0 | Felis | 46993.0 | 0 | Felis catus | 46993.0 | 0 |
| scf7180000023002 | 16819 | 0.5406 | 0 | 242.1555 | Eukaryota | 12152.0 | 0 | Chordata | 12152.0 | 0 | Carnivora | 12152.0 | 0 | Felidae | 12152.0 | 0 | Felis | 10947.0 | 1 | Felis catus | 10947.0 | 1 |
| scf7180000021103 | 27235 | 0.5093 | 0 | 453.2557 | no-hit | 0.0 | 0 | no-hit | 0.0 | 0 | no-hit | 0.0 | 0 | no-hit | 0.0 | 0 | no-hit | 0.0 | 0 | no-hit | 0.0 | 0 |
| scf7180000020289 | 20008 | 0.6079 | 0 | 29.762 | Eukaryota | 13053.0 | 0 | Chordata | 13053.0 | 0 | Carnivora | 13053.0 | 0 | Felidae | 7435.0 | 1 | Canis | 5618.0 | 5 | Canis lupus | 5618.0 | 5 |
| scf7180000018067 | 1161805 | 0.3815 | 0 | 51.3082 | Eukaryota | 84976.0 | 0 | Chordata | 84976.0 | 0 | Carnivora | 84976.0 | 0 | Felidae | 84976.0 | 0 | Felis | 54780.0 | 1 | Felis catus | 54780.0 | 1 |
| scf7180000020140 | 56043 | 0.4029 | 0 | 159.9769 | Eukaryota | 29430.0 | 0 | Chordata | 29430.0 | 0 | Carnivora | 29430.0 | 0 | Felidae | 29430.0 | 0 | Felis | 26514.0 | 1 | Felis catus | 26514.0 | 1 |
| scf7180000022007 | 63519 | 0.4457 | 0 | 24.8865 | Eukaryota | 19767.0 | 0 | Chordata | 19767.0 | 0 | Carnivora | 19767.0 | 0 | Felidae | 19767.0 | 0 | Felis | 15840.0 | 1 | Felis catus | 15840.0 | 1 |
| scf7180000021869 | 27976 | 0.4576 | 0 | 23.5147 | Eukaryota | 4298.0 | 0 | Chordata | 4298.0 | 0 | Carnivora | 1639.0 | 3 | Felidae | 1639.0 | 3 | Felis | 1639.0 | 3 | Felis catus | 1639.0 | 3 |
| scf7180000018864 | 562811 | 0.4622 | 0 | 50.3727 | Eukaryota | 14851.0 | 0 | Chordata | 14851.0 | 0 | Carnivora | 6415.0 | 2 | Felidae | 6415.0 | 2 | Felis | 6415.0 | 3 | Felis catus | 6415.0 | 3 |
| scf7180000018853 | 18829 | 0.4109 | 0 | 24.1201 | Eukaryota | 6612.0 | 0 | Chordata | 6612.0 | 0 | Carnivora | 6612.0 | 0 | Felidae | 3586.0 | 3 | Panthera | 1438.0 | 7 | Panthera tigri | 754.0 | 9 |
| scf7180000023670 | 63064 | 0.4487 | 0 | 388.9357 | Eukaryota | 7381.0 | 0 | Chordata | 7381.0 | 0 | Carnivora | 7381.0 | 0 | Otariidae | 5263.0 | 1 | Eumetopias | 2980.0 | 2 | netopias juba | 2980.0 | 2 |
| scf7180000019563 | 355887 | 0.4412 | 0 | 50.3869 | Eukaryota | 20885.0 | 0 | Chordata | 20885.0 | 0 | Carnivora | 20885.0 | 0 | Felidae | 20885.0 | 0 | Felis | 18779.0 | 1 | Felis catus | 18779.0 | 1 |
| scf7180000021244 | 44966 | 0.4226 | 0 | 26.2518 | Eukaryota | 12624.0 | 0 | Chordata | 12624.0 | 0 | Carnivora | 12624.0 | 0 | Felidae | 12624.0 | 0 | Felis | 11329.0 | 1 | Felis catus | 11329.0 | 1 |
| scf7180000020876 | 10982 | 0.4802 | 0 | 47.9917 | Eukaryota | 2756.0 | 0 | Chordata | 2756.0 | 0 | Carnivora | 2756.0 | 0 | Felidae | 2756.0 | 0 | Felis | 1926.0 | 1 | Felis catus | 1926.0 | 1 |
| scf7180000022480 | 15852 | 0.3574 | 0 | 22.7403 | Eukaryota | 29122.0 | 0 | Chordata | 29122.0 | 0 | Carnivora | 29122.0 | 0 | Felidae | 29122.0 | 0 | Felis | 23257.0 | 1 | Felis catus | 23257.0 | 1 |
| scf7180000019847 | 184086 | 0.4454 | 0 | 26.2729 | Eukaryota | 39419.0 | 0 | Chordata | 39419.0 | 0 | Carnivora | 39419.0 | 0 | Felidae | 39419.0 | 0 | Felis | 39419.0 | 0 | Felis catus | 39419.0 | 0 |
| scf7180000019410 | 374674 | 0.3724 | 0 | 50.1892 | Eukaryota | 41447.0 | 0 | Chordata | 41447.0 | 0 | Carnivora | 41447.0 | 0 | Felidae | 41447.0 | 0 | Felis | 37247.0 | 1 | Felis catus | 37247.0 | 1 |
| scf7180000018209 | 1640848 | 0.3624 | 0 | 49.0189 | Eukaryota | 47227.0 | 0 | Chordata | 47227.0 | 0 | Carnivora | 47227.0 | 0 | Felidae | 47227.0 | 0 | Felis | 47227.0 | 0 | Felis catus | 47227.0 | 0 |
| scf7180000018636 | 14653 | 0.3472 | 0 | 28.3846 | Eukaryota | 2551.0 | 0 | Chordata | 2551.0 | 0 | Carnivora | 2551.0 | 0 | Felidae | 2303.0 | 1 | Felis | 1796.0 | 2 | Felis catus | 1796.0 | 2 |
| scf7180000019716 | 214271 | 0.5285 | 0 | 51.2445 | Eukaryota | 73150.0 | 0 | Chordata | 73150.0 | 0 | Carnivora | 73150.0 | 0 | Felidae | 51436.0 | 1 | Felis | 51436.0 | 1 | Felis catus | 51436.0 | 1 |
| scf7180000018290 | 1477293 | 0.4267 | 100 | 51.8154 | Eukaryota | 30183.0 | 0 | Chordata | 30183.0 | 0 | Carnivora | 30183.0 | 0 | Felidae | 30183.0 | 0 | Felis | 27115.0 | 1 | Felis catus | 27115.0 | 1 |
| scf7180000024232 | 229871 | 0.4676 | 0 | 51.0809 | Eukaryota | 18737.0 | 0 | Chordata | 18737.0 | 0 | Carnivora | 18737.0 | 0 | Felidae | 17167.0 | 1 | Felis | 5759.0 | 4 | Felis catus | 5759.0 | 4 |
| scf7180000024068 | 1063682 | 0.369 | 0 | 50.3471 | Eukaryota | 59782.0 | 0 | Chordata | 59782.0 | 0 | Carnivora | 59782.0 | 0 | Felidae | 59782.0 | 0 | Panthera | 35922.0 | 1 | anthera pardu | 35922.0 | 1 |
| scf7180000019363 | 900204 | 0.4772 | 0 | 51.6588 | Eukaryota | 41128.0 | 0 | Chordata | 41128.0 | 0 | Carnivora | 41128.0 | 0 | Felidae | 41128.0 | 0 | Felis | 41128.0 | 0 | Felis catus | 41128.0 | 0 |
| scf7180000019279 | 10033 | 0.3971 | 0 | 26.8145 | Eukaryota | 41361.0 | 0 | Chordata | 41361.0 | 0 | Carnivora | 41361.0 | 0 | Felidae | 41361.0 | 0 | Felis | 41361.0 | 0 | Felis catus | 41361.0 | 0 |
| scf7180000020152 | 188684 | 0.5573 | 0 | 46.1004 | Eukaryota | 8715.0 | 0 | Chordata | 8715.0 | 0 | Carnivora | 8715.0 | 0 | Felidae | 8715.0 | 0 | Felis | 8715.0 | 0 | Felis catus | 8715.0 | 0 |
| scf7180000023687 | 65744 | 0.3861 | 0 | 691.9543 | Eukaryota | 12634.0 | 0 | Chordata | 12634.0 | 0 | Carnivora | 12634.0 | 0 | Felidae | 12634.0 | 0 | Felis | 7615.0 | 1 | Felis catus | 7615.0 | 1 |
| scf7180000023492 | 30252 | 0.4614 | 0 | 77.1486 | Eukaryota | 2175.0 | 0 | Chordata | 2175.0 | 0 | Carnivora | 2175.0 | 0 | Felidae | 1455.0 | 1 | Felis | 1240.0 | 4 | Felis catus | 1240.0 | 4 |
| scf7180000017939 | 169957 | 0.4499 | 0 | 48.8669 | Eukaryota | 46993.0 | 0 | Chordata | 46993.0 | 0 | Carnivora | 46993.0 | 0 | Felidae | 46993.0 | 0 | Felis | 46993.0 | 0 | Felis catus | 46993.0 | 0 |
| scf7180000017635 | 887813 | 0.3659 | 0 | 49.3766 | Eukaryota | 47214.0 | 0 | Chordata | 47214.0 | 0 | Carnivora | 47214.0 | 0 | Felidae | 47214.0 | 0 | Felis | 47214.0 | 0 | Felis catus | 47214.0 | 0 |
| scf7180000023485 | 15875 | 0.458 | 0 | 282.2014 | no-hit | 0.0 | 0 | no-hit | 0.0 | 0 | no-hit | 0.0 | 0 | no-hit | 0.0 | 0 | no-hit | 0.0 | 0 | no-hit | 0.0 | 0 |
| scf7180000018928 | 262601 | 0.5284 | 0 | 48.8249 | Eukaryota | 38580.0 | 0 | Chordata | 38580.0 | 0 | Carnivora | 38580.0 | 0 | Felidae | 38580.0 | 0 | Acinonyx | 21378.0 | 2 | cinonyx jubat | 21378.0 | 2 |
| scf7180000017491 | 79761 | 0.3903 | 0 | 44.4751 | Eukaryota | 23888.0 | 0 | Chordata | 23888.0 | 0 | Carnivora | 23888.0 | 0 | Felidae | 23888.0 | 0 | Felis | 23888.0 | 0 | Felis catus | 23888.0 | 0 |
| scf7180000021199 | 45488 | 0.3633 | 0 | 25.4663 | Eukaryota | 11071.0 | 0 | Chordata | 11071.0 | 0 | Carnivora | 9911.0 | 1 | Felidae | 7459.0 | 2 | Felis | 6572.0 | 3 | Felis catus | 6572.0 | 3 |
| scf7180000019148 | 423749 | 0.4712 | 0 | 51.4407 | Eukaryota | 46028.0 | 0 | Chordata | 46028.0 | 0 | Carnivora | 46028.0 | 0 | Felidae | 46028.0 | 0 | Felis | 46028.0 | 0 | Felis catus | 46028.0 | 0 |
| scf7180000019692 | 21530 | 0.5245 | 0 | 166.7359 | Eukaryota | 2632.0 | 0 | Chordata | 2632.0 | 0 | Carnivora | 2632.0 | 0 | Felidae | 2371.0 | 1 | Felis | 1910.0 | 2 | Felis catus | 1910.0 | 2 |
| scf7180000020112 | 30698 | 0.3957 | 0 | 289.4132 | Eukaryota | 46406.0 | 0 | Chordata | 46406.0 | 0 | Carnivora | 46406.0 | 0 | Felidae | 46406.0 | 0 | Felis | 46406.0 | 0 | Felis catus | 46406.0 | 0 |
| scf7180000021849 | 19067 | 0.3914 | 0 | 24.6368 | Eukaryota | 44245.0 | 0 | Chordata | 44245.0 | 0 | Carnivora | 44245.0 | 0 | Felidae | 44245.0 | 0 | Felis | 44245.0 | 0 | Felis catus | 44245.0 | 0 |
| scf7180000018829 | 475168 | 0.5345 | 0 | 55.556 | Eukaryota | 49927.0 | 0 | Chordata | 49927.0 | 0 | Carnivora | 49927.0 | 0 | Phocidae | 20242.0 | 5 | eptonychote | 20242.0 | 6 | nychotes wei | 20242.0 | 7 |
| scf7180000023915 | 961250 | 0.3485 | 0 | 48.4838 | Eukaryota | 47031.0 | 0 | Chordata | 47031.0 | 0 | Carnivora | 47031.0 | 0 | Felidae | 47031.0 | 0 | Felis | 47031.0 | 0 | Felis catus | 47031.0 | 0 |
| scf7180000022591 | 4173 | 0.5006 | 0 | 20.0465 | Eukaryota | 7144.0 | 0 | Chordata | 7144.0 | 0 | Carnivora | 7144.0 | 0 | Felidae | 6552.0 | 1 | Felis | 2585.0 | 3 | Felis catus | 2585.0 | 4 |
| scf7180000019508 | 261541 | 0.3795 | 0 | 51.0077 | Eukaryota | 37152.0 | 0 | Chordata | 37152.0 | 0 | Carnivora | 37152.0 | 0 | Felidae | 26374.0 | 2 | Panthera | 11238.0 | 4 | Felis catus | 7578.0 | 5 |
| scf7180000021341 | 40779 | 0.4188 | 0 | 23.9826 | Eukaryota | 8708.0 | 0 | Chordata | 8708.0 | 0 | Primates | 4168.0 | 2 | Hominidae | 4168.0 | 2 | Felis | 3740.0 | 3 | Felis catus | 3740.0 | 3 |
| scf7180000017426 | 3130513 | 0.426 | 0 | 50.7424 | Eukaryota | 73894.0 | 0 | Chordata | 73894.0 | 0 | Carnivora | 48099.0 | 1 | Felidae | 41137.0 | 2 | Equus | 25795.0 | 5 | cinonyx jubat | 17441.0 | 7 |
| scf7180000017605 | 2038450 | 0.4113 | 0 | 52.0165 | Eukaryota | 168998.0 | 0 | Chordata | 168998.0 | 0 | Carnivora | 168998.0 | 0 | Felidae | 104256.0 | 4 | Panthera | 50569.0 | 8 | anthera pardu | 27013.0 | 9 |
| scf7180000022395 | 13782 | 0.5745 | 0 | 42.2552 | Eukaryota | 1981.0 | 0 | Chordata | 1981.0 | 0 | Carnivora | 1609.0 | 1 | Felidae | 1363.0 | 2 | Felis | 954.0 | 3 | Felis catus | 954.0 | 3 |
| scf7180000021867 | 41003 | 0.3788 | 0 | 45.2072 | Eukaryota | 45875.0 | 0 | Chordata | 45875.0 | 0 | Carnivora | 45875.0 | 0 | Felidae | 45875.0 | 0 | Felis | 45875.0 | 0 | Felis catus | 45875.0 | 0 |
| scf7180000020499 | 26594 | 0.4264 | 0 | 414.0511 | Eukaryota | 256900.0 | 0 | Chordata | 256900.0 | 0 | Carnivora | 256900.0 | 0 | Hyaenidae | 256900.0 | 0 | Proteles | 155466.0 | 1 | roteles cristat | 155466.0 | 1 |
| scf7180000019350 | 43861 | 0.4655 | 0 | 160.2347 | Eukaryota | 16271.0 | 0 | Chordata | 16271.0 | 0 | Carnivora | 16271.0 | 0 | Felidae | 16271.0 | 0 | Felis | 14688.0 | 1 | Felis catus | 14688.0 | 1 |
| scf7180000019248 | 343425 | 0.4532 | 0 | 48.9139 | Eukaryota | 16091.0 | 0 | Chordata | 16091.0 | 0 | Carnivora | 16091.0 | 0 | Felidae | 16091.0 | 0 | Felis | 14497.0 | 1 | Felis catus | 14497.0 | 1 |
| scf7180000017940 | 239161 | 0.3741 | 0 | 51.2114 | Eukaryota | 74199.0 | 0 | Chordata | 74199.0 | 0 | Carnivora | 74199.0 | 0 | Herpestidae | 47116.0 | 1 | Suricata | 47116.0 | 2 | ricata suricat | 47116.0 | 2 |
| scf7180000020800 | 138195 | 0.3766 | 100 | 24.1914 | Eukaryota | 26686.0 | 0 | Chordata | 26686.0 | 0 | Carnivora | 26686.0 | 0 | Felidae | 26686.0 | 0 | Felis | 26686.0 | 0 | Felis catus | 26686.0 | 0 |
| scf7180000019887 | 187816 | 0.4039 | 0 | 51.8158 | Eukaryota | 31265.0 | 0 | Chordata | 31265.0 | 0 | Carnivora | 31265.0 | 0 | Felidae | 31265.0 | 0 | Felis | 31265.0 | 0 | Felis catus | 31265.0 | 0 |
| scf7180000018154 | 1182903 | 0.3799 | 0 | 50.8535 | Eukaryota | 144897.0 | 0 | Chordata | 144897.0 | 0 | Carnivora | 144897.0 | 0 | Felidae | 130467.0 | 1 | Panthera | 43595.0 | 5 | anthera pardu | 29100.0 | 6 |
| scf7180000022531 | 13319 | 0.5258 | 0 | 33.2901 | Eukaryota | 2511.0 | 0 | Chordata | 2511.0 | 0 | Carnivora | 2511.0 | 0 | Felidae | 2281.0 | 1 | Felis | 1265.0 | 3 | Felis catus | 1265.0 | 3 |
| scf7180000020502 | 102256 | 0.3639 | 0 | 25.0847 | Eukaryota | 25417.0 | 0 | Chordata | 25417.0 | 0 | Carnivora | 25417.0 | 0 | Felidae | 25417.0 | 0 | Felis | 20240.0 | 1 | Felis catus | 20240.0 | 1 |
| scf7180000019998 | 175770 | 0.4987 | 0 | 49.7275 | Eukaryota | 11899.0 | 1 | Chordata | 11899.0 | 1 | Primates | 6222.0 | 3 | Hominidae | 4654.0 | 4 | Homo | 4654.0 | 5 | Homo sapien | 4654.0 | 5 |
| scf7180000022487 | 21600 | 0.372 | 0 | 25.6034 | Eukaryota | 44359.0 | 0 | Chordata | 44359.0 | 0 | Carnivora | 44359.0 | 0 | Felidae | 44359.0 | 0 | Felis | 44359.0 | 0 | Felis catus | 44359.0 | 0 |
| scf7180000017536 | 2718788 | 0.3732 | 0 | 50.9725 | Eukaryota | 55073.0 | 0 | Chordata | 55073.0 | 0 | Carnivora | 55073.0 | 0 | Felidae | 55073.0 | 0 | Panthera | 33007.0 | 2 | ynx canadens | 16566.0 | 3 |
| scf7180000022029 | 58775 | 0.3802 | 0 | 25.0613 | Eukaryota | 23631.0 | 0 | Chordata | 23631.0 | 0 | Carnivora | 23631.0 | 0 | Felidae | 23631.0 | 0 | Felis | 21243.0 | 1 | Felis catus | 21243.0 | 1 |
| scf7180000018738 | 642238 | 0.3861 | 0 | 50.6389 | Eukaryota | 46027.0 | 0 | Chordata | 46027.0 | 0 | Carnivora | 46027.0 | 0 | Felidae | 46027.0 | 0 | Felis | 46027.0 | 0 | Felis catus | 46027.0 | 0 |
| scf7180000021575 | 58618 | 0.3727 | 0 | 48.4371 | Eukaryota | 10717.0 | 0 | Chordata | 10717.0 | 0 | Primates | 4002.0 | 6 | Hominidae | 4002.0 | 6 | Homo | 4002.0 | 6 | Homo sapien | 4002.0 | 6 |
| scf7180000023997 | 47457 | 0.3968 | 0 | 25.3041 | Eukaryota | 10072.0 | 0 | Chordata | 10072.0 | 0 | Carnivora | 8731.0 | 1 | Canidae | 5051.0 | 2 | Canis | 5051.0 | 2 | Canis lupus | 5051.0 | 2 |
| scf7180000017715 | 49462 | 0.3735 | 0 | 38.6993 | Eukaryota | 26536.0 | 0 | Chordata | 26536.0 | 0 | Carnivora | 26536.0 | 0 | Felidae | 26536.0 | 0 | Felis | 23828.0 | 1 | Felis catus | 23828.0 | 1 |
| scf7180000017543 | 2536410 | 0.3671 | 0 | 49.323 | Eukaryota | 46415.0 | 0 | Chordata | 46415.0 | 0 | Carnivora | 46415.0 | 0 | Felidae | 46415.0 | 0 | Felis | 46415.0 | 0 | Felis catus | 46415.0 | 0 |
| scf7180000020002 | 120885 | 0.5259 | 0 | 48.1584 | Eukaryota | 19649.0 | 0 | Chordata | 19649.0 | 0 | Primates | 6173.0 | 5 | Hominidae | 6173.0 | 5 | Homo | 6173.0 | 6 | Homo sapien | 6173.0 | 6 |
| scf7180000019292 | 1371455 | 0.5128 | 0 | 53.9505 | Eukaryota | 48985.0 | 0 | Chordata | 48985.0 | 0 | Carnivora | 48985.0 | 0 | Herpestidae | 25395.0 | 1 | Suricata | 25395.0 | 4 | ricata suricat | 25395.0 | 4 |
| scf7180000017981 | 1299757 | 0.388 | 0 | 49.878 | Eukaryota | 46258.0 | 0 | Chordata | 46258.0 | 0 | Carnivora | 46258.0 | 0 | Felidae | 46258.0 | 0 | Felis | 41676.0 | 1 | Felis catus | 41676.0 | 1 |
| scf7180000022870 | 18017 | 0.511 | 0 | 388.829 | no-hit | 0.0 | 0 | no-hit | 0.0 | 0 | no-hit | 0.0 | 0 | no-hit | 0.0 | 0 | no-hit | 0.0 | 0 | no-hit | 0.0 | 0 |
| scf7180000021202 | 64696 | 0.365 | 0 | 24.2157 | Eukaryota | 34436.0 | 0 | Chordata | 34436.0 | 0 | Carnivora | 34436.0 | 0 | Herpestidae | 19550.0 | 2 | Suricata | 19550.0 | 3 | ricata suricat | 19550.0 | 4 |
| scf7180000023296 | 42615 | 0.3987 | 0 | 309.01 | Eukaryota | 7003.0 | 0 | Chordata | 7003.0 | 0 | Artiodactyla | 4889.0 | 2 | Bovidae | 4889.0 | 2 | Ovis | 4083.0 | 3 | vis canadens | 4083.0 | 3 |
| scf7180000018710 | 575660 | 0.446 | 0 | 50.0111 | Eukaryota | 21334.0 | 0 | Chordata | 21334.0 | 0 | Carnivora | 21334.0 | 0 | Herpestidae | 19269.0 | 1 | Suricata | 19269.0 | 1 | ricata suricat | 19269.0 | 1 |
| scf7180000021719 | 16538 | 0.5267 | 0 | 46.3028 | Eukaryota | 2729.0 | 0 | Chordata | 2729.0 | 0 | Carnivora | 2729.0 | 0 | Felidae | 2729.0 | 0 | Felis | 1350.0 | 4 | Felis catus | 1350.0 | 4 |
| scf7180000017717 | 2357342 | 0.4381 | 0 | 52.9341 | Eukaryota | 46957.0 | 0 | Chordata | 46957.0 | 0 | Carnivora | 46957.0 | 0 | Felidae | 46957.0 | 0 | Felis | 46957.0 | 0 | Felis catus | 46957.0 | 0 |
| scf7180000019523 | 116116 | 0.3931 | 0 | 47.067 | Eukaryota | 42680.0 | 0 | Chordata | 42680.0 | 0 | Carnivora | 42680.0 | 0 | Felidae | 42680.0 | 0 | Felis | 42680.0 | 0 | Felis catus | 42680.0 | 0 |
| scf7180000021120 | 28368 | 0.521 | 0 | 375.5295 | no-hit | 0.0 | 0 | no-hit | 0.0 | 0 | no-hit | 0.0 | 0 | no-hit | 0.0 | 0 | no-hit | 0.0 | 0 | no-hit | 0.0 | 0 |
| scf7180000017656 | 677132 | 0.3593 | 0 | 49.0066 | Eukaryota | 44441.0 | 0 | Chordata | 44441.0 | 0 | Carnivora | 44441.0 | 0 | Felidae | 44441.0 | 0 | Felis | 44441.0 | 0 | Felis catus | 44441.0 | 0 |
| scf7180000017768 | 1094279 | 0.3603 | 100 | 48.5107 | Eukaryota | 45967.0 | 0 | Chordata | 45967.0 | 0 | Carnivora | 45967.0 | 0 | Felidae | 45967.0 | 0 | Felis | 45967.0 | 0 | Felis catus | 45967.0 | 0 |
| scf7180000017999 | 29604 | 0.3878 | 0 | 27.0998 | Eukaryota | 34595.0 | 0 | Chordata | 34595.0 | 0 | Carnivora | 34595.0 | 0 | Felidae | 34595.0 | 0 | Felis | 34595.0 | 0 | Felis catus | 34595.0 | 0 |
| scf7180000020941 | 77186 | 0.3882 | 0 | 25.4965 | Eukaryota | 18356.0 | 0 | Chordata | 18356.0 | 0 | Primates | 12300.0 | 3 | Hominidae | 8332.0 | 4 | Pan | 4872.0 | 5 | an troglodyte | 4872.0 | 5 |
| scf7180000023937 | 589333 | 0.445 | 0 | 51.6635 | Eukaryota | 46759.0 | 0 | Chordata | 46759.0 | 0 | Carnivora | 46759.0 | 0 | Felidae | 46759.0 | 0 | Felis | 46759.0 | 0 | Felis catus | 46759.0 | 0 |
| scf7180000022076 | 20218 | 0.6302 | 0 | 51.5548 | Eukaryota | 2249.0 | 0 | Chordata | 2249.0 | 0 | Carnivora | 2249.0 | 0 | Canidae | 917.0 | 5 | Canis | 680.0 | 7 | Canis lupus | 680.0 | 7 |
| scf7180000019859 | 226682 | 0.3913 | 0 | 29.9793 | Eukaryota | 45758.0 | 0 | Chordata | 45758.0 | 0 | Carnivora | 45758.0 | 0 | Felidae | 45758.0 | 0 | Felis | 45758.0 | 0 | Felis catus | 45758.0 | 0 |
| scf7180000020781 | 65529 | 0.5702 | 0 | 46.9583 | Eukaryota | 5209.0 | 0 | Chordata | 5209.0 | 0 | Carnivora | 3219.0 | 2 | Canidae | 2023.0 | 4 | Canis | 2023.0 | 5 | Canis lupus | 2023.0 | 5 |
| scf7180000022456 | 88710 | 0.3938 | 0 | 229.3276 | Eukaryota | 25442.0 | 0 | Chordata | 25442.0 | 0 | Carnivora | 25442.0 | 0 | Felidae | 25442.0 | 0 | Felis | 22841.0 | 1 | Felis catus | 22841.0 | 1 |
| scf7180000023562 | 24596 | 0.4805 | 0 | 421.2765 | no-hit | 0.0 | 0 | no-hit | 0.0 | 0 | no-hit | 0.0 | 0 | no-hit | 0.0 | 0 | no-hit | 0.0 | 0 | no-hit | 0.0 | 0 |
| scf7180000020198 | 162334 | 0.3977 | 0 | 25.766 | Eukaryota | 35181.0 | 0 | Chordata | 35181.0 | 0 | Carnivora | 35181.0 | 0 | Herpestidae | 24787.0 | 1 | Suricata | 24787.0 | 2 | ricata suricat | 24787.0 | 2 |
| scf7180000022187 | 25790 | 0.3803 | 0 | 23.0079 | Eukaryota | 26555.0 | 0 | Chordata | 26555.0 | 0 | Carnivora | 26555.0 | 0 | Felidae | 26555.0 | 0 | Felis | 23840.0 | 1 | Felis catus | 23840.0 | 1 |
| scf7180000019034 | 630152 | 0.4842 | 0 | 48.2152 | Eukaryota | 65359.0 | 1 | Chordata | 65359.0 | 1 | Carnivora | 25019.0 | 3 | Felidae | 25019.0 | 4 | Homo | 16704.0 | 6 | Homo sapien | 16704.0 | 8 |
| scf7180000023108 | 28510 | 0.3809 | 0 | 232.3227 | Eukaryota | 9543.0 | 0 | Chordata | 9543.0 | 0 | pterygiforme | 9543.0 | 0 | Apterygidae | 9543.0 | 0 | Apteryx | 9543.0 | 0 | pteryx mante | 9543.0 | 0 |
| scf7180000020078 | 151653 | 0.4229 | 0 | 50.7959 | Eukaryota | 13457.0 | 0 | Chordata | 13457.0 | 0 | Carnivora | 8369.0 | 1 | Felidae | 5754.0 | 3 | Globicephala | 3060.0 | 6 | obicephala me | 3060.0 | 6 |
| scf7180000022305 | 14065 | 0.6008 | 0 | 51.7078 | Eukaryota | 3658.0 | 0 | Chordata | 3658.0 | 0 | Carnivora | 3658.0 | 0 | Felidae | 3658.0 | 0 | Acinonyx | 1768.0 | 4 | cinonyx jubat | 1768.0 | 5 |
| scf7180000021140 | 45866 | 0.3641 | 0 | 23.7266 | Eukaryota | 17933.0 | 0 | Chordata | 17933.0 | 0 | Carnivora | 17933.0 | 0 | Felidae | 17933.0 | 0 | Felis | 16014.0 | 1 | Felis catus | 16014.0 | 1 |
| scf7180000019201 | 336075 | 0.5422 | 0 | 49.1923 | Eukaryota | 34164.0 | 0 | Chordata | 34164.0 | 0 | Carnivora | 34164.0 | 0 | Felidae | 14057.0 | 4 | Panthera | 3554.0 | 9 | anthera pardu | 3554.0 | 9 |
| scf7180000020988 | 30885 | 0.406 | 0 | 24.1207 | Eukaryota | 36391.0 | 0 | Chordata | 36391.0 | 0 | Carnivora | 36391.0 | 0 | Felidae | 36391.0 | 0 | Felis | 36391.0 | 0 | Felis catus | 36391.0 | 0 |
| scf7180000019487 | 274887 | 0.5448 | 0 | 50.2212 | Eukaryota | 32249.0 | 0 | Chordata | 32249.0 | 0 | Carnivora | 32249.0 | 0 | Felidae | 32249.0 | 0 | Lynx | 22757.0 | 1 | ynx canadens | 22757.0 | 1 |
| scf7180000022000 | 41985 | 0.3611 | 0 | 24.0659 | Eukaryota | 27246.0 | 0 | Chordata | 27246.0 | 0 | Carnivora | 27246.0 | 0 | Felidae | 27246.0 | 0 | Felis | 27246.0 | 0 | Felis catus | 27246.0 | 0 |
| scf7180000024286 | 100718 | 0.6265 | 0 | 54.4667 | Eukaryota | 33144.0 | 0 | Chordata | 33144.0 | 0 | Artiodactyla | 22233.0 | 2 | Bovidae | 22233.0 | 3 | Bos | 22233.0 | 3 | ndicus x Bos t | 12596.0 | 5 |
| scf7180000019384 | 418917 | 0.4351 | 0 | 49.0833 | Eukaryota | 23694.0 | 0 | Chordata | 23694.0 | 0 | Carnivora | 23694.0 | 0 | Felidae | 23694.0 | 0 | Felis | 23694.0 | 0 | Felis catus | 23694.0 | 0 |
| scf7180000020104 | 221921 | 0.4761 | 0 | 25.7779 | Eukaryota | 19266.0 | 0 | Chordata | 19266.0 | 0 | Carnivora | 19266.0 | 0 | Felidae | 19266.0 | 0 | Felis | 15919.0 | 1 | Felis catus | 15919.0 | 1 |
| scf7180000022768 | 7958 | 0.601 | 0 | 50.0095 | Eukaryota | 2257.0 | 0 | Chordata | 2257.0 | 0 | Carnivora | 2257.0 | 0 | Felidae | 2257.0 | 0 | Felis | 1819.0 | 1 | Felis catus | 1819.0 | 1 |

| | | | | | | | | | | | | | | | | | | | | | | |
|---|---|---|---|---|---|---|---|---|---|---|---|---|---|---|---|---|---|---|---|---|---|---|
| scf7180000020458 | 186445 | 0.4925 | 0 | 50.959 | Eukaryota | 44969.0 | 0 | Chordata | 44969.0 | 0 | Carnivora | 44969.0 | 0 | Felidae | 44969.0 | 0 | Felis | 44969.0 | 0 | Felis catus | 44969.0 | 0 |
| scf7180000018673 | 714164 | 0.491 | 0 | 52.9872 | Eukaryota | 34895.0 | 0 | Chordata | 34895.0 | 0 | Carnivora | 34895.0 | 0 | Felidae | 16161.0 | 2 | Zalophus | 7418.0 | 6 | phus californi | 7418.0 | 6 |
| scf7180000019040 | 515207 | 0.3526 | 0 | 49.5994 | Eukaryota | 45636.0 | 0 | Chordata | 45636.0 | 0 | Carnivora | 45636.0 | 0 | Felidae | 45636.0 | 0 | Felis | 45636.0 | 0 | Felis catus | 45636.0 | 0 |
| scf7180000018675 | 908499 | 0.4016 | 0 | 48.759 | Eukaryota | 45652.0 | 0 | Chordata | 45652.0 | 0 | Carnivora | 45652.0 | 0 | Felidae | 45652.0 | 0 | Felis | 45652.0 | 0 | Felis catus | 45652.0 | 0 |
| scf7180000018431 | 742661 | 0.3926 | 0 | 53.3332 | Eukaryota | 56115.0 | 0 | Chordata | 56115.0 | 0 | Carnivora | 56115.0 | 0 | Felidae | 44165.0 | 1 | Lynx | 16838.0 | 4 | ynx canadens | 16838.0 | 5 |
| scf7180000021423 | 10965 | 0.6694 | 0 | 181.2158 | Eukaryota | 2515.0 | 0 | Chordata | 2515.0 | 0 | Rodentia | 1022.0 | 4 | Caviidae | 765.0 | 5 | Cavia | 765.0 | 5 | avia porcellu | 765.0 | 6 |
| scf7180000019655 | 267276 | 0.5717 | 0 | 48.1515 | Eukaryota | 6976.0 | 1 | Chordata | 6976.0 | 1 | Carnivora | 5346.0 | 3 | Felidae | 4542.0 | 4 | Acinonyx | 2346.0 | 6 | cinonyx jubat | 2346.0 | 6 |
| scf7180000018669 | 261570 | 0.4031 | 0 | 50.8799 | Eukaryota | 44549.0 | 0 | Chordata | 44549.0 | 0 | Carnivora | 44549.0 | 0 | Felidae | 44549.0 | 0 | Felis | 40150.0 | 1 | Felis catus | 40150.0 | 1 |
| scf7180000023421 | 61689 | 0.4002 | 0 | 405.7672 | Eukaryota | 47746.0 | 0 | Chordata | 47746.0 | 0 | Carnivora | 47746.0 | 0 | Felidae | 47746.0 | 0 | Felis | 47746.0 | 0 | Felis catus | 47746.0 | 0 |
| scf7180000017818 | 2632095 | 0.3878 | 0 | 51.9331 | Eukaryota | 74115.0 | 0 | Chordata | 74115.0 | 0 | Carnivora | 74115.0 | 0 | Felidae | 46741.0 | 1 | Suricata | 27374.0 | 3 | ricata suricat | 27374.0 | 3 |
| scf7180000024045 | 298477 | 0.5043 | 0 | 50.2371 | Eukaryota | 29805.0 | 0 | Chordata | 29805.0 | 0 | Carnivora | 29805.0 | 0 | Felidae | 25266.0 | 2 | Lynx | 12509.0 | 5 | ynx canadens | 12509.0 | 5 |
| scf7180000023862 | 1701655 | 0.3905 | 0 | 51.6989 | Eukaryota | 132709.0 | 0 | Chordata | 132709.0 | 0 | Carnivora | 132709.0 | 0 | Felidae | 132709.0 | 0 | Felis | 53068.0 | 2 | Felis catus | 53068.0 | 2 |
| scf7180000018763 | 496833 | 0.4275 | 0 | 51.1552 | Eukaryota | 63752.0 | 0 | Chordata | 63752.0 | 0 | Carnivora | 57713.0 | 1 | Felidae | 26222.0 | 5 | Panthera | 13183.0 | 7 | Ursus arctos | 11998.0 | 8 |
| scf7180000018366 | 187294 | 0.3991 | 0 | 53.5611 | Eukaryota | 45285.0 | 0 | Chordata | 45285.0 | 0 | Carnivora | 45285.0 | 0 | Felidae | 45285.0 | 0 | Acinonyx | 22735.0 | 1 | cinonyx jubat | 22735.0 | 1 |
| scf7180000020812 | 70130 | 0.3688 | 0 | 26.6184 | Eukaryota | 11774.0 | 0 | Chordata | 11774.0 | 0 | Primates | 6706.0 | 3 | Hominidae | 6706.0 | 4 | Homo | 4818.0 | 5 | Homo sapiens | 4818.0 | 5 |
| scf7180000020825 | 68567 | 0.3672 | 0 | 25.5612 | Eukaryota | 15918.0 | 0 | Chordata | 15918.0 | 0 | Carnivora | 15918.0 | 0 | Felidae | 15918.0 | 0 | Felis | 12722.0 | 1 | Felis catus | 12722.0 | 1 |
| scf7180000018083 | 371462 | 0.4113 | 0 | 51.0413 | Eukaryota | 34077.0 | 0 | Chordata | 34077.0 | 0 | Carnivora | 34077.0 | 0 | Felidae | 34077.0 | 0 | Lynx | 27568.0 | 2 | ynx canadens | 27568.0 | 2 |
| scf7180000018078 | 1278487 | 0.3689 | 0 | 49.822 | Eukaryota | 73406.0 | 0 | Chordata | 73406.0 | 0 | Carnivora | 73406.0 | 0 | Phocidae | 54516.0 | 1 | eptonychote | 54516.0 | 1 | nychotes we | 54516.0 | 1 |
| scf7180000022497 | 21904 | 0.4283 | 0 | 44.5809 | Eukaryota | 14424.0 | 0 | Chordata | 14424.0 | 0 | Carnivora | 14424.0 | 0 | Felidae | 14424.0 | 0 | Felis | 14424.0 | 0 | Felis catus | 14424.0 | 0 |
| scf7180000018148 | 145667 | 0.3895 | 0 | 51.4748 | Eukaryota | 14755.0 | 0 | Chordata | 14755.0 | 0 | Carnivora | 14755.0 | 0 | Felidae | 14755.0 | 0 | Felis | 13338.0 | 1 | Felis catus | 13338.0 | 1 |
| scf7180000020057 | 52637 | 0.4217 | 0 | 72.5207 | Eukaryota | 44999.0 | 0 | Chordata | 44999.0 | 0 | Carnivora | 44999.0 | 0 | Felidae | 44999.0 | 0 | Felis | 44999.0 | 0 | Felis catus | 44999.0 | 0 |
| scf7180000020050 | 320371 | 0.4195 | 0 | 51.9745 | Eukaryota | 22843.0 | 0 | Chordata | 22843.0 | 0 | Carnivora | 22843.0 | 0 | Felidae | 22843.0 | 0 | Panthera | 13476.0 | 2 | anthera pardu | 13476.0 | 2 |
| scf7180000022283 | 28477 | 0.4115 | 0 | 23.6312 | Eukaryota | 5734.0 | 0 | Chordata | 5734.0 | 0 | Carnivora | 4893.0 | 1 | Felidae | 4893.0 | 1 | Felis | 4346.0 | 2 | Felis catus | 4346.0 | 2 |
| scf7180000017691 | 1928590 | 0.3821 | 0 | 49.2135 | Eukaryota | 46612.0 | 0 | Chordata | 46612.0 | 0 | Carnivora | 46612.0 | 0 | Felidae | 46612.0 | 0 | Felis | 46612.0 | 0 | Felis catus | 46612.0 | 0 |
| scf7180000021967 | 35387 | 0.5741 | 0 | 41.3941 | Eukaryota | 3103.0 | 0 | Chordata | 3103.0 | 0 | Carnivora | 3103.0 | 0 | Hyaenidae | 1282.0 | 3 | Suricata | 954.0 | 7 | ricata suricat | 954.0 | 7 |
| scf7180000022813 | 17457 | 0.4364 | 0 | 31.304 | Eukaryota | 26372.0 | 0 | Chordata | 26372.0 | 0 | Carnivora | 26372.0 | 0 | Felidae | 26372.0 | 0 | Felis | 23832.0 | 1 | Felis catus | 23832.0 | 1 |
| scf7180000017729 | 1389952 | 0.427 | 0 | 52.1449 | Eukaryota | 47096.0 | 0 | Chordata | 47096.0 | 0 | Carnivora | 47096.0 | 0 | Felidae | 41121.0 | 1 | Felis | 41121.0 | 1 | Felis catus | 41121.0 | 1 |
| scf7180000017522 | 2549028 | 0.3529 | 0 | 48.7551 | Eukaryota | 47395.0 | 0 | Chordata | 47395.0 | 0 | Carnivora | 47395.0 | 0 | Felidae | 47395.0 | 0 | Felis | 42643.0 | 1 | Felis catus | 42643.0 | 1 |
| scf7180000021481 | 81780 | 0.4616 | 0 | 27.0967 | Eukaryota | 8995.0 | 0 | Chordata | 8995.0 | 0 | Primates | 7489.0 | 1 | Hominidae | 7489.0 | 1 | Homo | 5414.0 | 2 | Homo sapiens | 5414.0 | 2 |
| scf7180000018563 | 662135 | 0.4405 | 0 | 51.3467 | Eukaryota | 32830.0 | 0 | Chordata | 32830.0 | 0 | Carnivora | 32830.0 | 0 | Felidae | 23023.0 | 1 | Panthera | 13081.0 | 2 | Felis catus | 9942.0 | 3 |
| scf7180000020561 | 104486 | 0.4024 | 0 | 50.1361 | Eukaryota | 9390.0 | 0 | Chordata | 9390.0 | 0 | Carnivora | 9390.0 | 0 | Felidae | 9390.0 | 0 | Felis | 8470.0 | 1 | Felis catus | 8470.0 | 1 |
| scf7180000017454 | 4374147 | 0.3822 | 0 | 50.8994 | Eukaryota | 71741.0 | 0 | Chordata | 71741.0 | 0 | Carnivora | 71741.0 | 0 | Felidae | 60005.0 | 1 | Acinonyx | 45064.0 | 3 | cinonyx jubat | 45064.0 | 3 |
| scf7180000022190 | 21701 | 0.3457 | 0 | 25.4044 | Eukaryota | 13719.0 | 0 | Chordata | 13719.0 | 0 | Carnivora | 13719.0 | 0 | Felidae | 13719.0 | 0 | Felis | 12291.0 | 1 | Felis catus | 12291.0 | 1 |
| scf7180000020764 | 45376 | 0.3874 | 0 | 25.3246 | Eukaryota | 43505.0 | 0 | Chordata | 43505.0 | 0 | Carnivora | 43505.0 | 0 | Felidae | 43505.0 | 0 | Felis | 43505.0 | 0 | Felis catus | 43505.0 | 0 |
| scf7180000021732 | 46249 | 0.3808 | 0 | 24.1643 | Eukaryota | 35979.0 | 0 | Chordata | 35979.0 | 0 | Carnivora | 35979.0 | 0 | Felidae | 35979.0 | 0 | Felis | 35979.0 | 0 | Felis catus | 35979.0 | 0 |
| scf7180000018456 | 779952 | 0.5228 | 0 | 53.6664 | Eukaryota | 46806.0 | 0 | Chordata | 46806.0 | 0 | Carnivora | 46806.0 | 0 | Felidae | 42495.0 | 1 | Lynx | 25600.0 | 4 | ynx canadens | 25600.0 | 4 |
| scf7180000017459 | 3502170 | 0.4011 | 0 | 51.1482 | Eukaryota | 52052.0 | 0 | Chordata | 52052.0 | 0 | Carnivora | 52052.0 | 0 | Felidae | 43745.0 | 1 | Felis | 24297.0 | 4 | Felis catus | 24297.0 | 4 |
| scf7180000023940 | 298440 | 0.3925 | 0 | 50.6393 | Eukaryota | 77464.0 | 0 | Chordata | 77464.0 | 0 | Carnivora | 77464.0 | 0 | Felidae | 77464.0 | 0 | Lynx | 63056.0 | 1 | ynx canadens | 63056.0 | 1 |
| scf7180000020760 | 87355 | 0.3917 | 0 | 53.7139 | Eukaryota | 44701.0 | 0 | Chordata | 44701.0 | 0 | Carnivora | 44701.0 | 0 | Felidae | 44701.0 | 0 | Felis | 44701.0 | 0 | Felis catus | 44701.0 | 0 |
| scf7180000021616 | 54782 | 0.3721 | 0 | 24.742 | Eukaryota | 47892.0 | 0 | Chordata | 47892.0 | 0 | Carnivora | 47892.0 | 0 | Felidae | 47892.0 | 0 | Felis | 28780.0 | 1 | Felis catus | 28780.0 | 1 |
| scf7180000023978 | 940316 | 0.4025 | 0 | 51.4845 | Eukaryota | 44652.0 | 0 | Chordata | 44652.0 | 0 | Carnivora | 44652.0 | 0 | Felidae | 44652.0 | 0 | Panthera | 18116.0 | 2 | anthera pardu | 18116.0 | 2 |
| scf7180000022446 | 202477 | 0.5473 | 0 | 48.3762 | Eukaryota | 11670.0 | 0 | Chordata | 11670.0 | 0 | Carnivora | 11670.0 | 0 | Felidae | 11670.0 | 0 | Felis | 4796.0 | 2 | Felis catus | 4796.0 | 2 |
| scf7180000019106 | 498915 | 0.4852 | 0 | 51.5557 | Eukaryota | 26038.0 | 0 | Chordata | 26038.0 | 0 | Carnivora | 26038.0 | 0 | Felidae | 23871.0 | 1 | Felis | 23871.0 | 1 | Felis catus | 23871.0 | 1 |
| scf7180000018307 | 950847 | 0.4169 | 0 | 51.1203 | Eukaryota | 46809.0 | 0 | Chordata | 46809.0 | 0 | Carnivora | 46809.0 | 0 | Felidae | 46809.0 | 0 | Felis | 46809.0 | 0 | Felis catus | 46809.0 | 0 |
| scf7180000023584 | 62032 | 0.4312 | 0 | 692.1379 | Eukaryota | 30363.0 | 0 | Chordata | 30363.0 | 0 | Carnivora | 30363.0 | 0 | Felidae | 30363.0 | 0 | Felis | 30363.0 | 0 | Felis catus | 30363.0 | 0 |
| scf7180000020143 | 198094 | 0.443 | 0 | 24.5137 | Eukaryota | 31514.0 | 0 | Chordata | 31514.0 | 0 | Carnivora | 18467.0 | 2 | Suidae | 7702.0 | 6 | Sus | 7702.0 | 7 | Sus scrofa | 7702.0 | 7 |
| scf7180000022616 | 17692 | 0.4255 | 0 | 26.6425 | Eukaryota | 3819.0 | 0 | Chordata | 3819.0 | 0 | Primates | 2085.0 | 1 | Hominidae | 2085.0 | 1 | Felis | 1488.0 | 3 | Felis catus | 1488.0 | 3 |
| scf7180000023000 | 26646 | 0.5328 | 0 | 150.3922 | Eukaryota | 12246.0 | 0 | Chordata | 12246.0 | 0 | Carnivora | 12246.0 | 0 | Felidae | 12246.0 | 0 | Felis | 12246.0 | 0 | Felis catus | 12246.0 | 0 |
| scf7180000022996 | 34146 | 0.4723 | 0 | 236.3241 | Eukaryota | 24287.0 | 0 | Chordata | 24287.0 | 0 | Carnivora | 24287.0 | 0 | Felidae | 24287.0 | 0 | Felis | 24287.0 | 0 | Felis catus | 24287.0 | 0 |
| scf7180000017449 | 4092222 | 0.359 | 0 | 50.0395 | Eukaryota | 54090.0 | 0 | Chordata | 54090.0 | 0 | Carnivora | 54090.0 | 0 | Felidae | 54090.0 | 0 | Panthera | 54090.0 | 0 | anthera pardu | 54090.0 | 0 |
| scf7180000023254 | 33227 | 0.3911 | 0 | 289.7945 | Eukaryota | 45519.0 | 0 | Chordata | 45519.0 | 0 | Carnivora | 45519.0 | 0 | Felidae | 45519.0 | 0 | Felis | 45519.0 | 0 | Felis catus | 45519.0 | 0 |
| scf7180000021187 | 91257 | 0.4219 | 0 | 35.8213 | Eukaryota | 13537.0 | 0 | Chordata | 13537.0 | 0 | Carnivora | 13537.0 | 0 | Felidae | 11403.0 | 2 | Felis | 4344.0 | 6 | Felis catus | 4344.0 | 7 |
| scf7180000018687 | 706699 | 0.3654 | 0 | 50.6915 | Eukaryota | 30645.0 | 0 | Chordata | 30645.0 | 0 | Carnivora | 30645.0 | 0 | Felidae | 30645.0 | 0 | Felis | 27542.0 | 1 | Felis catus | 27542.0 | 1 |
| scf7180000018989 | 502112 | 0.3725 | 0 | 49.8329 | Eukaryota | 46876.0 | 0 | Chordata | 46876.0 | 0 | Carnivora | 46876.0 | 0 | Felidae | 46876.0 | 0 | Felis | 46876.0 | 0 | Felis catus | 46876.0 | 0 |
| scf7180000018639 | 819050 | 0.4675 | 0 | 52.3195 | Eukaryota | 69284.0 | 0 | Chordata | 69284.0 | 0 | Carnivora | 60673.0 | 2 | Felidae | 47590.0 | 3 | Felis | 30725.0 | 5 | Felis catus | 30725.0 | 5 |
| scf7180000020983 | 94172 | 0.4238 | 0 | 51.9504 | Eukaryota | 16419.0 | 0 | Chordata | 16419.0 | 0 | Carnivora | 13634.0 | 1 | Felidae | 6950.0 | 2 | Canis | 6684.0 | 3 | Canis lupus | 6684.0 | 3 |
| scf7180000021137 | 92381 | 0.393 | 0 | 24.6677 | Eukaryota | 45031.0 | 0 | Chordata | 45031.0 | 0 | Carnivora | 45031.0 | 0 | Felidae | 45031.0 | 0 | Felis | 45031.0 | 0 | Felis catus | 45031.0 | 0 |
| scf7180000021396 | 52984 | 0.3938 | 0 | 28.0994 | Eukaryota | 42397.0 | 0 | Chordata | 42397.0 | 0 | Carnivora | 35405.0 | 1 | Felidae | 13961.0 | 5 | Canis | 8168.0 | 8 | Canis lupus | 8168.0 | 8 |
| scf7180000022749 | 7594 | 0.4259 | 0 | 23.596 | Eukaryota | 4404.0 | 0 | Chordata | 4404.0 | 0 | Carnivora | 4404.0 | 0 | Felidae | 3547.0 | 1 | Felis | 3547.0 | 1 | Felis catus | 3547.0 | 1 |
| scf7180000024067 | 19999 | 0.3962 | 0 | 24.6458 | Eukaryota | 15378.0 | 0 | Chordata | 15378.0 | 0 | Carnivora | 15378.0 | 0 | Felidae | 15378.0 | 0 | Felis | 13848.0 | 1 | Felis catus | 13848.0 | 1 |
| scf7180000021845 | 59251 | 0.5813 | 0 | 47.337 | Eukaryota | 2872.0 | 0 | Chordata | 2872.0 | 0 | Carnivora | 2872.0 | 0 | Felidae | 2006.0 | 2 | Felis | 883.0 | 5 | Felis catus | 883.0 | 5 |
| scf7180000019505 | 391019 | 0.3747 | 0 | 50.1414 | Eukaryota | 43827.0 | 0 | Chordata | 43827.0 | 0 | Carnivora | 43827.0 | 0 | Felidae | 43827.0 | 0 | Felis | 43827.0 | 0 | Felis catus | 43827.0 | 0 |
| scf7180000022968 | 31415 | 0.5469 | 0 | 365.6837 | Eukaryota | 5821.0 | 0 | Chordata | 5821.0 | 0 | Carnivora | 5821.0 | 0 | Felidae | 5248.0 | 1 | Felis | 3434.0 | 2 | Felis catus | 3434.0 | 2 |
| scf7180000023032 | 19767 | 0.5599 | 0 | 142.0226 | Eukaryota | 6413.0 | 0 | Chordata | 6413.0 | 0 | Carnivora | 6413.0 | 0 | Felidae | 6413.0 | 0 | Felis | 2721.0 | 2 | Felis catus | 2721.0 | 2 |
| scf7180000019643 | 17598 | 0.4456 | 0 | 26.4754 | Eukaryota | 29703.0 | 0 | Chordata | 29703.0 | 0 | Carnivora | 29703.0 | 0 | Felidae | 26550.0 | 1 | Panthera | 8866.0 | 3 | ynx canadens | 8850.0 | 4 |
| scf7180000019504 | 304431 | 0.3467 | 0 | 48.3635 | Eukaryota | 42828.0 | 0 | Chordata | 42828.0 | 0 | Carnivora | 42828.0 | 0 | Felidae | 42828.0 | 0 | Felis | 42828.0 | 0 | Felis catus | 42828.0 | 0 |
| scf7180000023948 | 955982 | 0.3814 | 0 | 50.2525 | Eukaryota | 46412.0 | 0 | Chordata | 46412.0 | 0 | Carnivora | 46412.0 | 0 | Felidae | 46412.0 | 0 | Felis | 46412.0 | 0 | Felis catus | 46412.0 | 0 |
| scf7180000020007 | 161653 | 0.5242 | 0 | 48.6809 | Eukaryota | 25359.0 | 0 | Chordata | 25359.0 | 0 | Carnivora | 25359.0 | 0 | Felidae | 25359.0 | 0 | Lynx | 12955.0 | 3 | ynx canadens | 12955.0 | 3 |
| scf7180000017894 | 1418676 | 0.3555 | 0 | 49.6268 | Eukaryota | 110154.0 | 0 | Chordata | 110154.0 | 0 | Carnivora | 110154.0 | 0 | Felidae | 43497.0 | 3 | Zalophus | 26014.0 | 6 | phus californi | 26014.0 | 6 |
| scf7180000018394 | 770401 | 0.4096 | 0 | 50.7879 | Eukaryota | 50765.0 | 0 | Chordata | 50765.0 | 0 | Carnivora | 50765.0 | 0 | Felidae | 50765.0 | 0 | Acinonyx | 35588.0 | 3 | cinonyx jubat | 35588.0 | 3 |
| scf7180000018618 | 907979 | 0.3669 | 100 | 48.8958 | Eukaryota | 74537.0 | 0 | Chordata | 74537.0 | 0 | Carnivora | 74537.0 | 0 | Ursidae | 23910.0 | 5 | Ursus | 15830.0 | 7 | Canis lupus | 14082.0 | 8 |
| scf7180000020997 | 51599 | 0.3958 | 0 | 26.6472 | Eukaryota | 24449.0 | 0 | Chordata | 24449.0 | 0 | Carnivora | 24449.0 | 0 | Felidae | 22046.0 | 1 | Felis | 19702.0 | 2 | Felis catus | 19702.0 | 2 |
| scf7180000024086 | 354420 | 0.4238 | 0 | 49.7746 | Eukaryota | 42841.0 | 0 | Chordata | 42841.0 | 0 | Carnivora | 42841.0 | 0 | Felidae | 42841.0 | 0 | Felis | 42841.0 | 0 | Felis catus | 42841.0 | 0 |
| scf7180000019153 | 343223 | 0.4537 | 0 | 53.359 | Eukaryota | 32825.0 | 0 | Chordata | 32825.0 | 0 | Primates | 18060.0 | 1 | Hominidae | 18060.0 | 2 | Homo | 18060.0 | 2 | Homo sapiens | 18060.0 | 2 |
| scf7180000023100 | 37103 | 0.5324 | 0 | 188.1494 | Eukaryota | 7053.0 | 0 | Chordata | 7053.0 | 0 | Carnivora | 7053.0 | 0 | Felidae | 7053.0 | 0 | Felis | 3003.0 | 2 | Felis catus | 3003.0 | 2 |
| scf7180000018901 | 498825 | 0.5338 | 0 | 50.366 | Eukaryota | 46135.0 | 0 | Chordata | 46135.0 | 0 | Carnivora | 46135.0 | 0 | Felidae | 46135.0 | 0 | Lynx | 32363.0 | 2 | ynx canadens | 32363.0 | 2 |
| scf7180000022477 | 32240 | 0.4278 | 0 | 56.2281 | Eukaryota | 5446.0 | 0 | Chordata | 5446.0 | 0 | Carnivora | 5446.0 | 0 | Felidae | 5446.0 | 0 | Felis | 4806.0 | 1 | Felis catus | 4806.0 | 1 |
| scf7180000017647 | 1703532 | 0.391 | 0 | 50.7761 | Eukaryota | 43940.0 | 0 | Chordata | 43940.0 | 0 | Carnivora | 43940.0 | 0 | Felidae | 43940.0 | 0 | Felis | 43940.0 | 0 | Felis catus | 43940.0 | 0 |
| scf7180000023220 | 32361 | 0.5646 | 0 | 191.2385 | Eukaryota | 3974.0 | 0 | Chordata | 3974.0 | 0 | Carnivora | 3974.0 | 0 | Felidae | 3707.0 | 1 | Acinonyx | 1896.0 | 3 | cinonyx jubat | 1896.0 | 3 |
| scf7180000020565 | 208366 | 0.368 | 0 | 25.2794 | Eukaryota | 13255.0 | 0 | Chordata | 13255.0 | 0 | Primates | 11476.0 | 1 | Hominidae | 11476.0 | 1 | Homo | 9885.0 | 2 | Homo sapiens | 9885.0 | 2 |
| scf7180000018916 | 108043 | 0.3868 | 0 | 43.5937 | Eukaryota | 45554.0 | 0 | Chordata | 45554.0 | 0 | Carnivora | 45554.0 | 0 | Felidae | 45554.0 | 0 | Felis | 45554.0 | 0 | Felis catus | 45554.0 | 0 |
| scf7180000021052 | 151708 | 0.4889 | 0 | 48.6983 | Eukaryota | 15076.0 | 1 | Chordata | 15076.0 | 1 | Primates | 8676.0 | 4 | Hominidae | 8676.0 | 4 | Homo | 8676.0 | 5 | Homo sapiens | 8676.0 | 5 |
| scf7180000018180 | 991026 | 0.3668 | 0 | 49.775 | Eukaryota | 46245.0 | 0 | Chordata | 46245.0 | 0 | Carnivora | 46245.0 | 0 | Felidae | 46245.0 | 0 | Felis | 46245.0 | 0 | Felis catus | 46245.0 | 0 |
| scf7180000022741 | 12178 | 0.39 | 0 | 26.6947 | Eukaryota | 3145.0 | 0 | Chordata | 3145.0 | 0 | Primates | 1394.0 | 2 | Hominidae | 1394.0 | 2 | Felis | 1019.0 | 3 | Felis catus | 1019.0 | 3 |
| scf7180000022645 | 3248 | 0.3658 | 0 | 21.2217 | Eukaryota | 2566.0 | 0 | Chordata | 2566.0 | 0 | Carnivora | 2566.0 | 0 | Felidae | 2566.0 | 0 | Felis | 1795.0 | 1 | Felis catus | 1795.0 | 1 |
| scf7180000022703 | 7245 | 0.6685 | 0 | 412.6832 | Eukaryota | 1418.0 | 0 | Chordata | 1277.0 | 1 | Carnivora | 1277.0 | 1 | Otariidae | 536.0 | 4 | Eumetopias | 402.0 | 5 | etopias juba | 402.0 | 5 |
| scf7180000021512 | 116154 | 0.3723 | 0 | 25.5181 | Eukaryota | 9992.0 | 0 | Chordata | 9992.0 | 0 | Carnivora | 6247.0 | 3 | Canidae | 3495.0 | 4 | Canis | 3495.0 | 5 | Canis lupus | 3495.0 | 5 |
| scf7180000022343 | 13795 | 0.3448 | 0 | 24.2474 | Eukaryota | 8072.0 | 0 | Chordata | 8072.0 | 0 | Carnivora | 8072.0 | 0 | Felidae | 4951.0 | 3 | Panthera | 1633.0 | 7 | unresolved | 833.0 | 9 |
| scf7180000019633 | 34317 | 0.3844 | 0 | 43.3591 | Eukaryota | 45067.0 | 0 | Chordata | 45067.0 | 0 | Carnivora | 45067.0 | 0 | Felidae | 45067.0 | 0 | Felis | 45067.0 | 0 | Felis catus | 45067.0 | 0 |
| scf7180000021810 | 21449 | 0.3974 | 0 | 27.7048 | Eukaryota | 12138.0 | 0 | Chordata | 12138.0 | 0 | Carnivora | 8885.0 | 3 | Felidae | 8885.0 | 5 | Felis | 8672.0 | 6 | Felis catus | 8672.0 | 6 |
| scf7180000018130 | 1427765 | 0.3736 | 0 | 50.4296 | Eukaryota | 63684.0 | 0 | Chordata | 63684.0 | 0 | Carnivora | 63684.0 | 0 | Felidae | 28473.0 | 5 | Suricata | 9631.0 | 8 | ricata suricat | 9631.0 | 8 |
| scf7180000019121 | 469635 | 0.3947 | 0 | 49.6794 | Eukaryota | 26162.0 | 0 | Chordata | 26162.0 | 0 | Carnivora | 26162.0 | 0 | Felidae | 26162.0 | 0 | Felis | 23521.0 | 1 | Felis catus | 23521.0 | 1 |
| scf7180000019252 | 479492 | 0.4053 | 0 | 50.6686 | Eukaryota | 13301.0 | 0 | Chordata | 13301.0 | 0 | Carnivora | 13301.0 | 0 | Felidae | 13301.0 | 0 | Felis | 10659.0 | 1 | Felis catus | 10659.0 | 1 |
| scf7180000021343 | 57664 | 0.6258 | 0 | 52.3073 | Eukaryota | 12742.0 | 0 | Chordata | 12742.0 | 0 | Carnivora | 12742.0 | 0 | Herpestidae | 8322.0 | 1 | Suricata | 8322.0 | 2 | ricata suricat | 8322.0 | 2 |
| scf7180000018827 | 17334 | 0.3477 | 0 | 23.9801 | Eukaryota | 5746.0 | 0 | Chordata | 5746.0 | 0 | Artiodactyla | 2678.0 | 2 | Bovidae | 2678.0 | 3 | Felis | 1787.0 | 4 | Felis catus | 1787.0 | 4 |
| scf7180000018576 | 632913 | 0.3918 | 0 | 48.9815 | Eukaryota | 56348.0 | 0 | Chordata | 56348.0 | 0 | Carnivora | 56348.0 | 0 | Felidae | 26459.0 | 3 | Suricata | 15090.0 | 6 | ricata suricat | 15090.0 | 8 |
| scf7180000019679 | 119773 | 0.4157 | 0 | 50.7194 | Eukaryota | 42400.0 | 0 | Chordata | 42400.0 | 0 | Carnivora | 42400.0 | 0 | Felidae | 42400.0 | 0 | Felis | 38200.0 | 1 | Felis catus | 38200.0 | 1 |
| scf7180000020528 | 11854 | 0.538 | 0 | 156.0272 | Eukaryota | 11987.0 | 0 | Chordata | 11987.0 | 0 | Carnivora | 11987.0 | 0 | Felidae | 11987.0 | 0 | Felis | 10795.0 | 1 | Felis catus | 10795.0 | 1 |
| scf7180000021238 | 72484 | 0.4701 | 0 | 24.9976 | Eukaryota | 14347.0 | 0 | Chordata | 14347.0 | 0 | Carnivora | 14347.0 | 0 | Felidae | 14347.0 | 0 | Felis | 9376.0 | 2 | Felis catus | 9376.0 | 2 |
| scf7180000021943 | 30066 | 0.5191 | 0 | 197.7525 | Eukaryota | 12915.0 | 0 | Chordata | 12915.0 | 0 | Carnivora | 12915.0 | 0 | Felidae | 12915.0 | 0 | Felis | 12915.0 | 0 | Felis catus | 12915.0 | 0 |
| scf7180000022549 | 12489 | 0.4322 | 0 | 27.7144 | Eukaryota | 10956.0 | 0 | Chordata | 10956.0 | 0 | Carnivora | 10956.0 | 0 | Felidae | 10956.0 | 0 | Felis | 9879.0 | 1 | Felis catus | 9879.0 | 1 |
| scf7180000019969 | 2336889 | 0.494 | 0 | 55.0524 | Eukaryota | 47188.0 | 0 | Chordata | 47188.0 | 0 | Carnivora | 47188.0 | 0 | Felidae | 47188.0 | 0 | Felis | 47188.0 | 0 | Felis catus | 47188.0 | 0 |
| scf7180000019206 | 407121 | 0.405 | 0 | 50.4848 | Eukaryota | 47500.0 | 0 | Chordata | 47500.0 | 0 | Carnivora | 47500.0 | 0 | Felidae | 47500.0 | 0 | Felis | 47500.0 | 0 | Felis catus | 47500.0 | 0 |
| scf7180000021077 | 86504 | 0.3578 | 0 | 23.4983 | Eukaryota | 45275.0 | 0 | Chordata | 45275.0 | 0 | Carnivora | 45275.0 | 0 | Felidae | 45275.0 | 0 | Felis | 45275.0 | 0 | Felis catus | 45275.0 | 0 |
| scf7180000017638 | 130911 | 0.3836 | 0 | 48.6029 | Eukaryota | 46819.0 | 0 | Chordata | 46819.0 | 0 | Carnivora | 46819.0 | 0 | Felidae | 46819.0 | 0 | Felis | 46819.0 | 0 | Felis catus | 46819.0 | 0 |
| scf7180000021637 | 20395 | 0.5634 | 0 | 47.5821 | Eukaryota | 2685.0 | 0 | Chordata | 2685.0 | 0 | Carnivora | 2685.0 | 0 | Felidae | 2420.0 | 1 | Felis | 2015.0 | 2 | Felis catus | 2015.0 | 2 |
| scf7180000018038 | 2377990 | 0.3793 | 0 | 50.9359 | Eukaryota | 59024.0 | 0 | Chordata | 59024.0 | 0 | Carnivora | 59024.0 | 0 | Felidae | 59024.0 | 0 | Lynx | 30511.0 | 1 | ynx canadens | 30511.0 | 1 |
| scf7180000021299 | 3901 | 0.5245 | 0 | 26.9005 | no-hit | 0.0 | 0 | no-hit | 0.0 | 0 | no-hit | 0.0 | 0 | no-hit | 0.0 | 0 | no-hit | 0.0 | 0 | no-hit | 0.0 | 0 |
| scf7180000017880 | 1506526 | 0.4172 | 0 | 50.263 | Eukaryota | 46878.0 | 0 | Chordata | 46878.0 | 0 | Carnivora | 46878.0 | 0 | Felidae | 46878.0 | 0 | Felis | 46878.0 | 0 | Felis catus | 46878.0 | 0 |
| scf7180000018890 | 474628 | 0.4471 | 0 | 47.8758 | Eukaryota | 51505.0 | 0 | Chordata | 51505.0 | 0 | Carnivora | 51505.0 | 0 | Felidae | 51505.0 | 0 | Felis | 51505.0 | 0 | Felis catus | 51505.0 | 0 |
| scf7180000023763 | 1234523 | 0.4015 | 0 | 50.1395 | Eukaryota | 71582.0 | 0 | Chordata | 71582.0 | 0 | Carnivora | 71582.0 | 0 | Felidae | 35101.0 | 2 | Panthera | 35101.0 | 2 | anthera pardu | 24267.0 | 3 |
| scf7180000021201 | 56914 | 0.5735 | 0 | 65.1814 | Eukaryota | 7320.0 | 0 | Chordata | 7320.0 | 0 | Carnivora | 7320.0 | 0 | Felidae | 7320.0 | 0 | Felis | 5313.0 | 2 | Felis catus | 5313.0 | 2 |
| scf7180000021301 | 40079 | 0.4415 | 0 | 47.6111 | Eukaryota | 10143.0 | 0 | Chordata | 10143.0 | 0 | Primates | 3945.0 | 4 | Hominidae | 3945.0 | 5 | Homo | 2955.0 | 6 | Homo sapiens | 2955.0 | 6 |
| scf7180000020180 | 11602 | 0.411 | 0 | 27.2093 | Eukaryota | 15148.0 | 0 | Chordata | 15148.0 | 0 | Carnivora | 15148.0 | 0 | Felidae | 10941.0 | 3 | Felis | 4331.0 | 6 | Felis catus | 4331.0 | 7 |
| scf7180000023805 | 1626043 | 0.4125 | 0 | 52.1164 | Eukaryota | 47534.0 | 0 | Chordata | 47534.0 | 0 | Carnivora | 47534.0 | 0 | Felidae | 47534.0 | 0 | Felis | 47534.0 | 0 | Felis catus | 47534.0 | 0 |
| scf7180000021552 | 64444 | 0.4802 | 0 | 53.3177 | Eukaryota | 29479.0 | 0 | Chordata | 29479.0 | 0 | Carnivora | 29479.0 | 0 | Felidae | 29479.0 | 0 | Felis | 26553.0 | 1 | Felis catus | 26553.0 | 1 |
| scf7180000022863 | 16716 | 0.4193 | 0 | 151.8726 | Eukaryota | 46080.0 | 0 | Chordata | 46080.0 | 0 | Carnivora | 46080.0 | 0 | Felidae | 46080.0 | 0 | Felis | 46080.0 | 0 | Felis catus | 46080.0 | 0 |
| scf7180000019397 | 45158 | 0.4078 | 0 | 122.4935 | Eukaryota | 45974.0 | 0 | Chordata | 45974.0 | 0 | Carnivora | 45974.0 | 0 | Felidae | 45974.0 | 0 | Felis | 45974.0 | 0 | Felis catus | 45974.0 | 0 |
| scf7180000022621 | 24321 | 0.5419 | 0 | 49.8383 | Eukaryota | 5838.0 | 0 | Chordata | 5838.0 | 0 | Carnivora | 5838.0 | 0 | Felidae | 5838.0 | 0 | Prionailurus | 5838.0 | 0 | ailurus benga | 5838.0 | 0 |
| scf7180000022268 | 8162 | 0.4296 | 0 | 26.5934 | Eukaryota | 2475.0 | 0 | Chordata | 2475.0 | 0 | Carnivora | 2475.0 | 0 | Felidae | 2205.0 | 1 | Felis | 1683.0 | 2 | Felis catus | 1683.0 | 2 |
| scf7180000017855 | 1648562 | 0.3861 | 0 | 49.8975 | Eukaryota | 47842.0 | 0 | Chordata | 47842.0 | 0 | Carnivora | 47842.0 | 0 | Felidae | 47842.0 | 0 | Felis | 47842.0 | 0 | Felis catus | 47842.0 | 0 |
| scf7180000017979 | 1337212 | 0.3821 | 0 | 51.4554 | Eukaryota | 47819.0 | 0 | Chordata | 47819.0 | 0 | Carnivora | 47819.0 | 0 | Felidae | 47819.0 | 0 | Felis | 47819.0 | 0 | Felis catus | 47819.0 | 0 |
| scf7180000019428 | 56515 | 0.5162 | 0 | 59.0026 | Eukaryota | 19128.0 | 0 | Chordata | 19128.0 | 0 | Carnivora | 19128.0 | 0 | Felidae | 15792.0 | 1 | Felis | 6828.0 | 5 | Felis catus | 6828.0 | 5 |
| scf7180000021602 | 31459 | 0.3703 | 0 | 23.059 | Eukaryota | 33254.0 | 0 | Chordata | 33254.0 | 0 | Carnivora | 33254.0 | 0 | Felidae | 33254.0 | 0 | Felis | 29964.0 | 1 | Felis catus | 29964.0 | 1 |
| scf7180000021099 | 56666 | 0.359 | 0 | 24.4435 | Eukaryota | 5578.0 | 0 | Chordata | 5578.0 | 0 | Carnivora | 5578.0 | 0 | Canidae | 2792.0 | 1 | Canis | 2792.0 | 1 | Canis lupus | 2792.0 | 1 |
| scf7180000024225 | 430871 | 0.3619 | 0 | 62.4494 | Eukaryota | 36699.0 | 0 | Chordata | 36699.0 | 0 | Carnivora | 36699.0 | 0 | Felidae | 36699.0 | 0 | Felis | 36699.0 | 0 | Felis catus | 36699.0 | 0 |
| scf7180000022051 | 65799 | 0.4837 | 0 | 50.1942 | Eukaryota | 17019.0 | 0 | Chordata | 17019.0 | 0 | Carnivora | 17019.0 | 0 | Felidae | 15270.0 | 1 | Felis | 5070.0 | 4 | Felis catus | 5070.0 | 4 |
| scf7180000022603 | 7987 | 0.4234 | 0 | 52.8492 | Eukaryota | 3670.0 | 0 | Chordata | 3670.0 | 0 | Carnivora | 2102.0 | 1 | Felidae | 2102.0 | 1 | Felis | 1576.0 | 2 | Felis catus | 1576.0 | 2 |
| scf7180000023927 | 1026655 | 0.3715 | 0 | 50.3677 | Eukaryota | 45948.0 | 0 | Chordata | 45948.0 | 0 | Carnivora | 45948.0 | 0 | Felidae | 45948.0 | 0 | Felis | 41344.0 | 1 | Felis catus | 41344.0 | 1 |
| scf7180000023512 | 21328 | 0.4642 | 0 | 386.3314 | Eukaryota | 2592.0 | 0 | Chordata | 2592.0 | 0 | Artiodactyla | 1699.0 | 1 | Bovidae | 1699.0 | 2 | Ovis | 1699.0 | 3 | vis canadens | 1699.0 | 3 |
| scf7180000024271 | 3746027 | 0.3991 | 100 | 51.3203 | Eukaryota | 47415.0 | 0 | Chordata | 47415.0 | 0 | Carnivora | 47415.0 | 0 | Felidae | 47415.0 | 0 | Felis | 40864.0 | 1 | Felis catus | 40864.0 | 1 |
| scf7180000023132 | 46952 | 0.5251 | 0 | 251.7828 | Eukaryota | 12822.0 | 0 | Chordata | 12822.0 | 0 | Carnivora | 12822.0 | 0 | Felidae | 12822.0 | 0 | Felis | 12822.0 | 0 | Felis catus | 12822.0 | 0 |
| scf7180000018821 | 422653 | 0.4658 | 0 | 53.5812 | Eukaryota | 68010.0 | 0 | Chordata | 68010.0 | 0 | Carnivora | 68010.0 | 0 | Felidae | 68010.0 | 0 | Panthera | 34030.0 | 1 | anthera pardu | 34030.0 | 1 |
| scf7180000021495 | 40429 | 0.3936 | 0 | 25.4268 | Eukaryota | 35145.0 | 0 | Chordata | 35145.0 | 0 | Carnivora | 35145.0 | 0 | Felidae | 20739.0 | 1 | Suricata | 14406.0 | 2 | ricata suricat | 14406.0 | 2 |
| scf7180000021207 | 65816 | 0.3728 | 0 | 24.503 | Eukaryota | 15335.0 | 0 | Chordata | 15335.0 | 0 | Carnivora | 7207.0 | 2 | Felidae | 7207.0 | 2 | Felis | 5779.0 | 4 | Felis catus | 5779.0 | 4 |
| scf7180000019660 | 599372 | 0.4185 | 100 | 51.7683 | Eukaryota | 48178.0 | 0 | Chordata | 48178.0 | 0 | Carnivora | 48178.0 | 0 | Felidae | 37910.0 | 1 | Felis | 37910.0 | 1 | Felis catus | 37910.0 | 1 |
| scf7180000019469 | 348439 | 0.3572 | 0 | 48.666 | Eukaryota | 45877.0 | 0 | Chordata | 45877.0 | 0 | Carnivora | 45877.0 | 0 | Felidae | 45877.0 | 0 | Felis | 45877.0 | 0 | Felis catus | 45877.0 | 0 |
| scf7180000018912 | 177234 | 0.3693 | 0 | 47.1474 | Eukaryota | 46128.0 | 0 | Chordata | 46128.0 | 0 | Carnivora | 46128.0 | 0 | Felidae | 46128.0 | 0 | Felis | 46128.0 | 0 | Felis catus | 46128.0 | 0 |
| scf7180000018547 | 763184 | 0.4597 | 0 | 50.9752 | Eukaryota | 38190.0 | 0 | Chordata | 38190.0 | 0 | Carnivora | 38190.0 | 0 | Felidae | 38190.0 | 0 | Panthera | 38190.0 | 0 | anthera pardu | 38190.0 | 0 |
| scf7180000022463 | 6248 | 0.3868 | 0 | 24.5398 | Eukaryota | 14434.0 | 0 | Chordata | 14434.0 | 0 | Carnivora | 14434.0 | 0 | Felidae | 14434.0 | 0 | Felis | 12999.0 | 1 | Felis catus | 12999.0 | 1 |
| scf7180000021664 | 36128 | 0.576 | 0 | 48.0533 | Eukaryota | 3199.0 | 0 | Chordata | 3199.0 | 0 | Carnivora | 3199.0 | 0 | Felidae | 2266.0 | 1 | Felis | 1878.0 | 2 | Felis catus | 1878.0 | 2 |
| scf7180000019788 | 226290 | 0.3493 | 0 | 47.9524 | Eukaryota | 41619.0 | 0 | Chordata | 41619.0 | 0 | Carnivora | 41619.0 | 0 | Felidae | 41619.0 | 0 | Felis | 41619.0 | 0 | Felis catus | 41619.0 | 0 |
| scf7180000017857 | 592161 | 0.3473 | 0 | 48.1949 | Eukaryota | 23791.0 | 0 | Chordata | 23791.0 | 0 | Carnivora | 23791.0 | 0 | Felidae | 23791.0 | 0 | Felis | 21406.0 | 1 | Felis catus | 21406.0 | 1 |
| scf7180000023384 | 27641 | 0.4397 | 0 | 102.8589 | Eukaryota | 5198.0 | 0 | Chordata | 5198.0 | 0 | Carnivora | 3148.0 | 1 | Bovidae | 2050.0 | 2 | Ovis | 2050.0 | 2 | vis canadens | 2050.0 | 2 |
| scf7180000021814 | 14128 | 0.3855 | 0 | 25.8481 | Eukaryota | 9353.0 | 0 | Chordata | 9353.0 | 0 | Carnivora | 9353.0 | 0 | Felidae | 8431.0 | 1 | Felis | 8431.0 | 1 | Felis catus | 8431.0 | 1 |
| scf7180000020955 | 82837 | 0.3743 | 0 | 25.4393 | Eukaryota | 9498.0 | 0 | Chordata | 9498.0 | 0 | Carnivora | 5911.0 | 3 | Canidae | 3762.0 | 5 | Canis | 3762.0 | 5 | Canis lupus | 3762.0 | 5 |
| scf7180000022506 | 15335 | 0.6104 | 0 | 50.8551 | Eukaryota | 4142.0 | 0 | Chordata | 4142.0 | 0 | Carnivora | 4142.0 | 0 | Felidae | 4142.0 | 0 | Felis | 2113.0 | 2 | Felis catus | 2113.0 | 2 |
| scf7180000021395 | 63487 | 0.384 | 0 | 23.5521 | Eukaryota | 39982.0 | 0 | Chordata | 39982.0 | 0 | Carnivora | 39982.0 | 0 | Felidae | 39982.0 | 0 | Felis | 39982.0 | 0 | Felis catus | 39982.0 | 0 |
| scf7180000017629 | 2085332 | 0.3573 | 0 | 49.8954 | Eukaryota | 49387.0 | 0 | Chordata | 49387.0 | 0 | Carnivora | 49387.0 | 0 | Felidae | 49387.0 | 0 | Felis | 43272.0 | 1 | Felis catus | 43272.0 | 1 |
| scf7180000021782 | 33098 | 0.4317 | 0 | 695.9391 | Eukaryota | 2368.0 | 0 | Chordata | 2368.0 | 0 | Carnivora | 2368.0 | 0 | Felidae | 2368.0 | 0 | Felis | 1918.0 | 1 | Felis catus | 1918.0 | 1 |
| scf7180000024274 | 637595 | 0.5826 | 100 | 56.2161 | Eukaryota | 46260.0 | 0 | Chordata | 46260.0 | 0 | Carnivora | 46260.0 | 0 | Felidae | 46260.0 | 0 | Felis | 41670.0 | 1 | Felis catus | 41670.0 | 1 |
| scf7180000022499 | 8893 | 0.3847 | 0 | 23.4751 | Eukaryota | 24672.0 | 0 | Chordata | 24672.0 | 0 | Carnivora | 24672.0 | 0 | Felidae | 24672.0 | 0 | Felis | 22413.0 | 1 | Felis catus | 22413.0 | 1 |
| scf7180000021484 | 52035 | 0.4702 | 0 | 30.0412 | Eukaryota | 35631.0 | 0 | Chordata | 35631.0 | 0 | Carnivora | 35631.0 | 0 | Felidae | 35631.0 | 0 | Felis | 35631.0 | 0 | Felis catus | 35631.0 | 0 |
| scf7180000018119 | 1071202 | 0.3692 | 0 | 50.5222 | Eukaryota | 46895.0 | 0 | Chordata | 46895.0 | 0 | Carnivora | 46895.0 | 0 | Felidae | 46895.0 | 0 | Felis | 46895.0 | 0 | Felis catus | 46895.0 | 0 |
| scf7180000022364 | 8969 | 0.4013 | 0 | 22.8208 | Eukaryota | 3862.0 | 0 | Chordata | 3862.0 | 0 | Carnivora | 3862.0 | 0 | Felidae | 3553.0 | 1 | Felis | 2513.0 | 2 | Felis catus | 2513.0 | 2 |
| scf7180000021551 | 6633 | 0.5197 | 0 | 77.6111 | no-hit | 0.0 | 0 | no-hit | 0.0 | 0 | no-hit | 0.0 | 0 | no-hit | 0.0 | 0 | no-hit | 0.0 | 0 | no-hit | 0.0 | 0 |
| scf7180000020556 | 12184 | 0.4673 | 0 | 23.0454 | Eukaryota | 3188.0 | 0 | Chordata | 3188.0 | 0 | Primates | 2222.0 | 1 | Hominidae | 1092.0 | 4 | Homo | 1092.0 | 5 | Homo sapiens | 1092.0 | 5 |
| scf7180000023962 | 781071 | 0.4743 | 0 | 51.0161 | Eukaryota | 50783.0 | 0 | Chordata | 50783.0 | 0 | Primates | 28513.0 | 2 | Hominidae | 28513.0 | 3 | Homo | 23080.0 | 5 | Homo sapiens | 23080.0 | 6 |
| scf7180000023012 | 50145 | 0.4098 | 0 | 268.4292 | Eukaryota | 6361.0 | 0 | Chordata | 6361.0 | 0 | Carnivora | 6361.0 | 0 | Felidae | 6361.0 | 0 | Felis | 5734.0 | 1 | Felis catus | 5734.0 | 1 |

| | | | | | | | | | | | | | | | | | | | |
|---|---|---|---|---|---|---|---|---|---|---|---|---|---|---|---|---|---|---|---|
| scf7180000024213 | 41150 | 0.4175 | 0 | 25.4203 | Eukaryota | 12496.0 | 0 | Chordata | 12496.0 | 0 | Carnivora | 9324.0 | 1 | Mustelidae | 4610.0 | 6 | Mustela | 3716.0 | 8 | ustela putori | 3716.0 | 8 |
| scf7180000023867 | 722233 | 0.3711 | 0 | 49.3738 | Eukaryota | 47205.0 | 0 | Chordata | 47205.0 | 0 | Carnivora | 47205.0 | 0 | Felidae | 47205.0 | 0 | Felis | 47205.0 | 0 | Felis catus | 47205.0 | 0 |
| scf7180000022199 | 16806 | 0.6268 | 0 | 46.6248 | Eukaryota | 2993.0 | 0 | Chordata | 2993.0 | 0 | Carnivora | 2993.0 | 0 | Felidae | 2084.0 | 1 | Acinonyx | 1192.0 | 3 | cinonyx jubat | 1192.0 | 4 |
| scf7180000022299 | 19993 | 0.3616 | 0 | 24.2694 | Eukaryota | 27909.0 | 0 | Chordata | 27909.0 | 0 | Carnivora | 27909.0 | 0 | Felidae | 27909.0 | 0 | Felis | 27909.0 | 0 | Felis catus | 27909.0 | 0 |
| scf7180000018241 | 1572428 | 0.4668 | 0 | 52.5454 | Eukaryota | 24956.0 | 0 | Chordata | 24956.0 | 0 | Carnivora | 24956.0 | 0 | Felidae | 17525.0 | 1 | Panthera | 12616.0 | 3 | Panthera tigri | 7536.0 | 4 |
| scf7180000017991 | 2539008 | 0.4006 | 0 | 51.6799 | Eukaryota | 61746.0 | 0 | Chordata | 61746.0 | 0 | Carnivora | 61746.0 | 0 | Felidae | 53454.0 | 1 | Panthera | 34278.0 | 2 | anthera pardu | 21406.0 | 3 |
| scf7180000017904 | 1337674 | 0.4278 | 0 | 51.4282 | Eukaryota | 46618.0 | 0 | Chordata | 46618.0 | 0 | Carnivora | 46618.0 | 0 | Felidae | 46618.0 | 0 | Felis | 46618.0 | 0 | Felis catus | 46618.0 | 0 |
| scf7180000023818 | 867631 | 0.4872 | 0 | 51.8767 | Eukaryota | 66149.0 | 0 | Chordata | 66149.0 | 0 | Carnivora | 66149.0 | 0 | Felidae | 50526.0 | 2 | Felis | 25136.0 | 5 | Felis catus | 25136.0 | 6 |
| scf7180000018616 | 759541 | 0.3677 | 0 | 49.8124 | Eukaryota | 46652.0 | 0 | Chordata | 46652.0 | 0 | Carnivora | 46652.0 | 0 | Felidae | 46652.0 | 0 | Felis | 36460.0 | 1 | Felis catus | 36460.0 | 1 |
| scf7180000019418 | 350597 | 0.3843 | 0 | 49.1374 | Eukaryota | 58120.0 | 0 | Chordata | 58120.0 | 0 | Carnivora | 58120.0 | 0 | Felidae | 33016.0 | 1 | Suricata | 25104.0 | 2 | ricata suricat | 25104.0 | 2 |
| scf7180000021424 | 21651 | 0.3894 | 0 | 22.4509 | Eukaryota | 32925.0 | 0 | Chordata | 32925.0 | 0 | Carnivora | 32925.0 | 0 | Felidae | 32925.0 | 0 | Felis | 32925.0 | 0 | Felis catus | 32925.0 | 0 |
| scf7180000023801 | 2183427 | 0.428 | 0 | 51.4267 | Eukaryota | 66894.0 | 0 | Chordata | 66894.0 | 0 | Carnivora | 66894.0 | 0 | Felidae | 61795.0 | 1 | Felis | 41539.0 | 3 | Felis catus | 41539.0 | 3 |
| scf7180000020322 | 111777 | 0.523 | 0 | 26.7177 | Eukaryota | 27470.0 | 0 | Chordata | 27470.0 | 0 | Carnivora | 23606.0 | 1 | Otariidae | 11690.0 | 3 | Eumetopias | 6930.0 | 5 | netopias juba | 6930.0 | 5 |
| scf7180000021000 | 74715 | 0.3872 | 0 | 25.1986 | Eukaryota | 43331.0 | 0 | Chordata | 43331.0 | 0 | Carnivora | 43331.0 | 0 | Felidae | 43331.0 | 0 | Felis | 43331.0 | 0 | Felis catus | 43331.0 | 0 |
| scf7180000019436 | 12397 | 0.3577 | 0 | 26.7022 | Eukaryota | 13001.0 | 0 | Chordata | 13001.0 | 0 | Carnivora | 13001.0 | 0 | Herpestidae | 11780.0 | 1 | Suricata | 11780.0 | 1 | ricata suricat | 11780.0 | 1 |
| scf7180000020784 | 139255 | 0.4487 | 0 | 51.2287 | Eukaryota | 36985.0 | 0 | Chordata | 36985.0 | 0 | Carnivora | 36985.0 | 0 | Felidae | 36985.0 | 0 | Panthera | 14776.0 | 2 | anthera pardu | 14776.0 | 2 |
| scf7180000022393 | 11225 | 0.3614 | 0 | 23.7686 | Eukaryota | 6853.0 | 0 | Chordata | 6853.0 | 0 | Primates | 3378.0 | 2 | Hominidae | 3378.0 | 5 | Pan | 1700.0 | 6 | an troglodyte | 1700.0 | 6 |
| scf7180000019169 | 1204360 | 0.3767 | 0 | 50.115 | Eukaryota | 48053.0 | 0 | Chordata | 48053.0 | 0 | Carnivora | 48053.0 | 0 | Felidae | 48053.0 | 0 | Felis | 48053.0 | 0 | Felis catus | 48053.0 | 0 |
| scf7180000023833 | 892531 | 0.3894 | 0 | 50.16 | Eukaryota | 46598.0 | 0 | Chordata | 46598.0 | 0 | Carnivora | 46598.0 | 0 | Felidae | 46598.0 | 0 | Felis | 46598.0 | 0 | Felis catus | 46598.0 | 0 |
| scf7180000020827 | 5550 | 0.6114 | 0 | 34.6719 | Eukaryota | 10010.0 | 0 | Chordata | 10010.0 | 0 | Carnivora | 9014.0 | 1 | Felidae | 6859.0 | 3 | Lynx | 4827.0 | 5 | ynx canadens | 4827.0 | 5 |
| scf7180000018050 | 409679 | 0.3931 | 0 | 51.5276 | Eukaryota | 47826.0 | 0 | Chordata | 47826.0 | 0 | Carnivora | 47826.0 | 0 | Felidae | 47826.0 | 0 | Felis | 47826.0 | 0 | Felis catus | 47826.0 | 0 |
| scf7180000019902 | 196006 | 0.4234 | 0 | 50.6135 | Eukaryota | 47699.0 | 0 | Chordata | 47699.0 | 0 | Carnivora | 47699.0 | 0 | Felidae | 47699.0 | 0 | Felis | 47699.0 | 0 | Felis catus | 47699.0 | 0 |
| scf7180000024220 | 152289 | 0.3761 | 0 | 51.0014 | Eukaryota | 46929.0 | 0 | Chordata | 46929.0 | 0 | Carnivora | 46929.0 | 0 | Felidae | 46929.0 | 0 | Felis | 46929.0 | 0 | Felis catus | 46929.0 | 0 |
| scf7180000018742 | 1341592 | 0.419 | 100 | 51.8699 | Eukaryota | 48187.0 | 0 | Chordata | 48187.0 | 0 | Carnivora | 48187.0 | 0 | Felidae | 48187.0 | 0 | Felis | 48187.0 | 0 | Felis catus | 48187.0 | 0 |
| scf7180000022606 | 21375 | 0.388 | 0 | 24.7034 | Eukaryota | 39533.0 | 0 | Chordata | 39533.0 | 0 | Carnivora | 39533.0 | 0 | Felidae | 39533.0 | 0 | Felis | 39533.0 | 0 | Felis catus | 39533.0 | 0 |
| scf7180000018418 | 848006 | 0.4184 | 0 | 50.1912 | Eukaryota | 68276.0 | 0 | Chordata | 68276.0 | 0 | Carnivora | 68276.0 | 0 | Felidae | 68276.0 | 0 | Acinonyx | 40980.0 | 1 | cinonyx jubat | 40980.0 | 1 |
| scf7180000021282 | 80081 | 0.3797 | 0 | 25.2748 | Eukaryota | 8980.0 | 0 | Chordata | 8980.0 | 0 | Primates | 4613.0 | 4 | Hominidae | 4613.0 | 4 | Homo | 3514.0 | 6 | Homo sapiens | 3514.0 | 6 |
| scf7180000024111 | 160523 | 0.6243 | 0 | 47.3269 | Eukaryota | 9310.0 | 0 | Chordata | 9310.0 | 0 | Carnivora | 7672.0 | 1 | Felidae | 4069.0 | 4 | Equus | 1638.0 | 8 | Equus caballu | 1638.0 | 8 |
| scf7180000022351 | 39152 | 0.3598 | 0 | 86.1392 | Eukaryota | 22899.0 | 0 | Chordata | 22899.0 | 0 | Carnivora | 22899.0 | 0 | Felidae | 22899.0 | 0 | Felis | 20599.0 | 1 | Felis catus | 20599.0 | 1 |
| scf7180000018120 | 414214 | 0.3881 | 0 | 50.4221 | Eukaryota | 47738.0 | 0 | Chordata | 47738.0 | 0 | Carnivora | 47738.0 | 0 | Felidae | 47738.0 | 0 | Felis | 47738.0 | 0 | Felis catus | 47738.0 | 0 |
| scf7180000018942 | 442538 | 0.4174 | 0 | 49.0892 | Eukaryota | 42744.0 | 0 | Chordata | 42744.0 | 0 | Carnivora | 42744.0 | 0 | Otariidae | 17138.0 | 3 | Callorhinus | 8592.0 | 8 | llorhinus ursir | 8592.0 | 8 |
| scf7180000023936 | 541827 | 0.4178 | 0 | 50.009 | Eukaryota | 19998.0 | 0 | Chordata | 19998.0 | 0 | Carnivora | 19998.0 | 0 | Felidae | 17763.0 | 1 | Felis | 17763.0 | 1 | Felis catus | 17763.0 | 1 |
| scf7180000017931 | 1485110 | 0.3722 | 0 | 51.0092 | Eukaryota | 58827.0 | 0 | Chordata | 58827.0 | 0 | Carnivora | 58827.0 | 0 | Felidae | 50542.0 | 1 | Felis | 32222.0 | 3 | Felis catus | 32222.0 | 3 |
| scf7180000020724 | 96384 | 0.5482 | 0 | 47.3032 | Eukaryota | 4966.0 | 0 | Chordata | 4966.0 | 0 | Carnivora | 4966.0 | 0 | Felidae | 4966.0 | 0 | Felis | 3972.0 | 1 | Felis catus | 3972.0 | 1 |
| scf7180000023895 | 627842 | 0.3488 | 0 | 49.0119 | Eukaryota | 46283.0 | 0 | Chordata | 46283.0 | 0 | Carnivora | 46283.0 | 0 | Felidae | 46283.0 | 0 | Felis | 46283.0 | 0 | Felis catus | 46283.0 | 0 |
| scf7180000018369 | 566776 | 0.3911 | 0 | 50.45 | Eukaryota | 47970.0 | 0 | Chordata | 47970.0 | 0 | Carnivora | 47970.0 | 0 | Felidae | 47970.0 | 0 | Felis | 47970.0 | 0 | Felis catus | 47970.0 | 0 |
| scf7180000018074 | 1235630 | 0.3496 | 0 | 49.1995 | Eukaryota | 56190.0 | 0 | Chordata | 56190.0 | 0 | Carnivora | 56190.0 | 0 | Ursidae | 35172.0 | 2 | Ursus | 35172.0 | 3 | Ursus arctos | 35172.0 | 3 |
| scf7180000021990 | 73723 | 0.3909 | 0 | 24.3727 | Eukaryota | 45519.0 | 0 | Chordata | 45519.0 | 0 | Carnivora | 45519.0 | 0 | Felidae | 45519.0 | 0 | Felis | 45519.0 | 0 | Felis catus | 45519.0 | 0 |
| scf7180000019251 | 357513 | 0.3951 | 0 | 52.4897 | Eukaryota | 36482.0 | 0 | Chordata | 36482.0 | 0 | Carnivora | 36482.0 | 0 | Felidae | 36482.0 | 0 | Felis | 36482.0 | 0 | Felis catus | 36482.0 | 0 |
| scf7180000022327 | 16787 | 0.4469 | 0 | 23.4641 | Eukaryota | 6154.0 | 0 | Chordata | 6154.0 | 0 | Carnivora | 6154.0 | 0 | Felidae | 6154.0 | 0 | Felis | 5544.0 | 1 | Felis catus | 5544.0 | 1 |
| scf7180000019657 | 312452 | 0.3836 | 0 | 52.0089 | Eukaryota | 47853.0 | 0 | Chordata | 47853.0 | 0 | Carnivora | 47853.0 | 0 | Otariidae | 15739.0 | 4 | Ursus | 10211.0 | 8 | phus californi | 6355.0 | 9 |
| scf7180000020919 | 67225 | 0.4028 | 0 | 49.5086 | Eukaryota | 15546.0 | 0 | Chordata | 15546.0 | 0 | Carnivora | 15546.0 | 0 | Felidae | 15546.0 | 0 | Felis | 14005.0 | 1 | Felis catus | 14005.0 | 1 |
| scf7180000022162 | 27463 | 0.406 | 0 | 2233.1614 | no-hit | 0.0 | 0 | no-hit | 0.0 | 0 | no-hit | 0.0 | 0 | no-hit | 0.0 | 0 | no-hit | 0.0 | 0 | no-hit | 0.0 | 0 |
| scf7180000023545 | 33882 | 0.427 | 0 | 190.5755 | Eukaryota | 5874.0 | 0 | Chordata | 5874.0 | 0 | Carnivora | 5874.0 | 0 | Phocidae | 2396.0 | 2 | eptonychote | 1797.0 | 4 | onychotes wec | 1797.0 | 4 |
| scf7180000018334 | 986748 | 0.3939 | 0 | 49.5841 | Eukaryota | 107281.0 | 0 | Chordata | 107281.0 | 0 | Carnivora | 107281.0 | 0 | Felidae | 107281.0 | 0 | Panthera | 51730.0 | 4 | anthera pardu | 41388.0 | 5 |
| scf7180000019851 | 213347 | 0.3889 | 0 | 50.1939 | Eukaryota | 43735.0 | 0 | Chordata | 43735.0 | 0 | Carnivora | 43735.0 | 0 | Felidae | 43735.0 | 0 | Felis | 43735.0 | 0 | Felis catus | 43735.0 | 0 |
| scf7180000017846 | 3258195 | 0.3803 | 0 | 50.9646 | Eukaryota | 57337.0 | 0 | Chordata | 57337.0 | 0 | Carnivora | 57337.0 | 0 | Felidae | 57337.0 | 0 | Felis | 38482.0 | 1 | Felis catus | 38482.0 | 1 |
| scf7180000018400 | 974134 | 0.4194 | 0 | 52.3285 | Eukaryota | 45910.0 | 0 | Chordata | 45910.0 | 0 | Carnivora | 45910.0 | 0 | Felidae | 45910.0 | 0 | Felis | 45910.0 | 0 | Felis catus | 45910.0 | 0 |
| scf7180000023776 | 412614 | 0.4596 | 0 | 48.2786 | Eukaryota | 46720.0 | 0 | Chordata | 46720.0 | 0 | Carnivora | 46720.0 | 0 | Felidae | 46720.0 | 0 | Felis | 46720.0 | 0 | Felis catus | 46720.0 | 0 |
| scf7180000017434 | 786576 | 0.5159 | 0 | 52.725 | Eukaryota | 45967.0 | 0 | Chordata | 45967.0 | 0 | Carnivora | 45967.0 | 0 | Felidae | 41547.0 | 1 | Felis | 25959.0 | 4 | Felis catus | 25959.0 | 4 |
| scf7180000019031 | 12841 | 0.4222 | 0 | 14.8217 | Eukaryota | 40855.0 | 0 | Chordata | 40855.0 | 0 | Carnivora | 40855.0 | 0 | Felidae | 40855.0 | 0 | Felis | 40855.0 | 0 | Felis catus | 40855.0 | 0 |
| scf7180000022432 | 4522 | 0.5228 | 0 | 17.8426 | no-hit | 0.0 | 0 | no-hit | 0.0 | 0 | no-hit | 0.0 | 0 | no-hit | 0.0 | 0 | no-hit | 0.0 | 0 | no-hit | 0.0 | 0 |
| scf7180000022390 | 47130 | 0.6345 | 0 | 1626.0508 | Eukaryota | 47420.0 | 0 | Chordata | 47420.0 | 0 | Carnivora | 47420.0 | 0 | Felidae | 17195.0 | 2 | Ursus | 15031.0 | 6 | Ursus arctos | 15031.0 | 6 |
| scf7180000022340 | 22066 | 0.3823 | 0 | 26.3525 | Eukaryota | 16684.0 | 0 | Chordata | 16684.0 | 0 | Carnivora | 16684.0 | 0 | Felidae | 16684.0 | 0 | Felis | 15008.0 | 1 | Felis catus | 15008.0 | 1 |
| scf7180000018201 | 232818 | 0.3555 | 0 | 47.217 | Eukaryota | 28175.0 | 0 | Chordata | 28175.0 | 0 | Carnivora | 28175.0 | 0 | Felidae | 28175.0 | 0 | Felis | 28175.0 | 0 | Felis catus | 28175.0 | 0 |
| scf7180000017854 | 446839 | 0.435 | 0 | 49.8438 | Eukaryota | 47634.0 | 0 | Chordata | 47634.0 | 0 | Carnivora | 47634.0 | 0 | Felidae | 47634.0 | 0 | Felis | 47634.0 | 0 | Felis catus | 47634.0 | 0 |
| scf7180000018780 | 1809034 | 0.3802 | 0 | 49.3601 | Eukaryota | 44701.0 | 0 | Chordata | 44701.0 | 0 | Carnivora | 44701.0 | 0 | Felidae | 44701.0 | 0 | Felis | 44701.0 | 0 | Felis catus | 44701.0 | 0 |
| scf7180000018242 | 1004651 | 0.3938 | 0 | 48.8303 | Eukaryota | 53062.0 | 0 | Chordata | 53062.0 | 0 | Carnivora | 53062.0 | 0 | Felidae | 53062.0 | 0 | Panthera | 29143.0 | 1 | Felis catus | 23919.0 | 2 |
| scf7180000018075 | 1110532 | 0.4994 | 0 | 51.0849 | Eukaryota | 47633.0 | 0 | Chordata | 47633.0 | 0 | Carnivora | 47633.0 | 0 | Felidae | 47633.0 | 0 | Felis | 47633.0 | 0 | Felis catus | 47633.0 | 0 |
| scf7180000017707 | 29334 | 0.5048 | 0 | 26.4296 | Eukaryota | 19659.0 | 0 | Chordata | 19659.0 | 0 | Carnivora | 19659.0 | 0 | Felidae | 11275.0 | 1 | Felis | 11275.0 | 2 | Felis catus | 11275.0 | 2 |
| scf7180000017499 | 3177301 | 0.4234 | 0 | 52.4071 | Eukaryota | 48727.0 | 0 | Chordata | 48727.0 | 0 | Carnivora | 48727.0 | 0 | Felidae | 29249.0 | 3 | Felis | 18683.0 | 5 | Felis catus | 18683.0 | 5 |
| scf7180000020925 | 63984 | 0.4097 | 0 | 24.6321 | Eukaryota | 44358.0 | 0 | Chordata | 44358.0 | 0 | Carnivora | 44358.0 | 0 | Felidae | 44358.0 | 0 | Felis | 44358.0 | 0 | Felis catus | 44358.0 | 0 |
| scf7180000023380 | 16933 | 0.5094 | 0 | 313.2192 | no-hit | 0.0 | 0 | no-hit | 0.0 | 0 | no-hit | 0.0 | 0 | no-hit | 0.0 | 0 | no-hit | 0.0 | 0 | no-hit | 0.0 | 0 |
| scf7180000022077 | 25584 | 0.412 | 0 | 38.8746 | Eukaryota | 4369.0 | 1 | Chordata | 4369.0 | 1 | Artiodactyla | 1794.0 | 3 | Otariidae | 1207.0 | 5 | Zalophus | 1207.0 | 5 | phus californi | 1207.0 | 5 |
| scf7180000020208 | 451420 | 0.387 | 100 | 35.7128 | Eukaryota | 42962.0 | 0 | Chordata | 42962.0 | 0 | Carnivora | 42962.0 | 0 | Felidae | 42962.0 | 0 | Felis | 42962.0 | 0 | Felis catus | 42962.0 | 0 |
| scf7180000019611 | 382073 | 0.3796 | 100 | 49.2651 | Eukaryota | 46434.0 | 0 | Chordata | 46434.0 | 0 | Carnivora | 46434.0 | 0 | Felidae | 46434.0 | 0 | Felis | 46434.0 | 0 | Felis catus | 46434.0 | 0 |
| scf7180000022827 | 21185 | 0.3879 | 0 | 282.2823 | Eukaryota | 45435.0 | 0 | Chordata | 45435.0 | 0 | Carnivora | 45435.0 | 0 | Felidae | 45435.0 | 0 | Felis | 45435.0 | 0 | Felis catus | 45435.0 | 0 |
| scf7180000019470 | 259381 | 0.4152 | 0 | 51.7654 | Eukaryota | 42946.0 | 0 | Chordata | 42946.0 | 0 | Carnivora | 42946.0 | 0 | Felidae | 42946.0 | 0 | Felis | 42946.0 | 0 | Felis catus | 42946.0 | 0 |
| scf7180000017860 | 2153131 | 0.3669 | 0 | 49.5229 | Eukaryota | 62870.0 | 0 | Chordata | 62870.0 | 0 | Carnivora | 62870.0 | 0 | Felidae | 62870.0 | 0 | Panthera | 43588.0 | 1 | anthera pardu | 43588.0 | 1 |
| scf7180000018973 | 521568 | 0.4206 | 0 | 50.7922 | Eukaryota | 42662.0 | 0 | Chordata | 42662.0 | 0 | Carnivora | 42662.0 | 0 | Felidae | 42662.0 | 0 | Felis | 38381.0 | 1 | Felis catus | 38381.0 | 1 |
| scf7180000024027 | 361103 | 0.4638 | 0 | 52.6457 | Eukaryota | 29939.0 | 0 | Chordata | 29939.0 | 0 | Carnivora | 29939.0 | 0 | Felidae | 29939.0 | 0 | Acinonyx | 12211.0 | 4 | cinonyx jubat | 12211.0 | 4 |
| scf7180000019125 | 119528 | 0.4465 | 0 | 47.3592 | Eukaryota | 29818.0 | 0 | Chordata | 29818.0 | 0 | Carnivora | 29818.0 | 0 | Felidae | 29818.0 | 0 | Lynx | 16190.0 | 3 | ynx canadens | 16190.0 | 4 |
| scf7180000024022 | 533520 | 0.4846 | 0 | 51.6433 | Eukaryota | 11992.0 | 0 | Chordata | 11992.0 | 0 | Carnivora | 11992.0 | 0 | Felidae | 11992.0 | 0 | Felis | 9607.0 | 1 | Felis catus | 9607.0 | 1 |
| scf7180000018695 | 12978 | 0.3944 | 0 | 18.9656 | Eukaryota | 28891.0 | 0 | Chordata | 28891.0 | 0 | Carnivora | 28891.0 | 0 | Felidae | 28891.0 | 0 | Felis | 26015.0 | 1 | Felis catus | 26015.0 | 1 |
| scf7180000019421 | 8065 | 0.4159 | 0 | 28.6164 | Eukaryota | 6201.0 | 0 | Chordata | 6201.0 | 0 | Carnivora | 6201.0 | 0 | Felidae | 6201.0 | 0 | Felis | 5578.0 | 1 | Felis catus | 5578.0 | 1 |
| scf7180000021016 | 75430 | 0.3642 | 0 | 24.8317 | Eukaryota | 42754.0 | 0 | Chordata | 42754.0 | 0 | Carnivora | 42754.0 | 0 | Felidae | 42754.0 | 0 | Felis | 42754.0 | 0 | Felis catus | 42754.0 | 0 |
| scf7180000023118 | 22326 | 0.5191 | 0 | 181.0944 | Eukaryota | 21930.0 | 0 | Chordata | 21930.0 | 0 | Carnivora | 21930.0 | 0 | Felidae | 21930.0 | 0 | Felis | 19717.0 | 1 | Felis catus | 19717.0 | 1 |
| scf7180000018698 | 37010 | 0.4369 | 0 | 44.9554 | Eukaryota | 8366.0 | 0 | Chordata | 8366.0 | 0 | Carnivora | 8366.0 | 0 | Felidae | 7586.0 | 1 | Felis | 7586.0 | 1 | Felis catus | 7586.0 | 1 |
| scf7180000022119 | 19019 | 0.5795 | 0 | 50.3154 | Eukaryota | 2345.0 | 0 | Chordata | 2345.0 | 0 | Carnivora | 2345.0 | 0 | Felidae | 2095.0 | 1 | Felis | 1633.0 | 2 | Felis catus | 1633.0 | 2 |
| scf7180000019911 | 188329 | 0.5911 | 0 | 47.7387 | Eukaryota | 5622.0 | 0 | Chordata | 5622.0 | 0 | Carnivora | 5622.0 | 0 | Canidae | 3204.0 | 1 | Canis | 3204.0 | 3 | Canis lupus | 3204.0 | 3 |
| scf7180000022932 | 31552 | 0.4013 | 0 | 174.8255 | Eukaryota | 6974.0 | 0 | Chordata | 6974.0 | 0 | Carnivora | 6974.0 | 0 | Felidae | 6974.0 | 0 | Felis | 6974.0 | 0 | Felis catus | 6974.0 | 0 |
| scf7180000020791 | 54856 | 0.4093 | 0 | 51.4318 | Eukaryota | 33659.0 | 0 | Chordata | 33659.0 | 0 | Carnivora | 33659.0 | 0 | Felidae | 33659.0 | 0 | Felis | 30246.0 | 1 | Felis catus | 30246.0 | 1 |
| scf7180000017790 | 562144 | 0.3756 | 0 | 47.3787 | Eukaryota | 47141.0 | 0 | Chordata | 47141.0 | 0 | Carnivora | 47141.0 | 0 | Felidae | 24329.0 | 1 | Suricata | 22812.0 | 2 | ricata suricat | 22812.0 | 2 |
| scf7180000022673 | 8679 | 0.3634 | 0 | 23.8686 | Eukaryota | 15255.0 | 0 | Chordata | 15255.0 | 0 | Carnivora | 15255.0 | 0 | Felidae | 15255.0 | 0 | Felis | 14082.0 | 1 | Felis catus | 14082.0 | 1 |
| scf7180000020310 | 103522 | 0.3611 | 0 | 49.1375 | Eukaryota | 25615.0 | 0 | Chordata | 25615.0 | 0 | Carnivora | 25615.0 | 0 | Felidae | 25615.0 | 0 | Felis | 25615.0 | 0 | Felis catus | 25615.0 | 0 |
| scf7180000017466 | 1281154 | 0.3726 | 0 | 49.1541 | Eukaryota | 47440.0 | 0 | Chordata | 47440.0 | 0 | Carnivora | 47440.0 | 0 | Felidae | 47440.0 | 0 | Felis | 47440.0 | 0 | Felis catus | 47440.0 | 0 |
| scf7180000018671 | 699196 | 0.4145 | 0 | 51.1544 | Eukaryota | 35903.0 | 0 | Chordata | 35903.0 | 0 | Carnivora | 35903.0 | 0 | Felidae | 14295.0 | 5 | Zalophus | 7114.0 | 8 | phus californi | 7114.0 | 8 |
| scf7180000023972 | 1906925 | 0.4713 | 0 | 51.2699 | Eukaryota | 91227.0 | 0 | Chordata | 91227.0 | 0 | Carnivora | 91227.0 | 0 | Felidae | 91227.0 | 0 | Lynx | 36516.0 | 2 | ynx canadens | 36516.0 | 2 |
| scf7180000020790 | 85073 | 0.3932 | 0 | 25.6085 | Eukaryota | 43829.0 | 0 | Chordata | 43829.0 | 0 | Carnivora | 43829.0 | 0 | Felidae | 43829.0 | 0 | Felis | 39478.0 | 1 | Felis catus | 39478.0 | 1 |
| scf7180000022578 | 66385 | 0.4092 | 0 | 514.5899 | no-hit | 0.0 | 0 | no-hit | 0.0 | 0 | no-hit | 0.0 | 0 | no-hit | 0.0 | 0 | no-hit | 0.0 | 0 | no-hit | 0.0 | 0 |
| scf7180000023777 | 2305191 | 0.3576 | 0 | 49.1576 | Eukaryota | 46150.0 | 0 | Chordata | 46150.0 | 0 | Carnivora | 46150.0 | 0 | Felidae | 46150.0 | 0 | Felis | 46150.0 | 0 | Felis catus | 46150.0 | 0 |
| scf7180000018329 | 790448 | 0.3815 | 0 | 50.563 | Eukaryota | 52760.0 | 0 | Chordata | 52760.0 | 0 | Carnivora | 52760.0 | 0 | Felidae | 52760.0 | 0 | Felis | 40333.0 | 2 | Felis catus | 40333.0 | 2 |
| scf7180000019139 | 484717 | 0.4102 | 0 | 50.8174 | Eukaryota | 97061.0 | 0 | Chordata | 97061.0 | 0 | Carnivora | 97061.0 | 0 | Felidae | 97061.0 | 0 | Acinonyx | 58379.0 | 1 | cinonyx jubat | 58379.0 | 1 |
| scf7180000019533 | 384357 | 0.3473 | 0 | 48.0162 | Eukaryota | 47567.0 | 0 | Chordata | 47567.0 | 0 | Carnivora | 47567.0 | 0 | Felidae | 47567.0 | 0 | Felis | 47567.0 | 0 | Felis catus | 47567.0 | 0 |
| scf7180000019989 | 420771 | 0.4471 | 0 | 50.3172 | Eukaryota | 36114.0 | 0 | Chordata | 36114.0 | 0 | Carnivora | 36114.0 | 0 | Felidae | 36114.0 | 0 | Felis | 36114.0 | 0 | Felis catus | 36114.0 | 0 |
| scf7180000019458 | 1444263 | 0.4393 | 0 | 49.8041 | Eukaryota | 42282.0 | 0 | Chordata | 42282.0 | 0 | Carnivora | 42282.0 | 0 | Felidae | 35274.0 | 1 | Felis | 11750.0 | 5 | Felis catus | 11750.0 | 5 |
| scf7180000022314 | 20638 | 0.4753 | 0 | 24.5063 | Eukaryota | 45274.0 | 0 | Chordata | 45274.0 | 0 | Carnivora | 45274.0 | 0 | Felidae | 45274.0 | 0 | Felis | 45274.0 | 0 | Felis catus | 45274.0 | 0 |
| scf7180000018090 | 279503 | 0.3839 | 0 | 49.4006 | Eukaryota | 46197.0 | 0 | Chordata | 46197.0 | 0 | Carnivora | 46197.0 | 0 | Felidae | 46197.0 | 0 | Felis | 46197.0 | 0 | Felis catus | 46197.0 | 0 |
| scf7180000021402 | 31872 | 0.3548 | 0 | 49.1158 | Eukaryota | 12100.0 | 0 | Chordata | 12100.0 | 0 | Carnivora | 12100.0 | 0 | Felidae | 12100.0 | 0 | Felis | 10905.0 | 1 | Felis catus | 10905.0 | 1 |
| scf7180000021044 | 54409 | 0.3846 | 0 | 23.1103 | Eukaryota | 20378.0 | 0 | Chordata | 20378.0 | 0 | Carnivora | 20378.0 | 0 | Felidae | 20378.0 | 0 | Felis | 16277.0 | 1 | Felis catus | 16277.0 | 1 |
| scf7180000020880 | 117149 | 0.4213 | 0 | 25.5126 | Eukaryota | 6920.0 | 0 | Chordata | 6920.0 | 0 | Primates | 3477.0 | 3 | Hominidae | 2152.0 | 6 | Sus | 1504.0 | 7 | Sus scrofa | 1504.0 | 7 |
| scf7180000018974 | 539173 | 0.4398 | 0 | 50.204 | Eukaryota | 46411.0 | 0 | Chordata | 46411.0 | 0 | Carnivora | 46411.0 | 0 | Felidae | 46411.0 | 0 | Felis | 46411.0 | 0 | Felis catus | 46411.0 | 0 |
| scf7180000017918 | 197377 | 0.4593 | 0 | 49.7453 | Eukaryota | 19142.0 | 0 | Chordata | 19142.0 | 0 | Carnivora | 19142.0 | 0 | Felidae | 19142.0 | 0 | Felis | 16829.0 | 1 | Felis catus | 16829.0 | 1 |
| scf7180000018195 | 13527 | 0.3761 | 0 | 19.3211 | Eukaryota | 35055.0 | 0 | Chordata | 35055.0 | 0 | Carnivora | 35055.0 | 0 | Felidae | 35055.0 | 0 | Felis | 35055.0 | 0 | Felis catus | 35055.0 | 0 |
| scf7180000018611 | 448129 | 0.4083 | 0 | 48.765 | Eukaryota | 46832.0 | 0 | Chordata | 46832.0 | 0 | Carnivora | 46832.0 | 0 | Felidae | 46832.0 | 0 | Felis | 46832.0 | 0 | Felis catus | 46832.0 | 0 |
| scf7180000017797 | 3043950 | 0.5089 | 100 | 51.4131 | Eukaryota | 46245.0 | 0 | Chordata | 46245.0 | 0 | Carnivora | 46245.0 | 0 | Felidae | 36623.0 | 1 | Felis | 29601.0 | 2 | Felis catus | 29601.0 | 2 |
| scf7180000021725 | 59086 | 0.3611 | 0 | 23.6644 | Eukaryota | 40485.0 | 0 | Chordata | 40485.0 | 0 | Carnivora | 40485.0 | 0 | Felidae | 40485.0 | 0 | Felis | 40485.0 | 0 | Felis catus | 40485.0 | 0 |
| scf7180000018545 | 96572 | 0.3646 | 0 | 44.0101 | Eukaryota | 34625.0 | 0 | Chordata | 34625.0 | 0 | Carnivora | 34625.0 | 0 | Felidae | 34625.0 | 0 | Felis | 34625.0 | 0 | Felis catus | 34625.0 | 0 |
| scf7180000020164 | 186737 | 0.3721 | 0 | 48.4545 | Eukaryota | 26762.0 | 0 | Chordata | 26762.0 | 0 | Carnivora | 26762.0 | 0 | Felidae | 26762.0 | 0 | Panthera | 26762.0 | 0 | anthera pardu | 26762.0 | 0 |
| scf7180000019524 | 408693 | 0.369 | 0 | 50.5252 | Eukaryota | 46440.0 | 0 | Chordata | 46440.0 | 0 | Carnivora | 46440.0 | 0 | Felidae | 46440.0 | 0 | Felis | 46440.0 | 0 | Felis catus | 46440.0 | 0 |
| scf7180000019089 | 426108 | 0.4138 | 0 | 52.441 | Eukaryota | 37217.0 | 0 | Chordata | 37217.0 | 0 | Carnivora | 37217.0 | 0 | Felidae | 32916.0 | 1 | Panthera | 19830.0 | 3 | anthera pardu | 19830.0 | 3 |
| scf7180000022078 | 20600 | 0.3795 | 0 | 23.7345 | Eukaryota | 19529.0 | 0 | Chordata | 19529.0 | 0 | Carnivora | 19529.0 | 0 | Felidae | 19529.0 | 0 | Felis | 17634.0 | 1 | Felis catus | 17634.0 | 1 |
| scf7180000021268 | 49247 | 0.5639 | 0 | 51.4197 | Eukaryota | 16952.0 | 0 | Chordata | 16952.0 | 0 | Carnivora | 16952.0 | 0 | Felidae | 16952.0 | 0 | Felis | 16952.0 | 0 | Felis catus | 16952.0 | 0 |
| scf7180000018487 | 717466 | 0.4559 | 0 | 50.8395 | Eukaryota | 17075.0 | 1 | Chordata | 17075.0 | 1 | Carnivora | 14938.0 | 2 | Felidae | 14938.0 | 2 | Felis | 13085.0 | 3 | Felis catus | 13085.0 | 3 |
| scf7180000018135 | 1175926 | 0.4412 | 0 | 52.4023 | Eukaryota | 46530.0 | 0 | Chordata | 46530.0 | 0 | Carnivora | 36307.0 | 1 | Felidae | 27453.0 | 2 | Panthera | 22225.0 | 4 | anthera pardu | 13349.0 | 5 |
| scf7180000022236 | 25037 | 0.4205 | 0 | 24.728 | Eukaryota | 10529.0 | 0 | Chordata | 10529.0 | 0 | Carnivora | 7637.0 | 1 | Felidae | 6791.0 | 2 | Felis | 6791.0 | 2 | Felis catus | 6791.0 | 2 |
| scf7180000021826 | 17577 | 0.375 | 0 | 23.895 | Eukaryota | 11911.0 | 0 | Chordata | 11911.0 | 0 | Carnivora | 11911.0 | 0 | Felidae | 11911.0 | 0 | Felis | 10736.0 | 1 | Felis catus | 10736.0 | 1 |
| scf7180000017750 | 1594112 | 0.4414 | 0 | 52.2621 | Eukaryota | 46928.0 | 0 | Chordata | 46928.0 | 0 | Carnivora | 46928.0 | 0 | Felidae | 46928.0 | 0 | Felis | 46928.0 | 0 | Felis catus | 46928.0 | 0 |
| scf7180000020388 | 33121 | 0.4102 | 0 | 22.4997 | Eukaryota | 9035.0 | 0 | Chordata | 9035.0 | 0 | Carnivora | 9035.0 | 0 | Felidae | 9035.0 | 0 | Felis | 9035.0 | 0 | Felis catus | 9035.0 | 0 |
| scf7180000019086 | 429945 | 0.3979 | 0 | 50.9613 | Eukaryota | 48276.0 | 0 | Chordata | 48276.0 | 0 | Carnivora | 48276.0 | 0 | Felidae | 48276.0 | 0 | Felis | 48276.0 | 0 | Felis catus | 48276.0 | 0 |
| scf7180000018982 | 246884 | 0.5722 | 0 | 49.3152 | Eukaryota | 37460.0 | 0 | Chordata | 37460.0 | 0 | Carnivora | 37460.0 | 0 | Felidae | 37460.0 | 0 | Acinonyx | 37460.0 | 0 | cinonyx jubat | 37460.0 | 0 |
| scf7180000024098 | 962759 | 0.3999 | 0 | 51.4498 | Eukaryota | 52060.0 | 0 | Chordata | 52060.0 | 0 | Carnivora | 52060.0 | 0 | Felidae | 52060.0 | 0 | Felis | 20190.0 | 3 | Felis catus | 20190.0 | 3 |
| scf7180000023765 | 358278 | 0.3736 | 0 | 49.8244 | Eukaryota | 30441.0 | 0 | Chordata | 30441.0 | 0 | Carnivora | 30441.0 | 0 | Felidae | 30441.0 | 0 | Felis | 27351.0 | 1 | Felis catus | 27351.0 | 1 |
| scf7180000022482 | 4025 | 0.3928 | 0 | 26.5637 | Eukaryota | 2466.0 | 0 | Chordata | 2466.0 | 0 | Carnivora | 1307.0 | 2 | Felidae | 1111.0 | 3 | Felis | 952.0 | 4 | Felis catus | 952.0 | 4 |
| scf7180000018536 | 788970 | 0.3556 | 0 | 51.0866 | Eukaryota | 87395.0 | 0 | Chordata | 87395.0 | 0 | Carnivora | 64273.0 | 1 | Felidae | 64273.0 | 1 | Felis | 41519.0 | 3 | Felis catus | 41519.0 | 3 |
| scf7180000023825 | 1383016 | 0.377 | 0 | 50.6138 | Eukaryota | 47483.0 | 0 | Chordata | 47483.0 | 0 | Carnivora | 47483.0 | 0 | Felidae | 47483.0 | 0 | Felis | 47483.0 | 0 | Felis catus | 47483.0 | 0 |
| scf7180000019027 | 501435 | 0.4398 | 0 | 50.2302 | Eukaryota | 48555.0 | 0 | Chordata | 48555.0 | 0 | Carnivora | 48555.0 | 0 | Felidae | 48555.0 | 0 | Felis | 43760.0 | 1 | Felis catus | 43760.0 | 1 |
| scf7180000022247 | 12158 | 0.4263 | 0 | 42.0977 | Eukaryota | 5198.0 | 0 | Chordata | 5198.0 | 0 | Carnivora | 5198.0 | 0 | Felidae | 5198.0 | 0 | Felis | 5198.0 | 0 | Felis catus | 5198.0 | 0 |
| scf7180000017538 | 2536175 | 0.4304 | 0 | 51.719 | Eukaryota | 46158.0 | 0 | Chordata | 46158.0 | 0 | Carnivora | 46158.0 | 0 | Felidae | 46158.0 | 0 | Felis | 46158.0 | 0 | Felis catus | 46158.0 | 0 |
| scf7180000022080 | 28503 | 0.4428 | 0 | 52.0645 | Eukaryota | 6987.0 | 0 | Chordata | 6987.0 | 0 | Carnivora | 3891.0 | 3 | Canidae | 1850.0 | 5 | Canis | 1850.0 | 6 | Canis lupus | 1850.0 | 6 |
| scf7180000020871 | 106355 | 0.4209 | 0 | 24.4902 | Eukaryota | 31349.0 | 0 | Chordata | 31349.0 | 0 | Carnivora | 31349.0 | 0 | Felidae | 31349.0 | 0 | Felis | 28132.0 | 1 | Felis catus | 28132.0 | 1 |
| scf7180000021079 | 59535 | 0.5998 | 0 | 43.8104 | Eukaryota | 20071.0 | 0 | Chordata | 20071.0 | 0 | Carnivora | 20071.0 | 0 | Felidae | 20071.0 | 0 | Felis | 18074.0 | 1 | Felis catus | 18074.0 | 1 |
| scf7180000021139 | 89232 | 0.4261 | 0 | 26.2455 | Eukaryota | 6364.0 | 0 | Chordata | 6364.0 | 0 | Primates | 2385.0 | 2 | Hominidae | 2385.0 | 3 | Homo | 2385.0 | 4 | Homo sapiens | 2385.0 | 4 |
| scf7180000018712 | 157017 | 0.3645 | 0 | 45.4392 | Eukaryota | 47370.0 | 0 | Chordata | 47370.0 | 0 | Carnivora | 47370.0 | 0 | Felidae | 47370.0 | 0 | Felis | 47370.0 | 0 | Felis catus | 47370.0 | 0 |
| scf7180000021491 | 15444 | 0.3618 | 0 | 39.6229 | Eukaryota | 4757.0 | 0 | Chordata | 4757.0 | 0 | Carnivora | 2949.0 | 2 | Felidae | 2949.0 | 2 | Felis | 1947.0 | 3 | Felis catus | 1947.0 | 3 |
| scf7180000023183 | 68722 | 0.496 | 0 | 1226.5143 | Eukaryota | 10878.0 | 0 | Chordata | 10878.0 | 0 | Carnivora | 10878.0 | 0 | Felidae | 10878.0 | 0 | Felis | 8681.0 | 1 | Felis catus | 8681.0 | 1 |
| scf7180000020525 | 118235 | 0.4181 | 0 | 25.7608 | Eukaryota | 8631.0 | 0 | Chordata | 8631.0 | 0 | Carnivora | 7629.0 | 1 | Felidae | 7629.0 | 1 | Felis | 6738.0 | 2 | Felis catus | 6738.0 | 2 |
| scf7180000020368 | 118377 | 0.365 | 0 | 25.0859 | Eukaryota | 38745.0 | 0 | Chordata | 38745.0 | 0 | Carnivora | 38745.0 | 0 | Felidae | 38745.0 | 0 | Felis | 34820.0 | 1 | Felis catus | 34820.0 | 1 |
| scf7180000021714 | 40072 | 0.3848 | 0 | 49.3611 | Eukaryota | 5482.0 | 0 | Chordata | 5482.0 | 0 | Carnivora | 2627.0 | 3 | Felidae | 2157.0 | 4 | Felis | 2157.0 | 6 | Felis catus | 2157.0 | 6 |
| scf7180000021627 | 60373 | 0.4178 | 0 | 25.8041 | Eukaryota | 9165.0 | 0 | Chordata | 9165.0 | 0 | Primates | 5909.0 | 2 | Hominidae | 4763.0 | 4 | Homo | 4763.0 | 4 | Homo sapiens | 4763.0 | 4 |
| scf7180000019017 | 393305 | 0.4154 | 0 | 52.0823 | Eukaryota | 106384.0 | 0 | Chordata | 106384.0 | 0 | Carnivora | 106384.0 | 0 | Felidae | 106384.0 | 0 | Panthera | 42598.0 | 3 | anthera pardu | 31974.0 | 4 |
| scf7180000020647 | 84738 | 0.5978 | 0 | 51.5553 | Eukaryota | 4595.0 | 0 | Chordata | 4595.0 | 0 | Carnivora | 4595.0 | 0 | Felidae | 4595.0 | 0 | Panthera | 3292.0 | 2 | anthera pardu | 3292.0 | 2 |
| scf7180000023995 | 472841 | 0.3924 | 0 | 49.9623 | Eukaryota | 29034.0 | 0 | Chordata | 29034.0 | 0 | Carnivora | 29034.0 | 0 | Felidae | 29034.0 | 0 | Felis | 22828.0 | 2 | Felis catus | 22828.0 | 2 |
| scf7180000021278 | 105140 | 0.3536 | 0 | 25.6737 | Eukaryota | 26650.0 | 0 | Chordata | 26650.0 | 0 | Primates | 18571.0 | 2 | Hominidae | 18571.0 | 2 | Homo | 13917.0 | 3 | Homo sapiens | 13917.0 | 3 |
| scf7180000022186 | 23941 | 0.365 | 0 | 25.4206 | Eukaryota | 28771.0 | 0 | Chordata | 28771.0 | 0 | Carnivora | 28771.0 | 0 | Felidae | 28771.0 | 0 | Felis | 25825.0 | 1 | Felis catus | 25825.0 | 1 |
| scf7180000022249 | 45275 | 0.4923 | 0 | 49.0676 | Eukaryota | 8388.0 | 0 | Chordata | 8388.0 | 0 | Carnivora | 8388.0 | 0 | Felidae | 4194.0 | 3 | Enhydra | 2529.0 | 5 | Enhydra lutris | 2529.0 | 6 |
| scf7180000021064 | 68883 | 0.4568 | 0 | 25.1547 | Eukaryota | 9263.0 | 0 | Chordata | 9263.0 | 0 | Carnivora | 9263.0 | 0 | Felidae | 9263.0 | 0 | Felis | 8337.0 | 1 | Felis catus | 8337.0 | 1 |
| scf7180000023162 | 17946 | 0.5908 | 0 | 156.8475 | Eukaryota | 2370.0 | 0 | Chordata | 2370.0 | 0 | Primates | 1074.0 | 3 | Muridae | 575.0 | 4 | Homo | 566.0 | 5 | Homo sapiens | 566.0 | 5 |
| scf7180000023199 | 36569 | 0.5388 | 0 | 263.5745 | Eukaryota | 7745.0 | 0 | Chordata | 7745.0 | 0 | Carnivora | 7745.0 | 0 | Felidae | 7255.0 | 1 | Felis | 4654.0 | 2 | Felis catus | 4654.0 | 2 |
| scf7180000022196 | 21042 | 0.5058 | 0 | 218.9801 | Eukaryota | 4372.0 | 0 | Chordata | 4372.0 | 0 | Carnivora | 4372.0 | 0 | Felidae | 3928.0 | 1 | Felis | 2636.0 | 2 | Felis catus | 2636.0 | 2 |
| scf7180000022049 | 6527 | 0.4655 | 0 | 19.7937 | Eukaryota | 2534.0 | 0 | Chordata | 2534.0 | 0 | Carnivora | 2534.0 | 0 | Felidae | 2534.0 | 0 | Felis | 2534.0 | 0 | Felis catus | 2534.0 | 0 |
| scf7180000018992 | 871788 | 0.4179 | 0 | 50.7733 | Eukaryota | 43857.0 | 0 | Chordata | 43857.0 | 0 | Carnivora | 43857.0 | 0 | Felidae | 43857.0 | 0 | Felis | 26325.0 | 2 | Felis catus | 26325.0 | 2 |
| scf7180000017692 | 2389182 | 0.4602 | 0 | 52.2821 | Eukaryota | 48622.0 | 0 | Chordata | 48622.0 | 0 | Carnivora | 48622.0 | 0 | Felidae | 44060.0 | 1 | Felis | 24181.0 | 3 | Felis catus | 24181.0 | 4 |
| scf7180000020673 | 118594 | 0.52 | 0 | 50.0487 | Eukaryota | 8313.0 | 0 | Chordata | 8313.0 | 0 | Carnivora | 8313.0 | 0 | Felidae | 8313.0 | 0 | Felis | 8313.0 | 0 | Felis catus | 8313.0 | 0 |
| scf7180000017924 | 80702 | 0.4625 | 0 | 41.9845 | Eukaryota | 37989.0 | 0 | Chordata | 37989.0 | 0 | Carnivora | 37989.0 | 0 | Felidae | 37989.0 | 0 | Felis | 37989.0 | 0 | Felis catus | 37989.0 | 0 |
| scf7180000018730 | 10817 | 0.3686 | 0 | 21.6677 | Eukaryota | 11535.0 | 0 | Chordata | 11535.0 | 0 | Carnivora | 11535.0 | 0 | Felidae | 11535.0 | 0 | Felis | 8094.0 | 1 | Felis catus | 8094.0 | 1 |
| scf7180000019898 | 194877 | 0.4312 | 0 | 25.1394 | Eukaryota | 45983.0 | 0 | Chordata | 45983.0 | 0 | Carnivora | 45983.0 | 0 | Felidae | 45983.0 | 0 | Felis | 45983.0 | 0 | Felis catus | 45983.0 | 0 |
| scf7180000022731 | 5024 | 0.5247 | 0 | 48.5733 | no-hit | 0.0 | 0 | no-hit | 0.0 | 0 | no-hit | 0.0 | 0 | no-hit | 0.0 | 0 | no-hit | 0.0 | 0 | no-hit | 0.0 | 0 |
| scf7180000020065 | 116102 | 0.3964 | 0 | 49.5348 | Eukaryota | 46154.0 | 0 | Chordata | 46154.0 | 0 | Carnivora | 46154.0 | 0 | Felidae | 46154.0 | 0 | Felis | 46154.0 | 0 | Felis catus | 46154.0 | 0 |

| | | | | | | | | | | | | | | | | | | | | | | |
|---|---|---|---|---|---|---|---|---|---|---|---|---|---|---|---|---|---|---|---|---|---|---|
| scf7180000020328 | 180592 | 0.4043 | 0 | 24.1903 | Eukaryota | 19780.0 | 0 | Chordata | 19780.0 | 0 | Carnivora | 19780.0 | 0 | Felidae | 19780.0 | 0 | Felis | 19780.0 | 0 | Felis catus | 19780.0 | 0 |
| scf7180000024016 | 526742 | 0.5681 | 0 | 52.4394 | Eukaryota | 42907.0 | 0 | Chordata | 42907.0 | 0 | Carnivora | 42907.0 | 0 | Otariidae | 33138.0 | 3 | Zalophus | 33138.0 | 3 | phus californi | 33138.0 | 3 |
| scf7180000021164 | 84584 | 0.3743 | 0 | 24.8861 | Eukaryota | 44701.0 | 0 | Chordata | 44701.0 | 0 | Carnivora | 44701.0 | 0 | Felidae | 44701.0 | 0 | Felis | 44701.0 | 0 | Felis catus | 44701.0 | 0 |
| scf7180000018327 | 1405925 | 0.4231 | 100 | 51.0807 | Eukaryota | 44544.0 | 0 | Chordata | 44544.0 | 0 | Carnivora | 44544.0 | 0 | Felidae | 44544.0 | 0 | Felis | 44544.0 | 0 | Felis catus | 44544.0 | 0 |
| scf7180000020796 | 84626 | 0.3693 | 0 | 23.4845 | Eukaryota | 34714.0 | 0 | Chordata | 34714.0 | 0 | Carnivora | 34714.0 | 0 | Felidae | 34714.0 | 0 | Felis | 34714.0 | 0 | Felis catus | 34714.0 | 0 |
| scf7180000018411 | 952876 | 0.3872 | 0 | 51.1564 | Eukaryota | 47254.0 | 0 | Chordata | 47254.0 | 0 | Carnivora | 47254.0 | 0 | Felidae | 47254.0 | 0 | Felis | 47254.0 | 0 | Felis catus | 47254.0 | 0 |
| scf7180000021721 | 33732 | 0.3809 | 0 | 27.4703 | Eukaryota | 46247.0 | 0 | Chordata | 46247.0 | 0 | Carnivora | 46247.0 | 0 | Felidae | 46247.0 | 0 | Felis | 46247.0 | 0 | Felis catus | 46247.0 | 0 |
| scf7180000019988 | 191876 | 0.4163 | 0 | 49.3173 | Eukaryota | 46516.0 | 0 | Chordata | 46516.0 | 0 | Carnivora | 46516.0 | 0 | Felidae | 46516.0 | 0 | Felis | 41884.0 | 1 | Felis catus | 41884.0 | 1 |
| scf7180000020257 | 28743 | 0.5047 | 0 | 266.1135 | Eukaryota | 3943.0 | 0 | Chordata | 3943.0 | 0 | Primates | 2136.0 | 1 | Hominidae | 1456.0 | 3 | Homo | 1456.0 | 5 | Homo sapiens | 1456.0 | 5 |
| scf7180000018371 | 838464 | 0.3599 | 0 | 49.9243 | Eukaryota | 45236.0 | 0 | Chordata | 45236.0 | 0 | Carnivora | 45236.0 | 0 | Felidae | 45236.0 | 0 | Felis | 45236.0 | 0 | Felis catus | 45236.0 | 0 |
| scf7180000024209 | 801460 | 0.5017 | 0 | 49.7329 | Eukaryota | 35027.0 | 0 | Chordata | 35027.0 | 0 | Carnivora | 29978.0 | 1 | Herpestidae | 17744.0 | 2 | Suricata | 17744.0 | 3 | ıricata suricat | 17744.0 | 3 |
| scf7180000020802 | 88482 | 0.4479 | 0 | 51.2493 | Eukaryota | 9085.0 | 0 | Chordata | 9085.0 | 0 | Carnivora | 9085.0 | 0 | Felidae | 9085.0 | 0 | Felis | 9085.0 | 0 | Felis catus | 9085.0 | 0 |
| scf7180000019122 | 369085 | 0.4451 | 0 | 60.4968 | Eukaryota | 42623.0 | 0 | Chordata | 42623.0 | 0 | Carnivora | 42623.0 | 0 | Felidae | 42623.0 | 0 | Felis | 38430.0 | 1 | Felis catus | 38430.0 | 1 |
| scf7180000019790 | 148533 | 0.5248 | 0 | 48.0979 | Eukaryota | 6954.0 | 1 | Chordata | 6954.0 | 1 | Primates | 2408.0 | 5 | Bovidae | 2358.0 | 6 | Homo | 1630.0 | 7 | Homo sapiens | 1630.0 | 7 |
| scf7180000020555 | 157227 | 0.4085 | 0 | 50.3638 | Eukaryota | 15614.0 | 0 | Chordata | 15614.0 | 0 | Carnivora | 10595.0 | 2 | Felidae | 5998.0 | 3 | Lynx | 4629.0 | 4 | ynx canadens | 4629.0 | 4 |
| scf7180000022867 | 712655 | 0.4775 | 0 | 109.2894 | Eukaryota | 37098.0 | 0 | Chordata | 37098.0 | 0 | Carnivora | 37098.0 | 0 | Felidae | 17545.0 | 2 | Ursus | 12627.0 | 4 | Ursus arctos | 12627.0 | 4 |
| scf7180000023645 | 21233 | 0.4917 | 0 | 86.4253 | no-hit | 0.0 | 0 | no-hit | 0.0 | 0 | no-hit | 0.0 | 0 | no-hit | 0.0 | 0 | no-hit | 0.0 | 0 | no-hit | 0.0 | 0 |
| scf7180000022832 | 61696 | 0.3925 | 0 | 242.3223 | Eukaryota | 44223.0 | 0 | Chordata | 44223.0 | 0 | Carnivora | 44223.0 | 0 | Felidae | 44223.0 | 0 | Felis | 44223.0 | 0 | Felis catus | 44223.0 | 0 |
| scf7180000019518 | 352466 | 0.5555 | 0 | 48.519 | Eukaryota | 12030.0 | 0 | Chordata | 12030.0 | 0 | Carnivora | 12030.0 | 0 | Felidae | 12030.0 | 0 | Felis | 4234.0 | 2 | Felis catus | 4234.0 | 2 |
| scf7180000021249 | 102149 | 0.4479 | 0 | 26.4981 | Eukaryota | 45023.0 | 0 | Chordata | 45023.0 | 0 | Carnivora | 45023.0 | 0 | Herpestidae | 45023.0 | 0 | Suricata | 45023.0 | 0 | ıricata suricat | 45023.0 | 0 |
| scf7180000024205 | 20286 | 0.6508 | 0 | 69.9815 | Eukaryota | 5877.0 | 0 | Chordata | 5877.0 | 0 | Carnivora | 5877.0 | 0 | Felidae | 4186.0 | 1 | Felis | 4186.0 | 1 | Felis catus | 4186.0 | 1 |
| scf7180000018881 | 551618 | 0.4005 | 0 | 52.0874 | Eukaryota | 32425.0 | 0 | Chordata | 32425.0 | 0 | Carnivora | 32425.0 | 0 | Felidae | 16302.0 | 3 | unresolved | 6508.0 | 7 | unresolved | 6508.0 | 7 |
| scf7180000020349 | 56163 | 0.5208 | 0 | 182.1493 | no-hit | 0.0 | 0 | no-hit | 0.0 | 0 | no-hit | 0.0 | 0 | no-hit | 0.0 | 0 | no-hit | 0.0 | 0 | no-hit | 0.0 | 0 |
| scf7180000022747 | 48972 | 0.3825 | 0 | 24.1802 | Eukaryota | 31292.0 | 0 | Chordata | 31292.0 | 0 | Carnivora | 31292.0 | 0 | Felidae | 31292.0 | 0 | Felis | 14645.0 | 2 | Felis catus | 14645.0 | 3 |
| scf7180000020191 | 204373 | 0.4034 | 0 | 37.893 | Eukaryota | 45837.0 | 0 | Chordata | 45837.0 | 0 | Carnivora | 45837.0 | 0 | Felidae | 45837.0 | 0 | Felis | 45837.0 | 0 | Felis catus | 45837.0 | 0 |
| scf7180000019378 | 492126 | 0.5605 | 0 | 51.0571 | Eukaryota | 28983.0 | 0 | Chordata | 28983.0 | 0 | Carnivora | 28983.0 | 0 | Felidae | 18198.0 | 1 | Panthera | 11116.0 | 2 | anthera pardı | 11116.0 | 2 |
| scf7180000022091 | 12964 | 0.4293 | 0 | 785.2461 | Eukaryota | 2214.0 | 0 | Chordata | 2214.0 | 0 | Carnivora | 2214.0 | 0 | Felidae | 2214.0 | 0 | Felis | 1128.0 | 2 | Felis catus | 1128.0 | 3 |
| scf7180000019473 | 269941 | 0.4385 | 0 | 51.3197 | Eukaryota | 27753.0 | 0 | Chordata | 27753.0 | 0 | Carnivora | 27753.0 | 0 | Felidae | 27753.0 | 0 | Felis | 24897.0 | 1 | Felis catus | 24897.0 | 1 |
| scf7180000018226 | 24097 | 0.407 | 0 | 25.5026 | Eukaryota | 36921.0 | 0 | Chordata | 36921.0 | 0 | Carnivora | 36921.0 | 0 | Felidae | 36921.0 | 0 | Felis | 36921.0 | 0 | Felis catus | 36921.0 | 0 |
| scf7180000019804 | 41497 | 0.5046 | 0 | 209.198 | Eukaryota | 15816.0 | 0 | Chordata | 15816.0 | 0 | Carnivora | 15816.0 | 0 | Felidae | 15816.0 | 0 | Felis | 15816.0 | 0 | Felis catus | 15816.0 | 0 |
| scf7180000024256 | 612365 | 0.4169 | 0 | 49.9915 | Eukaryota | 46031.0 | 0 | Chordata | 46031.0 | 0 | Carnivora | 46031.0 | 0 | Felidae | 46031.0 | 0 | Felis | 46031.0 | 0 | Felis catus | 46031.0 | 0 |
| scf7180000019253 | 401326 | 0.3979 | 0 | 50.3803 | Eukaryota | 46529.0 | 0 | Chordata | 46529.0 | 0 | Carnivora | 46529.0 | 0 | Felidae | 46529.0 | 0 | Felis | 46529.0 | 0 | Felis catus | 46529.0 | 0 |
| scf7180000021227 | 64567 | 0.5864 | 100 | 42.7065 | Eukaryota | 2824.0 | 1 | Chordata | 2824.0 | 1 | Carnivora | 1653.0 | 3 | Felidae | 1388.0 | 4 | Felis | 1388.0 | 4 | Felis catus | 1388.0 | 4 |
| scf7180000018110 | 222935 | 0.5853 | 0 | 51.9935 | Eukaryota | 25528.0 | 0 | Chordata | 25528.0 | 0 | Carnivora | 22896.0 | 1 | Felidae | 20367.0 | 2 | Acinonyx | 5190.0 | 6 | :inonyx jubatı | 5190.0 | 6 |
| scf7180000022008 | 21852 | 0.4441 | 0 | 15.5394 | Eukaryota | 7924.0 | 0 | Chordata | 7924.0 | 0 | Carnivora | 7924.0 | 0 | Felidae | 5515.0 | 1 | Panthera | 2381.0 | 4 | anthera pardı | 2381.0 | 4 |
| scf7180000018020 | 1214154 | 0.3557 | 0 | 47.5473 | Eukaryota | 50263.0 | 0 | Chordata | 50263.0 | 0 | Carnivora | 42266.0 | 1 | Felidae | 42266.0 | 1 | Felis | 42266.0 | 1 | Felis catus | 42266.0 | 1 |
| scf7180000023851 | 1076816 | 0.3993 | 0 | 50.7538 | Eukaryota | 59969.0 | 0 | Chordata | 59969.0 | 0 | Carnivora | 59969.0 | 0 | Felidae | 59969.0 | 0 | Acinonyx | 59969.0 | 0 | :inonyx jubatı | 59969.0 | 0 |
| scf7180000021305 | 40896 | 0.4985 | 0 | 27.7448 | Eukaryota | 7845.0 | 0 | Chordata | 7845.0 | 0 | Carnivora | 2479.0 | 4 | Felidae | 2479.0 | 4 | Homo | 2070.0 | 6 | Homo sapiens | 2070.0 | 6 |
| scf7180000018216 | 892346 | 0.3985 | 0 | 51.109 | Eukaryota | 45146.0 | 0 | Chordata | 45146.0 | 0 | Carnivora | 45146.0 | 0 | Felidae | 45146.0 | 0 | Felis | 45146.0 | 0 | Felis catus | 45146.0 | 0 |
| scf7180000023846 | 1100366 | 0.442 | 0 | 51.9839 | Eukaryota | 45260.0 | 0 | Chordata | 45260.0 | 0 | Carnivora | 45260.0 | 0 | Felidae | 45260.0 | 0 | Felis | 45260.0 | 0 | Felis catus | 45260.0 | 0 |
| scf7180000019986 | 146832 | 0.6135 | 0 | 45.4761 | Eukaryota | 10613.0 | 0 | Chordata | 10613.0 | 0 | Carnivora | 6606.0 | 3 | Felidae | 4586.0 | 4 | Homo | 2026.0 | 7 | Homo sapiens | 2026.0 | 7 |
| scf7180000019662 | 386400 | 0.4748 | 0 | 51.1552 | Eukaryota | 34018.0 | 0 | Chordata | 34018.0 | 0 | Carnivora | 14290.0 | 4 | Otariidae | 10384.0 | 5 | Equus | 4113.0 | 9 | Equus caballu | 4113.0 | 9 |
| scf7180000017582 | 2434737 | 0.3833 | 0 | 51.7713 | Eukaryota | 44358.0 | 0 | Chordata | 44358.0 | 0 | Carnivora | 44358.0 | 0 | Felidae | 44358.0 | 0 | Felis | 39912.0 | 1 | Felis catus | 39912.0 | 1 |
| scf7180000020643 | 99114 | 0.4373 | 0 | 48.8694 | Eukaryota | 17941.0 | 0 | Chordata | 17941.0 | 0 | Carnivora | 17941.0 | 0 | Felidae | 17941.0 | 0 | Felis | 16160.0 | 1 | Felis catus | 16160.0 | 1 |
| scf7180000020710 | 65475 | 0.3745 | 0 | 25.0937 | Eukaryota | 45762.0 | 0 | Chordata | 45762.0 | 0 | Carnivora | 45762.0 | 0 | Felidae | 45762.0 | 0 | Felis | 45762.0 | 0 | Felis catus | 45762.0 | 0 |
| scf7180000019726 | 219642 | 0.4756 | 0 | 50.2472 | Eukaryota | 9222.0 | 0 | Chordata | 9222.0 | 0 | Carnivora | 9222.0 | 0 | Felidae | 9222.0 | 0 | Felis | 8315.0 | 1 | Felis catus | 8315.0 | 1 |
| scf7180000020834 | 132576 | 0.4073 | 0 | 25.5149 | Eukaryota | 15572.0 | 0 | Chordata | 15572.0 | 0 | Carnivora | 15572.0 | 0 | Felidae | 15572.0 | 0 | Felis | 13848.0 | 1 | Felis catus | 13848.0 | 1 |
| scf7180000018836 | 529485 | 0.4064 | 0 | 49.7231 | Eukaryota | 25455.0 | 0 | Chordata | 25455.0 | 0 | Carnivora | 25455.0 | 0 | Herpestidae | 13610.0 | 1 | Suricata | 13610.0 | 2 | ıricata suricat | 13610.0 | 2 |
| scf7180000017636 | 265575 | 0.3443 | 0 | 46.3039 | Eukaryota | 45403.0 | 0 | Chordata | 45403.0 | 0 | Carnivora | 45403.0 | 0 | Felidae | 45403.0 | 0 | Felis | 45403.0 | 0 | Felis catus | 45403.0 | 0 |
| scf7180000019077 | 302654 | 0.5066 | 0 | 48.8851 | Eukaryota | 14950.0 | 0 | Chordata | 14950.0 | 0 | Carnivora | 14950.0 | 0 | Felidae | 14950.0 | 0 | Felis | 11949.0 | 1 | Felis catus | 11949.0 | 1 |
| scf7180000018331 | 822700 | 0.3405 | 0 | 48.8056 | Eukaryota | 47616.0 | 0 | Chordata | 47616.0 | 0 | Carnivora | 47616.0 | 0 | Felidae | 47616.0 | 0 | Felis | 47616.0 | 0 | Felis catus | 47616.0 | 0 |
| scf7180000017837 | 548385 | 0.3777 | 0 | 49.4408 | Eukaryota | 43545.0 | 0 | Chordata | 43545.0 | 0 | Carnivora | 43545.0 | 0 | Felidae | 43545.0 | 0 | Felis | 43545.0 | 0 | Felis catus | 43545.0 | 0 |
| scf7180000021527 | 21617 | 0.5516 | 0 | 48.1324 | Eukaryota | 11548.0 | 0 | Chordata | 11548.0 | 0 | Carnivora | 11548.0 | 0 | Otariidae | 7556.0 | 1 | Eumetopias | 5032.0 | 3 | netopias juba | 5032.0 | 3 |
| scf7180000019028 | 202575 | 0.397 | 0 | 26.0222 | Eukaryota | 43274.0 | 0 | Chordata | 43274.0 | 0 | Carnivora | 43274.0 | 0 | Felidae | 43274.0 | 0 | Felis | 43274.0 | 0 | Felis catus | 43274.0 | 0 |
| scf7180000021483 | 21719 | 0.3836 | 0 | 25.0414 | Eukaryota | 27102.0 | 0 | Chordata | 27102.0 | 0 | Carnivora | 27102.0 | 0 | Felidae | 27102.0 | 0 | Felis | 24372.0 | 1 | Felis catus | 24372.0 | 1 |
| scf7180000019008 | 428179 | 0.3461 | 0 | 49.7114 | Eukaryota | 47436.0 | 0 | Chordata | 47436.0 | 0 | Carnivora | 47436.0 | 0 | Felidae | 47436.0 | 0 | Felis | 47436.0 | 0 | Felis catus | 47436.0 | 0 |
| scf7180000017646 | 1813771 | 0.4646 | 0 | 52.2256 | Eukaryota | 40919.0 | 1 | Chordata | 40919.0 | 1 | Carnivora | 30959.0 | 3 | Felidae | 22736.0 | 4 | Felis | 9294.0 | 7 | Felis catus | 9294.0 | 7 |
| scf7180000018555 | 667780 | 0.3815 | 0 | 49.8971 | Eukaryota | 48355.0 | 0 | Chordata | 48355.0 | 0 | Carnivora | 48355.0 | 0 | Felidae | 37747.0 | 2 | Felis | 33030.0 | 3 | Felis catus | 33030.0 | 3 |
| scf7180000021442 | 71335 | 0.5787 | 0 | 56.1036 | Eukaryota | 9339.0 | 0 | Chordata | 9339.0 | 0 | Carnivora | 9339.0 | 0 | Felidae | 8380.0 | 1 | Felis | 3748.0 | 4 | Felis catus | 3748.0 | 4 |
| scf7180000017995 | 1717131 | 0.4195 | 0 | 52.6435 | Eukaryota | 90890.0 | 0 | Chordata | 90890.0 | 0 | Carnivora | 90890.0 | 0 | Felidae | 90890.0 | 0 | Felis | 45470.0 | 1 | Felis catus | 45470.0 | 1 |
| scf7180000023973 | 126381 | 0.3947 | 0 | 49.2278 | Eukaryota | 43505.0 | 0 | Chordata | 43505.0 | 0 | Carnivora | 43505.0 | 0 | Felidae | 43505.0 | 0 | Felis | 43505.0 | 0 | Felis catus | 43505.0 | 0 |
| scf7180000022387 | 6863 | 0.3922 | 0 | 22.3948 | Eukaryota | 5571.0 | 0 | Chordata | 5571.0 | 0 | Carnivora | 3951.0 | 1 | Felidae | 2845.0 | 2 | Panthera | 2033.0 | 4 | Sus scrofa | 1620.0 | 5 |
| scf7180000023552 | 14476 | 0.7258 | 0 | 2125.2343 | Eukaryota | 2459.0 | 0 | Chordata | 2459.0 | 0 | Carnivora | 2459.0 | 0 | Felidae | 1797.0 | 1 | Panthera | 1140.0 | 2 | anthera pardı | 1140.0 | 2 |
| scf7180000023823 | 2577994 | 0.3613 | 0 | 50.1297 | Eukaryota | 47116.0 | 0 | Chordata | 47116.0 | 0 | Carnivora | 47116.0 | 0 | Felidae | 47116.0 | 0 | Felis | 42462.0 | 1 | Felis catus | 42462.0 | 1 |
| scf7180000019313 | 520091 | 0.4083 | 0 | 47.0707 | Eukaryota | 38576.0 | 0 | Chordata | 38576.0 | 0 | Carnivora | 38576.0 | 0 | Felidae | 30409.0 | 1 | Lynx | 14883.0 | 4 | ynx canadens | 14883.0 | 4 |
| scf7180000018247 | 1094529 | 0.3855 | 0 | 49.0874 | Eukaryota | 46309.0 | 0 | Chordata | 46309.0 | 0 | Carnivora | 46309.0 | 0 | Felidae | 46309.0 | 0 | Felis | 46309.0 | 0 | Felis catus | 46309.0 | 0 |
| scf7180000023949 | 722453 | 0.4066 | 0 | 50.5624 | Eukaryota | 81132.0 | 0 | Chordata | 81132.0 | 0 | Carnivora | 81132.0 | 0 | Felidae | 81132.0 | 0 | Lynx | 48586.0 | 1 | ynx canadens | 48586.0 | 1 |
| scf7180000019291 | 391575 | 0.5191 | 0 | 50.6814 | Eukaryota | 14187.0 | 0 | Chordata | 14187.0 | 0 | Carnivora | 14187.0 | 0 | Felidae | 14187.0 | 0 | Felis | 11360.0 | 1 | Felis catus | 11360.0 | 1 |
| scf7180000021397 | 49388 | 0.3683 | 0 | 49.0128 | Eukaryota | 31915.0 | 0 | Chordata | 31915.0 | 0 | Carnivora | 31915.0 | 0 | Felidae | 31915.0 | 0 | Felis | 28694.0 | 1 | Felis catus | 28694.0 | 1 |
| scf7180000020987 | 68734 | 0.5452 | 0 | 45.4799 | Eukaryota | 4277.0 | 0 | Chordata | 4277.0 | 0 | Carnivora | 1828.0 | 3 | Felidae | 1828.0 | 4 | Felis | 1828.0 | 5 | Felis catus | 1828.0 | 5 |
| scf7180000018308 | 1440100 | 0.4442 | 0 | 51.7809 | Eukaryota | 31949.0 | 0 | Chordata | 31949.0 | 0 | Carnivora | 31949.0 | 0 | Felidae | 13402.0 | 3 | Canis | 8124.0 | 6 | Canis lupus | 8124.0 | 6 |
| scf7180000017724 | 2081084 | 0.349 | 0 | 48.0486 | Eukaryota | 46267.0 | 0 | Chordata | 46267.0 | 0 | Carnivora | 46267.0 | 0 | Felidae | 46267.0 | 0 | Felis | 46267.0 | 0 | Felis catus | 46267.0 | 0 |
| scf7180000021285 | 88874 | 0.56 | 0 | 28.2032 | Eukaryota | 45810.0 | 0 | Chordata | 45810.0 | 0 | Carnivora | 45810.0 | 0 | Felidae | 45810.0 | 0 | Felis | 45810.0 | 0 | Felis catus | 45810.0 | 0 |
| scf7180000021754 | 8119 | 0.5201 | 0 | 79.8063 | no-hit | 0.0 | 0 | no-hit | 0.0 | 0 | no-hit | 0.0 | 0 | no-hit | 0.0 | 0 | no-hit | 0.0 | 0 | no-hit | 0.0 | 0 |
| scf7180000018859 | 295644 | 0.4212 | 0 | 49.8596 | Eukaryota | 22126.0 | 0 | Chordata | 22126.0 | 0 | Carnivora | 22126.0 | 0 | Felidae | 17802.0 | 2 | Panthera | 8850.0 | 6 | anthera pardı | 6619.0 | 7 |
| scf7180000021874 | 39184 | 0.3934 | 0 | 22.6344 | Eukaryota | 6189.0 | 0 | Chordata | 6189.0 | 0 | Carnivora | 5387.0 | 1 | Phocidae | 2989.0 | 4 | .eptonychote | 1789.0 | 6 | ınychotes wet | 1789.0 | 6 |
| scf7180000021460 | 80412 | 0.4129 | 0 | 25.911 | Eukaryota | 11446.0 | 0 | Chordata | 11446.0 | 0 | Primates | 5708.0 | 3 | Hominidae | 5708.0 | 3 | Homo | 2856.0 | 5 | Homo sapiens | 2856.0 | 5 |
| scf7180000018635 | 771147 | 0.3676 | 0 | 49.9759 | Eukaryota | 47373.0 | 0 | Chordata | 47373.0 | 0 | Carnivora | 47373.0 | 0 | Felidae | 25572.0 | 1 | Felis | 25572.0 | 1 | Felis catus | 25572.0 | 1 |
| scf7180000022767 | 12874 | 0.5743 | 0 | 159.0353 | Eukaryota | 12114.0 | 0 | Chordata | 12114.0 | 0 | Carnivora | 12114.0 | 0 | Felidae | 12114.0 | 0 | Felis | 10915.0 | 1 | Felis catus | 10915.0 | 1 |
| scf7180000017933 | 701519 | 0.468 | 0 | 50.5613 | Eukaryota | 37357.0 | 0 | Chordata | 37357.0 | 0 | Carnivora | 28395.0 | 3 | Felidae | 24795.0 | 4 | Panthera | 6829.0 | 8 | ynx canadens | 4946.0 | 9 |
| scf7180000019503 | 817866 | 0.5064 | 100 | 55.0476 | Eukaryota | 46005.0 | 0 | Chordata | 46005.0 | 0 | Carnivora | 46005.0 | 0 | Felidae | 46005.0 | 0 | Felis | 46005.0 | 0 | Felis catus | 46005.0 | 0 |
| scf7180000023235 | 24250 | 0.4677 | 0 | 215.4144 | Eukaryota | 11882.0 | 0 | Chordata | 11882.0 | 0 | Carnivora | 11882.0 | 0 | Felidae | 11882.0 | 0 | Felis | 10707.0 | 1 | Felis catus | 10707.0 | 1 |
| scf7180000022101 | 81700 | 0.4062 | 0 | 23.7016 | Eukaryota | 45511.0 | 0 | Chordata | 45511.0 | 0 | Carnivora | 45511.0 | 0 | Felidae | 45511.0 | 0 | Felis | 45511.0 | 0 | Felis catus | 45511.0 | 0 |
| scf7180000022686 | 6883 | 0.4075 | 0 | 51.7252 | Eukaryota | 2605.0 | 0 | Chordata | 2605.0 | 0 | Carnivora | 1759.0 | 2 | Felidae | 1015.0 | 3 | Felis | 760.0 | 4 | Felis catus | 760.0 | 4 |
| scf7180000019673 | 21381 | 0.4446 | 0 | 33.5402 | Eukaryota | 3270.0 | 0 | Chordata | 3270.0 | 0 | Carnivora | 2124.0 | 2 | Felidae | 2124.0 | 2 | Felis | 1590.0 | 3 | Felis catus | 1590.0 | 3 |
| scf7180000020381 | 132274 | 0.3935 | 0 | 25.5779 | Eukaryota | 6103.0 | 0 | Chordata | 6103.0 | 0 | Carnivora | 4163.0 | 2 | Felidae | 4163.0 | 2 | Felis | 2964.0 | 3 | Felis catus | 2964.0 | 3 |
| scf7180000024043 | 296655 | 0.4564 | 0 | 52.6735 | Eukaryota | 43278.0 | 0 | Chordata | 43278.0 | 0 | Carnivora | 43278.0 | 0 | Felidae | 43278.0 | 0 | Lynx | 30313.0 | 2 | ynx canadens | 30313.0 | 2 |
| scf7180000019841 | 206467 | 0.5932 | 0 | 49.1585 | Eukaryota | 5584.0 | 0 | Chordata | 5584.0 | 0 | Carnivora | 5584.0 | 0 | Felidae | 5584.0 | 0 | Panthera | 2860.0 | 2 | anthera pardı | 2213.0 | 3 |
| scf7180000019522 | 71969 | 0.3464 | 0 | 47.7256 | Eukaryota | 13454.0 | 0 | Chordata | 13454.0 | 0 | Carnivora | 13454.0 | 0 | Felidae | 13454.0 | 0 | Felis | 12122.0 | 1 | Felis catus | 12122.0 | 1 |
| scf7180000020071 | 18635 | 0.4105 | 0 | 187.7114 | Eukaryota | 6141.0 | 0 | Chordata | 6141.0 | 0 | Carnivora | 6141.0 | 0 | Felidae | 6141.0 | 0 | Felis | 6141.0 | 0 | Felis catus | 6141.0 | 0 |
| scf7180000019674 | 277569 | 0.4458 | 0 | 51.9111 | Eukaryota | 44462.0 | 0 | Chordata | 44462.0 | 0 | Carnivora | 31922.0 | 1 | Herpestidae | 24636.0 | 3 | Suricata | 24636.0 | 4 | ıricata suricat | 24636.0 | 4 |
| scf7180000024105 | 277563 | 0.4876 | 0 | 50.2996 | Eukaryota | 5544.0 | 0 | Chordata | 5544.0 | 0 | Primates | 2527.0 | 2 | Felidae | 1816.0 | 3 | Homo | 1543.0 | 5 | Homo sapiens | 1543.0 | 5 |
| scf7180000017738 | 1802993 | 0.4512 | 0 | 50.3493 | Eukaryota | 47809.0 | 0 | Chordata | 47809.0 | 0 | Carnivora | 47809.0 | 0 | Felidae | 47809.0 | 0 | Felis | 42414.0 | 1 | Felis catus | 42414.0 | 1 |
| scf7180000021726 | 39358 | 0.3853 | 0 | 24.839 | Eukaryota | 13344.0 | 0 | Chordata | 13344.0 | 0 | Carnivora | 13344.0 | 0 | Felidae | 13344.0 | 0 | Felis | 12003.0 | 1 | Felis catus | 12003.0 | 1 |
| scf7180000019007 | 1147568 | 0.3911 | 0 | 49.558 | Eukaryota | 41498.0 | 0 | Chordata | 41498.0 | 0 | Carnivora | 41498.0 | 0 | Felidae | 41498.0 | 0 | Felis | 41498.0 | 0 | Felis catus | 41498.0 | 0 |
| scf7180000022513 | 5328 | 0.5199 | 0 | 80.6178 | no-hit | 0.0 | 0 | no-hit | 0.0 | 0 | no-hit | 0.0 | 0 | no-hit | 0.0 | 0 | no-hit | 0.0 | 0 | no-hit | 0.0 | 0 |
| scf7180000018628 | 1126111 | 0.4848 | 0 | 49.6403 | Eukaryota | 26479.0 | 1 | Chordata | 26479.0 | 1 | Carnivora | 20676.0 | 2 | Felidae | 20676.0 | 2 | Felis | 17649.0 | 3 | Felis catus | 17649.0 | 3 |
| scf7180000018770 | 606241 | 0.4172 | 0 | 50.244 | Eukaryota | 47565.0 | 0 | Chordata | 47565.0 | 0 | Carnivora | 47565.0 | 0 | Felidae | 47565.0 | 0 | Felis | 47565.0 | 0 | Felis catus | 47565.0 | 0 |
| scf7180000024212 | 56990 | 0.4006 | 0 | 23.7248 | Eukaryota | 13306.0 | 0 | Chordata | 13306.0 | 0 | Carnivora | 13306.0 | 0 | Felidae | 13306.0 | 0 | Felis | 11970.0 | 1 | Felis catus | 11970.0 | 1 |
| scf7180000018160 | 1001896 | 0.3933 | 0 | 50.5722 | Eukaryota | 29147.0 | 0 | Chordata | 29147.0 | 0 | Carnivora | 19124.0 | 1 | Felidae | 19124.0 | 1 | Felis | 16346.0 | 3 | Felis catus | 16346.0 | 3 |
| scf7180000018475 | 737999 | 0.5358 | 0 | 51.2825 | Eukaryota | 34025.0 | 0 | Chordata | 34025.0 | 0 | Carnivora | 34025.0 | 0 | Felidae | 27753.0 | 1 | Felis | 14860.0 | 3 | Felis catus | 14860.0 | 3 |
| scf7180000019425 | 350413 | 0.5069 | 0 | 52.1861 | Eukaryota | 21036.0 | 0 | Chordata | 21036.0 | 0 | Carnivora | 21036.0 | 0 | Felidae | 21036.0 | 0 | Felis | 18921.0 | 1 | Felis catus | 18921.0 | 1 |
| scf7180000020887 | 105307 | 0.3975 | 0 | 49.1119 | Eukaryota | 26548.0 | 0 | Chordata | 26548.0 | 0 | Carnivora | 26548.0 | 0 | Felidae | 26548.0 | 0 | Felis | 23851.0 | 1 | Felis catus | 23851.0 | 1 |
| scf7180000020691 | 102006 | 0.4729 | 0 | 27.7363 | Eukaryota | 10015.0 | 0 | Chordata | 10015.0 | 0 | Carnivora | 5348.0 | 2 | Felidae | 5348.0 | 2 | Felis | 4583.0 | 3 | Felis catus | 4583.0 | 3 |
| scf7180000018792 | 77040 | 0.3894 | 0 | 23.5579 | Eukaryota | 40183.0 | 0 | Chordata | 40183.0 | 0 | Carnivora | 40183.0 | 0 | Felidae | 40183.0 | 0 | Felis | 40183.0 | 0 | Felis catus | 40183.0 | 0 |
| scf7180000023123 | 45840 | 0.3991 | 0 | 261.2434 | Eukaryota | 7246.0 | 0 | Chordata | 7246.0 | 0 | Carnivora | 7246.0 | 0 | Felidae | 7246.0 | 0 | Felis | 7246.0 | 0 | Felis catus | 7246.0 | 0 |
| scf7180000017462 | 5077735 | 0.4513 | 0 | 51.0608 | Eukaryota | 48009.0 | 0 | Chordata | 48009.0 | 0 | Carnivora | 48009.0 | 0 | Felidae | 48009.0 | 0 | Felis | 42805.0 | 1 | Felis catus | 42805.0 | 1 |
| scf7180000018365 | 50505 | 0.3947 | 0 | 39.0737 | Eukaryota | 20444.0 | 0 | Chordata | 20444.0 | 0 | Carnivora | 20444.0 | 0 | Felidae | 20444.0 | 0 | Felis | 18386.0 | 1 | Felis catus | 18386.0 | 1 |
| scf7180000020779 | 22967 | 0.4485 | 0 | 392.9168 | Eukaryota | 6278.0 | 0 | Chordata | 6278.0 | 0 | Carnivora | 6278.0 | 0 | Felidae | 6278.0 | 0 | Felis | 6278.0 | 0 | Felis catus | 6278.0 | 0 |
| scf7180000024231 | 289159 | 0.4128 | 0 | 51.6163 | Eukaryota | 24796.0 | 0 | Chordata | 24796.0 | 0 | Carnivora | 24796.0 | 0 | Felidae | 15348.0 | 3 | Felis | 5534.0 | 6 | Felis catus | 5534.0 | 7 |
| scf7180000019324 | 319869 | 0.4509 | 0 | 50.3704 | Eukaryota | 8697.0 | 0 | Chordata | 8697.0 | 0 | Carnivora | 3865.0 | 2 | Felidae | 3865.0 | 2 | Felis | 3037.0 | 4 | Felis catus | 3037.0 | 4 |
| scf7180000024180 | 96600 | 0.3602 | 0 | 24.0029 | Eukaryota | 16029.0 | 0 | Chordata | 16029.0 | 0 | Carnivora | 16029.0 | 0 | Felidae | 16029.0 | 0 | Felis | 16029.0 | 0 | Felis catus | 16029.0 | 0 |
| scf7180000020684 | 104670 | 0.401 | 0 | 49.6508 | Eukaryota | 9916.0 | 0 | Chordata | 9916.0 | 0 | Carnivora | 9916.0 | 0 | Felidae | 9916.0 | 0 | Felis | 8937.0 | 1 | Felis catus | 8937.0 | 1 |
| scf7180000022176 | 20479 | 0.4065 | 0 | 24.2433 | Eukaryota | 35138.0 | 0 | Chordata | 35138.0 | 0 | Carnivora | 35138.0 | 0 | Felidae | 25229.0 | 3 | Panthera | 10848.0 | 6 | anthera pardı | 10848.0 | 6 |
| scf7180000018333 | 966727 | 0.3928 | 0 | 51.5164 | Eukaryota | 44679.0 | 0 | Chordata | 44679.0 | 0 | Carnivora | 44679.0 | 0 | Felidae | 44679.0 | 0 | Felis | 44679.0 | 0 | Felis catus | 44679.0 | 0 |
| scf7180000023795 | 1465344 | 0.4856 | 0 | 51.0188 | Eukaryota | 62985.0 | 0 | Chordata | 62985.0 | 0 | Carnivora | 62985.0 | 0 | Felidae | 33705.0 | 1 | Panthera | 33705.0 | 1 | anthera pardı | 33705.0 | 1 |
| scf7180000021226 | 63836 | 0.6354 | 0 | 49.1362 | Eukaryota | 19400.0 | 0 | Chordata | 19400.0 | 0 | Carnivora | 19400.0 | 0 | Felidae | 19400.0 | 0 | Felis | 19400.0 | 0 | Felis catus | 19400.0 | 0 |
| scf7180000020228 | 55686 | 0.4646 | 0 | 27.9227 | Eukaryota | 6156.0 | 0 | Chordata | 6156.0 | 0 | Carnivora | 6156.0 | 0 | Felidae | 6156.0 | 0 | Felis | 6156.0 | 0 | Felis catus | 6156.0 | 0 |
| scf7180000021267 | 75360 | 0.3675 | 0 | 25.221 | Eukaryota | 6801.0 | 0 | Chordata | 6801.0 | 0 | Carnivora | 2714.0 | 2 | Felidae | 2714.0 | 2 | Felis | 2714.0 | 3 | Felis catus | 2714.0 | 3 |
| scf7180000021017 | 62462 | 0.5474 | 0 | 43.6118 | Eukaryota | 3336.0 | 0 | Chordata | 3336.0 | 0 | Carnivora | 3336.0 | 0 | Felidae | 3336.0 | 0 | Panthera | 1461.0 | 3 | Felis catus | 1216.0 | 5 |
| scf7180000019445 | 716847 | 0.5086 | 0 | 50.4911 | Eukaryota | 26408.0 | 0 | Chordata | 26408.0 | 0 | Carnivora | 26408.0 | 0 | Felidae | 26408.0 | 0 | Felis | 23757.0 | 1 | Felis catus | 23757.0 | 1 |
| scf7180000020221 | 46111 | 0.4538 | 0 | 28.8472 | Eukaryota | 7706.0 | 0 | Chordata | 7706.0 | 0 | Carnivora | 7706.0 | 0 | Felidae | 7706.0 | 0 | Felis | 6946.0 | 1 | Felis catus | 6946.0 | 1 |
| scf7180000021012 | 68480 | 0.397 | 0 | 25.5376 | Eukaryota | 44851.0 | 0 | Chordata | 44851.0 | 0 | Carnivora | 44851.0 | 0 | Felidae | 44851.0 | 0 | Felis | 44851.0 | 0 | Felis catus | 44851.0 | 0 |
| scf7180000017836 | 967771 | 0.4023 | 0 | 51.1272 | Eukaryota | 47172.0 | 0 | Chordata | 47172.0 | 0 | Carnivora | 47172.0 | 0 | Felidae | 47172.0 | 0 | Felis | 47172.0 | 0 | Felis catus | 47172.0 | 0 |
| scf7180000018384 | 249585 | 0.346 | 0 | 47.0288 | Eukaryota | 10982.0 | 0 | Chordata | 10982.0 | 0 | Carnivora | 6964.0 | 2 | Felidae | 6964.0 | 2 | Felis | 5785.0 | 3 | Felis catus | 5785.0 | 3 |
| scf7180000020657 | 113170 | 0.3959 | 0 | 24.7596 | Eukaryota | 7531.0 | 0 | Chordata | 7531.0 | 0 | Carnivora | 6753.0 | 1 | Felidae | 6753.0 | 1 | Felis | 6753.0 | 1 | Felis catus | 6753.0 | 1 |
| scf7180000019719 | 159578 | 0.3669 | 0 | 52.2235 | Eukaryota | 15713.0 | 0 | Chordata | 15713.0 | 0 | Primates | 10435.0 | 3 | Hominidae | 10435.0 | 3 | Homo | 10435.0 | 4 | Homo sapiens | 10435.0 | 4 |
| scf7180000022661 | 13118 | 0.3721 | 0 | 24.2818 | Eukaryota | 20261.0 | 0 | Chordata | 20261.0 | 0 | Carnivora | 20261.0 | 0 | Felidae | 20261.0 | 0 | Felis | 18194.0 | 1 | Felis catus | 18194.0 | 1 |
| scf7180000019191 | 21679 | 0.3746 | 0 | 25.7726 | Eukaryota | 14117.0 | 0 | Chordata | 14117.0 | 0 | Carnivora | 14117.0 | 0 | Felidae | 14117.0 | 0 | Felis | 14117.0 | 0 | Felis catus | 14117.0 | 0 |
| scf7180000017777 | 1911938 | 0.3656 | 0 | 48.9632 | Eukaryota | 47121.0 | 0 | Chordata | 47121.0 | 0 | Carnivora | 47121.0 | 0 | Felidae | 47121.0 | 0 | Felis | 47121.0 | 0 | Felis catus | 47121.0 | 0 |
| scf7180000018422 | 910108 | 0.4218 | 0 | 51.1214 | Eukaryota | 45399.0 | 0 | Chordata | 45399.0 | 0 | Carnivora | 45399.0 | 0 | Felidae | 45399.0 | 0 | Felis | 45399.0 | 0 | Felis catus | 45399.0 | 0 |
| scf7180000021969 | 8046 | 0.5215 | 0 | 48.4782 | no-hit | 0.0 | 0 | no-hit | 0.0 | 0 | no-hit | 0.0 | 0 | no-hit | 0.0 | 0 | no-hit | 0.0 | 0 | no-hit | 0.0 | 0 |
| scf7180000020918 | 82851 | 0.5115 | 0 | 27.2916 | Eukaryota | 41156.0 | 0 | Chordata | 41156.0 | 0 | Carnivora | 41156.0 | 0 | Felidae | 41156.0 | 0 | Felis | 16463.0 | 3 | Felis catus | 16463.0 | 3 |
| scf7180000019918 | 276030 | 0.4425 | 0 | 50.0827 | Eukaryota | 30881.0 | 0 | Chordata | 30881.0 | 0 | Carnivora | 30881.0 | 0 | Felidae | 30881.0 | 0 | Felis | 27723.0 | 1 | Felis catus | 27723.0 | 1 |
| scf7180000021022 | 69733 | 0.3859 | 0 | 25.2757 | Eukaryota | 44246.0 | 0 | Chordata | 44246.0 | 0 | Carnivora | 44246.0 | 0 | Felidae | 44246.0 | 0 | Felis | 44246.0 | 0 | Felis catus | 44246.0 | 0 |
| scf7180000023838 | 1314066 | 0.4006 | 0 | 50.485 | Eukaryota | 27338.0 | 0 | Chordata | 27338.0 | 0 | Carnivora | 27338.0 | 0 | Felidae | 27338.0 | 0 | Felis | 24575.0 | 1 | Felis catus | 24575.0 | 1 |
| scf7180000024243 | 379662 | 0.4965 | 0 | 49.5468 | Eukaryota | 63589.0 | 0 | Chordata | 63589.0 | 0 | Carnivora | 63589.0 | 0 | Felidae | 63589.0 | 0 | Panthera | 38099.0 | 3 | anthera pardı | 31740.0 | 4 |
| scf7180000023079 | 36811 | 0.5747 | 0 | 264.2319 | Eukaryota | 3234.0 | 0 | Chordata | 3234.0 | 0 | Carnivora | 2630.0 | 1 | Felidae | 1528.0 | 3 | Felis | 1528.0 | 3 | Felis catus | 1528.0 | 3 |
| scf7180000019873 | 12311 | 0.394 | 0 | 22.7775 | Eukaryota | 17340.0 | 0 | Chordata | 17340.0 | 0 | Carnivora | 17340.0 | 0 | Felidae | 17340.0 | 0 | Felis | 13909.0 | 1 | Felis catus | 13909.0 | 1 |
| scf7180000019948 | 154835 | 0.5786 | 0 | 50.3346 | Eukaryota | 11026.0 | 0 | Chordata | 11026.0 | 0 | Primates | 4685.0 | 2 | Felidae | 4073.0 | 3 | Homo | 3752.0 | 6 | Homo sapiens | 3752.0 | 6 |
| scf7180000021197 | 69892 | 0.3593 | 0 | 24.8039 | Eukaryota | 45383.0 | 0 | Chordata | 45383.0 | 0 | Carnivora | 45383.0 | 0 | Felidae | 45383.0 | 0 | Felis | 40871.0 | 1 | Felis catus | 40871.0 | 1 |
| scf7180000018118 | 1067821 | 0.365 | 0 | 50.8751 | Eukaryota | 48257.0 | 0 | Chordata | 48257.0 | 0 | Carnivora | 36195.0 | 1 | Felidae | 36195.0 | 1 | Felis | 36195.0 | 1 | Felis catus | 36195.0 | 1 |
| scf7180000020652 | 4493 | 0.6359 | 0 | 19.1421 | Eukaryota | 2206.0 | 0 | Chordata | 2206.0 | 0 | Carnivora | 2206.0 | 0 | Felidae | 2206.0 | 0 | Felis | 1980.0 | 1 | Felis catus | 1980.0 | 1 |
| scf7180000024185 | 101723 | 0.4052 | 0 | 49.9213 | Eukaryota | 74130.0 | 0 | Chordata | 74130.0 | 0 | Carnivora | 74130.0 | 0 | Herpestidae | 38610.0 | 2 | Suricata | 38610.0 | 3 | ıricata suricat | 38610.0 | 4 |
| scf7180000019738 | 209997 | 0.5129 | 0 | 49.5738 | Eukaryota | 6014.0 | 0 | Chordata | 6014.0 | 0 | Carnivora | 6014.0 | 0 | Felidae | 6014.0 | 0 | Felis | 6014.0 | 0 | Felis catus | 6014.0 | 0 |
| scf7180000021607 | 24118 | 0.42 | 0 | 25.8344 | Eukaryota | 13988.0 | 0 | Chordata | 13988.0 | 0 | Primates | 6214.0 | 2 | Hominidae | 6214.0 | 3 | Homo | 6214.0 | 4 | Homo sapiens | 6214.0 | 4 |
| scf7180000017551 | 2650205 | 0.3669 | 0 | 49.8132 | Eukaryota | 76409.0 | 0 | Chordata | 76409.0 | 0 | Carnivora | 76409.0 | 0 | Felidae | 76409.0 | 0 | Panthera | 23032.0 | 4 | :inonyx jubatı | 22833.0 | 5 |
| scf7180000024082 | 1506581 | 0.4143 | 0 | 52.0059 | Eukaryota | 47357.0 | 0 | Chordata | 47357.0 | 0 | Carnivora | 47357.0 | 0 | Felidae | 47357.0 | 0 | Felis | 47357.0 | 0 | Felis catus | 47357.0 | 0 |
| scf7180000021375 | 79364 | 0.378 | 0 | 24.3275 | Eukaryota | 47125.0 | 0 | Chordata | 47125.0 | 0 | Carnivora | 47125.0 | 0 | Felidae | 47125.0 | 0 | Felis | 47125.0 | 0 | Felis catus | 47125.0 | 0 |
| scf7180000019597 | 294552 | 0.4174 | 0 | 50.7607 | Eukaryota | 56548.0 | 0 | Chordata | 56548.0 | 0 | Carnivora | 56548.0 | 0 | Herpestidae | 50895.0 | 1 | Suricata | 50895.0 | 1 | ıricata suricat | 50895.0 | 1 |
| scf7180000019782 | 331406 | 0.381 | 0 | 50.4033 | Eukaryota | 46938.0 | 0 | Chordata | 46938.0 | 0 | Carnivora | 46938.0 | 0 | Felidae | 46938.0 | 0 | Felis | 46938.0 | 0 | Felis catus | 46938.0 | 0 |
| scf7180000018443 | 2341310 | 0.3875 | 0 | 49.0525 | Eukaryota | 65204.0 | 0 | Chordata | 65204.0 | 0 | Carnivora | 53436.0 | 1 | Canidae | 25856.0 | 4 | Canis | 25856.0 | 5 | Canis lupus | 25856.0 | 5 |
| scf7180000021977 | 24601 | 0.5857 | 0 | 46.6201 | Eukaryota | 6880.0 | 0 | Chordata | 6880.0 | 0 | Carnivora | 6880.0 | 0 | Felidae | 3114.0 | 3 | Ursus | 2528.0 | 5 | Ursus arctos | 2528.0 | 5 |
| scf7180000022851 | 985684 | 0.4141 | 0 | 129.1761 | Eukaryota | 43994.0 | 0 | Chordata | 43994.0 | 0 | Carnivora | 43994.0 | 0 | Felidae | 43994.0 | 0 | Felis | 39589.0 | 1 | Felis catus | 39589.0 | 1 |
| scf7180000018059 | 1447346 | 0.4166 | 0 | 49.5555 | Eukaryota | 47789.0 | 0 | Chordata | 47789.0 | 0 | Carnivora | 47789.0 | 0 | Felidae | 47789.0 | 0 | Felis | 47789.0 | 0 | Felis catus | 47789.0 | 0 |
| scf7180000023004 | 65212 | 0.5391 | 0 | 230.5418 | Eukaryota | 6299.0 | 0 | Chordata | 6299.0 | 0 | Carnivora | 6299.0 | 0 | Felidae | 5687.0 | 1 | Panthera | 3075.0 | 3 | Panthera tigri: | 1692.0 | 4 |
| scf7180000023875 | 428390 | 0.5344 | 0 | 50.7416 | Eukaryota | 54522.0 | 0 | Chordata | 54522.0 | 0 | Carnivora | 54522.0 | 0 | Herpestidae | 33474.0 | 1 | Suricata | 33474.0 | 1 | ıricata suricat | 33474.0 | 1 |
| scf7180000024002 | 488770 | 0.3928 | 0 | 49.9985 | Eukaryota | 21060.0 | 0 | Chordata | 21060.0 | 0 | Carnivora | 21060.0 | 0 | Felidae | 21060.0 | 0 | Felis | 18982.0 | 1 | Felis catus | 18982.0 | 1 |
| scf7180000023173 | 489692 | 0.4757 | 0 | 46.6861 | Eukaryota | 26438.0 | 0 | Chordata | 26438.0 | 0 | Carnivora | 26438.0 | 0 | Felidae | 26438.0 | 0 | Felis | 21128.0 | 1 | Felis catus | 21128.0 | 1 |

| | | | | | | | | | | | | | | | | | | | | |
|---|---|---|---|---|---|---|---|---|---|---|---|---|---|---|---|---|---|---|---|---|
| scf7180000019482 | 372053 | 0.3909 | 0 | 52.2244 | Eukaryota | 47322.0 | 0 | Chordata | 47322.0 | 0 | Carnivora | 47322.0 | 0 | Felidae | 47322.0 | 0 | Felis | 47322.0 | 0 | Felis catus | 47322.0 | 0 |
| scf7180000021572 | 21277 | 0.3518 | 0 | 23.5086 | Eukaryota | 13255.0 | 0 | Chordata | 13255.0 | 0 | Carnivora | 13255.0 | 0 | Felidae | 13255.0 | 0 | Felis | 13255.0 | 0 | Felis catus | 13255.0 | 0 |
| scf7180000021051 | 95576 | 0.4917 | 0 | 47.0505 | Eukaryota | 44554.0 | 0 | Chordata | 44554.0 | 0 | Carnivora | 44554.0 | 0 | Felidae | 39495.0 | 1 | Panthera | 22979.0 | 3 | anthera pardı | 22979.0 | 3 |
| scf7180000018128 | 2005279 | 0.418 | 0 | 49.3365 | Eukaryota | 43010.0 | 0 | Chordata | 43010.0 | 0 | Carnivora | 43010.0 | 0 | Felidae | 43010.0 | 0 | Panthera | 29050.0 | 1 | anthera pardı | 20750.0 | 2 |
| scf7180000022344 | 7065 | 0.474 | 0 | 48.2558 | Eukaryota | 2423.0 | 0 | Chordata | 2423.0 | 0 | Carnivora | 2423.0 | 0 | Felidae | 2186.0 | 1 | Felis | 1218.0 | 4 | Felis catus | 983.0 | 6 |
| scf7180000023374 | 24404 | 0.4327 | 0 | 167.9003 | Eukaryota | 12172.0 | 0 | Chordata | 12172.0 | 0 | Carnivora | 12172.0 | 0 | Felidae | 12172.0 | 0 | Felis | 10973.0 | 1 | Felis catus | 10973.0 | 1 |
| scf7180000019762 | 345789 | 0.4144 | 0 | 51.6246 | Eukaryota | 47903.0 | 0 | Chordata | 47903.0 | 0 | Carnivora | 47903.0 | 0 | Felidae | 47903.0 | 0 | Felis | 47903.0 | 0 | Felis catus | 47903.0 | 0 |
| scf7180000018924 | 442792 | 0.3957 | 0 | 50.0624 | Eukaryota | 53899.0 | 1 | Chordata | 53899.0 | 1 | Carnivora | 21735.0 | 3 | Felidae | 21735.0 | 3 | Felis | 21735.0 | 3 | Felis catus | 21735.0 | 4 |
| scf7180000017813 | 1481213 | 0.4337 | 0 | 51.6646 | Eukaryota | 44010.0 | 0 | Chordata | 44010.0 | 0 | Carnivora | 44010.0 | 0 | Felidae | 44010.0 | 0 | Acinonyx | 44010.0 | 0 | :inonyx jubat | 44010.0 | 0 |
| scf7180000017542 | 2453719 | 0.3939 | 0 | 50.4655 | Eukaryota | 48500.0 | 0 | Chordata | 48500.0 | 0 | Carnivora | 48500.0 | 0 | Felidae | 48500.0 | 0 | Felis | 28257.0 | 3 | Felis catus | 28257.0 | 4 |
| scf7180000021508 | 7170 | 0.5787 | 0 | 33.9316 | no-hit | 0.0 | 0 | no-hit | 0.0 | 0 | no-hit | 0.0 | 0 | no-hit | 0.0 | 0 | no-hit | 0.0 | 0 | no-hit | 0.0 | 0 |
| scf7180000021704 | 33372 | 0.6135 | 0 | 46.1554 | Eukaryota | 4608.0 | 1 | Chordata | 4608.0 | 1 | Carnivora | 4074.0 | 2 | Felidae | 4074.0 | 2 | Felis | 4074.0 | 2 | Felis catus | 4074.0 | 2 |
| scf7180000020229 | 150437 | 0.4654 | 0 | 25.3277 | Eukaryota | 40833.0 | 0 | Chordata | 40833.0 | 0 | Carnivora | 40833.0 | 0 | Felidae | 40833.0 | 0 | Felis | 40833.0 | 0 | Felis catus | 40833.0 | 0 |
| scf7180000021753 | 49049 | 0.404 | 0 | 26.0495 | Eukaryota | 6165.0 | 0 | Chordata | 6165.0 | 0 | Primates | 3206.0 | 3 | Hominidae | 3206.0 | 3 | Sus | 1894.0 | 4 | Sus scrofa | 1894.0 | 4 |
| scf7180000018397 | 1655157 | 0.4057 | 0 | 50.6675 | Eukaryota | 77837.0 | 0 | Chordata | 77837.0 | 0 | Carnivora | 77837.0 | 0 | Felidae | 77837.0 | 0 | Panthera | 38962.0 | 2 | anthera pardı | 31178.0 | 3 |
| scf7180000019456 | 317602 | 0.3845 | 0 | 50.5496 | Eukaryota | 43296.0 | 0 | Chordata | 43296.0 | 0 | Carnivora | 43296.0 | 0 | Felidae | 43296.0 | 0 | Felis | 38959.0 | 1 | Felis catus | 38959.0 | 1 |
| scf7180000023993 | 577195 | 0.3752 | 0 | 49.6875 | Eukaryota | 42157.0 | 0 | Chordata | 42157.0 | 0 | Carnivora | 42157.0 | 0 | Felidae | 42157.0 | 0 | Felis | 42157.0 | 0 | Felis catus | 42157.0 | 0 |
| scf7180000018348 | 2210528 | 0.3662 | 0 | 49.5986 | Eukaryota | 47884.0 | 0 | Chordata | 47884.0 | 0 | Carnivora | 47884.0 | 0 | Felidae | 47884.0 | 0 | Felis | 47884.0 | 0 | Felis catus | 47884.0 | 0 |
| scf7180000024060 | 385417 | 0.417 | 0 | 52.245 | Eukaryota | 19995.0 | 0 | Chordata | 19995.0 | 0 | Carnivora | 19995.0 | 0 | Felidae | 19995.0 | 0 | Felis | 17308.0 | 1 | Felis catus | 17308.0 | 1 |
| scf7180000019885 | 20109 | 0.4238 | 0 | 36.2157 | Eukaryota | 7843.0 | 0 | Chordata | 7843.0 | 0 | Carnivora | 7843.0 | 0 | Felidae | 5777.0 | 2 | Felis | 3727.0 | 4 | Felis catus | 3727.0 | 4 |
| scf7180000020656 | 39717 | 0.3886 | 0 | 41.5455 | Eukaryota | 36013.0 | 0 | Chordata | 36013.0 | 0 | Carnivora | 36013.0 | 0 | Herpestidae | 14856.0 | 2 | Suricata | 14856.0 | 3 | ıricata suricat | 14856.0 | 3 |
| scf7180000021864 | 27556 | 0.4487 | 0 | 26.0107 | Eukaryota | 5367.0 | 0 | Chordata | 5367.0 | 0 | Carnivora | 4016.0 | 1 | Felidae | 4016.0 | 1 | Felis | 4016.0 | 2 | Felis catus | 4016.0 | 2 |
| scf7180000017489 | 1554567 | 0.4732 | 0 | 52.627 | Eukaryota | 62840.0 | 0 | Chordata | 62840.0 | 0 | Carnivora | 62840.0 | 0 | Felidae | 62840.0 | 0 | Acinonyx | 44010.0 | 1 | :inonyx jubat | 44010.0 | 1 |
| scf7180000024110 | 230204 | 0.4391 | 0 | 49.5308 | Eukaryota | 15746.0 | 1 | Chordata | 15746.0 | 1 | Artiodactyla | 8550.0 | 3 | Hominidae | 5428.0 | 4 | Homo | 5428.0 | 5 | Homo sapiens | 5428.0 | 5 |
| scf7180000022011 | 21293 | 0.4208 | 0 | 24.845 | Eukaryota | 4470.0 | 1 | Chordata | 4091.0 | 2 | Artiodactyla | 1819.0 | 4 | Equidae | 1629.0 | 5 | Equus | 1629.0 | 6 | :quus caballu | 1629.0 | 6 |
| scf7180000017843 | 220914 | 0.3653 | 0 | 49.3019 | Eukaryota | 43136.0 | 0 | Chordata | 43136.0 | 0 | Carnivora | 43136.0 | 0 | Felidae | 43136.0 | 0 | Felis | 43136.0 | 0 | Felis catus | 43136.0 | 0 |
| scf7180000022755 | 13146 | 0.3508 | 0 | 44.2336 | Eukaryota | 6079.0 | 0 | Chordata | 6079.0 | 0 | Carnivora | 6079.0 | 0 | Felidae | 6079.0 | 0 | Felis | 5532.0 | 1 | Felis catus | 5532.0 | 1 |
| scf7180000019995 | 160156 | 0.3858 | 0 | 48.5943 | Eukaryota | 22237.0 | 0 | Chordata | 22237.0 | 0 | Carnivora | 22237.0 | 0 | Felidae | 22237.0 | 0 | Acinonyx | 15610.0 | 1 | :inonyx jubat | 15610.0 | 1 |
| scf7180000019484 | 283951 | 0.4474 | 0 | 51.0364 | Eukaryota | 46330.0 | 0 | Chordata | 46330.0 | 0 | Carnivora | 46330.0 | 0 | Felidae | 33931.0 | 1 | Panthera | 19732.0 | 2 | anthera pardı | 19732.0 | 2 |
| scf7180000017839 | 1421202 | 0.4746 | 0 | 52.1325 | Eukaryota | 53905.0 | 0 | Chordata | 53905.0 | 0 | Carnivora | 53905.0 | 0 | Felidae | 37678.0 | 3 | Acinonyx | 11044.0 | 6 | :inonyx jubat | 11044.0 | 7 |
| scf7180000020703 | 31020 | 0.4618 | 0 | 146.5236 | Eukaryota | 2311.0 | 0 | Chordata | 2311.0 | 0 | Carnivora | 2311.0 | 0 | Felidae | 2311.0 | 0 | Felis | 2102.0 | 1 | Felis catus | 2102.0 | 1 |
| scf7180000023759 | 3984540 | 0.3818 | 0 | 50.3825 | Eukaryota | 48266.0 | 0 | Chordata | 48266.0 | 0 | Carnivora | 48266.0 | 0 | Felidae | 48266.0 | 0 | Felis | 48266.0 | 0 | Felis catus | 48266.0 | 0 |
| scf7180000017897 | 1554248 | 0.3583 | 0 | 49.037 | Eukaryota | 46659.0 | 0 | Chordata | 46659.0 | 0 | Carnivora | 46659.0 | 0 | Felidae | 46659.0 | 0 | Felis | 46659.0 | 0 | Felis catus | 46659.0 | 0 |
| scf7180000021320 | 58574 | 0.4715 | 0 | 283.8228 | Eukaryota | 45527.0 | 0 | Chordata | 45527.0 | 0 | Carnivora | 45527.0 | 0 | Felidae | 45527.0 | 0 | Felis | 40878.0 | 1 | Felis catus | 40878.0 | 1 |
| scf7180000022153 | 37469 | 0.3813 | 0 | 25.8172 | Eukaryota | 5285.0 | 0 | Chordata | 5285.0 | 0 | Carnivora | 5285.0 | 0 | Felidae | 5285.0 | 0 | Felis | 4767.0 | 1 | Felis catus | 4767.0 | 1 |
| scf7180000021434 | 60754 | 0.5877 | 0 | 40.7967 | Eukaryota | 27517.0 | 0 | Chordata | 27517.0 | 0 | Carnivora | 21543.0 | 1 | Felidae | 10942.0 | 6 | :eratotheriun | 3000.0 | 9 | ıtotherium sir | 3000.0 | 9 |
| scf7180000023406 | 22339 | 0.4257 | 0 | 429.6717 | Eukaryota | 246528.0 | 0 | Chordata | 246528.0 | 0 | Carnivora | 246528.0 | 0 | Hyaenidae | 246528.0 | 0 | Proteles | 147089.0 | 1 | roteles cristat | 147089.0 | 1 |
| scf7180000022066 | 28643 | 0.4443 | 0 | 25.6112 | Eukaryota | 6475.0 | 0 | Chordata | 6475.0 | 0 | Carnivora | 5983.0 | 1 | Felidae | 3857.0 | 2 | Felis | 3857.0 | 2 | Felis catus | 3857.0 | 2 |
| scf7180000018230 | 992238 | 0.4139 | 0 | 51.2083 | Eukaryota | 51817.0 | 0 | Chordata | 51817.0 | 0 | Carnivora | 51817.0 | 0 | Felidae | 51817.0 | 0 | Felis | 51817.0 | 0 | Felis catus | 51817.0 | 0 |
| scf7180000021516 | 47065 | 0.445 | 0 | 26.7874 | Eukaryota | 14000.0 | 0 | Chordata | 14000.0 | 0 | Carnivora | 14000.0 | 0 | Felidae | 13013.0 | 1 | Felis | 8959.0 | 3 | Felis catus | 8959.0 | 3 |
| scf7180000023045 | 8697 | 0.537 | 0 | 91.634 | Eukaryota | 12294.0 | 0 | Chordata | 12294.0 | 0 | Carnivora | 12294.0 | 0 | Felidae | 12294.0 | 0 | Felis | 11069.0 | 1 | Felis catus | 11069.0 | 1 |
| scf7180000021655 | 12816 | 0.4208 | 0 | 27.9728 | Eukaryota | 11053.0 | 0 | Chordata | 11053.0 | 0 | Carnivora | 11053.0 | 0 | Felidae | 11053.0 | 0 | Felis | 9955.0 | 1 | Felis catus | 9955.0 | 1 |
| scf7180000020399 | 106194 | 0.4651 | 0 | 47.1413 | Eukaryota | 4862.0 | 1 | Chordata | 4862.0 | 1 | Carnivora | 3563.0 | 3 | Felidae | 2971.0 | 4 | Felis | 2971.0 | 4 | Felis catus | 2971.0 | 4 |
| scf7180000019559 | 97871 | 0.4863 | 0 | 44.1442 | Eukaryota | 29400.0 | 0 | Chordata | 29400.0 | 0 | Carnivora | 29400.0 | 0 | Felidae | 26969.0 | 1 | Acinonyx | 7430.0 | 5 | :inonyx jubat | 7430.0 | 5 |
| scf7180000022111 | 11955 | 0.5202 | 0 | 62.0761 | no-hit | 0.0 | 0 | no-hit | 0.0 | 0 | no-hit | 0.0 | 0 | no-hit | 0.0 | 0 | no-hit | 0.0 | 0 | no-hit | 0.0 | 0 |
| scf7180000017664 | 2050316 | 0.3777 | 0 | 51.1325 | Eukaryota | 92020.0 | 0 | Chordata | 92020.0 | 0 | Carnivora | 61971.0 | 1 | Felidae | 61971.0 | 1 | Panthera | 35297.0 | 3 | Homo sapiens | 30049.0 | 4 |
| scf7180000019066 | 522624 | 0.3966 | 0 | 49.935 | Eukaryota | 47023.0 | 0 | Chordata | 47023.0 | 0 | Carnivora | 47023.0 | 0 | Felidae | 47023.0 | 0 | Felis | 47023.0 | 0 | Felis catus | 47023.0 | 0 |
| scf7180000022167 | 28740 | 0.6863 | 0 | 53.8483 | Eukaryota | 10267.0 | 1 | Chordata | 10267.0 | 1 | Carnivora | 7201.0 | 2 | Felidae | 3960.0 | 5 | Homo | 3066.0 | 7 | Homo sapiens | 3066.0 | 7 |
| scf7180000022170 | 30472 | 0.57 | 0 | 45.5529 | Eukaryota | 12319.0 | 0 | Chordata | 12319.0 | 0 | Carnivora | 6424.0 | 2 | Bovidae | 2479.0 | 7 | Felis | 1600.0 | 9 | Felis catus | 1600.0 | 9 |
| scf7180000022523 | 14612 | 0.4294 | 0 | 132.9628 | Eukaryota | 2313.0 | 0 | Chordata | 2313.0 | 0 | Carnivora | 2313.0 | 0 | Felidae | 2313.0 | 0 | Felis | 1443.0 | 2 | Felis catus | 1443.0 | 2 |
| scf7180000020885 | 71602 | 0.5298 | 0 | 45.3709 | Eukaryota | 15524.0 | 1 | Chordata | 15524.0 | 1 | Carnivora | 7137.0 | 3 | Canidae | 5598.0 | 5 | Canis | 5598.0 | 6 | Canis lupus | 5598.0 | 6 |
| scf7180000022936 | 522335 | 0.3971 | 0 | 82.7437 | Eukaryota | 44738.0 | 0 | Chordata | 44738.0 | 0 | Carnivora | 44738.0 | 0 | Felidae | 44738.0 | 0 | Felis | 40333.0 | 1 | Felis catus | 40333.0 | 1 |
| scf7180000019967 | 235383 | 0.4004 | 0 | 48.8763 | Eukaryota | 45332.0 | 0 | Chordata | 45332.0 | 0 | Carnivora | 45332.0 | 0 | Felidae | 45332.0 | 0 | Felis | 45332.0 | 0 | Felis catus | 45332.0 | 0 |
| scf7180000018385 | 56660 | 0.4185 | 0 | 47.1232 | Eukaryota | 42486.0 | 0 | Chordata | 42486.0 | 0 | Carnivora | 42486.0 | 0 | Felidae | 42486.0 | 0 | Felis | 38277.0 | 1 | Felis catus | 38277.0 | 1 |
| scf7180000019900 | 128709 | 0.3677 | 0 | 24.6442 | Eukaryota | 12897.0 | 0 | Chordata | 12897.0 | 0 | Carnivora | 12897.0 | 0 | Felidae | 11432.0 | 1 | Acinonyx | 8099.0 | 3 | :inonyx jubat | 8099.0 | 3 |
| scf7180000020931 | 62646 | 0.3922 | 0 | 24.3128 | Eukaryota | 11262.0 | 0 | Chordata | 11262.0 | 0 | Carnivora | 5392.0 | 3 | Hominidae | 3606.0 | 5 | Homo | 2387.0 | 7 | Homo sapiens | 2387.0 | 8 |
| scf7180000020823 | 38849 | 0.6401 | 0 | 49.0261 | Eukaryota | 20068.0 | 0 | Chordata | 20068.0 | 0 | Carnivora | 20068.0 | 0 | Felidae | 20068.0 | 0 | Felis | 10050.0 | 2 | Felis catus | 10050.0 | 2 |
| scf7180000017771 | 188148 | 0.4091 | 0 | 51.4008 | Eukaryota | 47311.0 | 0 | Chordata | 47311.0 | 0 | Carnivora | 47311.0 | 0 | Felidae | 47311.0 | 0 | Felis | 47311.0 | 0 | Felis catus | 47311.0 | 0 |
| scf7180000022454 | 9444 | 0.5188 | 0 | 58.2053 | no-hit | 0.0 | 0 | no-hit | 0.0 | 0 | no-hit | 0.0 | 0 | no-hit | 0.0 | 0 | no-hit | 0.0 | 0 | no-hit | 0.0 | 0 |
| scf7180000021326 | 55728 | 0.4632 | 0 | 26.7047 | Eukaryota | 6565.0 | 0 | Chordata | 6565.0 | 0 | Carnivora | 6565.0 | 0 | Felidae | 5633.0 | 1 | Felis | 2763.0 | 3 | Felis catus | 2763.0 | 4 |
| scf7180000020801 | 100439 | 0.4701 | 0 | 30.1181 | Eukaryota | 12495.0 | 0 | Chordata | 12495.0 | 0 | Carnivora | 12495.0 | 0 | Felidae | 8996.0 | 3 | Panthera | 3722.0 | 5 | anthera pardı | 3722.0 | 5 |
| scf7180000018765 | 5239325 | 0.3867 | 100 | 50.781 | Eukaryota | 56174.0 | 0 | Chordata | 56174.0 | 0 | Carnivora | 40477.0 | 2 | Felidae | 34580.0 | 3 | Panthera | 11545.0 | 6 | ynx canadens | 11528.0 | 7 |
| scf7180000020934 | 92801 | 0.5394 | 0 | 45.9972 | Eukaryota | 5307.0 | 0 | Chordata | 5307.0 | 0 | Carnivora | 5307.0 | 0 | Felidae | 4793.0 | 1 | Acinonyx | 2745.0 | 2 | :inonyx jubat | 2745.0 | 2 |
| scf7180000020054 | 186301 | 0.3876 | 0 | 26.1438 | Eukaryota | 40622.0 | 0 | Chordata | 40622.0 | 0 | Carnivora | 22880.0 | 2 | Felidae | 22880.0 | 2 | Felis | 22880.0 | 3 | Felis catus | 22880.0 | 3 |
| scf7180000018285 | 130344 | 0.4359 | 0 | 48.4583 | Eukaryota | 24036.0 | 0 | Chordata | 24036.0 | 0 | Carnivora | 24036.0 | 0 | Felidae | 24036.0 | 0 | Panthera | 14432.0 | 1 | anthera pardı | 9636.0 | 2 |
| scf7180000019322 | 310826 | 0.4833 | 0 | 52.0369 | Eukaryota | 14935.0 | 0 | Chordata | 14935.0 | 0 | Carnivora | 14935.0 | 0 | Felidae | 11544.0 | 2 | Panthera | 4888.0 | 4 | Felis catus | 3868.0 | 5 |
| scf7180000019870 | 218345 | 0.4555 | 0 | 34.4668 | Eukaryota | 24713.0 | 0 | Chordata | 24713.0 | 0 | Carnivora | 24713.0 | 0 | Felidae | 12000.0 | 3 | Zalophus | 5489.0 | 6 | phus californi | 5489.0 | 7 |
| scf7180000022291 | 24616 | 0.3551 | 0 | 24.6884 | Eukaryota | 20117.0 | 0 | Chordata | 20117.0 | 0 | Carnivora | 20117.0 | 0 | Felidae | 20117.0 | 0 | Felis | 18065.0 | 1 | Felis catus | 18065.0 | 1 |
| scf7180000020739 | 89950 | 0.3896 | 0 | 23.7606 | Eukaryota | 25965.0 | 0 | Chordata | 25965.0 | 0 | Carnivora | 25965.0 | 0 | Felidae | 25965.0 | 0 | Acinonyx | 23436.0 | 1 | :inonyx jubat | 23436.0 | 1 |
| scf7180000018280 | 366094 | 0.3608 | 0 | 50.5235 | Eukaryota | 46508.0 | 0 | Chordata | 46508.0 | 0 | Carnivora | 46508.0 | 0 | Felidae | 46508.0 | 0 | Felis | 41911.0 | 1 | Felis catus | 41911.0 | 1 |
| scf7180000022838 | 37766 | 0.6196 | 0 | 877.1746 | Eukaryota | 1283.0 | 0 | Chordata | 1283.0 | 0 | Carnivora | 1283.0 | 0 | Felidae | 1283.0 | 0 | Felis | 1283.0 | 0 | Felis catus | 1283.0 | 0 |
| scf7180000021940 | 169537 | 0.5935 | 0 | 49.5178 | Eukaryota | 36359.0 | 0 | Chordata | 36359.0 | 0 | Carnivora | 36359.0 | 0 | Felidae | 36359.0 | 0 | Felis | 32776.0 | 1 | Felis catus | 32776.0 | 1 |
| scf7180000022824 | 1447491 | 0.3575 | 0 | 49.5383 | Eukaryota | 96707.0 | 0 | Chordata | 96707.0 | 0 | Chiroptera | 64016.0 | 1 | Pteropodidae | 64016.0 | 1 | Rousettus | 64016.0 | 1 | ;ettus aegypti | 64016.0 | 1 |
| scf7180000019124 | 186688 | 0.4143 | 0 | 51.0199 | Eukaryota | 20302.0 | 0 | Chordata | 20302.0 | 0 | Carnivora | 20302.0 | 0 | Felidae | 20302.0 | 0 | Felis | 18191.0 | 1 | Felis catus | 18191.0 | 1 |
| scf7180000021165 | 168763 | 0.3798 | 0 | 25.9225 | Eukaryota | 84590.0 | 0 | Chordata | 84590.0 | 0 | Carnivora | 84590.0 | 0 | Felidae | 84590.0 | 0 | Acinonyx | 42345.0 | 1 | :inonyx jubat | 42345.0 | 1 |
| scf7180000023764 | 3563082 | 0.4234 | 0 | 51.2547 | Eukaryota | 84126.0 | 0 | Chordata | 84126.0 | 0 | Carnivora | 84126.0 | 0 | Felidae | 68107.0 | 2 | Felis | 25584.0 | 4 | Felis catus | 25584.0 | 4 |
| scf7180000018572 | 635124 | 0.4182 | 0 | 50.5597 | Eukaryota | 44164.0 | 0 | Chordata | 44164.0 | 0 | Carnivora | 44164.0 | 0 | Felidae | 44164.0 | 0 | Felis | 39783.0 | 1 | Felis catus | 39783.0 | 1 |
| scf7180000018595 | 190869 | 0.4247 | 0 | 55.2939 | Eukaryota | 44367.0 | 0 | Chordata | 44367.0 | 0 | Carnivora | 44367.0 | 0 | Felidae | 44367.0 | 0 | Felis | 44367.0 | 0 | Felis catus | 44367.0 | 0 |
| scf7180000019254 | 412589 | 0.5366 | 0 | 51.04 | Eukaryota | 21139.0 | 0 | Chordata | 21139.0 | 0 | Carnivora | 21139.0 | 0 | Felidae | 21139.0 | 0 | Felis | 19026.0 | 1 | Felis catus | 19026.0 | 1 |
| scf7180000022745 | 46841 | 0.5347 | 0 | 33.2323 | Eukaryota | 13326.0 | 0 | Chordata | 13326.0 | 0 | Carnivora | 13326.0 | 0 | Felidae | 8377.0 | 2 | Acinonyx | 2538.0 | 7 | :inonyx jubat | 2538.0 | 7 |
| scf7180000021621 | 69451 | 0.4067 | 0 | 25.9271 | Eukaryota | 24440.0 | 0 | Chordata | 24440.0 | 0 | Carnivora | 24440.0 | 0 | Felidae | 24440.0 | 0 | Felis | 22157.0 | 1 | Felis catus | 22157.0 | 1 |
| scf7180000020433 | 152857 | 0.3912 | 0 | 26.0415 | Eukaryota | 19848.0 | 0 | Chordata | 19848.0 | 0 | Primates | 11992.0 | 2 | Hominidae | 11992.0 | 2 | Homo | 9428.0 | 3 | Homo sapiens | 9428.0 | 3 |
| scf7180000017562 | 610588 | 0.4032 | 0 | 51.3005 | Eukaryota | 44061.0 | 0 | Chordata | 44061.0 | 0 | Carnivora | 44061.0 | 0 | Felidae | 44061.0 | 0 | Felis | 39680.0 | 1 | Felis catus | 39680.0 | 1 |
| scf7180000022215 | 28245 | 0.4333 | 0 | 24.6051 | Eukaryota | 27960.0 | 0 | Chordata | 27960.0 | 0 | Carnivora | 27960.0 | 0 | Felidae | 27960.0 | 0 | Felis | 25311.0 | 1 | Felis catus | 25311.0 | 1 |
| scf7180000021421 | 52847 | 0.3562 | 0 | 24.4186 | Eukaryota | 8427.0 | 0 | Chordata | 8427.0 | 0 | Primates | 6466.0 | 2 | Hominidae | 6466.0 | 2 | Homo | 5481.0 | 3 | Homo sapiens | 5481.0 | 3 |
| scf7180000020544 | 31984 | 0.4632 | 0 | 316.9433 | Eukaryota | 6272.0 | 0 | Chordata | 6272.0 | 0 | Artiodactyla | 6272.0 | 0 | Bovidae | 6272.0 | 0 | Ovis | 6272.0 | 0 | )vis canadens | 6272.0 | 0 |
| scf7180000017788 | 796439 | 0.3762 | 0 | 50.497 | Eukaryota | 44033.0 | 0 | Chordata | 44033.0 | 0 | Carnivora | 44033.0 | 0 | Felidae | 44033.0 | 0 | Felis | 44033.0 | 0 | Felis catus | 44033.0 | 0 |
| scf7180000018810 | 1693294 | 0.4246 | 0 | 51.8067 | Eukaryota | 48251.0 | 0 | Chordata | 48251.0 | 0 | Carnivora | 48251.0 | 0 | Felidae | 48251.0 | 0 | Felis | 48251.0 | 0 | Felis catus | 48251.0 | 0 |
| scf7180000020141 | 139816 | 0.4689 | 0 | 233.0801 | Eukaryota | 39079.0 | 0 | Chordata | 39079.0 | 0 | Carnivora | 39079.0 | 0 | Felidae | 30217.0 | 1 | Felis | 26494.0 | 2 | Felis catus | 26494.0 | 2 |
| scf7180000020513 | 203487 | 0.5468 | 0 | 46.1913 | Eukaryota | 19708.0 | 1 | Chordata | 19708.0 | 1 | Carnivora | 7750.0 | 3 | Felidae | 5421.0 | 5 | undef | 4304.0 | 7 | tic synthetic c | 4304.0 | 7 |
| scf7180000019561 | 3047244 | 0.373 | 100 | 50.5307 | Eukaryota | 46191.0 | 0 | Chordata | 46191.0 | 0 | Carnivora | 46191.0 | 0 | Felidae | 46191.0 | 0 | Felis | 46191.0 | 0 | Felis catus | 46191.0 | 0 |
| scf7180000023465 | 19475 | 0.4866 | 0 | 230.4234 | Eukaryota | 9292.0 | 0 | Chordata | 9292.0 | 0 | Carnivora | 9292.0 | 0 | Felidae | 9292.0 | 0 | Felis | 9292.0 | 0 | Felis catus | 9292.0 | 0 |
| scf7180000021387 | 31902 | 0.4215 | 0 | 24.4689 | Eukaryota | 5983.0 | 0 | Chordata | 5983.0 | 0 | Perissodactyla | 3995.0 | 3 | Equidae | 3995.0 | 4 | Equus | 3995.0 | 4 | juus przewals | 3636.0 | 5 |
| scf7180000023821 | 2228530 | 0.3862 | 0 | 51.1588 | Eukaryota | 73211.0 | 0 | Chordata | 73211.0 | 0 | Carnivora | 73211.0 | 0 | Felidae | 73211.0 | 0 | Lynx | 22449.0 | 3 | ynx canadens | 22449.0 | 4 |
| scf7180000022353 | 29783 | 0.3772 | 0 | 28.1419 | Eukaryota | 47258.0 | 0 | Chordata | 47258.0 | 0 | Carnivora | 47258.0 | 0 | Felidae | 47258.0 | 0 | Felis | 47258.0 | 0 | Felis catus | 47258.0 | 0 |
| scf7180000022871 | 30750 | 0.4632 | 0 | 163.0504 | Eukaryota | 40283.0 | 0 | Chordata | 40283.0 | 0 | Carnivora | 40283.0 | 0 | Felidae | 40283.0 | 0 | Felis | 36295.0 | 1 | Felis catus | 36295.0 | 1 |
| scf7180000020782 | 79814 | 0.4307 | 0 | 577.6387 | Eukaryota | 2374.0 | 0 | Chordata | 2374.0 | 0 | Carnivora | 2374.0 | 0 | Felidae | 2374.0 | 0 | Felis | 1718.0 | 2 | Felis catus | 1718.0 | 2 |
| scf7180000018429 | 698953 | 0.4218 | 0 | 52.8463 | Eukaryota | 45832.0 | 0 | Chordata | 45832.0 | 0 | Carnivora | 45832.0 | 0 | Felidae | 45832.0 | 0 | Felis | 45832.0 | 0 | Felis catus | 45832.0 | 0 |
| scf7180000022298 | 16560 | 0.4127 | 0 | 24.8543 | Eukaryota | 5456.0 | 0 | Chordata | 5456.0 | 0 | Artiodactyla | 3870.0 | 1 | Delphinidae | 2090.0 | 4 | Myotis | 1054.0 | 8 | Ayotis lucifugı | 1054.0 | 8 |
| scf7180000019338 | 12599 | 0.3945 | 0 | 28.6707 | Eukaryota | 10456.0 | 0 | Chordata | 10456.0 | 0 | Carnivora | 10456.0 | 0 | Felidae | 10456.0 | 0 | Felis | 9414.0 | 1 | Felis catus | 9414.0 | 1 |
| scf7180000019380 | 321069 | 0.4064 | 0 | 50.822 | Eukaryota | 39033.0 | 0 | Chordata | 39033.0 | 0 | Carnivora | 39033.0 | 0 | Felidae | 39033.0 | 0 | Felis | 39033.0 | 0 | Felis catus | 39033.0 | 0 |
| scf7180000021787 | 24894 | 0.5801 | 0 | 48.0996 | Eukaryota | 3321.0 | 1 | Chordata | 3321.0 | 1 | Carnivora | 1351.0 | 3 | Muridae | 1149.0 | 5 | Mus | 1149.0 | 6 | Mus musculus | 1149.0 | 6 |
| scf7180000021760 | 23027 | 0.5719 | 0 | 66.2676 | Eukaryota | 2966.0 | 0 | Chordata | 2966.0 | 0 | Carnivora | 2966.0 | 0 | Felidae | 2966.0 | 0 | Felis | 1487.0 | 2 | Felis catus | 1487.0 | 4 |
| scf7180000017878 | 11655 | 0.416 | 0 | 32.267 | Eukaryota | 11249.0 | 0 | Chordata | 11249.0 | 0 | Carnivora | 11249.0 | 0 | Felidae | 11249.0 | 0 | Felis | 10100.0 | 1 | Felis catus | 10100.0 | 1 |
| scf7180000024275 | 818152 | 0.3515 | 100 | 48.6528 | Eukaryota | 47925.0 | 0 | Chordata | 47925.0 | 0 | Carnivora | 47925.0 | 0 | Felidae | 38126.0 | 2 | Felis | 38126.0 | 2 | Felis catus | 38126.0 | 2 |
| scf7180000019774 | 181929 | 0.4847 | 0 | 50.1275 | Eukaryota | 51814.0 | 0 | Chordata | 51814.0 | 0 | Carnivora | 51814.0 | 0 | Felidae | 51814.0 | 0 | Acinonyx | 41544.0 | 2 | :inonyx jubat | 41544.0 | 2 |
| scf7180000018535 | 886547 | 0.4119 | 0 | 51.4471 | Eukaryota | 47123.0 | 0 | Chordata | 47123.0 | 0 | Carnivora | 47123.0 | 0 | Felidae | 47123.0 | 0 | Felis | 47123.0 | 0 | Felis catus | 47123.0 | 0 |
| scf7180000021015 | 49980 | 0.4324 | 0 | 23.8262 | Eukaryota | 6142.0 | 0 | Chordata | 6142.0 | 0 | Carnivora | 3842.0 | 2 | Felidae | 3842.0 | 2 | Felis | 3842.0 | 2 | Felis catus | 3842.0 | 2 |
| scf7180000018189 | 964128 | 0.4885 | 0 | 50.8016 | Eukaryota | 23722.0 | 0 | Chordata | 23722.0 | 0 | Carnivora | 23722.0 | 0 | Felidae | 18736.0 | 1 | Lynx | 18736.0 | 1 | ynx canadens | 18736.0 | 1 |
| scf7180000020371 | 116265 | 0.3783 | 0 | 24.3325 | Eukaryota | 21430.0 | 0 | Chordata | 21430.0 | 0 | Carnivora | 21430.0 | 0 | Felidae | 21430.0 | 0 | Acinonyx | 8175.0 | 2 | :inonyx jubat | 8175.0 | 2 |
| scf7180000021956 | 21135 | 0.4229 | 0 | 23.7309 | Eukaryota | 7199.0 | 0 | Chordata | 7199.0 | 0 | Carnivora | 7199.0 | 0 | Felidae | 4423.0 | 1 | Felis | 4423.0 | 1 | Felis catus | 4423.0 | 1 |
| scf7180000020842 | 90256 | 0.5909 | 0 | 47.4491 | Eukaryota | 41030.0 | 0 | Chordata | 41030.0 | 0 | Carnivora | 41030.0 | 0 | Felidae | 41030.0 | 0 | Felis | 36965.0 | 1 | Felis catus | 36965.0 | 1 |
| scf7180000020066 | 154683 | 0.3911 | 0 | 50.5257 | Eukaryota | 21064.0 | 0 | Chordata | 21064.0 | 0 | Carnivora | 21064.0 | 0 | Felidae | 19086.0 | 1 | Felis | 14237.0 | 3 | Felis catus | 14237.0 | 3 |
| scf7180000018034 | 1811004 | 0.4126 | 0 | 51.1079 | Eukaryota | 47237.0 | 0 | Chordata | 47237.0 | 0 | Carnivora | 47237.0 | 0 | Felidae | 47237.0 | 0 | Felis | 47237.0 | 0 | Felis catus | 47237.0 | 0 |
| scf7180000023157 | 458136 | 0.3597 | 0 | 48.0653 | Eukaryota | 55853.0 | 0 | Chordata | 55853.0 | 0 | Carnivora | 49345.0 | 1 | Felidae | 26933.0 | 4 | Acinonyx | 12135.0 | 7 | :inonyx jubat | 12135.0 | 7 |
| scf7180000020529 | 141231 | 0.5344 | 0 | 51.6265 | Eukaryota | 19469.0 | 0 | Chordata | 19469.0 | 0 | Carnivora | 19469.0 | 0 | Felidae | 19469.0 | 0 | Panthera | 10485.0 | 1 | anthera pardı | 10485.0 | 1 |
| scf7180000019560 | 12820 | 0.424 | 0 | 28.524 | Eukaryota | 10610.0 | 0 | Chordata | 10610.0 | 0 | Carnivora | 10610.0 | 0 | Felidae | 10610.0 | 0 | Felis | 10610.0 | 0 | Felis catus | 10610.0 | 0 |
| scf7180000023844 | 1837567 | 0.401 | 0 | 50.3002 | Eukaryota | 46196.0 | 0 | Chordata | 46196.0 | 0 | Carnivora | 46196.0 | 0 | Felidae | 46196.0 | 0 | Felis | 46196.0 | 0 | Felis catus | 46196.0 | 0 |
| scf7180000022636 | 2686 | 0.5149 | 0 | 318.9544 | no-hit | 0.0 | 0 | no-hit | 0.0 | 0 | no-hit | 0.0 | 0 | no-hit | 0.0 | 0 | no-hit | 0.0 | 0 | no-hit | 0.0 | 0 |
| scf7180000021824 | 57355 | 0.5328 | 0 | 46.4551 | Eukaryota | 10487.0 | 0 | Chordata | 10487.0 | 0 | Carnivora | 10487.0 | 0 | Felidae | 10487.0 | 0 | Felis | 6252.0 | 1 | Felis catus | 6252.0 | 2 |
| scf7180000019824 | 20106 | 0.4213 | 0 | 25.7394 | Eukaryota | 36597.0 | 0 | Chordata | 36597.0 | 0 | Carnivora | 36597.0 | 0 | Felidae | 36597.0 | 0 | Felis | 32875.0 | 1 | Felis catus | 32875.0 | 1 |
| scf7180000023376 | 16925 | 0.483 | 0 | 336.2452 | Eukaryota | 3166.0 | 0 | Chordata | 3166.0 | 0 | Carnivora | 3166.0 | 0 | Felidae | 2709.0 | 1 | Acinonyx | 867.0 | 5 | :inonyx jubat | 867.0 | 5 |
| scf7180000022056 | 102385 | 0.5342 | 0 | 47.568 | Eukaryota | 15212.0 | 0 | Chordata | 15212.0 | 0 | Carnivora | 15212.0 | 0 | Felidae | 13671.0 | 1 | Felis | 13671.0 | 1 | Felis catus | 13671.0 | 1 |
| scf7180000018666 | 620859 | 0.3858 | 0 | 50.834 | Eukaryota | 45062.0 | 0 | Chordata | 45062.0 | 0 | Carnivora | 45062.0 | 0 | Felidae | 45062.0 | 0 | Felis | 40592.0 | 1 | Felis catus | 40592.0 | 1 |
| scf7180000023827 | 391265 | 0.3917 | 0 | 49.2091 | Eukaryota | 26953.0 | 1 | Chordata | 26953.0 | 1 | Primates | 13573.0 | 4 | Hominidae | 13573.0 | 4 | Homo | 13573.0 | 5 | Homo sapiens | 13573.0 | 5 |
| scf7180000018380 | 303813 | 0.3788 | 0 | 50.7588 | Eukaryota | 46636.0 | 0 | Chordata | 46636.0 | 0 | Carnivora | 46636.0 | 0 | Felidae | 46636.0 | 0 | Felis | 46636.0 | 0 | Felis catus | 46636.0 | 0 |
| scf7180000019704 | 38885 | 0.401 | 0 | 48.3188 | Eukaryota | 22916.0 | 0 | Chordata | 22916.0 | 0 | Carnivora | 22916.0 | 0 | Felidae | 22916.0 | 0 | Felis | 20882.0 | 1 | Felis catus | 20882.0 | 1 |
| scf7180000017898 | 1377158 | 0.4523 | 0 | 51.1755 | Eukaryota | 44172.0 | 0 | Chordata | 44172.0 | 0 | Carnivora | 44172.0 | 0 | Felidae | 44172.0 | 0 | Felis | 39791.0 | 1 | Felis catus | 39791.0 | 1 |
| scf7180000021722 | 64737 | 0.5647 | 0 | 49.9815 | Eukaryota | 3488.0 | 1 | Chordata | 3488.0 | 1 | Carnivora | 1580.0 | 3 | Felidae | 1225.0 | 4 | Homo | 1016.0 | 6 | Homo sapiens | 1016.0 | 6 |
| scf7180000021384 | 41187 | 0.4702 | 0 | 27.2784 | Eukaryota | 26357.0 | 0 | Chordata | 26357.0 | 0 | Carnivora | 18041.0 | 1 | Felidae | 18041.0 | 1 | Homo | 8316.0 | 4 | Homo sapiens | 8316.0 | 4 |
| scf7180000017982 | 472815 | 0.4125 | 0 | 49.8112 | Eukaryota | 45722.0 | 0 | Chordata | 45722.0 | 0 | Carnivora | 45722.0 | 0 | Felidae | 45722.0 | 0 | Felis | 45722.0 | 0 | Felis catus | 45722.0 | 0 |
| scf7180000024282 | 95793 | 0.3987 | 100 | 49.5082 | Eukaryota | 30310.0 | 0 | Chordata | 30310.0 | 0 | Artiodactyla | 18209.0 | 2 | Delphinidae | 6097.0 | 6 | Balaenoptera | 6060.0 | 7 | optera acutor | 6060.0 | 8 |
| scf7180000018728 | 699070 | 0.3802 | 0 | 50.4468 | Eukaryota | 45934.0 | 0 | Chordata | 45934.0 | 0 | Carnivora | 45934.0 | 0 | Felidae | 45934.0 | 0 | Felis | 45934.0 | 0 | Felis catus | 45934.0 | 0 |
| scf7180000018373 | 806022 | 0.3793 | 0 | 49.2678 | Eukaryota | 26535.0 | 0 | Chordata | 26535.0 | 0 | Carnivora | 26535.0 | 0 | Felidae | 26535.0 | 0 | Felis | 21094.0 | 1 | Felis catus | 21094.0 | 1 |
| scf7180000022555 | 10880 | 0.4114 | 0 | 27.1496 | Eukaryota | 11292.0 | 0 | Chordata | 11292.0 | 0 | Primates | 7455.0 | 3 | Hominidae | 7455.0 | 3 | Pan | 5984.0 | 4 | an troglodyte | 5984.0 | 6 |
| scf7180000021141 | 60522 | 0.6104 | 0 | 48.1912 | Eukaryota | 8518.0 | 0 | Chordata | 8518.0 | 0 | Carnivora | 8518.0 | 0 | Felidae | 8518.0 | 0 | Felis | 8518.0 | 0 | Felis catus | 8518.0 | 0 |
| scf7180000022359 | 13293 | 0.3724 | 0 | 23.5246 | Eukaryota | 11253.0 | 0 | Chordata | 11253.0 | 0 | Carnivora | 11253.0 | 0 | Felidae | 10143.0 | 1 | Felis | 9040.0 | 2 | Felis catus | 9040.0 | 2 |
| scf7180000023904 | 1519942 | 0.3993 | 0 | 51.0197 | Eukaryota | 49794.0 | 0 | Chordata | 49794.0 | 0 | Carnivora | 49794.0 | 0 | Felidae | 49794.0 | 0 | Panthera | 35099.0 | 2 | anthera pardı | 18860.0 | 3 |
| scf7180000017784 | 1685305 | 0.3681 | 0 | 49.7614 | Eukaryota | 47564.0 | 0 | Chordata | 47564.0 | 0 | Carnivora | 47564.0 | 0 | Felidae | 47564.0 | 0 | Felis | 47564.0 | 0 | Felis catus | 47564.0 | 0 |
| scf7180000020680 | 58274 | 0.3589 | 0 | 25.2386 | Eukaryota | 43871.0 | 0 | Chordata | 43871.0 | 0 | Carnivora | 43871.0 | 0 | Felidae | 43871.0 | 0 | Felis | 43871.0 | 0 | Felis catus | 43871.0 | 0 |
| scf7180000019606 | 289980 | 0.392 | 0 | 51.9878 | Eukaryota | 29730.0 | 0 | Chordata | 29730.0 | 0 | Carnivora | 29730.0 | 0 | Felidae | 26932.0 | 1 | Panthera | 14940.0 | 2 | anthera pardı | 14940.0 | 2 |
| scf7180000019477 | 288876 | 0.3572 | 0 | 48.7667 | Eukaryota | 69750.0 | 0 | Chordata | 69750.0 | 0 | Carnivora | 69750.0 | 0 | Canidae | 69750.0 | 0 | Canis | 69750.0 | 0 | Canis lupus | 69750.0 | 0 |
| scf7180000018060 | 569767 | 0.4377 | 0 | 52.0643 | Eukaryota | 45922.0 | 0 | Chordata | 45922.0 | 0 | Carnivora | 45922.0 | 0 | Felidae | 45922.0 | 0 | Felis | 36138.0 | 1 | Felis catus | 36138.0 | 1 |
| scf7180000021272 | 38175 | 0.4407 | 0 | 45.9693 | Eukaryota | 23958.0 | 0 | Chordata | 23958.0 | 0 | Carnivora | 23958.0 | 0 | Felidae | 23958.0 | 0 | Felis | 21581.0 | 1 | Felis catus | 21581.0 | 1 |
| scf7180000019177 | 228824 | 0.4224 | 0 | 50.2379 | Eukaryota | 46615.0 | 0 | Chordata | 46615.0 | 0 | Carnivora | 46615.0 | 0 | Felidae | 46615.0 | 0 | Felis | 46615.0 | 0 | Felis catus | 46615.0 | 0 |
| scf7180000022046 | 8406 | 0.385 | 0 | 12.8666 | Eukaryota | 45102.0 | 0 | Chordata | 45102.0 | 0 | Carnivora | 45102.0 | 0 | Felidae | 45102.0 | 0 | Felis | 45102.0 | 0 | Felis catus | 45102.0 | 0 |
| scf7180000022352 | 21952 | 0.3716 | 0 | 24.2176 | Eukaryota | 6304.0 | 0 | Chordata | 6304.0 | 0 | Carnivora | 6304.0 | 0 | Felidae | 4238.0 | 2 | Felis | 4238.0 | 2 | Felis catus | 4238.0 | 2 |
| scf7180000021431 | 156131 | 0.5215 | 0 | 48.4283 | Eukaryota | 26820.0 | 0 | Chordata | 26820.0 | 0 | Carnivora | 26820.0 | 0 | Felidae | 26820.0 | 0 | Acinonyx | 26820.0 | 0 | :inonyx jubat | 26820.0 | 0 |
| scf7180000018769 | 112498 | 0.4451 | 0 | 48.0439 | Eukaryota | 58037.0 | 0 | Chordata | 58037.0 | 0 | Carnivora | 58037.0 | 0 | Felidae | 37847.0 | 1 | Suricata | 20190.0 | 4 | ıricata suricat | 20190.0 | 4 |
| scf7180000017713 | 1847378 | 0.3647 | 0 | 48.7477 | Eukaryota | 52572.0 | 0 | Chordata | 52572.0 | 0 | Carnivora | 52572.0 | 0 | Felidae | 52572.0 | 0 | Panthera | 27451.0 | 1 | Felis catus | 25121.0 | 2 |
| scf7180000021863 | 36750 | 0.3576 | 0 | 23.5409 | Eukaryota | 37038.0 | 0 | Chordata | 37038.0 | 0 | Carnivora | 37038.0 | 0 | Felidae | 37038.0 | 0 | Felis | 33374.0 | 1 | Felis catus | 33374.0 | 1 |
| scf7180000018767 | 618240 | 0.4003 | 0 | 51.6944 | Eukaryota | 47428.0 | 0 | Chordata | 47428.0 | 0 | Carnivora | 47428.0 | 0 | Felidae | 47428.0 | 0 | Felis | 47428.0 | 0 | Felis catus | 47428.0 | 0 |
| scf7180000020062 | 139924 | 0.384 | 0 | 49.7899 | Eukaryota | 20086.0 | 0 | Chordata | 20086.0 | 0 | Carnivora | 11264.0 | 2 | Otariidae | 7168.0 | 5 | Zalophus | 7168.0 | 6 | phus californi | 7168.0 | 6 |
| scf7180000023863 | 506805 | 0.3662 | 0 | 50.2373 | Eukaryota | 23439.0 | 0 | Chordata | 23439.0 | 0 | Primates | 19510.0 | 1 | Hominidae | 16390.0 | 2 | Homo | 16390.0 | 2 | Homo sapiens | 16390.0 | 2 |
| scf7180000020029 | 136425 | 0.4356 | 0 | 48.5824 | Eukaryota | 56970.0 | 0 | Chordata | 56970.0 | 0 | Carnivora | 56970.0 | 0 | Felidae | 35730.0 | 3 | Panthera | 13862.0 | 7 | Ursus arctos | 7588.0 | 8 |
| scf7180000020137 | 131756 | 0.5582 | 0 | 48.295 | Eukaryota | 22652.0 | 0 | Chordata | 22652.0 | 0 | Carnivora | 22652.0 | 0 | Felidae | 17154.0 | 2 | Lynx | 5386.0 | 5 | ynx canadens | 5386.0 | 5 |
| scf7180000020193 | 104191 | 0.5572 | 0 | 50.714 | Eukaryota | 6937.0 | 0 | Chordata | 6937.0 | 0 | Carnivora | 6937.0 | 0 | Felidae | 3744.0 | 4 | Felis | 754.0 | 9 | Felis catus | 754.0 | 9 |
| scf7180000021669 | 49017 | 0.3711 | 0 | 22.0726 | Eukaryota | 31493.0 | 0 | Chordata | 31493.0 | 0 | Carnivora | 31493.0 | 0 | Felidae | 31493.0 | 0 | Felis | 31493.0 | 0 | Felis catus | 31493.0 | 0 |
| scf7180000020239 | 32750 | 0.4139 | 0 | 35.3902 | Eukaryota | 24073.0 | 0 | Chordata | 24073.0 | 0 | Carnivora | 24073.0 | 0 | Felidae | 24073.0 | 0 | Felis | 21633.0 | 1 | Felis catus | 21633.0 | 1 |

| | | | | | | | | | | | | | | | | | | | | | | |
|---|---|---|---|---|---|---|---|---|---|---|---|---|---|---|---|---|---|---|---|---|---|---|
| scf7180000024097 | 395315 | 0.5261 | 0 | 54.7253 | Eukaryota | 45507.0 | 0 | Chordata | 45507.0 | 0 | Carnivora | 45507.0 | 0 | Felidae | 45507.0 | 0 | Felis | 40860.0 | 1 | Felis catus | 40860.0 | 1 |
| scf7180000018627 | 410578 | 0.3791 | 0 | 49.8267 | Eukaryota | 46248.0 | 0 | Chordata | 46248.0 | 0 | Carnivora | 46248.0 | 0 | Felidae | 46248.0 | 0 | Felis | 46248.0 | 0 | Felis catus | 46248.0 | 0 |
| scf7180000020185 | 58846 | 0.3899 | 0 | 365.8854 | Eukaryota | 37697.0 | 0 | Chordata | 37697.0 | 0 | Carnivora | 37697.0 | 0 | Felidae | 37697.0 | 0 | Felis | 33950.0 | 1 | Felis catus | 33950.0 | 1 |
| scf7180000020855 | 66233 | 0.3618 | 0 | 24.8196 | Eukaryota | 42828.0 | 0 | Chordata | 42828.0 | 0 | Carnivora | 42828.0 | 0 | Felidae | 42828.0 | 0 | Felis | 42828.0 | 0 | Felis catus | 42828.0 | 0 |
| scf7180000019467 | 187023 | 0.4235 | 0 | 46.3246 | Eukaryota | 40228.0 | 0 | Chordata | 40228.0 | 0 | Carnivora | 40228.0 | 0 | Felidae | 40228.0 | 0 | Felis | 36233.0 | 1 | Felis catus | 36233.0 | 1 |
| scf7180000019202 | 151997 | 0.407 | 0 | 50.8071 | Eukaryota | 33058.0 | 0 | Chordata | 33058.0 | 0 | Carnivora | 33058.0 | 0 | Felidae | 33058.0 | 0 | Felis | 29687.0 | 1 | Felis catus | 29687.0 | 1 |
| scf7180000021290 | 61088 | 0.4134 | 0 | 25.7938 | Eukaryota | 46863.0 | 0 | Chordata | 46863.0 | 0 | Carnivora | 46863.0 | 0 | Felidae | 46863.0 | 0 | Felis | 46863.0 | 0 | Felis catus | 46863.0 | 0 |
| scf7180000019994 | 200554 | 0.382 | 0 | 25.1515 | Eukaryota | 16525.0 | 0 | Chordata | 16525.0 | 0 | Carnivora | 16525.0 | 0 | Felidae | 16525.0 | 0 | Felis | 13218.0 | 1 | Felis catus | 13218.0 | 1 |
| scf7180000022165 | 5645 | 0.6377 | 0 | 31.8962 | Eukaryota | 2537.0 | 0 | Chordata | 2537.0 | 0 | Carnivora | 2537.0 | 0 | Felidae | 2037.0 | 1 | Felis | 1024.0 | 3 | Felis catus | 1024.0 | 3 |
| scf7180000017879 | 1701383 | 0.3797 | 100 | 50.7315 | Eukaryota | 45911.0 | 0 | Chordata | 45911.0 | 0 | Carnivora | 45911.0 | 0 | Felidae | 45911.0 | 0 | Felis | 41362.0 | 1 | Felis catus | 41362.0 | 1 |
| scf7180000024024 | 254028 | 0.3872 | 0 | 49.6591 | Eukaryota | 42370.0 | 0 | Chordata | 42370.0 | 0 | Carnivora | 42370.0 | 0 | Felidae | 42370.0 | 0 | Felis | 42370.0 | 0 | Felis catus | 42370.0 | 0 |
| scf7180000021411 | 53886 | 0.4323 | 0 | 26.9975 | Eukaryota | 11602.0 | 0 | Chordata | 11602.0 | 0 | Carnivora | 11602.0 | 0 | Felidae | 9709.0 | 1 | Felis | 8634.0 | 2 | Felis catus | 8634.0 | 2 |
| scf7180000019703 | 15842 | 0.552 | 0 | 249.4969 | Eukaryota | 2579.0 | 0 | Chordata | 2579.0 | 0 | Carnivora | 1429.0 | 2 | Felidae | 1429.0 | 2 | Felis | 1185.0 | 3 | Felis catus | 1185.0 | 3 |
| scf7180000021417 | 35334 | 0.3539 | 0 | 24.1901 | Eukaryota | 10269.0 | 0 | Chordata | 10269.0 | 0 | Carnivora | 6842.0 | 1 | Felidae | 6842.0 | 1 | Felis | 6842.0 | 2 | Felis catus | 6842.0 | 2 |
| scf7180000019729 | 310206 | 0.4838 | 0 | 51.7568 | Eukaryota | 39746.0 | 0 | Chordata | 39746.0 | 0 | Carnivora | 39746.0 | 0 | Felidae | 39746.0 | 0 | Felis | 39746.0 | 0 | Felis catus | 39746.0 | 0 |
| scf7180000021154 | 64688 | 0.6303 | 0 | 50.4823 | Eukaryota | 3513.0 | 0 | Chordata | 3513.0 | 0 | Carnivora | 2862.0 | 2 | Felidae | 2137.0 | 3 | Felis | 714.0 | 7 | Felis catus | 714.0 | 7 |
| scf7180000018764 | 89716 | 0.3733 | 0 | 46.5388 | Eukaryota | 9504.0 | 0 | Chordata | 9504.0 | 0 | Primates | 5620.0 | 2 | Hominidae | 3694.0 | 4 | Homo | 3694.0 | 5 | Homo sapiens | 3694.0 | 5 |
| scf7180000020484 | 61683 | 0.3799 | 0 | 40.4972 | Eukaryota | 21026.0 | 0 | Chordata | 21026.0 | 0 | Carnivora | 21026.0 | 0 | Felidae | 21026.0 | 0 | Felis | 18891.0 | 1 | Felis catus | 18891.0 | 1 |
| scf7180000018260 | 1048414 | 0.4313 | 0 | 48.6711 | Eukaryota | 17022.0 | 0 | Chordata | 17022.0 | 0 | Carnivora | 17022.0 | 0 | Felidae | 17022.0 | 0 | Felis | 8352.0 | 3 | Felis catus | 8352.0 | 3 |
| scf7180000019387 | 344176 | 0.477 | 0 | 51.031 | Eukaryota | 20425.0 | 0 | Chordata | 20425.0 | 0 | Carnivora | 20425.0 | 0 | Ursidae | 20425.0 | 0 | Ursus | 20425.0 | 0 | Ursus arctos | 14350.0 | 1 |
| scf7180000023783 | 2684425 | 0.4509 | 0 | 52.6427 | Eukaryota | 47696.0 | 0 | Chordata | 47696.0 | 0 | Carnivora | 47696.0 | 0 | Felidae | 47696.0 | 0 | Felis | 47696.0 | 0 | Felis catus | 47696.0 | 0 |
| scf7180000023059 | 59702 | 0.4005 | 0 | 387.9067 | Eukaryota | 47426.0 | 0 | Chordata | 47426.0 | 0 | Carnivora | 47426.0 | 0 | Felidae | 47426.0 | 0 | Felis | 47426.0 | 0 | Felis catus | 47426.0 | 0 |
| scf7180000022300 | 18674 | 0.4287 | 0 | 27.415 | Eukaryota | 65491.0 | 0 | Chordata | 65491.0 | 0 | Carnivora | 65491.0 | 0 | Felidae | 65491.0 | 0 | Panthera | 45963.0 | 3 | anthera pardu | 39462.0 | 4 |
| scf7180000021234 | 54910 | 0.601 | 0 | 48.3636 | Eukaryota | 17183.0 | 0 | Chordata | 17183.0 | 0 | Carnivora | 17183.0 | 0 | Canidae | 13452.0 | 1 | Canis | 13452.0 | 2 | Canis lupus | 13452.0 | 2 |
| scf7180000021497 | 36563 | 0.4394 | 0 | 46.4662 | Eukaryota | 15481.0 | 0 | Chordata | 15481.0 | 0 | Carnivora | 15481.0 | 0 | Felidae | 15481.0 | 0 | Felis | 15481.0 | 0 | Felis catus | 15481.0 | 0 |
| scf7180000017913 | 986721 | 0.3928 | 0 | 51.1871 | Eukaryota | 48012.0 | 0 | Chordata | 48012.0 | 0 | Carnivora | 48012.0 | 0 | Felidae | 41834.0 | 1 | Felis | 37448.0 | 2 | Felis catus | 37448.0 | 2 |
| scf7180000019442 | 465748 | 0.4067 | 0 | 50.7001 | Eukaryota | 46447.0 | 0 | Chordata | 46447.0 | 0 | Carnivora | 46447.0 | 0 | Felidae | 46447.0 | 0 | Felis | 46447.0 | 0 | Felis catus | 46447.0 | 0 |
| scf7180000020889 | 74836 | 0.3973 | 0 | 25.472 | Eukaryota | 29821.0 | 0 | Chordata | 29821.0 | 0 | Carnivora | 29821.0 | 0 | Felidae | 29821.0 | 0 | Felis | 29821.0 | 0 | Felis catus | 29821.0 | 0 |
| scf7180000019161 | 441486 | 0.5046 | 0 | 52.0164 | Eukaryota | 23597.0 | 0 | Chordata | 23597.0 | 0 | Carnivora | 23597.0 | 0 | Felidae | 23597.0 | 0 | Acinonyx | 11952.0 | 4 | cinonyx jubat | 11952.0 | 4 |
| scf7180000020831 | 68413 | 0.4182 | 0 | 25.3817 | Eukaryota | 25998.0 | 0 | Chordata | 25998.0 | 0 | Carnivora | 25998.0 | 0 | Felidae | 25998.0 | 0 | Acinonyx | 13013.0 | 1 | cinonyx jubat | 13013.0 | 1 |
| scf7180000018775 | 12493 | 0.3743 | 0 | 47.5164 | Eukaryota | 17640.0 | 0 | Chordata | 17640.0 | 0 | Carnivora | 17640.0 | 0 | Felidae | 13351.0 | 1 | Panthera | 6620.0 | 4 | Canis lupus | 4289.0 | 5 |
| scf7180000021937 | 23453 | 0.373 | 0 | 23.591 | Eukaryota | 36543.0 | 0 | Chordata | 36543.0 | 0 | Carnivora | 36543.0 | 0 | Felidae | 36543.0 | 0 | Felis | 36543.0 | 0 | Felis catus | 36543.0 | 0 |
| scf7180000021711 | 20727 | 0.5217 | 0 | 126.1726 | no-hit | 0.0 | 0 | no-hit | 0.0 | 0 | no-hit | 0.0 | 0 | no-hit | 0.0 | 0 | no-hit | 0.0 | 0 | no-hit | 0.0 | 0 |
| scf7180000018376 | 801116 | 0.3913 | 0 | 50.2798 | Eukaryota | 50688.0 | 0 | Chordata | 50688.0 | 0 | Carnivora | 50688.0 | 0 | Felidae | 44298.0 | 1 | Panthera | 24827.0 | 3 | anthera pardu | 14883.0 | 4 |
| scf7180000020181 | 144319 | 0.4226 | 0 | 50.2012 | Eukaryota | 46536.0 | 0 | Chordata | 46536.0 | 0 | Carnivora | 46536.0 | 0 | Felidae | 46536.0 | 0 | Felis | 46536.0 | 0 | Felis catus | 46536.0 | 0 |
| scf7180000018737 | 82383 | 0.3893 | 0 | 43.8467 | Eukaryota | 45528.0 | 0 | Chordata | 45528.0 | 0 | Carnivora | 45528.0 | 0 | Felidae | 45528.0 | 0 | Felis | 45528.0 | 0 | Felis catus | 45528.0 | 0 |
| scf7180000020059 | 162918 | 0.3782 | 0 | 47.8647 | Eukaryota | 41845.0 | 0 | Chordata | 41845.0 | 0 | Carnivora | 41845.0 | 0 | Felidae | 41845.0 | 0 | Felis | 37700.0 | 1 | Felis catus | 37700.0 | 1 |
| scf7180000021216 | 40347 | 0.4118 | 0 | 424.7055 | Eukaryota | 7028.0 | 0 | Chordata | 7028.0 | 0 | Carnivora | 7028.0 | 0 | Felidae | 7028.0 | 0 | Felis | 7028.0 | 0 | Felis catus | 7028.0 | 0 |
| scf7180000017556 | 2583930 | 0.3613 | 0 | 49.7545 | Eukaryota | 46840.0 | 0 | Chordata | 46840.0 | 0 | Carnivora | 46840.0 | 0 | Felidae | 46840.0 | 0 | Felis | 46840.0 | 0 | Felis catus | 46840.0 | 0 |
| scf7180000023693 | 3744 | 0.6282 | 0 | 4548.8187 | Eukaryota | 1257.0 | 0 | Chordata | 1257.0 | 0 | Carnivora | 1257.0 | 0 | Felidae | 1257.0 | 0 | Felis | 1257.0 | 0 | Felis catus | 1257.0 | 0 |
| scf7180000017451 | 4119637 | 0.3884 | 0 | 49.7716 | Eukaryota | 48962.0 | 0 | Chordata | 48962.0 | 0 | Carnivora | 48962.0 | 0 | Felidae | 48962.0 | 0 | Felis | 48962.0 | 0 | Felis catus | 48962.0 | 0 |
| scf7180000019051 | 207749 | 0.3893 | 0 | 51.0264 | Eukaryota | 44519.0 | 0 | Chordata | 44519.0 | 0 | Carnivora | 44519.0 | 0 | Felidae | 44519.0 | 0 | Felis | 44519.0 | 0 | Felis catus | 44519.0 | 0 |
| scf7180000021247 | 56946 | 0.5695 | 0 | 51.1088 | Eukaryota | 5230.0 | 0 | Chordata | 5230.0 | 0 | Carnivora | 2676.0 | 5 | Felidae | 1807.0 | 7 | Panthera | 1807.0 | 7 | anthera pardu | 1142.0 | 8 |
| scf7180000018483 | 634175 | 0.4263 | 0 | 50.1896 | Eukaryota | 44151.0 | 0 | Chordata | 44151.0 | 0 | Carnivora | 44151.0 | 0 | Felidae | 44151.0 | 0 | Felis | 44151.0 | 0 | Felis catus | 44151.0 | 0 |
| scf7180000018640 | 456921 | 0.3544 | 0 | 48.3954 | Eukaryota | 46736.0 | 0 | Chordata | 46736.0 | 0 | Carnivora | 46736.0 | 0 | Felidae | 46736.0 | 0 | Felis | 46736.0 | 0 | Felis catus | 46736.0 | 0 |
| scf7180000020674 | 73204 | 0.5707 | 0 | 47.1166 | Eukaryota | 2653.0 | 0 | Chordata | 2653.0 | 0 | Carnivora | 2653.0 | 0 | Felidae | 2388.0 | 1 | Felis | 2388.0 | 1 | Felis catus | 2388.0 | 1 |
| scf7180000022172 | 9913 | 0.42 | 0 | 25.9656 | Eukaryota | 8661.0 | 0 | Chordata | 8661.0 | 0 | Primates | 5617.0 | 3 | Cercopithecidae | 4056.0 | 4 | Papio | 2842.0 | 6 | Papio anubis | 2842.0 | 6 |
| scf7180000018294 | 288688 | 0.5557 | 0 | 46.9741 | Eukaryota | 19692.0 | 0 | Chordata | 19692.0 | 0 | Carnivora | 19692.0 | 0 | Felidae | 17725.0 | 1 | Panthera | 7821.0 | 4 | cinonyx jubat | 3960.0 | 5 |
| scf7180000020921 | 76208 | 0.4356 | 0 | 45.9283 | Eukaryota | 26745.0 | 0 | Chordata | 26745.0 | 0 | Carnivora | 26745.0 | 0 | Felidae | 26745.0 | 0 | Panthera | 10654.0 | 3 | ynx canadens | 5377.0 | 4 |
| scf7180000024052 | 318327 | 0.5208 | 0 | 48.0227 | Eukaryota | 18349.0 | 0 | Chordata | 18349.0 | 0 | Carnivora | 18349.0 | 0 | Felidae | 18349.0 | 0 | Felis | 14248.0 | 2 | Felis catus | 14248.0 | 2 |
| scf7180000017993 | 1661574 | 0.478 | 0 | 51.5017 | Eukaryota | 75947.0 | 0 | Chordata | 75947.0 | 0 | Carnivora | 75947.0 | 0 | Felidae | 41218.0 | 2 | Felis | 24812.0 | 5 | Felis catus | 24812.0 | 5 |
| scf7180000022589 | 3663 | 0.6563 | 0 | 45.8441 | Eukaryota | 2492.0 | 0 | Chordata | 2492.0 | 0 | Carnivora | 2244.0 | 1 | Otariidae | 1002.0 | 5 | Zalophus | 504.0 | 8 | phus californi | 504.0 | 8 |
| scf7180000018964 | 475111 | 0.4488 | 0 | 51.6745 | Eukaryota | 21919.0 | 1 | Chordata | 21919.0 | 1 | Carnivora | 12222.0 | 2 | Felidae | 12222.0 | 2 | Felis | 12222.0 | 2 | Felis catus | 12222.0 | 2 |
| scf7180000023178 | 90059 | 0.5211 | 0 | 437.5563 | Eukaryota | 21792.0 | 0 | Chordata | 21792.0 | 0 | Carnivora | 21792.0 | 0 | Felidae | 21792.0 | 0 | Felis | 19581.0 | 1 | Felis catus | 19581.0 | 1 |
| scf7180000017588 | 2793768 | 0.4045 | 100 | 51.2886 | Eukaryota | 52947.0 | 0 | Chordata | 52947.0 | 0 | Carnivora | 52947.0 | 0 | Felidae | 52947.0 | 0 | Felis | 52947.0 | 0 | Felis catus | 52947.0 | 0 |
| scf7180000023787 | 581956 | 0.4108 | 0 | 50.0033 | Eukaryota | 29306.0 | 0 | Chordata | 29306.0 | 0 | Carnivora | 21642.0 | 1 | Felidae | 13665.0 | 4 | Desmodus | 7664.0 | 8 | modus rotun | 7664.0 | 8 |
| scf7180000018179 | 979649 | 0.3776 | 0 | 49.6402 | Eukaryota | 56766.0 | 0 | Chordata | 56766.0 | 0 | Carnivora | 41654.0 | 2 | Felidae | 30148.0 | 3 | Acinonyx | 15152.0 | 4 | cinonyx jubat | 15152.0 | 5 |
| scf7180000023356 | 26112 | 0.5827 | 0 | 234.256 | no-hit | 0.0 | 0 | no-hit | 0.0 | 0 | no-hit | 0.0 | 0 | no-hit | 0.0 | 0 | no-hit | 0.0 | 0 | no-hit | 0.0 | 0 |
| scf7180000019819 | 189292 | 0.4031 | 0 | 25.1842 | Eukaryota | 10623.0 | 0 | Chordata | 10623.0 | 0 | Primates | 3286.0 | 3 | Hominidae | 3286.0 | 4 | Homo | 2150.0 | 6 | Homo sapiens | 2150.0 | 6 |
| scf7180000020441 | 63539 | 0.3945 | 0 | 50.6259 | Eukaryota | 18370.0 | 0 | Chordata | 18370.0 | 0 | Carnivora | 16179.0 | 1 | Felidae | 16179.0 | 1 | Lynx | 7180.0 | 4 | ynx canadens | 7180.0 | 4 |
| scf7180000019423 | 520036 | 0.5388 | 0 | 51.4549 | Eukaryota | 30673.0 | 0 | Chordata | 30673.0 | 0 | Carnivora | 30673.0 | 0 | Ursidae | 10816.0 | 3 | Ursus | 10816.0 | 3 | rsus maritimu | 10816.0 | 3 |
| scf7180000020398 | 86839 | 0.6122 | 0 | 43.7386 | Eukaryota | 25800.0 | 0 | Chordata | 25800.0 | 0 | Carnivora | 25800.0 | 0 | Mustelidae | 16485.0 | 1 | Mustela | 16485.0 | 1 | ustela putori | 16485.0 | 1 |
| scf7180000018626 | 1125848 | 0.485 | 0 | 51.7365 | Eukaryota | 39016.0 | 0 | Chordata | 39016.0 | 0 | Carnivora | 39016.0 | 0 | Felidae | 39016.0 | 0 | Felis | 27011.0 | 3 | Felis catus | 27011.0 | 3 |
| scf7180000018248 | 1786455 | 0.475 | 0 | 51.2313 | Eukaryota | 45573.0 | 0 | Chordata | 45573.0 | 0 | Carnivora | 34199.0 | 2 | Ursidae | 20478.0 | 3 | Ailuropoda | 20478.0 | 3 | poda melanc | 20478.0 | 3 |
| scf7180000018129 | 28904 | 0.3978 | 0 | 360.9063 | Eukaryota | 6783.0 | 0 | Chordata | 6783.0 | 0 | Carnivora | 6783.0 | 0 | Felidae | 6783.0 | 0 | Felis | 6783.0 | 0 | Felis catus | 6783.0 | 0 |
| scf7180000024151 | 117488 | 0.3783 | 0 | 24.7702 | Eukaryota | 36660.0 | 0 | Chordata | 36660.0 | 0 | Carnivora | 36660.0 | 0 | Felidae | 36660.0 | 0 | Felis | 33005.0 | 1 | Felis catus | 33005.0 | 1 |
| scf7180000023582 | 65537 | 0.4322 | 0 | 227.888 | Eukaryota | 29377.0 | 0 | Chordata | 29377.0 | 0 | Carnivora | 29377.0 | 0 | Felidae | 29377.0 | 0 | Felis | 29377.0 | 0 | Felis catus | 29377.0 | 0 |
| scf7180000018029 | 346696 | 0.4565 | 0 | 52.0093 | Eukaryota | 36801.0 | 0 | Chordata | 36801.0 | 0 | Carnivora | 36801.0 | 0 | Felidae | 22269.0 | 1 | Suricata | 14532.0 | 4 | ricata suricat | 14532.0 | 4 |
| scf7180000021150 | 103456 | 0.457 | 0 | 27.698 | Eukaryota | 45367.0 | 0 | Chordata | 45367.0 | 0 | Carnivora | 45367.0 | 0 | Felidae | 45367.0 | 0 | Felis | 45367.0 | 0 | Felis catus | 45367.0 | 0 |
| scf7180000019081 | 374828 | 0.5502 | 0 | 51.1957 | Eukaryota | 27503.0 | 0 | Chordata | 27503.0 | 0 | Carnivora | 27503.0 | 0 | Felidae | 16615.0 | 2 | Panthera | 8299.0 | 6 | ricata suricat | 5596.0 | 7 |
| scf7180000018197 | 1085791 | 0.3583 | 0 | 49.897 | Eukaryota | 61639.0 | 0 | Chordata | 61639.0 | 0 | Carnivora | 61639.0 | 0 | Felidae | 61639.0 | 0 | Panthera | 43125.0 | 2 | anthera pardu | 30785.0 | 3 |
| scf7180000023605 | 12400 | 0.6021 | 0 | 153.0591 | Eukaryota | 2366.0 | 0 | Chordata | 2366.0 | 0 | Carnivora | 1683.0 | 1 | Felidae | 1455.0 | 2 | Felis | 924.0 | 4 | Felis catus | 924.0 | 5 |
| scf7180000022626 | 20132 | 0.4636 | 0 | 27.1876 | Eukaryota | 8998.0 | 0 | Chordata | 8998.0 | 0 | Carnivora | 8998.0 | 0 | Felidae | 8998.0 | 0 | Felis | 7291.0 | 2 | Felis catus | 7291.0 | 2 |
| scf7180000020658 | 141691 | 0.3887 | 0 | 25.6928 | Eukaryota | 35747.0 | 0 | Chordata | 35747.0 | 0 | Carnivora | 35747.0 | 0 | Felidae | 30888.0 | 1 | Acinonyx | 17160.0 | 2 | cinonyx jubat | 17160.0 | 2 |
| scf7180000020411 | 143359 | 0.4076 | 0 | 49.3703 | Eukaryota | 46148.0 | 0 | Chordata | 46148.0 | 0 | Carnivora | 46148.0 | 0 | Felidae | 46148.0 | 0 | Felis | 46148.0 | 0 | Felis catus | 46148.0 | 0 |
| scf7180000020006 | 198511 | 0.3786 | 0 | 26.0269 | Eukaryota | 57219.0 | 0 | Chordata | 57219.0 | 0 | Carnivora | 57219.0 | 0 | Felidae | 57219.0 | 0 | Felis | 51501.0 | 1 | Felis catus | 51501.0 | 1 |
| scf7180000019752 | 171006 | 0.4506 | 0 | 54.132 | Eukaryota | 46306.0 | 0 | Chordata | 46306.0 | 0 | Carnivora | 46306.0 | 0 | Felidae | 46306.0 | 0 | Felis | 46306.0 | 0 | Felis catus | 46306.0 | 0 |
| scf7180000018057 | 2179717 | 0.4803 | 0 | 53.4008 | Eukaryota | 57290.0 | 0 | Chordata | 57290.0 | 0 | Carnivora | 57290.0 | 0 | Felidae | 51136.0 | 1 | Felis | 28440.0 | 2 | Felis catus | 28440.0 | 2 |
| scf7180000023557 | 21404 | 0.4628 | 0 | 297.051 | no-hit | 0.0 | 0 | no-hit | 0.0 | 0 | no-hit | 0.0 | 0 | no-hit | 0.0 | 0 | no-hit | 0.0 | 0 | no-hit | 0.0 | 0 |
| scf7180000023743 | 51443 | 0.433 | 0 | 477.3834 | Eukaryota | 29384.0 | 0 | Chordata | 29384.0 | 0 | Carnivora | 29384.0 | 0 | Felidae | 29384.0 | 0 | Felis | 29384.0 | 0 | Felis catus | 29384.0 | 0 |
| scf7180000019080 | 281933 | 0.4105 | 0 | 48.2177 | Eukaryota | 46932.0 | 0 | Chordata | 46932.0 | 0 | Carnivora | 46932.0 | 0 | Felidae | 46932.0 | 0 | Felis | 37577.0 | 2 | Felis catus | 37577.0 | 2 |
| scf7180000022108 | 40507 | 0.4504 | 0 | 25.6681 | Eukaryota | 76081.0 | 0 | Chordata | 76081.0 | 0 | Carnivora | 44500.0 | 1 | Herpestidae | 22545.0 | 4 | Suricata | 22545.0 | 6 | ricata suricat | 22545.0 | 6 |
| scf7180000021056 | 66447 | 0.3883 | 0 | 44.7806 | Eukaryota | 43712.0 | 0 | Chordata | 43712.0 | 0 | Carnivora | 43712.0 | 0 | Felidae | 43712.0 | 0 | Felis | 43712.0 | 0 | Felis catus | 43712.0 | 0 |
| scf7180000021225 | 67056 | 0.5924 | 0 | 96.2495 | Eukaryota | 2413.0 | 0 | Chordata | 2413.0 | 0 | Carnivora | 2413.0 | 0 | Felidae | 2413.0 | 0 | Felis | 2104.0 | 1 | Felis catus | 2104.0 | 1 |
| scf7180000023961 | 1038777 | 0.4656 | 0 | 50.1621 | Eukaryota | 22221.0 | 0 | Chordata | 22221.0 | 0 | Carnivora | 22221.0 | 0 | Felidae | 17774.0 | 1 | Felis | 15785.0 | 2 | Felis catus | 15785.0 | 2 |
| scf7180000021303 | 41085 | 0.5592 | 0 | 51.0958 | Eukaryota | 7078.0 | 0 | Chordata | 7078.0 | 0 | Carnivora | 5493.0 | 1 | Felidae | 5493.0 | 1 | Felis | 3432.0 | 3 | Felis catus | 3432.0 | 3 |
| scf7180000019756 | 256368 | 0.5403 | 0 | 60.5044 | Eukaryota | 22004.0 | 0 | Chordata | 22004.0 | 0 | Carnivora | 22004.0 | 0 | Felidae | 10475.0 | 2 | Suricata | 9508.0 | 5 | ricata suricat | 9508.0 | 5 |
| scf7180000019972 | 148147 | 0.4337 | 0 | 47.9651 | Eukaryota | 31872.0 | 0 | Chordata | 31872.0 | 0 | Carnivora | 31872.0 | 0 | Felidae | 31872.0 | 0 | Felis | 28616.0 | 1 | Felis catus | 28616.0 | 1 |
| scf7180000017447 | 58632 | 0.3928 | 0 | 24.5209 | Eukaryota | 46191.0 | 0 | Chordata | 46191.0 | 0 | Carnivora | 46191.0 | 0 | Felidae | 46191.0 | 0 | Felis | 46191.0 | 0 | Felis catus | 46191.0 | 0 |
| scf7180000022911 | 65505 | 0.5246 | 0 | 188.3367 | Eukaryota | 15713.0 | 0 | Chordata | 15713.0 | 0 | Carnivora | 15713.0 | 0 | Felidae | 15713.0 | 0 | Felis | 15713.0 | 0 | Felis catus | 15713.0 | 0 |
| scf7180000021158 | 42143 | 0.4355 | 0 | 48.4946 | Eukaryota | 4136.0 | 0 | Chordata | 4136.0 | 0 | Carnivora | 3256.0 | 1 | Felidae | 3256.0 | 1 | Lynx | 2849.0 | 2 | ynx canadens | 2849.0 | 2 |
| scf7180000020664 | 52676 | 0.4232 | 0 | 527.3223 | Eukaryota | 13291.0 | 0 | Chordata | 13291.0 | 0 | Carnivora | 13291.0 | 0 | Felidae | 13291.0 | 0 | Felis | 13291.0 | 0 | Felis catus | 13291.0 | 0 |
| scf7180000020571 | 18877 | 0.4041 | 0 | 26.1041 | Eukaryota | 13436.0 | 0 | Chordata | 13436.0 | 0 | Carnivora | 13436.0 | 0 | Felidae | 13436.0 | 0 | Felis | 12104.0 | 1 | Felis catus | 12104.0 | 1 |
| scf7180000020990 | 84408 | 0.4061 | 0 | 25.3309 | Eukaryota | 29200.0 | 0 | Chordata | 29200.0 | 0 | Carnivora | 29200.0 | 0 | Felidae | 29200.0 | 0 | Panthera | 11712.0 | 3 | Felis catus | 8691.0 | 4 |
| scf7180000019848 | 190679 | 0.4202 | 0 | 48.2478 | Eukaryota | 13615.0 | 0 | Chordata | 13615.0 | 0 | Carnivora | 13615.0 | 0 | Felidae | 13615.0 | 0 | Felis | 12250.0 | 1 | Felis catus | 12250.0 | 1 |
| scf7180000018175 | 984417 | 0.4026 | 0 | 51.52 | Eukaryota | 46046.0 | 0 | Chordata | 46046.0 | 0 | Carnivora | 46046.0 | 0 | Felidae | 46046.0 | 0 | Felis | 46046.0 | 0 | Felis catus | 46046.0 | 0 |
| scf7180000019896 | 129572 | 0.4884 | 0 | 49.1529 | Eukaryota | 40597.0 | 0 | Chordata | 40597.0 | 0 | Carnivora | 40597.0 | 0 | Felidae | 29032.0 | 3 | Panthera | 12518.0 | 6 | anthera pardu | 8344.0 | 7 |
| scf7180000018895 | 52581 | 0.3941 | 100 | 25.2539 | Eukaryota | 40928.0 | 0 | Chordata | 40928.0 | 0 | Carnivora | 40928.0 | 0 | Felidae | 40928.0 | 0 | Felis | 40928.0 | 0 | Felis catus | 40928.0 | 0 |
| scf7180000022048 | 23487 | 0.4332 | 0 | 453.513 | no-hit | 0.0 | 0 | no-hit | 0.0 | 0 | no-hit | 0.0 | 0 | no-hit | 0.0 | 0 | no-hit | 0.0 | 0 | no-hit | 0.0 | 0 |
| scf7180000018880 | 450812 | 0.4538 | 0 | 50.7943 | Eukaryota | 26331.0 | 1 | Chordata | 26331.0 | 1 | Primates | 15993.0 | 3 | Hominidae | 12116.0 | 4 | Homo | 12116.0 | 6 | Homo sapiens | 12116.0 | 6 |
| scf7180000020912 | 66290 | 0.3796 | 0 | 25.0036 | Eukaryota | 30389.0 | 0 | Chordata | 30389.0 | 0 | Carnivora | 30389.0 | 0 | Felidae | 30389.0 | 0 | Felis | 30389.0 | 0 | Felis catus | 30389.0 | 0 |
| scf7180000023946 | 1183164 | 0.398 | 0 | 51.2144 | Eukaryota | 45960.0 | 0 | Chordata | 45960.0 | 0 | Carnivora | 45960.0 | 0 | Felidae | 45960.0 | 0 | Felis | 41411.0 | 1 | Felis catus | 41411.0 | 1 |
| scf7180000022302 | 37190 | 0.6788 | 0 | 53.7885 | Eukaryota | 6989.0 | 0 | Chordata | 6989.0 | 0 | Carnivora | 6989.0 | 0 | Felidae | 6390.0 | 1 | Acinonyx | 5022.0 | 2 | cinonyx jubat | 5022.0 | 2 |
| scf7180000023145 | 34042 | 0.4341 | 0 | 795.9432 | Eukaryota | 18891.0 | 0 | Chordata | 18891.0 | 0 | Carnivora | 18891.0 | 0 | Felidae | 18891.0 | 0 | Felis | 15134.0 | 1 | Felis catus | 15134.0 | 1 |
| scf7180000024023 | 464228 | 0.5325 | 0 | 50.3204 | Eukaryota | 28113.0 | 0 | Chordata | 28113.0 | 0 | Carnivora | 28113.0 | 0 | Felidae | 25150.0 | 1 | Panthera | 11418.0 | 4 | ynx canadens | 7980.0 | 5 |
| scf7180000019513 | 206892 | 0.3658 | 0 | 47.8481 | Eukaryota | 45347.0 | 0 | Chordata | 45347.0 | 0 | Carnivora | 45347.0 | 0 | Felidae | 45347.0 | 0 | Felis | 45347.0 | 0 | Felis catus | 45347.0 | 0 |
| scf7180000017560 | 33400 | 0.4537 | 0 | 170.4026 | Eukaryota | 21636.0 | 0 | Chordata | 21636.0 | 0 | Carnivora | 21636.0 | 0 | Felidae | 21636.0 | 0 | Felis | 19425.0 | 1 | Felis catus | 19425.0 | 1 |
| scf7180000023503 | 44422 | 0.4919 | 0 | 291.8853 | Eukaryota | 12880.0 | 0 | Chordata | 12880.0 | 0 | Carnivora | 12880.0 | 0 | Felidae | 12880.0 | 0 | Felis | 12880.0 | 0 | Felis catus | 12880.0 | 0 |
| scf7180000022769 | 6928 | 0.4596 | 0 | 76.2176 | Eukaryota | 1989.0 | 0 | Chordata | 1989.0 | 0 | Carnivora | 1989.0 | 0 | Felidae | 1989.0 | 0 | Felis | 1595.0 | 1 | Felis catus | 1595.0 | 1 |
| scf7180000022451 | 8274 | 0.5904 | 0 | 49.3259 | Eukaryota | 885.0 | 0 | Chordata | 885.0 | 0 | Primates | 224.0 | 4 | Hominidae | 224.0 | 4 | Homo | 224.0 | 4 | Homo sapiens | 224.0 | 4 |
| scf7180000021659 | 38790 | 0.421 | 0 | 26.9893 | Eukaryota | 9031.0 | 0 | Chordata | 9031.0 | 0 | Carnivora | 3458.0 | 3 | Felidae | 2872.0 | 4 | Felis | 2872.0 | 4 | Felis catus | 2872.0 | 4 |
| scf7180000021034 | 78745 | 0.3702 | 0 | 26.2285 | Eukaryota | 8242.0 | 0 | Chordata | 8242.0 | 0 | Carnivora | 8242.0 | 0 | Felidae | 8242.0 | 0 | Felis | 8242.0 | 0 | Felis catus | 8242.0 | 0 |
| scf7180000020920 | 60420 | 0.4423 | 0 | 48.6249 | Eukaryota | 19085.0 | 0 | Chordata | 19085.0 | 0 | Carnivora | 19085.0 | 0 | Felidae | 19085.0 | 0 | Felis | 15285.0 | 1 | Felis catus | 15285.0 | 1 |
| scf7180000017714 | 1927414 | 0.3808 | 0 | 49.8742 | Eukaryota | 46980.0 | 0 | Chordata | 46980.0 | 0 | Carnivora | 46980.0 | 0 | Felidae | 46980.0 | 0 | Felis | 46980.0 | 0 | Felis catus | 46980.0 | 0 |
| scf7180000021966 | 26781 | 0.5858 | 0 | 49.2897 | Eukaryota | 2330.0 | 0 | Chordata | 2330.0 | 0 | Carnivora | 2330.0 | 0 | Otariidae | 1631.0 | 1 | Eumetopias | 1398.0 | 2 | netopias juba | 1398.0 | 2 |
| scf7180000019430 | 289454 | 0.4878 | 0 | 48.2341 | Eukaryota | 18793.0 | 0 | Chordata | 18793.0 | 0 | Carnivora | 18793.0 | 0 | Felidae | 18793.0 | 0 | Felis | 16909.0 | 1 | Felis catus | 16909.0 | 1 |
| scf7180000019876 | 221387 | 0.3782 | 0 | 50.537 | Eukaryota | 46932.0 | 0 | Chordata | 46932.0 | 0 | Carnivora | 46932.0 | 0 | Felidae | 46932.0 | 0 | Felis | 46932.0 | 0 | Felis catus | 46932.0 | 0 |
| scf7180000019311 | 365821 | 0.5492 | 0 | 49.4753 | Eukaryota | 20534.0 | 1 | Chordata | 20534.0 | 1 | Carnivora | 15057.0 | 2 | Felidae | 10140.0 | 4 | Acinonyx | 10140.0 | 4 | cinonyx jubat | 10140.0 | 4 |
| scf7180000020603 | 114063 | 0.4453 | 0 | 49.5556 | Eukaryota | 31360.0 | 0 | Chordata | 31360.0 | 0 | Carnivora | 31360.0 | 0 | Felidae | 31360.0 | 0 | Felis | 28148.0 | 1 | Felis catus | 28148.0 | 1 |
| scf7180000021886 | 34827 | 0.4024 | 0 | 24.7625 | Eukaryota | 16233.0 | 0 | Chordata | 16233.0 | 0 | Carnivora | 16233.0 | 0 | Felidae | 16233.0 | 0 | Felis | 12978.0 | 1 | Felis catus | 12978.0 | 1 |
| scf7180000019933 | 204989 | 0.3577 | 0 | 50.0466 | Eukaryota | 18247.0 | 0 | Chordata | 18247.0 | 0 | Carnivora | 18247.0 | 0 | Felidae | 18247.0 | 0 | Felis | 16542.0 | 1 | Felis catus | 16542.0 | 1 |
| scf7180000019594 | 9181 | 0.4347 | 0 | 25.7849 | Eukaryota | 3530.0 | 0 | Chordata | 3530.0 | 0 | Carnivora | 1449.0 | 3 | Felidae | 1092.0 | 5 | Equus | 1005.0 | 6 | Equus caballus | 1005.0 | 6 |
| scf7180000021576 | 53752 | 0.3759 | 0 | 26.0058 | Eukaryota | 45933.0 | 0 | Chordata | 45933.0 | 0 | Carnivora | 45933.0 | 0 | Felidae | 45933.0 | 0 | Felis | 45933.0 | 0 | Felis catus | 45933.0 | 0 |
| scf7180000023046 | 15387 | 0.5953 | 0 | 206.7463 | Eukaryota | 3148.0 | 0 | Chordata | 3148.0 | 0 | Carnivora | 2863.0 | 1 | Felidae | 2406.0 | 2 | Panthera | 1250.0 | 4 | anthera pardu | 1250.0 | 4 |
| scf7180000022113 | 26094 | 0.4259 | 0 | 24.874 | Eukaryota | 7653.0 | 0 | Chordata | 7653.0 | 0 | Carnivora | 6530.0 | 1 | Felidae | 6530.0 | 1 | Felis | 6530.0 | 1 | Felis catus | 6530.0 | 1 |
| scf7180000020121 | 176923 | 0.523 | 0 | 50.987 | Eukaryota | 19587.0 | 0 | Chordata | 19587.0 | 0 | Carnivora | 19587.0 | 0 | Felidae | 19587.0 | 0 | Panthera | 7820.0 | 3 | anthera pardu | 3934.0 | 4 |
| scf7180000020537 | 7904 | 0.556 | 0 | 172.8472 | Eukaryota | 12043.0 | 0 | Chordata | 12043.0 | 0 | Carnivora | 12043.0 | 0 | Felidae | 12043.0 | 0 | Felis | 10859.0 | 1 | Felis catus | 10859.0 | 1 |
| scf7180000020211 | 112188 | 0.5429 | 0 | 54.6747 | Eukaryota | 14223.0 | 0 | Chordata | 14223.0 | 0 | Carnivora | 9836.0 | 2 | Felidae | 9836.0 | 2 | Panthera | 6528.0 | 5 | anthera pardu | 6528.0 | 6 |
| scf7180000020758 | 112545 | 0.4107 | 0 | 25.57 | Eukaryota | 10193.0 | 0 | Chordata | 10193.0 | 0 | Carnivora | 6988.0 | 1 | Felidae | 6988.0 | 1 | Felis | 6007.0 | 3 | Felis catus | 6007.0 | 3 |
| scf7180000024089 | 339088 | 0.3518 | 0 | 49.2056 | Eukaryota | 11446.0 | 0 | Chordata | 11446.0 | 0 | Carnivora | 11446.0 | 0 | Felidae | 11446.0 | 0 | Felis | 11446.0 | 0 | Felis catus | 11446.0 | 0 |
| scf7180000021250 | 49384 | 0.6325 | 0 | 55.3641 | Eukaryota | 5564.0 | 0 | Chordata | 5564.0 | 0 | Carnivora | 5564.0 | 0 | Felidae | 5035.0 | 1 | Lynx | 2620.0 | 5 | ynx canadens | 2620.0 | 5 |
| scf7180000019850 | 21354 | 0.5113 | 0 | 209.1394 | Eukaryota | 12941.0 | 0 | Chordata | 12941.0 | 0 | Carnivora | 12941.0 | 0 | Felidae | 12941.0 | 0 | Felis | 12941.0 | 0 | Felis catus | 12941.0 | 0 |
| scf7180000021308 | 69915 | 0.3726 | 0 | 23.5727 | Eukaryota | 7443.0 | 0 | Chordata | 7443.0 | 0 | Carnivora | 5878.0 | 2 | Felidae | 5878.0 | 2 | Felis | 5878.0 | 2 | Felis catus | 5878.0 | 2 |
| scf7180000018668 | 267000 | 0.4681 | 0 | 51.2583 | Eukaryota | 46469.0 | 0 | Chordata | 46469.0 | 0 | Carnivora | 46469.0 | 0 | Herpestidae | 17376.0 | 2 | Suricata | 17376.0 | 3 | ricata suricat | 17376.0 | 3 |
| scf7180000017721 | 2045829 | 0.4065 | 100 | 51.1225 | Eukaryota | 73509.0 | 0 | Chordata | 73509.0 | 0 | Carnivora | 73509.0 | 0 | Felidae | 73509.0 | 0 | Panthera | 33128.0 | 3 | anthera pardu | 33128.0 | 3 |
| scf7180000021634 | 36347 | 0.4004 | 0 | 26.4813 | Eukaryota | 39892.0 | 0 | Chordata | 39892.0 | 0 | Carnivora | 39892.0 | 0 | Felidae | 39892.0 | 0 | Felis | 39892.0 | 0 | Felis catus | 39892.0 | 0 |
| scf7180000018058 | 826130 | 0.36 | 0 | 48.9525 | Eukaryota | 46393.0 | 0 | Chordata | 46393.0 | 0 | Carnivora | 46393.0 | 0 | Felidae | 46393.0 | 0 | Felis | 46393.0 | 0 | Felis catus | 46393.0 | 0 |
| scf7180000019112 | 447054 | 0.3848 | 0 | 49.6341 | Eukaryota | 44241.0 | 0 | Chordata | 44241.0 | 0 | Carnivora | 44241.0 | 0 | Felidae | 44241.0 | 0 | Felis | 44241.0 | 0 | Felis catus | 44241.0 | 0 |
| scf7180000023987 | 103474 | 0.3991 | 0 | 45.5521 | Eukaryota | 42697.0 | 0 | Chordata | 42697.0 | 0 | Carnivora | 42697.0 | 0 | Felidae | 42697.0 | 0 | Felis | 42697.0 | 0 | Felis catus | 42697.0 | 0 |
| scf7180000022886 | 13893 | 0.5487 | 0 | 196.9574 | Eukaryota | 2728.0 | 0 | Chordata | 2728.0 | 0 | Carnivora | 2728.0 | 0 | Felidae | 2461.0 | 1 | Felis | 1295.0 | 3 | Felis catus | 1295.0 | 3 |
| scf7180000021954 | 54171 | 0.3849 | 0 | 27.5911 | Eukaryota | 8012.0 | 0 | Chordata | 8012.0 | 0 | Carnivora | 3031.0 | 2 | Felidae | 3031.0 | 2 | Felis | 3031.0 | 3 | Felis catus | 3031.0 | 3 |
| scf7180000023969 | 674833 | 0.3799 | 0 | 50.5014 | Eukaryota | 48135.0 | 0 | Chordata | 48135.0 | 0 | Carnivora | 48135.0 | 0 | Felidae | 48135.0 | 0 | Felis | 32191.0 | 2 | Felis catus | 32191.0 | 2 |
| scf7180000019664 | 169556 | 0.5345 | 0 | 47.541 | Eukaryota | 28249.0 | 0 | Chordata | 28249.0 | 0 | Carnivora | 28249.0 | 0 | Felidae | 14950.0 | 4 | Lynx | 6097.0 | 8 | ynx canadens | 6097.0 | 8 |
| scf7180000021705 | 22045 | 0.3857 | 0 | 27.2814 | Eukaryota | 40848.0 | 0 | Chordata | 40848.0 | 0 | Carnivora | 40848.0 | 0 | Felidae | 40848.0 | 0 | Felis | 40848.0 | 0 | Felis catus | 40848.0 | 0 |
| scf7180000023563 | 30686 | 0.4793 | 0 | 430.0512 | no-hit | 0.0 | 0 | no-hit | 0.0 | 0 | no-hit | 0.0 | 0 | no-hit | 0.0 | 0 | no-hit | 0.0 | 0 | no-hit | 0.0 | 0 |
| scf7180000018925 | 461262 | 0.3659 | 0 | 49.1891 | Eukaryota | 34948.0 | 0 | Chordata | 34948.0 | 0 | Carnivora | 34948.0 | 0 | Felidae | 34948.0 | 0 | Felis | 31609.0 | 1 | Felis catus | 31609.0 | 1 |
| scf7180000021822 | 42463 | 0.3966 | 0 | 48.167 | Eukaryota | 6625.0 | 0 | Chordata | 6625.0 | 0 | Carnivora | 6625.0 | 0 | Felidae | 6625.0 | 0 | Felis | 6625.0 | 0 | Felis catus | 6625.0 | 0 |
| scf7180000019317 | 6276 | 0.3883 | 0 | 24.2528 | Eukaryota | 3937.0 | 0 | Chordata | 3937.0 | 0 | Carnivora | 3937.0 | 0 | Felidae | 3552.0 | 1 | Felis | 1104.0 | 5 | Felis catus | 1104.0 | 6 |
| scf7180000020956 | 54233 | 0.5075 | 0 | 88.7641 | Eukaryota | 18591.0 | 0 | Chordata | 18591.0 | 0 | Carnivora | 18591.0 | 0 | Felidae | 18591.0 | 0 | Felis | 13020.0 | 1 | Felis catus | 13020.0 | 1 |
| scf7180000020614 | 101851 | 0.5687 | 0 | 45.7458 | Eukaryota | 23175.0 | 0 | Chordata | 23175.0 | 0 | Carnivora | 23175.0 | 0 | Felidae | 23175.0 | 0 | Felis | 20825.0 | 1 | Felis catus | 20825.0 | 1 |
| scf7180000018012 | 14586 | 0.528 | 0 | 1000.2121 | no-hit | 0.0 | 0 | no-hit | 0.0 | 0 | no-hit | 0.0 | 0 | no-hit | 0.0 | 0 | no-hit | 0.0 | 0 | no-hit | 0.0 | 0 |
| scf7180000019041 | 403638 | 0.5196 | 0 | 51.8463 | Eukaryota | 35442.0 | 0 | Chordata | 35442.0 | 0 | Carnivora | 35442.0 | 0 | Felidae | 35442.0 | 0 | Panthera | 14252.0 | 3 | anthera pardu | 14252.0 | 3 |
| scf7180000020383 | 15230 | 0.5674 | 0 | 93.4005 | Eukaryota | 2434.0 | 0 | Chordata | 2434.0 | 0 | Carnivora | 1384.0 | 2 | Felidae | 1140.0 | 3 | Felis | 896.0 | 4 | Felis catus | 896.0 | 4 |
| scf7180000021335 | 76039 | 0.5814 | 0 | 47.6481 | Eukaryota | 2581.0 | 1 | Chordata | 2581.0 | 1 | Carnivora | 2024.0 | 2 | Felidae | 1421.0 | 3 | Felis | 1421.0 | 3 | Felis catus | 1421.0 | 3 |
| scf7180000020117 | 148580 | 0.5301 | 0 | 25.3844 | Eukaryota | 9769.0 | 0 | Chordata | 9769.0 | 0 | Primates | 4954.0 | 3 | Hominidae | 4954.0 | 4 | Pan | 2888.0 | 5 | an troglodyte | 2888.0 | 5 |
| scf7180000021111 | 118534 | 0.4676 | 0 | 76.8117 | Eukaryota | 7517.0 | 0 | Chordata | 7517.0 | 0 | Carnivora | 7517.0 | 0 | Felidae | 5233.0 | 2 | Panthera | 2288.0 | 5 | anthera pardu | 2288.0 | 5 |
| scf7180000021927 | 35291 | 0.3843 | 0 | 25.7659 | Eukaryota | 21407.0 | 0 | Chordata | 21407.0 | 0 | Carnivora | 21407.0 | 0 | Felidae | 21407.0 | 0 | Felis | 17123.0 | 1 | Felis catus | 17123.0 | 1 |
| scf7180000020015 | 198708 | 0.4809 | 0 | 49.9433 | Eukaryota | 12728.0 | 0 | Chordata | 12728.0 | 0 | Primates | 7030.0 | 5 | Hominidae | 7030.0 | 5 | Homo | 7030.0 | 5 | Homo sapiens | 7030.0 | 5 |
| scf7180000021470 | 48452 | 0.468 | 0 | 24.7819 | Eukaryota | 12891.0 | 0 | Chordata | 12891.0 | 0 | Primates | 5059.0 | 3 | Hominidae | 5059.0 | 4 | Homo | 5059.0 | 5 | Homo sapiens | 5059.0 | 5 |

| | | | | | | | | | | | | | | | | | | | | | | |
|---|---|---|---|---|---|---|---|---|---|---|---|---|---|---|---|---|---|---|---|---|---|---|
| scf7180000018654 | 20981 | 0.3914 | 0 | 26.1405 | Eukaryota | 45641.0 | 0 | Chordata | 45641.0 | 0 | Carnivora | 45641.0 | 0 | Felidae | 45641.0 | 0 | Felis | 45641.0 | 0 | Felis catus | 45641.0 | 0 |
| scf7180000022756 | 13692 | 0.4447 | 0 | 51.2817 | Eukaryota | 4051.0 | 0 | Chordata | 4051.0 | 0 | Carnivora | 2055.0 | 1 | Hominidae | 1996.0 | 2 | Homo | 1996.0 | 3 | Homo sapiens | 1996.0 | 3 |
| scf7180000021500 | 23723 | 0.5561 | 0 | 49.5088 | Eukaryota | 9155.0 | 0 | Chordata | 9155.0 | 0 | Carnivora | 8253.0 | 1 | Felidae | 3712.0 | 3 | Zalophus | 2700.0 | 6 | phus californi | 2700.0 | 6 |
| scf7180000018727 | 499570 | 0.5799 | 0 | 51.9486 | Eukaryota | 12905.0 | 0 | Chordata | 12905.0 | 0 | Carnivora | 12905.0 | 0 | Felidae | 12905.0 | 0 | Felis | 10465.0 | 1 | Felis catus | 10465.0 | 1 |
| scf7180000021852 | 42351 | 0.381 | 0 | 24.8134 | Eukaryota | 46223.0 | 0 | Chordata | 46223.0 | 0 | Carnivora | 46223.0 | 0 | Felidae | 46223.0 | 0 | Felis | 46223.0 | 0 | Felis catus | 46223.0 | 0 |
| scf7180000020865 | 371439 | 0.5887 | 0 | 49.4749 | Eukaryota | 20789.0 | 1 | Chordata | 20789.0 | 1 | Carnivora | 18367.0 | 2 | Canidae | 11696.0 | 4 | Canis | 11696.0 | 4 | Canis lupus | 11696.0 | 4 |
| scf7180000020017 | 149141 | 0.5947 | 0 | 51.8332 | Eukaryota | 12130.0 | 0 | Chordata | 12130.0 | 0 | Carnivora | 12130.0 | 0 | Felidae | 12130.0 | 0 | Felis | 8508.0 | 1 | Felis catus | 8508.0 | 1 |
| scf7180000022864 | 9280 | 0.6042 | 0 | 144.7512 | Eukaryota | 1821.0 | 0 | Chordata | 1821.0 | 0 | Carnivora | 1821.0 | 0 | Felidae | 1821.0 | 0 | Felis | 1281.0 | 1 | Felis catus | 1281.0 | 1 |
| scf7180000019316 | 15173 | 0.5437 | 0 | 52.018 | Eukaryota | 7437.0 | 0 | Chordata | 7437.0 | 0 | Carnivora | 7437.0 | 0 | Felidae | 6805.0 | 1 | Acinonyx | 3011.0 | 4 | cinonyx jubat | 3011.0 | 4 |
| scf7180000017645 | 2011261 | 0.4081 | 0 | 51.3776 | Eukaryota | 47863.0 | 0 | Chordata | 47863.0 | 0 | Carnivora | 47863.0 | 0 | Felidae | 42099.0 | 1 | Felis | 37465.0 | 2 | Felis catus | 37465.0 | 2 |
| scf7180000021083 | 14456 | 0.4976 | 0 | 222.5095 | Eukaryota | 17274.0 | 0 | Chordata | 17274.0 | 0 | Carnivora | 17274.0 | 0 | Felidae | 17274.0 | 0 | Felis | 13847.0 | 1 | Felis catus | 13847.0 | 1 |
| scf7180000022917 | 65090 | 0.3927 | 0 | 356.5287 | Eukaryota | 7410.0 | 0 | Chordata | 7410.0 | 0 | Carnivora | 5261.0 | 2 | Felidae | 5261.0 | 2 | Felis | 5261.0 | 2 | Felis catus | 5261.0 | 2 |
| scf7180000020362 | 147937 | 0.4748 | 0 | 25.3843 | Eukaryota | 26181.0 | 0 | Chordata | 26181.0 | 0 | Carnivora | 26181.0 | 0 | Felidae | 26181.0 | 0 | Felis | 23562.0 | 1 | Felis catus | 23562.0 | 1 |
| scf7180000020534 | 6396 | 0.5228 | 0 | 47.6924 | no-hit | 0.0 | 0 | no-hit | 0.0 | 0 | no-hit | 0.0 | 0 | no-hit | 0.0 | 0 | no-hit | 0.0 | 0 | no-hit | 0.0 | 0 |
| scf7180000021307 | 6713 | 0.5214 | 0 | 146.2014 | no-hit | 0.0 | 0 | no-hit | 0.0 | 0 | no-hit | 0.0 | 0 | no-hit | 0.0 | 0 | no-hit | 0.0 | 0 | no-hit | 0.0 | 0 |
| scf7180000020419 | 146368 | 0.5545 | 0 | 26.7809 | Eukaryota | 15314.0 | 0 | Chordata | 15314.0 | 0 | Carnivora | 15314.0 | 0 | Felidae | 10396.0 | 1 | Felis | 7568.0 | 3 | Felis catus | 7568.0 | 3 |
| scf7180000020814 | 16006 | 0.4077 | 0 | 15.8957 | Eukaryota | 46466.0 | 0 | Chordata | 46466.0 | 0 | Carnivora | 46466.0 | 0 | Felidae | 46466.0 | 0 | Felis | 46466.0 | 0 | Felis catus | 46466.0 | 0 |
| scf7180000017673 | 2005654 | 0.4327 | 0 | 51.7724 | Eukaryota | 42842.0 | 0 | Chordata | 42842.0 | 0 | Carnivora | 42842.0 | 0 | Felidae | 42842.0 | 0 | Felis | 42842.0 | 0 | Felis catus | 42842.0 | 0 |
| scf7180000019022 | 464037 | 0.3472 | 0 | 48.0245 | Eukaryota | 44452.0 | 0 | Chordata | 44452.0 | 0 | Carnivora | 44452.0 | 0 | Felidae | 44452.0 | 0 | Felis | 44452.0 | 0 | Felis catus | 44452.0 | 0 |
| scf7180000022226 | 22767 | 0.3887 | 0 | 23.8127 | Eukaryota | 3266.0 | 0 | Chordata | 3266.0 | 0 | Carnivora | 3266.0 | 0 | Felidae | 3266.0 | 0 | Felis | 2918.0 | 1 | Felis catus | 2918.0 | 1 |
| scf7180000024197 | 73770 | 0.3646 | 0 | 23.9596 | Eukaryota | 45210.0 | 0 | Chordata | 45210.0 | 0 | Carnivora | 45210.0 | 0 | Felidae | 45210.0 | 0 | Felis | 45210.0 | 0 | Felis catus | 45210.0 | 0 |
| scf7180000018998 | 398752 | 0.6108 | 0 | 51.2123 | Eukaryota | 19747.0 | 0 | Chordata | 19747.0 | 0 | Carnivora | 19747.0 | 0 | Felidae | 15367.0 | 2 | Felis | 3876.0 | 6 | Felis catus | 3876.0 | 6 |
| scf7180000018433 | 18666 | 0.391 | 0 | 36.211 | Eukaryota | 46691.0 | 0 | Chordata | 46691.0 | 0 | Carnivora | 46691.0 | 0 | Felidae | 46691.0 | 0 | Felis | 46691.0 | 0 | Felis catus | 46691.0 | 0 |
| scf7180000020615 | 104215 | 0.471 | 0 | 52.0809 | Eukaryota | 11106.0 | 0 | Chordata | 11106.0 | 0 | Carnivora | 11106.0 | 0 | Canidae | 6066.0 | 1 | Canis | 6066.0 | 1 | Canis lupus | 6066.0 | 1 |
| scf7180000020863 | 55447 | 0.424 | 0 | 25.9339 | Eukaryota | 13411.0 | 0 | Chordata | 12657.0 | 1 | Primates | 5849.0 | 4 | Hominidae | 5849.0 | 4 | Homo | 3902.0 | 5 | Homo sapiens | 3902.0 | 5 |
| scf7180000022306 | 25989 | 0.373 | 0 | 24.3221 | Eukaryota | 10440.0 | 0 | Chordata | 10440.0 | 0 | Carnivora | 10440.0 | 0 | Felidae | 10440.0 | 0 | Felis | 7297.0 | 1 | Felis catus | 7297.0 | 1 |
| scf7180000019102 | 56971 | 0.4277 | 0 | 39.6327 | Eukaryota | 27944.0 | 0 | Chordata | 27944.0 | 0 | Carnivora | 27944.0 | 0 | Felidae | 27944.0 | 0 | Felis | 25122.0 | 1 | Felis catus | 25122.0 | 1 |
| scf7180000017958 | 868162 | 0.4721 | 0 | 53.1187 | Eukaryota | 61006.0 | 0 | Chordata | 61006.0 | 0 | Carnivora | 61006.0 | 0 | Herpestidae | 24764.0 | 4 | Suricata | 24764.0 | 6 | ricata suricat | 24764.0 | 6 |
| scf7180000019171 | 428366 | 0.431 | 0 | 51.5972 | Eukaryota | 46287.0 | 0 | Chordata | 46287.0 | 0 | Carnivora | 46287.0 | 0 | Felidae | 46287.0 | 0 | Felis | 46287.0 | 0 | Felis catus | 46287.0 | 0 |
| scf7180000020522 | 132584 | 0.4207 | 0 | 49.2917 | Eukaryota | 18461.0 | 0 | Chordata | 18461.0 | 0 | Carnivora | 18461.0 | 0 | Otariidae | 8859.0 | 3 | Callorhinus | 3802.0 | 7 | llorhinus ursi | 3802.0 | 7 |
| scf7180000020032 | 18435 | 0.5059 | 0 | 264.9569 | Eukaryota | 12277.0 | 0 | Chordata | 12277.0 | 0 | Carnivora | 12277.0 | 0 | Felidae | 12277.0 | 0 | Felis | 11067.0 | 1 | Felis catus | 11067.0 | 1 |
| scf7180000019877 | 5807 | 0.4112 | 0 | 30.1966 | Eukaryota | 11883.0 | 0 | Chordata | 11883.0 | 0 | Carnivora | 7948.0 | 2 | Felidae | 7948.0 | 2 | Felis | 6826.0 | 4 | Felis catus | 6826.0 | 4 |
| scf7180000022629 | 10434 | 0.3646 | 0 | 26.2346 | Eukaryota | 4584.0 | 0 | Chordata | 4584.0 | 0 | Primates | 3425.0 | 2 | Hominidae | 3425.0 | 3 | Homo | 1741.0 | 4 | Homo sapiens | 1741.0 | 4 |
| scf7180000019480 | 277033 | 0.3961 | 0 | 49.846 | Eukaryota | 25813.0 | 0 | Chordata | 25813.0 | 0 | Carnivora | 25813.0 | 0 | Felidae | 25813.0 | 0 | Felis | 23198.0 | 1 | Felis catus | 23198.0 | 1 |
| scf7180000019176 | 539179 | 0.4579 | 0 | 49.4128 | Eukaryota | 46914.0 | 0 | Chordata | 46914.0 | 0 | Carnivora | 46914.0 | 0 | Felidae | 46914.0 | 0 | Felis | 46914.0 | 0 | Felis catus | 46914.0 | 0 |
| scf7180000024028 | 459782 | 0.4684 | 0 | 51.2401 | Eukaryota | 38197.0 | 0 | Chordata | 38197.0 | 0 | Artiodactyla | 25955.0 | 1 | Monodontidae | 22477.0 | 2 | elphinapteru | 18004.0 | 4 | hinapterus le | 18004.0 | 4 |
| scf7180000018256 | 1079513 | 0.3845 | 0 | 51.4383 | Eukaryota | 58793.0 | 0 | Chordata | 58793.0 | 0 | Carnivora | 58793.0 | 0 | Felidae | 58793.0 | 0 | Felis | 58793.0 | 0 | Felis catus | 58793.0 | 0 |
| scf7180000020914 | 120203 | 0.3984 | 0 | 25.1815 | Eukaryota | 21581.0 | 0 | Chordata | 21581.0 | 0 | Primates | 10071.0 | 3 | Hominidae | 10071.0 | 3 | Homo | 7960.0 | 6 | Homo sapiens | 7960.0 | 6 |
| scf7180000019478 | 210057 | 0.4067 | 0 | 50.547 | Eukaryota | 46496.0 | 0 | Chordata | 46496.0 | 0 | Carnivora | 46496.0 | 0 | Felidae | 46496.0 | 0 | Felis | 41857.0 | 1 | Felis catus | 41857.0 | 1 |
| scf7180000020662 | 19384 | 0.5632 | 0 | 248.9017 | Eukaryota | 2770.0 | 0 | Chordata | 2770.0 | 0 | Carnivora | 2770.0 | 0 | Felidae | 2507.0 | 1 | Felis | 1275.0 | 3 | Felis catus | 1275.0 | 4 |
| scf7180000020902 | 32658 | 0.5232 | 0 | 230.961 | no-hit | 0.0 | 0 | no-hit | 0.0 | 0 | no-hit | 0.0 | 0 | no-hit | 0.0 | 0 | no-hit | 0.0 | 0 | no-hit | 0.0 | 0 |
| scf7180000020382 | 144200 | 0.4471 | 0 | 25.0055 | Eukaryota | 19348.0 | 0 | Chordata | 19348.0 | 0 | Carnivora | 19348.0 | 0 | Canidae | 11332.0 | 2 | Canis | 11332.0 | 2 | Canis lupus | 11332.0 | 2 |
| scf7180000018779 | 652851 | 0.5335 | 0 | 51.0706 | Eukaryota | 50578.0 | 1 | Chordata | 50578.0 | 1 | Carnivora | 35014.0 | 2 | Hominidae | 15564.0 | 5 | Homo | 15564.0 | 7 | Homo sapiens | 15564.0 | 7 |
| scf7180000020123 | 179448 | 0.6536 | 0 | 54.2852 | Eukaryota | 15091.0 | 0 | Chordata | 15091.0 | 0 | Carnivora | 15091.0 | 0 | Felidae | 15091.0 | 0 | Felis | 9180.0 | 2 | Felis catus | 9180.0 | 2 |
| scf7180000019070 | 228196 | 0.4448 | 0 | 50.6792 | Eukaryota | 14850.0 | 1 | Chordata | 14850.0 | 1 | Carnivora | 13636.0 | 2 | Felidae | 9939.0 | 4 | Lynx | 5325.0 | 5 | ynx canadens | 5325.0 | 5 |
| scf7180000024041 | 425059 | 0.4379 | 0 | 50.6431 | Eukaryota | 27000.0 | 0 | Chordata | 27000.0 | 0 | Carnivora | 27000.0 | 0 | Felidae | 27000.0 | 0 | Felis | 24252.0 | 1 | Felis catus | 24252.0 | 1 |
| scf7180000023921 | 864184 | 0.3715 | 0 | 48.512 | Eukaryota | 48076.0 | 0 | Chordata | 48076.0 | 0 | Carnivora | 48076.0 | 0 | Felidae | 48076.0 | 0 | Felis | 48076.0 | 0 | Felis catus | 48076.0 | 0 |
| scf7180000021265 | 11188 | 0.5203 | 0 | 77.0829 | no-hit | 0.0 | 0 | no-hit | 0.0 | 0 | no-hit | 0.0 | 0 | no-hit | 0.0 | 0 | no-hit | 0.0 | 0 | no-hit | 0.0 | 0 |
| scf7180000017916 | 918872 | 0.3787 | 0 | 49.6344 | Eukaryota | 50977.0 | 0 | Chordata | 50977.0 | 0 | Carnivora | 50977.0 | 0 | Felidae | 50977.0 | 0 | Felis | 50977.0 | 0 | Felis catus | 50977.0 | 0 |
| scf7180000018278 | 834497 | 0.459 | 0 | 50.801 | Eukaryota | 12691.0 | 0 | Chordata | 12691.0 | 0 | Primates | 4785.0 | 2 | Hominidae | 4785.0 | 2 | Felis | 4547.0 | 4 | Felis catus | 4547.0 | 4 |
| scf7180000017547 | 3121025 | 0.361 | 0 | 49.6498 | Eukaryota | 47401.0 | 0 | Chordata | 47401.0 | 0 | Carnivora | 47401.0 | 0 | Felidae | 47401.0 | 0 | Felis | 47401.0 | 0 | Felis catus | 47401.0 | 0 |
| scf7180000020195 | 134562 | 0.4932 | 0 | 50.2546 | Eukaryota | 9613.0 | 0 | Chordata | 9613.0 | 0 | Carnivora | 9613.0 | 0 | Otariidae | 5670.0 | 1 | Eumetopias | 3786.0 | 4 | netopias juba | 3786.0 | 4 |
| scf7180000020632 | 105173 | 0.4208 | 0 | 25.6456 | Eukaryota | 28324.0 | 0 | Chordata | 28324.0 | 0 | Carnivora | 28324.0 | 0 | Felidae | 28324.0 | 0 | Acinonyx | 11236.0 | 2 | cinonyx jubat | 11236.0 | 2 |
| scf7180000021162 | 39554 | 0.3715 | 0 | 25.4574 | Eukaryota | 10048.0 | 0 | Chordata | 10048.0 | 0 | Primates | 7161.0 | 2 | Hominidae | 7161.0 | 2 | Homo | 5134.0 | 3 | Homo sapiens | 5134.0 | 3 |
| scf7180000019454 | 328083 | 0.4365 | 0 | 26.3103 | Eukaryota | 45476.0 | 0 | Chordata | 45476.0 | 0 | Carnivora | 45476.0 | 0 | Felidae | 45476.0 | 0 | Felis | 25679.0 | 1 | Felis catus | 25679.0 | 1 |
| scf7180000018621 | 15220 | 0.3897 | 0 | 24.4408 | Eukaryota | 10841.0 | 0 | Chordata | 10841.0 | 0 | Carnivora | 10841.0 | 0 | Felidae | 10841.0 | 0 | Felis | 8716.0 | 1 | Felis catus | 8716.0 | 1 |
| scf7180000019937 | 160847 | 0.4123 | 0 | 24.8448 | Eukaryota | 32191.0 | 0 | Chordata | 32191.0 | 0 | Carnivora | 18808.0 | 2 | Felidae | 16470.0 | 4 | Acinonyx | 5908.0 | 8 | cinonyx jubat | 5908.0 | 8 |
| scf7180000018870 | 433966 | 0.392 | 0 | 50.2604 | Eukaryota | 40478.0 | 0 | Chordata | 40478.0 | 0 | Carnivora | 40478.0 | 0 | Herpestidae | 23850.0 | 2 | Suricata | 23850.0 | 3 | ricata suricat | 23850.0 | 3 |
| scf7180000019917 | 333781 | 0.6206 | 0 | 47.5551 | Eukaryota | 19941.0 | 0 | Chordata | 19941.0 | 0 | Carnivora | 19941.0 | 0 | Felidae | 17894.0 | 1 | Acinonyx | 8032.0 | 3 | cinonyx jubat | 8032.0 | 3 |
| scf7180000024077 | 326438 | 0.4036 | 0 | 26.3026 | Eukaryota | 25163.0 | 0 | Chordata | 25163.0 | 0 | Carnivora | 22354.0 | 1 | Felidae | 19879.0 | 2 | Felis | 7458.0 | 4 | Felis catus | 7458.0 | 4 |
| scf7180000022366 | 19443 | 0.4027 | 0 | 25.4333 | Eukaryota | 5631.0 | 0 | Chordata | 5631.0 | 0 | Primates | 4917.0 | 1 | Hominidae | 4917.0 | 1 | Pan | 3010.0 | 2 | an troglodyte | 3010.0 | 2 |
| scf7180000019796 | 209482 | 0.4404 | 0 | 50.3449 | Eukaryota | 12812.0 | 0 | Chordata | 12812.0 | 0 | Carnivora | 10781.0 | 1 | Felidae | 10781.0 | 1 | Felis | 5110.0 | 4 | Felis catus | 5110.0 | 4 |
| scf7180000020845 | 68148 | 0.5219 | 0 | 48.3878 | Eukaryota | 7066.0 | 0 | Chordata | 7066.0 | 0 | Carnivora | 7066.0 | 0 | Felidae | 7066.0 | 0 | Felis | 7066.0 | 0 | Felis catus | 7066.0 | 0 |
| scf7180000019542 | 106775 | 0.4199 | 0 | 51.1139 | Eukaryota | 10075.0 | 0 | Chordata | 10075.0 | 0 | Carnivora | 5837.0 | 1 | Felidae | 5837.0 | 1 | Felis | 4185.0 | 3 | Felis catus | 4185.0 | 3 |
| scf7180000019082 | 529240 | 0.3917 | 0 | 51.295 | Eukaryota | 56360.0 | 0 | Chordata | 56360.0 | 0 | Carnivora | 56360.0 | 0 | Felidae | 56360.0 | 0 | Felis | 45120.0 | 1 | Felis catus | 45120.0 | 1 |
| scf7180000021696 | 35805 | 0.469 | 0 | 48.2725 | Eukaryota | 4729.0 | 0 | Chordata | 4729.0 | 0 | Carnivora | 3160.0 | 2 | Felidae | 3160.0 | 2 | Felis | 2458.0 | 3 | Felis catus | 2458.0 | 3 |
| scf7180000018863 | 315841 | 0.4263 | 0 | 50.3683 | Eukaryota | 39498.0 | 0 | Chordata | 39498.0 | 0 | Carnivora | 39498.0 | 0 | Felidae | 39498.0 | 0 | Felis | 39498.0 | 0 | Felis catus | 39498.0 | 0 |
| scf7180000024102 | 295371 | 0.3676 | 0 | 49.8068 | Eukaryota | 16330.0 | 0 | Chordata | 16330.0 | 0 | Carnivora | 16330.0 | 0 | Felidae | 16330.0 | 0 | Felis | 7816.0 | 2 | Felis catus | 7816.0 | 2 |
| scf7180000023068 | 40646 | 0.3857 | 0 | 325.7298 | Eukaryota | 8420.0 | 0 | Chordata | 8420.0 | 0 | pterygiforme | 5051.0 | 2 | Apterygidae | 5051.0 | 2 | Apteryx | 5051.0 | 2 | pteryx mante | 5051.0 | 2 |
| scf7180000023882 | 805390 | 0.3529 | 0 | 49.1318 | Eukaryota | 46044.0 | 0 | Chordata | 46044.0 | 0 | Carnivora | 46044.0 | 0 | Felidae | 46044.0 | 0 | Felis | 46044.0 | 0 | Felis catus | 46044.0 | 0 |
| scf7180000022367 | 17821 | 0.5033 | 0 | 28.4217 | Eukaryota | 4009.0 | 0 | Chordata | 4009.0 | 0 | Chiroptera | 1452.0 | 4 | Pteropodidae | 1452.0 | 5 | Pteropus | 1452.0 | 5 | ropus vampy | 1452.0 | 5 |
| scf7180000021965 | 32798 | 0.5562 | 0 | 51.8153 | Eukaryota | 3602.0 | 0 | Chordata | 3602.0 | 0 | Carnivora | 3602.0 | 0 | Felidae | 2977.0 | 1 | Felis | 1335.0 | 4 | Felis catus | 1335.0 | 5 |
| scf7180000021946 | 27598 | 0.5269 | 0 | 1420.0643 | no-hit | 0.0 | 0 | no-hit | 0.0 | 0 | no-hit | 0.0 | 0 | no-hit | 0.0 | 0 | no-hit | 0.0 | 0 | no-hit | 0.0 | 0 |
| scf7180000022691 | 3911 | 0.5277 | 0 | 32.7486 | no-hit | 0.0 | 0 | no-hit | 0.0 | 0 | no-hit | 0.0 | 0 | no-hit | 0.0 | 0 | no-hit | 0.0 | 0 | no-hit | 0.0 | 0 |
| scf7180000023502 | 34743 | 0.5159 | 0 | 216.3561 | Eukaryota | 27178.0 | 0 | Chordata | 27178.0 | 0 | Carnivora | 27178.0 | 0 | Felidae | 27178.0 | 0 | Acinonyx | 13312.0 | 2 | cinonyx jubat | 13312.0 | 2 |
| scf7180000019152 | 18111 | 0.4647 | 0 | 16.4055 | Eukaryota | 4885.0 | 0 | Chordata | 4885.0 | 0 | Carnivora | 4885.0 | 0 | Felidae | 4079.0 | 1 | Felis | 1337.0 | 4 | Felis catus | 1337.0 | 5 |
| scf7180000019262 | 384095 | 0.3924 | 0 | 45.8014 | Eukaryota | 52913.0 | 0 | Chordata | 52913.0 | 0 | Carnivora | 52913.0 | 0 | Otariidae | 15657.0 | 4 | Canis | 10537.0 | 8 | Canis lupus | 10537.0 | 8 |
| scf7180000023644 | 25826 | 0.4522 | 0 | 216.3756 | Eukaryota | 2629.0 | 0 | Chordata | 2629.0 | 0 | Carnivora | 2223.0 | 1 | Hyaenidae | 774.0 | 4 | Crocuta | 774.0 | 5 | rocuta crocut | 774.0 | 5 |
| scf7180000019521 | 37379 | 0.4411 | 0 | 365.1939 | Eukaryota | 4781.0 | 0 | Chordata | 4781.0 | 0 | Carnivora | 3878.0 | 1 | Felidae | 2519.0 | 2 | Felis | 2519.0 | 2 | Felis catus | 2519.0 | 2 |
| scf7180000020962 | 75894 | 0.4435 | 0 | 25.9452 | Eukaryota | 13029.0 | 0 | Chordata | 13029.0 | 0 | Primates | 8176.0 | 2 | Hominidae | 6300.0 | 4 | Homo | 6300.0 | 5 | Homo sapiens | 6300.0 | 5 |
| scf7180000023436 | 33850 | 0.4072 | 0 | 315.016 | Eukaryota | 47455.0 | 0 | Chordata | 47455.0 | 0 | Carnivora | 47455.0 | 0 | Felidae | 47455.0 | 0 | Felis | 47455.0 | 0 | Felis catus | 47455.0 | 0 |
| scf7180000022739 | 14331 | 0.3896 | 0 | 24.8084 | Eukaryota | 31137.0 | 0 | Chordata | 31137.0 | 0 | Carnivora | 31137.0 | 0 | Felidae | 31137.0 | 0 | Felis | 31137.0 | 0 | Felis catus | 31137.0 | 0 |
| scf7180000017572 | 1461305 | 0.3792 | 0 | 49.7604 | Eukaryota | 72137.0 | 0 | Chordata | 72137.0 | 0 | Artiodactyla | 40750.0 | 1 | Canidae | 23817.0 | 5 | Canis | 16570.0 | 7 | Canis lupus | 16570.0 | 7 |
| scf7180000017915 | 2949099 | 0.3908 | 0 | 49.552 | Eukaryota | 46871.0 | 0 | Chordata | 46871.0 | 0 | Carnivora | 46871.0 | 0 | Felidae | 46871.0 | 0 | Felis | 46871.0 | 0 | Felis catus | 46871.0 | 0 |
| scf7180000019021 | 504327 | 0.5462 | 0 | 50.9838 | Eukaryota | 34623.0 | 0 | Chordata | 34623.0 | 0 | Carnivora | 34623.0 | 0 | Felidae | 32096.0 | 1 | Felis | 12971.0 | 4 | Felis catus | 12971.0 | 4 |
| scf7180000019568 | 299765 | 0.3869 | 0 | 50.0819 | Eukaryota | 79229.0 | 0 | Chordata | 79229.0 | 0 | Carnivora | 79229.0 | 0 | Felidae | 57102.0 | 3 | Panthera | 24183.0 | 7 | anthera pardu | 16958.0 | 8 |
| scf7180000021524 | 66248 | 0.3627 | 0 | 25.1652 | Eukaryota | 12088.0 | 0 | Chordata | 12088.0 | 0 | Primates | 6693.0 | 2 | Hominidae | 6693.0 | 3 | Sus | 3763.0 | 4 | Sus scrofa | 3763.0 | 4 |
| scf7180000020092 | 125093 | 0.3608 | 0 | 50.8931 | Eukaryota | 13911.0 | 1 | Chordata | 13911.0 | 1 | Primates | 5220.0 | 3 | Hominidae | 5220.0 | 4 | Homo | 5220.0 | 5 | Homo sapiens | 5220.0 | 5 |
| scf7180000019642 | 294741 | 0.4563 | 0 | 26.8183 | Eukaryota | 59059.0 | 0 | Chordata | 59059.0 | 0 | Carnivora | 59059.0 | 0 | Felidae | 53524.0 | 1 | Felis | 23824.0 | 3 | Felis catus | 23824.0 | 3 |
| scf7180000021404 | 71480 | 0.4507 | 0 | 24.1784 | Eukaryota | 10105.0 | 0 | Chordata | 10105.0 | 0 | Carnivora | 10105.0 | 0 | Felidae | 10105.0 | 0 | Felis | 10105.0 | 0 | Felis catus | 10105.0 | 0 |
| scf7180000021699 | 42344 | 0.5337 | 0 | 46.7184 | Eukaryota | 3442.0 | 0 | Chordata | 3442.0 | 0 | Carnivora | 3442.0 | 0 | Felidae | 3442.0 | 0 | Felis | 3085.0 | 1 | Felis catus | 3085.0 | 1 |
| scf7180000018507 | 733317 | 0.4348 | 0 | 51.6261 | Eukaryota | 39108.0 | 0 | Chordata | 39108.0 | 0 | Carnivora | 39108.0 | 0 | Felidae | 39108.0 | 0 | Felis | 35192.0 | 1 | Felis catus | 35192.0 | 1 |
| scf7180000020085 | 107630 | 0.4165 | 0 | 49.4213 | Eukaryota | 20923.0 | 0 | Chordata | 20923.0 | 0 | Carnivora | 17083.0 | 2 | Felidae | 17083.0 | 2 | Felis | 6930.0 | 5 | Felis catus | 6930.0 | 6 |
| scf7180000019295 | 392955 | 0.5913 | 0 | 49.3785 | Eukaryota | 33958.0 | 1 | Chordata | 33958.0 | 1 | Carnivora | 29978.0 | 2 | Felidae | 23628.0 | 3 | Felis | 23628.0 | 3 | Felis catus | 23628.0 | 3 |
| scf7180000024239 | 1176842 | 0.4017 | 0 | 50.3217 | Eukaryota | 46795.0 | 0 | Chordata | 46795.0 | 0 | Carnivora | 46795.0 | 0 | Felidae | 46795.0 | 0 | Felis | 46795.0 | 0 | Felis catus | 46795.0 | 0 |
| scf7180000019185 | 640530 | 0.462 | 0 | 50.7212 | Eukaryota | 47391.0 | 0 | Chordata | 47391.0 | 0 | Carnivora | 47391.0 | 0 | Felidae | 47391.0 | 0 | Felis | 42691.0 | 1 | Felis catus | 42691.0 | 1 |
| scf7180000021671 | 46623 | 0.3753 | 0 | 25.1011 | Eukaryota | 35900.0 | 0 | Chordata | 35900.0 | 0 | Carnivora | 35900.0 | 0 | Felidae | 35900.0 | 0 | Felis | 32308.0 | 1 | Felis catus | 32308.0 | 1 |
| scf7180000023912 | 1037444 | 0.3879 | 0 | 50.242 | Eukaryota | 46984.0 | 0 | Chordata | 46984.0 | 0 | Carnivora | 46984.0 | 0 | Felidae | 46984.0 | 0 | Felis | 46984.0 | 0 | Felis catus | 46984.0 | 0 |
| scf7180000018962 | 399916 | 0.4833 | 0 | 49.8824 | Eukaryota | 39228.0 | 0 | Chordata | 39228.0 | 0 | Carnivora | 39228.0 | 0 | Felidae | 39228.0 | 0 | Felis | 39228.0 | 0 | Felis catus | 39228.0 | 0 |
| scf7180000023144 | 62235 | 0.3722 | 0 | 72.4218 | Eukaryota | 30217.0 | 0 | Chordata | 30217.0 | 0 | Artiodactyla | 19770.0 | 1 | Bovidae | 19770.0 | 1 | Ovis | 19770.0 | 2 | vis canadensi | 19770.0 | 2 |
| scf7180000021598 | 38735 | 0.4374 | 0 | 26.3663 | Eukaryota | 8869.0 | 0 | Chordata | 8869.0 | 0 | Carnivora | 3744.0 | 2 | Felidae | 3744.0 | 2 | Felis | 3744.0 | 3 | Felis catus | 3744.0 | 3 |
| scf7180000020403 | 76258 | 0.3592 | 0 | 24.6887 | Eukaryota | 31337.0 | 0 | Chordata | 31337.0 | 0 | Carnivora | 31337.0 | 0 | Felidae | 31337.0 | 0 | Felis | 28190.0 | 1 | Felis catus | 28190.0 | 1 |
| scf7180000020488 | 79967 | 0.3757 | 0 | 23.3223 | Eukaryota | 45188.0 | 0 | Chordata | 45188.0 | 0 | Carnivora | 45188.0 | 0 | Felidae | 45188.0 | 0 | Felis | 45188.0 | 0 | Felis catus | 45188.0 | 0 |
| scf7180000018283 | 565495 | 0.4026 | 0 | 50.447 | Eukaryota | 35939.0 | 1 | Chordata | 35939.0 | 1 | Carnivora | 32733.0 | 2 | Felidae | 29056.0 | 3 | Felis | 12341.0 | 6 | Felis catus | 12341.0 | 6 |
| scf7180000018317 | 717065 | 0.4228 | 0 | 52.3209 | Eukaryota | 29422.0 | 0 | Chordata | 29422.0 | 0 | Carnivora | 17147.0 | 2 | Felidae | 14476.0 | 4 | unresolved | 6568.0 | 7 | unresolved | 6568.0 | 7 |
| scf7180000021520 | 61270 | 0.4046 | 0 | 26.0565 | Eukaryota | 16200.0 | 0 | Chordata | 16200.0 | 0 | Carnivora | 7943.0 | 2 | Felidae | 7943.0 | 2 | Felis | 7943.0 | 3 | Felis catus | 7943.0 | 3 |
| scf7180000019393 | 124040 | 0.5727 | 0 | 49.4516 | Eukaryota | 42339.0 | 0 | Chordata | 42339.0 | 0 | Carnivora | 22117.0 | 1 | Cercopithecidae | 20222.0 | 2 | Acinonyx | 8618.0 | 6 | cinonyx jubat | 8618.0 | 6 |
| scf7180000019149 | 370240 | 0.3465 | 0 | 47.9726 | Eukaryota | 45967.0 | 0 | Chordata | 45967.0 | 0 | Carnivora | 45967.0 | 0 | Felidae | 45967.0 | 0 | Felis | 45967.0 | 0 | Felis catus | 45967.0 | 0 |
| scf7180000020507 | 364913 | 0.4328 | 0 | 49.7127 | Eukaryota | 43079.0 | 0 | Chordata | 43079.0 | 0 | Carnivora | 43079.0 | 0 | Felidae | 43079.0 | 0 | Felis | 43079.0 | 0 | Felis catus | 43079.0 | 0 |
| scf7180000023006 | 45917 | 0.4105 | 0 | 34.9548 | Eukaryota | 47117.0 | 0 | Chordata | 47117.0 | 0 | Carnivora | 47117.0 | 0 | Felidae | 47117.0 | 0 | Felis | 47117.0 | 0 | Felis catus | 47117.0 | 0 |
| scf7180000020742 | 89125 | 0.5891 | 0 | 49.4883 | Eukaryota | 5665.0 | 1 | Chordata | 5665.0 | 1 | Carnivora | 2845.0 | 3 | Hominidae | 2238.0 | 5 | Homo | 2238.0 | 6 | Homo sapiens | 2238.0 | 6 |
| scf7180000023892 | 1128960 | 0.416 | 0 | 52.4077 | Eukaryota | 35874.0 | 0 | Chordata | 35874.0 | 0 | Carnivora | 35874.0 | 0 | Felidae | 27302.0 | 2 | Felis | 17763.0 | 4 | Felis catus | 17763.0 | 4 |
| scf7180000022312 | 34241 | 0.5069 | 0 | 99.5417 | Eukaryota | 9921.0 | 0 | Chordata | 9921.0 | 0 | Carnivora | 8893.0 | 1 | Felidae | 7120.0 | 3 | Lynx | 6336.0 | 5 | ynx canadens | 6336.0 | 5 |
| scf7180000020437 | 87818 | 0.548 | 0 | 78.8898 | Eukaryota | 5211.0 | 0 | Chordata | 5211.0 | 0 | Carnivora | 5211.0 | 0 | Felidae | 4069.0 | 1 | Acinonyx | 1428.0 | 5 | cinonyx jubat | 1428.0 | 5 |
| scf7180000021930 | 16661 | 0.3828 | 0 | 16.2958 | Eukaryota | 27560.0 | 0 | Chordata | 27560.0 | 0 | Carnivora | 27560.0 | 0 | Felidae | 27560.0 | 0 | Felis | 27560.0 | 0 | Felis catus | 27560.0 | 0 |
| scf7180000019607 | 238160 | 0.3935 | 0 | 49.2739 | Eukaryota | 25811.0 | 1 | Chordata | 25811.0 | 1 | Carnivora | 17467.0 | 2 | Hominidae | 8344.0 | 5 | Homo | 8344.0 | 6 | Homo sapiens | 8344.0 | 6 |
| scf7180000022156 | 34154 | 0.4355 | 0 | 377.9742 | Eukaryota | 6211.0 | 0 | Chordata | 6211.0 | 0 | Carnivora | 6211.0 | 0 | Felidae | 6211.0 | 0 | Felis | 6211.0 | 0 | Felis catus | 6211.0 | 0 |
| scf7180000020945 | 50005 | 0.3746 | 0 | 48.9995 | Eukaryota | 43358.0 | 0 | Chordata | 43358.0 | 0 | Carnivora | 43358.0 | 0 | Felidae | 43358.0 | 0 | Felis | 43358.0 | 0 | Felis catus | 43358.0 | 0 |
| scf7180000019565 | 14803 | 0.6465 | 0 | 79.3837 | no-hit | 0.0 | 0 | no-hit | 0.0 | 0 | no-hit | 0.0 | 0 | no-hit | 0.0 | 0 | no-hit | 0.0 | 0 | no-hit | 0.0 | 0 |
| scf7180000021063 | 28731 | 0.4011 | 0 | 25.5241 | Eukaryota | 18026.0 | 0 | Chordata | 18026.0 | 0 | Carnivora | 16513.0 | 1 | Felidae | 12396.0 | 4 | Felis | 7550.0 | 5 | Felis catus | 7550.0 | 5 |
| scf7180000018796 | 89126 | 0.4394 | 0 | 48.0202 | Eukaryota | 44134.0 | 0 | Chordata | 44134.0 | 0 | Carnivora | 44134.0 | 0 | Felidae | 44134.0 | 0 | Felis | 39794.0 | 1 | Felis catus | 39794.0 | 1 |
| scf7180000019715 | 219460 | 0.3708 | 0 | 47.434 | Eukaryota | 33868.0 | 0 | Chordata | 33868.0 | 0 | Carnivora | 33868.0 | 0 | Felidae | 33868.0 | 0 | Felis | 33868.0 | 0 | Felis catus | 33868.0 | 0 |
| scf7180000018791 | 484582 | 0.442 | 0 | 51.794 | Eukaryota | 66840.0 | 0 | Chordata | 66840.0 | 0 | Carnivora | 66840.0 | 0 | Felidae | 52212.0 | 2 | Lynx | 21840.0 | 4 | ynx canadens | 21840.0 | 4 |
| scf7180000017791 | 2182412 | 0.4072 | 0 | 51.5216 | Eukaryota | 46905.0 | 0 | Chordata | 46905.0 | 0 | Carnivora | 46905.0 | 0 | Felidae | 46905.0 | 0 | Felis | 46905.0 | 0 | Felis catus | 46905.0 | 0 |
| scf7180000024245 | 723078 | 0.5534 | 100 | 49.5795 | Eukaryota | 28712.0 | 0 | Chordata | 28712.0 | 0 | Carnivora | 28712.0 | 0 | Felidae | 28712.0 | 0 | Felis | 22903.0 | 1 | Felis catus | 22903.0 | 1 |
| scf7180000021168 | 63142 | 0.3651 | 0 | 24.7254 | Eukaryota | 9582.0 | 0 | Chordata | 9582.0 | 0 | Carnivora | 9582.0 | 0 | Felidae | 9582.0 | 0 | Felis | 9582.0 | 0 | Felis catus | 9582.0 | 0 |
| scf7180000022929 | 89210 | 0.4477 | 0 | 698.1746 | Eukaryota | 23039.0 | 0 | Chordata | 23039.0 | 0 | Carnivora | 23039.0 | 0 | Felidae | 23039.0 | 0 | Felis | 20710.0 | 1 | Felis catus | 20710.0 | 1 |
| scf7180000024158 | 99292 | 0.3993 | 0 | 26.023 | Eukaryota | 13429.0 | 0 | Chordata | 13429.0 | 0 | Carnivora | 13429.0 | 0 | Felidae | 13429.0 | 0 | Felis | 12104.0 | 1 | Felis catus | 12104.0 | 1 |
| scf7180000017992 | 1460527 | 0.4048 | 0 | 51.8148 | Eukaryota | 53467.0 | 0 | Chordata | 53467.0 | 0 | Carnivora | 53467.0 | 0 | Felidae | 36235.0 | 1 | Acinonyx | 25190.0 | 3 | cinonyx jubat | 25190.0 | 3 |
| scf7180000021647 | 24619 | 0.3727 | 0 | 25.0473 | Eukaryota | 7832.0 | 0 | Chordata | 7832.0 | 0 | Carnivora | 4290.0 | 3 | Felidae | 4290.0 | 3 | Felis | 4290.0 | 3 | Felis catus | 4290.0 | 3 |
| scf7180000020515 | 13537 | 0.4419 | 0 | 57.2076 | Eukaryota | 10907.0 | 0 | Chordata | 10907.0 | 0 | Carnivora | 10907.0 | 0 | Ursidae | 6440.0 | 2 | Ursus | 5320.0 | 4 | Ursus arctos | 3195.0 | 5 |
| scf7180000021703 | 19282 | 0.4306 | 0 | 402.3915 | Eukaryota | 2188.0 | 0 | Chordata | 2188.0 | 0 | Carnivora | 2188.0 | 0 | Felidae | 2188.0 | 0 | Felis | 1549.0 | 2 | Felis catus | 1549.0 | 2 |
| scf7180000017512 | 2774830 | 0.4055 | 0 | 50.482 | Eukaryota | 48400.0 | 0 | Chordata | 48400.0 | 0 | Carnivora | 48400.0 | 0 | Felidae | 48400.0 | 0 | Felis | 48400.0 | 0 | Felis catus | 48400.0 | 0 |
| scf7180000017471 | 4405218 | 0.4176 | 100 | 51.0701 | Eukaryota | 49838.0 | 0 | Chordata | 49838.0 | 0 | Carnivora | 49838.0 | 0 | Felidae | 43780.0 | 1 | Felis | 43780.0 | 1 | Felis catus | 43780.0 | 1 |
| scf7180000022697 | 9240 | 0.5223 | 0 | 653.0953 | no-hit | 0.0 | 0 | no-hit | 0.0 | 0 | no-hit | 0.0 | 0 | no-hit | 0.0 | 0 | no-hit | 0.0 | 0 | no-hit | 0.0 | 0 |
| scf7180000020853 | 72510 | 0.5211 | 0 | 48.6069 | Eukaryota | 15275.0 | 0 | Chordata | 15275.0 | 0 | Carnivora | 15275.0 | 0 | Felidae | 15275.0 | 0 | Acinonyx | 5814.0 | 4 | cinonyx jubat | 5814.0 | 5 |
| scf7180000019016 | 4985428 | 0.3987 | 100 | 50.58 | Eukaryota | 130910.0 | 0 | Chordata | 130910.0 | 0 | Carnivora | 130910.0 | 0 | Felidae | 104028.0 | 1 | Panthera | 51979.0 | 3 | anthera pardu | 51979.0 | 3 |
| scf7180000020609 | 97993 | 0.462 | 0 | 27.0675 | Eukaryota | 34562.0 | 0 | Chordata | 34562.0 | 0 | Carnivora | 34562.0 | 0 | Felidae | 24437.0 | 2 | Felis | 10470.0 | 4 | Felis catus | 10470.0 | 4 |
| scf7180000019369 | 368319 | 0.521 | 0 | 49.6094 | Eukaryota | 8270.0 | 0 | Chordata | 8270.0 | 0 | Carnivora | 7422.0 | 1 | Felidae | 6160.0 | 2 | Felis | 6160.0 | 2 | Felis catus | 6160.0 | 2 |
| scf7180000022145 | 24535 | 0.4141 | 0 | 23.2164 | Eukaryota | 14021.0 | 0 | Chordata | 14021.0 | 0 | Carnivora | 14021.0 | 0 | Felidae | 14021.0 | 0 | Felis | 12626.0 | 1 | Felis catus | 12626.0 | 1 |
| scf7180000019012 | 24240 | 0.3627 | 0 | 24.8416 | Eukaryota | 9963.0 | 0 | Chordata | 9963.0 | 0 | Carnivora | 9963.0 | 0 | Felidae | 9043.0 | 1 | Felis | 9043.0 | 1 | Felis catus | 9043.0 | 1 |
| scf7180000017683 | 1958306 | 0.3897 | 0 | 50.1244 | Eukaryota | 49087.0 | 0 | Chordata | 49087.0 | 0 | Carnivora | 49087.0 | 0 | Felidae | 49087.0 | 0 | Felis | 40908.0 | 1 | Felis catus | 40908.0 | 1 |
| scf7180000020008 | 175550 | 0.3765 | 0 | 52.3931 | Eukaryota | 47303.0 | 0 | Chordata | 47303.0 | 0 | Carnivora | 47303.0 | 0 | Felidae | 47303.0 | 0 | Felis | 47303.0 | 0 | Felis catus | 47303.0 | 0 |
| scf7180000017708 | 723883 | 0.4567 | 0 | 52.3427 | Eukaryota | 49711.0 | 0 | Chordata | 49711.0 | 0 | Carnivora | 49711.0 | 0 | Otariidae | 18141.0 | 5 | Callorhinus | 8159.0 | 8 | llorhinus ursi | 8159.0 | 8 |
| scf7180000022582 | 5373 | 0.3959 | 0 | 22.9164 | Eukaryota | 2029.0 | 0 | Chordata | 2029.0 | 0 | Carnivora | 2029.0 | 0 | Felidae | 1820.0 | 1 | Leopardus | 633.0 | 6 | opardus tigri | 213.0 | 9 |
| scf7180000021390 | 41813 | 0.37 | 0 | 24.5763 | Eukaryota | 20831.0 | 0 | Chordata | 20831.0 | 0 | Carnivora | 20831.0 | 0 | Felidae | 20831.0 | 0 | Felis | 18729.0 | 1 | Felis catus | 18729.0 | 1 |
| scf7180000020212 | 152634 | 0.4684 | 0 | 49.409 | Eukaryota | 7775.0 | 1 | Chordata | 7775.0 | 1 | Carnivora | 3907.0 | 2 | Felidae | 3907.0 | 2 | Felis | 3907.0 | 3 | Felis catus | 3907.0 | 3 |
| scf7180000023315 | 26628 | 0.4903 | 0 | 605.6283 | Eukaryota | 17883.0 | 0 | Chordata | 17883.0 | 0 | Carnivora | 17883.0 | 0 | Felidae | 14369.0 | 1 | Acinonyx | 3534.0 | 5 | cinonyx jubat | 3534.0 | 5 |
| scf7180000023567 | 15777 | 0.4813 | 0 | 516.18 | Eukaryota | 17585.0 | 0 | Chordata | 17585.0 | 0 | Carnivora | 17585.0 | 0 | Felidae | 14145.0 | 1 | Acinonyx | 3477.0 | 5 | cinonyx jubat | 3477.0 | 5 |
| scf7180000019591 | 229972 | 0.3662 | 0 | 50.0635 | Eukaryota | 45934.0 | 0 | Chordata | 45934.0 | 0 | Carnivora | 45934.0 | 0 | Felidae | 45934.0 | 0 | Felis | 45934.0 | 0 | Felis catus | 45934.0 | 0 |
| scf7180000017485 | 719698 | 0.3635 | 0 | 49.7797 | Eukaryota | 45591.0 | 0 | Chordata | 45591.0 | 0 | Carnivora | 45591.0 | 0 | Felidae | 45591.0 | 0 | Felis | 41397.0 | 1 | Felis catus | 41397.0 | 1 |
| scf7180000023103 | 39768 | 0.556 | 0 | 173.4155 | Eukaryota | 5174.0 | 0 | Chordata | 5174.0 | 0 | Carnivora | 4298.0 | 1 | Felidae | 4048.0 | 2 | Felis | 2042.0 | 4 | Felis catus | 2042.0 | 4 |
| scf7180000019275 | 12060 | 0.3545 | 0 | 23.1593 | Eukaryota | 8651.0 | 0 | Chordata | 8651.0 | 0 | Carnivora | 8651.0 | 0 | Felidae | 8651.0 | 0 | Panthera | 5172.0 | 2 | cinonyx jubat | 2616.0 | 3 |
| scf7180000023404 | 20293 | 0.424 | 0 | 445.6364 | Eukaryota | 257160.0 | 0 | Chordata | 257160.0 | 0 | Carnivora | 257160.0 | 0 | Hyaenidae | 257160.0 | 0 | Proteles | 155586.0 | 1 | roteles cristat | 155586.0 | 1 |
| scf7180000022057 | 11913 | 0.5893 | 0 | 42.2883 | Eukaryota | 2575.0 | 0 | Chordata | 2575.0 | 0 | Carnivora | 2575.0 | 0 | Felidae | 2294.0 | 1 | Felis | 1539.0 | 3 | Felis catus | 1539.0 | 3 |
| scf7180000022124 | 15408 | 0.4357 | 0 | 23.2671 | Eukaryota | 4904.0 | 0 | Chordata | 4904.0 | 0 | Carnivora | 3643.0 | 1 | Felidae | 3643.0 | 1 | Felis | 3643.0 | 2 | Felis catus | 3643.0 | 2 |
| scf7180000024269 | 2225959 | 0.3938 | 0 | 50.5874 | Eukaryota | 70707.0 | 0 | Chordata | 70707.0 | 0 | Carnivora | 70707.0 | 0 | Felidae | 70707.0 | 0 | Felis | 26706.0 | 2 | Felis catus | 26706.0 | 2 |
| scf7180000021968 | 19746 | 0.4054 | 0 | 25.4441 | Eukaryota | 9604.0 | 0 | Chordata | 9604.0 | 0 | Carnivora | 8350.0 | 1 | Felidae | 8350.0 | 1 | Felis | 6497.0 | 2 | Felis catus | 6497.0 | 2 |
| scf7180000019168 | 512182 | 0.4318 | 0 | 26.8181 | Eukaryota | 37396.0 | 0 | Chordata | 37396.0 | 0 | Primates | 32292.0 | 1 | Hominidae | 32292.0 | 1 | Homo | 19170.0 | 2 | Homo sapiens | 19170.0 | 2 |

| | | | | | | | | | | | | | | | | | | | | | | |
|---|---|---|---|---|---|---|---|---|---|---|---|---|---|---|---|---|---|---|---|---|---|---|
| scf7180000023896 | 1496193 | 0.3618 | 0 | 49.5977 | Eukaryota | 47932.0 | 0 | Chordata | 47932.0 | 0 | Carnivora | 47932.0 | 0 | Felidae | 47932.0 | 0 | Felis | 47932.0 | 0 | Felis catus | 47932.0 | 0 |
| scf7180000021801 | 16712 | 0.4269 | 0 | 26.9978 | Eukaryota | 6464.0 | 0 | Chordata | 6464.0 | 0 | Primates | 4965.0 | 3 | Hominidae | 4965.0 | 3 | Homo | 3711.0 | 4 | Homo sapiens | 3711.0 | 4 |
| scf7180000019691 | 143284 | 0.4775 | 0 | 50.8626 | Eukaryota | 12821.0 | 0 | Chordata | 12821.0 | 0 | Carnivora | 12821.0 | 0 | Otariidae | 5826.0 | 3 | Zalophus | 5826.0 | 3 | phus californi | 5826.0 | 3 |
| scf7180000018472 | 1558826 | 0.4835 | 0 | 52.3663 | Eukaryota | 86573.0 | 0 | Chordata | 86573.0 | 0 | Carnivora | 86573.0 | 0 | Felidae | 70661.0 | 2 | Panthera | 25637.0 | 6 | anthera pardu | 17212.0 | 7 |
| scf7180000019268 | 374781 | 0.3794 | 0 | 49.9385 | Eukaryota | 33147.0 | 0 | Chordata | 33147.0 | 0 | Carnivora | 33147.0 | 0 | Felidae | 33147.0 | 0 | Felis | 29769.0 | 1 | Felis catus | 29769.0 | 1 |
| scf7180000023301 | 28170 | 0.4296 | 0 | 101.7488 | Eukaryota | 13064.0 | 0 | Chordata | 13064.0 | 0 | Carnivora | 13064.0 | 0 | Felidae | 13064.0 | 0 | Felis | 10480.0 | 1 | Felis catus | 10480.0 | 1 |
| scf7180000019706 | 38087 | 0.3967 | 0 | 25.4535 | Eukaryota | 22047.0 | 0 | Chordata | 22047.0 | 0 | Carnivora | 22047.0 | 0 | Felidae | 22047.0 | 0 | Felis | 19864.0 | 1 | Felis catus | 19864.0 | 1 |
| scf7180000019820 | 204021 | 0.381 | 0 | 48.8672 | Eukaryota | 33270.0 | 0 | Chordata | 33270.0 | 0 | Carnivora | 33270.0 | 0 | Herpestidae | 17590.0 | 1 | Suricata | 17590.0 | 1 | ıricata suricat | 17590.0 | 1 |
| scf7180000023977 | 1016136 | 0.4779 | 0 | 53.4525 | Eukaryota | 25240.0 | 0 | Chordata | 25240.0 | 0 | Carnivora | 25240.0 | 0 | Felidae | 20763.0 | 1 | Acinonyx | 9720.0 | 3 | :inonyx jubat | 9720.0 | 3 |
| scf7180000022804 | 12324 | 0.5019 | 0 | 52.6765 | Eukaryota | 4928.0 | 0 | Chordata | 4928.0 | 0 | Carnivora | 4928.0 | 0 | Felidae | 4682.0 | 1 | Felis | 1787.0 | 4 | Felis catus | 1787.0 | 4 |
| scf7180000021169 | 18788 | 0.5201 | 0 | 198.5326 | no-hit | 0.0 | 0 | no-hit | 0.0 | 0 | no-hit | 0.0 | 0 | no-hit | 0.0 | 0 | no-hit | 0.0 | 0 | no-hit | 0.0 | 0 |
| scf7180000017786 | 1248858 | 0.3688 | 0 | 49.9917 | Eukaryota | 45457.0 | 0 | Chordata | 45457.0 | 0 | Carnivora | 45457.0 | 0 | Felidae | 45457.0 | 0 | Felis | 45457.0 | 0 | Felis catus | 45457.0 | 0 |
| scf7180000018463 | 701622 | 0.3705 | 0 | 49.8773 | Eukaryota | 58530.0 | 0 | Chordata | 58530.0 | 0 | Carnivora | 58530.0 | 0 | Felidae | 58530.0 | 0 | Acinonyx | 58530.0 | 0 | :inonyx jubat | 58530.0 | 0 |
| scf7180000021220 | 79131 | 0.3722 | 0 | 24.4601 | Eukaryota | 17539.0 | 0 | Chordata | 17539.0 | 0 | Carnivora | 17539.0 | 0 | Felidae | 17539.0 | 0 | Felis | 15795.0 | 1 | Felis catus | 15795.0 | 1 |
| scf7180000018706 | 29323 | 0.3629 | 0 | 27.7182 | Eukaryota | 9494.0 | 0 | Chordata | 9494.0 | 0 | Carnivora | 9494.0 | 0 | Felidae | 9494.0 | 0 | Felis | 9494.0 | 0 | Felis catus | 9494.0 | 0 |
| scf7180000024228 | 306032 | 0.3883 | 0 | 50.3608 | Eukaryota | 48976.0 | 0 | Chordata | 48976.0 | 0 | Carnivora | 48976.0 | 0 | Felidae | 33589.0 | 1 | Suricata | 15387.0 | 3 | ıricata suricat | 15387.0 | 3 |
| scf7180000022203 | 19313 | 0.4363 | 0 | 29.8014 | Eukaryota | 28009.0 | 0 | Chordata | 28009.0 | 0 | Carnivora | 28009.0 | 0 | Felidae | 28009.0 | 0 | Felis | 25159.0 | 1 | Felis catus | 25159.0 | 1 |
| scf7180000021269 | 67220 | 0.3841 | 0 | 22.6584 | Eukaryota | 13593.0 | 0 | Chordata | 13593.0 | 0 | Carnivora | 13593.0 | 0 | Felidae | 13593.0 | 0 | Felis | 12230.0 | 1 | Felis catus | 12230.0 | 1 |
| scf7180000017630 | 2009031 | 0.3924 | 0 | 50.4911 | Eukaryota | 47802.0 | 0 | Chordata | 47802.0 | 0 | Carnivora | 47802.0 | 0 | Felidae | 47802.0 | 0 | Felis | 47802.0 | 0 | Felis catus | 47802.0 | 0 |
| scf7180000020258 | 21369 | 0.4611 | 0 | 42.8131 | Eukaryota | 5983.0 | 0 | Chordata | 5983.0 | 0 | Carnivora | 5983.0 | 0 | Felidae | 4165.0 | 1 | Panthera | 2396.0 | 3 | ıricata suricat | 1818.0 | 4 |
| scf7180000022389 | 4984903 | 0.4446 | 0 | 51.2751 | Eukaryota | 50142.0 | 0 | Chordata | 50142.0 | 0 | Carnivora | 50142.0 | 0 | Felidae | 26553.0 | 2 | Felis | 18257.0 | 3 | Felis catus | 18257.0 | 3 |
| scf7180000018808 | 6859 | 0.4807 | 0 | 44.0761 | Eukaryota | 3034.0 | 0 | Chordata | 3034.0 | 0 | Carnivora | 1819.0 | 1 | Felidae | 1155.0 | 5 | Panthera | 583.0 | 8 | rsus maritimu | 385.0 | 9 |
| scf7180000019298 | 295792 | 0.3537 | 0 | 48.8092 | Eukaryota | 16267.0 | 0 | Chordata | 16267.0 | 0 | Carnivora | 16267.0 | 0 | Felidae | 16267.0 | 0 | Felis | 14645.0 | 1 | Felis catus | 14645.0 | 1 |
| scf7180000020977 | 108237 | 0.6108 | 0 | 49.5408 | Eukaryota | 19462.0 | 0 | Chordata | 19462.0 | 0 | Carnivora | 19462.0 | 0 | Felidae | 8566.0 | 2 | Panthera | 8566.0 | 2 | anthera pardu | 8566.0 | 2 |
| scf7180000023685 | 79714 | 0.4078 | 0 | 634.1289 | Eukaryota | 23515.0 | 0 | Chordata | 23515.0 | 0 | Carnivora | 23515.0 | 0 | Felidae | 23515.0 | 0 | Felis | 23515.0 | 0 | Felis catus | 23515.0 | 0 |
| scf7180000020993 | 36342 | 0.4392 | 0 | 24.2921 | Eukaryota | 10616.0 | 0 | Chordata | 10616.0 | 0 | Primates | 3886.0 | 3 | Hominidae | 3886.0 | 6 | Sus | 2285.0 | 8 | Sus scrofa | 2285.0 | 8 |
| scf7180000017690 | 1703833 | 0.3719 | 0 | 49.5526 | Eukaryota | 47680.0 | 0 | Chordata | 47680.0 | 0 | Carnivora | 47680.0 | 0 | Felidae | 47680.0 | 0 | Felis | 47680.0 | 0 | Felis catus | 47680.0 | 0 |
| scf7180000020650 | 104321 | 0.4018 | 0 | 50.1668 | Eukaryota | 7599.0 | 0 | Chordata | 7599.0 | 0 | Primates | 4897.0 | 2 | Hominidae | 2216.0 | 4 | Homo | 2216.0 | 5 | Homo sapiens | 2216.0 | 5 |
| scf7180000017765 | 120826 | 0.3904 | 0 | 47.1485 | Eukaryota | 45440.0 | 0 | Chordata | 45440.0 | 0 | Carnivora | 45440.0 | 0 | Felidae | 45440.0 | 0 | Felis | 45440.0 | 0 | Felis catus | 45440.0 | 0 |
| scf7180000023263 | 18484 | 0.4277 | 0 | 114.4427 | Eukaryota | 4226.0 | 0 | Chordata | 4226.0 | 0 | Artiodactyla | 2976.0 | 1 | Bovidae | 2976.0 | 1 | Ovis | 2976.0 | 2 | Ovis canadensi | 2976.0 | 2 |
| scf7180000021445 | 60054 | 0.5817 | 0 | 48.4184 | Eukaryota | 5992.0 | 0 | Chordata | 5992.0 | 0 | Rodentia | 2275.0 | 4 | Muridae | 1931.0 | 5 | Mus | 1931.0 | 5 | Mus musculus | 1931.0 | 5 |
| scf7180000017731 | 1636625 | 0.4134 | 0 | 51.3227 | Eukaryota | 46976.0 | 0 | Chordata | 46976.0 | 0 | Carnivora | 46976.0 | 0 | Felidae | 42255.0 | 1 | Felis | 42255.0 | 1 | Felis catus | 42255.0 | 1 |
| scf7180000017624 | 1205364 | 0.3604 | 0 | 51.3462 | Eukaryota | 38755.0 | 0 | Chordata | 38755.0 | 0 | Carnivora | 38755.0 | 0 | Felidae | 21675.0 | 1 | Felis | 21675.0 | 1 | Felis catus | 21675.0 | 1 |
| scf7180000018813 | 650508 | 0.4913 | 0 | 53.0476 | Eukaryota | 73963.0 | 0 | Chordata | 73963.0 | 0 | Carnivora | 52489.0 | 1 | Felidae | 24618.0 | 3 | Equus | 21474.0 | 4 | Canis lupus | 20417.0 | 6 |
| scf7180000022534 | 7610 | 0.395 | 0 | 25.449 | Eukaryota | 4805.0 | 0 | Chordata | 4805.0 | 0 | Carnivora | 4182.0 | 1 | Felidae | 3205.0 | 2 | Felis | 3205.0 | 2 | Felis catus | 3205.0 | 2 |
| scf7180000020248 | 922583 | 0.4161 | 0 | 52.2863 | Eukaryota | 50660.0 | 0 | Chordata | 50660.0 | 0 | Carnivora | 50660.0 | 0 | Felidae | 50660.0 | 0 | Felis | 44918.0 | 1 | Felis catus | 44918.0 | 1 |
| scf7180000021913 | 29288 | 0.393 | 0 | 24.6035 | Eukaryota | 25821.0 | 0 | Chordata | 25821.0 | 0 | Carnivora | 25821.0 | 0 | Felidae | 25821.0 | 0 | Felis | 23202.0 | 1 | Felis catus | 23202.0 | 1 |
| scf7180000018914 | 500627 | 0.3995 | 0 | 50.0673 | Eukaryota | 41674.0 | 0 | Chordata | 41674.0 | 0 | Carnivora | 41674.0 | 0 | Felidae | 41674.0 | 0 | Felis | 37529.0 | 1 | Felis catus | 37529.0 | 1 |
| scf7180000018522 | 788116 | 0.4049 | 0 | 49.5055 | Eukaryota | 15165.0 | 1 | Chordata | 15165.0 | 1 | Primates | 8418.0 | 3 | Hominidae | 8418.0 | 4 | Homo | 8418.0 | 5 | Homo sapiens | 8418.0 | 5 |
| scf7180000023847 | 1361615 | 0.4853 | 0 | 51.4247 | Eukaryota | 70761.0 | 0 | Chordata | 70761.0 | 0 | Carnivora | 70761.0 | 0 | Felidae | 70761.0 | 0 | Lynx | 35450.0 | 2 | ynx canadens | 35450.0 | 2 |
| scf7180000023088 | 69936 | 0.5445 | 0 | 383.9505 | Eukaryota | 21907.0 | 0 | Chordata | 21907.0 | 0 | Carnivora | 21907.0 | 0 | Felidae | 21907.0 | 0 | Felis | 19690.0 | 1 | Felis catus | 19690.0 | 1 |
| scf7180000018407 | 832219 | 0.3807 | 0 | 50.1355 | Eukaryota | 41852.0 | 0 | Chordata | 41852.0 | 0 | Carnivora | 41852.0 | 0 | Felidae | 41852.0 | 0 | Felis | 37724.0 | 1 | Felis catus | 37724.0 | 1 |
| scf7180000023054 | 25823 | 0.5319 | 0 | 146.0131 | Eukaryota | 4622.0 | 0 | Chordata | 4622.0 | 0 | Carnivora | 4622.0 | 0 | Felidae | 4622.0 | 0 | Felis | 4350.0 | 1 | Felis catus | 4350.0 | 1 |
| scf7180000023561 | 19713 | 0.4794 | 0 | 513.9274 | no-hit | 0.0 | 0 | no-hit | 0.0 | 0 | no-hit | 0.0 | 0 | no-hit | 0.0 | 0 | no-hit | 0.0 | 0 | no-hit | 0.0 | 0 |
| scf7180000021537 | 44818 | 0.4398 | 0 | 214.7308 | Eukaryota | 2437.0 | 0 | Chordata | 2437.0 | 0 | Carnivora | 2437.0 | 0 | Felidae | 2230.0 | 1 | Felis | 1582.0 | 2 | Felis catus | 1582.0 | 2 |
| scf7180000019257 | 343693 | 0.5212 | 0 | 51.5142 | Eukaryota | 28531.0 | 0 | Chordata | 28531.0 | 0 | Carnivora | 28531.0 | 0 | Felidae | 28531.0 | 0 | Felis | 17611.0 | 1 | Felis catus | 17611.0 | 1 |
| scf7180000017465 | 937417 | 0.3988 | 0 | 50.8202 | Eukaryota | 44662.0 | 0 | Chordata | 44662.0 | 0 | Carnivora | 44662.0 | 0 | Felidae | 44662.0 | 0 | Felis | 44662.0 | 0 | Felis catus | 44662.0 | 0 |
| scf7180000021045 | 9969 | 0.3942 | 0 | 15.6408 | Eukaryota | 17470.0 | 0 | Chordata | 17470.0 | 0 | Carnivora | 17470.0 | 0 | Felidae | 17470.0 | 0 | Felis | 17470.0 | 0 | Felis catus | 17470.0 | 0 |
| scf7180000018785 | 552313 | 0.4915 | 0 | 51.7075 | Eukaryota | 24955.0 | 0 | Chordata | 24955.0 | 0 | Carnivora | 24955.0 | 0 | Felidae | 22585.0 | 1 | Panthera | 10679.0 | 5 | anthera pardu | 7873.0 | 6 |
| scf7180000020682 | 125899 | 0.4851 | 0 | 48.2781 | Eukaryota | 14709.0 | 0 | Chordata | 14709.0 | 0 | Carnivora | 14709.0 | 0 | Felidae | 14709.0 | 0 | Felis | 11780.0 | 1 | Felis catus | 11780.0 | 1 |
| scf7180000023792 | 1985373 | 0.4655 | 0 | 52.1245 | Eukaryota | 26799.0 | 0 | Chordata | 26799.0 | 0 | Carnivora | 26799.0 | 0 | Felidae | 26799.0 | 0 | Felis | 26799.0 | 0 | Felis catus | 26799.0 | 0 |
| scf7180000018473 | 695498 | 0.3926 | 0 | 51.1548 | Eukaryota | 54331.0 | 0 | Chordata | 54331.0 | 0 | Carnivora | 54331.0 | 0 | Felidae | 36701.0 | 3 | Acinonyx | 36701.0 | 3 | :inonyx jubat | 36701.0 | 3 |
| scf7180000021107 | 78470 | 0.4713 | 0 | 25.7874 | Eukaryota | 11792.0 | 0 | Chordata | 11792.0 | 0 | Carnivora | 6871.0 | 2 | Felidae | 6871.0 | 2 | Panthera | 2933.0 | 4 | anthera pardu | 2933.0 | 4 |
| scf7180000019924 | 1149872 | 0.3661 | 100 | 48.9625 | Eukaryota | 47637.0 | 0 | Chordata | 47637.0 | 0 | Carnivora | 47637.0 | 0 | Felidae | 47637.0 | 0 | Felis | 47637.0 | 0 | Felis catus | 47637.0 | 0 |
| scf7180000022026 | 29161 | 0.4998 | 0 | 26.6755 | Eukaryota | 7664.0 | 0 | Chordata | 7664.0 | 0 | Chiroptera | 3063.0 | 3 | Felidae | 2122.0 | 4 | Lynx | 2122.0 | 4 | ynx canadens | 2122.0 | 4 |
| scf7180000021976 | 55563 | 0.3631 | 0 | 23.9154 | Eukaryota | 15413.0 | 0 | Chordata | 15413.0 | 0 | Carnivora | 15413.0 | 0 | Felidae | 15413.0 | 0 | Felis | 14027.0 | 1 | Felis catus | 14027.0 | 1 |
| scf7180000022438 | 13791 | 0.4667 | 0 | 25.1517 | Eukaryota | 3111.0 | 0 | Chordata | 3111.0 | 0 | Carnivora | 3111.0 | 0 | Felidae | 3111.0 | 0 | Felis | 2530.0 | 1 | Felis catus | 2530.0 | 1 |
| scf7180000018174 | 17154 | 0.4351 | 0 | 36.8851 | Eukaryota | 3507.0 | 0 | Chordata | 3507.0 | 0 | Artiodactyla | 1828.0 | 1 | Felidae | 1679.0 | 2 | Felis | 1679.0 | 3 | Felis catus | 1442.0 | 4 |
| scf7180000022837 | 59441 | 0.6012 | 0 | 1105.0123 | Eukaryota | 4547.0 | 0 | Chordata | 4547.0 | 0 | Carnivora | 2412.0 | 1 | Camelidae | 2135.0 | 3 | Camelus | 2135.0 | 4 | ıelus dromeda | 2135.0 | 4 |
| scf7180000021533 | 62632 | 0.5203 | 0 | 1250.7634 | no-hit | 0.0 | 0 | no-hit | 0.0 | 0 | no-hit | 0.0 | 0 | no-hit | 0.0 | 0 | no-hit | 0.0 | 0 | no-hit | 0.0 | 0 |
| scf7180000021596 | 12436 | 0.5224 | 0 | 66.5611 | no-hit | 0.0 | 0 | no-hit | 0.0 | 0 | no-hit | 0.0 | 0 | no-hit | 0.0 | 0 | no-hit | 0.0 | 0 | no-hit | 0.0 | 0 |
| scf7180000022588 | 9363 | 0.3706 | 0 | 24.2131 | Eukaryota | 5352.0 | 0 | Chordata | 5352.0 | 0 | Carnivora | 5352.0 | 0 | Felidae | 5352.0 | 0 | Felis | 4783.0 | 1 | Felis catus | 4783.0 | 1 |
| scf7180000018449 | 171067 | 0.4199 | 0 | 51.6155 | Eukaryota | 34252.0 | 0 | Chordata | 34252.0 | 0 | Carnivora | 34252.0 | 0 | Felidae | 34252.0 | 0 | Lynx | 24651.0 | 1 | ynx canadens | 24651.0 | 1 |
| scf7180000022652 | 5414 | 0.5565 | 0 | 47.52 | Eukaryota | 2637.0 | 0 | Chordata | 2637.0 | 0 | Carnivora | 2637.0 | 0 | Felidae | 2637.0 | 0 | Felis | 2637.0 | 0 | Felis catus | 2637.0 | 0 |
| scf7180000019535 | 289327 | 0.5347 | 0 | 51.1821 | Eukaryota | 24019.0 | 0 | Chordata | 24019.0 | 0 | Carnivora | 24019.0 | 0 | Felidae | 24019.0 | 0 | Panthera | 9448.0 | 4 | anthera pardu | 7174.0 | 5 |
| scf7180000018597 | 486826 | 0.3596 | 0 | 49.3758 | Eukaryota | 52459.0 | 0 | Chordata | 52459.0 | 0 | Carnivora | 42245.0 | 1 | Canidae | 20272.0 | 3 | Vulpes | 20272.0 | 3 | Vulpes vulpes | 20272.0 | 3 |
| scf7180000021455 | 49905 | 0.4856 | 0 | 46.1147 | Eukaryota | 4583.0 | 0 | Chordata | 4583.0 | 0 | Carnivora | 4583.0 | 0 | Felidae | 4583.0 | 0 | Felis | 2051.0 | 3 | Felis catus | 2051.0 | 3 |
| scf7180000022821 | 698407 | 0.3575 | 0 | 48.7213 | Eukaryota | 45949.0 | 0 | Chordata | 45949.0 | 0 | Carnivora | 45949.0 | 0 | Felidae | 45949.0 | 0 | Felis | 45949.0 | 0 | Felis catus | 45949.0 | 0 |
| scf7180000020923 | 88076 | 0.5977 | 0 | 47.8367 | Eukaryota | 4119.0 | 1 | Chordata | 4119.0 | 1 | Carnivora | 3223.0 | 2 | Felidae | 1555.0 | 4 | Ursus | 1276.0 | 5 | Ursus arctos | 1276.0 | 5 |
| scf7180000022517 | 11390 | 0.6478 | 0 | 49.456 | Eukaryota | 12411.0 | 0 | Chordata | 12411.0 | 0 | Carnivora | 12411.0 | 0 | Herpestidae | 6536.0 | 1 | Suricata | 6536.0 | 1 | ıricata suricat | 6536.0 | 1 |
| scf7180000021131 | 15070 | 0.517 | 0 | 996.2132 | no-hit | 0.0 | 0 | no-hit | 0.0 | 0 | no-hit | 0.0 | 0 | no-hit | 0.0 | 0 | no-hit | 0.0 | 0 | no-hit | 0.0 | 0 |
| scf7180000020747 | 74022 | 0.5527 | 0 | 52.8216 | Eukaryota | 27359.0 | 0 | Chordata | 27359.0 | 0 | Carnivora | 27359.0 | 0 | Felidae | 27359.0 | 0 | Felis | 27359.0 | 0 | Felis catus | 27359.0 | 0 |
| scf7180000022625 | 9249 | 0.5663 | 0 | 55.7397 | Eukaryota | 1271.0 | 0 | Chordata | 1271.0 | 0 | Primates | 638.0 | 2 | Hominidae | 638.0 | 2 | Homo | 355.0 | 4 | Homo sapiens | 355.0 | 4 |
| scf7180000021755 | 16872 | 0.4066 | 0 | 24.8715 | Eukaryota | 2335.0 | 0 | Chordata | 2335.0 | 0 | Carnivora | 2335.0 | 0 | Felidae | 1862.0 | 2 | Felis | 1408.0 | 4 | Felis catus | 1408.0 | 4 |
| scf7180000018086 | 2933354 | 0.4149 | 0 | 51.1242 | Eukaryota | 46818.0 | 0 | Chordata | 46818.0 | 0 | Carnivora | 46818.0 | 0 | Felidae | 32829.0 | 1 | Felis | 32829.0 | 1 | Felis catus | 32829.0 | 1 |
| scf7180000023920 | 1063261 | 0.4829 | 0 | 52.0735 | Eukaryota | 42169.0 | 0 | Chordata | 42169.0 | 0 | Carnivora | 42169.0 | 0 | Felidae | 20394.0 | 2 | Acinonyx | 10738.0 | 6 | :inonyx jubat | 10738.0 | 6 |
| scf7180000021186 | 51659 | 0.4247 | 0 | 51.9701 | Eukaryota | 85554.0 | 0 | Chordata | 85554.0 | 0 | Carnivora | 85554.0 | 0 | Felidae | 54464.0 | 3 | Panthera | 27255.0 | 5 | anthera pardu | 18228.0 | 6 |
| scf7180000019305 | 68436 | 0.3496 | 0 | 294.2329 | Eukaryota | 47944.0 | 0 | Chordata | 47944.0 | 0 | Carnivora | 47944.0 | 0 | Felidae | 47944.0 | 0 | Felis | 47944.0 | 0 | Felis catus | 47944.0 | 0 |
| scf7180000021830 | 77188 | 0.3811 | 0 | 26.1468 | Eukaryota | 8429.0 | 0 | Chordata | 8429.0 | 0 | Carnivora | 4589.0 | 2 | Felidae | 3783.0 | 3 | Felis | 3783.0 | 4 | Felis catus | 3783.0 | 4 |
| scf7180000023097 | 41881 | 0.5681 | 0 | 266.588 | Eukaryota | 7165.0 | 0 | Chordata | 7165.0 | 0 | Carnivora | 7165.0 | 0 | Felidae | 5402.0 | 1 | Felis | 2052.0 | 3 | Felis catus | 2052.0 | 3 |
| scf7180000023788 | 931715 | 0.3784 | 0 | 50.5665 | Eukaryota | 43411.0 | 0 | Chordata | 43411.0 | 0 | Carnivora | 43411.0 | 0 | Felidae | 43411.0 | 0 | Felis | 43411.0 | 0 | Felis catus | 43411.0 | 0 |
| scf7180000017987 | 2191571 | 0.3773 | 0 | 50.9161 | Eukaryota | 51450.0 | 0 | Chordata | 51450.0 | 0 | Carnivora | 51450.0 | 0 | Felidae | 51450.0 | 0 | Felis | 51450.0 | 0 | Felis catus | 51450.0 | 0 |
| scf7180000024123 | 174579 | 0.4942 | 0 | 52.3174 | Eukaryota | 22176.0 | 0 | Chordata | 22176.0 | 0 | Carnivora | 22176.0 | 0 | Felidae | 22176.0 | 0 | Felis | 18279.0 | 1 | Felis catus | 18279.0 | 1 |
| scf7180000022128 | 38123 | 0.3834 | 0 | 25.556 | Eukaryota | 15261.0 | 0 | Chordata | 15261.0 | 0 | Carnivora | 15261.0 | 0 | Felidae | 15261.0 | 0 | Felis | 13746.0 | 1 | Felis catus | 13746.0 | 1 |
| scf7180000022585 | 3799 | 0.5596 | 0 | 58.0143 | Eukaryota | 2468.0 | 0 | Chordata | 2468.0 | 0 | Carnivora | 2468.0 | 0 | Felidae | 2468.0 | 0 | Felis | 1619.0 | 3 | Felis catus | 1382.0 | 4 |
| scf7180000018125 | 156669 | 0.3839 | 0 | 49.4318 | Eukaryota | 46644.0 | 0 | Chordata | 46644.0 | 0 | Carnivora | 46644.0 | 0 | Felidae | 46644.0 | 0 | Felis | 46644.0 | 0 | Felis catus | 46644.0 | 0 |
| scf7180000024222 | 31136 | 0.4362 | 0 | 25.0351 | Eukaryota | 10743.0 | 0 | Chordata | 10743.0 | 0 | Primates | 7103.0 | 1 | Hominidae | 7103.0 | 1 | Homo | 5863.0 | 3 | Homo sapiens | 5863.0 | 4 |
| scf7180000021865 | 22421 | 0.461 | 0 | 613.2237 | Eukaryota | 2630.0 | 0 | Chordata | 2630.0 | 0 | Artiodactyla | 1723.0 | 1 | Bovidae | 1723.0 | 2 | Ovis | 1723.0 | 3 | Ovis canadensi | 1723.0 | 3 |
| scf7180000018903 | 101484 | 0.4211 | 0 | 46.1868 | Eukaryota | 45077.0 | 0 | Chordata | 45077.0 | 0 | Carnivora | 45077.0 | 0 | Felidae | 45077.0 | 0 | Felis | 45077.0 | 0 | Felis catus | 45077.0 | 0 |
| scf7180000018506 | 176339 | 0.3733 | 0 | 48.3965 | Eukaryota | 13654.0 | 0 | Chordata | 13654.0 | 0 | Carnivora | 13654.0 | 0 | Felidae | 9645.0 | 1 | Felis | 8442.0 | 2 | Felis catus | 8442.0 | 2 |
| scf7180000018869 | 675933 | 0.3966 | 0 | 48.4621 | Eukaryota | 45499.0 | 0 | Chordata | 45499.0 | 0 | Carnivora | 45499.0 | 0 | Felidae | 45499.0 | 0 | Felis | 45499.0 | 0 | Felis catus | 45499.0 | 0 |
| scf7180000023911 | 1298084 | 0.426 | 0 | 52.1552 | Eukaryota | 37842.0 | 0 | Chordata | 37842.0 | 0 | Artiodactyla | 18081.0 | 2 | Delphinidae | 14461.0 | 3 | Pan | 11619.0 | 4 | an troglodyte | 11619.0 | 4 |
| scf7180000018106 | 1212032 | 0.4048 | 0 | 50.4279 | Eukaryota | 46054.0 | 0 | Chordata | 46054.0 | 0 | Carnivora | 46054.0 | 0 | Felidae | 46054.0 | 0 | Felis | 46054.0 | 0 | Felis catus | 46054.0 | 0 |
| scf7180000019025 | 222958 | 0.4387 | 0 | 49.4455 | Eukaryota | 37913.0 | 0 | Chordata | 37913.0 | 0 | Carnivora | 37913.0 | 0 | Felidae | 37913.0 | 0 | Felis | 37913.0 | 0 | Felis catus | 37913.0 | 0 |
| scf7180000019990 | 309626 | 0.3877 | 0 | 47.6184 | Eukaryota | 37218.0 | 0 | Chordata | 37218.0 | 0 | Carnivora | 37218.0 | 0 | Felidae | 37218.0 | 0 | Panthera | 22350.0 | 2 | anthera pardu | 18625.0 | 3 |
| scf7180000020466 | 63929 | 0.3678 | 0 | 22.7021 | Eukaryota | 29347.0 | 0 | Chordata | 29347.0 | 0 | Carnivora | 29347.0 | 0 | Felidae | 29347.0 | 0 | Felis | 26355.0 | 1 | Felis catus | 26355.0 | 1 |
| scf7180000023513 | 22802 | 0.4597 | 0 | 474.2303 | Eukaryota | 2595.0 | 0 | Chordata | 2595.0 | 0 | Artiodactyla | 1707.0 | 1 | Bovidae | 1707.0 | 2 | Ovis | 1707.0 | 3 | Ovis canadensi | 1707.0 | 3 |
| scf7180000021206 | 40469 | 0.588 | 0 | 51.9622 | Eukaryota | 9984.0 | 0 | Chordata | 9984.0 | 0 | Carnivora | 9984.0 | 0 | Felidae | 9984.0 | 0 | Felis | 9984.0 | 0 | Felis catus | 9984.0 | 0 |
| scf7180000022562 | 9284 | 0.4762 | 0 | 26.6851 | Eukaryota | 2709.0 | 0 | Chordata | 2709.0 | 0 | Carnivora | 2709.0 | 0 | Felidae | 1379.0 | 1 | Canis | 1330.0 | 3 | Canis lupus | 1330.0 | 4 |
| scf7180000020133 | 106137 | 0.4709 | 0 | 51.7231 | Eukaryota | 4175.0 | 0 | Chordata | 4175.0 | 0 | Carnivora | 3681.0 | 1 | Felidae | 2849.0 | 3 | Felis | 2849.0 | 3 | Felis catus | 2849.0 | 3 |
| scf7180000019136 | 483046 | 0.3692 | 0 | 50.5674 | Eukaryota | 43406.0 | 0 | Chordata | 43406.0 | 0 | Carnivora | 43406.0 | 0 | Felidae | 43406.0 | 0 | Felis | 43406.0 | 0 | Felis catus | 43406.0 | 0 |
| scf7180000023077 | 44360 | 0.5777 | 0 | 196.2012 | Eukaryota | 2793.0 | 0 | Chordata | 2793.0 | 0 | Carnivora | 2189.0 | 1 | Felidae | 2189.0 | 1 | Neofelis | 1098.0 | 2 | eofelis nebulo | 1098.0 | 2 |
| scf7180000022200 | 28584 | 0.3885 | 0 | 23.8155 | Eukaryota | 41960.0 | 0 | Chordata | 41960.0 | 0 | Carnivora | 41960.0 | 0 | Felidae | 41960.0 | 0 | Felis | 41960.0 | 0 | Felis catus | 41960.0 | 0 |
| scf7180000019151 | 414597 | 0.3667 | 0 | 49.6281 | Eukaryota | 46327.0 | 0 | Chordata | 46327.0 | 0 | Carnivora | 46327.0 | 0 | Felidae | 46327.0 | 0 | Felis | 46327.0 | 0 | Felis catus | 46327.0 | 0 |
| scf7180000021324 | 17295 | 0.5414 | 0 | 24.3705 | Eukaryota | 20295.0 | 0 | Chordata | 20295.0 | 0 | Carnivora | 20295.0 | 0 | Felidae | 20295.0 | 0 | Felis | 18241.0 | 1 | Felis catus | 18241.0 | 1 |
| scf7180000020975 | 16234 | 0.4247 | 0 | 79.9861 | Eukaryota | 2138.0 | 0 | Chordata | 2138.0 | 0 | Carnivora | 2138.0 | 0 | Felidae | 2138.0 | 0 | Felis | 1932.0 | 1 | Felis catus | 1932.0 | 1 |
| scf7180000019072 | 410737 | 0.5173 | 0 | 50.3235 | Eukaryota | 16140.0 | 0 | Chordata | 16140.0 | 0 | Carnivora | 9902.0 | 2 | Mustelidae | 8365.0 | 3 | Enhydra | 6880.0 | 4 | Enhydra lutris | 6880.0 | 4 |
| scf7180000021604 | 135399 | 0.3647 | 100 | 24.17 | Eukaryota | 35059.0 | 0 | Chordata | 35059.0 | 0 | Carnivora | 35059.0 | 0 | Felidae | 35059.0 | 0 | Felis | 35059.0 | 0 | Felis catus | 35059.0 | 0 |
| scf7180000017862 | 2433428 | 0.3688 | 0 | 50.9498 | Eukaryota | 75704.0 | 0 | Chordata | 75704.0 | 0 | Carnivora | 75704.0 | 0 | Herpestidae | 38641.0 | 1 | Suricata | 38641.0 | 3 | ıricata suricat | 38641.0 | 3 |
| scf7180000018783 | 600288 | 0.4281 | 0 | 52.431 | Eukaryota | 47996.0 | 0 | Chordata | 47996.0 | 0 | Carnivora | 47996.0 | 0 | Felidae | 47996.0 | 0 | Felis | 47996.0 | 0 | Felis catus | 47996.0 | 0 |
| scf7180000020254 | 53888 | 0.3851 | 0 | 40.9518 | Eukaryota | 45084.0 | 0 | Chordata | 45084.0 | 0 | Carnivora | 45084.0 | 0 | Felidae | 45084.0 | 0 | Felis | 40609.0 | 1 | Felis catus | 40609.0 | 1 |
| scf7180000021560 | 981047 | 0.4996 | 0 | 50.8464 | Eukaryota | 18082.0 | 0 | Chordata | 18082.0 | 0 | Carnivora | 18082.0 | 0 | Felidae | 18082.0 | 0 | Acinonyx | 8980.0 | 2 | :inonyx jubat | 8980.0 | 2 |
| scf7180000023834 | 2035808 | 0.4019 | 0 | 50.3055 | Eukaryota | 62440.0 | 0 | Chordata | 62440.0 | 0 | Carnivora | 62440.0 | 0 | Felidae | 55815.0 | 1 | Felis | 21333.0 | 4 | Felis catus | 21333.0 | 5 |
| scf7180000020795 | 52011 | 0.3589 | 0 | 23.5644 | Eukaryota | 11379.0 | 0 | Chordata | 11379.0 | 0 | Carnivora | 11379.0 | 0 | Felidae | 11379.0 | 0 | Felis | 10289.0 | 1 | Felis catus | 10289.0 | 1 |
| scf7180000023633 | 82824 | 0.5598 | 0 | 240.6317 | Eukaryota | 15749.0 | 0 | Chordata | 15749.0 | 0 | Carnivora | 15749.0 | 0 | Felidae | 15749.0 | 0 | Felis | 15749.0 | 0 | Felis catus | 15749.0 | 0 |
| scf7180000018814 | 499662 | 0.4005 | 0 | 50.3017 | Eukaryota | 39753.0 | 0 | Chordata | 39753.0 | 0 | Carnivora | 39753.0 | 0 | Felidae | 39753.0 | 0 | Lynx | 19865.0 | 2 | ynx canadens | 19865.0 | 2 |
| scf7180000020930 | 62674 | 0.3996 | 0 | 24.762 | Eukaryota | 11785.0 | 0 | Chordata | 11785.0 | 0 | Carnivora | 11785.0 | 0 | Felidae | 8876.0 | 2 | Felis | 5631.0 | 4 | Felis catus | 5631.0 | 4 |
| scf7180000023647 | 59263 | 0.4943 | 0 | 657.4736 | no-hit | 0.0 | 0 | no-hit | 0.0 | 0 | no-hit | 0.0 | 0 | no-hit | 0.0 | 0 | no-hit | 0.0 | 0 | no-hit | 0.0 | 0 |
| scf7180000021219 | 50864 | 0.4726 | 0 | 1110.1026 | Eukaryota | 17705.0 | 0 | Chordata | 17705.0 | 0 | Carnivora | 17705.0 | 0 | Felidae | 14247.0 | 1 | Acinonyx | 3501.0 | 5 | :inonyx jubat | 3501.0 | 5 |
| scf7180000023891 | 3510018 | 0.3847 | 0 | 50.883 | Eukaryota | 71077.0 | 0 | Chordata | 71077.0 | 0 | Carnivora | 71077.0 | 0 | Felidae | 54676.0 | 2 | Felis | 19073.0 | 5 | Felis catus | 19073.0 | 5 |
| scf7180000018952 | 29098 | 0.381 | 0 | 47.6444 | Eukaryota | 9553.0 | 0 | Chordata | 9553.0 | 0 | Artiodactyla | 4729.0 | 3 | Camelidae | 4729.0 | 3 | Canis | 2954.0 | 4 | Canis lupus | 2954.0 | 4 |
| scf7180000023794 | 1742785 | 0.3708 | 0 | 50.5126 | Eukaryota | 92166.0 | 0 | Chordata | 92166.0 | 0 | Primates | 68472.0 | 1 | Hominidae | 68472.0 | 2 | Homo | 35356.0 | 4 | Homo sapiens | 35356.0 | 4 |
| scf7180000023278 | 124650 | 0.4356 | 0 | 83.5266 | Eukaryota | 15661.0 | 0 | Chordata | 15661.0 | 0 | Carnivora | 14133.0 | 1 | Felidae | 4850.0 | 5 | unresolved | 3111.0 | 7 | nachus schau | 3111.0 | 8 |
| scf7180000020494 | 20078 | 0.4266 | 0 | 392.867 | Eukaryota | 151644.0 | 0 | Chordata | 151644.0 | 0 | Carnivora | 151644.0 | 0 | Hyaenidae | 151644.0 | 0 | Proteles | 88978.0 | 1 | roteles cristat | 88978.0 | 1 |
| scf7180000021639 | 24433 | 0.4212 | 0 | 23.9971 | Eukaryota | 5038.0 | 0 | Chordata | 5038.0 | 0 | Primates | 2119.0 | 2 | Hominidae | 2119.0 | 3 | Pan | 1420.0 | 5 | an troglodyte | 1420.0 | 5 |
| scf7180000020552 | 127912 | 0.411 | 0 | 26.0718 | Eukaryota | 26379.0 | 0 | Chordata | 26379.0 | 0 | Carnivora | 26379.0 | 0 | Felidae | 26379.0 | 0 | Felis | 23721.0 | 1 | Felis catus | 23721.0 | 1 |
| scf7180000019119 | 348866 | 0.443 | 0 | 50.1052 | Eukaryota | 46899.0 | 0 | Chordata | 46899.0 | 0 | Carnivora | 46899.0 | 0 | Felidae | 46899.0 | 0 | Felis | 46899.0 | 0 | Felis catus | 46899.0 | 0 |
| scf7180000021588 | 35969 | 0.3839 | 0 | 25.9786 | Eukaryota | 15340.0 | 0 | Chordata | 15340.0 | 0 | Carnivora | 6816.0 | 2 | Felidae | 6816.0 | 2 | Panthera | 3635.0 | 4 | anthera pardu | 3635.0 | 4 |
| scf7180000019170 | 34160 | 0.5251 | 0 | 259.6149 | Eukaryota | 12936.0 | 0 | Chordata | 12936.0 | 0 | Carnivora | 12936.0 | 0 | Felidae | 12936.0 | 0 | Felis | 12936.0 | 0 | Felis catus | 12936.0 | 0 |
| scf7180000020094 | 226631 | 0.4437 | 0 | 51.0674 | Eukaryota | 32422.0 | 0 | Chordata | 32422.0 | 0 | Carnivora | 32422.0 | 0 | Otariidae | 28311.0 | 1 | Zalophus | 18852.0 | 2 | phus californi | 18852.0 | 2 |
| scf7180000022450 | 24497 | 0.3777 | 0 | 25.9853 | Eukaryota | 7496.0 | 0 | Chordata | 7496.0 | 0 | Primates | 3753.0 | 3 | Hominidae | 3753.0 | 4 | Pan | 2484.0 | 6 | an troglodyte | 2484.0 | 6 |
| scf7180000020679 | 74597 | 0.6168 | 0 | 50.9729 | Eukaryota | 20985.0 | 0 | Chordata | 20985.0 | 0 | Carnivora | 20985.0 | 0 | Felidae | 10893.0 | 3 | Odobenus | 3972.0 | 8 | ›benus rosma | 3972.0 | 8 |
| scf7180000019010 | 523617 | 0.3905 | 0 | 49.846 | Eukaryota | 68229.0 | 0 | Chordata | 68229.0 | 0 | Carnivora | 68229.0 | 0 | Felidae | 61693.0 | 1 | Felis | 27470.0 | 4 | Felis catus | 27470.0 | 4 |
| scf7180000022956 | 58197 | 0.457 | 0 | 334.081 | Eukaryota | 5780.0 | 0 | Chordata | 5780.0 | 0 | Carnivora | 5780.0 | 0 | Phocidae | 2352.0 | 3 | .eptonychote | 1764.0 | 4 | ınychotes we | 1764.0 | 4 |
| scf7180000020096 | 146833 | 0.3725 | 0 | 46.8302 | Eukaryota | 46608.0 | 0 | Chordata | 46608.0 | 0 | Carnivora | 46608.0 | 0 | Felidae | 46608.0 | 0 | Felis | 46608.0 | 0 | Felis catus | 46608.0 | 0 |
| scf7180000024038 | 271949 | 0.3948 | 0 | 50.1785 | Eukaryota | 46620.0 | 0 | Chordata | 46620.0 | 0 | Carnivora | 46620.0 | 0 | Felidae | 46620.0 | 0 | Felis | 46620.0 | 0 | Felis catus | 46620.0 | 0 |
| scf7180000017574 | 1115789 | 0.4228 | 0 | 52.5276 | Eukaryota | 47463.0 | 0 | Chordata | 47463.0 | 0 | Carnivora | 47463.0 | 0 | Felidae | 47463.0 | 0 | Felis | 47463.0 | 0 | Felis catus | 47463.0 | 0 |
| scf7180000019763 | 273763 | 0.3817 | 0 | 50.5042 | Eukaryota | 33021.0 | 0 | Chordata | 33021.0 | 0 | Carnivora | 33021.0 | 0 | Felidae | 33021.0 | 0 | Felis | 33021.0 | 0 | Felis catus | 33021.0 | 0 |
| scf7180000021777 | 39280 | 0.3754 | 0 | 23.603 | Eukaryota | 43243.0 | 0 | Chordata | 43243.0 | 0 | Carnivora | 43243.0 | 0 | Felidae | 43243.0 | 0 | Felis | 38969.0 | 1 | Felis catus | 38969.0 | 1 |
| scf7180000019656 | 264510 | 0.4475 | 0 | 25.5048 | Eukaryota | 56690.0 | 0 | Chordata | 56690.0 | 0 | Carnivora | 56690.0 | 0 | Canidae | 17719.0 | 4 | Canis | 11822.0 | 7 | Canis lupus | 11822.0 | 8 |
| scf7180000024154 | 138271 | 0.4169 | 0 | 24.7057 | Eukaryota | 12946.0 | 0 | Chordata | 12946.0 | 0 | Primates | 6264.0 | 2 | Hominidae | 6264.0 | 3 | Pan | 3709.0 | 4 | an troglodyte | 3709.0 | 4 |
| scf7180000019982 | 159989 | 0.3374 | 0 | 46.9426 | Eukaryota | 47245.0 | 0 | Chordata | 47245.0 | 0 | Carnivora | 47245.0 | 0 | Felidae | 47245.0 | 0 | Felis | 47245.0 | 0 | Felis catus | 47245.0 | 0 |
| scf7180000019284 | 527280 | 0.5076 | 0 | 50.6058 | Eukaryota | 13445.0 | 0 | Chordata | 13445.0 | 0 | Carnivora | 13445.0 | 0 | Felidae | 10770.0 | 1 | Panthera | 4033.0 | 5 | Felis catus | 2721.0 | 6 |
| scf7180000023810 | 1759253 | 0.4045 | 0 | 50.3449 | Eukaryota | 47275.0 | 0 | Chordata | 47275.0 | 0 | Carnivora | 47275.0 | 0 | Felidae | 47275.0 | 0 | Felis | 47275.0 | 0 | Felis catus | 47275.0 | 0 |
| scf7180000018747 | 647395 | 0.378 | 0 | 51.2207 | Eukaryota | 47048.0 | 0 | Chordata | 47048.0 | 0 | Carnivora | 47048.0 | 0 | Felidae | 47048.0 | 0 | Felis | 47048.0 | 0 | Felis catus | 47048.0 | 0 |
| scf7180000020704 | 111114 | 0.4552 | 0 | 26.758 | Eukaryota | 15897.0 | 0 | Chordata | 15897.0 | 0 | Primates | 6621.0 | 3 | Hominidae | 6621.0 | 3 | Canis | 4904.0 | 4 | Canis lupus | 4904.0 | 4 |
| scf7180000021363 | 125540 | 0.3762 | 0 | 24.3249 | Eukaryota | 44458.0 | 0 | Chordata | 44458.0 | 0 | Carnivora | 44458.0 | 0 | Felidae | 44458.0 | 0 | Felis | 40046.0 | 1 | Felis catus | 40046.0 | 1 |
| scf7180000022850 | 24238 | 0.5425 | 0 | 463.4063 | Eukaryota | 2515.0 | 0 | Chordata | 2515.0 | 0 | Rodentia | 1022.0 | 4 | Caviidae | 765.0 | 5 | Cavia | 765.0 | 5 | :avia porcellu | 765.0 | 6 |
| scf7180000021068 | 41392 | 0.4297 | 0 | 51.527 | Eukaryota | 10645.0 | 0 | Chordata | 10645.0 | 0 | Carnivora | 9795.0 | 1 | Herpestidae | 6280.0 | 2 | Suricata | 6280.0 | 2 | ıricata suricat | 6280.0 | 2 |
| scf7180000018516 | 5836 | 0.3487 | 0 | 23.3248 | Eukaryota | 2821.0 | 0 | Chordata | 2821.0 | 0 | Carnivora | 2119.0 | 1 | Felidae | 2119.0 | 1 | Felis | 1318.0 | 2 | Felis catus | 1318.0 | 2 |
| scf7180000020566 | 106790 | 0.4726 | 0 | 45.5108 | Eukaryota | 36357.0 | 0 | Chordata | 36357.0 | 0 | Carnivora | 23104.0 | 1 | Felidae | 19369.0 | 2 | Equus | 13253.0 | 3 | anthera pardu | 11619.0 | 4 |
| scf7180000019019 | 480088 | 0.3905 | 0 | 48.225 | Eukaryota | 47262.0 | 0 | Chordata | 47262.0 | 0 | Carnivora | 47262.0 | 0 | Felidae | 47262.0 | 0 | Felis | 47262.0 | 0 | Felis catus | 47262.0 | 0 |
| scf7180000019596 | 296821 | 0.403 | 0 | 51.3274 | Eukaryota | 39508.0 | 0 | Chordata | 39508.0 | 0 | Carnivora | 39508.0 | 0 | Felidae | 39508.0 | 0 | Felis | 35476.0 | 1 | Felis catus | 35476.0 | 1 |
| scf7180000019299 | 309372 | 0.3909 | 0 | 51.0775 | Eukaryota | 91535.0 | 0 | Chordata | 91535.0 | 0 | Carnivora | 91535.0 | 0 | Felidae | 74580.0 | 2 | Lynx | 18814.0 | 5 | ynx canadens | 18814.0 | 5 |
| scf7180000017649 | 144143 | 0.5669 | 0 | 46.4258 | Eukaryota | 10104.0 | 0 | Chordata | 10104.0 | 0 | Carnivora | 10104.0 | 0 | Felidae | 10104.0 | 0 | Felis | 10104.0 | 0 | Felis catus | 10104.0 | 0 |
| scf7180000019622 | 250518 | 0.3947 | 0 | 52.0902 | Eukaryota | 60828.0 | 0 | Chordata | 60828.0 | 0 | Carnivora | 60828.0 | 0 | Mustelidae | 44058.0 | 1 | Mustela | 44058.0 | 1 | lustela putori | 44058.0 | 1 |
| scf7180000022809 | 21746 | 0.4226 | 0 | 25.164 | Eukaryota | 5140.0 | 0 | Chordata | 5140.0 | 0 | Artiodactyla | 2398.0 | 3 | Bovidae | 2043.0 | 4 | Homo | 1582.0 | 6 | Homo sapiens | 1582.0 | 6 |
| scf7180000021242 | 28255 | 0.5633 | 0 | 41.5109 | Eukaryota | 4759.0 | 1 | Chordata | 4759.0 | 1 | Carnivora | 1644.0 | 4 | Felidae | 1644.0 | 4 | Felis | 1644.0 | 5 | Felis catus | 1644.0 | 5 |
| scf7180000019063 | 488703 | 0.4811 | 0 | 52.2258 | Eukaryota | 42394.0 | 0 | Chordata | 42394.0 | 0 | Carnivora | 42394.0 | 0 | Felidae | 42394.0 | 0 | Felis | 42394.0 | 0 | Felis catus | 42394.0 | 0 |

| | | | | | | | | | | | | | | | | | | | | | |
|---|---|---|---|---|---|---|---|---|---|---|---|---|---|---|---|---|---|---|---|---|---|
| scf7180000024272 | 846964 | 0.4159 | 0 | 50.3944 | Eukaryota | 46876.0 | 0 | Chordata | 46876.0 | 0 | Carnivora | 46876.0 | 0 | Felidae | 46876.0 | 0 | Felis | 46876.0 | 0 | Felis catus | 46876.0 | 0 |
| scf7180000019810 | 159738 | 0.4384 | 0 | 50.016 | Eukaryota | 29762.0 | 0 | Chordata | 29762.0 | 0 | Carnivora | 29762.0 | 0 | Felidae | 23451.0 | 1 | Acinonyx | 10806.0 | 3 | cinonyx jubat | 10806.0 | 4 |
| scf7180000022653 | 9959 | 0.3816 | 0 | 24.8019 | Eukaryota | 4810.0 | 0 | Chordata | 4810.0 | 0 | Carnivora | 4810.0 | 0 | Felidae | 4810.0 | 0 | Felis | 4810.0 | 0 | Felis catus | 4810.0 | 0 |
| scf7180000021433 | 10713 | 0.4828 | 0 | 214.1603 | no-hit | 0.0 | 0 | no-hit | 0.0 | 0 | no-hit | 0.0 | 0 | no-hit | 0.0 | 0 | no-hit | 0.0 | 0 | no-hit | 0.0 | 0 |
| scf7180000021172 | 53205 | 0.4742 | 0 | 27.7966 | Eukaryota | 38265.0 | 0 | Chordata | 38265.0 | 0 | Carnivora | 38265.0 | 0 | Felidae | 38265.0 | 0 | Felis | 34486.0 | 1 | Felis catus | 34486.0 | 1 |
| scf7180000017430 | 6557508 | 0.3918 | 0 | 51.2482 | Eukaryota | 62266.0 | 0 | Chordata | 62266.0 | 0 | Carnivora | 41098.0 | 1 | Felidae | 41098.0 | 1 | Lynx | 31824.0 | 2 | ynx canadens | 31824.0 | 2 |
| scf7180000023998 | 511591 | 0.4178 | 0 | 51.0999 | Eukaryota | 40993.0 | 0 | Chordata | 40993.0 | 0 | Carnivora | 40993.0 | 0 | Felidae | 40993.0 | 0 | Felis | 40993.0 | 0 | Felis catus | 40993.0 | 0 |
| scf7180000021670 | 5059 | 0.6485 | 0 | 28.6928 | no-hit | 0.0 | 0 | no-hit | 0.0 | 0 | no-hit | 0.0 | 0 | no-hit | 0.0 | 0 | no-hit | 0.0 | 0 | no-hit | 0.0 | 0 |
| scf7180000022229 | 16545 | 0.3822 | 0 | 23.5306 | Eukaryota | 9935.0 | 0 | Chordata | 9935.0 | 0 | Carnivora | 9935.0 | 0 | Felidae | 9935.0 | 0 | Felis | 9935.0 | 0 | Felis catus | 9935.0 | 0 |
| scf7180000017881 | 92907 | 0.3875 | 0 | 47.9839 | Eukaryota | 47384.0 | 0 | Chordata | 47384.0 | 0 | Carnivora | 47384.0 | 0 | Felidae | 47384.0 | 0 | Felis | 47384.0 | 0 | Felis catus | 47384.0 | 0 |
| scf7180000020151 | 173460 | 0.5848 | 0 | 46.3198 | Eukaryota | 7230.0 | 1 | Chordata | 7230.0 | 1 | Carnivora | 4614.0 | 3 | Felidae | 4614.0 | 3 | Felis | 2949.0 | 6 | Felis catus | 2949.0 | 6 |
| scf7180000019991 | 202196 | 0.4269 | 0 | 26.6089 | Eukaryota | 24270.0 | 0 | Chordata | 24270.0 | 0 | Carnivora | 24270.0 | 0 | Canidae | 24270.0 | 0 | Canis | 24270.0 | 0 | Canis lupus | 24270.0 | 0 |
| scf7180000022544 | 19781 | 0.3852 | 0 | 257.7256 | Eukaryota | 4499.0 | 0 | Chordata | 4499.0 | 0 | Carnivora | 2261.0 | 1 | Hominidae | 2238.0 | 3 | Homo | 2238.0 | 3 | Homo sapiens | 2238.0 | 3 |
| scf7180000018851 | 451409 | 0.3415 | 0 | 47.673 | Eukaryota | 45176.0 | 0 | Chordata | 45176.0 | 0 | Carnivora | 45176.0 | 0 | Felidae | 45176.0 | 0 | Felis | 45176.0 | 0 | Felis catus | 45176.0 | 0 |
| scf7180000019922 | 205128 | 0.3499 | 0 | 49.042 | Eukaryota | 41856.0 | 0 | Chordata | 41856.0 | 0 | Carnivora | 41856.0 | 0 | Felidae | 41856.0 | 0 | Felis | 41856.0 | 0 | Felis catus | 41856.0 | 0 |
| scf7180000018700 | 1339429 | 0.4186 | 0 | 50.6592 | Eukaryota | 57813.0 | 0 | Chordata | 57813.0 | 0 | Carnivora | 57813.0 | 0 | Felidae | 52559.0 | 1 | Panthera | 18179.0 | 4 | ynx canadens | 16974.0 | 5 |
| scf7180000018490 | 19519 | 0.3912 | 0 | 21.5747 | Eukaryota | 46432.0 | 0 | Chordata | 46432.0 | 0 | Carnivora | 46432.0 | 0 | Felidae | 46432.0 | 0 | Felis | 46432.0 | 0 | Felis catus | 46432.0 | 0 |
| scf7180000023215 | 46157 | 0.5156 | 0 | 394.166 | Eukaryota | 5013.0 | 0 | Chordata | 5013.0 | 0 | Carnivora | 4417.0 | 2 | Felidae | 2960.0 | 5 | Felis | 2228.0 | 7 | Felis catus | 2228.0 | 7 |
| scf7180000018347 | 227785 | 0.3559 | 0 | 48.5525 | Eukaryota | 45324.0 | 0 | Chordata | 45324.0 | 0 | Carnivora | 45324.0 | 0 | Felidae | 45324.0 | 0 | Felis | 45324.0 | 0 | Felis catus | 45324.0 | 0 |
| scf7180000023570 | 22758 | 0.4867 | 0 | 337.1793 | no-hit | 0.0 | 0 | no-hit | 0.0 | 0 | no-hit | 0.0 | 0 | no-hit | 0.0 | 0 | no-hit | 0.0 | 0 | no-hit | 0.0 | 0 |
| scf7180000021818 | 34898 | 0.4227 | 0 | 50.3427 | Eukaryota | 14813.0 | 0 | Chordata | 14813.0 | 0 | Carnivora | 14813.0 | 0 | Felidae | 5964.0 | 5 | Felis | 1506.0 | 9 | Felis catus | 1506.0 | 9 |
| scf7180000024078 | 260535 | 0.5523 | 0 | 51.0882 | Eukaryota | 11618.0 | 0 | Chordata | 11618.0 | 0 | Carnivora | 8651.0 | 1 | Felidae | 7348.0 | 2 | Sus | 2967.0 | 6 | Sus scrofa | 2967.0 | 6 |
| scf7180000023167 | 24307 | 0.3925 | 0 | 297.13 | Eukaryota | 10125.0 | 0 | Chordata | 10125.0 | 0 | pterygiforme | 9055.0 | 1 | Apterygidae | 9055.0 | 1 | Apteryx | 9055.0 | 1 | pteryx mante | 9055.0 | 1 |
| scf7180000017755 | 2257037 | 0.4131 | 0 | 51.5517 | Eukaryota | 47694.0 | 0 | Chordata | 47694.0 | 0 | Carnivora | 47694.0 | 0 | Felidae | 47694.0 | 0 | Felis | 47694.0 | 0 | Felis catus | 47694.0 | 0 |
| scf7180000019088 | 329810 | 0.4275 | 0 | 50.7823 | Eukaryota | 12641.0 | 1 | Chordata | 12641.0 | 1 | Primates | 9762.0 | 2 | Hominidae | 8332.0 | 3 | Homo | 8332.0 | 4 | Homo sapiens | 8332.0 | 4 |
| scf7180000023592 | 48733 | 0.3972 | 0 | 247.9797 | Eukaryota | 21693.0 | 0 | Chordata | 21693.0 | 0 | Carnivora | 21693.0 | 0 | Felidae | 21693.0 | 0 | Felis | 19559.0 | 1 | Felis catus | 19559.0 | 1 |
| scf7180000019052 | 238710 | 0.4199 | 0 | 50.8727 | Eukaryota | 24525.0 | 0 | Chordata | 24525.0 | 0 | Carnivora | 24525.0 | 0 | Felidae | 24525.0 | 0 | Felis | 24525.0 | 0 | Felis catus | 24525.0 | 0 |
| scf7180000023931 | 720664 | 0.4437 | 0 | 51.0482 | Eukaryota | 24670.0 | 0 | Chordata | 24670.0 | 0 | Carnivora | 24670.0 | 0 | Felidae | 24670.0 | 0 | Felis | 22180.0 | 1 | Felis catus | 22180.0 | 1 |
| scf7180000021510 | 51656 | 0.3791 | 0 | 21.7835 | Eukaryota | 45043.0 | 0 | Chordata | 45043.0 | 0 | Carnivora | 45043.0 | 0 | Felidae | 45043.0 | 0 | Felis | 45043.0 | 0 | Felis catus | 45043.0 | 0 |
| scf7180000021693 | 7994 | 0.5188 | 0 | 61.9053 | no-hit | 0.0 | 0 | no-hit | 0.0 | 0 | no-hit | 0.0 | 0 | no-hit | 0.0 | 0 | no-hit | 0.0 | 0 | no-hit | 0.0 | 0 |
| scf7180000021574 | 47066 | 0.4096 | 0 | 24.7571 | Eukaryota | 45524.0 | 0 | Chordata | 45524.0 | 0 | Carnivora | 17652.0 | 3 | Canidae | 17652.0 | 4 | Canis | 17652.0 | 4 | Canis lupus | 17652.0 | 5 |
| scf7180000018893 | 446211 | 0.4129 | 0 | 48.7369 | Eukaryota | 44885.0 | 0 | Chordata | 44885.0 | 0 | Carnivora | 44885.0 | 0 | Felidae | 44885.0 | 0 | Felis | 44885.0 | 0 | Felis catus | 44885.0 | 0 |
| scf7180000019036 | 421215 | 0.4561 | 0 | 51.1588 | Eukaryota | 19812.0 | 0 | Chordata | 19812.0 | 0 | Carnivora | 19812.0 | 0 | Felidae | 15970.0 | 1 | Felis | 7973.0 | 4 | Felis catus | 7973.0 | 4 |
| scf7180000019075 | 872168 | 0.4083 | 0 | 50.0025 | Eukaryota | 46451.0 | 0 | Chordata | 46451.0 | 0 | Carnivora | 46451.0 | 0 | Felidae | 46451.0 | 0 | Felis | 46451.0 | 0 | Felis catus | 46451.0 | 0 |
| scf7180000019957 | 130334 | 0.3655 | 0 | 49.5055 | Eukaryota | 23665.0 | 0 | Chordata | 23665.0 | 0 | Carnivora | 23665.0 | 0 | Felidae | 23665.0 | 0 | Felis | 18962.0 | 1 | Felis catus | 18962.0 | 1 |
| scf7180000023070 | 51318 | 0.4428 | 0 | 228.3892 | Eukaryota | 11945.0 | 0 | Chordata | 11945.0 | 0 | Carnivora | 11945.0 | 0 | Felidae | 11945.0 | 0 | Felis | 10764.0 | 1 | Felis catus | 10764.0 | 1 |
| scf7180000019550 | 302585 | 0.3932 | 0 | 51.6527 | Eukaryota | 40334.0 | 0 | Chordata | 40334.0 | 0 | Carnivora | 40334.0 | 0 | Felidae | 40334.0 | 0 | Felis | 40334.0 | 0 | Felis catus | 40334.0 | 0 |
| scf7180000020224 | 63949 | 0.5281 | 0 | 314.0767 | Eukaryota | 15547.0 | 0 | Chordata | 15547.0 | 0 | Carnivora | 15547.0 | 0 | Felidae | 15547.0 | 0 | Felis | 15547.0 | 0 | Felis catus | 15547.0 | 0 |
| scf7180000021003 | 55529 | 0.4489 | 0 | 26.3436 | Eukaryota | 12442.0 | 0 | Chordata | 12442.0 | 0 | Carnivora | 12442.0 | 0 | Felidae | 12442.0 | 0 | Lynx | 7194.0 | 2 | ynx canadens | 7194.0 | 2 |
| scf7180000024066 | 193615 | 0.5041 | 0 | 51.895 | Eukaryota | 18459.0 | 0 | Chordata | 18459.0 | 0 | Carnivora | 17341.0 | 1 | Herpestidae | 11350.0 | 5 | Suricata | 11350.0 | 5 | ıricata suricat | 11350.0 | 5 |
| scf7180000021872 | 22912 | 0.3876 | 0 | 26.3733 | Eukaryota | 36603.0 | 0 | Chordata | 36603.0 | 0 | Carnivora | 36603.0 | 0 | Felidae | 36603.0 | 0 | Felis | 36603.0 | 0 | Felis catus | 36603.0 | 0 |
| scf7180000023289 | 18640 | 0.4274 | 0 | 445.3688 | Eukaryota | 13340.0 | 0 | Chordata | 13340.0 | 0 | Carnivora | 13340.0 | 0 | Felidae | 13340.0 | 0 | Felis | 12021.0 | 1 | Felis catus | 12021.0 | 1 |
| scf7180000021683 | 25381 | 0.4311 | 0 | 24.6512 | Eukaryota | 3750.0 | 0 | Chordata | 3750.0 | 0 | Carnivora | 3027.0 | 1 | Felidae | 2740.0 | 2 | Felis | 2740.0 | 2 | Felis catus | 2740.0 | 2 |
| scf7180000019770 | 257784 | 0.348 | 0 | 48.4018 | Eukaryota | 45633.0 | 0 | Chordata | 45633.0 | 0 | Carnivora | 45633.0 | 0 | Felidae | 45633.0 | 0 | Felis | 41125.0 | 1 | Felis catus | 41125.0 | 1 |
| scf7180000019426 | 420206 | 0.3648 | 0 | 49.9369 | Eukaryota | 45866.0 | 0 | Chordata | 45866.0 | 0 | Carnivora | 45866.0 | 0 | Felidae | 45866.0 | 0 | Felis | 41317.0 | 1 | Felis catus | 41317.0 | 1 |
| scf7180000021570 | 17094 | 0.6157 | 0 | 44.9054 | Eukaryota | 11315.0 | 0 | Chordata | 11315.0 | 0 | Carnivora | 11315.0 | 0 | Felidae | 11315.0 | 0 | Felis | 10192.0 | 1 | Felis catus | 10192.0 | 1 |
| scf7180000022403 | 19517 | 0.4667 | 0 | 25.0071 | Eukaryota | 8285.0 | 0 | Chordata | 8285.0 | 0 | Carnivora | 4354.0 | 2 | Canidae | 2602.0 | 5 | Canis | 1728.0 | 8 | Canis lupus | 1728.0 | 8 |
| scf7180000021492 | 58433 | 0.3599 | 0 | 23.6853 | Eukaryota | 43652.0 | 0 | Chordata | 43652.0 | 0 | Carnivora | 43652.0 | 0 | Felidae | 43652.0 | 0 | Felis | 43652.0 | 0 | Felis catus | 43652.0 | 0 |
| scf7180000019858 | 20226 | 0.5585 | 0 | 189.7344 | Eukaryota | 2795.0 | 0 | Chordata | 2795.0 | 0 | Carnivora | 2189.0 | 1 | Felidae | 2189.0 | 1 | Neofelis | 1098.0 | 2 | eofelis nebulo | 1098.0 | 2 |
| scf7180000019936 | 1469364 | 0.4577 | 0 | 52.6116 | Eukaryota | 49527.0 | 0 | Chordata | 49527.0 | 0 | Carnivora | 49527.0 | 0 | Felidae | 49527.0 | 0 | Felis | 40002.0 | 2 | Felis catus | 40002.0 | 2 |
| scf7180000022592 | 12263 | 0.7024 | 0 | 90.5788 | no-hit | 0.0 | 0 | no-hit | 0.0 | 0 | no-hit | 0.0 | 0 | no-hit | 0.0 | 0 | no-hit | 0.0 | 0 | no-hit | 0.0 | 0 |
| scf7180000021152 | 50818 | 0.6183 | 0 | 52.7255 | Eukaryota | 9266.0 | 0 | Chordata | 9266.0 | 0 | Carnivora | 9266.0 | 0 | Felidae | 9266.0 | 0 | Panthera | 4916.0 | 1 | Panthera tigri | 4916.0 | 1 |
| scf7180000024036 | 291491 | 0.4053 | 0 | 50.5912 | Eukaryota | 5519.0 | 1 | Chordata | 5519.0 | 1 | Carnivora | 2103.0 | 4 | Ursidae | 1565.0 | 5 | Ursus | 1565.0 | 6 | Homo sapiens | 1416.0 | 7 |
| scf7180000019739 | 746623 | 0.5284 | 0 | 52.4394 | Eukaryota | 50774.0 | 0 | Chordata | 50774.0 | 0 | Carnivora | 50774.0 | 0 | Felidae | 36094.0 | 2 | Acinonyx | 15570.0 | 6 | cinonyx jubat | 15570.0 | 6 |
| scf7180000018152 | 147229 | 0.4482 | 0 | 50.2588 | Eukaryota | 22818.0 | 0 | Chordata | 22818.0 | 0 | Carnivora | 22818.0 | 0 | Felidae | 14714.0 | 1 | Vulpes | 8104.0 | 5 | Vulpes vulpes | 8104.0 | 5 |
| scf7180000022030 | 22771 | 0.4759 | 0 | 249.1765 | no-hit | 0.0 | 0 | no-hit | 0.0 | 0 | no-hit | 0.0 | 0 | no-hit | 0.0 | 0 | no-hit | 0.0 | 0 | no-hit | 0.0 | 0 |
| scf7180000019925 | 351134 | 0.4882 | 0 | 54.5252 | Eukaryota | 68200.0 | 0 | Chordata | 68200.0 | 0 | Carnivora | 68200.0 | 0 | Felidae | 68200.0 | 0 | Acinonyx | 34540.0 | 1 | cinonyx jubat | 34540.0 | 1 |
| scf7180000020799 | 98050 | 0.3946 | 0 | 48.0026 | Eukaryota | 45898.0 | 0 | Chordata | 45898.0 | 0 | Carnivora | 45898.0 | 0 | Felidae | 45898.0 | 0 | Felis | 45898.0 | 0 | Felis catus | 45898.0 | 0 |
| scf7180000020701 | 11412 | 0.3842 | 0 | 22.3164 | Eukaryota | 8014.0 | 0 | Chordata | 8014.0 | 0 | Carnivora | 8014.0 | 0 | Felidae | 6406.0 | 1 | Felis | 5628.0 | 2 | Felis catus | 5628.0 | 2 |
| scf7180000023839 | 873460 | 0.387 | 0 | 51.5535 | Eukaryota | 39841.0 | 0 | Chordata | 39841.0 | 0 | Carnivora | 39841.0 | 0 | Felidae | 21382.0 | 3 | Felis | 12946.0 | 7 | Felis catus | 12946.0 | 7 |
| scf7180000019476 | 318892 | 0.4377 | 0 | 29.1071 | Eukaryota | 45235.0 | 0 | Chordata | 45235.0 | 0 | Carnivora | 45235.0 | 0 | Felidae | 45235.0 | 0 | Felis | 45235.0 | 0 | Felis catus | 45235.0 | 0 |
| scf7180000021862 | 69827 | 0.3686 | 0 | 24.7455 | Eukaryota | 42004.0 | 0 | Chordata | 42004.0 | 0 | Carnivora | 42004.0 | 0 | Felidae | 42004.0 | 0 | Felis | 42004.0 | 0 | Felis catus | 42004.0 | 0 |
| scf7180000018956 | 15354 | 0.3732 | 0 | 23.5907 | Eukaryota | 43286.0 | 0 | Chordata | 43286.0 | 0 | Carnivora | 43286.0 | 0 | Felidae | 43286.0 | 0 | Felis | 43286.0 | 0 | Felis catus | 43286.0 | 0 |
| scf7180000017970 | 1543751 | 0.3931 | 0 | 50.949 | Eukaryota | 62452.0 | 0 | Chordata | 62452.0 | 0 | Carnivora | 62452.0 | 0 | Felidae | 55975.0 | 1 | Lynx | 18765.0 | 5 | ynx canadens | 18765.0 | 5 |
| scf7180000019856 | 179589 | 0.4802 | 0 | 51.5497 | Eukaryota | 19845.0 | 0 | Chordata | 19845.0 | 0 | Carnivora | 19845.0 | 0 | Felidae | 19845.0 | 0 | Panthera | 9552.0 | 4 | anthera pard | 9552.0 | 4 |
| scf7180000019230 | 480535 | 0.3848 | 0 | 45.7883 | Eukaryota | 44917.0 | 0 | Chordata | 44917.0 | 0 | Carnivora | 44917.0 | 0 | Felidae | 44917.0 | 0 | Felis | 44917.0 | 0 | Felis catus | 44917.0 | 0 |
| scf7180000019208 | 318495 | 0.486 | 0 | 50.0562 | Eukaryota | 18986.0 | 0 | Chordata | 18986.0 | 0 | Carnivora | 18986.0 | 0 | Felidae | 18986.0 | 0 | Felis | 15192.0 | 1 | Felis catus | 15192.0 | 1 |
| scf7180000019450 | 205344 | 0.4666 | 0 | 52.2574 | Eukaryota | 16177.0 | 0 | Chordata | 16177.0 | 0 | Carnivora | 10707.0 | 1 | Hominidae | 5470.0 | 3 | Homo | 5470.0 | 3 | Homo sapiens | 5470.0 | 3 |
| scf7180000023873 | 1079022 | 0.4063 | 0 | 50.0902 | Eukaryota | 47544.0 | 0 | Chordata | 47544.0 | 0 | Carnivora | 47544.0 | 0 | Felidae | 47544.0 | 0 | Felis | 47544.0 | 0 | Felis catus | 47544.0 | 0 |
| scf7180000018672 | 947202 | 0.4913 | 0 | 51.7734 | Eukaryota | 19389.0 | 0 | Chordata | 19389.0 | 0 | Carnivora | 19389.0 | 0 | Felidae | 19389.0 | 0 | Felis | 19389.0 | 0 | Felis catus | 19389.0 | 0 |
| scf7180000019784 | 192097 | 0.4051 | 0 | 50.9232 | Eukaryota | 46541.0 | 0 | Chordata | 46541.0 | 0 | Carnivora | 46541.0 | 0 | Felidae | 46541.0 | 0 | Felis | 46541.0 | 0 | Felis catus | 46541.0 | 0 |
| scf7180000019333 | 9345 | 0.3899 | 0 | 17.1487 | Eukaryota | 36084.0 | 0 | Chordata | 36084.0 | 0 | Carnivora | 36084.0 | 0 | Felidae | 36084.0 | 0 | Felis | 32479.0 | 1 | Felis catus | 32479.0 | 1 |
| scf7180000021565 | 37832 | 0.3899 | 0 | 26.3185 | Eukaryota | 11710.0 | 0 | Chordata | 11710.0 | 0 | Carnivora | 6560.0 | 2 | Felidae | 4495.0 | 3 | Homo | 2840.0 | 7 | Homo sapiens | 2840.0 | 7 |
| scf7180000018596 | 625153 | 0.4545 | 0 | 51.7443 | Eukaryota | 47765.0 | 0 | Chordata | 47765.0 | 0 | Carnivora | 47765.0 | 0 | Felidae | 47765.0 | 0 | Panthera | 23926.0 | 2 | anthera pard | 19148.0 | 3 |
| scf7180000019817 | 270408 | 0.384 | 0 | 47.6285 | Eukaryota | 23100.0 | 0 | Chordata | 23100.0 | 0 | Carnivora | 23100.0 | 0 | Felidae | 20608.0 | 1 | Felis | 18318.0 | 2 | Felis catus | 18318.0 | 2 |
| scf7180000018968 | 476949 | 0.4626 | 0 | 52.5819 | Eukaryota | 47405.0 | 0 | Chordata | 47405.0 | 0 | Carnivora | 47405.0 | 0 | Felidae | 47405.0 | 0 | Felis | 47405.0 | 0 | Felis catus | 47405.0 | 0 |
| scf7180000018768 | 585274 | 0.4071 | 0 | 51.1513 | Eukaryota | 37806.0 | 0 | Chordata | 37806.0 | 0 | Carnivora | 37806.0 | 0 | Canidae | 18110.0 | 2 | Canis | 18110.0 | 2 | Canis lupus | 18110.0 | 2 |
| scf7180000019065 | 351809 | 0.4697 | 0 | 51.0082 | Eukaryota | 38349.0 | 0 | Chordata | 38349.0 | 0 | Carnivora | 38349.0 | 0 | Felidae | 34483.0 | 1 | Felis | 15396.0 | 4 | Felis catus | 15396.0 | 4 |
| scf7180000022805 | 8736 | 0.3883 | 0 | 21.8713 | Eukaryota | 27169.0 | 0 | Chordata | 27169.0 | 0 | Carnivora | 27169.0 | 0 | Felidae | 27169.0 | 0 | Felis | 24380.0 | 1 | Felis catus | 24380.0 | 1 |
| scf7180000020166 | 131536 | 0.3662 | 0 | 23.9922 | Eukaryota | 12696.0 | 0 | Chordata | 12696.0 | 0 | Primates | 10487.0 | 2 | Hominidae | 10487.0 | 2 | Pan | 6567.0 | 3 | an troglodyte | 6567.0 | 3 |
| scf7180000021088 | 582013 | 0.4276 | 0 | 52.1738 | Eukaryota | 31383.0 | 0 | Chordata | 31383.0 | 0 | Carnivora | 31383.0 | 0 | Felidae | 31383.0 | 0 | Felis | 28208.0 | 1 | Felis catus | 28208.0 | 1 |
| scf7180000021895 | 17530 | 0.501 | 0 | 54.1656 | Eukaryota | 2707.0 | 0 | Chordata | 2707.0 | 0 | Carnivora | 2072.0 | 1 | Felidae | 1809.0 | 2 | Felis | 1546.0 | 4 | Felis catus | 1546.0 | 4 |
| scf7180000023959 | 815882 | 0.5024 | 0 | 57.476 | Eukaryota | 43400.0 | 0 | Chordata | 43400.0 | 0 | Carnivora | 25707.0 | 3 | Ursidae | 9225.0 | 8 | Ursus | 9225.0 | 8 | posideros arm | 5293.0 | 9 |
| scf7180000018795 | 92088 | 0.3825 | 0 | 24.2114 | Eukaryota | 36223.0 | 0 | Chordata | 36223.0 | 0 | Carnivora | 36223.0 | 0 | Felidae | 36223.0 | 0 | Felis | 36223.0 | 0 | Felis catus | 36223.0 | 0 |
| scf7180000023914 | 1116590 | 0.3934 | 0 | 50.4446 | Eukaryota | 44414.0 | 0 | Chordata | 44414.0 | 0 | Carnivora | 44414.0 | 0 | Felidae | 44414.0 | 0 | Felis | 44414.0 | 0 | Felis catus | 44414.0 | 0 |
| scf7180000019961 | 279827 | 0.399 | 0 | 52.4846 | Eukaryota | 22964.0 | 0 | Chordata | 22964.0 | 0 | Carnivora | 16312.0 | 1 | Mustelidae | 11240.0 | 3 | Mustela | 11240.0 | 3 | ustela putori | 11240.0 | 4 |
| scf7180000019143 | 15397 | 0.4322 | 0 | 23.7338 | Eukaryota | 17008.0 | 0 | Chordata | 17008.0 | 0 | Carnivora | 17008.0 | 0 | Felidae | 9524.0 | 3 | Lynx | 3279.0 | 8 | ynx canadens | 3279.0 | 8 |
| scf7180000022459 | 29317 | 0.4392 | 0 | 49.8802 | Eukaryota | 9927.0 | 0 | Chordata | 9927.0 | 0 | Carnivora | 9927.0 | 0 | Herpestidae | 5431.0 | 1 | Suricata | 5431.0 | 1 | ıricata suricat | 5431.0 | 1 |
| scf7180000019705 | 72597 | 0.3745 | 0 | 45.7428 | Eukaryota | 46266.0 | 0 | Chordata | 46266.0 | 0 | Carnivora | 46266.0 | 0 | Felidae | 46266.0 | 0 | Felis | 41686.0 | 1 | Felis catus | 41686.0 | 1 |
| scf7180000021676 | 25623 | 0.4292 | 0 | 25.7417 | Eukaryota | 10326.0 | 0 | Chordata | 10326.0 | 0 | Carnivora | 10326.0 | 0 | Felidae | 10326.0 | 0 | Felis | 9312.0 | 1 | Felis catus | 9312.0 | 1 |
| scf7180000021277 | 68185 | 0.3449 | 0 | 43.7662 | Eukaryota | 19842.0 | 0 | Chordata | 19842.0 | 0 | Carnivora | 19842.0 | 0 | Felidae | 19842.0 | 0 | Felis | 18314.0 | 1 | Felis catus | 18314.0 | 1 |
| scf7180000020234 | 160708 | 0.6224 | 100 | 49.0848 | Eukaryota | 8691.0 | 0 | Chordata | 8691.0 | 0 | Carnivora | 8691.0 | 0 | Felidae | 7806.0 | 1 | Lynx | 4338.0 | 2 | ynx canadens | 4338.0 | 2 |
| scf7180000020153 | 158705 | 0.3781 | 0 | 50.2542 | Eukaryota | 28015.0 | 0 | Chordata | 28015.0 | 0 | Carnivora | 28015.0 | 0 | Felidae | 28015.0 | 0 | Felis | 25145.0 | 1 | Felis catus | 25145.0 | 1 |
| scf7180000022641 | 11657 | 0.3766 | 0 | 21.0512 | Eukaryota | 39252.0 | 0 | Chordata | 39252.0 | 0 | Carnivora | 39252.0 | 0 | Felidae | 39252.0 | 0 | Felis | 39252.0 | 0 | Felis catus | 39252.0 | 0 |
| scf7180000022152 | 8232 | 0.5248 | 0 | 41.6253 | no-hit | 0.0 | 0 | no-hit | 0.0 | 0 | no-hit | 0.0 | 0 | no-hit | 0.0 | 0 | no-hit | 0.0 | 0 | no-hit | 0.0 | 0 |
| scf7180000024087 | 192837 | 0.4363 | 0 | 51.2322 | Eukaryota | 30499.0 | 0 | Chordata | 30499.0 | 0 | Carnivora | 30499.0 | 0 | Felidae | 21720.0 | 2 | Panthera | 6198.0 | 7 | ynx canadens | 6154.0 | 8 |
| scf7180000020459 | 101152 | 0.5555 | 0 | 47.831 | Eukaryota | 6027.0 | 0 | Chordata | 6027.0 | 0 | Carnivora | 6027.0 | 0 | Felidae | 6027.0 | 0 | Felis | 5419.0 | 1 | Felis catus | 5419.0 | 1 |
| scf7180000021702 | 39972 | 0.4019 | 0 | 26.6215 | Eukaryota | 13280.0 | 0 | Chordata | 13280.0 | 0 | Primates | 6443.0 | 2 | Hominidae | 6443.0 | 2 | Felis | 4292.0 | 4 | Felis catus | 4292.0 | 4 |
| scf7180000023874 | 1199250 | 0.4391 | 0 | 50.5038 | Eukaryota | 46443.0 | 0 | Chordata | 46443.0 | 0 | Carnivora | 46443.0 | 0 | Felidae | 46443.0 | 0 | Felis | 46443.0 | 0 | Felis catus | 46443.0 | 0 |
| scf7180000017474 | 444352 | 0.4151 | 0 | 48.603 | Eukaryota | 29549.0 | 0 | Chordata | 29549.0 | 0 | Carnivora | 29549.0 | 0 | Felidae | 29549.0 | 0 | Felis | 26562.0 | 1 | Felis catus | 26562.0 | 1 |
| scf7180000018926 | 804164 | 0.475 | 0 | 52.0063 | Eukaryota | 63415.0 | 0 | Chordata | 63415.0 | 0 | Carnivora | 34895.0 | 1 | Felidae | 34895.0 | 3 | agenorhynchu | 16842.0 | 6 | rhynchus obli | 16842.0 | 7 |
| scf7180000020570 | 119289 | 0.4426 | 0 | 50.8858 | Eukaryota | 18310.0 | 1 | Chordata | 18310.0 | 1 | Carnivora | 16376.0 | 2 | Herpestidae | 16376.0 | 2 | Suricata | 16376.0 | 2 | ıricata suricat | 16376.0 | 2 |
| scf7180000019187 | 362977 | 0.5486 | 0 | 53.3087 | Eukaryota | 18121.0 | 1 | Chordata | 18121.0 | 1 | Carnivora | 9127.0 | 2 | Felidae | 9127.0 | 2 | Homo | 6761.0 | 6 | Homo sapiens | 6761.0 | 7 |
| scf7180000018193 | 91545 | 0.3956 | 0 | 47.9638 | Eukaryota | 33582.0 | 0 | Chordata | 33582.0 | 0 | Carnivora | 33582.0 | 0 | Felidae | 33582.0 | 0 | Felis | 30160.0 | 1 | Felis catus | 30160.0 | 1 |
| scf7180000021809 | 25798 | 0.4048 | 0 | 24.8284 | Eukaryota | 44197.0 | 0 | Chordata | 44197.0 | 0 | Carnivora | 44197.0 | 0 | Felidae | 44197.0 | 0 | Felis | 39825.0 | 1 | Felis catus | 39825.0 | 1 |
| scf7180000020084 | 376043 | 0.4398 | 0 | 53.3404 | Eukaryota | 19505.0 | 0 | Chordata | 19505.0 | 0 | Carnivora | 19505.0 | 0 | Felidae | 19505.0 | 0 | Felis | 17549.0 | 1 | Felis catus | 17549.0 | 1 |
| scf7180000018625 | 580231 | 0.4476 | 0 | 50.8428 | Eukaryota | 29760.0 | 0 | Chordata | 29760.0 | 0 | Carnivora | 29760.0 | 0 | Felidae | 12631.0 | 3 | Ursus | 7566.0 | 5 | Ursus arctos | 7566.0 | 5 |
| scf7180000018378 | 881716 | 0.3766 | 0 | 51.3023 | Eukaryota | 47733.0 | 0 | Chordata | 47733.0 | 0 | Carnivora | 47733.0 | 0 | Felidae | 47733.0 | 0 | Felis | 47733.0 | 0 | Felis catus | 47733.0 | 0 |
| scf7180000021309 | 58275 | 0.4372 | 0 | 26.2128 | Eukaryota | 8180.0 | 0 | Chordata | 8180.0 | 0 | Primates | 5765.0 | 1 | Cercopithecidae | 2723.0 | 3 | Homo | 2454.0 | 5 | Homo sapiens | 2454.0 | 6 |
| scf7180000017781 | 1501170 | 0.3927 | 0 | 49.4866 | Eukaryota | 72379.0 | 0 | Chordata | 72379.0 | 0 | Carnivora | 72379.0 | 0 | Felidae | 72379.0 | 0 | Acinonyx | 28996.0 | 2 | cinonyx jubat | 28996.0 | 2 |
| scf7180000018113 | 1051223 | 0.3948 | 0 | 49.4815 | Eukaryota | 45486.0 | 0 | Chordata | 45486.0 | 0 | Carnivora | 45486.0 | 0 | Felidae | 45486.0 | 0 | Felis | 45486.0 | 0 | Felis catus | 45486.0 | 0 |
| scf7180000019447 | 58377 | 0.4115 | 0 | 117.3531 | Eukaryota | 45259.0 | 0 | Chordata | 45259.0 | 0 | Carnivora | 45259.0 | 0 | Felidae | 45259.0 | 0 | Felis | 45259.0 | 0 | Felis catus | 45259.0 | 0 |
| scf7180000019495 | 217169 | 0.509 | 0 | 49.9607 | Eukaryota | 17242.0 | 0 | Chordata | 17242.0 | 0 | Carnivora | 17242.0 | 0 | Felidae | 17242.0 | 0 | Panthera | 6864.0 | 3 | anthera pard | 6864.0 | 3 |
| scf7180000018232 | 1909866 | 0.4507 | 100 | 52.537 | Eukaryota | 41895.0 | 0 | Chordata | 41895.0 | 0 | Carnivora | 41895.0 | 0 | Felidae | 21311.0 | 2 | Suricata | 12998.0 | 5 | ıricata suricat | 12998.0 | 6 |
| scf7180000023674 | 153696 | 0.5281 | 0 | 558.2568 | Eukaryota | 22347.0 | 0 | Chordata | 22347.0 | 0 | Carnivora | 22347.0 | 0 | Felidae | 22347.0 | 0 | Felis | 20090.0 | 1 | Felis catus | 20090.0 | 1 |
| scf7180000019336 | 390461 | 0.3612 | 0 | 44.1011 | Eukaryota | 47780.0 | 0 | Chordata | 47780.0 | 0 | Carnivora | 47780.0 | 0 | Mustelidae | 25515.0 | 1 | Mustela | 25515.0 | 1 | ustela putori | 25515.0 | 1 |
| scf7180000018121 | 1486853 | 0.402 | 0 | 50.7427 | Eukaryota | 45661.0 | 0 | Chordata | 45661.0 | 0 | Carnivora | 45661.0 | 0 | Felidae | 38811.0 | 1 | Acinonyx | 18757.0 | 3 | cinonyx jubat | 18757.0 | 3 |
| scf7180000018031 | 2236799 | 0.3505 | 0 | 48.146 | Eukaryota | 48762.0 | 0 | Chordata | 48762.0 | 0 | Carnivora | 48762.0 | 0 | Felidae | 41855.0 | 1 | Felis | 41855.0 | 1 | Felis catus | 41855.0 | 1 |
| scf7180000019854 | 12776 | 0.3887 | 0 | 23.9342 | Eukaryota | 8022.0 | 0 | Chordata | 8022.0 | 0 | Primates | 4742.0 | 1 | Hominidae | 4742.0 | 2 | Homo | 2736.0 | 4 | Homo sapiens | 2736.0 | 4 |
| scf7180000021373 | 61824 | 0.3747 | 0 | 48.8577 | Eukaryota | 7922.0 | 0 | Chordata | 7922.0 | 0 | Carnivora | 6217.0 | 1 | Felidae | 6217.0 | 1 | Felis | 6217.0 | 1 | Felis catus | 6217.0 | 1 |
| scf7180000024135 | 170166 | 0.4194 | 0 | 50.1377 | Eukaryota | 28400.0 | 0 | Chordata | 28400.0 | 0 | Carnivora | 28400.0 | 0 | Felidae | 28400.0 | 0 | Acinonyx | 11504.0 | 2 | cinonyx jubat | 11504.0 | 2 |
| scf7180000020553 | 74362 | 0.4109 | 0 | 46.729 | Eukaryota | 44172.0 | 0 | Chordata | 44172.0 | 0 | Carnivora | 44172.0 | 0 | Felidae | 44172.0 | 0 | Felis | 39824.0 | 1 | Felis catus | 39824.0 | 1 |
| scf7180000021116 | 55723 | 0.3845 | 0 | 25.992 | Eukaryota | 29450.0 | 0 | Chordata | 29450.0 | 0 | Carnivora | 29450.0 | 0 | Felidae | 29450.0 | 0 | Felis | 26439.0 | 1 | Felis catus | 26439.0 | 1 |
| scf7180000017891 | 643627 | 0.3676 | 0 | 48.7293 | Eukaryota | 42152.0 | 0 | Chordata | 42152.0 | 0 | Carnivora | 42152.0 | 0 | Felidae | 42152.0 | 0 | Felis | 37961.0 | 1 | Felis catus | 37961.0 | 1 |
| scf7180000022220 | 18184 | 0.6181 | 0 | 50.2371 | Eukaryota | 2692.0 | 0 | Chordata | 2692.0 | 0 | Carnivora | 2692.0 | 0 | Felidae | 1632.0 | 2 | Suricata | 801.0 | 5 | ıricata suricat | 801.0 | 5 |
| scf7180000021319 | 23009 | 0.4258 | 0 | 405.7635 | Eukaryota | 203424.0 | 0 | Chordata | 203424.0 | 0 | Carnivora | 203424.0 | 0 | Hyaenidae | 203424.0 | 0 | Proteles | 120464.0 | 1 | roteles cristat | 120464.0 | 1 |
| scf7180000017475 | 482833 | 0.4493 | 0 | 51.0649 | Eukaryota | 35286.0 | 1 | Chordata | 35286.0 | 1 | undef | 15206.0 | 6 | undef | 15206.0 | 6 | undef | 15206.0 | 7 | tic synthetic c | 15206.0 | 7 |
| scf7180000018679 | 55303 | 0.48 | 0 | 58.5883 | Eukaryota | 5144.0 | 0 | Chordata | 5144.0 | 0 | Carnivora | 5144.0 | 0 | Felidae | 3798.0 | 1 | Felis | 1985.0 | 3 | Felis catus | 1985.0 | 3 |
| scf7180000021273 | 50536 | 0.3724 | 0 | 25.6626 | Eukaryota | 42261.0 | 0 | Chordata | 42261.0 | 0 | Carnivora | 42261.0 | 0 | Felidae | 42261.0 | 0 | Felis | 42261.0 | 0 | Felis catus | 42261.0 | 0 |
| scf7180000021795 | 22128 | 0.5351 | 0 | 49.269 | Eukaryota | 2945.0 | 0 | Chordata | 2945.0 | 0 | Carnivora | 2945.0 | 0 | Felidae | 2945.0 | 0 | Felis | 2653.0 | 1 | Felis catus | 2653.0 | 1 |
| scf7180000021599 | 52713 | 0.4081 | 0 | 26.7062 | Eukaryota | 24376.0 | 0 | Chordata | 24376.0 | 0 | Carnivora | 24376.0 | 0 | Felidae | 24376.0 | 0 | Felis | 21890.0 | 1 | Felis catus | 21890.0 | 1 |
| scf7180000018024 | 258444 | 0.4063 | 0 | 49.2583 | Eukaryota | 43603.0 | 0 | Chordata | 43603.0 | 0 | Carnivora | 43603.0 | 0 | Felidae | 43603.0 | 0 | Felis | 43603.0 | 0 | Felis catus | 43603.0 | 0 |
| scf7180000019636 | 152899 | 0.4257 | 0 | 65.8899 | Eukaryota | 46335.0 | 0 | Chordata | 46335.0 | 0 | Carnivora | 46335.0 | 0 | Felidae | 46335.0 | 0 | Felis | 46335.0 | 0 | Felis catus | 46335.0 | 0 |
| scf7180000021986 | 13610 | 0.4069 | 0 | 26.2227 | Eukaryota | 4620.0 | 0 | Chordata | 4620.0 | 0 | Carnivora | 4160.0 | 1 | Felidae | 3692.0 | 2 | Felis | 2672.0 | 3 | Felis catus | 2672.0 | 3 |
| scf7180000018495 | 669340 | 0.4587 | 0 | 52.8561 | Eukaryota | 47312.0 | 0 | Chordata | 47312.0 | 0 | Carnivora | 47312.0 | 0 | Felidae | 47312.0 | 0 | Felis | 47312.0 | 0 | Felis catus | 47312.0 | 0 |
| scf7180000023926 | 84444 | 0.3781 | 0 | 23.5058 | Eukaryota | 26994.0 | 0 | Chordata | 26994.0 | 0 | Carnivora | 26994.0 | 0 | Felidae | 26994.0 | 0 | Felis | 21560.0 | 1 | Felis catus | 21560.0 | 1 |
| scf7180000019507 | 269989 | 0.4179 | 0 | 50.965 | Eukaryota | 30464.0 | 0 | Chordata | 30464.0 | 0 | Carnivora | 30464.0 | 0 | Felidae | 30464.0 | 0 | Felis | 27640.0 | 1 | Felis catus | 27640.0 | 1 |
| scf7180000018258 | 724428 | 0.3764 | 0 | 50.3513 | Eukaryota | 47063.0 | 0 | Chordata | 47063.0 | 0 | Carnivora | 47063.0 | 0 | Felidae | 47063.0 | 0 | Felis | 47063.0 | 0 | Felis catus | 47063.0 | 0 |
| scf7180000018009 | 1202345 | 0.4211 | 0 | 50.6787 | Eukaryota | 45897.0 | 0 | Chordata | 45897.0 | 0 | Carnivora | 45897.0 | 0 | Felidae | 45897.0 | 0 | Felis | 45897.0 | 0 | Felis catus | 45897.0 | 0 |
| scf7180000022277 | 18973 | 0.4633 | 0 | 23.5326 | Eukaryota | 7517.0 | 0 | Chordata | 7517.0 | 0 | Rodentia | 2118.0 | 7 | Muridae | 2118.0 | 8 | Mus | 2118.0 | 8 | Mus musculus | 2118.0 | 8 |
| scf7180000020455 | 101789 | 0.498 | 0 | 27.0161 | Eukaryota | 6571.0 | 0 | Chordata | 6571.0 | 0 | Carnivora | 6571.0 | 0 | Felidae | 4590.0 | 1 | Canis | 1981.0 | 4 | Canis lupus | 1981.0 | 5 |
| scf7180000018042 | 1183236 | 0.384 | 0 | 51.6588 | Eukaryota | 46087.0 | 0 | Chordata | 46087.0 | 0 | Carnivora | 46087.0 | 0 | Felidae | 46087.0 | 0 | Felis | 41488.0 | 1 | Felis catus | 41488.0 | 1 |
| scf7180000018055 | 406889 | 0.3922 | 0 | 51.8849 | Eukaryota | 45968.0 | 0 | Chordata | 45968.0 | 0 | Carnivora | 45968.0 | 0 | Felidae | 45968.0 | 0 | Felis | 45968.0 | 0 | Felis catus | 45968.0 | 0 |
| scf7180000020690 | 66692 | 0.6544 | 0 | 54.2945 | Eukaryota | 7711.0 | 0 | Chordata | 7711.0 | 0 | Carnivora | 5521.0 | 1 | Felidae | 4580.0 | 3 | Felis | 3737.0 | 4 | Felis catus | 3737.0 | 4 |
| scf7180000018771 | 19856 | 0.4575 | 0 | 78.1531 | Eukaryota | 6898.0 | 0 | Chordata | 6898.0 | 0 | Carnivora | 6898.0 | 0 | Felidae | 6898.0 | 0 | Panthera | 2884.0 | 2 | anthera pard | 2884.0 | 2 |
| scf7180000018501 | 674056 | 0.4551 | 0 | 49.8788 | Eukaryota | 30202.0 | 0 | Chordata | 30202.0 | 0 | Carnivora | 30202.0 | 0 | Felidae | 30202.0 | 0 | Felis | 27119.0 | 1 | Felis catus | 27119.0 | 1 |
| scf7180000018444 | 493321 | 0.4384 | 0 | 52.4435 | Eukaryota | 46697.0 | 0 | Chordata | 46697.0 | 0 | Carnivora | 46697.0 | 0 | Felidae | 46697.0 | 0 | Felis | 46697.0 | 0 | Felis catus | 46697.0 | 0 |
| scf7180000019472 | 233947 | 0.3548 | 0 | 59.1261 | Eukaryota | 45896.0 | 0 | Chordata | 45896.0 | 0 | Carnivora | 45896.0 | 0 | Felidae | 45896.0 | 0 | Felis | 45896.0 | 0 | Felis catus | 45896.0 | 0 |
| scf7180000021900 | 43109 | 0.4433 | 0 | 25.9732 | Eukaryota | 5318.0 | 0 | Chordata | 5318.0 | 0 | Carnivora | 3247.0 | 2 | Felidae | 2452.0 | 4 | Sus | 1123.0 | 8 | Sus scrofa | 1123.0 | 9 |
| scf7180000019912 | 167399 | 0.6138 | 0 | 50.5864 | Eukaryota | 13216.0 | 0 | Chordata | 13216.0 | 0 | Carnivora | 13216.0 | 0 | Felidae | 13216.0 | 0 | Panthera | 5312.0 | 3 | anthera pard | 3984.0 | 4 |
| scf7180000019227 | 356405 | 0.3511 | 0 | 48.0738 | Eukaryota | 47004.0 | 0 | Chordata | 47004.0 | 0 | Carnivora | 47004.0 | 0 | Felidae | 47004.0 | 0 | Felis | 47004.0 | 0 | Felis catus | 47004.0 | 0 |
| scf7180000020832 | 86262 | 0.3621 | 0 | 24.6573 | Eukaryota | 40963.0 | 0 | Chordata | 40963.0 | 0 | Carnivora | 40963.0 | 0 | Felidae | 40963.0 | 0 | Felis | 40963.0 | 0 | Felis catus | 40963.0 | 0 |
| scf7180000018019 | 1237375 | 0.3583 | 0 | 49.3263 | Eukaryota | 47947.0 | 0 | Chordata | 47947.0 | 0 | Carnivora | 47947.0 | 0 | Felidae | 47947.0 | 0 | Felis | 47947.0 | 0 | Felis catus | 47947.0 | 0 |
| scf7180000018143 | 73834 | 0.5334 | 0 | 49.1146 | Eukaryota | 3516.0 | 0 | Chordata | 3516.0 | 0 | Carnivora | 3122.0 | 1 | Felidae | 1382.0 | 3 | Felis | 1123.0 | 4 | Felis catus | 1123.0 | 4 |
| scf7180000017450 | 3580614 | 0.3496 | 0 | 48.8862 | Eukaryota | 121172.0 | 0 | Chordata | 121172.0 | 0 | Carnivora | 121172.0 | 0 | Felidae | 84695.0 | 1 | Felis | 36543.0 | 3 | Felis catus | 36543.0 | 3 |
| scf7180000017938 | 1277888 | 0.398 | 0 | 50.5301 | Eukaryota | 46765.0 | 0 | Chordata | 46765.0 | 0 | Carnivora | 46765.0 | 0 | Felidae | 37179.0 | 1 | Felis | 37179.0 | 1 | Felis catus | 37179.0 | 1 |
| scf7180000017816 | 358698 | 0.3861 | 0 | 50.5795 | Eukaryota | 59633.0 | 0 | Chordata | 59633.0 | 0 | Carnivora | 59633.0 | 0 | Felidae | 59633.0 | 0 | Neofelis | 24682.0 | 2 | eofelis nebulo | 24682.0 | 2 |
| scf7180000022259 | 41589 | 0.4857 | 0 | 2759.0769 | Eukaryota | 2333.0 | 0 | Chordata | 2333.0 | 0 | Carnivora | 1581.0 | 1 | Felidae | 1581.0 | 3 | Felis | 1581.0 | 3 | Felis catus | 1581.0 | 3 |
| scf7180000018082 | 185008 | 0.3953 | 0 | 48.094 | Eukaryota | 56677.0 | 0 | Chordata | 56677.0 | 0 | Carnivora | 56677.0 | 0 | Felidae | 22928.0 | 5 | Acinonyx | 11339.0 | 8 | cinonyx jubat | 11339.0 | 8 |

| | | | | | | | | | | | | | | | | | | | | | | |
|---|---|---|---|---|---|---|---|---|---|---|---|---|---|---|---|---|---|---|---|---|---|---|
| scf7180000022126 | 15861 | 0.4053 | 0 | 26.7689 | Eukaryota | 45473.0 | 0 | Chordata | 45473.0 | 0 | Carnivora | 45473.0 | 0 | Felidae | 45473.0 | 0 | Felis | 40871.0 | 1 | Felis catus | 40871.0 | 1 |
| scf7180000020377 | 107763 | 0.6281 | 0 | 50.7716 | Eukaryota | 12852.0 | 0 | Chordata | 12852.0 | 0 | Carnivora | 12852.0 | 0 | Herpestidae | 6714.0 | 1 | Suricata | 6714.0 | 5 | ricata suricat | 6714.0 | 5 |
| scf7180000021346 | 12558 | 0.5177 | 0 | 229.6809 | no-hit | 0.0 | 0 | no-hit | 0.0 | 0 | no-hit | 0.0 | 0 | no-hit | 0.0 | 0 | no-hit | 0.0 | 0 | no-hit | 0.0 | 0 |
| scf7180000021701 | 35428 | 0.3838 | 0 | 25.0375 | Eukaryota | 5031.0 | 0 | Chordata | 5031.0 | 0 | Primates | 3004.0 | 3 | Hominidae | 3004.0 | 3 | Pan | 2020.0 | 4 | an troglodyte | 2020.0 | 4 |
| scf7180000021156 | 77998 | 0.3867 | 0 | 24.5711 | Eukaryota | 45491.0 | 0 | Chordata | 45491.0 | 0 | Carnivora | 45491.0 | 0 | Felidae | 45491.0 | 0 | Felis | 45491.0 | 0 | Felis catus | 45491.0 | 0 |
| scf7180000017687 | 1789039 | 0.4591 | 0 | 51.2164 | Eukaryota | 50296.0 | 0 | Chordata | 50296.0 | 0 | Carnivora | 45321.0 | 1 | Felidae | 37483.0 | 2 | Felis | 37483.0 | 2 | Felis catus | 37483.0 | 2 |
| scf7180000019199 | 373209 | 0.3732 | 0 | 49.3769 | Eukaryota | 48085.0 | 0 | Chordata | 48085.0 | 0 | Carnivora | 48085.0 | 0 | Felidae | 42914.0 | 1 | Felis | 42914.0 | 1 | Felis catus | 42914.0 | 1 |
| scf7180000023450 | 72994 | 0.5212 | 0 | 2275.5356 | no-hit | 0.0 | 0 | no-hit | 0.0 | 0 | no-hit | 0.0 | 0 | no-hit | 0.0 | 0 | no-hit | 0.0 | 0 | no-hit | 0.0 | 0 |
| scf7180000017601 | 2232082 | 0.5202 | 0 | 51.6617 | Eukaryota | 32720.0 | 0 | Chordata | 32720.0 | 0 | Carnivora | 32720.0 | 0 | Herpestidae | 18667.0 | 1 | Suricata | 18667.0 | 3 | ricata suricat | 18667.0 | 3 |
| scf7180000021229 | 37965 | 0.6109 | 0 | 44.9589 | Eukaryota | 7946.0 | 1 | Chordata | 7946.0 | 1 | Carnivora | 4282.0 | 4 | Felidae | 2660.0 | 6 | Sus | 2523.0 | 7 | Sus scrofa | 2523.0 | 7 |
| scf7180000019514 | 298020 | 0.3808 | 0 | 50.675 | Eukaryota | 34373.0 | 0 | Chordata | 34373.0 | 0 | Carnivora | 34373.0 | 0 | Felidae | 34373.0 | 0 | Felis | 34373.0 | 0 | Felis catus | 34373.0 | 0 |
| scf7180000022751 | 10142 | 0.4185 | 0 | 26.0138 | Eukaryota | 5679.0 | 0 | Chordata | 5679.0 | 0 | Primates | 4422.0 | 2 | Hominidae | 4422.0 | 2 | Pan | 2639.0 | 4 | an troglodyte | 2639.0 | 4 |
| scf7180000019255 | 3573026 | 0.4584 | 0 | 52.718 | Eukaryota | 90408.0 | 0 | Chordata | 90408.0 | 0 | Carnivora | 90408.0 | 0 | Felidae | 72040.0 | 1 | Acinonyx | 26970.0 | 4 | inonyx jubat | 26970.0 | 4 |
| scf7180000023185 | 14283 | 0.387 | 0 | 263.1965 | Eukaryota | 5150.0 | 0 | Chordata | 5150.0 | 0 | Carnivora | 5150.0 | 0 | Felidae | 5150.0 | 0 | Felis | 4120.0 | 1 | Felis catus | 4120.0 | 1 |
| scf7180000021980 | 26511 | 0.404 | 0 | 24.2353 | Eukaryota | 25672.0 | 0 | Chordata | 25672.0 | 0 | Artiodactyla | 16419.0 | 1 | Bovidae | 16419.0 | 1 | Ovis | 16419.0 | 1 | vis canadens | 16419.0 | 1 |
| scf7180000021462 | 79455 | 0.4419 | 0 | 52.0085 | Eukaryota | 37581.0 | 0 | Chordata | 37581.0 | 0 | Carnivora | 37581.0 | 0 | Felidae | 37581.0 | 0 | Felis | 37581.0 | 0 | Felis catus | 37581.0 | 0 |
| scf7180000018286 | 788811 | 0.4133 | 0 | 58.2883 | Eukaryota | 38929.0 | 0 | Chordata | 38929.0 | 0 | Carnivora | 38929.0 | 0 | Felidae | 38929.0 | 0 | Acinonyx | 22180.0 | 2 | inonyx jubat | 22180.0 | 2 |
| scf7180000022425 | 12546 | 0.4198 | 0 | 23.0216 | Eukaryota | 3254.0 | 0 | Chordata | 3254.0 | 0 | Carnivora | 2642.0 | 1 | Felidae | 2642.0 | 1 | Felis | 2340.0 | 2 | Felis catus | 2340.0 | 2 |
| scf7180000020116 | 137647 | 0.3654 | 0 | 24.1911 | Eukaryota | 38648.0 | 0 | Chordata | 38648.0 | 0 | Carnivora | 38648.0 | 0 | Felidae | 38648.0 | 0 | Felis | 38648.0 | 0 | Felis catus | 38648.0 | 0 |
| scf7180000021181 | 59132 | 0.4973 | 0 | 25.3328 | Eukaryota | 10975.0 | 0 | Chordata | 10975.0 | 0 | Carnivora | 5656.0 | 3 | Felidae | 5656.0 | 3 | Felis | 5656.0 | 4 | Felis catus | 5656.0 | 4 |
| scf7180000018435 | 757864 | 0.4855 | 0 | 52.0218 | Eukaryota | 22711.0 | 1 | Chordata | 22711.0 | 1 | Carnivora | 12386.0 | 3 | Felidae | 5638.0 | 5 | Panthera | 5638.0 | 6 | anthera pardu | 5638.0 | 6 |
| scf7180000017908 | 2540446 | 0.4246 | 100 | 51.5973 | Eukaryota | 46379.0 | 0 | Chordata | 46379.0 | 0 | Carnivora | 46379.0 | 0 | Felidae | 46379.0 | 0 | Felis | 46379.0 | 0 | Felis catus | 46379.0 | 0 |
| scf7180000018840 | 454683 | 0.5242 | 0 | 51.7894 | Eukaryota | 37953.0 | 0 | Chordata | 37953.0 | 0 | Carnivora | 37953.0 | 0 | Otariidae | 11895.0 | 4 | Felis | 11816.0 | 6 | Felis catus | 11816.0 | 6 |
| scf7180000023778 | 830757 | 0.3999 | 0 | 51.0693 | Eukaryota | 32913.0 | 0 | Chordata | 32913.0 | 0 | Carnivora | 32913.0 | 0 | Felidae | 32913.0 | 0 | Felis | 32913.0 | 0 | Felis catus | 32913.0 | 0 |
| scf7180000020926 | 42636 | 0.5393 | 0 | 47.5798 | Eukaryota | 7909.0 | 0 | Chordata | 7909.0 | 0 | Carnivora | 7909.0 | 0 | Felidae | 7909.0 | 0 | Felis | 5931.0 | 1 | Felis catus | 5931.0 | 1 |
| scf7180000022355 | 14395 | 0.4163 | 0 | 23.8385 | Eukaryota | 11820.0 | 0 | Chordata | 11820.0 | 0 | Carnivora | 11820.0 | 0 | Felidae | 11820.0 | 0 | Felis | 10630.0 | 1 | Felis catus | 10630.0 | 1 |
| scf7180000018049 | 1570695 | 0.399 | 0 | 51.8305 | Eukaryota | 47128.0 | 0 | Chordata | 47128.0 | 0 | Carnivora | 37040.0 | 1 | Otariidae | 18283.0 | 3 | Homo | 10088.0 | 6 | Homo sapiens | 10088.0 | 6 |
| scf7180000020132 | 255457 | 0.3939 | 0 | 51.1936 | Eukaryota | 24562.0 | 0 | Chordata | 24562.0 | 0 | Carnivora | 24562.0 | 0 | Felidae | 24562.0 | 0 | Felis | 24562.0 | 0 | Felis catus | 24562.0 | 0 |
| scf7180000020822 | 197384 | 0.6336 | 0 | 43.8904 | Eukaryota | 6553.0 | 0 | Chordata | 6553.0 | 0 | Carnivora | 6553.0 | 0 | Felidae | 6553.0 | 0 | Felis | 3380.0 | 2 | Felis catus | 3380.0 | 2 |
| scf7180000021613 | 28437 | 0.3769 | 0 | 29.6784 | Eukaryota | 46557.0 | 0 | Chordata | 46557.0 | 0 | Carnivora | 46557.0 | 0 | Felidae | 46557.0 | 0 | Felis | 46557.0 | 0 | Felis catus | 46557.0 | 0 |
| scf7180000017841 | 1550239 | 0.3957 | 0 | 52.1284 | Eukaryota | 47731.0 | 0 | Chordata | 47731.0 | 0 | Carnivora | 47731.0 | 0 | Felidae | 47731.0 | 0 | Felis | 47731.0 | 0 | Felis catus | 47731.0 | 0 |
| scf7180000019123 | 273513 | 0.4658 | 0 | 47.0756 | Eukaryota | 30545.0 | 0 | Chordata | 30545.0 | 0 | Carnivora | 30545.0 | 0 | Ursidae | 14198.0 | 2 | Ursus | 14198.0 | 2 | Felis catus | 10373.0 | 3 |
| scf7180000019814 | 331797 | 0.3595 | 0 | 50.2969 | Eukaryota | 54048.0 | 0 | Chordata | 54048.0 | 0 | Carnivora | 42194.0 | 1 | Felidae | 42194.0 | 1 | Felis | 42194.0 | 1 | Felis catus | 42194.0 | 1 |
| scf7180000018723 | 519373 | 0.5255 | 0 | 52.7801 | Eukaryota | 35160.0 | 0 | Chordata | 35160.0 | 0 | Carnivora | 22043.0 | 2 | Felidae | 15887.0 | 3 | Homo | 7842.0 | 5 | Homo sapiens | 7842.0 | 5 |
| scf7180000021626 | 24638 | 0.3833 | 0 | 36.118 | Eukaryota | 32302.0 | 0 | Chordata | 32302.0 | 0 | Carnivora | 32302.0 | 0 | Felidae | 32302.0 | 0 | Felis | 29707.0 | 1 | Felis catus | 29707.0 | 1 |
| scf7180000018046 | 1073606 | 0.3904 | 0 | 50.5072 | Eukaryota | 99734.0 | 0 | Chordata | 99734.0 | 0 | Carnivora | 99734.0 | 0 | Felidae | 61641.0 | 2 | Lynx | 33276.0 | 3 | ynx canadens | 33276.0 | 4 |
| scf7180000019718 | 41149 | 0.4605 | 0 | 57.6565 | Eukaryota | 2484.0 | 0 | Chordata | 2484.0 | 0 | Carnivora | 2484.0 | 0 | Felidae | 2484.0 | 0 | Felis | 2484.0 | 0 | Felis catus | 2484.0 | 0 |
| scf7180000020991 | 75050 | 0.4075 | 0 | 26.4712 | Eukaryota | 42061.0 | 0 | Chordata | 42061.0 | 0 | Carnivora | 42061.0 | 0 | Felidae | 42061.0 | 0 | Felis | 42061.0 | 0 | Felis catus | 42061.0 | 0 |
| scf7180000017896 | 626424 | 0.4193 | 0 | 49.7698 | Eukaryota | 36245.0 | 0 | Chordata | 36245.0 | 0 | Carnivora | 36245.0 | 0 | Felidae | 36245.0 | 0 | Felis | 36245.0 | 0 | Felis catus | 36245.0 | 0 |
| scf7180000020780 | 94761 | 0.4772 | 0 | 49.0758 | Eukaryota | 3580.0 | 0 | Chordata | 3580.0 | 0 | Carnivora | 2463.0 | 2 | Felidae | 2198.0 | 3 | Felis | 1930.0 | 5 | Felis catus | 1930.0 | 5 |
| scf7180000022240 | 14247 | 0.4143 | 0 | 34.0315 | Eukaryota | 31415.0 | 0 | Chordata | 31415.0 | 0 | Carnivora | 31415.0 | 0 | Felidae | 31415.0 | 0 | Felis | 28229.0 | 1 | Felis catus | 28229.0 | 1 |
| scf7180000018805 | 607229 | 0.403 | 0 | 52.3375 | Eukaryota | 92937.0 | 0 | Chordata | 92937.0 | 0 | Carnivora | 92937.0 | 0 | Felidae | 92937.0 | 0 | Felis | 65063.0 | 1 | Felis catus | 65063.0 | 1 |
| scf7180000018386 | 881143 | 0.4336 | 0 | 50.7527 | Eukaryota | 92506.0 | 0 | Chordata | 92506.0 | 0 | Carnivora | 92506.0 | 0 | Felidae | 83826.0 | 1 | Acinonyx | 37611.0 | 4 | inonyx jubat | 37611.0 | 4 |
| scf7180000022325 | 4648 | 0.5148 | 0 | 40.6549 | no-hit | 0.0 | 0 | no-hit | 0.0 | 0 | no-hit | 0.0 | 0 | no-hit | 0.0 | 0 | no-hit | 0.0 | 0 | no-hit | 0.0 | 0 |
| scf7180000020890 | 71593 | 0.4022 | 0 | 25.224 | Eukaryota | 47206.0 | 0 | Chordata | 47206.0 | 0 | Carnivora | 47206.0 | 0 | Felidae | 47206.0 | 0 | Felis | 47206.0 | 0 | Felis catus | 47206.0 | 0 |
| scf7180000020129 | 158673 | 0.3556 | 0 | 23.1973 | Eukaryota | 45327.0 | 0 | Chordata | 45327.0 | 0 | Carnivora | 45327.0 | 0 | Felidae | 45327.0 | 0 | Felis | 45327.0 | 0 | Felis catus | 45327.0 | 0 |
| scf7180000021680 | 40021 | 0.3993 | 0 | 25.7819 | Eukaryota | 19479.0 | 0 | Chordata | 19479.0 | 0 | Carnivora | 19479.0 | 0 | Felidae | 19479.0 | 0 | Felis | 17525.0 | 1 | Felis catus | 17525.0 | 1 |
| scf7180000018943 | 442287 | 0.3661 | 0 | 47.7905 | Eukaryota | 46227.0 | 0 | Chordata | 46227.0 | 0 | Carnivora | 46227.0 | 0 | Felidae | 46227.0 | 0 | Felis | 46227.0 | 0 | Felis catus | 46227.0 | 0 |
| scf7180000021360 | 61692 | 0.4346 | 0 | 24.3297 | Eukaryota | 9786.0 | 0 | Chordata | 9786.0 | 0 | Carnivora | 5862.0 | 1 | unresolved | 2898.0 | 4 | Equus | 2898.0 | 7 | nychotes we | 1382.0 | 9 |
| scf7180000020075 | 154884 | 0.5121 | 0 | 55.886 | Eukaryota | 25473.0 | 0 | Chordata | 25473.0 | 0 | Carnivora | 25473.0 | 0 | Felidae | 25473.0 | 0 | Felis | 22902.0 | 1 | Felis catus | 22902.0 | 1 |
| scf7180000023798 | 1694913 | 0.4669 | 0 | 51.9479 | Eukaryota | 87291.0 | 0 | Chordata | 87291.0 | 0 | Carnivora | 87291.0 | 0 | Felidae | 60741.0 | 1 | Panthera | 34835.0 | 4 | ricata suricat | 26550.0 | 5 |
| scf7180000019813 | 128005 | 0.3891 | 0 | 46.8564 | Eukaryota | 46862.0 | 0 | Chordata | 46862.0 | 0 | Carnivora | 46862.0 | 0 | Felidae | 46862.0 | 0 | Felis | 46862.0 | 0 | Felis catus | 46862.0 | 0 |
| scf7180000018752 | 1470725 | 0.4263 | 0 | 52.1945 | Eukaryota | 48854.0 | 0 | Chordata | 48854.0 | 0 | Carnivora | 48854.0 | 0 | Felidae | 34010.0 | 1 | Felis | 24340.0 | 2 | Felis catus | 24340.0 | 2 |
| scf7180000018633 | 565465 | 0.3652 | 0 | 49.0224 | Eukaryota | 47535.0 | 0 | Chordata | 47535.0 | 0 | Carnivora | 47535.0 | 0 | Felidae | 47535.0 | 0 | Felis | 47535.0 | 0 | Felis catus | 47535.0 | 0 |
| scf7180000017598 | 1169876 | 0.4478 | 0 | 51.2383 | Eukaryota | 42337.0 | 0 | Chordata | 42337.0 | 0 | Carnivora | 42337.0 | 0 | Felidae | 42337.0 | 0 | Felis | 42337.0 | 0 | Felis catus | 42337.0 | 0 |
| scf7180000020641 | 136783 | 0.4229 | 0 | 55.2351 | Eukaryota | 39576.0 | 0 | Chordata | 39576.0 | 0 | Carnivora | 39576.0 | 0 | Felidae | 39576.0 | 0 | Felis | 35640.0 | 1 | Felis catus | 35640.0 | 1 |
| scf7180000021142 | 20904 | 0.5021 | 0 | 273.1006 | Eukaryota | 15601.0 | 0 | Chordata | 15601.0 | 0 | Carnivora | 15601.0 | 0 | Felidae | 15601.0 | 0 | Felis | 15601.0 | 0 | Felis catus | 15601.0 | 0 |
| scf7180000017501 | 12996 | 0.3891 | 0 | 22.6911 | Eukaryota | 3946.0 | 0 | Chordata | 3946.0 | 0 | Carnivora | 3946.0 | 0 | Felidae | 3946.0 | 0 | Felis | 3558.0 | 1 | Felis catus | 3558.0 | 1 |
| scf7180000021712 | 55006 | 0.4151 | 0 | 54.0565 | Eukaryota | 17516.0 | 0 | Chordata | 17516.0 | 0 | Carnivora | 17516.0 | 0 | Otariidae | 7109.0 | 3 | Callorhinus | 3558.0 | 8 | llorhinus ursi | 3558.0 | 8 |
| scf7180000018016 | 1283866 | 0.3705 | 0 | 49.6375 | Eukaryota | 138711.0 | 0 | Chordata | 138711.0 | 0 | Carnivora | 138711.0 | 0 | Felidae | 129084.0 | 1 | Panthera | 33208.0 | 4 | ynx canadens | 32180.0 | 5 |
| scf7180000022461 | 10719 | 0.4216 | 0 | 21.0692 | Eukaryota | 4641.0 | 0 | Chordata | 4641.0 | 0 | Primates | 2156.0 | 2 | Hominidae | 2156.0 | 2 | Felis | 1696.0 | 3 | Felis catus | 1696.0 | 3 |
| scf7180000018645 | 2296148 | 0.4131 | 0 | 51.4694 | Eukaryota | 46133.0 | 0 | Chordata | 46133.0 | 0 | Carnivora | 46133.0 | 0 | Felidae | 46133.0 | 0 | Felis | 46133.0 | 0 | Felis catus | 46133.0 | 0 |
| scf7180000020693 | 159486 | 0.366 | 0 | 24.7922 | Eukaryota | 16108.0 | 0 | Chordata | 16108.0 | 0 | Carnivora | 16108.0 | 0 | Felidae | 12568.0 | 2 | Felis | 12568.0 | 2 | Felis catus | 12568.0 | 2 |
| scf7180000017798 | 77836 | 0.3687 | 0 | 23.5252 | Eukaryota | 39250.0 | 0 | Chordata | 39250.0 | 0 | Carnivora | 39250.0 | 0 | Felidae | 39250.0 | 0 | Felis | 39250.0 | 0 | Felis catus | 39250.0 | 0 |
| scf7180000017486 | 3247415 | 0.3893 | 0 | 50.1678 | Eukaryota | 157556.0 | 0 | Chordata | 157556.0 | 0 | Carnivora | 157556.0 | 0 | Felidae | 127060.0 | 2 | Panthera | 46729.0 | 5 | inonyx jubat | 33382.0 | 6 |
| scf7180000017570 | 587779 | 0.3862 | 0 | 50.5516 | Eukaryota | 45647.0 | 0 | Chordata | 45647.0 | 0 | Carnivora | 45647.0 | 0 | Felidae | 45647.0 | 0 | Felis | 45647.0 | 0 | Felis catus | 45647.0 | 0 |
| scf7180000020385 | 167997 | 0.4351 | 0 | 25.788 | Eukaryota | 21256.0 | 0 | Chordata | 21256.0 | 0 | Artiodactyla | 11310.0 | 2 | Bovidae | 11310.0 | 2 | Bos | 7512.0 | 4 | Bos indicus | 3804.0 | 6 |
| scf7180000022548 | 16084 | 0.3915 | 0 | 25.9053 | Eukaryota | 18115.0 | 0 | Chordata | 18115.0 | 0 | Carnivora | 18115.0 | 0 | Felidae | 18115.0 | 0 | Felis | 13884.0 | 1 | Felis catus | 13884.0 | 1 |
| scf7180000019471 | 249341 | 0.4632 | 0 | 52.7535 | Eukaryota | 15860.0 | 0 | Chordata | 15860.0 | 0 | Carnivora | 15860.0 | 0 | Felidae | 15860.0 | 0 | Felis | 14303.0 | 1 | Felis catus | 14303.0 | 1 |
| scf7180000022001 | 87979 | 0.6037 | 0 | 49.3157 | Eukaryota | 14921.0 | 0 | Chordata | 14921.0 | 0 | Carnivora | 14921.0 | 0 | Felidae | 9527.0 | 3 | Lynx | 2385.0 | 8 | ynx canadens | 2385.0 | 8 |
| scf7180000020904 | 66376 | 0.419 | 0 | 50.7105 | Eukaryota | 45844.0 | 0 | Chordata | 45844.0 | 0 | Carnivora | 45844.0 | 0 | Felidae | 45844.0 | 0 | Felis | 45844.0 | 0 | Felis catus | 45844.0 | 0 |
| scf7180000024073 | 273122 | 0.4377 | 0 | 50.3903 | Eukaryota | 47113.0 | 0 | Chordata | 47113.0 | 0 | Carnivora | 47113.0 | 0 | Felidae | 37971.0 | 1 | Lynx | 14571.0 | 4 | ynx canadens | 14571.0 | 4 |
| scf7180000019343 | 314760 | 0.3706 | 0 | 50.4053 | Eukaryota | 30053.0 | 0 | Chordata | 30053.0 | 0 | Carnivora | 30053.0 | 0 | Felidae | 30053.0 | 0 | Felis | 26954.0 | 1 | Felis catus | 26954.0 | 1 |
| scf7180000019247 | 13250 | 0.4497 | 0 | 24.4228 | Eukaryota | 13116.0 | 0 | Chordata | 13116.0 | 0 | Carnivora | 13116.0 | 0 | Felidae | 11603.0 | 1 | Felis | 5786.0 | 4 | Felis catus | 5786.0 | 5 |
| scf7180000021894 | 48729 | 0.402 | 0 | 52.8679 | Eukaryota | 42642.0 | 0 | Chordata | 42642.0 | 0 | Carnivora | 42642.0 | 0 | Felidae | 42642.0 | 0 | Felis | 38442.0 | 1 | Felis catus | 38442.0 | 1 |
| scf7180000017929 | 630546 | 0.3756 | 0 | 49.7625 | Eukaryota | 43808.0 | 0 | Chordata | 43808.0 | 0 | Carnivora | 43808.0 | 0 | Felidae | 43808.0 | 0 | Felis | 39468.0 | 1 | Felis catus | 39468.0 | 1 |
| scf7180000022256 | 24582 | 0.4284 | 0 | 53.5579 | Eukaryota | 6284.0 | 0 | Chordata | 6284.0 | 0 | Primates | 4002.0 | 2 | Hominidae | 4002.0 | 2 | Homo | 2032.0 | 4 | Homo sapiens | 2032.0 | 4 |
| scf7180000019915 | 366454 | 0.4149 | 0 | 52.9748 | Eukaryota | 43614.0 | 0 | Chordata | 43614.0 | 0 | Carnivora | 43614.0 | 0 | Felidae | 26822.0 | 1 | Suricata | 16792.0 | 3 | ricata suricat | 16792.0 | 3 |
| scf7180000019688 | 217805 | 0.3677 | 0 | 24.1088 | Eukaryota | 41718.0 | 0 | Chordata | 41718.0 | 0 | Carnivora | 41718.0 | 0 | Felidae | 41718.0 | 0 | Felis | 41718.0 | 0 | Felis catus | 41718.0 | 0 |
| scf7180000023963 | 722026 | 0.4054 | 0 | 51.9276 | Eukaryota | 46000.0 | 0 | Chordata | 46000.0 | 0 | Carnivora | 46000.0 | 0 | Felidae | 46000.0 | 0 | Felis | 41401.0 | 1 | Felis catus | 41401.0 | 1 |
| scf7180000022435 | 11002 | 0.5376 | 0 | 51.3126 | Eukaryota | 2492.0 | 0 | Chordata | 2492.0 | 0 | Carnivora | 2492.0 | 0 | Felidae | 2242.0 | 1 | Felis | 1492.0 | 2 | Felis catus | 1492.0 | 2 |
| scf7180000018562 | 873281 | 0.3698 | 0 | 49.4879 | Eukaryota | 46449.0 | 0 | Chordata | 46449.0 | 0 | Carnivora | 46449.0 | 0 | Felidae | 46449.0 | 0 | Felis | 46449.0 | 0 | Felis catus | 46449.0 | 0 |
| scf7180000019921 | 1458422 | 0.4163 | 0 | 52.351 | Eukaryota | 46995.0 | 0 | Chordata | 46995.0 | 0 | Carnivora | 46995.0 | 0 | Felidae | 46995.0 | 0 | Felis | 46995.0 | 0 | Felis catus | 46995.0 | 0 |
| scf7180000021674 | 48901 | 0.4656 | 0 | 25.9032 | Eukaryota | 6188.0 | 0 | Chordata | 6188.0 | 0 | Primates | 2080.0 | 3 | Hominidae | 2080.0 | 5 | Homo | 2080.0 | 5 | Homo sapiens | 2080.0 | 5 |
| scf7180000017576 | 1926135 | 0.3479 | 0 | 48.5685 | Eukaryota | 45908.0 | 0 | Chordata | 45908.0 | 0 | Carnivora | 45908.0 | 0 | Felidae | 45908.0 | 0 | Felis | 45908.0 | 0 | Felis catus | 45908.0 | 0 |
| scf7180000023922 | 685958 | 0.4811 | 0 | 50.6764 | Eukaryota | 12772.0 | 0 | Chordata | 12772.0 | 0 | Carnivora | 12772.0 | 0 | Felidae | 12772.0 | 0 | Felis | 11506.0 | 1 | Felis catus | 11506.0 | 1 |
| scf7180000019725 | 272239 | 0.4286 | 0 | 49.1905 | Eukaryota | 31932.0 | 0 | Chordata | 31932.0 | 0 | Carnivora | 31932.0 | 0 | Felidae | 31932.0 | 0 | Felis | 28685.0 | 1 | Felis catus | 28685.0 | 1 |
| scf7180000023479 | 15044 | 0.6064 | 0 | 200.2237 | Eukaryota | 230.0 | 0 | Chordata | 230.0 | 0 | Carnivora | 230.0 | 0 | Felidae | 230.0 | 0 | Panthera | 230.0 | 0 | anthera pardu | 230.0 | 0 |
| scf7180000022539 | 14174 | 0.6344 | 0 | 52.1047 | Eukaryota | 2544.0 | 0 | Chordata | 2544.0 | 0 | Carnivora | 2255.0 | 1 | Felidae | 2255.0 | 1 | Felis | 1493.0 | 2 | Felis catus | 1493.0 | 2 |
| scf7180000018356 | 1056725 | 0.5781 | 0 | 53.3221 | Eukaryota | 104845.0 | 0 | Chordata | 104845.0 | 0 | Carnivora | 104845.0 | 0 | Felidae | 104845.0 | 0 | Felis | 62945.0 | 2 | Felis catus | 62945.0 | 2 |
| scf7180000020086 | 135414 | 0.417 | 0 | 24.7133 | Eukaryota | 15721.0 | 0 | Chordata | 15721.0 | 0 | Carnivora | 15721.0 | 0 | Otariidae | 6323.0 | 4 | Canis | 3178.0 | 6 | Canis lupus | 3178.0 | 6 |
| scf7180000020992 | 66991 | 0.3691 | 0 | 23.1145 | Eukaryota | 46352.0 | 0 | Chordata | 46352.0 | 0 | Carnivora | 46352.0 | 0 | Felidae | 46352.0 | 0 | Felis | 46352.0 | 0 | Felis catus | 46352.0 | 0 |
| scf7180000021736 | 5562 | 0.3585 | 0 | 18.9561 | Eukaryota | 1477.0 | 0 | Chordata | 1477.0 | 0 | Carnivora | 1477.0 | 0 | Felidae | 1477.0 | 0 | Felis | 1477.0 | 0 | Felis catus | 1477.0 | 0 |
| scf7180000021695 | 72978 | 0.3719 | 0 | 26.0256 | Eukaryota | 45945.0 | 0 | Chordata | 45945.0 | 0 | Carnivora | 45945.0 | 0 | Felidae | 45945.0 | 0 | Felis | 45945.0 | 0 | Felis catus | 45945.0 | 0 |
| scf7180000021646 | 55663 | 0.4292 | 0 | 30.3152 | Eukaryota | 42749.0 | 0 | Chordata | 42749.0 | 0 | Carnivora | 42749.0 | 0 | Felidae | 42749.0 | 0 | Felis | 42749.0 | 0 | Felis catus | 42749.0 | 0 |
| scf7180000022707 | 17807 | 0.5061 | 0 | 45.6995 | Eukaryota | 2796.0 | 0 | Chordata | 2796.0 | 0 | Carnivora | 2796.0 | 0 | Felidae | 2517.0 | 1 | Lynx | 889.0 | 5 | ynx canadens | 889.0 | 5 |
| scf7180000020101 | 156488 | 0.4088 | 0 | 25.8057 | Eukaryota | 57421.0 | 0 | Chordata | 57421.0 | 0 | Carnivora | 57421.0 | 0 | Felidae | 57421.0 | 0 | Felis | 34464.0 | 2 | Felis catus | 34464.0 | 2 |
| scf7180000021593 | 51762 | 0.4954 | 0 | 26.3362 | Eukaryota | 16979.0 | 0 | Chordata | 16979.0 | 0 | Carnivora | 16979.0 | 0 | Felidae | 16979.0 | 0 | Felis | 15292.0 | 1 | Felis catus | 15292.0 | 1 |
| scf7180000019695 | 257293 | 0.4552 | 0 | 49.5835 | Eukaryota | 45980.0 | 0 | Chordata | 45980.0 | 0 | Carnivora | 45980.0 | 0 | Felidae | 45980.0 | 0 | Felis | 41337.0 | 1 | Felis catus | 41337.0 | 1 |
| scf7180000018295 | 945539 | 0.3797 | 0 | 49.3181 | Eukaryota | 45833.0 | 0 | Chordata | 45833.0 | 0 | Carnivora | 45833.0 | 0 | Felidae | 45833.0 | 0 | Felis | 45833.0 | 0 | Felis catus | 45833.0 | 0 |
| scf7180000017964 | 1379650 | 0.451 | 0 | 51.3408 | Eukaryota | 54877.0 | 0 | Chordata | 54877.0 | 0 | Carnivora | 54877.0 | 0 | Felidae | 49981.0 | 1 | Acinonyx | 49981.0 | 1 | inonyx jubat | 49981.0 | 1 |
| scf7180000019481 | 436247 | 0.5484 | 0 | 49.2794 | Eukaryota | 36985.0 | 0 | Chordata | 36985.0 | 0 | Carnivora | 36985.0 | 0 | Felidae | 36985.0 | 0 | Felis | 36985.0 | 0 | Felis catus | 36985.0 | 0 |
| scf7180000021601 | 29743 | 0.6005 | 0 | 46.0578 | Eukaryota | 3107.0 | 0 | Chordata | 3107.0 | 0 | Carnivora | 3107.0 | 0 | Felidae | 3107.0 | 0 | Felis | 2826.0 | 1 | Felis catus | 2826.0 | 1 |
| scf7180000022680 | 5137 | 0.3693 | 0 | 24.1138 | Eukaryota | 1596.0 | 0 | Chordata | 1596.0 | 0 | Carnivora | 1158.0 | 1 | Felidae | 1158.0 | 1 | Felis | 598.0 | 3 | Felis catus | 598.0 | 3 |
| scf7180000022058 | 32338 | 0.3691 | 0 | 23.9783 | Eukaryota | 11284.0 | 0 | Chordata | 11284.0 | 0 | Carnivora | 11284.0 | 0 | Felidae | 6957.0 | 3 | Panthera | 2284.0 | 7 | anthera pardu | 2284.0 | 8 |
| scf7180000022779 | 13526 | 0.4444 | 0 | 24.9217 | Eukaryota | 6530.0 | 0 | Chordata | 6530.0 | 0 | Artiodactyla | 2805.0 | 2 | Bovidae | 2112.0 | 4 | Bos | 2112.0 | 4 | Bos mutus | 2112.0 | 4 |
| scf7180000020249 | 207555 | 0.3532 | 0 | 47.4574 | Eukaryota | 11969.0 | 0 | Chordata | 11969.0 | 0 | Carnivora | 11969.0 | 0 | Felidae | 11969.0 | 0 | Felis | 9571.0 | 1 | Felis catus | 9571.0 | 1 |
| scf7180000017533 | 113789 | 0.3807 | 0 | 25.1306 | Eukaryota | 24640.0 | 0 | Chordata | 24640.0 | 0 | Carnivora | 24640.0 | 0 | Felidae | 24640.0 | 0 | Felis | 22215.0 | 1 | Felis catus | 22215.0 | 1 |
| scf7180000018171 | 13147 | 0.4103 | 0 | 24.6989 | Eukaryota | 14022.0 | 0 | Chordata | 14022.0 | 0 | Carnivora | 14022.0 | 0 | Felidae | 14022.0 | 0 | Felis | 11240.0 | 1 | Felis catus | 11240.0 | 1 |
| scf7180000022070 | 25889 | 0.3574 | 0 | 23.1542 | Eukaryota | 23901.0 | 0 | Chordata | 23901.0 | 0 | Carnivora | 23901.0 | 0 | Felidae | 23901.0 | 0 | Felis | 21466.0 | 1 | Felis catus | 21466.0 | 1 |
| scf7180000020462 | 7362 | 0.733 | 0 | 33.0299 | Eukaryota | 300.0 | 0 | Chordata | 300.0 | 0 | Carnivora | 300.0 | 0 | Felidae | 300.0 | 0 | Panthera | 300.0 | 0 | anthera pardu | 300.0 | 0 |
| scf7180000021487 | 30352 | 0.5891 | 0 | 48.6155 | Eukaryota | 3241.0 | 0 | Chordata | 3241.0 | 0 | Primates | 1271.0 | 3 | Hominidae | 1271.0 | 4 | Homo | 1271.0 | 5 | Homo sapiens | 1271.0 | 5 |
| scf7180000018601 | 789723 | 0.4498 | 0 | 52.3686 | Eukaryota | 71438.0 | 0 | Chordata | 71438.0 | 0 | Carnivora | 71438.0 | 0 | Felidae | 71438.0 | 0 | Felis | 43127.0 | 1 | Felis catus | 43127.0 | 1 |
| scf7180000019844 | 205785 | 0.4138 | 0 | 26.929 | Eukaryota | 35110.0 | 0 | Chordata | 35110.0 | 0 | Carnivora | 35110.0 | 0 | Felidae | 30221.0 | 1 | Felis | 30221.0 | 1 | Felis catus | 30221.0 | 1 |
| scf7180000021447 | 41288 | 0.3885 | 0 | 24.9517 | Eukaryota | 17630.0 | 0 | Chordata | 17630.0 | 0 | Carnivora | 17630.0 | 0 | Felidae | 17630.0 | 0 | Felis | 15855.0 | 1 | Felis catus | 15855.0 | 1 |
| scf7180000020891 | 26108 | 0.5177 | 0 | 201.1175 | Eukaryota | 12270.0 | 0 | Chordata | 12270.0 | 0 | Carnivora | 12270.0 | 0 | Felidae | 12270.0 | 0 | Felis | 11060.0 | 1 | Felis catus | 11060.0 | 1 |
| scf7180000018733 | 486624 | 0.3872 | 0 | 49.6463 | Eukaryota | 45200.0 | 0 | Chordata | 45200.0 | 0 | Carnivora | 45200.0 | 0 | Felidae | 45200.0 | 0 | Felis | 45200.0 | 0 | Felis catus | 45200.0 | 0 |
| scf7180000018644 | 356874 | 0.3715 | 0 | 49.7714 | Eukaryota | 34758.0 | 0 | Chordata | 34758.0 | 0 | Carnivora | 34758.0 | 0 | Felidae | 34758.0 | 0 | Lynx | 27854.0 | 1 | ynx canadens | 27854.0 | 1 |
| scf7180000022519 | 19754 | 0.4011 | 0 | 34.179 | Eukaryota | 46456.0 | 0 | Chordata | 46456.0 | 0 | Carnivora | 46456.0 | 0 | Felidae | 46456.0 | 0 | Felis | 46456.0 | 0 | Felis catus | 46456.0 | 0 |
| scf7180000018993 | 561855 | 0.4303 | 0 | 51.3636 | Eukaryota | 53052.0 | 0 | Chordata | 53052.0 | 0 | Carnivora | 53052.0 | 0 | Mustelidae | 29625.0 | 1 | Enhydra | 29625.0 | 1 | Enhydra lutris | 29625.0 | 1 |
| scf7180000020715 | 67818 | 0.3672 | 0 | 24.8891 | Eukaryota | 31061.0 | 0 | Chordata | 31061.0 | 0 | Carnivora | 31061.0 | 0 | Felidae | 31061.0 | 0 | Felis | 27899.0 | 1 | Felis catus | 27899.0 | 1 |
| scf7180000019039 | 444190 | 0.3569 | 0 | 48.845 | Eukaryota | 74803.0 | 0 | Chordata | 74803.0 | 0 | Carnivora | 61987.0 | 1 | Felidae | 51479.0 | 2 | Felis | 27673.0 | 3 | Felis catus | 27673.0 | 3 |
| scf7180000019717 | 253112 | 0.3484 | 0 | 49.0199 | Eukaryota | 18346.0 | 0 | Chordata | 18346.0 | 0 | Carnivora | 14667.0 | 1 | Felidae | 14667.0 | 1 | Felis | 11439.0 | 2 | Felis catus | 11439.0 | 2 |
| scf7180000021075 | 81435 | 0.557 | 0 | 51.9202 | Eukaryota | 17619.0 | 0 | Chordata | 17619.0 | 0 | Carnivora | 15984.0 | 1 | Otariidae | 5370.0 | 5 | Panthera | 3591.0 | 8 | anthera pardu | 3591.0 | 8 |
| scf7180000022682 | 27326 | 0.4177 | 0 | 24.3367 | Eukaryota | 20333.0 | 0 | Chordata | 20333.0 | 0 | Carnivora | 20333.0 | 0 | Felidae | 20333.0 | 0 | Felis | 20333.0 | 0 | Felis catus | 20333.0 | 0 |
| scf7180000021180 | 43831 | 0.3819 | 0 | 25.1444 | Eukaryota | 5826.0 | 0 | Chordata | 5826.0 | 0 | Carnivora | 4324.0 | 2 | Felidae | 2583.0 | 3 | Felis | 2583.0 | 3 | Felis catus | 2583.0 | 3 |
| scf7180000022036 | 29842 | 0.3673 | 0 | 25.395 | Eukaryota | 32855.0 | 0 | Chordata | 32855.0 | 0 | Carnivora | 32855.0 | 0 | Felidae | 32855.0 | 0 | Felis | 29612.0 | 1 | Felis catus | 29612.0 | 1 |
| scf7180000018523 | 85548 | 0.3624 | 0 | 50.4871 | Eukaryota | 46523.0 | 0 | Chordata | 46523.0 | 0 | Carnivora | 46523.0 | 0 | Felidae | 46523.0 | 0 | Felis | 46523.0 | 0 | Felis catus | 46523.0 | 0 |
| scf7180000019420 | 206555 | 0.4133 | 0 | 52.646 | Eukaryota | 59558.0 | 0 | Chordata | 59558.0 | 0 | Carnivora | 59558.0 | 0 | Felidae | 59558.0 | 0 | Felis | 35724.0 | 1 | Felis catus | 35724.0 | 2 |
| scf7180000021868 | 7213 | 0.5211 | 0 | 162.6077 | no-hit | 0.0 | 0 | no-hit | 0.0 | 0 | no-hit | 0.0 | 0 | no-hit | 0.0 | 0 | no-hit | 0.0 | 0 | no-hit | 0.0 | 0 |
| scf7180000023876 | 776665 | 0.4747 | 0 | 51.7436 | Eukaryota | 46967.0 | 0 | Chordata | 46967.0 | 0 | Carnivora | 46967.0 | 0 | Felidae | 46967.0 | 0 | Felis | 46967.0 | 0 | Felis catus | 46967.0 | 0 |
| scf7180000023762 | 2117783 | 0.3773 | 0 | 49.5939 | Eukaryota | 49231.0 | 0 | Chordata | 49231.0 | 0 | Carnivora | 49231.0 | 0 | Felidae | 49231.0 | 0 | Felis | 49231.0 | 0 | Felis catus | 49231.0 | 0 |
| scf7180000018321 | 619954 | 0.4216 | 0 | 50.2964 | Eukaryota | 40999.0 | 0 | Chordata | 40999.0 | 0 | Carnivora | 40999.0 | 0 | Felidae | 40999.0 | 0 | Felis | 40999.0 | 0 | Felis catus | 40999.0 | 0 |
| scf7180000023504 | 19271 | 0.4913 | 0 | 316.6503 | Eukaryota | 12744.0 | 0 | Chordata | 12744.0 | 0 | Carnivora | 12744.0 | 0 | Felidae | 12744.0 | 0 | Felis | 12744.0 | 0 | Felis catus | 12744.0 | 0 |
| scf7180000017693 | 333175 | 0.4154 | 0 | 49.7223 | Eukaryota | 52053.0 | 0 | Chordata | 52053.0 | 0 | Carnivora | 52053.0 | 0 | Felidae | 26307.0 | 4 | Panthera | 10499.0 | 8 | llorhinus ursi | 5323.0 | 9 |
| scf7180000019781 | 14341 | 0.4073 | 0 | 25.4901 | Eukaryota | 9320.0 | 0 | Chordata | 9320.0 | 0 | Carnivora | 9320.0 | 0 | Felidae | 9320.0 | 0 | Felis | 9320.0 | 0 | Felis catus | 9320.0 | 0 |
| scf7180000019346 | 2260975 | 0.4374 | 100 | 51.7695 | Eukaryota | 61038.0 | 0 | Chordata | 61038.0 | 0 | Carnivora | 61038.0 | 0 | Felidae | 61038.0 | 0 | Lynx | 36690.0 | 1 | ynx canadens | 36690.0 | 1 |
| scf7180000019736 | 212044 | 0.4149 | 0 | 25.7974 | Eukaryota | 9835.0 | 0 | Chordata | 9835.0 | 0 | Carnivora | 9835.0 | 0 | Felidae | 9835.0 | 0 | Felis | 7895.0 | 1 | Felis catus | 7895.0 | 1 |
| scf7180000020356 | 184446 | 0.3808 | 0 | 50.7209 | Eukaryota | 35378.0 | 0 | Chordata | 35378.0 | 0 | Carnivora | 35378.0 | 0 | Felidae | 35378.0 | 0 | Felis | 31789.0 | 1 | Felis catus | 31789.0 | 1 |
| scf7180000021300 | 78928 | 0.4284 | 0 | 24.6578 | Eukaryota | 21584.0 | 0 | Chordata | 21584.0 | 0 | Carnivora | 21584.0 | 0 | Felidae | 21584.0 | 0 | Felis | 19414.0 | 1 | Felis catus | 19414.0 | 1 |
| scf7180000022658 | 12816 | 0.4361 | 0 | 26.7658 | Eukaryota | 4531.0 | 0 | Chordata | 4531.0 | 0 | Carnivora | 4531.0 | 0 | Felidae | 2771.0 | 1 | Felis | 2771.0 | 1 | Felis catus | 2771.0 | 1 |
| scf7180000017737 | 1828884 | 0.4289 | 0 | 51.7007 | Eukaryota | 48568.0 | 0 | Chordata | 48568.0 | 0 | Carnivora | 48568.0 | 0 | Canidae | 34016.0 | 1 | Canis | 34016.0 | 2 | Canis lupus | 34016.0 | 2 |
| scf7180000019372 | 262234 | 0.3911 | 0 | 51.9092 | Eukaryota | 34941.0 | 0 | Chordata | 34941.0 | 0 | Primates | 23858.0 | 2 | Hominidae | 23858.0 | 3 | Homo | 23858.0 | 4 | Homo sapiens | 23858.0 | 4 |
| scf7180000020288 | 114223 | 0.6046 | 0 | 47.2956 | Eukaryota | 2857.0 | 1 | Chordata | 2857.0 | 1 | Carnivora | 1348.0 | 3 | Felidae | 1348.0 | 3 | Felis | 1348.0 | 4 | Felis catus | 1348.0 | 4 |
| scf7180000022608 | 13569 | 0.5274 | 0 | 45.5906 | Eukaryota | 2603.0 | 0 | Chordata | 2603.0 | 0 | Carnivora | 2603.0 | 0 | Felidae | 2344.0 | 1 | Felis | 1557.0 | 2 | Felis catus | 1557.0 | 2 |
| scf7180000019250 | 41845 | 0.3993 | 0 | 48.8902 | Eukaryota | 38843.0 | 0 | Chordata | 38843.0 | 0 | Carnivora | 38843.0 | 0 | Felidae | 38843.0 | 0 | Felis | 38843.0 | 0 | Felis catus | 38843.0 | 0 |
| scf7180000020022 | 233694 | 0.4105 | 0 | 51.7131 | Eukaryota | 46860.0 | 0 | Chordata | 46860.0 | 0 | Carnivora | 46860.0 | 0 | Felidae | 46860.0 | 0 | Felis | 46860.0 | 0 | Felis catus | 46860.0 | 0 |
| scf7180000020591 | 111387 | 0.4226 | 0 | 25.5047 | Eukaryota | 26246.0 | 0 | Chordata | 26246.0 | 0 | Carnivora | 26246.0 | 0 | Felidae | 13601.0 | 4 | Panthera | 5345.0 | 8 | ynx canadens | 2767.0 | 9 |
| scf7180000019361 | 331703 | 0.4869 | 0 | 51.1976 | Eukaryota | 28416.0 | 0 | Chordata | 28416.0 | 0 | Carnivora | 28416.0 | 0 | Felidae | 28416.0 | 0 | Acinonyx | 16956.0 | 1 | inonyx jubat | 16956.0 | 1 |
| scf7180000021379 | 42976 | 0.4404 | 0 | 26.7245 | Eukaryota | 38892.0 | 0 | Chordata | 38892.0 | 0 | Carnivora | 38892.0 | 0 | Felidae | 38892.0 | 0 | Felis | 35026.0 | 1 | Felis catus | 35026.0 | 1 |
| scf7180000024009 | 174898 | 0.4372 | 0 | 55.7017 | Eukaryota | 50943.0 | 0 | Chordata | 50943.0 | 0 | Carnivora | 50943.0 | 0 | Felidae | 47064.0 | 1 | Panthera | 26200.0 | 2 | anthera pardu | 26200.0 | 2 |
| scf7180000019996 | 24772 | 0.4931 | 0 | 397.3273 | Eukaryota | 6667.0 | 0 | Chordata | 6667.0 | 0 | Carnivora | 6667.0 | 0 | Felidae | 6667.0 | 0 | Felis | 3565.0 | 2 | Felis catus | 3565.0 | 2 |
| scf7180000019274 | 305710 | 0.523 | 0 | 54.6654 | Eukaryota | 69245.0 | 0 | Chordata | 69245.0 | 0 | Carnivora | 69245.0 | 0 | Felidae | 43420.0 | 1 | Panthera | 43420.0 | 1 | anthera pardu | 43420.0 | 1 |
| scf7180000021042 | 27987 | 0.3692 | 0 | 246.52 | Eukaryota | 46855.0 | 0 | Chordata | 46855.0 | 0 | Carnivora | 46855.0 | 0 | Felidae | 46855.0 | 0 | Felis | 46855.0 | 0 | Felis catus | 46855.0 | 0 |
| scf7180000020786 | 72781 | 0.394 | 0 | 49.2369 | Eukaryota | 3813.0 | 1 | Chordata | 3813.0 | 1 | Primates | 2863.0 | 2 | Hominidae | 2489.0 | 3 | Homo | 2489.0 | 4 | Homo sapiens | 2489.0 | 4 |
| scf7180000017711 | 1902908 | 0.4207 | 0 | 51.8172 | Eukaryota | 74649.0 | 0 | Chordata | 74649.0 | 0 | Carnivora | 74649.0 | 0 | Felidae | 74649.0 | 0 | Panthera | 55450.0 | 1 | anthera pardu | 55450.0 | 1 |
| scf7180000019886 | 177626 | 0.4561 | 0 | 51.3628 | Eukaryota | 21691.0 | 0 | Chordata | 21691.0 | 0 | Carnivora | 21691.0 | 0 | Felidae | 19028.0 | 1 | Felis | 16876.0 | 2 | Felis catus | 16876.0 | 2 |
| scf7180000018985 | 474842 | 0.4642 | 0 | 52.6693 | Eukaryota | 62195.0 | 0 | Chordata | 62195.0 | 0 | Carnivora | 62195.0 | 0 | Felidae | 62195.0 | 0 | Panthera | 43679.0 | 1 | anthera pardu | 37464.0 | 2 |
| scf7180000019294 | 692398 | 0.5321 | 0 | 50.9918 | Eukaryota | 17475.0 | 0 | Chordata | 17475.0 | 0 | Carnivora | 17475.0 | 0 | Felidae | 17475.0 | 0 | Felis | 10195.0 | 3 | Felis catus | 10195.0 | 3 |

| | | | | | | | | | | | | | | | | | | | | | | |
|---|---|---|---|---|---|---|---|---|---|---|---|---|---|---|---|---|---|---|---|---|---|---|
| scf7180000020642 | 142113 | 0.5673 | 0 | 49.8345 | Eukaryota | 19304.0 | 0 | Chordata | 19304.0 | 0 | Carnivora | 19304.0 | 0 | Felidae | 19304.0 | 0 | Acinonyx | 7796.0 | 2 | cinonyx jubat | 7796.0 | 2 |
| scf7180000020001 | 997080 | 0.4796 | 0 | 52.2375 | Eukaryota | 52457.0 | 0 | Chordata | 52457.0 | 0 | Carnivora | 52457.0 | 0 | Felidae | 52457.0 | 0 | Lynx | 20876.0 | 3 | ynx canadens | 20876.0 | 3 |
| scf7180000021718 | 37697 | 0.5116 | 0 | 50.9346 | Eukaryota | 6480.0 | 1 | Chordata | 6480.0 | 1 | Primates | 2329.0 | 5 | Hominidae | 2329.0 | 5 | Homo | 2329.0 | 6 | Homo sapiens | 2329.0 | 6 |
| scf7180000018133 | 256073 | 0.3612 | 0 | 47.8215 | Eukaryota | 41186.0 | 0 | Chordata | 41186.0 | 0 | Carnivora | 41186.0 | 0 | Felidae | 41186.0 | 0 | Felis | 37023.0 | 1 | Felis catus | 37023.0 | 1 |
| scf7180000022445 | 28228 | 0.5241 | 0 | 51.4187 | Eukaryota | 5657.0 | 0 | Chordata | 5657.0 | 0 | Carnivora | 5657.0 | 0 | Felidae | 5657.0 | 0 | Felis | 5657.0 | 0 | Felis catus | 5657.0 | 0 |
| scf7180000017910 | 987050 | 0.418 | 0 | 49.9224 | Eukaryota | 46395.0 | 0 | Chordata | 46395.0 | 0 | Carnivora | 46395.0 | 0 | Felidae | 46395.0 | 0 | Felis | 46395.0 | 0 | Felis catus | 46395.0 | 0 |
| scf7180000023988 | 223290 | 0.4175 | 0 | 47.1258 | Eukaryota | 26820.0 | 0 | Chordata | 26820.0 | 0 | Carnivora | 26820.0 | 0 | Felidae | 26820.0 | 0 | Panthera | 9154.0 | 4 | anthera pardu | 7039.0 | 5 |
| scf7180000024240 | 617510 | 0.4394 | 0 | 51.5225 | Eukaryota | 29546.0 | 0 | Chordata | 29546.0 | 0 | Carnivora | 29546.0 | 0 | Felidae | 29546.0 | 0 | Felis | 29546.0 | 0 | Felis catus | 29546.0 | 0 |
| scf7180000021564 | 47900 | 0.546 | 0 | 42.8573 | Eukaryota | 14469.0 | 0 | Chordata | 14469.0 | 0 | Carnivora | 14469.0 | 0 | Felidae | 13150.0 | 1 | Felis | 5037.0 | 5 | Felis catus | 5037.0 | 6 |
| scf7180000023740 | 39503 | 0.4846 | 0 | 271.6874 | Eukaryota | 9316.0 | 0 | Chordata | 9316.0 | 0 | Carnivora | 9316.0 | 0 | Felidae | 9316.0 | 0 | Felis | 9316.0 | 0 | Felis catus | 9316.0 | 0 |
| scf7180000020238 | 189972 | 0.5198 | 0 | 48.2632 | Eukaryota | 12924.0 | 1 | Chordata | 12924.0 | 1 | Primates | 10072.0 | 2 | Hominidae | 8047.0 | 3 | Homo | 5485.0 | 4 | Homo sapiens | 5485.0 | 4 |
| scf7180000020241 | 107872 | 0.5313 | 0 | 50.4277 | Eukaryota | 19183.0 | 0 | Chordata | 19183.0 | 0 | Carnivora | 19183.0 | 0 | Felidae | 17267.0 | 1 | Panthera | 8475.0 | 4 | Panthera tigri | 6811.0 | 5 |
| scf7180000019617 | 196506 | 0.4042 | 0 | 50.0981 | Eukaryota | 13099.0 | 0 | Chordata | 13099.0 | 0 | Carnivora | 13099.0 | 0 | Felidae | 6570.0 | 4 | Panthera | 2642.0 | 7 | anthera pardu | 2642.0 | 7 |
| scf7180000017438 | 177693 | 0.3802 | 0 | 50.0272 | Eukaryota | 45185.0 | 0 | Chordata | 45185.0 | 0 | Carnivora | 45185.0 | 0 | Felidae | 45185.0 | 0 | Felis | 40714.0 | 1 | Felis catus | 40714.0 | 1 |
| scf7180000018326 | 939523 | 0.4417 | 0 | 50.5198 | Eukaryota | 24549.0 | 0 | Chordata | 24549.0 | 0 | Carnivora | 24549.0 | 0 | Felidae | 8752.0 | 4 | Callorhinus | 3743.0 | 8 | llorhinus ursir | 3743.0 | 8 |
| scf7180000019500 | 62537 | 0.3789 | 0 | 24.6381 | Eukaryota | 9936.0 | 0 | Chordata | 9936.0 | 0 | Artiodactyla | 3770.0 | 3 | Suidae | 3770.0 | 3 | Sus | 3770.0 | 3 | Sus scrofa | 3770.0 | 3 |
| scf7180000024093 | 209827 | 0.5245 | 0 | 48.3519 | Eukaryota | 38328.0 | 0 | Chordata | 38328.0 | 0 | Carnivora | 38328.0 | 0 | Felidae | 38328.0 | 0 | Felis | 23106.0 | 1 | Felis catus | 23106.0 | 1 |
| scf7180000018729 | 909122 | 0.473 | 0 | 52.051 | Eukaryota | 31578.0 | 0 | Chordata | 31578.0 | 0 | Carnivora | 31578.0 | 0 | Felidae | 31578.0 | 0 | Felis | 12165.0 | 3 | Felis catus | 12165.0 | 4 |
| scf7180000019015 | 378896 | 0.3577 | 0 | 46.3113 | Eukaryota | 48275.0 | 0 | Chordata | 48275.0 | 0 | Carnivora | 48275.0 | 0 | Felidae | 48275.0 | 0 | Felis | 48275.0 | 0 | Felis catus | 48275.0 | 0 |
| scf7180000019104 | 509396 | 0.4086 | 0 | 52.3038 | Eukaryota | 49208.0 | 0 | Chordata | 49208.0 | 0 | Carnivora | 49208.0 | 0 | Felidae | 49208.0 | 0 | Felis | 49208.0 | 0 | Felis catus | 49208.0 | 0 |
| scf7180000023209 | 29988 | 0.4738 | 0 | 116.2341 | Eukaryota | 12018.0 | 0 | Chordata | 12018.0 | 0 | Carnivora | 12018.0 | 0 | Felidae | 12018.0 | 0 | Felis | 10832.0 | 1 | Felis catus | 10832.0 | 1 |
| scf7180000018845 | 36777 | 0.4813 | 0 | 45.3042 | Eukaryota | 8319.0 | 0 | Chordata | 8319.0 | 0 | Carnivora | 8319.0 | 0 | Felidae | 8319.0 | 0 | Felis | 7515.0 | 1 | Felis catus | 7515.0 | 1 |
| scf7180000022207 | 24732 | 0.4329 | 0 | 38.9879 | Eukaryota | 6125.0 | 0 | Chordata | 6125.0 | 0 | Carnivora | 6125.0 | 0 | Felidae | 6125.0 | 0 | Felis | 4628.0 | 2 | Felis catus | 4628.0 | 2 |
| scf7180000021428 | 52557 | 0.5337 | 0 | 49.0761 | Eukaryota | 9830.0 | 0 | Chordata | 9830.0 | 0 | Carnivora | 9830.0 | 0 | Felidae | 9830.0 | 0 | Felis | 9830.0 | 0 | Felis catus | 9830.0 | 0 |
| scf7180000018293 | 505671 | 0.4192 | 0 | 50.07 | Eukaryota | 44566.0 | 0 | Chordata | 44566.0 | 0 | Carnivora | 44566.0 | 0 | Felidae | 44566.0 | 0 | Felis | 40095.0 | 1 | Felis catus | 40095.0 | 1 |
| scf7180000020769 | 25384 | 0.4181 | 0 | 175.9995 | Eukaryota | 46746.0 | 0 | Chordata | 46746.0 | 0 | Carnivora | 46746.0 | 0 | Felidae | 46746.0 | 0 | Felis | 46746.0 | 0 | Felis catus | 46746.0 | 0 |
| scf7180000019285 | 142610 | 0.3771 | 0 | 394.0648 | Eukaryota | 36232.0 | 0 | Chordata | 36232.0 | 0 | Carnivora | 36232.0 | 0 | Felidae | 36232.0 | 0 | Felis | 32523.0 | 1 | Felis catus | 32523.0 | 1 |
| scf7180000019511 | 347180 | 0.4625 | 0 | 50.3652 | Eukaryota | 14487.0 | 1 | Chordata | 14487.0 | 1 | Primates | 5270.0 | 5 | Hominidae | 5270.0 | 5 | Homo | 5270.0 | 6 | Homo sapiens | 5270.0 | 6 |
| scf7180000017760 | 1767277 | 0.4276 | 0 | 52.9805 | Eukaryota | 45748.0 | 0 | Chordata | 45748.0 | 0 | Carnivora | 45748.0 | 0 | Felidae | 45748.0 | 0 | Felis | 45748.0 | 0 | Felis catus | 45748.0 | 0 |
| scf7180000017936 | 1230951 | 0.4115 | 0 | 51.5522 | Eukaryota | 46950.0 | 0 | Chordata | 46950.0 | 0 | Carnivora | 46950.0 | 0 | Felidae | 46950.0 | 0 | Felis | 46950.0 | 0 | Felis catus | 46950.0 | 0 |
| scf7180000019203 | 28309 | 0.4852 | 0 | 186.9933 | Eukaryota | 5205.0 | 0 | Chordata | 5205.0 | 0 | Carnivora | 5205.0 | 0 | Felidae | 5205.0 | 0 | Felis | 5205.0 | 0 | Felis catus | 5205.0 | 0 |
| scf7180000019964 | 123770 | 0.4436 | 0 | 83.7585 | Eukaryota | 46776.0 | 0 | Chordata | 46776.0 | 0 | Carnivora | 46776.0 | 0 | Felidae | 46776.0 | 0 | Felis | 46776.0 | 0 | Felis catus | 46776.0 | 0 |
| scf7180000021292 | 47284 | 0.389 | 0 | 50.3118 | Eukaryota | 10173.0 | 0 | Chordata | 10173.0 | 0 | Carnivora | 5272.0 | 2 | Felidae | 4695.0 | 4 | Felis | 4055.0 | 5 | Felis catus | 4055.0 | 5 |
| scf7180000019076 | 8088 | 0.3813 | 0 | 26.5307 | Eukaryota | 33268.0 | 0 | Chordata | 33268.0 | 0 | Carnivora | 33268.0 | 0 | Felidae | 33268.0 | 0 | Felis | 29949.0 | 1 | Felis catus | 29949.0 | 1 |
| scf7180000021948 | 45978 | 0.3842 | 0 | 25.5071 | Eukaryota | 28671.0 | 0 | Chordata | 28671.0 | 0 | Carnivora | 28671.0 | 0 | Felidae | 28671.0 | 0 | Felis | 28671.0 | 0 | Felis catus | 28671.0 | 0 |
| scf7180000019785 | 266986 | 0.3845 | 0 | 50.1978 | Eukaryota | 30347.0 | 1 | Chordata | 30347.0 | 1 | Carnivora | 23543.0 | 2 | Otariidae | 23543.0 | 2 | Callorhinus | 10119.0 | 4 | llorhinus ursir | 10119.0 | 4 |
| scf7180000022363 | 16357 | 0.3808 | 0 | 25.1477 | Eukaryota | 24008.0 | 0 | Chordata | 24008.0 | 0 | Carnivora | 24008.0 | 0 | Felidae | 24008.0 | 0 | Felis | 21609.0 | 1 | Felis catus | 21609.0 | 1 |
| scf7180000019787 | 265184 | 0.3815 | 0 | 49.2962 | Eukaryota | 45055.0 | 0 | Chordata | 45055.0 | 0 | Carnivora | 45055.0 | 0 | Felidae | 45055.0 | 0 | Felis | 45055.0 | 0 | Felis catus | 45055.0 | 0 |
| scf7180000022728 | 5699 | 0.3773 | 0 | 24.8843 | Eukaryota | 8954.0 | 0 | Chordata | 8954.0 | 0 | Carnivora | 8954.0 | 0 | Felidae | 8954.0 | 0 | Felis | 7176.0 | 1 | Felis catus | 7176.0 | 1 |
| scf7180000017476 | 3473208 | 0.3937 | 0 | 50.5671 | Eukaryota | 122213.0 | 0 | Chordata | 122213.0 | 0 | Artiodactyla | 61084.0 | 2 | Delphinidae | 61084.0 | 2 | Globicephala | 42140.0 | 3 | bicephala me | 42140.0 | 3 |
| scf7180000021918 | 40433 | 0.3672 | 0 | 24.6346 | Eukaryota | 43759.0 | 0 | Chordata | 43759.0 | 0 | Carnivora | 43759.0 | 0 | Felidae | 43759.0 | 0 | Felis | 43759.0 | 0 | Felis catus | 43759.0 | 0 |
| scf7180000022923 | 4367440 | 0.3797 | 0 | 50.4503 | Eukaryota | 50211.0 | 0 | Chordata | 50211.0 | 0 | Carnivora | 50211.0 | 0 | Felidae | 50211.0 | 0 | Felis | 33801.0 | 3 | Felis catus | 33801.0 | 3 |
| scf7180000019113 | 127165 | 0.3541 | 0 | 46.8467 | Eukaryota | 39972.0 | 0 | Chordata | 39972.0 | 0 | Carnivora | 39972.0 | 0 | Felidae | 39972.0 | 0 | Felis | 39972.0 | 0 | Felis catus | 39972.0 | 0 |
| scf7180000024173 | 109216 | 0.5705 | 0 | 50.095 | Eukaryota | 22369.0 | 0 | Chordata | 22369.0 | 0 | Carnivora | 22369.0 | 0 | Otariidae | 15451.0 | 3 | Zalophus | 15451.0 | 3 | phus californi | 15451.0 | 3 |
| scf7180000021657 | 35489 | 0.37 | 0 | 25.6745 | Eukaryota | 14363.0 | 0 | Chordata | 14363.0 | 0 | Carnivora | 14363.0 | 0 | Felidae | 14363.0 | 0 | Felis | 12940.0 | 1 | Felis catus | 12940.0 | 1 |
| scf7180000021174 | 59627 | 0.4023 | 0 | 23.2442 | Eukaryota | 22715.0 | 0 | Chordata | 22715.0 | 0 | Carnivora | 22715.0 | 0 | Felidae | 13940.0 | 3 | Suricata | 5112.0 | 6 | ricata suricat | 5112.0 | 7 |
| scf7180000019392 | 110512 | 0.4836 | 0 | 51.3529 | Eukaryota | 15369.0 | 0 | Chordata | 15369.0 | 0 | Carnivora | 15369.0 | 0 | Felidae | 7094.0 | 2 | Suricata | 5520.0 | 6 | ricata suricat | 5520.0 | 7 |
| scf7180000024175 | 123789 | 0.5179 | 0 | 27.7329 | Eukaryota | 16325.0 | 0 | Chordata | 16325.0 | 0 | Carnivora | 16325.0 | 0 | Felidae | 12421.0 | 1 | Felis | 12421.0 | 1 | Felis catus | 12421.0 | 1 |
| scf7180000020329 | 94685 | 0.5086 | 0 | 55.1206 | Eukaryota | 25719.0 | 0 | Chordata | 25719.0 | 0 | Carnivora | 25719.0 | 0 | Felidae | 25719.0 | 0 | Felis | 23530.0 | 1 | Felis catus | 23530.0 | 1 |
| scf7180000018233 | 1027118 | 0.4326 | 0 | 52.0223 | Eukaryota | 42159.0 | 0 | Chordata | 42159.0 | 0 | Carnivora | 42159.0 | 0 | Felidae | 42159.0 | 0 | Felis | 42159.0 | 0 | Felis catus | 42159.0 | 0 |
| scf7180000022244 | 16765 | 0.4201 | 0 | 23.9525 | Eukaryota | 2877.0 | 0 | Chordata | 2877.0 | 0 | Carnivora | 2439.0 | 1 | Felidae | 2439.0 | 1 | Felis | 2176.0 | 2 | Felis catus | 2176.0 | 2 |
| scf7180000021632 | 69914 | 0.4864 | 0 | 49.8789 | Eukaryota | 26280.0 | 0 | Chordata | 26280.0 | 0 | Carnivora | 26280.0 | 0 | Felidae | 26280.0 | 0 | Felis | 23617.0 | 1 | Felis catus | 23617.0 | 1 |
| scf7180000019310 | 396559 | 0.3851 | 0 | 50.6679 | Eukaryota | 47222.0 | 0 | Chordata | 47222.0 | 0 | Carnivora | 47222.0 | 0 | Felidae | 47222.0 | 0 | Felis | 47222.0 | 0 | Felis catus | 47222.0 | 0 |
| scf7180000023367 | 18198 | 0.425 | 0 | 94.3313 | Eukaryota | 2147.0 | 0 | Chordata | 2147.0 | 0 | Carnivora | 2147.0 | 0 | Felidae | 2147.0 | 0 | Felis | 1941.0 | 1 | Felis catus | 1941.0 | 1 |
| scf7180000019811 | 264023 | 0.4312 | 0 | 47.8279 | Eukaryota | 29960.0 | 0 | Chordata | 29960.0 | 0 | Carnivora | 29960.0 | 0 | Felidae | 29960.0 | 0 | Felis | 27010.0 | 1 | Felis catus | 27010.0 | 1 |
| scf7180000019188 | 468983 | 0.4361 | 0 | 50.2609 | Eukaryota | 20614.0 | 0 | Chordata | 20614.0 | 0 | Carnivora | 20614.0 | 0 | Felidae | 17160.0 | 2 | Panthera | 6584.0 | 4 | anthera pardu | 6584.0 | 4 |
| scf7180000021793 | 37292 | 0.5691 | 0 | 53.9666 | Eukaryota | 4009.0 | 0 | Chordata | 4009.0 | 0 | Carnivora | 4009.0 | 0 | Felidae | 1591.0 | 3 | Suricata | 1236.0 | 6 | ricata suricat | 1236.0 | 6 |
| scf7180000023913 | 426492 | 0.3455 | 0 | 48.2933 | Eukaryota | 45158.0 | 0 | Chordata | 45158.0 | 0 | Carnivora | 45158.0 | 0 | Felidae | 45158.0 | 0 | Felis | 45158.0 | 0 | Felis catus | 45158.0 | 0 |
| scf7180000024264 | 2888018 | 0.3968 | 0 | 50.5486 | Eukaryota | 47734.0 | 0 | Chordata | 47734.0 | 0 | Carnivora | 47734.0 | 0 | Felidae | 47734.0 | 0 | Felis | 47734.0 | 0 | Felis catus | 47734.0 | 0 |
| scf7180000017778 | 3484760 | 0.4066 | 100 | 51.0637 | Eukaryota | 47706.0 | 0 | Chordata | 47706.0 | 0 | Carnivora | 47706.0 | 0 | Felidae | 47706.0 | 0 | Felis | 42978.0 | 1 | Felis catus | 42978.0 | 1 |
| scf7180000022796 | 13174 | 0.4354 | 0 | 4736.573 | Eukaryota | 8369.0 | 0 | Chordata | 8369.0 | 0 | Artiodactyla | 8369.0 | 0 | Bovidae | 8369.0 | 0 | Ovis | 8369.0 | 0 | vis canadens | 8369.0 | 0 |
| scf7180000021204 | 50639 | 0.4574 | 0 | 50.4437 | Eukaryota | 9210.0 | 0 | Chordata | 9210.0 | 0 | Carnivora | 9210.0 | 0 | Felidae | 9210.0 | 0 | Felis | 9210.0 | 0 | Felis catus | 9210.0 | 0 |
| scf7180000019765 | 500924 | 0.557 | 0 | 47.9499 | Eukaryota | 9927.0 | 0 | Chordata | 9393.0 | 1 | Carnivora | 4862.0 | 3 | Felidae | 4862.0 | 3 | Homo | 3975.0 | 4 | Homo sapiens | 3975.0 | 4 |
| scf7180000018566 | 590143 | 0.3824 | 0 | 51.3694 | Eukaryota | 43807.0 | 0 | Chordata | 43807.0 | 0 | Carnivora | 43807.0 | 0 | Felidae | 43807.0 | 0 | Felis | 43807.0 | 0 | Felis catus | 43807.0 | 0 |
| scf7180000023083 | 78651 | 0.5677 | 0 | 130.4276 | Eukaryota | 4053.0 | 0 | Chordata | 4053.0 | 0 | Carnivora | 3487.0 | 1 | Felidae | 3224.0 | 2 | Acinonyx | 1792.0 | 4 | cinonyx jubat | 1792.0 | 4 |
| scf7180000017505 | 855966 | 0.3703 | 0 | 50.6823 | Eukaryota | 46804.0 | 0 | Chordata | 46804.0 | 0 | Carnivora | 46804.0 | 0 | Felidae | 46804.0 | 0 | Felis | 42150.0 | 1 | Felis catus | 42150.0 | 1 |
| scf7180000018318 | 903106 | 0.4348 | 0 | 52.1107 | Eukaryota | 80177.0 | 0 | Chordata | 80177.0 | 0 | Carnivora | 73303.0 | 1 | Felidae | 66822.0 | 2 | Felis | 60452.0 | 3 | Felis catus | 60452.0 | 3 |
| scf7180000018137 | 562232 | 0.4531 | 0 | 71.0772 | Eukaryota | 45739.0 | 0 | Chordata | 45739.0 | 0 | Carnivora | 45739.0 | 0 | Felidae | 45739.0 | 0 | Felis | 41229.0 | 1 | Felis catus | 41229.0 | 1 |
| scf7180000020510 | 80906 | 0.4048 | 0 | 49.3551 | Eukaryota | 40120.0 | 0 | Chordata | 40120.0 | 0 | Carnivora | 40120.0 | 0 | Felidae | 40120.0 | 0 | Felis | 36118.0 | 1 | Felis catus | 36118.0 | 1 |
| scf7180000020579 | 118521 | 0.3899 | 0 | 25.6631 | Eukaryota | 46254.0 | 0 | Chordata | 46254.0 | 0 | Carnivora | 46254.0 | 0 | Herpestidae | 37472.0 | 1 | Suricata | 37472.0 | 2 | ricata suricat | 37472.0 | 2 |
| scf7180000017789 | 2916880 | 0.4767 | 0 | 50.0821 | Eukaryota | 47060.0 | 0 | Chordata | 47060.0 | 0 | Carnivora | 47060.0 | 0 | Felidae | 47060.0 | 0 | Felis | 47060.0 | 0 | Felis catus | 47060.0 | 0 |
| scf7180000020763 | 17758 | 0.3911 | 0 | 156.8147 | Eukaryota | 6631.0 | 0 | Chordata | 6631.0 | 0 | Carnivora | 6631.0 | 0 | Felidae | 6631.0 | 0 | Felis | 6631.0 | 0 | Felis catus | 6631.0 | 0 |
| scf7180000018448 | 196203 | 0.3772 | 0 | 48.1452 | Eukaryota | 46686.0 | 0 | Chordata | 46686.0 | 0 | Carnivora | 46686.0 | 0 | Felidae | 46686.0 | 0 | Felis | 46686.0 | 0 | Felis catus | 46686.0 | 0 |
| scf7180000020207 | 127684 | 0.3786 | 0 | 23.9969 | Eukaryota | 39499.0 | 0 | Chordata | 39499.0 | 0 | Carnivora | 39499.0 | 0 | Felidae | 39499.0 | 0 | Felis | 39499.0 | 0 | Felis catus | 39499.0 | 0 |
| scf7180000022053 | 31161 | 0.4187 | 0 | 24.8919 | Eukaryota | 35610.0 | 0 | Chordata | 35610.0 | 0 | Carnivora | 35610.0 | 0 | Felidae | 35610.0 | 0 | Felis | 35610.0 | 0 | Felis catus | 35610.0 | 0 |
| scf7180000017659 | 2017168 | 0.3845 | 0 | 50.4888 | Eukaryota | 48225.0 | 0 | Chordata | 48225.0 | 0 | Carnivora | 48225.0 | 0 | Felidae | 48225.0 | 0 | Felis | 48225.0 | 0 | Felis catus | 48225.0 | 0 |
| scf7180000022031 | 46681 | 0.43 | 0 | 28.7707 | Eukaryota | 21551.0 | 0 | Chordata | 21551.0 | 0 | Carnivora | 21551.0 | 0 | Felidae | 21551.0 | 0 | Felis | 17550.0 | 1 | Felis catus | 17550.0 | 1 |
| scf7180000020154 | 158196 | 0.469 | 0 | 53.455 | Eukaryota | 38875.0 | 0 | Chordata | 38875.0 | 0 | Carnivora | 38875.0 | 0 | Felidae | 38875.0 | 0 | Felis | 38875.0 | 0 | Felis catus | 38875.0 | 0 |
| scf7180000017963 | 151192 | 0.4005 | 0 | 47.4157 | Eukaryota | 46011.0 | 0 | Chordata | 46011.0 | 0 | Carnivora | 46011.0 | 0 | Herpestidae | 23205.0 | 1 | Suricata | 23205.0 | 2 | ricata suricat | 23205.0 | 3 |
| scf7180000024010 | 763623 | 0.4024 | 0 | 50.4724 | Eukaryota | 44600.0 | 0 | Chordata | 44600.0 | 0 | Carnivora | 44600.0 | 0 | Felidae | 44600.0 | 0 | Felis | 44600.0 | 0 | Felis catus | 44600.0 | 0 |
| scf7180000019649 | 320653 | 0.4155 | 0 | 52.6813 | Eukaryota | 46260.0 | 0 | Chordata | 46260.0 | 0 | Carnivora | 46260.0 | 0 | Felidae | 46260.0 | 0 | Felis | 41456.0 | 1 | Felis catus | 41456.0 | 1 |
| scf7180000018098 | 1403534 | 0.3854 | 100 | 50.4807 | Eukaryota | 29255.0 | 0 | Chordata | 29255.0 | 0 | Carnivora | 29255.0 | 0 | Felidae | 29255.0 | 0 | Felis | 26276.0 | 1 | Felis catus | 26276.0 | 1 |
| scf7180000017478 | 3194998 | 0.36 | 0 | 49.3893 | Eukaryota | 47223.0 | 0 | Chordata | 47223.0 | 0 | Carnivora | 47223.0 | 0 | Felidae | 47223.0 | 0 | Felis | 47223.0 | 0 | Felis catus | 47223.0 | 0 |
| scf7180000020163 | 301599 | 0.4219 | 0 | 24.9861 | Eukaryota | 36775.0 | 0 | Chordata | 36775.0 | 0 | Carnivora | 25850.0 | 2 | Felidae | 25850.0 | 2 | Panthera | 18994.0 | 3 | anthera pardu | 15586.0 | 4 |
| scf7180000024088 | 189243 | 0.4014 | 0 | 50.1565 | Eukaryota | 39631.0 | 0 | Chordata | 39631.0 | 0 | Carnivora | 35438.0 | 1 | Felidae | 35438.0 | 1 | Panthera | 35438.0 | 1 | anthera pardu | 27566.0 | 2 |
| scf7180000017513 | 688816 | 0.429 | 0 | 50.9393 | Eukaryota | 50303.0 | 0 | Chordata | 50303.0 | 0 | Carnivora | 50303.0 | 0 | Felidae | 50303.0 | 0 | Felis | 34504.0 | 3 | Felis catus | 34504.0 | 3 |
| scf7180000023815 | 583676 | 0.3434 | 0 | 46.5337 | Eukaryota | 44078.0 | 0 | Chordata | 44078.0 | 0 | Carnivora | 44078.0 | 0 | Felidae | 44078.0 | 0 | Felis | 44078.0 | 0 | Felis catus | 44078.0 | 0 |
| scf7180000022460 | 8393 | 0.3791 | 0 | 22.2343 | Eukaryota | 4854.0 | 0 | Chordata | 4854.0 | 0 | Carnivora | 3239.0 | 1 | Felidae | 2808.0 | 3 | Lynx | 1692.0 | 4 | ynx canadens | 1692.0 | 4 |
| scf7180000019906 | 162698 | 0.4608 | 0 | 51.7948 | Eukaryota | 28755.0 | 0 | Chordata | 28755.0 | 0 | Carnivora | 28755.0 | 0 | Felidae | 28755.0 | 0 | Panthera | 9216.0 | 3 | anthera pardu | 9216.0 | 3 |
| scf7180000023396 | 8837 | 0.5099 | 0 | 126.3813 | no-hit | 0.0 | 0 | no-hit | 0.0 | 0 | no-hit | 0.0 | 0 | no-hit | 0.0 | 0 | no-hit | 0.0 | 0 | no-hit | 0.0 | 0 |
| scf7180000021157 | 130135 | 0.582 | 0 | 50.515 | Eukaryota | 6355.0 | 0 | Chordata | 6355.0 | 0 | Artiodactyla | 4153.0 | 1 | Delphinidae | 4153.0 | 1 | Globicephala | 2985.0 | 2 | bicephala me | 2985.0 | 2 |
| scf7180000020994 | 88767 | 0.4092 | 0 | 25.4838 | Eukaryota | 27141.0 | 0 | Chordata | 27141.0 | 0 | Carnivora | 24230.0 | 1 | Felidae | 24230.0 | 1 | Acinonyx | 16128.0 | 2 | cinonyx jubat | 16128.0 | 2 |
| scf7180000019043 | 440685 | 0.4136 | 0 | 49.9142 | Eukaryota | 38354.0 | 0 | Chordata | 38354.0 | 0 | Carnivora | 38354.0 | 0 | Felidae | 38354.0 | 0 | Felis | 34512.0 | 1 | Felis catus | 34512.0 | 1 |
| scf7180000020417 | 104998 | 0.3786 | 0 | 24.63 | Eukaryota | 44309.0 | 0 | Chordata | 44309.0 | 0 | Carnivora | 44309.0 | 0 | Felidae | 44309.0 | 0 | Felis | 44309.0 | 0 | Felis catus | 44309.0 | 0 |
| scf7180000023010 | 41491 | 0.5596 | 0 | 369.4682 | Eukaryota | 20534.0 | 0 | Chordata | 20534.0 | 0 | Carnivora | 20534.0 | 0 | Felidae | 20534.0 | 0 | Felis | 18441.0 | 1 | Felis catus | 18441.0 | 1 |
| scf7180000024091 | 263353 | 0.5177 | 0 | 50.4344 | Eukaryota | 24040.0 | 0 | Chordata | 24040.0 | 0 | Carnivora | 24040.0 | 0 | Felidae | 20678.0 | 1 | Felis | 13764.0 | 4 | Felis catus | 13764.0 | 4 |
| scf7180000017852 | 2427826 | 0.4534 | 0 | 51.5111 | Eukaryota | 63629.0 | 0 | Chordata | 63629.0 | 0 | Carnivora | 63629.0 | 0 | Felidae | 63629.0 | 0 | Panthera | 36245.0 | 1 | anthera pardu | 36245.0 | 1 |
| scf7180000020083 | 262900 | 0.3749 | 0 | 51.1125 | Eukaryota | 8881.0 | 1 | Chordata | 8881.0 | 1 | Primates | 8121.0 | 2 | Hominidae | 4815.0 | 3 | Homo | 4815.0 | 4 | Homo sapiens | 4815.0 | 4 |
| scf7180000017984 | 89074 | 0.49 | 0 | 478.1647 | Eukaryota | 16563.0 | 0 | Chordata | 16563.0 | 0 | Carnivora | 15185.0 | 1 | Felidae | 11175.0 | 3 | Felis | 6519.0 | 4 | Felis catus | 6519.0 | 4 |
| scf7180000018838 | 633648 | 0.3914 | 0 | 51.6246 | Eukaryota | 46941.0 | 0 | Chordata | 46941.0 | 0 | Carnivora | 46941.0 | 0 | Felidae | 46941.0 | 0 | Felis | 46941.0 | 0 | Felis catus | 46941.0 | 0 |
| scf7180000018585 | 686205 | 0.3591 | 0 | 49.5404 | Eukaryota | 47424.0 | 0 | Chordata | 47424.0 | 0 | Carnivora | 47424.0 | 0 | Felidae | 47424.0 | 0 | Felis | 47424.0 | 0 | Felis catus | 47424.0 | 0 |
| scf7180000023883 | 1061213 | 0.4404 | 0 | 51.0538 | Eukaryota | 97075.0 | 0 | Chordata | 97075.0 | 0 | Carnivora | 97075.0 | 0 | Felidae | 97075.0 | 0 | Lynx | 48550.0 | 1 | ynx canadens | 48550.0 | 1 |
| scf7180000020023 | 162335 | 0.401 | 0 | 25.0325 | Eukaryota | 17751.0 | 0 | Chordata | 17751.0 | 0 | Carnivora | 17751.0 | 0 | Felidae | 17751.0 | 0 | Felis | 15946.0 | 1 | Felis catus | 15946.0 | 1 |
| scf7180000020263 | 2953 | 0.5066 | 0 | 29.1666 | Eukaryota | 2423.0 | 0 | Chordata | 2423.0 | 0 | Carnivora | 2423.0 | 0 | Felidae | 724.0 | 5 | Felis | 496.0 | 7 | Felis catus | 496.0 | 8 |
| scf7180000024126 | 210993 | 0.4864 | 0 | 52.2225 | Eukaryota | 38730.0 | 0 | Chordata | 38730.0 | 0 | Carnivora | 38730.0 | 0 | Felidae | 38730.0 | 0 | Felis | 38730.0 | 0 | Felis catus | 38730.0 | 0 |
| scf7180000019145 | 195772 | 0.3891 | 0 | 49.839 | Eukaryota | 27484.0 | 0 | Chordata | 27484.0 | 0 | Carnivora | 27484.0 | 0 | Felidae | 27484.0 | 0 | Felis | 24700.0 | 1 | Felis catus | 24700.0 | 1 |
| scf7180000023616 | 130147 | 0.3923 | 0 | 66.8828 | Eukaryota | 36864.0 | 0 | Chordata | 36864.0 | 0 | Carnivora | 36864.0 | 0 | Felidae | 36864.0 | 0 | Felis | 36864.0 | 0 | Felis catus | 36864.0 | 0 |
| scf7180000024216 | 45735 | 0.5361 | 0 | 40.5244 | Eukaryota | 5020.0 | 0 | Chordata | 5020.0 | 0 | Carnivora | 5020.0 | 0 | Felidae | 4761.0 | 1 | Panthera | 2131.0 | 4 | anthera pardu | 1074.0 | 5 |
| scf7180000020639 | 101040 | 0.4002 | 0 | 49.4002 | Eukaryota | 11857.0 | 0 | Chordata | 11857.0 | 0 | Carnivora | 9057.0 | 1 | Felidae | 9057.0 | 1 | Felis | 6775.0 | 2 | Felis catus | 6775.0 | 2 |
| scf7180000024235 | 25918 | 0.4169 | 0 | 25.4535 | Eukaryota | 9431.0 | 0 | Chordata | 9431.0 | 0 | Primates | 5314.0 | 3 | Hominidae | 5314.0 | 3 | Homo | 2751.0 | 4 | Homo sapiens | 2751.0 | 4 |
| scf7180000018052 | 1327379 | 0.4872 | 0 | 52.7969 | Eukaryota | 53772.0 | 0 | Chordata | 53772.0 | 0 | Carnivora | 53772.0 | 0 | Felidae | 43750.0 | 1 | Felis | 37870.0 | 2 | Felis catus | 37870.0 | 2 |
| scf7180000021338 | 36874 | 0.391 | 0 | 25.2941 | Eukaryota | 6124.0 | 0 | Chordata | 6124.0 | 0 | Carnivora | 6124.0 | 0 | Felidae | 6124.0 | 0 | Felis | 6124.0 | 0 | Felis catus | 6124.0 | 0 |
| scf7180000023218 | 33653 | 0.5637 | 0 | 121.6747 | Eukaryota | 6748.0 | 0 | Chordata | 6748.0 | 0 | Carnivora | 6748.0 | 0 | Felidae | 6748.0 | 0 | Felis | 2500.0 | 3 | Felis catus | 2500.0 | 3 |
| scf7180000021807 | 38032 | 0.3592 | 0 | 23.4111 | Eukaryota | 40588.0 | 0 | Chordata | 40588.0 | 0 | Carnivora | 40588.0 | 0 | Felidae | 40588.0 | 0 | Felis | 36560.0 | 1 | Felis catus | 36560.0 | 1 |
| scf7180000020179 | 243661 | 0.6067 | 0 | 51.6404 | Eukaryota | 15031.0 | 0 | Chordata | 15031.0 | 0 | Carnivora | 15031.0 | 0 | Felidae | 15031.0 | 0 | Acinonyx | 7606.0 | 1 | cinonyx jubat | 7606.0 | 1 |
| scf7180000019437 | 2213621 | 0.381 | 0 | 50.5904 | Eukaryota | 47879.0 | 0 | Chordata | 47879.0 | 0 | Carnivora | 47879.0 | 0 | Felidae | 47879.0 | 0 | Felis | 47879.0 | 0 | Felis catus | 47879.0 | 0 |
| scf7180000019580 | 236884 | 0.4152 | 0 | 50.5157 | Eukaryota | 20178.0 | 0 | Chordata | 20178.0 | 0 | Carnivora | 20178.0 | 0 | Felidae | 20178.0 | 0 | Felis | 12399.0 | 3 | Felis catus | 12399.0 | 3 |
| scf7180000022024 | 17199 | 0.5265 | 0 | 1991.5401 | no-hit | 0.0 | 0 | no-hit | 0.0 | 0 | no-hit | 0.0 | 0 | no-hit | 0.0 | 0 | no-hit | 0.0 | 0 | no-hit | 0.0 | 0 |
| scf7180000020540 | 91218 | 0.3881 | 0 | 24.8493 | Eukaryota | 57498.0 | 0 | Chordata | 57498.0 | 0 | Carnivora | 57498.0 | 0 | Felidae | 57498.0 | 0 | Panthera | 47456.0 | 2 | anthera pardu | 47456.0 | 2 |
| scf7180000022408 | 24489 | 0.3838 | 0 | 25.4733 | Eukaryota | 13348.0 | 0 | Chordata | 13348.0 | 0 | Carnivora | 13348.0 | 0 | Felidae | 13348.0 | 0 | Felis | 11874.0 | 1 | Felis catus | 11874.0 | 1 |
| scf7180000023796 | 3140948 | 0.4153 | 0 | 50.8661 | Eukaryota | 54097.0 | 0 | Chordata | 54097.0 | 0 | Carnivora | 54097.0 | 0 | Herpestidae | 34860.0 | 1 | Suricata | 34860.0 | 1 | ricata suricat | 34860.0 | 1 |
| scf7180000019825 | 9153 | 0.6232 | 0 | 130.6573 | Eukaryota | 2543.0 | 0 | Chordata | 2543.0 | 0 | Carnivora | 2289.0 | 1 | Felidae | 2289.0 | 1 | Felis | 1675.0 | 3 | Felis catus | 1675.0 | 3 |
| scf7180000018424 | 16879 | 0.5954 | 0 | 42.2735 | Eukaryota | 3904.0 | 0 | Chordata | 3904.0 | 0 | Carnivora | 3904.0 | 0 | Felidae | 3510.0 | 1 | Lynx | 1950.0 | 2 | ynx canadens | 1950.0 | 2 |
| scf7180000022318 | 13287 | 0.4203 | 0 | 23.3862 | Eukaryota | 31732.0 | 0 | Chordata | 31732.0 | 0 | Carnivora | 31732.0 | 0 | Felidae | 31732.0 | 0 | Felis | 28500.0 | 1 | Felis catus | 28500.0 | 1 |
| scf7180000023971 | 508272 | 0.4152 | 0 | 50.9666 | Eukaryota | 32892.0 | 0 | Chordata | 32892.0 | 0 | Carnivora | 32892.0 | 0 | Felidae | 18690.0 | 3 | Felis | 7672.0 | 7 | Felis catus | 7672.0 | 7 |
| scf7180000020706 | 3329 | 0.6858 | 0 | 117.3549 | Eukaryota | 357.0 | 0 | Chordata | 357.0 | 0 | Carnivora | 357.0 | 0 | Felidae | 357.0 | 0 | Panthera | 357.0 | 0 | anthera pardu | 357.0 | 0 |
| scf7180000021554 | 35586 | 0.363 | 0 | 23.9805 | Eukaryota | 38873.0 | 0 | Chordata | 38873.0 | 0 | Carnivora | 38873.0 | 0 | Felidae | 38873.0 | 0 | Felis | 38873.0 | 0 | Felis catus | 38873.0 | 0 |
| scf7180000017826 | 448534 | 0.3667 | 0 | 46.2627 | Eukaryota | 43421.0 | 0 | Chordata | 43421.0 | 0 | Carnivora | 43421.0 | 0 | Felidae | 43421.0 | 0 | Felis | 43421.0 | 0 | Felis catus | 43421.0 | 0 |
| scf7180000021955 | 14715 | 0.3811 | 0 | 24.2199 | Eukaryota | 41642.0 | 0 | Chordata | 41642.0 | 0 | Carnivora | 41642.0 | 0 | Felidae | 41642.0 | 0 | Felis | 41642.0 | 0 | Felis catus | 41642.0 | 0 |
| scf7180000017590 | 697938 | 0.3825 | 0 | 49.4983 | Eukaryota | 44792.0 | 0 | Chordata | 44792.0 | 0 | Carnivora | 44792.0 | 0 | Felidae | 44792.0 | 0 | Felis | 44792.0 | 0 | Felis catus | 44792.0 | 0 |
| scf7180000018647 | 729701 | 0.3858 | 0 | 51.7764 | Eukaryota | 37617.0 | 0 | Chordata | 37617.0 | 0 | Carnivora | 37617.0 | 0 | Felidae | 37617.0 | 0 | Felis | 37617.0 | 0 | Felis catus | 37617.0 | 0 |
| scf7180000020498 | 104336 | 0.3802 | 0 | 23.1471 | Eukaryota | 43971.0 | 0 | Chordata | 43971.0 | 0 | Carnivora | 43971.0 | 0 | Felidae | 43971.0 | 0 | Felis | 43971.0 | 0 | Felis catus | 43971.0 | 0 |
| scf7180000018833 | 548244 | 0.4361 | 0 | 49.0691 | Eukaryota | 44151.0 | 0 | Chordata | 44151.0 | 0 | Carnivora | 44151.0 | 0 | Felidae | 44151.0 | 0 | Felis | 44151.0 | 0 | Felis catus | 44151.0 | 0 |
| scf7180000020718 | 97123 | 0.3699 | 0 | 25.158 | Eukaryota | 37055.0 | 0 | Chordata | 37055.0 | 0 | Carnivora | 37055.0 | 0 | Felidae | 37055.0 | 0 | Felis | 37055.0 | 0 | Felis catus | 37055.0 | 0 |
| scf7180000023724 | 71040 | 0.5007 | 0 | 518.8453 | Eukaryota | 21863.0 | 0 | Chordata | 21863.0 | 0 | Carnivora | 21863.0 | 0 | Felidae | 21863.0 | 0 | Felis | 19641.0 | 1 | Felis catus | 19641.0 | 1 |
| scf7180000022786 | 8575 | 0.5237 | 0 | 56.6429 | no-hit | 0.0 | 0 | no-hit | 0.0 | 0 | no-hit | 0.0 | 0 | no-hit | 0.0 | 0 | no-hit | 0.0 | 0 | no-hit | 0.0 | 0 |
| scf7180000022538 | 11571 | 0.4347 | 0 | 23.9455 | Eukaryota | 3333.0 | 0 | Chordata | 3333.0 | 0 | Carnivora | 2196.0 | 2 | Felidae | 2196.0 | 2 | Felis | 1356.0 | 3 | Felis catus | 1356.0 | 3 |
| scf7180000021030 | 48589 | 0.3774 | 0 | 24.6025 | Eukaryota | 25487.0 | 0 | Chordata | 25487.0 | 0 | Carnivora | 25487.0 | 0 | Felidae | 25487.0 | 0 | Felis | 22905.0 | 1 | Felis catus | 22905.0 | 1 |
| scf7180000021128 | 81517 | 0.3565 | 0 | 24.0584 | Eukaryota | 20391.0 | 0 | Chordata | 20391.0 | 0 | Carnivora | 20391.0 | 0 | Felidae | 20391.0 | 0 | Felis | 18368.0 | 1 | Felis catus | 18368.0 | 1 |
| scf7180000019130 | 32358 | 0.3376 | 0 | 33.2031 | Eukaryota | 41370.0 | 0 | Chordata | 41370.0 | 0 | Carnivora | 41370.0 | 0 | Felidae | 41370.0 | 0 | Felis | 37272.0 | 1 | Felis catus | 37272.0 | 1 |
| scf7180000020052 | 245804 | 0.5213 | 0 | 49.9558 | Eukaryota | 10608.0 | 0 | Chordata | 10608.0 | 0 | Primates | 5788.0 | 4 | Hominidae | 3816.0 | 6 | Homo | 3816.0 | 6 | Homo sapiens | 3816.0 | 6 |
| scf7180000024258 | 462868 | 0.3491 | 0 | 47.5311 | Eukaryota | 45256.0 | 0 | Chordata | 45256.0 | 0 | Carnivora | 45256.0 | 0 | Felidae | 45256.0 | 0 | Felis | 45256.0 | 0 | Felis catus | 45256.0 | 0 |
| scf7180000018569 | 753558 | 0.4338 | 0 | 51.2938 | Eukaryota | 33919.0 | 1 | Chordata | 33919.0 | 1 | Carnivora | 26068.0 | 3 | Felidae | 26068.0 | 3 | Felis | 17702.0 | 5 | Felis catus | 17702.0 | 5 |
| scf7180000020901 | 67478 | 0.4738 | 0 | 50.9974 | Eukaryota | 37595.0 | 0 | Chordata | 37595.0 | 0 | Carnivora | 37595.0 | 0 | Felidae | 16211.0 | 3 | Canis | 10590.0 | 7 | Canis lupus | 10590.0 | 7 |
| scf7180000023533 | 83540 | 0.4161 | 0 | 212.0835 | Eukaryota | 17134.0 | 0 | Chordata | 17134.0 | 0 | Carnivora | 17134.0 | 0 | Felidae | 17134.0 | 0 | Felis | 17134.0 | 0 | Felis catus | 17134.0 | 0 |
| scf7180000018553 | 1168976 | 0.4693 | 0 | 51.139 | Eukaryota | 32815.0 | 1 | Chordata | 32815.0 | 1 | Artiodactyla | 15911.0 | 4 | Suidae | 15911.0 | 4 | Sus | 15911.0 | 5 | Sus scrofa | 15911.0 | 5 |
| scf7180000018253 | 524046 | 0.4591 | 0 | 51.2085 | Eukaryota | 45668.0 | 0 | Chordata | 45668.0 | 0 | Carnivora | 45668.0 | 0 | Felidae | 45668.0 | 0 | Felis | 45668.0 | 0 | Felis catus | 45668.0 | 0 |
| scf7180000018976 | 387378 | 0.4215 | 0 | 50.7727 | Eukaryota | 35459.0 | 0 | Chordata | 35459.0 | 0 | Carnivora | 35459.0 | 0 | Felidae | 29105.0 | 1 | Acinonyx | 14020.0 | 3 | cinonyx jubat | 14020.0 | 3 |
| scf7180000024157 | 141542 | 0.3714 | 0 | 24.0377 | Eukaryota | 17493.0 | 0 | Chordata | 17493.0 | 0 | Carnivora | 17493.0 | 0 | Felidae | 17493.0 | 0 | Felis | 17493.0 | 0 | Felis catus | 17493.0 | 0 |
| scf7180000023114 | 50642 | 0.4341 | 0 | 219.9652 | Eukaryota | 23919.0 | 0 | Chordata | 23919.0 | 0 | Carnivora | 23919.0 | 0 | Felidae | 23919.0 | 0 | Felis | 21505.0 | 1 | Felis catus | 21505.0 | 1 |
| scf7180000020251 | 160279 | 0.5054 | 0 | 51.23 | Eukaryota | 15943.0 | 0 | Chordata | 15943.0 | 0 | Carnivora | 15943.0 | 0 | Felidae | 15943.0 | 0 | Felis | 11627.0 | 1 | Felis catus | 11627.0 | 1 |
| scf7180000017549 | 383441 | 0.4006 | 0 | 50.2313 | Eukaryota | 46885.0 | 0 | Chordata | 46885.0 | 0 | Carnivora | 46885.0 | 0 | Felidae | 46885.0 | 0 | Felis | 46885.0 | 0 | Felis catus | 46885.0 | 0 |
| scf7180000022511 | 7824 | 0.4049 | 0 | 1418.3956 | no-hit | 0.0 | 0 | no-hit | 0.0 | 0 | no-hit | 0.0 | 0 | no-hit | 0.0 | 0 | no-hit | 0.0 | 0 | no-hit | 0.0 | 0 |
| scf7180000020231 | 32471 | 0.5388 | 0 | 53.9721 | Eukaryota | 8045.0 | 0 | Chordata | 8045.0 | 0 | Carnivora | 8045.0 | 0 | Felidae | 8045.0 | 0 | Felis | 5610.0 | 2 | Felis catus | 5610.0 | 2 |
| scf7180000019304 | 737220 | 0.508 | 0 | 188.1131 | Eukaryota | 62961.0 | 0 | Chordata | 62961.0 | 0 | Carnivora | 62961.0 | 0 | Herpestidae | 28600.0 | 3 | Suricata | 28600.0 | 3 | ricata suricat | 28600.0 | 3 |
| scf7180000018977 | 558861 | 0.4789 | 0 | 51.4867 | Eukaryota | 28829.0 | 0 | Chordata | 28829.0 | 0 | Carnivora | 20972.0 | 2 | Felidae | 15842.0 | 4 | Panthera | 5872.0 | 7 | Homo sapiens | 5092.0 | 8 |
| scf7180000021037 | 58920 | 0.4511 | 0 | 26.4425 | Eukaryota | 12566.0 | 0 | Chordata | 12566.0 | 0 | Carnivora | 12566.0 | 0 | Felidae | 11360.0 | 1 | Felis | 11360.0 | 1 | Felis catus | 11360.0 | 1 |
| scf7180000024074 | 182433 | 0.357 | 0 | 109.5235 | Eukaryota | 33568.0 | 0 | Chordata | 33568.0 | 0 | Carnivora | 33568.0 | 0 | Felidae | 33568.0 | 0 | Felis | 30253.0 | 1 | Felis catus | 30253.0 | 1 |
| scf7180000022533 | 14956 | 0.3762 | 0 | 23.3084 | Eukaryota | 7360.0 | 0 | Chordata | 7360.0 | 0 | Carnivora | 7360.0 | 0 | Felidae | 4442.0 | 1 | Felis | 3745.0 | 2 | Felis catus | 3745.0 | 2 |

| | | | | | | | | | | | | | | | | | | | | | | |
|---|---|---|---|---|---|---|---|---|---|---|---|---|---|---|---|---|---|---|---|---|---|---|
| scf7180000020805 | 83812 | 0.3827 | 0 | 50.6042 | Eukaryota | 17965.0 | 0 | Chordata | 17965.0 | 0 | Carnivora | 17965.0 | 0 | Felidae | 17965.0 | 0 | Felis | 17965.0 | 0 | Felis catus | 17965.0 | 0 |
| scf7180000022230 | 16120 | 0.3784 | 0 | 47.4421 | Eukaryota | 6082.0 | 0 | Chordata | 6082.0 | 0 | Primates | 2765.0 | 2 | Hominidae | 2765.0 | 2 | Homo | 2765.0 | 3 | Homo sapiens | 2765.0 | 3 |
| scf7180000020431 | 131567 | 0.4207 | 0 | 25.2413 | Eukaryota | 10210.0 | 0 | Chordata | 10210.0 | 0 | Carnivora | 10210.0 | 0 | Felidae | 10210.0 | 0 | Felis | 7507.0 | 2 | Felis catus | 7507.0 | 2 |
| scf7180000022914 | 701386 | 0.3644 | 0 | 49.9932 | Eukaryota | 47341.0 | 0 | Chordata | 47341.0 | 0 | Carnivora | 47341.0 | 0 | Felidae | 47341.0 | 0 | Felis | 47341.0 | 0 | Felis catus | 47341.0 | 0 |
| scf7180000018830 | 25983 | 0.3697 | 0 | 46.1313 | Eukaryota | 33159.0 | 0 | Chordata | 33159.0 | 0 | Carnivora | 33159.0 | 0 | Felidae | 33159.0 | 0 | Felis | 33159.0 | 0 | Felis catus | 33159.0 | 0 |
| scf7180000018502 | 835052 | 0.362 | 0 | 50.1214 | Eukaryota | 37113.0 | 0 | Chordata | 37113.0 | 0 | Carnivora | 37113.0 | 0 | Felidae | 37113.0 | 0 | Felis | 33382.0 | 1 | Felis catus | 33382.0 | 1 |
| scf7180000021193 | 111791 | 0.4136 | 0 | 25.4535 | Eukaryota | 33002.0 | 0 | Chordata | 33002.0 | 0 | Carnivora | 33002.0 | 0 | Felidae | 33002.0 | 0 | Felis | 33002.0 | 0 | Felis catus | 33002.0 | 0 |
| scf7180000018446 | 1582745 | 0.3848 | 0 | 50.4875 | Eukaryota | 47493.0 | 0 | Chordata | 47493.0 | 0 | Carnivora | 47493.0 | 0 | Felidae | 47493.0 | 0 | Felis | 47493.0 | 0 | Felis catus | 47493.0 | 0 |
| scf7180000017955 | 112005 | 0.435 | 0 | 48.9183 | Eukaryota | 23427.0 | 0 | Chordata | 23427.0 | 0 | Carnivora | 23427.0 | 0 | Felidae | 23427.0 | 0 | Felis | 18726.0 | 1 | Felis catus | 18726.0 | 1 |
| scf7180000019441 | 352713 | 0.3504 | 0 | 48.9612 | Eukaryota | 46340.0 | 0 | Chordata | 46340.0 | 0 | Carnivora | 46340.0 | 0 | Felidae | 46340.0 | 0 | Felis | 46340.0 | 0 | Felis catus | 46340.0 | 0 |
| scf7180000021271 | 54336 | 0.3772 | 0 | 24.9611 | Eukaryota | 43221.0 | 0 | Chordata | 43221.0 | 0 | Carnivora | 43221.0 | 0 | Felidae | 43221.0 | 0 | Felis | 43221.0 | 0 | Felis catus | 43221.0 | 0 |
| scf7180000019913 | 256347 | 0.4275 | 0 | 48.6639 | Eukaryota | 30564.0 | 0 | Chordata | 30564.0 | 0 | Carnivora | 30564.0 | 0 | Felidae | 30564.0 | 0 | Felis | 24449.0 | 1 | Felis catus | 24449.0 | 1 |
| scf7180000021611 | 31991 | 0.4385 | 0 | 24.3118 | Eukaryota | 2853.0 | 0 | Chordata | 2853.0 | 0 | Carnivora | 2526.0 | 1 | Felidae | 2252.0 | 2 | Felis | 1708.0 | 4 | Felis catus | 1708.0 | 4 |
| scf7180000021767 | 6486 | 0.5678 | 0 | 184.0012 | no-hit | 0.0 | 0 | no-hit | 0.0 | 0 | no-hit | 0.0 | 0 | no-hit | 0.0 | 0 | no-hit | 0.0 | 0 | no-hit | 0.0 | 0 |
| scf7180000019839 | 296481 | 0.3628 | 0 | 50.3666 | Eukaryota | 8705.0 | 0 | Chordata | 8705.0 | 0 | Carnivora | 5729.0 | 1 | Felidae | 5729.0 | 1 | Felis | 5729.0 | 1 | Felis catus | 5729.0 | 1 |
| scf7180000018145 | 1026835 | 0.3837 | 0 | 50.7045 | Eukaryota | 45802.0 | 0 | Chordata | 45802.0 | 0 | Carnivora | 45802.0 | 0 | Felidae | 45802.0 | 0 | Felis | 45802.0 | 0 | Felis catus | 45802.0 | 0 |
| scf7180000022434 | 13911 | 0.6318 | 0 | 50.0741 | Eukaryota | 2193.0 | 1 | Chordata | 2193.0 | 1 | Carnivora | 1697.0 | 2 | Felidae | 1460.0 | 3 | Felis | 1214.0 | 4 | Felis catus | 1214.0 | 4 |
| scf7180000020376 | 102599 | 0.5896 | 0 | 49.4276 | Eukaryota | 5904.0 | 0 | Chordata | 5904.0 | 0 | Carnivora | 3024.0 | 2 | Canidae | 1568.0 | 4 | Homo | 1234.0 | 6 | Homo sapiens | 1234.0 | 6 |
| scf7180000021897 | 32800 | 0.6523 | 0 | 45.006 | Eukaryota | 9026.0 | 0 | Chordata | 9026.0 | 0 | Carnivora | 5448.0 | 2 | Felidae | 4712.0 | 4 | Acinonyx | 2706.0 | 5 | cinonyx jubat | 2706.0 | 5 |
| scf7180000017504 | 901960 | 0.4172 | 0 | 51.5094 | Eukaryota | 113894.0 | 0 | Chordata | 113894.0 | 0 | Carnivora | 113894.0 | 0 | Felidae | 113894.0 | 0 | Panthera | 79772.0 | 1 | anthera pardu | 79772.0 | 1 |
| scf7180000022683 | 13249 | 0.67 | 0 | 976.4175 | Eukaryota | 630.0 | 0 | Chordata | 483.0 | 1 | Carnivora | 483.0 | 1 | Felidae | 344.0 | 2 | Felis | 344.0 | 2 | Felis catus | 344.0 | 2 |
| scf7180000018875 | 6304 | 0.4091 | 0 | 22.6336 | Eukaryota | 2694.0 | 0 | Chordata | 2694.0 | 0 | Carnivora | 2694.0 | 0 | Felidae | 2296.0 | 1 | Felis | 1828.0 | 2 | Felis catus | 1828.0 | 2 |
| scf7180000018815 | 65995 | 0.3995 | 0 | 138.322 | Eukaryota | 24138.0 | 0 | Chordata | 24138.0 | 0 | Carnivora | 24138.0 | 0 | Felidae | 24138.0 | 0 | Felis | 21713.0 | 1 | Felis catus | 21713.0 | 1 |
| scf7180000020622 | 114883 | 0.482 | 0 | 49.817 | Eukaryota | 10792.0 | 0 | Chordata | 10792.0 | 0 | Carnivora | 10792.0 | 0 | Felidae | 10792.0 | 0 | Lynx | 6432.0 | 2 | ynx canadens | 6432.0 | 2 |
| scf7180000021648 | 17440 | 0.4248 | 0 | 407.0293 | Eukaryota | 214907.0 | 0 | Chordata | 214907.0 | 0 | Carnivora | 214907.0 | 0 | Hyaenidae | 214907.0 | 0 | Proteles | 127931.0 | 1 | roteles cristat | 127931.0 | 1 |
| scf7180000019979 | 191683 | 0.5651 | 0 | 51.4038 | Eukaryota | 22072.0 | 0 | Chordata | 22072.0 | 0 | Carnivora | 22072.0 | 0 | Felidae | 8949.0 | 2 | eptonychote | 6855.0 | 6 | nychotes wed | 6855.0 | 6 |
| scf7180000021078 | 81032 | 0.556 | 0 | 53.5773 | Eukaryota | 9042.0 | 0 | Chordata | 9042.0 | 0 | Carnivora | 9042.0 | 0 | Felidae | 9042.0 | 0 | Panthera | 3576.0 | 2 | anthera pardu | 3576.0 | 2 |
| scf7180000018105 | 386641 | 0.3454 | 0 | 48.0206 | Eukaryota | 42704.0 | 0 | Chordata | 42704.0 | 0 | Carnivora | 42704.0 | 0 | Felidae | 42704.0 | 0 | Felis | 42704.0 | 0 | Felis catus | 42704.0 | 0 |
| scf7180000020378 | 166140 | 0.4041 | 0 | 49.448 | Eukaryota | 46667.0 | 0 | Chordata | 46667.0 | 0 | Carnivora | 46667.0 | 0 | Felidae | 46667.0 | 0 | Felis | 46667.0 | 0 | Felis catus | 46667.0 | 0 |
| scf7180000019539 | 278604 | 0.5696 | 0 | 50.7531 | Eukaryota | 9584.0 | 0 | Chordata | 9584.0 | 0 | Carnivora | 9584.0 | 0 | Felidae | 9584.0 | 0 | Felis | 9584.0 | 0 | Felis catus | 9584.0 | 0 |
| scf7180000022012 | 43161 | 0.4291 | 0 | 24.1528 | Eukaryota | 6525.0 | 0 | Chordata | 6525.0 | 0 | Carnivora | 6525.0 | 0 | Felidae | 5889.0 | 1 | Felis | 5222.0 | 2 | Felis catus | 5222.0 | 2 |
| scf7180000019492 | 247557 | 0.3837 | 0 | 49.3566 | Eukaryota | 20867.0 | 0 | Chordata | 20867.0 | 0 | Carnivora | 20867.0 | 0 | Felidae | 20867.0 | 0 | Felis | 18763.0 | 1 | Felis catus | 18763.0 | 1 |
| scf7180000021804 | 17356 | 0.3782 | 0 | 42.7231 | Eukaryota | 27057.0 | 0 | Chordata | 27057.0 | 0 | Carnivora | 27057.0 | 0 | Felidae | 27057.0 | 0 | Felis | 24360.0 | 1 | Felis catus | 24360.0 | 1 |
| scf7180000018937 | 575434 | 0.3874 | 0 | 50.406 | Eukaryota | 56118.0 | 0 | Chordata | 56118.0 | 0 | Carnivora | 56118.0 | 0 | Felidae | 56118.0 | 0 | Felis | 56118.0 | 0 | Felis catus | 56118.0 | 0 |
| scf7180000018651 | 610238 | 0.4054 | 0 | 51.8108 | Eukaryota | 18514.0 | 1 | Chordata | 18514.0 | 1 | Primates | 13983.0 | 3 | Hominidae | 13983.0 | 3 | Homo | 13983.0 | 3 | Homo sapiens | 13983.0 | 3 |
| scf7180000020516 | 164770 | 0.5292 | 0 | 47.0856 | Eukaryota | 4640.0 | 0 | Chordata | 4640.0 | 0 | Carnivora | 3654.0 | 2 | Mustelidae | 1512.0 | 5 | Mustela | 1512.0 | 6 | ustela putori | 1512.0 | 6 |
| scf7180000022402 | 8881 | 0.3958 | 0 | 23.7994 | Eukaryota | 10174.0 | 0 | Chordata | 10174.0 | 0 | Carnivora | 10174.0 | 0 | Felidae | 10174.0 | 0 | Felis | 9174.0 | 1 | Felis catus | 9174.0 | 1 |
| scf7180000020968 | 95655 | 0.3823 | 0 | 49.1786 | Eukaryota | 15342.0 | 0 | Chordata | 15342.0 | 0 | Carnivora | 15342.0 | 0 | Felidae | 15342.0 | 0 | Felis | 12290.0 | 1 | Felis catus | 12290.0 | 1 |
| scf7180000023055 | 60219 | 0.5142 | 0 | 171.0151 | Eukaryota | 4626.0 | 0 | Chordata | 4626.0 | 0 | Carnivora | 4626.0 | 0 | Felidae | 4365.0 | 1 | Felis | 4093.0 | 2 | Felis catus | 4093.0 | 2 |
| scf7180000021302 | 56113 | 0.36 | 0 | 23.4715 | Eukaryota | 43810.0 | 0 | Chordata | 43810.0 | 0 | Carnivora | 43810.0 | 0 | Felidae | 43810.0 | 0 | Felis | 43810.0 | 0 | Felis catus | 43810.0 | 0 |
| scf7180000022142 | 16984 | 0.5264 | 0 | 565.277 | Eukaryota | 163.0 | 0 | latyhelminthe | 163.0 | 0 | olystomatide | 163.0 | 0 | Polystomatidae | 163.0 | 0 | rotopolystom | 163.0 | 0 | olystoma xen | 163.0 | 0 |
| scf7180000020423 | 69788 | 0.4319 | 0 | 127.944 | Eukaryota | 12535.0 | 0 | Chordata | 12535.0 | 0 | Carnivora | 9385.0 | 1 | Felidae | 9385.0 | 1 | Felis | 7752.0 | 2 | Felis catus | 7752.0 | 2 |
| scf7180000018754 | 669470 | 0.3826 | 0 | 51.729 | Eukaryota | 46590.0 | 0 | Chordata | 46590.0 | 0 | Carnivora | 46590.0 | 0 | Felidae | 46590.0 | 0 | Felis | 46590.0 | 0 | Felis catus | 46590.0 | 0 |
| scf7180000019468 | 293732 | 0.3883 | 0 | 49.1254 | Eukaryota | 45346.0 | 0 | Chordata | 45346.0 | 0 | Carnivora | 45346.0 | 0 | Felidae | 45346.0 | 0 | Felis | 40860.0 | 1 | Felis catus | 40860.0 | 1 |
| scf7180000017586 | 2115317 | 0.425 | 0 | 50.3323 | Eukaryota | 46099.0 | 0 | Chordata | 46099.0 | 0 | Carnivora | 46099.0 | 0 | Felidae | 46099.0 | 0 | Felis | 46099.0 | 0 | Felis catus | 46099.0 | 0 |
| scf7180000022073 | 13896 | 0.3487 | 0 | 23.617 | Eukaryota | 3938.0 | 0 | Chordata | 3938.0 | 0 | Carnivora | 3938.0 | 0 | Felidae | 3938.0 | 0 | Felis | 3542.0 | 1 | Felis catus | 3542.0 | 1 |
| scf7180000018445 | 742180 | 0.5062 | 0 | 50.6399 | Eukaryota | 37086.0 | 0 | Chordata | 37086.0 | 0 | Carnivora | 37086.0 | 0 | Herpestidae | 23616.0 | 1 | Suricata | 23616.0 | 2 | ricata suricat | 23616.0 | 2 |
| scf7180000020147 | 160269 | 0.3856 | 0 | 26.0942 | Eukaryota | 12551.0 | 0 | Chordata | 12551.0 | 0 | Carnivora | 6762.0 | 2 | Felidae | 5703.0 | 3 | Pan | 2789.0 | 6 | an troglodyte | 2789.0 | 7 |
| scf7180000024255 | 824074 | 0.4534 | 0 | 52.623 | Eukaryota | 76225.0 | 0 | Chordata | 76225.0 | 0 | Carnivora | 76225.0 | 0 | Herpestidae | 40140.0 | 1 | Suricata | 40140.0 | 1 | ricata suricat | 40140.0 | 1 |
| scf7180000018950 | 644189 | 0.4434 | 0 | 50.9524 | Eukaryota | 47264.0 | 0 | Chordata | 47264.0 | 0 | Carnivora | 47264.0 | 0 | Felidae | 47264.0 | 0 | Felis | 47264.0 | 0 | Felis catus | 47264.0 | 0 |
| scf7180000021858 | 31639 | 0.5622 | 0 | 46.0278 | Eukaryota | 3892.0 | 1 | Chordata | 3892.0 | 1 | Artiodactyla | 1111.0 | 7 | Suidae | 1111.0 | 7 | Sus | 1111.0 | 7 | Sus scrofa | 1111.0 | 7 |
| scf7180000019003 | 72421 | 0.3874 | 0 | 47.3047 | Eukaryota | 46718.0 | 0 | Chordata | 46718.0 | 0 | Carnivora | 46718.0 | 0 | Felidae | 46718.0 | 0 | Felis | 46718.0 | 0 | Felis catus | 46718.0 | 0 |
| scf7180000019867 | 265722 | 0.397 | 0 | 26.0572 | Eukaryota | 26296.0 | 0 | Chordata | 26296.0 | 0 | Primates | 18568.0 | 1 | Hominidae | 17046.0 | 3 | Homo | 10218.0 | 4 | Homo sapiens | 10218.0 | 4 |
| scf7180000018624 | 1345162 | 0.467 | 100 | 50.4204 | Eukaryota | 37418.0 | 0 | Chordata | 37418.0 | 0 | Carnivora | 37418.0 | 0 | Felidae | 22752.0 | 3 | Lynx | 7784.0 | 5 | ynx canadens | 7784.0 | 5 |
| scf7180000019018 | 1106169 | 0.4135 | 0 | 50.3359 | Eukaryota | 44536.0 | 0 | Chordata | 44536.0 | 0 | Carnivora | 44536.0 | 0 | Felidae | 23701.0 | 3 | Panthera | 10228.0 | 7 | nychotes wed | 8580.0 | 8 |
| scf7180000020789 | 100272 | 0.3625 | 0 | 49.57 | Eukaryota | 6357.0 | 0 | Chordata | 6357.0 | 0 | Carnivora | 4801.0 | 1 | Felidae | 4801.0 | 1 | Felis | 4071.0 | 2 | Felis catus | 4071.0 | 2 |
| scf7180000022104 | 10278 | 0.3847 | 0 | 24.1525 | Eukaryota | 30034.0 | 0 | Chordata | 30034.0 | 0 | Carnivora | 30034.0 | 0 | Felidae | 30034.0 | 0 | Felis | 30034.0 | 0 | Felis catus | 30034.0 | 0 |
| scf7180000023930 | 1433526 | 0.3976 | 0 | 55.1865 | Eukaryota | 56125.0 | 0 | Chordata | 56125.0 | 0 | Carnivora | 56125.0 | 0 | Felidae | 56125.0 | 0 | Lynx | 28065.0 | 2 | ynx canadens | 28065.0 | 2 |
| scf7180000021953 | 15327 | 0.6562 | 0 | 52.8921 | Eukaryota | 2013.0 | 0 | Chordata | 2013.0 | 0 | Carnivora | 2013.0 | 0 | Felidae | 2013.0 | 0 | Felis | 2013.0 | 0 | Felis catus | 2013.0 | 0 |
| scf7180000017568 | 599940 | 0.3954 | 0 | 51.5603 | Eukaryota | 46502.0 | 0 | Chordata | 46502.0 | 0 | Carnivora | 46502.0 | 0 | Felidae | 46502.0 | 0 | Felis | 46502.0 | 0 | Felis catus | 46502.0 | 0 |
| scf7180000023546 | 23739 | 0.4053 | 0 | 441.6145 | Eukaryota | 5866.0 | 0 | Chordata | 5866.0 | 0 | Carnivora | 5866.0 | 0 | Phocidae | 2372.0 | 2 | Eumetopias | 2308.0 | 3 | netopias juba | 2308.0 | 3 |
| scf7180000021591 | 48978 | 0.5785 | 0 | 47.5181 | Eukaryota | 17788.0 | 0 | Chordata | 17788.0 | 0 | Carnivora | 17788.0 | 0 | Felidae | 17788.0 | 0 | Felis | 17788.0 | 0 | Felis catus | 17788.0 | 0 |
| scf7180000018447 | 878989 | 0.3564 | 0 | 49.4719 | Eukaryota | 36546.0 | 0 | Chordata | 36546.0 | 0 | Carnivora | 36546.0 | 0 | Felidae | 36546.0 | 0 | Felis | 36546.0 | 0 | Felis catus | 36546.0 | 0 |
| scf7180000021009 | 81204 | 0.5045 | 0 | 52.8633 | Eukaryota | 20414.0 | 0 | Chordata | 20414.0 | 0 | Carnivora | 20414.0 | 0 | Felidae | 8564.0 | 4 | Panthera | 4291.0 | 8 | anthera pardu | 4291.0 | 8 |
| scf7180000018528 | 626170 | 0.3882 | 0 | 52.055 | Eukaryota | 75742.0 | 0 | Chordata | 75742.0 | 0 | Carnivora | 75742.0 | 0 | Felidae | 75742.0 | 0 | Panthera | 75742.0 | 0 | anthera pardu | 45366.0 | 1 |
| scf7180000021211 | 21516 | 0.4741 | 0 | 38.3565 | Eukaryota | 22341.0 | 0 | Chordata | 22341.0 | 0 | Carnivora | 22341.0 | 0 | Felidae | 22341.0 | 0 | Felis | 20163.0 | 1 | Felis catus | 20163.0 | 1 |
| scf7180000021006 | 132534 | 0.5326 | 0 | 51.7921 | Eukaryota | 30682.0 | 0 | Chordata | 30682.0 | 0 | Carnivora | 30682.0 | 0 | Felidae | 18658.0 | 2 | Suricata | 7376.0 | 6 | ricata suricat | 7376.0 | 7 |
| scf7180000020541 | 97521 | 0.429 | 0 | 51.0826 | Eukaryota | 7351.0 | 0 | Chordata | 7351.0 | 0 | Carnivora | 5387.0 | 2 | Felidae | 5387.0 | 2 | Felis | 5387.0 | 2 | Felis catus | 5387.0 | 2 |
| scf7180000022365 | 15290 | 0.4104 | 0 | 23.5357 | Eukaryota | 2469.0 | 0 | Chordata | 2469.0 | 0 | Carnivora | 2469.0 | 0 | Felidae | 2469.0 | 0 | Felis | 2226.0 | 1 | Felis catus | 1982.0 | 2 |
| scf7180000020777 | 136342 | 0.5808 | 0 | 51.0154 | Eukaryota | 12997.0 | 0 | Chordata | 12997.0 | 0 | Carnivora | 12997.0 | 0 | Felidae | 12997.0 | 0 | Felis | 7831.0 | 1 | Felis catus | 7831.0 | 1 |
| scf7180000023842 | 1190881 | 0.3885 | 0 | 50.5839 | Eukaryota | 46686.0 | 0 | Chordata | 46686.0 | 0 | Carnivora | 46686.0 | 0 | Felidae | 41565.0 | 1 | Felis | 41565.0 | 1 | Felis catus | 41565.0 | 1 |
| scf7180000023529 | 39752 | 0.456 | 0 | 421.2813 | Eukaryota | 6039.0 | 0 | Chordata | 6039.0 | 0 | Carnivora | 6039.0 | 0 | Phocidae | 2464.0 | 2 | eptonychote | 1848.0 | 4 | nychotes wed | 1848.0 | 4 |
| scf7180000018720 | 626702 | 0.3995 | 0 | 51.5775 | Eukaryota | 46756.0 | 0 | Chordata | 46756.0 | 0 | Carnivora | 46756.0 | 0 | Felidae | 46756.0 | 0 | Felis | 46756.0 | 0 | Felis catus | 46756.0 | 0 |
| scf7180000023175 | 80781 | 0.4014 | 0 | 287.584 | Eukaryota | 27398.0 | 0 | Chordata | 27398.0 | 0 | Carnivora | 27398.0 | 0 | Felidae | 27398.0 | 0 | Felis | 24605.0 | 1 | Felis catus | 24605.0 | 1 |
| scf7180000020421 | 149899 | 0.4266 | 0 | 50.8875 | Eukaryota | 33863.0 | 0 | Chordata | 33863.0 | 0 | Carnivora | 33863.0 | 0 | Felidae | 26779.0 | 1 | Felis | 23382.0 | 2 | Felis catus | 23382.0 | 2 |
| scf7180000024079 | 300246 | 0.563 | 0 | 48.8753 | Eukaryota | 16034.0 | 0 | Chordata | 16034.0 | 0 | Carnivora | 16034.0 | 0 | Felidae | 6259.0 | 4 | Suricata | 4450.0 | 6 | ricata suricat | 4450.0 | 7 |
| scf7180000017893 | 1472418 | 0.3518 | 0 | 49.6734 | Eukaryota | 46432.0 | 0 | Chordata | 46432.0 | 0 | Carnivora | 46432.0 | 0 | Felidae | 46432.0 | 0 | Felis | 46432.0 | 0 | Felis catus | 46432.0 | 0 |
| scf7180000018127 | 349494 | 0.3839 | 0 | 49.6582 | Eukaryota | 47628.0 | 0 | Chordata | 47628.0 | 0 | Carnivora | 47628.0 | 0 | Felidae | 47628.0 | 0 | Felis | 47628.0 | 0 | Felis catus | 47628.0 | 0 |
| scf7180000023564 | 29348 | 0.4783 | 0 | 384.3721 | no-hit | 0.0 | 0 | no-hit | 0.0 | 0 | no-hit | 0.0 | 0 | no-hit | 0.0 | 0 | no-hit | 0.0 | 0 | no-hit | 0.0 | 0 |
| scf7180000023156 | 123103 | 0.4915 | 0 | 206.5134 | Eukaryota | 21950.0 | 0 | Chordata | 21950.0 | 0 | Carnivora | 21950.0 | 0 | Felidae | 21950.0 | 0 | Felis | 21950.0 | 0 | Felis catus | 21950.0 | 0 |
| scf7180000020275 | 119369 | 0.3912 | 0 | 51.0932 | Eukaryota | 22563.0 | 0 | Chordata | 22563.0 | 0 | Carnivora | 22563.0 | 0 | Felidae | 22563.0 | 0 | Felis | 18004.0 | 1 | Felis catus | 18004.0 | 1 |
| scf7180000018715 | 608538 | 0.4524 | 0 | 51.2934 | Eukaryota | 44179.0 | 0 | Chordata | 44179.0 | 0 | Carnivora | 44179.0 | 0 | Felidae | 44179.0 | 0 | Felis | 44179.0 | 0 | Felis catus | 44179.0 | 0 |
| scf7180000018760 | 382919 | 0.3941 | 0 | 50.9251 | Eukaryota | 45272.0 | 0 | Chordata | 45272.0 | 0 | Carnivora | 45272.0 | 0 | Felidae | 45272.0 | 0 | Felis | 45272.0 | 0 | Felis catus | 45272.0 | 0 |
| scf7180000023975 | 967938 | 0.5509 | 0 | 50.2945 | Eukaryota | 40316.0 | 0 | Chordata | 40316.0 | 0 | Carnivora | 36469.0 | 1 | Felidae | 12022.0 | 5 | Canis | 8183.0 | 8 | Canis lupus | 8183.0 | 8 |
| scf7180000024266 | 2257350 | 0.3901 | 0 | 51.6357 | Eukaryota | 61734.0 | 0 | Chordata | 61734.0 | 0 | Carnivora | 61734.0 | 0 | Felidae | 55506.0 | 1 | Lynx | 55506.0 | 1 | ynx canadens | 55506.0 | 1 |
| scf7180000020702 | 113896 | 0.455 | 0 | 45.97 | Eukaryota | 23861.0 | 0 | Chordata | 23861.0 | 0 | Carnivora | 23861.0 | 0 | Felidae | 21652.0 | 1 | Panthera | 7340.0 | 4 | Felis catus | 7071.0 | 5 |
| scf7180000021762 | 14915 | 0.3906 | 0 | 26.575 | Eukaryota | 46662.0 | 0 | Chordata | 46662.0 | 0 | Carnivora | 46662.0 | 0 | Felidae | 46662.0 | 0 | Felis | 46662.0 | 0 | Felis catus | 46662.0 | 0 |
| scf7180000021123 | 50950 | 0.4588 | 0 | 3220.0552 | no-hit | 0.0 | 0 | no-hit | 0.0 | 0 | no-hit | 0.0 | 0 | no-hit | 0.0 | 0 | no-hit | 0.0 | 0 | no-hit | 0.0 | 0 |
| scf7180000019609 | 21020 | 0.5498 | 0 | 276.5563 | Eukaryota | 5931.0 | 0 | Chordata | 5931.0 | 0 | Carnivora | 5931.0 | 0 | Phocidae | 2416.0 | 3 | eptonychote | 1812.0 | 4 | nychotes wed | 1812.0 | 4 |
| scf7180000019676 | 312106 | 0.3715 | 0 | 54.444 | Eukaryota | 43679.0 | 0 | Chordata | 43679.0 | 0 | Carnivora | 43679.0 | 0 | Felidae | 39631.0 | 1 | Panthera | 13165.0 | 5 | ynx canadens | 8906.0 | 6 |
| scf7180000017575 | 802664 | 0.371 | 0 | 50.08 | Eukaryota | 39594.0 | 0 | Chordata | 39594.0 | 0 | Carnivora | 39594.0 | 0 | Felidae | 39594.0 | 0 | Felis | 20323.0 | 2 | Felis catus | 20323.0 | 2 |
| scf7180000021061 | 95382 | 0.4027 | 0 | 25.8857 | Eukaryota | 44313.0 | 0 | Chordata | 44313.0 | 0 | Carnivora | 44313.0 | 0 | Felidae | 44313.0 | 0 | Felis | 44313.0 | 0 | Felis catus | 44313.0 | 0 |
| scf7180000021479 | 22845 | 0.5281 | 0 | 2003.0752 | no-hit | 0.0 | 0 | no-hit | 0.0 | 0 | no-hit | 0.0 | 0 | no-hit | 0.0 | 0 | no-hit | 0.0 | 0 | no-hit | 0.0 | 0 |
| scf7180000018270 | 589977 | 0.477 | 0 | 52.5642 | Eukaryota | 48112.0 | 0 | Chordata | 48112.0 | 0 | Carnivora | 48112.0 | 0 | Felidae | 38064.0 | 1 | Panthera | 21702.0 | 3 | anthera pardu | 16509.0 | 4 |
| scf7180000022649 | 18032 | 0.3764 | 0 | 25.8999 | Eukaryota | 10194.0 | 0 | Chordata | 10194.0 | 0 | Carnivora | 10194.0 | 0 | Felidae | 10194.0 | 0 | Felis | 8159.0 | 1 | Felis catus | 8159.0 | 1 |
| scf7180000020418 | 2377153 | 0.4909 | 0 | 50.7691 | Eukaryota | 25047.0 | 0 | Chordata | 25047.0 | 0 | Carnivora | 25047.0 | 0 | Felidae | 25047.0 | 0 | Felis | 19562.0 | 2 | Felis catus | 19562.0 | 2 |
| scf7180000023163 | 19596 | 0.6027 | 0 | 233.6737 | Eukaryota | 2467.0 | 0 | Chordata | 2467.0 | 0 | Primates | 1520.0 | 3 | Cercopithecidae | 942.0 | 4 | Macaca | 942.0 | 5 | caca nemestr | 733.0 | 6 |
| scf7180000019557 | 268099 | 0.3881 | 0 | 51.0099 | Eukaryota | 45136.0 | 0 | Chordata | 45136.0 | 0 | Carnivora | 45136.0 | 0 | Felidae | 45136.0 | 0 | Felis | 45136.0 | 0 | Felis catus | 45136.0 | 0 |
| scf7180000022173 | 19856 | 0.4229 | 0 | 24.8731 | Eukaryota | 5386.0 | 0 | Chordata | 5386.0 | 0 | Carnivora | 4678.0 | 1 | Felidae | 4678.0 | 1 | Felis | 3647.0 | 2 | Felis catus | 3647.0 | 2 |
| scf7180000022814 | 5151 | 0.3584 | 0 | 22.7749 | Eukaryota | 6353.0 | 0 | Chordata | 6353.0 | 0 | Carnivora | 3003.0 | 2 | Felidae | 3003.0 | 2 | Felis | 3003.0 | 3 | Felis catus | 3003.0 | 3 |
| scf7180000023976 | 789408 | 0.4021 | 0 | 51.8604 | Eukaryota | 45321.0 | 0 | Chordata | 45321.0 | 0 | Carnivora | 45321.0 | 0 | Felidae | 45321.0 | 0 | Panthera | 35796.0 | 1 | anthera pardu | 26784.0 | 2 |
| scf7180000018907 | 624668 | 0.3719 | 0 | 48.7012 | Eukaryota | 46323.0 | 0 | Chordata | 46323.0 | 0 | Carnivora | 46323.0 | 0 | Felidae | 46323.0 | 0 | Felis | 46323.0 | 0 | Felis catus | 46323.0 | 0 |
| scf7180000017725 | 1750291 | 0.4823 | 0 | 50.7144 | Eukaryota | 112974.0 | 0 | Chordata | 112974.0 | 0 | Carnivora | 112974.0 | 0 | Felidae | 112974.0 | 0 | Felis | 112974.0 | 0 | Felis catus | 112974.0 | 0 |
| scf7180000021239 | 82031 | 0.459 | 0 | 47.242 | Eukaryota | 4469.0 | 0 | Chordata | 4469.0 | 0 | Carnivora | 3597.0 | 1 | Felidae | 3597.0 | 1 | Felis | 2769.0 | 3 | Felis catus | 2769.0 | 3 |
| scf7180000022541 | 14533 | 0.3718 | 0 | 23.7228 | Eukaryota | 32771.0 | 0 | Chordata | 32771.0 | 0 | Carnivora | 32771.0 | 0 | Felidae | 32771.0 | 0 | Felis | 29404.0 | 1 | Felis catus | 29404.0 | 1 |
| scf7180000018352 | 928853 | 0.3855 | 0 | 49.9051 | Eukaryota | 82473.0 | 0 | Chordata | 82473.0 | 0 | Carnivora | 82473.0 | 0 | Felidae | 82473.0 | 0 | Acinonyx | 32931.0 | 3 | cinonyx jubat | 32931.0 | 3 |
| scf7180000020685 | 98406 | 0.5529 | 0 | 48.967 | Eukaryota | 14232.0 | 0 | Chordata | 14232.0 | 0 | Carnivora | 14232.0 | 0 | Felidae | 14232.0 | 0 | Felis | 11398.0 | 1 | Felis catus | 11398.0 | 1 |
| scf7180000018213 | 447190 | 0.3977 | 0 | 51.1933 | Eukaryota | 43370.0 | 0 | Chordata | 43370.0 | 0 | Carnivora | 43370.0 | 0 | Felidae | 43370.0 | 0 | Felis | 43370.0 | 0 | Felis catus | 43370.0 | 0 |
| scf7180000018609 | 585982 | 0.4135 | 0 | 50.3613 | Eukaryota | 46090.0 | 0 | Chordata | 46090.0 | 0 | Carnivora | 46090.0 | 0 | Felidae | 46090.0 | 0 | Felis | 46090.0 | 0 | Felis catus | 46090.0 | 0 |
| scf7180000018455 | 30154 | 0.3891 | 0 | 38.3385 | Eukaryota | 45026.0 | 0 | Chordata | 45026.0 | 0 | Carnivora | 45026.0 | 0 | Felidae | 45026.0 | 0 | Felis | 45026.0 | 0 | Felis catus | 45026.0 | 0 |
| scf7180000023115 | 43834 | 0.4269 | 0 | 480.0657 | Eukaryota | 24951.0 | 0 | Chordata | 24951.0 | 0 | Carnivora | 24951.0 | 0 | Felidae | 24951.0 | 0 | Felis | 22422.0 | 1 | Felis catus | 22422.0 | 1 |
| scf7180000019549 | 392523 | 0.4311 | 0 | 51.3746 | Eukaryota | 28822.0 | 0 | Chordata | 28822.0 | 0 | Carnivora | 28822.0 | 0 | Felidae | 28822.0 | 0 | Felis | 28822.0 | 0 | Felis catus | 28822.0 | 0 |
| scf7180000022546 | 5013 | 0.3792 | 0 | 27.475 | Eukaryota | 2422.0 | 0 | Chordata | 2422.0 | 0 | Carnivora | 2422.0 | 0 | Felidae | 2422.0 | 0 | Felis | 1934.0 | 1 | Felis catus | 1934.0 | 1 |
| scf7180000018296 | 583176 | 0.494 | 0 | 53.804 | Eukaryota | 46448.0 | 0 | Chordata | 46448.0 | 0 | Carnivora | 41991.0 | 1 | Felidae | 24951.0 | 4 | Felis | 10960.0 | 7 | Felis catus | 10960.0 | 7 |
| scf7180000017763 | 1846358 | 0.4219 | 0 | 51.9559 | Eukaryota | 55130.0 | 0 | Chordata | 55130.0 | 0 | Carnivora | 55130.0 | 0 | Felidae | 55130.0 | 0 | Panthera | 33210.0 | 2 | anthera pardu | 33210.0 | 2 |
| scf7180000018196 | 968973 | 0.3921 | 0 | 49.7339 | Eukaryota | 46995.0 | 0 | Chordata | 46995.0 | 0 | Carnivora | 46995.0 | 0 | Felidae | 46995.0 | 0 | Felis | 46995.0 | 0 | Felis catus | 46995.0 | 0 |
| scf7180000019180 | 159128 | 0.4491 | 0 | 45.2322 | Eukaryota | 29087.0 | 0 | Chordata | 29087.0 | 0 | Carnivora | 24281.0 | 1 | Felidae | 17742.0 | 3 | Felis | 5944.0 | 5 | Felis catus | 5944.0 | 5 |
| scf7180000021466 | 51923 | 0.3951 | 0 | 24.765 | Eukaryota | 12949.0 | 0 | Chordata | 12949.0 | 0 | Carnivora | 5957.0 | 2 | Felidae | 5957.0 | 2 | Felis | 4817.0 | 4 | Felis catus | 4817.0 | 4 |
| scf7180000022924 | 50679 | 0.3861 | 0 | 105.1347 | Eukaryota | 37487.0 | 0 | Chordata | 37487.0 | 0 | Carnivora | 37487.0 | 0 | Felidae | 37487.0 | 0 | Felis | 37487.0 | 0 | Felis catus | 37487.0 | 0 |
| scf7180000021114 | 121091 | 0.4005 | 0 | 24.8773 | Eukaryota | 28494.0 | 0 | Chordata | 28494.0 | 0 | Carnivora | 28494.0 | 0 | Felidae | 28494.0 | 0 | Felis | 23955.0 | 1 | Felis catus | 23955.0 | 1 |
| scf7180000021892 | 24813 | 0.3864 | 0 | 23.5498 | Eukaryota | 39731.0 | 0 | Chordata | 39731.0 | 0 | Carnivora | 39731.0 | 0 | Felidae | 39731.0 | 0 | Felis | 39731.0 | 0 | Felis catus | 39731.0 | 0 |
| scf7180000020583 | 116474 | 0.3789 | 0 | 25.7104 | Eukaryota | 17275.0 | 0 | Chordata | 17275.0 | 0 | Primates | 12591.0 | 2 | Hominidae | 12591.0 | 2 | Pan | 7472.0 | 3 | an troglodyte | 7472.0 | 3 |
| scf7180000023769 | 1157195 | 0.3992 | 0 | 50.2794 | Eukaryota | 45360.0 | 0 | Chordata | 45360.0 | 0 | Carnivora | 45360.0 | 0 | Felidae | 45360.0 | 0 | Felis | 45360.0 | 0 | Felis catus | 45360.0 | 0 |
| scf7180000020559 | 22406 | 0.3864 | 0 | 25.207 | Eukaryota | 46075.0 | 0 | Chordata | 46075.0 | 0 | Carnivora | 46075.0 | 0 | Felidae | 46075.0 | 0 | Felis | 46075.0 | 0 | Felis catus | 46075.0 | 0 |
| scf7180000022441 | 26698 | 0.3892 | 0 | 24.1166 | Eukaryota | 43065.0 | 0 | Chordata | 43065.0 | 0 | Carnivora | 43065.0 | 0 | Felidae | 43065.0 | 0 | Felis | 39222.0 | 1 | Felis catus | 39222.0 | 1 |
| scf7180000019590 | 240632 | 0.5544 | 0 | 48.2864 | Eukaryota | 12756.0 | 0 | Chordata | 12756.0 | 0 | Carnivora | 12756.0 | 0 | Felidae | 12756.0 | 0 | Felis | 11527.0 | 1 | Felis catus | 11527.0 | 1 |
| scf7180000019213 | 401137 | 0.4094 | 0 | 24.7413 | Eukaryota | 24668.0 | 0 | Chordata | 24668.0 | 0 | Carnivora | 24668.0 | 0 | Felidae | 24668.0 | 0 | Felis | 22228.0 | 1 | Felis catus | 22228.0 | 1 |
| scf7180000018503 | 814773 | 0.3679 | 0 | 49.7534 | Eukaryota | 43590.0 | 0 | Chordata | 43590.0 | 0 | Carnivora | 43590.0 | 0 | Felidae | 43590.0 | 0 | Felis | 43590.0 | 0 | Felis catus | 43590.0 | 0 |
| scf7180000017436 | 6902714 | 0.4198 | 0 | 51.2889 | Eukaryota | 48516.0 | 0 | Chordata | 48516.0 | 0 | Carnivora | 23921.0 | 4 | Felidae | 23921.0 | 4 | Felis | 23921.0 | 4 | Felis catus | 23921.0 | 4 |
| scf7180000019090 | 383920 | 0.6039 | 0 | 52.8949 | Eukaryota | 30838.0 | 0 | Chordata | 30838.0 | 0 | Carnivora | 19758.0 | 2 | Felidae | 12490.0 | 4 | Homo | 5952.0 | 7 | Homo sapiens | 5952.0 | 7 |
| scf7180000022100 | 24633 | 0.4028 | 0 | 25.9072 | Eukaryota | 4294.0 | 0 | Chordata | 4294.0 | 0 | Carnivora | 4294.0 | 0 | Felidae | 4294.0 | 0 | Felis | 3669.0 | 2 | Felis catus | 3669.0 | 2 |
| scf7180000023107 | 60526 | 0.3937 | 0 | 259.5948 | Eukaryota | 7078.0 | 0 | Chordata | 7078.0 | 0 | Carnivora | 7078.0 | 0 | Felidae | 7078.0 | 0 | Felis | 7078.0 | 0 | Felis catus | 7078.0 | 0 |
| scf7180000019641 | 280826 | 0.3667 | 0 | 50.6141 | Eukaryota | 14025.0 | 0 | Chordata | 14025.0 | 0 | Carnivora | 14025.0 | 0 | Felidae | 14025.0 | 0 | Felis | 12636.0 | 1 | Felis catus | 12636.0 | 1 |
| scf7180000021195 | 71026 | 0.5548 | 0 | 48.5161 | Eukaryota | 66020.0 | 0 | Chordata | 66020.0 | 0 | Carnivora | 66020.0 | 0 | Felidae | 55194.0 | 1 | Felis | 20978.0 | 5 | Felis catus | 20978.0 | 5 |
| scf7180000019780 | 26154 | 0.4635 | 0 | 294.4198 | Eukaryota | 3703.0 | 0 | Chordata | 3703.0 | 0 | Carnivora | 3703.0 | 0 | Felidae | 2785.0 | 2 | Panthera | 1768.0 | 3 | anthera pardu | 1424.0 | 4 |
| scf7180000018184 | 12484 | 0.6753 | 0 | 266.232 | Eukaryota | 2539.0 | 0 | Chordata | 2539.0 | 0 | Rodentia | 1040.0 | 4 | Caviidae | 783.0 | 5 | Cavia | 783.0 | 5 | avia porcellu | 783.0 | 6 |
| scf7180000020215 | 220461 | 0.3777 | 0 | 24.8226 | Eukaryota | 46137.0 | 0 | Chordata | 46137.0 | 0 | Carnivora | 46137.0 | 0 | Felidae | 46137.0 | 0 | Felis | 46137.0 | 0 | Felis catus | 46137.0 | 0 |
| scf7180000024011 | 1202208 | 0.4666 | 0 | 51.452 | Eukaryota | 43171.0 | 0 | Chordata | 43171.0 | 0 | Carnivora | 43171.0 | 0 | Felidae | 21784.0 | 4 | Acinonyx | 13038.0 | 5 | cinonyx jubat | 13038.0 | 5 |
| scf7180000019120 | 2771069 | 0.4167 | 0 | 51.1418 | Eukaryota | 47053.0 | 0 | Chordata | 47053.0 | 0 | Carnivora | 47053.0 | 0 | Felidae | 47053.0 | 0 | Felis | 42098.0 | 1 | Felis catus | 42098.0 | 1 |
| scf7180000020809 | 88598 | 0.3787 | 0 | 24.2083 | Eukaryota | 23141.0 | 0 | Chordata | 23141.0 | 0 | Carnivora | 23141.0 | 0 | Felidae | 23141.0 | 0 | Felis | 20827.0 | 1 | Felis catus | 20827.0 | 1 |
| scf7180000018847 | 172608 | 0.4198 | 0 | 50.2449 | Eukaryota | 44176.0 | 0 | Chordata | 44176.0 | 0 | Carnivora | 44176.0 | 0 | Felidae | 44176.0 | 0 | Felis | 39806.0 | 1 | Felis catus | 39806.0 | 1 |
| scf7180000022945 | 64726 | 0.4967 | 0 | 288.6792 | Eukaryota | 5722.0 | 0 | Chordata | 5722.0 | 0 | Carnivora | 5722.0 | 0 | Phocidae | 2328.0 | 3 | eptonychote | 1746.0 | 4 | nychotes wed | 1746.0 | 4 |
| scf7180000018080 | 1017878 | 0.4003 | 0 | 49.6603 | Eukaryota | 44909.0 | 0 | Chordata | 44909.0 | 0 | Carnivora | 44909.0 | 0 | Felidae | 44909.0 | 0 | Felis | 44909.0 | 0 | Felis catus | 44909.0 | 0 |
| scf7180000017719 | 434741 | 0.4036 | 0 | 50.265 | Eukaryota | 46969.0 | 0 | Chordata | 46969.0 | 0 | Carnivora | 46969.0 | 0 | Felidae | 46969.0 | 0 | Felis | 46969.0 | 0 | Felis catus | 46969.0 | 0 |
| scf7180000023942 | 524118 | 0.3985 | 0 | 47.3723 | Eukaryota | 25689.0 | 0 | Chordata | 25689.0 | 0 | Carnivora | 25689.0 | 0 | Herpestidae | 10420.0 | 3 | Suricata | 10420.0 | 4 | ricata suricat | 10420.0 | 5 |
| scf7180000023935 | 715117 | 0.393 | 0 | 52.4357 | Eukaryota | 58686.0 | 0 | Chordata | 58686.0 | 0 | Carnivora | 32715.0 | 2 | Herpestidae | 32715.0 | 2 | Suricata | 32715.0 | 2 | ricata suricat | 32715.0 | 2 |
| scf7180000021336 | 214277 | 0.604 | 100 | 48.6375 | Eukaryota | 8853.0 | 1 | Chordata | 8853.0 | 1 | Carnivora | 6215.0 | 3 | Felidae | 2780.0 | 6 | Acinonyx | 2780.0 | 6 | cinonyx jubat | 2780.0 | 6 |
| scf7180000017761 | 2943272 | 0.4278 | 0 | 51.3163 | Eukaryota | 47375.0 | 0 | Chordata | 47375.0 | 0 | Carnivora | 47375.0 | 0 | Felidae | 47375.0 | 0 | Felis | 47375.0 | 0 | Felis catus | 47375.0 | 0 |
| scf7180000018185 | 878148 | 0.4988 | 0 | 51.0445 | Eukaryota | 31667.0 | 0 | Chordata | 31667.0 | 0 | Carnivora | 31667.0 | 0 | Felidae | 25185.0 | 1 | Felis | 16205.0 | 3 | Felis catus | 16205.0 | 3 |
| scf7180000024277 | 491937 | 0.4076 | 0 | 47.9517 | Eukaryota | 43159.0 | 0 | Chordata | 43159.0 | 0 | Carnivora | 43159.0 | 0 | Felidae | 43159.0 | 0 | Felis | 43159.0 | 0 | Felis catus | 43159.0 | 0 |
| scf7180000020913 | 85979 | 0.4502 | 0 | 24.0606 | Eukaryota | 16776.0 | 0 | Chordata | 16776.0 | 0 | Primates | 11669.0 | 2 | Hominidae | 11669.0 | 2 | Pan | 6718.0 | 3 | an troglodyte | 6718.0 | 3 |
| scf7180000017428 | 7561702 | 0.39 | 0 | 50.5537 | Eukaryota | 57692.0 | 0 | Chordata | 57692.0 | 0 | Carnivora | 57692.0 | 0 | Canidae | 38939.0 | 1 | Canis | 38939.0 | 3 | Canis lupus | 38939.0 | 3 |
| scf7180000023279 | 58013 | 0.4953 | 0 | 185.1931 | Eukaryota | 5223.0 | 0 | Chordata | 5223.0 | 0 | Carnivora | 5223.0 | 0 | Felidae | 5223.0 | 0 | Felis | 4945.0 | 1 | Felis catus | 4945.0 | 1 |
| scf7180000017511 | 1177618 | 0.4167 | 0 | 49.9612 | Eukaryota | 61109.0 | 0 | Chordata | 61109.0 | 0 | Carnivora | 61109.0 | 0 | Felidae | 61109.0 | 0 | Panthera | 24749.0 | 3 | Panthera tigri | 20106.0 | 4 |
| scf7180000021246 | 48223 | 0.4822 | 0 | 48.0028 | Eukaryota | 4221.0 | 0 | Chordata | 4221.0 | 0 | Carnivora | 3820.0 | 1 | Felidae | 2071.0 | 2 | Felis | 2071.0 | 2 | Felis catus | 2071.0 | 2 |
| scf7180000021999 | 12777 | 0.6292 | 0 | 49.7822 | Eukaryota | 1972.0 | 0 | Chordata | 1972.0 | 0 | Carnivora | 1972.0 | 0 | Felidae | 1746.0 | 1 | Felis | 943.0 | 2 | Felis catus | 943.0 | 2 |
| scf7180000022310 | 24353 | 0.4008 | 0 | 50.7739 | Eukaryota | 15685.0 | 0 | Chordata | 15685.0 | 0 | Carnivora | 15685.0 | 0 | Felidae | 15685.0 | 0 | Felis | 14094.0 | 1 | Felis catus | 14094.0 | 1 |
| scf7180000020415 | 9982 | 0.4271 | 0 | 55.9485 | Eukaryota | 4981.0 | 0 | Chordata | 4981.0 | 0 | Carnivora | 4981.0 | 0 | Felidae | 4981.0 | 0 | Felis | 3266.0 | 2 | Felis catus | 3266.0 | 2 |
| scf7180000023443 | 15799 | 0.4579 | 0 | 357.3663 | no-hit | 0.0 | 0 | no-hit | 0.0 | 0 | no-hit | 0.0 | 0 | no-hit | 0.0 | 0 | no-hit | 0.0 | 0 | no-hit | 0.0 | 0 |

| | | | | | | | | | | | | | | | | | | | | | | |
|---|---|---|---|---|---|---|---|---|---|---|---|---|---|---|---|---|---|---|---|---|---|---|
| scf7180000023811 | 2136956 | 0.3694 | 0 | 50.5576 | Eukaryota | 47008.0 | 0 | Chordata | 47008.0 | 0 | Carnivora | 47008.0 | 0 | Felidae | 47008.0 | 0 | Felis | 47008.0 | 0 | Felis catus | 47008.0 | 0 |
| scf7180000020272 | 157853 | 0.6125 | 0 | 48.2434 | Eukaryota | 19081.0 | 0 | Chordata | 19081.0 | 0 | Carnivora | 19081.0 | 0 | Felidae | 17029.0 | 1 | Acinonyx | 5753.0 | 4 | cinonyx jubat | 5753.0 | 4 |
| scf7180000020868 | 1786058 | 0.3775 | 0 | 49.1536 | Eukaryota | 39031.0 | 0 | Chordata | 39031.0 | 0 | Carnivora | 39031.0 | 0 | Felidae | 39031.0 | 0 | Felis | 39031.0 | 0 | Felis catus | 39031.0 | 0 |
| scf7180000023750 | 16530 | 0.4621 | 0 | 337.1681 | no-hit | 0.0 | 0 | no-hit | 0.0 | 0 | no-hit | 0.0 | 0 | no-hit | 0.0 | 0 | no-hit | 0.0 | 0 | no-hit | 0.0 | 0 |
| scf7180000019665 | 249432 | 0.5635 | 0 | 48.609 | Eukaryota | 45430.0 | 0 | Chordata | 45430.0 | 0 | Carnivora | 37041.0 | 1 | Otariidae | 27827.0 | 3 | Zalophus | 23191.0 | 4 | phus californi | 23191.0 | 4 |
| scf7180000020392 | 108413 | 0.4015 | 0 | 43.2315 | Eukaryota | 17796.0 | 0 | Chordata | 17796.0 | 0 | Carnivora | 16006.0 | 1 | Felidae | 16006.0 | 1 | Felis | 14255.0 | 2 | Felis catus | 14255.0 | 2 |
| scf7180000021910 | 39680 | 0.3547 | 0 | 25.1863 | Eukaryota | 41185.0 | 0 | Chordata | 41185.0 | 0 | Carnivora | 41185.0 | 0 | Felidae | 41185.0 | 0 | Felis | 41185.0 | 0 | Felis catus | 41185.0 | 0 |
| scf7180000020004 | 138734 | 0.4649 | 0 | 49.283 | Eukaryota | 11723.0 | 0 | Chordata | 11723.0 | 0 | Carnivora | 11723.0 | 0 | Felidae | 11723.0 | 0 | Felis | 11723.0 | 0 | Felis catus | 11723.0 | 0 |
| scf7180000017433 | 6018991 | 0.3625 | 100 | 50.1937 | Eukaryota | 48990.0 | 0 | Chordata | 48990.0 | 0 | Carnivora | 48990.0 | 0 | Felidae | 33276.0 | 1 | Felis | 33276.0 | 1 | Felis catus | 33276.0 | 1 |
| scf7180000019126 | 400109 | 0.5227 | 0 | 50.7015 | Eukaryota | 31425.0 | 0 | Chordata | 31425.0 | 0 | Carnivora | 31425.0 | 0 | Felidae | 12802.0 | 4 | Lynx | 6742.0 | 7 | ynx canadens | 6742.0 | 7 |
| scf7180000021489 | 40699 | 0.4073 | 0 | 32.8444 | Eukaryota | 27335.0 | 0 | Chordata | 27335.0 | 0 | Carnivora | 27335.0 | 0 | Felidae | 27335.0 | 0 | Felis | 24551.0 | 1 | Felis catus | 24551.0 | 1 |
| scf7180000020457 | 152548 | 0.4048 | 0 | 51.2539 | Eukaryota | 9539.0 | 0 | Chordata | 9539.0 | 0 | Carnivora | 9539.0 | 0 | Felidae | 9539.0 | 0 | Felis | 9539.0 | 0 | Felis catus | 9539.0 | 0 |
| scf7180000021974 | 51794 | 0.3898 | 0 | 48.3368 | Eukaryota | 14315.0 | 0 | Chordata | 14315.0 | 0 | Carnivora | 14315.0 | 0 | Felidae | 14315.0 | 0 | Felis | 11676.0 | 1 | Felis catus | 11676.0 | 1 |
| scf7180000018169 | 1110028 | 0.4393 | 0 | 68.1702 | Eukaryota | 46487.0 | 0 | Chordata | 46487.0 | 0 | Carnivora | 46487.0 | 0 | Felidae | 46487.0 | 0 | Felis | 46487.0 | 0 | Felis catus | 46487.0 | 0 |
| scf7180000022442 | 16213 | 0.3724 | 0 | 24.249 | Eukaryota | 24758.0 | 0 | Chordata | 24758.0 | 0 | Carnivora | 24758.0 | 0 | Felidae | 24758.0 | 0 | Felis | 22277.0 | 1 | Felis catus | 22277.0 | 1 |
| scf7180000020064 | 139086 | 0.4552 | 0 | 25.4987 | Eukaryota | 33780.0 | 0 | Chordata | 33780.0 | 0 | Carnivora | 33780.0 | 0 | Felidae | 33780.0 | 0 | Felis | 33780.0 | 0 | Felis catus | 33780.0 | 0 |
| scf7180000021715 | 42676 | 0.4869 | 0 | 27.5561 | Eukaryota | 5195.0 | 0 | Chordata | 5195.0 | 0 | Primates | 2596.0 | 3 | Hominidae | 2596.0 | 5 | Homo | 2596.0 | 5 | Homo sapiens | 2596.0 | 5 |
| scf7180000024012 | 1458895 | 0.42 | 0 | 50.4321 | Eukaryota | 49511.0 | 0 | Chordata | 49511.0 | 0 | Carnivora | 49511.0 | 0 | Felidae | 37494.0 | 1 | Felis | 37494.0 | 1 | Felis catus | 37494.0 | 2 |
| scf7180000020819 | 63543 | 0.4253 | 0 | 35.0371 | Eukaryota | 46524.0 | 0 | Chordata | 46524.0 | 0 | Carnivora | 46524.0 | 0 | Felidae | 46524.0 | 0 | Felis | 46524.0 | 0 | Felis catus | 46524.0 | 0 |
| scf7180000024155 | 143923 | 0.4323 | 0 | 24.511 | Eukaryota | 39636.0 | 0 | Chordata | 39636.0 | 0 | Carnivora | 39636.0 | 0 | Herpestidae | 32710.0 | 1 | Suricata | 32710.0 | 1 | ricata suricat | 32710.0 | 1 |
| scf7180000023481 | 52348 | 0.3913 | 0 | 224.0207 | Eukaryota | 12335.0 | 0 | Chordata | 12335.0 | 0 | Carnivora | 12335.0 | 0 | Felidae | 12335.0 | 0 | Felis | 11114.0 | 1 | Felis catus | 11114.0 | 1 |
| scf7180000018462 | 561353 | 0.3524 | 0 | 48.5098 | Eukaryota | 46943.0 | 0 | Chordata | 46943.0 | 0 | Carnivora | 46943.0 | 0 | Felidae | 46943.0 | 0 | Felis | 46943.0 | 0 | Felis catus | 46943.0 | 0 |
| scf7180000018122 | 1146482 | 0.35 | 0 | 47.85 | Eukaryota | 45937.0 | 0 | Chordata | 45937.0 | 0 | Carnivora | 45937.0 | 0 | Felidae | 45937.0 | 0 | Felis | 45937.0 | 0 | Felis catus | 45937.0 | 0 |
| scf7180000020202 | 132366 | 0.6223 | 0 | 48.3024 | Eukaryota | 27798.0 | 0 | Chordata | 27798.0 | 0 | Primates | 22766.0 | 1 | Hominidae | 14993.0 | 3 | Homo | 14993.0 | 5 | Homo sapiens | 14993.0 | 5 |
| scf7180000020097 | 172892 | 0.3686 | 0 | 27.8071 | Eukaryota | 47678.0 | 0 | Chordata | 47678.0 | 0 | Carnivora | 47678.0 | 0 | Felidae | 47678.0 | 0 | Felis | 47678.0 | 0 | Felis catus | 47678.0 | 0 |
| scf7180000018164 | 106808 | 0.3809 | 0 | 47.7335 | Eukaryota | 42551.0 | 0 | Chordata | 42551.0 | 0 | Carnivora | 42551.0 | 0 | Felidae | 42551.0 | 0 | Felis | 42551.0 | 0 | Felis catus | 42551.0 | 0 |
| scf7180000020035 | 203438 | 0.4052 | 100 | 27.1995 | Eukaryota | 28776.0 | 0 | Chordata | 28776.0 | 0 | Carnivora | 25686.0 | 1 | Felidae | 25686.0 | 1 | Felis | 22808.0 | 2 | Felis catus | 22808.0 | 2 |
| scf7180000019240 | 374154 | 0.404 | 0 | 25.7152 | Eukaryota | 67427.0 | 0 | Chordata | 67427.0 | 0 | Carnivora | 47347.0 | 1 | Felidae | 26560.0 | 3 | Suricata | 20787.0 | 4 | ricata suricat | 20787.0 | 4 |
| scf7180000017594 | 2254894 | 0.4786 | 0 | 51.1904 | Eukaryota | 55728.0 | 0 | Chordata | 55728.0 | 0 | Carnivora | 55728.0 | 0 | Felidae | 55728.0 | 0 | Panthera | 39016.0 | 1 | Panthera tigri | 27883.0 | 2 |
| scf7180000019197 | 351094 | 0.3847 | 0 | 50.9218 | Eukaryota | 47891.0 | 0 | Chordata | 47891.0 | 0 | Carnivora | 47891.0 | 0 | Felidae | 47891.0 | 0 | Felis | 47891.0 | 0 | Felis catus | 47891.0 | 0 |
| scf7180000018504 | 116614 | 0.3987 | 0 | 51.6732 | Eukaryota | 40084.0 | 0 | Chordata | 40084.0 | 0 | Carnivora | 40084.0 | 0 | Felidae | 40084.0 | 0 | Felis | 40084.0 | 0 | Felis catus | 40084.0 | 0 |
| scf7180000021996 | 15999 | 0.44 | 0 | 27.6363 | Eukaryota | 9329.0 | 0 | Chordata | 9329.0 | 0 | Carnivora | 8298.0 | 1 | Felidae | 8298.0 | 1 | Felis | 4446.0 | 4 | Felis catus | 4446.0 | 4 |
| scf7180000024167 | 125688 | 0.4458 | 0 | 27.8801 | Eukaryota | 59646.0 | 0 | Chordata | 59646.0 | 0 | Carnivora | 59646.0 | 0 | Canidae | 34504.0 | 1 | Canis | 34504.0 | 1 | Canis lupus | 34504.0 | 1 |
| scf7180000021108 | 13575 | 0.5194 | 0 | 71.4688 | no-hit | 0.0 | 0 | no-hit | 0.0 | 0 | no-hit | 0.0 | 0 | no-hit | 0.0 | 0 | no-hit | 0.0 | 0 | no-hit | 0.0 | 0 |
| scf7180000020396 | 175867 | 0.5215 | 0 | 48.4842 | Eukaryota | 45980.0 | 0 | Chordata | 45980.0 | 0 | Carnivora | 45980.0 | 0 | Felidae | 45980.0 | 0 | Felis | 16827.0 | 3 | Felis catus | 16827.0 | 3 |
| scf7180000020177 | 233825 | 0.3517 | 0 | 49.772 | Eukaryota | 19508.0 | 0 | Chordata | 19508.0 | 0 | Carnivora | 19508.0 | 0 | Felidae | 19508.0 | 0 | Felis | 17541.0 | 1 | Felis catus | 17541.0 | 1 |
| scf7180000020916 | 113452 | 0.374 | 0 | 23.5887 | Eukaryota | 42053.0 | 0 | Chordata | 42053.0 | 0 | Carnivora | 42053.0 | 0 | Felidae | 42053.0 | 0 | Felis | 42053.0 | 0 | Felis catus | 42053.0 | 0 |
| scf7180000019592 | 249482 | 0.4449 | 0 | 50.0635 | Eukaryota | 24620.0 | 0 | Chordata | 24620.0 | 0 | Carnivora | 13310.0 | 1 | Hominidae | 11310.0 | 2 | Homo | 11310.0 | 4 | Homo sapiens | 11310.0 | 4 |
| scf7180000019434 | 313742 | 0.523 | 0 | 49.9436 | Eukaryota | 34393.0 | 0 | Chordata | 34393.0 | 0 | Carnivora | 34393.0 | 0 | Felidae | 34393.0 | 0 | Felis | 34393.0 | 0 | Felis catus | 34393.0 | 0 |
| scf7180000019920 | 167357 | 0.4323 | 0 | 52.6148 | Eukaryota | 9204.0 | 0 | Chordata | 9204.0 | 0 | Carnivora | 5202.0 | 2 | Canidae | 3305.0 | 4 | Canis | 3305.0 | 5 | Canis lupus | 3305.0 | 5 |
| scf7180000019256 | 466969 | 0.3732 | 0 | 50.5341 | Eukaryota | 46884.0 | 0 | Chordata | 46884.0 | 0 | Carnivora | 46884.0 | 0 | Felidae | 46884.0 | 0 | Felis | 46884.0 | 0 | Felis catus | 46884.0 | 0 |
| scf7180000022510 | 8098 | 0.3729 | 0 | 25.092 | Eukaryota | 2759.0 | 0 | Chordata | 2759.0 | 0 | Carnivora | 2759.0 | 0 | Felidae | 2759.0 | 0 | Felis | 1917.0 | 1 | Felis catus | 1917.0 | 1 |
| scf7180000021359 | 55275 | 0.3933 | 0 | 25.8534 | Eukaryota | 7889.0 | 0 | Chordata | 7889.0 | 0 | Carnivora | 3464.0 | 2 | Felidae | 3464.0 | 2 | Felis | 3464.0 | 2 | Felis catus | 3464.0 | 2 |
| scf7180000022462 | 20517 | 0.3669 | 0 | 23.7762 | Eukaryota | 8651.0 | 0 | Chordata | 8651.0 | 0 | Carnivora | 8651.0 | 0 | Felidae | 7014.0 | 2 | Felis | 6142.0 | 3 | Felis catus | 6142.0 | 3 |
| scf7180000017796 | 692901 | 0.3415 | 0 | 47.797 | Eukaryota | 45223.0 | 0 | Chordata | 45223.0 | 0 | Carnivora | 45223.0 | 0 | Felidae | 45223.0 | 0 | Felis | 45223.0 | 0 | Felis catus | 45223.0 | 0 |
| scf7180000021465 | 30807 | 0.3668 | 0 | 25.8728 | Eukaryota | 18011.0 | 0 | Chordata | 18011.0 | 0 | Carnivora | 18011.0 | 0 | Felidae | 18011.0 | 0 | Felis | 18011.0 | 0 | Felis catus | 18011.0 | 0 |
| scf7180000021919 | 22280 | 0.3588 | 0 | 22.1948 | Eukaryota | 34046.0 | 0 | Chordata | 34046.0 | 0 | Carnivora | 34046.0 | 0 | Felidae | 34046.0 | 0 | Felis | 34046.0 | 0 | Felis catus | 34046.0 | 0 |
| scf7180000018825 | 491646 | 0.3896 | 0 | 50.9119 | Eukaryota | 91748.0 | 0 | Chordata | 91748.0 | 0 | Carnivora | 91748.0 | 0 | Felidae | 61320.0 | 1 | Canis | 30428.0 | 3 | Canis lupus | 30428.0 | 3 |
| scf7180000024075 | 237532 | 0.6332 | 0 | 48.8259 | Eukaryota | 21890.0 | 0 | Chordata | 21890.0 | 0 | Carnivora | 21890.0 | 0 | Felidae | 21890.0 | 0 | Felis | 10959.0 | 1 | Felis catus | 10959.0 | 1 |
| scf7180000018311 | 890361 | 0.3763 | 0 | 50.2428 | Eukaryota | 46435.0 | 0 | Chordata | 46435.0 | 0 | Carnivora | 46435.0 | 0 | Felidae | 46435.0 | 0 | Felis | 46435.0 | 0 | Felis catus | 46435.0 | 0 |
| scf7180000020316 | 183660 | 0.3839 | 0 | 25.1931 | Eukaryota | 41753.0 | 0 | Chordata | 41753.0 | 0 | Carnivora | 41753.0 | 0 | Felidae | 41753.0 | 0 | Felis | 37636.0 | 1 | Felis catus | 37636.0 | 1 |
| scf7180000018066 | 1206376 | 0.3558 | 0 | 49.4397 | Eukaryota | 46424.0 | 0 | Chordata | 46424.0 | 0 | Carnivora | 46424.0 | 0 | Felidae | 46424.0 | 0 | Felis | 46424.0 | 0 | Felis catus | 46424.0 | 0 |
| scf7180000021617 | 49949 | 0.395 | 0 | 24.941 | Eukaryota | 4578.0 | 0 | Chordata | 4578.0 | 0 | Carnivora | 2265.0 | 2 | Canidae | 1869.0 | 3 | Canis | 1869.0 | 4 | Canis lupus | 1869.0 | 4 |
| scf7180000017966 | 552961 | 0.444 | 0 | 51.2928 | Eukaryota | 44830.0 | 0 | Chordata | 44830.0 | 0 | Carnivora | 44830.0 | 0 | Felidae | 44830.0 | 0 | Acinonyx | 22645.0 | 2 | cinonyx jubat | 22645.0 | 2 |
| scf7180000020369 | 152819 | 0.4153 | 0 | 51.7477 | Eukaryota | 12005.0 | 0 | Chordata | 12005.0 | 0 | Carnivora | 12005.0 | 0 | Felidae | 12005.0 | 0 | Felis | 7247.0 | 1 | Felis catus | 7247.0 | 1 |
| scf7180000020068 | 316185 | 0.3855 | 0 | 25.5638 | Eukaryota | 44532.0 | 0 | Chordata | 44532.0 | 0 | Carnivora | 44532.0 | 0 | Felidae | 44532.0 | 0 | Felis | 40105.0 | 1 | Felis catus | 40105.0 | 1 |
| scf7180000018794 | 1114881 | 0.3885 | 0 | 50.4819 | Eukaryota | 27012.0 | 0 | Chordata | 27012.0 | 0 | Carnivora | 27012.0 | 0 | Felidae | 27012.0 | 0 | Felis | 24310.0 | 1 | Felis catus | 24310.0 | 1 |
| scf7180000023755 | 5320541 | 0.399 | 0 | 51.8514 | Eukaryota | 54295.0 | 0 | Chordata | 54295.0 | 0 | Carnivora | 54295.0 | 0 | Felidae | 54295.0 | 0 | Felis | 22208.0 | 2 | Felis catus | 22208.0 | 3 |
| scf7180000023939 | 602459 | 0.4682 | 0 | 52.9343 | Eukaryota | 32873.0 | 0 | Chordata | 32873.0 | 0 | Carnivora | 32873.0 | 0 | Felidae | 32873.0 | 0 | Felis | 32873.0 | 0 | Felis catus | 32873.0 | 0 |
| scf7180000023953 | 736724 | 0.4273 | 0 | 50.5164 | Eukaryota | 24983.0 | 0 | Chordata | 24983.0 | 0 | Carnivora | 24983.0 | 0 | Felidae | 24983.0 | 0 | Felis | 22443.0 | 1 | Felis catus | 22443.0 | 1 |
| scf7180000018684 | 698830 | 0.4174 | 0 | 62.0622 | Eukaryota | 47464.0 | 0 | Chordata | 47464.0 | 0 | Carnivora | 47464.0 | 0 | Felidae | 47464.0 | 0 | Felis | 47464.0 | 0 | Felis catus | 47464.0 | 0 |
| scf7180000022723 | 4912 | 0.4049 | 0 | 21.263 | Eukaryota | 383.0 | 0 | Chordata | 383.0 | 0 | Artiodactyla | 383.0 | 0 | Bovidae | 383.0 | 0 | Bos | 383.0 | 0 | Bos mutus | 198.0 | 1 |
| scf7180000023551 | 19785 | 0.5644 | 0 | 605.3955 | Eukaryota | 17603.0 | 0 | Chordata | 17603.0 | 0 | Carnivora | 17603.0 | 0 | Felidae | 17603.0 | 0 | Felis | 14462.0 | 1 | Felis catus | 14462.0 | 1 |
| scf7180000019488 | 296001 | 0.3971 | 0 | 50.3956 | Eukaryota | 13804.0 | 0 | Chordata | 13804.0 | 0 | Carnivora | 9896.0 | 2 | Phocidae | 5272.0 | 4 | Neomonachu | 4008.0 | 5 | nachus schau | 4008.0 | 6 |
| scf7180000017678 | 1874195 | 0.3932 | 0 | 50.6899 | Eukaryota | 89350.0 | 0 | Chordata | 89350.0 | 0 | Carnivora | 89350.0 | 0 | Felidae | 89350.0 | 0 | Acinonyx | 89350.0 | 0 | cinonyx jubat | 89350.0 | 0 |
| scf7180000023944 | 789393 | 0.3596 | 0 | 50.0893 | Eukaryota | 45163.0 | 0 | Chordata | 45163.0 | 0 | Carnivora | 45163.0 | 0 | Felidae | 45163.0 | 0 | Felis | 40710.0 | 1 | Felis catus | 40710.0 | 1 |
| scf7180000021350 | 42802 | 0.3829 | 0 | 25.668 | Eukaryota | 11368.0 | 0 | Chordata | 11368.0 | 0 | Primates | 6748.0 | 2 | Hominidae | 5525.0 | 4 | Pan | 4139.0 | 7 | an troglodyte | 4139.0 | 7 |
| scf7180000019975 | 177669 | 0.5208 | 0 | 49.9977 | Eukaryota | 18459.0 | 0 | Chordata | 18459.0 | 0 | Carnivora | 18459.0 | 0 | Felidae | 14785.0 | 2 | Acinonyx | 3762.0 | 6 | cinonyx jubat | 3762.0 | 6 |
| scf7180000020689 | 102805 | 0.3671 | 0 | 25.0362 | Eukaryota | 47162.0 | 0 | Chordata | 47162.0 | 0 | Carnivora | 47162.0 | 0 | Felidae | 47162.0 | 0 | Felis | 42465.0 | 1 | Felis catus | 42465.0 | 1 |
| scf7180000022674 | 11355 | 0.3782 | 0 | 24.1939 | Eukaryota | 4070.0 | 0 | Chordata | 4070.0 | 0 | Carnivora | 3486.0 | 1 | Canidae | 1841.0 | 2 | Canis | 1841.0 | 2 | Canis lupus | 1841.0 | 2 |
| scf7180000017563 | 2559474 | 0.4094 | 0 | 51.2705 | Eukaryota | 47620.0 | 0 | Chordata | 47620.0 | 0 | Carnivora | 47620.0 | 0 | Felidae | 47620.0 | 0 | Felis | 47620.0 | 0 | Felis catus | 47620.0 | 0 |
| scf7180000018099 | 1030061 | 0.4557 | 0 | 53.5179 | Eukaryota | 28193.0 | 0 | Chordata | 28193.0 | 0 | Carnivora | 28193.0 | 0 | Felidae | 28193.0 | 0 | Felis | 25341.0 | 1 | Felis catus | 25341.0 | 1 |
| scf7180000021566 | 36584 | 0.3576 | 0 | 49.5684 | Eukaryota | 14329.0 | 0 | Chordata | 14329.0 | 0 | Carnivora | 14329.0 | 0 | Ursidae | 5642.0 | 3 | Ursus | 4271.0 | 7 | rsus maritimu | 2841.0 | 8 |
| scf7180000020854 | 118839 | 0.3811 | 0 | 24.6208 | Eukaryota | 46106.0 | 0 | Chordata | 46106.0 | 0 | Carnivora | 46106.0 | 0 | Felidae | 46106.0 | 0 | Felis | 46106.0 | 0 | Felis catus | 46106.0 | 0 |
| scf7180000018774 | 130202 | 0.4088 | 0 | 47.0114 | Eukaryota | 10684.0 | 0 | Chordata | 10684.0 | 0 | Carnivora | 8101.0 | 1 | Felidae | 7194.0 | 2 | Panthera | 4390.0 | 4 | Panthera tigri | 3298.0 | 6 |
| scf7180000020618 | 174303 | 0.4005 | 0 | 49.9203 | Eukaryota | 25675.0 | 0 | Chordata | 25675.0 | 0 | Carnivora | 25675.0 | 0 | Felidae | 25675.0 | 0 | Felis | 20449.0 | 1 | Felis catus | 20449.0 | 1 |
| scf7180000021213 | 36137 | 0.4646 | 0 | 290.6484 | Eukaryota | 5984.0 | 0 | Chordata | 5984.0 | 0 | Carnivora | 5984.0 | 0 | Phocidae | 2440.0 | 2 | eptonychote | 1830.0 | 4 | nychotes we | 1830.0 | 4 |
| scf7180000017876 | 65610 | 0.4951 | 0 | 48.9355 | Eukaryota | 47502.0 | 0 | Chordata | 47502.0 | 0 | Carnivora | 47502.0 | 0 | Felidae | 47502.0 | 0 | Felis | 47502.0 | 0 | Felis catus | 47502.0 | 0 |
| scf7180000020343 | 146915 | 0.5241 | 0 | 51.2413 | Eukaryota | 10411.0 | 1 | Chordata | 10411.0 | 1 | Carnivora | 6571.0 | 3 | Felidae | 6571.0 | 3 | Felis | 5485.0 | 5 | Felis catus | 5485.0 | 5 |
| scf7180000022525 | 8181 | 0.4281 | 0 | 583.907 | Eukaryota | 2158.0 | 0 | Chordata | 2158.0 | 0 | Carnivora | 2158.0 | 0 | Felidae | 2158.0 | 0 | Felis | 1525.0 | 2 | Felis catus | 1525.0 | 3 |
| scf7180000019329 | 424817 | 0.3982 | 0 | 51.4023 | Eukaryota | 45576.0 | 0 | Chordata | 45576.0 | 0 | Carnivora | 45576.0 | 0 | Felidae | 45576.0 | 0 | Felis | 45576.0 | 0 | Felis catus | 45576.0 | 0 |
| scf7180000020142 | 88363 | 0.4427 | 0 | 26.3986 | Eukaryota | 28804.0 | 0 | Chordata | 28804.0 | 0 | Carnivora | 28804.0 | 0 | Felidae | 28804.0 | 0 | Felis | 25900.0 | 1 | Felis catus | 25900.0 | 1 |
| scf7180000022374 | 28287 | 0.5899 | 0 | 45.1431 | Eukaryota | 14430.0 | 0 | Chordata | 14430.0 | 0 | Carnivora | 14430.0 | 0 | Felidae | 5993.0 | 4 | Panthera | 2969.0 | 6 | Canis lupus | 2782.0 | 8 |
| scf7180000017639 | 1926326 | 0.4293 | 0 | 52.3685 | Eukaryota | 47806.0 | 0 | Chordata | 47806.0 | 0 | Carnivora | 47806.0 | 0 | Felidae | 47806.0 | 0 | Felis | 47806.0 | 0 | Felis catus | 47806.0 | 0 |
| scf7180000019865 | 269294 | 0.4291 | 0 | 49.3723 | Eukaryota | 46141.0 | 0 | Chordata | 46141.0 | 0 | Carnivora | 46141.0 | 0 | Felidae | 46141.0 | 0 | Felis | 46141.0 | 0 | Felis catus | 46141.0 | 0 |
| scf7180000020106 | 159161 | 0.4188 | 0 | 26.7991 | Eukaryota | 30257.0 | 0 | Chordata | 30257.0 | 0 | Carnivora | 19721.0 | 2 | Felidae | 19721.0 | 2 | Felis | 17139.0 | 3 | Felis catus | 17139.0 | 3 |
| scf7180000019707 | 31342 | 0.3983 | 0 | 48.7935 | Eukaryota | 46183.0 | 0 | Chordata | 46183.0 | 0 | Carnivora | 46183.0 | 0 | Felidae | 46183.0 | 0 | Felis | 46183.0 | 0 | Felis catus | 46183.0 | 0 |
| scf7180000018238 | 980101 | 0.3753 | 0 | 50.0381 | Eukaryota | 46721.0 | 0 | Chordata | 46721.0 | 0 | Carnivora | 46721.0 | 0 | Felidae | 46721.0 | 0 | Felis | 46721.0 | 0 | Felis catus | 46721.0 | 0 |
| scf7180000018313 | 595599 | 0.4883 | 0 | 51.6045 | Eukaryota | 74458.0 | 0 | Chordata | 74458.0 | 0 | Carnivora | 74458.0 | 0 | Felidae | 47276.0 | 3 | Felis | 17570.0 | 7 | Felis catus | 17570.0 | 7 |
| scf7180000019836 | 366067 | 0.3959 | 0 | 51.8554 | Eukaryota | 46226.0 | 0 | Chordata | 46226.0 | 0 | Carnivora | 46226.0 | 0 | Felidae | 46226.0 | 0 | Felis | 46226.0 | 0 | Felis catus | 46226.0 | 0 |
| scf7180000019117 | 380993 | 0.4096 | 0 | 50.1051 | Eukaryota | 49007.0 | 0 | Chordata | 49007.0 | 0 | Carnivora | 49007.0 | 0 | Felidae | 39211.0 | 1 | Panthera | 19600.0 | 3 | Panthera tigri | 19600.0 | 3 |
| scf7180000023133 | 27776 | 0.5297 | 0 | 224.5661 | Eukaryota | 5282.0 | 0 | Chordata | 5282.0 | 0 | Carnivora | 5282.0 | 0 | Felidae | 5041.0 | 1 | Felis | 2877.0 | 3 | Felis catus | 2877.0 | 3 |
| scf7180000021522 | 10032 | 0.5224 | 0 | 160.323 | no-hit | 0.0 | 0 | no-hit | 0.0 | 0 | no-hit | 0.0 | 0 | no-hit | 0.0 | 0 | no-hit | 0.0 | 0 | no-hit | 0.0 | 0 |
| scf7180000024054 | 244105 | 0.5584 | 0 | 51.0512 | Eukaryota | 24458.0 | 0 | Chordata | 24458.0 | 0 | Carnivora | 20678.0 | 1 | Felidae | 5051.0 | 6 | Suricata | 4080.0 | 8 | ricata suricat | 4080.0 | 9 |
| scf7180000019091 | 356045 | 0.4306 | 0 | 51.0822 | Eukaryota | 37079.0 | 0 | Chordata | 37079.0 | 0 | Carnivora | 37079.0 | 0 | Felidae | 24483.0 | 2 | Felis | 11323.0 | 4 | Felis catus | 11323.0 | 5 |
| scf7180000022206 | 46588 | 0.42 | 0 | 51.2467 | Eukaryota | 4239.0 | 0 | Chordata | 4239.0 | 0 | Carnivora | 1746.0 | 2 | Hominidae | 1535.0 | 4 | Homo | 1535.0 | 5 | Homo sapiens | 1535.0 | 5 |
| scf7180000018987 | 389182 | 0.4997 | 0 | 51.0251 | Eukaryota | 96513.0 | 0 | Chordata | 96513.0 | 0 | Carnivora | 96513.0 | 0 | Felidae | 96513.0 | 0 | Acinonyx | 86832.0 | 1 | cinonyx jubat | 86832.0 | 1 |
| scf7180000020943 | 89804 | 0.4216 | 0 | 26.3099 | Eukaryota | 15363.0 | 0 | Chordata | 15363.0 | 0 | Carnivora | 15363.0 | 0 | Felidae | 15363.0 | 0 | Felis | 13732.0 | 1 | Felis catus | 13732.0 | 1 |
| scf7180000019234 | 398779 | 0.3631 | 0 | 48.3902 | Eukaryota | 78815.0 | 0 | Chordata | 78815.0 | 0 | Carnivora | 78815.0 | 0 | Felidae | 70781.0 | 1 | Acinonyx | 62912.0 | 2 | cinonyx jubat | 62912.0 | 2 |
| scf7180000018314 | 451761 | 0.3603 | 0 | 50.6219 | Eukaryota | 59570.0 | 0 | Chordata | 59570.0 | 0 | Carnivora | 59570.0 | 0 | Felidae | 59570.0 | 0 | Acinonyx | 35826.0 | 1 | cinonyx jubat | 35826.0 | 1 |
| scf7180000020938 | 66926 | 0.3593 | 0 | 43.7353 | Eukaryota | 44018.0 | 0 | Chordata | 44018.0 | 0 | Carnivora | 44018.0 | 0 | Felidae | 44018.0 | 0 | Felis | 44018.0 | 0 | Felis catus | 44018.0 | 0 |
| scf7180000023049 | 37418 | 0.5421 | 0 | 141.5248 | Eukaryota | 5327.0 | 0 | Chordata | 5327.0 | 0 | Carnivora | 5327.0 | 0 | Felidae | 5075.0 | 1 | Felis | 3249.0 | 2 | Felis catus | 3249.0 | 2 |
| scf7180000023775 | 2558068 | 0.3884 | 0 | 49.9748 | Eukaryota | 71864.0 | 0 | Chordata | 71864.0 | 0 | Carnivora | 71864.0 | 0 | Felidae | 71864.0 | 0 | Panthera | 50649.0 | 2 | anthera pardu | 28248.0 | 3 |
| scf7180000019651 | 43815 | 0.3784 | 0 | 38.7472 | Eukaryota | 39779.0 | 0 | Chordata | 39779.0 | 0 | Carnivora | 39779.0 | 0 | Felidae | 39779.0 | 0 | Felis | 39779.0 | 0 | Felis catus | 39779.0 | 0 |
| scf7180000021853 | 7922 | 0.5206 | 0 | 41.2235 | no-hit | 0.0 | 0 | no-hit | 0.0 | 0 | no-hit | 0.0 | 0 | no-hit | 0.0 | 0 | no-hit | 0.0 | 0 | no-hit | 0.0 | 0 |
| scf7180000023416 | 25169 | 0.4232 | 0 | 420.6685 | Eukaryota | 256563.0 | 0 | Chordata | 256563.0 | 0 | Carnivora | 256563.0 | 0 | Hyaenidae | 256563.0 | 0 | Proteles | 154798.0 | 1 | roteles cristat | 154798.0 | 1 |
| scf7180000021914 | 18440 | 0.3677 | 0 | 27.1052 | Eukaryota | 8850.0 | 0 | Chordata | 8850.0 | 0 | Carnivora | 4469.0 | 3 | Odobenidae | 1837.0 | 7 | Odobenus | 1837.0 | 8 | obenus rosma | 1837.0 | 8 |
| scf7180000022620 | 18273 | 0.5862 | 0 | 49.5116 | Eukaryota | 6894.0 | 0 | Chordata | 6894.0 | 0 | Artiodactyla | 5046.0 | 1 | Delphinidae | 4345.0 | 2 | Pseudorca | 4345.0 | 2 | udorca crassic | 4345.0 | 2 |
| scf7180000018588 | 9399 | 0.392 | 0 | 15.4307 | Eukaryota | 25577.0 | 0 | Chordata | 25577.0 | 0 | Carnivora | 25577.0 | 0 | Felidae | 25577.0 | 0 | Felis | 25577.0 | 0 | Felis catus | 25577.0 | 0 |
| scf7180000024065 | 314720 | 0.4442 | 0 | 51.425 | Eukaryota | 35255.0 | 0 | Chordata | 35255.0 | 0 | Carnivora | 35255.0 | 0 | Felidae | 35255.0 | 0 | Felis | 35255.0 | 0 | Felis catus | 35255.0 | 0 |
| scf7180000019733 | 20596 | 0.4139 | 0 | 194.9659 | Eukaryota | 6354.0 | 0 | Chordata | 6354.0 | 0 | Carnivora | 6354.0 | 0 | Felidae | 6354.0 | 0 | Felis | 6354.0 | 0 | Felis catus | 6354.0 | 0 |
| scf7180000021348 | 7163 | 0.5157 | 0 | 57.7021 | no-hit | 0.0 | 0 | no-hit | 0.0 | 0 | no-hit | 0.0 | 0 | no-hit | 0.0 | 0 | no-hit | 0.0 | 0 | no-hit | 0.0 | 0 |
| scf7180000024248 | 1792978 | 0.4548 | 0 | 50.0558 | Eukaryota | 46081.0 | 0 | Chordata | 46081.0 | 0 | Carnivora | 46081.0 | 0 | Felidae | 46081.0 | 0 | Felis | 46081.0 | 0 | Felis catus | 46081.0 | 0 |
| scf7180000023613 | 31265 | 0.4601 | 0 | 312.9555 | no-hit | 0.0 | 0 | no-hit | 0.0 | 0 | no-hit | 0.0 | 0 | no-hit | 0.0 | 0 | no-hit | 0.0 | 0 | no-hit | 0.0 | 0 |
| scf7180000022315 | 13333 | 0.3511 | 0 | 22.9714 | Eukaryota | 3363.0 | 0 | Chordata | 3363.0 | 0 | Carnivora | 2075.0 | 1 | Felidae | 2075.0 | 1 | Felis | 1856.0 | 2 | Felis catus | 1856.0 | 2 |
| scf7180000018383 | 965139 | 0.3926 | 0 | 52.578 | Eukaryota | 88280.0 | 0 | Chordata | 88280.0 | 0 | Carnivora | 88280.0 | 0 | Herpestidae | 44460.0 | 1 | Suricata | 44460.0 | 3 | ricata suricat | 44460.0 | 4 |
| scf7180000023201 | 71583 | 0.3852 | 0 | 356.2974 | Eukaryota | 19063.0 | 0 | Chordata | 19063.0 | 0 | Carnivora | 19063.0 | 0 | Felidae | 19063.0 | 0 | Felis | 19063.0 | 0 | Felis catus | 19063.0 | 0 |
| scf7180000018176 | 1025921 | 0.4526 | 0 | 51.2253 | Eukaryota | 46046.0 | 0 | Chordata | 46046.0 | 0 | Carnivora | 46046.0 | 0 | Felidae | 46046.0 | 0 | Felis | 46046.0 | 0 | Felis catus | 46046.0 | 0 |
| scf7180000018337 | 828560 | 0.4473 | 0 | 52.1136 | Eukaryota | 61555.0 | 0 | Chordata | 61555.0 | 0 | Carnivora | 61555.0 | 0 | Felidae | 55396.0 | 1 | Lynx | 30780.0 | 2 | ynx canadens | 30780.0 | 2 |
| scf7180000020511 | 99759 | 0.4009 | 0 | 50.5705 | Eukaryota | 20035.0 | 0 | Chordata | 20035.0 | 0 | Carnivora | 20035.0 | 0 | Felidae | 20035.0 | 0 | Panthera | 17630.0 | 1 | anthera pardu | 13706.0 | 2 |
| scf7180000019709 | 43918 | 0.3818 | 0 | 45.3906 | Eukaryota | 43831.0 | 0 | Chordata | 43831.0 | 0 | Carnivora | 43831.0 | 0 | Felidae | 43831.0 | 0 | Felis | 43831.0 | 0 | Felis catus | 43831.0 | 0 |
| scf7180000022504 | 16727 | 0.4199 | 0 | 22.949 | Eukaryota | 9104.0 | 0 | Chordata | 9104.0 | 0 | Carnivora | 9104.0 | 0 | Felidae | 9104.0 | 0 | Felis | 9104.0 | 0 | Felis catus | 9104.0 | 0 |
| scf7180000020792 | 23446 | 0.4343 | 0 | 38.4029 | Eukaryota | 20612.0 | 0 | Chordata | 20612.0 | 0 | Carnivora | 20612.0 | 0 | Felidae | 20612.0 | 0 | Felis | 20612.0 | 0 | Felis catus | 20612.0 | 0 |
| scf7180000023945 | 2135450 | 0.4089 | 0 | 51.8419 | Eukaryota | 49310.0 | 0 | Chordata | 49310.0 | 0 | Carnivora | 49310.0 | 0 | Felidae | 43557.0 | 1 | Felis | 37841.0 | 2 | Felis catus | 37841.0 | 2 |
| scf7180000022092 | 12177 | 0.6622 | 0 | 47.2861 | Eukaryota | 9157.0 | 0 | Chordata | 9157.0 | 0 | Carnivora | 9157.0 | 0 | Felidae | 8010.0 | 1 | Felis | 3244.0 | 4 | Felis catus | 3244.0 | 4 |
| scf7180000019190 | 48648 | 0.4189 | 0 | 39.9853 | Eukaryota | 30480.0 | 0 | Chordata | 30480.0 | 0 | Carnivora | 30480.0 | 0 | Felidae | 30480.0 | 0 | Felis | 27456.0 | 1 | Felis catus | 27456.0 | 1 |
| scf7180000020214 | 125101 | 0.5399 | 0 | 51.4695 | Eukaryota | 19779.0 | 0 | Chordata | 19779.0 | 0 | Carnivora | 19779.0 | 0 | Felidae | 12417.0 | 2 | Felis | 5468.0 | 6 | Felis catus | 5468.0 | 6 |
| scf7180000019551 | 612481 | 0.6044 | 0 | 50.4427 | Eukaryota | 16694.0 | 0 | Chordata | 16694.0 | 0 | Carnivora | 16694.0 | 0 | Felidae | 16694.0 | 0 | Felis | 15030.0 | 1 | Felis catus | 15030.0 | 1 |
| scf7180000021597 | 51285 | 0.4949 | 0 | 53.4422 | Eukaryota | 11168.0 | 1 | Chordata | 11168.0 | 1 | Carnivora | 9675.0 | 2 | Felidae | 9675.0 | 2 | Panthera | 3527.0 | 5 | anthera pardu | 3527.0 | 5 |
| scf7180000019282 | 389066 | 0.3809 | 0 | 40.1558 | Eukaryota | 41909.0 | 0 | Chordata | 41909.0 | 0 | Carnivora | 41909.0 | 0 | Felidae | 41909.0 | 0 | Felis | 41909.0 | 0 | Felis catus | 41909.0 | 0 |
| scf7180000021475 | 8524 | 0.5219 | 0 | 94.4519 | no-hit | 0.0 | 0 | no-hit | 0.0 | 0 | no-hit | 0.0 | 0 | no-hit | 0.0 | 0 | no-hit | 0.0 | 0 | no-hit | 0.0 | 0 |
| scf7180000019258 | 338546 | 0.4408 | 0 | 53.7446 | Eukaryota | 22018.0 | 0 | Chordata | 22018.0 | 0 | Carnivora | 22018.0 | 0 | Felidae | 12792.0 | 1 | Panthera | 12792.0 | 1 | anthera pardu | 12792.0 | 1 |
| scf7180000017833 | 1205871 | 0.4468 | 0 | 51.4754 | Eukaryota | 44466.0 | 0 | Chordata | 44466.0 | 0 | Carnivora | 44466.0 | 0 | Felidae | 44466.0 | 0 | Felis | 40080.0 | 1 | Felis catus | 40080.0 | 1 |
| scf7180000022736 | 7196 | 0.6729 | 0 | 637.2468 | Eukaryota | 1418.0 | 0 | Chordata | 1277.0 | 1 | Carnivora | 1277.0 | 1 | Otariidae | 536.0 | 4 | Eumetopias | 402.0 | 5 | netopias juba | 402.0 | 5 |
| scf7180000018556 | 840873 | 0.4525 | 0 | 51.3787 | Eukaryota | 49316.0 | 0 | Chordata | 49316.0 | 0 | Carnivora | 49316.0 | 0 | Felidae | 18365.0 | 2 | Zalophus | 16156.0 | 4 | phus californi | 16156.0 | 4 |
| scf7180000017726 | 40889 | 0.3344 | 0 | 24.2237 | Eukaryota | 7498.0 | 0 | Chordata | 7498.0 | 0 | Carnivora | 7498.0 | 0 | Felidae | 7498.0 | 0 | Felis | 6738.0 | 1 | Felis catus | 6738.0 | 1 |
| scf7180000022139 | 9454 | 0.4286 | 0 | 23.8622 | Eukaryota | 4395.0 | 0 | Chordata | 4395.0 | 0 | Carnivora | 4395.0 | 0 | Felidae | 4151.0 | 1 | Felis | 1464.0 | 4 | Felis catus | 1464.0 | 5 |
| scf7180000024156 | 125342 | 0.4738 | 0 | 25.0504 | Eukaryota | 16803.0 | 0 | Chordata | 16803.0 | 0 | Carnivora | 13362.0 | 2 | Felidae | 11895.0 | 3 | Acinonyx | 3284.0 | 6 | cinonyx jubat | 3284.0 | 6 |
| scf7180000017727 | 1732996 | 0.363 | 0 | 49.9841 | Eukaryota | 46660.0 | 0 | Chordata | 46660.0 | 0 | Carnivora | 46660.0 | 0 | Felidae | 46660.0 | 0 | Felis | 46660.0 | 0 | Felis catus | 46660.0 | 0 |
| scf7180000017757 | 1850084 | 0.3886 | 0 | 50.4295 | Eukaryota | 64040.0 | 0 | Chordata | 64040.0 | 0 | Carnivora | 64040.0 | 0 | Felidae | 64040.0 | 0 | Panthera | 25661.0 | 2 | anthera pardu | 19299.0 | 3 |
| scf7180000018649 | 560381 | 0.4082 | 0 | 51.3698 | Eukaryota | 47589.0 | 0 | Chordata | 47589.0 | 0 | Carnivora | 47589.0 | 0 | Herpestidae | 23064.0 | 2 | Suricata | 23064.0 | 4 | ricata suricat | 23064.0 | 4 |
| scf7180000023808 | 1936943 | 0.3679 | 0 | 50.4011 | Eukaryota | 47140.0 | 0 | Chordata | 47140.0 | 0 | Carnivora | 47140.0 | 0 | Felidae | 47140.0 | 0 | Felis | 47140.0 | 0 | Felis catus | 47140.0 | 0 |
| scf7180000023056 | 17614 | 0.5172 | 0 | 197.9809 | Eukaryota | 4583.0 | 0 | Chordata | 4583.0 | 0 | Carnivora | 4583.0 | 0 | Felidae | 4583.0 | 0 | Felis | 4311.0 | 1 | Felis catus | 4311.0 | 1 |
| scf7180000019366 | 320662 | 0.5149 | 0 | 50.7267 | Eukaryota | 21476.0 | 0 | Chordata | 21476.0 | 0 | Carnivora | 21476.0 | 0 | Felidae | 16314.0 | 2 | Panthera | 10009.0 | 5 | anthera pardu | 8125.0 | 6 |
| scf7180000017520 | 583371 | 0.3799 | 0 | 48.6956 | Eukaryota | 45557.0 | 0 | Chordata | 45557.0 | 0 | Carnivora | 45557.0 | 0 | Felidae | 45557.0 | 0 | Felis | 45557.0 | 0 | Felis catus | 45557.0 | 0 |
| scf7180000022704 | 14009 | 0.4927 | 0 | 27.4634 | Eukaryota | 2771.0 | 0 | Chordata | 2771.0 | 0 | Carnivora | 2771.0 | 0 | Felidae | 2771.0 | 0 | Felis | 2499.0 | 1 | Felis catus | 2499.0 | 1 |
| scf7180000018667 | 343645 | 0.421 | 0 | 52.2082 | Eukaryota | 72597.0 | 0 | Chordata | 72597.0 | 0 | Carnivora | 72597.0 | 0 | Felidae | 72597.0 | 0 | Acinonyx | 43028.0 | 2 | cinonyx jubat | 43028.0 | 2 |
| scf7180000021294 | 11085 | 0.4119 | 0 | 356.6996 | Eukaryota | 5094.0 | 0 | Chordata | 5094.0 | 0 | Carnivora | 5094.0 | 0 | Felidae | 5094.0 | 0 | Felis | 4075.0 | 1 | Felis catus | 4075.0 | 1 |
| scf7180000018532 | 1022733 | 0.3763 | 0 | 50.7733 | Eukaryota | 49896.0 | 0 | Chordata | 49896.0 | 0 | Carnivora | 49896.0 | 0 | Felidae | 49896.0 | 0 | Felis | 28418.0 | 1 | Felis catus | 28418.0 | 2 |
| scf7180000024100 | 69033 | 0.3885 | 0 | 25.921 | Eukaryota | 40031.0 | 0 | Chordata | 40031.0 | 0 | Carnivora | 40031.0 | 0 | Felidae | 40031.0 | 0 | Felis | 40031.0 | 0 | Felis catus | 40031.0 | 0 |
| scf7180000020607 | 30862 | 0.3847 | 0 | 23.1202 | Eukaryota | 41056.0 | 0 | Chordata | 41056.0 | 0 | Carnivora | 41056.0 | 0 | Felidae | 41056.0 | 0 | Felis | 41056.0 | 0 | Felis catus | 41056.0 | 0 |
| scf7180000021878 | 27293 | 0.4081 | 0 | 1153.9955 | no-hit | 0.0 | 0 | no-hit | 0.0 | 0 | no-hit | 0.0 | 0 | no-hit | 0.0 | 0 | no-hit | 0.0 | 0 | no-hit | 0.0 | 0 |
| scf7180000017495 | 3632334 | 0.4114 | 100 | 51.6723 | Eukaryota | 79500.0 | 0 | Chordata | 79500.0 | 0 | Carnivora | 79500.0 | 0 | Felidae | 79500.0 | 0 | Panthera | 79500.0 | 0 | anthera pardu | 79500.0 | 0 |
| scf7180000021933 | 38236 | 0.4315 | 0 | 45.2034 | Eukaryota | 4627.0 | 0 | Chordata | 4627.0 | 0 | Carnivora | 4627.0 | 0 | Felidae | 4627.0 | 0 | Felis | 2390.0 | 2 | Felis catus | 2390.0 | 2 |
| scf7180000018388 | 45656 | 0.5235 | 0 | 40.1148 | Eukaryota | 7277.0 | 1 | Chordata | 7277.0 | 1 | Artiodactyla | 3060.0 | 3 | Delphinidae | 3060.0 | 5 | agenorhynchu | 3060.0 | 5 | rhynchus obli | 3060.0 | 5 |
| scf7180000020900 | 59471 | 0.5605 | 0 | 54.2845 | Eukaryota | 8803.0 | 0 | Chordata | 8803.0 | 0 | Carnivora | 8803.0 | 0 | Felidae | 8803.0 | 0 | Panthera | 3556.0 | 2 | Felis catus | 3052.0 | 3 |
| scf7180000021177 | 88756 | 0.4549 | 0 | 49.5993 | Eukaryota | 3480.0 | 0 | Chordata | 3480.0 | 0 | Carnivora | 3480.0 | 0 | Felidae | 3480.0 | 0 | Felis | 2784.0 | 1 | Felis catus | 2784.0 | 1 |
| scf7180000019777 | 181340 | 0.5191 | 0 | 66.9671 | Eukaryota | 33363.0 | 0 | Chordata | 33363.0 | 0 | Carnivora | 33363.0 | 0 | Felidae | 33363.0 | 0 | Felis | 29985.0 | 1 | Felis catus | 29985.0 | 1 |
| scf7180000018480 | 631335 | 0.4196 | 0 | 52.1746 | Eukaryota | 62426.0 | 0 | Chordata | 62426.0 | 0 | Carnivora | 40593.0 | 3 | Otariidae | 11530.0 | 8 | Suricata | 9180.0 | 9 | ricata suricat | 9180.0 | 9 |

| | | | | | | | | | | | | | | | | | | | | | | |
|---|---|---|---|---|---|---|---|---|---|---|---|---|---|---|---|---|---|---|---|---|---|---|
| scf7180000021408 | 35412 | 0.3803 | 0 | 24.7134 | Eukaryota | 23891.0 | 0 | Chordata | 23891.0 | 0 | Carnivora | 23891.0 | 0 | Felidae | 23891.0 | 0 | Felis | 23891.0 | 0 | Felis catus | 23891.0 | 0 |
| scf7180000022065 | 16647 | 0.3668 | 0 | 45.2712 | Eukaryota | 12945.0 | 0 | Chordata | 12945.0 | 0 | Carnivora | 12945.0 | 0 | Felidae | 12945.0 | 0 | Felis | 12945.0 | 0 | Felis catus | 12945.0 | 0 |
| scf7180000019211 | 59006 | 0.4318 | 0 | 41.9696 | Eukaryota | 7652.0 | 1 | Chordata | 7652.0 | 1 | Primates | 3044.0 | 4 | Hominidae | 3044.0 | 4 | Pan | 1530.0 | 7 | an troglodyte | 1530.0 | 7 |
| scf7180000024226 | 1654467 | 0.5055 | 0 | 50.1623 | Eukaryota | 54644.0 | 0 | Chordata | 54644.0 | 0 | Carnivora | 54644.0 | 0 | Felidae | 31490.0 | 3 | Acinonyx | 14756.0 | 7 | cinonyx jubat | 14756.0 | 7 |
| scf7180000020756 | 97585 | 0.4314 | 0 | 25.5575 | Eukaryota | 21585.0 | 0 | Chordata | 21585.0 | 0 | Carnivora | 21585.0 | 0 | Felidae | 21585.0 | 0 | Felis | 17350.0 | 1 | Felis catus | 17350.0 | 1 |
| scf7180000017770 | 1855771 | 0.3488 | 0 | 47.9163 | Eukaryota | 47909.0 | 0 | Chordata | 47909.0 | 0 | Carnivora | 47909.0 | 0 | Felidae | 47909.0 | 0 | Felis | 38124.0 | 2 | Felis catus | 38124.0 | 2 |
| scf7180000020125 | 134310 | 0.3921 | 0 | 42.861 | Eukaryota | 42565.0 | 0 | Chordata | 42565.0 | 0 | Carnivora | 42565.0 | 0 | Felidae | 42565.0 | 0 | Felis | 42565.0 | 0 | Felis catus | 42565.0 | 0 |
| scf7180000019084 | 458982 | 0.4916 | 0 | 49.5553 | Eukaryota | 47672.0 | 0 | Chordata | 47672.0 | 0 | Carnivora | 47672.0 | 0 | Felidae | 47672.0 | 0 | Felis | 47672.0 | 0 | Felis catus | 47672.0 | 0 |
| scf7180000023820 | 2417913 | 0.3867 | 0 | 50.8533 | Eukaryota | 47572.0 | 0 | Chordata | 47572.0 | 0 | Carnivora | 47572.0 | 0 | Felidae | 47572.0 | 0 | Felis | 47572.0 | 0 | Felis catus | 47572.0 | 0 |
| scf7180000017617 | 2019310 | 0.4017 | 0 | 51.4696 | Eukaryota | 77635.0 | 0 | Chordata | 77635.0 | 0 | Carnivora | 77635.0 | 0 | Felidae | 77635.0 | 0 | Panthera | 46595.0 | 2 | anthera pardu | 38822.0 | 3 |
| scf7180000023950 | 860238 | 0.3866 | 0 | 51.2293 | Eukaryota | 47026.0 | 0 | Chordata | 47026.0 | 0 | Carnivora | 47026.0 | 0 | Felidae | 47026.0 | 0 | Felis | 42316.0 | 1 | Felis catus | 42316.0 | 1 |
| scf7180000018007 | 1308408 | 0.3858 | 0 | 50.6223 | Eukaryota | 49818.0 | 0 | Chordata | 49818.0 | 0 | Carnivora | 49818.0 | 0 | Felidae | 49818.0 | 0 | Felis | 38192.0 | 2 | Felis catus | 38192.0 | 2 |
| scf7180000019210 | 358186 | 0.4243 | 0 | 52.2235 | Eukaryota | 96516.0 | 0 | Chordata | 96516.0 | 0 | Carnivora | 96516.0 | 0 | Felidae | 43597.0 | 2 | Callorhinus | 16938.0 | 7 | llorhinus ursir | 16938.0 | 7 |
| scf7180000022913 | 39598 | 0.5302 | 0 | 335.5475 | Eukaryota | 15802.0 | 0 | Chordata | 15802.0 | 0 | Carnivora | 15802.0 | 0 | Felidae | 15802.0 | 0 | Felis | 15802.0 | 0 | Felis catus | 15802.0 | 0 |
| scf7180000020315 | 135687 | 0.504 | 0 | 50.1453 | Eukaryota | 15405.0 | 0 | Chordata | 15405.0 | 0 | Carnivora | 15405.0 | 0 | Felidae | 15405.0 | 0 | Felis | 6172.0 | 3 | Felis catus | 6172.0 | 3 |
| scf7180000017824 | 1071187 | 0.377 | 0 | 49.3026 | Eukaryota | 45769.0 | 0 | Chordata | 45769.0 | 0 | Carnivora | 45769.0 | 0 | Felidae | 45769.0 | 0 | Felis | 45769.0 | 0 | Felis catus | 45769.0 | 0 |
| scf7180000021942 | 35663 | 0.6452 | 0 | 52.986 | Eukaryota | 8999.0 | 0 | Chordata | 8999.0 | 0 | Carnivora | 8999.0 | 0 | Felidae | 4169.0 | 4 | Callorhinus | 1552.0 | 8 | llorhinus ursir | 1552.0 | 8 |
| scf7180000021536 | 37614 | 0.3837 | 0 | 24.8293 | Eukaryota | 45350.0 | 0 | Chordata | 45350.0 | 0 | Carnivora | 45350.0 | 0 | Felidae | 45350.0 | 0 | Felis | 45350.0 | 0 | Felis catus | 45350.0 | 0 |
| scf7180000022514 | 6933 | 0.6691 | 0 | 564.1423 | Eukaryota | 1414.0 | 0 | Chordata | 1273.0 | 1 | Carnivora | 1273.0 | 1 | Otariidae | 670.0 | 4 | Eumetopias | 402.0 | 5 | netopias juba | 402.0 | 5 |
| scf7180000020500 | 11374 | 0.6497 | 0 | 30.3224 | Eukaryota | 1851.0 | 0 | Chordata | 1851.0 | 0 | Carnivora | 1601.0 | 1 | Herpestidae | 1419.0 | 2 | Suricata | 1419.0 | 2 | ricata suricat | 1419.0 | 2 |
| scf7180000020840 | 72812 | 0.4175 | 0 | 26.4571 | Eukaryota | 44706.0 | 0 | Chordata | 44706.0 | 0 | Carnivora | 44706.0 | 0 | Felidae | 44706.0 | 0 | Felis | 40236.0 | 1 | Felis catus | 40236.0 | 1 |
| scf7180000024004 | 486027 | 0.4332 | 0 | 24.9207 | Eukaryota | 23311.0 | 0 | Chordata | 23311.0 | 0 | Carnivora | 23311.0 | 0 | Herpestidae | 13998.0 | 1 | Suricata | 13998.0 | 2 | ricata suricat | 13998.0 | 2 |
| scf7180000020005 | 8931 | 0.5364 | 0 | 22.3637 | Eukaryota | 3858.0 | 0 | Chordata | 3858.0 | 0 | Carnivora | 3858.0 | 0 | Felidae | 2932.0 | 2 | Panthera | 1123.0 | 5 | Felis catus | 1071.0 | 6 |
| scf7180000019604 | 803934 | 0.5236 | 0 | 51.0289 | Eukaryota | 18798.0 | 0 | Chordata | 18798.0 | 0 | Carnivora | 18798.0 | 0 | Felidae | 18798.0 | 0 | Lynx | 7577.0 | 2 | ynx canadens | 7577.0 | 2 |
| scf7180000019457 | 278125 | 0.5806 | 0 | 50.4133 | Eukaryota | 4656.0 | 1 | Chordata | 4656.0 | 1 | Carnivora | 2162.0 | 3 | Felidae | 2162.0 | 3 | Felis | 1494.0 | 6 | Felis catus | 1494.0 | 6 |
| scf7180000021218 | 98398 | 0.6356 | 0 | 40.6631 | Eukaryota | 3795.0 | 1 | Chordata | 3795.0 | 1 | Carnivora | 1639.0 | 4 | Felidae | 1639.0 | 4 | Felis | 1348.0 | 6 | Felis catus | 1348.0 | 6 |
| scf7180000022935 | 16997 | 0.4688 | 0 | 7212.9846 | Eukaryota | 9928.0 | 0 | Chordata | 9928.0 | 0 | Carnivora | 9928.0 | 0 | Felidae | 9928.0 | 0 | Felis | 8945.0 | 1 | Felis catus | 8945.0 | 1 |
| scf7180000019564 | 240398 | 0.3974 | 0 | 50.428 | Eukaryota | 47537.0 | 0 | Chordata | 47537.0 | 0 | Carnivora | 47537.0 | 0 | Felidae | 47537.0 | 0 | Felis | 47537.0 | 0 | Felis catus | 47537.0 | 0 |
| scf7180000022601 | 23779 | 0.5699 | 0 | 53.7689 | Eukaryota | 3837.0 | 0 | Chordata | 3837.0 | 0 | Carnivora | 2973.0 | 1 | Felidae | 1469.0 | 4 | Callorhinus | 802.0 | 6 | llorhinus ursir | 802.0 | 6 |
| scf7180000020523 | 130522 | 0.379 | 0 | 25.8548 | Eukaryota | 17080.0 | 0 | Chordata | 17080.0 | 0 | Carnivora | 8037.0 | 2 | Felidae | 8037.0 | 2 | Felis | 8037.0 | 3 | Felis catus | 8037.0 | 3 |
| scf7180000019455 | 39090 | 0.3565 | 0 | 45.0642 | Eukaryota | 14359.0 | 0 | Chordata | 14359.0 | 0 | Carnivora | 14359.0 | 0 | Felidae | 10174.0 | 2 | Panthera | 2971.0 | 6 | ynx canadens | 2791.0 | 8 |
| scf7180000022334 | 20722 | 0.4616 | 0 | 117.4423 | Eukaryota | 2139.0 | 0 | Chordata | 2139.0 | 0 | Carnivora | 2139.0 | 0 | Felidae | 1441.0 | 1 | Felis | 1232.0 | 4 | Felis catus | 1232.0 | 4 |
| scf7180000022829 | 539442 | 0.438 | 0 | 56.8133 | Eukaryota | 46016.0 | 0 | Chordata | 46016.0 | 0 | Carnivora | 46016.0 | 0 | Felidae | 46016.0 | 0 | Felis | 46016.0 | 0 | Felis catus | 46016.0 | 0 |
| scf7180000019209 | 28816 | 0.414 | 0 | 25.0108 | Eukaryota | 17402.0 | 0 | Chordata | 17402.0 | 0 | Carnivora | 17402.0 | 0 | Felidae | 16068.0 | 1 | Lynx | 7228.0 | 3 | ynx canadens | 7228.0 | 3 |
| scf7180000020460 | 118050 | 0.5129 | 0 | 48.5266 | Eukaryota | 19030.0 | 0 | Chordata | 19030.0 | 0 | Carnivora | 19030.0 | 0 | Felidae | 19030.0 | 0 | Panthera | 11418.0 | 1 | unresolved | 7612.0 | 2 |
| scf7180000019196 | 404668 | 0.4769 | 0 | 55.1687 | Eukaryota | 24322.0 | 0 | Chordata | 24322.0 | 0 | Carnivora | 21797.0 | 1 | Felidae | 14347.0 | 2 | Suricata | 7450.0 | 5 | ricata suricat | 7450.0 | 6 |
| scf7180000019585 | 73765 | 0.662 | 0 | 49.3636 | Eukaryota | 19223.0 | 0 | Chordata | 19223.0 | 0 | Carnivora | 19223.0 | 0 | Felidae | 11728.0 | 3 | Felis | 3911.0 | 7 | Felis catus | 3911.0 | 7 |
| scf7180000019466 | 32270 | 0.4562 | 0 | 183.5061 | Eukaryota | 8583.0 | 0 | Chordata | 8583.0 | 0 | Carnivora | 8583.0 | 0 | Felidae | 8583.0 | 0 | Felis | 6872.0 | 1 | Felis catus | 6872.0 | 1 |
| scf7180000018025 | 1188796 | 0.4242 | 0 | 51.7964 | Eukaryota | 30713.0 | 0 | Chordata | 30713.0 | 0 | Carnivora | 30713.0 | 0 | Felidae | 30713.0 | 0 | Felis | 27593.0 | 1 | Felis catus | 27593.0 | 1 |
| scf7180000021258 | 14668 | 0.3921 | 0 | 23.1248 | Eukaryota | 21638.0 | 0 | Chordata | 21638.0 | 0 | Carnivora | 21638.0 | 0 | Felidae | 21638.0 | 0 | Felis | 19471.0 | 1 | Felis catus | 19471.0 | 1 |
| scf7180000018301 | 906591 | 0.3887 | 0 | 50.1498 | Eukaryota | 48320.0 | 0 | Chordata | 48320.0 | 0 | Carnivora | 48320.0 | 0 | Felidae | 48320.0 | 0 | Felis | 48320.0 | 0 | Felis catus | 48320.0 | 0 |
| scf7180000024249 | 4427068 | 0.3705 | 0 | 49.9555 | Eukaryota | 47691.0 | 0 | Chordata | 47691.0 | 0 | Carnivora | 47691.0 | 0 | Felidae | 47691.0 | 0 | Felis | 47691.0 | 0 | Felis catus | 47691.0 | 0 |
| scf7180000019103 | 1121805 | 0.5327 | 0 | 50.2835 | Eukaryota | 20044.0 | 0 | Chordata | 20044.0 | 0 | Carnivora | 10883.0 | 3 | Ursidae | 6487.0 | 5 | Equus | 5528.0 | 6 | Equus caballu | 3710.0 | 8 |
| scf7180000020341 | 165959 | 0.4361 | 0 | 27.2583 | Eukaryota | 21842.0 | 0 | Chordata | 21842.0 | 0 | Carnivora | 21842.0 | 0 | Felidae | 21842.0 | 0 | Felis | 17466.0 | 1 | Felis catus | 17466.0 | 1 |
| scf7180000022955 | 29734 | 0.3949 | 0 | 370.7713 | Eukaryota | 6482.0 | 0 | Chordata | 6482.0 | 0 | Carnivora | 6482.0 | 0 | Felidae | 6482.0 | 0 | Felis | 6482.0 | 0 | Felis catus | 6482.0 | 0 |
| scf7180000023928 | 1052797 | 0.4312 | 0 | 52.8471 | Eukaryota | 29712.0 | 0 | Chordata | 29712.0 | 0 | Carnivora | 29712.0 | 0 | Felidae | 29712.0 | 0 | Felis | 26690.0 | 1 | Felis catus | 26690.0 | 1 |
| scf7180000021819 | 1406443 | 0.4611 | 0 | 51.3975 | Eukaryota | 61844.0 | 0 | Chordata | 61844.0 | 0 | Carnivora | 56614.0 | 1 | Felidae | 45806.0 | 2 | Panthera | 14374.0 | 6 | Felis catus | 12894.0 | 7 |
| scf7180000020041 | 25613 | 0.543 | 0 | 90.7641 | Eukaryota | 3677.0 | 0 | Chordata | 3677.0 | 0 | Carnivora | 3677.0 | 0 | Felidae | 3436.0 | 1 | Acinonyx | 1822.0 | 3 | cinonyx jubat | 1822.0 | 3 |
| scf7180000023767 | 3079841 | 0.3804 | 0 | 50.3614 | Eukaryota | 48513.0 | 0 | Chordata | 48513.0 | 0 | Carnivora | 48513.0 | 0 | Felidae | 48513.0 | 0 | Felis | 43475.0 | 1 | Felis catus | 43475.0 | 1 |
| scf7180000021076 | 84441 | 0.4281 | 0 | 26.5754 | Eukaryota | 18727.0 | 0 | Chordata | 18727.0 | 0 | Carnivora | 18727.0 | 0 | Felidae | 7046.0 | 4 | Suricata | 2169.0 | 9 | ricata suricat | 2169.0 | 9 |
| scf7180000018551 | 920016 | 0.4004 | 0 | 51.5752 | Eukaryota | 47334.0 | 0 | Chordata | 47334.0 | 0 | Carnivora | 47334.0 | 0 | Felidae | 47334.0 | 0 | Felis | 47334.0 | 0 | Felis catus | 47334.0 | 0 |
| scf7180000017952 | 1221712 | 0.3479 | 0 | 48.3807 | Eukaryota | 45405.0 | 0 | Chordata | 45405.0 | 0 | Carnivora | 45405.0 | 0 | Felidae | 45405.0 | 0 | Felis | 45405.0 | 0 | Felis catus | 45405.0 | 0 |
| scf7180000022114 | 13425 | 0.4261 | 0 | 24.3073 | Eukaryota | 3159.0 | 0 | Chordata | 3159.0 | 0 | Carnivora | 3159.0 | 0 | Felidae | 3159.0 | 0 | Felis | 1746.0 | 2 | Felis catus | 1746.0 | 2 |
| scf7180000020218 | 51545 | 0.4319 | 0 | 49.4106 | Eukaryota | 17178.0 | 0 | Chordata | 17178.0 | 0 | Carnivora | 17178.0 | 0 | Felidae | 17178.0 | 0 | Felis | 13744.0 | 1 | Felis catus | 13744.0 | 1 |
| scf7180000018619 | 600561 | 0.4214 | 0 | 51.835 | Eukaryota | 43537.0 | 0 | Chordata | 43537.0 | 0 | Carnivora | 43537.0 | 0 | Herpestidae | 29065.0 | 1 | Suricata | 29065.0 | 2 | ricata suricat | 29065.0 | 2 |
| scf7180000018299 | 26741 | 0.3448 | 0 | 36.4013 | Eukaryota | 10987.0 | 0 | Chordata | 10987.0 | 0 | Carnivora | 10987.0 | 0 | Felidae | 10987.0 | 0 | Felis | 8803.0 | 1 | Felis catus | 8803.0 | 1 |
| scf7180000024000 | 744374 | 0.4604 | 0 | 49.434 | Eukaryota | 47393.0 | 0 | Chordata | 47393.0 | 0 | Carnivora | 47393.0 | 0 | Felidae | 47393.0 | 0 | Felis | 42685.0 | 1 | Felis catus | 42685.0 | 1 |
| scf7180000021347 | 38434 | 0.3794 | 0 | 47.9263 | Eukaryota | 10920.0 | 0 | Chordata | 10920.0 | 0 | Carnivora | 10920.0 | 0 | Felidae | 7748.0 | 1 | Mustela | 3172.0 | 4 | ustela putori | 3172.0 | 4 |
| scf7180000017728 | 1710079 | 0.4329 | 0 | 51.3586 | Eukaryota | 44745.0 | 0 | Chordata | 44745.0 | 0 | Carnivora | 44745.0 | 0 | Felidae | 44745.0 | 0 | Felis | 40324.0 | 1 | Felis catus | 40324.0 | 1 |
| scf7180000018191 | 979043 | 0.384 | 0 | 50.955 | Eukaryota | 31622.0 | 0 | Chordata | 31622.0 | 0 | Carnivora | 31622.0 | 0 | Felidae | 31622.0 | 0 | Felis | 28399.0 | 1 | Felis catus | 28399.0 | 1 |
| scf7180000023216 | 47698 | 0.5314 | 0 | 311.6338 | Eukaryota | 12747.0 | 0 | Chordata | 12747.0 | 0 | Carnivora | 12747.0 | 0 | Felidae | 12747.0 | 0 | Felis | 12747.0 | 0 | Felis catus | 12747.0 | 0 |
| scf7180000017490 | 2764480 | 0.3798 | 0 | 50.0717 | Eukaryota | 73983.0 | 0 | Chordata | 73983.0 | 0 | Carnivora | 73983.0 | 0 | Felidae | 31128.0 | 2 | Lynx | 15428.0 | 6 | ynx canadens | 15428.0 | 6 |
| scf7180000021848 | 71659 | 0.5092 | 100 | 25.6925 | Eukaryota | 6662.0 | 0 | Chordata | 6662.0 | 0 | Primates | 4414.0 | 2 | Hominidae | 4414.0 | 3 | Homo | 3080.0 | 4 | Homo sapiens | 3080.0 | 4 |
| scf7180000021021 | 64442 | 0.4956 | 0 | 31.3279 | Eukaryota | 10414.0 | 0 | Chordata | 10414.0 | 0 | Carnivora | 10414.0 | 0 | Felidae | 9603.0 | 1 | Felis | 5111.0 | 3 | Felis catus | 5111.0 | 4 |
| scf7180000021667 | 59305 | 0.6173 | 0 | 50.9288 | Eukaryota | 6239.0 | 0 | Chordata | 6239.0 | 0 | Carnivora | 6239.0 | 0 | Felidae | 6239.0 | 0 | Felis | 6239.0 | 0 | Felis catus | 6239.0 | 0 |
| scf7180000020982 | 86289 | 0.3997 | 0 | 51.0116 | Eukaryota | 32712.0 | 0 | Chordata | 32712.0 | 0 | Carnivora | 32712.0 | 0 | Felidae | 32712.0 | 0 | Felis | 16357.0 | 1 | Felis catus | 16357.0 | 1 |
| scf7180000020587 | 97985 | 0.3923 | 0 | 25.6893 | Eukaryota | 22775.0 | 0 | Chordata | 22775.0 | 0 | Carnivora | 22775.0 | 0 | Felidae | 20185.0 | 1 | Felis | 12259.0 | 4 | Felis catus | 12259.0 | 4 |
| scf7180000020576 | 111939 | 0.3757 | 0 | 25.3602 | Eukaryota | 28914.0 | 0 | Chordata | 28914.0 | 0 | Carnivora | 28914.0 | 0 | Felidae | 22596.0 | 1 | Lynx | 7066.0 | 4 | ynx canadens | 7066.0 | 4 |
| scf7180000023841 | 2091635 | 0.4806 | 0 | 52.4749 | Eukaryota | 50903.0 | 0 | Chordata | 50903.0 | 0 | Carnivora | 50903.0 | 0 | Felidae | 35770.0 | 2 | Felis | 15530.0 | 6 | Felis catus | 15530.0 | 6 |
| scf7180000021231 | 82707 | 0.5476 | 0 | 49.0105 | Eukaryota | 13255.0 | 0 | Chordata | 13255.0 | 0 | Carnivora | 13255.0 | 0 | Felidae | 13255.0 | 0 | Felis | 9276.0 | 1 | Felis catus | 9276.0 | 1 |
| scf7180000022984 | 44121 | 0.5035 | 0 | 448.8057 | Eukaryota | 17085.0 | 0 | Chordata | 17085.0 | 0 | Carnivora | 17085.0 | 0 | Felidae | 17085.0 | 0 | Felis | 17085.0 | 0 | Felis catus | 17085.0 | 0 |
| scf7180000020886 | 65217 | 0.3675 | 0 | 25.1683 | Eukaryota | 41430.0 | 0 | Chordata | 41430.0 | 0 | Carnivora | 41430.0 | 0 | Felidae | 41430.0 | 0 | Felis | 41430.0 | 0 | Felis catus | 41430.0 | 0 |
| scf7180000021614 | 34338 | 0.3904 | 0 | 24.1133 | Eukaryota | 13292.0 | 0 | Chordata | 13292.0 | 0 | Carnivora | 13292.0 | 0 | Felidae | 13292.0 | 0 | Felis | 10643.0 | 1 | Felis catus | 10643.0 | 1 |
| scf7180000019767 | 13753 | 0.4198 | 0 | 34.4891 | Eukaryota | 2467.0 | 0 | Chordata | 2467.0 | 0 | Carnivora | 2467.0 | 0 | Felidae | 2193.0 | 1 | Felis | 1707.0 | 2 | Felis catus | 1707.0 | 2 |
| scf7180000023983 | 545477 | 0.5537 | 0 | 50.9802 | Eukaryota | 17855.0 | 0 | Chordata | 17855.0 | 0 | Carnivora | 16048.0 | 1 | Felidae | 10717.0 | 2 | Ailuropoda | 5331.0 | 5 | opoda melanc | 5331.0 | 5 |
| scf7180000019949 | 25501 | 0.5449 | 0 | 29.8198 | Eukaryota | 6703.0 | 0 | Chordata | 6703.0 | 0 | Carnivora | 6703.0 | 0 | Herpestidae | 4829.0 | 1 | Suricata | 4829.0 | 1 | ricata suricat | 4829.0 | 1 |
| scf7180000021104 | 95394 | 0.381 | 0 | 49.9488 | Eukaryota | 16146.0 | 0 | Chordata | 16146.0 | 0 | Carnivora | 16146.0 | 0 | Felidae | 16146.0 | 0 | Felis | 16146.0 | 0 | Felis catus | 16146.0 | 0 |
| scf7180000017783 | 1662581 | 0.3982 | 0 | 51.2324 | Eukaryota | 72226.0 | 0 | Chordata | 72226.0 | 0 | Carnivora | 72226.0 | 0 | Ursidae | 57108.0 | 1 | Ursus | 28784.0 | 2 | opoda melanc | 28324.0 | 3 |
| scf7180000022753 | 3709 | 0.5317 | 0 | 362.3375 | no-hit | 0.0 | 0 | no-hit | 0.0 | 0 | no-hit | 0.0 | 0 | no-hit | 0.0 | 0 | no-hit | 0.0 | 0 | no-hit | 0.0 | 0 |
| scf7180000023101 | 31696 | 0.5156 | 0 | 275.5843 | Eukaryota | 4810.0 | 0 | Chordata | 4810.0 | 0 | Carnivora | 4214.0 | 2 | Felidae | 3235.0 | 5 | Felis | 2235.0 | 7 | Felis catus | 2235.0 | 7 |
| scf7180000023925 | 1182863 | 0.408 | 0 | 50.9054 | Eukaryota | 46323.0 | 0 | Chordata | 46323.0 | 0 | Carnivora | 46323.0 | 0 | Felidae | 46323.0 | 0 | Felis | 46323.0 | 0 | Felis catus | 46323.0 | 0 |
| scf7180000019272 | 343058 | 0.3771 | 0 | 49.7854 | Eukaryota | 45166.0 | 0 | Chordata | 45166.0 | 0 | Carnivora | 27535.0 | 2 | Felidae | 27535.0 | 2 | Homo | 13191.0 | 4 | Homo sapiens | 13191.0 | 4 |
| scf7180000024165 | 150943 | 0.4285 | 0 | 26.4307 | Eukaryota | 32352.0 | 0 | Chordata | 32352.0 | 0 | Carnivora | 32352.0 | 0 | Felidae | 32352.0 | 0 | Acinonyx | 32352.0 | 0 | cinonyx jubat | 32352.0 | 0 |
| scf7180000022854 | 13371 | 0.6731 | 0 | 160.9352 | Eukaryota | 2565.0 | 0 | Chordata | 2565.0 | 0 | Primates | 1260.0 | 2 | Cercopithecidae | 1260.0 | 3 | Macaca | 1260.0 | 3 | acaca fascicul | 1260.0 | 3 |
| scf7180000021820 | 39928 | 0.4493 | 0 | 25.9225 | Eukaryota | 33554.0 | 0 | Chordata | 33554.0 | 0 | Carnivora | 33554.0 | 0 | Felidae | 33554.0 | 0 | Lynx | 16796.0 | 1 | ynx canadens | 16796.0 | 1 |
| scf7180000019331 | 271432 | 0.3801 | 0 | 50.7615 | Eukaryota | 46932.0 | 0 | Chordata | 46932.0 | 0 | Carnivora | 46932.0 | 0 | Felidae | 46932.0 | 0 | Felis | 46932.0 | 0 | Felis catus | 46932.0 | 0 |
| scf7180000018414 | 8611 | 0.3946 | 0 | 28.239 | Eukaryota | 5859.0 | 0 | Chordata | 5859.0 | 0 | Carnivora | 5859.0 | 0 | Felidae | 5859.0 | 0 | Felis | 5859.0 | 0 | Felis catus | 5859.0 | 0 |
| scf7180000022516 | 15543 | 0.371 | 0 | 23.4392 | Eukaryota | 20369.0 | 0 | Chordata | 20369.0 | 0 | Carnivora | 20369.0 | 0 | Felidae | 20369.0 | 0 | Felis | 20369.0 | 0 | Felis catus | 20369.0 | 0 |
| scf7180000020160 | 133773 | 0.3791 | 0 | 25.2283 | Eukaryota | 44107.0 | 0 | Chordata | 44107.0 | 0 | Carnivora | 44107.0 | 0 | Felidae | 44107.0 | 0 | Felis | 39608.0 | 1 | Felis catus | 39608.0 | 1 |
| scf7180000020762 | 23656 | 0.581 | 0 | 150.4119 | Eukaryota | 1960.0 | 0 | Chordata | 1960.0 | 0 | Primates | 884.0 | 2 | Felidae | 526.0 | 4 | Homo | 456.0 | 6 | Homo sapiens | 456.0 | 7 |
| scf7180000022604 | 23237 | 0.6738 | 0 | 48.2011 | Eukaryota | 15802.0 | 0 | Chordata | 15802.0 | 0 | Carnivora | 15802.0 | 0 | Felidae | 12811.0 | 2 | Lynx | 4924.0 | 6 | ynx canadens | 4924.0 | 6 |
| scf7180000019451 | 304750 | 0.4761 | 0 | 50.6068 | Eukaryota | 16066.0 | 0 | Chordata | 16066.0 | 0 | Carnivora | 16066.0 | 0 | Felidae | 16066.0 | 0 | Felis | 10975.0 | 2 | Felis catus | 10975.0 | 2 |
| scf7180000024133 | 283874 | 0.4182 | 0 | 51.7333 | Eukaryota | 42997.0 | 0 | Chordata | 42997.0 | 0 | Carnivora | 42997.0 | 0 | Felidae | 42997.0 | 0 | Felis | 42997.0 | 0 | Felis catus | 42997.0 | 0 |
| scf7180000022133 | 24749 | 0.3886 | 0 | 23.6157 | Eukaryota | 29670.0 | 0 | Chordata | 29670.0 | 0 | Carnivora | 29670.0 | 0 | Felidae | 29670.0 | 0 | Felis | 29670.0 | 0 | Felis catus | 29670.0 | 0 |
| scf7180000023877 | 1305082 | 0.3647 | 0 | 49.2032 | Eukaryota | 44831.0 | 0 | Chordata | 44831.0 | 0 | Carnivora | 44831.0 | 0 | Felidae | 44831.0 | 0 | Felis | 40282.0 | 1 | Felis catus | 40282.0 | 1 |
| scf7180000019235 | 313507 | 0.5203 | 0 | 51.5742 | Eukaryota | 25915.0 | 0 | Chordata | 25915.0 | 0 | Artiodactyla | 25915.0 | 0 | Delphinidae | 25915.0 | 0 | Globicephala | 18151.0 | 1 | obicephala me | 18151.0 | 1 |
| scf7180000020804 | 57298 | 0.5857 | 0 | 50.6156 | Eukaryota | 8752.0 | 0 | Chordata | 8752.0 | 0 | Carnivora | 8752.0 | 0 | Felidae | 5792.0 | 1 | Ursus | 2960.0 | 3 | Ursus arctos | 2960.0 | 3 |
| scf7180000023182 | 165332 | 0.4117 | 0 | 61.9259 | Eukaryota | 41209.0 | 0 | Chordata | 41209.0 | 0 | Carnivora | 41209.0 | 0 | Felidae | 41209.0 | 0 | Felis | 37103.0 | 1 | Felis catus | 37103.0 | 1 |
| scf7180000020184 | 122368 | 0.3769 | 0 | 25.7988 | Eukaryota | 44058.0 | 0 | Chordata | 44058.0 | 0 | Carnivora | 44058.0 | 0 | Felidae | 44058.0 | 0 | Felis | 44058.0 | 0 | Felis catus | 44058.0 | 0 |
| scf7180000023830 | 1406233 | 0.3709 | 0 | 49.9169 | Eukaryota | 44975.0 | 0 | Chordata | 44975.0 | 0 | Carnivora | 44975.0 | 0 | Felidae | 44975.0 | 0 | Felis | 44975.0 | 0 | Felis catus | 44975.0 | 0 |
| scf7180000021439 | 73795 | 0.3618 | 0 | 46.3401 | Eukaryota | 18446.0 | 0 | Chordata | 18446.0 | 0 | Carnivora | 16368.0 | 1 | Felidae | 16368.0 | 1 | Felis | 14499.0 | 2 | Felis catus | 14499.0 | 2 |
| scf7180000020366 | 141698 | 0.4921 | 0 | 27.536 | Eukaryota | 39294.0 | 0 | Chordata | 39294.0 | 0 | Carnivora | 35310.0 | 1 | Felidae | 23819.0 | 2 | Panthera | 11754.0 | 5 | ricata suricat | 11491.0 | 6 |
| scf7180000017487 | 10498 | 0.3904 | 0 | 35.7053 | Eukaryota | 19905.0 | 0 | Chordata | 19905.0 | 0 | Carnivora | 19905.0 | 0 | Felidae | 19905.0 | 0 | Felis | 17941.0 | 1 | Felis catus | 17941.0 | 1 |
| scf7180000023791 | 4407444 | 0.3856 | 100 | 50.8263 | Eukaryota | 57073.0 | 0 | Chordata | 57073.0 | 0 | Carnivora | 57073.0 | 0 | Felidae | 57073.0 | 0 | Felis | 57073.0 | 0 | Felis catus | 57073.0 | 0 |
| scf7180000017828 | 1970838 | 0.4761 | 0 | 52.9614 | Eukaryota | 70700.0 | 0 | Chordata | 70700.0 | 0 | Carnivora | 70700.0 | 0 | Felidae | 43853.0 | 3 | Panthera | 28782.0 | 6 | anthera pardu | 21247.0 | 7 |
| scf7180000022607 | 19113 | 0.3745 | 0 | 25.2509 | Eukaryota | 8664.0 | 0 | Chordata | 8664.0 | 0 | Carnivora | 8664.0 | 0 | Felidae | 8664.0 | 0 | Felis | 8664.0 | 0 | Felis catus | 8664.0 | 0 |
