## Supplementary material for "High-quality carnivore genomes from roadkill samples enable species delimitation in aardwolf and bat-eared fox": Table S7

| **Node** | **upper limit (My)** | **lower limit (My)** |
|---|---|---|
| Felis_catus,Lynx_lynx | NA | 5.3 |
| Felis_catus,Panthera_onca | 16.4 | NA |
| Felis_catus,Prionodon_pardicolor | NA | 28.5 |
| Proteles_cristatus_cristatus_TS307,Hyaena_hyaena | 16.4 | NA |
| Urva_javanica,Fossa_fossana_FMNH156648_SRR6053060 | NA | 16.4 |
| Proteles_cristatus_cristatus_TS307,Urva_javanica | NA | 16.4 |
| Civettictis_civetta,Genetta_servalina | NA | 11.2 |
| Canis_lupus,Urocyon_cinereoargenteus | NA | 5.3 |
| Mephitis_mephitis,Spilogale_putorius | NA | 1.8 |
| Procyon_lotor,Nasua_nasua | NA | 11.2 |
| Nasua_nasua,Ailurus_fulgens | NA | 16.4 |
| Arctocephalus_pusillus,Zalophus_californianus | NA | 1.8 |
| Mirounga_leonina,Phoca_fasciata | NA | 11.2 |
| Arctocephalus_pusillus,Phoca_fasciata | NA | 16.4 |
| Arctocephalus_pusillus,Nasua_nasua | NA | 28.5 |
| Ursus_arctos,Ailuropoda_melanoleuca | NA | 3.5 |
| Canis_lupus,Ursus_arctos | NA | 37 |
| Felis_catus,Canis_lupus | 65.8 | 50 |
