## Supplementary material for "High-quality carnivore genomes from roadkill samples enable species delimitation in aardwolf and bat-eared fox": Table S5

| **Species** | **Maker annotated genes** | **Number of orthologous genes after cleaning** | **Missing data** |
|---|---|---|---|
| *Acinonyx_jubatus* | NA | 12751 | 16.54 |
| *Ailuropoda_melanoleuca* | NA | 13411 | 9.57 |
| *Ailurus_fulgens* | 17619 | 11675 | 26.14 |
| *Arctocephalus_gazella* | 13714 | 8390 | 60.29 |
| *Bassariscus_sumichrasti* | 17573 | 11713 | 26.97 |
| *Callorhinus_ursinus* | 46164 | 7398 | 53.35 |
| *Canis_lupus_familiaris* | NA | 13808 | 6.04 |
| *Crocuta_Crocuta* | 16828 | 11525 | 29.17 |
| *Cryptoprocta_ferox* | 18199 | 12244 | 20.82 |
| *Enhydra_lutris* | NA | 13746 | 4.67 |
| *Eumetopias_jubatus* | 39926 | 8515 | 46.70 |
| *Felis_catus* | NA | 13047 | 11.98 |
| *Gulo_gulo* | 19696 | 8939 | 48.42 |
| *Helogale_parvula* | 18484 | 12005 | 23.92 |
| *Hyaena_hyaena* | 17663 | 12169 | 22.27 |
| *Leptonychotes_weddellii* | NA | 12439 | 22.57 |
| *Lutra_lutra* | 32278 | 12304 | 19.89 |
| *Lycaon_pictus* | 18068 | 11405 | 27.75 |
| *Lynx_canadensis* | 44864 | 7203 | 56.76 |
| *Lynx_pardinus* | 31025 | 9961 | 39.55 |
| *Manis_javanica* | NA | 12441 | 18.81 |
| *Mellivora_capensis* | 18194 | 12146 | 22.63 |
| *Mirounga_angustirostris* | 32777 | 12305 | 19.37 |
| *Mungos_mungo* | 18832 | 12241 | 21.30 |
| *Mustela_putorius* | NA | 13172 | 8.70 |
| *Nasua_narica* | 16858 | 11442 | 29.08 |
| *Neofelis_nebulosa* | 29785 | 12614 | 18.28 |
| *Neomonachus_schauinslandi* | NA | 13690 | 5.80 |
| *Neovison_vison* | 17294 | 11848 | 26.03 |
| *Odobenus_rosmarus* | NA | 13705 | 5.14 |
| *Otocyon_megalotis* | 18996 | 11981 | 22.02 |
| *Panthera_leo* | 17539 | 12162 | 22.91 |
| *Panthera_onca* | 16942 | 11758 | 27.22 |
| *Panthera_pardus* | NA | 13731 | 5.49 |
| *Panthera_tigris* | NA | 12841 | 15.37 |
| *Paradoxurus_hermaphroditus* | 17278 | 11387 | 30.18 |
| *Phoca_vitulina* | 18387 | 12057 | 22.73 |
| *Potos_flavus* | 27890 | 12234 | 22.80 |
| *Prionailurus_bengalensis* | 18057 | 12172 | 22.88 |
| *Procyon_lotor* | 16572 | 11165 | 32.46 |
| *Proteles_septentrionalis* | NA | 12050 | 22.43 |
| *Proteles_cristatus* | 17570 | 12062 | 22.96 |
| *Pteronura_brasiliensis* | 18445 | 12332 | 20.62 |
| *Puma_concolor* | 23358 | 10853 | 28.62 |
| *Spilogale_gracilis* | 17606 | 11744 | 26.61 |
| *Suricata_suricatta* | 17398 | 11707 | 26.32 |
| *Taxidea_taxus* | 15586 | 10141 | 42.19 |
| *Ursus_americanus* | 17694 | 11670 | 27.53 |
| *Ursus_arctos* | 18003 | 12145 | 22.21 |
| *Ursus_maritimus* | NA | 13111 | 13.56 |
| *Ursus_thibetanus* | 18365 | 12092 | 22.87 |
| *Vulpes_vulpes* | 18511 | 11688 | 26.02 |
| *Zalophus_californianus* | NA | 6305 | 64.65 |
