## Supplementary material for "High-quality carnivore genomes from roadkill samples enable species delimitation in aardwolf and bat-eared fox": Table S4

| NCBI Accession | Assembly_Name | Species | Order | Family | Assembly | Complete_All | Complete_Single | Complete_Duplicated | Fragmented | Missing |
|---|---|---|---|---|---|---|---|---|---|---|
| GCA_000181415.1 | ASM18141v1 | *Canis lupus* | Carnivora | Canidae | Canis lupus [ASM18141v1] | 3.4 | 3.4 | 0.0 | 20.1 | 76.5 |
| GCA_004023925.1 | FelNig_v1_BIUU | *Felis nigripes* | Carnivora | Felidae | Felis nigripes [FelNig_v1_BIUU] | 41.4 | 41.2 | 0.2 | 37.2 | 21.4 |
| GCA_000331495.1 | Beagle | *Canis lupus* | Carnivora | Canidae | Canis lupus [Beagle] | 41.8 | 41.2 | 0.6 | 38.1 | 20.1 |
| ISEM_TS307_SOAPdenovo | TS307_SOAPdenovo | *Proteles cristata* | Carnivora | Hyaenidae | Proteles cristata [ISEM_TS307_SOAPdenovo] | 44.9 | 44.6 | 0.3 | 31.7 | 23.4 |
| ISEM_TS305_SOAPdenovo | TS305_SOAPdenovo | *Otocyon megalotis* | Carnivora | Canidae | Otocyon megalotis [ISEM_TS305_SOAPdenovo] | 49.2 | 48.9 | 0.3 | 30.8 | 20.0 |
| GCA_003697995.1 | ASM369799v1 | *Taxidea taxus* | Carnivora | Mustelidae | Taxidea taxus [ASM369799v1] | 62.0 | 61.0 | 1.0 | 25.6 | 12.4 |
| GCA_000003115.1 | catChrV17e | *Felis catus* | Carnivora | Felidae | Felis catus [catChrV17e] | 68.7 | 68.3 | 0.4 | 13.8 | 17.5 |
| GCA_004023945.1 | HyaHya_v1_BIUU | *Hyaena hyaena* | Carnivora | Hyaenidae | Hyaena hyaena [HyaHya_v1_BIUU] | 74.7 | 73.7 | 1.0 | 16.2 | 9.1 |
| GCA_004023865.1 | MirAng_v1_BIUU | *Mirounga angustirostris* | Carnivora | Phocidae | Mirounga angustirostris [MirAng_v1_BIUU] | 75.4 | 74.7 | 0.7 | 16.5 | 8.1 |
| GCA_004024585.1 | ParHer_v1_BIUU | *Paradoxurus hermaphroditus* | Carnivora | Viverridae | Paradoxurus hermaphroditus [ParHer_v1_BIUU] | 78.3 | 77.7 | 0.6 | 15.6 | 6.1 |
| GCA_004023965.1 | SpiGra_v1_BIUU | *Spilogale gracilis* | Carnivora | Mephitidae | Spilogale gracilis [SpiGra_v1_BIUU] | 79.4 | 78.8 | 0.6 | 13.9 | 6.7 |
| GCA_004024625.1 | MelCap_v1_BIUU | *Mellivora capensis* | Carnivora | Mustelidae | Mellivora capensis [MelCap_v1_BIUU] | 81.9 | 81.3 | 0.6 | 12.8 | 5.3 |
| GCA_004023805.1 | PanOnc_v1_BIUU | *Panthera onca* | Carnivora | Felidae | Panthera onca [PanOnc_v1_BIUU] | 82.1 | 81.6 | 0.5 | 12.3 | 5.6 |
| GCF_000349705.1 | LepWed1.0 | *Leptonychotes weddellii* | Carnivora | Phocidae | Leptonychotes weddellii [LepWed1.0] | 82.4 | 81.4 | 1.0 | 11.9 | 5.7 |
| GCA_005406085.1 | Prionailurus_bengalensis_euptilurus_v01 | *Prionailurus bengalensis* | Carnivora | Felidae | Prionailurus bengalensis [Prionailurus_bengalensis_euptilurus_v01] | 84.2 | 83.7 | 0.5 | 11.0 | 4.8 |
| GCA_004023825.1 | VulLag_v1_BIUU | *Vulpes lagopus* | Carnivora | Canidae | Vulpes lagopus [VulLag_v1_BIUU] | 85.0 | 83.7 | 1.3 | 11.1 | 3.9 |
| GCA_004024605.1 | PteBra_v1_BIUU | *Pteronura brasiliensis* | Carnivora | Mustelidae | Pteronura brasiliensis [PteBra_v1_BIUU] | 85.2 | 84.6 | 0.6 | 10.3 | 4.5 |
| GCA_004123975.1 | Pco_k61 | *Puma concolor* | Carnivora | Felidae | Puma concolor [Pco_k61] | 86.2 | 85.6 | 0.6 | 8.8 | 5.0 |
| GCA_003344425.1 | ASM334442v1 | *Ursus americanus* | Carnivora | Ursidae | Ursus americanus [ASM334442v1] | 86.3 | 85.7 | 0.6 | 9.0 | 4.7 |
| GCA_004023845.1 | HelPar_v1_BIUU | *Helogale parvula* | Carnivora | Herpestidae | Helogale parvula [HelPar_v1_BIUU] | 87.0 | 86.3 | 0.7 | 9.0 | 4.0 |
| GCA_004216515.1 | sis1-161031-pseudohap | *Lycaon pictus* | Carnivora | Canidae | Lycaon pictus [sis1-161031-pseudohap] | 87.0 | 87.1 | 0.9 | 2.6 | 9.4 |
| GCA_004024565.1 | ZalCal_v1_BIUU | *Zalophus californianus* | Carnivora | Otariidae | Zalophus californianus [ZalCal_v1_BIUU] | 88.1 | 86.4 | 1.7 | 7.7 | 4.2 |
| GCA_004027395.1 | CanFam_VD_v1_BIUU | *Canis lupus* | Carnivora | Canidae | Canis lupus [CanFam_VD_v1_BIUU] | 88.4 | 87.4 | 1.0 | 7.9 | 3.7 |
| GCA_900500725.1 | ArcGazv1.4 | *Arctocephalus gazella* | Carnivora | Otariidae | Arctocephalus gazella [ArcGazv1.4] | 88.4 | 87.6 | 0.8 | 6.8 | 4.8 |
| GCA_900642305.1 | arcGaz3 | *Arctocephalus gazella* | Carnivora | Otariidae | Arctocephalus gazella [arcGaz3] | 88.6 | 87.9 | 0.7 | 6.7 | 4.7 |
| GCA_900006375.2 | Gulo_2.2_annotated | *Gulo gulo* | Carnivora | Mustelidae | Gulo gulo [Gulo_2.2] | 88.7 | 88.3 | 0.4 | 7.7 | 3.6 |
| GCA_004023905.1 | SurSur_v1_BIUU | *Suricata suricatta* | Carnivora | Herpestidae | Suricata suricatta [SurSur_v1_BIUU] | 88.9 | 87.8 | 1.1 | 7.4 | 3.7 |
| GCA_004023885.1 | CryFer_v1_BIUU | *Cryptoprocta ferox* | Carnivora | Eupleridae | Cryptoprocta ferox [CryFer_v1_BIUU] | 89.0 | 88.0 | 1.0 | 7.7 | 3.3 |
| GCA_004023785.1 | MunMun_v1_BIUU | *Mungos mungo* | Carnivora | Herpestidae | Mungos mungo [MunMun_v1_BIUU] | 90.1 | 89.3 | 0.8 | 6.7 | 3.2 |
| GCA_007922845.1 | UniMelb_Wolf_Refassem_1 | *Canis lupus* | Carnivora | Canidae | Canis lupus [UniMelb_Wolf_Refassem_1] | 90.4 | 89.4 | 1.0 | 5.5 | 4.1 |
| ISEM_TS307_MaSuRCA | TS307_MaSuRCA | *Proteles cristata* | Carnivora | Hyaenidae | Proteles cristata [ISEM_TS307_MaSuRCA] | 92.8 | 92.3 | 0.5 | 3.8 | 3.4 |
| ISEM_TS305_MaSuRCA | TS305_MaSuRCA | *Otocyon megalotis* | Carnivora | Canidae | Otocyon megalotis [ISEM_TS305_MaSuRCA] | 92.9 | 91.7 | 1.2 | 4.2 | 2.9 |
| GCA_006410715.1 | ASM641071v1 | *Enhydra lutris* | Carnivora | Mustelidae | Enhydra lutris [ASM641071v1] | 93.2 | 92.3 | 0.9 | 3.7 | 3.1 |
| GCF_003327715.1 | PumCon1.0 | *Puma concolor* | Carnivora | Felidae | Puma concolor [PumCon1] | 93.4 | 93.0 | 0.4 | 3.4 | 3.2 |
| GCF_001443585.1 | aciJub1 | *Acinonyx jubatus* | Carnivora | Felidae | Acinonyx jubatus [aciJub1] | 93.7 | 93.4 | 0.3 | 3.6 | 2.7 |
| GCA_001883655.1 | LycPicKen1.0 | *Lycaon pictus* | Carnivora | Canidae | Lycaon pictus [LycPicKen1.0] | 93.8 | 93.2 | 0.6 | 3.3 | 2.9 |
| GCA_002007445.1 | ASM200744v1 | *Ailuropoda melanoleuca* | Carnivora | Ursidae | Ailuropoda melanoleuca [ASM200744v1] | 94.4 | 93.1 | 1.3 | 2.6 | 3.0 |
| GCA_900661375.1 | LYPA1.0 | *Lynx pardinus* | Carnivora | Felidae | Lynx pardinus [LYPA1.0] | 94.4 | 93.9 | 0.5 | 3.1 | 2.5 |
| GCF_000464555.1 | PanTig1.0 | *Panthera tigris* | Carnivora | Felidae | Panthera tigris [PanTig1.0] | 94.4 | 94.0 | 0.4 | 3.2 | 2.4 |
| GCF_002201575.1 | ASM220157v1 | *Neomonachus schauinslandi* | Carnivora | Phocidae | Neomonachus schauinslandi [ASM220157v1] | 94.6 | 93.5 | 1.1 | 2.9 | 2.5 |
| GCF_003265705.1 | ASM326570v1 | *Callorhinus ursinus* | Carnivora | Otariidae | Callorhinus ursinus [ASM326570v1] | 94.7 | 82.7 | 12.0 | 2.3 | 3.0 |
| GCF_000687225.1 | UrsMar_1.0 | *Ursus maritimus* | Carnivora | Ursidae | Ursus maritimus [UrsMar_1] | 94.7 | 94.3 | 0.4 | 3.0 | 2.3 |
| GCA_001887905.1 | LycPicSAfr1.0 | *Lycaon pictus* | Carnivora | Canidae | Lycaon pictus [LycPicSAfr1.0] | 94.8 | 94.2 | 0.6 | 2.5 | 2.7 |
| GCF_000321225.1 | Oros_1.0 | *Odobenus rosmarus* | Carnivora | Odobenidae | Odobenus rosmarus [Oros_1.0] | 95.1 | 93.6 | 1.5 | 2.6 | 2.3 |
| GCF_004028035.1 | ASM402803v1 | *Eumetopias jubatus* | Carnivora | Otariidae | Eumetopias jubatus [ASM402803v1] | 95.2 | 92.9 | 2.3 | 2.4 | 2.4 |
| GCF_003160815.1 | VulVul2.2 | *Vulpes vulpes* | Carnivora | Canidae | Vulpes vulpes [VulVul2] | 95.2 | 93.8 | 0.6 | 2.8 | 2.0 |
| GCA_005444595.1 | UMICH_Zoey_3.1 | *Canis lupus* | Carnivora | Canidae | Canis lupus [UMICH_Zoey_3.1] | 95.2 | 93.9 | 1.3 | 2.6 | 2.2 |
| GCA_002007465.1 | ASM200746v1 | *Ailurus fulgens* | Carnivora | Ailuridae | Ailurus fulgens [ASM200746v1] | 95.2 | 94.3 | 0.9 | 2.6 | 2.2 |
| GCF_000215625.1 | MusPutFur1.0 | *Mustela putorius* | Carnivora | Mustelidae | Mustela putorius [MusPutFur1.0] | 95.2 | 94.6 | 0.6 | 2.5 | 2.3 |
| GCF_000004335.2 | AilMel_1.0 | *Ailuropoda melanoleuca* | Carnivora | Ursidae | Ailuropoda melanoleuca [AilMel_1.0] | 95.2 | 94.9 | 0.3 | 2.6 | 2.2 |
| GCF_900631625.1 | zalCal2.2 | *Zalophus californianus* | Carnivora | Otariidae | Zalophus californianus [zalCal2.2] | 95.3 | 93.8 | 1.5 | 2.2 | 2.5 |
| GCF_000002285.3 | CanFam3.1 | *Canis lupus* | Carnivora | Canidae | Canis lupus [CanFam3.1] | 95.3 | 94.0 | 1.3 | 2.4 | 2.3 |
| GCF_000181335.3 | Felis_catus_9.0 | *Felis catus* | Carnivora | Felidae | Felis catus [Felis_catus_9] | 95.3 | 94.9 | 0.4 | 2.4 | 2.3 |
| GCA_004886185.1 | Basenji_breed-1.1 | *Canis lupus* | Carnivora | Canidae | Canis lupus [Basenji_breed-1.1] | 95.5 | 94.1 | 1.4 | 2.1 | 2.4 |
| GCA_008641055.1 | ASM864105v1 | *Canis lupus* | Carnivora | Canidae | Canis lupus [ASM864105v1] | 95.5 | 94.2 | 1.3 | 2.1 | 2.4 |
| GCF_001857705.1 | PanPar1.0 | *Panthera pardus* | Carnivora | Felidae | Panthera pardus [PanPar1.0] | 95.5 | 94.7 | 0.8 | 2.3 | 2.2 |
| GCA_008692635.1 | BGI_CrCroc_1.0 | *Crocuta crocuta* | Carnivora | Hyaenidae | Crocuta crocuta [BGI_CrCroc_1.0] | 95.5 | 95.0 | 0.5 | 2.5 | 2.0 |
| GCA_003009895.1 | ASM300989v1 | *Hyaena hyaena* | Carnivora | Hyaenidae | Hyaena hyaena [ASM300989v1] | 95.6 | 94.9 | 0.7 | 2.4 | 2.0 |
| GCA_004348235.1 | GSC_HSeal_1.0 | *Phoca vitulina* | Carnivora | Phocidae | Phoca vitulina [GSC_HSeal_1.0] | 95.7 | 94.5 | 1.2 | 2.0 | 2.3 |
| GCF_003254725.1 | ASM325472v1 | *Canis lupus dingo* | Carnivora | Canidae | Canis lupus [ASM325472v1] | 95.8 | 94.4 | 1.4 | 1.9 | 2.3 |
| GCA_009660055.1 | ASM966005v1 | *Ursus thibetanus* | Carnivora | Ursidae | Ursus thibetanus [ASM966005v1] | 95.8 | 94.4 | 1.4 | 1.8 | 2.4 |
| GCA_900108605.1 | NNQGG.v01 | *Neovison vison* | Carnivora | Mustelidae | Neovison vison [NNQGG.v01] | 95.8 | 95.2 | 0.6 | 2.2 | 2.0 |
| GCF_000239315.1 | MusPutFurMale1.0 | *Mustela putorius* | Carnivora | Mustelidae | Mustela putorius [MusPutFurMale1] | 96.0 | 95.3 | 0.7 | 1.9 | 2.1 |
| GCF_002288905.2 | ASM228890v2 | *Enhydra lutris* | Carnivora | Mustelidae | Enhydra lutris [ASM228890v2] | 96.1 | 95.1 | 1.0 | 1.9 | 2.0 |
| GCF_003584765.1 | ASM358476v1 | *Ursus arctos* | Carnivora | Ursidae | Ursus arctos [ASM358476v1] | 96.1 | 95.2 | 0.9 | 1.9 | 2.0 |
| GCA_008795835.1 | PanLeo1.0 | *Panthera leo* | Carnivora | Felidae | Panthera leo [PanLeo1.0] | 96.2 | 95.6 | 0.6 | 1.9 | 1.9 |
| GCF_003709585.1 | Aci_jub_2 | *Acinonyx jubatus* | Carnivora | Felidae | Acinonyx jubatus [Aci_jub_2] | 96.3 | 95.8 | 0.5 | 1.8 | 1.9 |
