## Supplementary material for "High-quality carnivore genomes from roadkill samples enable species delimitation in aardwolf and bat-eared fox": Table S3

| Accession | Name | Species | Clade | Order | Status | ReleaseDate | LastUpdateDate | ScaffoldN50 | ContigN50 | ScaffoldCount | ContigCount | Total length | Submitter |
|---|---|---|---|---|---|---|---|---|---|---|---|---|---|
| GCF_000296735.1 | ChrAsi1.0 | *Chrysochloris asiatica* | Afrotheria | Afrosoricida | Scaffold | 20/09/12 | 19/02/14 | 13470186 | 19631 | 20500 | 391344 | 4210110458 | Broad Institute |
| GCA_004027935.1 | ChrAsi_v1_BIUU | *Chrysochloris asiatica* | Afrotheria | Afrosoricida | Scaffold | 16/01/19 | 16/01/19 | 6492 | 6444 | 3428078 | 3432954 | 4750461421 | Broad Institute |
| GCA_000313985.2 | ASM31398v2 | *Echinops telfairi* | Afrotheria | Afrosoricida | Scaffold | 16/07/19 | 17/07/19 | 54422506 | 20425 | 7489 | 277895 | 2947103070 | Broad Institute |
| GCA_004026705.1 | MicTal_v1_BIUU | *Microgale talazaci* | Afrotheria | Afrosoricida | Scaffold | 16/01/19 | 16/01/19 | 65143 | 56946 | 788895 | 795514 | 3236147583 | Broad Institute |
| GCA_004026845.1 | HetBruBak_v1_BIUU | *Heterohyrax brucei* | Afrotheria | Hyracoidea | Scaffold | 16/01/19 | 16/01/19 | 67904 | 62841 | 967933 | 971748 | 3618151652 | Broad Institute |
| GCA_004026925.2 | ProCapCap_v2_BIUU_UCD | *Procavia capensis* | Afrotheria | Hyracoidea | Scaffold | 15/07/19 | 17/07/19 | 9071062 | 38095 | 826553 | 1044315 | 3897621025 | Broad Institute |
| GCA_000152225.2 | Pcap_2.0 | *Procavia capensis* | Afrotheria | Hyracoidea | Scaffold | 16/12/14 | 16/12/14 | 905827 | 35792 | 65694 | 325568 | 3602182911 | Baylor College of Medicine |
| GCF_000299155.1 | EleEdw1.0 | *Elephantulus edwardii* | Afrotheria | Macroscelidea | Scaffold | 12/10/12 | 19/02/14 | 15011382 | 24219 | 8768 | 288162 | 3843982861 | Broad Institute |
| GCA_004027355.1 | EleEdw_v1_BIUU | *Elephantulus edwardii* | Afrotheria | Macroscelidea | Scaffold | 16/01/19 | 16/01/19 | 17713 | 16488 | 2475866 | 2486023 | 4288896762 | Broad Institute |
| GCF_000001905.1 | Loxafr3.0 | *Loxodonta africana* | Afrotheria | Proboscidea | Scaffold | 12/08/09 | 16/12/13 | 46401353 | 69023 | 2352 | 95866 | 3196738102 | Broad Institute |
| GCF_000243295.1 | TriManLat1.0 | *Trichechus manatus* | Afrotheria | Sirenia | Scaffold | 23/01/12 | 26/05/15 | 14442683 | 37750 | 6323 | 166490 | 3103808406 | Broad Institute |
| GCF_000298275.1 | OryAfe1.0 | *Orycteropus afer* | Afrotheria | Tubulidentata | Scaffold | 20/09/12 | 14/05/14 | 7875832 | 17683 | 22509 | 433653 | 4444080527 | Broad Institute |
| GCA_004365145.1 | OryAfe_v1_BIUU | *Orycteropus afer* | Afrotheria | Tubulidentata | Scaffold | 22/03/19 | 25/03/19 | 22965 | 20354 | 2675444 | 2690873 | 4687958587 | Broad Institute |
| GCA_004027255.2 | GalVar_v2_BIUU_UCD | *Galeopterus variegatus* | Euarchontoglires | Dermoptera | Scaffold | 15/07/19 | 17/07/19 | 7885395 | 34666 | 1523781 | 1653516 | 3349451543 | Broad Institute |
| GCF_000696425.1 | G_variegatus-3.0.2 | *Galeopterus variegatus* | Euarchontoglires | Dermoptera | Scaffold | 06/06/14 | 30/11/17 | 245189 | 20752 | 179514 | 511878 | 3187660572 | Washington University (WashU) |
| GCA_004026855.1 | LepAme_v1_BIUU | *Lepus americanus* | Euarchontoglires | Lagomorpha | Scaffold | 16/01/19 | 16/01/19 | 16725 | 15057 | 1592829 | 1602982 | 3123271264 | Broad Institute |
| GCF_000292845.1 | OchPri3.0 | *Ochotona princeps* | Euarchontoglires | Lagomorpha | Scaffold | 27/08/12 | 07/11/18 | 26863993 | 42347 | 10421 | 132963 | 2229840584 | Broad Institute |
| GCA_000164825.1 | ASM16482v1 | *Ochotona princeps* | Euarchontoglires | Lagomorpha | Scaffold | 16/10/10 | 29/11/16 | 88760 | 3265 | 187558 | 717779 | 3445784354 | Broad Institute |
| GCF_000260255.1 | OctDeg1.0 | *Octodon degus* | Euarchontoglires | Lagomorpha | Scaffold | 01/05/12 | 29/05/13 | 12091372 | 19847 | 7135 | 259905 | 2995889303 | Broad Institute |
| GCF_000003625.3 | OryCun2.0 | *Oryctolagus cuniculus* | Euarchontoglires | Lagomorpha | Chromosome | 29/10/09 | 05/06/14 | 35972871 | 64648 | 3241 | 84024 | 2737462810 | The Genome Sequencing Platform, The Assembly Computation and Development Core Team |
| GCA_004027835.1 | AloPal_v1_BIUU | *Alouatta palliata* | Euarchontoglires | Primates | Scaffold | 16/01/19 | 16/01/19 | 72427 | 51304 | 1136565 | 1152695 | 3033617595 | Broad Institute |
| GCF_000952055.2 | Anan_2.0 | *Aotus nancymaae* | Euarchontoglires | Primates | Scaffold | 12/06/17 | 28/06/17 | 8268663 | 126456 | 28922 | 112851 | 2861684817 | Baylor College of Medicine |
| GCA_004024785.1 | AteGeo_v1_BIUU | *Ateles geoffroyi* | Euarchontoglires | Primates | Scaffold | 15/01/19 | 15/01/19 | 73111 | 59033 | 869075 | 878490 | 2897033911 | Broad Institute |
| GCA_002754865.1 | ASM275486v1 | *Callithrix jacchus* | Euarchontoglires | Primates | Scaffold | 06/11/17 | 06/11/17 | 129239660 | 155284 | 39944 | 88439 | 2845375248 | Broad Institute |
| GCF_000004665.1 | Callithrix jacchus-3.2 | *Callithrix jacchus* | Euarchontoglires | Primates | Chromosome | 25/01/10 | 29/08/16 | 5167444 | 29273 | 14205 | 201371 | 2914958544 | Washington University (WashU) |
| GCA_001269965.1 | CIEA01 | *Callithrix jacchus* | Euarchontoglires | Primates | Contig | 18/08/15 | 18/08/15 | 61041 | 61041 | 109198 | 109198 | 2620625188 | Department of Biosciences and Informatics, Keio University |
| GCA_000832365.1 | ASM83236v1 | *Callithrix jacchus* | Euarchontoglires | Primates | Contig | 05/02/15 | 05/02/15 | 859 | 859 | 1992178 | 1992178 | 1504610980 | UN-NHPGC |
| GCF_000164805.1 | Tarsius_syrichta-2.0.1 | *Carlito syrichta* | Euarchontoglires | Primates | Scaffold | 18/09/13 | 15/07/17 | 401181 | 38165 | 337189 | 492903 | 3453864774 | Washington University (WashU) |
| GCA_004027755.1 | CebAlb_v1_BIUU | *Cebus albifrons* | Euarchontoglires | Primates | Scaffold | 16/01/19 | 16/01/19 | 31156 | 25652 | 1539166 | 1552889 | 3058667321 | Broad Institute |
| GCF_001604975.1 | Cebus_imitator-1.0 | *Cebus capucinus* | Euarchontoglires | Primates | Scaffold | 05/04/16 | 18/07/16 | 5274112 | 41196 | 7156 | 140597 | 2717703182 | McDonnell Genome Institute - Washington University School of Medicine |
| GCF_000955945.1 | Caty_1.0 | *Cercocebus atys* | Euarchontoglires | Primates | Scaffold | 19/03/15 | 30/03/15 | 12849131 | 112942 | 11433 | 76752 | 2848246356 | Human Genome Sequencing Center - BCM |
| GCA_004027615.1 | CertNeg_v1_BIUU | *Cercopithecus neglectus* | Euarchontoglires | Primates | Scaffold | 16/01/19 | 16/01/19 | 10270 | 9789 | 2119200 | 2126734 | 3278860052 | Broad Institute |
| GCA_004024725.1 | CheMed_v1_BIUU | *Cheirogaleus medius* | Euarchontoglires | Primates | Scaffold | 15/01/19 | 15/01/19 | 118572 | 90923 | 547751 | 555280 | 2393092913 | Broad Institute |
| GCF_000409795.2 | Chlorocebus_sabeus 1.1 | *Chlorocebus sabaeus* | Euarchontoglires | Primates | Chromosome | 25/03/14 | 08/12/17 | 81825804 | 90449 | 2004 | 162724 | 2789656328 | Vervet Genomics Consortium |
| GCF_000951035.1 | Cang.pa_1.0 | *Colobus angolensis* | Euarchontoglires | Primates | Scaffold | 12/03/15 | 30/03/15 | 7840981 | 38363 | 13124 | 197124 | 2970124662 | Baylor College of Medicine |
| GCA_004027145.1 | DauMad_v1_BIUU | *Daubentonia madagascariensis* | Euarchontoglires | Primates | Scaffold | 16/01/19 | 16/01/19 | 379919 | 298834 | 342451 | 344978 | 2498418007 | Broad Institute |
| GCA_000241425.1 | DauMad_1.0 | *Daubentonia madagascariensis* | Euarchontoglires | Primates | Scaffold | 05/01/12 | 30/11/16 | 3653 | 3358 | 3231305 | 3527892 | 2855365987 | Weill Cornell Medical College |
| GCA_004027335.1 | EryPat_v1_BIUU | *Erythrocebus patas* | Euarchontoglires | Primates | Scaffold | 16/01/19 | 16/01/19 | 34535 | 28523 | 2563960 | 2577035 | 3490485286 | Broad Institute |
| GCA_001262665.1 | Eflavifronsk33QCA | *Eulemur flavifrons* | Euarchontoglires | Primates | Scaffold | 06/08/15 | 06/08/15 | 413352 | 27331 | 38367 | 170563 | 2115568035 | University of Chicago |
| GCA_004027275.1 | EulFul_v1_BIUU | *Eulemur fulvus* | Euarchontoglires | Primates | Scaffold | 16/01/19 | 16/01/19 | 22561 | 21310 | 1693913 | 1697619 | 2748885701 | Broad Institute |
| GCA_001262655.1 | Emacaco_refEf_BWA_oneround | *Eulemur macaco* | Euarchontoglires | Primates | Scaffold | 06/08/15 | 06/08/15 | 405987 | 20038 | 26772 | 259238 | 2119875225 | University of Chicago |
| GCF_000151905.2 | gorGor4 | *Gorilla gorilla* | Euarchontoglires | Primates | Chromosome | 15/10/15 | 27/12/17 | 81227029 | 52934 | 40711 | 170105 | 3063362794 | Wellcome Trust Sanger Institute |
| GCA_900006655.3 | Susie3 | *Gorilla gorilla* | Euarchontoglires | Primates | Chromosome | 02/10/17 | 02/10/17 | 20634945 | 9406846 | 15297 | 16065 | 3084595669 | UNIVERSITY OF WASHINGTON |
| GCF_000001405.39 | GRCh38.p13 | *Homo sapiens* | Euarchontoglires | Primates | Chromosome | 01/03/19 | 14/06/19 | 59364414 | 56413054 | 639 | 1580 | 3272089205 | Genome Reference Consortium |
| GCA_004363605.1 | IndInd_v1_BIUU | *Indri indri* | Euarchontoglires | Primates | Scaffold | 22/03/19 | 22/03/19 | 28620 | 26447 | 1243540 | 1248425 | 2701849413 | Broad Institute |
| GCA_004024665.1 | LemCat_v1_BIUU | *Lemur catta* | Euarchontoglires | Primates | Scaffold | 15/01/19 | 15/01/19 | 215715 | 158439 | 575427 | 580026 | 2298449872 | Broad Institute |
| GCA_000222185.1 | MacFas_Jun2011 | *Macaca fascicularis* | Euarchontoglires | Primates | Chromosome | 09/08/11 | 30/11/16 | 155658674 | 3658 | 21 | 1628360 | 2878471044 | F. Hoffmann-La Roche Ltd |
| GCF_000364345.1 | Macaca_fascicularis_5.0 | *Macaca fascicularis* | Euarchontoglires | Primates | Chromosome | 12/06/13 | 08/12/17 | 88649475 | 86040 | 7601 | 87764 | 2946843737 | Washington University (WashU) |
| GCA_000230815.1 | CE_1.0 | *Macaca fascicularis* | Euarchontoglires | Primates | Chromosome | 20/10/11 | 30/11/16 | 1141162 | 13209 | 39921 | 439237 | 3126563191 | Beijing Genomics Institute |
| GCA_003118495.1 | macFus_1.0 | *Macaca fuscata* | Euarchontoglires | Primates | Contig | 11/05/18 | 11/05/18 | 94905 | 94905 | 90905 | 90905 | 2930706173 | National Institutes of Natural Sciences |
| GCF_003339765.1 | Mmul_10 | *Macaca mulatta* | Euarchontoglires | Primates | Chromosome | 13/03/19 | 26/04/19 | 82346004 | 46608966 | 2939 | 3182 | 2971331530 | The Genome Institute at Washington University School of Medicine |
| GCA_005453305.1 | ASM545330v1 | *Macaca mulatta* | Euarchontoglires | Primates | Contig | 17/05/19 | 17/05/19 | 8424208 | 8424208 | 1633 | 1633 | 2832421508 | None |
| GCF_000002255.3 | Mmul_051212 | *Macaca mulatta* | Euarchontoglires | Primates | Chromosome | 24/06/15 | 28/12/15 | 6094595 | 25707 | 122145 | 301015 | 3097387291 | Macaca mulatta Genome Sequencing Consortium |
| GCF_000772875.2 | Mmul_8.0.1 | *Macaca mulatta* | Euarchontoglires | Primates | Chromosome | 01/12/15 | 29/04/19 | 4193270 | 107156 | 284728 | 348494 | 3236224332 | Baylor College of Medicine Genome Sequencing Center |
| GCA_000230795.1 | CR_1.0 | *Macaca mulatta* | Euarchontoglires | Primates | Chromosome | 20/10/11 | 30/11/16 | 1660975 | 12841 | 34102 | 448689 | 2969971616 | Beijing Genomics Institute |
| GCA_005453525.2 | ASM545352v2 | *Macaca mulatta* | Euarchontoglires | Primates | Contig | 25/07/19 | 26/07/19 | 451490 | 451490 | 10217 | 10217 | 2179644428 | None |
| GCF_000956065.1 | Mnem_1.0 | *Macaca nemestrina* | Euarchontoglires | Primates | Scaffold | 19/03/15 | 30/03/15 | 15219753 | 106897 | 9733 | 94057 | 2948703511 | Baylor College of Medicine |
| GCF_000951045.1 | Mleu.le_1.0 | *Mandrillus leucophaeus* | Euarchontoglires | Primates | Scaffold | 12/03/15 | 30/03/15 | 3186748 | 31346 | 12821 | 246054 | 3061992840 | Baylor College of Medicine |
| GCA_004802615.1 | BGI_mandrill_1.0 | *Mandrillus sphinx* | Euarchontoglires | Primates | Scaffold | 22/04/19 | 22/04/19 | 3577924 | 21683 | 53550 | 282778 | 2860576561 | BGI |
| GCF_000165445.2 | Mmur_3.0 | *Microcebus murinus* | Euarchontoglires | Primates | Chromosome | 06/02/17 | 10/05/17 | 108171978 | 210702 | 7678 | 50984 | 2487409138 | Baylor College of Medicine |
| GCA_004024645.1 | MizCoq_v1_BIUU | *Mirza coquereli* | Euarchontoglires | Primates | Scaffold | 15/01/19 | 15/01/19 | 79947 | 60596 | 440879 | 451546 | 2340865962 | Broad Institute |
| GCF_000772465.1 | Charlie1.0 | *Nasalis larvatus* | Euarchontoglires | Primates | Chromosome | 12/11/14 | 01/12/17 | 137426304 | 13295 | 319549 | 553412 | 3011966170 | Proboscis Monkey Functional Genome Consortium |
| GCA_004027105.1 | NasLar_v1_BIUU | *Nasalis larvatus* | Euarchontoglires | Primates | Scaffold | 16/01/19 | 16/01/19 | 44943 | 34532 | 923815 | 942104 | 3050434315 | Broad Institute |
| GCF_000146795.2 | Nleu_3.0 | *Nomascus leucogenys* | Euarchontoglires | Primates | Chromosome | 10/12/12 | 05/12/13 | 52956880 | 35148 | 17484 | 197900 | 2962060179 | Gibbon Genome Sequencing Consortium |
| GCA_006542625.1 | Asia_NLE_v1 | *Nomascus leucogenys* | Euarchontoglires | Primates | Chromosome | 02/07/19 | 02/07/19 | 35886894 | 12782853 | 2251 | 2760 | 2843966406 | University of Washington |
| GCA_004027815.1 | NycCou_v1_BIUU | *Nycticebus coucang* | Euarchontoglires | Primates | Scaffold | 16/01/19 | 16/01/19 | 19724 | 17693 | 1937143 | 1946791 | 3294558825 | Broad Institute |
| GCF_000181295.1 | OtoGar3 | *Otolemur garnettii* | Euarchontoglires | Primates | Scaffold | 16/03/11 | 26/11/13 | 13852661 | 27100 | 7793 | 200240 | 2519724550 | Broad Institute of MIT and Harvard, USA, Cambridge |
| GCF_000258655.2 | panpan1.1 | *Pan paniscus* | Euarchontoglires | Primates | Chromosome | 18/08/15 | 12/01/18 | 8197324 | 66676 | 10274 | 121356 | 3286643896 | Max-Planck Institute for Evolutionary Anthropology |
| GCF_002880755.1 | Clint_PTRv2 | *Pan troglodytes* | Euarchontoglires | Primates | Chromosome | 19/01/18 | 03/04/18 | 53103722 | 12268567 | 4346 | 5061 | 3050398082 | University of Washington |
| GCF_000001515.7 | Pan_tro 3.0 | *Pan troglodytes* | Euarchontoglires | Primates | Chromosome | 03/05/16 | 27/03/18 | 26972556 | 384816 | 44449 | 72226 | 3231170666 | Chimpanzee Sequencing and Analysis Consortium |
| GCA_000325845.1 | ASM32584v1 | *Pan troglodytes* | Euarchontoglires | Primates | Contig | 19/12/12 | 19/12/12 | 15702 | 15702 | 361864 | 361864 | 2733948177 | Chimpanzee Sequencing and Analysis Consortium |
| GCF_000264685.3 | Panu_3.0 | *Papio anubis* | Euarchontoglires | Primates | Chromosome | 27/04/17 | 26/07/17 | 585721 | 149817 | 63235 | 118811 | 2959373024 | Human Genome Sequencing Center |
| GCF_002776525.2 | ASM277652v2 | *Piliocolobus tephrosceles* | Euarchontoglires | Primates | Scaffold | 11/09/18 | 25/09/18 | 13534873 | 98446 | 47449 | 111069 | 3038007327 | University of Oregon |
| GCA_004026645.1 | PitPit_v1_BIUU | *Pithecia pithecia* | Euarchontoglires | Primates | Scaffold | 16/01/19 | 16/01/19 | 83104 | 63292 | 892330 | 903646 | 2933562501 | Broad Institute |
| GCA_004027715.1 | CalDon_v1_BIUU | *Plecturocebus donacophilus* | Euarchontoglires | Primates | Scaffold | 16/01/19 | 16/01/19 | 46445 | 41149 | 1033983 | 1041365 | 2959764160 | Broad Institute |
| GCF_002880775.1 | Susie_PABv2 | *Pongo abelii* | Euarchontoglires | Primates | Chromosome | 19/01/18 | 05/12/18 | 98475126 | 11074009 | 5261 | 5814 | 3065052215 | University of Washington |
| GCA_000001545.3 | P_pygmaeus_2.0.2 | *Pongo abelii* | Euarchontoglires | Primates | Chromosome | 18/11/13 | 19/11/13 | 747460 | 15648 | 61538 | 408559 | 3441240245 | Orangutan Genome Sequencing Consortium |
| GCA_003258685.1 | Prosim_1.0 | *Prolemur simus* | Euarchontoglires | Primates | Scaffold | 20/06/18 | 20/06/18 | 2710671 | 47757 | 128596 | 208899 | 2411593676 | Henry Doorly Zoo & Aquarium |
| GCF_000956105.1 | Pcoq_1.0 | *Propithecus coquereli* | Euarchontoglires | Primates | Scaffold | 23/03/15 | 01/06/15 | 5604909 | 28129 | 22539 | 299070 | 2798152141 | Baylor College of Medicine |
| GCA_004024825.1 | PygNem_v1_BIUU | *Pygathrix nemaeus* | Euarchontoglires | Primates | Scaffold | 15/01/19 | 15/01/19 | 68569 | 51434 | 1431093 | 1445731 | 3217150031 | Broad Institute |
| GCF_001698545.1 | ASM169854v1 | *Rhinopithecus bieti* | Euarchontoglires | Primates | Scaffold | 03/08/16 | 25/08/16 | 2225337 | 36417 | 105032 | 226509 | 2977074946 | Laboratory for conservation and utilization of Bio-resource |
| GCA_007565055.1 | ASM756505v1 | *Rhinopithecus roxellana* | Euarchontoglires | Primates | Chromosome | 29/07/19 | 29/07/19 | 144559847 | 5723610 | 3256 | 6088 | 3038460517 | Northwest University |
| GCF_000769185.1 | Rrox_v1 | *Rhinopithecus roxellana* | Euarchontoglires | Primates | Scaffold | 29/10/14 | 08/12/17 | 1549224 | 77151 | 135512 | 196797 | 2899552139 | Novogene |
| GCA_004024885.1 | SagImp_v1_BIUU | *Saguinus imperator* | Euarchontoglires | Primates | Scaffold | 15/01/19 | 15/01/19 | 65636 | 53787 | 1666189 | 1675070 | 3493717886 | Broad Institute |
| GCF_000235385.1 | SaiBol1.0 | *Saimiri boliviensis* | Euarchontoglires | Primates | Scaffold | 18/11/11 | 16/12/13 | 18744880 | 38823 | 2686 | 151414 | 2608588537 | Broad Institute |
| GCA_004025065.1 | SemEnt_v1_BIUU | *Semnopithecus entellus* | Euarchontoglires | Primates | Scaffold | 15/01/19 | 15/01/19 | 29955 | 25064 | 1063749 | 1077535 | 3033044217 | Broad Institute |
| GCF_003255815.1 | Tgel_1.0 | *Theropithecus gelada* | Euarchontoglires | Primates | Chromosome | 19/06/18 | 29/06/18 | 130230028 | 310061 | 15308 | 35108 | 2889630685 | Kenneth Chiou's shared submissions |
| GCA_004027535.1 | AcoCah_v1_BIUU | *Acomys cahirinus* | Euarchontoglires | Rodentia | Scaffold | 16/01/19 | 16/01/19 | 65411 | 42476 | 371342 | 391811 | 2306070819 | Broad Institute |
| GCA_004027895.1 | AllBul_v1_BIUU | *Allactaga bullata* | Euarchontoglires | Rodentia | Scaffold | 16/01/19 | 16/01/19 | 36308 | 30651 | 1284382 | 1295369 | 3093575781 | Broad Institute |
| GCA_004027875.1 | AplRuf_v1_BIUU | *Aplodontia rufa* | Euarchontoglires | Rodentia | Scaffold | 16/01/19 | 16/01/19 | 37811 | 33245 | 1279526 | 1286779 | 3005535537 | Broad Institute |
| GCA_002335545.1 | Aspe_assembly01 | *Apodemus speciosus* | Euarchontoglires | Rodentia | Scaffold | 21/09/17 | 06/08/19 | 49031 | 4003 | 336124 | 609619 | 3494287372 | University of the Ryukyus |
| GCA_001305905.1 | ASM130590v1 | *Apodemus sylvaticus* | Euarchontoglires | Rodentia | Scaffold | 02/10/15 | 02/10/15 | 245982 | 1208 | 559629 | 3342720 | 3758143898 | University of Liverpool |
| GCA_004027915.1 | CapPil_v1_BIUU | *Capromys pilorides* | Euarchontoglires | Rodentia | Scaffold | 16/01/19 | 16/01/19 | 4081 | 4080 | 2352358 | 2355850 | 3023850356 | Broad Institute |
| GCF_001984765.1 | C.can genome v1.0 | *Castor canadensis* | Euarchontoglires | Rodentia | Scaffold | 04/02/17 | 17/02/17 | 317708 | 273954 | 21157 | 24477 | 2518306565 | Hospital for Sick Children |
| GCA_900168385.1 | bgp_v1 | *Castor canadensis* | Euarchontoglires | Rodentia | Scaffold | 03/05/17 | 06/08/19 | 295259 | 2701 | 629174 | 1785190 | 4167116470 | OREGON STATE UNIVERSITY |
| GCA_004027675.1 | CasCan_v1_BIUU | *Castor canadensis* | Euarchontoglires | Rodentia | Scaffold | 16/01/19 | 16/01/19 | 55723 | 49369 | 832147 | 837916 | 2761481163 | Broad Institute |
| GCA_000688575.1 | CavAp1.0 | *Cavia aperea* | Euarchontoglires | Rodentia | Scaffold | 09/05/14 | 09/05/14 | 27928671 | 1039 | 3131 | 2319938 | 2716396567 | Leibniz Institute for Zoo and Wildlife research |
| GCF_000151735.1 | Cavpor3.0 | *Cavia porcellus* | Euarchontoglires | Rodentia | Scaffold | 09/10/10 | 12/01/18 | 27942054 | 80583 | 3144 | 61604 | 2723219641 | The Genome Sequencing Platform, The Genome Assembly Team |
| GCA_004027695.1 | CavTsc_v1_BIUU | *Cavia tschudii* | Euarchontoglires | Rodentia | Scaffold | 16/01/19 | 16/01/19 | 91436 | 64807 | 1368179 | 1379500 | 3077352510 | Broad Institute |
| GCF_000276665.1 | ChiLan1.0 | *Chinchilla lanigera* | Euarchontoglires | Rodentia | Scaffold | 09/07/12 | 27/08/13 | 21893125 | 61105 | 2839 | 81656 | 2390868971 | Broad Institute |
| GCA_004027575.1 | CriGam_v1_BIUU | *Cricetomys gambianus* | Euarchontoglires | Rodentia | Scaffold | 16/01/19 | 16/01/19 | 110049 | 80761 | 1102952 | 1110425 | 2397721602 | Broad Institute |
| GCA_900186095.1 | CHOK1S_HZDv1 | *Cricetulus griseus* | Euarchontoglires | Rodentia | Scaffold | 31/10/17 | 31/10/17 | 62039716 | 97133 | 8264 | 71598 | 2358151106 | Eagle Genomics Ltd |
| GCF_003668045.1 | CriGri-PICR | *Cricetulus griseus* | Euarchontoglires | Rodentia | Scaffold | 23/10/18 | 18/12/18 | 19581774 | 2098530 | 1831 | 4826 | 2368923191 | CHO Genome Community |
| GCF_000419365.1 | C_griseus_v1.0 | *Cricetulus griseus* | Euarchontoglires | Rodentia | Scaffold | 12/07/13 | 26/12/18 | 1558295 | 27129 | 52710 | 218862 | 2360130144 | Beijing Genomics Institution-shenzhen |
| GCA_000448345.1 | Cgr1.0 | *Cricetulus griseus* | Euarchontoglires | Rodentia | Scaffold | 30/08/13 | 03/09/13 | 1236516 | 11899 | 28749 | 319219 | 2332774290 | CeBiTec, Bielefeld University |
| GCF_000223135.1 | CriGri_1.0 | *Cricetulus griseus* | Euarchontoglires | Rodentia | Scaffold | 26/08/11 | 27/12/17 | 1147233 | 39361 | 109152 | 265787 | 2399786748 | Beijing Genomics Institute |
| GCA_004027205.1 | CteGun_v1_BIUU | *Ctenodactylus gundi* | Euarchontoglires | Rodentia | Scaffold | 16/01/19 | 16/01/19 | 354548 | 218543 | 860073 | 865110 | 2322471743 | Broad Institute |
| GCA_004027165.1 | CteSoc_v1_BIUU | *Ctenomys sociabilis* | Euarchontoglires | Rodentia | Scaffold | 16/01/19 | 16/01/19 | 49073 | 37562 | 807571 | 823762 | 2834814551 | Broad Institute |
| GCA_004365215.1 | CunPac_v1_BIUU | *Cuniculus paca* | Euarchontoglires | Rodentia | Scaffold | 22/03/19 | 25/03/19 | 3892 | 3890 | 3768450 | 3771510 | 3399583187 | Broad Institute |
| GCA_004363535.1 | DasPun_v1_BIUU | *Dasyprocta punctata* | Euarchontoglires | Rodentia | Scaffold | 22/03/19 | 22/03/19 | 43703 | 38650 | 986664 | 993123 | 3012098298 | Broad Institute |
| GCA_004027595.1 | DinBra_v1_BIUU | *Dinomys branickii* | Euarchontoglires | Rodentia | Scaffold | 16/01/19 | 16/01/19 | 77918 | 64805 | 1158438 | 1164830 | 2688563719 | Broad Institute |
| GCF_000151885.1 | Dord_2.0 | *Dipodomys ordii* | Euarchontoglires | Rodentia | Scaffold | 12/12/14 | 07/07/15 | 11931245 | 48087 | 65193 | 148226 | 2236368823 | Baylor College of Medicine |
| GCA_004024685.1 | DipSte_v1_BIUU | *Dipodomys stephensi* | Euarchontoglires | Rodentia | Scaffold | 15/01/19 | 15/01/19 | 36811 | 30710 | 1269643 | 1278525 | 2346418196 | Broad Institute |
| GCA_004027295.1 | DolPat_v1_BIUU | *Dolichotis patagonum* | Euarchontoglires | Rodentia | Scaffold | 16/01/19 | 16/01/19 | 31026 | 27983 | 1905764 | 1911987 | 3289614431 | Broad Institute |
| GCA_001685075.1 | ASM168507v1 | *Ellobius lutescens* | Euarchontoglires | Rodentia | Scaffold | 14/07/16 | 06/08/19 | 242123 | 11648 | 58423 | 355225 | 2353188398 | Erasmus medical center (ErasmusMC) |
| GCA_001685115.1 | ASM168511v1 | *Ellobius lutescens* | Euarchontoglires | Rodentia | Scaffold | 14/07/16 | 06/08/19 | 13224 | 6461 | 413500 | 822944 | 2191766689 | Erasmus medical center (ErasmusMC) |
| GCA_001685095.1 | ETalpinus_0.1 | *Ellobius talpinus* | Euarchontoglires | Rodentia | Scaffold | 14/07/16 | 06/08/19 | 15246 | 8259 | 350460 | 644647 | 2265966243 | Erasmus medical center (ErasmusMC) |
| GCA_006547115.1 | GSC_porc_1.0 | *Erethizon dorsatum* | Euarchontoglires | Rodentia | Scaffold | 09/07/19 | 09/07/19 | 16680019 | 118744 | 15915 | 54415 | 2436547097 | Canada's Genomic Enterprise |
| GCF_000743615.1 | DMR_v1.0 | *Fukomys damarensis* | Euarchontoglires | Rodentia | Scaffold | 04/09/14 | 10/12/14 | 5314287 | 44830 | 74730 | 162545 | 2333892479 | BGI |
| GCA_004027185.1 | GliGli_v1_BIUU | *Glis glis* | Euarchontoglires | Rodentia | Scaffold | 16/01/19 | 16/01/19 | 30338 | 26062 | 433430 | 444115 | 2462087207 | Broad Institute |
| GCF_004785775.1 | NIH_TR_1.0 | *Grammomys surdaster* | Euarchontoglires | Rodentia | Scaffold | 12/04/19 | 21/04/19 | 8094907 | 51731 | 23656 | 100279 | 2412664998 | NIH Sequencing Center |
| GCA_004027655.1 | GraMur_v1_BIUU | *Graphiurus murinus* | Euarchontoglires | Rodentia | Scaffold | 16/01/19 | 16/01/19 | 28463 | 22138 | 1267939 | 1283630 | 2815014629 | Broad Institute |
| GCF_000247695.1 | HetGla_female_1.0 | *Heterocephalus glaber* | Euarchontoglires | Rodentia | Scaffold | 09/03/12 | 02/07/15 | 20532749 | 47778 | 4229 | 114653 | 2618204639 | Broad Institute |
| GCF_000230445.1 | HetGla_1.0 | *Heterocephalus glaber* | Euarchontoglires | Rodentia | Scaffold | 16/10/11 | 09/07/15 | 1603177 | 21750 | 39266 | 273990 | 2643961837 | Beijing Genomics Institute |
| GCA_004027455.1 | HydHyd_v1_BIUU | *Hydrochoerus hydrochaeris* | Euarchontoglires | Rodentia | Scaffold | 16/01/19 | 16/01/19 | 202224 | 148451 | 655342 | 661089 | 2734590088 | Broad Institute |
| GCA_004026905.1 | HysCri_v1_BIUU | *Hystrix cristata* | Euarchontoglires | Rodentia | Scaffold | 16/01/19 | 16/01/19 | 64768 | 57102 | 1136598 | 1140949 | 2470917678 | Broad Institute |
| GCA_004006685.1 | ASM400668v1 | *Ictidomys tridecemlineatus* | Euarchontoglires | Rodentia | Scaffold | 09/01/19 | 09/01/19 | 22643357 | 44127 | 10007 | 153524 | 2478641729 | 13-lined ground squirrel genome sequencing consortium |
| GCF_000236235.1 | SpeTri2.0 | *Ictidomys tridecemlineatus* | Euarchontoglires | Rodentia | Scaffold | 29/11/11 | 12/08/15 | 8192786 | 44137 | 12483 | 153488 | 2478393770 | Broad Institute |
| GCA_000181315.1 | ASM18131v1 | *Ictidomys tridecemlineatus* | Euarchontoglires | Rodentia | Contig | 19/11/10 | 29/11/16 | 2749 | 2749 | 797481 | 797481 | 1913367893 | The Genome Sequencing Platform, The Genome Assembly Team |
| GCF_000280705.1 | JacJac1.0 | *Jaculus jaculus* | Euarchontoglires | Rodentia | Scaffold | 07/08/12 | 29/05/13 | 22080993 | 15675 | 10898 | 337362 | 2835250225 | Broad Institute |
| GCA_003676075.2 | GSC_YBM_2.0 | *Marmota flaviventris* | Euarchontoglires | Rodentia | Scaffold | 29/05/19 | 29/05/19 | 11407078 | 143681 | 33623 | 68594 | 2581339348 | Canada's Genomic Enterprise |
| GCA_005280165.1 | ASM528016v1 | *Marmota himalayana* | Euarchontoglires | Rodentia | Scaffold | 10/05/19 | 11/05/19 | 1497034 | 102162 | 4363 | 46328 | 2471704417 | Xian Jiaotong University Health Science Center |
| GCF_001458135.1 | marMar2.1 | *Marmota marmota* | Euarchontoglires | Rodentia | Scaffold | 04/12/15 | 11/02/16 | 31340621 | 66492 | 14543 | 77915 | 2510587379 | MRC NATIONAL INSTITUTE FOR MEDICAL RESEARCH |
| GCF_002204375.1 | MunDraft-v1.0 | *Meriones unguiculatus* | Euarchontoglires | Rodentia | Scaffold | 21/06/17 | 27/06/17 | 374687 | 46546 | 68793 | 143857 | 2523124066 | Lucigen Corp. |
| GCA_004026785.1 | MerUng_v1_BIUU | *Meriones unguiculatus* | Euarchontoglires | Rodentia | Scaffold | 16/01/19 | 16/01/19 | 100883 | 69839 | 785187 | 796326 | 2637041433 | Broad Institute |
| GCF_000349665.1 | MesAur1.0 | *Mesocricetus auratus* | Euarchontoglires | Rodentia | Scaffold | 29/03/13 | 18/07/13 | 12753307 | 22511 | 21484 | 237700 | 2504925039 | Broad Institute |
| GCA_001305995.1 | ASM130599v1 | *Microtus agrestis* | Euarchontoglires | Rodentia | Scaffold | 02/10/15 | 02/10/15 | 896668 | 1380 | 230202 | 2813495 | 3124140565 | University of Liverpool |
| GCA_007455615.1 | ASM745561v1 | *Microtus arvalis* | Euarchontoglires | Rodentia | Scaffold | 25/07/19 | 26/07/19 | 52876 | 34797 | 1081432 | 1099517 | 2620579938 | Groningen Institute for Evolutionary Life Sciences |
| GCF_000317375.1 | MicOch1.0 | *Microtus ochrogaster* | Euarchontoglires | Rodentia | Chromosome | 07/12/12 | 21/08/13 | 17270019 | 21250 | 6335 | 187012 | 2287340943 | Broad Institute |
| GCA_007455595.1 | ASM745559v1 | *Microtus oeconomus* | Euarchontoglires | Rodentia | Scaffold | 25/07/19 | 26/07/19 | 114768 | 51221 | 562436 | 589027 | 2306554743 | Groningen Institute for Evolutionary Life Sciences |
| GCF_900094665.1 | CAROLI_EIJ_v1.1 | *Mus caroli* | Euarchontoglires | Rodentia | Chromosome | 02/05/17 | 21/05/17 | 122627250 | 30916 | 3163 | 136968 | 2553128900 | EBI |
| GCA_001624675.1 | NOD_ShiLtJ_v1 | *Mus musculus* | Euarchontoglires | Rodentia | Chromosome | 26/04/16 | 26/04/16 | 147352698 | 9439 | 5544 | 539650 | 2982063250 | Wellcome Trust Sanger Institute |
| GCA_001624475.1 | CBA_J_v1 | *Mus musculus* | Euarchontoglires | Rodentia | Chromosome | 26/04/16 | 26/04/16 | 144777665 | 13074 | 5466 | 370463 | 2921727955 | Wellcome Trust Sanger Institute |
| GCA_001632555.1 | C57BL_6NJ_v1 | *Mus musculus* | Euarchontoglires | Rodentia | Chromosome | 09/05/16 | 09/05/16 | 139118349 | 12707 | 3894 | 376733 | 2806800287 | Wellcome Trust Sanger Institute |
| GCA_001632615.1 | LP_J_v1 | *Mus musculus* | Euarchontoglires | Rodentia | Chromosome | 09/05/16 | 09/05/16 | 135164694 | 12608 | 3499 | 364701 | 2730798503 | Wellcome Trust Sanger Institute |
| GCA_001624185.1 | 129S1_SvImJ_v1 | *Mus musculus* | Euarchontoglires | Rodentia | Chromosome | 26/04/16 | 26/04/16 | 134541008 | 14243 | 7154 | 327878 | 2732673693 | Wellcome Trust Sanger Institute |
| GCA_001624445.1 | CAST_EiJ_v1 | *Mus musculus* | Euarchontoglires | Rodentia | Chromosome | 26/04/16 | 26/04/16 | 133752109 | 11642 | 2977 | 391250 | 2653990650 | Wellcome Trust Sanger Institute |
| GCA_001624835.1 | WSB_EiJ_v1 | *Mus musculus* | Euarchontoglires | Rodentia | Chromosome | 26/04/16 | 26/04/16 | 133184209 | 11777 | 2239 | 372890 | 2689657557 | Wellcome Trust Sanger Institute |
| GCA_001632575.1 | C3H_HeJ_v1 | *Mus musculus* | Euarchontoglires | Rodentia | Chromosome | 09/05/16 | 09/05/16 | 133067396 | 14540 | 4069 | 333188 | 2701131316 | Wellcome Trust Sanger Institute |
| GCA_001624295.1 | AKR_J_v1 | *Mus musculus* | Euarchontoglires | Rodentia | Chromosome | 26/04/16 | 26/04/16 | 132982296 | 23042 | 5953 | 242285 | 2712707555 | Wellcome Trust Sanger Institute |
| GCA_001624745.1 | NZO_HlLtJ_v1 | *Mus musculus* | Euarchontoglires | Rodentia | Chromosome | 26/04/16 | 26/04/16 | 132960827 | 16509 | 7022 | 294115 | 2699267296 | Wellcome Trust Sanger Institute |
| GCA_001632525.1 | BALB_cJ_v1 | *Mus musculus* | Euarchontoglires | Rodentia | Chromosome | 09/05/16 | 09/05/16 | 129642644 | 16196 | 3825 | 305391 | 2627344254 | Wellcome Trust Sanger Institute |
| GCA_001624215.1 | A_J_v1 | *Mus musculus* | Euarchontoglires | Rodentia | Chromosome | 26/04/16 | 26/04/16 | 129072804 | 19043 | 4688 | 306830 | 2629995394 | Wellcome Trust Sanger Institute |
| GCA_001624775.1 | PWK_PhJ_v1 | *Mus musculus* | Euarchontoglires | Rodentia | Chromosome | 26/04/16 | 26/04/16 | 128387505 | 5105 | 3140 | 832630 | 2559987392 | Wellcome Trust Sanger Institute |
| GCA_001624505.1 | DBA_2J_v1 | *Mus musculus* | Euarchontoglires | Rodentia | Chromosome | 26/04/16 | 26/04/16 | 128213663 | 13123 | 4105 | 368025 | 2606162158 | Wellcome Trust Sanger Institute |
| GCA_001624535.1 | FVB_NJ_v1 | *Mus musculus* | Euarchontoglires | Rodentia | Chromosome | 26/04/16 | 26/04/16 | 127063089 | 15797 | 5013 | 302598 | 2588619290 | Wellcome Trust Sanger Institute |
| GCF_000001635.26 | GRCm38.p6 | *Mus musculus* | Euarchontoglires | Rodentia | Chromosome | 15/09/17 | 02/10/17 | 52589046 | 32273079 | 239 | 885 | 2818974548 | Genome Reference Consortium |
| GCA_000002165.1 | Mm_Celera | *Mus musculus* | Euarchontoglires | Rodentia | Chromosome | 09/05/11 | 29/11/16 | 23250767 | 21690 | 668009 | 837963 | 3251246261 | Celera Genomics |
| GCA_000185105.2 | MmusALLPATHS2 | *Mus musculus* | Euarchontoglires | Rodentia | Scaffold | 29/08/12 | 29/08/12 | 18924318 | 19501 | 2314 | 191581 | 2505015129 | Broad Institute |
| GCA_000350445.1 | mm129svJae1.0 | *Mus musculus* | Euarchontoglires | Rodentia | Scaffold | 05/04/13 | 03/07/13 | 6876427 | 15255 | 9129 | 240489 | 2578286128 | Broad Institute |
| GCA_003774525.2 | ASM377452v2 | *Mus musculus* | Euarchontoglires | Rodentia | Scaffold | 19/11/18 | 19/11/18 | 1291016 | 391273 | 12661 | 36193 | 2801301292 | Jackson Laboratory |
| GCA_000185125.1 | MmusSOAP1 | *Mus musculus* | Euarchontoglires | Rodentia | Scaffold | 07/01/11 | 29/11/16 | 357132 | 15782 | 41268 | 391273 | 2561486811 | Broad Institute |
| GCA_000181195.1 | ASM18119v1 | *Mus musculus* | Euarchontoglires | Rodentia | Contig | 19/11/10 | 29/11/16 | 24821 | 24821 | 224713 | 224713 | 2477636994 | Mouse Genome Sequencing Consortium |
| GCA_000389885.1 | MSM_WGSv1 | *Mus musculus* | Euarchontoglires | Rodentia | Scaffold | 08/05/13 | 08/05/13 | 1170 | 1170 | 1237600 | 1237642 | 1195475039 | Research Organization of Information and Systems |
| GCF_900095145.1 | PAHARI_EIJ_v1.1 | *Mus pahari* | Euarchontoglires | Rodentia | Chromosome | 02/05/17 | 19/05/17 | 111406228 | 29465 | 2581 | 133720 | 2475012951 | EBI |
| GCA_003336285.1 | MUSP714 | *Mus spicilegus* | Euarchontoglires | Rodentia | Scaffold | 23/07/18 | 10/08/18 | 2198966 | 30918 | 37536 | 161217 | 2496083520 | Oklahoma State Universtiy |
| GCA_001624865.1 | SPRET_EiJ_v1 | *Mus spretus* | Euarchontoglires | Rodentia | Chromosome | 26/04/16 | 26/04/16 | 131945496 | 17887 | 5404 | 280393 | 2625587472 | Wellcome Trust Sanger Institute |
| GCA_004027005.1 | MusAve_v1_BIUU | *Muscardinus avellanarius* | Euarchontoglires | Rodentia | Scaffold | 16/01/19 | 16/01/19 | 59013 | 44294 | 466805 | 481881 | 2527147110 | Broad Institute |
| GCA_004027025.1 | MyoCoy_v1_BIUU | *Myocastor coypus* | Euarchontoglires | Rodentia | Scaffold | 16/01/19 | 16/01/19 | 35982 | 28446 | 1669667 | 1684916 | 2903175784 | Broad Institute |
| GCA_004368595.1 | ASM436859v1 | *Myodes glareolus* | Euarchontoglires | Rodentia | Scaffold | 26/03/19 | 26/03/19 | 1590265 | 164884 | 50926 | 72709 | 2396225068 | Lund University |
| GCA_001305785.1 | ASM130578v1 | *Myodes glareolus* | Euarchontoglires | Rodentia | Scaffold | 02/10/15 | 02/10/15 | 364535 | 1591 | 367242 | 2834384 | 3443072068 | University of Liverpool |
| GCF_000622305.1 | S.galili_v1.0 | *Nannospalax galili* | Euarchontoglires | Rodentia | Scaffold | 05/06/14 | 22/08/14 | 3618479 | 30353 | 154976 | 356097 | 3061424619 | BGI |
| GCA_001675575.1 | ASM167557v1 | *Neotoma lepida* | Euarchontoglires | Rodentia | Scaffold | 23/06/16 | 23/06/16 | 151693 | 9584 | 119373 | 398063 | 2353855022 | University of Utah |
| GCA_002564305.1 | Oct_mimax_AO248_v1 | *Octomys mimax* | Euarchontoglires | Rodentia | Scaffold | 17/10/17 | 06/08/19 | 4874 | 4478 | 1257804 | 1326410 | 2591288717 | McMaster University |
| GCA_004026605.1 | OndZib_v1_BIUU | *Ondatra zibethicus* | Euarchontoglires | Rodentia | Scaffold | 16/01/19 | 16/01/19 | 89093 | 73746 | 565356 | 571599 | 2562752769 | Broad Institute |
| GCA_004026725.1 | OnyTor_v1_BIUU | *Onychomys torridus* | Euarchontoglires | Rodentia | Scaffold | 16/01/19 | 16/01/19 | 20878 | 18287 | 2261735 | 2272285 | 3202219269 | Broad Institute |
| GCA_007922755.1 | PedCap_v1_BIUU | *Pedetes capensis* | Euarchontoglires | Rodentia | Scaffold | 08/08/19 | 08/08/19 | 74432 | 54009 | 673563 | 687430 | 2850076812 | Broad Institute |
| GCA_004363475.1 | PerLonPac_v1_BIUU | *Perognathus longimembris* | Euarchontoglires | Rodentia | Scaffold | 22/03/19 | 22/03/19 | 24714 | 17686 | 2409818 | 2428619 | 2601695796 | Broad Institute |
| GCA_007827085.1 | PCal_1.0 | *Peromyscus californicus* | Euarchontoglires | Rodentia | Chromosome | 02/08/19 | 02/08/19 | 97435531 | 32479 | 34400 | 155723 | 2401349449 | University of Maryland |
| GCF_004664715.1 | Pero_0.1 | *Peromyscus leucopus* | Euarchontoglires | Rodentia | Scaffold | 05/04/19 | 22/04/19 | 4476954 | 4102727 | 1763 | 1834 | 2474071336 | University of California Irvine |
| GCA_003704035.1 | HU_Pman_2.1 | *Peromyscus maniculatus* | Euarchontoglires | Rodentia | Chromosome | 01/11/18 | 01/11/18 | 115033041 | 30111 | 8523 | 155965 | 2512423440 | Harvard University / Howard Hughes Medical Institute |
| GCF_000500345.1 | Pman_1.0 | *Peromyscus maniculatus* | Euarchontoglires | Rodentia | Scaffold | 05/12/13 | 05/03/14 | 3760915 | 36367 | 30921 | 212962 | 2630541020 | Baylor College of Medicine |
| GCA_003704135.2 | HU_Ppol_1.3.3 | *Peromyscus polionotus* | Euarchontoglires | Rodentia | Chromosome | 29/04/19 | 29/04/19 | 117603569 | 13217 | 5292 | 318482 | 2645110392 | Harvard University / Howard Hughes Medical Institute |
| GCA_004026965.1 | PetTyp_v1_BIUU | *Petromus typicus* | Euarchontoglires | Rodentia | Scaffold | 16/01/19 | 16/01/19 | 35766 | 29916 | 910679 | 920324 | 2389366989 | Broad Institute |
| GCA_001707965.1 | Psun0.5 | *Phodopus sungorus* | Euarchontoglires | Rodentia | Contig | 18/08/16 | 06/08/19 | 2392 | 2392 | 1462128 | 1462128 | 2003225912 | University of Chicago |

| | | | | | | | | | | | | | |
|---|---|---|---|---|---|---|---|---|---|---|---|---|---|
| GCA_002215935.1 | ASM221593v1 | *Psammomys obesus* | Euarchontoglires | Rodentia | Contig | 07/07/17 | 20/07/17 | 76398 | 76398 | 96427 | 96427 | 2311471749 | Department of Zoology, University of Oxford, UK |
| GCF_000002265.2 | Rn_Celera | *Rattus norvegicus* | Euarchontoglires | Rodentia | Chromosome | 09/05/11 | 08/01/19 | 27430032 | 39779 | 7041 | 118432 | 2616422528 | Celera Genomics |
| GCF_000001895.5 | Rnor_6.0 | *Rattus norvegicus* | Euarchontoglires | Rodentia | Chromosome | 02/07/14 | 09/10/14 | 14986627 | 100461 | 955 | 75697 | 2870184193 | Rat Genome Sequencing Consortium |
| GCA_004026225.1 | RhiPru_v1_BIUU | *Rhizomys pruinosus* | Euarchontoglires | Rodentia | Scaffold | 16/01/19 | 16/01/19 | 2627 | 2627 | 4150536 | 4151498 | 3937265448 | Broad Institute |
| GCA_004025045.1 | SigHis_v1_BIUU | *Sigmodon hispidus* | Euarchontoglires | Rodentia | Scaffold | 15/01/19 | 15/01/19 | 101373 | 67983 | 883546 | 897290 | 2730600022 | Broad Institute |
| GCA_002406435.1 | ASM240643v1 | *Spermophilus dauricus* | Euarchontoglires | Rodentia | Scaffold | 04/10/17 | 04/10/17 | 1761345 | 34849 | 360804 | 502654 | 3106271744 | BGI-shenzhen |
| GCA_004025085.1 | ThrSwi_v1_BIUU | *Thryonomys swinderianus* | Euarchontoglires | Rodentia | Scaffold | 15/01/19 | 15/01/19 | 21523 | 18955 | 1889641 | 1898466 | 2662148533 | Broad Institute |
| GCA_002564285.1 | Tym_barrerae_AO245_v1 | *Tympanoctomys barrerae* | Euarchontoglires | Rodentia | Scaffold | 17/10/17 | 06/08/19 | 4698 | 4179 | 1558800 | 1647196 | 2651634209 | McMaster University |
| GCF_003426925.1 | ASM342692v1 | *Urocitellus parryii* | Euarchontoglires | Rodentia | Scaffold | 24/08/18 | 11/09/18 | 3964291 | 91013 | 15884 | 79883 | 2520505282 | UAF |
| GCA_004024805.1 | XerIna_v1_BIUU | *Xerus inauris* | Euarchontoglires | Rodentia | Scaffold | 15/01/19 | 15/01/19 | 83865 | 64054 | 577370 | 587115 | 2601418404 | Broad Institute |
| GCA_004024765.1 | ZapHud_v1_BIUU | *Zapus hudsonius* | Euarchontoglires | Rodentia | Scaffold | 15/01/19 | 15/01/19 | 26350 | 23350 | 1158100 | 1165710 | 2611189839 | Broad Institute |
| GCA_000181375.1 | ASM18137v1 | *Tupaia belangeri* | Euarchontoglires | Scandentia | Contig | 19/11/10 | 29/11/16 | 2974 | 2974 | 841006 | 841006 | 2137225476 | Broad Institute |
| GCF_000334495.1 | TupChi_1.0 | *Tupaia chinensis* | Euarchontoglires | Scandentia | Scaffold | 31/01/13 | 03/12/13 | 3670124 | 25938 | 50750 | 242445 | 2846580235 | BGI |
| GCA_004365275.1 | TupTan_v1_BIUU | *Tupaia tana* | Euarchontoglires | Scandentia | Scaffold | 22/03/19 | 25/03/19 | 2973 | 2971 | 3883620 | 3890086 | 3648908809 | Broad Institute |
| GCA_006408695.1 | IMP | *Aepyceros melampus* | Laurasiatheria | Artiodactyla | Scaffold | 24/06/19 | 24/06/19 | 344542 | 82459 | 24102 | 81980 | 2631303056 | Northwestern Polytechnical University |
| GCA_006408545.1 | HBT | *Alcelaphus buselaphus* | Laurasiatheria | Artiodactyla | Scaffold | 24/06/19 | 24/06/19 | 12034 | 889 | 539659 | 1241948 | 3882338275 | Northwestern Polytechnical University |
| GCA_007570765.1 | GSC_moose_1.0 | *Alces alces* | Laurasiatheria | Artiodactyla | Scaffold | 29/07/19 | 29/07/19 | 4131188 | 45090 | 48219 | 131786 | 2743737413 | Canada's Genomic Enterprise |
| GCA_002201775.1 | ALER1.0 | *Ammotragus lervia* | Laurasiatheria | Artiodactyla | Scaffold | 20/06/17 | 03/05/18 | 1301762 | 52017 | 77320 | 160040 | 2650836958 | Northwest A&F University |
| GCA_006408585.1 | SGB | *Antidorcas marsupialis* | Laurasiatheria | Artiodactyla | Scaffold | 24/06/19 | 24/06/19 | 694905 | 42749 | 94522 | 199605 | 3024612899 | Northwestern Polytechnical University |
| GCA_004027515.2 | AntAmePen_v2_BIUU_UCD | *Antilocapra americana* | Laurasiatheria | Artiodactyla | Scaffold | 15/07/19 | 17/07/19 | 18845065 | 61698 | 1585152 | 1647241 | 2955306661 | Broad Institute |
| GCA_007570785.1 | GSC_phorn_1.0 | *Antilocapra americana* | Laurasiatheria | Artiodactyla | Scaffold | 29/07/19 | 29/07/19 | 18739554 | 220431 | 22443 | 48446 | 2509063228 | Canada's Genomic Enterprise |
| GCA_003798545.1 | ASM379854v1 | *Axis porcinus* | Laurasiatheria | Artiodactyla | Scaffold | 20/11/18 | 20/11/18 | 20764858 | 172761 | 136093 | 168266 | 2676163295 | Novogene Bioinformatics Technology Corporation |
| GCA_004027495.1 | BeaHun_v1_BIUU | *Beatragus hunteri* | Laurasiatheria | Artiodactyla | Scaffold | 16/01/19 | 16/01/19 | 69303 | 57444 | 535889 | 545176 | 2697255800 | Broad Institute |
| GCF_000754665.1 | Bison_UMD1.0 | *Bison bison* | Laurasiatheria | Artiodactyla | Scaffold | 09/10/14 | 08/12/17 | 7192658 | 19971 | 128431 | 470415 | 2828031685 | University of Maryland |
| GCA_007844835.1 | NRC_Mithun_1 | *Bos frontalis* | Laurasiatheria | Artiodactyla | Scaffold | 05/08/19 | 05/08/19 | 1003348 | 28709 | 5015 | 179865 | 3002445034 | NRC on Mithun ICAR |
| GCA_005887515.1 | LU_Bosgru_v3.0 | *Bos grunniens* | Laurasiatheria | Artiodactyla | Chromosome | 10/06/19 | 10/06/19 | 114386978 | 44716738 | 414 | 1060 | 2832776395 | Lanzhou University |
| GCF_000247795.1 | Bos_indicus_1.0 | *Bos indicus* | Laurasiatheria | Artiodactyla | Chromosome | 16/12/14 | 12/01/17 | 106310653 | 28375 | 32 | 253770 | 2673965444 | Genoa Biotecnologia SA |
| GCA_002933975.1 | ASM293397v1 | *Bos indicus* | Laurasiatheria | Artiodactyla | Chromosome | 27/02/18 | 27/02/18 | 104295553 | 64498 | 216409 | 337292 | 2740330345 | Anand Agricultural University |
| GCF_003369695.1 | UOA_Brahman_1 | *Bos indicus x Bos taurus* | Laurasiatheria | Artiodactyla | Chromosome | 30/11/18 | 27/12/18 | 104466507 | 26764281 | 1250 | 1552 | 2680953056 | University of Adelaide |
| GCF_003369685.2 | UOA_Angus_1 | *Bos indicus x Bos taurus* | Laurasiatheria | Artiodactyla | Chromosome | 30/11/18 | 11/12/18 | 102763607 | 37061043 | 1435 | 1712 | 2580764822 | University of Adelaide |
| GCA_007646595.1 | ASM764659v1 | *Bos mutus* | Laurasiatheria | Artiodactyla | Scaffold | 30/07/19 | 30/07/19 | 16589160 | 90625 | 158691 | 209207 | 2761379179 | Lanzhou University |
| GCF_000298355.1 | BosGru_v2.0 | *Bos mutus* | Laurasiatheria | Artiodactyla | Scaffold | 09/01/13 | 01/11/13 | 1407960 | 22822 | 41192 | 233198 | 2645161911 | BGI-shenzhen |
| GCF_002263795.1 | ARS-UCD1.2 | *Bos taurus* | Laurasiatheria | Artiodactyla | Chromosome | 11/04/18 | 17/05/18 | 103308737 | 25896116 | 2211 | 2597 | 2715853792 | USDA ARS |
| GCF_000003205.7 | Btau_5.0.1 | *Bos taurus* | Laurasiatheria | Artiodactyla | Chromosome | 19/11/15 | 11/05/18 | 6806220 | 276285 | 3143 | 42267 | 2724980740 | Cattle Genome Sequencing International Consortium |
| GCF_000003055.6 | Bos_taurus_UMD_3.1.1 | *Bos taurus* | Laurasiatheria | Artiodactyla | Chromosome | 03/12/14 | 11/05/18 | 6380747 | 96955 | 3144 | 75618 | 2670139648 | Center for Bioinformatics and Computational Biology, University of Maryland |
| GCF_003121395.1 | UOA_WB_1 | *Bubalus bubalis* | Laurasiatheria | Artiodactyla | Chromosome | 14/05/18 | 30/08/18 | 117219835 | 22441509 | 509 | 919 | 2655780776 | University of Adelaide |
| GCA_004794615.1 | Bubbub1.0 | *Bubalus bubalis* | Laurasiatheria | Artiodactyla | Scaffold | 16/04/19 | 16/04/19 | 6957949 | 25038 | 33821 | 235948 | 2770260352 | BGI-Shenzhen |
| GCA_002993835.1 | ASM299383v1 | *Bubalus bubalis* | Laurasiatheria | Artiodactyla | Scaffold | 12/03/18 | 12/03/18 | 3579700 | 14568 | 99165 | 447289 | 3005952034 | Egyptian Water Buffalo Genome Consortium (Agriculture Genetic Engineering Research Institute and Nile University) |
| GCF_000471725.1 | UMD_CASPUR_WB_2.0 | *Bubalus bubalis* | Laurasiatheria | Artiodactyla | Scaffold | 18/10/13 | 07/07/18 | 1412388 | 21938 | 366983 | 630368 | 2836166969 | University of Maryland |
| GCA_000180995.3 | Bubalus_bubalis_Jaffrabadi_v3.0 | *Bubalus bubalis* | Laurasiatheria | Artiodactyla | Scaffold | 14/05/18 | 14/05/18 | 102345 | 13977 | 117845 | 432941 | 3759980894 | Anand Agricultural University, Anand, Gujarat, India |
| GCF_000767855.1 | Ca_bactrianus_MBC_1.0 | *Camelus bactrianus* | Laurasiatheria | Artiodactyla | Scaffold | 23/10/14 | 06/01/15 | 8812066 | 139019 | 35455 | 67435 | 1992663268 | Inner Mongolia Agricultural University, P.R.China |
| GCA_000604445.1 | CB1 | *Camelus bactrianus* | Laurasiatheria | Artiodactyla | Scaffold | 28/03/14 | 07/09/17 | 2814 | 1717 | 781462 | 1358656 | 1568773545 | VETMEDUNI_POPGEN |
| GCA_000803125.2 | CamDro2 | *Camelus dromedarius* | Laurasiatheria | Artiodactyla | Chromosome | 20/05/19 | 20/05/19 | 75021453 | 204356 | 23439 | 59913 | 2154386959 | Vetmeduni Vienna |
| GCF_000767585.1 | PRJNA234474_Ca_dromedarius_V1.0 | *Camelus dromedarius* | Laurasiatheria | Artiodactyla | Scaffold | 21/10/14 | 07/01/15 | 4188677 | 69131 | 32573 | 105348 | 2004063690 | King Abdulaziz City for Science and Technology |
| GCA_001640815.1 | ASM164081v1 | *Camelus dromedarius* | Laurasiatheria | Artiodactyla | Contig | 09/05/16 | 09/05/16 | 31503 | 31503 | 207871 | 207871 | 2084536768 | University of Bristol |
| GCF_000311805.1 | CB1 | *Camelus ferus* | Laurasiatheria | Artiodactyla | Scaffold | 05/08/13 | 29/11/13 | 2005940 | 90263 | 13334 | 68872 | 2009194609 | The Bactrian Camels Genome Sequencing and Analysis Consortium |
| GCA_000978405.1 | CapAeg_1.0 | *Capra aegagrus* | Laurasiatheria | Artiodactyla | Chromosome | 24/04/15 | 24/04/15 | 91317560 | 19347 | 89498 | 368693 | 2828873048 | BGI |
| GCA_000765075.1 | Caeg1 | *Capra aegagrus* | Laurasiatheria | Artiodactyla | Scaffold | 09/10/14 | 16/10/14 | 1750063 | 51868 | 6616 | 103827 | 2583320943 | EBI |
| GCA_004361675.1 | CVASU_BBG_1.0 | *Capra hircus* | Laurasiatheria | Artiodactyla | Chromosome | 25/03/19 | 25/03/19 | 102339471 | 819 | 3972 | 3696314 | 3041623660 | Bangladesh Goat Genome Consortium (BGGC) |
| GCF_001704415.1 | ARS1 | *Capra hircus* | Laurasiatheria | Artiodactyla | Chromosome | 24/08/16 | 03/02/17 | 87277232 | 26244591 | 29907 | 30399 | 2922813246 | USDA ARS |
| GCA_000317765.2 | CHIR_2.0 | *Capra hircus* | Laurasiatheria | Artiodactyla | Chromosome | 16/09/15 | 16/09/15 | 9094169 | 73533 | 102895 | 173141 | 2809418462 | Beijing Genomics Institute |
| GCA_006410555.1 | IBX | *Capra ibex* | Laurasiatheria | Artiodactyla | Scaffold | 24/06/19 | 24/06/19 | 61905114 | 380983 | 55914 | 75375 | 2701356656 | Northwestern Polytechnical University |
| GCA_003182615.2 | ASM318261v2 | *Capra sibirica* | Laurasiatheria | Artiodactyla | Scaffold | 30/10/18 | 30/10/18 | 15190720 | 376582 | 85609 | 113068 | 2733031394 | Northwest A&F University |
| GCA_000751575.1 | kmer631 | *Capreolus capreolus* | Laurasiatheria | Artiodactyla | Scaffold | 09/09/14 | 09/09/14 | 10458 | 4167 | 3088511 | 3781281 | 2785377831 | UNIMELB |
| GCA_004024745.2 | CatWag_v2_BIUU_UCD | *Catagonus wagneri* | Laurasiatheria | Artiodactyla | Scaffold | 15/07/19 | 17/07/19 | 19204659 | 65965 | 1183730 | 1240743 | 2640067814 | Broad Institute |
| GCA_006410635.1 | HVD | *Cephalophus harveyi* | Laurasiatheria | Artiodactyla | Scaffold | 24/06/19 | 24/06/19 | 365466 | 61368 | 85160 | 157791 | 2816643069 | Northwestern Polytechnical University |
| GCA_002197005.1 | CerEla1.0 | *Cervus elaphus* | Laurasiatheria | Artiodactyla | Chromosome | 19/06/17 | 07/03/18 | 107358006 | 7944 | 11479 | 406637 | 3438623608 | National Agricultural Research and Innovation Centre |
| GCA_006408615.1 | BWD | *Connochaetes taurinus* | Laurasiatheria | Artiodactyla | Scaffold | 24/06/19 | 24/06/19 | 366224 | 70608 | 78350 | 131840 | 2644042816 | Northwestern Polytechnical University |
| GCA_006408505.1 | CTB | *Damaliscus lunatus* | Laurasiatheria | Artiodactyla | Scaffold | 24/06/19 | 24/06/19 | 1166796 | 38843 | 92109 | 208023 | 3127764384 | Northwestern Polytechnical University |
| GCA_002443075.1 | Milu1.0 | *Elaphurus davidianus* | Laurasiatheria | Artiodactyla | Scaffold | 04/10/17 | 06/08/19 | 2844142 | 59950 | 194812 | 276603 | 2584693296 | Nanjing Normal University |
| GCA_006408755.1 | TMG | *Eudorcas thomsonii* | Laurasiatheria | Artiodactyla | Scaffold | 24/06/19 | 24/06/19 | 1581717 | 137476 | 123308 | 190522 | 2904272043 | Northwestern Polytechnical University |
| GCA_006408565.1 | GFE | *Giraffa camelopardalis* | Laurasiatheria | Artiodactyla | Scaffold | 24/06/19 | 24/06/19 | 3153258 | 25056 | 78945 | 253969 | 2473180627 | Northwestern Polytechnical University |
| GCA_001651235.1 | ASM165123v1 | *Giraffa tippelskirchi* | Laurasiatheria | Artiodactyla | Scaffold | 25/05/16 | 25/05/16 | 212164 | 47894 | 513177 | 595021 | 2705072464 | Pennsylvania State University |
| GCA_006547405.1 | ASM654740v1 | *Globicephala melas* | Laurasiatheria | Artiodactyla | Scaffold | 03/07/19 | 03/07/19 | 18102937 | 332801 | 6637 | 21236 | 2333871032 | Canada's Genomic Enterprise |
| GCA_004026825.1 | HemHyl_v1_BIUU | *Hemitragus hylocrius* | Laurasiatheria | Artiodactyla | Scaffold | 16/01/19 | 16/01/19 | 85340 | 66552 | 469163 | 479550 | 2715042536 | Broad Institute |
| GCA_004027065.2 | HipAmp_v2_BIUU_UCD | *Hippopotamus amphibius* | Laurasiatheria | Artiodactyla | Scaffold | 15/07/19 | 17/07/19 | 4444377 | 76609 | 910835 | 971145 | 2733493772 | Broad Institute |
| GCA_002995585.1 | ASM299558v1 | *Hippopotamus amphibius* | Laurasiatheria | Artiodactyla | Scaffold | 13/03/18 | 09/08/19 | 110752 | 34064 | 128772 | 208977 | 2425746322 | Senckenberg Gesellschaft fuer Naturforschung |
| GCA_006942125.1 | SCBI_HNig_1.0 | *Hippotragus niger* | Laurasiatheria | Artiodactyla | Scaffold | 12/07/19 | 12/07/19 | 4586323 | 45501 | 15557 | 135158 | 2595326149 | Saint-Petersburg State University |
| GCA_006459105.1 | NPU_HINE_1.0 | *Hydropotes inermis* | Laurasiatheria | Artiodactyla | Scaffold | 25/06/19 | 25/06/19 | 13818975 | 131446 | 22246 | 56894 | 2530176423 | Northwestern Polytechnical University |
| GCA_006410655.1 | DFW | *Kobus ellipsiprymnus* | Laurasiatheria | Artiodactyla | Scaffold | 24/06/19 | 24/06/19 | 779552 | 26169 | 88848 | 279061 | 2895340485 | Northwestern Polytechnical University |
| GCA_006410535.1 | GRK | *Litocranius walleri* | Laurasiatheria | Artiodactyla | Scaffold | 24/06/19 | 24/06/19 | 3126223 | 62351 | 79667 | 178573 | 2979336167 | Northwestern Polytechnical University |
| GCA_006408675.1 | KDD | *Madoqua kirkii* | Laurasiatheria | Artiodactyla | Scaffold | 24/06/19 | 24/06/19 | 27730 | 27722 | 216116 | 221029 | 2648126871 | Northwestern Polytechnical University |
| GCA_006459085.1 | FMD | *Moschus berezovskii* | Laurasiatheria | Artiodactyla | Scaffold | 25/06/19 | 25/06/19 | 2509225 | 57706 | 77520 | 154049 | 2818010997 | Northwestern Polytechnical University |
| GCA_006461545.1 | NPU_MBRE_1.0 | *Moschus berezovskii* | Laurasiatheria | Artiodactyla | Scaffold | 25/06/19 | 25/06/19 | 213462 | 24701 | 125813 | 275601 | 3835628252 | Northwestern Polytechnical University |
| GCA_006461725.1 | NPU_MCHR_1.0 | *Moschus chrysogaster* | Laurasiatheria | Artiodactyla | Scaffold | 25/06/19 | 25/06/19 | 100428 | 3769 | 127198 | 844280 | 4972482505 | Northwestern Polytechnical University |
| GCA_004024705.2 | MosMos_v2_BIUU_UCD | *Moschus moschiferus* | Laurasiatheria | Artiodactyla | Scaffold | 15/07/19 | 17/07/19 | 11728851 | 34785 | 1072499 | 1195517 | 3069608437 | Broad Institute |
| GCA_006408485.1 | BMJ | *Muntiacus crinifrons* | Laurasiatheria | Artiodactyla | Scaffold | 24/06/19 | 24/06/19 | 1305444 | 8265 | 21054 | 549998 | 2681196170 | Northwestern Polytechnical University |
| GCA_006409035.1 | CMJ | *Muntiacus muntjak* | Laurasiatheria | Artiodactyla | Scaffold | 24/06/19 | 24/06/19 | 1258210 | 23470 | 383561 | 556267 | 2703710594 | Northwestern Polytechnical University |
| GCA_006408525.1 | CIJ | *Muntiacus reevesi* | Laurasiatheria | Artiodactyla | Scaffold | 24/06/19 | 24/06/19 | 1221377 | 72382 | 200261 | 269408 | 2601888952 | Northwestern Polytechnical University |
| GCA_006408635.1 | GTG | *Nanger granti* | Laurasiatheria | Artiodactyla | Scaffold | 24/06/19 | 24/06/19 | 528456 | 7454 | 62189 | 671117 | 3032779593 | Northwestern Polytechnical University |
| GCA_006410615.1 | SUN | *Neotragus moschatus* | Laurasiatheria | Artiodactyla | Scaffold | 24/06/19 | 24/06/19 | 952090 | 59277 | 218204 | 366475 | 2675729750 | Northwestern Polytechnical University |
| GCA_006410875.1 | RAL | *Neotragus pygmaeus* | Laurasiatheria | Artiodactyla | Scaffold | 24/06/19 | 24/06/19 | 363895 | 30751 | 354089 | 471398 | 2895867692 | Northwestern Polytechnical University |
| GCA_004115125.1 | UofA_Ohem_1.0 | *Odocoileus hemionus* | Laurasiatheria | Artiodactyla | Scaffold | 30/01/19 | 02/02/19 | 838758 | 113295 | 16208 | 58896 | 2343701333 | University of Alberta |
| GCA_003697985.1 | ASM369798v1 | *Odocoileus hemionus* | Laurasiatheria | Artiodactyla | Scaffold | 31/10/18 | 31/10/18 | 9678 | 9488 | 358334 | 387819 | 2324921134 | Canada's Genomic Enterprise |
| GCF_002102435.1 | Ovir.te_1.0 | *Odocoileus virginianus* | Laurasiatheria | Artiodactyla | Scaffold | 20/04/17 | 28/04/17 | 850721 | 122019 | 17026 | 48653 | 2380505687 | Baylor College of Medicine |
| GCA_000191625.1 | RRL-RSL_1.0 | *Odocoileus virginianus* | Laurasiatheria | Artiodactyla | Contig | 07/03/11 | 29/11/16 | 451 | 451 | 93873 | 93873 | 37656191 | Texas A&M University |
| GCA_001660835.1 | ASM166083v1 | *Okapia johnstoni* | Laurasiatheria | Artiodactyla | Scaffold | 10/06/16 | 10/06/16 | 111538 | 39571 | 837179 | 925548 | 2878133036 | Pennsylvania State University |
| GCA_006410675.1 | KSP | *Oreotragus oreotragus* | Laurasiatheria | Artiodactyla | Scaffold | 24/06/19 | 24/06/19 | 339390 | 42127 | 24892 | 141857 | 3247632364 | Northwestern Polytechnical University |
| GCA_003945745.1 | UCDavis_Ogaz_1 | *Oryx gazella* | Laurasiatheria | Artiodactyla | Scaffold | 14/12/18 | 14/12/18 | 1579191 | 19498 | 31177 | 266309 | 2762920344 | University of California Davis |
| GCA_006410575.1 | GMB | *Oryx gazella* | Laurasiatheria | Artiodactyla | Scaffold | 24/06/19 | 24/06/19 | 1566830 | 19456 | 56922 | 292054 | 2768889188 | Northwestern Polytechnical University |
| GCA_006417275.1 | ORB | *Ourebia ourebi* | Laurasiatheria | Artiodactyla | Scaffold | 24/06/19 | 24/06/19 | 1259 | 1259 | 3508661 | 3508672 | 2524866914 | Northwestern Polytechnical University |
| GCA_003121645.1 | O_ammon_KGZ_v1.0 | *Ovis ammon* | Laurasiatheria | Artiodactyla | Scaffold | 11/05/18 | 17/05/18 | 72289505 | 151569 | 41955 | 75155 | 2643433589 | Northwest A&F University |
| GCA_006410595.1 | AGL | *Ovis ammon* | Laurasiatheria | Artiodactyla | Scaffold | 24/06/19 | 24/06/19 | 5620092 | 46562 | 240203 | 317544 | 2659657073 | Northwestern Polytechnical University |
| GCF_002742125.1 | Oar_rambouillet_v1.0 | *Ovis aries* | Laurasiatheria | Artiodactyla | Chromosome | 02/11/17 | 06/02/19 | 107697089 | 2572683 | 2641 | 7486 | 2869914396 | Baylor College of Medicine Human Genome Sequencing Center |
| GCF_000298735.2 | Oar_v4.0 | *Ovis aries* | Laurasiatheria | Artiodactyla | Chromosome | 22/11/15 | 25/02/19 | 100009711 | 150472 | 5466 | 48482 | 2615516299 | International Sheep Genome Consortium |
| GCA_000005525.1 | Ovis_aries_1.0 | *Ovis aries* | Laurasiatheria | Artiodactyla | Chromosome | 26/02/10 | 29/11/16 | 34092587 | 685 | 1217 | 2352347 | 2860496367 | International Sheep Genomics Consortium |
| GCF_000765115.1 | Oori1 | *Ovis aries* | Laurasiatheria | Artiodactyla | Scaffold | 09/10/14 | 25/02/19 | 2217029 | 39721 | 6173 | 128014 | 2589834840 | EBI |
| GCA_004026945.1 | OviCan_v1_BIUU | *Ovis canadensis* | Laurasiatheria | Artiodactyla | Scaffold | 16/01/19 | 16/01/19 | 69397 | 55973 | 1048136 | 1057548 | 2863658823 | Broad Institute |
| GCA_006410695.1 | MXD | *Philantomba maxwellii* | Laurasiatheria | Artiodactyla | Scaffold | 24/06/19 | 24/06/19 | 383899 | 11377 | 25633 | 334216 | 3145938691 | Northwestern Polytechnical University |
| GCA_006410515.1 | PLG | *Procapra przewalskii* | Laurasiatheria | Artiodactyla | Scaffold | 24/06/19 | 24/06/19 | 5522907 | 10405 | 274130 | 738871 | 2692986855 | Northwestern Polytechnical University |
| GCA_006408465.1 | WLD | *Przewalskium albirostris* | Laurasiatheria | Artiodactyla | Scaffold | 24/06/19 | 24/06/19 | 3769372 | 39627 | 171874 | 294078 | 2692225130 | Northwestern Polytechnical University |
| GCA_003182575.1 | ASM318257v1 | *Pseudois nayaur* | Laurasiatheria | Artiodactyla | Scaffold | 31/05/18 | 31/05/18 | 21385 | 20979 | 214772 | 220288 | 2584167189 | Northwest A&F University |
| GCA_004026565.1 | RanTarSib_v1_BIUU | *Rangifer tarandus* | Laurasiatheria | Artiodactyla | Scaffold | 16/01/19 | 16/01/19 | 89062 | 77671 | 1360739 | 1365508 | 2897299934 | Broad Institute |
| GCA_006410735.1 | SNB | *Raphicerus campestris* | Laurasiatheria | Artiodactyla | Scaffold | 24/06/19 | 24/06/19 | 537161 | 25050 | 57517 | 222189 | 3137860433 | Northwestern Polytechnical University |
| GCA_006410935.1 | BHR | *Redunca redunca* | Laurasiatheria | Artiodactyla | Scaffold | 24/06/19 | 24/06/19 | 423407 | 39282 | 534502 | 701851 | 2724978760 | Northwestern Polytechnical University |
| GCA_004024985.1 | SaiTat_v1_BIUU | *Saiga tatarica* | Laurasiatheria | Artiodactyla | Scaffold | 15/01/19 | 15/01/19 | 6453 | 6406 | 1772199 | 1775066 | 2876186568 | Broad Institute |
| GCA_001292865.1 | ss10.2_mar2013 | *Sus scrofa* | Laurasiatheria | Artiodactyla | Scaffold | 16/09/15 | 16/09/15 | 153651326 | 17259 | 5206 | 726487 | 2611360562 | F. Hoffmann - La Roche AG |
| GCA_007644095.1 | ASM764409v1 | *Sus scrofa* | Laurasiatheria | Artiodactyla | Chromosome | 01/08/19 | 01/08/19 | 140438739 | 1009524 | 6610 | 13644 | 2491053062 | Novogene |
| GCA_006511355.1 | NSME_pig_1 | *Sus scrofa* | Laurasiatheria | Artiodactyla | Chromosome | 03/07/19 | 03/07/19 | 138999934 | 52150866 | 613 | 1170 | 2502318605 | University of Messina |
| GCA_002844635.1 | USMARCv1.0 | *Sus scrofa* | Laurasiatheria | Artiodactyla | Chromosome | 20/12/17 | 20/12/17 | 131458098 | 6372407 | 14157 | 14818 | 2755438182 | USDA ARS |
| GCF_000003025.6 | Sscrofa11.1 | *Sus scrofa* | Laurasiatheria | Artiodactyla | Chromosome | 14/02/17 | 08/12/17 | 88231837 | 48231277 | 613 | 1118 | 2501912388 | The Swine Genome Sequencing Consortium (SGSC) |
| GCA_000325925.2 | minipig_v1.0 | *Sus scrofa* | Laurasiatheria | Artiodactyla | Scaffold | 18/03/15 | 18/03/15 | 5853977 | 31939 | 137577 | 272163 | 2508912457 | BGI-shenzhen |
| GCA_001700135.1 | Large_White_v1 | *Sus scrofa* | Laurasiatheria | Artiodactyla | Scaffold | 05/08/16 | 06/08/19 | 2441555 | 88831 | 102342 | 150742 | 2457907103 | Novogene |
| GCA_001700155.1 | Rongchang_pig_v1 | *Sus scrofa* | Laurasiatheria | Artiodactyla | Scaffold | 05/08/16 | 06/08/19 | 2325000 | 79093 | 120246 | 173508 | 2459027728 | Novogene |
| GCA_001700255.1 | Pietrain_pig_v1 | *Sus scrofa* | Laurasiatheria | Artiodactyla | Scaffold | 05/08/16 | 06/08/19 | 1663542 | 80611 | 88436 | 139497 | 2438316299 | Novogene |
| GCA_001700575.1 | Berkshire_pig_v1 | *Sus scrofa* | Laurasiatheria | Artiodactyla | Scaffold | 05/08/16 | 06/08/19 | 1655397 | 94651 | 94468 | 137661 | 2434706773 | Novogene |
| GCA_001700165.1 | Hampshire_pig_v1 | *Sus scrofa* | Laurasiatheria | Artiodactyla | Scaffold | 05/08/16 | 06/08/19 | 1550023 | 102417 | 82206 | 122452 | 2437109612 | Novogene |
| GCA_001700235.1 | Bamei_pig_v1 | *Sus scrofa* | Laurasiatheria | Artiodactyla | Scaffold | 05/08/16 | 06/08/19 | 1529027 | 70893 | 129335 | 187466 | 2460755741 | Novogene |
| GCA_001700295.1 | Jinhua_pig_v1 | *Sus scrofa* | Laurasiatheria | Artiodactyla | Scaffold | 05/08/16 | 06/08/19 | 1478908 | 95227 | 115554 | 158796 | 2453702738 | Novogene |
| GCA_001700215.1 | Landrace_pig_v1 | *Sus scrofa* | Laurasiatheria | Artiodactyla | Scaffold | 05/08/16 | 06/08/19 | 1407841 | 88142 | 94659 | 141909 | 2440981886 | Novogene |
| GCA_001700195.1 | Meishan_pig_v1 | *Sus scrofa* | Laurasiatheria | Artiodactyla | Scaffold | 05/08/16 | 06/08/19 | 1248180 | 63263 | 133833 | 201146 | 2467495771 | Novogene |
| GCA_000472085.2 | Tibetan_Pig_v2 | *Sus scrofa* | Laurasiatheria | Artiodactyla | Scaffold | 08/08/16 | 08/08/16 | 861885 | 57199 | 72068 | 148234 | 2437742524 | Novogene |
| GCA_000331475.1 | SscrofaMinipig | *Sus scrofa* | Laurasiatheria | Artiodactyla | Contig | 10/01/13 | 10/01/13 | 22008 | 22008 | 231585 | 231585 | 2358017222 | GlaxoSmithKline |
| GCA_006408735.1 | CMD | *Sylvicapra grimmia* | Laurasiatheria | Artiodactyla | Scaffold | 24/06/19 | 24/06/19 | 583330 | 9720 | 22324 | 468651 | 3145094493 | Northwestern Polytechnical University |
| GCA_006408785.1 | ABF | *Syncerus caffer* | Laurasiatheria | Artiodactyla | Scaffold | 24/06/19 | 24/06/19 | 2316376 | 52316 | 148371 | 248905 | 2929427825 | Northwestern Polytechnical University |
| GCA_006411685.1 | MTN | *Tragelaphus buxtoni* | Laurasiatheria | Artiodactyla | Scaffold | 24/06/19 | 24/06/19 | 7367 | 1286 | 2910462 | 4301196 | 3267488234 | Northwestern Polytechnical University |
| GCA_006410755.1 | BNG | *Tragelaphus eurycerus* | Laurasiatheria | Artiodactyla | Scaffold | 24/06/19 | 24/06/19 | 12574 | 1974 | 1539703 | 2829973 | 2962040965 | Northwestern Polytechnical University |
| GCA_006410775.1 | LEK | *Tragelaphus imberbis* | Laurasiatheria | Artiodactyla | Scaffold | 24/06/19 | 24/06/19 | 1775046 | 12858 | 272415 | 584429 | 4038235010 | Northwestern Polytechnical University |
| GCA_006416875.1 | CME | *Tragelaphus oryx* | Laurasiatheria | Artiodactyla | Scaffold | 24/06/19 | 24/06/19 | 1262 | 1262 | 4043025 | 4043033 | 2837493372 | Northwestern Polytechnical University |
| GCA_006410495.1 | BUB | *Tragelaphus scriptus* | Laurasiatheria | Artiodactyla | Scaffold | 24/06/19 | 24/06/19 | 890554 | 28350 | 237860 | 373810 | 2873326854 | Northwestern Polytechnical University |
| GCA_006411015.1 | STG | *Tragelaphus spekii* | Laurasiatheria | Artiodactyla | Scaffold | 24/06/19 | 24/06/19 | 46427 | 9477 | 1769205 | 2115196 | 3725951127 | Northwestern Polytechnical University |
| GCA_006410795.1 | GTK | *Tragelaphus strepsiceros* | Laurasiatheria | Artiodactyla | Scaffold | 24/06/19 | 24/06/19 | 511483 | 33649 | 320432 | 525298 | 2867754166 | Northwestern Polytechnical University |
| GCA_004024965.2 | ASM402496v2 | *Tragulus javanicus* | Laurasiatheria | Artiodactyla | Scaffold | 15/07/19 | 17/07/19 | 14082842 | 80230 | 1082489 | 1158497 | 2589955488 | Broad Institute |
| GCA_006408655.1 | LMD | *Tragulus kanchil* | Laurasiatheria | Artiodactyla | Scaffold | 24/06/19 | 24/06/19 | 243497 | 13850 | 185856 | 500458 | 3053423192 | Northwestern Polytechnical University |
| GCA_000164845.4 | VicPac3.1 | *Vicugna pacos* | Laurasiatheria | Artiodactyla | Scaffold | 16/07/19 | 17/07/19 | 24022313 | 29536 | 77389 | 243669 | 2119600897 | Deakin University |
| GCA_000767525.1 | Vi_pacos_V1.0 | *Vicugna pacos* | Laurasiatheria | Artiodactyla | Scaffold | 11/02/15 | 11/02/15 | 5303709 | 213649 | 52275 | 75733 | 2013701362 | BGI-Shenzhen |
| GCF_003709585.1 | Aci_jub_2 | *Acinonyx jubatus* | Laurasiatheria | Carnivora | Scaffold | 02/11/18 | 13/11/18 | 48500042 | 179924 | 3220 | 27393 | 2384851327 | Felidae consortium |
| GCF_001443585.1 | aciJub1 | *Acinonyx jubatus* | Laurasiatheria | Carnivora | Scaffold | 13/11/15 | 30/11/18 | 3122036 | 35120 | 14383 | 170171 | 2372553907 | Saint Petersburg State University |
| GCA_002007445.1 | ASM200744v1 | *Ailuropoda melanoleuca* | Laurasiatheria | Carnivora | Scaffold | 28/02/17 | 28/02/17 | 9947519 | 128604 | 57414 | 106175 | 2428263693 | Institute of Zoology, Chinese Academy of Sciences |
| GCF_000004335.2 | AilMel_1.0 | *Ailuropoda melanoleuca* | Laurasiatheria | Carnivora | Scaffold | 13/01/10 | 04/02/15 | 1281781 | 39886 | 81467 | 200593 | 2299509015 | Beijing Genomics Institute, Shenzhen |
| GCA_002007465.1 | ASM200746v1 | *Ailurus fulgens* | Laurasiatheria | Carnivora | Scaffold | 28/02/17 | 28/02/17 | 2983736 | 99577 | 11589 | 69825 | 2342939051 | Institute of Zoology, Chinese Academy of Sciences |
| GCA_900642305.1 | arcGaz3 | *Arctocephalus gazella* | Laurasiatheria | Carnivora | Scaffold | 13/02/19 | 13/02/19 | 139181869 | 537425 | 5180 | 13973 | 2313585684 | LMU |
| GCA_900500725.1 | ArcGazv1.4 | *Arctocephalus gazella* | Laurasiatheria | Carnivora | Scaffold | 25/08/18 | 31/08/18 | 6207322 | 538754 | 6169 | 13956 | 2313485084 | LMU |
| GCF_003265705.1 | ASM326570v1 | *Callorhinus ursinus* | Laurasiatheria | Carnivora | Scaffold | 27/06/18 | 21/07/18 | 31506801 | 133024 | 14230 | 52955 | 2706868872 | Canada's Genomics Enterprise |
| GCA_005444595.1 | UMICH_Zoey_3.1 | *Canis lupus* | Laurasiatheria | Carnivora | Chromosome | 30/05/19 | 30/05/19 | 64204256 | 4722218 | 794 | 1793 | 2343218756 | University of Michigan |
| GCA_007922845.1 | UniMelb_Wolf_Refassem_1 | *Canis lupus* | Laurasiatheria | Carnivora | Scaffold | 08/08/19 | 08/08/19 | 64175042 | 842 | 765 | 3050747 | 2342421267 | The University of Melbourne |
| GCA_004886185.1 | Basenji_breed-1.1 | *Canis lupus* | Laurasiatheria | Carnivora | Chromosome | 26/04/19 | 26/04/19 | 61087166 | 3131423 | 2243 | 3630 | 2410429933 | University of Missouri |
| GCF_000002285.3 | CanFam3.1 | *Canis lupus* | Laurasiatheria | Carnivora | Chromosome | 03/11/11 | 12/07/18 | 45876610 | 267478 | 3268 | 27106 | 2410976875 | Dog Genome Sequencing Consortium |
| GCF_003254725.1 | ASM325472v1 | *Canis lupus* | Laurasiatheria | Carnivora | Scaffold | 19/06/18 | 05/07/18 | 34446301 | 5390770 | 2444 | 3121 | 2439831507 | James Cook University |
| GCA_004027395.1 | CanFam_VD_v1_BIUU | *Canis lupus* | Laurasiatheria | Carnivora | Scaffold | 16/01/19 | 16/01/19 | 157754 | 100144 | 656742 | 667170 | 2544127025 | Broad Institute |
| GCA_000331495.1 | Beagle | *Canis lupus* | Laurasiatheria | Carnivora | Contig | 10/01/13 | 10/01/13 | 21834 | 21834 | 221474 | 221474 | 2254627151 | GlaxoSmithKline |
| GCA_000181415.1 | ASM18141v1 | *Canis lupus* | Laurasiatheria | Carnivora | Contig | 19/11/10 | 29/11/16 | 1516 | 1516 | 1089636 | 1089636 | 1517481071 | TIGR |
| GCA_004023885.1 | CryFer_v1_BIUU | *Cryptoprocta ferox* | Laurasiatheria | Carnivora | Scaffold | 15/01/19 | 15/01/19 | 173473 | 128639 | 464036 | 470371 | 2458846334 | Broad Institute |
| GCF_002288905.1 | ASM228890v2 | *Enhydra lutris* | Laurasiatheria | Carnivora | Scaffold | 15/09/17 | 20/09/17 | 38751465 | 244529 | 6771 | 29592 | 2455276666 | BC Cancer Agency |
| GCA_006410715.1 | ASM641071v1 | *Enhydra lutris* | Laurasiatheria | Carnivora | Scaffold | 24/06/19 | 24/06/19 | 9763901 | 23473 | 55496 | 212165 | 2425597608 | UCLA |
| GCF_004028035.1 | ASM402803v1 | *Eumetopias jubatus* | Laurasiatheria | Carnivora | Scaffold | 16/01/19 | 11/02/19 | 14018600 | 242372 | 7472 | 24747 | 2418263165 | Canada's Genomic Enterprise |
| GCF_000181335.3 | Felis_catus_9.0 | *Felis catus* | Laurasiatheria | Carnivora | Chromosome | 20/11/17 | 12/12/17 | 83967707 | 41915695 | 4508 | 4909 | 2521863845 | Genome Sequencing Center (GSC) at Washington University (WashU) School of Medicine |
| GCA_000003115.1 | catChrV17e | *Felis catus* | Laurasiatheria | Carnivora | Chromosome | 12/08/09 | 29/11/16 | 16304361 | 4577 | 104053 | 604920 | 3160286939 | NHGRI/Genome Technology Branch |
| GCA_004023925.1 | FelNig_v1_BIUU | *Felis nigripes* | Laurasiatheria | Carnivora | Scaffold | 15/01/19 | 15/01/19 | 18631 | 16007 | 590124 | 603600 | 2428424918 | Broad Institute |
| GCA_900006375.2 | Gulo_2.2_annotated | *Gulo gulo* | Laurasiatheria | Carnivora | Scaffold | 07/12/18 | 07/12/18 | 178272 | 3657 | 47417 | 1066768 | 2423180524 | Uppsala University |
| GCA_004023845.1 | HelPar_v1_BIUU | *Helogale parvula* | Laurasiatheria | Carnivora | Scaffold | 15/01/19 | 15/01/19 | 179119 | 113567 | 189370 | 200185 | 2392471390 | Broad Institute |
| GCA_003009895.1 | ASM300989v1 | *Hyaena hyaena* | Laurasiatheria | Carnivora | Scaffold | 19/03/18 | 05/06/18 | 2001327 | 311202 | 5760 | 18422 | 2374716107 | Potsdam University |
| GCA_004023945.1 | HyaHya_v1_BIUU | *Hyaena hyaena* | Laurasiatheria | Carnivora | Scaffold | 15/01/19 | 15/01/19 | 66490 | 51677 | 350223 | 362072 | 2445474026 | Broad Institute |
| GCF_000349705.1 | LepWed1.0 | *Leptonychotes weddellii* | Laurasiatheria | Carnivora | Scaffold | 29/03/13 | 14/02/14 | 904031 | 23664 | 16711 | 169547 | 3156902762 | Broad Institute |
| GCA_001883655.1 | LycPicKen1.0 | *Lycaon pictus* | Laurasiatheria | Carnivora | Chromosome | 01/12/16 | 01/12/16 | 64190785 | 2695 | 813 | 1430320 | 2356716622 | Smithsonian Institution |
| GCA_001887905.1 | LycPicSAfr1.0 | *Lycaon pictus* | Laurasiatheria | Carnivora | Chromosome | 02/12/16 | 02/12/16 | 63240551 | 5555 | 803 | 797755 | 2358143818 | Smithsonian Institution |
| GCA_004216515.1 | sis1-161031-pseudohap | *Lycaon pictus* | Laurasiatheria | Carnivora | Scaffold | 20/02/19 | 20/02/19 | 7494581 | 61225 | 1127 | 61776 | 2137698293 | Stanford University |
| GCA_007474595.1 | mLynCan4_v1.p | *Lynx canadensis* | Laurasiatheria | Carnivora | Chromosome | 26/07/19 | 06/08/19 | 146106016 | 7503561 | 67 | 902 | 2408900816 | Vertebrate Genomes Project |
| GCA_007474575.1 | mLynCan4_v1.alt | *Lynx canadensis* | Laurasiatheria | Carnivora | Scaffold | 25/07/19 | 26/07/19 | 130153 | 130121 | 15033 | 15036 | 1415234291 | Vertebrate Genomes Project |
| GCA_900661375.1 | LYPA1.0 | *Lynx pardinus* | Laurasiatheria | Carnivora | Scaffold | 10/04/19 | 10/04/19 | 1519745 | 99542 | 41700 | 91104 | 2413208909 | CNAG |
| GCA_004024625.1 | MelCap_v1_BIUU | *Mellivora capensis* | Laurasiatheria | Carnivora | Scaffold | 15/01/19 | 15/01/19 | 59143 | 50213 | 2068312 | 2074823 | 3091173380 | Broad Institute |
| GCA_004023865.1 | MirAng_v1_BIUU | *Mirounga angustirostris* | Laurasiatheria | Carnivora | Scaffold | 15/01/19 | 15/01/19 | 64640 | 52833 | 321256 | 330739 | 2407319723 | Broad Institute |
| GCA_004023785.1 | MunMun_v1_BIUU | *Mungos mungo* | Laurasiatheria | Carnivora | Scaffold | 15/01/19 | 15/01/19 | 236501 | 180702 | 312364 | 316874 | 2448211634 | Broad Institute |
| GCF_000239315.1 | MusPutFurMale1.0 | *Mustela putorius* | Laurasiatheria | Carnivora | Scaffold | 21/12/11 | 09/07/15 | 9567452 | 66755 | 4245 | 76915 | 2400181698 | Broad Institute |
| GCF_000215625.1 | MusPutFur1.0 | *Mustela putorius* | Laurasiatheria | Carnivora | Scaffold | 02/06/11 | 10/12/13 | 9335154 | 44823 | 7783 | 117483 | 2410879678 | Ferret Genome Sequencing Consortium |
| GCF_002201575.1 | ASM220157v1 | *Neomonachus schauinslandi* | Laurasiatheria | Carnivora | Scaffold | 20/06/17 | 28/06/17 | 29518589 | 112698 | 7873 | 46708 | 2400934290 | Johns Hopkins University |
| GCA_900108605.1 | NNQGG.v01 | *Neovison vison* | Laurasiatheria | Carnivora | Scaffold | 03/01/18 | 03/01/18 | 6814223 | 32336 | 7175 | 143887 | 2447189291 | Center for Quantitative Genetics and Genomics |
| GCF_000321225.1 | Oros_1.0 | *Odobenus rosmarus* | Laurasiatheria | Carnivora | Scaffold | 16/01/13 | 27/07/16 | 2616778 | 89951 | 3893 | 70656 | 2400150193 | Marine mammals |
| GCA_004023805.1 | PanOnc_v1_BIUU | *Panthera onca* | Laurasiatheria | Carnivora | Scaffold | 15/01/19 | 15/01/19 | 116574 | 62836 | 438792 | 460121 | 2504534411 | Broad Institute |
| GCF_001857705.1 | PanPar1.0 | *Panthera pardus* | Laurasiatheria | Carnivora | Scaffold | 16/11/16 | 06/12/16 | 21701857 | 20993 | 50377 | 265330 | 2578019207 | Ulsan National Institute of Science and Technology |
| GCF_000464555.1 | PanTig1.0 | *Panthera tigris* | Laurasiatheria | Carnivora | Scaffold | 06/09/13 | 07/03/14 | 8860407 | 30032 | 1479 | 157032 | 2391082183 | Personal Genomics Institute |
| GCA_004024585.1 | ParHer_v1_BIUU | *Paradoxurus hermaphroditus* | Laurasiatheria | Carnivora | Scaffold | 15/01/19 | 15/01/19 | 71823 | 62870 | 438125 | 442658 | 2477229103 | Broad Institute |
| GCA_004348235.1 | GSC_HSeal_1.0 | *Phoca vitulina* | Laurasiatheria | Carnivora | Scaffold | 13/03/19 | 13/03/19 | 41024070 | 283887 | 5542 | 21603 | 2363587482 | Canada's Genome Enterprise (CGEN) |
| GCA_005406085.1 | Prionailurus_bengalensis_euptilurus_v01 | *Prionailurus bengalensis* | Laurasiatheria | Carnivora | Scaffold | 15/05/19 | 15/05/19 | 152598 | 21454 | 65356 | 318468 | 2435348619 | Kyoto City Zoo |
| GCA_004024605.1 | PteBra_v1_BIUU | *Pteronura brasiliensis* | Laurasiatheria | Carnivora | Scaffold | 15/01/19 | 15/01/19 | 119023 | 90879 | 644728 | 652159 | 2602275310 | Broad Institute |
| GCF_003327715.1 | PumCon1.0 | *Puma concolor* | Laurasiatheria | Carnivora | Scaffold | 19/07/18 | 26/07/18 | 100532876 | 27187 | 2175 | 181166 | 2432966684 | Broad Institute |

| | | | | | | | | | | | | |
|---|---|---|---|---|---|---|---|---|---|---|---|---|
| GCA_004123975.1 | Pco_k61 | *Puma concolor* | Laurasiatheria | Carnivora | Scaffold | 31/01/19 | 31/01/19 | 193865 | 6267 | 152272 | 634810 | 2591702724 | Duke University |
| GCA_004023965.1 | SpiGra_v1_BIUU | *Spilogale gracilis* | Laurasiatheria | Carnivora | Scaffold | 15/01/19 | 15/01/19 | 85970 | 63054 | 422009 | 433831 | 2463318012 | Broad Institute |
| GCF_006229205.1 | meerkat_22Aug2017_6uvM2_HiC | *Suricata suricatta* | Laurasiatheria | Carnivora | Chromosome | 03/07/19 | 15/07/19 | 141453419 | 75409 | 63791 | 129036 | 2353578744 | DNA Zoo |
| GCA_004023905.1 | SurSur_v1_BIUU | *Suricata suricatta* | Laurasiatheria | Carnivora | Scaffold | 15/01/19 | 15/01/19 | 186735 | 148487 | 349868 | 354021 | 2471813441 | Broad Institute |
| GCA_003697995.1 | ASM369799v1 | *Taxidea taxus* | Laurasiatheria | Carnivora | Scaffold | 31/10/18 | 31/10/18 | 54556 | 37750 | 145161 | 173916 | 2416005247 | Canada's Genomic Enterprise |
| GCA_003344425.1 | ASM334442v1 | *Ursus americanus* | Laurasiatheria | Carnivora | Scaffold | 27/07/18 | 27/07/18 | 189900 | 27247 | 111495 | 231673 | 2588393916 | Jackson Laboratory |
| GCF_003584765.1 | ASM358476v1 | *Ursus arctos* | Laurasiatheria | Carnivora | Scaffold | 21/09/18 | 30/09/18 | 36708181 | 314844 | 6673 | 22245 | 2328659882 | Canada's Genomic Enterprise |
| GCF_000687225.1 | UrsMar_1.0 | *Ursus maritimus* | Laurasiatheria | Carnivora | Scaffold | 12/05/14 | 04/08/14 | 15940661 | 46506 | 23819 | 134162 | 2301379344 | BGI-Shenzhen |
| GCA_004023825.1 | VulLag_v1_BIUU | *Vulpes lagopus* | Laurasiatheria | Carnivora | Scaffold | 15/01/19 | 15/01/19 | 102053 | 73976 | 1263345 | 1272949 | 2688561625 | Broad Institute |
| GCF_003160815.1 | VulVul2.2 | *Vulpes vulpes* | Laurasiatheria | Carnivora | Scaffold | 25/05/18 | 31/07/18 | 12472085 | 55450 | 82424 | 183898 | 2421568072 | University of Illinois at Urbana-Champaign |
| GCF_900631625.1 | zalCal2.2 | *Zalophus californianus* | Laurasiatheria | Carnivora | Scaffold | 19/12/18 | 27/12/18 | 143424588 | 97662 | 10423 | 57871 | 2372369564 | LMU |
| GCA_004024565.1 | ZalCal_v1_BIUU | *Zalophus californianus* | Laurasiatheria | Carnivora | Scaffold | 15/01/19 | 15/01/19 | 132377 | 90567 | 571996 | 582272 | 2519860747 | Broad Institute |
| GCF_000493695.1 | BalAcu1.0 | *Balaenoptera acutorostrata* | Laurasiatheria | Cetacea | Scaffold | 31/10/13 | 09/03/16 | 12843668 | 22690 | 10776 | 184072 | 2431687698 | Korea Ocean Research & Development Institute |
| GCA_000978805.1 | ASM97880v1 | *Balaenoptera bonaerensis* | Laurasiatheria | Cetacea | Scaffold | 23/04/15 | 23/04/15 | 20082 | 8410 | 421444 | 720900 | 2234641557 | Division of Biological Science, Graduate School of Science, Kyoto University |
| GCF_002288925.1 | ASM228892v2 | *Delphinapterus leucas* | Laurasiatheria | Cetacea | Scaffold | 15/09/17 | 23/09/17 | 19885328 | 159142 | 6972 | 35370 | 2358522448 | BC Cancer Agency |
| GCA_002189225.1 | ASM218922v1 | *Eschrichtius robustus* | Laurasiatheria | Cetacea | Scaffold | 09/06/17 | 09/06/17 | 187455 | 10066 | 57203 | 375256 | 2849453582 | Purdue University |
| GCA_004363415.1 | EscRob_v1_BIUU | *Eschrichtius robustus* | Laurasiatheria | Cetacea | Scaffold | 22/03/19 | 22/03/19 | 94414 | 68559 | 1036148 | 1046770 | 2617572198 | Broad Institute |
| GCA_002738545.1 | EschrichtiusRobustus1.0 | *Eschrichtius robustus* | Laurasiatheria | Cetacea | Scaffold | 31/10/17 | 06/08/19 | 10674 | 2656 | 1213011 | 1595257 | 2922979301 | Laboratory of postgenomic studies, Engelhardt Institute of Molecular Biology, Russian Academy of Sciences |
| GCA_004363455.1 | EubJap_v1_BIUU | *Eubalaena japonica* | Laurasiatheria | Cetacea | Scaffold | 22/03/19 | 22/03/19 | 39813 | 34866 | 1353963 | 1361057 | 2693575300 | Broad Institute |
| GCA_004363515.1 | IniGeo_v1_BIUU | *Inia geoffrensis* | Laurasiatheria | Cetacea | Scaffold | 22/03/19 | 22/03/19 | 26707 | 24570 | 1213610 | 1218682 | 2596281358 | Broad Institute |
| GCA_004363705.1 | KogBre_v1_BIUU | *Kogia breviceps* | Laurasiatheria | Cetacea | Scaffold | 22/03/19 | 22/03/19 | 28812 | 26201 | 1252072 | 1258125 | 2759698159 | Broad Institute |
| GCF_003676395.1 | ASM367639v1 | *Lagenorhynchus obliquidens* | Laurasiatheria | Cetacea | Scaffold | 31/10/18 | 16/11/18 | 28371583 | 255779 | 5422 | 21793 | 2334467171 | Canada's Genomic Enterprise |
| GCF_000442215.1 | Lipotes_vexillifer_v1 | *Lipotes vexillifer* | Laurasiatheria | Cetacea | Scaffold | 09/08/13 | 01/04/14 | 2419148 | 31902 | 30713 | 155510 | 2429212129 | BGI |
| GCA_004329385.1 | megNov1 | *Megaptera novaeangliae* | Laurasiatheria | Cetacea | Scaffold | 06/03/19 | 06/03/19 | 9138802 | 12321 | 2558 | 387694 | 2265788366 | Arizona Cancer Evolution Center (ACE) |
| GCA_004027085.1 | MesBid_v1_BIUU | *Mesoplodon bidens* | Laurasiatheria | Cetacea | Scaffold | 16/01/19 | 16/01/19 | 33532 | 28959 | 1801720 | 1810317 | 2797694783 | Broad Institute |
| GCF_005190385.1 | NGI_Narwhal_1 | *Monodon monoceros* | Laurasiatheria | Cetacea | Scaffold | 07/05/19 | 21/05/19 | 107566389 | 255327 | 6972 | 25295 | 2355574979 | The Narwhal Genome Initiative |
| GCA_005125345.1 | ASM512534v1 | *Monodon monoceros* | Laurasiatheria | Cetacea | Scaffold | 03/05/19 | 04/05/19 | 1483363 | 10044 | 21006 | 813468 | 2350881526 | University of Copenhagen |
| GCA_004027045.1 | MonMon_v1_BIUU | *Monodon monoceros* | Laurasiatheria | Cetacea | Scaffold | 16/01/19 | 16/01/19 | 88921 | 70965 | 882704 | 890705 | 2546711041 | Broad Institute |
| GCA_004026685.1 | MonMon_M_v1_BIUU | *Monodon monoceros* | Laurasiatheria | Cetacea | Scaffold | 16/01/19 | 16/01/19 | 86766 | 67024 | 644873 | 653473 | 2472545502 | Broad Institute |
| GCF_003031525.1 | Neophocaena_asiaeorientalis_V1 | *Neophocaena asiaeorientalis* | Laurasiatheria | Cetacea | Scaffold | 03/04/18 | 20/04/18 | 6341296 | 86003 | 13699 | 66346 | 2284628084 | Nanjing normal university |
| GCF_000331955.2 | Oorc_1.1 | *Orcinus orca* | Laurasiatheria | Cetacea | Scaffold | 04/04/14 | 27/07/16 | 12735091 | 70300 | 1668 | 80100 | 2372919875 | Marine mammals |
| GCA_003071005.1 | ASM307100v1 | *Phocoena phocoena* | Laurasiatheria | Cetacea | Scaffold | 24/04/18 | 24/04/18 | 27499337 | 2773 | 142029 | 2347235 | 2441140962 | University of Potsdam |
| GCA_004363495.1 | PhoPho_v1_BIUU | *Phocoena phocoena* | Laurasiatheria | Cetacea | Scaffold | 22/03/19 | 22/03/19 | 115969 | 89111 | 1331158 | 1338272 | 2701010810 | Broad Institute |
| GCA_900411695.1 | PhyCat2.0 | *Physeter catodon* | Laurasiatheria | Cetacea | Scaffold | 07/07/18 | 07/07/18 | 122182240 | 43829 | 14676 | 140250 | 2512140680 | CNGB |
| GCF_002837175.2 | ASM283717v2 | *Physeter catodon* | Laurasiatheria | Cetacea | Chromosome | 08/03/19 | 21/03/19 | 122182240 | 42542 | 14677 | 143605 | 2512149402 | BGI |
| GCF_000472045.1 | Physeter_macrocephalus-2.0.2 | *Physeter catodon* | Laurasiatheria | Cetacea | Scaffold | 30/09/13 | 28/02/18 | 427290 | 35257 | 11711 | 110444 | 2280727784 | Aquatic Genome Models |
| GCA_004363435.1 | PlaMin_v1_BIUU | *Platanista minor* | Laurasiatheria | Cetacea | Scaffold | 22/03/19 | 22/03/19 | 23933 | 20879 | 1098790 | 1110492 | 2671072510 | Broad Institute |
| GCA_004363935.1 | PonBla_v1_BIUU | *Pontoporia blainvillei* | Laurasiatheria | Cetacea | Scaffold | 22/03/19 | 22/03/19 | 2541 | 2541 | 1885058 | 1885701 | 1685099017 | Broad Institute |
| GCA_007760645.1 | ASM776064v1 | *Sousa chinensis* | Laurasiatheria | Cetacea | Scaffold | 31/07/19 | 31/07/19 | 19436979 | 113766 | 23368 | 62803 | 2338882461 | BGI-SZ |
| GCA_003521335.2 | S_chinensis_fine_genome_map | *Sousa chinensis* | Laurasiatheria | Cetacea | Scaffold | 01/02/19 | 01/02/19 | 9008636 | 182701 | 20903 | 46900 | 2339085850 | Shantou university |
| GCA_003227395.1 | ASM322739v1 | *Tursiops aduncus* | Laurasiatheria | Cetacea | Scaffold | 14/06/18 | 14/06/18 | 1235788 | 206065 | 16249 | 44281 | 2503928010 | Ewha Womans University |
| GCA_003314715.1 | Tur_tru_Illumina_hap_v1 | *Tursiops truncatus* | Laurasiatheria | Cetacea | Scaffold | 11/07/18 | 11/07/18 | 27166507 | 30985 | 481 | 139544 | 2404363272 | Johns Hopkins University |
| GCF_001922835.1 | NIST Tur_tru v1 | *Tursiops truncatus* | Laurasiatheria | Cetacea | Scaffold | 28/12/16 | 12/01/17 | 26555543 | 44299 | 2648 | 116651 | 2132516271 | National Institute of Standards and Technology |
| GCA_003435595.3 | Tur_tru_Illumina_phased_v1 | *Tursiops truncatus* | Laurasiatheria | Cetacea | Scaffold | 02/11/18 | 02/11/18 | 781592 | 26636 | 97787 | 348040 | 4816480726 | Johns Hopkins University |
| GCF_000151865.2 | Ttru_1.4 | *Tursiops truncatus* | Laurasiatheria | Cetacea | Scaffold | 08/06/15 | 05/12/18 | 116287 | 11821 | 240558 | 554228 | 2551418184 | Baylor College of Medicine |
| GCA_004364475.1 | ZipCav_v1_BIUU | *Ziphius cavirostris* | Laurasiatheria | Cetacea | Scaffold | 22/03/19 | 25/03/19 | 3608 | 3606 | 3758276 | 3761505 | 3150417206 | Broad Institute |
| GCA_004027475.1 | AnoCau_v1_BIUU | *Anoura caudifer* | Laurasiatheria | Chiroptera | Scaffold | 16/01/19 | 16/01/19 | 185021 | 143417 | 337255 | 341895 | 2206589520 | Broad Institute |
| GCA_007922775.1 | AntPal_v1_BIUU | *Antrozous pallidus* | Laurasiatheria | Chiroptera | Scaffold | 08/08/19 | 08/08/19 | 13958 | 13442 | 902551 | 905784 | 2597469403 | Broad Institute |
| GCA_004027435.1 | ArtJam_v1_BIUU | *Artibeus jamaicensis* | Laurasiatheria | Chiroptera | Scaffold | 16/01/19 | 16/01/19 | 35347 | 32365 | 794376 | 798700 | 2424784351 | Broad Institute |
| GCA_004027735.1 | CarPer_v1_BIUU | *Carollia perspicillata* | Laurasiatheria | Chiroptera | Scaffold | 16/01/19 | 16/01/19 | 10739 | 10341 | 1925339 | 1929203 | 2689411905 | Broad Institute |
| GCA_004027555.1 | CraTho_v1_BIUU | *Craseonycteris thonglongyai* | Laurasiatheria | Chiroptera | Scaffold | 16/01/19 | 16/01/19 | 25762 | 23218 | 1224256 | 1230480 | 2272093077 | Broad Institute |
| GCF_002940915.1 | ASM294091v2 | *Desmodus rotundus* | Laurasiatheria | Chiroptera | Scaffold | 28/03/18 | 20/04/18 | 26869735 | 80250 | 29801 | 84109 | 2063808397 | University of Copenhagen |
| GCA_000465285.1 | ASM46528v1 | *Eidolon helvum* | Laurasiatheria | Chiroptera | Scaffold | 20/09/13 | 20/09/13 | 27684 | 12668 | 133538 | 288446 | 1837754460 | School of Biological & Chemical Sciences, Queen Mary University of London |
| GCA_003508835.1 | Espe.v1 | *Eonycteris spelaea* | Laurasiatheria | Chiroptera | Contig | 07/09/18 | 07/09/18 | 8002591 | 8002591 | 4469 | 4469 | 1966861576 | Duke-NUS Medical School |
| GCF_000308155.1 | EptFus1.0 | *Eptesicus fuscus* | Laurasiatheria | Chiroptera | Scaffold | 08/11/12 | 30/05/14 | 13454942 | 21392 | 6789 | 167058 | 2026629342 | Broad Institute |
| GCF_001890085.1 | ASM189008v1 | *Hipposideros armiger* | Laurasiatheria | Chiroptera | Scaffold | 06/12/16 | 20/12/16 | 2328177 | 39863 | 7571 | 128956 | 2236581172 | SKLEC & IECR |
| GCA_004027415.1 | HipGal_v1_BIUU | *Hipposideros galeritus* | Laurasiatheria | Chiroptera | Scaffold | 16/01/19 | 16/01/19 | 37985 | 33704 | 840200 | 846566 | 2440750851 | Broad Institute |
| GCA_004026805.1 | LasBor_v1_BIUU | *Lasiurus borealis* | Laurasiatheria | Chiroptera | Scaffold | 16/01/19 | 16/01/19 | 38543 | 35165 | 518900 | 525280 | 2857590849 | Broad Institute |
| GCA_004027375.1 | MacSob_v1_BIUU | *Macroglossus sobrinus* | Laurasiatheria | Chiroptera | Scaffold | 16/01/19 | 16/01/19 | 453401 | 338389 | 171985 | 174041 | 1897644983 | Broad Institute |
| GCA_007922815.1 | MacCal_v1_BIUU | *Macrotus californicus* | Laurasiatheria | Chiroptera | Scaffold | 08/08/19 | 08/08/19 | 16925 | 16386 | 1128787 | 1130929 | 2161582626 | Broad Institute |
| GCA_004026885.1 | MegLyr_v1_BIUU | *Megaderma lyra* | Laurasiatheria | Chiroptera | Scaffold | 16/01/19 | 16/01/19 | 96489 | 71674 | 1902801 | 1909067 | 2620948449 | Broad Institute |
| GCA_000465345.1 | ASM46534v1 | *Megaderma lyra* | Laurasiatheria | Chiroptera | Scaffold | 19/09/13 | 19/09/13 | 16881 | 7043 | 192872 | 518327 | 1735931796 | School of Biological & Chemical Sciences, Queen Mary University of London |
| GCA_004026765.1 | MicHir_v1_BIUU | *Micronycteris hirsuta* | Laurasiatheria | Chiroptera | Scaffold | 16/01/19 | 16/01/19 | 68868 | 62961 | 550090 | 553308 | 2314654405 | Broad Institute |
| GCF_001595765.1 | Mnat.v1 | *Miniopterus natalensis* | Laurasiatheria | Chiroptera | Scaffold | 22/03/16 | 11/04/16 | 4315193 | 29777 | 1269 | 114632 | 1803099001 | University of California, San Francisco |
| GCA_004026525.1 | MinSch_v1_BIUU | *Miniopterus schreibersii* | Laurasiatheria | Chiroptera | Scaffold | 16/01/19 | 16/01/19 | 108707 | 82539 | 177620 | 184538 | 1775846775 | Broad Institute |
| GCA_004026545.1 | MorMeg_v1_BIUU | *Mormoops blainvillei* | Laurasiatheria | Chiroptera | Scaffold | 16/01/19 | 16/01/19 | 156292 | 142682 | 205259 | 207094 | 2111750309 | Broad Institute |
| GCA_004026665.1 | MurFea_v1_BIUU | *Murina aurata* | Laurasiatheria | Chiroptera | Scaffold | 16/01/19 | 16/01/19 | 26051 | 23400 | 880177 | 886972 | 2331551382 | Broad Institute |
| GCF_000412655.1 | ASM41265v1 | *Myotis brandtii* | Laurasiatheria | Chiroptera | Scaffold | 05/07/13 | 30/10/13 | 3225832 | 23289 | 169750 | 325414 | 2107242811 | BGI |
| GCF_000327345.1 | ASM32734v1 | *Myotis davidii* | Laurasiatheria | Chiroptera | Scaffold | 20/12/12 | 12/02/14 | 3454484 | 15182 | 101769 | 325280 | 2059799708 | BGI |
| GCF_000147115.1 | Myoluc2.0 | *Myotis lucifugus* | Laurasiatheria | Chiroptera | Scaffold | 14/09/10 | 26/11/13 | 4293315 | 64330 | 11654 | 72785 | 2034575300 | Broad Institute |
| GCA_004026985.1 | MyoMyo_v1_BIUU | *Myotis myotis* | Laurasiatheria | Chiroptera | Scaffold | 16/01/19 | 16/01/19 | 21824 | 19322 | 1045197 | 1053330 | 2294472406 | Broad Institute |
| GCA_004026585.1 | NocLep_v1_BIUU | *Noctilio leporinus* | Laurasiatheria | Chiroptera | Scaffold | 16/01/19 | 16/01/19 | 191494 | 135651 | 298222 | 304393 | 2098501394 | Broad Institute |
| GCA_007922795.1 | NycHum_v1_BIUU | *Nycticeius humeralis* | Laurasiatheria | Chiroptera | Scaffold | 08/08/19 | 08/08/19 | 15095 | 14267 | 1676240 | 1680573 | 2779565143 | Broad Institute |
| GCF_004126475.1 | mPhyDis1_v1.p | *Phyllostomus discolor* | Laurasiatheria | Chiroptera | Chromosome | 17/05/19 | 17/05/19 | 110241909 | 6892556 | 141 | 964 | 2117764065 | Vertebrate Genomes Project |
| GCA_004115245.1 | mPhyDis1_v1.h | *Phyllostomus discolor* | Laurasiatheria | Chiroptera | Scaffold | 28/01/19 | 29/01/19 | 933674 | 933674 | 10835 | 10873 | 2050602545 | Vertebrate Genomes Project |
| GCA_004026625.1 | PipPip_v1_BIUU | *Pipistrellus pipistrellus* | Laurasiatheria | Chiroptera | Scaffold | 16/01/19 | 16/01/19 | 33992 | 26707 | 726104 | 736974 | 2090670443 | Broad Institute |
| GCA_000465405.1 | ASM46540v1 | *Pteronotus parnellii* | Laurasiatheria | Chiroptera | Scaffold | 19/09/13 | 19/09/13 | 22675 | 9502 | 177401 | 443121 | 1960317893 | School of Biological & Chemical Sciences, Queen Mary University of London |
| GCF_000325575.1 | ASM32557v1 | *Pteropus alecto* | Laurasiatheria | Chiroptera | Scaffold | 20/12/12 | 10/02/16 | 15954802 | 31841 | 65598 | 170164 | 1985975446 | BGI |
| GCF_000151845.1 | Pvam_2.0 | *Pteropus vampyrus* | Laurasiatheria | Chiroptera | Scaffold | 05/12/14 | 23/02/15 | 5954017 | 21866 | 36094 | 225433 | 2198284804 | Baylor College of Medicine |
| GCA_004115265.2 | mRhiFer1_v1.p | *Rhinolophus ferrumequinum* | Laurasiatheria | Chiroptera | Chromosome | 17/05/19 | 17/05/19 | 88025743 | 31900497 | 135 | 290 | 2075785400 | Vertebrate Genomes Project |
| GCA_004115295.1 | mRhiFer1_v1.h | *Rhinolophus ferrumequinum* | Laurasiatheria | Chiroptera | Scaffold | 28/01/19 | 29/01/19 | 301176 | 301176 | 13424 | 13426 | 1514615512 | Vertebrate Genomes Project |
| GCA_007922735.1 | RhiFer_v1_BIUU | *Rhinolophus ferrumequinum* | Laurasiatheria | Chiroptera | Scaffold | 08/08/19 | 08/08/19 | 156956 | 127195 | 345321 | 349275 | 2183991669 | Broad Institute |
| GCA_000465495.1 | ASM46549v1 | *Rhinolophus ferrumequinum* | Laurasiatheria | Chiroptera | Scaffold | 19/09/13 | 19/09/13 | 21151 | 11659 | 160500 | 290685 | 1926439238 | School of Biological & Chemical Sciences, Queen Mary University of London |
| GCF_001466805.2 | Raegyp2.0 | *Rousettus aegyptiacus* | Laurasiatheria | Chiroptera | Scaffold | 17/03/16 | 29/03/16 | 2007187 | 1488988 | 2490 | 3049 | 1910250568 | Boston University School of Medicine |
| GCA_004024865.1 | RouAeg_v1_BIUU | *Rousettus aegyptiacus* | Laurasiatheria | Chiroptera | Scaffold | 15/01/19 | 15/01/19 | 125332 | 100260 | 221754 | 226086 | 1972131931 | Broad Institute |
| GCA_004025005.1 | TadBra_v1_BIUU | *Tadarida brasiliensis* | Laurasiatheria | Chiroptera | Scaffold | 15/01/19 | 15/01/19 | 24311 | 22617 | 1067615 | 1071858 | 2709928648 | Broad Institute |
| GCA_004024845.1 | TonSau_v1_BIUU | *Tonatia saurophila* | Laurasiatheria | Chiroptera | Scaffold | 15/01/19 | 15/01/19 | 165561 | 141649 | 249810 | 252670 | 2105886965 | Broad Institute |
| GCF_000260355.1 | ConCri1.0 | *Condylura cristata* | Laurasiatheria | Eulipotyphla | Scaffold | 01/05/12 | 30/05/13 | 55520359 | 46163 | 2040 | 84597 | 1769662895 | Broad Institute |
| GCA_004027635.1 | CroInd_v1_BIUU | *Crocidura indochinensis* | Laurasiatheria | Eulipotyphla | Scaffold | 16/01/19 | 16/01/19 | 4929 | 4927 | 2415412 | 2420679 | 3082217326 | Broad Institute |
| GCF_000296755.1 | EriEur2.0 | *Erinaceus europaeus* | Laurasiatheria | Eulipotyphla | Scaffold | 20/09/12 | 09/03/16 | 3264618 | 21359 | 5803 | 225567 | 2715720925 | Broad Institute |
| GCA_000181395.1 | ASM18139v1 | *Erinaceus europaeus* | Laurasiatheria | Eulipotyphla | Contig | 19/11/10 | 29/11/16 | 2753 | 2753 | 847680 | 847680 | 2133117389 | Broad Institute |
| GCA_004024925.1 | ScaAqu_v1_BIUU | *Scalopus aquaticus* | Laurasiatheria | Eulipotyphla | Scaffold | 15/01/19 | 15/01/19 | 94879 | 72421 | 640296 | 647535 | 2068871057 | Broad Institute |
| GCA_004363575.1 | SolPar_v1_BIUU | *Solenodon paradoxus* | Laurasiatheria | Eulipotyphla | Scaffold | 22/03/19 | 22/03/19 | 407682 | 236847 | 176618 | 181804 | 2109877870 | Broad Institute |
| GCA_002901085.1 | ASM290108v1 | *Solenodon paradoxus* | Laurasiatheria | Eulipotyphla | Scaffold | 26/01/18 | 26/01/18 | 110915 | 89121 | 40482 | 48846 | 2049415269 | University of Puerto Rico - Mayaguez Campus |
| GCF_000181275.1 | SorAra2.0 | *Sorex araneus* | Laurasiatheria | Eulipotyphla | Scaffold | 27/08/12 | 10/03/16 | 22794405 | 22623 | 12845 | 201798 | 2423158183 | Broad Institute |
| GCA_004024945.1 | UroGra_v1_BIUU | *Uropsilus gracilis* | Laurasiatheria | Eulipotyphla | Scaffold | 15/01/19 | 15/01/19 | 55035 | 43618 | 723835 | 732083 | 2112710047 | Broad Institute |
| GCF_000283155.1 | CerSimSim1.0 | *Ceratotherium simum* | Laurasiatheria | Perissodactyla | Scaffold | 07/08/12 | 02/12/13 | 26277727 | 92960 | 3087 | 57824 | 2464367180 | Broad Institute |
| GCA_004027795.1 | CerCot_v1_BIUU | *Ceratotherium simum* | Laurasiatheria | Perissodactyla | Scaffold | 16/01/19 | 16/01/19 | 23005 | 21199 | 942426 | 949480 | 2666617437 | Broad Institute |
| GCA_002844835.1 | ASM284483v1 | *Dicerorhinus sumatrensis* | Laurasiatheria | Perissodactyla | Scaffold | 19/12/17 | 06/08/19 | 614498 | 89455 | 1061589 | 1110756 | 2949863658 | Marshall University Genomic Core Facility |
| GCA_004027315.2 | DicBicMic_v2_BIUU_UCD | *Diceros bicornis* | Laurasiatheria | Perissodactyla | Scaffold | 15/07/19 | 17/07/19 | 14664201 | 91002 | 1981226 | 2027983 | 3060207893 | Broad Institute |
| GCA_003033725.1 | ASM303372v1 | *Equus asinus* | Laurasiatheria | Perissodactyla | Scaffold | 04/04/18 | 17/05/18 | 15144613 | 134070 | 9021 | 49709 | 2321045832 | Centre for GeoGenetics |
| GCF_001305755.1 | ASM130575v1 | *Equus asinus* | Laurasiatheria | Perissodactyla | Scaffold | 02/10/15 | 02/12/15 | 3776412 | 66737 | 2167 | 71760 | 2391051217 | Inner Mongolian Agricultural University |
| GCF_002863925.1 | EquCab3.0 | *Equus caballus* | Laurasiatheria | Perissodactyla | Chromosome | 05/01/18 | 14/02/18 | 87230776 | 1502753 | 4701 | 10987 | 2506966135 | University of Louisville |
| GCF_000002305.2 | EquCab2.0 | *Equus caballus* | Laurasiatheria | Perissodactyla | Chromosome | 03/06/11 | 30/01/18 | 46749900 | 112381 | 9637 | 55317 | 2474929062 | The Genome Assembly Team |
| GCA_000696655.1 | Ajinai1.0 | *Equus caballus* | Laurasiatheria | Perissodactyla | Scaffold | 06/06/14 | 06/06/14 | 56256 | 40738 | 119776 | 155158 | 2377506909 | College of Animal Science, Inner Mongolian Agricultural University, China |
| GCF_000696695.1 | Burgud | *Equus przewalskii* | Laurasiatheria | Perissodactyla | Scaffold | 06/06/14 | 14/07/14 | 513800 | 57610 | 53097 | 134060 | 2395954271 | College of Animal Science, Inner Mongolian Agricultural University, China |
| GCA_004024905.1 | TapInd_v1_BIUU | *Tapirus indicus* | Laurasiatheria | Perissodactyla | Scaffold | 15/01/19 | 15/01/19 | 308930 | 225792 | 280135 | 284468 | 2425203111 | Broad Institute |
| GCA_004025025.1 | TapTer_v1_BIUU | *Tapirus terrestris* | Laurasiatheria | Perissodactyla | Scaffold | 15/01/19 | 15/01/19 | 186384 | 164193 | 803506 | 805779 | 2571458752 | Broad Institute |
| GCF_001685135.1 | ManJav1.0 | *Manis javanica* | Laurasiatheria | Pholidota | Scaffold | 14/07/16 | 08/08/16 | 204728 | 38922 | 80670 | 398998 | 2547395906 | The International Pangolin Research Consortium |
| GCA_000738955.1 | M_pentadactyla-1.1.1 | *Manis pentadactyla* | Laurasiatheria | Pholidota | Scaffold | 12/08/14 | 27/12/17 | 117920 | 28718 | 92772 | 230930 | 2204732179 | Washington University (WashU) |
| GCA_004765945.1 | ManTri_v1_BIUU | *Manis tricuspis* | Laurasiatheria | Pholidota | Scaffold | 09/04/19 | 09/04/19 | 25241 | 23755 | 2116242 | 2120018 | 3013399831 | Broad Institute |
| GCF_000189315.1 | Devil_ref v7.0 | *Sarcophilus harrisii* | Marsupialia | Dasyuromorphia | Scaffold | 20/01/12 | 26/11/13 | 1847106 | 20139 | 35974 | 237291 | 3174693010 | Wellcome Trust Sanger Institute |
| GCA_000219685.1 | Devil_CABOG_asm.06 | *Sarcophilus harrisii* | Marsupialia | Dasyuromorphia | Scaffold | 01/07/11 | 30/11/16 | 151762 | 11025 | 148774 | 457735 | 3232424150 | J. Craig Venter Institute |
| GCA_007646695.1 | UniMelb_Thylacine_Refassem_1 | *Thylacinus cynocephalus* | Marsupialia | Dasyuromorphia | Scaffold | 30/07/19 | 31/07/19 | 1923833 | 461 | 19549 | 5628809 | 3095956865 | The University of Melbourne |
| GCA_000002295.1 | ASM229v1 | *Monodelphis domestica* | Marsupialia | Didelphimorphia | Chromosome | 04/05/11 | 29/11/16 | 527952102 | 107990 | 5137 | 72811 | 3600487649 | The Genome Sequencing Platform, The Genome Assembly Team |
| GCF_000002295.2 | MonDom5 | *Monodelphis domestica* | Marsupialia | Didelphimorphia | Chromosome | 01/01/01 | 13/06/14 | 59809810 | 108014 | 5016 | 72674 | 3598443077 | The Genome Sequencing Platform, The Genome Assembly Team |
| GCA_000004035.1 | Meug_1.1 | *Notamacropus eugenii* | Marsupialia | Diprotodontia | Scaffold | 06/01/10 | 29/11/16 | 36602 | 2602 | 277711 | 1174382 | 3075184024 | Tammar Wallaby Genome Sequencing Consortium |
| GCF_002099425.1 | phaCin_unsw_v4.1 | *Phascolarctos cinereus* | Marsupialia | Diprotodontia | Contig | 18/04/17 | 10/05/17 | 11587828 | 11587828 | 1907 | 1907 | 3192581492 | The Earlham Institute |
| GCA_900166895.1 | phaCin_tgac_v2.0 | *Phascolarctos cinereus* | Marsupialia | Diprotodontia | Scaffold | 11/05/18 | 11/05/18 | 798273 | 126882 | 796464 | 819636 | 3603908365 | KOALA GENOME CONSORTIUM |
| GCF_900497805.2 | bare-nosed wombat genome assembly | *Vombatus ursinus* | Marsupialia | Diprotodontia | Scaffold | 19/12/18 | 24/01/19 | 28503419 | 113054 | 15416 | 81204 | 3486597218 | MRC Institute of Genetics and Molecular Medicine, University of Edinburgh |
| GCF_004115215.1 | mOrnAna1.p.v1 | *Ornithorhynchus anatinus* | Monotremata | Monotremata | Chromosome | 17/05/19 | 17/05/19 | 83338043 | 15146802 | 305 | 824 | 1858552590 | Vertebrate Genomes Project |
| GCA_002966995.1 | ASM296699v1 | *Ornithorhynchus anatinus* | Monotremata | Monotremata | Scaffold | 02/03/18 | 02/03/18 | 18718360 | 7578676 | 4568 | 5044 | 1990409179 | BGI |
| GCA_000002275.2 | ASM227v2 | *Ornithorhynchus anatinus* | Monotremata | Monotremata | Chromosome | 26/12/12 | 01/12/17 | 991605 | 11544 | 201524 | 445367 | 1996809494 | Washington University (WashU) |
| GCA_004115175.1 | mOrnAna1.h.v1 | *Ornithorhynchus anatinus* | Monotremata | Monotremata | Scaffold | 28/01/19 | 29/01/19 | 882175 | 882175 | 5850 | 5878 | 1575984168 | Vertebrate Genomes Project |
| GCA_004027955.1 | ChaVel_v1_BIUU | *Chaetophractus vellerosus* | Xenarthra | Cingulata | Scaffold | 16/01/19 | 16/01/19 | 1606 | 1606 | 7677333 | 7680080 | 5335596729 | Broad Institute |
| GCF_000208655.1 | Dasnov3.0 | *Dasypus novemcinctus* | Xenarthra | Cingulata | Scaffold | 06/01/12 | 03/12/13 | 1687935 | 26277 | 46559 | 314972 | 3631522711 | Baylor College of Medicine |
| GCA_004025125.1 | TolMat_v1_BIUU | *Tolypeutes matacus* | Xenarthra | Cingulata | Scaffold | 15/01/19 | 15/01/19 | 10217 | 9441 | 2925507 | 2941308 | 4115028163 | Broad Institute |
| GCA_004027775.1 | BraVar_v1_BIUU | *Bradypus variegatus* | Xenarthra | Pilosa | Scaffold | 16/01/19 | 16/01/19 | 1900 | 1900 | 1771805 | 1772267 | 1502475299 | Broad Institute |
| GCA_004027855.1 | ChoDid_v1_BIUU | *Choloepus didactylus* | Xenarthra | Pilosa | Scaffold | 16/01/19 | 16/01/19 | 7289 | 7167 | 2532333 | 2537432 | 3390661301 | Broad Institute |
| GCA_000164785.2 | C_hoffmanni-2.0.1 | *Choloepus hoffmanni* | Xenarthra | Pilosa | Scaffold | 02/10/14 | 14/01/15 | 366442 | 64490 | 269084 | 370041 | 3286013499 | The Genome Institute, Washington University School of Medicine |
| GCA_004026745.1 | MyrTri_v1_BIUU | *Myrmecophaga tridactyla* | Xenarthra | Pilosa | Scaffold | 16/01/19 | 16/01/19 | 41255 | 36434 | 1621408 | 1629691 | 3547273050 | Broad Institute |
| GCA_004025105.1 | TamTet_v1_BIUU | *Tamandua tetradactyla* | Xenarthra | Pilosa | Scaffold | 15/01/19 | 15/01/19 | 19789 | 18654 | 2144204 | 2150023 | 3849374830 | Broad Institute |
