## Supplementary material for "High-quality carnivore genomes from roadkill samples enable species delimitation in aardwolf and bat-eared fox": Table S2

| Species pair | LN | | | UGAM | | |
|---|---|---|---|---|---|---|
| | Chain 1 | Chain 2 | Chain 3 | Chain 1 | Chain 2 | Chain 3 |
| Proteles cristata/ Proteles septentrionalis | 1.228 [IC 95%: 5.34 – 0.58] | 1.014 [IC 95%: 3.01 – 0.56] | 0.841 [IC 95%: 1.26 – 0.55] | 1.327 [IC 95%: 1.86 – 0.93] | 1.335 [IC 95%: 1.86 – 0.94] | 1.34 [IC 95%: 1.88 – 0.94] |
| Otocyon megalotis megalotis/ Otocyon megalotis virgatus | 1.295 [IC 95%: 6.72 – 0.43] | 1.007 [IC 95%: 2.61 – 0.52] | 0.865 [IC 95%: 1.45– 0.48] | 0.569 [IC 95%: 0.81 – 0.39] | 0.573 [IC 95%: 0.83 – 0.39] | 0.57 [IC 95%: 0.82 – 0.40] |
