## Supplementary material for "High-quality carnivore genomes from roadkill samples enable species delimitation in aardwolf and bat-eared fox": Table S1

| Species 1 | Species 2 | Mitochondrial genetic distance (substitution per site) |
|---|---|---|
| Acinonyx_jubatus | Ailuropoda_melanoleuca | 0.23977690953504 |
| Acinonyx_jubatus | Ailurus_fulgens | 0.231539252415341 |
| Acinonyx_jubatus | Aonyx_cinereus | 0.240060980582738 |
| Acinonyx_jubatus | Arctocephalus_forsteri | 0.238382260513751 |
| Acinonyx_jubatus | Arctocephalus_pusillus | 0.229983370411641 |
| Acinonyx_jubatus | Arctocephalus_townsendi | 0.235337827600393 |
| Acinonyx_jubatus | Arctodus_simus | 0.225908176593236 |
| Acinonyx_jubatus | Arctonyx_collaris | 0.228430194933086 |
| Acinonyx_jubatus | Arctotherium_sp_ | 0.23420424346352 |
| Acinonyx_jubatus | Bdeogale_nigripes_FMNH167685_SRR6053065 | 0.194930813426156 |
| Acinonyx_jubatus | Callorhinus_ursinus | 0.22631498047223 |
| Acinonyx_jubatus | Canis_anthus | 0.24145785830046 |
| Acinonyx_jubatus | Canis_latrans | 0.237796921820336 |
| Acinonyx_jubatus | Canis_lupus | 0.234276505586901 |
| Acinonyx_jubatus | Caracal_caracal | 0.117833840620456 |
| Acinonyx_jubatus | Catopuma_badia | 0.104009476538531 |
| Acinonyx_jubatus | Catopuma_temminckii | 0.107868989398182 |
| Acinonyx_jubatus | Chrysocyon_brachyurus | 0.230998578614542 |
| Acinonyx_jubatus | Civettictis_civetta | 0.17116373666409 |
| Acinonyx_jubatus | Conepatus_chinga | 0.23923193902576 |
| Acinonyx_jubatus | Crocuta_crocuta | 0.199213643673961 |
| Acinonyx_jubatus | Cuon_alpinus | 0.240297705907397 |
| Acinonyx_jubatus | Cystophora_cristata | 0.216283630460222 |
| Acinonyx_jubatus | Enhydra_lutris | 0.229261360810057 |
| Acinonyx_jubatus | Erignathus_barbatus | 0.215943586223516 |
| Acinonyx_jubatus | Eumetopias_jubatus | 0.227199567677153 |
| Acinonyx_jubatus | Felis_catus | 0.116890392002287 |
| Acinonyx_jubatus | Felis_chaus | 0.115965887532984 |
| Acinonyx_jubatus | Felis_margarita | 0.117943689865935 |
| Acinonyx_jubatus | Felis_nigripes | 0.117198262510035 |
| Acinonyx_jubatus | Felis_silvestris | 0.116787117768395 |
| Acinonyx_jubatus | Fossa_fossana_FMNH156648_SRR6053060 | 0.182558137529981 |
| Acinonyx_jubatus | Genetta_servalina | 0.173173547105571 |
| Acinonyx_jubatus | Gulo_gulo | 0.236500479257428 |
| Acinonyx_jubatus | Halichoerus_grypus | 0.216370698840397 |
| Acinonyx_jubatus | Helarctos_malayanus | 0.235914052474836 |
| Acinonyx_jubatus | Hyaena_hyaena | 0.197077921305915 |
| Acinonyx_jubatus | Hydrurga_leptonyx | 0.208237883874568 |
| Acinonyx_jubatus | Leopardus_colocolo | 0.120168531290351 |
| Acinonyx_jubatus | Leopardus_geoffroyi | 0.12228306490072 |
| Acinonyx_jubatus | Leopardus_guigna | 0.122776934475837 |
| Acinonyx_jubatus | Leopardus_jacobita | 0.116385177407212 |
| Acinonyx_jubatus | Leopardus_pardalis | 0.116564592437902 |
| Acinonyx_jubatus | Leopardus_tigrinus | 0.120882775767795 |
| Acinonyx_jubatus | Leopardus_wiedii | 0.116941382207051 |
| Acinonyx_jubatus | Leptailurus_serval | 0.111798551698206 |
| Acinonyx_jubatus | Leptonychotes_weddellii | 0.209045727071242 |
| Acinonyx_jubatus | Lobodon_carcinophaga | 0.210718000676245 |
| Acinonyx_jubatus | Lutra_lutra | 0.232137189707761 |
| Acinonyx_jubatus | Lutra_sumatrana | 0.232630650001991 |
| Acinonyx_jubatus | Lutrogale_perspicillata | 0.24515010615481 |
| Acinonyx_jubatus | Lycaon_pictus | 0.233540351895116 |
| Acinonyx_jubatus | Lynx_canadensis | 0.106437786574005 |
| Acinonyx_jubatus | Lynx_lynx | 0.107823739916211 |
| Acinonyx_jubatus | Lynx_pardinus | 0.107299750060536 |
| Acinonyx_jubatus | Lynx_rufus | 0.107152109934135 |
| Acinonyx_jubatus | Manis_javanica | 0.2859691581718 |
| Acinonyx_jubatus | Martes_americana | 0.232179554881041 |
| Acinonyx_jubatus | Martes_flavigula | 0.234626420287232 |
| Acinonyx_jubatus | Martes_foina | 0.233092742851829 |
| Acinonyx_jubatus | Martes_martes | 0.231282330793169 |
| Acinonyx_jubatus | Martes_melampus | 0.229494739816668 |
| Acinonyx_jubatus | Martes_pennanti | 0.2353046464812 |
| Acinonyx_jubatus | Martes_zibellina | 0.233345143883257 |
| Acinonyx_jubatus | Meles_anakuma | 0.231406901686067 |
| Acinonyx_jubatus | Meles_leucurus | 0.224595093350034 |
| Acinonyx_jubatus | Meles_meles | 0.225960138321445 |
| Acinonyx_jubatus | Melogale_moschata | 0.235208474941846 |
| Acinonyx_jubatus | Melursus_ursinus | 0.241403688618916 |
| Acinonyx_jubatus | Mephitis_mephitis | 0.239156080733751 |
| Acinonyx_jubatus | Mirounga_leonina | 0.218861110535861 |
| Acinonyx_jubatus | Mungotictis_decemlineata | 0.185637276541506 |
| Acinonyx_jubatus | Mustela_altaica | 0.230167012482636 |
| Acinonyx_jubatus | Mustela_erminea | 0.224666904023612 |
| Acinonyx_jubatus | Mustela_eversmannii | 0.228512750880255 |
| Acinonyx_jubatus | Mustela_frenata | 0.224973532515469 |
| Acinonyx_jubatus | Mustela_itatsi | 0.230610439702013 |
| Acinonyx_jubatus | Mustela_kathiah | 0.226563079181231 |
| Acinonyx_jubatus | Mustela_nigripes | 0.225657569943898 |
| Acinonyx_jubatus | Mustela_nivalis | 0.230053973506493 |
| Acinonyx_jubatus | Mustela_putorius | 0.22678039962364 |
| Acinonyx_jubatus | Mustela_sibirica | 0.228532007112152 |
| Acinonyx_jubatus | Nandinia_binotata | 0.170968183851628 |
| Acinonyx_jubatus | Nasua_nasua | 0.24769135723093 |
| Acinonyx_jubatus | Neofelis_nebulosa | 0.150073514244868 |

| | | |
|---|---|---|
| Acinonyx_jubatus | Neomonachus_schauinslandi | 0.215641011428203 |
| Acinonyx_jubatus | Neophoca_cinerea | 0.233113503565574 |
| Acinonyx_jubatus | Neovison_vison | 0.228725224137654 |
| Acinonyx_jubatus | Nyctereutes_procyonoides | 0.237374626096488 |
| Acinonyx_jubatus | Odobenus_rosmarus_rosmarus | 0.252256975157905 |
| Acinonyx_jubatus | Otocolobus_manul | 0.114486022617344 |
| Acinonyx_jubatus | Otocyon_megalotis_megalotis_TS305 | 0.246574791037441 |
| Acinonyx_jubatus | Otocyon_megalotis_virgatus_FMNH158128 | 0.243939417105553 |
| Acinonyx_jubatus | Paguma_larvata | 0.177281692748496 |
| Acinonyx_jubatus | Panthera_leo | 0.139361527019643 |
| Acinonyx_jubatus | Panthera_onca | 0.140664399614199 |
| Acinonyx_jubatus | Panthera_pardus | 0.141481651491483 |
| Acinonyx_jubatus | Panthera_tigris | 0.138932170296036 |
| Acinonyx_jubatus | Panthera_uncia | 0.146015346381987 |
| Acinonyx_jubatus | Paradoxurus_hermaphroditus | 0.181193800214311 |
| Acinonyx_jubatus | Parahyaena_brunnea_FD126_Fecal | 0.193699696938982 |
| Acinonyx_jubatus | Pardofelis_marmorata | 0.114735538058673 |
| Acinonyx_jubatus | Phoca_fasciata | 0.216646710653187 |
| Acinonyx_jubatus | Phoca_groenlandica | 0.217012753503262 |
| Acinonyx_jubatus | Phoca_largha | 0.217833648833497 |
| Acinonyx_jubatus | Phoca_vitulina | 0.217485005801878 |
| Acinonyx_jubatus | Phocarctos_hookeri | 0.232064652855975 |
| Acinonyx_jubatus | Prionailurus_bengalensis | 0.107945871077883 |
| Acinonyx_jubatus | Prionailurus_bengalensis_euptilurus | 0.110671307137767 |
| Acinonyx_jubatus | Prionailurus_planiceps | 0.111356666478108 |
| Acinonyx_jubatus | Prionailurus_rubiginosus | 0.110834710397032 |
| Acinonyx_jubatus | Prionailurus_viverrinus | 0.113346296017142 |
| Acinonyx_jubatus | Prionodon_pardicolor | 0.180696359596869 |
| Acinonyx_jubatus | Procyon_lotor | 0.231533684648182 |
| Acinonyx_jubatus | Profelis_aurata | 0.116668380530676 |
| Acinonyx_jubatus | Proteles_cristatus_cristatus_TS307 | 0.203028546485901 |
| Acinonyx_jubatus | Proteles_cristatus_septentrionalis_NMS_Z_2018_54 | 0.20132460447473 |
| Acinonyx_jubatus | Puma_concolor | 0.094404735547944 |
| Acinonyx_jubatus | Puma_yagouaroundi | 0.0967657393086093 |
| Acinonyx_jubatus | Pusa_caspica | 0.219252407293145 |
| Acinonyx_jubatus | Pusa_hispida | 0.214822272375614 |
| Acinonyx_jubatus | Pusa_sibirica | 0.21542792375869 |
| Acinonyx_jubatus | Speothos_venaticus_T1624 | 0.241791526539076 |
| Acinonyx_jubatus | Spilogale_putorius | 0.243462624692334 |
| Acinonyx_jubatus | Taxidea_taxus | 0.221968223807158 |
| Acinonyx_jubatus | Tremarctos_ornatus | 0.235452035720581 |
| Acinonyx_jubatus | Urocyon_cinereoargenteus | 0.245086946433205 |
| Acinonyx_jubatus | Ursus_americanus | 0.23304765434824 |
| Acinonyx_jubatus | Ursus_arctos | 0.239687525468056 |
| Acinonyx_jubatus | Ursus_maritimus | 0.237769793021466 |
| Acinonyx_jubatus | Ursus_spelaeus | 0.232272632049575 |
| Acinonyx_jubatus | Ursus_thibetanus | 0.233959155787868 |
| Acinonyx_jubatus | Urva_javanica | 0.191776515806415 |
| Acinonyx_jubatus | Viverra_tangalunga_FMNH146957_SRR6053069 | 0.170091327334037 |
| Acinonyx_jubatus | Viverricula_indica | 0.17006284633793 |
| Acinonyx_jubatus | Vulpes_corsac | 0.2371849473692 |
| Acinonyx_jubatus | Vulpes_ferrilata | 0.236420628000453 |
| Acinonyx_jubatus | Vulpes_lagopus | 0.239833772974317 |
| Acinonyx_jubatus | Vulpes_vulpes_T3611 | 0.23150826401804 |
| Acinonyx_jubatus | Vulpes_zerda | 0.231496405493461 |
| Acinonyx_jubatus | Zalophus_californianus | 0.22974547352249 |
| Ailuropoda_melanoleuca | Ailurus_fulgens | 0.232263278615956 |
| Ailuropoda_melanoleuca | Aonyx_cinereus | 0.248833793553744 |
| Ailuropoda_melanoleuca | Arctocephalus_forsteri | 0.242221345251571 |
| Ailuropoda_melanoleuca | Arctocephalus_pusillus | 0.239379459688018 |
| Ailuropoda_melanoleuca | Arctocephalus_townsendi | 0.242354459296652 |
| Ailuropoda_melanoleuca | Arctodus_simus | 0.191101689538088 |
| Ailuropoda_melanoleuca | Arctonyx_collaris | 0.243835781159635 |
| Ailuropoda_melanoleuca | Arctotherium_sp_ | 0.192065689872458 |
| Ailuropoda_melanoleuca | Bdeogale_nigripes_FMNH167685_SRR6053065 | 0.249494456583247 |
| Ailuropoda_melanoleuca | Callorhinus_ursinus | 0.23371846549306 |
| Ailuropoda_melanoleuca | Canis_anthus | 0.248782567232256 |
| Ailuropoda_melanoleuca | Canis_latrans | 0.243431317645097 |
| Ailuropoda_melanoleuca | Canis_lupus | 0.250039713500152 |
| Ailuropoda_melanoleuca | Caracal_caracal | 0.238931495696184 |
| Ailuropoda_melanoleuca | Catopuma_badia | 0.236435097967748 |
| Ailuropoda_melanoleuca | Catopuma_temminckii | 0.241818373364514 |
| Ailuropoda_melanoleuca | Chrysocyon_brachyurus | 0.243009333508702 |
| Ailuropoda_melanoleuca | Civettictis_civetta | 0.235503789002343 |
| Ailuropoda_melanoleuca | Conepatus_chinga | 0.240485469225531 |
| Ailuropoda_melanoleuca | Crocuta_crocuta | 0.256739442963942 |
| Ailuropoda_melanoleuca | Cuon_alpinus | 0.250750113886205 |
| Ailuropoda_melanoleuca | Cystophora_cristata | 0.218111360471178 |
| Ailuropoda_melanoleuca | Enhydra_lutris | 0.238999907474521 |
| Ailuropoda_melanoleuca | Erignathus_barbatus | 0.224503609464742 |
| Ailuropoda_melanoleuca | Eumetopias_jubatus | 0.241149412362349 |
| Ailuropoda_melanoleuca | Felis_catus | 0.240810788864521 |
| Ailuropoda_melanoleuca | Felis_chaus | 0.238807699760246 |
| Ailuropoda_melanoleuca | Felis_margarita | 0.241645817880483 |
| Ailuropoda_melanoleuca | Felis_nigripes | 0.246903684029237 |
| Ailuropoda_melanoleuca | Felis_silvestris | 0.239286262939754 |
| Ailuropoda_melanoleuca | Fossa_fossana_FMNH156648_SRR6053060 | 0.248956902973566 |

| | | |
|---|---|---|
| Ailuropoda_melanoleuca | Genetta_servalina | 0.231487199740776 |
| Ailuropoda_melanoleuca | Gulo_gulo | 0.244492701667125 |
| Ailuropoda_melanoleuca | Halichoerus_grypus | 0.226862502497547 |
| Ailuropoda_melanoleuca | Helarctos_malayanus | 0.196275571485914 |
| Ailuropoda_melanoleuca | Hyaena_hyaena | 0.254412755130234 |
| Ailuropoda_melanoleuca | Hydrurga_leptonyx | 0.221143141166593 |
| Ailuropoda_melanoleuca | Leopardus_colocolo | 0.243075856819195 |
| Ailuropoda_melanoleuca | Leopardus_geoffroyi | 0.240994269256331 |
| Ailuropoda_melanoleuca | Leopardus_guigna | 0.240680615145148 |
| Ailuropoda_melanoleuca | Leopardus_jacobita | 0.243261467795497 |
| Ailuropoda_melanoleuca | Leopardus_pardalis | 0.24112589230978 |
| Ailuropoda_melanoleuca | Leopardus_tigrinus | 0.241396826631388 |
| Ailuropoda_melanoleuca | Leopardus_wiedii | 0.241013267013662 |
| Ailuropoda_melanoleuca | Leptailurus_serval | 0.233492272128156 |
| Ailuropoda_melanoleuca | Leptonychotes_weddellii | 0.21632319025665 |
| Ailuropoda_melanoleuca | Lobodon_carcinophaga | 0.216517675389875 |
| Ailuropoda_melanoleuca | Lutra_lutra | 0.247779671932659 |
| Ailuropoda_melanoleuca | Lutra_sumatrana | 0.242984149592209 |
| Ailuropoda_melanoleuca | Lutrogale_perspicillata | 0.25364319455647 |
| Ailuropoda_melanoleuca | Lycaon_pictus | 0.250354455073444 |
| Ailuropoda_melanoleuca | Lynx_canadensis | 0.246650525561469 |
| Ailuropoda_melanoleuca | Lynx_lynx | 0.244477852058704 |
| Ailuropoda_melanoleuca | Lynx_pardinus | 0.240611054340491 |
| Ailuropoda_melanoleuca | Lynx_rufus | 0.242629026001701 |
| Ailuropoda_melanoleuca | Manis_javanica | 0.314931370564602 |
| Ailuropoda_melanoleuca | Martes_americana | 0.244895517364148 |
| Ailuropoda_melanoleuca | Martes_flavigula | 0.245807680356008 |
| Ailuropoda_melanoleuca | Martes_foina | 0.244336811939874 |
| Ailuropoda_melanoleuca | Martes_martes | 0.247319400144277 |
| Ailuropoda_melanoleuca | Martes_melampus | 0.245010091150167 |
| Ailuropoda_melanoleuca | Martes_pennanti | 0.246426392012765 |
| Ailuropoda_melanoleuca | Martes_zibellina | 0.246701298166529 |
| Ailuropoda_melanoleuca | Meles_anakuma | 0.241885372289886 |
| Ailuropoda_melanoleuca | Meles_leucurus | 0.240254364222382 |
| Ailuropoda_melanoleuca | Meles_meles | 0.240521649699895 |
| Ailuropoda_melanoleuca | Melogale_moschata | 0.24753459895044 |
| Ailuropoda_melanoleuca | Melursus_ursinus | 0.198484745186704 |
| Ailuropoda_melanoleuca | Mephitis_mephitis | 0.241444974282113 |
| Ailuropoda_melanoleuca | Mirounga_leonina | 0.221750110228656 |
| Ailuropoda_melanoleuca | Mungotictis_decemlineata | 0.253837201374806 |
| Ailuropoda_melanoleuca | Mustela_altaica | 0.243601134520161 |
| Ailuropoda_melanoleuca | Mustela_erminea | 0.241335862379377 |
| Ailuropoda_melanoleuca | Mustela_eversmannii | 0.242798998681817 |
| Ailuropoda_melanoleuca | Mustela_frenata | 0.235685272405733 |
| Ailuropoda_melanoleuca | Mustela_itatsi | 0.24191274509629 |
| Ailuropoda_melanoleuca | Mustela_kathiah | 0.237607874029091 |
| Ailuropoda_melanoleuca | Mustela_nigripes | 0.242314890645653 |
| Ailuropoda_melanoleuca | Mustela_nivalis | 0.246255704919345 |
| Ailuropoda_melanoleuca | Mustela_putorius | 0.240568540071868 |
| Ailuropoda_melanoleuca | Mustela_sibirica | 0.244876793421007 |
| Ailuropoda_melanoleuca | Nandinia_binotata | 0.236657193690522 |
| Ailuropoda_melanoleuca | Nasua_nasua | 0.256023484461948 |
| Ailuropoda_melanoleuca | Neofelis_nebulosa | 0.254101926653267 |
| Ailuropoda_melanoleuca | Neomonachus_schauinslandi | 0.219245091042183 |
| Ailuropoda_melanoleuca | Neophoca_cinerea | 0.243120488112879 |
| Ailuropoda_melanoleuca | Neovison_vison | 0.241977595498175 |
| Ailuropoda_melanoleuca | Nyctereutes_procyonoides | 0.250258562443175 |
| Ailuropoda_melanoleuca | Odobenus_rosmarus_rosmarus | 0.253981116713936 |
| Ailuropoda_melanoleuca | Otocolobus_manul | 0.237977551279592 |
| Ailuropoda_melanoleuca | Otocyon_megalotis_megalotis_TS305 | 0.251411615609621 |
| Ailuropoda_melanoleuca | Otocyon_megalotis_virgatus_FMNH158128 | 0.252111871850559 |
| Ailuropoda_melanoleuca | Paguma_larvata | 0.237146778903835 |
| Ailuropoda_melanoleuca | Panthera_leo | 0.245487390263668 |
| Ailuropoda_melanoleuca | Panthera_onca | 0.247930261326098 |
| Ailuropoda_melanoleuca | Panthera_pardus | 0.245891799022661 |
| Ailuropoda_melanoleuca | Panthera_tigris | 0.251203698095122 |
| Ailuropoda_melanoleuca | Panthera_uncia | 0.248212439189137 |
| Ailuropoda_melanoleuca | Paradoxurus_hermaphroditus | 0.234229415142809 |
| Ailuropoda_melanoleuca | Parahyaena_brunnea_FD126_Fecal | 0.252167264921004 |
| Ailuropoda_melanoleuca | Pardofelis_marmorata | 0.24356149999499 |
| Ailuropoda_melanoleuca | Phoca_fasciata | 0.22152491516908 |
| Ailuropoda_melanoleuca | Phoca_groenlandica | 0.21935769563476 |
| Ailuropoda_melanoleuca | Phoca_largha | 0.225786394957653 |
| Ailuropoda_melanoleuca | Phoca_vitulina | 0.22574717825794 |
| Ailuropoda_melanoleuca | Phocarctos_hookeri | 0.247560806992175 |
| Ailuropoda_melanoleuca | Prionailurus_bengalensis | 0.238789657662244 |
| Ailuropoda_melanoleuca | Prionailurus_bengalensis_euptilurus | 0.241488112604143 |
| Ailuropoda_melanoleuca | Prionailurus_planiceps | 0.242156630227116 |
| Ailuropoda_melanoleuca | Prionailurus_rubiginosus | 0.24419822808357 |
| Ailuropoda_melanoleuca | Prionailurus_viverrinus | 0.241463920491668 |
| Ailuropoda_melanoleuca | Prionodon_pardicolor | 0.243194282352688 |
| Ailuropoda_melanoleuca | Procyon_lotor | 0.245549050597116 |
| Ailuropoda_melanoleuca | Profelis_aurata | 0.244314575755537 |
| Ailuropoda_melanoleuca | Proteles_cristatus_cristatus_TS307 | 0.256085850326363 |
| Ailuropoda_melanoleuca | Proteles_cristatus_septentrionalis_NMS_Z_2018_54 | 0.256867571544404 |
| Ailuropoda_melanoleuca | Puma_concolor | 0.241461925733242 |
| Ailuropoda_melanoleuca | Puma_yagouaroundi | 0.243936111798392 |

| | | |
|---|---|---|
| Ailuropoda_melanoleuca | Pusa_caspica | 0.230799877000653 |
| Ailuropoda_melanoleuca | Pusa_hispida | 0.223632250553136 |
| Ailuropoda_melanoleuca | Pusa_sibirica | 0.228274479778495 |
| Ailuropoda_melanoleuca | Speothos_venaticus_T1624 | 0.247749665046275 |
| Ailuropoda_melanoleuca | Spilogale_putorius | 0.244054592832145 |
| Ailuropoda_melanoleuca | Taxidea_taxus | 0.23613582536361 |
| Ailuropoda_melanoleuca | Tremarctos_ornatus | 0.193587643643118 |
| Ailuropoda_melanoleuca | Urocyon_cinereoargenteus | 0.255017214174943 |
| Ailuropoda_melanoleuca | Ursus_americanus | 0.199687720395171 |
| Ailuropoda_melanoleuca | Ursus_arctos | 0.199325095084287 |
| Ailuropoda_melanoleuca | Ursus_maritimus | 0.199151397449409 |
| Ailuropoda_melanoleuca | Ursus_spelaeus | 0.197850378023712 |
| Ailuropoda_melanoleuca | Ursus_thibetanus | 0.194909833717334 |
| Ailuropoda_melanoleuca | Urva_javanica | 0.247575963064872 |
| Ailuropoda_melanoleuca | Viverra_tangalunga_FMNH146957_SRR6053069 | 0.234277204979536 |
| Ailuropoda_melanoleuca | Viverricula_indica | 0.233088139945995 |
| Ailuropoda_melanoleuca | Vulpes_corsac | 0.247963564646517 |
| Ailuropoda_melanoleuca | Vulpes_ferrilata | 0.249990202098951 |
| Ailuropoda_melanoleuca | Vulpes_lagopus | 0.256368926798153 |
| Ailuropoda_melanoleuca | Vulpes_vulpes_T3611 | 0.249712872572395 |
| Ailuropoda_melanoleuca | Vulpes_zerda | 0.253438395379207 |
| Ailuropoda_melanoleuca | Zalophus_californianus | 0.237300627227397 |
| Ailurus_fulgens | Aonyx_cinereus | 0.230652205658395 |
| Ailurus_fulgens | Arctocephalus_forsteri | 0.232849390610146 |
| Ailurus_fulgens | Arctocephalus_pusillus | 0.230012051606305 |
| Ailurus_fulgens | Arctocephalus_townsendi | 0.234302254819627 |
| Ailurus_fulgens | Arctodus_simus | 0.231236092266674 |
| Ailurus_fulgens | Arctonyx_collaris | 0.217766812898886 |
| Ailurus_fulgens | Arctotherium_sp_ | 0.229893261345921 |
| Ailurus_fulgens | Bdeogale_nigripes_FMNH167685_SRR6053065 | 0.24493016208697 |
| Ailurus_fulgens | Callorhinus_ursinus | 0.231066658916549 |
| Ailurus_fulgens | Canis_anthus | 0.249848735901223 |
| Ailurus_fulgens | Canis_latrans | 0.246553823406005 |
| Ailurus_fulgens | Canis_lupus | 0.243354671535355 |
| Ailurus_fulgens | Caracal_caracal | 0.237147572463336 |
| Ailurus_fulgens | Catopuma_badia | 0.234689393722043 |
| Ailurus_fulgens | Catopuma_temminckii | 0.233713645432943 |
| Ailurus_fulgens | Chrysocyon_brachyurus | 0.240279604479558 |
| Ailurus_fulgens | Civettictis_civetta | 0.234969189794133 |
| Ailurus_fulgens | Conepatus_chinga | 0.234209343331612 |
| Ailurus_fulgens | Crocuta_crocuta | 0.252915115851182 |
| Ailurus_fulgens | Cuon_alpinus | 0.245224556971083 |
| Ailurus_fulgens | Cystophora_cristata | 0.220717185172626 |
| Ailurus_fulgens | Enhydra_lutris | 0.221653131772315 |
| Ailurus_fulgens | Erignathus_barbatus | 0.216263325582035 |
| Ailurus_fulgens | Eumetopias_jubatus | 0.224786652601139 |
| Ailurus_fulgens | Felis_catus | 0.232663971583998 |
| Ailurus_fulgens | Felis_chaus | 0.232033638889217 |
| Ailurus_fulgens | Felis_margarita | 0.231073219289615 |
| Ailurus_fulgens | Felis_nigripes | 0.233735515145842 |
| Ailurus_fulgens | Felis_silvestris | 0.231845079625939 |
| Ailurus_fulgens | Fossa_fossana_FMNH156648_SRR6053060 | 0.240530137474573 |
| Ailurus_fulgens | Genetta_servalina | 0.2340679619666 |
| Ailurus_fulgens | Gulo_gulo | 0.222879899114054 |
| Ailurus_fulgens | Halichoerus_grypus | 0.221203555376435 |
| Ailurus_fulgens | Helarctos_malayanus | 0.234996752708677 |
| Ailurus_fulgens | Hyaena_hyaena | 0.248960470198965 |
| Ailurus_fulgens | Hydrurga_leptonyx | 0.210891834555134 |
| Ailurus_fulgens | Leopardus_colocolo | 0.234490977325743 |
| Ailurus_fulgens | Leopardus_geoffroyi | 0.235411381513984 |
| Ailurus_fulgens | Leopardus_guigna | 0.237169128526589 |
| Ailurus_fulgens | Leopardus_jacobita | 0.233648373269666 |
| Ailurus_fulgens | Leopardus_pardalis | 0.236691963866762 |
| Ailurus_fulgens | Leopardus_tigrinus | 0.233815653991127 |
| Ailurus_fulgens | Leopardus_wiedii | 0.235333425794588 |
| Ailurus_fulgens | Leptailurus_serval | 0.22998980651688 |
| Ailurus_fulgens | Leptonychotes_weddellii | 0.211072529585694 |
| Ailurus_fulgens | Lobodon_carcinophaga | 0.210169199355087 |
| Ailurus_fulgens | Lutra_lutra | 0.230493287351274 |
| Ailurus_fulgens | Lutra_sumatrana | 0.22677343131084 |
| Ailurus_fulgens | Lutrogale_perspicillata | 0.238629148062772 |
| Ailurus_fulgens | Lycaon_pictus | 0.242726268972241 |
| Ailurus_fulgens | Lynx_canadensis | 0.238590983128999 |
| Ailurus_fulgens | Lynx_lynx | 0.239012362367425 |
| Ailurus_fulgens | Lynx_pardinus | 0.239908319601229 |
| Ailurus_fulgens | Lynx_rufus | 0.237102527140662 |
| Ailurus_fulgens | Manis_javanica | 0.304640193113998 |
| Ailurus_fulgens | Martes_americana | 0.227946981917736 |
| Ailurus_fulgens | Martes_flavigula | 0.226552829052738 |
| Ailurus_fulgens | Martes_foina | 0.223981864588202 |
| Ailurus_fulgens | Martes_martes | 0.230905556969352 |
| Ailurus_fulgens | Martes_melampus | 0.226245226253714 |
| Ailurus_fulgens | Martes_pennanti | 0.231852700253445 |
| Ailurus_fulgens | Martes_zibellina | 0.229794694616574 |
| Ailurus_fulgens | Meles_anakuma | 0.221238961973733 |
| Ailurus_fulgens | Meles_leucurus | 0.217632831721645 |
| Ailurus_fulgens | Meles_meles | 0.217093682006107 |

| | | |
|---|---|---|
| Ailurus_fulgens | Melogale_moschata | 0.229534137213971 |
| Ailurus_fulgens | Melursus_ursinus | 0.239627950319028 |
| Ailurus_fulgens | Mephitis_mephitis | 0.233022786293018 |
| Ailurus_fulgens | Mirounga_leonina | 0.211233269953164 |
| Ailurus_fulgens | Mungotictis_decemlineata | 0.244661055377943 |
| Ailurus_fulgens | Mustela_altaica | 0.223029478786132 |
| Ailurus_fulgens | Mustela_erminea | 0.21690428644236 |
| Ailurus_fulgens | Mustela_eversmannii | 0.225726454050963 |
| Ailurus_fulgens | Mustela_frenata | 0.220691316793025 |
| Ailurus_fulgens | Mustela_itatsi | 0.223358534408407 |
| Ailurus_fulgens | Mustela_kathiah | 0.225659161024507 |
| Ailurus_fulgens | Mustela_nigripes | 0.224590023109803 |
| Ailurus_fulgens | Mustela_nivalis | 0.221681814597339 |
| Ailurus_fulgens | Mustela_putorius | 0.224219494441957 |
| Ailurus_fulgens | Mustela_sibirica | 0.225970605595031 |
| Ailurus_fulgens | Nandinia_binotata | 0.227700098196426 |
| Ailurus_fulgens | Nasua_nasua | 0.248523110732208 |
| Ailurus_fulgens | Neofelis_nebulosa | 0.248542351117833 |
| Ailurus_fulgens | Neomonachus_schauinslandi | 0.211523894315403 |
| Ailurus_fulgens | Neophoca_cinerea | 0.232338803840724 |
| Ailurus_fulgens | Neovison_vison | 0.220659199302519 |
| Ailurus_fulgens | Nyctereutes_procyonoides | 0.25083272013193 |
| Ailurus_fulgens | Odobenus_rosmarus_rosmarus | 0.2420621731372 |
| Ailurus_fulgens | Otocolobus_manul | 0.233030876270038 |
| Ailurus_fulgens | Otocyon_megalotis_megalotis_TS305 | 0.257265226046281 |
| Ailurus_fulgens | Otocyon_megalotis_virgatus_FMNH158128 | 0.256649538358509 |
| Ailurus_fulgens | Paguma_larvata | 0.231346235097548 |
| Ailurus_fulgens | Panthera_leo | 0.240026935228721 |
| Ailurus_fulgens | Panthera_onca | 0.245143131491488 |
| Ailurus_fulgens | Panthera_pardus | 0.240720210146084 |
| Ailurus_fulgens | Panthera_tigris | 0.244417333640413 |
| Ailurus_fulgens | Panthera_uncia | 0.246918869497689 |
| Ailurus_fulgens | Paradoxurus_hermaphroditus | 0.23319356914037 |
| Ailurus_fulgens | Parahyaena_brunnea_FD126_Fecal | 0.250363064027515 |
| Ailurus_fulgens | Pardofelis_marmorata | 0.241878881276906 |
| Ailurus_fulgens | Phoca_fasciata | 0.215622704916077 |
| Ailurus_fulgens | Phoca_groenlandica | 0.217278948339794 |
| Ailurus_fulgens | Phoca_largha | 0.21979387529157 |
| Ailurus_fulgens | Phoca_vitulina | 0.219626448904736 |
| Ailurus_fulgens | Phocarctos_hookeri | 0.236595955536631 |
| Ailurus_fulgens | Prionailurus_bengalensis | 0.231041356314369 |
| Ailurus_fulgens | Prionailurus_bengalensis_euptilurus | 0.229590669274212 |
| Ailurus_fulgens | Prionailurus_planiceps | 0.236935155062588 |
| Ailurus_fulgens | Prionailurus_rubiginosus | 0.233893988850468 |
| Ailurus_fulgens | Prionailurus_viverrinus | 0.231020184213207 |
| Ailurus_fulgens | Prionodon_pardicolor | 0.245469430624016 |
| Ailurus_fulgens | Procyon_lotor | 0.234769127460683 |
| Ailurus_fulgens | Profelis_aurata | 0.235067114184851 |
| Ailurus_fulgens | Proteles_cristatus_cristatus_TS307 | 0.256314770774091 |
| Ailurus_fulgens | Proteles_cristatus_septentrionalis_NMS_Z_2018_54 | 0.258964546560961 |
| Ailurus_fulgens | Puma_concolor | 0.235433448034273 |
| Ailurus_fulgens | Puma_yagouaroundi | 0.229724144650564 |
| Ailurus_fulgens | Pusa_caspica | 0.218949376746173 |
| Ailurus_fulgens | Pusa_hispida | 0.21630876843134 |
| Ailurus_fulgens | Pusa_sibirica | 0.220287151523975 |
| Ailurus_fulgens | Speothos_venaticus_T1624 | 0.245390332819439 |
| Ailurus_fulgens | Spilogale_putorius | 0.229767527782166 |
| Ailurus_fulgens | Taxidea_taxus | 0.215863936375449 |
| Ailurus_fulgens | Tremarctos_ornatus | 0.237234169957692 |
| Ailurus_fulgens | Urocyon_cinereoargenteus | 0.258305759068051 |
| Ailurus_fulgens | Ursus_americanus | 0.231213445842017 |
| Ailurus_fulgens | Ursus_arctos | 0.235999695410885 |
| Ailurus_fulgens | Ursus_maritimus | 0.237831812161089 |
| Ailurus_fulgens | Ursus_spelaeus | 0.233948652548405 |
| Ailurus_fulgens | Ursus_thibetanus | 0.234113819782412 |
| Ailurus_fulgens | Urva_javanica | 0.245909331165258 |
| Ailurus_fulgens | Viverra_tangalunga_FMNH146957_SRR6053069 | 0.231208746729626 |
| Ailurus_fulgens | Viverricula_indica | 0.230936055124238 |
| Ailurus_fulgens | Vulpes_corsac | 0.254196500098939 |
| Ailurus_fulgens | Vulpes_ferrilata | 0.256027429699717 |
| Ailurus_fulgens | Vulpes_lagopus | 0.257729989059454 |
| Ailurus_fulgens | Vulpes_vulpes_T3611 | 0.255264393232293 |
| Ailurus_fulgens | Vulpes_zerda | 0.25533801214823 |
| Ailurus_fulgens | Zalophus_californianus | 0.22719479425592 |
| Aonyx_cinereus | Arctocephalus_forsteri | 0.237834505573667 |
| Aonyx_cinereus | Arctocephalus_pusillus | 0.231709025242604 |
| Aonyx_cinereus | Arctocephalus_townsendi | 0.235294418626602 |
| Aonyx_cinereus | Arctodus_simus | 0.23254550179107 |
| Aonyx_cinereus | Arctonyx_collaris | 0.181365921830473 |
| Aonyx_cinereus | Arctotherium_sp_ | 0.238452794449654 |
| Aonyx_cinereus | Bdeogale_nigripes_FMNH167685_SRR6053065 | 0.245112314603442 |
| Aonyx_cinereus | Callorhinus_ursinus | 0.229302782357131 |
| Aonyx_cinereus | Canis_anthus | 0.253305441303959 |
| Aonyx_cinereus | Canis_latrans | 0.253066979671233 |
| Aonyx_cinereus | Canis_lupus | 0.246106647785009 |
| Aonyx_cinereus | Caracal_caracal | 0.234490977325743 |
| Aonyx_cinereus | Catopuma_badia | 0.234261485584984 |

| | | |
|---|---|---|
| Aonyx_cinereus | Catopuma_temminckii | 0.238641176922012 |
| Aonyx_cinereus | Chrysocyon_brachyurus | 0.252466042245563 |
| Aonyx_cinereus | Civettictis_civetta | 0.234804368648929 |
| Aonyx_cinereus | Conepatus_chinga | 0.245803977303309 |
| Aonyx_cinereus | Crocuta_crocuta | 0.241638578072702 |
| Aonyx_cinereus | Cuon_alpinus | 0.258023239327359 |
| Aonyx_cinereus | Cystophora_cristata | 0.214838420346331 |
| Aonyx_cinereus | Enhydra_lutris | 0.141807496382441 |
| Aonyx_cinereus | Erignathus_barbatus | 0.219761546408901 |
| Aonyx_cinereus | Eumetopias_jubatus | 0.230583464552691 |
| Aonyx_cinereus | Felis_catus | 0.232861616410153 |
| Aonyx_cinereus | Felis_chaus | 0.239825754004412 |
| Aonyx_cinereus | Felis_margarita | 0.23200083983456 |
| Aonyx_cinereus | Felis_nigripes | 0.236456631788638 |
| Aonyx_cinereus | Felis_silvestris | 0.231251246179058 |
| Aonyx_cinereus | Fossa_fossana_FMNH156648_SRR6053060 | 0.232993609477282 |
| Aonyx_cinereus | Genetta_servalina | 0.236793545741667 |
| Aonyx_cinereus | Gulo_gulo | 0.178098811215985 |
| Aonyx_cinereus | Halichoerus_grypus | 0.216873193342062 |
| Aonyx_cinereus | Helarctos_malayanus | 0.243193434992212 |
| Aonyx_cinereus | Hyaena_hyaena | 0.23803862451977 |
| Aonyx_cinereus | Hydrurga_leptonyx | 0.212934704118024 |
| Aonyx_cinereus | Leopardus_colocolo | 0.231303123951947 |
| Aonyx_cinereus | Leopardus_geoffroyi | 0.23173926866657 |
| Aonyx_cinereus | Leopardus_guigna | 0.233129832797078 |
| Aonyx_cinereus | Leopardus_jacobita | 0.231591663025078 |
| Aonyx_cinereus | Leopardus_pardalis | 0.227308354985242 |
| Aonyx_cinereus | Leopardus_tigrinus | 0.234897595106942 |
| Aonyx_cinereus | Leopardus_wiedii | 0.229895243758958 |
| Aonyx_cinereus | Leptailurus_serval | 0.233559381544956 |
| Aonyx_cinereus | Leptonychotes_weddellii | 0.210225569346893 |
| Aonyx_cinereus | Lobodon_carcinophaga | 0.208141427762798 |
| Aonyx_cinereus | Lutra_lutra | 0.122405163679129 |
| Aonyx_cinereus | Lutra_sumatrana | 0.11968163285306 |
| Aonyx_cinereus | Lutrogale_perspicillata | 0.0676202558717779 |
| Aonyx_cinereus | Lycaon_pictus | 0.248395507701558 |
| Aonyx_cinereus | Lynx_canadensis | 0.231806465409516 |
| Aonyx_cinereus | Lynx_lynx | 0.233644802088657 |
| Aonyx_cinereus | Lynx_pardinus | 0.235437050382124 |
| Aonyx_cinereus | Lynx_rufus | 0.236757225369924 |
| Aonyx_cinereus | Manis_javanica | 0.29227508623858 |
| Aonyx_cinereus | Martes_americana | 0.172925244058687 |
| Aonyx_cinereus | Martes_flavigula | 0.174001008152506 |
| Aonyx_cinereus | Martes_foina | 0.170837912776295 |
| Aonyx_cinereus | Martes_martes | 0.17394002286992 |
| Aonyx_cinereus | Martes_melampus | 0.173697062017651 |
| Aonyx_cinereus | Martes_pennanti | 0.178713787892323 |
| Aonyx_cinereus | Martes_zibellina | 0.175301890440439 |
| Aonyx_cinereus | Meles_anakuma | 0.184385955147595 |
| Aonyx_cinereus | Meles_leucurus | 0.180854194909167 |
| Aonyx_cinereus | Meles_meles | 0.180049664265676 |
| Aonyx_cinereus | Melogale_moschata | 0.170554639950877 |
| Aonyx_cinereus | Melursus_ursinus | 0.23976589971758 |
| Aonyx_cinereus | Mephitis_mephitis | 0.245324778414503 |
| Aonyx_cinereus | Mirounga_leonina | 0.220425352941973 |
| Aonyx_cinereus | Mungotictis_decemlineata | 0.236585600566486 |
| Aonyx_cinereus | Mustela_altaica | 0.172131440350322 |
| Aonyx_cinereus | Mustela_erminea | 0.157848212977013 |
| Aonyx_cinereus | Mustela_eversmannii | 0.171300208854939 |
| Aonyx_cinereus | Mustela_frenata | 0.170998699909481 |
| Aonyx_cinereus | Mustela_itatsi | 0.174024869234759 |
| Aonyx_cinereus | Mustela_kathiah | 0.174657225245821 |
| Aonyx_cinereus | Mustela_nigripes | 0.173038435720038 |
| Aonyx_cinereus | Mustela_nivalis | 0.172042250713556 |
| Aonyx_cinereus | Mustela_putorius | 0.171597364212966 |
| Aonyx_cinereus | Mustela_sibirica | 0.171116990862799 |
| Aonyx_cinereus | Nandinia_binotata | 0.224469293232412 |
| Aonyx_cinereus | Nasua_nasua | 0.239810999801982 |
| Aonyx_cinereus | Neofelis_nebulosa | 0.242883367409512 |
| Aonyx_cinereus | Neomonachus_schauinslandi | 0.22109016934242 |
| Aonyx_cinereus | Neophoca_cinerea | 0.235117662101343 |
| Aonyx_cinereus | Neovison_vison | 0.176074984520443 |
| Aonyx_cinereus | Nyctereutes_procyonoides | 0.250218623638802 |
| Aonyx_cinereus | Odobenus_rosmarus_rosmarus | 0.250032613889804 |
| Aonyx_cinereus | Otocolobus_manul | 0.236324952946199 |
| Aonyx_cinereus | Otocyon_megalotis_megalotis_TS305 | 0.252258143219789 |
| Aonyx_cinereus | Otocyon_megalotis_virgatus_FMNH158128 | 0.252899238536227 |
| Aonyx_cinereus | Paguma_larvata | 0.247087630511177 |
| Aonyx_cinereus | Panthera_leo | 0.2386298445545 |
| Aonyx_cinereus | Panthera_onca | 0.235666852083877 |
| Aonyx_cinereus | Panthera_pardus | 0.240663884216326 |
| Aonyx_cinereus | Panthera_tigris | 0.24311267184833 |
| Aonyx_cinereus | Panthera_uncia | 0.238630951765334 |
| Aonyx_cinereus | Paradoxurus_hermaphroditus | 0.252489474807817 |
| Aonyx_cinereus | Parahyaena_brunnea_FD126_Fecal | 0.238616089994326 |
| Aonyx_cinereus | Pardofelis_marmorata | 0.23957940778779 |
| Aonyx_cinereus | Phoca_fasciata | 0.213866353446847 |

| | | |
|---|---|---|
| Aonyx_cinereus | Phoca_groenlandica | 0.217092070261481 |
| Aonyx_cinereus | Phoca_largha | 0.215338930329933 |
| Aonyx_cinereus | Phoca_vitulina | 0.216712006459752 |
| Aonyx_cinereus | Phocarctos_hookeri | 0.234948807146014 |
| Aonyx_cinereus | Prionailurus_bengalensis | 0.233619219413519 |
| Aonyx_cinereus | Prionailurus_bengalensis_euptilurus | 0.234326006820866 |
| Aonyx_cinereus | Prionailurus_planiceps | 0.2310565696254 |
| Aonyx_cinereus | Prionailurus_rubiginosus | 0.232747338794275 |
| Aonyx_cinereus | Prionailurus_viverrinus | 0.237147176169791 |
| Aonyx_cinereus | Prionodon_pardicolor | 0.247295040722384 |
| Aonyx_cinereus | Procyon_lotor | 0.224084717575246 |
| Aonyx_cinereus | Profelis_aurata | 0.240032006478527 |
| Aonyx_cinereus | Proteles_cristatus_cristatus_TS307 | 0.242095943456831 |
| Aonyx_cinereus | Proteles_cristatus_septentrionalis_NMS_Z_2018_54 | 0.245009443408288 |
| Aonyx_cinereus | Puma_concolor | 0.234804368648929 |
| Aonyx_cinereus | Puma_yagouaroundi | 0.237002885755315 |
| Aonyx_cinereus | Pusa_caspica | 0.216568338819747 |
| Aonyx_cinereus | Pusa_hispida | 0.213718584515162 |
| Aonyx_cinereus | Pusa_sibirica | 0.214991073938477 |
| Aonyx_cinereus | Speothos_venaticus_T1624 | 0.262075395278812 |
| Aonyx_cinereus | Spilogale_putorius | 0.248716067345673 |
| Aonyx_cinereus | Taxidea_taxus | 0.181661019990774 |
| Aonyx_cinereus | Tremarctos_ornatus | 0.23856643081378 |
| Aonyx_cinereus | Urocyon_cinereoargenteus | 0.25980803133657 |
| Aonyx_cinereus | Ursus_americanus | 0.24190283612445 |
| Aonyx_cinereus | Ursus_arctos | 0.242033167205761 |
| Aonyx_cinereus | Ursus_maritimus | 0.243191743883846 |
| Aonyx_cinereus | Ursus_spelaeus | 0.236842592756277 |
| Aonyx_cinereus | Ursus_thibetanus | 0.241187220770341 |
| Aonyx_cinereus | Urva_javanica | 0.245101714520134 |
| Aonyx_cinereus | Viverra_tangalunga_FMNH146957_SRR6053069 | 0.237739983715789 |
| Aonyx_cinereus | Viverricula_indica | 0.236195311175556 |
| Aonyx_cinereus | Vulpes_corsac | 0.250261367925842 |
| Aonyx_cinereus | Vulpes_ferrilata | 0.252291973155621 |
| Aonyx_cinereus | Vulpes_lagopus | 0.253955320812408 |
| Aonyx_cinereus | Vulpes_vulpes_T3611 | 0.248483789212883 |
| Aonyx_cinereus | Vulpes_zerda | 0.248641393866553 |
| Aonyx_cinereus | Zalophus_californianus | 0.231576004182548 |
| Arctocephalus_forsteri | Arctocephalus_pusillus | 0.075282829749243 |
| Arctocephalus_forsteri | Arctocephalus_townsendi | 0.0585238972905729 |
| Arctocephalus_forsteri | Arctodus_simus | 0.22998106461803 |
| Arctocephalus_forsteri | Arctonyx_collaris | 0.229649104514212 |
| Arctocephalus_forsteri | Arctotherium_sp_ | 0.231832652379089 |
| Arctocephalus_forsteri | Bdeogale_nigripes_FMNH167685_SRR6053065 | 0.239059333081604 |
| Arctocephalus_forsteri | Callorhinus_ursinus | 0.11994573688442 |
| Arctocephalus_forsteri | Canis_anthus | 0.251746613594979 |
| Arctocephalus_forsteri | Canis_latrans | 0.252145760697421 |
| Arctocephalus_forsteri | Canis_lupus | 0.252780966904772 |
| Arctocephalus_forsteri | Caracal_caracal | 0.238070067993772 |
| Arctocephalus_forsteri | Catopuma_badia | 0.238752661084428 |
| Arctocephalus_forsteri | Catopuma_temminckii | 0.239775890689084 |
| Arctocephalus_forsteri | Chrysocyon_brachyurus | 0.250907494076302 |
| Arctocephalus_forsteri | Civettictis_civetta | 0.238540896101673 |
| Arctocephalus_forsteri | Conepatus_chinga | 0.241925394101626 |
| Arctocephalus_forsteri | Crocuta_crocuta | 0.246186661919678 |
| Arctocephalus_forsteri | Cuon_alpinus | 0.255153329588481 |
| Arctocephalus_forsteri | Cystophora_cristata | 0.188930046307461 |
| Arctocephalus_forsteri | Enhydra_lutris | 0.237441331859527 |
| Arctocephalus_forsteri | Erignathus_barbatus | 0.181352694409384 |
| Arctocephalus_forsteri | Eumetopias_jubatus | 0.0857646053418883 |
| Arctocephalus_forsteri | Felis_catus | 0.240236802992497 |
| Arctocephalus_forsteri | Felis_chaus | 0.240805144207583 |
| Arctocephalus_forsteri | Felis_margarita | 0.238324706313287 |
| Arctocephalus_forsteri | Felis_nigripes | 0.241677032322216 |
| Arctocephalus_forsteri | Felis_silvestris | 0.238720930315952 |
| Arctocephalus_forsteri | Fossa_fossana_FMNH156648_SRR6053060 | 0.232601501507952 |
| Arctocephalus_forsteri | Genetta_servalina | 0.239584917065133 |
| Arctocephalus_forsteri | Gulo_gulo | 0.231492177881856 |
| Arctocephalus_forsteri | Halichoerus_grypus | 0.191205887979883 |
| Arctocephalus_forsteri | Helarctos_malayanus | 0.232384048934041 |
| Arctocephalus_forsteri | Hyaena_hyaena | 0.246973975095512 |
| Arctocephalus_forsteri | Hydrurga_leptonyx | 0.180086460826262 |
| Arctocephalus_forsteri | Leopardus_colocolo | 0.240221871318497 |
| Arctocephalus_forsteri | Leopardus_geoffroyi | 0.233734152670872 |
| Arctocephalus_forsteri | Leopardus_guigna | 0.235962933900384 |
| Arctocephalus_forsteri | Leopardus_jacobita | 0.237879756497846 |
| Arctocephalus_forsteri | Leopardus_pardalis | 0.234688150067633 |
| Arctocephalus_forsteri | Leopardus_tigrinus | 0.234796021515742 |
| Arctocephalus_forsteri | Leopardus_wiedii | 0.233315826629821 |
| Arctocephalus_forsteri | Leptailurus_serval | 0.239358974396438 |
| Arctocephalus_forsteri | Leptonychotes_weddellii | 0.17896277614654 |
| Arctocephalus_forsteri | Lobodon_carcinophaga | 0.178300304431157 |
| Arctocephalus_forsteri | Lutra_lutra | 0.235710055684387 |
| Arctocephalus_forsteri | Lutra_sumatrana | 0.232387473961735 |
| Arctocephalus_forsteri | Lutrogale_perspicillata | 0.24131847268946 |
| Arctocephalus_forsteri | Lycaon_pictus | 0.250398929731599 |
| Arctocephalus_forsteri | Lynx_canadensis | 0.235745965070948 |

| | | |
|---|---|---|
| Arctocephalus_forsteri | Lynx_lynx | 0.234881795306332 |
| Arctocephalus_forsteri | Lynx_pardinus | 0.232936756420691 |
| Arctocephalus_forsteri | Lynx_rufus | 0.236025628717343 |
| Arctocephalus_forsteri | Manis_javanica | 0.286855776048252 |
| Arctocephalus_forsteri | Martes_americana | 0.233690593586072 |
| Arctocephalus_forsteri | Martes_flavigula | 0.231274346822713 |
| Arctocephalus_forsteri | Martes_foina | 0.232701354829643 |
| Arctocephalus_forsteri | Martes_martes | 0.231660392164191 |
| Arctocephalus_forsteri | Martes_melampus | 0.23315755079929 |
| Arctocephalus_forsteri | Martes_pennanti | 0.236350039567666 |
| Arctocephalus_forsteri | Martes_zibellina | 0.233872106010338 |
| Arctocephalus_forsteri | Meles_anakuma | 0.23774240760337 |
| Arctocephalus_forsteri | Meles_leucurus | 0.233569471546458 |
| Arctocephalus_forsteri | Meles_meles | 0.231553641357977 |
| Arctocephalus_forsteri | Melogale_moschata | 0.239679079594717 |
| Arctocephalus_forsteri | Melursus_ursinus | 0.237320549455458 |
| Arctocephalus_forsteri | Mephitis_mephitis | 0.24326832632573 |
| Arctocephalus_forsteri | Mirounga_leonina | 0.182550555069499 |
| Arctocephalus_forsteri | Mungotictis_decemlineata | 0.242897058908415 |
| Arctocephalus_forsteri | Mustela_altaica | 0.239989584496938 |
| Arctocephalus_forsteri | Mustela_erminea | 0.231113236525694 |
| Arctocephalus_forsteri | Mustela_eversmannii | 0.242766194896679 |
| Arctocephalus_forsteri | Mustela_frenata | 0.23024213709183 |
| Arctocephalus_forsteri | Mustela_itatsi | 0.239765268656477 |
| Arctocephalus_forsteri | Mustela_kathiah | 0.237705421455711 |
| Arctocephalus_forsteri | Mustela_nigripes | 0.240883239586471 |
| Arctocephalus_forsteri | Mustela_nivalis | 0.240621141967789 |
| Arctocephalus_forsteri | Mustela_putorius | 0.239606777063122 |
| Arctocephalus_forsteri | Mustela_sibirica | 0.239693163417387 |
| Arctocephalus_forsteri | Nandinia_binotata | 0.22837022099111 |
| Arctocephalus_forsteri | Nasua_nasua | 0.243846820676561 |
| Arctocephalus_forsteri | Neofelis_nebulosa | 0.251214545992476 |
| Arctocephalus_forsteri | Neomonachus_schauinslandi | 0.184718762653607 |
| Arctocephalus_forsteri | Neophoca_cinerea | 0.0731676040288147 |
| Arctocephalus_forsteri | Neovison_vison | 0.232960664426236 |
| Arctocephalus_forsteri | Nyctereutes_procyonoides | 0.242855609667584 |
| Arctocephalus_forsteri | Odobenus_rosmarus_rosmarus | 0.197109973756599 |
| Arctocephalus_forsteri | Otocolobus_manul | 0.234965831118517 |
| Arctocephalus_forsteri | Otocyon_megalotis_megalotis_TS305 | 0.248392140445258 |
| Arctocephalus_forsteri | Otocyon_megalotis_virgatus_FMNH158128 | 0.249704952163157 |
| Arctocephalus_forsteri | Paguma_larvata | 0.24342768878181 |
| Arctocephalus_forsteri | Panthera_leo | 0.246835190760218 |
| Arctocephalus_forsteri | Panthera_onca | 0.245853318642676 |
| Arctocephalus_forsteri | Panthera_pardus | 0.249441428955758 |
| Arctocephalus_forsteri | Panthera_tigris | 0.245664472288459 |
| Arctocephalus_forsteri | Panthera_uncia | 0.24876022994031 |
| Arctocephalus_forsteri | Paradoxurus_hermaphroditus | 0.243533241018101 |
| Arctocephalus_forsteri | Parahyaena_brunnea_FD126_Fecal | 0.242984535546528 |
| Arctocephalus_forsteri | Pardofelis_marmorata | 0.242861902689016 |
| Arctocephalus_forsteri | Phoca_fasciata | 0.183625968728622 |
| Arctocephalus_forsteri | Phoca_groenlandica | 0.188707028971742 |
| Arctocephalus_forsteri | Phoca_largha | 0.190173655407983 |
| Arctocephalus_forsteri | Phoca_vitulina | 0.191302249889837 |
| Arctocephalus_forsteri | Phocarctos_hookeri | 0.0782294239287658 |
| Arctocephalus_forsteri | Prionailurus_bengalensis | 0.238234657234004 |
| Arctocephalus_forsteri | Prionailurus_bengalensis_euptilurus | 0.237303282946862 |
| Arctocephalus_forsteri | Prionailurus_planiceps | 0.234234439753299 |
| Arctocephalus_forsteri | Prionailurus_rubiginosus | 0.235579366999005 |
| Arctocephalus_forsteri | Prionailurus_viverrinus | 0.240777346241023 |
| Arctocephalus_forsteri | Prionodon_pardicolor | 0.247824541133215 |
| Arctocephalus_forsteri | Procyon_lotor | 0.240403567218753 |
| Arctocephalus_forsteri | Profelis_aurata | 0.24436681762997 |
| Arctocephalus_forsteri | Proteles_cristatus_cristatus_TS307 | 0.248028186974438 |
| Arctocephalus_forsteri | Proteles_cristatus_septentrionalis_NMS_Z_2018_54 | 0.249177569357676 |
| Arctocephalus_forsteri | Puma_concolor | 0.238292389029005 |
| Arctocephalus_forsteri | Puma_yagouaroundi | 0.241511421024629 |
| Arctocephalus_forsteri | Pusa_caspica | 0.190020418131331 |
| Arctocephalus_forsteri | Pusa_hispida | 0.187969953972515 |
| Arctocephalus_forsteri | Pusa_sibirica | 0.188835793828858 |
| Arctocephalus_forsteri | Speothos_venaticus_T1624 | 0.257452721199566 |
| Arctocephalus_forsteri | Spilogale_putorius | 0.246405897332844 |
| Arctocephalus_forsteri | Taxidea_taxus | 0.223706905683243 |
| Arctocephalus_forsteri | Tremarctos_ornatus | 0.23557460144274 |
| Arctocephalus_forsteri | Urocyon_cinereoargenteus | 0.253320284636788 |
| Arctocephalus_forsteri | Ursus_americanus | 0.233178193346933 |
| Arctocephalus_forsteri | Ursus_arctos | 0.24367992581801 |
| Arctocephalus_forsteri | Ursus_maritimus | 0.242663260816803 |
| Arctocephalus_forsteri | Ursus_spelaeus | 0.2362042966547 |
| Arctocephalus_forsteri | Ursus_thibetanus | 0.235752319222793 |
| Arctocephalus_forsteri | Urva_javanica | 0.240599995830182 |
| Arctocephalus_forsteri | Viverra_tangalunga_FMNH146957_SRR6053069 | 0.236301958556644 |
| Arctocephalus_forsteri | Viverricula_indica | 0.240629405435598 |
| Arctocephalus_forsteri | Vulpes_corsac | 0.249756866438811 |
| Arctocephalus_forsteri | Vulpes_ferrilata | 0.253213252596682 |
| Arctocephalus_forsteri | Vulpes_lagopus | 0.251874077574962 |
| Arctocephalus_forsteri | Vulpes_vulpes_T3611 | 0.247711953877617 |
| Arctocephalus_forsteri | Vulpes_zerda | 0.249400789384897 |

| | | |
|---|---|---|
| Arctocephalus_forsteri | Zalophus_californianus | 0.0816899532945693 |
| Arctocephalus_pusillus | Arctocephalus_townsendi | 0.0758766917298973 |
| Arctocephalus_pusillus | Arctodus_simus | 0.22680010375149 |
| Arctocephalus_pusillus | Arctonyx_collaris | 0.228352136288586 |
| Arctocephalus_pusillus | Arctotherium_sp_ | 0.231884975530894 |
| Arctocephalus_pusillus | Bdeogale_nigripes_FMNH167685_SRR6053065 | 0.236302378588888 |
| Arctocephalus_pusillus | Callorhinus_ursinus | 0.118278926137065 |
| Arctocephalus_pusillus | Canis_anthus | 0.247229739002938 |
| Arctocephalus_pusillus | Canis_latrans | 0.245299105250385 |
| Arctocephalus_pusillus | Canis_lupus | 0.245087580028085 |
| Arctocephalus_pusillus | Caracal_caracal | 0.23193509821737 |
| Arctocephalus_pusillus | Catopuma_badia | 0.232834201568839 |
| Arctocephalus_pusillus | Catopuma_temminckii | 0.236713707099199 |
| Arctocephalus_pusillus | Chrysocyon_brachyurus | 0.252679634401514 |
| Arctocephalus_pusillus | Civettictis_civetta | 0.234284445543932 |
| Arctocephalus_pusillus | Conepatus_chinga | 0.233251895170753 |
| Arctocephalus_pusillus | Crocuta_crocuta | 0.240306167699748 |
| Arctocephalus_pusillus | Cuon_alpinus | 0.25100964855941 |
| Arctocephalus_pusillus | Cystophora_cristata | 0.185861599787296 |
| Arctocephalus_pusillus | Enhydra_lutris | 0.232620867422667 |
| Arctocephalus_pusillus | Erignathus_barbatus | 0.180636729108402 |
| Arctocephalus_pusillus | Eumetopias_jubatus | 0.0816390246622461 |
| Arctocephalus_pusillus | Felis_catus | 0.235246188199005 |
| Arctocephalus_pusillus | Felis_chaus | 0.23766944101316 |
| Arctocephalus_pusillus | Felis_margarita | 0.233773952213167 |
| Arctocephalus_pusillus | Felis_nigripes | 0.234354145410135 |
| Arctocephalus_pusillus | Felis_silvestris | 0.233943700991801 |
| Arctocephalus_pusillus | Fossa_fossana_FMNH156648_SRR6053060 | 0.230396921440147 |
| Arctocephalus_pusillus | Genetta_servalina | 0.235647160548463 |
| Arctocephalus_pusillus | Gulo_gulo | 0.23193573524073 |
| Arctocephalus_pusillus | Halichoerus_grypus | 0.185353699874781 |
| Arctocephalus_pusillus | Helarctos_malayanus | 0.228191798203351 |
| Arctocephalus_pusillus | Hyaena_hyaena | 0.241084551043873 |
| Arctocephalus_pusillus | Hydrurga_leptonyx | 0.17789608643734 |
| Arctocephalus_pusillus | Leopardus_colocolo | 0.23366355862655 |
| Arctocephalus_pusillus | Leopardus_geoffroyi | 0.232064652855975 |
| Arctocephalus_pusillus | Leopardus_guigna | 0.232064652855975 |
| Arctocephalus_pusillus | Leopardus_jacobita | 0.234536564953351 |
| Arctocephalus_pusillus | Leopardus_pardalis | 0.228195289553435 |
| Arctocephalus_pusillus | Leopardus_tigrinus | 0.230381949903953 |
| Arctocephalus_pusillus | Leopardus_wiedii | 0.227163652589839 |
| Arctocephalus_pusillus | Leptailurus_serval | 0.229874068848194 |
| Arctocephalus_pusillus | Leptonychotes_weddellii | 0.172389139964522 |
| Arctocephalus_pusillus | Lobodon_carcinophaga | 0.176219387183703 |
| Arctocephalus_pusillus | Lutra_lutra | 0.230836569871257 |
| Arctocephalus_pusillus | Lutra_sumatrana | 0.23017336650222 |
| Arctocephalus_pusillus | Lutrogale_perspicillata | 0.23628591206685 |
| Arctocephalus_pusillus | Lycaon_pictus | 0.245696924657954 |
| Arctocephalus_pusillus | Lynx_canadensis | 0.232614480614995 |
| Arctocephalus_pusillus | Lynx_lynx | 0.232044680973746 |
| Arctocephalus_pusillus | Lynx_pardinus | 0.229821802721719 |
| Arctocephalus_pusillus | Lynx_rufus | 0.231884877567329 |
| Arctocephalus_pusillus | Manis_javanica | 0.283793918246429 |
| Arctocephalus_pusillus | Martes_americana | 0.232888949555598 |
| Arctocephalus_pusillus | Martes_flavigula | 0.229302663386898 |
| Arctocephalus_pusillus | Martes_foina | 0.230904254366668 |
| Arctocephalus_pusillus | Martes_martes | 0.229424278344033 |
| Arctocephalus_pusillus | Martes_melampus | 0.22784776638556 |
| Arctocephalus_pusillus | Martes_pennanti | 0.234536244475112 |
| Arctocephalus_pusillus | Martes_zibellina | 0.230739604798136 |
| Arctocephalus_pusillus | Meles_anakuma | 0.231853450639448 |
| Arctocephalus_pusillus | Meles_leucurus | 0.227700269869262 |
| Arctocephalus_pusillus | Meles_meles | 0.228709702782675 |
| Arctocephalus_pusillus | Melogale_moschata | 0.236398245110533 |
| Arctocephalus_pusillus | Melursus_ursinus | 0.234717577148485 |
| Arctocephalus_pusillus | Mephitis_mephitis | 0.239992668095835 |
| Arctocephalus_pusillus | Mirounga_leonina | 0.178925849882857 |
| Arctocephalus_pusillus | Mungotictis_decemlineata | 0.236842645439472 |
| Arctocephalus_pusillus | Mustela_altaica | 0.238332633452024 |
| Arctocephalus_pusillus | Mustela_erminea | 0.226152489140326 |
| Arctocephalus_pusillus | Mustela_eversmannii | 0.235088851973937 |
| Arctocephalus_pusillus | Mustela_frenata | 0.229624496101107 |
| Arctocephalus_pusillus | Mustela_itatsi | 0.231799281955352 |
| Arctocephalus_pusillus | Mustela_kathiah | 0.23694716471306 |
| Arctocephalus_pusillus | Mustela_nigripes | 0.233503089324732 |
| Arctocephalus_pusillus | Mustela_nivalis | 0.235911770666649 |
| Arctocephalus_pusillus | Mustela_putorius | 0.233968102266018 |
| Arctocephalus_pusillus | Mustela_sibirica | 0.234267978894543 |
| Arctocephalus_pusillus | Nandinia_binotata | 0.221035104926346 |
| Arctocephalus_pusillus | Nasua_nasua | 0.239594468729615 |
| Arctocephalus_pusillus | Neofelis_nebulosa | 0.245931116987921 |
| Arctocephalus_pusillus | Neomonachus_schauinslandi | 0.177000553529774 |
| Arctocephalus_pusillus | Neophoca_cinerea | 0.0782488798342201 |
| Arctocephalus_pusillus | Neovison_vison | 0.234046018589432 |
| Arctocephalus_pusillus | Nyctereutes_procyonoides | 0.240025541713931 |
| Arctocephalus_pusillus | Odobenus_rosmarus_rosmarus | 0.197327537690182 |
| Arctocephalus_pusillus | Otocolobus_manul | 0.230751778567479 |

| | | |
|---|---|---|
| Arctocephalus_pusillus | Otocyon_megalotis_megalotis_TS305 | 0.246789849947891 |
| Arctocephalus_pusillus | Otocyon_megalotis_virgatus_FMNH158128 | 0.246858342297476 |
| Arctocephalus_pusillus | Paguma_larvata | 0.238118808494364 |
| Arctocephalus_pusillus | Panthera_leo | 0.238964489809424 |
| Arctocephalus_pusillus | Panthera_onca | 0.238203292918984 |
| Arctocephalus_pusillus | Panthera_pardus | 0.240679095269961 |
| Arctocephalus_pusillus | Panthera_tigris | 0.238467907107806 |
| Arctocephalus_pusillus | Panthera_uncia | 0.24287294075523 |
| Arctocephalus_pusillus | Paradoxurus_hermaphroditus | 0.242380105244979 |
| Arctocephalus_pusillus | Parahyaena_brunnea_FD126_Fecal | 0.240551321181662 |
| Arctocephalus_pusillus | Pardofelis_marmorata | 0.235898548182429 |
| Arctocephalus_pusillus | Phoca_fasciata | 0.180315053424778 |
| Arctocephalus_pusillus | Phoca_groenlandica | 0.185802002823548 |
| Arctocephalus_pusillus | Phoca_largha | 0.182989134247085 |
| Arctocephalus_pusillus | Phoca_vitulina | 0.184857141732459 |
| Arctocephalus_pusillus | Phocarctos_hookeri | 0.081111273711675 |
| Arctocephalus_pusillus | Prionailurus_bengalensis | 0.232814250167005 |
| Arctocephalus_pusillus | Prionailurus_bengalensis_euptilurus | 0.229704017724613 |
| Arctocephalus_pusillus | Prionailurus_planiceps | 0.235468845336309 |
| Arctocephalus_pusillus | Prionailurus_rubiginosus | 0.234324424456871 |
| Arctocephalus_pusillus | Prionailurus_viverrinus | 0.235399559385392 |
| Arctocephalus_pusillus | Prionodon_pardicolor | 0.243634415580215 |
| Arctocephalus_pusillus | Procyon_lotor | 0.234441942285165 |
| Arctocephalus_pusillus | Profelis_aurata | 0.235669319325242 |
| Arctocephalus_pusillus | Proteles_cristatus_cristatus_TS307 | 0.239620285301158 |
| Arctocephalus_pusillus | Proteles_cristatus_septentrionalis_NMS_Z_2018_54 | 0.243982269297119 |
| Arctocephalus_pusillus | Puma_concolor | 0.235016772071204 |
| Arctocephalus_pusillus | Puma_yagouaroundi | 0.235496567349685 |
| Arctocephalus_pusillus | Pusa_caspica | 0.18611623767206 |
| Arctocephalus_pusillus | Pusa_hispida | 0.184271546955667 |
| Arctocephalus_pusillus | Pusa_sibirica | 0.184882898892577 |
| Arctocephalus_pusillus | Speothos_venaticus_T1624 | 0.251680598584436 |
| Arctocephalus_pusillus | Spilogale_putorius | 0.245887393421638 |
| Arctocephalus_pusillus | Taxidea_taxus | 0.220995696350693 |
| Arctocephalus_pusillus | Tremarctos_ornatus | 0.232386754050553 |
| Arctocephalus_pusillus | Urocyon_cinereoargenteus | 0.2484394253611 |
| Arctocephalus_pusillus | Ursus_americanus | 0.232047839126461 |
| Arctocephalus_pusillus | Ursus_arctos | 0.238969902278595 |
| Arctocephalus_pusillus | Ursus_maritimus | 0.235360379776555 |
| Arctocephalus_pusillus | Ursus_spelaeus | 0.234763403608741 |
| Arctocephalus_pusillus | Ursus_thibetanus | 0.234392301044217 |
| Arctocephalus_pusillus | Urva_javanica | 0.237964344902317 |
| Arctocephalus_pusillus | Viverra_tangalunga_FMNH146957_SRR6053069 | 0.233934662365976 |
| Arctocephalus_pusillus | Viverricula_indica | 0.235477418048275 |
| Arctocephalus_pusillus | Vulpes_corsac | 0.248473153640649 |
| Arctocephalus_pusillus | Vulpes_ferrilata | 0.250672228474313 |
| Arctocephalus_pusillus | Vulpes_lagopus | 0.245808956923715 |
| Arctocephalus_pusillus | Vulpes_vulpes_T3611 | 0.244884581693837 |
| Arctocephalus_pusillus | Vulpes_zerda | 0.246426906818943 |
| Arctocephalus_pusillus | Zalophus_californianus | 0.0821369129108798 |
| Arctocephalus_townsendi | Arctodus_simus | 0.230389618409102 |
| Arctocephalus_townsendi | Arctonyx_collaris | 0.232108947851035 |
| Arctocephalus_townsendi | Arctotherium_sp_ | 0.233365705575991 |
| Arctocephalus_townsendi | Bdeogale_nigripes_FMNH167685_SRR6053065 | 0.239934954721275 |
| Arctocephalus_townsendi | Callorhinus_ursinus | 0.117806694116463 |
| Arctocephalus_townsendi | Canis_anthus | 0.252667351187887 |
| Arctocephalus_townsendi | Canis_latrans | 0.250452990051263 |
| Arctocephalus_townsendi | Canis_lupus | 0.254272770139112 |
| Arctocephalus_townsendi | Caracal_caracal | 0.238705248752981 |
| Arctocephalus_townsendi | Catopuma_badia | 0.239876604821226 |
| Arctocephalus_townsendi | Catopuma_temminckii | 0.239898106602863 |
| Arctocephalus_townsendi | Chrysocyon_brachyurus | 0.255654932672022 |
| Arctocephalus_townsendi | Civettictis_civetta | 0.240715062232012 |
| Arctocephalus_townsendi | Conepatus_chinga | 0.239893993091109 |
| Arctocephalus_townsendi | Crocuta_crocuta | 0.246293197328591 |
| Arctocephalus_townsendi | Cuon_alpinus | 0.25771087012621 |
| Arctocephalus_townsendi | Cystophora_cristata | 0.189422167806268 |
| Arctocephalus_townsendi | Enhydra_lutris | 0.23528054979783 |
| Arctocephalus_townsendi | Erignathus_barbatus | 0.181111695436735 |
| Arctocephalus_townsendi | Eumetopias_jubatus | 0.0875124563346057 |
| Arctocephalus_townsendi | Felis_catus | 0.241097012737713 |
| Arctocephalus_townsendi | Felis_chaus | 0.240884533829979 |
| Arctocephalus_townsendi | Felis_margarita | 0.2395156952079 |
| Arctocephalus_townsendi | Felis_nigripes | 0.24330153258098 |
| Arctocephalus_townsendi | Felis_silvestris | 0.240136024000664 |
| Arctocephalus_townsendi | Fossa_fossana_FMNH156648_SRR6053060 | 0.23390806171203 |
| Arctocephalus_townsendi | Genetta_servalina | 0.236490493684712 |
| Arctocephalus_townsendi | Gulo_gulo | 0.235574587444363 |
| Arctocephalus_townsendi | Halichoerus_grypus | 0.188556403427421 |
| Arctocephalus_townsendi | Helarctos_malayanus | 0.23041860490029 |
| Arctocephalus_townsendi | Hyaena_hyaena | 0.246988957237839 |
| Arctocephalus_townsendi | Hydrurga_leptonyx | 0.181029917210402 |
| Arctocephalus_townsendi | Leopardus_colocolo | 0.240905997828059 |
| Arctocephalus_townsendi | Leopardus_geoffroyi | 0.238885121051414 |
| Arctocephalus_townsendi | Leopardus_guigna | 0.237773886012393 |
| Arctocephalus_townsendi | Leopardus_jacobita | 0.243051666899444 |
| Arctocephalus_townsendi | Leopardus_pardalis | 0.238181216154728 |

| | | |
|---|---|---|
| Arctocephalus_townsendi | Leopardus_tigrinus | 0.236845116992168 |
| Arctocephalus_townsendi | Leopardus_wiedii | 0.236136382954145 |
| Arctocephalus_townsendi | Leptailurus_serval | 0.236769754931966 |
| Arctocephalus_townsendi | Leptonychotes_weddellii | 0.179697835421269 |
| Arctocephalus_townsendi | Lobodon_carcinophaga | 0.18120977249475 |
| Arctocephalus_townsendi | Lutra_lutra | 0.233556061062166 |
| Arctocephalus_townsendi | Lutra_sumatrana | 0.233908515081245 |
| Arctocephalus_townsendi | Lutrogale_perspicillata | 0.236626901021875 |
| Arctocephalus_townsendi | Lycaon_pictus | 0.253156345808094 |
| Arctocephalus_townsendi | Lynx_canadensis | 0.237933225817369 |
| Arctocephalus_townsendi | Lynx_lynx | 0.236178512376875 |
| Arctocephalus_townsendi | Lynx_pardinus | 0.235556122374644 |
| Arctocephalus_townsendi | Lynx_rufus | 0.235683065275412 |
| Arctocephalus_townsendi | Manis_javanica | 0.287785983431861 |
| Arctocephalus_townsendi | Martes_americana | 0.234945877763916 |
| Arctocephalus_townsendi | Martes_flavigula | 0.230640286018342 |
| Arctocephalus_townsendi | Martes_foina | 0.233820354981592 |
| Arctocephalus_townsendi | Martes_martes | 0.230804903001264 |
| Arctocephalus_townsendi | Martes_melampus | 0.232077507998899 |
| Arctocephalus_townsendi | Martes_pennanti | 0.238084768068046 |
| Arctocephalus_townsendi | Martes_zibellina | 0.230351156965135 |
| Arctocephalus_townsendi | Meles_anakuma | 0.236926390260693 |
| Arctocephalus_townsendi | Meles_leucurus | 0.232184050243351 |
| Arctocephalus_townsendi | Meles_meles | 0.232870522111621 |
| Arctocephalus_townsendi | Melogale_moschata | 0.23728635528659 |
| Arctocephalus_townsendi | Melursus_ursinus | 0.235673771842899 |
| Arctocephalus_townsendi | Mephitis_mephitis | 0.243144261681552 |
| Arctocephalus_townsendi | Mirounga_leonina | 0.181272776240519 |
| Arctocephalus_townsendi | Mungotictis_decemlineata | 0.243648873345979 |
| Arctocephalus_townsendi | Mustela_altaica | 0.235755741074271 |
| Arctocephalus_townsendi | Mustela_erminea | 0.229517942485615 |
| Arctocephalus_townsendi | Mustela_eversmannii | 0.239514715314529 |
| Arctocephalus_townsendi | Mustela_frenata | 0.230476238184483 |
| Arctocephalus_townsendi | Mustela_itatsi | 0.235000304732401 |
| Arctocephalus_townsendi | Mustela_kathiah | 0.23865397681142 |
| Arctocephalus_townsendi | Mustela_nigripes | 0.238652433882032 |
| Arctocephalus_townsendi | Mustela_nivalis | 0.23792694685251 |
| Arctocephalus_townsendi | Mustela_putorius | 0.237046538444649 |
| Arctocephalus_townsendi | Mustela_sibirica | 0.237573274612976 |
| Arctocephalus_townsendi | Nandinia_binotata | 0.22969240897601 |
| Arctocephalus_townsendi | Nasua_nasua | 0.243582343017436 |
| Arctocephalus_townsendi | Neofelis_nebulosa | 0.250723197339094 |
| Arctocephalus_townsendi | Neomonachus_schauinslandi | 0.183976373906375 |
| Arctocephalus_townsendi | Neophoca_cinerea | 0.0721106288170296 |
| Arctocephalus_townsendi | Neovison_vison | 0.234391034925633 |
| Arctocephalus_townsendi | Nyctereutes_procyonoides | 0.243644753222642 |
| Arctocephalus_townsendi | Odobenus_rosmarus_rosmarus | 0.197852899976415 |
| Arctocephalus_townsendi | Otocolobus_manul | 0.235970136843327 |
| Arctocephalus_townsendi | Otocyon_megalotis_megalotis_TS305 | 0.251418936840076 |
| Arctocephalus_townsendi | Otocyon_megalotis_virgatus_FMNH158128 | 0.251941331982656 |
| Arctocephalus_townsendi | Paguma_larvata | 0.24312841536325 |
| Arctocephalus_townsendi | Panthera_leo | 0.247931797411507 |
| Arctocephalus_townsendi | Panthera_onca | 0.242356761534676 |
| Arctocephalus_townsendi | Panthera_pardus | 0.247479052615434 |
| Arctocephalus_townsendi | Panthera_tigris | 0.24520146742177 |
| Arctocephalus_townsendi | Panthera_uncia | 0.248951974596077 |
| Arctocephalus_townsendi | Paradoxurus_hermaphroditus | 0.247275960027673 |
| Arctocephalus_townsendi | Parahyaena_brunnea_FD126_Fecal | 0.24613932382583 |
| Arctocephalus_townsendi | Pardofelis_marmorata | 0.243053039817498 |
| Arctocephalus_townsendi | Phoca_fasciata | 0.184554734106757 |
| Arctocephalus_townsendi | Phoca_groenlandica | 0.191018743847245 |
| Arctocephalus_townsendi | Phoca_largha | 0.187419996635006 |
| Arctocephalus_townsendi | Phoca_vitulina | 0.189175644963111 |
| Arctocephalus_townsendi | Phocarctos_hookeri | 0.0776116197987364 |
| Arctocephalus_townsendi | Prionailurus_bengalensis | 0.23467782636793 |
| Arctocephalus_townsendi | Prionailurus_bengalensis_euptilurus | 0.234726544660565 |
| Arctocephalus_townsendi | Prionailurus_planiceps | 0.236855484573544 |
| Arctocephalus_townsendi | Prionailurus_rubiginosus | 0.234224322529321 |
| Arctocephalus_townsendi | Prionailurus_viverrinus | 0.237292721918264 |
| Arctocephalus_townsendi | Prionodon_pardicolor | 0.24918852088172 |
| Arctocephalus_townsendi | Procyon_lotor | 0.240455060920738 |
| Arctocephalus_townsendi | Profelis_aurata | 0.245233835042519 |
| Arctocephalus_townsendi | Proteles_cristatus_cristatus_TS307 | 0.248333765489616 |
| Arctocephalus_townsendi | Proteles_cristatus_septentrionalis_NMS_Z_2018_54 | 0.249010203022326 |
| Arctocephalus_townsendi | Puma_concolor | 0.241136832238077 |
| Arctocephalus_townsendi | Puma_yagouaroundi | 0.239717166664588 |
| Arctocephalus_townsendi | Pusa_caspica | 0.187245481510229 |
| Arctocephalus_townsendi | Pusa_hispida | 0.186569695602297 |
| Arctocephalus_townsendi | Pusa_sibirica | 0.187245481510229 |
| Arctocephalus_townsendi | Speothos_venaticus_T1624 | 0.255816215041425 |
| Arctocephalus_townsendi | Spilogale_putorius | 0.246257148032477 |
| Arctocephalus_townsendi | Taxidea_taxus | 0.228482926920779 |
| Arctocephalus_townsendi | Tremarctos_ornatus | 0.238366713705066 |
| Arctocephalus_townsendi | Urocyon_cinereoargenteus | 0.254908361115931 |
| Arctocephalus_townsendi | Ursus_americanus | 0.233277523307477 |
| Arctocephalus_townsendi | Ursus_arctos | 0.240552908161087 |
| Arctocephalus_townsendi | Ursus_maritimus | 0.237613782464678 |

| | | |
|---|---|---|
| Arctocephalus_townsendi | Ursus_spelaeus | 0.236250348831826 |
| Arctocephalus_townsendi | Ursus_thibetanus | 0.23579852456071 |
| Arctocephalus_townsendi | Urva_javanica | 0.24240810504353 |
| Arctocephalus_townsendi | Viverra_tangalunga_FMNH146957_SRR6053069 | 0.237489244591267 |
| Arctocephalus_townsendi | Viverricula_indica | 0.238172138799788 |
| Arctocephalus_townsendi | Vulpes_corsac | 0.254124404391842 |
| Arctocephalus_townsendi | Vulpes_ferrilata | 0.255093581119635 |
| Arctocephalus_townsendi | Vulpes_lagopus | 0.25274852767849 |
| Arctocephalus_townsendi | Vulpes_vulpes_T3611 | 0.249957503744342 |
| Arctocephalus_townsendi | Vulpes_zerda | 0.25095233225241 |
| Arctocephalus_townsendi | Zalophus_californianus | 0.0851097526984944 |
| Arctodus_simus | Arctonyx_collaris | 0.232327408335888 |
| Arctodus_simus | Arctotherium_sp_ | 0.107877315464513 |
| Arctodus_simus | Bdeogale_nigripes_FMNH167685_SRR6053065 | 0.235611494868114 |
| Arctodus_simus | Callorhinus_ursinus | 0.227973029790585 |
| Arctodus_simus | Canis_anthus | 0.248426029155959 |
| Arctodus_simus | Canis_latrans | 0.242802056756656 |
| Arctodus_simus | Canis_lupus | 0.244950166335442 |
| Arctodus_simus | Caracal_caracal | 0.228255327383799 |
| Arctodus_simus | Catopuma_badia | 0.226618790199666 |
| Arctodus_simus | Catopuma_temminckii | 0.227628325381731 |
| Arctodus_simus | Chrysocyon_brachyurus | 0.24277334119063 |
| Arctodus_simus | Civettictis_civetta | 0.231916904555018 |
| Arctodus_simus | Conepatus_chinga | 0.234899518943436 |
| Arctodus_simus | Crocuta_crocuta | 0.24278387977926 |
| Arctodus_simus | Cuon_alpinus | 0.245882072647729 |
| Arctodus_simus | Cystophora_cristata | 0.208081104695904 |
| Arctodus_simus | Enhydra_lutris | 0.227920641893543 |
| Arctodus_simus | Erignathus_barbatus | 0.206272591252728 |
| Arctodus_simus | Eumetopias_jubatus | 0.223418273369412 |
| Arctodus_simus | Felis_catus | 0.23127718377831 |
| Arctodus_simus | Felis_chaus | 0.235868256395128 |
| Arctodus_simus | Felis_margarita | 0.229467258516269 |
| Arctodus_simus | Felis_nigripes | 0.233107421919977 |
| Arctodus_simus | Felis_silvestris | 0.227268727987131 |
| Arctodus_simus | Fossa_fossana_FMNH156648_SRR6053060 | 0.231779071186441 |
| Arctodus_simus | Genetta_servalina | 0.226814843281276 |
| Arctodus_simus | Gulo_gulo | 0.232995091570715 |
| Arctodus_simus | Halichoerus_grypus | 0.206258589012011 |
| Arctodus_simus | Helarctos_malayanus | 0.166437521572924 |
| Arctodus_simus | Hyaena_hyaena | 0.239437149940887 |
| Arctodus_simus | Hydrurga_leptonyx | 0.201502462789301 |
| Arctodus_simus | Leopardus_colocolo | 0.228972640384094 |
| Arctodus_simus | Leopardus_geoffroyi | 0.231454235410899 |
| Arctodus_simus | Leopardus_guigna | 0.229819795635394 |
| Arctodus_simus | Leopardus_jacobita | 0.231709025242604 |
| Arctodus_simus | Leopardus_pardalis | 0.226268577162332 |
| Arctodus_simus | Leopardus_tigrinus | 0.228631966542845 |
| Arctodus_simus | Leopardus_wiedii | 0.229257318750088 |
| Arctodus_simus | Leptailurus_serval | 0.224853575467349 |
| Arctodus_simus | Leptonychotes_weddellii | 0.198726685243374 |
| Arctodus_simus | Lobodon_carcinophaga | 0.206563786671997 |
| Arctodus_simus | Lutra_lutra | 0.234816653473749 |
| Arctodus_simus | Lutra_sumatrana | 0.232248033005806 |
| Arctodus_simus | Lutrogale_perspicillata | 0.239469819304267 |
| Arctodus_simus | Lycaon_pictus | 0.238224467505097 |
| Arctodus_simus | Lynx_canadensis | 0.232330593442773 |
| Arctodus_simus | Lynx_lynx | 0.231221971177308 |
| Arctodus_simus | Lynx_pardinus | 0.232107122990687 |
| Arctodus_simus | Lynx_rufus | 0.229467258516269 |
| Arctodus_simus | Manis_javanica | 0.303647838410552 |
| Arctodus_simus | Martes_americana | 0.229624386154636 |
| Arctodus_simus | Martes_flavigula | 0.23185369663864 |
| Arctodus_simus | Martes_foina | 0.228692569552976 |
| Arctodus_simus | Martes_martes | 0.229629531973824 |
| Arctodus_simus | Martes_melampus | 0.228199076382383 |
| Arctodus_simus | Martes_pennanti | 0.235601627400291 |
| Arctodus_simus | Martes_zibellina | 0.230960424636925 |
| Arctodus_simus | Meles_anakuma | 0.238439455051213 |
| Arctodus_simus | Meles_leucurus | 0.231124596469978 |
| Arctodus_simus | Meles_meles | 0.227389879466989 |
| Arctodus_simus | Melogale_moschata | 0.23324655748369 |
| Arctodus_simus | Melursus_ursinus | 0.171586808479576 |
| Arctodus_simus | Mephitis_mephitis | 0.235786216409766 |
| Arctodus_simus | Mirounga_leonina | 0.203971485059191 |
| Arctodus_simus | Mungotictis_decemlineata | 0.235744921793757 |
| Arctodus_simus | Mustela_altaica | 0.235037780762957 |
| Arctodus_simus | Mustela_erminea | 0.225048561410094 |
| Arctodus_simus | Mustela_eversmannii | 0.22960026464837 |
| Arctodus_simus | Mustela_frenata | 0.2261009547356 |
| Arctodus_simus | Mustela_itatsi | 0.230195805471298 |
| Arctodus_simus | Mustela_kathiah | 0.231209498664567 |
| Arctodus_simus | Mustela_nigripes | 0.227359762309717 |
| Arctodus_simus | Mustela_nivalis | 0.233768974141147 |
| Arctodus_simus | Mustela_putorius | 0.228192671360816 |
| Arctodus_simus | Mustela_sibirica | 0.231296502322242 |
| Arctodus_simus | Nandinia_binotata | 0.21820952725463 |

| | | |
|---|---|---|
| Arctodus_simus | Nasua_nasua | 0.248711555486315 |
| Arctodus_simus | Neofelis_nebulosa | 0.24595564967918 |
| Arctodus_simus | Neomonachus_schauinslandi | 0.210658739078072 |
| Arctodus_simus | Neophoca_cinerea | 0.228243772056497 |
| Arctodus_simus | Neovison_vison | 0.228961852539245 |
| Arctodus_simus | Nyctereutes_procyonoides | 0.239400916035451 |
| Arctodus_simus | Odobenus_rosmarus_rosmarus | 0.23867463567648 |
| Arctodus_simus | Otocolobus_manul | 0.23702716508438 |
| Arctodus_simus | Otocyon_megalotis_megalotis_TS305 | 0.247473428780776 |
| Arctodus_simus | Otocyon_megalotis_virgatus_FMNH158128 | 0.245702451031661 |
| Arctodus_simus | Paguma_larvata | 0.230760816692788 |
| Arctodus_simus | Panthera_leo | 0.238688197180389 |
| Arctodus_simus | Panthera_onca | 0.239566581000475 |
| Arctodus_simus | Panthera_pardus | 0.242021467537038 |
| Arctodus_simus | Panthera_tigris | 0.244655375059352 |
| Arctodus_simus | Panthera_uncia | 0.242739005587416 |
| Arctodus_simus | Paradoxurus_hermaphroditus | 0.237828437339276 |
| Arctodus_simus | Parahyaena_brunnea_FD126_Fecal | 0.239676419255276 |
| Arctodus_simus | Pardofelis_marmorata | 0.236351893863162 |
| Arctodus_simus | Phoca_fasciata | 0.204381214217953 |
| Arctodus_simus | Phoca_groenlandica | 0.207172034622362 |
| Arctodus_simus | Phoca_largha | 0.207444957126843 |
| Arctodus_simus | Phoca_vitulina | 0.207433079000891 |
| Arctodus_simus | Phocarctos_hookeri | 0.231728663484342 |
| Arctodus_simus | Prionailurus_bengalensis | 0.226859685722288 |
| Arctodus_simus | Prionailurus_bengalensis_euptilurus | 0.232285339735145 |
| Arctodus_simus | Prionailurus_planiceps | 0.229213322304584 |
| Arctodus_simus | Prionailurus_rubiginosus | 0.2275417851275 |
| Arctodus_simus | Prionailurus_viverrinus | 0.231112296240177 |
| Arctodus_simus | Prionodon_pardicolor | 0.238901803003076 |
| Arctodus_simus | Procyon_lotor | 0.233467384247784 |
| Arctodus_simus | Profelis_aurata | 0.233778629505636 |
| Arctodus_simus | Proteles_cristatus_cristatus_TS307 | 0.243729582634851 |
| Arctodus_simus | Proteles_cristatus_septentrionalis_NMS_Z_2018_54 | 0.243996225625198 |
| Arctodus_simus | Puma_concolor | 0.231233069864885 |
| Arctodus_simus | Puma_yagouaroundi | 0.232518386342593 |
| Arctodus_simus | Pusa_caspica | 0.205669809195751 |
| Arctodus_simus | Pusa_hispida | 0.203902612360595 |
| Arctodus_simus | Pusa_sibirica | 0.20461723545366 |
| Arctodus_simus | Speothos_venaticus_T1624 | 0.249898475663568 |
| Arctodus_simus | Spilogale_putorius | 0.239791603506524 |
| Arctodus_simus | Taxidea_taxus | 0.224793415425225 |
| Arctodus_simus | Tremarctos_ornatus | 0.104123842585008 |
| Arctodus_simus | Urocyon_cinereoargenteus | 0.254272037980765 |
| Arctodus_simus | Ursus_americanus | 0.164904256834109 |
| Arctodus_simus | Ursus_arctos | 0.174186537006755 |
| Arctodus_simus | Ursus_maritimus | 0.174123915914674 |
| Arctodus_simus | Ursus_spelaeus | 0.171565967085526 |
| Arctodus_simus | Ursus_thibetanus | 0.164468467678562 |
| Arctodus_simus | Urva_javanica | 0.242844742913156 |
| Arctodus_simus | Viverra_tangalunga_FMNH146957_SRR6053069 | 0.225970336325855 |
| Arctodus_simus | Viverricula_indica | 0.227146847220936 |
| Arctodus_simus | Vulpes_corsac | 0.242619644428769 |
| Arctodus_simus | Vulpes_ferrilata | 0.246253270084108 |
| Arctodus_simus | Vulpes_lagopus | 0.243650570985511 |
| Arctodus_simus | Vulpes_vulpes_T3611 | 0.239830942052306 |
| Arctodus_simus | Vulpes_zerda | 0.238444477765207 |
| Arctodus_simus | Zalophus_californianus | 0.224447562784566 |
| Arctonyx_collaris | Arctotherium_sp_ | 0.23391740885146 |
| Arctonyx_collaris | Bdeogale_nigripes_FMNH167685_SRR6053065 | 0.242135005334988 |
| Arctonyx_collaris | Callorhinus_ursinus | 0.222679554445671 |
| Arctonyx_collaris | Canis_anthus | 0.241337205293932 |
| Arctonyx_collaris | Canis_latrans | 0.236412365390915 |
| Arctonyx_collaris | Canis_lupus | 0.237369898347434 |
| Arctonyx_collaris | Caracal_caracal | 0.232824861765389 |
| Arctonyx_collaris | Catopuma_badia | 0.228132310986806 |
| Arctonyx_collaris | Catopuma_temminckii | 0.230303491640771 |
| Arctonyx_collaris | Chrysocyon_brachyurus | 0.240659599431818 |
| Arctonyx_collaris | Civettictis_civetta | 0.227665083842702 |
| Arctonyx_collaris | Conepatus_chinga | 0.232055085752038 |
| Arctonyx_collaris | Crocuta_crocuta | 0.245620857134351 |
| Arctonyx_collaris | Cuon_alpinus | 0.242266780312128 |
| Arctonyx_collaris | Cystophora_cristata | 0.214154537504438 |
| Arctonyx_collaris | Enhydra_lutris | 0.173561219299169 |
| Arctonyx_collaris | Erignathus_barbatus | 0.212370639981085 |
| Arctonyx_collaris | Eumetopias_jubatus | 0.229653409506242 |
| Arctonyx_collaris | Felis_catus | 0.233865796567393 |
| Arctonyx_collaris | Felis_chaus | 0.234276225355715 |
| Arctonyx_collaris | Felis_margarita | 0.230499987988932 |
| Arctonyx_collaris | Felis_nigripes | 0.23323471985454 |
| Arctonyx_collaris | Felis_silvestris | 0.231110361400396 |
| Arctonyx_collaris | Fossa_fossana_FMNH156648_SRR6053060 | 0.229929681011993 |
| Arctonyx_collaris | Genetta_servalina | 0.222328627260376 |
| Arctonyx_collaris | Gulo_gulo | 0.171595157221552 |
| Arctonyx_collaris | Halichoerus_grypus | 0.216478298517905 |
| Arctonyx_collaris | Helarctos_malayanus | 0.240033389308155 |
| Arctonyx_collaris | Hyaena_hyaena | 0.243245658050171 |

| | | |
|---|---|---|
| Arctonyx_collaris | Hydrurga_leptonyx | 0.208194704273137 |
| Arctonyx_collaris | Leopardus_colocolo | 0.228132310986806 |
| Arctonyx_collaris | Leopardus_geoffroyi | 0.230430910422248 |
| Arctonyx_collaris | Leopardus_guigna | 0.230939065953482 |
| Arctonyx_collaris | Leopardus_jacobita | 0.230633017136003 |
| Arctonyx_collaris | Leopardus_pardalis | 0.23080517593849 |
| Arctonyx_collaris | Leopardus_tigrinus | 0.228568710876757 |
| Arctonyx_collaris | Leopardus_wiedii | 0.232415227576261 |
| Arctonyx_collaris | Leptailurus_serval | 0.224802630163764 |
| Arctonyx_collaris | Leptonychotes_weddellii | 0.208129861910002 |
| Arctonyx_collaris | Lobodon_carcinophaga | 0.206472635974124 |
| Arctonyx_collaris | Lutra_lutra | 0.175772149443916 |
| Arctonyx_collaris | Lutra_sumatrana | 0.175058461334092 |
| Arctonyx_collaris | Lutrogale_perspicillata | 0.184253848197172 |
| Arctonyx_collaris | Lycaon_pictus | 0.238464091760049 |
| Arctonyx_collaris | Lynx_canadensis | 0.22909708614214 |
| Arctonyx_collaris | Lynx_lynx | 0.228846916207623 |
| Arctonyx_collaris | Lynx_pardinus | 0.22952351087247 |
| Arctonyx_collaris | Lynx_rufus | 0.228916894403762 |
| Arctonyx_collaris | Manis_javanica | 0.296963133427027 |
| Arctonyx_collaris | Martes_americana | 0.171844996962337 |
| Arctonyx_collaris | Martes_flavigula | 0.170999567321824 |
| Arctonyx_collaris | Martes_foina | 0.164072329815196 |
| Arctonyx_collaris | Martes_martes | 0.169445793468122 |
| Arctonyx_collaris | Martes_melampus | 0.170313454614591 |
| Arctonyx_collaris | Martes_pennanti | 0.178747079252808 |
| Arctonyx_collaris | Martes_zibellina | 0.171512930411686 |
| Arctonyx_collaris | Meles_anakuma | 0.104596944488525 |
| Arctonyx_collaris | Meles_leucurus | 0.0980919744506725 |
| Arctonyx_collaris | Meles_meles | 0.104174365670318 |
| Arctonyx_collaris | Melogale_moschata | 0.174820704112348 |
| Arctonyx_collaris | Melursus_ursinus | 0.242700406384488 |
| Arctonyx_collaris | Mephitis_mephitis | 0.226511185949526 |
| Arctonyx_collaris | Mirounga_leonina | 0.211147508460373 |
| Arctonyx_collaris | Mungotictis_decemlineata | 0.2373522568232 |
| Arctonyx_collaris | Mustela_altaica | 0.178298000930741 |
| Arctonyx_collaris | Mustela_erminea | 0.165884580417402 |
| Arctonyx_collaris | Mustela_eversmannii | 0.1752449755261 |
| Arctonyx_collaris | Mustela_frenata | 0.169116055464446 |
| Arctonyx_collaris | Mustela_itatsi | 0.173170565645986 |
| Arctonyx_collaris | Mustela_kathiah | 0.170798738199939 |
| Arctonyx_collaris | Mustela_nigripes | 0.173723631452434 |
| Arctonyx_collaris | Mustela_nivalis | 0.173974929323802 |
| Arctonyx_collaris | Mustela_putorius | 0.175542287630834 |
| Arctonyx_collaris | Mustela_sibirica | 0.174937724389845 |
| Arctonyx_collaris | Nandinia_binotata | 0.215712665979983 |
| Arctonyx_collaris | Nasua_nasua | 0.236223126933071 |
| Arctonyx_collaris | Neofelis_nebulosa | 0.250063608608277 |
| Arctonyx_collaris | Neomonachus_schauinslandi | 0.21054395255206 |
| Arctonyx_collaris | Neophoca_cinerea | 0.235589491068492 |
| Arctonyx_collaris | Neovison_vison | 0.161808480100257 |
| Arctonyx_collaris | Nyctereutes_procyonoides | 0.241281518035909 |
| Arctonyx_collaris | Odobenus_rosmarus_rosmarus | 0.247461544922482 |
| Arctonyx_collaris | Otocolobus_manul | 0.226952553738214 |
| Arctonyx_collaris | Otocyon_megalotis_megalotis_TS305 | 0.250521427532801 |
| Arctonyx_collaris | Otocyon_megalotis_virgatus_FMNH158128 | 0.251655729289697 |
| Arctonyx_collaris | Paguma_larvata | 0.227286235879052 |
| Arctonyx_collaris | Panthera_leo | 0.238419019532889 |
| Arctonyx_collaris | Panthera_onca | 0.23770020715895 |
| Arctonyx_collaris | Panthera_pardus | 0.238399751036655 |
| Arctonyx_collaris | Panthera_tigris | 0.242311687361657 |
| Arctonyx_collaris | Panthera_uncia | 0.244221581252687 |
| Arctonyx_collaris | Paradoxurus_hermaphroditus | 0.228654246550725 |
| Arctonyx_collaris | Parahyaena_brunnea_FD126_Fecal | 0.24280355644171 |
| Arctonyx_collaris | Pardofelis_marmorata | 0.235869972561955 |
| Arctonyx_collaris | Phoca_fasciata | 0.215852497610798 |
| Arctonyx_collaris | Phoca_groenlandica | 0.216591002606063 |
| Arctonyx_collaris | Phoca_largha | 0.213912875015436 |
| Arctonyx_collaris | Phoca_vitulina | 0.214487112345254 |
| Arctonyx_collaris | Phocarctos_hookeri | 0.236469023276735 |
| Arctonyx_collaris | Prionailurus_bengalensis | 0.230201312933504 |
| Arctonyx_collaris | Prionailurus_bengalensis_euptilurus | 0.232415227576261 |
| Arctonyx_collaris | Prionailurus_planiceps | 0.235491176146476 |
| Arctonyx_collaris | Prionailurus_rubiginosus | 0.23287524944524 |
| Arctonyx_collaris | Prionailurus_viverrinus | 0.232882577084998 |
| Arctonyx_collaris | Prionodon_pardicolor | 0.231067892729335 |
| Arctonyx_collaris | Procyon_lotor | 0.214957462120387 |
| Arctonyx_collaris | Profelis_aurata | 0.23644558080906 |
| Arctonyx_collaris | Proteles_cristatus_cristatus_TS307 | 0.241087211841275 |
| Arctonyx_collaris | Proteles_cristatus_septentrionalis_NMS_Z_2018_54 | 0.24372505508218 |
| Arctonyx_collaris | Puma_concolor | 0.227655341366311 |
| Arctonyx_collaris | Puma_yagouaroundi | 0.229971873820807 |
| Arctonyx_collaris | Pusa_caspica | 0.215826433893886 |
| Arctonyx_collaris | Pusa_hispida | 0.212343192067207 |
| Arctonyx_collaris | Pusa_sibirica | 0.215656186066606 |
| Arctonyx_collaris | Speothos_venaticus_T1624 | 0.248945310910416 |
| Arctonyx_collaris | Spilogale_putorius | 0.225849673677687 |

| | | |
|---|---|---|
| Arctonyx_collaris | Taxidea_taxus | 0.175849362377692 |
| Arctonyx_collaris | Tremarctos_ornatus | 0.239993849397068 |
| Arctonyx_collaris | Urocyon_cinereoargenteus | 0.257587789325522 |
| Arctonyx_collaris | Ursus_americanus | 0.232963457472828 |
| Arctonyx_collaris | Ursus_arctos | 0.242517628259509 |
| Arctonyx_collaris | Ursus_maritimus | 0.242704852741252 |
| Arctonyx_collaris | Ursus_spelaeus | 0.239913211928191 |
| Arctonyx_collaris | Ursus_thibetanus | 0.232627323006387 |
| Arctonyx_collaris | Urva_javanica | 0.239785097617416 |
| Arctonyx_collaris | Viverra_tangalunga_FMNH146957_SRR6053069 | 0.22905569789544 |
| Arctonyx_collaris | Viverricula_indica | 0.225666619233656 |
| Arctonyx_collaris | Vulpes_corsac | 0.252701283205579 |
| Arctonyx_collaris | Vulpes_ferrilata | 0.252644486372819 |
| Arctonyx_collaris | Vulpes_lagopus | 0.24976445887352 |
| Arctonyx_collaris | Vulpes_vulpes_T3611 | 0.247846153654299 |
| Arctonyx_collaris | Vulpes_zerda | 0.247460739340957 |
| Arctonyx_collaris | Zalophus_californianus | 0.228681372472563 |
| Arctotherium_sp_ | Bdeogale_nigripes_FMNH167685_SRR6053065 | 0.243353958670655 |
| Arctotherium_sp_ | Callorhinus_ursinus | 0.231582357715214 |
| Arctotherium_sp_ | Canis_anthus | 0.250659735987674 |
| Arctotherium_sp_ | Canis_latrans | 0.244448611993991 |
| Arctotherium_sp_ | Canis_lupus | 0.245694789142986 |
| Arctotherium_sp_ | Caracal_caracal | 0.236679989005785 |
| Arctotherium_sp_ | Catopuma_badia | 0.232151993156406 |
| Arctotherium_sp_ | Catopuma_temminckii | 0.234795018520038 |
| Arctotherium_sp_ | Chrysocyon_brachyurus | 0.240609601997332 |
| Arctotherium_sp_ | Civettictis_civetta | 0.232958752420309 |
| Arctotherium_sp_ | Conepatus_chinga | 0.241066571068733 |
| Arctotherium_sp_ | Crocuta_crocuta | 0.248490287505388 |
| Arctotherium_sp_ | Cuon_alpinus | 0.249159998919307 |
| Arctotherium_sp_ | Cystophora_cristata | 0.209910396427472 |
| Arctotherium_sp_ | Enhydra_lutris | 0.237182189780878 |
| Arctotherium_sp_ | Erignathus_barbatus | 0.20943029424252 |
| Arctotherium_sp_ | Eumetopias_jubatus | 0.227167207328022 |
| Arctotherium_sp_ | Felis_catus | 0.239380077087717 |
| Arctotherium_sp_ | Felis_chaus | 0.240321592221929 |
| Arctotherium_sp_ | Felis_margarita | 0.237657911168236 |
| Arctotherium_sp_ | Felis_nigripes | 0.237787103885656 |
| Arctotherium_sp_ | Felis_silvestris | 0.236540467413373 |
| Arctotherium_sp_ | Fossa_fossana_FMNH156648_SRR6053060 | 0.235315811262932 |
| Arctotherium_sp_ | Genetta_servalina | 0.234588373632727 |
| Arctotherium_sp_ | Gulo_gulo | 0.229033856201772 |
| Arctotherium_sp_ | Halichoerus_grypus | 0.211744376461521 |
| Arctotherium_sp_ | Helarctos_malayanus | 0.168698372863198 |
| Arctotherium_sp_ | Hyaena_hyaena | 0.246744072466476 |
| Arctotherium_sp_ | Hydrurga_leptonyx | 0.206268201400843 |
| Arctotherium_sp_ | Leopardus_colocolo | 0.238518428527693 |
| Arctotherium_sp_ | Leopardus_geoffroyi | 0.23768127394468 |
| Arctotherium_sp_ | Leopardus_guigna | 0.237480904346078 |
| Arctotherium_sp_ | Leopardus_jacobita | 0.238164352165062 |
| Arctotherium_sp_ | Leopardus_pardalis | 0.234883365548079 |
| Arctotherium_sp_ | Leopardus_tigrinus | 0.235714031070437 |
| Arctotherium_sp_ | Leopardus_wiedii | 0.237362848212128 |
| Arctotherium_sp_ | Leptailurus_serval | 0.23392006396874 |
| Arctotherium_sp_ | Leptonychotes_weddellii | 0.202336897065885 |
| Arctotherium_sp_ | Lobodon_carcinophaga | 0.208063858132323 |
| Arctotherium_sp_ | Lutra_lutra | 0.234537975291126 |
| Arctotherium_sp_ | Lutra_sumatrana | 0.231160157321423 |
| Arctotherium_sp_ | Lutrogale_perspicillata | 0.245244554896876 |
| Arctotherium_sp_ | Lycaon_pictus | 0.241296165973174 |
| Arctotherium_sp_ | Lynx_canadensis | 0.237227381449504 |
| Arctotherium_sp_ | Lynx_lynx | 0.235259965088731 |
| Arctotherium_sp_ | Lynx_pardinus | 0.236017184489958 |
| Arctotherium_sp_ | Lynx_rufus | 0.232937579371022 |
| Arctotherium_sp_ | Manis_javanica | 0.300180677473062 |
| Arctotherium_sp_ | Martes_americana | 0.234067558357812 |
| Arctotherium_sp_ | Martes_flavigula | 0.238488396255057 |
| Arctotherium_sp_ | Martes_foina | 0.230187880718728 |
| Arctotherium_sp_ | Martes_martes | 0.236788091578226 |
| Arctotherium_sp_ | Martes_melampus | 0.233756256647509 |
| Arctotherium_sp_ | Martes_pennanti | 0.238866110515853 |
| Arctotherium_sp_ | Martes_zibellina | 0.236547759493473 |
| Arctotherium_sp_ | Meles_anakuma | 0.238586617750457 |
| Arctotherium_sp_ | Meles_leucurus | 0.233149371992832 |
| Arctotherium_sp_ | Meles_meles | 0.233407051708956 |
| Arctotherium_sp_ | Melogale_moschata | 0.234896099047832 |
| Arctotherium_sp_ | Melursus_ursinus | 0.175163174241398 |
| Arctotherium_sp_ | Mephitis_mephitis | 0.241055647053571 |
| Arctotherium_sp_ | Mirounga_leonina | 0.210721926331189 |
| Arctotherium_sp_ | Mungotictis_decemlineata | 0.241420985335527 |
| Arctotherium_sp_ | Mustela_altaica | 0.241571608180465 |
| Arctotherium_sp_ | Mustela_erminea | 0.231755347785004 |
| Arctotherium_sp_ | Mustela_eversmannii | 0.237726061344629 |
| Arctotherium_sp_ | Mustela_frenata | 0.231531833943686 |
| Arctotherium_sp_ | Mustela_itatsi | 0.239504601397638 |
| Arctotherium_sp_ | Mustela_kathiah | 0.235798750778953 |
| Arctotherium_sp_ | Mustela_nigripes | 0.238600339819645 |

| | | |
|---|---|---|
| Arctotherium_sp_ | Mustela_nivalis | 0.239822193725921 |
| Arctotherium_sp_ | Mustela_putorius | 0.237426302384213 |
| Arctotherium_sp_ | Mustela_sibirica | 0.23981397791681 |
| Arctotherium_sp_ | Nandinia_binotata | 0.221581280523554 |
| Arctotherium_sp_ | Nasua_nasua | 0.251486036080698 |
| Arctotherium_sp_ | Neofelis_nebulosa | 0.253998090973661 |
| Arctotherium_sp_ | Neomonachus_schauinslandi | 0.213945181099846 |
| Arctotherium_sp_ | Neophoca_cinerea | 0.230561750268901 |
| Arctotherium_sp_ | Neovison_vison | 0.235186466719802 |
| Arctotherium_sp_ | Nyctereutes_procyonoides | 0.242472342330595 |
| Arctotherium_sp_ | Odobenus_rosmarus_rosmarus | 0.23972899859314 |
| Arctotherium_sp_ | Otocolobus_manul | 0.241335448601755 |
| Arctotherium_sp_ | Otocyon_megalotis_megalotis_TS305 | 0.250686251770296 |
| Arctotherium_sp_ | Otocyon_megalotis_virgatus_FMNH158128 | 0.248223063336872 |
| Arctotherium_sp_ | Paguma_larvata | 0.237017988677135 |
| Arctotherium_sp_ | Panthera_leo | 0.243078387692164 |
| Arctotherium_sp_ | Panthera_onca | 0.247285354230984 |
| Arctotherium_sp_ | Panthera_pardus | 0.247147538331434 |
| Arctotherium_sp_ | Panthera_tigris | 0.245463129891682 |
| Arctotherium_sp_ | Panthera_uncia | 0.245452097285748 |
| Arctotherium_sp_ | Paradoxurus_hermaphroditus | 0.247981070419649 |
| Arctotherium_sp_ | Parahyaena_brunnea_FD126_Fecal | 0.243818432036052 |
| Arctotherium_sp_ | Pardofelis_marmorata | 0.239728605683955 |
| Arctotherium_sp_ | Phoca_fasciata | 0.207986187777562 |
| Arctotherium_sp_ | Phoca_groenlandica | 0.210574115230263 |
| Arctotherium_sp_ | Phoca_largha | 0.209464737254261 |
| Arctotherium_sp_ | Phoca_vitulina | 0.209994551259055 |
| Arctotherium_sp_ | Phocarctos_hookeri | 0.235386964318794 |
| Arctotherium_sp_ | Prionailurus_bengalensis | 0.237074328344797 |
| Arctotherium_sp_ | Prionailurus_bengalensis_euptilurus | 0.235194510574832 |
| Arctotherium_sp_ | Prionailurus_planiceps | 0.232993844570905 |
| Arctotherium_sp_ | Prionailurus_rubiginosus | 0.231089306109085 |
| Arctotherium_sp_ | Prionailurus_viverrinus | 0.237139034421462 |
| Arctotherium_sp_ | Prionodon_pardicolor | 0.24675233619735 |
| Arctotherium_sp_ | Procyon_lotor | 0.234798955744883 |
| Arctotherium_sp_ | Profelis_aurata | 0.233931626819149 |
| Arctotherium_sp_ | Proteles_cristatus_cristatus_TS307 | 0.247057806797605 |
| Arctotherium_sp_ | Proteles_cristatus_septentrionalis_NMS_Z_2018_54 | 0.247552265775045 |
| Arctotherium_sp_ | Puma_concolor | 0.23958710584854 |
| Arctotherium_sp_ | Puma_yagouaroundi | 0.23620981718318 |
| Arctotherium_sp_ | Pusa_caspica | 0.209826171715145 |
| Arctotherium_sp_ | Pusa_hispida | 0.20696597612129 |
| Arctotherium_sp_ | Pusa_sibirica | 0.20854995052993 |
| Arctotherium_sp_ | Speothos_venaticus_T1624 | 0.245982810591223 |
| Arctotherium_sp_ | Spilogale_putorius | 0.242213868113627 |
| Arctotherium_sp_ | Taxidea_taxus | 0.229132423062049 |
| Arctotherium_sp_ | Tremarctos_ornatus | 0.107259946326475 |
| Arctotherium_sp_ | Urocyon_cinereoargenteus | 0.252749519900611 |
| Arctotherium_sp_ | Ursus_americanus | 0.169007324783263 |
| Arctotherium_sp_ | Ursus_arctos | 0.173832676339985 |
| Arctotherium_sp_ | Ursus_maritimus | 0.173031713863919 |
| Arctotherium_sp_ | Ursus_spelaeus | 0.170062569127418 |
| Arctotherium_sp_ | Ursus_thibetanus | 0.165683435723708 |
| Arctotherium_sp_ | Urva_javanica | 0.246979047129644 |
| Arctotherium_sp_ | Viverra_tangalunga_FMNH146957_SRR6053069 | 0.228413266569683 |
| Arctotherium_sp_ | Viverricula_indica | 0.231969165248992 |
| Arctotherium_sp_ | Vulpes_corsac | 0.245393002976572 |
| Arctotherium_sp_ | Vulpes_ferrilata | 0.248299657660984 |
| Arctotherium_sp_ | Vulpes_lagopus | 0.24770570018186 |
| Arctotherium_sp_ | Vulpes_vulpes_T3611 | 0.245508202060626 |
| Arctotherium_sp_ | Vulpes_zerda | 0.246148715638076 |
| Arctotherium_sp_ | Zalophus_californianus | 0.226760985825056 |
| Bdeogale_nigripes_FMNH167685_SRR6053065 | Callorhinus_ursinus | 0.231978143109546 |
| Bdeogale_nigripes_FMNH167685_SRR6053065 | Canis_anthus | 0.246418282854776 |
| Bdeogale_nigripes_FMNH167685_SRR6053065 | Canis_latrans | 0.242993830287979 |
| Bdeogale_nigripes_FMNH167685_SRR6053065 | Canis_lupus | 0.244426737912035 |
| Bdeogale_nigripes_FMNH167685_SRR6053065 | Caracal_caracal | 0.191105506685969 |
| Bdeogale_nigripes_FMNH167685_SRR6053065 | Catopuma_badia | 0.189104634655914 |
| Bdeogale_nigripes_FMNH167685_SRR6053065 | Catopuma_temminckii | 0.197671091654524 |
| Bdeogale_nigripes_FMNH167685_SRR6053065 | Chrysocyon_brachyurus | 0.243971185761292 |
| Bdeogale_nigripes_FMNH167685_SRR6053065 | Civettictis_civetta | 0.187818036076885 |
| Bdeogale_nigripes_FMNH167685_SRR6053065 | Conepatus_chinga | 0.249418783010842 |
| Bdeogale_nigripes_FMNH167685_SRR6053065 | Crocuta_crocuta | 0.19454224068299 |
| Bdeogale_nigripes_FMNH167685_SRR6053065 | Cuon_alpinus | 0.24889797927512 |
| Bdeogale_nigripes_FMNH167685_SRR6053065 | Cystophora_cristata | 0.221712082928222 |
| Bdeogale_nigripes_FMNH167685_SRR6053065 | Enhydra_lutris | 0.241225107437254 |
| Bdeogale_nigripes_FMNH167685_SRR6053065 | Erignathus_barbatus | 0.226548808366202 |
| Bdeogale_nigripes_FMNH167685_SRR6053065 | Eumetopias_jubatus | 0.236555791579116 |
| Bdeogale_nigripes_FMNH167685_SRR6053065 | Felis_catus | 0.195460997235349 |
| Bdeogale_nigripes_FMNH167685_SRR6053065 | Felis_chaus | 0.197969697755713 |
| Bdeogale_nigripes_FMNH167685_SRR6053065 | Felis_margarita | 0.193786304440934 |
| Bdeogale_nigripes_FMNH167685_SRR6053065 | Felis_nigripes | 0.194586867006732 |
| Bdeogale_nigripes_FMNH167685_SRR6053065 | Felis_silvestris | 0.193825107284353 |
| Bdeogale_nigripes_FMNH167685_SRR6053065 | Fossa_fossana_FMNH156648_SRR6053060 | 0.172155095579022 |
| Bdeogale_nigripes_FMNH167685_SRR6053065 | Genetta_servalina | 0.190306216870581 |
| Bdeogale_nigripes_FMNH167685_SRR6053065 | Gulo_gulo | 0.243839162804944 |
| Bdeogale_nigripes_FMNH167685_SRR6053065 | Halichoerus_grypus | 0.224979335559311 |

| | | |
|---|---|---|
| Bdeogale_nigripes_FMNH167685_SRR6053065 | Helarctos_malayanus | 0.241596443243929 |
| Bdeogale_nigripes_FMNH167685_SRR6053065 | Hyaena_hyaena | 0.192632025732202 |
| Bdeogale_nigripes_FMNH167685_SRR6053065 | Hydrurga_leptonyx | 0.217247356172231 |
| Bdeogale_nigripes_FMNH167685_SRR6053065 | Leopardus_colocolo | 0.190028564303053 |
| Bdeogale_nigripes_FMNH167685_SRR6053065 | Leopardus_geoffroyi | 0.192278658189748 |
| Bdeogale_nigripes_FMNH167685_SRR6053065 | Leopardus_guigna | 0.192553070192119 |
| Bdeogale_nigripes_FMNH167685_SRR6053065 | Leopardus_jacobita | 0.196055142838988 |
| Bdeogale_nigripes_FMNH167685_SRR6053065 | Leopardus_pardalis | 0.192052572352557 |
| Bdeogale_nigripes_FMNH167685_SRR6053065 | Leopardus_tigrinus | 0.193415296360248 |
| Bdeogale_nigripes_FMNH167685_SRR6053065 | Leopardus_wiedii | 0.19342949777322 |
| Bdeogale_nigripes_FMNH167685_SRR6053065 | Leptailurus_serval | 0.19088932630281 |
| Bdeogale_nigripes_FMNH167685_SRR6053065 | Leptonychotes_weddellii | 0.213814037595973 |
| Bdeogale_nigripes_FMNH167685_SRR6053065 | Lobodon_carcinophaga | 0.219743734321551 |
| Bdeogale_nigripes_FMNH167685_SRR6053065 | Lutra_lutra | 0.241376407233214 |
| Bdeogale_nigripes_FMNH167685_SRR6053065 | Lutra_sumatrana | 0.242023671555859 |
| Bdeogale_nigripes_FMNH167685_SRR6053065 | Lutrogale_perspicillata | 0.253236630933223 |
| Bdeogale_nigripes_FMNH167685_SRR6053065 | Lycaon_pictus | 0.242299807031428 |
| Bdeogale_nigripes_FMNH167685_SRR6053065 | Lynx_canadensis | 0.192065130837271 |
| Bdeogale_nigripes_FMNH167685_SRR6053065 | Lynx_lynx | 0.188520090126885 |
| Bdeogale_nigripes_FMNH167685_SRR6053065 | Lynx_pardinus | 0.190718050936551 |
| Bdeogale_nigripes_FMNH167685_SRR6053065 | Lynx_rufus | 0.190878383830778 |
| Bdeogale_nigripes_FMNH167685_SRR6053065 | Manis_javanica | 0.284456677271915 |
| Bdeogale_nigripes_FMNH167685_SRR6053065 | Martes_americana | 0.2399026547343 |
| Bdeogale_nigripes_FMNH167685_SRR6053065 | Martes_flavigula | 0.240397978025344 |
| Bdeogale_nigripes_FMNH167685_SRR6053065 | Martes_foina | 0.237907556489402 |
| Bdeogale_nigripes_FMNH167685_SRR6053065 | Martes_martes | 0.236911980939234 |
| Bdeogale_nigripes_FMNH167685_SRR6053065 | Martes_melampus | 0.239923108068027 |
| Bdeogale_nigripes_FMNH167685_SRR6053065 | Martes_pennanti | 0.244471933067347 |
| Bdeogale_nigripes_FMNH167685_SRR6053065 | Martes_zibellina | 0.238783952250143 |
| Bdeogale_nigripes_FMNH167685_SRR6053065 | Meles_anakuma | 0.245156278227923 |
| Bdeogale_nigripes_FMNH167685_SRR6053065 | Meles_leucurus | 0.242115069657954 |
| Bdeogale_nigripes_FMNH167685_SRR6053065 | Meles_meles | 0.239236727887805 |
| Bdeogale_nigripes_FMNH167685_SRR6053065 | Melogale_moschata | 0.243720289205708 |
| Bdeogale_nigripes_FMNH167685_SRR6053065 | Melursus_ursinus | 0.251615295963004 |
| Bdeogale_nigripes_FMNH167685_SRR6053065 | Mephitis_mephitis | 0.24974323488184 |
| Bdeogale_nigripes_FMNH167685_SRR6053065 | Mirounga_leonina | 0.225458656635242 |
| Bdeogale_nigripes_FMNH167685_SRR6053065 | Mungotictis_decemlineata | 0.174741822789151 |
| Bdeogale_nigripes_FMNH167685_SRR6053065 | Mustela_altaica | 0.244613446872403 |
| Bdeogale_nigripes_FMNH167685_SRR6053065 | Mustela_erminea | 0.236217264015557 |
| Bdeogale_nigripes_FMNH167685_SRR6053065 | Mustela_eversmannii | 0.242217088010153 |
| Bdeogale_nigripes_FMNH167685_SRR6053065 | Mustela_frenata | 0.238051558452752 |
| Bdeogale_nigripes_FMNH167685_SRR6053065 | Mustela_itatsi | 0.24242932805727 |
| Bdeogale_nigripes_FMNH167685_SRR6053065 | Mustela_kathiah | 0.245054680118064 |
| Bdeogale_nigripes_FMNH167685_SRR6053065 | Mustela_nigripes | 0.24104393893104 |
| Bdeogale_nigripes_FMNH167685_SRR6053065 | Mustela_nivalis | 0.243889455173767 |
| Bdeogale_nigripes_FMNH167685_SRR6053065 | Mustela_putorius | 0.240824874585644 |
| Bdeogale_nigripes_FMNH167685_SRR6053065 | Mustela_sibirica | 0.241036686060114 |
| Bdeogale_nigripes_FMNH167685_SRR6053065 | Nandinia_binotata | 0.181393379514722 |
| Bdeogale_nigripes_FMNH167685_SRR6053065 | Nasua_nasua | 0.253717938358095 |
| Bdeogale_nigripes_FMNH167685_SRR6053065 | Neofelis_nebulosa | 0.202940393754396 |
| Bdeogale_nigripes_FMNH167685_SRR6053065 | Neomonachus_schauinslandi | 0.229861227852126 |
| Bdeogale_nigripes_FMNH167685_SRR6053065 | Neophoca_cinerea | 0.239140756385924 |
| Bdeogale_nigripes_FMNH167685_SRR6053065 | Neovison_vison | 0.240486983594023 |
| Bdeogale_nigripes_FMNH167685_SRR6053065 | Nyctereutes_procyonoides | 0.239747978336439 |
| Bdeogale_nigripes_FMNH167685_SRR6053065 | Odobenus_rosmarus_rosmarus | 0.259899197350891 |
| Bdeogale_nigripes_FMNH167685_SRR6053065 | Otocolobus_manul | 0.19563519530774 |
| Bdeogale_nigripes_FMNH167685_SRR6053065 | Otocyon_megalotis_megalotis_TS305 | 0.248599480352935 |
| Bdeogale_nigripes_FMNH167685_SRR6053065 | Otocyon_megalotis_virgatus_FMNH158128 | 0.250008866162515 |
| Bdeogale_nigripes_FMNH167685_SRR6053065 | Paguma_larvata | 0.196250918013017 |
| Bdeogale_nigripes_FMNH167685_SRR6053065 | Panthera_leo | 0.199546175684588 |
| Bdeogale_nigripes_FMNH167685_SRR6053065 | Panthera_onca | 0.199982587817508 |
| Bdeogale_nigripes_FMNH167685_SRR6053065 | Panthera_pardus | 0.200348768851358 |
| Bdeogale_nigripes_FMNH167685_SRR6053065 | Panthera_tigris | 0.20500885469376 |
| Bdeogale_nigripes_FMNH167685_SRR6053065 | Panthera_uncia | 0.204858592850127 |
| Bdeogale_nigripes_FMNH167685_SRR6053065 | Paradoxurus_hermaphroditus | 0.19982227405882 |
| Bdeogale_nigripes_FMNH167685_SRR6053065 | Parahyaena_brunnea_FD126_Fecal | 0.188032459557014 |
| Bdeogale_nigripes_FMNH167685_SRR6053065 | Pardofelis_marmorata | 0.19314973257782 |
| Bdeogale_nigripes_FMNH167685_SRR6053065 | Phoca_fasciata | 0.225511120151994 |
| Bdeogale_nigripes_FMNH167685_SRR6053065 | Phoca_groenlandica | 0.229135915627221 |
| Bdeogale_nigripes_FMNH167685_SRR6053065 | Phoca_largha | 0.227996938561487 |
| Bdeogale_nigripes_FMNH167685_SRR6053065 | Phoca_vitulina | 0.228929736687013 |
| Bdeogale_nigripes_FMNH167685_SRR6053065 | Phocarctos_hookeri | 0.240883391700344 |
| Bdeogale_nigripes_FMNH167685_SRR6053065 | Prionailurus_bengalensis | 0.18702461907528 |
| Bdeogale_nigripes_FMNH167685_SRR6053065 | Prionailurus_bengalensis_euptilurus | 0.188368983897586 |
| Bdeogale_nigripes_FMNH167685_SRR6053065 | Prionailurus_planiceps | 0.190756366469403 |
| Bdeogale_nigripes_FMNH167685_SRR6053065 | Prionailurus_rubiginosus | 0.183632765054126 |
| Bdeogale_nigripes_FMNH167685_SRR6053065 | Prionailurus_viverrinus | 0.190047646365319 |
| Bdeogale_nigripes_FMNH167685_SRR6053065 | Prionodon_pardicolor | 0.201618119804448 |
| Bdeogale_nigripes_FMNH167685_SRR6053065 | Procyon_lotor | 0.243496675606275 |
| Bdeogale_nigripes_FMNH167685_SRR6053065 | Profelis_aurata | 0.195585843007348 |
| Bdeogale_nigripes_FMNH167685_SRR6053065 | Proteles_cristatus_cristatus_TS307 | 0.199446356417948 |
| Bdeogale_nigripes_FMNH167685_SRR6053065 | Proteles_cristatus_septentrionalis_NMS_Z_2018_54 | 0.201485017400759 |
| Bdeogale_nigripes_FMNH167685_SRR6053065 | Puma_concolor | 0.191048379587434 |
| Bdeogale_nigripes_FMNH167685_SRR6053065 | Puma_yagouaroundi | 0.196215323668562 |
| Bdeogale_nigripes_FMNH167685_SRR6053065 | Pusa_caspica | 0.225701388892827 |
| Bdeogale_nigripes_FMNH167685_SRR6053065 | Pusa_hispida | 0.224096923333676 |
| Bdeogale_nigripes_FMNH167685_SRR6053065 | Pusa_sibirica | 0.223862952734888 |

| | | |
|---|---|---|
| Bdeogale_nigripes_FMNH167685_SRR6053065 | Speothos_venaticus_T1624 | 0.250590558072342 |
| Bdeogale_nigripes_FMNH167685_SRR6053065 | Spilogale_putorius | 0.254862080341055 |
| Bdeogale_nigripes_FMNH167685_SRR6053065 | Taxidea_taxus | 0.232822713931466 |
| Bdeogale_nigripes_FMNH167685_SRR6053065 | Tremarctos_ornatus | 0.242185451037147 |
| Bdeogale_nigripes_FMNH167685_SRR6053065 | Urocyon_cinereoargenteus | 0.2528426938781 |
| Bdeogale_nigripes_FMNH167685_SRR6053065 | Ursus_americanus | 0.243555909064313 |
| Bdeogale_nigripes_FMNH167685_SRR6053065 | Ursus_arctos | 0.249670139091499 |
| Bdeogale_nigripes_FMNH167685_SRR6053065 | Ursus_maritimus | 0.247583185951604 |
| Bdeogale_nigripes_FMNH167685_SRR6053065 | Ursus_spelaeus | 0.245541903914749 |
| Bdeogale_nigripes_FMNH167685_SRR6053065 | Ursus_thibetanus | 0.242022702012523 |
| Bdeogale_nigripes_FMNH167685_SRR6053065 | Urva_javanica | 0.1168216855764 |
| Bdeogale_nigripes_FMNH167685_SRR6053065 | Viverra_tangalunga_FMNH146957_SRR6053069 | 0.188204272034283 |
| Bdeogale_nigripes_FMNH167685_SRR6053065 | Viverricula_indica | 0.186548235940873 |
| Bdeogale_nigripes_FMNH167685_SRR6053065 | Vulpes_corsac | 0.245908287491292 |
| Bdeogale_nigripes_FMNH167685_SRR6053065 | Vulpes_ferrilata | 0.250235922835687 |
| Bdeogale_nigripes_FMNH167685_SRR6053065 | Vulpes_lagopus | 0.246692899749973 |
| Bdeogale_nigripes_FMNH167685_SRR6053065 | Vulpes_vulpes_T3611 | 0.240622381933046 |
| Bdeogale_nigripes_FMNH167685_SRR6053065 | Vulpes_zerda | 0.24171212831372 |
| Bdeogale_nigripes_FMNH167685_SRR6053065 | Zalophus_californianus | 0.234362957665786 |
| Callorhinus_ursinus | Canis_anthus | 0.248671869864046 |
| Callorhinus_ursinus | Canis_latrans | 0.244583915150682 |
| Callorhinus_ursinus | Canis_lupus | 0.247604895547815 |
| Callorhinus_ursinus | Caracal_caracal | 0.225955041599652 |
| Callorhinus_ursinus | Catopuma_badia | 0.232853961421477 |
| Callorhinus_ursinus | Catopuma_temminckii | 0.233793765493199 |
| Callorhinus_ursinus | Chrysocyon_brachyurus | 0.247749358030255 |
| Callorhinus_ursinus | Civettictis_civetta | 0.234955685501634 |
| Callorhinus_ursinus | Conepatus_chinga | 0.231439645216304 |
| Callorhinus_ursinus | Crocuta_crocuta | 0.235570457748426 |
| Callorhinus_ursinus | Cuon_alpinus | 0.253919271323568 |
| Callorhinus_ursinus | Cystophora_cristata | 0.176856187367282 |
| Callorhinus_ursinus | Enhydra_lutris | 0.230316345032968 |
| Callorhinus_ursinus | Erignathus_barbatus | 0.176283860267156 |
| Callorhinus_ursinus | Eumetopias_jubatus | 0.113540140601364 |
| Callorhinus_ursinus | Felis_catus | 0.233255809421345 |
| Callorhinus_ursinus | Felis_chaus | 0.232497509119274 |
| Callorhinus_ursinus | Felis_margarita | 0.233146027886159 |
| Callorhinus_ursinus | Felis_nigripes | 0.23517570876999 |
| Callorhinus_ursinus | Felis_silvestris | 0.23329489052055 |
| Callorhinus_ursinus | Fossa_fossana_FMNH156648_SRR6053060 | 0.229108418858615 |
| Callorhinus_ursinus | Genetta_servalina | 0.231459904088546 |
| Callorhinus_ursinus | Gulo_gulo | 0.224670061027408 |
| Callorhinus_ursinus | Halichoerus_grypus | 0.180093997489923 |
| Callorhinus_ursinus | Helarctos_malayanus | 0.231299653778228 |
| Callorhinus_ursinus | Hyaena_hyaena | 0.238128816953664 |
| Callorhinus_ursinus | Hydrurga_leptonyx | 0.171422030708966 |
| Callorhinus_ursinus | Leopardus_colocolo | 0.22690389269419 |
| Callorhinus_ursinus | Leopardus_geoffroyi | 0.230202045245654 |
| Callorhinus_ursinus | Leopardus_guigna | 0.230839087256812 |
| Callorhinus_ursinus | Leopardus_jacobita | 0.229267002510515 |
| Callorhinus_ursinus | Leopardus_pardalis | 0.226581936806962 |
| Callorhinus_ursinus | Leopardus_tigrinus | 0.227636610600721 |
| Callorhinus_ursinus | Leopardus_wiedii | 0.229364538368165 |
| Callorhinus_ursinus | Leptailurus_serval | 0.227603136504211 |
| Callorhinus_ursinus | Leptonychotes_weddellii | 0.168483338572331 |
| Callorhinus_ursinus | Lobodon_carcinophaga | 0.171797248951218 |
| Callorhinus_ursinus | Lutra_lutra | 0.226060175136558 |
| Callorhinus_ursinus | Lutra_sumatrana | 0.226696267112201 |
| Callorhinus_ursinus | Lutrogale_perspicillata | 0.236344793462505 |
| Callorhinus_ursinus | Lycaon_pictus | 0.24654498016307 |
| Callorhinus_ursinus | Lynx_canadensis | 0.225803200615523 |
| Callorhinus_ursinus | Lynx_lynx | 0.224682910765874 |
| Callorhinus_ursinus | Lynx_pardinus | 0.224107349887105 |
| Callorhinus_ursinus | Lynx_rufus | 0.227257973933042 |
| Callorhinus_ursinus | Manis_javanica | 0.288148829548972 |
| Callorhinus_ursinus | Martes_americana | 0.230603319351927 |
| Callorhinus_ursinus | Martes_flavigula | 0.224691185802613 |
| Callorhinus_ursinus | Martes_foina | 0.2280586490705 |
| Callorhinus_ursinus | Martes_martes | 0.226169704162187 |
| Callorhinus_ursinus | Martes_melampus | 0.224674396126535 |
| Callorhinus_ursinus | Martes_pennanti | 0.23265972627434 |
| Callorhinus_ursinus | Martes_zibellina | 0.22782638766888 |
| Callorhinus_ursinus | Meles_anakuma | 0.227631943579924 |
| Callorhinus_ursinus | Meles_leucurus | 0.225163552472235 |
| Callorhinus_ursinus | Meles_meles | 0.22510215554756 |
| Callorhinus_ursinus | Melogale_moschata | 0.234192840790745 |
| Callorhinus_ursinus | Melursus_ursinus | 0.236363600894264 |
| Callorhinus_ursinus | Mephitis_mephitis | 0.234829445151033 |
| Callorhinus_ursinus | Mirounga_leonina | 0.176476792742406 |
| Callorhinus_ursinus | Mungotictis_decemlineata | 0.235217152419794 |
| Callorhinus_ursinus | Mustela_altaica | 0.229162013160324 |
| Callorhinus_ursinus | Mustela_erminea | 0.224170364766514 |
| Callorhinus_ursinus | Mustela_eversmannii | 0.230668889723445 |
| Callorhinus_ursinus | Mustela_frenata | 0.228529582840264 |
| Callorhinus_ursinus | Mustela_itatsi | 0.229799896777656 |
| Callorhinus_ursinus | Mustela_kathiah | 0.227834552455046 |
| Callorhinus_ursinus | Mustela_nigripes | 0.229408397602286 |

| | | |
|---|---|---|
| Callorhinus_ursinus | Mustela_nivalis | 0.231141283565903 |
| Callorhinus_ursinus | Mustela_putorius | 0.227587530418295 |
| Callorhinus_ursinus | Mustela_sibirica | 0.228179708753026 |
| Callorhinus_ursinus | Nandinia_binotata | 0.220730405401971 |
| Callorhinus_ursinus | Nasua_nasua | 0.240077777263767 |
| Callorhinus_ursinus | Neofelis_nebulosa | 0.244423615637185 |
| Callorhinus_ursinus | Neomonachus_schauinslandi | 0.173451264314785 |
| Callorhinus_ursinus | Neophoca_cinerea | 0.121098769183702 |
| Callorhinus_ursinus | Neovison_vison | 0.2290486695761 |
| Callorhinus_ursinus | Nyctereutes_procyonoides | 0.2436547482295 |
| Callorhinus_ursinus | Odobenus_rosmarus_rosmarus | 0.188227176876366 |
| Callorhinus_ursinus | Otocolobus_manul | 0.230533079821511 |
| Callorhinus_ursinus | Otocyon_megalotis_megalotis_TS305 | 0.245819005087137 |
| Callorhinus_ursinus | Otocyon_megalotis_virgatus_FMNH158128 | 0.245549543562425 |
| Callorhinus_ursinus | Paguma_larvata | 0.234758182562948 |
| Callorhinus_ursinus | Panthera_leo | 0.235777917505071 |
| Callorhinus_ursinus | Panthera_onca | 0.234815055925909 |
| Callorhinus_ursinus | Panthera_pardus | 0.241485991299516 |
| Callorhinus_ursinus | Panthera_tigris | 0.233203849534657 |
| Callorhinus_ursinus | Panthera_uncia | 0.241197911315761 |
| Callorhinus_ursinus | Paradoxurus_hermaphroditus | 0.237244252370897 |
| Callorhinus_ursinus | Parahyaena_brunnea_FD126_Fecal | 0.236731132753393 |
| Callorhinus_ursinus | Pardofelis_marmorata | 0.235677626937257 |
| Callorhinus_ursinus | Phoca_fasciata | 0.173318491759481 |
| Callorhinus_ursinus | Phoca_groenlandica | 0.176815580695619 |
| Callorhinus_ursinus | Phoca_largha | 0.17979059090517 |
| Callorhinus_ursinus | Phoca_vitulina | 0.178560060830183 |
| Callorhinus_ursinus | Phocarctos_hookeri | 0.127423156496385 |
| Callorhinus_ursinus | Prionailurus_bengalensis | 0.225007586101957 |
| Callorhinus_ursinus | Prionailurus_bengalensis_euptilurus | 0.228990933030987 |
| Callorhinus_ursinus | Prionailurus_planiceps | 0.231517978367152 |
| Callorhinus_ursinus | Prionailurus_rubiginosus | 0.228843716286858 |
| Callorhinus_ursinus | Prionailurus_viverrinus | 0.231048316029943 |
| Callorhinus_ursinus | Prionodon_pardicolor | 0.24102278154192 |
| Callorhinus_ursinus | Procyon_lotor | 0.230934774451901 |
| Callorhinus_ursinus | Profelis_aurata | 0.234896180227083 |
| Callorhinus_ursinus | Proteles_cristatus_cristatus_TS307 | 0.240255327323549 |
| Callorhinus_ursinus | Proteles_cristatus_septentrionalis_NMS_Z_2018_54 | 0.24265543426042 |
| Callorhinus_ursinus | Puma_concolor | 0.232703737876813 |
| Callorhinus_ursinus | Puma_yagouaroundi | 0.232017252997244 |
| Callorhinus_ursinus | Pusa_caspica | 0.179312180550059 |
| Callorhinus_ursinus | Pusa_hispida | 0.176075293679016 |
| Callorhinus_ursinus | Pusa_sibirica | 0.17685730198659 |
| Callorhinus_ursinus | Speothos_venaticus_T1624 | 0.254774173332309 |
| Callorhinus_ursinus | Spilogale_putorius | 0.234229415142809 |
| Callorhinus_ursinus | Taxidea_taxus | 0.217420515765296 |
| Callorhinus_ursinus | Tremarctos_ornatus | 0.2305143985771 |
| Callorhinus_ursinus | Urocyon_cinereoargenteus | 0.247355629889499 |
| Callorhinus_ursinus | Ursus_americanus | 0.231360735274906 |
| Callorhinus_ursinus | Ursus_arctos | 0.237473766841705 |
| Callorhinus_ursinus | Ursus_maritimus | 0.234639032682539 |
| Callorhinus_ursinus | Ursus_spelaeus | 0.234319765596931 |
| Callorhinus_ursinus | Ursus_thibetanus | 0.234485047501521 |
| Callorhinus_ursinus | Urva_javanica | 0.234992722613077 |
| Callorhinus_ursinus | Viverra_tangalunga_FMNH146957_SRR6053069 | 0.231579040874541 |
| Callorhinus_ursinus | Viverricula_indica | 0.234714928371641 |
| Callorhinus_ursinus | Vulpes_corsac | 0.247797934091713 |
| Callorhinus_ursinus | Vulpes_ferrilata | 0.248258651134202 |
| Callorhinus_ursinus | Vulpes_lagopus | 0.248730641829966 |
| Callorhinus_ursinus | Vulpes_vulpes_T3611 | 0.247127995440633 |
| Callorhinus_ursinus | Vulpes_zerda | 0.246673160420242 |
| Callorhinus_ursinus | Zalophus_californianus | 0.112471076771718 |
| Canis_anthus | Canis_latrans | 0.0500092141412824 |
| Canis_anthus | Canis_lupus | 0.0326830310659574 |
| Canis_anthus | Caracal_caracal | 0.241338180184938 |
| Canis_anthus | Catopuma_badia | 0.234962721350201 |
| Canis_anthus | Catopuma_temminckii | 0.239812033981155 |
| Canis_anthus | Chrysocyon_brachyurus | 0.130845684654164 |
| Canis_anthus | Civettictis_civetta | 0.238944504666666 |
| Canis_anthus | Conepatus_chinga | 0.254353474699112 |
| Canis_anthus | Crocuta_crocuta | 0.256258823034714 |
| Canis_anthus | Cuon_alpinus | 0.105774264824254 |
| Canis_anthus | Cystophora_cristata | 0.236693096136528 |
| Canis_anthus | Enhydra_lutris | 0.247851187364077 |
| Canis_anthus | Erignathus_barbatus | 0.229036669503957 |
| Canis_anthus | Eumetopias_jubatus | 0.245868978525793 |
| Canis_anthus | Felis_catus | 0.240720114123427 |
| Canis_anthus | Felis_chaus | 0.246095909581808 |
| Canis_anthus | Felis_margarita | 0.240831299358486 |
| Canis_anthus | Felis_nigripes | 0.241125828923769 |
| Canis_anthus | Felis_silvestris | 0.239276951487424 |
| Canis_anthus | Fossa_fossana_FMNH156648_SRR6053060 | 0.243786008051646 |
| Canis_anthus | Genetta_servalina | 0.2430766111654 |
| Canis_anthus | Gulo_gulo | 0.246664057124575 |
| Canis_anthus | Halichoerus_grypus | 0.23521988100195 |
| Canis_anthus | Helarctos_malayanus | 0.247985482116967 |
| Canis_anthus | Hyaena_hyaena | 0.250469142751097 |

| | | |
|---|---|---|
| Canis_anthus | Hydrurga_leptonyx | 0.227071170993585 |
| Canis_anthus | Leopardus_colocolo | 0.240719723096643 |
| Canis_anthus | Leopardus_geoffroyi | 0.241730461689579 |
| Canis_anthus | Leopardus_guigna | 0.245732030041852 |
| Canis_anthus | Leopardus_jacobita | 0.247138689197689 |
| Canis_anthus | Leopardus_pardalis | 0.241497760377121 |
| Canis_anthus | Leopardus_tigrinus | 0.241751711443297 |
| Canis_anthus | Leopardus_wiedii | 0.240204847146355 |
| Canis_anthus | Leptailurus_serval | 0.235412091254814 |
| Canis_anthus | Leptonychotes_weddellii | 0.22413440422511 |
| Canis_anthus | Lobodon_carcinophaga | 0.221004143401333 |
| Canis_anthus | Lutra_lutra | 0.252663625555813 |
| Canis_anthus | Lutra_sumatrana | 0.251387674037336 |
| Canis_anthus | Lutrogale_perspicillata | 0.257679005234588 |
| Canis_anthus | Lycaon_pictus | 0.121485317385954 |
| Canis_anthus | Lynx_canadensis | 0.242826434219049 |
| Canis_anthus | Lynx_lynx | 0.238863805665453 |
| Canis_anthus | Lynx_pardinus | 0.239750920557734 |
| Canis_anthus | Lynx_rufus | 0.24137973025486 |
| Canis_anthus | Manis_javanica | 0.308251198239911 |
| Canis_anthus | Martes_americana | 0.243583208752462 |
| Canis_anthus | Martes_flavigula | 0.237336920495473 |
| Canis_anthus | Martes_foina | 0.24312841536325 |
| Canis_anthus | Martes_martes | 0.241043551243197 |
| Canis_anthus | Martes_melampus | 0.239112924138756 |
| Canis_anthus | Martes_pennanti | 0.2412193926437 |
| Canis_anthus | Martes_zibellina | 0.241187192195811 |
| Canis_anthus | Meles_anakuma | 0.243603676016944 |
| Canis_anthus | Meles_leucurus | 0.243006555974247 |
| Canis_anthus | Meles_meles | 0.240525379085049 |
| Canis_anthus | Melogale_moschata | 0.249290487154 |
| Canis_anthus | Melursus_ursinus | 0.250291377489183 |
| Canis_anthus | Mephitis_mephitis | 0.250140715486893 |
| Canis_anthus | Mirounga_leonina | 0.229713755934175 |
| Canis_anthus | Mungotictis_decemlineata | 0.248457243461336 |
| Canis_anthus | Mustela_altaica | 0.243623724871294 |
| Canis_anthus | Mustela_erminea | 0.235732746105167 |
| Canis_anthus | Mustela_eversmannii | 0.242134889665083 |
| Canis_anthus | Mustela_frenata | 0.2343228201378 |
| Canis_anthus | Mustela_itatsi | 0.242404909198193 |
| Canis_anthus | Mustela_kathiah | 0.238114525341214 |
| Canis_anthus | Mustela_nigripes | 0.238901792407129 |
| Canis_anthus | Mustela_nivalis | 0.240622583110973 |
| Canis_anthus | Mustela_putorius | 0.240392009942631 |
| Canis_anthus | Mustela_sibirica | 0.240512000339318 |
| Canis_anthus | Nandinia_binotata | 0.232897849316906 |
| Canis_anthus | Nasua_nasua | 0.258756434671469 |
| Canis_anthus | Neofelis_nebulosa | 0.255769322972223 |
| Canis_anthus | Neomonachus_schauinslandi | 0.232585873976126 |
| Canis_anthus | Neophoca_cinerea | 0.250307896503465 |
| Canis_anthus | Neovison_vison | 0.240972674107501 |
| Canis_anthus | Nyctereutes_procyonoides | 0.175857369959711 |
| Canis_anthus | Odobenus_rosmarus_rosmarus | 0.270276112747476 |
| Canis_anthus | Otocolobus_manul | 0.237749506531347 |
| Canis_anthus | Otocyon_megalotis_megalotis_TS305 | 0.178883656552334 |
| Canis_anthus | Otocyon_megalotis_virgatus_FMNH158128 | 0.181498992558463 |
| Canis_anthus | Paguma_larvata | 0.24154649038345 |
| Canis_anthus | Panthera_leo | 0.246812167334036 |
| Canis_anthus | Panthera_onca | 0.249374867140224 |
| Canis_anthus | Panthera_pardus | 0.247915574361349 |
| Canis_anthus | Panthera_tigris | 0.247551640169156 |
| Canis_anthus | Panthera_uncia | 0.250310956950693 |
| Canis_anthus | Paradoxurus_hermaphroditus | 0.24228018680999 |
| Canis_anthus | Parahyaena_brunnea_FD126_Fecal | 0.251269213675505 |
| Canis_anthus | Pardofelis_marmorata | 0.245245678576376 |
| Canis_anthus | Phoca_fasciata | 0.233539850187311 |
| Canis_anthus | Phoca_groenlandica | 0.235635247621425 |
| Canis_anthus | Phoca_largha | 0.235894714870153 |
| Canis_anthus | Phoca_vitulina | 0.236263452009317 |
| Canis_anthus | Phocarctos_hookeri | 0.2600183903888 |
| Canis_anthus | Prionailurus_bengalensis | 0.23703545154253 |
| Canis_anthus | Prionailurus_bengalensis_euptilurus | 0.238582414102505 |
| Canis_anthus | Prionailurus_planiceps | 0.241143969842413 |
| Canis_anthus | Prionailurus_rubiginosus | 0.236497918714023 |
| Canis_anthus | Prionailurus_viverrinus | 0.243433897451666 |
| Canis_anthus | Prionodon_pardicolor | 0.241559328798252 |
| Canis_anthus | Procyon_lotor | 0.247723994733117 |
| Canis_anthus | Profelis_aurata | 0.241641281977687 |
| Canis_anthus | Proteles_cristatus_cristatus_TS307 | 0.258961430742302 |
| Canis_anthus | Proteles_cristatus_septentrionalis_NMS_Z_2018_54 | 0.258481338439491 |
| Canis_anthus | Puma_concolor | 0.241012869952032 |
| Canis_anthus | Puma_yagouaroundi | 0.238916348271257 |
| Canis_anthus | Pusa_caspica | 0.233556852497245 |
| Canis_anthus | Pusa_hispida | 0.233805439918918 |
| Canis_anthus | Pusa_sibirica | 0.237414270738661 |
| Canis_anthus | Speothos_venaticus_T1624 | 0.136156311003246 |
| Canis_anthus | Spilogale_putorius | 0.246874924098869 |

| | | |
|---|---|---|
| Canis_anthus | Taxidea_taxus | 0.243710227991591 |
| Canis_anthus | Tremarctos_ornatus | 0.252634603788867 |
| Canis_anthus | Urocyon_cinereoargenteus | 0.185289183847499 |
| Canis_anthus | Ursus_americanus | 0.24613642044535 |
| Canis_anthus | Ursus_arctos | 0.251627017590636 |
| Canis_anthus | Ursus_maritimus | 0.252353472289664 |
| Canis_anthus | Ursus_spelaeus | 0.249377613671185 |
| Canis_anthus | Ursus_thibetanus | 0.243888278214526 |
| Canis_anthus | Urva_javanica | 0.243212111769169 |
| Canis_anthus | Viverra_tangalunga_FMNH146957_SRR6053069 | 0.234494881052361 |
| Canis_anthus | Viverricula_indica | 0.231695268656282 |
| Canis_anthus | Vulpes_corsac | 0.177888670522871 |
| Canis_anthus | Vulpes_ferrilata | 0.176725179322134 |
| Canis_anthus | Vulpes_lagopus | 0.17877136450713 |
| Canis_anthus | Vulpes_vulpes_T3611 | 0.178002955726656 |
| Canis_anthus | Vulpes_zerda | 0.178106379038639 |
| Canis_anthus | Zalophus_californianus | 0.248893681868382 |
| Canis_latrans | Canis_lupus | 0.0492600041258489 |
| Canis_latrans | Caracal_caracal | 0.239268998198813 |
| Canis_latrans | Catopuma_badia | 0.23334528518842 |
| Canis_latrans | Catopuma_temminckii | 0.236178890594316 |
| Canis_latrans | Chrysocyon_brachyurus | 0.125185606836687 |
| Canis_latrans | Civettictis_civetta | 0.233883934878109 |
| Canis_latrans | Conepatus_chinga | 0.246428263197356 |
| Canis_latrans | Crocuta_crocuta | 0.253970201546472 |
| Canis_latrans | Cuon_alpinus | 0.10093056423654 |
| Canis_latrans | Cystophora_cristata | 0.22916748689699 |
| Canis_latrans | Enhydra_lutris | 0.242256564872033 |
| Canis_latrans | Erignathus_barbatus | 0.226386730542888 |
| Canis_latrans | Eumetopias_jubatus | 0.244122482488669 |
| Canis_latrans | Felis_catus | 0.237655874673781 |
| Canis_latrans | Felis_chaus | 0.242356479179331 |
| Canis_latrans | Felis_margarita | 0.236219568305473 |
| Canis_latrans | Felis_nigripes | 0.23675127935862 |
| Canis_latrans | Felis_silvestris | 0.236219568305473 |
| Canis_latrans | Fossa_fossana_FMNH156648_SRR6053060 | 0.241710111361286 |
| Canis_latrans | Genetta_servalina | 0.238987762121918 |
| Canis_latrans | Gulo_gulo | 0.240412636081848 |
| Canis_latrans | Halichoerus_grypus | 0.230449053474839 |
| Canis_latrans | Helarctos_malayanus | 0.243520261878002 |
| Canis_latrans | Hyaena_hyaena | 0.249887275012006 |
| Canis_latrans | Hydrurga_leptonyx | 0.219853451176773 |
| Canis_latrans | Leopardus_colocolo | 0.23915738661555 |
| Canis_latrans | Leopardus_geoffroyi | 0.240396887105624 |
| Canis_latrans | Leopardus_guigna | 0.241802143225513 |
| Canis_latrans | Leopardus_jacobita | 0.243332617364762 |
| Canis_latrans | Leopardus_pardalis | 0.237857752532161 |
| Canis_latrans | Leopardus_tigrinus | 0.237873261924718 |
| Canis_latrans | Leopardus_wiedii | 0.23776704958332 |
| Canis_latrans | Leptailurus_serval | 0.234877898388371 |
| Canis_latrans | Leptonychotes_weddellii | 0.216628841209122 |
| Canis_latrans | Lobodon_carcinophaga | 0.216967669260549 |
| Canis_latrans | Lutra_lutra | 0.244140953982867 |
| Canis_latrans | Lutra_sumatrana | 0.246531034862856 |
| Canis_latrans | Lutrogale_perspicillata | 0.255059163991127 |
| Canis_latrans | Lycaon_pictus | 0.116410647162374 |
| Canis_latrans | Lynx_canadensis | 0.240753136105015 |
| Canis_latrans | Lynx_lynx | 0.236470138427327 |
| Canis_latrans | Lynx_pardinus | 0.237575477663823 |
| Canis_latrans | Lynx_rufus | 0.236540330744296 |
| Canis_latrans | Manis_javanica | 0.305631657577595 |
| Canis_latrans | Martes_americana | 0.239123464745811 |
| Canis_latrans | Martes_flavigula | 0.233273577502511 |
| Canis_latrans | Martes_foina | 0.237566423055614 |
| Canis_latrans | Martes_martes | 0.234194055882594 |
| Canis_latrans | Martes_melampus | 0.234775651287355 |
| Canis_latrans | Martes_pennanti | 0.240131918992259 |
| Canis_latrans | Martes_zibellina | 0.234093962034052 |
| Canis_latrans | Meles_anakuma | 0.240154727371763 |
| Canis_latrans | Meles_leucurus | 0.238276211631908 |
| Canis_latrans | Meles_meles | 0.234975143303294 |
| Canis_latrans | Melogale_moschata | 0.24432225354838 |
| Canis_latrans | Melursus_ursinus | 0.249820504527851 |
| Canis_latrans | Mephitis_mephitis | 0.245272233454671 |
| Canis_latrans | Mirounga_leonina | 0.227586868321412 |
| Canis_latrans | Mungotictis_decemlineata | 0.242136405076402 |
| Canis_latrans | Mustela_altaica | 0.238215336991927 |
| Canis_latrans | Mustela_erminea | 0.232826679000508 |
| Canis_latrans | Mustela_eversmannii | 0.240006586758604 |
| Canis_latrans | Mustela_frenata | 0.234347419677758 |
| Canis_latrans | Mustela_itatsi | 0.23815928014907 |
| Canis_latrans | Mustela_kathiah | 0.235120642200516 |
| Canis_latrans | Mustela_nigripes | 0.235815768623946 |
| Canis_latrans | Mustela_nivalis | 0.236941068467296 |
| Canis_latrans | Mustela_putorius | 0.237498923460523 |
| Canis_latrans | Mustela_sibirica | 0.237306671001272 |
| Canis_latrans | Nandinia_binotata | 0.228854539034724 |

| | | |
|---|---|---|
| Canis_latrans | Nasua_nasua | 0.256636111922792 |
| Canis_latrans | Neofelis_nebulosa | 0.25286172428809 |
| Canis_latrans | Neomonachus_schauinslandi | 0.225881570108342 |
| Canis_latrans | Neophoca_cinerea | 0.24947887241946 |
| Canis_latrans | Neovison_vison | 0.236374714771524 |
| Canis_latrans | Nyctereutes_procyonoides | 0.167652058551223 |
| Canis_latrans | Odobenus_rosmarus_rosmarus | 0.266250768135261 |
| Canis_latrans | Otocolobus_manul | 0.235780949653104 |
| Canis_latrans | Otocyon_megalotis_megalotis_TS305 | 0.172864866835507 |
| Canis_latrans | Otocyon_megalotis_virgatus_FMNH158128 | 0.177120422312505 |
| Canis_latrans | Paguma_larvata | 0.236978372870911 |
| Canis_latrans | Panthera_leo | 0.243454765535636 |
| Canis_latrans | Panthera_onca | 0.243452315963678 |
| Canis_latrans | Panthera_pardus | 0.246234397659647 |
| Canis_latrans | Panthera_tigris | 0.246454931604951 |
| Canis_latrans | Panthera_uncia | 0.246170260354343 |
| Canis_latrans | Paradoxurus_hermaphroditus | 0.237355764216485 |
| Canis_latrans | Parahyaena_brunnea_FD126_Fecal | 0.249109669403075 |
| Canis_latrans | Pardofelis_marmorata | 0.242258223069575 |
| Canis_latrans | Phoca_fasciata | 0.228201433469825 |
| Canis_latrans | Phoca_groenlandica | 0.229992121451569 |
| Canis_latrans | Phoca_largha | 0.231199718920782 |
| Canis_latrans | Phoca_vitulina | 0.230380155827655 |
| Canis_latrans | Phocarctos_hookeri | 0.256132413565616 |
| Canis_latrans | Prionailurus_bengalensis | 0.234409048644422 |
| Canis_latrans | Prionailurus_bengalensis_euptilurus | 0.235398930181993 |
| Canis_latrans | Prionailurus_planiceps | 0.240608953607184 |
| Canis_latrans | Prionailurus_rubiginosus | 0.233984146523292 |
| Canis_latrans | Prionailurus_viverrinus | 0.239892449640314 |
| Canis_latrans | Prionodon_pardicolor | 0.237811253335819 |
| Canis_latrans | Procyon_lotor | 0.243248695558433 |
| Canis_latrans | Profelis_aurata | 0.239550653736254 |
| Canis_latrans | Proteles_cristatus_cristatus_TS307 | 0.254975768381563 |
| Canis_latrans | Proteles_cristatus_septentrionalis_NMS_Z_2018_54 | 0.254385897110943 |
| Canis_latrans | Puma_concolor | 0.235939904882512 |
| Canis_latrans | Puma_yagouaroundi | 0.235512474016909 |
| Canis_latrans | Pusa_caspica | 0.228559139332894 |
| Canis_latrans | Pusa_hispida | 0.228480977459102 |
| Canis_latrans | Pusa_sibirica | 0.228757950916965 |
| Canis_latrans | Speothos_venaticus_T1624 | 0.131163068917726 |
| Canis_latrans | Spilogale_putorius | 0.240616845289307 |
| Canis_latrans | Taxidea_taxus | 0.241596975155988 |
| Canis_latrans | Tremarctos_ornatus | 0.251539309933083 |
| Canis_latrans | Urocyon_cinereoargenteus | 0.181971759211421 |
| Canis_latrans | Ursus_americanus | 0.24551699555384 |
| Canis_latrans | Ursus_arctos | 0.24946475103355 |
| Canis_latrans | Ursus_maritimus | 0.248677908052559 |
| Canis_latrans | Ursus_spelaeus | 0.248995131158662 |
| Canis_latrans | Ursus_thibetanus | 0.24316467465475 |
| Canis_latrans | Urva_javanica | 0.240345067316752 |
| Canis_latrans | Viverra_tangalunga_FMNH146957_SRR6053069 | 0.230005696467937 |
| Canis_latrans | Viverricula_indica | 0.229289463494028 |
| Canis_latrans | Vulpes_corsac | 0.17177673152066 |
| Canis_latrans | Vulpes_ferrilata | 0.171440778629868 |
| Canis_latrans | Vulpes_lagopus | 0.173160632069307 |
| Canis_latrans | Vulpes_vulpes_T3611 | 0.172708336465431 |
| Canis_latrans | Vulpes_zerda | 0.173631224340243 |
| Canis_latrans | Zalophus_californianus | 0.246690878203642 |
| Canis_lupus | Caracal_caracal | 0.233546299869036 |
| Canis_lupus | Catopuma_badia | 0.22887269598104 |
| Canis_lupus | Catopuma_temminckii | 0.234765617620384 |
| Canis_lupus | Chrysocyon_brachyurus | 0.124422393561232 |
| Canis_lupus | Civettictis_civetta | 0.233139928125205 |
| Canis_lupus | Conepatus_chinga | 0.248802004613542 |
| Canis_lupus | Crocuta_crocuta | 0.253651507721581 |
| Canis_lupus | Cuon_alpinus | 0.10106355340069 |
| Canis_lupus | Cystophora_cristata | 0.230518965128351 |
| Canis_lupus | Enhydra_lutris | 0.242702297016551 |
| Canis_lupus | Erignathus_barbatus | 0.229206341316065 |
| Canis_lupus | Eumetopias_jubatus | 0.24309795529891 |
| Canis_lupus | Felis_catus | 0.236972488597419 |
| Canis_lupus | Felis_chaus | 0.242558684199063 |
| Canis_lupus | Felis_margarita | 0.235318136145185 |
| Canis_lupus | Felis_nigripes | 0.236550415297254 |
| Canis_lupus | Felis_silvestris | 0.235208040361409 |
| Canis_lupus | Fossa_fossana_FMNH156648_SRR6053060 | 0.239256880131326 |
| Canis_lupus | Genetta_servalina | 0.237000785227287 |
| Canis_lupus | Gulo_gulo | 0.239007039314231 |
| Canis_lupus | Halichoerus_grypus | 0.228817135229517 |
| Canis_lupus | Helarctos_malayanus | 0.245216904157492 |
| Canis_lupus | Hyaena_hyaena | 0.245979668711975 |
| Canis_lupus | Hydrurga_leptonyx | 0.222478233122159 |
| Canis_lupus | Leopardus_colocolo | 0.23382173488896 |
| Canis_lupus | Leopardus_geoffroyi | 0.23625098419302 |
| Canis_lupus | Leopardus_guigna | 0.238108814375538 |
| Canis_lupus | Leopardus_jacobita | 0.240942509323357 |
| Canis_lupus | Leopardus_pardalis | 0.236020748878816 |

| | | |
|---|---|---|
| Canis_lupus | Leopardus_tigrinus | 0.23495877941597 |
| Canis_lupus | Leopardus_wiedii | 0.234738791456988 |
| Canis_lupus | Leptailurus_serval | 0.231060923050874 |
| Canis_lupus | Leptonychotes_weddellii | 0.222374999165645 |
| Canis_lupus | Lobodon_carcinophaga | 0.218091875731245 |
| Canis_lupus | Lutra_lutra | 0.244973312342892 |
| Canis_lupus | Lutra_sumatrana | 0.243705040791205 |
| Canis_lupus | Lutrogale_perspicillata | 0.25401197998215 |
| Canis_lupus | Lycaon_pictus | 0.11771229763962 |
| Canis_lupus | Lynx_canadensis | 0.239217542464772 |
| Canis_lupus | Lynx_lynx | 0.237727420640177 |
| Canis_lupus | Lynx_pardinus | 0.23820947854704 |
| Canis_lupus | Lynx_rufus | 0.239680580557875 |
| Canis_lupus | Manis_javanica | 0.30502508222721 |
| Canis_lupus | Martes_americana | 0.239791170164424 |
| Canis_lupus | Martes_flavigula | 0.235086116308195 |
| Canis_lupus | Martes_foina | 0.237898796212103 |
| Canis_lupus | Martes_martes | 0.23640229619493 |
| Canis_lupus | Martes_melampus | 0.233913992621078 |
| Canis_lupus | Martes_pennanti | 0.241234736530673 |
| Canis_lupus | Martes_zibellina | 0.235528050165543 |
| Canis_lupus | Meles_anakuma | 0.241046275368794 |
| Canis_lupus | Meles_leucurus | 0.237614252135397 |
| Canis_lupus | Meles_meles | 0.236475288532213 |
| Canis_lupus | Melogale_moschata | 0.243475691591441 |
| Canis_lupus | Melursus_ursinus | 0.249109325517957 |
| Canis_lupus | Mephitis_mephitis | 0.245121700270227 |
| Canis_lupus | Mirounga_leonina | 0.224506213504948 |
| Canis_lupus | Mungotictis_decemlineata | 0.243762531112228 |
| Canis_lupus | Mustela_altaica | 0.235577388468058 |
| Canis_lupus | Mustela_erminea | 0.228420590766989 |
| Canis_lupus | Mustela_eversmannii | 0.2351300393518 |
| Canis_lupus | Mustela_frenata | 0.227877340026608 |
| Canis_lupus | Mustela_itatsi | 0.235376359366351 |
| Canis_lupus | Mustela_kathiah | 0.231642512354083 |
| Canis_lupus | Mustela_nigripes | 0.231411635060501 |
| Canis_lupus | Mustela_nivalis | 0.231655052975863 |
| Canis_lupus | Mustela_putorius | 0.232752829079593 |
| Canis_lupus | Mustela_sibirica | 0.232779335636598 |
| Canis_lupus | Nandinia_binotata | 0.227732372116886 |
| Canis_lupus | Nasua_nasua | 0.257550273285124 |
| Canis_lupus | Neofelis_nebulosa | 0.251633315031653 |
| Canis_lupus | Neomonachus_schauinslandi | 0.226795453848892 |
| Canis_lupus | Neophoca_cinerea | 0.250570080844645 |
| Canis_lupus | Neovison_vison | 0.233780642015868 |
| Canis_lupus | Nyctereutes_procyonoides | 0.171236342360026 |
| Canis_lupus | Odobenus_rosmarus_rosmarus | 0.264763161591024 |
| Canis_lupus | Otocolobus_manul | 0.232378829415805 |
| Canis_lupus | Otocyon_megalotis_megalotis_TS305 | 0.178439492071846 |
| Canis_lupus | Otocyon_megalotis_virgatus_FMNH158128 | 0.178207481288034 |
| Canis_lupus | Paguma_larvata | 0.234107838036534 |
| Canis_lupus | Panthera_leo | 0.243182409630352 |
| Canis_lupus | Panthera_onca | 0.244835470348529 |
| Canis_lupus | Panthera_pardus | 0.243720289205708 |
| Canis_lupus | Panthera_tigris | 0.244722787741587 |
| Canis_lupus | Panthera_uncia | 0.246908858193572 |
| Canis_lupus | Paradoxurus_hermaphroditus | 0.238218670279836 |
| Canis_lupus | Parahyaena_brunnea_FD126_Fecal | 0.248568243569293 |
| Canis_lupus | Pardofelis_marmorata | 0.241714103874098 |
| Canis_lupus | Phoca_fasciata | 0.231087593703835 |
| Canis_lupus | Phoca_groenlandica | 0.232514616996478 |
| Canis_lupus | Phoca_largha | 0.230719478773844 |
| Canis_lupus | Phoca_vitulina | 0.230888290688046 |
| Canis_lupus | Phocarctos_hookeri | 0.257138973454682 |
| Canis_lupus | Prionailurus_bengalensis | 0.232054096154391 |
| Canis_lupus | Prionailurus_bengalensis_euptilurus | 0.234227938430941 |
| Canis_lupus | Prionailurus_planiceps | 0.236883364386988 |
| Canis_lupus | Prionailurus_rubiginosus | 0.233655199143006 |
| Canis_lupus | Prionailurus_viverrinus | 0.240448005253025 |
| Canis_lupus | Prionodon_pardicolor | 0.241133892653488 |
| Canis_lupus | Procyon_lotor | 0.239583234290907 |
| Canis_lupus | Profelis_aurata | 0.235127314039385 |
| Canis_lupus | Proteles_cristatus_cristatus_TS307 | 0.255225151886443 |
| Canis_lupus | Proteles_cristatus_septentrionalis_NMS_Z_2018_54 | 0.255544596656794 |
| Canis_lupus | Puma_concolor | 0.236219568305473 |
| Canis_lupus | Puma_yagouaroundi | 0.234891151793951 |
| Canis_lupus | Pusa_caspica | 0.226604500581174 |
| Canis_lupus | Pusa_hispida | 0.22794339101416 |
| Canis_lupus | Pusa_sibirica | 0.230021032057179 |
| Canis_lupus | Speothos_venaticus_T1624 | 0.137471861144668 |
| Canis_lupus | Spilogale_putorius | 0.241672777257357 |
| Canis_lupus | Taxidea_taxus | 0.238214107658694 |
| Canis_lupus | Tremarctos_ornatus | 0.250278275216728 |
| Canis_lupus | Urocyon_cinereoargenteus | 0.183887426593 |
| Canis_lupus | Ursus_americanus | 0.245211283918499 |
| Canis_lupus | Ursus_arctos | 0.247475665310199 |
| Canis_lupus | Ursus_maritimus | 0.248563791881803 |

| | | |
|---|---|---|
| Canis_lupus | Ursus_spelaeus | 0.248766820652412 |
| Canis_lupus | Ursus_thibetanus | 0.241408715213942 |
| Canis_lupus | Urva_javanica | 0.239903371174954 |
| Canis_lupus | Viverra_tangalunga_FMNH146957_SRR6053069 | 0.22893606963676 |
| Canis_lupus | Viverricula_indica | 0.22791328573853 |
| Canis_lupus | Vulpes_corsac | 0.169573112192282 |
| Canis_lupus | Vulpes_ferrilata | 0.169237552843758 |
| Canis_lupus | Vulpes_lagopus | 0.173656673431357 |
| Canis_lupus | Vulpes_vulpes_T3611 | 0.171296876257885 |
| Canis_lupus | Vulpes_zerda | 0.172217310144324 |
| Canis_lupus | Zalophus_californianus | 0.24926860258581 |
| Caracal_caracal | Catopuma_badia | 0.112511038627409 |
| Caracal_caracal | Catopuma_temminckii | 0.11561837139657 |
| Caracal_caracal | Chrysocyon_brachyurus | 0.239320812455847 |
| Caracal_caracal | Civettictis_civetta | 0.174444749125347 |
| Caracal_caracal | Conepatus_chinga | 0.237801558557444 |
| Caracal_caracal | Crocuta_crocuta | 0.201654465852557 |
| Caracal_caracal | Cuon_alpinus | 0.237384504226512 |
| Caracal_caracal | Cystophora_cristata | 0.217444097895185 |
| Caracal_caracal | Enhydra_lutris | 0.229632440334038 |
| Caracal_caracal | Erignathus_barbatus | 0.220071627434989 |
| Caracal_caracal | Eumetopias_jubatus | 0.236048504705654 |
| Caracal_caracal | Felis_catus | 0.112255638967459 |
| Caracal_caracal | Felis_chaus | 0.11127718756343 |
| Caracal_caracal | Felis_margarita | 0.114163918337425 |
| Caracal_caracal | Felis_nigripes | 0.115865863916165 |
| Caracal_caracal | Felis_silvestris | 0.110844696700542 |
| Caracal_caracal | Fossa_fossana_FMNH156648_SRR6053060 | 0.181416288900348 |
| Caracal_caracal | Genetta_servalina | 0.176551269048488 |
| Caracal_caracal | Gulo_gulo | 0.239077031866295 |
| Caracal_caracal | Halichoerus_grypus | 0.223835214784326 |
| Caracal_caracal | Helarctos_malayanus | 0.235425515451757 |
| Caracal_caracal | Hyaena_hyaena | 0.195101153565685 |
| Caracal_caracal | Hydrurga_leptonyx | 0.210155318726216 |
| Caracal_caracal | Leopardus_colocolo | 0.118610837648856 |
| Caracal_caracal | Leopardus_geoffroyi | 0.123299999979581 |
| Caracal_caracal | Leopardus_guigna | 0.121791426480044 |
| Caracal_caracal | Leopardus_jacobita | 0.119821088459452 |
| Caracal_caracal | Leopardus_pardalis | 0.117731489681177 |
| Caracal_caracal | Leopardus_tigrinus | 0.124295656968005 |
| Caracal_caracal | Leopardus_wiedii | 0.118773650172163 |
| Caracal_caracal | Leptailurus_serval | 0.101966783189911 |
| Caracal_caracal | Leptonychotes_weddellii | 0.210028421522237 |
| Caracal_caracal | Lobodon_carcinophaga | 0.212960262512494 |
| Caracal_caracal | Lutra_lutra | 0.236901768697629 |
| Caracal_caracal | Lutra_sumatrana | 0.234304443407718 |
| Caracal_caracal | Lutrogale_perspicillata | 0.24074284039115 |
| Caracal_caracal | Lycaon_pictus | 0.238180048832229 |
| Caracal_caracal | Lynx_canadensis | 0.11530682664122 |
| Caracal_caracal | Lynx_lynx | 0.118697098077865 |
| Caracal_caracal | Lynx_pardinus | 0.118354418810634 |
| Caracal_caracal | Lynx_rufus | 0.119220574854616 |
| Caracal_caracal | Manis_javanica | 0.290175702121086 |
| Caracal_caracal | Martes_americana | 0.236046238370442 |
| Caracal_caracal | Martes_flavigula | 0.239592883656308 |
| Caracal_caracal | Martes_foina | 0.232493918565221 |
| Caracal_caracal | Martes_martes | 0.236033936245471 |
| Caracal_caracal | Martes_melampus | 0.234447576152369 |
| Caracal_caracal | Martes_pennanti | 0.240698978340648 |
| Caracal_caracal | Martes_zibellina | 0.238139193884834 |
| Caracal_caracal | Meles_anakuma | 0.234533996101923 |
| Caracal_caracal | Meles_leucurus | 0.230318014285898 |
| Caracal_caracal | Meles_meles | 0.229507796627242 |
| Caracal_caracal | Melogale_moschata | 0.235891650716294 |
| Caracal_caracal | Melursus_ursinus | 0.238017069793433 |
| Caracal_caracal | Mephitis_mephitis | 0.238321605748617 |
| Caracal_caracal | Mirounga_leonina | 0.216041543683691 |
| Caracal_caracal | Mungotictis_decemlineata | 0.190296438429125 |
| Caracal_caracal | Mustela_altaica | 0.232364250554204 |
| Caracal_caracal | Mustela_erminea | 0.228003587703117 |
| Caracal_caracal | Mustela_eversmannii | 0.233085263448125 |
| Caracal_caracal | Mustela_frenata | 0.225357304666187 |
| Caracal_caracal | Mustela_itatsi | 0.229942206932607 |
| Caracal_caracal | Mustela_kathiah | 0.232585873976126 |
| Caracal_caracal | Mustela_nigripes | 0.230728168967951 |
| Caracal_caracal | Mustela_nivalis | 0.230280260641517 |
| Caracal_caracal | Mustela_putorius | 0.231775501943703 |
| Caracal_caracal | Mustela_sibirica | 0.232665298029379 |
| Caracal_caracal | Nandinia_binotata | 0.172944423486371 |
| Caracal_caracal | Nasua_nasua | 0.250413064587759 |
| Caracal_caracal | Neofelis_nebulosa | 0.145118527994115 |
| Caracal_caracal | Neomonachus_schauinslandi | 0.216548713979205 |
| Caracal_caracal | Neophoca_cinerea | 0.235247875274155 |
| Caracal_caracal | Neovison_vison | 0.240031185625285 |
| Caracal_caracal | Nyctereutes_procyonoides | 0.240227916377116 |
| Caracal_caracal | Odobenus_rosmarus_rosmarus | 0.255204451404185 |
| Caracal_caracal | Otocolobus_manul | 0.113651869625735 |

| | | |
|---|---|---|
| Caracal_caracal | Otocyon_megalotis_megalotis_TS305 | 0.243669451179804 |
| Caracal_caracal | Otocyon_megalotis_virgatus_FMNH158128 | 0.243107567348446 |
| Caracal_caracal | Paguma_larvata | 0.182429036180833 |
| Caracal_caracal | Panthera_leo | 0.133203979538325 |
| Caracal_caracal | Panthera_onca | 0.129749834770352 |
| Caracal_caracal | Panthera_pardus | 0.132970430535333 |
| Caracal_caracal | Panthera_tigris | 0.141076895558496 |
| Caracal_caracal | Panthera_uncia | 0.134619811149005 |
| Caracal_caracal | Paradoxurus_hermaphroditus | 0.181615700056669 |
| Caracal_caracal | Parahyaena_brunnea_FD126_Fecal | 0.194457602510366 |
| Caracal_caracal | Pardofelis_marmorata | 0.121724040328291 |
| Caracal_caracal | Phoca_fasciata | 0.220994797550679 |
| Caracal_caracal | Phoca_groenlandica | 0.223223374407651 |
| Caracal_caracal | Phoca_largha | 0.225771159976833 |
| Caracal_caracal | Phoca_vitulina | 0.226375896955024 |
| Caracal_caracal | Phocarctos_hookeri | 0.238519748841698 |
| Caracal_caracal | Prionailurus_bengalensis | 0.107923970822684 |
| Caracal_caracal | Prionailurus_bengalensis_euptilurus | 0.109605027359314 |
| Caracal_caracal | Prionailurus_planiceps | 0.113372237244151 |
| Caracal_caracal | Prionailurus_rubiginosus | 0.111576458916057 |
| Caracal_caracal | Prionailurus_viverrinus | 0.110996892311765 |
| Caracal_caracal | Prionodon_pardicolor | 0.186060853710953 |
| Caracal_caracal | Procyon_lotor | 0.232799072035292 |
| Caracal_caracal | Profelis_aurata | 0.0815738465751477 |
| Caracal_caracal | Proteles_cristatus_cristatus_TS307 | 0.19661078140728 |
| Caracal_caracal | Proteles_cristatus_septentrionalis_NMS_Z_2018_54 | 0.19652579797823 |
| Caracal_caracal | Puma_concolor | 0.114814893879699 |
| Caracal_caracal | Puma_yagouaroundi | 0.114906154942816 |
| Caracal_caracal | Pusa_caspica | 0.226456783579078 |
| Caracal_caracal | Pusa_hispida | 0.220108950082864 |
| Caracal_caracal | Pusa_sibirica | 0.222581342969691 |
| Caracal_caracal | Speothos_venaticus_T1624 | 0.245577598805576 |
| Caracal_caracal | Spilogale_putorius | 0.243706649547091 |
| Caracal_caracal | Taxidea_taxus | 0.219261996507968 |
| Caracal_caracal | Tremarctos_ornatus | 0.234718238877105 |
| Caracal_caracal | Urocyon_cinereoargenteus | 0.247409040403829 |
| Caracal_caracal | Ursus_americanus | 0.234914507281175 |
| Caracal_caracal | Ursus_arctos | 0.240659315976748 |
| Caracal_caracal | Ursus_maritimus | 0.240623098505154 |
| Caracal_caracal | Ursus_spelaeus | 0.234537947591591 |
| Caracal_caracal | Ursus_thibetanus | 0.232407497009692 |
| Caracal_caracal | Urva_javanica | 0.190472058656367 |
| Caracal_caracal | Viverra_tangalunga_FMNH146957_SRR6053069 | 0.174281740542265 |
| Caracal_caracal | Viverricula_indica | 0.172983438461676 |
| Caracal_caracal | Vulpes_corsac | 0.24371717199008 |
| Caracal_caracal | Vulpes_ferrilata | 0.243012216533136 |
| Caracal_caracal | Vulpes_lagopus | 0.243591769729014 |
| Caracal_caracal | Vulpes_vulpes_T3611 | 0.233334198446913 |
| Caracal_caracal | Vulpes_zerda | 0.23617075292828 |
| Caracal_caracal | Zalophus_californianus | 0.23296365169286 |
| Catopuma_badia | Catopuma_temminckii | 0.070543149601641 |
| Catopuma_badia | Chrysocyon_brachyurus | 0.235130316874934 |
| Catopuma_badia | Civettictis_civetta | 0.16974309267364 |
| Catopuma_badia | Conepatus_chinga | 0.241295881250462 |
| Catopuma_badia | Crocuta_crocuta | 0.195806198229886 |
| Catopuma_badia | Cuon_alpinus | 0.23543806003374 |
| Catopuma_badia | Cystophora_cristata | 0.217667992218986 |
| Catopuma_badia | Enhydra_lutris | 0.228045509012816 |
| Catopuma_badia | Erignathus_barbatus | 0.219094276582777 |
| Catopuma_badia | Eumetopias_jubatus | 0.233595083920676 |
| Catopuma_badia | Felis_catus | 0.112048814886078 |
| Catopuma_badia | Felis_chaus | 0.111875397929504 |
| Catopuma_badia | Felis_margarita | 0.111785247642125 |
| Catopuma_badia | Felis_nigripes | 0.111497419591792 |
| Catopuma_badia | Felis_silvestris | 0.109801505050406 |
| Catopuma_badia | Fossa_fossana_FMNH156648_SRR6053060 | 0.181595807896792 |
| Catopuma_badia | Genetta_servalina | 0.175599809333969 |
| Catopuma_badia | Gulo_gulo | 0.234498309261213 |
| Catopuma_badia | Halichoerus_grypus | 0.216702844466221 |
| Catopuma_badia | Helarctos_malayanus | 0.233324651010696 |
| Catopuma_badia | Hyaena_hyaena | 0.191449034427575 |
| Catopuma_badia | Hydrurga_leptonyx | 0.211489219124704 |
| Catopuma_badia | Leopardus_colocolo | 0.112760039682617 |
| Catopuma_badia | Leopardus_geoffroyi | 0.120155611464722 |
| Catopuma_badia | Leopardus_guigna | 0.119379773280036 |
| Catopuma_badia | Leopardus_jacobita | 0.113491798195846 |
| Catopuma_badia | Leopardus_pardalis | 0.113090578935195 |
| Catopuma_badia | Leopardus_tigrinus | 0.116667424098308 |
| Catopuma_badia | Leopardus_wiedii | 0.113294578994749 |
| Catopuma_badia | Leptailurus_serval | 0.103839913107972 |
| Catopuma_badia | Leptonychotes_weddellii | 0.212650315287034 |
| Catopuma_badia | Lobodon_carcinophaga | 0.211798753979263 |
| Catopuma_badia | Lutra_lutra | 0.232966982211815 |
| Catopuma_badia | Lutra_sumatrana | 0.230303491640771 |
| Catopuma_badia | Lutrogale_perspicillata | 0.240216306699586 |
| Catopuma_badia | Lycaon_pictus | 0.234571094245374 |
| Catopuma_badia | Lynx_canadensis | 0.101230469508855 |

| | | |
|---|---|---|
| Catopuma_badia | Lynx_lynx | 0.103648523034621 |
| Catopuma_badia | Lynx_pardinus | 0.105030757604549 |
| Catopuma_badia | Lynx_rufus | 0.102675846600316 |
| Catopuma_badia | Manis_javanica | 0.28492096827937 |
| Catopuma_badia | Martes_americana | 0.235150983487917 |
| Catopuma_badia | Martes_flavigula | 0.237007925145113 |
| Catopuma_badia | Martes_foina | 0.232950989902927 |
| Catopuma_badia | Martes_martes | 0.234340059091036 |
| Catopuma_badia | Martes_melampus | 0.233334579651156 |
| Catopuma_badia | Martes_pennanti | 0.238095767806841 |
| Catopuma_badia | Martes_zibellina | 0.236324952946199 |
| Catopuma_badia | Meles_anakuma | 0.23403965749653 |
| Catopuma_badia | Meles_leucurus | 0.226326357942912 |
| Catopuma_badia | Meles_meles | 0.225425415681038 |
| Catopuma_badia | Melogale_moschata | 0.233292463467245 |
| Catopuma_badia | Melursus_ursinus | 0.236706095640291 |
| Catopuma_badia | Mephitis_mephitis | 0.237045221101186 |
| Catopuma_badia | Mirounga_leonina | 0.214858151719855 |
| Catopuma_badia | Mungotictis_decemlineata | 0.183277298268705 |
| Catopuma_badia | Mustela_altaica | 0.231927089800883 |
| Catopuma_badia | Mustela_erminea | 0.223974265808653 |
| Catopuma_badia | Mustela_eversmannii | 0.231189463243639 |
| Catopuma_badia | Mustela_frenata | 0.230262452185085 |
| Catopuma_badia | Mustela_itatsi | 0.230466129818291 |
| Catopuma_badia | Mustela_kathiah | 0.230808087734555 |
| Catopuma_badia | Mustela_nigripes | 0.227978370649449 |
| Catopuma_badia | Mustela_nivalis | 0.232364250554204 |
| Catopuma_badia | Mustela_putorius | 0.230214795794736 |
| Catopuma_badia | Mustela_sibirica | 0.229479108663301 |
| Catopuma_badia | Nandinia_binotata | 0.167162629406704 |
| Catopuma_badia | Nasua_nasua | 0.250581855583016 |
| Catopuma_badia | Neofelis_nebulosa | 0.141963652282757 |
| Catopuma_badia | Neomonachus_schauinslandi | 0.216504131669436 |
| Catopuma_badia | Neophoca_cinerea | 0.237257165735606 |
| Catopuma_badia | Neovison_vison | 0.232254456965708 |
| Catopuma_badia | Nyctereutes_procyonoides | 0.232472973373779 |
| Catopuma_badia | Odobenus_rosmarus_rosmarus | 0.251893569679704 |
| Catopuma_badia | Otocolobus_manul | 0.108060660751673 |
| Catopuma_badia | Otocyon_megalotis_megalotis_TS305 | 0.242492815669253 |
| Catopuma_badia | Otocyon_megalotis_virgatus_FMNH158128 | 0.240788334593016 |
| Catopuma_badia | Paguma_larvata | 0.176326426425488 |
| Catopuma_badia | Panthera_leo | 0.136400750178501 |
| Catopuma_badia | Panthera_onca | 0.135475112006229 |
| Catopuma_badia | Panthera_pardus | 0.137727677037286 |
| Catopuma_badia | Panthera_tigris | 0.14008311482929 |
| Catopuma_badia | Panthera_uncia | 0.137159754324906 |
| Catopuma_badia | Paradoxurus_hermaphroditus | 0.182872003719224 |
| Catopuma_badia | Parahyaena_brunnea_FD126_Fecal | 0.193798035826099 |
| Catopuma_badia | Pardofelis_marmorata | 0.109217054652096 |
| Catopuma_badia | Phoca_fasciata | 0.216869638252097 |
| Catopuma_badia | Phoca_groenlandica | 0.218523313936682 |
| Catopuma_badia | Phoca_largha | 0.216522789705477 |
| Catopuma_badia | Phoca_vitulina | 0.217870110909774 |
| Catopuma_badia | Phocarctos_hookeri | 0.236353703436954 |
| Catopuma_badia | Prionailurus_bengalensis | 0.106089919699107 |
| Catopuma_badia | Prionailurus_bengalensis_euptilurus | 0.104752825358812 |
| Catopuma_badia | Prionailurus_planiceps | 0.109457907150524 |
| Catopuma_badia | Prionailurus_rubiginosus | 0.107166720825003 |
| Catopuma_badia | Prionailurus_viverrinus | 0.109939497782796 |
| Catopuma_badia | Prionodon_pardicolor | 0.182989968578211 |
| Catopuma_badia | Procyon_lotor | 0.233429511446932 |
| Catopuma_badia | Profelis_aurata | 0.110049053628049 |
| Catopuma_badia | Proteles_cristatus_cristatus_TS307 | 0.200315994311039 |
| Catopuma_badia | Proteles_cristatus_septentrionalis_NMS_Z_2018_54 | 0.199574782824177 |
| Catopuma_badia | Puma_concolor | 0.0986387478210769 |
| Catopuma_badia | Puma_yagouaroundi | 0.101958999466103 |
| Catopuma_badia | Pusa_caspica | 0.219699519344828 |
| Catopuma_badia | Pusa_hispida | 0.21429854863829 |
| Catopuma_badia | Pusa_sibirica | 0.216470328401987 |
| Catopuma_badia | Speothos_venaticus_T1624 | 0.245512432982233 |
| Catopuma_badia | Spilogale_putorius | 0.243654981253003 |
| Catopuma_badia | Taxidea_taxus | 0.220685111675588 |
| Catopuma_badia | Tremarctos_ornatus | 0.236226861628278 |
| Catopuma_badia | Urocyon_cinereoargenteus | 0.242785039192477 |
| Catopuma_badia | Ursus_americanus | 0.235548336374717 |
| Catopuma_badia | Ursus_arctos | 0.242177201730585 |
| Catopuma_badia | Ursus_maritimus | 0.240113176761588 |
| Catopuma_badia | Ursus_spelaeus | 0.240225599471449 |
| Catopuma_badia | Ursus_thibetanus | 0.236105687898263 |
| Catopuma_badia | Urva_javanica | 0.187148569601808 |
| Catopuma_badia | Viverra_tangalunga_FMNH146957_SRR6053069 | 0.170602683671343 |
| Catopuma_badia | Viverricula_indica | 0.169578703904822 |
| Catopuma_badia | Vulpes_corsac | 0.238874441799588 |
| Catopuma_badia | Vulpes_ferrilata | 0.240662630175889 |
| Catopuma_badia | Vulpes_lagopus | 0.237324580459656 |
| Catopuma_badia | Vulpes_vulpes_T3611 | 0.23517570876999 |
| Catopuma_badia | Vulpes_zerda | 0.23287377785723 |

| | | |
|---|---|---|
| Catopuma_badia | Zalophus_californianus | 0.233093582234877 |
| Catopuma_temminckii | Chrysocyon_brachyurus | 0.233036270449652 |
| Catopuma_temminckii | Civettictis_civetta | 0.174782604585837 |
| Catopuma_temminckii | Conepatus_chinga | 0.239177170173009 |
| Catopuma_temminckii | Crocuta_crocuta | 0.205471057734214 |
| Catopuma_temminckii | Cuon_alpinus | 0.239987998143634 |
| Catopuma_temminckii | Cystophora_cristata | 0.21756548891121 |
| Catopuma_temminckii | Enhydra_lutris | 0.230433738090514 |
| Catopuma_temminckii | Erignathus_barbatus | 0.223074523864886 |
| Catopuma_temminckii | Eumetopias_jubatus | 0.236480350044649 |
| Catopuma_temminckii | Felis_catus | 0.111181463984773 |
| Catopuma_temminckii | Felis_chaus | 0.112222115860231 |
| Catopuma_temminckii | Felis_margarita | 0.111307345109373 |
| Catopuma_temminckii | Felis_nigripes | 0.11139240956168 |
| Catopuma_temminckii | Felis_silvestris | 0.110331407474078 |
| Catopuma_temminckii | Fossa_fossana_FMNH156648_SRR6053060 | 0.188491466906767 |
| Catopuma_temminckii | Genetta_servalina | 0.179131165112381 |
| Catopuma_temminckii | Gulo_gulo | 0.236997524858634 |
| Catopuma_temminckii | Halichoerus_grypus | 0.224783161570562 |
| Catopuma_temminckii | Helarctos_malayanus | 0.233679449676651 |
| Catopuma_temminckii | Hyaena_hyaena | 0.196645735581804 |
| Catopuma_temminckii | Hydrurga_leptonyx | 0.213135347853926 |
| Catopuma_temminckii | Leopardus_colocolo | 0.115427448246375 |
| Catopuma_temminckii | Leopardus_geoffroyi | 0.11718191102327 |
| Catopuma_temminckii | Leopardus_guigna | 0.116111110027151 |
| Catopuma_temminckii | Leopardus_jacobita | 0.115495542049025 |
| Catopuma_temminckii | Leopardus_pardalis | 0.114078252601193 |
| Catopuma_temminckii | Leopardus_tigrinus | 0.116889834681572 |
| Catopuma_temminckii | Leopardus_wiedii | 0.115769824555755 |
| Catopuma_temminckii | Leptailurus_serval | 0.108246469010461 |
| Catopuma_temminckii | Leptonychotes_weddellii | 0.21399613336353 |
| Catopuma_temminckii | Lobodon_carcinophaga | 0.215917160273034 |
| Catopuma_temminckii | Lutra_lutra | 0.23529102358058 |
| Catopuma_temminckii | Lutra_sumatrana | 0.231072722561724 |
| Catopuma_temminckii | Lutrogale_perspicillata | 0.243066352311403 |
| Catopuma_temminckii | Lycaon_pictus | 0.239967081084792 |
| Catopuma_temminckii | Lynx_canadensis | 0.103757799169655 |
| Catopuma_temminckii | Lynx_lynx | 0.103511002359142 |
| Catopuma_temminckii | Lynx_pardinus | 0.103326958922813 |
| Catopuma_temminckii | Lynx_rufus | 0.103272875667127 |
| Catopuma_temminckii | Manis_javanica | 0.285687522183671 |
| Catopuma_temminckii | Martes_americana | 0.233224167814559 |
| Catopuma_temminckii | Martes_flavigula | 0.24054290850963 |
| Catopuma_temminckii | Martes_foina | 0.232152721731481 |
| Catopuma_temminckii | Martes_martes | 0.231643792554375 |
| Catopuma_temminckii | Martes_melampus | 0.231523242819689 |
| Catopuma_temminckii | Martes_pennanti | 0.235848350954976 |
| Catopuma_temminckii | Martes_zibellina | 0.234947535448243 |
| Catopuma_temminckii | Meles_anakuma | 0.234233951607235 |
| Catopuma_temminckii | Meles_leucurus | 0.228264471135869 |
| Catopuma_temminckii | Meles_meles | 0.226972388465445 |
| Catopuma_temminckii | Melogale_moschata | 0.236279389731855 |
| Catopuma_temminckii | Melursus_ursinus | 0.235479921800937 |
| Catopuma_temminckii | Mephitis_mephitis | 0.238501585802579 |
| Catopuma_temminckii | Mirounga_leonina | 0.223548612927358 |
| Catopuma_temminckii | Mungotictis_decemlineata | 0.186530215618053 |
| Catopuma_temminckii | Mustela_altaica | 0.231476760939137 |
| Catopuma_temminckii | Mustela_erminea | 0.222877889167373 |
| Catopuma_temminckii | Mustela_eversmannii | 0.226581936806962 |
| Catopuma_temminckii | Mustela_frenata | 0.228768571018627 |
| Catopuma_temminckii | Mustela_itatsi | 0.229148143860243 |
| Catopuma_temminckii | Mustela_kathiah | 0.232176386283361 |
| Catopuma_temminckii | Mustela_nigripes | 0.224043287960859 |
| Catopuma_temminckii | Mustela_nivalis | 0.229048638056777 |
| Catopuma_temminckii | Mustela_putorius | 0.226917508981322 |
| Catopuma_temminckii | Mustela_sibirica | 0.227272490204548 |
| Catopuma_temminckii | Nandinia_binotata | 0.172995945092603 |
| Catopuma_temminckii | Nasua_nasua | 0.252091031316511 |
| Catopuma_temminckii | Neofelis_nebulosa | 0.141004563681888 |
| Catopuma_temminckii | Neomonachus_schauinslandi | 0.225514423247621 |
| Catopuma_temminckii | Neophoca_cinerea | 0.241395925550545 |
| Catopuma_temminckii | Neovison_vison | 0.234295440575234 |
| Catopuma_temminckii | Nyctereutes_procyonoides | 0.23293624996129 |
| Catopuma_temminckii | Odobenus_rosmarus_rosmarus | 0.258022868738316 |
| Catopuma_temminckii | Otocolobus_manul | 0.111080150094921 |
| Catopuma_temminckii | Otocyon_megalotis_megalotis_TS305 | 0.24518818201355 |
| Catopuma_temminckii | Otocyon_megalotis_virgatus_FMNH158128 | 0.244626309896616 |
| Catopuma_temminckii | Paguma_larvata | 0.179596108282966 |
| Catopuma_temminckii | Panthera_leo | 0.136637869788229 |
| Catopuma_temminckii | Panthera_onca | 0.138054044363945 |
| Catopuma_temminckii | Panthera_pardus | 0.136988391526232 |
| Catopuma_temminckii | Panthera_tigris | 0.139638380208154 |
| Catopuma_temminckii | Panthera_uncia | 0.140508648499435 |
| Catopuma_temminckii | Paradoxurus_hermaphroditus | 0.185604231824481 |
| Catopuma_temminckii | Parahyaena_brunnea_FD126_Fecal | 0.199001756818369 |
| Catopuma_temminckii | Pardofelis_marmorata | 0.110225127667938 |
| Catopuma_temminckii | Phoca_fasciata | 0.221784867797863 |

| | | |
|---|---|---|
| Catopuma_temminckii | Phoca_groenlandica | 0.225654811388176 |
| Catopuma_temminckii | Phoca_largha | 0.223942579638474 |
| Catopuma_temminckii | Phoca_vitulina | 0.224110701922147 |
| Catopuma_temminckii | Phocarctos_hookeri | 0.243132148639122 |
| Catopuma_temminckii | Prionailurus_bengalensis | 0.108048888239833 |
| Catopuma_temminckii | Prionailurus_bengalensis_euptilurus | 0.106755074495888 |
| Catopuma_temminckii | Prionailurus_planiceps | 0.111788261853599 |
| Catopuma_temminckii | Prionailurus_rubiginosus | 0.108925981899231 |
| Catopuma_temminckii | Prionailurus_viverrinus | 0.110421736026744 |
| Catopuma_temminckii | Prionodon_pardicolor | 0.185156631567779 |
| Catopuma_temminckii | Procyon_lotor | 0.237729113649871 |
| Catopuma_temminckii | Profelis_aurata | 0.110598227209548 |
| Catopuma_temminckii | Proteles_cristatus_cristatus_TS307 | 0.199108389401713 |
| Catopuma_temminckii | Proteles_cristatus_septentrionalis_NMS_Z_2018_54 | 0.199743377316457 |
| Catopuma_temminckii | Puma_concolor | 0.105160438075307 |
| Catopuma_temminckii | Puma_yagouaroundi | 0.106082134284778 |
| Catopuma_temminckii | Pusa_caspica | 0.226542245887455 |
| Catopuma_temminckii | Pusa_hispida | 0.222866866309373 |
| Catopuma_temminckii | Pusa_sibirica | 0.223252115347864 |
| Catopuma_temminckii | Speothos_venaticus_T1624 | 0.244622073153922 |
| Catopuma_temminckii | Spilogale_putorius | 0.244941292671685 |
| Catopuma_temminckii | Taxidea_taxus | 0.220219823415084 |
| Catopuma_temminckii | Tremarctos_ornatus | 0.234641535583512 |
| Catopuma_temminckii | Urocyon_cinereoargenteus | 0.242446730046668 |
| Catopuma_temminckii | Ursus_americanus | 0.233635534230703 |
| Catopuma_temminckii | Ursus_arctos | 0.240261490574807 |
| Catopuma_temminckii | Ursus_maritimus | 0.238429862412162 |
| Catopuma_temminckii | Ursus_spelaeus | 0.234480888692934 |
| Catopuma_temminckii | Ursus_thibetanus | 0.231863385662124 |
| Catopuma_temminckii | Urva_javanica | 0.193295519572932 |
| Catopuma_temminckii | Viverra_tangalunga_FMNH146957_SRR6053069 | 0.172617930853578 |
| Catopuma_temminckii | Viverricula_indica | 0.173728563203839 |
| Catopuma_temminckii | Vulpes_corsac | 0.239505176692252 |
| Catopuma_temminckii | Vulpes_ferrilata | 0.238938869865232 |
| Catopuma_temminckii | Vulpes_lagopus | 0.238731492953591 |
| Catopuma_temminckii | Vulpes_vulpes_T3611 | 0.234574482529488 |
| Catopuma_temminckii | Vulpes_zerda | 0.233593705260768 |
| Catopuma_temminckii | Zalophus_californianus | 0.235669319325242 |
| Chrysocyon_brachyurus | Civettictis_civetta | 0.236604060821105 |
| Chrysocyon_brachyurus | Conepatus_chinga | 0.247783942315981 |
| Chrysocyon_brachyurus | Crocuta_crocuta | 0.257277372302547 |
| Chrysocyon_brachyurus | Cuon_alpinus | 0.138084503405805 |
| Chrysocyon_brachyurus | Cystophora_cristata | 0.230458989937908 |
| Chrysocyon_brachyurus | Enhydra_lutris | 0.243961363326999 |
| Chrysocyon_brachyurus | Erignathus_barbatus | 0.228285112602996 |
| Chrysocyon_brachyurus | Eumetopias_jubatus | 0.245095161469431 |
| Chrysocyon_brachyurus | Felis_catus | 0.235678096205678 |
| Chrysocyon_brachyurus | Felis_chaus | 0.235507086228357 |
| Chrysocyon_brachyurus | Felis_margarita | 0.236229485763101 |
| Chrysocyon_brachyurus | Felis_nigripes | 0.239026348000895 |
| Chrysocyon_brachyurus | Felis_silvestris | 0.234247322612147 |
| Chrysocyon_brachyurus | Fossa_fossana_FMNH156648_SRR6053060 | 0.23989206954479 |
| Chrysocyon_brachyurus | Genetta_servalina | 0.240941895950776 |
| Chrysocyon_brachyurus | Gulo_gulo | 0.242189784294988 |
| Chrysocyon_brachyurus | Halichoerus_grypus | 0.232757657281662 |
| Chrysocyon_brachyurus | Helarctos_malayanus | 0.248107027718979 |
| Chrysocyon_brachyurus | Hyaena_hyaena | 0.253945236918664 |
| Chrysocyon_brachyurus | Hydrurga_leptonyx | 0.225940261989544 |
| Chrysocyon_brachyurus | Leopardus_colocolo | 0.236718183034766 |
| Chrysocyon_brachyurus | Leopardus_geoffroyi | 0.236623545591959 |
| Chrysocyon_brachyurus | Leopardus_guigna | 0.23949870556579 |
| Chrysocyon_brachyurus | Leopardus_jacobita | 0.236393337490717 |
| Chrysocyon_brachyurus | Leopardus_pardalis | 0.231932239049136 |
| Chrysocyon_brachyurus | Leopardus_tigrinus | 0.236268212322789 |
| Chrysocyon_brachyurus | Leopardus_wiedii | 0.234160542155796 |
| Chrysocyon_brachyurus | Leptailurus_serval | 0.235717665445064 |
| Chrysocyon_brachyurus | Leptonychotes_weddellii | 0.224332158615321 |
| Chrysocyon_brachyurus | Lobodon_carcinophaga | 0.221394823049365 |
| Chrysocyon_brachyurus | Lutra_lutra | 0.248050682913126 |
| Chrysocyon_brachyurus | Lutra_sumatrana | 0.247488821970756 |
| Chrysocyon_brachyurus | Lutrogale_perspicillata | 0.263337443032816 |
| Chrysocyon_brachyurus | Lycaon_pictus | 0.134085652480143 |
| Chrysocyon_brachyurus | Lynx_canadensis | 0.240763542835552 |
| Chrysocyon_brachyurus | Lynx_lynx | 0.240195068932487 |
| Chrysocyon_brachyurus | Lynx_pardinus | 0.242533342874438 |
| Chrysocyon_brachyurus | Lynx_rufus | 0.237736351526699 |
| Chrysocyon_brachyurus | Manis_javanica | 0.307657954022597 |
| Chrysocyon_brachyurus | Martes_americana | 0.245153941801638 |
| Chrysocyon_brachyurus | Martes_flavigula | 0.239127069289081 |
| Chrysocyon_brachyurus | Martes_foina | 0.240778075193111 |
| Chrysocyon_brachyurus | Martes_martes | 0.242536730947649 |
| Chrysocyon_brachyurus | Martes_melampus | 0.243467182622179 |
| Chrysocyon_brachyurus | Martes_pennanti | 0.243569704749772 |
| Chrysocyon_brachyurus | Martes_zibellina | 0.245559447728239 |
| Chrysocyon_brachyurus | Meles_anakuma | 0.246583374641158 |
| Chrysocyon_brachyurus | Meles_leucurus | 0.242580588521434 |
| Chrysocyon_brachyurus | Meles_meles | 0.241447189070151 |

| | | |
|---|---|---|
| Chrysocyon_brachyurus | Melogale_moschata | 0.241074267617672 |
| Chrysocyon_brachyurus | Melursus_ursinus | 0.252869159649302 |
| Chrysocyon_brachyurus | Mephitis_mephitis | 0.244691871335549 |
| Chrysocyon_brachyurus | Mirounga_leonina | 0.229234700856636 |
| Chrysocyon_brachyurus | Mungotictis_decemlineata | 0.24835839679223 |
| Chrysocyon_brachyurus | Mustela_altaica | 0.244375028068977 |
| Chrysocyon_brachyurus | Mustela_erminea | 0.239150465863075 |
| Chrysocyon_brachyurus | Mustela_eversmannii | 0.237998065820047 |
| Chrysocyon_brachyurus | Mustela_frenata | 0.23324515008797 |
| Chrysocyon_brachyurus | Mustela_itatsi | 0.241478113604772 |
| Chrysocyon_brachyurus | Mustela_kathiah | 0.23090476334585 |
| Chrysocyon_brachyurus | Mustela_nigripes | 0.236117537819195 |
| Chrysocyon_brachyurus | Mustela_nivalis | 0.241421539244075 |
| Chrysocyon_brachyurus | Mustela_putorius | 0.236246061340707 |
| Chrysocyon_brachyurus | Mustela_sibirica | 0.238185095457779 |
| Chrysocyon_brachyurus | Nandinia_binotata | 0.230898758675895 |
| Chrysocyon_brachyurus | Nasua_nasua | 0.263647420529492 |
| Chrysocyon_brachyurus | Neofelis_nebulosa | 0.255638404079184 |
| Chrysocyon_brachyurus | Neomonachus_schauinslandi | 0.232276181624619 |
| Chrysocyon_brachyurus | Neophoca_cinerea | 0.247802987105751 |
| Chrysocyon_brachyurus | Neovison_vison | 0.23284091792322 |
| Chrysocyon_brachyurus | Nyctereutes_procyonoides | 0.175222374227279 |
| Chrysocyon_brachyurus | Odobenus_rosmarus_rosmarus | 0.266393876244629 |
| Chrysocyon_brachyurus | Otocolobus_manul | 0.234509288741996 |
| Chrysocyon_brachyurus | Otocyon_megalotis_megalotis_TS305 | 0.180942928146668 |
| Chrysocyon_brachyurus | Otocyon_megalotis_virgatus_FMNH158128 | 0.180163202347188 |
| Chrysocyon_brachyurus | Paguma_larvata | 0.234682334041832 |
| Chrysocyon_brachyurus | Panthera_leo | 0.24438113128634 |
| Chrysocyon_brachyurus | Panthera_onca | 0.248351399100329 |
| Chrysocyon_brachyurus | Panthera_pardus | 0.251576247444137 |
| Chrysocyon_brachyurus | Panthera_tigris | 0.244285090983024 |
| Chrysocyon_brachyurus | Panthera_uncia | 0.246448365541778 |
| Chrysocyon_brachyurus | Paradoxurus_hermaphroditus | 0.235294971985251 |
| Chrysocyon_brachyurus | Parahyaena_brunnea_FD126_Fecal | 0.252140615830206 |
| Chrysocyon_brachyurus | Pardofelis_marmorata | 0.243336311663711 |
| Chrysocyon_brachyurus | Phoca_fasciata | 0.229274516279676 |
| Chrysocyon_brachyurus | Phoca_groenlandica | 0.231047242143199 |
| Chrysocyon_brachyurus | Phoca_largha | 0.234060515917483 |
| Chrysocyon_brachyurus | Phoca_vitulina | 0.235113448596282 |
| Chrysocyon_brachyurus | Phocarctos_hookeri | 0.257562387312037 |
| Chrysocyon_brachyurus | Prionailurus_bengalensis | 0.235818017488377 |
| Chrysocyon_brachyurus | Prionailurus_bengalensis_euptilurus | 0.237273872519013 |
| Chrysocyon_brachyurus | Prionailurus_planiceps | 0.236822322936387 |
| Chrysocyon_brachyurus | Prionailurus_rubiginosus | 0.232225714853805 |
| Chrysocyon_brachyurus | Prionailurus_viverrinus | 0.238673050619434 |
| Chrysocyon_brachyurus | Prionodon_pardicolor | 0.235505177957852 |
| Chrysocyon_brachyurus | Procyon_lotor | 0.245402618693236 |
| Chrysocyon_brachyurus | Profelis_aurata | 0.238180048832229 |
| Chrysocyon_brachyurus | Proteles_cristatus_cristatus_TS307 | 0.257035927362418 |
| Chrysocyon_brachyurus | Proteles_cristatus_septentrionalis_NMS_Z_2018_54 | 0.254186916805099 |
| Chrysocyon_brachyurus | Puma_concolor | 0.239752246091076 |
| Chrysocyon_brachyurus | Puma_yagouaroundi | 0.23951888362815 |
| Chrysocyon_brachyurus | Pusa_caspica | 0.233350461592715 |
| Chrysocyon_brachyurus | Pusa_hispida | 0.230339804454896 |
| Chrysocyon_brachyurus | Pusa_sibirica | 0.232489248917327 |
| Chrysocyon_brachyurus | Speothos_venaticus_T1624 | 0.139807146798374 |
| Chrysocyon_brachyurus | Spilogale_putorius | 0.238120196151193 |
| Chrysocyon_brachyurus | Taxidea_taxus | 0.238296264540366 |
| Chrysocyon_brachyurus | Tremarctos_ornatus | 0.247397804462659 |
| Chrysocyon_brachyurus | Urocyon_cinereoargenteus | 0.189000239416137 |
| Chrysocyon_brachyurus | Ursus_americanus | 0.252053788305436 |
| Chrysocyon_brachyurus | Ursus_arctos | 0.252652072926573 |
| Chrysocyon_brachyurus | Ursus_maritimus | 0.25476843324339 |
| Chrysocyon_brachyurus | Ursus_spelaeus | 0.25117097581012 |
| Chrysocyon_brachyurus | Ursus_thibetanus | 0.246876687066569 |
| Chrysocyon_brachyurus | Urva_javanica | 0.240721702600087 |
| Chrysocyon_brachyurus | Viverra_tangalunga_FMNH146957_SRR6053069 | 0.230796157952212 |
| Chrysocyon_brachyurus | Viverricula_indica | 0.232682505179626 |
| Chrysocyon_brachyurus | Vulpes_corsac | 0.180578363765926 |
| Chrysocyon_brachyurus | Vulpes_ferrilata | 0.184503563127008 |
| Chrysocyon_brachyurus | Vulpes_lagopus | 0.182699288231745 |
| Chrysocyon_brachyurus | Vulpes_vulpes_T3611 | 0.182815814797271 |
| Chrysocyon_brachyurus | Vulpes_zerda | 0.18085663436452 |
| Chrysocyon_brachyurus | Zalophus_californianus | 0.248242682512017 |
| Civettictis_civetta | Conepatus_chinga | 0.237010351783144 |
| Civettictis_civetta | Crocuta_crocuta | 0.203212318595631 |
| Civettictis_civetta | Cuon_alpinus | 0.240812590179414 |
| Civettictis_civetta | Cystophora_cristata | 0.21709451894574 |
| Civettictis_civetta | Enhydra_lutris | 0.229864461868299 |
| Civettictis_civetta | Erignathus_barbatus | 0.220750252653193 |
| Civettictis_civetta | Eumetopias_jubatus | 0.23587778647774 |
| Civettictis_civetta | Felis_catus | 0.175798137751212 |
| Civettictis_civetta | Felis_chaus | 0.17369710552119 |
| Civettictis_civetta | Felis_margarita | 0.174485452933565 |
| Civettictis_civetta | Felis_nigripes | 0.175685172239617 |
| Civettictis_civetta | Felis_silvestris | 0.175696335556991 |
| Civettictis_civetta | Fossa_fossana_FMNH156648_SRR6053060 | 0.177823584338194 |

| | | |
|---|---|---|
| Civettictis_civetta | Genetta_servalina | 0.146618263848663 |
| Civettictis_civetta | Gulo_gulo | 0.234518710282203 |
| Civettictis_civetta | Halichoerus_grypus | 0.224690196428926 |
| Civettictis_civetta | Helarctos_malayanus | 0.234992771624092 |
| Civettictis_civetta | Hyaena_hyaena | 0.200082421382345 |
| Civettictis_civetta | Hydrurga_leptonyx | 0.214105479887927 |
| Civettictis_civetta | Leopardus_colocolo | 0.169907067427407 |
| Civettictis_civetta | Leopardus_geoffroyi | 0.174977372203206 |
| Civettictis_civetta | Leopardus_guigna | 0.176455311538995 |
| Civettictis_civetta | Leopardus_jacobita | 0.178567617941274 |
| Civettictis_civetta | Leopardus_pardalis | 0.174536602105963 |
| Civettictis_civetta | Leopardus_tigrinus | 0.178142614520429 |
| Civettictis_civetta | Leopardus_wiedii | 0.174741604158855 |
| Civettictis_civetta | Leptailurus_serval | 0.172419314097917 |
| Civettictis_civetta | Leptonychotes_weddellii | 0.211435208079633 |
| Civettictis_civetta | Lobodon_carcinophaga | 0.214770845262909 |
| Civettictis_civetta | Lutra_lutra | 0.236081130275635 |
| Civettictis_civetta | Lutra_sumatrana | 0.235989188893976 |
| Civettictis_civetta | Lutrogale_perspicillata | 0.241369981307135 |
| Civettictis_civetta | Lycaon_pictus | 0.234361001413787 |
| Civettictis_civetta | Lynx_canadensis | 0.180451015290245 |
| Civettictis_civetta | Lynx_lynx | 0.177885629773345 |
| Civettictis_civetta | Lynx_pardinus | 0.176013660420244 |
| Civettictis_civetta | Lynx_rufus | 0.174895928356213 |
| Civettictis_civetta | Manis_javanica | 0.293596396410142 |
| Civettictis_civetta | Martes_americana | 0.23412342994454 |
| Civettictis_civetta | Martes_flavigula | 0.235931778140475 |
| Civettictis_civetta | Martes_foina | 0.23223797572856 |
| Civettictis_civetta | Martes_martes | 0.231990449636422 |
| Civettictis_civetta | Martes_melampus | 0.232289687040076 |
| Civettictis_civetta | Martes_pennanti | 0.239266532382534 |
| Civettictis_civetta | Martes_zibellina | 0.233744716319595 |
| Civettictis_civetta | Meles_anakuma | 0.233374860414653 |
| Civettictis_civetta | Meles_leucurus | 0.228062355059641 |
| Civettictis_civetta | Meles_meles | 0.226069968554126 |
| Civettictis_civetta | Melogale_moschata | 0.232283910117209 |
| Civettictis_civetta | Melursus_ursinus | 0.241933354045951 |
| Civettictis_civetta | Mephitis_mephitis | 0.236574941058132 |
| Civettictis_civetta | Mirounga_leonina | 0.221126891019221 |
| Civettictis_civetta | Mungotictis_decemlineata | 0.183969665413985 |
| Civettictis_civetta | Mustela_altaica | 0.231298944303853 |
| Civettictis_civetta | Mustela_erminea | 0.224614532396187 |
| Civettictis_civetta | Mustela_eversmannii | 0.232464144693912 |
| Civettictis_civetta | Mustela_frenata | 0.22768508382562 |
| Civettictis_civetta | Mustela_itatsi | 0.234965831118517 |
| Civettictis_civetta | Mustela_kathiah | 0.229126363647892 |
| Civettictis_civetta | Mustela_nigripes | 0.233673876173546 |
| Civettictis_civetta | Mustela_nivalis | 0.229733274480412 |
| Civettictis_civetta | Mustela_putorius | 0.232144687481184 |
| Civettictis_civetta | Mustela_sibirica | 0.231516106448789 |
| Civettictis_civetta | Nandinia_binotata | 0.174586097602185 |
| Civettictis_civetta | Nasua_nasua | 0.257571922204967 |
| Civettictis_civetta | Neofelis_nebulosa | 0.18644213346916 |
| Civettictis_civetta | Neomonachus_schauinslandi | 0.220977848453299 |
| Civettictis_civetta | Neophoca_cinerea | 0.239801799389288 |
| Civettictis_civetta | Neovison_vison | 0.228966244455001 |
| Civettictis_civetta | Nyctereutes_procyonoides | 0.236402931203771 |
| Civettictis_civetta | Odobenus_rosmarus_rosmarus | 0.256996517753982 |
| Civettictis_civetta | Otocolobus_manul | 0.17204599777629 |
| Civettictis_civetta | Otocyon_megalotis_megalotis_TS305 | 0.245898215986641 |
| Civettictis_civetta | Otocyon_megalotis_virgatus_FMNH158128 | 0.244456864108071 |
| Civettictis_civetta | Paguma_larvata | 0.161251929676871 |
| Civettictis_civetta | Panthera_leo | 0.179773153022309 |
| Civettictis_civetta | Panthera_onca | 0.181312578288636 |
| Civettictis_civetta | Panthera_pardus | 0.182797488993601 |
| Civettictis_civetta | Panthera_tigris | 0.186411175251115 |
| Civettictis_civetta | Panthera_uncia | 0.18045486446088 |
| Civettictis_civetta | Paradoxurus_hermaphroditus | 0.163985301429217 |
| Civettictis_civetta | Parahyaena_brunnea_FD126_Fecal | 0.199968818874404 |
| Civettictis_civetta | Pardofelis_marmorata | 0.179333576135935 |
| Civettictis_civetta | Phoca_fasciata | 0.221103862927643 |
| Civettictis_civetta | Phoca_groenlandica | 0.217593829173986 |
| Civettictis_civetta | Phoca_largha | 0.224083914887549 |
| Civettictis_civetta | Phoca_vitulina | 0.225975137000229 |
| Civettictis_civetta | Phocarctos_hookeri | 0.23814447741479 |
| Civettictis_civetta | Prionailurus_bengalensis | 0.171859243537688 |
| Civettictis_civetta | Prionailurus_bengalensis_euptilurus | 0.172225272691142 |
| Civettictis_civetta | Prionailurus_planiceps | 0.17332981714158 |
| Civettictis_civetta | Prionailurus_rubiginosus | 0.174538384777388 |
| Civettictis_civetta | Prionailurus_viverrinus | 0.176466627424178 |
| Civettictis_civetta | Prionodon_pardicolor | 0.187855155051543 |
| Civettictis_civetta | Procyon_lotor | 0.236299920368028 |
| Civettictis_civetta | Profelis_aurata | 0.177496281824297 |
| Civettictis_civetta | Proteles_cristatus_cristatus_TS307 | 0.203474535312315 |
| Civettictis_civetta | Proteles_cristatus_septentrionalis_NMS_Z_2018_54 | 0.200877804722454 |
| Civettictis_civetta | Puma_concolor | 0.172013340628565 |
| Civettictis_civetta | Puma_yagouaroundi | 0.176044887081527 |

| | | |
|---|---|---|
| Civettictis_civetta | Pusa_caspica | 0.22523091617982 |
| Civettictis_civetta | Pusa_hispida | 0.219236812732164 |
| Civettictis_civetta | Pusa_sibirica | 0.223274381934666 |
| Civettictis_civetta | Speothos_venaticus_T1624 | 0.245360978396716 |
| Civettictis_civetta | Spilogale_putorius | 0.243454021073203 |
| Civettictis_civetta | Taxidea_taxus | 0.218066982466281 |
| Civettictis_civetta | Tremarctos_ornatus | 0.239423207306259 |
| Civettictis_civetta | Urocyon_cinereoargenteus | 0.249061729697401 |
| Civettictis_civetta | Ursus_americanus | 0.235981384987581 |
| Civettictis_civetta | Ursus_arctos | 0.241631606726108 |
| Civettictis_civetta | Ursus_maritimus | 0.239563681364002 |
| Civettictis_civetta | Ursus_spelaeus | 0.237725961651825 |
| Civettictis_civetta | Ursus_thibetanus | 0.234536244475112 |
| Civettictis_civetta | Urva_javanica | 0.19322264936773 |
| Civettictis_civetta | Viverra_tangalunga_FMNH146957_SRR6053069 | 0.10937015884428 |
| Civettictis_civetta | Viverricula_indica | 0.127553514653705 |
| Civettictis_civetta | Vulpes_corsac | 0.231595310745365 |
| Civettictis_civetta | Vulpes_ferrilata | 0.233921923449237 |
| Civettictis_civetta | Vulpes_lagopus | 0.235518811280234 |
| Civettictis_civetta | Vulpes_vulpes_T3611 | 0.228809393559018 |
| Civettictis_civetta | Vulpes_zerda | 0.229861227852126 |
| Civettictis_civetta | Zalophus_californianus | 0.235990718714001 |
| Conepatus_chinga | Crocuta_crocuta | 0.256944276733585 |
| Conepatus_chinga | Cuon_alpinus | 0.250849397346161 |
| Conepatus_chinga | Cystophora_cristata | 0.227590608721767 |
| Conepatus_chinga | Enhydra_lutris | 0.232666776701465 |
| Conepatus_chinga | Erignathus_barbatus | 0.218177850102609 |
| Conepatus_chinga | Eumetopias_jubatus | 0.233550281956215 |
| Conepatus_chinga | Felis_catus | 0.237658622226065 |
| Conepatus_chinga | Felis_chaus | 0.237993286655957 |
| Conepatus_chinga | Felis_margarita | 0.237327459834005 |
| Conepatus_chinga | Felis_nigripes | 0.241801078341276 |
| Conepatus_chinga | Felis_silvestris | 0.235254227105112 |
| Conepatus_chinga | Fossa_fossana_FMNH156648_SRR6053060 | 0.238944504666666 |
| Conepatus_chinga | Genetta_servalina | 0.232601501507952 |
| Conepatus_chinga | Gulo_gulo | 0.228904864731009 |
| Conepatus_chinga | Halichoerus_grypus | 0.227809955073168 |
| Conepatus_chinga | Helarctos_malayanus | 0.244921465446913 |
| Conepatus_chinga | Hyaena_hyaena | 0.250987646945752 |
| Conepatus_chinga | Hydrurga_leptonyx | 0.21590131510317 |
| Conepatus_chinga | Leopardus_colocolo | 0.237960624560688 |
| Conepatus_chinga | Leopardus_geoffroyi | 0.235125298159253 |
| Conepatus_chinga | Leopardus_guigna | 0.237418344139287 |
| Conepatus_chinga | Leopardus_jacobita | 0.235923767494154 |
| Conepatus_chinga | Leopardus_pardalis | 0.238884684106918 |
| Conepatus_chinga | Leopardus_tigrinus | 0.236550801378585 |
| Conepatus_chinga | Leopardus_wiedii | 0.238019512896419 |
| Conepatus_chinga | Leptailurus_serval | 0.237607064225957 |
| Conepatus_chinga | Leptonychotes_weddellii | 0.214323247143363 |
| Conepatus_chinga | Lobodon_carcinophaga | 0.217180244276139 |
| Conepatus_chinga | Lutra_lutra | 0.235422350940078 |
| Conepatus_chinga | Lutra_sumatrana | 0.234987414422638 |
| Conepatus_chinga | Lutrogale_perspicillata | 0.253800720592718 |
| Conepatus_chinga | Lycaon_pictus | 0.241040570895924 |
| Conepatus_chinga | Lynx_canadensis | 0.237557983494365 |
| Conepatus_chinga | Lynx_lynx | 0.239600403589931 |
| Conepatus_chinga | Lynx_pardinus | 0.240245694351754 |
| Conepatus_chinga | Lynx_rufus | 0.239147516435697 |
| Conepatus_chinga | Manis_javanica | 0.296677440206958 |
| Conepatus_chinga | Martes_americana | 0.23307520865996 |
| Conepatus_chinga | Martes_flavigula | 0.23915697438026 |
| Conepatus_chinga | Martes_foina | 0.230708748452644 |
| Conepatus_chinga | Martes_martes | 0.230777709520491 |
| Conepatus_chinga | Martes_melampus | 0.232425892965137 |
| Conepatus_chinga | Martes_pennanti | 0.239940702315476 |
| Conepatus_chinga | Martes_zibellina | 0.232431184597832 |
| Conepatus_chinga | Meles_anakuma | 0.233848089622007 |
| Conepatus_chinga | Meles_leucurus | 0.231487884626827 |
| Conepatus_chinga | Meles_meles | 0.22969991104998 |
| Conepatus_chinga | Melogale_moschata | 0.235775028735793 |
| Conepatus_chinga | Melursus_ursinus | 0.246439321276524 |
| Conepatus_chinga | Mephitis_mephitis | 0.184267862555883 |
| Conepatus_chinga | Mirounga_leonina | 0.218916377889704 |
| Conepatus_chinga | Mungotictis_decemlineata | 0.244526695840459 |
| Conepatus_chinga | Mustela_altaica | 0.239046803883302 |
| Conepatus_chinga | Mustela_erminea | 0.231023246626168 |
| Conepatus_chinga | Mustela_eversmannii | 0.234888931932667 |
| Conepatus_chinga | Mustela_frenata | 0.225412753933882 |
| Conepatus_chinga | Mustela_itatsi | 0.234922379561923 |
| Conepatus_chinga | Mustela_kathiah | 0.22542310023913 |
| Conepatus_chinga | Mustela_nigripes | 0.234234840088068 |
| Conepatus_chinga | Mustela_nivalis | 0.239259198158305 |
| Conepatus_chinga | Mustela_putorius | 0.232815712735489 |
| Conepatus_chinga | Mustela_sibirica | 0.237006489540111 |
| Conepatus_chinga | Nandinia_binotata | 0.227153946810056 |
| Conepatus_chinga | Nasua_nasua | 0.241878646258777 |
| Conepatus_chinga | Neofelis_nebulosa | 0.254159356180584 |

| | | |
|---|---|---|
| Conepatus_chinga | Neomonachus_schauinslandi | 0.218825532103826 |
| Conepatus_chinga | Neophoca_cinerea | 0.239402770838521 |
| Conepatus_chinga | Neovison_vison | 0.227622366097553 |
| Conepatus_chinga | Nyctereutes_procyonoides | 0.245096007392887 |
| Conepatus_chinga | Odobenus_rosmarus_rosmarus | 0.253694557311583 |
| Conepatus_chinga | Otocolobus_manul | 0.240974624253131 |
| Conepatus_chinga | Otocyon_megalotis_megalotis_TS305 | 0.258135584669321 |
| Conepatus_chinga | Otocyon_megalotis_virgatus_FMNH158128 | 0.258918624257547 |
| Conepatus_chinga | Paguma_larvata | 0.232813633324699 |
| Conepatus_chinga | Panthera_leo | 0.244658910328747 |
| Conepatus_chinga | Panthera_onca | 0.249112168082646 |
| Conepatus_chinga | Panthera_pardus | 0.251340578170108 |
| Conepatus_chinga | Panthera_tigris | 0.255414583834681 |
| Conepatus_chinga | Panthera_uncia | 0.253566576294145 |
| Conepatus_chinga | Paradoxurus_hermaphroditus | 0.230628741168158 |
| Conepatus_chinga | Parahyaena_brunnea_FD126_Fecal | 0.25054656064021 |
| Conepatus_chinga | Pardofelis_marmorata | 0.247471249396669 |
| Conepatus_chinga | Phoca_fasciata | 0.221597383510012 |
| Conepatus_chinga | Phoca_groenlandica | 0.222660554536482 |
| Conepatus_chinga | Phoca_largha | 0.225470363898957 |
| Conepatus_chinga | Phoca_vitulina | 0.224152185813048 |
| Conepatus_chinga | Phocarctos_hookeri | 0.243652546554147 |
| Conepatus_chinga | Prionailurus_bengalensis | 0.236503498172386 |
| Conepatus_chinga | Prionailurus_bengalensis_euptilurus | 0.237604698925843 |
| Conepatus_chinga | Prionailurus_planiceps | 0.240701026414606 |
| Conepatus_chinga | Prionailurus_rubiginosus | 0.235276416691254 |
| Conepatus_chinga | Prionailurus_viverrinus | 0.239886050944131 |
| Conepatus_chinga | Prionodon_pardicolor | 0.236158177111077 |
| Conepatus_chinga | Procyon_lotor | 0.229241193814641 |
| Conepatus_chinga | Profelis_aurata | 0.24141701718351 |
| Conepatus_chinga | Proteles_cristatus_cristatus_TS307 | 0.249348480488162 |
| Conepatus_chinga | Proteles_cristatus_septentrionalis_NMS_Z_2018_54 | 0.252923206129016 |
| Conepatus_chinga | Puma_concolor | 0.239693914977579 |
| Conepatus_chinga | Puma_yagouaroundi | 0.239746225850727 |
| Conepatus_chinga | Pusa_caspica | 0.227763981406347 |
| Conepatus_chinga | Pusa_hispida | 0.22241516934518 |
| Conepatus_chinga | Pusa_sibirica | 0.226095317401983 |
| Conepatus_chinga | Speothos_venaticus_T1624 | 0.25206743760278 |
| Conepatus_chinga | Spilogale_putorius | 0.183448506309057 |
| Conepatus_chinga | Taxidea_taxus | 0.223235895135651 |
| Conepatus_chinga | Tremarctos_ornatus | 0.24255043330234 |
| Conepatus_chinga | Urocyon_cinereoargenteus | 0.261859115888321 |
| Conepatus_chinga | Ursus_americanus | 0.243532090685437 |
| Conepatus_chinga | Ursus_arctos | 0.246856912363183 |
| Conepatus_chinga | Ursus_maritimus | 0.24737917662597 |
| Conepatus_chinga | Ursus_spelaeus | 0.243747739219893 |
| Conepatus_chinga | Ursus_thibetanus | 0.240802556259368 |
| Conepatus_chinga | Urva_javanica | 0.24727266869386 |
| Conepatus_chinga | Viverra_tangalunga_FMNH146957_SRR6053069 | 0.232751123450607 |
| Conepatus_chinga | Viverricula_indica | 0.228966369791175 |
| Conepatus_chinga | Vulpes_corsac | 0.251522976719011 |
| Conepatus_chinga | Vulpes_ferrilata | 0.254602907543362 |
| Conepatus_chinga | Vulpes_lagopus | 0.249342558240292 |
| Conepatus_chinga | Vulpes_vulpes_T3611 | 0.24926860258581 |
| Conepatus_chinga | Vulpes_zerda | 0.24926860258581 |
| Conepatus_chinga | Zalophus_californianus | 0.234804368648929 |
| Crocuta_crocuta | Cuon_alpinus | 0.262572355138836 |
| Crocuta_crocuta | Cystophora_cristata | 0.226671727595392 |
| Crocuta_crocuta | Enhydra_lutris | 0.239501529911986 |
| Crocuta_crocuta | Erignathus_barbatus | 0.230919314635632 |
| Crocuta_crocuta | Eumetopias_jubatus | 0.240850978593651 |
| Crocuta_crocuta | Felis_catus | 0.203956455964891 |
| Crocuta_crocuta | Felis_chaus | 0.204439521151065 |
| Crocuta_crocuta | Felis_margarita | 0.202973930626576 |
| Crocuta_crocuta | Felis_nigripes | 0.202219410380603 |
| Crocuta_crocuta | Felis_silvestris | 0.202336897065885 |
| Crocuta_crocuta | Fossa_fossana_FMNH156648_SRR6053060 | 0.185457100879987 |
| Crocuta_crocuta | Genetta_servalina | 0.200966431172206 |
| Crocuta_crocuta | Gulo_gulo | 0.245299105250385 |
| Crocuta_crocuta | Halichoerus_grypus | 0.228043603995067 |
| Crocuta_crocuta | Helarctos_malayanus | 0.24712494361454 |
| Crocuta_crocuta | Hyaena_hyaena | 0.107872020900959 |
| Crocuta_crocuta | Hydrurga_leptonyx | 0.224025109742321 |
| Crocuta_crocuta | Leopardus_colocolo | 0.194854553095434 |
| Crocuta_crocuta | Leopardus_geoffroyi | 0.194864858847438 |
| Crocuta_crocuta | Leopardus_guigna | 0.195539127010644 |
| Crocuta_crocuta | Leopardus_jacobita | 0.198927838027881 |
| Crocuta_crocuta | Leopardus_pardalis | 0.192712775052078 |
| Crocuta_crocuta | Leopardus_tigrinus | 0.197901406366516 |
| Crocuta_crocuta | Leopardus_wiedii | 0.195273936613279 |
| Crocuta_crocuta | Leptailurus_serval | 0.199596143716166 |
| Crocuta_crocuta | Leptonychotes_weddellii | 0.21796931696487 |
| Crocuta_crocuta | Lobodon_carcinophaga | 0.226631173677289 |
| Crocuta_crocuta | Lutra_lutra | 0.243304870881128 |
| Crocuta_crocuta | Lutra_sumatrana | 0.239832520151464 |
| Crocuta_crocuta | Lutrogale_perspicillata | 0.249479158879682 |
| Crocuta_crocuta | Lycaon_pictus | 0.249887275012006 |

| | | |
|---|---|---|
| Crocuta_crocuta | Lynx_canadensis | 0.190687548095539 |
| Crocuta_crocuta | Lynx_lynx | 0.193139768988733 |
| Crocuta_crocuta | Lynx_pardinus | 0.194772563150702 |
| Crocuta_crocuta | Lynx_rufus | 0.199870309808478 |
| Crocuta_crocuta | Manis_javanica | 0.280101593916874 |
| Crocuta_crocuta | Martes_americana | 0.24217563099349 |
| Crocuta_crocuta | Martes_flavigula | 0.246496473383627 |
| Crocuta_crocuta | Martes_foina | 0.243619276617516 |
| Crocuta_crocuta | Martes_martes | 0.242654915007502 |
| Crocuta_crocuta | Martes_melampus | 0.243453438939572 |
| Crocuta_crocuta | Martes_pennanti | 0.246459116764554 |
| Crocuta_crocuta | Martes_zibellina | 0.244751801567079 |
| Crocuta_crocuta | Meles_anakuma | 0.247315682362027 |
| Crocuta_crocuta | Meles_leucurus | 0.242000274491703 |
| Crocuta_crocuta | Meles_meles | 0.247703141513603 |
| Crocuta_crocuta | Melogale_moschata | 0.246891458254097 |
| Crocuta_crocuta | Melursus_ursinus | 0.250520079608665 |
| Crocuta_crocuta | Mephitis_mephitis | 0.254363819944361 |
| Crocuta_crocuta | Mirounga_leonina | 0.225898894929809 |
| Crocuta_crocuta | Mungotictis_decemlineata | 0.189415558593938 |
| Crocuta_crocuta | Mustela_altaica | 0.248152590883257 |
| Crocuta_crocuta | Mustela_erminea | 0.235847212327516 |
| Crocuta_crocuta | Mustela_eversmannii | 0.249149286265656 |
| Crocuta_crocuta | Mustela_frenata | 0.24317626868094 |
| Crocuta_crocuta | Mustela_itatsi | 0.247499300874713 |
| Crocuta_crocuta | Mustela_kathiah | 0.247529986805622 |
| Crocuta_crocuta | Mustela_nigripes | 0.244029396641932 |
| Crocuta_crocuta | Mustela_nivalis | 0.246093043060128 |
| Crocuta_crocuta | Mustela_putorius | 0.245564134677109 |
| Crocuta_crocuta | Mustela_sibirica | 0.245798533130083 |
| Crocuta_crocuta | Nandinia_binotata | 0.19177940685426 |
| Crocuta_crocuta | Nasua_nasua | 0.258571234797483 |
| Crocuta_crocuta | Neofelis_nebulosa | 0.206245280917413 |
| Crocuta_crocuta | Neomonachus_schauinslandi | 0.231968875774979 |
| Crocuta_crocuta | Neophoca_cinerea | 0.244136137218594 |
| Crocuta_crocuta | Neovison_vison | 0.245770211751232 |
| Crocuta_crocuta | Nyctereutes_procyonoides | 0.250060246973605 |
| Crocuta_crocuta | Odobenus_rosmarus_rosmarus | 0.254889765667551 |
| Crocuta_crocuta | Otocolobus_manul | 0.19852811912821 |
| Crocuta_crocuta | Otocyon_megalotis_megalotis_TS305 | 0.256132424103117 |
| Crocuta_crocuta | Otocyon_megalotis_virgatus_FMNH158128 | 0.258332331583926 |
| Crocuta_crocuta | Paguma_larvata | 0.204836975607256 |
| Crocuta_crocuta | Panthera_leo | 0.205527770746697 |
| Crocuta_crocuta | Panthera_onca | 0.204760617169806 |
| Crocuta_crocuta | Panthera_pardus | 0.206241067211568 |
| Crocuta_crocuta | Panthera_tigris | 0.202616102311385 |
| Crocuta_crocuta | Panthera_uncia | 0.208895828250174 |
| Crocuta_crocuta | Paradoxurus_hermaphroditus | 0.213140020357374 |
| Crocuta_crocuta | Parahyaena_brunnea_FD126_Fecal | 0.107186867884028 |
| Crocuta_crocuta | Pardofelis_marmorata | 0.196203698482834 |
| Crocuta_crocuta | Phoca_fasciata | 0.227111827063648 |
| Crocuta_crocuta | Phoca_groenlandica | 0.235948165335017 |
| Crocuta_crocuta | Phoca_largha | 0.227192519312319 |
| Crocuta_crocuta | Phoca_vitulina | 0.229104413233988 |
| Crocuta_crocuta | Phocarctos_hookeri | 0.243036857082563 |
| Crocuta_crocuta | Prionailurus_bengalensis | 0.194034840931646 |
| Crocuta_crocuta | Prionailurus_bengalensis_euptilurus | 0.196222830972795 |
| Crocuta_crocuta | Prionailurus_planiceps | 0.19933187611311 |
| Crocuta_crocuta | Prionailurus_rubiginosus | 0.19401210629058 |
| Crocuta_crocuta | Prionailurus_viverrinus | 0.198864705448381 |
| Crocuta_crocuta | Prionodon_pardicolor | 0.210557640533871 |
| Crocuta_crocuta | Procyon_lotor | 0.248448671465945 |
| Crocuta_crocuta | Profelis_aurata | 0.204681151026994 |
| Crocuta_crocuta | Proteles_cristatus_cristatus_TS307 | 0.128734101742495 |
| Crocuta_crocuta | Proteles_cristatus_septentrionalis_NMS_Z_2018_54 | 0.131856318555283 |
| Crocuta_crocuta | Puma_concolor | 0.198409090483595 |
| Crocuta_crocuta | Puma_yagouaroundi | 0.197626676620182 |
| Crocuta_crocuta | Pusa_caspica | 0.226038930671924 |
| Crocuta_crocuta | Pusa_hispida | 0.226001507067549 |
| Crocuta_crocuta | Pusa_sibirica | 0.222955586836617 |
| Crocuta_crocuta | Speothos_venaticus_T1624 | 0.262603761787846 |
| Crocuta_crocuta | Spilogale_putorius | 0.260820004163012 |
| Crocuta_crocuta | Taxidea_taxus | 0.240922928115796 |
| Crocuta_crocuta | Tremarctos_ornatus | 0.248198160859888 |
| Crocuta_crocuta | Urocyon_cinereoargenteus | 0.260785284934493 |
| Crocuta_crocuta | Ursus_americanus | 0.25452463448357 |
| Crocuta_crocuta | Ursus_arctos | 0.255449482321504 |
| Crocuta_crocuta | Ursus_maritimus | 0.253336096629798 |
| Crocuta_crocuta | Ursus_spelaeus | 0.248391550987408 |
| Crocuta_crocuta | Ursus_thibetanus | 0.248843187530195 |
| Crocuta_crocuta | Urva_javanica | 0.193769316022806 |
| Crocuta_crocuta | Viverra_tangalunga_FMNH146957_SRR6053069 | 0.202293239494212 |
| Crocuta_crocuta | Viverricula_indica | 0.200710213430889 |
| Crocuta_crocuta | Vulpes_corsac | 0.259407050799765 |
| Crocuta_crocuta | Vulpes_ferrilata | 0.258618063440265 |
| Crocuta_crocuta | Vulpes_lagopus | 0.254396734362676 |
| Crocuta_crocuta | Vulpes_vulpes_T3611 | 0.254825071113869 |

| | | |
|---|---|---|
| Crocuta_crocuta | Vulpes_zerda | 0.253776428044444 |
| Crocuta_crocuta | Zalophus_californianus | 0.242896418185599 |
| Cuon_alpinus | Cystophora_cristata | 0.236485329093055 |
| Cuon_alpinus | Enhydra_lutris | 0.246416199322054 |
| Cuon_alpinus | Erignathus_barbatus | 0.232335228702836 |
| Cuon_alpinus | Eumetopias_jubatus | 0.248778987396136 |
| Cuon_alpinus | Felis_catus | 0.244541644721254 |
| Cuon_alpinus | Felis_chaus | 0.24466464465426 |
| Cuon_alpinus | Felis_margarita | 0.243959704767357 |
| Cuon_alpinus | Felis_nigripes | 0.24407847636152 |
| Cuon_alpinus | Felis_silvestris | 0.242862099977737 |
| Cuon_alpinus | Fossa_fossana_FMNH156648_SRR6053060 | 0.25155352327911 |
| Cuon_alpinus | Genetta_servalina | 0.245788123929372 |
| Cuon_alpinus | Gulo_gulo | 0.246722183640288 |
| Cuon_alpinus | Halichoerus_grypus | 0.236057167505975 |
| Cuon_alpinus | Helarctos_malayanus | 0.253991812763056 |
| Cuon_alpinus | Hyaena_hyaena | 0.25493411020541 |
| Cuon_alpinus | Hydrurga_leptonyx | 0.229377544834614 |
| Cuon_alpinus | Leopardus_colocolo | 0.239851773121139 |
| Cuon_alpinus | Leopardus_geoffroyi | 0.241975618332505 |
| Cuon_alpinus | Leopardus_guigna | 0.240508010564243 |
| Cuon_alpinus | Leopardus_jacobita | 0.248089846874423 |
| Cuon_alpinus | Leopardus_pardalis | 0.242320612819337 |
| Cuon_alpinus | Leopardus_tigrinus | 0.242458944194718 |
| Cuon_alpinus | Leopardus_wiedii | 0.241358926386634 |
| Cuon_alpinus | Leptailurus_serval | 0.239207294358202 |
| Cuon_alpinus | Leptonychotes_weddellii | 0.231791028259914 |
| Cuon_alpinus | Lobodon_carcinophaga | 0.226782570998381 |
| Cuon_alpinus | Lutra_lutra | 0.257962880328915 |
| Cuon_alpinus | Lutra_sumatrana | 0.257140886712321 |
| Cuon_alpinus | Lutrogale_perspicillata | 0.267308548165891 |
| Cuon_alpinus | Lycaon_pictus | 0.124339716610416 |
| Cuon_alpinus | Lynx_canadensis | 0.244418709592801 |
| Cuon_alpinus | Lynx_lynx | 0.244178405464824 |
| Cuon_alpinus | Lynx_pardinus | 0.24788628762574 |
| Cuon_alpinus | Lynx_rufus | 0.246364245107176 |
| Cuon_alpinus | Manis_javanica | 0.311962552754021 |
| Cuon_alpinus | Martes_americana | 0.246089144952026 |
| Cuon_alpinus | Martes_flavigula | 0.243840291950356 |
| Cuon_alpinus | Martes_foina | 0.242730568642855 |
| Cuon_alpinus | Martes_martes | 0.246348980939891 |
| Cuon_alpinus | Martes_melampus | 0.24390527478116 |
| Cuon_alpinus | Martes_pennanti | 0.247409040403829 |
| Cuon_alpinus | Martes_zibellina | 0.247151282553589 |
| Cuon_alpinus | Meles_anakuma | 0.249348480488162 |
| Cuon_alpinus | Meles_leucurus | 0.243208178365792 |
| Cuon_alpinus | Meles_meles | 0.24445595009024 |
| Cuon_alpinus | Melogale_moschata | 0.246973975095512 |
| Cuon_alpinus | Melursus_ursinus | 0.255341609625693 |
| Cuon_alpinus | Mephitis_mephitis | 0.249997788582542 |
| Cuon_alpinus | Mirounga_leonina | 0.228846916207623 |
| Cuon_alpinus | Mungotictis_decemlineata | 0.254344558727251 |
| Cuon_alpinus | Mustela_altaica | 0.243533357432943 |
| Cuon_alpinus | Mustela_erminea | 0.238477772863372 |
| Cuon_alpinus | Mustela_eversmannii | 0.242927365587743 |
| Cuon_alpinus | Mustela_frenata | 0.23578832548302 |
| Cuon_alpinus | Mustela_itatsi | 0.244182034203068 |
| Cuon_alpinus | Mustela_kathiah | 0.239674416930744 |
| Cuon_alpinus | Mustela_nigripes | 0.239261816428227 |
| Cuon_alpinus | Mustela_nivalis | 0.239770155993042 |
| Cuon_alpinus | Mustela_putorius | 0.24260369481796 |
| Cuon_alpinus | Mustela_sibirica | 0.242764176915353 |
| Cuon_alpinus | Nandinia_binotata | 0.237706398593926 |
| Cuon_alpinus | Nasua_nasua | 0.263070197562104 |
| Cuon_alpinus | Neofelis_nebulosa | 0.254734580930739 |
| Cuon_alpinus | Neomonachus_schauinslandi | 0.234091909424139 |
| Cuon_alpinus | Neophoca_cinerea | 0.257433112932735 |
| Cuon_alpinus | Neovison_vison | 0.236805116185506 |
| Cuon_alpinus | Nyctereutes_procyonoides | 0.183664259630191 |
| Cuon_alpinus | Odobenus_rosmarus_rosmarus | 0.267446948664626 |
| Cuon_alpinus | Otocolobus_manul | 0.240205267869475 |
| Cuon_alpinus | Otocyon_megalotis_megalotis_TS305 | 0.180061635463494 |
| Cuon_alpinus | Otocyon_megalotis_virgatus_FMNH158128 | 0.180641043883489 |
| Cuon_alpinus | Paguma_larvata | 0.240655405754 |
| Cuon_alpinus | Panthera_leo | 0.254251172565907 |
| Cuon_alpinus | Panthera_onca | 0.257005700877108 |
| Cuon_alpinus | Panthera_pardus | 0.256720850341616 |
| Cuon_alpinus | Panthera_tigris | 0.25619955681329 |
| Cuon_alpinus | Panthera_uncia | 0.254729182767791 |
| Cuon_alpinus | Paradoxurus_hermaphroditus | 0.240893254099394 |
| Cuon_alpinus | Parahyaena_brunnea_FD126_Fecal | 0.254558630701412 |
| Cuon_alpinus | Pardofelis_marmorata | 0.247432937243096 |
| Cuon_alpinus | Phoca_fasciata | 0.235004437441473 |
| Cuon_alpinus | Phoca_groenlandica | 0.236994122640726 |
| Cuon_alpinus | Phoca_largha | 0.238819604978475 |
| Cuon_alpinus | Phoca_vitulina | 0.23831741320345 |
| Cuon_alpinus | Phocarctos_hookeri | 0.26107736042816 |

| | | |
|---|---|---|
| Cuon_alpinus | Prionailurus_bengalensis | 0.236852080267148 |
| Cuon_alpinus | Prionailurus_bengalensis_euptilurus | 0.239398587100185 |
| Cuon_alpinus | Prionailurus_planiceps | 0.2436400920792 |
| Cuon_alpinus | Prionailurus_rubiginosus | 0.241519553186454 |
| Cuon_alpinus | Prionailurus_viverrinus | 0.246734720135726 |
| Cuon_alpinus | Prionodon_pardicolor | 0.239691370569922 |
| Cuon_alpinus | Procyon_lotor | 0.247369450041697 |
| Cuon_alpinus | Profelis_aurata | 0.243139126924105 |
| Cuon_alpinus | Proteles_cristatus_cristatus_TS307 | 0.260372944305183 |
| Cuon_alpinus | Proteles_cristatus_septentrionalis_NMS_Z_2018_54 | 0.257939643671423 |
| Cuon_alpinus | Puma_concolor | 0.239055519218097 |
| Cuon_alpinus | Puma_yagouaroundi | 0.241620605593863 |
| Cuon_alpinus | Pusa_caspica | 0.234901766030956 |
| Cuon_alpinus | Pusa_hispida | 0.233708449360037 |
| Cuon_alpinus | Pusa_sibirica | 0.234661426957889 |
| Cuon_alpinus | Speothos_venaticus_T1624 | 0.141705116798514 |
| Cuon_alpinus | Spilogale_putorius | 0.243269242240404 |
| Cuon_alpinus | Taxidea_taxus | 0.246937479866155 |
| Cuon_alpinus | Tremarctos_ornatus | 0.25609001981025 |
| Cuon_alpinus | Urocyon_cinereoargenteus | 0.19570235902672 |
| Cuon_alpinus | Ursus_americanus | 0.249465170009471 |
| Cuon_alpinus | Ursus_arctos | 0.253198515928161 |
| Cuon_alpinus | Ursus_maritimus | 0.253468151012136 |
| Cuon_alpinus | Ursus_spelaeus | 0.252866328384078 |
| Cuon_alpinus | Ursus_thibetanus | 0.248449701750104 |
| Cuon_alpinus | Urva_javanica | 0.246610430904237 |
| Cuon_alpinus | Viverra_tangalunga_FMNH146957_SRR6053069 | 0.23266354325306 |
| Cuon_alpinus | Viverricula_indica | 0.236962528688185 |
| Cuon_alpinus | Vulpes_corsac | 0.181227830560347 |
| Cuon_alpinus | Vulpes_ferrilata | 0.183001430352061 |
| Cuon_alpinus | Vulpes_lagopus | 0.183687545077813 |
| Cuon_alpinus | Vulpes_vulpes_T3611 | 0.181447578551107 |
| Cuon_alpinus | Vulpes_zerda | 0.180347589020006 |
| Cuon_alpinus | Zalophus_californianus | 0.251871534739756 |
| Cystophora_cristata | Enhydra_lutris | 0.21622132739941 |
| Cystophora_cristata | Erignathus_barbatus | 0.130149343958081 |
| Cystophora_cristata | Eumetopias_jubatus | 0.181001858444354 |
| Cystophora_cristata | Felis_catus | 0.217514460454133 |
| Cystophora_cristata | Felis_chaus | 0.216274487190075 |
| Cystophora_cristata | Felis_margarita | 0.21512454802477 |
| Cystophora_cristata | Felis_nigripes | 0.214742322607661 |
| Cystophora_cristata | Felis_silvestris | 0.216140141395613 |
| Cystophora_cristata | Fossa_fossana_FMNH156648_SRR6053060 | 0.210798690771591 |
| Cystophora_cristata | Genetta_servalina | 0.220320582569036 |
| Cystophora_cristata | Gulo_gulo | 0.213885295545717 |
| Cystophora_cristata | Halichoerus_grypus | 0.0986796419279991 |
| Cystophora_cristata | Helarctos_malayanus | 0.216362427855074 |
| Cystophora_cristata | Hyaena_hyaena | 0.218767591180987 |
| Cystophora_cristata | Hydrurga_leptonyx | 0.135334510500863 |
| Cystophora_cristata | Leopardus_colocolo | 0.214370990135026 |
| Cystophora_cristata | Leopardus_geoffroyi | 0.213756505334663 |
| Cystophora_cristata | Leopardus_guigna | 0.214823172405873 |
| Cystophora_cristata | Leopardus_jacobita | 0.216851504067584 |
| Cystophora_cristata | Leopardus_pardalis | 0.211001093105812 |
| Cystophora_cristata | Leopardus_tigrinus | 0.215623577571669 |
| Cystophora_cristata | Leopardus_wiedii | 0.213571918851273 |
| Cystophora_cristata | Leptailurus_serval | 0.216213250290109 |
| Cystophora_cristata | Leptonychotes_weddellii | 0.134881498399005 |
| Cystophora_cristata | Lobodon_carcinophaga | 0.135265711989647 |
| Cystophora_cristata | Lutra_lutra | 0.211059681410728 |
| Cystophora_cristata | Lutra_sumatrana | 0.208868610578431 |
| Cystophora_cristata | Lutrogale_perspicillata | 0.222719933399503 |
| Cystophora_cristata | Lycaon_pictus | 0.230867530846872 |
| Cystophora_cristata | Lynx_canadensis | 0.218014383881505 |
| Cystophora_cristata | Lynx_lynx | 0.219654877553024 |
| Cystophora_cristata | Lynx_pardinus | 0.215971691319414 |
| Cystophora_cristata | Lynx_rufus | 0.213933951886341 |
| Cystophora_cristata | Manis_javanica | 0.276212921941302 |
| Cystophora_cristata | Martes_americana | 0.21485956705717 |
| Cystophora_cristata | Martes_flavigula | 0.214265973289599 |
| Cystophora_cristata | Martes_foina | 0.212972173558741 |
| Cystophora_cristata | Martes_martes | 0.213208374017001 |
| Cystophora_cristata | Martes_melampus | 0.211932686813233 |
| Cystophora_cristata | Martes_pennanti | 0.215193045034459 |
| Cystophora_cristata | Martes_zibellina | 0.214059862885611 |
| Cystophora_cristata | Meles_anakuma | 0.215472211863331 |
| Cystophora_cristata | Meles_leucurus | 0.21059345826338 |
| Cystophora_cristata | Meles_meles | 0.210954054047075 |
| Cystophora_cristata | Melogale_moschata | 0.214083459029471 |
| Cystophora_cristata | Melursus_ursinus | 0.211961247141255 |
| Cystophora_cristata | Mephitis_mephitis | 0.222484904423157 |
| Cystophora_cristata | Mirounga_leonina | 0.143606942951353 |
| Cystophora_cristata | Mungotictis_decemlineata | 0.220279618017438 |
| Cystophora_cristata | Mustela_altaica | 0.21759924076535 |
| Cystophora_cristata | Mustela_erminea | 0.209302805506628 |
| Cystophora_cristata | Mustela_eversmannii | 0.213919627855411 |
| Cystophora_cristata | Mustela_frenata | 0.206433271893105 |

| | | |
|---|---|---|
| Cystophora_cristata | Mustela_itatsi | 0.213031574189258 |
| Cystophora_cristata | Mustela_kathiah | 0.213138755325789 |
| Cystophora_cristata | Mustela_nigripes | 0.21154371590771 |
| Cystophora_cristata | Mustela_nivalis | 0.216206105775577 |
| Cystophora_cristata | Mustela_putorius | 0.214231861765404 |
| Cystophora_cristata | Mustela_sibirica | 0.215117102456434 |
| Cystophora_cristata | Nandinia_binotata | 0.201662442232323 |
| Cystophora_cristata | Nasua_nasua | 0.233315375076524 |
| Cystophora_cristata | Neofelis_nebulosa | 0.227365726499128 |
| Cystophora_cristata | Neomonachus_schauinslandi | 0.145015985097564 |
| Cystophora_cristata | Neophoca_cinerea | 0.187715271907426 |
| Cystophora_cristata | Neovison_vison | 0.214114417376895 |
| Cystophora_cristata | Nyctereutes_procyonoides | 0.227835398672469 |
| Cystophora_cristata | Odobenus_rosmarus_rosmarus | 0.206662471522859 |
| Cystophora_cristata | Otocolobus_manul | 0.218845651972988 |
| Cystophora_cristata | Otocyon_megalotis_megalotis_TS305 | 0.231038700539252 |
| Cystophora_cristata | Otocyon_megalotis_virgatus_FMNH158128 | 0.229795786951047 |
| Cystophora_cristata | Paguma_larvata | 0.226950967886812 |
| Cystophora_cristata | Panthera_leo | 0.224229187535556 |
| Cystophora_cristata | Panthera_onca | 0.223055067013355 |
| Cystophora_cristata | Panthera_pardus | 0.222236979498615 |
| Cystophora_cristata | Panthera_tigris | 0.223302237751376 |
| Cystophora_cristata | Panthera_uncia | 0.223448928048013 |
| Cystophora_cristata | Paradoxurus_hermaphroditus | 0.231649895173796 |
| Cystophora_cristata | Parahyaena_brunnea_FD126_Fecal | 0.222694419447155 |
| Cystophora_cristata | Pardofelis_marmorata | 0.21816772671343 |
| Cystophora_cristata | Phoca_fasciata | 0.0983513837923968 |
| Cystophora_cristata | Phoca_groenlandica | 0.103280747747496 |
| Cystophora_cristata | Phoca_largha | 0.0985886501170109 |
| Cystophora_cristata | Phoca_vitulina | 0.0996456654767586 |
| Cystophora_cristata | Phocarctos_hookeri | 0.186404014340071 |
| Cystophora_cristata | Prionailurus_bengalensis | 0.211091109412772 |
| Cystophora_cristata | Prionailurus_bengalensis_euptilurus | 0.214493929862712 |
| Cystophora_cristata | Prionailurus_planiceps | 0.21550765830035 |
| Cystophora_cristata | Prionailurus_rubiginosus | 0.215560692971231 |
| Cystophora_cristata | Prionailurus_viverrinus | 0.213605536687163 |
| Cystophora_cristata | Prionodon_pardicolor | 0.22747545979989 |
| Cystophora_cristata | Procyon_lotor | 0.220109648513734 |
| Cystophora_cristata | Profelis_aurata | 0.21841995282908 |
| Cystophora_cristata | Proteles_cristatus_cristatus_TS307 | 0.22483714135222 |
| Cystophora_cristata | Proteles_cristatus_septentrionalis_NMS_Z_2018_54 | 0.227839813414657 |
| Cystophora_cristata | Puma_concolor | 0.216338775789797 |
| Cystophora_cristata | Puma_yagouaroundi | 0.217781150851901 |
| Cystophora_cristata | Pusa_caspica | 0.0972338638473842 |
| Cystophora_cristata | Pusa_hispida | 0.095392679740328 |
| Cystophora_cristata | Pusa_sibirica | 0.0940018764319347 |
| Cystophora_cristata | Speothos_venaticus_T1624 | 0.235045470472705 |
| Cystophora_cristata | Spilogale_putorius | 0.22655216222571 |
| Cystophora_cristata | Taxidea_taxus | 0.20103084104539 |
| Cystophora_cristata | Tremarctos_ornatus | 0.216928563864304 |
| Cystophora_cristata | Urocyon_cinereoargenteus | 0.236647404970914 |
| Cystophora_cristata | Ursus_americanus | 0.216766351284372 |
| Cystophora_cristata | Ursus_arctos | 0.217928397117188 |
| Cystophora_cristata | Ursus_maritimus | 0.216073824784829 |
| Cystophora_cristata | Ursus_spelaeus | 0.213850713030913 |
| Cystophora_cristata | Ursus_thibetanus | 0.215466421168126 |
| Cystophora_cristata | Urva_javanica | 0.215017106085347 |
| Cystophora_cristata | Viverra_tangalunga_FMNH146957_SRR6053069 | 0.218100433611082 |
| Cystophora_cristata | Viverricula_indica | 0.216008317348589 |
| Cystophora_cristata | Vulpes_corsac | 0.230210111850894 |
| Cystophora_cristata | Vulpes_ferrilata | 0.233498264118726 |
| Cystophora_cristata | Vulpes_lagopus | 0.2342656357577 |
| Cystophora_cristata | Vulpes_vulpes_T3611 | 0.227417605352387 |
| Cystophora_cristata | Vulpes_zerda | 0.229703801236815 |
| Cystophora_cristata | Zalophus_californianus | 0.181942590733678 |
| Enhydra_lutris | Erignathus_barbatus | 0.217176770236965 |
| Enhydra_lutris | Eumetopias_jubatus | 0.22879585261044 |
| Enhydra_lutris | Felis_catus | 0.230365470512769 |
| Enhydra_lutris | Felis_chaus | 0.233195065828365 |
| Enhydra_lutris | Felis_margarita | 0.230145762720592 |
| Enhydra_lutris | Felis_nigripes | 0.230108636274824 |
| Enhydra_lutris | Felis_silvestris | 0.22919933935616 |
| Enhydra_lutris | Fossa_fossana_FMNH156648_SRR6053060 | 0.229379531755564 |
| Enhydra_lutris | Genetta_servalina | 0.229801926006346 |
| Enhydra_lutris | Gulo_gulo | 0.170277725877101 |
| Enhydra_lutris | Halichoerus_grypus | 0.216785000073273 |
| Enhydra_lutris | Helarctos_malayanus | 0.236994593816894 |
| Enhydra_lutris | Hyaena_hyaena | 0.23747614590378 |
| Enhydra_lutris | Hydrurga_leptonyx | 0.204081849202434 |
| Enhydra_lutris | Leopardus_colocolo | 0.226168160538398 |
| Enhydra_lutris | Leopardus_geoffroyi | 0.22557378004789 |
| Enhydra_lutris | Leopardus_guigna | 0.226950829062208 |
| Enhydra_lutris | Leopardus_jacobita | 0.227300221500903 |
| Enhydra_lutris | Leopardus_pardalis | 0.226268874026845 |
| Enhydra_lutris | Leopardus_tigrinus | 0.228720010493276 |
| Enhydra_lutris | Leopardus_wiedii | 0.227778676291585 |
| Enhydra_lutris | Leptailurus_serval | 0.227516992894637 |

| | | |
|---|---|---|
| Enhydra_lutris | Leptonychotes_weddellii | 0.205642762494038 |
| Enhydra_lutris | Lobodon_carcinophaga | 0.205573694886109 |
| Enhydra_lutris | Lutra_lutra | 0.138057569919181 |
| Enhydra_lutris | Lutra_sumatrana | 0.134553564922442 |
| Enhydra_lutris | Lutrogale_perspicillata | 0.146708887200456 |
| Enhydra_lutris | Lycaon_pictus | 0.243806420100366 |
| Enhydra_lutris | Lynx_canadensis | 0.22400184649128 |
| Enhydra_lutris | Lynx_lynx | 0.222602389723519 |
| Enhydra_lutris | Lynx_pardinus | 0.224100917902619 |
| Enhydra_lutris | Lynx_rufus | 0.223863526150997 |
| Enhydra_lutris | Manis_javanica | 0.285008737732605 |
| Enhydra_lutris | Martes_americana | 0.170861044615389 |
| Enhydra_lutris | Martes_flavigula | 0.167989709801363 |
| Enhydra_lutris | Martes_foina | 0.166986961567452 |
| Enhydra_lutris | Martes_martes | 0.169090124149128 |
| Enhydra_lutris | Martes_melampus | 0.169284119719622 |
| Enhydra_lutris | Martes_pennanti | 0.16493352969054 |
| Enhydra_lutris | Martes_zibellina | 0.170953586816038 |
| Enhydra_lutris | Meles_anakuma | 0.178083070747915 |
| Enhydra_lutris | Meles_leucurus | 0.175555197363498 |
| Enhydra_lutris | Meles_meles | 0.174463828862233 |
| Enhydra_lutris | Melogale_moschata | 0.162860876228816 |
| Enhydra_lutris | Melursus_ursinus | 0.237155224092935 |
| Enhydra_lutris | Mephitis_mephitis | 0.233375515094599 |
| Enhydra_lutris | Mirounga_leonina | 0.212786333263643 |
| Enhydra_lutris | Mungotictis_decemlineata | 0.236231493308408 |
| Enhydra_lutris | Mustela_altaica | 0.163703636206741 |
| Enhydra_lutris | Mustela_erminea | 0.150462850725549 |
| Enhydra_lutris | Mustela_eversmannii | 0.161372699410385 |
| Enhydra_lutris | Mustela_frenata | 0.160489446082567 |
| Enhydra_lutris | Mustela_itatsi | 0.159874902132879 |
| Enhydra_lutris | Mustela_kathiah | 0.16356318508191 |
| Enhydra_lutris | Mustela_nigripes | 0.16035881227671 |
| Enhydra_lutris | Mustela_nivalis | 0.164096615810017 |
| Enhydra_lutris | Mustela_putorius | 0.159554164412573 |
| Enhydra_lutris | Mustela_sibirica | 0.160487458482381 |
| Enhydra_lutris | Nandinia_binotata | 0.216058935220204 |
| Enhydra_lutris | Nasua_nasua | 0.232134172546986 |
| Enhydra_lutris | Neofelis_nebulosa | 0.240978316776289 |
| Enhydra_lutris | Neomonachus_schauinslandi | 0.213482614654051 |
| Enhydra_lutris | Neophoca_cinerea | 0.233720659953241 |
| Enhydra_lutris | Neovison_vison | 0.163431113525098 |
| Enhydra_lutris | Nyctereutes_procyonoides | 0.242115069657954 |
| Enhydra_lutris | Odobenus_rosmarus_rosmarus | 0.246943005706825 |
| Enhydra_lutris | Otocolobus_manul | 0.229031594574019 |
| Enhydra_lutris | Otocyon_megalotis_megalotis_TS305 | 0.246754902908554 |
| Enhydra_lutris | Otocyon_megalotis_virgatus_FMNH158128 | 0.246258059346499 |
| Enhydra_lutris | Paguma_larvata | 0.237044534181503 |
| Enhydra_lutris | Panthera_leo | 0.233672725432763 |
| Enhydra_lutris | Panthera_onca | 0.233134874126673 |
| Enhydra_lutris | Panthera_pardus | 0.234382896849759 |
| Enhydra_lutris | Panthera_tigris | 0.239951249999073 |
| Enhydra_lutris | Panthera_uncia | 0.234980038116256 |
| Enhydra_lutris | Paradoxurus_hermaphroditus | 0.239794199547709 |
| Enhydra_lutris | Parahyaena_brunnea_FD126_Fecal | 0.236744730336741 |
| Enhydra_lutris | Pardofelis_marmorata | 0.234415173116952 |
| Enhydra_lutris | Phoca_fasciata | 0.216362501690981 |
| Enhydra_lutris | Phoca_groenlandica | 0.218805325859175 |
| Enhydra_lutris | Phoca_largha | 0.214278069047355 |
| Enhydra_lutris | Phoca_vitulina | 0.217772267524315 |
| Enhydra_lutris | Phocarctos_hookeri | 0.239068658561979 |
| Enhydra_lutris | Prionailurus_bengalensis | 0.224031523871379 |
| Enhydra_lutris | Prionailurus_bengalensis_euptilurus | 0.225921359614226 |
| Enhydra_lutris | Prionailurus_planiceps | 0.226466913405393 |
| Enhydra_lutris | Prionailurus_rubiginosus | 0.22701020490167 |
| Enhydra_lutris | Prionailurus_viverrinus | 0.22688344423254 |
| Enhydra_lutris | Prionodon_pardicolor | 0.237659352499961 |
| Enhydra_lutris | Procyon_lotor | 0.214598896044802 |
| Enhydra_lutris | Profelis_aurata | 0.234100962578424 |
| Enhydra_lutris | Proteles_cristatus_cristatus_TS307 | 0.243547697446504 |
| Enhydra_lutris | Proteles_cristatus_septentrionalis_NMS_Z_2018_54 | 0.242806928217939 |
| Enhydra_lutris | Puma_concolor | 0.226962254503815 |
| Enhydra_lutris | Puma_yagouaroundi | 0.227933791370928 |
| Enhydra_lutris | Pusa_caspica | 0.214966101909444 |
| Enhydra_lutris | Pusa_hispida | 0.21356312235737 |
| Enhydra_lutris | Pusa_sibirica | 0.215311963539111 |
| Enhydra_lutris | Speothos_venaticus_T1624 | 0.252147875174657 |
| Enhydra_lutris | Spilogale_putorius | 0.235868256395128 |
| Enhydra_lutris | Taxidea_taxus | 0.170724204865439 |
| Enhydra_lutris | Tremarctos_ornatus | 0.235566584987065 |
| Enhydra_lutris | Urocyon_cinereoargenteus | 0.250601118788292 |
| Enhydra_lutris | Ursus_americanus | 0.234963675984585 |
| Enhydra_lutris | Ursus_arctos | 0.239952249453206 |
| Enhydra_lutris | Ursus_maritimus | 0.240507853408542 |
| Enhydra_lutris | Ursus_spelaeus | 0.233070124472487 |
| Enhydra_lutris | Ursus_thibetanus | 0.236077860683544 |
| Enhydra_lutris | Urva_javanica | 0.238078918078175 |

| | | |
|---|---|---|
| Enhydra_lutris | Viverra_tangalunga_FMNH146957_SRR6053069 | 0.228141493540791 |
| Enhydra_lutris | Viverricula_indica | 0.230390278718058 |
| Enhydra_lutris | Vulpes_corsac | 0.24718532284731 |
| Enhydra_lutris | Vulpes_ferrilata | 0.245914503678157 |
| Enhydra_lutris | Vulpes_lagopus | 0.2484394253611 |
| Enhydra_lutris | Vulpes_vulpes_T3611 | 0.241390153809939 |
| Enhydra_lutris | Vulpes_zerda | 0.242883371961954 |
| Enhydra_lutris | Zalophus_californianus | 0.23259799633301 |
| Erignathus_barbatus | Eumetopias_jubatus | 0.173516307561306 |
| Erignathus_barbatus | Felis_catus | 0.217797405566613 |
| Erignathus_barbatus | Felis_chaus | 0.222563939476688 |
| Erignathus_barbatus | Felis_margarita | 0.216353075929845 |
| Erignathus_barbatus | Felis_nigripes | 0.221445095615083 |
| Erignathus_barbatus | Felis_silvestris | 0.218011790820192 |
| Erignathus_barbatus | Fossa_fossana_FMNH156648_SRR6053060 | 0.215401475322332 |
| Erignathus_barbatus | Genetta_servalina | 0.220043779617529 |
| Erignathus_barbatus | Gulo_gulo | 0.211090530777951 |
| Erignathus_barbatus | Halichoerus_grypus | 0.127285896010371 |
| Erignathus_barbatus | Helarctos_malayanus | 0.216397599760979 |
| Erignathus_barbatus | Hyaena_hyaena | 0.225006388885302 |
| Erignathus_barbatus | Hydrurga_leptonyx | 0.127808396079741 |
| Erignathus_barbatus | Leopardus_colocolo | 0.214032265778768 |
| Erignathus_barbatus | Leopardus_geoffroyi | 0.21845906959778 |
| Erignathus_barbatus | Leopardus_guigna | 0.216940782592332 |
| Erignathus_barbatus | Leopardus_jacobita | 0.21768305476151 |
| Erignathus_barbatus | Leopardus_pardalis | 0.214671379628037 |
| Erignathus_barbatus | Leopardus_tigrinus | 0.218511810168532 |
| Erignathus_barbatus | Leopardus_wiedii | 0.212914644960476 |
| Erignathus_barbatus | Leptailurus_serval | 0.214911169167376 |
| Erignathus_barbatus | Leptonychotes_weddellii | 0.122231976607316 |
| Erignathus_barbatus | Lobodon_carcinophaga | 0.124653205562894 |
| Erignathus_barbatus | Lutra_lutra | 0.21547448343178 |
| Erignathus_barbatus | Lutra_sumatrana | 0.214351721849499 |
| Erignathus_barbatus | Lutrogale_perspicillata | 0.225639221585286 |
| Erignathus_barbatus | Lycaon_pictus | 0.229951997143568 |
| Erignathus_barbatus | Lynx_canadensis | 0.216397707009959 |
| Erignathus_barbatus | Lynx_lynx | 0.216076885696418 |
| Erignathus_barbatus | Lynx_pardinus | 0.216755073484747 |
| Erignathus_barbatus | Lynx_rufus | 0.215178191547811 |
| Erignathus_barbatus | Manis_javanica | 0.279082961558289 |
| Erignathus_barbatus | Martes_americana | 0.212878713989712 |
| Erignathus_barbatus | Martes_flavigula | 0.211891804060575 |
| Erignathus_barbatus | Martes_foina | 0.216622241482627 |
| Erignathus_barbatus | Martes_martes | 0.210717741398007 |
| Erignathus_barbatus | Martes_melampus | 0.211483264393511 |
| Erignathus_barbatus | Martes_pennanti | 0.217557435968923 |
| Erignathus_barbatus | Martes_zibellina | 0.21229594722919 |
| Erignathus_barbatus | Meles_anakuma | 0.217022968075695 |
| Erignathus_barbatus | Meles_leucurus | 0.214594662356424 |
| Erignathus_barbatus | Meles_meles | 0.210914176683613 |
| Erignathus_barbatus | Melogale_moschata | 0.217024151468609 |
| Erignathus_barbatus | Melursus_ursinus | 0.217020394832357 |
| Erignathus_barbatus | Mephitis_mephitis | 0.221817356532996 |
| Erignathus_barbatus | Mirounga_leonina | 0.132613677006054 |
| Erignathus_barbatus | Mungotictis_decemlineata | 0.226118323774117 |
| Erignathus_barbatus | Mustela_altaica | 0.219119318685637 |
| Erignathus_barbatus | Mustela_erminea | 0.209819708407438 |
| Erignathus_barbatus | Mustela_eversmannii | 0.22017752023389 |
| Erignathus_barbatus | Mustela_frenata | 0.213284150426426 |
| Erignathus_barbatus | Mustela_itatsi | 0.218867836559616 |
| Erignathus_barbatus | Mustela_kathiah | 0.217564418501479 |
| Erignathus_barbatus | Mustela_nigripes | 0.220322611561646 |
| Erignathus_barbatus | Mustela_nivalis | 0.216161706677493 |
| Erignathus_barbatus | Mustela_putorius | 0.219320979310078 |
| Erignathus_barbatus | Mustela_sibirica | 0.219129658040146 |
| Erignathus_barbatus | Nandinia_binotata | 0.203285574659248 |
| Erignathus_barbatus | Nasua_nasua | 0.231110446625756 |
| Erignathus_barbatus | Neofelis_nebulosa | 0.230173174441078 |
| Erignathus_barbatus | Neomonachus_schauinslandi | 0.140316999455911 |
| Erignathus_barbatus | Neophoca_cinerea | 0.179533994995973 |
| Erignathus_barbatus | Neovison_vison | 0.217042713727092 |
| Erignathus_barbatus | Nyctereutes_procyonoides | 0.223791004462628 |
| Erignathus_barbatus | Odobenus_rosmarus_rosmarus | 0.202176778114937 |
| Erignathus_barbatus | Otocolobus_manul | 0.219201745685824 |
| Erignathus_barbatus | Otocyon_megalotis_megalotis_TS305 | 0.230042985485632 |
| Erignathus_barbatus | Otocyon_megalotis_virgatus_FMNH158128 | 0.2319115301971 |
| Erignathus_barbatus | Paguma_larvata | 0.223616131626179 |
| Erignathus_barbatus | Panthera_leo | 0.225436912288293 |
| Erignathus_barbatus | Panthera_onca | 0.226593856853855 |
| Erignathus_barbatus | Panthera_pardus | 0.228668073313472 |
| Erignathus_barbatus | Panthera_tigris | 0.232831316585272 |
| Erignathus_barbatus | Panthera_uncia | 0.228270707339706 |
| Erignathus_barbatus | Paradoxurus_hermaphroditus | 0.230579195718542 |
| Erignathus_barbatus | Parahyaena_brunnea_FD126_Fecal | 0.224871852267871 |
| Erignathus_barbatus | Pardofelis_marmorata | 0.221731352802119 |
| Erignathus_barbatus | Phoca_fasciata | 0.122090072127014 |
| Erignathus_barbatus | Phoca_groenlandica | 0.128526386208627 |

| | | |
|---|---|---|
| Erignathus_barbatus | Phoca_largha | 0.126835081306094 |
| Erignathus_barbatus | Phoca_vitulina | 0.127574795282957 |
| Erignathus_barbatus | Phocarctos_hookeri | 0.186085840995034 |
| Erignathus_barbatus | Prionailurus_bengalensis | 0.214192848905151 |
| Erignathus_barbatus | Prionailurus_bengalensis_euptilurus | 0.216432852651637 |
| Erignathus_barbatus | Prionailurus_planiceps | 0.21884286825522 |
| Erignathus_barbatus | Prionailurus_rubiginosus | 0.217538230711023 |
| Erignathus_barbatus | Prionailurus_viverrinus | 0.217074701263526 |
| Erignathus_barbatus | Prionodon_pardicolor | 0.226265476721999 |
| Erignathus_barbatus | Procyon_lotor | 0.217653047155951 |
| Erignathus_barbatus | Profelis_aurata | 0.218271836338924 |
| Erignathus_barbatus | Proteles_cristatus_cristatus_TS307 | 0.231007491950657 |
| Erignathus_barbatus | Proteles_cristatus_septentrionalis_NMS_Z_2018_54 | 0.230768740354234 |
| Erignathus_barbatus | Puma_concolor | 0.22121986986353 |
| Erignathus_barbatus | Puma_yagouaroundi | 0.21662081357257 |
| Erignathus_barbatus | Pusa_caspica | 0.127701830522799 |
| Erignathus_barbatus | Pusa_hispida | 0.12359169458934 |
| Erignathus_barbatus | Pusa_sibirica | 0.124926172552408 |
| Erignathus_barbatus | Speothos_venaticus_T1624 | 0.235587512490936 |
| Erignathus_barbatus | Spilogale_putorius | 0.220359780898105 |
| Erignathus_barbatus | Taxidea_taxus | 0.201676620324451 |
| Erignathus_barbatus | Tremarctos_ornatus | 0.211937585698555 |
| Erignathus_barbatus | Urocyon_cinereoargenteus | 0.237738081972777 |
| Erignathus_barbatus | Ursus_americanus | 0.214066338099533 |
| Erignathus_barbatus | Ursus_arctos | 0.221220719763837 |
| Erignathus_barbatus | Ursus_maritimus | 0.220128122052153 |
| Erignathus_barbatus | Ursus_spelaeus | 0.214625823132142 |
| Erignathus_barbatus | Ursus_thibetanus | 0.215123054450222 |
| Erignathus_barbatus | Urva_javanica | 0.221957198247608 |
| Erignathus_barbatus | Viverra_tangalunga_FMNH146957_SRR6053069 | 0.217429882892247 |
| Erignathus_barbatus | Viverricula_indica | 0.213552671418808 |
| Erignathus_barbatus | Vulpes_corsac | 0.231795705932351 |
| Erignathus_barbatus | Vulpes_ferrilata | 0.233346662305626 |
| Erignathus_barbatus | Vulpes_lagopus | 0.230180115752803 |
| Erignathus_barbatus | Vulpes_vulpes_T3611 | 0.226507520538185 |
| Erignathus_barbatus | Vulpes_zerda | 0.227546254423585 |
| Erignathus_barbatus | Zalophus_californianus | 0.177433245999615 |
| Eumetopias_jubatus | Felis_catus | 0.2378974271952 |
| Eumetopias_jubatus | Felis_chaus | 0.237464833715595 |
| Eumetopias_jubatus | Felis_margarita | 0.235668239547439 |
| Eumetopias_jubatus | Felis_nigripes | 0.235878831412147 |
| Eumetopias_jubatus | Felis_silvestris | 0.23661059583541 |
| Eumetopias_jubatus | Fossa_fossana_FMNH156648_SRR6053060 | 0.229852634907228 |
| Eumetopias_jubatus | Genetta_servalina | 0.234008230638765 |
| Eumetopias_jubatus | Gulo_gulo | 0.227233992690941 |
| Eumetopias_jubatus | Halichoerus_grypus | 0.178893966957886 |
| Eumetopias_jubatus | Helarctos_malayanus | 0.227636594834021 |
| Eumetopias_jubatus | Hyaena_hyaena | 0.241588710810139 |
| Eumetopias_jubatus | Hydrurga_leptonyx | 0.169768438289572 |
| Eumetopias_jubatus | Leopardus_colocolo | 0.230855651251664 |
| Eumetopias_jubatus | Leopardus_geoffroyi | 0.232965473576604 |
| Eumetopias_jubatus | Leopardus_guigna | 0.234064375117844 |
| Eumetopias_jubatus | Leopardus_jacobita | 0.235215897442026 |
| Eumetopias_jubatus | Leopardus_pardalis | 0.23071334120832 |
| Eumetopias_jubatus | Leopardus_tigrinus | 0.230191001226468 |
| Eumetopias_jubatus | Leopardus_wiedii | 0.231427896391581 |
| Eumetopias_jubatus | Leptailurus_serval | 0.230632798703546 |
| Eumetopias_jubatus | Leptonychotes_weddellii | 0.16904382398959 |
| Eumetopias_jubatus | Lobodon_carcinophaga | 0.167390154445448 |
| Eumetopias_jubatus | Lutra_lutra | 0.228741904977765 |
| Eumetopias_jubatus | Lutra_sumatrana | 0.227994211520933 |
| Eumetopias_jubatus | Lutrogale_perspicillata | 0.236340341193266 |
| Eumetopias_jubatus | Lycaon_pictus | 0.245349264409986 |
| Eumetopias_jubatus | Lynx_canadensis | 0.229222892370221 |
| Eumetopias_jubatus | Lynx_lynx | 0.230005887119816 |
| Eumetopias_jubatus | Lynx_pardinus | 0.22851677643214 |
| Eumetopias_jubatus | Lynx_rufus | 0.228450818369677 |
| Eumetopias_jubatus | Manis_javanica | 0.284634069557926 |
| Eumetopias_jubatus | Martes_americana | 0.227739875402515 |
| Eumetopias_jubatus | Martes_flavigula | 0.224758221127061 |
| Eumetopias_jubatus | Martes_foina | 0.230239033752192 |
| Eumetopias_jubatus | Martes_martes | 0.225929395796227 |
| Eumetopias_jubatus | Martes_melampus | 0.225111172743085 |
| Eumetopias_jubatus | Martes_pennanti | 0.235981384987581 |
| Eumetopias_jubatus | Martes_zibellina | 0.226356799483495 |
| Eumetopias_jubatus | Meles_anakuma | 0.234888931932667 |
| Eumetopias_jubatus | Meles_leucurus | 0.230406238138534 |
| Eumetopias_jubatus | Meles_meles | 0.228008199094856 |
| Eumetopias_jubatus | Melogale_moschata | 0.235479921800937 |
| Eumetopias_jubatus | Melursus_ursinus | 0.233085813111258 |
| Eumetopias_jubatus | Mephitis_mephitis | 0.239729071339261 |
| Eumetopias_jubatus | Mirounga_leonina | 0.17339280926975 |
| Eumetopias_jubatus | Mungotictis_decemlineata | 0.23688196628268 |
| Eumetopias_jubatus | Mustela_altaica | 0.232863464180309 |
| Eumetopias_jubatus | Mustela_erminea | 0.221292400173322 |
| Eumetopias_jubatus | Mustela_eversmannii | 0.230407069411962 |
| Eumetopias_jubatus | Mustela_frenata | 0.229282146143318 |

| | | |
|---|---|---|
| Eumetopias_jubatus | Mustela_itatsi | 0.228661576023597 |
| Eumetopias_jubatus | Mustela_kathiah | 0.233508685534436 |
| Eumetopias_jubatus | Mustela_nigripes | 0.229148038324242 |
| Eumetopias_jubatus | Mustela_nivalis | 0.231869989818045 |
| Eumetopias_jubatus | Mustela_putorius | 0.227984363339229 |
| Eumetopias_jubatus | Mustela_sibirica | 0.229632440334038 |
| Eumetopias_jubatus | Nandinia_binotata | 0.221608385148618 |
| Eumetopias_jubatus | Nasua_nasua | 0.242015670773358 |
| Eumetopias_jubatus | Neofelis_nebulosa | 0.246315306515134 |
| Eumetopias_jubatus | Neomonachus_schauinslandi | 0.172927357889429 |
| Eumetopias_jubatus | Neophoca_cinerea | 0.0874109292242457 |
| Eumetopias_jubatus | Neovison_vison | 0.228391567302136 |
| Eumetopias_jubatus | Nyctereutes_procyonoides | 0.243385366998244 |
| Eumetopias_jubatus | Odobenus_rosmarus_rosmarus | 0.194057189835638 |
| Eumetopias_jubatus | Otocolobus_manul | 0.229708083019453 |
| Eumetopias_jubatus | Otocyon_megalotis_megalotis_TS305 | 0.244167816646639 |
| Eumetopias_jubatus | Otocyon_megalotis_virgatus_FMNH158128 | 0.245133558383059 |
| Eumetopias_jubatus | Paguma_larvata | 0.233160901182529 |
| Eumetopias_jubatus | Panthera_leo | 0.239400779657222 |
| Eumetopias_jubatus | Panthera_onca | 0.235938245574372 |
| Eumetopias_jubatus | Panthera_pardus | 0.238845532515208 |
| Eumetopias_jubatus | Panthera_tigris | 0.236703394497251 |
| Eumetopias_jubatus | Panthera_uncia | 0.240847314548436 |
| Eumetopias_jubatus | Paradoxurus_hermaphroditus | 0.240699624112213 |
| Eumetopias_jubatus | Parahyaena_brunnea_FD126_Fecal | 0.238059058945503 |
| Eumetopias_jubatus | Pardofelis_marmorata | 0.237283985529754 |
| Eumetopias_jubatus | Phoca_fasciata | 0.175962629944978 |
| Eumetopias_jubatus | Phoca_groenlandica | 0.180423134376105 |
| Eumetopias_jubatus | Phoca_largha | 0.180218734576106 |
| Eumetopias_jubatus | Phoca_vitulina | 0.180229388232071 |
| Eumetopias_jubatus | Phocarctos_hookeri | 0.0937090386063269 |
| Eumetopias_jubatus | Prionailurus_bengalensis | 0.230932039954289 |
| Eumetopias_jubatus | Prionailurus_bengalensis_euptilurus | 0.229692596468824 |
| Eumetopias_jubatus | Prionailurus_planiceps | 0.233336416999086 |
| Eumetopias_jubatus | Prionailurus_rubiginosus | 0.229328099599808 |
| Eumetopias_jubatus | Prionailurus_viverrinus | 0.23277526163767 |
| Eumetopias_jubatus | Prionodon_pardicolor | 0.24438444288115 |
| Eumetopias_jubatus | Procyon_lotor | 0.229678294427007 |
| Eumetopias_jubatus | Profelis_aurata | 0.236963536981435 |
| Eumetopias_jubatus | Proteles_cristatus_cristatus_TS307 | 0.241811521040998 |
| Eumetopias_jubatus | Proteles_cristatus_septentrionalis_NMS_Z_2018_54 | 0.243461534771612 |
| Eumetopias_jubatus | Puma_concolor | 0.232455849516596 |
| Eumetopias_jubatus | Puma_yagouaroundi | 0.235700019949942 |
| Eumetopias_jubatus | Pusa_caspica | 0.179121795353523 |
| Eumetopias_jubatus | Pusa_hispida | 0.177329065096754 |
| Eumetopias_jubatus | Pusa_sibirica | 0.1792250264884 |
| Eumetopias_jubatus | Speothos_venaticus_T1624 | 0.252615833620125 |
| Eumetopias_jubatus | Spilogale_putorius | 0.23928011728128 |
| Eumetopias_jubatus | Taxidea_taxus | 0.219692681123308 |
| Eumetopias_jubatus | Tremarctos_ornatus | 0.23013680646838 |
| Eumetopias_jubatus | Urocyon_cinereoargenteus | 0.248667948230236 |
| Eumetopias_jubatus | Ursus_americanus | 0.228586710982477 |
| Eumetopias_jubatus | Ursus_arctos | 0.237028021626892 |
| Eumetopias_jubatus | Ursus_maritimus | 0.237041260812769 |
| Eumetopias_jubatus | Ursus_spelaeus | 0.233964268747472 |
| Eumetopias_jubatus | Ursus_thibetanus | 0.230182834015259 |
| Eumetopias_jubatus | Urva_javanica | 0.238877004490339 |
| Eumetopias_jubatus | Viverra_tangalunga_FMNH146957_SRR6053069 | 0.228647160908195 |
| Eumetopias_jubatus | Viverricula_indica | 0.227154231157704 |
| Eumetopias_jubatus | Vulpes_corsac | 0.245521918881626 |
| Eumetopias_jubatus | Vulpes_ferrilata | 0.247253815222755 |
| Eumetopias_jubatus | Vulpes_lagopus | 0.245227696186765 |
| Eumetopias_jubatus | Vulpes_vulpes_T3611 | 0.241379091001229 |
| Eumetopias_jubatus | Vulpes_zerda | 0.241569279619093 |
| Eumetopias_jubatus | Zalophus_californianus | 0.0641475358123821 |
| Felis_catus | Felis_chaus | 0.0709354149992822 |
| Felis_catus | Felis_margarita | 0.0215147155129939 |
| Felis_catus | Felis_nigripes | 0.0719770205949549 |
| Felis_catus | Felis_silvestris | 0.0114959487434367 |
| Felis_catus | Fossa_fossana_FMNH156648_SRR6053060 | 0.185956892799881 |
| Felis_catus | Genetta_servalina | 0.177086797201779 |
| Felis_catus | Gulo_gulo | 0.235413886245369 |
| Felis_catus | Halichoerus_grypus | 0.223815873531876 |
| Felis_catus | Helarctos_malayanus | 0.23986577472748 |
| Felis_catus | Hyaena_hyaena | 0.198855363080451 |
| Felis_catus | Hydrurga_leptonyx | 0.210756103846898 |
| Felis_catus | Leopardus_colocolo | 0.116103089952621 |
| Felis_catus | Leopardus_geoffroyi | 0.117155850669398 |
| Felis_catus | Leopardus_guigna | 0.117121248680529 |
| Felis_catus | Leopardus_jacobita | 0.114607914607086 |
| Felis_catus | Leopardus_pardalis | 0.110019719965594 |
| Felis_catus | Leopardus_tigrinus | 0.117429438872883 |
| Felis_catus | Leopardus_wiedii | 0.115444389711356 |
| Felis_catus | Leptailurus_serval | 0.108238342478995 |
| Felis_catus | Leptonychotes_weddellii | 0.208624363936644 |
| Felis_catus | Lobodon_carcinophaga | 0.215142840068523 |
| Felis_catus | Lutra_lutra | 0.236235659258314 |

| | | |
|---|---|---|
| Felis_catus | Lutra_sumatrana | 0.232410240035891 |
| Felis_catus | Lutrogale_perspicillata | 0.24038420147216 |
| Felis_catus | Lycaon_pictus | 0.232482465834551 |
| Felis_catus | Lynx_canadensis | 0.111252947337958 |
| Felis_catus | Lynx_lynx | 0.114103844246869 |
| Felis_catus | Lynx_pardinus | 0.112307244891406 |
| Felis_catus | Lynx_rufus | 0.108077950398654 |
| Felis_catus | Manis_javanica | 0.290621546923033 |
| Felis_catus | Martes_americana | 0.232468529934228 |
| Felis_catus | Martes_flavigula | 0.237240130261922 |
| Felis_catus | Martes_foina | 0.232279316658961 |
| Felis_catus | Martes_martes | 0.230142493960724 |
| Felis_catus | Martes_melampus | 0.230770484152327 |
| Felis_catus | Martes_pennanti | 0.23931127941723 |
| Felis_catus | Martes_zibellina | 0.233655374765888 |
| Felis_catus | Meles_anakuma | 0.233387974055413 |
| Felis_catus | Meles_leucurus | 0.230166485930979 |
| Felis_catus | Meles_meles | 0.227536919225934 |
| Felis_catus | Melogale_moschata | 0.232268960335726 |
| Felis_catus | Melursus_ursinus | 0.237673523669827 |
| Felis_catus | Mephitis_mephitis | 0.245315293965801 |
| Felis_catus | Mirounga_leonina | 0.223965393404483 |
| Felis_catus | Mungotictis_decemlineata | 0.18906156604837 |
| Felis_catus | Mustela_altaica | 0.232097761268428 |
| Felis_catus | Mustela_erminea | 0.228253096315331 |
| Felis_catus | Mustela_eversmannii | 0.232464144693912 |
| Felis_catus | Mustela_frenata | 0.226200469859317 |
| Felis_catus | Mustela_itatsi | 0.23206598368286 |
| Felis_catus | Mustela_kathiah | 0.229981452039823 |
| Felis_catus | Mustela_nigripes | 0.231318367365951 |
| Felis_catus | Mustela_nivalis | 0.234294163905689 |
| Felis_catus | Mustela_putorius | 0.232474068257259 |
| Felis_catus | Mustela_sibirica | 0.230620173067675 |
| Felis_catus | Nandinia_binotata | 0.170297555632715 |
| Felis_catus | Nasua_nasua | 0.247646286107851 |
| Felis_catus | Neofelis_nebulosa | 0.140523071328642 |
| Felis_catus | Neomonachus_schauinslandi | 0.220985465461292 |
| Felis_catus | Neophoca_cinerea | 0.238383684376256 |
| Felis_catus | Neovison_vison | 0.233233460155059 |
| Felis_catus | Nyctereutes_procyonoides | 0.239388371776712 |
| Felis_catus | Odobenus_rosmarus_rosmarus | 0.254137573970769 |
| Felis_catus | Otocolobus_manul | 0.104499627949771 |
| Felis_catus | Otocyon_megalotis_megalotis_TS305 | 0.242506729803078 |
| Felis_catus | Otocyon_megalotis_virgatus_FMNH158128 | 0.243512741867761 |
| Felis_catus | Paguma_larvata | 0.187414941829728 |
| Felis_catus | Panthera_leo | 0.134075878634645 |
| Felis_catus | Panthera_onca | 0.134240871732316 |
| Felis_catus | Panthera_pardus | 0.133474008441614 |
| Felis_catus | Panthera_tigris | 0.143971118047308 |
| Felis_catus | Panthera_uncia | 0.137111914461043 |
| Felis_catus | Paradoxurus_hermaphroditus | 0.187570734108085 |
| Felis_catus | Parahyaena_brunnea_FD126_Fecal | 0.197470456133624 |
| Felis_catus | Pardofelis_marmorata | 0.118616437034858 |
| Felis_catus | Phoca_fasciata | 0.223264335652441 |
| Felis_catus | Phoca_groenlandica | 0.221365428695979 |
| Felis_catus | Phoca_largha | 0.224536740905171 |
| Felis_catus | Phoca_vitulina | 0.223729522436304 |
| Felis_catus | Phocarctos_hookeri | 0.241713343707726 |
| Felis_catus | Prionailurus_bengalensis | 0.101209301094294 |
| Felis_catus | Prionailurus_bengalensis_euptilurus | 0.0994702098753893 |
| Felis_catus | Prionailurus_planiceps | 0.104746173234029 |
| Felis_catus | Prionailurus_rubiginosus | 0.0987977625331954 |
| Felis_catus | Prionailurus_viverrinus | 0.104914974175536 |
| Felis_catus | Prionodon_pardicolor | 0.186914733320901 |
| Felis_catus | Procyon_lotor | 0.232296651228566 |
| Felis_catus | Profelis_aurata | 0.112442297828585 |
| Felis_catus | Proteles_cristatus_cristatus_TS307 | 0.200941641905484 |
| Felis_catus | Proteles_cristatus_septentrionalis_NMS_Z_2018_54 | 0.204134500775578 |
| Felis_catus | Puma_concolor | 0.112248985319631 |
| Felis_catus | Puma_yagouaroundi | 0.113189297036966 |
| Felis_catus | Pusa_caspica | 0.222975860132318 |
| Felis_catus | Pusa_hispida | 0.222051747994011 |
| Felis_catus | Pusa_sibirica | 0.22221762775131 |
| Felis_catus | Speothos_venaticus_T1624 | 0.245556549540525 |
| Felis_catus | Spilogale_putorius | 0.248125869101016 |
| Felis_catus | Taxidea_taxus | 0.223332710599376 |
| Felis_catus | Tremarctos_ornatus | 0.23667523738797 |
| Felis_catus | Urocyon_cinereoargenteus | 0.246439092257152 |
| Felis_catus | Ursus_americanus | 0.241461925733242 |
| Felis_catus | Ursus_arctos | 0.244821612617158 |
| Felis_catus | Ursus_maritimus | 0.244898083314711 |
| Felis_catus | Ursus_spelaeus | 0.236737494357551 |
| Felis_catus | Ursus_thibetanus | 0.236159812889779 |
| Felis_catus | Urva_javanica | 0.194954053985235 |
| Felis_catus | Viverra_tangalunga_FMNH146957_SRR6053069 | 0.175203305415073 |
| Felis_catus | Viverricula_indica | 0.176609355181575 |
| Felis_catus | Vulpes_corsac | 0.240217542798106 |

| | | |
|---|---|---|
| Felis_catus | Vulpes_ferrilata | 0.241299598706169 |
| Felis_catus | Vulpes_lagopus | 0.239421544390723 |
| Felis_catus | Vulpes_vulpes_T3611 | 0.23592830755169 |
| Felis_catus | Vulpes_zerda | 0.236359711211468 |
| Felis_catus | Zalophus_californianus | 0.235135849458626 |
| Felis_chaus | Felis_margarita | 0.0715517992857886 |
| Felis_chaus | Felis_nigripes | 0.0724025612357196 |
| Felis_chaus | Felis_silvestris | 0.0704690795621125 |
| Felis_chaus | Fossa_fossana_FMNH156648_SRR6053060 | 0.187884055864906 |
| Felis_chaus | Genetta_servalina | 0.175190769532038 |
| Felis_chaus | Gulo_gulo | 0.236020402539588 |
| Felis_chaus | Halichoerus_grypus | 0.228114110732944 |
| Felis_chaus | Helarctos_malayanus | 0.240378969291276 |
| Felis_chaus | Hyaena_hyaena | 0.198864311738604 |
| Felis_chaus | Hydrurga_leptonyx | 0.213792336090101 |
| Felis_chaus | Leopardus_colocolo | 0.113491798195846 |
| Felis_chaus | Leopardus_geoffroyi | 0.115667038000061 |
| Felis_chaus | Leopardus_guigna | 0.114974420694708 |
| Felis_chaus | Leopardus_jacobita | 0.114530462241755 |
| Felis_chaus | Leopardus_pardalis | 0.109939497782796 |
| Felis_chaus | Leopardus_tigrinus | 0.116128257286846 |
| Felis_chaus | Leopardus_wiedii | 0.113302424986089 |
| Felis_chaus | Leptailurus_serval | 0.107618090230881 |
| Felis_chaus | Leptonychotes_weddellii | 0.213747969350079 |
| Felis_chaus | Lobodon_carcinophaga | 0.219042120068691 |
| Felis_chaus | Lutra_lutra | 0.236779359192617 |
| Felis_chaus | Lutra_sumatrana | 0.235491176146476 |
| Felis_chaus | Lutrogale_perspicillata | 0.247790686154953 |
| Felis_chaus | Lycaon_pictus | 0.237243619244561 |
| Felis_chaus | Lynx_canadensis | 0.110691545612208 |
| Felis_chaus | Lynx_lynx | 0.113916729674223 |
| Felis_chaus | Lynx_pardinus | 0.112587025076403 |
| Felis_chaus | Lynx_rufus | 0.113029812576517 |
| Felis_chaus | Manis_javanica | 0.289336674799392 |
| Felis_chaus | Martes_americana | 0.237059376919021 |
| Felis_chaus | Martes_flavigula | 0.240031185625285 |
| Felis_chaus | Martes_foina | 0.235780662924496 |
| Felis_chaus | Martes_martes | 0.232394163700897 |
| Felis_chaus | Martes_melampus | 0.231371592598991 |
| Felis_chaus | Martes_pennanti | 0.240820923590959 |
| Felis_chaus | Martes_zibellina | 0.23490439599128 |
| Felis_chaus | Meles_anakuma | 0.233708531656614 |
| Felis_chaus | Meles_leucurus | 0.231895599914427 |
| Felis_chaus | Meles_meles | 0.229120602140602 |
| Felis_chaus | Melogale_moschata | 0.233555476495924 |
| Felis_chaus | Melursus_ursinus | 0.239840902147088 |
| Felis_chaus | Mephitis_mephitis | 0.243361315871673 |
| Felis_chaus | Mirounga_leonina | 0.219908459121338 |
| Felis_chaus | Mungotictis_decemlineata | 0.194706468130349 |
| Felis_chaus | Mustela_altaica | 0.236482057618913 |
| Felis_chaus | Mustela_erminea | 0.227180115323464 |
| Felis_chaus | Mustela_eversmannii | 0.235793733713226 |
| Felis_chaus | Mustela_frenata | 0.225891317362959 |
| Felis_chaus | Mustela_itatsi | 0.232434026441861 |
| Felis_chaus | Mustela_kathiah | 0.232135608096915 |
| Felis_chaus | Mustela_nigripes | 0.234636673175819 |
| Felis_chaus | Mustela_nivalis | 0.236769754931966 |
| Felis_chaus | Mustela_putorius | 0.236136382954145 |
| Felis_chaus | Mustela_sibirica | 0.23448796748845 |
| Felis_chaus | Nandinia_binotata | 0.176863959568003 |
| Felis_chaus | Nasua_nasua | 0.250308013906338 |
| Felis_chaus | Neofelis_nebulosa | 0.136267437397016 |
| Felis_chaus | Neomonachus_schauinslandi | 0.217074701263526 |
| Felis_chaus | Neophoca_cinerea | 0.241176487440597 |
| Felis_chaus | Neovison_vison | 0.231785072207452 |
| Felis_chaus | Nyctereutes_procyonoides | 0.241289158747665 |
| Felis_chaus | Odobenus_rosmarus_rosmarus | 0.257214554474513 |
| Felis_chaus | Otocolobus_manul | 0.104691635311147 |
| Felis_chaus | Otocyon_megalotis_megalotis_TS305 | 0.245548222499061 |
| Felis_chaus | Otocyon_megalotis_virgatus_FMNH158128 | 0.244759921195823 |
| Felis_chaus | Paguma_larvata | 0.184138309438768 |
| Felis_chaus | Panthera_leo | 0.137872287320153 |
| Felis_chaus | Panthera_onca | 0.132872791193727 |
| Felis_chaus | Panthera_pardus | 0.136580460747639 |
| Felis_chaus | Panthera_tigris | 0.139013015648543 |
| Felis_chaus | Panthera_uncia | 0.13608862897343 |
| Felis_chaus | Paradoxurus_hermaphroditus | 0.18870929199072 |
| Felis_chaus | Parahyaena_brunnea_FD126_Fecal | 0.196424644093827 |
| Felis_chaus | Pardofelis_marmorata | 0.119775504532316 |
| Felis_chaus | Phoca_fasciata | 0.224812413792443 |
| Felis_chaus | Phoca_groenlandica | 0.223144042901829 |
| Felis_chaus | Phoca_largha | 0.226088588219114 |
| Felis_chaus | Phoca_vitulina | 0.225277774686102 |
| Felis_chaus | Phocarctos_hookeri | 0.243044752298911 |
| Felis_chaus | Prionailurus_bengalensis | 0.0965645420343742 |
| Felis_chaus | Prionailurus_bengalensis_euptilurus | 0.100940111510726 |
| Felis_chaus | Prionailurus_planiceps | 0.103957920182126 |

| | | |
|---|---|---|
| Felis_chaus | Prionailurus_rubiginosus | 0.0988280775038844 |
| Felis_chaus | Prionailurus_viverrinus | 0.103266055278543 |
| Felis_chaus | Prionodon_pardicolor | 0.185197669309866 |
| Felis_chaus | Procyon_lotor | 0.233429511446932 |
| Felis_chaus | Profelis_aurata | 0.110046035078058 |
| Felis_chaus | Proteles_cristatus_cristatus_TS307 | 0.202589259814226 |
| Felis_chaus | Proteles_cristatus_septentrionalis_NMS_Z_2018_54 | 0.204830458870875 |
| Felis_chaus | Puma_concolor | 0.107456022601184 |
| Felis_chaus | Puma_yagouaroundi | 0.112081475246551 |
| Felis_chaus | Pusa_caspica | 0.228127815479233 |
| Felis_chaus | Pusa_hispida | 0.223380340796919 |
| Felis_chaus | Pusa_sibirica | 0.224634351485893 |
| Felis_chaus | Speothos_venaticus_T1624 | 0.246352260544381 |
| Felis_chaus | Spilogale_putorius | 0.24904284346695 |
| Felis_chaus | Taxidea_taxus | 0.227320512884809 |
| Felis_chaus | Tremarctos_ornatus | 0.238571769442952 |
| Felis_chaus | Urocyon_cinereoargenteus | 0.252254306408167 |
| Felis_chaus | Ursus_americanus | 0.241552486607199 |
| Felis_chaus | Ursus_arctos | 0.243509221335263 |
| Felis_chaus | Ursus_maritimus | 0.244151656207555 |
| Felis_chaus | Ursus_spelaeus | 0.239037665764117 |
| Felis_chaus | Ursus_thibetanus | 0.233993289656599 |
| Felis_chaus | Urva_javanica | 0.197371879751478 |
| Felis_chaus | Viverra_tangalunga_FMNH146957_SRR6053069 | 0.175767253829454 |
| Felis_chaus | Viverricula_indica | 0.17505944032048 |
| Felis_chaus | Vulpes_corsac | 0.239449879096637 |
| Felis_chaus | Vulpes_ferrilata | 0.239530097914591 |
| Felis_chaus | Vulpes_lagopus | 0.238342304390645 |
| Felis_chaus | Vulpes_vulpes_T3611 | 0.234745909013824 |
| Felis_chaus | Vulpes_zerda | 0.234294360078639 |
| Felis_chaus | Zalophus_californianus | 0.235697960911973 |
| Felis_margarita | Felis_nigripes | 0.073548266454854 |
| Felis_margarita | Felis_silvestris | 0.0197279789097873 |
| Felis_margarita | Fossa_fossana_FMNH156648_SRR6053060 | 0.186164836242263 |
| Felis_margarita | Genetta_servalina | 0.174791838986818 |
| Felis_margarita | Gulo_gulo | 0.234177466229251 |
| Felis_margarita | Halichoerus_grypus | 0.221093293798997 |
| Felis_margarita | Helarctos_malayanus | 0.238455996442979 |
| Felis_margarita | Hyaena_hyaena | 0.199193300670254 |
| Felis_margarita | Hydrurga_leptonyx | 0.209559000923369 |
| Felis_margarita | Leopardus_colocolo | 0.115650125678583 |
| Felis_margarita | Leopardus_geoffroyi | 0.116796104398888 |
| Felis_margarita | Leopardus_guigna | 0.116667364181968 |
| Felis_margarita | Leopardus_jacobita | 0.114232820816051 |
| Felis_margarita | Leopardus_pardalis | 0.10938505729639 |
| Felis_margarita | Leopardus_tigrinus | 0.118295841213938 |
| Felis_margarita | Leopardus_wiedii | 0.114897586272283 |
| Felis_margarita | Leptailurus_serval | 0.107419571194871 |
| Felis_margarita | Leptonychotes_weddellii | 0.209539997755065 |
| Felis_margarita | Lobodon_carcinophaga | 0.214255968991296 |
| Felis_margarita | Lutra_lutra | 0.234129459491642 |
| Felis_margarita | Lutra_sumatrana | 0.230870084802701 |
| Felis_margarita | Lutrogale_perspicillata | 0.239959268997471 |
| Felis_margarita | Lycaon_pictus | 0.232154058597785 |
| Felis_margarita | Lynx_canadensis | 0.111346309302565 |
| Felis_margarita | Lynx_lynx | 0.114291567170043 |
| Felis_margarita | Lynx_pardinus | 0.112213760736698 |
| Felis_margarita | Lynx_rufus | 0.107831667533089 |
| Felis_margarita | Manis_javanica | 0.288299383056908 |
| Felis_margarita | Martes_americana | 0.233109024343461 |
| Felis_margarita | Martes_flavigula | 0.233993882355807 |
| Felis_margarita | Martes_foina | 0.229951682198706 |
| Felis_margarita | Martes_martes | 0.229772871283618 |
| Felis_margarita | Martes_melampus | 0.230750039942046 |
| Felis_margarita | Martes_pennanti | 0.238864950443161 |
| Felis_margarita | Martes_zibellina | 0.233061292131479 |
| Felis_margarita | Meles_anakuma | 0.231256680162391 |
| Felis_margarita | Meles_leucurus | 0.229555877432456 |
| Felis_margarita | Meles_meles | 0.22697080131884 |
| Felis_margarita | Melogale_moschata | 0.232310469969867 |
| Felis_margarita | Melursus_ursinus | 0.238031698954354 |
| Felis_margarita | Mephitis_mephitis | 0.241538477632494 |
| Felis_margarita | Mirounga_leonina | 0.221117566982515 |
| Felis_margarita | Mungotictis_decemlineata | 0.189900932523951 |
| Felis_margarita | Mustela_altaica | 0.230347403939908 |
| Felis_margarita | Mustela_erminea | 0.225878305702229 |
| Felis_margarita | Mustela_eversmannii | 0.229360454321021 |
| Felis_margarita | Mustela_frenata | 0.224265763592836 |
| Felis_margarita | Mustela_itatsi | 0.230167012482636 |
| Felis_margarita | Mustela_kathiah | 0.229942206932607 |
| Felis_margarita | Mustela_nigripes | 0.227698666911114 |
| Felis_margarita | Mustela_nivalis | 0.232973740603085 |
| Felis_margarita | Mustela_putorius | 0.229151812120957 |
| Felis_margarita | Mustela_sibirica | 0.227527879492322 |
| Felis_margarita | Nandinia_binotata | 0.169108158907413 |
| Felis_margarita | Nasua_nasua | 0.245664483728951 |
| Felis_margarita | Neofelis_nebulosa | 0.14074040849412 |

| | | |
|---|---|---|
| Felis_margarita | Neomonachus_schauinslandi | 0.219674991278056 |
| Felis_margarita | Neophoca_cinerea | 0.2379193592292 |
| Felis_margarita | Neovison_vison | 0.230381949903953 |
| Felis_margarita | Nyctereutes_procyonoides | 0.238209400690681 |
| Felis_margarita | Odobenus_rosmarus_rosmarus | 0.253793891977629 |
| Felis_margarita | Otocolobus_manul | 0.104729699797655 |
| Felis_margarita | Otocyon_megalotis_megalotis_TS305 | 0.243917430380079 |
| Felis_margarita | Otocyon_megalotis_virgatus_FMNH158128 | 0.24571766464811 |
| Felis_margarita | Paguma_larvata | 0.184875097742716 |
| Felis_margarita | Panthera_leo | 0.13477597382045 |
| Felis_margarita | Panthera_onca | 0.134843720661493 |
| Felis_margarita | Panthera_pardus | 0.135048168683372 |
| Felis_margarita | Panthera_tigris | 0.143475676072566 |
| Felis_margarita | Panthera_uncia | 0.138557865123681 |
| Felis_margarita | Paradoxurus_hermaphroditus | 0.186008467544911 |
| Felis_margarita | Parahyaena_brunnea_FD126_Fecal | 0.196332126541762 |
| Felis_margarita | Pardofelis_marmorata | 0.118161268974335 |
| Felis_margarita | Phoca_fasciata | 0.221173161956711 |
| Felis_margarita | Phoca_groenlandica | 0.218959370857181 |
| Felis_margarita | Phoca_largha | 0.222440500173492 |
| Felis_margarita | Phoca_vitulina | 0.221206892575248 |
| Felis_margarita | Phocarctos_hookeri | 0.239352073018851 |
| Felis_margarita | Prionailurus_bengalensis | 0.101315761734441 |
| Felis_margarita | Prionailurus_bengalensis_euptilurus | 0.10049042012577 |
| Felis_margarita | Prionailurus_planiceps | 0.104024519216311 |
| Felis_margarita | Prionailurus_rubiginosus | 0.0982663412988399 |
| Felis_margarita | Prionailurus_viverrinus | 0.105392108545789 |
| Felis_margarita | Prionodon_pardicolor | 0.184794440860243 |
| Felis_margarita | Procyon_lotor | 0.232655106348674 |
| Felis_margarita | Profelis_aurata | 0.115106079051044 |
| Felis_margarita | Proteles_cristatus_cristatus_TS307 | 0.199226376951966 |
| Felis_margarita | Proteles_cristatus_septentrionalis_NMS_Z_2018_54 | 0.203046429489471 |
| Felis_margarita | Puma_concolor | 0.111425243234761 |
| Felis_margarita | Puma_yagouaroundi | 0.111898823803805 |
| Felis_margarita | Pusa_caspica | 0.221422332524515 |
| Felis_margarita | Pusa_hispida | 0.219856319002336 |
| Felis_margarita | Pusa_sibirica | 0.220127693760424 |
| Felis_margarita | Speothos_venaticus_T1624 | 0.24443710431751 |
| Felis_margarita | Spilogale_putorius | 0.24578526221512 |
| Felis_margarita | Taxidea_taxus | 0.222937359546936 |
| Felis_margarita | Tremarctos_ornatus | 0.23649829320647 |
| Felis_margarita | Urocyon_cinereoargenteus | 0.246033073456657 |
| Felis_margarita | Ursus_americanus | 0.236603193337645 |
| Felis_margarita | Ursus_arctos | 0.242470953796837 |
| Felis_margarita | Ursus_maritimus | 0.242999151288179 |
| Felis_margarita | Ursus_spelaeus | 0.236783250335599 |
| Felis_margarita | Ursus_thibetanus | 0.234757735719775 |
| Felis_margarita | Urva_javanica | 0.194241245722174 |
| Felis_margarita | Viverra_tangalunga_FMNH146957_SRR6053069 | 0.174956939295111 |
| Felis_margarita | Viverricula_indica | 0.176650552982314 |
| Felis_margarita | Vulpes_corsac | 0.237283985529754 |
| Felis_margarita | Vulpes_ferrilata | 0.240047035479291 |
| Felis_margarita | Vulpes_lagopus | 0.237707981834686 |
| Felis_margarita | Vulpes_vulpes_T3611 | 0.234795287215491 |
| Felis_margarita | Vulpes_zerda | 0.234564458553156 |
| Felis_margarita | Zalophus_californianus | 0.232694886376021 |
| Felis_nigripes | Felis_silvestris | 0.0714231419837151 |
| Felis_nigripes | Fossa_fossana_FMNH156648_SRR6053060 | 0.186945382247538 |
| Felis_nigripes | Genetta_servalina | 0.175584634765772 |
| Felis_nigripes | Gulo_gulo | 0.236395779526413 |
| Felis_nigripes | Halichoerus_grypus | 0.223806223643056 |
| Felis_nigripes | Helarctos_malayanus | 0.240941335788531 |
| Felis_nigripes | Hyaena_hyaena | 0.197532600141912 |
| Felis_nigripes | Hydrurga_leptonyx | 0.208718530973467 |
| Felis_nigripes | Leopardus_colocolo | 0.116128480111996 |
| Felis_nigripes | Leopardus_geoffroyi | 0.118315055984183 |
| Felis_nigripes | Leopardus_guigna | 0.116581786895195 |
| Felis_nigripes | Leopardus_jacobita | 0.115085649161098 |
| Felis_nigripes | Leopardus_pardalis | 0.112265579567021 |
| Felis_nigripes | Leopardus_tigrinus | 0.120291851208172 |
| Felis_nigripes | Leopardus_wiedii | 0.113107330607577 |
| Felis_nigripes | Leptailurus_serval | 0.10993717277055 |
| Felis_nigripes | Leptonychotes_weddellii | 0.210264930227226 |
| Felis_nigripes | Lobodon_carcinophaga | 0.215631584766838 |
| Felis_nigripes | Lutra_lutra | 0.23622318560913 |
| Felis_nigripes | Lutra_sumatrana | 0.230962759948569 |
| Felis_nigripes | Lutrogale_perspicillata | 0.241763598517246 |
| Felis_nigripes | Lycaon_pictus | 0.237817116043448 |
| Felis_nigripes | Lynx_canadensis | 0.111948961017477 |
| Felis_nigripes | Lynx_lynx | 0.116540986013905 |
| Felis_nigripes | Lynx_pardinus | 0.11533397088467 |
| Felis_nigripes | Lynx_rufus | 0.113848294092546 |
| Felis_nigripes | Manis_javanica | 0.287863394337324 |
| Felis_nigripes | Martes_americana | 0.234529366671424 |
| Felis_nigripes | Martes_flavigula | 0.236627385825411 |
| Felis_nigripes | Martes_foina | 0.235980960353754 |
| Felis_nigripes | Martes_martes | 0.23358728462244 |

| | | |
|---|---|---|
| Felis_nigripes | Martes_melampus | 0.232252456072798 |
| Felis_nigripes | Martes_pennanti | 0.239009353724936 |
| Felis_nigripes | Martes_zibellina | 0.234682862979237 |
| Felis_nigripes | Meles_anakuma | 0.232578686356793 |
| Felis_nigripes | Meles_leucurus | 0.230197675542082 |
| Felis_nigripes | Meles_meles | 0.2281345165794 |
| Felis_nigripes | Melogale_moschata | 0.22906668235991 |
| Felis_nigripes | Melursus_ursinus | 0.242543611430542 |
| Felis_nigripes | Mephitis_mephitis | 0.244045335437881 |
| Felis_nigripes | Mirounga_leonina | 0.22003452448079 |
| Felis_nigripes | Mungotictis_decemlineata | 0.192575034509044 |
| Felis_nigripes | Mustela_altaica | 0.235438049264562 |
| Felis_nigripes | Mustela_erminea | 0.226398742119394 |
| Felis_nigripes | Mustela_eversmannii | 0.234467326391746 |
| Felis_nigripes | Mustela_frenata | 0.226762492580629 |
| Felis_nigripes | Mustela_itatsi | 0.23296365169286 |
| Felis_nigripes | Mustela_kathiah | 0.233477555710049 |
| Felis_nigripes | Mustela_nigripes | 0.232214294511275 |
| Felis_nigripes | Mustela_nivalis | 0.235420250149683 |
| Felis_nigripes | Mustela_putorius | 0.235382482789584 |
| Felis_nigripes | Mustela_sibirica | 0.230638410417486 |
| Felis_nigripes | Nandinia_binotata | 0.173996635684339 |
| Felis_nigripes | Nasua_nasua | 0.248449601684953 |
| Felis_nigripes | Neofelis_nebulosa | 0.138946107418068 |
| Felis_nigripes | Neomonachus_schauinslandi | 0.22040410890443 |
| Felis_nigripes | Neophoca_cinerea | 0.240618762898664 |
| Felis_nigripes | Neovison_vison | 0.232444050886036 |
| Felis_nigripes | Nyctereutes_procyonoides | 0.237193610472286 |
| Felis_nigripes | Odobenus_rosmarus_rosmarus | 0.255921922927178 |
| Felis_nigripes | Otocolobus_manul | 0.106505040145176 |
| Felis_nigripes | Otocyon_megalotis_megalotis_TS305 | 0.245165442103042 |
| Felis_nigripes | Otocyon_megalotis_virgatus_FMNH158128 | 0.24471608456317 |
| Felis_nigripes | Paguma_larvata | 0.185156631567779 |
| Felis_nigripes | Panthera_leo | 0.13889405850375 |
| Felis_nigripes | Panthera_onca | 0.136570385158995 |
| Felis_nigripes | Panthera_pardus | 0.135429683537129 |
| Felis_nigripes | Panthera_tigris | 0.142759089953757 |
| Felis_nigripes | Panthera_uncia | 0.138081189685424 |
| Felis_nigripes | Paradoxurus_hermaphroditus | 0.188687230224584 |
| Felis_nigripes | Parahyaena_brunnea_FD126_Fecal | 0.19709880494953 |
| Felis_nigripes | Pardofelis_marmorata | 0.12034937912152 |
| Felis_nigripes | Phoca_fasciata | 0.22010523762062 |
| Felis_nigripes | Phoca_groenlandica | 0.222328440510988 |
| Felis_nigripes | Phoca_largha | 0.224546431697791 |
| Felis_nigripes | Phoca_vitulina | 0.222548584912347 |
| Felis_nigripes | Phocarctos_hookeri | 0.242777561282125 |
| Felis_nigripes | Prionailurus_bengalensis | 0.0990392972456348 |
| Felis_nigripes | Prionailurus_bengalensis_euptilurus | 0.100955643558366 |
| Felis_nigripes | Prionailurus_planiceps | 0.106911914828626 |
| Felis_nigripes | Prionailurus_rubiginosus | 0.103557624407491 |
| Felis_nigripes | Prionailurus_viverrinus | 0.107867112763566 |
| Felis_nigripes | Prionodon_pardicolor | 0.185301533041025 |
| Felis_nigripes | Procyon_lotor | 0.235040229251114 |
| Felis_nigripes | Profelis_aurata | 0.110861445251307 |
| Felis_nigripes | Proteles_cristatus_cristatus_TS307 | 0.196115341469443 |
| Felis_nigripes | Proteles_cristatus_septentrionalis_NMS_Z_2018_54 | 0.199595633943255 |
| Felis_nigripes | Puma_concolor | 0.11031511708837 |
| Felis_nigripes | Puma_yagouaroundi | 0.113043301028381 |
| Felis_nigripes | Pusa_caspica | 0.22555425726497 |
| Felis_nigripes | Pusa_hispida | 0.219865605890611 |
| Felis_nigripes | Pusa_sibirica | 0.222660478491032 |
| Felis_nigripes | Speothos_venaticus_T1624 | 0.247294108883239 |
| Felis_nigripes | Spilogale_putorius | 0.242753767450593 |
| Felis_nigripes | Taxidea_taxus | 0.224280329794117 |
| Felis_nigripes | Tremarctos_ornatus | 0.23915697438026 |
| Felis_nigripes | Urocyon_cinereoargenteus | 0.247307422912411 |
| Felis_nigripes | Ursus_americanus | 0.243246063929207 |
| Felis_nigripes | Ursus_arctos | 0.246346913401563 |
| Felis_nigripes | Ursus_maritimus | 0.244491676621882 |
| Felis_nigripes | Ursus_spelaeus | 0.239262132659021 |
| Felis_nigripes | Ursus_thibetanus | 0.236268158353998 |
| Felis_nigripes | Urva_javanica | 0.19571032108035 |
| Felis_nigripes | Viverra_tangalunga_FMNH146957_SRR6053069 | 0.176424422406788 |
| Felis_nigripes | Viverricula_indica | 0.172279419834215 |
| Felis_nigripes | Vulpes_corsac | 0.238283089649017 |
| Felis_nigripes | Vulpes_ferrilata | 0.24016245223337 |
| Felis_nigripes | Vulpes_lagopus | 0.238308068413143 |
| Felis_nigripes | Vulpes_vulpes_T3611 | 0.236361164281009 |
| Felis_nigripes | Vulpes_zerda | 0.236907536014625 |
| Felis_nigripes | Zalophus_californianus | 0.235105595213474 |
| Felis_silvestris | Fossa_fossana_FMNH156648_SRR6053060 | 0.185063062684744 |
| Felis_silvestris | Genetta_servalina | 0.176468726744808 |
| Felis_silvestris | Gulo_gulo | 0.23335696908379 |
| Felis_silvestris | Halichoerus_grypus | 0.221905502198478 |
| Felis_silvestris | Helarctos_malayanus | 0.240810788864521 |
| Felis_silvestris | Hyaena_hyaena | 0.197164459347081 |
| Felis_silvestris | Hydrurga_leptonyx | 0.208853755325067 |

| | | |
|---|---|---|
| Felis_silvestris | Leopardus_colocolo | 0.114122355758164 |
| Felis_silvestris | Leopardus_geoffroyi | 0.11707188337592 |
| Felis_silvestris | Leopardus_guigna | 0.11637697574314 |
| Felis_silvestris | Leopardus_jacobita | 0.112240703603293 |
| Felis_silvestris | Leopardus_pardalis | 0.109077955530325 |
| Felis_silvestris | Leopardus_tigrinus | 0.116307913778783 |
| Felis_silvestris | Leopardus_wiedii | 0.114496543688656 |
| Felis_silvestris | Leptailurus_serval | 0.10692808252254 |
| Felis_silvestris | Leptonychotes_weddellii | 0.207467442932468 |
| Felis_silvestris | Lobodon_carcinophaga | 0.213227332777366 |
| Felis_silvestris | Lutra_lutra | 0.235280166572608 |
| Felis_silvestris | Lutra_sumatrana | 0.230911133620168 |
| Felis_silvestris | Lutrogale_perspicillata | 0.238864950443161 |
| Felis_silvestris | Lycaon_pictus | 0.229861227852126 |
| Felis_silvestris | Lynx_canadensis | 0.108060802465438 |
| Felis_silvestris | Lynx_lynx | 0.112017163790642 |
| Felis_silvestris | Lynx_pardinus | 0.110320285250705 |
| Felis_silvestris | Lynx_rufus | 0.106675742873018 |
| Felis_silvestris | Manis_javanica | 0.288437689437634 |
| Felis_silvestris | Martes_americana | 0.230201312933504 |
| Felis_silvestris | Martes_flavigula | 0.235178540484677 |
| Felis_silvestris | Martes_foina | 0.229026339517475 |
| Felis_silvestris | Martes_martes | 0.229176199390246 |
| Felis_silvestris | Martes_melampus | 0.229604340869277 |
| Felis_silvestris | Martes_pennanti | 0.237794547077004 |
| Felis_silvestris | Martes_zibellina | 0.231251246179058 |
| Felis_silvestris | Meles_anakuma | 0.230528938334934 |
| Felis_silvestris | Meles_leucurus | 0.229069152010886 |
| Felis_silvestris | Meles_meles | 0.225702439516232 |
| Felis_silvestris | Melogale_moschata | 0.230351761822545 |
| Felis_silvestris | Melursus_ursinus | 0.239734844331287 |
| Felis_silvestris | Mephitis_mephitis | 0.241427205794937 |
| Felis_silvestris | Mirounga_leonina | 0.221173161956711 |
| Felis_silvestris | Mungotictis_decemlineata | 0.188685983075243 |
| Felis_silvestris | Mustela_altaica | 0.230912858082117 |
| Felis_silvestris | Mustela_erminea | 0.225769751641881 |
| Felis_silvestris | Mustela_eversmannii | 0.228782365357722 |
| Felis_silvestris | Mustela_frenata | 0.224393041841963 |
| Felis_silvestris | Mustela_itatsi | 0.229241861649171 |
| Felis_silvestris | Mustela_kathiah | 0.227797804066379 |
| Felis_silvestris | Mustela_nigripes | 0.227301167726479 |
| Felis_silvestris | Mustela_nivalis | 0.232663975359201 |
| Felis_silvestris | Mustela_putorius | 0.229555963485375 |
| Felis_silvestris | Mustela_sibirica | 0.226606083358301 |
| Felis_silvestris | Nandinia_binotata | 0.169460614583224 |
| Felis_silvestris | Nasua_nasua | 0.245393015056905 |
| Felis_silvestris | Neofelis_nebulosa | 0.139643429811992 |
| Felis_silvestris | Neomonachus_schauinslandi | 0.22003452448079 |
| Felis_silvestris | Neophoca_cinerea | 0.237094767911103 |
| Felis_silvestris | Neovison_vison | 0.231296856280838 |
| Felis_silvestris | Nyctereutes_procyonoides | 0.236771505661442 |
| Felis_silvestris | Odobenus_rosmarus_rosmarus | 0.251670646366178 |
| Felis_silvestris | Otocolobus_manul | 0.103380227500527 |
| Felis_silvestris | Otocyon_megalotis_megalotis_TS305 | 0.241522163177522 |
| Felis_silvestris | Otocyon_megalotis_virgatus_FMNH158128 | 0.243422692339548 |
| Felis_silvestris | Paguma_larvata | 0.185290725364321 |
| Felis_silvestris | Panthera_leo | 0.133105723710494 |
| Felis_silvestris | Panthera_onca | 0.132397963837151 |
| Felis_silvestris | Panthera_pardus | 0.132213938739024 |
| Felis_silvestris | Panthera_tigris | 0.142769862278752 |
| Felis_silvestris | Panthera_uncia | 0.13717143288029 |
| Felis_silvestris | Paradoxurus_hermaphroditus | 0.185572043368598 |
| Felis_silvestris | Parahyaena_brunnea_FD126_Fecal | 0.19651965425954 |
| Felis_silvestris | Pardofelis_marmorata | 0.11672083350842 |
| Felis_silvestris | Phoca_fasciata | 0.221013503011677 |
| Felis_silvestris | Phoca_groenlandica | 0.218048126048582 |
| Felis_silvestris | Phoca_largha | 0.222172858074595 |
| Felis_silvestris | Phoca_vitulina | 0.221046781723086 |
| Felis_silvestris | Phocarctos_hookeri | 0.240082835028445 |
| Felis_silvestris | Prionailurus_bengalensis | 0.0998206627081437 |
| Felis_silvestris | Prionailurus_bengalensis_euptilurus | 0.0985405891487524 |
| Felis_silvestris | Prionailurus_planiceps | 0.103350288709897 |
| Felis_silvestris | Prionailurus_rubiginosus | 0.0977775204684452 |
| Felis_silvestris | Prionailurus_viverrinus | 0.103334725437943 |
| Felis_silvestris | Prionodon_pardicolor | 0.184109488448736 |
| Felis_silvestris | Procyon_lotor | 0.23315714209919 |
| Felis_silvestris | Profelis_aurata | 0.112996861851144 |
| Felis_silvestris | Proteles_cristatus_cristatus_TS307 | 0.198378949809221 |
| Felis_silvestris | Proteles_cristatus_septentrionalis_NMS_Z_2018_54 | 0.201238824145614 |
| Felis_silvestris | Puma_concolor | 0.111864990159273 |
| Felis_silvestris | Puma_yagouaroundi | 0.11055022260501 |
| Felis_silvestris | Pusa_caspica | 0.220939541209785 |
| Felis_silvestris | Pusa_hispida | 0.219912246164718 |
| Felis_silvestris | Pusa_sibirica | 0.220184254057102 |
| Felis_silvestris | Speothos_venaticus_T1624 | 0.244213515716803 |
| Felis_silvestris | Spilogale_putorius | 0.242875674774503 |
| Felis_silvestris | Taxidea_taxus | 0.22135714044236 |

| | | |
|---|---|---|
| Felis_silvestris | Tremarctos_ornatus | 0.233159382635167 |
| Felis_silvestris | Urocyon_cinereoargenteus | 0.244639517713025 |
| Felis_silvestris | Ursus_americanus | 0.240296137609811 |
| Felis_silvestris | Ursus_arctos | 0.246094978722222 |
| Felis_silvestris | Ursus_maritimus | 0.246398597810817 |
| Felis_silvestris | Ursus_spelaeus | 0.23722982846063 |
| Felis_silvestris | Ursus_thibetanus | 0.236652099474564 |
| Felis_silvestris | Urva_javanica | 0.19469836488087 |
| Felis_silvestris | Viverra_tangalunga_FMNH146957_SRR6053069 | 0.172767278085862 |
| Felis_silvestris | Viverricula_indica | 0.176095934226161 |
| Felis_silvestris | Vulpes_corsac | 0.237796839039879 |
| Felis_silvestris | Vulpes_ferrilata | 0.240339163457339 |
| Felis_silvestris | Vulpes_lagopus | 0.237465906352939 |
| Felis_silvestris | Vulpes_vulpes_T3611 | 0.234205640623922 |
| Felis_silvestris | Vulpes_zerda | 0.233993974218917 |
| Felis_silvestris | Zalophus_californianus | 0.233213486349779 |
| Fossa_fossana_FMNH156648_SRR6053060 | Genetta_servalina | 0.181671631651608 |
| Fossa_fossana_FMNH156648_SRR6053060 | Gulo_gulo | 0.233783696812328 |
| Fossa_fossana_FMNH156648_SRR6053060 | Halichoerus_grypus | 0.217252874990444 |
| Fossa_fossana_FMNH156648_SRR6053060 | Helarctos_malayanus | 0.23350355878012 |
| Fossa_fossana_FMNH156648_SRR6053060 | Hyaena_hyaena | 0.179278152740301 |
| Fossa_fossana_FMNH156648_SRR6053060 | Hydrurga_leptonyx | 0.207305721758748 |
| Fossa_fossana_FMNH156648_SRR6053060 | Leopardus_colocolo | 0.179268407468891 |
| Fossa_fossana_FMNH156648_SRR6053060 | Leopardus_geoffroyi | 0.183983504840828 |
| Fossa_fossana_FMNH156648_SRR6053060 | Leopardus_guigna | 0.185374556828346 |
| Fossa_fossana_FMNH156648_SRR6053060 | Leopardus_jacobita | 0.183061832662743 |
| Fossa_fossana_FMNH156648_SRR6053060 | Leopardus_pardalis | 0.179263633792936 |
| Fossa_fossana_FMNH156648_SRR6053060 | Leopardus_tigrinus | 0.185001866708478 |
| Fossa_fossana_FMNH156648_SRR6053060 | Leopardus_wiedii | 0.178872386133189 |
| Fossa_fossana_FMNH156648_SRR6053060 | Leptailurus_serval | 0.182552770431067 |
| Fossa_fossana_FMNH156648_SRR6053060 | Leptonychotes_weddellii | 0.207427417991176 |
| Fossa_fossana_FMNH156648_SRR6053060 | Lobodon_carcinophaga | 0.211945963215301 |
| Fossa_fossana_FMNH156648_SRR6053060 | Lutra_lutra | 0.230295325403624 |
| Fossa_fossana_FMNH156648_SRR6053060 | Lutra_sumatrana | 0.234074265434389 |
| Fossa_fossana_FMNH156648_SRR6053060 | Lutrogale_perspicillata | 0.240650733560478 |
| Fossa_fossana_FMNH156648_SRR6053060 | Lycaon_pictus | 0.238873856897477 |
| Fossa_fossana_FMNH156648_SRR6053060 | Lynx_canadensis | 0.182356108345449 |
| Fossa_fossana_FMNH156648_SRR6053060 | Lynx_lynx | 0.183518577556697 |
| Fossa_fossana_FMNH156648_SRR6053060 | Lynx_pardinus | 0.181925233702543 |
| Fossa_fossana_FMNH156648_SRR6053060 | Lynx_rufus | 0.180433324971149 |
| Fossa_fossana_FMNH156648_SRR6053060 | Manis_javanica | 0.279149203006688 |
| Fossa_fossana_FMNH156648_SRR6053060 | Martes_americana | 0.236405803725618 |
| Fossa_fossana_FMNH156648_SRR6053060 | Martes_flavigula | 0.234335255505519 |
| Fossa_fossana_FMNH156648_SRR6053060 | Martes_foina | 0.232105172285316 |
| Fossa_fossana_FMNH156648_SRR6053060 | Martes_martes | 0.233915693502606 |
| Fossa_fossana_FMNH156648_SRR6053060 | Martes_melampus | 0.23439552842868 |
| Fossa_fossana_FMNH156648_SRR6053060 | Martes_pennanti | 0.236243010498711 |
| Fossa_fossana_FMNH156648_SRR6053060 | Martes_zibellina | 0.234367252404346 |
| Fossa_fossana_FMNH156648_SRR6053060 | Meles_anakuma | 0.238753355636765 |
| Fossa_fossana_FMNH156648_SRR6053060 | Meles_leucurus | 0.231398746987704 |
| Fossa_fossana_FMNH156648_SRR6053060 | Meles_meles | 0.227948710289748 |
| Fossa_fossana_FMNH156648_SRR6053060 | Melogale_moschata | 0.234784868519256 |
| Fossa_fossana_FMNH156648_SRR6053060 | Melursus_ursinus | 0.23738074067589 |
| Fossa_fossana_FMNH156648_SRR6053060 | Mephitis_mephitis | 0.243144443293569 |
| Fossa_fossana_FMNH156648_SRR6053060 | Mirounga_leonina | 0.213138277651302 |
| Fossa_fossana_FMNH156648_SRR6053060 | Mungotictis_decemlineata | 0.152898838680414 |
| Fossa_fossana_FMNH156648_SRR6053060 | Mustela_altaica | 0.234576269884038 |
| Fossa_fossana_FMNH156648_SRR6053060 | Mustela_erminea | 0.225759809454763 |
| Fossa_fossana_FMNH156648_SRR6053060 | Mustela_eversmannii | 0.234623642344667 |
| Fossa_fossana_FMNH156648_SRR6053060 | Mustela_frenata | 0.232426636526942 |
| Fossa_fossana_FMNH156648_SRR6053060 | Mustela_itatsi | 0.237087400939119 |
| Fossa_fossana_FMNH156648_SRR6053060 | Mustela_kathiah | 0.231716517550394 |
| Fossa_fossana_FMNH156648_SRR6053060 | Mustela_nigripes | 0.231731000660854 |
| Fossa_fossana_FMNH156648_SRR6053060 | Mustela_nivalis | 0.235522102256358 |
| Fossa_fossana_FMNH156648_SRR6053060 | Mustela_putorius | 0.232388179915306 |
| Fossa_fossana_FMNH156648_SRR6053060 | Mustela_sibirica | 0.232944830137172 |
| Fossa_fossana_FMNH156648_SRR6053060 | Nandinia_binotata | 0.172268371041101 |
| Fossa_fossana_FMNH156648_SRR6053060 | Nasua_nasua | 0.249203300864116 |
| Fossa_fossana_FMNH156648_SRR6053060 | Neofelis_nebulosa | 0.19277421015658 |
| Fossa_fossana_FMNH156648_SRR6053060 | Neomonachus_schauinslandi | 0.215686524554165 |
| Fossa_fossana_FMNH156648_SRR6053060 | Neophoca_cinerea | 0.232611034338993 |
| Fossa_fossana_FMNH156648_SRR6053060 | Neovison_vison | 0.236351436721558 |
| Fossa_fossana_FMNH156648_SRR6053060 | Nyctereutes_procyonoides | 0.234673998788925 |
| Fossa_fossana_FMNH156648_SRR6053060 | Odobenus_rosmarus_rosmarus | 0.245311507157958 |
| Fossa_fossana_FMNH156648_SRR6053060 | Otocolobus_manul | 0.186654049747279 |
| Fossa_fossana_FMNH156648_SRR6053060 | Otocyon_megalotis_megalotis_TS305 | 0.247149327291718 |
| Fossa_fossana_FMNH156648_SRR6053060 | Otocyon_megalotis_virgatus_FMNH158128 | 0.244400144359344 |
| Fossa_fossana_FMNH156648_SRR6053060 | Paguma_larvata | 0.188300965825613 |
| Fossa_fossana_FMNH156648_SRR6053060 | Panthera_leo | 0.190051953600546 |
| Fossa_fossana_FMNH156648_SRR6053060 | Panthera_onca | 0.18713291872569 |
| Fossa_fossana_FMNH156648_SRR6053060 | Panthera_pardus | 0.193014684548795 |
| Fossa_fossana_FMNH156648_SRR6053060 | Panthera_tigris | 0.19422817200057 |
| Fossa_fossana_FMNH156648_SRR6053060 | Panthera_uncia | 0.194795322156747 |
| Fossa_fossana_FMNH156648_SRR6053060 | Paradoxurus_hermaphroditus | 0.196699266790559 |
| Fossa_fossana_FMNH156648_SRR6053060 | Parahyaena_brunnea_FD126_Fecal | 0.178627877412564 |
| Fossa_fossana_FMNH156648_SRR6053060 | Pardofelis_marmorata | 0.188321934125103 |
| Fossa_fossana_FMNH156648_SRR6053060 | Phoca_fasciata | 0.213350749767535 |

| | | |
|---|---|---|
| Fossa_fossana_FMNH156648_SRR6053060 | Phoca_groenlandica | 0.21615040707736 |
| Fossa_fossana_FMNH156648_SRR6053060 | Phoca_largha | 0.216183584591463 |
| Fossa_fossana_FMNH156648_SRR6053060 | Phoca_vitulina | 0.217297447833389 |
| Fossa_fossana_FMNH156648_SRR6053060 | Phocarctos_hookeri | 0.23624118313787 |
| Fossa_fossana_FMNH156648_SRR6053060 | Prionailurus_bengalensis | 0.178868524558628 |
| Fossa_fossana_FMNH156648_SRR6053060 | Prionailurus_bengalensis_euptilurus | 0.179157049668466 |
| Fossa_fossana_FMNH156648_SRR6053060 | Prionailurus_planiceps | 0.181240107255293 |
| Fossa_fossana_FMNH156648_SRR6053060 | Prionailurus_rubiginosus | 0.178173537306339 |
| Fossa_fossana_FMNH156648_SRR6053060 | Prionailurus_viverrinus | 0.179817430701255 |
| Fossa_fossana_FMNH156648_SRR6053060 | Prionodon_pardicolor | 0.195663650336846 |
| Fossa_fossana_FMNH156648_SRR6053060 | Procyon_lotor | 0.237869105346333 |
| Fossa_fossana_FMNH156648_SRR6053060 | Profelis_aurata | 0.184740026625939 |
| Fossa_fossana_FMNH156648_SRR6053060 | Proteles_cristatus_cristatus_TS307 | 0.187114236931673 |
| Fossa_fossana_FMNH156648_SRR6053060 | Proteles_cristatus_septentrionalis_NMS_Z_2018_54 | 0.187322331825947 |
| Fossa_fossana_FMNH156648_SRR6053060 | Puma_concolor | 0.186955831794057 |
| Fossa_fossana_FMNH156648_SRR6053060 | Puma_yagouaroundi | 0.185500400446265 |
| Fossa_fossana_FMNH156648_SRR6053060 | Pusa_caspica | 0.215881078387446 |
| Fossa_fossana_FMNH156648_SRR6053060 | Pusa_hispida | 0.215720560655311 |
| Fossa_fossana_FMNH156648_SRR6053060 | Pusa_sibirica | 0.214849012035143 |
| Fossa_fossana_FMNH156648_SRR6053060 | Speothos_venaticus_T1624 | 0.25000466180893 |
| Fossa_fossana_FMNH156648_SRR6053060 | Spilogale_putorius | 0.250987321361719 |
| Fossa_fossana_FMNH156648_SRR6053060 | Taxidea_taxus | 0.227130019549696 |
| Fossa_fossana_FMNH156648_SRR6053060 | Tremarctos_ornatus | 0.233000073014217 |
| Fossa_fossana_FMNH156648_SRR6053060 | Urocyon_cinereoargenteus | 0.248282466676554 |
| Fossa_fossana_FMNH156648_SRR6053060 | Ursus_americanus | 0.234284453904263 |
| Fossa_fossana_FMNH156648_SRR6053060 | Ursus_arctos | 0.236791770808173 |
| Fossa_fossana_FMNH156648_SRR6053060 | Ursus_maritimus | 0.23767510926093 |
| Fossa_fossana_FMNH156648_SRR6053060 | Ursus_spelaeus | 0.234438129543592 |
| Fossa_fossana_FMNH156648_SRR6053060 | Ursus_thibetanus | 0.229985182640909 |
| Fossa_fossana_FMNH156648_SRR6053060 | Urva_javanica | 0.174353866427594 |
| Fossa_fossana_FMNH156648_SRR6053060 | Viverra_tangalunga_FMNH146957_SRR6053069 | 0.1781336256352 |
| Fossa_fossana_FMNH156648_SRR6053060 | Viverricula_indica | 0.177573419344846 |
| Fossa_fossana_FMNH156648_SRR6053060 | Vulpes_corsac | 0.240800696356848 |
| Fossa_fossana_FMNH156648_SRR6053060 | Vulpes_ferrilata | 0.243374419537148 |
| Fossa_fossana_FMNH156648_SRR6053060 | Vulpes_lagopus | 0.23760570856286 |
| Fossa_fossana_FMNH156648_SRR6053060 | Vulpes_vulpes_T3611 | 0.237294020413034 |
| Fossa_fossana_FMNH156648_SRR6053060 | Vulpes_zerda | 0.235658397153093 |
| Fossa_fossana_FMNH156648_SRR6053060 | Zalophus_californianus | 0.225928652922106 |
| Genetta_servalina | Gulo_gulo | 0.235197859886104 |
| Genetta_servalina | Halichoerus_grypus | 0.222768832592659 |
| Genetta_servalina | Helarctos_malayanus | 0.233444813981943 |
| Genetta_servalina | Hyaena_hyaena | 0.199968818874404 |
| Genetta_servalina | Hydrurga_leptonyx | 0.20958947826515 |
| Genetta_servalina | Leopardus_colocolo | 0.173252387181865 |
| Genetta_servalina | Leopardus_geoffroyi | 0.177762854797389 |
| Genetta_servalina | Leopardus_guigna | 0.176716435459649 |
| Genetta_servalina | Leopardus_jacobita | 0.176726397274745 |
| Genetta_servalina | Leopardus_pardalis | 0.174165193084169 |
| Genetta_servalina | Leopardus_tigrinus | 0.177970646803508 |
| Genetta_servalina | Leopardus_wiedii | 0.172820405663607 |
| Genetta_servalina | Leptailurus_serval | 0.169189452169752 |
| Genetta_servalina | Leptonychotes_weddellii | 0.211183229611302 |
| Genetta_servalina | Lobodon_carcinophaga | 0.212791031258937 |
| Genetta_servalina | Lutra_lutra | 0.232814316270616 |
| Genetta_servalina | Lutra_sumatrana | 0.23109349533836 |
| Genetta_servalina | Lutrogale_perspicillata | 0.243245259198008 |
| Genetta_servalina | Lycaon_pictus | 0.240649629373082 |
| Genetta_servalina | Lynx_canadensis | 0.175636486784676 |
| Genetta_servalina | Lynx_lynx | 0.177402375042416 |
| Genetta_servalina | Lynx_pardinus | 0.173106552360921 |
| Genetta_servalina | Lynx_rufus | 0.177765265818899 |
| Genetta_servalina | Manis_javanica | 0.294040818165723 |
| Genetta_servalina | Martes_americana | 0.237837716905993 |
| Genetta_servalina | Martes_flavigula | 0.235983775398236 |
| Genetta_servalina | Martes_foina | 0.235380237605972 |
| Genetta_servalina | Martes_martes | 0.233761875830427 |
| Genetta_servalina | Martes_melampus | 0.236303204452059 |
| Genetta_servalina | Martes_pennanti | 0.232331309112948 |
| Genetta_servalina | Martes_zibellina | 0.235724019242678 |
| Genetta_servalina | Meles_anakuma | 0.228261026843482 |
| Genetta_servalina | Meles_leucurus | 0.224705555714465 |
| Genetta_servalina | Meles_meles | 0.229984441868905 |
| Genetta_servalina | Melogale_moschata | 0.232446928672461 |
| Genetta_servalina | Melursus_ursinus | 0.239302007380227 |
| Genetta_servalina | Mephitis_mephitis | 0.231627531391254 |
| Genetta_servalina | Mirounga_leonina | 0.217261844247199 |
| Genetta_servalina | Mungotictis_decemlineata | 0.188542712914245 |
| Genetta_servalina | Mustela_altaica | 0.237987916744645 |
| Genetta_servalina | Mustela_erminea | 0.228365414774474 |
| Genetta_servalina | Mustela_eversmannii | 0.23318363176309 |
| Genetta_servalina | Mustela_frenata | 0.226217830217401 |
| Genetta_servalina | Mustela_itatsi | 0.230324073469742 |
| Genetta_servalina | Mustela_kathiah | 0.234305656943934 |
| Genetta_servalina | Mustela_nigripes | 0.23329489052055 |
| Genetta_servalina | Mustela_nivalis | 0.235526754383569 |
| Genetta_servalina | Mustela_putorius | 0.231218583030713 |
| Genetta_servalina | Mustela_sibirica | 0.232414738817192 |

| | | |
|---|---|---|
| Genetta_servalina | Nandinia_binotata | 0.17300976035897 |
| Genetta_servalina | Nasua_nasua | 0.252417869567822 |
| Genetta_servalina | Neofelis_nebulosa | 0.186068717540091 |
| Genetta_servalina | Neomonachus_schauinslandi | 0.22017000849514 |
| Genetta_servalina | Neophoca_cinerea | 0.240200100882142 |
| Genetta_servalina | Neovison_vison | 0.226237275520703 |
| Genetta_servalina | Nyctereutes_procyonoides | 0.241059862263688 |
| Genetta_servalina | Odobenus_rosmarus_rosmarus | 0.253233758359449 |
| Genetta_servalina | Otocolobus_manul | 0.173398488339221 |
| Genetta_servalina | Otocyon_megalotis_megalotis_TS305 | 0.246277447213458 |
| Genetta_servalina | Otocyon_megalotis_virgatus_FMNH158128 | 0.244007602881127 |
| Genetta_servalina | Paguma_larvata | 0.150017767258086 |
| Genetta_servalina | Panthera_leo | 0.184799394769303 |
| Genetta_servalina | Panthera_onca | 0.187146821715131 |
| Genetta_servalina | Panthera_pardus | 0.183740341004098 |
| Genetta_servalina | Panthera_tigris | 0.186343065499441 |
| Genetta_servalina | Panthera_uncia | 0.192230433771708 |
| Genetta_servalina | Paradoxurus_hermaphroditus | 0.159711949461808 |
| Genetta_servalina | Parahyaena_brunnea_FD126_Fecal | 0.200577137121279 |
| Genetta_servalina | Pardofelis_marmorata | 0.177484903721881 |
| Genetta_servalina | Phoca_fasciata | 0.219177499772319 |
| Genetta_servalina | Phoca_groenlandica | 0.21845288002839 |
| Genetta_servalina | Phoca_largha | 0.22258313703202 |
| Genetta_servalina | Phoca_vitulina | 0.222732566857426 |
| Genetta_servalina | Phocarctos_hookeri | 0.239793513906242 |
| Genetta_servalina | Prionailurus_bengalensis | 0.16915035399722 |
| Genetta_servalina | Prionailurus_bengalensis_euptilurus | 0.170116892439233 |
| Genetta_servalina | Prionailurus_planiceps | 0.1758673698829 |
| Genetta_servalina | Prionailurus_rubiginosus | 0.17280707266224 |
| Genetta_servalina | Prionailurus_viverrinus | 0.171581230860715 |
| Genetta_servalina | Prionodon_pardicolor | 0.18718059411482 |
| Genetta_servalina | Procyon_lotor | 0.236295049488816 |
| Genetta_servalina | Profelis_aurata | 0.179626134719049 |
| Genetta_servalina | Proteles_cristatus_cristatus_TS307 | 0.205048382735281 |
| Genetta_servalina | Proteles_cristatus_septentrionalis_NMS_Z_2018_54 | 0.203485450947441 |
| Genetta_servalina | Puma_concolor | 0.174548646638914 |
| Genetta_servalina | Puma_yagouaroundi | 0.177031560666479 |
| Genetta_servalina | Pusa_caspica | 0.226002420497758 |
| Genetta_servalina | Pusa_hispida | 0.219173046032238 |
| Genetta_servalina | Pusa_sibirica | 0.22440804777206 |
| Genetta_servalina | Speothos_venaticus_T1624 | 0.246359008471811 |
| Genetta_servalina | Spilogale_putorius | 0.240891633457439 |
| Genetta_servalina | Taxidea_taxus | 0.221133020063788 |
| Genetta_servalina | Tremarctos_ornatus | 0.236337890486947 |
| Genetta_servalina | Urocyon_cinereoargenteus | 0.24684426629779 |
| Genetta_servalina | Ursus_americanus | 0.235674472080326 |
| Genetta_servalina | Ursus_arctos | 0.238375401400366 |
| Genetta_servalina | Ursus_maritimus | 0.237665597393674 |
| Genetta_servalina | Ursus_spelaeus | 0.238387563935096 |
| Genetta_servalina | Ursus_thibetanus | 0.23566599788864 |
| Genetta_servalina | Urva_javanica | 0.184228466574011 |
| Genetta_servalina | Viverra_tangalunga_FMNH146957_SRR6053069 | 0.146150126182802 |
| Genetta_servalina | Viverricula_indica | 0.142862679749811 |
| Genetta_servalina | Vulpes_corsac | 0.241164167570351 |
| Genetta_servalina | Vulpes_ferrilata | 0.240336694443758 |
| Genetta_servalina | Vulpes_lagopus | 0.240851773454472 |
| Genetta_servalina | Vulpes_vulpes_T3611 | 0.233655199143006 |
| Genetta_servalina | Vulpes_zerda | 0.233769388568976 |
| Genetta_servalina | Zalophus_californianus | 0.234325428310562 |
| Gulo_gulo | Halichoerus_grypus | 0.215376895778748 |
| Gulo_gulo | Helarctos_malayanus | 0.236164089881605 |
| Gulo_gulo | Hyaena_hyaena | 0.242118385425725 |
| Gulo_gulo | Hydrurga_leptonyx | 0.209897344407413 |
| Gulo_gulo | Leopardus_colocolo | 0.232604359436642 |
| Gulo_gulo | Leopardus_geoffroyi | 0.233536260888875 |
| Gulo_gulo | Leopardus_guigna | 0.232944586260788 |
| Gulo_gulo | Leopardus_jacobita | 0.233146831411804 |
| Gulo_gulo | Leopardus_pardalis | 0.233208906987143 |
| Gulo_gulo | Leopardus_tigrinus | 0.236146894026847 |
| Gulo_gulo | Leopardus_wiedii | 0.232437556971905 |
| Gulo_gulo | Leptailurus_serval | 0.235076566031811 |
| Gulo_gulo | Leptonychotes_weddellii | 0.20986655086409 |
| Gulo_gulo | Lobodon_carcinophaga | 0.20953082291628 |
| Gulo_gulo | Lutra_lutra | 0.170220997459419 |
| Gulo_gulo | Lutra_sumatrana | 0.169706647038585 |
| Gulo_gulo | Lutrogale_perspicillata | 0.183366604332578 |
| Gulo_gulo | Lycaon_pictus | 0.237929662175271 |
| Gulo_gulo | Lynx_canadensis | 0.232947056919257 |
| Gulo_gulo | Lynx_lynx | 0.2344570270709 |
| Gulo_gulo | Lynx_pardinus | 0.231176571536165 |
| Gulo_gulo | Lynx_rufus | 0.231555842412438 |
| Gulo_gulo | Manis_javanica | 0.290544718903008 |
| Gulo_gulo | Martes_americana | 0.132407643341931 |
| Gulo_gulo | Martes_flavigula | 0.13409575431271 |
| Gulo_gulo | Martes_foina | 0.135952911771824 |
| Gulo_gulo | Martes_martes | 0.128266053528798 |
| Gulo_gulo | Martes_melampus | 0.131865354469285 |

| | | |
|---|---|---|
| Gulo_gulo | Martes_pennanti | 0.152636523903584 |
| Gulo_gulo | Martes_zibellina | 0.130588708174826 |
| Gulo_gulo | Meles_anakuma | 0.178126959506196 |
| Gulo_gulo | Meles_leucurus | 0.174808328112843 |
| Gulo_gulo | Meles_meles | 0.172259333285024 |
| Gulo_gulo | Melogale_moschata | 0.170395239291934 |
| Gulo_gulo | Melursus_ursinus | 0.241816666369181 |
| Gulo_gulo | Mephitis_mephitis | 0.237504889254993 |
| Gulo_gulo | Mirounga_leonina | 0.215605171674939 |
| Gulo_gulo | Mungotictis_decemlineata | 0.23878190742233 |
| Gulo_gulo | Mustela_altaica | 0.170378133655572 |
| Gulo_gulo | Mustela_erminea | 0.163964895374698 |
| Gulo_gulo | Mustela_eversmannii | 0.170533977940039 |
| Gulo_gulo | Mustela_frenata | 0.169216970926628 |
| Gulo_gulo | Mustela_itatsi | 0.172626936383312 |
| Gulo_gulo | Mustela_kathiah | 0.169325567867933 |
| Gulo_gulo | Mustela_nigripes | 0.170379651972282 |
| Gulo_gulo | Mustela_nivalis | 0.16977668704643 |
| Gulo_gulo | Mustela_putorius | 0.169218201341913 |
| Gulo_gulo | Mustela_sibirica | 0.171740039594199 |
| Gulo_gulo | Nandinia_binotata | 0.218523313936682 |
| Gulo_gulo | Nasua_nasua | 0.228475597988235 |
| Gulo_gulo | Neofelis_nebulosa | 0.244973745379745 |
| Gulo_gulo | Neomonachus_schauinslandi | 0.216351311006232 |
| Gulo_gulo | Neophoca_cinerea | 0.230682941980627 |
| Gulo_gulo | Neovison_vison | 0.16860107247245 |
| Gulo_gulo | Nyctereutes_procyonoides | 0.242862099977737 |
| Gulo_gulo | Odobenus_rosmarus_rosmarus | 0.249724796074463 |
| Gulo_gulo | Otocolobus_manul | 0.236178512376875 |
| Gulo_gulo | Otocyon_megalotis_megalotis_TS305 | 0.24517719099695 |
| Gulo_gulo | Otocyon_megalotis_virgatus_FMNH158128 | 0.246212946233481 |
| Gulo_gulo | Paguma_larvata | 0.242965398114538 |
| Gulo_gulo | Panthera_leo | 0.238949213783659 |
| Gulo_gulo | Panthera_onca | 0.239345934159937 |
| Gulo_gulo | Panthera_pardus | 0.242211724199742 |
| Gulo_gulo | Panthera_tigris | 0.241851979556319 |
| Gulo_gulo | Panthera_uncia | 0.245694884353459 |
| Gulo_gulo | Paradoxurus_hermaphroditus | 0.250970032745853 |
| Gulo_gulo | Parahyaena_brunnea_FD126_Fecal | 0.241097655778155 |
| Gulo_gulo | Pardofelis_marmorata | 0.237869105346333 |
| Gulo_gulo | Phoca_fasciata | 0.214350094388567 |
| Gulo_gulo | Phoca_groenlandica | 0.215291579180385 |
| Gulo_gulo | Phoca_largha | 0.214598277756768 |
| Gulo_gulo | Phoca_vitulina | 0.215193045034459 |
| Gulo_gulo | Phocarctos_hookeri | 0.23434978251555 |
| Gulo_gulo | Prionailurus_bengalensis | 0.231606337235465 |
| Gulo_gulo | Prionailurus_bengalensis_euptilurus | 0.232348071512336 |
| Gulo_gulo | Prionailurus_planiceps | 0.233229711590887 |
| Gulo_gulo | Prionailurus_rubiginosus | 0.232176386283361 |
| Gulo_gulo | Prionailurus_viverrinus | 0.234229275067911 |
| Gulo_gulo | Prionodon_pardicolor | 0.237216443675958 |
| Gulo_gulo | Procyon_lotor | 0.219184328069777 |
| Gulo_gulo | Profelis_aurata | 0.240160587872726 |
| Gulo_gulo | Proteles_cristatus_cristatus_TS307 | 0.243912202333737 |
| Gulo_gulo | Proteles_cristatus_septentrionalis_NMS_Z_2018_54 | 0.243352229153917 |
| Gulo_gulo | Puma_concolor | 0.23990578178428 |
| Gulo_gulo | Puma_yagouaroundi | 0.234577726038916 |
| Gulo_gulo | Pusa_caspica | 0.215291492709525 |
| Gulo_gulo | Pusa_hispida | 0.211686951077845 |
| Gulo_gulo | Pusa_sibirica | 0.214772039353049 |
| Gulo_gulo | Speothos_venaticus_T1624 | 0.249467068198699 |
| Gulo_gulo | Spilogale_putorius | 0.232265143774396 |
| Gulo_gulo | Taxidea_taxus | 0.17119471034008 |
| Gulo_gulo | Tremarctos_ornatus | 0.234432313088534 |
| Gulo_gulo | Urocyon_cinereoargenteus | 0.252171776318742 |
| Gulo_gulo | Ursus_americanus | 0.236950701580145 |
| Gulo_gulo | Ursus_arctos | 0.240632621274938 |
| Gulo_gulo | Ursus_maritimus | 0.24016275802116 |
| Gulo_gulo | Ursus_spelaeus | 0.238143419917436 |
| Gulo_gulo | Ursus_thibetanus | 0.234147560092634 |
| Gulo_gulo | Urva_javanica | 0.241629288686688 |
| Gulo_gulo | Viverra_tangalunga_FMNH146957_SRR6053069 | 0.231319934757069 |
| Gulo_gulo | Viverricula_indica | 0.235973483310338 |
| Gulo_gulo | Vulpes_corsac | 0.244839884935136 |
| Gulo_gulo | Vulpes_ferrilata | 0.245323098677158 |
| Gulo_gulo | Vulpes_lagopus | 0.243908675760281 |
| Gulo_gulo | Vulpes_vulpes_T3611 | 0.244560106150458 |
| Gulo_gulo | Vulpes_zerda | 0.242763207764111 |
| Gulo_gulo | Zalophus_californianus | 0.229897913716824 |
| Halichoerus_grypus | Helarctos_malayanus | 0.216616729818173 |
| Halichoerus_grypus | Hyaena_hyaena | 0.225799671659227 |
| Halichoerus_grypus | Hydrurga_leptonyx | 0.133978446128374 |
| Halichoerus_grypus | Leopardus_colocolo | 0.2174018223775 |
| Halichoerus_grypus | Leopardus_geoffroyi | 0.221835508397171 |
| Halichoerus_grypus | Leopardus_guigna | 0.223893570228364 |
| Halichoerus_grypus | Leopardus_jacobita | 0.221203582337766 |
| Halichoerus_grypus | Leopardus_pardalis | 0.219584934112871 |

| | | |
|---|---|---|
| Halichoerus_grypus | Leopardus_tigrinus | 0.223726754914053 |
| Halichoerus_grypus | Leopardus_wiedii | 0.218742568175255 |
| Halichoerus_grypus | Leptailurus_serval | 0.218336206642417 |
| Halichoerus_grypus | Leptonychotes_weddellii | 0.130062183980121 |
| Halichoerus_grypus | Lobodon_carcinophaga | 0.132901765693086 |
| Halichoerus_grypus | Lutra_lutra | 0.216124604305355 |
| Halichoerus_grypus | Lutra_sumatrana | 0.212064326146038 |
| Halichoerus_grypus | Lutrogale_perspicillata | 0.222389563237557 |
| Halichoerus_grypus | Lycaon_pictus | 0.231956412676645 |
| Halichoerus_grypus | Lynx_canadensis | 0.220409431970238 |
| Halichoerus_grypus | Lynx_lynx | 0.21939834062945 |
| Halichoerus_grypus | Lynx_pardinus | 0.222672176299277 |
| Halichoerus_grypus | Lynx_rufus | 0.217870110909774 |
| Halichoerus_grypus | Manis_javanica | 0.275132688482184 |
| Halichoerus_grypus | Martes_americana | 0.21622132739941 |
| Halichoerus_grypus | Martes_flavigula | 0.218283836806204 |
| Halichoerus_grypus | Martes_foina | 0.215842255253346 |
| Halichoerus_grypus | Martes_martes | 0.21651413152559 |
| Halichoerus_grypus | Martes_melampus | 0.216416353999833 |
| Halichoerus_grypus | Martes_pennanti | 0.223935983936971 |
| Halichoerus_grypus | Martes_zibellina | 0.21726276258904 |
| Halichoerus_grypus | Meles_anakuma | 0.220908394835519 |
| Halichoerus_grypus | Meles_leucurus | 0.216314272833765 |
| Halichoerus_grypus | Meles_meles | 0.213979340731761 |
| Halichoerus_grypus | Melogale_moschata | 0.221614974208947 |
| Halichoerus_grypus | Melursus_ursinus | 0.215267736667937 |
| Halichoerus_grypus | Mephitis_mephitis | 0.225266983919842 |
| Halichoerus_grypus | Mirounga_leonina | 0.142937556492474 |
| Halichoerus_grypus | Mungotictis_decemlineata | 0.223173201382388 |
| Halichoerus_grypus | Mustela_altaica | 0.222095548441886 |
| Halichoerus_grypus | Mustela_erminea | 0.207564276517713 |
| Halichoerus_grypus | Mustela_eversmannii | 0.217203587142899 |
| Halichoerus_grypus | Mustela_frenata | 0.213181199844441 |
| Halichoerus_grypus | Mustela_itatsi | 0.216741706942895 |
| Halichoerus_grypus | Mustela_kathiah | 0.219569805876777 |
| Halichoerus_grypus | Mustela_nigripes | 0.215048282451881 |
| Halichoerus_grypus | Mustela_nivalis | 0.21752752625368 |
| Halichoerus_grypus | Mustela_putorius | 0.215577050501245 |
| Halichoerus_grypus | Mustela_sibirica | 0.216806938692339 |
| Halichoerus_grypus | Nandinia_binotata | 0.208142190259817 |
| Halichoerus_grypus | Nasua_nasua | 0.230586573702466 |
| Halichoerus_grypus | Neofelis_nebulosa | 0.231660079475358 |
| Halichoerus_grypus | Neomonachus_schauinslandi | 0.148350714145232 |
| Halichoerus_grypus | Neophoca_cinerea | 0.184051868456238 |
| Halichoerus_grypus | Neovison_vison | 0.219731240323858 |
| Halichoerus_grypus | Nyctereutes_procyonoides | 0.232314834292046 |
| Halichoerus_grypus | Odobenus_rosmarus_rosmarus | 0.205823633028615 |
| Halichoerus_grypus | Otocolobus_manul | 0.223175201147531 |
| Halichoerus_grypus | Otocyon_megalotis_megalotis_TS305 | 0.23660702275052 |
| Halichoerus_grypus | Otocyon_megalotis_virgatus_FMNH158128 | 0.23680237330562 |
| Halichoerus_grypus | Paguma_larvata | 0.229712440546778 |
| Halichoerus_grypus | Panthera_leo | 0.228193015547523 |
| Halichoerus_grypus | Panthera_onca | 0.228450362208316 |
| Halichoerus_grypus | Panthera_pardus | 0.231974261832763 |
| Halichoerus_grypus | Panthera_tigris | 0.229873213883889 |
| Halichoerus_grypus | Panthera_uncia | 0.227933791370928 |
| Halichoerus_grypus | Paradoxurus_hermaphroditus | 0.239769129475816 |
| Halichoerus_grypus | Parahyaena_brunnea_FD126_Fecal | 0.225971488856077 |
| Halichoerus_grypus | Pardofelis_marmorata | 0.225465313032065 |
| Halichoerus_grypus | Phoca_fasciata | 0.0829251031121342 |
| Halichoerus_grypus | Phoca_groenlandica | 0.0874443473918359 |
| Halichoerus_grypus | Phoca_largha | 0.0383213296500316 |
| Halichoerus_grypus | Phoca_vitulina | 0.0390438009879371 |
| Halichoerus_grypus | Phocarctos_hookeri | 0.187111041071437 |
| Halichoerus_grypus | Prionailurus_bengalensis | 0.220099598830131 |
| Halichoerus_grypus | Prionailurus_bengalensis_euptilurus | 0.21934618140203 |
| Halichoerus_grypus | Prionailurus_planiceps | 0.223156028492915 |
| Halichoerus_grypus | Prionailurus_rubiginosus | 0.221985110787999 |
| Halichoerus_grypus | Prionailurus_viverrinus | 0.220619672606842 |
| Halichoerus_grypus | Prionodon_pardicolor | 0.2314376410846 |
| Halichoerus_grypus | Procyon_lotor | 0.219151843119156 |
| Halichoerus_grypus | Profelis_aurata | 0.225712359389969 |
| Halichoerus_grypus | Proteles_cristatus_cristatus_TS307 | 0.233413494369339 |
| Halichoerus_grypus | Proteles_cristatus_septentrionalis_NMS_Z_2018_54 | 0.234143894983598 |
| Halichoerus_grypus | Puma_concolor | 0.220488830783191 |
| Halichoerus_grypus | Puma_yagouaroundi | 0.223254758312796 |
| Halichoerus_grypus | Pusa_caspica | 0.0306330755622393 |
| Halichoerus_grypus | Pusa_hispida | 0.03391303016489 |
| Halichoerus_grypus | Pusa_sibirica | 0.0296502993237379 |
| Halichoerus_grypus | Speothos_venaticus_T1624 | 0.241990767863412 |
| Halichoerus_grypus | Spilogale_putorius | 0.229852782968918 |
| Halichoerus_grypus | Taxidea_taxus | 0.207314800556743 |
| Halichoerus_grypus | Tremarctos_ornatus | 0.216521474844066 |
| Halichoerus_grypus | Urocyon_cinereoargenteus | 0.238274802815835 |
| Halichoerus_grypus | Ursus_americanus | 0.217526567011315 |
| Halichoerus_grypus | Ursus_arctos | 0.221605006007084 |
| Halichoerus_grypus | Ursus_maritimus | 0.220732440163752 |

| | | |
|---|---|---|
| Halichoerus_grypus | Ursus_spelaeus | 0.221142738721169 |
| Halichoerus_grypus | Ursus_thibetanus | 0.217670304208516 |
| Halichoerus_grypus | Urva_javanica | 0.222743521818421 |
| Halichoerus_grypus | Viverra_tangalunga_FMNH146957_SRR6053069 | 0.223302237751376 |
| Halichoerus_grypus | Viverricula_indica | 0.219046547068281 |
| Halichoerus_grypus | Vulpes_corsac | 0.237783606005503 |
| Halichoerus_grypus | Vulpes_ferrilata | 0.23832953407332 |
| Halichoerus_grypus | Vulpes_lagopus | 0.23455509341401 |
| Halichoerus_grypus | Vulpes_vulpes_T3611 | 0.233003417259097 |
| Halichoerus_grypus | Vulpes_zerda | 0.230904737540165 |
| Halichoerus_grypus | Zalophus_californianus | 0.181566025265901 |
| Helarctos_malayanus | Hyaena_hyaena | 0.241274351396788 |
| Helarctos_malayanus | Hydrurga_leptonyx | 0.215060324994928 |
| Helarctos_malayanus | Leopardus_colocolo | 0.238533327166408 |
| Helarctos_malayanus | Leopardus_geoffroyi | 0.23839030570595 |
| Helarctos_malayanus | Leopardus_guigna | 0.238077828009509 |
| Helarctos_malayanus | Leopardus_jacobita | 0.240355232196622 |
| Helarctos_malayanus | Leopardus_pardalis | 0.237487956628382 |
| Helarctos_malayanus | Leopardus_tigrinus | 0.240586044507439 |
| Helarctos_malayanus | Leopardus_wiedii | 0.238070500166269 |
| Helarctos_malayanus | Leptailurus_serval | 0.238896491060456 |
| Helarctos_malayanus | Leptonychotes_weddellii | 0.214213166887384 |
| Helarctos_malayanus | Lobodon_carcinophaga | 0.215654996427528 |
| Helarctos_malayanus | Lutra_lutra | 0.244613294964905 |
| Helarctos_malayanus | Lutra_sumatrana | 0.238275035403124 |
| Helarctos_malayanus | Lutrogale_perspicillata | 0.247780442131762 |
| Helarctos_malayanus | Lycaon_pictus | 0.244004364875759 |
| Helarctos_malayanus | Lynx_canadensis | 0.238453372479744 |
| Helarctos_malayanus | Lynx_lynx | 0.236214190059566 |
| Helarctos_malayanus | Lynx_pardinus | 0.239232645285905 |
| Helarctos_malayanus | Lynx_rufus | 0.238614144545727 |
| Helarctos_malayanus | Manis_javanica | 0.304258301891886 |
| Helarctos_malayanus | Martes_americana | 0.235028926661737 |
| Helarctos_malayanus | Martes_flavigula | 0.237209210866427 |
| Helarctos_malayanus | Martes_foina | 0.234398833171846 |
| Helarctos_malayanus | Martes_martes | 0.239004544346564 |
| Helarctos_malayanus | Martes_melampus | 0.23856643081378 |
| Helarctos_malayanus | Martes_pennanti | 0.244439497047968 |
| Helarctos_malayanus | Martes_zibellina | 0.238763081344946 |
| Helarctos_malayanus | Meles_anakuma | 0.240748232841366 |
| Helarctos_malayanus | Meles_leucurus | 0.235569579509769 |
| Helarctos_malayanus | Meles_meles | 0.232457509577728 |
| Helarctos_malayanus | Melogale_moschata | 0.236616322812377 |
| Helarctos_malayanus | Melursus_ursinus | 0.103484203451219 |
| Helarctos_malayanus | Mephitis_mephitis | 0.243872371695784 |
| Helarctos_malayanus | Mirounga_leonina | 0.217758368028611 |
| Helarctos_malayanus | Mungotictis_decemlineata | 0.240373976514538 |
| Helarctos_malayanus | Mustela_altaica | 0.242568614617513 |
| Helarctos_malayanus | Mustela_erminea | 0.232735652651723 |
| Helarctos_malayanus | Mustela_eversmannii | 0.238151754962652 |
| Helarctos_malayanus | Mustela_frenata | 0.234775644904008 |
| Helarctos_malayanus | Mustela_itatsi | 0.240018595958509 |
| Helarctos_malayanus | Mustela_kathiah | 0.239243117890839 |
| Helarctos_malayanus | Mustela_nigripes | 0.239139488167242 |
| Helarctos_malayanus | Mustela_nivalis | 0.240589427321173 |
| Helarctos_malayanus | Mustela_putorius | 0.237764826432351 |
| Helarctos_malayanus | Mustela_sibirica | 0.240154152552905 |
| Helarctos_malayanus | Nandinia_binotata | 0.226398917188819 |
| Helarctos_malayanus | Nasua_nasua | 0.250098751953735 |
| Helarctos_malayanus | Neofelis_nebulosa | 0.248780127443149 |
| Helarctos_malayanus | Neomonachus_schauinslandi | 0.220739615158688 |
| Helarctos_malayanus | Neophoca_cinerea | 0.229699290435597 |
| Helarctos_malayanus | Neovison_vison | 0.238823711465163 |
| Helarctos_malayanus | Nyctereutes_procyonoides | 0.243594509348177 |
| Helarctos_malayanus | Odobenus_rosmarus_rosmarus | 0.249716047648154 |
| Helarctos_malayanus | Otocolobus_manul | 0.239456961126365 |
| Helarctos_malayanus | Otocyon_megalotis_megalotis_TS305 | 0.250176586510841 |
| Helarctos_malayanus | Otocyon_megalotis_virgatus_FMNH158128 | 0.24862871998581 |
| Helarctos_malayanus | Paguma_larvata | 0.236701740958095 |
| Helarctos_malayanus | Panthera_leo | 0.248285784444246 |
| Helarctos_malayanus | Panthera_onca | 0.245665416556439 |
| Helarctos_malayanus | Panthera_pardus | 0.247512099107451 |
| Helarctos_malayanus | Panthera_tigris | 0.246600002172272 |
| Helarctos_malayanus | Panthera_uncia | 0.246992923564198 |
| Helarctos_malayanus | Paradoxurus_hermaphroditus | 0.243525846814425 |
| Helarctos_malayanus | Parahyaena_brunnea_FD126_Fecal | 0.24456177256933 |
| Helarctos_malayanus | Pardofelis_marmorata | 0.243640764799295 |
| Helarctos_malayanus | Phoca_fasciata | 0.213593838249837 |
| Helarctos_malayanus | Phoca_groenlandica | 0.218442912033028 |
| Helarctos_malayanus | Phoca_largha | 0.218043420865174 |
| Helarctos_malayanus | Phoca_vitulina | 0.220233528346143 |
| Helarctos_malayanus | Phocarctos_hookeri | 0.233800163541167 |
| Helarctos_malayanus | Prionailurus_bengalensis | 0.238854638335273 |
| Helarctos_malayanus | Prionailurus_bengalensis_euptilurus | 0.237017988677135 |
| Helarctos_malayanus | Prionailurus_planiceps | 0.238135807161939 |
| Helarctos_malayanus | Prionailurus_rubiginosus | 0.235012191469721 |
| Helarctos_malayanus | Prionailurus_viverrinus | 0.240515211082735 |

| | | |
|---|---|---|
| Helarctos_malayanus | Prionodon_pardicolor | 0.239185845105495 |
| Helarctos_malayanus | Procyon_lotor | 0.238812565921427 |
| Helarctos_malayanus | Profelis_aurata | 0.238860752886323 |
| Helarctos_malayanus | Proteles_cristatus_cristatus_TS307 | 0.245394887612549 |
| Helarctos_malayanus | Proteles_cristatus_septentrionalis_NMS_Z_2018_54 | 0.243876355590837 |
| Helarctos_malayanus | Puma_concolor | 0.240544477254784 |
| Helarctos_malayanus | Puma_yagouaroundi | 0.239356538629088 |
| Helarctos_malayanus | Pusa_caspica | 0.216242986594548 |
| Helarctos_malayanus | Pusa_hispida | 0.214527360657044 |
| Helarctos_malayanus | Pusa_sibirica | 0.216816647099504 |
| Helarctos_malayanus | Speothos_venaticus_T1624 | 0.246614719603797 |
| Helarctos_malayanus | Spilogale_putorius | 0.242966163926198 |
| Helarctos_malayanus | Taxidea_taxus | 0.230621967391961 |
| Helarctos_malayanus | Tremarctos_ornatus | 0.172371799641255 |
| Helarctos_malayanus | Urocyon_cinereoargenteus | 0.253707022979686 |
| Helarctos_malayanus | Ursus_americanus | 0.0939268005678426 |
| Helarctos_malayanus | Ursus_arctos | 0.0996051400635661 |
| Helarctos_malayanus | Ursus_maritimus | 0.0976182031471276 |
| Helarctos_malayanus | Ursus_spelaeus | 0.0980290338995658 |
| Helarctos_malayanus | Ursus_thibetanus | 0.089057152766268 |
| Helarctos_malayanus | Urva_javanica | 0.245294171486471 |
| Helarctos_malayanus | Viverra_tangalunga_FMNH146957_SRR6053069 | 0.230881637351795 |
| Helarctos_malayanus | Viverricula_indica | 0.233895614534794 |
| Helarctos_malayanus | Vulpes_corsac | 0.243746151608003 |
| Helarctos_malayanus | Vulpes_ferrilata | 0.247023571835953 |
| Helarctos_malayanus | Vulpes_lagopus | 0.249122787607631 |
| Helarctos_malayanus | Vulpes_vulpes_T3611 | 0.24847491089251 |
| Helarctos_malayanus | Vulpes_zerda | 0.249871607093296 |
| Helarctos_malayanus | Zalophus_californianus | 0.226047947737354 |
| Hyaena_hyaena | Hydrurga_leptonyx | 0.220493439934861 |
| Hyaena_hyaena | Leopardus_colocolo | 0.188011451333578 |
| Hyaena_hyaena | Leopardus_geoffroyi | 0.193270281794348 |
| Hyaena_hyaena | Leopardus_guigna | 0.191537362512746 |
| Hyaena_hyaena | Leopardus_jacobita | 0.192362919932727 |
| Hyaena_hyaena | Leopardus_pardalis | 0.187880596134756 |
| Hyaena_hyaena | Leopardus_tigrinus | 0.191452767300028 |
| Hyaena_hyaena | Leopardus_wiedii | 0.187711691475008 |
| Hyaena_hyaena | Leptailurus_serval | 0.194718647749256 |
| Hyaena_hyaena | Leptonychotes_weddellii | 0.215457101951618 |
| Hyaena_hyaena | Lobodon_carcinophaga | 0.22538488712589 |
| Hyaena_hyaena | Lutra_lutra | 0.241293245458708 |
| Hyaena_hyaena | Lutra_sumatrana | 0.240833298868764 |
| Hyaena_hyaena | Lutrogale_perspicillata | 0.243352229153917 |
| Hyaena_hyaena | Lycaon_pictus | 0.248634401293307 |
| Hyaena_hyaena | Lynx_canadensis | 0.187757794425032 |
| Hyaena_hyaena | Lynx_lynx | 0.189368299006949 |
| Hyaena_hyaena | Lynx_pardinus | 0.191507806340769 |
| Hyaena_hyaena | Lynx_rufus | 0.193488744410429 |
| Hyaena_hyaena | Manis_javanica | 0.283434740762321 |
| Hyaena_hyaena | Martes_americana | 0.240704408908443 |
| Hyaena_hyaena | Martes_flavigula | 0.240638893415971 |
| Hyaena_hyaena | Martes_foina | 0.240026237235351 |
| Hyaena_hyaena | Martes_martes | 0.239977532438672 |
| Hyaena_hyaena | Martes_melampus | 0.240440036961305 |
| Hyaena_hyaena | Martes_pennanti | 0.241913539015213 |
| Hyaena_hyaena | Martes_zibellina | 0.242958278738264 |
| Hyaena_hyaena | Meles_anakuma | 0.243545616813097 |
| Hyaena_hyaena | Meles_leucurus | 0.238572376531947 |
| Hyaena_hyaena | Meles_meles | 0.241087211841275 |
| Hyaena_hyaena | Melogale_moschata | 0.241665527770875 |
| Hyaena_hyaena | Melursus_ursinus | 0.238563283484368 |
| Hyaena_hyaena | Mephitis_mephitis | 0.249687556930388 |
| Hyaena_hyaena | Mirounga_leonina | 0.226061440695296 |
| Hyaena_hyaena | Mungotictis_decemlineata | 0.187722158006068 |
| Hyaena_hyaena | Mustela_altaica | 0.244192062183947 |
| Hyaena_hyaena | Mustela_erminea | 0.235297640937828 |
| Hyaena_hyaena | Mustela_eversmannii | 0.241833076430312 |
| Hyaena_hyaena | Mustela_frenata | 0.239449879096637 |
| Hyaena_hyaena | Mustela_itatsi | 0.239549088232173 |
| Hyaena_hyaena | Mustela_kathiah | 0.237869723732489 |
| Hyaena_hyaena | Mustela_nigripes | 0.238996846073096 |
| Hyaena_hyaena | Mustela_nivalis | 0.238290887606354 |
| Hyaena_hyaena | Mustela_putorius | 0.238962721192576 |
| Hyaena_hyaena | Mustela_sibirica | 0.237189080197077 |
| Hyaena_hyaena | Nandinia_binotata | 0.191004032840861 |
| Hyaena_hyaena | Nasua_nasua | 0.252114828116004 |
| Hyaena_hyaena | Neofelis_nebulosa | 0.203100321189543 |
| Hyaena_hyaena | Neomonachus_schauinslandi | 0.227768673877373 |
| Hyaena_hyaena | Neophoca_cinerea | 0.243954566625047 |
| Hyaena_hyaena | Neovison_vison | 0.240397978025344 |
| Hyaena_hyaena | Nyctereutes_procyonoides | 0.245107678917022 |
| Hyaena_hyaena | Odobenus_rosmarus_rosmarus | 0.257500223152682 |
| Hyaena_hyaena | Otocolobus_manul | 0.195350276538006 |
| Hyaena_hyaena | Otocyon_megalotis_megalotis_TS305 | 0.25040215622211 |
| Hyaena_hyaena | Otocyon_megalotis_virgatus_FMNH158128 | 0.251214001633084 |
| Hyaena_hyaena | Paguma_larvata | 0.205050605176838 |
| Hyaena_hyaena | Panthera_leo | 0.199806963301887 |

| | | |
|---|---|---|
| Hyaena_hyaena | Panthera_onca | 0.196943419086829 |
| Hyaena_hyaena | Panthera_pardus | 0.198409090483595 |
| Hyaena_hyaena | Panthera_tigris | 0.203724304572954 |
| Hyaena_hyaena | Panthera_uncia | 0.201780964370814 |
| Hyaena_hyaena | Paradoxurus_hermaphroditus | 0.210347456372394 |
| Hyaena_hyaena | Parahyaena_brunnea_FD126_Fecal | 0.0634561477226536 |
| Hyaena_hyaena | Pardofelis_marmorata | 0.198060527548154 |
| Hyaena_hyaena | Phoca_fasciata | 0.224223150906158 |
| Hyaena_hyaena | Phoca_groenlandica | 0.233866111140282 |
| Hyaena_hyaena | Phoca_largha | 0.227661560617074 |
| Hyaena_hyaena | Phoca_vitulina | 0.226422304955157 |
| Hyaena_hyaena | Phocarctos_hookeri | 0.246394828065281 |
| Hyaena_hyaena | Prionailurus_bengalensis | 0.189503104382449 |
| Hyaena_hyaena | Prionailurus_bengalensis_euptilurus | 0.18867202170617 |
| Hyaena_hyaena | Prionailurus_planiceps | 0.1964336679989 |
| Hyaena_hyaena | Prionailurus_rubiginosus | 0.188334098194383 |
| Hyaena_hyaena | Prionailurus_viverrinus | 0.19060885977213 |
| Hyaena_hyaena | Prionodon_pardicolor | 0.207422865883322 |
| Hyaena_hyaena | Procyon_lotor | 0.248516989595922 |
| Hyaena_hyaena | Profelis_aurata | 0.195297938842625 |
| Hyaena_hyaena | Proteles_cristatus_cristatus_TS307 | 0.127546104394469 |
| Hyaena_hyaena | Proteles_cristatus_septentrionalis_NMS_Z_2018_54 | 0.131335013107742 |
| Hyaena_hyaena | Puma_concolor | 0.196571489988189 |
| Hyaena_hyaena | Puma_yagouaroundi | 0.194806441227031 |
| Hyaena_hyaena | Pusa_caspica | 0.225314041153366 |
| Hyaena_hyaena | Pusa_hispida | 0.22182496421688 |
| Hyaena_hyaena | Pusa_sibirica | 0.22331455665509 |
| Hyaena_hyaena | Speothos_venaticus_T1624 | 0.256297845665384 |
| Hyaena_hyaena | Spilogale_putorius | 0.256514779395336 |
| Hyaena_hyaena | Taxidea_taxus | 0.233797670379694 |
| Hyaena_hyaena | Tremarctos_ornatus | 0.24574709283874 |
| Hyaena_hyaena | Urocyon_cinereoargenteus | 0.252766092804787 |
| Hyaena_hyaena | Ursus_americanus | 0.24394821978786 |
| Hyaena_hyaena | Ursus_arctos | 0.245500612064029 |
| Hyaena_hyaena | Ursus_maritimus | 0.240509059474605 |
| Hyaena_hyaena | Ursus_spelaeus | 0.240845382357241 |
| Hyaena_hyaena | Ursus_thibetanus | 0.242629618739954 |
| Hyaena_hyaena | Urva_javanica | 0.191323253615204 |
| Hyaena_hyaena | Viverra_tangalunga_FMNH146957_SRR6053069 | 0.19942516279645 |
| Hyaena_hyaena | Viverricula_indica | 0.196245008472302 |
| Hyaena_hyaena | Vulpes_corsac | 0.251973161057273 |
| Hyaena_hyaena | Vulpes_ferrilata | 0.254733353237027 |
| Hyaena_hyaena | Vulpes_lagopus | 0.249950528236443 |
| Hyaena_hyaena | Vulpes_vulpes_T3611 | 0.250057129268927 |
| Hyaena_hyaena | Vulpes_zerda | 0.247439542976224 |
| Hyaena_hyaena | Zalophus_californianus | 0.242136405076402 |
| Hydrurga_leptonyx | Leopardus_colocolo | 0.208119413270298 |
| Hydrurga_leptonyx | Leopardus_geoffroyi | 0.206482413643752 |
| Hydrurga_leptonyx | Leopardus_guigna | 0.208465414286968 |
| Hydrurga_leptonyx | Leopardus_jacobita | 0.206573408120283 |
| Hydrurga_leptonyx | Leopardus_pardalis | 0.20519176984592 |
| Hydrurga_leptonyx | Leopardus_tigrinus | 0.206820999136199 |
| Hydrurga_leptonyx | Leopardus_wiedii | 0.20678377942874 |
| Hydrurga_leptonyx | Leptailurus_serval | 0.207196382293116 |
| Hydrurga_leptonyx | Leptonychotes_weddellii | 0.0549110031672649 |
| Hydrurga_leptonyx | Lobodon_carcinophaga | 0.0774915371702852 |
| Hydrurga_leptonyx | Lutra_lutra | 0.207477504047417 |
| Hydrurga_leptonyx | Lutra_sumatrana | 0.205536605381113 |
| Hydrurga_leptonyx | Lutrogale_perspicillata | 0.217200295370129 |
| Hydrurga_leptonyx | Lycaon_pictus | 0.223343940512975 |
| Hydrurga_leptonyx | Lynx_canadensis | 0.21044475304969 |
| Hydrurga_leptonyx | Lynx_lynx | 0.211437626591813 |
| Hydrurga_leptonyx | Lynx_pardinus | 0.212037084965759 |
| Hydrurga_leptonyx | Lynx_rufus | 0.208271324317694 |
| Hydrurga_leptonyx | Manis_javanica | 0.267065571761415 |
| Hydrurga_leptonyx | Martes_americana | 0.21095771225502 |
| Hydrurga_leptonyx | Martes_flavigula | 0.211954614159354 |
| Hydrurga_leptonyx | Martes_foina | 0.209664302404547 |
| Hydrurga_leptonyx | Martes_martes | 0.207717680997651 |
| Hydrurga_leptonyx | Martes_melampus | 0.20628713398249 |
| Hydrurga_leptonyx | Martes_pennanti | 0.214688412405431 |
| Hydrurga_leptonyx | Martes_zibellina | 0.209948299553497 |
| Hydrurga_leptonyx | Meles_anakuma | 0.211341829874024 |
| Hydrurga_leptonyx | Meles_leucurus | 0.205488923950558 |
| Hydrurga_leptonyx | Meles_meles | 0.205879115362371 |
| Hydrurga_leptonyx | Melogale_moschata | 0.215189686608509 |
| Hydrurga_leptonyx | Melursus_ursinus | 0.214502434560595 |
| Hydrurga_leptonyx | Mephitis_mephitis | 0.221332482822975 |
| Hydrurga_leptonyx | Mirounga_leonina | 0.103651848200311 |
| Hydrurga_leptonyx | Mungotictis_decemlineata | 0.215346947367738 |
| Hydrurga_leptonyx | Mustela_altaica | 0.214248810046555 |
| Hydrurga_leptonyx | Mustela_erminea | 0.199681983692407 |
| Hydrurga_leptonyx | Mustela_eversmannii | 0.212593048851056 |
| Hydrurga_leptonyx | Mustela_frenata | 0.205906846362656 |
| Hydrurga_leptonyx | Mustela_itatsi | 0.210693139439815 |
| Hydrurga_leptonyx | Mustela_kathiah | 0.213732716496945 |
| Hydrurga_leptonyx | Mustela_nigripes | 0.211846768449316 |

| | | |
|---|---|---|
| Hydrurga_leptonyx | Mustela_nivalis | 0.213314005143745 |
| Hydrurga_leptonyx | Mustela_putorius | 0.210768491882632 |
| Hydrurga_leptonyx | Mustela_sibirica | 0.209469537641968 |
| Hydrurga_leptonyx | Nandinia_binotata | 0.193476796171663 |
| Hydrurga_leptonyx | Nasua_nasua | 0.226585285135784 |
| Hydrurga_leptonyx | Neofelis_nebulosa | 0.222463266876248 |
| Hydrurga_leptonyx | Neomonachus_schauinslandi | 0.115770819961699 |
| Hydrurga_leptonyx | Neophoca_cinerea | 0.182314745773612 |
| Hydrurga_leptonyx | Neovison_vison | 0.208001148625392 |
| Hydrurga_leptonyx | Nyctereutes_procyonoides | 0.222924621956051 |
| Hydrurga_leptonyx | Odobenus_rosmarus_rosmarus | 0.199023961797219 |
| Hydrurga_leptonyx | Otocolobus_manul | 0.208790259042611 |
| Hydrurga_leptonyx | Otocyon_megalotis_megalotis_TS305 | 0.228559783370805 |
| Hydrurga_leptonyx | Otocyon_megalotis_virgatus_FMNH158128 | 0.229621976656277 |
| Hydrurga_leptonyx | Paguma_larvata | 0.216104504736095 |
| Hydrurga_leptonyx | Panthera_leo | 0.214955724232753 |
| Hydrurga_leptonyx | Panthera_onca | 0.21404101800299 |
| Hydrurga_leptonyx | Panthera_pardus | 0.2197249705494 |
| Hydrurga_leptonyx | Panthera_tigris | 0.221359599769577 |
| Hydrurga_leptonyx | Panthera_uncia | 0.223636511769193 |
| Hydrurga_leptonyx | Paradoxurus_hermaphroditus | 0.224845860988794 |
| Hydrurga_leptonyx | Parahyaena_brunnea_FD126_Fecal | 0.217716402502469 |
| Hydrurga_leptonyx | Pardofelis_marmorata | 0.213579043063122 |
| Hydrurga_leptonyx | Phoca_fasciata | 0.133407605900809 |
| Hydrurga_leptonyx | Phoca_groenlandica | 0.137553184490587 |
| Hydrurga_leptonyx | Phoca_largha | 0.13225852125008 |
| Hydrurga_leptonyx | Phoca_vitulina | 0.13375735644569 |
| Hydrurga_leptonyx | Phocarctos_hookeri | 0.18381618439425 |
| Hydrurga_leptonyx | Prionailurus_bengalensis | 0.206645262470969 |
| Hydrurga_leptonyx | Prionailurus_bengalensis_euptilurus | 0.205893388289167 |
| Hydrurga_leptonyx | Prionailurus_planiceps | 0.209335127331364 |
| Hydrurga_leptonyx | Prionailurus_rubiginosus | 0.206756647251515 |
| Hydrurga_leptonyx | Prionailurus_viverrinus | 0.208718530973467 |
| Hydrurga_leptonyx | Prionodon_pardicolor | 0.221339206118575 |
| Hydrurga_leptonyx | Procyon_lotor | 0.215029926097779 |
| Hydrurga_leptonyx | Profelis_aurata | 0.212284555348314 |
| Hydrurga_leptonyx | Proteles_cristatus_cristatus_TS307 | 0.219375252067044 |
| Hydrurga_leptonyx | Proteles_cristatus_septentrionalis_NMS_Z_2018_54 | 0.220571093009102 |
| Hydrurga_leptonyx | Puma_concolor | 0.208043829101789 |
| Hydrurga_leptonyx | Puma_yagouaroundi | 0.206877551559078 |
| Hydrurga_leptonyx | Pusa_caspica | 0.132773448158474 |
| Hydrurga_leptonyx | Pusa_hispida | 0.131271509077149 |
| Hydrurga_leptonyx | Pusa_sibirica | 0.131459375972272 |
| Hydrurga_leptonyx | Speothos_venaticus_T1624 | 0.237049814688563 |
| Hydrurga_leptonyx | Spilogale_putorius | 0.212848200269807 |
| Hydrurga_leptonyx | Taxidea_taxus | 0.195743362146997 |
| Hydrurga_leptonyx | Tremarctos_ornatus | 0.207737199081754 |
| Hydrurga_leptonyx | Urocyon_cinereoargenteus | 0.230753250367439 |
| Hydrurga_leptonyx | Ursus_americanus | 0.211169763294198 |
| Hydrurga_leptonyx | Ursus_arctos | 0.217030429914212 |
| Hydrurga_leptonyx | Ursus_maritimus | 0.214854181256135 |
| Hydrurga_leptonyx | Ursus_spelaeus | 0.210166313062509 |
| Hydrurga_leptonyx | Ursus_thibetanus | 0.209409146882119 |
| Hydrurga_leptonyx | Urva_javanica | 0.21629837588892 |
| Hydrurga_leptonyx | Viverra_tangalunga_FMNH146957_SRR6053069 | 0.208689689442733 |
| Hydrurga_leptonyx | Viverricula_indica | 0.208752525099111 |
| Hydrurga_leptonyx | Vulpes_corsac | 0.232011244074969 |
| Hydrurga_leptonyx | Vulpes_ferrilata | 0.234272203911516 |
| Hydrurga_leptonyx | Vulpes_lagopus | 0.228022973571107 |
| Hydrurga_leptonyx | Vulpes_vulpes_T3611 | 0.227893584466991 |
| Hydrurga_leptonyx | Vulpes_zerda | 0.227885134240779 |
| Hydrurga_leptonyx | Zalophus_californianus | 0.172239879613768 |
| Leopardus_colocolo | Leopardus_geoffroyi | 0.0828649155987939 |
| Leopardus_colocolo | Leopardus_guigna | 0.0810978170992731 |
| Leopardus_colocolo | Leopardus_jacobita | 0.0752843493207397 |
| Leopardus_colocolo | Leopardus_pardalis | 0.0734013493242693 |
| Leopardus_colocolo | Leopardus_tigrinus | 0.0811380520979468 |
| Leopardus_colocolo | Leopardus_wiedii | 0.0740216105498313 |
| Leopardus_colocolo | Leptailurus_serval | 0.112400746527957 |
| Leopardus_colocolo | Leptonychotes_weddellii | 0.20547355893818 |
| Leopardus_colocolo | Lobodon_carcinophaga | 0.208781836615849 |
| Leopardus_colocolo | Lutra_lutra | 0.230262452185085 |
| Leopardus_colocolo | Lutra_sumatrana | 0.226930912418401 |
| Leopardus_colocolo | Lutrogale_perspicillata | 0.238095767806841 |
| Leopardus_colocolo | Lycaon_pictus | 0.240487616864752 |
| Leopardus_colocolo | Lynx_canadensis | 0.114172078835284 |
| Leopardus_colocolo | Lynx_lynx | 0.115948646568892 |
| Leopardus_colocolo | Lynx_pardinus | 0.113541945582601 |
| Leopardus_colocolo | Lynx_rufus | 0.116393462975831 |
| Leopardus_colocolo | Manis_javanica | 0.286464577924419 |
| Leopardus_colocolo | Martes_americana | 0.237018960510447 |
| Leopardus_colocolo | Martes_flavigula | 0.235005375689313 |
| Leopardus_colocolo | Martes_foina | 0.229990981793699 |
| Leopardus_colocolo | Martes_martes | 0.232888541482431 |
| Leopardus_colocolo | Martes_melampus | 0.234755186107742 |
| Leopardus_colocolo | Martes_pennanti | 0.237384627883923 |
| Leopardus_colocolo | Martes_zibellina | 0.233650213282341 |

| | | |
|---|---|---|
| Leopardus_colocolo | Meles_anakuma | 0.229861513338165 |
| Leopardus_colocolo | Meles_leucurus | 0.224713527427367 |
| Leopardus_colocolo | Meles_meles | 0.228162600580537 |
| Leopardus_colocolo | Melogale_moschata | 0.227417605352387 |
| Leopardus_colocolo | Melursus_ursinus | 0.238691343057399 |
| Leopardus_colocolo | Mephitis_mephitis | 0.236086311450735 |
| Leopardus_colocolo | Mirounga_leonina | 0.21315547900922 |
| Leopardus_colocolo | Mungotictis_decemlineata | 0.187758829052652 |
| Leopardus_colocolo | Mustela_altaica | 0.229597824708169 |
| Leopardus_colocolo | Mustela_erminea | 0.220344614100184 |
| Leopardus_colocolo | Mustela_eversmannii | 0.228990933030987 |
| Leopardus_colocolo | Mustela_frenata | 0.225386493973332 |
| Leopardus_colocolo | Mustela_itatsi | 0.228052979180689 |
| Leopardus_colocolo | Mustela_kathiah | 0.223684124749223 |
| Leopardus_colocolo | Mustela_nigripes | 0.226658880794326 |
| Leopardus_colocolo | Mustela_nivalis | 0.228573859284852 |
| Leopardus_colocolo | Mustela_putorius | 0.228455318578374 |
| Leopardus_colocolo | Mustela_sibirica | 0.228811125417375 |
| Leopardus_colocolo | Nandinia_binotata | 0.1739856934715 |
| Leopardus_colocolo | Nasua_nasua | 0.246619940442993 |
| Leopardus_colocolo | Neofelis_nebulosa | 0.146876590064393 |
| Leopardus_colocolo | Neomonachus_schauinslandi | 0.211234282279213 |
| Leopardus_colocolo | Neophoca_cinerea | 0.236301958556644 |
| Leopardus_colocolo | Neovison_vison | 0.22843258120144 |
| Leopardus_colocolo | Nyctereutes_procyonoides | 0.237837942302872 |
| Leopardus_colocolo | Odobenus_rosmarus_rosmarus | 0.251640945678693 |
| Leopardus_colocolo | Otocolobus_manul | 0.113079564275785 |
| Leopardus_colocolo | Otocyon_megalotis_megalotis_TS305 | 0.24379653041945 |
| Leopardus_colocolo | Otocyon_megalotis_virgatus_FMNH158128 | 0.242449130546991 |
| Leopardus_colocolo | Paguma_larvata | 0.177788283566471 |
| Leopardus_colocolo | Panthera_leo | 0.136172933582934 |
| Leopardus_colocolo | Panthera_onca | 0.132427034445924 |
| Leopardus_colocolo | Panthera_pardus | 0.133057658707616 |
| Leopardus_colocolo | Panthera_tigris | 0.137211935020255 |
| Leopardus_colocolo | Panthera_uncia | 0.137474445076208 |
| Leopardus_colocolo | Paradoxurus_hermaphroditus | 0.182474797250853 |
| Leopardus_colocolo | Parahyaena_brunnea_FD126_Fecal | 0.190176390235967 |
| Leopardus_colocolo | Pardofelis_marmorata | 0.119045319143711 |
| Leopardus_colocolo | Phoca_fasciata | 0.213348659813204 |
| Leopardus_colocolo | Phoca_groenlandica | 0.214369297041203 |
| Leopardus_colocolo | Phoca_largha | 0.215473386303663 |
| Leopardus_colocolo | Phoca_vitulina | 0.215321063318018 |
| Leopardus_colocolo | Phocarctos_hookeri | 0.237399195063054 |
| Leopardus_colocolo | Prionailurus_bengalensis | 0.108401238399188 |
| Leopardus_colocolo | Prionailurus_bengalensis_euptilurus | 0.110525839789321 |
| Leopardus_colocolo | Prionailurus_planiceps | 0.113261046698809 |
| Leopardus_colocolo | Prionailurus_rubiginosus | 0.111291914790529 |
| Leopardus_colocolo | Prionailurus_viverrinus | 0.113508472790835 |
| Leopardus_colocolo | Prionodon_pardicolor | 0.178275826347641 |
| Leopardus_colocolo | Procyon_lotor | 0.235251035223928 |
| Leopardus_colocolo | Profelis_aurata | 0.114146702860237 |
| Leopardus_colocolo | Proteles_cristatus_cristatus_TS307 | 0.199637049652041 |
| Leopardus_colocolo | Proteles_cristatus_septentrionalis_NMS_Z_2018_54 | 0.199552311069754 |
| Leopardus_colocolo | Puma_concolor | 0.116051328291701 |
| Leopardus_colocolo | Puma_yagouaroundi | 0.119299695185133 |
| Leopardus_colocolo | Pusa_caspica | 0.216977685076489 |
| Leopardus_colocolo | Pusa_hispida | 0.213730785182004 |
| Leopardus_colocolo | Pusa_sibirica | 0.213925995213904 |
| Leopardus_colocolo | Speothos_venaticus_T1624 | 0.247700805655828 |
| Leopardus_colocolo | Spilogale_putorius | 0.240951991781434 |
| Leopardus_colocolo | Taxidea_taxus | 0.22188643749189 |
| Leopardus_colocolo | Tremarctos_ornatus | 0.238748557216605 |
| Leopardus_colocolo | Urocyon_cinereoargenteus | 0.245149995515184 |
| Leopardus_colocolo | Ursus_americanus | 0.238135807161939 |
| Leopardus_colocolo | Ursus_arctos | 0.239894098129033 |
| Leopardus_colocolo | Ursus_maritimus | 0.239410060355889 |
| Leopardus_colocolo | Ursus_spelaeus | 0.236460015178765 |
| Leopardus_colocolo | Ursus_thibetanus | 0.23496886835416 |
| Leopardus_colocolo | Urva_javanica | 0.191001459239959 |
| Leopardus_colocolo | Viverra_tangalunga_FMNH146957_SRR6053069 | 0.172335712424407 |
| Leopardus_colocolo | Viverricula_indica | 0.169015431447773 |
| Leopardus_colocolo | Vulpes_corsac | 0.240821083937345 |
| Leopardus_colocolo | Vulpes_ferrilata | 0.241296860832255 |
| Leopardus_colocolo | Vulpes_lagopus | 0.24131005307059 |
| Leopardus_colocolo | Vulpes_vulpes_T3611 | 0.236040316680164 |
| Leopardus_colocolo | Vulpes_zerda | 0.236028686020325 |
| Leopardus_colocolo | Zalophus_californianus | 0.234785829310722 |
| Leopardus_geoffroyi | Leopardus_guigna | 0.0302637364079918 |
| Leopardus_geoffroyi | Leopardus_jacobita | 0.0805531784553565 |
| Leopardus_geoffroyi | Leopardus_pardalis | 0.0748116463762269 |
| Leopardus_geoffroyi | Leopardus_tigrinus | 0.058554200011052 |
| Leopardus_geoffroyi | Leopardus_wiedii | 0.0753467253101816 |
| Leopardus_geoffroyi | Leptailurus_serval | 0.115009783200814 |
| Leopardus_geoffroyi | Leptonychotes_weddellii | 0.208147030191545 |
| Leopardus_geoffroyi | Lobodon_carcinophaga | 0.210328654746131 |
| Leopardus_geoffroyi | Lutra_lutra | 0.232672474058929 |
| Leopardus_geoffroyi | Lutra_sumatrana | 0.228260726330587 |

| | | |
|---|---|---|
| Leopardus_geoffroyi | Lutrogale_perspicillata | 0.23978118643007 |
| Leopardus_geoffroyi | Lycaon_pictus | 0.24075080979219 |
| Leopardus_geoffroyi | Lynx_canadensis | 0.115925002328073 |
| Leopardus_geoffroyi | Lynx_lynx | 0.116385655049578 |
| Leopardus_geoffroyi | Lynx_pardinus | 0.116514735292684 |
| Leopardus_geoffroyi | Lynx_rufus | 0.119400702240772 |
| Leopardus_geoffroyi | Manis_javanica | 0.282885678811484 |
| Leopardus_geoffroyi | Martes_americana | 0.235144996733027 |
| Leopardus_geoffroyi | Martes_flavigula | 0.234133905344907 |
| Leopardus_geoffroyi | Martes_foina | 0.227398075222294 |
| Leopardus_geoffroyi | Martes_martes | 0.231687860148807 |
| Leopardus_geoffroyi | Martes_melampus | 0.229702417759075 |
| Leopardus_geoffroyi | Martes_pennanti | 0.237305929135527 |
| Leopardus_geoffroyi | Martes_zibellina | 0.234433967152547 |
| Leopardus_geoffroyi | Meles_anakuma | 0.230160188494886 |
| Leopardus_geoffroyi | Meles_leucurus | 0.225643211448668 |
| Leopardus_geoffroyi | Meles_meles | 0.227238347942074 |
| Leopardus_geoffroyi | Melogale_moschata | 0.231148542367462 |
| Leopardus_geoffroyi | Melursus_ursinus | 0.238861351787487 |
| Leopardus_geoffroyi | Mephitis_mephitis | 0.243098889063924 |
| Leopardus_geoffroyi | Mirounga_leonina | 0.217966679751807 |
| Leopardus_geoffroyi | Mungotictis_decemlineata | 0.186806665209541 |
| Leopardus_geoffroyi | Mustela_altaica | 0.232497509119274 |
| Leopardus_geoffroyi | Mustela_erminea | 0.221626222897254 |
| Leopardus_geoffroyi | Mustela_eversmannii | 0.230452634746555 |
| Leopardus_geoffroyi | Mustela_frenata | 0.22600225329135 |
| Leopardus_geoffroyi | Mustela_itatsi | 0.228730639669987 |
| Leopardus_geoffroyi | Mustela_kathiah | 0.226941293620252 |
| Leopardus_geoffroyi | Mustela_nigripes | 0.227661560617074 |
| Leopardus_geoffroyi | Mustela_nivalis | 0.227000853231547 |
| Leopardus_geoffroyi | Mustela_putorius | 0.228824729684522 |
| Leopardus_geoffroyi | Mustela_sibirica | 0.229469565068862 |
| Leopardus_geoffroyi | Nandinia_binotata | 0.17033505984933 |
| Leopardus_geoffroyi | Nasua_nasua | 0.248301650895813 |
| Leopardus_geoffroyi | Neofelis_nebulosa | 0.141830073383861 |
| Leopardus_geoffroyi | Neomonachus_schauinslandi | 0.213899684972679 |
| Leopardus_geoffroyi | Neophoca_cinerea | 0.236132342060897 |
| Leopardus_geoffroyi | Neovison_vison | 0.23152770683357 |
| Leopardus_geoffroyi | Nyctereutes_procyonoides | 0.234438047909159 |
| Leopardus_geoffroyi | Odobenus_rosmarus_rosmarus | 0.250692709326695 |
| Leopardus_geoffroyi | Otocolobus_manul | 0.119628532623254 |
| Leopardus_geoffroyi | Otocyon_megalotis_megalotis_TS305 | 0.241251722332615 |
| Leopardus_geoffroyi | Otocyon_megalotis_virgatus_FMNH158128 | 0.239129021397272 |
| Leopardus_geoffroyi | Paguma_larvata | 0.182098760803858 |
| Leopardus_geoffroyi | Panthera_leo | 0.133767163240875 |
| Leopardus_geoffroyi | Panthera_onca | 0.135782461091399 |
| Leopardus_geoffroyi | Panthera_pardus | 0.136594232577886 |
| Leopardus_geoffroyi | Panthera_tigris | 0.136578268230192 |
| Leopardus_geoffroyi | Panthera_uncia | 0.137622309151697 |
| Leopardus_geoffroyi | Paradoxurus_hermaphroditus | 0.187424758255263 |
| Leopardus_geoffroyi | Parahyaena_brunnea_FD126_Fecal | 0.192692512038757 |
| Leopardus_geoffroyi | Pardofelis_marmorata | 0.123772608010848 |
| Leopardus_geoffroyi | Phoca_fasciata | 0.217092716915851 |
| Leopardus_geoffroyi | Phoca_groenlandica | 0.222759104953894 |
| Leopardus_geoffroyi | Phoca_largha | 0.220755710979339 |
| Leopardus_geoffroyi | Phoca_vitulina | 0.221896008051601 |
| Leopardus_geoffroyi | Phocarctos_hookeri | 0.233983817744588 |
| Leopardus_geoffroyi | Prionailurus_bengalensis | 0.112087011137068 |
| Leopardus_geoffroyi | Prionailurus_bengalensis_euptilurus | 0.112280889013917 |
| Leopardus_geoffroyi | Prionailurus_planiceps | 0.115402068614299 |
| Leopardus_geoffroyi | Prionailurus_rubiginosus | 0.111379987346052 |
| Leopardus_geoffroyi | Prionailurus_viverrinus | 0.116968762270993 |
| Leopardus_geoffroyi | Prionodon_pardicolor | 0.178873588144828 |
| Leopardus_geoffroyi | Procyon_lotor | 0.236585600566486 |
| Leopardus_geoffroyi | Profelis_aurata | 0.123480946875647 |
| Leopardus_geoffroyi | Proteles_cristatus_cristatus_TS307 | 0.196203698482834 |
| Leopardus_geoffroyi | Proteles_cristatus_septentrionalis_NMS_Z_2018_54 | 0.197574284475474 |
| Leopardus_geoffroyi | Puma_concolor | 0.11703685517527 |
| Leopardus_geoffroyi | Puma_yagouaroundi | 0.123443982411631 |
| Leopardus_geoffroyi | Pusa_caspica | 0.223143056990073 |
| Leopardus_geoffroyi | Pusa_hispida | 0.21867709391064 |
| Leopardus_geoffroyi | Pusa_sibirica | 0.219290189503243 |
| Leopardus_geoffroyi | Speothos_venaticus_T1624 | 0.247766157880328 |
| Leopardus_geoffroyi | Spilogale_putorius | 0.245763925669296 |
| Leopardus_geoffroyi | Taxidea_taxus | 0.218549798605055 |
| Leopardus_geoffroyi | Tremarctos_ornatus | 0.234672178164979 |
| Leopardus_geoffroyi | Urocyon_cinereoargenteus | 0.249444806714052 |
| Leopardus_geoffroyi | Ursus_americanus | 0.23731850452789 |
| Leopardus_geoffroyi | Ursus_arctos | 0.238406408811755 |
| Leopardus_geoffroyi | Ursus_maritimus | 0.236806212854287 |
| Leopardus_geoffroyi | Ursus_spelaeus | 0.23462389095024 |
| Leopardus_geoffroyi | Ursus_thibetanus | 0.231056213716156 |
| Leopardus_geoffroyi | Urva_javanica | 0.191474058992708 |
| Leopardus_geoffroyi | Viverra_tangalunga_FMNH146957_SRR6053069 | 0.173812749482868 |
| Leopardus_geoffroyi | Viverricula_indica | 0.173351901472356 |
| Leopardus_geoffroyi | Vulpes_corsac | 0.241421510939977 |
| Leopardus_geoffroyi | Vulpes_ferrilata | 0.242303503114811 |

| | | |
|---|---|---|
| Leopardus_geoffroyi | Vulpes_lagopus | 0.241447270028673 |
| Leopardus_geoffroyi | Vulpes_vulpes_T3611 | 0.237052682513194 |
| Leopardus_geoffroyi | Vulpes_zerda | 0.235075468870494 |
| Leopardus_geoffroyi | Zalophus_californianus | 0.231845162071063 |
| Leopardus_guigna | Leopardus_jacobita | 0.0785172596215288 |
| Leopardus_guigna | Leopardus_pardalis | 0.0744516129292244 |
| Leopardus_guigna | Leopardus_tigrinus | 0.056109413484542 |
| Leopardus_guigna | Leopardus_wiedii | 0.0752479846533295 |
| Leopardus_guigna | Leptailurus_serval | 0.113413237892391 |
| Leopardus_guigna | Leptonychotes_weddellii | 0.209711631648915 |
| Leopardus_guigna | Lobodon_carcinophaga | 0.212871302454621 |
| Leopardus_guigna | Lutra_lutra | 0.235061594017044 |
| Leopardus_guigna | Lutra_sumatrana | 0.231289108391655 |
| Leopardus_guigna | Lutrogale_perspicillata | 0.239848865862172 |
| Leopardus_guigna | Lycaon_pictus | 0.240528439319323 |
| Leopardus_guigna | Lynx_canadensis | 0.112995886024942 |
| Leopardus_guigna | Lynx_lynx | 0.113082218528439 |
| Leopardus_guigna | Lynx_pardinus | 0.112741400311527 |
| Leopardus_guigna | Lynx_rufus | 0.116830812919017 |
| Leopardus_guigna | Manis_javanica | 0.2815672691381 |
| Leopardus_guigna | Martes_americana | 0.232626902231325 |
| Leopardus_guigna | Martes_flavigula | 0.232594935239085 |
| Leopardus_guigna | Martes_foina | 0.229513470076003 |
| Leopardus_guigna | Martes_martes | 0.230939065953482 |
| Leopardus_guigna | Martes_melampus | 0.228518975662417 |
| Leopardus_guigna | Martes_pennanti | 0.235619640109008 |
| Leopardus_guigna | Martes_zibellina | 0.233129832797078 |
| Leopardus_guigna | Meles_anakuma | 0.232706205694811 |
| Leopardus_guigna | Meles_leucurus | 0.226643504776496 |
| Leopardus_guigna | Meles_meles | 0.2278043704416 |
| Leopardus_guigna | Melogale_moschata | 0.230599574372043 |
| Leopardus_guigna | Melursus_ursinus | 0.238212859154272 |
| Leopardus_guigna | Mephitis_mephitis | 0.243657361142537 |
| Leopardus_guigna | Mirounga_leonina | 0.218610042482649 |
| Leopardus_guigna | Mungotictis_decemlineata | 0.187974705574603 |
| Leopardus_guigna | Mustela_altaica | 0.230181601835918 |
| Leopardus_guigna | Mustela_erminea | 0.220873534583069 |
| Leopardus_guigna | Mustela_eversmannii | 0.231019056669785 |
| Leopardus_guigna | Mustela_frenata | 0.22600225329135 |
| Leopardus_guigna | Mustela_itatsi | 0.229621280055614 |
| Leopardus_guigna | Mustela_kathiah | 0.227484890107803 |
| Leopardus_guigna | Mustela_nigripes | 0.228985995686507 |
| Leopardus_guigna | Mustela_nivalis | 0.226657209450637 |
| Leopardus_guigna | Mustela_putorius | 0.229389091559079 |
| Leopardus_guigna | Mustela_sibirica | 0.230034696732467 |
| Leopardus_guigna | Nandinia_binotata | 0.172015850309104 |
| Leopardus_guigna | Nasua_nasua | 0.248844526814051 |
| Leopardus_guigna | Neofelis_nebulosa | 0.142224901160769 |
| Leopardus_guigna | Neomonachus_schauinslandi | 0.214201542273146 |
| Leopardus_guigna | Neophoca_cinerea | 0.236111650438908 |
| Leopardus_guigna | Neovison_vison | 0.22803424284023 |
| Leopardus_guigna | Nyctereutes_procyonoides | 0.234987921493667 |
| Leopardus_guigna | Odobenus_rosmarus_rosmarus | 0.248396613209815 |
| Leopardus_guigna | Otocolobus_manul | 0.119895033733327 |
| Leopardus_guigna | Otocyon_megalotis_megalotis_TS305 | 0.245356650651652 |
| Leopardus_guigna | Otocyon_megalotis_virgatus_FMNH158128 | 0.244231248991718 |
| Leopardus_guigna | Paguma_larvata | 0.183589491321179 |
| Leopardus_guigna | Panthera_leo | 0.135597406805912 |
| Leopardus_guigna | Panthera_onca | 0.138207312589844 |
| Leopardus_guigna | Panthera_pardus | 0.137062824033074 |
| Leopardus_guigna | Panthera_tigris | 0.141274309439257 |
| Leopardus_guigna | Panthera_uncia | 0.138580789940835 |
| Leopardus_guigna | Paradoxurus_hermaphroditus | 0.188656366203198 |
| Leopardus_guigna | Parahyaena_brunnea_FD126_Fecal | 0.190648103321461 |
| Leopardus_guigna | Pardofelis_marmorata | 0.123963702987377 |
| Leopardus_guigna | Phoca_fasciata | 0.217735528232158 |
| Leopardus_guigna | Phoca_groenlandica | 0.223952422942759 |
| Leopardus_guigna | Phoca_largha | 0.22237628283881 |
| Leopardus_guigna | Phoca_vitulina | 0.221896008051601 |
| Leopardus_guigna | Phocarctos_hookeri | 0.236191326849146 |
| Leopardus_guigna | Prionailurus_bengalensis | 0.111526786988823 |
| Leopardus_guigna | Prionailurus_bengalensis_euptilurus | 0.111813380798463 |
| Leopardus_guigna | Prionailurus_planiceps | 0.115496150842444 |
| Leopardus_guigna | Prionailurus_rubiginosus | 0.11229619592113 |
| Leopardus_guigna | Prionailurus_viverrinus | 0.116780118616614 |
| Leopardus_guigna | Prionodon_pardicolor | 0.180316840581013 |
| Leopardus_guigna | Procyon_lotor | 0.238410565581744 |
| Leopardus_guigna | Profelis_aurata | 0.121398782597108 |
| Leopardus_guigna | Proteles_cristatus_cristatus_TS307 | 0.196563167582475 |
| Leopardus_guigna | Proteles_cristatus_septentrionalis_NMS_Z_2018_54 | 0.199202810499196 |
| Leopardus_guigna | Puma_concolor | 0.116094436651049 |
| Leopardus_guigna | Puma_yagouaroundi | 0.120875972201239 |
| Leopardus_guigna | Pusa_caspica | 0.224229187535556 |
| Leopardus_guigna | Pusa_hispida | 0.219645460901517 |
| Leopardus_guigna | Pusa_sibirica | 0.221773459561274 |
| Leopardus_guigna | Speothos_venaticus_T1624 | 0.245146091690138 |
| Leopardus_guigna | Spilogale_putorius | 0.246774821217508 |

| | | |
|---|---|---|
| Leopardus_guigna | Taxidea_taxus | 0.218729219084632 |
| Leopardus_guigna | Tremarctos_ornatus | 0.233027771241659 |
| Leopardus_guigna | Urocyon_cinereoargenteus | 0.250623662871007 |
| Leopardus_guigna | Ursus_americanus | 0.239805422061774 |
| Leopardus_guigna | Ursus_arctos | 0.241327109096059 |
| Leopardus_guigna | Ursus_maritimus | 0.241178438023824 |
| Leopardus_guigna | Ursus_spelaeus | 0.237769793021466 |
| Leopardus_guigna | Ursus_thibetanus | 0.233181896026637 |
| Leopardus_guigna | Urva_javanica | 0.190916036513513 |
| Leopardus_guigna | Viverra_tangalunga_FMNH146957_SRR6053069 | 0.174897536488363 |
| Leopardus_guigna | Viverricula_indica | 0.175030044987936 |
| Leopardus_guigna | Vulpes_corsac | 0.245225937932783 |
| Leopardus_guigna | Vulpes_ferrilata | 0.244878231741706 |
| Leopardus_guigna | Vulpes_lagopus | 0.243416865505854 |
| Leopardus_guigna | Vulpes_vulpes_T3611 | 0.239915039636761 |
| Leopardus_guigna | Vulpes_zerda | 0.237818809719435 |
| Leopardus_guigna | Zalophus_californianus | 0.23614270918903 |
| Leopardus_jacobita | Leopardus_pardalis | 0.0715565865898995 |
| Leopardus_jacobita | Leopardus_tigrinus | 0.079900240509121 |
| Leopardus_jacobita | Leopardus_wiedii | 0.0705169951212171 |
| Leopardus_jacobita | Leptailurus_serval | 0.113157632921173 |
| Leopardus_jacobita | Leptonychotes_weddellii | 0.206659765783849 |
| Leopardus_jacobita | Lobodon_carcinophaga | 0.209064770310262 |
| Leopardus_jacobita | Lutra_lutra | 0.232084466814475 |
| Leopardus_jacobita | Lutra_sumatrana | 0.228987557116511 |
| Leopardus_jacobita | Lutrogale_perspicillata | 0.239613043828833 |
| Leopardus_jacobita | Lycaon_pictus | 0.238280273416538 |
| Leopardus_jacobita | Lynx_canadensis | 0.114651011495418 |
| Leopardus_jacobita | Lynx_lynx | 0.115316649368994 |
| Leopardus_jacobita | Lynx_pardinus | 0.11646423415207 |
| Leopardus_jacobita | Lynx_rufus | 0.111806820600175 |
| Leopardus_jacobita | Manis_javanica | 0.287229284530234 |
| Leopardus_jacobita | Martes_americana | 0.234444510642355 |
| Leopardus_jacobita | Martes_flavigula | 0.235487252383044 |
| Leopardus_jacobita | Martes_foina | 0.2296330398981 |
| Leopardus_jacobita | Martes_martes | 0.230002266266348 |
| Leopardus_jacobita | Martes_melampus | 0.231859639826083 |
| Leopardus_jacobita | Martes_pennanti | 0.232709634595826 |
| Leopardus_jacobita | Martes_zibellina | 0.233292463467245 |
| Leopardus_jacobita | Meles_anakuma | 0.234655132698562 |
| Leopardus_jacobita | Meles_leucurus | 0.229250989048798 |
| Leopardus_jacobita | Meles_meles | 0.228333024922081 |
| Leopardus_jacobita | Melogale_moschata | 0.230650449393942 |
| Leopardus_jacobita | Melursus_ursinus | 0.239728605683955 |
| Leopardus_jacobita | Mephitis_mephitis | 0.240376629072954 |
| Leopardus_jacobita | Mirounga_leonina | 0.214147491597941 |
| Leopardus_jacobita | Mungotictis_decemlineata | 0.182033774311668 |
| Leopardus_jacobita | Mustela_altaica | 0.230832278765699 |
| Leopardus_jacobita | Mustela_erminea | 0.221940837213279 |
| Leopardus_jacobita | Mustela_eversmannii | 0.23027043451551 |
| Leopardus_jacobita | Mustela_frenata | 0.225041004464418 |
| Leopardus_jacobita | Mustela_itatsi | 0.22810952484961 |
| Leopardus_jacobita | Mustela_kathiah | 0.226757133005967 |
| Leopardus_jacobita | Mustela_nigripes | 0.227276801237541 |
| Leopardus_jacobita | Mustela_nivalis | 0.228971810313163 |
| Leopardus_jacobita | Mustela_putorius | 0.228204889010854 |
| Leopardus_jacobita | Mustela_sibirica | 0.228323529946918 |
| Leopardus_jacobita | Nandinia_binotata | 0.172435749643766 |
| Leopardus_jacobita | Nasua_nasua | 0.250876195785662 |
| Leopardus_jacobita | Neofelis_nebulosa | 0.1436630108063 |
| Leopardus_jacobita | Neomonachus_schauinslandi | 0.21463565594059 |
| Leopardus_jacobita | Neophoca_cinerea | 0.239606796359028 |
| Leopardus_jacobita | Neovison_vison | 0.230489372668981 |
| Leopardus_jacobita | Nyctereutes_procyonoides | 0.235269391417305 |
| Leopardus_jacobita | Odobenus_rosmarus_rosmarus | 0.252693752749436 |
| Leopardus_jacobita | Otocolobus_manul | 0.114157756840014 |
| Leopardus_jacobita | Otocyon_megalotis_megalotis_TS305 | 0.2410607674551 |
| Leopardus_jacobita | Otocyon_megalotis_virgatus_FMNH158128 | 0.240610924778622 |
| Leopardus_jacobita | Paguma_larvata | 0.182273058764086 |
| Leopardus_jacobita | Panthera_leo | 0.135745514964341 |
| Leopardus_jacobita | Panthera_onca | 0.137650195613105 |
| Leopardus_jacobita | Panthera_pardus | 0.134965179261669 |
| Leopardus_jacobita | Panthera_tigris | 0.138078984782893 |
| Leopardus_jacobita | Panthera_uncia | 0.139612541720044 |
| Leopardus_jacobita | Paradoxurus_hermaphroditus | 0.188312401550349 |
| Leopardus_jacobita | Parahyaena_brunnea_FD126_Fecal | 0.19308101645165 |
| Leopardus_jacobita | Pardofelis_marmorata | 0.119726169494778 |
| Leopardus_jacobita | Phoca_fasciata | 0.216568269432046 |
| Leopardus_jacobita | Phoca_groenlandica | 0.216418242693359 |
| Leopardus_jacobita | Phoca_largha | 0.21990824482572 |
| Leopardus_jacobita | Phoca_vitulina | 0.220545929627175 |
| Leopardus_jacobita | Phocarctos_hookeri | 0.237995935533085 |
| Leopardus_jacobita | Prionailurus_bengalensis | 0.108371944522302 |
| Leopardus_jacobita | Prionailurus_bengalensis_euptilurus | 0.108839750712968 |
| Leopardus_jacobita | Prionailurus_planiceps | 0.111678161572326 |
| Leopardus_jacobita | Prionailurus_rubiginosus | 0.111294893691184 |
| Leopardus_jacobita | Prionailurus_viverrinus | 0.112001893451447 |

| | | |
|---|---|---|
| Leopardus_jacobita | Prionodon_pardicolor | 0.178313666266358 |
| Leopardus_jacobita | Procyon_lotor | 0.235556122374644 |
| Leopardus_jacobita | Profelis_aurata | 0.117817098357507 |
| Leopardus_jacobita | Proteles_cristatus_cristatus_TS307 | 0.197352762475813 |
| Leopardus_jacobita | Proteles_cristatus_septentrionalis_NMS_Z_2018_54 | 0.200953328707527 |
| Leopardus_jacobita | Puma_concolor | 0.11266458949596 |
| Leopardus_jacobita | Puma_yagouaroundi | 0.119025696558019 |
| Leopardus_jacobita | Pusa_caspica | 0.22327212304715 |
| Leopardus_jacobita | Pusa_hispida | 0.219768945369822 |
| Leopardus_jacobita | Pusa_sibirica | 0.220602269753535 |
| Leopardus_jacobita | Speothos_venaticus_T1624 | 0.247488821970756 |
| Leopardus_jacobita | Spilogale_putorius | 0.246138388456107 |
| Leopardus_jacobita | Taxidea_taxus | 0.220756640298449 |
| Leopardus_jacobita | Tremarctos_ornatus | 0.236738720322352 |
| Leopardus_jacobita | Urocyon_cinereoargenteus | 0.249320096421635 |
| Leopardus_jacobita | Ursus_americanus | 0.237264442990328 |
| Leopardus_jacobita | Ursus_arctos | 0.241814304458008 |
| Leopardus_jacobita | Ursus_maritimus | 0.242455013928064 |
| Leopardus_jacobita | Ursus_spelaeus | 0.23746921276387 |
| Leopardus_jacobita | Ursus_thibetanus | 0.235751862098379 |
| Leopardus_jacobita | Urva_javanica | 0.193132584356024 |
| Leopardus_jacobita | Viverra_tangalunga_FMNH146957_SRR6053069 | 0.173025844398535 |
| Leopardus_jacobita | Viverricula_indica | 0.17152719368573 |
| Leopardus_jacobita | Vulpes_corsac | 0.245733229311473 |
| Leopardus_jacobita | Vulpes_ferrilata | 0.243277477565441 |
| Leopardus_jacobita | Vulpes_lagopus | 0.243315681612883 |
| Leopardus_jacobita | Vulpes_vulpes_T3611 | 0.24280355644171 |
| Leopardus_jacobita | Vulpes_zerda | 0.24047548837928 |
| Leopardus_jacobita | Zalophus_californianus | 0.233434150219467 |
| Leopardus_pardalis | Leopardus_tigrinus | 0.0757264626050351 |
| Leopardus_pardalis | Leopardus_wiedii | 0.0409992763934196 |
| Leopardus_pardalis | Leptailurus_serval | 0.109428127055965 |
| Leopardus_pardalis | Leptonychotes_weddellii | 0.205804003520515 |
| Leopardus_pardalis | Lobodon_carcinophaga | 0.209259065979253 |
| Leopardus_pardalis | Lutra_lutra | 0.229679660998921 |
| Leopardus_pardalis | Lutra_sumatrana | 0.227864859771141 |
| Leopardus_pardalis | Lutrogale_perspicillata | 0.237938838847611 |
| Leopardus_pardalis | Lycaon_pictus | 0.23655238467499 |
| Leopardus_pardalis | Lynx_canadensis | 0.109293595027741 |
| Leopardus_pardalis | Lynx_lynx | 0.106414109766675 |
| Leopardus_pardalis | Lynx_pardinus | 0.110997941667412 |
| Leopardus_pardalis | Lynx_rufus | 0.111219685222758 |
| Leopardus_pardalis | Manis_javanica | 0.280837408070892 |
| Leopardus_pardalis | Martes_americana | 0.232231556922772 |
| Leopardus_pardalis | Martes_flavigula | 0.234264876428386 |
| Leopardus_pardalis | Martes_foina | 0.225910853547564 |
| Leopardus_pardalis | Martes_martes | 0.22999471454236 |
| Leopardus_pardalis | Martes_melampus | 0.22921681368874 |
| Leopardus_pardalis | Martes_pennanti | 0.234361449784842 |
| Leopardus_pardalis | Martes_zibellina | 0.232294422877823 |
| Leopardus_pardalis | Meles_anakuma | 0.231319934757069 |
| Leopardus_pardalis | Meles_leucurus | 0.226248620452788 |
| Leopardus_pardalis | Meles_meles | 0.228394553301283 |
| Leopardus_pardalis | Melogale_moschata | 0.229086937584424 |
| Leopardus_pardalis | Melursus_ursinus | 0.23605977359588 |
| Leopardus_pardalis | Mephitis_mephitis | 0.243465221317861 |
| Leopardus_pardalis | Mirounga_leonina | 0.215925274096958 |
| Leopardus_pardalis | Mungotictis_decemlineata | 0.183819616819371 |
| Leopardus_pardalis | Mustela_altaica | 0.230929055768804 |
| Leopardus_pardalis | Mustela_erminea | 0.219753888915236 |
| Leopardus_pardalis | Mustela_eversmannii | 0.226076577479598 |
| Leopardus_pardalis | Mustela_frenata | 0.221595314631335 |
| Leopardus_pardalis | Mustela_itatsi | 0.228949146282167 |
| Leopardus_pardalis | Mustela_kathiah | 0.225689544819105 |
| Leopardus_pardalis | Mustela_nigripes | 0.223323626683024 |
| Leopardus_pardalis | Mustela_nivalis | 0.22719638054517 |
| Leopardus_pardalis | Mustela_putorius | 0.224892835497346 |
| Leopardus_pardalis | Mustela_sibirica | 0.224360689360975 |
| Leopardus_pardalis | Nandinia_binotata | 0.169104422965488 |
| Leopardus_pardalis | Nasua_nasua | 0.246731762955199 |
| Leopardus_pardalis | Neofelis_nebulosa | 0.13844478860116 |
| Leopardus_pardalis | Neomonachus_schauinslandi | 0.21550765830035 |
| Leopardus_pardalis | Neophoca_cinerea | 0.233864642582377 |
| Leopardus_pardalis | Neovison_vison | 0.226997814563472 |
| Leopardus_pardalis | Nyctereutes_procyonoides | 0.232420587886307 |
| Leopardus_pardalis | Odobenus_rosmarus_rosmarus | 0.25228525533803 |
| Leopardus_pardalis | Otocolobus_manul | 0.11285353257743 |
| Leopardus_pardalis | Otocyon_megalotis_megalotis_TS305 | 0.23617075292828 |
| Leopardus_pardalis | Otocyon_megalotis_virgatus_FMNH158128 | 0.235168240802363 |
| Leopardus_pardalis | Paguma_larvata | 0.178173537306339 |
| Leopardus_pardalis | Panthera_leo | 0.128649328419342 |
| Leopardus_pardalis | Panthera_onca | 0.127448095017707 |
| Leopardus_pardalis | Panthera_pardus | 0.12701486873871 |
| Leopardus_pardalis | Panthera_tigris | 0.134834746798567 |
| Leopardus_pardalis | Panthera_uncia | 0.132088192538724 |
| Leopardus_pardalis | Paradoxurus_hermaphroditus | 0.184833788935562 |
| Leopardus_pardalis | Parahyaena_brunnea_FD126_Fecal | 0.190672693572939 |

| | | |
|---|---|---|
| Leopardus_pardalis | Pardofelis_marmorata | 0.117054348566894 |
| Leopardus_pardalis | Phoca_fasciata | 0.215178191547811 |
| Leopardus_pardalis | Phoca_groenlandica | 0.217061329598731 |
| Leopardus_pardalis | Phoca_largha | 0.215614530161396 |
| Leopardus_pardalis | Phoca_vitulina | 0.216853057552957 |
| Leopardus_pardalis | Phocarctos_hookeri | 0.235579366999005 |
| Leopardus_pardalis | Prionailurus_bengalensis | 0.10707755263588 |
| Leopardus_pardalis | Prionailurus_bengalensis_euptilurus | 0.107823001704427 |
| Leopardus_pardalis | Prionailurus_planiceps | 0.112510390850958 |
| Leopardus_pardalis | Prionailurus_rubiginosus | 0.111189608856484 |
| Leopardus_pardalis | Prionailurus_viverrinus | 0.110608181447008 |
| Leopardus_pardalis | Prionodon_pardicolor | 0.177245356641263 |
| Leopardus_pardalis | Procyon_lotor | 0.234402421404503 |
| Leopardus_pardalis | Profelis_aurata | 0.118002749229854 |
| Leopardus_pardalis | Proteles_cristatus_cristatus_TS307 | 0.194992873648456 |
| Leopardus_pardalis | Proteles_cristatus_septentrionalis_NMS_Z_2018_54 | 0.197521057692877 |
| Leopardus_pardalis | Puma_concolor | 0.112077024631103 |
| Leopardus_pardalis | Puma_yagouaroundi | 0.118906602051385 |
| Leopardus_pardalis | Pusa_caspica | 0.219052688724218 |
| Leopardus_pardalis | Pusa_hispida | 0.217938821677319 |
| Leopardus_pardalis | Pusa_sibirica | 0.215012015863854 |
| Leopardus_pardalis | Speothos_venaticus_T1624 | 0.240814760235161 |
| Leopardus_pardalis | Spilogale_putorius | 0.244185001930749 |
| Leopardus_pardalis | Taxidea_taxus | 0.218324592269826 |
| Leopardus_pardalis | Tremarctos_ornatus | 0.2355567699488 |
| Leopardus_pardalis | Urocyon_cinereoargenteus | 0.243865095029373 |
| Leopardus_pardalis | Ursus_americanus | 0.233103343727621 |
| Leopardus_pardalis | Ursus_arctos | 0.239362512919441 |
| Leopardus_pardalis | Ursus_maritimus | 0.238341327661348 |
| Leopardus_pardalis | Ursus_spelaeus | 0.232616247100936 |
| Leopardus_pardalis | Ursus_thibetanus | 0.230713779973165 |
| Leopardus_pardalis | Urva_javanica | 0.192166534200163 |
| Leopardus_pardalis | Viverra_tangalunga_FMNH146957_SRR6053069 | 0.171372744192163 |
| Leopardus_pardalis | Viverricula_indica | 0.170385503195199 |
| Leopardus_pardalis | Vulpes_corsac | 0.239359422288776 |
| Leopardus_pardalis | Vulpes_ferrilata | 0.237464833715595 |
| Leopardus_pardalis | Vulpes_lagopus | 0.237496667844697 |
| Leopardus_pardalis | Vulpes_vulpes_T3611 | 0.23280714118037 |
| Leopardus_pardalis | Vulpes_zerda | 0.231150834400401 |
| Leopardus_pardalis | Zalophus_californianus | 0.231256412047988 |
| Leopardus_tigrinus | Leopardus_wiedii | 0.0738957570466379 |
| Leopardus_tigrinus | Leptailurus_serval | 0.113813877504207 |
| Leopardus_tigrinus | Leptonychotes_weddellii | 0.207540416054083 |
| Leopardus_tigrinus | Lobodon_carcinophaga | 0.211530798438426 |
| Leopardus_tigrinus | Lutra_lutra | 0.235615733495243 |
| Leopardus_tigrinus | Lutra_sumatrana | 0.231949029296956 |
| Leopardus_tigrinus | Lutrogale_perspicillata | 0.239848865862172 |
| Leopardus_tigrinus | Lycaon_pictus | 0.240568582774765 |
| Leopardus_tigrinus | Lynx_canadensis | 0.115855291014333 |
| Leopardus_tigrinus | Lynx_lynx | 0.116410663394326 |
| Leopardus_tigrinus | Lynx_pardinus | 0.119468629268424 |
| Leopardus_tigrinus | Lynx_rufus | 0.117721941870542 |
| Leopardus_tigrinus | Manis_javanica | 0.283412024018511 |
| Leopardus_tigrinus | Martes_americana | 0.234813138313764 |
| Leopardus_tigrinus | Martes_flavigula | 0.23556720290854 |
| Leopardus_tigrinus | Martes_foina | 0.229582915928381 |
| Leopardus_tigrinus | Martes_martes | 0.233560727822772 |
| Leopardus_tigrinus | Martes_melampus | 0.233219302244217 |
| Leopardus_tigrinus | Martes_pennanti | 0.23752834906802 |
| Leopardus_tigrinus | Martes_zibellina | 0.235762430238926 |
| Leopardus_tigrinus | Meles_anakuma | 0.231626633616652 |
| Leopardus_tigrinus | Meles_leucurus | 0.224269289689506 |
| Leopardus_tigrinus | Meles_meles | 0.229028242940672 |
| Leopardus_tigrinus | Melogale_moschata | 0.231787583204289 |
| Leopardus_tigrinus | Melursus_ursinus | 0.238926854167155 |
| Leopardus_tigrinus | Mephitis_mephitis | 0.240568582774765 |
| Leopardus_tigrinus | Mirounga_leonina | 0.215438731695457 |
| Leopardus_tigrinus | Mungotictis_decemlineata | 0.185448035682541 |
| Leopardus_tigrinus | Mustela_altaica | 0.234435439312106 |
| Leopardus_tigrinus | Mustela_erminea | 0.22322476586666 |
| Leopardus_tigrinus | Mustela_eversmannii | 0.231656795153416 |
| Leopardus_tigrinus | Mustela_frenata | 0.228688073492691 |
| Leopardus_tigrinus | Mustela_itatsi | 0.229384305739772 |
| Leopardus_tigrinus | Mustela_kathiah | 0.228445795602369 |
| Leopardus_tigrinus | Mustela_nigripes | 0.229294126281674 |
| Leopardus_tigrinus | Mustela_nivalis | 0.231688557904877 |
| Leopardus_tigrinus | Mustela_putorius | 0.231337846863367 |
| Leopardus_tigrinus | Mustela_sibirica | 0.231218583030713 |
| Leopardus_tigrinus | Nandinia_binotata | 0.175493308630205 |
| Leopardus_tigrinus | Nasua_nasua | 0.246721514140521 |
| Leopardus_tigrinus | Neofelis_nebulosa | 0.144734060187549 |
| Leopardus_tigrinus | Neomonachus_schauinslandi | 0.215125317120648 |
| Leopardus_tigrinus | Neophoca_cinerea | 0.234855504551432 |
| Leopardus_tigrinus | Neovison_vison | 0.231836959110407 |
| Leopardus_tigrinus | Nyctereutes_procyonoides | 0.23634163780656 |
| Leopardus_tigrinus | Odobenus_rosmarus_rosmarus | 0.250507435852275 |
| Leopardus_tigrinus | Otocolobus_manul | 0.120677445485277 |

| | | |
|---|---|---|
| Leopardus_tigrinus | Otocyon_megalotis_megalotis_TS305 | 0.24214959708611 |
| Leopardus_tigrinus | Otocyon_megalotis_virgatus_FMNH158128 | 0.242146076958997 |
| Leopardus_tigrinus | Paguma_larvata | 0.182061915311619 |
| Leopardus_tigrinus | Panthera_leo | 0.132427034445924 |
| Leopardus_tigrinus | Panthera_onca | 0.135389816498854 |
| Leopardus_tigrinus | Panthera_pardus | 0.135439677185034 |
| Leopardus_tigrinus | Panthera_tigris | 0.137748157995863 |
| Leopardus_tigrinus | Panthera_uncia | 0.140895844886785 |
| Leopardus_tigrinus | Paradoxurus_hermaphroditus | 0.188113292217468 |
| Leopardus_tigrinus | Parahyaena_brunnea_FD126_Fecal | 0.193006681072027 |
| Leopardus_tigrinus | Pardofelis_marmorata | 0.125228417635425 |
| Leopardus_tigrinus | Phoca_fasciata | 0.218628183838011 |
| Leopardus_tigrinus | Phoca_groenlandica | 0.21815539916635 |
| Leopardus_tigrinus | Phoca_largha | 0.221886527679825 |
| Leopardus_tigrinus | Phoca_vitulina | 0.222920414004602 |
| Leopardus_tigrinus | Phocarctos_hookeri | 0.233723582124059 |
| Leopardus_tigrinus | Prionailurus_bengalensis | 0.110622596564443 |
| Leopardus_tigrinus | Prionailurus_bengalensis_euptilurus | 0.112307244891406 |
| Leopardus_tigrinus | Prionailurus_planiceps | 0.117499955698903 |
| Leopardus_tigrinus | Prionailurus_rubiginosus | 0.112697950861391 |
| Leopardus_tigrinus | Prionailurus_viverrinus | 0.117748162499419 |
| Leopardus_tigrinus | Prionodon_pardicolor | 0.180194896829488 |
| Leopardus_tigrinus | Procyon_lotor | 0.23716618444841 |
| Leopardus_tigrinus | Profelis_aurata | 0.124783908359437 |
| Leopardus_tigrinus | Proteles_cristatus_cristatus_TS307 | 0.199965694967499 |
| Leopardus_tigrinus | Proteles_cristatus_septentrionalis_NMS_Z_2018_54 | 0.200580951942023 |
| Leopardus_tigrinus | Puma_concolor | 0.117910781453563 |
| Leopardus_tigrinus | Puma_yagouaroundi | 0.122229463457441 |
| Leopardus_tigrinus | Pusa_caspica | 0.225475321977137 |
| Leopardus_tigrinus | Pusa_hispida | 0.220235452381531 |
| Leopardus_tigrinus | Pusa_sibirica | 0.22076184240177 |
| Leopardus_tigrinus | Speothos_venaticus_T1624 | 0.247557688543056 |
| Leopardus_tigrinus | Spilogale_putorius | 0.245147502710336 |
| Leopardus_tigrinus | Taxidea_taxus | 0.22194915078792 |
| Leopardus_tigrinus | Tremarctos_ornatus | 0.233381284336358 |
| Leopardus_tigrinus | Urocyon_cinereoargenteus | 0.250963341513494 |
| Leopardus_tigrinus | Ursus_americanus | 0.241420618011796 |
| Leopardus_tigrinus | Ursus_arctos | 0.241818373364514 |
| Leopardus_tigrinus | Ursus_maritimus | 0.242007783116875 |
| Leopardus_tigrinus | Ursus_spelaeus | 0.237476379379278 |
| Leopardus_tigrinus | Ursus_thibetanus | 0.233535784433957 |
| Leopardus_tigrinus | Urva_javanica | 0.192505378909243 |
| Leopardus_tigrinus | Viverra_tangalunga_FMNH146957_SRR6053069 | 0.175554094100401 |
| Leopardus_tigrinus | Viverricula_indica | 0.171948071149453 |
| Leopardus_tigrinus | Vulpes_corsac | 0.242543802199229 |
| Leopardus_tigrinus | Vulpes_ferrilata | 0.241307169035084 |
| Leopardus_tigrinus | Vulpes_lagopus | 0.240631447159081 |
| Leopardus_tigrinus | Vulpes_vulpes_T3611 | 0.237374626096488 |
| Leopardus_tigrinus | Vulpes_zerda | 0.233288037390173 |
| Leopardus_tigrinus | Zalophus_californianus | 0.231496405493461 |
| Leopardus_wiedii | Leptailurus_serval | 0.108109671846174 |
| Leopardus_wiedii | Leptonychotes_weddellii | 0.206764818836027 |
| Leopardus_wiedii | Lobodon_carcinophaga | 0.210650227721569 |
| Leopardus_wiedii | Lutra_lutra | 0.228014873560828 |
| Leopardus_wiedii | Lutra_sumatrana | 0.22697080131884 |
| Leopardus_wiedii | Lutrogale_perspicillata | 0.239567928184637 |
| Leopardus_wiedii | Lycaon_pictus | 0.237143704527258 |
| Leopardus_wiedii | Lynx_canadensis | 0.110241836080328 |
| Leopardus_wiedii | Lynx_lynx | 0.109009351481623 |
| Leopardus_wiedii | Lynx_pardinus | 0.112961662164268 |
| Leopardus_wiedii | Lynx_rufus | 0.114308714986222 |
| Leopardus_wiedii | Manis_javanica | 0.281670343871684 |
| Leopardus_wiedii | Martes_americana | 0.233071761631836 |
| Leopardus_wiedii | Martes_flavigula | 0.23301353492609 |
| Leopardus_wiedii | Martes_foina | 0.23024201643141 |
| Leopardus_wiedii | Martes_martes | 0.2317121445469 |
| Leopardus_wiedii | Martes_melampus | 0.229285740413797 |
| Leopardus_wiedii | Martes_pennanti | 0.234650730104552 |
| Leopardus_wiedii | Martes_zibellina | 0.23424009119914 |
| Leopardus_wiedii | Meles_anakuma | 0.23138877526961 |
| Leopardus_wiedii | Meles_leucurus | 0.22686315853601 |
| Leopardus_wiedii | Meles_meles | 0.227368903392512 |
| Leopardus_wiedii | Melogale_moschata | 0.228519456738908 |
| Leopardus_wiedii | Melursus_ursinus | 0.237557716172175 |
| Leopardus_wiedii | Mephitis_mephitis | 0.238924506974837 |
| Leopardus_wiedii | Mirounga_leonina | 0.213917098866588 |
| Leopardus_wiedii | Mungotictis_decemlineata | 0.184361212482263 |
| Leopardus_wiedii | Mustela_altaica | 0.231656795153416 |
| Leopardus_wiedii | Mustela_erminea | 0.218342161729841 |
| Leopardus_wiedii | Mustela_eversmannii | 0.226889095276242 |
| Leopardus_wiedii | Mustela_frenata | 0.223719997046938 |
| Leopardus_wiedii | Mustela_itatsi | 0.228038581774494 |
| Leopardus_wiedii | Mustela_kathiah | 0.224459653750519 |
| Leopardus_wiedii | Mustela_nigripes | 0.224890161861212 |
| Leopardus_wiedii | Mustela_nivalis | 0.227792414261433 |
| Leopardus_wiedii | Mustela_putorius | 0.226029553841481 |
| Leopardus_wiedii | Mustela_sibirica | 0.225189952886646 |

| | | |
|---|---|---|
| Leopardus_wiedii | Nandinia_binotata | 0.169791685600184 |
| Leopardus_wiedii | Nasua_nasua | 0.247092902152728 |
| Leopardus_wiedii | Neofelis_nebulosa | 0.140115551826537 |
| Leopardus_wiedii | Neomonachus_schauinslandi | 0.216274487190075 |
| Leopardus_wiedii | Neophoca_cinerea | 0.234367252404346 |
| Leopardus_wiedii | Neovison_vison | 0.228830368927478 |
| Leopardus_wiedii | Nyctereutes_procyonoides | 0.232420587886307 |
| Leopardus_wiedii | Odobenus_rosmarus_rosmarus | 0.253340252479507 |
| Leopardus_wiedii | Otocolobus_manul | 0.11240274868267 |
| Leopardus_wiedii | Otocyon_megalotis_megalotis_TS305 | 0.238964489809424 |
| Leopardus_wiedii | Otocyon_megalotis_virgatus_FMNH158128 | 0.237181771069035 |
| Leopardus_wiedii | Paguma_larvata | 0.180023297919809 |
| Leopardus_wiedii | Panthera_leo | 0.131663290402701 |
| Leopardus_wiedii | Panthera_onca | 0.130530659448569 |
| Leopardus_wiedii | Panthera_pardus | 0.13021191057372 |
| Leopardus_wiedii | Panthera_tigris | 0.134270037082116 |
| Leopardus_wiedii | Panthera_uncia | 0.137831466042795 |
| Leopardus_wiedii | Paradoxurus_hermaphroditus | 0.184418678907567 |
| Leopardus_wiedii | Parahyaena_brunnea_FD126_Fecal | 0.189774270340643 |
| Leopardus_wiedii | Pardofelis_marmorata | 0.118282386214258 |
| Leopardus_wiedii | Phoca_fasciata | 0.217229574669755 |
| Leopardus_wiedii | Phoca_groenlandica | 0.216115372268171 |
| Leopardus_wiedii | Phoca_largha | 0.219173046032238 |
| Leopardus_wiedii | Phoca_vitulina | 0.217836348877267 |
| Leopardus_wiedii | Phocarctos_hookeri | 0.235752035165176 |
| Leopardus_wiedii | Prionailurus_bengalensis | 0.107726061157402 |
| Leopardus_wiedii | Prionailurus_bengalensis_euptilurus | 0.108380156558252 |
| Leopardus_wiedii | Prionailurus_planiceps | 0.113822850590207 |
| Leopardus_wiedii | Prionailurus_rubiginosus | 0.110444521636552 |
| Leopardus_wiedii | Prionailurus_viverrinus | 0.113132023036834 |
| Leopardus_wiedii | Prionodon_pardicolor | 0.177942242777172 |
| Leopardus_wiedii | Procyon_lotor | 0.232127904181611 |
| Leopardus_wiedii | Profelis_aurata | 0.117718550662142 |
| Leopardus_wiedii | Proteles_cristatus_cristatus_TS307 | 0.199129048461243 |
| Leopardus_wiedii | Proteles_cristatus_septentrionalis_NMS_Z_2018_54 | 0.198705364504104 |
| Leopardus_wiedii | Puma_concolor | 0.11331094837251 |
| Leopardus_wiedii | Puma_yagouaroundi | 0.118906602051385 |
| Leopardus_wiedii | Pusa_caspica | 0.22264057837437 |
| Leopardus_wiedii | Pusa_hispida | 0.217957630437207 |
| Leopardus_wiedii | Pusa_sibirica | 0.217388388044864 |
| Leopardus_wiedii | Speothos_venaticus_T1624 | 0.241852892677161 |
| Leopardus_wiedii | Spilogale_putorius | 0.245416805465305 |
| Leopardus_wiedii | Taxidea_taxus | 0.220129745166925 |
| Leopardus_wiedii | Tremarctos_ornatus | 0.235245414273446 |
| Leopardus_wiedii | Urocyon_cinereoargenteus | 0.24252161011896 |
| Leopardus_wiedii | Ursus_americanus | 0.236663661396633 |
| Leopardus_wiedii | Ursus_arctos | 0.241975821683989 |
| Leopardus_wiedii | Ursus_maritimus | 0.240161009305606 |
| Leopardus_wiedii | Ursus_spelaeus | 0.234526507803461 |
| Leopardus_wiedii | Ursus_thibetanus | 0.230846489821337 |
| Leopardus_wiedii | Urva_javanica | 0.192518526643852 |
| Leopardus_wiedii | Viverra_tangalunga_FMNH146957_SRR6053069 | 0.171372744192163 |
| Leopardus_wiedii | Viverricula_indica | 0.170975701437506 |
| Leopardus_wiedii | Vulpes_corsac | 0.239713442672521 |
| Leopardus_wiedii | Vulpes_ferrilata | 0.238592825418894 |
| Leopardus_wiedii | Vulpes_lagopus | 0.234975243187128 |
| Leopardus_wiedii | Vulpes_vulpes_T3611 | 0.233944512194563 |
| Leopardus_wiedii | Vulpes_zerda | 0.230751778567479 |
| Leopardus_wiedii | Zalophus_californianus | 0.23316477783101 |
| Leptailurus_serval | Leptonychotes_weddellii | 0.208756649918188 |
| Leptailurus_serval | Lobodon_carcinophaga | 0.214342483671294 |
| Leptailurus_serval | Lutra_lutra | 0.232750344551258 |
| Leptailurus_serval | Lutra_sumatrana | 0.231471486088472 |
| Leptailurus_serval | Lutrogale_perspicillata | 0.242429528772681 |
| Leptailurus_serval | Lycaon_pictus | 0.232501493315442 |
| Leptailurus_serval | Lynx_canadensis | 0.107061400620582 |
| Leptailurus_serval | Lynx_lynx | 0.110996892311765 |
| Leptailurus_serval | Lynx_pardinus | 0.109690921420041 |
| Leptailurus_serval | Lynx_rufus | 0.110521686833903 |
| Leptailurus_serval | Manis_javanica | 0.288797686634328 |
| Leptailurus_serval | Martes_americana | 0.232126617077663 |
| Leptailurus_serval | Martes_flavigula | 0.234025971403362 |
| Leptailurus_serval | Martes_foina | 0.229936481929656 |
| Leptailurus_serval | Martes_martes | 0.233063012896575 |
| Leptailurus_serval | Martes_melampus | 0.232500044488349 |
| Leptailurus_serval | Martes_pennanti | 0.239266532382534 |
| Leptailurus_serval | Martes_zibellina | 0.235378387431121 |
| Leptailurus_serval | Meles_anakuma | 0.230162933621244 |
| Leptailurus_serval | Meles_leucurus | 0.226954881158723 |
| Leptailurus_serval | Meles_meles | 0.228597458194454 |
| Leptailurus_serval | Melogale_moschata | 0.230606094707512 |
| Leptailurus_serval | Melursus_ursinus | 0.237015409411924 |
| Leptailurus_serval | Mephitis_mephitis | 0.239297837784116 |
| Leptailurus_serval | Mirounga_leonina | 0.217521165956635 |
| Leptailurus_serval | Mungotictis_decemlineata | 0.189815334157378 |
| Leptailurus_serval | Mustela_altaica | 0.230032029982168 |
| Leptailurus_serval | Mustela_erminea | 0.223140908322568 |

| | | |
|---|---|---|
| Leptailurus_serval | Mustela_eversmannii | 0.231406901686067 |
| Leptailurus_serval | Mustela_frenata | 0.223156028492915 |
| Leptailurus_serval | Mustela_itatsi | 0.231975062051604 |
| Leptailurus_serval | Mustela_kathiah | 0.229901448262435 |
| Leptailurus_serval | Mustela_nigripes | 0.228709015494853 |
| Leptailurus_serval | Mustela_nivalis | 0.228827980652825 |
| Leptailurus_serval | Mustela_putorius | 0.23042933798616 |
| Leptailurus_serval | Mustela_sibirica | 0.232286409388657 |
| Leptailurus_serval | Nandinia_binotata | 0.171968053925113 |
| Leptailurus_serval | Nasua_nasua | 0.248463372137857 |
| Leptailurus_serval | Neofelis_nebulosa | 0.137160671391508 |
| Leptailurus_serval | Neomonachus_schauinslandi | 0.217710598764341 |
| Leptailurus_serval | Neophoca_cinerea | 0.236194757938982 |
| Leptailurus_serval | Neovison_vison | 0.230429222688188 |
| Leptailurus_serval | Nyctereutes_procyonoides | 0.233707766518134 |
| Leptailurus_serval | Odobenus_rosmarus_rosmarus | 0.253482884226524 |
| Leptailurus_serval | Otocolobus_manul | 0.10923604419066 |
| Leptailurus_serval | Otocyon_megalotis_megalotis_TS305 | 0.24248187280775 |
| Leptailurus_serval | Otocyon_megalotis_virgatus_FMNH158128 | 0.241807898215111 |
| Leptailurus_serval | Paguma_larvata | 0.172978666873801 |
| Leptailurus_serval | Panthera_leo | 0.134276352273543 |
| Leptailurus_serval | Panthera_onca | 0.133642311265322 |
| Leptailurus_serval | Panthera_pardus | 0.13396415614486 |
| Leptailurus_serval | Panthera_tigris | 0.139310926945694 |
| Leptailurus_serval | Panthera_uncia | 0.140218014005843 |
| Leptailurus_serval | Paradoxurus_hermaphroditus | 0.176931348465246 |
| Leptailurus_serval | Parahyaena_brunnea_FD126_Fecal | 0.193341442360381 |
| Leptailurus_serval | Pardofelis_marmorata | 0.114959299650638 |
| Leptailurus_serval | Phoca_fasciata | 0.2152223672231 |
| Leptailurus_serval | Phoca_groenlandica | 0.217621733571901 |
| Leptailurus_serval | Phoca_largha | 0.219140374661597 |
| Leptailurus_serval | Phoca_vitulina | 0.218557277282958 |
| Leptailurus_serval | Phocarctos_hookeri | 0.237583472681785 |
| Leptailurus_serval | Prionailurus_bengalensis | 0.0993789273557382 |
| Leptailurus_serval | Prionailurus_bengalensis_euptilurus | 0.101574369533815 |
| Leptailurus_serval | Prionailurus_planiceps | 0.105857361879999 |
| Leptailurus_serval | Prionailurus_rubiginosus | 0.104668413711787 |
| Leptailurus_serval | Prionailurus_viverrinus | 0.105101926898535 |
| Leptailurus_serval | Prionodon_pardicolor | 0.177433132196574 |
| Leptailurus_serval | Procyon_lotor | 0.234586845281435 |
| Leptailurus_serval | Profelis_aurata | 0.101118814481041 |
| Leptailurus_serval | Proteles_cristatus_cristatus_TS307 | 0.198800563627851 |
| Leptailurus_serval | Proteles_cristatus_septentrionalis_NMS_Z_2018_54 | 0.200177829039412 |
| Leptailurus_serval | Puma_concolor | 0.108719117085631 |
| Leptailurus_serval | Puma_yagouaroundi | 0.110525839789321 |
| Leptailurus_serval | Pusa_caspica | 0.218663237299396 |
| Leptailurus_serval | Pusa_hispida | 0.214875873950169 |
| Leptailurus_serval | Pusa_sibirica | 0.21576685503151 |
| Leptailurus_serval | Speothos_venaticus_T1624 | 0.242651188812361 |
| Leptailurus_serval | Spilogale_putorius | 0.241973007885649 |
| Leptailurus_serval | Taxidea_taxus | 0.222436571993092 |
| Leptailurus_serval | Tremarctos_ornatus | 0.230504858227069 |
| Leptailurus_serval | Urocyon_cinereoargenteus | 0.246528695367516 |
| Leptailurus_serval | Ursus_americanus | 0.236615139166099 |
| Leptailurus_serval | Ursus_arctos | 0.242701216449312 |
| Leptailurus_serval | Ursus_maritimus | 0.240406328843014 |
| Leptailurus_serval | Ursus_spelaeus | 0.23811836505366 |
| Leptailurus_serval | Ursus_thibetanus | 0.237093154159172 |
| Leptailurus_serval | Urva_javanica | 0.192110816141173 |
| Leptailurus_serval | Viverra_tangalunga_FMNH146957_SRR6053069 | 0.169308112798186 |
| Leptailurus_serval | Viverricula_indica | 0.167919382445118 |
| Leptailurus_serval | Vulpes_corsac | 0.238301153288637 |
| Leptailurus_serval | Vulpes_ferrilata | 0.237495160418177 |
| Leptailurus_serval | Vulpes_lagopus | 0.237396171456371 |
| Leptailurus_serval | Vulpes_vulpes_T3611 | 0.231849725733611 |
| Leptailurus_serval | Vulpes_zerda | 0.232056209278358 |
| Leptailurus_serval | Zalophus_californianus | 0.232584159396935 |
| Leptonychotes_weddellii | Lobodon_carcinophaga | 0.0727410273987992 |
| Leptonychotes_weddellii | Lutra_lutra | 0.208027272643992 |
| Leptonychotes_weddellii | Lutra_sumatrana | 0.208233482000519 |
| Leptonychotes_weddellii | Lutrogale_perspicillata | 0.215649531962092 |
| Leptonychotes_weddellii | Lycaon_pictus | 0.221080779288593 |
| Leptonychotes_weddellii | Lynx_canadensis | 0.208659557945523 |
| Leptonychotes_weddellii | Lynx_lynx | 0.207678286919888 |
| Leptonychotes_weddellii | Lynx_pardinus | 0.207323456735244 |
| Leptonychotes_weddellii | Lynx_rufus | 0.206507069912609 |
| Leptonychotes_weddellii | Manis_javanica | 0.267850397485526 |
| Leptonychotes_weddellii | Martes_americana | 0.208423781539159 |
| Leptonychotes_weddellii | Martes_flavigula | 0.207737446883417 |
| Leptonychotes_weddellii | Martes_foina | 0.209053687114331 |
| Leptonychotes_weddellii | Martes_martes | 0.207215151576817 |
| Leptonychotes_weddellii | Martes_melampus | 0.205275025332436 |
| Leptonychotes_weddellii | Martes_pennanti | 0.214479504521325 |
| Leptonychotes_weddellii | Martes_zibellina | 0.208590052866418 |
| Leptonychotes_weddellii | Meles_anakuma | 0.208909856016542 |
| Leptonychotes_weddellii | Meles_leucurus | 0.207452549768771 |
| Leptonychotes_weddellii | Meles_meles | 0.205461729348154 |

| | | |
|---|---|---|
| Leptonychotes_weddellii | Melogale_moschata | 0.214251944919893 |
| Leptonychotes_weddellii | Melursus_ursinus | 0.212653028677782 |
| Leptonychotes_weddellii | Mephitis_mephitis | 0.218041014878227 |
| Leptonychotes_weddellii | Mirounga_leonina | 0.10234694900819 |
| Leptonychotes_weddellii | Mungotictis_decemlineata | 0.213597949919582 |
| Leptonychotes_weddellii | Mustela_altaica | 0.212505460166618 |
| Leptonychotes_weddellii | Mustela_erminea | 0.201571886348366 |
| Leptonychotes_weddellii | Mustela_eversmannii | 0.211439010648988 |
| Leptonychotes_weddellii | Mustela_frenata | 0.208104402494225 |
| Leptonychotes_weddellii | Mustela_itatsi | 0.208204860103502 |
| Leptonychotes_weddellii | Mustela_kathiah | 0.212972566909046 |
| Leptonychotes_weddellii | Mustela_nigripes | 0.210590926844361 |
| Leptonychotes_weddellii | Mustela_nivalis | 0.211345229317301 |
| Leptonychotes_weddellii | Mustela_putorius | 0.209301683106333 |
| Leptonychotes_weddellii | Mustela_sibirica | 0.209510051766109 |
| Leptonychotes_weddellii | Nandinia_binotata | 0.191814503697353 |
| Leptonychotes_weddellii | Nasua_nasua | 0.22427009280852 |
| Leptonychotes_weddellii | Neofelis_nebulosa | 0.220305708899232 |
| Leptonychotes_weddellii | Neomonachus_schauinslandi | 0.111404222946601 |
| Leptonychotes_weddellii | Neophoca_cinerea | 0.179116068455128 |
| Leptonychotes_weddellii | Neovison_vison | 0.209510051766109 |
| Leptonychotes_weddellii | Nyctereutes_procyonoides | 0.222066897826778 |
| Leptonychotes_weddellii | Odobenus_rosmarus_rosmarus | 0.197026656273186 |
| Leptonychotes_weddellii | Otocolobus_manul | 0.209208963625654 |
| Leptonychotes_weddellii | Otocyon_megalotis_megalotis_TS305 | 0.226445334904151 |
| Leptonychotes_weddellii | Otocyon_megalotis_virgatus_FMNH158128 | 0.230234913118256 |
| Leptonychotes_weddellii | Paguma_larvata | 0.218342161729841 |
| Leptonychotes_weddellii | Panthera_leo | 0.216441784979653 |
| Leptonychotes_weddellii | Panthera_onca | 0.215952377387605 |
| Leptonychotes_weddellii | Panthera_pardus | 0.219730190105851 |
| Leptonychotes_weddellii | Panthera_tigris | 0.222640104778787 |
| Leptonychotes_weddellii | Panthera_uncia | 0.220483850885927 |
| Leptonychotes_weddellii | Paradoxurus_hermaphroditus | 0.222470877746088 |
| Leptonychotes_weddellii | Parahyaena_brunnea_FD126_Fecal | 0.214925190437239 |
| Leptonychotes_weddellii | Pardofelis_marmorata | 0.21206880153077 |
| Leptonychotes_weddellii | Phoca_fasciata | 0.125691582707577 |
| Leptonychotes_weddellii | Phoca_groenlandica | 0.132189469494348 |
| Leptonychotes_weddellii | Phoca_largha | 0.128166002864321 |
| Leptonychotes_weddellii | Phoca_vitulina | 0.129673198836118 |
| Leptonychotes_weddellii | Phocarctos_hookeri | 0.181663494819751 |
| Leptonychotes_weddellii | Prionailurus_bengalensis | 0.205771900975405 |
| Leptonychotes_weddellii | Prionailurus_bengalensis_euptilurus | 0.205336994032795 |
| Leptonychotes_weddellii | Prionailurus_planiceps | 0.210654185016394 |
| Leptonychotes_weddellii | Prionailurus_rubiginosus | 0.207774636464599 |
| Leptonychotes_weddellii | Prionailurus_viverrinus | 0.208049970376755 |
| Leptonychotes_weddellii | Prionodon_pardicolor | 0.218523898341329 |
| Leptonychotes_weddellii | Procyon_lotor | 0.211884818991687 |
| Leptonychotes_weddellii | Profelis_aurata | 0.212260189293797 |
| Leptonychotes_weddellii | Proteles_cristatus_cristatus_TS307 | 0.21792768703079 |
| Leptonychotes_weddellii | Proteles_cristatus_septentrionalis_NMS_Z_2018_54 | 0.22213881309543 |
| Leptonychotes_weddellii | Puma_concolor | 0.210045165973208 |
| Leptonychotes_weddellii | Puma_yagouaroundi | 0.210773771929507 |
| Leptonychotes_weddellii | Pusa_caspica | 0.130108462338766 |
| Leptonychotes_weddellii | Pusa_hispida | 0.127851749847316 |
| Leptonychotes_weddellii | Pusa_sibirica | 0.128705175063553 |
| Leptonychotes_weddellii | Speothos_venaticus_T1624 | 0.232964943589175 |
| Leptonychotes_weddellii | Spilogale_putorius | 0.21207266398318 |
| Leptonychotes_weddellii | Taxidea_taxus | 0.195337176925476 |
| Leptonychotes_weddellii | Tremarctos_ornatus | 0.209228074910211 |
| Leptonychotes_weddellii | Urocyon_cinereoargenteus | 0.230318014285898 |
| Leptonychotes_weddellii | Ursus_americanus | 0.214104281869532 |
| Leptonychotes_weddellii | Ursus_arctos | 0.214078970411004 |
| Leptonychotes_weddellii | Ursus_maritimus | 0.212676684500978 |
| Leptonychotes_weddellii | Ursus_spelaeus | 0.211251909426681 |
| Leptonychotes_weddellii | Ursus_thibetanus | 0.210385451937025 |
| Leptonychotes_weddellii | Urva_javanica | 0.215923844335669 |
| Leptonychotes_weddellii | Viverra_tangalunga_FMNH146957_SRR6053069 | 0.210392769185271 |
| Leptonychotes_weddellii | Viverricula_indica | 0.209398344359617 |
| Leptonychotes_weddellii | Vulpes_corsac | 0.227774301698034 |
| Leptonychotes_weddellii | Vulpes_ferrilata | 0.23021146780297 |
| Leptonychotes_weddellii | Vulpes_lagopus | 0.226733007987981 |
| Leptonychotes_weddellii | Vulpes_vulpes_T3611 | 0.224845860988794 |
| Leptonychotes_weddellii | Vulpes_zerda | 0.226029491936061 |
| Leptonychotes_weddellii | Zalophus_californianus | 0.174803067379716 |
| Lobodon_carcinophaga | Lutra_lutra | 0.208122347430919 |
| Lobodon_carcinophaga | Lutra_sumatrana | 0.204793523979885 |
| Lobodon_carcinophaga | Lutrogale_perspicillata | 0.214938725739459 |
| Lobodon_carcinophaga | Lycaon_pictus | 0.217556457635996 |
| Lobodon_carcinophaga | Lynx_canadensis | 0.2141381882769 |
| Lobodon_carcinophaga | Lynx_lynx | 0.214422390848851 |
| Lobodon_carcinophaga | Lynx_pardinus | 0.213215156787153 |
| Lobodon_carcinophaga | Lynx_rufus | 0.212178308004098 |
| Lobodon_carcinophaga | Manis_javanica | 0.274308483083908 |
| Lobodon_carcinophaga | Martes_americana | 0.209497936901773 |
| Lobodon_carcinophaga | Martes_flavigula | 0.209284558044379 |
| Lobodon_carcinophaga | Martes_foina | 0.208354830594411 |
| Lobodon_carcinophaga | Martes_martes | 0.20854028297134 |

| | | |
|---|---|---|
| Lobodon_carcinophaga | Martes_melampus | 0.208877741668847 |
| Lobodon_carcinophaga | Martes_pennanti | 0.21035594234129 |
| Lobodon_carcinophaga | Martes_zibellina | 0.211503640342629 |
| Lobodon_carcinophaga | Meles_anakuma | 0.21085612662194 |
| Lobodon_carcinophaga | Meles_leucurus | 0.208526404880942 |
| Lobodon_carcinophaga | Meles_meles | 0.20412596833276 |
| Lobodon_carcinophaga | Melogale_moschata | 0.213567709775691 |
| Lobodon_carcinophaga | Melursus_ursinus | 0.213111419883219 |
| Lobodon_carcinophaga | Mephitis_mephitis | 0.216535637396999 |
| Lobodon_carcinophaga | Mirounga_leonina | 0.106512455358081 |
| Lobodon_carcinophaga | Mungotictis_decemlineata | 0.217029901210155 |
| Lobodon_carcinophaga | Mustela_altaica | 0.213907308511482 |
| Lobodon_carcinophaga | Mustela_erminea | 0.203582841071061 |
| Lobodon_carcinophaga | Mustela_eversmannii | 0.212480675289177 |
| Lobodon_carcinophaga | Mustela_frenata | 0.200847928149359 |
| Lobodon_carcinophaga | Mustela_itatsi | 0.208908404671238 |
| Lobodon_carcinophaga | Mustela_kathiah | 0.212295413420988 |
| Lobodon_carcinophaga | Mustela_nigripes | 0.210763752024317 |
| Lobodon_carcinophaga | Mustela_nivalis | 0.210438720576314 |
| Lobodon_carcinophaga | Mustela_putorius | 0.209916743604601 |
| Lobodon_carcinophaga | Mustela_sibirica | 0.208937588499476 |
| Lobodon_carcinophaga | Nandinia_binotata | 0.198157217215496 |
| Lobodon_carcinophaga | Nasua_nasua | 0.228691079641892 |
| Lobodon_carcinophaga | Neofelis_nebulosa | 0.229021260815997 |
| Lobodon_carcinophaga | Neomonachus_schauinslandi | 0.111376054276749 |
| Lobodon_carcinophaga | Neophoca_cinerea | 0.179176807900094 |
| Lobodon_carcinophaga | Neovison_vison | 0.20457295489228 |
| Lobodon_carcinophaga | Nyctereutes_procyonoides | 0.218986633423612 |
| Lobodon_carcinophaga | Odobenus_rosmarus_rosmarus | 0.197866992453281 |
| Lobodon_carcinophaga | Otocolobus_manul | 0.210320296243127 |
| Lobodon_carcinophaga | Otocyon_megalotis_megalotis_TS305 | 0.228514433774902 |
| Lobodon_carcinophaga | Otocyon_megalotis_virgatus_FMNH158128 | 0.231020184213207 |
| Lobodon_carcinophaga | Paguma_larvata | 0.219771867321074 |
| Lobodon_carcinophaga | Panthera_leo | 0.218173671890002 |
| Lobodon_carcinophaga | Panthera_onca | 0.218324457055128 |
| Lobodon_carcinophaga | Panthera_pardus | 0.219300036299344 |
| Lobodon_carcinophaga | Panthera_tigris | 0.226667995056878 |
| Lobodon_carcinophaga | Panthera_uncia | 0.22485168674558 |
| Lobodon_carcinophaga | Paradoxurus_hermaphroditus | 0.223894009132219 |
| Lobodon_carcinophaga | Parahyaena_brunnea_FD126_Fecal | 0.221493790430185 |
| Lobodon_carcinophaga | Pardofelis_marmorata | 0.2152223672231 |
| Lobodon_carcinophaga | Phoca_fasciata | 0.127002082701178 |
| Lobodon_carcinophaga | Phoca_groenlandica | 0.134040492626859 |
| Lobodon_carcinophaga | Phoca_largha | 0.131580025735026 |
| Lobodon_carcinophaga | Phoca_vitulina | 0.132614517073608 |
| Lobodon_carcinophaga | Phocarctos_hookeri | 0.185243157473442 |
| Lobodon_carcinophaga | Prionailurus_bengalensis | 0.210226947273009 |
| Lobodon_carcinophaga | Prionailurus_bengalensis_euptilurus | 0.210721915074832 |
| Lobodon_carcinophaga | Prionailurus_planiceps | 0.210125699606237 |
| Lobodon_carcinophaga | Prionailurus_rubiginosus | 0.209670173696945 |
| Lobodon_carcinophaga | Prionailurus_viverrinus | 0.213676678555516 |
| Lobodon_carcinophaga | Prionodon_pardicolor | 0.22747545979989 |
| Lobodon_carcinophaga | Procyon_lotor | 0.212769450967342 |
| Lobodon_carcinophaga | Profelis_aurata | 0.214582435395132 |
| Lobodon_carcinophaga | Proteles_cristatus_cristatus_TS307 | 0.228270654412686 |
| Lobodon_carcinophaga | Proteles_cristatus_septentrionalis_NMS_Z_2018_54 | 0.229707006933737 |
| Lobodon_carcinophaga | Puma_concolor | 0.212463835062348 |
| Lobodon_carcinophaga | Puma_yagouaroundi | 0.212874752643341 |
| Lobodon_carcinophaga | Pusa_caspica | 0.133825530828411 |
| Lobodon_carcinophaga | Pusa_hispida | 0.131064776671182 |
| Lobodon_carcinophaga | Pusa_sibirica | 0.132879389057179 |
| Lobodon_carcinophaga | Speothos_venaticus_T1624 | 0.230857588503187 |
| Lobodon_carcinophaga | Spilogale_putorius | 0.218277219385062 |
| Lobodon_carcinophaga | Taxidea_taxus | 0.197105317427457 |
| Lobodon_carcinophaga | Tremarctos_ornatus | 0.209924131388322 |
| Lobodon_carcinophaga | Urocyon_cinereoargenteus | 0.231482127692776 |
| Lobodon_carcinophaga | Ursus_americanus | 0.211313256158714 |
| Lobodon_carcinophaga | Ursus_arctos | 0.216505760795666 |
| Lobodon_carcinophaga | Ursus_maritimus | 0.216846624958094 |
| Lobodon_carcinophaga | Ursus_spelaeus | 0.210841987048444 |
| Lobodon_carcinophaga | Ursus_thibetanus | 0.210950329410731 |
| Lobodon_carcinophaga | Urva_javanica | 0.217321254303067 |
| Lobodon_carcinophaga | Viverra_tangalunga_FMNH146957_SRR6053069 | 0.213454768648435 |
| Lobodon_carcinophaga | Viverricula_indica | 0.208698351230233 |
| Lobodon_carcinophaga | Vulpes_corsac | 0.227416871559408 |
| Lobodon_carcinophaga | Vulpes_ferrilata | 0.228758458431035 |
| Lobodon_carcinophaga | Vulpes_lagopus | 0.228280702301302 |
| Lobodon_carcinophaga | Vulpes_vulpes_T3611 | 0.22083126447837 |
| Lobodon_carcinophaga | Vulpes_zerda | 0.221158988806256 |
| Lobodon_carcinophaga | Zalophus_californianus | 0.17402527030701 |
| Lutra_lutra | Lutra_sumatrana | 0.0593308722362363 |
| Lutra_lutra | Lutrogale_perspicillata | 0.124730016480128 |
| Lutra_lutra | Lycaon_pictus | 0.247985947163591 |
| Lutra_lutra | Lynx_canadensis | 0.231685748538133 |
| Lutra_lutra | Lynx_lynx | 0.228290711213832 |
| Lutra_lutra | Lynx_pardinus | 0.229912924712641 |
| Lutra_lutra | Lynx_rufus | 0.233109326649668 |

| | | |
|---|---|---|
| Lutra_lutra | Manis_javanica | 0.289253839180757 |
| Lutra_lutra | Martes_americana | 0.170450008806736 |
| Lutra_lutra | Martes_flavigula | 0.169140484492618 |
| Lutra_lutra | Martes_foina | 0.171513184139783 |
| Lutra_lutra | Martes_martes | 0.169193306902339 |
| Lutra_lutra | Martes_melampus | 0.169033144542618 |
| Lutra_lutra | Martes_pennanti | 0.168953291258548 |
| Lutra_lutra | Martes_zibellina | 0.170828904326033 |
| Lutra_lutra | Meles_anakuma | 0.182726402952261 |
| Lutra_lutra | Meles_leucurus | 0.177328236114333 |
| Lutra_lutra | Meles_meles | 0.173989507017601 |
| Lutra_lutra | Melogale_moschata | 0.16602717322021 |
| Lutra_lutra | Melursus_ursinus | 0.247897325490301 |
| Lutra_lutra | Mephitis_mephitis | 0.237310043250603 |
| Lutra_lutra | Mirounga_leonina | 0.221945440095722 |
| Lutra_lutra | Mungotictis_decemlineata | 0.236759226977176 |
| Lutra_lutra | Mustela_altaica | 0.167840975867149 |
| Lutra_lutra | Mustela_erminea | 0.154199509018916 |
| Lutra_lutra | Mustela_eversmannii | 0.168851549239022 |
| Lutra_lutra | Mustela_frenata | 0.163558181121722 |
| Lutra_lutra | Mustela_itatsi | 0.167819441316893 |
| Lutra_lutra | Mustela_kathiah | 0.167025315001106 |
| Lutra_lutra | Mustela_nigripes | 0.169308532853012 |
| Lutra_lutra | Mustela_nivalis | 0.168249719602977 |
| Lutra_lutra | Mustela_putorius | 0.16996646434807 |
| Lutra_lutra | Mustela_sibirica | 0.168362277355474 |
| Lutra_lutra | Nandinia_binotata | 0.220643566699323 |
| Lutra_lutra | Nasua_nasua | 0.23888810363189 |
| Lutra_lutra | Neofelis_nebulosa | 0.247916350215327 |
| Lutra_lutra | Neomonachus_schauinslandi | 0.213190942144006 |
| Lutra_lutra | Neophoca_cinerea | 0.234781415357084 |
| Lutra_lutra | Neovison_vison | 0.167690872560402 |
| Lutra_lutra | Nyctereutes_procyonoides | 0.247782079885317 |
| Lutra_lutra | Odobenus_rosmarus_rosmarus | 0.247268906725766 |
| Lutra_lutra | Otocolobus_manul | 0.234540037166309 |
| Lutra_lutra | Otocyon_megalotis_megalotis_TS305 | 0.251476083357545 |
| Lutra_lutra | Otocyon_megalotis_virgatus_FMNH158128 | 0.249275018043081 |
| Lutra_lutra | Paguma_larvata | 0.237563791198945 |
| Lutra_lutra | Panthera_leo | 0.241507181438107 |
| Lutra_lutra | Panthera_onca | 0.234782879888275 |
| Lutra_lutra | Panthera_pardus | 0.243190459949471 |
| Lutra_lutra | Panthera_tigris | 0.242135074189645 |
| Lutra_lutra | Panthera_uncia | 0.239042919073007 |
| Lutra_lutra | Paradoxurus_hermaphroditus | 0.247606989006018 |
| Lutra_lutra | Parahyaena_brunnea_FD126_Fecal | 0.23840446016888 |
| Lutra_lutra | Pardofelis_marmorata | 0.238139652502401 |
| Lutra_lutra | Phoca_fasciata | 0.210927380272498 |
| Lutra_lutra | Phoca_groenlandica | 0.216882590373639 |
| Lutra_lutra | Phoca_largha | 0.214076656560389 |
| Lutra_lutra | Phoca_vitulina | 0.21641919433446 |
| Lutra_lutra | Phocarctos_hookeri | 0.234658174243771 |
| Lutra_lutra | Prionailurus_bengalensis | 0.231486473497745 |
| Lutra_lutra | Prionailurus_bengalensis_euptilurus | 0.231334418783975 |
| Lutra_lutra | Prionailurus_planiceps | 0.232785216382554 |
| Lutra_lutra | Prionailurus_rubiginosus | 0.231780648264338 |
| Lutra_lutra | Prionailurus_viverrinus | 0.233052286647426 |
| Lutra_lutra | Prionodon_pardicolor | 0.241336002610353 |
| Lutra_lutra | Procyon_lotor | 0.222427747622671 |
| Lutra_lutra | Profelis_aurata | 0.235733519802533 |
| Lutra_lutra | Proteles_cristatus_cristatus_TS307 | 0.238321700222868 |
| Lutra_lutra | Proteles_cristatus_septentrionalis_NMS_Z_2018_54 | 0.241994913537351 |
| Lutra_lutra | Puma_concolor | 0.229683513763232 |
| Lutra_lutra | Puma_yagouaroundi | 0.232273411395164 |
| Lutra_lutra | Pusa_caspica | 0.218272385439667 |
| Lutra_lutra | Pusa_hispida | 0.212757027255938 |
| Lutra_lutra | Pusa_sibirica | 0.216318834007262 |
| Lutra_lutra | Speothos_venaticus_T1624 | 0.259946501857827 |
| Lutra_lutra | Spilogale_putorius | 0.234718238877105 |
| Lutra_lutra | Taxidea_taxus | 0.172737389929089 |
| Lutra_lutra | Tremarctos_ornatus | 0.237050402664865 |
| Lutra_lutra | Urocyon_cinereoargenteus | 0.253726671177603 |
| Lutra_lutra | Ursus_americanus | 0.250180983488064 |
| Lutra_lutra | Ursus_arctos | 0.249040456650132 |
| Lutra_lutra | Ursus_maritimus | 0.248823466065353 |
| Lutra_lutra | Ursus_spelaeus | 0.246078760456486 |
| Lutra_lutra | Ursus_thibetanus | 0.241119507746411 |
| Lutra_lutra | Urva_javanica | 0.238353739728971 |
| Lutra_lutra | Viverra_tangalunga_FMNH146957_SRR6053069 | 0.234776332355672 |
| Lutra_lutra | Viverricula_indica | 0.233529432517585 |
| Lutra_lutra | Vulpes_corsac | 0.25013704183995 |
| Lutra_lutra | Vulpes_ferrilata | 0.251712692635574 |
| Lutra_lutra | Vulpes_lagopus | 0.253258761951869 |
| Lutra_lutra | Vulpes_vulpes_T3611 | 0.248406305885639 |
| Lutra_lutra | Vulpes_zerda | 0.248359758542067 |
| Lutra_lutra | Zalophus_californianus | 0.232762964084762 |
| Lutra_sumatrana | Lutrogale_perspicillata | 0.122872172571363 |
| Lutra_sumatrana | Lycaon_pictus | 0.247914971718236 |

| | | |
|---|---|---|
| Lutra_sumatrana | Lynx_canadensis | 0.233851058030606 |
| Lutra_sumatrana | Lynx_lynx | 0.228797446406271 |
| Lutra_sumatrana | Lynx_pardinus | 0.230201312933504 |
| Lutra_sumatrana | Lynx_rufus | 0.230863176717228 |
| Lutra_sumatrana | Manis_javanica | 0.285858425080573 |
| Lutra_sumatrana | Martes_americana | 0.167619871561261 |
| Lutra_sumatrana | Martes_flavigula | 0.165348767585654 |
| Lutra_sumatrana | Martes_foina | 0.16942115881108З |
| Lutra_sumatrana | Martes_martes | 0.166470798741625 |
| Lutra_sumatrana | Martes_melampus | 0.165677327455475 |
| Lutra_sumatrana | Martes_pennanti | 0.168645472656583 |
| Lutra_sumatrana | Martes_zibellina | 0.167813370329557 |
| Lutra_sumatrana | Meles_anakuma | 0.179291449663915 |
| Lutra_sumatrana | Meles_leucurus | 0.174645911786158 |
| Lutra_sumatrana | Meles_meles | 0.173165821594269 |
| Lutra_sumatrana | Melogale_moschata | 0.16657278592485 |
| Lutra_sumatrana | Melursus_ursinus | 0.242718116843729 |
| Lutra_sumatrana | Mephitis_mephitis | 0.23449227246149 |
| Lutra_sumatrana | Mirounga_leonina | 0.218586645470344 |
| Lutra_sumatrana | Mungotictis_decemlineata | 0.234788262591461 |
| Lutra_sumatrana | Mustela_altaica | 0.166644768267317 |
| Lutra_sumatrana | Mustela_erminea | 0.148427321446303 |
| Lutra_sumatrana | Mustela_eversmannii | 0.165675243878817 |
| Lutra_sumatrana | Mustela_frenata | 0.157950764227491 |
| Lutra_sumatrana | Mustela_itatsi | 0.166537513057848 |
| Lutra_sumatrana | Mustela_kathiah | 0.168840683501988 |
| Lutra_sumatrana | Mustela_nigripes | 0.164687674804445 |
| Lutra_sumatrana | Mustela_nivalis | 0.165205444058349 |
| Lutra_sumatrana | Mustela_putorius | 0.165258195865179 |
| Lutra_sumatrana | Mustela_sibirica | 0.164964150707357 |
| Lutra_sumatrana | Nandinia_binotata | 0.219979282196752 |
| Lutra_sumatrana | Nasua_nasua | 0.234273008187967 |
| Lutra_sumatrana | Neofelis_nebulosa | 0.240988570490307 |
| Lutra_sumatrana | Neomonachus_schauinslandi | 0.212372401792618 |
| Lutra_sumatrana | Neophoca_cinerea | 0.233375515094599 |
| Lutra_sumatrana | Neovison_vison | 0.161769652590792 |
| Lutra_sumatrana | Nyctereutes_procyonoides | 0.246589943968194 |
| Lutra_sumatrana | Odobenus_rosmarus_rosmarus | 0.241029296844061 |
| Lutra_sumatrana | Otocolobus_manul | 0.232468529934228 |
| Lutra_sumatrana | Otocyon_megalotis_megalotis_TS305 | 0.250456945874143 |
| Lutra_sumatrana | Otocyon_megalotis_virgatus_FMNH158128 | 0.25043057625534 |
| Lutra_sumatrana | Paguma_larvata | 0.237965478203961 |
| Lutra_sumatrana | Panthera_leo | 0.23989241512594 |
| Lutra_sumatrana | Panthera_onca | 0.239166565397559 |
| Lutra_sumatrana | Panthera_pardus | 0.238552449814864 |
| Lutra_sumatrana | Panthera_tigris | 0.240854885209546 |
| Lutra_sumatrana | Panthera_uncia | 0.238419019532889 |
| Lutra_sumatrana | Paradoxurus_hermaphroditus | 0.248178639780682 |
| Lutra_sumatrana | Parahyaena_brunnea_FD126_Fecal | 0.239390418800417 |
| Lutra_sumatrana | Pardofelis_marmorata | 0.23396573522031 |
| Lutra_sumatrana | Phoca_fasciata | 0.208396719929208 |
| Lutra_sumatrana | Phoca_groenlandica | 0.214199530306453 |
| Lutra_sumatrana | Phoca_largha | 0.210478274654035 |
| Lutra_sumatrana | Phoca_vitulina | 0.211627011516115 |
| Lutra_sumatrana | Phocarctos_hookeri | 0.233341053927753 |
| Lutra_sumatrana | Prionailurus_bengalensis | 0.229892617143102 |
| Lutra_sumatrana | Prionailurus_bengalensis_euptilurus | 0.228290711213832 |
| Lutra_sumatrana | Prionailurus_planiceps | 0.231072722561724 |
| Lutra_sumatrana | Prionailurus_rubiginosus | 0.230628552717172 |
| Lutra_sumatrana | Prionailurus_viverrinus | 0.231230593203363 |
| Lutra_sumatrana | Prionodon_pardicolor | 0.237404080624351 |
| Lutra_sumatrana | Procyon_lotor | 0.216836533596719 |
| Lutra_sumatrana | Profelis_aurata | 0.233802573846508 |
| Lutra_sumatrana | Proteles_cristatus_cristatus_TS307 | 0.243767883961011 |
| Lutra_sumatrana | Proteles_cristatus_septentrionalis_NMS_Z_2018_54 | 0.247364617465465 |
| Lutra_sumatrana | Puma_concolor | 0.229115697037861 |
| Lutra_sumatrana | Puma_yagouaroundi | 0.226021530959309 |
| Lutra_sumatrana | Pusa_caspica | 0.214328366264989 |
| Lutra_sumatrana | Pusa_hispida | 0.20880268924334 |
| Lutra_sumatrana | Pusa_sibirica | 0.213464434761509 |
| Lutra_sumatrana | Speothos_venaticus_T1624 | 0.259625065917679 |
| Lutra_sumatrana | Spilogale_putorius | 0.229400807570983 |
| Lutra_sumatrana | Taxidea_taxus | 0.169675851954824 |
| Lutra_sumatrana | Tremarctos_ornatus | 0.235262606028957 |
| Lutra_sumatrana | Urocyon_cinereoargenteus | 0.254135728319176 |
| Lutra_sumatrana | Ursus_americanus | 0.240346243670064 |
| Lutra_sumatrana | Ursus_arctos | 0.244453682341762 |
| Lutra_sumatrana | Ursus_maritimus | 0.244439288939216 |
| Lutra_sumatrana | Ursus_spelaeus | 0.238979811482801 |
| Lutra_sumatrana | Ursus_thibetanus | 0.235461594322811 |
| Lutra_sumatrana | Urva_javanica | 0.239108179747397 |
| Lutra_sumatrana | Viverra_tangalunga_FMNH146957_SRR6053069 | 0.23190112404369 |
| Lutra_sumatrana | Viverricula_indica | 0.233566065128429 |
| Lutra_sumatrana | Vulpes_corsac | 0.251769301124371 |
| Lutra_sumatrana | Vulpes_ferrilata | 0.252667351187887 |
| Lutra_sumatrana | Vulpes_lagopus | 0.251485703640642 |
| Lutra_sumatrana | Vulpes_vulpes_T3611 | 0.246963085834201 |

| | | |
|---|---|---|
| Lutra_sumatrana | Vulpes_zerda | 0.24646583037718 |
| Lutra_sumatrana | Zalophus_californianus | 0.230924714429244 |
| Lutrogale_perspicillata | Lycaon_pictus | 0.262190735533072 |
| Lutrogale_perspicillata | Lynx_canadensis | 0.237270910439255 |
| Lutrogale_perspicillata | Lynx_lynx | 0.237092195392774 |
| Lutrogale_perspicillata | Lynx_pardinus | 0.236558095333455 |
| Lutrogale_perspicillata | Lynx_rufus | 0.240005987715836 |
| Lutrogale_perspicillata | Manis_javanica | 0.296054454144362 |
| Lutrogale_perspicillata | Martes_americana | 0.174499709679199 |
| Lutrogale_perspicillata | Martes_flavigula | 0.174301905724713 |
| Lutrogale_perspicillata | Martes_foina | 0.174888501287586 |
| Lutrogale_perspicillata | Martes_martes | 0.175725500289021 |
| Lutrogale_perspicillata | Martes_melampus | 0.174858397319964 |
| Lutrogale_perspicillata | Martes_pennanti | 0.178610410314488 |
| Lutrogale_perspicillata | Martes_zibellina | 0.173972395400519 |
| Lutrogale_perspicillata | Meles_anakuma | 0.187696034559507 |
| Lutrogale_perspicillata | Meles_leucurus | 0.186700085655924 |
| Lutrogale_perspicillata | Meles_meles | 0.185196749710034 |
| Lutrogale_perspicillata | Melogale_moschata | 0.17609804243511 |
| Lutrogale_perspicillata | Melursus_ursinus | 0.245735188169201 |
| Lutrogale_perspicillata | Mephitis_mephitis | 0.252111871850559 |
| Lutrogale_perspicillata | Mirounga_leonina | 0.229703417205728 |
| Lutrogale_perspicillata | Mungotictis_decemlineata | 0.241705609612312 |
| Lutrogale_perspicillata | Mustela_altaica | 0.174325365835613 |
| Lutrogale_perspicillata | Mustela_erminea | 0.163563445816038 |
| Lutrogale_perspicillata | Mustela_eversmannii | 0.177200575920326 |
| Lutrogale_perspicillata | Mustela_frenata | 0.176333647997259 |
| Lutrogale_perspicillata | Mustela_itatsi | 0.178956746728859 |
| Lutrogale_perspicillata | Mustela_kathiah | 0.181249436934732 |
| Lutrogale_perspicillata | Mustela_nigripes | 0.176900757058614 |
| Lutrogale_perspicillata | Mustela_nivalis | 0.174266707222179 |
| Lutrogale_perspicillata | Mustela_putorius | 0.17822811122987 |
| Lutrogale_perspicillata | Mustela_sibirica | 0.177327521913422 |
| Lutrogale_perspicillata | Nandinia_binotata | 0.231319934757069 |
| Lutrogale_perspicillata | Nasua_nasua | 0.243994496349858 |
| Lutrogale_perspicillata | Neofelis_nebulosa | 0.250764703315998 |
| Lutrogale_perspicillata | Neomonachus_schauinslandi | 0.227049380959441 |
| Lutrogale_perspicillata | Neophoca_cinerea | 0.240741824081355 |
| Lutrogale_perspicillata | Neovison_vison | 0.1807494828669 |
| Lutrogale_perspicillata | Nyctereutes_procyonoides | 0.256048019268469 |
| Lutrogale_perspicillata | Odobenus_rosmarus_rosmarus | 0.254189956084519 |
| Lutrogale_perspicillata | Otocolobus_manul | 0.24425602368687 |
| Lutrogale_perspicillata | Otocyon_megalotis_megalotis_TS305 | 0.256175643401315 |
| Lutrogale_perspicillata | Otocyon_megalotis_virgatus_FMNH158128 | 0.260105905974588 |
| Lutrogale_perspicillata | Paguma_larvata | 0.257316543674362 |
| Lutrogale_perspicillata | Panthera_leo | 0.245668117988065 |
| Lutrogale_perspicillata | Panthera_onca | 0.242770866999022 |
| Lutrogale_perspicillata | Panthera_pardus | 0.245913630244807 |
| Lutrogale_perspicillata | Panthera_tigris | 0.252266621034206 |
| Lutrogale_perspicillata | Panthera_uncia | 0.247247825794382 |
| Lutrogale_perspicillata | Paradoxurus_hermaphroditus | 0.262994200826631 |
| Lutrogale_perspicillata | Parahyaena_brunnea_FD126_Fecal | 0.241526970929381 |
| Lutrogale_perspicillata | Pardofelis_marmorata | 0.245566832442565 |
| Lutrogale_perspicillata | Phoca_fasciata | 0.220632655621864 |
| Lutrogale_perspicillata | Phoca_groenlandica | 0.221713135347098 |
| Lutrogale_perspicillata | Phoca_largha | 0.220513887785786 |
| Lutrogale_perspicillata | Phoca_vitulina | 0.221572501085754 |
| Lutrogale_perspicillata | Phocarctos_hookeri | 0.239315090336197 |
| Lutrogale_perspicillata | Prionailurus_bengalensis | 0.237026685988324 |
| Lutrogale_perspicillata | Prionailurus_bengalensis_euptilurus | 0.235737412414472 |
| Lutrogale_perspicillata | Prionailurus_planiceps | 0.2365362691625 |
| Lutrogale_perspicillata | Prionailurus_rubiginosus | 0.23881267841552 |
| Lutrogale_perspicillata | Prionailurus_viverrinus | 0.240574414769187 |
| Lutrogale_perspicillata | Prionodon_pardicolor | 0.251738237148601 |
| Lutrogale_perspicillata | Procyon_lotor | 0.230584714199764 |
| Lutrogale_perspicillata | Profelis_aurata | 0.245828348889173 |
| Lutrogale_perspicillata | Proteles_cristatus_cristatus_TS307 | 0.248019050687267 |
| Lutrogale_perspicillata | Proteles_cristatus_septentrionalis_NMS_Z_2018_54 | 0.253931569838514 |
| Lutrogale_perspicillata | Puma_concolor | 0.239646287804973 |
| Lutrogale_perspicillata | Puma_yagouaroundi | 0.23984752466803 |
| Lutrogale_perspicillata | Pusa_caspica | 0.224168334412442 |
| Lutrogale_perspicillata | Pusa_hispida | 0.22018575034962 |
| Lutrogale_perspicillata | Pusa_sibirica | 0.222675111466339 |
| Lutrogale_perspicillata | Speothos_venaticus_T1624 | 0.266281227035876 |
| Lutrogale_perspicillata | Spilogale_putorius | 0.254130577286879 |
| Lutrogale_perspicillata | Taxidea_taxus | 0.185296363197797 |
| Lutrogale_perspicillata | Tremarctos_ornatus | 0.2437804341849 |
| Lutrogale_perspicillata | Urocyon_cinereoargenteus | 0.26287564270939 |
| Lutrogale_perspicillata | Ursus_americanus | 0.246539706191659 |
| Lutrogale_perspicillata | Ursus_arctos | 0.248949453975264 |
| Lutrogale_perspicillata | Ursus_maritimus | 0.251399333818698 |
| Lutrogale_perspicillata | Ursus_spelaeus | 0.244500479457403 |
| Lutrogale_perspicillata | Ursus_thibetanus | 0.248329141816677 |
| Lutrogale_perspicillata | Urva_javanica | 0.24982633443952 |
| Lutrogale_perspicillata | Viverra_tangalunga_FMNH146957_SRR6053069 | 0.247408380856611 |
| Lutrogale_perspicillata | Viverricula_indica | 0.246919856967716 |
| Lutrogale_perspicillata | Vulpes_corsac | 0.262825877115599 |

| | | |
|---|---|---|
| Lutrogale_perspicillata | Vulpes_ferrilata | 0.263045411128352 |
| Lutrogale_perspicillata | Vulpes_lagopus | 0.266070434898755 |
| Lutrogale_perspicillata | Vulpes_vulpes_T3611 | 0.25820155337558 |
| Lutrogale_perspicillata | Vulpes_zerda | 0.25471957160399 |
| Lutrogale_perspicillata | Zalophus_californianus | 0.235879972954874 |
| Lycaon_pictus | Lynx_canadensis | 0.235327927874705 |
| Lycaon_pictus | Lynx_lynx | 0.235541401599185 |
| Lycaon_pictus | Lynx_pardinus | 0.236863716419468 |
| Lycaon_pictus | Lynx_rufus | 0.236641179916054 |
| Lycaon_pictus | Manis_javanica | 0.299831923808869 |
| Lycaon_pictus | Martes_americana | 0.240603518564001 |
| Lycaon_pictus | Martes_flavigula | 0.238058288970629 |
| Lycaon_pictus | Martes_foina | 0.241979173277509 |
| Lycaon_pictus | Martes_martes | 0.237787980608813 |
| Lycaon_pictus | Martes_melampus | 0.234115281898804 |
| Lycaon_pictus | Martes_pennanti | 0.242048701254419 |
| Lycaon_pictus | Martes_zibellina | 0.238442921387864 |
| Lycaon_pictus | Meles_anakuma | 0.247669871626318 |
| Lycaon_pictus | Meles_leucurus | 0.241460617672379 |
| Lycaon_pictus | Meles_meles | 0.243684820628397 |
| Lycaon_pictus | Melogale_moschata | 0.246058014310544 |
| Lycaon_pictus | Melursus_ursinus | 0.246499832850724 |
| Lycaon_pictus | Mephitis_mephitis | 0.244439560832603 |
| Lycaon_pictus | Mirounga_leonina | 0.223174199681586 |
| Lycaon_pictus | Mungotictis_decemlineata | 0.244722787741587 |
| Lycaon_pictus | Mustela_altaica | 0.241006769841264 |
| Lycaon_pictus | Mustela_erminea | 0.232739707129741 |
| Lycaon_pictus | Mustela_eversmannii | 0.238406348204395 |
| Lycaon_pictus | Mustela_frenata | 0.234436295193406 |
| Lycaon_pictus | Mustela_itatsi | 0.238213814796691 |
| Lycaon_pictus | Mustela_kathiah | 0.235321561822562 |
| Lycaon_pictus | Mustela_nigripes | 0.236858348715246 |
| Lycaon_pictus | Mustela_nivalis | 0.241145170917719 |
| Lycaon_pictus | Mustela_putorius | 0.238837678328889 |
| Lycaon_pictus | Mustela_sibirica | 0.236720472942892 |
| Lycaon_pictus | Nandinia_binotata | 0.227967911533802 |
| Lycaon_pictus | Nasua_nasua | 0.261477653603562 |
| Lycaon_pictus | Neofelis_nebulosa | 0.252817408383618 |
| Lycaon_pictus | Neomonachus_schauinslandi | 0.230379294545719 |
| Lycaon_pictus | Neophoca_cinerea | 0.249665717043945 |
| Lycaon_pictus | Neovison_vison | 0.23503860659957 |
| Lycaon_pictus | Nyctereutes_procyonoides | 0.177290422141369 |
| Lycaon_pictus | Odobenus_rosmarus_rosmarus | 0.265061980752811 |
| Lycaon_pictus | Otocolobus_manul | 0.23882360110478 |
| Lycaon_pictus | Otocyon_megalotis_megalotis_TS305 | 0.184613626678996 |
| Lycaon_pictus | Otocyon_megalotis_virgatus_FMNH158128 | 0.185854598242453 |
| Lycaon_pictus | Paguma_larvata | 0.240107510365313 |
| Lycaon_pictus | Panthera_leo | 0.245133558383059 |
| Lycaon_pictus | Panthera_onca | 0.244031482396664 |
| Lycaon_pictus | Panthera_pardus | 0.24733731424567 |
| Lycaon_pictus | Panthera_tigris | 0.250316834860732 |
| Lycaon_pictus | Panthera_uncia | 0.249662445328836 |
| Lycaon_pictus | Paradoxurus_hermaphroditus | 0.240861649348523 |
| Lycaon_pictus | Parahyaena_brunnea_FD126_Fecal | 0.248757648602584 |
| Lycaon_pictus | Pardofelis_marmorata | 0.241742344768619 |
| Lycaon_pictus | Phoca_fasciata | 0.231718663119711 |
| Lycaon_pictus | Phoca_groenlandica | 0.232726744071454 |
| Lycaon_pictus | Phoca_largha | 0.233036638573893 |
| Lycaon_pictus | Phoca_vitulina | 0.233467222667478 |
| Lycaon_pictus | Phocarctos_hookeri | 0.253875959074141 |
| Lycaon_pictus | Prionailurus_bengalensis | 0.233898412935709 |
| Lycaon_pictus | Prionailurus_bengalensis_euptilurus | 0.233569104471555 |
| Lycaon_pictus | Prionailurus_planiceps | 0.234900764586987 |
| Lycaon_pictus | Prionailurus_rubiginosus | 0.233016698101019 |
| Lycaon_pictus | Prionailurus_viverrinus | 0.238048235419935 |
| Lycaon_pictus | Prionodon_pardicolor | 0.236708595484584 |
| Lycaon_pictus | Procyon_lotor | 0.242802963413915 |
| Lycaon_pictus | Profelis_aurata | 0.240259123339661 |
| Lycaon_pictus | Proteles_cristatus_cristatus_TS307 | 0.255927977897384 |
| Lycaon_pictus | Proteles_cristatus_septentrionalis_NMS_Z_2018_54 | 0.253311362141846 |
| Lycaon_pictus | Puma_concolor | 0.236822322936387 |
| Lycaon_pictus | Puma_yagouaroundi | 0.235020342168673 |
| Lycaon_pictus | Pusa_caspica | 0.228767449719307 |
| Lycaon_pictus | Pusa_hispida | 0.228598555991064 |
| Lycaon_pictus | Pusa_sibirica | 0.231928403420379 |
| Lycaon_pictus | Speothos_venaticus_T1624 | 0.137023427912557 |
| Lycaon_pictus | Spilogale_putorius | 0.24264129246301 |
| Lycaon_pictus | Taxidea_taxus | 0.238510218591431 |
| Lycaon_pictus | Tremarctos_ornatus | 0.245551844721766 |
| Lycaon_pictus | Urocyon_cinereoargenteus | 0.195513714054509 |
| Lycaon_pictus | Ursus_americanus | 0.242334045297101 |
| Lycaon_pictus | Ursus_arctos | 0.250941486419975 |
| Lycaon_pictus | Ursus_maritimus | 0.248779298845681 |
| Lycaon_pictus | Ursus_spelaeus | 0.248208457363423 |
| Lycaon_pictus | Ursus_thibetanus | 0.239344790626736 |
| Lycaon_pictus | Urva_javanica | 0.239444949849108 |
| Lycaon_pictus | Viverra_tangalunga_FMNH146957_SRR6053069 | 0.232188153010288 |

| | | |
|---|---|---|
| Lycaon_pictus | Viverricula_indica | 0.229870448180946 |
| Lycaon_pictus | Vulpes_corsac | 0.181540428257306 |
| Lycaon_pictus | Vulpes_ferrilata | 0.182583151544859 |
| Lycaon_pictus | Vulpes_lagopus | 0.185552439782434 |
| Lycaon_pictus | Vulpes_vulpes_T3611 | 0.184851897432711 |
| Lycaon_pictus | Vulpes_zerda | 0.183408563411669 |
| Lycaon_pictus | Zalophus_californianus | 0.248522362480459 |
| Lynx_canadensis | Lynx_lynx | 0.0366828332395707 |
| Lynx_canadensis | Lynx_pardinus | 0.0369421642366548 |
| Lynx_canadensis | Lynx_rufus | 0.0792083597325853 |
| Lynx_canadensis | Manis_javanica | 0.282640875775841 |
| Lynx_canadensis | Martes_americana | 0.230142493960724 |
| Lynx_canadensis | Martes_flavigula | 0.23106691038867 |
| Lynx_canadensis | Martes_foina | 0.228638926661906 |
| Lynx_canadensis | Martes_martes | 0.23021427761538 |
| Lynx_canadensis | Martes_melampus | 0.228450362208316 |
| Lynx_canadensis | Martes_pennanti | 0.232152721731481 |
| Lynx_canadensis | Martes_zibellina | 0.232294422877823 |
| Lynx_canadensis | Meles_anakuma | 0.232930117456857 |
| Lynx_canadensis | Meles_leucurus | 0.227717245514867 |
| Lynx_canadensis | Meles_meles | 0.228025071465687 |
| Lynx_canadensis | Melogale_moschata | 0.228669228484518 |
| Lynx_canadensis | Melursus_ursinus | 0.239415490889873 |
| Lynx_canadensis | Mephitis_mephitis | 0.239206377415164 |
| Lynx_canadensis | Mirounga_leonina | 0.218762863922795 |
| Lynx_canadensis | Mungotictis_decemlineata | 0.183902268278922 |
| Lynx_canadensis | Mustela_altaica | 0.230620173067675 |
| Lynx_canadensis | Mustela_erminea | 0.221312668613617 |
| Lynx_canadensis | Mustela_eversmannii | 0.228907903786416 |
| Lynx_canadensis | Mustela_frenata | 0.229186332015466 |
| Lynx_canadensis | Mustela_itatsi | 0.229186886053184 |
| Lynx_canadensis | Mustela_kathiah | 0.231436004118793 |
| Lynx_canadensis | Mustela_nigripes | 0.225595620206087 |
| Lynx_canadensis | Mustela_nivalis | 0.228978194524896 |
| Lynx_canadensis | Mustela_putorius | 0.229354721674459 |
| Lynx_canadensis | Mustela_sibirica | 0.227400542389586 |
| Lynx_canadensis | Nandinia_binotata | 0.169118484370383 |
| Lynx_canadensis | Nasua_nasua | 0.247613226748635 |
| Lynx_canadensis | Neofelis_nebulosa | 0.138873347511336 |
| Lynx_canadensis | Neomonachus_schauinslandi | 0.216468663819346 |
| Lynx_canadensis | Neophoca_cinerea | 0.236226157620699 |
| Lynx_canadensis | Neovison_vison | 0.231216816225525 |
| Lynx_canadensis | Nyctereutes_procyonoides | 0.239501529911986 |
| Lynx_canadensis | Odobenus_rosmarus_rosmarus | 0.247779074597007 |
| Lynx_canadensis | Otocolobus_manul | 0.115581589259414 |
| Lynx_canadensis | Otocyon_megalotis_megalotis_TS305 | 0.240856211389321 |
| Lynx_canadensis | Otocyon_megalotis_virgatus_FMNH158128 | 0.240720611785255 |
| Lynx_canadensis | Paguma_larvata | 0.184657089315148 |
| Lynx_canadensis | Panthera_leo | 0.133291221455549 |
| Lynx_canadensis | Panthera_onca | 0.13446587222243 |
| Lynx_canadensis | Panthera_pardus | 0.13432344399691 |
| Lynx_canadensis | Panthera_tigris | 0.138879093571792 |
| Lynx_canadensis | Panthera_uncia | 0.143491965557013 |
| Lynx_canadensis | Paradoxurus_hermaphroditus | 0.190870091166816 |
| Lynx_canadensis | Parahyaena_brunnea_FD126_Fecal | 0.187222374716268 |
| Lynx_canadensis | Pardofelis_marmorata | 0.110785030774548 |
| Lynx_canadensis | Phoca_fasciata | 0.215694329964523 |
| Lynx_canadensis | Phoca_groenlandica | 0.216274487190075 |
| Lynx_canadensis | Phoca_largha | 0.220301568740394 |
| Lynx_canadensis | Phoca_vitulina | 0.220146492323658 |
| Lynx_canadensis | Phocarctos_hookeri | 0.233393701624545 |
| Lynx_canadensis | Prionailurus_bengalensis | 0.103188626019122 |
| Lynx_canadensis | Prionailurus_bengalensis_euptilurus | 0.101169241596268 |
| Lynx_canadensis | Prionailurus_planiceps | 0.109140537702994 |
| Lynx_canadensis | Prionailurus_rubiginosus | 0.106984977186624 |
| Lynx_canadensis | Prionailurus_viverrinus | 0.107528450125011 |
| Lynx_canadensis | Prionodon_pardicolor | 0.186071414034583 |
| Lynx_canadensis | Procyon_lotor | 0.233934034109644 |
| Lynx_canadensis | Profelis_aurata | 0.117507618707387 |
| Lynx_canadensis | Proteles_cristatus_cristatus_TS307 | 0.195040614206951 |
| Lynx_canadensis | Proteles_cristatus_septentrionalis_NMS_Z_2018_54 | 0.195460561738199 |
| Lynx_canadensis | Puma_concolor | 0.102340152006858 |
| Lynx_canadensis | Puma_yagouaroundi | 0.10362536402805 |
| Lynx_canadensis | Pusa_caspica | 0.222364106905168 |
| Lynx_canadensis | Pusa_hispida | 0.217471906761229 |
| Lynx_canadensis | Pusa_sibirica | 0.219915315636482 |
| Lynx_canadensis | Speothos_venaticus_T1624 | 0.246887269210785 |
| Lynx_canadensis | Spilogale_putorius | 0.246041969122454 |
| Lynx_canadensis | Taxidea_taxus | 0.223202178444766 |
| Lynx_canadensis | Tremarctos_ornatus | 0.234150219617046 |
| Lynx_canadensis | Urocyon_cinereoargenteus | 0.243299410102305 |
| Lynx_canadensis | Ursus_americanus | 0.241576647354957 |
| Lynx_canadensis | Ursus_arctos | 0.244100281024324 |
| Lynx_canadensis | Ursus_maritimus | 0.242253542765198 |
| Lynx_canadensis | Ursus_spelaeus | 0.23872482750679 |
| Lynx_canadensis | Ursus_thibetanus | 0.234906389497221 |
| Lynx_canadensis | Urva_javanica | 0.192433687413378 |

| | | |
|---|---|---|
| Lynx_canadensis | Viverra_tangalunga_FMNH146957_SRR6053069 | 0.173596183636401 |
| Lynx_canadensis | Viverricula_indica | 0.177609922703732 |
| Lynx_canadensis | Vulpes_corsac | 0.243389475851275 |
| Lynx_canadensis | Vulpes_ferrilata | 0.243919607060733 |
| Lynx_canadensis | Vulpes_lagopus | 0.242770866999022 |
| Lynx_canadensis | Vulpes_vulpes_T3611 | 0.235227379179356 |
| Lynx_canadensis | Vulpes_zerda | 0.23479859848867 |
| Lynx_canadensis | Zalophus_californianus | 0.232724491616539 |
| Lynx_lynx | Lynx_pardinus | 0.0387461373328366 |
| Lynx_lynx | Lynx_rufus | 0.0784009115486961 |
| Lynx_lynx | Manis_javanica | 0.282329499336888 |
| Lynx_lynx | Martes_americana | 0.230150748705462 |
| Lynx_lynx | Martes_flavigula | 0.230578529486914 |
| Lynx_lynx | Martes_foina | 0.229414005834801 |
| Lynx_lynx | Martes_martes | 0.230441103376252 |
| Lynx_lynx | Martes_melampus | 0.229663282242083 |
| Lynx_lynx | Martes_pennanti | 0.230523151172036 |
| Lynx_lynx | Martes_zibellina | 0.23296144723536 |
| Lynx_lynx | Meles_anakuma | 0.228886819899046 |
| Lynx_lynx | Meles_leucurus | 0.225722586757547 |
| Lynx_lynx | Meles_meles | 0.225851063879598 |
| Lynx_lynx | Melogale_moschata | 0.229205086508734 |
| Lynx_lynx | Melursus_ursinus | 0.239522205696721 |
| Lynx_lynx | Mephitis_mephitis | 0.24353519203474 |
| Lynx_lynx | Mirounga_leonina | 0.218189839991218 |
| Lynx_lynx | Mungotictis_decemlineata | 0.184543402712001 |
| Lynx_lynx | Mustela_altaica | 0.228891428505717 |
| Lynx_lynx | Mustela_erminea | 0.222083839345234 |
| Lynx_lynx | Mustela_eversmannii | 0.229080885357225 |
| Lynx_lynx | Mustela_frenata | 0.229414005834801 |
| Lynx_lynx | Mustela_itatsi | 0.229360454321021 |
| Lynx_lynx | Mustela_kathiah | 0.231117080992192 |
| Lynx_lynx | Mustela_nigripes | 0.226531900215329 |
| Lynx_lynx | Mustela_nivalis | 0.230481629959472 |
| Lynx_lynx | Mustela_putorius | 0.228763262140386 |
| Lynx_lynx | Mustela_sibirica | 0.22833683940233 |
| Lynx_lynx | Nandinia_binotata | 0.167996404281552 |
| Lynx_lynx | Nasua_nasua | 0.243871777359072 |
| Lynx_lynx | Neofelis_nebulosa | 0.139870956393623 |
| Lynx_lynx | Neomonachus_schauinslandi | 0.217181756183109 |
| Lynx_lynx | Neophoca_cinerea | 0.235693513128912 |
| Lynx_lynx | Neovison_vison | 0.23234444166928 |
| Lynx_lynx | Nyctereutes_procyonoides | 0.23948915860738 |
| Lynx_lynx | Odobenus_rosmarus_rosmarus | 0.247868865385437 |
| Lynx_lynx | Otocolobus_manul | 0.11786894088727 |
| Lynx_lynx | Otocyon_megalotis_megalotis_TS305 | 0.239650051129513 |
| Lynx_lynx | Otocyon_megalotis_virgatus_FMNH158128 | 0.241685019382585 |
| Lynx_lynx | Paguma_larvata | 0.183291537704817 |
| Lynx_lynx | Panthera_leo | 0.13152696777832 |
| Lynx_lynx | Panthera_onca | 0.134464567800075 |
| Lynx_lynx | Panthera_pardus | 0.130328538538658 |
| Lynx_lynx | Panthera_tigris | 0.137676439295714 |
| Lynx_lynx | Panthera_uncia | 0.142181384488374 |
| Lynx_lynx | Paradoxurus_hermaphroditus | 0.192000831923797 |
| Lynx_lynx | Parahyaena_brunnea_FD126_Fecal | 0.188520291352437 |
| Lynx_lynx | Pardofelis_marmorata | 0.10950339107429 |
| Lynx_lynx | Phoca_fasciata | 0.216228975011188 |
| Lynx_lynx | Phoca_groenlandica | 0.216917160800407 |
| Lynx_lynx | Phoca_largha | 0.217039967856138 |
| Lynx_lynx | Phoca_vitulina | 0.217208834540867 |
| Lynx_lynx | Phocarctos_hookeri | 0.234945554079645 |
| Lynx_lynx | Prionailurus_bengalensis | 0.108080932312261 |
| Lynx_lynx | Prionailurus_bengalensis_euptilurus | 0.105677110418474 |
| Lynx_lynx | Prionailurus_planiceps | 0.111558218129063 |
| Lynx_lynx | Prionailurus_rubiginosus | 0.109332473216342 |
| Lynx_lynx | Prionailurus_viverrinus | 0.111091277489972 |
| Lynx_lynx | Prionodon_pardicolor | 0.185904361555326 |
| Lynx_lynx | Procyon_lotor | 0.23484470360937 |
| Lynx_lynx | Profelis_aurata | 0.118571631437805 |
| Lynx_lynx | Proteles_cristatus_cristatus_TS307 | 0.195300655904689 |
| Lynx_lynx | Proteles_cristatus_septentrionalis_NMS_Z_2018_54 | 0.193521294596192 |
| Lynx_lynx | Puma_concolor | 0.103074274350141 |
| Lynx_lynx | Puma_yagouaroundi | 0.106298111453667 |
| Lynx_lynx | Pusa_caspica | 0.220371522363631 |
| Lynx_lynx | Pusa_hispida | 0.217860974882888 |
| Lynx_lynx | Pusa_sibirica | 0.219294400016872 |
| Lynx_lynx | Speothos_venaticus_T1624 | 0.243252227900571 |
| Lynx_lynx | Spilogale_putorius | 0.245017437420524 |
| Lynx_lynx | Taxidea_taxus | 0.219432720994256 |
| Lynx_lynx | Tremarctos_ornatus | 0.231067230593142 |
| Lynx_lynx | Urocyon_cinereoargenteus | 0.240992427645395 |
| Lynx_lynx | Ursus_americanus | 0.239882315487926 |
| Lynx_lynx | Ursus_arctos | 0.242620110872917 |
| Lynx_lynx | Ursus_maritimus | 0.240556321431373 |
| Lynx_lynx | Ursus_spelaeus | 0.237264442990328 |
| Lynx_lynx | Ursus_thibetanus | 0.237870014632252 |
| Lynx_lynx | Urva_javanica | 0.193674531358214 |

| | | |
|---|---|---|
| Lynx_lynx | Viverra_tangalunga_FMNH146957_SRR6053069 | 0.175152635468891 |
| Lynx_lynx | Viverricula_indica | 0.179486981607312 |
| Lynx_lynx | Vulpes_corsac | 0.242831662223582 |
| Lynx_lynx | Vulpes_ferrilata | 0.242021433377203 |
| Lynx_lynx | Vulpes_lagopus | 0.24248187280775 |
| Lynx_lynx | Vulpes_vulpes_T3611 | 0.236209714023631 |
| Lynx_lynx | Vulpes_zerda | 0.235779865423416 |
| Lynx_lynx | Zalophus_californianus | 0.232744800492608 |
| Lynx_pardinus | Lynx_rufus | 0.0806904541192048 |
| Lynx_pardinus | Manis_javanica | 0.282815810678974 |
| Lynx_pardinus | Martes_americana | 0.228489302194059 |
| Lynx_pardinus | Martes_flavigula | 0.228590112022451 |
| Lynx_pardinus | Martes_foina | 0.232028146772433 |
| Lynx_pardinus | Martes_martes | 0.226485854901342 |
| Lynx_pardinus | Martes_melampus | 0.225171487831954 |
| Lynx_pardinus | Martes_pennanti | 0.2276951047069 |
| Lynx_pardinus | Martes_zibellina | 0.229752671712956 |
| Lynx_pardinus | Meles_anakuma | 0.230441103376252 |
| Lynx_pardinus | Meles_leucurus | 0.226177754515199 |
| Lynx_pardinus | Meles_meles | 0.22823159169509 |
| Lynx_pardinus | Melogale_moschata | 0.229951682198706 |
| Lynx_pardinus | Melursus_ursinus | 0.241207411201673 |
| Lynx_pardinus | Mephitis_mephitis | 0.24123531462259 |
| Lynx_pardinus | Mirounga_leonina | 0.217689989645536 |
| Lynx_pardinus | Mungotictis_decemlineata | 0.184398751685787 |
| Lynx_pardinus | Mustela_altaica | 0.229128856955734 |
| Lynx_pardinus | Mustela_erminea | 0.220882393280398 |
| Lynx_pardinus | Mustela_eversmannii | 0.230082994183693 |
| Lynx_pardinus | Mustela_frenata | 0.227684896243348 |
| Lynx_pardinus | Mustela_itatsi | 0.228398391137673 |
| Lynx_pardinus | Mustela_kathiah | 0.233183673698611 |
| Lynx_pardinus | Mustela_nigripes | 0.226984578341398 |
| Lynx_pardinus | Mustela_nivalis | 0.229626617784247 |
| Lynx_pardinus | Mustela_putorius | 0.230202045245654 |
| Lynx_pardinus | Mustela_sibirica | 0.228682886760208 |
| Lynx_pardinus | Nandinia_binotata | 0.171146651527907 |
| Lynx_pardinus | Nasua_nasua | 0.246642004744901 |
| Lynx_pardinus | Neofelis_nebulosa | 0.143520967670303 |
| Lynx_pardinus | Neomonachus_schauinslandi | 0.215756270103515 |
| Lynx_pardinus | Neophoca_cinerea | 0.23154578565398 |
| Lynx_pardinus | Neovison_vison | 0.23166657797317 |
| Lynx_pardinus | Nyctereutes_procyonoides | 0.238843894343051 |
| Lynx_pardinus | Odobenus_rosmarus_rosmarus | 0.248304086793617 |
| Lynx_pardinus | Otocolobus_manul | 0.114948812014449 |
| Lynx_pardinus | Otocyon_megalotis_megalotis_TS305 | 0.239826956498207 |
| Lynx_pardinus | Otocyon_megalotis_virgatus_FMNH158128 | 0.241975030383785 |
| Lynx_pardinus | Paguma_larvata | 0.182071847189076 |
| Lynx_pardinus | Panthera_leo | 0.134962494565495 |
| Lynx_pardinus | Panthera_onca | 0.137311822423905 |
| Lynx_pardinus | Panthera_pardus | 0.133077312444367 |
| Lynx_pardinus | Panthera_tigris | 0.138599623320052 |
| Lynx_pardinus | Panthera_uncia | 0.145685851548063 |
| Lynx_pardinus | Paradoxurus_hermaphroditus | 0.188980570036733 |
| Lynx_pardinus | Parahyaena_brunnea_FD126_Fecal | 0.189616820862832 |
| Lynx_pardinus | Pardofelis_marmorata | 0.11211997493568 |
| Lynx_pardinus | Phoca_fasciata | 0.218088220367518 |
| Lynx_pardinus | Phoca_groenlandica | 0.218995567987691 |
| Lynx_pardinus | Phoca_largha | 0.221852511695566 |
| Lynx_pardinus | Phoca_vitulina | 0.22158783951779 |
| Lynx_pardinus | Phocarctos_hookeri | 0.233003565130654 |
| Lynx_pardinus | Prionailurus_bengalensis | 0.106815761150757 |
| Lynx_pardinus | Prionailurus_bengalensis_euptilurus | 0.106171242533198 |
| Lynx_pardinus | Prionailurus_planiceps | 0.112246566777756 |
| Lynx_pardinus | Prionailurus_rubiginosus | 0.108960829605438 |
| Lynx_pardinus | Prionailurus_viverrinus | 0.111091277489972 |
| Lynx_pardinus | Prionodon_pardicolor | 0.184947939829348 |
| Lynx_pardinus | Procyon_lotor | 0.235909574654839 |
| Lynx_pardinus | Profelis_aurata | 0.118097517994353 |
| Lynx_pardinus | Proteles_cristatus_cristatus_TS307 | 0.196095224285534 |
| Lynx_pardinus | Proteles_cristatus_septentrionalis_NMS_Z_2018_54 | 0.195465170647611 |
| Lynx_pardinus | Puma_concolor | 0.103381536192362 |
| Lynx_pardinus | Puma_yagouaroundi | 0.105867565656366 |
| Lynx_pardinus | Pusa_caspica | 0.224851979345694 |
| Lynx_pardinus | Pusa_hispida | 0.2206951120632 |
| Lynx_pardinus | Pusa_sibirica | 0.221636108316296 |
| Lynx_pardinus | Speothos_venaticus_T1624 | 0.245806656981458 |
| Lynx_pardinus | Spilogale_putorius | 0.245038419411854 |
| Lynx_pardinus | Taxidea_taxus | 0.220183713518231 |
| Lynx_pardinus | Tremarctos_ornatus | 0.23159775806074 |
| Lynx_pardinus | Urocyon_cinereoargenteus | 0.239097007826223 |
| Lynx_pardinus | Ursus_americanus | 0.239882315487926 |
| Lynx_pardinus | Ursus_arctos | 0.244996756361723 |
| Lynx_pardinus | Ursus_maritimus | 0.243826886831212 |
| Lynx_pardinus | Ursus_spelaeus | 0.239055356031068 |
| Lynx_pardinus | Ursus_thibetanus | 0.236351893863162 |
| Lynx_pardinus | Urva_javanica | 0.191398621105887 |
| Lynx_pardinus | Viverra_tangalunga_FMNH146957_SRR6053069 | 0.171764060728133 |

| | | |
|---|---|---|
| Lynx_pardinus | Viverricula_indica | 0.178742373995362 |
| Lynx_pardinus | Vulpes_corsac | 0.241266715719118 |
| Lynx_pardinus | Vulpes_ferrilata | 0.240683490779326 |
| Lynx_pardinus | Vulpes_lagopus | 0.238953888153521 |
| Lynx_pardinus | Vulpes_vulpes_T3611 | 0.233234170747824 |
| Lynx_pardinus | Vulpes_zerda | 0.234234439753299 |
| Lynx_pardinus | Zalophus_californianus | 0.2312466173641 |
| Lynx_rufus | Manis_javanica | 0.28073055098633 |
| Lynx_rufus | Martes_americana | 0.232510024159429 |
| Lynx_rufus | Martes_flavigula | 0.231296233119542 |
| Lynx_rufus | Martes_foina | 0.234397436292646 |
| Lynx_rufus | Martes_martes | 0.232693470434355 |
| Lynx_rufus | Martes_melampus | 0.233324029492 |
| Lynx_rufus | Martes_pennanti | 0.234369302591601 |
| Lynx_rufus | Martes_zibellina | 0.235115253044764 |
| Lynx_rufus | Meles_anakuma | 0.230053200272839 |
| Lynx_rufus | Meles_leucurus | 0.227536919225934 |
| Lynx_rufus | Meles_meles | 0.229549121339999 |
| Lynx_rufus | Melogale_moschata | 0.229612948311936 |
| Lynx_rufus | Melursus_ursinus | 0.239669606038608 |
| Lynx_rufus | Mephitis_mephitis | 0.244153676845531 |
| Lynx_rufus | Mirounga_leonina | 0.217877513455721 |
| Lynx_rufus | Mungotictis_decemlineata | 0.183847666888855 |
| Lynx_rufus | Mustela_altaica | 0.22852237198987 |
| Lynx_rufus | Mustela_erminea | 0.22340588171723 |
| Lynx_rufus | Mustela_eversmannii | 0.230022269957008 |
| Lynx_rufus | Mustela_frenata | 0.22782375885685 |
| Lynx_rufus | Mustela_itatsi | 0.230102411627197 |
| Lynx_rufus | Mustela_kathiah | 0.23184508059507 |
| Lynx_rufus | Mustela_nigripes | 0.227683600332821 |
| Lynx_rufus | Mustela_nivalis | 0.230768740354234 |
| Lynx_rufus | Mustela_putorius | 0.229485462371991 |
| Lynx_rufus | Mustela_sibirica | 0.229276692969282 |
| Lynx_rufus | Nandinia_binotata | 0.17332603248681 |
| Lynx_rufus | Nasua_nasua | 0.243568309700452 |
| Lynx_rufus | Neofelis_nebulosa | 0.14365801034098 |
| Lynx_rufus | Neomonachus_schauinslandi | 0.218247309850655 |
| Lynx_rufus | Neophoca_cinerea | 0.233303813983127 |
| Lynx_rufus | Neovison_vison | 0.228180621099388 |
| Lynx_rufus | Nyctereutes_procyonoides | 0.233817572162585 |
| Lynx_rufus | Odobenus_rosmarus_rosmarus | 0.250495185726333 |
| Lynx_rufus | Otocolobus_manul | 0.117370590270792 |
| Lynx_rufus | Otocyon_megalotis_megalotis_TS305 | 0.238281731205946 |
| Lynx_rufus | Otocyon_megalotis_virgatus_FMNH158128 | 0.239977333262044 |
| Lynx_rufus | Paguma_larvata | 0.181915249564938 |
| Lynx_rufus | Panthera_leo | 0.133609905854377 |
| Lynx_rufus | Panthera_onca | 0.135213256325059 |
| Lynx_rufus | Panthera_pardus | 0.136295027598106 |
| Lynx_rufus | Panthera_tigris | 0.14080873968251 |
| Lynx_rufus | Panthera_uncia | 0.140969382393042 |
| Lynx_rufus | Paradoxurus_hermaphroditus | 0.189869348541957 |
| Lynx_rufus | Parahyaena_brunnea_FD126_Fecal | 0.191592473177579 |
| Lynx_rufus | Pardofelis_marmorata | 0.11152373499771 |
| Lynx_rufus | Phoca_fasciata | 0.214511950263382 |
| Lynx_rufus | Phoca_groenlandica | 0.216268653165908 |
| Lynx_rufus | Phoca_largha | 0.217532899956326 |
| Lynx_rufus | Phoca_vitulina | 0.217290806955051 |
| Lynx_rufus | Phocarctos_hookeri | 0.236330464804093 |
| Lynx_rufus | Prionailurus_bengalensis | 0.107903474447418 |
| Lynx_rufus | Prionailurus_bengalensis_euptilurus | 0.104761846538868 |
| Lynx_rufus | Prionailurus_planiceps | 0.109964156053723 |
| Lynx_rufus | Prionailurus_rubiginosus | 0.10768380421088 |
| Lynx_rufus | Prionailurus_viverrinus | 0.111099631886622 |
| Lynx_rufus | Prionodon_pardicolor | 0.184483463344258 |
| Lynx_rufus | Procyon_lotor | 0.234502456063369 |
| Lynx_rufus | Profelis_aurata | 0.114294345515081 |
| Lynx_rufus | Proteles_cristatus_cristatus_TS307 | 0.197221943420395 |
| Lynx_rufus | Proteles_cristatus_septentrionalis_NMS_Z_2018_54 | 0.197136931092788 |
| Lynx_rufus | Puma_concolor | 0.0969277491692051 |
| Lynx_rufus | Puma_yagouaroundi | 0.104798763397371 |
| Lynx_rufus | Pusa_caspica | 0.219917753187945 |
| Lynx_rufus | Pusa_hispida | 0.216015048384756 |
| Lynx_rufus | Pusa_sibirica | 0.218303112387588 |
| Lynx_rufus | Speothos_venaticus_T1624 | 0.243361315871673 |
| Lynx_rufus | Spilogale_putorius | 0.250365829700014 |
| Lynx_rufus | Taxidea_taxus | 0.21762216370539 |
| Lynx_rufus | Tremarctos_ornatus | 0.232272759328477 |
| Lynx_rufus | Urocyon_cinereoargenteus | 0.242861902689016 |
| Lynx_rufus | Ursus_americanus | 0.236425746785966 |
| Lynx_rufus | Ursus_arctos | 0.244215593848144 |
| Lynx_rufus | Ursus_maritimus | 0.242031815156742 |
| Lynx_rufus | Ursus_spelaeus | 0.238058832152017 |
| Lynx_rufus | Ursus_thibetanus | 0.234424180003602 |
| Lynx_rufus | Urva_javanica | 0.191369979058943 |
| Lynx_rufus | Viverra_tangalunga_FMNH146957_SRR6053069 | 0.172772557168436 |
| Lynx_rufus | Viverricula_indica | 0.180776056581304 |
| Lynx_rufus | Vulpes_corsac | 0.239209701344278 |

| | | |
|---|---|---|
| Lynx_rufus | Vulpes_ferrilata | 0.238422321231476 |
| Lynx_rufus | Vulpes_lagopus | 0.23256555878747 |
| Lynx_rufus | Vulpes_vulpes_T3611 | 0.231778753630578 |
| Lynx_rufus | Vulpes_zerda | 0.231457172798583 |
| Lynx_rufus | Zalophus_californianus | 0.232604359436642 |
| Manis_javanica | Martes_americana | 0.283098521864539 |
| Manis_javanica | Martes_flavigula | 0.288478650816527 |
| Manis_javanica | Martes_foina | 0.281882400652972 |
| Manis_javanica | Martes_martes | 0.280270365439078 |
| Manis_javanica | Martes_melampus | 0.280706953523125 |
| Manis_javanica | Martes_pennanti | 0.291719978862532 |
| Manis_javanica | Martes_zibellina | 0.28200896707231 |
| Manis_javanica | Meles_anakuma | 0.296348999320283 |
| Manis_javanica | Meles_leucurus | 0.293808528087823 |
| Manis_javanica | Meles_meles | 0.298168272824112 |
| Manis_javanica | Melogale_moschata | 0.294327555779101 |
| Manis_javanica | Melursus_ursinus | 0.30428223333959 |
| Manis_javanica | Mephitis_mephitis | 0.311604866019023 |
| Manis_javanica | Mirounga_leonina | 0.272768998783732 |
| Manis_javanica | Mungotictis_decemlineata | 0.279491031576479 |
| Manis_javanica | Mustela_altaica | 0.297257736991473 |
| Manis_javanica | Mustela_erminea | 0.288493294130417 |
| Manis_javanica | Mustela_eversmannii | 0.299770894650632 |
| Manis_javanica | Mustela_frenata | 0.29147155837361 |
| Manis_javanica | Mustela_itatsi | 0.297714769551743 |
| Manis_javanica | Mustela_kathiah | 0.299530905855827 |
| Manis_javanica | Mustela_nigripes | 0.297081587104038 |
| Manis_javanica | Mustela_nivalis | 0.297803001923256 |
| Manis_javanica | Mustela_putorius | 0.29740527693171 |
| Manis_javanica | Mustela_sibirica | 0.296502059919128 |
| Manis_javanica | Nandinia_binotata | 0.266563692631458 |
| Manis_javanica | Nasua_nasua | 0.302059228164338 |
| Manis_javanica | Neofelis_nebulosa | 0.290976912965 |
| Manis_javanica | Neomonachus_schauinslandi | 0.279738296149687 |
| Manis_javanica | Neophoca_cinerea | 0.288276481144631 |
| Manis_javanica | Neovison_vison | 0.295175809236646 |
| Manis_javanica | Nyctereutes_procyonoides | 0.294627023369431 |
| Manis_javanica | Odobenus_rosmarus_rosmarus | 0.30437090475901 |
| Manis_javanica | Otocolobus_manul | 0.28835039166674 |
| Manis_javanica | Otocyon_megalotis_megalotis_TS305 | 0.300733302280175 |
| Manis_javanica | Otocyon_megalotis_virgatus_FMNH158128 | 0.305188897953076 |
| Manis_javanica | Paguma_larvata | 0.298269308929876 |
| Manis_javanica | Panthera_leo | 0.286711448222863 |
| Manis_javanica | Panthera_onca | 0.290771211974436 |
| Manis_javanica | Panthera_pardus | 0.290405500964806 |
| Manis_javanica | Panthera_tigris | 0.293102824159939 |
| Manis_javanica | Panthera_uncia | 0.291730537080076 |
| Manis_javanica | Paradoxurus_hermaphroditus | 0.311837400333838 |
| Manis_javanica | Parahyaena_brunnea_FD126_Fecal | 0.278313489165036 |
| Manis_javanica | Pardofelis_marmorata | 0.287269493029305 |
| Manis_javanica | Phoca_fasciata | 0.278129006524325 |
| Manis_javanica | Phoca_groenlandica | 0.282709598971689 |
| Manis_javanica | Phoca_largha | 0.273595561548085 |
| Manis_javanica | Phoca_vitulina | 0.275670705283975 |
| Manis_javanica | Phocarctos_hookeri | 0.287348090036438 |
| Manis_javanica | Prionailurus_bengalensis | 0.285584932566625 |
| Manis_javanica | Prionailurus_bengalensis_euptilurus | 0.283322141404241 |
| Manis_javanica | Prionailurus_planiceps | 0.286896456218905 |
| Manis_javanica | Prionailurus_rubiginosus | 0.284940619486862 |
| Manis_javanica | Prionailurus_viverrinus | 0.285038132896192 |
| Manis_javanica | Prionodon_pardicolor | 0.290497593988362 |
| Manis_javanica | Procyon_lotor | 0.294718371686731 |
| Manis_javanica | Profelis_aurata | 0.295015195773448 |
| Manis_javanica | Proteles_cristatus_cristatus_TS307 | 0.287034321418026 |
| Manis_javanica | Proteles_cristatus_septentrionalis_NMS_Z_2018_54 | 0.287208981615656 |
| Manis_javanica | Puma_concolor | 0.283253883437209 |
| Manis_javanica | Puma_yagouaroundi | 0.289855292277373 |
| Manis_javanica | Pusa_caspica | 0.2744327972035 |
| Manis_javanica | Pusa_hispida | 0.272532006473922 |
| Manis_javanica | Pusa_sibirica | 0.2746001700397 |
| Manis_javanica | Speothos_venaticus_T1624 | 0.31436343145877 |
| Manis_javanica | Spilogale_putorius | 0.309809709870876 |
| Manis_javanica | Taxidea_taxus | 0.28448491617756 |
| Manis_javanica | Tremarctos_ornatus | 0.30647810542834 |
| Manis_javanica | Urocyon_cinereoargenteus | 0.312119278385643 |
| Manis_javanica | Ursus_americanus | 0.305964390839219 |
| Manis_javanica | Ursus_arctos | 0.309163777396046 |
| Manis_javanica | Ursus_maritimus | 0.305353890152872 |
| Manis_javanica | Ursus_spelaeus | 0.30537839824016 |
| Manis_javanica | Ursus_thibetanus | 0.303526124680643 |
| Manis_javanica | Urva_javanica | 0.282656471151523 |
| Manis_javanica | Viverra_tangalunga_FMNH146957_SRR6053069 | 0.28933730191031 |
| Manis_javanica | Viverricula_indica | 0.297273098202859 |
| Manis_javanica | Vulpes_corsac | 0.306506255812507 |
| Manis_javanica | Vulpes_ferrilata | 0.306274357955613 |
| Manis_javanica | Vulpes_lagopus | 0.30377487683364 |
| Manis_javanica | Vulpes_vulpes_T3611 | 0.303025879021246 |

| | | |
|---|---|---|
| Manis_javanica | Vulpes_zerda | 0.30510934323296 |
| Manis_javanica | Zalophus_californianus | 0.282090420145848 |
| Martes_americana | Martes_flavigula | 0.123235890457274 |
| Martes_americana | Martes_foina | 0.0792373506226254 |
| Martes_americana | Martes_martes | 0.0420224636121351 |
| Martes_americana | Martes_melampus | 0.0415172717823236 |
| Martes_americana | Martes_pennanti | 0.139994838663331 |
| Martes_americana | Martes_zibellina | 0.0449784886927754 |
| Martes_americana | Meles_anakuma | 0.174589276989745 |
| Martes_americana | Meles_leucurus | 0.168941887156988 |
| Martes_americana | Meles_meles | 0.173645878715727 |
| Martes_americana | Melogale_moschata | 0.161354694549386 |
| Martes_americana | Melursus_ursinus | 0.234545078086757 |
| Martes_americana | Mephitis_mephitis | 0.241448226317356 |
| Martes_americana | Mirounga_leonina | 0.217773645355352 |
| Martes_americana | Mungotictis_decemlineata | 0.234540037166309 |
| Martes_americana | Mustela_altaica | 0.167529771483366 |
| Martes_americana | Mustela_erminea | 0.156972572126143 |
| Martes_americana | Mustela_eversmannii | 0.167560691333229 |
| Martes_americana | Mustela_frenata | 0.167084468524478 |
| Martes_americana | Mustela_itatsi | 0.170583837784206 |
| Martes_americana | Mustela_kathiah | 0.169840824898598 |
| Martes_americana | Mustela_nigripes | 0.16606970424313 |
| Martes_americana | Mustela_nivalis | 0.169227006464486 |
| Martes_americana | Mustela_putorius | 0.166231441884051 |
| Martes_americana | Mustela_sibirica | 0.166038928973606 |
| Martes_americana | Nandinia_binotata | 0.219420726335539 |
| Martes_americana | Nasua_nasua | 0.230162930227705 |
| Martes_americana | Neofelis_nebulosa | 0.248209078598835 |
| Martes_americana | Neomonachus_schauinslandi | 0.217935495723982 |
| Martes_americana | Neophoca_cinerea | 0.235909857908422 |
| Martes_americana | Neovison_vison | 0.166733003990711 |
| Martes_americana | Nyctereutes_procyonoides | 0.239499371888168 |
| Martes_americana | Odobenus_rosmarus_rosmarus | 0.245490876737243 |
| Martes_americana | Otocolobus_manul | 0.235615733495243 |
| Martes_americana | Otocyon_megalotis_megalotis_TS305 | 0.244864110642124 |
| Martes_americana | Otocyon_megalotis_virgatus_FMNH158128 | 0.247928675629798 |
| Martes_americana | Paguma_larvata | 0.246427630804962 |
| Martes_americana | Panthera_leo | 0.244909256506344 |
| Martes_americana | Panthera_onca | 0.245436458810976 |
| Martes_americana | Panthera_pardus | 0.246035173955914 |
| Martes_americana | Panthera_tigris | 0.244706829532803 |
| Martes_americana | Panthera_uncia | 0.245583655695182 |
| Martes_americana | Paradoxurus_hermaphroditus | 0.24917896359101 |
| Martes_americana | Parahyaena_brunnea_FD126_Fecal | 0.238373317927539 |
| Martes_americana | Pardofelis_marmorata | 0.236242132033244 |
| Martes_americana | Phoca_fasciata | 0.215821434654089 |
| Martes_americana | Phoca_groenlandica | 0.219848710854139 |
| Martes_americana | Phoca_largha | 0.214643700890838 |
| Martes_americana | Phoca_vitulina | 0.215475815286988 |
| Martes_americana | Phocarctos_hookeri | 0.235617256443435 |
| Martes_americana | Prionailurus_bengalensis | 0.233340025987364 |
| Martes_americana | Prionailurus_bengalensis_euptilurus | 0.233092742851829 |
| Martes_americana | Prionailurus_planiceps | 0.234640047358096 |
| Martes_americana | Prionailurus_rubiginosus | 0.230890581177077 |
| Martes_americana | Prionailurus_viverrinus | 0.23294054664292 |
| Martes_americana | Prionodon_pardicolor | 0.238507200628065 |
| Martes_americana | Procyon_lotor | 0.215838182858802 |
| Martes_americana | Profelis_aurata | 0.238887168721372 |
| Martes_americana | Proteles_cristatus_cristatus_TS307 | 0.238505065235349 |
| Martes_americana | Proteles_cristatus_septentrionalis_NMS_Z_2018_54 | 0.239213595436287 |
| Martes_americana | Puma_concolor | 0.232337759316138 |
| Martes_americana | Puma_yagouaroundi | 0.23485525364577 |
| Martes_americana | Pusa_caspica | 0.215376895778748 |
| Martes_americana | Pusa_hispida | 0.215117221076698 |
| Martes_americana | Pusa_sibirica | 0.214295697390097 |
| Martes_americana | Speothos_venaticus_T1624 | 0.247592083809792 |
| Martes_americana | Spilogale_putorius | 0.234182915085361 |
| Martes_americana | Taxidea_taxus | 0.171952235302943 |
| Martes_americana | Tremarctos_ornatus | 0.235077170198065 |
| Martes_americana | Urocyon_cinereoargenteus | 0.250192679303633 |
| Martes_americana | Ursus_americanus | 0.234346291582309 |
| Martes_americana | Ursus_arctos | 0.239576435565417 |
| Martes_americana | Ursus_maritimus | 0.238339183906107 |
| Martes_americana | Ursus_spelaeus | 0.236153590511533 |
| Martes_americana | Ursus_thibetanus | 0.233578484990216 |
| Martes_americana | Urva_javanica | 0.241103704724582 |
| Martes_americana | Viverra_tangalunga_FMNH146957_SRR6053069 | 0.23118945554761 |
| Martes_americana | Viverricula_indica | 0.238709532049074 |
| Martes_americana | Vulpes_corsac | 0.244456864108071 |
| Martes_americana | Vulpes_ferrilata | 0.243547697446504 |
| Martes_americana | Vulpes_lagopus | 0.241872529690826 |
| Martes_americana | Vulpes_vulpes_T3611 | 0.243840291950356 |
| Martes_americana | Vulpes_zerda | 0.241374246970075 |
| Martes_americana | Zalophus_californianus | 0.233954794403263 |
| Martes_flavigula | Martes_foina | 0.127737785439562 |
| Martes_flavigula | Martes_martes | 0.120425688645002 |

| | | |
|---|---|---|
| Martes_flavigula | Martes_melampus | 0.12052057358014 |
| Martes_flavigula | Martes_pennanti | 0.145530287829177 |
| Martes_flavigula | Martes_zibellina | 0.122425975435984 |
| Martes_flavigula | Meles_anakuma | 0.174691236697035 |
| Martes_flavigula | Meles_leucurus | 0.170014152542471 |
| Martes_flavigula | Meles_meles | 0.175510156986965 |
| Martes_flavigula | Melogale_moschata | 0.16716444820778 |
| Martes_flavigula | Melursus_ursinus | 0.238315848581915 |
| Martes_flavigula | Mephitis_mephitis | 0.238262103925751 |
| Martes_flavigula | Mirounga_leonina | 0.219601458117707 |
| Martes_flavigula | Mungotictis_decemlineata | 0.242098145740461 |
| Martes_flavigula | Mustela_altaica | 0.167854662739171 |
| Martes_flavigula | Mustela_erminea | 0.159064048036835 |
| Martes_flavigula | Mustela_eversmannii | 0.169674905875595 |
| Martes_flavigula | Mustela_frenata | 0.168481452296197 |
| Martes_flavigula | Mustela_itatsi | 0.170214832368247 |
| Martes_flavigula | Mustela_kathiah | 0.170319278109707 |
| Martes_flavigula | Mustela_nigripes | 0.167270661424897 |
| Martes_flavigula | Mustela_nivalis | 0.169432597205848 |
| Martes_flavigula | Mustela_putorius | 0.169970154100612 |
| Martes_flavigula | Mustela_sibirica | 0.170958705555589 |
| Martes_flavigula | Nandinia_binotata | 0.222717793789706 |
| Martes_flavigula | Nasua_nasua | 0.228525539455035 |
| Martes_flavigula | Neofelis_nebulosa | 0.249513642427401 |
| Martes_flavigula | Neomonachus_schauinslandi | 0.215189415593527 |
| Martes_flavigula | Neophoca_cinerea | 0.231794397230917 |
| Martes_flavigula | Neovison_vison | 0.167052922425976 |
| Martes_flavigula | Nyctereutes_procyonoides | 0.240003020394019 |
| Martes_flavigula | Odobenus_rosmarus_rosmarus | 0.24517499848063 |
| Martes_flavigula | Otocolobus_manul | 0.235066024242014 |
| Martes_flavigula | Otocyon_megalotis_megalotis_TS305 | 0.240668250100516 |
| Martes_flavigula | Otocyon_megalotis_virgatus_FMNH158128 | 0.246010803242239 |
| Martes_flavigula | Paguma_larvata | 0.242345855114068 |
| Martes_flavigula | Panthera_leo | 0.243761056429241 |
| Martes_flavigula | Panthera_onca | 0.240380851002984 |
| Martes_flavigula | Panthera_pardus | 0.242485092513593 |
| Martes_flavigula | Panthera_tigris | 0.246438450803771 |
| Martes_flavigula | Panthera_uncia | 0.247815655737816 |
| Martes_flavigula | Paradoxurus_hermaphroditus | 0.248789352547403 |
| Martes_flavigula | Parahyaena_brunnea_FD126_Fecal | 0.243517717567362 |
| Martes_flavigula | Pardofelis_marmorata | 0.235577350591065 |
| Martes_flavigula | Phoca_fasciata | 0.217381510312239 |
| Martes_flavigula | Phoca_groenlandica | 0.219652263150894 |
| Martes_flavigula | Phoca_largha | 0.214236275434127 |
| Martes_flavigula | Phoca_vitulina | 0.216580371846677 |
| Martes_flavigula | Phocarctos_hookeri | 0.236663661396633 |
| Martes_flavigula | Prionailurus_bengalensis | 0.235106740583196 |
| Martes_flavigula | Prionailurus_bengalensis_euptilurus | 0.233534136896662 |
| Martes_flavigula | Prionailurus_planiceps | 0.232743726463891 |
| Martes_flavigula | Prionailurus_rubiginosus | 0.230260622443902 |
| Martes_flavigula | Prionailurus_viverrinus | 0.235589672964083 |
| Martes_flavigula | Prionodon_pardicolor | 0.23770741613629 |
| Martes_flavigula | Procyon_lotor | 0.216839295275794 |
| Martes_flavigula | Profelis_aurata | 0.242535239013367 |
| Martes_flavigula | Proteles_cristatus_cristatus_TS307 | 0.246437628892713 |
| Martes_flavigula | Proteles_cristatus_septentrionalis_NMS_Z_2018_54 | 0.248599000647984 |
| Martes_flavigula | Puma_concolor | 0.235337827600393 |
| Martes_flavigula | Puma_yagouaroundi | 0.235628302232322 |
| Martes_flavigula | Pusa_caspica | 0.218977914944474 |
| Martes_flavigula | Pusa_hispida | 0.215359842278637 |
| Martes_flavigula | Pusa_sibirica | 0.217436197971157 |
| Martes_flavigula | Speothos_venaticus_T1624 | 0.246236053837332 |
| Martes_flavigula | Spilogale_putorius | 0.231576513538506 |
| Martes_flavigula | Taxidea_taxus | 0.173647071378391 |
| Martes_flavigula | Tremarctos_ornatus | 0.239052201264972 |
| Martes_flavigula | Urocyon_cinereoargenteus | 0.243066514904093 |
| Martes_flavigula | Ursus_americanus | 0.239565849232957 |
| Martes_flavigula | Ursus_arctos | 0.243348915997156 |
| Martes_flavigula | Ursus_maritimus | 0.242532878366232 |
| Martes_flavigula | Ursus_spelaeus | 0.238429355960382 |
| Martes_flavigula | Ursus_thibetanus | 0.236045416733993 |
| Martes_flavigula | Urva_javanica | 0.239202947058969 |
| Martes_flavigula | Viverra_tangalunga_FMNH146957_SRR6053069 | 0.233593705260768 |
| Martes_flavigula | Viverricula_indica | 0.236094480590762 |
| Martes_flavigula | Vulpes_corsac | 0.245023122928611 |
| Martes_flavigula | Vulpes_ferrilata | 0.2461143263144 |
| Martes_flavigula | Vulpes_lagopus | 0.248211828348423 |
| Martes_flavigula | Vulpes_vulpes_T3611 | 0.24355466317937 |
| Martes_flavigula | Vulpes_zerda | 0.241139575482038 |
| Martes_flavigula | Zalophus_californianus | 0.228839065311436 |
| Martes_foina | Martes_martes | 0.0782776106442571 |
| Martes_foina | Martes_melampus | 0.0776356515488546 |
| Martes_foina | Martes_pennanti | 0.146433283673728 |
| Martes_foina | Martes_zibellina | 0.0797765796529293 |
| Martes_foina | Meles_anakuma | 0.170512157817584 |
| Martes_foina | Meles_leucurus | 0.166432597935218 |
| Martes_foina | Meles_meles | 0.170442782497727 |

| | | |
|---|---|---|
| Martes_foina | Melogale_moschata | 0.160288270635825 |
| Martes_foina | Melursus_ursinus | 0.234505027907506 |
| Martes_foina | Mephitis_mephitis | 0.238577231178484 |
| Martes_foina | Mirounga_leonina | 0.213205827100192 |
| Martes_foina | Mungotictis_decemlineata | 0.232609822344675 |
| Martes_foina | Mustela_altaica | 0.167316736598737 |
| Martes_foina | Mustela_erminea | 0.15692467365027 |
| Martes_foina | Mustela_eversmannii | 0.169892599297734 |
| Martes_foina | Mustela_frenata | 0.170321529601332 |
| Martes_foina | Mustela_itatsi | 0.168942423867102 |
| Martes_foina | Mustela_kathiah | 0.170487219700397 |
| Martes_foina | Mustela_nigripes | 0.166971621903976 |
| Martes_foina | Mustela_nivalis | 0.168079172978597 |
| Martes_foina | Mustela_putorius | 0.169780066141218 |
| Martes_foina | Mustela_sibirica | 0.167631545735202 |
| Martes_foina | Nandinia_binotata | 0.215498635908277 |
| Martes_foina | Nasua_nasua | 0.225666880063993 |
| Martes_foina | Neofelis_nebulosa | 0.246093082921871 |
| Martes_foina | Neomonachus_schauinslandi | 0.216646116880392 |
| Martes_foina | Neophoca_cinerea | 0.236233443630996 |
| Martes_foina | Neovison_vison | 0.167982446232204 |
| Martes_foina | Nyctereutes_procyonoides | 0.243395407541945 |
| Martes_foina | Odobenus_rosmarus_rosmarus | 0.243645152658767 |
| Martes_foina | Otocolobus_manul | 0.232048768765505 |
| Martes_foina | Otocyon_megalotis_megalotis_TS305 | 0.242926089218307 |
| Martes_foina | Otocyon_megalotis_virgatus_FMNH158128 | 0.245639934958149 |
| Martes_foina | Paguma_larvata | 0.244838890882623 |
| Martes_foina | Panthera_leo | 0.24494397079586 |
| Martes_foina | Panthera_onca | 0.241672301172877 |
| Martes_foina | Panthera_pardus | 0.241437363130992 |
| Martes_foina | Panthera_tigris | 0.243863061876015 |
| Martes_foina | Panthera_uncia | 0.246455522289601 |
| Martes_foina | Paradoxurus_hermaphroditus | 0.249488797028645 |
| Martes_foina | Parahyaena_brunnea_FD126_Fecal | 0.238473928940001 |
| Martes_foina | Pardofelis_marmorata | 0.2354982519225 |
| Martes_foina | Phoca_fasciata | 0.21330555239772 |
| Martes_foina | Phoca_groenlandica | 0.218402018501895 |
| Martes_foina | Phoca_largha | 0.21269123353674 |
| Martes_foina | Phoca_vitulina | 0.214038742257091 |
| Martes_foina | Phocarctos_hookeri | 0.235605734674235 |
| Martes_foina | Prionailurus_bengalensis | 0.230290718388189 |
| Martes_foina | Prionailurus_bengalensis_euptilurus | 0.227744382488148 |
| Martes_foina | Prionailurus_planiceps | 0.229844010505555 |
| Martes_foina | Prionailurus_rubiginosus | 0.227655104744141 |
| Martes_foina | Prionailurus_viverrinus | 0.229016288733871 |
| Martes_foina | Prionodon_pardicolor | 0.237026685988324 |
| Martes_foina | Procyon_lotor | 0.216177779305863 |
| Martes_foina | Profelis_aurata | 0.23845160155404 |
| Martes_foina | Proteles_cristatus_cristatus_TS307 | 0.237759587953648 |
| Martes_foina | Proteles_cristatus_septentrionalis_NMS_Z_2018_54 | 0.241363429214223 |
| Martes_foina | Puma_concolor | 0.232720039472863 |
| Martes_foina | Puma_yagouaroundi | 0.233340025987364 |
| Martes_foina | Pusa_caspica | 0.213399422195718 |
| Martes_foina | Pusa_hispida | 0.211118212659304 |
| Martes_foina | Pusa_sibirica | 0.210817105287297 |
| Martes_foina | Speothos_venaticus_T1624 | 0.249638093354242 |
| Martes_foina | Spilogale_putorius | 0.232303258726479 |
| Martes_foina | Taxidea_taxus | 0.169756342389869 |
| Martes_foina | Tremarctos_ornatus | 0.232869455078617 |
| Martes_foina | Urocyon_cinereoargenteus | 0.245236844871494 |
| Martes_foina | Ursus_americanus | 0.236764918760316 |
| Martes_foina | Ursus_arctos | 0.242246007037029 |
| Martes_foina | Ursus_maritimus | 0.2428326775041 |
| Martes_foina | Ursus_spelaeus | 0.236964277437935 |
| Martes_foina | Ursus_thibetanus | 0.234724957683933 |
| Martes_foina | Urva_javanica | 0.239549258585083 |
| Martes_foina | Viverra_tangalunga_FMNH146957_SRR6053069 | 0.230329751666658 |
| Martes_foina | Viverricula_indica | 0.236521674446394 |
| Martes_foina | Vulpes_corsac | 0.247082015649588 |
| Martes_foina | Vulpes_ferrilata | 0.247747972506334 |
| Martes_foina | Vulpes_lagopus | 0.242606861616022 |
| Martes_foina | Vulpes_vulpes_T3611 | 0.240773963955017 |
| Martes_foina | Vulpes_zerda | 0.237857841722414 |
| Martes_foina | Zalophus_californianus | 0.234699574148898 |
| Martes_martes | Martes_melampus | 0.0310352488073757 |
| Martes_martes | Martes_pennanti | 0.141047282214503 |
| Martes_martes | Martes_zibellina | 0.0206844662550012 |
| Martes_martes | Meles_anakuma | 0.17556759705686 |
| Martes_martes | Meles_leucurus | 0.170632859097205 |
| Martes_martes | Meles_meles | 0.172337672866724 |
| Martes_martes | Melogale_moschata | 0.160968516323423 |
| Martes_martes | Melursus_ursinus | 0.235343784661491 |
| Martes_martes | Mephitis_mephitis | 0.240921013003109 |
| Martes_martes | Mirounga_leonina | 0.217089760752053 |
| Martes_martes | Mungotictis_decemlineata | 0.233266460380979 |
| Martes_martes | Mustela_altaica | 0.168312344658585 |
| Martes_martes | Mustela_erminea | 0.156004240726038 |

| | | |
|---|---|---|
| Martes_martes | Mustela_eversmannii | 0.167327866096259 |
| Martes_martes | Mustela_frenata | 0.169056069131433 |
| Martes_martes | Mustela_itatsi | 0.168719345088059 |
| Martes_martes | Mustela_kathiah | 0.167160895557172 |
| Martes_martes | Mustela_nigripes | 0.16626184102298 |
| Martes_martes | Mustela_nivalis | 0.168281371460509 |
| Martes_martes | Mustela_putorius | 0.166302537100832 |
| Martes_martes | Mustela_sibirica | 0.165684381117044 |
| Martes_martes | Nandinia_binotata | 0.219416898079472 |
| Martes_martes | Nasua_nasua | 0.232115837682559 |
| Martes_martes | Neofelis_nebulosa | 0.247724816286853 |
| Martes_martes | Neomonachus_schauinslandi | 0.213129107065292 |
| Martes_martes | Neophoca_cinerea | 0.231537990401062 |
| Martes_martes | Neovison_vison | 0.168109625827504 |
| Martes_martes | Nyctereutes_procyonoides | 0.237314027804921 |
| Martes_martes | Odobenus_rosmarus_rosmarus | 0.248360982344518 |
| Martes_martes | Otocolobus_manul | 0.233345143883257 |
| Martes_martes | Otocyon_megalotis_megalotis_TS305 | 0.242431251393841 |
| Martes_martes | Otocyon_megalotis_virgatus_FMNH158128 | 0.244694398225261 |
| Martes_martes | Paguma_larvata | 0.245131440525522 |
| Martes_martes | Panthera_leo | 0.243389897269784 |
| Martes_martes | Panthera_onca | 0.242815694866068 |
| Martes_martes | Panthera_pardus | 0.244851687407757 |
| Martes_martes | Panthera_tigris | 0.242513533391433 |
| Martes_martes | Panthera_uncia | 0.24779087742244 |
| Martes_martes | Paradoxurus_hermaphroditus | 0.254350704609021 |
| Martes_martes | Parahyaena_brunnea_FD126_Fecal | 0.239870702201027 |
| Martes_martes | Pardofelis_marmorata | 0.235319216913506 |
| Martes_martes | Phoca_fasciata | 0.214902020770885 |
| Martes_martes | Phoca_groenlandica | 0.220362378233768 |
| Martes_martes | Phoca_largha | 0.213208374017001 |
| Martes_martes | Phoca_vitulina | 0.214902020770885 |
| Martes_martes | Phocarctos_hookeri | 0.234249994072478 |
| Martes_martes | Prionailurus_bengalensis | 0.228708104852564 |
| Martes_martes | Prionailurus_bengalensis_euptilurus | 0.229645091457848 |
| Martes_martes | Prionailurus_planiceps | 0.233719106509983 |
| Martes_martes | Prionailurus_rubiginosus | 0.227893584466991 |
| Martes_martes | Prionailurus_viverrinus | 0.230372221669609 |
| Martes_martes | Prionodon_pardicolor | 0.238094422899832 |
| Martes_martes | Procyon_lotor | 0.211966601591603 |
| Martes_martes | Profelis_aurata | 0.239391339526228 |
| Martes_martes | Proteles_cristatus_cristatus_TS307 | 0.241543690601033 |
| Martes_martes | Proteles_cristatus_septentrionalis_NMS_Z_2018_54 | 0.244523750974301 |
| Martes_martes | Puma_concolor | 0.233292463467245 |
| Martes_martes | Puma_yagouaroundi | 0.234155077437736 |
| Martes_martes | Pusa_caspica | 0.215020169157803 |
| Martes_martes | Pusa_hispida | 0.212925467370742 |
| Martes_martes | Pusa_sibirica | 0.213399422195718 |
| Martes_martes | Speothos_venaticus_T1624 | 0.249116192149124 |
| Martes_martes | Spilogale_putorius | 0.234935222289305 |
| Martes_martes | Taxidea_taxus | 0.168881722296008 |
| Martes_martes | Tremarctos_ornatus | 0.234069832300839 |
| Martes_martes | Urocyon_cinereoargenteus | 0.245684207343355 |
| Martes_martes | Ursus_americanus | 0.234692988163841 |
| Martes_martes | Ursus_arctos | 0.240985658450457 |
| Martes_martes | Ursus_maritimus | 0.23985802731677 |
| Martes_martes | Ursus_spelaeus | 0.238233618296015 |
| Martes_martes | Ursus_thibetanus | 0.234941953835137 |
| Martes_martes | Urva_javanica | 0.236763194505005 |
| Martes_martes | Viverra_tangalunga_FMNH146957_SRR6053069 | 0.230561091873556 |
| Martes_martes | Viverricula_indica | 0.237051220447697 |
| Martes_martes | Vulpes_corsac | 0.242828874828314 |
| Martes_martes | Vulpes_ferrilata | 0.243939599787812 |
| Martes_martes | Vulpes_lagopus | 0.2390499809768 |
| Martes_martes | Vulpes_vulpes_T3611 | 0.239870702201027 |
| Martes_martes | Vulpes_zerda | 0.237420067291732 |
| Martes_martes | Zalophus_californianus | 0.230816233945021 |
| Martes_melampus | Martes_pennanti | 0.141172695509561 |
| Martes_melampus | Martes_zibellina | 0.0335249982822325 |
| Martes_melampus | Meles_anakuma | 0.174576128540657 |
| Martes_melampus | Meles_leucurus | 0.168850664000388 |
| Martes_melampus | Meles_meles | 0.172028390061416 |
| Martes_melampus | Melogale_moschata | 0.160139996108049 |
| Martes_melampus | Melursus_ursinus | 0.236153590511533 |
| Martes_melampus | Mephitis_mephitis | 0.24241529626218 |
| Martes_melampus | Mirounga_leonina | 0.216042416618627 |
| Martes_melampus | Mungotictis_decemlineata | 0.233523795074424 |
| Martes_melampus | Mustela_altaica | 0.165968469174957 |
| Martes_melampus | Mustela_erminea | 0.154980736009282 |
| Martes_melampus | Mustela_eversmannii | 0.165957858847237 |
| Martes_melampus | Mustela_frenata | 0.167143746569777 |
| Martes_melampus | Mustela_itatsi | 0.167570340214177 |
| Martes_melampus | Mustela_kathiah | 0.167392589873522 |
| Martes_melampus | Mustela_nigripes | 0.164189191346454 |
| Martes_melampus | Mustela_nivalis | 0.167621975900527 |
| Martes_melampus | Mustela_putorius | 0.164935906035212 |
| Martes_melampus | Mustela_sibirica | 0.164521784085506 |

| | | |
|---|---|---|
| Martes_melampus | Nandinia_binotata | 0.216809991150441 |
| Martes_melampus | Nasua_nasua | 0.230360699859099 |
| Martes_melampus | Neofelis_nebulosa | 0.248029529259939 |
| Martes_melampus | Neomonachus_schauinslandi | 0.21286314486245 |
| Martes_melampus | Neophoca_cinerea | 0.232590268403947 |
| Martes_melampus | Neovison_vison | 0.167343380465291 |
| Martes_melampus | Nyctereutes_procyonoides | 0.237615713150628 |
| Martes_melampus | Odobenus_rosmarus_rosmarus | 0.249009096614299 |
| Martes_melampus | Otocolobus_manul | 0.231220287750185 |
| Martes_melampus | Otocyon_megalotis_megalotis_TS305 | 0.24342768878181 |
| Martes_melampus | Otocyon_megalotis_virgatus_FMNH158128 | 0.24490808704439 |
| Martes_melampus | Paguma_larvata | 0.243703990580119 |
| Martes_melampus | Panthera_leo | 0.243490946270425 |
| Martes_melampus | Panthera_onca | 0.241997210910903 |
| Martes_melampus | Panthera_pardus | 0.244886185652056 |
| Martes_melampus | Panthera_tigris | 0.242254990584902 |
| Martes_melampus | Panthera_uncia | 0.246646699523488 |
| Martes_melampus | Paradoxurus_hermaphroditus | 0.249723300487215 |
| Martes_melampus | Parahyaena_brunnea_FD126_Fecal | 0.239553539347834 |
| Martes_melampus | Pardofelis_marmorata | 0.235087154693578 |
| Martes_melampus | Phoca_fasciata | 0.213083403698598 |
| Martes_melampus | Phoca_groenlandica | 0.219282420746467 |
| Martes_melampus | Phoca_largha | 0.213760252812658 |
| Martes_melampus | Phoca_vitulina | 0.214806800493138 |
| Martes_melampus | Phocarctos_hookeri | 0.233970519755512 |
| Martes_melampus | Prionailurus_bengalensis | 0.228896830761659 |
| Martes_melampus | Prionailurus_bengalensis_euptilurus | 0.229854229196143 |
| Martes_melampus | Prionailurus_planiceps | 0.233597915480953 |
| Martes_melampus | Prionailurus_rubiginosus | 0.227665083842702 |
| Martes_melampus | Prionailurus_viverrinus | 0.228280702301302 |
| Martes_melampus | Prionodon_pardicolor | 0.236769496207312 |
| Martes_melampus | Procyon_lotor | 0.21377776901509 |
| Martes_melampus | Profelis_aurata | 0.235500420781241 |
| Martes_melampus | Proteles_cristatus_cristatus_TS307 | 0.240799252032033 |
| Martes_melampus | Proteles_cristatus_septentrionalis_NMS_Z_2018_54 | 0.243413627638057 |
| Martes_melampus | Puma_concolor | 0.231299369814924 |
| Martes_melampus | Puma_yagouaroundi | 0.232268960335726 |
| Martes_melampus | Pusa_caspica | 0.213411141433932 |
| Martes_melampus | Pusa_hispida | 0.211969993356868 |
| Martes_melampus | Pusa_sibirica | 0.212872464372563 |
| Martes_melampus | Speothos_venaticus_T1624 | 0.247680813402254 |
| Martes_melampus | Spilogale_putorius | 0.234716919405618 |
| Martes_melampus | Taxidea_taxus | 0.168200050812479 |
| Martes_melampus | Tremarctos_ornatus | 0.233412348072941 |
| Martes_melampus | Urocyon_cinereoargenteus | 0.245157293135335 |
| Martes_melampus | Ursus_americanus | 0.236208008631023 |
| Martes_melampus | Ursus_arctos | 0.241115237301088 |
| Martes_melampus | Ursus_maritimus | 0.239076660028839 |
| Martes_melampus | Ursus_spelaeus | 0.235502115611235 |
| Martes_melampus | Ursus_thibetanus | 0.234620672774235 |
| Martes_melampus | Urva_javanica | 0.238149827758337 |
| Martes_melampus | Viverra_tangalunga_FMNH146957_SRR6053069 | 0.230311010169573 |
| Martes_melampus | Viverricula_indica | 0.236553573664096 |
| Martes_melampus | Vulpes_corsac | 0.242438245239738 |
| Martes_melampus | Vulpes_ferrilata | 0.244333419911484 |
| Martes_melampus | Vulpes_lagopus | 0.240290459607261 |
| Martes_melampus | Vulpes_vulpes_T3611 | 0.241404790355991 |
| Martes_melampus | Vulpes_zerda | 0.238949312265906 |
| Martes_melampus | Zalophus_californianus | 0.22974284309389 |
| Martes_pennanti | Martes_zibellina | 0.142119451929461 |
| Martes_pennanti | Meles_anakuma | 0.177198688100989 |
| Martes_pennanti | Meles_leucurus | 0.173520544056705 |
| Martes_pennanti | Meles_meles | 0.175923977983099 |
| Martes_pennanti | Melogale_moschata | 0.16933073851955 |
| Martes_pennanti | Melursus_ursinus | 0.245705258081405 |
| Martes_pennanti | Mephitis_mephitis | 0.239988869214126 |
| Martes_pennanti | Mirounga_leonina | 0.220545150757825 |
| Martes_pennanti | Mungotictis_decemlineata | 0.237417126734117 |
| Martes_pennanti | Mustela_altaica | 0.166709695789736 |
| Martes_pennanti | Mustela_erminea | 0.160148314943205 |
| Martes_pennanti | Mustela_eversmannii | 0.170790952904561 |
| Martes_pennanti | Mustela_frenata | 0.168233006349265 |
| Martes_pennanti | Mustela_itatsi | 0.171105155110462 |
| Martes_pennanti | Mustela_kathiah | 0.173105607443633 |
| Martes_pennanti | Mustela_nigripes | 0.171023926939818 |
| Martes_pennanti | Mustela_nivalis | 0.171991710626131 |
| Martes_pennanti | Mustela_putorius | 0.171006057764861 |
| Martes_pennanti | Mustela_sibirica | 0.172326895132248 |
| Martes_pennanti | Nandinia_binotata | 0.226089177778649 |
| Martes_pennanti | Nasua_nasua | 0.231509314319869 |
| Martes_pennanti | Neofelis_nebulosa | 0.248424179544394 |
| Martes_pennanti | Neomonachus_schauinslandi | 0.219928613919172 |
| Martes_pennanti | Neophoca_cinerea | 0.235418178472137 |
| Martes_pennanti | Neovison_vison | 0.169938375625165 |
| Martes_pennanti | Nyctereutes_procyonoides | 0.242337364002063 |
| Martes_pennanti | Odobenus_rosmarus_rosmarus | 0.254268652353223 |
| Martes_pennanti | Otocolobus_manul | 0.237807823397305 |

| | | |
|---|---|---|
| Martes_pennanti | Otocyon_megalotis_megalotis_TS305 | 0.245729076476784 |
| Martes_pennanti | Otocyon_megalotis_virgatus_FMNH158128 | 0.247940019877146 |
| Martes_pennanti | Paguma_larvata | 0.242503772915513 |
| Martes_pennanti | Panthera_leo | 0.24799931582428 |
| Martes_pennanti | Panthera_onca | 0.244842755137108 |
| Martes_pennanti | Panthera_pardus | 0.24912937563508 |
| Martes_pennanti | Panthera_tigris | 0.24764105333574 |
| Martes_pennanti | Panthera_uncia | 0.24647892142995 |
| Martes_pennanti | Paradoxurus_hermaphroditus | 0.248123505805523 |
| Martes_pennanti | Parahyaena_brunnea_FD126_Fecal | 0.241247543004574 |
| Martes_pennanti | Pardofelis_marmorata | 0.24111171237011 |
| Martes_pennanti | Phoca_fasciata | 0.222439013403638 |
| Martes_pennanti | Phoca_groenlandica | 0.218559329793913 |
| Martes_pennanti | Phoca_largha | 0.217666167077321 |
| Martes_pennanti | Phoca_vitulina | 0.218830073851061 |
| Martes_pennanti | Phocarctos_hookeri | 0.237941982649307 |
| Martes_pennanti | Prionailurus_bengalensis | 0.231849303878282 |
| Martes_pennanti | Prionailurus_bengalensis_euptilurus | 0.232483409574223 |
| Martes_pennanti | Prionailurus_planiceps | 0.234804368648929 |
| Martes_pennanti | Prionailurus_rubiginosus | 0.236342382641309 |
| Martes_pennanti | Prionailurus_viverrinus | 0.234055108713084 |
| Martes_pennanti | Prionodon_pardicolor | 0.241156423925396 |
| Martes_pennanti | Procyon_lotor | 0.223107820943944 |
| Martes_pennanti | Profelis_aurata | 0.241991291349321 |
| Martes_pennanti | Proteles_cristatus_cristatus_TS307 | 0.246418966638724 |
| Martes_pennanti | Proteles_cristatus_septentrionalis_NMS_Z_2018_54 | 0.250079249176105 |
| Martes_pennanti | Puma_concolor | 0.236438752151549 |
| Martes_pennanti | Puma_yagouaroundi | 0.235061594017044 |
| Martes_pennanti | Pusa_caspica | 0.222843329961552 |
| Martes_pennanti | Pusa_hispida | 0.2182077848995 |
| Martes_pennanti | Pusa_sibirica | 0.219909618708412 |
| Martes_pennanti | Speothos_venaticus_T1624 | 0.24663105733266 |
| Martes_pennanti | Spilogale_putorius | 0.241519591639203 |
| Martes_pennanti | Taxidea_taxus | 0.17143300857442 |
| Martes_pennanti | Tremarctos_ornatus | 0.237186773655485 |
| Martes_pennanti | Urocyon_cinereoargenteus | 0.243884284466956 |
| Martes_pennanti | Ursus_americanus | 0.246195605025054 |
| Martes_pennanti | Ursus_arctos | 0.244999886632071 |
| Martes_pennanti | Ursus_maritimus | 0.244554244681761 |
| Martes_pennanti | Ursus_spelaeus | 0.2406884359955 |
| Martes_pennanti | Ursus_thibetanus | 0.240600532933755 |
| Martes_pennanti | Urva_javanica | 0.242813792166974 |
| Martes_pennanti | Viverra_tangalunga_FMNH146957_SRR6053069 | 0.232914123571211 |
| Martes_pennanti | Viverricula_indica | 0.239992668095835 |
| Martes_pennanti | Vulpes_corsac | 0.251157011308948 |
| Martes_pennanti | Vulpes_ferrilata | 0.248742707260018 |
| Martes_pennanti | Vulpes_lagopus | 0.24682535731609 |
| Martes_pennanti | Vulpes_vulpes_T3611 | 0.245609986785513 |
| Martes_pennanti | Vulpes_zerda | 0.245494447564833 |
| Martes_pennanti | Zalophus_californianus | 0.23543019934505 |
| Martes_zibellina | Meles_anakuma | 0.17555582758459 |
| Martes_zibellina | Meles_leucurus | 0.17165232843027 |
| Martes_zibellina | Meles_meles | 0.173956244957847 |
| Martes_zibellina | Melogale_moschata | 0.161138264512652 |
| Martes_zibellina | Melursus_ursinus | 0.238277740616364 |
| Martes_zibellina | Mephitis_mephitis | 0.244187558759723 |
| Martes_zibellina | Mirounga_leonina | 0.21827178212521 |
| Martes_zibellina | Mungotictis_decemlineata | 0.235824139532809 |
| Martes_zibellina | Mustela_altaica | 0.167103482659747 |
| Martes_zibellina | Mustela_erminea | 0.155814566760961 |
| Martes_zibellina | Mustela_eversmannii | 0.168559185962858 |
| Martes_zibellina | Mustela_frenata | 0.170706313221641 |
| Martes_zibellina | Mustela_itatsi | 0.168200389285614 |
| Martes_zibellina | Mustela_kathiah | 0.168292289689502 |
| Martes_zibellina | Mustela_nigripes | 0.167083603628203 |
| Martes_zibellina | Mustela_nivalis | 0.168393123621426 |
| Martes_zibellina | Mustela_putorius | 0.168141453177021 |
| Martes_zibellina | Mustela_sibirica | 0.167114390333795 |
| Martes_zibellina | Nandinia_binotata | 0.221134137696365 |
| Martes_zibellina | Nasua_nasua | 0.231818590316681 |
| Martes_zibellina | Neofelis_nebulosa | 0.250303552508891 |
| Martes_zibellina | Neomonachus_schauinslandi | 0.217327808636244 |
| Martes_zibellina | Neophoca_cinerea | 0.23285881433919 |
| Martes_zibellina | Neovison_vison | 0.166592648576052 |
| Martes_zibellina | Nyctereutes_procyonoides | 0.239297816672941 |
| Martes_zibellina | Odobenus_rosmarus_rosmarus | 0.248947738456518 |
| Martes_zibellina | Otocolobus_manul | 0.235970130164179 |
| Martes_zibellina | Otocyon_megalotis_megalotis_TS305 | 0.244795727493355 |
| Martes_zibellina | Otocyon_megalotis_virgatus_FMNH158128 | 0.246731762955199 |
| Martes_zibellina | Paguma_larvata | 0.247580644010213 |
| Martes_zibellina | Panthera_leo | 0.245745995757587 |
| Martes_zibellina | Panthera_onca | 0.244017869251536 |
| Martes_zibellina | Panthera_pardus | 0.247552394325641 |
| Martes_zibellina | Panthera_tigris | 0.242951556561621 |
| Martes_zibellina | Panthera_uncia | 0.24902765640361 |
| Martes_zibellina | Paradoxurus_hermaphroditus | 0.255486840246803 |
| Martes_zibellina | Parahyaena_brunnea_FD126_Fecal | 0.238394065123893 |

| | | |
|---|---|---|
| Martes_zibellina | Pardofelis_marmorata | 0.238641935274432 |
| Martes_zibellina | Phoca_fasciata | 0.21553975567337 |
| Martes_zibellina | Phoca_groenlandica | 0.221006331181602 |
| Martes_zibellina | Phoca_largha | 0.215031365781925 |
| Martes_zibellina | Phoca_vitulina | 0.216947444746124 |
| Martes_zibellina | Phocarctos_hookeri | 0.233571572241721 |
| Martes_zibellina | Prionailurus_bengalensis | 0.230101782599064 |
| Martes_zibellina | Prionailurus_bengalensis_euptilurus | 0.232053001429065 |
| Martes_zibellina | Prionailurus_planiceps | 0.236035323365619 |
| Martes_zibellina | Prionailurus_rubiginosus | 0.229086937584424 |
| Martes_zibellina | Prionailurus_viverrinus | 0.232121388364433 |
| Martes_zibellina | Prionodon_pardicolor | 0.238864172081748 |
| Martes_zibellina | Procyon_lotor | 0.214128111175072 |
| Martes_zibellina | Profelis_aurata | 0.238820700788677 |
| Martes_zibellina | Proteles_cristatus_cristatus_TS307 | 0.239838992215945 |
| Martes_zibellina | Proteles_cristatus_septentrionalis_NMS_Z_2018_54 | 0.24267304456775 |
| Martes_zibellina | Puma_concolor | 0.235494117897636 |
| Martes_zibellina | Puma_yagouaroundi | 0.236360093288769 |
| Martes_zibellina | Pusa_caspica | 0.216418024166641 |
| Martes_zibellina | Pusa_hispida | 0.214425252401648 |
| Martes_zibellina | Pusa_sibirica | 0.215550627223527 |
| Martes_zibellina | Speothos_venaticus_T1624 | 0.250260359389586 |
| Martes_zibellina | Spilogale_putorius | 0.23671341877311 |
| Martes_zibellina | Taxidea_taxus | 0.170340192836797 |
| Martes_zibellina | Tremarctos_ornatus | 0.233830783194476 |
| Martes_zibellina | Urocyon_cinereoargenteus | 0.247025231002292 |
| Martes_zibellina | Ursus_americanus | 0.236063523427623 |
| Martes_zibellina | Ursus_arctos | 0.239943974889598 |
| Martes_zibellina | Ursus_maritimus | 0.239046345045031 |
| Martes_zibellina | Ursus_spelaeus | 0.237651366431572 |
| Martes_zibellina | Ursus_thibetanus | 0.236400209820835 |
| Martes_zibellina | Urva_javanica | 0.236773013470627 |
| Martes_zibellina | Viverra_tangalunga_FMNH146957_SRR6053069 | 0.231539884172252 |
| Martes_zibellina | Viverricula_indica | 0.238932497493125 |
| Martes_zibellina | Vulpes_corsac | 0.245041405226776 |
| Martes_zibellina | Vulpes_ferrilata | 0.24694657485698 |
| Martes_zibellina | Vulpes_lagopus | 0.241580240699132 |
| Martes_zibellina | Vulpes_vulpes_T3611 | 0.243862259341552 |
| Martes_zibellina | Vulpes_zerda | 0.241172461859675 |
| Martes_zibellina | Zalophus_californianus | 0.230915583687937 |
| Meles_anakuma | Meles_leucurus | 0.0361104243666735 |
| Meles_anakuma | Meles_meles | 0.0792013127508662 |
| Meles_anakuma | Melogale_moschata | 0.174665105828655 |
| Meles_anakuma | Melursus_ursinus | 0.247032378933602 |
| Meles_anakuma | Mephitis_mephitis | 0.228221473410335 |
| Meles_anakuma | Mirounga_leonina | 0.216171540120507 |
| Meles_anakuma | Mungotictis_decemlineata | 0.23895994587049 |
| Meles_anakuma | Mustela_altaica | 0.18397492374421 |
| Meles_anakuma | Mustela_erminea | 0.169747387214357 |
| Meles_anakuma | Mustela_eversmannii | 0.178780610471934 |
| Meles_anakuma | Mustela_frenata | 0.17180725648808 |
| Meles_anakuma | Mustela_itatsi | 0.175362040905009 |
| Meles_anakuma | Mustela_kathiah | 0.174805221781172 |
| Meles_anakuma | Mustela_nigripes | 0.179430158350838 |
| Meles_anakuma | Mustela_nivalis | 0.18115485052845 |
| Meles_anakuma | Mustela_putorius | 0.178152534503342 |
| Meles_anakuma | Mustela_sibirica | 0.175534069742122 |
| Meles_anakuma | Nandinia_binotata | 0.220210753604722 |
| Meles_anakuma | Nasua_nasua | 0.234215927630581 |
| Meles_anakuma | Neofelis_nebulosa | 0.247038224328121 |
| Meles_anakuma | Neomonachus_schauinslandi | 0.215950125977872 |
| Meles_anakuma | Neophoca_cinerea | 0.238826470785829 |
| Meles_anakuma | Neovison_vison | 0.172787394040106 |
| Meles_anakuma | Nyctereutes_procyonoides | 0.243588278018706 |
| Meles_anakuma | Odobenus_rosmarus_rosmarus | 0.251745160531007 |
| Meles_anakuma | Otocolobus_manul | 0.231116812639965 |
| Meles_anakuma | Otocyon_megalotis_megalotis_TS305 | 0.251882898762029 |
| Meles_anakuma | Otocyon_megalotis_virgatus_FMNH158128 | 0.256967962001422 |
| Meles_anakuma | Paguma_larvata | 0.232575336788439 |
| Meles_anakuma | Panthera_leo | 0.243883857497868 |
| Meles_anakuma | Panthera_onca | 0.241944113130195 |
| Meles_anakuma | Panthera_pardus | 0.244985602152835 |
| Meles_anakuma | Panthera_tigris | 0.238668978380273 |
| Meles_anakuma | Panthera_uncia | 0.247650199870846 |
| Meles_anakuma | Paradoxurus_hermaphroditus | 0.231740232685982 |
| Meles_anakuma | Parahyaena_brunnea_FD126_Fecal | 0.244130235686734 |
| Meles_anakuma | Pardofelis_marmorata | 0.240396313748853 |
| Meles_anakuma | Phoca_fasciata | 0.217295597927444 |
| Meles_anakuma | Phoca_groenlandica | 0.217821508211459 |
| Meles_anakuma | Phoca_largha | 0.218730540324054 |
| Meles_anakuma | Phoca_vitulina | 0.216817238634235 |
| Meles_anakuma | Phocarctos_hookeri | 0.242666283014439 |
| Meles_anakuma | Prionailurus_bengalensis | 0.229204986271522 |
| Meles_anakuma | Prionailurus_bengalensis_euptilurus | 0.23138877526961 |
| Meles_anakuma | Prionailurus_planiceps | 0.234233951607235 |
| Meles_anakuma | Prionailurus_rubiginosus | 0.234315887469905 |
| Meles_anakuma | Prionailurus_viverrinus | 0.232296651228566 |

| | | |
|---|---|---|
| Meles_anakuma | Prionodon_pardicolor | 0.237205509882763 |
| Meles_anakuma | Procyon_lotor | 0.217305998721664 |
| Meles_anakuma | Profelis_aurata | 0.237871532001234 |
| Meles_anakuma | Proteles_cristatus_cristatus_TS307 | 0.246111362608328 |
| Meles_anakuma | Proteles_cristatus_septentrionalis_NMS_Z_2018_54 | 0.245865487492049 |
| Meles_anakuma | Puma_concolor | 0.229593091743209 |
| Meles_anakuma | Puma_yagouaroundi | 0.230688142553231 |
| Meles_anakuma | Pusa_caspica | 0.217954677837089 |
| Meles_anakuma | Pusa_hispida | 0.217697051743851 |
| Meles_anakuma | Pusa_sibirica | 0.218862996129126 |
| Meles_anakuma | Speothos_venaticus_T1624 | 0.251052296115213 |
| Meles_anakuma | Spilogale_putorius | 0.228373181529645 |
| Meles_anakuma | Taxidea_taxus | 0.179575253522719 |
| Meles_anakuma | Tremarctos_ornatus | 0.242303110640072 |
| Meles_anakuma | Urocyon_cinereoargenteus | 0.251043249609535 |
| Meles_anakuma | Ursus_americanus | 0.239379459688018 |
| Meles_anakuma | Ursus_arctos | 0.246014546554239 |
| Meles_anakuma | Ursus_maritimus | 0.246602867998983 |
| Meles_anakuma | Ursus_spelaeus | 0.239606217061519 |
| Meles_anakuma | Ursus_thibetanus | 0.236284832838039 |
| Meles_anakuma | Urva_javanica | 0.237077555453544 |
| Meles_anakuma | Viverra_tangalunga_FMNH146957_SRR6053069 | 0.231346326062051 |
| Meles_anakuma | Viverricula_indica | 0.231715227853985 |
| Meles_anakuma | Vulpes_corsac | 0.253611760816992 |
| Meles_anakuma | Vulpes_ferrilata | 0.256867488343691 |
| Meles_anakuma | Vulpes_lagopus | 0.252056575986205 |
| Meles_anakuma | Vulpes_vulpes_T3611 | 0.254443486634237 |
| Meles_anakuma | Vulpes_zerda | 0.254671562774177 |
| Meles_anakuma | Zalophus_californianus | 0.23400113766019 |
| Meles_leucurus | Meles_meles | 0.0740810599495699 |
| Meles_leucurus | Melogale_moschata | 0.17205887812257 |
| Meles_leucurus | Melursus_ursinus | 0.239206760059223 |
| Meles_leucurus | Mephitis_mephitis | 0.228578425967662 |
| Meles_leucurus | Mirounga_leonina | 0.208678825889831 |
| Meles_leucurus | Mungotictis_decemlineata | 0.235651590069995 |
| Meles_leucurus | Mustela_altaica | 0.179380877458534 |
| Meles_leucurus | Mustela_erminea | 0.165595997326146 |
| Meles_leucurus | Mustela_eversmannii | 0.17386113406804 |
| Meles_leucurus | Mustela_frenata | 0.165236307633821 |
| Meles_leucurus | Mustela_itatsi | 0.171687952270199 |
| Meles_leucurus | Mustela_kathiah | 0.169729926321944 |
| Meles_leucurus | Mustela_nigripes | 0.173974929323802 |
| Meles_leucurus | Mustela_nivalis | 0.174353866427594 |
| Meles_leucurus | Mustela_putorius | 0.173339872018902 |
| Meles_leucurus | Mustela_sibirica | 0.172267073902876 |
| Meles_leucurus | Nandinia_binotata | 0.216415028933039 |
| Meles_leucurus | Nasua_nasua | 0.232152118273391 |
| Meles_leucurus | Neofelis_nebulosa | 0.243964158167841 |
| Meles_leucurus | Neomonachus_schauinslandi | 0.211281133826678 |
| Meles_leucurus | Neophoca_cinerea | 0.235225637799901 |
| Meles_leucurus | Neovison_vison | 0.168150669353241 |
| Meles_leucurus | Nyctereutes_procyonoides | 0.241108344540155 |
| Meles_leucurus | Odobenus_rosmarus_rosmarus | 0.251686045347932 |
| Meles_leucurus | Otocolobus_manul | 0.226683086800372 |
| Meles_leucurus | Otocyon_megalotis_megalotis_TS305 | 0.250012781659998 |
| Meles_leucurus | Otocyon_megalotis_virgatus_FMNH158128 | 0.252166980003566 |
| Meles_leucurus | Paguma_larvata | 0.231077702212021 |
| Meles_leucurus | Panthera_leo | 0.24091177320891 |
| Meles_leucurus | Panthera_onca | 0.236775499988201 |
| Meles_leucurus | Panthera_pardus | 0.239657397631776 |
| Meles_leucurus | Panthera_tigris | 0.236514499604638 |
| Meles_leucurus | Panthera_uncia | 0.242884867495008 |
| Meles_leucurus | Paradoxurus_hermaphroditus | 0.229697425444414 |
| Meles_leucurus | Parahyaena_brunnea_FD126_Fecal | 0.241489979844283 |
| Meles_leucurus | Pardofelis_marmorata | 0.237994032052589 |
| Meles_leucurus | Phoca_fasciata | 0.216611569320672 |
| Meles_leucurus | Phoca_groenlandica | 0.214547983130266 |
| Meles_leucurus | Phoca_largha | 0.216206258143347 |
| Meles_leucurus | Phoca_vitulina | 0.214625072141819 |
| Meles_leucurus | Phocarctos_hookeri | 0.236794724551969 |
| Meles_leucurus | Prionailurus_bengalensis | 0.22415102277354 |
| Meles_leucurus | Prionailurus_bengalensis_euptilurus | 0.227628269626393 |
| Meles_leucurus | Prionailurus_planiceps | 0.228264471135869 |
| Meles_leucurus | Prionailurus_rubiginosus | 0.229971873820807 |
| Meles_leucurus | Prionailurus_viverrinus | 0.22754701365029 |
| Meles_leucurus | Prionodon_pardicolor | 0.231286256740071 |
| Meles_leucurus | Procyon_lotor | 0.212745981497043 |
| Meles_leucurus | Profelis_aurata | 0.231866561731676 |
| Meles_leucurus | Proteles_cristatus_cristatus_TS307 | 0.239803866762484 |
| Meles_leucurus | Proteles_cristatus_septentrionalis_NMS_Z_2018_54 | 0.242457574475498 |
| Meles_leucurus | Puma_concolor | 0.22258636461047 |
| Meles_leucurus | Puma_yagouaroundi | 0.225060407005252 |
| Meles_leucurus | Pusa_caspica | 0.213917510823564 |
| Meles_leucurus | Pusa_hispida | 0.212587759272966 |
| Meles_leucurus | Pusa_sibirica | 0.214281674744964 |
| Meles_leucurus | Speothos_venaticus_T1624 | 0.24926404504932 |
| Meles_leucurus | Spilogale_putorius | 0.223740265176397 |

| | | |
|---|---|---|
| Meles_leucurus | Taxidea_taxus | 0.175285001417176 |
| Meles_leucurus | Tremarctos_ornatus | 0.237986346454656 |
| Meles_leucurus | Urocyon_cinereoargenteus | 0.248726153008313 |
| Meles_leucurus | Ursus_americanus | 0.234548025560747 |
| Meles_leucurus | Ursus_arctos | 0.237214376027858 |
| Meles_leucurus | Ursus_maritimus | 0.237201166816049 |
| Meles_leucurus | Ursus_spelaeus | 0.232976226803676 |
| Meles_leucurus | Ursus_thibetanus | 0.230580849387523 |
| Meles_leucurus | Urva_javanica | 0.234984420659393 |
| Meles_leucurus | Viverra_tangalunga_FMNH146957_SRR6053069 | 0.227844085930343 |
| Meles_leucurus | Viverricula_indica | 0.227269632825091 |
| Meles_leucurus | Vulpes_corsac | 0.2478895541266 |
| Meles_leucurus | Vulpes_ferrilata | 0.252497136144201 |
| Meles_leucurus | Vulpes_lagopus | 0.247631756616948 |
| Meles_leucurus | Vulpes_vulpes_T3611 | 0.249527389074969 |
| Meles_leucurus | Vulpes_zerda | 0.249321081907783 |
| Meles_leucurus | Zalophus_californianus | 0.231288225071878 |
| Meles_meles | Melogale_moschata | 0.176189045510794 |
| Meles_meles | Melursus_ursinus | 0.233537956034962 |
| Meles_meles | Mephitis_mephitis | 0.226229901745758 |
| Meles_meles | Mirounga_leonina | 0.206053742104824 |
| Meles_meles | Mungotictis_decemlineata | 0.23381302244292 |
| Meles_meles | Mustela_altaica | 0.178165371485508 |
| Meles_meles | Mustela_erminea | 0.167087860816458 |
| Meles_meles | Mustela_eversmannii | 0.177808702470208 |
| Meles_meles | Mustela_frenata | 0.172737389929089 |
| Meles_meles | Mustela_itatsi | 0.173625302977945 |
| Meles_meles | Mustela_kathiah | 0.174608713597168 |
| Meles_meles | Mustela_nigripes | 0.174772500352678 |
| Meles_meles | Mustela_nivalis | 0.175521089864377 |
| Meles_meles | Mustela_putorius | 0.175964675953729 |
| Meles_meles | Mustela_sibirica | 0.172265636050643 |
| Meles_meles | Nandinia_binotata | 0.215828096508624 |
| Meles_meles | Nasua_nasua | 0.234544357845749 |
| Meles_meles | Neofelis_nebulosa | 0.240999830477564 |
| Meles_meles | Neomonachus_schauinslandi | 0.214789579380296 |
| Meles_meles | Neophoca_cinerea | 0.234714766624689 |
| Meles_meles | Neovison_vison | 0.170775420266196 |
| Meles_meles | Nyctereutes_procyonoides | 0.236913133353304 |
| Meles_meles | Odobenus_rosmarus_rosmarus | 0.24812168508384 |
| Meles_meles | Otocolobus_manul | 0.225683498001666 |
| Meles_meles | Otocyon_megalotis_megalotis_TS305 | 0.249654819124848 |
| Meles_meles | Otocyon_megalotis_virgatus_FMNH158128 | 0.248767124323052 |
| Meles_meles | Paguma_larvata | 0.229366532158103 |
| Meles_meles | Panthera_leo | 0.23767081416367 |
| Meles_meles | Panthera_onca | 0.232196859822015 |
| Meles_meles | Panthera_pardus | 0.240216306699586 |
| Meles_meles | Panthera_tigris | 0.237114145212827 |
| Meles_meles | Panthera_uncia | 0.240216306699586 |
| Meles_meles | Paradoxurus_hermaphroditus | 0.229716602487819 |
| Meles_meles | Parahyaena_brunnea_FD126_Fecal | 0.242452744616198 |
| Meles_meles | Pardofelis_marmorata | 0.238306243337979 |
| Meles_meles | Phoca_fasciata | 0.214255775800711 |
| Meles_meles | Phoca_groenlandica | 0.211897686506268 |
| Meles_meles | Phoca_largha | 0.214542794100237 |
| Meles_meles | Phoca_vitulina | 0.214168599442056 |
| Meles_meles | Phocarctos_hookeri | 0.236105687898263 |
| Meles_meles | Prionailurus_bengalensis | 0.220638450051592 |
| Meles_meles | Prionailurus_bengalensis_euptilurus | 0.224772532019176 |
| Meles_meles | Prionailurus_planiceps | 0.22659642071396 |
| Meles_meles | Prionailurus_rubiginosus | 0.225396251490878 |
| Meles_meles | Prionailurus_viverrinus | 0.223833990214657 |
| Meles_meles | Prionodon_pardicolor | 0.232424906759578 |
| Meles_meles | Procyon_lotor | 0.215696064551012 |
| Meles_meles | Profelis_aurata | 0.231514136425771 |
| Meles_meles | Proteles_cristatus_cristatus_TS307 | 0.243059428837606 |
| Meles_meles | Proteles_cristatus_septentrionalis_NMS_Z_2018_54 | 0.243149852927569 |
| Meles_meles | Puma_concolor | 0.226128552473394 |
| Meles_meles | Puma_yagouaroundi | 0.225208097747762 |
| Meles_meles | Pusa_caspica | 0.213139379579406 |
| Meles_meles | Pusa_hispida | 0.210847079292709 |
| Meles_meles | Pusa_sibirica | 0.213354410224586 |
| Meles_meles | Speothos_venaticus_T1624 | 0.247965381149142 |
| Meles_meles | Spilogale_putorius | 0.221527365112848 |
| Meles_meles | Taxidea_taxus | 0.171317727133171 |
| Meles_meles | Tremarctos_ornatus | 0.234776695407306 |
| Meles_meles | Urocyon_cinereoargenteus | 0.246625552068007 |
| Meles_meles | Ursus_americanus | 0.225874049940358 |
| Meles_meles | Ursus_arctos | 0.232289264507303 |
| Meles_meles | Ursus_maritimus | 0.23283743198892 |
| Meles_meles | Ursus_spelaeus | 0.229824813168805 |
| Meles_meles | Ursus_thibetanus | 0.226255453853551 |
| Meles_meles | Urva_javanica | 0.232597623861917 |
| Meles_meles | Viverra_tangalunga_FMNH146957_SRR6053069 | 0.22890685559708 |
| Meles_meles | Viverricula_indica | 0.226476120013761 |
| Meles_meles | Vulpes_corsac | 0.245641096149788 |
| Meles_meles | Vulpes_ferrilata | 0.249660410051109 |

| | | |
|---|---|---|
| Meles_meles | Vulpes_lagopus | 0.247208399270677 |
| Meles_meles | Vulpes_vulpes_T3611 | 0.242554275977528 |
| Meles_meles | Vulpes_zerda | 0.244520054484192 |
| Meles_meles | Zalophus_californianus | 0.229214372843346 |
| Melogale_moschata | Melursus_ursinus | 0.242934389981556 |
| Melogale_moschata | Mephitis_mephitis | 0.238453372479744 |
| Melogale_moschata | Mirounga_leonina | 0.222106261029708 |
| Melogale_moschata | Mungotictis_decemlineata | 0.23383408981237 |
| Melogale_moschata | Mustela_altaica | 0.173559055115088 |
| Melogale_moschata | Mustela_erminea | 0.162633082134354 |
| Melogale_moschata | Mustela_eversmannii | 0.168298697022978 |
| Melogale_moschata | Mustela_frenata | 0.164890514563819 |
| Melogale_moschata | Mustela_itatsi | 0.170164161779818 |
| Melogale_moschata | Mustela_kathiah | 0.168228504851755 |
| Melogale_moschata | Mustela_nigripes | 0.168551737220016 |
| Melogale_moschata | Mustela_nivalis | 0.173137203876283 |
| Melogale_moschata | Mustela_putorius | 0.169534139139647 |
| Melogale_moschata | Mustela_sibirica | 0.16707694931674 |
| Melogale_moschata | Nandinia_binotata | 0.221877061084715 |
| Melogale_moschata | Nasua_nasua | 0.238073616778381 |
| Melogale_moschata | Neofelis_nebulosa | 0.243332856146379 |
| Melogale_moschata | Neomonachus_schauinslandi | 0.22052639569966 |
| Melogale_moschata | Neophoca_cinerea | 0.237622301547828 |
| Melogale_moschata | Neovison_vison | 0.162945215987903 |
| Melogale_moschata | Nyctereutes_procyonoides | 0.243574269216634 |
| Melogale_moschata | Odobenus_rosmarus_rosmarus | 0.251899356370808 |
| Melogale_moschata | Otocolobus_manul | 0.233423905243889 |
| Melogale_moschata | Otocyon_megalotis_megalotis_TS305 | 0.245338272240153 |
| Melogale_moschata | Otocyon_megalotis_virgatus_FMNH158128 | 0.245697356090672 |
| Melogale_moschata | Paguma_larvata | 0.234729404878482 |
| Melogale_moschata | Panthera_leo | 0.242142782325904 |
| Melogale_moschata | Panthera_onca | 0.233939667811043 |
| Melogale_moschata | Panthera_pardus | 0.236739538739947 |
| Melogale_moschata | Panthera_tigris | 0.239411853307378 |
| Melogale_moschata | Panthera_uncia | 0.242434282389987 |
| Melogale_moschata | Paradoxurus_hermaphroditus | 0.240462482823342 |
| Melogale_moschata | Parahyaena_brunnea_FD126_Fecal | 0.240910747182456 |
| Melogale_moschata | Pardofelis_marmorata | 0.23737123580402 |
| Melogale_moschata | Phoca_fasciata | 0.216129532221913 |
| Melogale_moschata | Phoca_groenlandica | 0.218326595941678 |
| Melogale_moschata | Phoca_largha | 0.219369731466922 |
| Melogale_moschata | Phoca_vitulina | 0.220338176583318 |
| Melogale_moschata | Phocarctos_hookeri | 0.235417376636021 |
| Melogale_moschata | Prionailurus_bengalensis | 0.23296144723536 |
| Melogale_moschata | Prionailurus_bengalensis_euptilurus | 0.232693470434355 |
| Melogale_moschata | Prionailurus_planiceps | 0.232011244074969 |
| Melogale_moschata | Prionailurus_rubiginosus | 0.229383961393389 |
| Melogale_moschata | Prionailurus_viverrinus | 0.232803784830832 |
| Melogale_moschata | Prionodon_pardicolor | 0.238644111518115 |
| Melogale_moschata | Procyon_lotor | 0.219836835745151 |
| Melogale_moschata | Profelis_aurata | 0.237860013946469 |
| Melogale_moschata | Proteles_cristatus_cristatus_TS307 | 0.24588935443209 |
| Melogale_moschata | Proteles_cristatus_septentrionalis_NMS_Z_2018_54 | 0.245708342567905 |
| Melogale_moschata | Puma_concolor | 0.23090855869484 |
| Melogale_moschata | Puma_yagouaroundi | 0.234150081525248 |
| Melogale_moschata | Pusa_caspica | 0.219783859799415 |
| Melogale_moschata | Pusa_hispida | 0.21583393233917 |
| Melogale_moschata | Pusa_sibirica | 0.219348177384309 |
| Melogale_moschata | Speothos_venaticus_T1624 | 0.249654819124848 |
| Melogale_moschata | Spilogale_putorius | 0.241369603012879 |
| Melogale_moschata | Taxidea_taxus | 0.174815415987895 |
| Melogale_moschata | Tremarctos_ornatus | 0.237740353715638 |
| Melogale_moschata | Urocyon_cinereoargenteus | 0.253560325851555 |
| Melogale_moschata | Ursus_americanus | 0.238801755301871 |
| Melogale_moschata | Ursus_arctos | 0.239156161487919 |
| Melogale_moschata | Ursus_maritimus | 0.239397177602593 |
| Melogale_moschata | Ursus_spelaeus | 0.238116137767388 |
| Melogale_moschata | Ursus_thibetanus | 0.235620246652122 |
| Melogale_moschata | Urva_javanica | 0.24185377779783 |
| Melogale_moschata | Viverra_tangalunga_FMNH146957_SRR6053069 | 0.231120654768443 |
| Melogale_moschata | Viverricula_indica | 0.235478442354315 |
| Melogale_moschata | Vulpes_corsac | 0.244639517713025 |
| Melogale_moschata | Vulpes_ferrilata | 0.24620488141817 |
| Melogale_moschata | Vulpes_lagopus | 0.247375765165861 |
| Melogale_moschata | Vulpes_vulpes_T3611 | 0.24218630218195 |
| Melogale_moschata | Vulpes_zerda | 0.241112560655468 |
| Melogale_moschata | Zalophus_californianus | 0.236710049763789 |
| Melursus_ursinus | Mephitis_mephitis | 0.242160474265782 |
| Melursus_ursinus | Mirounga_leonina | 0.219810513423083 |
| Melursus_ursinus | Mungotictis_decemlineata | 0.243735644977761 |
| Melursus_ursinus | Mustela_altaica | 0.248146446390991 |
| Melursus_ursinus | Mustela_erminea | 0.23507772294556 |
| Melursus_ursinus | Mustela_eversmannii | 0.241002169365989 |
| Melursus_ursinus | Mustela_frenata | 0.241055768398894 |
| Melursus_ursinus | Mustela_itatsi | 0.244730591703396 |
| Melursus_ursinus | Mustela_kathiah | 0.242300519175263 |
| Melursus_ursinus | Mustela_nigripes | 0.240969461590911 |

| | | |
|---|---|---|
| Melursus_ursinus | Mustela_nivalis | 0.243971174893465 |
| Melursus_ursinus | Mustela_putorius | 0.240728139051211 |
| Melursus_ursinus | Mustela_sibirica | 0.240313865604998 |
| Melursus_ursinus | Nandinia_binotata | 0.229773928942013 |
| Melursus_ursinus | Nasua_nasua | 0.251110569898663 |
| Melursus_ursinus | Neofelis_nebulosa | 0.248614590978749 |
| Melursus_ursinus | Neomonachus_schauinslandi | 0.220043395415308 |
| Melursus_ursinus | Neophoca_cinerea | 0.234098435781102 |
| Melursus_ursinus | Neovison_vison | 0.238839146345569 |
| Melursus_ursinus | Nyctereutes_procyonoides | 0.247324604640319 |
| Melursus_ursinus | Odobenus_rosmarus_rosmarus | 0.252706686225936 |
| Melursus_ursinus | Otocolobus_manul | 0.239456961126365 |
| Melursus_ursinus | Otocyon_megalotis_megalotis_TS305 | 0.253339837969925 |
| Melursus_ursinus | Otocyon_megalotis_virgatus_FMNH158128 | 0.251239343859545 |
| Melursus_ursinus | Paguma_larvata | 0.24239880864533 |
| Melursus_ursinus | Panthera_leo | 0.248083005769157 |
| Melursus_ursinus | Panthera_onca | 0.246258729494729 |
| Melursus_ursinus | Panthera_pardus | 0.249149023135009 |
| Melursus_ursinus | Panthera_tigris | 0.252789192946796 |
| Melursus_ursinus | Panthera_uncia | 0.250770414059119 |
| Melursus_ursinus | Paradoxurus_hermaphroditus | 0.249313473850442 |
| Melursus_ursinus | Parahyaena_brunnea_FD126_Fecal | 0.241491861530195 |
| Melursus_ursinus | Pardofelis_marmorata | 0.24486131693424 |
| Melursus_ursinus | Phoca_fasciata | 0.212819363757688 |
| Melursus_ursinus | Phoca_groenlandica | 0.215935019244944 |
| Melursus_ursinus | Phoca_largha | 0.214830530388031 |
| Melursus_ursinus | Phoca_vitulina | 0.215364780043927 |
| Melursus_ursinus | Phocarctos_hookeri | 0.241728297807424 |
| Melursus_ursinus | Prionailurus_bengalensis | 0.239350636067326 |
| Melursus_ursinus | Prionailurus_bengalensis_euptilurus | 0.238607026030171 |
| Melursus_ursinus | Prionailurus_planiceps | 0.239976973262185 |
| Melursus_ursinus | Prionailurus_rubiginosus | 0.235591267976656 |
| Melursus_ursinus | Prionailurus_viverrinus | 0.237087279372319 |
| Melursus_ursinus | Prionodon_pardicolor | 0.241319960442441 |
| Melursus_ursinus | Procyon_lotor | 0.24011574792462 |
| Melursus_ursinus | Profelis_aurata | 0.242037017810028 |
| Melursus_ursinus | Proteles_cristatus_cristatus_TS307 | 0.247765707115391 |
| Melursus_ursinus | Proteles_cristatus_septentrionalis_NMS_Z_2018_54 | 0.248538614349956 |
| Melursus_ursinus | Puma_concolor | 0.24169367267316 |
| Melursus_ursinus | Puma_yagouaroundi | 0.238082182291638 |
| Melursus_ursinus | Pusa_caspica | 0.215028448951356 |
| Melursus_ursinus | Pusa_hispida | 0.213618595659677 |
| Melursus_ursinus | Pusa_sibirica | 0.213498560313764 |
| Melursus_ursinus | Speothos_venaticus_T1624 | 0.253144212578329 |
| Melursus_ursinus | Spilogale_putorius | 0.24675213389659 |
| Melursus_ursinus | Taxidea_taxus | 0.232884924309907 |
| Melursus_ursinus | Tremarctos_ornatus | 0.17300344516162 |
| Melursus_ursinus | Urocyon_cinereoargenteus | 0.261484496308231 |
| Melursus_ursinus | Ursus_americanus | 0.105146101389676 |
| Melursus_ursinus | Ursus_arctos | 0.0985537667155264 |
| Melursus_ursinus | Ursus_maritimus | 0.0991100181166187 |
| Melursus_ursinus | Ursus_spelaeus | 0.102090359305777 |
| Melursus_ursinus | Ursus_thibetanus | 0.0981807916553123 |
| Melursus_ursinus | Urva_javanica | 0.250669874545989 |
| Melursus_ursinus | Viverra_tangalunga_FMNH146957_SRR6053069 | 0.236309630381194 |
| Melursus_ursinus | Viverricula_indica | 0.241845323813341 |
| Melursus_ursinus | Vulpes_corsac | 0.247749665046275 |
| Melursus_ursinus | Vulpes_ferrilata | 0.249415103556253 |
| Melursus_ursinus | Vulpes_lagopus | 0.251881737870955 |
| Melursus_ursinus | Vulpes_vulpes_T3611 | 0.244602404711451 |
| Melursus_ursinus | Vulpes_zerda | 0.247349479621101 |
| Melursus_ursinus | Zalophus_californianus | 0.235236204700719 |
| Mephitis_mephitis | Mirounga_leonina | 0.222364959583414 |
| Mephitis_mephitis | Mungotictis_decemlineata | 0.252045528856332 |
| Mephitis_mephitis | Mustela_altaica | 0.234129459491642 |
| Mephitis_mephitis | Mustela_erminea | 0.226589552105219 |
| Mephitis_mephitis | Mustela_eversmannii | 0.231219308096952 |
| Mephitis_mephitis | Mustela_frenata | 0.226053533291102 |
| Mephitis_mephitis | Mustela_itatsi | 0.22844251796971 |
| Mephitis_mephitis | Mustela_kathiah | 0.223873399061595 |
| Mephitis_mephitis | Mustela_nigripes | 0.231035625041005 |
| Mephitis_mephitis | Mustela_nivalis | 0.233697806285136 |
| Mephitis_mephitis | Mustela_putorius | 0.231691993258529 |
| Mephitis_mephitis | Mustela_sibirica | 0.232076588607445 |
| Mephitis_mephitis | Nandinia_binotata | 0.23307327500654 |
| Mephitis_mephitis | Nasua_nasua | 0.249071786118617 |
| Mephitis_mephitis | Neofelis_nebulosa | 0.253179192668316 |
| Mephitis_mephitis | Neomonachus_schauinslandi | 0.220671336949114 |
| Mephitis_mephitis | Neophoca_cinerea | 0.24494397079586 |
| Mephitis_mephitis | Neovison_vison | 0.222275848100591 |
| Mephitis_mephitis | Nyctereutes_procyonoides | 0.249364055032438 |
| Mephitis_mephitis | Odobenus_rosmarus_rosmarus | 0.258048464247926 |
| Mephitis_mephitis | Otocolobus_manul | 0.240649629373082 |
| Mephitis_mephitis | Otocyon_megalotis_megalotis_TS305 | 0.257287251481228 |
| Mephitis_mephitis | Otocyon_megalotis_virgatus_FMNH158128 | 0.254614833324357 |
| Mephitis_mephitis | Paguma_larvata | 0.229343766993993 |
| Mephitis_mephitis | Panthera_leo | 0.250614228460023 |

| | | |
|---|---|---|
| Mephitis_mephitis | Panthera_onca | 0.246630200262041 |
| Mephitis_mephitis | Panthera_pardus | 0.250432161997721 |
| Mephitis_mephitis | Panthera_tigris | 0.253362021470014 |
| Mephitis_mephitis | Panthera_uncia | 0.253179275648259 |
| Mephitis_mephitis | Paradoxurus_hermaphroditus | 0.237101401588393 |
| Mephitis_mephitis | Parahyaena_brunnea_FD126_Fecal | 0.248566067929209 |
| Mephitis_mephitis | Pardofelis_marmorata | 0.245829848283239 |
| Mephitis_mephitis | Phoca_fasciata | 0.223988952370076 |
| Mephitis_mephitis | Phoca_groenlandica | 0.222552095810282 |
| Mephitis_mephitis | Phoca_largha | 0.227520288222877 |
| Mephitis_mephitis | Phoca_vitulina | 0.2280586490705 |
| Mephitis_mephitis | Phocarctos_hookeri | 0.246429678169315 |
| Mephitis_mephitis | Prionailurus_bengalensis | 0.234184612572195 |
| Mephitis_mephitis | Prionailurus_bengalensis_euptilurus | 0.235536377387377 |
| Mephitis_mephitis | Prionailurus_planiceps | 0.238864858957045 |
| Mephitis_mephitis | Prionailurus_rubiginosus | 0.239255825747963 |
| Mephitis_mephitis | Prionailurus_viverrinus | 0.236623545591959 |
| Mephitis_mephitis | Prionodon_pardicolor | 0.246286011168672 |
| Mephitis_mephitis | Procyon_lotor | 0.234434036866304 |
| Mephitis_mephitis | Profelis_aurata | 0.240013091791771 |
| Mephitis_mephitis | Proteles_cristatus_cristatus_TS307 | 0.258637619043737 |
| Mephitis_mephitis | Proteles_cristatus_septentrionalis_NMS_Z_2018_54 | 0.259371276421522 |
| Mephitis_mephitis | Puma_concolor | 0.2418819580804 |
| Mephitis_mephitis | Puma_yagouaroundi | 0.237360252262247 |
| Mephitis_mephitis | Pusa_caspica | 0.227233992690941 |
| Mephitis_mephitis | Pusa_hispida | 0.222340141414722 |
| Mephitis_mephitis | Pusa_sibirica | 0.224384604110326 |
| Mephitis_mephitis | Speothos_venaticus_T1624 | 0.254191286641779 |
| Mephitis_mephitis | Spilogale_putorius | 0.164236902726097 |
| Mephitis_mephitis | Taxidea_taxus | 0.229663914997718 |
| Mephitis_mephitis | Tremarctos_ornatus | 0.243116051267186 |
| Mephitis_mephitis | Urocyon_cinereoargenteus | 0.262580657765678 |
| Mephitis_mephitis | Ursus_americanus | 0.241344976919752 |
| Mephitis_mephitis | Ursus_arctos | 0.249039963777522 |
| Mephitis_mephitis | Ursus_maritimus | 0.247362989843895 |
| Mephitis_mephitis | Ursus_spelaeus | 0.243884668972171 |
| Mephitis_mephitis | Ursus_thibetanus | 0.2453365285022 |
| Mephitis_mephitis | Urva_javanica | 0.24417420736731 |
| Mephitis_mephitis | Viverra_tangalunga_FMNH146957_SRR6053069 | 0.232825912488679 |
| Mephitis_mephitis | Viverricula_indica | 0.226685573481885 |
| Mephitis_mephitis | Vulpes_corsac | 0.247650773638243 |
| Mephitis_mephitis | Vulpes_ferrilata | 0.247885833180505 |
| Mephitis_mephitis | Vulpes_lagopus | 0.250202314553081 |
| Mephitis_mephitis | Vulpes_vulpes_T3611 | 0.249181650993481 |
| Mephitis_mephitis | Vulpes_zerda | 0.252713279046409 |
| Mephitis_mephitis | Zalophus_californianus | 0.239613043828833 |
| Mirounga_leonina | Mungotictis_decemlineata | 0.22449623059648 |
| Mirounga_leonina | Mustela_altaica | 0.218765989944991 |
| Mirounga_leonina | Mustela_erminea | 0.208743735760496 |
| Mirounga_leonina | Mustela_eversmannii | 0.22156323565885 |
| Mirounga_leonina | Mustela_frenata | 0.210972177516042 |
| Mirounga_leonina | Mustela_itatsi | 0.217981912170851 |
| Mirounga_leonina | Mustela_kathiah | 0.214716503029281 |
| Mirounga_leonina | Mustela_nigripes | 0.219286084458842 |
| Mirounga_leonina | Mustela_nivalis | 0.219571876451121 |
| Mirounga_leonina | Mustela_putorius | 0.220576512587671 |
| Mirounga_leonina | Mustela_sibirica | 0.218563731220724 |
| Mirounga_leonina | Nandinia_binotata | 0.201453163871646 |
| Mirounga_leonina | Nasua_nasua | 0.23330370370232 |
| Mirounga_leonina | Neofelis_nebulosa | 0.229436015796591 |
| Mirounga_leonina | Neomonachus_schauinslandi | 0.124812011042788 |
| Mirounga_leonina | Neophoca_cinerea | 0.182786701421994 |
| Mirounga_leonina | Neovison_vison | 0.216143051197085 |
| Mirounga_leonina | Nyctereutes_procyonoides | 0.228413320735348 |
| Mirounga_leonina | Odobenus_rosmarus_rosmarus | 0.196240539551587 |
| Mirounga_leonina | Otocolobus_manul | 0.216459890713417 |
| Mirounga_leonina | Otocyon_megalotis_megalotis_TS305 | 0.236173902442675 |
| Mirounga_leonina | Otocyon_megalotis_virgatus_FMNH158128 | 0.236123038981868 |
| Mirounga_leonina | Paguma_larvata | 0.220532682145829 |
| Mirounga_leonina | Panthera_leo | 0.227159109099124 |
| Mirounga_leonina | Panthera_onca | 0.226050551794078 |
| Mirounga_leonina | Panthera_pardus | 0.232011244074969 |
| Mirounga_leonina | Panthera_tigris | 0.231162131889467 |
| Mirounga_leonina | Panthera_uncia | 0.235369396519125 |
| Mirounga_leonina | Paradoxurus_hermaphroditus | 0.224332913767589 |
| Mirounga_leonina | Parahyaena_brunnea_FD126_Fecal | 0.223351522798318 |
| Mirounga_leonina | Pardofelis_marmorata | 0.222600367729382 |
| Mirounga_leonina | Phoca_fasciata | 0.139512919940181 |
| Mirounga_leonina | Phoca_groenlandica | 0.141324317044781 |
| Mirounga_leonina | Phoca_largha | 0.14427705361621 |
| Mirounga_leonina | Phoca_vitulina | 0.144859331219172 |
| Mirounga_leonina | Phocarctos_hookeri | 0.183295784680329 |
| Mirounga_leonina | Prionailurus_bengalensis | 0.211492430133989 |
| Mirounga_leonina | Prionailurus_bengalensis_euptilurus | 0.216317050685036 |
| Mirounga_leonina | Prionailurus_planiceps | 0.215036230260017 |
| Mirounga_leonina | Prionailurus_rubiginosus | 0.216227995558972 |
| Mirounga_leonina | Prionailurus_viverrinus | 0.214165059457408 |

| | | |
|---|---|---|
| Mirounga_leonina | Prionodon_pardicolor | 0.22570949220113 |
| Mirounga_leonina | Procyon_lotor | 0.214901631649393 |
| Mirounga_leonina | Profelis_aurata | 0.217797405566613 |
| Mirounga_leonina | Proteles_cristatus_cristatus_TS307 | 0.231041425209593 |
| Mirounga_leonina | Proteles_cristatus_septentrionalis_NMS_Z_2018_54 | 0.229929681011993 |
| Mirounga_leonina | Puma_concolor | 0.216878525289498 |
| Mirounga_leonina | Puma_yagouaroundi | 0.217502571844927 |
| Mirounga_leonina | Pusa_caspica | 0.14391107161822 |
| Mirounga_leonina | Pusa_hispida | 0.139699324369866 |
| Mirounga_leonina | Pusa_sibirica | 0.142193578413667 |
| Mirounga_leonina | Speothos_venaticus_T1624 | 0.235115665752625 |
| Mirounga_leonina | Spilogale_putorius | 0.216552999355661 |
| Mirounga_leonina | Taxidea_taxus | 0.204452267039034 |
| Mirounga_leonina | Tremarctos_ornatus | 0.213509085303861 |
| Mirounga_leonina | Urocyon_cinereoargenteus | 0.236882999182107 |
| Mirounga_leonina | Ursus_americanus | 0.216967292718526 |
| Mirounga_leonina | Ursus_arctos | 0.21951779144647 |
| Mirounga_leonina | Ursus_maritimus | 0.218540349394851 |
| Mirounga_leonina | Ursus_spelaeus | 0.215472855111093 |
| Mirounga_leonina | Ursus_thibetanus | 0.215315361997409 |
| Mirounga_leonina | Urva_javanica | 0.218765221482212 |
| Mirounga_leonina | Viverra_tangalunga_FMNH146957_SRR6053069 | 0.216880448594106 |
| Mirounga_leonina | Viverricula_indica | 0.211319135825465 |
| Mirounga_leonina | Vulpes_corsac | 0.234491970971145 |
| Mirounga_leonina | Vulpes_ferrilata | 0.235916191361932 |
| Mirounga_leonina | Vulpes_lagopus | 0.231107853761066 |
| Mirounga_leonina | Vulpes_vulpes_T3611 | 0.231938709332967 |
| Mirounga_leonina | Vulpes_zerda | 0.233187854661569 |
| Mirounga_leonina | Zalophus_californianus | 0.176517364377142 |
| Mungotictis_decemlineata | Mustela_altaica | 0.24279520446718 |
| Mungotictis_decemlineata | Mustela_erminea | 0.233976001250614 |
| Mungotictis_decemlineata | Mustela_eversmannii | 0.244429499380422 |
| Mungotictis_decemlineata | Mustela_frenata | 0.23736154635768 |
| Mungotictis_decemlineata | Mustela_itatsi | 0.242915388247142 |
| Mungotictis_decemlineata | Mustela_kathiah | 0.243290604543699 |
| Mungotictis_decemlineata | Mustela_nigripes | 0.242830300341671 |
| Mungotictis_decemlineata | Mustela_nivalis | 0.241014507538472 |
| Mungotictis_decemlineata | Mustela_putorius | 0.240927200490608 |
| Mungotictis_decemlineata | Mustela_sibirica | 0.24093770594416 |
| Mungotictis_decemlineata | Nandinia_binotata | 0.175420753247679 |
| Mungotictis_decemlineata | Nasua_nasua | 0.258855753646152 |
| Mungotictis_decemlineata | Neofelis_nebulosa | 0.200130972513066 |
| Mungotictis_decemlineata | Neomonachus_schauinslandi | 0.223420818868629 |
| Mungotictis_decemlineata | Neophoca_cinerea | 0.239960881659125 |
| Mungotictis_decemlineata | Neovison_vison | 0.241480741726502 |
| Mungotictis_decemlineata | Nyctereutes_procyonoides | 0.241286567055663 |
| Mungotictis_decemlineata | Odobenus_rosmarus_rosmarus | 0.258883994358586 |
| Mungotictis_decemlineata | Otocolobus_manul | 0.192846445399943 |
| Mungotictis_decemlineata | Otocyon_megalotis_megalotis_TS305 | 0.245581850520339 |
| Mungotictis_decemlineata | Otocyon_megalotis_virgatus_FMNH158128 | 0.245672161885614 |
| Mungotictis_decemlineata | Paguma_larvata | 0.198492512766018 |
| Mungotictis_decemlineata | Panthera_leo | 0.191191724775527 |
| Mungotictis_decemlineata | Panthera_onca | 0.198919282380916 |
| Mungotictis_decemlineata | Panthera_pardus | 0.191335003846543 |
| Mungotictis_decemlineata | Panthera_tigris | 0.195554780439929 |
| Mungotictis_decemlineata | Panthera_uncia | 0.196808570617381 |
| Mungotictis_decemlineata | Paradoxurus_hermaphroditus | 0.209019292858158 |
| Mungotictis_decemlineata | Parahyaena_brunnea_FD126_Fecal | 0.183899206563277 |
| Mungotictis_decemlineata | Pardofelis_marmorata | 0.192212597861011 |
| Mungotictis_decemlineata | Phoca_fasciata | 0.223024753573667 |
| Mungotictis_decemlineata | Phoca_groenlandica | 0.226009367574078 |
| Mungotictis_decemlineata | Phoca_largha | 0.221981951089635 |
| Mungotictis_decemlineata | Phoca_vitulina | 0.223627986401748 |
| Mungotictis_decemlineata | Phocarctos_hookeri | 0.240693942653964 |
| Mungotictis_decemlineata | Prionailurus_bengalensis | 0.183921329730539 |
| Mungotictis_decemlineata | Prionailurus_bengalensis_euptilurus | 0.182988534143091 |
| Mungotictis_decemlineata | Prionailurus_planiceps | 0.186565365331936 |
| Mungotictis_decemlineata | Prionailurus_rubiginosus | 0.183298276690416 |
| Mungotictis_decemlineata | Prionailurus_viverrinus | 0.183277298268705 |
| Mungotictis_decemlineata | Prionodon_pardicolor | 0.203500013252455 |
| Mungotictis_decemlineata | Procyon_lotor | 0.246627301844131 |
| Mungotictis_decemlineata | Profelis_aurata | 0.192994577179323 |
| Mungotictis_decemlineata | Proteles_cristatus_cristatus_TS307 | 0.189858719040325 |
| Mungotictis_decemlineata | Proteles_cristatus_septentrionalis_NMS_Z_2018_54 | 0.190403934791481 |
| Mungotictis_decemlineata | Puma_concolor | 0.191048714984422 |
| Mungotictis_decemlineata | Puma_yagouaroundi | 0.186760547449093 |
| Mungotictis_decemlineata | Pusa_caspica | 0.22078103170174 |
| Mungotictis_decemlineata | Pusa_hispida | 0.221518272198086 |
| Mungotictis_decemlineata | Pusa_sibirica | 0.219975090907052 |
| Mungotictis_decemlineata | Speothos_venaticus_T1624 | 0.255335230460535 |
| Mungotictis_decemlineata | Spilogale_putorius | 0.255253086649417 |
| Mungotictis_decemlineata | Taxidea_taxus | 0.225853978576965 |
| Mungotictis_decemlineata | Tremarctos_ornatus | 0.239715553150352 |
| Mungotictis_decemlineata | Urocyon_cinereoargenteus | 0.247747972506334 |
| Mungotictis_decemlineata | Ursus_americanus | 0.240205416008257 |
| Mungotictis_decemlineata | Ursus_arctos | 0.244729894768598 |
| Mungotictis_decemlineata | Ursus_maritimus | 0.2465115144269 |

| | | |
|---|---|---|
| Mungotictis_decemlineata | Ursus_spelaeus | 0.243948264675459 |
| Mungotictis_decemlineata | Ursus_thibetanus | 0.236485888430268 |
| Mungotictis_decemlineata | Urva_javanica | 0.178927759144224 |
| Mungotictis_decemlineata | Viverra_tangalunga_FMNH146957_SRR6053069 | 0.185910113776396 |
| Mungotictis_decemlineata | Viverricula_indica | 0.187627813478082 |
| Mungotictis_decemlineata | Vulpes_corsac | 0.237948216779882 |
| Mungotictis_decemlineata | Vulpes_ferrilata | 0.239310420932103 |
| Mungotictis_decemlineata | Vulpes_lagopus | 0.239213595436287 |
| Mungotictis_decemlineata | Vulpes_vulpes_T3611 | 0.240799252032033 |
| Mungotictis_decemlineata | Vulpes_zerda | 0.239213595436287 |
| Mungotictis_decemlineata | Zalophus_californianus | 0.238303869398192 |
| Mustela_altaica | Mustela_erminea | 0.0922012271738949 |
| Mustela_altaica | Mustela_eversmannii | 0.0918134822266983 |
| Mustela_altaica | Mustela_frenata | 0.133815092651245 |
| Mustela_altaica | Mustela_itatsi | 0.0917956049493742 |
| Mustela_altaica | Mustela_kathiah | 0.130258377871536 |
| Mustela_altaica | Mustela_nigripes | 0.0895213950732904 |
| Mustela_altaica | Mustela_nivalis | 0.0746124939234553 |
| Mustela_altaica | Mustela_putorius | 0.0897449601348745 |
| Mustela_altaica | Mustela_sibirica | 0.0892945617447271 |
| Mustela_altaica | Nandinia_binotata | 0.226572501011784 |
| Mustela_altaica | Nasua_nasua | 0.234103149846342 |
| Mustela_altaica | Neofelis_nebulosa | 0.245920460843569 |
| Mustela_altaica | Neomonachus_schauinslandi | 0.219300596513063 |
| Mustela_altaica | Neophoca_cinerea | 0.234311764783771 |
| Mustela_altaica | Neovison_vison | 0.141078017815705 |
| Mustela_altaica | Nyctereutes_procyonoides | 0.247279774961058 |
| Mustela_altaica | Odobenus_rosmarus_rosmarus | 0.251998068506367 |
| Mustela_altaica | Otocolobus_manul | 0.232197677001467 |
| Mustela_altaica | Otocyon_megalotis_megalotis_TS305 | 0.249654819124848 |
| Mustela_altaica | Otocyon_megalotis_virgatus_FMNH158128 | 0.253100513249059 |
| Mustela_altaica | Paguma_larvata | 0.237256153377939 |
| Mustela_altaica | Panthera_leo | 0.241944113130195 |
| Mustela_altaica | Panthera_onca | 0.243460245236949 |
| Mustela_altaica | Panthera_pardus | 0.244569248914118 |
| Mustela_altaica | Panthera_tigris | 0.252049219280573 |
| Mustela_altaica | Panthera_uncia | 0.246212946233481 |
| Mustela_altaica | Paradoxurus_hermaphroditus | 0.240457190855051 |
| Mustela_altaica | Parahyaena_brunnea_FD126_Fecal | 0.245665734698484 |
| Mustela_altaica | Pardofelis_marmorata | 0.236973786592109 |
| Mustela_altaica | Phoca_fasciata | 0.219238012141773 |
| Mustela_altaica | Phoca_groenlandica | 0.220659274187837 |
| Mustela_altaica | Phoca_largha | 0.218786144188498 |
| Mustela_altaica | Phoca_vitulina | 0.218558780782138 |
| Mustela_altaica | Phocarctos_hookeri | 0.23893129938305 |
| Mustela_altaica | Prionailurus_bengalensis | 0.230105541997732 |
| Mustela_altaica | Prionailurus_bengalensis_euptilurus | 0.229887105996005 |
| Mustela_altaica | Prionailurus_planiceps | 0.23200535835947 |
| Mustela_altaica | Prionailurus_rubiginosus | 0.229499466009132 |
| Mustela_altaica | Prionailurus_viverrinus | 0.233663977506906 |
| Mustela_altaica | Prionodon_pardicolor | 0.236842146856421 |
| Mustela_altaica | Procyon_lotor | 0.217440371102742 |
| Mustela_altaica | Profelis_aurata | 0.239718266652656 |
| Mustela_altaica | Proteles_cristatus_cristatus_TS307 | 0.250482491622203 |
| Mustela_altaica | Proteles_cristatus_septentrionalis_NMS_Z_2018_54 | 0.248317961608824 |
| Mustela_altaica | Puma_concolor | 0.229331943624742 |
| Mustela_altaica | Puma_yagouaroundi | 0.230565863801385 |
| Mustela_altaica | Pusa_caspica | 0.219786937570559 |
| Mustela_altaica | Pusa_hispida | 0.217748672412855 |
| Mustela_altaica | Pusa_sibirica | 0.223121013546512 |
| Mustela_altaica | Speothos_venaticus_T1624 | 0.244033365396489 |
| Mustela_altaica | Spilogale_putorius | 0.236634932589634 |
| Mustela_altaica | Taxidea_taxus | 0.177041333056452 |
| Mustela_altaica | Tremarctos_ornatus | 0.245215865995933 |
| Mustela_altaica | Urocyon_cinereoargenteus | 0.252443978039563 |
| Mustela_altaica | Ursus_americanus | 0.24030737353032 |
| Mustela_altaica | Ursus_arctos | 0.247041654290435 |
| Mustela_altaica | Ursus_maritimus | 0.246050167592008 |
| Mustela_altaica | Ursus_spelaeus | 0.241537514367303 |
| Mustela_altaica | Ursus_thibetanus | 0.237616871827031 |
| Mustela_altaica | Urva_javanica | 0.242825337235467 |
| Mustela_altaica | Viverra_tangalunga_FMNH146957_SRR6053069 | 0.231545026756365 |
| Mustela_altaica | Viverricula_indica | 0.232958981767899 |
| Mustela_altaica | Vulpes_corsac | 0.247752656002348 |
| Mustela_altaica | Vulpes_ferrilata | 0.246467759408231 |
| Mustela_altaica | Vulpes_lagopus | 0.252499249395634 |
| Mustela_altaica | Vulpes_vulpes_T3611 | 0.246184325580539 |
| Mustela_altaica | Vulpes_zerda | 0.245147502710336 |
| Mustela_altaica | Zalophus_californianus | 0.23392426464586 |
| Mustela_erminea | Mustela_eversmannii | 0.0979275666830796 |
| Mustela_erminea | Mustela_frenata | 0.122908436071702 |
| Mustela_erminea | Mustela_itatsi | 0.0939444129840234 |
| Mustela_erminea | Mustela_kathiah | 0.119213473077592 |
| Mustela_erminea | Mustela_nigripes | 0.0956021281596778 |
| Mustela_erminea | Mustela_nivalis | 0.0964313939488136 |
| Mustela_erminea | Mustela_putorius | 0.0971075987061482 |
| Mustela_erminea | Mustela_sibirica | 0.0983932460851375 |

| | | |
|---|---|---|
| Mustela_erminea | Nandinia_binotata | 0.21506796262525 |
| Mustela_erminea | Nasua_nasua | 0.22674799016659 |
| Mustela_erminea | Neofelis_nebulosa | 0.236015144100991 |
| Mustela_erminea | Neomonachus_schauinslandi | 0.20968964218941 |
| Mustela_erminea | Neophoca_cinerea | 0.229429203466137 |
| Mustela_erminea | Neovison_vison | 0.128671080404718 |
| Mustela_erminea | Nyctereutes_procyonoides | 0.235306589709753 |
| Mustela_erminea | Odobenus_rosmarus_rosmarus | 0.242457689830305 |
| Mustela_erminea | Otocolobus_manul | 0.228949146282167 |
| Mustela_erminea | Otocyon_megalotis_megalotis_TS305 | 0.247183375402813 |
| Mustela_erminea | Otocyon_megalotis_virgatus_FMNH158128 | 0.245689706429364 |
| Mustela_erminea | Paguma_larvata | 0.222773783009827 |
| Mustela_erminea | Panthera_leo | 0.231856655600128 |
| Mustela_erminea | Panthera_onca | 0.230757917780556 |
| Mustela_erminea | Panthera_pardus | 0.233246739806448 |
| Mustela_erminea | Panthera_tigris | 0.23594146524792 |
| Mustela_erminea | Panthera_uncia | 0.234550230307444 |
| Mustela_erminea | Paradoxurus_hermaphroditus | 0.230072502991933 |
| Mustela_erminea | Parahyaena_brunnea_FD126_Fecal | 0.233974687215978 |
| Mustela_erminea | Pardofelis_marmorata | 0.227519079381785 |
| Mustela_erminea | Phoca_fasciata | 0.210060776381566 |
| Mustela_erminea | Phoca_groenlandica | 0.208263638806126 |
| Mustela_erminea | Phoca_largha | 0.207296197847592 |
| Mustela_erminea | Phoca_vitulina | 0.208435007340633 |
| Mustela_erminea | Phocarctos_hookeri | 0.228890421729191 |
| Mustela_erminea | Prionailurus_bengalensis | 0.223072677668705 |
| Mustela_erminea | Prionailurus_bengalensis_euptilurus | 0.222622806736001 |
| Mustela_erminea | Prionailurus_planiceps | 0.225038644314945 |
| Mustela_erminea | Prionailurus_rubiginosus | 0.223467205206844 |
| Mustela_erminea | Prionailurus_viverrinus | 0.223018009146725 |
| Mustela_erminea | Prionodon_pardicolor | 0.233495005270161 |
| Mustela_erminea | Procyon_lotor | 0.207716246557932 |
| Mustela_erminea | Profelis_aurata | 0.234576247207734 |
| Mustela_erminea | Proteles_cristatus_cristatus_TS307 | 0.236153617098782 |
| Mustela_erminea | Proteles_cristatus_septentrionalis_NMS_Z_2018_54 | 0.238341337242812 |
| Mustela_erminea | Puma_concolor | 0.223261193755147 |
| Mustela_erminea | Puma_yagouaroundi | 0.225044475128507 |
| Mustela_erminea | Pusa_caspica | 0.207234256047395 |
| Mustela_erminea | Pusa_hispida | 0.204483217470404 |
| Mustela_erminea | Pusa_sibirica | 0.208307376442179 |
| Mustela_erminea | Speothos_venaticus_T1624 | 0.241185396398514 |
| Mustela_erminea | Spilogale_putorius | 0.225122565999523 |
| Mustela_erminea | Taxidea_taxus | 0.165962587485045 |
| Mustela_erminea | Tremarctos_ornatus | 0.231569109549837 |
| Mustela_erminea | Urocyon_cinereoargenteus | 0.250745606841384 |
| Mustela_erminea | Ursus_americanus | 0.231457477488645 |
| Mustela_erminea | Ursus_arctos | 0.235836827060177 |
| Mustela_erminea | Ursus_maritimus | 0.233071073057862 |
| Mustela_erminea | Ursus_spelaeus | 0.228356440380603 |
| Mustela_erminea | Ursus_thibetanus | 0.229205711055641 |
| Mustela_erminea | Urva_javanica | 0.23412753290654 |
| Mustela_erminea | Viverra_tangalunga_FMNH146957_SRR6053069 | 0.2189473987946 |
| Mustela_erminea | Viverricula_indica | 0.225286498892091 |
| Mustela_erminea | Vulpes_corsac | 0.239319693977437 |
| Mustela_erminea | Vulpes_ferrilata | 0.242175362700806 |
| Mustela_erminea | Vulpes_lagopus | 0.24684426629779 |
| Mustela_erminea | Vulpes_vulpes_T3611 | 0.235966730524192 |
| Mustela_erminea | Vulpes_zerda | 0.236043497067503 |
| Mustela_erminea | Zalophus_californianus | 0.222906465131684 |
| Mustela_eversmannii | Mustela_frenata | 0.135328843352096 |
| Mustela_eversmannii | Mustela_itatsi | 0.0538704947490548 |
| Mustela_eversmannii | Mustela_kathiah | 0.128384027870531 |
| Mustela_eversmannii | Mustela_nigripes | 0.0197213706670916 |
| Mustela_eversmannii | Mustela_nivalis | 0.0948151952469172 |
| Mustela_eversmannii | Mustela_putorius | 0.0126613509584173 |
| Mustela_eversmannii | Mustela_sibirica | 0.0340598911027018 |
| Mustela_eversmannii | Nandinia_binotata | 0.220535854505886 |
| Mustela_eversmannii | Nasua_nasua | 0.238749179310997 |
| Mustela_eversmannii | Neofelis_nebulosa | 0.244197681172562 |
| Mustela_eversmannii | Neomonachus_schauinslandi | 0.218797252132554 |
| Mustela_eversmannii | Neophoca_cinerea | 0.240190661155948 |
| Mustela_eversmannii | Neovison_vison | 0.138405969402341 |
| Mustela_eversmannii | Nyctereutes_procyonoides | 0.242806302013708 |
| Mustela_eversmannii | Odobenus_rosmarus_rosmarus | 0.251728931889367 |
| Mustela_eversmannii | Otocolobus_manul | 0.227608584935543 |
| Mustela_eversmannii | Otocyon_megalotis_megalotis_TS305 | 0.248698429607515 |
| Mustela_eversmannii | Otocyon_megalotis_virgatus_FMNH158128 | 0.249345455108966 |
| Mustela_eversmannii | Paguma_larvata | 0.228021816236026 |
| Mustela_eversmannii | Panthera_leo | 0.237625722263573 |
| Mustela_eversmannii | Panthera_onca | 0.235186557311264 |
| Mustela_eversmannii | Panthera_pardus | 0.239653738760037 |
| Mustela_eversmannii | Panthera_tigris | 0.243725814380815 |
| Mustela_eversmannii | Panthera_uncia | 0.239452374458345 |
| Mustela_eversmannii | Paradoxurus_hermaphroditus | 0.236219568305473 |
| Mustela_eversmannii | Parahyaena_brunnea_FD126_Fecal | 0.240407880129051 |
| Mustela_eversmannii | Pardofelis_marmorata | 0.232983667967562 |
| Mustela_eversmannii | Phoca_fasciata | 0.219037091877337 |

| | | |
|---|---|---|
| Mustela_eversmannii | Phoca_groenlandica | 0.219392746725305 |
| Mustela_eversmannii | Phoca_largha | 0.218284201352152 |
| Mustela_eversmannii | Phoca_vitulina | 0.218273928668405 |
| Mustela_eversmannii | Phocarctos_hookeri | 0.241185264536157 |
| Mustela_eversmannii | Prionailurus_bengalensis | 0.227740450239768 |
| Mustela_eversmannii | Prionailurus_bengalensis_euptilurus | 0.227087637613032 |
| Mustela_eversmannii | Prionailurus_planiceps | 0.226690608835209 |
| Mustela_eversmannii | Prionailurus_rubiginosus | 0.227028407541548 |
| Mustela_eversmannii | Prionailurus_viverrinus | 0.229864391054186 |
| Mustela_eversmannii | Prionodon_pardicolor | 0.237615990854038 |
| Mustela_eversmannii | Procyon_lotor | 0.217864425943386 |
| Mustela_eversmannii | Profelis_aurata | 0.24066413485337 |
| Mustela_eversmannii | Proteles_cristatus_cristatus_TS307 | 0.247604895547815 |
| Mustela_eversmannii | Proteles_cristatus_septentrionalis_NMS_Z_2018_54 | 0.247851187364077 |
| Mustela_eversmannii | Puma_concolor | 0.231206952476938 |
| Mustela_eversmannii | Puma_yagouaroundi | 0.227981770736782 |
| Mustela_eversmannii | Pusa_caspica | 0.217197839115439 |
| Mustela_eversmannii | Pusa_hispida | 0.216226441076492 |
| Mustela_eversmannii | Pusa_sibirica | 0.218342049092572 |
| Mustela_eversmannii | Speothos_venaticus_T1624 | 0.246174902524645 |
| Mustela_eversmannii | Spilogale_putorius | 0.236556726222646 |
| Mustela_eversmannii | Taxidea_taxus | 0.175052196362878 |
| Mustela_eversmannii | Tremarctos_ornatus | 0.240487521946095 |
| Mustela_eversmannii | Urocyon_cinereoargenteus | 0.249670227477034 |
| Mustela_eversmannii | Ursus_americanus | 0.236486344717187 |
| Mustela_eversmannii | Ursus_arctos | 0.238590662359965 |
| Mustela_eversmannii | Ursus_maritimus | 0.236373795476419 |
| Mustela_eversmannii | Ursus_spelaeus | 0.233531147684518 |
| Mustela_eversmannii | Ursus_thibetanus | 0.232450525574512 |
| Mustela_eversmannii | Urva_javanica | 0.24174895988108 |
| Mustela_eversmannii | Viverra_tangalunga_FMNH146957_SRR6053069 | 0.233893816843607 |
| Mustela_eversmannii | Viverricula_indica | 0.234565200515085 |
| Mustela_eversmannii | Vulpes_corsac | 0.244426726840819 |
| Mustela_eversmannii | Vulpes_ferrilata | 0.245423526289323 |
| Mustela_eversmannii | Vulpes_lagopus | 0.249403430456943 |
| Mustela_eversmannii | Vulpes_vulpes_T3611 | 0.241932704575869 |
| Mustela_eversmannii | Vulpes_zerda | 0.242704402686467 |
| Mustela_eversmannii | Zalophus_californianus | 0.234578455181539 |
| Mustela_frenata | Mustela_itatsi | 0.130702082094666 |
| Mustela_frenata | Mustela_kathiah | 0.131242624804744 |
| Mustela_frenata | Mustela_nigripes | 0.134483254261101 |
| Mustela_frenata | Mustela_nivalis | 0.132062251987677 |
| Mustela_frenata | Mustela_putorius | 0.136091490644555 |
| Mustela_frenata | Mustela_sibirica | 0.133998837849968 |
| Mustela_frenata | Nandinia_binotata | 0.215800690757834 |
| Mustela_frenata | Nasua_nasua | 0.230505248596335 |
| Mustela_frenata | Neofelis_nebulosa | 0.237779298430234 |
| Mustela_frenata | Neomonachus_schauinslandi | 0.211718037839114 |
| Mustela_frenata | Neophoca_cinerea | 0.229549762484077 |
| Mustela_frenata | Neovison_vison | 0.0980533383432167 |
| Mustela_frenata | Nyctereutes_procyonoides | 0.237274083970001 |
| Mustela_frenata | Odobenus_rosmarus_rosmarus | 0.243484389576814 |
| Mustela_frenata | Otocolobus_manul | 0.226421429805167 |
| Mustela_frenata | Otocyon_megalotis_megalotis_TS305 | 0.244274357218255 |
| Mustela_frenata | Otocyon_megalotis_virgatus_FMNH158128 | 0.244560106150458 |
| Mustela_frenata | Paguma_larvata | 0.229235629382593 |
| Mustela_frenata | Panthera_leo | 0.230939065953482 |
| Mustela_frenata | Panthera_onca | 0.233467222667478 |
| Mustela_frenata | Panthera_pardus | 0.232747338794275 |
| Mustela_frenata | Panthera_tigris | 0.237570476513814 |
| Mustela_frenata | Panthera_uncia | 0.233408714552084 |
| Mustela_frenata | Paradoxurus_hermaphroditus | 0.232434026441861 |
| Mustela_frenata | Parahyaena_brunnea_FD126_Fecal | 0.235797488098323 |
| Mustela_frenata | Pardofelis_marmorata | 0.234344497705408 |
| Mustela_frenata | Phoca_fasciata | 0.212642501470157 |
| Mustela_frenata | Phoca_groenlandica | 0.212632625857825 |
| Mustela_frenata | Phoca_largha | 0.212216052091676 |
| Mustela_frenata | Phoca_vitulina | 0.211437052691212 |
| Mustela_frenata | Phocarctos_hookeri | 0.23127986480217 |
| Mustela_frenata | Prionailurus_bengalensis | 0.22353606723662 |
| Mustela_frenata | Prionailurus_bengalensis_euptilurus | 0.22404509458765 |
| Mustela_frenata | Prionailurus_planiceps | 0.226930912418401 |
| Mustela_frenata | Prionailurus_rubiginosus | 0.226043868264576 |
| Mustela_frenata | Prionailurus_viverrinus | 0.22603119066143 |
| Mustela_frenata | Prionodon_pardicolor | 0.235482662776719 |
| Mustela_frenata | Procyon_lotor | 0.216690632953 |
| Mustela_frenata | Profelis_aurata | 0.231622898673574 |
| Mustela_frenata | Proteles_cristatus_cristatus_TS307 | 0.242670441016694 |
| Mustela_frenata | Proteles_cristatus_septentrionalis_NMS_Z_2018_54 | 0.244505433362488 |
| Mustela_frenata | Puma_concolor | 0.224214230926874 |
| Mustela_frenata | Puma_yagouaroundi | 0.22888690073434 |
| Mustela_frenata | Pusa_caspica | 0.212960352238737 |
| Mustela_frenata | Pusa_hispida | 0.209982217723754 |
| Mustela_frenata | Pusa_sibirica | 0.213436430806429 |
| Mustela_frenata | Speothos_venaticus_T1624 | 0.243380561332295 |
| Mustela_frenata | Spilogale_putorius | 0.233380174865927 |
| Mustela_frenata | Taxidea_taxus | 0.173924887767903 |

| | | |
|---|---|---|
| Mustela_frenata | Tremarctos_ornatus | 0.233422022918838 |
| Mustela_frenata | Urocyon_cinereoargenteus | 0.251976134810041 |
| Mustela_frenata | Ursus_americanus | 0.235591662223548 |
| Mustela_frenata | Ursus_arctos | 0.242669380449831 |
| Mustela_frenata | Ursus_maritimus | 0.241316479294164 |
| Mustela_frenata | Ursus_spelaeus | 0.234503483036844 |
| Mustela_frenata | Ursus_thibetanus | 0.235254166429432 |
| Mustela_frenata | Urva_javanica | 0.233862287748465 |
| Mustela_frenata | Viverra_tangalunga_FMNH146957_SRR6053069 | 0.222769839464089 |
| Mustela_frenata | Viverricula_indica | 0.225802039618952 |
| Mustela_frenata | Vulpes_corsac | 0.238168994163154 |
| Mustela_frenata | Vulpes_ferrilata | 0.241219547141463 |
| Mustela_frenata | Vulpes_lagopus | 0.243815659414059 |
| Mustela_frenata | Vulpes_vulpes_T3611 | 0.238260231171597 |
| Mustela_frenata | Vulpes_zerda | 0.238564206091835 |
| Mustela_frenata | Zalophus_californianus | 0.229780394785583 |
| Mustela_itatsi | Mustela_kathiah | 0.126469880015898 |
| Mustela_itatsi | Mustela_nigripes | 0.0530809323229164 |
| Mustela_itatsi | Mustela_nivalis | 0.0940402969356443 |
| Mustela_itatsi | Mustela_putorius | 0.05353305681388 |
| Mustela_itatsi | Mustela_sibirica | 0.05461505768165 |
| Mustela_itatsi | Nandinia_binotata | 0.22198037996625 |
| Mustela_itatsi | Nasua_nasua | 0.234639461884989 |
| Mustela_itatsi | Neofelis_nebulosa | 0.242416517803904 |
| Mustela_itatsi | Neomonachus_schauinslandi | 0.216269322764597 |
| Mustela_itatsi | Neophoca_cinerea | 0.238097552828681 |
| Mustela_itatsi | Neovison_vison | 0.136078185277991 |
| Mustela_itatsi | Nyctereutes_procyonoides | 0.244593987126093 |
| Mustela_itatsi | Odobenus_rosmarus_rosmarus | 0.251601660622477 |
| Mustela_itatsi | Otocolobus_manul | 0.229649729025854 |
| Mustela_itatsi | Otocyon_megalotis_megalotis_TS305 | 0.248977694543311 |
| Mustela_itatsi | Otocyon_megalotis_virgatus_FMNH158128 | 0.248497140658188 |
| Mustela_itatsi | Paguma_larvata | 0.229316861984527 |
| Mustela_itatsi | Panthera_leo | 0.238085920236657 |
| Mustela_itatsi | Panthera_onca | 0.235095538885674 |
| Mustela_itatsi | Panthera_pardus | 0.240226138446985 |
| Mustela_itatsi | Panthera_tigris | 0.247345499662782 |
| Mustela_itatsi | Panthera_uncia | 0.241944113130195 |
| Mustela_itatsi | Paradoxurus_hermaphroditus | 0.236430342608588 |
| Mustela_itatsi | Parahyaena_brunnea_FD126_Fecal | 0.243208178365792 |
| Mustela_itatsi | Pardofelis_marmorata | 0.238414869509648 |
| Mustela_itatsi | Phoca_fasciata | 0.220001850485774 |
| Mustela_itatsi | Phoca_groenlandica | 0.218732891885353 |
| Mustela_itatsi | Phoca_largha | 0.217065261742797 |
| Mustela_itatsi | Phoca_vitulina | 0.217250740703124 |
| Mustela_itatsi | Phocarctos_hookeri | 0.237972349143673 |
| Mustela_itatsi | Prionailurus_bengalensis | 0.228872301504029 |
| Mustela_itatsi | Prionailurus_bengalensis_euptilurus | 0.228763262140386 |
| Mustela_itatsi | Prionailurus_planiceps | 0.230440333117469 |
| Mustela_itatsi | Prionailurus_rubiginosus | 0.22826752576207 |
| Mustela_itatsi | Prionailurus_viverrinus | 0.232314834292046 |
| Mustela_itatsi | Prionodon_pardicolor | 0.235687967126182 |
| Mustela_itatsi | Procyon_lotor | 0.219379410497585 |
| Mustela_itatsi | Profelis_aurata | 0.238690525099017 |
| Mustela_itatsi | Proteles_cristatus_cristatus_TS307 | 0.246113863789652 |
| Mustela_itatsi | Proteles_cristatus_septentrionalis_NMS_Z_2018_54 | 0.24759427018937 |
| Mustela_itatsi | Puma_concolor | 0.230411078184966 |
| Mustela_itatsi | Puma_yagouaroundi | 0.229222892370221 |
| Mustela_itatsi | Pusa_caspica | 0.217167041926986 |
| Mustela_itatsi | Pusa_hispida | 0.212544347206178 |
| Mustela_itatsi | Pusa_sibirica | 0.216472818012923 |
| Mustela_itatsi | Speothos_venaticus_T1624 | 0.244642746290422 |
| Mustela_itatsi | Spilogale_putorius | 0.236100411227879 |
| Mustela_itatsi | Taxidea_taxus | 0.174224306082522 |
| Mustela_itatsi | Tremarctos_ornatus | 0.240254364222382 |
| Mustela_itatsi | Urocyon_cinereoargenteus | 0.25078902483088 |
| Mustela_itatsi | Ursus_americanus | 0.239844121555545 |
| Mustela_itatsi | Ursus_arctos | 0.24321195833598 |
| Mustela_itatsi | Ursus_maritimus | 0.23848742202332 |
| Mustela_itatsi | Ursus_spelaeus | 0.237748862515498 |
| Mustela_itatsi | Ursus_thibetanus | 0.232984124096026 |
| Mustela_itatsi | Urva_javanica | 0.238534920838034 |
| Mustela_itatsi | Viverra_tangalunga_FMNH146957_SRR6053069 | 0.230262931213503 |
| Mustela_itatsi | Viverricula_indica | 0.228469593324852 |
| Mustela_itatsi | Vulpes_corsac | 0.244884581693837 |
| Mustela_itatsi | Vulpes_ferrilata | 0.244874154163741 |
| Mustela_itatsi | Vulpes_lagopus | 0.250816543643285 |
| Mustela_itatsi | Vulpes_vulpes_T3611 | 0.243260021558484 |
| Mustela_itatsi | Vulpes_zerda | 0.243273161968221 |
| Mustela_itatsi | Zalophus_californianus | 0.233592001924585 |
| Mustela_kathiah | Mustela_nigripes | 0.127719460955734 |
| Mustela_kathiah | Mustela_nivalis | 0.130345620976934 |
| Mustela_kathiah | Mustela_putorius | 0.126486479872325 |
| Mustela_kathiah | Mustela_sibirica | 0.128861628441758 |
| Mustela_kathiah | Nandinia_binotata | 0.219217235434502 |
| Mustela_kathiah | Nasua_nasua | 0.23458856121169 |
| Mustela_kathiah | Neofelis_nebulosa | 0.238906428182689 |

| | | |
|---|---|---|
| Mustela_kathiah | Neomonachus_schauinslandi | 0.21895573319408 |
| Mustela_kathiah | Neophoca_cinerea | 0.239819359543356 |
| Mustela_kathiah | Neovison_vison | 0.134642123120209 |
| Mustela_kathiah | Nyctereutes_procyonoides | 0.242786761187697 |
| Mustela_kathiah | Odobenus_rosmarus_rosmarus | 0.255036116483248 |
| Mustela_kathiah | Otocolobus_manul | 0.227532251601016 |
| Mustela_kathiah | Otocyon_megalotis_megalotis_TS305 | 0.251172198699273 |
| Mustela_kathiah | Otocyon_megalotis_virgatus_FMNH158128 | 0.251782878303545 |
| Mustela_kathiah | Paguma_larvata | 0.229263166769255 |
| Mustela_kathiah | Panthera_leo | 0.232084952161936 |
| Mustela_kathiah | Panthera_onca | 0.230678868869488 |
| Mustela_kathiah | Panthera_pardus | 0.23442621403699 |
| Mustela_kathiah | Panthera_tigris | 0.235566673265863 |
| Mustela_kathiah | Panthera_uncia | 0.235372043307241 |
| Mustela_kathiah | Paradoxurus_hermaphroditus | 0.233807907006583 |
| Mustela_kathiah | Parahyaena_brunnea_FD126_Fecal | 0.243245658050171 |
| Mustela_kathiah | Pardofelis_marmorata | 0.235659043743298 |
| Mustela_kathiah | Phoca_fasciata | 0.213654040128593 |
| Mustela_kathiah | Phoca_groenlandica | 0.21344948841384 |
| Mustela_kathiah | Phoca_largha | 0.2188564605172 |
| Mustela_kathiah | Phoca_vitulina | 0.220343824860694 |
| Mustela_kathiah | Phocarctos_hookeri | 0.23725819951711 |
| Mustela_kathiah | Prionailurus_bengalensis | 0.224043287960859 |
| Mustela_kathiah | Prionailurus_bengalensis_euptilurus | 0.223647565793805 |
| Mustela_kathiah | Prionailurus_planiceps | 0.226086024044935 |
| Mustela_kathiah | Prionailurus_rubiginosus | 0.226640395387719 |
| Mustela_kathiah | Prionailurus_viverrinus | 0.22916748689699 |
| Mustela_kathiah | Prionodon_pardicolor | 0.23146043647111 |
| Mustela_kathiah | Procyon_lotor | 0.214405037373198 |
| Mustela_kathiah | Profelis_aurata | 0.233387974055413 |
| Mustela_kathiah | Proteles_cristatus_cristatus_TS307 | 0.247380753176979 |
| Mustela_kathiah | Proteles_cristatus_septentrionalis_NMS_Z_2018_54 | 0.246052728108552 |
| Mustela_kathiah | Puma_concolor | 0.229633717398208 |
| Mustela_kathiah | Puma_yagouaroundi | 0.230888085115722 |
| Mustela_kathiah | Pusa_caspica | 0.218503050597929 |
| Mustela_kathiah | Pusa_hispida | 0.216426560957339 |
| Mustela_kathiah | Pusa_sibirica | 0.218632663931778 |
| Mustela_kathiah | Speothos_venaticus_T1624 | 0.243163021246943 |
| Mustela_kathiah | Spilogale_putorius | 0.228554306740885 |
| Mustela_kathiah | Taxidea_taxus | 0.178309013741936 |
| Mustela_kathiah | Tremarctos_ornatus | 0.235701160671253 |
| Mustela_kathiah | Urocyon_cinereoargenteus | 0.253100513249059 |
| Mustela_kathiah | Ursus_americanus | 0.238590662359965 |
| Mustela_kathiah | Ursus_arctos | 0.238916806425566 |
| Mustela_kathiah | Ursus_maritimus | 0.238477668009202 |
| Mustela_kathiah | Ursus_spelaeus | 0.236361054032099 |
| Mustela_kathiah | Ursus_thibetanus | 0.234965363394525 |
| Mustela_kathiah | Urva_javanica | 0.23830039633582 |
| Mustela_kathiah | Viverra_tangalunga_FMNH146957_SRR6053069 | 0.225874419128692 |
| Mustela_kathiah | Viverricula_indica | 0.227782976454417 |
| Mustela_kathiah | Vulpes_corsac | 0.243109278444242 |
| Mustela_kathiah | Vulpes_ferrilata | 0.241245536243842 |
| Mustela_kathiah | Vulpes_lagopus | 0.243078153768341 |
| Mustela_kathiah | Vulpes_vulpes_T3611 | 0.24218609287159 |
| Mustela_kathiah | Vulpes_zerda | 0.244594510585905 |
| Mustela_kathiah | Zalophus_californianus | 0.236033936245471 |
| Mustela_nigripes | Mustela_nivalis | 0.0928507793669754 |
| Mustela_nigripes | Mustela_putorius | 0.0182354673131782 |
| Mustela_nigripes | Mustela_sibirica | 0.0333119226030871 |
| Mustela_nigripes | Nandinia_binotata | 0.217770452931367 |
| Mustela_nigripes | Nasua_nasua | 0.235735255259572 |
| Mustela_nigripes | Neofelis_nebulosa | 0.241767179262704 |
| Mustela_nigripes | Neomonachus_schauinslandi | 0.216015826979111 |
| Mustela_nigripes | Neophoca_cinerea | 0.238229714622651 |
| Mustela_nigripes | Neovison_vison | 0.138733092898919 |
| Mustela_nigripes | Nyctereutes_procyonoides | 0.240638598246543 |
| Mustela_nigripes | Odobenus_rosmarus_rosmarus | 0.249355250696481 |
| Mustela_nigripes | Otocolobus_manul | 0.226151717754574 |
| Mustela_nigripes | Otocyon_megalotis_megalotis_TS305 | 0.248553736801044 |
| Mustela_nigripes | Otocyon_megalotis_virgatus_FMNH158128 | 0.248296857910169 |
| Mustela_nigripes | Paguma_larvata | 0.230096350081477 |
| Mustela_nigripes | Panthera_leo | 0.235669319325242 |
| Mustela_nigripes | Panthera_onca | 0.233463863787837 |
| Mustela_nigripes | Panthera_pardus | 0.237357621652442 |
| Mustela_nigripes | Panthera_tigris | 0.243646709469831 |
| Mustela_nigripes | Panthera_uncia | 0.236271082305438 |
| Mustela_nigripes | Paradoxurus_hermaphroditus | 0.236001238234704 |
| Mustela_nigripes | Parahyaena_brunnea_FD126_Fecal | 0.239010846826712 |
| Mustela_nigripes | Pardofelis_marmorata | 0.229977204204026 |
| Mustela_nigripes | Phoca_fasciata | 0.217502194210182 |
| Mustela_nigripes | Phoca_groenlandica | 0.217012935282934 |
| Mustela_nigripes | Phoca_largha | 0.217072613908401 |
| Mustela_nigripes | Phoca_vitulina | 0.216974814213255 |
| Mustela_nigripes | Phocarctos_hookeri | 0.24020717372786 |
| Mustela_nigripes | Prionailurus_bengalensis | 0.225629838336155 |
| Mustela_nigripes | Prionailurus_bengalensis_euptilurus | 0.224763662285841 |
| Mustela_nigripes | Prionailurus_planiceps | 0.226426308069655 |

| | | |
|---|---|---|
| Mustela_nigripes | Prionailurus_rubiginosus | 0.226657209450637 |
| Mustela_nigripes | Prionailurus_viverrinus | 0.227419172023691 |
| Mustela_nigripes | Prionodon_pardicolor | 0.234897399178928 |
| Mustela_nigripes | Procyon_lotor | 0.216482422908615 |
| Mustela_nigripes | Profelis_aurata | 0.238600782658854 |
| Mustela_nigripes | Proteles_cristatus_cristatus_TS307 | 0.243228407904641 |
| Mustela_nigripes | Proteles_cristatus_septentrionalis_NMS_Z_2018_54 | 0.244120866865368 |
| Mustela_nigripes | Puma_concolor | 0.228535747928607 |
| Mustela_nigripes | Puma_yagouaroundi | 0.226631173677289 |
| Mustela_nigripes | Pusa_caspica | 0.216526120004123 |
| Mustela_nigripes | Pusa_hispida | 0.21534205055948 |
| Mustela_nigripes | Pusa_sibirica | 0.217126167885262 |
| Mustela_nigripes | Speothos_venaticus_T1624 | 0.245698734153736 |
| Mustela_nigripes | Spilogale_putorius | 0.234369302591601 |
| Mustela_nigripes | Taxidea_taxus | 0.173087248046144 |
| Mustela_nigripes | Tremarctos_ornatus | 0.239547902672488 |
| Mustela_nigripes | Urocyon_cinereoargenteus | 0.245481587117622 |
| Mustela_nigripes | Ursus_americanus | 0.23831306930071 |
| Mustela_nigripes | Ursus_arctos | 0.238161639466839 |
| Mustela_nigripes | Ursus_maritimus | 0.236622355336264 |
| Mustela_nigripes | Ursus_spelaeus | 0.232860262558171 |
| Mustela_nigripes | Ursus_thibetanus | 0.232648703689887 |
| Mustela_nigripes | Urva_javanica | 0.239691955687969 |
| Mustela_nigripes | Viverra_tangalunga_FMNH146957_SRR6053069 | 0.233473766619903 |
| Mustela_nigripes | Viverricula_indica | 0.232804667911208 |
| Mustela_nigripes | Vulpes_corsac | 0.241073733073976 |
| Mustela_nigripes | Vulpes_ferrilata | 0.241305896379081 |
| Mustela_nigripes | Vulpes_lagopus | 0.245973414922147 |
| Mustela_nigripes | Vulpes_vulpes_T3611 | 0.239238499968682 |
| Mustela_nigripes | Vulpes_zerda | 0.239448512185916 |
| Mustela_nigripes | Zalophus_californianus | 0.234301995658375 |
| Mustela_nivalis | Mustela_putorius | 0.0942570347706395 |
| Mustela_nivalis | Mustela_sibirica | 0.0940588057750174 |
| Mustela_nivalis | Nandinia_binotata | 0.224073172447242 |
| Mustela_nivalis | Nasua_nasua | 0.240805241131635 |
| Mustela_nivalis | Neofelis_nebulosa | 0.243099266923172 |
| Mustela_nivalis | Neomonachus_schauinslandi | 0.217012397701404 |
| Mustela_nivalis | Neophoca_cinerea | 0.236968748113328 |
| Mustela_nivalis | Neovison_vison | 0.139288214095817 |
| Mustela_nivalis | Nyctereutes_procyonoides | 0.241550244518004 |
| Mustela_nivalis | Odobenus_rosmarus_rosmarus | 0.254012662186423 |
| Mustela_nivalis | Otocolobus_manul | 0.227046847155575 |
| Mustela_nivalis | Otocyon_megalotis_megalotis_TS305 | 0.249740063954914 |
| Mustela_nivalis | Otocyon_megalotis_virgatus_FMNH158128 | 0.248017406545151 |
| Mustela_nivalis | Paguma_larvata | 0.2338791580535 |
| Mustela_nivalis | Panthera_leo | 0.239718266652656 |
| Mustela_nivalis | Panthera_onca | 0.23815494978225 |
| Mustela_nivalis | Panthera_pardus | 0.240567450745661 |
| Mustela_nivalis | Panthera_tigris | 0.246429297397507 |
| Mustela_nivalis | Panthera_uncia | 0.240790764728828 |
| Mustela_nivalis | Paradoxurus_hermaphroditus | 0.23850125688443 |
| Mustela_nivalis | Parahyaena_brunnea_FD126_Fecal | 0.241073733073976 |
| Mustela_nivalis | Pardofelis_marmorata | 0.232364250554204 |
| Mustela_nivalis | Phoca_fasciata | 0.220038336193622 |
| Mustela_nivalis | Phoca_groenlandica | 0.215194692300643 |
| Mustela_nivalis | Phoca_largha | 0.216536179132751 |
| Mustela_nivalis | Phoca_vitulina | 0.21620291378731 |
| Mustela_nivalis | Phocarctos_hookeri | 0.237938072365679 |
| Mustela_nivalis | Prionailurus_bengalensis | 0.228394116309553 |
| Mustela_nivalis | Prionailurus_bengalensis_euptilurus | 0.226525531537579 |
| Mustela_nivalis | Prionailurus_planiceps | 0.227215324082943 |
| Mustela_nivalis | Prionailurus_rubiginosus | 0.227389190503053 |
| Mustela_nivalis | Prionailurus_viverrinus | 0.231596201248034 |
| Mustela_nivalis | Prionodon_pardicolor | 0.233517199796443 |
| Mustela_nivalis | Procyon_lotor | 0.219559605374381 |
| Mustela_nivalis | Profelis_aurata | 0.238478677067094 |
| Mustela_nivalis | Proteles_cristatus_cristatus_TS307 | 0.246075038384405 |
| Mustela_nivalis | Proteles_cristatus_septentrionalis_NMS_Z_2018_54 | 0.246839496371434 |
| Mustela_nivalis | Puma_concolor | 0.228891428505717 |
| Mustela_nivalis | Puma_yagouaroundi | 0.233413494369339 |
| Mustela_nivalis | Pusa_caspica | 0.216873830031847 |
| Mustela_nivalis | Pusa_hispida | 0.214396711341104 |
| Mustela_nivalis | Pusa_sibirica | 0.21737283256033 |
| Mustela_nivalis | Speothos_venaticus_T1624 | 0.249343151258991 |
| Mustela_nivalis | Spilogale_putorius | 0.234386820357272 |
| Mustela_nivalis | Taxidea_taxus | 0.178258845701585 |
| Mustela_nivalis | Tremarctos_ornatus | 0.237942669734851 |
| Mustela_nivalis | Urocyon_cinereoargenteus | 0.253003800191908 |
| Mustela_nivalis | Ursus_americanus | 0.237404094896002 |
| Mustela_nivalis | Ursus_arctos | 0.244561730495574 |
| Mustela_nivalis | Ursus_maritimus | 0.242092860661984 |
| Mustela_nivalis | Ursus_spelaeus | 0.237951794982775 |
| Mustela_nivalis | Ursus_thibetanus | 0.235116078682992 |
| Mustela_nivalis | Urva_javanica | 0.239978263032701 |
| Mustela_nivalis | Viverra_tangalunga_FMNH146957_SRR6053069 | 0.228143044929659 |
| Mustela_nivalis | Viverricula_indica | 0.230951278269105 |
| Mustela_nivalis | Vulpes_corsac | 0.245268140822368 |

| | | |
|---|---|---|
| Mustela_nivalis | Vulpes_ferrilata | 0.247457473191805 |
| Mustela_nivalis | Vulpes_lagopus | 0.254329409476154 |
| Mustela_nivalis | Vulpes_vulpes_T3611 | 0.244192825287575 |
| Mustela_nivalis | Vulpes_zerda | 0.242692046390821 |
| Mustela_nivalis | Zalophus_californianus | 0.237581536419663 |
| Mustela_putorius | Mustela_sibirica | 0.0321304776458506 |
| Mustela_putorius | Nandinia_binotata | 0.219021232661252 |
| Mustela_putorius | Nasua_nasua | 0.236983817565586 |
| Mustela_putorius | Neofelis_nebulosa | 0.243288350525501 |
| Mustela_putorius | Neomonachus_schauinslandi | 0.215448851732973 |
| Mustela_putorius | Neophoca_cinerea | 0.237716780349084 |
| Mustela_putorius | Neovison_vison | 0.139189138925488 |
| Mustela_putorius | Nyctereutes_procyonoides | 0.241688895370278 |
| Mustela_putorius | Odobenus_rosmarus_rosmarus | 0.249315419785032 |
| Mustela_putorius | Otocolobus_manul | 0.226748914107624 |
| Mustela_putorius | Otocyon_megalotis_megalotis_TS305 | 0.247021861657172 |
| Mustela_putorius | Otocyon_megalotis_virgatus_FMNH158128 | 0.247664252782245 |
| Mustela_putorius | Paguma_larvata | 0.227596821834629 |
| Mustela_putorius | Panthera_leo | 0.236637873067131 |
| Mustela_putorius | Panthera_onca | 0.234093962034052 |
| Mustela_putorius | Panthera_pardus | 0.239107664729284 |
| Mustela_putorius | Panthera_tigris | 0.243512276625956 |
| Mustela_putorius | Panthera_uncia | 0.238795168725176 |
| Mustela_putorius | Paradoxurus_hermaphroditus | 0.235017230375356 |
| Mustela_putorius | Parahyaena_brunnea_FD126_Fecal | 0.237542318724549 |
| Mustela_putorius | Pardofelis_marmorata | 0.231347607771301 |
| Mustela_putorius | Phoca_fasciata | 0.217426225487025 |
| Mustela_putorius | Phoca_groenlandica | 0.218084173013942 |
| Mustela_putorius | Phoca_largha | 0.216438469902428 |
| Mustela_putorius | Phoca_vitulina | 0.216536179132751 |
| Mustela_putorius | Phocarctos_hookeri | 0.239044314388111 |
| Mustela_putorius | Prionailurus_bengalensis | 0.227423425830644 |
| Mustela_putorius | Prionailurus_bengalensis_euptilurus | 0.225577002531063 |
| Mustela_putorius | Prionailurus_planiceps | 0.226808777872924 |
| Mustela_putorius | Prionailurus_rubiginosus | 0.228341968908052 |
| Mustela_putorius | Prionailurus_viverrinus | 0.229327919148889 |
| Mustela_putorius | Prionodon_pardicolor | 0.236851732797506 |
| Mustela_putorius | Procyon_lotor | 0.214944706199247 |
| Mustela_putorius | Profelis_aurata | 0.2392232129317 |
| Mustela_putorius | Proteles_cristatus_cristatus_TS307 | 0.245035608338156 |
| Mustela_putorius | Proteles_cristatus_septentrionalis_NMS_Z_2018_54 | 0.245840315845883 |
| Mustela_putorius | Puma_concolor | 0.230012523985779 |
| Mustela_putorius | Puma_yagouaroundi | 0.22799120420238 |
| Mustela_putorius | Pusa_caspica | 0.216539821493327 |
| Mustela_putorius | Pusa_hispida | 0.214904107029776 |
| Mustela_putorius | Pusa_sibirica | 0.21768201022587 |
| Mustela_putorius | Speothos_venaticus_T1624 | 0.246296560375964 |
| Mustela_putorius | Spilogale_putorius | 0.235811268772597 |
| Mustela_putorius | Taxidea_taxus | 0.172835236659373 |
| Mustela_putorius | Tremarctos_ornatus | 0.240300479240908 |
| Mustela_putorius | Urocyon_cinereoargenteus | 0.247113094209958 |
| Mustela_putorius | Ursus_americanus | 0.237113767737295 |
| Mustela_putorius | Ursus_arctos | 0.240354589894532 |
| Mustela_putorius | Ursus_maritimus | 0.23779073952073 |
| Mustela_putorius | Ursus_spelaeus | 0.23393001275203 |
| Mustela_putorius | Ursus_thibetanus | 0.231619002968646 |
| Mustela_putorius | Urva_javanica | 0.239804522813216 |
| Mustela_putorius | Viverra_tangalunga_FMNH146957_SRR6053069 | 0.233683657587006 |
| Mustela_putorius | Viverricula_indica | 0.233583632516559 |
| Mustela_putorius | Vulpes_corsac | 0.241538222298284 |
| Mustela_putorius | Vulpes_ferrilata | 0.24264101065702 |
| Mustela_putorius | Vulpes_lagopus | 0.247722740149933 |
| Mustela_putorius | Vulpes_vulpes_T3611 | 0.239942894619439 |
| Mustela_putorius | Vulpes_zerda | 0.240598762902112 |
| Mustela_putorius | Zalophus_californianus | 0.231670703406121 |
| Mustela_sibirica | Nandinia_binotata | 0.218511810168532 |
| Mustela_sibirica | Nasua_nasua | 0.236608572165821 |
| Mustela_sibirica | Neofelis_nebulosa | 0.243658310601433 |
| Mustela_sibirica | Neomonachus_schauinslandi | 0.214306929579026 |
| Mustela_sibirica | Neophoca_cinerea | 0.239143984942746 |
| Mustela_sibirica | Neovison_vison | 0.139083607453788 |
| Mustela_sibirica | Nyctereutes_procyonoides | 0.24120134726956 |
| Mustela_sibirica | Odobenus_rosmarus_rosmarus | 0.250506918047881 |
| Mustela_sibirica | Otocolobus_manul | 0.22690389269419 |
| Mustela_sibirica | Otocyon_megalotis_megalotis_TS305 | 0.248875738321887 |
| Mustela_sibirica | Otocyon_megalotis_virgatus_FMNH158128 | 0.249411900181418 |
| Mustela_sibirica | Paguma_larvata | 0.229182025546878 |
| Mustela_sibirica | Panthera_leo | 0.236759226977176 |
| Mustela_sibirica | Panthera_onca | 0.236865866392608 |
| Mustela_sibirica | Panthera_pardus | 0.238895767476674 |
| Mustela_sibirica | Panthera_tigris | 0.244761944967594 |
| Mustela_sibirica | Panthera_uncia | 0.240387610795455 |
| Mustela_sibirica | Paradoxurus_hermaphroditus | 0.238734557040333 |
| Mustela_sibirica | Parahyaena_brunnea_FD126_Fecal | 0.239521453304083 |
| Mustela_sibirica | Pardofelis_marmorata | 0.235076182034877 |
| Mustela_sibirica | Phoca_fasciata | 0.2172137565697 |
| Mustela_sibirica | Phoca_groenlandica | 0.216381160487325 |

| | | |
|---|---|---|
| Mustela_sibirica | Phoca_largha | 0.217542727286304 |
| Mustela_sibirica | Phoca_vitulina | 0.217727983642424 |
| Mustela_sibirica | Phocarctos_hookeri | 0.240024625180545 |
| Mustela_sibirica | Prionailurus_bengalensis | 0.225477891560457 |
| Mustela_sibirica | Prionailurus_bengalensis_euptilurus | 0.225911684017988 |
| Mustela_sibirica | Prionailurus_planiceps | 0.228015740214539 |
| Mustela_sibirica | Prionailurus_rubiginosus | 0.225745751333023 |
| Mustela_sibirica | Prionailurus_viverrinus | 0.229228381032076 |
| Mustela_sibirica | Prionodon_pardicolor | 0.235085496770816 |
| Mustela_sibirica | Procyon_lotor | 0.216559793761579 |
| Mustela_sibirica | Profelis_aurata | 0.239011131332261 |
| Mustela_sibirica | Proteles_cristatus_cristatus_TS307 | 0.243837147214247 |
| Mustela_sibirica | Proteles_cristatus_septentrionalis_NMS_Z_2018_54 | 0.246005030341981 |
| Mustela_sibirica | Puma_concolor | 0.229038985470988 |
| Mustela_sibirica | Puma_yagouaroundi | 0.229417855295196 |
| Mustela_sibirica | Pusa_caspica | 0.214732680019131 |
| Mustela_sibirica | Pusa_hispida | 0.214947965275417 |
| Mustela_sibirica | Pusa_sibirica | 0.217168719631451 |
| Mustela_sibirica | Speothos_venaticus_T1624 | 0.24726199656099 |
| Mustela_sibirica | Spilogale_putorius | 0.231766948491449 |
| Mustela_sibirica | Taxidea_taxus | 0.17280591935632 |
| Mustela_sibirica | Tremarctos_ornatus | 0.24196099636056 |
| Mustela_sibirica | Urocyon_cinereoargenteus | 0.248607213698448 |
| Mustela_sibirica | Ursus_americanus | 0.237577932364979 |
| Mustela_sibirica | Ursus_arctos | 0.23618700476913 |
| Mustela_sibirica | Ursus_maritimus | 0.234427606911663 |
| Mustela_sibirica | Ursus_spelaeus | 0.232152118273391 |
| Mustela_sibirica | Ursus_thibetanus | 0.231854974400855 |
| Mustela_sibirica | Urva_javanica | 0.239665749146138 |
| Mustela_sibirica | Viverra_tangalunga_FMNH146957_SRR6053069 | 0.232464144693912 |
| Mustela_sibirica | Viverricula_indica | 0.230301692929662 |
| Mustela_sibirica | Vulpes_corsac | 0.242904775699959 |
| Mustela_sibirica | Vulpes_ferrilata | 0.243787286945389 |
| Mustela_sibirica | Vulpes_lagopus | 0.2478895541266 |
| Mustela_sibirica | Vulpes_vulpes_T3611 | 0.240992519489523 |
| Mustela_sibirica | Vulpes_zerda | 0.240851773454472 |
| Mustela_sibirica | Zalophus_californianus | 0.232870571270046 |
| Nandinia_binotata | Nasua_nasua | 0.241522163177522 |
| Nandinia_binotata | Neofelis_nebulosa | 0.183304999912754 |
| Nandinia_binotata | Neomonachus_schauinslandi | 0.200227694545793 |
| Nandinia_binotata | Neophoca_cinerea | 0.226219739158198 |
| Nandinia_binotata | Neovison_vison | 0.22184217039965 |
| Nandinia_binotata | Nyctereutes_procyonoides | 0.224384015376974 |
| Nandinia_binotata | Odobenus_rosmarus_rosmarus | 0.245006490741517 |
| Nandinia_binotata | Otocolobus_manul | 0.171028470994912 |
| Nandinia_binotata | Otocyon_megalotis_megalotis_TS305 | 0.233903811022557 |
| Nandinia_binotata | Otocyon_megalotis_virgatus_FMNH158128 | 0.23475533779186 |
| Nandinia_binotata | Paguma_larvata | 0.179192988289714 |
| Nandinia_binotata | Panthera_leo | 0.177607044080106 |
| Nandinia_binotata | Panthera_onca | 0.177931433139194 |
| Nandinia_binotata | Panthera_pardus | 0.180727940295847 |
| Nandinia_binotata | Panthera_tigris | 0.180967578379831 |
| Nandinia_binotata | Panthera_uncia | 0.176177888350053 |
| Nandinia_binotata | Paradoxurus_hermaphroditus | 0.188165307572729 |
| Nandinia_binotata | Parahyaena_brunnea_FD126_Fecal | 0.191084580857854 |
| Nandinia_binotata | Pardofelis_marmorata | 0.174044255458681 |
| Nandinia_binotata | Phoca_fasciata | 0.202231840773504 |
| Nandinia_binotata | Phoca_groenlandica | 0.206156770132338 |
| Nandinia_binotata | Phoca_largha | 0.206839771115102 |
| Nandinia_binotata | Phoca_vitulina | 0.206691556469353 |
| Nandinia_binotata | Phocarctos_hookeri | 0.228195289553435 |
| Nandinia_binotata | Prionailurus_bengalensis | 0.165778894786947 |
| Nandinia_binotata | Prionailurus_bengalensis_euptilurus | 0.165212996859205 |
| Nandinia_binotata | Prionailurus_planiceps | 0.171400879526504 |
| Nandinia_binotata | Prionailurus_rubiginosus | 0.168599005153419 |
| Nandinia_binotata | Prionailurus_viverrinus | 0.170139094511896 |
| Nandinia_binotata | Prionodon_pardicolor | 0.183548112551836 |
| Nandinia_binotata | Procyon_lotor | 0.223953401317237 |
| Nandinia_binotata | Profelis_aurata | 0.177022185269555 |
| Nandinia_binotata | Proteles_cristatus_cristatus_TS307 | 0.194811720890663 |
| Nandinia_binotata | Proteles_cristatus_septentrionalis_NMS_Z_2018_54 | 0.197297728308078 |
| Nandinia_binotata | Puma_concolor | 0.17194539713947 |
| Nandinia_binotata | Puma_yagouaroundi | 0.170246487630484 |
| Nandinia_binotata | Pusa_caspica | 0.204786849769322 |
| Nandinia_binotata | Pusa_hispida | 0.205644856913563 |
| Nandinia_binotata | Pusa_sibirica | 0.204627772973321 |
| Nandinia_binotata | Speothos_venaticus_T1624 | 0.240215845242495 |
| Nandinia_binotata | Spilogale_putorius | 0.231089019060495 |
| Nandinia_binotata | Taxidea_taxus | 0.202237854250315 |
| Nandinia_binotata | Tremarctos_ornatus | 0.223307730091809 |
| Nandinia_binotata | Urocyon_cinereoargenteus | 0.242995191262431 |
| Nandinia_binotata | Ursus_americanus | 0.228451455205209 |
| Nandinia_binotata | Ursus_arctos | 0.229278320604995 |
| Nandinia_binotata | Ursus_maritimus | 0.230706908049739 |
| Nandinia_binotata | Ursus_spelaeus | 0.228852617855408 |
| Nandinia_binotata | Ursus_thibetanus | 0.223484134602883 |
| Nandinia_binotata | Urva_javanica | 0.183632134951353 |

| | | |
|---|---|---|
| Nandinia_binotata | Viverra_tangalunga_FMNH146957_SRR6053069 | 0.174891130656892 |
| Nandinia_binotata | Viverricula_indica | 0.173156055518007 |
| Nandinia_binotata | Vulpes_corsac | 0.236048504705654 |
| Nandinia_binotata | Vulpes_ferrilata | 0.233524452315429 |
| Nandinia_binotata | Vulpes_lagopus | 0.2358377413063 |
| Nandinia_binotata | Vulpes_vulpes_T3611 | 0.227497744563402 |
| Nandinia_binotata | Vulpes_zerda | 0.227661560617074 |
| Nandinia_binotata | Zalophus_californianus | 0.221699815442823 |
| Nasua_nasua | Neofelis_nebulosa | 0.25718449184033 |
| Nasua_nasua | Neomonachus_schauinslandi | 0.233902448116061 |
| Nasua_nasua | Neophoca_cinerea | 0.242966163926198 |
| Nasua_nasua | Neovison_vison | 0.234339714951261 |
| Nasua_nasua | Nyctereutes_procyonoides | 0.256452364678782 |
| Nasua_nasua | Odobenus_rosmarus_rosmarus | 0.258188124508726 |
| Nasua_nasua | Otocolobus_manul | 0.252101608467346 |
| Nasua_nasua | Otocyon_megalotis_megalotis_TS305 | 0.265398696444086 |
| Nasua_nasua | Otocyon_megalotis_virgatus_FMNH158128 | 0.268424341403175 |
| Nasua_nasua | Paguma_larvata | 0.258883136835736 |
| Nasua_nasua | Panthera_leo | 0.255010906755268 |
| Nasua_nasua | Panthera_onca | 0.257015025345714 |
| Nasua_nasua | Panthera_pardus | 0.259420920924389 |
| Nasua_nasua | Panthera_tigris | 0.25731295902309 |
| Nasua_nasua | Panthera_uncia | 0.255192246866583 |
| Nasua_nasua | Paradoxurus_hermaphroditus | 0.262330760973392 |
| Nasua_nasua | Parahyaena_brunnea_FD126_Fecal | 0.257094144704165 |
| Nasua_nasua | Pardofelis_marmorata | 0.25253728027917 |
| Nasua_nasua | Phoca_fasciata | 0.230345176504758 |
| Nasua_nasua | Phoca_groenlandica | 0.234644160483585 |
| Nasua_nasua | Phoca_largha | 0.231191646487404 |
| Nasua_nasua | Phoca_vitulina | 0.230246852003094 |
| Nasua_nasua | Phocarctos_hookeri | 0.2464279053801 |
| Nasua_nasua | Prionailurus_bengalensis | 0.248246158116876 |
| Nasua_nasua | Prionailurus_bengalensis_euptilurus | 0.247975619719116 |
| Nasua_nasua | Prionailurus_planiceps | 0.248269145122034 |
| Nasua_nasua | Prionailurus_rubiginosus | 0.244705417729888 |
| Nasua_nasua | Prionailurus_viverrinus | 0.247523079808146 |
| Nasua_nasua | Prionodon_pardicolor | 0.254170235372789 |
| Nasua_nasua | Procyon_lotor | 0.216845660875893 |
| Nasua_nasua | Profelis_aurata | 0.249633250959002 |
| Nasua_nasua | Proteles_cristatus_cristatus_TS307 | 0.25638412833985 |
| Nasua_nasua | Proteles_cristatus_septentrionalis_NMS_Z_2018_54 | 0.25824106572547 |
| Nasua_nasua | Puma_concolor | 0.24710423150937 |
| Nasua_nasua | Puma_yagouaroundi | 0.25004874738043 |
| Nasua_nasua | Pusa_caspica | 0.233010983649912 |
| Nasua_nasua | Pusa_hispida | 0.229373208460363 |
| Nasua_nasua | Pusa_sibirica | 0.231168336481998 |
| Nasua_nasua | Speothos_venaticus_T1624 | 0.264288270463859 |
| Nasua_nasua | Spilogale_putorius | 0.249810111785401 |
| Nasua_nasua | Taxidea_taxus | 0.22354269164089 |
| Nasua_nasua | Tremarctos_ornatus | 0.249752531512896 |
| Nasua_nasua | Urocyon_cinereoargenteus | 0.266473492546099 |
| Nasua_nasua | Ursus_americanus | 0.251774633049522 |
| Nasua_nasua | Ursus_arctos | 0.256094166435026 |
| Nasua_nasua | Ursus_maritimus | 0.255029426501379 |
| Nasua_nasua | Ursus_spelaeus | 0.251833459372035 |
| Nasua_nasua | Ursus_thibetanus | 0.248278815567303 |
| Nasua_nasua | Urva_javanica | 0.256945932236404 |
| Nasua_nasua | Viverra_tangalunga_FMNH146957_SRR6053069 | 0.250861447947129 |
| Nasua_nasua | Viverricula_indica | 0.255338235095518 |
| Nasua_nasua | Vulpes_corsac | 0.267788464417371 |
| Nasua_nasua | Vulpes_ferrilata | 0.268254401688475 |
| Nasua_nasua | Vulpes_lagopus | 0.267674882144803 |
| Nasua_nasua | Vulpes_vulpes_T3611 | 0.265310856926465 |
| Nasua_nasua | Vulpes_zerda | 0.26322777641409 |
| Nasua_nasua | Zalophus_californianus | 0.243116668624777 |
| Neofelis_nebulosa | Neomonachus_schauinslandi | 0.234072321448311 |
| Neofelis_nebulosa | Neophoca_cinerea | 0.252800583964194 |
| Neofelis_nebulosa | Neovison_vison | 0.241915323924004 |
| Neofelis_nebulosa | Nyctereutes_procyonoides | 0.248891198601684 |
| Neofelis_nebulosa | Odobenus_rosmarus_rosmarus | 0.261568068483641 |
| Neofelis_nebulosa | Otocolobus_manul | 0.144390163700645 |
| Neofelis_nebulosa | Otocyon_megalotis_megalotis_TS305 | 0.249222220546184 |
| Neofelis_nebulosa | Otocyon_megalotis_virgatus_FMNH158128 | 0.249429623311232 |
| Neofelis_nebulosa | Paguma_larvata | 0.193642634513946 |
| Neofelis_nebulosa | Panthera_leo | 0.121714721716077 |
| Neofelis_nebulosa | Panthera_onca | 0.121496279648885 |
| Neofelis_nebulosa | Panthera_pardus | 0.124160851640155 |
| Neofelis_nebulosa | Panthera_tigris | 0.128665274932349 |
| Neofelis_nebulosa | Panthera_uncia | 0.12721394592118 |
| Neofelis_nebulosa | Paradoxurus_hermaphroditus | 0.198948838384657 |
| Neofelis_nebulosa | Parahyaena_brunnea_FD126_Fecal | 0.200623114149446 |
| Neofelis_nebulosa | Pardofelis_marmorata | 0.144723290297953 |
| Neofelis_nebulosa | Phoca_fasciata | 0.229010940993107 |
| Neofelis_nebulosa | Phoca_groenlandica | 0.230910076307014 |
| Neofelis_nebulosa | Phoca_largha | 0.23159662992031 |
| Neofelis_nebulosa | Phoca_vitulina | 0.231083101946197 |
| Neofelis_nebulosa | Phocarctos_hookeri | 0.249050665924666 |

| | | |
|---|---|---|
| Neofelis_nebulosa | Prionailurus_bengalensis | 0.137241362936723 |
| Neofelis_nebulosa | Prionailurus_bengalensis_euptilurus | 0.135994751973126 |
| Neofelis_nebulosa | Prionailurus_planiceps | 0.140756347672694 |
| Neofelis_nebulosa | Prionailurus_rubiginosus | 0.139020042490315 |
| Neofelis_nebulosa | Prionailurus_viverrinus | 0.139142301260099 |
| Neofelis_nebulosa | Prionodon_pardicolor | 0.193765749121037 |
| Neofelis_nebulosa | Procyon_lotor | 0.249896263733243 |
| Neofelis_nebulosa | Profelis_aurata | 0.143297441213056 |
| Neofelis_nebulosa | Proteles_cristatus_cristatus_TS307 | 0.206677749484998 |
| Neofelis_nebulosa | Proteles_cristatus_septentrionalis_NMS_Z_2018_54 | 0.208503548957673 |
| Neofelis_nebulosa | Puma_concolor | 0.140031719087842 |
| Neofelis_nebulosa | Puma_yagouaroundi | 0.14396314200978 |
| Neofelis_nebulosa | Pusa_caspica | 0.234112017236952 |
| Neofelis_nebulosa | Pusa_hispida | 0.231304302487448 |
| Neofelis_nebulosa | Pusa_sibirica | 0.233646649953402 |
| Neofelis_nebulosa | Speothos_venaticus_T1624 | 0.257623115256254 |
| Neofelis_nebulosa | Spilogale_putorius | 0.259474425760218 |
| Neofelis_nebulosa | Taxidea_taxus | 0.235849321133037 |
| Neofelis_nebulosa | Tremarctos_ornatus | 0.251859127224874 |
| Neofelis_nebulosa | Urocyon_cinereoargenteus | 0.256503468578629 |
| Neofelis_nebulosa | Ursus_americanus | 0.249034606658537 |
| Neofelis_nebulosa | Ursus_arctos | 0.254311548661816 |
| Neofelis_nebulosa | Ursus_maritimus | 0.251071869738768 |
| Neofelis_nebulosa | Ursus_spelaeus | 0.24623309596424 |
| Neofelis_nebulosa | Ursus_thibetanus | 0.245651650379253 |
| Neofelis_nebulosa | Urva_javanica | 0.205915302905314 |
| Neofelis_nebulosa | Viverra_tangalunga_FMNH146957_SRR6053069 | 0.184577108334189 |
| Neofelis_nebulosa | Viverricula_indica | 0.187103649471591 |
| Neofelis_nebulosa | Vulpes_corsac | 0.246754902908554 |
| Neofelis_nebulosa | Vulpes_ferrilata | 0.249682251306153 |
| Neofelis_nebulosa | Vulpes_lagopus | 0.244670536454202 |
| Neofelis_nebulosa | Vulpes_vulpes_T3611 | 0.245684207343355 |
| Neofelis_nebulosa | Vulpes_zerda | 0.245098076171109 |
| Neofelis_nebulosa | Zalophus_californianus | 0.247693804585552 |
| Neomonachus_schauinslandi | Neophoca_cinerea | 0.180397148491602 |
| Neomonachus_schauinslandi | Neovison_vison | 0.217878249678146 |
| Neomonachus_schauinslandi | Nyctereutes_procyonoides | 0.225289019863976 |
| Neomonachus_schauinslandi | Odobenus_rosmarus_rosmarus | 0.200945833190943 |
| Neomonachus_schauinslandi | Otocolobus_manul | 0.22180693706108 |
| Neomonachus_schauinslandi | Otocyon_megalotis_megalotis_TS305 | 0.235600486268754 |
| Neomonachus_schauinslandi | Otocyon_megalotis_virgatus_FMNH158128 | 0.238339944563849 |
| Neomonachus_schauinslandi | Paguma_larvata | 0.227045299718399 |
| Neomonachus_schauinslandi | Panthera_leo | 0.225208097747762 |
| Neomonachus_schauinslandi | Panthera_onca | 0.224271527909433 |
| Neomonachus_schauinslandi | Panthera_pardus | 0.228748359798696 |
| Neomonachus_schauinslandi | Panthera_tigris | 0.227658969950581 |
| Neomonachus_schauinslandi | Panthera_uncia | 0.229076802989837 |
| Neomonachus_schauinslandi | Paradoxurus_hermaphroditus | 0.232184378255771 |
| Neomonachus_schauinslandi | Parahyaena_brunnea_FD126_Fecal | 0.227472025234918 |
| Neomonachus_schauinslandi | Pardofelis_marmorata | 0.223098842061066 |
| Neomonachus_schauinslandi | Phoca_fasciata | 0.139387043901437 |
| Neomonachus_schauinslandi | Phoca_groenlandica | 0.14308677281343 |
| Neomonachus_schauinslandi | Phoca_largha | 0.143111034761855 |
| Neomonachus_schauinslandi | Phoca_vitulina | 0.144483322866974 |
| Neomonachus_schauinslandi | Phocarctos_hookeri | 0.191077998896205 |
| Neomonachus_schauinslandi | Prionailurus_bengalensis | 0.214564803905583 |
| Neomonachus_schauinslandi | Prionailurus_bengalensis_euptilurus | 0.21573838814223 |
| Neomonachus_schauinslandi | Prionailurus_planiceps | 0.219973260105689 |
| Neomonachus_schauinslandi | Prionailurus_rubiginosus | 0.216931847692097 |
| Neomonachus_schauinslandi | Prionailurus_viverrinus | 0.21947232013825 |
| Neomonachus_schauinslandi | Prionodon_pardicolor | 0.230157498090863 |
| Neomonachus_schauinslandi | Procyon_lotor | 0.222793287950335 |
| Neomonachus_schauinslandi | Profelis_aurata | 0.217949982449471 |
| Neomonachus_schauinslandi | Proteles_cristatus_cristatus_TS307 | 0.232745982791692 |
| Neomonachus_schauinslandi | Proteles_cristatus_septentrionalis_NMS_Z_2018_54 | 0.234284445543932 |
| Neomonachus_schauinslandi | Puma_concolor | 0.218422659770868 |
| Neomonachus_schauinslandi | Puma_yagouaroundi | 0.217904099487155 |
| Neomonachus_schauinslandi | Pusa_caspica | 0.145597925100271 |
| Neomonachus_schauinslandi | Pusa_hispida | 0.143776972094789 |
| Neomonachus_schauinslandi | Pusa_sibirica | 0.144727962030443 |
| Neomonachus_schauinslandi | Speothos_venaticus_T1624 | 0.237974883149814 |
| Neomonachus_schauinslandi | Spilogale_putorius | 0.221125714280003 |
| Neomonachus_schauinslandi | Taxidea_taxus | 0.202445374184165 |
| Neomonachus_schauinslandi | Tremarctos_ornatus | 0.217519209193334 |
| Neomonachus_schauinslandi | Urocyon_cinereoargenteus | 0.238898479027648 |
| Neomonachus_schauinslandi | Ursus_americanus | 0.219272639922448 |
| Neomonachus_schauinslandi | Ursus_arctos | 0.21667569877485 |
| Neomonachus_schauinslandi | Ursus_maritimus | 0.216469008822395 |
| Neomonachus_schauinslandi | Ursus_spelaeus | 0.215667262119809 |
| Neomonachus_schauinslandi | Ursus_thibetanus | 0.219398213262113 |
| Neomonachus_schauinslandi | Urva_javanica | 0.219642671010236 |
| Neomonachus_schauinslandi | Viverra_tangalunga_FMNH146957_SRR6053069 | 0.217929437752113 |
| Neomonachus_schauinslandi | Viverricula_indica | 0.215026742425638 |
| Neomonachus_schauinslandi | Vulpes_corsac | 0.233129832797078 |
| Neomonachus_schauinslandi | Vulpes_ferrilata | 0.233381852938118 |
| Neomonachus_schauinslandi | Vulpes_lagopus | 0.233650213282341 |
| Neomonachus_schauinslandi | Vulpes_vulpes_T3611 | 0.229314555584158 |

| | | |
|---|---|---|
| Neomonachus_schauinslandi | Vulpes_zerda | 0.231701734572028 |
| Neomonachus_schauinslandi | Zalophus_californianus | 0.178813943603701 |
| Neophoca_cinerea | Neovison_vison | 0.234900323771047 |
| Neophoca_cinerea | Nyctereutes_procyonoides | 0.242492571359341 |
| Neophoca_cinerea | Odobenus_rosmarus_rosmarus | 0.197987103333141 |
| Neophoca_cinerea | Otocolobus_manul | 0.235266453086013 |
| Neophoca_cinerea | Otocyon_megalotis_megalotis_TS305 | 0.249241064658731 |
| Neophoca_cinerea | Otocyon_megalotis_virgatus_FMNH158128 | 0.250669710325339 |
| Neophoca_cinerea | Paguma_larvata | 0.242276693577673 |
| Neophoca_cinerea | Panthera_leo | 0.242167901877357 |
| Neophoca_cinerea | Panthera_onca | 0.241955022108857 |
| Neophoca_cinerea | Panthera_pardus | 0.243288350525501 |
| Neophoca_cinerea | Panthera_tigris | 0.245042721754527 |
| Neophoca_cinerea | Panthera_uncia | 0.249269013119223 |
| Neophoca_cinerea | Paradoxurus_hermaphroditus | 0.244824176116358 |
| Neophoca_cinerea | Parahyaena_brunnea_FD126_Fecal | 0.24308571357717 |
| Neophoca_cinerea | Pardofelis_marmorata | 0.2411077481194 |
| Neophoca_cinerea | Phoca_fasciata | 0.184241221712746 |
| Neophoca_cinerea | Phoca_groenlandica | 0.187724915607893 |
| Neophoca_cinerea | Phoca_largha | 0.184229985802006 |
| Neophoca_cinerea | Phoca_vitulina | 0.185787965158745 |
| Neophoca_cinerea | Phocarctos_hookeri | 0.0775680658308659 |
| Neophoca_cinerea | Prionailurus_bengalensis | 0.231007491950657 |
| Neophoca_cinerea | Prionailurus_bengalensis_euptilurus | 0.232613801730506 |
| Neophoca_cinerea | Prionailurus_planiceps | 0.23475533779186 |
| Neophoca_cinerea | Prionailurus_rubiginosus | 0.234906384610771 |
| Neophoca_cinerea | Prionailurus_viverrinus | 0.234285238365367 |
| Neophoca_cinerea | Prionodon_pardicolor | 0.248333765489616 |
| Neophoca_cinerea | Procyon_lotor | 0.234987414422638 |
| Neophoca_cinerea | Profelis_aurata | 0.242064901868531 |
| Neophoca_cinerea | Proteles_cristatus_cristatus_TS307 | 0.245118937797738 |
| Neophoca_cinerea | Proteles_cristatus_septentrionalis_NMS_Z_2018_54 | 0.246127793649477 |
| Neophoca_cinerea | Puma_concolor | 0.239887348558472 |
| Neophoca_cinerea | Puma_yagouaroundi | 0.240523626111087 |
| Neophoca_cinerea | Pusa_caspica | 0.185103265306363 |
| Neophoca_cinerea | Pusa_hispida | 0.182347461593749 |
| Neophoca_cinerea | Pusa_sibirica | 0.182681442777276 |
| Neophoca_cinerea | Speothos_venaticus_T1624 | 0.254557485657552 |
| Neophoca_cinerea | Spilogale_putorius | 0.244854357030226 |
| Neophoca_cinerea | Taxidea_taxus | 0.224567454081226 |
| Neophoca_cinerea | Tremarctos_ornatus | 0.234631129201969 |
| Neophoca_cinerea | Urocyon_cinereoargenteus | 0.251962858729914 |
| Neophoca_cinerea | Ursus_americanus | 0.233001806781017 |
| Neophoca_cinerea | Ursus_arctos | 0.238453052770884 |
| Neophoca_cinerea | Ursus_maritimus | 0.238806805564007 |
| Neophoca_cinerea | Ursus_spelaeus | 0.235128744027271 |
| Neophoca_cinerea | Ursus_thibetanus | 0.235519715450447 |
| Neophoca_cinerea | Urva_javanica | 0.239482324282337 |
| Neophoca_cinerea | Viverra_tangalunga_FMNH146957_SRR6053069 | 0.239022809692435 |
| Neophoca_cinerea | Viverricula_indica | 0.238331995153354 |
| Neophoca_cinerea | Vulpes_corsac | 0.251238351723198 |
| Neophoca_cinerea | Vulpes_ferrilata | 0.25251978383034 |
| Neophoca_cinerea | Vulpes_lagopus | 0.250730493817912 |
| Neophoca_cinerea | Vulpes_vulpes_T3611 | 0.246394828065281 |
| Neophoca_cinerea | Vulpes_zerda | 0.246255893309971 |
| Neophoca_cinerea | Zalophus_californianus | 0.0857313672000139 |
| Neovison_vison | Nyctereutes_procyonoides | 0.241125081921915 |
| Neovison_vison | Odobenus_rosmarus_rosmarus | 0.246759806644213 |
| Neovison_vison | Otocolobus_manul | 0.232117150321863 |
| Neovison_vison | Otocyon_megalotis_megalotis_TS305 | 0.244685328859541 |
| Neovison_vison | Otocyon_megalotis_virgatus_FMNH158128 | 0.245918928229263 |
| Neovison_vison | Paguma_larvata | 0.22690151703967 |
| Neovison_vison | Panthera_leo | 0.243456038452647 |
| Neovison_vison | Panthera_onca | 0.239648059427672 |
| Neovison_vison | Panthera_pardus | 0.243165160010815 |
| Neovison_vison | Panthera_tigris | 0.246408897375276 |
| Neovison_vison | Panthera_uncia | 0.244467388387301 |
| Neovison_vison | Paradoxurus_hermaphroditus | 0.230827238484488 |
| Neovison_vison | Parahyaena_brunnea_FD126_Fecal | 0.24021491435612 |
| Neovison_vison | Pardofelis_marmorata | 0.237974883149814 |
| Neovison_vison | Phoca_fasciata | 0.22031237782003 |
| Neovison_vison | Phoca_groenlandica | 0.21793931011293 |
| Neovison_vison | Phoca_largha | 0.218454381759893 |
| Neovison_vison | Phoca_vitulina | 0.218206898879887 |
| Neovison_vison | Phocarctos_hookeri | 0.237314241723829 |
| Neovison_vison | Prionailurus_bengalensis | 0.229061795016735 |
| Neovison_vison | Prionailurus_bengalensis_euptilurus | 0.229318301004271 |
| Neovison_vison | Prionailurus_planiceps | 0.23016100137872 |
| Neovison_vison | Prionailurus_rubiginosus | 0.231389058750174 |
| Neovison_vison | Prionailurus_viverrinus | 0.229626617784247 |
| Neovison_vison | Prionodon_pardicolor | 0.235396288899299 |
| Neovison_vison | Procyon_lotor | 0.216559793761579 |
| Neovison_vison | Profelis_aurata | 0.240352359909312 |
| Neovison_vison | Proteles_cristatus_cristatus_TS307 | 0.247219061829037 |
| Neovison_vison | Proteles_cristatus_septentrionalis_NMS_Z_2018_54 | 0.250059563202734 |
| Neovison_vison | Puma_concolor | 0.227291861277927 |
| Neovison_vison | Puma_yagouaroundi | 0.230684661290301 |

| | | |
|---|---|---|
| Neovison_vison | Pusa_caspica | 0.220708717506245 |
| Neovison_vison | Pusa_hispida | 0.216544861268146 |
| Neovison_vison | Pusa_sibirica | 0.218321108138511 |
| Neovison_vison | Speothos_venaticus_T1624 | 0.242531910564278 |
| Neovison_vison | Spilogale_putorius | 0.223668882956321 |
| Neovison_vison | Taxidea_taxus | 0.17831908371586 |
| Neovison_vison | Tremarctos_ornatus | 0.237813116277171 |
| Neovison_vison | Urocyon_cinereoargenteus | 0.249891654689 |
| Neovison_vison | Ursus_americanus | 0.233456372477648 |
| Neovison_vison | Ursus_arctos | 0.242060572704738 |
| Neovison_vison | Ursus_maritimus | 0.241165105352396 |
| Neovison_vison | Ursus_spelaeus | 0.234812726200874 |
| Neovison_vison | Ursus_thibetanus | 0.236097991878914 |
| Neovison_vison | Urva_javanica | 0.236236224875227 |
| Neovison_vison | Viverra_tangalunga_FMNH146957_SRR6053069 | 0.223054399339407 |
| Neovison_vison | Viverricula_indica | 0.221663135390655 |
| Neovison_vison | Vulpes_corsac | 0.241662016093186 |
| Neovison_vison | Vulpes_ferrilata | 0.245048567458618 |
| Neovison_vison | Vulpes_lagopus | 0.246513091428757 |
| Neovison_vison | Vulpes_vulpes_T3611 | 0.240872305211318 |
| Neovison_vison | Vulpes_zerda | 0.24137973025486 |
| Neovison_vison | Zalophus_californianus | 0.233049561217999 |
| Nyctereutes_procyonoides | Odobenus_rosmarus_rosmarus | 0.26187854314355 |
| Nyctereutes_procyonoides | Otocolobus_manul | 0.235742196900608 |
| Nyctereutes_procyonoides | Otocyon_megalotis_megalotis_TS305 | 0.174888101197203 |
| Nyctereutes_procyonoides | Otocyon_megalotis_virgatus_FMNH158128 | 0.176578902221474 |
| Nyctereutes_procyonoides | Paguma_larvata | 0.240416794578338 |
| Nyctereutes_procyonoides | Panthera_leo | 0.245560845694297 |
| Nyctereutes_procyonoides | Panthera_onca | 0.240336694443758 |
| Nyctereutes_procyonoides | Panthera_pardus | 0.246416199322054 |
| Nyctereutes_procyonoides | Panthera_tigris | 0.250271534798133 |
| Nyctereutes_procyonoides | Panthera_uncia | 0.247878682083712 |
| Nyctereutes_procyonoides | Paradoxurus_hermaphroditus | 0.24264101065702 |
| Nyctereutes_procyonoides | Parahyaena_brunnea_FD126_Fecal | 0.243970413155 |
| Nyctereutes_procyonoides | Pardofelis_marmorata | 0.243123298671544 |
| Nyctereutes_procyonoides | Phoca_fasciata | 0.226889095276242 |
| Nyctereutes_procyonoides | Phoca_groenlandica | 0.232025066729362 |
| Nyctereutes_procyonoides | Phoca_largha | 0.233303519663078 |
| Nyctereutes_procyonoides | Phoca_vitulina | 0.231099381268068 |
| Nyctereutes_procyonoides | Phocarctos_hookeri | 0.249414666463704 |
| Nyctereutes_procyonoides | Prionailurus_bengalensis | 0.231978823436796 |
| Nyctereutes_procyonoides | Prionailurus_bengalensis_euptilurus | 0.233220902335826 |
| Nyctereutes_procyonoides | Prionailurus_planiceps | 0.237545981822451 |
| Nyctereutes_procyonoides | Prionailurus_rubiginosus | 0.234461252417164 |
| Nyctereutes_procyonoides | Prionailurus_viverrinus | 0.235379450849843 |
| Nyctereutes_procyonoides | Prionodon_pardicolor | 0.239953895908931 |
| Nyctereutes_procyonoides | Procyon_lotor | 0.237409247527803 |
| Nyctereutes_procyonoides | Profelis_aurata | 0.238773739040539 |
| Nyctereutes_procyonoides | Proteles_cristatus_cristatus_TS307 | 0.251860706990391 |
| Nyctereutes_procyonoides | Proteles_cristatus_septentrionalis_NMS_Z_2018_54 | 0.252065688758984 |
| Nyctereutes_procyonoides | Puma_concolor | 0.23146043647111 |
| Nyctereutes_procyonoides | Puma_yagouaroundi | 0.237023355526129 |
| Nyctereutes_procyonoides | Pusa_caspica | 0.230002792070077 |
| Nyctereutes_procyonoides | Pusa_hispida | 0.228654246550725 |
| Nyctereutes_procyonoides | Pusa_sibirica | 0.230440333117469 |
| Nyctereutes_procyonoides | Speothos_venaticus_T1624 | 0.183883166052195 |
| Nyctereutes_procyonoides | Spilogale_putorius | 0.244375057641886 |
| Nyctereutes_procyonoides | Taxidea_taxus | 0.235455075411525 |
| Nyctereutes_procyonoides | Tremarctos_ornatus | 0.248468567604776 |
| Nyctereutes_procyonoides | Urocyon_cinereoargenteus | 0.185423059921993 |
| Nyctereutes_procyonoides | Ursus_americanus | 0.250397335376066 |
| Nyctereutes_procyonoides | Ursus_arctos | 0.256650467271085 |
| Nyctereutes_procyonoides | Ursus_maritimus | 0.255739895415187 |
| Nyctereutes_procyonoides | Ursus_spelaeus | 0.246894899400427 |
| Nyctereutes_procyonoides | Ursus_thibetanus | 0.245423144686709 |
| Nyctereutes_procyonoides | Urva_javanica | 0.237554766903809 |
| Nyctereutes_procyonoides | Viverra_tangalunga_FMNH146957_SRR6053069 | 0.235216182464535 |
| Nyctereutes_procyonoides | Viverricula_indica | 0.22858302122362 |
| Nyctereutes_procyonoides | Vulpes_corsac | 0.172726526235126 |
| Nyctereutes_procyonoides | Vulpes_ferrilata | 0.172964176480677 |
| Nyctereutes_procyonoides | Vulpes_lagopus | 0.168548743699491 |
| Nyctereutes_procyonoides | Vulpes_vulpes_T3611 | 0.172057876711545 |
| Nyctereutes_procyonoides | Vulpes_zerda | 0.172489193279635 |
| Nyctereutes_procyonoides | Zalophus_californianus | 0.240733222101977 |
| Odobenus_rosmarus_rosmarus | Otocolobus_manul | 0.25291492194777 |
| Odobenus_rosmarus_rosmarus | Otocyon_megalotis_megalotis_TS305 | 0.267093917222448 |
| Odobenus_rosmarus_rosmarus | Otocyon_megalotis_virgatus_FMNH158128 | 0.265442312825411 |
| Odobenus_rosmarus_rosmarus | Paguma_larvata | 0.260765269536338 |
| Odobenus_rosmarus_rosmarus | Panthera_leo | 0.257945690592809 |
| Odobenus_rosmarus_rosmarus | Panthera_onca | 0.254149824114379 |
| Odobenus_rosmarus_rosmarus | Panthera_pardus | 0.255409369728761 |
| Odobenus_rosmarus_rosmarus | Panthera_tigris | 0.262464227441718 |
| Odobenus_rosmarus_rosmarus | Panthera_uncia | 0.258472520928469 |
| Odobenus_rosmarus_rosmarus | Paradoxurus_hermaphroditus | 0.260687519547564 |
| Odobenus_rosmarus_rosmarus | Parahyaena_brunnea_FD126_Fecal | 0.255409335239992 |
| Odobenus_rosmarus_rosmarus | Pardofelis_marmorata | 0.254723218672632 |
| Odobenus_rosmarus_rosmarus | Phoca_fasciata | 0.201935168515219 |

| | | |
|---|---|---|
| Odobenus_rosmarus_rosmarus | Phoca_groenlandica | 0.206801787888996 |
| Odobenus_rosmarus_rosmarus | Phoca_largha | 0.201541793858973 |
| Odobenus_rosmarus_rosmarus | Phoca_vitulina | 0.201852558022935 |
| Odobenus_rosmarus_rosmarus | Phocarctos_hookeri | 0.195320661255218 |
| Odobenus_rosmarus_rosmarus | Prionailurus_bengalensis | 0.252742868915334 |
| Odobenus_rosmarus_rosmarus | Prionailurus_bengalensis_euptilurus | 0.253127000727797 |
| Odobenus_rosmarus_rosmarus | Prionailurus_planiceps | 0.253634557181653 |
| Odobenus_rosmarus_rosmarus | Prionailurus_rubiginosus | 0.250956560947683 |
| Odobenus_rosmarus_rosmarus | Prionailurus_viverrinus | 0.250890752422572 |
| Odobenus_rosmarus_rosmarus | Prionodon_pardicolor | 0.261440083791528 |
| Odobenus_rosmarus_rosmarus | Procyon_lotor | 0.249136114299988 |
| Odobenus_rosmarus_rosmarus | Profelis_aurata | 0.253331448131247 |
| Odobenus_rosmarus_rosmarus | Proteles_cristatus_cristatus_TS307 | 0.263145277867881 |
| Odobenus_rosmarus_rosmarus | Proteles_cristatus_septentrionalis_NMS_Z_2018_54 | 0.26166297253032 |
| Odobenus_rosmarus_rosmarus | Puma_concolor | 0.251905735303657 |
| Odobenus_rosmarus_rosmarus | Puma_yagouaroundi | 0.246563349661319 |
| Odobenus_rosmarus_rosmarus | Pusa_caspica | 0.206741147469422 |
| Odobenus_rosmarus_rosmarus | Pusa_hispida | 0.202328749917775 |
| Odobenus_rosmarus_rosmarus | Pusa_sibirica | 0.205979532699161 |
| Odobenus_rosmarus_rosmarus | Speothos_venaticus_T1624 | 0.272382525396281 |
| Odobenus_rosmarus_rosmarus | Spilogale_putorius | 0.25681615428132 |
| Odobenus_rosmarus_rosmarus | Taxidea_taxus | 0.234635201341444 |
| Odobenus_rosmarus_rosmarus | Tremarctos_ornatus | 0.24038421500344 |
| Odobenus_rosmarus_rosmarus | Urocyon_cinereoargenteus | 0.267434174878856 |
| Odobenus_rosmarus_rosmarus | Ursus_americanus | 0.250759526287753 |
| Odobenus_rosmarus_rosmarus | Ursus_arctos | 0.252052701982409 |
| Odobenus_rosmarus_rosmarus | Ursus_maritimus | 0.254634664863067 |
| Odobenus_rosmarus_rosmarus | Ursus_spelaeus | 0.24841481438024 |
| Odobenus_rosmarus_rosmarus | Ursus_thibetanus | 0.251458716788819 |
| Odobenus_rosmarus_rosmarus | Urva_javanica | 0.257401649556313 |
| Odobenus_rosmarus_rosmarus | Viverra_tangalunga_FMNH146957_SRR6053069 | 0.249692408291958 |
| Odobenus_rosmarus_rosmarus | Viverricula_indica | 0.258716751120986 |
| Odobenus_rosmarus_rosmarus | Vulpes_corsac | 0.260882906972727 |
| Odobenus_rosmarus_rosmarus | Vulpes_ferrilata | 0.260870700843958 |
| Odobenus_rosmarus_rosmarus | Vulpes_lagopus | 0.26444946528735 |
| Odobenus_rosmarus_rosmarus | Vulpes_vulpes_T3611 | 0.260051997301403 |
| Odobenus_rosmarus_rosmarus | Vulpes_zerda | 0.259499355053723 |
| Odobenus_rosmarus_rosmarus | Zalophus_californianus | 0.1930860802204 |
| Otocolobus_manul | Otocyon_megalotis_megalotis_TS305 | 0.244853152734434 |
| Otocolobus_manul | Otocyon_megalotis_virgatus_FMNH158128 | 0.24330153258098 |
| Otocolobus_manul | Paguma_larvata | 0.177059871217098 |
| Otocolobus_manul | Panthera_leo | 0.141989591581244 |
| Otocolobus_manul | Panthera_onca | 0.136078797643561 |
| Otocolobus_manul | Panthera_pardus | 0.141258488100389 |
| Otocolobus_manul | Panthera_tigris | 0.142249752058459 |
| Otocolobus_manul | Panthera_uncia | 0.142176855674275 |
| Otocolobus_manul | Paradoxurus_hermaphroditus | 0.183118210543934 |
| Otocolobus_manul | Parahyaena_brunnea_FD126_Fecal | 0.196386370048088 |
| Otocolobus_manul | Pardofelis_marmorata | 0.120733964998456 |
| Otocolobus_manul | Phoca_fasciata | 0.221937926175266 |
| Otocolobus_manul | Phoca_groenlandica | 0.223500263462588 |
| Otocolobus_manul | Phoca_largha | 0.223984588049088 |
| Otocolobus_manul | Phoca_vitulina | 0.224587393990052 |
| Otocolobus_manul | Phocarctos_hookeri | 0.236119567083143 |
| Otocolobus_manul | Prionailurus_bengalensis | 0.102744929655757 |
| Otocolobus_manul | Prionailurus_bengalensis_euptilurus | 0.104861949282526 |
| Otocolobus_manul | Prionailurus_planiceps | 0.105506096359887 |
| Otocolobus_manul | Prionailurus_rubiginosus | 0.102167252712954 |
| Otocolobus_manul | Prionailurus_viverrinus | 0.106538188974865 |
| Otocolobus_manul | Prionodon_pardicolor | 0.185138394112716 |
| Otocolobus_manul | Procyon_lotor | 0.231366461697578 |
| Otocolobus_manul | Profelis_aurata | 0.113840118932797 |
| Otocolobus_manul | Proteles_cristatus_cristatus_TS307 | 0.203227037942624 |
| Otocolobus_manul | Proteles_cristatus_septentrionalis_NMS_Z_2018_54 | 0.203035409026525 |
| Otocolobus_manul | Puma_concolor | 0.113032272162886 |
| Otocolobus_manul | Puma_yagouaroundi | 0.110957576474684 |
| Otocolobus_manul | Pusa_caspica | 0.225643233153444 |
| Otocolobus_manul | Pusa_hispida | 0.219991898237362 |
| Otocolobus_manul | Pusa_sibirica | 0.222296916754373 |
| Otocolobus_manul | Speothos_venaticus_T1624 | 0.246093043060128 |
| Otocolobus_manul | Spilogale_putorius | 0.246994266950573 |
| Otocolobus_manul | Taxidea_taxus | 0.22233564281369 |
| Otocolobus_manul | Tremarctos_ornatus | 0.240933181681101 |
| Otocolobus_manul | Urocyon_cinereoargenteus | 0.249048209704129 |
| Otocolobus_manul | Ursus_americanus | 0.239480496452925 |
| Otocolobus_manul | Ursus_arctos | 0.247093521781748 |
| Otocolobus_manul | Ursus_maritimus | 0.245576756213281 |
| Otocolobus_manul | Ursus_spelaeus | 0.241351219037343 |
| Otocolobus_manul | Ursus_thibetanus | 0.239646828286862 |
| Otocolobus_manul | Urva_javanica | 0.193929655115366 |
| Otocolobus_manul | Viverra_tangalunga_FMNH146957_SRR6053069 | 0.17066555660937 |
| Otocolobus_manul | Viverricula_indica | 0.170573686105239 |
| Otocolobus_manul | Vulpes_corsac | 0.243688137799403 |
| Otocolobus_manul | Vulpes_ferrilata | 0.244708107254479 |
| Otocolobus_manul | Vulpes_lagopus | 0.241750210133894 |
| Otocolobus_manul | Vulpes_vulpes_T3611 | 0.235556818149793 |
| Otocolobus_manul | Vulpes_zerda | 0.236410439579902 |

| | | |
|---|---|---|
| Otocolobus_manul | Zalophus_californianus | 0.231099381268068 |
| Otocyon_megalotis_megalotis_TS305 | Otocyon_megalotis_virgatus_FMNH158128 | 0.0204876292273266 |
| Otocyon_megalotis_megalotis_TS305 | Paguma_larvata | 0.256525423817248 |
| Otocyon_megalotis_megalotis_TS305 | Panthera_leo | 0.243422692339548 |
| Otocyon_megalotis_megalotis_TS305 | Panthera_onca | 0.248505566821278 |
| Otocyon_megalotis_megalotis_TS305 | Panthera_pardus | 0.24695976104117 |
| Otocyon_megalotis_megalotis_TS305 | Panthera_tigris | 0.246733695704438 |
| Otocyon_megalotis_megalotis_TS305 | Panthera_uncia | 0.247793618205907 |
| Otocyon_megalotis_megalotis_TS305 | Paradoxurus_hermaphroditus | 0.257596798295945 |
| Otocyon_megalotis_megalotis_TS305 | Parahyaena_brunnea_FD126_Fecal | 0.250931760716856 |
| Otocyon_megalotis_megalotis_TS305 | Pardofelis_marmorata | 0.249664588345173 |
| Otocyon_megalotis_megalotis_TS305 | Phoca_fasciata | 0.234525927353754 |
| Otocyon_megalotis_megalotis_TS305 | Phoca_groenlandica | 0.233324029492 |
| Otocyon_megalotis_megalotis_TS305 | Phoca_largha | 0.233851058030606 |
| Otocyon_megalotis_megalotis_TS305 | Phoca_vitulina | 0.233797711165859 |
| Otocyon_megalotis_megalotis_TS305 | Phocarctos_hookeri | 0.254775545221595 |
| Otocyon_megalotis_megalotis_TS305 | Prionailurus_bengalensis | 0.246179243160047 |
| Otocyon_megalotis_megalotis_TS305 | Prionailurus_bengalensis_euptilurus | 0.244287746921532 |
| Otocyon_megalotis_megalotis_TS305 | Prionailurus_planiceps | 0.250436569829006 |
| Otocyon_megalotis_megalotis_TS305 | Prionailurus_rubiginosus | 0.244333419911484 |
| Otocyon_megalotis_megalotis_TS305 | Prionailurus_viverrinus | 0.246336955056984 |
| Otocyon_megalotis_megalotis_TS305 | Prionodon_pardicolor | 0.245774566068473 |
| Otocyon_megalotis_megalotis_TS305 | Procyon_lotor | 0.25091868157942 |
| Otocyon_megalotis_megalotis_TS305 | Profelis_aurata | 0.251988918417604 |
| Otocyon_megalotis_megalotis_TS305 | Proteles_cristatus_cristatus_TS307 | 0.25644100707132 |
| Otocyon_megalotis_megalotis_TS305 | Proteles_cristatus_septentrionalis_NMS_Z_2018_54 | 0.256326577550118 |
| Otocyon_megalotis_megalotis_TS305 | Puma_concolor | 0.242460030245095 |
| Otocyon_megalotis_megalotis_TS305 | Puma_yagouaroundi | 0.24466142650961 |
| Otocyon_megalotis_megalotis_TS305 | Pusa_caspica | 0.23381302244292 |
| Otocyon_megalotis_megalotis_TS305 | Pusa_hispida | 0.235851931716656 |
| Otocyon_megalotis_megalotis_TS305 | Pusa_sibirica | 0.232530855648366 |
| Otocyon_megalotis_megalotis_TS305 | Speothos_venaticus_T1624 | 0.186626601599288 |
| Otocyon_megalotis_megalotis_TS305 | Spilogale_putorius | 0.262976563010779 |
| Otocyon_megalotis_megalotis_TS305 | Taxidea_taxus | 0.247475944765284 |
| Otocyon_megalotis_megalotis_TS305 | Tremarctos_ornatus | 0.252195235358105 |
| Otocyon_megalotis_megalotis_TS305 | Urocyon_cinereoargenteus | 0.188052321556795 |
| Otocyon_megalotis_megalotis_TS305 | Ursus_americanus | 0.251348611486836 |
| Otocyon_megalotis_megalotis_TS305 | Ursus_arctos | 0.253646071133514 |
| Otocyon_megalotis_megalotis_TS305 | Ursus_maritimus | 0.252994454580645 |
| Otocyon_megalotis_megalotis_TS305 | Ursus_spelaeus | 0.25417310001951 |
| Otocyon_megalotis_megalotis_TS305 | Ursus_thibetanus | 0.244420505614339 |
| Otocyon_megalotis_megalotis_TS305 | Urva_javanica | 0.249334717229537 |
| Otocyon_megalotis_megalotis_TS305 | Viverra_tangalunga_FMNH146957_SRR6053069 | 0.243490615584334 |
| Otocyon_megalotis_megalotis_TS305 | Viverricula_indica | 0.241571420109388 |
| Otocyon_megalotis_megalotis_TS305 | Vulpes_corsac | 0.177142989039933 |
| Otocyon_megalotis_megalotis_TS305 | Vulpes_ferrilata | 0.176232883222296 |
| Otocyon_megalotis_megalotis_TS305 | Vulpes_lagopus | 0.172167626798465 |
| Otocyon_megalotis_megalotis_TS305 | Vulpes_vulpes_T3611 | 0.175276909955419 |
| Otocyon_megalotis_megalotis_TS305 | Vulpes_zerda | 0.174863002647014 |
| Otocyon_megalotis_megalotis_TS305 | Zalophus_californianus | 0.245752433552391 |
| Otocyon_megalotis_virgatus_FMNH158128 | Paguma_larvata | 0.254582156279334 |
| Otocyon_megalotis_virgatus_FMNH158128 | Panthera_leo | 0.244051311961908 |
| Otocyon_megalotis_virgatus_FMNH158128 | Panthera_onca | 0.248574321904603 |
| Otocyon_megalotis_virgatus_FMNH158128 | Panthera_pardus | 0.244771695882099 |
| Otocyon_megalotis_virgatus_FMNH158128 | Panthera_tigris | 0.244119030943544 |
| Otocyon_megalotis_virgatus_FMNH158128 | Panthera_uncia | 0.245086307289271 |
| Otocyon_megalotis_virgatus_FMNH158128 | Paradoxurus_hermaphroditus | 0.256684186034768 |
| Otocyon_megalotis_virgatus_FMNH158128 | Parahyaena_brunnea_FD126_Fecal | 0.25095233225241 |
| Otocyon_megalotis_virgatus_FMNH158128 | Pardofelis_marmorata | 0.250237335889026 |
| Otocyon_megalotis_virgatus_FMNH158128 | Phoca_fasciata | 0.235946093419102 |
| Otocyon_megalotis_virgatus_FMNH158128 | Phoca_groenlandica | 0.235161671076878 |
| Otocyon_megalotis_virgatus_FMNH158128 | Phoca_largha | 0.235824139532809 |
| Otocyon_megalotis_virgatus_FMNH158128 | Phoca_vitulina | 0.235437050382124 |
| Otocyon_megalotis_virgatus_FMNH158128 | Phocarctos_hookeri | 0.254348269008034 |
| Otocyon_megalotis_virgatus_FMNH158128 | Prionailurus_bengalensis | 0.245933796311274 |
| Otocyon_megalotis_virgatus_FMNH158128 | Prionailurus_bengalensis_euptilurus | 0.244175010440933 |
| Otocyon_megalotis_virgatus_FMNH158128 | Prionailurus_planiceps | 0.247923463712408 |
| Otocyon_megalotis_virgatus_FMNH158128 | Prionailurus_rubiginosus | 0.243635466318962 |
| Otocyon_megalotis_virgatus_FMNH158128 | Prionailurus_viverrinus | 0.244737630056953 |
| Otocyon_megalotis_virgatus_FMNH158128 | Prionodon_pardicolor | 0.244401522225916 |
| Otocyon_megalotis_virgatus_FMNH158128 | Procyon_lotor | 0.247298037004685 |
| Otocyon_megalotis_virgatus_FMNH158128 | Profelis_aurata | 0.249375147749828 |
| Otocyon_megalotis_virgatus_FMNH158128 | Proteles_cristatus_cristatus_TS307 | 0.255663995538659 |
| Otocyon_megalotis_virgatus_FMNH158128 | Proteles_cristatus_septentrionalis_NMS_Z_2018_54 | 0.256579806062101 |
| Otocyon_megalotis_virgatus_FMNH158128 | Puma_concolor | 0.241093280606216 |
| Otocyon_megalotis_virgatus_FMNH158128 | Puma_yagouaroundi | 0.243266274582253 |
| Otocyon_megalotis_virgatus_FMNH158128 | Pusa_caspica | 0.233562130431206 |
| Otocyon_megalotis_virgatus_FMNH158128 | Pusa_hispida | 0.232504441601653 |
| Otocyon_megalotis_virgatus_FMNH158128 | Pusa_sibirica | 0.231838158015521 |
| Otocyon_megalotis_virgatus_FMNH158128 | Speothos_venaticus_T1624 | 0.188792513268867 |
| Otocyon_megalotis_virgatus_FMNH158128 | Spilogale_putorius | 0.263318170070906 |
| Otocyon_megalotis_virgatus_FMNH158128 | Taxidea_taxus | 0.246755375864937 |
| Otocyon_megalotis_virgatus_FMNH158128 | Tremarctos_ornatus | 0.251221868939575 |
| Otocyon_megalotis_virgatus_FMNH158128 | Urocyon_cinereoargenteus | 0.189409536715772 |
| Otocyon_megalotis_virgatus_FMNH158128 | Ursus_americanus | 0.25052406697172 |
| Otocyon_megalotis_virgatus_FMNH158128 | Ursus_arctos | 0.250511000592523 |
| Otocyon_megalotis_virgatus_FMNH158128 | Ursus_maritimus | 0.250550243026914 |

| | | |
|---|---|---|
| Otocyon_megalotis_virgatus_FMNH158128 | Ursus_spelaeus | 0.252540618747386 |
| Otocyon_megalotis_virgatus_FMNH158128 | Ursus_thibetanus | 0.242015670773358 |
| Otocyon_megalotis_virgatus_FMNH158128 | Urva_javanica | 0.249059916432426 |
| Otocyon_megalotis_virgatus_FMNH158128 | Viverra_tangalunga_FMNH146957_SRR6053069 | 0.241248614040466 |
| Otocyon_megalotis_virgatus_FMNH158128 | Viverricula_indica | 0.241100570233021 |
| Otocyon_megalotis_virgatus_FMNH158128 | Vulpes_corsac | 0.178286440681731 |
| Otocyon_megalotis_virgatus_FMNH158128 | Vulpes_ferrilata | 0.176440094165999 |
| Otocyon_megalotis_virgatus_FMNH158128 | Vulpes_lagopus | 0.173507695634846 |
| Otocyon_megalotis_virgatus_FMNH158128 | Vulpes_vulpes_T3611 | 0.174035582368571 |
| Otocyon_megalotis_virgatus_FMNH158128 | Vulpes_zerda | 0.172486794346504 |
| Otocyon_megalotis_virgatus_FMNH158128 | Zalophus_californianus | 0.246045838362022 |
| Paguma_larvata | Panthera_leo | 0.194086998903629 |
| Paguma_larvata | Panthera_onca | 0.192437471062186 |
| Paguma_larvata | Panthera_pardus | 0.19705310929694 |
| Paguma_larvata | Panthera_tigris | 0.194855909919815 |
| Paguma_larvata | Panthera_uncia | 0.200987639709167 |
| Paguma_larvata | Paradoxurus_hermaphroditus | 0.092124645580435 |
| Paguma_larvata | Parahyaena_brunnea_FD126_Fecal | 0.204143731608473 |
| Paguma_larvata | Pardofelis_marmorata | 0.185633932972757 |
| Paguma_larvata | Phoca_fasciata | 0.220640162799725 |
| Paguma_larvata | Phoca_groenlandica | 0.221044212211982 |
| Paguma_larvata | Phoca_largha | 0.228867204713969 |
| Paguma_larvata | Phoca_vitulina | 0.227964688941557 |
| Paguma_larvata | Phocarctos_hookeri | 0.244397173476208 |
| Paguma_larvata | Prionailurus_bengalensis | 0.175067948100684 |
| Paguma_larvata | Prionailurus_bengalensis_euptilurus | 0.173985565937512 |
| Paguma_larvata | Prionailurus_planiceps | 0.176893357182136 |
| Paguma_larvata | Prionailurus_rubiginosus | 0.179684761965991 |
| Paguma_larvata | Prionailurus_viverrinus | 0.179552281948674 |
| Paguma_larvata | Prionodon_pardicolor | 0.189681932115668 |
| Paguma_larvata | Procyon_lotor | 0.234214093086659 |
| Paguma_larvata | Profelis_aurata | 0.181693605606295 |
| Paguma_larvata | Proteles_cristatus_cristatus_TS307 | 0.212276525969581 |
| Paguma_larvata | Proteles_cristatus_septentrionalis_NMS_Z_2018_54 | 0.208613111240556 |
| Paguma_larvata | Puma_concolor | 0.18257509442016 |
| Paguma_larvata | Puma_yagouaroundi | 0.181693335198831 |
| Paguma_larvata | Pusa_caspica | 0.228628853960655 |
| Paguma_larvata | Pusa_hispida | 0.224431056602432 |
| Paguma_larvata | Pusa_sibirica | 0.227228402592238 |
| Paguma_larvata | Speothos_venaticus_T1624 | 0.249978959252513 |
| Paguma_larvata | Spilogale_putorius | 0.235495883390706 |
| Paguma_larvata | Taxidea_taxus | 0.224295511711087 |
| Paguma_larvata | Tremarctos_ornatus | 0.240378969291276 |
| Paguma_larvata | Urocyon_cinereoargenteus | 0.253862719774082 |
| Paguma_larvata | Ursus_americanus | 0.238533327166408 |
| Paguma_larvata | Ursus_arctos | 0.243265557494012 |
| Paguma_larvata | Ursus_maritimus | 0.243455335593438 |
| Paguma_larvata | Ursus_spelaeus | 0.239788564596107 |
| Paguma_larvata | Ursus_thibetanus | 0.237129658136002 |
| Paguma_larvata | Urva_javanica | 0.195120658545844 |
| Paguma_larvata | Viverra_tangalunga_FMNH146957_SRR6053069 | 0.150501049152542 |
| Paguma_larvata | Viverricula_indica | 0.153363495215052 |
| Paguma_larvata | Vulpes_corsac | 0.245868978525793 |
| Paguma_larvata | Vulpes_ferrilata | 0.247267665319204 |
| Paguma_larvata | Vulpes_lagopus | 0.24384958152461 |
| Paguma_larvata | Vulpes_vulpes_T3611 | 0.239751089387355 |
| Paguma_larvata | Vulpes_zerda | 0.242390494296127 |
| Paguma_larvata | Zalophus_californianus | 0.235139982384413 |
| Panthera_leo | Panthera_onca | 0.0772894042421567 |
| Panthera_leo | Panthera_pardus | 0.0636506511683941 |
| Panthera_leo | Panthera_tigris | 0.101263153234747 |
| Panthera_leo | Panthera_uncia | 0.079682722660575 |
| Panthera_leo | Paradoxurus_hermaphroditus | 0.195612133467053 |
| Panthera_leo | Parahyaena_brunnea_FD126_Fecal | 0.19979758567126 |
| Panthera_leo | Pardofelis_marmorata | 0.137574901857413 |
| Panthera_leo | Phoca_fasciata | 0.225435420262086 |
| Panthera_leo | Phoca_groenlandica | 0.229309012530541 |
| Panthera_leo | Phoca_largha | 0.228391567302136 |
| Panthera_leo | Phoca_vitulina | 0.227228402592238 |
| Panthera_leo | Phocarctos_hookeri | 0.243479574059296 |
| Panthera_leo | Prionailurus_bengalensis | 0.130907912879267 |
| Panthera_leo | Prionailurus_bengalensis_euptilurus | 0.131488124027034 |
| Panthera_leo | Prionailurus_planiceps | 0.13677958968716 |
| Panthera_leo | Prionailurus_rubiginosus | 0.1355436085994 |
| Panthera_leo | Prionailurus_viverrinus | 0.134281229253285 |
| Panthera_leo | Prionodon_pardicolor | 0.196727467110231 |
| Panthera_leo | Procyon_lotor | 0.240462673218172 |
| Panthera_leo | Profelis_aurata | 0.138373477772657 |
| Panthera_leo | Proteles_cristatus_cristatus_TS307 | 0.201834827583815 |
| Panthera_leo | Proteles_cristatus_septentrionalis_NMS_Z_2018_54 | 0.203882700014146 |
| Panthera_leo | Puma_concolor | 0.133144258599855 |
| Panthera_leo | Puma_yagouaroundi | 0.137424622225282 |
| Panthera_leo | Pusa_caspica | 0.22623851444978 |
| Panthera_leo | Pusa_hispida | 0.222612036741451 |
| Panthera_leo | Pusa_sibirica | 0.22527749272307 |
| Panthera_leo | Speothos_venaticus_T1624 | 0.255150802932049 |
| Panthera_leo | Spilogale_putorius | 0.250592125245855 |

| | | |
|---|---|---|
| Panthera_leo | Taxidea_taxus | 0.230071210590289 |
| Panthera_leo | Tremarctos_ornatus | 0.242335882038925 |
| Panthera_leo | Urocyon_cinereoargenteus | 0.257917651902777 |
| Panthera_leo | Ursus_americanus | 0.241210546193583 |
| Panthera_leo | Ursus_arctos | 0.246208008409024 |
| Panthera_leo | Ursus_maritimus | 0.244578815340174 |
| Panthera_leo | Ursus_spelaeus | 0.242275262787103 |
| Panthera_leo | Ursus_thibetanus | 0.244099027647382 |
| Panthera_leo | Urva_javanica | 0.197552217151326 |
| Panthera_leo | Viverra_tangalunga_FMNH146957_SRR6053069 | 0.176333849827398 |
| Panthera_leo | Viverricula_indica | 0.182099437558172 |
| Panthera_leo | Vulpes_corsac | 0.248041894914094 |
| Panthera_leo | Vulpes_ferrilata | 0.24769197017457 |
| Panthera_leo | Vulpes_lagopus | 0.245593881265234 |
| Panthera_leo | Vulpes_vulpes_T3611 | 0.242792882412509 |
| Panthera_leo | Vulpes_zerda | 0.242207687786272 |
| Panthera_leo | Zalophus_californianus | 0.240279604479558 |
| Panthera_onca | Panthera_pardus | 0.0773539183845899 |
| Panthera_onca | Panthera_tigris | 0.100047529033308 |
| Panthera_onca | Panthera_uncia | 0.0836275181881455 |
| Panthera_onca | Paradoxurus_hermaphroditus | 0.199081774489792 |
| Panthera_onca | Parahyaena_brunnea_FD126_Fecal | 0.196910157383982 |
| Panthera_onca | Pardofelis_marmorata | 0.134133996886145 |
| Panthera_onca | Phoca_fasciata | 0.228221525012198 |
| Panthera_onca | Phoca_groenlandica | 0.231675238518176 |
| Panthera_onca | Phoca_largha | 0.229176199390246 |
| Panthera_onca | Phoca_vitulina | 0.230091639759122 |
| Panthera_onca | Phocarctos_hookeri | 0.245063827355317 |
| Panthera_onca | Prionailurus_bengalensis | 0.130203519567335 |
| Panthera_onca | Prionailurus_bengalensis_euptilurus | 0.132505026648355 |
| Panthera_onca | Prionailurus_planiceps | 0.132737278735973 |
| Panthera_onca | Prionailurus_rubiginosus | 0.13355231281989 |
| Panthera_onca | Prionailurus_viverrinus | 0.133280243912446 |
| Panthera_onca | Prionodon_pardicolor | 0.19413299873806 |
| Panthera_onca | Procyon_lotor | 0.241279657274515 |
| Panthera_onca | Profelis_aurata | 0.135015614355459 |
| Panthera_onca | Proteles_cristatus_cristatus_TS307 | 0.205044431271701 |
| Panthera_onca | Proteles_cristatus_septentrionalis_NMS_Z_2018_54 | 0.204553713291629 |
| Panthera_onca | Puma_concolor | 0.13334802031379 |
| Panthera_onca | Puma_yagouaroundi | 0.136127183371302 |
| Panthera_onca | Pusa_caspica | 0.229998758239002 |
| Panthera_onca | Pusa_hispida | 0.22449623059648 |
| Panthera_onca | Pusa_sibirica | 0.227368903392512 |
| Panthera_onca | Speothos_venaticus_T1624 | 0.256875374989926 |
| Panthera_onca | Spilogale_putorius | 0.253053944422541 |
| Panthera_onca | Taxidea_taxus | 0.227717701768405 |
| Panthera_onca | Tremarctos_ornatus | 0.246766342860502 |
| Panthera_onca | Urocyon_cinereoargenteus | 0.255578985873698 |
| Panthera_onca | Ursus_americanus | 0.243447770808934 |
| Panthera_onca | Ursus_arctos | 0.24860293243263 |
| Panthera_onca | Ursus_maritimus | 0.246739123063947 |
| Panthera_onca | Ursus_spelaeus | 0.244541384559607 |
| Panthera_onca | Ursus_thibetanus | 0.244868603132895 |
| Panthera_onca | Urva_javanica | 0.198580310718662 |
| Panthera_onca | Viverra_tangalunga_FMNH146957_SRR6053069 | 0.180730867729689 |
| Panthera_onca | Viverricula_indica | 0.181072267325122 |
| Panthera_onca | Vulpes_corsac | 0.247061613721369 |
| Panthera_onca | Vulpes_ferrilata | 0.246037778395374 |
| Panthera_onca | Vulpes_lagopus | 0.243229600099796 |
| Panthera_onca | Vulpes_vulpes_T3611 | 0.243389334389173 |
| Panthera_onca | Vulpes_zerda | 0.241017640936561 |
| Panthera_onca | Zalophus_californianus | 0.239446628266827 |
| Panthera_pardus | Panthera_tigris | 0.102702821596818 |
| Panthera_pardus | Panthera_uncia | 0.0840617948012861 |
| Panthera_pardus | Paradoxurus_hermaphroditus | 0.199078412731333 |
| Panthera_pardus | Parahyaena_brunnea_FD126_Fecal | 0.197108944538011 |
| Panthera_pardus | Pardofelis_marmorata | 0.138656447449515 |
| Panthera_pardus | Phoca_fasciata | 0.231832688581261 |
| Panthera_pardus | Phoca_groenlandica | 0.23265702508808 |
| Panthera_pardus | Phoca_largha | 0.232063475858155 |
| Panthera_pardus | Phoca_vitulina | 0.231220287750185 |
| Panthera_pardus | Phocarctos_hookeri | 0.244467975820029 |
| Panthera_pardus | Prionailurus_bengalensis | 0.130599042069092 |
| Panthera_pardus | Prionailurus_bengalensis_euptilurus | 0.131855958204797 |
| Panthera_pardus | Prionailurus_planiceps | 0.136563711250344 |
| Panthera_pardus | Prionailurus_rubiginosus | 0.132030637288079 |
| Panthera_pardus | Prionailurus_viverrinus | 0.132901970589997 |
| Panthera_pardus | Prionodon_pardicolor | 0.196416031274882 |
| Panthera_pardus | Procyon_lotor | 0.245925287562672 |
| Panthera_pardus | Profelis_aurata | 0.137686597083728 |
| Panthera_pardus | Proteles_cristatus_cristatus_TS307 | 0.202558479687474 |
| Panthera_pardus | Proteles_cristatus_septentrionalis_NMS_Z_2018_54 | 0.201728433286147 |
| Panthera_pardus | Puma_concolor | 0.136119876030544 |
| Panthera_pardus | Puma_yagouaroundi | 0.138635716428319 |
| Panthera_pardus | Pusa_caspica | 0.229021260815997 |
| Panthera_pardus | Pusa_hispida | 0.226792595840386 |
| Panthera_pardus | Pusa_sibirica | 0.229041942265669 |

| | | |
|---|---|---|
| Panthera_pardus | Speothos_venaticus_T1624 | 0.253896691761241 |
| Panthera_pardus | Spilogale_putorius | 0.255734309610072 |
| Panthera_pardus | Taxidea_taxus | 0.230172946509931 |
| Panthera_pardus | Tremarctos_ornatus | 0.250408714649845 |
| Panthera_pardus | Urocyon_cinereoargenteus | 0.254269571699554 |
| Panthera_pardus | Ursus_americanus | 0.240173002829435 |
| Panthera_pardus | Ursus_arctos | 0.249010797855307 |
| Panthera_pardus | Ursus_maritimus | 0.246346913401563 |
| Panthera_pardus | Ursus_spelaeus | 0.244036689824493 |
| Panthera_pardus | Ursus_thibetanus | 0.242915143764943 |
| Panthera_pardus | Urva_javanica | 0.20014739072634 |
| Panthera_pardus | Viverra_tangalunga_FMNH146957_SRR6053069 | 0.181589758573966 |
| Panthera_pardus | Viverricula_indica | 0.182059730684313 |
| Panthera_pardus | Vulpes_corsac | 0.250806432427895 |
| Panthera_pardus | Vulpes_ferrilata | 0.249886160832865 |
| Panthera_pardus | Vulpes_lagopus | 0.246111475909983 |
| Panthera_pardus | Vulpes_vulpes_T3611 | 0.243304870881128 |
| Panthera_pardus | Vulpes_zerda | 0.241825063069156 |
| Panthera_pardus | Zalophus_californianus | 0.243075094261845 |
| Panthera_tigris | Panthera_uncia | 0.108057773666168 |
| Panthera_tigris | Paradoxurus_hermaphroditus | 0.201714322641423 |
| Panthera_tigris | Parahyaena_brunnea_FD126_Fecal | 0.203054134194693 |
| Panthera_tigris | Pardofelis_marmorata | 0.143917164204504 |
| Panthera_tigris | Phoca_fasciata | 0.228196068044827 |
| Panthera_tigris | Phoca_groenlandica | 0.230685832301026 |
| Panthera_tigris | Phoca_largha | 0.229962110675651 |
| Panthera_tigris | Phoca_vitulina | 0.229449729963786 |
| Panthera_tigris | Phocarctos_hookeri | 0.243747838496084 |
| Panthera_tigris | Prionailurus_bengalensis | 0.133796954431159 |
| Panthera_tigris | Prionailurus_bengalensis_euptilurus | 0.13545671390923 |
| Panthera_tigris | Prionailurus_planiceps | 0.142471263443049 |
| Panthera_tigris | Prionailurus_rubiginosus | 0.13843560904638 |
| Panthera_tigris | Prionailurus_viverrinus | 0.138171213345527 |
| Panthera_tigris | Prionodon_pardicolor | 0.202369974006551 |
| Panthera_tigris | Procyon_lotor | 0.244686874175777 |
| Panthera_tigris | Profelis_aurata | 0.139933830018526 |
| Panthera_tigris | Proteles_cristatus_cristatus_TS307 | 0.210684219032801 |
| Panthera_tigris | Proteles_cristatus_septentrionalis_NMS_Z_2018_54 | 0.209193024121028 |
| Panthera_tigris | Puma_concolor | 0.136570385158995 |
| Panthera_tigris | Puma_yagouaroundi | 0.141937553873149 |
| Panthera_tigris | Pusa_caspica | 0.228571602988039 |
| Panthera_tigris | Pusa_hispida | 0.225903468914039 |
| Panthera_tigris | Pusa_sibirica | 0.225746492770104 |
| Panthera_tigris | Speothos_venaticus_T1624 | 0.252986876884934 |
| Panthera_tigris | Spilogale_putorius | 0.25420692921578 |
| Panthera_tigris | Taxidea_taxus | 0.23572494176536 |
| Panthera_tigris | Tremarctos_ornatus | 0.247911724984155 |
| Panthera_tigris | Urocyon_cinereoargenteus | 0.256684186034768 |
| Panthera_tigris | Ursus_americanus | 0.245993024994244 |
| Panthera_tigris | Ursus_arctos | 0.252191388482688 |
| Panthera_tigris | Ursus_maritimus | 0.248134606900744 |
| Panthera_tigris | Ursus_spelaeus | 0.245386562433588 |
| Panthera_tigris | Ursus_thibetanus | 0.244351497714275 |
| Panthera_tigris | Urva_javanica | 0.201122218823728 |
| Panthera_tigris | Viverra_tangalunga_FMNH146957_SRR6053069 | 0.181052637514735 |
| Panthera_tigris | Viverricula_indica | 0.184743301801309 |
| Panthera_tigris | Vulpes_corsac | 0.247589692565967 |
| Panthera_tigris | Vulpes_ferrilata | 0.245324622281079 |
| Panthera_tigris | Vulpes_lagopus | 0.241777989506626 |
| Panthera_tigris | Vulpes_vulpes_T3611 | 0.246743025064922 |
| Panthera_tigris | Vulpes_zerda | 0.243345388909312 |
| Panthera_tigris | Zalophus_californianus | 0.237113075813345 |
| Panthera_uncia | Paradoxurus_hermaphroditus | 0.203145651256133 |
| Panthera_uncia | Parahyaena_brunnea_FD126_Fecal | 0.201856462837825 |
| Panthera_uncia | Pardofelis_marmorata | 0.141037093935217 |
| Panthera_uncia | Phoca_fasciata | 0.229326402730274 |
| Panthera_uncia | Phoca_groenlandica | 0.235978928570568 |
| Panthera_uncia | Phoca_largha | 0.229227002173943 |
| Panthera_uncia | Phoca_vitulina | 0.228280702301302 |
| Panthera_uncia | Phocarctos_hookeri | 0.246720739261451 |
| Panthera_uncia | Prionailurus_bengalensis | 0.13702597822603 |
| Panthera_uncia | Prionailurus_bengalensis_euptilurus | 0.137811120228791 |
| Panthera_uncia | Prionailurus_planiceps | 0.139808966786409 |
| Panthera_uncia | Prionailurus_rubiginosus | 0.142394432515766 |
| Panthera_uncia | Prionailurus_viverrinus | 0.140533377042969 |
| Panthera_uncia | Prionodon_pardicolor | 0.196576509849189 |
| Panthera_uncia | Procyon_lotor | 0.245496719112562 |
| Panthera_uncia | Profelis_aurata | 0.137362474592291 |
| Panthera_uncia | Proteles_cristatus_cristatus_TS307 | 0.207455348548072 |
| Panthera_uncia | Proteles_cristatus_septentrionalis_NMS_Z_2018_54 | 0.208338627928338 |
| Panthera_uncia | Puma_concolor | 0.136487485411566 |
| Panthera_uncia | Puma_yagouaroundi | 0.142142827442516 |
| Panthera_uncia | Pusa_caspica | 0.229041942265669 |
| Panthera_uncia | Pusa_hispida | 0.225831225587383 |
| Panthera_uncia | Pusa_sibirica | 0.227419742587765 |
| Panthera_uncia | Speothos_venaticus_T1624 | 0.260074661894639 |
| Panthera_uncia | Spilogale_putorius | 0.254751232595477 |

| | | |
|---|---|---|
| Panthera_uncia | Taxidea_taxus | 0.234185162674479 |
| Panthera_uncia | Tremarctos_ornatus | 0.247700224174107 |
| Panthera_uncia | Urocyon_cinereoargenteus | 0.258930446697806 |
| Panthera_uncia | Ursus_americanus | 0.2468161045311 |
| Panthera_uncia | Ursus_arctos | 0.24845429441247 |
| Panthera_uncia | Ursus_maritimus | 0.24613234374078 |
| Panthera_uncia | Ursus_spelaeus | 0.246551110247831 |
| Panthera_uncia | Ursus_thibetanus | 0.243720993306575 |
| Panthera_uncia | Urva_javanica | 0.202737448807142 |
| Panthera_uncia | Viverra_tangalunga_FMNH146957_SRR6053069 | 0.185227272018927 |
| Panthera_uncia | Viverricula_indica | 0.184217616209235 |
| Panthera_uncia | Vulpes_corsac | 0.248901201230197 |
| Panthera_uncia | Vulpes_ferrilata | 0.249568988195464 |
| Panthera_uncia | Vulpes_lagopus | 0.244897071231166 |
| Panthera_uncia | Vulpes_vulpes_T3611 | 0.240784399461826 |
| Panthera_uncia | Vulpes_zerda | 0.240422768443159 |
| Panthera_uncia | Zalophus_californianus | 0.242312716606985 |
| Paradoxurus_hermaphroditus | Parahyaena_brunnea_FD126_Fecal | 0.214286359266121 |
| Paradoxurus_hermaphroditus | Pardofelis_marmorata | 0.190489957955703 |
| Paradoxurus_hermaphroditus | Phoca_fasciata | 0.229304458786436 |
| Paradoxurus_hermaphroditus | Phoca_groenlandica | 0.228280719513379 |
| Paradoxurus_hermaphroditus | Phoca_largha | 0.240574505023603 |
| Paradoxurus_hermaphroditus | Phoca_vitulina | 0.239982501788598 |
| Paradoxurus_hermaphroditus | Phocarctos_hookeri | 0.244951954815644 |
| Paradoxurus_hermaphroditus | Prionailurus_bengalensis | 0.178604479354611 |
| Paradoxurus_hermaphroditus | Prionailurus_bengalensis_euptilurus | 0.181419393877214 |
| Paradoxurus_hermaphroditus | Prionailurus_planiceps | 0.183304999912754 |
| Paradoxurus_hermaphroditus | Prionailurus_rubiginosus | 0.183158087938293 |
| Paradoxurus_hermaphroditus | Prionailurus_viverrinus | 0.186643299228406 |
| Paradoxurus_hermaphroditus | Prionodon_pardicolor | 0.190303558554297 |
| Paradoxurus_hermaphroditus | Procyon_lotor | 0.243742153478884 |
| Paradoxurus_hermaphroditus | Profelis_aurata | 0.182222296463188 |
| Paradoxurus_hermaphroditus | Proteles_cristatus_cristatus_TS307 | 0.220502984272051 |
| Paradoxurus_hermaphroditus | Proteles_cristatus_septentrionalis_NMS_Z_2018_54 | 0.217302784152073 |
| Paradoxurus_hermaphroditus | Puma_concolor | 0.187508140228854 |
| Paradoxurus_hermaphroditus | Puma_yagouaroundi | 0.184934453533274 |
| Paradoxurus_hermaphroditus | Pusa_caspica | 0.238696737060409 |
| Paradoxurus_hermaphroditus | Pusa_hispida | 0.236270687834084 |
| Paradoxurus_hermaphroditus | Pusa_sibirica | 0.237915835518365 |
| Paradoxurus_hermaphroditus | Speothos_venaticus_T1624 | 0.246722450221081 |
| Paradoxurus_hermaphroditus | Spilogale_putorius | 0.231143001263579 |
| Paradoxurus_hermaphroditus | Taxidea_taxus | 0.231840118863514 |
| Paradoxurus_hermaphroditus | Tremarctos_ornatus | 0.248918868033807 |
| Paradoxurus_hermaphroditus | Urocyon_cinereoargenteus | 0.254605338110233 |
| Paradoxurus_hermaphroditus | Ursus_americanus | 0.246038575779479 |
| Paradoxurus_hermaphroditus | Ursus_arctos | 0.251181344952775 |
| Paradoxurus_hermaphroditus | Ursus_maritimus | 0.253413615582716 |
| Paradoxurus_hermaphroditus | Ursus_spelaeus | 0.24590087955424 |
| Paradoxurus_hermaphroditus | Ursus_thibetanus | 0.244509339229913 |
| Paradoxurus_hermaphroditus | Urva_javanica | 0.196705874018062 |
| Paradoxurus_hermaphroditus | Viverra_tangalunga_FMNH146957_SRR6053069 | 0.156266354299463 |
| Paradoxurus_hermaphroditus | Viverricula_indica | 0.15321040327122 |
| Paradoxurus_hermaphroditus | Vulpes_corsac | 0.24731666914346 |
| Paradoxurus_hermaphroditus | Vulpes_ferrilata | 0.250008866162515 |
| Paradoxurus_hermaphroditus | Vulpes_lagopus | 0.244601104168453 |
| Paradoxurus_hermaphroditus | Vulpes_vulpes_T3611 | 0.243008037786464 |
| Paradoxurus_hermaphroditus | Vulpes_zerda | 0.245689784573251 |
| Paradoxurus_hermaphroditus | Zalophus_californianus | 0.242714814618183 |
| Parahyaena_brunnea_FD126_Fecal | Pardofelis_marmorata | 0.194992873648456 |
| Parahyaena_brunnea_FD126_Fecal | Phoca_fasciata | 0.228535747928607 |
| Parahyaena_brunnea_FD126_Fecal | Phoca_groenlandica | 0.233444100634409 |
| Parahyaena_brunnea_FD126_Fecal | Phoca_largha | 0.224691075179891 |
| Parahyaena_brunnea_FD126_Fecal | Phoca_vitulina | 0.226160822463226 |
| Parahyaena_brunnea_FD126_Fecal | Phocarctos_hookeri | 0.246016038375634 |
| Parahyaena_brunnea_FD126_Fecal | Prionailurus_bengalensis | 0.186478698207224 |
| Parahyaena_brunnea_FD126_Fecal | Prionailurus_bengalensis_euptilurus | 0.187721320779083 |
| Parahyaena_brunnea_FD126_Fecal | Prionailurus_planiceps | 0.193372129123316 |
| Parahyaena_brunnea_FD126_Fecal | Prionailurus_rubiginosus | 0.188441418843202 |
| Parahyaena_brunnea_FD126_Fecal | Prionailurus_viverrinus | 0.187391193833006 |
| Parahyaena_brunnea_FD126_Fecal | Prionodon_pardicolor | 0.206641743591826 |
| Parahyaena_brunnea_FD126_Fecal | Procyon_lotor | 0.244153126024647 |
| Parahyaena_brunnea_FD126_Fecal | Profelis_aurata | 0.196549602991687 |
| Parahyaena_brunnea_FD126_Fecal | Proteles_cristatus_cristatus_TS307 | 0.126000435155031 |
| Parahyaena_brunnea_FD126_Fecal | Proteles_cristatus_septentrionalis_NMS_Z_2018_54 | 0.12745470829822 |
| Parahyaena_brunnea_FD126_Fecal | Puma_concolor | 0.191626711472987 |
| Parahyaena_brunnea_FD126_Fecal | Puma_yagouaroundi | 0.191045383658912 |
| Parahyaena_brunnea_FD126_Fecal | Pusa_caspica | 0.227551294146543 |
| Parahyaena_brunnea_FD126_Fecal | Pusa_hispida | 0.222748230038487 |
| Parahyaena_brunnea_FD126_Fecal | Pusa_sibirica | 0.223054399339407 |
| Parahyaena_brunnea_FD126_Fecal | Speothos_venaticus_T1624 | 0.256650909356036 |
| Parahyaena_brunnea_FD126_Fecal | Spilogale_putorius | 0.257894167901587 |
| Parahyaena_brunnea_FD126_Fecal | Taxidea_taxus | 0.232823419710224 |
| Parahyaena_brunnea_FD126_Fecal | Tremarctos_ornatus | 0.242149445879559 |
| Parahyaena_brunnea_FD126_Fecal | Urocyon_cinereoargenteus | 0.255778649160841 |
| Parahyaena_brunnea_FD126_Fecal | Ursus_americanus | 0.244530366307335 |
| Parahyaena_brunnea_FD126_Fecal | Ursus_arctos | 0.247906187102832 |
| Parahyaena_brunnea_FD126_Fecal | Ursus_maritimus | 0.24480457894921 |

| | | |
|---|---|---|
| Parahyaena_brunnea_FD126_Fecal | Ursus_spelaeus | 0.239626684859281 |
| Parahyaena_brunnea_FD126_Fecal | Ursus_thibetanus | 0.243891229905257 |
| Parahyaena_brunnea_FD126_Fecal | Urva_javanica | 0.188830516076696 |
| Parahyaena_brunnea_FD126_Fecal | Viverra_tangalunga_FMNH146957_SRR6053069 | 0.199182670482944 |
| Parahyaena_brunnea_FD126_Fecal | Viverricula_indica | 0.197056356318864 |
| Parahyaena_brunnea_FD126_Fecal | Vulpes_corsac | 0.252685441446321 |
| Parahyaena_brunnea_FD126_Fecal | Vulpes_ferrilata | 0.25476647824159 |
| Parahyaena_brunnea_FD126_Fecal | Vulpes_lagopus | 0.251833518925569 |
| Parahyaena_brunnea_FD126_Fecal | Vulpes_vulpes_T3611 | 0.249525189971545 |
| Parahyaena_brunnea_FD126_Fecal | Vulpes_zerda | 0.247134170732217 |
| Parahyaena_brunnea_FD126_Fecal | Zalophus_californianus | 0.236611452822069 |
| Pardofelis_marmorata | Phoca_fasciata | 0.222624365169058 |
| Pardofelis_marmorata | Phoca_groenlandica | 0.222023106541318 |
| Pardofelis_marmorata | Phoca_largha | 0.222907988153355 |
| Pardofelis_marmorata | Phoca_vitulina | 0.223293150550835 |
| Pardofelis_marmorata | Phocarctos_hookeri | 0.242257983852929 |
| Pardofelis_marmorata | Prionailurus_bengalensis | 0.112188756949838 |
| Pardofelis_marmorata | Prionailurus_bengalensis_euptilurus | 0.112382889082668 |
| Pardofelis_marmorata | Prionailurus_planiceps | 0.113438889205014 |
| Pardofelis_marmorata | Prionailurus_rubiginosus | 0.114214911836581 |
| Pardofelis_marmorata | Prionailurus_viverrinus | 0.113643352198736 |
| Pardofelis_marmorata | Prionodon_pardicolor | 0.188948827948736 |
| Pardofelis_marmorata | Procyon_lotor | 0.241907403994805 |
| Pardofelis_marmorata | Profelis_aurata | 0.118195850387738 |
| Pardofelis_marmorata | Proteles_cristatus_cristatus_TS307 | 0.199618290147866 |
| Pardofelis_marmorata | Proteles_cristatus_septentrionalis_NMS_Z_2018_54 | 0.197402161572669 |
| Pardofelis_marmorata | Puma_concolor | 0.106807814342975 |
| Pardofelis_marmorata | Puma_yagouaroundi | 0.114581751048826 |
| Pardofelis_marmorata | Pusa_caspica | 0.224932311142332 |
| Pardofelis_marmorata | Pusa_hispida | 0.222935935738111 |
| Pardofelis_marmorata | Pusa_sibirica | 0.223853667038622 |
| Pardofelis_marmorata | Speothos_venaticus_T1624 | 0.250300026510554 |
| Pardofelis_marmorata | Spilogale_putorius | 0.252192144655854 |
| Pardofelis_marmorata | Taxidea_taxus | 0.225831260455022 |
| Pardofelis_marmorata | Tremarctos_ornatus | 0.243447770808934 |
| Pardofelis_marmorata | Urocyon_cinereoargenteus | 0.249025952413121 |
| Pardofelis_marmorata | Ursus_americanus | 0.241729713021356 |
| Pardofelis_marmorata | Ursus_arctos | 0.251259777871262 |
| Pardofelis_marmorata | Ursus_maritimus | 0.247335179375532 |
| Pardofelis_marmorata | Ursus_spelaeus | 0.246386131883776 |
| Pardofelis_marmorata | Ursus_thibetanus | 0.24229360165674 |
| Pardofelis_marmorata | Urva_javanica | 0.193628828954577 |
| Pardofelis_marmorata | Viverra_tangalunga_FMNH146957_SRR6053069 | 0.177349568005995 |
| Pardofelis_marmorata | Viverricula_indica | 0.178020128719139 |
| Pardofelis_marmorata | Vulpes_corsac | 0.245728711632431 |
| Pardofelis_marmorata | Vulpes_ferrilata | 0.245382037492417 |
| Pardofelis_marmorata | Vulpes_lagopus | 0.24000329721767 |
| Pardofelis_marmorata | Vulpes_vulpes_T3611 | 0.239494672501388 |
| Pardofelis_marmorata | Vulpes_zerda | 0.238620332437348 |
| Pardofelis_marmorata | Zalophus_californianus | 0.236111650438908 |
| Phoca_fasciata | Phoca_groenlandica | 0.0696295227161188 |
| Phoca_fasciata | Phoca_largha | 0.0839421608272918 |
| Phoca_fasciata | Phoca_vitulina | 0.0851406836783863 |
| Phoca_fasciata | Phocarctos_hookeri | 0.185358584373089 |
| Phoca_fasciata | Prionailurus_bengalensis | 0.216548713979205 |
| Phoca_fasciata | Prionailurus_bengalensis_euptilurus | 0.216548713979205 |
| Phoca_fasciata | Prionailurus_planiceps | 0.218119017927616 |
| Phoca_fasciata | Prionailurus_rubiginosus | 0.217547179644251 |
| Phoca_fasciata | Prionailurus_viverrinus | 0.218566411805148 |
| Phoca_fasciata | Prionodon_pardicolor | 0.227920193215067 |
| Phoca_fasciata | Procyon_lotor | 0.219315049215352 |
| Phoca_fasciata | Profelis_aurata | 0.220718109585175 |
| Phoca_fasciata | Proteles_cristatus_cristatus_TS307 | 0.230501030511204 |
| Phoca_fasciata | Proteles_cristatus_septentrionalis_NMS_Z_2018_54 | 0.230919314635632 |
| Phoca_fasciata | Puma_concolor | 0.217128912089042 |
| Phoca_fasciata | Puma_yagouaroundi | 0.21884286825522 |
| Phoca_fasciata | Pusa_caspica | 0.0827431882795146 |
| Phoca_fasciata | Pusa_hispida | 0.0825545823738626 |
| Phoca_fasciata | Pusa_sibirica | 0.0822014750145542 |
| Phoca_fasciata | Speothos_venaticus_T1624 | 0.238022968257532 |
| Phoca_fasciata | Spilogale_putorius | 0.228827581112048 |
| Phoca_fasciata | Taxidea_taxus | 0.208172838304985 |
| Phoca_fasciata | Tremarctos_ornatus | 0.212181632018958 |
| Phoca_fasciata | Urocyon_cinereoargenteus | 0.233904756727524 |
| Phoca_fasciata | Ursus_americanus | 0.214539622501139 |
| Phoca_fasciata | Ursus_arctos | 0.219627589421405 |
| Phoca_fasciata | Ursus_maritimus | 0.21644977764124 |
| Phoca_fasciata | Ursus_spelaeus | 0.212372133396759 |
| Phoca_fasciata | Ursus_thibetanus | 0.212239057833358 |
| Phoca_fasciata | Urva_javanica | 0.219372317650988 |
| Phoca_fasciata | Viverra_tangalunga_FMNH146957_SRR6053069 | 0.218552423317906 |
| Phoca_fasciata | Viverricula_indica | 0.21421808884188 |
| Phoca_fasciata | Vulpes_corsac | 0.231048895717789 |
| Phoca_fasciata | Vulpes_ferrilata | 0.233502744538481 |
| Phoca_fasciata | Vulpes_lagopus | 0.234255060265943 |
| Phoca_fasciata | Vulpes_vulpes_T3611 | 0.229823615075181 |
| Phoca_fasciata | Vulpes_zerda | 0.229158270628709 |

| | | |
|---|---|---|
| Phoca_fasciata | Zalophus_californianus | 0.17831091356199 |
| Phoca_groenlandica | Phoca_largha | 0.0862460463686915 |
| Phoca_groenlandica | Phoca_vitulina | 0.0865582604357981 |
| Phoca_groenlandica | Phocarctos_hookeri | 0.187822956775243 |
| Phoca_groenlandica | Prionailurus_bengalensis | 0.217217887977516 |
| Phoca_groenlandica | Prionailurus_bengalensis_euptilurus | 0.218308537301077 |
| Phoca_groenlandica | Prionailurus_planiceps | 0.218682936808943 |
| Phoca_groenlandica | Prionailurus_rubiginosus | 0.219986736254333 |
| Phoca_groenlandica | Prionailurus_viverrinus | 0.219490952127088 |
| Phoca_groenlandica | Prionodon_pardicolor | 0.226783886424064 |
| Phoca_groenlandica | Procyon_lotor | 0.219761546408901 |
| Phoca_groenlandica | Profelis_aurata | 0.222946689425416 |
| Phoca_groenlandica | Proteles_cristatus_cristatus_TS307 | 0.238636943686072 |
| Phoca_groenlandica | Proteles_cristatus_septentrionalis_NMS_Z_2018_54 | 0.23937298503828 |
| Phoca_groenlandica | Puma_concolor | 0.216960169807008 |
| Phoca_groenlandica | Puma_yagouaroundi | 0.219659644418259 |
| Phoca_groenlandica | Pusa_caspica | 0.0869079076120956 |
| Phoca_groenlandica | Pusa_hispida | 0.085917709534073 |
| Phoca_groenlandica | Pusa_sibirica | 0.0860953452051837 |
| Phoca_groenlandica | Speothos_venaticus_T1624 | 0.237927124985442 |
| Phoca_groenlandica | Spilogale_putorius | 0.227475114985924 |
| Phoca_groenlandica | Taxidea_taxus | 0.20463605281939 |
| Phoca_groenlandica | Tremarctos_ornatus | 0.214590860038183 |
| Phoca_groenlandica | Urocyon_cinereoargenteus | 0.235845962594261 |
| Phoca_groenlandica | Ursus_americanus | 0.218432952033241 |
| Phoca_groenlandica | Ursus_arctos | 0.222087498126417 |
| Phoca_groenlandica | Ursus_maritimus | 0.221325610512362 |
| Phoca_groenlandica | Ursus_spelaeus | 0.217983771751677 |
| Phoca_groenlandica | Ursus_thibetanus | 0.211865482802024 |
| Phoca_groenlandica | Urva_javanica | 0.227244256573524 |
| Phoca_groenlandica | Viverra_tangalunga_FMNH146957_SRR6053069 | 0.217846883662076 |
| Phoca_groenlandica | Viverricula_indica | 0.215330117236519 |
| Phoca_groenlandica | Vulpes_corsac | 0.23754226051247 |
| Phoca_groenlandica | Vulpes_ferrilata | 0.238331748465551 |
| Phoca_groenlandica | Vulpes_lagopus | 0.236754170412986 |
| Phoca_groenlandica | Vulpes_vulpes_T3611 | 0.233871012579371 |
| Phoca_groenlandica | Vulpes_zerda | 0.233755287118385 |
| Phoca_groenlandica | Zalophus_californianus | 0.181237429145317 |
| Phoca_largha | Phoca_vitulina | 0.0209865582787697 |
| Phoca_largha | Phocarctos_hookeri | 0.188288718784679 |
| Phoca_largha | Prionailurus_bengalensis | 0.218575560023392 |
| Phoca_largha | Prionailurus_bengalensis_euptilurus | 0.218897759583462 |
| Phoca_largha | Prionailurus_planiceps | 0.221623396416079 |
| Phoca_largha | Prionailurus_rubiginosus | 0.222252609698941 |
| Phoca_largha | Prionailurus_viverrinus | 0.220690052865627 |
| Phoca_largha | Prionodon_pardicolor | 0.231547206111882 |
| Phoca_largha | Procyon_lotor | 0.219521729061763 |
| Phoca_largha | Profelis_aurata | 0.226088588219114 |
| Phoca_largha | Proteles_cristatus_cristatus_TS307 | 0.232244557514274 |
| Phoca_largha | Proteles_cristatus_septentrionalis_NMS_Z_2018_54 | 0.2327540086725 |
| Phoca_largha | Puma_concolor | 0.217584972330674 |
| Phoca_largha | Puma_yagouaroundi | 0.220320499705529 |
| Phoca_largha | Pusa_caspica | 0.0354443015100179 |
| Phoca_largha | Pusa_hispida | 0.0385101082325893 |
| Phoca_largha | Pusa_sibirica | 0.0331711588751885 |
| Phoca_largha | Speothos_venaticus_T1624 | 0.239193434186037 |
| Phoca_largha | Spilogale_putorius | 0.229610270280281 |
| Phoca_largha | Taxidea_taxus | 0.205021769246686 |
| Phoca_largha | Tremarctos_ornatus | 0.216715699035458 |
| Phoca_largha | Urocyon_cinereoargenteus | 0.235158434341826 |
| Phoca_largha | Ursus_americanus | 0.21904282405188 |
| Phoca_largha | Ursus_arctos | 0.219920150979288 |
| Phoca_largha | Ursus_maritimus | 0.218059529973797 |
| Phoca_largha | Ursus_spelaeus | 0.217941138027235 |
| Phoca_largha | Ursus_thibetanus | 0.215416434529069 |
| Phoca_largha | Urva_javanica | 0.223837601667139 |
| Phoca_largha | Viverra_tangalunga_FMNH146957_SRR6053069 | 0.22282775459861 |
| Phoca_largha | Viverricula_indica | 0.221705608976103 |
| Phoca_largha | Vulpes_corsac | 0.237162331559265 |
| Phoca_largha | Vulpes_ferrilata | 0.237395446637155 |
| Phoca_largha | Vulpes_lagopus | 0.237417126734117 |
| Phoca_largha | Vulpes_vulpes_T3611 | 0.23537256558771 |
| Phoca_largha | Vulpes_zerda | 0.234593601173415 |
| Phoca_largha | Zalophus_californianus | 0.182260195244709 |
| Phoca_vitulina | Phocarctos_hookeri | 0.188780267301355 |
| Phoca_vitulina | Prionailurus_bengalensis | 0.218959370857181 |
| Phoca_vitulina | Prionailurus_bengalensis_euptilurus | 0.219066782243201 |
| Phoca_vitulina | Prionailurus_planiceps | 0.222764824509187 |
| Phoca_vitulina | Prionailurus_rubiginosus | 0.221342385304196 |
| Phoca_vitulina | Prionailurus_viverrinus | 0.221721966772275 |
| Phoca_vitulina | Prionodon_pardicolor | 0.231608092813474 |
| Phoca_vitulina | Procyon_lotor | 0.220033260574651 |
| Phoca_vitulina | Profelis_aurata | 0.227129225287778 |
| Phoca_vitulina | Proteles_cristatus_cristatus_TS307 | 0.232214943897844 |
| Phoca_vitulina | Proteles_cristatus_septentrionalis_NMS_Z_2018_54 | 0.234484368083167 |
| Phoca_vitulina | Puma_concolor | 0.218290159458596 |
| Phoca_vitulina | Puma_yagouaroundi | 0.219414501049196 |

| | | |
|---|---|---|
| Phoca_vitulina | Pusa_caspica | 0.0370482665888212 |
| Phoca_vitulina | Pusa_hispida | 0.0397155346156532 |
| Phoca_vitulina | Pusa_sibirica | 0.0363812386517211 |
| Phoca_vitulina | Speothos_venaticus_T1624 | 0.24034423645624 |
| Phoca_vitulina | Spilogale_putorius | 0.230239238285971 |
| Phoca_vitulina | Taxidea_taxus | 0.207655234699505 |
| Phoca_vitulina | Tremarctos_ornatus | 0.216702904508195 |
| Phoca_vitulina | Urocyon_cinereoargenteus | 0.234461428077478 |
| Phoca_vitulina | Ursus_americanus | 0.22115584633254 |
| Phoca_vitulina | Ursus_arctos | 0.223923726425896 |
| Phoca_vitulina | Ursus_maritimus | 0.221715809331637 |
| Phoca_vitulina | Ursus_spelaeus | 0.221018914682905 |
| Phoca_vitulina | Ursus_thibetanus | 0.21584208685398 |
| Phoca_vitulina | Urva_javanica | 0.224117173825449 |
| Phoca_vitulina | Viverra_tangalunga_FMNH146957_SRR6053069 | 0.222216389201151 |
| Phoca_vitulina | Viverricula_indica | 0.220235452381531 |
| Phoca_vitulina | Vulpes_corsac | 0.237284273762878 |
| Phoca_vitulina | Vulpes_ferrilata | 0.238296652666631 |
| Phoca_vitulina | Vulpes_lagopus | 0.237051220447697 |
| Phoca_vitulina | Vulpes_vulpes_T3611 | 0.232268960335726 |
| Phoca_vitulina | Vulpes_zerda | 0.231602060882871 |
| Phoca_vitulina | Zalophus_californianus | 0.182167178488861 |
| Phocarctos_hookeri | Prionailurus_bengalensis | 0.232594252363434 |
| Phocarctos_hookeri | Prionailurus_bengalensis_euptilurus | 0.231815523545662 |
| Phocarctos_hookeri | Prionailurus_planiceps | 0.233793765493199 |
| Phocarctos_hookeri | Prionailurus_rubiginosus | 0.235898548182429 |
| Phocarctos_hookeri | Prionailurus_viverrinus | 0.233785001430006 |
| Phocarctos_hookeri | Prionodon_pardicolor | 0.24644805158382 |
| Phocarctos_hookeri | Procyon_lotor | 0.239606513632839 |
| Phocarctos_hookeri | Profelis_aurata | 0.241897850541349 |
| Phocarctos_hookeri | Proteles_cristatus_cristatus_TS307 | 0.252486083768162 |
| Phocarctos_hookeri | Proteles_cristatus_septentrionalis_NMS_Z_2018_54 | 0.253190586526255 |
| Phocarctos_hookeri | Puma_concolor | 0.239187008727204 |
| Phocarctos_hookeri | Puma_yagouaroundi | 0.237638385715213 |
| Phocarctos_hookeri | Pusa_caspica | 0.188005805320894 |
| Phocarctos_hookeri | Pusa_hispida | 0.184815793089356 |
| Phocarctos_hookeri | Pusa_sibirica | 0.185293711572316 |
| Phocarctos_hookeri | Speothos_venaticus_T1624 | 0.262528100735624 |
| Phocarctos_hookeri | Spilogale_putorius | 0.248800409495937 |
| Phocarctos_hookeri | Taxidea_taxus | 0.226323571053443 |
| Phocarctos_hookeri | Tremarctos_ornatus | 0.239152150998458 |
| Phocarctos_hookeri | Urocyon_cinereoargenteus | 0.256512048680941 |
| Phocarctos_hookeri | Ursus_americanus | 0.23888576599012 |
| Phocarctos_hookeri | Ursus_arctos | 0.242139379179023 |
| Phocarctos_hookeri | Ursus_maritimus | 0.239761122460678 |
| Phocarctos_hookeri | Ursus_spelaeus | 0.242153199013155 |
| Phocarctos_hookeri | Ursus_thibetanus | 0.240926174894768 |
| Phocarctos_hookeri | Urva_javanica | 0.246305179643794 |
| Phocarctos_hookeri | Viverra_tangalunga_FMNH146957_SRR6053069 | 0.236883623395748 |
| Phocarctos_hookeri | Viverricula_indica | 0.237505297915795 |
| Phocarctos_hookeri | Vulpes_corsac | 0.254117298413288 |
| Phocarctos_hookeri | Vulpes_ferrilata | 0.254969361081897 |
| Phocarctos_hookeri | Vulpes_lagopus | 0.254469478508582 |
| Phocarctos_hookeri | Vulpes_vulpes_T3611 | 0.252964532199843 |
| Phocarctos_hookeri | Vulpes_zerda | 0.253167979543814 |
| Phocarctos_hookeri | Zalophus_californianus | 0.0890387026977033 |
| Prionailurus_bengalensis | Prionailurus_bengalensis_euptilurus | 0.0404398196253018 |
| Prionailurus_bengalensis | Prionailurus_planiceps | 0.0660468902434153 |
| Prionailurus_bengalensis | Prionailurus_rubiginosus | 0.0707365033790168 |
| Prionailurus_bengalensis | Prionailurus_viverrinus | 0.045107466478183 |
| Prionailurus_bengalensis | Prionodon_pardicolor | 0.18279841859651 |
| Prionailurus_bengalensis | Procyon_lotor | 0.230780727282315 |
| Prionailurus_bengalensis | Profelis_aurata | 0.10705325239834 |
| Prionailurus_bengalensis | Proteles_cristatus_cristatus_TS307 | 0.190833602761434 |
| Prionailurus_bengalensis | Proteles_cristatus_septentrionalis_NMS_Z_2018_54 | 0.190228493178002 |
| Prionailurus_bengalensis | Puma_concolor | 0.104192620466259 |
| Prionailurus_bengalensis | Puma_yagouaroundi | 0.105282617475623 |
| Prionailurus_bengalensis | Pusa_caspica | 0.219350650160123 |
| Prionailurus_bengalensis | Pusa_hispida | 0.216718734308307 |
| Prionailurus_bengalensis | Pusa_sibirica | 0.216274487190075 |
| Prionailurus_bengalensis | Speothos_venaticus_T1624 | 0.243066429989438 |
| Prionailurus_bengalensis | Spilogale_putorius | 0.23987340389006 |
| Prionailurus_bengalensis | Taxidea_taxus | 0.218895519605709 |
| Prionailurus_bengalensis | Tremarctos_ornatus | 0.229985182640909 |
| Prionailurus_bengalensis | Urocyon_cinereoargenteus | 0.245999689890376 |
| Prionailurus_bengalensis | Ursus_americanus | 0.239143984942746 |
| Prionailurus_bengalensis | Ursus_arctos | 0.241661657084677 |
| Prionailurus_bengalensis | Ursus_maritimus | 0.240273482593675 |
| Prionailurus_bengalensis | Ursus_spelaeus | 0.2367753308555 |
| Prionailurus_bengalensis | Ursus_thibetanus | 0.237003667799239 |
| Prionailurus_bengalensis | Urva_javanica | 0.188333454997082 |
| Prionailurus_bengalensis | Viverra_tangalunga_FMNH146957_SRR6053069 | 0.169361138526019 |
| Prionailurus_bengalensis | Viverricula_indica | 0.167448488471161 |
| Prionailurus_bengalensis | Vulpes_corsac | 0.239672051950771 |
| Prionailurus_bengalensis | Vulpes_ferrilata | 0.238219538141415 |
| Prionailurus_bengalensis | Vulpes_lagopus | 0.237143993544355 |
| Prionailurus_bengalensis | Vulpes_vulpes_T3611 | 0.234805205523475 |

| | | |
|---|---|---|
| Prionailurus_bengalensis | Vulpes_zerda | 0.23246561548012 |
| Prionailurus_bengalensis | Zalophus_californianus | 0.231037137842083 |
| Prionailurus_bengalensis_euptilurus | Prionailurus_planiceps | 0.0671838027335832 |
| Prionailurus_bengalensis_euptilurus | Prionailurus_rubiginosus | 0.070411561374979 |
| Prionailurus_bengalensis_euptilurus | Prionailurus_viverrinus | 0.0379214711294659 |
| Prionailurus_bengalensis_euptilurus | Prionodon_pardicolor | 0.182838118732794 |
| Prionailurus_bengalensis_euptilurus | Procyon_lotor | 0.231749592505291 |
| Prionailurus_bengalensis_euptilurus | Profelis_aurata | 0.109011648292039 |
| Prionailurus_bengalensis_euptilurus | Proteles_cristatus_cristatus_TS307 | 0.194386758063161 |
| Prionailurus_bengalensis_euptilurus | Proteles_cristatus_septentrionalis_NMS_Z_2018_54 | 0.196509325270011 |
| Prionailurus_bengalensis_euptilurus | Puma_concolor | 0.103748301549771 |
| Prionailurus_bengalensis_euptilurus | Puma_yagouaroundi | 0.105774264824254 |
| Prionailurus_bengalensis_euptilurus | Pusa_caspica | 0.220320582569036 |
| Prionailurus_bengalensis_euptilurus | Pusa_hispida | 0.215667357503662 |
| Prionailurus_bengalensis_euptilurus | Pusa_sibirica | 0.215846496579735 |
| Prionailurus_bengalensis_euptilurus | Speothos_venaticus_T1624 | 0.243826890306365 |
| Prionailurus_bengalensis_euptilurus | Spilogale_putorius | 0.241204814862058 |
| Prionailurus_bengalensis_euptilurus | Taxidea_taxus | 0.220452083203231 |
| Prionailurus_bengalensis_euptilurus | Tremarctos_ornatus | 0.232018775962645 |
| Prionailurus_bengalensis_euptilurus | Urocyon_cinereoargenteus | 0.242900272057128 |
| Prionailurus_bengalensis_euptilurus | Ursus_americanus | 0.23717584548785 |
| Prionailurus_bengalensis_euptilurus | Ursus_arctos | 0.242717545078487 |
| Prionailurus_bengalensis_euptilurus | Ursus_maritimus | 0.241327109096059 |
| Prionailurus_bengalensis_euptilurus | Ursus_spelaeus | 0.237804895891618 |
| Prionailurus_bengalensis_euptilurus | Ursus_thibetanus | 0.235352121641739 |
| Prionailurus_bengalensis_euptilurus | Urva_javanica | 0.192688291526023 |
| Prionailurus_bengalensis_euptilurus | Viverra_tangalunga_FMNH146957_SRR6053069 | 0.166606383003594 |
| Prionailurus_bengalensis_euptilurus | Viverricula_indica | 0.168314455803022 |
| Prionailurus_bengalensis_euptilurus | Vulpes_corsac | 0.239692718601528 |
| Prionailurus_bengalensis_euptilurus | Vulpes_ferrilata | 0.240147863882148 |
| Prionailurus_bengalensis_euptilurus | Vulpes_lagopus | 0.233214484241955 |
| Prionailurus_bengalensis_euptilurus | Vulpes_vulpes_T3611 | 0.235556818149793 |
| Prionailurus_bengalensis_euptilurus | Vulpes_zerda | 0.234224052285395 |
| Prionailurus_bengalensis_euptilurus | Zalophus_californianus | 0.230778556090092 |
| Prionailurus_planiceps | Prionailurus_rubiginosus | 0.0764959476237557 |
| Prionailurus_planiceps | Prionailurus_viverrinus | 0.0703546622156108 |
| Prionailurus_planiceps | Prionodon_pardicolor | 0.182488757149783 |
| Prionailurus_planiceps | Procyon_lotor | 0.233355722186801 |
| Prionailurus_planiceps | Profelis_aurata | 0.112588666383573 |
| Prionailurus_planiceps | Proteles_cristatus_cristatus_TS307 | 0.196067608474763 |
| Prionailurus_planiceps | Proteles_cristatus_septentrionalis_NMS_Z_2018_54 | 0.196699387015174 |
| Prionailurus_planiceps | Puma_concolor | 0.110929664870037 |
| Prionailurus_planiceps | Puma_yagouaroundi | 0.111559860596791 |
| Prionailurus_planiceps | Pusa_caspica | 0.222838335323789 |
| Prionailurus_planiceps | Pusa_hispida | 0.218790336658997 |
| Prionailurus_planiceps | Pusa_sibirica | 0.220498312959531 |
| Prionailurus_planiceps | Speothos_venaticus_T1624 | 0.2447296977223 |
| Prionailurus_planiceps | Spilogale_putorius | 0.243299668859388 |
| Prionailurus_planiceps | Taxidea_taxus | 0.224105853338935 |
| Prionailurus_planiceps | Tremarctos_ornatus | 0.23246194018399 |
| Prionailurus_planiceps | Urocyon_cinereoargenteus | 0.247465470622164 |
| Prionailurus_planiceps | Ursus_americanus | 0.240692500337572 |
| Prionailurus_planiceps | Ursus_arctos | 0.246030785353053 |
| Prionailurus_planiceps | Ursus_maritimus | 0.244629919310422 |
| Prionailurus_planiceps | Ursus_spelaeus | 0.242116375242583 |
| Prionailurus_planiceps | Ursus_thibetanus | 0.237068636272416 |
| Prionailurus_planiceps | Urva_javanica | 0.191881019246965 |
| Prionailurus_planiceps | Viverra_tangalunga_FMNH146957_SRR6053069 | 0.170825244372327 |
| Prionailurus_planiceps | Viverricula_indica | 0.17604442296939 |
| Prionailurus_planiceps | Vulpes_corsac | 0.241899227125461 |
| Prionailurus_planiceps | Vulpes_ferrilata | 0.243565395948549 |
| Prionailurus_planiceps | Vulpes_lagopus | 0.240342973948949 |
| Prionailurus_planiceps | Vulpes_vulpes_T3611 | 0.236701466687279 |
| Prionailurus_planiceps | Vulpes_zerda | 0.235697960911973 |
| Prionailurus_planiceps | Zalophus_californianus | 0.233003565130654 |
| Prionailurus_rubiginosus | Prionailurus_viverrinus | 0.0733575884404672 |
| Prionailurus_rubiginosus | Prionodon_pardicolor | 0.180712617194855 |
| Prionailurus_rubiginosus | Procyon_lotor | 0.231248203052991 |
| Prionailurus_rubiginosus | Profelis_aurata | 0.109430298975229 |
| Prionailurus_rubiginosus | Proteles_cristatus_cristatus_TS307 | 0.194556263959984 |
| Prionailurus_rubiginosus | Proteles_cristatus_septentrionalis_NMS_Z_2018_54 | 0.195816371736051 |
| Prionailurus_rubiginosus | Puma_concolor | 0.105682426577592 |
| Prionailurus_rubiginosus | Puma_yagouaroundi | 0.108530805868078 |
| Prionailurus_rubiginosus | Pusa_caspica | 0.222857745192511 |
| Prionailurus_rubiginosus | Pusa_hispida | 0.220010591263548 |
| Prionailurus_rubiginosus | Pusa_sibirica | 0.219186815293487 |
| Prionailurus_rubiginosus | Speothos_venaticus_T1624 | 0.240508879656052 |
| Prionailurus_rubiginosus | Spilogale_putorius | 0.243613908009735 |
| Prionailurus_rubiginosus | Taxidea_taxus | 0.220774947271604 |
| Prionailurus_rubiginosus | Tremarctos_ornatus | 0.22879585261044 |
| Prionailurus_rubiginosus | Urocyon_cinereoargenteus | 0.242766784344583 |
| Prionailurus_rubiginosus | Ursus_americanus | 0.238378742974403 |
| Prionailurus_rubiginosus | Ursus_arctos | 0.240527141888212 |
| Prionailurus_rubiginosus | Ursus_maritimus | 0.23959271730403 |
| Prionailurus_rubiginosus | Ursus_spelaeus | 0.236325660353195 |
| Prionailurus_rubiginosus | Ursus_thibetanus | 0.234834965055565 |
| Prionailurus_rubiginosus | Urva_javanica | 0.186499358892469 |

| | | |
|---|---|---|
| Prionailurus_rubiginosus | Viverra_tangalunga_FMNH146957_SRR6053069 | 0.172132129359705 |
| Prionailurus_rubiginosus | Viverricula_indica | 0.17012417415658 |
| Prionailurus_rubiginosus | Vulpes_corsac | 0.23437470639878 |
| Prionailurus_rubiginosus | Vulpes_ferrilata | 0.236119159196016 |
| Prionailurus_rubiginosus | Vulpes_lagopus | 0.235416403467209 |
| Prionailurus_rubiginosus | Vulpes_vulpes_T3611 | 0.230771081847212 |
| Prionailurus_rubiginosus | Vulpes_zerda | 0.228682886760208 |
| Prionailurus_rubiginosus | Zalophus_californianus | 0.231376975112767 |
| Prionailurus_viverrinus | Prionodon_pardicolor | 0.18441300538445 |
| Prionailurus_viverrinus | Procyon_lotor | 0.233445015874686 |
| Prionailurus_viverrinus | Profelis_aurata | 0.109969635046903 |
| Prionailurus_viverrinus | Proteles_cristatus_cristatus_TS307 | 0.195416279626536 |
| Prionailurus_viverrinus | Proteles_cristatus_septentrionalis_NMS_Z_2018_54 | 0.197121016638557 |
| Prionailurus_viverrinus | Puma_concolor | 0.109947706992034 |
| Prionailurus_viverrinus | Puma_yagouaroundi | 0.113175807672372 |
| Prionailurus_viverrinus | Pusa_caspica | 0.221833116133881 |
| Prionailurus_viverrinus | Pusa_hispida | 0.218990341114216 |
| Prionailurus_viverrinus | Pusa_sibirica | 0.22053637886069 |
| Prionailurus_viverrinus | Speothos_venaticus_T1624 | 0.245665734698484 |
| Prionailurus_viverrinus | Spilogale_putorius | 0.24155837430521 |
| Prionailurus_viverrinus | Taxidea_taxus | 0.218984973046148 |
| Prionailurus_viverrinus | Tremarctos_ornatus | 0.231045308330985 |
| Prionailurus_viverrinus | Urocyon_cinereoargenteus | 0.245571683431549 |
| Prionailurus_viverrinus | Ursus_americanus | 0.236148371098358 |
| Prionailurus_viverrinus | Ursus_arctos | 0.241411742201779 |
| Prionailurus_viverrinus | Ursus_maritimus | 0.240924732262847 |
| Prionailurus_viverrinus | Ursus_spelaeus | 0.238205848071498 |
| Prionailurus_viverrinus | Ursus_thibetanus | 0.238500561972643 |
| Prionailurus_viverrinus | Urva_javanica | 0.190528683043401 |
| Prionailurus_viverrinus | Viverra_tangalunga_FMNH146957_SRR6053069 | 0.16806426246535 |
| Prionailurus_viverrinus | Viverricula_indica | 0.171707717147167 |
| Prionailurus_viverrinus | Vulpes_corsac | 0.240227916377116 |
| Prionailurus_viverrinus | Vulpes_ferrilata | 0.241219547141463 |
| Prionailurus_viverrinus | Vulpes_lagopus | 0.238765503647101 |
| Prionailurus_viverrinus | Vulpes_vulpes_T3611 | 0.236349608295768 |
| Prionailurus_viverrinus | Vulpes_zerda | 0.233803727036629 |
| Prionailurus_viverrinus | Zalophus_californianus | 0.23145590827193 |
| Prionodon_pardicolor | Procyon_lotor | 0.243345388909312 |
| Prionodon_pardicolor | Profelis_aurata | 0.189138368219165 |
| Prionodon_pardicolor | Proteles_cristatus_cristatus_TS307 | 0.21226954130142 |
| Prionodon_pardicolor | Proteles_cristatus_septentrionalis_NMS_Z_2018_54 | 0.212119976485089 |
| Prionodon_pardicolor | Puma_concolor | 0.180381105174795 |
| Prionodon_pardicolor | Puma_yagouaroundi | 0.182509491015129 |
| Prionodon_pardicolor | Pusa_caspica | 0.229276692969282 |
| Prionodon_pardicolor | Pusa_hispida | 0.227910715540108 |
| Prionodon_pardicolor | Pusa_sibirica | 0.228730900970976 |
| Prionodon_pardicolor | Speothos_venaticus_T1624 | 0.249170848072191 |
| Prionodon_pardicolor | Spilogale_putorius | 0.244851678156498 |
| Prionodon_pardicolor | Taxidea_taxus | 0.2282184217247 |
| Prionodon_pardicolor | Tremarctos_ornatus | 0.247562310632924 |
| Prionodon_pardicolor | Urocyon_cinereoargenteus | 0.247331261709802 |
| Prionodon_pardicolor | Ursus_americanus | 0.242632240342221 |
| Prionodon_pardicolor | Ursus_arctos | 0.243120742651838 |
| Prionodon_pardicolor | Ursus_maritimus | 0.24489571184011 |
| Prionodon_pardicolor | Ursus_spelaeus | 0.243861266567642 |
| Prionodon_pardicolor | Ursus_thibetanus | 0.23903186474943 |
| Prionodon_pardicolor | Urva_javanica | 0.200621724315756 |
| Prionodon_pardicolor | Viverra_tangalunga_FMNH146957_SRR6053069 | 0.188502874780223 |
| Prionodon_pardicolor | Viverricula_indica | 0.185308993029814 |
| Prionodon_pardicolor | Vulpes_corsac | 0.24047884892773 |
| Prionodon_pardicolor | Vulpes_ferrilata | 0.244781192664914 |
| Prionodon_pardicolor | Vulpes_lagopus | 0.237143993544355 |
| Prionodon_pardicolor | Vulpes_vulpes_T3611 | 0.242016335617924 |
| Prionodon_pardicolor | Vulpes_zerda | 0.243315681612883 |
| Prionodon_pardicolor | Zalophus_californianus | 0.244039936985909 |
| Procyon_lotor | Profelis_aurata | 0.233092742851829 |
| Procyon_lotor | Proteles_cristatus_cristatus_TS307 | 0.250954923374798 |
| Procyon_lotor | Proteles_cristatus_septentrionalis_NMS_Z_2018_54 | 0.250363684100829 |
| Procyon_lotor | Puma_concolor | 0.236005628801397 |
| Procyon_lotor | Puma_yagouaroundi | 0.233166620265924 |
| Procyon_lotor | Pusa_caspica | 0.218456166312395 |
| Procyon_lotor | Pusa_hispida | 0.216287671099434 |
| Procyon_lotor | Pusa_sibirica | 0.219305486612334 |
| Procyon_lotor | Speothos_venaticus_T1624 | 0.249510316192642 |
| Procyon_lotor | Spilogale_putorius | 0.228136221907909 |
| Procyon_lotor | Taxidea_taxus | 0.204763236589905 |
| Procyon_lotor | Tremarctos_ornatus | 0.233191085412243 |
| Procyon_lotor | Urocyon_cinereoargenteus | 0.255447705367096 |
| Procyon_lotor | Ursus_americanus | 0.238299421713372 |
| Procyon_lotor | Ursus_arctos | 0.243123322515612 |
| Procyon_lotor | Ursus_maritimus | 0.242880535315002 |
| Procyon_lotor | Ursus_spelaeus | 0.241454838284877 |
| Procyon_lotor | Ursus_thibetanus | 0.24022505088848 |
| Procyon_lotor | Urva_javanica | 0.240447632984262 |
| Procyon_lotor | Viverra_tangalunga_FMNH146957_SRR6053069 | 0.231914812230911 |
| Procyon_lotor | Viverricula_indica | 0.23315459609391 |
| Procyon_lotor | Vulpes_corsac | 0.249290487154 |

| | | |
|---|---|---|
| Procyon_lotor | Vulpes_ferrilata | 0.253429495247996 |
| Procyon_lotor | Vulpes_lagopus | 0.252246634390438 |
| Procyon_lotor | Vulpes_vulpes_T3611 | 0.247315682362027 |
| Procyon_lotor | Vulpes_zerda | 0.246818006916112 |
| Procyon_lotor | Zalophus_californianus | 0.23609162025857 |
| Profelis_aurata | Proteles_cristatus_cristatus_TS307 | 0.200930895097944 |
| Profelis_aurata | Proteles_cristatus_septentrionalis_NMS_Z_2018_54 | 0.20337875013611 |
| Profelis_aurata | Puma_concolor | 0.112493619341483 |
| Profelis_aurata | Puma_yagouaroundi | 0.114291567170043 |
| Profelis_aurata | Pusa_caspica | 0.227142809141188 |
| Profelis_aurata | Pusa_hispida | 0.224140136883645 |
| Profelis_aurata | Pusa_sibirica | 0.22561442367163 |
| Profelis_aurata | Speothos_venaticus_T1624 | 0.248307736891159 |
| Profelis_aurata | Spilogale_putorius | 0.245038419411854 |
| Profelis_aurata | Taxidea_taxus | 0.228439473321847 |
| Profelis_aurata | Tremarctos_ornatus | 0.235380418649714 |
| Profelis_aurata | Urocyon_cinereoargenteus | 0.251456767216626 |
| Profelis_aurata | Ursus_americanus | 0.239470040513209 |
| Profelis_aurata | Ursus_arctos | 0.239731033298608 |
| Profelis_aurata | Ursus_maritimus | 0.238908432007573 |
| Profelis_aurata | Ursus_spelaeus | 0.234952605451351 |
| Profelis_aurata | Ursus_thibetanus | 0.239253437397967 |
| Profelis_aurata | Urva_javanica | 0.191325323747434 |
| Profelis_aurata | Viverra_tangalunga_FMNH146957_SRR6053069 | 0.173718490463047 |
| Profelis_aurata | Viverricula_indica | 0.170264463764235 |
| Profelis_aurata | Vulpes_corsac | 0.240910747182456 |
| Profelis_aurata | Vulpes_ferrilata | 0.240321925137346 |
| Profelis_aurata | Vulpes_lagopus | 0.242485092513593 |
| Profelis_aurata | Vulpes_vulpes_T3611 | 0.236832436594569 |
| Profelis_aurata | Vulpes_zerda | 0.23595986723121 |
| Profelis_aurata | Zalophus_californianus | 0.238213238057563 |
| Proteles_cristatus_cristatus_TS307 | Proteles_cristatus_septentrionalis_NMS_Z_2018_54 | 0.0427682848418366 |
| Proteles_cristatus_cristatus_TS307 | Puma_concolor | 0.196080989325412 |
| Proteles_cristatus_cristatus_TS307 | Puma_yagouaroundi | 0.194760875446972 |
| Proteles_cristatus_cristatus_TS307 | Pusa_caspica | 0.231395966010945 |
| Proteles_cristatus_cristatus_TS307 | Pusa_hispida | 0.230608739794905 |
| Proteles_cristatus_cristatus_TS307 | Pusa_sibirica | 0.229394072552069 |
| Proteles_cristatus_cristatus_TS307 | Speothos_venaticus_T1624 | 0.262361993409555 |
| Proteles_cristatus_cristatus_TS307 | Spilogale_putorius | 0.260671387591489 |
| Proteles_cristatus_cristatus_TS307 | Taxidea_taxus | 0.237639094694749 |
| Proteles_cristatus_cristatus_TS307 | Tremarctos_ornatus | 0.247399142571471 |
| Proteles_cristatus_cristatus_TS307 | Urocyon_cinereoargenteus | 0.257723308864739 |
| Proteles_cristatus_cristatus_TS307 | Ursus_americanus | 0.250164506646637 |
| Proteles_cristatus_cristatus_TS307 | Ursus_arctos | 0.248589947893152 |
| Proteles_cristatus_cristatus_TS307 | Ursus_maritimus | 0.249604720196414 |
| Proteles_cristatus_cristatus_TS307 | Ursus_spelaeus | 0.24356149999499 |
| Proteles_cristatus_cristatus_TS307 | Ursus_thibetanus | 0.244797027254432 |
| Proteles_cristatus_cristatus_TS307 | Urva_javanica | 0.198592529413035 |
| Proteles_cristatus_cristatus_TS307 | Viverra_tangalunga_FMNH146957_SRR6053069 | 0.202259167836758 |
| Proteles_cristatus_cristatus_TS307 | Viverricula_indica | 0.201695451892312 |
| Proteles_cristatus_cristatus_TS307 | Vulpes_corsac | 0.255126937233918 |
| Proteles_cristatus_cristatus_TS307 | Vulpes_ferrilata | 0.258559726799419 |
| Proteles_cristatus_cristatus_TS307 | Vulpes_lagopus | 0.254704944101964 |
| Proteles_cristatus_cristatus_TS307 | Vulpes_vulpes_T3611 | 0.253456928931668 |
| Proteles_cristatus_cristatus_TS307 | Vulpes_zerda | 0.251936412427073 |
| Proteles_cristatus_cristatus_TS307 | Zalophus_californianus | 0.244213515716803 |
| Proteles_cristatus_septentrionalis_NMS_Z_2018_54 | Puma_concolor | 0.198399711862033 |
| Proteles_cristatus_septentrionalis_NMS_Z_2018_54 | Puma_yagouaroundi | 0.19823201592629 |
| Proteles_cristatus_septentrionalis_NMS_Z_2018_54 | Pusa_caspica | 0.231904819913696 |
| Proteles_cristatus_septentrionalis_NMS_Z_2018_54 | Pusa_hispida | 0.229912920585331 |
| Proteles_cristatus_septentrionalis_NMS_Z_2018_54 | Pusa_sibirica | 0.230229995420952 |
| Proteles_cristatus_septentrionalis_NMS_Z_2018_54 | Speothos_venaticus_T1624 | 0.262707496129741 |
| Proteles_cristatus_septentrionalis_NMS_Z_2018_54 | Spilogale_putorius | 0.2598201696787 |
| Proteles_cristatus_septentrionalis_NMS_Z_2018_54 | Taxidea_taxus | 0.237308005398908 |
| Proteles_cristatus_septentrionalis_NMS_Z_2018_54 | Tremarctos_ornatus | 0.250290748202505 |
| Proteles_cristatus_septentrionalis_NMS_Z_2018_54 | Urocyon_cinereoargenteus | 0.259303769069013 |
| Proteles_cristatus_septentrionalis_NMS_Z_2018_54 | Ursus_americanus | 0.250000195057554 |
| Proteles_cristatus_septentrionalis_NMS_Z_2018_54 | Ursus_arctos | 0.250481530509221 |
| Proteles_cristatus_septentrionalis_NMS_Z_2018_54 | Ursus_maritimus | 0.250697743792731 |
| Proteles_cristatus_septentrionalis_NMS_Z_2018_54 | Ursus_spelaeus | 0.246455841560354 |
| Proteles_cristatus_septentrionalis_NMS_Z_2018_54 | Ursus_thibetanus | 0.242826035969269 |
| Proteles_cristatus_septentrionalis_NMS_Z_2018_54 | Urva_javanica | 0.196150185328047 |
| Proteles_cristatus_septentrionalis_NMS_Z_2018_54 | Viverra_tangalunga_FMNH146957_SRR6053069 | 0.201153866017329 |
| Proteles_cristatus_septentrionalis_NMS_Z_2018_54 | Viverricula_indica | 0.199448207651298 |
| Proteles_cristatus_septentrionalis_NMS_Z_2018_54 | Vulpes_corsac | 0.255875184205944 |
| Proteles_cristatus_septentrionalis_NMS_Z_2018_54 | Vulpes_ferrilata | 0.256114072800359 |
| Proteles_cristatus_septentrionalis_NMS_Z_2018_54 | Vulpes_lagopus | 0.254223948627854 |
| Proteles_cristatus_septentrionalis_NMS_Z_2018_54 | Vulpes_vulpes_T3611 | 0.252388747638292 |
| Proteles_cristatus_septentrionalis_NMS_Z_2018_54 | Vulpes_zerda | 0.251435093026654 |
| Proteles_cristatus_septentrionalis_NMS_Z_2018_54 | Zalophus_californianus | 0.248140824956441 |
| Puma_concolor | Puma_yagouaroundi | 0.0883602624344147 |
| Puma_concolor | Pusa_caspica | 0.219093761489164 |
| Puma_concolor | Pusa_hispida | 0.217980808981499 |
| Puma_concolor | Pusa_sibirica | 0.217625915833444 |
| Puma_concolor | Speothos_venaticus_T1624 | 0.243593458554634 |
| Puma_concolor | Spilogale_putorius | 0.248017406545151 |
| Puma_concolor | Taxidea_taxus | 0.221428900262509 |

| | | |
|---|---|---|
| Puma_concolor | Tremarctos_ornatus | 0.238064114864195 |
| Puma_concolor | Urocyon_cinereoargenteus | 0.247702661387053 |
| Puma_concolor | Ursus_americanus | 0.237453267186279 |
| Puma_concolor | Ursus_arctos | 0.244013701825776 |
| Puma_concolor | Ursus_maritimus | 0.242168694523554 |
| Puma_concolor | Ursus_spelaeus | 0.236963988825479 |
| Puma_concolor | Ursus_thibetanus | 0.23594025743426 |
| Puma_concolor | Urva_javanica | 0.192535372581833 |
| Puma_concolor | Viverra_tangalunga_FMNH146957_SRR6053069 | 0.172322055635545 |
| Puma_concolor | Viverricula_indica | 0.171497881510346 |
| Puma_concolor | Vulpes_corsac | 0.239279332374721 |
| Puma_concolor | Vulpes_ferrilata | 0.240714889541185 |
| Puma_concolor | Vulpes_lagopus | 0.238525894268625 |
| Puma_concolor | Vulpes_vulpes_T3611 | 0.233546299869036 |
| Puma_concolor | Vulpes_zerda | 0.232455849516596 |
| Puma_concolor | Zalophus_californianus | 0.2356518068124 |
| Puma_yagouaroundi | Pusa_caspica | 0.224564952319542 |
| Puma_yagouaroundi | Pusa_hispida | 0.218370014471716 |
| Puma_yagouaroundi | Pusa_sibirica | 0.218228818325853 |
| Puma_yagouaroundi | Speothos_venaticus_T1624 | 0.242843867315693 |
| Puma_yagouaroundi | Spilogale_putorius | 0.246674590550224 |
| Puma_yagouaroundi | Taxidea_taxus | 0.219262351712808 |
| Puma_yagouaroundi | Tremarctos_ornatus | 0.237686926020974 |
| Puma_yagouaroundi | Urocyon_cinereoargenteus | 0.24637249966124 |
| Puma_yagouaroundi | Ursus_americanus | 0.238348021926789 |
| Puma_yagouaroundi | Ursus_arctos | 0.240587008508028 |
| Puma_yagouaroundi | Ursus_maritimus | 0.240101254020173 |
| Puma_yagouaroundi | Ursus_spelaeus | 0.234490176431807 |
| Puma_yagouaroundi | Ursus_thibetanus | 0.233181896026637 |
| Puma_yagouaroundi | Urva_javanica | 0.195039341832544 |
| Puma_yagouaroundi | Viverra_tangalunga_FMNH146957_SRR6053069 | 0.170237086311509 |
| Puma_yagouaroundi | Viverricula_indica | 0.171584917528544 |
| Puma_yagouaroundi | Vulpes_corsac | 0.241129073965772 |
| Puma_yagouaroundi | Vulpes_ferrilata | 0.241745529616441 |
| Puma_yagouaroundi | Vulpes_lagopus | 0.241426001418238 |
| Puma_yagouaroundi | Vulpes_vulpes_T3611 | 0.233016863179368 |
| Puma_yagouaroundi | Vulpes_zerda | 0.232694886376021 |
| Puma_yagouaroundi | Zalophus_californianus | 0.235125792901206 |
| Pusa_caspica | Pusa_hispida | 0.0304992950294954 |
| Pusa_caspica | Pusa_sibirica | 0.0263405922062245 |
| Pusa_caspica | Speothos_venaticus_T1624 | 0.239865875392657 |
| Pusa_caspica | Spilogale_putorius | 0.229920338418497 |
| Pusa_caspica | Taxidea_taxus | 0.207177629636993 |
| Pusa_caspica | Tremarctos_ornatus | 0.21623606421642 |
| Pusa_caspica | Urocyon_cinereoargenteus | 0.236116567341901 |
| Pusa_caspica | Ursus_americanus | 0.21677448996223 |
| Pusa_caspica | Ursus_arctos | 0.221514585063213 |
| Pusa_caspica | Ursus_maritimus | 0.219646403712946 |
| Pusa_caspica | Ursus_spelaeus | 0.220440830187412 |
| Pusa_caspica | Ursus_thibetanus | 0.215432194223652 |
| Pusa_caspica | Urva_javanica | 0.222795605257833 |
| Pusa_caspica | Viverra_tangalunga_FMNH146957_SRR6053069 | 0.221685746151208 |
| Pusa_caspica | Viverricula_indica | 0.220042607601952 |
| Pusa_caspica | Vulpes_corsac | 0.235029568055241 |
| Pusa_caspica | Vulpes_ferrilata | 0.23583756136046 |
| Pusa_caspica | Vulpes_lagopus | 0.232300071477552 |
| Pusa_caspica | Vulpes_vulpes_T3611 | 0.23073257812646 |
| Pusa_caspica | Vulpes_zerda | 0.229429035882639 |
| Pusa_caspica | Zalophus_californianus | 0.182298205195908 |
| Pusa_hispida | Pusa_sibirica | 0.0285621783822475 |
| Pusa_hispida | Speothos_venaticus_T1624 | 0.238800463941133 |
| Pusa_hispida | Spilogale_putorius | 0.226814882587774 |
| Pusa_hispida | Taxidea_taxus | 0.203356961869811 |
| Pusa_hispida | Tremarctos_ornatus | 0.213991292729202 |
| Pusa_hispida | Urocyon_cinereoargenteus | 0.236517489219256 |
| Pusa_hispida | Ursus_americanus | 0.215820296494699 |
| Pusa_hispida | Ursus_arctos | 0.220550214572818 |
| Pusa_hispida | Ursus_maritimus | 0.218135764056773 |
| Pusa_hispida | Ursus_spelaeus | 0.220030078887074 |
| Pusa_hispida | Ursus_thibetanus | 0.213825999749173 |
| Pusa_hispida | Urva_javanica | 0.221737924151749 |
| Pusa_hispida | Viverra_tangalunga_FMNH146957_SRR6053069 | 0.219933912801043 |
| Pusa_hispida | Viverricula_indica | 0.217723952744059 |
| Pusa_hispida | Vulpes_corsac | 0.232730458418483 |
| Pusa_hispida | Vulpes_ferrilata | 0.232630650001991 |
| Pusa_hispida | Vulpes_lagopus | 0.232410240035891 |
| Pusa_hispida | Vulpes_vulpes_T3611 | 0.22821147226993 |
| Pusa_hispida | Vulpes_zerda | 0.226257199805655 |
| Pusa_hispida | Zalophus_californianus | 0.178649699531111 |
| Pusa_sibirica | Speothos_venaticus_T1624 | 0.24107159082883 |
| Pusa_sibirica | Spilogale_putorius | 0.22833249694171 |
| Pusa_sibirica | Taxidea_taxus | 0.208026616363309 |
| Pusa_sibirica | Tremarctos_ornatus | 0.213848306828983 |
| Pusa_sibirica | Urocyon_cinereoargenteus | 0.238362543400999 |
| Pusa_sibirica | Ursus_americanus | 0.217907752833672 |
| Pusa_sibirica | Ursus_arctos | 0.21999092143032 |
| Pusa_sibirica | Ursus_maritimus | 0.218568764277306 |

| | | |
|---|---|---|
| Pusa_sibirica | Ursus_spelaeus | 0.219581984031791 |
| Pusa_sibirica | Ursus_thibetanus | 0.214084402534006 |
| Pusa_sibirica | Urva_javanica | 0.222883524341719 |
| Pusa_sibirica | Viverra_tangalunga_FMNH146957_SRR6053069 | 0.223968358811233 |
| Pusa_sibirica | Viverricula_indica | 0.218602930591308 |
| Pusa_sibirica | Vulpes_corsac | 0.236403413178942 |
| Pusa_sibirica | Vulpes_ferrilata | 0.235748203840074 |
| Pusa_sibirica | Vulpes_lagopus | 0.233908269485823 |
| Pusa_sibirica | Vulpes_vulpes_T3611 | 0.229545652898225 |
| Pusa_sibirica | Vulpes_zerda | 0.228025071465687 |
| Pusa_sibirica | Zalophus_californianus | 0.18186048218544 |
| Speothos_venaticus_T1624 | Spilogale_putorius | 0.247577862849361 |
| Speothos_venaticus_T1624 | Taxidea_taxus | 0.245429398371515 |
| Speothos_venaticus_T1624 | Tremarctos_ornatus | 0.253744637502482 |
| Speothos_venaticus_T1624 | Urocyon_cinereoargenteus | 0.198163059232977 |
| Speothos_venaticus_T1624 | Ursus_americanus | 0.255164338383727 |
| Speothos_venaticus_T1624 | Ursus_arctos | 0.258426235147313 |
| Speothos_venaticus_T1624 | Ursus_maritimus | 0.257770370510277 |
| Speothos_venaticus_T1624 | Ursus_spelaeus | 0.254703349530051 |
| Speothos_venaticus_T1624 | Ursus_thibetanus | 0.247498284323225 |
| Speothos_venaticus_T1624 | Urva_javanica | 0.250632177745253 |
| Speothos_venaticus_T1624 | Viverra_tangalunga_FMNH146957_SRR6053069 | 0.242671588613439 |
| Speothos_venaticus_T1624 | Viverricula_indica | 0.239972945428167 |
| Speothos_venaticus_T1624 | Vulpes_corsac | 0.184911737518271 |
| Speothos_venaticus_T1624 | Vulpes_ferrilata | 0.188800753499891 |
| Speothos_venaticus_T1624 | Vulpes_lagopus | 0.188088230966587 |
| Speothos_venaticus_T1624 | Vulpes_vulpes_T3611 | 0.186099802336968 |
| Speothos_venaticus_T1624 | Vulpes_zerda | 0.18536535671126 |
| Speothos_venaticus_T1624 | Zalophus_californianus | 0.251045023827545 |
| Spilogale_putorius | Taxidea_taxus | 0.226990403559497 |
| Spilogale_putorius | Tremarctos_ornatus | 0.242701216449312 |
| Spilogale_putorius | Urocyon_cinereoargenteus | 0.257861536957582 |
| Spilogale_putorius | Ursus_americanus | 0.241320407470161 |
| Spilogale_putorius | Ursus_arctos | 0.243569823748125 |
| Spilogale_putorius | Ursus_maritimus | 0.245994063595226 |
| Spilogale_putorius | Ursus_spelaeus | 0.24009002960796 |
| Spilogale_putorius | Ursus_thibetanus | 0.241570761525079 |
| Spilogale_putorius | Urva_javanica | 0.244298926642033 |
| Spilogale_putorius | Viverra_tangalunga_FMNH146957_SRR6053069 | 0.239688351749517 |
| Spilogale_putorius | Viverricula_indica | 0.234290536500864 |
| Spilogale_putorius | Vulpes_corsac | 0.245897508258918 |
| Spilogale_putorius | Vulpes_ferrilata | 0.248970894924851 |
| Spilogale_putorius | Vulpes_lagopus | 0.249921847767934 |
| Spilogale_putorius | Vulpes_vulpes_T3611 | 0.248119253362841 |
| Spilogale_putorius | Vulpes_zerda | 0.248366177072932 |
| Spilogale_putorius | Zalophus_californianus | 0.239858630953546 |
| Taxidea_taxus | Tremarctos_ornatus | 0.22927414967136 |
| Taxidea_taxus | Urocyon_cinereoargenteus | 0.246722450221081 |
| Taxidea_taxus | Ursus_americanus | 0.230423634299569 |
| Taxidea_taxus | Ursus_arctos | 0.234850181537301 |
| Taxidea_taxus | Ursus_maritimus | 0.235810134580535 |
| Taxidea_taxus | Ursus_spelaeus | 0.231845822030591 |
| Taxidea_taxus | Ursus_thibetanus | 0.227660332531096 |
| Taxidea_taxus | Urva_javanica | 0.227214911169922 |
| Taxidea_taxus | Viverra_tangalunga_FMNH146957_SRR6053069 | 0.219334310910782 |
| Taxidea_taxus | Viverricula_indica | 0.217861810726207 |
| Taxidea_taxus | Vulpes_corsac | 0.24349148588513 |
| Taxidea_taxus | Vulpes_ferrilata | 0.246083049855292 |
| Taxidea_taxus | Vulpes_lagopus | 0.247233223441552 |
| Taxidea_taxus | Vulpes_vulpes_T3611 | 0.237975323452486 |
| Taxidea_taxus | Vulpes_zerda | 0.238691552904489 |
| Taxidea_taxus | Zalophus_californianus | 0.22040563858029 |
| Tremarctos_ornatus | Urocyon_cinereoargenteus | 0.25369586829267 |
| Tremarctos_ornatus | Ursus_americanus | 0.172996093243544 |
| Tremarctos_ornatus | Ursus_arctos | 0.175504213161323 |
| Tremarctos_ornatus | Ursus_maritimus | 0.1793761603056 |
| Tremarctos_ornatus | Ursus_spelaeus | 0.175942519522698 |
| Tremarctos_ornatus | Ursus_thibetanus | 0.171222654767319 |
| Tremarctos_ornatus | Urva_javanica | 0.243746151608003 |
| Tremarctos_ornatus | Viverra_tangalunga_FMNH146957_SRR6053069 | 0.231630633958692 |
| Tremarctos_ornatus | Viverricula_indica | 0.237345317421244 |
| Tremarctos_ornatus | Vulpes_corsac | 0.247092528746109 |
| Tremarctos_ornatus | Vulpes_ferrilata | 0.252532608237212 |
| Tremarctos_ornatus | Vulpes_lagopus | 0.247048092551925 |
| Tremarctos_ornatus | Vulpes_vulpes_T3611 | 0.246351205914389 |
| Tremarctos_ornatus | Vulpes_zerda | 0.246652585412862 |
| Tremarctos_ornatus | Zalophus_californianus | 0.230936193151945 |
| Urocyon_cinereoargenteus | Ursus_americanus | 0.254450243459218 |
| Urocyon_cinereoargenteus | Ursus_arctos | 0.263136440077018 |
| Urocyon_cinereoargenteus | Ursus_maritimus | 0.264027962482652 |
| Urocyon_cinereoargenteus | Ursus_spelaeus | 0.260337904859732 |
| Urocyon_cinereoargenteus | Ursus_thibetanus | 0.253413615582716 |
| Urocyon_cinereoargenteus | Urva_javanica | 0.250815419026295 |
| Urocyon_cinereoargenteus | Viverra_tangalunga_FMNH146957_SRR6053069 | 0.246666156909741 |
| Urocyon_cinereoargenteus | Viverricula_indica | 0.239734224151074 |
| Urocyon_cinereoargenteus | Vulpes_corsac | 0.188772518935795 |
| Urocyon_cinereoargenteus | Vulpes_ferrilata | 0.188817746856629 |

| | | |
|---|---|---|
| Urocyon_cinereoargenteus | Vulpes_lagopus | 0.184642701374744 |
| Urocyon_cinereoargenteus | Vulpes_vulpes_T3611 | 0.183036364421726 |
| Urocyon_cinereoargenteus | Vulpes_zerda | 0.185238780148462 |
| Urocyon_cinereoargenteus | Zalophus_californianus | 0.251384284040134 |
| Ursus_americanus | Ursus_arctos | 0.10257446554408 |
| Ursus_americanus | Ursus_maritimus | 0.100943032725693 |
| Ursus_americanus | Ursus_spelaeus | 0.0969953086998193 |
| Ursus_americanus | Ursus_thibetanus | 0.084899423669254 |
| Ursus_americanus | Urva_javanica | 0.247856098142246 |
| Ursus_americanus | Viverra_tangalunga_FMNH146957_SRR6053069 | 0.230240610866961 |
| Ursus_americanus | Viverricula_indica | 0.235322514257126 |
| Ursus_americanus | Vulpes_corsac | 0.246913941364642 |
| Ursus_americanus | Vulpes_ferrilata | 0.250176344815665 |
| Ursus_americanus | Vulpes_lagopus | 0.250837662959636 |
| Ursus_americanus | Vulpes_vulpes_T3611 | 0.24623765987098 |
| Ursus_americanus | Vulpes_zerda | 0.247374282319988 |
| Ursus_americanus | Zalophus_californianus | 0.23144721802246 |
| Ursus_arctos | Ursus_maritimus | 0.0211588392830005 |
| Ursus_arctos | Ursus_spelaeus | 0.0614312070045428 |
| Ursus_arctos | Ursus_thibetanus | 0.0955159124568741 |
| Ursus_arctos | Urva_javanica | 0.252406166546028 |
| Ursus_arctos | Viverra_tangalunga_FMNH146957_SRR6053069 | 0.234493854490029 |
| Ursus_arctos | Viverricula_indica | 0.241656870971537 |
| Ursus_arctos | Vulpes_corsac | 0.250585900654876 |
| Ursus_arctos | Vulpes_ferrilata | 0.253784955153301 |
| Ursus_arctos | Vulpes_lagopus | 0.253016971116277 |
| Ursus_arctos | Vulpes_vulpes_T3611 | 0.251704261942477 |
| Ursus_arctos | Vulpes_zerda | 0.254120281369541 |
| Ursus_arctos | Zalophus_californianus | 0.236316704209046 |
| Ursus_maritimus | Ursus_spelaeus | 0.0609352244631397 |
| Ursus_maritimus | Ursus_thibetanus | 0.0964481335497451 |
| Ursus_maritimus | Urva_javanica | 0.254460009536707 |
| Ursus_maritimus | Viverra_tangalunga_FMNH146957_SRR6053069 | 0.234904720201151 |
| Ursus_maritimus | Viverricula_indica | 0.241845323813341 |
| Ursus_maritimus | Vulpes_corsac | 0.24945692825749 |
| Ursus_maritimus | Vulpes_ferrilata | 0.252992378384551 |
| Ursus_maritimus | Vulpes_lagopus | 0.253374790548099 |
| Ursus_maritimus | Vulpes_vulpes_T3611 | 0.251806142654314 |
| Ursus_maritimus | Vulpes_zerda | 0.254337793632587 |
| Ursus_maritimus | Zalophus_californianus | 0.234526508770732 |
| Ursus_spelaeus | Ursus_thibetanus | 0.0950840002918061 |
| Ursus_spelaeus | Urva_javanica | 0.247677127217376 |
| Ursus_spelaeus | Viverra_tangalunga_FMNH146957_SRR6053069 | 0.233637630696032 |
| Ursus_spelaeus | Viverricula_indica | 0.236513498537016 |
| Ursus_spelaeus | Vulpes_corsac | 0.248001310093645 |
| Ursus_spelaeus | Vulpes_ferrilata | 0.250265384082923 |
| Ursus_spelaeus | Vulpes_lagopus | 0.248844652838357 |
| Ursus_spelaeus | Vulpes_vulpes_T3611 | 0.248121547403206 |
| Ursus_spelaeus | Vulpes_zerda | 0.248691477933397 |
| Ursus_spelaeus | Zalophus_californianus | 0.232582502547882 |
| Ursus_thibetanus | Urva_javanica | 0.246632423557129 |
| Ursus_thibetanus | Viverra_tangalunga_FMNH146957_SRR6053069 | 0.233221887257337 |
| Ursus_thibetanus | Viverricula_indica | 0.233082007907357 |
| Ursus_thibetanus | Vulpes_corsac | 0.246209388141265 |
| Ursus_thibetanus | Vulpes_ferrilata | 0.248552467025663 |
| Ursus_thibetanus | Vulpes_lagopus | 0.249833291628966 |
| Ursus_thibetanus | Vulpes_vulpes_T3611 | 0.245961716830947 |
| Ursus_thibetanus | Vulpes_zerda | 0.246667351275194 |
| Ursus_thibetanus | Zalophus_californianus | 0.230593536725188 |
| Urva_javanica | Viverra_tangalunga_FMNH146957_SRR6053069 | 0.185361137948073 |
| Urva_javanica | Viverricula_indica | 0.183057426617247 |
| Urva_javanica | Vulpes_corsac | 0.246366004875882 |
| Urva_javanica | Vulpes_ferrilata | 0.248200174097425 |
| Urva_javanica | Vulpes_lagopus | 0.246101760623857 |
| Urva_javanica | Vulpes_vulpes_T3611 | 0.242015741522008 |
| Urva_javanica | Vulpes_zerda | 0.244242420775403 |
| Urva_javanica | Zalophus_californianus | 0.238295507294376 |
| Viverra_tangalunga_FMNH146957_SRR6053069 | Viverricula_indica | 0.125872485525471 |
| Viverra_tangalunga_FMNH146957_SRR6053069 | Vulpes_corsac | 0.23702250260193 |
| Viverra_tangalunga_FMNH146957_SRR6053069 | Vulpes_ferrilata | 0.240513127168277 |
| Viverra_tangalunga_FMNH146957_SRR6053069 | Vulpes_lagopus | 0.235857806462479 |
| Viverra_tangalunga_FMNH146957_SRR6053069 | Vulpes_vulpes_T3611 | 0.23448477794573 |
| Viverra_tangalunga_FMNH146957_SRR6053069 | Vulpes_zerda | 0.236420628000453 |
| Viverra_tangalunga_FMNH146957_SRR6053069 | Zalophus_californianus | 0.230234007720772 |
| Viverricula_indica | Vulpes_corsac | 0.230690721318159 |
| Viverricula_indica | Vulpes_ferrilata | 0.232814498312105 |
| Viverricula_indica | Vulpes_lagopus | 0.233683657587006 |
| Viverricula_indica | Vulpes_vulpes_T3611 | 0.229626617784247 |
| Viverricula_indica | Vulpes_zerda | 0.230231270571353 |
| Viverricula_indica | Zalophus_californianus | 0.23317528127361 |
| Vulpes_corsac | Vulpes_ferrilata | 0.0402095029874956 |
| Vulpes_corsac | Vulpes_lagopus | 0.108424956838712 |
| Vulpes_corsac | Vulpes_vulpes_T3611 | 0.0840861988458248 |
| Vulpes_corsac | Vulpes_zerda | 0.0941569398401654 |
| Vulpes_corsac | Zalophus_californianus | 0.247117316293077 |
| Vulpes_ferrilata | Vulpes_lagopus | 0.107459636689297 |
| Vulpes_ferrilata | Vulpes_vulpes_T3611 | 0.0851957163344219 |

| | | |
|---|---|---|
| Vulpes_ferrilata | Vulpes_zerda | 0.0953735008309456 |
| Vulpes_ferrilata | Zalophus_californianus | 0.24976445887352 |
| Vulpes_lagopus | Vulpes_vulpes_T3611 | 0.102394828118212 |
| Vulpes_lagopus | Vulpes_zerda | 0.110521438797377 |
| Vulpes_lagopus | Zalophus_californianus | 0.244822234865656 |
| Vulpes_vulpes_T3611 | Vulpes_zerda | 0.0290924414399201 |
| Vulpes_vulpes_T3611 | Zalophus_californianus | 0.242844668278792 |
| Vulpes_zerda | Zalophus_californianus | 0.244514351726062 |
