## Supplementary figures and images for "High-quality carnivore genomes from roadkill samples enable species delimitation in aardwolf and bat-eared fox"

### Figure S1

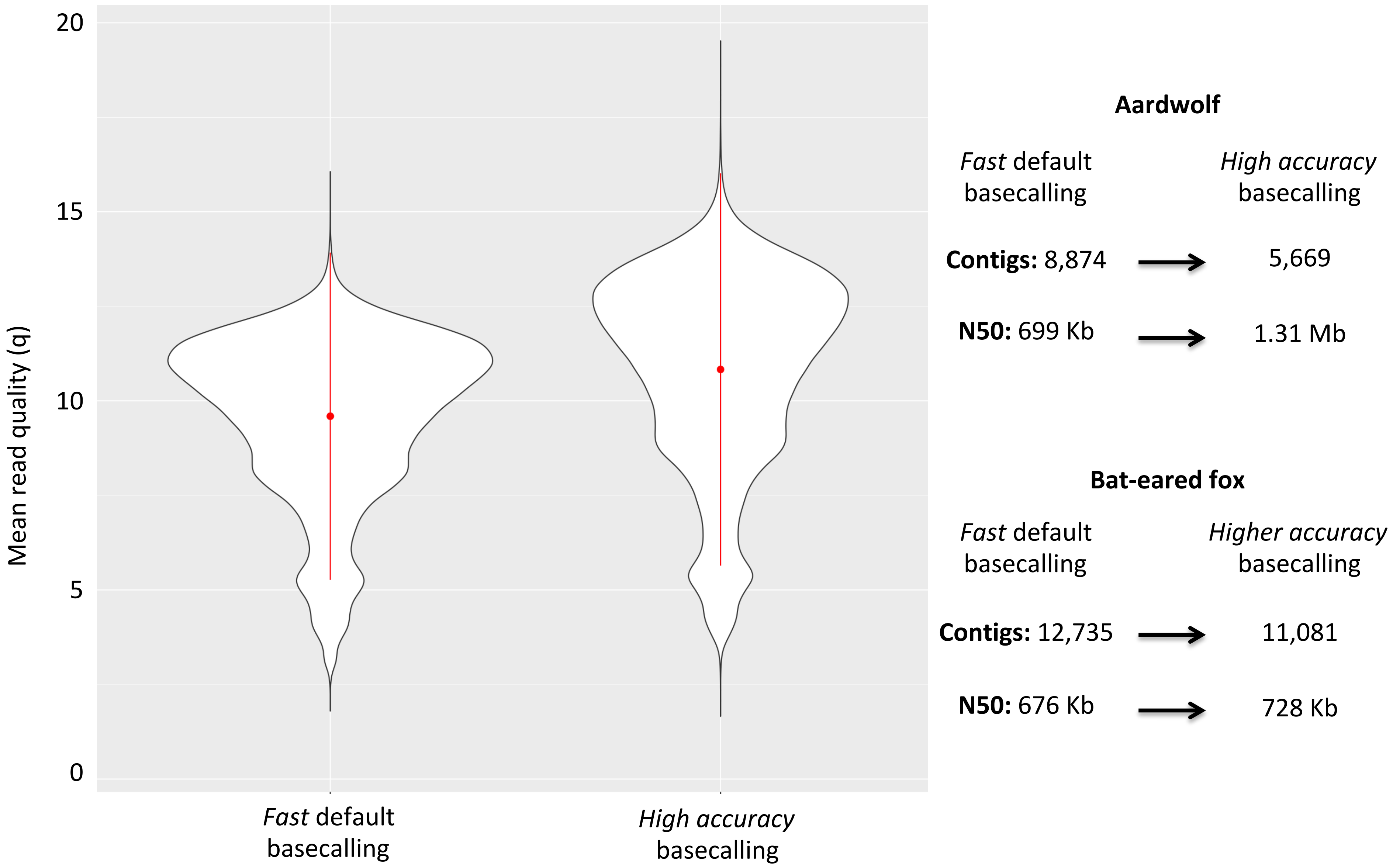

### Figure S3

#### a) BlobTools results for *Proteles cristatus*

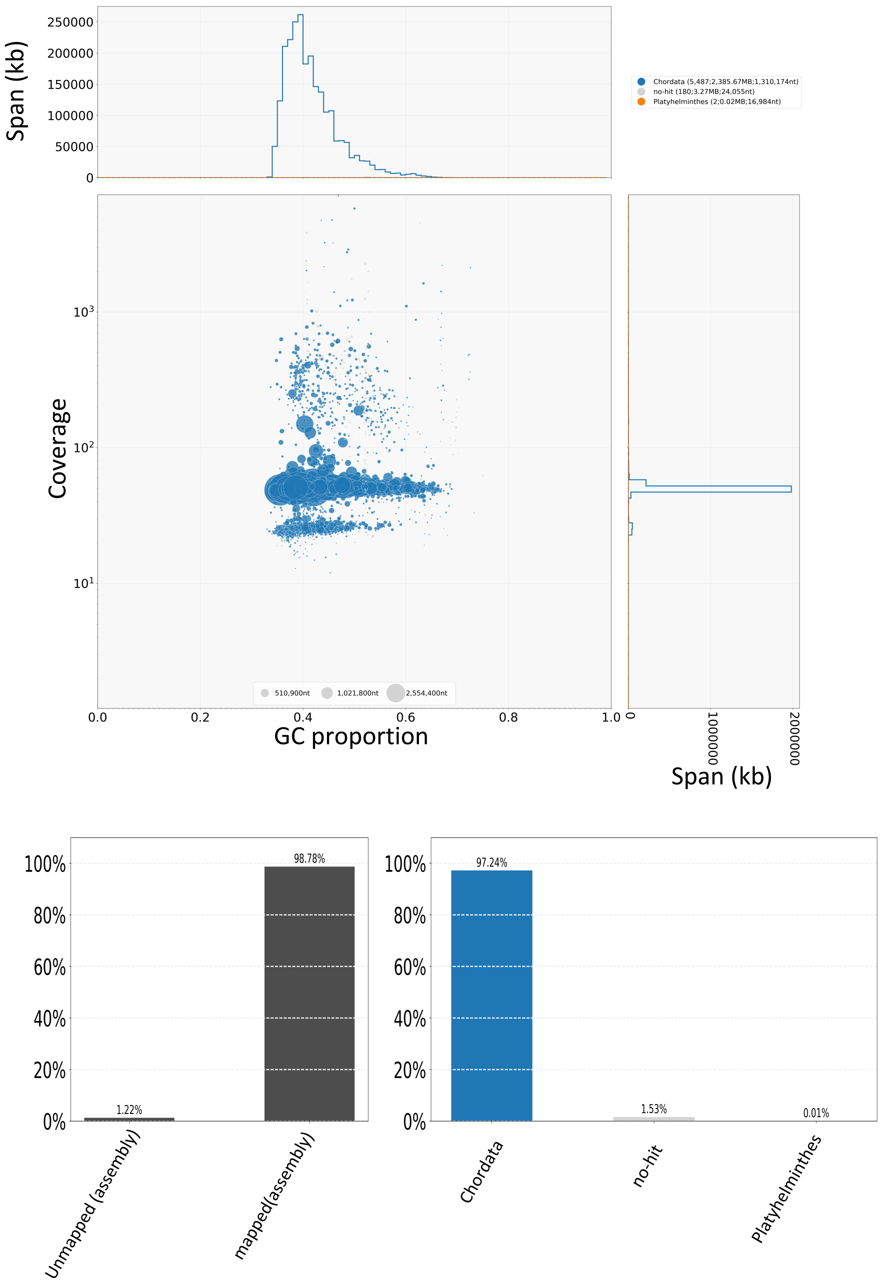


#### b) BlobTools results for *Otocyon megalotis*

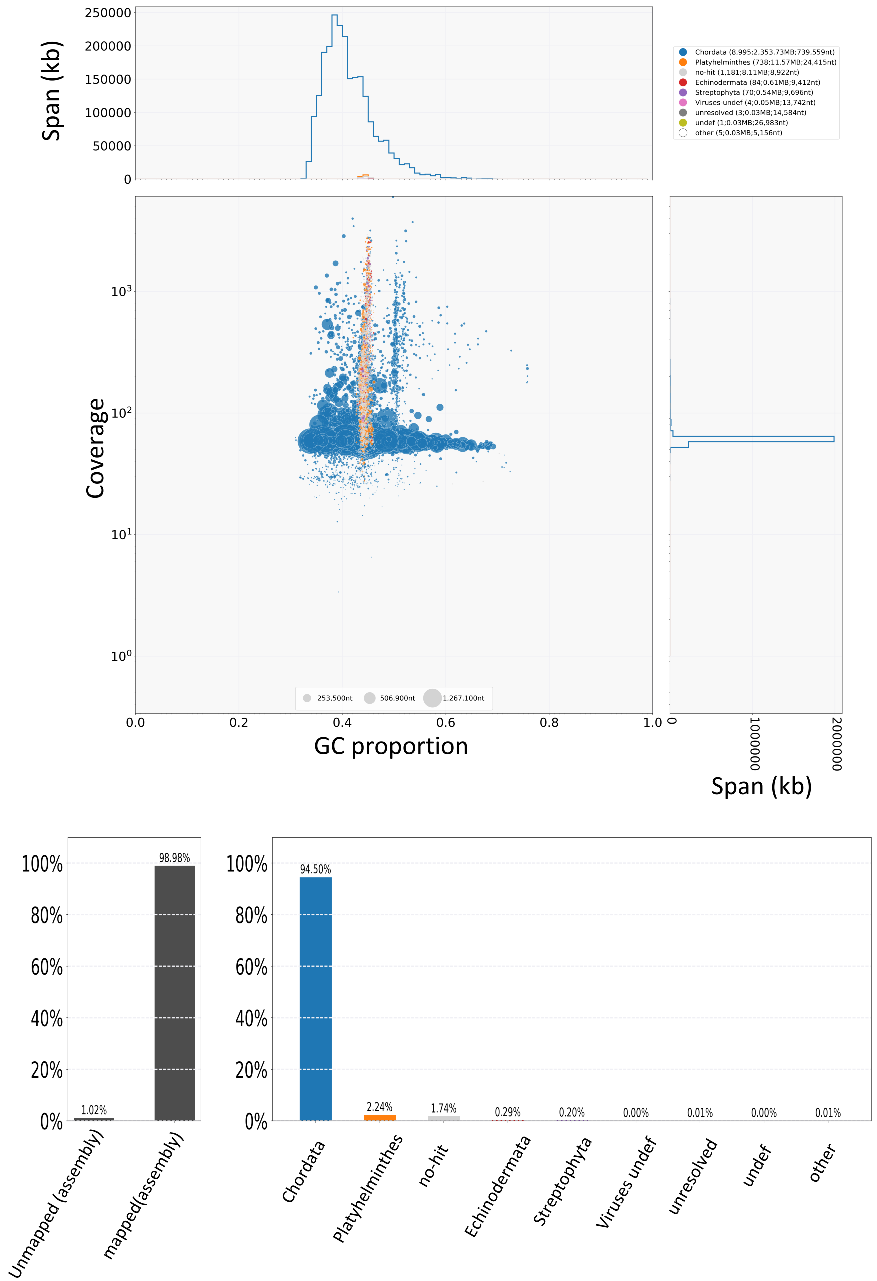
