## Supplementary material for "High-quality carnivore genomes from roadkill samples enable species delimitation in aardwolf and bat-eared fox": Figure S2

### F-Stastistic:

$$\mathbf{FST} = 1 - \frac{\pi_{within}}{\pi_{between}}$$

Hudson et al 1992

FST = 1 = Highly structured
FST = 0 = No structuration

### Genetic differentiation index (GDI, based on heterozygosity):

| Fixed | Private A | Private B | Shared |
|---|---|---|---|
| AA/TT | AT/AA | AA/AT | AT/AT |

$\pi_{AB}$ = fixed + private$_A$ + private$_B$ + shared$_{AB}$
$\pi_A$ = private$_A$ + shared$_{AB}$
$\pi_B$ = private$_B$ + shared$_{AB}$

$$1 - \frac{(\pi_A + \pi_B)/2}{\pi_{tot\ AB}}$$

### For different population level:

$$1 - \frac{(\pi_{A1} + \pi_{A2})/2}{\pi_{tot\ A}} \quad \text{vs} \quad 1 - \frac{(\pi_{A1} + \pi_{B1})/2}{\pi_{tot\ A1B1}}$$

GDI within pop A (control) | global GDI

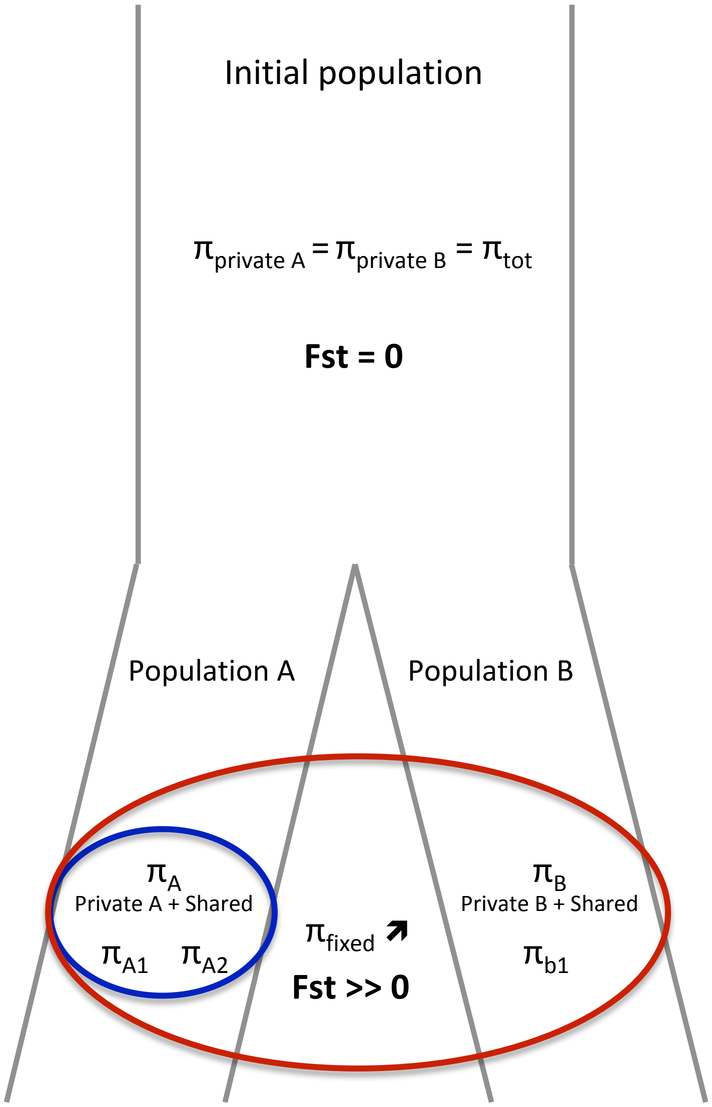
